# Transcriptional Determinism and Stochasticity Contribute to the Complexity of Autism Associated *SHANK* Family Genes

**DOI:** 10.1101/2024.03.18.585480

**Authors:** Xiaona Lu, Pengyu Ni, Paola Suarez-Meade, Yu Ma, Emily Niemitz Forrest, Guilin Wang, Yi Wang, Alfredo Quiñones-Hinojosa, Mark Gerstein, Yong-hui Jiang

## Abstract

Precision of transcription is critical because transcriptional dysregulation is disease causing. Traditional methods of transcriptional profiling are inadequate to elucidate the full spectrum of the transcriptome, particularly for longer and less abundant mRNAs. *SHANK3* is one of the most common autism causative genes. Twenty-four *Shank3* mutant animal lines have been developed for autism modeling. However, their preclinical validity has been questioned due to incomplete *Shank3* transcript structure. We applied an integrative approach combining cDNA-capture and long-read sequencing to profile the *SHANK3* transcriptome in human and mice. We unexpectedly discovered an extremely complex *SHANK3* transcriptome. Specific *SHANK3* transcripts were altered in *Shank3* mutant mice and postmortem brains tissues from individuals with ASD. The enhanced *SHANK3* transcriptome significantly improved the detection rate for potential deleterious variants from genomics studies of neuropsychiatric disorders. Our findings suggest the stochastic transcription of genome associated with *SHANK* family genes.

## Introduction

In the central dogma of molecular biology, RNA transcription acts as a rheostat, orchestrating the cellular functions of the genes in response to intrinsic and extrinsic signals. The complex functions in the organs such as brain require a diverse proteome from a relatively small gene pool. This diversity is facilitated by transcriptional regulations involving alternative promoter usage and splicing, occurring in >90% of neuronal genes in mammalian brains^1–4^. Disruption in transcript-specific regulatory elements due to DNA mutations can lead to diseases. Transcriptome-wide changes are implicated in neuropsychiatric conditions, including autism spectrum disorder (ASD)^5–8^. Accurate annotation and interpretation of these changes relied on a comprehensive transcriptomic profile, either for a given gene or on a genome-wide scale. However, popular short-read sequencing is suboptimal for delineating longer transcripts and discovering novel exons and splicing events^9^. Standard long-read sequencing techniques are not sufficiently sensitive to detect transcripts with lower abundance. A theoretical solution lies in the combination of mRNA/cDNA-capture methods^10^ and long-read sequencing, could identify both long and low abundant transcripts. However, this approach has been sparingly reported, probably due to the technical challenge of preserving the mRNA integrity. The current inability to construct a complete transcriptome fuels a continuing debate over the extent of pervasive transcription across the genome and the significance of transcriptional “dark matter” endogenously^11–15^. The incomplete transcriptome impedes accurate annotation of disease-linked variants and interpretation of transcriptomic data. This shortfall affects the validation of genetically modified disease models used in preclinical research to develop molecular therapies. Previous studies have indicated specific functions of *SHANK3* mRNA transcripts at synapses^16–29^. An incomplete human *SHANK3* transcriptome could underestimate the contribution of the genetic risk for ASD and other neuropsychiatric disorders. Similarly, the incomplete mouse transcriptome complicate interpretations of their relevance to human *SHANK3* disorders from studies of more than 24 lines of genetically modified animal models^30^ ^18,31^. To bridge these substantial gaps in knowledge, we performed standard Iso-Seq (**SIS**) for whole transcriptome analysis and paired with targeted cDNA capture and long-read sequencing techniques (Capture-Iso-Seq, **CIS**) to specifically investigate the *SHANK* family genes in humans and mouse brain. We discovered a drastically intricate *SHANK3* transcript structure and a broad transcriptomic diversity across the human and mouse genomes. We identified unexpected extensive fusion transcripts and atypical patterns of transcripts in *Shank3* mutant mice. The enhanced *SHANK3* transcriptome has significantly improved the discovery rate of deleterious variants in genomic and transcriptomic studies of neuropsychiatric disorders. Our study advocates for a paradigm shift in experimental design and evaluation of genetic disease models using genetically modified animals, emphasizing the need to carefully evaluate the molecular validity of these mutant animal models in preclinical research.

## Results

### Dataset overview and experimental strategy evolution and optimization

We sequenced 56 SMRT libraries of human and mouse brains using the PacBio Sequel II System (**Fig. 1A-B**). Sixteen libraries proceeded using the SIS method. Forty libraries were constructed following the CIS method, which employed targeted capture enrichment with specific oligonucleotide probe panels, that covered the full genomic regions of *SHANK/Shank* family genes (*SHANK1-3*, **Supplementary Table 1-2**). A non-neuronal gene, *TP53,* was included as a comparison. Twenty libraries were synthesized from cerebral cortex of neurotypical children aged 5-6 years, and young adults, aged 24-30 years. For mice, 35 libraries were derived from striatum (ST) and prefrontal cortices (PFC) of 21-day-old wild-type (WT) C57BL/6J and *Shank3* mutants (*Shank3*^Δe4–9^, *Shank3*^Δe21^ and *Shank3*^Δe4–22^)^19,20,32–34^. We processed only the RNA with an Integrity Number (RIN) above 7 for human and above 8 for mouse samples for subsequent sequencing. The quality and reproducibility of the SIS and CIS platforms were optimized (**Supplementary Fig.1A-I)**. For experimental validation, RT-PCR and Sanger sequencing confirmed novel *SHANK3* transcripts from CIS. We performed *in silico* transcriptome analyses using short-read bulk RNA-seq (**srRNA-seq**) and single-cell RNA-seq (**scRNA-seq**) data, and gene discovery analyses of exome sequencing (ES) and whole genome sequencing (WGS) data from PsychENCODE project along with other genomics studies^5,35–39^.

**Fig. 1.**
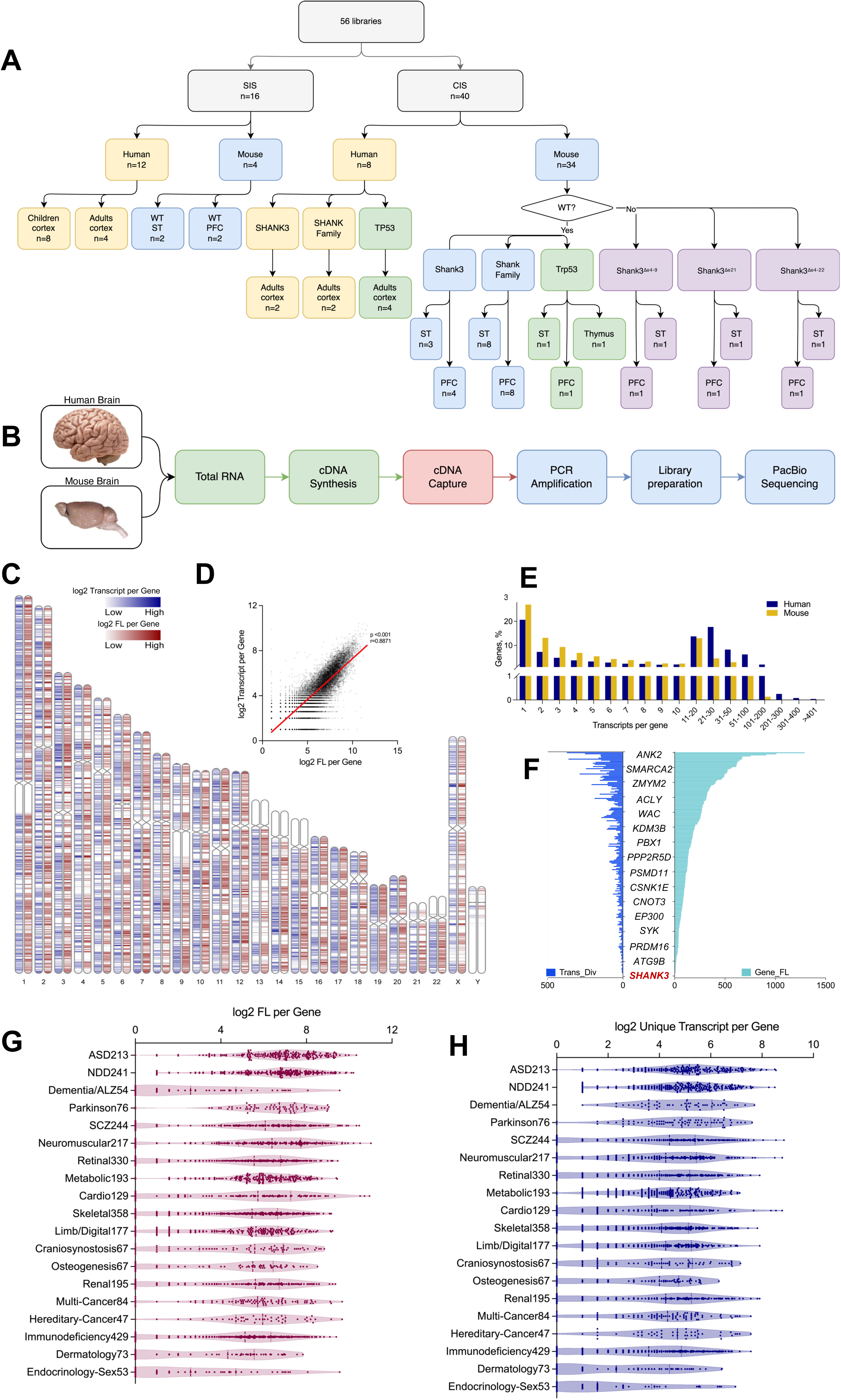
Genome wide transcript diversity and abundance in brains detected by SIS. **A.** Experimental design of SIS and CIS of human and mouse tissues. **B.** Schematic of experimental procedure of RNA capture and long read-sequencing. **C.** Number of unique transcripts (transcript diversity) for individual genes (blue) and the number of sequences reads (abundance) (red) for an individual transcript detected in human cerebral cortex by SIS with projected chromosome coordinates and ideograms. **D.** Transcript diversity was significantly correlated with the sequence reads (abundance) of the transcripts. **E.** Number of transcripts per gene genome-wide from SIS in human and mouse brains. **F.** Number of unique transcripts (Trans_Div) and abundance (Gene_FL) for 213 ASD risk genes, shown an average of 56 transcripts per gene and a median of 35. **G-H.** Human SIS data showed heightened transcript diversity in genes associated with brain disorders, especially ASD and NDD, compared to other diseases. We observed a strong correlation between transcript diversity and abundance in all gene clusters except for those related to dementia/Alzheimer’s.

### Standard Iso-Seq uncovered more diverse transcriptome genome-wide in mouse and human brains

From the SIS of 12 SMRT libraries of human brain, we uncovered 131,585 unique transcripts across 15,308 annotated genes, including 311 novel transcripts (**UCSC Track 1**). Distribution of unique transcripts and sequencing reads per gene are shown in **Fig. 1C**. The number of unique transcripts for a given gene was significantly correlated (Pearson r=0.8871, p<0.001) with its abundance (**Fig. 1D**). From 4 SIS of mouse ST and PFC, we uncovered 154,492 unique transcripts from 16,556 annotated genes, with 1,570 being novel (**Fig.1E** and **UCSC Track 2 and 3)**.

In human brains, the average number of isoforms per gene was 19, with an average sequence read count of 63. Notably, 595 genes exhibited over 100 isoforms (**Fig.1E** and **Supplementary Table 3a)**. *SEPTIN4* has the highest number of isoforms at 692, a gene encoding a presynaptic scaffold and GTP-binding protein, involved in exocytosis, and interacted with alpha-synuclein, implicated in Parkinson’s disease^40^. In mouse brains, the average number of unique transcripts per gene was found to be 8, with an average of 17 sequence reads per transcript. *Sorbs1* had the highest number of isoforms at 158; this gene encodes a Sorbin and SH3 domain containing protein involved in insulin signaling and stimulation^41^ **(Supplementary Table 3b)**. We identified 182 genes with more than 50 isoforms and 19 genes with over 100 isoforms in mouse brains. Of these, seven have human orthologs that also exhibit more than 100 isoforms. Our studies revealed a greater transcript diversity than other studies using the same sequencing platform and analytic algorithm^8,42^. We examined the transcript diversity of 213 highly confident ASD risk genes consolidated from 3 recent extensive ASD genomics studies using our SIS data^43–45^ (**Fig. 1F** and **Supplementary Table 4**). On average, individual ASD risk genes exhibited 56 transcripts, with a 90% CI ranging 8-140. *ANK2* was noted for having the highest number of transcripts of 372. Remarkably, the expression level of *SHANK3* was one of the lowest, ranking 212 of 213 ASD risk genes (**Fig. 1F**). Genes associated with brain disorders, especially ASD and neurodevelopmental disorders (NDD), have significantly greater numbers of transcripts compared to genes implicated in disorders not related to the brain (**Fig. 1G-H).**

### A complex mouse *Shank3* transcriptome from CIS

We noted that the longest annotated *SHANK3/Shank3* transcripts in humans (NM_001372044.2, 7,691 bp, hg38) and mice (NM_021423.4, 7,380 bp, mm39) have not been detected in any published long-read RNA-seq datasets^6,8,42^. From 4 SIS of mouse ST and PFC, we identified only 5 *Shank3* transcripts (ranging 5,625-6,463 bp) in ST, with none detected in PFC upon validation. The discrepancy in transcript number and the variation between ST and PFC were consistent with the highest expression level of *Shank3* in ST and lower expression in neocortex at P21 days^25^. The failure to detect longer *Shank3* mRNAs by SIS was most likely due to their low abundance, as transcripts up to 14.5 kb were successfully sequenced in our libraries (**Supplementary Fig. 1F-G**).

With CIS, we detected 545 *Shank3* transcripts in the mouse ST (**Fig. 2A**) and 345 in PFC (**Fig. 3A**), including the longest annotated transcript (NM_021423.4). We successfully validated 51 (85%) out of 60 representative novel transcripts by RT-PCR and sequencing (**Fig. 2E-H** and **Supplementary Table 5**). To evaluate the quality of each transcript, we employed a confidence metric that integrates the transcript abundance, the length of predicted open reading frame (ORF), and validations with srRNA-seq data (**Supplementary Fig. 2A**). In ST, 223 (41%) of *Shank3* transcripts were classified as high confidence, while 382 (59%) were in moderate confidence. In PFC, 168 (49%) transcripts were in high confidence, with the remaining and 176 (51%) of moderate confidence. Analysis revealed 36 and 26 potential transcription start sites (TSS) in ST and PFC, respectively. In the ST, 142 *Shank3* transcripts originated at exon 1 of the annotated referenced transcript (NM_021423.4) and terminated at 26 different sites **(Fig. 2B)**. Thirty-five transcripts terminated within exon 21, each presenting different ORFs. Exon 21, the largest coding exon of 2,257 bp, was spliced out in many transcripts. Over 90% of transcripts terminated within 100-500 bp of annotated transcription termination sites (TTS) and poly A signals (**Supplementary Fig. 3**). This indicates that the early terminations are not artifacts of RNA degradation or cDNA synthesis errors. Intron retentions were observed in introns 1, 2, 11, 12, and 19, leading to altered ORFs and earlier stop codons. While some transcript structure variations were subtle, they are predicted to encode different ORFs (**Fig. 2C-D)**.

**Fig. 2.**
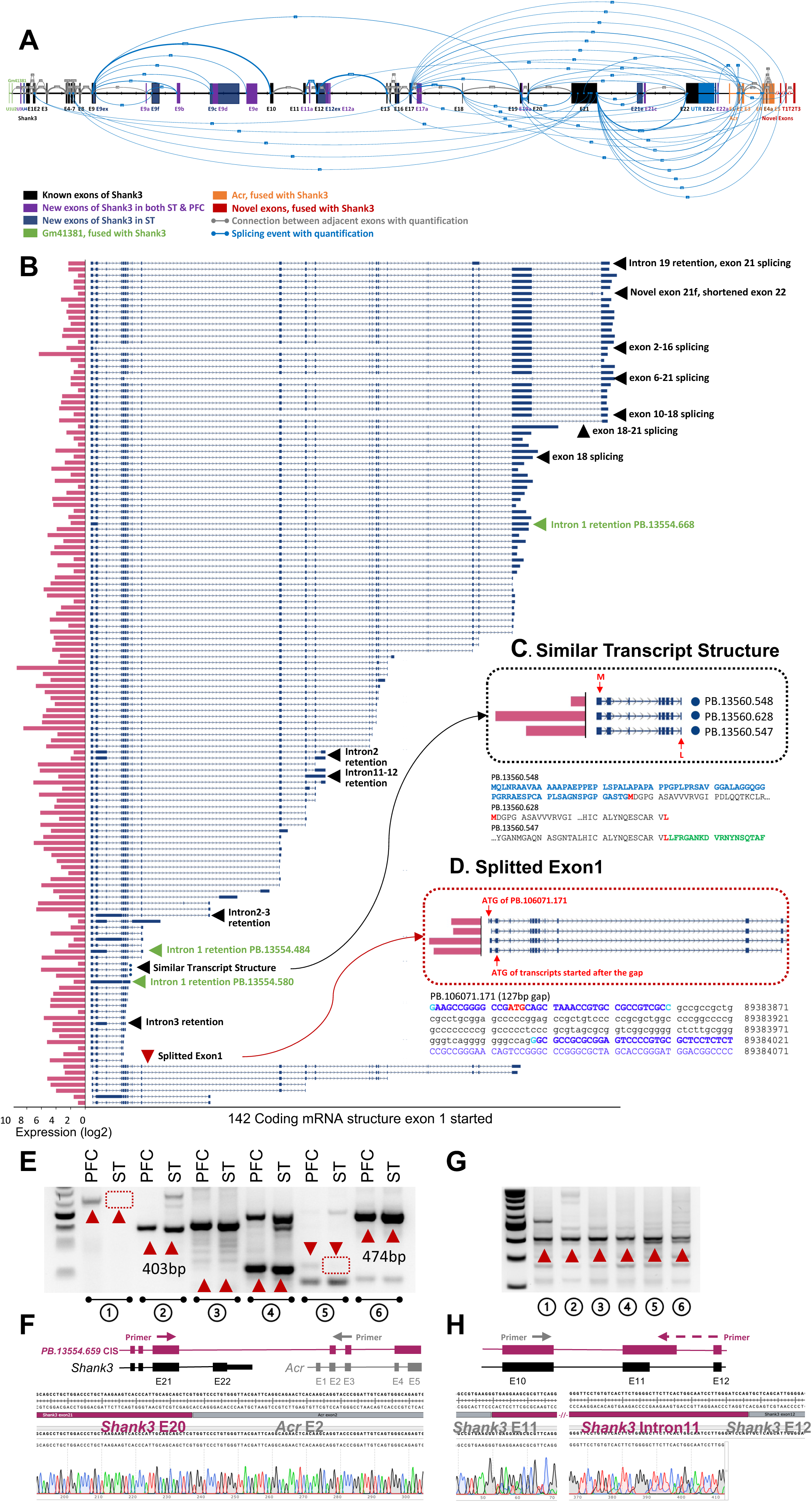
Novel *Shank3* transcriptome in mouse striatum (ST) by CIS. **A.** CIS revealed a refined *Shank3* gene structure and splicing patterns in WT mouse striatum. The established *Shank3* structure (NM_001034115, mm39) is expanded with newly detected exons shared between striatum and PFC, depicted in purple. Unique splicing events, represented by grey lines and thickness indicating read quantity, include novel striatum-specific exons in dark blue and alternative splices in light blue. Fusion transcript exons near Gm41381 and *Acr*, shown in green and orange, respectively, feature unique splicing with newly identified red exons (T1-T3) exclusive to *Shank3*. New exon U3 is shared between striatum and PFC. U4 is linked to Gm4138 and striatum specific. 21e is a new in-frame exon and 21c is a new exon harbor a stop codon. **B.** 142 unique transcripts started with the canonical exon 1 of annotated *Shank3* (NM_001034115) in ST and terminated at different positions. Pink bar plots on the left are the abundance (log2 counts). Arrows describe the features of given transcripts. **C.** Example of transcripts with similar structures in panorama but different at the sequence level with predicted ORFs and ATG codons. The transcripts of PB.13560.548, PB.13560.628, and PB.13560.547 are similar but the predicted ORFs show different ATG codons and protein domains. **D.** Details of the split exon 1. There is a cryptic splicing of 127 bp (non-capitalized sequence in black) within the annotated exon 1 of transcript PB.106071.171 which resulted in a predicted upstream ATG codon and additional 134 amino acids. Other transcripts have transcriptional starting sites (TSS) in exon 1 but predicted ATG codon in exon 2. Variability in TSS and intron 1 retention, as seen in transcripts PB.13554.484, PB.13554.580, and PB.13554.668, leads to ORFs of 304 aa, 106 aa, and 1,290 aa, respectively. **E.** Validations new transcripts from paired mouse PFC and ST samples. Pair 1, novel exon U1; Pair 2, fusion transcript between *Shank3* exon 21 and *Acr* exon2; Pair 3, splicing event between *Shank3* exon 9 and exon19; Pair 4, splicing event between *Shank3* exon 5 and exon 21; Pair 5, novel exon 9b of *Shank3*; Pair 6, *Shank3* exon11 extension/intron11 retention. The red arrows are the novel products confirmed by Sanger sequencing. Other bands are products from known transcripts. **F.** Sanger sequencing confirmation of a fusion transcript between *Shank3* exon21 and *Acr* exon2 in mouse brain (pair 2 of E) **G.** Fusion transcripts in other tissues. Forward and reverse primers were from exon 20 of *Shank3* and exon 5 of mouse *Acr* respectively. lane1, liver in P21 mouse; lane 2, thymus in P21 mouse; lane 3, ovary in P21 mouse; lane 4, ovary in 3 months old mouse; lane 5, testis in P21 mouse; lane 6, testis in 3-month-old mouse. The red arrows are the novel products confirmed by Sanger sequencing as indicated. Other bands are known products. **H.** Sanger sequencing of *Shank3* exon 11 extension/intron 11 retention in mouse brain (lane 6 of G).

**Fig. 3.**
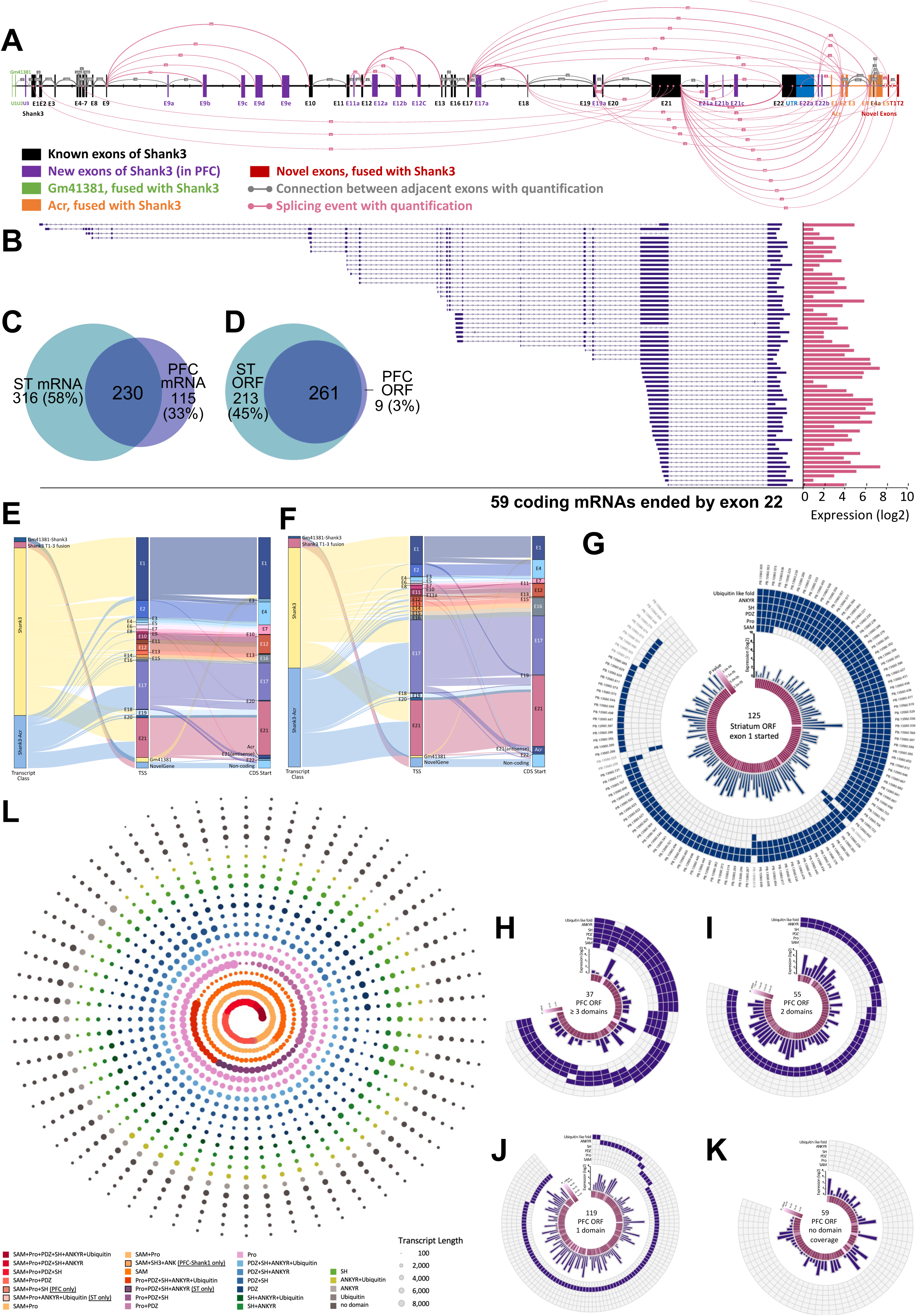
Novel *Shank3* transcriptome in mouse PFC by CIS and predicted domain structures of ORFs. **A.** New *Shank3* transcript structure and conch plot of splicing events discovered in WT mouse PFC by CIS. Color code is the same as Fig. 2A. The novel exon 9a (chr15:89394416-89394465, mm39) is shared between PFC and ST. Other novel exons such as exon 12e (chr15:89414330-89414640, mm39) were unique to PFC. Novel exons 21a, 21b and 21c are predicted to result in an early stop codon and shorter ORFs (chr15: 89394416-89394465, chr15: 89408698-89408784, chr15: 89418571-89418609, mm39). **B.** Structure of 59 transcripts with different TSSs but terminating at annotated exon 22 of *Shank3*. Pink bar plot represents the abundance (log2 counts) of each transcript. **C-D.** The comparison of transcripts and predicted ORFs between mouse ST and PFC. **E-F.** The pattern of deduced TSS and predicted starting sites of the coding sequence (CDS) for all *Shank3* transcripts including new 5′ and 3′ fusion transcripts from CIS in mouse ST (E) and PFC (F). Each filament represents an individual transcript in different classes of GM41381(U1-U2)-*Shank3*, *Shank3*-T1-3, *Shank3,* Shank3-Acr (first column), deduced TSS (middle column), and predicted starting sites of CDS (third column). **G.** A total of 125 unique ORFs are predicted from 142 transcripts starting with exon 1 in ST. The pattern of the combination of 6 protein domains is shown in the outermost ring of the windmill plot. The middle layer shows the abundance of each RNA transcript and the p value of its expression level compared to other transcripts. Only 4 ORFs of transcripts contained all 6 protein domains. **H-K.** Four windmill plots showing 270 predicted ORFs from all 345 transcripts detected in PFC classified by the combination of functional domains. **L.** Spiral plot showing an aggregated functional domain coverage of the transcripts captured by *Shank1-3* joint probe panel by CIS of mouse PFC and ST. Each dot represents a unique transcript. Each color represents a unique combination of functional domain. The dots are ordered from the longest to the shortest transcript, while the colors are arranged from the SAM to UBL domain.

In the PFC, we identified 59 *Shank3* transcripts initiating from 19 different exons and terminating within the last coding exon 22 (**Fig. 3A-B**). Notably, 28 of these transcripts started within exon 21 with different ATG codons. This finding aligns with our prior results obtained from 5′ RACE experiments^25^. We discovered 12 new exons in ST and 17 in PFC, with 11 being shared to both regions (**Fig. 2A** and **Fig. 3A**). Additionally, we discovered 4 new untranslated exons, U1-4, located 5′ upstream of the annotated *Shank3* exon 1 **(Fig. 2A)**. Six new and alternative spliced exons, E9a-f, were identified between exons 9 and 10. The spliced variants between exons 9 and 10 were the most abundant with 4,326 reads in ST and 641 in PFC, while exon 12e was exclusive to the PFC (**Fig. 2E**).

Surprisingly, we observed a considerable number of novel fusion transcripts, in which different *Shank3* exons were joined to downstream exons 2-5 of the *Acr* gene, which encodes the acrosin protein in the acrosome of spermatozoa^46^ (**Fig. 2A** and **Fig. 3A**). These fusion transcripts were validated by PCR and sequencing (**Fig.2E-F**). We noted that splice events linking *Shank3* exons 17 and 21 to *Acr* exons occurred more frequently than others. Specifically, fusions from *Shank3* exon 21 to *Acr* exon 2 (208 reads) and exon 3 (243 reads) were the most abundant. We also identified splice products from *Shank3* exons 17 and 21 to three novel exons/transcripts (T1-3) situated downstream of *Acr* (**Fig. 2A**). These transcripts in ST and PFC are predicted to yield five ORFs, extending the SHANK3 protein by an additional 64 amino acids (NP_001358973).

The transcriptomic architecture of *Shank3* revealed by CIS in ST and PFC displayed both shared and unique characteristics. Overall, 230 transcripts (42% of ST, 67% of PFC) were common to both brain regions **(Fig. 3C).** We analyzed the tissue specific usage of TSS and coding sequence starting sites (CDS). Transcripts were categorized as follows: overlapping with the annotated *Shank3* mRNA, U1-4 to *Shank3*, *Shank3*-*Acr* fusion, and *Shank3*-T1-3. In ST, 75% of transcripts belonged to the category overlapping with the annotated *Shank3*, and 24% fell within the *Shank3*-*Acr* fusion category (**Fig. 3E**). In PFC, 52% of the transcripts were overlapping with annotated *Shank3*, while 43% were classified as *Shank3-Acr* fusion transcripts **(Fig. 3F).**

### Protein domain specific mouse SHANK3 proteome

SHANK3 and its family encode proteins possess 6 domains of ubiquitin-like (Ubl), ankyrin repeats (ANKYR), postsynaptic density protein 95/discs large homologue 1/zonula occludens 1 (PDZ), an Src homology 3 (SH3), a proline-rich region containing Homer and Cortactin-binding sites (Pro), and a sterile alpha motif (SAM)^47–49^. As scaffold proteins in the postsynaptic density (PSD) of synapses, SHANK3 protein interacts with various synaptic proteins via these domains, contributing to synaptic architecture and function. There are 474 ORFs predicted from 545 *Shank3* transcripts in ST and 270 ORFs in PFC using GeneMarkS-T^50^, with 261 ORFs being common to both brain regions (**Fig. 3D).** ORFs of novel transcripts were further corroborated by proteome data derived from various *in silico* datasets, utilizing graded criteria for sequence identity and overlap **(Supplementary Fig. 2B**).

Among the 125 ORFs predicted from 140 *Shank3* transcripts starting from exon 1 in ST, only 4 ORFs encompassed all 6 protein domains (**Fig. 3G**). Among the 270 ORFs predicted from 345 *Shank3* transcripts in PFC, only one ORF contained the complete set of 6 protein domains, while 37 ORFs have more than 3 protein domains **(Fig. 3H-K).** One hundred nineteen SHANK3 ORFs (30%) in PFC comprised only a single protein domain, typically the Pro domain. Approximate 15% of the predicted ORFs lacked recognized protein domains. The protein domain combinations were found to be non-random and tissue-specific; for instance, no predicted ORFs included the SAM-SH3 combination. The SAM-Pro-SH3 and SAM-SH3-ANKYR domains combinations were exclusive to PFC, while the Ubl-ANKYR-Pro-SAM and ANKYR-SH3-PDZ-Pro combination were identified only in ST (**Fig. 3L**).

### Uniquely altered *Shank3* transcriptome in *Shank3* mutant mice

Sixteen *Shank3* mutant mouse lines and 8 mutant rat, dog, and non-human primate featuring various exonic deletions or point mutations, have been generated to model *SHANK3* associated ASD^30^ (**Fig. 4A**). Using the same *Shank3* probe design, we conducted CIS on *Shank3* mutant mice: those with deletions of exons 4-9 (*Shank3*^Δe4–9^), exons 4-22 (*Shank3*^Δe4–22^), and exon 21 (*Shank3*^Δe21^)^19,20,32,33^. In *Shank3*^Δe4–9^homozygous mice, we detected 69 *Shank3* transcripts in ST and 56 in PFC. Representative mutant and residual transcripts are diagramed in **Fig. 4B**, with details provided in **Supplementary Fig. 4A-B**. In ST and PFC of *Shank3*^Δe4–9^ mice, we identified 3 long transcripts (∼7.3 kb), harboring a deletion of exons 4-9. Interestingly, the first exon of these transcripts, with the exon 4-9 deletion, was in intron 1 of the annotated *Shank3*, a TSS was not utilized in WT mice, suggesting an alternative TSS due to the exon 4-9 deletion. These transcripts also lacked coding exon 22 and exhibited fusions between exon 21 of *Shank3* and exon 2 of *Acr*. ORF prediction suggests that the resultant SHANK3-ACR fusion proteins for these mutant transcripts is 1254 aa for PB.6361.147, 1073 aa for PB.6623.114, and 833aa for PB.6623.199. Approximately 70% of the residual transcripts are initiated from intron 16/exon 17 and terminate within exon 21/intron 21 of *Shank3* or exon 5 of *Acr*. Transcripts starting at exon 11 were exclusively detected in ST. The proportion of transcripts initiated from intron 16/exon 17 was increased in *Shank3*^Δe4–9^ mice compared to WT. A total of 54 ORFs (ranging 113 to 1,327 aa), were predicted in ST, with a similar pattern observed in PFC from residual transcripts of *Shank3*^Δe4–9^ mice.

**Fig. 4.**
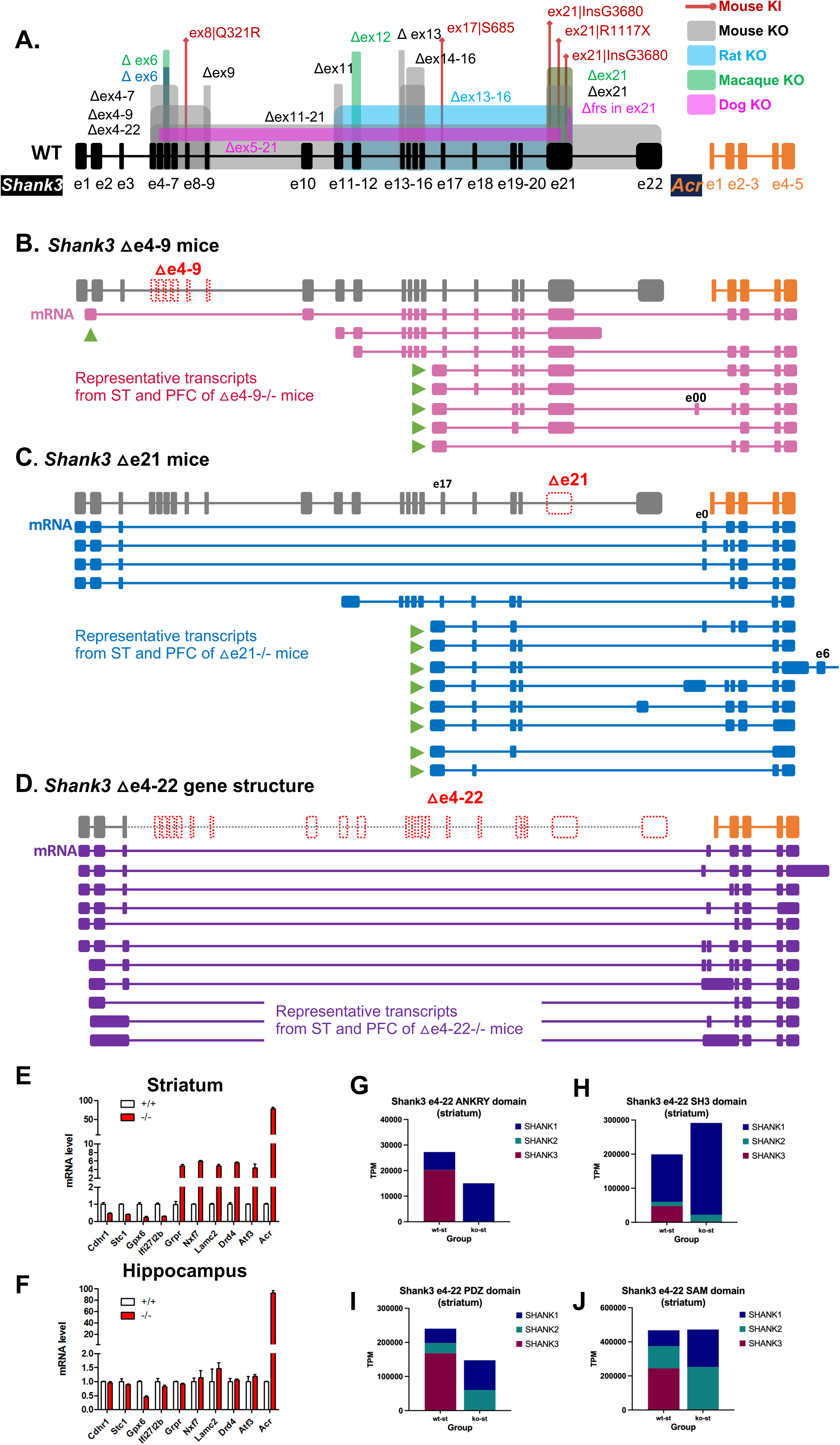
The summary and illustration of altered *Shank3* transcripts in *Shank3*^Δe4–9^, *Shank3*^Δe21^ and *Shank3*^Δe4–22^ mutant mice from CIS. **A.** Current annotated mouse *Shank3* and *Acr* (NM_013455, mm39) gene structure. The annotations of genetically targeted mutations in mice, rat, monkey, and dog are shown. (KO: exonic deletions; KI; knock-in mutation) **B.** The gene structure of *Shank3*^Δe4–9^ mutant mice in grey and representative mRNA transcripts from *Shank3*^Δe4–9-/-^ mice are in pink. No transcript using first annotated exon 1 was detected. Instead, the first exon, presumably a cryptic TSS (arrow), was detected in intron 1. The exon 4-9 deleted transcript missed exon 11, 12, and 22 but with fusion between *Shank3* and *Acr*. The transcripts starting at intron 16/exon 17 (arrows) as first exon were most abundant. Extensive fusion transcripts between *Shank3* exon 21 and *Acr* exon 2 were observed. The last coding exon 22 was not detected in any transcripts. **C.** The gene structure of *Shank3*^Δe21^ mutant mice and *Acr* gene in grey and representative3 mRNA transcripts from of *Shank3*^Δe21-/-^ mice in blue. The splicing between exon 4 of *Shank3* and exons of *Acr* that resulted in fusion transcripts were observed. The transcripts starting at intron 16/exon 17 (arrows) as first exon and fusion between *Shank3* and *Acr* were most common. The coding exon 22 were not detected in any transcript. **D.** The gene structure of *Shank3*^Δe4–22^ mutant mice and *Acr* gene in grey and representative mRNA transcripts in purple. The number of fusion transcripts between *Shank3* and *Acr* is significantly increased in *Shank3*^Δe4–22-/-^ mutant mice. **E-F.** Increased expression of *Acr* transcript in *Shank3*^Δe4–22-/-^ mutant mouse by RT-qPCR. The expression of *Acr* gene was significantly increased in both striatum and hippocampus by >100 folds. **G-J.** Compensatory expression of the functional domains of *SHANK* family proteins in striatum of *Shank3*^Δe4–22^ mutant mice. The bulk RNA-seq data of *Shank3*^Δe4–22^ were analyzed for the compensatory expression of other functional domains of *Shank1* and *Shank2* genes. The deficiency of ANKRY and SH3 domains of SHANK3 was compensated by SHANK1 but the deficiency of PDZ and SAM domains were compensated by both SHANK1 and SHANK2. The deficiency of SAM and SH3 domain was fully compensated but the deficiency of ANKRY and PDZ domains was partially compensated.

In *Shank3*^Δe21^ homozygous mice, we identified 401 *Shank3* transcripts in ST and 148 in PFC (**Fig. 4C**, **Supplementary Fig. 4C-D**). In *Shank3*^Δe4–22^ homozygous mice, the number were 436 in ST and 792 in PFC (**Fig. 4D**, **Supplementary Fig. 4E-F**). Remarkably, over 99% of these transcripts were *Shank3-ACR* fusion events in both brain regions of *Shank3*^Δe21^ and *Shank3*^Δe4–22^ mice. The predominant transcripts in *Shank3*^Δe21^ mice was from the intron 16/exon 17 region in both ST and PFC. Conversely, in *Shank3*^Δe4–22^ mice, transcription primarily initiated from intron 1/exon 2. We also detected multiple novel exons interposed between *Shank3* and *Acr* genes (**Supplementary Fig. 4C*-D\***), exclusive to these *Shank3* mutant lines and absent in WT. Fusion transcripts of *Shank3-Acr* were more prevalent in *Shank3*^Δe4–22^, *Shank3*^Δe4–9^, and *Shank3*^Δe21^ mutants. Moreover, a significant overexpression of *Acr* transcripts was found in neocortex and hippocampus of *Shank3*^Δe4–22-/-^ mice (**Fig. 4E-F**). Bulk RNA-seq data analysis from ST of *Shank3*^Δe4–22-/-^ mice also indicated a compensatory expression from *Shank1* and *Shank2*, which was protein domain-specific (**Fig. 4H-K**).

The *Shank3* transcriptomic findings from CIS prompted us to extend our approach to include all *Shank* family genes (*Shank1-3*) using a joint capture strategy. This joint CIS for the *Shank* family genes identified 664 *Shank1* and 495 *Shank2* transcripts in PFC, and 320 *Shank1* and 326 *Shank2* transcripts in ST (**UCSC Tracks 4 and 5**). The overall transcript structures and patterns of *Shank3* from both single-gene and joint CIS were similar. We discovered 7 novel exons upstream of the annotated exon 1 of *SHANK1* (**Supplementary Fig. 5A**). Fusion transcripts involving *Shank1* and *Shank2* with adjacent genes were also detected. The most upstream novel exon of *Shank1* overlapped with the last exon of *Clec11a* gene (NM_009131.3), which is transcribed in the reverse direction to *Shank1* **(Supplementary Fig. 5B)**. The fusion transcripts between *Shank1* and *Josd2*, a gene located approximately 100 kb downstream, were exclusively detected in PFC. Two new untranslated exons, U1 and U2, were found about 24kb upstream from the annotated 5′ exon 1 of *SHANK2* (**Supplementary Fig. 5C**)

### Transcript diversity of *SHANK* family genes in human brains

In current reference genome (hg38), an annotated human *SHANK3* mRNA (7,691 bp, NM_001372044) is displayed, yet it has not been experimentally validated. With CIS on *SHANK* family genes, we discovered 472 unique *SHANK3* transcripts (**Fig. 5A-C, UCSC Track 6**), with the longest was 6,824 bp. Notably, the annotated 7,691 bp *SHANK3* transcript (NM_001372044) was absent. The absence of the longest *SHANK3* transcript is unlikely a result of RNA degradation, because a 10.8 kb *SHANK2* transcript was detected in the same captured sample. Instead, it appeared due to extremely low or no expression of the full-length *SHANK3* transcript in adult frontal and temporal cortices. Most of the 472 unique *SHANK3* transcripts clustered within regions spanning exons 1-9 and 10-22. None incorporated splicing between exon 9 and 10, an area characterized by high GC content (77% of GC) and a CpG island (hg38). Accordingly, 43 unique transcripts initiated from this CpG island, implying a TSS within intron 9. *In silico* analysis using a parameter-free assembly approach (Cufflinks-Cuffmerge)^51^ applied to srRNA-seq data also failed to detect any transcripts connecting exon 9 and 10.

**Fig. 5.**
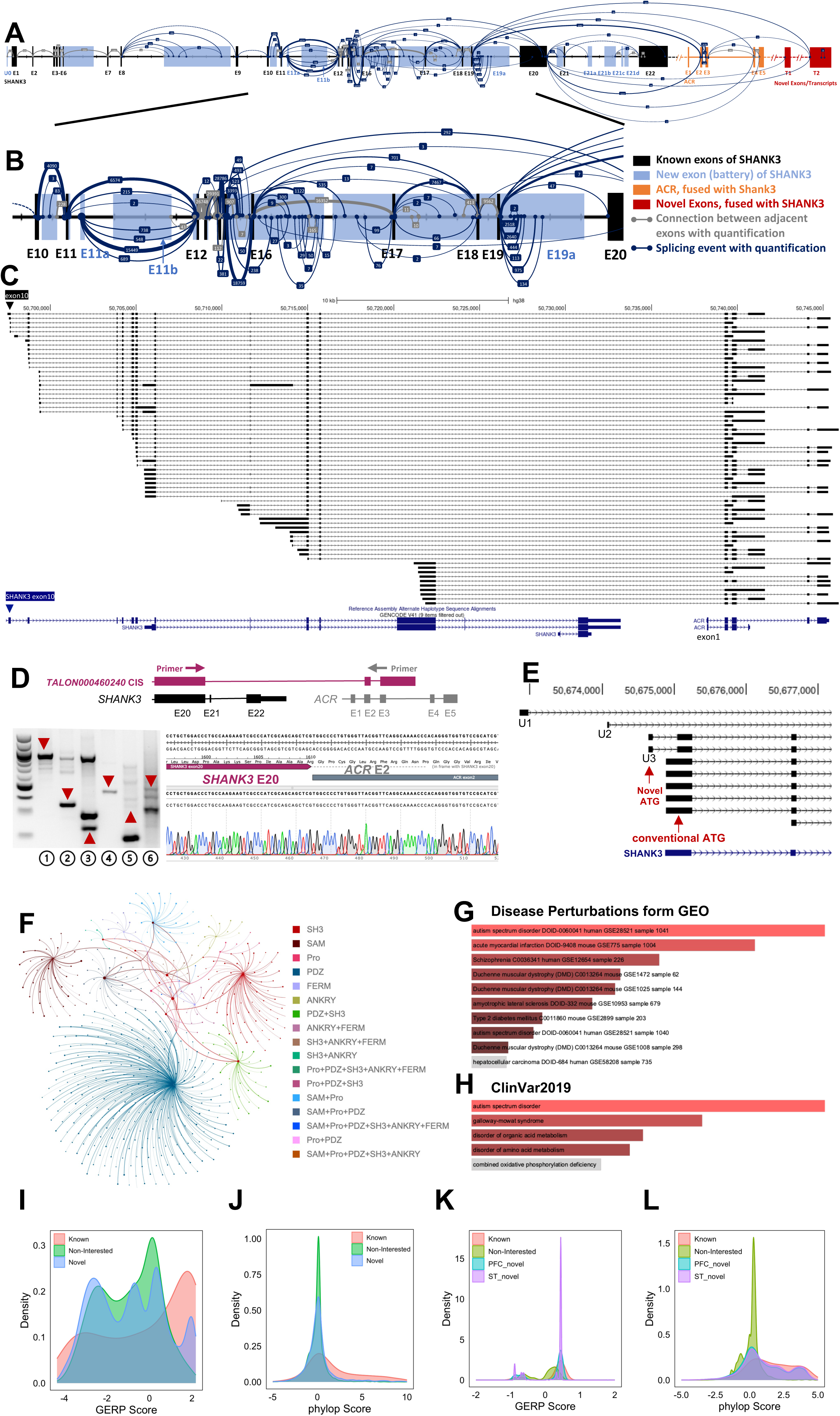
The novel transcripts of human *SHANK3* genes detected by CIS and predicted ORFs. **A.** New *SHANK3* transcript structure and Conch plot of *SHANK3* transcripts discovered by CIS in normal human cortex. Black backbone is the annotated *SHANK3* transcript of NM_001372044 (hg38). Blue rectangles represent novel exons of *SHANK3*. The exons of *ACR* are in orange rectangles. The new and uncharacterized exons distal to *ACR* are in red rectangles. The grey line connects adjacent exons while the light blue line illustrates alternative splicing events. The number of sequences reads for the splicing event is shown in the middle of connecting lines and reflected in the thickness of the connecting lines. **B.** Zoomed view of the splicing events between exons 10 and 20 in the human cortex. Exons 16 and 20 of *SHANK3* in humans corresponds to exons 17 and 21 of *Shank3* in mice. **C.** Structure and abundance of the fusion transcripts between *SHANK3* and *ACR* in the human cortex. Majority of fusion transcripts are initiated after exon 10, mainly from introns 16, 17, and exon 21. The fusion transcripts are notably skipping exon 20 (the largest exon) of *SHANK3* and exon 1 of ACR. **D.** Validations novel *SHANK3* transcripts in in human brain tissue by RT-PCR and Sanger sequencing. Diagram for the primer design of L1 is shown. RT-PCR gel: L1, fusion transcript between *SHANK3* exon 20 and *ACR* exon 2; L2, fusion transcript between *SHANK3* exon 20 and *ACR* exon 4; Lane 3, fusion transcript between SHANK3 exon 19 and ACR exon 2; L 3, novel exon U3; Lane 4; L5, intron14 retention; L6, intron 15 retention. M, DNA marker. Sanger sequence of RT-PCR product of *SHANK3* exon 20 and *ACR* exon 2 fusion from L1 **E.** Three new exons upstream of the annotated exon 1 of *SHANK3* mRNA (NM_001372044) (U1, chr22:50672853-50672979; U2, chr22:50674076-50674097; U3, chr22:50674642-50674705, hg38). A new ATG codon is in U2. **F.** Dandelion plot shows functional domain combinations of the *SHANK1*, *SHANK2*, and *SHANK3* transcripts from CIS. Each dot represents a unique transcript, and each color is a unique combination of functional domains. There are 17 combinations of functional domains of human *SHANK* family genes. The PDZ domain was significantly more present (∼70%) in predicted ORFs. **G-H.** Significant enrichment of fusion transcripts in transcriptome data of ASD and schizophrenia. For Gene Ontology enrichment analysis with Enrichr95 in 41 disease-related datasets. The fusion transcripts were significantly enriched in ASD and schizophrenia in Disease Perturbations form GEO dataset (G) and the ClinVar2019 dataset (H). **I-J.** Distribution of GERP (G) and PhyloP (H) scores across human *SHANK3* genomic regions of known coding exons, novel exons from CIS, and non-transcribed region in cerebral cortex. **I**. GERP score for novel exons from CIS in cerebral cortex is significantly high than non-transcribed region (D=0.097; p<0.001) but significantly lower than that of *SHANK3* known exons (D=0.299; p<0.001). **J.** PhyloP score for novel exons from CIS in cerebral cortex is significantly higher than non-transcribed region (D=0.133, p<0.001) but significantly lower than that of *SHANK3* known coding exons (D=0.296, p<0.001). **K-L.** Distribution of GERP and PhyloP scores across mouse *Shank3* genomic regions of known coding exons, novel exons from CIS, and non-transcribed region in PFC and ST. **K**. GERP score for novel exons from CIS in PFC and ST is significantly high than that of non-transcribed region (PFC: D=0.548, p<0.001; ST:D=0.602, p<0.001) but significantly lower than known *Shank3* coding exons (PFC:0.15, p<0.001; ST:D=0.0960; p<0.001). **L**. PhyloP score for novel exons from CIS in PFC and ST is significantly higher than that of non-transcribed region (PFC:D=0.385, p<0.001; ST:D=0.439, p<0.001) but significantly lower than known *Shank3* coding exons (PFC:D=0.184, p<0.001; ST:D=0.128, P<0.001).

Similar to mouse *Shank3*, we detected 66 fusion transcripts between *SHANK3* and *ACR* (**Fig. 5C**). These fusion transcripts, intron retention, and novel exons were validated by RT-PCR and sequencing **(Fig. 5D)**. Fifty-eight of them were fusion transcripts constituted exon 19/exon 20 of *SHANK3* (exon 20 is the largest exon in human equivalent to exon 21 in mice) to exons 2-5 of *ACR*. Nine transcripts started within *SHANK3* exon 20 were found to be fused with *ACR*. We observed splicing events connecting *SHANK3* exons 19-20 to uncharacterized downstream exons, T1-2, of *ACR*. We also detected 3 novel untranslated exons (U1-U3) upstream exon 1 of *SHANK3* mRNA (**Fig. 5E**). The sequence of U2 is highly conserved in mouse.

With the joint capture for *SHANK* family genes, we detected 86 *SHANK1* and 277 *SHANK2* transcripts (**UCSC Track 6**), from which 69 ORFs for *SHANK1* and 165 ORF for *SHANK2* were predicted. Across these *SHANK* family ORFs, we observed 17 different combinations of six functional domains, with the PDZ domain appearing most frequently (**Fig. 5F**). A complete set of all six functional domains (Ubl, ANKYR, SH3, PDZ, Pro, and SAM) was predicted only in one *SHANK2* transcript.

The unexpected discovery of extensive fusion transcripts between *SHANK3* and *ACR* in human brain tissue led to a comprehensive genome-wide analysis for fusion transcripts in SIS data. We detected 2,265 fusion transcripts (1.7% of the total transcripts) associated with 3,499 genes in the brains of children and adults, with 963 fusion transcripts common to both groups. About 98% of fusion transcripts are between two adject genes. A small number of fusion transcripts are among 3 adjacent genes. No fusion transcript is from distant genes or genes from two chromosomes. Gene Ontology enrichment analysis revealed a significantly enrichment of fusion transcripts in genes associated with ASD (**Fig. 5G-H**).

To access the functional constraint of novel *SHANK3/Shank3* exons in humans and mice identified by CIS, we utilized Evolutionary Rate Profiling (GERP)^52,53^ and PhyloP^54^ conservation scores. In mice, GERP and PhyloP scores for most *Shank3* novel exons were significantly higher than those of non-transcribed region, but they were lower than scores for known coding exons in both PFC and ST (**Fig. 5I-J, supplementary Table 6A-B**). A concordant pattern was observed in human *SHANK3* (**Fig. 5K-L, supplementary Table 6C-D**). These results suggest that the novel exons of *SHANK3/ Shank3* uncovered by CIS are evolutionarily constrained elements, underscoring their potential functional significance.

### Transcript diversity and novel transcripts of *TP53* gene in human and mice

To examine whether the transcriptional complexity is exclusively associated with synaptic genes, we applied SIS and CIS to *TP53* in human brain, and to *Trp53* in mouse brains and thymus, where *Trp53* expression is the highest. SIS detected only 5 *Trp53* transcripts in mouse ST and 3 in mouse PFC that is consistent with the data in literature ^55,56^,. In contrast, CIS identified a comprehensive set of 243 transcripts from thymus, 164 from PFC, and 188 from ST (**Supplementary Fig. 6A-C**, UCSC Track 7). The pattern of unique *Trp53* transcripts is similar amongst the 3 tissues, with 18 alternative TSS deduced from thymus transcripts. A significantly higher percentage of transcripts exhibited intron retention in *Trp53* compared to *Shank3*. Additionally, novel tissue-specific 5′ exons unique to brain (bU1) and thymus (tU1/tU2) were discovered.

In human brain, CIS detected 106 *TP53* transcripts, which predicted 60 ORFs, 18 TSSs, and three 3′ transcriptional ends (**Supplementary Fig. 6D, UCSC tracks)**. We also discovered 3 novel exons (hT1-3) at the 3′ end, which extended the C-terminus of TP53 ORF by 72aa and is conserved with the mouse *TRP53* (77% identical). These observations underscore the complexity of the *TP53/Trp53* transcriptome, which is complex but less heterogenous than *SHANK* family genes.

### Developmental, tissue, and tell type specificity of SHANK3/Shank3 transcripts from CIS

To investigate the developmental specificity of *Shank3* transcriptome, we aligned mouse srRNA-seq data of cerebral cortex at different ages from E14.5 to P180 day^57–59^ to *Shank3* transcripts from CIS (**Fig. 6A**). The E14.5 embryos exhibited the least diversity of *Shank3* transcripts of. As development, the number of unique *Shank3* transcripts increased, reaching a maximum at P56 day before declining at P180 day.

**Fig. 6.**
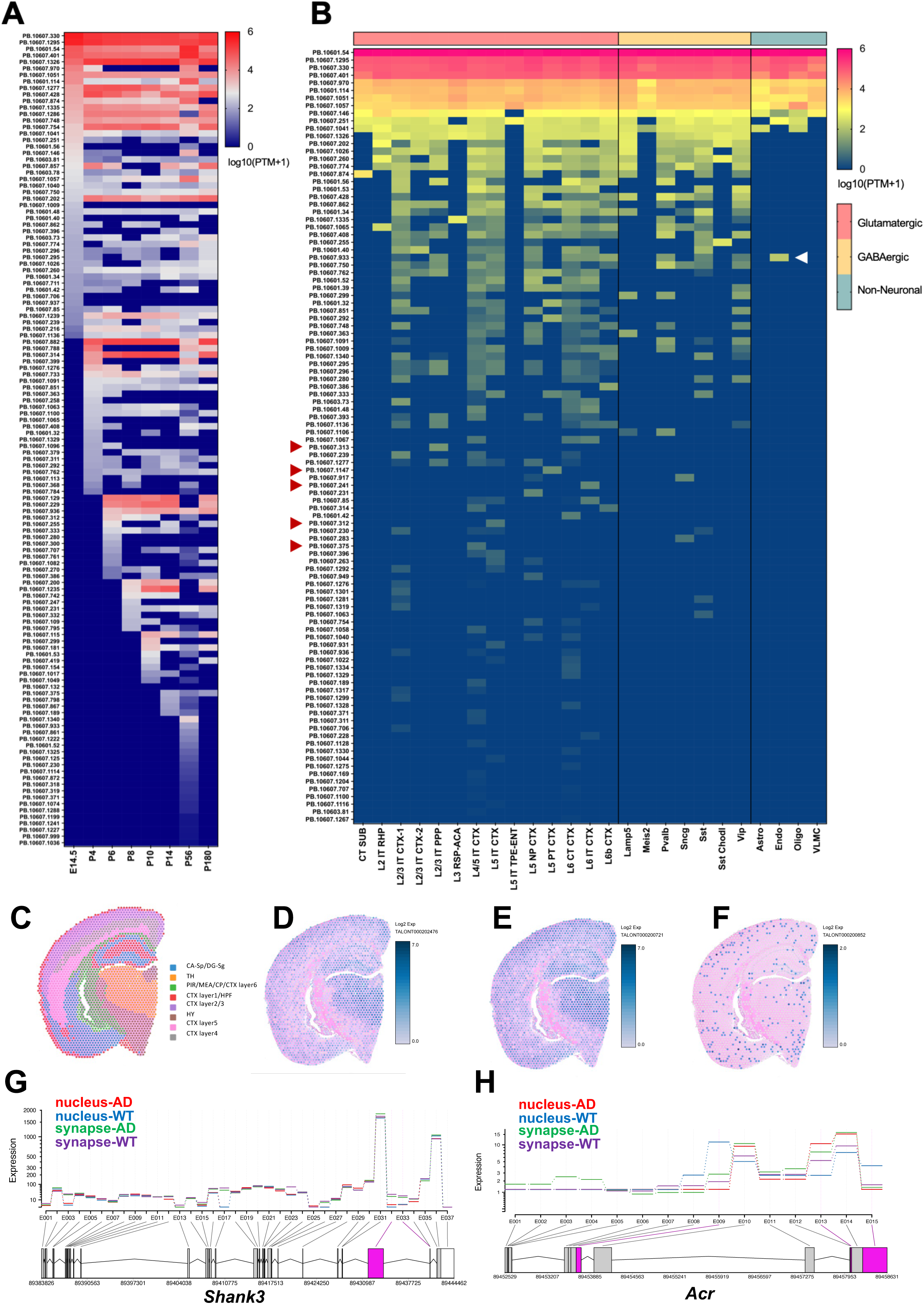
Developmental, cell type, cell compartment specific, and spatial transcriptome of *Shank3* in mouse brains. **A.** Developmental specific *Shank3* transcripts in mouse cerebral cortex. **B**. Cell type specific *Shank3* transcripts in mouse brains. The scRNA-seq of anterior cingulate cortex (ACA)^5^ was aligned to *Shank3* transcripts detected by CIS. Glutamatergic neurons, especially the L2/3, L4/5, and L6 CTX, have more diverse *Shank3* transcripts compared to GABAergic neuron and non-neuronal cells. Certain transcripts were cell type specific. *Shank3* transcript (PB.10607.933) including exon 18 was only detected in endothelial cells. **C-F.** Mouse *Shank3* transcripts in Visium spatial transcriptome. C. Visium spatial anatomy (CA: Cornu Ammonis, DG: Dentate Gyrus, TH: Thalamus, PIR: Piriform cortex, MEA: Medial Amygdala, CP: choroid plexus, CTX: Cortex, HPF: Hippocampal Formation, HY: Hypothalamus). **G.** Cellular compartment specific changes of *Shank3* exon usage in the hippocampus of Alzheimer’s disease (AD) mouse model from scRNA-seq data from different cellular compartment. The nucleus, compared to synapses, expressed significantly fewer splicing events of 32 and 33 that correspond to the exon 21, the largest exon of mouse *Shank3*. **H.** Different pattern of *Shank3-Acr fusion* transcripts in nucleus and synapse between WT and AD mice.

Further analysis on cell type specificity aligning scRNA-seq data from anterior cingulate area (ACA) of 8-week-old mice^39^ to *Shank3* transcripts identified by CIS, demonstrated a significantly higher abundance of *Shank3* transcripts in glutamatergic neurons compared to GABAergic neurons. The *Shank3* transcripts including exon 18 was exclusively found in endothelial cells (**Fig. 6B**).

To investigate tissue specificity, we analyzed the exon usage in mouse *Shank3* transcripts from CIS against scRNA-seq data from 5 cerebral cortex subregions^39^. The exon usage patterns of *Shank3* CIS transcripts within the same cell type exhibited unique variations across different brain subregions (**Supplementary Fig.7**). The pattern of human *SHANK3* transcripts in infants and children was distinct from adults, when aligned human srRNA-seq data to *SHANK3* transcripts from CIS. The *SHANK3* exon usages were also changes with age.

We mapped *Shank3* transcripts to 10x Genomics Visium spatial transcriptome of mouse to visualize the expressive pattern *in situ*^60^. Two probes targeting *Shank3* exons 11 and 22, and one for *Acr* exon 5, facilitated this analysis. Three *Shank3* transcripts identified by CIS were enriched to distinct anatomical regions (**Fig. 6C-F**). *Shank3* transcript TALONT000202476 containing exon 11, and TALONT000200721 incorporating exon 22, have a similar cell-specific expression pattern, albeit at different levels of abundance. Transcript TALONT000200852, a fusion transcript connecting *Shank3* exon 21 and *Acr* exon 5, displayed a cell type-specific expression pattern. Furthermore, we found a cellular compartment-specific preference for the *Shank3* transcripts. The inclusion of *Shank3* largest exon 21 is significantly more common in synapses than in nuclei from mouse brain scRNA-seq data^61^ (**Fig.6G**). The exon 2 of *Acr* gene, frequently fused with *Shank3* exons, was significantly less present in nucleus of AD models compared to WT. The splicing events involving 5′ segment of *Acr* exon 5 were more common across both nucleus and synapses in AD mice, while splicing involving the latter 2/3 of *Acr* exon 5 was more frequent in nucleus of WT (**Fig. 6H).**

### Applications of *SHANK3* transcriptome from CIS to genome sequencing and transcriptome analyses of ASD and other neuropsychiatric disorders

Human *SHANK3* transcripts identified through CIS exhibit expression patterns that are specific to developmental stages and brain regions, such as the cerebral cortex and cerebellum (**Fig. 7A-D** and **Supplementary Fig. 8**). We extended the *in-silico* transcriptome diversity analysis to 213 highly confident ASD risk genes consolidated from 3 recent extensive ASD genomics studies (**Supplementary Table 4)**^43–45^. The transcriptome diversity of ASD risk genes was significantly greater than of non-ASD associated genes (Pearson r=0.386, p<0.001). Specifically, ASD risk genes associated with gene expression regulation and neuronal communication showed significantly higher level of transcriptome complexity, compared to genes in other functional categories (Pearson r=0.825 and Pearson r=0.793, respectively, both p<0.001). *SHANK3*, consistently reported as one of the top 5 ASD causing genes in these studies^43–45^, is also implicated in schizophrenia (SCZ)^62^, bipolar disorder (BPD)^63^, and major depressive disorder (MDD)^64^. To investigate alterations in *SHANK3* transcriptomes across these disorders, we analyzed srRNA-seq data from the PsychENCODE project^35^. Principle component analysis (PCA) revealed unique transcript patterns for each disorder, especially for ASD and SCZ **(Fig. 7E).** The expressions of a subset of *SHANK3* transcripts varied across ASD, MDD, BD, SCZ, and controls **(Fig. 7E-I)**. Brain region and age-specific expression of *SHANK*3 transcripts formed distinct cluster in PCA analysis (**Fig. J**). The exons 12, 15, 20, and 22 of *SHANK3* transcripts in BA7 were significantly more represented in ASD brains than controls **(Fig. 7K)**, and exon 10 showed a higher expression in BA38 of ASD brains **(Fig. 7L).**

**Fig. 7.**
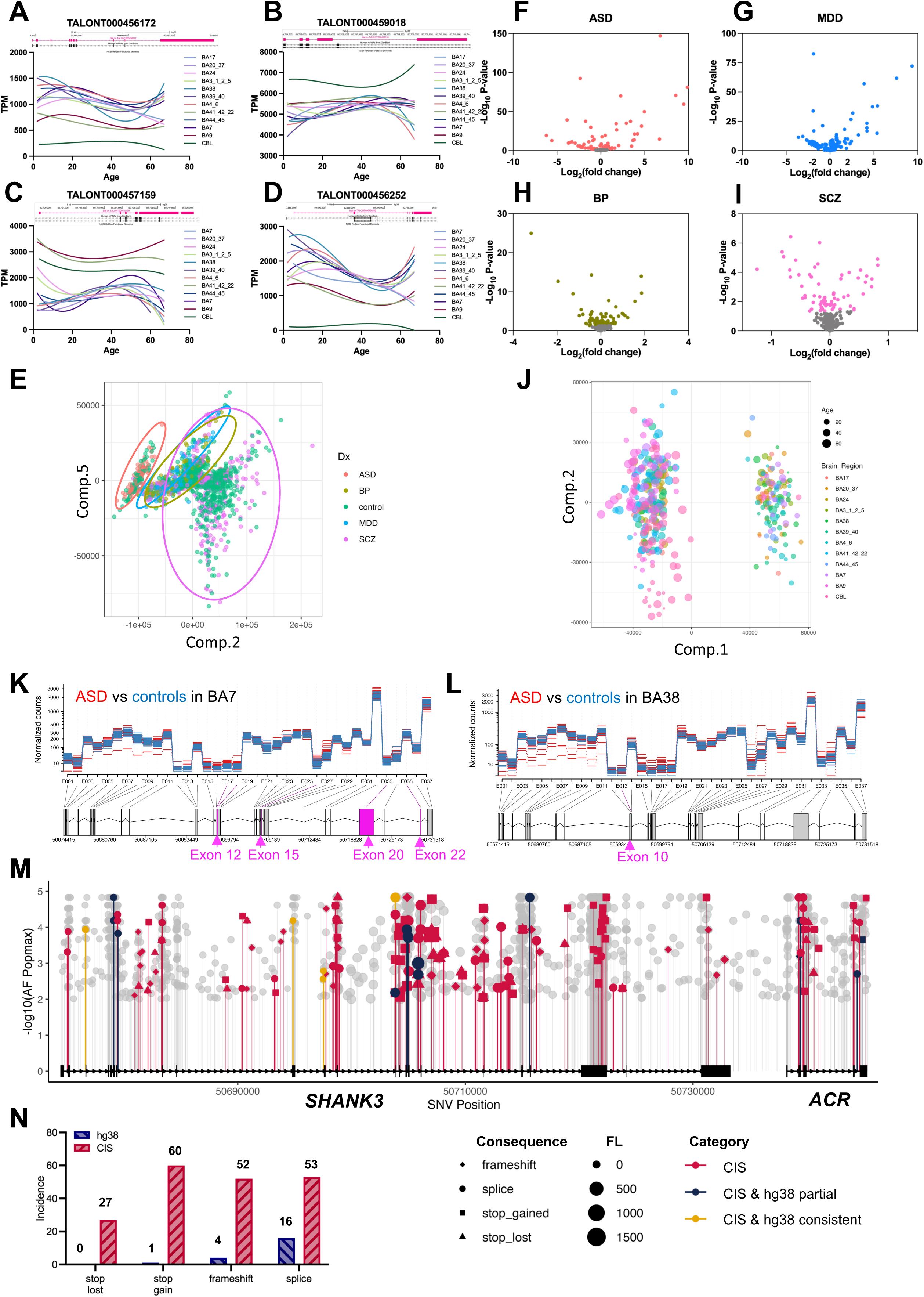
Improved transcriptome analysis of ASD transcriptome and sequence variant annotations of genome sequence data using *SHANK3* transcript structure from CIS A-D. The pattern of human *SHANK3* transcripts from CIS changed at different ages and brain regions. Bulk RNA-seq data of normal controls was aligned to *SHANK3* transcripts detected using CIS (BA, Brodmann area; CBL; cerebellum). **E-I.** PCA of human *SHANK3* transcripts from CIS and bulk RNA-seq data of 2,474 cases with ASD, BPD, MDD), or SCZ, and normal controls from PsychENCODE (only data from prefrontal cortex is included). The clusters of MDD and BPD overlapped but are separate from ASD and SCZ. The volcano plots for individual disorders ASD (n=68), MDD (n=87), BPD (n=297), and SCZ (n=736) compared to controls (n=1,286). **J.** PCA analysis of *SHANK3* transcripts in different brain regions and age (BA, Brodmann area; CBL, cerebellum) **K-L:** Brain region-specific change of *SHANK3* transcripts in ASD brains. Bulk RNA-seq data of subregions of the brain from ASD and controls were aligned to *SHANK3* transcripts from CIS. **K.** Exons 11, 15, 20, and 22 of *SHANK3* transcripts were significantly more represented in the BA7 region of ASD. **L.** Exon 10 of *SHANK3* transcripts is significantly more represented in BA38 of ASD brain. **M.** Utilizing the updated *SHANK3* transcript structure from CIS enhanced PTV detection in ASD, SCZ, and BPD exome and genome sequencing data. From 55,000 cases, we identified 1,530 new PTVs, a significant increase from previous annotations using the SHANK3 transcript NM_001372044.2 in hg38. Of these, 192 variants were likely deleterious, including 27 stop-loss, 60 stop-gain, 52 frameshift, and 53 splice variants, compared to the earlier finding of 22 such variants. **N.** The discovery rate of PTVs for *SHANK3* is increased from 1.3% using NM_001372044.2/hg38 as a reference to 12.5% using the transcript structure from CIS in this study.

While *SHANK3* genetic mutations are implicated in 1-2% of ASD cases and to a lesser extent in other neuropsychiatric disorders^43–45,65,66^, we sought to examine whether incorporating the enhanced *SHANK3* transcript structure from CIS into public available ES and WGS of ASD/SCZ/BPD datasets could uncover additional disease-associated single nucleotide variants (SNVs)^45,67–70^. We re-analyzed sequence variants on a large cohort of 177,000 samples of both controls and disease subjects, including ES data from the Autism Sequencing Consortium^45^, BPD Exomes^68^, SCZ Exome Meta-analysis Consortium^67^, as well as WGS of ASD, SCZ, and BP cohorts from BrainVar^69^ and BrainGVEX^70^. Variant identifications and annotations of were previously based on the mRNA reference NM_001372044.2 and hg38 genome assembly. We used Variant Effect Predictor (VEP, release 107)^71^ and Genome Aggregation Database (gnomAD, v3.1.2)^72^, for annotation and filtering, including variants with a population allele frequency of <= 0.01 for protein truncating variants (PTVs), and excluding missense and synonymous variants for further analysis. SpliceAI^73^ and SnpEff^74^ were used to analyze splice variants and evaluate the pathogenic potential of stop-loss, stop-gain, and frameshift variants. This re-annotation identified 1,530 new SNVs across 55,000 cases pooled from ASD (11,986 ES; 923 WGS), BP (14,210 ES), and SCZ (27,648 ES) cohorts **(Fig. 7M)**, resulting in the discovery of 27 stop-loss, 60 stop-gain, 52 frameshift, and 53 splice variants in *SHANK3* considered potentially deleterious or PTVs using CIS annotation in disease subjects but not in controls. This was a marked contrast to the variants analyzed using current reference (0 stop-loss, 1 stop-gain, 4 frameshift, and 16 splice variants). Accordingly, the detection rate for potential deleterious SNVs of *SHANK3* increased from 1.3% when using current reference (NM_001372044) to 12.5% when annotated with the *SHANK3* CIS transcripts, highlighting the significance of comprehensive transcriptome annotation in uncovering genetic contributions to neuropsychiatric disorders **(Fig. 7N).**

## Discussion

Diverse transcription is crucial for generating proteomic diversity and facilitating complex cellular functions. Precision of transcription is critical because mutations in the transcriptional regulatory DNA elements can cause numerous single gene disorders. Despite the recent report of the completed human genome^75^, the transcriptome remains largely uncharted. Our work applying SIS on human and mouse brains discovered unprecedented transcriptome diversity^8,42^. Glinos *et al*^14^ reported a maximum of 178 isoforms for a single gene, with only 5 genes exhibiting more than 100 isoforms, and a median of 2 isoforms per gene across various tissues and cell lines. Leung *et al*.’s study^42^ noted a peak of 40 isoforms per gene in the human cortex. Furthermore, Chau *et al*^76^ assembled an average of 4 isoforms per gene from bulk RNA-seq of human developing brains. Significantly, these studies uncovered only a few incomplete *SHANK3* mRNA isoforms. However, our study identified as many as 692 isoforms for a single gene, with 595 genes having more than 100 isoforms, and an average of 19 isoforms per gene in the human cerebral cortex. Our results suggest that the extent of transcript complexity described in existing literature is significantly underestimated, particularly for genes like *SHANK3*.

Our targeted capture and long-read sequencing have mapped the SHANK family transcriptomes in detail, with the majority of novel transcripts likely endogenously expressed. This is supported by our strict identification process, validation through RT-PCR and Sanger sequencing, consistency across experiments and brain regions, and conservation between species. Additionally, the specificity of these transcripts was confirmed in Shank3 mutant mice. Despite the high confidence, it remains a possibility that a small fraction might not be expressed endogenously. The discovery of a substantial number of fusion transcripts for *SHANK3/Shank3* in our study was unexpected, with a prevalence that surpassed the findings from other studies^8,42^. Until recently, fusion transcripts have been largely investigated in cancer-related studies because of their oncogenic properities^77,78^. Yet, their presence in normal cells has only recently been acknowledged ^8,42,79^. Two recent studies using the SIS method^8,42^ reported a mere 136 fusion transcripts (0.41% of total transcripts) in human brains. In contrast, our study identified 2,265 fusion transcripts in human brains, constituting 1.7% of total transcripts. Interestingly, these fusion transcripts were found to be particularly more enriched in the human ASD-associated transcriptome.

The enhanced *SHANK3* transcript structure from CIS has significantly increased the detection rate of PTVs or predicted LOF variants in ES and WGS data for neuropsychiatric disorders. Further functional validations are warranted to determine the pathogenicity of these new identified PTVs. Our findings highlight the significance of employing fully characterized transcript structures in genomics studies of disease gene discovery. Transcriptional dysregulations in the brain have been implicated in neuropsychiatric disorders^5,80^. By integrating the *SHANK3* transcriptome data from CIS and the transcriptome data from PsychENCODE, we discovered brain region-specific dysregulations in *SHANK3* transcriptome associated with ASD and other neuropsychiatric disorders. Notably, brain region-specific DNA methylation in intragenic CpG islands, which show altered methylation in ASD brains^81,82^, suggest that epigenetic changes could be instrumental in *SHANK3* transcript variations.

In *Shank3* mutant mice, stable transcripts with exonic deletions indicated truncated protein production or upregulated non-mutant isoforms^30,83^. Cryptic promoters, especially within intron 16/exon 17, suggest alternative initiation and potential novel protein isoforms. These could perturb the postsynaptic density protein interactome, indicating possible loss and gain of function in *Shank3* mutants. Such complexities question the molecular and phenotypic consistency of *Shank3* mouse models with SHANK3 disorders^16–18,30,84,85^. For example, differential behavioral phenotypes and receptor subunit alterations are noted across mutant lines^19,30,34^. Specifically, *Shank3*^Δe21^ mutants show unique upregulation of alternative transcripts and fusion transcripts, diverging behaviorally from *Shank3*^Δe4–22^ mutants^30,86^. These molecular nuances challenge the translational fidelity of *Shank3* mouse models for preclinical studies and necessitate re-evaluation, particularly for models in therapeutic development.

Our study’s detailed alignment of *SHANK3/Shank3* transcripts underscores its proteomic diversity at the PSD, essential for complex synaptic functions^48,49,87^. However, about 15% of the transcripts, possibly arising from cryptic promoters or alternative splicing, lack substantial ORFs or are lowly expressed, hinting at stochastic transcription events previously noted in other species^88–97^. Challenges to the ENCODE projects’ findings on genome transcription by subsequent short-read RNA-seq studies^11–13,15,98–100^, align with our discovery that *SHANK3/Shank3* and *TP53* transcription involves intragenic promoters and frequent intron retention. These regions, less conserved evolutionarily, affirm pervasive transcription and suggest a more deterministic transcriptional landscape for these genes in humans and mice.

### Limitations of the study

Several limitations of study that are warranted for discussion. We will not be able quantify the extent of stochasticity of transcription from current analysis. The extensive functional validation of transcripts at the protein level remains a challenge, as some transcripts may function uniquely at the RNA level, eluding protein-interaction analyses. Also, our capture-based method trades sensitivity for efficiency when scaling up, as increased gene targets reduce sequence depth, necessitating careful experimental design for quality data.

## Acknowledgement

We thank the valuable discussion and assistance of Antonio Giraldez, Hongyu Zhao, Gang Peng. And Antonio Jorge Forte. We thank Emily Qian and Rao Nivedita for editorial assistant. YHJ is supported by NIH Grants R01MH117289, R01HD088007, and R01HD088626. XL is the postdoctoral fellow of Foundation for Angelman Syndrome Therapeutics (FAST).

## Author contributions

XL and YHJ conceived and designed the project. XL performed most of data collection and data analysis. PSM, YM, YW and AQH prepared and process human brain tissues. GW assisted the long-read sequencing production. MG and NP assist the data analysis. XL and YHJ wrote the manuscript together with all co-authors.

## STAR Methods

### RESOURCE AVAILABILITY

#### Lead contact

- Further information and requests for resources and reagents should be directed to and will be fulfilled by the lead contact, Yong-Hui Jiang.

#### Materials availability

##### Materials availability statement

- Oligonucleotide probe panels were synthesized by Integrated DNA Technologies (IDT). The probe coverage and design are provided in Supplementary Tables.

##### Data and code availability

- Both human and mouse raw sequencing data have been deposited at SRA under BioProject: PRJNA1066952 and are publicly available as of the date of publication. Accession numbers are listed in the key resources table. All UCSC tracks described in manuscript have been deposited at Mendeley and are publicly available as of the date of publication. The DOI is listed in the key resources table.
- This paper does not report original code.
- Any additional information required to reanalyze the data reported in this paper is available from the lead contact upon request.

### EXPERIMENTAL MODEL AND STUDY PARTICIPANT DETAILS

#### Human brain tissues

Adult human cortex tissues (n=4, 24-33 years old; frontal cortex, n=2; temporal cortex, n=2) were obtained from Mayo Clinic Florida Biospecimen Bank and processed at Yale University School of Medicine. Children cortex tissues (n=4, 5-12 years old; temporal cortex, n=3; amygdala, n=1) were obtained and processed from the Children’s Hospital of Fudan University in Shanghai, followed the same RNA extraction, library preparation and sequencing protocols as Yale site. The IRB protocols were approved both at Mayo Clinic Florida and the Children’s Hospital of Fudan University in Shanghai.

#### Mice

Wild type C57BL/6J mice were obtained from the Jackson Laboratory. *Shank3* mutant mice of Shank*3* exons 4-9 deletion (*Shank3*^Δe4–9^) ^34^ and *Shank3* exons 4-22 (*Shank3*^Δe4–22^)^19^ were generated and maintained in Jiang’s lab. *Shank3* exon 21 deletion (*Shank3*^Δe21^) *was obtained from Jackson Laboratory (Shank3^tm^*^1^*^.1Pfw^*/J and Strain #:018398)^101^. Mice were housed of 4-5 per cage in pathogen-free mouse facility with free access to food and water on a 12-hour light: dark cycle at the ambient temperature of 20-22°C and humidity of 30-70%. An equal number of male and female mice were used for all experiments. All procedures were performed following the approved animal protocol by Yale University School of Medicine Animal Care and Use Committee.

## METHOD DETAILS

### RNA Isolation and Quality Control

Mouse brain tissues were snap-frozen in liquid nitrogen immediately after dissection. Human brain tissues were snap-frozen in liquid nitrogen within an hour after dissection. All tissues were stored in liquid nitrogen thereafter. Total RNA was isolated from 20 mg frozen tissues, using NucleoZOL™ (Takara Bio, 740404.200) and NucleoSpin® RNA set for NucleoZOL™ (Takara Bio, 740406.50) following the manufactures specifications, followed by rDNase Set (Takara Bio, 740963) to digest DNA, and NucleoSpin® RNA Clean-up XS (Takara Bio, 740903) for RNA repurification. RNA purity (260/280, 260/230) and concentration were measured on NanoDrop™ 2000/2000c Spectrophotometers. RNA integrity number (RIN) was assessed using Agilent 2100 Bioanalyzer system.

### Generation of standard and captured Iso-seq libraries

The Iso-seq libraries were prepared by following the manufacturer’s instructions for each step (Iso-Seq™ Express Template Preparation for Sequel® and Sequel II Systems for standard Iso-seq; Customer Collaboration – Iso-Seq® Express Capture Using IDT xGen® Lockdown® Probes for capture Iso-seq). The 600 ng of total RNA was used as input. Only the RNA with RIN higher than 7 of human samples, and 8 of mouse samples were processed for reverse transcription, amplification, enrichment, and library preparations.

### Hybridization Capture Panel Design

Hybridization capture panel design was assisted by IDT (Integrated DAN Technologies). Briefly, after extracted as 120-base-length sequence of interested gene, xGen Lockdown probes were aligned to the genome and calculated the number of possible enrichment sites. A “perfect” probe was considered as only has 1 hit (the target of interest) with genome, but most of the sequences returned more than 1 hit. Following IDT proprietary xGen Off-Target QC Method, any probes with more than 50 hits were removed because of non-specific targets in genome. The specifics and details of each probe panel are presented in supplementary table 3.

### Hybridization Protocol

300 ng of total RNA in less than 5.4 μL of volume mixed with 2 μL of NEBNext Single Cell RT Primer Mix. The final volume was brought up to 9 μL with nuclease-free water. The reaction was placed in a thermocycler and run for 5 minutes at 70°C, followed by holding at 4°C for primer annealing and first-strand synthesis. Reverse transcription template switching reaction was then performed by adding 5 μL of NEBNext Single Cell RT Buffer, 3 μL of nuclease-free water, and 2 μL of NEBNext Single cell RT Enzyme Mix to the first-strand cDNA. The reaction was incubated in a thermocycler at 42°C with the lid at 52°C for 75 minutes, followed by holding at 4°C. After adding 1 μL of Iso-Seq Express Template Switching oligo to the 19 μL reaction for a final volume of 20 μL, the reaction was incubated again in a thermocycler at 42°C with the lid at 52°C for 15 minutes, followed by holding at 4°C.

The Reverse Transcription and Template Switching reaction product was then purified using ProNex Beads before proceeding with cDNA amplification. For amplification, 50 μL of NEBNext Single Cell cDNA PCR master Mix, 2 μL of NEBNext Single Cell cDNA PCR Primer, 2 μL of Iso-Seq Express cDNA PCR primer, and 0.5 μL of NEBNext Cell Lysis Buffer were added to the purified product. The reaction was incubated in a thermocycler and run for 45 seconds at 98°C, followed by 14 cycles of the following steps: 10 seconds at 98°C, 15 seconds at 62°C, and 3 minutes at 72°C. The reaction was then held for 5 minutes at 72°C, followed by holding at 4°C. Finally, the product was purified again using ProNex Beads before proceeding with either the library preparation for standard Iso-Seq (SIS) or the capture steps for capture-based Iso-Seq (CIS).

As for the capture steps, first concentrate a total of 500ng cDNA in a 1.5 mL LoBind tube along with 7.5 μL of Cot DNA. To this mixture, add 1.8X volume of ProNex beads and gently pipette mix 10 times, followed by incubation for 10 min at room temperature. Place the tube on a magnet stand and wait until supernatant is clear. Remove the supernatant and wash twice with 200μL of freshly prepared 80% ethanol while on the magnet stand. Spin the tube strip briefly after removing the second wash, return to magnetic stand, and remove residual ethanol. Next, immediately add the hybridization reaction mix (which comprises 2X Hybridization Buffer, Hybridization Buffer Enhancer, xGen Asym TSO block, xGen RT-primer-barcode block, and 1X xGen Lockdown Panel) to elute the cDNA. Gently pipette mix 10 times and incubate for 5 min at room temperature. Then, place the tube on the magnetic stand to separate the beads from the supernatant. Transfer 17 μL of the supernatant to a new 0.2 mL PCR tube and briefly centrifuge it. Ensure that the tube is tightly sealed to prevent evaporation. Finally, place the sample tube in the thermal cycler and start the hybridization program: HYB program (lid set at 100°C), 95°C for 30 sec, 65°C for 4 hr, and lastly hold at 65°C.

During the incubation, prepare 1X working buffers and beads for capture. Preheat the wash buffers to +65°C in a heat block or water bath. To prepare the capture beads, allow the Dynabeads M-270 Streptavidin to warm to room temperature for 30 minutes prior to use. Thoroughly vortex the beads for 15 seconds to mix them, then aliquot 50 μL of beads into a 0.2 mL PCR tube, followed by adding 100 μL of 1X Bead Wash Buffer per capture, and pipette the mixture 10 times. Place the PCR tube on a magnetic rack. When the supernatant is clear, carefully remove and discard it without disturbing the beads. Note: Allow the Dynabeads to settle for at least 1 minute before removing the supernatant. Thereafter, two washes are performed as follows: Add 100 μL of 1X Bead Wash Buffer, pipette 10 times to mix, then place the PCR tube on a magnetic rack, allowing the beads to fully separate from the supernatant. Carefully remove and discard the clear supernatant. Repeat this process for a total of two washes. Finally, resuspend the beads in 17 μL of Bead Resuspension Mix per capture. The Bead Resuspension Mix includes xGen 2X Hybridization Buffer (8.5 μL), xGen Hybridization Buffer Enhancer (2.7 μL), and Nuclease-Free Water (5.8 μL). By following these steps carefully, you can ensure that the buffers and beads are prepared correctly for the capture step and obtain reliable results.

Then Bind cDNA to the capture beads, by incubating the samples in a thermocycler set to +65C for 45 minutes. Then Wash the captured cDNA with 1X wash buffers and elute the cDNA with 46ul elution buffer. To amplify the captured DNA sample, NEBNext High-Fidelity 2X PCR Master Mix is recommended, and the NEBNext Single Cell cDNA PCR Master Mix is alternative for post capture amplification. Assemble the following PCR reaction: 50 μL of NEBNext High-Fidelity 2X PCR Master Mix, 2 μL of NEBNext Single Cell cDNA PCR Primer, 2 μL of Iso-Seq Express cDNA PCR Primer, 0.5 μL of NEBNext Cell Lysis Buffer, and 45.5 μL of the captured library. Amplify the PCR reaction mix using the following PCR protocol: Denature the DNA at 98°C for 45 seconds. Perform 14 cycles of the following steps: a. Denature the DNA at 98°C for 10 seconds. b. Anneal the primers at 62°C for 15 seconds. c. Extend the DNA at 72°C for 3 minutes. Final extension at 72°C for 5 minutes, and hold at 4°C. Finally perform the post amplification clean up steps with ProNex brands and ethanol. Use 1 μL of sample to quantifiy with Qubit dsDNA HS kit and dilute 1 μL of sample to 1.5ng/μL and run 1 μL on an Agilent Bioanalyzer using the High Sensitivity DNA kit. We used 500ng cDNA for library construction as Sequel II sequence platform required. After DNA damage repair, end repair/A-Tailing, overhang adapter ligation, and purification with ProNex Beads, the cDNA library is ready for sequencing

### Sequencing Platform

To load the cDNA library onto the PacBio Sequel II System, the diffusion method was applied and followed by a 24-hour movie time and a 2-hour pre-extension time. The samples were cleaned up using ProNex beads and loaded onto the plate at a concentration of 50-100 pM.

### Sequence data filtering algorithm

The following pipeline was diagramed in Supplementary Fig.1. Sequencing reads were screened initially with Lima (v2.5.0) and IsoSeq (v3). A transcript with both cDNA primers and the poly(A) was identified and called Full-length reads^102^. The Full-length reads which had less than 100 base pairs 5’ end overhang, less than 30 bases pairs 3’ end overhang, and less than 10 base pairs gaps in the middle are considered as the same transcript. Clustering using hierarchical alignment, and iterative cluster merging, generate polished sequence, with quality scores. The output further filtered with SQANTI3 (v4.3) after cluster and collapse to generate unique transcripts. SQANTI3 filtered the transcripts as below: If a transcript is Full-Splice Match (FSM), then it was retained unless the 3’ end was unreliable (intrapriming). If a transcript was not Full-Splice Match, then it was retained only if all below were met: (1) 3’ end is reliable. (2) did not have a junction that was labeled as RT-Switching. (3) all intro-exon junctions were canonical ^102^. Further criteria included a transcript had to include at least 2 exons, and in the sense orientation and predicted open reading frame (ORF) had longer than 100 amino acids for the given transcript.

### Transcript Confidence Score

To assess the quality of individual transcript, transcripts after filtering steps were scored by the following scoring metrics: (1) Score of 3 point: If the exons of transcript were presented in the sequences of by either Illumina short read methods of the bulk RNAseq (human dataset: UCLA-2022, BrainGVEX, CMC, CommonMind and LIDB) and SMART scRNAseq. (2) Score of 2 points: If a transcript had predicted ORF longer than 100AA. (3) If the abundance of a transcript were higher than 20 percentage of the rank of the abundance of all transcripts. The summation of scores was confidence score to define each transcript: high confidence (≥ 4 points), moderate confidence (2-3 points), and low confidence (0-1 point).

### Iso-seq data analysis pipeline

The flow chart below described the analytic pipeline for ISO-Seq sequence dat. The subreads.bam file of an Iso-Seq SMRT cell was a raw input. The number of the SMRT cells, instead of the number of multiplex samples sequenced on a SMRT cell, regardless of the library preparation methods [Sta-Iso-Seq (SIS) or Cap-Iso-Seq (CIS)], dictated the direction of the analysis flow.

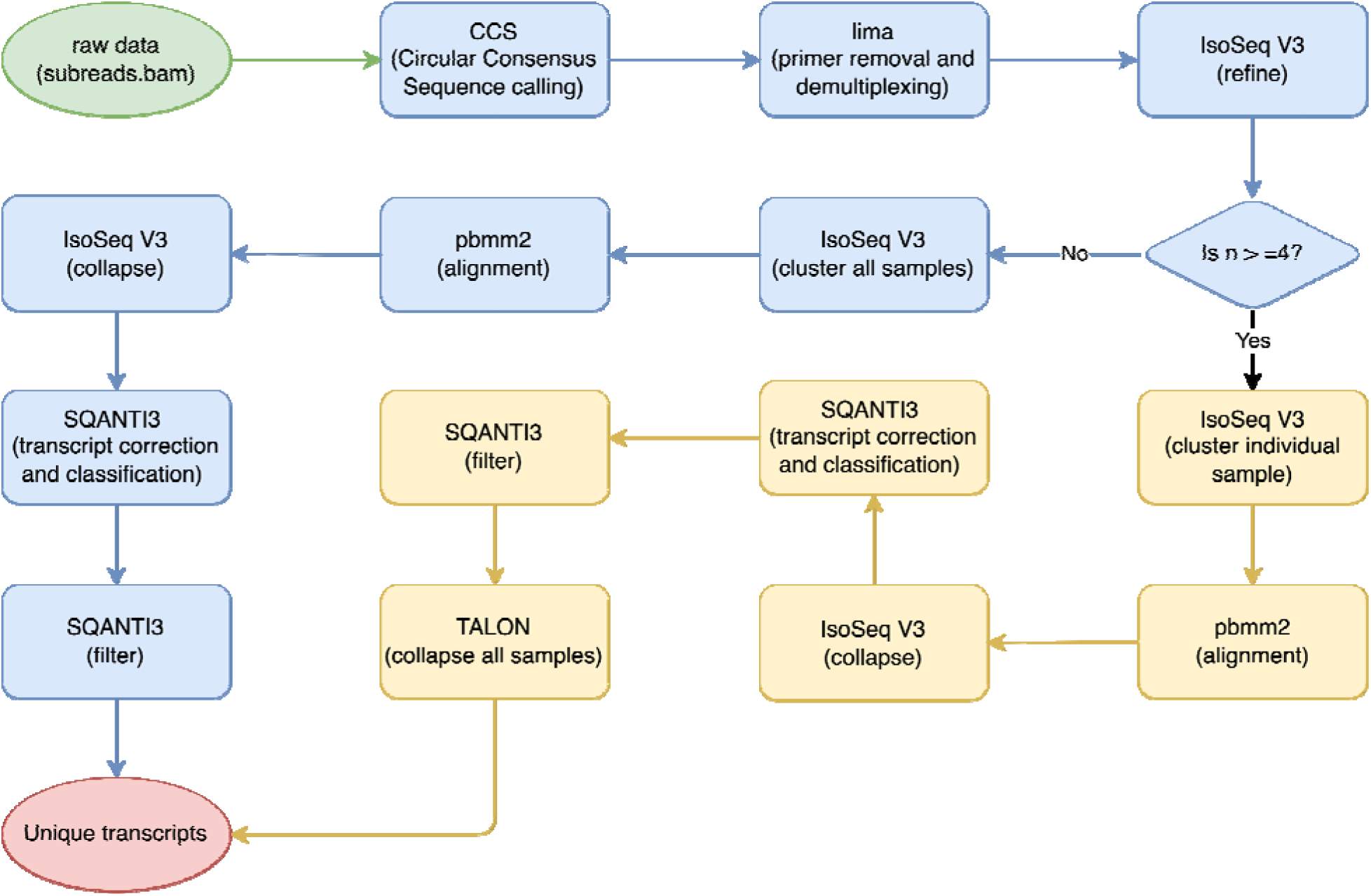

### Real-time quantitative polymerase chain reaction (RT-qPCR)

Two μg of total RNA was reverse transcribed into cDNA templates using RNA to cDNA EcoDry™ Premix kit including both random hexamer and oligo(dT)_18_ primers (Takara Bio, 639548). KAPA SYBR® FAST qPCR Master Mix (2X) Universal (Kapa Biosystems, KK4602) was used for qPCR reactions with 18 ng of cDNA as template input. The following program on CFX96 Touch Real-Time PCR Detection System (BIO-RAD) was used: 3 minutes at 95°C for enzyme activation, followed by 40 cycles of denaturation (95°C, 3 seconds) and annealing, extension, data acquisition (60°C, 30 seconds), followed by dissociation and holding at 4°C. The PCR primers are shown in supplementary Table 3.

### RNA-Seq Data Processing

Illumina bulk RNAseq raw data in FASTQ format after quality control and filtering with fastp ^103^, and SMART scRNAseq FASTQ data, were aligned to hg38 for human sequences and mm39 for mouse sequences using HISAT 2.2.1 ^104^. Aligned RNA-Seq data (aligned to hg37/38) in BAM format were converted to FASTQ format using SAMtools ^105^ when the raw FASTQ was not available, followed by the same process as above. Gene expression counts and DEXSeq-counts were calculated using FeatureCount ^106^ for further gene expression and exon usage analysis. Detailed RNAseq datasets information summarized in supplementary table 4.

### Differential Transcript Usage

Transcript-level quantification of the processed RNA-Seq data was performed using the software Salmon 1.4.0 ^107^. The transcriptome index used for quantification was built from the reference genome annotation (in GTF format), along with the reference genome FASTA file. Transcript abundances were estimated using the quasi-mapping algorithm (--quasiMAP) mode, which performs a lightweight alignment-free estimation of abundances based on k-mer matching. The output files were generated in TPM (transcripts per million) format.

### Differential Exon Usage (DEU)

DEXSeq-counts tables were imported into R, analysis with R package DEXSeq^108^. Normalization and filtering were performed to remove lowly expressed exons. DexSeq uses a binomial generalized linear model to estimate exon expression, accounting for the variability in exon-exon junction usage across samples. DEU was then tested using the DEXSeq function, which fits a statistical model to test for differences in exon usage between two or more groups of samples. Exons with an adjusted p-value ≤ 0.05 and a log2 fold change ≥ 1 or ≤ -1 were considered significantly differentially used, and visualized with built-in function of DEXSeq.

### Whole Genome Sequencing and Exome Analysis

DNA variation data post variation calling in VCF format were downloaded from Autism Sequencing Consortium (ASC), Bipolar Exomes (BipEx), whole-exome sequencing case-control study of epilepsy (Epi25), Schizophrenia exome meta-analysis consortium (SCHEMA), and PsychENCODE. VCFs initially aligned to hg38 (BipEx and Epi25) and the datasets (ASC, SCHEMA and PsychENCODE) after alignment lift over from hg37 to hg38 with UCSC LiftOver tool and chain file, were subsetted to the region of interest (SHANK3, chr22:50670000-50770000) using BCFtools (v 1.16) (Danecek et al. 2021). The data format was modified using HTSlib (v 1.16)^109^ and TAB-delimited file InderXer (Tabix, v 0.2.5)^110^. Then the data were annotated with Ensembl Variant Effect Predictor (VET, release 107)(McLaren et al. 2016) and filtered with Genome Aggregation Database (gnomAD, v3.1.2)^72^ by INFO/AF_popmax<=0.01. Filtered DNA variation were aligned to novel exons detected in SIS and CIS with SpliceAI ^73^ for splicing event analysis, and with SnpEff ^74^ to evaluate other deleterious SNV (stop lost, stop gain and frameshift).

### Data Visualization

Visualization was performed using ggplot2 (version 3.3.2) in R (version 4.2.2) for plotting gene expression, transcript and exon usage profiles and heatmaps.

### Spatial Transcriptional Analysis

An open access Visium dataset of mouse brain coronal section from 10x Genomics^111^ in FASTQ format was analyzed using customized references and annotation generated from mouse *Shank3* CIS transcripts using Cell Ranger ^112^, followed by quantitation with customized probe-set (probe-transcripts relation spreadsheet) using 10x Genomics Space Ranger v2.0. The output cloupe file was visualized using 10x Genomics Loupe Visualization Software v6.5.

## KEY RESOURCES TABLE

Submitted as a separate file

