## Supplemental Figures for "Transcriptional Determinism and Stochasticity Contribute to the Complexity of Autism Associated *SHANK* Family Genes"

#### Supplementary Fig.1

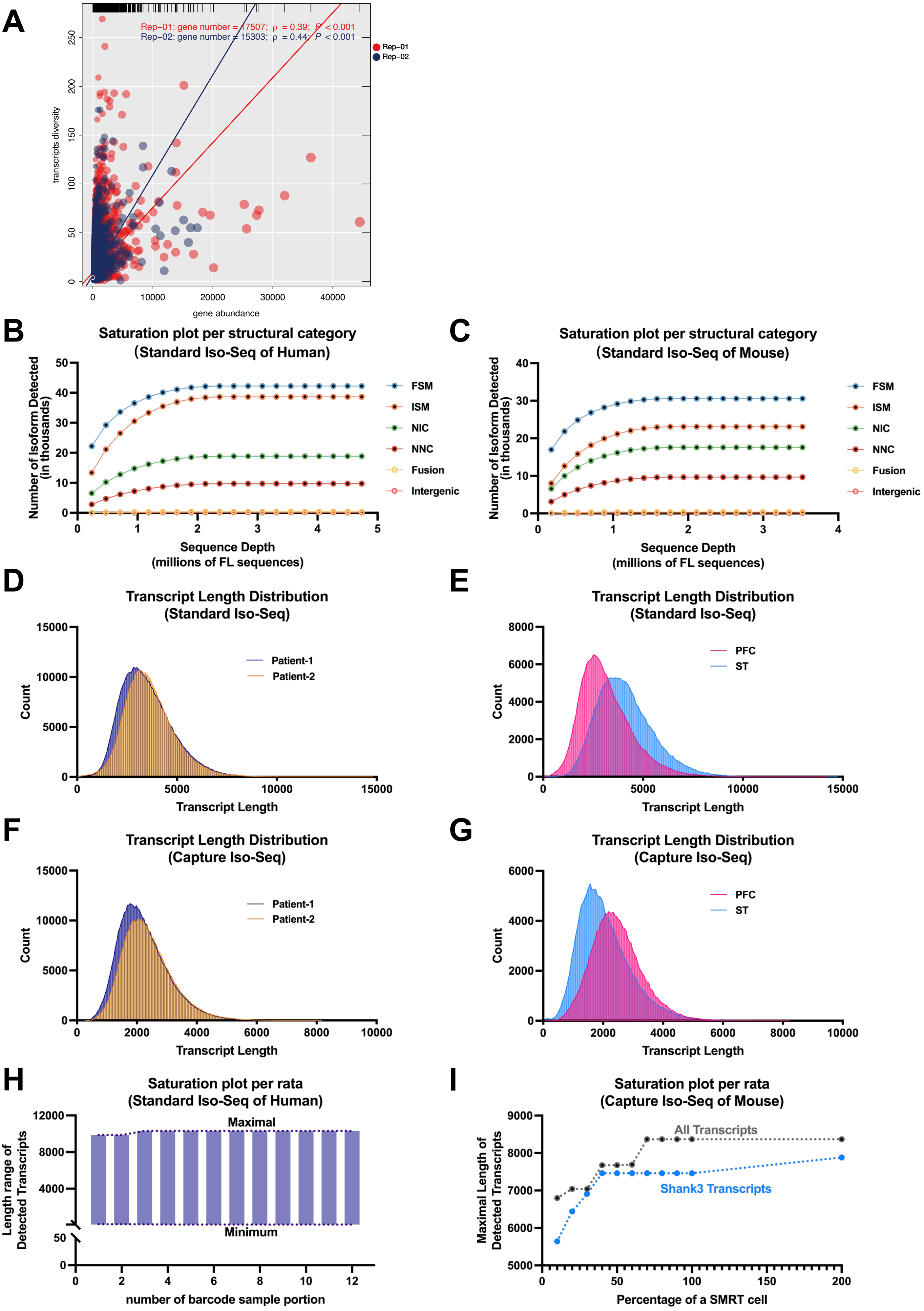

**Supplementary Fig.1. Reproducibility, QC, and sequence depth optimization of Iso-Seq**

**A.** The reproducibility of Iso-Seq were evaluated using unique transcript discovery rates of individual gene among repeated runs of a sample. The ratio of transcript diversity and the abundance between 2 runs of a sample was comparable [Pearson r (transcript length) = 0.999, $p<0.001$; Pearson r (transcript diversity of each gene) = 0.387, $p < 0.001$)]. The transcript to gene ratio is 0.39 in the first run and 0.44 in the second. The transcripts discover ratio was not significantly different between 2 batches.

**B-C.** Unique transcript discover rate saturating at about 2 million Full-Length (FL) reads/sequences per sample of both human and mouse, in each structural category (FSM: Full Splice Match; ISM: Incomplete Splice Match; NIC: Novel In Catalog; NNC: Novel Not in Catalog; Fusion: Fusion transcript, Intergenic: transcripts is in the intergenic region) of transcripts predicted by SQANTI3 [1].

**D-E.** The median of transcripts length detected was 3487 bp in human cortex, 3065 bp in mouse PFC, and 3963bp in mouse ST by using SIS, with the longest transcripts being 14809 bp, 14250 bp and 14748 bp respectively. The transcript length distributions between 2 human samples were significantly overlapped, while the transcript length distribution of ST shifted to right compared to PFC in mouse.

**F-G.** The median of transcript length detected was 2258 bp in human cortex, 2396 bp in mouse PFC and 1933bp mouse ST by CIS. The longest transcripts were 8177 bp, 8223bp and 8426 bp respectively.

**H.** Transcript length observed in accumulative 1/12 barcoded a human cortex sample using SIS, showed the maximum of transcript length detected in the 4/12 samples, indicating the sequencing depth saturation point.

**I**. The maximal length of transcripts in accumulating randomly divided (random seed method) tenth of 2 mouse ST samples indicating the sequencing depth saturated at about 40% of a SMRT cell in CIS. The maximal transcript length of targeting sequences was not increased after saturation point, but the off-targeting sequences increased.

#### Supplementary Fig.2

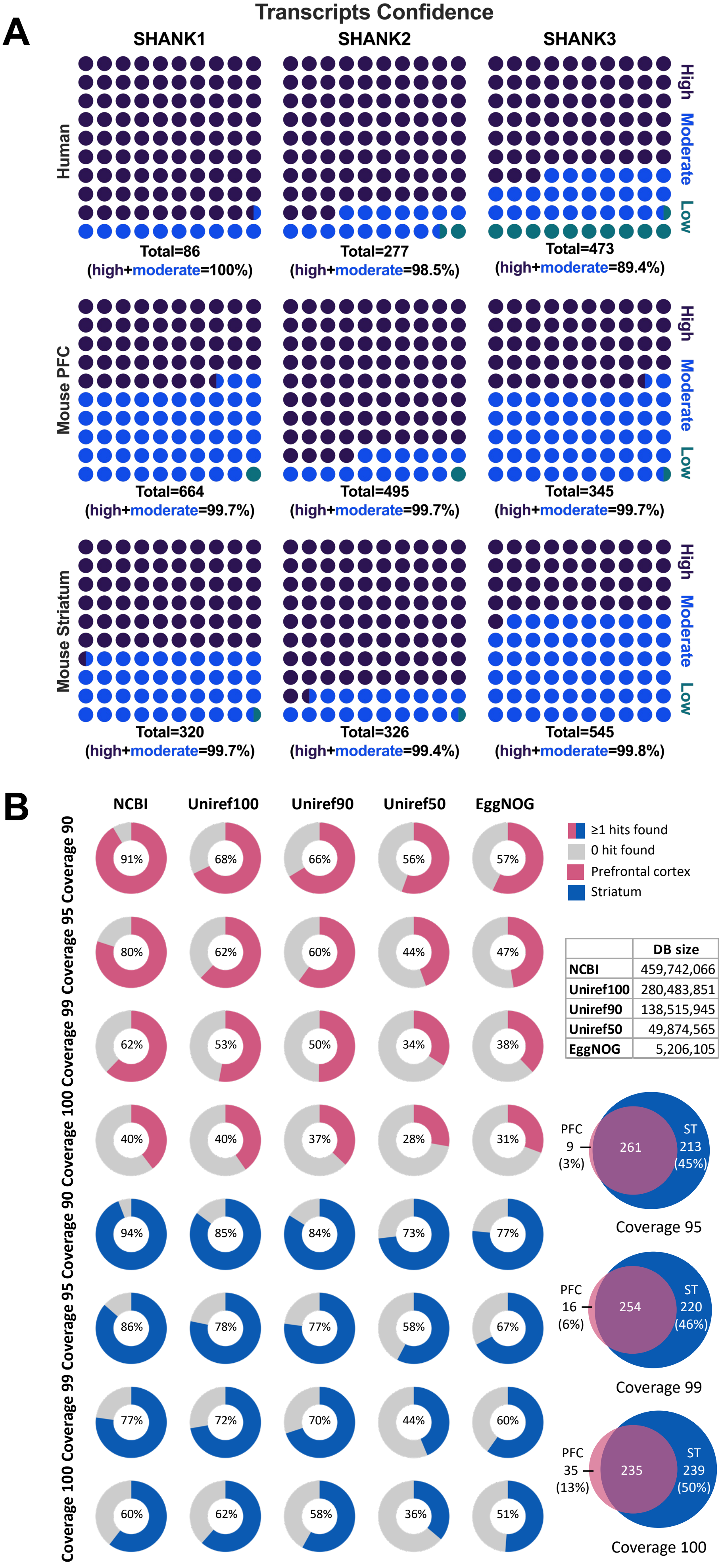


| | DB size |
|---|---|
| NCBI | 459,742,066 |
| Uniref100 | 280,483,851 |
| Uniref90 | 138,515,945 |
| Uniref50 | 49,874,565 |
| EggNOG | 5,206,105 |

**Supplementary Fig.2. In silico validation of SHANK family transcripts.**
**A. Confidence of CIS Transcripts of human and mouse *SHANK* family.** The scoring metrics (described in method section) were used to assess the confidence of individual transcripts of *SHANK* family genes detected by CIS in human and mouse brain samples. Black dots represent the transcripts scoring high confidence, while blue and green dots represent moderate and low confidence respectively. The percentage was shown. **Top:** The 100%, 98.5%, and 89.4% of *SHANK1, SHANK2*, and *SHANK3* transcripts of human cortex are in moderate to high confidence category respectively. **Middle:** The 99.7% of *Shank1*, *Shank2*, and *Shank3* transcripts in mouse PFC are in the category of moderate to high confidence. **Bottom:** The 99.7%, 99.4%, and 99.8% of *Shank1*, *Shank2*, and *Shank3* transcripts in mouse ST are in the category of moderate to high confidence.
**B. In silico validation of predicted SHANK3 ORFs in multiple protein databases.** The alignment of amino acid (AA) sequence of predicted ORFs of mouse PFC (pink) and striatum (blue) to NCBI, Uniref100, Uniref90, Uniref50 and EggNOG database. The higher alignment correlated significantly with the size of protein database (e.g., $R^2$ = 0.9161 and p=0.009, coverage=95 in PFC). The identity of ORFs between 2 brain regions varied with the parameters but 87% of ORFs detected in PFC were conserved with striatum even using the most rigorous criterion (coverage=100).

### Supplementary Fig. 3

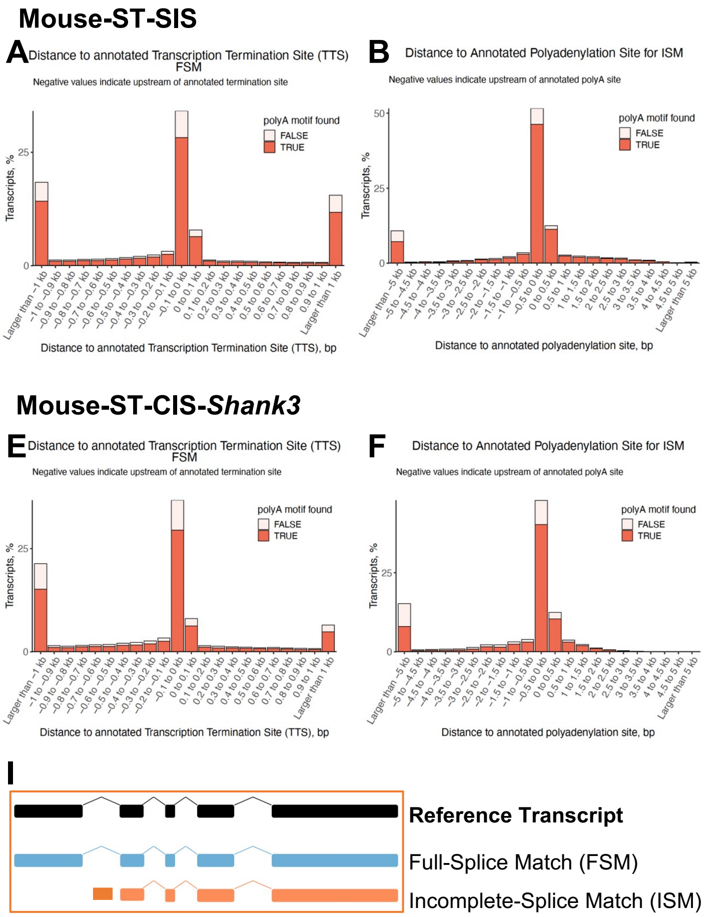

**Supplementary Fig. 3. Correlation of relative position of the end of transcript sequence reads from SIS and the annotated transcription termination site (TTS) and poly A signal for genome wide transcripts detected by SIS and *Shank3* transcripts by CIS.** The transcripts were classified as either Fully-Splice Match(FSM) or Incomplete-Splice Match (ISM) (i). The distance is between the end of transcript sequence read from SIS or CIS and annotated TTS and poly A site for individual transcript from SIS and CIS. The negative value indicates that the TTS or poly site is upstream of the end of transcript sequence read and the positive value indicates that the TTS or poly A site is downstream of the end of sequence read. The true is for then transcript with significant correlation and the false is no correlation.
**A.** ~25% of FSM transcripts have TTS within 0.1 kb upstream of the end of transcript sequence reads in mouse ST
**B.** ~50% of ISM transcripts have poly A signal within 0.5 kb upstream of the end of transcript sequence read in mouse ST
**C.** ~25% of FSM transcripts have TTS within 0.1 kb upstream of the end of sequence reads in mouse PFC
**D.** ~50% of ISM transcripts have poly A within 0.1 kb of the end of transcript sequence reads in mouse PFC
**E.** ~30% of FSM transcripts have TTS within 0.1 kb upstream of the end of transcript sequence reads in mouse ST
**F.** ~40% of ISM transcripts have poly A signal within 0.5 kb upstream of the end of transcript sequence read in mouse ST
**G.** ~30% of FSM transcripts have TTS within 0.1 kb upstream of the end of sequence reads in mouse PFC
**H.** ~40% of ISM transcripts have poly A within 0.1 kb of the end of transcript sequence reads in mouse PFC
**I.** Illustration of full-splice match (FSM) and incomplete-splice match (ISM) transcripts

### Supplementary Fig.4

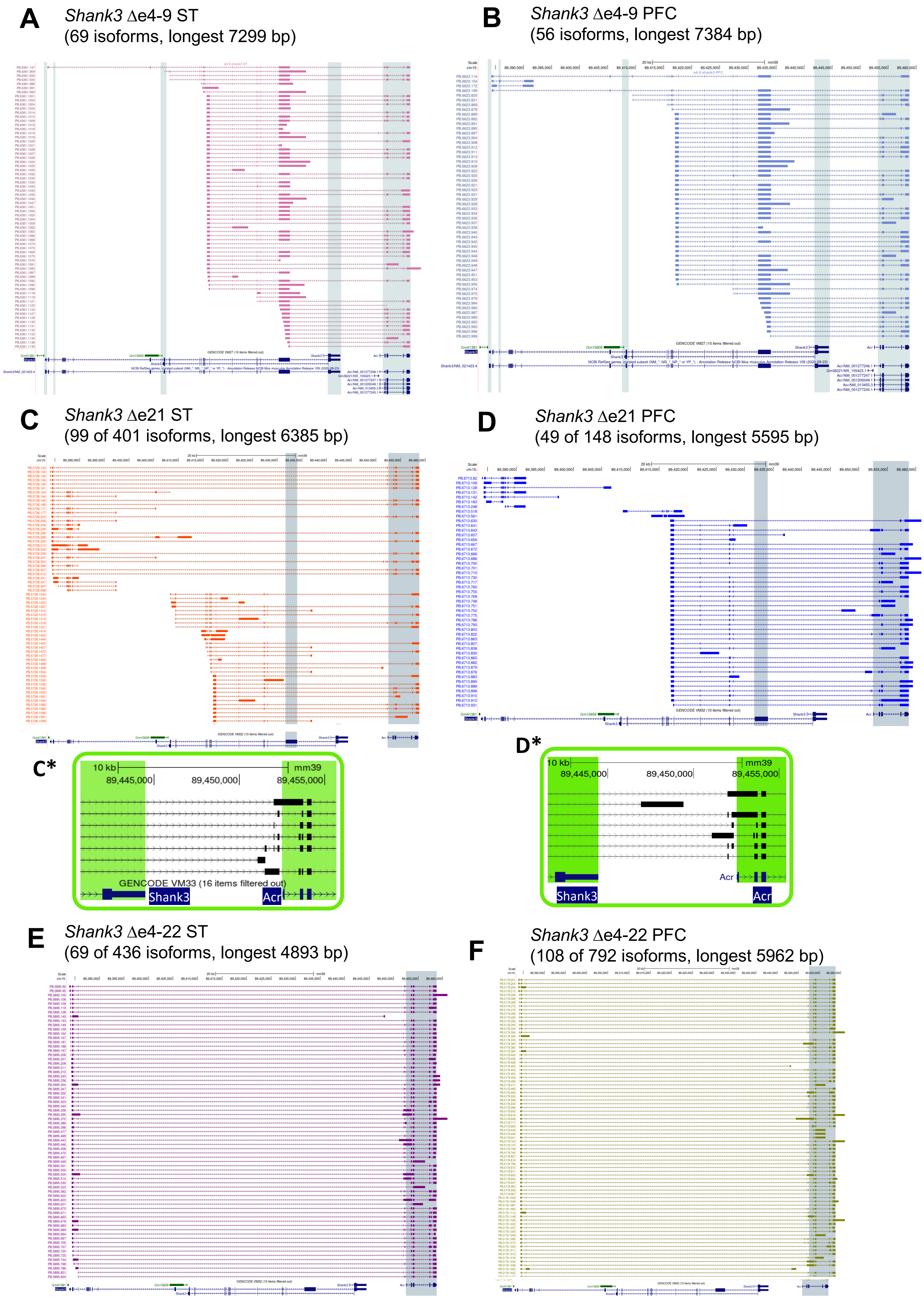

**Supplementary Fig. 4**
**Transcriptional map of *Shank3*$^{\Delta e4\text{-}9}$, *Shank3*$^{\Delta e21}$, and *Shank3*$^{\Delta e4\text{-}22}$ homozygous mice in ST and PFC by CIS**

**A**. Transcriptional map of *Shank3*$^{\Delta e4\text{-}9}$ homozygous mice in ST. A total of 69 different *Shank3* transcripts were detected by CIS. The deletion of exon 4-9 of *Shank3* was confirmed in the transcript PB 6361.147. The exon 3 was not detected in any of the transcripts. The long transcripts with exon 4-9 deletion did not include exons 1 and 22 (the last exon of annotated *Shank3*) indicating that a different TSS and poly A was used after exon 4-9 deletion. The transcripts starting exon 1 were most common in ST of WT but intron/exon 17 was the first exon for most of *Shank3* transcripts (48/69 transcripts) in *Shank3*$^{\Delta e4\text{-}9}$ mouse

**B.** Transcriptional map of *Shank3*$^{\Delta e4\text{-}9}$ homozygous mice in PFC. A total of 56 Shank3 transcripts were detected and that was 16% of that in WT PFC. In contrast with ST (a), the transcripts containing exon 3 were detected. Exon 11 was the first exon for many transcripts in ST but not in PFC. Same as in ST, a significantly higher percentage of fusion transcripts between *Shank3* and *Acr* genes was observed and the transcripts starting at intron16/exon 17 were predominant in *Shank3*$^{\Delta e4\text{-}9}$ mice (37/56 transcripts).

**C**. Transcriptional map of *Shank3*$^{\Delta e21}$ homozygous mutant mice in ST. A total of 401 *Shank3* transcripts with the longest transcript of 6385 bp were uncovered. About 85% (339/401) of *Shank3* transcripts were *Shank3* and *Acr* fusion transcripts in the ST of *Shank3*$^{\Delta e21}$. Similar to the *Shank3*$^{\Delta e4\text{-}9}$ mutant mouse, the intron 16/exon 17 became the predominant starting exon in the *Shank3*$^{\Delta e21}$ mouse but exon 1 is used as an alternative TSS. Among the 401 transcripts, 390 transcripts predicted 99 significant and unique ORFs

**D.** Transcriptional map of *Shank3*$^{\Delta e21}$ homozygous mutant in PFC. A total of 148 transcripts were detected. The longest transcript was 5595 bp. The transcripts with intron16/exon17 as the first exon accounted for 92% of residual transcripts and 87% (n=129) of these transcripts were fused with the *Acr* gene. Among 148 transcripts, 146 transcripts predicted 49 significant and unique ORFs.

**C*-D*.** Novel exons were found between *Shank3* and *Acr* in the fusion transcripts that were unique to *Shank3*$^{\Delta e21}$ mutant mouse. The expression of *Acr* exon 1 as part of the fusion transcripts, presented together with intron 1 retention of *Acr* in both ST and PFC was unique to *Shank3*$^{\Delta e21}$ mutant mouse.

**E.** Transcriptional map of *Shank3*$^{\Delta e4\text{-}22}$ homozygous mice in ST. A total of 436 transcripts were detected in the ST of *Shank3*$^{\Delta e4\text{-}22}$ homozygous mouse with the most extended transcript of 4893bp. Among the 69 ORFs detected in ST, 36 ORFs aligned to *SHANK3-ACR* fusion (n=35) or *SHANK3* only; 33 aligned to ACR only.

**F.** Transcriptional map of *Shank3*$^{\Delta e4\text{-}22}$ homozygous mice in PFC. A total of 792 transcripts were discovered in the PFC of *Shank3*$^{\Delta e4\text{-}22}$ homozygous mutant mouse compared to WT PFC. The most extended transcript was 5962bp. Among the 108 ORFs detected in PFC, 50 ORFs were aligned to *SHANK3-ACR* fusion transcripts (n=47) or *SHANK3* only; 58 aligned to ACR (n=56).

#### Supplementary Fig.5

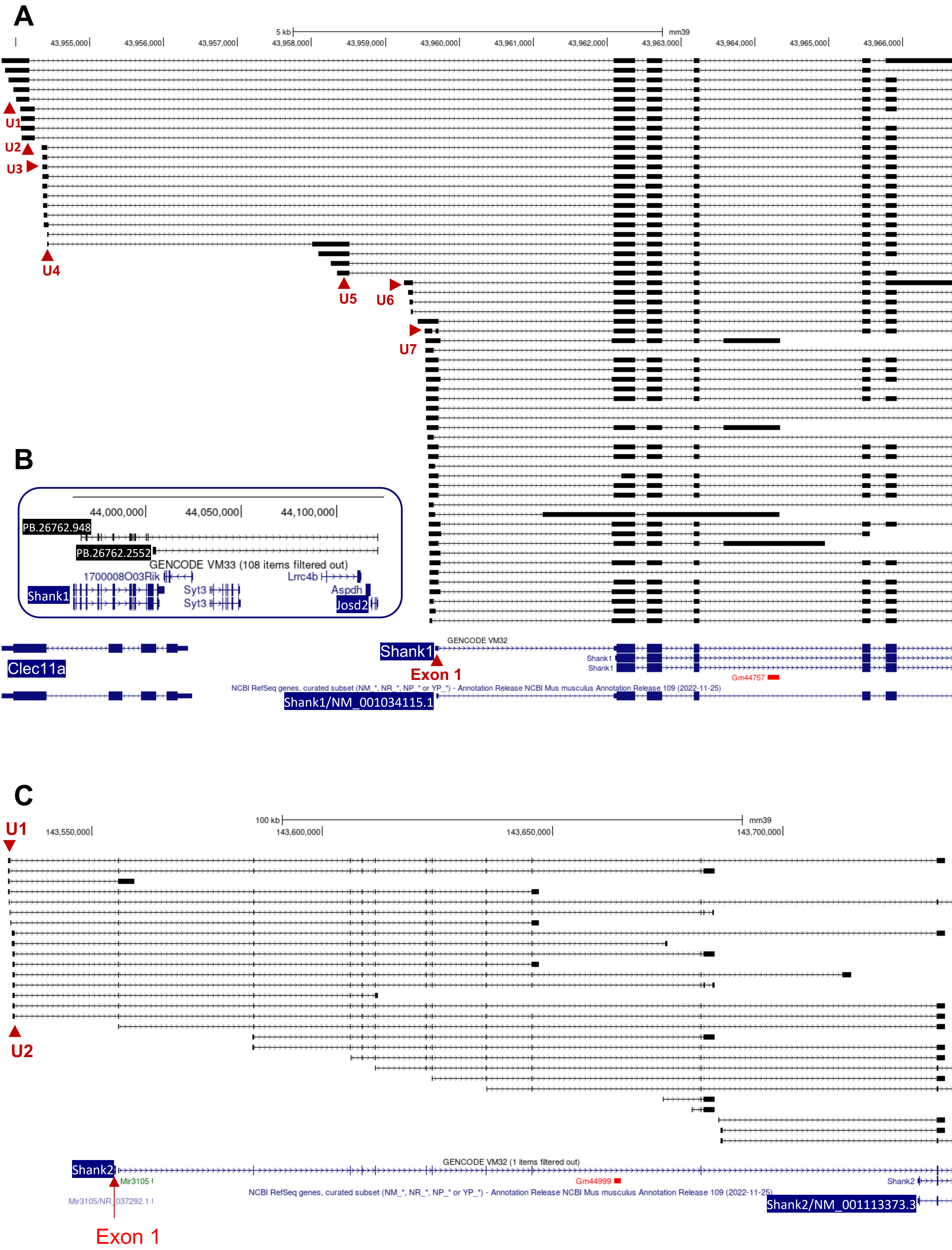

**Supplementary Fig. 5 The transcript structure of *Shank1 and Shank2* from CIS in mouse brains**

**A.** Seven novel exons (U1-7) found at upstream of annotated exon 1 of *Shank1* (NM_001034115.1) transcript. The most upstream novel exon (U1, chr7: 43953916-43954183, mm39) of *Shank1* overlapped with the last exon of annotated *Clec11a* gene (NM_009131.3) that was transcribed in an opposite direction of *Shank1* and extended the genome of *Shank1* by 5.76 kb**.**

**B.** Two novel fusion transcripts (PB.26762.948 and PB.26762.2552) between *Shank1* and *Josd2* gene, a gene ~100 kb downstream of *Shank1*, were detected in PFC only. Transcript PB.26762.948 aligned with exon6-24 of Shank1 and exon 5 (the last exon) of *Josd2* and encodes a 773 AA ORF. Transcript PB.26762.2552 aligned with exon 24 and a novel exon in intron 24 of *Shank1* and the last exon of Josd2 that encodes a 103AA ORF.

**C.** Two novel exons (U1 -2) found at upstream of exon 1 of annotated *Shank2* (NM_001113373.3) transcript. The most upstream novel exon (U1: chr7:143531825-143532147, mm39) of *Shank2* extended the genome of *Shank2* by 23.84 kb.

### Supplementary Fig.6

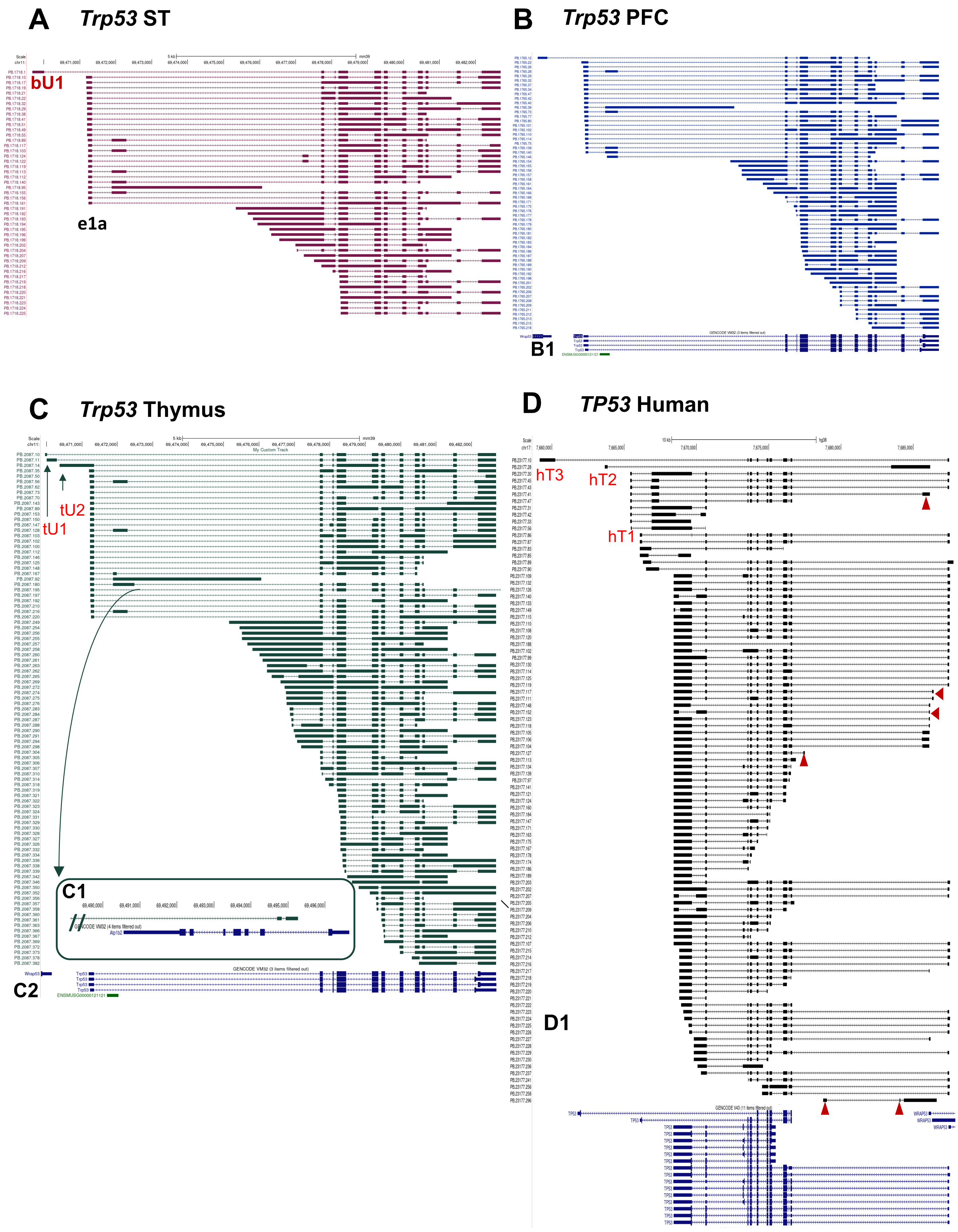

**Supplementary Fig.6 *Trp53* transcripts detected by CIS in mouse ST and PFC and human TP53 detected by CIS in human brain**

**A-B.** *Trp53* transcripts detected by CIS in mouse ST and PFC using mouse *Trp53* probe panel.

A brain specific novel exon (bU1, 451 bp/chr11:69469678-694700129, mm39) at 1458 bp upstream of current annotated exon1 of *Trp53* was detected in both ST (a. red) and PFC of mice (b. blue) The bU1 exon was predicted to have a new ATG that extend an extra 88AA to the existing ORF. A novel exon 1a (441bp/chr11: 69471880-69472291, mm39) was observed in both ST and PFC that were predicted as a new TSS. In addition to annotated exon 1 of *Trp53* as canonical TSS, additional TSS were predicted in the intron 1, exon 2 and exon 4 based on the structure of the transcripts. Intron retention was widely observed in both brain regions of mouse.

**B1**. Annotated *Trp53* transcript structure in genome browser

**C.** *Trp53* transcripts detected with CIS in thymus tissue using mouse *Trp53* probe panel. Two new exons (tU1: 71bp/chr11:69469941-69470012 and tU2, chr11: 285bp/69470001-69470286, mm39) at 5′ were identified in thymus but not in brain that suggest tissue specific transcription start site (TSS). The first TSS extended the gene structure of *Trp53* by 1.24 kb.

**C1.** Two 3′ new exons (tT1, chr11: 69494709-69494832 and tT2, chr11: 69494939-69495271, mm39) were found after the last exon of annotated *Trp53* in thymus but not in PFC and ST. The tT2 exon was 12kb downstream of the last exon and extended the ORF of Trp53 by 42 AA.

**C2**. Annotated *Trp53* transcript structure in genome browser

**D.** Human *TP53* transcripts detected in human brain using human *TP53* probe panel by CIS. Three novel exons (hT1-3) were detected on the 3′ of *TP53* in human brain. The farthest exon (hT3, chr17:7659158-7660233, hg38) is 1.5 kb from the last exon of current annotated *TP53* in human. We also detected 6 novel exons (red arrows). The intron retention was significantly less frequently in human brain compared to mice.

**D1.** Annotated human *TP53* in genome browser (NM_000546, hg19)

### Supplementary Fig. 7

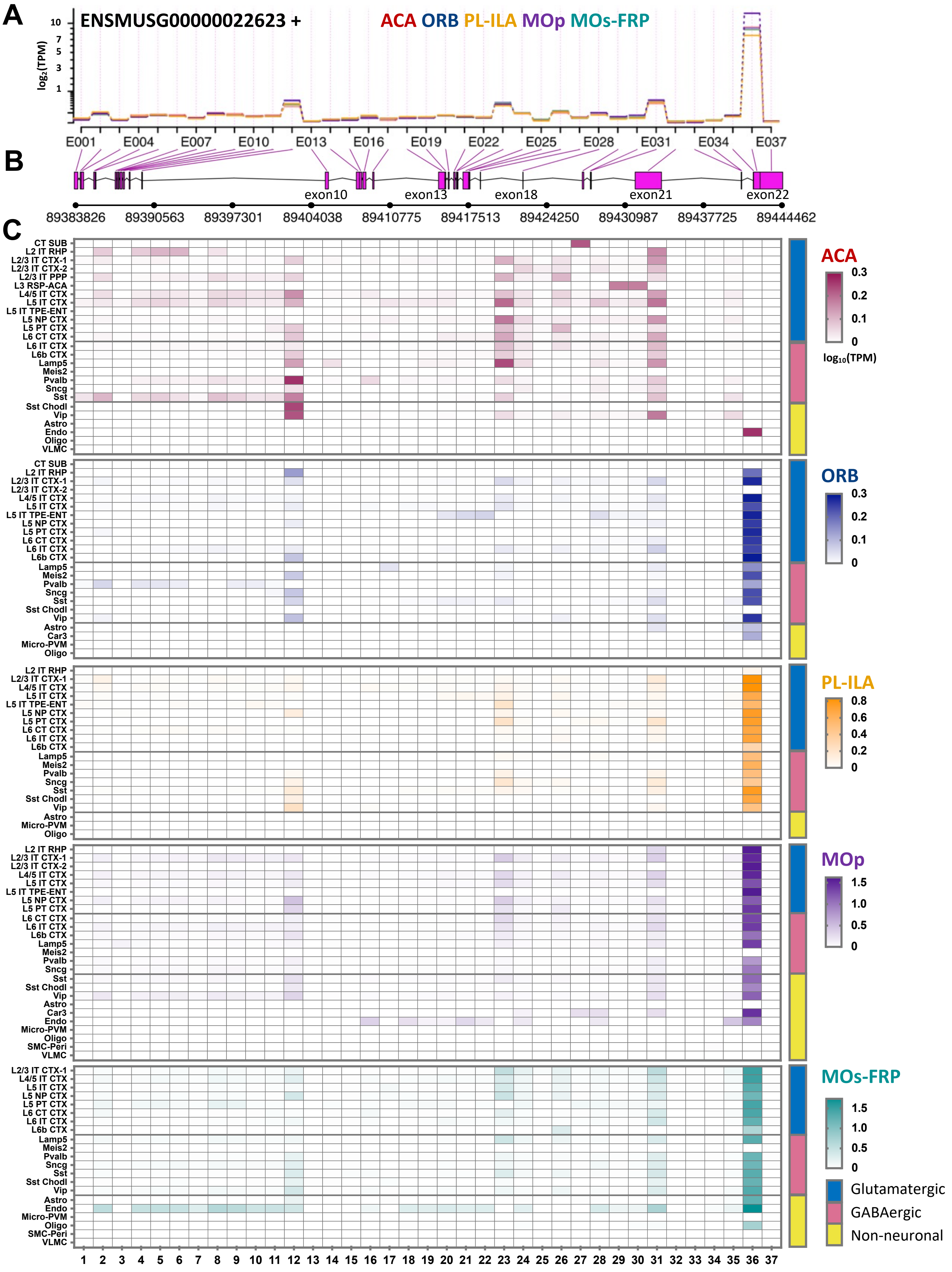

**Supplementary Fig.7 *Shank3* transcripts from CIS in mouse brain single cell data.**
A. The splicing events corresponding to the exons of *Shank3* transcripts from CIS**.** sc-RNA-seq dataset[5] of 5 sub-regions (ACA, ORB, PL-ILA, MOp, MOs-FRP) of WT mouse cortex were analyzed to evaluate the exon usage using DEXSeq[6]. The numbers listed (E001-E037) were splicing events rather annotated exons (exon 1-22). The splicing event could occur within an annotated exon. It is a more general concept than alternative splicing, since it also includes changes in the usage of alternative transcript start sites and polyadenylation sites, which can cause differential usage of exons at the 5′ and 3′ transcripts. A line connecting individual splicing event to corresponding annotated exons.
**B.** The exons of *Shank3* that expressed significantly different statistically among different brain regions in pink. The black boxes are annotated exons of mouse *Shank3* gene, and the coordinates were referred to mm39).
**C.** Cell type specificity for the single splicing event and exon usage. For example, the ACA tends to use the splicing event 12 (corresponding to annotated exon 9), 23 (exon15) and 31(exon 20). In ACA, only the endothelial cells expressed the splicing site 36 (exon 22). However, the splicing site 36 was preferred in most of other cell types of other 4 brain regions. The exon 22 of *Shank3* annotated transcript was significantly more represented in 4 sub-regions of cortex except in ACA.  The exon 22 fusion with *Acr* gene were found predominately in ACA. The exon 10, 17 and 21 were more represented in ACA than that other 4 sub-regions in the majority of cell types.

ACA: Anterior cingulate, ORB: Orbital area, PL-ILA: Prelimbic-infralimbic area, MOp: Primary motor area, MOs-FRP: Secondary motor-frontal pole.

#### Supplementary Fig. 8

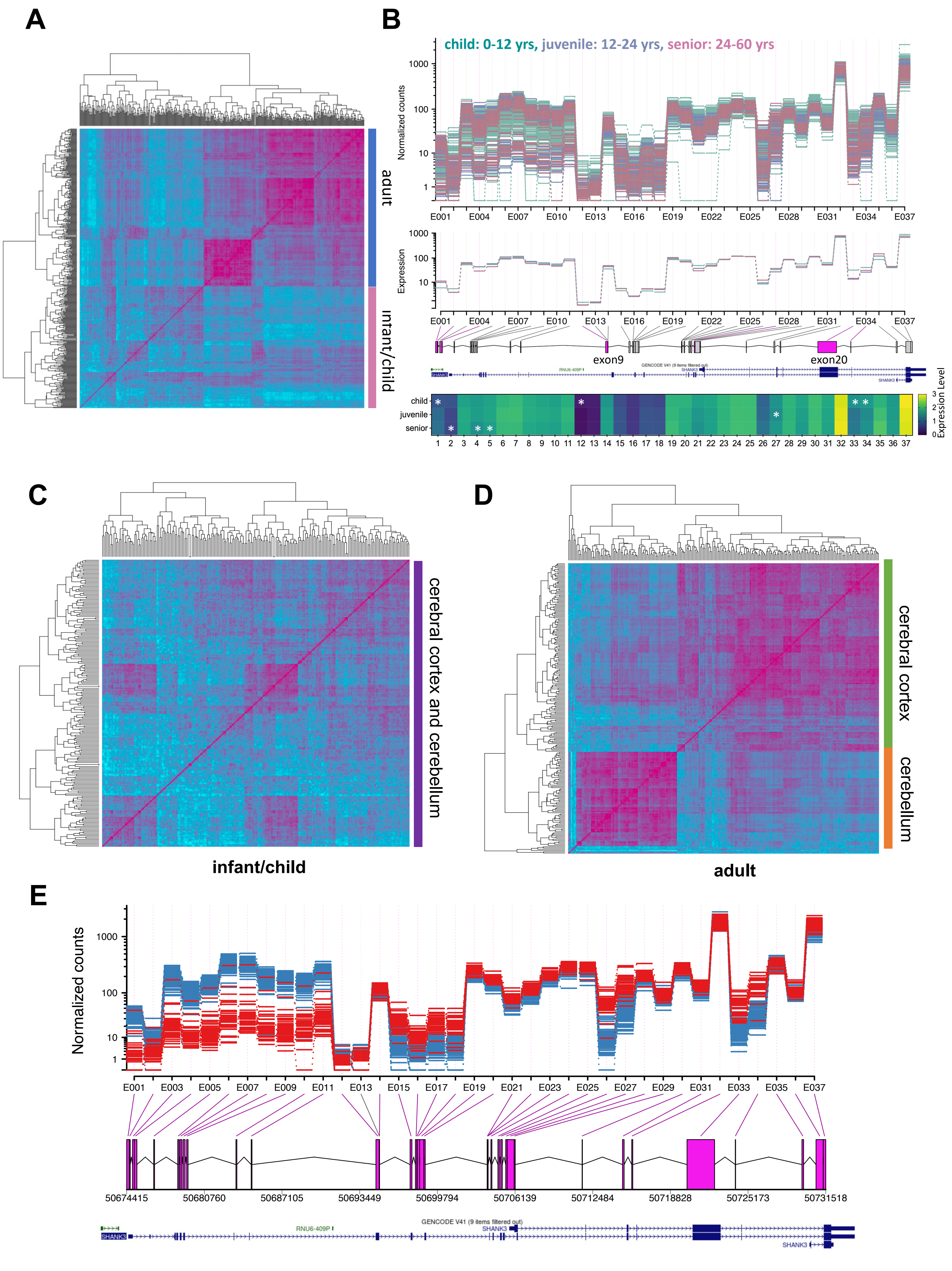

**Supplementary Fig. 8 Development and brain region-specific expression of *SHANK3* transcripts in human brains**

**A.** *SHANK3* transcripts and exon usage changed with age in human brains. Short-read RNA-seq data of normal human cortex samples from PsychENCODE [UCLA group (n=253) and YALE groups (n=227)] were aligned to human CIS *SHANK3* transcripts. The pattern of *SHANK3* transcripts in infants and children was different from adults.

**B.** The exon usage of *SHANK3* transcripts was significantly different among the child (0-12 years), juvenile (12 to 24 years), and adult/senior (24-60 years) groups. The 3′ exons were more abundant than that of 5′ exons of the transcripts. The transcripts contacting exon 1 of annotated *SHANK3* was more abundant in developing brains, while the 5′ transcripts terminated prior to exon 9 were more prevalent in aging brains.

**C-D.** Human *SHANK3* transcripts and exon usage in different brain regions**.** Short read RNA-seq data of normal brain tissues from PsychENCODE[7] were aligned to *SHANK3* transcripts from CIS.

**C**. The heatmap showed no brain region specificity was found among prefrontal cortex, temporal cortex and cerebellum of children (mean age 3.5 years, n=227). **D.** The heatmap showed no brain region specificity was found between cerebral cortex and cerebellum in adult brains (mean age 31 years, n=253).

**E.** The exon usages of *SHANK3* in cerebellum and cerebral cortex. In normal cerebellum, the 3′ exons were more transcribed than the 5′ exons of *SHANK3.* In normal cerebral cortex, the 5’ exons of *SHANK3* were more transcribed.
