## Supplementary Table 1 for "Transcriptional Determinism and Stochasticity Contribute to the Complexity of Autism Associated *SHANK* Family Genes"

**Supplementary Table 1a. The probe design for human SHANK3 capture**

| Chromosome | Start | Stop | Name | Score | Strand | Code | Seq | Len |
|---|---|---|---|---|---|---|---|---|
| chr22 | 50682617 | 50682737 | 785447_45746696_NM_001372044.2_83 | 44 | + | good | GGAGTGTGGATATACTTGCCTGTTCTGGGGGTGTACACGTGTGTGTGCACACAGGTGACCCTGTACAGTGATTGCATGCGTGCACCAGGGAGTGTGGATATACTTGCCTGTTCTGGGGGT | 120 |
| chr22 | 50679977 | 50680097 | 785447_45746696_NM_001372044.2_61 | 42 | + | good | TGCGTGCACCAGGTAGTGTGGATATACTTGCCTGTTCTGGGGGTGTACACGTGTGTTTGCACACAGATGACCTTGTACAGTGATTGCATGCGTGCACCAGGGAGTGTGGATATACTTGTC | 120 |
| chr22 | 50681297 | 50681417 | 785447_45746696_NM_001372044.2_72 | 42 | + | good | AGGTAGTGTGGATATACTTGCCTGTTCTGGGGGTGTACACGTGTGTTTGCACACAGATGACCCTGTACAGTGATTGCATGCGTGCACCAGGGAGTGTGGATATACTTGCCTGTTCTGGGG | 120 |
| chr22 | 50681657 | 50681777 | 785447_45746696_NM_001372044.2_75 | 42 | + | good | GGAGTGTGGATATACTTGCCTGTTCTGGGGGTGTACACGTGTGTGCACACAGATGACCCTGTACAGTGATTGTACACGTGTACCAGGGAGTGTGGATATACTTGCCTGTTCTGGGGGTGT | 120 |
| chr22 | 50682257 | 50682377 | 785447_45746696_NM_001372044.2_80 | 42 | + | good | CGTGTACCAGGGAGTGTGGATATACTTGCCTGTTCTGGGGGTGTACATGTGTGTGCACACAGATGACCCTGTACAGTGATTGCATGCGTGCACCAGGGAGTGTGGATATACTTGCCTGTT | 120 |
| chr22 | 50679617 | 50679737 | 785447_45746696_NM_001372044.2_58 | 40 | + | good | TGATTGCATGCGTGCACCAGGGAGTGTGGATATACTTGCCTGTTCTGGGGTTGTACATGTGTGTGCACACAGATGACCCTGTACAGTGATTGTATGCGTGCACCAGTGAGTGTGGATATA | 120 |
| chr22 | 50679737 | 50679857 | 785447_45746696_NM_001372044.2_59 | 40 | + | good | CTTGCCTGTTCTGGGGGTGTACGTGTGTTTGTGTGCACACAGGTGACCCTGTACAGTGATTGCATGTGTGCACCAGGGAGTGTGGATATACTTGCCTGTTCTGGGGGTGTACACGTGTGT | 120 |
| chr22 | 50680697 | 50680817 | 785447_45746696_NM_001372044.2_67 | 40 | + | good | TGGATATACTTGTCTGTTCTGGGGGTGTACACGTGTGTTTGCACACAGATGACCTTGTACAGTGATTGCATGCGTGCACCAGGGAGTGTGGATATACTTGCCTGTTCTGGGGGTGTACAT | 120 |
| chr22 | 50680937 | 50681057 | 785447_45746696_NM_001372044.2_69 | 40 | + | good | CATGCGTGCACCAGGGAGTGTGGATATACTTGCCTGTTCTGGGGGTGTACATGTGTGTGCACACAGATGACCCTGTACACTGATTGCATGCGTGCACCAGGGAGTGTGGATATACTTGCC | 120 |
| chr22 | 50681177 | 50681297 | 785447_45746696_NM_001372044.2_71 | 40 | + | good | AGATGACCCTGTACAGTGATTGTATGCGTGTACCAGGGAGTGTGGATATACTTGCCTGTTCTGGGGGTGTACATGTGTGTGCACACAGATGACCCTGTACAGTGATTGTATGCGTGCACC | 120 |
| chr22 | 50681537 | 50681657 | 785447_45746696_NM_001372044.2_74 | 40 | + | good | CCCTGTACAGTGATTGTACACGTGTACCAGGGAGTGTGGATATACTTGCCTGTTCTGGGGGTGTACGTGTGTTTGTGTGCACACAGGTGACCCTGTACAGTGATTGCATGCGTGCACCAG | 120 |
| chr22 | 50681897 | 50682017 | 785447_45746696_NM_001372044.2_77 | 40 | + | good | TGATTGTACACGTGTACCAGGGAGTGTGGATATACTTGCCTGTTCTGGGGGTGTACACGTGTGTTTGCACACAGATGACCCTGTACAGTGATTGTACACGTGTACCAGGGAGTGTGGATA | 120 |
| chr22 | 50682497 | 50682617 | 785447_45746696_NM_001372044.2_82 | 40 | + | good | GATCCTGTACAGTGATTGCATGCGTGCACCAGGGAGTGTGGATATACTTGCCTGTTCTGGGGGTGTACACATGTGTTTGCACACAGATGACCCTGTACAGTGATTGCATGCGTGCACCAG | 120 |
| chr22 | 50682857 | 50682977 | 785447_45746696_NM_001372044.2_85 | 40 | + | good | AGTGATTGCATGCGTGCACCAGGGAGTGTGGAAATACTTGCCTGTTCTGGGGGTGTACACGTGTGTGTGCACACAGATGACCCTGTACAGTGATTGTGTATGTGCATCCCTGCCTCTGTG | 120 |
| chr22 | 50679857 | 50679977 | 785447_45746696_NM_001372044.2_60 | 39 | + | good | TTGCACACAGATGACCCTGTACAGTGATTGCATGCGTGCACCAGGGAGTGTGGATATACTTGCCTGTTCTGGGGGTGTACACGTGTGTGTGCACACAGATGACCCTGTACAGTGATTGCA | 120 |
| chr22 | 50680097 | 50680217 | 785447_45746696_NM_001372044.2_62 | 39 | + | good | TGTTCTGGGGGTGTACACGTGTGTTTGCACACAGATGACCCTGTACAGTGATTGCATGCGTGCACCAGGGAGTGTGGATATACTTGCCTGTTCTGGGGGTGTACACGTGTGTTTGCACAC | 120 |
| chr22 | 50680217 | 50680337 | 785447_45746696_NM_001372044.2_63 | 39 | + | good | AGGTGACCCTGTACAGTGATCGTACACGTGTACCAGGGAGTGTGGATATACTTGCCTGTTCTGGGGTTGTACGTGTGTGTGTGCACACAGATGACTCTGTACAGTGATTGCATGCGTGCA | 120 |
| chr22 | 50680337 | 50680457 | 785447_45746696_NM_001372044.2_64 | 39 | + | good | CCAGGTAGTGTGGATATACTTGTCTGTTCTGGGGGTGTACACGTGTGTTTGCACACAGATGACCTTGTACAGTGATTGCATGCGTGCACCAGGGAGTGTGGATATACTTGCCTGTTCTGG | 120 |
| chr22 | 50680457 | 50680577 | 785447_45746696_NM_001372044.2_65 | 39 | + | good | GGGTGTACATGTGTGTGCACACAGATGACCCTGTACAGTGATTGCATGCGTGCACCAGGGAGTGTGGATATACTTGCCTGTTCTGGGGGTGTACACGTGTGTTTGCACACAGGTGACCCT | 120 |
| chr22 | 50680577 | 50680697 | 785447_45746696_NM_001372044.2_66 | 39 | + | good | GTACAGTGATTGTACACGTGTACCAGGGAGTGTGGATATACTTGCCTGTTCTGGGGTTGTACGTGTGTGTGTGCACACAGATGACTCTGTACAGTGATTGCATGCGTGCACCAGGTAGTG | 120 |
| chr22 | 50680817 | 50680937 | 785447_45746696_NM_001372044.2_68 | 39 | + | good | GTGTGTGCACACAGATGACCCTGTACACTGATTGCATGCGTGCACCAGGGAGTGTGGATATACTTGCCTGTTCTGGGGGTGTACATGTGTGTGCACACAGATGACCCTGTACAGTGATTG | 120 |
| chr22 | 50681417 | 50681537 | 785447_45746696_NM_001372044.2_73 | 39 | + | good | TTGTACATGTGTCTGTGTGCACACAGATGACTCTGTACAGTGATTGTATGCATGTACCAGTGAGTGTGGATATACTTGTCTGTTCTGGGGGTGTACGTGTGTTTGTGTGCACACAGGTGA | 120 |
| chr22 | 50682017 | 50682137 | 785447_45746696_NM_001372044.2_78 | 39 | + | good | TACTTGCCTGTTCTGGGGGTGTACATGTGTGTGCACACAGATGACCCTGTACAGTGATTGCATGCGTGCACTAGGGAGTGTGGATATACTTGCCTGTTCTGGGGGTGTACACATGTGTTT | 120 |
| chr22 | 50682137 | 50682257 | 785447_45746696_NM_001372044.2_79 | 39 | + | good | GCACACAGATGACCCTGTACAGTGATTGTACACGTGTATCAGGGAGTGTGGATATACTTGCCTGTTCTGGGGGTGTACACGTGTGTTTGCACACAGATGACCCTGTACAGTGATTGTACA | 120 |
| chr22 | 50682377 | 50682497 | 785447_45746696_NM_001372044.2_81 | 39 | + | good | CTGGGGGTGTACACGTGTGTGTGCACACAGGTGACCCTGTACAGTGATTGCATGCGTGCACCAGGGAGTGTGGATATACTTGCCTGTTCTGGGGGTGTACACGTGTGTGTGCACACAGGT | 120 |
| chr22 | 50682737 | 50682857 | 785447_45746696_NM_001372044.2_84 | 39 | + | good | GTACACGTGTGTGTGCACACAGGTGACCCTGTACAGTGATTGCATGCGTGCACCAGGGAGTGTGGATATACTTGCCTGTTCTGGGGGTGTACATGTGTGTGCACACAGGTGACCCTGTAC | 120 |
| chr22 | 50681057 | 50681177 | 785447_45746696_NM_001372044.2_70 | 38 | + | good | TGTTCTGGTGTTGTACATGTGTGTGTGTGCACACAGATGACTCTACAGTGATTGTATGCGTGCAGCAGGTAGTGTGGATATACTTGCCTGTTCTGGGGGTGTACACGTGTGTTTGCACAC | 120 |
| chr22 | 50681777 | 50681897 | 785447_45746696_NM_001372044.2_76 | 38 | + | good | ACACGTGTGTTTGCACACAGATGACCCTGTACAGTGATTGTACACGTGTACCAGGGAGTGTGGATATACTTGCCTGTTCTGGGGGTGTACACGTGTGTGCACACAGATGACCCTGTACAG | 120 |
| chr22 | 50679497 | 50679617 | 785447_45746696_NM_001372044.2_57 | 34 | + | good | TGTGTGTGTGGGCACACAGGTGACCCTGTACGGTGATTGCATGTGTGCACCGAGTGTGGATATACTTGCCTGTTCTGGGGGTGTACGTGTGTTTGTGTGCACACAGGTGACCCTGTACAG | 120 |
| chr22 | 50686217 | 50686337 | 785447_45746696_NM_001372044.2_113 | 33 | + | good | CCGGAGTCCCCTCTGGGAGAGTTGTAGCTGGCCAGGCTGGGACCGTCCTGTTGTAGCTGCCGGAGTCCCCTCTGGGAGAGTTGCTGCTGGCCAGGCTGGGACTGTCCTGTTGTAGCTGCC | 120 |
| chr22 | 50685497 | 50685617 | 785447_45746696_NM_001372044.2_107 | 32 | + | good | GTTGTAGCTGCCGGAGTCCCCTCTGGGAGAGTTGCTGCTGGCCAGGCTGGGACCGTCCTGTTGTAGCTGCCGGAGTCCCCTCTGGGAGAGTTGTAGCTGCCGGAGTCCCCTCTGGGAGAG | 120 |
| chr22 | 50685737 | 50685857 | 785447_45746696_NM_001372044.2_109 | 32 | + | good | GTAGCTGCCGGAGTCCCCTCTGGGAGAGTTGCTGCTGGCCAGGCTGGGATGGTCCTGTTGTAGCTGCCGGAGTCCCCTCTGGGAGAGTTGCTGCTGGCCAGGCTGGGACCGTCCTGTTGT | 120 |
| chr22 | 50685617 | 50685737 | 785447_45746696_NM_001372044.2_108 | 31 | + | good | TTGCTGCTGGCCAGGCTGGGACGGTCCTGTTGTAGCTGCCGGAGTCCCCTCTGGGAGAGTTGTAGCTGGCCAGGCTGGGACCGTCCTGTTGTAGCTGCCGGAGTCCCCTCTGGGAGAGTT | 120 |
| chr22 | 50685857 | 50685977 | 785447_45746696_NM_001372044.2_110 | 31 | + | good | AGCTGCTGGAGTCCCCTCTGGGAGAGTTGTAGCTGCCGGAGTCCCCTCTGGGAGAGTTGCTGCTGGCCAGGCTGGGACCGTCCTGTTGTAGCTGCCGGAGTCCCCTCTGGGAGAGTTGCT | 120 |
| chr22 | 50686457 | 50686577 | 785447_45746696_NM_001372044.2_115 | 31 | + | good | AGTCCCCTCTGGGAGAGTTGTAGCTGCCGGAGTCCCCTCTGGGAGAGTTGCTGCTGGCCAGGCTGGGACGGTCCTGTTGTAGCTGCCGGAGTCCCCTCTGGGAGAGTTGCTGCTGGCCAG | 120 |
| chr22 | 50686577 | 50686697 | 785447_45746696_NM_001372044.2_116 | 31 | + | good | GCTGGGACGGTCCTGTTGTAGCTGCCGGAGTCCCCTCTGGGAGAGTTGTAGCTGGCCAGGCTGGGACCGTCCTGTTGTAGCTGCCGGAGTCCCCTCTGGGAGAGTTGCTGCTGGCCAGGC | 120 |
| chr22 | 50685377 | 50685497 | 785447_45746696_NM_001372044.2_106 | 30 | + | good | GAGTTGCTGCTGGCCAGGCTGGGACCGTCCTGTTGTAGCTGCCGGAGTCCCCTCTGGGAGAGTTGTAGCTGCCGGAGTCCCCTCTGGGAGAGTTGCTGCTGGCCAGGCTGGGACGGTCCT | 120 |
| chr22 | 50686097 | 50686217 | 785447_45746696_NM_001372044.2_112 | 30 | + | good | TGCCGGAGTCCCCTCTGGGAGAGTTGCTGCTGGCCAGGCTGGGACCGTCCTGTTGTAGCTGCCGGAGTCCCCTCTGGGAGAGTTGCTGCTGGCCAGGCTGGGACCATCCTGTTGTAGCTG | 120 |
| chr22 | 50686697 | 50686817 | 785447_45746696_NM_001372044.2_117 | 30 | + | good | TGGGACCGTCCTGTTGTAGCTGCCGGAGTCCCCTCTGGGAGAGTTGTAGCTGCCGGAGTCCCCTCTGGGAGAGTTGCTGCTGGCCAGGCTGGGACCGTCCTGTTGTAGCTGCCGGAGTCC | 120 |
| chr22 | 50685977 | 50686097 | 785447_45746696_NM_001372044.2_111 | 26 | + | good | GCTGGCCAGGCTGGGACGGTCCTGTTGTAGCTGCCGGAGTCCCCTCTGGGAGAGTTGCTGTTGGCCAGGCTGGGACCGTCCTGTTGTAGCTGCCGGAGTCCCCTCTGGGAGAGTTGTAGC | 120 |
| chr22 | 50686337 | 50686457 | 785447_45746696_NM_001372044.2_114 | 26 | + | good | GGAGTCCCCTCTGGGAGAGTTGCTGCTGGCCAGGCTGGGACCGTCCTGTTGTAGCTGCCGGAGTCCCCTCTGGGAGAGTTGCTGCTGGCCAGGCTGGGACTGTCCTGTTGTAGCTGCCGG | 120 |
| chr22 | 50686817 | 50686937 | 785447_45746696_NM_001372044.2_118 | 24 | + | good | CCTCTGGGCGAGTTGCTGCTGGCCAGGCTGGGACGGTCCTGTTGTAGCTGCCGGAGTCCCCTCTGGGAGAGTTGCTGCTGGCCAGGCTGGGACCATCCTGTTGTAGCTGCCGGAGTCCCC | 120 |
| chr22 | 50685257 | 50685377 | 785447_45746696_NM_001372044.2_105 | 22 | + | good | GCCAGGATACCCGAGCCTCTTGGAGGTCTCTGGTGGGTGTGGGGCCAGGCTGGGACCGTCCTGTTGTAGCTGCCGGAGTCCCCTCTGGGAGAGTTGTAGCTGCCGGAGTCCCCTCTGGGA | 120 |
| chr22 | 50716937 | 50717057 | 785447_45746696_NM_001372044.2_369 | 22 | + | good | GCTGGACACTGAGCATTTAATACTTAGAATAACCATATTGAGGTTGAGACCACTAGTACCTCCATTTTACAGATGAGAAAACTGAAGCAAGGAGGGTAAATGACCTGCCCAAGATCACAC | 120 |
| chr22 | 50694257 | 50694377 | 785447_45746696_NM_001372044.2_180 | 17 | + | good | GTGGTGGTTTTCATGCTGCGGTGGCAGAGTTGAGTCATTGCAACACAGACTATATAGCTTGCAAAGCCTTAAGTAGTTACTGTCCAATTCTTTCCAGAAAAGGTTTGCTGATCCCTGGTC | 120 |
| chr22 | 50691857 | 50691977 | 785447_45746696_NM_001372044.2_160 | 16 | + | good | TTTCTATTTGTTGGCATGTAATTGTTCATAGTGTTATATTCCTTTTTAACTTCTGTAACACTGATAGTAAGTCCCCAAATTTCATTTGTGATTTTAGTTATTTCTGTCCATCTTCTCTCC | 120 |
| chr22 | 50675537 | 50675657 | 785447_45746696_NM_001372044.2_24 | 11 | + | good | CATGTGTAGTGAGCAGGCAGTGTGTGCAGTTAGCAGGCACAGTGTGTGCAGTGAGTGGGCAGTGTGCACAGCATGTACAGTGTGAGTGGTGTGTGCTGTGTGCCGTGAGCATGTGTAGTG | 120 |
| chr22 | 50690537 | 50690657 | 785447_45746696_NM_001372044.2_149 | 11 | + | good | AACTTTTGAAATCAGGAGGTTTGAGTGCTTCAACTTGTTCTTATTTGTTATCGTTTTGGCTAATCAGGGCCCCTGGCAATTCCATATAGACTTGAACATTGCTTTTCTATTTGTGCAAAA | 120 |
| chr22 | 50693057 | 50693177 | 785447_45746696_NM_001372044.2_170 | 10 | + | good | TAATTAGGTTATTGATTTGAGATTGTTTTCCTTTTTAAAAGTAAGTGTTTACAGCGTTGTTTCCCTTTAAGTACTGCTTTTGCTCCATATGTTGTATTTTTGTTCTTGTTTGTCTTAAAG | 120 |
| chr22 | 50710937 | 50711057 | 785447_45746696_NM_001372044.2_319 | 7 | + | good | GACACGTGCTGTGTGCACCTGCATGCGTGTGGCACACGTGTGTGCATGGGTGAGCGGATACGCACTGTGTGCGCCTGCATGCGTGTGGCACACATGTCTGTGCGTGGGTGAGCAGACGCG | 120 |
| chr22 | 50711057 | 50711177 | 785447_45746696_NM_001372044.2_320 | 7 | + | good | CTGTGTGCACCTGCATGCGTGTGGCACACGTGTGTGCGTGGGTGAGCGGATACGCGCTGTATGCACCTGCGTGCATGCAGCACACGTCTGTGCGTGGGTGAGCGGATACGCGCTGTGTGC | 120 |
| chr22 | 50711177 | 50711297 | 785447_45746696_NM_001372044.2_321 | 7 | + | good | GCCTGCATGCGTGTGGCACACGTCTGTGCGTGGGTGAGCGGATATGCAGTGTGCGCCTGCATGCGTGTGGCACACGTGTGTGCGTGGGTGAGCGGATACGCGCTGTGTGCCCCTGCGTGC | 120 |
| chr22 | 50694617 | 50694737 | 785447_45746696_NM_001372044.2_183 | 6 | + | good | TTCATTTGTAGGGTGGCCAACTCTTTTGGTTTGCCTGGAACCAAGGCAGTTCCTGGAACATAGGATTTAGTTTTAACATCTGGAAAGCCTGGGCAAACTGGACAAGTTGGTCACCCAGCT | 120 |
| chr22 | 50674337 | 50674457 | 785447_45746696_NM_001372044.2_14 | 5 | + | good | TTCCCTCCCGTCTCAGGCCCCCTCCCCCCGCCGCCCCCGCCCCCGGGGAAGGCAGGCGCCGAGCTGAGCCGGGGCCGATGCAGCTGAGCCGCGCCGCCGCCGCCGCCGCCGCCGCCCCTG | 120 |
| chr22 | 50683817 | 50683937 | 785447_45746696_NM_001372044.2_93 | 4 | + | good | GGGATGGTGGAGGGGGGTGGTGGAGAGGGGGTGGTGGAGAGGGGGGTGGTGGAGAGGGGTGTGATCCTAGCCGCTGATGTAATTCAGGGGAGGTTTCCGGGCCCTTTCTCTGGCCAGGT | 120 |
| chr22 | 50701937 | 50702057 | 785447_45746696_NM_001372044.2_244 | 3 | + | good | CACCTATGTTTACAGCCATCTATCCTTTGACTCACCCACCCATCCCACCATCATCTTCTTAACCGTTGACTCACCAGCCAACCCAATCCACCACCCCTCCCTTCCTCCATCCATCCACCT | 120 |
| chr22 | 50710817 | 50710937 | 785447_45746696_NM_001372044.2_318 | 3 | + | good | AGCCAAGCATGATGGAGTTGTGCATATCTGTGTGTGTGTGCGTGTCTGTGCGTTGGTGAGCGGACACGCGCTGTCTGCACCTGCGTGCATGCGGCACACATGTCTGTGCGTGGGTGAGTG | 120 |
| chr22 | 50716097 | 50716217 | 785447_45746696_NM_001372044.2_362 | 3 | + | good | CCTCCCATGGGCAGCCTCATCCCTGTCCCCAGCTGCCACTCCCTGTCCACTGGGCACCCCCACCTCCCCATCACCTCTCATCCTTCCCATGGGCAGCCTCATCCCTGTCCCCAGCTCAGC | 120 |
| chr22 | 50723657 | 50723777 | 785447_45746696_NM_001372044.2_425 | 3 | + | good | GGATGCTGCCTCCAGCTCCTGTCATCCTGGGGAATTGCTCTGGGGTCCTCTCCCCTTTTAGCCTTTTCCAACTCTCAGCCAAACTGGAAAGCCCTCTTCCCAGCAGTGCAGTGTTGAAGG | 120 |

| | | | | | | | | |
|---|---|---|---|---|---|---|---|---|
| chr22 | 50679257 | 50679377 | 785447_45746696_NM_001372044.2_55 | 2 | + | **good** | GTGGGGGCCCTAGCCTCTGCCCAGGGACCCTACAGCACCTTGCTCTTCCCCCAGGCCTGCCGCTTTGGGCACGTGCAGCATCTGGAGCACCTGCTGTTCTATGGGGCAGACATGGGGGCC | 120 |
| chr22 | 50683697 | 50683817 | 785447_45746696_NM_001372044.2_92 | 2 | + | **good** | CTGACCCCCCAACCCCGAGAGACCAGCAGGGGTGCAGAAGCCAAACTGCAGAGGGGGTGGAGAGGGGGGTGGTGGAGGGGGGTGGTGGAGAGGGGGGTGGTGGAGGGGGATGGTGG | 120 |
| chr22 | 50691497 | 50691617 | 785447_45746696_NM_001372044.2_157 | 2 | + | **good** | TTTTTCCCCCTTTGTTCTATTAATGTGATGTATTATATTGATTACATTAGTCTGTAGATTTCTTTTCCTGCAATGCCCTTATCTGGCTTTGGTAGCAGGGTAATACTGGCCTCGTGAAAT | 120 |
| chr22 | 50693897 | 50694017 | 785447_45746696_NM_001372044.2_177 | 2 | + | **good** | GAACTGTCTATTTCTCTCTTAAATTGTGTTGTTTGTATCATATTTTGATGTTCTGTTGTTCGGTGTGTATATGTTAGTAATTGTTAATCAACATATAATGTCCTTCTCTGTCTCTTGTTA | 120 |
| chr22 | 50702057 | 50702177 | 785447_45746696_NM_001372044.2_245 | 2 | + | **good** | CTCCACCTACCTGTCTACACACCCAGTCCATCCATCCATCTACCCATGTATCTGTCTGCTCTTTGACCTATCTACCCATGGCCCTCTCCTTCCCTCCCTCCATCCACTGGTCCAGTGCCT | 120 |
| chr22 | 50702177 | 50702297 | 785447_45746696_NM_001372044.2_246 | 2 | + | **good** | TTCAATCCACCCTTCTGTCCGTCTATCTCAGATGCACCTGTCCACCATTTACTCTCCTAGTCAGCATCCCTTCTGTCCGTCTATCTCAGATGCACCTGTCCACCATTTACTCTCCTAGTC | 120 |
| chr22 | 50702297 | 50702417 | 785447_45746696_NM_001372044.2_247 | 2 | + | **good** | AGCATGCAGAATGCCTGTGACGTGCCAGGTGCTGTGCTGGGTGCTAGGATCACAGAGATAAAGAACATGTGGCCCTTACCCTCAAGGAGCTCACAATTAGTGGGGGAAACAGGTCATCGA | 120 |
| chr22 | 50704697 | 50704817 | 785447_45746696_NM_001372044.2_267 | 2 | + | **good** | AACCTAGCTGGTGAAGCGCCTTCCTAATTGCCCCCCGCAGCAGAGACCCCCATCGAGGAGTTCACGCCCACGCCAGCCTTCCCGGCGCTGCAGTATCTCGAGTCGGTGGACGTGGAGGGT | 120 |
| chr22 | 50711297 | 50711417 | 785447_45746696_NM_001372044.2_322 | 2 | + | **good** | ATGCAGCACACATGTCTGTGCGTGGGTGAGTGGACATGCGCTGTGTGCACCTGCATGCATGTGGCACACACACACAAGTCCCTGCTTGTGGTTGTTTCCTCCTTGGCTTTCAAGGCATGT | 120 |
| chr22 | 50715977 | 50716097 | 785447_45746696_NM_001372044.2_361 | 2 | + | **good** | CTCCCTGTCCAGTGGGCACCGCCCCCCATCGCCTCATCCCTCCCATGGGCAGTCTCATCCCTGTCCCCAGCTGCCACTCCCTGTCCACTGGGCACCCCCACCTCCCCATCACCTCTCATC | 120 |
| chr22 | 50722937 | 50723057 | 785447_45746696_NM_001372044.2_419 | 2 | + | **good** | GCGGCTTGGAGCACACTGACTCCTTTTCTTTTGGGGGATCTGCCACTAACTCCCTATTTCCCATCCCAGAAACATTTGTCTCTTGGCACCACCTTAGAATCCCTTAAACATGGATTCCCG | 120 |
| chr22 | 50723297 | 50723417 | 785447_45746696_NM_001372044.2_422 | 2 | + | **good** | GGGCCCGTGGGCTGCATGGATGCTGGCGGCAGAGCTGGTCATCCCCCACCCGCCCCCTTGTCTGCCTTTTTAAAGCTGCTTTTGCCTTCTGTGCCCCTAGGTCTCCCCTCTCCTCTTTGG | 120 |
| chr22 | 50723417 | 50723537 | 785447_45746696_NM_001372044.2_423 | 2 | + | **good** | GTCTGGGGGGGTGGTATGTGGATGCCACCTCTTGACTCCTGCTTCTTGCTGCCTGGAAGACCAACCTAGTGGGCCCCGTACTGTCAGCCTTGGAGGACAGAGTTCACAGCGTAGCAACGT | 120 |
| chr22 | 50723537 | 50723657 | 785447_45746696_NM_001372044.2_424 | 2 | + | **good** | GTTCAGAACTTAAGGACTTTGCAGGTCTTACAAAGGCCTGGCCATTCTACCTTCTTTAGTTCAGGATTCAAAAGACAGGTAGGAGCTTGGGAAGCTCATGAGGCCTCTCCTAAGGTCCCG | 120 |
| chr22 | 50729177 | 50729297 | 785447_45746696_NM_001372044.2_471 | 2 | + | **good** | AAACCTGCTTTAAGGGGAGCCCCATGCGTGTGTGGGGCAGGCACTGGTGATGCTGTGTGGTGAGGACAGCCTTTAAGGGGAGCCCCATGCCTGTATGGGGCAGGGACTGGTGATGCTGTG | 120 |
| chr22 | 50732897 | 50733017 | 785447_45746696_NM_001372044.2_502 | 2 | + | **good** | CCACTCATCCTTCCGCCAACAGTCCAGGTCACACAGCAGCAGTCACTGTAACAGACTGCCACATACACACTCGGTCTCACACTCACCTGTGGGTTTTGGTTCCGTTCAATTTGGGTTTTT | 120 |
| chr22 | 50733017 | 50733137 | 785447_45746696_NM_001372044.2_503 | 2 | + | **good** | AACTTTACAGGGTCAGTTCCGCTTCACCTCCTTTTGTATGGAGTTCCATCCGGGGGGTTTCACCCCCTGCTCCAGTCCTGAGGCCTCCTGACCCTGACGTTGTGATACGCCCCACAGAGA | 120 |
| chr22 | 50733137 | 50733257 | 785447_45746696_NM_001372044.2_504 | 2 | + | **good** | TCTATGTTTCTTATATTATTATTATTGATAATAATTATTATAATATTATTATGTAATAAATTTATAAGAAATGAAGCCATGGCTCAGTTGCCTGCTTGAGGGGATATTTGTGTCTGTCCC | 120 |
| chr22 | 50672777 | 50672897 | 785447_45746696_NM_001372044.2_1 | 1 | + | **good** | CAGGGAGCAACCGTCTAGGCGTGGGGAGAACGCAGGACGTGACCCACACACCGCACTGGAGGCTCCGCTCTGCCCGGTCAGTGGCGTCCTGGGGAGGCGGGAGGAGGAAGCCAGACTGG | 120 |
| chr22 | 50672897 | 50673017 | 785447_45746696_NM_001372044.2_2 | 1 | + | **good** | CCCAGGGAAGTGAGGCCCTGCTCTGGGGTTGACCAGAGGCTGTTTTGAGGGGAGAGGGCTCATCAGAAACACTCTGTCAGGGGTGCTCTGGGGTGAGCAGGTGCTGGTCGAGGTGGATCT | 120 |
| chr22 | 50673017 | 50673137 | 785447_45746696_NM_001372044.2_3 | 1 | + | **good** | GCTGGGGAGTGGGGGCTGTCCTTGGAGGGAGTGCCAAGCTCTGTTCTGTGTGGGTCAGGGTGTTTAGGATGGTCCAGGTTGTCACGCGGGGGAGGGTAGGGAGGTTTTTCTGCGTGTATG | 120 |
| chr22 | 50673137 | 50673257 | 785447_45746696_NM_001372044.2_4 | 1 | + | **good** | GGGGCCGGAGCAGTTCAAGGTTATTCCTGGGAGGCTCAGGCAAAATCCGTTTTGTGGGTGATCTGGAGCTGCTCTTGGGGAGGCTGGGGGTTAGAGGCTGTTTTCCTCCTTAGGCAGAGG | 120 |
| chr22 | 50673257 | 50673377 | 785447_45746696_NM_001372044.2_5 | 1 | + | **good** | GGTTCCCCAGGCTGGGCTCTTCTGTGAACGTTCTCAAGCTCTGCCCTCAGGGGGCCAGCGGCTCCTCTAGGAGGATCTGGGGCTGCTTGGGGGCGGAGGAGGGGGGTGGGGAACGCCAGG | 120 |
| chr22 | 50673377 | 50673497 | 785447_45746696_NM_001372044.2_6 | 1 | + | **good** | CAAAGCGGGTTGGGAGAGTTCTGCAGGAGGCGGAACTATTTCGACTTAGGGGCTGCTCTTGGGGGCACAGGGGCTCGCAGTGCAGGACAGTTCCGAGAGGCCGCTCAGGGCTGGGGGGCC | 120 |
| chr22 | 50673497 | 50673617 | 785447_45746696_NM_001372044.2_7 | 1 | + | **good** | TGGTGCACCCCGATGGGGCGGCGGTGCCTGAGCTCGTTTCTTTACTGAGCTCCCCCCTCCTCCGCCCACTCATCACCCCCTCCCGCGTTGCCACGACAACGCGTAAAACTAAAATTCACT | 120 |
| chr22 | 50673617 | 50673737 | 785447_45746696_NM_001372044.2_8 | 1 | + | **good** | TCCCAGTCGCAGGTGGAGACTAGTAGCGCCCCCTCCCCTAGGCCCTTCCCCGTAGGTCGCGCCCCCATCCCGCCGTCGCTAAGGTGACGGGGAGGGGGCGACAGGCGTTGGATCCGGACC | 120 |
| chr22 | 50673737 | 50673857 | 785447_45746696_NM_001372044.2_9 | 1 | + | **good** | GGCCAGGGGTCCTTGGGGGGAAGTCCGGAGCAAACTCCCCAAACTAGGCTGGCGTGGGGGAGCCGCGCCGAACACGCCAGGGCGCGGTCGAAAGGGTTAATTCGGAGGGCCTCTCGCCG | 120 |
| chr22 | 50673857 | 50673977 | 785447_45746696_NM_001372044.2_10 | 1 | + | **good** | GGCGGTGGGAAGGGCCCCGGGCCCTCCCCGGTCTGGAGGTCCCCGTGGTCCGACCCCCAGGCCTGGGGCGGGGGGGGGTCCCCGCCATCTCCTCCAGGCCCGAAGCTGGGGGTCGGTGG | 120 |
| chr22 | 50673977 | 50674097 | 785447_45746696_NM_001372044.2_11 | 1 | + | **good** | GTGGGGGGGAGAAGCCGCCACCTCCGGAAATTAATTGTTTTTCTTTCCCCTTCTCGCCCTACCTTGGTCTTCGTGCCCCGACGCGGCCCCCACCCCAGCTCCGGGACCCCTTCCTCCGCC | 120 |
| chr22 | 50674097 | 50674217 | 785447_45746696_NM_001372044.2_12 | 1 | + | **good** | GCACCCGCCCCGGTGGTCCGCGGATGCCCGCCCTTGCCGCTCAGCCACTCCCCCCGCACCGAGGCCTAGGACTCCCCCCCCCAACCCCGTCACAGCCCCCCAGACCCCCGCCCCGTGGCT | 120 |
| chr22 | 50674217 | 50674337 | 785447_45746696_NM_001372044.2_13 | 1 | + | **good** | CGGCCCCCGCCCTCCGCACACACCTCCCGCCCCCACCCGGGACCCCGCAAGTAACCCCCCAGCACTGGCCCTGAGCCCTCCCGGCCCCCGCCTCCGGCGCAGCCCCCTCGCCACCCCCGC | 120 |
| chr22 | 50674457 | 50674577 | 785447_45746696_NM_001372044.2_15 | 1 | + | **good** | CGGAGCCCCCGGAGCCGCTGTCCCCCGCGCCGGCCCCGGCCCCGGCCCCCCCCGGCCCCCTCCCGCGCAGCGCGGCCGACGGGGCTCCGGCGGGGGGGAAGGGGGGGCCGGGGCGCC | 120 |
| chr22 | 50674577 | 50674697 | 785447_45746696_NM_001372044.2_16 | 1 | + | **good** | CGCGGAGTCCCCGGGCGCTCCGTTCCCCGGCGCGAGCGGCCCCGGCCCGGGCCCCGGCGCGGGGATGGACGGCCCCGGGGCCAGCGCCGTGGTCGTGCGCGTCGGCATCCCGGACCTG | 120 |
| chr22 | 50674697 | 50674817 | 785447_45746696_NM_001372044.2_17 | 1 | + | **good** | GCAGACGGTGAGCCCCGCCGCCCTGGGCCCGGCCGTGCCCCTGCGCTCCCCGCCCGGGATTCCCCCACCCCCGCCGGGCGCGCCCGGCGCCCGGGACCCCCGGCCCACGGCTACTCACCC | 120 |
| chr22 | 50674817 | 50674937 | 785447_45746696_NM_001372044.2_18 | 1 | + | **good** | CTCCCCCGCCGCCTCCGCCGGGACCCTCCCCATCGCCAGGGCGGGGCCCCGGGAAGCCCGGCCCCTGGGGCGGGGCTTCGGCCGCGGTTCGCGGAGGCGCGGGGTCCCGGGCGCCGGC | 120 |
| chr22 | 50674937 | 50675057 | 785447_45746696_NM_001372044.2_19 | 1 | + | **good** | CCGAGCCCCGGACTCCTTCGGCGGGGGCCCGGGGCTCGGCACCCCGCATGGGGCCGGCGGGGCGGGTCCGCGCTCCCGGGACCTGAGCTCACGAGCCCGCTCCGCTGCAGAAGTGCCTG | 120 |
| chr22 | 50675057 | 50675177 | 785447_45746696_NM_001372044.2_20 | 1 | + | **good** | GCCTGGACCCGGCCGCGCCCGTGTGGGCCGCCAAGCAGCGCGTGCTCTGCGCCCTCAACCACAGCCTCCAGGACGCGCTCAACTATGGGCTTTTCCAGCCGCCCTCCCGGGGCCGCGCCG | 120 |
| chr22 | 50675177 | 50675297 | 785447_45746696_NM_001372044.2_21 | 1 | + | **good** | GCAAGTTCCTGGATGAGGAGCGGCTCCTGCAGGAGTACCCGCCCAACCTGGACACGCCCCTGCCCTACCTGGAGGTAAGTGGCCGGCGCGGGGGTGAGCTGAGGAGCGCGCAGGGTGGAT | 120 |
| chr22 | 50675297 | 50675417 | 785447_45746696_NM_001372044.2_22 | 1 | + | **good** | CACCAAGCCCCGTGGCGGGACCAGTGAAGGGCACGGCAGTGGGGAAACACAAGTGGGAGGGGTGAGGGGTGGAGGCTGTGTGTGTGTGTGTGTGTGTGTGTGTGGTGCTGTGTGCAG | 120 |
| chr22 | 50675417 | 50675537 | 785447_45746696_NM_001372044.2_23 | 1 | + | **good** | GTGCAGTGAGCGTGTACAGGGTGCAGCGAGCGGACACAGTGTATGCGATGAGTAGGCGCGGTGTGTGCAGTGAGCGGGCAGGGCGAGCAGTAAGGATGTACAGTGTGGGCAGTGTGCGA | 120 |
| chr22 | 50675657 | 50675777 | 785447_45746696_NM_001372044.2_25 | 1 | + | **good** | AGTGGGCACAGTGCAGTCAGTGTGCACAAAGTATGCAGTGGGCACGTACCGTGTGTGCAGTGAGTGGTGTGCAGTGTGTAGTGTGCAGTGAGCAGTGTGTACAGCATTGCAGTGTGGGGC | 120 |
| chr22 | 50675777 | 50675897 | 785447_45746696_NM_001372044.2_26 | 1 | + | **good** | AGTGAGTACAGTGTGAGTGTTGTGATTGTGGTAAGCAGGATGCGCAGTATACAGTGAACAGTGTGCACAGTGTGTGCAGTGTGGGCTGTGTGCCACAGAGTCAGTGTGGTGTGTGTAGTT | 120 |
| chr22 | 50675897 | 50676017 | 785447_45746696_NM_001372044.2_27 | 1 | + | **good** | TGAACAGTGTGTGCATTGAGCAGCATGGATGGTGTGGACGCTGAGCATTGTTCTCCAGGGAGGAGTGTGAGCACGAGAGAGTGCCAGAGGGGTGTGTGGTGTGAGCAGGGCTATCTGTGT | 120 |
| chr22 | 50676017 | 50676137 | 785447_45746696_NM_001372044.2_28 | 1 | + | **good** | GCACGTTTGTTCCTTTCTCCAGCTGTGAAGTCTTGTGAAGGCCAACCAAGTCCCCTCCTTTACTCACCCATGCATGGTGTGAAGATGTATTGAGTGCCTTGCTAGGCATGGGGACGTAGA | 120 |
| chr22 | 50676137 | 50676257 | 785447_45746696_NM_001372044.2_29 | 1 | + | **good** | CGGGGTCAGTCCTGTGGAAGGTCTTGGAGTTTGGCAGGGTGGAGGGGGGTGCCCAACTGCAGTGGGCATTGAATAAAGATTTTGTGAGCTGAGCTCAAAGTTGGGCGGGCCTCCGTGGTG | 120 |
| chr22 | 50676257 | 50676377 | 785447_45746696_NM_001372044.2_30 | 1 | + | **good** | GCACAGGAAAGTGGGGTCAGTTCTACCTGGAGAGTGGAGGGGGATGTCTAGGATGGTAAGCCTGGAATCAGGCCTTCAGAGAGGAGTGGGATTTTGCCGAGAATCCTGGGGATGGAAGG | 120 |
| chr22 | 50676377 | 50676497 | 785447_45746696_NM_001372044.2_31 | 1 | + | **good** | CGACGGGACAGTGCAGGCTGCGGGCAGCTAGGCACGTGCCTTCCATCGGGCTGCATCATGCCTGAGTGTGGTGGGTGCATGGCAGCTGTTAGCTCTGTCCACTGTGGTAGTATGACTGAT | 120 |
| chr22 | 50676497 | 50676617 | 785447_45746696_NM_001372044.2_32 | 1 | + | **good** | GGTGTGTACAGGAGGGCAGTGAGGGGTGCGGTGTGGCCAGCATGAGCGGGACGGGGTTTGTGCATGGACTCACTTGCTCAGCCGGGGTGGGGGCATTTTCTCTACCTTTTCTTTATCTGA | 120 |
| chr22 | 50676617 | 50676737 | 785447_45746696_NM_001372044.2_33 | 1 | + | **good** | GCAGTTTCGATACAAGCGGCGAGTTTATGCCCAGAACCTCATCGATGATAAGCAGTTTGCAAAGCTTCACACAAAGGTAAAGGATCACGGGGAGGGGGCTCCTGAGGTTCCCTCCTGCCT | 120 |
| chr22 | 50676737 | 50676857 | 785447_45746696_NM_001372044.2_34 | 1 | + | **good** | CCCTGTGGCTGCTGTCCCCCACCCCAGCTTGGGGCTGACCACAGTCCCCCAGCTTTAGCTCAGTCCATTTCCCCATCATCAGGGGCCCAGAGCCTGTACTGGGTGTGGCTGAAGGGCTGG | 120 |
| chr22 | 50676857 | 50676977 | 785447_45746696_NM_001372044.2_35 | 1 | + | **good** | GCACAGATTCCTGGCCCCATGGTTGGGGTCAGGTGGTACAGTGATGTGTTCCAGTTAGATCAGACTCTTGCCGACCCCCCTGGCTTAGGGGCTGGAGTGTCCTGTGAGAAGCTGGGTGGA | 120 |
| chr22 | 50676977 | 50677097 | 785447_45746696_NM_001372044.2_36 | 1 | + | **good** | GAGGGAGTGGACAAGCATCTGATGTGATGGCTGTCGGGACAAGGCACCCAGCATCGAGTGGTCGACCAGTGCTAGGCATTTGTGATTAGACCTCTCATTAATCCTCTTCCAGAGGGACTA | 120 |
| chr22 | 50677097 | 50677217 | 785447_45746696_NM_001372044.2_37 | 1 | + | **good** | TTATAACCCTGTTTCACAGATGGGGACACAGAGTACCCGACTGTGTAGACAGTGAAGCTGGGGCTGAACCCAGATCTGTCTGATTCCAGGCCCTGTGCCCAGGCATGCCTTTGAGGTGTC | 120 |
| chr22 | 50677217 | 50677337 | 785447_45746696_NM_001372044.2_38 | 1 | + | **good** | TACTGCTGGGTGCCACCCCCAACTGGGCCTGACCCCAAATGCTCTTGAGGGGGGCACCTTACTATTTCCCGCTGAATGTCAGAGGGGCAGGGTGGGTGCCACAGCCCCTCCCCAGGGGCT | 120 |
| chr22 | 50677337 | 50677457 | 785447_45746696_NM_001372044.2_39 | 1 | + | **good** | TCCACCCGCAGCTCACAGTCCAGCAGGCACTTGTTTGCTGGATACTTTATGGGCGGTCTCGAGCTCAGGAGAGGGGTCAGAATGGAGGGTCTCTGGAGCCCAGGAGAGGAATCAGAACTG | 120 |
| chr22 | 50677457 | 50677577 | 785447_45746696_NM_001372044.2_40 | 1 | + | **good** | TCAGCCCTGTGTCCTAATGGTCCCATCAGTGACTTTGGAGCCTCAGGCTTCTGTGCCAGGCTTTTCTGCTGCCCCAGGGCAGGCGGTGGTCAACGGCCTTGGCACTATGGCTGGGCAGAG | 120 |
| chr22 | 50677577 | 50677697 | 785447_45746696_NM_001372044.2_41 | 1 | + | **good** | TCCACTGTGGAAAGGTCCCCCTCTCCTGCCACTGGCCCTCACTTGACCTTGTCAGCCTGGGTTCCTACCCCATGGCCACCTCTCCCTCCGGATCTCTCTTCAGTGACCAACAAGACATGA | 120 |
| chr22 | 50677697 | 50677817 | 785447_45746696_NM_001372044.2_42 | 1 | + | **good** | GTGACTCACTCTGAAGTAGGTCTCGTTTGTTTTTAGGATGAGCCTGAACTTCTTACATAGCCCGTGTTTCTGCTTGCCTGGGCTCCAGCTGGCCTTTCCCTCTCCCCTGGTAGTGAGCAC | 120 |
| chr22 | 50677817 | 50677937 | 785447_45746696_NM_001372044.2_43 | 1 | + | **good** | TTCCTCCTCTTTCACTGCCTGGCCCGCCTGAGGAGGTTGCCTCCCAAGGCAGGGGTCCCTATGGCACCCTCCCCTATTTCTGAGGTACACTGGGTGTTTATGGAAGGGCCCCGGCCTTTG | 120 |
| chr22 | 50677937 | 50678057 | 785447_45746696_NM_001372044.2_44 | 1 | + | **good** | GCCAGGGCACCTTGCCTTGTGTGTGTGGTGTCCGTAGTGCCGGCTGGGGAAGTGAGGCCTTGTGGGGTGAGTGAATAAACTGGGTGAATGAGTACAAGGCCATCAAAGTCATGTCACAGG | 120 |
| chr22 | 50678057 | 50678177 | 785447_45746696_NM_001372044.2_45 | 1 | + | **good** | ATGGGTCCCTGGGGTGGGGGTCTGCGGTGGGGTGAGCTGGAGAGGAAGATGGCCGAAGAAGGAGCAGGTTGTCTCAGGGTGCTGAGGATTCAGGGCTGCTGTGGGGCTGGTTCCTGCC | 120 |
| chr22 | 50678177 | 50678297 | 785447_45746696_NM_001372044.2_46 | 1 | + | **good** | GGGTGGACCTCGGGCTCTTCTAGGCCAGGCAGGTCGGGGCAGCAGGGCCGGAGAGACGGAGCCAGGGCAGTGACTGGGCCTGGAGTGGGGGACTTGCTTGGGGCCCCACCAGGGTGAC | 120 |

| | | | | | | | | |
|---|---|---|---|---|---|---|---|---|
| chr22 | 50678297 | 50678417 | 785447_45746696_NM_001372044.2_47 | 1 | + | good | GGCCTTGGTGAGGGGCTACTTGGGTACAAGCTGACAGTCGTGACTGGTTTGGCTATCAGGGCGCAAGGAAGGGCTTTGAGCCGTGCATGGCCCACCCGAGTGGGAATTGGGGCCGTGGT | 120 |
| chr22 | 50678417 | 50678537 | 785447_45746696_NM_001372044.2_48 | 1 | + | good | GGGAGTGCAGGACCGTGGTTGACAATTGTGATGTCAGGTGACAGGTCAGACTGGTAGGGATGTGGCGGGGGTTGCCTGAAGGTGGCCTGAGGCTTGCCGGAGGAAGGCGGGTGATGTTCA | 120 |
| chr22 | 50678537 | 50678657 | 785447_45746696_NM_001372044.2_49 | 1 | + | good | GATGATGGAGGCCTTGGTGCCAGGCTGACTGACGGCCGGTGTTCCAGGCGAACCTGAAGAAGTTCATGGACTACGTCCAGCTGCATAGCACGGACAAGGTGGCACGCCTGTTGGACAAGG | 120 |
| chr22 | 50678657 | 50678777 | 785447_45746696_NM_001372044.2_50 | 1 | + | good | GGCTGGACCCCAACTTCCATGACCCTGACTCAGGAGGTGAGGAGTGGAGTCGGGGAGGGGCATGGCCTTTGCGCGGCTGGGAGCCTGACCCTTATCTGTCTGTGAACCCAGAGTGCCCCC | 120 |
| chr22 | 50678777 | 50678897 | 785447_45746696_NM_001372044.2_51 | 1 | + | good | TGAGCCTCGCAGCCCAGCTGGACAACGCCACGGACCTGCTAAAGGTGCTGAAGAATGGTGGTGCCCACCTGGACTTCCGCACTCGCGATGGGCTCACTGCCGTGCACTGTGCCACACGCC | 120 |
| chr22 | 50678897 | 50679017 | 785447_45746696_NM_001372044.2_52 | 1 | + | good | AGCGGAATGCGGCAGCACTGACGGTCAGTGAGGGCGGGGCCTGGCCTGGAGGGGCTCTTGCCTGGTGATGGGGCTGGGGGCAGCTGGGCCTGGTGTGGATACTGAGGCTGCTCACCCTCA | 120 |
| chr22 | 50679017 | 50679137 | 785447_45746696_NM_001372044.2_53 | 1 | + | good | GACCCTGCTGGACCTGGGGGCTTCACCTGACTACAAGGACAGCCGCGGCTTGACACCCCTCTACCACAGCGCCCTGGGGGGTGGGGATGCCCTCTGCTGTGAGCTGCTTCTCCACGACCA | 120 |
| chr22 | 50679137 | 50679257 | 785447_45746696_NM_001372044.2_54 | 1 | + | good | CGCTCAGCTGGGGATCACCGACGAGAATGGCTGGCAGGAGATCCACCAGGTGTGCAGGGAGCCGAGGTGGGGTCCCGGCCTCTGTGTGCTGGGTTGGGGGTCCTGGCTCTGTCTGTAGGG | 120 |
| chr22 | 50679377 | 50679497 | 785447_45746696_NM_001372044.2_56 | 1 | + | good | CAGAACGCCTCGGGGAACACAGCCCTGCACATCTGTGCCCTCTACAACCAGGTGCGACTGTGTGTCCTGCACATGCCTGCACCAGCGAGTGTGCATATACTTGCCTCTTCTGGGGGTGTA | 120 |
| chr22 | 50682977 | 50683097 | 785447_45746696_NM_001372044.2_86 | 1 | + | good | CCATGGTATATATATGTGCTCTGTGTCCTGCAGTGAGTTGTGGCTGCAGCACAGCCTCATAGGCATATGTGTGCACATTTGTTCTCTGAACACACAGGGGCTTCACATGTGTGCACGTGT | 120 |
| chr22 | 50683097 | 50683217 | 785447_45746696_NM_001372044.2_87 | 1 | + | good | GTTCTGAATAACCAGGTATGAATTGGGTACATCTAGGCCCTCTGGGCCAGGTGAGACCTGAGCGTGTATACCTACTGGCTTGTCTCTGCAACTCAGGTGTACATGGAACAAATAGGTGTG | 120 |
| chr22 | 50683217 | 50683337 | 785447_45746696_NM_001372044.2_88 | 1 | + | good | AGTCCGTGTGTGTGAGCCTGTGCCCTGCGCACGCCATGTGTGCATTCCTGTGTGCGCATGTGCTGTTGTGCTCGGATGGTCTCTCCAGCCACCCAGCTGTGATTCCCTCTTCCCCGCAAC | 120 |
| chr22 | 50683337 | 50683457 | 785447_45746696_NM_001372044.2_89 | 1 | + | good | AGGAGAGCTGTGCTCGTGTCCTGCTCTTCCGTGGAGCTAACAGGGATGTCCGCAACTACAACAGCCAGACAGCCTTCCAGGTACACCGGTGGTTTACAGGAGCTCAAGGCTGCCCCAGAG | 120 |
| chr22 | 50683457 | 50683577 | 785447_45746696_NM_001372044.2_90 | 1 | + | good | GTGTCTGTCTCTGTGTCCATGTGACTTGACTTCTCTGAACCTTGGTTCTTCCCTGGAAGGCCCTAAGGGAGCACCTCCCCCAGGACTGCCCACAGGAGGTGTTGGGGGACGAGCCCAGCA | 120 |
| chr22 | 50683577 | 50683697 | 785447_45746696_NM_001372044.2_91 | 1 | + | good | CGCGAGGGGTATTTGGTGTTGATGTTCCCTTCGTCCCCTCGCCAGGGAGAGAGGAGGGTCAGCAGGGCTCTGGGGCAGGGGTATGGGGAAAATGAGAAGACTGGGGTGACAGGTGTGGGT | 120 |
| chr22 | 50683937 | 50684057 | 785447_45746696_NM_001372044.2_94 | 1 | + | good | GGTTGGCACTGATGACCAGGTGGACGTGGTCTCTGACTTGGAGTTTGTTGGGGAGTCGGGAATATTGTGGGGGTTTGCGGACCAGGAACAGAGCCAGAACTCCTCTATTTTCAGTAGGAG | 120 |
| chr22 | 50684057 | 50684177 | 785447_45746696_NM_001372044.2_95 | 1 | + | good | TGAGATGTGGGAAGAGTGGTCAGGCTGCTGAGGTTGTGTGGTCATAGAAGGTGGAGGCAGCAGTGGGCTGCCAGATGGGTTTGGGCACACTGGGAAGCAGATACGCCCGGGCCTGTTGGA | 120 |
| chr22 | 50684177 | 50684297 | 785447_45746696_NM_001372044.2_96 | 1 | + | good | TGACTGGGTGCAGGGTCACAGGAAGGGGAGGATGAGCTTGGCAGCTTGGGCAGCTGCGGGGCCTTGGCTGAGATGGGAAACATGGCTGCATGTTGGGACAGACCAGTGTGCCCGTTTGCT | 120 |
| chr22 | 50684297 | 50684417 | 785447_45746696_NM_001372044.2_97 | 1 | + | good | TGGGACTGCTGGTTTTAGCTCTGAACTTCTGTCCAGGCAGCCCCTCAGTCCCAGGCAGACAGAGGGTGGGTCACTCCAGTTATAGGGTGTGCCTGTGGCATCCCAGTTACAGACAAGAGT | 120 |
| chr22 | 50684417 | 50684537 | 785447_45746696_NM_001372044.2_98 | 1 | + | good | GTGTGTCAGTAGAGAAGAGGAATGGGCTGGGGTCTCCGGGGCCCCCAACCAAGGAAGACTCCACACACGAACTGGAGAAGGAGCAGCTGGGGCTGGCTGGCGAGGCAAAGCTGGTGCTGC | 120 |
| chr22 | 50684537 | 50684657 | 785447_45746696_NM_001372044.2_99 | 1 | + | good | TCAGCCCCTCCCTGCTCCTCAAGGCCTTGACCTCCCCTTTCCCTCAGGTGGCCATCATCGCAGGGAACTTTGAGCTTGCAGAGGTTATCAAGACCCACAAAGACTCGGATGTTGGTGAGT | 120 |
| chr22 | 50684657 | 50684777 | 785447_45746696_NM_001372044.2_100 | 1 | + | good | TCTGCCCACCTGGGCGACCCTGCTGAATGTAGATTCGTGTGGTTTTCTGGGGCCCAGCAGATGTGGGGGTCAGTGGTGACATAAAGGGGCCCCACACCCCACATCTACACTGTGTGGCCA | 120 |
| chr22 | 50684777 | 50684897 | 785447_45746696_NM_001372044.2_101 | 1 | + | good | GTGGTCTCTGAGTGGCCACTGCGGGCATGAGGAACGGGCCACAGGGCCTGTGTTGAGTGAACATTGTTTTTGGTCTGGGAGAAGAGCATGGTGAGGTTTGGGCTGTGGGCCGGGGCTAGC | 120 |
| chr22 | 50684897 | 50685017 | 785447_45746696_NM_001372044.2_102 | 1 | + | good | ATTCAGCTTAGGGGGCAGCATAGGTTGAGAGTGGGGTTGAAGTTAGGGTCAGAGTCCCTGTTTCCCGTTAAATCCTAGGAAGAGCCAGCAGGAGACATGGGGGTGGTGGACCTCACTTCA | 120 |
| chr22 | 50685017 | 50685137 | 785447_45746696_NM_001372044.2_103 | 1 | + | good | CCCTGGAGCAAGCATGAGTCTTGGTGGTCCGTAGCAAGCTCTCAGCATGGAAAGATCATGTAAACGACACATAAGGAAAACAACAGTGATGGGAAAGCAGAATGGGGCTGGGAATAGTGC | 120 |
| chr22 | 50685137 | 50685257 | 785447_45746696_NM_001372044.2_104 | 1 | + | good | CATAGCCATCCTCAGGGCAAGGAGCTCTGAGCCCCCTGCTGCCTGCGAGTAGAGGAAAGCCTGGTGTGTGCGTGACTCACGGTCCTGGAAGGAAAACCAGGAGACACACACGTGGCCTCA | 120 |
| chr22 | 50686937 | 50687057 | 785447_45746696_NM_001372044.2_119 | 1 | + | good | TCTGGTGCCTGTGCCTCTCTCTGGTATATGGACTCCCAGGACCCCCTTCCTGCCATGAGTGTTTACTCGTGGGTGTCCCCATTTTTTGTTTTTGTTTTAATTTGAAGTGACAGCGGGCAG | 120 |
| chr22 | 50687057 | 50687177 | 785447_45746696_NM_001372044.2_120 | 1 | + | good | CAAACGTCTGGAGCAGATGTTAGGGTTGGACTCTCACTTGGTTCTAATGTGAGGTCATGGTCAGGGCTGGAGTTGAATTTTGGACTGGGGTTAAGATTAAGGTCGGTTTCATGTTTAAGG | 120 |
| chr22 | 50687177 | 50687297 | 785447_45746696_NM_001372044.2_121 | 1 | + | good | GCCCTTCTTCTGTAGAGTTCCCCCTGAACATCCCCCTGTAAGGCTTGGTTATGAGCCCCTCCACCCCAGGCTGTGTCAGTTGAACCATCTGAGGCTGTGGCAGACCCTCGGTGAATGCCA | 120 |
| chr22 | 50687297 | 50687417 | 785447_45746696_NM_001372044.2_122 | 1 | + | good | CAGCTGCTCTCAAGGCCACTGCAGGGAAGTGTGTGTTGAGGAAGTAACTGGGCCAGTGGGATGGGTAGGATCGGAGTCCAGGGTCAGGTCAGGGGAGAACTGTGGTAAGGTTTGGGTTTG | 120 |
| chr22 | 50687417 | 50687537 | 785447_45746696_NM_001372044.2_123 | 1 | + | good | CATAAGGGGCAGCAGATAGTTACTTCTGGTTTGGGCTGGTGTTTTGGAGGTGGCCCTTAGTGTTGGAAGGCCCTGGTTTGTGCATGTGTGGGGCCTCTTCCTCTTTTTTTTTTTTTTTTT | 120 |
| chr22 | 50687897 | 50688017 | 785447_45746696_NM_001372044.2_127 | 1 | + | good | AAACATGGTCTCTTATGGTCATTTGCATCCCCTTCCCCCTGTGGTGTCCTGTGGCGTTTGATCCTGGGTGGGGCTGGACTCTTGCCGGTGGCCTTGGTCTAGTTGCTTGGAGGGAGGGCG | 120 |
| chr22 | 50688017 | 50688137 | 785447_45746696_NM_001372044.2_128 | 1 | + | good | GGAAGGCCCCATCTGCCCCTCATCTGTGCGTTGTGGACAGATACCCCAGGGCTCTTGGAGTCACCATGGTTGCGACTTACTGGCTATCCAACCTAGGTGTGCTTGCTTGGTTGGAGTTAG | 120 |
| chr22 | 50688137 | 50688257 | 785447_45746696_NM_001372044.2_129 | 1 | + | good | ACACACGTCCCTCAGTGGTAAGGAGCAGGCACCTGTGTCAGTTCATCCTGATGTGGTCGTCGGCTCCATGGGCCAACAACACCCTTACCTGATCAGGTGCCACACACAGACACTAGGGTG | 120 |
| chr22 | 50688257 | 50688377 | 785447_45746696_NM_001372044.2_130 | 1 | + | good | ACACCGCCAGATTTCCCTGAGTACAGGCAGCATATTTGTCAGGATTTAATACAAGCCCCTTATCTCAATTTGCCCTCTCAACAACCCCCCAAGGAGTGATTAGACTGTTTTACGGAGAAG | 120 |
| chr22 | 50688377 | 50688497 | 785447_45746696_NM_001372044.2_131 | 1 | + | good | GAAACTGAGGCACAGAGGGGGTTAATTCAGACCTGATTGATTCAGGGTCTGCCTTTGTCACTCCCATGCCCTCCGTACACATTTGCTCCAAGGACAGGACAGTCATGACTTGCTACATCC | 120 |
| chr22 | 50688497 | 50688617 | 785447_45746696_NM_001372044.2_132 | 1 | + | good | TATGCCCACTGGGGTCCCAGAGTGGGGCCTGTACACAGAAGAGGAACTGGAAGGAGGTGCCGCCTTCTCTGGTGGGCAGGTGCCACCCGGTGGGATCTGGTAGACTGGGACCTGTGTAAC | 120 |
| chr22 | 50688617 | 50688737 | 785447_45746696_NM_001372044.2_133 | 1 | + | good | CCTTCAGGCAACAGGATAGCTTGCTGCCTCTCGACAGCCTTGCTTGCCCCCTCTAGACTGACAGGCCTTGGGTGGGGTCCCTGGCCTGGGCGTGGAATGGGCATCCCACAGTGCTTTTGA | 120 |
| chr22 | 50688737 | 50688857 | 785447_45746696_NM_001372044.2_134 | 1 | + | good | TGAGTACCTTGGGGAAATGCTTCTCCAGGGTGCTGTCATGCCCATGAGAACACACCTGTGCCTGGGAGTGTGTGCCTGAGAGAGGGTAAGAGACAATGTGGACCCAAAGAGTGACCCGAC | 120 |
| chr22 | 50688857 | 50688977 | 785447_45746696_NM_001372044.2_135 | 1 | + | good | GTGACAGAGGCAAGGAAGCAGAGGTGACCACTTCTAGGCCAGAATGATTCAGTAGCGTTCTGTATATTCACGGTGTTGTGCAGCCGATTCTATTGAATTCAAAACATTTTCATCACCACA | 120 |
| chr22 | 50688977 | 50689097 | 785447_45746696_NM_001372044.2_136 | 1 | + | good | AAAGAAGACCCTGGACACATTAAGCAATAATTCTGCATCAATTACCCTCTCCCTCCAACTCTGTCAGCCATTAACACACTTTCGTCTCTGTGGACTATAGATTTACCTATTCTTGATATT | 120 |
| chr22 | 50694137 | 50694257 | 785447_45746696_NM_001372044.2_179 | 1 | + | good | GGATTTAAAGTGATTATTTTGTAAGTAGCATATAGTTGGATAATACTTTTTAATCACTATCTGCTTTTTAAGTAAAATTCTATTGGAACACAAACACACCTATCTGCTTGTATGTTGTTT | 120 |
| chr22 | 50694377 | 50694497 | 785447_45746696_NM_001372044.2_181 | 1 | + | good | TAGAATGTCTTTTATAGTTTCCCTGGTTATAGTTAACTGCAAATGGTCCTGTTCAGCAGTAGTCTGGTGAGATTCGTCACATGTAGCCTGTTTTGTCTCTCCTGAAGCTTGTGAAATGCA | 120 |
| chr22 | 50694497 | 50694617 | 785447_45746696_NM_001372044.2_182 | 1 | + | good | GTTCTTAGAGCATATCTGAGCACCCTCAAGGCCCCTTTCGGCTGAGAGGAGAAGCAGGCTTCTACCTCTGAAGCTAGGGTGAGAGGCAGTTTGACTGGGATGTGGCCCCTTAATGGTAGC | 120 |
| chr22 | 50694737 | 50694857 | 785447_45746696_NM_001372044.2_184 | 1 | + | good | GGCTGTTCAGAGTCTTACCTATGCCCCCTTACCCCAGTACCATTCAGGGAAACCCCCAGCTATGCGAAGCGGCGGCGACTGGCTGGCCCCAGTGGCTTGGCATCCCCTCGGCCTCTGCAG | 120 |
| chr22 | 50694857 | 50694977 | 785447_45746696_NM_001372044.2_185 | 1 | + | good | CGCTCAGCCAGCGATATCAACCTGAAGGGGGAGGCACAGCCAGCAGCTTCTCCTGGACCCTCGCTGAGAAGCCTCCCCCACCAGCTGCTGCTCCAGCGGCTGCAAGAGGAGAAAGATCGT | 120 |
| chr22 | 50694977 | 50695097 | 785447_45746696_NM_001372044.2_186 | 1 | + | good | GACCGGGATGCCGACCAGGAGAGCAACATCAGTGGCCCTTTAGCAGGCAGGGCCGGCCAAAGCAAGATCAGGTAGGAGGGGGCTGGCAGGCCCTGGAGGGGTTGGGAGGGTGGGGTGCC | 120 |
| chr22 | 50695097 | 50695217 | 785447_45746696_NM_001372044.2_187 | 1 | + | good | GGACCTGAGCCAGGAGGAGCCAGCACCGGGAGGCAGAGAGAGGCCTGGTGGTGCCTAGCACCTGTGTAGTGATGGGCTAGGGCTTCCTGGTTGGGGAAGGGAAGAAGGCATCTCTGGAGC | 120 |
| chr22 | 50695217 | 50695337 | 785447_45746696_NM_001372044.2_188 | 1 | + | good | TGGGGACAGGCACATGCACGCTGATTTCCGGTTGGAGCTGGGCTTGCCATGAGGACAGTGGGCAGTTGAGGGAGCGTGTTAGGTGGAGAGGGTATGCTCCTACCTGTGATTTAGAAAACA | 120 |
| chr22 | 50695337 | 50695457 | 785447_45746696_NM_001372044.2_189 | 1 | + | good | CACTGCATTGTCAGGCATGGCTGGGAAGAGTCAGGGCTGGTGGCCTGTGGTAAGGGGGACAGGGATGGAGTTGCCACTCCAGTAGGGCCCAAAACCTACAGGTAAGCCCCAGCCCTGTCC | 120 |
| chr22 | 50695457 | 50695577 | 785447_45746696_NM_001372044.2_190 | 1 | + | good | CCAGGGTATACTCATGGTTTCCCCCAGCCCTGGCACACCTACAGGACTCTGTGGCTTCTATTCTTTACCCTCACCCTGAGGCCTGGGTCCCTGAGGCAGGGATGCCTGGACCGGAGCCCA | 120 |
| chr22 | 50695577 | 50695697 | 785447_45746696_NM_001372044.2_191 | 1 | + | good | CTGGCCTCTCTGGGCTGTGCATTGGGTGCTGAGGTGGGCGCACTGGCTCTCAGCCACTTTGTGCCTGGGCTTCTCATCAGTAAGTGGGAATAGTCCTACCTAAGTCATTGTCTTTGGGAA | 120 |
| chr22 | 50695697 | 50695817 | 785447_45746696_NM_001372044.2_192 | 1 | + | good | GCCTTCCAGAATGCCCTGGGCAGGTGAGGGATCACCTGGCAGAGTCTGGCAACTGCCGGTAAAAGCACCTGCCCTCCCTGTGCCTTGAAGGCAGGCACCGTCTTTGTCGTTTCCTAGTTT | 120 |
| chr22 | 50695817 | 50695937 | 785447_45746696_NM_001372044.2_193 | 1 | + | good | CATCAGAGCCAGGCACTGAGGAGGGTGTTTGGCAGACATTGTTGAGTGGAAGAGTTACATAAACAGTATGTTGTTGTCAGGATGGCCGCCGGCCTTTACAGTTCTGTCAGTCCAGGCCCG | 120 |
| chr22 | 50695937 | 50696057 | 785447_45746696_NM_001372044.2_194 | 1 | + | good | CTGTTGGGCTGAATGGCATGTGTATACTAGCTCCTTTAACCTTGACGAACGTACATGTTCATGCTCTATTAATACCATGTTACAGATGAGCGTCCTGGTACGCAGGTGGGGCCTGTGACG | 120 |
| chr22 | 50696057 | 50696177 | 785447_45746696_NM_001372044.2_195 | 1 | + | good | GGTCCAGAGTCACAGAGCTACGGATCAGCTTCACTGGGCCCCAGGTAGACCCAATCCCCAACTTTCTAAGCCTGAGTGTTCAGCAGAGGGCTCAGGGTGCAGCAGGGCCTATGGAAGGCG | 120 |
| chr22 | 50696177 | 50696297 | 785447_45746696_NM_001372044.2_196 | 1 | + | good | CCCCTTCCTGTGCTGCCGCCCCTGCTCTGCAGCCTGGTCTCAGCGATGTCTCCGTGTCTGTCTTTGCCTGGGCTTGTTGGACTCGCAAGGCTGGCTGTACTGCCCCAAATCTCAATTCTC | 120 |
| chr22 | 50696297 | 50696417 | 785447_45746696_NM_001372044.2_197 | 1 | + | good | CCCTGAAGCTTTCTTATAGACCCGCGGATTTTCTAAGGGCTGGGCGGCCCAGTGGCTGAAAGAACACATTCTGCATTCCGGATGTTTCCATCCCGCAGAAAGGCTGCTCCTGAGTTGGAG | 120 |
| chr22 | 50696417 | 50696537 | 785447_45746696_NM_001372044.2_198 | 1 | + | good | GCACGCGTTGCCACAGGCTCCTCCCTGCATTCATTTTTTGTACGTGTTCATTCATCCTTTCAGCAGACACAAAACTAAGACCTGCTGTGTGAGGGGCTCCAGCTCACCTGCTCTGTGGGG | 120 |
| chr22 | 50696537 | 50696657 | 785447_45746696_NM_001372044.2_199 | 1 | + | good | ATGGCAGAATTCTGTTTTTTCCTTCCTCGCAAACATTTATTGGGTGCCTGGTGCGTCAGAGGTGGTGGGTCTCAGGAAGTTGACAGCCGAGGGCCCGGGCTAAGCAGCAGTTAGGTGGGC | 120 |
| chr22 | 50696657 | 50696777 | 785447_45746696_NM_001372044.2_200 | 1 | + | good | AGTGATCCCCTTATGGGGCTGCCCTGGTGAGTGGACAAGGGTCTGCAGCTCTCGGGAGGGGCTGAGGAAGTTCGGTGCCCCGAGGAGAGAAGTCACCCCCTGGGGGAGGCGAGGCTGGGC | 120 |
| chr22 | 50696777 | 50696897 | 785447_45746696_NM_001372044.2_201 | 1 | + | good | CCTGCACCGCGCGGGTCGCTGCGCCCTCTGTCGGTTGAGCCGGACCGGGACCCGACCCCCATCAGCCCCCCAGTATCACACGGGGCACGGCGGGGGAGTTTGGGCTGAGAGCCCGTCACT | 120 |

| | | | | | | | | |
|---|---|---|---|---|---|---|---|---|
| chr22 | 50696897 | 50697017 | 785447_45746696_NM_001372044.2_202 | 1 | + | good | TAACAGCTGGCCCAGGGGTCAGGATGTTAGAGTAGTTCAGGTGTTGGCAACTGCGTGACGGACAGCCCACAGGAGGGGAAGGAGAGGTGCCCCGTGTGACTAGAGCGTAGCAAACTGCCA | 120 |
| chr22 | 50697017 | 50697137 | 785447_45746696_NM_001372044.2_203 | 1 | + | good | AGCCCGAGGGAGCCTGGCTCTTGGAAAAGCGGGTGGCTCTGGCCTGCCTGGGTACTAGGGAGCCACTGCAGGCTTCTGAGCAGAGCCCCAGTGGAGGAGTGAGAGCTCAGGCTCTACCCC | 120 |
| chr22 | 50697137 | 50697257 | 785447_45746696_NM_001372044.2_204 | 1 | + | good | AGTCTGGGAGTGGTCTGGGGAAGGTGGGCATCACGCAGTGGGCGTGGGCAGGGGCCGGTGTCCAGGACGAGGGGACCCAGGCCTAGAGGGGGACTGGGCACCCAGCGATCCGGGCCCT | 120 |
| chr22 | 50697257 | 50697377 | 785447_45746696_NM_001372044.2_205 | 1 | + | good | ACCTGGAGGGGTGGGGGGGGCGCCCCTCCCTCCCGTTCACCGGCTCCAGGCGGCTTTGCTGGTGCCCGAAGCCCCCGCCCCATCCCCCGCTCCCACTAGGCTGCCCTGACCACCGTCCCG | 120 |
| chr22 | 50697377 | 50697497 | 785447_45746696_NM_001372044.2_206 | 1 | + | good | CGTCCGCGTCCGAACTCCCCCTCCCGGGGGTCGGCGGCGAGGGGAGGGCGGGAGGGAGGGCGCGAGGGCCGCCACCACCGCCCGCAGAGGGAGGAGCCCGGCCGTGGAGGAGGCGG | 120 |
| chr22 | 50697497 | 50697617 | 785447_45746696_NM_001372044.2_207 | 1 | + | good | CGGGGCGGCCGCGGCATGGAGCGAGCCTGGCGCGCCCAGGAGCTGTATTCGAATTCGAGCTCGGTTCCCCGCGCCCCCTGCGCCCCCCGCACCGCCGCCCCGGGGCCCGAAGCGGAAACT | 120 |
| chr22 | 50697617 | 50697737 | 785447_45746696_NM_001372044.2_208 | 1 | + | good | TTACAGCGCCGTCCCCGGCCGCAAGTTCATCGCCGTGAAGGCGCACAGCCCGCAGGGTGAAGGCGAGATCCCGCTGCACCGCGGCGAGGCCGTGAAGGGTGAGGGGCGCGGGGGGGCG | 120 |
| chr22 | 50697737 | 50697857 | 785447_45746696_NM_001372044.2_209 | 1 | + | good | GGGGGGCGGGCCCGGCGCGGGGAGGGGGCGGCGCCGCGCGCGGTGCTGGCCGGGCCGGGGCAGTGGCTCTGGGGTCTCCTCTGCCGGGGCGGCCCTGGGCCCTTGTGGGATCCCTGG | 120 |
| chr22 | 50697857 | 50697977 | 785447_45746696_NM_001372044.2_210 | 1 | + | good | CACGGTGAAGGGCTCTGCCTGGGGAAGGTTCCTGCCGTGCGGGTCCCTCCGGTGCTCTGTCGTTCCGGGCTCCCTGTGTCACCACGGAGGCTCCTCTCTCGCCACGGGCGTTTCTGTGTC | 120 |
| chr22 | 50697977 | 50698097 | 785447_45746696_NM_001372044.2_211 | 1 | + | good | CCCGGGGGTCTCTGCCTGAAGGACCCTGTCCCATTACAGAGCTTCCTTGCATTGCGGGGTTCCCGTGGCACTTCTGCAGCTTCTCCATTGGAGGCCCCTGCGATGTGGGGGACCCTTGCC | 120 |
| chr22 | 50698097 | 50698217 | 785447_45746696_NM_001372044.2_212 | 1 | + | good | GTCGTGGGGTGTCTGTGACCGTCATGTGGGTGTTTGTGTTATGCAGGCTTCTGTCACCGGGGCTTCCTGTGTTCTGGGGGGATCGCGTGCCATGACAAACCCCTCTCATTCTGGGGGTCT | 120 |
| chr22 | 50698217 | 50698337 | 785447_45746696_NM_001372044.2_213 | 1 | + | good | CGGGGCCATCACTGGGCTCCTAGCCTCAGGGCCGGCTGAGGTGGAAACAGCCCAGCTGGTGCATCACGTGGCCTCACCCACTGGCCACAGCACGATGACCCCGAGCTCTCGGCAGTGACC | 120 |
| chr22 | 50698337 | 50698457 | 785447_45746696_NM_001372044.2_214 | 1 | + | good | CCTGGGTGGGTGACAGAGCCAGGATGGGGGTCGGTTGAAGGGGCTGGGGGAGCATGGTCAGCTGGGGGTGGGGGCAGCAGCAGGAGTGTGGCCCCTGCCCCTGCCTGCGCCCCTCCCC | 120 |
| chr22 | 50698457 | 50698577 | 785447_45746696_NM_001372044.2_215 | 1 | + | good | GTGTGTCCATCTGTGTGTCTCTCTGTCCCCCACATGCCCACCCTGTGCCGAGCCCATCTGTTCCTTTCTCTCTTCTGCGTGGATCCCAAAATCTTCCCAGGGAAAAAGCTGGGAGAAAGT | 120 |
| chr22 | 50698577 | 50698697 | 785447_45746696_NM_001372044.2_216 | 1 | + | good | GGGAAGGGAAGGAGGGAAAGGGCAGGGGGTGGGTGGGCAGAACCTGCTCCTGAGGTGGGGTAGGCGCCCAGCTCTGCTCCCCACTGACGGCCTGTCTGGCTTCTTCCTCCAGTGCTCAGC | 120 |
| chr22 | 50698697 | 50698817 | 785447_45746696_NM_001372044.2_217 | 1 | + | good | ATTGGGGAGGGCGGTTTCTGGGAGGGAACCGTGAAAGGCCGCACGGGCTGGTTCCCGGCCGACTGCGTGGAGGAAGTGCAGATGAGGCAGCATGACACACGGCCTGGTGAGTGACCCCAC | 120 |
| chr22 | 50698817 | 50698937 | 785447_45746696_NM_001372044.2_218 | 1 | + | good | GGCTCCCCGGGCAGCTCCCAAGGGGACCACCCCTTCCAGTTTCCCTTTGTTCTCTCTTGGTGCTAAATCCACATGGATATTCATAGAGAAAAGACTAGAGGTAAACTCAAACAAACACTC | 120 |
| chr22 | 50698937 | 50699057 | 785447_45746696_NM_001372044.2_219 | 1 | + | good | AAAGTAGATGCAAACTTGTGTACATGACACAGACACGTGTGCACACACTCTGCATATACTTGGGAACATGCGTGTATGTAACCTGACGCTTCACGTGCAGTGGAGACACATAGGACGTGT | 120 |
| chr22 | 50699057 | 50699177 | 785447_45746696_NM_001372044.2_220 | 1 | + | good | GTGACGGGGCCTCTGTCTGTGTGATCCCACATTGACTCCCAGTGACTTGCACCCCACCAGCACAGTCCTTCAGAAACACCAGGTGTGTGGAACGCATGATTCCTGCGTAGCTGGCAGACA | 120 |
| chr22 | 50699177 | 50699297 | 785447_45746696_NM_001372044.2_221 | 1 | + | good | TGTAAGGAGGTCAGTGTGAGAAAAGAGGCGTTTTTCCAAAGTGGACAGATTGTCCAAGTGGGCAGAGCAGGCAGGCCCGGAGCAGCCAAGAGGAAATGAGGCCAGTTGGGCCTGGAGGTG | 120 |
| chr22 | 50699297 | 50699417 | 785447_45746696_NM_001372044.2_222 | 1 | + | good | CCCTGGTGAGTGCCCTGGGGCAGGGATGGTCTGAGGGCAGCAGGTGAGGCTGGGCTGGTCCTCAGCCTGCAGGGAGGATGTGAAGGTGAAGGCTGCAGCTCCTGGGTTGGTGAGTGGCT | 120 |
| chr22 | 50699417 | 50699537 | 785447_45746696_NM_001372044.2_223 | 1 | + | good | CTGTCCAGCCCTGCTGACCATCTGGTCCTTTGGGGGCCCCCGGGCTGGAGCTGGGTGCGTGTCTTGGGGGTGCCCTTGCAAGGAACCCTCAGGGGTCCCGGGAGGCCCCCAGATCCATGC | 120 |
| chr22 | 50699537 | 50699657 | 785447_45746696_NM_001372044.2_224 | 1 | + | good | ATGTGCTTCTGTCCTGGAGAGCTGGTGGGCCAAGCAAACCTCTCCTGAGTGATGGTCACTGGGGGCCATCGGTGGTGTGTCTGGATCAAGGGTGCATGCACCCTCCCTCTGCATGTGAAG | 120 |
| chr22 | 50699657 | 50699777 | 785447_45746696_NM_001372044.2_225 | 1 | + | good | GGCTCAGGCCTGGGGTTACTGTGTCCCCATCTCTGTGTCCCCACCTCTGAGAGTTTCCCAGCGACTCCACCCCTGTACGGCCTGGACCCCTGCCCTGTGCTGAGCTCAGCAGAGGCCCAG | 120 |
| chr22 | 50699777 | 50699897 | 785447_45746696_NM_001372044.2_226 | 1 | + | good | GGAGGCAGGAGCTTCGCCACTGACCTTTTCCTGGGCCGGTGCCCTTTCCTCCTTCCTTGGCCTTGTTCTGCCTTGCTCTGACTGGTGGCTTAGAGTGTGGAAGGGACTTGGCCCCCTGTT | 120 |
| chr22 | 50699897 | 50700017 | 785447_45746696_NM_001372044.2_227 | 1 | + | good | CTCAGCCCGCGGAGGGTGGATAGGGCAGGGTCCAGATGGGAAATGGTTTCAGAGATTTGGGCCCTGGTTCAGACAGTATGGGAGAGGGGAGGGGAAGGAGAGGTAGGGGGATCCACACC | 120 |
| chr22 | 50700017 | 50700137 | 785447_45746696_NM_001372044.2_228 | 1 | + | good | GTGGCGGGAGCGGGTCTTGAGCTCCCCCACGTGGTGGTGAGGAAGGTTCTGGTTTGAGGACTGTGTGGAGTGAGGGGCATTTGGACCTACGTCTGAAGCTGACGGGGAGGGAGGGGGGTT | 120 |
| chr22 | 50700137 | 50700257 | 785447_45746696_NM_001372044.2_229 | 1 | + | good | TAGGCTGGAGTCAGAATTGGAGAGCAGCAAATGTTTTTTAGGTGTTTGTCATGTGCTGGGCCCTGTGCTGGGAGCCGGGATCACGGAGGCTCAGCTGGAAGCCTTGTGGGAGCAGTGGCT | 120 |
| chr22 | 50700257 | 50700377 | 785447_45746696_NM_001372044.2_230 | 1 | + | good | GTGACGGGGGCCGTGTGAGCGTGGGAGCTCCCTGGGCTGGCCTGAAGGAGTCAGAGAGGGCCTCAAAACAGAGGCATTGTCTGAGCTGAGTCCTCTAGGTGGTCAGCGGCTGGGGCTGAG | 120 |
| chr22 | 50700377 | 50700497 | 785447_45746696_NM_001372044.2_231 | 1 | + | good | GGGACAGCCGTGTGTGCAGGGTCCGTGTGGCGCTCTACAGCTGTGTGTCAGCACGGCCTGCTCTGAGTGCCTGGCACAGTTTTAGCACTTCTACACGTATAAGTCGTTGACTCATCCCAG | 120 |
| chr22 | 50700497 | 50700617 | 785447_45746696_NM_001372044.2_232 | 1 | + | good | CAGTCCCAAGAGGTGGGGCCACTTGTTGTTATTCCATTTCTAGGGGGTCACATGGCCCTGAGGGGCAAAGTCTTGTTTGCAACACAGGCAGCAAGTTCTATGTGAGTGAAACGCCTGCAT | 120 |
| chr22 | 50700617 | 50700737 | 785447_45746696_NM_001372044.2_233 | 1 | + | good | GGGGTGAGGGGCGAGGCTGGTGTGGGGACCACGGGGTGGTCAGGTGGCATGGAGTCAAGTGACAGTGTGGCATCATGGCGGAGGGAACCCAGCCGCAGTGTCTGATTGTGCAGGTTGTG | 120 |
| chr22 | 50700737 | 50700857 | 785447_45746696_NM_001372044.2_234 | 1 | + | good | AGGATCAAAGCCCGGGCTGGGCAGAGCAGAGCAATGGAGAGGAGGGTTCAGGTCTGGGCTCCAGCCAGCAGCCATGGTGTGTGTGACCTGGGGAGTGAGTGTCAGAGGTCCCGGGGTAG | 120 |
| chr22 | 50700857 | 50700977 | 785447_45746696_NM_001372044.2_235 | 1 | + | good | CGGGAGTATTTCATGAGTGGGCAGCGGGAGAGGGACCTGCGGAGGCTGCCTGCGAAGGGCAGGGAGTGGGAGCAGTGACAGTGCCCTCTGACCTTGGCTGCAGGCCCTGCTGGCCACACT | 120 |
| chr22 | 50700977 | 50701097 | 785447_45746696_NM_001372044.2_236 | 1 | + | good | GCGCAGAGCCTCTGCTCTGGGATCCCGGGGCCGAGTTCTGCCAGAGGCCAGAGTGCAGCCCAGGAACAGGAACCTGCTTCCAACAGGCCCAGATCTGCTCCCCCCACCCCCCAAAAAAAT | 120 |
| chr22 | 50701097 | 50701217 | 785447_45746696_NM_001372044.2_237 | 1 | + | good | CCATTTCCGGGACTGGAGTGGGGACAGAAAAGGACTTAAAGGTTAGGGAGGGGACATAGTCCCTGTGCGTGCAGGGGCTGGTGCCCAGAGCCGGTGCCTGCTGTGGGATCGGGAGGCGAG | 120 |
| chr22 | 50701217 | 50701337 | 785447_45746696_NM_001372044.2_238 | 1 | + | good | TCCTGGTCACTTACCCAGCAGAGAGCAGTCTCCCACGGCACCTCCTGGCAGGCAGAGGGTGGAGCTGTGAGGGAGGGAATGGAGAAGCGAGGTCAGGTGTGTTCTGGAGGCCCAGCAGGA | 120 |
| chr22 | 50701337 | 50701457 | 785447_45746696_NM_001372044.2_239 | 1 | + | good | GCAGGCCCTGTGCCTTCCCTCCCCTCCCTCCTTCCCTTTCTTTGCAGTAAGGTCAGCACTTGGGTGGGTAGTAGGAGATGGCTGTGTTGGGCATTTTCTCTTCAGCTGGCTTTTATGTGT | 120 |
| chr22 | 50701457 | 50701577 | 785447_45746696_NM_001372044.2_240 | 1 | + | good | CTTATTAACAAAACTGGAAAAATGAGTTTGTGTTATGTAATCCATACAATTTGTTTGGTTGACAACATCTGAAAACAGGAGGCTGGTCAAGTGCAAGTGTGGCGGGTTCTTCAGCAGGCA | 120 |
| chr22 | 50701577 | 50701697 | 785447_45746696_NM_001372044.2_241 | 1 | + | good | GCACCATGACACTTCCGTGCCTCTGTCGACGTGGAGAAGGGTGTGATGGAGGAAAGGAGGGGTCTGATGCCCCACCTGAGTGGTGATGAGGCTTGGCCTGAGCTGGCCTTAGGTTCAGCG | 120 |
| chr22 | 50701697 | 50701817 | 785447_45746696_NM_001372044.2_242 | 1 | + | good | GGATGAGGGGTGCAGGTGTGGTTGGGGAATGCCAGGATAAGATATAGGTCATCAGAACCAAGGGGATCCTCCAAGTTGGATTTCTCTATTCTATATGGGCGGGCTTAGAGGGGGCTGTTG | 120 |
| chr22 | 50701817 | 50701937 | 785447_45746696_NM_001372044.2_243 | 1 | + | good | ACTTACTGAGAGTCACACAACGTAGTAGCTGGGACCTGGAATCCAGCCCGATACTTTTTTGATATCTGTGCCTTCTGATAGAAATAGGCACTCTTTCCTCTTTCCGTCCGGCCACTGCTC | 120 |
| chr22 | 50702417 | 50702537 | 785447_45746696_NM_001372044.2_248 | 1 | + | good | AAGCTGCAGGTTATCTTTCTGATCTCATTTCCTGCCACTCTCTCTAGCTCCCTCTGCTCCAGCTGCACAGGCTCTGAACTCTCTCAAATGCTGCAGGCTGATGCCTGCCGTGGGCTTCCC | 120 |
| chr22 | 50702537 | 50702657 | 785447_45746696_NM_001372044.2_249 | 1 | + | good | CTGACCACCCCCGCTCCTGGGAAGCCCTTTGCCCACATATCCACAGGCTGCTCCCCATCCTCAGCCTCTGCTCAGCCCTCCCGTGGGGGTGAGGCCTTTGCGTGCCTTTCTCTGCCCTCC | 120 |
| chr22 | 50702657 | 50702777 | 785447_45746696_NM_001372044.2_250 | 1 | + | good | CTGATCTGCTCGCTGGTCTCCTGAGCACTTGCCAGCATTCGCATGCTATGTCTTTATCCTCTGTTTATCCTGTCTCCCAGAACAGATTATAAACCCCACGAGTGCAGAGACTTTGTTCTC | 120 |
| chr22 | 50702777 | 50702897 | 785447_45746696_NM_001372044.2_251 | 1 | + | good | TTCTCTGCTGTATTTGCAGAGCTGGTACACAGTAGGAGCTCAATAAATACTTGCTGAAGGAAAACATGGATCTTGAGAAATGCTACCAAGAGAGGAAATGAGGAAGGTGCTGAGGGATGA | 120 |
| chr22 | 50702897 | 50703017 | 785447_45746696_NM_001372044.2_252 | 1 | + | good | AAGCGAGGCAGTGGTGGTGCTGCCTGTGGGGCCTTTGCGGTCCTGGAGAGTGGAGCTGCGGGGCTGGGAAGGCTGGTGGGTGGGGCTCGCAGGCAGCTGGAGGAGCTCGTGGGAGCAG | 120 |
| chr22 | 50703017 | 50703137 | 785447_45746696_NM_001372044.2_253 | 1 | + | good | AGGTCATGGCTGGAGGTGTGGCTCGAACCTTCAGGTCAGTGAGGGTGGAGTGGTGGGCAGACAGCTGGCTAGCTTGGTGCTGGCCATGTGGGACTTGAGATCAAGAGGACAGTTGGGGAC | 120 |
| chr22 | 50703137 | 50703257 | 785447_45746696_NM_001372044.2_254 | 1 | + | good | CAAGCTCTGTGGGATGCATCGAGGTCAGGGCAGGGGACCCAGCCCACACACCTAGATCCCCTGCCCCACTCACAGTTAGTCTGAAGCAAGGGACCCCATGAGGTTGAGCATCAAGGAGCT | 120 |
| chr22 | 50703257 | 50703377 | 785447_45746696_NM_001372044.2_255 | 1 | + | good | ATTGTGAAGAAGTTGGTGAAGACAGAACGCTGCTTCCATGCCAGCGCCTATAGGCACATAGCACGGCATTTGTACATGAGTGTGGGTAATGCAGATTCAGATGTGCACACACACATGCAT | 120 |
| chr22 | 50703377 | 50703497 | 785447_45746696_NM_001372044.2_256 | 1 | + | good | GAACACACCTCACACGCACATGACGGCAGGTCACACAGACACATGAGCACGGCACTTGCACACAGTGGGCTCATATAGAGACACGTGCATGAGCACAGCATCTACACATGACCACGGATC | 120 |
| chr22 | 50703497 | 50703617 | 785447_45746696_NM_001372044.2_257 | 1 | + | good | TCTCACACACACACGTATGTGCACACCCCCCTCTTTGCTCGCAGCCTTGGGGCTCACTTACGCTCTGTGGACGGCTCTGGCCTTGTTTGGACTCCTGGTTCAGCAGCTACTTAGGAAAGA | 120 |
| chr22 | 50703617 | 50703737 | 785447_45746696_NM_001372044.2_258 | 1 | + | good | GATGAGCTGCTGCCCCCTCTTGGGCCCTGACCCACGGAGGTCAGAATTCCCTACATCATCCCCTCAGCCTTCTTACCTCTGGACCCAGGACTCCAGGATTTGGGGCCCCCTACTTCCATC | 120 |
| chr22 | 50703737 | 50703857 | 785447_45746696_NM_001372044.2_259 | 1 | + | good | TGGACACAACCAGGCCAAGGGGGTGTGTGTGGGCAGAGACTGGTGACCAGCATGGGTGAGGGCAGTTGAGGCAGGCCAGCTGGAGGCCACGACTGTCCATCAGCTCCCGATACTCCCTTC | 120 |
| chr22 | 50703857 | 50703977 | 785447_45746696_NM_001372044.2_260 | 1 | + | good | AGAAACGCGGGAGGACCGGACGAAGCGGCTCTTTCGGCACTACACAGTGGGCTCCTACGACAGCCTCACCTCACACAGGTACGTGCAGGGACCCTGGCTGGCGGGAGCATGCATGTGGGA | 120 |
| chr22 | 50703977 | 50704097 | 785447_45746696_NM_001372044.2_261 | 1 | + | good | CGTGGGGCAGGACCTGCAGTGTAGAGGGAGGCCAGGTATGAAGAGGTCAGAGGAGGGCAAGGGAATGTGCCTGGCCGACCTCTCTTCTGCTGCCCGGCCTGTCAGTCCCGGGGTGACTGC | 120 |
| chr22 | 50704097 | 50704217 | 785447_45746696_NM_001372044.2_262 | 1 | + | good | AGGGTGGACTGTGGCCGGCCAACCTCGAGGCAGGGCTTACCTCACTCCTCCCTGCTTTCCTTCATCAGCGATTATGTCATTGATGACAAAGTGGCTGTCCTGCAGAAACGGGACCACGAG | 120 |
| chr22 | 50704217 | 50704337 | 785447_45746696_NM_001372044.2_263 | 1 | + | good | GGCTTTGGTTTTGTGCTCCGGGGAGCCAAAGGTAATGGGGAGTGGGTGCCCGGGGGTCAGGCAGGCAGGGGCTGCACACCGCTCAGGGATGTCTTCAGCTTCTTGCTTCTGCCCCCAACT | 120 |
| chr22 | 50704337 | 50704457 | 785447_45746696_NM_001372044.2_264 | 1 | + | good | CTGCTCTGCCCCACACTCCCTCCCTGCCTTGCCCCTGCGGCTCCCGCTTTTGGCCCCGGCTTTCCTGGTTCCTCTGCTCGGTCGCGACCGTCGTGGTTTGCAACGTCCCGGCCCAGGCGG | 120 |
| chr22 | 50704457 | 50704577 | 785447_45746696_NM_001372044.2_265 | 1 | + | good | CAATTCCTTCCCTTCGTTGTTTTCCACTGTCTTTGCTCTAGAGTCTAACCGAGTGGGGGCTTTTCTGGCATGGGCAAAGGGAGTTTTGCAGAGGGTCATGTCTAGGATCCTTGGGCCGGC | 120 |
| chr22 | 50704577 | 50704697 | 785447_45746696_NM_001372044.2_266 | 1 | + | good | TCGTCCCTGCGCTGTGGCTCCCCGTGGCCCTCCCCCAGCCCCGGGTCACTGCCCCCCGCAGTCATCTCTTTCCTGTGCTGAGCCACTCGGAGGTTGCTGTGTGGCAGCCGCCCCCACCCG | 120 |
| chr22 | 50704817 | 50704937 | 785447_45746696_NM_001372044.2_268 | 1 | + | good | GTGGCCTGGAGGGCCGGGCTGCGCACGGGAGACTTCCTCATCGAGGTGAGGTCGTTCTGGCCGGTGCTGCCCAGTGGGATGCTGAGCCCTGAGCCCTGTGGTGGACGTGCCTGGGGCGTC | 120 |

| | | | | | | | | |
|---|---|---|---|---|---|---|---|---|
| chr22 | 50704937 | 50705057 | 785447_45746696_NM_001372044.2_269 | 1 | + | good | CCCACCCAGCTGCCTGTCTATCCCAGGTGAACGGGGTGAACGTGGTGAAGGTCGGACACAAGCAGGTGGTGGCTCTGATTCGCCAGGGTGGCAACCGCCTCGTCATGAAGGTTGTGTCTG | 120 |
| chr22 | 50705057 | 50705177 | 785447_45746696_NM_001372044.2_270 | 1 | + | good | TGACAAGGAAGCCAGAAGAGGACGGGGCTCGGCGCAGAGGTGAGGGGTCACGCTTCAGGCCTCTGTGCCCAAACTTTCCCCTGACCTTAGACCTTTGACTTCAGGCCCACATTCCTCTTC | 120 |
| chr22 | 50705177 | 50705297 | 785447_45746696_NM_001372044.2_271 | 1 | + | good | CCATCCCTGACCCCCAACTGCTGACATCAGATTCCTGAACCTCAATGAACCACCCACACGTGTGGGCTGGGACCCCTCCCGATCCCTGGTGCTAGACTGCCAGTCCTGTCCCCTGGCCTG | 120 |
| chr22 | 50705297 | 50705417 | 785447_45746696_NM_001372044.2_272 | 1 | + | good | AGGCCTTTGACCCCTGACCCCTGACCTCAGGCTGCCCCTCCTCAGATCCCTGCCCCCTCCTTCCCGGCGTTCCCCCAGCCCCTGGCGTGGCAGAGCCCTCCTTCCCTACACCGCCAGCAG | 120 |
| chr22 | 50705417 | 50705537 | 785447_45746696_NM_001372044.2_273 | 1 | + | good | CTGCCTGGAGGCCGGGGTTGCCATAGCAACTGTGAGGTTGACGCTGCCGCGCTGCTTATTGTCGGAGGGAAGGGGGAGGCGGCACCTGAGGGAAGAGGAAAAGCGTCTTCTCCCCGCGAC | 120 |
| chr22 | 50705537 | 50705657 | 785447_45746696_NM_001372044.2_274 | 1 | + | good | GACGCGGCACAGCCTGGGCCCAGGAGGCTTTGCCGGCCAGGCCCAGCCCAGGCCCGTGGCTTGGCTGACCACACCCTGGCCCCACAAGGGGCAGTGGCCACCCGGGCCTGGGGGGAGCC | 120 |
| chr22 | 50705657 | 50705777 | 785447_45746696_NM_001372044.2_275 | 1 | + | good | TGGAGCAGCAGGAGCCCCGCCCGCAAGGAGACCCCTCCCAGCCCCCACCCCATGGCTCTGGTGCCCCCTTGCTTAGCCCTCGAAGCCTGCAGGGTGGGTGCTCAGGCCAGAGCCAGCCGG | 120 |
| chr22 | 50705777 | 50705897 | 785447_45746696_NM_001372044.2_276 | 1 | + | good | CACCGTGACCCCCGCCAGCACCCTTGATCGTCTCCTGAGGCCCTGATTCTGGGGGTTCCCAGGGCCCCTGAGGCCCTGACCGCCCGCCCCCCCCCCCCGCACTCCCTGAGCTCCTTGAG | 120 |
| chr22 | 50705897 | 50706017 | 785447_45746696_NM_001372044.2_277 | 1 | + | good | CCATGAAGAAGTTTGCGTCCAGCCGTAACCTGAACAAGATCCTGGCACAGTGCGACTCATCCTCGCGGGAGTACGAGGAGATCCAGGCGGTGGAGCGCAAGTGGCACCTGCACCTGGCCA | 120 |
| chr22 | 50706017 | 50706137 | 785447_45746696_NM_001372044.2_278 | 1 | + | good | CGCCGCGCCGCCTGCTGCTGGACAGGAGGTCCAAGGCCTCCCTCTTCTTTGCAGCCCCACCGCCCCCCAAGAGGGCCCCCAGCACCACACTGACCCTGCGCTCCAAGTCCATGACAGCTG | 120 |
| chr22 | 50706137 | 50706257 | 785447_45746696_NM_001372044.2_279 | 1 | + | good | AGCTCGAGGAACTTGGTGAGTGGCGGGGGTGGCGGTGGAGGTGGACGCAGGTGGACGGTCCATGATGGGCAGACAGGGCCGGGAGACACAGGTGGTACGGGAGGGCAGGCGGTCCAG | 120 |
| chr22 | 50706257 | 50706377 | 785447_45746696_NM_001372044.2_280 | 1 | + | good | GAGGGTAGGAAGGACAGGTGGGTACAAGAGCAGGTCAGAGGTGAATGGGTGGGTGCTGGCGCGATGGGGAAATTTGGTGCTTGTGGGGTGAGGAGAGGGTGGCATGTGGAAGGGGTGA | 120 |
| chr22 | 50706377 | 50706497 | 785447_45746696_NM_001372044.2_281 | 1 | + | good | TGTGAGAGGCTGGGGATGGACAGTGGGGTGTCCAGGACAGATGACGGGGAGCAGGGCCTGGCCCACCAGTATGTTTGCATGTGTGTGCATGTGTGTGCGTGTGCTGCCTCTGGTTTTCCA | 120 |
| chr22 | 50706497 | 50706617 | 785447_45746696_NM_001372044.2_282 | 1 | + | good | CGTCTCCACTTTGACTTTACTCCTGCATTTCATCCAAGTGAGGAGAGAGCGGTAGTGAGGTGACCAGGGTCTGCTGCTGGCTCTGCCAGGCACTGGCTGTGTCAGCTCTGTGTTTATTTT | 120 |
| chr22 | 50706617 | 50706737 | 785447_45746696_NM_001372044.2_283 | 1 | + | good | ACCCTCTTGGTGACTGTGCACTCATTGACAGGTGGCCCTGCTGACCCCACAGGCCACAGGAGTACTGTCGGTGGGATGATGGGGCAGGGCCTCGAGCGCGGCAGGAGGGGAGCAGGAGTG | 120 |
| chr22 | 50706737 | 50706857 | 785447_45746696_NM_001372044.2_284 | 1 | + | good | TGTTGGGGTTTGATGGGGCATGTTCTGGGCACAAGCCCTGGTTGACCATCAACAGCCTCTTCATGGGCAGGTCTTATATATGTGTCCATTAACAACCTTGTCATGTGCACGTCCAGATAT | 120 |
| chr22 | 50706857 | 50706977 | 785447_45746696_NM_001372044.2_285 | 1 | + | good | ACTTGATCATGTCCTGACGTGACAGTCAGTAGCCTCTTCATGGGACTGTCCCATTACGTGTGACCGTTGTCCTCACTGTGGGTATGTGCTATACATGTGGCCATCAACAGCCTCGTCATG | 120 |
| chr22 | 50707337 | 50707457 | 785447_45746696_NM_001372044.2_289 | 1 | + | good | TCTGTGATCTGAGCCCAGTTCTGTTCCCAGACTTAACTCTGCCTCTGGTCATAGAAGGAACCCCAACTTTGGTCCTAAACTGAACCCTGAAGTCCTGGCCACAGGAAGGGACCCCATTCT | 120 |
| chr22 | 50707457 | 50707577 | 785447_45746696_NM_001372044.2_290 | 1 | + | good | GGTTCTAGGCTGAGCCCTATCCCTGGCTTCAGTCTTGGTTCTACCCCAGCCATAGAAGGAACCCTTTTCCTGGTCCCCAGAATTATCCCTGACCTCACTCATGAACCAAGCCCTGACCCT | 120 |
| chr22 | 50707577 | 50707697 | 785447_45746696_NM_001372044.2_291 | 1 | + | good | GTCCTTGCCACAGGCTGAGCTGCCCCATGCCAAGCAGGTGGTCTTAACTGCAGTTTAGTTCAGGGTGGAGAACAGCTACCAACGTCCATTGGGCTGACGTGGTCAGAATTTTCACCATAG | 120 |
| chr22 | 50707697 | 50707817 | 785447_45746696_NM_001372044.2_292 | 1 | + | good | CTATTCCAAATAAAATAAAACCCAAAGAAAAAAACAAAAAACCCCATGTAGATAATTCAGAATGAAAAGGAGAGAAACCAAAGCCAGAGGCTTCCATAGCTCGTTATCTAGTACGCAGGG | 120 |
| chr22 | 50707817 | 50707937 | 785447_45746696_NM_001372044.2_293 | 1 | + | good | GCAAGGGGTTGGGGTCCTGAGGCCCCCACAAGGCTGGGAGACGTGGCTGGACCTGTGAGAAGCGGGAAGGTTAAAAACCAAGCCCAACCACTGGAATTAGCTTCGAAAGAACTGGAAATC | 120 |
| chr22 | 50707937 | 50708057 | 785447_45746696_NM_001372044.2_294 | 1 | + | good | AGAGAAGAATATGAAAAACATTAAAATAAAGTAAATTGTCACAAAGATTAAAAGAACTAACAGAACTAATGAGAGAATAGGACAGAACAAACAAGCAAAAAAAGAATAAGCACATTTTAG | 120 |
| chr22 | 50708417 | 50708537 | 785447_45746696_NM_001372044.2_298 | 1 | + | good | AAAAAAGAAAGCCATGCCTGGGATAAACAGTACACTAGACCCAGGTGAAGAAAGAAACAGCAAGTTAGAAGATAAATCTGAGGTTACCCAGAATATAGCACAGAAATATAAGGAGATAGA | 120 |
| chr22 | 50708537 | 50708657 | 785447_45746696_NM_001372044.2_299 | 1 | + | good | AAATATCAAAGAGCAGTTAGGACACACACACACAAAAATCACATAGATAATTCAATGGGACTATTGGGAAAATCCCAGCATCTACCTAATGCAAGGCGTGGAAGCAAAAATAAAATGTTG | 120 |
| chr22 | 50708657 | 50708777 | 785447_45746696_NM_001372044.2_300 | 1 | + | good | CCTGGAGGCAATATTTGGAGATGAGATGGCTTAAGAATTTCCGAAAGTTGAACAAAGACATGAGTCTTCAGTTTAAAGAAATATAAGTCTCAGCAAGGAAATAAAAACAAATCTGTATCC | 120 |
| chr22 | 50709137 | 50709257 | 785447_45746696_NM_001372044.2_304 | 1 | + | good | CAAAAAATGATGAGAGAGAGAGAGAGAGATGGGCTGCTGGATGTCTTAGACGGGAGCAGTGGTCCAACTGACAACAGACTTCTCATCAGCAGTGGCAGATGACAGTAGATGATGGAAGAAGA | 120 |
| chr22 | 50709257 | 50709377 | 785447_45746696_NM_001372044.2_305 | 1 | + | good | GCTTCAAAGTGTGGAGTCAAAATTCTTTTCCACCTAGAATTCTATACCCCGCTAATCCTGCTCATTCAGGAGTGAGGCAGAGAGGCTAATCCTAGACCTCTGGCACTCGGGTCATGAGGG | 120 |
| chr22 | 50709377 | 50709497 | 785447_45746696_NM_001372044.2_306 | 1 | + | good | TCCAAGGTCAGTCACGGGGAGCTGACCTGTCAATTGAAGATCCAAACTGATCAATAAGCCAGCACCAGTAGGCAGCTGGGTCCCAGGATTCTCCTGCCAGAGGCTGTCACATAGCCACAG | 120 |
| chr22 | 50709497 | 50709617 | 785447_45746696_NM_001372044.2_307 | 1 | + | good | TCAGCTGCATGTTCAGAAGGGACTCTTCTATTCAGCACCACGGGGCTTTTGGGCAGCCTGAGATCCGTAATCCTGGGACCCAGGGACTCTGCCCTATGGAAACGGAAGGACTGCGCTTTG | 120 |
| chr22 | 50709617 | 50709737 | 785447_45746696_NM_001372044.2_308 | 1 | + | good | CTCTCTGGGCTTTTCTGCCTGAAGAAACTGGTGAATGAAGACGCTCCTTTCCCTCCCCTCTGGGCTGAGTGTTCCCTCTTCTGCAACAAGCCTCAGGACAGGTTGCTGGAGTCACTTTGC | 120 |
| chr22 | 50709737 | 50709857 | 785447_45746696_NM_001372044.2_309 | 1 | + | good | AGAGCCCGGTCTTCCAGAGGAGTTGCTCACAGCTTCAGGCCACCATAATTTTCCAGGCTTTTTTCTGAGCAGAGTCTGGGGGAGGATGAAGGGACAGCTTAGCGCAGAGCCTCTACCCAC | 120 |
| chr22 | 50709857 | 50709977 | 785447_45746696_NM_001372044.2_310 | 1 | + | good | ACTTCTGGAGCTCAGGAAGCCACCCTGTCCCCAGAAGACAGCTGTCCTGGGAAGCCAGCAGGGCTTCTGGAAGTCCCCTCTGGAAATGCCTGATTTGAGGACGCCATGCCAGCATCAGGG | 120 |
| chr22 | 50709977 | 50710097 | 785447_45746696_NM_001372044.2_311 | 1 | + | good | TTTCCCCAGGTACCTAAGGGACCATCCCACAACCCAGTGTTTTTGCTTTCAGAAGATAGAAATGAGGCCTGTTGGTCACTGCTGGCATGAACACATCAGAACTGCCTCACACTGTCAAGA | 120 |
| chr22 | 50710097 | 50710217 | 785447_45746696_NM_001372044.2_312 | 1 | + | good | GCTATTAGAGCCCTTTAGGATCCATAGTCTATCGAGAAGGTGCCTGGACACCTAAAAAGAAAAGAAAGTGAGGTGTCAGAGAAAGCAGGTGCAAGTCGGAGGCTCCTGTCCAGGTGGCTG | 120 |
| chr22 | 50710217 | 50710337 | 785447_45746696_NM_001372044.2_313 | 1 | + | good | TTTGCTGCTCGGAGGATGTGGGAGAGGCATGGCCCTGTTGGAGGAAGGGGAGGGAAGGGGTATCCTGTGGGGGAATGGGGTGAGCAAAGGCTGGTGCCTGGAAAATGGAGGGTTGCAGA | 120 |
| chr22 | 50710337 | 50710457 | 785447_45746696_NM_001372044.2_314 | 1 | + | good | CTGGGAGTGAGAGGAAGGGGCTGGGTGGGGACCCTCCTGAGATAGGTGGTGAGAGAGAGAAGAGGGCAGAACACCCCTCCTCCCCTCAGCACTGGGGGCTCCTCATGGTCGCCACATGCA | 120 |
| chr22 | 50710457 | 50710577 | 785447_45746696_NM_001372044.2_315 | 1 | + | good | TGTCTGCTCGGTTTGTACCAGGAATGTGCTGGGGCCACAGGTCAGTGGAGACCCAGGCCAGGGAAGGCCCTGTGGCGGGTCCAAGCCCTTCCTGTGTATTCACGTTCCCGGGAGAACTCT | 120 |
| chr22 | 50710577 | 50710697 | 785447_45746696_NM_001372044.2_316 | 1 | + | good | GGACTTCCCTGTAATTTCTGGGCCCCTTTTTCTTTTGTGATGAGAACATCAGGGTAGGTGCCACACCCAGCATCTATGTGAGTGGCCTTGTCCTGGAAAAGGTTTCTGTGCCAAACACAC | 120 |
| chr22 | 50710697 | 50710817 | 785447_45746696_NM_001372044.2_317 | 1 | + | good | TTGCCCTGCCTTTTAAGACCGACTTTGAACAGAACACAGAGGTGACCACCTTAATTTTCAGCGACACCTGAACTTGTTTTCTATGGGGTTTTGCCTTTTTTCTGAACAAACGTTTGACTG | 120 |
| chr22 | 50711417 | 50711537 | 785447_45746696_NM_001372044.2_323 | 1 | + | good | ACCAACTGACTCCAGAGTTGGGCAGATTTGCTATTCACGGTCTGATAAAGAGGTGTAGGCTGGGGGCAGATTTGGGAGGTCATGGTTTTTGGGGGAGGTGGTGGTGAGAGGGATGGGGTG | 120 |
| chr22 | 50711537 | 50711657 | 785447_45746696_NM_001372044.2_324 | 1 | + | good | GGGAAATGAGTGTGGATAATGGGGCCCGGGTGCCGAGCCTGCCTCTTACTCCCTTTACTCTGTTTCTTGATTCCAAGCCTCCATTCGGAGAAGAAAAGGGGGTGAGTCATCTGCCTGTGT | 120 |
| chr22 | 50711657 | 50711777 | 785447_45746696_NM_001372044.2_325 | 1 | + | good | CCCCAGGGCCTTGGCTTTGCCTGACCCCTGTCTGGGGGAGTGCTGGTGTGTGAGGAGGTTCCAGCGCAAGTCAGGGTGGCCTGGAAGCCTGGGCTGCACCCTCCTGTTGGCCTTCTCCCT | 120 |
| chr22 | 50711777 | 50711897 | 785447_45746696_NM_001372044.2_326 | 1 | + | good | GGGCATCTGGTAGGGGCATGAGAGAAGGGCCTGCAAGGATGGATGGGGAGTTCCCAGGGGTGTGACCCTCTGAGACTGTGATGGCCTTGGGGGCTCAGGTGAGCCTGGGGCCTGTGAGCT | 120 |
| chr22 | 50711897 | 50712017 | 785447_45746696_NM_001372044.2_327 | 1 | + | good | GTGCAGATAGCCTGATGGAGGCAGCCCGTGGGGAGGAGCTGGGAGAGCCCCGTGGAGGCCACTGCATGCACCATGGCCCGGACAGCAGGAGGGAAAGGGCTCCGAGGGTCCCAGAGGG | 120 |
| chr22 | 50712017 | 50712137 | 785447_45746696_NM_001372044.2_328 | 1 | + | good | AGCTCTAGGGAGCAGGGAGACTGGGCCGTGTTGGCGTGCACGTTGCTTCTTTCTGCTGGTCCTCTCGGGGCCTGTGCTTGGGTTGTGGGACCTACTCCCGTGCTGGTTGGTTGTAGGGCC | 120 |
| chr22 | 50712137 | 50712257 | 785447_45746696_NM_001372044.2_329 | 1 | + | good | AAGCCCAGAGCTGATGGTCCCTTCAGAGCAGGGGTCTGAGTCCCACAGCACTGGGGAAGGAGCTTGGAGCTCCCCTGGCCAAAGACCCCAGGCTGTTGACTACCCTCCTGAACCCCCGTT | 120 |
| chr22 | 50712257 | 50712377 | 785447_45746696_NM_001372044.2_330 | 1 | + | good | CAGGCCTCCCTCTACCCTTTGGGGCTGCTCCCTGTATGTCCTTTATGTTCAGCCTGGTTGCAGGGGAGGGCGCATTCAGGTGGGGACAGTCACCTGGGCCTGGGTACTGAGGTCCACTGT | 120 |
| chr22 | 50712377 | 50712497 | 785447_45746696_NM_001372044.2_331 | 1 | + | good | CTGGTGAAGTCCTGGGACCCGGCTTCCGGCAGCCTCCTCTGCCTCTCGGGTGGATGGGAATCAGGGCCCAGACAGAGGCTTGAGGTGGCTTTCTCCACCCAGGCCTGTATATTTGCTCCA | 120 |
| chr22 | 50712497 | 50712617 | 785447_45746696_NM_001372044.2_332 | 1 | + | good | CGTCCGTGTACCTGATGCCCATCCCTCCCTCTCTCTCGACCCACTGGGCTTTCCCTGGGTGAGGACACATCTGCCAGGCCCTTGGCCCAGCCCTCTTGAGGTCCCCTTGCTGCCTCTTGC | 120 |
| chr22 | 50712617 | 50712737 | 785447_45746696_NM_001372044.2_333 | 1 | + | good | CCCAGCCCAGGTGAAGGGCTTCCCCAGTGTCTTCTCAGGTATTAAAAACAGGTGACAAGGTCTCCTCCAGTGACCTTTTCTGCATGCCTTTGGGGCAGACATACAAGCTATTTCTGGAAT | 120 |
| chr22 | 50712737 | 50712857 | 785447_45746696_NM_001372044.2_334 | 1 | + | good | CTTCTGGCTCTGCCCTCTAGCTCGTTAGTGGGGCTCCCAGGGTGCTGTGTTGCAGCCCAGGTGCTGCTCCAGTTGTGGAAGAAGTGATGGCTGGTTGATCATCCCGCAGGCTGGGGCCCT | 120 |
| chr22 | 50712857 | 50712977 | 785447_45746696_NM_001372044.2_335 | 1 | + | good | GTTGATCAAACAGCTGCAAACCCAGCTGTGTCCTGCTGTGCACCCAGAGGGCCCTGGGAGTGAGCGGGTCCACTTGGTTCCTGATGGGCTTTGTTTCCCTTAATTATCTAAAATTATTTA | 120 |
| chr22 | 50712977 | 50713097 | 785447_45746696_NM_001372044.2_336 | 1 | + | good | TTTAAGAATATATCCTCTAATATTGGGGGGAGATTTATGTCAACCTCATCAGTTTGTAATTTGCAGAATTCTTGATGAATCCTTTCCCTTTTGAAAACACTTCTCACCCGGCTCCAGCAC | 120 |
| chr22 | 50713097 | 50713217 | 785447_45746696_NM_001372044.2_337 | 1 | + | good | CATCGCCGGATGACCAGAAACAGCTGCTCACTAAACGTTATGTGCGTCCCGCTGTGGGTCTCTGGGATGGGTCTTCCAGACCTAGAGAGACAGCCTCTCCTAGCCCGTGGCTCAGGCGGC | 120 |
| chr22 | 50713217 | 50713337 | 785447_45746696_NM_001372044.2_338 | 1 | + | good | AACGTGTCTCCCATGATGCTGAGCCGTGTGTGCCTCACGGAGTTTTCTCTCCATTCATATCTTTACTTCTGCAGTTTCCGGGGAGTATGGTGCTTAAGTCTCATCTCTTACTTCCTAGTC | 120 |
| chr22 | 50713337 | 50713457 | 785447_45746696_NM_001372044.2_339 | 1 | + | good | AGAGATTTATGGCAATTTCGCTTCAGTTTGCTCCTGGGTTGGATAAGGGTGATGATAAAATCCTCGTGTAGTTGTGGGATGAGAGGAGTGGTTTGCATAAAGTCCCGCATACTCTAGAAA | 120 |
| chr22 | 50713457 | 50713577 | 785447_45746696_NM_001372044.2_340 | 1 | + | good | TCACCAAGAACTTAACAGATGGTGGCAATGGTGACTTCAACTCCCAAAAAGAAATCATTCACTTCATGACAAAAGCAAATGAGAAGTCCCCTCAAACCAGCAGCCCCTGGTCTGTACCTG | 120 |
| chr22 | 50713577 | 50713697 | 785447_45746696_NM_001372044.2_341 | 1 | + | good | GGCCCCTGTCACCTGGTGGCACTGGGAGCTGTGCACAGGCCCAGATATCTGCAGTTGGAGCCTGTTTTGCTGGTAAACTCTTGTATCGTTCAGGACACAGGCTGGGCGCTGATGGAACAC | 120 |
| chr22 | 50713697 | 50713817 | 785447_45746696_NM_001372044.2_342 | 1 | + | good | AGAGAACAGTATTTGCTCTGTTCTGCAGACCCGGAAGCTGTCTGGGGAGATGGGGCACACAGACATGAATTGTACTGAATTGTGTCCAGTGCCAGCACAGTTGTAAGCCTTTCTTCTGTG | 120 |
| chr22 | 50713817 | 50713937 | 785447_45746696_NM_001372044.2_343 | 1 | + | good | CTAATTCTCACTAGAGACACGTGAGCGAGCACTGTTTTTATGCCTGTTTTATCAATGAGGACACAGAGGTCACAAAGAGGTTATGTCAGGACAGAGTCAGTGTACAGGGTGGGTCTGGTC | 120 |
| chr22 | 50713937 | 50714057 | 785447_45746696_NM_001372044.2_344 | 1 | + | good | TGGGTTGGGTACAGGACAGTCTTGGGGAGTTCCTTCCTTCCAGCAATGCTAAGCTTGGGGAGTTCCTTCCTTCCTTCCAGCAATGCTAAGCAGATTGCAGAGGTAGAATCAGCAGGTGCT | 120 |

| | | | | | | | | |
|---|---|---|---|---|---|---|---|---|
| chr22 | 50714057 | 50714177 | 785447_45746696_NM_001372044.2_345 | 1 | + | good | GTGAACAGCTGCAGATTGGAGTTGGAAGGAGTAGGAGGTGTCCAGTCTGACCGTTGAAATTCTCCAACAAGGGGCCAAGACAAGAGTGAAAGGTGCCAAGACAAGTGTGGCCAGACAGTG | 120 |
| chr22 | 50714177 | 50714297 | 785447_45746696_NM_001372044.2_346 | 1 | + | good | GTCCCTCTAGGGTTGGCATAGCCCCAAAGAGTGGAGAAGGGCTTGGGGTTGGGTGCAGCCTGAGTGATGGGTGGGGTGCAGTGGGGCTGCTGAATGAGAGGGCCGGTGCAGCAGCCAGG | 120 |
| chr22 | 50714297 | 50714417 | 785447_45746696_NM_001372044.2_347 | 1 | + | good | GGACATGAGTGCTGCCCCAGCCCTTGCTGCATGCTGCACGCACTGGTCCTCAGTGGACCCACAGGGGGCGAGAGCTGCTTATGTCCTGGGGGAGCCATCCAACCAGGGCTCTGATGGTCC | 120 |
| chr22 | 50714417 | 50714537 | 785447_45746696_NM_001372044.2_348 | 1 | + | good | TGAGATACACGTATTAGTTTAACAGTTTCCAAGTCACTTTGATCTTCTGTGGAAACCTGCTTGTACATCAGGGTCTGTCCTCGGGCTGTGACCCGCTGTGAACAGCTTCCCTTCCAGTGC | 120 |
| chr22 | 50714537 | 50714657 | 785447_45746696_NM_001372044.2_349 | 1 | + | good | GGTACAGATAGCCAAGAGTGGCCTGTATGTGCTGAGAACAGTGACAGGACGAAGGGAAATGGAGGGAAATCCAGGGAGACTGTGGTCTCTGAATCGTTTGTCTGTCCTGTGCCCTGGTCT | 120 |
| chr22 | 50714657 | 50714777 | 785447_45746696_NM_001372044.2_350 | 1 | + | good | TCTCCCTCATGGTCAGAGTGGCCTTTCCTTAAGTGAGGTCCACAGGGTCACCTGGAGTCTCCTTGAGGCTAGAGAGGCTGCCTGGGGTGGGGTCTGAGTTCTGGGGTGAGGGCCTTGTCT | 120 |
| chr22 | 50714777 | 50714897 | 785447_45746696_NM_001372044.2_351 | 1 | + | good | TGGGAGCCTGGGGTGGGAGCTCAAGGAGGGAGGGGAGCGGTGGCCTCTGTCAGCATCACGGGTGGCCGGGTTGGGGAGGACATGGCAGTGGGGCAGGAGGTGGCCCAGGCAGCTGAG | 120 |
| chr22 | 50714897 | 50715017 | 785447_45746696_NM_001372044.2_352 | 1 | + | good | GAGCCTCCTTGCTGTGCAGAGAAGCTGGACGAGATGCTGGCAGCCGCCGCAGAGCCAACGCTGCGGCCAGACATCGCAGACGCAGACTCCAGAGCCGCCACCGTCAAACAGAGGCCCACC | 120 |
| chr22 | 50715017 | 50715137 | 785447_45746696_NM_001372044.2_353 | 1 | + | good | AGTCGGAGGATCACACCCGCCGAGATTAGCGTAAGGGCCACGGGCGGCTGGAGCGCTGGGTCGGGCAGGCATGGGGGTCAGACTGTCCTGGGCCCTCTTGACGGAGGTGAGAGCCTATC | 120 |
| chr22 | 50715137 | 50715257 | 785447_45746696_NM_001372044.2_354 | 1 | + | good | ATGTGGACCCCTCTCCAGAGGGGTGTGTGCACCCCATGGCCTCTCTCAGAGACTCTTGGGGCTGTCTCTGCCCCCTAAATGGCCACAGCTCAGCACCTGCTGGGTTTGCTCTTGTAGCTT | 120 |
| chr22 | 50715257 | 50715377 | 785447_45746696_NM_001372044.2_355 | 1 | + | good | TTGGTGCCTCTGCCTTCAGGCAGTCCCTGTCTCACGCCTGAGACCACCAAGGTACCCCAGCTCCGCTGAGAACACGACAAAGCATGTCTCTGTTCCCTGCCCACTTGCCAACCCGGCGCC | 120 |
| chr22 | 50715377 | 50715497 | 785447_45746696_NM_001372044.2_356 | 1 | + | good | TTCTAGGCTCCTCCCCCACCCTCACCCAGGGTCGGCCAGGGTCTGCACCCACCCTGCCCAACACCGTCTACACTGTACCACCATCTGCACTGCCCACACAGCCCAGGCTCCAATGGGCTG | 120 |
| chr22 | 50715497 | 50715617 | 785447_45746696_NM_001372044.2_357 | 1 | + | good | CATCCTCTGCTCCCTGAGGGCAGAGTCCAGACGTGATTCCTGGGTCCAGGCACCCACAGGCACTAGTGCCATTGGAGTGAGAGCGTGGGGTGTCTCACCTCTGGCTTAGGAGGAGGACTG | 120 |
| chr22 | 50715617 | 50715737 | 785447_45746696_NM_001372044.2_358 | 1 | + | good | GGGGCCCCCAGTGCCCTGGAACCTCCATATTCCCCTCCCTGACCCCCACAGTCATTGTTTGAACGCCAGGGCCTCCCAGGCCCAGAGAAGCTGCCGGGCTCCTTGCGGAAGGGGATTCCA | 120 |
| chr22 | 50715737 | 50715857 | 785447_45746696_NM_001372044.2_359 | 1 | + | good | CGGACCAAGTCTGTAGGTATGGCTGCGCTGTGGGGCTGCATGGGGTGGGGAGGAACGGGGCTGGGGCCGGCAGGGTGGACTTGGGTTTGAAGGACACTGCCTCTCTCTGCCCATAACTGG | 120 |
| chr22 | 50715857 | 50715977 | 785447_45746696_NM_001372044.2_360 | 1 | + | good | GGTTTCCAGTTTCCCTCTCAGTCCTGTCTCTTCCTGGCCCCAGCTAAAGGATCTCATACGTTGATGGACATGTGGGGATTAGGCCTTCCCCAACCCAGAGCTTTCCCCCGGCAGCCGACA | 120 |
| chr22 | 50716217 | 50716337 | 785447_45746696_NM_001372044.2_363 | 1 | + | good | TGCCTCCATCGCGGTTGCTCCCTTGCAGCCCAAGTGCATGTGAAGTTTCTGACCCTCAAACCCCCTGAACTTGCCTCTCCCCTACTTCTCCATGTATTGCTGTCTCCCCCCTTCACAGCC | 120 |
| chr22 | 50716337 | 50716457 | 785447_45746696_NM_001372044.2_364 | 1 | + | good | AGTTTCCCCAAAAAGTCACCTCTACTGTCCATCTTGTCTCCCGTCCCCCACCGGCTCCTCAGCCTGCGGCAGACTTCCTCTCTCCATACCCAAATTGAAACTGCTCCCACCAGGGTCAC | 120 |
| chr22 | 50716457 | 50716577 | 785447_45746696_NM_001372044.2_365 | 1 | + | good | CGGCGGCCTCCAGGGCCTTCCTCCCCATGGATGCATGGCCAGCACTCCTGCTGACCTCAGCTGGCATTTGGTTTCTTTGCTCCTTCTTCCGTGAAACGCTCCTCCCCAAAGCTTTAGGAC | 120 |
| chr22 | 50716577 | 50716697 | 785447_45746696_NM_001372044.2_366 | 1 | + | good | AACACTTACTGGTTTTCCCCTCCTTTCTCAGATGGCACCTATTTCAACCCTGCCGCCCCAGCCTCCCAAATATGGGACCTTTTCAAGGCTGTGTCTTCAGCCTCCTTCCCCTCTGTGCCT | 120 |
| chr22 | 50716697 | 50716817 | 785447_45746696_NM_001372044.2_367 | 1 | + | good | CACCACTCCCCAGCCACCATCACACTCCACACTCACCGTCTCCTTTCCTGACCCAGATGGGGACATGTAACAAGCCCTGCCCACTGTGCATCCCTCCAGACATCCACTCTCCCGTCCCCT | 120 |
| chr22 | 50716817 | 50716937 | 785447_45746696_NM_001372044.2_368 | 1 | + | good | GCCCACCTGTGCATCTGTCCACCCACACACCTGCCCACTTGACCGTCCGTTGTCCATACAAACCCCAGCACACCTCCCATCTGTTACACATAGGAGTAAACCTCTCAGCACTTACTACGG | 120 |
| chr22 | 50717057 | 50717177 | 785447_45746696_NM_001372044.2_370 | 1 | + | good | AGCTGCCCAAACTCGGATTCCTCACCTGCCTCAGGACCCTGCCAATTTGTGTTCACACAAGGCCGATCCTGGGCCAAGTCTGGGGCTTGGTTAGGGAATGTGACGTGTGCAGAGAGCTCA | 120 |
| chr22 | 50717177 | 50717297 | 785447_45746696_NM_001372044.2_371 | 1 | + | good | GCGGGCACATTGTGGGTGCCAGGGCTCACGGACCAATGGAGCCCACTGGGGGGCTCCAAGCACACCTGGTGTCACAGAAGGCTGTGTGTGTCGGGGTGGGGGTACTTTTGAGCTAAGATG | 120 |
| chr22 | 50717297 | 50717417 | 785447_45746696_NM_001372044.2_372 | 1 | + | good | TGCAAGATAAATGGGAGTTTCTGGGCGATAATGGGGTGAACTGACAGAGCAGTGTGGAAAAGGCATATGGGTGTGTGACCTGGTCCCTCAGAGACTGCCAAGAGTTATTTCTGTAGGTGG | 120 |
| chr22 | 50717417 | 50717537 | 785447_45746696_NM_001372044.2_373 | 1 | + | good | GATTGTGGGGAAGAAGGCTGGTGAGGCTGCAGCTACTTTGGAGCTGAGAAGCGCTCTCCCCTGCTCTCCCTGGCCTAAACACTGAGTGTCTTCCATGGTTTCTTTCTTATCCTCTGCCTC | 120 |
| chr22 | 50717537 | 50717657 | 785447_45746696_NM_001372044.2_374 | 1 | + | good | ACCTGATGTGTCGGGGTGGGTGGCCTCACAGCCCTGCCCTGCACAGGTCTCATCACCTTGCAGCTGGACCATGGCTTTGTTCTCATGACTGCCTGCCCTGGCTCTGCCCTCTGGTCCATC | 120 |
| chr22 | 50717657 | 50717777 | 785447_45746696_NM_001372044.2_375 | 1 | + | good | CCTCTGGCCAAATCAGGCTTTCCGCAGAGCTGCTATCACCCCATATCCGTCCAGCTTGACCCTCTGTGCACTCCTCATTGCCTCAGGGCCCTGGAGATCGGCCCCAGTGCTCTCCTGCTT | 120 |
| chr22 | 50717777 | 50717897 | 785447_45746696_NM_001372044.2_376 | 1 | + | good | CATCTGGAAACATTTGTCTCTGCCTCCCTTGACCTGGGTCTTCTGACTCTTGCTGTTCCTGGGCCCGACCCTGCAGGTCTAGGTCCATGCACCTGCCCTGCAGGCCTTGAGCCACTCTCC | 120 |
| chr22 | 50717897 | 50718017 | 785447_45746696_NM_001372044.2_377 | 1 | + | good | TAGGACTGCCAGTTCTGCACTGGTCCCTCAGGCACCCTCTTGTACTCGGTTGATGGATGGGTGGACAGGTGCTCCTTTTTGGCTCCTCTTGGCCTGCTGGGCCTCAGCTGCCAGCATGCA | 120 |
| chr22 | 50718017 | 50718137 | 785447_45746696_NM_001372044.2_378 | 1 | + | good | CATTGGTTGGGTTGTCAGACTCCCCAGGCCCCTGCCCCATGCCCTGGCTCTGTTGCTCTGGGCACACGATGCAGCCCTTGAAACATCAGGATTTTGATCCATAAACATGGAGACCACCTA | 120 |
| chr22 | 50718137 | 50718257 | 785447_45746696_NM_001372044.2_379 | 1 | + | good | TTACCCACCTCCAAGGACCACTCTGAGTCTCCCCCGGTAGATGGCACTCACAGACCGCGTGGCATGCTGGGAGTGCTCAGCAGCTGCCAGTGTTAGTAGACGTAGATGCTGCCTCTTGAG | 120 |
| chr22 | 50718257 | 50718377 | 785447_45746696_NM_001372044.2_380 | 1 | + | good | GTTGGCACACAGTGGGTGCTCGAGAAATGCTCTCTGCTATGACCTAGGCAGACAAAGAGGGTCTGCCTCTGCCCCCAAGGGGTCCCACCCCCGCCCCAGATGGTATCAGAACAGATTCCC | 120 |
| chr22 | 50718377 | 50718497 | 785447_45746696_NM_001372044.2_381 | 1 | + | good | CATTCCATGCTGAGGCAAGTGACTTTTCTTTACCCTAAGTCAGAACTTGGCATGCTATTAGCATTAAGTTTCCCCTTATTTCATCTTTGCACGTCACTCCCGCCTTCTGCATCCTCAAGG | 120 |
| chr22 | 50718497 | 50718617 | 785447_45746696_NM_001372044.2_382 | 1 | + | good | TGCGCTCATTCTGCGTCGCTCCATCTAACACATAGGCTGTGGGTGTTCACTGCGTGTTCATACACAGTGCAGGGAATGCAGCAGGAGACCCAGGACGAACTGGGCACAATTTCTCTGTCC | 120 |
| chr22 | 50718617 | 50718737 | 785447_45746696_NM_001372044.2_383 | 1 | + | good | AGCGGTTGGGGATGGGGGGAAGGCAGACCTAAAAATCACGATGATGATGACCTCGGCCCCCTTCCCCCATTCCCAGGATACTGCTGCAGTTCAGCCCTGCACTGTAAAGGTGAAATCGCC | 120 |
| chr22 | 50718737 | 50718857 | 785447_45746696_NM_001372044.2_384 | 1 | + | good | CCACAATAGTTGCAGGCTGTCCTCTTGACATGAGGAAACTGAAGTTCCTAGAGGCTTCTGAAGTTCCTGGCAGATCTAAAATTCAAGCCTAGGTCTGTACACAAAAGCCCACCCTTGGTA | 120 |
| chr22 | 50718857 | 50718977 | 785447_45746696_NM_001372044.2_385 | 1 | + | good | CCTTGGCTACAAAGAGATATTATGCCCAGCAAGAAGCAGTGTAGTACTGGAGGCCTCATTAGCACTGCGGTGGCTGGAGGAGGATGCGATCAGGAACAGGCAGTCTGTATTTTAGCGAGT | 120 |
| chr22 | 50718977 | 50719097 | 785447_45746696_NM_001372044.2_386 | 1 | + | good | TCATAAAAATGTTGACCAATCTCAGAGTGAGGCTGGGAACAGGAGCACCACGAGACCTGGGTGTGTGGGTGTGGGGCCGTCCGCCCCCTGCAGCCTCACTGAACACACCTGCAGCACGTG | 120 |
| chr22 | 50719097 | 50719217 | 785447_45746696_NM_001372044.2_387 | 1 | + | good | TGTCCGCATCATGATGTCCACAGGGGTGTTCAGGAGGTTGTGTCAGGTTCTGCTGGACAGCAGGAGCATGAGGGCCCTGAGCACTCCAGCCCCTGCACAGGACCCTAACAATGATGTCTG | 120 |
| chr22 | 50719217 | 50719337 | 785447_45746696_NM_001372044.2_388 | 1 | + | good | AGCAAGGCATGCAGAGGGGGGTTACTGAAGCTCAAAGGGGCTCAGGGCCTGGCCTGGGGTCACGCTGCCAGTAGGGGTTAAAATGAAACATGCTGGGTGCTCAGCCTGCAGTTCCATTTT | 120 |
| chr22 | 50719337 | 50719457 | 785447_45746696_NM_001372044.2_389 | 1 | + | good | GATTGATCCGTCCGTTCTTCCCTTCACCCACTCACCCATGCGGGTCTCTGGGAGGAGCCTCTCAAGGCTTCCTCCACAGGGCACCTCCTTACCTGGGTGGGCATTAGGTCTTTCTCTGCT | 120 |
| chr22 | 50719457 | 50719577 | 785447_45746696_NM_001372044.2_390 | 1 | + | good | GGAGGCCTGGCTCCAGTGGCCACAGCCCTGCTGCCCCTCACCCTGCCAGTGTGTCACTCCTGCTCATTTCCGACTATCTTCCTCTCACCTCTGGTTGCCTCCAGGGCTGGTACTTCTGCG | 120 |
| chr22 | 50719577 | 50719697 | 785447_45746696_NM_001372044.2_391 | 1 | + | good | TCGGCCTTGTTGATTCTGGGGTGGGGAGCCCTGTACTGGCCCCTCCAAGCCCCTCAGCAGTTCTGTCCCCATGTCTTGCGGGGCTGTCCCTGCCTTTCTGGGATATCTTCTCAGGGCCTG | 120 |
| chr22 | 50719697 | 50719817 | 785447_45746696_NM_001372044.2_392 | 1 | + | good | CTTGATGACCCTGGGTTTGGGCAGGTCTTGGCCCCAACCCAGGCCGTCAGAGTTTGTGTCCTTTCTCAGGGGTCCCCGGTGGGGCCCTCCTCCATCCTGTAACTGAACGCACACCTCTCT | 120 |
| chr22 | 50719817 | 50719937 | 785447_45746696_NM_001372044.2_393 | 1 | + | good | CCTGTCCTCTTCACAAGAGCCCTCCCCGTGCAGCCCTGGGCCTGGGGCACAGAGCTTGGGCATCCAGGGACCAGCCCAGACCAGGGTCTTGCTCGGAGCCCGGGCTCTGGGCTCCCTGTT | 120 |
| chr22 | 50719937 | 50720057 | 785447_45746696_NM_001372044.2_394 | 1 | + | good | TCTCCCCTGCCCTCCATTCCCCGCCCACCACGGGTCCCAACCCCATCTTCCCGAGCATTCTAGCTCCTCGCGCCGGGTTCTGCCGCGGGCGTCCATTGTGTCCGGACGGTGGCTTCCCCG | 120 |
| chr22 | 50720057 | 50720177 | 785447_45746696_NM_001372044.2_395 | 1 | + | good | GGGTGGAGTCGGGTCAAGGCTGGCCTCTGTGGGAGGGGGTTGCCGGGGTCCCCAGGAACCTCTCCGAAGGCAGCACCACCCCCCGCCCAGCGCCCTGGCTGGTCTCACCGGCCCTTCCGT | 120 |
| chr22 | 50720177 | 50720297 | 785447_45746696_NM_001372044.2_396 | 1 | + | good | CCGCAGGGGAGGACGAGAAGCTGGCGTCCCTGCTGGAAGGGCGCTTCCCGCGGAGCACCTCGATGCAAGACCCGGTGCGCGAGGGTCGCGGCATCCCGCCCCCGCCGCAGACCGCGCCG | 120 |
| chr22 | 50720297 | 50720417 | 785447_45746696_NM_001372044.2_397 | 1 | + | good | CTCCCCCGCCCGCGCCCTACTACTTCGACTCGGGGCCGCCCCCGGCCTTCTCGCCGCCGCCCCCGCCGGGCCGCGCCTACGACACGGTGCGCTCCAGCTTCAAGCCCGGCCTGGAGGCGC | 120 |
| chr22 | 50720417 | 50720537 | 785447_45746696_NM_001372044.2_398 | 1 | + | good | GCCTGGGCGCGGGCGCTGCCGGCCTGTACGAGCCGGGCGCGGCCCTCGGCCCGCTGCCGTATCCCGAGCGGCAGAAGCGCGCGCTCCATGATCATCCTGCAGGACTCGGCGCCCGAGT | 120 |
| chr22 | 50720537 | 50720657 | 785447_45746696_NM_001372044.2_399 | 1 | + | good | CGGGCGACGCCCCTCGACCCCCGCCCGCGGCCACCCCGCCCGAGCGACCCAAGCGCCGGCCGCGGCCGCCCGGCCCCGACAGCCCCTACGCCAACCTGGGCGCCTTCAGCGCCAGCCTCT | 120 |
| chr22 | 50720657 | 50720777 | 785447_45746696_NM_001372044.2_400 | 1 | + | good | TCGCTCCGTCCAAGCCGCAGCGCCGCAAGAGCCCCCTGGTGAAGCAGCTGCAGGTGGAGGACGCGCAGGAGCGCGCGGCCCTGGCCGTGGGCAGCCCCGGTCCCGGCGGCGGCAGCTTC | 120 |
| chr22 | 50720777 | 50720897 | 785447_45746696_NM_001372044.2_401 | 1 | + | good | CCCGCGAGCCCTCCCCGACCCACCGCGGTCCGCGCCCGGGTGGCCTCGACTACGGCGCGGGCGATGGCCCGGGGCTCGCGTTCGGCGGCCCGGGCCCGGCCAAGGACCGGCGGCTGGA | 120 |
| chr22 | 50720897 | 50721017 | 785447_45746696_NM_001372044.2_402 | 1 | + | good | AGCGGCGCCGCTCCACTGTGTTCCTGTCCGTGGGGGCCATCGAGGGCAGCGCCCCCGGCGCGGATCTGCCATCCCTACAGCCCTCCCGCTCCATCGACGAGCGCCTCCTGGGGACCGGCC | 120 |
| chr22 | 50721017 | 50721137 | 785447_45746696_NM_001372044.2_403 | 1 | + | good | CCACCGCCGGCCGCGACCTGCTGCTGCCCTCCCCGGTGTCTGCCCTGAAGCCGTTGGTCAGCGGCCCGAGCCTGGGGCCCTCGGGTTCCACCTTCATCCACCCACTCACCGGCAAACCCC | 120 |
| chr22 | 50721137 | 50721257 | 785447_45746696_NM_001372044.2_404 | 1 | + | good | TGGACCCCAGCTCACCCCTGGCCCTTGCCCTGGCTGCCCGAGAGCGAGCTCTGGCCTCCCAGGCGCCCTCCCGGTCCCCCACACCCGTGCACAGTCCCGACGCCGACCGCCCCGGACCCC | 120 |
| chr22 | 50721257 | 50721377 | 785447_45746696_NM_001372044.2_405 | 1 | + | good | TGTTTGTGGATGTACAGGCCCGGGACCCAGAGCGAGGGTCCCTGGCTTCCCCGGCTTTCTCCCCACGGAGCCCAGCCTGGATTCCTGTGCCTGCTCGCAGGGAGGCAGAGAAGGTCCCCC | 120 |
| chr22 | 50721377 | 50721497 | 785447_45746696_NM_001372044.2_406 | 1 | + | good | GGGAGGAGCGGAAGTCACCCGAGGACAAGAAGTCCATGATCCTCAGCGTCCTGGACACATCCCTGCAGCGGCCAGCTGGCCTCATCGTTGTGCACGCCACCAGCAACGGGCAGGAGCCCA | 120 |
| chr22 | 50721497 | 50721617 | 785447_45746696_NM_001372044.2_407 | 1 | + | good | GCAGGCTGGGGGGGGCCGAAGAGGAGCGCCCGGGCACCCCGGAGTTGGCCCCGGCCCCCATGCAGTCAGCGGCTGTGGCAGAGCCCCTGCCCAGCCCCCGGGCCCAGCCCCCTGGTGG | 120 |
| chr22 | 50721617 | 50721737 | 785447_45746696_NM_001372044.2_408 | 1 | + | good | CCCCGGCAGACGCCGGGCCAGGCCAGGGCAGCTCAGAGGAAGAGCCAGAGCTGGTGTTTGCTGTGAACCTGCCACCTGCCCAGCTGTCGTCCAGCGATGAGGAGACCAGGGAGGAGCTGG | 120 |
| chr22 | 50721737 | 50721857 | 785447_45746696_NM_001372044.2_409 | 1 | + | good | CCCGAATTGGGTTGGTGCCACCCCCTGAAGAGTTTGCCAACGGGGTCCTGCTGGCCACCCCACTCGCTGGCCCGGGCCCCTCGCCCACCACGGTGCCCAGCCCGGCCTCAGGGAAGCCCA | 120 |

| | | | | | | | | |
|---|---|---|---|---|---|---|---|---|
| chr22 | 50721857 | 50721977 | 785447_45746696_NM_001372044.2_410 | 1 | + | good | GCAGTGAGCCACCCCCTGCCCCTGAGTCTGCAGCCGACTCTGGGGTGGAGGAGGCTGACACACGCAGCTCCAGCGACCCCCACCTGGAGACCACAAGCACCATCTCCACGGTGTCCAGCA | 120 |
| chr22 | 50721977 | 50722097 | 785447_45746696_NM_001372044.2_411 | 1 | + | good | TGTCCACCTTGAGCTCGGAGAGCGGGGAACTCACTGACACCCACACCTCCTTCGCTGACGGACACACTTTTCTACTCGAGAAGCCACCAGTGCCTCCCAAGCCCAAGCTCAAGTCCCCGC | 120 |
| chr22 | 50722097 | 50722217 | 785447_45746696_NM_001372044.2_412 | 1 | + | good | TGGGGAAGGGGCCGGTGACCTTCAGGGACCCGCTGCTGAAGCAGTCCTCGGACAGCGAGCTCATGGCCCAGCAGCACCACGCCGCCTCTGCCGGGCTGGCCTCTGCCGCCGGGCCTGCCC | 120 |
| chr22 | 50722217 | 50722337 | 785447_45746696_NM_001372044.2_413 | 1 | + | good | GCCCTCGCTACCTCTTCCAGAGAAGGTCCAAGCTATGGGGGGACCCCGTGGAGAGCCGGGGGCTCCCTGGGCCTGAAGACGACAAACCAACTGTGATCAGTGAGCTCAGCTCCCGCCTGC | 120 |
| chr22 | 50722337 | 50722457 | 785447_45746696_NM_001372044.2_414 | 1 | + | good | AGCAGCTGAACAAGGACACGCGTTCCCTGGGGGAGGAACCAGTTGGTGGCCTGGGCAGCCTGCTGGACCCTGCCAAGAAGTCGCCCATCGCAGCAGCTCGGTGAGCAGGGCGGTGCGGGG | 120 |
| chr22 | 50722457 | 50722577 | 785447_45746696_NM_001372044.2_415 | 1 | + | good | AGGGATCCGTGCCTTGTCCGTGGCCCCGTCTGTCATTCCTCTTGTCTGTTCTCTGGCTCTCCTGTTCCTCCTTGTTCTGTTCCATTCCTTCTGTGGCAACCCCCAACCCGCCCCCGCCAT | 120 |
| chr22 | 50722577 | 50722697 | 785447_45746696_NM_001372044.2_416 | 1 | + | good | CCACCTCTGAATCCGGTCTTGCTTGGCCTGCCCGAGAGAGGAGGTTCTTCTGGGCGTCTGACACGTCAGGGTTGCTGCTCACTGTGTCCCTGTTGGTGCCAGAAGTGGGAGCTGGGCTCC | 120 |
| chr22 | 50722697 | 50722817 | 785447_45746696_NM_001372044.2_417 | 1 | + | good | CCTCAGAGACTCAGGTTGGGCATGAGGCGCCTCCATGGCCCTCCTGGAGGCTCTTGGCCCCAGGGATTCCTGTGAGTTCTCTCTCTCCCTCCCGACACAGGCTGTCTTTACCATTAAGGG | 120 |
| chr22 | 50722817 | 50722937 | 785447_45746696_NM_001372044.2_418 | 1 | + | good | TTGTTTTGTGTTTTACTTTGGAGGTACGCCTCCCTCCTCCCATCCTCTCTGTGGCCATGTGGCTGCCCAGTGTGGCTGATACCCTCTGCCTTACAGCTGCCTCCTGCCTTGCTTCCTCTG | 120 |
| chr22 | 50723057 | 50723177 | 785447_45746696_NM_001372044.2_420 | 1 | + | good | TGTGTCATTTCTAAAGTGCAGTAAGAAGATAGATGGAAGTCACGCCATCTCCCTGCACTCCATCTGCCCTCTTGCCTTCTCCCCACCACTTCCCCGTCTGTCCTGCCCCTGCCCAGAAGG | 120 |
| chr22 | 50723177 | 50723297 | 785447_45746696_NM_001372044.2_421 | 1 | + | good | ATCTGAGCAGGGGCCTGGCTTGCTGCTGAGGCTGTCACAGCTGCCAGAGCCCTGGGCTGATCCCATAGGTCTTTCCTTGAGGACCCGACTCCCAAGGCTCCTTCCAAAGAGGAGCCCTTC | 120 |
| chr22 | 50723777 | 50723897 | 785447_45746696_NM_001372044.2_426 | 1 | + | good | TGCCCGTAGAATGGGTGTTATAATCAGAGTGAGCAGCCTGGTCCTAGGCCTCTGTACAGGACCAGACCCCTGAGGCTGGGGTCTCCTGACCCACACCTGACCAGCCCCCATCTTCCCTCT | 120 |
| chr22 | 50723897 | 50724017 | 785447_45746696_NM_001372044.2_427 | 1 | + | good | CTGCTTCTCCCTCCGCTCTTCTCTGCCTCTTGGTCTTGATGAAAATCAAAGCCATTTTAAAAAGTGCATAGCACAGTGCCTGGCCTGGTTCGGGCCCTCAATAAACATTTCTTAAATGGA | 120 |
| chr22 | 50724017 | 50724137 | 785447_45746696_NM_001372044.2_428 | 1 | + | good | TGAAAGAACAAAGCAAAATGCAAATGCTGTGTTTTGTGATTTGAGATCTAGGGAGGTGGCTTAGGACAAAAACCCACAGAAGGACTTACTCAGCGTTCAGACTCATCAGGTGTCGATCCC | 120 |
| chr22 | 50724137 | 50724257 | 785447_45746696_NM_001372044.2_429 | 1 | + | good | CCATGGTCGGGCTCAGGTGGGCCCAGGTCTCGTCATTGGCTCTGTCCTCTCCTGTGAGGCAGCTGCAGCAGTTGCAGGGCAGGGTCAGGGGATTCCTCAGCCGGAACCTCTGGCACTTCC | 120 |
| chr22 | 50724257 | 50724377 | 785447_45746696_NM_001372044.2_430 | 1 | + | good | CCTTCTCTGAGTTCATCCTCCCACGGTGCTCTTTAGCATTCTGTCTTGATCAGGGTACGGTTCTCTTAGCCCTTGGCTCTGGCTTTCACCAGGACCCTGTTTGTTTTTCTTTCTCAGATG | 120 |
| chr22 | 50724377 | 50724497 | 785447_45746696_NM_001372044.2_431 | 1 | + | good | TGGGATGGGGAAGGACAAGACCAGCAGCCCTGACCCTTACGTGACGTCTGTTCTTTTACTCAGTAGTTCCTGCAAAATTCTGATCTCTGATTGGGTTCATGTGGGTCCTCTGCTTTTCCC | 120 |
| chr22 | 50724497 | 50724617 | 785447_45746696_NM_001372044.2_432 | 1 | + | good | TGACTGATCACTGGGGTCTGGGGACATAGGGCTGTCATCGGCCAAGCTTGAGTCCTCTGTTCTCCCAGGAGCCAGGGGGATGGGTCCGCCCACCTAGCAAGCAGGTGCTGAGAGTGAGGG | 120 |
| chr22 | 50724617 | 50724737 | 785447_45746696_NM_001372044.2_433 | 1 | + | good | GAGGGGGCTCTCCAGATGGAAGTCCAGTGCTGTCCCTTGAGTAAGTAGATGCTGGACAGCCTGTAGCAACCAATGTTCTGTGACACGGCCCCCACTGGCATCAGCAACTCACTTCCTTGC | 120 |
| chr22 | 50724737 | 50724857 | 785447_45746696_NM_001372044.2_434 | 1 | + | good | CGGTCATTGGCTTGGAGCCATCAGAGGGCCCTGATGTCGGTGCTCAGGAGGTCACAGCTTAGTGCTGTATCCCTCCCCTGTGAAGTGTATTTACAGCGAGCCAACTGCACAGGCTATCGC | 120 |
| chr22 | 50724857 | 50724977 | 785447_45746696_NM_001372044.2_435 | 1 | + | good | GAGGCTGCACACGGCATCACCTGTGAAGCACCTGGTAGGCCCAGGGCTGGGGCCTGTGGACCCCTCACTGGTGTGAGCTGCCATTGTCTTTGCTGTCACTGACGACACCTAGGTGAGGTC | 120 |
| chr22 | 50724977 | 50725097 | 785447_45746696_NM_001372044.2_436 | 1 | + | good | TTAGAGTCCTTTGTGGGAAAACAAGCCTGTTGGATGCAGCACTGGAGACGGTCCTGCCGTATCAGTTTGTCTTCACACCGTTCTGTTAGACGGACACTAACAGAATCCTCATGTTCCAGA | 120 |
| chr22 | 50725097 | 50725217 | 785447_45746696_NM_001372044.2_437 | 1 | + | good | CAATGAAACTGAGGTACCAGAGAGCTTCCTTGCAGACAGAAAACGGTGGTCTGCCTGACCATGAGCCCTGCTCTCTGGCTGCCGTAACCAGGGCCGCTGTTGTTAAGCCCCTTCTGATGA | 120 |
| chr22 | 50725217 | 50725337 | 785447_45746696_NM_001372044.2_438 | 1 | + | good | CACACAATGGGCTTTTGCCCAGTAACGCCCCTCCACAGAAGTGGCAGGTGTGTGCATGGCACAGACTCACCCCCGCTGTTTACTGGTCATGTTTGTCATAAGTAGCACCTGGTGCCACCT | 120 |
| chr22 | 50725337 | 50725457 | 785447_45746696_NM_001372044.2_439 | 1 | + | good | GAGTGGCCTACCCGGGCCTGCTTTTCCTTCTCAGTTTGCCTCTGTGTTACCTTCTGTTCCAGCTCACTCTTCCTTTCACCCTCTTGCGCTCCTTCTCGCCCTCAGACCTGGGCTGAACTA | 120 |
| chr22 | 50725457 | 50725577 | 785447_45746696_NM_001372044.2_440 | 1 | + | good | CAGCCATCTGGAGGGGCCCCTGCCCACCCTGGGCCCTTAACTTCCGGTGCTTCTGGCTGCAGGCAGCGGAGCGGAGCACAGGACACTTGGTGATGTCTGAGTCCAGTCCTGTTTCAGATC | 120 |
| chr22 | 50725577 | 50725697 | 785447_45746696_NM_001372044.2_441 | 1 | + | good | TGGTCCTTGTGTGCCCACCGTGGCTGTTGTCAGCTCTTGTCCTGCGGCCCCTAATGGCTGGAGTCCGGTACTCACTGCTCACTTAGGGATAAAACTGCTGGTGGAACTGCCTAAAGTGTG | 120 |
| chr22 | 50725697 | 50725817 | 785447_45746696_NM_001372044.2_442 | 1 | + | good | GGAGGTTCTGACATCCAGGTATGGCTGCTTCCTTCCCAGGGCCAGTCGTGGCAGAGGCGCTAGGTCACCCTCAGGGCCATACCCTGAGCTGTGTCTCCAGCCCCTTGTGTTCCACAGTGT | 120 |
| chr22 | 50725817 | 50725937 | 785447_45746696_NM_001372044.2_443 | 1 | + | good | TTGTCTCCCAGGCCCATCCTTCTCCTCCCTGACCGACACCATGAGGAAGGGAATGCCCACCCACCTTTTCCAGAGATGGCTCATTCCTGATGTGCTTTAGCCACAAGGTGTTCTGTGGCC | 120 |
| chr22 | 50725937 | 50726057 | 785447_45746696_NM_001372044.2_444 | 1 | + | good | TGAGAGCTTTCTGGACAAGTTAAGCTGTCCTGTATGTCGTGGAACCAAAGCTCGGTAACCTGTGGAAAAATGATGAGCTCTGGGGCCACCTTTGATTTTCAGGGGAACCTGGAGGATGTG | 120 |
| chr22 | 50726057 | 50726177 | 785447_45746696_NM_001372044.2_445 | 1 | + | good | GCACTCTCATGCTGCCTCTGGAAGGTCCAGGACCTTCGGCCTTTTGAATCCTCGATCCCAGATTGTCCACCTGCTCTGTGCTCCCAGCTCCAAGTCTAGCTCTGCTGGACAGACCATTCA | 120 |
| chr22 | 50726177 | 50726297 | 785447_45746696_NM_001372044.2_446 | 1 | + | good | TCTGCCCAGAGCCCCTCCTCGGTCTCTGGGCCCCACACCTGACTCTGAGGCTACATTTGGGCATGATTCAGACCTGACTCCGAGCCCCACAACTGGCCACAGGCACCCCATCTGTCTGAG | 120 |
| chr22 | 50726297 | 50726417 | 785447_45746696_NM_001372044.2_447 | 1 | + | good | GCTCTCACTTGTCTCTGAGACCCTGGCTTCACACCTGGCTCTAAACTCCATTTGTCTTCCAGGCCCCGTGCTCGGATCTGAATCCCATGTGGGGTCTGAGGTCCATAGTAAGGCTCACGT | 120 |
| chr22 | 50726417 | 50726537 | 785447_45746696_NM_001372044.2_448 | 1 | + | good | CTGCCTTGGCACTTAGCTTTACATGGGACTCTCACATCTGGTTTTGTAGGTAAAGTTCATGGCAAAGTAGCTTTGGAAACACAAAAAGAGGTTCTTTGCTGCAGTACTTCTGGTGCCTTT | 120 |
| chr22 | 50726537 | 50726657 | 785447_45746696_NM_001372044.2_449 | 1 | + | good | GACAAGCTTGTGTGGAGGAGGGCACTGTGTGCCGTGGTCTCCAGACCTGCTAGAGTCAGTTCCTAGGTCTCCCAGAAGACCCATGTTCCTCAAAATGGTCAAGAAGCCCTGCCCTGGTGG | 120 |
| chr22 | 50726657 | 50726777 | 785447_45746696_NM_001372044.2_450 | 1 | + | good | CCTGCTGCAGACCCTGCCTGCTAATATATTTGTCAGGGCTCTGGACAAAGCACTGTGTGCTTGGTGGATCACACTGGTAAGCAGGGCCAAGGCCAGCAGGGCCTGCTCTTTTGAAACCCA | 120 |
| chr22 | 50726777 | 50726897 | 785447_45746696_NM_001372044.2_451 | 1 | + | good | GACTGAATGTGTCACATACCTGCCAGTGGGGGCCCCTGCAGGCCACATGCAGCTGGGGCAGGTGTGTTCCCAGGATCCCTGAACCCTGTTCGGGGAGTAGGAAGGAGGTACAGAGGAGGG | 120 |
| chr22 | 50726897 | 50727017 | 785447_45746696_NM_001372044.2_452 | 1 | + | good | GTCATAGGACACTGGGGCCCATCAGAGCTGGAGCTGGGAACAGGCGGGGTGCCGAGGAGCAAGACGTGTTACCGCAGAGGATGCACTGGAAGCTTCCGAGCCCATGTCTGGTTTTGTGCT | 120 |
| chr22 | 50727017 | 50727137 | 785447_45746696_NM_001372044.2_453 | 1 | + | good | CCTCTGAGAATTAGTTCCCTTAATGAGGCCCCTAGTTTGCAAAAGGCCACAGGCCTAGACCGGCCCCACGTGAGTGCAGTGGCAGAAGGCCCGAGGCAGTATCCCGGGCCCGAGTGTGGA | 120 |
| chr22 | 50727137 | 50727257 | 785447_45746696_NM_001372044.2_454 | 1 | + | good | CATTTTCCTGCCTGTTGGCCCTAGTGGGCTGACTGAGTTGGGCACGCCTGGCAGGGACATGTCATACATACTATTTTTGCATTTTGTTAGTGCATCTCGTATTCCTGGCTTTGTGCTAGG | 120 |
| chr22 | 50727257 | 50727377 | 785447_45746696_NM_001372044.2_455 | 1 | + | good | TACCAGGAGGCAAGGAGAACTGGTAAGAAGTGGGGTGAAGGGCCTGAGACCATGAACGGTCTGGATCCAGAAATGGCAAGGGGTAAAGAGAAGGTCTGGGTTGATATTAGGAGTTAGCCA | 120 |
| chr22 | 50727377 | 50727497 | 785447_45746696_NM_001372044.2_456 | 1 | + | good | TTGGCTGTAGGGTAAAAAGAGGAGCTGATGGTGGCTCCCAGGCTCTGTGTGTGGCAATGACAGGGTGCCGTTCACTGAGTTGGGGATGGAGGGACATGTGTGGATACCCCAGTAGAGGTG | 120 |
| chr22 | 50727497 | 50727617 | 785447_45746696_NM_001372044.2_457 | 1 | + | good | TTGGACTCATCAGAGGCTGTCTGGGTCTCACATGTAGCATTCCTGGGCACACTGGCCATAGCAAAGGGTGTGAGTGGATAATGTTGCCCAGGAGAAACTGCAAACTAAGAGGGGAGCTCT | 120 |
| chr22 | 50727617 | 50727737 | 785447_45746696_NM_001372044.2_458 | 1 | + | good | CCGAAAATCCATCTAGGAAGGAAGAGGCCTGGGAAAGAATGTCTCGAAGGGAAACATGAACAGGTCAGCAGCGGGGTCTTGCTGACCGCCTGGCCAACTGTGGGGCAGAGGGCGGAGTGG | 120 |
| chr22 | 50727737 | 50727857 | 785447_45746696_NM_001372044.2_459 | 1 | + | good | GGAGGGTGTGGCAAGGCCAGACCTCTCTCACTGGCTGAGAGTCACTGGGAAACAGTGAGACGGTGCAGGATTCAAATGTAGGTGGAAAGACGTGGGGCCGTGGGGGCCTCAGTGTGATCT | 120 |
| chr22 | 50727857 | 50727977 | 785447_45746696_NM_001372044.2_460 | 1 | + | good | GGACTCAGCCTCTTCAGCGTGGCTGCTGGAGGTGTTCGTGGGTGACGGTGCCTGGTGAAGTATCATGTGTTAGAGGGTGGGTGGCCCAGAAGCTCTCTGGGAGGCTGGCAGGTGCTGTGA | 120 |
| chr22 | 50727977 | 50728097 | 785447_45746696_NM_001372044.2_461 | 1 | + | good | TGTGGCTGTTTGGCGGCCGGAGCCTGGCCAGGGAGGTGGCCAGACGGCTGAGAGTGACAGTGGGTGTGCAGAGAGGCCATGGAAACATGGAAGTCAGACATGGACACTGCTCGGGTTTGT | 120 |
| chr22 | 50728097 | 50728217 | 785447_45746696_NM_001372044.2_462 | 1 | + | good | GCCCTGGGGATGGTGTGGGCAGTACCCTCTTCCCCAAGAGAAAGAGGAGAAGCAGACACCTGGGCCATGGGTGACCCCAGGCCCCTCCCTCCTGTGCCCTGGGGCCTTTGGCAGGCACAG | 120 |
| chr22 | 50728217 | 50728337 | 785447_45746696_NM_001372044.2_463 | 1 | + | good | CCTTCCTTGGGCGTTTTCTCTTGGCCTCCTGACATCTCCAGCATGCTCTGGGAGGCGGGAGCAATGTGAATGGCTCTCCTCACATCTCTTGCCCTCGGGCCCCTGGCTCCGTGGGTGCCT | 120 |
| chr22 | 50728337 | 50728457 | 785447_45746696_NM_001372044.2_464 | 1 | + | good | CTGGAGCCAGCCTGCCCTCTGCCCAGGTCTCCAGCTCAGCCCCTGCCAGTACTACCCAGACAGGCACAACTCTGCTCTGCCCAGCTCGGACCTCCTGGAGCCCAGACCCTGCTCTGGGCT | 120 |
| chr22 | 50728457 | 50728577 | 785447_45746696_NM_001372044.2_465 | 1 | + | good | CCGGCCGAGCCTCCTGGCTGCGCTGCCCGCGCCTGCCCTGTGCCTCTGCCTTCACTGCTGTGGTCCTGCAACTCATCAGTGTGCTTCAGTCACACCTGAGACTCCAGGTACACCTGCCAT | 120 |
| chr22 | 50728577 | 50728697 | 785447_45746696_NM_001372044.2_466 | 1 | + | good | CACAACTCCACTGAAAGCTAAGCCTTGGGGCAGGGGCAGGGCCAGGGTAGGTGTCCATCCTGTGATACAGGTCCCCAGGCCAGGGCCACTTTGTTGGGGCTGGTGGGTGGAGGCCAGGTG | 120 |
| chr22 | 50728697 | 50728817 | 785447_45746696_NM_001372044.2_467 | 1 | + | good | CATATTCCTGGTCCTGCCCCTCTCTGGGTGTCCACAGTCAGGTCCCTCTGAGTGATTCTCTCGGAGTCTCCTCTGGCAAGCCCCTCCAGCAGTCACCACCACCTGGGCACCCACCCCCAG | 120 |
| chr22 | 50728817 | 50728937 | 785447_45746696_NM_001372044.2_468 | 1 | + | good | GGGACCCCCCGACCCAACACCACCTGGCACCTCCCCTTGGGAATCCCCCCAACCTTGGCTGCCTCCCCACTTTGGCCCACTGTGCAGGTGGATCCATGGAGGCCAGGTCCAATGGGAGCA | 120 |
| chr22 | 50728937 | 50729057 | 785447_45746696_NM_001372044.2_469 | 1 | + | good | AAGACAGCTTATTCTGGTGGTTGGTGCCTGTGCTGAGGCCCACCTCAGGTGTTCTTGAAGCCCCTTTCTGTTTGGGGGCACAGTGTGGCCTCTTCAGGAAGTTCTGCTCTGGATCTGAGC | 120 |
| chr22 | 50729057 | 50729177 | 785447_45746696_NM_001372044.2_470 | 1 | + | good | AGCTGGGAGGAGGCAGTAGGGACAGAGGCTGAGGAGAGGACAGAAATGAGGTCCCATGGGAGAGACAGAGCTCAGATCTCCACAGAGGGCTGACCTCTCACAGGTGGGAGACAAAAGACC | 120 |
| chr22 | 50729297 | 50729417 | 785447_45746696_NM_001372044.2_472 | 1 | + | good | TGGTGAGGACAGCACAGAGGTTGGGGTGGGGGGACTCAGCCTGGGCAGGGGCAGGGAGAGGGGGTGGGAGGGCCACTGGAGAGGGGCCCGGCCAAAGAGGATGTCAGGGGCTCAGCA | 120 |
| chr22 | 50729417 | 50729537 | 785447_45746696_NM_001372044.2_473 | 1 | + | good | CCAGGCGGCCAGGCACCTGCCGAGCACTGCAGCCACGCTTGTGGTTCACCTGCACTTGGAGACTCTCGCACAGGCGCCTCTCCTCCACCAGCAGCTCCAGCTTTGTCCCTCCCACAGCAG | 120 |
| chr22 | 50729537 | 50729657 | 785447_45746696_NM_001372044.2_474 | 1 | + | good | CTGGGCCCTCTGCACCTGAGTTCCTCCCTCCACACCTGGCTGCCCATTGTGTGGTTGCAGCTGTGGGTCGCGTGGGCCTGGCCGGCTAACTCTCTCTCTCATTGCTCTCTCCTGCCCATC | 120 |
| chr22 | 50729657 | 50729777 | 785447_45746696_NM_001372044.2_475 | 1 | + | good | TGCATGTGGCTGCTCTGTCGAGCTGCGCCGTGGTCCCGAGCGCCGGCTGTGTAAGTGAGCATGCCCATCCCATGCCTCTTGCCGTCCACACGCTGTGCCTGTCTGCCTGGGTCTCTGCTC | 120 |
| chr22 | 50729777 | 50729897 | 785447_45746696_NM_001372044.2_476 | 1 | + | good | TGCGTGGCTGTGAGAGGCTCTGGTGCCACTGACAGCCCCTTGAGGCTTCCCCTGACACAGTGGGGATAGGGATGGGAATGGGAGGACATGGGAGTGGGTTTTCTCTGGAGCTATCACCCC | 120 |
| chr22 | 50729897 | 50730017 | 785447_45746696_NM_001372044.2_477 | 1 | + | good | AGGTAGGCCTCCCACCACGGCAGAGCCAAGGAGGGGCTAGAGCTCTAGGGTCCTGTCAGGTGAGGCTGGGAAGTGAGCTGCCATCGGTTCTTGTGTGTGTGCGTGCGTGCATGTGTGCGT | 120 |

| | | | | | | | | |
|---|---|---|---|---|---|---|---|---|
| chr22 | 50730017 | 50730137 | 785447_45746696_NM_001372044.2_478 | 1 | + | good | GTTGCGTGTCTGCGTGTGTGTGCTGCTGCCACCGCAGCATGTCTGTAACGCGTGTGGGCACCGTTGCTGCTGTTGTGTGCCGTCTGTGCAGGAGGCTGCCTTTGTGTTGAGGGTGTGCAC | 120 |
| chr22 | 50730137 | 50730257 | 785447_45746696_NM_001372044.2_479 | 1 | + | good | GGCCTCACACCTGCCCTGCATGTGCTGCTGCTCCATACGGGTACGAGCCCTGCCTAGTGTCTGTCTCCTGTGTCACGGACCTGTTCAACCTCGTGCTGCTGCCAGCCTTTATCGCGACTC | 120 |
| chr22 | 50730257 | 50730377 | 785447_45746696_NM_001372044.2_480 | 1 | + | good | AGCTGTCCCTGGAACCTGCCCAGGATCCCCTGGGTCTTCTCATGAGAGCAGAGCTGTGTGGGGGGTGGGCGGTGAAGGGTACTGCCCAAGTCTCAGCGTCCCGGGTATCTGTGGATCCCG | 120 |
| chr22 | 50730377 | 50730497 | 785447_45746696_NM_001372044.2_481 | 1 | + | good | CCATGCCCAGAGCCGGTGTCGGAGGCTGGCAGGAGGGAGAAGCCCGCCCTTTGCCATGAGAGGCTGTCTTTTCTTTGGTTGGGCTGCACTTGGAGCCTGGATGGAGTGGAGGGGGCCACC | 120 |
| chr22 | 50730497 | 50730617 | 785447_45746696_NM_001372044.2_482 | 1 | + | good | AGTCATTCCTCATATTCCAGCCAGTCGCTGGCTCTGGTCCCAGGGGCCAAAGAAAAGGGCCAGGGTAACCGTAGGATCCCACCCTTTATTTCTTCCTCTGGCCGGGCTACTCCCGCCAGC | 120 |
| chr22 | 50730617 | 50730737 | 785447_45746696_NM_001372044.2_483 | 1 | + | good | CGCAGCCCCAGCCCGTTTCCTCCTGGACCCTGCCCGCTCCCCTCCGCCCGTCCCCCCTTGGCTGTGCGCCCCTCACCTGGCGCTGACCCCTCTCCCTCCGCAGGCTCTTCAGCAGCCTCG | 120 |
| chr22 | 50730737 | 50730857 | 785447_45746696_NM_001372044.2_484 | 1 | + | good | GTGAGCTGAGCTCCATTTCAGCGCAGCGCAGCCCCGGGGGCCCGGGCGGCGGGGCCTCGTACTCGGTGAGGCCCAGTGGCCGCTACCCCGTGGCGAGACGCGCCCCGAGCCCGGTGAAG( | 120 |
| chr22 | 50730857 | 50730977 | 785447_45746696_NM_001372044.2_485 | 1 | + | good | CCGCGTCGCTGGAGCGGGTGGAGGGGCTGGGGGCGGGCGCGGGGGGCGCAGGGCGGCCCTTCGGCCTCACGCCCCCCACCATCCTCAAGTCGTCCAGCCTCTCCATCCCGCACGAGCCCA | 120 |
| chr22 | 50730977 | 50731097 | 785447_45746696_NM_001372044.2_486 | 1 | + | good | AGGAGGTGCGCTTCGTGGTGCGCAGCGTGAGCGCGCGCAGTCGCTCCCCCTCGCCGTCGCCGCTGCCCTCGCCCGCGTCCGGCCCCGGCCCCGGCGCCCCCGGCCCACGCCGACCCTTCC | 120 |
| chr22 | 50731097 | 50731217 | 785447_45746696_NM_001372044.2_487 | 1 | + | good | AGCAGAAGCCGCTGCAGCTCTGGAGCAAGTTCGACGTGGGCGACTGGCTGGAGAGCATCCACCTAGGCGAGCACCGCGACCGCTTCGAGGACCATGAGATAGAAGGCGCGCACCTACCCG | 120 |
| chr22 | 50731217 | 50731337 | 785447_45746696_NM_001372044.2_488 | 1 | + | good | CGCTTACCAAGGACGACTTCGTGGAGCTGGGCGTCACGCGCGTGGGCCACCGCATGAACATCGAGCGCGCGCTCAGGCAGCTGGACGGCAGCTGACGCCCCACCCCCACTCCCGCCCCGG | 120 |
| chr22 | 50731337 | 50731457 | 785447_45746696_NM_001372044.2_489 | 1 | + | good | CCGTGCCCTGCCGGCAGGGCCCCCACCCCCACCCCGGGCCGCGGGCTCGGCCTGCCCCTTACGACGGCGCCCGGGCCAGGAATGTTGCATGAATCGTCCTGTTTGCTGTTGCTCGGAGA | 120 |
| chr22 | 50731457 | 50731577 | 785447_45746696_NM_001372044.2_490 | 1 | + | good | CTCGCCCTGTACATTGCTTAGTGCCCTCACCGGCCGCCCAGCCCACCCAGCGCACAGTCAGGAAGGGCGTGGACCAGGGAGGCTGGGGCGGGAGGTGCCGGGGGTGGGGTGCCCTAGCGT | 120 |
| chr22 | 50731577 | 50731697 | 785447_45746696_NM_001372044.2_491 | 1 | + | good | GACCACCTCCTTCGCAGCTCCTGGTGGCCATTCTCCCAGAGGGGGAACCTAGTCCAGCATGCGAGGTCAGGACCCGCCTTGGTGACTCGGGGGGAGGGGGGAGACATTGGGATTCTCGAT | 120 |
| chr22 | 50731697 | 50731817 | 785447_45746696_NM_001372044.2_492 | 1 | + | good | GGGGGCCAAGGAGCCCCCCTGTTTTGCATATTTTAATCCACTCTATATTTGGAACGAGAAAAGGAACAAATATCTCTGTCCGTAATAGTTTCCTCTCCCCTCCCTTCTACTTCCACTGGT | 120 |
| chr22 | 50731817 | 50731937 | 785447_45746696_NM_001372044.2_493 | 1 | + | good | CCCACTGCAGCTGCCCAGTCTTCCATCTCCGGCCCCTCACTGCCACTGCCACCCCACAACGGGGCAGGGGACGCTCCAGCTGGTCTGGGGTTGGCCAGGGCCCTAGTGGCCCGCCCTGGG | 120 |
| chr22 | 50731937 | 50732057 | 785447_45746696_NM_001372044.2_494 | 1 | + | good | GCCCCAGCTCGGCCCCTCGCCTCGCTGAGCTCTAGTGTGCCCCACCGACCCTTCAGGTGCTGCTCGTGGTGGGAGGGGCGGCAGGCCGCGGGTCCTGCTGTGCACCCGCGGGACCAGCCG | 120 |
| chr22 | 50732057 | 50732177 | 785447_45746696_NM_001372044.2_495 | 1 | + | good | GCCTGGGAGACCATCGGCCGGGGGGGATGAGGGCAGGGCCCTGCCGCTCCACCGCAGCCATCTTCCTCACAGGGTCTCTCCCCAAGGAGGGGGCTAGCTTGGTCCCCATGCTCTTGGGCA | 120 |
| chr22 | 50732177 | 50732297 | 785447_45746696_NM_001372044.2_496 | 1 | + | good | ACTACAGCAGAGAAGCCTCCCTGCCTTGGACCCCAAAGTCTCCTGTCCTGCCCTTTATGTGTGTGGGTGAAACTGGGTGCGTCTGAGCACGTGGGAGCCGTGTGTGTGCCTGATTACTGA | 120 |
| chr22 | 50732297 | 50732417 | 785447_45746696_NM_001372044.2_497 | 1 | + | good | GTGGCCACCAGGGGCCGCTCTGGACTAGCGCGGGGCCGTGGAGGCGTGCACCGTGTGCATGCGTGGGGTGTACCTGTGAGAGCACCCTGTCTCCTCTTCCAAAGAAAGTCAGAGGCCATC | 120 |
| chr22 | 50732417 | 50732537 | 785447_45746696_NM_001372044.2_498 | 1 | + | good | CTGCACCCTGGGTCCAGCTGTTTGCCCAGCCTGTCCTTCCAGAGCCTCACCCAGCCTGAGCGGGGTTCCCTGGTGAATCCCTGCTGCTTGGGGAGGCCCCAAGGGCCCCTTGGAGGCAGC | 120 |
| chr22 | 50732537 | 50732657 | 785447_45746696_NM_001372044.2_499 | 1 | + | good | GCCCCCACCTTGGGCTTCTGAGGGCATCATAGGGGGACCCCTAGAGTCAGTTCACCACAGGCCCTGGGGAGAGTCAAAGACCCCCGAGGGTGCCCAGCCCCCCACACTGTGACTCCTCAC | 120 |
| chr22 | 50732657 | 50732777 | 785447_45746696_NM_001372044.2_500 | 1 | + | good | ACTCAGCGATGACCTGTGGGGTGGGGGGCCCTGGGACGTTTTTAAACCTAGGGTTTGGAGTCTGGACTAAGCTCCATCCACGTCACTCACAAGTTTCTGTTTATATTTCTAGCTTTTTTT | 120 |
| chr22 | 50732777 | 50732897 | 785447_45746696_NM_001372044.2_501 | 1 | + | good | AATAAAATAAAAAAAAAAAGAAAACAGAAGTTTTCACAACCCAGGGGCCTGGCACGCCGGTCTGTGCCTGCCCGCCCCGCCCTGGCCCACCGGCCCCACTCCCTGGGCACAGAGTCACAC | 120 |

**Supplementary Table 1b. The probe design for human SHANK1-2 capture**

| Chromosome | Start | Stop | Name | Score | Strand | Code | Seq | Len |
|---|---|---|---|---|---|---|---|---|
| chr11 | 70515455 | 70515575 | 788855_47188552_ENST00000601538.6,E | 50 | - | good | CCCACCCCAGCTTCCCAGAGTGCTGGGATTATAGACATGAGCCACCGCTCCGGGCTGGCATTAAAAATTTGAAAGATAGATTGAAAGATAAACTTGAAAAGAATTTCTCAGAAAAGAAAA | 120 |
| chr11 | 70895855 | 70895975 | 788855_47188552_ENST00000601538.6,E | 50 | - | good | TGGCCCACTCCAGGGGGGTGTTGATGACTGGGGAGACTGCACGTGAGGGGCAGGGGCTCTCTGGGTAAGCTCTATTCCTTCTGCTTGGTTTTACTGTGAACCTAAAACTGCTCTAAAAAT | 120 |
| chr11 | 71182535 | 71182655 | 788855_47188552_ENST00000601538.6,E | 50 | - | good | ATGTTAGATGGGTCCAGATCTAGAATGACTGGCATCCTTATCAGAAGGAGGAATTCAGACACAGACATGGAGGGAAGACCGTGTGAAGACACAGGGAGAAGATGGTCACCTACCAGCCAA | 120 |
| chr11 | 70843535 | 70843655 | 788855_47188552_ENST00000601538.6,E | 49 | - | good | TGGGGCCTCCTGCAGCCTGTCTCTGCGGCTTGTAGGTGCCGCCTTCTCCTTGTGACCTCACAGGGTCGTCTCTCTGTGCGTCTTTGTCCTCATCTCCTCTTCTTCAAGGACATCAATGAG | 120 |
| chr11 | 71133335 | 71133455 | 788855_47188552_ENST00000601538.6,E | 49 | - | good | GTCTGTCCGTCCGTCCCTCCCTCCCTCCCTCCATCTGTCCATCCATCCTTCCCTCCATCTGTGCATCCACCCAGCCACCCACCTACCCAGCCAGCCAGCCAGCCAGCCATGCATCCTCCC | 120 |
| chr11 | 71227415 | 71227535 | 788855_47188552_ENST00000601538.6,E | 49 | - | good | GTACACGTTTAGAATAACTATGTCTTCTTGGTGGAGTCCTCTCTTACATTATGTAATGTTCTCTTTGTGGTATTTTCTTTGCCCTGAAGTCTACTTTATCTAATATTAACATAGCCATGA | 120 |
| chr11 | 70516535 | 70516655 | 788855_47188552_ENST00000601538.6,E | 48 | - | good | TTTATTCTCTTCAGAGTGCATTTTGCAGAGTTAATTTAAGGAAGTCCAACATCAATTTTGACTTTTATAGATTATGCTTTTGGTGTTGTATCTAAGAAATCATCACCAAACTCAAGGTCA | 120 |
| chr11 | 70823135 | 70823255 | 788855_47188552_ENST00000601538.6,E | 48 | - | good | CAGCGCCACCTCTGTCCCCCGTGAACTCTGCCAGCGCCACCTCTGTCCCCCGTGAGCTCTGCCAGCGCCACCTCTGTCCCCCGTGAGCTCTGCCAGCGCCACCTCTGTCCCCCGTGAGCT | 120 |
| chr11 | 70836935 | 70837055 | 788855_47188552_ENST00000601538.6,E | 48 | - | good | CTGATGAGGGCTCCCTTCCTGGCTTGCAGACGGCAGCCTTCTGGTTGAGCCCTCCCATGGTGGAGAGACAGAGAGACAGACAGCGACCAACTCTCAAGTCTCTTCTTACAAGGACACCAG | 120 |
| chr11 | 70822655 | 70822775 | 788855_47188552_ENST00000601538.6,E | 47 | - | good | CCCATGACCTCTGCCAGCGCCACCTCTGTCCCCCATGACCTCTGCCAGCGCCACCTCTGTCCCCCATGACCTCTGCCAGCGCCACCTCTGTCCCCCATGGGCTCTGCCAACGCCACCTCT | 120 |
| chr11 | 70822775 | 70822895 | 788855_47188552_ENST00000601538.6,E | 47 | - | good | TGACCTCTGCCAGTGCCACCTCTGCCCCCCGTGACCTCTGCCAACGCCACCTCTGACCCCCATGAGCTCTGCCAGCGCCACCTCTGTCCCCTATGAGCTCTGCCAGCGCCACCTCTGTCC | 120 |
| chr19 | 50678908 | 50679028 | 788855_47188554_ENST00000293441.6,E | 47 | - | good | CCTGACCCCTCCCCATCACCCTGACCCCTCCCCACACCCTGACCCCTCCCCACACCCTGACCCCTCCCCATCACCCTGACCCCTCCCCATCATCTTGACCCCTCCCCACACCCTGACCCC | 120 |
| chr11 | 70823015 | 70823135 | 788855_47188552_ENST00000601538.6,E | 46 | - | good | CTGCCAGCGCCACCTCTGTCCCCCGTGAGCTCTGCCAGCGCCACCTCTGTCCCCCGTGAGCTCTGCCAGCGCCACCTCTGTCCCCCGTGGGCTCTGCCAACGCCACCTGTCCCCCATGAA | 120 |
| chr11 | 70823255 | 70823375 | 788855_47188552_ENST00000601538.6,E | 46 | - | good | GCCACCTCTGTCCCCCATGAGTTCTGCCAACACCACCTCTGTCCCCCATTAACTCTGCCACCGCCACCTCTGTCCCCCATGAACTCTGCCAGCGCCACCTCTGTCCCCCGTGAACTCTGC | 120 |
| chr11 | 70942175 | 70942295 | 788855_47188552_ENST00000601538.6,E | 46 | - | good | GTTGCAGCACAGAAACCCTCACCAGAAGCCAAGGCGATGCCCTTGAACTTCCCACACCACAGAACCTTGAGCTAAATAAACCTCTTTTCTTTCCTTTTCCTTTTTTTTCGAGACAGAGTC | 120 |
| chr11 | 70942535 | 70942655 | 788855_47188552_ENST00000601538.6,E | 46 | - | good | AGTTCATCTGTTAGAAGTGAAGTGGAAATGACACTCTGCTAGTCTGCTAGGTCTGTGCTATGGTTTGGATATGGCTCGCTTGTCCCCACCAAATCTCATGTTGAAATTTGATTCCTCAAT | 120 |
| chr11 | 70636775 | 70636895 | 788855_47188552_ENST00000601538.6,E | 45 | - | good | GCACACATGCTACCATGTGCACACACATGTTCACACATGCTTACATACATTCTTATAAATACACAAGTTCTCACACATGCCCACACACATGTTCACACACACGCTCACACATGCTCACAC | 120 |
| chr11 | 70760015 | 70760135 | 788855_47188552_ENST00000601538.6,E | 45 | - | good | TGTTCACCTGGGGTGGAGGCTTCACATTTAAAGGCATTACAGTCAGGCTCTGTATGTCAAAAGGTGAAGCAGGAACGTAAAGCCGCTCAGTGAGCAGCCTCTGTAAACCGGCAGAGCCGG | 120 |
| chr11 | 70796855 | 70796975 | 788855_47188552_ENST00000601538.6,E | 45 | - | good | GGTGTGCCGCCTTCCTTTCTGTCCTTTTACGGTTTCTTTCTGTTGTTGTCATGGTGATTGTCAACTATCCTGGTGCTGGGGGGAGGGTCATTTAGCATGGAAATCAGACTGTAATGAGGT | 120 |
| chr11 | 70823495 | 70823615 | 788855_47188552_ENST00000601538.6,E | 44 | - | good | TCTGTCCCCCGTGAACGCTGCCAACGCCACCTCTGTCCCCCGTGAACGCTGCCAGCGCCACCTCTGTCCCTCATGAACTCTGCCAGCGCCACCTCTGTCCCTCATGAGTTCTGCCAATGC | 120 |
| chr11 | 71070695 | 71070815 | 788855_47188552_ENST00000601538.6,E | 44 | - | good | AGTTGGGTAACATTTTAGGCTTTGCGTGTCGCATGGACTCCGTCATAACGATTCAACTGCCACTGTAGTGCAAAAGCAGCCATAGACAGTATGTAAGTCAGAGTGTAGCCGTGTTCCAAT | 120 |
| chr11 | 71143655 | 71143775 | 788855_47188552_ENST00000601538.6,E | 44 | - | good | TGTGAAGCTCATCACAACCTTCTTGCCTTAGGAACAAGTAGGCAGCACTTCAGCCCTTTGCTGGGACCCCTCTTAAATGGCAAAATCACCAACAAAAAGCACAAGCAATGTGAGAATGAC | 120 |
| chr19 | 50678788 | 50678908 | 788855_47188554_ENST00000293441.6,E | 44 | - | good | TCCCCACACCCTGACCCCTCCCCATCACCCTGACCCCTCCCCACACCCTGACCCCTCCCCATCATCCTGACCCCTCCCCACACCCTGACCCCTCCCCATCATCTTGACCCCTCCCCATCA | 120 |
| chr11 | 70774655 | 70774775 | 788855_47188552_ENST00000601538.6,E | 43 | - | good | AATTGATATATAATTTCAATGCCATAAAATTTACTGTTTTAAAGTGCACCATTAAGTGGCTTTTTGCATATTCACGAGATTGTGCAAACATCACCACTCTCTAATCTCAGAGCATTTGCA | 120 |
| chr11 | 70927775 | 70927895 | 788855_47188552_ENST00000601538.6,E | 43 | - | good | CAGCTGGGGAGAGCTAATGCCCCCTTCCTCTTCCCTTTGTTCTATTCAGACCCTCAACGGATTGGATGGGGCCCACCCACTTTGGGGAGGGCATCTGCTTTTCTGGGTTTACCGATTCAA | 120 |
| chr11 | 71086535 | 71086655 | 788855_47188552_ENST00000601538.6,E | 43 | - | good | CAGGGGACCTCAGTCTTAAGGCCTTCAACTGATTGGATGGGGCCCACTCAATATTATGGAGAATCATCTGCTTAACTCCAAGTCTAATGATATATATATTATTATATATAATATATTTAT | 120 |
| chr11 | 70733495 | 70733615 | 788855_47188552_ENST00000601538.6,E | 42 | - | good | GTGGCAGCCTCTCAGCTCTGCCTTTGTAGCAGGCAGCAGCCACAGACAGTACCCCCATGAATGGAGTGGCAGCATGCCAATAAAACTTTATTGACAAAAGCAGGCAGCAGCCCTGGGTTC | 120 |
| chr11 | 70739735 | 70739855 | 788855_47188552_ENST00000601538.6,E | 42 | - | good | GGGTGCCTCCCTGGTGCCTCGTGTGTCTGGCTCTCCTTGTCTTACAAAGACACTGTCATGCTGGATGAGGGCCCATCACAGTACCTCGTTTTAACTTAATCACCTCTTTAAAGACCTTAT | 120 |
| chr11 | 70636175 | 70636295 | 788855_47188552_ENST00000601538.6,E | 41 | - | good | CATACTCATACATGTACACACACATGCATGCTCGCATACACGCATGCTCATACACTCAGACTCACATTCACACACACACACGTGCTGACATGCATTCACACATGCTCACGCATATACTCT | 120 |
| chr11 | 70668455 | 70668575 | 788855_47188552_ENST00000601538.6,E | 41 | - | good | CACCACAGACTAGGGGGCTCAAAACAAGAGAAGTCACTTACCTGTCACAGCCCTGGAGGCCAGAGGTCCAAAACCAAGATGACAGCAGGGCCACGCTCCCTCCGGGGGCTCCAGGGGAGA | 120 |
| chr11 | 70822535 | 70822655 | 788855_47188552_ENST00000601538.6,E | 41 | - | good | GTCCCCCATGAGCTCTGCCAGTGCCACCTCTGTCCCCCGTGAACTCTGCCAGTGCCACCTCTGTCCCCCGTGAACTCTGCCAGCGCCACCTCTGTCCCCCGTGAACTCTTCCAGCGCCAC | 120 |
| chr11 | 70822895 | 70823015 | 788855_47188552_ENST00000601538.6,E | 41 | - | good | CTCTGCCAACATCACCTCTGTCCCCCATGGGCTCTGCCAGCGCCACCTCTGTCCCCCGTGACCTCTGCCAGTGCCACCTCTGTCCCCCGTGACCTCTGCCAGTGCCACCTCTGTCCCCCG | 120 |
| chr11 | 70823735 | 70823855 | 788855_47188552_ENST00000601538.6,E | 41 | - | good | CCAGGAGCTCTGTCAACGCCACCTCTGTCCCTCATGAACTCGGCCACCGCCACCTCTGCCCCCCATGAATGCTGCCAACGCCACCTCTGTCCCCCATGAACGCTGCCAACGCCACCTCTG | 120 |
| chr11 | 70860815 | 70860935 | 788855_47188552_ENST00000601538.6,E | 41 | - | good | AGAATTTGGAATGAAGGGGCTTTGTGGTATGATTCACATACACATTTTCTTTTATTTCTTTTTTGTTTGTTTGTTTAAATAGAGATGGGATTTTGCCATGTTGCCCAGGTTGGTCTTGAA | 120 |
| chr11 | 70730975 | 70731095 | 788855_47188552_ENST00000601538.6,E | 40 | - | good | TTAAAGGTCCCATCTCCAAATACAGTCAAATTAGGCGTTGGGGCTTCAACATATGTGTGTTGCGGGCACACAGGTTAGTCCATAACATGGAGTGACTGCTTCATGGGTATGAGATTTTCT | 120 |
| chr11 | 70823615 | 70823735 | 788855_47188552_ENST00000601538.6,E | 40 | - | good | TCCCTCATGAGCTCTACTAACACCACCTCTGTCCCTCATGAGTTCTGCCAGCGCCACCTCTGTCCCTCATGAGTTCTGCCAATGCCACCTCTGTCCCCCGTGAATGCTGCCAACGCCACC | 120 |
| chr11 | 70917935 | 70918055 | 788855_47188552_ENST00000601538.6,E | 40 | - | good | TGGCCTTATTTAAAAAAAAAAAAAAAAAAAAAAAAAAAACTTTTAAGTTCAGGGGTATATGTGCAGAACGGGCAGGTTTGTTACCTAGGTAAACACATGTCTTGGGGGTTTATTGTACACA | 120 |
| chr11 | 71133455 | 71133575 | 788855_47188552_ENST00000601538.6,E | 40 | - | good | CATCCCCCCTCCACTCACTCATCCGTCTATCTATCCATCCAGCCAGCCAGCCTGCCTGCCTGTCTGCCTGCCATCCATCCGTCCGTCCATCCCTCTGTCTCTCTCTCCCTCCCTCCCTCC | 120 |
| chr19 | 50679028 | 50679148 | 788855_47188554_ENST00000293441.6,E | 40 | - | good | TTGTAACCCTGACCCCTCCCCATCACCCTGACCCCTCCCCACACCCTGACCCCTCCCCATCACCCTGACCCCTCCCCATCATCTTGACCCCTCCCCATCACCCTGACCCCTCCCCATCAC | 120 |
| chr11 | 70729175 | 70729295 | 788855_47188552_ENST00000601538.6,E | 39 | - | good | TGGCTTCATTCACTGAGCAGGAAGTTCCTGAGACGCATCTGTGTTGCCACGTGTATCAGTATTTCTTTCTTTCTTCTTTTTTTTTTTTTAATTTTTTTTTAGATGAATTCTTGCTCTGTT | 120 |
| chr11 | 70751375 | 70751495 | 788855_47188552_ENST00000601538.6,E | 38 | - | good | ACCTGCTGGCCACAAATGTTCTTGGTTTTGCTTCATCTAAAAATGTCTTGATTTAACCTTTATTCCTGAAGGATATTTTCACTGGGTATAGAATCCTGCATTTGCAGTATTTTAAAAATA | 120 |
| chr11 | 70759775 | 70759895 | 788855_47188552_ENST00000601538.6,E | 38 | - | good | AAGAGAAAGCTACTGAAATCAGCCTCTTATCCAATCACAGTTATAGTTATGGCTGCTGGCATGGGGTCTGTTATCAGCATCTGGTGGAGCTGCAATGGTTTTAATGTTGTCCATCCCAAG | 120 |
| chr11 | 70793255 | 70793375 | 788855_47188552_ENST00000601538.6,E | 38 | - | good | GTGGGGATTGGTTCCAGGACACCACCACCTCACATCATATACCAAAATCCACCTCAAGTCCCACAGTCATCTCCATGGAACCTGTGTATGTGAAGTCAGCCCTCTGTATATGCAGGTTTT | 120 |
| chr11 | 71155175 | 71155295 | 788855_47188552_ENST00000601538.6,E | 38 | - | good | TAGGTCCACCCCTCTGGGAGCGTGGGCTGGGGTAGGTCCACTCCTCCTCCGGGAGCGTGGGCTGGGGTAGGTCCACCCCTCTGGGAGCTTGGGCTGGGGTAGGTCCACCCCTCTGGGAGC | 120 |
| chr11 | 71155295 | 71155415 | 788855_47188552_ENST00000601538.6,E | 38 | - | good | CTCTTCTGGGAGCGTGGGCGGGGGTAGGTCCACCCCTCTGGGAGCGTGGGCTGGGGTAGGTCCACCTCTCTGGGAGCGTGGGCTGGGGTAGGTCCACCCCTCTGGGAGCTTGGGCTGGGG | 120 |
| chr11 | 70823375 | 70823495 | 788855_47188552_ENST00000601538.6,E | 37 | - | good | CACCTCTGTCCCCCGTGAACTCTGCCAACGCCACCTCTGTCCCCTATGACCTCTGCCAATGACACCTCTGTCCCCCGAGTTCTGCCAACGCCACCTCTGTCCCCTATGAACTCTGCCAGT | 120 |
| chr11 | 70929575 | 70929695 | 788855_47188552_ENST00000601538.6,E | 37 | - | good | TGGACTATCTGAGTCAGGCCCATGTTATCACAAGGGTCCTTATAAGAGGAAGGCAGAGGGTCAGAGTCAGCAGTCATGTGACAACAGGCACGGAGGTCGGAGAGAGATTTGAAGATGCTG | 120 |
| chr11 | 71088095 | 71088215 | 788855_47188552_ENST00000601538.6,E | 37 | - | good | CCCAGGAAGAGTTGATGCCGCTGCGGGGATCCTGTTTGTCTCTGGCAGAATCTCTTCCTCCTCAGGGGACCTCAGTCTTTTTTCTCTTGAGGCCTCCAACTGATTGGATGGGGCCCACCC | 120 |
| chr11 | 71191415 | 71191535 | 788855_47188552_ENST00000601538.6,E | 37 | - | good | AAACAGATGACCCTGTGGATGCGTGGAGGGATGACCCTATGAGGGATGCGTGGAGGGATGACCCTGTGAGGACACAGTGAGAAGGTGGTGTCTGCAAGCCAAGCAGAGAGGCCTCAGGAG | 120 |
| chr11 | 70587575 | 70587695 | 788855_47188552_ENST00000601538.6,E | 35 | - | good | GACGTGCTCAACAATGGGTTGCCATAAACCTTCAGTTTGTAACAAATGCAGTATCCCCAAAGCACAATAAAGCAAGGGGTAATAAAACAAGATATGCCTCTGCATCTTTTTTATACCTTT | 120 |
| chr11 | 70697015 | 70697135 | 788855_47188552_ENST00000601538.6,E | 35 | - | good | TAATGCAAACTCCCCACTCTACCTCCCTCAGACTCCAGCTCCCAACCTGTCTACTTTCTGTCTCTATGAATGTGATGACTCTAGGAACCTCAGATGAGTGGAATCCCGAAGTATTCATCT | 120 |
| chr11 | 70727015 | 70727135 | 788855_47188552_ENST00000601538.6,E | 35 | - | good | CTTGCCCAGAGTCATAATGTATTAGTCAGCTGTTGCTGCATAACAAATTATCCCAAAAGTTAGTGGCTTAAAACAACCGTCATTTGTCTCATGATTCTGTAAGTCAAATTAGGGTGGGTT | 120 |
| chr11 | 70774055 | 70774175 | 788855_47188552_ENST00000601538.6,E | 35 | - | good | TTTTGAGATTCTCCCATGCCATAGCATGTATCAGAATCTCACTGCTTCTTATGGCTGAGTTGTATTTCATTGTATGGTTATACCATACTTTGTTTCTATTAGAAGATGGACACTTTGGTT | 120 |
| chr11 | 71155055 | 71155175 | 788855_47188552_ENST00000601538.6,E | 35 | - | good | TTGGGCTGGGGTAGGTCCACTCCTCCTCCGGGAGCGTGGGCTGGGGTAGGTCCACCCCTCCTCCGGGAGCGTGGGCTGGGGTAGGTCCACCCCTCCTCCGGGAGCGTGGGCTGGGGTAGG | 120 |
| chr11 | 71228015 | 71228135 | 788855_47188552_ENST00000601538.6,E | 35 | - | good | GTGTGTAGAGCTGATTTAGTATTCCCTTAGTATCTTTTTCATGTCTGCAGGGTCTTTAATGGTGTCTCCTATTTAATTCCTGGTATTGGTAATTTGTGTTGTTTTTTTTTCTTGGTCAGT | 120 |
| chr11 | 70543775 | 70543895 | 788855_47188552_ENST00000601538.6,E | 34 | - | good | GCAGGTGCCCCTAATTCAATTCAGTTCTGCACTGCTCACCTGGAGAGTGTGTTGATCCCATAGGGTGAGGCTCAGTCCCACAAGATGGGCCCCACGTCCTTTGCAGGTGCCAAGAACAAG | 120 |
| chr11 | 70620815 | 70620935 | 788855_47188552_ENST00000601538.6,E | 34 | - | good | ACAGACCAGGGGGCTTAAACAAGCATTTATTTGGAGACTAGAAGTCCCAGATCAAGGTGCCAGCAATTTGGTTCCTGGTGAGGGGTCGCTTCCTGCTTATAGGCAGCTGGCTTCCTCACA | 120 |
| chr11 | 70953215 | 70953335 | 788855_47188552_ENST00000601538.6,E | 34 | - | good | ATTCTCCCTTAAAAAACTCCCTTTCCTGTGTAAATGTATCCTATCAGTTCTGTCCCTCTGGAGATCCCTGATTTAATACACTCACTTCGGAGGGGAGGGCGTGGATCTGTTCATGAGGGA | 120 |
| chr11 | 71100215 | 71100335 | 788855_47188552_ENST00000601538.6,E | 34 | - | good | ATAAAGCTGCTGTAAACACTAACATTCAGATTTCCTTGTGGACGTAAGTTTTCAGCTCATTCAGGGAGACACCTAGGAGCTGGGTTGCTGGATTGTGGAGTAAGTGCGTTTTTAGTTTTG | 120 |

| | | | | | | | | |
|---|---|---|---|---|---|---|---|---|
| chr11 | 71154935 | 71155055 | 788855_47188552_ENST00000601538.6,E | 34 | - | **good** | TCCACCTCTCTGGGAGCGTGGGCTGGGGTAGGTCCATCCCTCCTCTGGGAGCGTGGGCTGGGGTAGGTCCATCCCTCTTCTGGGAGCGTGGGCGGGGGTAGGTCCATCCCTCCTCTGGGA | 120 |
| chr11 | 71192135 | 71192255 | 788855_47188552_ENST00000601538.6,E | 34 | - | **good** | AACTGTATTTGGAGACAAGTAATCAAGTTAAGTTGGGGTCATATGGATGGACCCTAATCCAAAGGACTGGTGTCCTGATAAGAAGATGAGATGAAGGCCGGGTGTGGTGGCTCACGCCTG | 120 |
| chr11 | 71223335 | 71223455 | 788855_47188552_ENST00000601538.6,E | 34 | - | **good** | ATGACAAGAAAAATGTCTGTACATGTTCGGTACAGATGCAACTAAAAAATATTTCCAATGTGCGCATGGTTGGTTGAGTTCAGGGTAACTGAAGCCACGCATATGGAAGGCTGTTGCTGT | 120 |
| chr11 | 71228255 | 71228375 | 788855_47188552_ENST00000601538.6,E | 34 | - | **good** | GGAAGAAATTGTGTAGAATTAGTATTAATTATTCTTTAAGCATTTCATAGAATTCTCCAGTGAACTCATTTAGGTCTAAAGATTTCTTTTTCTCAAAAAAACTGGAGTTTAAAAATTATG | 120 |
| chr11 | 70542215 | 70542335 | 788855_47188552_ENST00000601538.6,E | 33 | - | **good** | ACACAGCCTCATCCTCCTCTCTCTTCTCATCCCTCATAAGGGCACCCGTCACTGGATTCAGGGCCCACTTCAGTAACCGAGGATGATCTCATCTCGAAATCCTTAACTTAATTATATCTA | 120 |
| chr11 | 70635095 | 70635215 | 788855_47188552_ENST00000601538.6,E | 33 | - | **good** | CTTCGGGCATAGATGCAGAAGTGGCATTGCTGGCTCACGTGCTCGTTCTGTGTTTAAGGTTTTGAGGAGCAGCCATGCCATGTTCCACAGGGGCTGCACCACTTTACCTTCCTGCCAGCG | 120 |
| chr11 | 70946855 | 70946975 | 788855_47188552_ENST00000601538.6,E | 33 | - | **good** | TAGTGGAGAGGGAGGGTGAGCCTGGGGAAGAGTTGGTTAGTGGAGAGGGAGGGTGAGCCTGGGGAAGAGTTGGTTAGTGGAGAGGGAGGTTGAGCCTGGGAAGGGTTAGTTAGTGGAG | 120 |
| chr11 | 71154695 | 71154815 | 788855_47188552_ENST00000601538.6,E | 33 | - | **good** | CCATCCCTCTTCTGGGAGCATGGGCGGGGGTAGGTCCATCCCTCTGGGAGCGTGGGCGGGGGTAGGTCCATCCCTCCTCTGGGAGCGTGGGTGGGGGTAGGTCCATCCCTCCTCTGGGAG | 120 |
| chr11 | 71154815 | 71154935 | 788855_47188552_ENST00000601538.6,E | 33 | - | **good** | GCGTGGGCGGGGGTAGGTCCATCCCTCTGGGAGTGTGGGCGGGGGTAGGTCCATCCCTCCTCTGGGAGCGTGGGCGGGGGTAGGTCCATCCCTCCTCTGGGAGCGTGGGCTGGGGTAGG | 120 |
| chr11 | 71155895 | 71156015 | 788855_47188552_ENST00000601538.6,E | 33 | - | **good** | CCCTTGTGTGTCCAGAGTCCCTCTTCTGATAAAAAGACACCAGCCTTGCATTAGGGCCCACCTGTAAGACCTTGTGGAACCTTAAGTACCTTTTTAAAGGCCCTGTCTCCAATACTGTCA | 120 |
| chr11 | 70526615 | 70526735 | 788855_47188552_ENST00000601538.6,E | 32 | - | **good** | CAAAATCATAAAATTTCGAAAGGAGACTTTATTTGTAAAGGGTTACGCCTGCGAGGTGGCATTGTGACCAGCTGGGAAGCATAGCCTCTGGCCAAAGCCGGAAGCAGGCACTTCAGGGG | 120 |
| chr11 | 70562135 | 70562255 | 788855_47188552_ENST00000601538.6,E | 32 | - | **good** | TTTTCAAGTGTACTTGGAAGGGTTATTAAGATAAACCATATTTTAGTCCATAAAACAAGTCTCAATAAATTTGGAACTATTCAAGTCATACAGATGTGAGCTCTAACCAAAACGGAATCG | 120 |
| chr11 | 70690655 | 70690775 | 788855_47188552_ENST00000601538.6,E | 32 | - | **good** | TAAACTTCTCTCTCTTGTTTAATACATTTTAGACACTATCACTCGACCCTTTGGTATATTTATATTTATATTTATATTTATATTTATATTTATATTTATATTTATGGGTTCTAAGTTTTG | 120 |
| chr11 | 71105015 | 71105135 | 788855_47188552_ENST00000601538.6,E | 32 | - | **good** | TGTTCCATTTTCCAATGTACCCCAGTTTGCTTATCCACTTGCCTCCTAAAGGACATCTGGGCTGCTTCCAGCTTTTGGCAATTGTGGGTCAAGCTGCCATAAACACCCGTGTGTGGGTTT | 120 |
| chr19 | 50663188 | 50663308 | 788855_47188554_ENST00000293441.6,E | 32 | - | **good** | CTCCCTCTCTTCATCTCTCTCTCTCTGTGTCCCCTGTCCCTTCCATCTCTCTCCCCTGGATGCTCACCCTCTTTTTGTCTCTCTCTCACTGTCTCCCCATCTTTCACTCTGCTACGACTG | 120 |
| chr11 | 70697135 | 70697255 | 788855_47188552_ENST00000601538.6,E | 31 | - | **good** | ACCATTTGTTAGTGAATCACTAGTTTAGTGGTATTAAGAACATTCACAGTGTTGTGTAGCCAACACCACCATCCAGCTCCAGAGCTCTTTTGTCTTGCAAAGTTGGAACTCTGTCTGCAT | 120 |
| chr11 | 70754375 | 70754495 | 788855_47188552_ENST00000601538.6,E | 31 | - | **good** | CATTAGTTTGTTTATATGGTGGCTTACATCAATAAATTTTTCAAATATTGAACCAGCCTTGCACTCCCAGGATAAAGCACACTTGCTCATGATTCCCTTTCTACTAATATATGTCTGGAT | 120 |
| chr11 | 71160575 | 71160695 | 788855_47188552_ENST00000601538.6,E | 31 | - | **good** | GTGGGTGTCTTGGGCAGCTCAGGCTATAACAGAATAGATAGGATGACTTAGCAGAAATTTATTTCTCACCTTTCTGGAGGCTGGACATCTGAGATCAAGAAGCTGGCTGATTCAGCTCC | 120 |
| chr11 | 71243735 | 71243855 | 788855_47188552_ENST00000601538.6,E | 31 | - | **good** | GCTTGTCATAGGTCTTTTTAGATTTTCTGTTTCTTCTTGAGTCAAGTTTGTTAGTTTGTGTCCCACCAGAAATGTGTCCACTTCATGTGAGTTTCTCTAATGTGGTTGCTATGTAGTTGT | 120 |
| chr11 | 70517135 | 70517255 | 788855_47188552_ENST00000601538.6,E | 30 | - | **good** | ATAAGCATTCCGGTTGCTGCACATCCTCGCCGGCATTTGCTGTTGTCGGTGTTTGAGATTCTGGCCATTCCATAGGTGTGTAGTGGTGTCTCACTGTTGGCTTGATTTGCTAATTCCACA | 120 |
| chr11 | 70701815 | 70701935 | 788855_47188552_ENST00000601538.6,E | 30 | - | **good** | TGATGGTGGTGGTGATGATGGTGGTGATGATGGAGGGTGGTGGTGATGATGATGGGAGTGGTGATGGTGATAATGATCTTGTCCCTTAGGGTCCCTTTCCTGTCCCTTAGGGACCATATC | 120 |
| chr11 | 70723415 | 70723535 | 788855_47188552_ENST00000601538.6,E | 30 | - | **good** | TGCATTCTCATCTGGAGGTTCAATGGGGAAGGGTCCACTTCCAAGCTCTCACAGGTTGTTGGCAGAAATTCAGTTCCTTGTATTTTTATCAGTAAGGGCTTCATTGCTGGCTGGGGGCTG | 120 |
| chr11 | 70946255 | 70946375 | 788855_47188552_ENST00000601538.6,E | 30 | - | **good** | CTGATCCTGGGAAGTGTTGGTTAGTGGAGAGGGAGGCTGATCCTGGGAAGTGTTGGTTAGTGGAGAGAGAGGTTGAGCCTGGGAAGCGTTGTTTAGTGGAGAGGGAGGTTGAGCCTGGAA | 120 |
| chr19 | 50699908 | 50700028 | 788855_47188554_ENST00000293441.6,E | 30 | - | **good** | TCCAATGCCCCGAGCCTTCCAACCCTCCAATGCCCCGATCCCTCCAATCCTCCAATGCCCCGAGCCCTCCAATCCTCCAATGCCCCGAGCCCTCCAATCCTCCAATGCCCCGAGCCCTCC | 120 |
| chr11 | 70619135 | 70619255 | 788855_47188552_ENST00000601538.6,E | 29 | - | **good** | CTGACCTACAGAACTGGCACGTAAGAACATTTTTGTTGTTTTAAGCCACTAAATTTGTAGTAATTAATTACAGCGAGCAATAAGAATCTAACCCAAGGGTGGTTGTCAGGCTGAAATGAT | 120 |
| chr11 | 71155415 | 71155535 | 788855_47188552_ENST00000601538.6,E | 29 | - | **good** | GTCTCCATGTTGAGGGTGTGGCTGGGGGCATGGGAGGGGCAGGTCCATCCCTCCTCTGGGAGCGTGGGCAGGGGTAGGTCCACCCCTCCTCTGGGAGCGTGGGCGGGGGTAGGTCCATCC | 120 |
| chr11 | 71227655 | 71227775 | 788855_47188552_ENST00000601538.6,E | 29 | - | **good** | CACTTGAAAAGAATGTGAATTCTGCTGTAGTACAGTGGAGGAAGGTTCTACAACTATCACTTTGATCTTGTTGATTGGGTATATTACTGAGTTCTGTATCCTTGCTGATTTTCCTGTCCT | 120 |
| chr11 | 70783055 | 70783175 | 788855_47188552_ENST00000601538.6,E | 28 | - | **good** | GAGGCTGGACAGTCAGAGATGAGGGTGCGAGCATGGCCATTTCTGGGAAGGGCCCTCTTCCTGGCTTGCAGACAGCCTCTGTCTCTCTGTGTCCTCATCCGCAGAAAGACAGAGACAGAG | 120 |
| chr11 | 70822415 | 70822535 | 788855_47188552_ENST00000601538.6,E | 28 | - | **good** | CTCTGTCCCTCATGAACTCTGCCAACTCCACCTCTGTCCCCTATGAACTCTGCCAGTGCCATCTTTGTCCCTGTGACCTCTGCCAGCACCACCTCTATCCCCCTTGAGTTCTGCCAGCAC | 120 |
| chr11 | 70946375 | 70946495 | 788855_47188552_ENST00000601538.6,E | 28 | - | **good** | GACAGGGAGGCTGATGCTGGGGAAGAGTTGGTTAGTGGAGAGGGAGGTTGAGCCTGGGAAGGGTTGGTTAGTGGAGAGGGAGGGTGAACCTGGGGAAGGGTTGGTTAGTAGAGAGGGA | 120 |
| chr11 | 70946615 | 70946735 | 788855_47188552_ENST00000601538.6,E | 28 | - | **good** | TAGGGAAGGGTTGGTTAGTGGAGAGGGAGGGTGAGTCTGGGAAGGGTTGGTTAGTGGAGAGGGAGGTTGACCCTGGGGAAGAGTTGGTTAGCAGAGAGGGAGGGTGAGCCTGGGGAAG | 120 |
| chr11 | 70946735 | 70946855 | 788855_47188552_ENST00000601538.6,E | 28 | - | **good** | GGAGGATGAGCCTGGGAAGGGTTGGTTAGTGGAGAGGGAGGGTGAGTCTGGGAAGGGTTGGTTAGTGGAGAGAGAGGTTGAGCCTGGGAAGGGTTGGTTAGTGGAGAGGGAGGTTGAG | 120 |
| chr11 | 70543535 | 70543655 | 788855_47188552_ENST00000601538.6,E | 27 | - | **good** | GGTTTCTTATAAAGGATATTACAAATGGAGTGATGTGTGGGGTGAGGTACAGGGGAAGGCGTGCAGCTTTGATGCCCTGCCCAGGCACGCCACCCCCAAGGAACCTCCACGTGTTCATCT | 120 |
| chr11 | 70560935 | 70561055 | 788855_47188552_ENST00000601538.6,E | 27 | - | **good** | CTTTGCCCTTTATCAAAAACCCGTTCTTCGTGTGTGTGTGGGGCTATCTCTGCGCTCTCTTTTCTCTTCCATTATCTATTTGTCTAGCTTGGTGCCACTGTCTTGATTACTGTATCTTTA | 120 |
| chr11 | 70625015 | 70625135 | 788855_47188552_ENST00000601538.6,E | 27 | - | **good** | CCTTGCCTGATCTCAGCCTGGATCCCTTTGCTGTAATAAACCATAACTGTGAGCATAGGTGCTTTTAGTGAGTTCTACGAGTCTTTCCCACAAATCACCAAACCTGAGGGTGGTCTTGGA | 120 |
| chr11 | 70946015 | 70946135 | 788855_47188552_ENST00000601538.6,E | 27 | - | **good** | GCGGAGAGGCAGGTTGTGTCTGGGAAGGGTTGGTTAGTGGAGAGAGAGGTTGAGCCTGAGAAGGGTTGGTTAGCGGAGAGGGAGGTTGAGTCTGGGAAGGGGTGGTTAGTGGAGAGGGA | 120 |
| chr11 | 70946135 | 70946255 | 788855_47188552_ENST00000601538.6,E | 27 | - | **good** | AGGGTTGGTTAGCGGAGAGGGAGGTTGAGCTTGGGGAAGAGTTGGTTAGCGGAGAGGGAGGTTGAGCCTGGGAAGGGTTGGTTAGTGGAGAGGGAGGTTGAGCCTGGGAAGGGTTAGTT | 120 |
| chr19 | 50699788 | 50699908 | 788855_47188554_ENST00000293441.6,E | 27 | - | **good** | AATCCTCCAATGCCCCGAGCCCTCCAATCCTCCAATGCCCCGAGCCCTCCAATCCTCCAATGCCCCAAACCCTCTAATCCTCTAATGCCCCAATCCCTCCAATCTCCCAAAGCCCCAAGC | 120 |
| chr19 | 50700028 | 50700148 | 788855_47188554_ENST00000293441.6,E | 27 | - | **good** | CAACCCGAGCCTTCCAACCCTCCAATGCCCCGAGCCCTCCAATCCCCCAATGCCCCGATCCCTCCAATCCTCCAATGCCCTGAGCCTTCCAACCCTCCAATGCCCCGAGCCCTCCAATCC | 120 |
| chr19 | 50700148 | 50700268 | 788855_47188554_ENST00000293441.6,E | 27 | - | **good** | CTCCCAATGCCCCGAGCCCTCCAATTCTCCAATGCCCCAAACCCTCTAATCCTCCAATGCCCCGAGTCCTCCAATCTCCCAATGCCCCAAGCCCTCCAATCTTCCAGTGCCCGAGCCTTC | 120 |
| chr11 | 70636295 | 70636415 | 788855_47188552_ENST00000601538.6,E | 26 | - | **good** | CTCACACATGCTCGCATACACACTCATACACACAGACTCACATTCATACACACATATGCACATGTACTAACACGCATTCACACATGCTAACATTCACACGCTCACACATCTGCTCACACA | 120 |
| chr11 | 71076455 | 71076575 | 788855_47188552_ENST00000601538.6,E | 26 | - | **good** | TAAATGGAAATTTCTCGTGTGTGTGTGTGGTTTTTTTTTTTGTTTTTCGTTTTTTGTTTTTGTTTGTTTTTTTTTTTGTTTTGTTTTGTTTTTATAGCTCATCCAATTAATAGTTTCAATA | 120 |
| chr11 | 71099375 | 71099495 | 788855_47188552_ENST00000601538.6,E | 26 | - | **good** | CATTTGTAAACAAACTTTTCATTTTAGAACAGTTCTAGATTTGCAGAATAAGTGCAAAGATAGTACACAGAGTTACCATATATCCTATACCCAGCTTCCCCCCTTATCAGCATCTTACTC | 120 |
| chr11 | 70620095 | 70620215 | 788855_47188552_ENST00000601538.6,E | 25 | - | **good** | GCCGTGGACCGAATGTCTGTGTGACCCCCACACTCAAATGTTGGAACCCTAACCCCCACCAGGATGGTGTCAGGAGCAGGGCCTTTGGGAGGGGGCTAGGGCTTGATGAGGTCAGAGGGT | 120 |
| chr11 | 70946495 | 70946615 | 788855_47188552_ENST00000601538.6,E | 25 | - | **good** | TTGGTTAGTGGAGAGGGAGGCTGATCCTGGGAAGTGTTGGTTAGTGGAGAGGAAGGCTGACCCTGGGGAAGAGTTGGTTAGTGGAGAGGGATGGTGAGCCTGGGGAAGGGTTGGTTAGTG | 120 |
| chr11 | 71226215 | 71226335 | 788855_47188552_ENST00000601538.6,E | 25 | - | **good** | ATTTTATTATTGTCCCATAACTCCCTGAGACTCTGTTTATTTATTTTCAGTCTGTTTCACTCTATTGTTCAGTTGGGATAATGTCTGTTGTTCTGTCTTCCAGTTCACTGATTATTTCCT | 120 |
| chr11 | 70542335 | 70542455 | 788855_47188552_ENST00000601538.6,E | 24 | - | **good** | GCAAAGCCATGAGCCCCCTCTAGGCTCTAGGGGAGGGTCCTTCCTTGCCTCCTCCAGCTCTGGGGCTCCCGGTATTCCTTGGCTGCTAGCCACATGGTGCCAACCTGTGTCTCTGTCGTC | 120 |
| chr11 | 70556975 | 70557095 | 788855_47188552_ENST00000601538.6,E | 24 | - | **good** | GGGTCCTTCTGAGTCACACCAGCAATGCCACTACATTTATGGCCCTGGCTTTGTAAAAATACAGTCAGGCTCGCCTAGGTGTGGTGACTCATGCCTATAATCCCAGCACCTTGGGAGGCC | 120 |
| chr11 | 70792895 | 70793015 | 788855_47188552_ENST00000601538.6,E | 24 | - | **good** | GGTGCCATCAGGGCTCATTACATGCCTTGACTTCCCAGTCTCAAATGATTCTCCTGCCTTATTTTAAAATTTATTTTATTAGTGTATTTTTTTGTAGAGACGAGTTCTCACTATGTTGCC | 120 |
| chr11 | 70927895 | 70928015 | 788855_47188552_ENST00000601538.6,E | 24 | - | **good** | TGTGATTACGGAGGCTGAGAAGTCCCACAATCTGCCATCTGCAAACCAAAGACCTGGGAAAGCTGGTGGTGTAAATGCCAGGACAAGGGCAGAAAACGATTTGTGTTCAGCTCCAGCAGT | 120 |
| chr11 | 70945895 | 70946015 | 788855_47188552_ENST00000601538.6,E | 24 | - | **good** | GGCGAGCCTGGGAAGGGTTGGTTAGTGGAGAGGGAGGTAGAGCCTGGGAAGGGTTGGTTAGTGGAGAGGGAGGTTGAACTGGGAAGGTTTGGTTAGTGGAGAGGGAGGCTGTGAAAGA | 120 |
| chr11 | 71129615 | 71129735 | 788855_47188552_ENST00000601538.6,E | 24 | - | **good** | AGCCTCTCACCTTGCAGAATACTTGTAACTCATGTGAACCAAGTGTGAGTGACCTTCCCTCACTGTCTCCATTTTTTCTTTTGTTTTGTTTTGAGTCAGGGTCTTGCTCTGTCACCCAGG | 120 |
| chr11 | 71170895 | 71171015 | 788855_47188552_ENST00000601538.6,E | 24 | - | **good** | ATTGTACCGCACCTTGATCTTGGACGCCCAGTCCCCAGAATGTGAGCACACACACATTTCTGCTGTCATAGCCGCAGCCGTGGTATTCTGTGATGGCGGCAGAGCTGACTAATGCACCCC | 120 |
| chr11 | 71176535 | 71176655 | 788855_47188552_ENST00000601538.6,E | 24 | - | **good** | ATTTTAGGCAGTTTCTATTGCTCTGACTTCAAGTTCACAAATCTTTTTGTGTGATGTGTAATTTGCCATTAATTCTATTCAGGGTATTTTTCACTTCAGATATTGTAGTTTTTATCTTTG | 120 |
| chr11 | 71227895 | 71228015 | 788855_47188552_ENST00000601538.6,E | 24 | - | **good** | CTTGCTAGAGTTTTGTCAATTGCATTGACTTTTACAAAGAGTCAGCTTATTGTTTCACTGATTTTCTCTGTTGTTTTTCTGTTTCCTGTTCATTTTTTCCCACTCATATGTTTATTATTT | 120 |
| chr19 | 50700268 | 50700388 | 788855_47188554_ENST00000293441.6,E | 24 | - | **good** | AATGCCCTGAGCTCTCCAATCCTCCTATGCCCCGAGCCCTCCAATCCTCCAATGCCCCGAGCCCTTCAATCCTCCAATGCCCCGAGCCCTCCAATCCTCCAATGCCCCGAGTCCTCCACT | 120 |
| chr11 | 70736375 | 70736495 | 788855_47188552_ENST00000601538.6,E | 23 | - | **good** | TGTGTCTGTGTCTTCTCTTCTGACTCTCAGAAGAACACGTATCATTGGATTTAAGACCCCCCTGGGAAATCCAGGGTGATCTCCTCATCCCAAGATCCTTAACTCAATCACACCTACAAG | 120 |
| chr11 | 70781495 | 70781615 | 788855_47188552_ENST00000601538.6,E | 23 | - | **good** | TAGAGCTTAAAGTATAATAAATAAATAAGTAAATATATATATATATATATATATATAATAAGTAAGTAAATTGCAAGCTGCTTTTAGAAATAAATGTGTAGGAATGACTGTGTTTGGAAA | 120 |
| chr11 | 70796735 | 70796855 | 788855_47188552_ENST00000601538.6,E | 23 | - | **good** | TAGAGGTTCTTCAGAAGCCAAGTGAGCTGCCATCTTGGATCCCACTAGTCTTAGCCTTTTTGGTCACCAGGGGGGAGCTTCTGACCTCAGGCATCCTGTTGCCTCAAGATAAGCAGAGTTG | 120 |
| chr11 | 70891535 | 70891655 | 788855_47188552_ENST00000601538.6,E | 23 | - | **good** | ATGAAAGAATTCCAACTATGAGACCTTGGAGGCTGTCCACCCAGAGCACCAGCAGAAGGAACCAGCCCTGCCAACACATCGATTTTGGACTTCTGGCCTTCAGAACTATGAGATAGTAAA | 120 |
| chr11 | 71090375 | 71090495 | 788855_47188552_ENST00000601538.6,E | 23 | - | **good** | TAGAAGCAGCAGGACACACACACACACACACACACACACACACACACACACACGTAGTGTTTCTCTGGAGAACCCTGGCTGATACACCCTGAATGTAGAAGCAGCAGGGGACACACACAC | 120 |
| chr11 | 71226815 | 71226935 | 788855_47188552_ENST00000601538.6,E | 23 | - | **good** | GAACCACATAGATTATGTTGTAATTTTTGCTTCAACCATCAAATATAATCTATAAAACTTGAGAAGGAAAGCCTATTATATTTGCCAAACATTTTGCTTACCATACTTTTTTCCTTCCTA | 120 |

| | | | | | | | | |
|---|---|---|---|---|---|---|---|---|
| chr11 | 71227295 | 71227415 | 788855_47188552_ENST00000601538.6,E | 23 | - | good | CTGCTTTCTTTTGACTTCTGTTTGTATAAGGTATCTTATTCCATCCTTTTACTTTCATCCTACCTATGGCATTATATTTGAAGCAAATTTTTATAGACCTCATTTAATTGTATCTTTTAA | 120 |
| chr11 | 70533695 | 70533815 | 788855_47188552_ENST00000601538.6,E | 22 | - | good | GCTTGCCTATGGCTGGGGAAGGGGAGAGTGAGGACTGACTGCTAATGGGTAGGGGGTTTCCTTGTAGGGGGATGAACTAGAGGTGGTGCTTGCACAACACTGTGAATGTACTAAATGCCA | 120 |
| chr11 | 70576295 | 70576415 | 788855_47188552_ENST00000601538.6,E | 22 | - | good | TGATCTGCCCGCCTCGGCCTCCCAAAGTGCTGGGATTACAGGCATGAGTGTGGGTTTTTTGACTCTAAGTCCAGTGCTCCCCACTGGCACGGCTCTCTCCCCAGGAGGTGCATGGGGTCT | 120 |
| chr11 | 70772495 | 70772615 | 788855_47188552_ENST00000601538.6,E | 22 | - | good | GAGGCATCTCTATCACTGGCTGCATGTAGGAAGCGACAGGTCAGCTCACCCTTTCTGAAACTATAGTTTCTCCAATGTTTCAACATGATATAGTCAGTATAATCAGGCATATGTGAGGAT | 120 |
| chr11 | 70870655 | 70870775 | 788855_47188552_ENST00000601538.6,E | 22 | - | good | TGAATGAATGAAATGTGGCCACAGCCATGCACTCGGAGCAAACCTCACACAAAGATGAAGGCAGAGCTCTGGGTGATGCTTCTGTCAGTCAAGGAATGCCAAAGATTGCCTACAGGCCAC | 120 |
| chr11 | 71065295 | 71065415 | 788855_47188552_ENST00000601538.6,E | 22 | - | good | CACCCCCCCACAACTTCCCCAGTCACTGCTCATCTCCCTGGAAGAGTTCTGCACACACACACCCCCACAACTTCCCCACTCACTGCTCATCTCCCTGGGAGAGTTCTGCACACACCTCCC | 120 |
| chr11 | 71065415 | 71065535 | 788855_47188552_ENST00000601538.6,E | 22 | - | good | CTGGGAGAGTTCTGCACATACACCCCAACTTCCCCACTCACTGCTCATCTCCCTGGAAGAGTTCTGCACACACCCCCCAACTTCCCCACTCACTGCTCATCTCCCTGGAAGAGTTCTGCA | 120 |
| chr11 | 71065775 | 71065895 | 788855_47188552_ENST00000601538.6,E | 22 | - | good | GTTCTGTATCCCCCCCAACTTCCCCACTCACCGCTCATCTCCCTGGGAGAGTTCTGTATGCCCCGCCCCCAACTTCCCCACTCACCACTCATCTCCCTGGGAGAGTTCTGTACCCCTCCT | 120 |
| chr11 | 71065895 | 71066015 | 788855_47188552_ENST00000601538.6,E | 22 | - | good | TCTCCCTGGGAGAGTTCTGTACTCCCCTCCCAACTTCCCCACTCACTGCTCATCTCCCTGGGAGACTTCTGTACCCCGCCCCCACCAACTTCCCCACTCACCGCTCATCTCCCTGGGAGA | 120 |
| chr11 | 71066255 | 71066375 | 788855_47188552_ENST00000601538.6,E | 22 | - | good | CTGGGAGACTTCTGCACCCCCCCAACTTCCCCACTCACTGCTCATCTCCCTGGGAGACTTCTGCACCCCCCACAACTTCCCCACTCTCTGCTCATCTTCGTGGGAGAGTTCTGTACCCTC | 120 |
| chr11 | 71098535 | 71098655 | 788855_47188552_ENST00000601538.6,E | 22 | - | good | CTGTTCCTTTTGCCTTTAGTCTTACAGACTCCACTCATGTCTGGAGCTACTGAGGTCAGCGGTTCATTCCCCATCCCTTTCAGCGAGGTTGATTCCTACACCTGTGATATGTTCTGTGGT | 120 |
| chr11 | 71220455 | 71220575 | 788855_47188552_ENST00000601538.6,E | 22 | - | good | TGAATGGACCACACCTTGTTTATGCGTTCATCCAGTGGTGGACACTGGGTTGCTCCAACCTTTTGGCTGTTGTGAATAGTGTTCTGTGAACATAGGTGTGCGGATATATCTCCTCAAGAT | 120 |
| chr19 | 50672188 | 50672308 | 788855_47188554_ENST00000293441.6,E | 22 | - | good | CTCCCAAAGTGCTGGGATTACAGGCGTGAGCCACGTTGCCCGGCCTCCTTCCACCGTCTTCACACTGCTCTCCCTTACGGCAGGGGCTTAGGGAAGGAATGGGGGCAGCCGTATAGTGAA | 120 |
| chr11 | 70769735 | 70769855 | 788855_47188552_ENST00000601538.6,E | 21 | - | good | AGATGAAAGATACCCATACACACACACCTACACACGTGCAAATGTGCACATACGTGCATGCCTGCACACAGACACACATGCCTGCACATGCCCACATAGAACATACACACATGCCTACAC | 120 |
| chr11 | 70772735 | 70772855 | 788855_47188552_ENST00000601538.6,E | 21 | - | good | TTTTGTACTTTTTGACAAAGAATGATAAATTTGAAAAGAAACGACAGGGCAAAGAAAATCTGGCTAGGACAGTAAATTTTCTAGTGGAGTCCCTAGGACATATACAGGTGGTGTAAAGCA | 120 |
| chr11 | 70870535 | 70870655 | 788855_47188552_ENST00000601538.6,E | 21 | - | good | TGGGAGCTGGAGAGACAGGCCTGGAACAGAGTCTCCCTGCAGTCCCCAGTGTGAACCCCCCCACTGCCAACACCTTGACCTCAGACTTCCAACCTCCAGAACCATGAGACAATTCATCCC | 120 |
| chr11 | 71065655 | 71065775 | 788855_47188552_ENST00000601538.6,E | 21 | - | good | CCCCAACTTCGCCACTCACTGCTCATCTCCCTGGGAGAGTTCTGTACCACCCCCAACTTCCCCACTCACTGTTCATCTCCCTGGAAGAGTTCCGCACACCCCCCACAAACTTCCCCAGTC | 120 |
| chr11 | 71066015 | 71066135 | 788855_47188552_ENST00000601538.6,E | 21 | - | good | ACTCACTGCTCATCTCCCTGGGAGAGTTCTGCACACACCCCCCCCACCAACTTCCCCACTCACTGCTCATCTCCCTGGGAGAGTTCTGTACCCTCCCCCAACTTCCCCACTCACCACTCA | 120 |
| chr11 | 71066135 | 71066255 | 788855_47188552_ENST00000601538.6,E | 21 | - | good | CCCCGCCAACTTCCCCACTCACTGCTCATCTCCGGGGGAGAGTTCTATACCCCCCCCAACTTCCCCACTCACTGCTCATCTCCCCAGGAGAGTTCTGCACCCCCCCCACCCAACTTCCCC | 120 |
| chr19 | 50676388 | 50676508 | 788855_47188554_ENST00000293441.6,E | 21 | - | good | ACCCTTCAGCTCCTGGCCTCCTTCCTATTCCTCCTGAGCCTGCCTGGCTCACCTTCACCTCAGGGCCTTTGCACCTGCTGTTCCACCTGCCTGGAACACTGTTCCTCTACACATCATTAT | 120 |
| chr11 | 70640375 | 70640495 | 788855_47188552_ENST00000601538.6,E | 20 | - | good | ACTCACAGTGCTGGAGGCCACAGGTTTGAAATCCAGGGCCTGGCAGGGCCCTGCTCCCCATGGTGGCTCTAGGGGAGGATCCTTCCCAGCCTCTTCCGGCTTCCCGTGTCTCCAGGCGTC | 120 |
| chr11 | 70946975 | 70947095 | 788855_47188552_ENST00000601538.6,E | 20 | - | good | AGCACAGGGAGATTTTCCAAGTCAGTTGGTTAATTGAGAAGGGTTGGTTAGTGCAGAGGGAGGTTGAGCCTGGGAAGGGTTGGTTAGCGGAGAGGGAGGTTGAGCCTGGGAAGGGTTGGT | 120 |
| chr11 | 71176895 | 71177015 | 788855_47188552_ENST00000601538.6,E | 20 | - | good | CTGTCATTCTTAAGTTTGTTTTCTGGATGTAACATGTCTCCCTTCCCTAGTCTGCTTTTAAGATTTTTTCTTTATCTTTGGTTTTGAGCCATTTTGTTATGTTGTGCCTTGGTGTAATTT | 120 |
| chr11 | 71227535 | 71227655 | 788855_47188552_ENST00000601538.6,E | 20 | - | good | ATTCTATCAATTGTGGAGAAAGGTTTGTTAAAGTTGTATTCAGTTATAAGTGTGGATTTATGTCTTCTTTCTTCAGTTCTATCCATTTTTGCTTCATGTATTTTGCTCCTCTGTTGTTTG | 120 |
| chr19 | 50714668 | 50714788 | 788855_47188554_ENST00000293441.6,E | 20 | - | good | GTAGCTATATCAGTTAAATAACATATATCATTAACACGAATTTCAACACAACTAACACTTCACCCCCAACCACCCTGATTTTTTTTTTTTTTTTTTTTTGAGACGGGGTCTTGCTCTGTCA | 120 |
| chr11 | 70509815 | 70509935 | 788855_47188552_ENST00000601538.6,E | 19 | - | good | ACAGAATACACAGATCAGGTGGCTTAAACAGCACTTATTTCTCCCAGCTCGGGAGACCAGAAGTCCAAGATCATGGCACTGGCCGGCAAACTGGTTTGCAGAAAGCCATCTTCTCTTCAC | 120 |
| chr11 | 70526375 | 70526495 | 788855_47188552_ENST00000601538.6,E | 19 | - | good | CATGTTCACCTCGGGGTAGAGACTTAACATTGAAAGGCATGGCAATCAGGCCCTGCATGTGGAAAGGTGAAGCAGGGACTCAAAGGGGCTCAGTGCAGCCTCTGTGACCCAGCGAGAGCC | 120 |
| chr11 | 70556255 | 70556375 | 788855_47188552_ENST00000601538.6,E | 19 | - | good | GAGAAAATAAATGTGATCAGGCTGCACGTTAAGTGCTTATAGCAGCTTTATTGCCAAAAACTGGATGTAGCCTGGATGTCCTTCAGTAGGTGAGTGGTCAAACAAACTGTGATATATTCA | 120 |
| chr11 | 70695335 | 70695455 | 788855_47188552_ENST00000601538.6,E | 19 | - | good | CACTACGTGGTCTTTTGTGTCTGGCTTCTTCCACTCTGCATCATGTTTTCAAAGTGCATCCACACAGTGACACACACCAGTACTTCATTCTTTCCCCACCCCCCCATTTTTTCTTACTGT | 120 |
| chr11 | 70739615 | 70739735 | 788855_47188552_ENST00000601538.6,E | 19 | - | good | CTCCAAATACAGTCACATTCTGAGGAGCAGGGGTTAGGCTTTCAGCATATGAATTTGGAGGGACTCCGTTCAGCCCATCACAGCCTCCCTGGGGCCCAGCTGACCCTGGCTGGTGGTGGG | 120 |
| chr11 | 70797815 | 70797935 | 788855_47188552_ENST00000601538.6,E | 19 | - | good | TGATTGCCGAGAGATTGAACTGATTAAAATGTGAAGAGGAGACCACTTATATAAAATGGTACCGAGTGTGTGTGTGTGTGTGTGTGTGTGTGTGTGTGTGTGTATGAGAGTGGAGGAAAA | 120 |
| chr11 | 70929455 | 70929575 | 788855_47188552_ENST00000601538.6,E | 19 | - | good | CCATGCTGGTCTTGAAGATGGAGGAGGAGCCATGAGCCCAAAAATGCAGATGGCCTCTAGAACCTGGCAGAGGGGGAGGAAATGGATTCTTCCTTACTGCCTCCAGGAGGAATGCTGCCCT | 120 |
| chr11 | 71210135 | 71210255 | 788855_47188552_ENST00000601538.6,E | 19 | - | good | AAATATATATATATATATATATATATATATATATATATATATATATACCTGTGGATTTCCAATGAAGAGGACCAGGTGGCAAGGGCTGAGACCAGGTATGCACCACTAAATACTGGGGATGG | 120 |
| chr11 | 71219375 | 71219495 | 788855_47188552_ENST00000601538.6,E | 19 | - | good | TTTTTATGAGCTTATTGGTTATTTGAATATCTTCAGAAAAATGTCTATTCAAGTTCTTTGCCTATTTTTGAGTTGTTGGATTGTATGAGTTCTTGATATATGCTAGATGCTAATCCCTGG | 120 |
| chr11 | 71220335 | 71220455 | 788855_47188552_ENST00000601538.6,E | 19 | - | good | ATCCACCCGTTCTTTCAGGCATATACCTACCCCCAGAAATGGAATGGCAGGTTCATATGGTTTTTTATGTTTAGTTTTTGGAGAAACTGCCGTACTGTTTTCCACAGCGGTTATACTATT | 120 |
| chr11 | 71243135 | 71243255 | 788855_47188552_ENST00000601538.6,E | 19 | - | good | TTTATTGCTTTTCTAGCTTAGATTTCATTAACTTCAGCCCTAACCTTTATTATTTCCTTCCTTCTGCTAGCTCAGTTTCAGTTTCCCGTTCTTTTTCTGACTCTCACAAGGTGGAAGGTT | 120 |
| chr11 | 70541735 | 70541855 | 788855_47188552_ENST00000601538.6,E | 18 | - | good | CAGGGTCAGCCCCGGCATTGGTGGCACCTGGGACTCAGCAATTCTCTGTTGTGGGGGACTGTCCTGTGCTTTGTAGGATGTATAGCAGCTTCCTTGGTCTCCACCCACAAGATGCCAGTA | 120 |
| chr11 | 70557095 | 70557215 | 788855_47188552_ENST00000601538.6,E | 18 | - | good | GCACTTGGTAAATATTGAGTGAAGAGGTGAATGAATGAATGAATGAATGAATGAATGAATGAATGAGGCTTTTAGTGCAATGCCTGGTATGTGGTGAGCACTTGAAATATGGAAGTGGTT | 120 |
| chr11 | 70615295 | 70615415 | 788855_47188552_ENST00000601538.6,E | 18 | - | good | ATTCCAGTCAGCCCTCCCTCTCTCGGGGTGTTGCATTTCTAGCACATTCTATGGTTATTCTTAAGAACTGAAAGCAAAAGAGAGGGGAGAACTGGGCTCGTCCAAGGCCACCTAGAGGAC | 120 |
| chr11 | 70796975 | 70797095 | 788855_47188552_ENST00000601538.6,E | 18 | - | good | AGCTCCCGAGGGCTGGGAAGTCACAGAGCTGGGGCCTCTGTAATAAAGGGATTGGGCATTAAATCAAGGGGAGGAATATTCATGTCTTTTCTGGAAATGGGCAGAGAGCTTCTCAGAACC | 120 |
| chr11 | 70928015 | 70928135 | 788855_47188552_ENST00000601538.6,E | 18 | - | good | AGCCAGGGTTCTCTAGAAAAATGGAACCAATAAGATACATGTATATATGTGTACATATATAAATATATACATATATATAGAAAGAGAATGAGGAGAGATTTATTATAAGGAATTGGCGCA | 120 |
| chr11 | 70950455 | 70950575 | 788855_47188552_ENST00000601538.6,E | 18 | - | good | AGGTGGGACCTTTTGGAGGTGATTACATCATGAGGGTAGAGCCCCCGTGATGGGATTCGTGCCCTTGTACAAAAGGCTGGAGGTGGACCTGGTGCAGTGGCTCACGCCTGTAATCCCAGC | 120 |
| chr11 | 71065535 | 71065655 | 788855_47188552_ENST00000601538.6,E | 18 | - | good | ACTGCTCATCTCCCTGGGAGAGTTCTGCACACACCCCCCTGCAACGTCCCCACTCACTGCTCATCTCTCTGGAAGAGTTCTGCACACACCCCTCAACTTCCCCACTCACTGCTCATCTCC | 120 |
| chr11 | 71226575 | 71226695 | 788855_47188552_ENST00000601538.6,E | 18 | - | good | TCTTAATTTTCCTTTCATTCTTACAGGATATTTTTGTTGGTATAAAATTATGGATTGAGAATACTTTTCTTTCATCACTTGAAAATTACTGTGCCACTTTCTTTTGGTCTCCACAGTTTC | 120 |
| chr11 | 71233175 | 71233295 | 788855_47188552_ENST00000601538.6,E | 18 | - | good | TCTTTTTGATGGCTGAATCACGTTCCATTTTAAGGCTGGACCACAAGATGTTTATGCATTCCTCAGCTTAGGGATGTTGGGGTTACCTCCACTTTTGGGCTGTTATGAGGTATGCTGCCG | 120 |
| chr11 | 70520015 | 70520135 | 788855_47188552_ENST00000601538.6,E | 17 | - | good | TATGGGAACTCCAGATACTGAGTTCACAATGTTTTTTGTAAATCTAAAACTATTCTAAAATAAAAGATTTGTTTAAAAACAAAGACTAGACCAGGCGTGGTGGCTTACGCCTGTCATCCC | 120 |
| chr11 | 70608215 | 70608335 | 788855_47188552_ENST00000601538.6,E | 17 | - | good | GTAGCCCAGTCAGTAGCTGAAAATAGTAGCTCTTTTCACAAAGAGGAAAGAAATATCCCTTCTGGCTGGGTGAGGTGGTTCGCACCCACAATCCAAGCACTTTGGGAGGCCAAGGCAGGG | 120 |
| chr11 | 70795775 | 70795895 | 788855_47188552_ENST00000601538.6,E | 17 | - | good | GCAGGGCTGCATTCCTTTCTGCAATCTGTGGGGGAGAATTTGTTTCCTTGCCTTTCCGACATCCTTGGCTCATGGCCCCTTCCTCCGTCTCCAAAGCCAGCACCTGTTGGTAGGGTCCTT | 120 |
| chr11 | 71118215 | 71118335 | 788855_47188552_ENST00000601538.6,E | 17 | - | good | TCTCAGGTATTTCTTTATAGCAGTGTGAAAATGGACTAATAAAAGGTTGCTTCTGATTTTGCCATGAATCTCACACTGTTGTACAACTTTTATACATTTTTTTTTCACTACAGATTCAAA | 120 |
| chr11 | 71169935 | 71170055 | 788855_47188552_ENST00000601538.6,E | 17 | - | good | ACAAGTTAAAATTAATGAATATTGTATCTTTTTTTAAAAAAAGATGAGGTCTTGCTATTATGTCCAGGCTGGAAAGGATATTCACAGGTGGGGTCACAGAGCACTGCAGCCTTGAACTCC | 120 |
| chr11 | 70478135 | 70478255 | 788855_47188552_ENST00000601538.6,E | 16 | - | good | GCAGGAGGATCACTTGAGGAGGTTGAGGCTGCAGTGAGCTATGATTGCTGCATGACCCCGTCTCCAAAAAAAAAAAAAGCCAAATTGAATTTGTAAACAGTAAAAAAGCCACTTGCTTCTT | 120 |
| chr11 | 70572095 | 70572215 | 788855_47188552_ENST00000601538.6,E | 16 | - | good | TGCACTCCAGCCTGGGCGACAGAGTGAGACTCCGTCTCAAAACCCAACAGGGGTTGATTTTCTCCCAGTTCTGGAGGCTGAAGCCTGAGAATAGGATGCCGGCAGGGCCGGATTCTGGCG | 120 |
| chr11 | 70610135 | 70610255 | 788855_47188552_ENST00000601538.6,E | 16 | - | good | GCACCAAAGCTCCCATTGGTCCCATAACTGTTTTCTTTTCATTGTGGTCAAATACATGTAATGTAAAATGTGCCATTTTAACCATTTAAAGGGGGAATAATTCAGTGGCATTTAGTACAT | 120 |
| chr11 | 70790615 | 70790735 | 788855_47188552_ENST00000601538.6,E | 16 | - | good | GGCAGTCCAGCAGCCTGGAACGCCCAGACAGGAGCTGGTGCTGCAAACTTGATGCAGAATCTCTTCTTCCTCAAAGAAACCTCAATTTTGCTCTTGAGGCCTTTTAAATGATTGGAGGAG | 120 |
| chr11 | 71178455 | 71178575 | 788855_47188552_ENST00000601538.6,E | 16 | - | good | TCATCTTAACCATTCAGGAGTGTACGACTCAGTGGTATTAAATACATTCACAACATTGTGTAACCACCACTCTGTCTATACCCAAAACTTTCTTTTGTCATTTTTCACAAAATCTCTGTG | 120 |
| chr11 | 71225615 | 71225735 | 788855_47188552_ENST00000601538.6,E | 16 | - | good | GTGGAAGCATTCATCATGGTTGCTTTCAACTCTTTGTCAGATCATCCTGACATCTCTGTCACCTCCATGTTGGCATCCATTGATTTTCTTTTTTTCAGTTTGAAATCTTCCTGATCCTTG | 120 |
| chr11 | 71242175 | 71242295 | 788855_47188552_ENST00000601538.6,E | 16 | - | good | GTAGAACTCTTCAGCTTGCAGAACTGAAACTACGTCCCCACTAAACACTAGCAAACCAGCCCCCTTCTCCCAGCCCCTGGCGACCACCAGTCTGTTTTCTGTCTCTTTGACTTCTGTAGG | 120 |
| chr19 | 50699668 | 50699788 | 788855_47188554_ENST00000293441.6,E | 16 | - | good | CCTCCATTCCCCCAATGGCCCAAGCCATCCAATCCTCCCAGTGCCCAAGCCTTCCAGTTCCCCAGTGCTACCAACCCCTCCATTCCTCCAATGCCCCAAGCCTTCCAATCCCCCAATGCC | 120 |
| chr11 | 70516055 | 70516175 | 788855_47188552_ENST00000601538.6,E | 15 | - | good | TGTGTTGGCTATTTTGGGCCTCTTAGCTTTCCATATAAACTTTAGGATCCATTTGTCAGTATTCACAAAATAATTGGCTGTGATCGAACTTAGTCTTTTAGGAGCTACTCTGAAATATGA | 120 |
| chr11 | 70532735 | 70532855 | 788855_47188552_ENST00000601538.6,E | 15 | - | good | GTTGCGTCCATCTTTTGGCTACTGAATGGTGCTGCTGTGGACACTCATGCGCACGTGTCGGTCTAAGACCCTGCTCTGAATTATTCTGGGGTTATAAGTAGGCATGGAATTGCTGGGTCA | 120 |
| chr11 | 70577735 | 70577855 | 788855_47188552_ENST00000601538.6,E | 15 | - | good | CTCATTTTAATTTAATCAGCTCTTTCAAGGCCCTATCTGCAAATACAGTCACGTTCTGAGGTTGTGGGGGTCGGGACTGTAACACAGGAATGGTGACCCACAGTTTACCTCCGTTTATCA | 120 |
| chr11 | 70627535 | 70627655 | 788855_47188552_ENST00000601538.6,E | 15 | - | good | ATGTGATTATATGTGCAGATATACCAAATGTACATGCAAATATTCCAAAATCCAAAAATATATGCAATCCAGAATACTTCTAGTCCAAGCGTTTAGGATAAGGGCTATTCTGCTGTAGCC | 120 |

| | | | | | | | | |
|---|---|---|---|---|---|---|---|---|
| chr11 | 70647935 | 70648055 | 788855_47188552_ENST00000601538.6,E | 15 | - | good | CATGTGGTTAGACCACAATTAGTTTATCCATTCACTAGTTCAGGGACACTTGGGCTGTTTCTAACTTATGGTTGTGAATAATGTCGACATAAACGTTTTTGTACCAGTTTTCGTTTTCAG | 120 |
| chr11 | 70730855 | 70730975 | 788855_47188552_ENST00000601538.6,E | 15 | - | good | TTTTGGGTGATTTAAATGTTCTGGAACTAGATAAAAGTGCTGGTTGCACAGCATTGTGGATGTGTTAACTGCCAACAAATTGTATTTTTTTAATGATTAGTATTCAGTTTTGTAAATTTC | 120 |
| chr11 | 70821335 | 70821455 | 788855_47188552_ENST00000601538.6,E | 15 | - | good | AAACCCTGTCTCAAAAACAACAACAACAACAACAAAACAACCAGATATTTACACCTCCCAGTCCTGGGGACTACAAGTCTGAGATCAGGGCATCGCAGGGCTGGTTCCTCCTGAGGCCTC | 120 |
| chr11 | 70863815 | 70863935 | 788855_47188552_ENST00000601538.6,E | 15 | - | good | ACCGTTGTGGGGGCGGACGCCTGTGATTAGGGTGTCCCAGGGCTGGTTTCTCCTGAGGCCTCTCCTCCTGGCTTGCAGGTGGCTGTCGTCATCCTCACTCGGTTTCTTCCTGTCTGTCCA | 120 |
| chr11 | 70868495 | 70868615 | 788855_47188552_ENST00000601538.6,E | 15 | - | good | GTCCATAGGCTGAGCCTGGACTCTGGAACCCACCTTCTTGGCTTGTGTCTTGACTCTGCCCTTCACAGGCTGTGTGACTTTGGACAAGTTACTTAACCTCTCTGTGCCACAGTTGCCTCC | 120 |
| chr11 | 70919135 | 70919255 | 788855_47188552_ENST00000601538.6,E | 15 | - | good | GCATGTGAGGCGGTGTGCGTGTTCATCAGCTCGATTTAGCCATTGCACATTGTATGCATGTTTCTCTTTCTGTTTGTTTTTATTTTTTTGAGACAGGGTCTTGCTCTGTTGTTGCCTGGG | 120 |
| chr11 | 70951175 | 70951295 | 788855_47188552_ENST00000601538.6,E | 15 | - | good | ACAGCAACCAGGGGAAACGAATTTATGACATCACGGTGCAGCAGCCAGGGGAAATGAATTTATGACGTCACGGTGCAGCAGCCAGGGGAAATGAATTTATGATGTCACGGCACAGCAGCC | 120 |
| chr11 | 71065175 | 71065295 | 788855_47188552_ENST00000601538.6,E | 15 | - | good | ACCAACTTCCCCACTCACTGTTTATCTCCCTGGAAGAGTTCTGCACACAGACCCCTAACTTCCCCACTCACTGCTTATCTCCCCGGGAGAGTTCTGCACACGTGGCTTCCACCTTCTCAT | 120 |
| chr11 | 71088215 | 71088335 | 788855_47188552_ENST00000601538.6,E | 15 | - | good | GTGCACATGCATGCATGTGTGTGTGTGTACAGATACAGAATTTTTTAAAATAGTATTTTGAAGAATTGGCTCATGCAGTGGTGGAGCTGCAAGTCCAAAATCTGCAGGGCAGGCTGGAGA | 120 |
| chr11 | 71169215 | 71169335 | 788855_47188552_ENST00000601538.6,E | 15 | - | good | TGACCTCAGCCTTGCTGGATGTGCAACACCTGAGTGAAATCTAACTTGAGCTATTTCTGATAAATGTCTATATTAAAGAAAAATGAAACACGAGCTTAGTCAATCAGAAGCTGTCAATTA | 120 |
| chr11 | 70614095 | 70614215 | 788855_47188552_ENST00000601538.6,E | 14 | - | good | AGGCTCCAGGAGGCCCTTCCTGGCCTCTTCCAGCTTCTGGTGGTGCTGACACTCCTTGGCCTTCCCTGGCTTGCGCTGCCTGGCTCCAGTCTCGGCCTTTATTGTCACATGGTGGGCTCC | 120 |
| chr11 | 70747655 | 70747775 | 788855_47188552_ENST00000601538.6,E | 14 | - | good | GTGGCATACATGCACACACGTGCCTGTTCACACATATGCACACTATACATATGTGCAGCAAGCATGCACACACACATACATACTCATGTGCACACACATGCTGTTGTCTCTGCTATTCAG | 120 |
| chr11 | 70940015 | 70940135 | 788855_47188552_ENST00000601538.6,E | 14 | - | good | GGGGTTCAGAAGTCTGAAATCAAGGTGTGGGTGGGGCTGCCTTCCTTCTGGAAGCCCTGGGGAAAAACCATTCTTCATCTCCTCCAGCTCTGGGTGGTGATGGGCAGTCTTGGTGCTCCT | 120 |
| chr11 | 70951295 | 70951415 | 788855_47188552_ENST00000601538.6,E | 14 | - | good | TGTCACAATGCAACAGCCAGGGGAAATGAATTTATGACATCACCACCCAGCCGCCAGGGGAAATGAATTTATGACGTCATGGCACAGCAGCCAGGGGAAATGAATTTATGATGTCACGGC | 120 |
| chr11 | 70951535 | 70951655 | 788855_47188552_ENST00000601538.6,E | 14 | - | good | GGGGAAATGAATTTATGATGTCACAGTGCAGCAGCCAGAGGAAATGAATTTATGATGTCACAGCACAGCAGCCAGAGGAAATGAATTTATGACGTCACCACGCAGCAGCCAGGGGAAATG | 120 |
| chr11 | 71056775 | 71056895 | 788855_47188552_ENST00000601538.6,E | 14 | - | good | AGCTCCAGATTTTCTTTTCTTTTCTTCTTTTCTTTTCTTTCTTCTTTCTTCCTTTCTGTATTTTACTGAAGATATTTCATTTATTGTATATTTTATATACATGTGCAAGACTTTCTATAG | 120 |
| chr11 | 71097815 | 71097935 | 788855_47188552_ENST00000601538.6,E | 14 | - | good | ACAAACAGGCATGCACACATAGGTGTGCATACACACAGCCATGCACACACAGGCACACAGGCATGCACATAGGTGTGCACACATACACTGGCGTACACATGCTCCAGCCTGCACTGCCCT | 120 |
| chr11 | 71117615 | 71117735 | 788855_47188552_ENST00000601538.6,E | 14 | - | good | CTTCCATCCCTGTGGAGTTGGGCGTGCACCACCCTCCCAGTATGTGGATGTGTTTGCTGGCTCAGAAGCCCCCAGGCCCCATTGTTGAGGTATTTTTATGGAAGTTTCATTACATAGGCA | 120 |
| chr11 | 71230655 | 71230775 | 788855_47188552_ENST00000601538.6,E | 14 | - | good | TATTTGTGTGGGTTTTTTCTGGGCTCTGTATTGAGTTCTATTGCTATGTGCCTGTCCCTCCACCAGAACCACACAGCTTTGATTACTATACTGTATAATAAATCTTGAAGCTGAGTATAC | 120 |
| chr11 | 70526135 | 70526255 | 788855_47188552_ENST00000601538.6,E | 13 | - | good | GCTACAATTATATTAATGTTGCTTATCCCAAGGCCAGCGCTTGCTTAGCTGCTAGAGAATCAGAAAAGCCTTGTGGCGGCCAGAATCCACCTTATTCTTAGGTTTAGGGTGCAGGATCAC | 120 |
| chr11 | 70530695 | 70530815 | 788855_47188552_ENST00000601538.6,E | 13 | - | good | CATCTCAGAACTTTCCATCTTCCCAATAAACACTGTCCCCAGTAAACTCCGTCCCCAGTAAACACTAGCTCTTCATTCTCTCTTCTCCCAGCCCCGGGCACCCACTGTTCTACTTTCTGT | 120 |
| chr11 | 70615415 | 70615535 | 788855_47188552_ENST00000601538.6,E | 13 | - | good | CACTAAGGGACCCTGGAAAGCAGCTCCTCCTGGGTTTTATACCCCGGGGCCATATGACTTTTGGACTAAAGTGTTGATGGACATCATGTTTCCGGGGAAGACTGGGACAGAGCTCAGGCT | 120 |
| chr11 | 70674695 | 70674815 | 788855_47188552_ENST00000601538.6,E | 13 | - | good | GCCCCACCCATCATTTACATATTGTCATGGCTGCCTCTGTGCCACAACTAGCAGAGTGAGTAGTTGCAACAGGGACCACCCATCCACCCTCAAAGCCTGGAATGTTCACTCTCTGGGCCG | 120 |
| chr11 | 70767575 | 70767695 | 788855_47188552_ENST00000601538.6,E | 13 | - | good | CACCCCTGGCAGCCACTAATCTACTTCTCATCACTGCAGATTCCCTTTCCTGTGCATTTCCTATCAATGAAATCCTCCAGCATGGCCTTTGTGACTGGCATCCTTCACTCAGTGCCATGT | 120 |
| chr11 | 70793615 | 70793735 | 788855_47188552_ENST00000601538.6,E | 13 | - | good | CTCCTACCGGCAGTGTGTGAGAGTTCCAGTTGCTCCACTTTCTCATCAACATTTGGCAATATCAGTCTGTTTAACTTCAGCTGTTCTAATAGATATGAAATGATATCTTATTGTGGGTGT | 120 |
| chr11 | 70951415 | 70951535 | 788855_47188552_ENST00000601538.6,E | 13 | - | good | AATTTATGACATCACAGCACAGCAGCCAGGGGAAATGAATTTATGATGTCACCATGCAGCAACCAGGGGAAATGAATTTATGACATCACCACACAGCAGCCAGGGGAAATGAATTTATGA | 120 |
| chr11 | 71100335 | 71100455 | 788855_47188552_ENST00000601538.6,E | 13 | - | good | TTGACAGCTTGTCTCTCTTTAGTGTTGAACAACATCTTGTTGTCTGCATGTGCAATGGATAATGTATCCACTCACCTACTGAAGGACCTCTGGATTGTTTGCCATTTTTGGCAACGATGA | 120 |
| chr11 | 71193935 | 71194055 | 788855_47188552_ENST00000601538.6,E | 13 | - | good | TCTTGACTCAGCCCTTCCCGCCAGCAGCACTGCTGTGGGCTGAACTGTGTCCCCGCAAATTCACAGGTTGAAGCTCTAACCCCAAGTACTCAGAACGTGACCTTATTTGGAGCGAGGGTC | 120 |
| chr19 | 50680588 | 50680708 | 788855_47188554_ENST00000293441.6,E | 13 | - | good | ACTCCAGCCTGGGTGACAGGGTGCGATTCTGTCTCGGGGAAAAAAAAAAAAAAGATGCCACTATCTAAGGACTTGGAAACTGGGAGATTTCAACCCTCTCCAAAGATGCACTTTTGCATAT | 120 |
| chr19 | 50694988 | 50695108 | 788855_47188554_ENST00000293441.6,E | 13 | - | good | GCGGCGGCGGCGGCGGCGGCGGGGGCGGCCGGCACCGGGACCCGGGCCGGGGCTGGGGCCGCGGCGGCGATGGCTTTGTCTGCGGTCGGCGGCGGGGGTCTCGGGGGCGGCCCCGG | 120 |
| chr11 | 70512095 | 70512215 | 788855_47188552_ENST00000601538.6,E | 12 | - | good | GCAGCTGGCAGGTAGAATTCAGGGACACTGCTAAACATCCTACAGTGCACACCCACAACAAAGAATTAATTGGTCCAAAATCACCATAGTGCCACATTTCACAAACTCTGAGCCAAGTCA | 120 |
| chr11 | 70539815 | 70539935 | 788855_47188552_ENST00000601538.6,E | 12 | - | good | CAGCCACCACCAGGGGCACAAAAATGCTTGGAACAGAAACTGCCTCGCAGTCCCCAGAGGAACCCACCCTGCCCACACCTTGATCTTGGACTTCACCTCCAGGACTGTAGGAACATGAAC | 120 |
| chr11 | 70553015 | 70553135 | 788855_47188552_ENST00000601538.6,E | 12 | - | good | GGTGACAGAGCAAGACTCTGTCTCAAAAAAAAATAAAGGAGATTATTAAAGTCAAATGAGGCCATTAGAGGGGGCCTTAATCCGATATCACTGTGTCCTTAAGAGGGGATTTGGACACAG | 120 |
| chr11 | 70571975 | 70572095 | 788855_47188552_ENST00000601538.6,E | 12 | - | good | AGGGCCTGTCCCTGGTGTGGAGATGGCCGCCTTCTCACGCTGTGCTCAAGTGGCAGAGGGAGAAAGTGGAAGCGAGCTCTCTGGGGTCCCTTTTATAAGGCTACCGATCCCATCACAAGG | 120 |
| chr11 | 70778495 | 70778615 | 788855_47188552_ENST00000601538.6,E | 12 | - | good | CCTCACAGGGCCTTTCCTTGTGCATGCACACCTCTGCTCTCTCTTCCTCTTAAGAATGCCAGTCCTCTTGGGTCAAGGCCCTGCCTTGTACCTTCATTTAATCTTAATTACCTCTCTAAA | 120 |
| chr11 | 70871135 | 70871255 | 788855_47188552_ENST00000601538.6,E | 12 | - | good | CTGACTCCCCTTTGAAGTGTGTCCCTGCAAATGCATATGTTGAAGTATCACCCTGAGGATCTCAGAGTGTGACCTTGTTTAGAGGTAGGGTCTTTACAGAAGTCATCAGCTGAAAATGAG | 120 |
| chr11 | 71090735 | 71090855 | 788855_47188552_ENST00000601538.6,E | 12 | - | good | ACTCCAGATTCCCGCTGAACACTGCCCTGGGTCTCCAGCCTGCCCTGCAGATTTCAGACTTACAGCCTCCGTGATTGCATGAGCCAACTCTTTAAAATATTTCTATCTTTAATTTTATGT | 120 |
| chr11 | 71136215 | 71136335 | 788855_47188552_ENST00000601538.6,E | 12 | - | good | CAAAAGACATGCTCTCCTTTTTTATGGCTGAGTAGTATTCCGTTGCACATATGCACCACGTTTCCTTGATCCAGTCTTGGTGGACCCTGAGGCTGAAGCATGACAGCTTTGAGGTCCATC | 120 |
| chr11 | 71177495 | 71177615 | 788855_47188552_ENST00000601538.6,E | 12 | - | good | ATTCTGCTTTTCTTCTTTTTCTGTCTTCTTTTAGATTAATAGAATTTTATGATTCCATTTATCTTTATTGGTTTATCTGCGATTACTCTTTGTTTTGTTCTTTCAATGGTTGGCTTAGGG | 120 |
| chr11 | 71216135 | 71216255 | 788855_47188552_ENST00000601538.6,E | 12 | - | good | CATCTTTTTAATCCAATATTATGTTTATCACATGCTGGACATTTGGGTTGTTCCCACTTTTTGGCTTTTACACACTTGGACATTCTTGTACAGGCTTTTGTGTGGACGCGATTTCGTTTC | 120 |
| chr11 | 71216375 | 71216495 | 788855_47188552_ENST00000601538.6,E | 12 | - | good | TCACTCCCTCCTCCTGAATCCCTCTAACCAATCACCTCCGGCCCTGGCGACCACAAATCTCCATTTCATCTCTACGGATTTGCCCATCTGGGACAGTTCATATAAATGGAATCGTATAAT | 120 |
| chr11 | 71226455 | 71226575 | 788855_47188552_ENST00000601538.6,E | 12 | - | good | TGATGAGAAATGTAAGTATTGGTTTCTCTGCATGTTTTCAATATTTTTTCTTAGTGTTCAGAAGTTTAATTCTGATGTATATTGGTGTGAATTTTCTTGGGTTTATCCTATTTGGGATTC | 120 |
| chr11 | 71236775 | 71236895 | 788855_47188552_ENST00000601538.6,E | 12 | - | good | TGCCACATTGGGTTTTGTTTATTTGTTTGTTTTGAGATGGAGTCTCACTGTGTCACCCAAGCTAAAGTAAGTTTGTTTTGAGACAAGAGTCTCGCTCTATCAACCAGGCTGGAGTAAAAA | 120 |
| chr11 | 71242895 | 71243015 | 788855_47188552_ENST00000601538.6,E | 12 | - | good | TCGTTTCAAAGTGTTTTCCAATTTCCCCGTGGTTTATTCTCTGACCCATTGGCTATTTAGATGTCTGTTTAATTTCCACATATTTGTGCATTTGTAAAATAAATTTCGAATTTCATTCTA | 120 |
| chr11 | 71243975 | 71244095 | 788855_47188552_ENST00000601538.6,E | 12 | - | good | TTTGTAGGTGTCTGGCTGCCCAAGGAAGCCTATAGTTTTCTTTTGAAGTCTGGTTTTGGTATCAGAGTAATACTGGCCTCATAAGATGACTTGGGAAGCATTGTCTTCCATTTTTTTCTG | 120 |
| chr11 | 70508855 | 70508975 | 788855_47188552_ENST00000601538.6,E | 11 | - | good | AAAGCATCACCCTGATCTCACCCTTCATGGCGCCTTCTCCCTGTGTGCATATCTCTATATCCAGACTCCCCCGACTCTCTTTCTTTTGAGACAGGGTCTCACTCTGTTGCCCAGACTGGA | 120 |
| chr11 | 70562015 | 70562135 | 788855_47188552_ENST00000601538.6,E | 11 | - | good | TATTAGAAACCAGTAAGAGAAAGATCTCTAGAAAATCCCCCAGATATTAGGAAAGTAAATAGTATTATTTTAAATAACACCGGGATCAAAGTAGAAATCAAATGGGAAATTAGAAAATAT | 120 |
| chr11 | 70565015 | 70565135 | 788855_47188552_ENST00000601538.6,E | 11 | - | good | TACAATTCATGATGTCTGGCATCCATTCAGAAATTCCCAGTCTTGCAAAGAAGCAGGAAAATACAGCCCGTAAGGAAGAGAAAAATCCATTATCAAAATTGACTGGGGATTAACACCGAT | 120 |
| chr11 | 70615535 | 70615655 | 788855_47188552_ENST00000601538.6,E | 11 | - | good | TGGTTCATTGCTTACAGATATGCAGCAAGGGACAATAGAAGCCTGGGATTCGGCCGAGCCAGCCTGTAAGGCTCAGGAAAGCTGACCGGAGTCTTGACTGCATGTGCTCCCTTGCGCCGC | 120 |
| chr11 | 70624655 | 70624775 | 788855_47188552_ENST00000601538.6,E | 11 | - | good | GGATCACTCTGCTGCGGGCCCCTCCTGCACACTGTGGGGTGTCCAGCAGCAGCCTGGCCTCTCCCCACTCCATGCCAGCACCAGCCCCCAGTTGTGAAAACCAAGAATGTCTCCAGACGT | 120 |
| chr11 | 70760135 | 70760255 | 788855_47188552_ENST00000601538.6,E | 11 | - | good | GCATTGGCTAAGGTTATGGCAGGTTAAGCATACTCAGCAGGTTACAGGAGGAGCTGTGAATATTCACAAAGCAGGTCCTGACATGTGCACTGAGCACACATGCGTGTTACATGCGTCCCA | 120 |
| chr11 | 70795175 | 70795295 | 788855_47188552_ENST00000601538.6,E | 11 | - | good | TTAATTCCCTTTTGCCTTAAGGGAGCATATTCACAGGTTCTAGGGATTAGGACCTGGGCATCCTAGGGAGGAGCCATCTTTCTCCCTAACACAGATGGCAAAAATGAACATCAGAACTCT | 120 |
| chr11 | 70821095 | 70821215 | 788855_47188552_ENST00000601538.6,E | 11 | - | good | CACTAATGGCCCTCCACCCAGATGATCTCTGTAAAGACCTTACCTGCAAATATTGGCACATTTTGAAAACTGGGGTTTAGGACGTCAACACACGAACTTGGGGGACACAGTTCAGCCCAC | 120 |
| chr11 | 70853975 | 70854095 | 788855_47188552_ENST00000601538.6,E | 11 | - | good | TGAATCACTGTTAACACTGCAGGGTCTTTCTTCTGGGTCAAATAATGTCCCCTCAATTTCATGTCCAACTTGAACCTCAGAATGTGACCTTATTTAGAAATAGCGTCTTTGTGGGTTTAA | 120 |
| chr11 | 70865015 | 70865135 | 788855_47188552_ENST00000601538.6,E | 11 | - | good | AGTACCATGAACTAGATGGCTCGTAAAAAATACATATTTATTTCTCACCATTCTGGAGGCTAGAAGTCTGAAACCAGGGTACCAGAGAACTCTCTAGGGCCCCTTTTTCTTTAAGACACT | 120 |
| chr11 | 70905455 | 70905575 | 788855_47188552_ENST00000601538.6,E | 11 | - | good | TTACCTCCCAAAGGCCCCAGCTCCTAACACCCTCACACTGGGGCTTCCATTTCAACACATGAATTCGGAGGGGGGCCTCAAATATCCAGACCATGGCAGAAATCCGCACGGCCACTGGGCT | 120 |
| chr11 | 70920335 | 70920455 | 788855_47188552_ENST00000601538.6,E | 11 | - | good | CCCTCCGCCTCTCCTTTCCACTCTTGTCTCTTCCTTCATCATAGGTAATCATTTTTTAAAATAGACTGGGTGTGGCAGTTCACACTTGTAAACCCAGCCCTTTGGGAGGCCAAGGCGGGA | 120 |
| chr11 | 71099615 | 71099735 | 788855_47188552_ENST00000601538.6,E | 11 | - | good | CATACGGTGTGGAGCATCTTCTCGTGTGTGTTGGCTATCTGAATATCTTCTGTTGTGAGGTATCCATTTAAATATTTTGTTCCCTTTTTAATTGGGTGGTTTGCTTTCTTGTGGAGTTTT | 120 |
| chr11 | 71231135 | 71231255 | 788855_47188552_ENST00000601538.6,E | 11 | - | good | TGCTCTTTTAAGTGGATTGTCATTTTTTTAACTGTTGAGCTTTGAGAGTTCTTTATTCTAGACACATATTATTTCGAATTATTTGTTAAATATGTGGTTTGCAGACATTTTTCTCCCACT | 120 |
| chr11 | 71232455 | 71232575 | 788855_47188552_ENST00000601538.6,E | 11 | - | good | TGAAGAGGAGGAGGAGGAGGAGGAGAGGAGGGGGGAAAGAGGAGGAGGAGGAGAAGGGGTTGGTCTTGCCTTCTCAGAGGCGGCAGAGGTGGGAGAATATCCACGTGTAAGAGGACTT | 120 |
| chr11 | 70561415 | 70561535 | 788855_47188552_ENST00000601538.6,E | 10 | - | good | CAACAAGCAAGCAGAAGAAGTGATTAAGCTCACAGCAATCAATGACATAGAAAATAGAAAAACAAGGAAAATGGAGCAACCAAAAGTTGGTTCTTTGGCAACGTCGAGAAAATCGATGCA | 120 |
| chr11 | 70562495 | 70562615 | 788855_47188552_ENST00000601538.6,E | 10 | - | good | ATACCACATTAACACTAATCCAAAGAAAACTGGGATGACGATATTAGTATCAAAATAGATTTTAGAACAAAGAATGTTAACAGTATTAAAGAAGGTTATTTCATTATGGTAAAAGGGTCA | 120 |

| | | | | | | | | |
|---|---|---|---|---|---|---|---|---|
| chr11 | 70599935 | 70600055 | 788855_47188552_ENST00000601538.6,E | 10 | - | good | TTGCATTGACCCAGAATAGACCAATTTGGGAAAAATTAACCCCTTCACAATGTTGTCTTCCTGTTCAAGAACATGATACATTTTCTTTCTTTCTTTCTTTCTTTCTTTCTTTCTTTCTTT | 120 |
| chr11 | 70636415 | 70636535 | 788855_47188552_ENST00000601538.6,E | 10 | - | good | CTCACACATATTCACACATTCTCATGCATCCTCGTACATATGTTCACACACGTGCTTACACATACACAGATGCTCACACGTGCTCACACATATTCATATGCTCACACACTTGCACACATG | 120 |
| chr11 | 70743455 | 70743575 | 788855_47188552_ENST00000601538.6,E | 10 | - | good | ATGAATAAAGTCCCAGTAGGAATCACGTCTCCTCCTGAGTCAGCTCAGGCTGCTGTAACAAAAATCCTCTAGACTGTGTGGCTCAAACAACAGAAGTGTTTCTCAGCCCTCTGGAGGCTG | 120 |
| chr11 | 70815095 | 70815215 | 788855_47188552_ENST00000601538.6,E | 10 | - | good | GTGTGTGTGTGTGTGTGTGTGTGTGTGTGTGTGTGTGTGTTTCTTCTCCCATCCTCCTAAAGGTGCTGGTCCGGGAGGGTAGGAGCCAGTGCACCTGCACACCTCTGCTCCCTCAGGTGT | 120 |
| chr11 | 70859135 | 70859255 | 788855_47188552_ENST00000601538.6,E | 10 | - | good | TCCATCTACTTATCTACACATCTATTCACCCACTTGCCTACTTACCCTCATCCACCCATCCATCCAGACATCTGTTCATCTCTCTATTATCTATACATTCATTCTGTCCATCCAATTCAT | 120 |
| chr11 | 70868975 | 70869095 | 788855_47188552_ENST00000601538.6,E | 10 | - | good | CTGTGTCCTGATCTCTTCTTGTAAGGACGCCAGTTGGATTGGATTTAGGACCTGCCCCATCTGGCCTCACTTTAATTTAGTTGCCTCTTTAAAGCCCATCTCCCTATACAGTCGCACACA | 120 |
| chr11 | 71072735 | 71072855 | 788855_47188552_ENST00000601538.6,E | 10 | - | good | TAGATTTTCCTGCTGTAGGGACTTGATATGAGTAGAAACATAAAGTGTTTGTCCTTTGGTGTCTGGCTGATTTCACTGAGTGCAATGTTTTCGAGGTTTGTCCACGCTGTAGGCTGTGTC | 120 |
| chr11 | 71074175 | 71074295 | 788855_47188552_ENST00000601538.6,E | 10 | - | good | TGTTCTCCGGGTCTCTAAGTCCATTCTTTGGGTTTAAACGGGTAAGTTTGTTTCTCACACGCTGCAGTGGCACGCACCCCAGCCAGCCCTGACCCTGGGGGGCACCGGGAAGGCCTTGCA | 120 |
| chr11 | 71187455 | 71187575 | 788855_47188552_ENST00000601538.6,E | 10 | - | good | TGAGAAACCCATGAACTGTGCTGAATAAAACATGAGAAACCCATGAACTGTCCTGAATAAAACATGAAAAACCCATGAACTGTCCTGAATAAAACATGAGAAACCCATGAAGTGTCCTGA | 120 |
| chr11 | 71231015 | 71231135 | 788855_47188552_ENST00000601538.6,E | 10 | - | good | CTATGGTTTGTGTTTTTCTCCTTTCAGCAGGTCTTTCAGAGCAAAAGTTTTTAATTTGGATGAGAACCTATTTGTCAATATTTTTGTATGTGGATTGTGCTTTTTACATTGGGTCTATGA | 120 |
| chr11 | 70508495 | 70508615 | 788855_47188552_ENST00000601538.6,E | 9 | - | good | TGGCCCAAATTTCCCCTTTTCATAAGAACACTAGTCACCTGGGTTAGGCTCACCCTAATGGCCCCTTTTTAACTTGACTGTATCTGCAAAGTAAGGTCATATTTTGAGATACTAGGGATT | 120 |
| chr11 | 70584335 | 70584455 | 788855_47188552_ENST00000601538.6,E | 9 | - | good | TTAGTTGGTTAGCATTTGAAAATCCAGCTCTCAGGTGAGCTGTTTACTTTCTCTAGGAATTGCCTGATTCTTGAGGGGCCATCCCTCCAGGGTCAGCAAAACCCTGGATGCCAGAGCATC | 120 |
| chr11 | 70754255 | 70754375 | 788855_47188552_ENST00000601538.6,E | 9 | - | good | TTGACTTCCTAATATTTTAAGGATCTGTGTCTATGTTCCTGAGGGATGTTGGTGTATTGATGTCTTATGCTGTCTTTGTCTGGTTTGTGTATCAGAATAATGCTGCCCTTATAAATTAAT | 120 |
| chr11 | 70772615 | 70772735 | 788855_47188552_ENST00000601538.6,E | 9 | - | good | GTGGAAGATGAGGGCTACTCCATTATGTATGTTTACTCAGGCCCACTGCAGCCTCCAGTGCCCAGCCTCTGGTGACAAGGGCCATTTTCTTGCCCTGGTGCAGAGAGGGGACCCCTTCCA | 120 |
| chr11 | 70890935 | 70891055 | 788855_47188552_ENST00000601538.6,E | 9 | - | good | AATGGCAGGTCCTGAAGATGCCTGCACTCTCACCAGCTCAAACACCTATGTGGTCACCATCTTCCCGTGACAACTCTTTTTTTTTTTTTTGAGGCGGAGTCTATCACTCTGTCACCCAGG | 120 |
| chr11 | 70953095 | 70953215 | 788855_47188552_ENST00000601538.6,E | 9 | - | good | TCCACCCCCATGGCCCGGACACCTCCTGCTAGGCCCAGCTCCCAACACCACCACACTGGGGATTAAATTTTAATGTGAGGATTGGAAAGGGCCAGTGTTCAAAGTGTAGCACTGGGTAAG | 120 |
| chr11 | 71143655 | 71143775 | 788855_47188552_ENST00000601538.6,E | 9 | - | good | ATGTATGTTGCTGTTTAAAGATGCCATATTTACTATGTAATGTTGATTCGTTAACATGGAACTCACAGCCAACAGTGCTCTCAGGCAGGCCTCAGTGGAGCTTCTGTGGCATGTTTTCTC | 120 |
| chr11 | 71157095 | 71157215 | 788855_47188552_ENST00000601538.6,E | 9 | - | good | GTTATTTCCCTAGAGCCCATTTCTTTTAGATAGAGAACCATTGCAGAGTTCCTGCTGTCAGTGTCTTTATATATATGTGTGTGTATATATATATATATTTATTATACTTTAAGTTCTAGG | 120 |
| chr11 | 71179175 | 71179295 | 788855_47188552_ENST00000601538.6,E | 9 | - | good | ACAGTTAATTTCAAATGTTGAACCAAATGTTAAATTTGATCAGTTCCTGATCAAAATCACACTTGGTTATGAAATATCCTTTTTACATATTGTTGGATTTGATTTGTTGTTTTATTTTGA | 120 |
| chr11 | 71212895 | 71213015 | 788855_47188552_ENST00000601538.6,E | 9 | - | good | TCCCTGCTCCTCTCCCTGTGCCTCCCCCTGGGGTCCCTGCTCCTCTCCCTGTGCCTCCGCCTGGGGTCCCTGCTCCTCTCCCTGTGCCTCCGCCTGGGGTCCCTGCTCCTCTCCCTGTGC | 120 |
| chr11 | 71224895 | 71225015 | 788855_47188552_ENST00000601538.6,E | 9 | - | good | ACCTTGCAGAGTGTTAGGTTTGTTTTATATACAGCATCCTGAGTTTTAGTTGTGCTTTGTGTGAGGAATAGGGAAAAATGTGTCTACTCCAATTGCCCGGAAGCAGAAGTTCGATCCCTG | 120 |
| chr11 | 71229335 | 71229455 | 788855_47188552_ENST00000601538.6,E | 9 | - | good | TATTGTATCTTTAATTTTGCTATCTGCATGATTATTGCTAGAGAATACAATTGATTTTAATATGTTGATCTGGTGTCTTGTGACCTTGCTCAACTCATTTATTATTTCCAGGAGGTTTTG | 120 |
| chr11 | 71229935 | 71230055 | 788855_47188552_ENST00000601538.6,E | 9 | - | good | TTGGCCTCTCAAAGTGCTGAATAATCTTGTCCATATCTACAAAAAAATTTTGCTGAGATTTTGACTGGAATTGTGTTCATCATCATCACCTGCACATCAGTTTGGGAAAAATTGACCTTC | 120 |
| chr11 | 71244575 | 71244695 | 788855_47188552_ENST00000601538.6,E | 9 | - | good | AGAAGAGTGGCATTGTTTAACATTTTTGTGGTCTTTTTGCTGTGAGGTTTAATGGAAGACAGCTGGGTTCCCGTATCTGCTCCTGCAGTCAGCTTGTTGCAATATCACAAGTTTTGTAGC | 120 |
| chr19 | 50679148 | 50679268 | 788855_47188554_ENST00000293441.6,E | 9 | - | good | TCTAACCAGTTCCCTGCCACTCTAAGCCCTCTCCCGTCACCCTGACCCCTTGCCTGTAACCCTGACCCCTCCCCGTCACACTGACACCCCCCCATCACCCTCACCCCTTACCTGTAATGC | 120 |
| chr19 | 50694148 | 50694268 | 788855_47188554_ENST00000293441.6,E | 9 | - | good | CGGTGCGTGTGTCAGTGTGGCTGTGTCGATGTGCGTGTGTGTGACTGCCTGCAGCCGCGTGCGTGTGTGTGCGTGTGTGTGTGAGAGAGAGAAATGGAGAGATGGTGGGCTCCCAGCCCA | 120 |
| chr11 | 70508975 | 70509095 | 788855_47188552_ENST00000601538.6,E | 8 | - | good | GGAGACAAGAAGTTTGAGATCAAGGTGGCAGTAGCATGGGTTCCCTCTAAGGCTGTGAGGGAAAATCTTCCAGGCCTCTCCCATAGGAGCTGACGGTTTCCTGGCAACCTTTAGCATCTC | 120 |
| chr11 | 70518215 | 70518335 | 788855_47188552_ENST00000601538.6,E | 8 | - | good | TGGAGTACAGTGGTGTGATCCCAGCTAACTGCAGCTAATCTCCCAGGCTCGAGCAATCCTCCCACTTCAGTTGTGTTTCTTCTTAATAAGAGTCTCTGTCTTTTCATAGAAAGCATGCTG | 120 |
| chr11 | 70579655 | 70579775 | 788855_47188552_ENST00000601538.6,E | 8 | - | good | CTCCAGGTCTTCATCACATCTGCAAAGTCCCTTTTGCGGCTTATGGTCACACAGGTCCCAGGGATTAGGGCGTAGACATCTTGCAGCCGTGGCTTTGCTGACCACAGCAGGCTGACCTGC | 120 |
| chr11 | 70630775 | 70630895 | 788855_47188552_ENST00000601538.6,E | 8 | - | good | GCTCACCCTTTGTGCTGGTGTCCTGGGGCTGCTCTCACAAAGGGTTCAAAGCCACCAGCTGGGGCTTGAACAACAGAAGTGTCTCCCCTCACAGTTCTGGAGGCAGAGACCAATGTCAAG | 120 |
| chr11 | 70637015 | 70637135 | 788855_47188552_ENST00000601538.6,E | 8 | - | good | CACACAGGCACCCACACACGTGCACATACACACAGGTACCCATACACATGCACACACACGTGCACACACATGAATGCACCCACATACGTGCCCACCAGAGGAGTGTGGCCCTGCTTACCC | 120 |
| chr11 | 70646735 | 70646855 | 788855_47188552_ENST00000601538.6,E | 8 | - | good | AATAAATAAATAAATAAATAAATAAAATAAATAAAAGTTAGATTCCTTTTAATAACTGGAGAAGTACTATTAAAGATCAGAAACCCTGTTTTCTATCTGTTTGGTATCTGGACTGTGATT | 120 |
| chr11 | 70648055 | 70648175 | 788855_47188552_ENST00000601538.6,E | 8 | - | good | ACAGAATCATGCTCTAGGCAACCTTTTGTCTGGCTTCTTTCAGCATCGTGTGCTTTAGGTTCATCCGTATGGTAGCATGTGTCCGTACTACCTTCTGTTTGTGGGTGAATAACGTTCCAC | 120 |
| chr11 | 70668215 | 70668335 | 788855_47188552_ENST00000601538.6,E | 8 | - | good | ACTGCCTCTGGGTGTCAACCTCCCTCTGCTTTTCTCGCGTAAGGGCACAGTCATTGGTTTAGGACCCTCCCTAAATCCAGGAAGATCTCAACTCAAGACCCTTAACTAATTATACCTGCA | 120 |
| chr11 | 70673855 | 70673975 | 788855_47188552_ENST00000601538.6,E | 8 | - | good | GGCCCCACCAACAACATTGGGATCAAATTTCACCATGGGATTTGGAGGGGACAGATAGCCAAATTATATGAGCCTCCTACTTTTAACTGTTTTTACAACTGCCCCTCTGGTGACACCAGT | 120 |
| chr11 | 70761335 | 70761455 | 788855_47188552_ENST00000601538.6,E | 8 | - | good | TGATGTCTCAGCTCAGGCTGCTGAAACAAACACAATCGAGACTGGGGCTCACACACCAGAAATCGATTTTCTCACAGTGCTGGAGAGTGAAAGTCCCAGATCAGGGAGCCTGCCTGGCTC | 120 |
| chr11 | 70774175 | 70774295 | 788855_47188552_ENST00000601538.6,E | 8 | - | good | GAGCCCTGGCAACCATGACTCCACTTTCTCAATCTAGATTTGCCTGTTCTGTCCATTTCATGTAATGGGATCATCCATGTGGGGTCCTTTATGACCAGCTTCTTTTAGTTAGGACAGTGT | 120 |
| chr11 | 70776695 | 70776815 | 788855_47188552_ENST00000601538.6,E | 8 | - | good | GGACAGGTGGGAGGATAGATGGATGGGTGAATGGATGGGTGGCTGAGTGGGTGAGTAGATGGATAAATAGGTGAGTAGGTGGGAATCTAGTAAGACCCACCCAGTTAGGTGGGTAGGTGG | 120 |
| chr11 | 70849775 | 70849895 | 788855_47188552_ENST00000601538.6,E | 8 | - | good | TCACTGACCTGTACACTTAAAAATGATGAACATGGTAAATTTTATATGTATTTTGCCATGAGTAAAAACATTTTTTTTAAAAAAAGAGATAACGTAGGGAGATCATGGACCCCAGATCAC | 120 |
| chr11 | 70862615 | 70862735 | 788855_47188552_ENST00000601538.6,E | 8 | - | good | CCCTCTTTGTTCATGATCCACTCAAGCCGGAAGCCTGAGTATCATCCACAGTCCTGTCCATCAGCCAGTCCAGTCTATCAGCCAGTCCAGTCCATCAGCCAGTCCAGTCCAGTCCATCAG | 120 |
| chr11 | 70864655 | 70864775 | 788855_47188552_ENST00000601538.6,E | 8 | - | good | GGGATTGCAGGTTCACACCACCATATCCAGGTCCTGGGGGTCCCTTTTATAAGGACATTAATCCTATTTATGAAGGTTCTACACTTATGACTTGCTTACCTCCCAAAGACCCCACCTCAT | 120 |
| chr11 | 70897415 | 70897535 | 788855_47188552_ENST00000601538.6,E | 8 | - | good | TGAGGTTGGCCAATTTGAAATCAGTAGGTAAGGCTGGAAACTCAGGCAGGAGTTGATGCCTTAGTCCTAAGGAAAGACTTCTTCCATAGGAAACCTCAATTTCTGCTCATAAGGCCCCCA | 120 |
| chr11 | 70926575 | 70926695 | 788855_47188552_ENST00000601538.6,E | 8 | - | good | TCTGGTCTCAGCTCTTGTACCCTAAACAAGTGTCCTTTTTGCCATCTCTTTAGTGCTATGGTTTTTGAAATTGTGTGCCGTTTGTTGGTGGTTTTGCTGTATGGAATGGCCCCTATCGTA | 120 |
| chr11 | 70931115 | 70931235 | 788855_47188552_ENST00000601538.6,E | 8 | - | good | CATGTAACATGTGGCTGTGTGTGTCTGGCTGCTTTCACTGAGCATGGTGTGGTCAAGTCTCATCCATGTGTTAGCGTGTGTTGGGGCTTCATTCCCTTTCATGGCTGAATTTTGTGCATC | 120 |
| chr11 | 70933415 | 70933535 | 788855_47188552_ENST00000601538.6,E | 8 | - | good | TGTAGTCAAATACACATAACCTAAAATTTATCAATTTTGATCATTTCTCAGCATGCCCTTCAGTGGCATTAAGTGCATTCATGGGGTGGTGTAGCCATGACTACCAGAATTTTGTCATTT | 120 |
| chr11 | 71088815 | 71088935 | 788855_47188552_ENST00000601538.6,E | 8 | - | good | GTTCAGGACGAGGGGAGGCCTCTGGTGGGGTGGTGCAGTCACCCACAAGGTCAAGTTCAGGACGAGGGGAGGCCTCTGGTGGGGTGGTGCAGTCACCCACAAGGTCAAGTTCAGGACGAG | 120 |
| chr11 | 71090135 | 71090255 | 788855_47188552_ENST00000601538.6,E | 8 | - | good | GTTTCTCTGGAGAACCCTGGCTGATACACCCTGAATGTAGAAGCAGCAGGACACATACACACACACACACACACACACACACAGGTTGTGTTTCTCTGGAGAACCCTGGCTGATACACCC | 120 |
| chr11 | 71090495 | 71090615 | 788855_47188552_ENST00000601538.6,E | 8 | - | good | TTCTCTGGAGAACCCTGGCTGATACACCCTGAATGTAGAAGCAGCAGGACACACACACACACACACACACACACACAGAGGTTGTGTTTCTCTGGAGAGCCCTGGCTGATACACCCTGAATG | 120 |
| chr11 | 71143295 | 71143415 | 788855_47188552_ENST00000601538.6,E | 8 | - | good | TGTTTTGTGCGAAAAGCATGGATTTTGGGGTTACAAATAAATTTTAGCAAGTAGGCATACCCCAATACAAAATTCATGAATAGTGAGGATTGGCTGTGTTTTGAAAATGTCAGCTTGTTC | 120 |
| chr11 | 71175575 | 71175695 | 788855_47188552_ENST00000601538.6,E | 8 | - | good | GTGGTTTGGATATATTTTGGGTGGTTTCAGGTAGGACGCTAAATCCAGTTTCCCTTCCTCCATCTTGGCTGGAAGCGATGTCTCTGTCTCTGTCTCTCTCTCTCTCTCTCTCTCTCTCTC | 120 |
| chr11 | 71177015 | 71177135 | 788855_47188552_ENST00000601538.6,E | 8 | - | good | TTCGGCTACTTCATTTCTGAAAGATATATTCACTGCTTATAGAATTCTAGTTTGACTTTTTTCGTTTAAAGATGCCAGTCCACTGTCTGCTCACTTGAATCATTTCCATCAAGAAATCTG | 120 |
| chr11 | 71200055 | 71200175 | 788855_47188552_ENST00000601538.6,E | 8 | - | good | TCCACAGAGGCAGAAAATAGACTAGCAGTTGCCAGAGGCTGGGAGGTGGCTGCTAATGGGTTTAGGGTTTCCATGTGGGGTGCTGAGAATGTCCTGACGTTAGATAGTGGTGATGGTTAC | 120 |
| chr11 | 71216255 | 71216375 | 788855_47188552_ENST00000601538.6,E | 8 | - | good | ACATGACCTTTGTGTCCATCTTCTCACTCAGCATCATGTTTTCAAGGCTTGACCACGTTTAGCATGATTCAGTGATTCCTTCCTTTTAAAGGCTGAGTGATACTGCACCATATGATACAG | 120 |
| chr19 | 50662948 | 50663068 | 788855_47188554_ENST00000293441.6,E | 8 | - | good | AATAATAGTAACCTGCATCAGGGCTGTCGTGGGGATTAAACTGAGATGCAATCATGTAAAGCACTTAGAACAATGCCTGGCACACGTGAGCATCAGTAAGCGGTGACGATTATTATTATT | 120 |
| chr19 | 50675308 | 50675428 | 788855_47188554_ENST00000293441.6,E | 8 | - | good | CTGGACTAGAACCAATAGAGCTCTCTGTGATGATGAAATGTTCCATATCTATACTGTCCAATTTGGTAGCCACCAGCCACAGGTAACTGTTGAATACTTCAAATGTGGCCAGTGCAACAG | 120 |
| chr11 | 70533095 | 70533215 | 788855_47188552_ENST00000601538.6,E | 7 | - | good | AAAAGTGTACAGTGATATTCGACACAATCACAGGGTGTGCAACCATCACCTTCTTCCGGTTCCTGGACATTTCCATCACCCCAGAAGGAAACCCCGTCCCCAAGGGCCGTCCCTCCCATT | 120 |
| chr11 | 70563455 | 70563575 | 788855_47188552_ENST00000601538.6,E | 7 | - | good | CAAAAGCAAAACTCCATCTCAAAAAAAAAAAAAAAAAGTGACCACAGATGGAAATACACCAAGGAAGGAAGGACTTAGCTGCATGGCTATATATGTAAAATTTGTTTTGTTTAATTCTTTC | 120 |
| chr11 | 70588175 | 70588295 | 788855_47188552_ENST00000601538.6,E | 7 | - | good | GAGTGACTCTTCCTTTCACTTGAGCAGTTAGAGGCCATTGTGTGATCATTAATTGGCCTTATTTCTTTTTTTATTCCCTTGGCATGTGAAATAAGAGCTATAATCAGCCTAATTTCTATA | 120 |
| chr11 | 70600055 | 70600175 | 788855_47188552_ENST00000601538.6,E | 7 | - | good | AATGGCTGGGATTACAGGTGTGAGCCACCGTGCCTGCTCCATATGTATTTTCCAATCAGCTTATCAAGTTACACCAAAACAAAACAGCAACAACAAACCCTATTGGGATTTTGGTTAGAA | 120 |
| chr11 | 70609775 | 70609895 | 788855_47188552_ENST00000601538.6,E | 7 | - | good | GGATCTGTGTCTCATTCCTTTTCATGCCCGAATTATATTCCATAATATACATATTCCATATTGTATATTATGGAATATACAATATAGATATATTATGTATATCTATATACATATACAATA | 120 |
| chr11 | 70630535 | 70630655 | 788855_47188552_ENST00000601538.6,E | 7 | - | good | TCTGCACGTGGCCATGTCTCTGTGTGTCTGTATTTTCCCTCCTTATAAGGGCACCAACCAGTGGACTTAGGGCCCACTTATTCCAGGATGACCTCATCCCAAGTGATTGCATCTCCAGCA | 120 |
| chr11 | 70683575 | 70683695 | 788855_47188552_ENST00000601538.6,E | 7 | - | good | GCAGCCACCAGAAGCGGCAAGAAGCAAGGAGGGTTCCTCCCCTGGGGCCTCCTGAGGGAGCGTGGCCCTGCCGATGCCTTGATTTCACCACAGTGGTCATGCTGATACTGCACCTCTGGC | 120 |

| | | | | | | | | |
|---|---|---|---|---|---|---|---|---|
| chr11 | 70736615 | 70736735 | 788855_47188552_ENST00000601538.6,E | 7 | - | good | ATGAAGCCGTGTGACTCTCCTGAGGACCACAAACTGGGTGGCTCAGAACAGCAGAGAACGGATTGTCTCAGTTCAGGAGTCCAGAAGTCCAATATCACGGTGGCAGCAGGGCCGTGCTCC | 120 |
| chr11 | 70751015 | 70751135 | 788855_47188552_ENST00000601538.6,E | 7 | - | good | GATTGGATAATTTCTATGGATCTGTCTTCAAGGTCACTATTTGTTTCCTCTGTCTTCTATATTCTGCTTTTGACACATCTAGTAATTTTTTTTTCATTTGTATTGTTTCCGTTGAGAGTT | 120 |
| chr11 | 70753295 | 70753415 | 788855_47188552_ENST00000601538.6,E | 7 | - | good | GTCTATCTTGGCGAATATTCTGTGGACATTGAAAAGAATGTATATTATGCTCGTTATTGTTGAGTGCGGTGTTCTTTAAATGTCAGTTCGGTCCTTTTGATGGTATTTTTAAGTGTTTAT | 120 |
| chr11 | 70797095 | 70797215 | 788855_47188552_ENST00000601538.6,E | 7 | - | good | TTCTCTGTTGTGTGCTACACAGCCAGTATCCAAAAGCAAAAGTGAGCTTCAAGCAGCACAGGCTTTATTTGTTGGCCATGGAACCGAGAAGCGGGAGCTTGGCTCACAGATCCATTTATC | 120 |
| chr11 | 70843655 | 70843775 | 788855_47188552_ENST00000601538.6,E | 7 | - | good | CCCTGTATCAGTCTCCCGTGTGCAGAGAGCCCTGCAGGTGGGCTGGGGGGCATTCAACAGCAGGCCTCTATGTTCTCACAGTCCTGGAGGCTGGAGCTCGAGATCAAGGTGTCGCAGGGC | 120 |
| chr11 | 71059415 | 71059535 | 788855_47188552_ENST00000601538.6,E | 7 | - | good | CTGTCTGTCTTCTTAAATATGTATGTGACCCTCTATTGAGATGCAATTCACACACCATACAGTTCACCCATCTAAAGTGTACAGCTCCATGATCGCTGGTCTGTCCACAGAGTTGTGCAG | 120 |
| chr11 | 71088695 | 71088815 | 788855_47188552_ENST00000601538.6,E | 7 | - | good | GGGCAGCCCCTGGTGGGGTGGTGCAGTCACCCACAAGGTCAAGTTCAGGATGAGGGGCGGCCCCTGGTGGGGTGGTGCAGTCACCCACAAGGTCAAGTTCAGGATGAGGGGAGGCCCCTG | 120 |
| chr11 | 71088935 | 71089055 | 788855_47188552_ENST00000601538.6,E | 7 | - | good | GGTCAAGGTCAAGTTCAGGACGAGGGGCGGCCCCTGGTGGGGTGGTGCAGTCACCCACAAGGTCAAGTTCAGGATGAGGGGAGGCCCCTGGTGGGGTGGTGCAGTCACCCACAAGGTCAA | 120 |
| chr11 | 71133935 | 71134055 | 788855_47188552_ENST00000601538.6,E | 7 | - | good | AGCTCTTTTATCCACGAATGTAGCCGGGTGCAGTGGTGCACAACTGTAATCCCAGCTACTCAGAAGGCTCAGGCAGGGGGATCACCTGGGCAACGTAATGAGACCCCTTCCCTAAAAAAA | 120 |
| chr11 | 71156135 | 71156255 | 788855_47188552_ENST00000601538.6,E | 7 | - | good | CCTATCAGGAGGGTCCTGGGGACTGGAGGAGTCAGCTCAGGCTGCCATCACAGAGCACCACAGAACTGGCAGTCTAGACGACAGACATTGATGGTCTCACAGTCCTGGAGGCTGTTTGTC | 120 |
| chr11 | 71159135 | 71159255 | 788855_47188552_ENST00000601538.6,E | 7 | - | good | AGGTCACCACCAACATGCGAGAAAGGCAAGGAAGTGGATTGTTCCCCAGAATCTTGAGAAGGAGCCAGCTCTGACAACACCTTGGCTTTAGCCCCATAAACTCATTTAGGACTTCTGACC | 120 |
| chr11 | 71179655 | 71179775 | 788855_47188552_ENST00000601538.6,E | 7 | - | good | AGAAATATAATTGATTGCTGTATATTCCTGTTGCCTTCTTCAGTCTTGCTAAAAATCACTTGTTCTAGTAGTTTTTTCAAGGATTCCAATGGATTTTCTATGGGGACAATCATGTCTTCT | 120 |
| chr11 | 71196215 | 71196335 | 788855_47188552_ENST00000601538.6,E | 7 | - | good | CCAGCCTGGGTAACAGAGCAAGACACTGTCTCGGGAAAAAGAAAAAAACAAAAAACCAGAGGTGGGTTGAGAGCCTTGGGTAGATTAAAGCATGTAGCTAGAATTTAAGAACCATCCATT | 120 |
| chr11 | 71200295 | 71200415 | 788855_47188552_ENST00000601538.6,E | 7 | - | good | GGCAGCGTTATTTATAATAGCCCAAAACTGGAATCAGTCAGATGTCTGTCAGCTGAGTAATAAATAAACAGTGTGGTGTACCCATGTGATGGGATGTGACTTGGCCTGAAAAGGAATGGA | 120 |
| chr11 | 71225015 | 71225135 | 788855_47188552_ENST00000601538.6,E | 7 | - | good | TCCTGCTCCTGTTGGCTTTTCTGGGTTGCTGGCTTATTCAGCACCACGTCTGGGATGTATGAGGCAAGAAGAAAACCCAGAGAGCTCACACCATGTCCCTGGCTGGTCTGCCTTCTCTCC | 120 |
| chr11 | 71226695 | 71226815 | 788855_47188552_ENST00000601538.6,E | 7 | - | good | ATGTTCCAAAACTCCTTCTTTTATTATTATTTTTTTATTTAGAAAGGCTTCTTTTTAGCTATTCTTTTAGAATAGGTCTGCCAGTGGCAAATTTCCTTTGTTTCCTTTCATCTGAGACTA | 120 |
| chr11 | 71228135 | 71228255 | 788855_47188552_ENST00000601538.6,E | 7 | - | good | AATTCACTTCCCTTAATAGTTGTAGGAAATGATTCAAATTACCTTTTTCAAATTGGGCAAGGTGAAGTAGTTTGTATTTCTTGAATAATTGCTTTATTTCATCTACATTGTCAAATTTAT | 120 |
| chr11 | 71236535 | 71236655 | 788855_47188552_ENST00000601538.6,E | 7 | - | good | TTGGCCCATAGGCGGTCTCTTGCTGACCCTGATTTATTTAGGTCATGGTTCTGTAAGTTGCCTACTGGAACCAGGCTCAAGCTGAGCGTGATCTCGGCTCACTGCAACCTCCACCTCCCA | 120 |
| chr19 | 50680108 | 50680228 | 788855_47188554_ENST00000293441.6,E | 7 | - | good | GTCCTTGTCTCTCCCCCTTCTAGGCCCCATCTCTGTCTGTCGCTCTTCGCCTCTCTCACTCTCCCTATCTTTCTCTTCCCATCCCTCCCTGTTTCCCCACCTCCCTCTCTCCCCCATCTG | 120 |
| chr11 | 70483655 | 70483775 | 788855_47188552_ENST00000601538.6,E | 6 | - | good | AGAATTATGTACGTATACACCTACAAGTACACTCAACAGCCTCCAGGAATTTTGAGAAAGGGTGTTGCTTTGCCGCTCAGGCTGGAGTGCTGTGACGTGATCACGGCTCATTGCAGCCTT | 120 |
| chr11 | 70484375 | 70484495 | 788855_47188552_ENST00000601538.6,E | 6 | - | good | TATGGGAAGCATGATGTTGGCTTCTGCTTGGCTTCTGGAGAAGCCTCAAGAAACTTTCAGTGATCACCAAAGGGGGAGCAAGAGGGAGTCGGGGGAGGTACCACAGTCTATTTAAACAAC | 120 |
| chr11 | 70540055 | 70540175 | 788855_47188552_ENST00000601538.6,E | 6 | - | good | TTCTTACGTTGAAGCCCTAACCCCCATTACCTCAGAGTGAGGTCTTGCTTGGAAATTGGGACATTGCGATGTGACTGGTTAAGATGGGGTCACAAGGGTAGGGCAGCCCCTGATCTACTG | 120 |
| chr11 | 70564895 | 70565015 | 788855_47188552_ENST00000601538.6,E | 6 | - | good | GTTAGAGTTAGCAGACAAGGATATTAAAAGAATCATTAAAACTGTATTCCATATGTCCAAAAAGTTAAATGTAGACACAGAAGATATTTTGAAAGCCCACTGGAATGTCCATTTTTAAAT | 120 |
| chr11 | 70644335 | 70644455 | 788855_47188552_ENST00000601538.6,E | 6 | - | good | GCCAAATACAACATTCTCTTCCTTTGAACTGTATGATTCCGCTCAAGTGAGGTTCCTAGAGTAGTCACATCCGTCACATTCATAGAGACAGAAAGTAGGATGGTGGGTGTCGGGGGCTGG | 120 |
| chr11 | 70683455 | 70683575 | 788855_47188552_ENST00000601538.6,E | 6 | - | good | CTCCAGCGCTGTGAGAGAACCCATTTCTGCTGTTTAAACCACCCGGTTTGTGGCAATTTGTTACAGTGTCCACAGGAAACTCTGAGAGCATGTAGCAGCATATCAGTGAGCTAACAGACA | 120 |
| chr11 | 70751255 | 70751375 | 788855_47188552_ENST00000601538.6,E | 6 | - | good | CTGTTCCACTTCCTTCTGACCTCCATAGTTCTGATGAGGAATCTGCAGTAATTCCTGTGGTTTTTCTTCTGTATGAAATATGTCATTTTTCTCTAGCTGCTTTCAGTATATATTTTTTCG | 120 |
| chr11 | 70794815 | 70794935 | 788855_47188552_ENST00000601538.6,E | 6 | - | good | AAATCTAGTTCAGTGCATTTTGACAAATGTATATACCCACGTCTCCAACTGCAAATAAGATATTAAACCCTTCCTTCACAGGCCAGGTGTGGTGGCTCATGCCTGTAATCTTTGCACCTT | 120 |
| chr11 | 70926455 | 70926575 | 788855_47188552_ENST00000601538.6,E | 6 | - | good | GTGCTGCAGTGCTGTCTGGGGTTCCCAAGTGCGAGAAGGCTGTGGTGTGCTCATGAAGAAAACACACTTTAGAGAAACTTCATTCATGTCTGAGTCATAGTGCCATTGGCCGTGAATGCA | 120 |
| chr11 | 70926815 | 70926935 | 788855_47188552_ENST00000601538.6,E | 6 | - | good | TGCCTCCTCCAGCTCGAGTGGGTCAGACCATGGACGCCTCACACTTCTGCACAGTTCGTCCTCCTTATCCACAGATTCCATATTTGTGATTTTGCCTACTCACTAAAATTTATTTGTAAC | 120 |
| chr11 | 71078015 | 71078135 | 788855_47188552_ENST00000601538.6,E | 6 | - | good | AGCTCACCTTGCTCTTTCCTTGCCTCAGGAATCAGCTGCTTCTCCAAGGAGCCTGATTCTTTTTAGTGTGGAGTGGTATTTAGAAGCCAAGGTGCAGGTGCTTGGGGTGCTGATGGCTGC | 120 |
| chr11 | 71090015 | 71090135 | 788855_47188552_ENST00000601538.6,E | 6 | - | good | TGAATGTAGAAGCAGCAGGATACACACACACACACACACACACACACGTTCTGTGTCTCTGTAGAACCCTGACTGATATATCCTGAATGTGGAAGCAGCAGGTGAATTCTTGCACAGCAC | 120 |
| chr11 | 71096015 | 71096135 | 788855_47188552_ENST00000601538.6,E | 6 | - | good | GCTTTTCACATGGAGACAGGATGAATGAAGATGCTGGCATGTCTACGCACGGCATCCTAGTTGGGGAAGACAGGAAGCCTGCTTTTCACATGGAGACAGGATGAATGAAGATGCTGGCAT | 120 |
| chr11 | 71171255 | 71171375 | 788855_47188552_ENST00000601538.6,E | 6 | - | good | GGCCTTTGGCTTCAGAGTGACTTACGCCATCACCTTCTCTGGTTCTGAGGCTTAGGACCTGGATTGAGTCTCGCTCTCAGCCTCGAGGGACTCCAGCCTGCAGACAGCCCGTCGCAGGTC | 120 |
| chr11 | 71176175 | 71176295 | 788855_47188552_ENST00000601538.6,E | 6 | - | good | TGTGTGTTTTGGATTTCTGTACATATTCTTGAGCTTTTTGGGGGCGGGATTTAGTTAATTTACTTAGAAACAGCTTGATCCTTTTGGGTCTTGCTTTGAATATTCGTTGGGCGGGATGAT | 120 |
| chr11 | 71176775 | 71176895 | 788855_47188552_ENST00000601538.6,E | 6 | - | good | TCTTTATGATTCTTGTGCTTAGGTTACATTGAGCTTCATGAAGCTGGGAGTTTATAGTTTTCATAAAATTTGAATAACTTTTGGCCAAAAATAGAAATATTTGTTCTGTTCCCCCCTCTT | 120 |
| chr11 | 71177135 | 71177255 | 788855_47188552_ENST00000601538.6,E | 6 | - | good | CCATTTCTGATCATCTTCACTCTTTTGTGTAATTCATATTTTTATCTGGTATCATTTTCTTCTGCCTCAAGAATTTCTATTAACATTTATTGTAGGACAGATCTCCTGGTCATGAATTCT | 120 |
| chr11 | 71211695 | 71211815 | 788855_47188552_ENST00000601538.6,E | 6 | - | good | AAGAGAATTGACTCACCATACATCATGTGATTCCATTTATATGAAATGCCTCAAAGTGGCCAAGTTATATATAGAGAAAATAATTAGTGGTTGGGGAGAGACTGAGGGAGAGGATGGGGG | 120 |
| chr11 | 71242295 | 71242415 | 788855_47188552_ENST00000601538.6,E | 6 | - | good | TTATAAATTGTGCAAAATACCTATAACATAAAAAGTTACCCTTTTCACCATTTTTAGGTGTGTAGTTCTGAGGCATTAAGGATGCTCATCGTGCTCTGCAGCCATCGTCACCATGTATCT | 120 |
| chr19 | 50694028 | 50694148 | 788855_47188554_ENST00000293441.6,E | 6 | - | good | CCCCCCAACTGTGTCCATCTGTGTGTCCCTGTGTGACAGCGTCTGTCTCTGTATGACTTGGAGCGACAGCGAGGCTGTCTGGGACTGTGTGTGTGTGTGTGTGTGTGTGTGTGTGTGTGT | 120 |
| chr19 | 50701588 | 50701708 | 788855_47188554_ENST00000293441.6,E | 6 | - | good | GAGCTGAGCGGTATAGAGGTTGAGAGCCTGGCAGCCTGGGTTCAAATCCAGACCCCACCACTTCCTAGCAGGGTGACCTCAGCCAAGTGATTTCTCTGGGCCTCAGTTTCTTCATCTGTA | 120 |
| chr19 | 50704708 | 50704828 | 788855_47188554_ENST00000293441.6,E | 6 | - | good | CCCACAGTCTTAGGATTACAGGCTTGAGCCACCACGCCTGGCTATAGACATTTAAATTCTTTCTTGTTTTCATATGGCATATCCTTGAAATCCAGTGTGTAGTTCGCATGCATGGTGCAT | 120 |
| chr11 | 70513535 | 70513655 | 788855_47188552_ENST00000601538.6,E | 5 | - | good | TTTCCTTTACCTAGTTGGGTATTGTTACAGTAGCTGTTTTTTAAAGTCCTTCTCTGATAATTCCAACATCTGGGTCATCTCTGGGTTTGCTAAATAAGTTTGAGTTGTATCCCAAATTTC | 120 |
| chr11 | 70517855 | 70517975 | 788855_47188552_ENST00000601538.6,E | 5 | - | good | TTGCCACATACCCCCTTATATCTCTCCCTGTTTTCCCTATTTTTCAATTGGCTTGTCAATTGTAGCAAATGTCCCACACAGTTGAGAAGCCAATATTGATACACCATTATTAACTAAAGC | 120 |
| chr11 | 70539935 | 70540055 | 788855_47188552_ENST00000601538.6,E | 5 | - | good | GCCCACTTCCTTGTAAGAAGAGGGAGGTTTGGGCGCAGACAGGCACACAGTGGGAAGGCCAGGCGAAGATGGAGGCAGAGGGGGGTGATGCTCCTCCAAGCCAGGGACACCAAGGCTTGC | 120 |
| chr11 | 70584575 | 70584695 | 788855_47188552_ENST00000601538.6,E | 5 | - | good | TTAGGAGGCAGAGGGAGAGGGGGGAAATGTGGACAGGAGCTTTACCATGATTTCTGCAGGAAGGAGCAGACTAAGTAGGCAGTGTAAGTGGGTTTAGCGTTGGCAGCTGGAATCATTTCA | 120 |
| chr11 | 70595855 | 70595975 | 788855_47188552_ENST00000601538.6,E | 5 | - | good | CTCACGATGTTCTCCTCTGTGTTTGCGACTGAAACCCCGCTCTGTTTTCTCTCACAAGGACACCAGTCGCTGGAATAAAGTCCTACCCTAATCCAGCATGACCTTACCTTAACTAAATCT | 120 |
| chr11 | 70631375 | 70631495 | 788855_47188552_ENST00000601538.6,E | 5 | - | good | CCATGATGGAGAAGGTGGATACCAAGAGAAAAGGACCAGGGACATGGTTTTATTTGGAGAAGGCAGGTGGGAGGGAGGTTGTGTGGGTGTGTGTGTGTGTGTGTGTGTGTGTGTGTGTGT | 120 |
| chr11 | 70747775 | 70747895 | 788855_47188552_ENST00000601538.6,E | 5 | - | good | TGCTATATGTACATCATGTGTGCATGTATGCACACACAAACACTGCACATACATGCACCGTGCAGCATGCATGCACACGTGGACACTCACAAACATGCACACTGCACATACATGTACCAT | 120 |
| chr11 | 70751135 | 70751255 | 788855_47188552_ENST00000601538.6,E | 5 | - | good | TTTTTAGTCTTCAGCAATTTGATTTGGACGTGTCTGGGAATGGATTTTTTGAGCTAATCCTGTTTAGGATGCATTGAGCTTCTTGAATCTGTAAGTTTATGTGCTTTTCTCTCTTGTTCA | 120 |
| chr11 | 70751975 | 70752095 | 788855_47188552_ENST00000601538.6,E | 5 | - | good | AAAAATAAAACTCCATCTGGTAATCTATTTTATTTGATGTGTTTAGAGCATTTAAATTTAATGTTATTATTAATGTGTTAGTATTTATATCTACCATTTTATTTCTTATTTTCTATTTGT | 120 |
| chr11 | 70771175 | 70771295 | 788855_47188552_ENST00000601538.6,E | 5 | - | good | GAGCCCGTCACTCCTTCGCGATGGTGTGCTGAGATGGCAAGAAGCCGTCTTTTATAGAAACTGTCTCTGGGCCGAAGGCCGTGGGTCATGCCTGTAATCCCAGCACTTTCAGAGGCTGAG | 120 |
| chr11 | 70772855 | 70772975 | 788855_47188552_ENST00000601538.6,E | 5 | - | good | GAAACCAGATTGACAGCAGGCAGATTAATAAGAATAGCATACCAATTGTATTGGTTTTACATATACATGGGGATCTTCTGCAAGAGAGCGAAATGCAGAGAAGTGACCCAAGCAAGACGC | 120 |
| chr11 | 70788455 | 70788575 | 788855_47188552_ENST00000601538.6,E | 5 | - | good | GCCCTGTACTGCATACACCATGTTTTTGGTCTGATATCAGAGCCAGCTCCTAAGTGACTAACCAGTGGGTGGTGTGGACAGCAGGAGTGCGCTGGACAAAGGGATGAGTCACTTCCCCCG | 120 |
| chr11 | 70824095 | 70824215 | 788855_47188552_ENST00000601538.6,E | 5 | - | good | CTCCAGGGCTGCCCTGTGTCCTGGGGAGGTGGGGAAGGTGCGCCGGGTCACAACTGTCTCCCGTGAGCTCTGCCAGCACCCTCCATCCCCCATGAGCTCTGCCAGCACCTCCTCTGCCCT | 120 |
| chr11 | 70840895 | 70841015 | 788855_47188552_ENST00000601538.6,E | 5 | - | good | GCTGGGATTACAGACATGAGCCACCACGCCTGGCCTATTCCTATATGTTTTATACACCTCTGGAAGAGATCAGAGCCAGCCAGAGTGTTTCTCAGGTGAAGTCTTATCAAGCTCCAGTGA | 120 |
| chr11 | 70862495 | 70862615 | 788855_47188552_ENST00000601538.6,E | 5 | - | good | CCAGTCCAGTCCATCAGCAGTCCAGTCCATCAGCCCGTCCAGTCCATCAGCCAGTCCAGTCATGTCCATCAGCCAGTCCAGTCTATCAGCCAGTCCAGTCCATCAGCCAGTCTAGTCCTG | 120 |
| chr11 | 70918775 | 70918895 | 788855_47188552_ENST00000601538.6,E | 5 | - | good | ACTGTGCCTGGCCTGTATGCGTTTTTCAAAACATGTTGTACACCATCAGTGTATACTATTTTTATTTGTCAACTTAAAAAGTCAATTAAGGCTGGGCATGGTGGGTCATGCCCTGTAATC | 120 |
| chr11 | 71137895 | 71138015 | 788855_47188552_ENST00000601538.6,E | 5 | - | good | CCATTAGAAGCTGCATTCGGGTGCTTTGCTGTGTGTGTGCGCCCGCCCATTGTGAGAGAGGGCTCTGGGCGGCTGGTGCAGGGGTGGTTCGTTACCGTTAGAAGCTGCGTTTCGATGCTT | 120 |
| chr11 | 71138135 | 71138255 | 788855_47188552_ENST00000601538.6,E | 5 | - | good | CCATTGTGAGAGGGGGCTCTGGACGGCCGGTGCAGGGGTGGTTAGTTACCGTTAGAAGCTGTGTTCAGGTGCTTTGCTGTGTGAGTGCGCCCGCCCATTGTGAGAGAGGGCTCTGGGCGG | 120 |
| chr11 | 71187335 | 71187455 | 788855_47188552_ENST00000601538.6,E | 5 | - | good | ATAAAACACGAGAAACCCATGAAGAACTGTCCCTGAATAAAACGTGAGAAACCCATGAAAAACTGTCTCCAAACAAAACATGAGAAACCCATGAAAAACTGTCCCCAAACAAAACATGAG | 120 |
| chr11 | 71200535 | 71200655 | 788855_47188552_ENST00000601538.6,E | 5 | - | good | CTGAGGAAGACGGCCTGGCAAGTCCTAAAAAAAGTTAAATGTAGAGTGAAGTTAAATGTAGAGTTATGTTAAATGTAGAGTGAAGTCAAATGTAGAGTGAAGTCAAATGTAGAGTGAAGT | 120 |
| chr11 | 70498895 | 70499015 | 788855_47188552_ENST00000601538.6,E | 4 | - | good | AAAGTCAGGCCTTTACAGTTGTATTGGGTTGAAAAGTGACCCCCAAAGTTCTGTGAATGGGACCTTATTTGGAAATAGGATCCTTGTAGATCTAATCAAGATGAGGTCATGCTGGACTCG | 120 |

| | | | | | | | | |
|---|---|---|---|---|---|---|---|---|
| chr11 | 70514975 | 70515095 | 788855_47188552_ENST00000601538.6,E | 4 | - | good | GCACTCCAGCCTGGGCGACAGAGCAAGACCCTGTGTCAAAAATTTATTTTGTCTGATATTAACAGCCATTCTGCCCAGGAACTGTCTTTTCTCATTTATTAACTCTCAGCCTCTCAGTGT | 120 |
| chr11 | 70542095 | 70542215 | 788855_47188552_ENST00000601538.6,E | 4 | - | good | CAAAGAGGCTGTTTCCAAATAAGGTCACATTGGCAGGTGCCAGGGGTTAAAGGCATGGACATATCTTTCGAGGGCCACCGGTCAGCTCACTATGTAGTCTGTTTTTGTAACTTCTCTGCA | 120 |
| chr11 | 70548935 | 70549055 | 788855_47188552_ENST00000601538.6,E | 4 | - | good | GTCTCTGACAGTTTGGGGGCTAGCAGTCTGAAGTCGGGATGTCAGCAGGGCCATGCTTTCCCGGAAGGCCCTGGGGGAGGAGCCTTGCCTACTCCTCCAGCTTCTTCCTGCTCCCGGTGG | 120 |
| chr11 | 70554455 | 70554575 | 788855_47188552_ENST00000601538.6,E | 4 | - | good | AACAGAGGAACCTGGGGGAGGGGGGAGGAGAGGGAATCTGGGAACTCTGAACTTTCTGTGCAGTGCAATTTTTCTGTAAATCCAAAAATGTTCCAAAATATAAAGTCTTTGTTAAAATAA | 120 |
| chr11 | 70560455 | 70560575 | 788855_47188552_ENST00000601538.6,E | 4 | - | good | TGTTTTGCTTACTCTAGATATTTTGCATTTCCCATGAATTTTACAATCAGCTTGTCCTTTTCTACCAAAAAAAAAAAAAAAAAAAAAAAAAAACCCTACTGGGGATGTGACAGGAATTACA | 120 |
| chr11 | 70594055 | 70594175 | 788855_47188552_ENST00000601538.6,E | 4 | - | good | AACTGTTGACTCATTCATCTGTGGATGGGCATTGTTTATAAAGTCAGCTGAGGCACAATAAAATTTTAAAGAATTGATGTGAGCAAGCAGCGATTTACGAATCGGGCAGCTCCAACACGG | 120 |
| chr11 | 70635335 | 70635455 | 788855_47188552_ENST00000601538.6,E | 4 | - | good | GGTACATTTCACTCAGTGTAATACCTTCGAGGTTCCTCCGTGTTGTAGCATGCATCAGAATCTCCTTCTGGTTTAAGCCTGAATGATAACTATCCCAACTCTACGGAAACACCACCTTTT | 120 |
| chr11 | 70636895 | 70637015 | 788855_47188552_ENST00000601538.6,E | 4 | - | good | CTTGATCTTAGACTTCCCGCCGCCAGACTGAGAGAAAAATTCCTGTTGTTTGAAGCCCCTTGGTTTGCGGTGATTTGTCGTGACAACCCCAGGAAACGGTACACATGTTCACACACACTT | 120 |
| chr11 | 70740095 | 70740215 | 788855_47188552_ENST00000601538.6,E | 4 | - | good | GTGTCCAGTGAGCACCTGCTGTGTGGCAGGCACTGTCTGTCCTAGCTGCTGTGGACGGAGCCATGAGCACCTGCTGTGTGGCAGGCACTGTCTGTCCTAGGTGCTGTGGACGGAGCCATG | 120 |
| chr11 | 70746215 | 70746335 | 788855_47188552_ENST00000601538.6,E | 4 | - | good | GGAGATGGGGGCTGTGAGCATGGGGCAGCCCGCATCCCCCACCCTGAAGGGAGGCCCAGCCTTCCCTGTCCTGGAGAGGCGGGAGAGTGTGGTGGGAGAGGCTGCTTCTGTGCTACAAGT | 120 |
| chr11 | 70746335 | 70746455 | 788855_47188552_ENST00000601538.6,E | 4 | - | good | GCGGGAGAGTGTGGTGGGAGAGGAGATGGGGGCTGTGAGCATGGGGCACCCTGCATCCCCCACCCTGAAGGGAGGCCAAGCCTTCCCTGTCCTGGAGAGGCGGGAGAGTGTGGTGGGAG | 120 |
| chr11 | 70759655 | 70759775 | 788855_47188552_ENST00000601538.6,E | 4 | - | good | GCCGGGGCTTGCTGGCTGCTAGAGGAAGAGAAAAGCACTATGACAGCCAGAACTCGTTTATTTAAGTGTGGGGTGCATGACTTCACCCTTGCCTCACACAGTCTTAGGTCTTGCTTACAA | 120 |
| chr11 | 70792415 | 70792535 | 788855_47188552_ENST00000601538.6,E | 4 | - | good | GCTGCTCAGGACCCAGTGAATTCTCATAGCCTCATGACAGTTGCTGGAGGTTGGGGATAACAAACAGCAAAATCAAATGGTGGTCCTCGTTGGTTGGTTGGTTGGTTGGTTGGTTGGTTG | 120 |
| chr11 | 70793015 | 70793135 | 788855_47188552_ENST00000601538.6,E | 4 | - | good | AGGGTTTTTAAAATTTTGAATCTAATCCAAATTATTCTTTTTTTCTTCCACGTTAGTGCTTTTTGTCACTTTTTTCTTTTGAGTCAGGGTCTTGCTGTGTCATCGAGGTGGGAATGCAGT | 120 |
| chr11 | 70793735 | 70793855 | 788855_47188552_ENST00000601538.6,E | 4 | - | good | AGTCTTTTTAAGGACATACATTTTTATTTCTCTAGGTAGAAAACTGCTGGGTTGTATGATAACATTTTCATTTTATAAGAAACTACCACACAGTTCTCCAAAGTGCTGTCCCATTTTGTT | 120 |
| chr11 | 70843175 | 70843295 | 788855_47188552_ENST00000601538.6,E | 4 | - | good | CTTTAGCAGCCCAGCCCACCCCATAGCAGCCCAGCCCACCCCATAGGAGCCCAGCCCACCCCATAGCAGCCCAGCCCACCCTAATAGCTGTCTTTGCAACCTCCCAGTCACAGTCACCAC | 120 |
| chr11 | 70898735 | 70898855 | 788855_47188552_ENST00000601538.6,E | 4 | - | good | TCAAGTGATGTTCCAGCCATACCCTCAGTGTTCGTTTTGTTGCAATGTGGTTAGGCTTTTTCACTTTTTGCCGTATGGATAGACTGAGAATCTTCCAAATCTTCAAGCTCTGCTTCCTTT | 120 |
| chr11 | 70929935 | 70930055 | 788855_47188552_ENST00000601538.6,E | 4 | - | good | CCTGTGTTTGCTGCTGTGCTAGGAGGGCAGAGTTGAGGAGCTGTGACAGAGACCGTGTGGTCCTCAAAGCTGAAGGTGTTTACTCTCTGGCTTTCTGCAGGGCTTGCTCACACCAGATGT | 120 |
| chr11 | 70937855 | 70937975 | 788855_47188552_ENST00000601538.6,E | 4 | - | good | CGCTCACACATGCACACACACACACAGGCATGCACACAGGCATCCACACACACATGCAAATGAAGAGAGAGGCCCCAAAGGCCCTCTGGCTTCTGAGAGGTCACACACCCTGCACTAGGT | 120 |
| chr11 | 71055815 | 71055935 | 788855_47188552_ENST00000601538.6,E | 4 | - | good | TTTAAAGTGTACACTTTGGTGGCATTAAGAATATTCATGATGCTGTGCAAAACCATCCCCATTGTCTAGCTAAGAACTTAGCTTTGGGAACTTTCCTTTCCCAAAAGGAAACCCTGTACC | 120 |
| chr11 | 71095895 | 71096015 | 788855_47188552_ENST00000601538.6,E | 4 | - | good | GTCTACACACGGCATTCTAGTTGGGGAAGACAGGAAGCATGCTTTTCACATGGAGACAGGATGAATGAAGATGTTGGCATGTCTACACACAGCATCCTAGTTGGGGAAGACAGGAAGCCT | 120 |
| chr11 | 71099015 | 71099135 | 788855_47188552_ENST00000601538.6,E | 4 | - | good | GTTTCAGAGGAGGACTGGTCAGGTGCTTGTAGGGTGCCCCTTATTGGGATCCATCTGGTGTTTTCACGATTAGGGGCTGTGGGTGTGAGGAGGAAGAGCACAGAGCCTTTCTCATCACCT | 120 |
| chr11 | 71105855 | 71105975 | 788855_47188552_ENST00000601538.6,E | 4 | - | good | CTTGGAGTCGTAACCCAGTAGTGATCCTCAAGTACAGAGCTTACATGAGGGTTTACGCTTGGTGCTGTACCCAACAGTGATCCTCAAGTCCAGAGCTTACATGAGGGTTTATGCTTGGTG | 120 |
| chr11 | 71138015 | 71138135 | 788855_47188552_ENST00000601538.6,E | 4 | - | good | CTGGTGCAGGGGTGTGGTTCGTTACTGTTAGAAGCTGTGTTCGGGTGCTTTGCTGTATGTGTGCGCCCACCCATTGTGAGAGAGGGCTCTGGGCGGCTGGTGCAGGAGTGTGGTTCGTTA | 120 |
| chr11 | 71159255 | 71159375 | 788855_47188552_ENST00000601538.6,E | 4 | - | good | GTCATTATTTTTAAGAGGGAGATTTGGCTGCAGAAGAGGAGATGGTGATGTGAAGGTGGAAGCTGAGATAGGAGTGACCCACTTTGAAATTGGAGGAAGGGGCCATGAGTGTCAGAGTAC | 120 |
| chr11 | 71195375 | 71195495 | 788855_47188552_ENST00000601538.6,E | 4 | - | good | AAAATTATGCTTTCAGTATAAAATTTCATGTCTCCATCAATCATTCACCTGCATATTTTGGCTGTGATGACAAGATTAATTGGTTAGGCACTTTTTTTTTTTTTGAGACGGAGTCTCGCT | 120 |
| chr11 | 71214095 | 71214215 | 788855_47188552_ENST00000601538.6,E | 4 | - | good | ACATTTCCCAAGAGCAGGGGGCATGAGCCCTGGGGCCACAGGGGAAGCCCCAGGGTTTGGTCAGGAGGCAGGAGGTGGGAGGAAACTGAGGCCAGACCCTTTCTGGGGGTTTCTGTGAGA | 120 |
| chr11 | 71233535 | 71233655 | 788855_47188552_ENST00000601538.6,E | 4 | - | good | TCTTAGAGAATTTGTTGAGATATAATTTACATAACATTGTGATTCACCCTTTGAAAACCTGCAGTGCCGTGTTTTTTAGTAAATTAGGGAGATTGTGCAACCATCGGAGTCGATTTAGGA | 120 |
| chr11 | 71234375 | 71234495 | 788855_47188552_ENST00000601538.6,E | 4 | - | good | TCAGAACTCCTCACAATGATGGTTTCCACCTCTGTAAGAGTGATGAGCACCTCCAAAAATAGAGGTGCATCTGCAGGCTAACATTTTGTTGTTATTTATTTATTTATTTATTTATTTATT | 120 |
| chr11 | 71236655 | 71236775 | 788855_47188552_ENST00000601538.6,E | 4 | - | good | TGTTGTGCTTATGTGCTGTACAGTTTGTCTCCAAAAAAGCAGTCAAAGGCAATACATGAGGGAATGGACACAGCTGTATTCCAGTAAAACTTTACTTATAAAAACAGGCCTGGGCTAGAA | 120 |
| chr19 | 50683588 | 50683708 | 788855_47188554_ENST00000293441.6,E | 4 | - | good | GGTGCCTTCTCATTTCGGCTCTTATACTGTGAACGAGTGTCTTTTCTCGGTCTGTTTAGTGCCACGTGCTTTGCATCTTTGGGCCTTTTAGTGGTGATATCAGCGTTTAAAAGGTCCCCC | 120 |
| chr19 | 50706508 | 50706628 | 788855_47188554_ENST00000293441.6,E | 4 | - | good | GATTCCAAGCAGAGGGAGCAGCCGTGCAAAGGTCCTGGGGCATTAGTGTGCTGTCAGAGGAAGGAGGTGGCCAGTGTGGCTGCAGCCTGGGAGCAAGGGCAGGTGGTAGAAGATGAAGTT | 120 |
| chr19 | 50719708 | 50719828 | 788855_47188554_ENST00000293441.6,E | 4 | - | good | AGGGAGGGAGAGAGACAGGGTGAGGGAGAGACAGCGAGAGCGAGAGAGCGAGAAGGGTGGCAGAGGAGGCGCGGAGCGGGCGGCGGCGGCGGCCCGAGGAAGAGGAGGAGG | 120 |
| chr11 | 70520255 | 70520375 | 788855_47188552_ENST00000601538.6,E | 3 | - | good | AGGTCAACAAAACAAAGTCTAAGAAACCATCCCAGCAGGCAGAGCCTAGAGCAGCAACTAAACGTAATGTGGTGCTCTGGACAGATCCTGGGACAGACAAGGGACGTCAGGTAAAAATTA | 120 |
| chr11 | 70533815 | 70533935 | 788855_47188552_ENST00000601538.6,E | 3 | - | good | GAGACGTGCCACCTTGTGGGTGAGCATGGAAGGGGTGACACTGAGTAAAAGAAGCAGACACAAAAGGAGACATGGTGCACAGTTCCATTTTCATGAAATGTCCAGGACAGGCCGATTGGT | 120 |
| chr11 | 70536695 | 70536815 | 788855_47188552_ENST00000601538.6,E | 3 | - | good | ATAAAAAGGAAGAAAGTGATGAGACATGCCACCTTGTGGGTGAGCATGGAAGGGGTGACACTGAGTGAAAGAAGCAGATACAAAAGGTGGCACGGTGCACGGTTCCATTTTCACGAAATG | 120 |
| chr11 | 70556135 | 70556255 | 788855_47188552_ENST00000601538.6,E | 3 | - | good | GACAATGGAAAAGTATTCAGCAATGAAAAGAAATGGCTATCGAGGCATAAAAAAGAAATGAGGTGTATTGAGTCACAAAATATATGGAGGAAACTTAAATCCGTATCATTAAGCAAAAGA | 120 |
| chr11 | 70561055 | 70561175 | 788855_47188552_ENST00000601538.6,E | 3 | - | good | TCTCAAAAAAATAGAAACCTCTAGCCAGGCTGATCCAGAAGGAGAAAAGAGAAGACACACATAGCCAACAGCAGAAATGAGAGAGGAAGAATCCTGCAGATTCCTACTCTGTTGAATTGC | 120 |
| chr11 | 70564055 | 70564175 | 788855_47188552_ENST00000601538.6,E | 3 | - | good | TCAATAGACTCCAAGCACAAGACACATGAGAGAAAAAGAAACAAAAACACACCCAGGCACATAGGATTCAAATTGCTCCAAATTAGAGATAAGGACAAATATGTTAAAAGCAGCCACATG | 120 |
| chr11 | 70584455 | 70584575 | 788855_47188552_ENST00000601538.6,E | 3 | - | good | GAGCTCTGGGGCCTAGGGCCTGTCCCTGGCTGTCCGGTACCCAGCCCTGGGGTGATGGGGGCAGGGTTAGTGGCTGGGAGCATGGGACCCGGATAGAGGAGGGGTGGGGTGGCCTCTGT | 120 |
| chr11 | 70594415 | 70594535 | 788855_47188552_ENST00000601538.6,E | 3 | - | good | ATACTGTTGATACTCAAGCTCCTTGGGTTGACTTCATGCAGATTTTTTCCAACCAAACATGGGTCAAAAATACAGTATTCGTGGAATGCAGAACCCACCCCACGTAGAAAGGAGGCTGGC | 120 |
| chr11 | 70608455 | 70608575 | 788855_47188552_ENST00000601538.6,E | 3 | - | good | TGCTGTGTCCTGCCCACCTGCAGAGCCTTTACATGACCCCATTTTACAGATGAAGAAGCTGAGGCTCAGGGAAAGGCTGGGTGACTTAGCCCAGGTCCCAGAACTCAGAGGTGCAGACCT | 120 |
| chr11 | 70623695 | 70623815 | 788855_47188552_ENST00000601538.6,E | 3 | - | good | TCTGGTAGAACTTTCTGGAAATGTCCCATGTGCTCGATATGGCACCACTGGCCACACACAGCTACTGAGCACTCAAAATGCAGCCAGTGTGGCTGAGGAGCTAATTCTTAGTTTTACGTA | 120 |
| chr11 | 70655375 | 70655495 | 788855_47188552_ENST00000601538.6,E | 3 | - | good | TAACCCTCACAGTCATCCATATAGCATGTGTGTTCCTCCCCATTTTACAGATGGGGAAACTGAGGTACATAGGTGCAGTAACTACATGGCCATGTAGCTGGTGAGTGGAGAAGGTGGAAT | 120 |
| chr11 | 70686455 | 70686575 | 788855_47188552_ENST00000601538.6,E | 3 | - | good | CTGGCAAATGTCCAGCTCTCAGAGAATGGAGAGTTGAGTGGAGAGAGAGGTGAGAAGATGGATGGATAGACAGATGATGGATGGCTGGAAGGAAGGAAGGATAGATGGGGAGCTGGATGG | 120 |
| chr11 | 70689695 | 70689815 | 788855_47188552_ENST00000601538.6,E | 3 | - | good | TTTTTCTTTTCAGTTCTCTCGGATTTCGCTTCATGTATTTTGAAGTTCTGATGTTAGGTTTGTACTCATTCAGGACTGTTTTGTCTTCTTGGTGATGGTCCTAGGCCTGGAAGTTCAGGT | 120 |
| chr11 | 70723295 | 70723415 | 788855_47188552_ENST00000601538.6,E | 3 | - | good | GAGACTGCTCTCAGCAACCAGAGGCCACCTGTGTCTCCTTGCTAGATGGGGTTCCCCAAGGCAACTGCTTGCTTCATCAGGCCAGCGAGAGAGAGAGAGAGAGAGAGAGAGAGAGACAGA | 120 |
| chr11 | 70746455 | 70746575 | 788855_47188552_ENST00000601538.6,E | 3 | - | good | CCCAGCCTTCCCGTCCTGGAGAAGCGGGAGAGTGGTGGGAGAGGAGATGCGTGCTGTGAGCACGGGGCACCCCGCATCCCCCGCCCTGAAGGAAGGTCCAGCCTTCCCTGTCCTGGAGAG | 120 |
| chr11 | 70752335 | 70752455 | 788855_47188552_ENST00000601538.6,E | 3 | - | good | TGAGCCACCGCGCCCGGCCGAATTGATTTTTTTTAATCATTATTCAATGTTCCTATTTATCCCTGGCAATTTTCCTTGCACTTGAGACTACTTTGATATTAATATAGCCACTCTGGCTTT | 120 |
| chr11 | 70758695 | 70758815 | 788855_47188552_ENST00000601538.6,E | 3 | - | good | TTATTTTTAGTTCTCACATGTGAAGATGGAGGCAGAAGTTGGAGGGATACGGCCACAAGCCAGGGACTGCTGAGGGGTGCCAGCAGCCCCAGAGGCTGGGAGAGAGGCTTGGGGCAGATT | 120 |
| chr11 | 70760495 | 70760615 | 788855_47188552_ENST00000601538.6,E | 3 | - | good | GTCCTGTCAGGCTGAATGCTGGCCCCACCCATGACCAGTATGCGTCCTAACTTCCAGGTGCTATGAGTCAGACCTTATTTGGAAAATAAAAGCATCTTTGCAAATGTAGTTAAGGATCTT | 120 |
| chr11 | 70821695 | 70821815 | 788855_47188552_ENST00000601538.6,E | 3 | - | good | AGATTGAATCATCCAGCCAGTGGACTCGGAGACTGTATGGTTGGCTCAGGCGTGGCAGAGCGCCACAGACAGGTGGCTTAAACTCCAGATATTGGCCGGGCACGGTGGCTCATGCCTGTA | 120 |
| chr11 | 70823855 | 70823975 | 788855_47188552_ENST00000601538.6,E | 3 | - | good | TTGGGAGCTGTCAATACCACCTCTGTACCTGTGAGCTCTGCTAGTGCCACCTTTGTCCTCAGTGAAATCTACCAGCACCACCTTTGTCCTCATGTGCTTTGCCAGTGCCACCTCTGTCCC | 120 |
| chr11 | 70843295 | 70843415 | 788855_47188552_ENST00000601538.6,E | 3 | - | good | GCGCCAGCCATCCATAAACTCTTCCCAAAAGCCACCAAGACTTGCTTCCCTTCCTCCTCCTGGGGCCCACCAAGCCCACCCCATAGGAGCCCAGCCCACCCCATAGCAGCCCAGCCCACC | 120 |
| chr11 | 70863935 | 70864055 | 788855_47188552_ENST00000601538.6,E | 3 | - | good | TGCCCAGGTCGTAGAGACTGAGGGGCCAGGACAGGGGAGGCCATGGAGCCAGTTTTTGTTTGCTGGGGCTGCCATAAGAAAGTATCAAAAACTGGGTGGTTTTATCAGAAACATTCTCTC | 120 |
| chr11 | 70889015 | 70889135 | 788855_47188552_ENST00000601538.6,E | 3 | - | good | AACTCATCACACCTGAAAAGACCCTGTTTCCAAATGAGGGTGCAGTTTAGGTCCCACGGATTAGGACTTGGGCCCATCTTTTTGAGGACCACCATTCCACCCACTCTAGGGAACTCACAT | 120 |
| chr11 | 70921895 | 70922015 | 788855_47188552_ENST00000601538.6,E | 3 | - | good | TACTAATCATTTTTAAGTTCTTCGCTGCTCACTCCAATATCTGGATCATCCATGGGTCTGTTTCTATTGACTGTTTTTTCTCTTTGTTTTCCAACACATGGTACTGTCTGTGGGAATGAC | 120 |
| chr11 | 70941455 | 70941575 | 788855_47188552_ENST00000601538.6,E | 3 | - | good | ACACCCCTAAAAGGTCCCAATCCAAATGCCATCCCCTTGGGGATTAGAGCTCTGACATATGAATTTGGGAGAGTTGGGAGGGGACACACAATTTAGTCCATAGCGACCCCAACAGCAAGT | 120 |
| chr11 | 70951895 | 70952015 | 788855_47188552_ENST00000601538.6,E | 3 | - | good | AATGTGATATTGTTTGGAACTAGGGCCTTTGCAGATGTCACTGAGTTAAGACGACCTCATGCTGGATTGGGGAGGCCCTAATCTTGAAAGATGTCCTCTTAAGAAAAGCGAAAAGAGACA | 120 |
| chr11 | 71075135 | 71075255 | 788855_47188552_ENST00000601538.6,E | 3 | - | good | GCAGCACCTGGAGCACCTGCTGTTCTACGGGGCAGACATGAGTGCCCAGAATGCCTCGGGGAACACGGCCTTGCACATCTGCGCCCTCTACAACCAGGTGAGTGCTGAGCGCATTTAAAG | 120 |
| chr11 | 71078135 | 71078255 | 788855_47188552_ENST00000601538.6,E | 3 | - | good | TCCCAGTTTTAGCCAGTGGGTGTCCCTTCAGGATGGCTCCTGTGTCCTTGGGACACAGCCCCATTGATTTTTTTTTTTTTTTTTACCATTTCCTTTATTTCTGGCACAACAAGATATTTCA | 120 |
| chr11 | 71099135 | 71099255 | 788855_47188552_ENST00000601538.6,E | 3 | - | good | GTCCTTTCCTGTCCCTGGGCCCCACCCGGGTTCCCACATGACATTCCCTCATTCGTCTCCCTGGGCACCTCTTGGTGGTGATTGTTTCCCAGGCTCTCCCTGTTGGTGATGACCTGGACA | 120 |

| | | | | | | | | |
|---|---|---|---|---|---|---|---|---|
| chr11 | 71105735 | 71105855 | 788855_47188552_ENST00000601538.6,E | 3 | - | good | CTGTACCCAGTAGTGATCCTCAAGTCCGGAGCTTACATGAGGGTTTACGCCTGGAGTTGTACCCAACAGTGATCCTCCAGTCCGGAGCTTACATGAGAGTTCACTCTTGGTGTTGTACGT | 120 |
| chr11 | 71110775 | 71110895 | 788855_47188552_ENST00000601538.6,E | 3 | - | good | CCCAAAGGCCTCACCTCTCCATACTATTACATTGGAGATTATTCAACCTATGAATTTTGCGGGGACACACACGTTAAGTCCGTAGCACTAGGATGGCGAACGGTGAGGGGATTTTAAACA | 120 |
| chr11 | 71111015 | 71111135 | 788855_47188552_ENST00000601538.6,E | 3 | - | good | AAAATACCTTGGGTAACTTATAAACAACAAAAGTGTGACTCTCACCATTTTGGAGGCTGCAAGTCTGAGATCGTGGCATCGGCAGGTTCAGCATCTGGCCAAGGCCCAGTTCCTGGTTTG | 120 |
| chr11 | 71118815 | 71118935 | 788855_47188552_ENST00000601538.6,E | 3 | - | good | GCCTGTTCCAGCCGGCCAGCAATGGGCGTGACGGCAAGTTCCTGGATGAGGAGCGGCTCCTGCGCGAGTACCCACAGCCCGTGGGTGAGGGCGTTCCTTCCCTGGAGGTATATTTCAGCC | 120 |
| chr11 | 71134295 | 71134415 | 788855_47188552_ENST00000601538.6,E | 3 | - | good | TAATTCACCCATTGGAAGTGTGCAAGTCGTTGGATTTTAGGAAATTCACGGATGAGTGAGCCCATCACCACTGTGAATCTCAGGAAGCTTCCATCCCCCCAAGAGGAAACCCCACCCCTG | 120 |
| chr11 | 71146295 | 71146415 | 788855_47188552_ENST00000601538.6,E | 3 | - | good | AGGAAGGGGTGTGGCTTCATCTTGTCTGTAAAGGGGAGCTTCGTGTGCAGAGGAAGGGGTGTGGCTTCATCTTGTCTGTAAAGGGGAGCTTCGTGTGCAGAGGAAGGGGTGTGGCTTCAT | 120 |
| chr11 | 71160935 | 71161055 | 788855_47188552_ENST00000601538.6,E | 3 | - | good | AAAGAGCAGCTGGGGGATCCATTCAGTTGGTGGTGGACTTAGAATTTTATTTCTGGTTAACAAAAGTAAACTTCTTAAAGAAGGCTGTCTGCAAACTTAGTTTTCAGTTTTGAGGCAACT | 120 |
| chr11 | 71163095 | 71163215 | 788855_47188552_ENST00000601538.6,E | 3 | - | good | TATAAACACAGAAAAGTAGACAAACACAATAATGACTCTATACCCACCATCAGTATTAAACATTTTAAGATTTTGCCCTATTTGCATATTTGTTTCTATTCTGTATATATATATATATGT | 120 |
| chr11 | 71163215 | 71163335 | 788855_47188552_ENST00000601538.6,E | 3 | - | good | CCCTGTGCCATGGTCATCTAATTTCTAAAGTTTTTATTTTTAGTGAAAAACATTATAAACACAGAAAAGTAGACAAACACAATAATGACTCTATACCCACCATCAGTATTAAACAAACAT | 120 |
| chr11 | 71187575 | 71187695 | 788855_47188552_ENST00000601538.6,E | 3 | - | good | CACCAAGTACTGCCTCCCACTGAGGCTTCTGCCTGGGTCTGTACATTCCTGACATGAAAACCCATGAAGAACGGTTCCTGGGCAAACATGAGAAACCCTTGAACTGTGCTGAATAAAACA | 120 |
| chr11 | 71187695 | 71187815 | 788855_47188552_ENST00000601538.6,E | 3 | - | good | CAGAGGAAGACACTGTCTCTAACTTCCAAGGCTGCTCGCAGTACAAACATCTTAGAGGATGGGCCAGAAAAAGGCAGAGTCAGTGCCTCTGCCCACGCAACACACAGAACAAGAGAAACC | 120 |
| chr11 | 71203415 | 71203535 | 788855_47188552_ENST00000601538.6,E | 3 | - | good | CTCTGCCTCCCCAGTTCAAGCAATTGTCCTGGGGACCCAGTCTTTCTTTGGAGTGTGTGGGGTTTGGACAACCCAGGCCTGCTGAGTTAACCCTTTATTGCACTACAGTATTAAAATGTC | 120 |
| chr11 | 71211575 | 71211695 | 788855_47188552_ENST00000601538.6,E | 3 | - | good | AGCTTGAGGAGTGATGGCTAAGGGATACAGGTTAATTTTTTTTTTTTTTTTTTTGCAAATGCTCAAAAATGGATTGTGGTGATAGATACCCTACTCTGTGAACTCACTAAAACCCACTGAA | 120 |
| chr11 | 71225135 | 71225255 | 788855_47188552_ENST00000601538.6,E | 3 | - | good | GCCACAGATTTTTCTGCAGTGTCTGGCTGGAGTAGACCTGTTGTCTAAATATATTGCCCTGTTAGCTGCCGCTTTCCTAGTCTTTCGACTAAAGAGAATATGCTTTTGTTGTGGTACTGT | 120 |
| chr11 | 71225495 | 71225615 | 788855_47188552_ENST00000601538.6,E | 3 | - | good | GTTTGATGGGTGATTTTCTAATGAAACTTGGACATATTTGTATTATGTTATGAGACTTTTAGTCTTAGTTAAACTTTTTTTGTAAGCTAACTTTTCCTGACGCCACTTCCACAAGGGAAG | 120 |
| chr11 | 71228375 | 71228495 | 788855_47188552_ENST00000601538.6,E | 3 | - | good | ATGAGCCACCATACCCAGCCTGTAATCTTCTTTTTGTGCTCATTTTCTCTGGCTTTGGTATCAAGGTAGTAGTTGCTCCATAAAATAAATTAAGAAATGTTCTCTCCTATTCTACTTTCT | 120 |
| chr11 | 71235095 | 71235215 | 788855_47188552_ENST00000601538.6,E | 3 | - | good | GTGACCAGTCCCAGTCTGAGACTGGGGCCCCCCACCCTAAGTCACCTCATTAGCATAAGCTCCAGTGGGGTCAAAAGGGACTCACTTCTGCTACTCAGGAAATTCTAAGGGTATTAGGAT | 120 |
| chr19 | 50692348 | 50692468 | 788855_47188554_ENST00000293441.6,E | 3 | - | good | ATATATATATATCTGGTGAATTCAAATCTTGGCTTTGACATTTTTCTAGCTGGTGTCATCTTGAGCAAGTTATTTCTCTGTGCCTCGGTTTCCCCATTTATAAAATGGGAAGAATAACAG | 120 |
| chr19 | 50700388 | 50700508 | 788855_47188554_ENST00000293441.6,E | 3 | - | good | ACCTCATCTCATGCGACTCTTCCTTTCACCCACAAGGTTCCAGGGACCTCATCCTACTCTCCAGTCCTCCAATGCCCCAAGCCCTCCAATTCTCCAATGCCCCAAGTCCTCCAATCTCCC | 120 |
| chr19 | 50712748 | 50712868 | 788855_47188554_ENST00000293441.6,E | 3 | - | good | AGTAAAATTAAAATTTCAGTTCCTCAGGCTCACTAGCCACGTTCAAGTGCTCAGGGGCCCACACGTGCCTGCCATATAGGGTGGTGCAGTTGCAGACCATCCCCATGATGGGGGAAACGT | 120 |
| chr11 | 70473575 | 70473695 | 788855_47188552_ENST00000601538.6,E | 2 | - | good | GGTCCCCCAGCCCCTCTGTCCCAGTTCAGGTGAGTGGGCTGACGTGGTGTGCTCTGGTGTCTCCAACTTTGCTCTGGCCCCTTTCTCCCCCTCCCCCACCCTGGCATGGCAAATCATTCG | 120 |
| chr11 | 70473695 | 70473815 | 788855_47188552_ENST00000601538.6,E | 2 | - | good | TCAGTGGCCACCCTCCCGAGGGTCCCCCGGCCCTGCCCCCCAGCACAGGCAGGTCCCCCAGCCCCTCTGTCCCAGTTCAGGTGAGTGGGCTGACGTGGTGTGCTCCCCCCAGCACAGGCA | 120 |
| chr11 | 70484135 | 70484255 | 788855_47188552_ENST00000601538.6,E | 2 | - | good | CTACAATTTCACATATGAGATTTGGGTGGGAACACAGATCCAAACCGTATCAGGGCTCAGCAGTCTGTGGCTTTGGGGAGCCCCATTGGCTCCCGTGATGGCTTATGGTTCTTGGTTCTC | 120 |
| chr11 | 70512215 | 70512335 | 788855_47188552_ENST00000601538.6,E | 2 | - | good | TTTTTTAAAAAGAGCCAAATTAGTGGTTCTCAACTGATGGTGTCTTGACCTTCCATAGGACATTTGGCAGTATCTGGAAACATTCTTGATTTTCACGCTTGGGGGTGGAGATTATTGCTG | 120 |
| chr11 | 70512815 | 70512935 | 788855_47188552_ENST00000601538.6,E | 2 | - | good | TTTATACCCAGCCAAACTACCAATTAAGTGTACAGATAAAGCCATTTTCAAATAGTCAATGTCTCAAAAAAATTATGTCCTAACTCTATTTGGGGAGAAGATACTGGAGGATGTCCTTTA | 120 |
| chr11 | 70513775 | 70513895 | 788855_47188552_ENST00000601538.6,E | 2 | - | good | CTCGTTCACGTTATGTTAGACCTTTTGATCCTTAATACTCTCTGTTCTACTTTTTAAAATCTTTTTTTCTCTCTTCTTTGGATTGGATATGTTCTCTTGGCATGTCTTCAAATTCACTAA | 120 |
| chr11 | 70520495 | 70520615 | 788855_47188552_ENST00000601538.6,E | 2 | - | good | AAAGTAGGGAAAGGAGAGTAACTTCACAGCGGGGGAGCCCAGTGAGCCCTCCCTTGGCCAGGCGAGCAGGGCCAGCACCAACAGCGAAAAGTCTTGGTGACAGTGTGTGCCCTCGATACT | 120 |
| chr11 | 70520615 | 70520735 | 788855_47188552_ENST00000601538.6,E | 2 | - | good | AATAGACAAATCTTCCCTAACATGTGCAAAGATACTCTGTCCTCAAGGAGGTGGTGTGTAACTCCCCACCCCTTAGGTGTGGGCAGTATGCAGTGACTTGCTTTCGAGAGCACTGTGTGG | 120 |
| chr11 | 70520735 | 70520855 | 788855_47188552_ENST00000601538.6,E | 2 | - | good | AGACCACGGCCGGGACAATTTGAGCTAGAAAATAAAGTCACATTTGGTTATCATTCAAAGTACAACATGAGTATTCATGGGTCCACACTGATAAATAAACAACTGAAAGATAAAGGGAGG | 120 |
| chr11 | 70526495 | 70526615 | 788855_47188552_ENST00000601538.6,E | 2 | - | good | AGTGGGGCTGGAATGAGGATACGCTGGCTAAGCACACTCAGCAGGTTATAGAAGCCATGAGTATTCACACAGGGGCCCTGACGCGTGTGCGCTGAGCACACATGCATGGTACACGCCTCC | 120 |
| chr11 | 70530215 | 70530335 | 788855_47188552_ENST00000601538.6,E | 2 | - | good | TCGAACTCCTGGGTGCGAGGATTGTTTCAGCCCGGAAGTTTGAGACCAGCCTGGGCAACATAGTGCAGAGATTACAGGCATGAGCCACGGTGCCCGGCCTCCTCCTTTTCACGGCTGAGT | 120 |
| chr11 | 70532975 | 70533095 | 788855_47188552_ENST00000601538.6,E | 2 | - | good | GCCCCTCCCCCCAGCCTCCGGCAACGTCTGCTTTCCGCCGCTGTGAATTTGGCTACTCTGGACATTACATAGAAATGCAATCATGCAAAGTGGGGTGCTTCGTGGCTTCTTTCATTCAGC | 120 |
| chr11 | 70533935 | 70534055 | 788855_47188552_ENST00000601538.6,E | 2 | - | good | CCTCCTTCACCAAGAGACACTGGGCTGGGATGAGCTCAGACCAGCCATGTACCACAGCTCTTGGAGAATGGGCACCCATGCAGTGGAACACTACTTGGCCATAAAGAGGAAGAAAGTGAT | 120 |
| chr11 | 70535495 | 70535615 | 788855_47188552_ENST00000601538.6,E | 2 | - | good | CAGAGGCAACCTGACAGACATGAGATGCACCAGCAGACAGAAGGACGGGGAGATGGACGAACAGACAGATGAATAGACTGATGGATGATGGTGAATGGATGAATAGATGGACGGACGGAC | 120 |
| chr11 | 70536815 | 70536935 | 788855_47188552_ENST00000601538.6,E | 2 | - | good | CTGTCCTCACCCCATGGTTCTCTCCTTCACCAAGAGACACTGGGCTGGGATGAGCTCAGACCAGCCATGTACCACAGCTCTTGGAGAATGGGCACCCATGCAGTGGAACACTACTTGGCC | 120 |
| chr11 | 70542455 | 70542575 | 788855_47188552_ENST00000601538.6,E | 2 | - | good | GGCCGTGTCCCATAAGCAGGGCTGAGCACAGCCCACATGCACCTCCAGGGGCTGTGCTGCAAACGACACACATGGATTTTCTCACAGTTCGGAGGCCAGAATTCCAAAATCAAGGTGTGG | 120 |
| chr11 | 70545815 | 70545935 | 788855_47188552_ENST00000601538.6,E | 2 | - | good | ATGCATGTTCCCGAGTATTGTGAATGGTTCTAGCAAATATCAGACCTGAGCAAGGGTTATGGGAGCTCTGACTCGTAGTCAGCCAGGCAGAAGTGTGGGTCATCTGGGGAGCCACTTCTT | 120 |
| chr11 | 70554575 | 70554695 | 788855_47188552_ENST00000601538.6,E | 2 | - | good | CAACGAATATACAATGCCCAGAAGGAACCCTCTGATGACTGTGGATTTAGTTAATTATAATGTGTCAGTATCGGTTCATTAATTGCAACAAATAACAGCCGATTCATGCAAGGTGTTACT | 120 |
| chr11 | 70560215 | 70560335 | 788855_47188552_ENST00000601538.6,E | 2 | - | good | TGGCACATCTTTTGTTCTATTTATTCCTAGTTTGTATTTTTTGAAGATATGGTAGAGGTATTTTCGAATTTCTGTTTGTGATCATTTTTTGCTAGTATATAGAATTACTCACCGGGGTGT | 120 |
| chr11 | 70565135 | 70565255 | 788855_47188552_ENST00000601538.6,E | 2 | - | good | AATAAATGGATGGATGGATGGATGGATGGATGGATGGATGAAACTGCTTCCAGGTGACTTAACTACACCCAGAACAAAGCTCAAGACCAGGATACAGAACTACCCAGCGCCCCCAACAAG | 120 |
| chr11 | 70571855 | 70571975 | 788855_47188552_ENST00000601538.6,E | 2 | - | good | GCTCAGTCTCCTCCCAGAGGCCTCACCCCCAACACCATGCCATTGGAGGTTAGGGTCCAACCCAGGAATGTTGCAGAGACATGGACATTCAGCCCACAGCACCCAGCCCTACCCCTCCTC | 120 |
| chr11 | 70573295 | 70573415 | 788855_47188552_ENST00000601538.6,E | 2 | - | good | AGTTTTCACGTGTGTATTCTTGGGAAACCATCGCCAAGGTCAAGGTGACAAGTCCATCCACCAGCCGGCTTGCTCCTGCCCCCGCCCCCCCCCCCCACCGCTGGAGTGACCACCCAGCTC | 120 |
| chr11 | 70579535 | 70579655 | 788855_47188552_ENST00000601538.6,E | 2 | - | good | ACAGGACAGGTCCCGGGGACACCGGGGGTCCTGGGATCCGGCCCAGGCCCGGGAGACACCACCTGGGGCCCAGCTCCATCCATGCCTGTCTTACATTCATGCCATCACCGCCCAGGCCTG | 120 |
| chr11 | 70579775 | 70579895 | 788855_47188552_ENST00000601538.6,E | 2 | - | good | CTCCAGCCAACAGCGTATCCCTCCAACTTCTGCTTCCATCCTCACCTCTCCAGCTCTGCTCCTCCCGGCTCCTCCTCCACTTATCAGGACGCTTGTGATGACGCTGAACATCAGGGCCAT | 120 |
| chr11 | 70580015 | 70580135 | 788855_47188552_ENST00000601538.6,E | 2 | - | good | CAGCTGTGCACTTCCCGTGGCTGAGGCATTGGCAGCAAGTCACCACGAACCTGGTGGCTTCCAGCAACACGTGTTTATTATCTTAGCGTTGTGAAAGCCAGAAGTCCAATGCAGGGCTCA | 120 |
| chr11 | 70592735 | 70592855 | 788855_47188552_ENST00000601538.6,E | 2 | - | good | TCCCTCTGCTGGACATCGGGTCCCTGAAGGTGCATGCAGAGGGACATGGCTGCAGGGCTGAAGGAGCATGGATGGGAATCCGCAGCTGCCTTGGGCCATAGCAAAAGGGCAGACGCAGCC | 120 |
| chr11 | 70593815 | 70593935 | 788855_47188552_ENST00000601538.6,E | 2 | - | good | GCAGGTCCCACATGTGCATCAGTCACTATTGTATCACCAGAGCCTAGAACAGCATCTGGACTGTAGCAATTCATTGAATGAATGAATGAATGAATGAACGAACGAACGAAGGAACTAATG | 120 |
| chr11 | 70594775 | 70594895 | 788855_47188552_ENST00000601538.6,E | 2 | - | good | GAAGGTGCATGCAGAGGGGCATGGGCTGCAGGGCTGGAGGAGCATGGGTGGGAATCCGCAGCTGCCTTGGGCCATAGCGAAAGGGCAGACGCAGCCACTGTTGCTTAGGTGATGGGCAAC | 120 |
| chr11 | 70596935 | 70597055 | 788855_47188552_ENST00000601538.6,E | 2 | - | good | TAGGGTCTGAGGATGTCTGCGCTTGCAAAACAGATATAGACCAGTCACCCCCGAGCACCGGTGCACAAGTATGCAGCTTTGTTCATAGGATCTGAGGATGTCTGCGCTTGCAAAATAGAT | 120 |
| chr11 | 70597055 | 70597175 | 788855_47188552_ENST00000601538.6,E | 2 | - | good | GGGCTCAGTGACCTTGTGCCCATTTTGTGCGAAACCTGCTCCTCTGTGCTGCTGCGCCCCTGAGTCTGGAGGGAACAGTCACCCCCTGAGCACCGGTGCACAATTATTCAGCTTTGTTCA | 120 |
| chr11 | 70600775 | 70600895 | 788855_47188552_ENST00000601538.6,E | 2 | - | good | AAATTATCTTCTCCCAAGAGTTTGTAAAGCTGTTTCCCAATCTTATTTTCCAGGAGTTTAAAGTTTGTCTTTCATATGTAAGTCCTTAATCCAAGAATTCAGATTGTGTGTGTATGTGTG | 120 |
| chr11 | 70601735 | 70601855 | 788855_47188552_ENST00000601538.6,E | 2 | - | good | GTCAATCTGGGAGGTACTTAATAATTTTTCATTGCCATTTGAATGTGCGTTTCCTGATTACTAATGGACTTGAGCATCTTTTATGTTTCTGGACTGTTTGTGGTGCCTTTGCTGGGAAAT | 120 |
| chr11 | 70601855 | 70601975 | 788855_47188552_ENST00000601538.6,E | 2 | - | good | TGAGATTTGGGCAGGGTTAAATATCCAAACTGTATCAGCTGCGTTCCAGCCACCTGGGTGGCAGGTCTCCTGGCTCCATGCTGCAGCTGCCCCTGTTCTGGTTCAGACCTGGCAGCTTTT | 120 |
| chr11 | 70614215 | 70614335 | 788855_47188552_ENST00000601538.6,E | 2 | - | good | CTGTCTCAAAAAAAAAAACAAAACCCACAGACTGTCTGGCTTAAAACAGCAAACTTTTACTGTGTCATTGGATGCTGGAAGACAAAGGTCAGGGTGTCAGCAGGGCAGCGCTCCCTCCAG | 120 |
| chr11 | 70619375 | 70619495 | 788855_47188552_ENST00000601538.6,E | 2 | - | good | GGGCCTGAGTGTGACCACAGGCGTCCTTATAAGGGGGAGGCAGAGGTAGATTTGATGCATGCAGGGGAGAAGTCCATGGGACCCCCCAAGGCAGAACTTGGAGTAATGAACCCTCAGCTA | 120 |
| chr11 | 70621415 | 70621535 | 788855_47188552_ENST00000601538.6,E | 2 | - | good | CCTGCCATGGGCCACGTCCTCGCCAAGGGTGCTGTAGGGGCACAGCCTTTGGTGTGCATCTGGCAGTGTGGTGGGGCAGAGGCAGGAGCCAGGCCAGAAAGAAATGGGCAGATGGTGTGC | 120 |
| chr11 | 70621535 | 70621655 | 788855_47188552_ENST00000601538.6,E | 2 | - | good | GTATCAGACCTCCCATCTGATGAAATCACCTTCCATCTGGGAAATGAAGGAGCTGGGAAGAATTTCCAGTCCCCTCTCTCCTTCCTTCCTTCCACCCAGTCCACCCGTATTTCCCAAGCA | 120 |
| chr11 | 70623455 | 70623575 | 788855_47188552_ENST00000601538.6,E | 2 | - | good | GTCCCCTCTCTCCTTCCTTCCTTCCTTCCAGCCAGTCCACCCATATTTCCCAAGCACCTGCCATGGGCCGTGTCCTCATCAAGGGTGCTGTGGGGGCGCAGCCTTCGGGGTGCATCTGGC | 120 |
| chr11 | 70623575 | 70623695 | 788855_47188552_ENST00000601538.6,E | 2 | - | good | TTTAATATCAATTAATTGAAACTTAAGCAGCCGCGGGTGGCTGGGGCTACTGTATCAGACCTCCCATTTGATGAAAACCACCTTCCATCTGGGAAATGAAGGAGCTGGGAAGAATTTCCA | 120 |
| chr11 | 70625255 | 70625375 | 788855_47188552_ENST00000601538.6,E | 2 | - | good | AAGGATTAGCCCTAACCTCCAGGAGGAGCTGGAAACTAAAGCCCTGCCACCAAGGCAGTGTGTGACAGAGCCCCAGCAAAGACTCGGGACACCAAAGGTCTGGGAGCATCCCTGGCGGCA | 120 |
| chr11 | 70625375 | 70625495 | 788855_47188552_ENST00000601538.6,E | 2 | - | good | TGTATGTGTCTCTGTTTGCCTGGGGGCTTTGACTGCTGGACAGTCTGACAGTGTGATTTATGGTGGGAGCTCTGGGTCATGCCACATCAGCTTGGACCTCCAGAAGAAACTGGAGACCAG | 120 |
| chr11 | 70626455 | 70626575 | 788855_47188552_ENST00000601538.6,E | 2 | - | good | ACGAGCTCTGCCTCTGGCCGAGGACTAGAGCTGTCCATCTTTCCTGTTTCCTGCAGTCCAATGGGCTTTGTGGCGTAAAGCTTGCCACCTGGGGGTTGTCGAACTTGACCAGATGAGACT | 120 |

| | | | | | | | | |
|---|---|---|---|---|---|---|---|---|
| chr11 | 70626575 | 70626695 | 788855_47188552_ENST00000601538.6,E | 2 | - | good | AGTAGCCTCTGGTCCAGGGCTGGAGCCATCGGTCTCTCCTGTTTCTTGCAGTCCAGTGGGCTCTGTGGCATAAAGCTTGCCACTCGGCGGTTGTGGGATTAGACCAGATGAGACTGACTG | 120 |
| chr11 | 70626695 | 70626815 | 788855_47188552_ENST00000601538.6,E | 2 | - | good | AGGCCAGCTCTGCCTCCTGTGGGTGGGCCACCTTGGACTCCTCCTCTGCAGAATAAGGGGCTGCATCTGGACCTTTCAGGGCTGCTACAAGGATTGGCCTTTCAGAGGTACTACAGGGAT | 120 |
| chr11 | 70627415 | 70627535 | 788855_47188552_ENST00000601538.6,E | 2 | - | good | ACCTGACCTGTAACAGTCAGGACAGCGGGGTTAGGCTGCAGTAGCAAATAATCTCCAGGTTGCCATGGCTTAAAAAAACAAAGGCTTATTTCCTGCTCACTCTCACTGCCCTGTCAGGTC | 120 |
| chr11 | 70628495 | 70628615 | 788855_47188552_ENST00000601538.6,E | 2 | - | good | GTCGAGGCCAGCTCTGCCTCCCATGGGTGGGCCGCCTTAGGCTCCTCCTCTGCAAAATAGGGAGCTGTTGCAGGACATTTCAGAGCTACTATAAGGACTGAAGGAGGCCCCGGGGAAAAG | 120 |
| chr11 | 70652855 | 70652975 | 788855_47188552_ENST00000601538.6,E | 2 | - | good | GGAAAGAAAATGAAGTCCAGATGCAGCATGATCAGTCATTATGCTTGCTGGGTCCCAGTATTCGTGTTGATAAAATGGGTGGTTGTGCTGAACTGGGCTCTTAGCACTGGACTGCTGGTC | 120 |
| chr11 | 70653575 | 70653695 | 788855_47188552_ENST00000601538.6,E | 2 | - | good | TCAGGTGATTCTGCTACTGTCACAACCACCCAATTTATCAACAGGAATACTGGGACCCAGCAAGCCTAATGACTGATCATGCTGAATATGGACTTCATTTTCTTTCTTTTTTTTTTTTTT | 120 |
| chr11 | 70655015 | 70655135 | 788855_47188552_ENST00000601538.6,E | 2 | - | good | AACAAGAGCCATAAAGCTGAACCAGGAATTTTACATCTAGGAATCCATGCTAAGGGAATAATTTTTTAAAAAACAGGTAGAAATCACGCCTGTAATCCCAGCACTTTGGGAGGCCAAGAC | 120 |
| chr11 | 70659815 | 70659935 | 788855_47188552_ENST00000601538.6,E | 2 | - | good | GAATTCACACCAACACCGGCTTTCCCAGCCCTACAGTACCTGGAGTCCGTGGATGAAGGTGGGGTGGCGTGGCAAGCCGGACTAAGGACCGGGGACTTCTTGATTGAGGTAGGGACACAG | 120 |
| chr11 | 70668335 | 70668455 | 788855_47188552_ENST00000601538.6,E | 2 | - | good | GTCTTCCTCCTCCACCTCTGCTGTTGGCCTGCAGTCCTTGGTGCTCCTCAGCTGGGCCACATCACTCCAGCCTCCACATCACTCCAGCCTCTGCCTCCACCTTCCCATGGCCTCCTCTGC | 120 |
| chr11 | 70683695 | 70683815 | 788855_47188552_ENST00000601538.6,E | 2 | - | good | GGAGACTCTGTCTGAAAAAAAAAAAACAAAAAGGAGATCATGAAGCTAGAAGAGGAGAAAGCCACATGACCACAGAGGCAGAGATTAGAGAGATTGACCCCAAGACAAGGAATGCCTGCT | 120 |
| chr11 | 70684055 | 70684175 | 788855_47188552_ENST00000601538.6,E | 2 | - | good | GGGGCCGTGGATATGAAGAGTGAGTAGCAGGGACAGGGACCCAGGTCTCTCTCTCTCTTTTCATTCATTTATTCAAAACTATGCATCCAGGGCCAGGTGCAGTGGCTCACGCCTGTCA | 120 |
| chr11 | 70687415 | 70687535 | 788855_47188552_ENST00000601538.6,E | 2 | - | good | CATTTACTGATTTCAGTGGTGTAAATTGCCCCACAATGGCCCATTCCGAGCTGCCAACGTGAGGTCACTGAGCAAGCTGGAAAGAGGCGCCCACAGTCAGCCCTGGCAGCTCCTAAGAGC | 120 |
| chr11 | 70697255 | 70697375 | 788855_47188552_ENST00000601538.6,E | 2 | - | good | GCTGCATCATGTTTTCAAAGTCTATCCACATGGTGGCACATACCAGTACTTCATTCCTTCCCCGCCCCCAATTTTTTCTTATTGTGGTAAAATACATGTAATATGAAGTTTCCTATTCCG | 120 |
| chr11 | 70699775 | 70699895 | 788855_47188552_ENST00000601538.6,E | 2 | - | good | ACATCTCAGTGGTGAGAACTAGTCACGTGGCCACATCTGGGTTCAAGGGAGGCTGGGAAATGTAGTCCTGGGCTCAACACCTGCCTCCTTTCCTGCCCTCAAGGGCAACACCACACTGCT | 120 |
| chr11 | 70701095 | 70701215 | 788855_47188552_ENST00000601538.6,E | 2 | - | good | CTTTACTGAAATAGTCTTTTTTTCTGGTAAATACACACACACACACACGCGTACACACACACGCACACATTCCACTACACACTCCTCTCTGCTTCTGGCCCCGGCCTCCGATTTCAGCCA | 120 |
| chr11 | 70711655 | 70711775 | 788855_47188552_ENST00000601538.6,E | 2 | - | good | AGCCAGTTGGCGGCATTTTTGTGATACGTGTTGCTTTTCAAATCAAAGCTCTGTGCCTCTCAGTTGGGTAGTTCTGAGAGCCAGGGTCCCCGCCCTTCTGTCCCCCTCGCGGTGGGTCCC | 120 |
| chr11 | 70711775 | 70711895 | 788855_47188552_ENST00000601538.6,E | 2 | - | good | TGGGCATTCTGCACAGGGTCCTTATTTTCTCTTCCCCTCCCCTGCAAGGTCAGTCCCGGGAGCCCAGAAGGTGCTTGATGAATGATCTGCTGCTGATTGAGAAAAGGAACAAGTGAGCCG | 120 |
| chr11 | 70711895 | 70712015 | 788855_47188552_ENST00000601538.6,E | 2 | - | good | GCAGCTCTGTAAGGGGAGACGTTTGTGTTGTTTGAGGCCACCGAGCCTGGTCGTCTGTGCCGGCAGCTGCAGGAGCCCAGGCCACCATCCCCACGAGGTCCGTGTGCTGTGACGTGTGCC | 120 |
| chr11 | 70712015 | 70712135 | 788855_47188552_ENST00000601538.6,E | 2 | - | good | GCCTCTGGAAGCCGGGCGGGGGAAGCAAGTGGATCCCCCACACTCCAGGAGCCTCCAGCAGGAAGGAAGCTCCACAAACGCCCTAACTTCAGCCCAGAGACCTGGTCAGACTTCCAGCCT | 120 |
| chr11 | 70726655 | 70726775 | 788855_47188552_ENST00000601538.6,E | 2 | - | good | CACAGGACCAGACCAAGTTTAAGGTCTGGGCAGACAGGCTTTGTCTCTTGATAAGAGGAGCTGCAAAGTCATATCGCATACGGGTGTAGGTTCAGGGAGGGGAATAATTTTAGGTCTTGT | 120 |
| chr11 | 70731335 | 70731455 | 788855_47188552_ENST00000601538.6,E | 2 | - | good | GAAACAGAAAGTAGATGAGTGGTTGCCAGGAGATGGGGGAGAAGGAGCGGGGAGTATGTCAGTCAGCCTGGACTTTCATAACAAAATGCCACAGACTGTTGCTTAAACAACAGACACTTA | 120 |
| chr11 | 70739975 | 70740095 | 788855_47188552_ENST00000601538.6,E | 2 | - | good | AGCACCTGCTGTGTGGCAGGCACTGTCTGTCCTAGGTGCTGTGGACGGAGCCATTAACAATCCCAAAATTCCTGCCCCTGCCGGGCCCCTGGTCTGGGTCGTGTGACTGTCCAAGGTCTC | 120 |
| chr11 | 70743215 | 70743335 | 788855_47188552_ENST00000601538.6,E | 2 | - | good | CCTCCAACCCTAATCACCTCCCAAAGCCCCCACCTCCAGACACCCTCACACCGGGGATTTCGGCCCCAATATATGAACTGGGGGGGGCACGCAGACGTGCAGCTCGAGCGCTGCCCCTGG | 120 |
| chr11 | 70750895 | 70751015 | 788855_47188552_ENST00000601538.6,E | 2 | - | good | GGCATATTTCTAGTTCTTGGTATGCCAAGTAAATGAAACCTGTATCATGGACATTTGGAATATTTTGTTAAGATACTCTGGGTCTTGTTTAAACCCGATGGAATAGGTTGACATGAGCAC | 120 |
| chr11 | 70753175 | 70753295 | 788855_47188552_ENST00000601538.6,E | 2 | - | good | AGCCTTGCTGACTTGTCCTCTGTTGTTGATTGCTGAGAACGTAGGAATGTGGAAATTTTCCAGTATCATCGTGTGGCTGTCCATTTTGTTTTTTTTTGTTTGTTTTTTGTTTTTTTTTTT | 120 |
| chr11 | 70753655 | 70753775 | 788855_47188552_ENST00000601538.6,E | 2 | - | good | TTCCTAGGTGAGTTCTGGAAGTTTGCAACTTTTAGAGAATTAGTTTATTTTACCTAAATTGTCAAATTTATCTTAGACTTGGTCACAGTATTCCCTCATTATCTGTTTAAAGTCAGCCCT | 120 |
| chr11 | 70754135 | 70754255 | 788855_47188552_ENST00000601538.6,E | 2 | - | good | TTGGGAAGCGTTCCCTCCACTTCTCTTTTCTGGGAGGAATTGTGTAGAGTTGGTGTTGCTGCTTCTTGAAATGATTGGCAAAATTCACCAGCGAAATCACTGGAGTCAAGAAGTCTTTTT | 120 |
| chr11 | 70755335 | 70755455 | 788855_47188552_ENST00000601538.6,E | 2 | - | good | TGAGGGGCCGGAGTGCACCCCTGCACACAGTTTCTGCAGGAAGCCCCTGGAGGGTGGACACCCACAGGAAGAAGGCTGCCTGCGCGCTGGGCGCAGTGGCTCACACCTGTAATCCCAGCG | 120 |
| chr11 | 70757375 | 70757495 | 788855_47188552_ENST00000601538.6,E | 2 | - | good | GTGTTGGCTCAACACGCCATGGGAGCTGCTGCCGCAGGGCTGGGTGTCCCGGCCTCTCCATGTAGGGGACATTTCAGCTCCCAGATGCCAGCTAGGGGCCAGCCCTGCGGGCAGGACTTT | 120 |
| chr11 | 70757495 | 70757615 | 788855_47188552_ENST00000601538.6,E | 2 | - | good | GCCACATGTGTGAAATATTTCTCCCTTGGGAATCCTGTCACGTAGGCACCCTCAGCCTGGCACGTGCCACAATTCCCGACTTCTGGAAGGAAAGCTGGGGTTTTGCTTGCATACACCACA | 120 |
| chr11 | 70759535 | 70759655 | 788855_47188552_ENST00000601538.6,E | 2 | - | good | TCCAGTCTCTTATTGCCACCAGGAGCCCATTCTATCAGTCTCACAATCTCTCCTTTAGTGTTAACGCCGGCCAGTTGTGGCTAAACCTCAACAGGGAGCAGCATAACAAGACGTGCCCAA | 120 |
| chr11 | 70761215 | 70761335 | 788855_47188552_ENST00000601538.6,E | 2 | - | good | ATGGACACCTCCTCGCTATGCCCCCGTGTGGCCAAGGGAGAAAAAGACTGGGACTCTCTTATCTCTCCTAGTAAGGGCACAAATCCCATCATAGGGGCCCCACCATCATGACCTTATCTC | 120 |
| chr11 | 70772975 | 70773095 | 788855_47188552_ENST00000601538.6,E | 2 | - | good | GCTCAGTAAAGATAAGACCTGCCCACAGCCCTGGGGATGGGCGGAGACTGGGGGATTAGAGGAGCCAGAGGTCACCTCCAACTGCCATTTAGCTCTTAGTGGGCTCTTCAGCTGGATCTG | 120 |
| chr11 | 70793855 | 70793975 | 788855_47188552_ENST00000601538.6,E | 2 | - | good | TGGCTAGTGTTTTATGGTGTATTTACACCACAGTTTGATCATTCTCCTGATGATGGACATGTGGATTGCTTCTCAGTTTTGGCTAATATGAATGAACCTGCTAAAAAAGGTTTTTGTGTG | 120 |
| chr11 | 70823975 | 70824095 | 788855_47188552_ENST00000601538.6,E | 2 | - | good | CAGGAGCTCCATCAACGCCACCTCTGTCCTGTGAGCCCTGCCAATGCCATCTTTGTCTTCAATGAAACCTGCCAGCACCACATGTGTCCCTTGTGAGCGCTGCCAGCACCACCTCTGTCC | 120 |
| chr11 | 70827695 | 70827815 | 788855_47188552_ENST00000601538.6,E | 2 | - | good | TATTGACACAAAGTGTGTCAACCATATGGAATTAAACTTTAGTGCTCTGGTTGCTTTTTCCTCGTCTTGTCTCGTCTTTCTCTGGTTGTGTGTGTGTGTGTGTGTGTGTGTGTGTGTG | 120 |
| chr11 | 70831175 | 70831295 | 788855_47188552_ENST00000601538.6,E | 2 | - | good | CATGGACAACAAACAGACTCAAAAGGTGCTCGCCTGCGTTTTGAGATGGGTGGGCGTGCGGGCAGGCAACAAACAGACTCAAAAGGTGCTTGCCTGAGTTTTGGGACGGGTGGGTGTGCG | 120 |
| chr11 | 70834175 | 70834295 | 788855_47188552_ENST00000601538.6,E | 2 | - | good | TAGGCTCAGGTGCTCCCCTGTGTCTTCCCCTGAAATATGGGGTGGGACCAGAGTTAGAGCCACCTCCTGTGTATTTTGTGTTAAGGTGGAAATAGGCTCAGGTGCTCCCCTGCGTCTCCC | 120 |
| chr11 | 70835375 | 70835495 | 788855_47188552_ENST00000601538.6,E | 2 | - | good | GACTGAGCGGACCATGTCCCCAAGAACTGGTGGAGCACATCTCAGCACATGAGCCTCTTGACTGAGCGTCCTGTGTCCTCGAGAACTGGCGGAGCATGTCTCAGCACATGAGCCTCTTGA | 120 |
| chr11 | 70862015 | 70862135 | 788855_47188552_ENST00000601538.6,E | 2 | - | good | AGATCAGCTGAAACAGAGACAGCCGCATCGTGGGGAGAGGTGTCACCTCATGGAGTCCATCCAGGTTGGGCTCCACACCAGATTCCCCCTGGCCTGGGAGACCCAGTGCGACAGGATCCT | 120 |
| chr11 | 70862135 | 70862255 | 788855_47188552_ENST00000601538.6,E | 2 | - | good | ACTTAATCGCAGCAGGTTTGCAGCAGGAATCTGGATGAACTGGGATGAGGTCAGGCTGTGTTCGGGGAGCTGGCACCTGGGCACTGGGCAGGTGTGCCATCAGTTCAGTGTCTGGTGCTT | 120 |
| chr11 | 70863215 | 70863335 | 788855_47188552_ENST00000601538.6,E | 2 | - | good | CTGGGCACTGGGCAGTTGTGCCATCAGCTCAGGTTCTGGTGCTTAGATCAGCTGAAACAGAGACAGCCATGTCATGGGGAGAGGTGTCACCTCATGGAGCCCACACCAGTGGGCCACTCA | 120 |
| chr11 | 70870055 | 70870175 | 788855_47188552_ENST00000601538.6,E | 2 | - | good | AGATTCTGCTGCTGTGCTCATCCTCAAACCTCTTCCATAGTGCCCTGGGCCACACTCTCTTTATAAGGACACTAATTATTGGATTTAGGACCCTCCCTAATCCAGTATTACCTCGTCGCA | 120 |
| chr11 | 70872095 | 70872215 | 788855_47188552_ENST00000601538.6,E | 2 | - | good | CTTGACCTGAACTGATTTGAGGCTATTTATAGTTTTTATTTATCCTACTTAGCATGATTATTCATCCGTGCCCTGAAGGACTCCTCCCGCAATTGGCAATGGGACACTGCCCAGACCCAC | 120 |
| chr11 | 70874495 | 70874615 | 788855_47188552_ENST00000601538.6,E | 2 | - | good | ATACCTTTAAAATTGACATTCATTCTTTCAAAATATTTGGTTTTAAAACTGATGCACAAATCAATAAAATGATGGTTATTACATGACTGTTAATGAAAATAATTTTGCCATACAGAGGAA | 120 |
| chr11 | 70874975 | 70875095 | 788855_47188552_ENST00000601538.6,E | 2 | - | good | AGATCTAAGCAATTGCTACAGTTGCTGTTGAAATGTATCTGTGAACTTCAAATTAGCATCTTTTTATAATGTATCCTTTAGTGAATACCATATTCTGTATGAACATTAACTTGGGGAAGC | 120 |
| chr11 | 70886615 | 70886735 | 788855_47188552_ENST00000601538.6,E | 2 | - | good | TAATATGTTAGGGGTGTGTGTGTGTGTGTGTGTGTGTGATTGTGTGATTGTGTGTGTGAAAGTAAAATTCAGGCTGATTTTGAGCCATGATCCTTTAAAAATATACCTGTTTGCTAGATA | 120 |
| chr11 | 70889375 | 70889495 | 788855_47188552_ENST00000601538.6,E | 2 | - | good | TGGGCCTGCTCAGGACCCAGGGATGGGCTGTGTACCCGGTTCCCGGGGCTGCCACAACAAATGACCACCAACTGTGTGGCTTTAAGCAACTGGAATTGACTGTCTCCCTGTTTTGAAGGC | 120 |
| chr11 | 70897655 | 70897775 | 788855_47188552_ENST00000601538.6,E | 2 | - | good | GTCCTTTGAGGCCTACAGATGCCCTAAGTAAGGAAGAAAAAACACAAAACCAAAAATCACCCACCCTACCTTACCAGATCACCTCATACACACAAGAAGAAAGGTTGATTTAAATTAAAC | 120 |
| chr11 | 70897775 | 70897895 | 788855_47188552_ENST00000601538.6,E | 2 | - | good | GCCAATAATTCTAAATAAAGTCCCAAATTATTACACAAATTTGAGCTTTAAAAATATTCTTTAAGAAAAAATAAAAACTGAAGAGCGAAGACTTTCTGAGGAATATGACCTAAAGGAAAG | 120 |
| chr11 | 70898975 | 70899095 | 788855_47188552_ENST00000601538.6,E | 2 | - | good | TAAGTCTTTGGGCAGAGACCATCTAACTGTTGAAACCAAGGGGTGGATCTGATGATCTCTGAATTCCTTTCAGGGTTATTTTGCCTTCTTCCTGAAGAGTAGCACACATTTGTGACCAGA | 120 |
| chr11 | 70901015 | 70901135 | 788855_47188552_ENST00000601538.6,E | 2 | - | good | CGCTCTGTGCCTCAGTTTCCCCCTCTGTCGAGTAGAGATAATGGTGCCCACTTCATAGAGCTGTTAAGATCAAAAGAGGTCGCTCCTGGGAACTCACTCAGAGCAGCGTGTGGCACCTGA | 120 |
| chr11 | 70904975 | 70905095 | 788855_47188552_ENST00000601538.6,E | 2 | - | good | CCTGGCACCACCCCACTCCACCTTCCACCATCCTGAGACCATGCCATGCCTAAGAGCACCCAGGCTGTTCCTCCCATTTCCCCACTGTATTAGTCCATTTTCACGCTGCCGATAAAGACA | 120 |
| chr11 | 70910975 | 70911095 | 788855_47188552_ENST00000601538.6,E | 2 | - | good | TGTGGAGTGATTTTGGAGGAAGTCTGGGGGTGAATGCCGATCTTCTGAGTGTGACATTACTTACAGGAGCACCCCCTTCGTTTCTGCAAATGCTGAGGTGTTTAGAGAGGGATGCATCAT | 120 |
| chr11 | 70912655 | 70912775 | 788855_47188552_ENST00000601538.6,E | 2 | - | good | AAACTGGAAACAAAAAGAGGTTTAAGTGGTTTTGAAGGAAGTCTGGGGGTGAGTGCCGATCTTCTGAGTGTGACGTTTCTTGCAGGAGCGCCCCCTTCATTTCTGCAAATGCTGAGGTGT | 120 |
| chr11 | 70919975 | 70920095 | 788855_47188552_ENST00000601538.6,E | 2 | - | good | TTATTACCTCTGTTTACACATGAAAACACTGAGGCTCAGAGATGTTAAGTAACTTGGCCAGAGTGCTACAGCTGGTAAACTTGAACTTGAGACTAGAAGCCAGGTCTTATCCCAACATGG | 120 |
| chr11 | 70920455 | 70920575 | 788855_47188552_ENST00000601538.6,E | 2 | - | good | AATTTTATTTTTGATTTTTAATGACTTGCTTTTTAAATTTTAATTTTGGTTTGTAATTCTATAGAATATTTGCATTTTAAAGTCAAATCTATAAACCAAGTTATACACAAAGAAGTTCCA | 120 |
| chr11 | 70922855 | 70922975 | 788855_47188552_ENST00000601538.6,E | 2 | - | good | TGTGGATTTTTTTTTCAAATTTATCTCCTTTCAACTTTGCTTATGGGTTTTTTGTTTGTTTGTTTGTTTGTTTGTTTAAAGTATAGGGGATGTTGTATTTTGTTTATGTCACCAGGTCAG | 120 |
| chr11 | 70939895 | 70940015 | 788855_47188552_ENST00000601538.6,E | 2 | - | good | TGGCTTGTGGCAGCATCACACCAATCTCTGGACCCCCACCCCAGCTCAACCCCTGTGGTCACATTACCTTCTCTTCTGCGTCAAATCACCCTCTTAGAAGGACACTTGGGATACATTTAG | 120 |
| chr11 | 70951055 | 70951175 | 788855_47188552_ENST00000601538.6,E | 2 | - | good | AGGGGAAATGAATGTATGACATCACAGTGCAGCAGCCAGGGGAAATGAATTTATGACGTCATGGTGCAGCAGTCAGGGGAGATGAATGCAACTCCTCTCTACTTCTTTGAAATGGGGCTA | 120 |
| chr11 | 70953815 | 70953935 | 788855_47188552_ENST00000601538.6,E | 2 | - | good | CAGTCCACAGGCGGGAGCTGGCCCAGCTCCCTGGCAGGGAGGTGCCAGGCCCTTCCTAACAACCAGCTCTCAGTGGGAACTGGCAGAGCAAGAACCCACTCACTGTGATGGTTAATGTTA | 120 |
| chr11 | 71063855 | 71063975 | 788855_47188552_ENST00000601538.6,E | 2 | - | good | GTCTTCATCTGTAAATGGAAATGATGGGGTCTGGGGGTCCAGCAGCTGATGGGAACTGCAGGCACACTGTGTTCTCAGCGGGCACAGTGCGAGCCCCAGGAACGCTCCTTCGTGGGGAGG | 120 |

| | | | | | | | | |
|---|---|---|---|---|---|---|---|---|
| chr11 | 71063975 | 71064095 | 788855_47188552_ENST00000601538.6,E | 2 | - | good | TTCATCTGTAAATGGAAATGATGGGGTCTGGGGAATCCAGCAGCCAGTGGGAACTGCAGGTCTTCATATGTAAATGGAAATGATGGGGTCTGGGGGTCCAGCAGCTGGTGGGAACTGCAG | 120 |
| chr11 | 71084975 | 71085095 | 788855_47188552_ENST00000601538.6,E | 2 | - | good | ATTTCCTTTAATGAACAAACCCTTCCAGGCACATGACAGAGGCTTGTCTTTGTAAAAATGGTCAAATGTGTATAGGACCCGTAGCAAACCCAGGCACATGACAGAGGCTTGTCTTTGTAA | 120 |
| chr11 | 71098895 | 71099015 | 788855_47188552_ENST00000601538.6,E | 2 | - | good | GGTGCCGAGGGCCCGTCCTGTTGGTGTGGCTCATCTTGGGCGATGCTGCCCCCCCATCTCCTGGCTGAGCTCGTGTCCATCAGGTTTCCCCACGGAGAGATGGTCCCGTTTTTCTGTTTC | 120 |
| chr11 | 71103335 | 71103455 | 788855_47188552_ENST00000601538.6,E | 2 | - | good | GTGACCAGTGTATCCGGTTTTCCCGGGACTGCCCTGGTTTTAGCACTGAATGCCTCATATCCTGGGAAGCCTGTTGGTCCCAGGGAGACTGGCACTGTTACCCACCCTACCAGTAACAAG | 120 |
| chr11 | 71104655 | 71104775 | 788855_47188552_ENST00000601538.6,E | 2 | - | good | CCAAATCAGGAGAGAAACCAGCAGTGCTGTTAGAGGGGCAGGGAAGTGGGGACCCAGGGTGATGGGGGAAAAGAGAAAAGTGAGGACCCAGGGTGACGGGGGAAAAGAGAAAAGTGAAC | 120 |
| chr11 | 71106455 | 71106575 | 788855_47188552_ENST00000601538.6,E | 2 | - | good | CTGACCATGTTTCCCAGAGCTGAGTAGAAATGGCTTGGGAGCCTGATCCATGGAGACAGTAGCACAGAGTTACCTTATGAATGTTCTGAAATATTCTCATGATGACAAGCCTCTGGGCTA | 120 |
| chr11 | 71106575 | 71106695 | 788855_47188552_ENST00000601538.6,E | 2 | - | good | AGGCACGTGCCACCATACTTGGAATTCTGCCGCTGGAATTCTGGATTAGGCCGGTACCAAAGCAGAAGGTCCATAGTTGTTGTGAGCCGTTGTGAAGCAGCATAATTAAAGGAGAAAGAC | 120 |
| chr11 | 71107535 | 71107655 | 788855_47188552_ENST00000601538.6,E | 2 | - | good | TTAGGCTGGTGCCAGAGCAAAAAGTTCATAGTTGTTGTGAGCCGTTGTGGAGCAGCATAATTAAAGGAGAAAGACCTGACCACTTTCCCCGGAACTGAGTAGAAAAGGCTCGGGAACCTG | 120 |
| chr11 | 71115095 | 71115215 | 788855_47188552_ENST00000601538.6,E | 2 | - | good | ATTACAGGCGTGAGCCACCACGCCCGGCCAAAATTGATGCCGATACTTAGGTTTATTAATCTCCTTTGCATGACCCACTGCAAAGGCTGAGATTGGCTCAGGGATTGACACATTAGGGGC | 120 |
| chr11 | 71117375 | 71117495 | 788855_47188552_ENST00000601538.6,E | 2 | - | good | GGAAGCTACCTGGGAGTTGAGAGTCATCTCATTAGTGCTCCAGAGATACTCTTATCACTGAGGACAGTACAAAGGTTTTAGGAACTCTGCCAGGAGTCAGGCACAAAGAAGGAATTTTTA | 120 |
| chr11 | 71117495 | 71117615 | 788855_47188552_ENST00000601538.6,E | 2 | - | good | TTACATAGGATGAAATCACTGACCATGGTGATTGGACTCCATCCTGCCAGTCAGCAGGGCGACTGGGAGTCCTGACCCTCGAGTCATAGTCTGATCCTTCCGGCCACCAGCCCCCATCCT | 120 |
| chr11 | 71117735 | 71117855 | 788855_47188552_ENST00000601538.6,E | 2 | - | good | CAGCTCAGAGAACCTAAGAAGGCATTTTAACTGCTACCCCTGGCTTATTACAAAGTGTACAAACGCACAGCTGATGAAGAGGTGCACTAGGCGACGTCGGGAGGGCCCAGCGTGCAGGAG | 120 |
| chr11 | 71117855 | 71117975 | 788855_47188552_ENST00000601538.6,E | 2 | - | good | AGTCCCATAAGACTGCCCTCCATAAACGTCAAGTCCCCAGGCCACCCACATTTCTGTCCCATTTGGCTGTGAACTAGGGGTTCCCACCACCCCCTTTTTGGGTTCCATCCCTTGCTGGAG | 120 |
| chr11 | 71117975 | 71118095 | 788855_47188552_ENST00000601538.6,E | 2 | - | good | TTTAAATTTTTCCTTCCCAGCGCCTAATGGGTGGTGTATTTTCCTTCTCCACCTCTCCAGACACCAGCTGGATGTCAATAATTAATTCAATTCTGACACTAACTACAAGTTTAAGGGCTC | 120 |
| chr11 | 71118095 | 71118215 | 788855_47188552_ENST00000601538.6,E | 2 | - | good | GAATCCCTGCAGAGAAATCTTGCTGTGAATAATAAACCGTGGGTCTGATTCAGGCTTTCATAACCAGGACCATGGGCCCTGGGCATGGAGAACGGGGCTCCCCATGCTTCTCTGAGGGGT | 120 |
| chr11 | 71118695 | 71118815 | 788855_47188552_ENST00000601538.6,E | 2 | - | good | CACAGTGCATGGACTGTGGTTGTGTCACAGCCTGGATGCCATGGTGGGTGGGGGAGACATGGTGGTGGGGCGAGACCAATTCCATTTGCATTGGGAAATTATCCCCACATTTAAAAATCA | 120 |
| chr11 | 71118935 | 71119055 | 788855_47188552_ENST00000601538.6,E | 2 | - | good | GCTGTCTTTGTCCTTCCTGGCAGAAATGCATTCGATTTAACCCGGATGCCACAGTGTGGGTTGCAAAGCAGCGGATCCTGTGTACATTAACCCAGAGTTTGAAAGATGTCCTGAACTACG | 120 |
| chr11 | 71119055 | 71119175 | 788855_47188552_ENST00000601538.6,E | 2 | - | good | CTGTGAAAAGCCGTGGGTTCACTGCTCAGAGGAAGTGTCGGGAAGGGGTGGAAGCAGCTTTGCCTCTCTGACCACGCGCCCCACATGGGTGACTCAGGTAGCGTCCTCTGTCACTCATCA | 120 |
| chr11 | 71119175 | 71119295 | 788855_47188552_ENST00000601538.6,E | 2 | - | good | CTGGAGATTTGAGTCCATGGTGAGATCAAGGCCAAAGAGACCAATGCCTCGGTCTTTCCTTTTTCCCTTTTATATTTACCGAATGTCTGTGTCCCAGCAACCAGTAGGCTTTATTTTTCA | 120 |
| chr11 | 71119655 | 71119775 | 788855_47188552_ENST00000601538.6,E | 2 | - | good | AGGTTCTCAGGTCCCACCCAGACCTGCTGAATCAGACATCATGTTAGGGAGACATAAGTGGGGTCTCTGACCTGTGTTTTTCAAACTGCTGGAGTTTGAGAAGTGCCATGCAAGTGAGAG | 120 |
| chr11 | 71119775 | 71119895 | 788855_47188552_ENST00000601538.6,E | 2 | - | good | ATGCTAAAGAACGATCCATGTCAAAATTAAGTCACATTGAAATATGCGCGGGCTGTGGCTCTGAATGTGTGTTCCCCAGGACGGCAGCAGCAGTAGCACCTTGGGATTTGTCTGAGATGT | 120 |
| chr11 | 71119895 | 71120015 | 788855_47188552_ENST00000601538.6,E | 2 | - | good | TCCTTTGGGAGGAGAGGTGATGACTGCATTTTTAAAGCTTTTGTGTTCAGAGTCAGCGTAGGTAGTATGAAAACTTTCTCGTCAGTGTGGTTATAAGACAAGGCTTCCACCGAATCTGAA | 120 |
| chr11 | 71120015 | 71120135 | 788855_47188552_ENST00000601538.6,E | 2 | - | good | TTGAAGTGTTGTGATTGGTTTGCAGGCCCTGTTGACACTATTTAACTTTGCAAAAATGAACTCTCATGGCAACTGATTTGATTTTCCATCAAATCGTTTTTCCGTAATTGCTTTCCTGTG | 120 |
| chr11 | 71120135 | 71120255 | 788855_47188552_ENST00000601538.6,E | 2 | - | good | GGCTAGAAGTGGATTTGCCTGGCCTTTGCCTTTCCTGGCTCACTGGTGAGTCTCACAGAGGGGACCTGGGAACATGACCAAGTTGTTGTTATTTCAGTCCTCAGAGGTGCTATTATATGT | 120 |
| chr11 | 71120255 | 71120375 | 788855_47188552_ENST00000601538.6,E | 2 | - | good | ATATTCAATCAAACCTCCTTTTGTTCTGAACACTTTGACAGTGACAAAGTGAATTATTGAGCCCCTGCTGACAGCTGTTCAGTAGAATATAATTTTAATTCAACAAAAATTAGAAGTCAT | 120 |
| chr11 | 71120375 | 71120495 | 788855_47188552_ENST00000601538.6,E | 2 | - | good | GGATCATTTTCCCAGCGATGAGGAACTTACCCTCGGGCTGTCTTGGGAAGATGATAGGGCCTTTTGCTTTCTTGGGCACAAACCCACAGAATCCTCAGATAGAGAATTGGTTTGAGTCTT | 120 |
| chr11 | 71120495 | 71120615 | 788855_47188552_ENST00000601538.6,E | 2 | - | good | ACGTGAGCTCAGCCTTTGACATTGGAAGGGAGAAACAGGAGCTTTTGTGCACAGGGTCAGGTTTCCTGGCAGGGAGGACTCTGGCCGCCCACTGAACCCGACCTCCTGTTGGCCGTCCCA | 120 |
| chr11 | 71120615 | 71120735 | 788855_47188552_ENST00000601538.6,E | 2 | - | good | AGTTTCCAGAGAGTCCTTGCTTAGCCAAGGATGGGGGAGGGTGCGGGTCTGGAAATCTTTCTGGAATAAACAACTGCCCCCACCCAGCCTCACAGCCCTGTGGGGAGCTTTGAGAGCCAC | 120 |
| chr11 | 71120735 | 71120855 | 788855_47188552_ENST00000601538.6,E | 2 | - | good | TGGGCTCTGTGGATGTTGGAGCTTGAATTGGAGGGGCCTAGAAGGATTCCTGCAACTGAAGGGTGAGTAGTGAGCGGCCCCACCTGGTCACACCTGATACAGCTTGTTTCTTGTGCCAGG | 120 |
| chr11 | 71120855 | 71120975 | 788855_47188552_ENST00000601538.6,E | 2 | - | good | TATCTTTGCAATCACCTGCAGTGGCACAGAAGTCCCCTGAGAGGCAGTTTGGAGCCCTGTGCTCCTCCAAGGAATTGCCCCTTGGCTGAAGCTTCTCTGCAGGTGTCAGGGAGGGAGCCA | 120 |
| chr11 | 71120975 | 71121095 | 788855_47188552_ENST00000601538.6,E | 2 | - | good | TGATTCTCAAGGTGGTGCTCTTAGCATTTTCTGGAGCAGATGGTTCTTGCTTTGTGGGGTTGGCCCCCGTGCTGCACAGACACAGCATCCCCACCCTGTCCACAAATGACGGTGGACACC | 120 |
| chr11 | 71121095 | 71121215 | 788855_47188552_ENST00000601538.6,E | 2 | - | good | TTTTCATGGGGGAATCAGAGGACTGCTGGGTGGGTGGGTTTCCCTAGGAATCAAGGTTGGTGTGCCCCACTGGGTTGCAAACCCAGGCTTTCCGACCTTTGAGCTCTTGCTCCTTCTAGA | 120 |
| chr11 | 71121215 | 71121335 | 788855_47188552_ENST00000601538.6,E | 2 | - | good | GGTATGGGCATTAGGACTTCAACATTTATTTAGAGGGGCACAGTTCAGCCCATAACAACATGTCTTATTTATATTCTGTTTGATCCTGACAGCAGCCCTGAAGTTAAGTAGTGGCTCTGA | 120 |
| chr11 | 71122655 | 71122775 | 788855_47188552_ENST00000601538.6,E | 2 | - | good | GGGGGCTTACCAGAAATCTTAGCATTGCTGGGTTTGCAGATGGGACACACCAGTCTCTGCCTTCCCATTCACGTGGTCGTCTCCCCGTGTGCATGATGCCGTGTCCAGATTTTCTCACTT | 120 |
| chr11 | 71122895 | 71123015 | 788855_47188552_ENST00000601538.6,E | 2 | - | good | GCTCTCAGAGATGATGATGCTGCTGTTTTTTTGCTGCCACAATTAGTGGCAGGTGATTCACACGTTTATGAGTTCCCATGCCTGCCATAACCAACCACAGACTGGTGCCTTAAATCCCAG | 120 |
| chr11 | 71123015 | 71123135 | 788855_47188552_ENST00000601538.6,E | 2 | - | good | AAAGTGTCTTCTGACGTCGTGGGCTTGCCGGGAGCTGGTGTGAACACGTGGGTGGCTCCCATCACGTATGGAGGGAGGGGAGCTCCCTGGGAGCTGGGAATGCTCGACCTGCTGAGTTCC | 120 |
| chr11 | 71123135 | 71123255 | 788855_47188552_ENST00000601538.6,E | 2 | - | good | ACATCATTCAGCCAGTGTCAGTCACCTGTACTGTGTCAGGCCTGCTCTTGGACCTGGGGTTTGCCTGATTGGACAGTGAGTCAATGATGGGTAGGTGAAGAATTGAAGTAGGATTCATCC | 120 |
| chr11 | 71123255 | 71123375 | 788855_47188552_ENST00000601538.6,E | 2 | - | good | TCGTGTTAGCATGGATTTTTACCTTTTCCATATTGTGTGACAGTTGAGTAGGTGTTTTTCCCCACTACAAATGGTGACCTCAGGCTGGTTTGCTTCTTATTTATTGTCTTTGTGGCTCCC | 120 |
| chr11 | 71123375 | 71123495 | 788855_47188552_ENST00000601538.6,E | 2 | - | good | TCAACCTGGAAGTCCTACCCCAGACTCGCATATCCCTCTGTATTCCTGGGCAGCAGCCTATGATGTGTGCTCCTTACCAAGCAGCGTCACTTACTCCTTCCTCTGGGTCCCCAAGGGAGC | 120 |
| chr11 | 71123495 | 71123615 | 788855_47188552_ENST00000601538.6,E | 2 | - | good | CACACTGGTAACAGTGGGACCAGGAAGGGTCACTTAAGGCAGAAACTGTCTTGGGCCATCCACCTGCTAGACTCATCGCTGTTCTACCTTTGCCTCCCTCCCTCTCCTTATGATGGGCCC | 120 |
| chr11 | 71123615 | 71123735 | 788855_47188552_ENST00000601538.6,E | 2 | - | good | ATCACTCACTCACTCAGAACATCTCTAACAAATTATAAAGGAATAGCCACCTCGCAGGGCTGTGGGGAGACCAGAATCAGATACTGTTCTCCTCAAGGAGAACTTGACATGACCAAGGCT | 120 |
| chr11 | 71123735 | 71123855 | 788855_47188552_ENST00000601538.6,E | 2 | - | good | CTCAAGGAGAAGAGAGTTGTTTCCTCCAGTAATCCTTTAGAGGAGTTGGCCAGGGTTGGGACTTCCTATACACTGTTGTCTTTGCCATCCTTTCTGTACTATCTGTGAGTCATCTAACAA | 120 |
| chr11 | 71123855 | 71123975 | 788855_47188552_ENST00000601538.6,E | 2 | - | good | TCCCCTCCCTGTAGACACTGAACTCTTTAAGGGGCAAGAGCACATCTTGGTCATCTTCGTGTCCTCTCACCCACTCCTGTCCTGGTGCTGGTATAGGACCCTGTGCACAGAAGATAGAAG | 120 |
| chr11 | 71124455 | 71124575 | 788855_47188552_ENST00000601538.6,E | 2 | - | good | GAGAATGAGAATAATATTATTATATAAACAGTAATAATAATATTATACAAAAAGTAATAATAATATCCACTTAGGGTTGTGGTGAAGGTGATAAGGTTAGCGTAATTTCTGGCTTATTAG | 120 |
| chr11 | 71124575 | 71124695 | 788855_47188552_ENST00000601538.6,E | 2 | - | good | CAGCAAGAACTCGGGAGATGGGGAGATTTGACTTGGGTTTAAACCAAACCTCTACTGCTTTCTAGAAGTGTGACCTTGGGGTAGTTACTTGGAGTCTCTGAGGCTCAGCTTCTTTATCTA | 120 |
| chr11 | 71124695 | 71124815 | 788855_47188552_ENST00000601538.6,E | 2 | - | good | TTCTCTGTGAAGCTCAATAAAGCAAGTGAAATAGAATGGGATATGCCTCTAATTTATGATTTGCCTGCACTAAGATTGGCAGTGGGTGTGTATAGAGAGGAACATCATCTATCACTTGAA | 120 |
| chr11 | 71127815 | 71127935 | 788855_47188552_ENST00000601538.6,E | 2 | - | good | GGGGCGCGGGGTTGTATGCAGCCTGGAAGGTCTGAACAAAGGGAATATGTCTTCTGATCCAGAAATGACCTGAACATCAACATCTTCTTCAGTATTCTCAATGACTTACATAACAGGCAT | 120 |
| chr11 | 71127935 | 71128055 | 788855_47188552_ENST00000601538.6,E | 2 | - | good | GTGGTCAGCTCAGCCTGCAGGAGGCCAGGGAGATGACATCGTGAGACCTCGGCCCCGGTGGAAAGGAGAGTGGGGCTCCAGCCGGCAGACGCGGAGAGGATGGCTGGTCCACCAGAGGC | 120 |
| chr11 | 71128055 | 71128175 | 788855_47188552_ENST00000601538.6,E | 2 | - | good | AATGCAAGGTGAAGAGGCCGAGACTCATGGGCCAGACAGGAAAGGGAGCAGGGCCAGTCAGCTGGCCTGAGGGCCGTGGGCAGTCGGAGCAGCTCCTTGTGTTCCCACCCAAGGTGCCCA | 120 |
| chr11 | 71128175 | 71128295 | 788855_47188552_ENST00000601538.6,E | 2 | - | good | TTTAGGAGATTGAGTGACAGAGGGACCAACTCCTGCCTACCAGTTAGGAGAGAGTGAGGAGAAACATCCTAAGGCAAAATATGAAACTCAGCACTGCTCCCACCTTCCTGGGAGACAGGA | 120 |
| chr11 | 71128295 | 71128415 | 788855_47188552_ENST00000601538.6,E | 2 | - | good | GAGAAGACAGCCTGGTTTGTGTCTCGGAGATCTCGTTACATCCATTTGCAGGTTTTCTGTGTCTTGAGCCCCTTGGGCCAGGCACGCAGCAGGTTTCGGAAGGATCTGGACATCACAGCC | 120 |
| chr11 | 71128415 | 71128535 | 788855_47188552_ENST00000601538.6,E | 2 | - | good | ATCGCAAGGCTGGTGTCAGCCACAAACTCCGTCTCCACAGCTCCTGACCATGTCAGCCCCTGTCCCACTCCATCACCACCCAGGTCTCATGCTGTATGTACTGCCTTTCTGGAGATAAGG | 120 |
| chr11 | 71128535 | 71128655 | 788855_47188552_ENST00000601538.6,E | 2 | - | good | CTGTGCTAGGACACAGCTCCCTTTTGTGGCTGAGGGGCTTTGTGGTATGAAGGGAACTTAGAGAGAACTGAGGCTGGCCTGAAGAAGTCCTGAGAACTTTCCGGGATCGGTGTGTACATC | 120 |
| chr11 | 71128655 | 71128775 | 788855_47188552_ENST00000601538.6,E | 2 | - | good | CTCGGGCTGGGGCAGGGAGGAGCACTTAACTGGCTGGACGTTGCAGTGGTGCTGAGACAGATGTCGGGGGAGAGAGAGATGGGGAGGAGTAGGAGGGACGGGAAGATGTGGATGGTTCA | 120 |
| chr11 | 71129135 | 71129255 | 788855_47188552_ENST00000601538.6,E | 2 | - | good | AAGACTCAGTAGGCAGAAACTCCCGTCAGCCCCGCTGTGAAGTCCAGACCTTTCTTGACTGTTGGATGATACTACCCAGGAGTGTTCCATGTCTGGAATGTTGATCCAATGCTGTTCAAT | 120 |
| chr11 | 71129255 | 71129375 | 788855_47188552_ENST00000601538.6,E | 2 | - | good | TCAACAGCATGGCCAACCCTAGCTCTGTAACTTACAGCTCTGCAGACCTCGGTGGCGTAGCCTCCCCTCGCCAGAATCATTGCTGTAGTTAATGGTCTGCACAGTCCACTGAGTACAAGC | 120 |
| chr11 | 71129375 | 71129495 | 788855_47188552_ENST00000601538.6,E | 2 | - | good | TTTTTAAAAATTTTCATTTTTCGTAGAGACGGTTTCCCTATGTTGCCTCGCCTGCTCTTTGGTTTTGATTTGAGCTTGACCATGAACTGTACTAAGAGGGGTGTAGATGTATCACTCTGT | 120 |
| chr11 | 71129735 | 71129855 | 788855_47188552_ENST00000601538.6,E | 2 | - | good | TGCCACAGCAGCCCGAGTGACTCATAACTCAGCGTAAGTGGCAGACTCAGGCCAAACACAGCTGCCTCTGCCTGTCTCCAGTGCCCGCATTCTGCCCACAGTCCAGTAGTCTCAGTTCTT | 120 |
| chr11 | 71130215 | 71130335 | 788855_47188552_ENST00000601538.6,E | 2 | - | good | CGGCGCTGGTTTGCTGCTCCACCATTGTTTATGTGTCTTACTCAGCAGTGCGGACGTGTCCCAACACAGGCAGGTGGTGGCCCTTGGTAGGTTGCAGAGACTCGCCCCATGGTCATTCAG | 120 |
| chr11 | 71130335 | 71130455 | 788855_47188552_ENST00000601538.6,E | 2 | - | good | CCCAAAACAGATTTGGAAAGCAATTTAGTCTCTCACAGGAAACTCTTCTTGCTACATTGCTGTAAATCAAAACAGCCTTGCCCAAGAAGCTCCTGTGACCCTGGCAGGCCGGATACTTTT | 120 |
| chr11 | 71130455 | 71130575 | 788855_47188552_ENST00000601538.6,E | 2 | - | good | GGCAATAAGACCTATTTTATATGGTATAATATTTACCATTTTCTCAAGCTTTGTCTAGACACTTCCTTGCCCTCACTACAATCTTCCAAAATCACCCGTCATCATCGCCATTTTACAGGG | 120 |
| chr11 | 71130575 | 71130695 | 788855_47188552_ENST00000601538.6,E | 2 | - | good | AAGTCTCACCTGAGGAAGGTGAGTTAGCAGATGCCGTAGAGGTGGAGGTCCCTGAATCCATCTGTGTAATCCAAGCCTCATGTTGTAGATGGCCCTTTCCACAGCTTTCAGATAACCCAT | 120 |

| | | | | | | | | |
|---|---|---|---|---|---|---|---|---|
| chr11 | 71130695 | 71130815 | 788855_47188552_ENST00000601538.6,E | 2 | - | good | GATCCTGAATAGGGAATATCCCTGTGGGGCCACGACCTGAGATGCTCTGGCATTTCCTCCCCCTTGTTCTCTTGGCTTTTCCTGGGCTCCTTGGGTAGAAGTCAAGAGGAAGGTGGATGT | 120 |
| chr11 | 71130815 | 71130935 | 788855_47188552_ENST00000601538.6,E | 2 | - | good | ATGACGTGCTCAGAAACTTGGAGTTTTTTCCGGCTTTCCTAAGTTCTTGCTGAACGCTGCTTACATGCGTTTGCAGAGCTGGCATCTTGATCGTCCCCGGGGTCTGCGTTTCTGGCTCCA | 120 |
| chr11 | 71130935 | 71131055 | 788855_47188552_ENST00000601538.6,E | 2 | - | good | GCCCCTGCCAAGCTTGTGTTTATGACGGGTCCTTAAAAATGCATCATGCAGAACTTGATGATGACAGAAACCGTAGCTGCAGAGGAAATTCTGAATGTGAAATGCGCTTCTTTGTAATGA | 120 |
| chr11 | 71131055 | 71131175 | 788855_47188552_ENST00000601538.6,E | 2 | - | good | TAAACCAATTGTAGTTTGGGGTGACCTTGATGTTGGTTATTTGTTGGAGGTGTGCAAGTACACGGATTGCACGTGAACAGCTATCAAAGACCCGAAGACGCCTGCGCGCACCTTTCCGTG | 120 |
| chr11 | 71131175 | 71131295 | 788855_47188552_ENST00000601538.6,E | 2 | - | good | ATTTCAGCTTCGTGTGTTACCCGTGCCGGGGTTTTAGTGTTTGCCACCACGAGCACCAGCCCAATGGGCCTCTGTCCGCAGTAAGAAGTATTATTTCATTTGGATAATTGCATTTGGGAA | 120 |
| chr11 | 71131295 | 71131415 | 788855_47188552_ENST00000601538.6,E | 2 | - | good | TTATTGGCGTTTAAACATGCAGAAGCACTAAGCCTGACTCTTGGACCTCAGCACCCCCCGACTAGGCTGTCCTTCTGAGTCGAGGTCACCCCACATTTCCCTGTTCCATTGACATGTCTG | 120 |
| chr11 | 71131415 | 71131535 | 788855_47188552_ENST00000601538.6,E | 2 | - | good | TAATCGCCTGGAGATACAGTCTGCCCACCGCTTCGCCACAGCATCCAGCGGCTGCCATGAAGCTTTCACTGTGCGAAAAATGTTCTGATTGGAAAAACGAATCCACAGCTCCGTTACCCT | 120 |
| chr11 | 71131535 | 71131655 | 788855_47188552_ENST00000601538.6,E | 2 | - | good | TATGAAAAGAGGAAGATTAAGTCACCCCCAAGGAGTCCATGGACCACTGGCAGAACCAAGCAGTGTCTCCCAGTTTAGCATCACCTTGACAATAGCAGAGACAGACAGGCAGCCTCTTAC | 120 |
| chr11 | 71131655 | 71131775 | 788855_47188552_ENST00000601538.6,E | 2 | - | good | TTTTAATTATAACCCCCCTTCCATGGCTGACTTACTTCACTTCAATTTATTGCTTCTAAAGGGACTTCTCTGGCCAGCATCTGATCTCTGCTTTACAATGGCTCTGCCTGAAATCATGGC | 120 |
| chr11 | 71131775 | 71131895 | 788855_47188552_ENST00000601538.6,E | 2 | - | good | TTGGGCCTGGGGGTGCTAATGCGGCCTCCCGGAGAACATAAGGAAGAGATGATGTGTTTAACATGCTTACGAGCCAGGCTATGGGCACATAGTAAGTGTTTTGTGTCTGCAGTCAGGACT | 120 |
| chr11 | 71131895 | 71132015 | 788855_47188552_ENST00000601538.6,E | 2 | - | good | TGTGCTGCCAGGATCGAGTTGGACTCATTAGCTCGGCCTCGGGGACCCCCTTAATCCCTGCTCCTCCTCTTGCAGCTGTGTGGCTCTGGGAATGATCTGGCATTTTTGGAGCATTCCTCT | 120 |
| chr11 | 71132015 | 71132135 | 788855_47188552_ENST00000601538.6,E | 2 | - | good | CCACAGCTTCTGAGCAGCATCTGCGTCCCGGCAGCAGCCTCACCTGCCTCTGTGGCCTCGTCGCTAATCCACTTTCTGACCTCCTGCTGGCTTAATCTTCCTGACTCATGGCTCCGTCTG | 120 |
| chr11 | 71132135 | 71132255 | 788855_47188552_ENST00000601538.6,E | 2 | - | good | GCATCTTTGCTCTGAGGTTGGACCTGGGCGATTCCTCCAGCCTGGCTCCGACTCTGCCTCCCCGCCAGCCTCCACCCTTCCCTCGCTGCTCTCCCTTCTGGGCGGACAAGGCCTCCCACA | 120 |
| chr11 | 71132255 | 71132375 | 788855_47188552_ENST00000601538.6,E | 2 | - | good | CGTCCCCTCTTTGCTGGCATCCCAAGCTGGTAAGGCTTATCCAGAGGGTGGCAATACCAGGGCCGCAGATCATCAGTTTTTACTTTTCTGTGTTAAATCCAAGGGTGAGGCTTTTTTCCA | 120 |
| chr11 | 71132375 | 71132495 | 788855_47188552_ENST00000601538.6,E | 2 | - | good | AACAATGTACGGGAAAAGTGACTTTGGTCTGGAATTACTACAAGTTCTGTAGCAAAGCTCTGTAAATCCCACAGTGTAGGGGGGAGTATTGTGGTTTGGGGCCATGCTGGTGAGGAGCAGC | 120 |
| chr11 | 71132495 | 71132615 | 788855_47188552_ENST00000601538.6,E | 2 | - | good | GTATATGGGAGAGATTGGCCCAGTTTGCAGGGTCTCCGTAGCCTGCACATAAAACGTCTTGACTTGAGGTGCTCTTCATCGTTGGCTTAGAGTTAATGCCTACTTAAATCTCATGGAGAA | 120 |
| chr11 | 71132615 | 71132735 | 788855_47188552_ENST00000601538.6,E | 2 | - | good | GTCACAGGGCATCCTCGATAAAGGCAGAGAAGGACTTTGTTTTCTTACACGTCATTTAAAGCAAATATTGTAAGGATTATATGTGGACCCTTCCTCTGTTACGTATTGGGTTCGTGCTTT | 120 |
| chr11 | 71132735 | 71132855 | 788855_47188552_ENST00000601538.6,E | 2 | - | good | CTATTTTAGTCAAGAAACTGGTGAGTTACAAATTGTTTAGAAAAAAAGGGAAGGCTGTGGGGCCTTCCAATGAAAGTTGCCATGGAAAGTCAAGCCTTGATCATATTGGTTATTATTCAC | 120 |
| chr11 | 71132855 | 71132975 | 788855_47188552_ENST00000601538.6,E | 2 | - | good | GCTAAGACCAAAGAGAGTGCTGCTCCCCAGCCCCTCAGCCTTCCCTCTGCCTCCACTGTCTCTGTAATACAACAATGAAAACTAGCCTACCTGTCCTTAAGAGGAGTGGTTTACACTTGA | 120 |
| chr11 | 71132975 | 71133095 | 788855_47188552_ENST00000601538.6,E | 2 | - | good | TGGTTAGGATCTTTGGTCTCCACTTTGAGTAGCCCCGCCATGTCCTTTGTAGTATGCATTTGGATTACCTCTTGGGGAGGTAGAAGGCATGGAATCTGATTGTGGTTTCATTAACTGGTT | 120 |
| chr11 | 71133575 | 71133695 | 788855_47188552_ENST00000601538.6,E | 2 | - | good | AGTCCATGGAGTGTGGCCCAACACTGACTCCAGCCTAACCAGGGAGGCCCCCTAGACCACCCCTCCACTCTACGCTTCCACCCACCTACCCACCCATCCCCTGACCCTCCAATCCATCCA | 120 |
| chr11 | 71133695 | 71133815 | 788855_47188552_ENST00000601538.6,E | 2 | - | good | TAGAAGTTTAGGTATATGTTGCCTTGACATGAGCACATTTTCTGCCCTAAAAGAAAAGTCATCAAAAAGGACTTGGGCAGAAGATGGTAACATGATAATGGTCTTTATTCAAAAACTTTG | 120 |
| chr11 | 71133815 | 71133935 | 788855_47188552_ENST00000601538.6,E | 2 | - | good | AAAAAAAAATACATAGTTTTCCTTTGGGCTGCCCTTTCAGTTTCAGATGTTTTGTTCCAGAATGTTCTCTCCCAGTTGCTTGTTAGCTAAATGAAATTCTTCTGGGATTATAGACTTAAT | 120 |
| chr11 | 71134055 | 71134175 | 788855_47188552_ENST00000601538.6,E | 2 | - | good | TCCTGCTGGAGATGAGCCCCTCCCCTCCCACCTGTGCTGTGTCCAGTGCCCACTCCGGGCCTCCTGCCACGGAGGAGCTTTCTGCACTGTCAAGTCACCCAGTCCACCGAGAAAGGTAAA | 120 |
| chr11 | 71134175 | 71134295 | 788855_47188552_ENST00000601538.6,E | 2 | - | good | TTGGCAGTCACTTCCCCTTTCCCCCTCTCCCCTCCCCAGATGCCTGCTCTGGCCATCGTCAGTGATGGGGTGGCACCGCACGTGGACTGAGAGGGAAGCCTTGTCTCAGTTCTTTGCCTC | 120 |
| chr11 | 71134775 | 71134895 | 788855_47188552_ENST00000601538.6,E | 2 | - | good | GGAGAAGGATTAGAAGAGATGGGGGCTCCAGGTAAATGGTGCTGGAGGCTGGGACCCTGGGAAGTGCAGGGAAAGCTGGAGACCCACAGACGCTTCCCTATGACAAAGCCAGTGGGGCTG | 120 |
| chr11 | 71134895 | 71135015 | 788855_47188552_ENST00000601538.6,E | 2 | - | good | GCGTCCAGGTATAGGGTCATGACAAGGAGGCCATGGTTGTATTTGATGGGCAGAGGTTTGGTACAGAGCATGTGACATCGTTTATATTTGAAAAGGGTAATCTTGGCCTCTCTGGTGAGA | 120 |
| chr11 | 71135015 | 71135135 | 788855_47188552_ENST00000601538.6,E | 2 | - | good | AGGGTGAGGGTCTTCAGGGACAGCCAGCCTGAGGTATGCACAAACTTGGTGTGTTGGCTCAGCAGAGAAGGCTGAGGGCTGCGGGAGCAGTGTGTGTGAGGCAGCAGAGGTGGGCGGTG[illegible] | 120 |
| chr11 | 71135135 | 71135255 | 788855_47188552_ENST00000601538.6,E | 2 | - | good | TGCTGGGTATCTGGCTATGGAGTGAGTGAATTCATTTAAACCCTTGCCTCTGTGGGGTGAGGCGCCAATGGCTGTACAGAGCCTCAGAGGATGGGAGAGGGTCTTTGCTTGGAATCTTCT | 120 |
| chr11 | 71135255 | 71135375 | 788855_47188552_ENST00000601538.6,E | 2 | - | good | ACAGGGGGATTTCACAGCTAAGCGCCTGGGGGGCAGCTGGGCAGCAATTCCGCTCCACAAGGACACCTAGTGCAGGCTCTGCTCATCCTTGTACCCCATCAGCCGGTACAACCGAGTGAA | 120 |
| chr11 | 71135375 | 71135495 | 788855_47188552_ENST00000601538.6,E | 2 | - | good | AAAAAAAAAAAAACATATCCCCCCTAGTCTGGGCACACACCTATTGGAAGTGTGCAAGTCAGTGGATTCTTTTTCCTGAGGATTCCATGGCTCCATGGCTTCTCCTCGGTCTCTTTGTTT | 120 |
| chr11 | 71135735 | 71135855 | 788855_47188552_ENST00000601538.6,E | 2 | - | good | GGCCTTGGGTGGTTGTGTACATGTGAACACCGTGGCCACCGCTGAGCCCAGCCCTCCTGTGACTTCTTCCCGGTCTGAGGATAAAAATCAGGTCCTCCCCGGCTGGGCACGGTGGCTCAT | 120 |
| chr11 | 71135855 | 71135975 | 788855_47188552_ENST00000601538.6,E | 2 | - | good | TAGACCCCCGCAAGCCTGTGTTGTTCCCTCAGCCTTCATCACCACCCCACCTTTCCTGTCATCTTCCTCTTGGGCAACCAGAACAGCTTCAGACCAGACTGTCAGAAGTTCTCTAGGTGA | 120 |
| chr11 | 71135975 | 71136095 | 788855_47188552_ENST00000601538.6,E | 2 | - | good | TTTCGCCGTGGGGACCATCTTTGATTCAGCCGTCGCAACCAGAATGTAGGCGTCCTCCTGAGCTTCCATCTCTACAGCTCCTATAGCCAGTCCCTCCCTAAGTCTGGTCACTGTACCTTC | 120 |
| chr11 | 71136095 | 71136215 | 788855_47188552_ENST00000601538.6,E | 2 | - | good | CCCAGGGTGGCGTTTAGCAGAGCCCTGTTGAGCCTTGTTCTGATTCTCATCATCTTGCATGGAGGGACCACATCTTGTGGATCCATTCCCACTCAATGGTCAGCTGGATTCTTTCCACTT | 120 |
| chr11 | 71136575 | 71136695 | 788855_47188552_ENST00000601538.6,E | 2 | - | good | GACATGGAGTCAAACAACATGTGGCCTTCTGTGCCTGATTTCTTTGACTCAGCATAATCAGCTTTTTTGGGTATAAATTCAGTGGGTACAAATGCAGTTTTATTACATGGGAATGTTGCA | 120 |
| chr11 | 71136695 | 71136815 | 788855_47188552_ENST00000601538.6,E | 2 | - | good | CACCATCCTGGTGTGGGAGGTTTCCATCCCTTATGCTGGGTTGCAGTGGATCCCCACCCCCACTTCCAGCCCCTGACGACCAGGAAGCCACTTTCTGTCTGTGTGGATTTGCTTATTCCA | 120 |
| chr11 | 71137175 | 71137295 | 788855_47188552_ENST00000601538.6,E | 2 | - | good | AAGGATTTTAATACTTATTTTAAAAAAGGATTTAAAAAAAAAGAAAAGACTAAAACTTTCTTTTTTTCAAAAAGTTTTCTTGAGTTATAATTTATAAATTATAGGATTTACTTACTTTAA | 120 |
| chr11 | 71137295 | 71137415 | 788855_47188552_ENST00000601538.6,E | 2 | - | good | CTCAAGTTGTGCAAAACCAATGCTATGACCGTAACTTTCTCTAGGAACCTCCAGGTCTGGCCCCACTGTTATTTGCTCTCTTAGATAGCAGTTACCTTTTTTTTTTTTTTTTTTTTAAGT | 120 |
| chr11 | 71137415 | 71137535 | 788855_47188552_ENST00000601538.6,E | 2 | - | good | GATAGAATTTCCATTCACACAGCATCCACGGTCACATCTCCTGCTGCAGCCTCCTTTGGGGGGGCTGGGCCTTATTTCCCCCTCCATACCTTCTCTGTTGAAAGCCCAGATGCCCCTCAA | 120 |
| chr11 | 71137535 | 71137655 | 788855_47188552_ENST00000601538.6,E | 2 | - | good | GGGCCTTCCAGGGAGTGGGTGGCGGGAGGGCTTCTCGGCCAGGCCCAGCTCCCTGCAGCCAGCACCCTGGCACCGATAGAACTTCTGTTAGCGGCTCCCTGCACCCAGCACCCTGGCGGC | 120 |
| chr11 | 71137655 | 71137775 | 788855_47188552_ENST00000601538.6,E | 2 | - | good | AATGGCAGCCTCTTAATTCCACTTCAAGAAGGTTTGTCCTTGTTTCCCTCGGGTGGATTTTTCCAGGTAGAAAGCCAGTTCTGGTCAGATTTGCCTTCTGCTGCATTCTGTTCAACACCT | 120 |
| chr11 | 71137775 | 71137895 | 788855_47188552_ENST00000601538.6,E | 2 | - | good | TGCTGTGTGTGTGTCTGATTCCTTGCTCCTGGACCCTGCAAGGTGAGAGCTGGGTGTCTTCTAGGAGTTGATGGAGTCCATGTGCCTGAGAGATTGGCAGCTTGGGTGGTCGTGATCCTG | 120 |
| chr11 | 71138255 | 71138375 | 788855_47188552_ENST00000601538.6,E | 2 | - | good | GAGGATTCCTCTACTTTTGAATATTGGGAGGACTCTACCTGTGCTGGCAACGTCTGAGCCTATCTTCCCCTCCTGTACCTGGGGCTGGCTCCTCATCCTCAGAGACCATGAGGCGCCCGG | 120 |
| chr11 | 71138375 | 71138495 | 788855_47188552_ENST00000601538.6,E | 2 | - | good | TATTATTGTGGATTAAAATAGAACCCCAAAAGTAAGGAGGTTTCACCTCCTCCACCTTCAGGATTTAATCCCCTCTTTTTGGTCACAGGCACATAATTACCGCATTTGCATTTTGTATTT | 120 |
| chr11 | 71138855 | 71138975 | 788855_47188552_ENST00000601538.6,E | 2 | - | good | CACTCAAAATCCCCACTACCTTTGCCATGCTGGGAGGGCCCTGCCCTCTCATTCCCGCTGTGCTGCCTCACCCTGTGGCATGCATCTGCTGACGGGGATGTCCTCTCGTCTGTCTGCTGC | 120 |
| chr11 | 71138975 | 71139095 | 788855_47188552_ENST00000601538.6,E | 2 | - | good | CACGTGCATCTTCCAGTGGGGTACTTCATTTAGGAAAAAGCATTTGCACTGCCAACCTGACCCCCTCAAAGACCATGCATTGTCACTCAGCTGTCCTCCCTCTCTCCCCGCCTGTACAGT | 120 |
| chr11 | 71139095 | 71139215 | 788855_47188552_ENST00000601538.6,E | 2 | - | good | AGCTCAGCTGAAGCTGTCACTCTGAGGGCTGATCCTTGCGACCTGGCCCACCTCCCAGCAGACGCTCAGGGAAAATGAGTGTCCCATTCTGTGCTTGAGTCAGTGCTTCCATTTATAAAG | 120 |
| chr11 | 71139215 | 71139335 | 788855_47188552_ENST00000601538.6,E | 2 | - | good | GCCATGTTGCAAGGAGAGCAGAGAAGAAAATAGAACGGGGCTGTGGCGGCCTTTGTCGGGCTCTGCTGGGGTGTCTACGTCGGCCCCGCATCCGCCTCTCCCTGAGGCCTGGCACCGAGT | 120 |
| chr11 | 71139575 | 71139695 | 788855_47188552_ENST00000601538.6,E | 2 | - | good | AATGCACAGTGGTGGAGTGCTGGCTCTGGACATTCAGCATGGCCTTTAAAACAAGTGGCATGTGCCAAAATCAATACAAACTGGTGAGCTCTGTTATTTCTGGGGGAGTGTGAATGAGGA | 120 |
| chr11 | 71139695 | 71139815 | 788855_47188552_ENST00000601538.6,E | 2 | - | good | CTTCAGGCACAACCGGAGCCCTCACTTGAATAGTCGTGTGTGGGTGGGCTTCCCGGGCCACGCAGGGAGTGGTGGGGTGGGCTCTGGGCACAGGGAGGGCAGCCTCTCTGCATCCTCTCA | 120 |
| chr11 | 71139815 | 71139935 | 788855_47188552_ENST00000601538.6,E | 2 | - | good | CCTCCGCTGCTGTCCGGGCCCACTGCTGCTGGCCGAGTGAGTGCTGTTTCTGAGCCTCACCCTCCAGTGGAACTGAAGCTTTAAGTCATTGAGTGAGAGCCTCTAGCAGTGCTCACAACC | 120 |
| chr11 | 71139935 | 71140055 | 788855_47188552_ENST00000601538.6,E | 2 | - | good | CCGGAGGAGCCAGCCAGGAGCTGCCGAGTGAGTCTGCACCGAGGCCCCAGTGCCCGTCCACTGGGAGTTGGTGCTGGAGAAGCACCTGGGCGGCCCCTTTTCTCCCCCAATGCCACCCCT | 120 |
| chr11 | 71140055 | 71140175 | 788855_47188552_ENST00000601538.6,E | 2 | - | good | AGAAATCCCCCGGGTGTGAGCTCTCAGCAACAAAGCAGACATTGTTCCCGTGTGAGAAATCCCTTGTGGGGCTGAGTCCTCTGTGGAGCCAGGGACACGAGACAGTTCCTGGAACCCTGC | 120 |
| chr11 | 71140175 | 71140295 | 788855_47188552_ENST00000601538.6,E | 2 | - | good | AGCCAGGGCCTCTTCAGGGTTTAGTCTCTACTCGTGGGAGCCTGGTTGTTTCTGGAACCTTCTGAAGACCACTTTGCTTCACTCTGTTCAAGACAAACAAAGCCCCTGGAGGCTGAAACC | 120 |
| chr11 | 71140295 | 71140415 | 788855_47188552_ENST00000601538.6,E | 2 | - | good | GCTGTTTCCCCAGCAGGTGTTCTGCCCACTTTGCTAGAGTCACTGGAGACCTGTTGGAGTGATGGATGGAGCCACAGGTCTGCTCCAGCCCAGATTTAACCTACAGCACCTCCTCGTGAA | 120 |
| chr11 | 71140415 | 71140535 | 788855_47188552_ENST00000601538.6,E | 2 | - | good | CCATCGGCACTGGGGCAAGGGTGTTGGGAGATCAGGGGCCTTCTACACCCGGCCGGACCCTGCCTCCCTGCCTGTTTGACGCTTTCCTTGGCAGGGTAATCTTCCCCTTGCAGGGCAGGG | 120 |
| chr11 | 71140535 | 71140655 | 788855_47188552_ENST00000601538.6,E | 2 | - | good | AGGGGCCCATTACTGTTTCTGTTCTCCTGGGTCCACCCTCCTGCTGGTGCTACCCACACTCGGAGGCTCTCATCAGCCACGTAGAAGGGAGGCCCCGGTCATCTCAGCCACTGAGGTCCT | 120 |
| chr11 | 71143775 | 71143895 | 788855_47188552_ENST00000601538.6,E | 2 | - | good | GTTGCCACAGATGCTGAATGAGTGACGCCTTGGCCATTGCTCCCAGAGGAAGTGCAGTGTTCGATTCCTACAAGCCCCTGGTCACATGTTTGACCATAGGTCAATACGGAATCTTACTTT | 120 |
| chr11 | 71148695 | 71148815 | 788855_47188552_ENST00000601538.6,E | 2 | - | good | TTGAATCCTTCCGTGAATTGCCTTCCTTTTGCACAGTGTTATGTTTCAGGGATCATCCACATGTAGCTTGGGTTCATTCATTGTCACAGCTGTGGAGTTCATGGGAATGAAGAGACCACA | 120 |
| chr11 | 71148815 | 71148935 | 788855_47188552_ENST00000601538.6,E | 2 | - | good | GGCATGTGCCTTTCCAGTTGACTGCTTCTGTGCCGTTACGCACGGTGAAAATGCCCATGTACCATCTGTAGGACTGTGTGGCATCTTTCCAGCCTTTATCTGGAAATGATAGTATGTGGT | 120 |
| chr11 | 71148935 | 71149055 | 788855_47188552_ENST00000601538.6,E | 2 | - | good | CCAACTGTACCAAGTTTGACCTTGTCCTGTAGTTGCTGGAATTGCTTTCAGGAAGTTCATGGCTCCAAGCAGCGCGGCCCACCCTGTCTTCCTCCGAGAGGGAACTGCTCTCCTGCATTG | 120 |

| | | | | | | | | |
|---|---|---|---|---|---|---|---|---|
| chr11 | 71149055 | 71149175 | 788855_47188552_ENST00000601538.6,E | 2 | - | good | CTTATGCCCAGGCAGCATATTTTGGGGTGGCCTCTTCTGCCCCCTGCACTCCTGCCCTCCTTCCTTGCTCCTTGTGAATGATTTCTAACAAAGAGAATAATACGGCAAACATCTGTGTCC | 120 |
| chr11 | 71149175 | 71149295 | 788855_47188552_ENST00000601538.6,E | 2 | - | good | TGTCTCTAGTGATGAACTTCTGAAATTTACGTCTTGCTTTGAGGCACACCAGTGGCGGGGGAAGGCAGCGAGCTGTTCTCGCACCTGCTTTTCCTCCATCGTCTTCAGCTCAGGACTGTC | 120 |
| chr11 | 71149295 | 71149415 | 788855_47188552_ENST00000601538.6,E | 2 | - | good | TCTAGAGGGCAAATTATGGGGCGGCCAATCTTTAGGAAAGAAACAGCAGCTGAAGGCTAGTCTGTTGGTGAAGTCAGCTCTGTAGGTTCCTCCAGGGCGGGGCTGACAAGGGTCTAAAGC | 120 |
| chr11 | 71149415 | 71149535 | 788855_47188552_ENST00000601538.6,E | 2 | - | good | AGCACCCGTCAGTGAAGAGTGTCTGAAAAAGTGCCTACAGCTTGGGCTGGGTTGGTGCCTCAGAGCAGGAACAGTACATTTGTAGAGAAGCGACAAGACAAAGGGAAAGGACTGTGAGTT | 120 |
| chr11 | 71149535 | 71149655 | 788855_47188552_ENST00000601538.6,E | 2 | - | good | CTAATCCCTCCCCCTGCAGAGGAGAAATTTCTTTTCCTACTACCCTCCTCTCTACCTGGGGCCCTGAAATTGAGCTGACATAAGACAGTCGCAAGAGAAAAATGGTTTGTGAACACATAG | 120 |
| chr11 | 71150495 | 71150615 | 788855_47188552_ENST00000601538.6,E | 2 | - | good | TGTATAATTGGTGCACAGTAAATATATTTTGGTTTATAGCTAGAGACTAAATAGAAGATTAAACTATTGATTTGGGATTATCAATGTCATTGTTACCCTGACATTCCACTAATGTGAATA | 120 |
| chr11 | 71150615 | 71150735 | 788855_47188552_ENST00000601538.6,E | 2 | - | good | TGTGTTAGGTGGTTGGACACAGGGTGCTCCGAAAACTGTGGACCATGGCATGGCGCCTTTCTTACAAAGAAAGAACATGCGCTGGACATGAAGCCACACACACCCTAGGGGGATCCCTCA | 120 |
| chr11 | 71150735 | 71150855 | 788855_47188552_ENST00000601538.6,E | 2 | - | good | CCCCTTTTCTTGGCTCGTGCTGTCATCTGTGCCTTGTGAAGTGTCTAAACACAGGGTGAGAAGTTCCTGCCCCCACCCACTCCACGCAGCCGAGAGTCCCCAGGATATCCATGCAGGGAT | 120 |
| chr11 | 71150855 | 71150975 | 788855_47188552_ENST00000601538.6,E | 2 | - | good | CGGGAGCCTGTGATTTTTCTAGAGATGACTGAAATCCAGATGTTGTGTGAAAACTCCTAAGTTCAGGATTCTTATAAAGCCCATTCTCATTTTAGGAACTCGAGGGTAAGAAACAGGTGC | 120 |
| chr11 | 71150975 | 71151095 | 788855_47188552_ENST00000601538.6,E | 2 | - | good | CAGGGTGCAGAGGACATAGTGAGGTGGGGCTGGGTGAATCATGTCGCCCTTTCCCTGTTGAGAGGGCCACACGGCTCGGTGCCCACATGGCTGCCAGGTGGGTGGAGTGGGCTCAGGAGG | 120 |
| chr11 | 71151095 | 71151215 | 788855_47188552_ENST00000601538.6,E | 2 | - | good | AGTTATAAAACACTAGCACAGGGTCCACAATTCTGCTGCCAGGGCCCAGGGCTGATGGGGCGGGTGAGCTGAGTGCAGGGTGAGCCTCAGTGTACCTGGATGGTGAGGGGCAGTGGCTTC | 120 |
| chr11 | 71151215 | 71151335 | 788855_47188552_ENST00000601538.6,E | 2 | - | good | CGGAAGAGCGCTGGGAGTCAACAGTGGGTATCAGCTTTCTCCTAATTGGACATAGTGTTGGTTCCTGGTGAACTGCCCGTGGCCTCCCCGGGAGGTTCCTGAACACTCACAGGGCCCCCC | 120 |
| chr11 | 71151335 | 71151455 | 788855_47188552_ENST00000601538.6,E | 2 | - | good | TATCTGATGAGGAGTGGCTGCTGTCAGGTTTCTGGGCAGCAAAGCTGAGGGGTGAGGCCGGAGTGCCTGCTGAGGACATGGTTCCAGTGCACCCCGCTACCCGGCAGGCTCTTCCCACAC | 120 |
| chr11 | 71151455 | 71151575 | 788855_47188552_ENST00000601538.6,E | 2 | - | good | CTCTTTCCCTTCACCCTCATGCCTGCACTGCTTTCCAACCCAGCAGGGCCCAGGAGAATGAGGACGAAGAATAAAAACGGGTGAAATGGGGGTGGCATCGCAGCTACAGGTGGAGGTTGA | 120 |
| chr11 | 71151575 | 71151695 | 788855_47188552_ENST00000601538.6,E | 2 | - | good | CCCCCCGCCCTCCGTTTTTCAGATGTTGACGCTGAGGCTGGCCTCGCTACCACACTAGGCTTCTTTCTGCATTAATGAGTGACATGAGCTCTGAGTAGCTTTGGCTCCACTTAAGAAGGG | 120 |
| chr11 | 71151695 | 71151815 | 788855_47188552_ENST00000601538.6,E | 2 | - | good | GCAGCTTGGAGCCTGGACGGTCTCTGCCTGCTCTTGCCAGTCAGACGTTATCTGACCTGAGCTGGGGCAGAACCAGTGTGGGCCGTGTCTGCTCTGTGCCAGGGCTCTGAGCTGCGGGCT | 120 |
| chr11 | 71151815 | 71151935 | 788855_47188552_ENST00000601538.6,E | 2 | - | good | GCCCCCCTGACTGCTCTTGTGGGCGTCTGCGATCGGGAGCCCTGGCCTCCCACCCAGCGTGTGTTCACTGGGTGCCCACTGTCCACTGAGGGGGGGCTTCCTACACATGCACCCCTGGAA | 120 |
| chr11 | 71151935 | 71152055 | 788855_47188552_ENST00000601538.6,E | 2 | - | good | AGATATCTGCAGGCCTGGCAACACCAGACCTGCGTTCCTACCTACCAGCTGTGGCCAGAGTGGGAATGCTGCTGCCTGGCTGATTTGGGGACCACATTCTGCCAGCATCGCCCCCTTCGT | 120 |
| chr11 | 71152415 | 71152535 | 788855_47188552_ENST00000601538.6,E | 2 | - | good | CAGGAGCCCCTCCTTGCCGCCCTTCCTGCTGTGGGCTGTTGGTTTTGTTCCTGGACTCAGGTCCAGCTGTAGATGTGTACTATGTGGCCTGCAAAGTAATCCGTTAAAAACAGTGAATTT | 120 |
| chr11 | 71152535 | 71152655 | 788855_47188552_ENST00000601538.6,E | 2 | - | good | CACCATGGTTTGGGAGCAGATGTGCCTGTTTGTGACCATCACCCCCAGAGCCGGTACCAAGGGGACACAGGAGTCCTAGGGGGTGAAGATGTCCGCCGGCAGCAGCGGGGGCTGTGGCCT | 120 |
| chr11 | 71152655 | 71152775 | 788855_47188552_ENST00000601538.6,E | 2 | - | good | CTCCCAGGATAGTTCTGGAAAGAGCCCGGGAGGTGTGACCCAGCCTTCTCCTCCTCTCTGCTCACCTGCTTCTGCACGAAATCCCGGTTCCTCTTCAGACCCAGCACTGGAGCCCTTCTG | 120 |
| chr11 | 71152775 | 71152895 | 788855_47188552_ENST00000601538.6,E | 2 | - | good | ATAATTAAATGTTTTCCCAGGCAGTGATCAGGTGTCTGCTGTTCCCAGGAAGCTGCTAATCAGGTTACTCAAGACCCTTACCAAGGTCAAGACCCTCCCAGTCAGATCTCTGGCCATGAG | 120 |
| chr11 | 71152895 | 71153015 | 788855_47188552_ENST00000601538.6,E | 2 | - | good | TAGTATAAGAATGTCCCACGCAATTATTTGGGACATGCAGTAAAGTGTCTTTGTTGTTTGTCTGAAATGCAGATGGGACTTGCGCCTTCTGCCTTGCCTGGTGGCTCTCCCCGGGAGTTA | 120 |
| chr11 | 71153015 | 71153135 | 788855_47188552_ENST00000601538.6,E | 2 | - | good | GCTGGTCCTCACGGGGTTAGAGAACTCAGCCCAGCCCCCCTGGCTCTCCCATGGTGTTGCCAGATAAAACGCAGCGCATCCAGTTCTGCCTGAATTTCAGATGAACAGCAGAGAGTTTTT | 120 |
| chr11 | 71153135 | 71153255 | 788855_47188552_ENST00000601538.6,E | 2 | - | good | CCCGGGGTGGTGAGTAACCTTCCCCTCCTCTGATCTGGCACTGGCTTTTCCTGGGGAAGCCAGGATTTCAGATTGCGTCTTCGGCTTTGCTTTCATTCTCTCTGCGGAGCCTCCCGGCGT | 120 |
| chr11 | 71153255 | 71153375 | 788855_47188552_ENST00000601538.6,E | 2 | - | good | TAGGAGGGCCAAGGAGAGAAACACACACTAAGAGTCATTGAAGTTCATGGACCTGGGGTTTTATGGCCTCTCCCCCGCCCCCGACCCGCCCGAGAGATTAGTTACACAGAAAACGGTTGC | 120 |
| chr11 | 71153375 | 71153495 | 788855_47188552_ENST00000601538.6,E | 2 | - | good | GGACGAATGAGCCAGTTGAAGATTTTAGGTTTTCTTGGAGCAAGTTAGCAGCAAAATGCTGTTTTTGTTGAACTTGCTTTGACTGTGATGATTTTCTTTTCTCCTCGTTAGTAGCTACTC | 120 |
| chr11 | 71153495 | 71153615 | 788855_47188552_ENST00000601538.6,E | 2 | - | good | CGTGAGGTTTTTTGTTTGTTTTTATTATGTATCTTCTCTCCCAGGTCAGCTTTCATATTTTCACAGTGGCTGAAATAATTAGTGGAAAAGTCTGTCTACCTTTTCATCAGAGGTACTGGA | 120 |
| chr11 | 71153975 | 71154095 | 788855_47188552_ENST00000601538.6,E | 2 | - | good | TCACATACAGGGTGTCTCTCAGAGATTCCATATCAATAAAAACCTTCTTTTCTTTTGGTGAAAAAGTAAATCAGAGTTCAGATGAGGAAACTCTTGCACTTTTTTTGTTGTTGTTGTTTA | 120 |
| chr11 | 71154095 | 71154215 | 788855_47188552_ENST00000601538.6,E | 2 | - | good | TGTAAACGGCACCAGGGGAGCCTCGGGGTCCTCACTGCCTTGAGGCCGAGCACATTTTACACACAGGAATGCACAAAATAAATGAAATGTGTAGTCACAAAATCTCGGTTTCCATGTGAT | 120 |
| chr11 | 71154215 | 71154335 | 788855_47188552_ENST00000601538.6,E | 2 | - | good | AGAACGGCAATTTTCTAACATCCGATGGAACCTGGATTCCTTGAATATTCTGTTCTAGTTCCACAGGTTCCAATGGAGTGCTGTCAGTCCTGCAGCCGTTTGACCCCGAGTTTAAATGTC | 120 |
| chr11 | 71154335 | 71154455 | 788855_47188552_ENST00000601538.6,E | 2 | - | good | CTGCCCGGCTGGCCTCCCCTTCCCTCACATCTGGCCAACTCTGGACTGGCTGGCAGCCCACCCTTCCCCAAAGCCCACACCCTCGCACTTGCTCTCAGCCCTATGTCTATGTCAGCTCTT | 120 |
| chr11 | 71154455 | 71154575 | 788855_47188552_ENST00000601538.6,E | 2 | - | good | CTTGGCCAGGGCCCCCATGTGTTGTGTTCTAGCTCCTTCTCAGCTGATGTGTTTTGCAGGTGCAGGGAGGAGAACAGTGGGTTAAATGAACAGAAGGAAGCCAGCGTGCCTTTCTCTCTT | 120 |
| chr11 | 71154575 | 71154695 | 788855_47188552_ENST00000601538.6,E | 2 | - | good | CGTGGGCTGGGGTAGGTCCATCCCTCTGGGAGCGTGGGTGGGGTAAGTTCACCCCTCCAGTATTTGGGGCAGTGTGTCAGAAGGTCTGCTTTTGCAGAAACTTCCAGAGCAGTCCACCCC | 120 |
| chr11 | 71155535 | 71155655 | 788855_47188552_ENST00000601538.6,E | 2 | - | good | GGGACAGCTTCTAGGCAAGACTGCTGGAAGAGGTGGTGCTAGACTGAATCCCGGCATTGGTGTGGTGGGATCCTCAGACCTCCCTGCCAGGTTCTGCTTCAGTTACTGGGGGCTGGTGGT | 120 |
| chr11 | 71155655 | 71155775 | 788855_47188552_ENST00000601538.6,E | 2 | - | good | CCTGACATGGGGTTGCCACGTGGTTTTGTGTGGCCACGGCCATAGTCACTCTACTTGGGGTGGGTGGGACGGGCCACCTCTCTGGGGAGCTTTGAGGGGTATGGGGCAGGGCGGGTTTGG | 120 |
| chr11 | 71155775 | 71155895 | 788855_47188552_ENST00000601538.6,E | 2 | - | good | CTTTTCCAGGCCCCGGGGATTCGGACTTCCACATACGGATTTGAGCAGGGGACACAGTTCAGCTCACAGTGTGTCTAATGCCATGCTGGTGCCTGGGGGCTACGTGGCAGATGGCACAGC | 120 |
| chr11 | 71156255 | 71156375 | 788855_47188552_ENST00000601538.6,E | 2 | - | good | GATGCCAGCAGGGCTGGTAACAGAGTGTTTGTTTGATGGTGATTAGGGAAATCCCAGGAAACCTGGATAGATGGAGGTTCTTTTCAGAAATCTGTTTGTTCCATGCATCTGTTTCACAAA | 120 |
| chr11 | 71156375 | 71156495 | 788855_47188552_ENST00000601538.6,E | 2 | - | good | GAGCGAGTCTGAAAGGGAAGGATTTGGCTGAACGGGGAGAGAAACTCAGGACAGGAGACGGTGACACCAGTGGAGGGCGGAAACTCAGAGGCAGGATCCCGTTTCTTGAGTTCATCCAGA | 120 |
| chr11 | 71156495 | 71156615 | 788855_47188552_ENST00000601538.6,E | 2 | - | good | GAGTCAGGAGGCACAGCTTAGGAAGGAATTAGATGTGGAATTAGGCGTGGACTCCAGGCCCTCTGAGGTGAGAGGAAGGGGAGCTTGTGTACCATAGGCAGTTGGTTCTGACCTGGGGTA | 120 |
| chr11 | 71156615 | 71156735 | 788855_47188552_ENST00000601538.6,E | 2 | - | good | TAAAAATTCAGCAGCAAAATGACCTGTGTGTTTTTGTTCTATCCAGTTACATTTCACAAATGTATGTGTCCAGAGGAGGGTTGGAGGGTGTATGTGTAATGGAATTAGTTCTCAGAGCTG | 120 |
| chr11 | 71156735 | 71156855 | 788855_47188552_ENST00000601538.6,E | 2 | - | good | TCTCTTCCTGTTGTGGGAGATTGTTTATGAGCCCCAGGGGAAAAGAATGTCTGGGGCAAGTGCACTTGGGAAATGCTGCGTTTACCCCTTTCTCTGAGACACTGCCATATAAAGCCCTCT | 120 |
| chr11 | 71156855 | 71156975 | 788855_47188552_ENST00000601538.6,E | 2 | - | good | GATAATACTTGGCAGGGTGCTGGCCTCAGTGGGCCATGCATTGTAGGTATGTGATTGGCTTTTGCTTCTAAGATCTTGAGTGTAGCAAGGAAGGGACTATGGGCAACTCAGACCATAGCT | 120 |
| chr11 | 71157215 | 71157335 | 788855_47188552_ENST00000601538.6,E | 2 | - | good | CATCTAAGGGAGTGCTCTTAAGTGACGTATCATTTTAATTCAGGCATCTGATTTATTAGTAGCTTGTTTTCTGATTTGGGGATATTGGTTTTGGCATTAAGAGGAAGGTGTGAAGTAGCC | 120 |
| chr11 | 71157335 | 71157455 | 788855_47188552_ENST00000601538.6,E | 2 | - | good | TGAATGCTGAGGCATGGAAGAGAAATAGACTTACCTAATGAATGTACCATCTCACCAGTCCTTCTCAAGACCTGGACACCCCAATGTCTTAATAGAACACACCAGTGAGGCATTGCAATT | 120 |
| chr11 | 71157455 | 71157575 | 788855_47188552_ENST00000601538.6,E | 2 | - | good | TTTTCTAGCAGAAGCTGAGAGACTGTTGGAGGCTGTGTTGAGTCTGGCCAAAGAAAGACGGTGCAGACCACATTAGTGGTGTGGTTACTGGTGCCACCAAGAAAACGGCACACTTCCACA | 120 |
| chr11 | 71157575 | 71157695 | 788855_47188552_ENST00000601538.6,E | 2 | - | good | CTGAAATGGAAACCTGTCATAAAATGAAGACTTCTTTATAAATCATGTGCTCTTTACAGCAGCCTGGACAGAAACCCACTGAGGCCCAATTAGAAAGCCAGACAGACTCTTCTCTCTGGC | 120 |
| chr11 | 71157695 | 71157815 | 788855_47188552_ENST00000601538.6,E | 2 | - | good | CCTGGTCACGGTGAATGGTTTCGGCAGAACTGTGAGATGGGTAGGCCAGCATATCACAAACAGGCTAAGAGTGGGTTCATGCTAGCTGGCCAGAACCCCTTGAATTCTGCTCTAAGAAAA | 120 |
| chr11 | 71157815 | 71157935 | 788855_47188552_ENST00000601538.6,E | 2 | - | good | TAAGAGCTGCACACAGGGGCAGTGTCCCAGGAGGGGCTTGTTGACCTGTTGTCTTGATTGGTTTCCTTCTCTGTAGGTTGTCTCCCAGGAAGCCTTTTACCTTGGGCAAGTCGCTCCCTT | 120 |
| chr11 | 71157935 | 71158055 | 788855_47188552_ENST00000601538.6,E | 2 | - | good | TCTTCTATTCCCAGTTTTCCAAGACTTTGGTCATGACTAGGGGTGGTTTGTGTCCCCCCAGGCTGTCCGTGGCTGGGCCTCTTGTTCCTGCTGGACACAGGGCCTTGTTGGACCCACACT | 120 |
| chr11 | 71158055 | 71158175 | 788855_47188552_ENST00000601538.6,E | 2 | - | good | AAAATAATGATTTCCCTGCTTTTAGTCCATAAACAATGCATTCGGTTATCTGTTCGCTTACTTATTTTTGATGTTTGCTACAGGATTTGAGTAGGTGCTCTGTTCGAGTTTGGGAAATTC | 120 |
| chr11 | 71158175 | 71158295 | 788855_47188552_ENST00000601538.6,E | 2 | - | good | TTTTGTATATTCTTATCTGGCATTTTGCTGGACTTTTTTTTAACTAGTTTTATGGTTACACATTATTTAATTTGTCCCTAGATTTTCTTGGATTTTCTATTCACAGAAGCATATTATCTG | 120 |
| chr11 | 71158295 | 71158415 | 788855_47188552_ENST00000601538.6,E | 2 | - | good | AAGGTCTCATACATATTGTGTTAGATTTTTTTTTAGTTGACTTTGGTTATTGTTGGTGGTATTATAGTGTTTTTTGAATGTTTTTCAATTGTTTATTGCTAATGTAACAATTCACTTAAC | 120 |
| chr11 | 71158415 | 71158535 | 788855_47188552_ENST00000601538.6,E | 2 | - | good | TTGAAGATATTAAAGCCTAATGTCCATGATCATTGTATAGCTCTTCATTTATTTGGGTCTACCTTCAATTCTTTCTTTATTTCAAAAAGATAATGCTATGTAGTATTACTTTTCTCCACA | 120 |
| chr11 | 71158535 | 71158655 | 788855_47188552_ENST00000601538.6,E | 2 | - | good | TTTTATGTGGCTCTCCATCTTCTTTGAGCTTTAAGATCAGAGTGTTAAGTTCTACCAACAGTCCTATCAGGGTTTTATTAGAAGTTACATCAAGTTTATATTTTGTGGAGAATTTTCATC | 120 |
| chr11 | 71158655 | 71158775 | 788855_47188552_ENST00000601538.6,E | 2 | - | good | TTTTTAAATCCATCTATTTGTCTTTGTCTAATACCATACTGTTTTCATATTATAACTTTTTCTGGTAAGCCTTGGCATCTGGTAGAATGAGTCTCCTCTTAGTCTCCAGAGTTATCTTGG | 120 |
| chr11 | 71158775 | 71158895 | 788855_47188552_ENST00000601538.6,E | 2 | - | good | TTTCTCATCTTCTCCATGTGAATACCTGCTAGTGTCAGCCTCATAGGTTTATTAATTCTTCTCTTTCCCCCAACACATATGGGAGTCTGATATTTACAAGCCCAGCCTAAGGTTCTGTGC | 120 |
| chr11 | 71158895 | 71159015 | 788855_47188552_ENST00000601538.6,E | 2 | - | good | CCTTACATGAAGGCATTGCCTCTTACTTTTTCTTCTAGAGATTTGTTGTGTTGCTTTTCACGTGACAGTAGTCAGTGCATCTGGAATGCGTTTGTGTGTCTGGAGTGGCATGGGGCTCTA | 120 |
| chr11 | 71159015 | 71159135 | 788855_47188552_ENST00000601538.6,E | 2 | - | good | TCAGCGTGATGAGAGTACATTGCTGCACATTTCTGTTGCTCTAACCCACTAAGTCTGTGCTATCTTCTTACACAGCAATAAAGCAGCATTTATTGAGAAGCCAGAATCCCGTCTTAGGGA | 120 |
| chr11 | 71159495 | 71159615 | 788855_47188552_ENST00000601538.6,E | 2 | - | good | TGCCTTGTAGAAAGAAACACTTTTTTTCTCGCCTTGTTTTATTGAGCAAATAACAATTGAACATATTTGTGGCTGGCTAATTAATGGCCCCCAAAGATATTCAGGTTCTAGTCTCTGGTA | 120 |
| chr11 | 71159615 | 71159735 | 788855_47188552_ENST00000601538.6,E | 2 | - | good | TTGGGATTACGGGCATGAGCCACCATGCCCGGCCTGAGGTGTCTTCTTGCTGTTGTGTCTGATGGAAGTCTTAACTTTTACATTGTCTAATTTGTCTGTCCCTTCATGCCTTGTACCCGA | 120 |
| chr11 | 71159975 | 71160095 | 788855_47188552_ENST00000601538.6,E | 2 | - | good | TGTACCCATTTGCATCTGGAATTGTTTCTTTTTTGCCCTGTAGGAATTCTTCCCAGATTTTCCACAGCGCTCTGTCCTATGTTGTAAAGGTCCCTTTCAGGTGTCTTTTTTTTTTTTTTT | 120 |

| | | | | | | | | |
|---|---|---|---|---|---|---|---|---|
| chr11 | 71160095 | 71160215 | 788855_47188552_ENST00000601538.6,E | 2 | - | good | TCCCTTCCCCATCCTGGGACTGCATTTTCACTGTGGGCCTTGGCATTGCTTGTTTTCCTGGTACCTTTGGGATGTGTTCTCCCATGCCAACTAGCCCTTCAGGTGTCCCCTGCGATGGAC | 120 |
| chr11 | 71160215 | 71160335 | 788855_47188552_ENST00000601538.6,E | 2 | - | good | GGCCTTAGCATATGAATTTTAGAGGGAGACAGTTCAGTTCGTAGCAGTGAGAATTTGAGGGCTGCTTTAAGGGGGGATGACAGAGTCCCCAGGAGCCTCCGGAGCAGTCCCACTTTGCAT | 120 |
| chr11 | 71160455 | 71160575 | 788855_47188552_ENST00000601538.6,E | 2 | - | good | CATTGAGGGACTGCTTCCTGGCTTGCAGAGGGTTACCTTCTGACTGTGTCCACACATGGTTGGTTGCAGGGAGATGGGAATGCATGACCACACGTGAACACGCATGCTGGCTCTCTGATG | 120 |
| chr11 | 71160695 | 71160815 | 788855_47188552_ENST00000601538.6,E | 2 | - | good | CCCAGCAGGGATTTTCCTGGTTTCTGCAGGGCTCTGGGGACAAAATGGCATACGTTGGAGTGGCCTCCTCAACTTTTTCCCCATCAACTCTGTTATGTTTTACACATGATGTTGTAGAAG | 120 |
| chr11 | 71160815 | 71160935 | 788855_47188552_ENST00000601538.6,E | 2 | - | good | GGTTTCTATTGGAAGCCATCTTCTTTGGAGACAGACATGTCTCAGGAGGCTCAGCAGCCCAGGGATGAAGGCTGCCAACGTGGACTGTCATTTAAGGTGTCATTGCTGGAGCTGAAATAA | 120 |
| chr11 | 71161895 | 71162015 | 788855_47188552_ENST00000601538.6,E | 2 | - | good | AGGAGAGGCTGTTGCAATGGCTTAGGGAGCCTCCACACTGTCTACCAGATTTGAGGATGCTGCCCTGCCTGTGGACAGACTGTGAAGACATTTCCTCCTGTGTCTAAGAAAATGCCAATG | 120 |
| chr11 | 71162015 | 71162135 | 788855_47188552_ENST00000601538.6,E | 2 | - | good | AAGACAGTGGCATTTAAGCTGTTTCCAGTTCATTGGTTAATTCACCAAGTGTCCATTGAGATCATGTGATGCTCCAGAACAAAAAGCAGAAAATGCTACAGGGGTGTCTCATTCAGTGTA | 120 |
| chr11 | 71162495 | 71162615 | 788855_47188552_ENST00000601538.6,E | 2 | - | good | TGACATTTCCCCATTTCACTGTCACATGCCAACGTATGTTACTTGATTTGTCTTCACTGTTCTATAATTTTACCTGATATTACTATACAATCATTTACTTAGTTTCCTGTTGGTGGCTGC | 120 |
| chr11 | 71162615 | 71162735 | 788855_47188552_ENST00000601538.6,E | 2 | - | good | TTTGAAGCAGTAATCACTGTTTTGAAGTTGGGCTATAGATAGGTGAAGATAGCCATTGCCATCTATATTTTTAATTCATGTATGAAAGTGTTTAGATAAATGTATCTATTCTTTTGCAAT | 120 |
| chr11 | 71163335 | 71163455 | 788855_47188552_ENST00000601538.6,E | 2 | - | good | CAGCACGGCCTGCCGTGTTCTCACCCACTGAGATCCCCCCACAAGCACATCTGTCCAGCTTCCAAAATGAATAGATTTGATTTATCAAGGGCTTGCACTGGAAAGACCATTACAGGCAGA | 120 |
| chr11 | 71163455 | 71163575 | 788855_47188552_ENST00000601538.6,E | 2 | - | good | AGTCACAATGCCCTCCTATCACATTCTCCTGGGACTGACAAGAGATACTAAAATGTAGGCCTAGCTGGTAGGCTGATGTGCAGGAAGCCCTCAAGAAGCATCAGCTCTGCCTCCCTGCAG | 120 |
| chr11 | 71163575 | 71163695 | 788855_47188552_ENST00000601538.6,E | 2 | - | good | TATCTTTTGAAAATCGTCTGGAGAATCTTGAATTTGGATTGAGTGTGAGCTGCCTCTCTTTCTGGCTGTGATGGACCAGGAGCTTATCTTCTAAGGCTCCTTATCTATCAGATAAGGGGA | 120 |
| chr11 | 71163695 | 71163815 | 788855_47188552_ENST00000601538.6,E | 2 | - | good | AGGGTGTGTGATTTGCGTGAAAGGATGGGGAGTGAATTGATCCGAATTTCACTCCTCTAAGAGCTCACACTGCAATCTTTTGGGGCAGTTTCCACTGAGCTCCTTTTTGCACAAGCAAAC | 120 |
| chr11 | 71163815 | 71163935 | 788855_47188552_ENST00000601538.6,E | 2 | - | good | TACACCTAAACCTGCTCTAGAAAAATAAAGCCTATTAATTTAGAAGTGCACAATAAAAAGCACGCATGTATAGATATTTTGCATTCTTCCTTCTGTCTTCCTCCAGCCTCCCTGTTGAAG | 120 |
| chr11 | 71163935 | 71164055 | 788855_47188552_ENST00000601538.6,E | 2 | - | good | TGTATTTTGATTTTACATTTCCTTTTAGATCCTAAGTGCAACAGGTTTGTTAGAGCAGACAGACTACAAACTAGGAACTAGATCAGTGGGAACTCTCTGTACTATCAGCTGAGTTTCCTA | 120 |
| chr11 | 71164055 | 71164175 | 788855_47188552_ENST00000601538.6,E | 2 | - | good | CCCTTGCATTCACAACCCAGGAGCCAGAACTCTCCAGATGGACCCAAGTCAACTGTGGGGAGGCTGAGGAACAAACACAGGGTACCTGAAAATTTGGAGACCACTGGCATCTGGGAGGGT | 120 |
| chr11 | 71164175 | 71164295 | 788855_47188552_ENST00000601538.6,E | 2 | - | good | ATATTTAATCTTCAAAGGCCAGTTAATCCCCCAGCCCTCCACTGCCAGGTACTGGGATTTTAAGAACTTGGCTTGTACCTGGGGATGATTTTAGTGGCTAGTCTTGACTGCCACACCCCT | 120 |
| chr11 | 71164295 | 71164415 | 788855_47188552_ENST00000601538.6,E | 2 | - | good | GGAGCTGGAATAGAACCCGGTCATGCTTCTGTCTTCTCTGCAGTGGCCTAGGTTGCTGGGTCTGTTTAATGCCAAAGCCTCTCTGCCTAGACAGACACTGCTGAGATGCAATGTGCTGTG | 120 |
| chr11 | 71164415 | 71164535 | 788855_47188552_ENST00000601538.6,E | 2 | - | good | GCTGTCATGAGGCCCCATTGTGAATTGGAGGTGTCTTAGGTGGGTCACCCTGCGGTCAGCCCCTCACTGGGGACACTGGGGCCCGCTGGGTTATTCTGTGGCTGCTCTGGTGTTGGGTCT | 120 |
| chr11 | 71164535 | 71164655 | 788855_47188552_ENST00000601538.6,E | 2 | - | good | GACATTTCATCGCTCCATAGAGGTCTCAGAGCTGTGTGATTTTGTTGTGTGATGTTCCCAGTTCCTAGACACAGTAATTTAACTTAGTTTCTGGTAACCCATAGTTTAGAAGGTGCGGCA | 120 |
| chr11 | 71164655 | 71164775 | 788855_47188552_ENST00000601538.6,E | 2 | - | good | AATTGGGTATGGCACAGAAAGCTGTCGAGAGTTCTTACCCTTGATTTGAAATCTGTTTAGTGTACAAGAACATCTATACACTAAATATGTTTAGTAACAGGCACATCTCCGTTTACCAGT | 120 |
| chr11 | 71164775 | 71164895 | 788855_47188552_ENST00000601538.6,E | 2 | - | good | ACTCATGGTGGAGTCATAGCCGTCCCCAAGCAGGTGACTGGAAAGTGCCCTTCAGACTCAAGGACTTGGCGTCTCTCCTAGGCCCCCAACAGATAGATGCCTGAGGCCGGCGTTTGATGG | 120 |
| chr11 | 71164895 | 71165015 | 788855_47188552_ENST00000601538.6,E | 2 | - | good | AAGGATTGCAAAGGACAGCTGGGTTCCCACAAACATTGGAATAATAGGAATTAAAGGAACCCACCTCCAGCTTAAATACTGTCATTACAGATGATGGCAGGATGGAGTAGCCACAGCCCC | 120 |
| chr11 | 71165375 | 71165495 | 788855_47188552_ENST00000601538.6,E | 2 | - | good | CACAGCATAGCGTCCTAGGAGCAGGATACACTGAGGACACGCACACACACCTGGGGGCAGCTCCCAGGGAGGTCTGCAAGAGATTCTCTCGCCATGAGGTTAGCGCAATAGTTTAAAATG | 120 |
| chr11 | 71165495 | 71165615 | 788855_47188552_ENST00000601538.6,E | 2 | - | good | TGAATGATCTCGCTGTAGGGTTTGACCAGGCTTCGCAGAGAGAAGCGACAACAGCCTTCCCGCTTGGTGTTCGGGCTTCATTCTTCATGCTGTAGGGGCCTTGAGGCTGTGGGGCTGCTG | 120 |
| chr11 | 71165615 | 71165735 | 788855_47188552_ENST00000601538.6,E | 2 | - | good | TTGGCTGAGCCCCTGTTTCACTTGGGGGTATTTCAAGGTGAATGATTACCAGGTCCCCCGGCAACCCCTGGCATCCAAATCTCAAGGGCTGAGAAGTTCCCAGGAGGAAGATTTCTGGGA | 120 |
| chr11 | 71165735 | 71165855 | 788855_47188552_ENST00000601538.6,E | 2 | - | good | TGGACTGTGAGATCAGCCGGGAAGCGTTCCAGGAGGGAGGCCCGAGGATGAGACGTGTCTCTGCTCCTGAGAAAAGGTTACATGACTACACCACGTAAAAGATGTTGCCAGTTCTCAACC | 120 |
| chr11 | 71165855 | 71165975 | 788855_47188552_ENST00000601538.6,E | 2 | - | good | TGGAGAGTTGTCCTCCACTCACGCTCAGCAGAGGATGGAGGAGATGGCTTCCTGACAGAGTGCTCATGGAGGGTGCTCCCCGGACATGGTCTACACTGGGTCCTGAGGAGTTGGTGGGGC | 120 |
| chr11 | 71176415 | 71176535 | 788855_47188552_ENST00000601538.6,E | 2 | - | good | GAAGTTTAATTTTTTTCCATGTTGCCACTTACCTTTTTGAACATACAGAATACAGTTAATATGATTTGAATGCTTTGTCTGCTAATTATAACATTGGTATTTTTGCTTACTTTTGATTGG | 120 |
| chr11 | 71177375 | 71177495 | 788855_47188552_ENST00000601538.6,E | 2 | - | good | TTCATAGCATAAATCTTTATCTTACTACACTTTTATCATCAAGTAGTATTATACCAGTTCAGGTAAGCATAAGAACTTTACAATAATATGCTTCCATTTTCCCACTCCAACCTTTGTGCT | 120 |
| chr11 | 71182895 | 71183015 | 788855_47188552_ENST00000601538.6,E | 2 | - | good | CAGGTCAGAGATGAAGGAGTTTATTACTCACACCAGTAGCACTAGACAGAGCCTCAGGGTTTTCGTGCTGGTTTACTGAGCCCTGATTCCCACACTGTGATGCAGAGAGGCATGGATGAT | 120 |
| chr11 | 71183135 | 71183255 | 788855_47188552_ENST00000601538.6,E | 2 | - | good | TTTGCCCCTTGTCCCTCCATCACTTGGACGAGAACATTTGCTACCTGATCCAGGGAGGGTTCTCATATCCTTGGAGCCCAGCCTGCAGAGCCTCTTTATTTCGCCTTCCAGAAAGCCTTG | 120 |
| chr11 | 71185415 | 71185535 | 788855_47188552_ENST00000601538.6,E | 2 | - | good | AGAGGGAAGAAAGAAGCTTCATGCCCAAGAATGGTCAGGGTGGAACCTTGAGTTTTCACAGTTGGTCTATATAATCCCCATCCTTCTCTTCCTGGCCATTACTCAGAATGTGGCCAAGAG | 120 |
| chr11 | 71200175 | 71200295 | 788855_47188552_ENST00000601538.6,E | 2 | - | good | TTACGGATGAGGACTACTGCATAGATACACCTTGAGGACATTACGTGTGGTGAAAGGAGCCAGACACAGAGGCCACGTATGATGTGACTCCATTTATCCGCATTGTCCAGATTGGTCAAA | 120 |
| chr11 | 71203895 | 71204015 | 788855_47188552_ENST00000601538.6,E | 2 | - | good | GGGTGGAATCAGCAAAGGCAAAAGGCATGTGGGGCAGAGTCCAGGAGAGACCAGGCGCCCGCTTTCCGTGTTCTCTCCCGGGGCGGCACGTAGATGGTTCCCCCCAAAGATGTGTGACAG | 120 |
| chr11 | 71223815 | 71223935 | 788855_47188552_ENST00000601538.6,E | 2 | - | good | AGCCCTGTGGGCAATTTCTCAATCCTTCTGAGCTATCCTCACCAACGTGCCCATGCCAGACCCAGACAGTGTGGCTGCAGACAGATGAGTGGTGCAGGCGCTTCCAGTTCCTCTCCGTCA | 120 |
| chr11 | 71224295 | 71224415 | 788855_47188552_ENST00000601538.6,E | 2 | - | good | AGGGTGGCTGCAGACAGATGAGTGGTGCAGGTGCTTCCAGCTCCTCTCCGTCATTTTCACATGAACGGGGGTGCGCCTCGTGATGCCCACGTGCAGGGAAGGGTGCAGTGAAGCCCAGGT | 120 |
| chr11 | 71226335 | 71226455 | 788855_47188552_ENST00000601538.6,E | 2 | - | good | GTTCAGCTTCTGGAATCTGTAGGTTTATATCTCTCAGCAAATTTGGGACATTTTGAGTACTTTTTCAGACCCTCCCTCTCTTCTCTCTCCTTCTGGGATTTCAAACATTTAAAGTTAGGT | 120 |
| chr11 | 71228735 | 71228855 | 788855_47188552_ENST00000601538.6,E | 2 | - | good | TCCTTTGTTAAAGATTTCTGTGTCTATATTCATGAAGACCTTTGATTTACGTTTGTATAAGGTATCTTCTTTTTTAAAAAATTATTAATTAATTTTTGAGATGGAGTCTTACTCTTGTCG | 120 |
| chr11 | 71229095 | 71229215 | 788855_47188552_ENST00000601538.6,E | 2 | - | good | CAGCTCTATGTTGAGTAAGAATGGTGAGAGCAGGCATCTTTGCCTTGGTCCTGATCATATGGGGAAAGCATTTTGTTTTTATTTGGTGAATCTTGGCTCATGTTTCTTCGCAGATGTTCT | 120 |
| chr11 | 71232935 | 71233055 | 788855_47188552_ENST00000601538.6,E | 2 | - | good | ACAGGTGTGAGTCACTGTGGCTGGCCTACATGTTTTTGTGTGGATGTATGTTTTATTATTTTTAGTTGTATGCCTAGGAGTGAAATTGCTGCCTCTCATTGAAGCTGTGTGAAAGTTTTG | 120 |
| chr11 | 71235215 | 71235335 | 788855_47188552_ENST00000601538.6,E | 2 | - | good | AGGTGTGAGCCACCGTGCCCGTCCTGTGCAAGAGTTTTTATAGAGCTTAATCTCCGGTACCTTCCGGAGCTCAGCTAGTGGGGCCAGAAGTTCCCACCTTCTTAGACACCTGTGTTCCTG | 120 |
| chr11 | 71245295 | 71245415 | 788855_47188552_ENST00000601538.6,E | 2 | - | good | TATTTACGCTATTAGAATTTACAAATATCTAATTATTAATGCATTTAAAAATAGTAATAAACTCATGACATGTTGGTATAAATAATGTGTTTTTATGGATAACAACTATTTTCAAAATAA | 120 |
| chr19 | 50661988 | 50662108 | 788855_47188554_ENST00000293441.6,E | 2 | - | good | CACCGAGCCCAGTTCCTGGACCACGAGATCGATGGCTCCCACCTGCCCGCCTTGACCAAGGAGGACTACGTCGATCTAGGTGTGACCAGGGTGGGCCACCGCATGAACATCGACCGGGCT | 120 |
| chr19 | 50663068 | 50663188 | 788855_47188554_ENST00000293441.6,E | 2 | - | good | GCCATAACCTGAAAGGCAGTATAGAATAGTGGTGAAGAGCTCAGACTTTGGTGCCATATAACCGGGCTCTGCCACTTCCTGTGTGTCCTTAGGCAAGTTATTTAACCTCTCTGTGCTGAT | 120 |
| chr19 | 50668348 | 50668468 | 788855_47188554_ENST00000293441.6,E | 2 | - | good | CACCAGTAGCAGCGGCCGCAGCAGCCAGGGCAGCAGCACCGAGGCGGAGCCCCCACCCAGCCGGAGCCCACGGGAGGCGGCGGCGGCGGCGGCTCCTCGCCCAGCCCCGCCCCGGCC | 120 |
| chr19 | 50670268 | 50670388 | 788855_47188554_ENST00000293441.6,E | 2 | - | good | AATGATAGTGGCTGGCCCAGGGTGTTGACTCTGGAGGGGTGAGAAGGGATTGGATTCCGGATATGTTTTGAAACAACAGGATTTGCTGGTGGATTGGATGTGGGTGACTTGAGGAAGAAA | 120 |
| chr19 | 50673748 | 50673868 | 788855_47188554_ENST00000293441.6,E | 2 | - | good | TGAGCCGTGTTAACGCCACTGCACTCCAGCCCTGTCTCAAAAAAAAAAAAAAAAAGTTTAAATTTTATGACTTTGTTGCTGCAAAGACCGATATTATCCAGGGTACAATTTTCTCTTTTCT | 120 |
| chr19 | 50676268 | 50676388 | 788855_47188554_ENST00000293441.6,E | 2 | - | good | GCCCCCTCCTTCACCTCCTCCAGGAGTTTGCTCAAATAATACCTTCTCAGTGAGGCCTTCCTTGACCTCCTTATTTTATTTATTTTGAGACAAGGTCTTGTTCCGTTACCCAAGCAGGAG | 120 |
| chr19 | 50678668 | 50678788 | 788855_47188554_ENST00000293441.6,E | 2 | - | good | CCCTGACCCCTCCCCATCACCCTGACCTTTCCCCATCACCCTGACCCCTCCGTTTTCCCTTTGACTCCTCCACTGTCTCTCTTTTACCTCTTCTCTCACCCTGACCCCCACCCTGTCACC | 120 |
| chr19 | 50688868 | 50688988 | 788855_47188554_ENST00000293441.6,E | 2 | - | good | CGGCTCCTCCCTGTCTGCAGCGCAGACCCCCATCGAGGAGTTCACCCCCACCCCGGCCTTCCCGGCGCTGCAGTACCTGGAGTCGGTGGACGAGGGTGGCGTGGCATGGCGAGCTGGACT | 120 |
| chr19 | 50689948 | 50690068 | 788855_47188554_ENST00000293441.6,E | 2 | - | good | CCAATGTGGTGGCCACCAGCCATGTGCAGCTTCTGCACACTTGGAAATATGGCCACTGTGACTGAGGAATTGAATTTTTCCATTTGATTTTGATTAGCTTTCATAAAAACAGCCACGTGC | 120 |
| chr19 | 50691988 | 50692108 | 788855_47188554_ENST00000293441.6,E | 2 | - | good | TTACCACTGCAGTTACCCGAGTATTCTCTTGGGTATTGTTGAAGAATGATACTCACTGTGTGACCTTGGACAAGTTACTTAACCTTTCTGTGCCTCTGTTTCCTCACCTATAAAGTGAAA | 120 |
| chr19 | 50707468 | 50707588 | 788855_47188554_ENST00000293441.6,E | 2 | - | good | CTCTGTCTCAAAAAAAAAAAAAAAAGGAAGAAGAAGGCCAGGTTGTTGGAATCCTAACTGCTCCACTAACTGGTTGTGTGGCCCTGGGCGTAGCATCTGCCCTCTCTGAACCTGTCCTGTA | 120 |
| chr19 | 50714188 | 50714308 | 788855_47188554_ENST00000293441.6,E | 2 | - | good | TCTTTGCCCATGGCTCAGACGGGGTTGAAGAAGTTCCTGGAGTATGTGCAGCTCGGGACATCTGACAAGGTGGCGCGGCTGCTGGACAAGGGGCTGGACCCCAATTACCATGACTCGGAT | 120 |
| chr11 | 70467815 | 70467935 | 788855_47188552_ENST00000601538.6,E | 1 | - | good | CACGTGATTCTGGAGCCTTTCTCATCATAAGACTGTGGAAAATACACAGTCGTTTAAGTTAAAAATAAAAAGTAAATTGAAATAAGATTGTGTTATCAATTAATTTTCCATTAAAAGTGC | 120 |
| chr11 | 70467935 | 70468055 | 788855_47188552_ENST00000601538.6,E | 1 | - | good | CAATGTATTAATGTAGCTGATAAAATAGTTATAGTGCCTTGTTTTCTGAACTGAGAAGCCCATATCAATGCACAACGTAAAGAAAGGACGGTTCGAGCTATGAAATGTTGAGGACACCTA | 120 |
| chr11 | 70468055 | 70468175 | 788855_47188552_ENST00000601538.6,E | 1 | - | good | GAAAAGAAAAAAAAAAAGGAAAAAAAATAAAGTGTGTGCCTTGGGTGTTTTGAACCTGATCTGCATAAAACCATATTGTATTCAAATGCTGTATCTTTCATGAAAGGATAAATATATCTT | 120 |
| chr11 | 70468175 | 70468295 | 788855_47188552_ENST00000601538.6,E | 1 | - | good | TTCCACAGTCTTTACCGTTTTATGTTCAAAATTACAACAATCCCTGTCCATTGATTCCACTCTGGAACTCTTTGTTCATGCCAATTTTGAAATTTTAATACGAGCCTTCAAATAAACACA | 120 |
| chr11 | 70468295 | 70468415 | 788855_47188552_ENST00000601538.6,E | 1 | - | good | TTGTGGGGAATCAGAGAATTTCCAAACTTGTTTCTCAGACTTCCGCAGATCTCATCACTTTGATTTCTAATCCATGCTGTATTGGTGATTTTGTTTATCGTTCCTGTAACTTGTTCTACA | 120 |
| chr11 | 70468415 | 70468535 | 788855_47188552_ENST00000601538.6,E | 1 | - | good | ACTGGCGAACTGGAATGTGTTTCTGTATTTGTAGACAACCATGAACCCATGCAAGTAGGTGAACATTCCACAGTGGCTGGGTGACCACAGCAGCTGCATGCAGACAGGACTGCCCGTGCT | 120 |
| chr11 | 70468535 | 70468655 | 788855_47188552_ENST00000601538.6,E | 1 | - | good | AGACTTTGCTTCCAATGGTTTCTTCCAGACCACTTTTCCTAGATGAATATATTCGTTTACCTTACTAGGAAAATTATTGGAAGATTTTTTCTTTTACTTGAAATTGGAGGCATTTTAATA | 120 |

| | | | | | | | | |
|---|---|---|---|---|---|---|---|---|
| chr11 | 70468655 | 70468775 | 788855_47188552_ENST00000601538.6,E | 1 | - | good | GCAGCAGCCTCTATGCTTCCACGTTCAGGAAAAATTCCAGTCCTCATATCTTTTGTAGTTCACCCTCAAGCTCTCCCGCTTCACCATCCAATAGTTTCTCCCAAACCTTGGCACCCCCCT | 120 |
| chr11 | 70468775 | 70468895 | 788855_47188552_ENST00000601538.6,E | 1 | - | good | TCCGCTTTAGGGAAGCATATTTTAAACCTAAACGTTGAACTTCTTCTTTGGCCTCACCAGTGAAAACTTGTTGTCTTTAGTTCCTAAAGTTTCTTCTACTTTGGCACATTCCCCAGTTGA | 120 |
| chr11 | 70468895 | 70469015 | 788855_47188552_ENST00000601538.6,E | 1 | - | good | GGGAGTGGCAGAACTGGGCCGCCTCTCTGGTTGACAAGCAAACCACATGCTAAGGCTTGGAGCAAGAGAGAATTTGTGTCTATTGGCAAAGAACTAAGCCAGGAAGACATGGGCCATCCC | 120 |
| chr11 | 70469015 | 70469135 | 788855_47188552_ENST00000601538.6,E | 1 | - | good | CTTTTCCTTTTGTTCTGGGTGGAAGCAGCGGTGCCGCGGAGGGCCAGCCAGATCCGGACCCTTCCCTTAGGGTCCAGTCTCCCCACACCCCAGCAGGGTGTCTTCTAGCCATAAGGCCAA | 120 |
| chr11 | 70469135 | 70469255 | 788855_47188552_ENST00000601538.6,E | 1 | - | good | CTTGGTTCCATTTCCTTATTCTCAATTTATCAATGGATACGTGCATGTTATTTCAGAATTATGCAAAACGTCAAAATCTGCTTCTGTGACCGCTGCTATAGGCGTGGAGCTGAGGCTCGG | 120 |
| chr11 | 70469255 | 70469375 | 788855_47188552_ENST00000601538.6,E | 1 | - | good | GCCCGACTTGCAATCTAATGTAGATTATCTCAGGCATTGTGGCCAGCTCTGCCTCTCTAAAACTGACCAGAAAAATCTCTCTCATCGAGTAAACAGGCTCCTGTCACTGAGCTAATCTGC | 120 |
| chr11 | 70469375 | 70469495 | 788855_47188552_ENST00000601538.6,E | 1 | - | good | TCCTTAAATCTCTTGTATGGCAACCAAAATTACTGTAAAAAAATAAATATACTATTGCACTAAGGTTGTGGTTCTGATTGCAAACAAACAGTGAACACTGTCTGAATTAAACAAAAAGCT | 120 |
| chr11 | 70469495 | 70469615 | 788855_47188552_ENST00000601538.6,E | 1 | - | good | CTATAGTTTGTGTAACAACACGTGTTCGCATTATCTATGTTGCTGTTACTTTTGTGCTTTTATTCTTTTTAGACTTTATAAAAAAAAAAAAAAAGCTCCTGTAATTTGCACTTTCTCCCAA | 120 |
| chr11 | 70469615 | 70469735 | 788855_47188552_ENST00000601538.6,E | 1 | - | good | GTTATTGAACAAGCAAGCATTATCCAGTTGATCTGGCAATGACTTTTTGTGTGTGGGCCACAATATTGATTTTCCCATTAACAATTTTTTTTTGTTTTTTAAATACTAATATGTTTCACA | 120 |
| chr11 | 70469735 | 70469855 | 788855_47188552_ENST00000601538.6,E | 1 | - | good | TAAGTTAATTCACATAATATAAAAATTACTCAATGTGACTGTTGATTTGCTGAACTTTACATATCACAAAGTGAATTATTTGTGATACTTTAGTTAATAAAATGGTAAATTTTTTTCTCA | 120 |
| chr11 | 70469855 | 70469975 | 788855_47188552_ENST00000601538.6,E | 1 | - | good | AAATAAACTGCATTTACATGCCTAGTTTCTGTAATATTGTGTATACAAAACCCAAATCTCTCAAAATGTAAATTATGTATACCTGCCAAGATACCTTTTCCAGGGTGTCTGCGCACATTT | 120 |
| chr11 | 70469975 | 70470095 | 788855_47188552_ENST00000601538.6,E | 1 | - | good | GGACGAGGCCCTATTCCATGCCCCTCTTTAATGGTGGAACAAATGTTAAACTGCTCATCTAAAGATCATGTTGATATTATTCCAGGTTTTAAGATCAACTTTTGTTACATACTGTAATTT | 120 |
| chr11 | 70470095 | 70470215 | 788855_47188552_ENST00000601538.6,E | 1 | - | good | AAATAAACACATGTTGCCAAGATGAATCGGTCTCTATCTCTATCTGCTTATTTTGGTACTGAAAGCAATAGTTCCTCATTCAAATCACCACCCACTGTTCTCCCCCTTTGGGACATGTTA | 120 |
| chr11 | 70470215 | 70470335 | 788855_47188552_ENST00000601538.6,E | 1 | - | good | CCACATGCCACTAACCACAAGCAAGCTGGCTGCAAGACTAAAGAGCTGATAACATAGTTTATTTTTACACTGTCTTATTATAGAGAAGTAATAGACCTATCAGAACCTGCACTGACCAAC | 120 |
| chr11 | 70470335 | 70470455 | 788855_47188552_ENST00000601538.6,E | 1 | - | good | AGAGTCAAGGAAACCGCTGGCAGACGATAACCAGTCGAAACTCGTGACTTCGGTTTGTTGAACTTTGGCAGCCAGTTGGTGAGGGCCAGATGTTATTCCCTTTCTTAAAGATACTCCAAG | 120 |
| chr11 | 70470455 | 70470575 | 788855_47188552_ENST00000601538.6,E | 1 | - | good | TCTTCCATTTGATGATACCGCAAAATTCCGTGAGTCCATTCCTTTGGCATGGCACTTTCCCTGGGCCTACAGTTGGTATTACCTCTGTGCTCAGTGCCAGGCAAAACACTAGCTCAAAGG | 120 |
| chr11 | 70470575 | 70470695 | 788855_47188552_ENST00000601538.6,E | 1 | - | good | CCATCATGAGATGAATGAGCATTACTCTAGATAAATTTCAAGACAGGATACTACAGGTGGCCTGCTGAGGCTGCCCCATATTTTAGAAAATGTAAAAATGGTGGTTTGGCCATTAATTTG | 120 |
| chr11 | 70470695 | 70470815 | 788855_47188552_ENST00000601538.6,E | 1 | - | good | GTTTTTGGACCTTCTGTTCCAAAACCTTTTGCAGGTCAATCTTTGTATTTGAAATGATCCAATCCAACTTGAAGTCAATTGAATATTAAGGCGCTTTACTTCCGTGTGCTTTCAGTTTTT | 120 |
| chr11 | 70470815 | 70470935 | 788855_47188552_ENST00000601538.6,E | 1 | - | good | AAGGACAATTTCTTCAAAGTGGTTACAATTCATAATGCAGCAGTTTCTCCAAAAACAAAAACAAAACACACACCACACACACGCGCTTTTCCAGTCACACACCCCTGATGTTGGAACCAA | 120 |
| chr11 | 70470935 | 70471055 | 788855_47188552_ENST00000601538.6,E | 1 | - | good | TAATACTATTTCTGTGGAATAATTATAAAAGTATGACCTTTTTAAATCAACCTTATTTGGATGCATCTGAACCAGCAGAGCTGTGTTATATTTTCTATCTTTGCTAGAACTTCGTCATTG | 120 |
| chr11 | 70471055 | 70471175 | 788855_47188552_ENST00000601538.6,E | 1 | - | good | AGTTGTACAGATAGTTTATAAGCACAATATTTTAAGAAAAAAAAAGTGGCTGGTCTACTAGGCAGCCTTTGTGCCACTTCAGTGCTAGAAAGTTAAAGAAAAAAAAACTTTTGTGATTTAA | 120 |
| chr11 | 70471175 | 70471295 | 788855_47188552_ENST00000601538.6,E | 1 | - | good | ATTATAAAACATGGTTTTGTTTTTTTTTTTCCTTTTTTTCTTGATTAGGTCTGGAACAGCTCTAGAATGAACACATAAAATTTAGCAATTTAAAATCTTTCTTTACTGCAAGTTTAAAT | 120 |
| chr11 | 70471295 | 70471415 | 788855_47188552_ENST00000601538.6,E | 1 | - | good | TGCACATGGGCCGCAAAACACACAGTCAGGCTTCCAGCACATTCTCCCCCATTTGGAGGATACTCGAATGTCAGGTTTTTGGTTTTATTATTATTTCAGAACTAGCTCAGCCCATCTCTA | 120 |
| chr11 | 70471415 | 70471535 | 788855_47188552_ENST00000601538.6,E | 1 | - | good | GGTCTCCTGGGCTGCCAGGTCAGCCTCTCTGGCAAGGCTTTCTTGAGCCCCGCCCCTTTCTTTCCCCGGAGTCCCTCCACCCCATAACAATACCTCGAATTTCCAAAAGAGGTCACCAGA | 120 |
| chr11 | 70471535 | 70471655 | 788855_47188552_ENST00000601538.6,E | 1 | - | good | TACAACAGCTTCTTTGAGGACCCAGTGGGTATGGAGTATAGACAGAACCCAGGGTTGAGAACAGAAGGTGGGCGGCAGGATCAGAGTGAAAGCAGAGGCGTGAGGAGAGGAAAGCAGGGA | 120 |
| chr11 | 70471655 | 70471775 | 788855_47188552_ENST00000601538.6,E | 1 | - | good | AAGGGTGAGCATCTCTGCTGAGACAGTCCTTTTGCTCTCGGAGGCCAGGGAAGATGGTACTTAGAGGCTTTTCCCCTATCGCTCTGGGTGTCTAGGAATCCCACCAGCTTGTCTTAACAG | 120 |
| chr11 | 70471775 | 70471895 | 788855_47188552_ENST00000601538.6,E | 1 | - | good | CCACTTTGACATGAAGGACAAGAAGACGCATGGCTCATGGCGGGCACATGCGGCTGCCAGTGAGACAGCGTCTCCTCTGGGAGCTGGGCGGGCACAGCATCCTCAGTTCTGTGCCCAGCC | 120 |
| chr11 | 70471895 | 70472015 | 788855_47188552_ENST00000601538.6,E | 1 | - | good | AAGAGCAGTTGTCTGCCATGAGCACCCATCCAGGATGGAGAATAAGGGCTTCTCTGCCTCTCAGAATTCTTTTTAATTGAAGATGTCTTGAGCTCTGCAAAGATCAGAGCAGGTGAGCAT | 120 |
| chr11 | 70472015 | 70472135 | 788855_47188552_ENST00000601538.6,E | 1 | - | good | GGCCAAGCAGATGTCACATGGAGTTAGTCAAAGCACAAAGTCACGATTCCACAATGGAGGGGAGACCTGGCCAAGGGAGCCAGCCAGCGTGCAACTGCCCAAGCTCCAGGTCTCCAGGAC | 120 |
| chr11 | 70472135 | 70472255 | 788855_47188552_ENST00000601538.6,E | 1 | - | good | TTTGCATCCATGGCTCTTTGGCCATGAGGTTCCTGGCAGTGCTGGGAGTTTGGATGGGATCGTGCCCAGCTTTGCTTAGCTTTCTTTATTTCTGCAAATCTGTTAGCATAATTCCAAGGT | 120 |
| chr11 | 70472255 | 70472375 | 788855_47188552_ENST00000601538.6,E | 1 | - | good | CTCCTGCACTTGGCTTGCAGTGAGATGCTCCCAGATGCATTTGTCCAGTGCCCCATCATGGGCCTGAAAGGCAGAGAAACTTTTTCCTACACAGATTCTTTTCCCCATCTCCTCCTGTGG | 120 |
| chr11 | 70472375 | 70472495 | 788855_47188552_ENST00000601538.6,E | 1 | - | good | CTCCTGAGACCTCCGTCCTCTGCTTTCCGGGCAGCTCTCACCACCCCAGGCCCCGGCATGAGGCCTTTCCTCAGTCCTGTGGCCTCTCAGAGGACACCTGATGCTCACCTGCCCCTCTTT | 120 |
| chr11 | 70472495 | 70472615 | 788855_47188552_ENST00000601538.6,E | 1 | - | good | AGATTCTCAGCCTGCCGCCCAGCATGCAGGTGGCCTCGCTTTGCCGGGTCCGAGAGGCTCCCCGTCAATTTTGCACGGGATCCTAGCTCTTGTAGGCAGACACCAGTGCACTCTAGATAC | 120 |
| chr11 | 70472615 | 70472735 | 788855_47188552_ENST00000601538.6,E | 1 | - | good | AATACTGAGTTGTGTCCACAACATGGCTGGGTCTTCAGACCCCTGGCTCACCATGTGGGTGTCTTGGGCAGTTTCTATCACACATGGGACAAGGGGAGGGAGTTTTTCTAACATGGAAAA | 120 |
| chr11 | 70472735 | 70472855 | 788855_47188552_ENST00000601538.6,E | 1 | - | good | CCACCTCGCAGACTGCTCTTGTTATAAGTAGAGATGGGCTCGTGCTGAAACATCTGAATGCCAAGCGAAGTCTGTGAGCATCAACCCCACTCCATGGGTTTGTCTCCTGGTACCCAAAGA | 120 |
| chr11 | 70472855 | 70472975 | 788855_47188552_ENST00000601538.6,E | 1 | - | good | GTCACTTACCAAACCTGCAGAAGGAGGACCTCATCGATCTTGGGGTAACTCGAGTCGGGCACAGAATGAACATAGAAAGGGCTTTGAAACAGCTGCTGGACAGATAAGGACGGCTGCTCT | 120 |
| chr11 | 70472975 | 70473095 | 788855_47188552_ENST00000601538.6,E | 1 | - | good | ATAAGCCTTTTACAACTAAACCTGTCCACCTGTGGACTAAACCAGATGTGGCCGATTGGCTGGAAAGTCTAAACTTGGGTGAACATAAAGAGGCCTTCATGGACAATGAGATCGATGGCA | 120 |
| chr11 | 70473095 | 70473215 | 788855_47188552_ENST00000601538.6,E | 1 | - | good | CCGCTCTCTCAGATGTCTTTAGCCTTCCAAGCCAGCCCCCTTCTGGGGATCTATTTGGCTTGAACCCAGCGGGACGCAGTAGGTCGCCATCCCCCTCGATACTGCAACAGCCAATCTCAA | 120 |
| chr11 | 70473215 | 70473335 | 788855_47188552_ENST00000601538.6,E | 1 | - | good | CCGACTATGAAAGCAGGACCTCAGGAACAAGACGTGCCCCAAGCCCTGTGGTCTCGCCAACAGAGATGAACAAAGAGACCCTGCCCGCCCCCCTGTCTGCTGCCACCGCCTCTCCTTCTC | 120 |
| chr11 | 70473335 | 70473455 | 788855_47188552_ENST00000601538.6,E | 1 | - | good | TGTGTTGCTGTTTCAGAAGCCCGGAGATCATGAGCACCATCTCAGGTACACGGAGCACGACGGTCACCTTCACTGTTCGCCCCGGCACCTCCCAGCCCATCACCCTGCAGAGCCGGCCCC | 120 |
| chr11 | 70473455 | 70473575 | 788855_47188552_ENST00000601538.6,E | 1 | - | good | TTCACTGCCAGTGGATCTGCCACTAGACACTCTGGCATGGTTTGGGCGTCTCCCTCGCCTTCCTGGGTGATCTGCAGCCAGCTGCCCTGACTCGGTGACCTGCCTTGTGATGGTTTTCTC | 120 |
| chr11 | 70473815 | 70473935 | 788855_47188552_ENST00000601538.6,E | 1 | - | good | TCCTTCCCACCTGCTGAAGCTGAGGCGCTGGAGATGTTGCCTGGCACGTTCTACCCAGGTGACAGGACCCCACTCTGTCCCTGTCTCCGTGCCCTCTTCAAGGGACCCCTTCCAAATGCC | 120 |
| chr11 | 70473935 | 70474055 | 788855_47188552_ENST00000601538.6,E | 1 | - | good | GCTACAGGGGTTCCAGTTACTGAAAGGGGCCACCTGGGAAGGATCCTGCCCTGCTCTTGTCCATTTACCCAAAAAGCCTAAGGCTGCCAGGCGGCTGAGGGGCCTGTCCTCCAGGTGTGC | 120 |
| chr11 | 70474055 | 70474175 | 788855_47188552_ENST00000601538.6,E | 1 | - | good | CACTGGGTGGGCGCTGGACAGTGCTGCCTCAGGAAGGTGAGCGCAGCCCACACTGGCCACACAAGCCACGCCCTGCTGTCTGGGTTTCCGGGTGATCTCAACCCCTGAGTGCCCTGCGTA | 120 |
| chr11 | 70474175 | 70474295 | 788855_47188552_ENST00000601538.6,E | 1 | - | good | CAGGGCTCCTCACGCGTTCTGCTTTTCAGTAAGGCCCGTGTGTCCTACCAGTGAGCCTCTCACATGAAAGGGGCTAAAAATAACACAGGGTACCCCAGCTCCTTCTGCAACAAAACTGTC | 120 |
| chr11 | 70474295 | 70474415 | 788855_47188552_ENST00000601538.6,E | 1 | - | good | TAAAGGTTGGAGCTAGAAGGGTCGTCCTCCCTGACTTGAGCTCTGAGGGCTTTGCCTGCCCAGCCAGAGCGGCAAGGCACAGGGGACCCTCGGGGACGCCCATGGCCACCCTGGGGAAGA | 120 |
| chr11 | 70474415 | 70474535 | 788855_47188552_ENST00000601538.6,E | 1 | - | good | TGAGGGAGTGGTGTGTGAAAGGCCACCTCCACGTGGGTAAGCGTGAGGACTTGGACTTCTCTGGCACTGAGATGGGACCTCCTGCCTGTGGGAGTCATCTGGCCACCACCCTGGGGCCAG | 120 |
| chr11 | 70474535 | 70474655 | 788855_47188552_ENST00000601538.6,E | 1 | - | good | TCCTAGGGACAATAGGATGAATGGGCTTTCAGTGTGGGGACAGCAAAACATGCACTAGGGCCCAGAGTGGCAGTTCTCTTGGTGTGGAGAGTGCCTGCCACAGGCCTTGGCCAGAGCCCG | 120 |
| chr11 | 70474655 | 70474775 | 788855_47188552_ENST00000601538.6,E | 1 | - | good | AGAGAGCCGTGGTAGGATGGATCTTTTCAGAAGCGGTGGGTGGGCCGTTCATATCCACGTGAATGGGCCCAAAAGGCAAGAGACCCACACCTCTCCTCCCCTCCCCAGTGGCTGTGGCTT | 120 |
| chr11 | 70474775 | 70474895 | 788855_47188552_ENST00000601538.6,E | 1 | - | good | TAGGAGGGAGAGGAGCCCACTTCATGTGGGAAGATGTGGAGACAGGAGAAAAGTGCACCCAAGCTGGCATCTGACAGAGCCAGGGCTCCAGCAGACATCTCTCAAGTCCAGGTCTTGAAG | 120 |
| chr11 | 70474895 | 70475015 | 788855_47188552_ENST00000601538.6,E | 1 | - | good | CCTCTTGCCCATGACCCTTTCACTGAGACACTGCAGCCCACCACCATTCACGGTCACGATGTCACACCAGTGCACTCCTCATGCTCTCCAGAGCCCCTCCACCCAGGGTAACCCACCACA | 120 |
| chr11 | 70475015 | 70475135 | 788855_47188552_ENST00000601538.6,E | 1 | - | good | TCCAAAGACAGCCCCAGTGCAAGTCGATGAACTGGCCCCCAGCAGACGGGCAGGGGCTTCAGTCTTGCCCTCCAGGGAGTCTCCCCACACGGCAGCATCTCTACTGTTTGCAGCTCTCCC | 120 |
| chr11 | 70475135 | 70475255 | 788855_47188552_ENST00000601538.6,E | 1 | - | good | TCCACAGGTTTGAGACGAGCTCAGGAGTCCTCACGGCCCACCAGTGTCCCAGGTGGTTTCTATGAAGGGTCTGTGGGCTCCTCTTTGGGAAGTGCTGCTCTAACTGGTCAGCCATTGAGC | 120 |
| chr11 | 70475255 | 70475375 | 788855_47188552_ENST00000601538.6,E | 1 | - | good | GTGCATACCCTGGAATGAAAACCAAGGCCCAACCAAATGGCCTCTAAGGTTCAGGGTCTCTCAGTGTCTGGGGAGTTTGTTAATTGCAGACACCTGGTCCCCATGCACAGCACTCCTGCT | 120 |
| chr11 | 70475375 | 70475495 | 788855_47188552_ENST00000601538.6,E | 1 | - | good | CCATGGTAAAGAAAGAGAAAAGAAAAAAACGGAGAGAGACAGACTAGACCAGCAGCGGGGCCCGTCTCCAGGTCACTCCCTTTGCTTTTAGCTCCTGGAGGGTTCTGGGCCATTAGCTTT | 120 |
| chr11 | 70475495 | 70475615 | 788855_47188552_ENST00000601538.6,E | 1 | - | good | TCCTTTCTCACTATGGACTAGTCCACACTTTAAGCAACAGTGACTAACATCAAGTAGGAAACTGACCCACAGCTTAGGGGAACCTAGGTAACTGATGGGTGGTTTCACCCAAATATACAT | 120 |
| chr11 | 70475615 | 70475735 | 788855_47188552_ENST00000601538.6,E | 1 | - | good | AACACCCTATGGGTGGATGGGGTCCCTGGCCCAGCAGTGCACCCACCAGCCCCTGCAAACACTTCTCACAGTAGGATTGAAGCCTCAGTCCACAGCACAGGCCCCTTTCAGATTTGCCCC | 120 |
| chr11 | 70475735 | 70475855 | 788855_47188552_ENST00000601538.6,E | 1 | - | good | AGCAGTATAACCAATGACCAGCAAGCCACCATTGAGGGTGGGGGCAGGTTCTAGATGGGGATGCCTGGTTTGTAACATCTCCCACTCAGGGGCACCCAGCCCTCTGGAAAATAGCACGTG | 120 |
| chr11 | 70475855 | 70475975 | 788855_47188552_ENST00000601538.6,E | 1 | - | good | TGCTGGGATTACAGGCGTGAGCCACCGTGTGCAGCCTCTGATTTCTTGATTGTGAGTAACTGGGCCCAGAAACAGTGGAGGAATCCTCAGAGTCTCACCTGCCTTGTGCCTGTTGGTGTG | 120 |
| chr11 | 70476215 | 70476335 | 788855_47188552_ENST00000601538.6,E | 1 | - | good | GAGATACTAGAGGGTGCTGGCCCCTCTCCCCTGCTGTGAAGCCCCACTGGGCTCTGATTTGATTTCGGGTTTTTTGGGGGTTTTTTTATTGTTGTTGTTGTTGTTTTTGAGAAGAAGTCT | 120 |
| chr11 | 70476335 | 70476455 | 788855_47188552_ENST00000601538.6,E | 1 | - | good | GTTTTAAGTTGGAACTGGAGCTGGTCTTTCGACCTTCTTTGAGTACCAGAGAAGGGGGGGTTGCAGGTTTTTTCGTGTATCCTATGTCCATAGGCCACCTCTCTCTTGAGAGTATAAACT | 120 |
| chr11 | 70476455 | 70476575 | 788855_47188552_ENST00000601538.6,E | 1 | - | good | TTATTTGATTTCAATCATCTTGGCTTAATTGTTTTATCAACGTGGACAAATAAGCCTTCCCCAAAACTGTGCTCTTGTGACTCTGGAATAAACACAGAAGAACAGGCCACAGCACCCCCA | 120 |

| | | | | | | | | |
|---|---|---|---|---|---|---|---|---|
| chr11 | 70476575 | 70476695 | 788855_47188552_ENST00000601538.6,E | 1 | - | good | CATATGTCTTCTTGTAAGATGTTTTCCCATTTTATAGGAAACATTCCCTGTGCCCCAAGGTCCTTGGCCCTAATTAACACCCCAAATCACCTCTGAATGCAGTGGCTTTCAACTTACCCA | 120 |
| chr11 | 70476695 | 70476815 | 788855_47188552_ENST00000601538.6,E | 1 | - | good | ACGGTGGAGCTGCCCAGAGTCAGGGCTGCAGGCACAGTCACCATGTAGTCAGCTTTAGCAAAAGCACATGGAGACCAGAACTTTACCCCAGCTACCCCTCCTCCTCTAGTCCTCTGTGGG | 120 |
| chr11 | 70476815 | 70476935 | 788855_47188552_ENST00000601538.6,E | 1 | - | good | CCAGAGATACCCGTGGTGAATACTGCCCAGGCGTGAGGTAGGTGCTCTCACTCTGGGGCTTTACTTGCCTCTGCTTCACAGGCCTGGTGAGCACCAAGGCCAGGCACCAGTCATGCTGCC | 120 |
| chr11 | 70476935 | 70477055 | 788855_47188552_ENST00000601538.6,E | 1 | - | good | CCTAACCACACACACAGCTGCCTGCAGAGCCTTTCCTTACAAAAACCAAAATCAATCCCAACTACCTGGGAACAGCTGGGTATTTTCCCAGTGAGGGCAGCACTGGGGCAACCCTCTGGG | 120 |
| chr11 | 70477055 | 70477175 | 788855_47188552_ENST00000601538.6,E | 1 | - | good | AGAAGAGGGTCGTGCTGAACAGAAGTAAGTGCCACCTGCTGTGCTGCTGCGTGACCCCGTGTGTGCCACTCTCGCGCTCCCGCTGCCCACCGAGGTGTTTGCAGAGCCTCCTCTTCGCTT | 120 |
| chr11 | 70477175 | 70477295 | 788855_47188552_ENST00000601538.6,E | 1 | - | good | ATTTCATCTGTCGAAAAGCAGCTTTTTTCTGTTTCTTCCTTTCTTGTTCATTAATTTTTGCAGTGTTGTCTCAAGTCATTCCATTTGTCATTTTCATTTGCTCCCATGACCGGTGTTAAC | 120 |
| chr11 | 70477295 | 70477415 | 788855_47188552_ENST00000601538.6,E | 1 | - | good | AGTTTCCCACGCTCCCTATGAAAGTGATGCCATATTGATGTTTCAGCTCTGGGCAGGCTGTGGCAGGCCGAAGGGACCACCACGGATGGGCTTTCAGCCAGGGTCTGCCCCACTGCCTTC | 120 |
| chr11 | 70477415 | 70477535 | 788855_47188552_ENST00000601538.6,E | 1 | - | good | CTTCAGGCAGAGGCGAACTTCACTCCAATGCTACTATTTTCTTTCTTTTTTCCTGGTTATTTTTCTTCTTTACAAGTGGTTACATTATCCCAAGCCAAAGAAGGAGAAGACAAGCTTGTG | 120 |
| chr11 | 70477535 | 70477655 | 788855_47188552_ENST00000601538.6,E | 1 | - | good | CACTTGTCCTGTTGGGGTGCCCTCCCCCACCAGGAATGTTCGTTTATCCTCCCTAAGCCACAGGAAAGACAGTTCAGGCACCTGAGGTAGGAAATTGGCAATAAGCTGCTTCTCAGTGGG | 120 |
| chr11 | 70477655 | 70477775 | 788855_47188552_ENST00000601538.6,E | 1 | - | good | CACTCACAGGCGGTTACTTTCTATAGCAATGTGGGGCAGGACACTGGCAGGCCCCAAGCACTTGACGCCCATGTGATTAACGGCTGGGCTGTCATCTCATGATGTAGACTTGAAGCAAAG | 120 |
| chr11 | 70477775 | 70477895 | 788855_47188552_ENST00000601538.6,E | 1 | - | good | CGTAACCATCTGTGCAACCGGGAAGATCCAGTGAATATCATCAATGTGATTAACCAAGAGAGTCAAGCTGCTGAGGAGGGCAGGGCGGGCAGCCCAGGGTGGCCTTCACCCTGCGTCTCA | 120 |
| chr11 | 70477895 | 70478015 | 788855_47188552_ENST00000601538.6,E | 1 | - | good | AGGGTAGACAGTGGTGACCTGGTGGGGAGTTGGGTGTGAGTGTGTATATGTGTCTGTGTTCTAAATCAGGGCAGCAGATTCCTGTGTGTTCCGACTGGGAACCCAGCGGTTGCTTAGCAA | 120 |
| chr11 | 70478015 | 70478135 | 788855_47188552_ENST00000601538.6,E | 1 | - | good | TACTTACCTCACCCCAATTAACTTGTACTGACTGATGAAATAAGAGCACCTCTCTGAGTGTCTCAATTACCTTGATTTTTTTTAATTCTCACATTTCTACATGAATGTGATTTAAAAGGC | 120 |
| chr11 | 70478255 | 70478375 | 788855_47188552_ENST00000601538.6,E | 1 | - | good | TGACAGGCGGCTCATGGGAGGGACAGAACCAGCATGCCTAATCAGCATCACTGACCAGCCCCTGGCCCATGTTCAGCTACAGTTAAAGAGCAGATTACTCTAGCTATGCCAGAGGCTGAT | 120 |
| chr11 | 70478375 | 70478495 | 788855_47188552_ENST00000601538.6,E | 1 | - | good | CTATTTTCATGCAAGATGGGGGCTCATCACTAAAATGCAGAGGCGTGCGGTGTGCTGGAGAGAACATATGTTGACAGGCTCCAAGTCCAGCCTCAGTGCTGGCCAGGCCCAGCAGGCCCT | 120 |
| chr11 | 70478495 | 70478615 | 788855_47188552_ENST00000601538.6,E | 1 | - | good | CATCCAGGCCTCGGGGAAGCTAGGTAAGGACTTGCTGCCTCCGGTTCTATCTTTTCAATGCCCTGCTTATCAGGAGGCTGTAAAGATGGAAAAAATAAGAAAGCAGTTTGGAAATAAGTG | 120 |
| chr11 | 70478615 | 70478735 | 788855_47188552_ENST00000601538.6,E | 1 | - | good | CTAAAACTAGACCAGGACCCCCTCATGGCACTCAGAGCTGGGGGTGAGGCGGAGGCCTGGCAGCTGCTGAGACACAACATAACACATGGCACAGCTGGGCAATATGGGCTCCAGTCCCCA | 120 |
| chr11 | 70478735 | 70478855 | 788855_47188552_ENST00000601538.6,E | 1 | - | good | CTGGCTCTGTAACTCGGTCAAGTCACTTAACTTTCTGATCTGGGAAAAGGATATATTATCCATCAACACTTCAGTCAACAATTATTTATCGCACCCCTGTTCTGGGCCAGACCTCAAGGA | 120 |
| chr11 | 70478855 | 70478975 | 788855_47188552_ENST00000601538.6,E | 1 | - | good | AATCTCTCTTCCCAAGAGGTGGGCACCTTCTTGATCTTAGCTGCGAACCTCCTGGCCTGAAGACCTGGATTCCTGGTGGAACTGCCGCGGAGCCTCAGGCTGCCTGCCCCACTCCCAGGT | 120 |
| chr11 | 70478975 | 70479095 | 788855_47188552_ENST00000601538.6,E | 1 | - | good | CGTTGGGCAAGGATGAATGGGAGTGCCTCGTCCTTGCCCATCAACAGGCAAATCTCAAGTCAGAATCTTCCTCCCCATTTTGGACATGAGGCCCTTTTGGCAAAAGCATATCACAGGGTT | 120 |
| chr11 | 70479095 | 70479215 | 788855_47188552_ENST00000601538.6,E | 1 | - | good | TAGGTGGTTTGAGGGTTTCCAGGCCCTGATGGAAAAGTGAATGTGACATCAGTGAATGCGACTGTGGGTGGCTGGGAGCACTAGTCAGGATGTGCGTCCATCGGGGCTGGTACTTGCGGG | 120 |
| chr11 | 70479215 | 70479335 | 788855_47188552_ENST00000601538.6,E | 1 | - | good | CAAATAACTTTGGGTTGTTAAAAAGTTATCTCTGCCCAGCTGGTCCCAGAGCACAGACTGCTTCAAAACTGACTTGGTACTAGAAGCTGCCTCACGGTGTTTTCAGCATCTCAGGACCAC | 120 |
| chr11 | 70479335 | 70479455 | 788855_47188552_ENST00000601538.6,E | 1 | - | good | AGGCGAGTTTTCCTCCCGAGGCCCCTGCGCCTGCTGCCTCACGGTACCTGCCTCCCCTTCCCAAGGCCTCCAGTCCCTACTGGGCTCTCTGCAGTATTATCCTGACAGCCTCTTGATTTT | 120 |
| chr11 | 70479455 | 70479575 | 788855_47188552_ENST00000601538.6,E | 1 | - | good | GGACCTCGAGATTGGCCTTCGCTTCCCCATGGGATGGATGGGCTTCAGGGAGATTTGTCCTGTGAAACTCCAGGTCCCAGAGGCCCCTGAGGGAGTTCAGGCTACCTGGGATGTGAGCTC | 120 |
| chr11 | 70479575 | 70479695 | 788855_47188552_ENST00000601538.6,E | 1 | - | good | GAGCAGTGAGTGTGTGCTTGTGGGGGTCCCCCATGGCTTCTGTTAGGGTGCATGGCCTATAAGAGTTCCCCAGAGTCCCCTCTAGGTCGCCAGCCCTTTGCAGTGGGTGGGGTCCAGAAG | 120 |
| chr11 | 70479695 | 70479815 | 788855_47188552_ENST00000601538.6,E | 1 | - | good | TATTGAGAGATACTGACACATACCAGGCGCCGATAGTAAAGCCAGATAAATCTGATGTTCTTTACCATCGCTTCCATGGCAGAGGTCAGCTTCACATTTGCTCTGTATGTGGGGGCAACA | 120 |
| chr11 | 70479815 | 70479935 | 788855_47188552_ENST00000601538.6,E | 1 | - | good | TCAAGGGCTAAGTGCCTAAGGAACCCAGGAGAACACAGCAGCTTGCCTGACTTATATGCTTTGACCCAAAACAAACTTGCTATGACCCAGGAGCCCCAGGTGTCATGTGCCCTCCAGCTT | 120 |
| chr11 | 70479935 | 70480055 | 788855_47188552_ENST00000601538.6,E | 1 | - | good | GACCAGACTTTGTTTCCTGAAGATGGCCCCAGGCAGCTGTGTGATCAGGTGAATGCACTGGTGGCACAGGGAGGCTGCTGTGGGAGTGTAGGGCCTGCATTCTGGAGACAAGCAGGGAGAC | 120 |
| chr11 | 70480055 | 70480175 | 788855_47188552_ENST00000601538.6,E | 1 | - | good | ATGGCCAGATGAGGAGGGCTGGGCAGGGCCAGGTCACAAGAGGCTTGCAGGTAATGGGCATGAGCTGAATGGAGTTCCAGATATGGCTGCCTTCAGAGGGCTTCAGCAAAGTCTTGGCAT | 120 |
| chr11 | 70480175 | 70480295 | 788855_47188552_ENST00000601538.6,E | 1 | - | good | AATGACCAGAAATTTAAAAAGGAGCCAACCTTGCAGAAAACCAGGAGTGGAGCATCCCAGGCACGGGACAGAGAGTACAAGGCCCTGAGGCCAGAAAGTGCCTTCTGTGCTCATGAAACC | 120 |
| chr11 | 70480295 | 70480415 | 788855_47188552_ENST00000601538.6,E | 1 | - | good | CCACGGGACACCAGCAGGTTAACCCTGACCACATGCAGAAAGGGCGGACTAGGATGCCCTAGCAGGTTACAGTGAGCATGTGCAGAAAGGGACTGACACTCTGCAAGGAAATGACATTTG | 120 |
| chr11 | 70480415 | 70480535 | 788855_47188552_ENST00000601538.6,E | 1 | - | good | GACAAACCCAAACTGTGACAAATGCCACAAAGGAGCCACAACATGCTAATAGGTTAGTGACCAGTGCAGAAAGGGGCTGTGATGCACCAGCAGGTAAAAATAACCACGTGCAGAAGAGAG | 120 |
| chr11 | 70480535 | 70480655 | 788855_47188552_ENST00000601538.6,E | 1 | - | good | CTTTATGTGCCAGGTGCCATGCTGTGCCCAGGATTTATCACTGAACAAGAAAACATTCCAGATGCCCCTCCCTTTGGGGAGCTCAAATTCTCGAGGGGAGAAATGGACAACAAATAATAA | 120 |
| chr11 | 70480655 | 70480775 | 788855_47188552_ENST00000601538.6,E | 1 | - | good | AGCCTAGTGCCAGGCTGAGGCTAGAGAGTAACGAGGAATACACCAACAAGGATGCCTGGACCCTCAGAATGGGGCAGCAATTCCGTCACTAATCATTCACTCAACAAAGTTAGGGATTGC | 120 |
| chr11 | 70480775 | 70480895 | 788855_47188552_ENST00000601538.6,E | 1 | - | good | GCCGGAAGCTCCTCAGGTGCAGGCCTAGGAGACTGACTGGACACTCAGACTTGGAATCATGGCCCAGCCCCACTGTCTCCTGAATTCACCAGTCAGGAGTTCACTGAGCTACTTTCTATA | 120 |
| chr11 | 70480895 | 70481015 | 788855_47188552_ENST00000601538.6,E | 1 | - | good | GTGAAGAGTCAGTGGTCCCTCCTCAGCAGAATGCCAAGACAACAGCTGCCAGCCCAGAGGCCTGACAGTGCTGCACGCTGGGGGTCCCAGCACAGGAGCCTGGGGCTGCAAAAGCTCAGA | 120 |
| chr11 | 70481015 | 70481135 | 788855_47188552_ENST00000601538.6,E | 1 | - | good | TAAGAAAAGTGTTCCAGGACCACCTGGCCTGATTATTCCAGGAGGTAGTCTCATCTTTGATCTCAAACTGGATCAAAGTGAAGGAAACTGGTTTGGGGAAGGAAGAGTGAGTGTGTCTCT | 120 |
| chr11 | 70481135 | 70481255 | 788855_47188552_ENST00000601538.6,E | 1 | - | good | GGAAGATGATGAAGTAGTTCTCTTTAGTACATAATGTTAAAAGAGAAAAAAGTACATCATATTAGAGGCATCTCCCAGGGTGATCTGAAAACACCCTGCCAAGAATGGAGGCGTCAGTTC | 120 |
| chr11 | 70481255 | 70481375 | 788855_47188552_ENST00000601538.6,E | 1 | - | good | CCTTGGGTATGTTTGATTTAACTTTAGCCGTGAATGAGTCTCTGATGCTCAAAGCTCTTCCCTGAAATCAAGGATGACATGGAATTTGCAGGCGTCTTAATTACAAATACGCCGACGAAA | 120 |
| chr11 | 70481375 | 70481495 | 788855_47188552_ENST00000601538.6,E | 1 | - | good | AACTTGACCCTTTTAAAGTTCCTCTCATCCTTGGATTCTTCCTTTCTTTTCACTTCTTGCAAGAATCCAGGATGGGATCAAAGGAAGCAGCTGAAGATTAAACAACTAGATCCTTGTTTT | 120 |
| chr11 | 70481495 | 70481615 | 788855_47188552_ENST00000601538.6,E | 1 | - | good | ATAAGCAGTGGTGTAAAAATCCCATTTCCATAGAAACAAGTTTGACTGCCCAAGAAAGGGACAATTTTTAGAGCTGTTCCTCAAATGTTCATTCATGAAATGGGCAGAAACCTCAGCTG | 120 |
| chr11 | 70481615 | 70481735 | 788855_47188552_ENST00000601538.6,E | 1 | - | good | CTGTGAATGGCCCTGGGCAAAATGAGGCACGAAATTGGATGAAGGGAATTCCTACCGAGGAAAAAGAAATGAGTTTTTCTAGGGAAAATTGAAAACCAGCCATGCTTTTTGAAATGACAG | 120 |
| chr11 | 70481735 | 70481855 | 788855_47188552_ENST00000601538.6,E | 1 | - | good | TCACTGATGGGCCCTGGGCATTTGATCATCTGAGGATCAGAGGGAGCCTTAAGAACTCTCACCTGAGTTCACCGCCTGGGCGACTCACCACCACCTTTAAATGCATCTAAAAATTAAGAG | 120 |
| chr11 | 70481855 | 70481975 | 788855_47188552_ENST00000601538.6,E | 1 | - | good | CTGGGGGGGGGTCCAGGACGGAGTTGCCAGGGCACCAGAGCTGTCCCCCTTGCCGTGAGGGCTGTCTCCTCTCAGCCACAGACCTATCTCTCAGCATGTCTCCTTTGCTGGGCGTGGTCT | 120 |
| chr11 | 70481975 | 70482095 | 788855_47188552_ENST00000601538.6,E | 1 | - | good | CTTGCTTCTCGGATCTAATAACCTCACCAGGCTGAGCACTTGGGACAAAGCCCCCATGGAGGTGCGCCACGGCCACGGGACACACTGACAAGGACAAGTGGGGGGCAGTCAGAACCACCA | 120 |
| chr11 | 70482095 | 70482215 | 788855_47188552_ENST00000601538.6,E | 1 | - | good | GGCCAAGGGTTACACTTTACACTCAGAGTTGGACTAGAAACAGGATGGCCTGAGGCCAGAGGACAAAGATGAAGAGCATTCACTGTAGGCACTAAACTAGGCTCAATCCTCTCCCTCGCC | 120 |
| chr11 | 70482215 | 70482335 | 788855_47188552_ENST00000601538.6,E | 1 | - | good | CTAATAGCAGGCTTAGTCACTAGGAGGTCACCCTGCGGCCGATAACAGCAGGCTTAGTCACTAGGGGTTCACTGGCACTGCCAAGCTCCCAGAGGCCATGTCTGTTCCCTTCTTGCTCCC | 120 |
| chr11 | 70482335 | 70482455 | 788855_47188552_ENST00000601538.6,E | 1 | - | good | CTGCCTTGGAAGGAATGACCAACTGGCAAACCGAACACCTGCAAGGCTGAGACTGCCGCGGGCGGGGCTGCTCTCAGGCCTCTGACTGTCCTTAGGCCAAGCTCTTGGTCAGCAAGAGG | 120 |
| chr11 | 70482455 | 70482575 | 788855_47188552_ENST00000601538.6,E | 1 | - | good | GCAGAAGGCAGGGCGGTGAAGCTACTGAGCCGAGGCTACTTCACAGGAAGACGCGGAACTAGACCTCCAACCCATGGAGCTGGCTCAGGGTTTGACTCCTCACCTTGTGGGAGCCAGGGA | 120 |
| chr11 | 70482575 | 70482695 | 788855_47188552_ENST00000601538.6,E | 1 | - | good | CTTACAGTAAGAACAGGAGCCCACACCTATGACACAAAGCATTCACCAAACGCCAGGCACTACATTCCACGGCACCCCTGAGCCAGGGCCTGTAATTTTTCCCACTGTGTGAACAAGGAA | 120 |
| chr11 | 70482695 | 70482815 | 788855_47188552_ENST00000601538.6,E | 1 | - | good | CTTCATAGGGCAGAATCCTGCTGTACTCTGAAGGAGAGCAGCTGCATACCACCGCCGACCCACGCGTGAGTGCCAAGAACGGCCGCCTCCCAGCTGTCCTGTTATTTATAGCCCGGCTCT | 120 |
| chr11 | 70482815 | 70482935 | 788855_47188552_ENST00000601538.6,E | 1 | - | good | GGCTGGGCCAGCTCCTGGGGTGCCCACAGGGGAGTGGGACCCCAACTTGCAGCGTCTCTAGGGTTTCTCTGCCCTGCATTTCTCCAGGGGGACCCTGAGAGGGGGCTGGCTCAAAGCAGA | 120 |
| chr11 | 70482935 | 70483055 | 788855_47188552_ENST00000601538.6,E | 1 | - | good | GGGCATCCTGCACCAAGCTGTGCTCCCCTGAGCACTCCACTCGGGGAGTCCAGGGACCCCGTCCATGGAGACCTCTCCCTGAGTGGCAGCTGGACAGCTCATCCCCTCTACTCGTCTTCA | 120 |
| chr11 | 70483055 | 70483175 | 788855_47188552_ENST00000601538.6,E | 1 | - | good | GGACACAGGCTGGGGTTTCCTGACTCTCACTTCCCCTTGTAAGCAATGTCTGCAGTAAGGAAGGAGCTGGAAGGGAATGGCTCTAGGGCTGGAGGAAGGAATCCACTGTTCAGGTCAGCA | 120 |
| chr11 | 70483175 | 70483295 | 788855_47188552_ENST00000601538.6,E | 1 | - | good | CAAAACACTTTCTCATGGGTGGGTAATTTTTATGACAACTGGGGAGCAGGTATGAGTCACACCCATTTTTCAGAGGATAAGTGGCTTGTCCACATACACAGCCAGCAGGAGGCAGAGCTG | 120 |
| chr11 | 70483295 | 70483415 | 788855_47188552_ENST00000601538.6,E | 1 | - | good | AAAATCTGAAAAACTGACTAGTTTAACCCTTCCCCATTACATACTAATGTGGCTCTTCCGTCATTACTCTATGTGGAAGCAATAATTAAAGAGGCCTTATCTTAGTGTTTCACATTTTTA | 120 |
| chr11 | 70483775 | 70483895 | 788855_47188552_ENST00000601538.6,E | 1 | - | good | ATAAATGACTTGGATGTCATCTATTACTCTAAAGGAGTATCGATAGGCTGGCATTGGTATAGAGAAGCACAGTTACACTGCAGACATTCAGACAGTGCCTCCAGTCAAGGAGAAAGTCTA | 120 |
| chr11 | 70483895 | 70484015 | 788855_47188552_ENST00000601538.6,E | 1 | - | good | CTGGGCAGGCATTTGATTTAAGCCACTCAGTGCATTAATCAGTGACATCACATACAGCTGGGGCTCAAGATGCAGACTGAGAGCCCCAATTTTCCCAGTTATTTTCAGTGTGGATGTAA | 120 |
| chr11 | 70484015 | 70484135 | 788855_47188552_ENST00000601538.6,E | 1 | - | good | CTTCCCTGTGGTGTGCACCATTTGCTGTTCCCCAGGTGGCAGCATCTTCAGTCTCTCCCACTGATATTTGATCTTGGGTTTCATCATTTTGACTAATGAAGCCTCTCTCCATACCCCTG | 120 |
| chr11 | 70484615 | 70484735 | 788855_47188552_ENST00000601538.6,E | 1 | - | good | AGAAGCCTCTAGCTGTGTCCATGTCAGCCCTTCACAGTATTGTGCAGGGTTTACATTATTTCCATTGGAACTGAACGTAGAATAAGTCAGGAGTCATTTCTAGACGCAACCAGAGGGCTC | 120 |
| chr11 | 70484735 | 70484855 | 788855_47188552_ENST00000601538.6,E | 1 | - | good | TTCTTCTTCCCCTCCCATCACTACCAGGTTTGCGGGCTTGGGAGGCACAGGGTCATGGCTGTACTGGTGAACTCTAAGTGCAGTACATTCTTCCCTGCCAGCCCATCCTGTCCCAGGGGC | 120 |
| chr11 | 70484855 | 70484975 | 788855_47188552_ENST00000601538.6,E | 1 | - | good | GCCGTTTAGAGCCTCGCCAAAGCCCAAACCAAGCTTCGAGTTGGTTGCAGGAGACTAGCTGCCCCAAGAGGGCAGCGAGTGGGTGGGCGGTGGGTGCAGGTGTCTGAATACACGTGTGGG | 120 |

| | | | | | | | | |
|---|---|---|---|---|---|---|---|---|
| chr11 | 70484975 | 70485095 | 788855_47188552_ENST00000601538.6,E | 1 | - | good | TCCTTCATAAAGACTTGTAATGAAATGAGGTTGCTGGAAGCCCTCCTCACCTTGGCTTCCAAAGAGAAGCTCTACTCTCTCTTCTGTGCCCTCCAGATCTCTGCCAGGACCACCTAGAAG | 120 |
| chr11 | 70485095 | 70485215 | 788855_47188552_ENST00000601538.6,E | 1 | - | good | GGGAATTCGTAAAGCCTGTGTGGGCCACGTCTGTTAATGCAGTAACACTTCTTTTTCTCATGCTCCTGGACTCCGGGATAGCCGAGTAATGATCACAGCCATCCTGGTGGGACACAGCAC | 120 |
| chr11 | 70485215 | 70485335 | 788855_47188552_ENST00000601538.6,E | 1 | - | good | AGTCCGCCAGCCTCGCTCCAAGGTAAGTCCGCGGTTGGTGCACATGCACACCGCTCTGCGTGCACTGTCCCTGCTGACCAGGAAAGGCCCTCGGGTAGCAGCCCCAGCGGACACGAAGAG | 120 |
| chr11 | 70485335 | 70485455 | 788855_47188552_ENST00000601538.6,E | 1 | - | good | GCCCGATTCTCTCAGGCCCAAAGGCAAACGTTATTAGTGAATTGAACTCTATCCTACAGCAAATGAACCGAGAGAAATTGGCAAAGCCGGGGGAAGGACTGGATTCACCAATGGGAGCCA | 120 |
| chr11 | 70485455 | 70485575 | 788855_47188552_ENST00000601538.6,E | 1 | - | good | ATAGCTTTGTTATCCCCCCGCCCGCTCCCCCGCCCCCGCCGGGCAGTGCCCAGCCTGGGATGGCCAAGGTTCTCCAGCCAAGGACCTCCAAGTTGTGGGGCGACGTCACAGAGATCAAAA | 120 |
| chr11 | 70485575 | 70485695 | 788855_47188552_ENST00000601538.6,E | 1 | - | good | ATGCAGATGGGCAAGCATTTATGGTTGACAAACCCCCAGTACCTCCTAAGCCAAAAATGAAGCCCATCATTCACAAAAGCAATGCACTTTATCAAGACGCGCTCGTGGAAGAAGATGTAG | 120 |
| chr11 | 70485695 | 70485815 | 788855_47188552_ENST00000601538.6,E | 1 | - | good | AGGTGGACAGCCGGAGTAGCAGCGACCACCACCTCGAGACGACCAGCACTATCTCCACCGTGTCTAGCATCTCCACCCTGTCTTCCGAAGGTGGAGAGAATGTGGACACCTGCACAGTCT | 120 |
| chr11 | 70485815 | 70485935 | 788855_47188552_ENST00000601538.6,E | 1 | - | good | CCAGCCAACCAACCAACTCTGCAGACAGCAAGAAGCCAGCCAGTCTTTCAAACTGTCTGCCTGCCTCATTCCTGCCACCCCCTGAAAGCTTTGACGCCGTCGCCGACTCTGGGATCGAGG | 120 |
| chr11 | 70485935 | 70486055 | 788855_47188552_ENST00000601538.6,E | 1 | - | good | CTCCCCTGGAATTTGCAAATAGTTTTGATATCCCCGATGACCGGGCAGCTTCTGTCCCGGCTCTCTCAGACTTAGTGAAGCAGAAGAAAAGCGACACCCCTCAGTCCCCTTCGTTGAACT | 120 |
| chr11 | 70486055 | 70486175 | 788855_47188552_ENST00000601538.6,E | 1 | - | good | GAACCATCGTCGCGGTGGGCTCCATGGAAGAGGCGGTGATTTTGCCATTCCGCATCCCTCCTCCCCCTCTGGCATCCGTGGACTTGGATGAGGATTTTATTTTTACAGAGCCATTGCCTC | 120 |
| chr11 | 70486175 | 70486295 | 788855_47188552_ENST00000601538.6,E | 1 | - | good | AGAAGGCAGAGGTGGAGATGAAGCCAGACAGCTCGCCGTCCGAGGTGCCAGAAGGTGTTTCCGAAACCGAAGGTGCTTTACAGATCTCCGCTGCCCCCGAGCCCACCACCGTGCCCGGCA | 120 |
| chr11 | 70486295 | 70486415 | 788855_47188552_ENST00000601538.6,E | 1 | - | good | ATGACAAGAAGAACATGCTGATCGACATCATGGACACGTCCCAGCAGAAGTCGGCTGGCCTGCTGATGGTGCACACCGTGGACGCCACTAAGCTGGACAACGCCCTGCAGGAAGAGGACG | 120 |
| chr11 | 70486415 | 70486535 | 788855_47188552_ENST00000601538.6,E | 1 | - | good | TGCGGCCCAGCCTGGATGCCGGCTTCCCTACGGTCACCAGGCAGAACACCCGGGGACCCCTGAGGCGGCAGGAGACGGAGAACAAGTACGAGACCGACCTGGGCCGAGACCGGAAAGGC | 120 |
| chr11 | 70486535 | 70486655 | 788855_47188552_ENST00000601538.6,E | 1 | - | good | GCTCCCCGCTGGCCCTGGCACTCTCCGCAAGGGACCGAGCCATGAAGGAGTCTCAACAGGGACCCAAAGGGGAGGCCCCCAAGGCCGACCTCAACAAACCTCTTTACATTGATACCAAAA | 120 |
| chr11 | 70486655 | 70486775 | 788855_47188552_ENST00000601538.6,E | 1 | - | good | TGAATTCCACGTCCAAAGCCCAGGGGCCCGAGAGCAGCCCAGCAGTGCCCTCCGCGAGCAGCGGCACAGCCGGCCCCGGGAATTATGTCCACCCACTCACAGGGCGGCTGCTTGATCCCA | 120 |
| chr11 | 70486775 | 70486895 | 788855_47188552_ENST00000601538.6,E | 1 | - | good | CTGACGAGGACAGCGCTGAGCAGCTGTCATCCCCCATGCCGAGTGCCACGCCCAGGGAGCCCGAAAACCATTTCGTGGGTGGCGCCGAGGCCAGTGCTCCGGGTGAGGCTGGGAGGCCGC | 120 |
| chr11 | 70486895 | 70487015 | 788855_47188552_ENST00000601538.6,E | 1 | - | good | TGGAAGCCAGGAGGAACTCCCCGGCCTTCCTCTCCACAGACCTGGGGGATGAGGATGTGGGCCTGGGGCCACCCGCCCCCAGGACGCGGCCCTCCATGTTCCCCGAGGAGGGGGATTTTG | 120 |
| chr11 | 70487015 | 70487135 | 788855_47188552_ENST00000601538.6,E | 1 | - | good | TGGAGATCGACCCCCAGGCCCCGGAGCCACCGAGCCAGCTGCGGCCTGACGAAAGCCTGACCGTCAGCAGCCCCTTTGCCGCCGCCATCGCCGGAGCCGTCCGCGACCGTGAGAAGCGGC | 120 |
| chr11 | 70487135 | 70487255 | 788855_47188552_ENST00000601538.6,E | 1 | - | good | TGGTGAAGCAGTCCAACGTGGAGGACAGCCCCGAGAAGACGTGCTCCATCCCTATCCCGACCATCATCGTGAAGGAGCCGTCCACCAGCAGCAGCGGCAAGAGCAGCCAGGGCAGCAGCA | 120 |
| chr11 | 70487255 | 70487375 | 788855_47188552_ENST00000601538.6,E | 1 | - | good | CGCAAGCCAACTTCCGCAACAAGAGAGGCCAGATGCCAGAAAACCCATACTCAGAGGTGGGGAAGATCGCCAGCAAAGCCGTCTACGTCCCCGCCAAGCCCGCCAGGCGGAAGGGGATGC | 120 |
| chr11 | 70487375 | 70487495 | 788855_47188552_ENST00000601538.6,E | 1 | - | good | CCCCCGCCACCAGGTCCGACACCGTGGCCACCATGATGAGGGAGAAGGGGATGTACTTCAGGAGAGAGCTGGACCGCTACTCCTTGGACTCTGAAGACCTCTACAGTCGGAATGCCGGCC | 120 |
| chr11 | 70487495 | 70487615 | 788855_47188552_ENST00000601538.6,E | 1 | - | good | CTGTGCCCCCGTCCCCACCACCACCTTCCCCAACCACTTACAACTGCCCCAAGTCCCCAACTCCAAGAGTCTACGGGACGATTAAGCCTGCGTTCAATCAGAATTCTGCCGCCAAGGTGT | 120 |
| chr11 | 70487615 | 70487735 | 788855_47188552_ENST00000601538.6,E | 1 | - | good | GTTTTCTGCTTGTAGGAATAACAGAGGAAGAGCGGCAGTTTCTGGCTCCTCCAATGCTGAAGTTCACCAGAAGCCTGTCCATGCCGGACACCTCTGAGGACATCCCCCCTCCACCGCAGT | 120 |
| chr11 | 70487735 | 70487855 | 788855_47188552_ENST00000601538.6,E | 1 | - | good | TGTGCTGTGTTTCCTGAACATCCGGTTCATCATCTGAATTTGTGAGTTTGTGGAGATGTAGCGTATCGCACGTTCCTAGGAACTTAGGCTTTGTTGCTATTGTGACTTAAAGCTAAACCT | 120 |
| chr11 | 70487855 | 70487975 | 788855_47188552_ENST00000601538.6,E | 1 | - | good | TGTGCCTGGAGGCCCCACAGCAGCAGCGTACTCACAGCATCCCCAGTGCCTTTCTGTTCTTCCCTTTCTTGTCCTTTGGCTCACGACGGGCCATGTGACTAACCACTCGGCATCGTGGCT | 120 |
| chr11 | 70487975 | 70488095 | 788855_47188552_ENST00000601538.6,E | 1 | - | good | CGCACACACACACGCACAAGCAGCCCCAAGGCGAGCTGCCCACTTGGCCTGCAGACTGGCACACGGCTCAGCGTCAGGGGCCCTCCTGCCCTGGCCTCACACAGCTCCCCTGCTGCAGCC | 120 |
| chr11 | 70488095 | 70488215 | 788855_47188552_ENST00000601538.6,E | 1 | - | good | CCTAAAAACACCCCCAGGAGCCCTTACCATGCCTGAAGGGGCATGGTGGAAGGATACCTCCTGGTCTAGCCAACCCCGCCCCAAGCACAATGCCACAATGCCAGCACAAGCACACGTAAG | 120 |
| chr11 | 70488215 | 70488335 | 788855_47188552_ENST00000601538.6,E | 1 | - | good | GTGGGGCTCAAGTGCCATCTGGACGGTGGACTGCACACCATCCTTGCCACATAAACACAGCAATACCTGCAACCCTTTGCTCTCATTTCAGATGCACAGGTGACAGCCTTAGACCCCTGC | 120 |
| chr11 | 70488335 | 70488455 | 788855_47188552_ENST00000601538.6,E | 1 | - | good | CAATGAAAGGATTCACATGACCCTGGAAGAAAAGCACCTCCTCCCAGCACGCACAGAGCCAACCGTGGGCAGATGGCTTCGCTCAGGGTGGCAGAGGCCTCCCTGTACATCTGGGCAGCC | 120 |
| chr11 | 70488455 | 70488575 | 788855_47188552_ENST00000601538.6,E | 1 | - | good | CAGGCTCTAGAGCCCACATGGCCCCTCCTCTGAGCTGCAATTCCCCAGCCATCCATCTCTCTGAAGGTCCAGAGAGCTCGAGGACACATAGAACCCTTAGTGGGGCCCAGTGAAGTCAGG | 120 |
| chr11 | 70488575 | 70488695 | 788855_47188552_ENST00000601538.6,E | 1 | - | good | CAGGACCACTTTCAGGGCTGCCGCTGGGACCCAGAAGGGGCTCACTGAGGCCTGGCTCAGGATCCTATTCCTGGCTCAGTCCAGGACTTTAAAGGAACCTGGGCTCCCTCCCCTCAGGGG | 120 |
| chr11 | 70488695 | 70488815 | 788855_47188552_ENST00000601538.6,E | 1 | - | good | ATCCACATATTTTCATTAGGTCAACTCATACTTTGTGCCAGGTCAGCCACACTTGGACCCTGGTTAGTTGCGATTATGGTGGTCCACTGGGCTCTCAGCCCCTGCTCCAAAGGGTCTTGG | 120 |
| chr11 | 70488815 | 70488935 | 788855_47188552_ENST00000601538.6,E | 1 | - | good | TGTCTGTGTCTTCAGCTGTTGAACAGTCTGTAAAGTCACACCTAGGCAACACACCCAGAGAACCCTCAGGCCACCCTGCTGGCTGTCGTAGGACCTCAGCTGCATCCCCACACATTCAAC | 120 |
| chr11 | 70488935 | 70489055 | 788855_47188552_ENST00000601538.6,E | 1 | - | good | AGTAATTTATTTTGCTACCTCTTTGAAAACCTGCATGCTCCCAACATGAAATCCCTGTGCCACGCGTCTGTCGGTGTGTGTGCGTGTCTCGCTTCCCTTCTCCAAGTGCTCCTGTTGACG | 120 |
| chr11 | 70489055 | 70489175 | 788855_47188552_ENST00000601538.6,E | 1 | - | good | CTTTGCAATGCCAAAATGTCCTCCCACAAGGAGTATGTTAAATTTGGAATACCCTTATCGAATGGAAGTTTTCATTGGAATAAATTATTGTAGTTTGTCCTAGGACTTTTTTTCATTTCA | 120 |
| chr11 | 70489175 | 70489295 | 788855_47188552_ENST00000601538.6,E | 1 | - | good | CTGAATGTAACCCAGTTTACTTTATAAGGTGTATTCTTAAATTAGTACTCCTTGGGAACAGAATTGCCAGACAGCCAACTCAGAGTGACTGGTCAATGGAGAACACTTTGAACGAGCCAT | 120 |
| chr11 | 70489295 | 70489415 | 788855_47188552_ENST00000601538.6,E | 1 | - | good | AACTCAGGGAAAGCTGCGTATTTGTCTTGCGGTTACTGGTTAATAACAACTGATTTCCTCCTGTCAGACAGCAGAATCTTTCTATCAGGTTAGTGAACCTTACGGTTTGGAATCTGTAAT | 120 |
| chr11 | 70489415 | 70489535 | 788855_47188552_ENST00000601538.6,E | 1 | - | good | CTGCTGTGCAAAAGCGCAGCGTGCTTGTGAGTCTCAGGTGGATAACTGTAGGCAGTCACTAGGCAGTAAGTGGCCGTGGAGGTGGTGTCTGCTGTGCTTAAAGCTCCCCCTGCACCAGCA | 120 |
| chr11 | 70489535 | 70489655 | 788855_47188552_ENST00000601538.6,E | 1 | - | good | TCAATTCTGGTAGAAGCTAAGGCCCTTGCCCTGTGAGAGTTGTGGAGGCAACCCTGGGACTCTGGTCTGCCCAGCCCACCCCAGGGGCACTGTCCTGTGTGTTCTGGAAATGCCTCTGGG | 120 |
| chr11 | 70489655 | 70489775 | 788855_47188552_ENST00000601538.6,E | 1 | - | good | GTCACAATTATTCCTATTGACCCTGTTGCAGGCAAAGGTACACTTGACCTTGAGACAAGGTCACACTTGACCCTTTGAGACAGTCCCTTTGTGAGATAATTCAGCATTGCCTTTACTGTG | 120 |
| chr11 | 70489775 | 70489895 | 788855_47188552_ENST00000601538.6,E | 1 | - | good | AGCCACCACCAGGCCTGGAGTATGAGCTCAGCAGATGTACTTCCAAGGGTGGTTGGGTCAGGTCTCGAAATGAAATAATGTACTGTGTGTTGCAGAAAACTGCCTTCCTTTCTTCTTAAT | 120 |
| chr11 | 70489895 | 70490015 | 788855_47188552_ENST00000601538.6,E | 1 | - | good | TATCTGTGATAATTCCAGCAGGGTAAAGCGGGAGGGTGGGCGGGGTGGTGCGAGCCAACCCCCCTCCCCCACCCTGCCCTGCTCAGGGTGGGAAGGGTAAGTGGGTTCCCTGCATTCCCC | 120 |
| chr11 | 70490015 | 70490135 | 788855_47188552_ENST00000601538.6,E | 1 | - | good | CAGAGGGTGGGAACCTGGCCTTGCCCACTAGGGAGCCATTGGAACATCTGAGTGGCATGAAACTCATCCCCCCTGAAGGACACAAAATCTAGGGTGGCTTCTGTCATCTTGAAGCCCAGT | 120 |
| chr11 | 70490135 | 70490255 | 788855_47188552_ENST00000601538.6,E | 1 | - | good | CCACAGAAGTTGCCCCTGGGCTTTCAAACACAGCCTTAGGCCCCTCAGGAATTCTGAGCCCTGGGCCTCTCACAGTTGCCTTGGCGTCCCACCCCAGCAAGGCCCTGCCAGCCCACCCAC | 120 |
| chr11 | 70490255 | 70490375 | 788855_47188552_ENST00000601538.6,E | 1 | - | good | CGCCAAGGAATCGCCGTGATGACGCCCACTGTTCCTGGGAGCCCAAAAGCCCCGTTTCTGGGCATCCCTCGAGGTACGATGCGAAGGCAGAAATCAATAGGTAAGAAGTGTGGCACTCCC | 120 |
| chr11 | 70490375 | 70490495 | 788855_47188552_ENST00000601538.6,E | 1 | - | good | GCTTGGGGAAGCCACAGTGACGGAAGTTCCCTTGTGGTCTCTGGGCCCTGTGACCGTGGGTGGGGGCTGGCCAGGGCTGCGTCTCACCCTAGGCTCTCTGGTTTGCAGTCTGTGTACGAA | 120 |
| chr11 | 70490495 | 70490615 | 788855_47188552_ENST00000601538.6,E | 1 | - | good | GGCCAAAGGCCAATAGCAGCTGCAGCGAATAGGAAAGGCACCTAGAGAGGGACCTTCCTGGGTCCTCAATGCTGTGGTTCTCATCAGCCCCACTGGAGACTTTAGGTGGCTCCAGAAGCA | 120 |
| chr11 | 70490615 | 70490735 | 788855_47188552_ENST00000601538.6,E | 1 | - | good | TCCCACAGGACATCCAGCCTGGCCCCCCTCTCAGAGAGACGTATCTGTCCAGCTGCTCTGGCAAATGGTGTTGCAGCAGACCACATCTCCATGGAACAGGGCATGCTGCCATCCTAACAG | 120 |
| chr11 | 70490735 | 70490855 | 788855_47188552_ENST00000601538.6,E | 1 | - | good | GAGCCCATTTTATGAAAGAGGAGATGGTGGCTCCTATGTAAGGACTTGATGGTCACATGGCACATGGGGAGTGGGGTATGGACCCAGGCCAGACCCCACTGCCCACCACGCTGTGCCCTT | 120 |
| chr11 | 70490855 | 70490975 | 788855_47188552_ENST00000601538.6,E | 1 | - | good | CTCCAGGCTGTGCCCCTGAGCAGCTCCAGGGCTTGCTTAAGCTAGCTGGCACATCATGCTCACCCACACACCACCACAATACCTGGTTGGCCTTTGCACACACTGTGGGGCTGGTGTCAG | 120 |
| chr11 | 70490975 | 70491095 | 788855_47188552_ENST00000601538.6,E | 1 | - | good | AGCGAAAGAGTTGGCAGCACCAGGACTCTGCTTCCAAGGTGGAGGGGAGGAGGGGAGCTCTAGGCTCCTCCTGGGACAGGCTGAAGAGGGAGGAAGCTGGGGACAGCTCACTGCTGCAG | 120 |
| chr11 | 70491095 | 70491215 | 788855_47188552_ENST00000601538.6,E | 1 | - | good | TTGCACACGAATATTTCCATTGAGCGTCCGCATCTGATATTTTCCCGGGTTGGCCTTCAACACTACAGGTTTGGGGCCCTGAGCCATCTCAGTTTTGCAGACAAAACAACAGCCTGGGGA | 120 |
| chr11 | 70491215 | 70491335 | 788855_47188552_ENST00000601538.6,E | 1 | - | good | AAGGTGCCCTTGATCTCCCAAAGGGTCCCACTTTGCTCCTCAGCCTCAGGGCCATAGAGTGAGACAAACAGAGGCTGTGGACACATGGAAACTGCATCTCAAAGCTGGAGCAATAAAAGC | 120 |
| chr11 | 70491335 | 70491455 | 788855_47188552_ENST00000601538.6,E | 1 | - | good | CTCTGAACTCCACAAGGGGACTTCCCCTCCAGGCCCTGCCTGCCGGTCCTTGCCTCCCACTTTGTTCCCAGTCCTTCGTGGCACCCACATGCCCCACCAAGTTGGGGGGGTGTCTGAGAC | 120 |
| chr11 | 70491455 | 70491575 | 788855_47188552_ENST00000601538.6,E | 1 | - | good | TTCTTGAACAATAACACCTATAAATATTATAAATTTACCACAAGGCTTTTCTGTAATGTGCATTAAACAACTGTTAATAATTACAGTTACAATATCTGCCCTCACACCTCCTCCCATCAC | 120 |
| chr11 | 70491575 | 70491695 | 788855_47188552_ENST00000601538.6,E | 1 | - | good | CTCCATTGTGCAGAGGCAGCCATCTGAACCTGATGTCAGATTCTAGCACTGCCTACCTGTCCTGATTGCTTACCTTAGTATTCTCTTTGAACCAGTATCAAACTGCCTATGTCTAGCTTT | 120 |
| chr11 | 70491695 | 70491815 | 788855_47188552_ENST00000601538.6,E | 1 | - | good | GGTCACCTTCCCTAGGCAGTCAGGAGGGCACACAGCTGGAAAAGGGCCTGGGGCAGAGGCTGCTGATTCCCCCAGGCACGTCCTGAACAAGGCGTGGCCTCCTGGACCCTTGGATTGCTT | 120 |
| chr11 | 70491815 | 70491935 | 788855_47188552_ENST00000601538.6,E | 1 | - | good | TCCCCCACACTCTCCGAGATCCTGGGCAACACTTGTCCTCGGAGCCAGAAGTGCCATGAAGGGCTTCAGTGGCTGCTGGGCGCAGTGGGTGCCTGGCTCAGAGCAGGCTGCCCGTGCCAT | 120 |
| chr11 | 70491935 | 70492055 | 788855_47188552_ENST00000601538.6,E | 1 | - | good | ATGGAAGTAAGGGGCCTTGACACACTCTTCAGACAGAGGGGCTCTTGCCTCCTGGTCCACCCAGAGCACAGAAATGCTCCGGTGTGGAAGAGGCCGTCTGTCCCACATCCCATGTGGCAG | 120 |
| chr11 | 70492055 | 70492175 | 788855_47188552_ENST00000601538.6,E | 1 | - | good | CTGTGCAGACTCACAGTGGTGGGACAGAGCAGGCTGCATCCAAGCCCTGCCCTTACCGCTCACTCAGTGCCCTGAGCCTCGAAGGTGTCCTGTCATACTTGGCTGAGGACCCGTGAGCAC | 120 |
| chr11 | 70492175 | 70492295 | 788855_47188552_ENST00000601538.6,E | 1 | - | good | ACGGTGCCCAGGAAGTGGGTGGGCTGTCTCAGTAGCTGAGGTGCACACGCTTTCATGTCTGGTTCGTGCTGTCCAGAGGTGGGGCCCAAATCTAAGTGGCCACAGGACAAAGAGCCGGGT | 120 |
| chr11 | 70492295 | 70492415 | 788855_47188552_ENST00000601538.6,E | 1 | - | good | AACATGGCTGTGGAACCGAGGGTGGCGACCATCAAGCAGCGGCCCAGCAGCCGGTGCTTCCCGGCGGGCTCAGACATGAACGTGAGTGGCCGTACCTTGGGACCACGAGGGCGGGGGCC | 120 |

| | | | | | | | | |
|---|---|---|---|---|---|---|---|---|
| chr11 | 70492415 | 70492535 | 788855_47188552_ENST00000601538.6,E | 1 | - | good | TGCGCTTTCCTATCTACACCCAGCTTCCCCCACTGGTGTTGCTGCTCCAATCCTCACGGTTCTATTGCAGATAAACCCGAGGAGATAGTCCCGGCCTCCAAGCCCTCCCGCGCTGCTGAG | 120 |
| chr11 | 70492535 | 70492655 | 788855_47188552_ENST00000601538.6,E | 1 | - | good | TCCTGGGAAGGAGCCCATGCCCCAGTGGGAATGCATGTGGCAGAGGCACACGCATGCTCCTCACAGCAGCAACAAATGCCCCCTGCACAGTTTTGGATGTTGGAGTGGCTGCTGCACCTG | 120 |
| chr11 | 70492655 | 70492775 | 788855_47188552_ENST00000601538.6,E | 1 | - | good | CCAAAAAACAGAAATGTCCCAAAAAGCTTCAGAGCCCTTGGGGGCTGGGCACTTAAGAAGGTGGCCGAGCTGTGGAGGGTCTTCTATGTCCATGCCTCGTGCATGCCTCCGAGGGCAAGG | 120 |
| chr11 | 70493135 | 70493255 | 788855_47188552_ENST00000601538.6,E | 1 | - | good | TGGCAGATCCTCATGTCTTGGCAATCACAGTTACAAAGGAACCATATGTCTGAGTCTCTTCAAAAAAAAAAACAAAAACAAACAAAAAAAAAAAACTTCAAAAAGTCAGTTTAGTTAACCC | 120 |
| chr11 | 70493255 | 70493375 | 788855_47188552_ENST00000601538.6,E | 1 | - | good | TTAAAGGAAATGGGAAGGTTCTGCATTGGGGAAAATAAACAGAGACCTTCCCTCTGCAACACTTACTAACGCCTGGGGCCACGAGCACCCTGGCCTCTGAAACACCCACGCTGCCTTGGG | 120 |
| chr11 | 70493375 | 70493495 | 788855_47188552_ENST00000601538.6,E | 1 | - | good | TTCAGTTTTCAAGTAAAAAAAAAAAAAAAAAATGTGCTGCAAATGTCCTCTTTAAAAAACTTTTGTTAATTGCTAAGGATCCCTTATAACTAAACTTTGGTTTTCTTTTTTTTTTTTTTTT | 120 |
| chr11 | 70493495 | 70493615 | 788855_47188552_ENST00000601538.6,E | 1 | - | good | AGCTCTCCAGGTCTCAGTGGCACAGGGCCCTTGCATTTAAACAGCCTTAGACACACCCTTATGCCTGATCCCAAACACCCTGAGATGAAGCTAAGGCGTCAGGAGAGCTCTGGGATGTGC | 120 |
| chr11 | 70493615 | 70493735 | 788855_47188552_ENST00000601538.6,E | 1 | - | good | CAAAGGCTCAGAATGGGGTACACCCACCAGATACCACACTGGCTTTGTCCCAACCCAGTGACATCTTAGAAGTACAGCTGGGGTCACCTTTTCCAGAATGCACATCGCGACTGGAAGACC | 120 |
| chr11 | 70493735 | 70493855 | 788855_47188552_ENST00000601538.6,E | 1 | - | good | CAAGGCCCTAGCATACACTTCATGGGAGCCACCAGACCCCAAGAGTGGGCTTAGGGTGGCCAGAAGGCAGAATGTTCCAGATGGCCTGCTTCTCCCCAGGGAGTAAGGCCAGTGCTGTCT | 120 |
| chr11 | 70493855 | 70493975 | 788855_47188552_ENST00000601538.6,E | 1 | - | good | AAGCTGCCATGACAGGTAACTTCCGCCTCAGCTTGAAGAACAATTTGGAGATGGGGGCCTTCCTCCCTGCACTCTGCTCAGCTCTAACTTGCACAATAGCCTCATGGCTCAAAGTCTGGG | 120 |
| chr11 | 70493975 | 70494095 | 788855_47188552_ENST00000601538.6,E | 1 | - | good | CCTCTTGTGGGGACCTCTGGTCTGGGCTGCCTTTGTCAGGAGCCTGATGCCTTTGTGTTGGACAACCCTGATCCGTGCCGTGGCTCTGGGAGCTTTGCCCGTGGGTCAGCTCCCCGTGAG | 120 |
| chr11 | 70494095 | 70494215 | 788855_47188552_ENST00000601538.6,E | 1 | - | good | TGAGAGTGTCCATAGGTGTCAGAGTTGTGTACGTCTCAGCTGCTCTGCAAGGTTGAGGGGCCATCACAGGGCGCCTGGCTCAGTGTCCCCAGCACACCCCCATTTCCTCTCCCCAAACTT | 120 |
| chr11 | 70494215 | 70494335 | 788855_47188552_ENST00000601538.6,E | 1 | - | good | ATCTTAAAGAAAAGAAAAAAAAGTATAGACTGCCCTGAGGTTCTGGCCCGGTGGGGGCAGGACTGGCCCAAGGGTCTGGAACACAGGCCTGGAACCAGGCCCCTCATGCAGACTCTACTG | 120 |
| chr11 | 70494575 | 70494695 | 788855_47188552_ENST00000601538.6,E | 1 | - | good | AGGGCCCCATGGTAAGTGTGCTGGGCGTGACAGCCATATGGCCTCTGCTGCACATTCTCCTGCCTTTTCTTCCCCATCCTTGTTTGAAGACTAAAAGTGTATGGCTGGGTGTGGTGGCTC | 120 |
| chr11 | 70494695 | 70494815 | 788855_47188552_ENST00000601538.6,E | 1 | - | good | GGGAGATTTGGCTGCCCGGACACTATGGGGCAAGGGCTTTCTTGCTTTGCGTGAACACTTGGCATTTTGCTGACAACTGGAAACGTGGTCCAGGCCAGGGGTCGGCAAACACCTCCCTGA | 120 |
| chr11 | 70494815 | 70494935 | 788855_47188552_ENST00000601538.6,E | 1 | - | good | GGGCTTGTGGGTCCAGGAATAACAGGAGAGAATGCATGAGCAGGGACGCAGAATTGGGGCCCAGTGAGCCTGCAGGCCTGGTCAGCTGGACTTGGTCATGAAGGGAGAGAGTGCTGTGGG | 120 |
| chr11 | 70494935 | 70495055 | 788855_47188552_ENST00000601538.6,E | 1 | - | good | AGATGAGAATCGGGTGTTTCCTACCGTGCCTGACCAGTTCCCTGAGAAGCAGATGCTTAGCATCCACTGGGGGAATGACAAGGTCCCCAAGGAGGCATCAGGCTCATCGAAGAGCCAGCA | 120 |
| chr11 | 70495055 | 70495175 | 788855_47188552_ENST00000601538.6,E | 1 | - | good | AGCCATGCAGCACCTGCTGCAGCCGGAGGAGCTGGGATGAGGGCCTTGAACGCAGGTGAGGACAGCACGGCTCTGGACTGGGGTCACAGGTCACAGCCAGGATGTCCCCACTGCCAAGTG | 120 |
| chr11 | 70495175 | 70495295 | 788855_47188552_ENST00000601538.6,E | 1 | - | good | CCCTGGGAGGGGAAAGGCTCCTTGGAGGAAGCAAATTGCTGGGACCATGGCAGGTGGACCCAGCCCGTGCAGCCCAGGGAGGTTGACAGGGACCCCAGCAAAGGGAATGTGAGATGAGG( | 120 |
| chr11 | 70495295 | 70495415 | 788855_47188552_ENST00000601538.6,E | 1 | - | good | ACCCAAGCATCACCTGGTCCCAAGTCAAGGGGGAGAAGCCCAGCTGATCGGCGAGGCCAGGGAACAGAGGCCGGCACAGAAACCAATTCGACCCCAGCCATCCAAGGAGTCCCAGCCCAG | 120 |
| chr11 | 70495415 | 70495535 | 788855_47188552_ENST00000601538.6,E | 1 | - | good | ACTCTGGGGGCCACCAGCGGCCACATCTCACTTGAGGCAAAGAACAGCCACTGCCTAGCGCCCCCATTTCCAGGGATAGACCAAGCTGGAGTTGCACATCACAGCCGGCCAGAGGAGATC | 120 |
| chr11 | 70495535 | 70495655 | 788855_47188552_ENST00000601538.6,E | 1 | - | good | CTGTCAGGAAGCAGGTCCCAGGGTATCCCCACCCCGCAGTGCCCTGGCTCACCACAGCTATGGGACATAACTCCCTGGTGCAGGAGACACATCTCAACCCCCGTTTCATTGGAGAAAAAT | 120 |
| chr11 | 70495655 | 70495775 | 788855_47188552_ENST00000601538.6,E | 1 | - | good | AGCTGCTCTTCCCCTAACTTGGTGGCATCCCTTCTGGGTCATGTTCATGGAGGTACATGCTGCTGCACCCCGCTCAGGCCTCACGCTCCACTGGGGTCAGAGGAGGGGCCCGGGCCAGCA | 120 |
| chr11 | 70495775 | 70495895 | 788855_47188552_ENST00000601538.6,E | 1 | - | good | CAGCCAGGAAGGCACCCTTCCCTGTCTGCACCTGTGTACAGTATCAGGTGGGCTGGATTCTGCGGCTGCCCCCAGAGCTCGCCACCTGGAATGCCCCCGTGGGATGTTTTGTTGGGCTCC | 120 |
| chr11 | 70495895 | 70496015 | 788855_47188552_ENST00000601538.6,E | 1 | - | good | ACCTTAGCCTCTGAGCTTACCATAGCCTGGAAGTCAGGCCTGGAGGGATGCCCAGGCGTTTCTGGCTCGGATGGGCCTTCCTGCATCTCCACAGGTGCAGGTCTGAGCACAGGCAGGTGG | 120 |
| chr11 | 70496015 | 70496135 | 788855_47188552_ENST00000601538.6,E | 1 | - | good | CACCTGCGCCCAGGCCTTCCCCTGCAGGGCCTCGGCCGTCCTGCATCTTCTGGGTCTGACACCTGTCAATGTTCAACGCCGATGACTTTCCCTCACCTCCGCTATTCAGCCCCAGCTGCC | 120 |
| chr11 | 70496135 | 70496255 | 788855_47188552_ENST00000601538.6,E | 1 | - | good | GATTTGGCTCATTTAAATTGTAATCCCAGGAAAGCTGTTATGCCCTAGAGTCCCACCTTCCTTGATGTGCCCAGACATGGCCAACAGCACAGCAGAGCCTGGCCACAGCCTTCCCACACC | 120 |
| chr11 | 70496255 | 70496375 | 788855_47188552_ENST00000601538.6,E | 1 | - | good | GACCCTCAGTGTGCAGTGGATGGAAATGAATCAGCCTCATCAGCTGTGGGGTTTGTTAATACATAAAAAACAGACACATGGCTTCTGGAAGGGCCCAGTTGGAAACGGGAGCTTCTTGAA | 120 |
| chr11 | 70496375 | 70496495 | 788855_47188552_ENST00000601538.6,E | 1 | - | good | ATGAGGTGTATTTAACAAGATGCCTGCCGCAGGCCCTGAGGTCCTAAAATCGCAGCCTGCAGCCTGTTCATTTTGGGAATGTGAGAAGTCGAGGCGGTGCCCCAGGGAGGCCGGGCAGCA | 120 |
| chr11 | 70496495 | 70496615 | 788855_47188552_ENST00000601538.6,E | 1 | - | good | TACAGCCAGCTTCTCATTAGGATGAGAAACAGGATGATCGGCCCAGAACTTCCACCAGATTCAGGGACGGCTGTCATTATTGACCAGGACACCTTTCCTTCACCCTGGGCACACAGTTAC | 120 |
| chr11 | 70496615 | 70496735 | 788855_47188552_ENST00000601538.6,E | 1 | - | good | GGCTTGGGGGACCCCAGCGCCAGCCCCAGCTCCCGCCTGGTCCCATGGTTTGCCCCCATATGGCCTTACGCATCTCAGACCTGAAATGTTTTCCCTTCGACATGGGAACAATCCCTTTAC | 120 |
| chr11 | 70496735 | 70496855 | 788855_47188552_ENST00000601538.6,E | 1 | - | good | TGGTTCTCCTGGACGAGGCAAACCATGCCAATGGTGACCCCAACCATTTCCAAACAAACTTTCTGCAGCCGTTGCCCACCAGGGCCCAGGGCTGGGCAGCCCATCACATGGCAGGCACAA | 120 |
| chr11 | 70496855 | 70496975 | 788855_47188552_ENST00000601538.6,E | 1 | - | good | ACCAGATGTCCACAGGCAAGGTGGCCACAGTTTCAGGGATGCACCTTCAATGACATGACCAGCCCCACCCTCTGATGTGGCGACTGTGATGATGGGTAGGGTGCCCCCACCCCCGCCTCG | 120 |
| chr11 | 70496975 | 70497095 | 788855_47188552_ENST00000601538.6,E | 1 | - | good | TTAAAAACGAATGCTAGCCCCTGACCCTACCACCCCAACACGCCCGGCTGCTCTCACATGCTCTAGTTCAAAAGAACATTTGTTTTGGGGTTATTAGGAAGGGAAAGGTTTAAAACAGAG | 120 |
| chr11 | 70497095 | 70497215 | 788855_47188552_ENST00000601538.6,E | 1 | - | good | TCAGGCCAGCATGATCCCACTGGCCACGTCCCCAAGAGACAAGGGCCAGGAGGGACCCACACCCCTGTGGGGAGCCAATGCAGGGTTTTCCTTTTTCATTTGAATTACACTAACTCTAAC | 120 |
| chr11 | 70497215 | 70497335 | 788855_47188552_ENST00000601538.6,E | 1 | - | good | TATAAATTAAGCATAATCCTTAGGAGCCTAGAATGAGAAGCAGGAATCTCAGTCCTGTGGCTTTTCTGAAAAGTCTCCATATCCCGGTAAACAAGATCTCCAAGATAAGGACTGAGCAAG | 120 |
| chr11 | 70497335 | 70497455 | 788855_47188552_ENST00000601538.6,E | 1 | - | good | AATCTGTCTGAGGCTTGTTTGTTTTCTGTCACCGAACAATTCATTCTAAGACTGCCAGGGGCCAGATTCACTAATCCCCTTCATCCAGAGTCCCTGAATAAAGACCTAGTTCTGAGCTTT | 120 |
| chr11 | 70497455 | 70497575 | 788855_47188552_ENST00000601538.6,E | 1 | - | good | GGTCAGCATCACGTATCTAGTAGAGAGAACACACGTTTGCTCAGAGGGCTGAAATGCGGGTGACTGCTGGGGCATCTTCATCAGCTCCTCACTGGAGTGTTTTGGGATTTGACAGTTTCA | 120 |
| chr11 | 70497575 | 70497695 | 788855_47188552_ENST00000601538.6,E | 1 | - | good | GCCCCCTATCAGGGGAACCTGCACTGAAGTCAGCCCGTGGTCCAGAGTCAGCTCATGGGCTGGGGGGCTCTGGTAACAGGCAGGTGACAAACCTACCTCCCCTCCCTTTTCCCTGTGCCT | 120 |
| chr11 | 70497695 | 70497815 | 788855_47188552_ENST00000601538.6,E | 1 | - | good | CTCCCAGCCTCGGCATGGACGCTGGGGTCGTGTCACTCCTCGATTTGGTGACTCCCTTGTGATTTGCTGAGGATGCAGAGCAGTGGCAAGATTTCTCTAGCACAGGGCACACTGCTTCCT | 120 |
| chr11 | 70497815 | 70497935 | 788855_47188552_ENST00000601538.6,E | 1 | - | good | TCTGGGCCTATTGTGTGGCCACAGGCAAAGGAACTCGCTGGACAGTGGACCTAGTGTCACTTCAGTGCAGAGCCTATTTTCCAGGAGACTCAGCCACCAGGTGCCTGGGTGGTCCTGGGC | 120 |
| chr11 | 70497935 | 70498055 | 788855_47188552_ENST00000601538.6,E | 1 | - | good | GTGGAGCAGCCCCATGCAGCCCTCCACAGGTCTGGTCTGCTGCTTTCCCCCAGAAGGTTCTGAAGAGTCAACAGGCAAACAGCTGGGCACTCTGCCCCTCAATGGGGCTGGCTCCCAGGC | 120 |
| chr11 | 70498055 | 70498175 | 788855_47188552_ENST00000601538.6,E | 1 | - | good | GACAGACCACTTCAAGCCCACATAAGGGGAGAGAGCAGTGGTTGAGAACTGACAGGGTCCCTGGACCATGCCAGAGGATGTGAAGCAGACCATGAATCAGGTCCCAGGAAAGCTGAAGAG | 120 |
| chr11 | 70498175 | 70498295 | 788855_47188552_ENST00000601538.6,E | 1 | - | good | GCTTCTGTCCTCTAACATCTGGCGCTGCAGATGTGTGTGTTTTCCTTTTGGAAATGGAGTGACATTTGTACTCTCTCACCCAGAGACAGGATGTAGTAGCACTGCAACTCCAGAGAAGTA | 120 |
| chr11 | 70498295 | 70498415 | 788855_47188552_ENST00000601538.6,E | 1 | - | good | TGCAGGCCAGGACCAGTCCACCAGGGTCCAAGCACACCCTTCCCACCCTGCACTGCCGCTTTGGAAGATTAAACCAAGAGGCCGGCCAACTGGCAACAGCCGCTGCTTCTAACACACGCG | 120 |
| chr11 | 70498415 | 70498535 | 788855_47188552_ENST00000601538.6,E | 1 | - | good | AGGATTTGGGGTGGGGCCTCTGCCCCTGGGCACATGAAGGGGACTGTGGCCTTCACCCTACCAGGCACCTATGGCTGTACCTTCAGACTCTAGCCCTTGGAAGGTGTGTGACTCAGAGCC | 120 |
| chr11 | 70498535 | 70498655 | 788855_47188552_ENST00000601538.6,E | 1 | - | good | TGATCCCAGGTTTTCAGTTCTCCATCTGTAAAATCCTTGTTTAGTCTTTAAGTGTTCTCTGAGCACCTGCCCTAGTGCCCAGACCCTGTGCTGGGCTCAGGAAGAAGCCGGCTGTCAGGC | 120 |
| chr11 | 70498655 | 70498775 | 788855_47188552_ENST00000601538.6,E | 1 | - | good | GAACTGTGCAGGGATACGTTTCCATTGTTTTAAGCCCCCCGTGCATGGTCATTTATCATAACATCCCTGGCAAAGCACCTCAAATCCCGGGTCTGTCCCCGCTGGCTTTACCCTGCTTAG | 120 |
| chr11 | 70498775 | 70498895 | 788855_47188552_ENST00000601538.6,E | 1 | - | good | GGTGGGCCCTAAATCCAGTGGTGTGTCCTTGTAAAGAAGGAAGGAACAAGCCAAGAAACACCATGGATTTGATCCTCTCCAGACCCCTGAAGGAGCCTGACCTTGGACTTCCAGCCTCCA | 120 |
| chr11 | 70499015 | 70499135 | 788855_47188552_ENST00000601538.6,E | 1 | - | good | CACATAGTGGATGCACGGCAAATGCTCCAGGCCCCTCGTGCTGCCTCCACCTGTCTCTGGGAGGACTGGCCTCGCCTCGGAAAAAGCACTCGTCACACTCCAGAAAGGCCATGTGGAACA | 120 |
| chr11 | 70499135 | 70499255 | 788855_47188552_ENST00000601538.6,E | 1 | - | good | CGTGGCCCAGCCTGGTCACACTGGGGGATCCAGCTGGGGACTGAAGACACCGAGCGTTGCTGACAGCCCCTGCCTCTAGGGCCATGAGGGCATCCCATGGCGCACATGCGAAGTGCGGGG | 120 |
| chr11 | 70499255 | 70499375 | 788855_47188552_ENST00000601538.6,E | 1 | - | good | AAGATCCACCTCCCGTCTGAGCAAGAGAAACTGGCTACAAATTTGAGCTTCCAGAGCCTCAACCCCAGATATAACAGGGGGCAAGGGTGTGGGCAGGAAGACCCCAAAGGGACCAGCAGC | 120 |
| chr11 | 70499375 | 70499495 | 788855_47188552_ENST00000601538.6,E | 1 | - | good | CAGAAGTCCCTTGTCAGGGAGAACAGTTCGGGACTCAGGTGGCTGAGCCACCGTGGGCGCCTGTGGCCTGCTGGGACCTGGCCCCATCCACACCACCCTCCTCCCCCAGGGTGGCATCAG | 120 |
| chr11 | 70499495 | 70499615 | 788855_47188552_ENST00000601538.6,E | 1 | - | good | ACCCCTTCAATCCAAGGCCAAACCCCAGCTCACGGGAATGGCCCTCTATTCATCACCTAACTGAGGAAGATGCCCCACCATGGCTGCCAGGGACCCAGTTGCCAGAACAGCGTGGGACAA | 120 |
| chr11 | 70499615 | 70499735 | 788855_47188552_ENST00000601538.6,E | 1 | - | good | GCTGCCCAGAAAACTCCCAGCGGGCTTTCTAGGCCCCACTGCTGCGCCGAGGATGTGGTGGACACCCGGGGTACAGAATACCCTTCTTTCTATGTCGTGCAATTACGTTTGTGTGTGAAC | 120 |
| chr11 | 70499735 | 70499855 | 788855_47188552_ENST00000601538.6,E | 1 | - | good | CTCTGTCCCAATCATGTTAGATGAGATTGGGTGAGGTGGGCACAGAACCACCACTGTCCCGACCTGTGTCATCCCAGCGGCCTCTCTGCAGCAGCCCCGGGCTTGCACCAGCATCCTACA | 120 |
| chr11 | 70499855 | 70499975 | 788855_47188552_ENST00000601538.6,E | 1 | - | good | GGTATTGCTCGCCTGCTGCTGGACACAGCTCCCCCAACAATTAAAAGCAGCACTCCCAGCTTTTAATTAATCCCTTGGAGTTGGGAGGAGCCTAGCACCTGGCCCTAAGGCACATCTTAA | 120 |
| chr11 | 70499975 | 70500095 | 788855_47188552_ENST00000601538.6,E | 1 | - | good | CCTCCCCACCCAGGGCTGGTGTGGAGTGGGAGACTGGCAGTCACTGTGGCTGTGGGCAGCCGGACCGCAGGCCCCAGATAGACTCAGGTTCCCCCTTTCACAGCAGCAAGCACATCGGGG | 120 |
| chr11 | 70500095 | 70500215 | 788855_47188552_ENST00000601538.6,E | 1 | - | good | TTGTGGTGGGCAGGAACTGCCTCCCTTTAAGGCCCGGAAATCACCCATTCCCCAAGAGGCTGCCCTGTGCCCTGTGGGGCACTAGAAGGAGAAGGGAGCAGTGTGAGCCCTTTGGCTACA | 120 |
| chr11 | 70500215 | 70500335 | 788855_47188552_ENST00000601538.6,E | 1 | - | good | TTCACATCGTCATGACCGTGAAGCTGGCCGGACCCCTGTTTGGTCTCAGCTGGGGCTGCCTGGATGGGGGGTTGCAGATGAGCAGAGCCTCTGGCTGCATTCCTGACCCTCAGGACCCCT | 120 |
| chr11 | 70500335 | 70500455 | 788855_47188552_ENST00000601538.6,E | 1 | - | good | TCCTGCTGGGTCCTGCCCGAGGAGCCCTGCCGCCCTGGAGCACATTCATCTGTCTGTTCTTTCTGCTTGGCTTCTGTCTGTGATTCCCCAGCGTGTCCTGTGTTTCTTCCCTGACCTGTG | 120 |
| chr11 | 70500455 | 70500575 | 788855_47188552_ENST00000601538.6,E | 1 | - | good | AGAAGGGTAAGGGAGGCCCAGTGTCTGTCTCAGCCCCCTGCCCATCACCCCCCTGTGGAACAGAAACATCCTTTGTGATGCCGTGAAGTCTCAAATGTGTCGCAAAGCCTTGTTGGCTTC | 120 |

| | | | | | | | | |
|---|---|---|---|---|---|---|---|---|
| chr11 | 70500575 | 70500695 | 788855_47188552_ENST00000601538.6,E | 1 | - | good | GCCCAGCCTGCTCCTGAGGCGCCCTGAGCTGACAAGGCCCGAGTGTAGGCTGCGGGCGGGCGCTGCATCCTGGTGGCTCATGGCGGTCCCCTTTCTGTTTGCAAGCCTCGGTCCGGAAGA | 120 |
| chr11 | 70500695 | 70500815 | 788855_47188552_ENST00000601538.6,E | 1 | - | good | GAGCAGGGCAGACAGGACAGAGTTGTGCGGGCAGCCAAGGTTTGGGGAAAGCCACTCCATGGATAGCGCAGCCTGCGTTCGCAGCGTTCCAGACAGCCCCTGCCTGCCGCCCCATTGCCG | 120 |
| chr11 | 70500815 | 70500935 | 788855_47188552_ENST00000601538.6,E | 1 | - | good | CCCGCCACCTCTGCTCATGTCTGCTCTGAGAACTCCTCCCTACTTTGGCAGGGTGCTGGTGGGACAAGGGGTGCTGAGGAAGAAGGAAAGCCCCTTTCCACTGCCAGGCAGTAGGTTAAC | 120 |
| chr11 | 70500935 | 70501055 | 788855_47188552_ENST00000601538.6,E | 1 | - | good | TCCCCAGAGGGACAAGCCTAGCAGCTCTCATCTTTGCAAGGGGTGCTCCTGAGAGCTCAGCTCAGTCAAGAAACAACAGAATGAGGCAGGAAAGGCCACAGGTGCACTCTAAAGGCTGGC | 120 |
| chr11 | 70501055 | 70501175 | 788855_47188552_ENST00000601538.6,E | 1 | - | good | CCCTGCAGTGGCGCTGCCCCACCCACCCAGAACCAGTAGTGCCCCGCGTGCCCCAGCCTCACAGGCGACGGAGGAGGTGACCAGCCTCCAGGCCCTCTCCACAGAGGCCTGTCCTGGGGC | 120 |
| chr11 | 70501175 | 70501295 | 788855_47188552_ENST00000601538.6,E | 1 | - | good | GCCACAGCAGTGATGTCATCCTTCCTCGGCCCAGGGCGCTGAGGCCGGCCAGGCCGCTTCTCACGAGGTTCTGGGGCTAGAGTCTGAGGATGGGCATGAGGCCTCAGTCCAAAACAGCAG | 120 |
| chr11 | 70501295 | 70501415 | 788855_47188552_ENST00000601538.6,E | 1 | - | good | CAGCTGGCCTGGGGGGTTGAGAGCCAAGCCCTGGCTGGTGCTCTCACTCACAGGCCCCGTCCTCCCTGCACTCTGGCACAGGCAGCTTTCCCGAGAGGAGTCTAGCAAGTGCTAAGTGCT | 120 |
| chr11 | 70501415 | 70501535 | 788855_47188552_ENST00000601538.6,E | 1 | - | good | AACACACAGAGATGGCATGTAGGTGGAAACGCTTGCTCCACTGGAGCACAAATTGGAGCTGGTGAAGGCAGCTTACATGGGCGCAGAAACGAGCCCCCGCCCAAGGCCCACGGTCCAGTC | 120 |
| chr11 | 70501535 | 70501655 | 788855_47188552_ENST00000601538.6,E | 1 | - | good | CTTGTAGGCAGGGGAAGGGCGACTAGCAGCCACGTGGGCCATGCTGGCCATGCAGGATCCTGTTTGTTCCGGGGAGTCACCATGGAGCCAGGGACACCTTTGGAAAGAGGGTGGCTCTGA | 120 |
| chr11 | 70501655 | 70501775 | 788855_47188552_ENST00000601538.6,E | 1 | - | good | TGTGTAGATAGCGTGGGCTTGGCAGAGCAGTCTGGTCTGCACAGCAGGCTTAGGGAACGCCCGGCACCTTTGTGTAAGGTCTGCAAGCAACAGAGAGAGGCTCTGGCCCTCTGACTCCAT | 120 |
| chr11 | 70501775 | 70501895 | 788855_47188552_ENST00000601538.6,E | 1 | - | good | CAGCTCCCCCTGCTGTCTAGGGCACCCCCACTGACATCCACCTCAGCCCAGGCCTCGGAGGCCTGTGTGAGCTGCTCCCCACGTGGCTCAGACCAGAAGGCTCCATCCTCCGGGGATCTG | 120 |
| chr11 | 70501895 | 70502015 | 788855_47188552_ENST00000601538.6,E | 1 | - | good | GCGGGGCGTTGTTGCTAGTCCTGCCTTTCCTCTTTTCTGTTTATCTAACATTGTTTTGTTTTGAATTTTTGGGCCTCTCACTAGTGGATAAAGGTAAGCAGCCCCACTGGGTCCCCAAAG | 120 |
| chr11 | 70502015 | 70502135 | 788855_47188552_ENST00000601538.6,E | 1 | - | good | CAGGCTGTGCATGACCCCACGCTTGCTCGCTACCCCCCTCCTGCCCCTGTACCCAGACGTGCATCAGGAAATGCCCCTGCATGCCCCGCAGCCCCATGTGTGCACCTCCCTCGGAGCCTC | 120 |
| chr11 | 70502135 | 70502255 | 788855_47188552_ENST00000601538.6,E | 1 | - | good | AGCCCTCACCCTGCGCTCCAAGTCCATGACCTCGGAGCTGGAGGAGCTCGGTAAGTCGCACACCCGGCCCAGCCCTGTACAGTCACCATGCCGGTCCCTGAGCTGCCCTTTGCAAAGCAC | 120 |
| chr11 | 70502255 | 70502375 | 788855_47188552_ENST00000601538.6,E | 1 | - | good | CCTGCCAGCCACAGGGCCACTGCTAGCCTTATTCCCATATCTGGAAACATGGGCTGGGGTCTCACACCCATCTCTCCTCTGCCCTCCAGCTCCCCCGCCTCCAAAGCGGGCACCGACCAC | 120 |
| chr11 | 70502375 | 70502495 | 788855_47188552_ENST00000601538.6,E | 1 | - | good | CCATGAGGGCCTGCACTGCAGTTCAGGGATATGGCAAGATCCTCCTGAGCACCTGTCCTTTCCCATCATAAGCACACCTGCACAATCCCTGACCACCGCAGCCAAACAGCCTGAGACCCA | 120 |
| chr11 | 70502495 | 70502615 | 788855_47188552_ENST00000601538.6,E | 1 | - | good | CACACGCTGCCGGAATCATCCATGTCCCACCAGGAGTGCCCTTTTAGTCACTAGCAGGCGTCCCTTCCGCAGGCTTCCCACTCAGTGCCCTGTCCCCTCCCTCCCCCTGCCCAGGCCCAC | 120 |
| chr11 | 70502615 | 70502735 | 788855_47188552_ENST00000601538.6,E | 1 | - | good | CGGGCAGGACAGCTGGGGGGCAGTGGGCCCTCTCTCCAGCTGATCCCACACTTCTGAGAGCCCAGGCACAGGGGTGAGGCTGGAATTTACCGCCAGAGCCCACGACAATGACTTGGGGCT | 120 |
| chr11 | 70502735 | 70502855 | 788855_47188552_ENST00000601538.6,E | 1 | - | good | TCCTTAAGGTGGTCACGGTGACCAGGAATCTGGACCCCGACGACACCGCCAGGAAGAAAGGTACATGCCCCACATCGCCCAGCTCCAGCCTGGGGCCCTACTGGGGGGGGGGGTGGGGG | 120 |
| chr11 | 70502855 | 70502975 | 788855_47188552_ENST00000601538.6,E | 1 | - | good | CTCCGCTGGCTCTCACTGATGCCATCCTCTCCGCCTTCCCACAGGTTAACAATGAGAATGTTGTCAAAGTCGGCCACAGGCAGGTGGTGAACATGATCCGGCAGGGAGGGAATCACCTGG | 120 |
| chr11 | 70502975 | 70503095 | 788855_47188552_ENST00000601538.6,E | 1 | - | good | TCCCCAAAAGCACCTTTGCTTCCAAAAAGATGACGGTCTCACTCCCTTTGCTTCCCTGCATTTGCCCCCTTTTTAGGAGCCCACATGTCTCTGCCTTGGGTGGGTGCCAACTGTCTTCCT | 120 |
| chr11 | 70503095 | 70503215 | 788855_47188552_ENST00000601538.6,E | 1 | - | good | TTCCTCTGCAGCAGCTGTGGAGGACTCAGCAGAGAGCTGCAGTGTCAGAGGAAGGCCTCCCCCAACCTCCTGAGCCACAGTTAAGTGACCTCCACAGTCACAGCAGCATTCATTCTGGTT | 120 |
| chr11 | 70503215 | 70503335 | 788855_47188552_ENST00000601538.6,E | 1 | - | good | CCCAGCCCTGGAACTCGCCTGTCAGGCTCTTCACATGCAGTCTCTTGGTGGCCCTGAAAATTGCCCCAGGCCTGTAAAAGTAAGCACCTAAGGATGCTCAGGTTAAATAGAAATCAGCCC | 120 |
| chr11 | 70503335 | 70503455 | 788855_47188552_ENST00000601538.6,E | 1 | - | good | GGAGGGAGGGAGTGGGAGGAAAGGGGCTGCTTCAGTGTGATGTGCATTCCCAGAATACATGAGAAGCAATTAAGCGCCACGTTTCCTCCTCCTGTGAGGATGCAAAGCCTGGCACTGCAG | 120 |
| chr11 | 70503455 | 70503575 | 788855_47188552_ENST00000601538.6,E | 1 | - | good | CCCAGGAGCACTGGGGCAGCTAGTTGTACAGACCAGCACAGGGACAGGGCCCCCACTCTTGTAAGCACCCTGCTGGGCCAGGCTTTTCTGCCTTCCCAGGCCTCTGGAGCGGGAGAGCAG | 120 |
| chr11 | 70503575 | 70503695 | 788855_47188552_ENST00000601538.6,E | 1 | - | good | TGCATTTGCCATGGTGAGGACAAATAGGGCAGGAGGTAACCGGCTCCTAAGAGTTCCTCAACTGCAGGGCAGTGGCCATAGCATGGCAGCCTTGGGGGTCCCCAGCCTGCCACTCACCGT | 120 |
| chr11 | 70503695 | 70503815 | 788855_47188552_ENST00000601538.6,E | 1 | - | good | AGCGTGGGGCTGGGAGGCAGAGGCCAAGGCCATCTGCTGCTGAGCATTGCTCAGGGAGGTGAGAATAGCAGAGGGGCCAGGGGGATGGCACCCTGAGTCCGTTCCTACTTCCGTGTCAGG | 120 |
| chr11 | 70503815 | 70503935 | 788855_47188552_ENST00000601538.6,E | 1 | - | good | CTTTTCACAGCAACATTTGAAAACCACTAAGTGTTACTCCTGCAGTAGCCAGGGTGGGCAGTGTCAGCCCAGACGCCCACCCAGTTCCAGTCTCCATCACTCGGGTTTTCCTCTGGGAGT | 120 |
| chr11 | 70503935 | 70504055 | 788855_47188552_ENST00000601538.6,E | 1 | - | good | CCCTGCTGCCACCTAGAGACAGCTGCCTCTGGGCTGCGCTCCCATTCAGGAGCTGCTGGGATGAAGGTGCCAGAGGCCCCCTGTGGAGCTCAGTTCCATGGGCAGGGGTGAGGCCGGGGC | 120 |
| chr11 | 70504055 | 70504175 | 788855_47188552_ENST00000601538.6,E | 1 | - | good | GCACAGTGTGGCTGGTGAGGGGCATGACGTCTTGGCTGTTGTTTACGATCAGGGTGCTTTTCTTTGTTCGAAGGTGGCGTGTGAAGCAGCTGTAAACTTCACCTTCTGGGGCTTCCCCCT | 120 |
| chr11 | 70504175 | 70504295 | 788855_47188552_ENST00000601538.6,E | 1 | - | good | TTACAGGGATGTTGGATTTGTGAAAATCCACCAAGCAGCACACAGGCTTTAAGTTTTCTGTTTGTGGGTATACCAGATGTTTAGTTTTTGAAGAATCTATAGCAGGAGGATATGCGTACA | 120 |
| chr11 | 70504295 | 70504415 | 788855_47188552_ENST00000601538.6,E | 1 | - | good | GCAGCCGTCTCGTAGGTGACCAAAACTGGCACATCCTGGGCCTGTCAACAGTATGGGGAGCCAGGCACTGGGCCTGTCAACAGTATGGTGAGCCAGGCAGTACCTGCTTCTATCCAGCTC | 120 |
| chr11 | 70504415 | 70504535 | 788855_47188552_ENST00000601538.6,E | 1 | - | good | TCTGAGTACATGGGAATGAGCCGGTCCCACAGCAGAGACTGCAGAGCAGCTGGCACGGGGACTGCCCTGGCCCCACTGAGTCCTCCCTGGGCTCCTCGCAGCCCTGCCTGGAGGACCTCT | 120 |
| chr11 | 70504535 | 70504655 | 788855_47188552_ENST00000601538.6,E | 1 | - | good | CTCTCCCAAGGCCCCATTAGGATAGTCCTGCTCCCAAGACTCTATCTTCAGGGCAGTGCAGGGTGACCAGCTGTCCAGTGGACCCAGGACACTGGATACTCACTGCTGAAACCGGGTGGG | 120 |
| chr11 | 70504655 | 70504775 | 788855_47188552_ENST00000601538.6,E | 1 | - | good | GATCCGGCTCCTTTACTTCCTAAATCCATCTGCTTTCCTAATTACATTCTGAGAACCTCAGTGAAAGGAACAGCCCCCAGGAGTCACCTTGCAAAGACCGCTGCCGCTGTAGGTGGGCCT | 120 |
| chr11 | 70504775 | 70504895 | 788855_47188552_ENST00000601538.6,E | 1 | - | good | ATGGGAGGGGAGAGAACCCGATAACGCGTGGCTGGTGTTCGGTTGCCTGTCTGTACCAGAGCAGCGAGGGTCCTGCAAGCTGAGGTCTGCCCTCCGAGAACTCCCGGTGCTGTCTTCCTC | 120 |
| chr11 | 70504895 | 70505015 | 788855_47188552_ENST00000601538.6,E | 1 | - | good | ATGGAGGCAGAGACAGCTGAACTGCCTGTGCCTCATGGCTTTTAGTGGTACGGGGAACAGAAACACTGGAATGAGTGAATGAATGAATGAATGAGCAACAGCCACTACCCGGGTGTGCTC | 120 |
| chr11 | 70505015 | 70505135 | 788855_47188552_ENST00000601538.6,E | 1 | - | good | TGTTCAGTCTCCTTTTACTTGGTTCTGCCTTCTTAGGTGGTACTCAGCCACCTGAAGTTATGTGTTCACTTCTTTGTGTGTCGGCCAAAACTCCCTGATGAGGACGCAAACTTCCCACAG | 120 |
| chr11 | 70505135 | 70505255 | 788855_47188552_ENST00000601538.6,E | 1 | - | good | GCATCCAAGGCCCTGCTCTCAGGCCTTGGCCTTTTCTGCGCCCCCCTGCCTACCGCTTCCTTGCCCATTCCCACTCCATGACTCCGCCCACACTGCTCTTTCCTGGTGGGTCCAGGCCTC | 120 |
| chr11 | 70505255 | 70505375 | 788855_47188552_ENST00000601538.6,E | 1 | - | good | GACCCCAGTGTCTCGGTGCCCCCCCACGTGTGTTTGGCACCTCATACTCACAGCTGCTCGGGTCAGATGCCTCATCTCTGCATCGGGCCTCAGCACTCCCTGGCCCATAACTTCACCCTG | 120 |
| chr11 | 70505375 | 70505495 | 788855_47188552_ENST00000601538.6,E | 1 | - | good | TGGGGATCTAGTGGCCAGGCAGCTGGGGGCTCCCTACTGCCAGGTCTTCTCACCAAGGCCTCACACATTGGCCTCGCGGCTCCTGAGCCTGGGCAGCCTGCCTCCTCCATCTCAGCTTGA | 120 |
| chr11 | 70505495 | 70505615 | 788855_47188552_ENST00000601538.6,E | 1 | - | good | GGAGCCCCCTCAAGGCCCGGCACTGGGGGCCCGCCCAGAGCAGGAAAGCTGGTCCCAGCCCATCTCAAGGTGGGGCCGAGTCCTCCCCTCGCTCCCCACAGCCTTGCAAGGGATGACCCC | 120 |
| chr11 | 70505615 | 70505735 | 788855_47188552_ENST00000601538.6,E | 1 | - | good | CTCCAGTGACTGGCAGCCCATCCCCGTGCCAGTCTCGTACTCTCCCCAAGCATCAGATCCAGAGAATGCCCCACAGCGGGTGGCAGGAGCAAGCAGGCAGTAATATGCACTTTCTTGGTT | 120 |
| chr11 | 70505735 | 70505855 | 788855_47188552_ENST00000601538.6,E | 1 | - | good | ATGGTTAGCAGGCTGCAGTCAGAGGCCACATCCCCAGCCCCTCCCTGCCTAAGTGGCCAGGCTGTTAAAGTCCCCATCAGCCCGACGAGCCCTCAGGAGGCCGCAACTTGAGCTTGGCTT | 120 |
| chr11 | 70505855 | 70505975 | 788855_47188552_ENST00000601538.6,E | 1 | - | good | CCCAAAGCGGCCAAGGGCCAGCTCTGGCCAAGATCACAGAAGGACAGCATCCAGCCACCACAAATGAACCACGGGAGATTCTGGAGGATGAGGATTCTAGGGAGTGAGCCCAAGGCCGTG | 120 |
| chr11 | 70505975 | 70506095 | 788855_47188552_ENST00000601538.6,E | 1 | - | good | CCTTTCTTTGGTCCTTTGCCTCCCCTGCCCAGAAAAGAGTAATGTCTGTCTCTGACACTTCTGAGGAAAAGGGCCCTCCAGCCCCTTGACCCTTCCAAGGCACAGGAACAAACTGCTTCT | 120 |
| chr11 | 70506095 | 70506215 | 788855_47188552_ENST00000601538.6,E | 1 | - | good | ACCAGAACGTGCACCCAGGCTGTCGGCTACAGACAAGGAAGGTGCTGGGGCCGCCAAGGTCCTTGGCCCCGCAGCCAAGCTTCCTGAGAATCCCCCATGAGTGGCTCATTCCCCGAGAGG | 120 |
| chr11 | 70506215 | 70506335 | 788855_47188552_ENST00000601538.6,E | 1 | - | good | GGAGTTGGGTGGGAACAGCAGCTTCTGCAGACTCAGATATGAGACATCAAATCTGGGCAGCCAGCCTGGCCCTGTGCAGGAGCAGCTCTGGCTGGGGCAGCATGAGTGTGTGGAGGTGCC | 120 |
| chr11 | 70506335 | 70506455 | 788855_47188552_ENST00000601538.6,E | 1 | - | good | GGGAGCCTCATTTCCTCTGAGCTCTCCTGAGCAGGTCCTGCCATTTGCTGCAGTCTGCTGAGGGCAGCAGGTCTACGAGGGGTACAGGGGATGGAATCAGGAACTGGGGAGCCCTGATTA | 120 |
| chr11 | 70506455 | 70506575 | 788855_47188552_ENST00000601538.6,E | 1 | - | good | CGTCAGAGGGGCATCAGATGCCCAACAACTGATTCCAAACCTAGCTCCACCCCTGCAAGCCCTGTATGCCTCTTTGTCATCAGCTTTCAGAGAAGGCCAAGTGATGTTTCCAGGGCCCAT | 120 |
| chr11 | 70506575 | 70506695 | 788855_47188552_ENST00000601538.6,E | 1 | - | good | CCATGAAGCAGGAGGGAGGGCCCTCCAAGTGGAGCTGGGACTGCAGGGAAGCTCGTCCAAAGCAACTTTGTTTGGAAGTGGCTCATGACCCCGAAAGACCTGGGCGAGACAGATTCTGGA | 120 |
| chr11 | 70506695 | 70506815 | 788855_47188552_ENST00000601538.6,E | 1 | - | good | GTGTCCTGCCCACGGGCACCCAGCAAGCAGCAGGGCCCCCACAGGCTCCGGACCCAATAATATGCTGATTCTGGAGGGGTGCAGGCCCACCAGGACCAGGGCTCACCGGATACCATCCCT | 120 |
| chr11 | 70506815 | 70506935 | 788855_47188552_ENST00000601538.6,E | 1 | - | good | GAACATCCTAGGATGTGTGTGGATGGTTTGAGGCCCTGTTGGCACCACCTCCCTGCCCCCAAGTGCTTGCCAGGGCAGGAAGCCATGCTGTTTACTGAGGACACCCCGCTAGAGAGGCAG | 120 |
| chr11 | 70506935 | 70507055 | 788855_47188552_ENST00000601538.6,E | 1 | - | good | TTCTAGACTTTCTATGCAAAGGTCTTATACTCTTTAAATTGTGACAGCTAAGTTAAAACTGACTGAGGCACTGTGTGGGTGGGATTTGACCCTTGGTAATCGCTGGATGGGGACCAAAGG | 120 |
| chr11 | 70507055 | 70507175 | 788855_47188552_ENST00000601538.6,E | 1 | - | good | GGGGCGGGCGGGCACAGCAGCAGTCACAGCCTCAGCTAACAGCAGGGGAGGATGCCGCATGCAAGTGGGAGGCCCAACTCCCCAAGTGCGAGGGTTCTAATTTTTCAATGAAAACTGAAA | 120 |
| chr11 | 70507175 | 70507295 | 788855_47188552_ENST00000601538.6,E | 1 | - | good | TCTAAACGCAGATGGCATCAAAGGCAGTGATAGCAGTTTTCTGTTTCACTGGTGGCTGGCAAGAGCCACAAATCCAATGCCCCCTCAAGCTGAGAAGGTGACGTCAGGGCCCAGGAGGCT | 120 |
| chr11 | 70507295 | 70507415 | 788855_47188552_ENST00000601538.6,E | 1 | - | good | ATACAGGGCTCACCAGTCACCTGTGGCTCCCGTAGGAAATAGAAATCAGTTTCATGAATGTCACCCAGGCTGATTTTGAGACCAGCAATGTGCAGAGTTCAATCTGTCCCTTCCCAAAAT | 120 |
| chr11 | 70507415 | 70507535 | 788855_47188552_ENST00000601538.6,E | 1 | - | good | TGTTGCAGGCCCCTCACAGACATCCCTCTCCCTCTGCAGGGGAAAGATTCTGGGCAGAAGGTCAATGTGTTTTGAGATCTGAGCTCTGACACAGGAGACTGGCCTGGGAAGCTTATATTG | 120 |
| chr11 | 70507535 | 70507655 | 788855_47188552_ENST00000601538.6,E | 1 | - | good | GGGCCTGCATCTAGGGTTCAGGGTAATGCGGAGTGGTGACAGACGGGATGGCCACCTGCCTCTGCCCCAGCGCCCACATGACATTCCCCCATTCTGCCTCCTCCTCATGCTCAGAAAGGC | 120 |
| chr11 | 70507655 | 70507775 | 788855_47188552_ENST00000601538.6,E | 1 | - | good | ACGATGTGGTTTCTATATTAAGTGCAAGATTCGGGGCCACTTTCCTGGCTTTTGCAAGCCCAAGTTTGAGTTGGGGTCCCCCACAGGCCCCTCCAGACCTCATGTCCGTGTCTGTCCACT | 120 |
| chr11 | 70507775 | 70507895 | 788855_47188552_ENST00000601538.6,E | 1 | - | good | CCCCAGCTGCTTTGGGGAAGTCGGCATTTTTAGTGTCTTGATGGTATTTAGGGTCTAATTACAAACGCGCGTCACTGGGACCACATGCTTGGTGCTGGAACTGGCTCTTCATTGATGGAA | 120 |
| chr11 | 70507895 | 70508015 | 788855_47188552_ENST00000601538.6,E | 1 | - | good | TACCCTGGAGGGGTGCAAAGCAAATGTGCGTTGCAGGAGTTACTACCGGTCTGAGTTCATTTGCAGAGCCTCTAAACACTCCAAAAAATCACAAGCCCCGCTCTTGCGGGTTCCTCCCAG | 120 |

| | | | | | | | | |
|---|---|---|---|---|---|---|---|---|
| chr11 | 70508015 | 70508135 | 788855_47188552_ENST00000601538.6,E | 1 | - | good | GGCACAGGCTGTCAGTGCGGCCAGGTTCTCACCAAGGGGTCTTCCTGTCCTGGGCCTGTATGATAGAGGTTCCTGTGGCACACTGTCACGGCCTGTGCCCTCTCGGCAGAATCTAAACCA | 120 |
| chr11 | 70508135 | 70508255 | 788855_47188552_ENST00000601538.6,E | 1 | - | good | AGCTGAGGGGGACAGGGCAGGTGTGTGCCCAGCAGCACGTGGGCACCATCCACCCCCATGAGCCCTGGTGCCCTGTGGCGTTCCCAGTAGGCAGGCTCTCATTGCCTGAGGTCAGAGCCG | 120 |
| chr11 | 70508255 | 70508375 | 788855_47188552_ENST00000601538.6,E | 1 | - | good | GACTTAGCCCTGATGGTGGCCTTAGGGGAAGCTGGAGGGATGGGATGTGGGGAGGGGTGCTGAGCATTCTGTCGGGAAAGGTCAGTGTGGGCGGAAGGTTGGGATGGGTTCTCTGTGCTC | 120 |
| chr11 | 70508375 | 70508495 | 788855_47188552_ENST00000601538.6,E | 1 | - | good | AGGACTCCAGCATATAAATTTGGGCAGGACACAGTTCAGTCCAGAACAAGTGTCTGGGGACACAGAGATGATGCACCAGTAGGCAAAGTAGGGCAGGGGGCAGACTCTGCCCCGCCCCAG | 120 |
| chr11 | 70509095 | 70509215 | 788855_47188552_ENST00000601538.6,E | 1 | - | good | GAGGGCTGCAATAGAACAGTCCTCAGGGTGGTAACCAAGAAGGCCGGGCAAGGCACAGGCCTGGTGACCAAGCACTGCAGGTGGGCAGCTTCAACCAGACAGGGACTGTCCCACAGCTCT | 120 |
| chr11 | 70509215 | 70509335 | 788855_47188552_ENST00000601538.6,E | 1 | - | good | CCCAATCAGTGTCTGCTCACATTGGGCAGCGCTGGAGACCTGTGCCACTGCATGGGCAACAGCTGGTGCTGCAGGGGCAGGTGAGCCTCAGTGGGCAGCAGACAGAGATGCACTGTGGCT | 120 |
| chr11 | 70509335 | 70509455 | 788855_47188552_ENST00000601538.6,E | 1 | - | good | GGGAGCCCCCCCTCAATGGGATAAATGGGACTTGAGGCCATTTTTGCTCTGAGACCAATAGCCAGTGTCCCCACACTGGGCCCCTGGGGGAGGTCGTGGGACCAAGAGCCAAGACTGTGG | 120 |
| chr11 | 70509455 | 70509575 | 788855_47188552_ENST00000601538.6,E | 1 | - | good | AGGGGGCAGGGGAGCTGAGGAAAGCTGACCCGAAGGCTACACCAATGCTGAAGACTCAGAGCTACATGCCAGCTGGCCGTGCTCTGAGCTCCTGCAAGGAAGAGTGCAGAGCTCCATGCT | 120 |
| chr11 | 70509575 | 70509695 | 788855_47188552_ENST00000601538.6,E | 1 | - | good | CACTGGCGGTGAGGGTTTCAACACATAGACTGGGGGGACATAGACATGCAGCCACTGCAAGTCCCCAGCTTCAAGTCGTGGCCAGCAGGGTGGGCTGAGTCAGAGCTGAAAGACAGGGTG | 120 |
| chr11 | 70509935 | 70510055 | 788855_47188552_ENST00000601538.6,E | 1 | - | good | CTGGTACAATATATGTAATTGCAGGGGATGGGAGACGGAAACAGCTGTGGCCAGGCTCCTGGGTAAGTAAGCACAGCTATGGTTTGTCACTTGAGTTCTTAGTCCGTTTGGGCTGCCGCG | 120 |
| chr11 | 70510055 | 70510175 | 788855_47188552_ENST00000601538.6,E | 1 | - | good | CTGGGTGAACTGCGGCCCCAGGGTCTGCTCCTCCAGTCCCTGCCTCCCAGCTTCCCTTTAAGACCAGTCACTCCTGGGTCACCTGCTGGCCAAGACACCTCCTCCCTCCCCATCTCAAGC | 120 |
| chr11 | 70510175 | 70510295 | 788855_47188552_ENST00000601538.6,E | 1 | - | good | TTCCCCCAGGACCTTTAAGATGCCCCAGCCACCAACCTTGGCCCCAGGTTTGCTAACAGCAGCAGCATCTGGGGGCTGTGGCCTTTGCATGCCAGAGCTGCCTCTGCCATCTCTGCAGGT | 120 |
| chr11 | 70510295 | 70510415 | 788855_47188552_ENST00000601538.6,E | 1 | - | good | CTGCTACAAGCATGAGCGCGGGAGGGCTGTGGCCTCCAGCCTGCACTGCCCACTCCCTCCCAGGGCTGCGGCAGTCATTTTCCAATGACAGTGCCTCTGAGTTCCCACTCACAACACCCC | 120 |
| chr11 | 70510415 | 70510535 | 788855_47188552_ENST00000601538.6,E | 1 | - | good | GCGCTGGCAGAGGCAGGAAAGACAATCTCTTCTCAGGGACCCACATTACATGCCCCAGTTAGCCCTGCTAAAGGGGACAGTTTCAATAGGACCTCTGTGGGGTGGGGAAGGGCAAAAGCT | 120 |
| chr11 | 70510535 | 70510655 | 788855_47188552_ENST00000601538.6,E | 1 | - | good | TTCCTGCAGACAGGTTCCCCCAGCTCCTGGACAGAGGAGGAGCAGGGAAAGGAGGGACAGGATCTGTATCCAGAGGCCTCACAGACAGCTGGGTCCAGTGGCAGAAACTTGGGGGTGGTG | 120 |
| chr11 | 70510655 | 70510775 | 788855_47188552_ENST00000601538.6,E | 1 | - | good | CCAGCCTAGGGAGGACTCCAGGAACCCCTAGGACTCACTGTGCTATGACTCGCACAAGGCAGAGACACTTCCTCTAAAGGTAGAAGACAGTGACATGGGCCCATCCTGGAGGACACCCTC | 120 |
| chr11 | 70510775 | 70510895 | 788855_47188552_ENST00000601538.6,E | 1 | - | good | AGGAGTAGGTGCTGCCACTTCATGTGGCTGCGCAGCTTCTGTCCGGAAAATGGAGTCACACTTCCTCTTCACCAGGAAAGGGGAAGGCAGCTGGGAGCATGGGCTCCACCCTTTGTGTGC | 120 |
| chr11 | 70510895 | 70511015 | 788855_47188552_ENST00000601538.6,E | 1 | - | good | TTGGGGAAGGACTTCATCCAAAGCTTGTTTATTTGCAGAATAGGATGAGAACTTAAGACCAGCATGGTCGACGGGAGGTTGGCGCAGCCTCTTTGGTTGGGGCTGGCTCTGAGGCAGCA | 120 |
| chr11 | 70511015 | 70511135 | 788855_47188552_ENST00000601538.6,E | 1 | - | good | CACACTCCGTTGGCATCCTTGAGCCTTGTTGTCAAAGTGCTCACCTTCACACGGGGAGCAGGTGCACAAAAACCCAAAGCCACACGACGCACAGCTTTGTCCTTGGTATGTTTCTTACTT | 120 |
| chr11 | 70511135 | 70511255 | 788855_47188552_ENST00000601538.6,E | 1 | - | good | GAAGTATCCAGTCCTGCCTTGTTGCGTGCTGGTCTTGTCAATTAATTTCACGTTGAGTTGAGTTGCCAGCCATGTGCCTTTGCCCCACTCTTCCCATAGAAGTGTTGATGGGCGCTGGGG | 120 |
| chr11 | 70511255 | 70511375 | 788855_47188552_ENST00000601538.6,E | 1 | - | good | GGTTCCAAAGTCTCATTGCAGGTTAAACTGATTTTAGGTTTGTTTAAAGGCTGGGAAATTCTAAAGATCCTAAAGCTTTTTTTCCTCTGTCCCTGCTGAACTGGGTAGGGAAATGTAACT | 120 |
| chr11 | 70511375 | 70511495 | 788855_47188552_ENST00000601538.6,E | 1 | - | good | ACCCCATGGAAGCTCAAGATGCCAACCTGAGCATATGAGGAAACAGCAAGCATTTCACCTATTCTGCAGTTTCTGTCACCACATGAGCAATTTCTGGCATCTGTTTTAAGGACTGTAGCT | 120 |
| chr11 | 70511495 | 70511615 | 788855_47188552_ENST00000601538.6,E | 1 | - | good | TCCCCAGGAGCTGCAGTACTTACCTGAGCCTTCTATAAATCTGGCAAGTGATCAAGCCATACCTTCCCCTGGTGGCAGCATTCCAGAATTGCAGGTCCAGGTTTTTTGAGGGGCACCGAA | 120 |
| chr11 | 70511615 | 70511735 | 788855_47188552_ENST00000601538.6,E | 1 | - | good | CTCTGTAAATGTCTCCATGTCTGTATGTAAGGGTGATGCATGGACAGGCACATTACTCTTTTTTTGGAGTTATCTGGATCCATGATTTCTCAACAGTTTCTCTCTACTCCAAGGCAGTTA | 120 |
| chr11 | 70511735 | 70511855 | 788855_47188552_ENST00000601538.6,E | 1 | - | good | GAAGAGAATTATTTACAAAGGAAGGCAGGTATGCCCATTCTTCACAGAAATTCCCTGGACTGTGGCTGATGATAGGCTTGGGTTCTTGCCATCTACCTGGATGTGGCTGTGATTCACTCA | 120 |
| chr11 | 70511855 | 70511975 | 788855_47188552_ENST00000601538.6,E | 1 | - | good | GAAGTAAGGCTGGGTTGGTGGGGTTTCATCATGACAATCCTTGTGGAACTCTTTGTTTCTGTAAACTCTGTGTATGTGTATGTACAACTTTTGATATTTTAAATAACGATTTAAGTGGAA | 120 |
| chr11 | 70511975 | 70512095 | 788855_47188552_ENST00000601538.6,E | 1 | - | good | TCCTTTTCCATCACAGTGTGTCAGCGGATAATATCGAAAATGTAAATATCAAGTAGCAAAAATAGGTGTTCAGAGATTGAGTGATAGATATAAAATTAAATAGCTGAAGGCTCAGAGTGA | 120 |
| chr11 | 70512335 | 70512455 | 788855_47188552_ENST00000601538.6,E | 1 | - | good | TTAAGCTATGAATAGATTTTACATAGTCATGATGTAAATCCTGAATGGTGATCTAGTTAAAATTTTGATATCAATACATCTAGAGGATGGTGGTACTGGGAAAACAGTGGGGAGGGAGGA | 120 |
| chr11 | 70512455 | 70512575 | 788855_47188552_ENST00000601538.6,E | 1 | - | good | CATTTGGATTTGAATTACTGTTAAGTACATAGAAAAACCGACCAACAAACAAAATTATTTTGCTCTAGGGAGAACAAAATGTACAGGAAAAGATGAAGTAACCATAGTATACTATAGAGT | 120 |
| chr11 | 70512575 | 70512695 | 788855_47188552_ENST00000601538.6,E | 1 | - | good | CTTAATCCTGGTTGTTCAGTGCAGATCAAAAGTCTTCAGGAAATATTTCTTCAGGAAGACAAACTTAAAATGGATAATGTATTTGAAGACTCTGAAAAGAGATTTACATAATTAGGGAAG | 120 |
| chr11 | 70512695 | 70512815 | 788855_47188552_ENST00000601538.6,E | 1 | - | good | CAAAAAATGAGGTTTTTGAACCAAATAAACAAATACAGGGATTCAGGAGATCCAACACAGGAAAGAATCAGTCAAAGGAATCCTGAGGATTAGAGTAATGGGGACAGCTGCTAGAAAACA | 120 |
| chr11 | 70512935 | 70513055 | 788855_47188552_ENST00000601538.6,E | 1 | - | good | GAGAATTTATTTGTTGGTTTGGGGGTGGGGATTGTACTCTGTCTAGATTTTATAGCTATTACCTGCAGAAGAGTTAGTTAGTCTTAGGAATTTACTTTACCTTATTCCCCAACCTCAAAT | 120 |
| chr11 | 70513055 | 70513175 | 788855_47188552_ENST00000601538.6,E | 1 | - | good | AAAAAAAAAAAACAAAAAACTGCAGGTGACTCATACCATAATGCATCCATTGCACAAAGCTTCTAATTTTCAACTCTCATCTAAAATCTGCCTGTCTTTGTTCACTCTTCAGAACCTTCA | 120 |
| chr11 | 70513175 | 70513295 | 788855_47188552_ENST00000601538.6,E | 1 | - | good | ACTACCCTTCTCTGGCTCTCTCTTTTCTGGAGTTCTCCCCTCACTTTGGCTGCTATGGCCACTGTGGGCTGTATCCTCTGATTTCTATCAGAACTGCAGGTTTGTGCAGGATCTGGATTT | 120 |
| chr11 | 70513295 | 70513415 | 788855_47188552_ENST00000601538.6,E | 1 | - | good | AGTGGCAGCAGCTCAAATTTCAGCCTTATCATAGCAAGCCTATTTGAGTCTTCCTTACACACACAGCTCAGCTCAGACATCAACCAAAGACCTATGAGAAGTTTACACACAGAATATGAA | 120 |
| chr11 | 70513415 | 70513535 | 788855_47188552_ENST00000601538.6,E | 1 | - | good | GTGAATGTTATGCTTTGGCAACTCTGGATTATATTACATTCCTTTGATGGGTGTTAATGTTTTTGTTTTAGCTGATAATGAGTTTGGTTAAACTCAAATTGCAACCTTTGTCTGCCCTGC | 120 |
| chr11 | 70513655 | 70513775 | 788855_47188552_ENST00000601538.6,E | 1 | - | good | TAATTAGCTCTGTCATATCCATGCTGATGTTAAGCCCAGCTGATGCATTTCTAACAATTCCATATAATTCTTTTTAAAATTTCCTTGGGATTTCCTGTCTTTTATTCATTATAAGCCTAT | 120 |
| chr11 | 70513895 | 70514015 | 788855_47188552_ENST00000601538.6,E | 1 | - | good | ATCTATAAATTTATGTCCTTCACCTAATTGGGGAATTCCTTGGGTACCATTTTTTCAAGTATTTTTTATGACTCATTTTCTCTGTCTTCTTTGACTCTAATTTTCCCCCCTTCTGAGACT | 120 |
| chr11 | 70514015 | 70514135 | 788855_47188552_ENST00000601538.6,E | 1 | - | good | ATAATTTCTCTGGCTGCTTTAAGATTCTCACCTTGTCTTTGGCTTTGAGCAGAATTGACTATAATGTCCCTATATAATTTTTTAAATACTTATCTTTTTGGGGATTCACAGGCTTCTTGA | 120 |
| chr11 | 70514135 | 70514255 | 788855_47188552_ENST00000601538.6,E | 1 | - | good | TTTACTTTCACCTGTTAAAGATGTTGTTCCGCCGTCTTCTACAAGAGGTTCTTCTCCATGGTTTCTGATGCAAAGTCTGCAATCATTCAGTCATTTTTATTTAGAGTGTGAATAATGTAA | 120 |
| chr11 | 70514255 | 70514375 | 788855_47188552_ENST00000601538.6,E | 1 | - | good | TTCTTTAGTGCAGGCTTTCTGGCAACAAATAACCTGGTTTTCATTTTTTTATAAAGGTCTTTATTTTGCCTGTATTCTTTAAAGAATTTTCACCAGAAATGGAATTCTAGTTGTGTTTTT | 120 |
| chr11 | 70514375 | 70514495 | 788855_47188552_ENST00000601538.6,E | 1 | - | good | ATCTGCTTTTAAAATTAAGAGGGAGAATCATCTTTTATTTTGTATCCAGGTGTTCACCATTTGTGGCACCCTTTGTGCCATCCATAGATCCAGTCCCGCTTCACTTCAGCTTGAAGAACG | 120 |
| chr11 | 70514495 | 70514615 | 788855_47188552_ENST00000601538.6,E | 1 | - | good | GCAACCATATAGATGCATCTTCTTCACTCCTGCCTCACTTTATTCTATAATTGTCATGTGTGTTACATCTACACACTTGTAAACCCCAAAATCAGTATTATTTTTGCCTAAAACAGTGAT | 120 |
| chr11 | 70514615 | 70514735 | 788855_47188552_ENST00000601538.6,E | 1 | - | good | TTACACTGTAGCTCTTCACATTCTTTTAGCAGTGGATCAAGGATTACAGTATACATTCTTAGCTTTCCATAGTCTACTTAGAGTTCAAATTGTACACTCCATGTAAAATATAAAAACCCA | 120 |
| chr11 | 70514735 | 70514855 | 788855_47188552_ENST00000601538.6,E | 1 | - | good | CCATCTAGGTTTACAAGTTTGTCTTCTGTTTATCTCCTCAGTTTTTATTTTTCCCTTTTTCTGCCTTATTTTAGATTATTTAACTATTCTTAGAACCTTATTTTAATTTTTCCATTGGCG | 120 |
| chr11 | 70514855 | 70514975 | 788855_47188552_ENST00000601538.6,E | 1 | - | good | CTTTGTATTGAAAGTGTACGGACAGCATAATTAGGTCTTCTTTTTAAACCATTTTATAAGGCCTGATGTTTAGTTGGAGTTTTATTCTATTTAACATTTAATAGAATTGTTCATGTGGTT | 120 |
| chr11 | 70515335 | 70515455 | 788855_47188552_ENST00000601538.6,E | 1 | - | good | AGATATGGCCTTCGAAGAAGATAATAAAATTAGAAGGTCAGTCCAGACGGTCTACTGTGTATAAAATAAGGATACCTCAAAGAGAAAATAAAAGAAATATTTAAAAAGTCATTGAATGGC | 120 |
| chr11 | 70515815 | 70515935 | 788855_47188552_ENST00000601538.6,E | 1 | - | good | ATTAATATGCTCAGTGAGATCAAAGAAAAAAAAAGTAAAATGTTCTTTCTGAAATGATCAGAGAACAAAATAAGAGTTCCTGAAAATTAAAAATATTATATAGCAGGTATAAAAAAAATT | 120 |
| chr11 | 70515935 | 70516055 | 788855_47188552_ENST00000601538.6,E | 1 | - | good | GCTGCCTTCCAAGGAACATCAGACATTTGATGAACTAATATGAAATACGGCCCAAAACAAACACACTGAAAACAACTGAAGAAACCAGTATGCAGGAAGAGTAAGATTAAGAAATGTATT | 120 |
| chr11 | 70517975 | 70518095 | 788855_47188552_ENST00000601538.6,E | 1 | - | good | TACTGTCATGTAACATTGGGCTTCTTCCATAATAAAAACTTGTAGAAATGAGCCTCTTTTTAATTAGTAGACTTTATATTTTGAGGAGTTTATGGGGAAAAATGAGTGAAAAGTACCGAC | 120 |
| chr11 | 70518095 | 70518215 | 788855_47188552_ENST00000601538.6,E | 1 | - | good | AGCCATTTCCAGGGAACGCAGAGGGATATGATCGCCCCAGGGGCTGGGTGGGTGCTCTCGGAGCCAGCAGTGGCTGACGCTGGCTAAAGGGACAGCACTGGCTCAGGGTACTCATCTCTC | 120 |
| chr11 | 70518335 | 70518455 | 788855_47188552_ENST00000601538.6,E | 1 | - | good | AGCCTGAGCCATGACGCGGTGCCTCATGATACCAGGAATGCAGTCGCTTTTTAGGCATGAGAATGGCATTGTGGTTTTTGGTTTTTGTTTTGTTTGAGACAGGTTCTCTGTTGCTCAGGC | 120 |
| chr11 | 70518455 | 70518575 | 788855_47188552_ENST00000601538.6,E | 1 | - | good | AAAGGGATAAAAGAGGAATGGGGAAAGCCTAGAGATGTAAAAAGGCTTCAGAGACGTGCCAGCCAGATGTGACGTGTGCATCTTGCTCATGAGTCAAGTAAACCAGCTGTACACTCAGGG | 120 |
| chr11 | 70518575 | 70518695 | 788855_47188552_ENST00000601538.6,E | 1 | - | good | CGTGGGTATGCCCGGCACCGCCTGCAAAGGGCTCCTGGATGCAATCCCGCCTCTGACCACCACTAATAGAAGCAATGCCGCAGGATGCAATTGACTCGGTTTCTTCAGCAAATGAACTGC | 120 |
| chr11 | 70518695 | 70518815 | 788855_47188552_ENST00000601538.6,E | 1 | - | good | CAGTGGCTGCTAAAAACACTAAGTGGCAAGCTGACGGGGAGTGGGACAGGCTGAGGGTGCTTGACCCACCGGTGCTGTCTCGTCGCAGAAAGACCCAAGCAGCTGTGCCGGGGCTCCTGG | 120 |
| chr11 | 70518815 | 70518935 | 788855_47188552_ENST00000601538.6,E | 1 | - | good | CCTGTCTCAAGAAGTTAAAATAAAAATTATTCCAACTAGTAAGTGGAGAAGGAGCAACAGAAGTTGAACATCAGCGTTCTGCAGCTCCCAGGAGGTGAAGGATCTGGGCATGGCATTTAC | 120 |
| chr11 | 70519175 | 70519295 | 788855_47188552_ENST00000601538.6,E | 1 | - | good | TCCGGCCCGTATCTGGGCCTTCTCACATACACTGTGCCTCAGAGTCATCCAAGTGTTGAAAAGGGAAAGTTTCTTCTTAAAAATTAGTGTGGCTGGGTGCAGTGCTCATGCCTATGATCC | 120 |
| chr11 | 70519655 | 70519775 | 788855_47188552_ENST00000601538.6,E | 1 | - | good | TAACAGAATAAGACCCCGTTTCAATAAAATAAAATAAAAACAATAGTAATTGTAGTGGATTCAAACTTTTATCAGCTATGTTCTTATAAACATTCTTGAGTTCATAATGGCACTTTTTGT | 120 |
| chr11 | 70520375 | 70520495 | 788855_47188552_ENST00000601538.6,E | 1 | - | good | ATGTGACAGGAATGGCTCTTTACCTCCTGAACACCCATATATACCTCAGTCTAATAATGAGGAAAAAAAATCTTAATTCAGGGGTAGAACAAAAGACCTGACCAGCAGCCAAAACTGTCA | 120 |
| chr11 | 70520855 | 70520975 | 788855_47188552_ENST00000601538.6,E | 1 | - | good | AAAGCACGGCAAAAGGTAGCAGCCTAGAGCATCCTGTAGAACCGGAAAGGAAGGAAGTGGTCAAAAAACAAAAGGCTGGAGCAAATCCCAGGGAGCTCCCAGAGGCCAAGGAGCTCCCAA | 120 |
| chr11 | 70520975 | 70521095 | 788855_47188552_ENST00000601538.6,E | 1 | - | good | AGGGTAGAATATTTTGTCACATCCAAAAGCAAGGAAGGTATTGAAGACACAAAGTCACATGTGATGGGCTCAGGAGCCAGTGTGAGGAAGGGCTGCTCATACAGCACAGTGGAGGATTCG | 120 |
| chr11 | 70521095 | 70521215 | 788855_47188552_ENST00000601538.6,E | 1 | - | good | AATGGCAGTCATTAGGAACCTAGATTTTCAGCAAAGAGAAAGAGACACCAATATAAAAGAAAATAAGTAAAAACAGCATCATCCTAAATTTCTGTTGAAGCTATCAGTATAATCATGAAG | 120 |

| | | | | | | | | |
|---|---|---|---|---|---|---|---|---|
| chr11 | 70521215 | 70521335 | 788855_47188552_ENST00000601538.6,E | 1 | - | good | GAAATCATAACCTCTTTTCGGTAATCCTGTTATGAGTAATGACATTGGTGGTGTTAGTCTGAAAGTAATGATAGATGGATAGATGATTAATAAATGATAGAGCAAATAGATCATGACCTT | 120 |
| chr11 | 70521335 | 70521455 | 788855_47188552_ENST00000601538.6,E | 1 | - | good | AAATTTCAAAAATCTCAAAATCCAAAACAAAATAATAGAAATGAACATAACTGCTAATTAAATTAATGGCATTAACAAATTATGTCAAGTGGCAATTTGCATGTCATTGCAAACTGGAAA | 120 |
| chr11 | 70521455 | 70521575 | 788855_47188552_ENST00000601538.6,E | 1 | - | good | ACTTGTTTTAATTTAGCATTGCCTCAAGGGGGAGAATACAAGAAGAGGAAGTAGAAGAGAGAGAGTAGCAGGTTCAATTTTGGAATTGAGTAAATATTTCACGTAATTGTAAAACAGTCA | 120 |
| chr11 | 70521575 | 70521695 | 788855_47188552_ENST00000601538.6,E | 1 | - | good | CCACACACATGCTCACACTCACCCACACACATGCTCACGTGCACACACACATACACATACAGAAACACACAGGTATCTGCTCATTTCTCCTTTTTAATGGAAGGATAATCCACCAAAATA | 120 |
| chr11 | 70521695 | 70521815 | 788855_47188552_ENST00000601538.6,E | 1 | - | good | GCACCGTCTCCAGGGGGTGTTGTTACGTGGGAAAGGCAAGGCGAGAAGAATGTGACCAGCGTGCTGCTGTTTACCCAAGGAAGGTGGGGAGGGAATGTGCACACACACGTGCACACTCAC | 120 |
| chr11 | 70521815 | 70521935 | 788855_47188552_ENST00000601538.6,E | 1 | - | good | AGCAGAACACTAAGCACCAGACCCACCAGTAGGAGCCTGCAGTGTGTGCCGCGTAACAGCCTTGCAGCGAGGACATGTGGCTGCAGTCAGGTGCAGGGAGCGTCTCTGGGAGCTGCCGTG | 120 |
| chr11 | 70521935 | 70522055 | 788855_47188552_ENST00000601538.6,E | 1 | - | good | TGGGTAGTTTGGATAAGAAACCGTTAGAGCCAGCAACCCCACAGATGTGAACATAGCCCAAAGGTGCACTCGGAAAATTAGAACAAGGACTCACGTGGCTGTTTTCAGCTCTGTTTGGAG | 120 |
| chr11 | 70522055 | 70522175 | 788855_47188552_ENST00000601538.6,E | 1 | - | good | CTCAGAAAGCTGGAGCGGGGCTGTGGTGGGGAAAGCTGGTACAGTCAGAATCACGTGGCGGAACTGCAAGAGGCTGTGGCCTCTGCCTAGGAGGTGGAGGTAGCCTGGGCTAAGGGCTGA | 120 |
| chr11 | 70522175 | 70522295 | 788855_47188552_ENST00000601538.6,E | 1 | - | good | AGGTTAAGCCGTGCATTCAGCCATGCACGCAGCAGGATAAGCCATCCTTAAAGAGGCAGAGACCAACAAGGACAGAGCAGAAACGGCGATGGCGGCTGCACAGCAAGTGGACTCAAAGGC | 120 |
| chr11 | 70522295 | 70522415 | 788855_47188552_ENST00000601538.6,E | 1 | - | good | TGCTGTGGCCACCCCTTTGGTACTTTCTTGTATCTTTTTCAAGAATAAATGAATTTTTAGAGGTTTGATTGGCAACGTCCAGCAAAAGAGGGACACTGCCAATCAATCAAATCTAGATAG | 120 |
| chr11 | 70522415 | 70522535 | 788855_47188552_ENST00000601538.6,E | 1 | - | good | AGGAGGAAATAAGATCGGGGTTTGAATAAGCCATGGCTGGAGACAAGCTAAGAATTAATGAGAACCCTGCACGTGCAGAAGTGTGGCTGCCCTGGGAGGGCGCCCGTGCCTGGACAGCCA | 120 |
| chr11 | 70522535 | 70522655 | 788855_47188552_ENST00000601538.6,E | 1 | - | good | CGCGGAGCTGGAGCCGGGTAAGGGGCCTGGGCTTTCAAAGCCTGAGCCAGCAGGGACTCGGAGGCCCTGGGTAGGATGAGAGACGATAGAGGGGGTACCACCAGAGAGCTTTCTAGTCAA | 120 |
| chr11 | 70522655 | 70522775 | 788855_47188552_ENST00000601538.6,E | 1 | - | good | TCCTTCTTTCCTCTCCCCCTCTGATGGCTGCAATAACTGAGGGACTGGGACACGGTCCGAATGGATGCCAAGGACAGCCTTGAAAAAGCTTAGACAGTGTGAGCAGGTGGCCGGCCAGCC | 120 |
| chr11 | 70522775 | 70522895 | 788855_47188552_ENST00000601538.6,E | 1 | - | good | GCTAAGAGAGGAGCTGTGGCATTTAGGGGCCCAAATGCCTGGCTGGCTTGGGGCGGCTTCCCTTCTGCACGAGTCTCAAATGATGCCGCCTGGTTGGAGGTGGGGGCAGTGCCCTTCTCT | 120 |
| chr11 | 70522895 | 70523015 | 788855_47188552_ENST00000601538.6,E | 1 | - | good | CTCGTGGCACTGTGTGTAAACATTTTTCAAAGAGGGTGTGGATTGGAGCTGCGGGTCTGGAGGGAGGCCTCAGTGAAGGTAGCCTGCCCGCGGCCGCACGTTGCCGGGGCAACTGTGGCC | 120 |
| chr11 | 70523015 | 70523135 | 788855_47188552_ENST00000601538.6,E | 1 | - | good | CCGCGTGCTTTGATGCTCGGGGAGGAGCAGGAGACAAGCAGCTGCCCCTCCACACCCACACCAGCATTGCAGGTGTGGGCCTGTGGGTTTCCTTGCACCCGCAAGCCCTCTGCCGGAGAC | 120 |
| chr11 | 70523135 | 70523255 | 788855_47188552_ENST00000601538.6,E | 1 | - | good | CCAGTCGCATTGAGGTGTGAGGCCAGCCTCCTCCTGGTGCTCTGGACAGAGAGGCCTCCCGGTGTGCACACAGCAGCCAGCTGGCCCGGGCAGATGCAGAAGCACACCCTGCTGTGCCAC | 120 |
| chr11 | 70523255 | 70523375 | 788855_47188552_ENST00000601538.6,E | 1 | - | good | GCTGAGACTTGAACCCAGGCATTCCTACCCTGCTGTGTTTTACCACAGTCCCAAAGAGAACTGCTCAATAGTATTTATAGAGCACCTGCTGCTGTGAGGGCTGGGAATCTGTATTTGTAA | 120 |
| chr11 | 70523375 | 70523495 | 788855_47188552_ENST00000601538.6,E | 1 | - | good | CACCCGTCCTCAGGCATTCTCATGGATAGACTGCCAGAACTGCTGCTCCCCTGCTGCAGATAGAGCTGATGAGGGCCAGAGCCCTGCTCTGAGGTCACAAACTCACCTGACAGTGGGGCA | 120 |
| chr11 | 70523495 | 70523615 | 788855_47188552_ENST00000601538.6,E | 1 | - | good | GCAGGACCATAGACAAAGCCTGCTTAGGTGGGAAAGCCGGGCTGCAGTACAGGGCTTCGGGGCAGCACCGGGGATCCTCCCCAGCAGAGCAGGGCCTCTGCATCCCCCCTTCCAGTCACA | 120 |
| chr11 | 70523615 | 70523735 | 788855_47188552_ENST00000601538.6,E | 1 | - | good | CCCTGATTCGTGCACAGCCCAGCCCCAGCAGCGAGCGCCTCCTGACACAGGAAGCGGTGAACACTGCCCTCCAGGACCGGCTGCTGCACAGGGTACATGGGAAGGACAGCCAGGGAGGGG | 120 |
| chr11 | 70523735 | 70523855 | 788855_47188552_ENST00000601538.6,E | 1 | - | good | CAGAGGAAGCTACCCCAGCTCTCCGACTCTGCCTCCATATGATGCTCTTCCACTGCTTTGAACTGAAATTATGGATATTCTTGTTGAGAACAAGATAGTGGACTGAGAATCCCGTACAGC | 120 |
| chr11 | 70523855 | 70523975 | 788855_47188552_ENST00000601538.6,E | 1 | - | good | AGGTATTCAGCCTGGGCCCTGACCCCTGACAGGGGCCTTGAAGCCAGCATGCAGTGGCCGTGCGGAGCAGCACAGGGGAGGGAGGACAGCTGAAGGACACAGCAGGATGAAGGGTGCTGT | 120 |
| chr11 | 70523975 | 70524095 | 788855_47188552_ENST00000601538.6,E | 1 | - | good | GTGTAACTATGTGAAGTCCTTAGCCCAGGGGCAGCCATGTCGTCAGTGCTCAGTTGACACTGGCTGTTAATATTCCCTTGCTAGTCACGCTGCAAATATGGCCAGGCCCCCTCATGTGCC | 120 |
| chr11 | 70524095 | 70524215 | 788855_47188552_ENST00000601538.6,E | 1 | - | good | CTTGGTTACTCTCCTGGTAGAGAGGGGAGGACCCAGGGCTCCGCTGTAGCTTCTGTCCCTCATCTGTAAAATGGGATCCCGTCGCCACCTGGTCCATAGGGTGCTAGTGAGGACTGAATG | 120 |
| chr11 | 70524215 | 70524335 | 788855_47188552_ENST00000601538.6,E | 1 | - | good | CTCTGTGCATCCACGCTGGTAAGGCGAGCGTGGATGTGAGCGACACAGCTGCCCCTGGTGCTTTCCCATCTCCGATGGTGGCATTCACAGCTGCACTGCCCCCCAGGGGGTGGGACTGCT | 120 |
| chr11 | 70524335 | 70524455 | 788855_47188552_ENST00000601538.6,E | 1 | - | good | TGTCTGGGGTTGTACCGATTTGCTCCGGTAAAGATAGCACATTGGGAACAGCAACCTCAGTGGGTGTCACCACCAGATTAAGTTGAAGATAGTCCATACCTTTCTCCAGGCTCCCTGGGA | 120 |
| chr11 | 70524455 | 70524575 | 788855_47188552_ENST00000601538.6,E | 1 | - | good | CCTGAGGACGGGCTGTGCATGTGGACCACTGGTGTTGTCAGGTGAAAGCAGACTCCTACTAGTACTCCTTGCAGATCAATGTGGAGCAAAGGAATTCACAAGGTCTTAGCTGCATCCGAG | 120 |
| chr11 | 70524575 | 70524695 | 788855_47188552_ENST00000601538.6,E | 1 | - | good | CTCATGATCTAGTTACCGTAATGTCATCAGTATCGTGGACAGCATGATGTTTTCTGGAAAGCTGAGCTGATGGGATCTCTGAGGGCCCCGGCATGGCAGAGAGCAGGAGGCTCGGGGTAG | 120 |
| chr11 | 70524695 | 70524815 | 788855_47188552_ENST00000601538.6,E | 1 | - | good | AACATGTTCCAGAGCACTGAGGACGCAGAACCCTCACTGAACTGGCAGGCCCCTGCATCTCTGTGGGCTTCCCCCACCCTTCTTTGCATAGAAGGCATCCGAAGCATTTGCTACTTCCTG | 120 |
| chr11 | 70524815 | 70524935 | 788855_47188552_ENST00000601538.6,E | 1 | - | good | TCCACACTCCACAAAGACCTGCCAAGCCTCGTGCCTGTGTGCGTGTGAGCATGTATGTGTCCATGTGTGCATGAGATGGTGAGAGGCAGCGGAATGTTCCCCCCTTGGAGGGGCTATCTC | 120 |
| chr11 | 70524935 | 70525055 | 788855_47188552_ENST00000601538.6,E | 1 | - | good | TGCTGCAGCTGGGACCTGCACAGAGCCTTATCTTACTCTGGTCCCCACTAGAAGCGGGCAGGCTTGCAGAGGACTTTGTAGATGGGCTGGAGAAAATGTTCAAATGTGGCCTGTGTTGCC | 120 |
| chr11 | 70525055 | 70525175 | 788855_47188552_ENST00000601538.6,E | 1 | - | good | CACTGGGATTCTGTTGCTGGTGCTGGCCAGAGCCTCCACACGCGTCCCTGTTTGCCATGGGTACTGTAAGTGTCGCTGCAGCTGCCGGGCTGTCAAGCCGATTGGCAGCAAGGCCTGTGC | 120 |
| chr11 | 70525175 | 70525295 | 788855_47188552_ENST00000601538.6,E | 1 | - | good | GCTGGGCAAGCCTGGGCAAGGTATACACCGTGGCCCAGGAGCAAGCCAGAGAGGTGTTTCCCAGAAGAAGAGTTACCTGCGGAGAAGGGCCTGGATGCGCTCCCAAGTCCTCGAGGCCTG | 120 |
| chr11 | 70525295 | 70525415 | 788855_47188552_ENST00000601538.6,E | 1 | - | good | TGAGCTGTCCTGGGTGTTAGGGGCTGCTGTCCTTAACTCTTGGAGAGGGGCTGCCACCCACAGAGTCTGTTCCTTGGGTGAGCCTGGGGACTGGTACACAAAGCTAACAGAAGTGACCTA | 120 |
| chr11 | 70525415 | 70525535 | 788855_47188552_ENST00000601538.6,E | 1 | - | good | GCCCTTTTCTCTCACCTCTCTCTTAGTCTGTCTCATGCTCCTAGCAGAAGGCAGAAAAAAAAAAATCCACCATCCCCTGCCCATCCCTGGGCCTGCTGGCTGGGGTTCACAGACCCGGGA | 120 |
| chr11 | 70525535 | 70525655 | 788855_47188552_ENST00000601538.6,E | 1 | - | good | GTTGGGAGGTGGGGCTGCACAGTACCAGGACCCTGTTCGCTCCTGTTCTGATGCTGCCAACCCTGGCATTTCAAAAGAAAGAGCCTGTTTTCTCCCCTACCTCGAAGCTCGTTTCCAAAT | 120 |
| chr11 | 70525655 | 70525775 | 788855_47188552_ENST00000601538.6,E | 1 | - | good | TTAGCTGGAGAGCAGAGATCCCCTTCTGAGTGTGCTGTTCAGGGGGCAAAGTGGTGGCTTGGTCCCATCCAGGACCTGGTGTGTGTGTTTACGGGGGTGCCAGGGCTCCGTTCCCCTGGC | 120 |
| chr11 | 70525775 | 70525895 | 788855_47188552_ENST00000601538.6,E | 1 | - | good | CAGCAGAGGAGACAGGAGTCACAGAGCAGGGAGGGGGAAGGAAGCTTCTGAGAACAGGACTTTCCCGCCATCACCTCCACAAAACTAAAGGAGCCACATCCCTGGGGGCTAGGAGGGGGG | 120 |
| chr11 | 70525895 | 70526015 | 788855_47188552_ENST00000601538.6,E | 1 | - | good | GAAAAGGTGGGTGGGGGGTGGGGGCAGAGTATGCCAGGGCCTGTCCTCCTGAGAGAGGCCTGCTGGCTGAGAATGATGGGCAGTTGCCCCCCAGGCCACCAAGACTAAGGTGGAGAAGTC | 120 |
| chr11 | 70526735 | 70526855 | 788855_47188552_ENST00000601538.6,E | 1 | - | good | CGAGGGGGCCAGGCCTGCAGAGGTCGGCAGCACATGATAGGGGGTGTTCAGGAGGGAGGGAGGGGGTCCTTGCCTGAGAGTAGGGGAGGGGGAGGAGTGGTGTCAACTGAAAAGAAAT | 120 |
| chr11 | 70526855 | 70526975 | 788855_47188552_ENST00000601538.6,E | 1 | - | good | CTGGGCGGGGCGGTTGGCATACCTCTGAGACGAGGACAGGGATGCTCCCAGGGTGGTGGCCAGATGGGGCCGGTACAGGGAGGAGAGTGGGCTCAGAGGGGCTCCCCCGGGATGTGATT | 120 |
| chr11 | 70526975 | 70527095 | 788855_47188552_ENST00000601538.6,E | 1 | - | good | ACGATGTGGAGGAGGCCTGGAGGCCAGGATCAGGGAAGGGGACGGTGTGAGGAGGGTCTGGGATGTGGTCGTGTAGGAAGCGTTGAGGACTGGGAGCGTCTGGTCAGCAGCCAAGGTGA | 120 |
| chr11 | 70527095 | 70527215 | 788855_47188552_ENST00000601538.6,E | 1 | - | good | CAGAATGTTCAGGAGCATCTGGTCAATAAAGGACCAGGTCTCTGGGCTCTCCCTCCCCACACAGGGCCCGCCCAGACCACAGCTGTGGAATCAGAGCCGTTTGTGGGTCAGAACACCAGC | 120 |
| chr11 | 70527215 | 70527335 | 788855_47188552_ENST00000601538.6,E | 1 | - | good | TGAGTCCAGCCCGGCCCCAGAAGGGGTCAGCACAAGTCCACAGCCAGGCCGGCACCACACCCTCCCCAGGAGGCTTTAACTACAGGAGTCCTGGGGGTGGGGACAGGGACAGGACCCAGG | 120 |
| chr11 | 70527335 | 70527455 | 788855_47188552_ENST00000601538.6,E | 1 | - | good | CAGGCCCCACCTGAGCCCACACCATTCCCCAGGGCCCACTGGAAGCGGGGACCATGGGGGTCGTTCTAGAATGAGCTGCAGGTGACTGCAAGCTCTGCTCAAGAGGAACAAGGTCAGGG | 120 |
| chr11 | 70527455 | 70527575 | 788855_47188552_ENST00000601538.6,E | 1 | - | good | TCAGTGCCATCTCCAGGCTGCTGGAGGCCATTTATAGCAACTGCCCTTCTGAGTGAGGAGCAAATGAGTATTTTAAACACACCAGGAAGCCATGCCACTTTAGGATTCCCCAACTTGAAT | 120 |
| chr11 | 70527575 | 70527695 | 788855_47188552_ENST00000601538.6,E | 1 | - | good | GAGGCCACATGAATGGCCAGAGAGATTTGGGGGCCCGGCCTCAGGGCCCCTGCTCCTTTCACCACCACGGGGGCTCCAGGGAGGAGGCGCCTCTCAGACCCGAGCTGGCTGATGACACCA | 120 |
| chr11 | 70527695 | 70527815 | 788855_47188552_ENST00000601538.6,E | 1 | - | good | TGGGTGTCCTCTGTCCTCTGTCCTAGCCTCCTGAAGGAGAGTCAGGCACATTCGTAGCTCAGCTTATCAGAGGGGCCTTCTCCATGACCTCGGAGTCTGAGTCACCACCACATCCCCTGA | 120 |
| chr11 | 70527815 | 70527935 | 788855_47188552_ENST00000601538.6,E | 1 | - | good | AGACACCTGAGTCCCTGGCAAGGTCTGGGTCGCCCGGTGAAGGAGCATGGGCGGGAGTCTGGCTGCCTGGCTGCCTCCGCAGCCTCCTCTCTGCCCTTTGCCCTGGTCTGCCATTGTCCC | 120 |
| chr11 | 70527935 | 70528055 | 788855_47188552_ENST00000601538.6,E | 1 | - | good | ATGGGAGGACGGTCATGAGAGTGTGGCCTGCCCATTTGCTGCAGATGCTCCTGGCAGCTCTCCAGGTCCTTAACATGCACTCAGGCCCCTCCCTCTCACCACCATCCCATTTTGTGGCTG | 120 |
| chr11 | 70528055 | 70528175 | 788855_47188552_ENST00000601538.6,E | 1 | - | good | GATCCTCAGCTGGGGCTGGCAGCATCACGCCGAGGTGCACCTGATGCTCGGTTGGTTTCTGCGCCGCCATCCTGGTTCCTACCTTCTGCCTTGTCGTCTGGGTCCGGATCATCTGCAGGG | 120 |
| chr11 | 70528175 | 70528295 | 788855_47188552_ENST00000601538.6,E | 1 | - | good | TTCCACCGGATACTAGGGCACCACAGCAGCCCGAGAGCCACGTTCGTCCCACACATGGACAACCTGACCCTCAGACCTCAGGGACGCCTGGGGACGGGGTCTCGGCCTCTCGGCCTGCCA | 120 |
| chr11 | 70528295 | 70528415 | 788855_47188552_ENST00000601538.6,E | 1 | - | good | CCTTTCCAGGGTCCCATGGCCACCAGGTGGCACCTGCATTGCGCGTCGAGGCACTGGCAAGGAGCCTCGCGTCAGCCCAGGAGGCTGCATCCCTGCTGAGGACCTGCCAAACCTTCCTGC | 120 |
| chr11 | 70528415 | 70528535 | 788855_47188552_ENST00000601538.6,E | 1 | - | good | GCTAATGGAGCTCTGGGCAACTTGAACACTGCGCAGTAAACGGGAAAGAGCCCCATCCAGCCCCTCCCATGGGTGCCATAAAGGGCCATGAACTGGGGTGTTCTTGTGGGACCTGGAGCC | 120 |
| chr11 | 70528535 | 70528655 | 788855_47188552_ENST00000601538.6,E | 1 | - | good | CCCCCTTCTGCCCTCAGCCTCAGGCTCTGCCGGGCCTGCTTCTGCCTGGGCCTTCATCAGGATGCAGGAGCCCCCCTCCCCATGGAGCTCTGCTGCCCCCACGCTGGCTGACCGGGGCAG | 120 |
| chr11 | 70528655 | 70528775 | 788855_47188552_ENST00000601538.6,E | 1 | - | good | CCACTTGGCGTAGGTGATCTCCCCCACGGATGCCGAGGACGCTTCCTTCCTTCCATGCCCCTGGGGAGCCGGCGAGCTGACCCGCTTGTCAGGGCATAGTAAAGAATCGCCCAGGAGAGA | 120 |
| chr11 | 70528775 | 70528895 | 788855_47188552_ENST00000601538.6,E | 1 | - | good | CCCTCCTGAGAGGCGGGTGTGGGGACCCTGGAGCAGGAGTCGCCACCTGGGTTCCTCTGGGTGCTGTGCTTCTCCCAGCTCGCCGGCTGCCACGGACCCCCCTTCACCTTTCACCCTCCC | 120 |
| chr11 | 70528895 | 70529015 | 788855_47188552_ENST00000601538.6,E | 1 | - | good | CCCCCATCATCACATTGAGGAGATCCCGTCTCCTCCCCTAAGGCTCCCACAAGGCCACCAGTGGGGGGAAATCGTCACGGGGGCCTCCCCAGGACCTTTTATTTCTTCAAACAGGGGAGC | 120 |
| chr11 | 70529015 | 70529135 | 788855_47188552_ENST00000601538.6,E | 1 | - | good | CCACTGGCTCCCCAGGCACCCGTTTCTTACTCCCTGGTCCCAAAGGGGAGTAAGTGACCAGGGTGTCTGAACCCCACTGAGTTCCGCCCCAAGCCAGGCTCCAGGATGCCCCATGATCTG | 120 |
| chr11 | 70529135 | 70529255 | 788855_47188552_ENST00000601538.6,E | 1 | - | good | GTGAGATTCTGCAGTGGGTACAGACCCTGCTGTTGCAAGTAGTGGGTGCCATGATGGTTTCCTAGCATGACATGGGTCCTGAGCTGGTGCCACATGAGACACTGGTCCTGGCCTGGTGTG | 120 |
| chr11 | 70529255 | 70529375 | 788855_47188552_ENST00000601538.6,E | 1 | - | good | CCAGCCACCATTACCAGCTGTGTGGCCTTGGCCAAGTCACCCGCCTTCTCTGGGCCCAGCCTCCCATCTGGAAGATGGGGATGCAGTAGTGGCTGCTGACAGGGCTGCTGGGAGGACTGG | 120 |

| | | | | | | | | |
|---|---|---|---|---|---|---|---|---|
| chr11 | 70529375 | 70529495 | 788855_47188552_ENST00000601538.6,E | 1 | - | good | AAAATAACCCAGAATCTGCATCCGCCGCACTGCAGCCTGCATCCCACGGCACCTTGTGGGGGGTTTTGGGTGGTAATGGGGAACCCGTGTGTTGAACCCCAACTGCCCGGTTCCACTGCA | 120 |
| chr11 | 70529495 | 70529615 | 788855_47188552_ENST00000601538.6,E | 1 | - | good | GCTATGACTGTACTAAATGTTATCAATGGTCAGTGTTAGGCCACGTGTATTTTACCACAATGCGAAGGTGTATAGATCTTGGAATCCCTGTCAGGAGAGGAGGACCTTCTAGGCTATTCT | 120 |
| chr11 | 70529855 | 70529975 | 788855_47188552_ENST00000601538.6,E | 1 | - | good | TCCTACATCCCCACAACCCAAACCATAAATGGTCGTAACTGGTAAGCAAAAAAACCAACTGTGGTCTGTCTGTGCAATGGAATATTATTCAGCCATAAAGAGGAACAGAAGACTGATGCC | 120 |
| chr11 | 70531175 | 70531295 | 788855_47188552_ENST00000601538.6,E | 1 | - | good | TCATGGGGCTGGCAGGTGAGGCGCCTGGGCCTGGGCCATGCCTCTCCCTTTTAAAAAGTTATTATTTTAAATCATAGTAAAATACATGTAAACTCTATAATCTTAGCCTTTTTGTTTTTT | 120 |
| chr11 | 70531295 | 70531415 | 788855_47188552_ENST00000601538.6,E | 1 | - | good | GCACCCCAGGCTCAGGCCCCCGGGCGCGTGGGTGCATTTGCATTTGATCCTCCAAGACCCCAAAGAGTGGCCCCACCACCAGCCCCCAGCACAGATGGATGCAAAGGCCCAGAGAGGCAC | 120 |
| chr11 | 70531415 | 70531535 | 788855_47188552_ENST00000601538.6,E | 1 | - | good | CATGATCCCAGGCCTCCCTGGGGCCTCACCCTCAACCGTGACCTGGGAACAGTCGTCCACCCGCGAGGCAGTCAGTCTAGTGTGGGACAAAGAAAACCCCAGCTCACCTCTGCCAAGCAT | 120 |
| chr11 | 70531535 | 70531655 | 788855_47188552_ENST00000601538.6,E | 1 | - | good | AGAGGCGCAGCAAAATGCCTGTGGCCACACAGCAAGGCCTGGGGACACTGAGATGGGACCTTAGCCCTAGACTTGTCCTCACAGCCCTGCTGCATGCCACAGTCACCCTGCCTGCCCTGG | 120 |
| chr11 | 70531655 | 70531775 | 788855_47188552_ENST00000601538.6,E | 1 | - | good | CCACTTCCAGCCCCGAGTGACATGCAGGCCCAGGACTTCCCAAGGCGCTCAGTGTCTCTGCGCTCCTACCCAGTCCCCATGCCAGCCCCGAGGTGCTGCTTATCCCCACTCCTAGGACGC | 120 |
| chr11 | 70531775 | 70531895 | 788855_47188552_ENST00000601538.6,E | 1 | - | good | AGACAAGCCCCAGCACCCAGAGCTGCGAGGCAGGAAAGAAATGTATGGGGAGAAGTGGAGCTGCAAGGACCCCGTCTCATCAGTGGGGGGCCAGCCAGCAAGTGACCAGGTGTGGAGAAG | 120 |
| chr11 | 70531895 | 70532015 | 788855_47188552_ENST00000601538.6,E | 1 | - | good | TCCGTACCCAGCCTCTGTCCTAGAGAGGGGCATATATCCCAGAACTGTCACTACACACCACGGTCCAGGCAGACAAGGAGAGATCTGGTCCCTACTGTCAGCTCCGGGAAGGTTTGGCAG | 120 |
| chr11 | 70532015 | 70532135 | 788855_47188552_ENST00000601538.6,E | 1 | - | good | CAGCATTGAGTTTCTCAATGACAAGCAAAAGAATCAAGGAGATTTGTTTCCTTGGGCCTCTCTGCCCAGGTGCAGTGATCCAGAGGGAAGAAGCCCAGGTTGGAGGAAAATGCTCAGAGC | 120 |
| chr11 | 70532135 | 70532255 | 788855_47188552_ENST00000601538.6,E | 1 | - | good | CGGAAACACATTTATCTTGCACAGTATGGGGGCACCTGCCTTATGGAACTGACATGTTCCTGAGAACAGCTGCTGCCACAGGAATTTCAGCTCAGCATGCCTGAAATTTCAACTCATGCT | 120 |
| chr11 | 70532255 | 70532375 | 788855_47188552_ENST00000601538.6,E | 1 | - | good | GCAGCTGTGCACAGCATGGGGGGCAGAGCACAGAGTGGACTTGAACACCACCCCCAGCAGACCCCCTTGTGGAGTGAACAACCTACACAGTTGTCCAGGGCAGTCCTGCCTGCTGACACC | 120 |
| chr11 | 70532375 | 70532495 | 788855_47188552_ENST00000601538.6,E | 1 | - | good | CGCACACAGACCCACAATTCTGCCCATAGACTTGAGGCGTTTGCTGATGGGCCACCTCTGTGAGCCCACAATACCTTTGTGCAAATGTGAGAAGGCCCCCCACTCTAGGCAGGCATGCCA | 120 |
| chr11 | 70532495 | 70532615 | 788855_47188552_ENST00000601538.6,E | 1 | - | good | GAGCTGGCTTCACTGTGCTGTGCCTGCTTCCCTGGAAGAGGGGCTCTGGGCCTCTTTCTGGGTTGTGACTCCATTTTAGAGTCTGGCGAGGACCAGGGGCCCTCCCTCTAGGAAAATGCA | 120 |
| chr11 | 70532615 | 70532735 | 788855_47188552_ENST00000601538.6,E | 1 | - | good | TTGGGTAATTCTGTGTTTAACTCTTTGAGGAACCTTCAAACTCTTTCACTGCAGCTGCACCATTCTGCATTCCCACTAAATGCTTTTTTTTTTTTTTCCATGGACAATGGACTGTATTGG | 120 |
| chr11 | 70532855 | 70532975 | 788855_47188552_ENST00000601538.6,E | 1 | - | good | ATCATGTCTCTGAGGCTCTCCTTGTTGGAACACGTGTCAGCATTTCATTCCTTTCTGTGAGTGTCGCACTGTATGGCTCGACCACAGAGTGTTATTCATCCCTCTGTCCGTAGACACTGG | 120 |
| chr11 | 70533455 | 70533575 | 788855_47188552_ENST00000601538.6,E | 1 | - | good | AGCATGGCTGCGGCAGAGGGAACGCCAGGCAAGCAGGGGGAGGGGCCAGATGACATGGGCCCCCGGGTTGTGATTGGAGCTAGGTTGGACATTCTCAGCAGTTCTGACACTGGAATAAGG | 120 |
| chr11 | 70533575 | 70533695 | 788855_47188552_ENST00000601538.6,E | 1 | - | good | TGGAGTCGCACACTTTAAAAGTGTGAATTTTACATTATATAAGCTCATCTCAATTAAAGAAATATGCATGTCCCCCAAAAAGCTGATGGGAGCCAGTCATTGGGGAACAGAGGGGCACCC | 120 |
| chr11 | 70534535 | 70534655 | 788855_47188552_ENST00000601538.6,E | 1 | - | good | TCCAGCTGGCATTTGTTGAAGCCAGCCTGGCTCAGGGGCAAGACCACAGACTGCAGACCAGGACACCTGGGGCCTTGTCCCAGCCCTGTGTCCTGCCCTGGGCCCTCCATGGTGGCTTCT | 120 |
| chr11 | 70534655 | 70534775 | 788855_47188552_ENST00000601538.6,E | 1 | - | good | TCTGAAGCGTCACCTGAACTGATGAACTGTGTCACTGTCATTTATGGGAAGAAGTGACATGCTAGAGTGAAGCCACCAGTTTGGACTCCCCAGCAGTCTTAGGACACTGTCATTCTTTCA | 120 |
| chr11 | 70534775 | 70534895 | 788855_47188552_ENST00000601538.6,E | 1 | - | good | GCAGCTTCATGGCCTGACTGCACTCTCGGTGAGGCAGACAGTCAAGGGAAAGACCGTCATGCAGATCGTGAGTCCCCACCTCGCCATTGTCCTCTACATCCCTCGGTCTGTGCAGTAGTG | 120 |
| chr11 | 70534895 | 70535015 | 788855_47188552_ENST00000601538.6,E | 1 | - | good | TGACCAGGGTCCTGGAGCCATGGCTCCTTTGTCAGTGTTTCCCCAGCCAGAGGCACTATCCACATCTCCTGCCCCTGCATAGCTAACGAACTTGACTATCCTCATCTCTGTTGGGCCGAA | 120 |
| chr11 | 70535015 | 70535135 | 788855_47188552_ENST00000601538.6,E | 1 | - | good | TGGTCATGAGCACTACTTCATGTGCCTGCCTCACCTGCCCCAAAGCCAGTCTTCCCTCCAGGTCAGCTCTGCCCTGTTCAACATGGGTGGGAAGCAGAGTGTGAGAGCTGAGGACCACCC | 120 |
| chr11 | 70535135 | 70535255 | 788855_47188552_ENST00000601538.6,E | 1 | - | good | AATGGATGGATAGAGCGATGGATGGACTGACTCATGTGGGAAGATGGATTAGGTGAGCAGATGGATGGAAAATGGATAATAGAGGGTGGGAGGGAGGAAGGTGTTCGTGGGAACCATCCC | 120 |
| chr11 | 70535615 | 70535735 | 788855_47188552_ENST00000601538.6,E | 1 | - | good | CCTGCTGCAGGGCTGGCCTCGGTCCCCCTGGGTGGCCATCCTCCTGCCCAACTGTGCAGCCCTTAAAGGCGGGGCCAGGAGTTATTCATTCCCCAGCGTCTAGCACAACACTTGACCTCA | 120 |
| chr11 | 70535735 | 70535855 | 788855_47188552_ENST00000601538.6,E | 1 | - | good | TTCTTCTCAGGCTCCCAAGCTTCCCAAGTGCCTGGGCCCAGCTCCCTGCCAGCGGATGGCCCACCCGAACCTCTCCTGCCCCGTGGAGCACAGGGACTTCCCACAGTGCTGCTGGGTGGC | 120 |
| chr11 | 70535855 | 70535975 | 788855_47188552_ENST00000601538.6,E | 1 | - | good | CACGCAGCCCCCTCAGTCAACCAGCCTCGGGTGAGCCACATGTCCATTCTGGGCTATGTGCCTTTCCTGGGGTCTCCGGCCAGTCACTGCTTCCCCCACAGCTTACCAGGCCCTGACACC | 120 |
| chr11 | 70535975 | 70536095 | 788855_47188552_ENST00000601538.6,E | 1 | - | good | CATGGTGCTGTCGGTGGCTGGGGCCTGGGGGTGCAGGTGTGTCCACGTGCCTTGGCGACAGTGTGTGCACCTGCAGAGAGTTGGGCCCTGTAGGGCTGGTCTGGCTTCAGGCCCAGCCCC | 120 |
| chr11 | 70536095 | 70536215 | 788855_47188552_ENST00000601538.6,E | 1 | - | good | CACAGGGCAGGCTCAGCACAGAGCCGGGCAGTGAGGCCAGGGTAAGGAGGTCTCCTGGGTCGGTGCCTCCAGCAGACCTGGGTGGGCTTCAGCCCAGGGTGAGGGCCCCCAGGGGGCAG | 120 |
| chr11 | 70536215 | 70536335 | 788855_47188552_ENST00000601538.6,E | 1 | - | good | CCTCATCTCAGCTTGGTCCAACGGAGTTGAACTTTAACAAACAACAAGCGTGATGAGCCCGCCACACCCCGCCACTTACAGCTGGCCCAGCCCCTGGCACTGTGAGCTCCGCTCAGACCA | 120 |
| chr11 | 70536335 | 70536455 | 788855_47188552_ENST00000601538.6,E | 1 | - | good | GCCAGCCTGGGTGGGGACAGCCCTGGGGTGCACAGCCACACTTTGCTGGCTGTGGAGTGGCCTCCAGCCTACAGAGCCGGCCCACACACTGACGCAGCTCCGGTTTTATTTTAGACCAAG | 120 |
| chr11 | 70536455 | 70536575 | 788855_47188552_ENST00000601538.6,E | 1 | - | good | CTGGGAGTGGGCCCTGTGCCCACCGCAGCCATGTGGGTGCAGACACAGCCTCAGGATCGGCCGGCCATTCTGCCTCTCGGGTTCACCAGAGGCCACAGCTGGGCCTCATGGGGACAGCTC | 120 |
| chr11 | 70536575 | 70536695 | 788855_47188552_ENST00000601538.6,E | 1 | - | good | TCCAAGACATTGGCTCAAGAAGTGAGAGCCAAGCCTGAGGTGTCTGACCCCAGCTGAAGGGGAGCCACAAGCCTGAAGCCCAGGAGCTCATGGGAGACACTGAGTGTGGGAATCGGGTCC | 120 |
| chr11 | 70536935 | 70537055 | 788855_47188552_ENST00000601538.6,E | 1 | - | good | CCACCCCTCCCCAGCCTCATCTGGGGAACCCTCTTCTAGCTGACGTACGACACAAAGCCCAACACTCAACACGGCTCAGAAAATGCTGTTTATTTCCTGGAGTAAGAAGGGTCCTCTGCT | 120 |
| chr11 | 70537055 | 70537175 | 788855_47188552_ENST00000601538.6,E | 1 | - | good | CACCCCTGCAGGGGACTCTGAGAGGGGGGCGCTCAGGGAACTTGGCAGATCCTTTCAAACTAGGATCTCAGTGTTTGGGAGGCCCTGAGGGAGCAAAAGGAGCATGGGCAGACCGTGCCC | 120 |
| chr11 | 70537175 | 70537295 | 788855_47188552_ENST00000601538.6,E | 1 | - | good | AGGACATATCTTTTGGGGGCTGCCATGCTGGCCCTGGGGGCTTCTTTCCTAATGAGCATCCTGGGGGGAGGTTTCCCTCCTCCGCTTCTCAGGCCTAAATTCTGGAGGGGCTGTTGGGGA | 120 |
| chr11 | 70537535 | 70537655 | 788855_47188552_ENST00000601538.6,E | 1 | - | good | AAGCAAGGGGAATGTGTTCCCCTACAGTCCTGGGAGCCAGAAGTCAGACAGCAGCCTCACAGGGCTGCAGCCAGGTGTGGCAGGGCCCCGTGCCATCCCAAGACTCCGGGGAGGAGGCCT | 120 |
| chr11 | 70537655 | 70537775 | 788855_47188552_ENST00000601538.6,E | 1 | - | good | GCAGAGCTGACTCGTGCCTCTGCAGGACTGAGCCAGGGCCCAGCCGCCTATTTCTTCACTTTGTTGGGTGTGTTTGAGTGGCCTGGGCTTGCCACAAGTGGCTACAAACGGGGCAGCTTC | 120 |
| chr11 | 70537775 | 70537895 | 788855_47188552_ENST00000601538.6,E | 1 | - | good | TGCCTCCTGGAAGACCCCCCACGGGCCTCTGTGACCTGCTGCAGGGTTGACGTCGGCTACACCAGGCCTGGCCCCGCACACACGGTCCCTGTTGCCAGCAGAATGTCGAGGGTCCCCTCC | 120 |
| chr11 | 70537895 | 70538015 | 788855_47188552_ENST00000601538.6,E | 1 | - | good | GAGCTACTTTGAGATCACTGTTTTGTGACACTCATGGGGTCTCTCGCTGCCCTCTCAATGTAAATCCAGCTAGGGCAGGTGACTGGCCAGAGAGTTCCTCAGGGCACGTGCCCCATAAAA | 120 |
| chr11 | 70538015 | 70538135 | 788855_47188552_ENST00000601538.6,E | 1 | - | good | GACCGCCTGAATCCCCACTGGTCTCTGCCCCTCATCTCCACCGCTCGCTACTGCCCCGTGGGTGTGCTGGGCCCAGGATGGTCACGAGTGTTCAAGGGTAACGCCCCTCTGCAAGTGAGT | 120 |
| chr11 | 70538135 | 70538255 | 788855_47188552_ENST00000601538.6,E | 1 | - | good | AGCGCCCGGGGACTCCGTGTGGGCTGCTGGCTATGTCCACAGGCAGGGCTGCAGGAGCCCCCGGGTGGGGTGACTCAGCTTTCTGGCCTTACCCAGGCCTCAGGCTGGGGAGGTGACAGT | 120 |
| chr11 | 70538255 | 70538375 | 788855_47188552_ENST00000601538.6,E | 1 | - | good | ACTGTGCATCCTTGGCCAAGCCCCTCGGGACAAAGGCGGTGGCCTCAGATCCCCTGGTCCCTCCGGGTCACTGTCGGCTAATTCTGGGGGAAGGACTGGCAAGGCTGTTTGGAAGAGGTC | 120 |
| chr11 | 70538375 | 70538495 | 788855_47188552_ENST00000601538.6,E | 1 | - | good | AGGACGCATCCCCAGAATACCATATGAGCCATGTGCCACCTGTGATTTTGCCCCATCCAGAGCCTGGCACAGAGGCCTAGCGTGCAGTCCACAAGAGGCCTGGCAAGCAGTGTCCACTCC | 120 |
| chr11 | 70538495 | 70538615 | 788855_47188552_ENST00000601538.6,E | 1 | - | good | AGCAGACAGGTTAGCCCCACGTTATCCTGCTCCTTCCTCCTGACCAGAGCTTTACTCTGACCAAGGGCCAGGACACTGCATGGGACAGTGGGAGCCACCTCCTTAGCACTGGCTCAGGGG | 120 |
| chr11 | 70538615 | 70538735 | 788855_47188552_ENST00000601538.6,E | 1 | - | good | GCCCCCTGGCTGGTGGATCTTATCTCCCCCTGGAGCAGGCAGAGCCCCAGGGAACCAGCACAGACTCCACCACTGACGGAGCAAGAGTGACAGCCACCTCCTCCTCCATGCATTTCCTGC | 120 |
| chr11 | 70538735 | 70538855 | 788855_47188552_ENST00000601538.6,E | 1 | - | good | TCAGGGCTGCCCCCAACACTCAGCCCTCCCCACCGTCTGTGCGTGCCTCCCGCCCCACCCAGTGCTTTCTCTGTGTGGAATGGGGCTGTTCCCTGGAGGCTCTGCCCCGGGGGCTGTTCT | 120 |
| chr11 | 70538855 | 70538975 | 788855_47188552_ENST00000601538.6,E | 1 | - | good | GGCCGACGGGGTAACCCAGGAGGCTTCCTAGCCCACGTCCTCTTCTAGGCTGCATCTCAGTAGCCAAGGCTAATGGAGGGGTGGTTGATGGTGAGGCTGGGGGTGCCCCTGAGACAGCAC | 120 |
| chr11 | 70538975 | 70539095 | 788855_47188552_ENST00000601538.6,E | 1 | - | good | CAGATCCAGTTAGCCACCCTGTTAGAACTCTGCATAAGGTAAAAGTAAACCCGTACATCGGCCCCGAGCGAGAGGCAGAGTAGTTCTGCAAGCCTAGTGCCTGGGTGCCAGGCTTGTGGC | 120 |
| chr11 | 70539095 | 70539215 | 788855_47188552_ENST00000601538.6,E | 1 | - | good | TTGCATCCTGGGGTCGGCCGTGCACGCTCTGCCATCCGGCCTCTCCCTCTGTGTGCATGGGCCTGCCTCTGCAGCCCTTCTTCCTTGTCTGATTTCCTGATCCTGATCCTCTGTGTGCCG | 120 |
| chr11 | 70539215 | 70539335 | 788855_47188552_ENST00000601538.6,E | 1 | - | good | AACATCATTTAATGTCTCCAGTCCATCCAAGGCTGTCCTGCCAGCTGCGTTTGGGCACCAGGTCATGCCAGGCCCGCGTGTGGCACAGCCTGCTCTCTGTGGCACTGGCATTCGGTGCTG | 120 |
| chr11 | 70539335 | 70539455 | 788855_47188552_ENST00000601538.6,E | 1 | - | good | GCGGTGTCACAGCACCTTTCCCCCGCAGATCTGTTGGGCACCCCGACTCGGAGTCTGGCCCCCATTTGGGATGGAATGTTCTAAAATGTTCTTCATTTACTCTACTTTCTTTAAATGGAG | 120 |
| chr11 | 70539455 | 70539575 | 788855_47188552_ENST00000601538.6,E | 1 | - | good | CTCATGAGTGAGCGTGGTGAGTATGGTGAGTGAGCGTGGCGAGTGAGCGTGGTGAGCGTGGCGAGTGAGCGTGGTGAGCGTGGCGAGTGAGCTTTCTGTGTGCCCTAAACCCCCCAGACC | 120 |
| chr11 | 70539575 | 70539695 | 788855_47188552_ENST00000601538.6,E | 1 | - | good | TGGGTCAGCGCCCAGATTACATCCTGGTCCCCACAGCAGAGTCTGGGCAGGGAGAAGCTGGGTAGATAAGAGCGCCGGAGTTGCCGCCAGCCTGTGGTGCTGTCTGTGATTGCTCATCCA | 120 |
| chr11 | 70539695 | 70539815 | 788855_47188552_ENST00000601538.6,E | 1 | - | good | GTCTGTGGTTAAAGCACCCAGGCCCTCTGCTACGGCCCCCCGGGAGAGAAGCGCACTCCCTGGAAACCCCAGCCTGCTGGGTCATGGCTCATCCCAGCTGAACGAGGAGAGGGTGGACA | 120 |
| chr11 | 70540175 | 70540295 | 788855_47188552_ENST00000601538.6,E | 1 | - | good | CTTGCAGAGATGGGTGAGCCCTGGCCCCTGCTCTTGGGGACACTGGCCTGGGGGAGGCATGTGAGATGAGACAAGCATGCCCATAACTCACGGCTGTGGAATGAATCGTGTTCCCCTGGA | 120 |
| chr11 | 70540295 | 70540415 | 788855_47188552_ENST00000601538.6,E | 1 | - | good | CCAGCCTGCCTTCATCACCTCGACTCCACAGGAGGGAGGGGCCCCAGCCTGCCTGGGTTTCTCAGACGCCGTCCCCTTGACCTGCCCGTGGGCGCCCCGCTTGCTGGTTCTGTTCTCCTT | 120 |
| chr11 | 70540415 | 70540535 | 788855_47188552_ENST00000601538.6,E | 1 | - | good | GGCCATGGGTATCGTGTGTGTGTCAGGCTGGACATGAGACTCCCAGGCTCCCGCGGGCAGGGAGGAGGTGGTAGGATGAGAATGGTCCTCATCACTCCGACTCCGCAGGAGAGAGGAGCC | 120 |
| chr11 | 70540535 | 70540655 | 788855_47188552_ENST00000601538.6,E | 1 | - | good | TTTACCGCAGTACAGTTAAAAATTAAAACTATAAAGATGCAGCCCTGGGTGGGGCTGACGGGGTGGGTTTCAAGGCCTCCATGGAGTCAGCTCACCCCCCGTCGCTAATCGTGAAGCTGC | 120 |
| chr11 | 70541135 | 70541255 | 788855_47188552_ENST00000601538.6,E | 1 | - | good | AGCTTAAGTGTCCATCACCGGTGCAGGAGGAAGGCAAATGTGGTGTATCCCGATCAGGGAGCATCGTTCCATCTTAGGCAGGAGGGAAAACCGACACTGGCTACAAAGGCCTCCTATGGA | 120 |
| chr11 | 70541255 | 70541375 | 788855_47188552_ENST00000601538.6,E | 1 | - | good | AAGGATGGTTGGTGGGGGGCCGGTGGCTATTCCTACTGTTCCCGCCAGGCTGATGAGTAGGGAAGGCTTAGAGGTGCAGCCAGGCTGACGGAAGGACAGGTGTTCTCAGGTGAAAGGGCC | 120 |

| | | | | | | | | |
|---|---|---|---|---|---|---|---|---|
| chr11 | 70541375 | 70541495 | 788855_47188552_ENST00000601538.6,E | 1 | - | good | GTGTCCTAGCTGGCTCGTGCCTGCCAGCCCACCTGCTACCTGCAGAGGTGACTGGCCAGGAGGGCACGCCAGGAAGCTGCCCAGCCAGCACAGCCAGCTGTCCATTTTCTTGATGAAGCC | 120 |
| chr11 | 70541495 | 70541615 | 788855_47188552_ENST00000601538.6,E | 1 | - | good | GCCAAGTTGGACACCAGGCTCCACGCCCCACATCTGTCAGTGCCTGCTGGCCTCACCCCTGCCTGCCCAGAGGTCTCTGGTGAGGCCCACAGTCACCTACACATCTAGTCATCACTCACC | 120 |
| chr11 | 70541615 | 70541735 | 788855_47188552_ENST00000601538.6,E | 1 | - | good | GCAACACAAGTTGTTGTCACAACCAAAAATATCTCCCAACATGGCTGAATGTGTTTTGTGGGGGGCACCGTTTCCCCCGCTTGAAAACCACTGCCTTAGGCCATAAACCGGGCTCTCAGA | 120 |
| chr11 | 70541855 | 70541975 | 788855_47188552_ENST00000601538.6,E | 1 | - | good | TCACGGGGTGTGCAGAAGCTTCTGAGTGAGGAGCTCTCTGGTTTGCATGGGTCCTGCTCTGTGCCAGCTGCCACGAGAACTGCAGGCTGCCTTCCAAGATGGCTGCCCTCAGCCTTAGGG | 120 |
| chr11 | 70541975 | 70542095 | 788855_47188552_ENST00000601538.6,E | 1 | - | good | ATAACTCAACCATGCCATGCAGCAGCCACTCAGCATTTGCCCAGGCACTACTGGGCACTCAGTGGCCCGTGCACAGAGGAGTAGCGTGGATCCTGGGCAGTAAACCCCCTCTAGCCACTG | 120 |
| chr11 | 70542575 | 70542695 | 788855_47188552_ENST00000601538.6,E | 1 | - | good | CTGAGTCGTTTGCCAACCTTCCTGGGCTGTTGGGTTTGTGCTCCAGGAAGGACCTGATGGTTAGAGTGGAGGTGTTGGAGAGAGTCCCAGAGCCCTGCCCAGTGAGGGTCAGGAGGCTAT | 120 |
| chr11 | 70542695 | 70542815 | 788855_47188552_ENST00000601538.6,E | 1 | - | good | AGGAGGCCCATTGGAAACCCCGCCAGGGTACAGGCAGGAAGCTGCGTGATCAGGTTGTCTTGTGAAAATGCTTCTTTGGGAACCAGCTTCCAGGAGTAGCCACTCAGGCCCACATGACTT | 120 |
| chr11 | 70542815 | 70542935 | 788855_47188552_ENST00000601538.6,E | 1 | - | good | CTGCTGAGATCTCCAGATGGGGGGACAGTCAGCCCTTGAGATACTAGAGAAATCCCCTGGGAACATTTCCTGCCTCTCTTCATGGCAGTGGGGAAACTGAGGCTATGGTGCCCGGGAGTT | 120 |
| chr11 | 70542935 | 70543055 | 788855_47188552_ENST00000601538.6,E | 1 | - | good | CCGGTTGAGGGCTCTGCCTCCTCACCCAGCGCATGCCGCCCCTGCCCTTCTACCCGCCTGCTTGCTCTGTTCAAGTTCTCATTTGTGGGATTGGTCAGCGGTCCCGGGGGGGCCCCCTTC | 120 |
| chr11 | 70543055 | 70543175 | 788855_47188552_ENST00000601538.6,E | 1 | - | good | GCCTTTCTCTAACAGAGCATCCTCTGATCTGAAAGGTGCCCGGCCATCAAACTGCAGTTCCAGTTCCATGAAAAGCACATGCAGTTACCCTGGGGAGCTCAGCCGCCACCCCACGCCAGG | 120 |
| chr11 | 70543175 | 70543295 | 788855_47188552_ENST00000601538.6,E | 1 | - | good | CTAGCATACAAAGACACTCAGCACTCTGGAGGGTCCTGAGATTTTAGGAGCCATGTGTCAGTAGAGGAGAGGAAAACCAAATACCCATTTCACAACGCAGACTGTGTGTGGCTCCTCCGT | 120 |
| chr11 | 70543895 | 70544015 | 788855_47188552_ENST00000601538.6,E | 1 | - | good | AAAGCAAACTCAGTTTCTTCCCACTATACTCTCTCAACGCAATCCACACAGAGCCCTCAGTGGCCAGATGAGTGGGGTTCCCACACACAGCAAGCAAGCAATCGGTTCCCTCGGACACCA | 120 |
| chr11 | 70544015 | 70544135 | 788855_47188552_ENST00000601538.6,E | 1 | - | good | GTTGGGTTTTCCAAATAGGCTGTTTCCTGACTTCTGGAAAGTTGCCTTAAGAAATGGGAACGTGTCTGCTAACAGCCTCTCGTTGAGGCCAAATTTGGGGGGACTGTGAGGCTATAGGAA | 120 |
| chr11 | 70544135 | 70544255 | 788855_47188552_ENST00000601538.6,E | 1 | - | good | GAGTGGGCAGGCCGGGGCTGGAGAGGTTCTGCGGCCCCAGGACTCTCTCCTGCATGTGGAGAGTGGAGATTTGAACTGGGCCTGTCATCCCCAGGGAGAGGCACAGGATGGGAGGTCAGG | 120 |
| chr11 | 70544255 | 70544375 | 788855_47188552_ENST00000601538.6,E | 1 | - | good | GGTCTGGAAGGTCCTACGCGTGGGGCTCCACCTTTGGGGGCTCATTTTATCTTCACAACAGCTATGTTCCCCGTAGCTGCTGGGACTGGCCCATTTTCCAGATGAGGACACTGAGGCACA | 120 |
| chr11 | 70544375 | 70544495 | 788855_47188552_ENST00000601538.6,E | 1 | - | good | AGTGAGATCCGTCAGTCACTCTCCCACGGGCAGCTGCTCCGGGTCTCACAGGGACAGACAGGGGCAGTAGCCTTGCCCCCCTCCAAACCCCAGGCTAAACAATGACAATTGCTAACAGCT | 120 |
| chr11 | 70544495 | 70544615 | 788855_47188552_ENST00000601538.6,E | 1 | - | good | TTGGGATGGGATGGTGTCTACTGTGACCTGAGGGCGGGGTGGCCCCTCCTGACCCTGCTGAGTGCCTCTCCCTGGGCAGGGATTTGGGCCCTGGTGAACCATACACACTGCACGGCCCAC | 120 |
| chr11 | 70544615 | 70544735 | 788855_47188552_ENST00000601538.6,E | 1 | - | good | AGCAGCAGCAGGGGCCTCCCCTTTTGTCGCCTACCGGATGAAAGACCTGTGTCCCTGCAGAGGTGGCCACCCCGCTCCTGGTCCCAAGTCCATCTGCTGGTCCAGGCCAGGGAGGGGGTG | 120 |
| chr11 | 70544735 | 70544855 | 788855_47188552_ENST00000601538.6,E | 1 | - | good | GACACCGGTGCCTTGCCCAGGACAGCCCCGGTGACATTTGCTCCTTTGTGGCATCAGAAGCCTGAGGGTGGGCTGAGGTCTCGCCTCTGTCGTCGCAGTGAGACTGCGGTTTCAGGTCAC | 120 |
| chr11 | 70544855 | 70544975 | 788855_47188552_ENST00000601538.6,E | 1 | - | good | AGCAGGAGACACAGCAGCTGGCCCATCCCCAGAGGCCGCAGCCCCACGTGGTCCCGCGACGGTGTCCTTGGTGCTCACGTGGGCTCAGAGGATGCAATCCCTGAAGGACCTGTGGGTGGG | 120 |
| chr11 | 70544975 | 70545095 | 788855_47188552_ENST00000601538.6,E | 1 | - | good | CCCCAGTGGCCCTTGCCTGCCCTGGCCTGGGCTCTTGCCACCGGCCCATCCTCTTCCAAGAGGCCATCCTGTGCCCGTGCTTGGCATGTGAGAGAAGCTCCCGAGGTCCACCGCCACCCC | 120 |
| chr11 | 70545095 | 70545215 | 788855_47188552_ENST00000601538.6,E | 1 | - | good | GCTCAGACAGCGCGAGCCCAGGGACTGCAGGAGCCACTTCCATCCATTCTCAGACAAAAAGAGACGCCCACGGCAGGACTGGGGTGGTGATCGGGGAATCAGAGCGCAACCCCTCCCAGG | 120 |
| chr11 | 70545215 | 70545335 | 788855_47188552_ENST00000601538.6,E | 1 | - | good | AATGTCTCTGGAGCAGGAGCTGCAGACAGGCAGGAGCTGGGAGCTGGTCAGGAAGACGAGACTGGCACACAAAGGAGCTGATTAGCAATGCGACGCTGCCTCCGGCCAGCCGGATTAGGA | 120 |
| chr11 | 70545335 | 70545455 | 788855_47188552_ENST00000601538.6,E | 1 | - | good | GGAGAGAGCGTGCGGTGGCTCGGGGCCTCCCAGAGCTGCCCTGACTCCCCTGCTTGGCCAGCCCCTCCCTGGGAGCCCCACCCCAGGGCCAGCAAAGGCTGCAGTGAGTCAGGGGCCGTG | 120 |
| chr11 | 70545455 | 70545575 | 788855_47188552_ENST00000601538.6,E | 1 | - | good | TCCCGTCCCAGTGGCCTTAGGGAGACGGTGCTGTCACTGAGCTGCTCTCAGGGGAGAAAACCAAGACCCCGAGGTGAGGACCCTCAGCTCTGCAGTGGGCGCTGAGGTGGGCGCCATCCA | 120 |
| chr11 | 70545575 | 70545695 | 788855_47188552_ENST00000601538.6,E | 1 | - | good | GGGATTAGAGAATTGGTTGATGTTGGAATGATGTATTTGGTGTCAGAAAAAAATACAACACCAGCTGTTTAATGAGCAATTACCGTGTGCAGGCGCCGTACGGAGGGTGTGCGTGAATTA | 120 |
| chr11 | 70545935 | 70546055 | 788855_47188552_ENST00000601538.6,E | 1 | - | good | TCCAAGCTGGTGGTACCCAAGGGGGTGTGGAAGCTCTGCACCCCCCATTTGCCCCGAGCTTCTCTTCCCTGTGGGCTGTTCAGAGGTGCAGCCTCCATAATCAACTGGTTGCCATAAGTG | 120 |
| chr11 | 70546295 | 70546415 | 788855_47188552_ENST00000601538.6,E | 1 | - | good | TACTTAGGGGAGGCGGAGGCTAGATAGCTTCAGGATGGCAGGGGCTGGTCACCAGAAAAACCAACCACATGATTCTAGAGGTGCAGAAGGCTCAGGAAGAGGGGCTGGAGGTTGAGCTCA | 120 |
| chr11 | 70546415 | 70546535 | 788855_47188552_ENST00000601538.6,E | 1 | - | good | TCGACAACTTGACAACTTGGAATAATAAAAAAGCATGCATTGGGGTCTCTGTCCCTAGTTCCTGGCACAGAACTCTTAAAACCTCTTGAATTTCCTGTCTTTTATATGTTAATGAGATGA | 120 |
| chr11 | 70546535 | 70546655 | 788855_47188552_ENST00000601538.6,E | 1 | - | good | ACCAGCCCAGTCGGCAGGGGGAGGCTTCAATGGGACTGGGCTGTAGCATGAGGGTCAGGGAAGCTGGGAGGGTCTCGATCTTGAGGAAAGACTGACCAATGAGGCAAGGTGTCTGGACTC | 120 |
| chr11 | 70546655 | 70546775 | 788855_47188552_ENST00000601538.6,E | 1 | - | good | AGCAGGGAGGTGTATGCATTGTTCCCCCTTGTTCCCAGGGAGGGGACTGGGGCAGCCCAGCATCTGCCAAGGCCAGCAAGGCAGTGGACACCCTCCCACCAAGGCCCACCAGGGGTCTAC | 120 |
| chr11 | 70546775 | 70546895 | 788855_47188552_ENST00000601538.6,E | 1 | - | good | AAGCAAAGAATCTTAATCATAGCTTGGCAAGCACAGAGCTGGCCAAGGCAGGATGCCCCCCAGAAGGTTCTGCAGCTGGAGGAACCTCGAGGGCCTGACACGCTGGAGACACCCAGAGCC | 120 |
| chr11 | 70546895 | 70547015 | 788855_47188552_ENST00000601538.6,E | 1 | - | good | CCAGGTGGGGGTGGGAGGTGAGCGGCCTGATCAGGTATCCAGGGTCGGGAGTTCTTGCTAAAGTTGGATTTGATACGGAAGCGCACAGATGGACTCAGGAGAAGTCCAGGGAGTCTGGTG | 120 |
| chr11 | 70547135 | 70547255 | 788855_47188552_ENST00000601538.6,E | 1 | - | good | AGTTTTTATCCAGGACTATTGCGGTAGGGGAGTAGCAGAATATTGAATAGGGAGGAGACTTAGTACGGGATAGGACTCAGTTCCTAATACTGCATGGGCTAGTGGGGATTTAGGGCCCAG | 120 |
| chr11 | 70547615 | 70547735 | 788855_47188552_ENST00000601538.6,E | 1 | - | good | CTATTCAAGCGGGGGTCCCAGAGAAGTTGCTTTTCTTTTCAAATCAGATCTGTACCTTCCTCAAGACTGAGGAATAGCTGTTTGATACTCAGACACGCTGTCCAGCTGGGATTACAGGCA | 120 |
| chr11 | 70547735 | 70547855 | 788855_47188552_ENST00000601538.6,E | 1 | - | good | ATTGAAATTCTCTGTCTCGATTGTGTCTAAATGGTTATGGGGACCAAGGATTCCTTTAGCAGAGTTAATTTGCACTTGGAAGACAATTACGTAGAGTAAGAGCTCAGCCTTTCAAGTCAG | 120 |
| chr11 | 70547855 | 70547975 | 788855_47188552_ENST00000601538.6,E | 1 | - | good | GGCTGGCAGGGAGACAGGATGTTTCCGGTTCTAGTTCAAACTCAGTGAGCTGATAACAAGCTTATCTTCACTCTGCTGCCATGAAGCTCAGCCTCGGGCTTGCAGGGGCTTGCCAAGAAC | 120 |
| chr11 | 70547975 | 70548095 | 788855_47188552_ENST00000601538.6,E | 1 | - | good | TCCCACATCTGTCCAGTACAAGCCACATATGTCAGCGCCTCCTTTGCCCAGAAGGCTGCACCAGGCCAGGCCAGGATGCTGATGCCAGCCACACCCACAGCCTTCTAGGGCATTCTGTGT | 120 |
| chr11 | 70548095 | 70548215 | 788855_47188552_ENST00000601538.6,E | 1 | - | good | TGAGTGGGGAGAGCAGCCGCCTGCTCCAGGGGTGGCCAGTGTCTGGGCACAGCCCACCCGGATGCTGGCAACCCCTGCCGGCCATCCTTGGGAAGGAAGGCGGAGTGAGAGGGCAGTGG | 120 |
| chr11 | 70548215 | 70548335 | 788855_47188552_ENST00000601538.6,E | 1 | - | good | TCCAGCCATCCATCGATTGTTAGAGATGGCTGTGTGTGCACGTCTGTGTCAGAGGCTCTTTTCAGAAAGATCAAAATCACAGGAGAAAACTGGAAGCATCTATTTCAGGCAGCGGGCCGC | 120 |
| chr11 | 70548335 | 70548455 | 788855_47188552_ENST00000601538.6,E | 1 | - | good | GAGACGGGCTGAGGTCAAGAGCAGGGAAGTGGCGGGTGGACGGGATGGGACTTGGGCTAGGCAGCCCTAGAAGCAGAGTCTCCCCACCTGGGATCAGGGCAGTCCCAGTAGGAAGGGAA | 120 |
| chr11 | 70548455 | 70548575 | 788855_47188552_ENST00000601538.6,E | 1 | - | good | GTTCTCATCCTTGGGGACTCTGGAAGGGGAAGGACAGAGAGTGGGCTCCATGTCCGGGCACAGGCGCCGGCCAGCCTCCCACTAAACTCTAGATGCTGCCCTGAGAACGGAGCCCAGCGG | 120 |
| chr11 | 70548575 | 70548695 | 788855_47188552_ENST00000601538.6,E | 1 | - | good | GGGGTAACACTTTTCAACCCACAATAACCCGTGATCCCAACAGCCAGGGTTCAGCACTTGCGCAGGGCAGCCTCTCCCATAGACCACACCTGCCTCCTAGGGGAGAAAGTGACAGCCTCT | 120 |
| chr11 | 70548815 | 70548935 | 788855_47188552_ENST00000601538.6,E | 1 | - | good | CTGCTGGCAACCCCTAGCACCCCTTGGCTTGCAGCAGCAGCCCCCATCTCTGCATCAGTGTTCACCTGGCCTCCTTCCCACCATGCGTGTCCACATTGTCCTCTTGTTTGAGAACACCAA | 120 |
| chr11 | 70549055 | 70549175 | 788855_47188552_ENST00000601538.6,E | 1 | - | good | TGGTCAGCATGAGCTTTCACGTCGAGCGCCCCCGTGTGCACAGCTGTGCCTGGGAATCCATTTCCTGGGGCTGCTGTCACAGCGTCCCACAGACCCTGTGGCTAGAACAACAGAGGCGTG | 120 |
| chr11 | 70549175 | 70549295 | 788855_47188552_ENST00000601538.6,E | 1 | - | good | ATGCCTTTGGCTCAGTCGGCGTGGAATCCTTTGTTCCGCTGCACAGGTCACTGGGGGCTGCGGTGCTTGCAGCCTGTGATAAGTACCTACGTAAGTCGGACCATGGTCTCAGGGCCTGTG | 120 |
| chr11 | 70549295 | 70549415 | 788855_47188552_ENST00000601538.6,E | 1 | - | good | GGCCATCGGGAGCTTATTTTTAAGGGCCCTTCACTAGGGGGACCAGTGTGACTGTGGGCTTGGGGTGAGAGAGGCAGAAGAAACTTGAAATGACCCGCACCTCATCCATGTCTGGAAGGG | 120 |
| chr11 | 70549415 | 70549535 | 788855_47188552_ENST00000601538.6,E | 1 | - | good | GGAGAGGACAGCGGAAGCAACCTCGCTTCGCAAATGTCATGAGGCCGTCCTAGTCGATGTCCCATCACCAAAGAGGAAGCAGGCAGCTCCATTTCCTAAATACAGAGGAGCACCTTTCTC | 120 |
| chr11 | 70549535 | 70549655 | 788855_47188552_ENST00000601538.6,E | 1 | - | good | CCCACTCAAGCTCCCCAGTCCCTGGCCGGGGACCTCGGGTCACTTCCCAGCTGAGCCCAGCTTCTGAGAGGGGAGCCACGGCCCCCGCCTGCCCCTGGCCACACTGCCTCGAGCACCAAA | 120 |
| chr11 | 70549655 | 70549775 | 788855_47188552_ENST00000601538.6,E | 1 | - | good | TAACTGTCTGTGGAAAAGGTCTCACCTCCAAGAGGCATGAATTCCAACAGGGCTGAAGATTCCTTCGAGGTCCTAATACATTCGTCCCATCCCAGGGACAGAGCTCTGGGACCCTGGTGT | 120 |
| chr11 | 70549775 | 70549895 | 788855_47188552_ENST00000601538.6,E | 1 | - | good | AACCGGAGCTTCGCTGTGTCTTTCGCAAGCCCCTTGCCTGGTGAAAAACCCAAACATGGAGGTATGGTGAGAATCACGCCTAAAATGCAGCTCTGTTTCCTCGAGCACCTGTGTCGGTGT | 120 |
| chr11 | 70549895 | 70550015 | 788855_47188552_ENST00000601538.6,E | 1 | - | good | GGGGAGGCTACAGGGCAGTGCCCTGCACATGGGGAGGCCTCGGCCAGCACTGGACAGAGCCCTGACAGAGCCCAGCAGGAGAGAGCCCACCAAGTCCCCCAGAGGCGGTGGCTGCGAGT | 120 |
| chr11 | 70550015 | 70550135 | 788855_47188552_ENST00000601538.6,E | 1 | - | good | CAGGAACCCCAGGCAGGAGTAGGAAGAGGGGAGCCTGGCGGGAGCTGGTAGGCCTGAGCTGCTCTTGGCTTCCTGGCTTCTGGGGTGGTGGCCAAGCCGGTGCGGTTTTTCTGGGGGACC | 120 |
| chr11 | 70550135 | 70550255 | 788855_47188552_ENST00000601538.6,E | 1 | - | good | TCATGGCAGGGATGTGTCCCCTTCCAGAGGTGCTACCGGGAGTGGGTCTGCAGCAGAACACCTGGGCTGCTGGGTGGGCATAGAGGGGGTGGGTTGATGGTGCCTCCCTTTCCAGGGATA | 120 |
| chr11 | 70550255 | 70550375 | 788855_47188552_ENST00000601538.6,E | 1 | - | good | CAGCAGGGGAGGGGTGGCGTCTGCTCCAAGACCAGCTGTGCTCCTGCCCACCCTCCAGGTAGGGAGGGTCTCCTCTGCCTAACTGACATGGGGAAAGGGCTGTAAGGAGGCTGAGGTCGT | 120 |
| chr11 | 70550375 | 70550495 | 788855_47188552_ENST00000601538.6,E | 1 | - | good | TGAATAGAAAGACCCTGGTATAGTGAGAAGCAGGGTCCAGGGTGTAGAACGCAGCATTTGCCAAAAACCGAAGAGCATCCCCTGCCCCAGCCCACCTGGGGGTACGCATGCAGCAGGCTC | 120 |
| chr11 | 70550495 | 70550615 | 788855_47188552_ENST00000601538.6,E | 1 | - | good | GGACCAAATGCGAGCCGACCCCTTGGATGTGGTCTTTCCTCCTGAACGGAGGCCTCGCCAGCAGATCCAGCAGATGAGATTTTCACTGAGCAGCAGTAGCAGTGGGCTAGACTTTAGTCT | 120 |
| chr11 | 70550615 | 70550735 | 788855_47188552_ENST00000601538.6,E | 1 | - | good | GCGCTGAGCCTACAGGGTATACCCCCTTCACCACCGGGCCCCTCCCCAGGGTTCTCCTTCACGTGGGACCTTCGCTAACAACATCGCTCAGTTCTTAGCCCACCAACCCACGCGAATCCG | 120 |
| chr11 | 70550735 | 70550855 | 788855_47188552_ENST00000601538.6,E | 1 | - | good | CCCTGAGGCTGGACTCACATTTCCACCACCTCTGGGCCCTCCTGGAGCTTGTCCTGCCCTGAGATTCACCCGCTGACTGCCTGGGAGAGCCCCGTGGCCCAGGACTGAAGGGAGGTTTGG | 120 |
| chr11 | 70550855 | 70550975 | 788855_47188552_ENST00000601538.6,E | 1 | - | good | GCGTTTACCACTGCAGCGTGGCTGTTTGGGTCCACACGTGTCTGCTCATGAGGCTAGACAAGTTCCTCTCCATAACCATCCATCCACACCTCGATGTTTCTCAGACCTGCTCCATGGAGG | 120 |
| chr11 | 70550975 | 70551095 | 788855_47188552_ENST00000601538.6,E | 1 | - | good | GTACAGGGATGAGGTCTCGGAGGGGGTGGGAGGGGGTCCGCTGGCACTCCAGCCCCTTGGCCCTCCCCGCTTGGCCCTCAGGGGCATCGCCTTCTCAGTGTCCCTGGACCTTCCTGGAGA | 120 |
| chr11 | 70551095 | 70551215 | 788855_47188552_ENST00000601538.6,E | 1 | - | good | AGGCAACACCAGGGGAAGTTTCTGTATGTGTTTTCTCCCTCAAAAAGAACATTTGCTGCAGCAGACACAGGCAGAGTGAGCACATGTGAAAATGCAGACCCTGGTGAAGCCCCCAGCAAA | 120 |

| | | | | | | | | |
|---|---|---|---|---|---|---|---|---|
| chr11 | 70551215 | 70551335 | 788855_47188552_ENST00000601538.6,E | 1 | - | good | GGCGGTGGGAGGGAGGGGGAGGGGCAGCGGGCAGGGCACAGAGCAGAGGCTCCAGCCTCCGTCTCAAGGCCTCGGGAAAGAGCCTCAGACACATCTTTGAAGCTGAAAGAGCCACATCA | 120 |
| chr11 | 70551335 | 70551455 | 788855_47188552_ENST00000601538.6,E | 1 | - | good | AGGTGTGACATTCTGCATGTGGTAGGAATGTTTTGAGAATTCTGCCCACAAATTAAGAGGTAGAGCTCCACTCCCCGTCCACGCAGCTTAGGGTCAGACTGGCCGGGTGGAGATGGGGGA | 120 |
| chr11 | 70551455 | 70551575 | 788855_47188552_ENST00000601538.6,E | 1 | - | good | TGAGGAAGCAGCCATGCCTGGAATGCAGGCCATGGCAAGCATCACACCGGGGCTGCCGAGGCCGGGGGTAGATAACAGGCATGCGGTCGCCATCAGAGCCCAGATGCTGGGGCGAAAGTC | 120 |
| chr11 | 70551575 | 70551695 | 788855_47188552_ENST00000601538.6,E | 1 | - | good | TTATTTTAGTGTCTTAGTACAAAGACGTAACTAGCGCATCGTTGTAGAATTAAAGCTCTATTCATGCTCCTGCTTAGTGGAGGCTTGGGAATTTGAGTGAAAAAAACCACTGAGTCAGCC | 120 |
| chr11 | 70551695 | 70551815 | 788855_47188552_ENST00000601538.6,E | 1 | - | good | GTTTCACATCACTCCAGGCCAGGGACCAGGTGGACGAGCAGAGACACCACATCCCCAGGTGGGGCCCACAGCTGAGGCTAAGTGGGATATAGGGCTATGCTTCATGTTGATGGAGCATAT | 120 |
| chr11 | 70551815 | 70551935 | 788855_47188552_ENST00000601538.6,E | 1 | - | good | AGGCTTGGCGCATAGAGAGTCTCGCTCCCCAGCCCTCCAGCCTCAAGGCCATACGGCAGGGTCCTAGCCCAGCACTCCTGACTGTGCCTGCTACAGAACCCAGAGTCCAGGCCTGCGGGA | 120 |
| chr11 | 70551935 | 70552055 | 788855_47188552_ENST00000601538.6,E | 1 | - | good | AATTACTCATCGCATTTAAGTTTCATCCATGCTCTCCAATAAAATGTACAGACCCCAGGAGGGAGGAAGCTCACACCCCCACGGGCTCCCCCGTGCCTCGGGCTGGGAAGCTGGCAGCAC | 120 |
| chr11 | 70552055 | 70552175 | 788855_47188552_ENST00000601538.6,E | 1 | - | good | CACCTACTCATATTATTCTTCAAAGCCGAGGTCAAAGCCTCCTCCTCCATGCAGCCCTCCTGGGAATCACCCTGACCTCAGGCATTGTCACTGGTAAACGTGGTCTTGGGTACTGCCTGG | 120 |
| chr11 | 70552175 | 70552295 | 788855_47188552_ENST00000601538.6,E | 1 | - | good | ATTCTCTCAGCTTCTCTCCTCCCAGGAGAGGCCCACTCTGTCTGCTCGCCTGTGTGCGTTTGTTCGCAGGCAGCCTGGCTGTACCTGGGAAATGCCGGTTGCCCCTTCCAACTGTCACCT | 120 |
| chr11 | 70552295 | 70552415 | 788855_47188552_ENST00000601538.6,E | 1 | - | good | TACAGGAACTAAATTGAGCTATAAATCTCCCTGAGCTTGGTGCACGAGTGTTCCAGGGCCCTGCTGGGCACCAGGAGTCCCAACCTCCCGAGGCTTGGCCTGGGCTGGAGGCGCCCAGGC | 120 |
| chr11 | 70552415 | 70552535 | 788855_47188552_ENST00000601538.6,E | 1 | - | good | AGAGCCTGGGAAATGAGACATCTGATCCCCTTGCAGAGTTAATCGCTTGCTGTTGGGATTAATTTTTCAGTTGTTGTCAAGTTGAAGCACTCGATAAATGACTCGCTGTGACATGGGACA | 120 |
| chr11 | 70552535 | 70552655 | 788855_47188552_ENST00000601538.6,E | 1 | - | good | CCCAGCGACGAGCCCCAGAGGCAGCCCATGGTGGCCCGCGGCTTGCCCAGGATCAGCCTCTGCCTGTGTCATCTTGGGTAGAAAGTGACGAATCGCTTCATAGACGAATGCCTGGAGATT | 120 |
| chr11 | 70552655 | 70552775 | 788855_47188552_ENST00000601538.6,E | 1 | - | good | CACGTGTGTAAGGAAAACGTCCCGATGCAGCTGGACACGGTGACTCCCTTGCACCCCAGGCTGTGTCCGTCACTGGCCCCCTGCGCCACGGAGAACCCTCGAGCCACAGGTTTAATGACG | 120 |
| chr11 | 70552775 | 70552895 | 788855_47188552_ENST00000601538.6,E | 1 | - | good | GTGGTTTAGGCTGCCCTGTCTGTGCTCTGTTAGGGCAGCCCAAGCTGACGACTAGAGGCTTCCGGAACTGGGTCCAGCTTAGAGGGCAGGTGGGCACCAGAGTCACCCTCCGTCCTGAGA | 120 |
| chr11 | 70553495 | 70553615 | 788855_47188552_ENST00000601538.6,E | 1 | - | good | CGTGCGTCCCACTGCACTCCTGCACCTCACGTGGCTGAGTGTTCCAGTGACTGAGCTGTCAGGACAGGAGCTGGGACCGGCAGTCAGGATGGCAGGGGGCTGCTGGGGATCGTCACAGCT | 120 |
| chr11 | 70553615 | 70553735 | 788855_47188552_ENST00000601538.6,E | 1 | - | good | CAAACCTCTGACATTGTCTTGTTCTGCTGCATGTTCCCCCAGAGAGGAGAAGCGTCTTGCTGATGATCGCATGGCCCTCGGTGAAAGAGCCCGGGTGCCAGGCTCCTGACCCTGAATTGA | 120 |
| chr11 | 70553735 | 70553855 | 788855_47188552_ENST00000601538.6,E | 1 | - | good | ATGTGACTGGGCTCTTATTCACTGAGGGGGTGTTAACTAATATAGGGATGTATCACACATCCTGACTTCAACCCTATGTCACGCAGATCATCACTGGAAGCTCAAGTGGCAGGGCAGAAG | 120 |
| chr11 | 70553855 | 70553975 | 788855_47188552_ENST00000601538.6,E | 1 | - | good | TTGCTGCTGACTTCCTGTCCACCAGGCCTGTAGCCTCTTCTGCGGCACACGCAGGGACCCCTGAATCACCCATTGCAAGTGAGCTTATCCCAACCCCACCTGGATGAGGAGTGACTCCTG | 120 |
| chr11 | 70553975 | 70554095 | 788855_47188552_ENST00000601538.6,E | 1 | - | good | ATCTGCTCCCCACAAAGGAGAAGGCAAAGCAAGGTTGTCATCCCCAGCGGGGACTTCCTGCAGCCAGTCCGGTCAGCGACCCCAGGCCATTTCGAGTTCTCTTTCATGACTTTATTCAGA | 120 |
| chr11 | 70554095 | 70554215 | 788855_47188552_ENST00000601538.6,E | 1 | - | good | GAGGGTCAAGGAGGTGCCTGTGATTTCTTATCAGGAGGGGAGGGTTGCAGATGTTGTGTTTTCTGGGGACAGGGGCTGGACTCGGGAGTTCCTCCCCTATCAGCACAGGCTGTTCCTACC | 120 |
| chr11 | 70554215 | 70554335 | 788855_47188552_ENST00000601538.6,E | 1 | - | good | GCACTGTGGGTGATCATCTGAGCACGCATTGACTTTTGAGGCCATCTGAGACTGGACCCCCTTGGGGGCTCATCAAAGCCAGGCTCTTAGGAGGGGAATGGAATGTTCTAGAAGTGGTTG | 120 |
| chr11 | 70554335 | 70554455 | 788855_47188552_ENST00000601538.6,E | 1 | - | good | GATCAGGTAGGTCCTTGCAAGCTTTGACTCATGGTGGAATCCAGGATTACAAGGGGAACCCATTATCCGAGAGGACAGAACAGGTGGAAAAGAGGCCGCTCTGGAACCCTCCCTGAAGCT | 120 |
| chr11 | 70554695 | 70554815 | 788855_47188552_ENST00000601538.6,E | 1 | - | good | AGCAGTTGCCAGGGATTGGGGGCAGGGAGGGTGAAAGATGGGGCACAGCCATTGTTGGGCAGTGGCGCTCTTCTGTATGGCACTGGAGTGGGGGAACGGGGCGTTATAGATAGTTACAAC | 120 |
| chr11 | 70554815 | 70554935 | 788855_47188552_ENST00000601538.6,E | 1 | - | good | CTGCAGGCCTTCGATTTGTTAAAAAAAAAAAAAAATGCAATGTTCGTGAAGTGCAATGAAGTGAGGTGTGCCTGTATATGACATTCTCAAAAAAAAAAAAAAAATGGAGATAGTGAAAAATC | 120 |
| chr11 | 70557215 | 70557335 | 788855_47188552_ENST00000601538.6,E | 1 | - | good | CCCTGGAACTCACATGTGATCATCTGTTTGCTTGCTTGCTTTTTGTCTGTAATTCGTGAGATCCTGAGGCCAGTTGATAGCTTCATTCCCAGCACCCAGGACACAGCCTGCATCTAGGAG | 120 |
| chr11 | 70557335 | 70557455 | 788855_47188552_ENST00000601538.6,E | 1 | - | good | ATCCAGAACATCTGTCTGGAAGCTTCCATATGGAACGCCCTCCATGTGAGTCCCTTCTTAGGAGGAGGTGGTGGTTAGCAGCAAGAGCCTCAGGGTGGGCTGGCCTGGGTTCAGTCCCCG | 120 |
| chr11 | 70557455 | 70557575 | 788855_47188552_ENST00000601538.6,E | 1 | - | good | CTGCTCCCGGCTTTAGCAGCAAGACCGACATCAAGCTCTGGAATGAGCCTCCCAAGGGTCACATCCAGCCTCAGAGCCAGGACTGCCAACTCCATCTGGGCTCCTCCCCCACTGCTGAGC | 120 |
| chr11 | 70557575 | 70557695 | 788855_47188552_ENST00000601538.6,E | 1 | - | good | CTTGTCCTTCCAGACTGTGGGAATCTTGGAGGACACAGCTGTGAGGGCCCCCCACCCACAGCTCCCCCAGGCCCCACTGTCAGCTCTTGCCCTGGCCCCGCGGTGCTGCCAGCCTGCTCC | 120 |
| chr11 | 70557695 | 70557815 | 788855_47188552_ENST00000601538.6,E | 1 | - | good | TGCATGAAGAAAGCCGCTTGTCCGTTGGAAACGTGCAGGCCCAGCCTCCTCGGGATCACTGGGGATCTCCGGAACCACTTGGGCCTTTGCACAGGAAAGCCCAGGTCAGGGGGCTTTCTG | 120 |
| chr11 | 70557815 | 70557935 | 788855_47188552_ENST00000601538.6,E | 1 | - | good | AAGGGGAAAGTGTTCAAATAGTAGCAGTGACTCAGCCCCGGCTCGGCACGCCTGGGAGCTGTGGCTGCAGAAAGCCCAGCTGCAGGGTCCAGGTGGGCTCAGATGCACTGGGCAGCTGTG | 120 |
| chr11 | 70557935 | 70558055 | 788855_47188552_ENST00000601538.6,E | 1 | - | good | TCGCCTCCTCTCCCTTCTCCCCTGGCATTTCCTTGCTCAGCAGATGAAGGATGATCAGAGTTCCTGGTGGCATTTGTTCTCAATTGGTATTTAACATATTTGCCAAAGACTGTCATTTTG | 120 |
| chr11 | 70558055 | 70558175 | 788855_47188552_ENST00000601538.6,E | 1 | - | good | ACCTCCAGATAGAAAGAACCTCTCCGAGAATCCCCAGAAAGACCCAGGCCTCCTTGGTGGGCAGCTGGACCCCTGCCACGGAACAGCCAGAGAGCATGGACCCCGGGCCCCCTCACGGTC | 120 |
| chr11 | 70558175 | 70558295 | 788855_47188552_ENST00000601538.6,E | 1 | - | good | GTACAGGTGTCCCCCACATGACATCAAGCCCACAAGCCCATGGCCTGACGCTCCTACCGAAGCCCCAGTGGGCAGGTGAATGAGCCTCCAGTCGGGGGGCCAGGGGGCTCCCCCTCGCAC | 120 |
| chr11 | 70558295 | 70558415 | 788855_47188552_ENST00000601538.6,E | 1 | - | good | GTGACAGGCACGACTGGGCTGGGTCGTGCACGCCGAATGGGCTGAGCCCGATGCTGACGCAGGGGGAAGCCGGCTGCTCCCTTCCTGCTGAGAGCTCCCGTGGCATTGTCCAGAGGGCAG | 120 |
| chr11 | 70558415 | 70558535 | 788855_47188552_ENST00000601538.6,E | 1 | - | good | CTTGTCTAAGACCATCCAAGTGAGAGCTGGCAGATCCAGGCTTGGAACCCTGGCAGGCTGGCTCCAGCCTGAGCTCCGAAGCTCTGTCCAGGCACTGTCTTGAAGGCCAGCTCTGTGGCG | 120 |
| chr11 | 70558535 | 70558655 | 788855_47188552_ENST00000601538.6,E | 1 | - | good | GCATCAGGCCCTGGACTGACTACCTCCCCACGTGGCCTCCCAGCTAGCATCCCCATGGGGGAGCACTGATCTTGCCTCTGTGTTACAGACGAGGAAACTGAGGCACAGAGAGGTAAGCCA | 120 |
| chr11 | 70558655 | 70558775 | 788855_47188552_ENST00000601538.6,E | 1 | - | good | CGTCAGTGTGTCTTTCTGGTCGCCTCCCGCTCAGGTGTGCACAGTCCTGTCTCTTTCGGTTCACTCACCCAGTGCCCAAGTGTAGAATCTGGCTCTCTGCAAAACTCAGGGGCCCAAGGT | 120 |
| chr11 | 70558775 | 70558895 | 788855_47188552_ENST00000601538.6,E | 1 | - | good | CTCCAACGGGGGCTGCTTTCTCTCCATGCAGCAGCTGTGCTGCTGTTTTCTTTCACGTCCTCCAGGAAGCCGGAGCCCTCCTGTGCTCTCCTGTAGCTGCCTCATCCCTGCGCTCTGGGC | 120 |
| chr11 | 70558895 | 70559015 | 788855_47188552_ENST00000601538.6,E | 1 | - | good | GTGAAATGAGAAAATTGTTATTAAGAGTCTCCGAAATGAGGAAATCGTTATTAAGAGGCGTGTGTGTGCACATTGCCCCGAGCACATCTCTCTTCCGTGTGCAGACGCGCTGCAGGCCGA | 120 |
| chr11 | 70559375 | 70559495 | 788855_47188552_ENST00000601538.6,E | 1 | - | good | GAATGATCTCCCGAGTAAGTGATTTGAGGTGAAAAAAGCCAGGAGCAGACAGACTGTTAACTGTGTTGCTATTTGTGTGAGAAAAAGAAGCGTGGTGTGTGGAAACTCTCTGAACGCTCT | 120 |
| chr11 | 70559495 | 70559615 | 788855_47188552_ENST00000601538.6,E | 1 | - | good | GCTCTACTTGTCCAGGTGTGTCCCCAGGAAAGCGCGGGCCAGGTAAGCTATGGAGCGCCCAGTGTGAGGAGTGCCGTGCCACTGTCAGAAAGCTGAGGCGGCTCTGGGGGTCCTGCTAGG | 120 |
| chr11 | 70559615 | 70559735 | 788855_47188552_ENST00000601538.6,E | 1 | - | good | GTTTTGGGGAGAGGTTACGATGCACGTCCTCCTTGGAGGACAGCTCATCAATTCCTATGAAAATGATCGATGCAGATGCCCTCTGGCCCAGCCTCTGTGCTAACAGGACTGGATCCCACA | 120 |
| chr11 | 70559735 | 70559855 | 788855_47188552_ENST00000601538.6,E | 1 | - | good | AGAATATGGGTGGTCTGGGGCAGGAAGAGGAAAGGGCTGTTTTGTGTGAGGGCTCTTTTTTTGGAGTCTGATCATCTACCATTTGGGCAGGAGGGGACTGGGCAGGGCGCACCGTCTTCT | 120 |
| chr11 | 70560335 | 70560455 | 788855_47188552_ENST00000601538.6,E | 1 | - | good | TTGAGTCCATAGATCCAGTTGGGAGAGATTCAGTCTCCTGCCACATGATCTAAAACCTCTCCATTTATTGAGATCTTCCATTTCTCTCAGCCATGTTTAATGCTTTTGCTATACAGGTCT | 120 |
| chr11 | 70562255 | 70562375 | 788855_47188552_ENST00000601538.6,E | 1 | - | good | TTCTCCTTCTGTAACTGATGGAAGTGTACAGAAAGTCAGCCGGGATATAGGAAACCATGAACAAACCCATCAACCAGCTTGACTTAACTGACAGTTGTGGAAAGAATCCTCTGTGCTTTG | 120 |
| chr11 | 70562615 | 70562735 | 788855_47188552_ENST00000601538.6,E | 1 | - | good | ATATAAACACCCTTATTTTAAGGGCAGATTCTTCAAGTTGAATAAATGCTCACCTCCAAGGAGCACTCGCTTTAATTATAAAGACACAAAAGAGTTAAAAGTAAGAAGGTGGAAAGGCAT | 120 |
| chr11 | 70562735 | 70562855 | 788855_47188552_ENST00000601538.6,E | 1 | - | good | AAGGCAGAAAATGAGAGAATTGGCGGGGGTGGGCAAAATATACAAGGGACAAATGAAAAGCAAAATGGAAAGAGGACAGAGTTAAACTTCCTATCAATAATCACATATTAAATGCAGACG | 120 |
| chr11 | 70563215 | 70563335 | 788855_47188552_ENST00000601538.6,E | 1 | - | good | GTAAGTTCTTGTATTGTACACGAAGTGGTAGAACATCAGCAGTGATAAGTTAAGGGTGTCTACTATAAATATAAATTCTAAAGCAACTGCTGAAATAACAAAACAAAGAATAATAGCTGA | 120 |
| chr11 | 70563335 | 70563455 | 788855_47188552_ENST00000601538.6,E | 1 | - | good | GAAAGATAACTGTCTTTAAACAGAAGAATAAATGTAGCTTGGGGATTACGACATGGAAAGATCGTAGGTGTGACAACAATGGCACAAAGTCTAGGAGGGGAAGAGTGGAAGTGTACTGTT | 120 |
| chr11 | 70563815 | 70563935 | 788855_47188552_ENST00000601538.6,E | 1 | - | good | AAAAACAGAATTATTCAACAGAGTTATTTCAACATAATTCAAACAGAATTATTTCAACAAAGGTAAAATACTTTTGAAAAATACAAAAGGTTTAATCATTCATCACCAGCAGGTACACTA | 120 |
| chr11 | 70563935 | 70564055 | 788855_47188552_ENST00000601538.6,E | 1 | - | good | GGGATAAAAGGACACGTGACACATGAAGACGAAAGAGAAAGATGACAGCAGGCATAGTGTCAGAAGCAATGCCAGTGAGCAGATCGTGGACAAATACCTTTACAGTATTGAAAGAGAAGA | 120 |
| chr11 | 70564535 | 70564655 | 788855_47188552_ENST00000601538.6,E | 1 | - | good | TGAGCTGCAAGATAGCTTCCAACAGCCCAATATACATGTAAGTAGAGTCCCTGAAGAGGGGGCGATTGTAGGGGGAGGGGAGAAAAAATACTTGAAGGCTGGGTGCGGTGGCTCATGCCT | 120 |
| chr11 | 70564655 | 70564775 | 788855_47188552_ENST00000601538.6,E | 1 | - | good | AGATCGAGGGTAGATTAAATGTTATAAAGTAATAGTGAACTCAGAGAAGTCTCAATAGAAACTATCCTGAGTGAAACACAGAGAAAAAAAAAGAATTCTTTTTAAAAGAACATCAGTCAG | 120 |
| chr11 | 70564775 | 70564895 | 788855_47188552_ENST00000601538.6,E | 1 | - | good | GGTTAATATGATAAATTTTGTTGTATATATTTTAGCATAAAGCAGTTCTTTACATTTTAACCCAAATCAAACTTCCAGTGATGAAAACCACAGTGTCTGAGATGACGGATACACAGGATG | 120 |
| chr11 | 70565615 | 70565735 | 788855_47188552_ENST00000601538.6,E | 1 | - | good | AAATATATAAAGTAATAGGTTTTACAACCAGACACCAGGTAATAAATGATAGTGATCACTAAAGCATGAGAAACAGATGGGATGAGCCCTATACTTGCCCCAGCTTCTTGCACTGAGCGT | 120 |
| chr11 | 70565735 | 70565855 | 788855_47188552_ENST00000601538.6,E | 1 | - | good | GAAAATTAAAAAAAATAATAATAATAATGCCTACTTCTGCTCCAACTTCTACTTGTGGCCAAGATGGAGAAACAGAAACTGGATTTACCTTTCCTCATGAAACAACTGAAAGAAAAGATA | 120 |
| chr11 | 70566335 | 70566455 | 788855_47188552_ENST00000601538.6,E | 1 | - | good | CCCTGTCCTGCGAGGGTGCAGTGTCACTTTTCCACTCCCATTTGGGGTCAACTCTTAACAGCATCCCTGAGAAGTGAGGTGGTGTAAAGTCAGGGAAGTGTGGTCTTTGTCTTATTAAAA | 120 |
| chr11 | 70566455 | 70566575 | 788855_47188552_ENST00000601538.6,E | 1 | - | good | ACTCAGAGTGGAAAGTCTTGGCAACCATGTTGCGGTTCAGACAGTGGGAGCCATGCTTCTGCGAATGAGCCTGGGGACTGGGGACTGTGGTCCCAGCCCTGGATTTGATGAGGGGAGTAT | 120 |
| chr11 | 70566575 | 70566695 | 788855_47188552_ENST00000601538.6,E | 1 | - | good | AAACACGGGAGGGGAAGGGCAGGGTGGGGTAAATTTCACTCTTAGAAACACCAGAATTTCTTCCTTTTCAGGATCAATGTCACCCATATCTTGTCAATTGTGACTGCAAACTTCAGAAGG | 120 |
| chr11 | 70566695 | 70566815 | 788855_47188552_ENST00000601538.6,E | 1 | - | good | GAGGAAATGGATTGCTAAGTGGTTATGATGGAGACTAAATGAATTACTTGTTAATCACCCTTGTCTGCTCTGGTAGTATCTGTGCAAGCCAGGATAGCTGCTCGGGGCACACTCACCATC | 120 |
| chr11 | 70566815 | 70566935 | 788855_47188552_ENST00000601538.6,E | 1 | - | good | TAGAGATTCTTCCTTCTTCCTGTCCAGGTAATTAATGAGGGACTTCAAAGGCATGAGGCTTTTCTGCTCCAATCAATTATTCAGATTGGCATATCCAAAGGTCTCTGAACAGGAAATAGA | 120 |
| chr11 | 70566935 | 70567055 | 788855_47188552_ENST00000601538.6,E | 1 | - | good | TCAGAGGGTTCCAGAAGCATTCCTCCAGCCTCGAGGGGAAATGGTGCCTCTGGCTAACGGAGGGAACCATCAAAGCCGGGTATGGTTTCTTTGATCGTGTTTAGCCGCTTTGTGGCTGAG | 120 |

| | | | | | | | | |
|---|---|---|---|---|---|---|---|---|
| chr11 | 70567055 | 70567175 | 788855_47188552_ENST00000601538.6,E | 1 | - | good | ATCACAGCTTCATAACAGATGCTCTGCAGACACCTCTCCATCAATCATAGCCCGGTGTGTTTAACTTAGAGTAGATGGTGGCACCACCCGATAACAATGGTGGTGGCATTCAGCAGCAGC | 120 |
| chr11 | 70567775 | 70567895 | 788855_47188552_ENST00000601538.6,E | 1 | - | good | CTTCCCTCCCAGCCCTGCAGCTATTCCAGGGTGTGGCATCTGTCTTAGTTCGGGCTGCCCCAGCAAAGCACCACAGACAGGGTGGCTGGGACAGTAGACGTTGACTCTCACAGTTCTGGA | 120 |
| chr11 | 70567895 | 70568015 | 788855_47188552_ENST00000601538.6,E | 1 | - | good | CATTTCACATCCTGTCTCCCCATTAACTCTTTTGTTGCCCGTCTCCTCGCTGAGTTTTGGTTTGAATAGCTCAGTCCATTTGACGTGTTTGGGTTGGACAAATACATAGGGTGTCTGCAG | 120 |
| chr11 | 70568015 | 70568135 | 788855_47188552_ENST00000601538.6,E | 1 | - | good | GACCCTTTCACTGGATCCATTGCAGGCAGGATCGGAGCCAGACTCCACCACCCCGCTGCTGGCCTCACGGCAGTTTTGCACCCAGCCTCCAGTACTGGCCATCAGCTGATGGCCCAGACC | 120 |
| chr11 | 70568135 | 70568255 | 788855_47188552_ENST00000601538.6,E | 1 | - | good | CGCCGCCCCTCGAAGGATCCCTGGCTCCAGCCTCCTCAGCTTAAGTCTCGCGGAAGAGGCTGCAGCTACAGCTGCCTGCGGGCCTGCTTTCTAGGTTATCCTTGTTGGAAGCTTTGCCCT | 120 |
| chr11 | 70568255 | 70568375 | 788855_47188552_ENST00000601538.6,E | 1 | - | good | GAGAGGCAAGGAGGCTTCACCCACCGAGTGAGGCACTGCCTTGCCTGAGCTCTGTCCCTAAGCTCACTTCCAGGGCGGTTTCGAAGAGGAAGTGAAGATTCTTCCTGGTGCTGGAGACGG | 120 |
| chr11 | 70568375 | 70568495 | 788855_47188552_ENST00000601538.6,E | 1 | - | good | TCCGAGGATTGCAGCCGCCTCCAAGAATCCATTAAATGAGTTCTTCAGACTCGCTTTTAATGCTTTTCCGAGGTTATTAAACAGAGAAGTTCTCTGCAAGGAGCTTTCAGGGAGCTGTGA | 120 |
| chr11 | 70568495 | 70568615 | 788855_47188552_ENST00000601538.6,E | 1 | - | good | GGTGCTGGGAGATGTATCCTCCCAGAAGGAACAACAACTCGAATTGGCTTTGGGGAAATTCAAACAGTTTCTCTCCAGCTGGAGGCTCCACCAAGAATAAGTTCCCCAGCTTCACGGAGC | 120 |
| chr11 | 70568615 | 70568735 | 788855_47188552_ENST00000601538.6,E | 1 | - | good | ACCATGCCTGGCCTGGGGCGCCCCTGCCTGCTACTCGTGGCTTTGCTGAAAAGAACAGCTCCCGGCCGGGAGGAAGTGGGCGGGGGGGGCAGGAATCCGCTGCCCACATGGGAGAGGGGA | 120 |
| chr11 | 70568735 | 70568855 | 788855_47188552_ENST00000601538.6,E | 1 | - | good | AAAGGAAGGAGGGGGCTGTCTAGCCATGGTGGCTCAAAGAATCAAAGTCTTGGAGCCACACCCTCCAGAGGGTGGTTTATCTTGCAGAATTGCTCCCTGAACCTGGCTTCTTCCAGACGG | 120 |
| chr11 | 70568855 | 70568975 | 788855_47188552_ENST00000601538.6,E | 1 | - | good | CTCGGACGTGGATGCTGAGTCCACTGGCCACACAGCTGCACTCTTGCTGGACACAGGCAGGCTTCTCAGCCATCTATAAACAGCTGGGTTGCGGTGCCAAGGGCTCAAGTCCTCTGTCAG | 120 |
| chr11 | 70568975 | 70569095 | 788855_47188552_ENST00000601538.6,E | 1 | - | good | TTTCCTGGCTCCGTTCACTGTAAGTCCAGCGGCAGGAGTGGGGCCTGCATGCTGCATGCCTGAAAACGGGGAAAGGGCACATGGGCAGCCTTGAACATCTTCAGAGAACGTGAGCAGTCC | 120 |
| chr11 | 70569095 | 70569215 | 788855_47188552_ENST00000601538.6,E | 1 | - | good | TTTGGGACTGTCCTCAGTCCGATTGGCCCTGAGAGGACAGAGGATCCCCCTCTCTGTGGGACTGAGGTGGTTCAGGTGGGGCCTTCCTGCAGAATCCAGGACAAGTCAAGGCAGATTCGC | 120 |
| chr11 | 70569215 | 70569335 | 788855_47188552_ENST00000601538.6,E | 1 | - | good | CGCATGAGCAGGATCACCCCAGACGCTCCCGAGCCTGACCCCCCAGGACCCCAGCGGCATGGCTTAGCCACCAGCCTCCCCTGGGGTCACTACACACTTAGGCTCGGGATGGCCCGGGGG | 120 |
| chr11 | 70569335 | 70569455 | 788855_47188552_ENST00000601538.6,E | 1 | - | good | CCCAGCACAGCAGCCTACGAGCCGCCTTCTCCCTCAGGACCCAGGCCCTGCTCACTCTGCCAGCCCCGGAGCTCCCCGCTCCACCGCACCTGGAAACCGGGGCAAGATGCGTCCATGAGG | 120 |
| chr11 | 70569455 | 70569575 | 788855_47188552_ENST00000601538.6,E | 1 | - | good | GCCACAGAAAGGAGAGGGCTGCGGCACCAGCAGAGGTAAGATGGGGAAGCGGCCGGCAGCGCCACCCATGCCTCTCTTCCTGGGGCCTCATCGTTGGTCAACCCCGCCATCCACCCTCTG | 120 |
| chr11 | 70569575 | 70569695 | 788855_47188552_ENST00000601538.6,E | 1 | - | good | CCTCAACCAATCACTGTGGCCTGGGAAACGGGAACCCCTGATTGGCCAGGACTGGGTCACTGGGCACTCCTGAGCCCGAGGGCTGTCCTAACCTCCTCGGTATCCTCCACGTGGGGAGGG | 120 |
| chr11 | 70569695 | 70569815 | 788855_47188552_ENST00000601538.6,E | 1 | - | good | AACCCCCTCCGTGCTCACACATCTGCCAGCATCACAGCCCCCCAGGGACAGGAGAGCATCTTCCCCAGTAGATGTAGAGGAAAAACCCAGGCAGGATTCTGATTGGCCTGGATGACCATC | 120 |
| chr11 | 70569815 | 70569935 | 788855_47188552_ENST00000601538.6,E | 1 | - | good | CTCTGTGTCTCTGTCTCGGTGAGTGTCTCTGTCTTGCTCTGTACCGTCGTCCCCGTCTCCTCGTACTGACCTCCTGCCCTTGCTTCTTTGTGCCGGCCTCAGCCTTTTCCCGGGCTGAGG | 120 |
| chr11 | 70569935 | 70570055 | 788855_47188552_ENST00000601538.6,E | 1 | - | good | AGCTCAAGTTTCATAGGTGAATAATGTGCAGGGGGAGACTCAACAGCTGGAGGCAGGAGTCCACAATGGTGCTGGGGGCCTGTCTCTGTCTCTCACTCCCTTCTCTCTCTCTCTCTATGT | 120 |
| chr11 | 70570055 | 70570175 | 788855_47188552_ENST00000601538.6,E | 1 | - | good | GGACTCTCGGCGGCAAGCCACAGAAGTTCCCTCTTGTCATTCGGTCAAGCCTTGTGAAGCTGCCATTTTTATAGGTCAAAAAACAGTCAGGTAGCGACAGCTTATGAGGCCCAACCGAAC | 120 |
| chr11 | 70570175 | 70570295 | 788855_47188552_ENST00000601538.6,E | 1 | - | good | ATCAAGAACGGGAGCGACCTGGGCCCCTTGTAAACCCGGCAGATCTAAAAGAGGGAAGGGCTTCAAAGGGGCAGGACTCCTGCGGGTCGGCGTGGCCGCCAGCCAAACCTACTTTCTCCA | 120 |
| chr11 | 70570295 | 70570415 | 788855_47188552_ENST00000601538.6,E | 1 | - | good | AAAGAGAGCCAGACAGTATCCATCCTGGGTCAGGGGCACCCACGCGTGCACCCTAGGGAGGCGCTGGGGCCTGAAGTTCTACCATCAGGAATTAGAAATACCAGTTTTAGAAAATTAATC | 120 |
| chr11 | 70570415 | 70570535 | 788855_47188552_ENST00000601538.6,E | 1 | - | good | GGAGTTCACAGCTCGGGCTGCCCAGGGCCCCTTTCAGGGCAAGGCCTCCCTGGGGAGCCTGGCAGTGGGACCCCGGTCCTGGTCCTGCAGCAAGCAGGGGTACCTTTGACCACAGCTCCT | 120 |
| chr11 | 70570535 | 70570655 | 788855_47188552_ENST00000601538.6,E | 1 | - | good | TGATCCTGGAGGGTGACCCTGTCTGAAGCTGCAGCCCACACACACCACTGGAGGCCGCAGCTGTGGAGGCAGACCCGGGCCTGCAGCAGAGCAGGTGAGGAGCTAGCCCTGTCCAAAGGA | 120 |
| chr11 | 70570655 | 70570775 | 788855_47188552_ENST00000601538.6,E | 1 | - | good | TTGATTCCCAGAGGGCCTCGGTCTATGACACCCCATTCCCCACTCCCCACTCCCCACTCCAGTACACCCACATCCTCCAGCCCCTTGGCACTCCCTCTCATGGAGCCAGCTGGGCTGCAG | 120 |
| chr11 | 70570775 | 70570895 | 788855_47188552_ENST00000601538.6,E | 1 | - | good | GCCCCACCCATATTTCCAGTTGGGTGACAAGCAGTGTATGTCCCTGGAACTGCACCCATAGCTCCACAGGGAGCTCAGTGTGAGGCCCTGGTGGGTGTGGGCCCCGGGTGTGAGTGGGCA | 120 |
| chr11 | 70570895 | 70571015 | 788855_47188552_ENST00000601538.6,E | 1 | - | good | AGAAAGAAATGCTAACACCCATGAACACCACCGCACAGGAGTGCTTTGACCCCCAAGATGCAGCATAGTTATGTTATGCACAAGATAACTACAAGATTTATGAAACTGTCCACTCTCAGA | 120 |
| chr11 | 70571015 | 70571135 | 788855_47188552_ENST00000601538.6,E | 1 | - | good | TGGGGGAGAGGCTGGACAGGGTCAGAGAGGCCTTGGGGGAAGCCAGGGAGAGTCATTCCCTAGAATCCGCCTCCAGGACCCAGCTGCTGTGGTTTTCTGAGCCTCCTGGATTGCTTCAGA | 120 |
| chr11 | 70571135 | 70571255 | 788855_47188552_ENST00000601538.6,E | 1 | - | good | TGGCCCGCAGCGGGGAGGATTAAATGCCATCTTCTGCAGGGCAATGGAGAGTCACAAACACGGGTGCAGATGCAGAGCCCAGCCTTGACCCCAGAGTGGACACTCCTGACCTCCAGGGAG | 120 |
| chr11 | 70571255 | 70571375 | 788855_47188552_ENST00000601538.6,E | 1 | - | good | GCCTGGCGTTGATTCTCTGCCCTTCCATTTTCGGGCTGTGGATCACCGGAAGTCACTCAGCCTCTAAGGTGAATACCCACAAGGTATTTGATGCTTACTTCATGCTCACAACACTGTCCT | 120 |
| chr11 | 70571375 | 70571495 | 788855_47188552_ENST00000601538.6,E | 1 | - | good | CTGGAAAATGAACATTTTATTTCAAAGAAAAAGTAGCACGATGGTATCCAAGTATCCAGGACCTGCTGCCATGCCCTTTCTCTGCAGAAGGCCTGGCTGGGCCCAGTGGCCTAGCCGAGA | 120 |
| chr11 | 70571495 | 70571615 | 788855_47188552_ENST00000601538.6,E | 1 | - | good | CACCTCTCCTCCCTAGACCCAAGGCCACTGTATTAGCCTGTGGGGTCAGGGGACCACTCCAGCTCAGCCAGCTATTGCTGGGGTGGGGCACATGGTGGGGGTAGCATCTAGACTTTTATA | 120 |
| chr11 | 70571615 | 70571735 | 788855_47188552_ENST00000601538.6,E | 1 | - | good | TTGCCGACTCCTCCAGTATCGACCTCAGCCCTTGCCTGTAGGGGCCTCCAACTGAGCTCCCTGCACAGACACCTGGATCCCCTGGTTCTGCCCCAAGGACAGAATTTGCTAGGCTCAGCT | 120 |
| chr11 | 70571735 | 70571855 | 788855_47188552_ENST00000601538.6,E | 1 | - | good | CTCCTCTCTCGCTTTCCAGTTCCCTCTCCCCTCGTCTCCACTCCTCCTTGGGGCCCCCTGATGCCGCACCCCTCCCTCTCCTCAGGGTTCATCACTTCTCCCTGCACGCACCACCCTCCT | 120 |
| chr11 | 70572455 | 70572575 | 788855_47188552_ENST00000601538.6,E | 1 | - | good | CCTTGTTTCCCTCTTCTCTCTCCTCCCTTAGTTTCCTTCCCTTCCCTTCTGTCTGAGTCAGCTAAATCCACTAGGGCAAAATACCCTAGATGGAGGGGCTTCAAAACAGCAGGGGTGACC | 120 |
| chr11 | 70572575 | 70572695 | 788855_47188552_ENST00000601538.6,E | 1 | - | good | TAGCTCACATTTCCCTGGAGACCCCTGATGTTGGGCCTCTTTTCTTATTCTTGCCAACCATCCCTGTGTCTTCTGTGAAGCCGGAAGTGGAATGTTGTGGGGAACAGAGGGGCTTTCTCT | 120 |
| chr11 | 70572695 | 70572815 | 788855_47188552_ENST00000601538.6,E | 1 | - | good | ACCATCTTGCATCCCTGCCAGCCACGTGTGGCGGGTCAGCCTCTCCTGGGCCTCGCCGGCACTTGGTGTGTGCCGTCTTTTTCATTTGGAACAGTGTGAGGTGGCGTCTCATTGTGACTT | 120 |
| chr11 | 70572815 | 70572935 | 788855_47188552_ENST00000601538.6,E | 1 | - | good | GACTTGGGGTAGTTAGGTGCCTTCATTTCTCTTGGATACTCAGAAGAGGAAAGCCTGGGTCACGTGATAGATGGATGTGTAAGTTTGAGAGACGGCCAAGCGGTCTTCCAAAGCAGCCTC | 120 |
| chr11 | 70572935 | 70573055 | 788855_47188552_ENST00000601538.6,E | 1 | - | good | GTGATGGGCCGTCATCATGTCGATGGCCCACAATTTGGTGACCAGTCACGATCGATGGGCGTTTGGTTGCAGCCACTCCGACCTACCACGCATAAAGTTGCTGTGAATGTCCATGCACTA | 120 |
| chr11 | 70573055 | 70573175 | 788855_47188552_ENST00000601538.6,E | 1 | - | good | TAGAACATCACATAAAATGGAGTCACGCGACTGAAGCTATTTTTTGGGTTGGTTCCTCTCAGGCAGTGTGATTATTCTGAGGCGTGTGTGTTCCGGCAGTCCGTTCCTTCGCGTTGCCGA | 120 |
| chr11 | 70573175 | 70573295 | 788855_47188552_ENST00000601538.6,E | 1 | - | good | CATCACAGCCCCCCCGCCCCCACCGCTGCAGTGACCACCCAGCTCACATCACAGCCCTGCCCCGCCCCCACCGCTGCAGTGACCGCCCAGCTCCGTCACATCACAGCAGCCTGCACCCTC | 120 |
| chr11 | 70573415 | 70573535 | 788855_47188552_ENST00000601538.6,E | 1 | - | good | AGACACATGGGGCAACCTCTCCTGTGGTATCGGTCATATCATGATACACTTTTTAAAAACTTTATTGGGGCATAATCAATGTCCAGTCAAGTTCACATATTTAAAGTGTGCAATTCTATA | 120 |
| chr11 | 70573535 | 70573655 | 788855_47188552_ENST00000601538.6,E | 1 | - | good | AGCTTGATCTGGTCAATGTAGGAGGGGCCACTGGGGTCATCTTCATGTCTTGGTCCCAGTAACTCCTATTTTCTTTATATCAAATGCCATGCCCCACCCCCCCCCCGCCCCCACACACAC | 120 |
| chr11 | 70573655 | 70573775 | 788855_47188552_ENST00000601538.6,E | 1 | - | good | AAGTGAGGTGTGGGGTTGGGGGCTTGGGGCATCGGCTGGCGCCCCGTTTCCCCTCCTCACCCTCAGGGGCAGCCAGGGGCTTGCGTGGGATGTGGACTCCAGCTCTCTCCCAGAACCTGA | 120 |
| chr11 | 70573775 | 70573895 | 788855_47188552_ENST00000601538.6,E | 1 | - | good | TTTCAGGGCCCCCAGCGGGTTCCGCTGTCACCAGGATGGTAAGTGTGCCTTGAAGTCTTCGGTGTGAGAAGTACTTTGCATACTCTAAGTCAGACAGAACATAAAAGTGGCCGTTGGCTG | 120 |
| chr11 | 70573895 | 70574015 | 788855_47188552_ENST00000601538.6,E | 1 | - | good | GAGAAGGACCCGGGCTGGGGGCCAGGTGCATGGGTCTTGGTGGCTGGAGAGAAACTGGCATAGCGCCATCCGCCCCTGGGGCAGGGTGTGGAGCCCAAGACCACATCATCAGCCCAGGCC | 120 |
| chr11 | 70574015 | 70574135 | 788855_47188552_ENST00000601538.6,E | 1 | - | good | GCCCTGTGGTCATAGACAAGAGGAACTGGCCGAGTTCTCCATCAGCACCATGCTGGGGGCTTGGCCATCCCTCGGCGCTGGTAGGAGGAGATCAGACTTCCTGAGAAGTGTCCATGTTTG | 120 |
| chr11 | 70574135 | 70574255 | 788855_47188552_ENST00000601538.6,E | 1 | - | good | ATGGGCTGGCTTTGGGCCACACCTCCGCAGTCTGTGTCGGCCCATCTGGAGATTGATCAGCAGGGTAAACTGAGGCAGGGTCTAAGAGAGATTGGGCAGGGGCCGCCATGAGTGCTGTGA | 120 |
| chr11 | 70574255 | 70574375 | 788855_47188552_ENST00000601538.6,E | 1 | - | good | CCCCTACCCTGATGTTGCCTCTCCCAGTGGGGAAGGTGGGGAATTTAGTGCCCTGTGGTATGAGTTCCCAGCAGGCTGTGGGGAGTGTGGGGAAGCTCAGAAGAAAGGCCCCAAACAAGC | 120 |
| chr11 | 70574375 | 70574495 | 788855_47188552_ENST00000601538.6,E | 1 | - | good | TGAAAGAATGACTGAATGAATGAATGGGAGACATTAGGTGATCAGTCACAGTTGCAGTGCTGGGATGGAGCAGAAGCAGGCATCAAGCCAGCCTGTTCTCCTGCCTCCAAAGACCCCCAC | 120 |
| chr11 | 70574495 | 70574615 | 788855_47188552_ENST00000601538.6,E | 1 | - | good | GAGGGGACCCTTCAGGGCACGCCACCCCCATGGGCAAAGGGACAGTCACTAGAAGTCCCCTGGCAAAGTGCAAATGTGTGAAAATGGGAGGAGCCATGGCCCCAGAGTCAGATGAATGAG | 120 |
| chr11 | 70574615 | 70574735 | 788855_47188552_ENST00000601538.6,E | 1 | - | good | TGCTCCATGCTGCAGAACGGCCTGGTCCCCAGTGCACAGGGTGGATCCCATCAGAGCCAGCCCCTCCTTCTCTTCCTGCTCCGTGGTGGAGCCAGGGCTGCATGCAGAGTCAGAGAAGGG | 120 |
| chr11 | 70574735 | 70574855 | 788855_47188552_ENST00000601538.6,E | 1 | - | good | GAGTTTTGTAAGGCTGCCAACCATAATGGTTTCAGAGGGGCTTCCATGTCCTGGGTGTCCTCGCACTACAGCAAACATGTGACCTAATCCCCCAGTTCATAGACTGAGACCCCCGGTCTG | 120 |
| chr11 | 70574855 | 70574975 | 788855_47188552_ENST00000601538.6,E | 1 | - | good | TGAAGGACAGATGGGACTCATAGCGGGGCCTCTCTGCTCAGCCTCCGGGTATCATGGCAAAATGAAGAGGATGGGGAAGGAAAGCAAAGAAAGCTTGCCGGGCAGCGAGGTTTGGGGTCA | 120 |
| chr11 | 70574975 | 70575095 | 788855_47188552_ENST00000601538.6,E | 1 | - | good | GCCCAGCTTGAATGCACCGCAGAGGATCAGCTGCAATGAAGGCTTTGAAACAGAAATGTGCCCTTTGTGTGTGTGGCTTCCTGTGTGTGAGTGTGGGGATGGGGATGGTTGGGCAGGGCC | 120 |
| chr11 | 70575095 | 70575215 | 788855_47188552_ENST00000601538.6,E | 1 | - | good | GACAGTCACCGTTGGGTGACCATTCCTCTGCTGCCCGCCTGGGTGGCTTGGCCTGGACCCCAGTGTCCTTCCCACTCAGTGTGATGTCCTTGTCCCCACCCAGAGCAGGGAGCAAGGAGG | 120 |
| chr11 | 70575575 | 70575695 | 788855_47188552_ENST00000601538.6,E | 1 | - | good | GTCGACTTACTTGCAGTCTGTAAAGCGCTCACACATTGCAAAACACATGTGGAAACACCAAAGGTGGAGGCAGAGGGAATGATAAAGAAAAGCCTGGTGTCTCCTCCCCACCCCAGGGGA | 120 |
| chr11 | 70575695 | 70575815 | 788855_47188552_ENST00000601538.6,E | 1 | - | good | CAAGTTTGCCACCCCATCACCCACCACGGGCGGGGGCTATGCAGAAGCCAGTCAGCAGCCACCTCACAGCCCTAGAGTCTCCACACATCTCATCCTGCGCCCCCCACCCACCACCTGAGT | 120 |
| chr11 | 70575815 | 70575935 | 788855_47188552_ENST00000601538.6,E | 1 | - | good | GGCTGCAGCACGGCCCAGGGGATTCTGCCGTAAGGAATTAAAACCTACCAGGGCCCCAGCCTAGCTCCGAGGAACCTCACAGGAGCTCAGCATCCTCCACCCTATCCAGCCCACGCCCCC | 120 |
| chr11 | 70575935 | 70576055 | 788855_47188552_ENST00000601538.6,E | 1 | - | good | GAGGGGGGAACACATTTCCACTCTGGGAAATGGGACATGCAGCACACCGAAGCTCATTCCTGACTCTCAATGCTCAGGTGTCTCCACCAACAAATAATTACTGAGCCCCTGACACTGCCA | 120 |
| chr11 | 70576055 | 70576175 | 788855_47188552_ENST00000601538.6,E | 1 | - | good | AGGGCTTGTAAGAAGTGCTGAGGGACAGGTAGTTCCCCGTAGACACAAATGGCCTGGTGCTCAGATTCCAAGAGTCTGTCTGCCAAAATTTAAGCCCGTGAAATGCTCTCCTTGTGGGGA | 120 |

| | | | | | | | | |
|---|---|---|---|---|---|---|---|---|
| chr11 | 70576175 | 70576295 | 788855_47188552_ENST00000601538.6,E | 1 | - | good | GCCGCAGCCGCTTGCTGTCCCTCGCAGGAGGGCTCTGCACCACATCTTCACCTGCAGCTTCCGTGTCTGTACCCCCAGCCTCTCCTTTCCTCGTCTGCATCCCAGCACCTGTGACAACAA | 120 |
| chr11 | 70576655 | 70576775 | 788855_47188552_ENST00000601538.6,E | 1 | - | good | GCCTCACAGCAGCCCTGGGCATCCCAGTCTTACTGATGGGGAAACGGAGGCCCTGGGAGAGCCAGGTAGCTTCTTCAGGCTGGTAAGGGCAGAGTTAGGCCTGGGAGTGTGGGGTTTTTT | 120 |
| chr11 | 70576775 | 70576895 | 788855_47188552_ENST00000601538.6,E | 1 | - | good | GTTGCTGACTCATGGCCCCCCTCACAGTCCCTCAGAGTTGGCAGCACTACTGCCCTATTTAGTAGGGGAGAAAAGTGAGGGATAAGGGTATGATGAAACCTGTTTGCAGGGGCTTTCGTT | 120 |
| chr11 | 70576895 | 70577015 | 788855_47188552_ENST00000601538.6,E | 1 | - | good | GGCGCTCTCCTGGGCTCCCACAGTGTCCTGGCCACCCCTTCCGCTAACTGCGTGATACTCATTATTTCTAGAGAGCCACCAGCACTCACCAGGCATTTACCACCTGCCTGCTGTGTCCTT | 120 |
| chr11 | 70577015 | 70577135 | 788855_47188552_ENST00000601538.6,E | 1 | - | good | GTCACTGTATCCAGACTGAGGGAAGAGGCTGGCGTCTGGTGATGTGGGGCTGGGGACACTGGGGGAGCTCATCCCCAGGGAGGGAGACAGCGGCGTGGCCTCCGTGGCAGGTGCAGGAC | 120 |
| chr11 | 70577135 | 70577255 | 788855_47188552_ENST00000601538.6,E | 1 | - | good | TGATGATCCCTTATCATGCTTTCCAGAAAAATGCAGCTCTGAGTCACTCTTCCTTCCTCTCACCCCACGGGGATCACGAGTCCTCCTGGGGCTGTCTGGGCTCCACGGTGTCACGTCTGT | 120 |
| chr11 | 70577255 | 70577375 | 788855_47188552_ENST00000601538.6,E | 1 | - | good | CACCAAGGAGACAGGCGTGGCCACGGCCCTCCTCACTCTCCACATGGTTGGGTCATTCCTGGTGGTCTTGGGGTCACTGGAGAAATCATGGCCCCTGCCTGGAGCTCGGGGTTTTGACAC | 120 |
| chr11 | 70577375 | 70577495 | 788855_47188552_ENST00000601538.6,E | 1 | - | good | TTCTGCAAACCCGCCTGTGGCTTTTGGTCCCCATTTGCAGTGGAATTGAGGCTCAGGGAATCATGTGACTTTCCCCAGCCCAGAGATGTCTCGTGGTCCGTGGGTGCCATGAAGTGGCTC | 120 |
| chr11 | 70577495 | 70577615 | 788855_47188552_ENST00000601538.6,E | 1 | - | good | TGCCTCATCTATTTTGTGCACATCAGGAAAGGATGATTCCCCTGCAAAAGGAGTAGCAGCAGTGCCTGTGGGCCTGGGGGTGGTCACTGGCCAGGACCCCACGCCTGTTACCTGGCTCAA | 120 |
| chr11 | 70577615 | 70577735 | 788855_47188552_ENST00000601538.6,E | 1 | - | good | GCCCACCTGGCTGTGGGGTACAGTGCAGCCAGGGCAGGGAGCCACGAGTCCAGTGCCTGAATGCTCAGGTGGGGGTGGGCCTGGGTGGGAAACCAGTCCTGGCCCCTCAAATCAATCTGT | 120 |
| chr11 | 70578095 | 70578215 | 788855_47188552_ENST00000601538.6,E | 1 | - | good | CTTGGACAGGCGCCGTATCTGTAGGCCGCTTCCCTGTCTGTGCAATGGCGAGAGTACAGTGTATGAGATGCCATGGCACCGCACTAAGTCAGTGTGTCTTCGTGTGTCTGAGTCAGCTCA | 120 |
| chr11 | 70578215 | 70578335 | 788855_47188552_ENST00000601538.6,E | 1 | - | good | CAAAAGCAATCTGCGTTCCTCCTGTCTTGGGGGAGCAGGAGTCTCTGTTTCTTGCAGCCTGTTGTGGTTCTGGGAAGTACAGTGTGCCCTGCAGGCATCCTGCCGGGATTCGAGTCCTAC | 120 |
| chr11 | 70578335 | 70578455 | 788855_47188552_ENST00000601538.6,E | 1 | - | good | CGTCGTGATGCGCCTGTCCCCCTGAGACCTCGCCTTCTGTGCCAACAACATCACCATCCTCAGGTGAGCAGGGAAGGTGCCATTCCTGAAACCCACTCTTGCGGGGCTGGAATAAAAGTT | 120 |
| chr11 | 70578455 | 70578575 | 788855_47188552_ENST00000601538.6,E | 1 | - | good | AGGGGAGGAACCACTGTCAGTACCAGAATGGTGCAAATGAAGGATGTGGGCCTCCGGAGCCCGTCGTGGCCACGAGCTCTTGGGCCGTGCAATGCTGTGGTCCAGGATGGGAAGGCTGTG | 120 |
| chr11 | 70578575 | 70578695 | 788855_47188552_ENST00000601538.6,E | 1 | - | good | GATGACACCAAGCATGAGGTCACGCACCTGGCTGGTTCATGATGGTGTGAGATGAAGGTTCTGCCTGTGAGACGTGGCCCAGGGCCCCCCACCTTCTCGGGGACACGCTGTCTACTCCAC | 120 |
| chr11 | 70578695 | 70578815 | 788855_47188552_ENST00000601538.6,E | 1 | - | good | CCAAGGAAAGAGCAATCATCCCCCTTTGAGAAGAAAACCAATCTCTTGGTTATCTCTGAACTGGGCTAAAATCTTCCATGGGCCATCCTGTTGCTCCATTTCATGTAAATTAGCAGATTC | 120 |
| chr11 | 70578815 | 70578935 | 788855_47188552_ENST00000601538.6,E | 1 | - | good | CTGTAAGGGGAGGCCGCCCATCCTGTGGGGAGGCCAGGGAAGAGCTGAGAGTGACCCGGCAGATGGTCTCTGGGCTCATTGCATCCCCGACATCTAAACCATCATGAATAAGTAACATTT | 120 |
| chr11 | 70578935 | 70579055 | 788855_47188552_ENST00000601538.6,E | 1 | - | good | CTCGTTCTCACCTGCATTCATGCTCCTCTGTGTTCAGGTTCCCACGTCCCTTTATTCAGCAACAGCTGGGCAGAGGGTGGATGGGGTCGCCCGGCTGGTGCTGAGGTGTGTGGGAATGCG | 120 |
| chr11 | 70579055 | 70579175 | 788855_47188552_ENST00000601538.6,E | 1 | - | good | TTGGCCACCTTGCCAGGGGTGAGCTGGACCTGGAGACTGAGCAAGACCTAATTTAATAAATCAGAGTAGGGCCTAGGGTGTGCAGCCCTGTGTTTCTACTCCCCTGGATCCAGCACAGGC | 120 |
| chr11 | 70579175 | 70579295 | 788855_47188552_ENST00000601538.6,E | 1 | - | good | TGTTTCATGTAGTTGCTAAGGTAGCCCAGTGAGCTCACTGGCTGGAAATGATTTTCTTGAGAAATATGATGCCCCACGCCCAGCACAGGGCAGGGTTGGACTGGGGATGCATCCATGCCC | 120 |
| chr11 | 70579295 | 70579415 | 788855_47188552_ENST00000601538.6,E | 1 | - | good | AAGGGATGTAGTGCCAGCTGCAGCCTCTTCTGGCTGTCCTCAGAGGCAGCCACCATTGTCCTGTCCCACAGCTGGGCCTTTTCAGGGACACTTCATGCTCCTATGAGGGACTTTGATCTG | 120 |
| chr11 | 70579415 | 70579535 | 788855_47188552_ENST00000601538.6,E | 1 | - | good | GGAGATGCCACCTGGGGCCCAGCCTCATCCATGCCTGTCTTACATTCATACCATCACCCAAGACTGGGCTTTGCCTTATGGGGTGGCCTGCAAGCGCCACTTTCTGGGCAGAAGGAAGGG | 120 |
| chr11 | 70580135 | 70580255 | 788855_47188552_ENST00000601538.6,E | 1 | - | good | CAGCCACCCCACGCCCCTGTTTATTAATGACCAAAAGGGAGGGGCTGGTCTGCAAAGTGTCCCTTCGCAGCAGCCTGGTGGATGTCCACGGCTTTCTGATGTTCTCAGCCTTGTAGAAGT | 120 |
| chr11 | 70580255 | 70580375 | 788855_47188552_ENST00000601538.6,E | 1 | - | good | GCTGGTGGCCCCCTGGACACTGGGGCCATTTGGGGTACAGGGCTGAGCTCTCACCTTGTGGCTCATTACAGGGTCTTCAGTTGCAAAATGTAGCCAGTAAGGCTGCGCTGTGCCTCAGCC | 120 |
| chr11 | 70580375 | 70580495 | 788855_47188552_ENST00000601538.6,E | 1 | - | good | TGCAGGCCCACGCCGCTCATTGGCGGTCCAATTTCATCCTGCAGGAGGGTAAGGGACGCGATTGAGCATTCTGGAGGCTGGCACGGTCTGCAGTGACAAAGGGATGTTCAGAGACACCCT | 120 |
| chr11 | 70580495 | 70580615 | 788855_47188552_ENST00000601538.6,E | 1 | - | good | CCACTCCCTCCTCCTTCATCTTGAGCCCCCGTGGGTGCCACCAGGTGCCCACAATTTCTTCTGGCAACACCGAGCAGACCCAGACTGCCCAGACAAGGCCGGGTCCCCAGCAGCCCAGGC | 120 |
| chr11 | 70580615 | 70580735 | 788855_47188552_ENST00000601538.6,E | 1 | - | good | CCTGGCTCTGGCTGGGCAGCCAGCAGCGGGGAAGCGGCCTCAGGCCTGAAGAGTGGGAGGAAGTGGCTCCTGTAACCGCTGAGCCAGTGCCACGGCGCTGTCCTTGGTCCTGGCAAAACA | 120 |
| chr11 | 70580735 | 70580855 | 788855_47188552_ENST00000601538.6,E | 1 | - | good | CAGGGTCCCCCAGCCAAACTGGCTCCAGACTTTGAGAGCAGCACAGGCTTTGTGGTCTCAGGGTGGCATGAAGATGTGGCCCGTGGCCTGCTGGCCTCGCTGTTCCCGCCTGCCCCAGC | 120 |
| chr11 | 70580855 | 70580975 | 788855_47188552_ENST00000601538.6,E | 1 | - | good | AGCCCCTGGACCTGGTGGGCTCCAGGTCAGGGCAGAGTGACAGAGGAGTGTGGGGTGTATACCCACACGTGTGTGCTGCAGCATCCTGGGACCCACAGAGGGAACAGGAGGTGAGCCCCC | 120 |
| chr11 | 70580975 | 70581095 | 788855_47188552_ENST00000601538.6,E | 1 | - | good | TCTTGGTCCAGGTGCCTCTTGCCCCTAGATCTGGCATCCCCCCTCCCTGGTGGCTCCTCAGAAGGCCTCGCAGTGACGAACACTTAGGTGCTGATGACGGGGAAGAGCCAGAACTTTGAG | 120 |
| chr11 | 70581095 | 70581215 | 788855_47188552_ENST00000601538.6,E | 1 | - | good | GTGACTACAAGAGCTTTTCTTCCCTTGCCCTTCAAACGCTAGACTTACTTTTGCCGTCGTCCCTGATGGAGATCCTCATGGTGATTTCTGGAGAAGGGAGTGCCCTGAACCTTAATCTAT | 120 |
| chr11 | 70581215 | 70581335 | 788855_47188552_ENST00000601538.6,E | 1 | - | good | ACCAACAACCTTAATGTAACACAAAATATACCATTCAGATAGGAAGATGCCATCTTTGAGGAAGGCAGAGGCCTTTCCTAAGGTACTGGAACAGCCAGTCAGGTTTGTGGTTTTATTTCA | 120 |
| chr11 | 70581695 | 70581815 | 788855_47188552_ENST00000601538.6,E | 1 | - | good | GGGGTGAGTTCCTCCAGTTCCAGGGCTGGACTCTCTATACTCATGGAATGCTGCTGTGTGCACCAGGCCCCAACTTCAGTCTTTGCCTTCAGCTAGCTAGCTTGATTGATTGATTGATTG | 120 |
| chr11 | 70581815 | 70581935 | 788855_47188552_ENST00000601538.6,E | 1 | - | good | GCCCAAGAGAGGAGATCAGGAGCTTTGCTCTACAGGACATTAAATCATAGTCCAAAGAGATTAAGTGGCTTCCCAGGGGCCATACCGCTCACACAGCTCAGAGCTGGCAGAGTGAGGCTT | 120 |
| chr11 | 70581935 | 70582055 | 788855_47188552_ENST00000601538.6,E | 1 | - | good | ACCTGGCCCGGGATCTGTCCCCTCCCATTGACCAGCAGCTCCTAGGGGAGGCAGGATGGCCTCTGTGGGCATGAGGCTCGGGGCAGAGCTGGCAAACATTTACGGCCCCAACCCTCGTGT | 120 |
| chr11 | 70582055 | 70582175 | 788855_47188552_ENST00000601538.6,E | 1 | - | good | TTCTTTGCAAGGCCTTCTCTGACTGATGGCACCTTGGACTCTCCGCCCTGGGTCTGTCTGCTGTCTCCCCAGGCCAACCCCTCCTCTGGCCTCAGCTCTTCGGCAGGGAAAGGAGACTGG | 120 |
| chr11 | 70582175 | 70582295 | 788855_47188552_ENST00000601538.6,E | 1 | - | good | CTTGGAGGCCCCGGATGTGTGGGGACAGTTAGGTGGTCCTTTCAGCTGGTGCCAATGCCAGGCCCCTTCTCATCATGAGGATGGCGGCATTGGCCCCCTGGGGTGGAGCCCCTCTCTGCT | 120 |
| chr11 | 70582295 | 70582415 | 788855_47188552_ENST00000601538.6,E | 1 | - | good | CTAGCAGAGAGCATCTCAACCCAGCCCTCCCGTCACTTGCAGATGTGTGCATGAGCTAATTGGATCCGGCCGGGGCTGGCGGCCTCCACGGCATTCAGTCAGCAGTAGTAGAGTTAGCGG | 120 |
| chr11 | 70582415 | 70582535 | 788855_47188552_ENST00000601538.6,E | 1 | - | good | TGGTCAGATATGGACCTGAGCTTATATCTGCTGCCTGCCGCCTCTATAGGGAGCTCAGAGGGACACACTGGCTCAGGGACTCCACAGGGAAGCTGAGTGGGTCCCATCTGCCGAGCAGGC | 120 |
| chr11 | 70582535 | 70582655 | 788855_47188552_ENST00000601538.6,E | 1 | - | good | TGCATCCCCACACCCGACTGGGTGCCCACCAGGACCTGTGAAGGCCCATCGCTCCAGGGCCACCCTCAGTGAACGAGGGGAGAGGTTTGCGTATCCTCCCGCTGCCTAGCCTGCACGCAC | 120 |
| chr11 | 70582655 | 70582775 | 788855_47188552_ENST00000601538.6,E | 1 | - | good | CCAGCTTCTAGAACACCACCATAGCACACACAGATGCTGCCTGTGACACGCTGTGTTTTCGGCTGTCATCTCTGGGATAAGGCCCCTTCAGGGACCCCGACAGCATCGGAAGTGAGACCC | 120 |
| chr11 | 70582775 | 70582895 | 788855_47188552_ENST00000601538.6,E | 1 | - | good | GCTCATCAGACACGCACACCCCACCCCCATCCCCAGCCCAGACTGACTGACTCAGATCACCAGGGAGAGGGCCCAGGGTCTACATTTGTCTAAAGCCCCCCAGGTGGTTCTGAAGCTCCG | 120 |
| chr11 | 70582895 | 70583015 | 788855_47188552_ENST00000601538.6,E | 1 | - | good | GCCACAGGGCAGCACCCTGGCCCCACGAGCCTGCTGCAAGCCTCAGGAGAGTGAGCTGCACAGCAGACCCTGCCAGTGATTCCTGGGGCTTCTCAGACCTCACCAACACCTGAGGGTCCT | 120 |
| chr11 | 70583015 | 70583135 | 788855_47188552_ENST00000601538.6,E | 1 | - | good | CATGTTCTCAGCCACAGCCCAGTGCCCCTGCCCCAGGGCCAGGCACCATGCAGATTCTCCCCCAATCTTCCCTCTGTCCCCCGCTCCATGTGGCTGTCAGAGTGCTGAGCTGCACCCCCA | 120 |
| chr11 | 70583135 | 70583255 | 788855_47188552_ENST00000601538.6,E | 1 | - | good | CAGTTGGGGACATGCTGCTAGGCCTTCTGGAGTCAGCCTGGGAATCCCCCATCTTTGGAGCCATACCTGAGCCCGTAGATAGATGAAGGCATCCCCGCTCTGGCCTCCGTAACTCCCTTA | 120 |
| chr11 | 70583255 | 70583375 | 788855_47188552_ENST00000601538.6,E | 1 | - | good | CTGGCCCCTGTTCTGCGGCTGGATCTTCTGGCTCTATCCTGGCCCTGTACCCTTGCCCTCCAGGTCACCTTCTCGCCACATGCTCCCTCTGCCAGCAAGGCCTCCTCCCACCTCTTGCAT | 120 |
| chr11 | 70583375 | 70583495 | 788855_47188552_ENST00000601538.6,E | 1 | - | good | ATCGCACCTGCCAGTATTCACCGAGCACTTGCTGTGCTGTAGGCACAGGGCAGGACTGGACTCCCTCTGCCTGTGCCACAGCCCCGAAGATCAGGGCACTGTGGATGCTGCTGCCACACG | 120 |
| chr11 | 70583495 | 70583615 | 788855_47188552_ENST00000601538.6,E | 1 | - | good | ACCCGGAGTGCATGTTTTCCGTGCCATCTGGGCAGCTGGCATTTAGCTGCGCTTTCCTGGCACTGCCCTCCTTGGAGCCAGAAGTCAGCCAGCCCCTTGCAAGAATCAGAGCTCATGAAA | 120 |
| chr11 | 70583615 | 70583735 | 788855_47188552_ENST00000601538.6,E | 1 | - | good | CAGCCTCGATGGTTCATTAGGAGGCTCCTGACAGTGAGCGTGGAGTGGACAGGCTCCAGATTTTATCCTGAGCCACTCAAGATGGAGGGGGCTCAACCCCAGTGAATTTGGGATTCGATG | 120 |
| chr11 | 70583735 | 70583855 | 788855_47188552_ENST00000601538.6,E | 1 | - | good | TTTACCTGTGAGGAGCGGGCAGCCCGCAGACGTTATACCGTGGGGTGATGGTGAGATCTTGTTGGGTTGCGGAGGATCGTTCTGGCCACTGGGTGGACAAAAGACTGCAGAAGGGACACA | 120 |
| chr11 | 70583855 | 70583975 | 788855_47188552_ENST00000601538.6,E | 1 | - | good | GAGGGAAGCAGGAGGGCTGTGCAGCTGGGCTGAGCAAGCTGCAGAGAGTGGCAAGGGAGAGAGGGGAGGCATGGCTGGGCCAGGCCGTGGGAACCTGTGGGCCACTGGGCCATGCTGG | 120 |
| chr11 | 70583975 | 70584095 | 788855_47188552_ENST00000601538.6,E | 1 | - | good | GGCAGCCATCACGGCCGATAATTGAATGGCTCTGGGGGTCGGCCGGGGATGAGACCTTGAGGAGCATGCTCGCGGTGGCTGCAGAGTCATGCACGGTGGTCAGGCCATGCCCAGAGGTTA | 120 |
| chr11 | 70584095 | 70584215 | 788855_47188552_ENST00000601538.6,E | 1 | - | good | GACTCACCAGACCCGCAAGTGTGCCCTTGATCCAGGCCCCACTCTCAGAGCTGGGGACACATCAGGGGCCAACCAAGGGCAGAGTGGAGCCAAAGCGCTGGCAGGTAGACCCGAGTCACA | 120 |
| chr11 | 70584215 | 70584335 | 788855_47188552_ENST00000601538.6,E | 1 | - | good | AGAAAGAGAAAATAAAAGACACAGATGATAGGCCTGTCACACGGGATCGGAACACAGACTCAGCCCTGCATGTGATTCGAGCGCCAGGATGCCTGGCTTAGTAACTACCTCCTACCTTCG | 120 |
| chr11 | 70584695 | 70584815 | 788855_47188552_ENST00000601538.6,E | 1 | - | good | GTTTTATTATTATTCAGGTATGGCGAGACCAACGGATCAGGGGGACAGCTGCTAACTCACTCTTCCCAGGAGGTAGGGGAGGCCATGCCTTGGGGGCCATTCAGGGAAGCACAAGGTCCA | 120 |
| chr11 | 70584815 | 70584935 | 788855_47188552_ENST00000601538.6,E | 1 | - | good | AGCACTGCCACTCTTTTCAGAACATGTAGTGGAATGACACAAAACAATTGCCTTCAGGGACTTGCTTTCCTGGATCCTTTGTCAGGCAGGTCCTGTGTACCGCAGTGTTTTAAAATGTAG | 120 |
| chr11 | 70584935 | 70585055 | 788855_47188552_ENST00000601538.6,E | 1 | - | good | AATCCAGTGCATCCAAGCCCCTGGGGTGGTAGCCACGGAGCTCAGGGGCTTGTCCTCCAGCCCTTAGGCCTCTGCGTCTCCGCTGACGATCTCAGGCACCAGCATGCACCAGACCGTGCC | 120 |
| chr11 | 70585055 | 70585175 | 788855_47188552_ENST00000601538.6,E | 1 | - | good | GGTCTCCCAGGGCCCAGAGCCTGGCACCATTTCCCTTGAGCCAGAGTGGGCGTGCTGTGGCCTGAGCTGTGCCAGCTCAGAGGGGCTGCGAGGCGGTATTCACCATGCCCCCACACACAA | 120 |
| chr11 | 70585175 | 70585295 | 788855_47188552_ENST00000601538.6,E | 1 | - | good | GCTGGCTCTACTGGTCATCCCTGCCCCATTGGCAGTGCCCTTTGCATCAGCTCTTGGCTTTAGCCCCTGCCAGCCAGTGGATGGACCACCTGTGATCAGGAGCCCACCCTCGAGTCAGGC | 120 |
| chr11 | 70585295 | 70585415 | 788855_47188552_ENST00000601538.6,E | 1 | - | good | GGGCAGCTTTACCGACTGGCTGCTTCCCCATTCCTCCCTCATGCTGTCCCTGCCCCACTCTGCCAGTCTGTTCTATTGTCACTGGTCCCAAATTCTCTGCTTCAAAAATCAGCCCATCCA | 120 |
| chr11 | 70585415 | 70585535 | 788855_47188552_ENST00000601538.6,E | 1 | - | good | CCTGTCCTTATTAGGAACAAAGAACAGGGAATGATTCTGGTCAATTTGTGTTAAATGCCAGGTGAAATTGGAGAGTATTGCAGGGCAGTGTGGCCTGGGACTTAGAGCTGTGAGGACCTG | 120 |
| chr11 | 70586015 | 70586135 | 788855_47188552_ENST00000601538.6,E | 1 | - | good | ATGTTACACAACCCAGGGCAGGCACCTGTGCGATTCTCCAAGAAACTTCACCACACTCGTGGCTACTTGTTCAGTTGTCACTTGCAGGTTGCAAACTCCACAAGGTCAGGGTTGAATCGG | 120 |

| | | | | | | | | |
|---|---|---|---|---|---|---|---|---|
| chr11 | 70586135 | 70586255 | 788855_47188552_ENST00000601538.6,E | 1 | - | good | AGCTCTCTTTCCTCAAGCACTCCTCTCTCCCTTGCTGACTCCTGGGCGACATTCAGTCCTGACAGTTACATCCTAGAGAAGCCTTCCCTGACCTTCCCCCTAATTAGAATAGCCTCCACC | 120 |
| chr11 | 70586255 | 70586375 | 788855_47188552_ENST00000601538.6,E | 1 | - | good | CCAGCAAGCCTTCTAAGGCGCAAGTGATCCTTGTGCCCAGCTTTTAAAGATTCTTCCAGAGGGTCCCTGAGGAGGCACCCCATTCCCATCTTCCTGATCTGCCCCCACTGCCCACCTCAC | 120 |
| chr11 | 70586375 | 70586495 | 788855_47188552_ENST00000601538.6,E | 1 | - | good | TGACAAGCTATGCAGCACCTTGGGGCCCTCCTTGCCACTAGCCCCGGCCACGCCTTAAGAGAAGCCAATAGGTGACCCCGGCCCCAGAACCTCCCTCATTCGCACCTGCCCCTCCTGCCC | 120 |
| chr11 | 70586495 | 70586615 | 788855_47188552_ENST00000601538.6,E | 1 | - | good | ATCTGTACTGAGAGTCTGCGTCCCAGCATGTGCCCCTTGGCCAGCCATGCAGGGGCCGCCTCGGAGTTAGCTTCCGAGGTCTCTGTGAAGAGTAGTGTGTCCAGGCTGCCCAGCTCTGAG | 120 |
| chr11 | 70586615 | 70586735 | 788855_47188552_ENST00000601538.6,E | 1 | - | good | CTCGCCATTCCCCTAGGATAAGGGTCAAGAGATATAAATAAGGGAAGGAAATGCTTTTTTCAAATTGCTAAATTTCTCATTGCCAGTTGGTTTCCACAGCAACGGTGCCAGAGGCTGGCG | 120 |
| chr11 | 70586735 | 70586855 | 788855_47188552_ENST00000601538.6,E | 1 | - | good | TGGCGTCTCTAGGCTCTGCCCACTCTGGGGGGGTGAGGTACTGGTGTCCTTCCCCGTTCCGGGGAGCGGTACCCTCGAGACACCTGCCCACTCCCGGCTCCAGCTGCGAGCCGTGCTTCC | 120 |
| chr11 | 70586855 | 70586975 | 788855_47188552_ENST00000601538.6,E | 1 | - | good | CAATCTTCGGAGTGACGTTTGAGGTCGTCTCTGGGAGAGGAGTGAGGTGGGGGAGGGAAGAGGGAGACTTTGCCTTTTCTGCATTATTTTCATCTTTCCCAGGAGTGACGTATTCTGGGA | 120 |
| chr11 | 70586975 | 70587095 | 788855_47188552_ENST00000601538.6,E | 1 | - | good | GGAGTGGGCTGCAGAGGAGCCATGGGCTAGAGTGTGACCCCATTTTGTTCAACAAGAGCACTGGCATGTGTCTGTATTCCTGCATGTGCCCAGCGGTGCATGGAGAATGACCTCGAATAA | 120 |
| chr11 | 70587095 | 70587215 | 788855_47188552_ENST00000601538.6,E | 1 | - | good | TCTGAGAGAACTGCTTTCATAAAGCCCGGCCCAGCTCGCCGTGGAACTCTGTACGGCAGTGGAAGTGACGCAACAACCCTAGTGTTGTGGTCGGGAGAGCAACACAGGCCCTCAGCAAAG | 120 |
| chr11 | 70587215 | 70587335 | 788855_47188552_ENST00000601538.6,E | 1 | - | good | GGCCTGTCTAGTTAGGGGTCCTGTTGGCCAGGCTGCCTTTAGAGAAAGCATTTAGAGGAGGGACTGGCAAGTGCATTGCTTCATTTCTTGGTGATTCATGTGGCGTGGCATCCCTGGACA | 120 |
| chr11 | 70587335 | 70587455 | 788855_47188552_ENST00000601538.6,E | 1 | - | good | AGAATGCCCCAGACCAATTTCGTTTTTCCCTGACTCTCATGTTTACAGATGTACCCGTGGTGCCGCGACAGGCTCCTTTCTCCAGATCGCTCTCCCCGCTTCTCCCCTCTGCCCCATTAG | 120 |
| chr11 | 70587455 | 70587575 | 788855_47188552_ENST00000601538.6,E | 1 | - | good | TCTCTGAATGTCGGACTCGCTGTTCCATACCATTTTGTAACCTACCCTTTTTTTCTGAACATGTGGGGAATATTTTTTCCTGTCGGTAAGCACTGTCATCCATGCCTTTGATGGCTGCGT | 120 |
| chr11 | 70590815 | 70590935 | 788855_47188552_ENST00000601538.6,E | 1 | - | good | AGGCATGAAAGCCTTGGCCCCAAGAGGAGCCTCGTTAACTCCAGGATGACTCTCAGCTCTTGTTTTCTGCACAGGCCGACCTGGGGATATTTGTGGGTTCAGCCCCAGACCACCACAGGA | 120 |
| chr11 | 70590935 | 70591055 | 788855_47188552_ENST00000601538.6,E | 1 | - | good | CCTTCCTTTTTCTGACCCATTTGTAGACCTTCCCTTTCCCTCCATTTCTTTGTCTTTTCCTTTTTTAAAAGATAACATCTACTGGGGAGAGGGATACTAATTATAAGCAATGAATCTGCC | 120 |
| chr11 | 70591415 | 70591535 | 788855_47188552_ENST00000601538.6,E | 1 | - | good | GAAAAGCAAATGTTGTATCTCCAGAAGTTGACAGCGTTTGAGTCCAGGGCAATGCGCTTACTCCTGTAATTTATCTGCACAGATTGGCTCTGTGGATTCCGGCTCCTTCTTCTATCTTTT | 120 |
| chr11 | 70591535 | 70591655 | 788855_47188552_ENST00000601538.6,E | 1 | - | good | ATGTTTTCACATGGTCTTCCTGTCCCTTTTTAAGTTGTTGAAATGTGTTATGTTTCCTTGGAGAAGACAAAAATGGAGCTATGATGTTGCTTTTTTTTTCTTTTTCTTTTTCTTTTTTAA | 120 |
| chr11 | 70592135 | 70592255 | 788855_47188552_ENST00000601538.6,E | 1 | - | good | GCTGCCTCTGTCTTTCCCCTGTCCCCTGCGTGCAGAGGGAAGACGGAGCAGGTAGAGTGAGGAGGCCGGCGGTCAGCACACATCAATTCACAGCCTGAGCCACAGGAGAGTATTAGAGGC | 120 |
| chr11 | 70592255 | 70592375 | 788855_47188552_ENST00000601538.6,E | 1 | - | good | TCTCACTCCGAGTGGGAGCAGACTGCATCTCCTCTCGGACTGACCCAAAGACAGTGCCAGCCTCCCCAGGCAGGACCCAGTCCTCACCTGCCCTGGCTGTCCCCTGGGGGACCAGCGACA | 120 |
| chr11 | 70592375 | 70592495 | 788855_47188552_ENST00000601538.6,E | 1 | - | good | TGCTGGGATGCAGAGCTGGGTGTGTGCTGCTTGAGTCTGTGTTGGCTGCATCAGATTGGTGGCCACGTCCCACCCAAAAGCCTGCGTGGAAGCTCCACCCGAGTCAGCAGAAGTCCTGCC | 120 |
| chr11 | 70592495 | 70592615 | 788855_47188552_ENST00000601538.6,E | 1 | - | good | CAACGGGGGGTGCAGACAGAGGGACAGAGGCTGCCCAAGACCCGCCTTGGGCAGAAGGGGGGCCACCGAGATGCCGCACGGCCTTCAGAAAGACTCCCCTGGCCGTGGCTGGAGGGGGT | 120 |
| chr11 | 70592615 | 70592735 | 788855_47188552_ENST00000601538.6,E | 1 | - | good | ACTGTTGCTTAGATGATGGGCCGAAATGGGAGGGGGCTGGAGGAGGGAGTTGGCCTAACCACACAGCCCTTGTTCTAAGAGGTGAAGCAGGCCAGGACCAGGGCTGCACACATTTGCCCT | 120 |
| chr11 | 70592855 | 70592975 | 788855_47188552_ENST00000601538.6,E | 1 | - | good | TTCGGCGAGGGTTTGCTGGACATCTTGGGGTTGCATCACGTAGGTGACAGGCCTGGGGGCAGGGCCGAGGAGGCTGGAAACCCTGAGTGCTACCCCAGGCGAGGTGGGGATGTCTGGGTG | 120 |
| chr11 | 70592975 | 70593095 | 788855_47188552_ENST00000601538.6,E | 1 | - | good | GGGCAGCCACAGGCCTTCCCCGAACTATGGGGCCAGACAGACGTGGGAAGTTTTTCATGCGAGGCCAGGGCTGGGACCCACCCCTCCGATAGCTGCAGGCTCCCTTCGTGAGTCCCTCCT | 120 |
| chr11 | 70593095 | 70593215 | 788855_47188552_ENST00000601538.6,E | 1 | - | good | ATAGAATATAATAAGTAGAGGAGCTACACCTAGCCAAAAATGCCACCAAAGTCCTACGCGTGCTGGGACAGTGTGCGTTCATGTTGGAAATGGTGCTGGGAGGACTTTGACAGGCAGCAG | 120 |
| chr11 | 70593215 | 70593335 | 788855_47188552_ENST00000601538.6,E | 1 | - | good | TTACTATATCCTACCTTCATTCACTGTTAAATGCAACATTCATTTAAAAACAGGCTGTAGGTTAGATTTTTCTCAATGCAGCAAGAGTTCTCAGAACTGCGTGGAGACTGCTGAGAAATT | 120 |
| chr11 | 70593335 | 70593455 | 788855_47188552_ENST00000601538.6,E | 1 | - | good | CTCCAGGACCACTTTGGTGTAAGAGCTCTTGAGCATTCCAGGATCCAGATCTAGAAACAACATTTCACAGCCCCAGAATAAAAAAAGGCAGCAAGAATGTTCAGATTAGTAAATATTTAA | 120 |
| chr11 | 70593455 | 70593575 | 788855_47188552_ENST00000601538.6,E | 1 | - | good | CTGCCACTCTAGTATTTTCTGACACACCTGCTCAGACGTCCCGTGGGGCAGGCAGGCGAGGTAGCCACCATGCCCAGGTGGGGAAACTGAGGCTGAGTCTGATGAAACAGCACCAGCCTC | 120 |
| chr11 | 70593575 | 70593695 | 788855_47188552_ENST00000601538.6,E | 1 | - | good | GCCCCAGGCCTCCTGACACTGAATGCCCTTTAGCTCACTCTTCACTCCGTTCATTCACTCAACAGGAGTTGGTTGTGGACAGAGCTCCCTGGGGGTGTTGTAGGTGTGGGAGGTGCCTTC | 120 |
| chr11 | 70593695 | 70593815 | 788855_47188552_ENST00000601538.6,E | 1 | - | good | ATAACAGTTGCATAGGTCCCTCCAGCTGAGGCCAAAGGGATCCAGATTGGAGTCCTTGTCACTCGAGAAGGTGGACACTCCGTTTGCCCACAAAGTCAGGAATGTAACCTGCCCAACCCT | 120 |
| chr11 | 70593935 | 70594055 | 788855_47188552_ENST00000601538.6,E | 1 | - | good | CAGTGCCTAGGAAGAGGCCCTGGAGGACATCTGGTCCACCCTTCTTTCTGCAAGCCCCTCCCTGGCACTTGTATCCCTTCCTCACACCCTCCACGTGGGTTCACTCAGCATCTGACACCA | 120 |
| chr11 | 70594175 | 70594295 | 788855_47188552_ENST00000601538.6,E | 1 | - | good | CTCACGTAACGCGATCTTTCGTGTTTGGCATGAAGTCGCTGAGCATCCCCCACGCCCCGGCATGTCAAAGCTGCGGTCCTTTTTGTGGCCGAATCATATCCAATGGTGTGGGTAGACCAC | 120 |
| chr11 | 70594295 | 70594415 | 788855_47188552_ENST00000601538.6,E | 1 | - | good | TTTTCATATACGTGGGTTCCTCAGGGCCAACTGTGTGACTTTTGAATATTCTTGGATTTGGGTCTGTGCTGGGGTCCTGGGACCGATCTCCAGAGTGCACCAAGGGCAACCGTGTGTGGA | 120 |
| chr11 | 70594535 | 70594655 | 788855_47188552_ENST00000601538.6,E | 1 | - | good | ACCTTAGAATGAAACCCTGGACCCAGTAAGCACCTGAGCCCCATTGCCCCTGCCCGCCCTCAGCCTGGCAACCACCAAGCTACTTCTCCTCTAAAAGTGTGTCTATTCCAGGCTTTTCAT | 120 |
| chr11 | 70594655 | 70594775 | 788855_47188552_ENST00000601538.6,E | 1 | - | good | GAATGCCCATTAGCTCACTCTTCACTCCATTCATTCACTCAGTGTGTATAATCAGTGACATTTCGAGGTTCACGGTGTTGTTCCACCACAACTTTTATCCAGTTCCCAGTCATTTTCCTC | 120 |
| chr11 | 70594895 | 70595015 | 788855_47188552_ENST00000601538.6,E | 1 | - | good | TGGGCTCAGAGAGACCCGGCTCCCGGCTCCCGGCCCTGAAGCCCTTCTCTCATCATTTGTTCAGGGTCACTTTTTTTGAAATTTTACTGAGGTAAAATGAACCACGAACATGGGGTCCCT | 120 |
| chr11 | 70595015 | 70595135 | 788855_47188552_ENST00000601538.6,E | 1 | - | good | GCAAGCCTTGGGCCCATTGGCAGGTGAGGGCTGAGAGGTGGGGGGCCCTGCAGGCTGCAGAAACCACCTTGGGGAATGAGGAAGAATGAGGCCCACCCAGATGTGAAGCCTGCCAGGAAA | 120 |
| chr11 | 70595135 | 70595255 | 788855_47188552_ENST00000601538.6,E | 1 | - | good | GGCCGGGCTGCACCCTTCAAACTGCCCTCCTGGCCAAAGCTGCCTGGAGGGCCACCCCCCACTCCATCCGGTGGCCGATGTGAGGGGCCAGCCTGGGTTTTTCTTGTACCCTGGCAGAGA | 120 |
| chr11 | 70595255 | 70595375 | 788855_47188552_ENST00000601538.6,E | 1 | - | good | CAAGTTGCTGCAGATGGTGCTGCGTATTTTTAACAAACGCAGCGGCCTCGGCTGGCTGCGAGGGGGCGCTTGGAGGGTGGCACTGGCACCGGGGCTTCCTGAGAGGGGGTCGGGGTGGAC | 120 |
| chr11 | 70595375 | 70595495 | 788855_47188552_ENST00000601538.6,E | 1 | - | good | TTTTCAGAGAGAGTGAGCCAAGCTGGGTGAGCCTCATAGGCCTCAGGAGCCTCATCTCAGGGATAAAAACTCGACTTTAAAAGAAGCCTCAGTCGCTTTATAAATAAATTTATTAAATTC | 120 |
| chr11 | 70595495 | 70595615 | 788855_47188552_ENST00000601538.6,E | 1 | - | good | CCGTGGAAGCCCTGTGGTTCTCAACTTTCCCTCACCTCTCCTGTGACAGGGGCTGTAGGCTGGGCTCCAGGCCACAGTTCACCGTCGCCTTCGCAGAGGGTTGTCTGGGCCAGCGTGTGG | 120 |
| chr11 | 70595615 | 70595735 | 788855_47188552_ENST00000601538.6,E | 1 | - | good | CTTCCCCCGCTACCCGGGGCGGTGGGGCCTCCCAGCAGCTGGGGAAGGGCTGGCAAGCCCCTTGTGGGTAGACACGAATTTTGGGCAGATGCCATTCAACACGGTCCAGGGGAACCTGAG | 120 |
| chr11 | 70595735 | 70595855 | 788855_47188552_ENST00000601538.6,E | 1 | - | good | GCAAAGACCCCGTGTCCAAATCAGGCCACAATCCGCAGTTTGAGAGAAGGGTTTTGTGGCTGGGAGCTGGGTCTCTCTGAGTCATTTCCTGGTGAAGTCCACTTCTGGGTGAGCCTCAAT | 120 |
| chr11 | 70595975 | 70596095 | 788855_47188552_ENST00000601538.6,E | 1 | - | good | TCAGCTGGGTGGACTCCCTCTGGAGGCTCTTGGGGAGGGGCCTTCTTGCCTTTCCCGTATCCTGGCGAGGTGCCCCTGATGCCTTGGCGTGGAGGTGGTCACCGCAATCTCTGCTTCATC | 120 |
| chr11 | 70596095 | 70596215 | 788855_47188552_ENST00000601538.6,E | 1 | - | good | AGCAGGCGGGGAAGGGGGCTCGCCTGGGCTGCCAAACAAATCACCACACGCAGGACAGCTCAGCACCGCAGAGATTGATCCTCCTGCAGCTCTAGAGGCCAGAATTCTGAAACCAAGGTG | 120 |
| chr11 | 70596215 | 70596335 | 788855_47188552_ENST00000601538.6,E | 1 | - | good | CCCCTCCCCTCTTGGTTCCCAGGCGCCAGGTCTGCCTGTGGGAATTGAAGTCAGCAGGAGGCACCATGGCACTGGGAAGCGTCCGGGCTCTGGCCCAGCCCTGGCCCAGGTGCCCCACAG | 120 |
| chr11 | 70596335 | 70596455 | 788855_47188552_ENST00000601538.6,E | 1 | - | good | GGGTAGGGGGTCCTCAGAGCTCTTTCCCTTTGCCCCACCCACCCTGGCCTGTCCCTTGGAAATACCCTGGACAGTTTCTCTGCCTGGTGCCTGTGGCTCCTTCCGGAACTGGTGCCCATG | 120 |
| chr11 | 70596455 | 70596575 | 788855_47188552_ENST00000601538.6,E | 1 | - | good | GCCCTCAGTGGTTTCTACTTCAGGAAAGTCCTGCAGTCCTCTTGGCATAGCAAGGCCTGGTGGGGGTGGGTCCCAGACCAGGACCCTGATGCCCCTTCCAACCCCAGAACCCTAGAACCC | 120 |
| chr11 | 70596575 | 70596695 | 788855_47188552_ENST00000601538.6,E | 1 | - | good | GGAAGGCAGGAAGGAAGGAAGGCCTGAGGGAAGGGGCAGCCACCAGTCCCTGCGCTGCACTGGCCCTGCTGTCCTGGCAGTTGCCCTGCTGGGCCTGCCGCCCCCCCGCATCAGGATTCT | 120 |
| chr11 | 70596695 | 70596815 | 788855_47188552_ENST00000601538.6,E | 1 | - | good | TCCTCACACTGGGGGAGAGCACAGCTCCAGACAGCCCCTGAGTGTACCCCACCAAAGTGCCCGTGACCCCGGAGAGAGTGGACTGTTGTGGTCTAGAAGGTGAGCCTTGCAGCAGCCCAG | 120 |
| chr11 | 70596815 | 70596935 | 788855_47188552_ENST00000601538.6,E | 1 | - | good | ATAGACCATAGGAGACCTCAGTCATGGGAAAGGCCCCTTGCCTCCAAGCCTGTGCCCATGTTGTGCCACAGCCCGACGCACTGTCCTCTGTTCTCTCCTCCTCCGAGAAGCCTCGATGAT | 120 |
| chr11 | 70597175 | 70597295 | 788855_47188552_ENST00000601538.6,E | 1 | - | good | GGGTAGTGGAAGGGAGTAGCTCAAAGGCGGGGGGCAGTGGGTGGAGAGAGAATGAGGGACAGGGAGGAGTTTCCCCTGGCGGCGGGTGCTGAGCCTGGCATGAGTGCCAGGAGTAATC | 120 |
| chr11 | 70597295 | 70597415 | 788855_47188552_ENST00000601538.6,E | 1 | - | good | GTGAGCCCCAAATCCATGGTGTGGCAAAGAGGCAACCCCTTCCCAGGGCTCCTCCAGGTTTCCTGTGTCTACTAAGTACTTGCTGCTTCTTGGCATCTCCACAGTTATTATGGGGTGCCT | 120 |
| chr11 | 70597415 | 70597535 | 788855_47188552_ENST00000601538.6,E | 1 | - | good | AGGTAAAGAAAAATCTCTTATCCTCTCTACCCTCTCTATCCTCTCTACCCTCTCTATCCTCTCTACCCTCTTATCTTACCGCCGCCAGCTCCACTGGCCGCCTCTCAGGGGGCTTTCCCT | 120 |
| chr11 | 70597895 | 70598015 | 788855_47188552_ENST00000601538.6,E | 1 | - | good | GAAGCACAGCTTCCAGGAGCTGCTGTCAGACTTGGTTGGTGCAGGGGTGCCCGCAAGGCCTCTCTCAGGGGTTGCATCTCAGGTGGGGTCTGAAGGGGTCTGGATCCTGCCACTCTGGTG | 120 |
| chr11 | 70598015 | 70598135 | 788855_47188552_ENST00000601538.6,E | 1 | - | good | GCACGTTCCCCTGCTGGCTCTGGACTCCAGCGCATGGCCAGAGTTCTGCAGCCCACTCCACCCCTGAGTGTCCAGAGGGGCAGTTAGCGGACAGCATTGCCCTGTGTGCTGTGCAGAGAC | 120 |
| chr11 | 70598135 | 70598255 | 788855_47188552_ENST00000601538.6,E | 1 | - | good | CTCCAGCCCAACTTGTGTGGCCCTGAGGTCATTCGACAAATGACCTCAGTCATTCGTCCCGCCCCATCGCCTGCAGCTGCAAACGGGCTCCATCTGCCCCTCTATGGGACCCTGTCTACG | 120 |
| chr11 | 70598255 | 70598375 | 788855_47188552_ENST00000601538.6,E | 1 | - | good | GTGTCATTTGTGCTAACTTCAGATCCATTTTCAGCTCCTTCCCAGATAGTCAAGGGCTCCCTGCCAAGCCTCCATTTCCAGCCTCTTCCTCTATTCTTCTCCACCTTCCAATACCCTTCT | 120 |
| chr11 | 70598375 | 70598495 | 788855_47188552_ENST00000601538.6,E | 1 | - | good | GATGGCTTGACCCTGCTCAGAAAGCCATCTGGGCCTCATGTTTTCTTTATGAAAGGATTTGCAACTTCTGGTTCAATCTCTTTCAAGGTTACAGAACTCTTCGAGTTTTTATTTTTTCTT | 120 |
| chr11 | 70598495 | 70598615 | 788855_47188552_ENST00000601538.6,E | 1 | - | good | TCTGAGGCCATTCTTTTCCAACCGGAGTCAAGCTCTTGCCAGTCTCCCAAGAAGCTGGGGAGTGTTCTCTCTTTTTCTCCAGAAGACATTGTTCGGGTTAGAATTATGTGTTCCTTCTGC | 120 |
| chr11 | 70598735 | 70598855 | 788855_47188552_ENST00000601538.6,E | 1 | - | good | GCGGATAATTAATTTCCTCAAGTCCTTTTTTGTATCACTTGCAAAAATTACATGGCTTTTCTTTGCTAATGAATTGTATTAATTGATGTTTCTGTTAGATCAGTCTTAGCTTCCTAAGAT | 120 |
| chr11 | 70598855 | 70598975 | 788855_47188552_ENST00000601538.6,E | 1 | - | good | AAAAAGCTTTCAAAATTTCTGTTGACTCTAGTCTTTTTTTTTTTTTTTTGACATGCAGCTTATCAGGTTAAAGTATTTTATTCTTTTCCTAGTGTGCAAGAGATTTTTATTTAATTGCAA | 120 |
| chr11 | 70598975 | 70599095 | 788855_47188552_ENST00000601538.6,E | 1 | - | good | GACTGATTATCTTTTTTCTTCCTTTCTCTTTTTTTCTCCGCTAGCTAGGACCTGCAATAAAATGCTGAATAGAAGTGGGAATAGTGGGAATACCTGCTTTGTTCTTGATCTTTAAAAAAA | 120 |

| | | | | | | | | |
|---|---|---|---|---|---|---|---|---|
| chr11 | 70599095 | 70599215 | 788855_47188552_ENST00000601538.6,E | 1 | - | good | TTTCAGAATTGCATTTGCTGATTCTTGCTGAAATGTAGAAATATGGCTGGCTTTGGTATATTGATTTTTATTTTATCCAGCCACTTTGCTGAATTCTCTTATTAATTCTCATCTTTTATC | 120 |
| chr11 | 70599215 | 70599335 | 788855_47188552_ENST00000601538.6,E | 1 | - | good | GCGCCCTGCCTACATTTCTCTTTCAAACTGGTTTTTTAAGTATCTCCACAAAGAGCTTCTGCAAAACCTTTTTTAGATTTATTCCTAGGTAGTTTCACTGATGTCACTTCATGATATATT | 120 |
| chr11 | 70600535 | 70600655 | 788855_47188552_ENST00000601538.6,E | 1 | - | good | ACCAAACTTCCACATGGATGCAGGTCTGTGGTGTGGCTGTCCGTTGGTCCATTTGTCTATCTGAGCACCACCTTCATTACCTCTGAATATTTCTCCCCTTCACCCCCAACCCACCCACCT | 120 |
| chr11 | 70600655 | 70600775 | 788855_47188552_ENST00000601538.6,E | 1 | - | good | TGTGTTATGTGAGGCATGGATCTAATTTCACTTTTTCCCCTGAGGACACCCCATCCTTCCAGCAGCATTGGTGGAATAAAACACTTTTCTCCAAGATCTTCCGTGCTGGAGGCGTCATAA | 120 |
| chr11 | 70600895 | 70601015 | 788855_47188552_ENST00000601538.6,E | 1 | - | good | TTCTCTTTACTAATCTTTAGCTTGTCTTTTCATTCTTTTATGACAAGTCATTTCTAATTTAATGTAGATGAATTTATCAAACTTTCCTTCAGAGTTTGTGATTTTTCTGTCTTATTTAAG | 120 |
| chr11 | 70602335 | 70602455 | 788855_47188552_ENST00000601538.6,E | 1 | - | good | TGAGCACTCCTGCGTATTCTCCTGATGTCTGAGTGTAGGTGTTAGGAGAGCAGCTGCTAGATCACTAGGAGGCAAGTGTTCAGCTCTACCACGTGGAGGCCGAGACCTCTCTGCAGGGAG | 120 |
| chr11 | 70602455 | 70602575 | 788855_47188552_ENST00000601538.6,E | 1 | - | good | TCGTGGAGGTGGGAATCTGGGGTCTGGTGTTGTGACCCGTCCCATCACTGATGTGTCTGTTCTCTTGCTGAGGGACGTGTGACCCCCTCCTGGTTTGGGGGGCTGTAAACAGAGGAACTC | 120 |
| chr11 | 70602575 | 70602695 | 788855_47188552_ENST00000601538.6,E | 1 | - | good | GATTTTTCTGCATCAAAATGTCCAACTATTCAGTGGGATAAATGCAGAGAGTAAATACAGGCGTTGCCACCAGTATAATTACCGAAAAGAAGAGGATGAAATGTTTTTGTTTCCAGAGCC | 120 |
| chr11 | 70602695 | 70602815 | 788855_47188552_ENST00000601538.6,E | 1 | - | good | TTAATATTGTAAAATGTAATGTTATACAATTTATACTATAAAATAAATTTCAGAGGATGTAACATTTTATGTATTTTATATATAATTCCAGTCCAATCATGTCATGTAATCAAGCACACT | 120 |
| chr11 | 70602815 | 70602935 | 788855_47188552_ENST00000601538.6,E | 1 | - | good | TTTATTCATTTTCCCACCTGTGTAGGACTCTATCATATGATAGACAGCTTCTTAGTCGCTATCCAAAGCCTGTGGGTCCACGTGCATTTCAGAATAAACATACATCATGCATGTATTTGT | 120 |
| chr11 | 70602935 | 70603055 | 788855_47188552_ENST00000601538.6,E | 1 | - | good | AACGTTTACCTGTTTTCCAGCTTTCTGTAAATGGACATTTGCCACCTGCATTCTTTGCCGACCTCTGCCACGGGCCAGCAGTATGCTCTGGGGTTCTGCCTCCCCGGTGTGTGTGTCTCG | 120 |
| chr11 | 70603055 | 70603175 | 788855_47188552_ENST00000601538.6,E | 1 | - | good | CTGGCCTGAGCCAAGGGACACCGTGCAGAGCCAGTAGGGCTGTGAGGGGTCCCCCACCACCGCTTCCTCAGGATTTTCCTTGGGGTTTTACCACCTGTGTTGATATCACTAAACAAAGCT | 120 |
| chr11 | 70603175 | 70603295 | 788855_47188552_ENST00000601538.6,E | 1 | - | good | CCACAACTGCTCATGGGGCCTGAGCCTCGGGCTCAACTGCCATCTGGGCCTGGACCCATCTGGTGTACATGGCCCATGGTTGGCGGTTTCTCCCCATGTCATGGAAGCATCACGGAGGTG | 120 |
| chr11 | 70603295 | 70603415 | 788855_47188552_ENST00000601538.6,E | 1 | - | good | TTCTCCTTCGGGCAGCTGGGAAGATGGGCTGCTGCCCCAGAGGAGGAAGGGAAGCCTGGAGGAGGAGCGGTGATGGCTGTGGCTTGGTGTTGGTCTCAGGACGGCAGCCAAGCTTCCTGC | 120 |
| chr11 | 70603415 | 70603535 | 788855_47188552_ENST00000601538.6,E | 1 | - | good | AAGAATGATTTGTGTGCCTACTGAGCAAAGGTTCTGCTGGACTCTGGGTTTACTGCAGCTGTGGACCAGCTGCTGGACAGGGTCAGGGTCAGGATGTGGAGGGACACTGGGGATATGCCT | 120 |
| chr11 | 70603535 | 70603655 | 788855_47188552_ENST00000601538.6,E | 1 | - | good | GGAGGAGGCACCTTCTCCGAGGTCCCAGCTCAGTACCTCACTAGCTGTGTGGCCTGAAGCAAGTCACTCAACCCTCTGTTCCTTGGTGTCAGTGACACATTATTAGTGGCTCACCTGCTC | 120 |
| chr11 | 70603655 | 70603775 | 788855_47188552_ENST00000601538.6,E | 1 | - | good | GCAAATTAGGCTGAGAAGGGGGACTGTGGGATTCTCCTGGCGCCTCTTGAGTTTTTCTGAAAGCACCAAGGCTGTTGGGAGTGGGGAGCAGGACTGGCCCAGGAGAGGGCTGCCTAGCAT | 120 |
| chr11 | 70603775 | 70603895 | 788855_47188552_ENST00000601538.6,E | 1 | - | good | AAGCACCATCTGCTCAGCATTCCTGCGGCCACCACCAGGGCATCGGGCAGGTCTCTGCCTGTCCCCTGGGCAGGGGATGGATCCTGATGAGTGGAGAGCTCCACAGACATTTCCTTGCCT | 120 |
| chr11 | 70603895 | 70604015 | 788855_47188552_ENST00000601538.6,E | 1 | - | good | AGCTAACATTTCATGGGCCCCTCGATGTTACTTTTGCATTGCTGCAGTGCCTGTGGTTAAATATTTGCATAAATAATTTTATCGGGTCGTTCTGCTCTCCTCACCCCACTTCTGGGTGCC | 120 |
| chr11 | 70604015 | 70604135 | 788855_47188552_ENST00000601538.6,E | 1 | - | good | AGACAGAGGGGCTGTCCCTCCCTCCAGGCTGCTCCCTCCAAGGCACCGTCTCCTCCCCAAACCCTCCCGCCCTCTCTGGCTTGCTGCTGGCTCTGCACACAGGCACTTGGGGCGGCAGGC | 120 |
| chr11 | 70604135 | 70604255 | 788855_47188552_ENST00000601538.6,E | 1 | - | good | GTGTCTCCCAGTTTTGCTCACCATGGAAATGCCCCACCATGGGTGAGAGAGCCCTTCATGGGTGTTTATTAAAGGAGGGAGGAGAAGGGAGACTCTGCCTGAGCAGCTGGGCCTGCGCAG | 120 |
| chr11 | 70604255 | 70604375 | 788855_47188552_ENST00000601538.6,E | 1 | - | good | TGCCCCACAGTTCCCATCTGCCATGTGCAGATCTGCCAGAGAGGCTGGGAGGAGCAGGGAGCGCAGGGTAGCCCCCCACGGGCATCCAGCTTCACTCTCGTGCTCTTGCACCGCTCACTT | 120 |
| chr11 | 70604375 | 70604495 | 788855_47188552_ENST00000601538.6,E | 1 | - | good | GCCCACTGCAGCCCAGAGCACCAGCACCACCTCTCCCAGGCCACCCCCTGGCCCTCCTGGACTCACTGTCCCAACCATTCAGAGGGCAGAGGGTTCTGGAGGGTCCTCCACGTAGCTGTG | 120 |
| chr11 | 70604495 | 70604615 | 788855_47188552_ENST00000601538.6,E | 1 | - | good | ACTGCTTTCAGTGCGGTCCTCATTGTCTCTACACAGCAACCACTCACCAGGAGGTCTCGGGTCAGGCGCCTGCCCTCCCCAAGCCGAGCGTTTATGTCTGTAAACTGGCGTGATCATAGT | 120 |
| chr11 | 70604615 | 70604735 | 788855_47188552_ENST00000601538.6,E | 1 | - | good | TATGAAAAGCACTGCGTATGCTTGCGGCATGCCGCTATGCTGTATGTAGAATGCATGTTGTATGTATACATGTGTAGTATATCAACGCATCATGTATGACATTGGGGAAGCTTTGTTGCC | 120 |
| chr11 | 70604735 | 70604855 | 788855_47188552_ENST00000601538.6,E | 1 | - | good | GCTTCCGTCCCATTGCCACCCTCAGCCCTTCTGTGCCTCAGTTTGTCATCTGTGAAATGGGGTGAGGATCATATGACTGACCCCCTTCAGGGGCCATTGTGAGACTTCAGTGGTTTTGTT | 120 |
| chr11 | 70604855 | 70604975 | 788855_47188552_ENST00000601538.6,E | 1 | - | good | CTGAAACCGCGGAACCTGCCTTGTGACTCACACGCTGCCCACACACCACTGCCCTGTGTCCAGCACCCTTGTGTGGCCAGTGCAGATGGATTGGGAGGGGGCCAGAGCCAGGCCACTGGG | 120 |
| chr11 | 70604975 | 70605095 | 788855_47188552_ENST00000601538.6,E | 1 | - | good | GAGCCTCCCATCCTCGGAGGGCTGCTGCAGGCCCCTGGCCCTGCCCCAGGCTGCCTGACCAAGGCCCATCTGGTCCTGACTGTAGAGCCTGGTGCTGCTGCTGGTGAATCCCGATGGGCC | 120 |
| chr11 | 70605095 | 70605215 | 788855_47188552_ENST00000601538.6,E | 1 | - | good | AGAGGCATGTTTACTTTAAGATTTGGCATGACAGGGGGCTTTTTTATGTAGAAAGACACGTGGCCCTGTTAAAAAGCGCCATCCCAGGCTCAAGCCCTGCCCTTCACTGAGGATTCTCA | 120 |
| chr11 | 70605215 | 70605335 | 788855_47188552_ENST00000601538.6,E | 1 | - | good | GCCAGCCAGGCTGTCTCCGGGCTGGGCTGCAGGAGGCTTCACTTCTCACTCTGGGTGTTCCCACCCCCGGGGCTCACAACGATGGAGTGGCCAGGCCTAGCAGGGAGAGCATCTTCCAGA | 120 |
| chr11 | 70605335 | 70605455 | 788855_47188552_ENST00000601538.6,E | 1 | - | good | AGTGTGGAGTGTCCAGCCTAAACATGAGTGGTCACAGGGGGGCTTCCTGCCAAGGGCCCTTTCCTAGCCACTGAGGGTGCGCCTGTCGCGGGCTCACCCGCTTGCTGCCTTCTGGGCGTG | 120 |
| chr11 | 70605455 | 70605575 | 788855_47188552_ENST00000601538.6,E | 1 | - | good | TAAAAATAAGGCCACCGGAGGCAGCCGGCCATTAAGAAATCTGCAGGAGCTCACATGGCTCTGTGGTCCCAGCGTGCTAATCGCTGGAGGGTGAGAGACCAGCTGCATTAGGAAAATGCA | 120 |
| chr11 | 70605575 | 70605695 | 788855_47188552_ENST00000601538.6,E | 1 | - | good | CAGTTGAGCTCTGCGTTCCTCGTCCCCCTCCGCCTCCTTGGAGAGAGTTCGATATGGACATGCTCCCCGTGCGCCTGATACATCTGAGCTTCACTGAGCGGTCAGTAGAGTGGAAATGGC | 120 |
| chr11 | 70605695 | 70605815 | 788855_47188552_ENST00000601538.6,E | 1 | - | good | CCTGTTGCCGTGCAGTTTCCTCGGGTTTTCTTTGCTCTGGTTCCAGGAGGAAAACATCAAGAAATGGGTTGGAAATCTGCATCTGTAACTCAATTTGTGCAATGAGCCCAGGGAAACGTG | 120 |
| chr11 | 70605815 | 70605935 | 788855_47188552_ENST00000601538.6,E | 1 | - | good | TCTAGAAGGGTCATTTAAATTTGTGCTTCCTCCTTCCTCCTTCCTCGCTGGCTTCAGGCCCCTAAATCCAGCTCTAGATAAGAAAGTGGTCTTGCTGCCAGCCCACCTGCCGTCCTGGGG | 120 |
| chr11 | 70605935 | 70606055 | 788855_47188552_ENST00000601538.6,E | 1 | - | good | ATCTGCTGTCGTCTGTCTGGGGAGCCCGACCCTCGGATGGTTATCTTTCCCCAGGATTCTTAACTGAGGGTACTTCATGCCATCTTTTTACCCCAAGGAGGGGGGGGAATAATAGGTTGT | 120 |
| chr11 | 70606055 | 70606175 | 788855_47188552_ENST00000601538.6,E | 1 | - | good | TGGGTTGGGCTGATTTTTGAGTGGGGTCTTTGCTGGTATTGGGGTGGCTCTGGATGACCTTCCAGCTGCCCCCCTGCCATGCACACTTCCTGGCAGCAGAGCACTCATGAAGCAGCCCCG | 120 |
| chr11 | 70606175 | 70606295 | 788855_47188552_ENST00000601538.6,E | 1 | - | good | AAAGGCAGGAGCAACCAAGCCTGTTCATGCTAGGATCTTTAAAGAGTGAGTAGGAGTCTCTTCAGTGGAAAAGGTGGGGGAAGGATGGGCAATGGAGCCAGAAGAGCTGATTTCCAGACA | 120 |
| chr11 | 70606535 | 70606655 | 788855_47188552_ENST00000601538.6,E | 1 | - | good | AGACGCCCAGGAGGGGAGTCTGAGGGGTAATTGTACTGCAGGACTTGGCCCCCCACAAAGGACCTGCTGATTCAAACTCTGAAGTCTGCACAGTAAGGCTGAGGATACAAGCTAGGGCCT | 120 |
| chr11 | 70606655 | 70606775 | 788855_47188552_ENST00000601538.6,E | 1 | - | good | GCGCCTGAGGCCCAAGGAGCAGACGGTCCATTCCATCTTGCACCTGGGCAGCTCCTAACCCCAAGCCAGGACCTAGAAGTTACAGAAATGGTGAGGATGACCACAGGCCCTCAAAGACAC | 120 |
| chr11 | 70606775 | 70606895 | 788855_47188552_ENST00000601538.6,E | 1 | - | good | TTGCACTGAAGGATGAAAACAGCCGAAGAACAGGGTTGCTCAGAGACATCACCTAGCGCACCTCCCTTCCGTATCTTCCTGCTGCGGTTTGACCCAGTAACAGGAATCAGGACTGCAGAA | 120 |
| chr11 | 70606895 | 70607015 | 788855_47188552_ENST00000601538.6,E | 1 | - | good | ACCCCTTGCTGGCCTGCCCCTCCAGGCACCTGTCTAGGAGCTGCAGACACAGACGACATGACTGGGAACTCAGGTCCCGAGCCTCATCCTTCTCCAGGGCTGTGCAGAGATTTAAGGGCT | 120 |
| chr11 | 70607015 | 70607135 | 788855_47188552_ENST00000601538.6,E | 1 | - | good | TGTTTCCAGGAAGGCCGCCCCCGTCTGAGGGGCCACAGAAGAGGGCTGAGGCCTCAGATCCGAGTGTGGATTTTGGACTTCCACTGACCCCGGCTATGTGGAGGGGCCAAGTGGGTGGGC | 120 |
| chr11 | 70607135 | 70607255 | 788855_47188552_ENST00000601538.6,E | 1 | - | good | ATTCCCCTAGGACCTAGGGTACGTGACCTGGGCAATCTGCAGTACCCAGACCCCGGAGCCACATTCTTCCCAGACCTGAAAAGGCACATATTGCTGTTGGTGCCCGGGGAGGAAGGCGGA | 120 |
| chr11 | 70607255 | 70607375 | 788855_47188552_ENST00000601538.6,E | 1 | - | good | CACATAGGGAGAGAGGAGCCGGAAAGTGGCCAGGTTCTTAGAATCTGCCAGCTGGTCACCTCTGACCTTCCACGGAGGCTGGGCTCAGAGTTCAGCCTGGAAACCAGTTCATCTGTTAAA | 120 |
| chr11 | 70607375 | 70607495 | 788855_47188552_ENST00000601538.6,E | 1 | - | good | ATGGGCACCTGAGGGCAGGGAGTGGGGTGTGATGGTGACCAGGACCCAGTTGGTCACCTCTGACTTCCGGGAAGACCTGGCCAAGCATGTACTTGGTTGGGGAGCTGGGTGTGGGGCAGG | 120 |
| chr11 | 70607495 | 70607615 | 788855_47188552_ENST00000601538.6,E | 1 | - | good | GCTCTGTTTGCTGACGGCCACAGGCATTTTCCAAGAAGTCCCTGGACCCGTGGGCCCAGGGAGTAAATGCTGGCAAACAGGCCCTAGAGCCAGTGTGTGCGTGCAGGTGCTGGGCACAGA | 120 |
| chr11 | 70607615 | 70607735 | 788855_47188552_ENST00000601538.6,E | 1 | - | good | GCTCACAGGATGGTGGCATGTCCTGGGGGCCCTGTGGTGTGGGGAGTTGGGAGTTGCCCTTGTTCTCCTTGGACAAGCCAGGGGTATACAGGGAGGCCATGCCTTCTTCCAAGCTGATCT | 120 |
| chr11 | 70607735 | 70607855 | 788855_47188552_ENST00000601538.6,E | 1 | - | good | GTCTCTTTCTGGCCACCACATGGGGCCTCTCCCGCTTAACCCCAACTCTCGGAACTGGCCAGCCAGGCTGGCATTAGGCGGTGCTGCTGCTTCAACACAGAGCAGAGCTGGCCACTGCCT | 120 |
| chr11 | 70607855 | 70607975 | 788855_47188552_ENST00000601538.6,E | 1 | - | good | CAACTGTGGAAATGAGCAGCTGCTCATGCTGCCCCACAGGAACGCAAGGATGGAGCTGGAACCAGATTCTTCTTTCTGAAAGGGGGGCTATATGTTGTCCAAGAGACAGAATACTCAGTG | 120 |
| chr11 | 70607975 | 70608095 | 788855_47188552_ENST00000601538.6,E | 1 | - | good | ACATTTACCTTTATCTGGCAAAGGCAGCAGGAATCGGACATGGCGGGCAGTGGTCAGGAGCCCTGGCACTGCCTTTTCTGTGTGACTCTGGGCAAGCCACTCAGCCTCTCTGGTCAGAGT | 120 |
| chr11 | 70608095 | 70608215 | 788855_47188552_ENST00000601538.6,E | 1 | - | good | AGATCGCTTGAGCTCAGGCTGGGCTACGTGGCAAAACCCTGTCTCTAAAAGAAAACAAAGAAAAAGAAAAATCTCTTGCAAATTAGGGTCAGATTCCGATTTCCCACAAGAGCAGCTGGC | 120 |
| chr11 | 70608335 | 70608455 | 788855_47188552_ENST00000601538.6,E | 1 | - | good | ACCCTCCAGACCCCATGATCCCCCATCAGGAGCTGGGGTGTTCGAGAGAAAAAGTTCCCATCGCCAGGTGGCCTTGGAAGCCTGCAGGCCCCCTACGTGTTCAACTAGGGAGCTCCTCCT | 120 |
| chr11 | 70608575 | 70608695 | 788855_47188552_ENST00000601538.6,E | 1 | - | good | GTCGAGGTGGGTGAAGCAGGGAGAGAGGCCCTGAGGAAGCAGCGCTCCTGCGGCTGATGGGGCCCAGGCTCGGCCAAGAGGAGGAGGGACAGTAGCTCTCAGTGGCTCCCTGCTGCCGA | 120 |
| chr11 | 70608695 | 70608815 | 788855_47188552_ENST00000601538.6,E | 1 | - | good | TTAGCTGAGAGGTTTGTAATTTGAGATCAATTCTCTGATCGGCTGCACACAGATGAGACTAAATAGCACGAGCTTTCGGCTGTGAATGGGTTTCCGGGAGCCCGTGGCCCTCACCCCCGT | 120 |
| chr11 | 70608815 | 70608935 | 788855_47188552_ENST00000601538.6,E | 1 | - | good | TTGTGTTTCCATTATCTCGTTATTCAAAACCGTGCCACATCTTTCCAGAGAGTTCCTGGGGACTGTGTTGTAAGGGTTATTTATAGCCGCTGGAATGAGGTTTTGCGTAATAGACCGCGT | 120 |
| chr11 | 70608935 | 70609055 | 788855_47188552_ENST00000601538.6,E | 1 | - | good | CAACAGGTTTCCCTTGAGACGTCTCCCGTTTCATCCCAGGGTTTTCTTGATTGAGTCCCACTGACCATGGCTAAGCGAGGAGCGAGCTCTCTCTGAGCAGTGAAATTAATTAGCATGGGT | 120 |
| chr11 | 70609055 | 70609175 | 788855_47188552_ENST00000601538.6,E | 1 | - | good | CCGAGCAGGGCTTTGAGAGGTGGGAGCCGGCCAGGCCCTCTCTGGGCCGCCCGGTGCCACAGGTCCGTGAACAAACAACGCGACCACTTGCTGCAGCTCCACGTGGCTGGGCCCTCCTGC | 120 |
| chr11 | 70609175 | 70609295 | 788855_47188552_ENST00000601538.6,E | 1 | - | good | CCTCGGATTCTTCCTCTTTCGCTTTGAAACCATCTGCCGTCAGACCTAGCCCAGGGCTGCCTTTGTGTCTGATTCCTCTGTTGGGTGGGCCGTCCCGGAGCCTTGCCTCAGCAGCATTGT | 120 |
| chr11 | 70609295 | 70609415 | 788855_47188552_ENST00000601538.6,E | 1 | - | good | GAATGCTTCCTGTGACCCATGCTGGTTTCTTCACACGCCAGGTCATCATCTTTCTGGGGGTCTCTAATTTGAACAGATAAAGCCACCTCCTGGGGCCTCCTGTGACCCTGCAACAGGAGT | 120 |
| chr11 | 70609415 | 70609535 | 788855_47188552_ENST00000601538.6,E | 1 | - | good | CTCGGCCATCACGGATGGGGCTGCTCCGAGCATGGCTGTGCGTGCTTGTTTTTGCCTCCCTGTGTGGTGGAATTACTGGGTCACATGGTGGTTGGGTGTTTAACTTACTGAGGAACCTCT | 120 |
| chr11 | 70609535 | 70609655 | 788855_47188552_ENST00000601538.6,E | 1 | - | good | TACAATATAGATATATTATGTATCTATTCCATTCTATCATATATCCCGCATCATTTGTACGTGCCACGTTTTGTTTGTTCATTCACCAGTGGATGTGCGTTGGGGCTGTTTCCACCGCCT | 120 |

| | | | | | | | | |
|---|---|---|---|---|---|---|---|---|
| chr11 | 70610015 | 70610135 | 788855_47188552_ENST00000601538.6,E | 1 | - | good | TCACAAGTCTGTGTGACCACCACCGAGACCTAGTTAACCGCCCCAAAAGGAACCCCGAACCTCCTGGCAGCCGCTCCCCTCACCCCTCCCTGGTCCCTGGCCATCCCCAGTGAGCTTTCT | 120 |
| chr11 | 70610255 | 70610375 | 788855_47188552_ENST00000601538.6,E | 1 | - | good | CAGGCAATAATTCATTTTTTTGCAGCTGAGCAGCAAGAGGGGACTGGGAGCTGAGTTTGCACGCCTCGTATTAAGAGGTGAACAAGCCTAAGAACAGGCAATGCTGATGGAATTTTAGGG | 120 |
| chr11 | 70610375 | 70610495 | 788855_47188552_ENST00000601538.6,E | 1 | - | good | GAGATCAGGGACAGTGGGACTGGCCGGGGGCACAGGCGTGTCATGTGCCACCCTGACTGACAGACCTCCTGGCCTTGGCATCAGTGGCAGGAGCAGGTCACAACATTAGACTCATAAAAC | 120 |
| chr11 | 70610495 | 70610615 | 788855_47188552_ENST00000601538.6,E | 1 | - | good | AGAGTCGAGGATTACTTTACAAATGTAGCCTGCTTGGAAAACCATCATCCTTGGATTCAGGGAGAGAAGAACAAAGGCCAGCAGCCTGTTTCTAGCTTCTGAGTTTGCATGGGAGAAGCT | 120 |
| chr11 | 70610615 | 70610735 | 788855_47188552_ENST00000601538.6,E | 1 | - | good | CATGCCGTGTTCTGTGCTCTGGGCTGGCATTGTGCCGGCTGCCCCACGGAATTCTCCGGGCCTGTCCCATCCGCTGGAGCCGATCAGCAGCTCTGGCTGGCTTTCATTAATATAAAACGG | 120 |
| chr11 | 70610735 | 70610855 | 788855_47188552_ENST00000601538.6,E | 1 | - | good | TGAGTCCTGTGGATCCCCACAGGCCCTGGAATGGAGTCTGTGGGCTCTGAGAGCTGCTTAAAAGATGCAGAAAGCCCAGAGGGGCCGGCAGGGGTGCCTGGTTTTCCTGCACGCCAGCCC | 120 |
| chr11 | 70610855 | 70610975 | 788855_47188552_ENST00000601538.6,E | 1 | - | good | TGGAGCTCTTCTGATCACCGTGCCCCATAGTTCCATATGGAAGCAGTGTTCTTCAAAGGACCATTTGTTTTTCAAATTCAGAGAGTTTTGAGCTCAGAAAGATTTCCGTCATTCTTGGCA | 120 |
| chr11 | 70610975 | 70611095 | 788855_47188552_ENST00000601538.6,E | 1 | - | good | TTGGGTTGTGGAGATTTTGAAAGGCTGCATTTGCCTCTTAGTCACTATGGCTCTAGGTTCAGCCTTTTTCTGGACGTTCAAAGGGATCAGGGTGCATCATTTGCTGGTTTTTAAAGAGGC | 120 |
| chr11 | 70611095 | 70611215 | 788855_47188552_ENST00000601538.6,E | 1 | - | good | TTAGCACCAGCCTCAGGAAGTGGCATTTGCGCTTAAATCTGAGAACCTCCAAACTCCTTTTGAAAGCTGCCACTTTTCTTGCATTCTGAAGCGTTGGCTTCACATGTGAAAAGGCAATTC | 120 |
| chr11 | 70611215 | 70611335 | 788855_47188552_ENST00000601538.6,E | 1 | - | good | CCGCCTCACCATCCCCACAGAGAACAAGGCCAACACCAATAGGTGTCCGTGAACATGCCCTTTGCAACACTGATAAACTCCCCTTTGTGAGCTTCTGCCAACCAGCCTTTTCCTGCAGTT | 120 |
| chr11 | 70611335 | 70611455 | 788855_47188552_ENST00000601538.6,E | 1 | - | good | TGCAGGGAAGATGGGGAAGCAGGGCAGGTGGGGAGCAGCCCAGGGCTCTGGCTGGAGTGACGATGAGTTGAGAAAACTGTCGAGGAACGGGTAATGCCACAGCCCTGCAGGCTGCTGGC | 120 |
| chr11 | 70611455 | 70611575 | 788855_47188552_ENST00000601538.6,E | 1 | - | good | CCTCTGTTTCTGTCCCATGTCACTCTGTCCTTCCAAACAGCCCAGAGGTGAGGCAGTCCCTGAAGGATAGGACGTGGGTCGTGGGGTGGTAACCAAGTACCCAGGGCTGCCATGGGGCAT | 120 |
| chr11 | 70611575 | 70611695 | 788855_47188552_ENST00000601538.6,E | 1 | - | good | CCTGGGGCCAGTGGGTCCCTTCCCTTTGTACATTCTTAGCCTTTTTGAAATCATGGTTTGAATTAGACCCAAAATAAAGAATGGAATGATAAATGCAGAAGTAGCCCTTAAACCAGACAC | 120 |
| chr11 | 70611695 | 70611815 | 788855_47188552_ENST00000601538.6,E | 1 | - | good | CCCCCATTCATCCTGGAAGGCCCATCCTGGGTGCTTGACTACCTAGAAATTCTTCCTAAATGGCTAGAGGAGTGAAGGAGTCTCAGACGACACCTCCACCAGGACACCCTCCCATCTTGC | 120 |
| chr11 | 70611815 | 70611935 | 788855_47188552_ENST00000601538.6,E | 1 | - | good | GGAACGAGGCTTCTCGTTCCCTTTCAGCAGCAGCCCAGTTTCACACTGCTGAGTGCTGCAAGAACTGAAAATTGCCTTTTTTCTCCATTTTCAGTAGAAACGAAGGCAGTTTCCTAAACT | 120 |
| chr11 | 70611935 | 70612055 | 788855_47188552_ENST00000601538.6,E | 1 | - | good | CGCCTCCTTCCTGGATCTTGGGCAGAACGCTCAAGACGCTCCCCTTCTGTGATGCCTTCCCAGTGTTCACGCCCCTCGCCACCCGCCCCCCCTCCCCCACCTCCCATGGCAAGTAAATTT | 120 |
| chr11 | 70612055 | 70612175 | 788855_47188552_ENST00000601538.6,E | 1 | - | good | ACACATCCCCACAGGGGCCCCACCACCAGCACAGACCCCTGGTTCCCAGGCAGGAGTCGCAGGCAACTTCCCCACCACCCCATCCAGCTCCCCTCCTGGGCCTGTGTCACGCCCGGACGC | 120 |
| chr11 | 70612175 | 70612295 | 788855_47188552_ENST00000601538.6,E | 1 | - | good | TCCCCTCTGCGTTTCCCTGATCAGCCCAGCCCTGATTCTCTCCTGCCCAGCACGTGTGAGTTCCCATACTCAGCAGCAGGACGCCTTCAAAACCCTTATTAAAGTCCTCTCTGTGAGAAC | 120 |
| chr11 | 70612295 | 70612415 | 788855_47188552_ENST00000601538.6,E | 1 | - | good | AACCTTTCATGCCTTCGCCTGCAGCTGTACCTGGCGAGAGATGCAGCTGCTATCAGGTAAATGGCAGGTGCTTGTTCATCTTGCCTCTGTCAGAGATGGGGGCTACAGGTAGGCTTCCTT | 120 |
| chr11 | 70612415 | 70612535 | 788855_47188552_ENST00000601538.6,E | 1 | - | good | AAGTATGCATGGATCTAAAGATATTCGTAATTGTACAAAGAAGGCAAAACACCCCCAAGAATATGTATGCACGCATGCATATGTGTATGTATACGTGTACACACGTATGTAGAGAAACCC | 120 |
| chr11 | 70612535 | 70612655 | 788855_47188552_ENST00000601538.6,E | 1 | - | good | GTGGCTGCCGGGGGCCGCTGGGGGCTTCTGGAGCGTTGAGCGTTTTGCATTTTTCTTCGTGGAGTTTCGTTTAACCTTGCCTGTACAGTATGCATACTTCAGGGATGAAGTTCCATCAGA | 120 |
| chr11 | 70612655 | 70612775 | 788855_47188552_ENST00000601538.6,E | 1 | - | good | ATTTGAAAAAAGGCAAAGGGAATCCATGTATTGTTTAAGGAAGTGAGCAGGTGACAGTGAGTGCTAGATAGAGCGCGGGAATTGCACGCGCAGACCCCGGGTGCCTCTGCAGGGAGGAAG | 120 |
| chr11 | 70612775 | 70612895 | 788855_47188552_ENST00000601538.6,E | 1 | - | good | AGGCGCGTGAATCTCAGACATGCTGTGGAGGGGAACGTGGAGGCCACAGAGGAATACATAGAGTCTGATATTGCGTATGGAAAGTTCCAAAGAAAAAAAGGCCAAAATGATATCTGTAAA | 120 |
| chr11 | 70612895 | 70613015 | 788855_47188552_ENST00000601538.6,E | 1 | - | good | AAGACGCTGGAAATGATGCAAGTGCCTGTGAACAAGAAAGGATCCGGATGCCGTGAGTTGCTCACACGTGGAGTACAGTGCGCTGGTGGGAGAATGAATGAATGCCAACTGCATTCATCA | 120 |
| chr11 | 70613015 | 70613135 | 788855_47188552_ENST00000601538.6,E | 1 | - | good | AAAGACACACCTGCTGTGCAAACCAGCATGTGACTCCATGGCACACACCCCTGACACATTCTCCCACGTGGGCCAGGAGACACACACGAGAAGTTTCCAAGCACCTCTGTTCATAACAGC | 120 |
| chr11 | 70613495 | 70613615 | 788855_47188552_ENST00000601538.6,E | 1 | - | good | CTTCCACACCTCTTTTTGGGGGACACAGTTTAACTCATAACACTGCTGCACATGGCCAGGTCTTCCAGAGCCTTTTGAGTTTCTACACTTTGTTTTTAAAAAAAGAAAAACATGGCCGGG | 120 |
| chr11 | 70614575 | 70614695 | 788855_47188552_ENST00000601538.6,E | 1 | - | good | CAAAGACTGGAGGATGGGATGAAGGAAGGGCCTGGAATGCTGCTTGTACTTATTTCTTAGGGCTGCTAAGAAACCACAGGTTGGGCTGGACATGGTGGCTCATGCCTGCAGTCCCAGCAT | 120 |
| chr11 | 70614695 | 70614815 | 788855_47188552_ENST00000601538.6,E | 1 | - | good | GAAGGGCTCAGAGGCTCACCCCAGGCTCAGAGGAGCACACAGACTCTTGGAGATCTGCAGAAGGGCTTGGTCATCCTTGCTCTCACAGCAACAGACCCAAGGCCTATTTGGTCACTGAGA | 120 |
| chr11 | 70614815 | 70614935 | 788855_47188552_ENST00000601538.6,E | 1 | - | good | AGCAGCATAGCTGCCCACTTCCTCTTCCCTCTCCAGAAACCTCTAACACGGCCATGGGGACATCCAGTACTCCCGTGTTGTTAAGTTAGTGGGGAGGGGAGGCCTGCAGCCCTGTGTTGA | 120 |
| chr11 | 70614935 | 70615055 | 788855_47188552_ENST00000601538.6,E | 1 | - | good | ACGAATGGGTTCTGTGCATGGGGGGAATCCAATCTAAGCAGGCTTCTCTTCTTTCCTTCCTTTTGCCTTTCCAGTTAACCCATTAAAGCAAGGCCCTGGGACCACAGCCCCCCAGCCAGG | 120 |
| chr11 | 70615055 | 70615175 | 788855_47188552_ENST00000601538.6,E | 1 | - | good | TCTCCACCCCCTACTCATTCCCCAACAGCTCTTGGTTCAGAGGGAGGCTGAGTGTTGAGTTCATGATCCCCCTCTGCCATCGGAAGATGTCTTCCATGAAGCATTGCAGATGGCATGTCC | 120 |
| chr11 | 70615175 | 70615295 | 788855_47188552_ENST00000601538.6,E | 1 | - | good | AGTCCCACACCAGGTATCAGGCCCTCATCACAGAGTCCACCAGGAGGAGCGGGGTGTGGGCTGCATTAGGCAGAGGTGTATAGGCAATGTTCAGCTTCTCCTTCTAGGCCCACCCACTGT | 120 |
| chr11 | 70615655 | 70615775 | 788855_47188552_ENST00000601538.6,E | 1 | - | good | AGGATTGGACTGAATGGCCTGACAGCTCTGCCCCAAGGTGTGGAGCAGGTTCACCACGCGCCGCTTACCACCTTGTCCTAGCCTGGTGAGCCAGCCGCCCACACAGCGAGTTCCATGAAC | 120 |
| chr11 | 70615775 | 70615895 | 788855_47188552_ENST00000601538.6,E | 1 | - | good | ACAGACCGAGGATGTAAGAGCCACAGAGAGCAAGTTCCTCCCTTGTTGCCAACAAAGCCAATACCGGCCAAGGTGGCTTAGACCTGTGGGTGGGTGGGCTTCCTTCTACCACCAAACCCC | 120 |
| chr11 | 70615895 | 70616015 | 788855_47188552_ENST00000601538.6,E | 1 | - | good | TCTCGGCATCCACGCTGTCTGCCACTCCTTCCTCAGCAGCCCTCTCAGGGACAGCTGGGGTCTGAGTGGCCCCAGTAAATGCTGGTGCCCACCCTCACCTCCCCACTAACCAACAAAGAG | 120 |
| chr11 | 70616015 | 70616135 | 788855_47188552_ENST00000601538.6,E | 1 | - | good | GCCCCCCATCCTGATGAGCAGCTGAGGCCCTGAATGCAGGGAAGGGCATCCTAACGGCAAGGACAGTGAAGGGAGGATGGGTTCCAAGGCCTCTGTCCATTTCCCAGAGGCAGGCTCTGG | 120 |
| chr11 | 70616135 | 70616255 | 788855_47188552_ENST00000601538.6,E | 1 | - | good | TCAGCTCAGTAGCTACTTATCGGAGCAAGGAGCAGTCGGCCGTGGGGTCCGTTCAGTTGCATCATGTCAAACTAGCAAATATTGAACTCTTATCAACCTACAAGAATGGCAATCTCATGT | 120 |
| chr11 | 70616255 | 70616375 | 788855_47188552_ENST00000601538.6,E | 1 | - | good | CTCCCAGGCTGGGGAGGGCCTCCTCTTGGGGTCCCTGGAGCCAGTTCCGACAGCCCAGTGTGGGGTCGCCAACAGACGCAGGGCCTGGGCCTTCTGCTCATTGAGACTTCTCGGGGACCT | 120 |
| chr11 | 70616375 | 70616495 | 788855_47188552_ENST00000601538.6,E | 1 | - | good | GCCTCCCCAGTACCACCCAGCGCCCTCCTCGGGCGGTCCTAGCACTCGAATTCACATCCTCTGCTCAGCTCACCCTCTGATCTCATTGTAGGCCCCAACTCAGAGGTGCCATCCAGGCCC | 120 |
| chr11 | 70616495 | 70616615 | 788855_47188552_ENST00000601538.6,E | 1 | - | good | ACCCAGCACTGGGCACCAGGGACCTGGTTCCAGGGTCTGTGGGGAACAGTGCCTCCTCTGTCCTCGTGGCCTGGCCCTGCCTCCCTCATCCCTGGACTTTCAGCCTCCTGAGTTGTCTCT | 120 |
| chr11 | 70616615 | 70616735 | 788855_47188552_ENST00000601538.6,E | 1 | - | good | TCCCTTCTGGCACTCCCAGCCTTCACCCAGCAGCCTCCGGAGGCCTCAGATCCCCCCACACCCCACCCTGCCTTCCTTCTCAGGTGTGAATGTTTTAACCCTCAGGAGAACTTGCTCAGG | 120 |
| chr11 | 70616735 | 70616855 | 788855_47188552_ENST00000601538.6,E | 1 | - | good | AGCGGCCCCTGTCGGGGGACGCCCCGCAGGCTCCCCTCGCCTGGGGACTGGTCAGGAGCCAGCCTGGCAGGAGTCCCCTGGGCCAGGGAGAGGCTCTGGGCCACCCCAGCCTGGAAGTA | 120 |
| chr11 | 70616855 | 70616975 | 788855_47188552_ENST00000601538.6,E | 1 | - | good | ACACTGTCTCACACACATACACACACACTATCATAGACACACACAGGAGCACACACGGCTCTGCCCCTGGCTGCTGCACCCTGTGGCCCTTTGCGGGGCATTTGTCATCTCCTCTTCCTC | 120 |
| chr11 | 70617335 | 70617455 | 788855_47188552_ENST00000601538.6,E | 1 | - | good | GCGTTTCTTTCTTTTATGGCAGGGCTGGGTGAACTCTGGCCTGGGGGCCACATTCCATCCATGACTGAGTTTCTGAGGAAGATTTTTACATTTTTAAAGGGTTTTAACATGCATGTACAT | 120 |
| chr11 | 70617455 | 70617575 | 788855_47188552_ENST00000601538.6,E | 1 | - | good | GCTTCCCGTGGCGACAGCTGCGTCTGCCCCCTAGCTCTGCTCTGAAGCCCACGATCAGCTCTGCAGAGGCTCTGACTGCCCGTTTTGACAAATCTGCTCCACCTCCCCACTCCGTTCCTC | 120 |
| chr11 | 70617575 | 70617695 | 788855_47188552_ENST00000601538.6,E | 1 | - | good | AACTCCACCTTGATCATGAAAGCCTCTTTCCCTCATTCCGGCCACCTTCTTCTCCCGTTAACTCTGAACTCGAGCTTGCTCTAGGCTTTGTAGGTCCCGTGGGGTTCTGAGGGTTCCTAT | 120 |
| chr11 | 70617695 | 70617815 | 788855_47188552_ENST00000601538.6,E | 1 | - | good | TTCCCCTTCCTGTGTCCATGTGTTCTCTTTGAATCAAACATTTCTAACATGCTTGGCATGGTAGACGTTTATTCAGAAAAATACTGTTTTTCAGAGTTTTGGTCATATGCACGTGGGGTA | 120 |
| chr11 | 70618295 | 70618415 | 788855_47188552_ENST00000601538.6,E | 1 | - | good | GGAGACTTTTGTCCTTTTGTTCAAAGACTTTTCCCTCTCAGTGGGTTCATTCTGACCTTCTGAAATCACCCCTGCACGAATCACATCAGAATCAAACATTTCTTTTTTTTTTTTTTTTTC | 120 |
| chr11 | 70618415 | 70618535 | 788855_47188552_ENST00000601538.6,E | 1 | - | good | CAGGCCTGGCTGTGGGCAGAGCCAGGGGCCATTCATTGGAACCTAGGGCGGGGTGGCCCTGGGATTCTAGAACGTTCCTTCCTCAGATGGTTAGGCTAGACATAAATGGCGGTTAACTCT | 120 |
| chr11 | 70618535 | 70618655 | 788855_47188552_ENST00000601538.6,E | 1 | - | good | TAAGCTGGATTACCGTATTCACAAGCTGAAATGCCTCGTAATTCTGCTCTAAGTAGCCAGCCTTGTGTTTTATGACTTAATTACTGAATCCGAGCCATGCACATCACCTGCAGGTGAGGC | 120 |
| chr11 | 70618655 | 70618775 | 788855_47188552_ENST00000601538.6,E | 1 | - | good | TTTATTCCACACAATGACAAATGCTCAGCATAGCGTGATGAGGGTAAGCGGCGGCCAGGGTTTCCGCATGGGCTGATTTAACCCAGAAGCCTGCGCCTTGCTCTCCCAGTCGGTGTTAAT | 120 |
| chr11 | 70618775 | 70618895 | 788855_47188552_ENST00000601538.6,E | 1 | - | good | ACGCCTCTCAGGGTTGCTGGATTTGTTCCATTTTAACACAAAGGCCCTGGAGAGCCAGCCTTGGGTGGTTTCCGGGCCTTTCCAGGGTCACCTGCCCAGTTTTCCAAGCTCAGGTTTATT | 120 |
| chr11 | 70618895 | 70619015 | 788855_47188552_ENST00000601538.6,E | 1 | - | good | GGGTAAGGGACTGGGGCGGCTGTGCACGGCACCTGCAATGATTCACCCCGTCCATCTCCTGGCCATCTGCCGGGTCAGCCTGTGTGTCCCCTGTCCCAGAGAAGCTCGCCTGCCCTGGAT | 120 |
| chr11 | 70619015 | 70619135 | 788855_47188552_ENST00000601538.6,E | 1 | - | good | GAACCCTCAGGAAGTCCTTAGAGTGTCACAGGCTGGGCTGGGGTCCGCCTCCCCTGGGCAGGGCCATGGGCAGAAAGCGGGGGTCCTGAAGGGAGATCGAGCCTGCAGGACGAAGGGTGT | 120 |
| chr11 | 70619255 | 70619375 | 788855_47188552_ENST00000601538.6,E | 1 | - | good | AGGAACGCCAGGGGTCCCCAGAGCTGGAAGAGACAGGAAAGGACCCTCGCTGAGAGCTTTCGGAAGGAGCCAGCCCTGCCAACGCCGCTCTCTCAGCCTCCTGAGCCCTGGGTTGGCCCT | 120 |
| chr11 | 70619495 | 70619615 | 788855_47188552_ENST00000601538.6,E | 1 | - | good | TCCACGTGCTTATCCCCAGAGCCTGTCGACCGACTGCCTTCCATGGGAAAGGCGGCTTTGCAGATGCAATGAACTGAGGGATCTTGCTGCAATCAGGAGGTCATCACGGATCGCTTAGGG | 120 |
| chr11 | 70619615 | 70619735 | 788855_47188552_ENST00000601538.6,E | 1 | - | good | GGAGATGGGGAAGAGAGGGGCCCTGAGGCGTTATCACTGCCTGGAAGGGTCCCTGGGCAGCCTCTCTGCAGAGCCCAGTTGTGCCTGCCAGTGGCCAGCTGAATAAGGGTCCCCAAGATG | 120 |
| chr11 | 70619735 | 70619855 | 788855_47188552_ENST00000601538.6,E | 1 | - | good | GGGCAGCGTCTCACCCTCCAGCATCCAAGGGCCTTGCTGTTATGGAGGCCTCCTCCCTGTGAGCCGGCACTGACTAGCACGCCTCATGGAGGCAGCCAGGACAGCCCCAAAGAGAATGAG | 120 |
| chr11 | 70619855 | 70619975 | 788855_47188552_ENST00000601538.6,E | 1 | - | good | CGGGGTACTTTGTGGCTCTTTTAAAACTCTCCCCTCCTGCCTCCCTGAGCATCAGTTAGCAGAGCTGAGCCCCAAACGTGCTCCTCCAAGTCCCATGGTAACAGAGGTAGATTTCACACT | 120 |
| chr11 | 70619975 | 70620095 | 788855_47188552_ENST00000601538.6,E | 1 | - | good | GGGGACTCATGATGGGATCCGTGTCCTTATGAGAAGAGAGAGGGAGACCAGCACACCCTCTCCCAGGATAGACATTCTGTTCCAGTCTTAAACAAGGTCCTCAAGTGTCCTTCTAAGTTC | 120 |
| chr11 | 70620215 | 70620335 | 788855_47188552_ENST00000601538.6,E | 1 | - | good | ATTTTTGTTCTGATGAGGCCCCGGCAAGGTAATGATTCTTCTGCCTTGGGAAAGAGAAGAGCCAGCCGGGCTATGCAGAGGCCTCTCCCCATCTCAGACACCAGGAAGCATCAGCTTGGT | 120 |
| chr11 | 70620335 | 70620455 | 788855_47188552_ENST00000601538.6,E | 1 | - | good | TGTGTTTCTTGTAATAACGATGGGATTACCTGTTCTGCTTAATTAAGTACTTCATGGACAGCCACCAAAATCTCCATTTCTCCCAACTCAGGAGGTTTCTAAAAGGCACAAAACAATTAC | 120 |

| | | | | | | | | |
|---|---|---|---|---|---|---|---|---|
| chr11 | 70620455 | 70620575 | 788855_47188552_ENST00000601538.6,E | 1 | - | good | AATGCAACAACCCAGATAGTTGTGAAGGAAGGCGCCCATGGTGGGGGTGGGGGAGACCAGGCGGGGCAGGCAGCCTCCAGGCCCACAGGCAACTGCATTTCCACAGATATTGGAGCTCTG | 120 |
| chr11 | 70620575 | 70620695 | 788855_47188552_ENST00000601538.6,E | 1 | - | good | CCGAGTTACCTCCAATAAAGGCCCTGTCTCCAAATGTGTCAACATAGGAACTTGGGGGGATACAGACATTCAGACGGTAACAACAGACTCCACATTTTGTCTGGGGGCAAGACAAGCCTT | 120 |
| chr11 | 70620935 | 70621055 | 788855_47188552_ENST00000601538.6,E | 1 | - | good | GCTGTTACCATCCATCCCCATTTCAGATGGAGAAACTGAGGCCTGTTGTGGTCACACAGATATTACGTCATACCGGACCCCACATGTCTTATTCAGTTTGGGTTGCTGTAATAAGGTTCG | 120 |
| chr11 | 70621055 | 70621175 | 788855_47188552_ENST00000601538.6,E | 1 | - | good | CCCAGGGGCTGCTGTCATTGATGTGAGCCTGTAGAGGTGAAGTCGGGATCACACCCTGCCCGATCCCGGTGCCTTGGTGCCTGAGCTTTCATTTCAGCCCCCACAGCCTTTGAGGCAGGT | 120 |
| chr11 | 70621175 | 70621295 | 788855_47188552_ENST00000601538.6,E | 1 | - | good | CAAAATAGTGATCAGGACAAGAGGGGCTGATGTCTTATGACACTTAATCTCCTCAAGTGTCTAGGACTGCCCCTTGGACACCCCAGAAATTGGGTCTTCCTGGGAGGTCAGAGCAGGCTG | 120 |
| chr11 | 70621295 | 70621415 | 788855_47188552_ENST00000601538.6,E | 1 | - | good | AGGGCATAAGCAATAGCCATGCAGTGCTGGGCTCCAAGCTGGTCTGGGTTCTGCTTCGCATGTTCCACCTGTGACCTCAAATGAGTACCCTGGCCGCTCCTGCCTCGGTTTCCTCATCTG | 120 |
| chr11 | 70621655 | 70621775 | 788855_47188552_ENST00000601538.6,E | 1 | - | good | ATCCCCATGTATCTCCATCATTCTTCCCCATACCCACGTCCTCAGGCAGAGTCTCCCCTTCTACACTCACTTGCCTGGGTTGTTTTGTAATGTAAGCAGCCACGCATGGCTGGGGCTACC | 120 |
| chr11 | 70621775 | 70621895 | 788855_47188552_ENST00000601538.6,E | 1 | - | good | GCTGGGATGTCTGTCGGCAACACCTGTCAGTCACACTTGATCACCTCTGCCATTAAGCCCTCAATCATGCTTTTCTGGGATCATTTATGGGTGTCTCCAGAAGAGTTACAGCAATGGATC | 120 |
| chr11 | 70621895 | 70622015 | 788855_47188552_ENST00000601538.6,E | 1 | - | good | GCTCAGCTCCAGAACCTCCCGGCACCGTGTGGACAGAGCCGGGAAGAGAAACGAGCTCAGAAGTTGGCCCTCACCCCTCCTGTGTGGACAGAAGAGCTGGCCAGGCCTGGGGGTGTGCAG | 120 |
| chr11 | 70622015 | 70622135 | 788855_47188552_ENST00000601538.6,E | 1 | - | good | GACCAGCTGTTCTTGTCAGGCTCACCCACCCCACTAGTGACAGGCAAACACCCCTTTGCCCCAATGTGCCTGGGGCAGCCAGCCCAGCAAGGTGGGCCAGGCCCCTCCACATGGCGCTGG | 120 |
| chr11 | 70622135 | 70622255 | 788855_47188552_ENST00000601538.6,E | 1 | - | good | TGAGACGGGGAAGCCTTTGGCAGGCAGGGGAAGTTGCGGTGCTGTGTTGGATTGGAGACGTCCATTCCATTGGACATCCTGCTGGGTTTGGATGGTTCTAGAAGGAGCCGAGGAGTGGTT | 120 |
| chr11 | 70622255 | 70622375 | 788855_47188552_ENST00000601538.6,E | 1 | - | good | GGGTGGAGAGGACTGAGGCTGGGGGTGGTTTTGGGGCACAGCACAGGTGGGGGCGGGGGTTGACATGAGTTGAGTCCTGGGCTCTTCCATACACTGCCGGGCAGCTGTGGGGCCCTTTGC | 120 |
| chr11 | 70622375 | 70622495 | 788855_47188552_ENST00000601538.6,E | 1 | - | good | GAGTGAATCAATTTCTGTTGTTTTAAGCCCCCCAGTTTGTGTTGCTTTGTAACAAAAGCCCAGGACACTCGCACAGAGGGTCAGGAAGCCCCATTGGCGGCTGTTCCACATCAAACACCG | 120 |
| chr11 | 70622495 | 70622615 | 788855_47188552_ENST00000601538.6,E | 1 | - | good | CAGTTACAAGCCAAGGGGAGGGCACCAGGGTGGCCGCAGCCCCTGGAGCTGAGGGAGGGCCCCGGAACGGATGGCCCCTCCCAGAAGCTTTGGTCTTGGACTGGAGGCCGCCTGAACTGT | 120 |
| chr11 | 70622615 | 70622735 | 788855_47188552_ENST00000601538.6,E | 1 | - | good | GGGAATCTCATTTGGAAACACGGTCTTCGCAGATGTCCTTAGATGTCATTTATAGTTAAGGATCTTGAGATGAGATCTTGAGATCATGAGCAGAAGGCCACGTGCAGAGGCTGGCAGGTG | 120 |
| chr11 | 70622735 | 70622855 | 788855_47188552_ENST00000601538.6,E | 1 | - | good | TTTGTGGCAGGGAGCAGTGGTGCTTGCGCGTGTGTGTTGTTCAGCTTGTTTGACTTCCGTTCGGAGACTGGATTGGAGACAGTAGGGGGTTGGATGGTGTTCCACAGAACTGTGTCCACT | 120 |
| chr11 | 70623215 | 70623335 | 788855_47188552_ENST00000601538.6,E | 1 | - | good | GAAGGTCGGCAACCTCGGAGGAGGTGGGCTTAGAGCCCCAGAGGCAAAAACCGCCCGGGGACCTGGAGGGGCAGAGGCCATGCACCCAAGTGCGGGAAGGGTGGGAGAGGCAGGCACA | 120 |
| chr11 | 70623335 | 70623455 | 788855_47188552_ENST00000601538.6,E | 1 | - | good | TGCATGATGGGACAGAGGCAGGAGCCAGGCCAGAAAGAAATGAATAAATAACGATGATCCTCCAAAAGGGCAAATGGTGCGTGGAGCATAAGCAATAGCTGTGCAGTGCTGGCCGGCAGG | 120 |
| chr11 | 70623815 | 70623935 | 788855_47188552_ENST00000601538.6,E | 1 | - | good | GTTGCTGTCCACACCCTGCCCTGGCCTGGGAGGGGCCATAGGCTGGAAGAGATGCCCTGGGACCTTCATAATGCTGACAGCCCTGAATTCTCTCTCCTTCCTTGGTCTAAAGCAACTCTG | 120 |
| chr11 | 70623935 | 70624055 | 788855_47188552_ENST00000601538.6,E | 1 | - | good | CCTGCACAGGGACGAAGGTGCCCTGCCTCCCATGGCTCCAGTTCTTGGTGGCAGCTCTGGGCGCACCCTTAGCCCTGAGTTGTGACCCATCCTATACCCCAGCTCCCCAGTGGGGTCTCT | 120 |
| chr11 | 70624055 | 70624175 | 788855_47188552_ENST00000601538.6,E | 1 | - | good | ACACTCCCCCGGCCTCAGCCCCTCCTGAGGCTGCAGGCTCCCAGGAAGCTCCGGGCCCACCCCTCCCCTAGGGAAACCAGGGCCTCTGCTCCCAGAAGTCCACACCCAAGGCTGCCCCTC | 120 |
| chr11 | 70624175 | 70624295 | 788855_47188552_ENST00000601538.6,E | 1 | - | good | CTCCATGGGGGAGAGGGGTAGGATGCAAGCCTGGGAATAAGACCATGCAGGTGACAGCCCGCCCCCATCAGCTCTGCCCTCAGCTCAGGAAGTGGCAGGCATGGTGTCCGTTAGTCCTGC | 120 |
| chr11 | 70624295 | 70624415 | 788855_47188552_ENST00000601538.6,E | 1 | - | good | GGGTGCAGTGTGTCCAACACCCCCTGGCCTCAAGTCTGAGGGCCGCAGAGCTGGCAGGAATCCCTTGGGAGCACCAGGGAACCAGGGGATTGGCTGGGGGCGTTGCTCCATTCCCACCTC | 120 |
| chr11 | 70624415 | 70624535 | 788855_47188552_ENST00000601538.6,E | 1 | - | good | TTTGAAGAAGGTTGTGGCCATGGGAGCATCTCTTTGGAAAGATGGAAATGGATTCTGAGGGATGTCTGGGGGTGAATCCTGGCTCAGGCAGGAGGGGATGGTTGTTCCTCTCTCTGTGAG | 120 |
| chr11 | 70624535 | 70624655 | 788855_47188552_ENST00000601538.6,E | 1 | - | good | TGACACGTGGTTCCCTGGTTCCTCCTCTGGTTCCCTGGGGTTTGGGGGGGCACCATCACCCCTGGCTGAGGACTGCTGCCTTAAGGGAAAGACGGTTTGAGGCTTTTTCTTTTTTTTTTT | 120 |
| chr11 | 70624775 | 70624895 | 788855_47188552_ENST00000601538.6,E | 1 | - | good | CACCAGGTGAGGGAGCCGATAGAGACCAGGCTCTGGGTGGAGGCATTTCGCAGAGCCGCCCTGCCAGGTGTGGAACTGGGACAGTGTTGCCCCAGGACGACACCGGGGACGTTTCTAGCA | 120 |
| chr11 | 70624895 | 70625015 | 788855_47188552_ENST00000601538.6,E | 1 | - | good | GACCCCCAGAACTTGCAGCTGGTGTCTGAATTGAGGAAGGGTGGTCTTGTGGGGGCTGTTCTCTCAAACTGTACATTGGCTAAACTCCTGGTGAGAACCCTTGGGACACCCCCCGCAGGC | 120 |
| chr11 | 70625135 | 70625255 | 788855_47188552_ENST00000601538.6,E | 1 | - | good | ATGCCCCATGCGTGTTGTCACACGTCATCTCTGGGAGGGACTTATCCCCACTTTCCCAGGGAGGACGGCAGACACCCCATGTCTGGACCCTCCTGGACTCTGCCCCCCACGCCTGCCTGT | 120 |
| chr11 | 70625495 | 70625615 | 788855_47188552_ENST00000601538.6,E | 1 | - | good | ATCTAAAATGGAGGGAGGGAGGTCTGGGTAGTGAAGCACCAACTTGATCCAAAGAGAGAGCTAACCTTTGCCTTCACCTCCTGGGAGGTGGTCTCTGGGCTCCCAGAATATCCTGCCTGA | 120 |
| chr11 | 70625615 | 70625735 | 788855_47188552_ENST00000601538.6,E | 1 | - | good | GGACTTGGGGAGCGGAGGGGCCTGAGTAAAAGAGGAGAGTGCAGGGACTGGCTCAGAGCCCACATCCTCCACTGCTCAGAGCCTGCTCAGATTTCCTAAGGTCTGGGCCTCAGTTTTCTG | 120 |
| chr11 | 70625735 | 70625855 | 788855_47188552_ENST00000601538.6,E | 1 | - | good | TTATGCACAAACGTGTCTTTTTTTTTTTTTTTTTTTTTTTTTTTTTCATTTTTGCACAGAGATCCAAGAGGCTGCACTGAGATTTCCCCCAGCTTGGGCTTGTGGACCTCTGGGAGGCAGG | 120 |
| chr11 | 70625855 | 70625975 | 788855_47188552_ENST00000601538.6,E | 1 | - | good | GGAGGGGCCCCAGGTTTAGAGGATTAATACTCCAGGTCCCACGCTGGCCTGGTGGTTGCCATGGGGTGAATTACTGGAAACTTCTCTTCCTGCATTTTTCCTTCCTCTACAGTGTGTATT | 120 |
| chr11 | 70625975 | 70626095 | 788855_47188552_ENST00000601538.6,E | 1 | - | good | GGGGTACCATGCTCAGCCGGGATGGGAGGCTGGGACGGAGGCTGCCTGGGAGTGAGAACAAGGCAGCGGAACTGGAGGCGGGGAGGCCCCAGAGGCCTGCAGCCCTCAGCCCTCCGAG | 120 |
| chr11 | 70626095 | 70626215 | 788855_47188552_ENST00000601538.6,E | 1 | - | good | TTGAGTGGACCAGAAAACATGTCTTCCTAAATGGGCCCAGCAAACGGTGCTGCTGGGGGTGTTTTCCGTTGCCAGCTGAGTTGTGACTCTTGGCAACGGTTTCTTAGCAACCGTGTGTGC | 120 |
| chr11 | 70626215 | 70626335 | 788855_47188552_ENST00000601538.6,E | 1 | - | good | CAGATATTGTGGGTGATATGTTTGTTTCATAACCGTTCCTAAATATAGAGTGCATGCTAAGTGATATCTAGAAGCCATGTAAAAGAAAACGTTTGAAATGTTTGCAAAGAGGATGAGGCA | 120 |
| chr11 | 70626335 | 70626455 | 788855_47188552_ENST00000601538.6,E | 1 | - | good | TACCGACAAGCTCCCAAATCAGCCAGGAGCACCAGGCTTCGCTGTGATATGAGGCAGGATTTTTCCCTGGTGTCTCCCTGCCTCTTGATCCCAGTACCAGGCTTCATGCCATAGTCAGTG | 120 |
| chr11 | 70626815 | 70626935 | 788855_47188552_ENST00000601538.6,E | 1 | - | good | CTGGGAACCGCAGCCTTCATCCTTGCTGGGATGGAGTCTTTCCTCTTCAGGCTGCCCACTCTGGGCTGAAGAGTGTCATGATGTGGCTTGGGGTGCAGTATTTCTCCACTACAGCAACAG | 120 |
| chr11 | 70626935 | 70627055 | 788855_47188552_ENST00000601538.6,E | 1 | - | good | CCACGCCCCCCACCTCCAGGCTCCCATCTGACCTTCACGGCTGCGCCAAAGCCAACCAAAGGGCCTCCCGGGGTCCCTGTCTTGGGTGCCCAGGAAATGAGATACGTGCTGACGGAGGAC | 120 |
| chr11 | 70627055 | 70627175 | 788855_47188552_ENST00000601538.6,E | 1 | - | good | AATCGGAATTTTGGGGAACAACTCTAATATCACAAACCTCAAAAACAGGAACCTTTAGTGCTAGAGCGAGCTGTGAGAGGAGACACTTCAGGGACAAAGTCCCCCAACACCATGCCACCC | 120 |
| chr11 | 70627175 | 70627295 | 788855_47188552_ENST00000601538.6,E | 1 | - | good | TGAGTTGGTCTTCATCTTTCCTGGAGTGACACACGTCCCTGCAGCCGCCGGCTCACTTGGCCGCACCTAACTTCAGGAGGGTGGAGACCTGCAGCCCTGTGTGTGCCCAAGAAGGAGGAA | 120 |
| chr11 | 70627295 | 70627415 | 788855_47188552_ENST00000601538.6,E | 1 | - | good | TGCTGGGTGCTGTCCTTTGCCCTCCTCACTTAGGAACCGGGTTGGGTGACGCCCCCACCAGCTGGACCATCATCCATTTATAGTTGCCACATCTGGGTTCAAGGGCAGTGTGGTGGGTGG | 120 |
| chr11 | 70628255 | 70628375 | 788855_47188552_ENST00000601538.6,E | 1 | - | good | TTTGGTGCCATCAGCCTCCACCCCTGCTGCGCCCGTCAGAATTTGGCTTCCACGTTCTGCTCCCGGACCCTCCCAGCCTAACCTGTGGATCCTGCCACACAAAGATGGGCTTACCTGGGA | 120 |
| chr11 | 70628375 | 70628495 | 788855_47188552_ENST00000601538.6,E | 1 | - | good | AGCTCTTGATATATTAAGGCACTGCTTAGTAGTGACTATGCTTACTTTGCGAGCAGGGAAACCGAGGCCTGGGTAGGACAGAGGGGGGCACATGTGTTTACTGCCCTCTCCGCCCCCGAC | 120 |
| chr11 | 70628615 | 70628735 | 788855_47188552_ENST00000601538.6,E | 1 | - | good | AGAGGCGGTGCTAACAGCCTGCTGTATGGTGGGTCTCCACTGCCTGGAGCTCGGGAAGCCGGAAGTTGGGGGATGACAGCAGCATCATGATGCTGGGTGTGGAGTGAGCATGGGGCTGGC | 120 |
| chr11 | 70628735 | 70628855 | 788855_47188552_ENST00000601538.6,E | 1 | - | good | GTACTGTGATCCTCTGGGACCTGTTCACTCCCCTAGTCAAGAGACTCTTCCTAGACACTCAGTTTGAGCCTGTCCTGGGGAGGGCAGGTGGGCCGAAACTTCGGCAGCCCCTCACCACCC | 120 |
| chr11 | 70628855 | 70628975 | 788855_47188552_ENST00000601538.6,E | 1 | - | good | ACAACAATGGTACAGCCTCTCCCCAAGTACATCCTGTCTTTGTGCAGAACACACTGCCTTGCCGCGCTGCAGCCCAGAGCCCCAGAGCTGGGATGATTTACCCAGCCAGAGCCCACAGCA | 120 |
| chr11 | 70628975 | 70629095 | 788855_47188552_ENST00000601538.6,E | 1 | - | good | GCTGCAGTGAGCACGTGAGCTGTGGCGCGCACATGCCTGAAACCATCCCTCTCCAAGTGTTGGCGTTTCCCTCCCAGGCTGGAATCAACTCCAGGCTTAAAATTAATTTCTCTTCCGCTC | 120 |
| chr11 | 70629095 | 70629215 | 788855_47188552_ENST00000601538.6,E | 1 | - | good | ACCCAAGGCTCTCTAGATGGAAGGAGGGGCCAGGCTCTTCCGGTGCTGGGATGTGGGGGTGGCTTGGAACACTGACAGAGGAGGGAGGGTGCCCGGGGCGCGTTTCCCCACTGAATTCAG | 120 |
| chr11 | 70629215 | 70629335 | 788855_47188552_ENST00000601538.6,E | 1 | - | good | GTTCGAAAGAGGCGGTTTCAGTCTGGCTCGCTGCCCCTCTCCCAGCCTTTTCGGCAGCAGCCTGGGTGGTGTGTTTTAGTTGGTGGGGCAGGGGACAGGGTGTGGCTCAGCGGAGAACAC | 120 |
| chr11 | 70629335 | 70629455 | 788855_47188552_ENST00000601538.6,E | 1 | - | good | GTGTCCCACTCCATCAGGCCTGGAGACCCGGGAGGCCAGCAGCAGAGCCACACTCTATTCTGTGTCCTCATCACCAAGCACTAGGTTAAATGACAGAGAACTGACCAAACCCTTCAGAGA | 120 |
| chr11 | 70629455 | 70629575 | 788855_47188552_ENST00000601538.6,E | 1 | - | good | GTGTTTGCCAGCTCATCCCGCTGCCCTTCCCACCACATGCGGCCCCCAGGGTCACGGTGGCCCGCAGCCTGTGATCCAGACCTCCCTCTCCCGCCAGGCCGGAACCTGCTGGACGTCCGT | 120 |
| chr11 | 70629575 | 70629695 | 788855_47188552_ENST00000601538.6,E | 1 | - | good | AGGTGGCACCTTCCTGACGGCAGCACCCCCTCTCCCCGGCTTGGAGCTGATGTCAGAGTGCACCTAGCAGTAGCGGCTGTGGCAGCAGTTCGGAGGCCCTTAGGGGAGCTGGAGTCCCCA | 120 |
| chr11 | 70629695 | 70629815 | 788855_47188552_ENST00000601538.6,E | 1 | - | good | CATTGCACACTTGGGGAAACTGAGGCCCAGAGATAGGCTTATTGGAGTCAGGGCATTAGAGGGCACCAGGATCATCCAGCTCTGCAGCAGGTGCTCCCTGGTGTCCCGTGCCTCTGGCCC | 120 |
| chr11 | 70629815 | 70629935 | 788855_47188552_ENST00000601538.6,E | 1 | - | good | GGGTTTTGTAGCTGGGTCTGTTAACTGCCAAGACAATATTAGGGTCCTCATAGTGAGGAAGGGGACTAGAATGGATGTCAGGGGGCCGGATCTGCTGGGGGTCTGGATGGCCAGGCCCTT | 120 |
| chr11 | 70629935 | 70630055 | 788855_47188552_ENST00000601538.6,E | 1 | - | good | GCAGTGTGCATGCTGTCTGCCCCTCTCTGGAAGCCCACAGACGATGTCCTCTCCTGTCTCATTGATAAGCACCCTGGGGTCTCCTGGTCACTCCTAGCAGCCAGGGGAAACTGAGAAAAG | 120 |
| chr11 | 70630055 | 70630175 | 788855_47188552_ENST00000601538.6,E | 1 | - | good | GCGTGGGCTCTGCAGCCCACCTCACCCCACTGCCATCAGGGAACCTTCTAGCATAGGGCAAGCTGAGTGGGGGCCAAAGCCCACGGTAGTGCATCTCCCGGACCAGGGCCAGGAGCAAAA | 120 |
| chr11 | 70630175 | 70630295 | 788855_47188552_ENST00000601538.6,E | 1 | - | good | GGCTGACCAGCCTTCTGGGGCCTGGGACCTGGGGCATCGCTGCTGCCTGCCGTCTGGCCTCAGGGCACCTGCAAGGTCAGTGCCAGCCCAGCAGGATGGCTGACCTTTCTCTTGCTCTAA | 120 |
| chr11 | 70630295 | 70630415 | 788855_47188552_ENST00000601538.6,E | 1 | - | good | GATGCCACTTTGTCCGCAGATCTCTTCCTGGCCCCGCCTCTGCCTGGGCAGCCTGGGTCTGATTGTCCTTCTGTCTGCCACCCTCACAGTCCTCAGCCGTGGCCTGGTTCCTGTCCTGGG | 120 |
| chr11 | 70630415 | 70630535 | 788855_47188552_ENST00000601538.6,E | 1 | - | good | ACCTTACTTCCAAAGAAGGTCACACTCTGAGGTTCTACATAGACGTGGATTTTAGGGGGACAAAATTCAACCACTACCGTCTCTTTGCTTGAAATCACACACAATTTCCAGAGGCCTAGA | 120 |
| chr11 | 70630895 | 70631015 | 788855_47188552_ENST00000601538.6,E | 1 | - | good | CACCCTCTGCTGCCAGTTGCTCCCTGTCCGGTCCCCAGGCCCCGGCTAGTCCTGAGGATGCGTTGGGCGCCGGAGGGAGGGACCACAGGATGCGCAGGTCACCACACTGGCCTCCTACAT | 120 |
| chr11 | 70631015 | 70631135 | 788855_47188552_ENST00000601538.6,E | 1 | - | good | TTCGCGGTCCTCTGAAGGGCTCTCGCTGCTGCTTCTGTCATTCCTGCCGGTCTTCCGTGGCTTCTCCTATGAATGCCACAGCCCTTTGTGGGTCAGATTCCGGAATGCAGGCTCTGTCAT | 120 |
| chr11 | 70631135 | 70631255 | 788855_47188552_ENST00000601538.6,E | 1 | - | good | CCTGTTTTAGGCAACCTGGACCTTATTGCTTTAGAAAAGTCAGCTCTCCTGTTCCTAAGGCCTCCTGGATGGGGGCCCGCCTCTTCCCCTTGGTGGTTTGCTGGGGTGGGAGTTCGCGT | 120 |

| | | | | | | | | |
|---|---|---|---|---|---|---|---|---|
| chr11 | 70631255 | 70631375 | 788855_47188552_ENST00000601538.6,E | 1 | - | good | AACTCCGGGAACCCTCAGAATGGCACGGCCCCCAGCAGGCTCCTCCCAGATTCTAGCTAAATGTGCCACTTGTACAATTCTAAGTTGTCATGAGTCTGGGAGAGCCTTAGGCTACCCTCC | 120 |
| chr11 | 70631495 | 70631615 | 788855_47188552_ENST00000601538.6,E | 1 | - | good | CCAGCGCCCTCTGGGTCACCTTGCCTTGGAGGCCCTGAGCTCAGCTTGTGCCTGGCCCAGCACTCAGTGACTCACTGTCCGCTGGGACTTTGCCGAGTGGATTAGGAGCTGAGACCTCCT | 120 |
| chr11 | 70631615 | 70631735 | 788855_47188552_ENST00000601538.6,E | 1 | - | good | GGGCCAGGCTTCATCCTACAGGCTTTTGTGTTCTTATAACCCCAGCTTGTCATGCAGGAAGCATCCTCACAGACACCCAGCCACATCCCCCACTGCGGGGGCACCTACGCTGTCCTATTC | 120 |
| chr11 | 70631735 | 70631855 | 788855_47188552_ENST00000601538.6,E | 1 | - | good | GCGTGATCCAATTACACAGCAGATTTCAGACGCCCAGACTTGGAGCTGGGGAAGCCTGGCGCCCTGGGGAGAACACTGCCTGGAGAGAGGCCGGGGAGGTCAGGGGACCCCGACTGCTTT | 120 |
| chr11 | 70631855 | 70631975 | 788855_47188552_ENST00000601538.6,E | 1 | - | good | CTGCGTGTGATTCCAAGAAATGTAATATATTTGATAGAAAATTACCTGTGGCAACTGCTTTCCTTCTCTGAGACTTGTGTTTTCCAAGACGGGCGCAGATGGGCGCTCAGCACGGGCCGG | 120 |
| chr11 | 70631975 | 70632095 | 788855_47188552_ENST00000601538.6,E | 1 | - | good | AAGGACAAATCGCATTCTAATGTTTAGCGTCTAAATGAAGCGTTTCCCAGAACAGAGTGAACCTTTCCATTGTCCTCTGAAAGCATTTCAAAAATAAGGGGGAATCTAATTAGTTAATTT | 120 |
| chr11 | 70632455 | 70632575 | 788855_47188552_ENST00000601538.6,E | 1 | - | good | AAGGCGGTTCTATAAAACAATCACAATTTGGATTGTTATCAAGGAGAGAAAGGAGCCTCTGGTTGAACCAGAGAGGTGGGTCTGACTCAGAAGTGTTTCCCAGACTGCCTTCACACACCT | 120 |
| chr11 | 70632575 | 70632695 | 788855_47188552_ENST00000601538.6,E | 1 | - | good | GAGTGCTCAGGGCGGGTGGGAGGTAGAGCAGCATGTGGCTGTGTGCCCACATCCCTGGGAGACAAGGATTGTCAGGCAGCATCAGGGAAAATGGTATGTGGGGAGATGATTCTGTTGTAA | 120 |
| chr11 | 70632695 | 70632815 | 788855_47188552_ENST00000601538.6,E | 1 | - | good | TTCCAGATCAACATGACCTTGTTTCAAAAAATAAACCAATTAGTATGTTCTTCTGCAGCCCGGATGTACCCTCATGCGCCCTTTTCTGGTTGGCTGGCCGGCAGAGTCCCAGCATCCACC | 120 |
| chr11 | 70632815 | 70632935 | 788855_47188552_ENST00000601538.6,E | 1 | - | good | ACTGGGATTAAAGGGGCCCAGGCTTCTCAGAGACCCAGCTTCCCAAGCCCTCATCTTGCTGAATGATGTCCTTGTCCTCTGTCCTCCCTCTGGTCCTGTCTACTCTTCCCCCGAGTTACC | 120 |
| chr11 | 70632935 | 70633055 | 788855_47188552_ENST00000601538.6,E | 1 | - | good | CCCAGTGGCAGCCCCCTCATGCCCAGGTCTGGGGATCCCAGGACCTCTCTCCTCAATGAGCACCTGCGCCTGTCCTTGTCCCCCTGCCCCGTATCCATCAACGCTCCATGCAAGGGGCTC | 120 |
| chr11 | 70633055 | 70633175 | 788855_47188552_ENST00000601538.6,E | 1 | - | good | AATATATGTATAAATATGTTATATATAACATACATAATATATATAACGTATTTATATATTCAACGTGTCTAAATCAGCTGCAGACACTATTCTGCATGGTGCCCAGGTTGCCCATCTGTG | 120 |
| chr11 | 70633295 | 70633415 | 788855_47188552_ENST00000601538.6,E | 1 | - | good | CATGTTGTTGAGGGATGAAGTAGGGTCTTTTTCACTCTCCCTGTGTCTTCTCTGCCCTGGTCTGTCTCTCTCTTCCTCACCAGTATGTCATTATGAACTCAGGATTTTTGTATATTCAAC | 120 |
| chr11 | 70633415 | 70633535 | 788855_47188552_ENST00000601538.6,E | 1 | - | good | CACTGTTTGTACAAGAAGGCAAGATAAATGCTTAATTCTTTCTTTTTCATTGTCAAGTCTCTCTATAGGAAGCTGATGCTTCAGTGATCTCTACTGATGTCCAGTGAGTTGGTTTTCTAT | 120 |
| chr11 | 70633535 | 70633655 | 788855_47188552_ENST00000601538.6,E | 1 | - | good | GGCTTGCAGGCTTGCATCTGATGACTTCCTAATCCTGGCATGCCTTCGCAGTGATGAGCTGAGATTCTTCCATCTCGAAGAACTTTCACATATCAAATAAGATGACTTGGTTAAGCTGAA | 120 |
| chr11 | 70633655 | 70633775 | 788855_47188552_ENST00000601538.6,E | 1 | - | good | AAAGCTGCACACAGGATCCAGGTGGTGATGGTGAATCCTTCCTCTTGAAGCTCCTCACCGGCTGGTCACCCAGGGGCCTCTCTCTGGGGTGGGCCCTGCAGACATTGAGGAGTACACTGA | 120 |
| chr11 | 70633775 | 70633895 | 788855_47188552_ENST00000601538.6,E | 1 | - | good | TATAAATGGAGGTGAGCTCTTGCAGCGCTGTTAATGTTGGGCCCAGCCACACGCAGGTGCTGTGAGCTTCACACCCCATCTCCCCAGGAGGCAGAAACTGTGCTCGCCACCGCTGTGACA | 120 |
| chr11 | 70633895 | 70634015 | 788855_47188552_ENST00000601538.6,E | 1 | - | good | TAGGGACAGGGTAGGTTGTTAGAGGGACAGGGTAGGTTGGCCATAGGCTACCCCCAACCCCCGATGTGCCTTCTGCCTCCTGTGGTGTCATCGGCCAGTCCTTGGGTCCTTCCCACTTCC | 120 |
| chr11 | 70634015 | 70634135 | 788855_47188552_ENST00000601538.6,E | 1 | - | good | ATCTCAGAAATGTCTTTTCACGGTGGTGACTTTGACCCAGGACCCACGTGGCACTTGGTAGATATGTCCCCAGGTCTCCAGTCTGAGCCATCTCCCGTGTTTCCTGCCAGCAACGTGTTA | 120 |
| chr11 | 70634135 | 70634255 | 788855_47188552_ENST00000601538.6,E | 1 | - | good | GTCTGTCTCTCACAGCATGGGATCTTTCTTGCGGGCCCCAGTGTCACTGTCCCATCCACGCACCCAGCAGTGGTCTCTGGGTGGAATCTGAAACCCCTCCATTAGCACATGTCCCAAATT | 120 |
| chr11 | 70634255 | 70634375 | 788855_47188552_ENST00000601538.6,E | 1 | - | good | TTTATTGGCTGTGCTTTTGTTACCAAGAAATTCTTGCCATCCATATACAGACGGACCTTTTTATTCTTTTTTTTTCTTCAGCAAATCCCAGGCATCGCGTCACTCCCTCTTCCCCTTCA | 120 |
| chr11 | 70635695 | 70635815 | 788855_47188552_ENST00000601538.6,E | 1 | - | good | CCCCCAAGAGATCACAGCCGCCATTTGCTCCATTCACTTCCTCCCCCCTTTTTTTTTTTACAGTTATACTTTGTATTATGTTTAAAATTGTAGTACAGTGCACATACCGTAAAATTTACC | 120 |
| chr11 | 70635815 | 70635935 | 788855_47188552_ENST00000601538.6,E | 1 | - | good | CTCATGCCCTACGGGGCCTTTCAGGCTGTGTCAAGGCCTAGGGCAATGGGGAGCCATTGGAGGCCCTTGAGCAGCAGATGAGCGTGACCGAGGGTTTATGAAGACCTTGTGGAGAGCTGG | 120 |
| chr11 | 70635935 | 70636055 | 788855_47188552_ENST00000601538.6,E | 1 | - | good | TGAGGTGAGGGTGGGCCGGGCACCGGGGTGGTGAGTGCAAGGCCCCACCTGGACCAGGGAAAGACTCTGCGTATTCTGGAACATTCGGGAGGGGTGGATGTGGCAGAGAAAGGGGCGG[illegible] | 120 |
| chr11 | 70636055 | 70636175 | 788855_47188552_ENST00000601538.6,E | 1 | - | good | CACACACACTAGGATGCTGTGCAGCTACAGGGCAGAGACCACGCCTTTGTCGACAGAGTGGTCAGGGAAGGCTCCCCAAGGGGAAGGATTTAGGCTGAGACCTGAAGAGGAGGGTAGAGC | 120 |
| chr11 | 70637135 | 70637255 | 788855_47188552_ENST00000601538.6,E | 1 | - | good | CTGGGGAAACAGGCACTGCACAGATTGACAGGCACTATGTGGGTCAACAGGCTCCATAACTGGAGGCTGTGGCTGTGCTGTGAATGGAAACACACACATACTAATGCACACACACATGCA | 120 |
| chr11 | 70637255 | 70637375 | 788855_47188552_ENST00000601538.6,E | 1 | - | good | ACCCTGGGTCTAATCAGTTGTGTGCGGGCCACACACTGAGTCCGTACTCAGCACCAGCCTGGGCCTTTTTGGGATCCTGCAGAGAGGAAGAGCAGGCCCTGCCCTCACAGAGCCTGAGTG | 120 |
| chr11 | 70637375 | 70637495 | 788855_47188552_ENST00000601538.6,E | 1 | - | good | GGGCACCTGGATTCTGCACCGCAGGAGGAGAGCTGGCCGTGGCCCGGCCAGGAGGGGAGTAAACACCCAGCCCAGCCCCACCTGCCCCCATCTGTGAATAGCAGGAGGGTCTCTATAGGC | 120 |
| chr11 | 70637495 | 70637615 | 788855_47188552_ENST00000601538.6,E | 1 | - | good | GCAGCGGCCTCCCACGGGTCACACCCTCTCTGCAGTGGCTGCTTCCCGATGGGTTGGTGGGGGGAGGTCTGAGGAGCTGCCTATCATGGGCTGGCTCTCCGGCCGCTCATTTGTCTCTGT | 120 |
| chr11 | 70637615 | 70637735 | 788855_47188552_ENST00000601538.6,E | 1 | - | good | TGCCAGGATCTGGGCCCCAGGGCTTTCGAGGGTCTGGAGCAGCCAGTCTCCCCTCGCCAGGGCGCCAGCCTCACTCCCAGTGCCTAGCATCCTTTGTGCCCCGGCCTTCCTGAGCGCTGG | 120 |
| chr11 | 70637735 | 70637855 | 788855_47188552_ENST00000601538.6,E | 1 | - | good | ACTCACTGGTCAGCCTGAGGCTGGCTGGGATACCAAGGCTCCTTGGGATTCCCCTGGGCAGCCTAGAGTGTGGGGTGCGGGAAGGAAAGGGGTCTGACACCTAGGTCTGTTTTGCAAGAC | 120 |
| chr11 | 70637855 | 70637975 | 788855_47188552_ENST00000601538.6,E | 1 | - | good | CATTCCCCGAGAACCTGTGCTCAGTGGGCCTGAATTGATGGACAGAGCGGAAGCCGGCTTGGCGTGAGCTGTGGGCAGCTCGCAGACCCCAAACAGGGCTCCAGCCCCGTCACTGGGTAG | 120 |
| chr11 | 70637975 | 70638095 | 788855_47188552_ENST00000601538.6,E | 1 | - | good | TTGCTGTGTGCCTCACAGCTCACTGTGGGCTGCTAGCTGTCCTGGGGGTACTTTCCAGCTTCCTGCTGGTTGGTGAGCAGACATCTGTGAATGGGACCCACCTGGCATGGGGCTCTTGAG | 120 |
| chr11 | 70638095 | 70638215 | 788855_47188552_ENST00000601538.6,E | 1 | - | good | CCGAGCCTGTTCCCCCCGGACTCTGTAGCTCTAAGCTCCTGCTCTGAGCAGTGCAGTGACGATTAGTGTATTTGCCAGATTATGGGATCAACTTCTTTGCAGGAGAAAATTAGCCCCAAA | 120 |
| chr11 | 70638215 | 70638335 | 788855_47188552_ENST00000601538.6,E | 1 | - | good | CAGCTGGTCTGTGGCCTGGGTCCTCTGTCACTGGGAGTAGCTTGTGGTCCATTGCCAGCTTTGGGTTTCATATCCTGGCAGCACATTCTTCAAGCCGGCAGTCCGTGGTCTCATTCTATG | 120 |
| chr11 | 70638335 | 70638455 | 788855_47188552_ENST00000601538.6,E | 1 | - | good | GGTCATCAGCAGGGCGCCGAGCACATCCTGCAGCCCCAGACCCTCAGCTTTCCCATCTGCAAGCGGGGCCGGGCGTGGATGCAGGGTTCCGTGACAATAGAAGCATTGTGCCCAGCACAG | 120 |
| chr11 | 70638455 | 70638575 | 788855_47188552_ENST00000601538.6,E | 1 | - | good | TTTCCGGTGCATCTGTTGACAGAAGAGCTGCTCCTTGGTCCTCGGCTGTTACAACTCATTCTTTGCTTCCGTTAAAGCAGAGAGGATCAGAGGAGCCGGGGGTCAGGCTGGCCCAGGGAA | 120 |
| chr11 | 70638575 | 70638695 | 788855_47188552_ENST00000601538.6,E | 1 | - | good | AATAGATCATTTTCTAGGAAATCTTTCTATTATATTTGTTGGAAATTCAGATGTTTTTATTATTGTACAAGAAATGCTGAGCAGACTAAAGAAAATAGGGCTTCCTCGGCCACCGCGGAT | 120 |
| chr11 | 70639055 | 70639175 | 788855_47188552_ENST00000601538.6,E | 1 | - | good | CCAACACAGAGGCCTCTAGATGCCATCAGGGAGTGGCAAGTCCTGGCTGCAGAGACCCTCAGCCAGCAGGGGAAGCCCTTCCCAGCACCAGCCCAGGCTGGCTCTGTGAGACTCAGGGCA | 120 |
| chr11 | 70639175 | 70639295 | 788855_47188552_ENST00000601538.6,E | 1 | - | good | TGTGGCATTTTGCTACGGCACCCTGAGCAAAGGCCTGATGGACTGAGACTCCCTTTCAGGCACTTGCGGAGCCCACTTCCACCCCAAACTGGCCATTTCCTGAGGCTGCCATCTGGGTCC | 120 |
| chr11 | 70639295 | 70639415 | 788855_47188552_ENST00000601538.6,E | 1 | - | good | CCTTCTGCCCTTTGAAGACACCAAGAGAAGAAACCAGGACAGGGGCCCTCACCAGACACTGACTCTTCCAGCCCACAGCATGGTGAGAAATACATTTCTGTTGTCTAGAAGCCACCAGTT | 120 |
| chr11 | 70639535 | 70639655 | 788855_47188552_ENST00000601538.6,E | 1 | - | good | TTGCTGGGAAAATCAGATGGAAATTGACTCTCAAAAAGAAAAGAAAAAGAAAAACCAGGCCCTCTTTGCTGTGGTGTGATGGTCTGTGTCACCTGAAAAATTCATACAGTAAATCCTCAT | 120 |
| chr11 | 70639655 | 70639775 | 788855_47188552_ENST00000601538.6,E | 1 | - | good | CACCCTGTGGGCCTGCAGGGCACCTTCATGTCCCTGGCCAGCCAGGGGAGCCTCAGGCCACCCTCCTTCCTGCACCCACCCCACCTACTCATGTGGATTTGGCTTTACCCAATTAATGTG | 120 |
| chr11 | 70639775 | 70639895 | 788855_47188552_ENST00000601538.6,E | 1 | - | good | GATGTCCCAATCCCAGCCACGTAGAACTCCACCACTGAGAGGCCGAGAGTTTAGAGAGGAGCCAGGTAAAGACCCCTTGGGGCTCAAGGGGGGTCCGCTCAGGCCCTGGAGGCCAGAACC | 120 |
| chr11 | 70639895 | 70640015 | 788855_47188552_ENST00000601538.6,E | 1 | - | good | CCCTGGACGGCCTGGGCAAAGCCCTCTGAGCCCCAGACGGGCCTGTGGGCCTGAAGACCTGCCCCCTCCTCCTTATGTCTGAGACCTAGCCCCTTCCTTGGGGCCGTCCCTGGGTCCACT | 120 |
| chr11 | 70640015 | 70640135 | 788855_47188552_ENST00000601538.6,E | 1 | - | good | GGCACATCTTTCTGGGAGACACCACTCAGCCCACTGCACTAGCTGTGCGTGGCCCCAGGTTGCCTGCTTACCATTGGCGACACATCCCAGCTGCACCTGCACGTTCTGCAGCCTCCAGGC | 120 |
| chr11 | 70640495 | 70640615 | 788855_47188552_ENST00000601538.6,E | 1 | - | good | GGTCAGGGCTCGGGCCTCAGCTTCGTGCAGTCCTCTCTGAAGCATAGCTGTATTCATGTCCTGAGGCCGCTGTCACAAATGACCAGAAACTGAGTGGCCTGAGACAACAGCGATGCGTTC | 120 |
| chr11 | 70640615 | 70640735 | 788855_47188552_ENST00000601538.6,E | 1 | - | good | GAGGGACTGTTCTAGAGAATGGGAGACAGGGAGGGATCCTGACTGGGCATGCGCCCCTCCTACCCCAGCCCTCCCCTGCTCGTAGTGGGCCCCCAGCTCCAGCACATGCCTGTGATCCCA | 120 |
| chr11 | 70640735 | 70640855 | 788855_47188552_ENST00000601538.6,E | 1 | - | good | CGCCTCGATTCTTCCTAGGAGCTCTGCAAGGTGGACTTGCCTGACTCTCCGTTTTACAGATGAGTAAACTGAGGCTTTCTGAGATGGAGCAACTTACAGAGGATTCCAGAGCCCAGCCTT | 120 |
| chr11 | 70640855 | 70640975 | 788855_47188552_ENST00000601538.6,E | 1 | - | good | TAGCTGTCTGAACCTTCAGGCCTCAAGCGAAGAGGCTCTTTGCATCCCTCTAAGTCCCCAGGCTCTCTGTGTAGACGCTCATTGCCTGCACTCATGGTGCTAAGCAGGTTACCCAGCAGT | 120 |
| chr11 | 70640975 | 70641095 | 788855_47188552_ENST00000601538.6,E | 1 | - | good | AAAAAAGAAAAAAAAGCATAGTAAAGAAAAAAAAAAATACTTAGAATTTTAAAAATTCTGTACACGATCTATTGTTATTTTTCATCAGGGAGGGATGCCAGGCAGTAGCCAGCCCATGTTT | 120 |
| chr11 | 70641455 | 70641575 | 788855_47188552_ENST00000601538.6,E | 1 | - | good | CCGCTTCTGTTCCGTTGACTGCAATCTCAACTGGCGGTCATTCCTCCCTGCAGGGGAGGCTGGGGCAGGCGGTCTTGCTGTGCCTCTGGAAGGAGAGCACATGCCATTCTCTGCTGTTGA | 120 |
| chr11 | 70641575 | 70641695 | 788855_47188552_ENST00000601538.6,E | 1 | - | good | TCTCCTCCAAGCAGCTTCAGGCTCCTCCCAGCTCCCAGCTCCTCCATCCTGAATACGCAGATGCCCAGGTCACCGGGCTCGGCCGCCTCTGGCTGGCAGAAGAGAAAGTGCATATCACTT | 120 |
| chr11 | 70641695 | 70641815 | 788855_47188552_ENST00000601538.6,E | 1 | - | good | TAGACATTGTGTGTCCGTGCCCGCAATGCTGGACACCGTAACAAGCCACATACTGAAAACTGGCTGTTCATTTGAGCCGTGGAAAGATATTTATTGCCCCTCTCAAGTCGGGCAGGCAGC | 120 |
| chr11 | 70641815 | 70641935 | 788855_47188552_ENST00000601538.6,E | 1 | - | good | CCCAGAGAGATTGGCCCTTTTGTGCTAGCACAGCTGCAAGCTGGGGCAGGGGCTTGCCTGCCCGCTGGACTGCCCACAGCCCAGGGATCTCGGGTGTGCTTTGTAGAACTGCACCCATTT | 120 |
| chr11 | 70641935 | 70642055 | 788855_47188552_ENST00000601538.6,E | 1 | - | good | CCATCCTGGTACCCAGTCCCCCGTAGCCTTCTCTCCAGACCCAGGAACTCCCAGGAAAGAGCCTGAACCTTGGTCACCCAGAAGAGTCATCCCCAGTGGGATTGCCTAGTGCAACCCTTG | 120 |
| chr11 | 70642055 | 70642175 | 788855_47188552_ENST00000601538.6,E | 1 | - | good | GTCTGGTTTCCTTTTAAAGAGACCCTGCGGGCAAAGGCTTCACCAACTGGAAAAGCCTGTGAAATGCGGCCTCCCTCAGGGGAATGCCAGTGACCCCAGCCCCTGTCCTTCTGACCTTAC | 120 |
| chr11 | 70642175 | 70642295 | 788855_47188552_ENST00000601538.6,E | 1 | - | good | TTTTAAGACCCATGGGCATTTTTAGGCGAGAAAATTTATCTCAAAGGCAGCCTAATTGCCAGTAGCTAGGCAGGATGGTTTGAGGTCAGGGCGCAGCGCAGCAAGTATTGGGTTGGCTTT | 120 |
| chr11 | 70642295 | 70642415 | 788855_47188552_ENST00000601538.6,E | 1 | - | good | CTCTGTCACTCACTGGTTGGTTTGTTAGCTAGGACATTGTCTTTATCCTGGGCATTTCCCCCCACCCCCCACAGCACTGCGATTCTGGATTTAAATTTGAATCGAACGGAAAATCAAAG | 120 |
| chr11 | 70642415 | 70642535 | 788855_47188552_ENST00000601538.6,E | 1 | - | good | ATTAGACAGGCTGAGCCGCCGTCCCCATAAATCTAACAAAGCTACCACTCTGTTCTCCAAAGCAGTTCACATGGGGTGACCTAGCAAGCTCCGTTCTGCCCAGACTTCCTGCTGAGGCCC | 120 |
| chr11 | 70642535 | 70642655 | 788855_47188552_ENST00000601538.6,E | 1 | - | good | GATGACGGGAGGTGTCAGGGGAAGGATGGAAGTGGCAGCCCGGCCTCTGTGTCCGTGGGAGGGAAGCACCCAGCTAGCGTGGACACAGGGGAGACACCTCTCGGCCCCGCCTCAACTGC[illegible] | 120 |
| chr11 | 70642655 | 70642775 | 788855_47188552_ENST00000601538.6,E | 1 | - | good | ATTCTCAGTCGCATGGAGGGCCTTTGTCATCTTCACCTCTTCAGCAGGTGCAACTCCTGAGCCAGATATAAATTTCCAGGAAATCAGTGTCCTGTTTGTGTCACTTGGTGTTTCCCAGAA | 120 |

| | | | | | | | | |
|---|---|---|---|---|---|---|---|---|
| chr11 | 70642775 | 70642895 | 788855_47188552_ENST00000601538.6,E | 1 | - | good | GTAGTCACCTTGGCTCTTCTGTAAAGTAACTTTTTTTAAAAACTTGTATTCTTTATCTAATTAGAAATAAACGAGCTGGTGAAATTATGGCTCTCAAGCCTGAGGGACCATCTCTGCGTT | 120 |
| chr11 | 70643015 | 70643135 | 788855_47188552_ENST00000601538.6,E | 1 | - | good | CCGTTGGAGGAAGAGATGGGTGGAGGGTGAGTCACAGGGGGCTCAGCTATTTGTTAACTGATACCTAGTAATTGTACATATGTACGGGCCCATGTGACGTTTTGATACATGCACACGATG | 120 |
| chr11 | 70643135 | 70643255 | 788855_47188552_ENST00000601538.6,E | 1 | - | good | ATATTATATGTATTCTCTGTACTCCACATTTTCCAAGGGAACATAAATGTTATGCAAAAGTTCTGGAAAGCTCCACACCAATGTGATAATCCTTCAAGATAAGCCTTCCTCCAAGGTAAC | 120 |
| chr11 | 70643615 | 70643735 | 788855_47188552_ENST00000601538.6,E | 1 | - | good | AACCCTCATTCAAATGACCAAAAACAATGCCAACGTATAAAAGAAAGAGGAATATTTGCATACCAACCACGACGTGATTCCCAAGACACCATCCCAAATACTAACCACAGGTCACAGACC | 120 |
| chr11 | 70643735 | 70643855 | 788855_47188552_ENST00000601538.6,E | 1 | - | good | GTTTCTGTTCACCTTTTGTTGTTGGTCAGTCTTCCCCCATGGGACCATAACTCTATGAGAGCAAAGACTGTCTTTCTTGCTCACCTACAAGAGAAAGCACCCAGCACATGGCAGGAGCTC | 120 |
| chr11 | 70643855 | 70643975 | 788855_47188552_ENST00000601538.6,E | 1 | - | good | GCTCCAGCTCAGCCCCCGACCCCAGATCTCCTCTGCCCCTCTGTGTTCCTCCCCCCCCTCCACCTGCAGTCCCTTAGTCGGCTTCCTGTTTATTCATAGCTCTGGCCAGCACACGGTATG | 120 |
| chr11 | 70643975 | 70644095 | 788855_47188552_ENST00000601538.6,E | 1 | - | good | GCTTTCCTGTTAAATCTGGGGCAAGTCCCAGCATGTTTCCTTAGCTCTGAAACAGGACCAGTGATAACCCCTGCTCGCCGCCACGTACAAACCCTTGGGTCAGCTGAGACCCTGGGCAGA | 120 |
| chr11 | 70644095 | 70644215 | 788855_47188552_ENST00000601538.6,E | 1 | - | good | ACTGGCCACTGTAAAATGGCAAATTTTATGCTTTGTGTCTTCTACTACATTTTTTTAAATTTGGAATTTTTAAAAAAGCAATGTTCTTCACTTTTTGTTGTTTAACTTTTGAATAGTTGG | 120 |
| chr11 | 70644455 | 70644575 | 788855_47188552_ENST00000601538.6,E | 1 | - | good | GAGGGCTGCTGGTCCTGCCCTTGGCCGGGGGCCCCCTCAGTTCCTGTCGCTGCGTCCACCCCACATGGAGGTATCCAGCCTGGGACCCATCGGGAAGCAGATGGACAGATGCCAGGAGTG | 120 |
| chr11 | 70644575 | 70644695 | 788855_47188552_ENST00000601538.6,E | 1 | - | good | TGTCCTCACCCCTGACTTTGCTGAGATCTGCATTGTCTCTGGGAATGCATGTTGGAAAGGTGTGTTCTTCCAAGATCAACCTCCCTCTTGCATTTGCAGCAGGCGGGAGCTGCCATGCGG | 120 |
| chr11 | 70644695 | 70644815 | 788855_47188552_ENST00000601538.6,E | 1 | - | good | AGGCATGGGCTGTTGTTTGCTGAATGCATAAAGGAAGGAATCAATGAAATGAGCACAGCAAAGTGGGACTCAGTGACACTGTGGGGTTTAGCAAGCAGGTCTGGAGATGGATTTCAGCAC | 120 |
| chr11 | 70644815 | 70644935 | 788855_47188552_ENST00000601538.6,E | 1 | - | good | ACGAGATCTTTTGATGAATTTCCTGCTTGTTCATAGTCCCTGCCAGGCTGGGGTCATCTTAAAGGCAGTCACCTTCTTCTCCACGTGTTTGTCTGAGGCACAGGTGTCGGTCTTGGCAAA | 120 |
| chr11 | 70645295 | 70645415 | 788855_47188552_ENST00000601538.6,E | 1 | - | good | GTTGTTGTTGTTCCTGGAGAACAGGCTGTGTGCCCAGCAGCCCCATGTCCAGCCCTATCCCCAGCCTATAGCCCCCCTAGACAGCCCCTGCCTCCGACCAGCAACACCGGGATTCTGGCA | 120 |
| chr11 | 70645415 | 70645535 | 788855_47188552_ENST00000601538.6,E | 1 | - | good | CTCATCTTTTCCGGGAAGACATTTTTTGTTGTCAGTTAATTTTTCCAGAGTGTTTTTCGATGAGCCTTCTGATTACAGTGGACTGATTTGTTGTTCAGTTTTTTGTTTTTTGTTTTTGTT | 120 |
| chr11 | 70645535 | 70645655 | 788855_47188552_ENST00000601538.6,E | 1 | - | good | GCGTTGCCTCCAATTCGTGACATATGCATCATTCTCTTGTTGTAAATATAGCAGCTGAACCTCGGCAGCACCCGTGTGGGGTCAGCAGTGATTGGTTTTAATTCCTGTATTGGGCCCAGC | 120 |
| chr11 | 70645655 | 70645775 | 788855_47188552_ENST00000601538.6,E | 1 | - | good | CAAAGCCCAGGGCACTCTGCCTTGCCCCAGCTCCCCAGTCTGCTGCTGGACTGAAGCGAAACTGTGAGGTGTCGGCCACTGAATTTGAGGGGACTCAGCCCTGAGGGCCTGTGGCAGGTG | 120 |
| chr11 | 70645775 | 70645895 | 788855_47188552_ENST00000601538.6,E | 1 | - | good | GGCAGGATTCAAACCCAAGTCTGGCTCCGGCTCCAGAGTTCTTGAACACTGTATGCCCTGCTGTGGCAGGGCAGGGGGTCAGAGGGTTCTCGGGGGTCCTGGCCACCCCGATGCTGGACT | 120 |
| chr11 | 70645895 | 70646015 | 788855_47188552_ENST00000601538.6,E | 1 | - | good | CCCCTTCCACGCTCAGCAGTGTGAGGAAGGAGATGGCATTAGTGTCAGTTTGCACATGGAACAGCTGAGGTTTGGTGAACATAGGATGACCCATCACAGGTGATGATGAGGGAGTGACAG | 120 |
| chr11 | 70646015 | 70646135 | 788855_47188552_ENST00000601538.6,E | 1 | - | good | TTGCCGCCACAGGCTACCAGCATGCGAGTTCTGGGGGTCCACACAGCTCTGGGAACAATAACAAGGACTGTTTCAATAATGACCTCTGCCACTTCTTGAGCACTTACTGTATGCCAGGGT | 120 |
| chr11 | 70646135 | 70646255 | 788855_47188552_ENST00000601538.6,E | 1 | - | good | ATGGATGAGTCTTTCTAGCTGGGAAGGGAAACGATCTTTCCAGCCTGAGGCTCACAGTCATTTACACTGAATGTCCTGGGTTTGAGGGGGCACCAAGCACAGGAGATCCAAACGCTGCCC | 120 |
| chr11 | 70646255 | 70646375 | 788855_47188552_ENST00000601538.6,E | 1 | - | good | TGGAGAGTAGAGAGCCTGATTGGGCACTTTGCCTTGGGAAGCCCTAAATGCTGCACTTGGGACACAAGCTTCCTTACCTTTGACCACCCTGGAGGGCTCAGGTGCGTCTGACACAGCCTG | 120 |
| chr11 | 70646375 | 70646495 | 788855_47188552_ENST00000601538.6,E | 1 | - | good | ATTGTCAAACTGAAGGAAGGAGGGAAAACATGGGCTGAGGAACTTGAGGGATGGCAGGGTGGCCCTTGGAGATCCTGTGTATTGGGAGAGGTGAGCTGTGGCTACCATGCACCACGAACT | 120 |
| chr11 | 70646495 | 70646615 | 788855_47188552_ENST00000601538.6,E | 1 | - | good | GTTGGTGAAACAGATTTGAATCCTATCTGATCCTTCTTCCCAAAGAGTGCAGAGTTCAGGTCCCATGCTTCTCCATAGCCCCTAAGCAGAGACATCCTGCAAAGTGGTTGCTCCACCCCT | 120 |
| chr11 | 70646615 | 70646735 | 788855_47188552_ENST00000601538.6,E | 1 | - | good | CCTTGAAAATCTTTCCAACAAGGACTGGACCACGTGGGGTTAGGGTATCTATCTGGGTTACACACATCTACCAAGTTATTTTGAGCAACTGCTTGTTAGGCTCAAGCCTGGAGCCTAGTG | 120 |
| chr11 | 70647215 | 70647335 | 788855_47188552_ENST00000601538.6,E | 1 | - | good | GGAAAGAGGAAAACAAAGAAACAGAGGGAGCTTAGGCAGAAAGCCGGAAGGAACAAAAGTTATGCACAGGCAAGACTTGAAGGAAAACTGGGGGCTTACTTCTACATACAGATGATAAAA | 120 |
| chr11 | 70647335 | 70647455 | 788855_47188552_ENST00000601538.6,E | 1 | - | good | CTGTTCTTCCAAGATCCACGTATCACTTCCCACCCTCCTGCTCCAACCACCCTCCTGTGCCACCTGTCAGGGAGGCGGGGTCTTGCCCAACTGGATATCTTCTGTGATGCAGATTCTAGA | 120 |
| chr11 | 70647455 | 70647575 | 788855_47188552_ENST00000601538.6,E | 1 | - | good | CTCACTTTGTGTTAAGTTTAACTCGGTGACTTTTATCAGCTCCTTTCTCTGCATGAGGAGGCCACAGCACAGAGTGGGGACTCACATACCCACAGACACACGGCGAGTGGTAGAGCCAGG | 120 |
| chr11 | 70647575 | 70647695 | 788855_47188552_ENST00000601538.6,E | 1 | - | good | CATGATACAGACTGCAAATAAGACTCCGAACAGCCACGGCAGTGACTGTGTAAAATGCTTGCGCTGCTTGAGCACCTACTGTGTGCCAGCACCGTTTTTGAGAGAGTTCACACACATGAA | 120 |
| chr11 | 70647695 | 70647815 | 788855_47188552_ENST00000601538.6,E | 1 | - | good | GAGTGCTGAGTTCACCACCTTCGTCACATGCCTGGGCAGAAGGACCGCAGAGGGAAGGGGGTCCTGTCCCCTTTTGTTAAGACAGAGCTCAGATCCAACAAAAGCTTAACTGCTGAGCAG | 120 |
| chr11 | 70647815 | 70647935 | 788855_47188552_ENST00000601538.6,E | 1 | - | good | TTCCTTAGGGTACATACCTAGGAGTGGAATTGCTGGGTCAGGTGGTCATTCTGTGTTTCTTGAAACTGCTGATGCACACCCCAGAGTCCCCATCCGGGGTGGGATGGGCGGGTGTTCAGT | 120 |
| chr11 | 70648175 | 70648295 | 788855_47188552_ENST00000601538.6,E | 1 | - | good | ACAGACCATAAAATTTACCATCAAACCCATTAGCACTCACTCTCCATCTCCCTCCCAGCCCCAGGCAAATACTCATCCCCTCTGGGTCTTGGGTCTGCCTGTTCCGGGCATTTTCCACCC | 120 |
| chr11 | 70648295 | 70648415 | 788855_47188552_ENST00000601538.6,E | 1 | - | good | ACAAGGCTTATCTTTGTGGAATTTTACTTCTTTATATGTGTCTGTTAGTAATACATTTTGTCATGTGTTTATACACTTTTTTTTTCACTTGGGTATGTCTTTTTTATTGTGGTAAAATAC | 120 |
| chr11 | 70648415 | 70648535 | 788855_47188552_ENST00000601538.6,E | 1 | - | good | TACATTAGCATTATTTCACTGTGTTTCGAGTAAATCATGTCAGACACAAATAAAATTGGTAATGCAGAGAAGCCAAAGAAAAAACCAGGGATGCTGCCACCAGGAGACAGTGTCCGCATA | 120 |
| chr11 | 70648535 | 70648655 | 788855_47188552_ENST00000601538.6,E | 1 | - | good | TGCCAGCCTTCTCTCCCCAGGCCCACTGGAATCCTGAGGGGTTTACAGGGGGAATTGTACCCCTGGTCCTAGGAAATTTTGAGTAAACTTTACCCCCATATCTGCCTGATAGCCAGCTTT | 120 |
| chr11 | 70648655 | 70648775 | 788855_47188552_ENST00000601538.6,E | 1 | - | good | CTTCCTCCCAAACCCCTTCCCCCTCTCCATCAGGGCCTGAACTTGAAAAGCTGATGGAGGTGGAGGGGTGGATTCATTTACATCAGAGAGAGCGTGAGAAGCCAGCCCTTCCCTGGAAGG | 120 |
| chr11 | 70648775 | 70648895 | 788855_47188552_ENST00000601538.6,E | 1 | - | good | ATAAGGGACCACCCAGAGCCGAGCACATGGCAGGCCACCAATAAATATTCCTTAAGGAAAGGAAGGCACCACTGAGGAGAGCACAGCCAGGATAGAACTTCCGTCTTGGACAAGGGTCTC | 120 |
| chr11 | 70648895 | 70649015 | 788855_47188552_ENST00000601538.6,E | 1 | - | good | ACAGTTTTGCCTCCTCCACCTGCTTTACCAGGGCACCTCCTGCATCTGCTCCCTGAAAGTTGCCTGTGATGCCCGCTCATAACCCTCTGCCATAGTCACTTCCTGCACATGCCCCATCAG | 120 |
| chr11 | 70649015 | 70649135 | 788855_47188552_ENST00000601538.6,E | 1 | - | good | TATGGGTGAGAAGGGGAGATGGCACGTGTGCCAGGCCAGTGAGGAGATGAACGTCGCCATTGGACACTCTCTCTGGTCTCCGGAATGTGCCACACCCCTCCTGTCCCCAGACACTTGCCC | 120 |
| chr11 | 70649135 | 70649255 | 788855_47188552_ENST00000601538.6,E | 1 | - | good | TTTGTGTTCAGTCTCCATGGCAGGACCTAGGGGGTCATGGAGATAGACAGGGACGGTCCTGGGCATGTGCACCTCCTTCAGCACCCTCAGGGCTGGGGTTGCAGATGGAGGAGTGGGTTA | 120 |
| chr11 | 70649255 | 70649375 | 788855_47188552_ENST00000601538.6,E | 1 | - | good | CTGAGAATGGTAGCCCCGCTGTGGTCTCATGAAATCAACCACCCACCAGCAAGTTCTCTACTTCCCTCTCCCCAGGATCTTCAGATGGGTCCCATGGAGCAGACTTTCCCATGCATCTCC | 120 |
| chr11 | 70649375 | 70649495 | 788855_47188552_ENST00000601538.6,E | 1 | - | good | AAGTGCCCCACAGGATCTTTTAATTGATGGAGTTTGTCAGCCCTGGGAATTAGGGTGTGTGTGCGCCAACAGTAAGCAGAGTGGGATGAGAAGGGGAGTTAGAGCTGATGCTGTGGAGCC | 120 |
| chr11 | 70649495 | 70649615 | 788855_47188552_ENST00000601538.6,E | 1 | - | good | CAGTGTTCTCAGCAGGCTCTTGTTAGTACTGATTTCTCTTCGCCTTTGGTGCCCTTTCGTGGGGAGTTTCATGATACAAGTGGTTAGACTGGCAAATGAACTGGAGGAAGGCCCTGTGCT | 120 |
| chr11 | 70649615 | 70649735 | 788855_47188552_ENST00000601538.6,E | 1 | - | good | TGGGCTGACATCTCCATGTTGGAAGCTGCTGGCGCCAGGTCCTCTGATGCACCCATTGGTGCCTCTGGGTCCTCTCCACCTGCCCCCCTCCTCCCTCCCATACACCCACTGCCCACCTCC | 120 |
| chr11 | 70649735 | 70649855 | 788855_47188552_ENST00000601538.6,E | 1 | - | good | CTGATGCTCCAGGTAAGTCCCAGATCATTGGGATGTAATCTTAACAGCCATCTTTTTCTGTGTGTGGATGCTGTACACCCAGCCCCAGGCTACATGGAGCGAAGGCTGCAGGAGGCATAG | 120 |
| chr11 | 70649855 | 70649975 | 788855_47188552_ENST00000601538.6,E | 1 | - | good | CTCTGCTACATCTCCAGCATGCAGCACAGTTCAGAGAGATTAAGAAGCCTGCCCAGGGCCCTCCAGGGAGGTCTGAATCTAAGCTGACTCCAAAGCTTGTTTCTTTTGCACTGTGTCTGT | 120 |
| chr11 | 70649975 | 70650095 | 788855_47188552_ENST00000601538.6,E | 1 | - | good | ACTCACACCGTGTGCTTCCATGACCATAGGAATCATACATCTCCCGTGTTAGGTGACTGACTTACTATCCACCTCCCTCAGGGAACATTCCCTGTGGGTCAGAGAGTTTGCTGACCAGGT | 120 |
| chr11 | 70650095 | 70650215 | 788855_47188552_ENST00000601538.6,E | 1 | - | good | CCTGCCATGTCCATCTGCATCTGCTTTGAGCCCCACTCCCCTCACCACTGAGGTTTCAGCTCACATGATGCCTTCTCAGAGAGGCCCCTCTTGGCACCCTCAGGTTGCCCTGACTCTCAC | 120 |
| chr11 | 70650215 | 70650335 | 788855_47188552_ENST00000601538.6,E | 1 | - | good | TGTCCTTGGAGTTTCTGGGCATAGAGGCTCATTACTTAGTGACAGTCTCATCTACTCGATGGAGAAATCTCTTCCACAAGGAGGGGCCAAGGAGCCAACATGGTTGAGCATCTGCTATGT | 120 |
| chr11 | 70650335 | 70650455 | 788855_47188552_ENST00000601538.6,E | 1 | - | good | CTCTGCAGGGTGCTGAACTAGACAGCTGGGAGGATTGGATGCAGCCTTCCTCAGGAGTCTGGAGAAAGCCCTCAGCATGTCCCCTCCCTCCATCCCTGCACCTTTCATTTCTGTCCCCTG | 120 |
| chr11 | 70650455 | 70650575 | 788855_47188552_ENST00000601538.6,E | 1 | - | good | CAAATGTGTGTGGCTGTACAGCTGCACAGAGCAAGAGCATTGCTGGATCTCTCCAGGGTGCTGAACCCTTGTGTGGCATGGTCTCTCTCCAGGGTGCTGAACCACTCCAGGGCACTGGCT | 120 |
| chr11 | 70650575 | 70650695 | 788855_47188552_ENST00000601538.6,E | 1 | - | good | TTGAGCAAAAGCTTAGGGGGTCAGGAATGTTCAAGGTGGATTTGGGGAGTGATCAGATGCTATGAAGCTGGGCATGACCGGCTGGTCTAGTGGGGGCCTGGGTTGTTAGTGAAGTAGGTA | 120 |
| chr11 | 70650695 | 70650815 | 788855_47188552_ENST00000601538.6,E | 1 | - | good | AAGCAGAGTGTGGTGAAGGCCCCAAAGTAAATCAGGAAAAGGGCTTCATGAAAGATGGTGTTGGCTCCAGGCTTTGATGGGTAAGTGAGGTTTTTGCAGGCAGAACTGAGGAGGAAGGTC | 120 |
| chr11 | 70650815 | 70650935 | 788855_47188552_ENST00000601538.6,E | 1 | - | good | TGAAGCTTCACTCATTCATTTCTGAAGCATACATTCTGCCTCCCAGGTACTGGGCTGGAAGAAAGACGTGAATCAGACACAGTTCCTTGAGGAACACACAGTCTAGAAGGGGTAGGCAAA | 120 |
| chr11 | 70650935 | 70651055 | 788855_47188552_ENST00000601538.6,E | 1 | - | good | CAGGTTGCTCATTTCTTGGCCTCAGTTTCTTGAACTGTAAATAAAAGTGTGTGTGAAGCATGTAGCAGTTTCTGGCATCATGAGTTCTCAACAAATGGTGGCAATTGTAAAAATTACTTC | 120 |
| chr11 | 70651055 | 70651175 | 788855_47188552_ENST00000601538.6,E | 1 | - | good | AGATGGCAGCACACAGGGTTGGAGAGAGGCAGAGAGGAGCCGCGGTCAAGGGCACAGGCTGTGCCATATGATCCAGGTTTGAATCCAGGCTCTGCCTCTTCCTAACTGTGTGAACTTGGG | 120 |
| chr11 | 70651175 | 70651295 | 788855_47188552_ENST00000601538.6,E | 1 | - | good | TGGCTCGTTCATTTTGGGGTTCTAATTTGAAGCATCAGTGATTTCAATTTCAGGCTATAAAAAGATTCAGAATCACTTCAGGGGACCATGGCTGCTGTGCTCAGATGCCATGGGACTGGC | 120 |
| chr11 | 70651295 | 70651415 | 788855_47188552_ENST00000601538.6,E | 1 | - | good | GAGGTCGTTCTCTTTGTCCTCCATATTGACTGGGAACAGGTGGTTCGGGCTGAGGAGTGGCCCAGGAATCTTGCTGAGAAGGTGGATGCGGGCAGCTCGGCAACTGCAAATCTGAAGTCT | 120 |
| chr11 | 70651415 | 70651535 | 788855_47188552_ENST00000601538.6,E | 1 | - | good | TCTCTGAGGCTTAATGCACTGTGGTCTTTCTTCTCCCAGACAGAGGCCACTGCTCTGTGGGGAGCCCTGGATGGCGGGGAGGTTGGATCCCTTTTCTGCATGTGGCCTTCTGGGCCTGGG | 120 |
| chr11 | 70651535 | 70651655 | 788855_47188552_ENST00000601538.6,E | 1 | - | good | GCTTCTCACCGTGTCGTCAGGATCCATTGACCAAGGGTAATAGCATTGGGGGCCTTTACTCTCTAAGGCTAATAGCACTGGGGATCTTCAGTCTCTGAGGCCAACAGCATTGGGGTCTGC | 120 |
| chr11 | 70651655 | 70651775 | 788855_47188552_ENST00000601538.6,E | 1 | - | good | GGGTCACATTGGTGCTTTGGTCATCAAGGCCAATATTTCATTGCATTCACACCAAAGACAAACAGGGTCTGTCCTGCATCCCTCTTGAGTGTGGGGTGTTTCTGTCCTGGCAGCTCAGTG | 120 |
| chr11 | 70651775 | 70651895 | 788855_47188552_ENST00000601538.6,E | 1 | - | good | CCCAAAATTTCTTCCACTCTTATTTAGATGGTTTTGAATGGCTACTTAATAACCAAGTTAAAGACCAAACACCAGCTGATGTTAGCAGGAACCCAGGGTGACTGCTGAATTCTTCAGCCA | 120 |
| chr11 | 70651895 | 70652015 | 788855_47188552_ENST00000601538.6,E | 1 | - | good | GGTTCAGCCACATCACACAGCTCACATTTCTCTAGCTTTCCTTTTGGGGTATTGTGAGAGACTTTGTCACATGTCCTGCTGGTGGACCTGATGACCAATCTAGGAACTCTCCACCCCATC | 120 |

| | | | | | | | | |
|---|---|---|---|---|---|---|---|---|
| chr11 | 70652015 | 70652135 | 788855_47188552_ENST00000601538.6,E | 1 | - | good | TTTTAGATCACCTTGGAATTCCTATTCCTGATGCCCACTTTGCCCTCTAGCAGCAAATCCCAGGGCCAAGGCCAGAGCCCTGTTGTGGTTTGCCTGGTGCCAGCATCCCTTGGGTTTAGT | 120 |
| chr11 | 70652135 | 70652255 | 788855_47188552_ENST00000601538.6,E | 1 | - | good | TGTCCCACCCCACCCGCCAGCCTCCAAAGTCCCTGGCAGGCATATCTCCTCCTTTGGCTCAAACCACTGCTCCCAGAGCACTTACCAGCCTGTACAATTCTCTGTTGGCATGGGTCTGGC | 120 |
| chr11 | 70652255 | 70652375 | 788855_47188552_ENST00000601538.6,E | 1 | - | good | TGTAGCAGTCACAGGGCACACCCACCATGTGCCAAGTACGTCGGACACTTCCAGCCAAGAGTACATGAGCCATGGTTTCCTTGTGCCTCTGTTGTGAGGAACATCACTAATAGAGGTACC | 120 |
| chr11 | 70652375 | 70652495 | 788855_47188552_ENST00000601538.6,E | 1 | - | good | CTTTCATCATCATTAATGATGCCTGACTCAGAAGAGGAAACCAGTGGCTATTAGTTGAACAAACGGGGATAATAATATTATAATAATGACTGATGTTTGTTACCACTATTATTTGAAGCA | 120 |
| chr11 | 70653335 | 70653455 | 788855_47188552_ENST00000601538.6,E | 1 | - | good | AAATATATATGTTTCTTGGGCTGGTATTTTAAAATGGGACTTGAGAAATGTGAGGTGGCCTCTAGCATGTTTGACTTTGCAACCAAGCTGGTGTGAAGAACAGTTGTTAGAGCTCGCTGC | 120 |
| chr11 | 70653935 | 70654055 | 788855_47188552_ENST00000601538.6,E | 1 | - | good | GTATTAAATACTTGCACGAGAGAGGGCGTGGATCTTCTTTATACTCGTGGTTTGTATTTAAGTCAGGATGCTAAGAGGGATGCCCTGGTGTTTTCCTCGGAGACCCTGGGAGTATCTCAT | 120 |
| chr11 | 70654055 | 70654175 | 788855_47188552_ENST00000601538.6,E | 1 | - | good | GTGGCCAATCTCATTTTTGTGTTAATTTCTGCAAATAGGTTTTCCCTTGCAGTTAAAGCAGGTGGCAGTATCCTTCATCCTTGAAAAGTGGTGCTCGAGTTTGCTAATTACATTTTCCTC | 120 |
| chr11 | 70654175 | 70654295 | 788855_47188552_ENST00000601538.6,E | 1 | - | good | TGCTGTCAGAGGAAAAAGCCTTCCTGGGCATTAATTAATTTCCCGGGGCCATTCCTTACCCGCCTGCTGCAGGGCTCTGATGACGCCTATGGTTTTTGTGATTGTTTTCTTGGCCCTGAA | 120 |
| chr11 | 70654295 | 70654415 | 788855_47188552_ENST00000601538.6,E | 1 | - | good | GAATCGAAGTATTCATATTTTGAAAAGTATCCATTTCATACATAGGAAACTGTGTCTTAGAAGAAAGTTATCTCACAGAGGAGACAAATGTTCAGCCCCCTTGGCACAGATAGTGCTGTT | 120 |
| chr11 | 70654415 | 70654535 | 788855_47188552_ENST00000601538.6,E | 1 | - | good | TACCATCTTTCCCCAGTATAGTCATAGAATAGAAAACAAAAAGGAAAGGAAATATTAACCCCTAGGTGGTGAGATTGTGTTATTTTCTCTTTCTTTCAGATGTTTGTCTCAGATTTCCTA | 120 |
| chr11 | 70654535 | 70654655 | 788855_47188552_ENST00000601538.6,E | 1 | - | good | GAATTCATAAGAAAGCATTAAAAATGATGTTTACAAGGAGCTCTTAATACCAACCAGAAAAGGTTACATTCTACTTAGTGAGAAAGGCAGAATCAAAATGATCAGCACAATATCATCTAA | 120 |
| chr11 | 70654655 | 70654775 | 788855_47188552_ENST00000601538.6,E | 1 | - | good | AAACAAAAAAAACAAAAACAAAACCAGGTAGCAATTACAGTGTGGCAAAAACAAGGAGTTGGGGCAGGGAGGGGGCGGGGCTAATAGGAGAATGGTTCAGTGGCAATGGACATCTGTTGT | 120 |
| chr11 | 70655135 | 70655255 | 788855_47188552_ENST00000601538.6,E | 1 | - | good | ATTTTCAAATTAGCAAATACTTTTTTGTGACTATACTCCTTGCTGGTGACAGCACTCATGTGTTGCTGGGGGAAAGCCTAAGTTGCCATAAATGTTTTCACAATCAATGTGGTAGTATGT | 120 |
| chr11 | 70655255 | 70655375 | 788855_47188552_ENST00000601538.6,E | 1 | - | good | TCCAACCCACATTGTCTGACTCTGTAATGTCATGGCTCAAAGATGCCTGAACAACACTTTGATGTGCTTTATGTATAAATGGGCAAAAGGTAAAAAGAAAAGCAGTGCGGTGACTGTTTT | 120 |
| chr11 | 70655495 | 70655615 | 788855_47188552_ENST00000601538.6,E | 1 | - | good | TCAACAGCAGGCAGGCAGAGGGCAAACAACATTTGCAACAAATGTCCCAAGAAGGCTGTGTCCTTGTTTATAGCAGCTGAGCTGGATCTCCAAACCCTACAAGCCCTGCAGTGCTCCGCT | 120 |
| chr11 | 70655615 | 70655735 | 788855_47188552_ENST00000601538.6,E | 1 | - | good | CCTAGTCTTAAGACTAGGACTGTCTTCCACAAACCCCTTGTTTGACAACCTAAATGTTGTTTCTTGAATGACTTTTGTGATCTAAAATATCCTGTAACTGAGACATCCTTACAGTTCAAC | 120 |
| chr11 | 70655735 | 70655855 | 788855_47188552_ENST00000601538.6,E | 1 | - | good | CCAACAGGCAGGCAAGGTGATGACCCACTGGCCCAGTGCAAGTTGACATTCCAGCCTCTTGATGTGGCCTGCCAACTAAAGTCTCCTCGACCCAAACACAAGCAGGTGCTGCGTTAATCT | 120 |
| chr11 | 70655855 | 70655975 | 788855_47188552_ENST00000601538.6,E | 1 | - | good | GGCTGGGCGTCAGTCACCTGAGGGTTATAGAAACCTCGTCTCCAGCCCACACGCCCTGAATCAGAATCATGAGGCGAGAGCTGGGCATCCATGAAAGCTGCAGTGGTGATGCCGGGGGTC | 120 |
| chr11 | 70655975 | 70656095 | 788855_47188552_ENST00000601538.6,E | 1 | - | good | AGGAGGATGTGGGTGGTTCCTGGATGACGGTAGTGCATCAAGTGTGTAAGGAGATGTGGAGGTGGGGACGCCACTTGGATGTCATGCAGGGCTCACTGTAGGTCTCACTCAGTGAAACCT | 120 |
| chr11 | 70656095 | 70656215 | 788855_47188552_ENST00000601538.6,E | 1 | - | good | AGAGTTTGCTGAAGACTTTGTCTGCTGTCTCTTTGAGATGGCCCTTATTAGGCCCCTTCACTCCCACAACATGGGGCTGGAGAGATCACAGGTTGCTTCTGAAGTTCACGACATGTGCTA | 120 |
| chr11 | 70656215 | 70656335 | 788855_47188552_ENST00000601538.6,E | 1 | - | good | ACAGAGGCCCATGTCCCTCTGGTGACAACAGGAAACCCAGTCCAAACTGGTTTTCATAATGATTAGAGCTTGTCAGGAAGCTGAGACCAGAGAGTTTCAAAGGAGAGTGAGCCTGCCATA | 120 |
| chr11 | 70656335 | 70656455 | 788855_47188552_ENST00000601538.6,E | 1 | - | good | ACCTGCTGAGGGAGGCTGAAGCCGAGGCTTGGAGCAGTGACAGCCCCCGAGAGAAGCAGGGCAGGTGTTTCTTTCTATGCAGGTCAGAGGTGAGGATGGGGCCTCCACCAGCCAGGAGTG | 120 |
| chr11 | 70656455 | 70656575 | 788855_47188552_ENST00000601538.6,E | 1 | - | good | GCCGTCCCAGATTGTGAAATGTCAGCCAGCCCAGGGGAGAAGGAGCCAAAGAGATTTCAGAGGCCGTGGGAGGGGCAGGGCGGCGGGAAGTTGAATGCCCCTGCGGAGAATGCCCGTGG | 120 |
| chr11 | 70656575 | 70656695 | 788855_47188552_ENST00000601538.6,E | 1 | - | good | AAAGATAAGTGAATGTCCCACGGGTAGGGACATTTGATCACAGTGGCCCCATGGATGGTGCGATTTGCCATTTCAGATCAGCTAGTGTGCCGGAATTCCCAGGATAGACAGTGACACTAA | 120 |
| chr11 | 70656695 | 70656815 | 788855_47188552_ENST00000601538.6,E | 1 | - | good | GACCCTACCAACCACAATCAAAGGCTTTGAGCTATCTCCAGATCGGGAGTCTCATGAGACAGATAGATGTTATCACTCAAGCTGTGTGGTAATAAAAGGTTCACCTCAGAGGTGGACGTG | 120 |
| chr11 | 70656815 | 70656935 | 788855_47188552_ENST00000601538.6,E | 1 | - | good | TCCTGCTGTGCATGACTTTCTTGGGTACTTAGCTTGTAGAATTTATTTCTCTCCTACCTAAGCTTTTGGGAAGAGTTTGTTCTTTTGCTGTTGTTTTTGTTCCTTGGTTGACCACCCCCT | 120 |
| chr11 | 70656935 | 70657055 | 788855_47188552_ENST00000601538.6,E | 1 | - | good | TGAAGTGAATGTTGGCTTTTAATGTCTTTGGTATCTCTAGAACTTCATTTTCCCTCGTTAACTTTGTGAGAATGCCCCGACGTGGCCATCCGGCTCTCGGCCTTGGTGCCTGTCATGATT | 120 |
| chr11 | 70657055 | 70657175 | 788855_47188552_ENST00000601538.6,E | 1 | - | good | CCTTGGCAAGCCTTCCTTTCTCTCTGCTATTTTGAAAGAACCTCGAAAGTGTGACAGATTTTTATTTTCTCTTTAAAAATGCATGCTGCGGTTTGAACTTTTGGACCAATGGGCACCCTT | 120 |
| chr11 | 70657175 | 70657295 | 788855_47188552_ENST00000601538.6,E | 1 | - | good | TTTTTGGTGAGCAGCCAGTCCTGGAGTCAGTGGGGTTTCTGAGCTCCCGGTGAGGTGCAAGGTCTCCACGTGAAAAGCTGCCCAAAATGCAGGCATTCTGTCTTCCTTACCCCAAGCAGC | 120 |
| chr11 | 70657295 | 70657415 | 788855_47188552_ENST00000601538.6,E | 1 | - | good | CTTGTATAGCATAATAGGTGCGTGTTTGTTCCAACAGACCATGTTTCTCTATTTTTAGTGAAGAGCTTTGACCTTGGAGGGATGGTGTGAAATAGCCTCACATGTCTCACGTGAATTTTC | 120 |
| chr11 | 70657415 | 70657535 | 788855_47188552_ENST00000601538.6,E | 1 | - | good | TCCCAGTTCATGTCCACAGCTGCCTCTGGGACTCAGAAATTCTCCTCTAGGGGCTTAAGACTGAGAATTCCATTGAGTTGTGTTCCTAGCGACTGGATTTTGCCACCGGAATATTAAAAT | 120 |
| chr11 | 70657535 | 70657655 | 788855_47188552_ENST00000601538.6,E | 1 | - | good | ATCACAGATAACCCAAGGAGCACTGGGCCCTCCACAGCCCTACGAGGCTGCTCCACACCAGTCATACACTAATGAGGGTGCGGGGTGCAGGCAGCTGGACCCTTGCACGATCCTGGCTCC | 120 |
| chr11 | 70657655 | 70657775 | 788855_47188552_ENST00000601538.6,E | 1 | - | good | TGGTATGCAAATAGATTTGCCATTTTGGGGCAGGTTTTATATTCTGGGTAACCTCTGACTTCCAGTATTCTAGGAATGGCTGGTCCTGAGATGCATTAGGGCTGCGCCTGCATTATGATT | 120 |
| chr11 | 70657775 | 70657895 | 788855_47188552_ENST00000601538.6,E | 1 | - | good | TGGGTGGGGCTGAGAAGGTGGACCTGTCCTGTCGGTTTGAAGCCTTTGTGGAAATTTGGAGCTTTGAGAGTTTCCACACTTCTGTTTCTAAGCACCTCCTCTGACATTGCTTTTAACCTC | 120 |
| chr11 | 70657895 | 70658015 | 788855_47188552_ENST00000601538.6,E | 1 | - | good | TAACCTCATCAATAAGAGAAACGTGGCCTGCTGAGGCACGTGTGTGGGCCCTGCACTTGCCCAGTGAGCGGAGCCTCAGTGGGTTAGAAGATGTAGCTGTCTGTGGCTGCAGCGCTCACC | 120 |
| chr11 | 70658015 | 70658135 | 788855_47188552_ENST00000601538.6,E | 1 | - | good | CAGGCAGGGTGGGGGGGGTGTTGAAATAGGTCACTCTTGGTTCTTGATTTTCAATAATAAAAGAGGGCTGCAGATGCAAGAACAACCCCCGTTGTTCAGCTCATTGGACTGTTTCCTGGA | 120 |
| chr11 | 70658375 | 70658495 | 788855_47188552_ENST00000601538.6,E | 1 | - | good | AATGAGCTTATTTGGGTTCTTGTGTGTTGAAACTGTCGGGTGCACCACCAGAATGACAGAGATTGGAGCCCGCTGTGGCTTTGTGGGCTAACTCGATTTGTAACCTTCATGGAAATGCAG | 120 |
| chr11 | 70658495 | 70658615 | 788855_47188552_ENST00000601538.6,E | 1 | - | good | AGCCAGGTAAGATGGAGCAGGTGCTTAAACTTATTCTGAGTGCCTCATGTTGATCTCTTTGATTTGCTGGAAGATGCATTTAAATTATTATTGACCAATCAGAACCTGGTTTGAATGAGA | 120 |
| chr11 | 70658615 | 70658735 | 788855_47188552_ENST00000601538.6,E | 1 | - | good | CCAATCAGGCCTAAGCTCCCAAGCTTCTTGTTCATGCTATGATTCTGCTAGTGATCTCCTTAGAGATCCTCAGGACTATTACGGGGCAATGGAAAGTCAGAAGGGAAGAGACTATTGGAG | 120 |
| chr11 | 70658735 | 70658855 | 788855_47188552_ENST00000601538.6,E | 1 | - | good | TGGCATCCAGCCAGCCATTTGGTGGTCCCTGCCATTGGGCTGGACAGACAGACATTCATTCTGTAAATGAGGAGCCCACATCTCCACCCGTGGTGAGAGGGCTACCCGTGATCATTTCTC | 120 |
| chr11 | 70658855 | 70658975 | 788855_47188552_ENST00000601538.6,E | 1 | - | good | AGACTCGAATGGTCCTCGTTGGGAGTGCGCATTTCTTGGATGCACGCGCTCTTCCATGTCACCCTGAGCCCAGCCTCGTGTACATCAGAACTCCACATGCAGGTGGAAGGGGTGGGTACG | 120 |
| chr11 | 70658975 | 70659095 | 788855_47188552_ENST00000601538.6,E | 1 | - | good | TTTTGTGAGTTTAGGTGTTCAGTTTTAATCTTGTTATGCAAAGGAACATGTGACTCTTTGGAAGTAGGAAGAAGAAAAGAATGTTACAGGTTGAAAAAGATGATATTTTAAAGGTTCAGT | 120 |
| chr11 | 70659095 | 70659215 | 788855_47188552_ENST00000601538.6,E | 1 | - | good | CATTCTCTTCCTTTAGAGAATAGAGGTTTGAGTGCTCCGGCTCTGCAAACAGTCCCTACGGTTGGCACGTTGGTGTGTGCGAGGGGTTCGGAGGCCCCAGTACATCTGATCACCTCGCCT | 120 |
| chr11 | 70659215 | 70659335 | 788855_47188552_ENST00000601538.6,E | 1 | - | good | GGCCTTCCAAGACCAAGGAGCCCGGGCAAGAGGGATGGCTGGTGTCGATGGTGCAAAAAGGAAACTTAGGAATTCAGATGCTTTGAAAATGAACTTTTGCTGGTACTGAAAAGGAAAGAG | 120 |
| chr11 | 70659335 | 70659455 | 788855_47188552_ENST00000601538.6,E | 1 | - | good | TCCTATGGGTGAGTCGGCAACCACTCTTGTGTGGCAGTTGCTGGCGTGAGGTCTGTAACATTGATGGCTAAGAGCTTGTAGATTTGCAGGTTGTGGTAACCACCCCATCAGATGGACGAT | 120 |
| chr11 | 70659455 | 70659575 | 788855_47188552_ENST00000601538.6,E | 1 | - | good | GAAGTAGAAGGTAGAAAGAGGAGGCATGTGGTCACCATGATGTTGGGGACATGTGCAGACCTGTGGGTGGTTTAGTTGTGACTAAATAGGCACCAAGAGGAGTCATCGTCCCTTCTTGTG | 120 |
| chr11 | 70659575 | 70659695 | 788855_47188552_ENST00000601538.6,E | 1 | - | good | GGTCAGTTTCTCCCACCGTGGGTGCCACTGCCTGTGGACCTGGTGTTGGGCTTGTCTTGGAGGCCTTGGCCGTAACCCGTTGGAAGGAGGAAAAGTCTGTGGAATTTGGTCATTGGTCTT | 120 |
| chr11 | 70659695 | 70659815 | 788855_47188552_ENST00000601538.6,E | 1 | - | good | GTGTCTGTATCTTTCTGCTTGGGGAGAGCGCCGACTCTCGGGGAGAGGGTCGTAGTCCTGGCAGCACAGCCACGAGGCCCAGTCTGGGGGTGCTTGTGGAGGCTGCCATGAACTTTCATT | 120 |
| chr11 | 70659935 | 70660055 | 788855_47188552_ENST00000601538.6,E | 1 | - | good | ATGAGTGGGCCCACTTCCTCCCGGGGTCCTGTGGGGAGGTCAGGCAATGTGAACTTGGAAGACATCTCCCAGGCTTGTAACCAGGTGCTGCTTGTCTTCCCAGCTGACACACCCATTGAA | 120 |
| chr11 | 70660055 | 70660175 | 788855_47188552_ENST00000601538.6,E | 1 | - | good | ATATGATGGCGGCCCCAAGGTGCAATTGACTTCATTGCTTGAGTTGCATCAGCCCATGAGTATGAATCTCAAGCCCTTCCTGGGGGAGAGTTGGCCTGGACTAGCCACCTGCAATGGGAG | 120 |
| chr11 | 70660175 | 70660295 | 788855_47188552_ENST00000601538.6,E | 1 | - | good | AACTTTTCCATTTTTAGAACCGAAAATGTTTAGTCACTGATGCTTTTAAAGAAAATGACAGGCGTGTAGACATGTACCACAGAGCACATTATCCTGTGGTTTCAGCAGAGAGGCCAAGAA | 120 |
| chr11 | 70660295 | 70660415 | 788855_47188552_ENST00000601538.6,E | 1 | - | good | GGTCAGCGTGATGTTCCGTCCGCGTTCCAAGTTGAATTCCCTTGGTTGCGAGTCTACGTAGATGTAAATTTGGCAAGTGGTTTAGTGTGGTGTTTCTGTAGCTCTGTGTGGCAGGGGAGG | 120 |
| chr11 | 70660415 | 70660535 | 788855_47188552_ENST00000601538.6,E | 1 | - | good | CACCCACACACACACATGCCAGTGCCCAGGGACACTGGTCACACACAGAGTGGTCACAGAGGCCTGGACTCCCTGTAGAGAGGACTCATGAGCACATTCCAAATCTGTAGCCACCCACTG | 120 |
| chr11 | 70660535 | 70660655 | 788855_47188552_ENST00000601538.6,E | 1 | - | good | ACCCACCCTTGGCCACTCATGCCTCCCTAGAAAAGTCCATCTCCTGCCCTGGAGTTGCACCTGGCAGCTGTGTGGGGCCCCAGAAGCCAAGCCTCACTGTGCTGGGCAGTAGTGGTTCAG | 120 |
| chr11 | 70660655 | 70660775 | 788855_47188552_ENST00000601538.6,E | 1 | - | good | CTCCATAGCGAGCTGTCCAGAGAGAGAAAGAGCAGATGCATCTGTCTCTGTGAGAGCATCCACGCCTGTCTGTGTGATTTGCTCTCCCCTCCCCCTGCTGGAATCTTTATCTCCACACCC | 120 |
| chr11 | 70660775 | 70660895 | 788855_47188552_ENST00000601538.6,E | 1 | - | good | GTGGCTCAGCCATGTGGAAGTTTCATCTTGGAGCTTACGCGCTGTGCCAGGGGTCCCGCACCTGTGCTGCAGGAGGCCTCCTCCCCAGCTGGCATGCCCACCTGCCTCAGACTCAGGCAG | 120 |
| chr11 | 70660895 | 70661015 | 788855_47188552_ENST00000601538.6,E | 1 | - | good | CTCAGGAATGCAGTCCATGGCGGTGCACCTGCCAGTCACCCTGCACGCGCAGCTTCCCCTCCAAAAGAAAACAAAATGGCATCACCTCCCCATGAGAGGGGACATCCGCCCACTGTGTGT | 120 |
| chr11 | 70661015 | 70661135 | 788855_47188552_ENST00000601538.6,E | 1 | - | good | GGCGGCAGGCATTTGTGTGCTCACCTCTAAGGTGACTTTGCAAAGGAAATGTGACAAGATGGTCCCCTCTGCCGTGGACTGAGCCGGCAGCACACACGTTACGTGGCCCGTCGTGGGAGG | 120 |
| chr11 | 70661135 | 70661255 | 788855_47188552_ENST00000601538.6,E | 1 | - | good | TAGTTGAGCCAGACTGTGGCTACCCTCAGAGCTCTGGGTCTCAAATCCCCAGGATTGCCCTGGCCTGAGTTTCAGGCACCTAGAAACTGCCCCATTCTGGGAGTGGGCTTTGGCTCCAAG | 120 |
| chr11 | 70661255 | 70661375 | 788855_47188552_ENST00000601538.6,E | 1 | - | good | GCTTCCTTGAGGAATGTTTTTAATTTTCTGTAGATGATGTTGTTCTGAATGGGTTCTCTCTCCCTCCTGCCCCTCCTACCTTTTGTTTTTAAATGATGGGGTGGTCAGTATTTTGTGTTC | 120 |
| chr11 | 70661375 | 70661495 | 788855_47188552_ENST00000601538.6,E | 1 | - | good | CCTGCATGAAAAACGTTCCCCAGCTCAGGAATAAGCATTAAATAAGCATTTTTCCTGGGGGCACTCAGTTGCTAGACAACCATGTTCCTTCACATTGTGCCTTACTCAGAAAACGTTAGG | 120 |
| chr11 | 70661615 | 70661735 | 788855_47188552_ENST00000601538.6,E | 1 | - | good | CGCGGTGATGATGACGGGCTACAATAATGGTCGCTGTCCCCGGAATTCTCTCTACAGTGACTGCATTATTGAGGAGAAGACGGTGGTCCTGCAGAAAAAAGACAATGAGGGCTTTGGATT | 120 |

| | | | | | | | | |
|---|---|---|---|---|---|---|---|---|
| chr11 | 70661735 | 70661855 | 788855_47188552_ENST00000601538.6,E | 1 | - | good | CCCCCTCTTCCTTCGCTCCTCCTGCACCCTCGCTGCCTCCTTTCCTCCATGCTGCCTGGATCTGGCGAGCTGGGGTGATTAATTGGCTATGATGATGAACGTCCCCGGCGGAGGAGCGGC | 120 |
| chr11 | 70661855 | 70661975 | 788855_47188552_ENST00000601538.6,E | 1 | - | good | TCAGCCATCCCTCAGGCAGCCGCCCCACCATGTATGACTCCTTCCTCCGGCTGGCTCCGGCTCTGCCTGCCCCCACCCTGCAAGTCTCATTGTGCCACGCTCGCCTATCCCGGTAGCCAC | 120 |
| chr11 | 70661975 | 70662095 | 788855_47188552_ENST00000601538.6,E | 1 | - | good | CGGAGGAGCTCATTCCCGACGCCGCCGCTGCTGAGGCCGCCGCTGCCGCCGCCGCCGTTCGGGCCACTTACTTTCTTTGCCCTGGGCGGCGTCCTGGCCAGCCCCCCTTGAGCCTCCAGC | 120 |
| chr11 | 70662095 | 70662215 | 788855_47188552_ENST00000601538.6,E | 1 | - | good | ATAGGATTTGATTTTATTTTAAACCCCCTTCAAGGAAAAGCAAAACAAAACTGCTGCCTCATTTCTCCTCTGGCTGATTTTCCCGGGCTTCCCTGGAAGATGCGTGTGGTCGGGACAAGC | 120 |
| chr11 | 70662215 | 70662335 | 788855_47188552_ENST00000601538.6,E | 1 | - | good | TGTGGGGGCTCCTGCCTGCCGCACACCGCCGGCGCCGGGACGGGAGAGGGAGGGGGCGCTGGATGCGACGCGCAGGTCCCTGCGTCTCTGGTGCCTGCTGCCGGCTCGTGCGGGGAGCA | 120 |
| chr11 | 70662335 | 70662455 | 788855_47188552_ENST00000601538.6,E | 1 | - | good | GGGATGCAGAGAGAGCCCCAGCCGCTCCCCCTCCCCGCCCTCTCCCAGCCCAGCCTGCGGATGCTCGCGCGGGATGACGCCCAGGCAGCCGCCGGGCGCACTGTGGGCTCGTCAGATGAT | 120 |
| chr11 | 70662455 | 70662575 | 788855_47188552_ENST00000601538.6,E | 1 | - | good | GTCCTCAGGGGCTTTAACACCGAAATGTCCTGAGTCCTCTGCCAGGGAGCCCCCACCCCTCCCCGACCGGCCACCGGCCACCTTCCAAGGCTGCAAGGGACCGGGGGGACGCTGGAGGGG | 120 |
| chr11 | 70662575 | 70662695 | 788855_47188552_ENST00000601538.6,E | 1 | - | good | AAAAAAATGCCATGGCTAGAGGCAAGCAGTTGTGGCTTTAGCTTTGTCATCTCAAAGGGGCAGCCACATACCCCCTCCAGTTCCCCAGCAATGAATTCATCATCCCCAGAGGGGGCTGGT | 120 |
| chr11 | 70662695 | 70662815 | 788855_47188552_ENST00000601538.6,E | 1 | - | good | CCACACACTCTGGGTGCCTCGGGGGCTGGATGTCATGTGGCGAGTGCCAGCAGGAAAAGCGAGAGGACACTGTGGCCACTAGAGCAAAGTTGGAGCTTTTTTGGTTTCTTTCCTTAAAAA | 120 |
| chr11 | 70662815 | 70662935 | 788855_47188552_ENST00000601538.6,E | 1 | - | good | GGGCGGGGGGGTGCCGGTAGCCTTCATCTCACTCCCCTTCTGCCCCCGCAGGCCACCCAGAAGGTGGGGGTGGAGGACAGTTCAGCCGCAGCCTGGAGGCTCACTGAGCAGGGCACCGTC | 120 |
| chr11 | 70662935 | 70663055 | 788855_47188552_ENST00000601538.6,E | 1 | - | good | CACAGCCCCGCAAGACCCTGTAGAGGCCTCCCTCCACGCTGCCTCCCTCCCTTCCCCGCTCCCCTCCACAGAGGCCTTCCCAGCTCCCCACTAACTTCCGGGCAGCAGCAGAGTCCTACG | 120 |
| chr11 | 70663055 | 70663175 | 788855_47188552_ENST00000601538.6,E | 1 | - | good | TGCATCCCTGGGACTGCCCCTGTGCGACGGGGGCCTGGTGGGTTCCTCATGACCCATCTGCAGCTTTTTCCGGCTGGGCCGCTGGCCTGGTGTCCTTTCTCCTCACCCCCGAGGCTCACA | 120 |
| chr11 | 70663175 | 70663295 | 788855_47188552_ENST00000601538.6,E | 1 | - | good | TGGCTGCACCTGGGGCCCTGTGCTTCCTGTAGACCGTGAACCTGGTGCTGGTGCCAGGCAGAGCTGACCCGCACTCCTCTGGGACCAAGAGTCCGTCAGCACATGGCCTTGTTTGAAATT | 120 |
| chr11 | 70663295 | 70663415 | 788855_47188552_ENST00000601538.6,E | 1 | - | good | GAGGATCGAGGGGCCGCTGCTCGTCCCTTGTCCAGGATACTGAGCCCTGTCATCCACGTTTGCCCCGTCCCTCCTCCTCCCCAGGGCCCCCCAATCCGGGGTGGCTGTGCTGTCACCAGC | 120 |
| chr11 | 70663415 | 70663535 | 788855_47188552_ENST00000601538.6,E | 1 | - | good | CTCCATGCACGACCCTCTGATGCTGTCGGGGGTGGGTGCGTGGAGTGGTGAGAAGAGCACGCGTTCCCTAGCTGGGTCCTCCGATCTCTCTGAGCCTCACCAGCCTCATCTGTAAAATGG | 120 |
| chr11 | 70663535 | 70663655 | 788855_47188552_ENST00000601538.6,E | 1 | - | good | AACCCCCCTCCCAGCTGCTGACAACGGGAGTGACCAGTCACCTGGATTGTCCTGGTCTTCGGCTGTGAAGCTATCATTCATGCTGGGCTATTCCTGGAAGACAGGGTCACCCCAGCATGC | 120 |
| chr11 | 70663655 | 70663775 | 788855_47188552_ENST00000601538.6,E | 1 | - | good | GGCTTCCAGGGATGAGCCAAGAGGGGCTCACAGTTGTGACCGAGTTCCCCAGAGTGGACCTCCCTGCACTTAGCAGGCTGTCTCTGGGGAGGGGGCAAGGTAAGGAAGGTCAGGGGCCAG | 120 |
| chr11 | 70663775 | 70663895 | 788855_47188552_ENST00000601538.6,E | 1 | - | good | GAGAGCCCCATCATTTTCTAAAGGTCACAATAACCATGTGACCTCACCCCCAGGCTCCCAGGGTCCTGGACCTCTGACCCTGTGTCAGCCCTTGTTTGGTGGCCCTGAGCAGGGTAAGTG | 120 |
| chr11 | 70663895 | 70664015 | 788855_47188552_ENST00000601538.6,E | 1 | - | good | CCTGAGGCATCCTATTACACCACACCAGCATGGCATGGCCTCATCTGACCCCATGGTGATCCCAAGAGGCCTGTAGGTTTGTTCTTACCCACTCTTGCTGTGCAGGGAAACTGAGGTTTA | 120 |
| chr11 | 70664015 | 70664135 | 788855_47188552_ENST00000601538.6,E | 1 | - | good | ACCCCTAAATCCATGGCAGTAGAGCCCCAGCTCTCCCTGCTCAGCCTCCTTAGGGCTGCAGAGTCCTGGCCACCTACCAGGCCTGCTGGAGGTTAATGGCAGCCGTCACAGGGACAGCGG | 120 |
| chr11 | 70664135 | 70664255 | 788855_47188552_ENST00000601538.6,E | 1 | - | good | AAATGGGTGGTGACTAGAAGGCATTTCCCATGAGGGACACAAGTCTGTGTTCTCGTTGTGGGGGGCCGTGGAGACAGGTCGAGGGACCTGGTAGAGGGTCTGCTCATCAATGCCTCATGG | 120 |
| chr11 | 70664255 | 70664375 | 788855_47188552_ENST00000601538.6,E | 1 | - | good | AACCTTCCATGAAGGAGACACAGGAGACACACAGGCTGGTCACACAGCTTGCGCCTCCCCAGCCGTGAGTGAGCCATGGGGCCGGGGTGGGCTGACTCCCCTTTCAGGAGTTCAGAGGGA | 120 |
| chr11 | 70664375 | 70664495 | 788855_47188552_ENST00000601538.6,E | 1 | - | good | CGGCATCATCCTTCCAGGGTTTCAGGGAGGCGGTCACCCTCAGCAGCTCCTCTTGCAGACAGCAGGTGCACAGCAGGGTTGAGCAGAGGCTCTTGGCTTTGGCAGCCTTGGCCTTTAGAG | 120 |
| chr11 | 70664495 | 70664615 | 788855_47188552_ENST00000601538.6,E | 1 | - | good | TGCAAGTGAGCACGCCTGGAGCGCATGACAGGTTCCCCTTGGAGCTGGGTGGAGATGGACTTGATCAGCCCATGTGTCCCCAGGGAGAGTGCAGGGCCCTGAGCTCCAGACCCCAGGGTT | 120 |
| chr11 | 70664615 | 70664735 | 788855_47188552_ENST00000601538.6,E | 1 | - | good | CAGCCCCCCTGGCCCTAAGAAGTTCCCGTCGCAGGGTATGCTGGCAGGAAGTGGTCTGGGTCATCTTGGAGATGCAAGGTTTGGATGGCGTTGGGCACTGCAGGAGGAGCTGCCCAGCAG | 120 |
| chr11 | 70664735 | 70664855 | 788855_47188552_ENST00000601538.6,E | 1 | - | good | AAAGCAAAGCTGGGATTTGAACCACGCTCCGTGTGATGCCAAAGCCTGTGCCTCCACCAGCCACACTGGAATGCCCCTGCCCTTCTCTAGTCTATGGGAGACAGCTAGCAAGTGTCTGGG | 120 |
| chr11 | 70664855 | 70664975 | 788855_47188552_ENST00000601538.6,E | 1 | - | good | GGTTCTGAGGCATGGATGCACGACTCCACGGCCTCCAGCTCAGGACCCATCAGGTGCATCTTGTTGTTCTCATTTTGCAGAGCAGGCTCTGGGGCCTGATAGGTCAGTGACCTTGGGGTC | 120 |
| chr11 | 70664975 | 70665095 | 788855_47188552_ENST00000601538.6,E | 1 | - | good | GTCGAGATCAGAGGGGTCCCTGTAGGAGGAAGGGGGTCCGGGGAAGAGAGGGTGGCGTGTGCAGACAGGGATCCCACTTGTAACTGTTGGAGTTATGAACATGTGCTGCTTGCCTGGGCT | 120 |
| chr11 | 70665455 | 70665575 | 788855_47188552_ENST00000601538.6,E | 1 | - | good | AAGTTAGAAACAGTCCTTCTTAATGTCAGAGACTAAGCAGTTTGGTGTTTTACTGAATAAACAGGACAGGGCCATTACAGTCTTGGGGCAAGATCTTTGCACAGTTACAAGTGTGGGCTA | 120 |
| chr11 | 70665575 | 70665695 | 788855_47188552_ENST00000601538.6,E | 1 | - | good | AAGGTGGTCAATGTCTGGGTTGAAATTCAAACCCATGTCTGTCTGCCAGTGAAAGTGAGATGGGTTCATGGTGGGTGGGGAGTGGGGGCTGTTTAGATCCTGAAACAATGCCGGAAAGAG | 120 |
| chr11 | 70665695 | 70665815 | 788855_47188552_ENST00000601538.6,E | 1 | - | good | AAGGAGCCTCCTGCTCATGGAGCATTCACTGGGCATCGCCTCACTTGCTCCTCCTAAAACTTACAAGCCAGAGGTTTCTCCCCAACTTTGCAGGTCAGGAAATTGAGGCTGTAAGTAGTC | 120 |
| chr11 | 70665815 | 70665935 | 788855_47188552_ENST00000601538.6,E | 1 | - | good | ACCTTGTAGAAAATGTCCCCTCCCCCACAAGAGACAAGCTCAGAGGGGACATTTTCTCCAGGGTCACCCAGCTCAAGAAGGTGGACGGGGATGCTAAGAAGAGCAGCTTCCCCTGGTTAC | 120 |
| chr11 | 70665935 | 70666055 | 788855_47188552_ENST00000601538.6,E | 1 | - | good | CTCCTGACTTGAGTTCATACCCATTTTGCATCCCCTCTGGGCACACATGGGGCACCTGGCCTATGCGGGGTGCTGTGGGCATTTGGGGTGGTCTGCAGGATGAGGTCACTGCCCCCATTG | 120 |
| chr11 | 70666055 | 70666175 | 788855_47188552_ENST00000601538.6,E | 1 | - | good | ACTGAGTCTACCTTCTGGGTTTAACTAGGAATCCAGGGATTGTTCTGGTGCCTTTTTACTCGAAACCCTGTAGTCATCCTTCGCCCACCTCCCACCCAACATCCGTTTCTCCCTGCTCTG | 120 |
| chr11 | 70666175 | 70666295 | 788855_47188552_ENST00000601538.6,E | 1 | - | good | CCATTGCTTTTCATGAATTCCCCAAAACATTGAAAGTTTCTAAGGGCCTGGGGCTGCCTTTTCTGTTTCCTCTCTGACTTCCTGGGCCAAGCCAGGTGCATGCCGTAGGTCCTCTGCAGC | 120 |
| chr11 | 70666295 | 70666415 | 788855_47188552_ENST00000601538.6,E | 1 | - | good | CTAAGGCCTGCGTGTGAGACCCCCCTGTCTGATTGCATGGGGATGGGGGGTGCTGCCAAGTTCCGCCTTTCAGACACAGTGACTGCAGCTCCTTTCAAGGAATGTCTCTTCTAGAGTGTC | 120 |
| chr11 | 70666415 | 70666535 | 788855_47188552_ENST00000601538.6,E | 1 | - | good | ACAGAACTCAATGCCACAACTTCCGGACCCAGCCGCAGCTGTCATCCTGGGATAGGTTTTAGATTCATGAACCAGCAGAAGGAGCCTCTGAGGGTTTCTTGTCTCCAGAAAGGCCCCTTC | 120 |
| chr11 | 70666535 | 70666655 | 788855_47188552_ENST00000601538.6,E | 1 | - | good | TGCATCTCATCTGCATGTGGAGAGTGGGTCTGCTGGTGCAGGGGCAGGGCAGGCTGGCAGGGGCCAAGGTCAGGGCAGACTCTGGTTCACAGGGACTTCAGCACAGTTCTAGGGCAGCAC | 120 |
| chr11 | 70666655 | 70666775 | 788855_47188552_ENST00000601538.6,E | 1 | - | good | CTTCAGGCACGTGATCAGTGAGCAGGTTAGGGGGTGAGACGGGCTTGGAGAAGGCTGTCCCAACACAGCGTGGGGAACGTGTAGGCGCCTCGGTCATTTCCCCATGAAACTGGATTGCAG | 120 |
| chr11 | 70666775 | 70666895 | 788855_47188552_ENST00000601538.6,E | 1 | - | good | GAGTGCGCCGCTGGATTACATTTTTTGTTCAATCAATAAGCACGGGTCAATGTTGTGTTGCATGGAGATACTCAGAGGATCAGGGCATGGCATTTGCTTACAAGGAGCCCACAGTGCCTA | 120 |
| chr11 | 70666895 | 70667015 | 788855_47188552_ENST00000601538.6,E | 1 | - | good | GGGGCTGGTTGCCCACCCCCTTCACCTTCCACAGCTCCTGCTGCAAGCAGATGTTGGCAAAAGAGGGTACATTTGGGGCATGTCAAAGTGCCCTTCCACCTAGCTCAGAAAACATTTGGG | 120 |
| chr11 | 70667015 | 70667135 | 788855_47188552_ENST00000601538.6,E | 1 | - | good | TTAAGACGTTTTTCTTCCCTCTGTGACGAAATTACCATTAGTACCTGTGCCGTGATGGCCATGATGGCAATGGTTTCCTCCATAGTTGTACAGTGCACAACCTGCACAGCTGTACTTGGT | 120 |
| chr11 | 70667135 | 70667255 | 788855_47188552_ENST00000601538.6,E | 1 | - | good | TGTCTCCGGGAAGCCAATATGGCTCCCATAGGGATACCTCTTGCATTCTCCATGGCAGCATGGCCAGGGCAGCCTAGACCCCATCTTTGTTTGAACTAGGAAAAGGTGCTTGCCCCTTCC | 120 |
| chr11 | 70667255 | 70667375 | 788855_47188552_ENST00000601538.6,E | 1 | - | good | GTGTATCCCATAAGCTCTCTTGCAGCATACAGTGGGACCGTGGGTGAGCAGAAACAGGCAAACCTTTGATGCTCTCTCCTCCAAGTGGCTCCCAGGAGGGCATCAACATGGAGGTAGTTG | 120 |
| chr11 | 70667375 | 70667495 | 788855_47188552_ENST00000601538.6,E | 1 | - | good | GGCAGACACCAGGAGCATCTTTAGGCCTGCCAGCGGGGAGAGACCTCTGGGGCCTGCTGCGTGGCCATCCCTGGGTCTCCACAACCCTCCTGTCTTGCTTCAAGGTGCTAAGTGCTCCAA | 120 |
| chr11 | 70667495 | 70667615 | 788855_47188552_ENST00000601538.6,E | 1 | - | good | CCCGCTGGACCAGGGAAAGTGCAGCCTTCCCTGTGAGTCTGTCATGGCGCCCAGTGGGAGAAAGGGAACCCGGGCCACTGCTAAAAGCCACAGGGTGTGTGGCCACCTGTGGTCTTGCGA | 120 |
| chr11 | 70667615 | 70667735 | 788855_47188552_ENST00000601538.6,E | 1 | - | good | GGCCTGGCTTCCTGTGCCTCCAAATGGACATCACCTCGAGGATCAGGTGCTGCTGCCAAGGGAGCTTGTTAAAGTGACTGACAGATCCTAAACGCCAACTGTCATCACTGGCACCTGCTG | 120 |
| chr11 | 70667735 | 70667855 | 788855_47188552_ENST00000601538.6,E | 1 | - | good | TGGTCTACCCCAATAACACTGGTGTTCAGAGTTAAAATGTCCACATCTATCCACCCCATGTGTGGGGAGGCACACAGAGCCACCCAGAGCCCCTCAGGCCCATACGTCTGGGCCAGCTCA | 120 |
| chr11 | 70667855 | 70667975 | 788855_47188552_ENST00000601538.6,E | 1 | - | good | TGGCCCCAGTTACTTCTGAACTTTGAAACCCAGTCCCAGCTGGTCACGGAAGAACTAAAGCTGTCACCAGACGGGGCCACCAACCCAGCTGGGATCCTGAGAGCTATGAGTAAAGCCATC | 120 |
| chr11 | 70667975 | 70668095 | 788855_47188552_ENST00000601538.6,E | 1 | - | good | AGACTGGGCCTGGTTTGTTCCTTCCTAAATGCCCAGCACCCAGTTCATCCCCTGACTCTCAGTGGGTATTCAGGATGTAACAGCTGGATTGATTTATTGATTTACACGGCGGCTTGACAC | 120 |
| chr11 | 70668095 | 70668215 | 788855_47188552_ENST00000601538.6,E | 1 | - | good | AAGACCCAAATTCCAAGTGTCGTCACTTTCACAGACGCTGGGGGTTAGGAATTGGACACATTTTTTTGGGTGGGTTGAGTCACCATTCAACTGACCACACCAGACAACTCTATAAGGACC | 120 |
| chr11 | 70668575 | 70668695 | 788855_47188552_ENST00000601538.6,E | 1 | - | good | GTCTTCCAGGTGCCCGTATGGTTCTCACAGAACCACGTCCTGCACTGTTGAAGGCACACATTTCTGTGACTGTGGATTGATGTTTATCTTCCTCACTTGCCCGGCCTTTGCTCTAACGAT | 120 |
| chr11 | 70668695 | 70668815 | 788855_47188552_ENST00000601538.6,E | 1 | - | good | TCAGGGCCGGCTGCCTACCTTCCAACCCCACTCTGTCTGGAATGTTCTTTTCTCTTCTGTCCCCTCCCCAGTCCTCCAACCCATTCTCCTATGTATACTTCAGTCCTCCTCCTCCAGGAA | 120 |
| chr11 | 70668815 | 70668935 | 788855_47188552_ENST00000601538.6,E | 1 | - | good | ACTTGTGCGTCACCGCCAAGCTGTGTCCCTAGACTGCCCTGGGTGAGTAGGCAGCCAACGTCTGCAGCCCCGTGTGGTCAGCTGCTCCTGCCCTCCCGCCTCATCACGGCCCTTCAGACA | 120 |
| chr11 | 70668935 | 70669055 | 788855_47188552_ENST00000601538.6,E | 1 | - | good | TCGGGCCTCACCGGAAAGAACGCCAGGAAAAGCCATAGGGCAACAGCTCTTCTGCCATTTCAGTCCTCATGGGTGAAACTGGGGTAAAGCGTGAGTGGCTGGGGCCCTGCTGACACCTGC | 120 |
| chr11 | 70669055 | 70669175 | 788855_47188552_ENST00000601538.6,E | 1 | - | good | GGAGCCTCCTCAGCCAGCAGCAGCCGACTCGTGCTTGTGCATTGTTCAGATGGAAGCATGATCATTGGATGGAAGTTCCTGGCACAGGTGCAAGTTGACCCTGAGTCATTCCAGAAGGAT | 120 |
| chr11 | 70669175 | 70669295 | 788855_47188552_ENST00000601538.6,E | 1 | - | good | CATGGAGGCAGAGGTGGGAACTCTGTCATCCACCGTTCCCCACAGACCACTCACAGCCGCTTTAAGAGAACAGCCAGGGCTTCTAGGCAGGAGCACGACAGTGTTCACGCTGATGGCCAG | 120 |
| chr11 | 70669295 | 70669415 | 788855_47188552_ENST00000601538.6,E | 1 | - | good | CTGGACCTTGGACCTCACACGTGCACATGTTCTGACATCAGTGCCTCATCCGATGTCCAACCCCGGCCAGAGGAACCACCGTCGCTTCAGCCCCACTCTGACCTTGGAAGCCCCACCCTG | 120 |
| chr11 | 70669415 | 70669535 | 788855_47188552_ENST00000601538.6,E | 1 | - | good | GAGGCCCCTTCCGCACATCCAGGGCGGGTCGGCTGCTCCAGTTCTGCCTTCCCACCCAGTGCTGGGATGTGCAACGCCCCCCACCGAATGCTCAGATGTGCAGCCCCCTCCTCTGCTGGC | 120 |
| chr11 | 70669535 | 70669655 | 788855_47188552_ENST00000601538.6,E | 1 | - | good | TTTATTTTTCAGGTTGCAGACACATTTGGGGGTCCCTGCAACCCTCGGAGACAGCTGGAGTCTGGACCTGTAGGGCCTGGTTGTGGAAACAGCCCCGCTGCCTCCTGCCCGGTGGCTCTG | 120 |
| chr11 | 70669655 | 70669775 | 788855_47188552_ENST00000601538.6,E | 1 | - | good | TTTATGGATACCAGCTTAAATGGGTCGTGACGTTGACCCCGTGTTAAATATCAGAGACACTGAATTCAACAGAGATTAACCGCACTGTTGTTTTCTCTTTGGATTTTGGAGCCAGTGAAT | 120 |
| chr11 | 70669775 | 70669895 | 788855_47188552_ENST00000601538.6,E | 1 | - | good | TACACTGATCGTTCCCCAGCAGCTCCTGTGATTCCCCAGCCGCCATCAGCACACCCAAGGATTCTTGCAAGGTGGGAACACCTGCCCTCCGAGTGGTTCTACACGCAATGAAGTGATGCG | 120 |

| | | | | | | | | |
|---|---|---|---|---|---|---|---|---|
| chr11 | 70669895 | 70670015 | 788855_47188552_ENST00000601538.6,E | 1 | - | good | CCCGCCTGGGTCTGATCAGACATCACACCCACAGCCCATGTTCAATGTACCTTTCTGCCATGAAAAACATTCAGAAGTGATTTAATCATTCCGCCTTTGAACTTGTTTGGCCGTGAGTGA | 120 |
| chr11 | 70670015 | 70670135 | 788855_47188552_ENST00000601538.6,E | 1 | - | good | GGACTTTCAGGGGCAATAAACGCACCACCCAGTCTCTCTGTGGGGAGTCCTTCCTGGCTTTCCCAGGTCCCACCACCCACCTCTATGGACCTTGCTCCACCCCCCTGCATTCCAGCGGTC | 120 |
| chr11 | 70670135 | 70670255 | 788855_47188552_ENST00000601538.6,E | 1 | - | good | TGCCTTCACCAGGCCCTGCCACTGAAATGCGATTATACCAGGAATGATTTAGCGAAGCTGCTCTCAGGCCCTTCCCTGCCTTGGAGAAGGCAGATCTCCTGGCTCCAGCTGCTCTGGGAA | 120 |
| chr11 | 70670255 | 70670375 | 788855_47188552_ENST00000601538.6,E | 1 | - | good | GCTGAAAGCTCTCTGCTTAGCCTCATCTTTCCTGATTGGCAGAGCCCGGAAATTTCTAGGACATCCACTGCCATCACCGTTCCCTGGAGGATCCTCTTTCTAATATAGTGCTCCAGTTCC | 120 |
| chr11 | 70670375 | 70670495 | 788855_47188552_ENST00000601538.6,E | 1 | - | good | CAGCTCTGCCACGGTGAGCTGGGCTGGCTGCCCGCACCCGGCGGCCAAAGCGCCCGAGCTGGCATCTCCGCTGAGCTGGGACACCTGTTGGGTCCTCACCTCAACACCCCCCACGTAACT | 120 |
| chr11 | 70670495 | 70670615 | 788855_47188552_ENST00000601538.6,E | 1 | - | good | TGTGGTCATGGGCTGAGAAGCTCCCCCATGCCCCTTGGGAGTGACAGATGACGGGGAAACCCTCCTGTTCTCCCGCAGCCTGCCCCGCCTGTCGAGCGGCCCTCTGTGCTGCCCTGCAGC | 120 |
| chr11 | 70670615 | 70670735 | 788855_47188552_ENST00000601538.6,E | 1 | - | good | CAGGATGGCACTGGGCGTTGGGCATCATTTCCCCTGTGAGTTATGCAGGAAGGGCAAGAGAGCAGCATCACACACTGTAACTCTCCGACCAGGAAGCAGGATGCTGGGAGCAGCTGCACG | 120 |
| chr11 | 70670735 | 70670855 | 788855_47188552_ENST00000601538.6,E | 1 | - | good | GCTCCTGGGAGCCAGGAGGAGTCCCCTCCGTGCAGGAATTGCCCAAAGTCATCACCTCCTGCCCTGGAAACAGCAATGGCCTCCCCAGGAGCAGGCTGGCCGGGGCTGTCACTCAGGCAC | 120 |
| chr11 | 70670855 | 70670975 | 788855_47188552_ENST00000601538.6,E | 1 | - | good | AGAAATGGCTTCCGCCAGCGCACGTCTCTGGGCACGGGTGTGCGGGCTGCATGAGTACAGAACCACTGCCCTCCTGATGCCACGTGAAGCGACGCAGCCCCCTGCCTGTGGGTGAGAGTG | 120 |
| chr11 | 70670975 | 70671095 | 788855_47188552_ENST00000601538.6,E | 1 | - | good | AAGTTCTTTGTACATTATTGTTTCTATTAGGAAAAACTTGCATTGCTTATCATGTTAGCAATGAGGAGGAAAATGGTCATAATAAGCGCCGGGCAGGGCGTTTTCATAATTGCCGATTTG | 120 |
| chr11 | 70671095 | 70671215 | 788855_47188552_ENST00000601538.6,E | 1 | - | good | GCAGTTAAGCCGTATCTGTCTCCTCCGCCAGCCGTTCCTGGCAGGTAACATGCCTGGTAGTGCTCAGGAAATGCTTGTGGGTGATTTATTACAAAGTAATGAGACAATTTAATTTGAATC | 120 |
| chr11 | 70671215 | 70671335 | 788855_47188552_ENST00000601538.6,E | 1 | - | good | TTCCACCGGGGTGCTCAGCTTCCAAGGCCGGCATTCTCCCTCCTAGCATTCTCCCTCCTAGACGACGTTTCCGTTCGTCTCGGCAGGGAGCGCTGATCACACCCCGTCATGATCATGAGC | 120 |
| chr11 | 70671335 | 70671455 | 788855_47188552_ENST00000601538.6,E | 1 | - | good | CCACCCCACTGTGGTCTCCAGGCTGTGGTTGACATCTGTAGTCGTCAAGTGAGGATACTGGCGCTCGACGAGGTTGAGGAACTTGGCCAGGGGCGTGGCGCTGGTTTGGGGAAGCTGGGG | 120 |
| chr11 | 70671455 | 70671575 | 788855_47188552_ENST00000601538.6,E | 1 | - | good | GTTTATGCATAAGGTCTCCAGAATGAAGCAAGTTCTAAGGTGCATTCTCACATGCTAATAGCCTTCCAGCCCCTGCTGAGTGCTGAATCCTGGCCAGGGGCCTTTTAAACAAGGAGCTTC | 120 |
| chr11 | 70671575 | 70671695 | 788855_47188552_ENST00000601538.6,E | 1 | - | good | TTTCTCCTTGTTGGCTAACACACCCGTGTGTCTTGCTCTCCATAAGGAAGGACACCCAAGGGCAGGCCTCTGCTGATCTCAGTAGGAGAGTCTTTTACCTGGGGACAAGTGAAAATATCT | 120 |
| chr11 | 70671695 | 70671815 | 788855_47188552_ENST00000601538.6,E | 1 | - | good | TGGAAGCACAGGGAGGCATCCTGGCATCCCTTGCTTGGGGGCCCCCTGCCCTCATCCCATCTGCCCACAGCCCTGTGTGCAGAGACCCGCCCACCTCTGAGACACAAACAGTGTGGGGAC | 120 |
| chr11 | 70671815 | 70671935 | 788855_47188552_ENST00000601538.6,E | 1 | - | good | TCATGGTACTTGCAGACAGGTTGGCTGTGGGGGAAGGAAAGAGCTCACTTCAGGACAGCTCCTCGGGTTTGGGCCAGCACTTGGGTGTTCTGCCTGGCACCGCAGAGTCCGTGCCAGCTT | 120 |
| chr11 | 70671935 | 70672055 | 788855_47188552_ENST00000601538.6,E | 1 | - | good | GAAAAGTATGGTGGGGATCTGTTGGGGTGTTTTAAGTGGGGGAGATACAGTGTGATTTAAGGTTTTTACATAAAGAGTGAATTCAGAGGAGACTTTTCAGTGGGCTCAGAATAGAGATAC | 120 |
| chr11 | 70672415 | 70672535 | 788855_47188552_ENST00000601538.6,E | 1 | - | good | GCACAGGCAACGCAAAGGCACTGTGTGGAGATGAGTGTGACCTGCTGGAGGAAGAGAGGATGGCAGCATGGCTGGAATGGGAGGAGTCAGGAGGAGAAGGGGCATGAATCAGATCACAA | 120 |
| chr11 | 70672535 | 70672655 | 788855_47188552_ENST00000601538.6,E | 1 | - | good | GCTGGGGACACAGGAGGACGGCAGCCACCAGGCGGCAAGGCCCATCTCTGGAGGTGACCCAGGGGACCTGCAGGGACAGCAGCCTGGCTCGGGGAGACCTGGAGCAGTGTCCTGAGAAT | 120 |
| chr11 | 70672655 | 70672775 | 788855_47188552_ENST00000601538.6,E | 1 | - | good | CCTTCGGGGGCACTCATGTGCTACTGGGGGAGTCAGACAGCAAATGAGTGAACAAGCAAATCACAATTTCAAACATAGTAACTGGCCAGAAGAAAGCAAAGCAGAGAGAAAAGCTGGAGG | 120 |
| chr11 | 70672775 | 70672895 | 788855_47188552_ENST00000601538.6,E | 1 | - | good | GAGCCAGGACTAGAAAGTGGGCCTTCTCCACCTAGAACAGCACTTTTCCTTCCTTCCAAAAATAACACCGGTGGCTGGCACCATCCAAGGCTGCAGAGTGTGGTGGGCAGAACCGTCCTG | 120 |
| chr11 | 70672895 | 70673015 | 788855_47188552_ENST00000601538.6,E | 1 | - | good | CGTCAGGAGCACTGATGAAGCTTCTGCTCAGATATCCTGGTCTTTCATTTTGCAGACAGGAAAACCCAGGCTCAGGGTGGTTGAGGGTCCTGTCCAGGGTCCCACTGCCAGCTCTTGCCA | 120 |
| chr11 | 70673015 | 70673135 | 788855_47188552_ENST00000601538.6,E | 1 | - | good | TTTTAAAACTGACAGCCCCATATCCCAGAAAACCCCTTTGTCAGGAAACATCAGGACAGGGGGACACCCAGCCTGCATCAAAGACCACATAGGAGGTCCCAGCTCAGGCTTCCTTCACTG | 120 |
| chr11 | 70673135 | 70673255 | 788855_47188552_ENST00000601538.6,E | 1 | - | good | AGTGTGACTTGGCCCTGCTGCTAGTGGAGGCATGTGCCCTGGGCAAACCTGTCGCTGCAGGCTGCCTGGACCTTCAGTGGCGGGCGACCAACTCACCCCATTTGTCCAGGACCATCTTGG | 120 |
| chr11 | 70673255 | 70673375 | 788855_47188552_ENST00000601538.6,E | 1 | - | good | GGAAAAGTCATCAACAGTCCCTTCCTGCACTCACAGCAAAATGCCAGGCAGTTGTACTGTTCCCCTGATTTTTTTTTTTTTTTAAAGAACATTCAGTTCAGAGTCAAAGGCTGAGGCTTC | 120 |
| chr11 | 70673375 | 70673495 | 788855_47188552_ENST00000601538.6,E | 1 | - | good | CTTGCCTCACAAGGAGACCCCTGTGGCCTGTCCCATGGCCCGCCAGCATGGTCTTTGTGGGCGAGCTTTGAGACAGACAGATGGGGCAGGCTTGGCCTGTCATTTTAATTAACGAATGAA | 120 |
| chr11 | 70673495 | 70673615 | 788855_47188552_ENST00000601538.6,E | 1 | - | good | ACAAAGCCCAGGACATCAGGTGACTGTCACACTGAGTCTGGTTCCCGGGAGAGGAGGACATTTCTGGTTTTGTAGCTGTGGGGCCATCTACTCCCCACCCCCACCAGAGGGACTTGGGAA | 120 |
| chr11 | 70673615 | 70673735 | 788855_47188552_ENST00000601538.6,E | 1 | - | good | GGCCTGGGTTTCTCATGGCCTCCCACTCAACAGCTCTGTGCCATCCAGGTTGGGCAGGTGATCCTGGTGGGGGGACCACGGGCTGCAGGTGGGCCCCCAGCCCTAGAATCAGCCTTCCCT | 120 |
| chr11 | 70673735 | 70673855 | 788855_47188552_ENST00000601538.6,E | 1 | - | good | CTCTTAATGAAGCATCAGCACTTGGAAGAGAAGCAAACTTTTCAGTGGCCCTAGTGAGTGGAGGGAGGTTGGTGGCCAGGGGTGGACGGCAGCAGTGGCCTTCTCCTCCGGCATGGGCCT | 120 |
| chr11 | 70674935 | 70675055 | 788855_47188552_ENST00000601538.6,E | 1 | - | good | AGAGCTCAGGTGTTACATCAAGCCAGGCACACCTGTACACAGTAATGTCAGCATGGCTTCTGCAGCAAGATTGAGACTTTTCTTAGTTTTTATTATACATAGTTATATGCCATGTACATG | 120 |
| chr11 | 70675055 | 70675175 | 788855_47188552_ENST00000601538.6,E | 1 | - | good | ACAAAGTAAAGAGGTGTTTCTTTGTTTTCCAGAAAGTACTCCCCATGTGATCCTTCTGAACCTCTCAGGCATCTCACAGATGATCTCATTTCTAAAACGCGTTCTCCTTGCCTCTTTTCC | 120 |
| chr11 | 70675175 | 70675295 | 788855_47188552_ENST00000601538.6,E | 1 | - | good | AAGGTGCTCAAAGGCCGTAGCATAATAATTACACTAATTACTCTAATTATGAATTATTACTCATTAAGCCAGAATTCAGGTCTATCCCCACTTGGAGCCAACCAGAGCATAAAAACCTAG | 120 |
| chr11 | 70675295 | 70675415 | 788855_47188552_ENST00000601538.6,E | 1 | - | good | TCTCTGCTTTATTTGCTCCCCTGTAAAGTGAGCGTCATCACAGCTCCTGCCTCATACGGTCATTATGAGGATTCAATTAGTTAAACCCACAGAACACTTAGAACAACGCCTGGTGTGTGG | 120 |
| chr11 | 70675415 | 70675535 | 788855_47188552_ENST00000601538.6,E | 1 | - | good | TGGCCCCCACCTGGCCAGGAATCAACAGTGCTTCAGTATCACTATCGTTCAGTAACCCACAGTGGTTCAGACAGCCTGGCTTTGCCTCCAGCCCTGTGTTCTGGTGCAAATCACTCAACC | 120 |
| chr11 | 70675535 | 70675655 | 788855_47188552_ENST00000601538.6,E | 1 | - | good | CCAATTCCTTCTCCAGTTCTTTCTCCAGAACATGCTGCTAGGACCAACGCCTCCCCTGAGGGTTCTTTCCTTTCTCTCTGTCATTGTCTGGGTTGGGAGGGCTGTGAGCTGTCGGGGAAC | 120 |
| chr11 | 70675655 | 70675775 | 788855_47188552_ENST00000601538.6,E | 1 | - | good | TTGGTCCACCATTTCCAATTTCAAATTTTTTGATACAGAGCAGGATCTCCTGGCAAAATCTTCTCAGAGCATCCCAGGGTGCCCTCCCAGCAGCTCAGATACTGGCAGGACAGTAAATCA | 120 |
| chr11 | 70675775 | 70675895 | 788855_47188552_ENST00000601538.6,E | 1 | - | good | TGAACTGTAAACGTTTATCCAGATGTTCCTGCACCCATTTTTAGGTGTGGGGCCTTGCTTTGCCTTTGGCCTTGCTCACCCCACCGTGTCCTTCCTCCCTGGCATTACCCAGCCCAGCAT | 120 |
| chr11 | 70675895 | 70676015 | 788855_47188552_ENST00000601538.6,E | 1 | - | good | AAAAGTGTGAACCACAGACCAGCACACACCTGCTGTGCCCAGGAGATGCCTTGTCTTCTGACACATTGCCAGTTCATTGAGATGGGAATCATAAAACACAGCCATAACCAGCATGGGGTG | 120 |
| chr11 | 70676015 | 70676135 | 788855_47188552_ENST00000601538.6,E | 1 | - | good | GTCACGTGAGGGACAGCATTAAGGTCCTCTTGGAAGTGGTGGCTATTGCAGTGCGCTTTGAAGAATGAATAGGAGTTCAACTGATAGAGATAGCAGGTATAGAGTGTTGTAGGTGGGAGT | 120 |
| chr11 | 70676135 | 70676255 | 788855_47188552_ENST00000601538.6,E | 1 | - | good | AGTGTCTGGCTTTTCCAGTCGCTGCAGCAGGCAGCAAGCTGGGAAGAGTGCCAGAGTCTGTGACAGGAGCTGGGTGTGGTGGAGAAAGAAGACTTAGTCCCCTCCCTCTAAGCACACAGT | 120 |
| chr11 | 70676255 | 70676375 | 788855_47188552_ENST00000601538.6,E | 1 | - | good | AGTCCAGCCTTTCCCAGAGGTCCCAGCAGACCTCGGGTCCGCTTGGCCTTGGGTCTCCTTGGCCAAAACTATTCCATAGGGTCATGACCATCCATAACTGCAGGAAAGGCTGCAATAGCA | 120 |
| chr11 | 70676375 | 70676495 | 788855_47188552_ENST00000601538.6,E | 1 | - | good | CATGGTGGCAAGATGGCAGCCATAGCTCCAAGTATGATGTCCCCATAGCACGTCCTACTCAGGAACGTAAAGGAGGGGGTGCCAGGGAGCAGTTCCCCAACCCCCACTCCCTTTCCAAAT | 120 |
| chr11 | 70676495 | 70676615 | 788855_47188552_ENST00000601538.6,E | 1 | - | good | GCAAGTAATAAAAAACCTCACTCACAGTGGCTGAGACCACAATGGCGAGTCCAGAGGAGGGCACAGCTAGCATCCATTGCACGGGCAGTGACATCTCTGAAGGCCTCTTGGCCTCTCCCT | 120 |
| chr11 | 70676615 | 70676735 | 788855_47188552_ENST00000601538.6,E | 1 | - | good | CTCTTAAGTTGTCAGGAAGCCATCCCAGTGTTTTGCCCGTATCTTCTAATTAATAACTACCCTGCCCCCTCACTCCAAGCTCCAAATATTCACATTATGTTCATTAGAAAGCATTTGGCT | 120 |
| chr11 | 70676735 | 70676855 | 788855_47188552_ENST00000601538.6,E | 1 | - | good | ATAGAGCAGGGCCCCCATCTGTGTCACTGGGGCTCATGAATCCTCCTTGCCAATGACTAGAGTCCTTGAAGGAATAAGGGCTGGGTCAGTGTTTCCCAACCCAGAATATAACTCGGAGAC | 120 |
| chr11 | 70676855 | 70676975 | 788855_47188552_ENST00000601538.6,E | 1 | - | good | GTTAGTGGGAAATTCAGACCGGTTGAGTCCAAGGAAACCACTTTTCCTTAGTTCTGTTTTCTGAACCAAAGTTCATTTGAGATGAGGGGTTGTGACTAGATCTTCTCTGGACAGTTGAAT | 120 |
| chr11 | 70676975 | 70677095 | 788855_47188552_ENST00000601538.6,E | 1 | - | good | CTTCACTTGGCTTTTTAGAGATTTCTCAGCCTTGGTGTGTGTTGTCCCCAGCCCTGTGATGTACAAGTAATTGCCTGTATACGTGGGATTAGCCAGCTTGCTCAGGAAGTCAGTGTTGTG | 120 |
| chr11 | 70677095 | 70677215 | 788855_47188552_ENST00000601538.6,E | 1 | - | good | GCAGCGTCTGGCCTCAGACATCCACTCATCAGGTCCAGACCAGACCGTGGGCCTTGGCTGTATCTTCTGATCCACTCCGTTGCTAAGTTCCCCTCTGTACCTAAACTCTGTCTCCCTTTA | 120 |
| chr11 | 70677215 | 70677335 | 788855_47188552_ENST00000601538.6,E | 1 | - | good | CTCAGCTGGATGCATTGAATGACAAAGACTTTGCAACTGTGTGGCTTGTCTGGGGTGTCTGGAAGCCCAGTCTGTGGACAGGCTGGTCCTCAGGTCTAGCAGAGGGACCCATTAGCAGAG | 120 |
| chr11 | 70677335 | 70677455 | 788855_47188552_ENST00000601538.6,E | 1 | - | good | GGTTTCATTCGAGCGAGGCTAATTTTCAGAATCGTGGGTGGTGGCCCGGGTTCTAGGAACACTTAAAGGATAGAGGAGACTCGGCACCACGTGCTGAATCCAGGAGGAGGATGTTGGCCG | 120 |
| chr11 | 70677455 | 70677575 | 788855_47188552_ENST00000601538.6,E | 1 | - | good | AGGGGCAACCAGCCTTACCTTTACCTCATTGGGAGACGCTTCAGCCCCCATGCTGTGGCCAAGGGTCCGGTCACAAAGACAGGGCATCAGTGGTGTCATGGAAGGTCAGAGGGAAGGCAG | 120 |
| chr11 | 70677575 | 70677695 | 788855_47188552_ENST00000601538.6,E | 1 | - | good | TGTTGGACCTGGTGCTCTGAGGGTTCCCTGCCTGCCCGCTTCTTGGAGGTCATCTCAGCAGTGGACGTGGTCATATTAATGGTGGACTTCCTTGGGTGGGTGCAATGAGCAGGGTTGTCC | 120 |
| chr11 | 70677695 | 70677815 | 788855_47188552_ENST00000601538.6,E | 1 | - | good | ACGCAAGGACCCTGGCTTTGGTGGTCGAGTGCAGAGCAGTGTGAGAGGGAGTGGGCATCTCGGGCTCAGGGGGAGCCTGGCTTCCGGAGCCCTCAACAGGGCTTCTGGTGGGACGTGCTG | 120 |
| chr11 | 70677815 | 70677935 | 788855_47188552_ENST00000601538.6,E | 1 | - | good | GGGAACAGAGGGCTGGAACCTTCTGCAAAAAGAAGGAGCGAGCCTACAACTGGGACAAGCAGCTCGTTCTTAAGATGTGTCTTCCTCAGCGAGAAGTTTGCACACATCTGGGAAATGGGT | 120 |
| chr11 | 70677935 | 70678055 | 788855_47188552_ENST00000601538.6,E | 1 | - | good | AAGACCTGGGAAAATTGGTGCAAGGGGTTGTCTTTTGTCTTTGGAGGGGGCCCCCTAACAAAATAAATGAACTGGACACCATGGGAAATGGCAGTGAGGATGCCAGTGGAGATGGTGGAA | 120 |
| chr11 | 70678055 | 70678175 | 788855_47188552_ENST00000601538.6,E | 1 | - | good | GACTCTGTTTCAAAAACAAACAAACAACAACACAACAAAAGAACAAGAAGGAAAATGAGAAGGAAATCAGGAAGTCCTGGAGAATGGGGAAGATTTAAAGTGAGAAACTGGCTGGGGCTG | 120 |
| chr11 | 70678415 | 70678535 | 788855_47188552_ENST00000601538.6,E | 1 | - | good | AAAGCCTGTTCTTAGCAGACTGGAAATGAGCAGGTCGAGAGGCAGGGAGAGGCAGGGAGAGGCAGGACAGATAACTCCAGCTTTAATAAAGAACAGGCAGGAGGCTGGGCGTGGTGGCTC | 120 |
| chr11 | 70678895 | 70679015 | 788855_47188552_ENST00000601538.6,E | 1 | - | good | GTCCTAGCGAGCATGCTGAGATCCCCTGGCTAAGGAGAGTGAGTGTGTTGGAGCTTGGAGGTAACCTGGGGATATCTGGAGGATGTATGTCAAGGCAAATGAATACCTGTGCAGCCTGGA | 120 |
| chr11 | 70679015 | 70679135 | 788855_47188552_ENST00000601538.6,E | 1 | - | good | CCAGCGTTGCGGTCAGCCGTGTTTTACTACCTGCATTTGCACGGATGACCCCGTGCCGTCACCTGTTACGGGCCTGGTGTCTTCTTCTTCCCAGTCATGTGAGGGCGGGGGAGGAAGGTT | 120 |
| chr11 | 70679135 | 70679255 | 788855_47188552_ENST00000601538.6,E | 1 | - | good | TGTTTGTAAAATGGGAAAGAAAATGGTGCCTCCCTCCACAGTGTGTTGTGTGAGAGAGAGAATGAGGGCAGGGGCTCCCCATGGGGCTTGGCATGGTGTGCGTACCCGGCCTGCCAGAGC | 120 |
| chr11 | 70679255 | 70679375 | 788855_47188552_ENST00000601538.6,E | 1 | - | good | GGCTGCTGGTGGTGGGAACAGGCTCTGCAGTTGGCCTCGGGGGTTGGATGCCCAGCTCCGCCATTTGCCAGCTGTGTGACCTTGAGCTCCTCACTGGACCTTTCTGAGCCGCAGCTTCCT | 120 |

| | | | | | | | | |
|---|---|---|---|---|---|---|---|---|
| chr11 | 70679375 | 70679495 | 788855_47188552_ENST00000601538.6,E | 1 | - | good | AGAGGGACTGCCCCAGGCAGGTGCACACCCCAAAAGAGGGGGCCGAGGGTGCAGGGAGATCTCCCACCTCCCTGATGGTCCGGGGATAGGCCTGCCCTGCTCCTCAAAAGGGGGCTGGCT | 120 |
| chr11 | 70679495 | 70679615 | 788855_47188552_ENST00000601538.6,E | 1 | - | good | GGCAGGACCCAGGCCCAGCCCTGACCCACTTCTCCCCAGCACAGGGGAATGGGACACAGCCCTTTGTACTGGTCATGGGGAGACAGCTGATGGCTGGAAGCCTCCGCTGCTGCCAGGCAC | 120 |
| chr11 | 70679615 | 70679735 | 788855_47188552_ENST00000601538.6,E | 1 | - | good | TCCTCATGATACCCCAAAATGCGCCAGACACCGTTCCAGGCCCTGGGGCGTATAGTCAGCACGACAGATGGCACCGGGCCTCCCAAGCCTCCCGTTCAGTGGAGGTGCCCTCTGTTCCTC | 120 |
| chr11 | 70679735 | 70679855 | 788855_47188552_ENST00000601538.6,E | 1 | - | good | GGTGGACGTCCAGGGTCCCGAGGGAGGGGCACCGGTTCTAGTTGTCACTCCATGGGCCAGAGGGCCTGGCTGGGGTGTGGCAGCGGGCAGCGGTGGCCTCTGTTGCTCAGGCCACAGACA | 120 |
| chr11 | 70679855 | 70679975 | 788855_47188552_ENST00000601538.6,E | 1 | - | good | GACCCCAAGTCATTTCCAGTCACAGTGATCACAGGGGTCTGGGAAGGGACTGTGCAGCCCCAAAGGCCACAGTCACCCTGGGCTCTGTGGATTTGGGAATGGATCACACAACTGTTGGCA | 120 |
| chr11 | 70679975 | 70680095 | 788855_47188552_ENST00000601538.6,E | 1 | - | good | TCCCTTCCCCAGGGCCAAGACATGCAGTAGAAAAGGACAGGAAATGCCGCCTCTCTCAGACTGTGCAGACTGCATGCTGCCTGCTCCACCGACCCCTGTCCCACCCGTCGGCCTCCGTGG | 120 |
| chr11 | 70680095 | 70680215 | 788855_47188552_ENST00000601538.6,E | 1 | - | good | GAGAACCCGCTCTGACAGAGGCTGTTTTGGCCGACATTAGCTGTGAGTGTGTAACTCCCTGGGATGGGGTGCGATGGCCCCACGTTACAGATGGAGAAACCGAGGCTGCAGGAGTTCTGC | 120 |
| chr11 | 70680215 | 70680335 | 788855_47188552_ENST00000601538.6,E | 1 | - | good | ACCCCCAGAACTGTAAGAGGCTAAATGTGTCTAGCTGCCAAGTTTGTGTTACGTTGTTACAGCCGTCACAGGACACTCACAGAGTGCTTGCAAGCATTCAATGCATAGACTGAAGAGGTT | 120 |
| chr11 | 70680335 | 70680455 | 788855_47188552_ENST00000601538.6,E | 1 | - | good | ATCATGGGTCAGGGAACGTGGTGGCCACTAGAAACCAGAGAACACAAGCAGCCCCTCCTCTGGAGCCTGGGTGGGGAGCCCGCCCTCCAACACCTCAGTCTTAGCCCAGTGAGACCTCGG | 120 |
| chr11 | 70680455 | 70680575 | 788855_47188552_ENST00000601538.6,E | 1 | - | good | CTGGCGAATGCAGGTGGGCCCTGTGCCATCACAGGGATCCTCATCAGAGGGCAGGGGGTCAGAGGCAGAGAGTGGGGAGCTGGGACGATGGGAGCGGAGGCTGGAGTGATCTGAGGAAT | 120 |
| chr11 | 70680575 | 70680695 | 788855_47188552_ENST00000601538.6,E | 1 | - | good | GAGACATCCATGTCCTAACCCCTGGGGCCTGTGAGTGTGTGACTTCACCCGGCCAAGGGGAATGAAGCTTCAGAGGGGGACTTGTCAGCTAATCAGCTGGTGGTGGATTAGGAGATGGTC | 120 |
| chr11 | 70680695 | 70680815 | 788855_47188552_ENST00000601538.6,E | 1 | - | good | CAATCTGCGAAAGATTGTTGAAAACACCCCCACTCCATATTAAAGCCTCATCCACCTTGAGCCTCACCAGGGAAAGAGAGAGAGGCTGGGGTGGCATGGCCAAGTGAACTGTGGAGCCCA | 120 |
| chr11 | 70680815 | 70680935 | 788855_47188552_ENST00000601538.6,E | 1 | - | good | GGAGTATCTCGGCCGCAGGACCTGCCTGCTTGCTTGCTGGTGCCCAATCTGAGGTGTGCCTGCATCCTGTTGAGAGTCCCCACAAGGAGGACGGTGCCTGCAGTTGGATGAGTCAGCCCA | 120 |
| chr11 | 70680935 | 70681055 | 788855_47188552_ENST00000601538.6,E | 1 | - | good | TCAGCTGGGCCACCAGCACCCCAGAGCCAGTTGCCTTTTAAGTGGGTACTGGCCTGTGGTCTGAGCATTGACGGGGATGGTTCACACCCAGGGAGAGTGTGATTTACTAGGAAGAGTGTG | 120 |
| chr11 | 70681055 | 70681175 | 788855_47188552_ENST00000601538.6,E | 1 | - | good | CGCTGTGTGCTTCAAGAGGCAGGACCCTCCCAGCAGGTGCCGGCAGCCCCAAGGCACAGATGCAGAGGCATCCCCGAGCAGCCAGTGTCCTCTGAGTCCTCCTGCAAGTCAGGGCACCTG | 120 |
| chr11 | 70681175 | 70681295 | 788855_47188552_ENST00000601538.6,E | 1 | - | good | GAAACATCTCTGTGCAGGAGCTGTTTATTCAGGGGGTTTCCGCCGTCAGACTTTGCCAAGAGAAAAGGGACCCCACGCTGAGCAGCAGAGAGAGAACGGGGGACTTGGCAGAGCCGGTGC | 120 |
| chr11 | 70681295 | 70681415 | 788855_47188552_ENST00000601538.6,E | 1 | - | good | CTGATAGACCCTTTAACCTTGCCAGAGAAGGTTACCGCTGAGGCTGGCGGAAATTTGGCAGCATCAAATGATGGGCTTGGACTTCTGAATTCTCCTCCAGTGCGAATCCTGAGCCGCGGA | 120 |
| chr11 | 70681415 | 70681535 | 788855_47188552_ENST00000601538.6,E | 1 | - | good | TGGGTCCCATTTCTGGGCATGGTGGGGTGGATGTGGTGGATGTGCACCTTTCGGCCTTTGAGGAGATACCCCAGGGGGCCCAGGTTTCCTCTTTATCTAATTAGTTTCTGGGCCGGGAAT | 120 |
| chr11 | 70681535 | 70681655 | 788855_47188552_ENST00000601538.6,E | 1 | - | good | AGCGTCCGTTCAGTCTCCCTTCATGCGAATGAGCTGGCCTCTCCCCTGTGGAGGTACAGCAGGGATCCGGCTGAGAGCCTGGATGGTTACAGTCCAGGGTGGCTGAGGCAGTGCCGCTGG | 120 |
| chr11 | 70681655 | 70681775 | 788855_47188552_ENST00000601538.6,E | 1 | - | good | CCCTTGCCTCGTGCTCCCACCCCAGTCACTCCATTAGGTACTGGGCTTGGGTCGCCTCCCAAACCAAAAGGCATGGGCAGGAGGGTTGCTGTGAGTAATCTGGGGAGCTCAGCGTGCCCC | 120 |
| chr11 | 70681775 | 70681895 | 788855_47188552_ENST00000601538.6,E | 1 | - | good | CATCTTTGACCTGAAGCCCGTGCAGAACCCCACAGTGTTCAAGATCTCTGGAGGTGGTTTCTGTTGCTAAAACCTAGAGCTCTGGTATCCAGAAAGGGGTTTCTCTGTGGAGCCTGGATA | 120 |
| chr11 | 70681895 | 70682015 | 788855_47188552_ENST00000601538.6,E | 1 | - | good | AGGGAAAAGCCGCAGCCCCTTGCCTGGGGGAAGGCCACACCCAGATGCGTTTCCCTTAGATTTCACCCCGAGCATGCCCAGACCCTGTTATTATAAAGCCCGGTGATGTCCGCCCCTCCA | 120 |
| chr11 | 70682015 | 70682135 | 788855_47188552_ENST00000601538.6,E | 1 | - | good | TCTGTCGGATCAGTCCTGGAACCTGGCCTGTGGGTCAAAGGCCCTCTCACAAAGCACCCTCCAGCAGGTTTGGGCTGCCTTATACTCTTCTCCAGGTTGTCGAACTCCTGTTTCTCTGAA | 120 |
| chr11 | 70682135 | 70682255 | 788855_47188552_ENST00000601538.6,E | 1 | - | good | GTCCGGGGGTGGGGTGGACCCAGCATGTCCCCTGGAGATGGCTCAAATATTTCCAGTTCAGACAGTGGCAGTAGCAGCAGCAGCCCTGGCCCGGCCTTGTGGTGCTCTTGAGGGTGCGTT | 120 |
| chr11 | 70682255 | 70682375 | 788855_47188552_ENST00000601538.6,E | 1 | - | good | ATGTATTTGTGGGGCACAGCGGGCTCCGCCACACAGCATCCTGCAGTTTGTTTCTGCACTCTGTTGGTGCCCAGCTTCCATGCCCATGGGTTCGTGTAGACAGTGTCCTTCCTAGTAACC | 120 |
| chr11 | 70682495 | 70682615 | 788855_47188552_ENST00000601538.6,E | 1 | - | good | AGTGCCCACTTTTCCATCTCAAGATATTCAAAGAATGCAGAAATATAGAAAGTCAATATCATCCTCCTGGTCCCACTTCCCAGAAGGGGCCACTGTGCATGGCTGGCTGTGCCCACCAGC | 120 |
| chr11 | 70682615 | 70682735 | 788855_47188552_ENST00000601538.6,E | 1 | - | good | GACAAGGGCTCCAGGCCTCCTTCTCTGTGGCCTTGGGCAAACCGGAAACTTCCTCCTACCTGCCTCCCCCATTTCCTCCCTCTCTCCCTCCCATCTTCATTGTCTGTTTTAGAAAACAAC | 120 |
| chr11 | 70682735 | 70682855 | 788855_47188552_ENST00000601538.6,E | 1 | - | good | GTCATCCTCCTGTGCTTAGGACTTGCGTTTTGTCTTTTCATTTTCTTCCATCCACGGCTGCTCTTGGGTCCTGAGAGAGTGAGTTCAACAAAGGAACCAGGATGCTCCCCGACCTGGGAG | 120 |
| chr11 | 70682855 | 70682975 | 788855_47188552_ENST00000601538.6,E | 1 | - | good | CCAGCCAATCTGCCTTAGACTTGTCACCTAACCCCTGGCCTCGGTTTCCTCACCTTCAGGGTAACACCTGACCCATGCCTCGGTGGTGCGGTGAAGAGCAGTGTCATTCCCCATCCTCAC | 120 |
| chr11 | 70682975 | 70683095 | 788855_47188552_ENST00000601538.6,E | 1 | - | good | GGTGAGCCTGGTGGAGAGACCAGCAGGGAAACTGAGGCCATCAGGGAGACAGCGTCTTACAGGGGATGTGGCTCAGCGGTCACACCAACCCTCTCGGCCACGTCCCAGCTTTGCCCTTCT | 120 |
| chr11 | 70683215 | 70683335 | 788855_47188552_ENST00000601538.6,E | 1 | - | good | AGCTCGCTGAGGTGAGAGCTGGCCAGAGACAGGAAGAAGGTTAGGACTAAGGAAAAGAAAGGAGGAAGTGGCCAGGCACGGTGGAATCCCAGAGCTTCAGGAGGCCAAGGTGGGAGGAT | 120 |
| chr11 | 70683335 | 70683455 | 788855_47188552_ENST00000601538.6,E | 1 | - | good | CAGAGATGCACCATCGGTAGAAGGCGGCAAGTTCCAGGGGTGGAAGGAAAGTGTTAAAAGCAGCACGGATTGGGTGGGGAAGAGGGCATGTTGCTGAATTACACAGAAACGGCAGAGTGG | 120 |
| chr11 | 70684175 | 70684295 | 788855_47188552_ENST00000601538.6,E | 1 | - | good | GCCGTGCCCAGCCAGGAGTGTGCCTTTGAACCTTCATGGTTTCAGGCTCTGGGAGCCATTTTTCACACTTCAAGGGTGATGACTGGGTAGGGGGACGGGGACAGTGAAGACCCTGGGAGA | 120 |
| chr11 | 70684535 | 70684655 | 788855_47188552_ENST00000601538.6,E | 1 | - | good | AGCCTGTGCTTCCTGGGTGAAGGGAGTGTGAGATACCAAACCGGATTCTCCTAGGAAACCAGAGAGTATGCCTTTGGGGTGGGGGTAGGGGGCTAAGAGACAGGGTTTCCCTCTGTCGCC | 120 |
| chr11 | 70684655 | 70684775 | 788855_47188552_ENST00000601538.6,E | 1 | - | good | ATTGGCAGGGCCAGGGGCACTGTCCCTGGCCCTGAGTGTGACTCAGGCACTCACTGAGTCACACTGAGTGACGGACAGGGGCCAAGGGTTTACCTGCTCCCACCTTGACACTCTTTCCAC | 120 |
| chr11 | 70684775 | 70684895 | 788855_47188552_ENST00000601538.6,E | 1 | - | good | GATCTCTGGAGTCCCTTCTTCTTCCTAAAGGGTACACTGAGGCAGAGTGAGGCATGTGACCTATCAGAAGGGCTGCAGCCGGTCAGACAGGGCAGGGCTGGGACCAACAACGCTCACTGG | 120 |
| chr11 | 70684895 | 70685015 | 788855_47188552_ENST00000601538.6,E | 1 | - | good | TTGCCCAGGGTCCTTTGCCTCACCTTTCTGCAGCACAGCATTCCCACTGCCAAAATGCACAAAGTAATTGTCCTGCCTCTTTTCCCAAATTATCCTCACCTTTGCATTTGAGCCAAGAGG | 120 |
| chr11 | 70685015 | 70685135 | 788855_47188552_ENST00000601538.6,E | 1 | - | good | TGTCAGATGGGAAGGCAGGGATTAAGGAACTGGAGTCCTCTGGCCACACCATCTGGGGGAATGGCTTGGACTGAGGTCAGGGACCTCAGGCAGGGACTAAGGTGGTGTTGGGGTCAGGCC | 120 |
| chr11 | 70685135 | 70685255 | 788855_47188552_ENST00000601538.6,E | 1 | - | good | GAGTTCAAAAGCAGATTGGAAGAGAGGGGCCCAGACTGGAGGTTGGGAGCTTTTTCTTTCCTGGGTTTCATTTGCCAGTTTTTTGTTTTTTGGGGGTTTTTTCGCCAAGCCTCTGGGAC | 120 |
| chr11 | 70685255 | 70685375 | 788855_47188552_ENST00000601538.6,E | 1 | - | good | CCCAGGTCCCCTGGCCTGGAGTTGGCCCCGTGGAGCTCTGAGGCCTCCAAGATGAGTGAAGGGGAGGAGCCAGGAAGATGGCAGCTCTTTTCTGGGCGGCAGGAGGAGGGTGGAAAAAGC | 120 |
| chr11 | 70685375 | 70685495 | 788855_47188552_ENST00000601538.6,E | 1 | - | good | CAAGCAGGGCTTAGCTCTTCAGGGCATGTGCTGATTAAAACCTGACTCTACTGCCCCCAAGGTCAAGGTGGAGGCTGGCTCGGTGGTCTAATGCCTGGTATAACAGCTGGCACCATGTGT | 120 |
| chr11 | 70685495 | 70685615 | 788855_47188552_ENST00000601538.6,E | 1 | - | good | CAGCCCATCCAAGTCTTCGGGAGGGTCCAGCTTGGAATCTGCTTCCCTCTCTTTCCCTGTCAGGGAGCTTGGTCAGATGGGTCAGGACTAGAGGCTGCACTCCCAGGCCCTGCCAACCAA | 120 |
| chr11 | 70685615 | 70685735 | 788855_47188552_ENST00000601538.6,E | 1 | - | good | AGGGCAGTCTAAGGAGTTGAGACTGCCCCCTGTCCTCCCTAGGGCTCTGCTGCCCCGTTCTCTCCAGGACAGAATGAAGCACTGGGCACCTGTCCCCATGCTTAGGATTACAAAGAGATG | 120 |
| chr11 | 70685735 | 70685855 | 788855_47188552_ENST00000601538.6,E | 1 | - | good | ACTTGGTACATTGTCAGCACTCAATAAATTTTCACTTCATGGGAAGGAGAGGATAGCATGGCCCCAGTGAGAAGGAGACTTGTGTCCCCAGTGCCAAGGAGTATGTCATGAAAAGTGCTC | 120 |
| chr11 | 70685855 | 70685975 | 788855_47188552_ENST00000601538.6,E | 1 | - | good | CATGGGTGGACCCATGGATGCATGGATGGGTGAGTGGGTGGGTGGCTGACTGAATCTGTGACTTGGGGTAAGGGTGGGGACCAAAGTTGGGCAGTGTGCTCAGACCTCTCTATAGGCAGT | 120 |
| chr11 | 70686575 | 70686695 | 788855_47188552_ENST00000601538.6,E | 1 | - | good | CCTGGGCCATGCAGGACTGCCTTTTCTCTGCCCACCCCTGTCCTTCTACACCTGTGTCTCCTCCAGATAGGACCAGGACTGACTGACTTGTCTCCTTCTCCCTGGCATCTGGCACAGGGT | 120 |
| chr11 | 70686695 | 70686815 | 788855_47188552_ENST00000601538.6,E | 1 | - | good | CCCCGCCTCTGCCCAAAGCCTGGTGCTCTGGGCCTTCCTCCAGGAAGCAGGGAGGTGTCTTTGGTGCCTGACTTGTAGAGCTAGAGTGAGAAGCAATCAGACCTCCCTGACTCAACATTA | 120 |
| chr11 | 70686815 | 70686935 | 788855_47188552_ENST00000601538.6,E | 1 | - | good | TCAGTGGCATACAAGGCCAAGGGGTGCCTCTCCAGGGAGCAGATGAGGAGACTGGTTCTGGAGGGGTACCCAAGGCTGTGCCTGGGAAGCAGCAGGTTTTCGGGAAATGCTCACACTGAT | 120 |
| chr11 | 70686935 | 70687055 | 788855_47188552_ENST00000601538.6,E | 1 | - | good | ACCCTGAGACAGGCTGGGCAGGGTTGGGATTCCAGGGCCTGACCCAGAGTGGGCAGAGGTATCCAGAGGGCAGCCCAAGATGGGCAGGAGTGCTCTTAGTTGGGGTCCAGACAGGGGCCA | 120 |
| chr11 | 70687055 | 70687175 | 788855_47188552_ENST00000601538.6,E | 1 | - | good | AATGCCTACATCATTGCTGAGATCTCTTGTGAATATAAAATTCTACACATTGGGCCTGAAACTGCCATAACCACTGGCCAGTGGTCACCATGAATGAATGCGCTTTAGACAAAACTGCAA | 120 |
| chr11 | 70687175 | 70687295 | 788855_47188552_ENST00000601538.6,E | 1 | - | good | GTTTCAAGCCTGATAGGCCTAGGCCTGGGAGTTCAGGTGCCAGGGGTCCTGGTCCCAGTTCTGACATCACCTTCCTGCACGGGCCTTAGTTTCCCCATTTGTAAGATGGCTACATTGTAA | 120 |
| chr11 | 70687295 | 70687415 | 788855_47188552_ENST00000601538.6,E | 1 | - | good | CAGCTTCGGCTCATCCCTGGGCAATATCCGTGCCTCTGCACCCCTGCTCCAACCTTGTGCTCTGAGTCGGGGTCTTACACAGGGTAATGAAGTGTGCTGATCTGAAGCTCCAGCAAGCAG | 120 |
| chr11 | 70687535 | 70687655 | 788855_47188552_ENST00000601538.6,E | 1 | - | good | GTTCTGACAATAGCTGCCAAAGCTGACCTCTGCCTCCACACCGCCCCCGCCCACCCCCCTACCCCCACACTGTACCTGGCCCCTCTCGTGCTCCTGTCTTATATCCACCTTGATTGGTAG | 120 |
| chr11 | 70687655 | 70687775 | 788855_47188552_ENST00000601538.6,E | 1 | - | good | GGGGCAGGGGCCCCCAGAGCTCAGGAGCTCTGTAGCTGCGCAGGAATCTTCTGTCTCCTAAAATCCCTGAGCCCCGCCGGCAGGCAAAGGAGAGGAGTTAGTCATGTTCTCTCTGGAGCT | 120 |
| chr11 | 70687775 | 70687895 | 788855_47188552_ENST00000601538.6,E | 1 | - | good | GTGGTGGGTGGGGCCATCGGAGCAGCAGCAGAGGAGGGCCTGGGGGGCTGGGATTCAGCACCTTCAGGCAGGGCCCCACTGAGGGCACTCCCAGAGGCCTGCCTTGCTCCCCACCTGGAT | 120 |
| chr11 | 70687895 | 70688015 | 788855_47188552_ENST00000601538.6,E | 1 | - | good | CCCCTGCTTTGGGGCTGGGAGCTGAGTTCCTCACCACAGGGAGGCTCCTGGCTCAGAGCAGCCTCAGACCACGTTTAGTACATTTCCCAGGTGGCGAGTGGGTGGATGGTGGGGACACCG | 120 |
| chr11 | 70688015 | 70688135 | 788855_47188552_ENST00000601538.6,E | 1 | - | good | CCAGGACCACTCTGGCTGGAGAGCAGGGAGGCCAAGCTCTCAGCAGCCTGGACACCCTCATGACGGCCTTTTCTGTGGGGCCAGGCAGGAAGGTCTGGCTGCTTTGTTCTGGACCCAAGT | 120 |
| chr11 | 70688135 | 70688255 | 788855_47188552_ENST00000601538.6,E | 1 | - | good | ATGGGTTCTGAGGGTGGAGAAGGGGAGGGGCCTCCTCATGCCCTGCAGGGGGGATCCAGCTCAGCAACTGTTCTGCATTGGGGCTGGGCGCACTTGGTCTTCAGGTTGGGGTAAGTGGCT | 120 |
| chr11 | 70688255 | 70688375 | 788855_47188552_ENST00000601538.6,E | 1 | - | good | CCCCTTGCAGTTTTATGACGTCAAGCACCTGACTTGGGGACTTTGGACCTTGGCCCCAGACCAAGGTGAGGGTAGGTTTTTCTGGCTGGAGGTACGCTCAGACCCTCTGAAAGCAGTTGG | 120 |
| chr11 | 70688375 | 70688495 | 788855_47188552_ENST00000601538.6,E | 1 | - | good | GGGATGGCCGGGGCTGCTGGCTTGGCTGGGCCAGGCCTGGACTCCTCCTTCTCCAGTACTCAGTGATGCGAAATTAGGTGCACGAAATTAGGTGTGCTCCGACCTGGCCACGGCCAACTT | 120 |
| chr11 | 70688495 | 70688615 | 788855_47188552_ENST00000601538.6,E | 1 | - | good | ACGGAGCTTTGAGATTCAGATTTCTTAGATATCATCTGTGAAATATATAATTTCAAAGAGCCAAGCATAAAGCTGACACTTGCTATGTTTCCTGGGTGCACAGTGGGTCATGAGTCTCTT | 120 |
| chr11 | 70688615 | 70688735 | 788855_47188552_ENST00000601538.6,E | 1 | - | good | GAGTTAGACAAGCCCGTGTTCCCTCCCAGCCTCTCCACCTGCCAGCCCCATGACAAGATCATCCACTCTCAGCTCTGCCTGTGCCATCTGCAAATGAGCATTTGTGATTCTGCCTCTGTC | 120 |

| | | | | | | | | |
|---|---|---|---|---|---|---|---|---|
| chr11 | 70688735 | 70688855 | 788855_47188552_ENST00000601538.6,E | 1 | - | **good** | TGATAGAGCTGTGGTGGGGAGACATGGTCCCAGGAGCGCAGTGTCCCTGCCGAGTCATGAAAGGGGAGGAGGCTCTCTGATGGGAAAGGGGGCAGTGAGGCACAAAGGGCCTAGACTCAT | 120 |
| chr11 | 70688855 | 70688975 | 788855_47188552_ENST00000601538.6,E | 1 | - | **good** | AGTTCTAAGCAAAGATAGACATAGGTTCTCATGCATACCTATGGGACAAGTAGTAGGATCCAAATTGAAAATTGAAGTGCTGGTCTGTTTATCTTGAAGTGAGTTGCACAGGTGGCCAGG | 120 |
| chr11 | 70688975 | 70689095 | 788855_47188552_ENST00000601538.6,E | 1 | - | **good** | ATGGGCAGATCTCTGCACGTAACTGCTCTCACGGCAGAGCTCAGGTGAACTGCACGTTCCTCAGGTCAAACACCAAAAAGAGAGGCTTAGAGACATGCTCCTGGCCCCACTCCCACCAGA | 120 |
| chr11 | 70689095 | 70689215 | 788855_47188552_ENST00000601538.6,E | 1 | - | **good** | TTATTTTGTCAATCACACTGTTACTGTCCAAGAGCTTTTCCTTTCAACAATGGGATAAGCTTCTAAAGCTTGAGAGAAGTGGTTTCCGGATCTCTTTGTGAACAAACCAGCCAGCCAGGT | 120 |
| chr11 | 70689215 | 70689335 | 788855_47188552_ENST00000601538.6,E | 1 | - | **good** | CTATTAGACCATTTACGTTCTCCCCATTTGTCACTTATTCTTTCTCAAGATTTTTTTCTTTGCCTTCTTGCTCTGTATTCTGGGAAATGTCCTTAACTTCCAACCTGTCTATGTAAGTTA | 120 |
| chr11 | 70689335 | 70689455 | 788855_47188552_ENST00000601538.6,E | 1 | - | **good** | TGAGCTCATTGCTCTGGAGACCCCTCCAGACTGAAGCAGCTTCCACTTCAAGAAGGTATTTTATTACCTCCTTGACAATTCCATCGCTTCCTCTTTCAGTTCAGCCTTTCTGGATTTCCT | 120 |
| chr11 | 70689455 | 70689575 | 788855_47188552_ENST00000601538.6,E | 1 | - | **good** | ACTGTAGGTAGCGACCCTTTATTTTCCTCCCTAGAAGTTTGTAGGAATGTCATATTTTCTACTTTTTCTTAGTGTTCTGAAATCTGATGGTGGATGTGAGTGTGCGCTTCCTTTGCACAT | 120 |
| chr11 | 70689575 | 70689695 | 788855_47188552_ENST00000601538.6,E | 1 | - | **good** | GTCTGGGGGTTCTGTTCCCAGTTCTGAATTGATCTTTTTCTCATTATTCAATGTTCCTGTTTATCCCTGGCAGGTGAGAATTGCCAGTAAGAAGTTATAGCACTGTCTGATTCTCATTCC | 120 |
| chr11 | 70689815 | 70689935 | 788855_47188552_ENST00000601538.6,E | 1 | - | **good** | GGCATTGCTTCATCACCATCCCGTCTCTACTCATGGCTTAGAGCCTTGCTGACTTGTCCTCTGTTGTCAATTTTTGAGAGAGGAATGCTGAAATTCTCCAATATTATTGTGTTTGTCTAT | 120 |
| chr11 | 70689935 | 70690055 | 788855_47188552_ENST00000601538.6,E | 1 | - | **good** | TCTGTCAGGATACATCCTCCACTTCAACAAAAGGTATCTGGAATGTAACCTCCCTGTCTGGAAATTCCCTAGTTATCAAATGTTCTCCAGGTTGTAGAACTCCAGTTTCAAAACTCTCCA | 120 |
| chr11 | 70690055 | 70690175 | 788855_47188552_ENST00000601538.6,E | 1 | - | **good** | TGGAGTCCTCTGACATTCTGCTCCAATTTGAACTCCAGCTCTCAGTACACGACGATCATCCTGCAACTTCACTTCACTGAGCTAGTGGATACGTGAGTTGCTTCTTTTCTTTGCCCTTAT | 120 |
| chr11 | 70690175 | 70690295 | 788855_47188552_ENST00000601538.6,E | 1 | - | **good** | TTTCCTGGAGTTTCTGATTGCCTTTTTTTTTCTCGTGGGAGAAAAATTATGGCTTCTTGAGCATACCACTAAAATATCATTCTTTTTCTGATGTATCACTGGAATCTGTCCTCTTCTTGT | 120 |
| chr11 | 70690295 | 70690415 | 788855_47188552_ENST00000601538.6,E | 1 | - | **good** | AAGAAGGGTACTAATAGTATCACAACCCAATGAGTTTTTTTTCCACTCTATCAGTGTTTCAGAGTAATTCTACATTTCAGCTTAATATTAGGATCATGATTTTCACATGAAGATTTTGTT | 120 |
| chr11 | 70690415 | 70690535 | 788855_47188552_ENST00000601538.6,E | 1 | - | **good** | CCAGCATAGACACCAAAAAAAAAAAAAAAAAAAAAAAAAAAAAAAAAACATGGGAAGTATTACATTATGATGATAATACAAATGGGGTTAGTATTCCAATTCCTCTTTACAGACCTATGAG | 120 |
| chr11 | 70690775 | 70690895 | 788855_47188552_ENST00000601538.6,E | 1 | - | **good** | AAATGTGAAACATCCTTGACACATTGCTGGAAAAAATGAGGCTTTCATAGTTTGACCCTCACCTCCAATTTTGAATAGTTCTTGCCAGTATGCTAATGGATTACCTTTAAATAACATTCT | 120 |
| chr11 | 70690895 | 70691015 | 788855_47188552_ENST00000601538.6,E | 1 | - | **good** | ACCCAGCCTCATCCACACAGCCCCACTTGGCAGAAAAAAAAGTTTTTAAATTATGTCTGCTTCATTCTTCTGGTGGGTATCTTCCCAACGTGATGATTACTCCCGTATTTCTTAATTTTA | 120 |
| chr11 | 70691015 | 70691135 | 788855_47188552_ENST00000601538.6,E | 1 | - | **good** | TCACATTTTTCCTCCCTCCATAAATTTTCATCCAAGTACTGCATGCATGTAATGAAAGTAATTAATTAAACTACAGAGATAGGGCCAAGAGTAGAAATAAATTATTATGTGCCCACTCTG | 120 |
| chr11 | 70691135 | 70691255 | 788855_47188552_ENST00000601538.6,E | 1 | - | **good** | CTGCTGTTTCCCTGGGCTGGCCCCTTGGCTCTTGTCCATGTTCTCTGTTCCCCTTAGCACCCCACAGTCCGGCTGCCCCAGGGACCCCTCCCCTCATCCATTCTCAGTGAACTCTCAGTT | 120 |
| chr11 | 70691255 | 70691375 | 788855_47188552_ENST00000601538.6,E | 1 | - | **good** | CTCCCAGTGCCTAAACGGGGGACCTGAGTGTCCCCCTCCTCATTCTCCCTTGCCTTTCTCCCTCCTTCCCTCCTTTAGTCCATCACAAGGGTCAAGGAGTCCTGTCTTCTCCTCGTCCTC | 120 |
| chr11 | 70691375 | 70691495 | 788855_47188552_ENST00000601538.6,E | 1 | - | **good** | TCTTCCTTTGCGCTCTCCCAGCATGGTAAAGATGGTGGCCGGAACCAGGTCTCCATTTGGGTGAGCCATTTCTCCAGGGAGTACTGCTCTTCTCTGTGGTCTCTATTGTGATGGACAGCC | 120 |
| chr11 | 70691495 | 70691615 | 788855_47188552_ENST00000601538.6,E | 1 | - | **good** | CTGGCCCATATTTTTATTTTTATTTTTGCAAGTATATAGTTCTTTTGTAGTTTTTTGAAACAGAGAGGTATTTCTCTGGGCATAGGCAAAGGAAGGCATCACAACCACAGATGGCCCAGC | 120 |
| chr11 | 70691975 | 70692095 | 788855_47188552_ENST00000601538.6,E | 1 | - | **good** | TATGTCTACTGTCTGGACACAGAAGGAAGCACACCCCATTTGTTCCTTACTCTCAACTGAAGGCTGGGCCTCGCAGTTAATTTGGAAGCCCCCATGGCGGCTTCTTTGAGGCATAGTTTG | 120 |
| chr11 | 70692095 | 70692215 | 788855_47188552_ENST00000601538.6,E | 1 | - | **good** | GGTTGTAAGAACTGAATGGCAAGAAACATTGATTTCCCTCATTCTTCCTCCGCCATTATTTGTATTTTTATATATTTTTTCTTATTTCTTAGACTTGAAAAGAAGGCTTTATCCCTGAG | 120 |
| chr11 | 70692215 | 70692335 | 788855_47188552_ENST00000601538.6,E | 1 | - | **good** | TCCTAAGAGGCATGTGCTATATAAATCCAGAGGGGTGCTGAGTTGTGAGCAGTGTGTGTGGGTAGCTGGCTTTCCTTTGAAACCCTAACTCTGCGCTCTAGAAGACTGCCTCAGTGTGTA | 120 |
| chr11 | 70692335 | 70692455 | 788855_47188552_ENST00000601538.6,E | 1 | - | **good** | TCCCCTCCCTGAGTCTTCAGTTCCTGTTGTTTGCAAAGTAGAGCTGCTTCCAACTCATTCTGTATCTCCAGTTGTGTTCGTGTTAAGGATGAATGAGAAAATATGTTTAGTACTTGCAGC | 120 |
| chr11 | 70692455 | 70692575 | 788855_47188552_ENST00000601538.6,E | 1 | - | **good** | GCAAATAGAAGCATATATGCATGTTATCTGAAGCGTTCTAAAGACGCTGAGAAATCACCGGTCCTAAGTTATCTCTCTGGCTTTCCAGCTAAGTCACTATGCAACCTTGGGTAGCTCGCC | 120 |
| chr11 | 70692575 | 70692695 | 788855_47188552_ENST00000601538.6,E | 1 | - | **good** | AGGAAATTGATAGAGTGTGAAGAGTAGGTCTCCCATCTTTGCCTTCTGGTCCCCAAGTTCTTCTCCCTAGAGAGAATCTCTGTTGCTGGGTTCCTGAGGCTCTTTCTAGAGAAGGGCCCT | 120 |
| chr11 | 70692695 | 70692815 | 788855_47188552_ENST00000601538.6,E | 1 | - | **good** | GAATATCCACTGTCCACAGCTCCCTATTTTCATGGCTTTCGAGCCTCACAGCCCAATTTTTATCTTTTCTAAATCAGTAATAAGAAATACATATACCTGGTTTAAAAATTCAAGATATGC | 120 |
| chr11 | 70692815 | 70692935 | 788855_47188552_ENST00000601538.6,E | 1 | - | **good** | CCACTCAAGGAAAAGGTTTAAGAAGGTGCTGAGCATCTGTTTTTTGTTCTCCCTTAGAGAAAACCAGTTTTCTCTTAGAGAGCGAATTAGAAGGATGGCAAAGTATGCCTGCTCACCATT | 120 |
| chr11 | 70692935 | 70693055 | 788855_47188552_ENST00000601538.6,E | 1 | - | **good** | GGTGAGGAAGAGCATGGAGGACTTAGTTCTCACCTTCTCTTCGTTTTTGCCGGGAGGAGCTCCCAGAGGCACATCCTTACTGGCAGCAAGGTCTGCTCAAAAGGAACATGTCCATTAATG | 120 |
| chr11 | 70693055 | 70693175 | 788855_47188552_ENST00000601538.6,E | 1 | - | **good** | ATTGCCATTATTCAGGCCAAGCAACAATGGTGGTTGGGGCTGGTGGAGGAGGTGGGGTAGAGAGGTGGATGGAACTGAGGGGTAGGATGGAGTTAGATTGGATAGGGGCATGGACATGGA | 120 |
| chr11 | 70693175 | 70693295 | 788855_47188552_ENST00000601538.6,E | 1 | - | **good** | TTGGGTCTTCTCGCACTTGCAGTGGAGGCATTGGAGGGCGTTAGGCAGAAGAGACATCATCTGAATCACTTTTCTGGAGGAAAGCAGATGGCAGGATGGATGGAAACCAGTGAGGAGCCC | 120 |
| chr11 | 70693295 | 70693415 | 788855_47188552_ENST00000601538.6,E | 1 | - | **good** | GTTCAGCAAGTGCAGAAGCACTGGGGTGATGAGAAGATTCGACACTGCAGGAATAGCCATGGCCTAGAAGAGATTAGAGGCAGTAATGTCAGAGAAAGACCTCGTGCTTCATGAAAAGAG | 120 |
| chr11 | 70693415 | 70693535 | 788855_47188552_ENST00000601538.6,E | 1 | - | **good** | TCCTGGGAGGTTGGGTCATGCCGGCTTTATGTAGGGATGCCAGGGAGGGAGTGTCTCCCTGAAGAAGTGACTTTTGAGTTGAAACACAAAGATTGAGGAGGAATCACCCCTCAAGAGAAG | 120 |
| chr11 | 70693535 | 70693655 | 788855_47188552_ENST00000601538.6,E | 1 | - | **good** | TTCTTTGACTTCAAGTAGATTACAGTCTAATTAGTAAAGGTACACAATAAAGTTAACAAATGAAGCAGAGGATCTCACTTTGTAGTTAATGCTATGAAGCAGATGAGTGCTGTCATAGAA | 120 |
| chr11 | 70693655 | 70693775 | 788855_47188552_ENST00000601538.6,E | 1 | - | **good** | CTACCCACCCATCCATCCACCCATCCACCCAACCACTGCATTCATCTGTCCATCTGTTTAATGAATATTTATCAAGATCTGACTATTCTGGGAGCTGAGGATATAAACAAGACAAGCACA | 120 |
| chr11 | 70693895 | 70694015 | 788855_47188552_ENST00000601538.6,E | 1 | - | **good** | CTGAACGGACTCCATTTATACATCCATCCTTCCCGCCATCCACTCACTCAGCCATTTATCTTTCTATCACTCATCTATCCACCTATTTATCTATCCACCCAGTCACCTGTCCACCTGTCC | 120 |
| chr11 | 70694015 | 70694135 | 788855_47188552_ENST00000601538.6,E | 1 | - | **good** | CTCAGGGAATGGGAATCTTAAGTTCCCACTCCTCGTTAACCAAATCCCCAGCCTCATCATGGGATACTCCAGAGAAATCTACCTGGTGGGATGGGGGAGAACTCAGGGTCTCACTTTAGG | 120 |
| chr11 | 70694135 | 70694255 | 788855_47188552_ENST00000601538.6,E | 1 | - | **good** | AGTGAGAGAAGGATCTAGAAATGGCCACATTAATTAAGGTCAAACAGCAGAGGCCCTAAGGGCCATTTCTACTTGGGACTTTACAGATATTTCAATGTCATTTTTAAAAACATGTCTCCA | 120 |
| chr11 | 70694255 | 70694375 | 788855_47188552_ENST00000601538.6,E | 1 | - | **good** | ACTGCTCAGAGTCTCAGAGCTGGAGAGCCAAATGTTGAAACACACCAATTCATTAGTGAACATATTCAGCTGTTCCAGCCAATTCCCTCCCTGGGATTCCCCTCCACTCACCCGGAAAAC | 120 |
| chr11 | 70694375 | 70694495 | 788855_47188552_ENST00000601538.6,E | 1 | - | **good** | CCACCCCACTGCCGATGCAAGTGTAATGAGAGAAAATGCCATGGGAAGTGAGACTCAGTGGAGACATTTTCTGAATCAGTCAGGAGGTTTCTCCCCTTCCAACCTCCTCTTATGTCAAAA | 120 |
| chr11 | 70694495 | 70694615 | 788855_47188552_ENST00000601538.6,E | 1 | - | **good** | CATTCATGTTGCAAGTAGACTAATGAGCAGCAGAAACCTGGGTGCAGTGCCAGCCCTGTGGGCCAATTATCCACAGTAGGCTGTGGGGTCACAATCTGGGCAGACAGCACAGGGCTGAGG | 120 |
| chr11 | 70694615 | 70694735 | 788855_47188552_ENST00000601538.6,E | 1 | - | **good** | AGTGGGTGTGGAGTGAGAGGCTGACTGCCAGCTTCTCCAATGGAGCAGGAGGAGCATCTGGAGGCCATGGTCACCAGGGCATGGGTGCAGAACAGCTTGTAGCTCACAAACCCTTGTTGG | 120 |
| chr11 | 70694735 | 70694855 | 788855_47188552_ENST00000601538.6,E | 1 | - | **good** | GAGCGGGCAGTGTGGAAGGAGAGCAGGCAGCATGGAAGGAGGGCAGGCAGCATGGGAGCTTGTGGACAGAGATGAGTAGGTCTTGTGCACCCCTGTAACCCTGGAATGGCATAACTCTGG | 120 |
| chr11 | 70694855 | 70694975 | 788855_47188552_ENST00000601538.6,E | 1 | - | **good** | GACCAGCAGAACTGGCCTCAGCTAGACTGTGAACCTATTAGCATGGCAGGGTGGACAGGGGGACAGGCAGTGCTGACAGAGGGTGGGCAGCACGGATGGAGGGTGGGCAGTGCAGGCAG | 120 |
| chr11 | 70694975 | 70695095 | 788855_47188552_ENST00000601538.6,E | 1 | - | **good** | CCTCAGATGAGTGGAATCCTGACGTATTCAGGATGATGGTATTATCGTTGTTCATCCCTGACTGTCTGAGCTGGTTCATAACGGGTGCAGGGTTAGCCTGAGGGAGGCCACCCTGCCCAG | 120 |
| chr11 | 70695455 | 70695575 | 788855_47188552_ENST00000601538.6,E | 1 | - | **good** | ACTGCCCAAAGCCAGCCTGGGTTGAGGGAAACTGACGAGGAGGAGGGCCAGTGTTTCTTGTGGTAACAGTCAGCACCTCCCAAAGCATTGTTGGGAGAGATGCACATAGATGGAATCAGC | 120 |
| chr11 | 70695575 | 70695695 | 788855_47188552_ENST00000601538.6,E | 1 | - | **good** | GGACACTGATGGAGGGAAAGGAAGGCCAAGTTAATGAGCATCTGAGCCTGAGTATGCGGCCTGATCTGTGGGCAGTAGAGCCCTGGCATTCAGGCCCCACCCTCCCGCGGTGAGGATGGT | 120 |
| chr11 | 70695695 | 70695815 | 788855_47188552_ENST00000601538.6,E | 1 | - | **good** | CAACAGCCCTGCTTATCCACTCCTCCCGTGCGCCACCCACCCTGCATGGAGCACAGCCCTGTTCATAGCTGAGCCTCCCAGCAACCTGGTAACGCAGGTGTTACATTCCACCTGCTAGAG | 120 |
| chr11 | 70695815 | 70695935 | 788855_47188552_ENST00000601538.6,E | 1 | - | **good** | GGTGAAAAAGCTTCAGCCTTGATCAGAGTTAGGAGTCAGCTCCAGTACTCCCATGGCGAGGGATCAGACAAGCTGTGGGCCTCATGTGACCCTGTGGTTCATTGAAAGAAACAAGAGTCA | 120 |
| chr11 | 70695935 | 70696055 | 788855_47188552_ENST00000601538.6,E | 1 | - | **good** | GTCAAGGGCTTTTGTTTTACATGTGTACTGAACACCTGTTCTGCCAGGTCCTGGGACCTCACCTCAGAGAGGAGGAACATGCAGACCACACTGGGGTCAACTGGTAGCTTCTAAGCAAAT | 120 |
| chr11 | 70696055 | 70696175 | 788855_47188552_ENST00000601538.6,E | 1 | - | **good** | GCTCCCCATACTAGAAGGAGCCAAGCAGGGCTGAGGATTCTCTTGCCAGAATGTCACAGAAGATTGGACTGGATGACCTCCAAGATCCCCACCAACTCTGCATCCCTGTTTTGGTGTCCA | 120 |
| chr11 | 70696175 | 70696295 | 788855_47188552_ENST00000601538.6,E | 1 | - | **good** | AAAGGTGTGACACATGGCTAACAGCAGGGCCAGAAAATACCAGGGACATGGGCTGGAGTTCAGCCAAGAGAAGACTTGCAGAACTATTGTGGCTGTTCAGGCTGCCTCAGGAGGTAGTGA | 120 |
| chr11 | 70696295 | 70696415 | 788855_47188552_ENST00000601538.6,E | 1 | - | **good** | GATAGCAAGCAGAGTCTGCACAATGATACATAGTGCCTGAGTCTCCCCACAGAGGAGCAGAGGAAGGACCTGCCTGGGAAGAGAGACATCTGTGGGCAAAGCTGGCTAGGGTGTTGTAGT | 120 |
| chr11 | 70696415 | 70696535 | 788855_47188552_ENST00000601538.6,E | 1 | - | **good** | TGATGAGCCTTTATTTGAGCCAGATGAATTGTTTCTGCAGCAGCAGAAGGCTGCCTGTAGAAGTGGACCCTGGGCAGGTGGTCATAGTCAGTGGGCCTATCACACTGTGATAGGATCTGT | 120 |
| chr11 | 70696535 | 70696655 | 788855_47188552_ENST00000601538.6,E | 1 | - | **good** | ACAGACTTTCCAGGTCTTGCTACACTCGCCCACTTTGTGACCTTGAAGCCCAAGTGTGATCCATGGCCTGGCCACAAACGCAGCCCTAAAGGCAGGTCAGGGACTGACCACAGCTTCTCC | 120 |
| chr11 | 70696655 | 70696775 | 788855_47188552_ENST00000601538.6,E | 1 | - | **good** | ATTGCTCATTGGACTCAGTTATAATGCTGACCTGTCCCAACCGCTTCTCTTGTAAATGAGGTGTGCTGTAAAAGGAATGCCATAGCCAGGAGAGGAGCCAAGACCCCTCCAGGGAATGTC | 120 |
| chr11 | 70696775 | 70696895 | 788855_47188552_ENST00000601538.6,E | 1 | - | **good** | ATTTTGCTCATCCCTTTATCCATCAGTGAGCACAGGTTATTTCCACCTTTTGGTTGTTGTGAATGGTATTGATTGCTGTTGATATTTGCATGCAAGTATGTGTGTGCACTTAGGAGCAGA | 120 |
| chr11 | 70697375 | 70697495 | 788855_47188552_ENST00000601538.6,E | 1 | - | **good** | CCCACTCCCATTGCCCCTCGCACCCACCCACCTGCTTCTGTCTGTGGGCTCTGCCTGCCTCGGTGCTGCGCGTAGATGGAATCAGCCGCTACGTGATCTTTCATGTCTGGCCTCTTCCAC | 120 |
| chr11 | 70697615 | 70697735 | 788855_47188552_ENST00000601538.6,E | 1 | - | **good** | CTGTGATGAGCAGCCCAGACCCCTGCACTCGACTCTTAGACCTGCCTGTGGAGGGATCCAACCTGGGTTAGCTGATTGATTTGTTTTTGTTTGTCTGTTTTGATTGAAGTAAAATGTACG | 120 |
| chr11 | 70697735 | 70697855 | 788855_47188552_ENST00000601538.6,E | 1 | - | **good** | TCCTGTGCCCGGCCCTGCTGGCTGAGGAGCCCCTTCTCAGAAGCTGACCCCACTCATCTCCTGCAGAGGTGGGAGGGTCCGGAAGCCCTACGCTGGGTTCTTCCCCGACCTCCAGGATAT | 120 |
| chr11 | 70697855 | 70697975 | 788855_47188552_ENST00000601538.6,E | 1 | - | **good** | CCCCTCCACGTGGGGAGCCCCAGATAGAGGGTTCTTCCTGGATGTTGAAGCTCAACAAGGGCCAGTGTGTGAGGTAGAAAAGAGAAGGAGGTGCAGGTGCAGGCCCCGGGGATGTGAGGG | 120 |

| | | | | | | | | |
|---|---|---|---|---|---|---|---|---|
| chr11 | 70697975 | 70698095 | 788855_47188552_ENST00000601538.6,E | 1 | - | good | GTCACCAGCTGTGGGGAAGTCCTTGGGTCTCTGTGGGGAGGTTGAGGCTGGGGGGTGCCTGCAGGGGAGGTTGAGGCTGGGCAGGGGAGGTGTCCTGCTGGCCCCGAGTGTGGGGAAAG | 120 |
| chr11 | 70698095 | 70698215 | 788855_47188552_ENST00000601538.6,E | 1 | - | good | CAGATCTTCTAAGCACAGCAGGTTTGAGTTCGTCTGATGTGTGGAAAGAGCCCTGCATCTTTGACAATGTGACTGGCACAGGCACGTGGTTGGATTTTTCTGCAGAAACCACACCTGTGT | 120 |
| chr11 | 70698215 | 70698335 | 788855_47188552_ENST00000601538.6,E | 1 | - | good | CGGATGCCTCAGGTCTGGGTTTCTGCCTTCTCTGTGGTCCTCGGGTTCTCATCCATTCCTGCCACTTCTGCTTAGGTGGAATGGCTGTGAAGGAAGGGGGCAGGCGAGCCACCAGGTAGC | 120 |
| chr11 | 70698335 | 70698455 | 788855_47188552_ENST00000601538.6,E | 1 | - | good | CAGGGAGAGGGCTGGGTAGCATTGTTTTTGCGAAAGCATGCAAATGTCAAGGATGACGGCAGTGGGCCACAGGAGCCTTGAGGCAGTATTGGGGCAGCAGCATCCCCCGGGGGAGTATTT | 120 |
| chr11 | 70698455 | 70698575 | 788855_47188552_ENST00000601538.6,E | 1 | - | good | CTGTGCCTGTTACCGGGCTAGGACTATGGAAGCCACCATCTCCTGGAGCCCCTGCCTGAGAGGTGTACTTGGTGCCACCTTTGGTTTGTGTCCAGCCAGAGCACACCTGTGTGGTGGCCC | 120 |
| chr11 | 70698575 | 70698695 | 788855_47188552_ENST00000601538.6,E | 1 | - | good | ACTTCCAGGTAAGACGCGCTGCTTTCTCTTCCAGGACCCTCTGGTCAAAGCTGTTCGTCTGTGCTGCAAATGCTTTGGACCATCTTGCATGCAGCTTTCAGGCCCCTCGTGTCTGCCTCC | 120 |
| chr11 | 70698695 | 70698815 | 788855_47188552_ENST00000601538.6,E | 1 | - | good | GTTCGGGGACCATGACTTTTCTGAATGGTGTTTCTCTTCCCTCTCTGTACAGAAACCCGCGCTGACCGCAGCAAGAAGCTTTTCAGGCACTACACCGTGGGCTCCTATGACAGCTTCGAC | 120 |
| chr11 | 70698815 | 70698935 | 788855_47188552_ENST00000601538.6,E | 1 | - | good | ACAAAGTTGAGCCCCAGAACCAGGCCTCATTTTCCTCCAGACTCTTCCCCAGGACAGTGCATCTCACATTTTGGGGAGTAGGTGAGAGGGCCCACCAGCCTCATGTGCTGTGGGTTCCGC | 120 |
| chr11 | 70698935 | 70699055 | 788855_47188552_ENST00000601538.6,E | 1 | - | good | GGCTGCTTCCTCCTGCCTCCTCTGTGAGCTGAGGCCTGTGGGACTCTGCCTCCCTTAGGGAGTAGTGCTCCTGACTCTTCAGTAGCCAGCATTGAGGTGCACATGCATTCCCCAAATTTC | 120 |
| chr11 | 70699055 | 70699175 | 788855_47188552_ENST00000601538.6,E | 1 | - | good | GTTTTCAGTTTTGCTGGAAGGCTCAGCAGCACCACGGCAGAATGCTGTTCTAGTGGATTCCATGAGCCAAGCGGGCTGCCCTGTTGGGGTCTGAGCTGGCCTGGCCCTCTCATCTTTGCA | 120 |
| chr11 | 70699175 | 70699295 | 788855_47188552_ENST00000601538.6,E | 1 | - | good | CTTAATTAGAAATGTTAATCATGTGGGTTTGGTGCTTTTTTTTTTTTTTAGGAGAAAGGAACTATAATCAGGAAAATGACAATAAATGTTCTTTCTTTCCCAAATGGCAGTTGTCTGCAG | 120 |
| chr11 | 70699295 | 70699415 | 788855_47188552_ENST00000601538.6,E | 1 | - | good | TGTGTGTGTTTTTTTTCTCCCTCCAGCACTGAGCTATCCATTAAATCTAATTACATCACTCATATCATTTCCCATGAAATTTATGTCTCAATCTGCTGAGTAAGGGGAGCAAGCAGAGGA | 120 |
| chr11 | 70699415 | 70699535 | 788855_47188552_ENST00000601538.6,E | 1 | - | good | CAGTTGCATAATTTCTTTGGATGTTTTGCAGGGCTTAGGTAGGAGATCATGCAGGAAATCCTGTTTGGGAGGAAAGTAAACAGGAACTCGTGAGCTGCTATGTTTTATGTCTTTTTGTTG | 120 |
| chr11 | 70699535 | 70699655 | 788855_47188552_ENST00000601538.6,E | 1 | - | good | ATTTTGGGGTTTGAGTGGAAGAAAAAATGATTTCACCTCAGAATTTGGGATGGAGGAGGCTAACAACAGAGAAGGTTGTTTACAGAGTTTAGATTAAACTTTGGGCCATTTTCCTGAAGT | 120 |
| chr11 | 70699655 | 70699775 | 788855_47188552_ENST00000601538.6,E | 1 | - | good | GGAAAAGGGAAAGGGTTCTTGGAACATGTGCTTTTGGTGAACAGCTGGCTGCCTCTGGGCACATGAAAGGTGTGGAGACCTTTCTCAATTCCCTTATTTACTTTTGAACTGCAAATGTGC | 120 |
| chr11 | 70699895 | 70700015 | 788855_47188552_ENST00000601538.6,E | 1 | - | good | CCCAGGGTCTCAGAGGCCTGAGCGTTGAGCTGGCAGAAGCAAGAAGACAGAGTCGGGAGAGCACACTGGCTTCTAAAAAGCCTAAGCTTGGAAGGGACCCTTCCCACCCTGCTTGTGCTC | 120 |
| chr11 | 70700015 | 70700135 | 788855_47188552_ENST00000601538.6,E | 1 | - | good | GAGCTTCCATGTGAAAGACTTAGCCACTGTGTCAAATTAATCCGAGGCTGCAGCCTAATGTGGTAGAAATGAAACTCTTGCTTACGTAAGAGTCCAGGGCACAGAGACCCAGCTGTTGAC | 120 |
| chr11 | 70700135 | 70700255 | 788855_47188552_ENST00000601538.6,E | 1 | - | good | ATAAGTGTCAGTGACGATGATGGAGATGGTGATGATGTGGCTGTTTGGGGAGTCCTGTTGTGTTTTGTAGGAGCCTCGGGGGTTTGTGGGGAGCAGTGTGGAAGGGGTGGAGACAGCTGG | 120 |
| chr11 | 70700255 | 70700375 | 788855_47188552_ENST00000601538.6,E | 1 | - | good | GCTGCACTGGAGGGGCCTGCGAAGTGGAGTGTGCCCCACTCCCTGGGTGGTCATGGCAGTTAAACTATTGAATACTCGGGCAACACTTTATAGCAGGGCCTGGTACAGAGTAAGTGCTCA | 120 |
| chr11 | 70700375 | 70700495 | 788855_47188552_ENST00000601538.6,E | 1 | - | good | GTGTTTTTATCTGGTAATCGTGCAATGCGTCTTTTTGGTTGTACAAGGAGGGCTCCCACGGCCCTGAGCAAGGCTTTGAGGAGTGGCATTCAATGGAGTGGAGGGCAGAGGGTGCCCCTG | 120 |
| chr11 | 70700495 | 70700615 | 788855_47188552_ENST00000601538.6,E | 1 | - | good | CCTGCACCTGGGCATATTTGTTGCAGATGCCAAGCAAAGGGGAGGTGTGTGCAGCTTGGCTGGACCGGTGCTGGCTTGGTGAGGGCCCTGTGCACTCTGACCAATCTGATTGTGTATACT | 120 |
| chr11 | 70700615 | 70700735 | 788855_47188552_ENST00000601538.6,E | 1 | - | good | TCTTCTAATCCCTGCAGTCGAGGCACGGGCCTGGCTGCCAGGCAGACTTCATAAATGATGATTGGAATTGGACCTGTGCAAGATGCTGGATGCCAGGGAGGAGGGGGGCACAGGTGTCCT | 120 |
| chr11 | 70700735 | 70700855 | 788855_47188552_ENST00000601538.6,E | 1 | - | good | TGTGTTCCCACCTCACAGCTACCCACGTGCTGGGCCCCTCATCCAGCCTGTGGAGGCTCAGTACCCTGGGGCTCCTCCTTCGCCCCATCCCCGGGATCCTGAGGCCCTGCCCTGCTGAAT | 120 |
| chr11 | 70700855 | 70700975 | 788855_47188552_ENST00000601538.6,E | 1 | - | good | TCCCTGTGGCCCTTCATTGGCATTGTGAGCTACTCTGTTTAACCCTAGGGAAAGGTGCAAGTCTCACTTGTCCCTGCACATGCATGGCCCAACATAAATGTCACAGGGATTGCCTGACCA | 120 |
| chr11 | 70700975 | 70701095 | 788855_47188552_ENST00000601538.6,E | 1 | - | good | GGATCCTGCACCAATCAGCTGGTAGCACCCAGCAGTTCCCTAGACAGCCAGCACAAAGCTGCCACATGAGGGTGTAAGGAGCAGAGACAATTAAAAGTAATAATAGCAATTAACAGCCAG | 120 |
| chr11 | 70701215 | 70701335 | 788855_47188552_ENST00000601538.6,E | 1 | - | good | GTAAAACTTGGTGAATGTTATGTTAGTCTACACCCTTTTTCTCTTTGTTTAAAGTTTTGTTGGTCCATGAAACTCCAGTCTGGGACACACTGGCTACCTTACCTAGCCTAGAAATGGTTC | 120 |
| chr11 | 70701695 | 70701815 | 788855_47188552_ENST00000601538.6,E | 1 | - | good | CCAGAGCCCAAGTATAACTGTTTCTGGGAACCAGGAGACAGGGAAGGCCTGTGCAGGGGGATTCCTGCATGAAGCAGTCAGAGATAATGTCCTTTTAAAGGCTTCTTTTTCTGGCCGGGT | 120 |
| chr11 | 70702655 | 70702775 | 788855_47188552_ENST00000601538.6,E | 1 | - | good | GTAAACCAGGGACAAGAAATCTGCTTACTTTGGGATATTGTGTGAGTACCAATTAAGGGTAAAGACTTACTTTTAATGAAATGAGATAATTTAGCAGAGCATTCCACACTTGACAGTAGA | 120 |
| chr11 | 70702775 | 70702895 | 788855_47188552_ENST00000601538.6,E | 1 | - | good | TCATCCTTTTAAAGATGAAATAATGAGGACTTGCAGCCAGTAAGTGGCAGAGCTGGGTTCTGAATCGAGCCAGCCTGGCCTTGGAGCCCACATCCCTTATCATTGCACTCTGCTACTTCT | 120 |
| chr11 | 70702895 | 70703015 | 788855_47188552_ENST00000601538.6,E | 1 | - | good | ATGACCACGACAGCATTAGTTAACATTACATAGCACTTTCAACATGCCATGCACCATTCTAGTGCCTTACATGTATTGATTCCTCAATCCTTCCCCTTTAAGGAGGGACTGTTACTATCA | 120 |
| chr11 | 70703015 | 70703135 | 788855_47188552_ENST00000601538.6,E | 1 | - | good | TGGTTGCTGTCAGCACTGATTGTGGAGGCACAGATCTGAGGTCGGAGGCCCACACTGCTGCTGACAGACTCCATGACCTTGAGAGGCCACCTTGGGAGGCAGAGGTTTCTAATCTAATGT | 120 |
| chr11 | 70703135 | 70703255 | 788855_47188552_ENST00000601538.6,E | 1 | - | good | TGAGCCTCCCCCTCCTCACCCGTAAACCAGACACAGCAACGTTACTGAGCTCAGACAGGTGAAGCACACACACCTAGAATACGTGTGGCCCCTGGTAAGTGGATCACCAACTATGGCCAT | 120 |
| chr11 | 70703255 | 70703375 | 788855_47188552_ENST00000601538.6,E | 1 | - | good | GATTTACACGGGTCGCTCATGAGATGCCACCTCCCAGGAGCCCTATATGGGTACTACCCCCGGAGGCTGCACCACAATGGGCTCTGCCACTTCCTGGCTGGGTGACTCACTCAAACTCTG | 120 |
| chr11 | 70703375 | 70703495 | 788855_47188552_ENST00000601538.6,E | 1 | - | good | ACTGTAACACTGGGGGCAGGGTCCCAAGGGCCGTCTAGCCCACCCCGCTTTTTCCAGGTTAGAAACCGAGGCTGCTGAGGGCACCTCCACAGGATGTGGCAGAGTTAAGAAGACCACCTG | 120 |
| chr11 | 70703495 | 70703615 | 788855_47188552_ENST00000601538.6,E | 1 | - | good | CACGTTTTCACCAGTGAGATCCAGGGAGCCTTGGGAAGGGGCAGCCCAGCAGCCCTTGCCTCTCAGCGCAGTGCACAGCAAGTCCCAGGGGGCCAAGCCTGGACATGCCATGAGGTGTGC | 120 |
| chr11 | 70703615 | 70703735 | 788855_47188552_ENST00000601538.6,E | 1 | - | good | ACCTCTCAGCGGCAGGAAACAGCCAGCATCCTTTGTTGTCACTGGTGCACAGGGAGGCTGGTGTGGCCCCTGACCTCCTGCTCCTCCCCTCGTCTGCCTCAGGGTGTTTTATTTGTGACC | 120 |
| chr11 | 70703735 | 70703855 | 788855_47188552_ENST00000601538.6,E | 1 | - | good | GGCTGGTCCCACAGAAGACAGGGAGGATGTGGCCAGCCAGGTGAGCAGGTGCATGCGCGGCTGAAGGTGTGGTCAGCTCTGAGGCAGAATTGACCTCGGGCCCCGGCACCTGGGAAGCTG | 120 |
| chr11 | 70703855 | 70703975 | 788855_47188552_ENST00000601538.6,E | 1 | - | good | TGGGATCAAAGGGACATTCCAGCCAGGCTCCCATGGGAACTGCAGGAGCCCAGGGCTGGGCAGTCAGGTGGGGAGGGGGCAGGGTGGCATCACCTGACCACATGGCTGGAGCTGAGAAA[illegible] | 120 |
| chr11 | 70703975 | 70704095 | 788855_47188552_ENST00000601538.6,E | 1 | - | good | TATCCCCCTCAGGAGGCTGGCCTCCCTCTGTGGCCTGAGGCGTGGGAACCCCAGCAGCAGGAGTAGGGCGGCATTGTCCCCGCTGTGGTCCCAGGCCCCTCCCCTGAGCCCGTCACTCAC | 120 |
| chr11 | 70704095 | 70704215 | 788855_47188552_ENST00000601538.6,E | 1 | - | good | GGCATGGCTGTGTCCAGGGTTTCCAGCTGTGTCTCTAGGGCAGTCGCCCGTCTCTCTGGGTCTCTCTGTCTCTGTTCCTGCCCTTCTTCCTCCCGAGCCCTGTGCCATCCTGCACATGGG | 120 |
| chr11 | 70704215 | 70704335 | 788855_47188552_ENST00000601538.6,E | 1 | - | good | CAGCACAACTATTAAATGCATTACTGCATCATTCCTAGTTCTTTCCATTGTAAACAACAGAAAGCCAAGTTTTTTTATACCACCAAGGATATGCATTTAGTTTGTACAAGTGAGAAGCCA | 120 |
| chr11 | 70704335 | 70704455 | 788855_47188552_ENST00000601538.6,E | 1 | - | good | GGGTGGGCTGCTTGGCCTCCTGAGGGTCAGCCTCACCAGGACCTTCCCATGGGGAAGGGAAGCGGCTGTGTCCCCAGAGAAGCTATGTAGCCACGGCAGTTTCTTAGCCACGCAGTGCAG | 120 |
| chr11 | 70704455 | 70704575 | 788855_47188552_ENST00000601538.6,E | 1 | - | good | ATCTCAGAAAAAAAGCCACCTGTCCTTAAGTGGGCCCTGGGTCACTTACTGCTAATGACAGGGCCCAGGAGAGTCCAGCAGGATCCCTAGACCACACCCACATATGCACCCCTGTGTGCC | 120 |
| chr11 | 70704575 | 70704695 | 788855_47188552_ENST00000601538.6,E | 1 | - | good | GGCTTCATTGCTGGACTCACTCCCTCCCTGTGCTTTGAATGGTGGCCATAGCCCGTGTCCTTCCATCGCATCTCTAAAAGGAAGCCAAGATCACCTTTCTCAGCAGTCAGACTACAGAAC | 120 |
| chr11 | 70704695 | 70704815 | 788855_47188552_ENST00000601538.6,E | 1 | - | good | ATTAGTAGCCTCAGTGACCACCACAAGCCACAATCCAGGGACTCCTAGCATCAAGGTCTTTTGTGTCCATCTCTCCCTGACCCCAGACCTTGAACTTGGTTCTGTCTTCCGCCATGAGCT | 120 |
| chr11 | 70704815 | 70704935 | 788855_47188552_ENST00000601538.6,E | 1 | - | good | GAATTCACTTCCTCTCAAAATTCACTGGCCAAAATTAATTTCAAGGGCCTACTCAACCACAGGAGAGCTAAGAAGTAGTGTCTGACCACATGCGCCCTGGTGGAAAGCCAGAAGTATTTG | 120 |
| chr11 | 70704935 | 70705055 | 788855_47188552_ENST00000601538.6,E | 1 | - | good | CAGACTAATAGAACCACCAGCTTCTCAAATGTTTTGGTGGCCATGCCCAAGGACAAAAAGACAATGCTGGTGGGTGGGGGTCTTGCATTTGGAATTAAAACCTCCAGCCAGGAAATAACA | 120 |
| chr11 | 70705055 | 70705175 | 788855_47188552_ENST00000601538.6,E | 1 | - | good | TAGCCTTAAAAAAATAAAAAATAAAAAATTCCCCAATCTCTGTGGTTTGGAACAACAAAAGTTTCTTTCTCATTCACACCTTATGCCCATCATACATCAACAGGGGGTGACTTGAGGACC | 120 |
| chr11 | 70705295 | 70705415 | 788855_47188552_ENST00000601538.6,E | 1 | - | good | CCTATACTTGCTAGGACAATTTTGGTTGCTGGTGACAGAAACCCATCTCTGCCTAGGATGTACTGGCCCACAGGACTAAGAGGGCCATGGAGATGTGTATCAGTCAGCTTGCCCTATATG | 120 |
| chr11 | 70705415 | 70705535 | 788855_47188552_ENST00000601538.6,E | 1 | - | good | TTAATGGGATCCATTTCCATTGTCTAGTAGGTGGTTTTTTCCTTTTCTCCTAGAAAGTACAGCCATTCTCTCAGAGGAGGCCAAACTCAGCCTTTGCCTTGATTTTGTCTGGTCTTCCCT | 120 |
| chr11 | 70705535 | 70705655 | 788855_47188552_ENST00000601538.6,E | 1 | - | good | AGCAGACAAGTCAATACTTCCCTCTTGGATGGTTGGCTTCCCTGGCTTCTTTATCACAGGCAGAGGGGTCTCCTCTGAGACACATAAAGTGCCTGCTGCTATGCTGTGTTCTGGCCATGC | 120 |
| chr11 | 70705655 | 70705775 | 788855_47188552_ENST00000601538.6,E | 1 | - | good | GGAGCACTACAAACATTGCTGAGCCCTTGCTATATGATGCACTGATCCCAGCACTGACTGGGAGGAAGGCCTGGTTGGAAGCTTGGTTTTGACGTCAAGAAGTTTCTCCATGGTTAAGGA | 120 |
| chr11 | 70705775 | 70705895 | 788855_47188552_ENST00000601538.6,E | 1 | - | good | TATCAGTAAAGGGAAGGGCAAGCATGGAAGCAAGGAGGCAGGAGGCTGCTGCAACAGTCCAGGTGCAAGATCATGGTGGTTCAGACTAGAGTGGTAGTAACTGCTAGCCTCCATGTATGT | 120 |
| chr11 | 70705895 | 70706015 | 788855_47188552_ENST00000601538.6,E | 1 | - | good | GGCAGCTTGAACGTGACTCTGGTTGCATTGGGAAGCTGTTGGAAAACGTTAAGCAGCAGAGTTGAAAATCACCTGATTTACATTTTTTAAAAGCTCATTCTATTGTCTCTATAGAATGTG | 120 |
| chr11 | 70706015 | 70706135 | 788855_47188552_ENST00000601538.6,E | 1 | - | good | GACCGAGGAGGGGAGAGCGGTGGGAGATGAGCCAGGGAGCAGGCAGGGCCACGTGGTGAGCGGTGGGAGATGAGCCAGGGAGCAGGCAGGGCCACGTGGTGTCTGGCTTGTGCCATA[illegible] | 120 |
| chr11 | 70706135 | 70706255 | 788855_47188552_ENST00000601538.6,E | 1 | - | good | CAGTGAAATGTGAGCTGGGGAAGGAATGTTCCAGGCAGAGTTAAGCACACACAAAGGCCCTGAGGTGTGAATGAGCTCGACTTGTTCAAGAGACAAAGAGGAGGCTAACACGGCTGGGGG | 120 |
| chr11 | 70706255 | 70706375 | 788855_47188552_ENST00000601538.6,E | 1 | - | good | TGGAATATGAGCAAAGGGTCCAAGGGTGTGTGGCTTTGACCCATTCGCTCACTTTCTCCAGAGTTACTGAGTGCCCGCTATGCACCAGGCTGTGTTGTAGGATCTGGAGAGCTAGCATTG | 120 |
| chr11 | 70706375 | 70706495 | 788855_47188552_ENST00000601538.6,E | 1 | - | good | ACTGCCCCAGCCAGGGCTGTGTCCCCTGCCTCGGCAATCCCCAGCTGGAGCTGTAATCATCCAGCTTCCTGTATCTCCAACTTGCCCCATTCAGACCCAAGGAGATGACGTGGCCAACGG | 120 |
| chr11 | 70706495 | 70706615 | 788855_47188552_ENST00000601538.6,E | 1 | - | good | AGAGGGATGTGTCGGGTTGACAAGTTCAAGAGCCTAAGGAGCTGTTTAGTCCCACCGGCCCACTTCACAGCTGAGAAGGGGAAGGCCCAGAGATGCAAAGTGAGGAGGTCACAGCCAGCC | 120 |
| chr11 | 70706615 | 70706735 | 788855_47188552_ENST00000601538.6,E | 1 | - | good | GATTTTTCTAGGCCATACTCAGAAAACAGCTTTCAGGCCCTGCATGCTCAAATGCTGCATGTGTGAGGTGTCTGGACCGATGGCCCTCTCCTGGGACATGGGGTGAGTCGCCTTACACCC | 120 |
| chr11 | 70706735 | 70706855 | 788855_47188552_ENST00000601538.6,E | 1 | - | good | ATTTTAAATGCTGACAGTCTGAAGCTTGTCATGCAATCTGTCACTGCCTCTCGTGAGTCCCAAAAGGTATATTAAGTGGGATTTCTAGGTTTGGCCTTTTTTTTTTTTCCTTTCAGTGTA | 120 |
| chr11 | 70706855 | 70706975 | 788855_47188552_ENST00000601538.6,E | 1 | - | good | TTTTCCAGAAACAGTCAGAGCGCAGGATCAGAGGCCACTGGAATCTCCCCAGATGCTCCATCTTGGCTTCCTTGCCAAGCTGTACAGCATCCTCTTTCCAGCAGCCCTGCTTAAAGGTCT | 120 |

| | | | | | | | | |
|---|---|---|---|---|---|---|---|---|
| chr11 | 70706975 | 70707095 | 788855_47188552_ENST00000601538.6,E | 1 | - | good | CTTGCCCCACCTTCTATCTCTTAGTTCTATAGCCCACCTCCTCCCAACCCATCCTTAAAGATTAGCTTTTGGCCTTTCAGTGTCTTAGTAAATTTCTTTGAGAAAAATCCTCTTATTTCC | 120 |
| chr11 | 70707455 | 70707575 | 788855_47188552_ENST00000601538.6,E | 1 | - | good | TTTAAGCTCGAGCTTGTCTTGTCTAAAGATGGACTGTTGTGTATCCTTCCTGCCTCTGTCTGTCCACGTGTGTCTCATGAACAGAGGGGACCACTGCCCATGGAATGTGGCTGGAGTGTA | 120 |
| chr11 | 70707575 | 70707695 | 788855_47188552_ENST00000601538.6,E | 1 | - | good | GCATTCATCTTGTCCTCAGTGCAGCCCTGTAAACTCTGCTCTTGGCCCAGGGCACAGTTGAGAAAACTGAGGCTTGGAGGGAGATGGTTTGCTGCTTGGGGTTGTGTAGCTAGCGAGAGA | 120 |
| chr11 | 70707695 | 70707815 | 788855_47188552_ENST00000601538.6,E | 1 | - | good | TTTCAGAGAGACTTGTAGACCAGAACGTCCAAGCTGGATTCCTGGACAGACAACAGTAGCCAACGTGACTCGACGGCTTCCCCAGGCCGGATGCTGGGGAAAGGATTTATTTAGAAGTGT | 120 |
| chr11 | 70707815 | 70707935 | 788855_47188552_ENST00000601538.6,E | 1 | - | good | GGCCTGGCTTGACACAGGTGCCATCTCATGGCCCAGCCCTCTGTAACTGGGGCGTGGGGTGAGGTGGTACAGAGGTGGTCAGGGAGCAGAGCTTCCTGTCTGCAAAGGGCAGGGGGATGT | 120 |
| chr11 | 70707935 | 70708055 | 788855_47188552_ENST00000601538.6,E | 1 | - | good | CCTCCCTCCACTGGTGCTGTGTGGTGACAGCCTTGTCGCCCCAGCTGTGATGTCCTCGGTGCACACCGGGAGCAGACTGTCCACATCTACTCCAGATTCCCAGGGGAGGTTCTGATGATT | 120 |
| chr11 | 70708055 | 70708175 | 788855_47188552_ENST00000601538.6,E | 1 | - | good | AAAGCTTGCCTGAGCTGGGTTGCATGAAGAAATGGACCTTGGTCTAGGGACGCCAGGGCTTTTCCTGGTCATCGAGGGCAGAAGGCAGGGCCCTGCCTCCTGAGGCCCTGGAAGCACATG | 120 |
| chr11 | 70708175 | 70708295 | 788855_47188552_ENST00000601538.6,E | 1 | - | good | AGTCCATCGGGAGAAATGGGCCACCGAGCCAGGACCTGAGATGACGTCTTGGGTATAACAGGTTCCACCCAGCTTGACAGACAGACTGAAAGATGGCACTCCTCCCATTGCAAGAAATAC | 120 |
| chr11 | 70708295 | 70708415 | 788855_47188552_ENST00000601538.6,E | 1 | - | good | AATCCCAATGTTTAGAGATTGGAGGCAAGGATGCCCAAAGCCCTGCAGTGTCCAGGGCAGTCCATCGGGAGAAATGCAGTGTCCAGGGCCGTCCATCAGGAGAAATGCAGTGTCCAGGGC | 120 |
| chr11 | 70708415 | 70708535 | 788855_47188552_ENST00000601538.6,E | 1 | - | good | ATTGTCACTGCCCTTTGACAAAAAGTAACCCAGAAATGTCTGCGGCATGTGGCTCTCGTGGTGGGGGAGTGCGGGGACTGATTGGAAACGTGGAGGGGACGTTACTTGGTTGTCATCTTG | 120 |
| chr11 | 70708535 | 70708655 | 788855_47188552_ENST00000601538.6,E | 1 | - | good | TGCTCTGTGGTCATTTGAAGTCAGCCTTAAGGTCCTTGGCCAAACCTCGGGGCCTCACTGTCTTTGTCATGGACACAGGAGACATCTGACCCCTGCCTGTTCAGTGCCCTCTGTCTGGTC | 120 |
| chr11 | 70708655 | 70708775 | 788855_47188552_ENST00000601538.6,E | 1 | - | good | ACGTAGCACTTGGGGAAAAGAATGAAAAAGCTCAGTATATTATCATCTGGGACCCCTCAAAGGGCTTGGAGGAGCGGCTCCCAGCCAAGGCAGACGATCCAAGGCTGCCTGACCCGGAAG | 120 |
| chr11 | 70708775 | 70708895 | 788855_47188552_ENST00000601538.6,E | 1 | - | good | CAGCAGGGGGGGTCCACACATCAGAAGGCCTTTCAATCCCCTCTTCAGGACGCATGGCATCATCTTCCCACTCTCCCGTGGCATCTGTGCTTATTGTCAATTTACAAAAGTAAATTTGGG | 120 |
| chr11 | 70709255 | 70709375 | 788855_47188552_ENST00000601538.6,E | 1 | - | good | GCAGAGGACACAGTCAAATAACCTGCCCAGGCTCCTGGGCCCGTGTTCATGTGTGTTCAGGGCTTTAGCTGATGTGACTCAGCTCTCTGCAGTCACCTTTTGAACTGTCCCATTCAGAAT | 120 |
| chr11 | 70709375 | 70709495 | 788855_47188552_ENST00000601538.6,E | 1 | - | good | ACTTCCTCCTCCTTCCTCAGCCAGGTCAGACCTGACCTCAACCTTCAGGGTTCTCCCCTGCTCCCTATTCCCAGCAGGAGACTTGGCAATAGCCAGCATTAGCTAAGGTCCTGCCATGTG | 120 |
| chr11 | 70709495 | 70709615 | 788855_47188552_ENST00000601538.6,E | 1 | - | good | ACAGAATTTTTCTAGGGCTTGTCCTGAGCATTTACCCATGGGATGTCAGGATGGCTGTGGTCACCGAGTAGGCTCTGTCAAGCTATAGCTGACATATGATATTTCCCCATCTCTGCTGAG | 120 |
| chr11 | 70709615 | 70709735 | 788855_47188552_ENST00000601538.6,E | 1 | - | good | CCTTTGCCAGCTGTCTCTGCCACGGCCACACGCTCCATCGGCCCAACACTGAGTGTGTGCGCTAAGCCTTCCCTTAACTATGCTCTGCCTCAGTGTCCGTGTCTGTGAAATGGGCATCCT | 120 |
| chr11 | 70709735 | 70709855 | 788855_47188552_ENST00000601538.6,E | 1 | - | good | TGCGGCTGGCTCCTCTTCTCTCCTCTCTCCCCAGCCTCTGTGCTCAGCCGTCACATGGACTCGCTCGCCAGGGCTCCCCCAAGTCTCTTCCCCAACCCCAAATCTCATGATACTTGCGTA | 120 |
| chr11 | 70709855 | 70709975 | 788855_47188552_ENST00000601538.6,E | 1 | - | good | TCTCCGTGGGAACCGTTCTCACCACCTGTCCCCCTCCGTCCCTGAAAGCATAGACTCAGCTCCGAGGACATCCTACCCTGGACCCCCGGAGCCCACCCGCCCCACTGGTGACCTGAGGCC | 120 |
| chr11 | 70709975 | 70710095 | 788855_47188552_ENST00000601538.6,E | 1 | - | good | TGGTGGTGGGTTTTGATCTGGGAGAGCGCCTAGAAAACCTCTTCTGCTGCCCCACCCTTTCCTGGCCTCTGAATTTCAAGTCCAGCGGCCAGGGAATCTCAGTGTTCATCCTCCTTTACT | 120 |
| chr11 | 70710095 | 70710215 | 788855_47188552_ENST00000601538.6,E | 1 | - | good | GCCCCCAGGTGTGGGGAGAGCCAGTAGGGGTCAAGGCCCTGTCTTCCCACGGCCACTGAGGAGGCGCTCAGCAGCCCTAAAATCGCGTTCATCTCAGGACAGGGGTTTTCAGTGCAAAGG | 120 |
| chr11 | 70710215 | 70710335 | 788855_47188552_ENST00000601538.6,E | 1 | - | good | CAGGTCAAAGGTTGTGTGCAGTGGTTGCTCACCCTTGACTTCAGAGCCTGGTCACTTGTCGTGCGGCAGCTCTTAGGCCGTTGTGAAGAGGGAGCCAGGCCTGCCTGAGGGGACGGCACA | 120 |
| chr11 | 70710335 | 70710455 | 788855_47188552_ENST00000601538.6,E | 1 | - | good | TCCGGCACAGTCCGCATTCAGCAGGGTGTCTGCAGCCCTCCTTGGGTTTTAAGCTTCCCTCCTACCCCATTCCATAGCCGAGAGGGGGCAGTGGGACACAGTCTCCTACAGTCCTCTCTG | 120 |
| chr11 | 70710455 | 70710575 | 788855_47188552_ENST00000601538.6,E | 1 | - | good | TCGGTCCCACCTGATAATCCCCCTTTCACCATCAACCCTGACATTACTGTGGGGGTGATGCCAGGGGGCGAAGGTCTAGGGGGCATTGTCAGTTCTGCCTCCCTCGCCGGCCTCATATGC | 120 |
| chr11 | 70710575 | 70710695 | 788855_47188552_ENST00000601538.6,E | 1 | - | good | GGCTCCCTGAGTCTATGGGACTGGCCTCCCCAACTCTTCTAGTCTCTCCGACTTCCCCTTCTGCCACCCGCCAGAGAAAAGTACTTGCTTTCAAAGGGCTCATGAGATGATGAGATTAGT | 120 |
| chr11 | 70710695 | 70710815 | 788855_47188552_ENST00000601538.6,E | 1 | - | good | ACTGGGTTCTCTGCCCAGGCTCTCACAGGGCTCCGGTGGAGGGGTCTGAGGCCTGGGCCCTTAGCTGCAGGCTCGGGGAGGAGTCTGCCTCTGGTCACATTCAGGTTGTGGGCAGGATTT | 120 |
| chr11 | 70710815 | 70710935 | 788855_47188552_ENST00000601538.6,E | 1 | - | good | TGTGAAGGAACCTGCAGCCCCAGTTCCCTCCGTTTCCTGTGCTGCAGAACCCGTGACCACAGGCGTGTCTGCTCACACCCCCGACCCCGAGGCTCTGTGAGTCCAAAGTCAGGCAGCTGG | 120 |
| chr11 | 70710935 | 70711055 | 788855_47188552_ENST00000601538.6,E | 1 | - | good | GGAAGATGTTCTGGGAATTCCCTTCACCCACATGCCTCTCCCTCCCTCTCTCTCTCTCTTTTTCAATCTCTGTCTTGGGTGAGATTGCAGATAAATCTTCAGGAAAAGAAAAAAAAGATC | 120 |
| chr11 | 70711055 | 70711175 | 788855_47188552_ENST00000601538.6,E | 1 | - | good | CGGAAGTCCTGGGGACCCCACCCCGCCCCAGAGCCCACTGCTGTGATTTTTGCCTCTTTGGGATGAAGAGTGAAGAGTGAGAGTTCAGAGAGGATGATGGATTTCAAGGGAGGGAAGAAG | 120 |
| chr11 | 70711175 | 70711295 | 788855_47188552_ENST00000601538.6,E | 1 | - | good | GGACAGCCCAGCCACGTTGCCACCCGGGTGACCCTGAGCCCTCTAAACGGCAGGAAGGGTGGCCAGAGCGGACTCAAGGCTGCACGTTGCAGGAGCTCAGCAATCACAACCAGGATGACA | 120 |
| chr11 | 70711295 | 70711415 | 788855_47188552_ENST00000601538.6,E | 1 | - | good | CGCTTTTGGCTCAGCAGCCTGGGCACAGCCTGAGCGGCGTCTGATTTAAAACCTGCGGAGACGGCAGGAGGCTGCGTGGGAGGGAGGCGTTCGGGTGCAGCACTGGGGGGATTCGGAGCT | 120 |
| chr11 | 70711415 | 70711535 | 788855_47188552_ENST00000601538.6,E | 1 | - | good | GCTGCCTGTGTGTTCCTCTGCCTTGCAAGGAGCCCATCAGCATCTTTGGTTTTAAAAAAGAAAAGTTATTTCGGTCTTTCATCCCCATCAGTAATTTCTTCCTCCTTGTGGTTCTGATCC | 120 |
| chr11 | 70711535 | 70711655 | 788855_47188552_ENST00000601538.6,E | 1 | - | good | ATACCCAACAGCTTGCCAGAGCAGAGACTGGGCCCCAAGAGTGGAGTCCTGAGCCTGGAGGCCGGGCGGGCGTGCTGGCTCCAAGCCCCGCACACTCCTAGCATTCCTGCCTGCGGCACT | 120 |
| chr11 | 70712135 | 70712255 | 788855_47188552_ENST00000601538.6,E | 1 | - | good | GTGGGAAAGGAGGCATAGGGCTGAGAGTCAGAGGGATGCAATGCAGTGGGGGGAAGGACTTGTCCCGCTTTTGCTGGCTTTGAAGGTGGAGGCAGTGGCCAGGATCCGAGGAACACCGCA | 120 |
| chr11 | 70712255 | 70712375 | 788855_47188552_ENST00000601538.6,E | 1 | - | good | AGAAGCTCTGAATACATCAGCCTCATGGTCACGTGGAGTGGAGGTCGGAGAGGGAACTCAGGTTGCTGTCAGCTGAGGGTTATCTGGATGGGTTCAGTGTGACCCCAAGGGTCCTTAAAT | 120 |
| chr11 | 70712375 | 70712495 | 788855_47188552_ENST00000601538.6,E | 1 | - | good | GGGGAGGTAAACAGTTTTCAGAGCACAGCCTGCCCAGCTGTGCTGTCTGACAGGGATGGTCTCTTTGTGGTTATGGTGGGCACAATAGCGGCCCCCAGAGATGTTCACGTCCTAATGTCC | 120 |
| chr11 | 70712495 | 70712615 | 788855_47188552_ENST00000601538.6,E | 1 | - | good | TCCGAATTGGAACTGAGTGTTGTTTGTATCTGGGAACAGGGAAGTGCATTTCCTGGAAGCCGCTTCTCTGCCACCCTCATAGTTTCTGTCCTGTTGAAAGTTCTGAGCTCTGTTGCTGAT | 120 |
| chr11 | 70712615 | 70712735 | 788855_47188552_ENST00000601538.6,E | 1 | - | good | TTAGCCGAGCGAGGAACCATCGTGCTGGCGGCTCAGTCTGAGTGAGGGCCACGCCCACCCATTGCAGGAGGAGTCTCCAGGAACTGGGCCCCGCCCTAGGGCTCACCATAAGTGGCAGAA | 120 |
| chr11 | 70712735 | 70712855 | 788855_47188552_ENST00000601538.6,E | 1 | - | good | TTCCAGCAGATAGGCAGCTTCCACTACTGGAAACCAGTGTGATTCCAGGAAGGGGAGCGTGGCCCCCTGAGCATGTGGTGCAGACCAGGAAAAGGGCTGTTGGTGCAGAGCTGGGCGCTT | 120 |
| chr11 | 70712855 | 70712975 | 788855_47188552_ENST00000601538.6,E | 1 | - | good | CAAAGAGGAGCAGGGCGGATGACACACGCCATACACATGTGCGCACATGGGAGGAACCACGATCTTCCAGGGAGCTGGGAACAAGTGGAACTATGGACTCGGGTGCTTGTTGGTGCTGAA | 120 |
| chr11 | 70712975 | 70713095 | 788855_47188552_ENST00000601538.6,E | 1 | - | good | AACCCATGCCAATGTTAAGGGAATTCAAAGCACGTGATTCTTTTGGCAAACTTCCCCTGCACCTCGCTGCTGGGGTCTTACCCAGGCGTGTGACACGAGGCTGTTCTTCCCAGCATCGAG | 120 |
| chr11 | 70713095 | 70713215 | 788855_47188552_ENST00000601538.6,E | 1 | - | good | CACCTGCTTCCTTGGCCCCTGGATTTGGAGTCCATAGATCTGGCCCTTGTGTGGTGTCAGAGTGCTGAGGACCCACTGGCACGTGCCCGACAGGGCTGGTGCCTCCCCTGCCCACACACA | 120 |
| chr11 | 70713215 | 70713335 | 788855_47188552_ENST00000601538.6,E | 1 | - | good | GTACAACCAACAATTTCCTGAGATGACTAAGAAAAACCAACTGTGCCAGGAGCAGACGGAGCCGGCTCTCTGAAGTGATTTCTGTTTATTGGAGGATGGAAACACAGCCCAGCATGCTGG | 120 |
| chr11 | 70713335 | 70713455 | 788855_47188552_ENST00000601538.6,E | 1 | - | good | TGCTGCTACGGGGGTTCTGTGAAAGCGGACACAGCTACGCTTGGCAAAAAGACTGCTTCGTAAAGTGGCAAGTCAGTACTGTAACCCGTGTTCTTCTTAAGGTGCTTCACTCATCTTGCA | 120 |
| chr11 | 70713455 | 70713575 | 788855_47188552_ENST00000601538.6,E | 1 | - | good | TGGAACATGCAACTGAGCCACTGCCTTGCTGGACACCCTCAGCCAACCTGTGAGCCTCCCCCTCGGCTCCGTACATGAAGCCACTGCTACGTGGGCCTTGGCTGCTCCACGCCCAAGCCC | 120 |
| chr11 | 70713575 | 70713695 | 788855_47188552_ENST00000601538.6,E | 1 | - | good | CCCGGTGCTGTGACGGACACCCGGCGCTGTGACAGCTCTGCCGCCTGGAGGCATCTCCCTCCCATTCTCTTCCTCAGACTCTTGGAGGCTGACACCTCACCCTGGCTTTCCGATGATCAG | 120 |
| chr11 | 70713695 | 70713815 | 788855_47188552_ENST00000601538.6,E | 1 | - | good | GAGAGTGTGTTTTACACTTAAATGGTGTGTGTGGCACACGGCAAGGCATGACGGATGGTGGAAAAGGACAGATCAAGATCACATGGTGGCAGCAATGGACACCCGGCGCTGTGACGGACA | 120 |
| chr11 | 70713815 | 70713935 | 788855_47188552_ENST00000601538.6,E | 1 | - | good | CAGTGCCCAGAGCTTGGACACTGGCACAGATCGCTGGTGTGCACCTGAGCTTCTTCACCTGTACAGCAGAGGCCAACGCTGCTAGCACTTTTCTTGTAGTGTTGCTGGGAGCATTCAGAA | 120 |
| chr11 | 70713935 | 70714055 | 788855_47188552_ENST00000601538.6,E | 1 | - | good | GGAGAAGCGCTGAGCTTCCTGGCCCCTGGGGTAGAGGGAGGAGCTGCTGCTTGCAAATAGCTGTTGAGTGAGCCAAAGCAGTTTGTTGGAAGGAAGAAGGCACTTAGCATGGCCACAAGG | 120 |
| chr11 | 70714055 | 70714175 | 788855_47188552_ENST00000601538.6,E | 1 | - | good | AGGAAGGGCTGAAAAGCTTCAGGCTCTGGGGAGGTTTGGTTTGGGGAAAGGAAGCCCCTCCTGGCAGGCGCCTGCTCCTGTCTCATTGGCCAGGACCTAGGCACTCCTGGAGGGAGGCTG | 120 |
| chr11 | 70714175 | 70714295 | 788855_47188552_ENST00000601538.6,E | 1 | - | good | GCATAGCTGCGCCACGGTGATCTGGGCATGTGGCTCTTTCTCTCCTTCTCCTCCGTGTCCTCAGCATGCTCTGGGGGCAGCATAGATGTAGAGCCACTTCCACATTCCAGACAGGGAAGA | 120 |
| chr11 | 70714295 | 70714415 | 788855_47188552_ENST00000601538.6,E | 1 | - | good | TAAGGCCTTGCAGTTAGGACCCTGCCTGGCTGGGAGAAACAGAAACCGTTGCCTAAACACACAGGGCTTTACTCATCTCACCCAACAGACATCAGGGGCCAGGCTGCAGGGGCCAGGTGG | 120 |
| chr11 | 70714415 | 70714535 | 788855_47188552_ENST00000601538.6,E | 1 | - | good | TTGCTGATTGTGTTATAGAGGAATACCATTTAACCTTCCGGTGCCTCGATTTCCACATCTGATAAGTGATTATTATAATGACAGTGCCTGTCTCATGGGGCTCTTTGTGGCTAAACAAGA | 120 |
| chr11 | 70714535 | 70714655 | 788855_47188552_ENST00000601538.6,E | 1 | - | good | GTGACCTTGGAAATGATCAGATCACACCTGATTTTCTCCTGGGTCGAGTCCTCTCGTGGAGTCTCTGTGTGCAATACTGCAGAAATGTTAAGATATTAATGTAATCACCAGCTCCATCAG | 120 |
| chr11 | 70714655 | 70714775 | 788855_47188552_ENST00000601538.6,E | 1 | - | good | AAAAAAAGGAAGGAAGAAAAATGCCGTAGGGAGGATGTAGCCAGAGGCCATGGGGTGACCTTGACTCTCAGAGCCAGGAGGACCTGGGTTTGGATCCCAGCTCTGATGTTCAGCAGCCAG | 120 |
| chr11 | 70715135 | 70715255 | 788855_47188552_ENST00000601538.6,E | 1 | - | good | ATTCTCCAATGAGGGTCCTTCCTCCAAGAACCTAGGATTCCAGGTTTTTCCCACATTCCCGGATAATTCTCAAGTGTAGCATTGTAAAAAATATTATCTACAGGGCACCACTCACCTCTG | 120 |
| chr11 | 70715255 | 70715375 | 788855_47188552_ENST00000601538.6,E | 1 | - | good | GGGTGGGAGGCATCCACTGAGCTTACAAAGCAGGCAAAGGAGCCCAGTGGCGGGAGGGCAGCAGGGCAGAGCTGAGCATCCTCCCTTCCCGGCTGGGTGACCCTGGGCACAGCCCGTCCC | 120 |
| chr11 | 70715375 | 70715495 | 788855_47188552_ENST00000601538.6,E | 1 | - | good | CTGAGTGTAGCCTCCCCTCCTACGGTGTCCCCAGCCTGGCTTGGCCCGAGAGCTGCAAGAAGGGCTAGTGTGGGCAAAAGAGAAGTCCACACTGCCCTCCCCATAGGCACTGGGTGCCCA | 120 |
| chr11 | 70715495 | 70715615 | 788855_47188552_ENST00000601538.6,E | 1 | - | good | TGATTTTTTTAAAACATTCTAATTACTATAGAGTAGAAACCAACTCAGTGACAGCCAACAGCTTGATTCATTTGTCCAGCCAATTTGGGAGGGTGGTAATTGGGAAGCATCACCAGAGA | 120 |
| chr11 | 70715615 | 70715735 | 788855_47188552_ENST00000601538.6,E | 1 | - | good | CAGCCGAGCTCATCAGCAAATGCAAATCAGGCACAGATTCATCCATTTACTCTAATTTCCATTTAAACTGTGAGTTTGGCTCGATGTATTGGTATACATGCTTCCACTGAGCTCCTGGGC | 120 |
| chr11 | 70715735 | 70715855 | 788855_47188552_ENST00000601538.6,E | 1 | - | good | CTCCAGCCCTGGGCTGCCGGGTCCAGGGGCATGTGGGTCACATTCGCCAGGCATCCTTCTGGACAGATAGAGGAGAGAGAGCCATTCTGGAAACGGAGCCTGCTTCGGCATGGACACTCC | 120 |
| chr11 | 70715855 | 70715975 | 788855_47188552_ENST00000601538.6,E | 1 | - | good | GCCCTGTCCACAGCCAGAGGCAGGCTCTGCTCCCCTCTGGGCCCAGGCCGGATTAGCTCAGCCGTGGACAAGTCTTTTGGAGTCTGGCCACAATACCCCACCCCTCCTCCCCACTGGCGA | 120 |

| | | | | | | | | |
|---|---|---|---|---|---|---|---|---|
| chr11 | 70715975 | 70716095 | 788855_47188552_ENST00000601538.6,E | 1 | - | good | GCGGGTGTTGGAGGGGCAGGCATGGGGGGCAGAGCCCTAAGACAGTTTTGGGGTTAGGCAGTGGGTGTTGGAGGGGCAGGCATAGGGGGCAGGGGTACACAGAGTGAGGCCCCTCCAG/ | 120 |
| chr11 | 70716095 | 70716215 | 788855_47188552_ENST00000601538.6,E | 1 | - | good | TTTGCTCTGAGAATGGTGGCTCTGATTATCATCGGCCTTCTTCGAGGTGCTCTGATTGGCCGGGGAAAGAGGCCGACCACAAAGAGGGCCCTAAGACAGGATGTTCTGTGGGGTTAGGCA | 120 |
| chr11 | 70716215 | 70716335 | 788855_47188552_ENST00000601538.6,E | 1 | - | good | TTAAATCCAGGCTCTGCTGCGTGCCCTTGGACATGGCCCCCTCCTGGGCCCTCTCTTGGGGGTAACAAAGCCTGGCCGGGAGCAGTGTTGAGCACCTCGTGCCAGGAGATGCTTGGGGTG | 120 |
| chr11 | 70716335 | 70716455 | 788855_47188552_ENST00000601538.6,E | 1 | - | good | TTCCTTGAGCTCAACAGGTCTTTTTTCTAGATGTCCTATTTCCGAAGCCTCAGCTGCCGGCATCCTTAAAAAAGATTATAAAGCTAAAAAGAATCCTCCCGCCCCCACCCATTCCTAGAT | 120 |
| chr11 | 70716455 | 70716575 | 788855_47188552_ENST00000601538.6,E | 1 | - | good | CCCAAGAAGGCACCCTGATGGAGATGGGTCCCCTTGACCAAGAACTTCGGTTGGCCAAGGTCCCAGCACCGAAACCCGAGCAATAAAGAAGCAGCTGTCTTCCCACACGATCAGGGCGGT | 120 |
| chr11 | 70716575 | 70716695 | 788855_47188552_ENST00000601538.6,E | 1 | - | good | CCCGTGTTGGTGAGTGGCAGGGAATGAGCCCTGAGCATCTGGCCCTGGCTTGCTAAGGTCACGGCCAGGGGCGGAGGGCGGGTGGTCTTTGTTAGCACCAGTGGGACCCACCTCATCCTC | 120 |
| chr11 | 70716695 | 70716815 | 788855_47188552_ENST00000601538.6,E | 1 | - | good | GATCTATGTGAGCCATGAGCTGTCGAGACTACGGTGACCCCTGCCCAGTCTCCCCGCTGGTCTTCACCCGCCTGGCAACTCTGGGTTTTGTCAGCGACGTGTAGAATCTTTTGGGGGTGC | 120 |
| chr11 | 70716815 | 70716935 | 788855_47188552_ENST00000601538.6,E | 1 | - | good | ATGGCCAGGCTGGGAGGGGTCCAGTTGGGCGGGGGGGGGGCTCCGGGACCCGGTCCAGTGAGATGTGAGGCGCCTGCCACCGAGGAGCATCCCTCCCAGATGTTGCTTTGCGGTTGAAGA | 120 |
| chr11 | 70716935 | 70717055 | 788855_47188552_ENST00000601538.6,E | 1 | - | good | AACGGTGGAGAGGGAGAAACACCCTGTGCGTTTGCAGGCTGCATAATGGTTTATATCAGCTTCTCCAGTAGCTTCTCACCAACTGCTCCCTCTTGCCCACAAAGGCCGAGGAGAGCAGGA | 120 |
| chr11 | 70717055 | 70717175 | 788855_47188552_ENST00000601538.6,E | 1 | - | good | GCTGGCCTGGGGCAGCGTGCCTGCTCCCGAGTGCAGCCTGCCTTTCTCCAAAGACAGCATGGCTGGGTGGATTTCCGTTGTTGTTTTAACCAAGTGGACGGTGTGCTCCTTCCGCGTGGT | 120 |
| chr11 | 70717175 | 70717295 | 788855_47188552_ENST00000601538.6,E | 1 | - | good | GTGGTTAACCTCAAAGGGAGGTCGGTTGCATGATGAGGTGCTGTGGCTACAGATGTGGCTTGTTCATTTTCTGGCGTGTTCTTTCCTAACCCGACAAACTGTATTGAAAGTCCCTGCAAG | 120 |
| chr11 | 70717295 | 70717415 | 788855_47188552_ENST00000601538.6,E | 1 | - | good | GGAAACAGGCTTCTCAGTTGCATCCAGAATCCCAGGGATCCTGTTTATAGTCTCAAGGGTCCCTTTGAGCGGCTAGTGGGTTTTTAAGACAAATTCATACATCTCTTCCTAGTAAGCACT | 120 |
| chr11 | 70717415 | 70717535 | 788855_47188552_ENST00000601538.6,E | 1 | - | good | TGTGTTTTAGGGGGACTGTGTCCTGCTTGCTGGGCGCCCCGAGGCTATTGTTTCCATCACGGCTTTCGCCTGCTTCTGATTTAGAATGGAAATAAAGATATAGCCTCAGATCATTCGGGA | 120 |
| chr11 | 70717535 | 70717655 | 788855_47188552_ENST00000601538.6,E | 1 | - | good | CTGTGGGCTGGTGTGCGGAGCCTTTGTTCTTTACGAGAGCTGCGGGGACGGGAGGACCGTGGGACTTGGGCAGTCCTGCCGGCGCTTGGTGAGACGGGGTGGCAATGGCTTTGCACTTGG | 120 |
| chr11 | 70717655 | 70717775 | 788855_47188552_ENST00000601538.6,E | 1 | - | good | ACGGACTCAGTCTCCGTGGAGACCCCGAGGAGCAGTGGTCCGAGTGAGATGGTCACACCCAGGTTAGAAAGGATGGCCGAGAATTCACGGTTCAAGAAGAAAGTTCATTTTGGAGGTACG | 120 |
| chr11 | 70717775 | 70717895 | 788855_47188552_ENST00000601538.6,E | 1 | - | good | AGCCATCTTTCTTCTTCATCTGATTTTTTTTTTCTTTTCTTCTTCAATTCTGCCCCCAGAGGGCTGCAACTTTTTTTTTTTCCTGTTGAATTAACAGGCAGTAGTAGATGAGGCCGCCCC | 120 |
| chr11 | 70717895 | 70718015 | 788855_47188552_ENST00000601538.6,E | 1 | - | good | GGACGCGGCTCCAGTCTGGGGCTTACAACTTGTGTCAAAGAAGCGAATTTTCAGCTGATTTGGATGTTCAGTCCCCAGAAGTCAAATTTGTGAACTTCCTCATGGATGAAACATTCCCGG | 120 |
| chr11 | 70718015 | 70718135 | 788855_47188552_ENST00000601538.6,E | 1 | - | good | GTAATTGCACTGGTTTTTCAGGCGTCCTTCACCCGTACTTATTTAGACTAGTGACTGATGCATTGCCATGGAAACTGGCACTAGGGGAAAAAAAAGTCCCTGGTACAGTGGGAAGTGGAA | 120 |
| chr11 | 70718135 | 70718255 | 788855_47188552_ENST00000601538.6,E | 1 | - | good | AAGCCTTGCAAAATGAGAAAGCAATTACTGGGGACGAGGACAGCTGCCGCCTCTAAGTGACACCTCCGGCCCATCCTAATAGCCGGGTTGAATCACTGCCTCTCCCTGCTAATGGCATGC | 120 |
| chr11 | 70718255 | 70718375 | 788855_47188552_ENST00000601538.6,E | 1 | - | good | GGCGCAGTTCCTTCCCTCCAGCGTAGGCTCAGGGCGTCTGTGTAGCCAAAACGGGCTTGGCTGCCCGGCTGCCCCTTCACAGAAAGCAGGGCTCTGAACGGCCAAGACCGAGGAAACACC | 120 |
| chr11 | 70718375 | 70718495 | 788855_47188552_ENST00000601538.6,E | 1 | - | good | GCATGTCCATGGATTTTTCCCCAAATCCCTCCTTGCCTCCTGATGCCCCAAGCCCCGCTAAGGACCCACTGCCCAGAGGAAGGCACAGTGCACGCCTTGGGCCAGGGCCACCTCCACCGA | 120 |
| chr11 | 70718495 | 70718615 | 788855_47188552_ENST00000601538.6,E | 1 | - | good | AAGTATTTGAAAGCCGAGAAGTCCTTGCAGGTATTTGGAAGGAGGCTCAGCCCCCAGCCCTGTGCCCTGACCCTGATGCAGGGTGGACTGCTGGCCGTGACCTTGCAGATGGAATCTGCA | 120 |
| chr11 | 70718615 | 70718735 | 788855_47188552_ENST00000601538.6,E | 1 | - | good | TGGACATTCAGGCCAGCAATTCAAAAAAAAAAAAAGAGAATGAGCTCTATCAAAAAGGGAGGAGCACTCCCTCCCTGCCCTGCTGTGACATTCGCCTTCTCACGGGACGTACCGATTGGG | 120 |
| chr11 | 70718735 | 70718855 | 788855_47188552_ENST00000601538.6,E | 1 | - | good | TCTGCCTCTCCAGGGGGATGAAGGACATCCATTGAGTATCCCCAACCCCTTCTAATCCGTTTTCTAACCCACTGAAAGCCTGCAGCTCAGGAAGGTGAGGGCTTCTGTCCCGGGCCATGA | 120 |
| chr11 | 70718855 | 70718975 | 788855_47188552_ENST00000601538.6,E | 1 | - | good | TACCGATAATTCGTGATTATCCCCAGCTGACAAATCAAGATAACTGGAAACAAGCCCAAGCTCATAGCGCCAGTGACCGGCAAAGTGGGATTCAGTCCAGGCTGCCTCCCTGCCATACAC | 120 |
| chr11 | 70718975 | 70719095 | 788855_47188552_ENST00000601538.6,E | 1 | - | good | TGCTCTTCACATGGGCAGTGAACGTAAGAACAGGAACAAAGCTGATCTTCCTTGCGTGCCTACTGTATGCCAGGCACCCTTCTTTAATTCATTGTATTCTCATGACCCTGATTTCCATCT | 120 |
| chr11 | 70719095 | 70719215 | 788855_47188552_ENST00000601538.6,E | 1 | - | good | GTCTATTCCAACGTGGGTGGGCTAACGGGAATCAAAGGAGTGAGTTACACCCCACGCTATCTGGAACCCCAGGCAGCGTTCACTGCGCCACGGCGAGCTTGCAGGTGATGGCACCTGACG | 120 |
| chr11 | 70719215 | 70719335 | 788855_47188552_ENST00000601538.6,E | 1 | - | good | TCTAGCCCCAAGTCCACCCTTACCTGGCTGTGTGACTTTGAGCCAGTCGCTAACCTTCTCCAGGTCCACGCTTCTCATCTTTGAACTGTTACAAAGTGTCTATTTGAACCGTCACAAAGT | 120 |
| chr11 | 70719335 | 70719455 | 788855_47188552_ENST00000601538.6,E | 1 | - | good | CCAGCTCCATCCTCATCCCCAGCAACCCACTGGGTGCCCATGCACACAGCACCATGACGTCCATGTCCCCAGGAGTGCTGGGGCAGCAGGCATGGATAGTGGGTGGGCACAGCGCTGGGT | 120 |
| chr11 | 70719455 | 70719575 | 788855_47188552_ENST00000601538.6,E | 1 | - | good | AAAGGGCCACAGAAACCAGGATCCCACGTAGGTCAGGGTTCTGCACCCCTCCACCATGCTGCATGCGGGCAGGATTAGGGTTAGGTTCAGCATCTGCCAGGCCACCCATGAGAGTGAAGC | 120 |
| chr11 | 70719575 | 70719695 | 788855_47188552_ENST00000601538.6,E | 1 | - | good | TGCCTCCCACTGAAATCCCAGTGAGCAGCAGCAGCCCCCTGGGCCCCTGGCGTGCCAGGTAGGGCTATTTTTATGACCCTCCGATTTGCCCATAGATCACATCAGACCACGGGCAGAAGG | 120 |
| chr11 | 70719695 | 70719815 | 788855_47188552_ENST00000601538.6,E | 1 | - | good | AAGATTCCAGCGCAGTGGGACAAGAGGATCTCGTGTGCTGCCGTGTCAAATGCTGGAGCAGGGGAGCGCGGGGCTGTTGTGGAAACCCTTGCTATTTTTACCTCCTGTCTCCGCAGCTTC | 120 |
| chr11 | 70719815 | 70719935 | 788855_47188552_ENST00000601538.6,E | 1 | - | good | CCTAGTCAGCCTGATGTGGGGGATTTAAGGACTTCAGGGAAGGTGGAGCCTCTGCTGTGTGCAGCTGCGAGCTCCTCATTTTCCTGGCAAGATAGGGTGTCTCCAAGGACAGGAACCTTT | 120 |
| chr11 | 70719935 | 70720055 | 788855_47188552_ENST00000601538.6,E | 1 | - | good | TGCTTATTCTTGATGCAGAAGCTGAATCTCAGAGAGGTTGAGGAACCTGCCTGGGGTCACCCACTGGGAAGGGACCAACCCTTTTGCTTTGAAGCTCAGCCCTCACCCACTTCTGGCCTT | 120 |
| chr11 | 70720055 | 70720175 | 788855_47188552_ENST00000601538.6,E | 1 | - | good | TACCAGGCCCTGGGGGTGCAGCTGTGTCCAGGGGCAACGTGGACCCTGCCCTCATGGAGCTGGTTCCAGCGGTCCGATGGTTCTGGCTCTGCCTTCCAGCAGGAGCTGTTCTGGTCTGCC | 120 |
| chr11 | 70720175 | 70720295 | 788855_47188552_ENST00000601538.6,E | 1 | - | good | TAAACCAGTTAGACGAGGTGATGACCCCTACATCAGAAATGACCGTGGCACTTTGTACAGTACTCTACAGTTTATAAGCACGTGTGTGCAGGTAGTCGGTGTTTATGGGCATCTCCCAGG | 120 |
| chr11 | 70720295 | 70720415 | 788855_47188552_ENST00000601538.6,E | 1 | - | good | TCCAGTCCCATCACCCATGGCATCAGGGAGCAGATTGACAGGCGGGGGCTGTGGACACCAGACCCCAGGCTCATTCTCTGTAGCCTCTGGAGATGTCCCTGGATCACATCCATTTTGTGA | 120 |
| chr11 | 70720415 | 70720535 | 788855_47188552_ENST00000601538.6,E | 1 | - | good | AGACACTTTCCACACACCCTCATGGGTTCTCCTCATGGCCCCCTGGGAGCTGGCACTTTCTCCATCCCCACTTTTTCACCAACGAAGGAGTCACAGGGCCAAGAATGCCAGGGACATTCC | 120 |
| chr11 | 70720535 | 70720655 | 788855_47188552_ENST00000601538.6,E | 1 | - | good | AATTAGAGACTGGTGTTGGGCCAAGAACAGCGTAAACAACTCTATCTCCAGCCCTCCCTCCTGGGGACCTCTTAGGGCCTCCCTGCCATGGGGGTGTGACGGATGAGTGGATTTCAACTG | 120 |
| chr11 | 70720655 | 70720775 | 788855_47188552_ENST00000601538.6,E | 1 | - | good | CGGTGTGCTACATGAATGTGTGTGTGTATGTTCACATGCGTGTTCACGTGTGTCAGCACATGTATACCGCAAGAACACACAGCTATCAGTCCCACCTGTCATTCCTGCATGTTTCTGGGT | 120 |
| chr11 | 70720895 | 70721015 | 788855_47188552_ENST00000601538.6,E | 1 | - | good | TGGGGGAATGAGGACTGGACTCCTTTGGGTTCCAGAGGGAAAGAAAGCCTTGAGATTATTTTTCCAAAGAAAAGAATTCCCAATTCCCAGTCTCTCCTCTGTTCCTCCCCAACATGTGTG | 120 |
| chr11 | 70721015 | 70721135 | 788855_47188552_ENST00000601538.6,E | 1 | - | good | GATGTCCAGGTGTTAACTGGAGGAAAGGGAGGGGCACAGAGAGAAGGTGCCCGGACTGCCGGTCAGTCAGGGATCTCACCTCAGGCCTGCCTGGCACACGGAAGCCCGGGCTCAGGTCAC | 120 |
| chr11 | 70721135 | 70721255 | 788855_47188552_ENST00000601538.6,E | 1 | - | good | CCTCACAGCCCCACTAGGTGTGATGCGCCTGAAATGAAACTCAAGAGCTGCTTGCCTCTTTCGGGAAGGAGCAGCAGAGCGTGCCAGGGAGACCTTTGTCTCCCTAGACGTCGGTGGAGA | 120 |
| chr11 | 70721255 | 70721375 | 788855_47188552_ENST00000601538.6,E | 1 | - | good | TGTATTAAATCCAAATGGGCAAGAGTTCATTGAGAACCGACCTTCAGGCATGCCCATGTGAGACACTGCGGAGGAAGCCTCAGAGTGTGCCCAGGAGCGGCTGCAGTCCCTGAGGGAAAC | 120 |
| chr11 | 70721375 | 70721495 | 788855_47188552_ENST00000601538.6,E | 1 | - | good | ACCTCCAATATTCAGTGGAGCCCAGCCCCTACAATTCCCTGTGGTGGAGCACCAGCTCCTGTACACACGGCCTGCCCCAAGATGTTGGCATCTCTGACCCCTGCATTTAATCCTGACTCA | 120 |
| chr11 | 70721495 | 70721615 | 788855_47188552_ENST00000601538.6,E | 1 | - | good | ACACCCTGGGGGCAGAACTGCCTCAGTAAGAACGCTGCTCACAAAGTGGTGCCCTGAGCCCAGCCCCACTCACAGCTGGCCTCAGAAAGAACAGGCCTTCCCCAACAGTGGGGTGTACAC | 120 |
| chr11 | 70721615 | 70721735 | 788855_47188552_ENST00000601538.6,E | 1 | - | good | AAGTCCTGCATGCAGGGTAACTGGCTGCTGACGGTCCCACATCCCAGCCCTGCCTGTGGCATGCCAGTGGTACCTCCCCAGCACTGGGACAGCACTAGCTGGCCCACGAACCTCCAGACC | 120 |
| chr11 | 70721735 | 70721855 | 788855_47188552_ENST00000601538.6,E | 1 | - | good | TTGTATGTATTCAAGCCGTGCTCAAAACACGCTGTGCTTCTTGCGAAAAGAGTTTAGGCTCTGAGCTCAGATGGTTAGGAAAAGACACTGCACCTGTCTAGGGTTGGGTGGGACCTTTGG | 120 |
| chr11 | 70721855 | 70721975 | 788855_47188552_ENST00000601538.6,E | 1 | - | good | TCGACGGGCCCTGCCACCAGTGGCATCATTTGGGGGAATTGGTGAGCAGTTCTTTCAAAGCCTCAGCTAAACAAAGGACATTCTTCCCTTGATGTTGAAGTCATTGGCATTCCTGAAGAA | 120 |
| chr11 | 70721975 | 70722095 | 788855_47188552_ENST00000601538.6,E | 1 | - | good | TGACAAAGCGGGAAGATGATTTCCAGATCCATGCACTTGGGTGCCACTGGACGGGAAAACAAAGGATGGGATGGATGATCCTGCAGTCACCTTGAGTGCCACATGAGGTGCACTCACACC | 120 |
| chr11 | 70722095 | 70722215 | 788855_47188552_ENST00000601538.6,E | 1 | - | good | AGTGGGAAATGCAGTGTGCATACTTAGCCCCTGAGAGCACTGAAGGGCAGGAATGGGGCTGAGTACCAAGAGACAGCACACATCCCAGCAGTGACAGTGAGGACAGAGGAGAGGATGGCA | 120 |
| chr11 | 70722215 | 70722335 | 788855_47188552_ENST00000601538.6,E | 1 | - | good | TGCTGAGAGCGTGAGGAGCATCCTATTGACATTCCTCAGTGGCCACCTGTGCCAGCCACTAAGGCCAACTTTGCACAAATGCCTCCCATCAGCAGAGCCTGACTGCTGCCCTGGCAAGAG | 120 |
| chr11 | 70722335 | 70722455 | 788855_47188552_ENST00000601538.6,E | 1 | - | good | CAAAACTGTTGGAAAGGCTGGAGGAGCAGAAAGGGACAGGTGATGTCACCCAGAGATTGGTCACTGCAGGAAGCCTTCATGTCCCTTAGGGTTACAGGAGTAAAAAGAACCAAGAGGTAT | 120 |
| chr11 | 70722455 | 70722575 | 788855_47188552_ENST00000601538.6,E | 1 | - | good | AACAATATACGTAATTTCAAAAAACTCAAGCTAATACCTTTTATGAATCCATGCAGTCAGAAAAAACAATCGCCTATGTAGGTATTTCAAACAGAGGGGATTGAATCAGGAAATTGTTTC | 120 |
| chr11 | 70722575 | 70722695 | 788855_47188552_ENST00000601538.6,E | 1 | - | good | TCTCTCATTTTTGCTCTGTGGTTCTCACCTCTAATAGACACATTGCACCCATTTTTTCAAACAATGATGTGCAAGGATTCCCCTTGTTATCTTTAAATGAAATTCTTATAGAATAAAATT | 120 |
| chr11 | 70722695 | 70722815 | 788855_47188552_ENST00000601538.6,E | 1 | - | good | AGATTCTTTTAATAAGTCTCCAGGTTAAGTCACCAGCAGCCTTCGAGGTACACTATGTTAGGAGGTAGTGGTTTTGACCTTTATTCCTAGCTCCACCATCCTCAACCCTTTAAGAAACCA | 120 |
| chr11 | 70722815 | 70722935 | 788855_47188552_ENST00000601538.6,E | 1 | - | good | GTTTCTTTTGAAAGCATTTGGTTATTTCAAAACCCTCTTTTTCAGGAGGCTTAGAATCTTTCTGAGCACATTATTCTCCTGCCACTTCTCATAAGTCGTAAGTCCTGCTTTTAAGCTAAG | 120 |
| chr11 | 70722935 | 70723055 | 788855_47188552_ENST00000601538.6,E | 1 | - | good | GGGGTCTGACGGTGCTTTGTAAAAAAGTCCCATAAAGGTAAGGAGATGAGGGACCAGGCAGAAAAGTTAAAACTGAGGCTTATTCATTATTAACAGTGAGATTTTTATCTAACAAAGTAG | 120 |
| chr11 | 70723055 | 70723175 | 788855_47188552_ENST00000601538.6,E | 1 | - | good | CCCAACCCATGCTCATTGGGAGAGAATCACATCAGGGCCTGAAGACCAGAAGGTAGAATCCAGGGCCACCTTATGAGTTTGTCTACCACAAGGACCATGAGCATATTTCTAGCCCCACAT | 120 |
| chr11 | 70723175 | 70723295 | 788855_47188552_ENST00000601538.6,E | 1 | - | good | CCCTCCAGCAAGAAAATTGCTACTATCTAATATAATCGTATAATCCTATAATCATGTGCATGAAGTCATGTATATCTTGTCCCCTTTGCTGTATTCTACTTGGAAGAAGCAAGTCTGAGA | 120 |
| chr11 | 70723535 | 70723655 | 788855_47188552_ENST00000601538.6,E | 1 | - | good | TACCATAAAATCAGTGGCTCCAACAACATGCATTTACTCTCTCATGGTGTCTTTGGGTTAGGAGAGAGGCATGCTTTAGCTGAGCCTTCTGCAAGGCTTCACTCAGCCTGTGAGTTGGGC | 120 |
| chr11 | 70723655 | 70723775 | 788855_47188552_ENST00000601538.6,E | 1 | - | good | GAGAGACAGTTTCACCCATAGAGTATTTTATAGTGTTACACCAAGTAAGGCTATATGGCATGGCTCATAAGTCTAAACAGTGTGGGACTTGATTAGTCACCTGTTGCTGCATGACAATAT | 120 |

| | | | | | | | | |
|---|---|---|---|---|---|---|---|---|
| chr11 | 70723775 | 70723895 | 788855_47188552_ENST00000601538.6,E | 1 | - | **good** | CCTAGGAGAGTGCTGGCTCCTTGGAGCTGCTTCATTACTATTTGAGGACACCTGCTCTTGTTTCTTTCTTCAGTTTGCTTCTCCCATATTGCCTCTCTACATCCTCTTCCTCAGGTCTAG | 120 |
| chr11 | 70723895 | 70724015 | 788855_47188552_ENST00000601538.6,E | 1 | - | **good** | CACACCTCTTGAGGAGTCAGGGTTTGATGCGGGATTTTTTTTTTTAAGTGGATTTGGCAAGGTCTTGAGCTGTGAAAAGGAACTGCTCTTATTTCTGTCCCTAGAGTACTCAGAAGAGCA | 120 |
| chr11 | 70724375 | 70724495 | 788855_47188552_ENST00000601538.6,E | 1 | - | **good** | ATGCTTAGTGACATGGAGTCCTCTGACAGCAATAGTGCTTGGAGACATTTTTACACCTATGGCCACAGGCACATGCATACACACACACCCCACCACCACAACAAAACCTGCTGCACAGAT | 120 |
| chr11 | 70724495 | 70724615 | 788855_47188552_ENST00000601538.6,E | 1 | - | **good** | TTGCATCCTCTAAATTGGATGCTGTCTGATATTTCCAAAAGGATACCATGCTTGATGAGATCAAAGGAGGAGAGGATGACTCAGAAAAGACCCAGAAGAGCTCACAGTGCTCTTAGCAGG | 120 |
| chr11 | 70724615 | 70724735 | 788855_47188552_ENST00000601538.6,E | 1 | - | **good** | AAGGCTGCCCAGAGAGCCTGGCTGCAGGTGTCGACACAGCTCTCCACAGGACGTGGTTGGCGGCAGTCACCTGGTCACCACAACTCTTCTAAGCCTCCCAGCTCGGGAATTCTGGACCTC | 120 |
| chr11 | 70724735 | 70724855 | 788855_47188552_ENST00000601538.6,E | 1 | - | **good** | CTGCTACCTGATCACCCCGTTAGATCTTGAGAGGTTTCCCTTCCCAAACACTGAGCAGGTGACATTTCCGGAACGCAGAGTTAGCGTCTTCCTGCTGCCTCTGAGCTGGTGTTTGGACAC | 120 |
| chr11 | 70724855 | 70724975 | 788855_47188552_ENST00000601538.6,E | 1 | - | **good** | CATTTGGTTGGGACTGGTTTTAACTACCTTGAGGACTTTTCTTATCTCACCCATCTCAAATGTCTCCTCCCCTGTAGGGTGTATGGGCATCAGCTGCAAGTTGCTTCTTCTGACTAGAGT | 120 |
| chr11 | 70724975 | 70725095 | 788855_47188552_ENST00000601538.6,E | 1 | - | **good** | AATCAAGCTTTGGAATATTCCTTTTTTAAAACATACTTTAAAAAATCTTTTCTTACCTGTGAAAGCTTATGGCACATAGTAGGCATTTAGCAAATGTTGGTTTCCTTCCTTTTAGACCGA | 120 |
| chr11 | 70725095 | 70725215 | 788855_47188552_ENST00000601538.6,E | 1 | - | **good** | CCACACAGTAGAAAAACATCCCTCCCTTTAGTACACTGTACGGTGCTTATTTTTGCCCATGTTATTTACGCTAAAGATAGTCTTATAATGGCCTTAGCTTAAGTGTTTTCCCCCCCAAAG | 120 |
| chr11 | 70725215 | 70725335 | 788855_47188552_ENST00000601538.6,E | 1 | - | **good** | TTGAAAAATGATTCATTTTAAATATGACACAGAACCAGCTGCAAATAAGTTAAATGTTGCTGCTTTGGGGTGTTAGTGGCATTTATGAATATTGTAGGGAAACATTTACTCACTAAAATT | 120 |
| chr11 | 70725335 | 70725455 | 788855_47188552_ENST00000601538.6,E | 1 | - | **good** | TTTGGGAGGAGCGGGTCTGGGCCACCGCACCCTGGCAAGCCCTCAGCGCCATGTGCCTTCCTGTCCCAGTACCCAAGGAAATTCCTCCTGCCACGCTCAGCTTTTAAGAGTCCGCAAATG | 120 |
| chr11 | 70725455 | 70725575 | 788855_47188552_ENST00000601538.6,E | 1 | - | **good** | TCCAAGTGACTCGGTCCATAGTGAGCCTCTTTGGGCACCAGGCCCTTGCTCTGACCTCATTAGCCTCGGGCTCCAGCCAGCAGTTGCAGCTGACATGCAGCTGTCAAAGGCCCCAGGCCC | 120 |
| chr11 | 70725575 | 70725695 | 788855_47188552_ENST00000601538.6,E | 1 | - | **good** | ATTCTCAGTGTTGCTGTGGATAGCAGTGTTTCTGTGTTGATATGGATGCATCTTTTCTCCTCGCATCCTCTTGGGATCGTGCGGTATTGGGGTGAGGTTCATCTGAACAACACCCACTGG | 120 |
| chr11 | 70725695 | 70725815 | 788855_47188552_ENST00000601538.6,E | 1 | - | **good** | CATTGTACACCATCTTTCAGTTGTCTTCAGAGGTCTACCGGGTCATCCATGAAGACAGGGAATAAAATACGTAGAGTTACCGACAATGTGCTGACAAGAAGTGGGCCTTTGATAAATGAC | 120 |
| chr11 | 70725815 | 70725935 | 788855_47188552_ENST00000601538.6,E | 1 | - | **good** | GCAGAGCAGTTAGAACAACACGTCTGCCTTGCTTTGTCTAGGTATTTGTGAGGAACAGAGATGATGCAGCTTAGAAGGTGGCTGTTCTTCCCCCCATGCTTCTGTACACATTTTTTCATT | 120 |
| chr11 | 70725935 | 70726055 | 788855_47188552_ENST00000601538.6,E | 1 | - | **good** | ACATGGAGGAGAGAGCTGGTGCTCCGAGTTGAGCTCTGTCCCTCCACTGCCTGTGTGTGTGAGTTCACGCATACCCTCCTGGGCATCCAGCTGCTTCGACATGTTTTTATGGACTGTCAG | 120 |
| chr11 | 70726055 | 70726175 | 788855_47188552_ENST00000601538.6,E | 1 | - | **good** | CCGGGTAAGATTTCACTGAAAAAGAGGCACCCATCCCTAAAGCGGGGAAGTAGGTGTAAAAGTCCTCAATGCACTTACCAAGTACAGCTCTGAGAAACTCAGCTGAGCATGGAGAAGCTT | 120 |
| chr11 | 70726175 | 70726295 | 788855_47188552_ENST00000601538.6,E | 1 | - | **good** | CAGGCAAGAGCTCTTTATCCCCAGCTTCCAAACCCAGCCCCAGGGAGCTCAGGTCCCTCCAAGATGCTGGTGGGAGGTGAGGGGGCAGGGCCTCTGTACTCGCTGCTTCTGTGTTAGGTT | 120 |
| chr11 | 70726295 | 70726415 | 788855_47188552_ENST00000601538.6,E | 1 | - | **good** | CCAGAGCCTGGCATGTGAGCTCCATAAGGCATTGGTGAATTAACAAAGTGCACCTTGACCAGAGGTTTCTGGTGCTCCTTTTAGTGGCCAGACCTCTGCACTTCCTGCTTTTGGTACAGA | 120 |
| chr11 | 70726415 | 70726535 | 788855_47188552_ENST00000601538.6,E | 1 | - | **good** | TTTGTCCACCTCCGTTCCAGCCCTGATCCCAAGGGGGTCTAATTTATCTGCCCACCCCACCCACCCACACCAGACTCAGAACAGCTTCAGGGCAAAGACCCCACATCTCTAGCCTCCTGA | 120 |
| chr11 | 70726535 | 70726655 | 788855_47188552_ENST00000601538.6,E | 1 | - | **good** | TGCCATCTACCACAGCCAACCCCCACCTGCCTTCCCAAATGAGCTTACTTTCTCCCTCTCTGAAAATCTTTGATCTCCACAGACAGAGACACTTGCCCCTGCCTTTTGGCTTCCACCAGC | 120 |
| chr11 | 70726775 | 70726895 | 788855_47188552_ENST00000601538.6,E | 1 | - | **good** | TAGCCCAGGCTTACTGACATGGCAGCTGTACAGCCTTCCAGGGGTGAGCAGAAGCACATGAGGCCTTTTGGAGTCTAGACTCAGAACTGGTCAATTGCACCACATGCTATTGGCCAGTAT | 120 |
| chr11 | 70726895 | 70727015 | 788855_47188552_ENST00000601538.6,E | 1 | - | **good** | CTGCAGGGAGGGTCTTCTGTTCCCAGGTAGATTTCCATTTGTGTCTGTAGTGGCAGTTGGTGAGCTCTCAGCAAGTTGGCAGGGATGACCAGGGCATCTGGGTCTCATCCTTTAGTGGGC | 120 |
| chr11 | 70727135 | 70727255 | 788855_47188552_ENST00000601538.6,E | 1 | - | **good** | AAATTCGCAATGCATTTTTACATAGTTTCCCATGTGACTCTCACAACTGCTCCGTGCCAGAGGCAAGATTTGTCATCTCCATTTTGTAGATGGTAAAACTGTTGCTAAATTATCAAGTGG | 120 |
| chr11 | 70727255 | 70727375 | 788855_47188552_ENST00000601538.6,E | 1 | - | **good** | GTCTCTGGAGAAAGACCCCAAATATAGCAGTTCATATGTGAAGTATTGGCCAACCTCCCACATTTTGTCTTCCAGGTTATTCTAATAGTAATACTTCCACCCCGGCTACCTTGCACTTTA | 120 |
| chr11 | 70727375 | 70727495 | 788855_47188552_ENST00000601538.6,E | 1 | - | **good** | GGTGCCTCTTACATCAGGGAACAAATTACCTACCACCTTAGTTGCCAGAAAGCCCTGGAAACTACACAAGAGTCAGTGGAGACTTGGCATCCCCGAAGGAACCCCTCAGGGAATTCTGCT | 120 |
| chr11 | 70727495 | 70727615 | 788855_47188552_ENST00000601538.6,E | 1 | - | **good** | AAAGAAACCACCTAGCAAGAGGAACAGTCTTAGCTTAGAGCGTAAGCTACAAAGCAATAGGGCCAGAATACCAACACCTTGCCTGGGATGCCAGCTCCTAGGATACCCCTGCTCTACCAT | 120 |
| chr11 | 70727615 | 70727735 | 788855_47188552_ENST00000601538.6,E | 1 | - | **good** | TCCTCTATCCTGAGTCTTAACCCAGGGTGGTACCAATTTTTGACATGGGCCACCTCATTTTTACCGTATGCCTGCCCCATGTGGAATGCAGAGCAGGGGTAGCAGCATCCATTTACCCAC | 120 |
| chr11 | 70727735 | 70727855 | 788855_47188552_ENST00000601538.6,E | 1 | - | **good** | AACTGAGGACAGCCCATGGCCCCCATCCAGCGTCTTGGGGGAACAGAGGCTGTGTTGCTGACCACACCATCTGAATGCTTGCGGGTAGGGCTGGGGCCCCCAACCCGGGACCTTGGGAGG | 120 |
| chr11 | 70727855 | 70727975 | 788855_47188552_ENST00000601538.6,E | 1 | - | **good** | TTCTGAAAAATGTCAGCCAACAAAACTCATCAGAACACCTCTCGCCTCCTTTGTCCCTTGAGGAAAATTACTTACTGCTCTGGAAGTTTTCTTTTGTTGATGTAAATGTTTGTGCACAGA | 120 |
| chr11 | 70727975 | 70728095 | 788855_47188552_ENST00000601538.6,E | 1 | - | **good** | CACCTGCCTGTTCTTGCCCTGACACTGGGTGCTTGCATATTCATTTACTCTCCTTTTCCAAAGCATTTATAGCATAGAACACTAAACACAATAAATGCCTTTGCTGAGCCCGTAAATAAC | 120 |
| chr11 | 70728095 | 70728215 | 788855_47188552_ENST00000601538.6,E | 1 | - | **good** | AGTACTTTTCCCTGTGTTCCTAACCAAGCACTGCCCTTTGGGAGTCGGTTTAACCCCATGGCTCGGTTTCTATGGTGACAGTCAACAGGGAGCCGCCAGGGAAGCTGTAGGCTCCGCATA | 120 |
| chr11 | 70728215 | 70728335 | 788855_47188552_ENST00000601538.6,E | 1 | - | **good** | CTGCAGACAAACCTCTGTCCCTTCTCACATATCCCAGAGGGATCCTCTTTGGATCCCACAAGGACCCATCTGCATCTGGGGAGGTGTGGGACACTCTCATTTGTTGGATTCCAACTGGGG | 120 |
| chr11 | 70728335 | 70728455 | 788855_47188552_ENST00000601538.6,E | 1 | - | **good** | GATGAACCAGTCCAGCTGCCAGGCTGTCTTGAGTTATATTGGCAGTAAGGGGGCTGCCTTCTACCAAGTAAACCGAGTGTGACCACTCCCAAGTCACGGTGCCTCGGATCCGGCCTCTGG | 120 |
| chr11 | 70728455 | 70728575 | 788855_47188552_ENST00000601538.6,E | 1 | - | **good** | CGGACTCAAAAAGCCCAGCCCCGTTCCACTCAGAAGCTCTGGACCTGTGGCTTTCATCTGGGGCTCCTTGAAGCCCTGGGATGACTTTGCTTCCGCCCAGGACGCTTCCTGAGTCTGCGT | 120 |
| chr11 | 70728575 | 70728695 | 788855_47188552_ENST00000601538.6,E | 1 | - | **good** | GCACCTTGGGCAGAAGAAAGACCACCCATGGTCAGTGTTCTGGGCGTGCGTATTTGTTTGTGCCTTGTTTGTTTCCAGGAAGATGTGTTGTTTGGGTAAGAATCCATTATCTCTGTCCAT | 120 |
| chr11 | 70728695 | 70728815 | 788855_47188552_ENST00000601538.6,E | 1 | - | **good** | CTTACGTGATCAGAGGGCATGCGTAGGAGAAGGTGTGAGTGTCACTGAAAGTGTTTGGAACTGGTTTTTTGTTTTTCTTTAAAGCAAACCTGAATTGGGGGTAACACAGTAATTCAAACA | 120 |
| chr11 | 70728815 | 70728935 | 788855_47188552_ENST00000601538.6,E | 1 | - | **good** | ACCACGCCCGGCCCTTCATTCCTTCTCAGGTGCCAACACCAGTCAGTCCGTGCTGCACCATCTCCCACTGGGAGGATCATTTTCTTGAACCCCTTATGGGTATGTTTAGCTGCAGGCCTC | 120 |
| chr11 | 70729295 | 70729415 | 788855_47188552_ENST00000601538.6,E | 1 | - | **good** | CCATGAAGCAGTTACTTCCTCCCTATTTCCCCCAGCCCCCAGTCCCTGGAGCCCACCAGTCTGTTTTCTGCCTCTGTGGACATGTGTATCTGGACATGTGTAGATAGAGCCCTTGGTGTC | 120 |
| chr11 | 70729415 | 70729535 | 788855_47188552_ENST00000601538.6,E | 1 | - | **good** | AAAAAAAAATGAAGAAAGTACACAGTGCAGTGGCATGAGTACATTCACAGGTATGTACAACAGCCACCTCTCTCTCGCTCACTCCAGAATACTTTTAGATCCCAAAAGGAAGCCCCGTGC | 120 |
| chr11 | 70729895 | 70730015 | 788855_47188552_ENST00000601538.6,E | 1 | - | **good** | GAGGCAGGGAGAGGCGGTTTTGCTTGCCCCACATCATGAACCCCTAGGTGGTGGAGCAGGTTTGGGACTTAGACCACCTGGCTCAGGCACCTATCGGCTTTTTTTTTTTTTTTTTAGTAA | 120 |
| chr11 | 70730015 | 70730135 | 788855_47188552_ENST00000601538.6,E | 1 | - | **good** | GCAGCTGGCACACAGGTCCCACTTCCTCCTCTCCAAGGCCCTTCTAAGAGCTTCACGTGCATTGACACATCCAGGCCCGACACATCTGCAATGGTAGGTATCATGACAAAGGAGAAAACC | 120 |
| chr11 | 70730135 | 70730255 | 788855_47188552_ENST00000601538.6,E | 1 | - | **good** | CCAATGTACACATCCTGCTTCAAAGAGCTCGGGGAACTCCACAGAAGGAGATTCACCTGTGCCAGCTCCTGGGGCCTCATGCTGCAAATAGCATTGATGATAAGATTGAAACACATAATA | 120 |
| chr11 | 70730255 | 70730375 | 788855_47188552_ENST00000601538.6,E | 1 | - | **good** | TTGGGCGTGAGTGGGGAGGTTGCAATCTCAGGGCCTGTTGCACTAGGCAGGTGCAAGGGCAGGAGTGGAGAAGGCAAGGAACTGACAGGCCAGGAACTCACAGTCTAGGTGTTCGTTGAA | 120 |
| chr11 | 70730375 | 70730495 | 788855_47188552_ENST00000601538.6,E | 1 | - | **good** | GGGGTTGAGTCTGGGATAGTCGGGTGAGATGCAGTCCAGTTAGGGGTGTGCACAGCTTTATAGGGATGGGCTTCGGGGGTGAGTCCTGGGCATCTATTTGTGATGAGATCATGGGGACAA | 120 |
| chr11 | 70730495 | 70730615 | 788855_47188552_ENST00000601538.6,E | 1 | - | **good** | CCAGGGTAGGAATGGATGTGAGGAGACATCAGCTCATCAGGCAGCTCTGAGGACAGTGCTGGAAGGGCTGGGAGATGCGGTCCAGTTAGGGGCACACTCAGCGTTACAGGGGTGGGCATT | 120 |
| chr11 | 70730615 | 70730735 | 788855_47188552_ENST00000601538.6,E | 1 | - | **good** | GATGTCTTGGATGATTTGGGAGTTGGTGGCCCTGCCGCAGCAGCCTCATCCTTCAGCAGGGAGAGGAGGAGATGCTGGCCTGGCCAGGAGCCTGGCTGCTCATCTGCAGGGACCAGGTGC | 120 |
| chr11 | 70730735 | 70730855 | 788855_47188552_ENST00000601538.6,E | 1 | - | **good** | ACCTAAATTACAAAAAAAAAAAAAAAAAAAGAAATAAAAAAGAGAGGTCCCAGGCATAGCAGTGAGCAGACGCCACTTTGGGAAGGAAAGTTGCTGAAATGAAGACATGGAGTCCAGAGAA | 120 |
| chr11 | 70731695 | 70731815 | 788855_47188552_ENST00000601538.6,E | 1 | - | **good** | CACCTTGCTGGTAGCTGGTGTGAATCTGGTTTTGAGGGCTCAAGAAAGAAGAGAGCCCCAGAGAATTAAAAACAGGTGTCCAGACAAGTCCCTGTGCACAGATGTTCACAGCAGCACTAT | 120 |
| chr11 | 70731815 | 70731935 | 788855_47188552_ENST00000601538.6,E | 1 | - | **good** | TCTGGAGAGCCTGGCCACACCCCAGCCCGGCAGAGCACGTGCAGGAGGAGGAGGCGGCTGTGTCAGCTGAGACGAGGGCTGCTCATCAGCCGTTCCACCTCGCAGCGTGGAGGCCACAGG | 120 |
| chr11 | 70731935 | 70732055 | 788855_47188552_ENST00000601538.6,E | 1 | - | **good** | TCACCCTGGGGGTGTGTGTGGCCAGAGAGGAAGACACAGAGGACTGCGTTCAGAGGATGGGGAGCTGAGGAGGAGGCCACAGGAGTGGCCAGGAGCATCCAGGAAGTGAGAAAGGGGA | 120 |
| chr11 | 70732055 | 70732175 | 788855_47188552_ENST00000601538.6,E | 1 | - | **good** | TGGGCGAGTTACATTTGAGCAGCCTGTAGTGTGTCAGGGTGGAGCCTGCATGGGTTGGGCTGGAGGGTGAGGCTGGGGCCACAGGAGTCTGATGGCATCTGAAGCCAAGACTTGAGCAGG | 120 |
| chr11 | 70732175 | 70732295 | 788855_47188552_ENST00000601538.6,E | 1 | - | **good** | GTGACCTGCAGGCACTTGCTGTCTGATTGGCTGTGGAGGGTGAGAGCAGAGGCGAGGCTTGCGGCTTCAGCAACTGGATGAGGCGCCAGGGCAGAGATGGGGTGATGTTACGTGTGATGT | 120 |
| chr11 | 70732295 | 70732415 | 788855_47188552_ENST00000601538.6,E | 1 | - | **good** | AGCAGTAGAGAAGTGGGGAGACTGGTCCAGAGCCAGGCACAGGGCCCAGGAGAGAGAGGAAGGTGGCTCAGGCCAGAGCAGAGATTGTGAATGGGCAGCAAGAAGGGTCCGGAGTTGTC | 120 |
| chr11 | 70732415 | 70732535 | 788855_47188552_ENST00000601538.6,E | 1 | - | **good** | CTCAGTGAGGCTGGGGGACACAGAAAAAAATCCTGGGCCATGCAAGGGTCTGAGCAGGCAAGAGATGTGATGGGTTCGGCTTTCTAAGGATGGCTTTGGGGCCAGGTGGAGAAAGGACAG | 120 |
| chr11 | 70732535 | 70732655 | 788855_47188552_ENST00000601538.6,E | 1 | - | **good** | GGTGCAAAGGCCCCATGGCGGGAATGGACTTGTGCTTGGGGAACAGCAAGAAGGGCAGCGCGGCTGTAGCTTCTAGGGAGGGGAGGGTCGGGGAGGTGTGGTCACATAGGCTGAGCCCA | 120 |
| chr11 | 70732655 | 70732775 | 788855_47188552_ENST00000601538.6,E | 1 | - | **good** | CTTCCTAGGCAGGACAAAAATGGCCACCTAGAGGTGGGATGTGTCAGCAGAGACCGGACAGAAGCAAGGGCATGGGCTGCGTGGGCATGGGCTGGGGCACTGCAGGCAGCAGGAACAGC | 120 |
| chr11 | 70732775 | 70732895 | 788855_47188552_ENST00000601538.6,E | 1 | - | **good** | GCAAAACAGAGAGTCCCTGCCCTCACAGAGCCGGCAGGCCTGTCCCCTGCAGGGAAATGGCAGCTGCTGGGGATGTGAAGCAGCTGCAGGGCTGGGGAGTGCTGGAGAGGGGAGGGCAG | 120 |
| chr11 | 70732895 | 70733015 | 788855_47188552_ENST00000601538.6,E | 1 | - | **good** | CACTCCATCGGCAGCATCTGACATGCAGGAGAAGCTTAGAATTCATTCATTCCACAGAGCACTATTCGCGCGGGCTGTGTGTGCCAGACACCATTTCCAGAGCTGGGGTTACAGCCGGAA | 120 |
| chr11 | 70733015 | 70733135 | 788855_47188552_ENST00000601538.6,E | 1 | - | **good** | GGCTCCGGTAGCATGTGACACGTCTTTCTGTCGCGGACGATCCCACTCAGTTGAGATGATCTGCTCCGCGTCTGTCTCTCCCACCCCGCCCAGCCTGTGCCCTCGCCACTGCTTAGTCAT | 120 |
| chr11 | 70733135 | 70733255 | 788855_47188552_ENST00000601538.6,E | 1 | - | **good** | TCCTGGGATGCCCTTCCCTCACATGTCCATCTACCCAACTCCTACTCATCCTGCAAAACCCAGCTGTGACAGTCCCTCCTCTCCACGCTTCAAGCAAAGTCCTCTCCCGGCACTCTCAGT | 120 |
| chr11 | 70733255 | 70733375 | 788855_47188552_ENST00000601538.6,E | 1 | - | **good** | TGCATTGAATGAGACATTCTGTATCACACATAAATTGTTAATATGAATCAACTCCTTGCTTCTGTGTAAACATACTAAACTTTTTCACACCTCTGGGCCTTTGCTTATGTTCTTCCTCCT | 120 |

| | | | | | | | | |
|---|---|---|---|---|---|---|---|---|
| chr11 | 70733375 | 70733495 | 788855_47188552_ENST00000601538.6,E | 1 | - | good | CAATCCCAGCTCAGTCACCCATAGCTGTGAAGCCTCCAGAAAGGGATGGAGCCTCTCACATTCTTTTTTCAGATCCTAAAATGGCGATGATCACTGTACCCACCACACAGGGCTAATATG | 120 |
| chr11 | 70733615 | 70733735 | 788855_47188552_ENST00000601538.6,E | 1 | - | good | GCACGGCCCACCCTAACTTACCTGCCACCCCCCTCTCCATGACCTAGGCCAGGGGTCCACAGAGTTTATCCGTAAAGGCCAGAGAGTAAATATTTTAGCCTTTGGAGCCATGTGCTCCCT | 120 |
| chr11 | 70733735 | 70733855 | 788855_47188552_ENST00000601538.6,E | 1 | - | good | TCCTCACCGTTCCCTGTGCCTGCACAGACACCTCATGGGAGCTGAGCCTGCCCCTCTTCTTGGGCCTCCCTCTTCCCCAGACCTGCCAGGCCCTCCTTCCCGGGTCCCTCTTCTAGTGGG | 120 |
| chr11 | 70733855 | 70733975 | 788855_47188552_ENST00000601538.6,E | 1 | - | good | ATGGCTTCGCTTTGTCTCCAGGTCCCACCTTGGCCAGGCACGAGGTGGGCCTAGCAAAGGCACCCAGGGGTAGCAAGTGCTTGGGGGGTCACCCTCCCCGCCTTGGCAGCTGAGTCCCTG | 120 |
| chr11 | 70733975 | 70734095 | 788855_47188552_ENST00000601538.6,E | 1 | - | good | GCCCCCTCCCTGCTGCCAGGGACCTCCCAGCCCCTGGTGACTCTTCCCCTCTCCCCAGTCACTGCACACCCTCCCCTGTCTTCCATGAGTGTGTGAGTGCGCGAGCAAGTACACAGAGCC | 120 |
| chr11 | 70734095 | 70734215 | 788855_47188552_ENST00000601538.6,E | 1 | - | good | CTGGAAGGGCCTCCACACCCTTGCCCCTGCCGCACCTGCTGCCCGGCACTCCCTCCCGCGGGCTCTGCTCTGCATACTCCACCTGCTCCTCCCAGATGCGCGGCTCAGGGGGTGCCTCCT | 120 |
| chr11 | 70734215 | 70734335 | 788855_47188552_ENST00000601538.6,E | 1 | - | good | CTCCCTCGGGTTTTAACTATTCTGCCCGTTCAATCCCTGCTCAGAAGCATCAGGCAAAACATTCTCGAACAACCCAGTTAAAACATCTCCTCGCATAGCCAGTTCCCCCGTCACGGTGCC | 120 |
| chr11 | 70734335 | 70734455 | 788855_47188552_ENST00000601538.6,E | 1 | - | good | TGAGCTGTCAGTGTGGAGCGTGTGGACTGGGTATGGGAGTCAGGCGGCCCACTGGCGCCCTCACGCTCATCATGAAATGCGCCTCACCTTTGAAGTCACCTTCACACACCTTCACTCGAC | 120 |
| chr11 | 70734455 | 70734575 | 788855_47188552_ENST00000601538.6,E | 1 | - | good | CCTCGCCCATTGGATCCTTTGAAGCACACAGCAGGAGGGCTCTGTGTCCGCTGGGCTCCTGGACTCTGCCTGTGGTTGGTGAAGCCTGGCTGAATTTTGCCAGCCCAGCTACAGGCACCC | 120 |
| chr11 | 70734575 | 70734695 | 788855_47188552_ENST00000601538.6,E | 1 | - | good | GCACAGGCTGTGGGGCCCCTGCTGAAAGCCTGGGCCAGGGCAGCCTCTGGCTGGGAGGAGAGTTGGTATTGGTCGGGGGGCCCTGGCTGGGCCTGTGTCCTCCCCACCCCACCCCTTGGT | 120 |
| chr11 | 70734695 | 70734815 | 788855_47188552_ENST00000601538.6,E | 1 | - | good | AGGTGGTTGCTGGGTAGGCCCTCGGGGCCAGCAGCCATTCCTGTGGCTATGTAACACTGGCTGGTCCCCAGGCCACCCCTCAGCACCCTTCGGTCTTGAGGACACCCCTTGGCCTCCAGG | 120 |
| chr11 | 70734815 | 70734935 | 788855_47188552_ENST00000601538.6,E | 1 | - | good | TATCCTGCCATTCCCAGCTTCATGGGCCTTGTACTCTGTTCTTGGCAGAATTTCTCTTTCCTTCCCAAGCCTGGGGGAGGGTGGTGGCAGGAGGGCCGTCCCCAAGGGTCCTCATCGCTC | 120 |
| chr11 | 70734935 | 70735055 | 788855_47188552_ENST00000601538.6,E | 1 | - | good | CCTTCGCCCCTGAGCCCAGACACGGGACTGGCCTCACAGAGCACAGATGTGCTACAGCCGCCGAGGCCTGGGCCCGCTCGTCTCCAGGGCAACCCAGAATCCCCCTTGTACTGCTTAAAA | 120 |
| chr11 | 70735055 | 70735175 | 788855_47188552_ENST00000601538.6,E | 1 | - | good | CCTTGAACACAGGGCAGTGGGTTCCCTGACAGATTGGGCAGAGTCCCGAAAACCTGCCCACCTGGGGGGAGGCCACGGCCTCGCCCACCTTCCCACACTGACCCCGCGATGGAGGGGTCC | 120 |
| chr11 | 70735175 | 70735295 | 788855_47188552_ENST00000601538.6,E | 1 | - | good | GGAAGCCAGGCCTTCCCAAGACCCTGCCTGGTTGCCCAGCTCCACCTGGGACACTTGCAGAACACTCACAAGGTGCGCCTCTGTCCTCAGTCCCCGCCGCCTCTCTTGGGGAAGATAGTC | 120 |
| chr11 | 70735295 | 70735415 | 788855_47188552_ENST00000601538.6,E | 1 | - | good | ATTTTCTATCATGCTGGGTGAGGCTGGCTTGCGGGACCCCAGGAAGCCTCCACAGGTGTGGCTTCCTCTTCCTCTCTGGAGTTCCCTGCACATCCCTCCTCCCCGAGGAAGGATCTCTCG | 120 |
| chr11 | 70735775 | 70735895 | 788855_47188552_ENST00000601538.6,E | 1 | - | good | TAGGGTTTGGAGAGCATGACCACGCAGCCACCTGCCTGAAGGGTGGACACGCTGAAGAGAGAGCTACAAGAACCTCAAGTTGGGCCAGTGCATGGTGCTCTGGGGAGGTGGACCTTGGGG | 120 |
| chr11 | 70735895 | 70736015 | 788855_47188552_ENST00000601538.6,E | 1 | - | good | CCTGGATCAGGAAAACATGGCAAGTGGCAGGATCTGAGGTCACATCATAAGGAGTGTGTACCCAAGGCAAGGGAGGTGAATGGAGCTTGTGGAGGTGAGCAGCTAGGATGGTGATGGGTG | 120 |
| chr11 | 70736015 | 70736135 | 788855_47188552_ENST00000601538.6,E | 1 | - | good | ACTGGCTCCTCTGCCACTGGGCTTGGCTGGGTTCTTGTTCCCGAGCCTCAGACCTGGGTCAGGCTTGGATCTAGTTTGCCCATCCTTGTCTACTCTAGTGGCTGTGGAGGAAGGCGATGG | 120 |
| chr11 | 70736135 | 70736255 | 788855_47188552_ENST00000601538.6,E | 1 | - | good | CACTGTAACTATTTGCTGAATGAGTGAGCTCTGAAACGAGCGGTTGGCCACCAGGAGGCCATGCATGGCCTGAATGTGAACACAGGTGTGTGTGGTTTCAGAACCCAGCCAGGGGCCTTG | 120 |
| chr11 | 70736255 | 70736375 | 788855_47188552_ENST00000601538.6,E | 1 | - | good | GAGCTGTTTTCCAAATCAGGTCACACTCGTGGGTTTCGGGGGCTAGGGTACAGTCACATCTTTTGGGGACCAGCACTCACCCCGCCCACACAGTTACATCCTCACAACTAGAGAGGGGCA | 120 |
| chr11 | 70736735 | 70736855 | 788855_47188552_ENST00000601538.6,E | 1 | - | good | TGGAGAAGCCACTTCACCCCCAGGGAGATGGTGCTGTCCTTTGCTTCATCCCTGGGATCTGAAGGGTGTGAGGGTGGGGTGGGGGTTGTGTTGGGGTGCGTGGCGGGGAGCCGCCATGGA | 120 |
| chr11 | 70736855 | 70736975 | 788855_47188552_ENST00000601538.6,E | 1 | - | good | CTGGGATTCTAGAGGCCACGTTTCAGAACCCCCAGGTTCTCGCAAGGAAGAGGCTGAATTCCGTTCCCATTTCACGTTCAGTCCCTGGCCCTGACCACATCTTGACAAACCGTTTGGTTC | 120 |
| chr11 | 70736975 | 70737095 | 788855_47188552_ENST00000601538.6,E | 1 | - | good | CGGCTCGGGACTTTGGCTCAGAAAGGATTTTATCCCCACTTTGGCCGCGGGTGAAGGAGAGTAGGGACAGGTTCCCCAGGCCCCTGGGAACACACATGGTACAAGAAATTCACACAGAAA | 120 |
| chr11 | 70737095 | 70737215 | 788855_47188552_ENST00000601538.6,E | 1 | - | good | TGCACCCAGAAGCCAGTCCCCTACCCCTGGGGGGCTGCACCCAGCCTGTGAGGGACCTGACTCCACCAGGGTGGCTGGGCAGGGCTGGCAGTCCCAGGAGTCTCTGAGGGGTGTAGCCCG | 120 |
| chr11 | 70737215 | 70737335 | 788855_47188552_ENST00000601538.6,E | 1 | - | good | CCTCACAGGAAGCTGGCAGGCAGATCAGGCAGGTGAATGCATCTTCCTTGAACTTGTTTCTATTTTAACATTGAAACGGGAGGATGCGCCACATGGCTGGGAAATGATAGTGAATGCAGC | 120 |
| chr11 | 70737335 | 70737455 | 788855_47188552_ENST00000601538.6,E | 1 | - | good | GACAGATCTCAAAGAAGGCCAGGCCACCTGGGGGCTGGCGTGTCCCAGGTGCAGAGGGCACAGAGCTGAGCCCGGGACCCTGATGGGTGAGGATCATTTTCAGAGCCGTTGGGGCTGCCA | 120 |
| chr11 | 70737455 | 70737575 | 788855_47188552_ENST00000601538.6,E | 1 | - | good | CATGTGGGTTTGGACTCAATGCGATGGAGGTGGCCAGGGATGTCTGGAGCGAGATGCATGCAGGGCTGGGGAACTCGGACAGAGTCTCATTGAGCCACCTGTGCCTGGCACGTGTTTCTG | 120 |
| chr11 | 70737575 | 70737695 | 788855_47188552_ENST00000601538.6,E | 1 | - | good | CCCTGCCCCGGGCTGCCCGTCCTATACTGGCTGCCCACTGGGGCCTGGGCTGCCCTCTCTAGAGGGTCAGGGAGCGCGGAAGCCTTTGCAGACAGTCTCCTGGAGAGAAATGGGGTGTAT | 120 |
| chr11 | 70737695 | 70737815 | 788855_47188552_ENST00000601538.6,E | 1 | - | good | CAGGGAAAACCGGCCCCAAGAAGCCTTCATTCTGGAATCATTTGTCCTCAAAATGTTTGCGATGTAAGGGTCTGTGGGAGGAGCACCACCCTCGCATTCCAGATGGGGCTGAGGCTGCAG | 120 |
| chr11 | 70737815 | 70737935 | 788855_47188552_ENST00000601538.6,E | 1 | - | good | ACCTTCACACTTGTCATTGTCACACATGCTCACCCTGGAACCTTATTAGCAAATATACTTAGTGTTTTTTCATTTAAGTGCCAGCTCTTGCAAATGACGGGCCTCCTTGCCATCTGTGTG | 120 |
| chr11 | 70737935 | 70738055 | 788855_47188552_ENST00000601538.6,E | 1 | - | good | TTGTCACCGTGCAGCCAGCCGGTGCAGTAGTTGCCGAGCCCGGCACCCGGCTCCTGGCATCATGAGGCCTGGGCCCAGATTCTGGTGTGATGCCCACGTCTCCGCGGCTCACCGTAGGGC | 120 |
| chr11 | 70738055 | 70738175 | 788855_47188552_ENST00000601538.6,E | 1 | - | good | CCTGGAATGTGTGCTGATTCTCTTGTGCGATGCATGGTTGCTTTGGTGTCAGGACAGATTTCGGGGCTTTCAGCCAACCATGCGATATGCAAGGGAGCTCTTGCCACCGTCTTGCGGGCT | 120 |
| chr11 | 70738175 | 70738295 | 788855_47188552_ENST00000601538.6,E | 1 | - | good | GAGTCCGGCTTTCTGCCAGGCCAGACGCGGTTTCCTCGGAAGCTTCGGTTGTTGTAAGGCTCGAGCATCTACCAGGTCTCTTAGCAGTTTTGAAACTCGTGGCATTGCTGCCAAGCCCCT | 120 |
| chr11 | 70738295 | 70738415 | 788855_47188552_ENST00000601538.6,E | 1 | - | good | TGACCTGCAAACAAGCACAGCGCCCCTCAGGGCTGTGGTGCAGCCTCCACTGAGTGTTTATGGTTGGGGTTGGTGCACAGAAGACAGAGGGGAGCGCCTTTCCCTGCACATTGTTGATGG | 120 |
| chr11 | 70738415 | 70738535 | 788855_47188552_ENST00000601538.6,E | 1 | - | good | ACTTCCCTGCAATGGGCCCTGGGAATGCACTTGCTCCCCGCACACGAGGGGCAGTGATGGCGGCCTCTTCCTGTGCAGCCTCAGCCACGGTACTGCTCGGAGCCTTGGGCGTCCAACTTG | 120 |
| chr11 | 70738535 | 70738655 | 788855_47188552_ENST00000601538.6,E | 1 | - | good | CAGGACGAGGGGTCCAGCGGCTGGCTGTGCTTCCATTGGTCCCTTGGGTAGCTGAAACAGTGGATAAGTTTGGGCCTAAACAGACCAGAACACCATGGGGCAGCTTTGTAGATTCCGGGG | 120 |
| chr11 | 70738655 | 70738775 | 788855_47188552_ENST00000601538.6,E | 1 | - | good | GCGGCTTCTGGCACAGTGGGCGTTCTGGCTAGTTGGCTCTATTGTCAGGGGACATAGCCTGGAGACGGGTCAGCACTGATGGGGGTTATAAGAATGAGTACGGTGGCCACTGCAGAAAGA | 120 |
| chr11 | 70738775 | 70738895 | 788855_47188552_ENST00000601538.6,E | 1 | - | good | GGCAGCAGAACCCTCTGGGGCCTGGGCAGGCTGGTGGCAGTGCATGGACGTGGCTGGAGGCCGTGAGGTGGTTTGCTTACTGCTTTTGTTGTAAACTACCACTTCCCTCCAAAGAGGCGG | 120 |
| chr11 | 70738895 | 70739015 | 788855_47188552_ENST00000601538.6,E | 1 | - | good | AGAGGGGTGTCTGAACACCACAGGTCCTTCCGTCGGGCCACAGACCCCCTCCCTACATGGGAAAGTTGAGTCCAGTGACCCCGCTGTCTTTCTCCATGGGCCGGATCTCCATTCGAGGGG | 120 |
| chr11 | 70739015 | 70739135 | 788855_47188552_ENST00000601538.6,E | 1 | - | good | TGCCTAACTCACTGGCCAGGAGATAAAGTGGAGTGTGTCCTGTCCCCAGACAATGGGGCCAGATGGTGGGATATGCAGCATCTTGGCTGGAGAAACTCACCAGCCCTACTCACTGAATCC | 120 |
| chr11 | 70739135 | 70739255 | 788855_47188552_ENST00000601538.6,E | 1 | - | good | GCGTCTGGGCTTTGTGTGCCCCTGAAGTTCACATGCCGCATCCCCACCATCAGGCCCTCTGGCTGAGGTTCAGTTGGTTGCTGGTGGGTCAGGCTCCTGATGGGGCTCAGCCATGCCACC | 120 |
| chr11 | 70739255 | 70739375 | 788855_47188552_ENST00000601538.6,E | 1 | - | good | TGTCTTCATTGTCCACCTCCTGGCCTCCTCCCTTCTGTGGGGAGGTGGAGATGTGTGGAGATGTGTGGAGATGGGTGGAGATGAGTGGAGATGGGTGGAGATGGGAGATGCCCCGGGTCT | 120 |
| chr11 | 70739375 | 70739495 | 788855_47188552_ENST00000601538.6,E | 1 | - | good | CAAACACCCTCCCCACGTGGGCACAGGTAGACTTCAGGGGCAAGAGGAGAGGACGGGGCAGCCTGGGCGTCAAGGCTGGAGTCTGTGCCTGGCCTCACCGCAGCAGCTGCTCCACACGAT | 120 |
| chr11 | 70739495 | 70739615 | 788855_47188552_ENST00000601538.6,E | 1 | - | good | ACCCTGGACCAACCTCACCAGGCTCAGAATCTGAAACCAGAACAAGACAAATCACATGATCTGGTGGAGCCCAGCAGCCACAGGTCATGATCCAGGTGTTCCCCAGACAAGGCACAGGCA | 120 |
| chr11 | 70739855 | 70739975 | 788855_47188552_ENST00000601538.6,E | 1 | - | good | CTGATGGCGAAGGAGCCAAGCTGGACTTCTGTGCCCTACCTGCGTGCCACCGAATGCCAGGATTCGGGATCATCATCCTCTGCGGCGGCTCTGGCCTGTCTTCACGTGGCCGTCCTCTGT | 120 |
| chr11 | 70740215 | 70740335 | 788855_47188552_ENST00000601538.6,E | 1 | - | good | GGAGCCCTTCTCATGGGTTGTCACGAGAACCCATTGCGGACAACCTTTATGGTTTTCATTCATACAGATGTGGGCTGCAGCGGCAGAGAAGCCCTTTCAGTCTTTCATTCAGCACCTCCT | 120 |
| chr11 | 70740335 | 70740455 | 788855_47188552_ENST00000601538.6,E | 1 | - | good | GGTGGAGGGTGGGTGGTCCCAGCCTAGGGGAATCTCAGGCCACAGAGACACAGAGCAGAGGTCAGGATAAACGGGCTGGCATTTATTTGTCGAACACATACAGTGTTTCTATCCCCAGAG | 120 |
| chr11 | 70740455 | 70740575 | 788855_47188552_ENST00000601538.6,E | 1 | - | good | ACTTCAGTCCCAGTTCCTCTGACGCATCACCCCAGGGGATGAGGGCGCTTCAGCTACTCCCTGTTTACAACCGAGGATTTGAGAGACCCAGAAAAGACCGGGGAAGCTTCAGGAACTCAA | 120 |
| chr11 | 70740575 | 70740695 | 788855_47188552_ENST00000601538.6,E | 1 | - | good | AACTGCTTAAGTGACAAAAGACCCCGGGTAATCATGTGATTAAAAATAAACTTGGGCTGACTGATGCTGCTTGCTGGGGCCTATCCAACCGGGCAGGGGGTGAGGGGGAGGAACAGGAAG | 120 |
| chr11 | 70740695 | 70740815 | 788855_47188552_ENST00000601538.6,E | 1 | - | good | GAACTGCCTCTGGGGAGAGGCTGGCTCCAAGGACTGGCTCCCTCTGTCCACTGCTACCAGGCTGTGCCAGCTGACGCTTCTTCTGGCTGCTCTGCTGTGGGAGTGCTGCACTCCCAGTTA | 120 |
| chr11 | 70740815 | 70740935 | 788855_47188552_ENST00000601538.6,E | 1 | - | good | TTGATCAACACACACATGGGCACCACCTGGCCAGGGGCTGCAGGTACTCAGGATGTTCCCTCAGTGAGGGATTCTGCCCTTTCCCATCTGCTGGATAGGACCAGACTGGCCTTGGACATCCA | 120 |
| chr11 | 70740935 | 70741055 | 788855_47188552_ENST00000601538.6,E | 1 | - | good | ATAGACAGATGGGATGAGCGGAAGTTGAATGGATGAACAGTTAAGCAAGAGATAACACAAATAACCTCATCCCCTCCACCTTGCTGCCCATCAAATCAGTCTCAGTCATAGTTTGTCCCC | 120 |
| chr11 | 70741055 | 70741175 | 788855_47188552_ENST00000601538.6,E | 1 | - | good | GATGGATGGGTAGGCGACACCTCTGAGAATTTGAGGGAGTTCAAAAGTGGGAGAAGGTATATCAGGTAGGTGCTCAATGAGCAATAGCTCAGTTTATGAGATGGATGATTGGATGGATGG | 120 |
| chr11 | 70741415 | 70741535 | 788855_47188552_ENST00000601538.6,E | 1 | - | good | AGGTACTCAGTGAGGTACAGACAGATACATGGGGTGGATGGGTGGGCAGATGGATGAGTGACTATGCAATGCCTCTGGGAAGCTAAGTGGGTCTGAAGGTAGAAGAAGATACCTGTTGGT | 120 |
| chr11 | 70741535 | 70741655 | 788855_47188552_ENST00000601538.6,E | 1 | - | good | AGGAGACTGGAAACCTGGCAGGGGACATGTCAGGGGCCTGGGGAGGACACACAGGGTACCCAGGCAGCTGGGAACAATGAGTAGGCAGTACCTCCAAGAAAGTGGGGGAAGGTACCTGG | 120 |
| chr11 | 70741655 | 70741775 | 788855_47188552_ENST00000601538.6,E | 1 | - | good | TTAATGAAAAAACAGAGGCAGCCTGTGTGCTCAACACATAGTAGGTTTTCTGCTAGTTTCCTTCCTTCTCCGAGGAGGTAAGCATCAACCACTTAAGTTATGGTGAGGCATAGGGTGCAG | 120 |
| chr11 | 70741775 | 70741895 | 788855_47188552_ENST00000601538.6,E | 1 | - | good | GCTGGGGGCAGCAGAAGGAATCCTGGATCTAGAGTTAAATATGCCTTCTGTTCCTTACTCCTTGCCAGTTGTGACTTCTTAGAAAAGTAGGATGGTCCCTTATGGACGTGTGATGACTGT | 120 |
| chr11 | 70741895 | 70742015 | 788855_47188552_ENST00000601538.6,E | 1 | - | good | AGACCAGTCCTGTAGCTGGGGCAGCAGGGGAACCAGGTGGGAATCTGGAATTCTGGACACTGACTCTTTCCTTAACAGTTCTCAGGGTCCACCAGCATGGGGGCTACTTCCTGTGGGACT | 120 |
| chr11 | 70742015 | 70742135 | 788855_47188552_ENST00000601538.6,E | 1 | - | good | GGGTTTGGAAGTTGTCAGAATTTGGAGTTTCCTAGGTGGACAGCCAGTGTATCTAATGTTTTGCCTTGTGAGTGGTTTTCATTACTCCATTTTACAGATGAGGAGGTGGGGACTCGGAAG | 120 |
| chr11 | 70742135 | 70742255 | 788855_47188552_ENST00000601538.6,E | 1 | - | good | GAGATTGCCAGCAGAAGGAGGAGGAAGGGGCTTTGCTTCATAGAGGACTGCAAGCTGTCTGCTTCCCTGGCCCATGAGGCCACAGGGGATGTGGGCTCAGGGACAGGGCAGTGGCCTCCA | 120 |
| chr11 | 70742255 | 70742375 | 788855_47188552_ENST00000601538.6,E | 1 | - | good | CCTTTTTCTTCACCCTGTGGGTTCAAGAGCAGTCGGTAGAAACCAAGAGACAATAGGGTCTGGAGAGCTGTGACTTCCCAGAGTGCGCAACAGTCCACCTGGGTTTTGAAGGATGAGTAG | 120 |

| | | | | | | | | |
|---|---|---|---|---|---|---|---|---|
| chr11 | 70742375 | 70742495 | 788855_47188552_ENST00000601538.6,E | 1 | - | good | TAGTTACTAGTAAGAGAAACCTGCCAAGAATTGTCCTGGACCTAGGCAGAGAACTAATATGCCTACCTTTTGAAAAAGCATTCCAAACTTCTACTTTGTGCAGTGGCCTGAGGGACAGGC | 120 |
| chr11 | 70742495 | 70742615 | 788855_47188552_ENST00000601538.6,E | 1 | - | good | GGATTATGGGATGGCAGAGCCGGGTGGGGGGTGGCGTGAACTGGACTGGGACTGTGGATCTGGAGTTAAATATGTGATGACTGCTTAGGATTCCCCGTCCCTCTCAGAGCTTGGCCTGGG | 120 |
| chr11 | 70742615 | 70742735 | 788855_47188552_ENST00000601538.6,E | 1 | - | good | GCCCAGTGGGGTGGCTGTGCATGGAGAGCTCTGAATCCGCCTCCTTTTGAGCCTAAACAGATTAGTGCTGACATTCATTCCGCGTCCTGGGCCTGCCTCTCTGTCATGCGGGGAAGTGCT | 120 |
| chr11 | 70742735 | 70742855 | 788855_47188552_ENST00000601538.6,E | 1 | - | good | ATCAGTGCTGGTCACACCTGTGCAGGGAGACTCCAGCCCCCGCCCCCAGCAGGGAGTGGCCATGCCTGCGCTCCAGGCCGCCCCCTTGCCTCTTATCTCCGGCAGCCAGTCAGCCACGCT | 120 |
| chr11 | 70742855 | 70742975 | 788855_47188552_ENST00000601538.6,E | 1 | - | good | TCCAGATGCCAGACATCCAAGGACGCCAGGCCAGGAGCCGGCTCATCAAATATGTGCAAGGCGTCAGAGGTGTGAATTTTCTACTTCCAGCCCAAGGCAAACTTTCTTTCCATTTCCTTT | 120 |
| chr11 | 70742975 | 70743095 | 788855_47188552_ENST00000601538.6,E | 1 | - | good | CAGATGCACGCTGGTCCTCAGGCACCTCCCATCTCCCAGGTTCTGATCCCCTCTCAGGTCCCACCAGCACTGCAGCCACATGGTGTGTTCAGCAAAATATGCACGAAGCTTCTGCAGAAT | 120 |
| chr11 | 70743095 | 70743215 | 788855_47188552_ENST00000601538.6,E | 1 | - | good | CATCTGGCAGCTCTAGCTCCTGCAGCTCCTGCGTTCCTGGCTGTGCTGTTATCGATCATTCTGCAGGTTTCCCCTCCAGCCCCCGCCCTGTGCCTTTGTGTCCCCACCGCTCAGCACCGG | 120 |
| chr11 | 70743335 | 70743455 | 788855_47188552_ENST00000601538.6,E | 1 | - | good | GAAAGTCCTTTATGGTCTGCAGACAGCAGCCTTTGTGCCATGTGCTCGCATTGGGGGGCGAGGGGTACTGGCGTCTATTCTCGCAAAGACACTAATCACCAGAGCCCACCCTCATGACCT | 120 |
| chr11 | 70743575 | 70743695 | 788855_47188552_ENST00000601538.6,E | 1 | - | good | TCCCGTGGGGGCTCTCCCGGCCCCTCTGTGTAGGGTAACTGCTGTCGTCGCTCTATAGAGAAGCTTTTCTGTAGCTGTGTGCTTATCCCAAAGCATCCAGTGTGTGTGCAGTAAATGTTG | 120 |
| chr11 | 70743695 | 70743815 | 788855_47188552_ENST00000601538.6,E | 1 | - | good | TTGGAGCCTGAATCCACAGGAATTGGTGGACAGGGAATGTAGGAGCTGGGCAGAGTTCACCAGGATCTGAGGATCCCGCCTGGAGGAGGGAGGAGAAGGTGAGGGGGACATGGAGCAACT | 120 |
| chr11 | 70743815 | 70743935 | 788855_47188552_ENST00000601538.6,E | 1 | - | good | GTCTTAGGAAGAGGACCCTGAGGGTTTGGCAGGGCCTCACAGAGCCACAAAGAGCCATGTGGGGGTGTCGCAGCAGCCGGGTAGGAAAGGAGGGAAGGAGACCTGAGAAAGAAAGACGT | 120 |
| chr11 | 70743935 | 70744055 | 788855_47188552_ENST00000601538.6,E | 1 | - | good | GGCCCGGGATGCCCAGCTGGAGTTAGATCCGGAGGGAAGGCTGAGGCCAGGTGTGGGGCTGAGGGCTTGGGCCCGGGAGCAGCTGAGTCTCTGCAGGCAGAAGGAGGTGCAGTGGACG | 120 |
| chr11 | 70744055 | 70744175 | 788855_47188552_ENST00000601538.6,E | 1 | - | good | TGAATGTGGCTCTGGGGCCCTGTGTGTTCCTGGTTCTTCCACTTTCCTCCCCTTGGGTAGGAGGCAGGGGCCTGCAGCCACACATTGGTGGCAGCTCAGGGTGGCTGCTGCGTGAAGTGT | 120 |
| chr11 | 70744175 | 70744295 | 788855_47188552_ENST00000601538.6,E | 1 | - | good | CCTCCCTGGGATCTGATCGCCCGAGCCCACAAGGGCAGGAGCTTCCTGGGGGAGGCCAGAGACCCACCCTGTGCCTTGGGCTAAAGCCTCGGCCATGGGGGTGGCGACTTGCCAGGTGCG | 120 |
| chr11 | 70744295 | 70744415 | 788855_47188552_ENST00000601538.6,E | 1 | - | good | TTGCAGAGCGCAGAGAGGGTGCTTAGAGGTGAGGAAGGAGGGCCCGGCAGGGTTCCCCACATGGGTCTCAGTGATGCTTGCTGCAGGGTGTGACCAGCCTCTACTGCAGTCATGGAAAGA | 120 |
| chr11 | 70744415 | 70744535 | 788855_47188552_ENST00000601538.6,E | 1 | - | good | GGAAGCTGCCTGCCCTCTTCCAACCTTCTGGGCTCATTCATTCCTCCCGTAGGCAGATCCTAGCAGGAACCCCCCAGGGGGCCGGGGGCATAGCCTGCCAGGCCCCACTGCAGGCTCACA | 120 |
| chr11 | 70744535 | 70744655 | 788855_47188552_ENST00000601538.6,E | 1 | - | good | CCACGGTCACCGCTGCAGGGGCACAGCCAGGTGGGCAGTGGAAGCACAGAGAGATGCCAGGACCACTGGAGCTGCTGTGGGCTGCCCTAGAGGGACTTGGAAAGGAGCAAGGAGGAGGG | 120 |
| chr11 | 70744655 | 70744775 | 788855_47188552_ENST00000601538.6,E | 1 | - | good | CCCAGCCAGGGCTCCCACAGCCCAGGGCTATTGGCCCCAAGAAACTTCGTCTGGCTGTGCGATGTCCTAGTGCTTGGAGAAACGCTGGCACTTCGGTCCCCAGCCATCACAATTCCACCA | 120 |
| chr11 | 70744775 | 70744895 | 788855_47188552_ENST00000601538.6,E | 1 | - | good | AGCCTTGGTGGGGAGCAAGACAGTGCTTATCGCTCAAGAGGTGGCCTGTGTGCTTGACCAAGCTGAGACCTTGCAGGGGTGCAGCCAAGGGGCCCCTCAGATGCCTGAGGTGGGGTGTGG | 120 |
| chr11 | 70744895 | 70745015 | 788855_47188552_ENST00000601538.6,E | 1 | - | good | GCTGGGTCAGGGCGCCGGTGTCCCCCGTGGTGTCTGTCCTCTTGTGCCTTCACTCTCACTTCAGAGCTGCAGGGTGCAGCTGCCACTGCCTGGGCTGGGGGAAGTGGCTACCTGGACTCA | 120 |
| chr11 | 70745015 | 70745135 | 788855_47188552_ENST00000601538.6,E | 1 | - | good | CTGAGTTGGAGGTCGGGTAGAAATTGGGGTGAAAATGACAATCCTTGATGAGGAGGTCCTGGCCGCTTGGGGCGGCTTTTTGCTGAGAGGTTTGCTTGGCTTCCTGGTGCTGCCGCAGAT | 120 |
| chr11 | 70745135 | 70745255 | 788855_47188552_ENST00000601538.6,E | 1 | - | good | AGAGGCTCAAAGGCAGGCAGTGAGGGAAACGCTTTGTAGCAGAGAAAGGCGGGGCTCACTAGCACTGGGGCCTTCTAAGTCATTGGTTGGGGTAGCAGATTTGGCTTTCTCTGGTTGACC | 120 |
| chr11 | 70745255 | 70745375 | 788855_47188552_ENST00000601538.6,E | 1 | - | good | AGATGTGGGCACCCAGGGCTGGCATGGAAATACGTGGAGACCACTCAACCGAGAGTAAGCAGAGGCTGGGTATCACGAGCTTGCTGTGGCAACAGCCAGGCCCTGCTACTTACATGAGGC | 120 |
| chr11 | 70745375 | 70745495 | 788855_47188552_ENST00000601538.6,E | 1 | - | good | CAAGACTGTGTTTTCAAGGCAGCAGGAGGAGGGGACGGAGGCCCGGAGGTTTGTTTTGAGGGACTGTGAGCCCCAGGGCTGAGACACGGGTGTTTCAACCCAAGGAGGGTGATAGATGAG | 120 |
| chr11 | 70745495 | 70745615 | 788855_47188552_ENST00000601538.6,E | 1 | - | good | GGGGCTCTCTGAGTTCCAGGCTTACCTCAGGCCAGGCCAGCGCAGGGCAGGACACTCCAAGGGTAGTCACCACAGTGGCAGGACCACTCATAGGGGCGGGACAGAGCCAAGGGACACAGC | 120 |
| chr11 | 70745615 | 70745735 | 788855_47188552_ENST00000601538.6,E | 1 | - | good | GTGGGACGGCTCACAGAAGTAGGAGGCAGACCCTGGGCCAGGTGGGGGCCAGGCCAGCTCCAAGCAGCTCTACCCATCCAAGCTTACTCTACACGAGACCCACGTTCCCCGGAGAGGATT | 120 |
| chr11 | 70745735 | 70745855 | 788855_47188552_ENST00000601538.6,E | 1 | - | good | GGCATGAGGCCACCTTTTGAGCCTCAGTTGCATTTGCATCCTCAGCAATGCGTTTCCCACCAAGGCTCTGCAACTTCCAAGGCGGCGGGTGACCTAGGGACAGCTCAGGTGAGGCAGGGA | 120 |
| chr11 | 70745855 | 70745975 | 788855_47188552_ENST00000601538.6,E | 1 | - | good | GCTGCCTTCTTTTCCGTAAATGCGCCCAAATGGTTTCACATCCGAGGCCTTTTTTGACAACCGTGAAGGACACGGATGAAGGAATAACGCTCGCTCGAGGTCTCAGGTGTGGCGGATCAG | 120 |
| chr11 | 70745975 | 70746095 | 788855_47188552_ENST00000601538.6,E | 1 | - | good | GTCCTCAGCCCTGTCTCGGGTCAGCACCTGTGTGATTTTCCCTTCAGTAACTCCCATGAAAGCCCCCAGGCCCTGGGTTGGCTGGAGGCTGGAGAGGAGGGTGCAGCATGCATGTGCTCA | 120 |
| chr11 | 70746095 | 70746215 | 788855_47188552_ENST00000601538.6,E | 1 | - | good | GTGGCTGGGCACGCTACCCAGCCCTGGGCTCTGCCCCCAGATCTGCTGGGTGAGTTCACAGCCTCCTGGACCGTGTCAGAGTCACCTTTGAAGCGTGGGTGTGAGATTCTGACCCAGCAG | 120 |
| chr11 | 70746575 | 70746695 | 788855_47188552_ENST00000601538.6,E | 1 | - | good | GGAGGGCAGGGACCAGACCCAGAGGCTCCCATCGGCCCCTCCTAGGGGTGACCCTGAAAGCAGCGGGGAGATGGGGGCTGTGAGCACGGGGCACCCTGCATCCCCCACCCTGGAGGGAG | 120 |
| chr11 | 70746695 | 70746815 | 788855_47188552_ENST00000601538.6,E | 1 | - | good | GTGTGGGGGCAGCGGGGCATAGAGGAGCTGAGGTGGGAGGTCGTGTGACATGCCTGGGGACACAGAAGAGTCACCCAAGGCCACCTGCAAGTGCTGATGACCAAAGGGGAGCAACAAGA | 120 |
| chr11 | 70746815 | 70746935 | 788855_47188552_ENST00000601538.6,E | 1 | - | good | GGAGTACAGCGGTGGAGGGAAGCAGACAAGCCACCCTCATCCCCAGGGGCTGTGGCAGAGAGATCTGGAGAGCCCAGACCTGAGGCTGGGATGGAAAGGGTAGGGCAGAGCCCTCCCCC | 120 |
| chr11 | 70747055 | 70747175 | 788855_47188552_ENST00000601538.6,E | 1 | - | good | CAGTCTCCTACTTCTGTGAACCCAAGCGGAGTCTTACCACCTCGGGCTTGTCCCCATGCAGCCCAGTCCCCAGTCACTCGTGGGGACCATGGATGAGCCAGGGGAAGAGACTGAGCAGGG | 120 |
| chr11 | 70747175 | 70747295 | 788855_47188552_ENST00000601538.6,E | 1 | - | good | AGCAGGGCAATCCTAAGCATGGCACCGAGCCCCACCTCCCCGCCTGCCACCTCCCTGTGCACTTGCTGCAAACTCTGCCACCGGTTTCCACTTCAGGGTGGTGTACCCCACCCAGCCCTT | 120 |
| chr11 | 70747295 | 70747415 | 788855_47188552_ENST00000601538.6,E | 1 | - | good | AGAATGTCAGGCCATGAAAAGACTTTTCCACCAGCAGAAAACAGGACTGTGTTCTCCGGGTAGCCAGCTGGTAACTGCCCGGGTCCCTCCCATCCCTGCTGCCCAGGGCTAGTTCTGGAG | 120 |
| chr11 | 70747415 | 70747535 | 788855_47188552_ENST00000601538.6,E | 1 | - | good | CAGGCTTCCAAATCCCAACACTAGTCTGTAAGCCAATCTTCAGTTACTCAGCTGTCCTTCCAGGATGTGACAAGGAAACCCGGCCCCCATGGCTCCTGGCCTGGGGTGATTAGGAGCCTG | 120 |
| chr11 | 70747535 | 70747655 | 788855_47188552_ENST00000601538.6,E | 1 | - | good | CATGGAATTGGGGCCTGGGGATATGGAGGGAGGGCCAGGTCAGCCCCACCCAAACAGTGCTATCTCCAACGCAAGCCTGGTGGCCACCAGCAGTGCCAGGTCTAAAGGCTGCCCTGGAAA | 120 |
| chr11 | 70747895 | 70748015 | 788855_47188552_ENST00000601538.6,E | 1 | - | good | TCCATATAGCAGTGCAGTCTCCTGCCTCCTTCCCTCTCCTCTTTTTTCGTCTAATATTTATTATTGTCCATGTATACACACACATTCACATATGTGCACATACACGTATGCTCATGTGTG | 120 |
| chr11 | 70748015 | 70748135 | 788855_47188552_ENST00000601538.6,E | 1 | - | good | AGTCAAGACCCAGGGTCCTTCTCTGCTGTGTGATCTCAGACAAGTCATGTGACTTCTTTAGTTTTCCTGACTAGGAGGTGTACAGAGTGGACAAGATAACCCCCGTTTCAGCTCCACTGT | 120 |
| chr11 | 70748135 | 70748255 | 788855_47188552_ENST00000601538.6,E | 1 | - | good | GGCGAGGCCTCCTCAGTTCAGGGTTTGTCTGTGGTGGGTATCCTGGGAGCAGCTCAGAATTCCTGGGAACAGGAAGAGCCTCCAGGCATTGTGGTGCAGTGGGCTGAGAAATGAACTGGG | 120 |
| chr11 | 70748255 | 70748375 | 788855_47188552_ENST00000601538.6,E | 1 | - | good | GTTCACCCTTGACTTCCTCACTCTTGCCAAAAGGACCTTTTGCCCCAGGGATTCCATCCATGGGGGACCTGCGGCACTGGTGGCTTTCAGAACAGCGTGTCTCTTGGTGATTTATGTCTG | 120 |
| chr11 | 70748375 | 70748495 | 788855_47188552_ENST00000601538.6,E | 1 | - | good | GGGAGTGTCTTTTCCTGCCCTCCTGCCAGCAGTCCCTCCAGGGAGCACCAGGCATTAGCCAGGCACCCACCATGGTTGTGGGGGTGGGCAGGGGACGGTCCCTTTTTCTGGTTAGTGGAT | 120 |
| chr11 | 70748495 | 70748615 | 788855_47188552_ENST00000601538.6,E | 1 | - | good | ATCGGGGTGGCATTAGAAAATATTTATTTTAAAGGTGGACTGTTGGATCTGAGGTCATTAGGAGCTGATGGGTTAGGTGCTGATGGGGCCCCCTTTGGGCACTGGCCCTCCCTGATTCTA | 120 |
| chr11 | 70748615 | 70748735 | 788855_47188552_ENST00000601538.6,E | 1 | - | good | GTTGTGTTGTAAACAATGTCCTTCTTCTAGTTTGTCTGCACCTTAACAGTTAGAAGGAATTATATTGGGGGTGGGTATACTGAAAGTTTCTAGTATCTGTGAATTTTATTTCCTGGCAAT | 120 |
| chr11 | 70748735 | 70748855 | 788855_47188552_ENST00000601538.6,E | 1 | - | good | TCACTGTTAAGCAGCAGGGCTGGTGGGAGGGGGTCGCCAGCTTATTCGGCACTAGTGGTCTTCACATTCACGGTCACTCCATCTGTGTAGGTATGAGCATTTCACATGGAGATGCAGACG | 120 |
| chr11 | 70748855 | 70748975 | 788855_47188552_ENST00000601538.6,E | 1 | - | good | TGTTTCTGTTGTGTAGTGTTCTGATTGCTGTCAGTGGAGGGTTGTTCTACAGTGTCCTAAGGGCAGCACTCTCCAGTGCTGGCTTTTAATCTGGAACTTTTCCAAGAGCTTTTCTCCACT | 120 |
| chr11 | 70748975 | 70749095 | 788855_47188552_ENST00000601538.6,E | 1 | - | good | CCTCAGGACAAAGCCCCCATGAAATGCAAACTAACACCTGAGCCCATCTGACTGCTCTTATTTACTCTTCAGAACCCTCAGGTCGGTCGCTGTGTGTGTGTGATGGGGGGGATTCGTTAT | 120 |
| chr11 | 70749095 | 70749215 | 788855_47188552_ENST00000601538.6,E | 1 | - | good | CACTCCCCAGTTCCCCTACAGTTGCCTGCACTTCTTCCCCTGAATCCTTCAGCCTGAAAGATGTCAGCTGTACTGGGGTTTTAGTCGCCTGTGCCACATGGTTGTGTGTCTGCAGCAGGT | 120 |
| chr11 | 70749215 | 70749335 | 788855_47188552_ENST00000601538.6,E | 1 | - | good | CACTCCTTTGTGCCTGTCCTACACATGCGTGCTTCGGGGTCAGCCTGGGGCTTTTGCAGATGCACACCCAGAACTGGGGACTCCCTTCCTCTGGCTCTTCTCCCTCCAGGGCTCTCCTCT | 120 |
| chr11 | 70749335 | 70749455 | 788855_47188552_ENST00000601538.6,E | 1 | - | good | TCCTCTGGAGGATGTTTATGATTTGTTTGAGCTGACAGCTACTATGATTAGACTCAAACTGGAATCTCCATCTTGCCTTCAGTGGTTAAAGTCTCAGATCACTCATTTAAGCTTTTGCTC | 120 |
| chr11 | 70749455 | 70749575 | 788855_47188552_ENST00000601538.6,E | 1 | - | good | CTCAAGCCTGCTATCTCTTGTCTTTTCTCTTGAGAATGGGGTCATATTTTCCCAGGAATTTTGGATTGTTTGCTTGATATTGTGAGTATTATGTTGTGTAGATTGTGGGTTGTGTTATAT | 120 |
| chr11 | 70749575 | 70749695 | 788855_47188552_ENST00000601538.6,E | 1 | - | good | TTTAACATTTTCATTTGGTTCCTTTTTATGAAAATGTTCACCACATTTACCTTTCCTTGTTGAAATAGTTACAATAGTGGCATTAAACTCATTGCCTCCTAGGTCCAGCATCTGCATCAT | 120 |
| chr11 | 70749695 | 70749815 | 788855_47188552_ENST00000601538.6,E | 1 | - | good | TCTTTCTTCTTCAGATTAGACAAATGTCATTGATTTGTCAAGTTCCCTGAGTCTTGTCTTCGTCACTTTTATCCTACTAAGGCTGTCCATTGGATTGTTTCAGATGTTGTGTTTTTCAAC | 120 |
| chr11 | 70749815 | 70749935 | 788855_47188552_ENST00000601538.6,E | 1 | - | good | ATTTCTTAAAATACTTTTTTCTGCCTCGTTCTCTGTGACTGTGATTATCCAATCCTTAGATTTTTTGTAATACTGTCCTTTGGAACCCTGACATTGTCTATTTTTTTTTCAGTCTTTTTT | 120 |
| chr11 | 70749935 | 70750055 | 788855_47188552_ENST00000601538.6,E | 1 | - | good | TTAACTAGGGAAACAGAGTGTGGTGTGTGTACTCCATCTTTCCTGGAACTAGAAGTACTTGCTGAGTTTCTTGAATCTGTGCATTCGTTTCTTTCATCAAATTTGAGAAGTTTGTTTATT | 120 |
| chr11 | 70750055 | 70750175 | 788855_47188552_ENST00000601538.6,E | 1 | - | good | TGGGAAACTCACAGCTTCTTTAGTGGTACTTGGAAATCTGCCTTTCTTCTCCAATTGCCTGCTACTCTTTATTTTTCAGAGTCCATAAATAACAGCATCTGCCCAGGTTTGCTAGCTGCG | 120 |
| chr11 | 70750175 | 70750295 | 788855_47188552_ENST00000601538.6,E | 1 | - | good | CATGCACTCTGAACCTTGGGGCTCCCTTTTCTTGAAGCTCTTTTTATCCACACTTGTTATACACATGTGGCTTTCAGGCTACCTTTGAGTCCCAGCCAGGTGACAGCCCAAGGGAAAAAA | 120 |
| chr11 | 70750295 | 70750415 | 788855_47188552_ENST00000601538.6,E | 1 | - | good | GCCAGAAAGAAGGATCCCCTCCTCCAGAGTTTTAGGTGCCTGCCCAACCTCCGCCACCACAGGATTGCTTGAGGCTGTGGGTGGGAGAAAAGGCAATAAAGAATGAAAGGGAAGATCTCC | 120 |
| chr11 | 70750415 | 70750535 | 788855_47188552_ENST00000601538.6,E | 1 | - | good | CCTGCTTTGCCATAAACTTACTGCAAATGGACCTTCATCCACGGTCAAATGGCAGAAAAAGAGAGAGAGAGGCCAAAAGCAATGGGGATTCCTCCACATGTGTGGACCACAGCTCCTCTG | 120 |
| chr11 | 70750535 | 70750655 | 788855_47188552_ENST00000601538.6,E | 1 | - | good | ACCACTTCACACGATCCCTGTTCTGCTCCCTCCTCACTACAGTCTCCCTGACATTTTCCGTGTCCATGGAGACCCCTTTATCAGGCTTCTGGATGAAAAACTGGGACCTTAATTAATTTC | 120 |
| chr11 | 70750655 | 70750775 | 788855_47188552_ENST00000601538.6,E | 1 | - | good | CGCCAAGCTTGGAGATGGTGGTGGTTTTTATCTCTTAGTTCTTTTCTGAATATCATTGATATGCTGTTCGAGGTCAGATCCACCTGTGCACAACTTGGAGCTGAGCCTAGAAGTTCACAC | 120 |

| | | | | | | | | |
|---|---|---|---|---|---|---|---|---|
| chr11 | 70750775 | 70750895 | 788855_47188552_ENST00000601538.6,E | 1 | - | good | TTGGTTGAGATCAAGCCCTAAATTCCTTTTATAGAACTTTTATGCATGGTTTCCAACGTCAAATGAGCTTCTAACCCCTTGGCTTGCGATTGAGATCTGTCCCACGTATGCACAGACACT | 120 |
| chr11 | 70751495 | 70751615 | 788855_47188552_ENST00000601538.6,E | 1 | - | good | TAGTCTATTAAAAGATATTGACCATTTTGTCTGTACTTCTTTATTCCTGACTTTCAAAGGCTCTCTGGTATCATATCCATTCCATTCCAAGCATTATCTTTGGCATTCCTCTTAAAGTTG | 120 |
| chr11 | 70751615 | 70751735 | 788855_47188552_ENST00000601538.6,E | 1 | - | good | TCTCCCTTTTTTGGTGTAGCTATCACGTGTATTAAACCCACATATATTGAAACTACTATCATAAAATATAATAATTTTTACTTCAACCATCGTATTTTCAAGAATGTAAGAAGTGAAGTA | 120 |
| chr11 | 70751735 | 70751855 | 788855_47188552_ENST00000601538.6,E | 1 | - | good | TTTATTAAAGCAATTAGTCTGTGATTTCAATACACATATGTAGCTTTTCCCAGTTTACTTAATGCTTTACCACTGCAAGTAGAACATGGAAACCTTACCACCATATAGATCGCTTTATTC | 120 |
| chr11 | 70751855 | 70751975 | 788855_47188552_ENST00000601538.6,E | 1 | - | good | TTCCTTTGCACGTTATTCTCTATTTTTCTTTCCCGATAATTTTTGGATTATTATTTGACTTGTTACTTTTCCATTTTAATTTAATTTTGGAGGCTTTAAAACAATTATATAGCTTTTTTT | 120 |
| chr11 | 70752215 | 70752335 | 788855_47188552_ENST00000601538.6,E | 1 | - | good | CTTCTTATTAGTGTGTGTATGTCTTTTTCTATCCCTTTGTTTTTAACCTCCCTATGTTGTTATATGTAAATTTGAGTCCTTATAGACTGCATGTCATTGGTTAGTTTTCTGAAAGTCCAT | 120 |
| chr11 | 70752695 | 70752815 | 788855_47188552_ENST00000601538.6,E | 1 | - | good | AAGTTCCAATGTTAGGTTGTAGGTTTATGCTGGCATTAGGTATTATTATACTGGTTAGGTTATTCACGACTGGCGTGTCTTCTTGGTGAATTGATTTTTTTTTTTTTTTTGAGACGGAGT | 120 |
| chr11 | 70753415 | 70753535 | 788855_47188552_ENST00000601538.6,E | 1 | - | good | TTTTCTCCTTTCTCCCTGTGCTTTTAGCTTATTGATTGCTGTCCTTATTTTGTTATTTGCTTCCTTCTCCATGCTTTGAGTTGATTTTGTTCTTTTTCTATTTTTTTGACTCAGGCTGTG | 120 |
| chr11 | 70754495 | 70754615 | 788855_47188552_ENST00000601538.6,E | 1 | - | good | GCATGGGCCACAAGTTCCAGGCTGGGCTCTGCTCGAGCTCACCGGGGGGAGGCTGTCACCTATGTAGAATTTCCTCAAAGGCTTTTTCTGCACCAGTTGCTATGATTATCTGTTTTTCTT | 120 |
| chr11 | 70754615 | 70754735 | 788855_47188552_ENST00000601538.6,E | 1 | - | good | AACTCTTATGGGTCCATGGTATTCAAGTAATGTCAGAGACCATGGGGGCCAGAAAGCTCTTTATTTCCACCTCCTCCTACCGCTGCCTTTGCCAGGCTACTCAGAACCCAAATGTGGAAG | 120 |
| chr11 | 70754735 | 70754855 | 788855_47188552_ENST00000601538.6,E | 1 | - | good | AGCTCTTCGGTTTTGTCAGCATTTCTTGGAAACGTGCTGTCTGGAAGGTGTTCAGCATTGCCCAGACACCTGGACTTTTCATCCTTGTCCTCCCTCTCCTGCCACTGAGGCCCCGCCACT | 120 |
| chr11 | 70754855 | 70754975 | 788855_47188552_ENST00000601538.6,E | 1 | - | good | ATCATAGCCGTTATTATTGCTGTATTAGTATCTTCTTAGGGACCCAGGTGCCTCCCTGTAATCGATGTCTCTCTGCATCTGGGGCCGTGTTAAATCTAGATATTTGAAGCAGCACTTCCC | 120 |
| chr11 | 70754975 | 70755095 | 788855_47188552_ENST00000601538.6,E | 1 | - | good | CATCTCAGAAAAAAAAAAAAAAAGATTATGAACAGGGTTTTCATACATAGGAATTGTGCGGGACATTTCGAAGTACTTCTGGGATCTTGCTTTTACTAGTATCGTTACTTATTATTTTGTA | 120 |
| chr11 | 70755455 | 70755575 | 788855_47188552_ENST00000601538.6,E | 1 | - | good | CCACCACTGCAGGCCGGGGCCGGGGCTGGGGACGTGCCTGGAAAGCCACTGGCTCTGCCCGGGTGCCCACAGGGCACCAGAGAGGCCAAGAGGTTGGACGCTCACAGGGCAGGGATGGG | 120 |
| chr11 | 70755575 | 70755695 | 788855_47188552_ENST00000601538.6,E | 1 | - | good | AGCCGCGGGGTGCGGAGCCGAGGGCGGCCTCCAGGAGTGAGTCCCTTCCGCACGAGAAAGGCCGGTTGCCAGGAGAACCTGCTGGGCCTTGTCCCCAGAATCTCCACTGTGGTCCCACTG | 120 |
| chr11 | 70755695 | 70755815 | 788855_47188552_ENST00000601538.6,E | 1 | - | good | TCCGGAAGCCGCGGCGTTCCAGGAATCTGCTGGCGCGGTGTCTTCCCTCTTGTGGCCTGTCGCTCGCCTGGTAGGAGCGGACTCACTGCCTGGCGTCCCTGCTTGTGGCCACTAGATGGC | 120 |
| chr11 | 70755815 | 70755935 | 788855_47188552_ENST00000601538.6,E | 1 | - | good | GGGTATTCAAGAGCTGGGCAACGGAAAGCCATCCTGTAAGATGGAAATAAACTTGTATTTCCAACAATCGGAGGGTGATGCAAAAGTGGCCCTACGATGGCCCCAGCACTGGGAGGCCCC | 120 |
| chr11 | 70755935 | 70756055 | 788855_47188552_ENST00000601538.6,E | 1 | - | good | CAGAGCCATCTCCGCAAGATCACAGAGGCCTTTGGCCCCAGGACGTCAGTGGGGTCCGGGGCTCGGTGAAGCGGCTAAGTCATAGGCTTGGCTTTCTGCACAGGCACAGCTGGGAGGGCT | 120 |
| chr11 | 70756055 | 70756175 | 788855_47188552_ENST00000601538.6,E | 1 | - | good | GCCTCGCTCTGGGAACCTGCCTCTGGGGGAGCTGGGGATGTCTCCTGCACCTGCTCGCACGGGGGCTGCAGCTGTCACCCTGAACCCGCTGCGGCCAGTGAGGAAACGAGCTCAGAGCAG | 120 |
| chr11 | 70756175 | 70756295 | 788855_47188552_ENST00000601538.6,E | 1 | - | good | GCATTTGAGCGTTTTCAGGTAAAGGATCTTTCCCATCAGAGCCTGGACATTCAGGAAGGGAGCTTTCCCTGGCTGGAGGGTGGAAGAGAGACCTCCAGCCTGGAGTCTGGAAGGTTCTCT | 120 |
| chr11 | 70756295 | 70756415 | 788855_47188552_ENST00000601538.6,E | 1 | - | good | CCTCCTGTCTATCCCTAGGCCCAGCACCGTGGCCAGCCCCCAGGATCAGTGAGTGCTCCTGGAACCCTTATTATCCCTGTTTCAGAAAATGTGCCTCTAAGGCTCAGAGAACCCAGGCCA | 120 |
| chr11 | 70756415 | 70756535 | 788855_47188552_ENST00000601538.6,E | 1 | - | good | AGCTGGAGGGAAGGAGCGTGCAGGCGGCGGTGGATTCAAATGCCTGTTCTGGGAGAGGGTGTGCTCTGTGCCAGGCAGCGTGGGGGCTTCACCTGCTAACCTGGGCTCTTCTCCGGTTCA | 120 |
| chr11 | 70756535 | 70756655 | 788855_47188552_ENST00000601538.6,E | 1 | - | good | TCCCCTGCTAGTGGCAGAAAGCAGGGTCCAGTATGATGCCATGGAACCTGCCTGGACCCCAGAGAGGGCAGCAGGGGTTGTGAGACTAGTTAGAGAGGTAGCCACAGCCCCTGCCCACAC | 120 |
| chr11 | 70756655 | 70756775 | 788855_47188552_ENST00000601538.6,E | 1 | - | good | CGTATGTCAGAATATGTTCAACGCTGCGTGGCCACTGAGATGATAGGGGTGTGGGGCTGCCTGGTGCCCCAGAGTGTGCAGCCGAGGCCCTGCCCTCAGCAGCTCCAAGCCTCCCCGTCT | 120 |
| chr11 | 70756775 | 70756895 | 788855_47188552_ENST00000601538.6,E | 1 | - | good | TATCCGTTCTCTTTTGTAACATTCTGTGCTGATCTACATAAGACAAGCTGCAGATCTGTGTAAAGGTGAGCTTAGAGGACACGTTCCCAGAAAGACAAAGCAGCTTACAAAGCAAAATTC | 120 |
| chr11 | 70756895 | 70757015 | 788855_47188552_ENST00000601538.6,E | 1 | - | good | CCAATATCATGGAAACTGCTAGTGCTCCATTTTGCAGACGGGGGCGCAAGGACCTGGGGAAGTGGAGTGGTTGGCCCAGGAGGTTTGGAGCCTGGGTCTGAGCCCAGGGTCTGCTCTATG | 120 |
| chr11 | 70757015 | 70757135 | 788855_47188552_ENST00000601538.6,E | 1 | - | good | CTTGTCTACTAAAGGGATGCATGGAGGAATAAAAATAAGAATAGTCACATTTATTAAGCATTTGTTACCTGCCAGCCTTTGACCTTCACACGGCTCCCCTCCATAATCCTTGCCAAACGT | 120 |
| chr11 | 70757135 | 70757255 | 788855_47188552_ENST00000601538.6,E | 1 | - | good | TCAGCCAGCTCATAAATGGCAAAGCTGGGTGGGATTCAAACCCAGGAGTCCAGCTCCTAACACTGCTATTAACCTTCCTGCTTCCTGTCTCTCCTGCTCAGGGCAGGGGCTCAGTAGATA | 120 |
| chr11 | 70757255 | 70757375 | 788855_47188552_ENST00000601538.6,E | 1 | - | good | CTCAGGGCAGCAGCCACAGGCCTGCTATGCTCACTCTTATCTGCATGGGTTGATATGGTTGTTATCCCTTTTTACAGATAAGGAAACCGAGGCTAGAGAGTGTAAGCCCCTTGCTCAAGG | 120 |
| chr11 | 70757615 | 70757735 | 788855_47188552_ENST00000601538.6,E | 1 | - | good | AGGTTTATAACAGTGCCACAGCTTAGGCGGAGTCTGCAGGCATCCATGAGCAGACTGAAGCCCCCTCCCTCTCTCCTTGAGGGGACACAGCACATTCCCCACCCACTAGGAAAAAATGCA | 120 |
| chr11 | 70757735 | 70757855 | 788855_47188552_ENST00000601538.6,E | 1 | - | good | TAGCAACCCTGGGACATTCATGCAGGCCTATGGGATGGTGTGTCGGGGGACCCCACCCACACATTCTGGGGTCCTCTGGAAGGACTCGCAGGACTCGATATAGAGTCATCCTGTTGTCTA | 120 |
| chr11 | 70760255 | 70760375 | 788855_47188552_ENST00000601538.6,E | 1 | - | good | AATTTAGAAGGGAGACTATTATAAAGGGTTACAACCTGTGAGCTGGGAATTGGGCCTCTAGCCAAAACCAGAGGCAGGCGCTTCGAGGAGGTGAGGGTGGAACGGGGCTTTATGCTGAAT | 120 |
| chr11 | 70760375 | 70760495 | 788855_47188552_ENST00000601538.6,E | 1 | - | good | GAGATGAGATCTTAGATTACGATGGCCTCCAAATCCAATGGCAGGTGACCTTAGAAGAGAGGAGGAACAAACGTGGGGAGCAGAAAGTCACATGTCAGCCCAAGAATTGCACGATTTATA | 120 |
| chr11 | 70760615 | 70760735 | 788855_47188552_ENST00000601538.6,E | 1 | - | good | AGGCAGGGACAGGTGTGCAATGCTCAGGGGAGCAGCCACCATGGGGCCTTCCCGAGCCGCTCCCTGTGCTGCACACTCCCACAGCACTGTAGGGTAGGGATTCATCGAGTCCCCTCAAGG | 120 |
| chr11 | 70760735 | 70760855 | 788855_47188552_ENST00000601538.6,E | 1 | - | good | ACTCCTCCCGGGAGCCTTCCTACACTGACTACCACCCTCCAGGTTAATCCCTTCATTTGCTCCTGGGGTACCTGCAGCCTCAGCAACTGCCTTCAGCTGGAAGCCTCCAGGCCACTGCCT | 120 |
| chr11 | 70760855 | 70760975 | 788855_47188552_ENST00000601538.6,E | 1 | - | good | CAGCTCCCTGAGCACATCCTCCAGGGGACCATCACCTCCCTGCATTGAAGCCCTGCAAACATGGCTCTGGACCCGCCTCCTGCTGGCTCTTCTCATCCTCCAGCCTCTGCCCCACAGGTC | 120 |
| chr11 | 70760975 | 70761095 | 788855_47188552_ENST00000601538.6,E | 1 | - | good | TTTATCCAGAGACCCAGCACACTGATAGAGGAGGGTCCCACGCTCTCCTCCCAGTGGGGCCTCTCCCACCCGACCTGCAGCCTGGAAAGCCGGCACCTCCACCACTGCCCCTGGGCATTG | 120 |
| chr11 | 70761455 | 70761575 | 788855_47188552_ENST00000601538.6,E | 1 | - | good | CACTGGTGCACCCGGGATGAGCCAGAGGACCCATAGAGGTGCGTCACTCACTGGACACCCTTCACCATCATCATCATCCCACGCAGACATGGTGCCTCGTTCTGTGGGTCTGGGGAAGGC | 120 |
| chr11 | 70761575 | 70761695 | 788855_47188552_ENST00000601538.6,E | 1 | - | good | GAACCTACTGAATGAGAGCCTGCCTGCTGGCAAGACCCCCAGGGACTGGAACACATGGCAAAGCCGGGGAAGCCCTGCACCTTGGAGTGTTTCAGAGTCTTCTCCAGAATGTGAGGGGAA | 120 |
| chr11 | 70761695 | 70761815 | 788855_47188552_ENST00000601538.6,E | 1 | - | good | CAAGGATGGCTGAGGTCAAGGGCCACTCGTGAAGATCTTGCCACTCAACGTGGCCCATGGAGCACAGCATGGGCATGGTCACCTGGGTGCTTGTTAGGAATGCAGGCTCCCAGGCCCCCA | 120 |
| chr11 | 70761815 | 70761935 | 788855_47188552_ENST00000601538.6,E | 1 | - | good | GCAGCAGCAGCAGCAGCCGGGAACTTGCTAGGAATGCAGAATCCCAGACTCCACCCTGGAATCTGAAGTTCTATGCTTTAAAAAGCCCTCCGGGGCAGCTCAGGTTTGGGGGTATCTGCA | 120 |
| chr11 | 70761935 | 70762055 | 788855_47188552_ENST00000601538.6,E | 1 | - | good | ACCCCACGCCCACAGATGCTGATGGAGTGGGCGGGGCCAGGGCAGCAGTCATTGAGAAAAGCCGCTGGGGCTGAGAACCACCTGGGCAGTGTTGCTCCAGCTGGATGCCCCAGACCAGCA | 120 |
| chr11 | 70762055 | 70762175 | 788855_47188552_ENST00000601538.6,E | 1 | - | good | TTGCTCTATTCTGTCTGCTTTAAAGATGAAATATTCATGAGATGAGCATTGGCAGATTGAGGAAAAGAGAGCAGGAGCGAAGTCAGGGGTTCTCTGAATCTCTGGGGCGCTGCAAAAAGA | 120 |
| chr11 | 70762175 | 70762295 | 788855_47188552_ENST00000601538.6,E | 1 | - | good | GCGATTGACATGGGCTTCATTTTCCTTCGGCTTAAAAATAAGCTAATTTCAGGCCACGTTTGTAAAACCACAGAATTTTGAGAACTACCGCGTTGGCTTAGGTTTTGCTTTGTTGGAGCT | 120 |
| chr11 | 70762295 | 70762415 | 788855_47188552_ENST00000601538.6,E | 1 | - | good | TTCCTTCTCATTTCCTCTGCATCCTGCTGGTCTGGGGGGCCCTGGATGCCACTAACCGACAGCCTGACAGTACCCACCCAGGGAGAACTTCTGTCTCCAGCCAGGACACTTGGGAGGTCT | 120 |
| chr11 | 70762415 | 70762535 | 788855_47188552_ENST00000601538.6,E | 1 | - | good | CAAGCACCCGGACACTTGGCACCACAAAAACCCTGGTGGAAGACTCTCACCATGGCCCTCAGACCAGCTGCCCACCAACTCCAGAATACAGACTCAAAAGAGGCGCCCCCACCACTGACA | 120 |
| chr11 | 70762535 | 70762655 | 788855_47188552_ENST00000601538.6,E | 1 | - | good | CACTGCTTGTGGGATCTGTGGACCTGGGGCCTGGGGTCTGTGCACCTCTGTACTGGACCATCAGCCCTGTCATCCAGTGGCTGGGCTTAGACTGGAGGGGCAGCACCCTGGGGTGGTCTC | 120 |
| chr11 | 70762655 | 70762775 | 788855_47188552_ENST00000601538.6,E | 1 | - | good | CCCCTCACCTCTTGCCTCTCCAGGTCAGTGTGGCTTACCTGAGAGGACCCAGGGGCACCTGCCCTAAGGTAGACACTAGGCCATCCAGATGGCTCATACTGAGGAAGGAGCTATGACCTC | 120 |
| chr11 | 70762775 | 70762895 | 788855_47188552_ENST00000601538.6,E | 1 | - | good | CGCCTCCATTCTGAGCTTCTCTAACAAGGCCACCACCTGTCTGTGCTTCCCTCCTCCTCCGGGCTGAAATGAGGGCACACATTTCAGGTGGCCGCTGTAGAGGGCCAGGTGGCCTTAATG | 120 |
| chr11 | 70762895 | 70763015 | 788855_47188552_ENST00000601538.6,E | 1 | - | good | GAGGGATGTGCCCAGGTGGAATGCAGTATGGCCGGGCCTCATGGTGGTGTGCAGGTGAGCGTGCTCCCATCACGTGGCCTGGCCACCAGCCATGACTTTGAAAAGTGACAGCATCTCTCC | 120 |
| chr11 | 70763015 | 70763135 | 788855_47188552_ENST00000601538.6,E | 1 | - | good | AGAGGTTAAGTGACTTGTTCCAGGTCGCACAGCCAATGAGTTCCCCTGTCACACCCTGCTCCTTCAGATCAGGCTCCTTGGCAGGGAAAGGGGGTGGTTAGGAAAGAGTTGAGTGGCTGG | 120 |
| chr11 | 70763135 | 70763255 | 788855_47188552_ENST00000601538.6,E | 1 | - | good | GCTTCATCACTGGTGTTCATTAGGATGCATCATTTGGGTTGCATTTGTTGTAACTTAGACCCACCCTCCAAGGCAGATTGGTTTTCGCTCTATCCTCCACATGAGGAAACTAAGGCTCAG | 120 |
| chr11 | 70763255 | 70763375 | 788855_47188552_ENST00000601538.6,E | 1 | - | good | ACCCCTGCCAGCAAACCCTCTGTGCCCCTGTGTCCCCTGCTGTCCCATGGGGATGGGAGTTCCCATCCCATGGCCCTGTCTCAAAGATGGTTATGTACGTGGCTGCTGCACGTGGCAGGT | 120 |
| chr11 | 70763375 | 70763495 | 788855_47188552_ENST00000601538.6,E | 1 | - | good | CATGGGAGTCATGAGAACGTCCAACTTCTGCTACACTTCCCTGCAGAGCCAGGCCCTGATGCTTCACCCACATCCACTCACCCAATCCTCAGGGCAGCACAGTTATCCCGGAGCAGCTCC | 120 |
| chr11 | 70763495 | 70763615 | 788855_47188552_ENST00000601538.6,E | 1 | - | good | GTGCTGTTGGCCCGAAAGCACATGGCCCTCAGCCCCTTCATGAGGCTGGCCCCAGTAGGTGGCCTTTCACTGACCACTGCGAGCCCAGCCTGGTCCCGAGCAGCTCAGAGCACAGAAGGG | 120 |
| chr11 | 70763615 | 70763735 | 788855_47188552_ENST00000601538.6,E | 1 | - | good | TTCTGCCTGCCTGCTCTTTATTGCAGTCAGCTGCAGACCATTATGCATCGGGCATCCATATGGCCCAACAGGCTCAGAGCCAGGCATGCTGCTGTCCAGGTCACAGGGTGCTTTCTGTGG | 120 |
| chr11 | 70763735 | 70763855 | 788855_47188552_ENST00000601538.6,E | 1 | - | good | CCCAGGACAGCTGAGCCTCTCTCTCTAGTAAGCCTGCAATGGGGCTGTGCTCCCTTGAAGCTTGGGCCTCTCCGTCTCCCTGGGTGTGGTTCTAGGGGTGACACCCTCCTGTCAAATGCA | 120 |
| chr11 | 70763855 | 70763975 | 788855_47188552_ENST00000601538.6,E | 1 | - | good | GCAGTTTAATGTGGTATGATTTCATGTCAGCATCTAACTTTCATACATTCCACCCTTTTAAAAAGCGGTTTAATAGTGTGGTTGATGCTGAGTCACATCCACTCACAGTGTGCCCAGAGG | 120 |
| chr11 | 70764935 | 70765055 | 788855_47188552_ENST00000601538.6,E | 1 | - | good | ATCAAAACAGAACTGGTGTAGTCTTCTTCCCTTGGGCTGTTGTTCAGTGGCTTGTCTTTCATTAATGGATTACTTCCCTCCAGGCCTAAGAATCTGTTGGGAGAGGGGCCATATGTAACA | 120 |
| chr11 | 70765055 | 70765175 | 788855_47188552_ENST00000601538.6,E | 1 | - | good | ACGTGGGCTATAATGTTTCCCCAGGAAACAGGAAATGGATGGAAGGCAGGCCCAACTCCCTCTTCCCTTCCACCTCAGTTCTGCACCTCCCCACTTCCTGCCCTGACACTTCTCTCCCCG | 120 |
| chr11 | 70765175 | 70765295 | 788855_47188552_ENST00000601538.6,E | 1 | - | good | TCCCTGACCCAATCATGCTGGCCAAGGGAGTTGGATGAACCCTTTGACCAGACCTGAATTGCATGCATATCTTTGGAGGCGGCCAGCCATGCCACTTGGGTCACAAAGACTGGCACACAG | 120 |
| chr11 | 70765295 | 70765415 | 788855_47188552_ENST00000601538.6,E | 1 | - | good | GCACCAGGCTAGCATTCTGCTGTTTTAGTAGTCACATAAGAGCAAGACTGCCTCTTTCCCAGTCATTGTAAAAGAAGGTTCAGGATCTAATTTCACTGGACTGATTTGAATCACTGCCCA | 120 |

| | | | | | | | | |
|---|---|---|---|---|---|---|---|---|
| chr11 | 70765415 | 70765535 | 788855_47188552_ENST00000601538.6,E | 1 | - | good | ATAACAGCTTCAGTCCTGGCTGGATCCAGGTGCTTAGGGAGCCTTAAGAAGCTCTCATTCTCTTTTCTCTGGTGTTGTCTGTATGAACTCCCCCATGCCATTCAGAGGTGGTCACCGGCA | 120 |
| chr11 | 70765535 | 70765655 | 788855_47188552_ENST00000601538.6,E | 1 | - | good | GGATCTCTTTGCTGAGGCTATACCCTGCCATGTTATTAAGGACACTTAGTTACAGATGACAGAAAACAAGCCTATCCTGGCTGAAACCAAACAGAAATTTAACGGCTTTTAAGTTTAGGA | 120 |
| chr11 | 70765655 | 70765775 | 788855_47188552_ENST00000601538.6,E | 1 | - | good | GAGCTGGGGAGGGACTTGCCCCAGATGCATTTGTGCCCACTTAATGTTACACCAGCCACCCTGGGTCTCTCGCAGGGGATTTCAGAGCCTCATGGGAGTTCTTGCTTCTTGGGAGCTCCT | 120 |
| chr11 | 70765775 | 70765895 | 788855_47188552_ENST00000601538.6,E | 1 | - | good | ACTTCCTCCCAATTTCAGAGCAGTTCAGGATATAAGGTACAAAATGTCATTCAAGAAAAGTGTTGGGAAGCTTTAAGTGAGAAAACAAAACTTATTTCATGTATGGGGAAACTGAAGCCT | 120 |
| chr11 | 70765895 | 70766015 | 788855_47188552_ENST00000601538.6,E | 1 | - | good | ATGGATGAGTAAGTTGAGAACTGATGTTAGAGTAGGCAGCTGGCCACCTCTGTCACAGTACCTCCTAGGTAATGGCTTTCTTCCCGTTTCCAAGGTGCTCACCCCGTACAGATGTCCTCT | 120 |
| chr11 | 70766015 | 70766135 | 788855_47188552_ENST00000601538.6,E | 1 | - | good | GAAAACATTCCCTCTTTTAAGGACATGCACATTGTCCACTCTGCTTCCAATTATGGATGATCAGCCATAACATCATCTCAAAGGAGGCTGAGCTGTTGGCTTTATTCCAGATAACCTGCT | 120 |
| chr11 | 70766135 | 70766255 | 788855_47188552_ENST00000601538.6,E | 1 | - | good | CTGGCCTCTGCCTTTCTTGCTGCTCCAACATCCTTAATAGGCTGCCTCGCAGTTCAAGGTGGCTGCTTAGACTCCAGCCATTGTATCTGCGTTCCAATAGAGGAAGAGCGAAAGGTTCAA | 120 |
| chr11 | 70766255 | 70766375 | 788855_47188552_ENST00000601538.6,E | 1 | - | good | GACAGAAAAACAACTAAAGAACATGGCTTACATGACACAGAGGCATATTTCTCTGTCATATTAAAGAAGACAGAGGGCAGCAGTCCAGAGCTGATATGAGGGCTTCAAACATACAGGGAC | 120 |
| chr11 | 70766375 | 70766495 | 788855_47188552_ENST00000601538.6,E | 1 | - | good | ACAAAAATCCAAATATCCATATCTTGAAAGAAGAGACCCCTTCCTCCCCTCTGACTTCTCCTTCCTCCCAGCCAGATTATCCAAGACGTGCTGTTATTCAGAGTCCCATCAGCCACATGT | 120 |
| chr11 | 70766495 | 70766615 | 788855_47188552_ENST00000601538.6,E | 1 | - | good | ACCCCTGATGCGGTTGCTTAGGAGTGGAAACTATTTTAGTGACTTAAACCAGGGCCTGGAGGTGTTTGAGGGAAATGTTCTGTAATTCGAAAGCATCAAATAAGATGCTTTCAAATTAAC | 120 |
| chr11 | 70766615 | 70766735 | 788855_47188552_ENST00000601538.6,E | 1 | - | good | GTACATTTTAACAAGCGAGGAGCAATCCCTGCCTACCTGTTAGAGCGGCAAGGAGCATTCCGGCCCCGTCTGTTCTGAGGGGCATTGATATCCACGTACACTGCTCTATTGTTTGATTAA | 120 |
| chr11 | 70766735 | 70766855 | 788855_47188552_ENST00000601538.6,E | 1 | - | good | TGCCAAGAGCTCTTCCAGCAGCCTACTTTTCCAAAATGATCGCTGATTGGAACAAAAACATTGCCACTAAAATTAGAGTGACAAAATGGAGTTGTTATGAAACAAGAGGAAAATCGACAG | 120 |
| chr11 | 70766855 | 70766975 | 788855_47188552_ENST00000601538.6,E | 1 | - | good | CATCTCCCGGACAGCTAGTTATTTTCTAAGGAGCGCTAACCAGCTCAAAACAGATTTAGAATTCCGTGTGGTTAAAAATCACCCGTCTTATGCCGTTTAAAAATTGCTTGTCATTTTGTA | 120 |
| chr11 | 70766975 | 70767095 | 788855_47188552_ENST00000601538.6,E | 1 | - | good | TCTCTCATCTGTGTCCCCACTCTGCAACTCCTTGGGGCAGGGACCCTGTCTGATGCCTCCCCGTGTCCGTGCCTGGCACACAGTAGGCCTGCCACACCAGAGCTAGCAGAGTGGACATCA | 120 |
| chr11 | 70767095 | 70767215 | 788855_47188552_ENST00000601538.6,E | 1 | - | good | GGGGCTGGGCAGGACCCTGGGCTGGAAGGCAGAGGTGAGGCTCTCCCAGCTGCTTGGCCTAACATCATCTCTACAGCCAGCCTTCAGTTCAAAAGCCCAGCGCCCATGTCCATTTTCACT | 120 |
| chr11 | 70767215 | 70767335 | 788855_47188552_ENST00000601538.6,E | 1 | - | good | ATTGGTGGGTCATATGGTAACTCTGTGTTTAACTTTTCGAAAAACCGCCAAACTTTCTTACATTCTGGGATCTTGAGCCCAGAGAATGAGGTCAGTCTGGCTGCTGCCTGGCCCTCTCCT | 120 |
| chr11 | 70767335 | 70767455 | 788855_47188552_ENST00000601538.6,E | 1 | - | good | GGGTTGCTCCCACCCTTCGGGCTGCATCTGTGAGTGGTGCGGCTGTGAACGTCCATATGTAAGGATATGCTTACCTGTTTTTCCTTCTTTTTCTTTTGGGTGTGCCTACCTGGGAATGGA | 120 |
| chr11 | 70767455 | 70767575 | 788855_47188552_ENST00000601538.6,E | 1 | - | good | TCTCAAGGCTCGTCCACATTGTGGCGTTGTGGTGGTACGTCGCTCCTTTTCAAGGCTGAGTAGTGCTTCCTTTTGTGTATGGACCATGTTCGTTATGTGCTCATCTGTTGAGGGACACTT | 120 |
| chr11 | 70767815 | 70767935 | 788855_47188552_ENST00000601538.6,E | 1 | - | good | ATCTGAGTGCAAGAAGCAGAGGGAAAGGGAGCACTGGCATATCTGAGCCCCAGCAGACCATTCTGGAAGGTTTTTGTGTTTTGATTCAGGCGAAATTGACATTAACCATTTTAAAGTACA | 120 |
| chr11 | 70767935 | 70768055 | 788855_47188552_ENST00000601538.6,E | 1 | - | good | AGAAAGTGGCCAGCTTGGATCTAAAGGGGATCTGGGTGTTGTTTCTAGCACCCTGCCCCGCCTTTCTGCACGGATCCCTGCAGAGTCAGGCAGGGGAGACCCAGAGGACCCCTGGTGGTA | 120 |
| chr11 | 70768055 | 70768175 | 788855_47188552_ENST00000601538.6,E | 1 | - | good | ATGCTGCACCTTCTCTTTGATGCCACTCCTGGCCCCAGGATGTGCAGCTGTCGCCACTTTACAGAGAAGGAAACAGGCTCCAAGAAATTCAGCAACATCCTCAGCCCCCCCACCCCAGCT | 120 |
| chr11 | 70768175 | 70768295 | 788855_47188552_ENST00000601538.6,E | 1 | - | good | TACTTCACTGTCTGGTGCAGAGGCCTGCTCAGTAAATCGCCACTGGCAGAACTACCCTGTCCATCAGACATGGCCTCTGCCCCAGTAGACAGGCAGGGAGGGAACTCACATCTACACTGC | 120 |
| chr11 | 70768295 | 70768415 | 788855_47188552_ENST00000601538.6,E | 1 | - | good | TTGGTGTGGTGACACTCTTGTGGCCAGAATTACTTGGCTGATTTGGGGAGCTCTATTCTTGCCTATCTCTCAGGTGATGCTGGGGACTTTATCCTCCCTGTCCACGACTTCCTCCATCAC | 120 |
| chr11 | 70768415 | 70768535 | 788855_47188552_ENST00000601538.6,E | 1 | - | good | CCAGCTCTCTCTGCTGCCAACAAGCTTCCCCACAGGGCCAGGCCACCTCCCCACTGTGCCGCCACTCCAAGGGCCTTCCCTGAACTGCCTCCGGGTCACATTCCTCACCTCATCCTTTTC | 120 |
| chr11 | 70768535 | 70768655 | 788855_47188552_ENST00000601538.6,E | 1 | - | good | CCCTTCCTTTGCAAGATAAATAACTCAGGGGTGCAGGCTGTGGTGTGTGCCCCACAGACTCCTGAGATGCCAAAGTGGTTAGAGATGGCCCCTCCTTCCGGCCCTCACTCCTGCAGGCAC | 120 |
| chr11 | 70768655 | 70768775 | 788855_47188552_ENST00000601538.6,E | 1 | - | good | CAGCCACCACTGCCCGCCTCCTCCCTCCATCTCCAGGCTCCCCATGGCCTGAGGCTTCCCCCAGCCTTGTTTCTCCTGATGAGGGAAACCTGCCTGGAAAACCTCCTGCAAATGGCTACT | 120 |
| chr11 | 70768775 | 70768895 | 788855_47188552_ENST00000601538.6,E | 1 | - | good | GACACTGCCACTGGTGCCCACCTGGCCCCTCTTCCCATGTGCAGGGTGAGAACCTTGGAAGCCAGGAGCCAGAGCCAAGGCCTCAGCAGCCAAGTCCTTGGCCTCACCCTCACACCCCTG | 120 |
| chr11 | 70768895 | 70769015 | 788855_47188552_ENST00000601538.6,E | 1 | - | good | AGACCCAAGATGCGGATGTGAGACCAAAGCCTGGGAAGCTTCGGCCACCTGTGCAGACACCTTTGGGGAAAGCTGGAGACACCTGCCTTCCCCACCCCCCTGGAGTGGGTCTCTGGACCT | 120 |
| chr11 | 70769015 | 70769135 | 788855_47188552_ENST00000601538.6,E | 1 | - | good | CCGCCAACCTCTTACTAGAACAAGTAGGATGCCAGGCCCTGCGTTCCAGGGAATAAAGCCTCTCAGCGAATTACACCTGGGGAAGAGTGGTGGCCTAGCGTGGACAGACCCCGACAAGGC | 120 |
| chr11 | 70769135 | 70769255 | 788855_47188552_ENST00000601538.6,E | 1 | - | good | ACTTACTGCCAGGAGCACGGGCCCTGATGATGGGCTGCACTCGCTCCGCTGAAGGAGACCGGAACTGGAACTTCTCAGGCGAGTCATTCACTCAACCACCCACCCACCCATTCATTCACC | 120 |
| chr11 | 70769255 | 70769375 | 788855_47188552_ENST00000601538.6,E | 1 | - | good | AGAAGGAGTCTATGTGCCAGGGTGACCGCATCATTTATCATCCAAACGAGGACATGTCTGCGAGCAAAGGAGGGCAGTGGGCACCGTCCCTGGGGCCACCGCGTGAAGCAGGGCTGCCCT | 120 |
| chr11 | 70769375 | 70769495 | 788855_47188552_ENST00000601538.6,E | 1 | - | good | GCTCCAGCGTTGTTCAGGGACATGCTCTGTCCTTGAGCTCTGGCACAGATAGAAAACCCCAGTGGCGACCTGACCTATTTGTTGTTCCTGAAGCTCCTGGCATTTCTCCTGGATCCTCCT | 120 |
| chr11 | 70769495 | 70769615 | 788855_47188552_ENST00000601538.6,E | 1 | - | good | CACGCACATGCCATTTCATGTGCTCTCGCAGAAGGCTCAGAGAGCCCCCCTGCCCCACCCCCTGCCGTGTGCACCAGGTATGCATCACGTCTGTTGAGCACAAGCTCAGGTCACCCTGCA | 120 |
| chr11 | 70769615 | 70769735 | 788855_47188552_ENST00000601538.6,E | 1 | - | good | ATGCCCACATAGAACACACGCACATGTGTGCTTATGCACATACTACACACATGCTGCACATGCTTCACTTGTGCTCAGCACACGTACACCCTCACATTTACACATGTGTACATGTGCATG | 120 |
| chr11 | 70769855 | 70769975 | 788855_47188552_ENST00000601538.6,E | 1 | - | good | GATGGCGGAGGAAGATGATGGATGCCCACCCCCTTCAGCCTCCAGCACCTGCAGGACCCAGGGTGTGCTGGCTGGTGCCTCTCTCACAAACACATGCAACCACAGACCTCATTGAGTAGC | 120 |
| chr11 | 70769975 | 70770095 | 788855_47188552_ENST00000601538.6,E | 1 | - | good | CGACCAGAATCAGAGGGAGCCAGAGGGCCCAGGTCACACTGTCCGTGGGCCACTGGGAAGGTACATGCTCAACACAAGGTTACGTAAGGCCTTCCTCCATGGCCCAGGGATAGGGTAGGA | 120 |
| chr11 | 70770095 | 70770215 | 788855_47188552_ENST00000601538.6,E | 1 | - | good | CTCCAAGTCACACATCAAGTGGGCGGCAGCAGAGACTAGAACTTTCTCCCCTAGACCAGAGCTTTATCTGACCCACATACTCAGCTGAGAAGGGTATTCATCTTCCCCTCAGCTGCTTTG | 120 |
| chr11 | 70770215 | 70770335 | 788855_47188552_ENST00000601538.6,E | 1 | - | good | CTCAGGGGCCTTCCCAAGTCTGCTGTGTATACAAGATTCCTACTATCAGAGACCAGGCTGGTCTGGTGGTGCTGATTCCCAGGAGGACACCAGAGGCAAAGAGCAGGAGACACAGCCGAG | 120 |
| chr11 | 70770335 | 70770455 | 788855_47188552_ENST00000601538.6,E | 1 | - | good | GCCTCAGTGTTAACACAGCTGGACAGCTGGTCTCTACAGTGCTGGGCACAGAAAGTTCCCACCCAGGCGGATGAGCAGAGGCATGAGGACCACAGGCCCATGGCCATGACCCCAGTCACC | 120 |
| chr11 | 70770455 | 70770575 | 788855_47188552_ENST00000601538.6,E | 1 | - | good | TTTAGAGAGAAGGCCCCTGGGCCTAAGTAACTTCCCAGCCATCACATGGTGACCCCGGCTCATTGGCCGCTCCAGGCACAGTGGCTTCTGTCCAGGTGGGCATTTTCAGGCCAGCTGGGG | 120 |
| chr11 | 70770575 | 70770695 | 788855_47188552_ENST00000601538.6,E | 1 | - | good | CTTTCATCATCACTGTGGAGCTTTTTATTTGTAAACGGTTACAACTTCTCAAGGTGCCCAAGCCCTTTCCCAGAGCCGTCCATTGGATAAACAGGGCCAGAATTGCATCTGAGGACCCGT | 120 |
| chr11 | 70770695 | 70770815 | 788855_47188552_ENST00000601538.6,E | 1 | - | good | GCACTCTGTGTGTCGGAGAGGGACTGATCCCAGAGCAGGAACCTGGGGGAAAGGTTACTTTTCCTCGGAGAGAGGAGGCCCATGCTCCAGTTGATTCTCAGAGAACCGAAATGGGTGTCA | 120 |
| chr11 | 70770815 | 70770935 | 788855_47188552_ENST00000601538.6,E | 1 | - | good | AAAAAAAAAAAAAAAAAAAAGGAAGTAAGAGGGAGGGAAGGAGAGAAGGAGGGATCTCTTATTGACTACCAGGGTCCTGCAGATCTGTGCACAGATTAACTGTGCCTGTTAACTTCTGGA | 120 |
| chr11 | 70771295 | 70771415 | 788855_47188552_ENST00000601538.6,E | 1 | - | good | TGCCCTGGGATGGGCCCCACCTGGGTCAGGCGCCCAGGGACCCAGCTGCCCCCACCACTTCTGTGTGCTGGTCACTTGTGCATGTGGTCACAAGGTGCCTGGCCCTCCATGTCTCTGAAA | 120 |
| chr11 | 70771415 | 70771535 | 788855_47188552_ENST00000601538.6,E | 1 | - | good | CACCCAGCCCAGCTGCTCCCTTCACCTTTTTAACCCTGAGAATGTGCTCAGAAGGTTCCAGCACTGCAGTCCTCATGCAGGAAGATGAGCTGCCACTGGGCATCTCCATCCAGATGCCAC | 120 |
| chr11 | 70771535 | 70771655 | 788855_47188552_ENST00000601538.6,E | 1 | - | good | CGCGGCTCCACGCCCCTCCTCCAGGTGGCTTGGGTGAGAGCCTCTGTCCTGTGAAGTAACTGCCAAAAACTGCCAGATCCCCTGGAGCCATCGTCCGACCCGATGGATACCCACCCACTC | 120 |
| chr11 | 70771655 | 70771775 | 788855_47188552_ENST00000601538.6,E | 1 | - | good | TAAATCCATCAGTGGCCGTGTGTGGCCACAGTGGGGCAGGCTGGCCGGCAGCCCAGCTGCTCCTGGAATGAGCTTAGAGTCACCCCTCCCGGTGGAGGCAGAAGCTCGCTGGGCCCCCAG | 120 |
| chr11 | 70771775 | 70771895 | 788855_47188552_ENST00000601538.6,E | 1 | - | good | TTTCCTGGCTTCCTCACCTTGATGGCACAGACCAGGGGCAGATTCTGGAGTCCTGGCAGCTCAACTGGGAACGGGGATTTGGCTCCAGGAAGTGTTCATTATTTATGAACCTCCAAATCC | 120 |
| chr11 | 70771895 | 70772015 | 788855_47188552_ENST00000601538.6,E | 1 | - | good | AGTGGGTTCCACTGACTTGAGCCGTCCTCATCCTTAGAGACCTTCACCTTTTAGACCTCAGAGCTTTGCTGCCTTTAAACAAAGACACCTTGTGGTGAAAACCCCCAGGCCCAGTAGCGA | 120 |
| chr11 | 70772015 | 70772135 | 788855_47188552_ENST00000601538.6,E | 1 | - | good | GTACCCACTGTTACACGGGCCCCATGCTTGTGTAATGTTCTGCTGTCAGCACCCGGAGACTCTGAATCATTTTAAACAAGGGGCTCTTTGTCATTTTGCCCGGGGTCTCACAGCTTACAC | 120 |
| chr11 | 70773095 | 70773215 | 788855_47188552_ENST00000601538.6,E | 1 | - | good | GACATCACCTTGTAGCTGTTCTTGAGCGGTGCACTGGCATCCAGGCCAGTAGTGTGGTTGCCTCACCCCCAGCTCCTCCTACAAGTTCTAGAAGGCCTCTCTCTGGTTCAGCTCTTGGCA | 120 |
| chr11 | 70773215 | 70773335 | 788855_47188552_ENST00000601538.6,E | 1 | - | good | TGTTCGCATGTTTCCCTCTTGTGAATAACGCTTCAGGAACGGCCTCATCATCTCAAAGCTTTGCTCTTGTCTTTAGGACGCTGCCCTGTGAGCCGCTCGCCCAGCTACTTTGCACAGCAA | 120 |
| chr11 | 70773335 | 70773455 | 788855_47188552_ENST00000601538.6,E | 1 | - | good | AGAGATGTCATGAATTGAAAAGATCCTTCCAAAGCACCATTGCTAATGGCTCTTCCAGTTGTCACAGATGTGTGACATTTGTCGCCTGTTCCCCCCACTTCTGACTTATGATACTGCAGT | 120 |
| chr11 | 70773455 | 70773575 | 788855_47188552_ENST00000601538.6,E | 1 | - | good | CCTGAGTCTATCCCACGTTCATAGTCACCACTGAGGCTGTAGTCCTTTGAGCATAAATGTATATTCAGCAACATTAAGCTCATTCTCTATGATTATGAATCTGCATTTTTCCAAGTGAAA | 120 |
| chr11 | 70773575 | 70773695 | 788855_47188552_ENST00000601538.6,E | 1 | - | good | TGCATCTTCTTTGGAGAGAAGGTCACTTATTAAACTAATACTTTATTATACAAGTAATATCTAGTCCTGTAGCAAATTTGGAAAATAAAGAAAGGTATAAGGAAGAAAATTAAAATCACT | 120 |
| chr11 | 70774775 | 70774895 | 788855_47188552_ENST00000601538.6,E | 1 | - | good | TGAGCCACCACACCTGGCCTAAAATCAAATTTTGACGGTTAACCCTCGCCTGTTTAGTTTTATATTATTCTGTGATCCGAAGGAATTCACACAGGACACCAACCTTTTTTTCTTTTCTTA | 120 |
| chr11 | 70775495 | 70775615 | 788855_47188552_ENST00000601538.6,E | 1 | - | good | CAGGAAGCCAAGAGGACATGGCCGCGAGTGGTGGATTGGGCACCATCTGCAAATGTGGGTCTAGAGTCAACACTCACCTTCAGCAGACGCAACCTCCAGATACCTCAAAGTGGTTAATTG | 120 |
| chr11 | 70775615 | 70775735 | 788855_47188552_ENST00000601538.6,E | 1 | - | good | AATCATGTAACCAAGGAGAGCATCTTGGATGACACTCTGTAGCCCCCTGTCCACAGCCAGGTTGTGTGGCTTAGGAGCACCGGGTCCAGGGAGGAACCCGTGTTTCTAGGAGACCTGGAA | 120 |
| chr11 | 70775735 | 70775855 | 788855_47188552_ENST00000601538.6,E | 1 | - | good | AATAAACATTGGCTTATGGGGGCCGCGTGCTGGTAACGATGAAACCTCCACATTTGCAGATTAAATTTAAGACAGGCATATGTAGAGTGGATGTGTAAAGGGCAGGCTCCTGCTCAAACA | 120 |
| chr11 | 70775855 | 70775975 | 788855_47188552_ENST00000601538.6,E | 1 | - | good | AGAGCATGCCATGGCTCATTCTGGGAAATGAAAGCTGTCATATGGTGCTCTGTTATTGAATGAAGATGTTGTTTTTGGCTCCTCTGAAATAGACGATGGGAAACAGATTCGGGGATTATG | 120 |
| chr11 | 70775975 | 70776095 | 788855_47188552_ENST00000601538.6,E | 1 | - | good | CGTTCTCTTGCTTGGCACTGGGACACCAAGTCTGGGTCTGGTCAGCCAGGGCTGGGAGCCCCCTTCTAGCTGGGGCAAAGATGGAGGCCAAAGGCCACCCCTGCCCCCGCTGCAACCTCC | 120 |
| chr11 | 70776095 | 70776215 | 788855_47188552_ENST00000601538.6,E | 1 | - | good | TTCCAGAAGTCTGATGTTGTCATGCCCTGAAAAGGTGGAAGGAATGGGCTGCTCCCAGTACATGCAGTGGAAACACTGATGTGCCATCTGTCAGATGGGAACCTAGTGAGGACAGACCTG | 120 |

| | | | | | | | | |
|---|---|---|---|---|---|---|---|---|
| chr11 | 70776215 | 70776335 | 788855_47188552_ENST00000601538.6,E | 1 | - | good | GGCCTCAGCCCTACCCCTGTGGAGCCAGAGGAATGGGGTTGGGGCTTGGACATCCACATGGGAGGGGTGGGTGACCTGCCCCCACTCACTCATGCCATCAGTGTCTTGGATTCTTGGGTG | 120 |
| chr11 | 70776335 | 70776455 | 788855_47188552_ENST00000601538.6,E | 1 | - | good | GGTTGTCTTCGAGTCTTTGGCCCCTAGAGAGTGTTAAGGTATCAGTTACCTTCAGCCAACCCAGATGCACTGTGGCCTTAGACTTTGCATGCCTGTCCTATCCCCCAAGTCCAGTTTGCA | 120 |
| chr11 | 70776455 | 70776575 | 788855_47188552_ENST00000601538.6,E | 1 | - | good | ACCTTTACATTACCCAATGAGATGTCTTTAAAACCATAGAGTTTGTGAGCAACAGGGAAGGCAGTGAGGCGTGCATGAGGCTCTTGGCGATACCAGCAACATCACTGGCATCCCAGGCCA | 120 |
| chr11 | 70776575 | 70776695 | 788855_47188552_ENST00000601538.6,E | 1 | - | good | GAGGATGGGTGACTAGGTAGATGGATGGATGGGCTTTGCTTGGGGATGTTTATAGATACTCACTCTACCTTTCATACACAGCTCACCACAATTCAAGTTGTGCATCCCTTCCTATTGTAA | 120 |
| chr11 | 70777535 | 70777655 | 788855_47188552_ENST00000601538.6,E | 1 | - | good | CCTCCATGAGAGCCAAGAATCAGGAGGATACAGTTGACTCACATCAGTATTCTCTGCACCTCACACAGTGCCTGGCATGGAGTAGGGGCTGAGAAAGTACTTCTGATGGATGGATGGATG | 120 |
| chr11 | 70777655 | 70777775 | 788855_47188552_ENST00000601538.6,E | 1 | - | good | GTGTTGGAGCAGATTTAATAAGTGCAAATGGTGCAAGCCAAGAGGACTGAAGGCAAGGACCTGGAGGTTAATTAATTTGTAAACGCCTGTTTGTGGGGTCAATTGATCAAGGTTTACTAC | 120 |
| chr11 | 70777775 | 70777895 | 788855_47188552_ENST00000601538.6,E | 1 | - | good | TGGATGCCAGGGCTCTATCTCCACCTCATGAAACCTGAATGTCTGGGGTGGGACCTGGGAATCTGTATTTTTAAGCAGCTTTCCCAGTGATGTGTTTGATGAGCTGGGATCAGGAACCTC | 120 |
| chr11 | 70777895 | 70778015 | 788855_47188552_ENST00000601538.6,E | 1 | - | good | TAGCTCTCCTGTCATTATTCTCCTTCCTCAGCTGCAGATACCCAGGAAACATGGGTTGTGTGTGTCTGGACAGCCATGTGTTAGGGTATGGCCCTCAACCTGGACTATAGAAAAGTAATG | 120 |
| chr11 | 70778015 | 70778135 | 788855_47188552_ENST00000601538.6,E | 1 | - | good | GTGACCTTGGGCAAGTTACTCAGCTTCTCTGAGCCTCTGTTGGCATTGACAGTAATGTGTTTGCCACAGGGCTGCTATGAGAGGGGAATGAATTCAAACTGGAAAGCACCAGGAGCAAGC | 120 |
| chr11 | 70778135 | 70778255 | 788855_47188552_ENST00000601538.6,E | 1 | - | good | CATCAGCATTTAGAAGCCTCTTTAGGTCAGTCTCAGGGCTGTTCTGGGGGTTTGACCTGGATTCTTTTCCATTGAAAAGGCAAACATCAGAGTCCATGACTCCTGATTTATGTCTCCTGT | 120 |
| chr11 | 70778255 | 70778375 | 788855_47188552_ENST00000601538.6,E | 1 | - | good | ATTCTGTGGATGACTTCTCAGTTTCACCCTTTCTTACCACACAATCCCATCCTCTGTGGGGCCATTTTGTTGTGTGAGCCGAAGGCAGAGACTGCTACTTGACAAGGGTGTACCCTGGCC | 120 |
| chr11 | 70778375 | 70778495 | 788855_47188552_ENST00000601538.6,E | 1 | - | good | GTCCTTTCTCCAAATCCAGTCACACTGAGGGGGTCATGGCTTCAGCATAGGAGTTTTGAGGGGGACACACACATTTAGTCTATTTCAGAGGCAAAGAGTCACAGCTGTAAACCAAGCTGC | 120 |
| chr11 | 70778735 | 70778855 | 788855_47188552_ENST00000601538.6,E | 1 | - | good | GGGTTAGAAACCTTCTTAGTGCTACACTGGGCTGCCAGTGTGGAGGGCAGTAGGCAGGGAAGAGTCTATATTAGTCAGTTTGGGCTGCTATCATGAAATACTGTAGAATGGGCAGCTTAA | 120 |
| chr11 | 70778855 | 70778975 | 788855_47188552_ENST00000601538.6,E | 1 | - | good | TTCTCCACAGCCCCTCCTGCCGCTACAGGATGTGGTGTCTGCAAAAGTCACCTTGTACAGTGGCCACTGTGGGCAGAGGGTTCAGGAGAGAAGGGCAGTGACATGGCCCGGACATTCACG | 120 |
| chr11 | 70778975 | 70779095 | 788855_47188552_ENST00000601538.6,E | 1 | - | good | ATCCAGTCTCTGGCTTTGCTGTGGGGCCCAGGCCTTCCTCGTCCCCTCCCAGCAGTGGTGGTGCCTGCTCTGGGTTACTATGGCAACCTGGTAATGCTGGGTACCAACCCTGTCCTTTTT | 120 |
| chr11 | 70779095 | 70779215 | 788855_47188552_ENST00000601538.6,E | 1 | - | good | GTATGTGCTGCTGTTTCGTAGCAGCCGCTCTGTGTTTTAAGGACCAGCTTCCATGCTTGGGTTCACCCTCAAAATAGAGTTCTGCCAGGTCTGTGTGTTATTTCAGGTTCCTTTGGGAGG | 120 |
| chr11 | 70779215 | 70779335 | 788855_47188552_ENST00000601538.6,E | 1 | - | good | TGTCCTGTCTGCTAATGATTATGTTTTTGTTAAACAACCGTGGCAGACTCCTGCATCCCCCACCCTTTCTGTCTCGTATGCCACTTTATTAAAACACACTCAGAGCCTCGCTGCCGTGAA | 120 |
| chr11 | 70779335 | 70779455 | 788855_47188552_ENST00000601538.6,E | 1 | - | good | CAGGCAAATGACTCTATTAGCACAATTGTAATCTAAAATATTAATAAGCCACAAAAGCCAGCTTCCAAAGAAGTTACTAGCTGTTCCCTCACAGAACATAAAATAGCTATGAGGACGCAT | 120 |
| chr11 | 70779455 | 70779575 | 788855_47188552_ENST00000601538.6,E | 1 | - | good | CTATTAAACAGAAGTGTCGTTAGTCACAGTGTGTATAAACCCTCGGCAAAACATCAGCATACCCATGTTTGGTTCTTCCAAAATAGAATGTCTTTGTTATCTTTAGAGCATTTAAAAATA | 120 |
| chr11 | 70779575 | 70779695 | 788855_47188552_ENST00000601538.6,E | 1 | - | good | CTGTGACCTGCTTTCTCAGGGAGTAGGAATGCCCTCCCAAGGATGTGAAAGGGGTGGGAGCTGTCACCCACATCTCTGGGCGGCTCTGGTTGGAGCCATGGAACAGGGCATGGTACAGTC | 120 |
| chr11 | 70779695 | 70779815 | 788855_47188552_ENST00000601538.6,E | 1 | - | good | GTGCTGGAACCAGCAGAAGGAGGGGGTGGCAGGCAGGACCTCTGCAGGCCATAGTTAGAGATATGCCTTCAGCAGGGACGCCAAGCGCCTCAGCTCCTTGAAGATCAGAGGAGGATATTG | 120 |
| chr11 | 70779815 | 70779935 | 788855_47188552_ENST00000601538.6,E | 1 | - | good | ATGATGTCCATGCCCACCCAGAACCAGTCACTCTGGCTAGGGGGTGGGGGCACCAGCAAGGGTGACTGGCCGGGTCTGGGTCAGGGGTGTAGGGAAGATGGCTTGCCAAGGACAGTTGGG | 120 |
| chr11 | 70779935 | 70780055 | 788855_47188552_ENST00000601538.6,E | 1 | - | good | CTCCCTCTCAGGATGGTGCCTGGCAGCTCCAGGGCTTATGTCCTTATAGGTCCAAGTCCTGCAGAAAGAAAATGGCTCTGTGCTAGCTTCTTAGCGAGAGTCCTGATGGGACCGACCTGA | 120 |
| chr11 | 70780055 | 70780175 | 788855_47188552_ENST00000601538.6,E | 1 | - | good | AAGCCTAAAATTGTCTGGCATCAGGCATAGTTGGATCCAGCTGCTCAAATGGTATCCCCAGGGAGCTCTCATTCCATCTCTTGGCCTGATTTCTCCATGGGGGCTTTATTCTTTGAGCAG | 120 |
| chr11 | 70780175 | 70780295 | 788855_47188552_ENST00000601538.6,E | 1 | - | good | CTGTACCAACTCAAGGGCTTCTCTGGTCATTATTGCTTTTTAAATTGGAGGCAACAAAACCCCAGGGCTCCCTCCAACCCCTGTCCCTACGAAAGGAATTTATTAGTGTACACAATGGAG | 120 |
| chr11 | 70780295 | 70780415 | 788855_47188552_ENST00000601538.6,E | 1 | - | good | CCTCTATTTTTCAGAGAAGGAAATCTGGGCTTCTGGTTACTGGGGATCGAATGCCTGGCTCATGGCAGAGCTGGCCTGGAGCCCTGGGGTTAAACTCTGACCCAGGGCCTCTCCCTTCCC | 120 |
| chr11 | 70780415 | 70780535 | 788855_47188552_ENST00000601538.6,E | 1 | - | good | ATAGGGTCATGTATTTAGGTTATCATCTACATGTTGGCACCAAATAAACAGACATGAGATGCCCCTCAGATGACTTGGAGTTTCTGGACTCCAATTTGGAAAAACTCATTTAGAGCCAGG | 120 |
| chr11 | 70780895 | 70781015 | 788855_47188552_ENST00000601538.6,E | 1 | - | good | ATGATAGTTACTGCTTAGTTCAAAATCTCTCTGACTTAGTAACGTGTAGGTCACTGTTAAGAGGGAAATCTCCTGCGTTAAAACAACAACAACAAAAACACAAAACAAAAAAACTATTGT | 120 |
| chr11 | 70781015 | 70781135 | 788855_47188552_ENST00000601538.6,E | 1 | - | good | ATGAGTTTCAGCCTGAGGATTTGCATGTGGGTCAGCTGACGAATGGAACCCCTCACTCCTTGGGCTCATAAACAGAGAACCGACACAAACACCATCCAGAGAAGGTGAGAACCGGAAGGG | 120 |
| chr11 | 70781135 | 70781255 | 788855_47188552_ENST00000601538.6,E | 1 | - | good | AGAACAATCGTGAACTGTGTGTTCAGTATTGTCGTTCCTATTTTGCATGTAAGGATGTGAGCTCCTTACATGCAACCGGTCCGGGGTCTCACGGCAACCGGTCCGGGGTCTCACAGCCCT | 120 |
| chr11 | 70781255 | 70781375 | 788855_47188552_ENST00000601538.6,E | 1 | - | good | GGGTACCATGGGGCTCTGAGGGTCCTGACAATAATCCTGACCCCGATTCCAACACCAGCCAGTGCCTGTTGGGCAGCTGGCTTCAAACCGGACGTTTCTCATGGTTTTCCTCCAATCTCC | 120 |
| chr11 | 70781375 | 70781495 | 788855_47188552_ENST00000601538.6,E | 1 | - | good | ACTCTGCCCTGCAGCTTTGAGGATGACCAAGCAATGGTCCCTTTCCCAGACAGGCTCCTTAACTTAAGGCTCTCAGAGATGCCTGGGTTGGGGGAGCTTCAGAGGTTTGCACCAGGCAGT | 120 |
| chr11 | 70782335 | 70782455 | 788855_47188552_ENST00000601538.6,E | 1 | - | good | TAGCCTGCTTTCCCCCTTGTTGGATGTGAAGGCTTCCTTCCATCTATGCCAGGGAGGCTGGTTTTCCACTTAAAATGGTGACTTAAAGAACTGCTTTAAGGCTGGGCGCGGTGGCTCACG | 120 |
| chr11 | 70782455 | 70782575 | 788855_47188552_ENST00000601538.6,E | 1 | - | good | TCTGGGGGCTGGGGACTCTTCCACACAGCTCTGTGGTCTGTAGTCCTCCTTATAACTTGGAGAAGGCGGGCCTGGCCCAGTGCCTCCCACAGACAGCAGGGCAGCTCGACTGGGACCCCA | 120 |
| chr11 | 70782575 | 70782695 | 788855_47188552_ENST00000601538.6,E | 1 | - | good | TTCAGGCTTGTCTGTTTCGTATTGATTTGTGGTCAGTCCTTGTATACTTCAGAAGCAAGACCTTTGTCAGTGGTGCAGGTGTGCGGATGACTGCTCCCACAGCGTGGTTTGCCTTTTTTT | 120 |
| chr11 | 70782695 | 70782815 | 788855_47188552_ENST00000601538.6,E | 1 | - | good | ATTGCCATTTGCGGTTCCCTTATTACTTCTGAGGCCGAGCACCTTTCTGTGCATGTATTGGGTGGCTGTTTGTGTGTCGTCTTTGGCCATGCGCCAGTTTGGGTCTTCTGCCTGTTTTTC | 120 |
| chr11 | 70782815 | 70782935 | 788855_47188552_ENST00000601538.6,E | 1 | - | good | TCTAGAGGGGGTCACAGACATTTAGTCTGTAAGAGTAATGGTGCCCACGTGTCACCAGCTCACTACTGACGGGCCCTTCCGCTGCGGACGTGCTGCGTGTGCGATGATGATATGGTTGTG | 120 |
| chr11 | 70782935 | 70783055 | 788855_47188552_ENST00000601538.6,E | 1 | - | good | AAGAGCACTGTGCTATCTCTTCTTAGGACGGCAGGAATCCCATCTTGGGGACCCCAGCTTCGTGACTTATCTAAACCCAATCACACCCCAAAGTCCCCATCACATTGCGGGTCCGTGAAT | 120 |
| chr11 | 70783055 | 70783175 | 788855_47188552_ENST00000601538.6,E | 1 | - | good | CAGGCTGGACACGTCCTCTGCTTTTCTGAATTCTGCAACCCTGTCCTGCCTGGACCTCATTGTCAGTCCAGGCTTCTGTAACAGAACACTGAGATCTGGAGGGCTAAAATGCTGGTTCTG | 120 |
| chr11 | 70783175 | 70783295 | 788855_47188552_ENST00000601538.6,E | 1 | - | good | TTTGTCCCTCTGTCCCGCCGTGGGCAGTCGTAGTCCGCTTGCTGTGGGAGTCCCTACCCTTGTGTCTCGATGTAAACGTGGGTGCCCTTCCATCGCCCATACCCCAGGGGTGACCGTGGT | 120 |
| chr11 | 70783295 | 70783415 | 788855_47188552_ENST00000601538.6,E | 1 | - | good | GAGACAGGATTGGCGTGTGCTGTGCTGTGCTGGCTTCTCTCCCAACAGCTGTCCCTGGTATCTGGGCCATGTGAGTAGATCTGGGTCACTTGTTTTCACTGTGAGGTCGTTCACTGGGGA | 120 |
| chr11 | 70783415 | 70783535 | 788855_47188552_ENST00000601538.6,E | 1 | - | good | CTGTCTCTGCCCAGGGCTGGTCTCGCACACAGTTGCTTCCTGATTTTATTACTAGTCCCAGGCTCTTATTACACAGCCCCAGACAGTGTCGTGGGACTCTGCCCACTCGTGAGCCATATG | 120 |
| chr11 | 70783535 | 70783655 | 788855_47188552_ENST00000601538.6,E | 1 | - | good | GAGAGGTTCACAAAACAACCGCACGTCCAGTGGGCAGTTATAAAACAAACCCTGTGGACCCAGCACCCCCAACCCCACCCAGACGCCCCCTCTGCCTCCTCAGCTCCCTCCCCAGGATAA | 120 |
| chr11 | 70783655 | 70783775 | 788855_47188552_ENST00000601538.6,E | 1 | - | good | CAGCAGTCCCCCTTCACTCCTTCCTCTTCCGGGCCTTCCGAATAGATGGGAGGATCCTGTTTGGTGCCTGTGTGATGCTTTTATTTTTAATTTTTCTGTGTGGCGCGTAACAAGCATACA | 120 |
| chr11 | 70783775 | 70783895 | 788855_47188552_ENST00000601538.6,E | 1 | - | good | TCCCCAGTGGCCCCTGCCCTGTGTCTCACACCCCCTGAGTCTGTGTCACCTCCAAGGCTCACACAGTGGGTGCCCAGGGCAGGCATCTGGGAGATGACCTTGCAGGTCAGACCCAGAATC | 120 |
| chr11 | 70783895 | 70784015 | 788855_47188552_ENST00000601538.6,E | 1 | - | good | AGACCCTGGCTGGGTGAACAGGGTGCCCGGCTTCTTCCTGCAGCACCTGAAACATCCCGGCTTTGCCCCGGAGACACCCTTGGGGCCTTCAGGCAGCCACTTGTTCCCACTTCCAGCCTG | 120 |
| chr11 | 70784015 | 70784135 | 788855_47188552_ENST00000601538.6,E | 1 | - | good | TGGACCTCAGCCACAGGCTCCCAGGGAGGCAGACAGAGCTCAGCCCATGCCACAGAGGGGGACAGTAACACCTGGGGACATCACCTCGCTGGTCAATGGCAAATGCGGATTAGAACTCAG | 120 |
| chr11 | 70784615 | 70784735 | 788855_47188552_ENST00000601538.6,E | 1 | - | good | AGCAGTGGCGGTTGGAGCGTGGGCCCTGGCCCCGGGGCTGAAGGACTCTGGGCAGAGGCCCTGACATTTACCGAGATGGGGAAGCCAGAGGAGTCCTGACTTCCCCACCCAGCATCTCCC | 120 |
| chr11 | 70784735 | 70784855 | 788855_47188552_ENST00000601538.6,E | 1 | - | good | GCTGAAAGTGGCACCTCGTGTCCCTGGCCCTGACAGCTCAGCCTGCAGGGGACCACCTGTGGCTTCCTGTGTCCATGGATGGGAGGCAGGACAGCAGCCCCCAATAGCGGGAGGGCTCTT | 120 |
| chr11 | 70784855 | 70784975 | 788855_47188552_ENST00000601538.6,E | 1 | - | good | TCCTACGGGCCTAGCAGCTGCAGGCACACAGGCCTCAGGACACCCCATGGAAACCCAGAGACCCACTGCAGATGGCCCTCCTGGGAAGCCACCACTTGTGCTCAGGACCTGGGACCAAAA | 120 |
| chr11 | 70784975 | 70785095 | 788855_47188552_ENST00000601538.6,E | 1 | - | good | TGTCCTGTGTGGTGTGCAGGTCTCCTGTCCCCTCCTGGCCATGCCCATTGGGCCCATAGCCTGGGTCCTCCCGGAGGCTCACAGGCTACCCAGAAATCAGGTCTGTGCTGGACGGAGGCA | 120 |
| chr11 | 70785095 | 70785215 | 788855_47188552_ENST00000601538.6,E | 1 | - | good | GGGCCCCTCCCCCTCGGGTCACCCCAACACTTTTAGCAGCAAACCTCATGCCTCCTCAAGTCTGGGGGTAAGATGGTGTGTGGACGAGGGTAGGAGGGGGCCCAGGCGCCCCCAGGGGTC | 120 |
| chr11 | 70785215 | 70785335 | 788855_47188552_ENST00000601538.6,E | 1 | - | good | AGGAGACTTTTGGAGAGAGGAGAGTTAGGTAAGGGGATGGGCCACCACCCTGGACAGGAGAGCTTTGGGGGGCATCTGGAGCAATGCAGGTTTAAAGCAGAGATCTGCTTTGATGAGTGA | 120 |
| chr11 | 70785335 | 70785455 | 788855_47188552_ENST00000601538.6,E | 1 | - | good | GCATTCACTGGGAACAGACTCCTCTCCCTGCAACCGGCAGGCTGCCTCTGCTCCACTGAGCGGGTGATGGAGCTTGGCGGGGTTGTGGGGAGCAGGGCCAGCTGGCAAAAAAAAGGAGAG | 120 |
| chr11 | 70785455 | 70785575 | 788855_47188552_ENST00000601538.6,E | 1 | - | good | CCTGGAAGCCTGAGACTAACGGCAGCTCCGGCCACCTGGAACACCCTCTAGAGACCTCCCTTCACCCCACTGACAGGCCCTGCCCAGCCCCTGGGGGGTGCTGGAGAGACAGAGGTAGTG | 120 |
| chr11 | 70785575 | 70785695 | 788855_47188552_ENST00000601538.6,E | 1 | - | good | AGCCTGGTGCCCGAGATGGGGGATGATGCCCAAGGCCGCTGTTACCTCTCTTGCTTTGTCACTGCACCCCCTGTCCTGGAACATGTGAGCCCCTGGAAGGTGTGGCAGGGTCCAGCACCA | 120 |
| chr11 | 70785695 | 70785815 | 788855_47188552_ENST00000601538.6,E | 1 | - | good | TAGTGGAACCCCTCCTCTTTGAGCCCCTGCTTAGAAACCAAAGAGTCCCTGAAGCGGACAGAGCATCCCCATGTCCACTCCCTCGACCTGGTGCAAATTAATCTCGGCTTTTGTTTGGGC | 120 |
| chr11 | 70785815 | 70785935 | 788855_47188552_ENST00000601538.6,E | 1 | - | good | GGAGACTTGGGCTGCAGTGTGGGAACCTCTACACTGCTAGAGGTCTGGTCCCGGGCCCATCTGTGCATAGCAGGTAGGACACCAGCCCATCAGTGGTCCCAGGCGACCACCCTCAGCTGA | 120 |
| chr11 | 70785935 | 70786055 | 788855_47188552_ENST00000601538.6,E | 1 | - | good | ATTCCACTTTATTGCCCCGTCATTTATAAGCCCTGTGCTGCCCGTGCCAGCTGTGTTTCCACTTGAAAGTTACTCCTGGTGTTTGTTGGAAATTCTGGCACGGTCCCGCTTACCCTGAGA | 120 |
| chr11 | 70786055 | 70786175 | 788855_47188552_ENST00000601538.6,E | 1 | - | good | CTTGGTCTAATAGACTGTCCTACCCCCAGCTCCATTGTAAAAGGAGGCCAGGCCAGAGACAGCCCAGGGTGGCTGCAGAGGAGAAAGGCACAGAAGCAAAGGACAGGTGCAAAGGTCATA | 120 |
| chr11 | 70786175 | 70786295 | 788855_47188552_ENST00000601538.6,E | 1 | - | good | CAGCACCACCAAGGCTCTGGCCATGCAGGGCACTGTGGCTACAGTAGCAGGTCATTGTCACTCCTGCTCTGCCCTCACCTCCACCCAGCCCCTGCCCATCTGCTTCCCTGCTTTGTGAGG | 120 |
| chr11 | 70786295 | 70786415 | 788855_47188552_ENST00000601538.6,E | 1 | - | good | CAGGGTTGTTGCAGTGGCTCACAGAGAAGAATGTAAAGGTATAAAAGTCCATACGCATACAAGAATTGTTACCAACAGCACTGATGGAATGGGCTTCCTGATAGTAATACCTGGGAACAC | 120 |

| | | | | | | | | |
|---|---|---|---|---|---|---|---|---|
| chr11 | 70786535 | 70786655 | 788855_47188552_ENST00000601538.6,E | 1 | - | good | TTGATGTAAAGAAATCAACACCACACTTGGTGCCCTTGAGAAAAATTATTTTACCCTTGTATGTCTCAGCTTTCTCATCTGTAAGTTGGATTTATATGCCCTGCAGATAGACACGCCCTG | 120 |
| chr11 | 70786655 | 70786775 | 788855_47188552_ENST00000601538.6,E | 1 | - | good | TGGCCATGAACAAATATTCTTCCTCATCCCAGCCCTGTGCAGCCTTCCCTTTGTTTGGAGGAGGTTCCTGGCTTTCCCCTTCTAGTTCTTGAAATTTCATTTCTTTCCTCACATCCCCTC | 120 |
| chr11 | 70786775 | 70786895 | 788855_47188552_ENST00000601538.6,E | 1 | - | good | TGATGGTGGTAGTGATGTGACTAGCAATGATGCTGGTGGTGATGGTGTCAGAGGTTTTAAGATGATAAAGGGATGATGCTAGTCAAGTCTGGCACTCCACCTTGGTGAACGCTACTGCTC | 120 |
| chr11 | 70787735 | 70787855 | 788855_47188552_ENST00000601538.6,E | 1 | - | good | CCGGCCCAGGACCTAAACATAGATACATCTGAGCTGACATCTGTTTCCTTTACCTGGAGGCTGGAGCATCCCCAGGGTCAGCCAGCCTCTGAGATGTTAAGATTTTGTGGCCTTGGAGTC | 120 |
| chr11 | 70787855 | 70787975 | 788855_47188552_ENST00000601538.6,E | 1 | - | good | GATGTTTATAGGTAACTCATGTGATCCTCAGAGTAAACTTGGGAGGGGGGCATGCAGTGATGTTAGTCCATTTGACACGAGGTCAGAGATGGTGAGTGACTTGCTCAGAAGCAGGGGTGG | 120 |
| chr11 | 70787975 | 70788095 | 788855_47188552_ENST00000601538.6,E | 1 | - | good | TGCACTGGAGTAAGTCATGTTCCGCCATCCTGCAGGCCATGTAACGTTATCGTTCCAGCCTTTTCAGAGCACACAGTTCCAAACCTTGTCCCTGTGGTTGCATCCTCTCACCTCTCATTT | 120 |
| chr11 | 70788095 | 70788215 | 788855_47188552_ENST00000601538.6,E | 1 | - | good | AAGTCCTCATCCACAATGGCCCTGGTGGGGGACCTGTCCTCTTGGCTGTCACACCTCCCAGTTCAACACTAGGTCAGCTTCTGGGATCCACCTGCTCTCCCTCGGGAGCCATATCATGTA | 120 |
| chr11 | 70788215 | 70788335 | 788855_47188552_ENST00000601538.6,E | 1 | - | good | AAACCTCAGAAAGTGAAACCATGGATAAAGGGAAGCTGCTGAAGTCACAAGTGACTTGGGGCCACTGTCCTTTGAGATTTCTTGTCTGTGGCCATTGATAAGAAGACGACACAATGAGGA | 120 |
| chr11 | 70788575 | 70788695 | 788855_47188552_ENST00000601538.6,E | 1 | - | good | GGGACCTCTTGGTTTTACCAACTCTTTTGTGACTGCAGCAGTCCCCTGTTATCTTCAGGGGCTACCTCCCAAGACCCCCATTGGGTGCCTGAAATGGGGGACAGCGCTGAACCCTGTACT | 120 |
| chr11 | 70788695 | 70788815 | 788855_47188552_ENST00000601538.6,E | 1 | - | good | ACCTCCTGTCTCCTATAGGACACCCTGTGTGGCCTCGTCGGGGTTTCAGGACACACTTGGAGTTTAAATCTCCTCCTCCCAAATCCAACCCTGGGGTCAGGGCTGGGCAGGAGAAGGATG | 120 |
| chr11 | 70788815 | 70788935 | 788855_47188552_ENST00000601538.6,E | 1 | - | good | TTCAGATCGTGTGGACCCGACAGGGTGGTGGTGAGGCTGGCATGTGTTTCCTCCTCCCAGAAATCGACCTGAGAGCAGTGCGTTGCTCCTCTATCATCTGAAGCCACACCTGGAGCCGTC | 120 |
| chr11 | 70788935 | 70789055 | 788855_47188552_ENST00000601538.6,E | 1 | - | good | GGGGCCCTTACTGGGAGCATCTTACAGCTTAGCATTGCCATTTGCTCTGAGAAGTGTCTCCGGACCGCCACCAGCCCCACACCCCAAGGACTAATGGGCATAACATTCTCGGATCTCTCC | 120 |
| chr11 | 70789055 | 70789175 | 788855_47188552_ENST00000601538.6,E | 1 | - | good | TCTGGGAAGCATGTGTGCCAGGCTGTCCAGATTCCCAAGCTCTCTTCCCATGGACTGTTCCCATTCAGAATCCCTGCCGTGGACTCTCCTTTCCAGCCACTCTCTAATTTACCAGCGTAG | 120 |
| chr11 | 70789175 | 70789295 | 788855_47188552_ENST00000601538.6,E | 1 | - | good | TACTTTTTTGTGGCACTCAATGGAGAAGAGACAGGACCTCATTTGAGCCTCGTAATTGCCCTTAATGGGAGGTGCCTTTATGATCCCCGCTTATTCATAAGGAAACACGTGTTGTGTATG | 120 |
| chr11 | 70789295 | 70789415 | 788855_47188552_ENST00000601538.6,E | 1 | - | good | TTTAAGAATAGGCCCCATGGCTCTCTAGAAGTTTCTAGAAGCTAGCCATAAGCATAGGATCTAGATCAGAAAGTCAGAGCTCTGACCAGGAAACAGATGTGGTTTGCAGTGTGTCTTTTC | 120 |
| chr11 | 70789415 | 70789535 | 788855_47188552_ENST00000601538.6,E | 1 | - | good | TTCCGGAATCTTCTGGCTGACTCCCTGGTGCAGGGGTCTCGACTGGGTAGAAGGTGTGAGTGTAGGATGGGATTAGCCGTGCCTTCCCTTGACTGGCCACTCTTGTGGGGTAAGTCTGCA | 120 |
| chr11 | 70789535 | 70789655 | 788855_47188552_ENST00000601538.6,E | 1 | - | good | AGTGTAGTTTTGGCATGTCCTGTGCCGTATGCACCACCAGAGCCTGTTGGAGTCGGGATCCTCAATCAGCTGCAGAGACCACCGAGGAAGTGACTTGATGGGATTCTCTGCAGAATGGGC | 120 |
| chr11 | 70789655 | 70789775 | 788855_47188552_ENST00000601538.6,E | 1 | - | good | AAGCCACTGAGGTAGGAGTGGGGGCCTCACAGCCAGGCAGCTCAGACTGGGGCATAAGATGCCACATTCTCAACTCCTACTTCACTCCTGGGCACCTACCCAGATGAGCAGCACCTACCC | 120 |
| chr11 | 70789775 | 70789895 | 788855_47188552_ENST00000601538.6,E | 1 | - | good | AAGACTTGACTATCAGTGAGCACCATTCATGTAGAACATGATGAGAAAATGGGTGTGATGAGTCACCGCTGTACATCCCTGTTTCCATCCTCCAGAGACAGGTAGGATCTGGTCACCTGA | 120 |
| chr11 | 70789895 | 70790015 | 788855_47188552_ENST00000601538.6,E | 1 | - | good | AGCGACACATCTTATGTGTACATGGAAACTGTCAGGCCAGGGCTCTCTGTTTATCCCTGAGAAAATGACAAGGCCAAGCAGCACCAGAGCAAATCGTATTTTAGTTGCTGAAACAAACAA | 120 |
| chr11 | 70790015 | 70790135 | 788855_47188552_ENST00000601538.6,E | 1 | - | good | GCCCCTGTTGATGTATCTTGACTAAGAAACAAAGATAAAATATTTCCGAAGTAATCAGTTGGCACCCATGGATGAGGCAGTTTATGAAAAATGAATGCAATGCACGTGTGTGCAGTTTTC | 120 |
| chr11 | 70790135 | 70790255 | 788855_47188552_ENST00000601538.6,E | 1 | - | good | GACTTGACTGCCTTCCGCATCTTCCAAGACATCAATTTCCTCATCTGTGAATTCAGGCTGGTGGATTGTAGAATCACTTGATCCTTCCTCCTCAGAATTCTACACATTGAAATCAGGGGT | 120 |
| chr11 | 70790255 | 70790375 | 788855_47188552_ENST00000601538.6,E | 1 | - | good | CTATAGCCTGGTCCGTTGGCCCATAAAGCTGACCCTCTTTATGCACCCTGGGTGGAGAGACAGTGCCCTGGGTGGACATGCACCCTGGGTGGAGAGACAGTGCCCACCTCCCCTACTGGG | 120 |
| chr11 | 70790615 | 70790735 | 788855_47188552_ENST00000601538.6,E | 1 | - | good | AAACAGAACCAGTAGGACGTAGACACAGATGTAGATACAGATAGAGCGTATGTATTAAGAGACTCATTTCAAGGACTTGGCTCACCCGATCATGGGAACAGGCAAGTCTAAAATCCTCTG | 120 |
| chr11 | 70790735 | 70790855 | 788855_47188552_ENST00000601538.6,E | 1 | - | good | GACACACATCATCGCCCGTGACACCCTGTGGAGGGAACCTGGATGCTGCAGATGCTGGGACCTGTGGTCCCGGGCCAGGAGCCACCTCCTGGTGACTGTTTGAGTCAAGATTCTCCAGAG | 120 |
| chr11 | 70790855 | 70790975 | 788855_47188552_ENST00000601538.6,E | 1 | - | good | CTTCCATTCTGTGGCCTGGTCATCCTCGAGGACCTTAGGGTCCCCTGCCCCCAGCCTGCAGAGACGGGGAGAGAGAATGGAGAAACGCTGCCCTCTTGTGTGAGGCTCTGGCACATCAGG | 120 |
| chr11 | 70790975 | 70791095 | 788855_47188552_ENST00000601538.6,E | 1 | - | good | ACAACTTCAGTGGCTTAATGGAGTGCAGGTTTTTTTCCTGCCCAGTAGCCACCCAGTGTGGTCTCTTCTGGTCAGCAGACGGGCTGATTTTCTCCATGATGTGACCAAGCAGTCCAGGCC | 120 |
| chr11 | 70791095 | 70791215 | 788855_47188552_ENST00000601538.6,E | 1 | - | good | TTGGGTGGTGTTGCACTTGATCAGTTAGGAGTTGGAAGGTCTGGCTTTTGTCAAGACTCTAAAGCTTTCCACCCATTTGTATTAGGTAGCTCAATGTCAGCTGCTACGACAAGTGAACCC | 120 |
| chr11 | 70791215 | 70791335 | 788855_47188552_ENST00000601538.6,E | 1 | - | good | GTCCTCACCCCATCCTGCTGGAGTTTAGGCTCTCTGAGGGCAGGGACACCTGTGCCCTGCATGGTGCCTATCCAGGACAGAGTGCATGTCAGTTGGCCCACCGCGTTTGTGAAAAGGAGG | 120 |
| chr11 | 70791335 | 70791455 | 788855_47188552_ENST00000601538.6,E | 1 | - | good | ATGCATCAGCCCAGAGACCTGAGAGCTAAGTGCCACATATGACTATGTTGTTATTATATGCAAGAGGAGATGCATCTTGGAAGATGGTGCTCTAGAAAATTGGTGCTTAATTACATGCAT | 120 |
| chr11 | 70791455 | 70791575 | 788855_47188552_ENST00000601538.6,E | 1 | - | good | ATGTGAAGCCTGGAGATGATGCGTATGAAGAGTTGGGTGGAAGTTTATGATCATAGCTTTTTTGTTTCTGTGCGTAAGAGACAGAGGCATGCCCAGATTTATAAAATCTCACTTCTTTGC | 120 |
| chr11 | 70791575 | 70791695 | 788855_47188552_ENST00000601538.6,E | 1 | - | good | ACTGCCAGTGATTAATCCTCCTGCACATCCACCTGAGGCAATGGGGAAACAGAAACATATCCGGGTCTATTCTCAGCTCAGCTGTCTCTGGCTAGCTCTGAGTAGACTTCTTTATCTGAA | 120 |
| chr11 | 70791695 | 70791815 | 788855_47188552_ENST00000601538.6,E | 1 | - | good | GGGAGTTACTGGGCAGGGACTATGACCCATTATTTGTAACTACTGATGCTTGAGCTGTGACCTGTACCTGCATCAGGACAGTGTTTCTCATGCAGTGATCAGGGTGGTGATCCTGGTGTG | 120 |
| chr11 | 70791815 | 70791935 | 788855_47188552_ENST00000601538.6,E | 1 | - | good | TCTCCAGTGGGCTTTCCTGCATGCTGTCTCATCCAACCTTCTGGAATAAGTAGGGCAGGAATCATCACCTGCATTGTACACATGAGCATCCCCATCTCAGCATGTCGGAGCACACAGATA | 120 |
| chr11 | 70791935 | 70792055 | 788855_47188552_ENST00000601538.6,E | 1 | - | good | ATACAAGTTCAGCTTAGTTCCATGAATAGTTTGGTAGTTATGCCTTCAAGATATAGTGGGCTTACTCACCTGAGCCTTAGTTGTAATTTCTTAAGGCTGCATGTTTATGGGACTTCACAG | 120 |
| chr11 | 70792535 | 70792655 | 788855_47188552_ENST00000601538.6,E | 1 | - | good | CTTTTATGGAGCACCATTGCTTTATGTTGTAAAATGAACATTTAGTTGGTAAACTCCATTCATGAACATTAAATGAAATAAGTGAGTGACTCACAGAATGAATAACTCAGTAAAGAAGTG | 120 |
| chr11 | 70792655 | 70792775 | 788855_47188552_ENST00000601538.6,E | 1 | - | good | ATTAGAAGACTTTGGATATACATAGAATCTGGAAGATGGTACAAAATGTTTTTAGATGTTCCCATCCACCAAGCTCTTTCTACACAGTAGACCTTGCTTGCACCCTTGGAAGCTGTTTCT | 120 |
| chr11 | 70793495 | 70793615 | 788855_47188552_ENST00000601538.6,E | 1 | - | good | AATTTGCATTTTTCTGGGGATGATGATTAGGCATCTCTTTGTGTGCTCATTTGCTGTATGTATAGCTTCTTTAAGTCTTTGGTTCATTTTTAATGGCAGTGTTCATCTTTTCCTTATTGG | 120 |
| chr11 | 70794335 | 70794455 | 788855_47188552_ENST00000601538.6,E | 1 | - | good | GTCACCATAGACCATAGATTTCTGTCACCATAGATTTTGCCTATCACTGGATTTCATGTAGATGAGACCCTGCAACTTGTACTCTCGGGCCCAACTTCTTTCCCTCAGCCTAATGCCTGT | 120 |
| chr11 | 70794455 | 70794575 | 788855_47188552_ENST00000601538.6,E | 1 | - | good | CAAAAAATATATAAAATAATTTTTAAAAACACCTTTCTTTCACCCCAGGAAGTGATCTCAGGCTGTCTTCCAGCCATCCCTCCCCATCCCACACCGGAAGCAAGTGTGGTTCTGATTTCT | 120 |
| chr11 | 70794935 | 70795055 | 788855_47188552_ENST00000601538.6,E | 1 | - | good | TTCCTGCTGGGTTCTGCTTGCCCTTGGGTGTTCTAAGATCGTGACTTAGCACCTTCTTTTGTATTATCTTCTTTTATAGAGGTGTGATTTACATACAGGAAAACACACAGATTATAAGAA | 120 |
| chr11 | 70795055 | 70795175 | 788855_47188552_ENST00000601538.6,E | 1 | - | good | CCTCTAACACAAAGGCTGTGGGTCAGGGAAGGAAGGCATCTCTGTGACCTGGGATGGAAGGAGGTACACCAAGATCACAGCACAGAGGAGTCGAGGAGGGCGGGGAGGATTCCACAGTGG | 120 |
| chr11 | 70795295 | 70795415 | 788855_47188552_ENST00000601538.6,E | 1 | - | good | AAAAAAAAGAAAAAGAAAGAAAAGAAAAAAGAAAAGAAAAGAACCCATGTGATTAGATGGGCCCACCTTGCTAATCCAGGTCAAGTAAACTGATGGGCCGCCTTAATTCTGCCTGCAACC | 120 |
| chr11 | 70795895 | 70796015 | 788855_47188552_ENST00000601538.6,E | 1 | - | good | GTTTCTAGAAGGGTGATCTTGCTCCAGCTCACACTCCCAGGACAATGTCCTTGATAAAACCGTATTTTCTACCTGGGCTGATGCGGGTGTGAGTCTCACTTGGCTAAAGTCAGGGCATCA | 120 |
| chr11 | 70796015 | 70796135 | 788855_47188552_ENST00000601538.6,E | 1 | - | good | GCAGGTGGGCGAGGTGGAGTCTGCCAGAAAATGTACATCCTGTGCAGGCAGCTTCCAGAGGGGCGGTGGTTTGTGTTTGTGTTTACTGTGGGAGGAGGTTATTTAAGGGGTAAGACAGAG | 120 |
| chr11 | 70796135 | 70796255 | 788855_47188552_ENST00000601538.6,E | 1 | - | good | CAACCTGTCTTCCTCCTCTCTCCTGTGGGCTTTTCCCGTCCACCTGCCAGCATCACCCTGTGTGCCCTAGATGAGAACGTATGGCTTTCTCTAGCCCCGTTGACAGGGTCCCCACCCAGG | 120 |
| chr11 | 70796255 | 70796375 | 788855_47188552_ENST00000601538.6,E | 1 | - | good | GGATGGGGCAGTCCTGGAGGGCTGCTGCGTAGGGGGATCATTTGAGCTTCCTGCTCTCTCCATCCTCCAGAGAGTAGCGGCTTCCAGGATAGTGTGCAGGAGGAAGCAGGGCAGCTGGAG | 120 |
| chr11 | 70796375 | 70796495 | 788855_47188552_ENST00000601538.6,E | 1 | - | good | CCACCACTCCTCCCTCTCCTGGCACTCACCTTCAGAGCCACACTGGCTGCCCTCCCTCTACTCTTTGTTGGAAGGACTCAACTGTGCCAGCACCGGGCTTTGTTGGCTATGCCACACCT | 120 |
| chr11 | 70796495 | 70796615 | 788855_47188552_ENST00000601538.6,E | 1 | - | good | AAGCAACCTTCCCCTGCCTCACTTGTCTGAGAAGGGCCAGGCCCTGGAGCTCAGCTTGCCTCCCAGGGTTGCTCAATGCCTTCCTTCTCCCTGTCCTCCAGTCCTCCTCTTCCTGAGCAG | 120 |
| chr11 | 70796615 | 70796735 | 788855_47188552_ENST00000601538.6,E | 1 | - | good | AACCAGGTAGACCACACGGGATTTACACTGGGCAGCAATCACACTGAGCACTGGCACAGGCCAGACCTCCAAGCCCTGCCATTGCCCGTCCACCCCTGGTCCACACCCACCCTCTCCTGC | 120 |
| chr11 | 70797215 | 70797335 | 788855_47188552_ENST00000601538.6,E | 1 | - | good | CTATGAAGGAGAAGCTGGGAATGTGCTCCACTCGGAAGGCTGGGTGGGAGGGAGTGTGTTTTTCTGGATGTGGCGATGACAATGTTACCAACAAAGCTGGTCTTCCCAAGAATGGGGTCT | 120 |
| chr11 | 70797335 | 70797455 | 788855_47188552_ENST00000601538.6,E | 1 | - | good | GATCTTGGCTTCAACGGGGACTCAGTTCGTGCATCAGCTTCCCTGCTGATGAGTCCCTGCTCAGGCTTAGACACCAGACGGTATTGTGGGGTATTGAGTTAAGACATGTCCCAAACTTAA | 120 |
| chr11 | 70797455 | 70797575 | 788855_47188552_ENST00000601538.6,E | 1 | - | good | GATCTGCCAGGGCCAGCCAGGCCCTACAAACGTGTGTTCTGGAAAGCAAGGAGGAGTCTGTCCTGGACCTGGCTGGCTTTGCAGTCTCTAGCAGGGAGGCTCAGTAGGTTTTCTGTCTGA | 120 |
| chr11 | 70797575 | 70797695 | 788855_47188552_ENST00000601538.6,E | 1 | - | good | CCATGCACGCCTTCTTGTGCGGAGCCTGCCGCCTCTGAGCTCTGTCCAAACGGGCCCATTGCCCTGTTTTTGAGGTTCCTAGTCTGTGGGCTCACTAGGTAGCCCTCTGCCGCCTGATCA | 120 |
| chr11 | 70797695 | 70797815 | 788855_47188552_ENST00000601538.6,E | 1 | - | good | GTGAAGCCTTAAGTGTCTTATAAGAAGAAAGCAGTCCGCATGTGAACGGCATCTGGCTTTTAGTAGAATCCCAGGCTCACAGAACACGTGTCAAGTGGCCACGGACGCTGCGGCCACCAA | 120 |
| chr11 | 70797935 | 70798055 | 788855_47188552_ENST00000601538.6,E | 1 | - | good | ACACGTAGCAAACGCCTCCGTAAAAACTGCTCGAAAGAGTTCAGTGTTCCGACGGGAAGCCATTCCAGCATCTCTCTTAGAACCTCCAGATATAGCACTTGCTGAAAATCGTTTCAAAAT | 120 |
| chr11 | 70798055 | 70798175 | 788855_47188552_ENST00000601538.6,E | 1 | - | good | CTAGCCATCAGAGAGAGACTTTTGGGGGTTGATTAGATTTCTTTTTCGATCTCTTCTATCGATGGCATAAAACAGGCAGCTTCCCATGTGCAGGCGAGATGCTCCTGTGCCGCCCTGAGA | 120 |
| chr11 | 70798175 | 70798295 | 788855_47188552_ENST00000601538.6,E | 1 | - | good | GGTTGGTTGCTTTCATTCCTCTGGGCTTCATCCAAACCAGAAATTCCTTACAAGATGGTGTGGAAGCTTTTAGTTAAAAGGCTGCTCAGTACACGTCCCAGAAAACCTGGACCGTATGTT | 120 |
| chr11 | 70798295 | 70798415 | 788855_47188552_ENST00000601538.6,E | 1 | - | good | CCTCGATCCTGCAGGCCTATCTCAGGCCCGTGGTCGGTGTTGTCCGTGGCAAGATTCGGCCGTTGCGTCGTAGAGGGAGGCTGAGTTGCGAATTCAGGACTTTCTAGATCTTTGGAAACA | 120 |
| chr11 | 70798415 | 70798535 | 788855_47188552_ENST00000601538.6,E | 1 | - | good | GCTTCTGGGAAGGCAGCGCCCGCGGCCACATCGGATGGTTTCCGGCGGAGTGCGTGGAGGAGGTCCAGTGTAAGCCCAGGGACAGCCAGGCAGGTAAGGCGAGCGCTCGTGGCCGCCCAC | 120 |
| chr11 | 70798535 | 70798655 | 788855_47188552_ENST00000601538.6,E | 1 | - | good | CGCTGATTCTCCTGCCTCTGCCCAGCCCCACTCGTCTTGTTCAACCTCAGGCCCTCCACGTCCCCCCTGCTAACACCTCTCCCTTTTTGTCGTCTCTAGTTCTGAGCATCGGTGAAGGGG | 120 |
| chr11 | 70798655 | 70798775 | 788855_47188552_ENST00000601538.6,E | 1 | - | good | AGGGGCACAGGGCACCTGTGGTCTGCAGTGGTGATGCCCCGGTGCCTCTTAGCACTGTCAGCAGCTGAAGGAAGTGCAGTCACACCAGAGCCGCTTGCTCTGCAGGCCTGGCCAGGGGTG | 120 |

| | | | | | | | | |
|---|---|---|---|---|---|---|---|---|
| chr11 | 70798775 | 70798895 | 788855_47188552_ENST00000601538.6,E | 1 | - | good | TGTCTCATGTGCAACACTGAGAGATGCTACAGGGCCGCCGTGCACATCCATTCATATGACGTGGTGTCAGATACGTAAAGATGCAAACATGTCAGTGCTTCATTCCTGTCTCATCACCTA | 120 |
| chr11 | 70798895 | 70799015 | 788855_47188552_ENST00000601538.6,E | 1 | - | good | CCCAGCTTAGGAGGGGAAGGTACGACCTCATCATTTCCTCCCTGCTTTGTGGCGGTGCCCCCCGACTCCACTTTTATATCTGAAATCATTGATGGTAAGGACAGCCAAGCACCACTGTCC | 120 |
| chr11 | 70799015 | 70799135 | 788855_47188552_ENST00000601538.6,E | 1 | - | good | ATGAGCCACCGCGCCCAGACCTCTGCTGTGATTTCTCCTTCCGCACGCTTGTTTAGAAGATCCAGACAGAGCCGGGCAGGTGGATTTTCTTTCCTCCAGGCCATCTGCCTCCGGTTCTTG | 120 |
| chr11 | 70799495 | 70799615 | 788855_47188552_ENST00000601538.6,E | 1 | - | good | TTGCCAGCAACCACGTAATGAATAGGTTCCTGATTAATAAGCCAGCATGCCTGTGGGCCAGGCCTCCAAGGGGCTGGTGCTGACTGTGTGAGGTTTTTAGTGCAGTTTCTTCCATTTTAA | 120 |
| chr11 | 70799615 | 70799735 | 788855_47188552_ENST00000601538.6,E | 1 | - | good | GGCCGTGTCTGTCCAGTGATGTGCGTGGTCCCCAGGACAGGGTCACTGTGTGGCCACCATCCCTTCTCTCCTGGGGAATTGGCTTCTTTCAACCACAAGTCAGAAAGAAAACTGCAGTCC | 120 |
| chr11 | 70799735 | 70799855 | 788855_47188552_ENST00000601538.6,E | 1 | - | good | AACAAACATGCTTTATTCTTCCAGCGGATGTTCATTGAGGGCCTACTGTGTACCCAGTACATGGGCTGGGGAGGGATGCCTGTGTCTGGACTCAGCAACTTCCACCCTGTAGGCTGGGGT | 120 |
| chr11 | 70799855 | 70799975 | 788855_47188552_ENST00000601538.6,E | 1 | - | good | TGACAGAAATAGTTATCACAGTAGGGCCATGGGGACCGGCGGGGTCCTCCTGACCTTGCCCCCACCCCATGGTGTCTTTGGGCTTATGTCCCTCACCCATGCTCAGCAGTTGAGTTTAAA | 120 |
| chr11 | 70800335 | 70800455 | 788855_47188552_ENST00000601538.6,E | 1 | - | good | CCTGAGTAATTTAAAAGAAGGTAAAGATGTGGAGAAATCAGATTGTTTTCCTCCTTGCTTAGACACATTAGGAGGGAGTGAAGGTGTTGAGGGCATCCCTGGCCCAGCACCAAAGTCCTA | 120 |
| chr11 | 70800455 | 70800575 | 788855_47188552_ENST00000601538.6,E | 1 | - | good | AACCTTCAAAAATCGTGGGCTGAGTGGTTCTCTGTCTGGCCTTTGAGATTCAAATGAGGGCTAGAGGAAAACACTTATGGGGCCATAATTCCTCTTTCAGCAGGGCTGTTGCCTTTGGAT | 120 |
| chr11 | 70800575 | 70800695 | 788855_47188552_ENST00000601538.6,E | 1 | - | good | GGAGGAGATGCTTCAGGCAGTGATGATTGTCCTGGTCACCTGGGATGTGGCTGATCTGTTTCAGTTAGGTCATTTCTTCAGGTTATTAGGAACTTATAAAGTGGGGTTTTCAGTTCCATT | 120 |
| chr11 | 70800695 | 70800815 | 788855_47188552_ENST00000601538.6,E | 1 | - | good | TGCAGCCCGGGGTTACTGCTCTGAGATGACCCTCTCAGGTCCAGGGTTCTGCTGGGAAACTGTGAGGCACATCCCGGCCAGCTGGGCTGAGAGGAGCCCACAGGCCCTGCATCGAGTTTT | 120 |
| chr11 | 70800815 | 70800935 | 788855_47188552_ENST00000601538.6,E | 1 | - | good | CCCTTCTCCCCAGGCCCATATAAAACAGCAGAATGACCTCACAGCCGTGAAGGAAGGAGAGTGTGGGTATCTGTGGGCACTGGCTCCTGGTGGCGCAGCAGCAAGTCCTTGGCCTCCCTT | 120 |
| chr11 | 70800935 | 70801055 | 788855_47188552_ENST00000601538.6,E | 1 | - | good | GTCATGCTCTGATAAATGAGGAAACCGAGGTACAGAGAGACCGAGTTTTCCCAAGCATCACATAGCTAAGATAGGGCAGAGCTTCCATGGAAGATGATACCAATCTCTACCTATGCAGTT | 120 |
| chr11 | 70801055 | 70801175 | 788855_47188552_ENST00000601538.6,E | 1 | - | good | GATTCTCCTGGAGAGGGGAGTTTGGAAGGAAAGTCAGGCAGGAGTGTTGAGTCCTCATTCCACACTGGGCTCACGTGCCTCCGATTCACGCATAGAAACACTTATGAGCTAGATAGTATT | 120 |
| chr11 | 70801175 | 70801295 | 788855_47188552_ENST00000601538.6,E | 1 | - | good | CTGCCTCCTTGTACAAGCCCATTGGAAACAGCCTTTCATATTTTTGGTTGGTCTTGTAATAACAATAACACATCCTGAAGTGCATACTTCATAGAGATTAAATATGGAACTAAAAGGTGT | 120 |
| chr11 | 70801295 | 70801415 | 788855_47188552_ENST00000601538.6,E | 1 | - | good | AGAAACCCAGAAGGGGCCCCCTCCTCCATTCCCCTGCACCCTCGGCACACGTCCTGCTCTTTTTACGCCCCTGACATGAAGGCTCCCCGTGTTGGGGGGACCCCTTGCAGCACGTCACCT | 120 |
| chr11 | 70801415 | 70801535 | 788855_47188552_ENST00000601538.6,E | 1 | - | good | TATGAGGAACACTCATTCCTACCTGGACGCAGATGTTCTGTCAGGGAGGCACTCAAGGCACAAACCCCTCTCTTGCTCCCCATTCTCTGAAGAGGACCAGCAGGTACCATTTGCACAACC | 120 |
| chr11 | 70801535 | 70801655 | 788855_47188552_ENST00000601538.6,E | 1 | - | good | GGGTGGATGGGGACTTCCTCTTCTCCTCCACCTCTTCTTACCCAGGGTGTGGAGTCTTTACTTTTGAATGAATTGCATCAGCAAACAATGGGGTAGCCAAGCCCAGCCCCTTCCTCACCA | 120 |
| chr11 | 70801655 | 70801775 | 788855_47188552_ENST00000601538.6,E | 1 | - | good | AGTGTTCCACATGGAGTAGGTGGGGCCACAGAGAGAACAGGAGTCCCCGGGATCCCTCAGCATCCCAGTGATGGGGCCAAGTCTCTCACCTCCCAGCCCAGTGCTCGCTCTTGTACTGCT | 120 |
| chr11 | 70801775 | 70801895 | 788855_47188552_ENST00000601538.6,E | 1 | - | good | GGCTCCAGGGACAGGGTCCTTGACAATTTTTCTAGCCAGAGCACCCAAGAGCAAGGCAGGACTCCAGACTCCAGAATTCCCCTGGTTGCCTGGTCAGAATTCACTCTCAGAATTCACCCT | 120 |
| chr11 | 70801895 | 70802015 | 788855_47188552_ENST00000601538.6,E | 1 | - | good | CCGCACCCCCTCTGTGCTTCCGTAGTCTTCCTCCTGCCCCTCGGTTTGGCTGGGACGATTCTCATCTGATCCTCTGTGGTAGAGTCAGTCCTATCACACCGTATTCCCCTGCAAGTCTGG | 120 |
| chr11 | 70802015 | 70802135 | 788855_47188552_ENST00000601538.6,E | 1 | - | good | CCCCAGCCGTGCCGTATGTGCCCCTCTCCCGGTCCTCATGCCCTGGGTGCAGTGCCTTCACCTGCTCTCTACCCACCATTCAAAGCACTGCCTCTTCCTGCAGGGTTGCCCCAAGTCTCG | 120 |
| chr11 | 70802135 | 70802255 | 788855_47188552_ENST00000601538.6,E | 1 | - | good | GCATGTGGGGCCCATAGAGAGATCCGCAGTTGGTTTTCATAGGTGCCTTAAAGCATCGTCTGTGTTGGAGCAGACATGGTGTTGAGGATGAACCTTCTTTCTAGCCAGGTTTCCCAGGGC | 120 |
| chr11 | 70802255 | 70802375 | 788855_47188552_ENST00000601538.6,E | 1 | - | good | GCTGCTTTCCAGTTTGCAAGAGAGCTGCTGAGGCAGGGAGCGGCCCACTGTGGTGGCCATAGTGAGGACAGCAGAGCTGGCCCTTCTCCAACAAGCTTATCCCCACTGTGGCCTCAGGAA | 120 |
| chr11 | 70802375 | 70802495 | 788855_47188552_ENST00000601538.6,E | 1 | - | good | CCTCCCTCCTCCGCAGTGACCTGGCAGGTTTCTGTACCCAGGCTGGGGTCAAGGGGGAGAAGGTGGAGTGTAGCAGCGGGGTCGGAGCCACCCCTGACCCATCCAGTACAGGGAATCACC | 120 |
| chr11 | 70802495 | 70802615 | 788855_47188552_ENST00000601538.6,E | 1 | - | good | TAAGCAGAAGGATATGTGGATTCAAGAACACCATGCATGGTTTGAAATTGAAGATGGCCTAGAGGTACATTAGAAACCAGCATGTTGGGTACAGAACAGGAGAAACACTGGGTCTCGGTT | 120 |
| chr11 | 70802615 | 70802735 | 788855_47188552_ENST00000601538.6,E | 1 | - | good | GGACAGTAAGGCCATCCTCCTTTTCTGAAGCTCCTCCAGGTGTCTGCCCCAAATCCTGCACCATCTTTGTGACCCCAGACACCAGCACAGTCCTTGACCCAAGTAAGTCCTGTGTCAATT | 120 |
| chr11 | 70802735 | 70802855 | 788855_47188552_ENST00000601538.6,E | 1 | - | good | GCTTACTGCAAGCCCCCATTCTGGCAAGCAAAAAACTAAGCCCATGACATAGCATTGCCAATGCTGGGGGGTGGGGGCACAGGGGGCGCCAGCGTTTTTGCTCTGTGACATTAGGTAGGT | 120 |
| chr11 | 70802855 | 70802975 | 788855_47188552_ENST00000601538.6,E | 1 | - | good | CAGCACGTAGCACAGTGAGCGTGGTCATTTTAGGGGTTCATTTATTTAACAGACACACAGTGAACCCCTGAGGGTTCGTGGCCATGGGTGAGGCCTAGGGAAGCAGTGATGATTGACCCT | 120 |
| chr11 | 70802975 | 70803095 | 788855_47188552_ENST00000601538.6,E | 1 | - | good | CTTCCCATCAGCTCCTCCTTCAGGATTCGGCTGCCATGTCCCTTCCTCTGGGAGTGTCCTCGCCCCCACCTGCCACGCTCCCTCTCTACTTTTTGGCCCAGGCCACACAGGCCTCTGATG | 120 |
| chr11 | 70803095 | 70803215 | 788855_47188552_ENST00000601538.6,E | 1 | - | good | GCATGGCAGCGTGAAAGAATGACAGAGTTTTCTAAATAGCAGTGTGTGTTTTGTCCCACGCTCTCACCACCATGCTTCCATCCTGCTTTTCCTCTGCCCAAACAGCAGCTATCCCTGTGT | 120 |
| chr11 | 70803455 | 70803575 | 788855_47188552_ENST00000601538.6,E | 1 | - | good | TGTCTCCAAGGACACAGCCTGGGAATCTGAATAGTTAACAAGCTCCTCAGGTGATTTTGCAGCATTCCTGACTGTGACGGATTGTAGACCCTCTGCAGGCAGGACTTTGTCTTCTTCTTG | 120 |
| chr11 | 70803575 | 70803695 | 788855_47188552_ENST00000601538.6,E | 1 | - | good | GCTGTGTACCTGAGTGCAGTTTCTCAGCCTCTCTTCCCGAAGTCCCTGCATCTGAGACATGAGAGGAGTTTGTTAAATGTGCAGATCTCCTGGCCATCCATCCCTACATCTGCAGAATTG | 120 |
| chr11 | 70803695 | 70803815 | 788855_47188552_ENST00000601538.6,E | 1 | - | good | TTTTTAATTCAAGGGCTCTTCTTTGTCCTTCACTGTGCCTGGCACATTCTTCCCTCCACTCCCGATAGTTCTTTATGCGAGATCCAGCATCAGGCTCCTGAGCCAGAGGCTCCTGTCCAA | 120 |
| chr11 | 70803815 | 70803935 | 788855_47188552_ENST00000601538.6,E | 1 | - | good | ACAGTTGCATACCATGTACCTGGGAAAACATACCCCAACATTTTTCTTCTAAAATAAGGGGTGCCCAAAAATAGGTGTGTACATGTTCCGATTCCCCCACTGACGCTCCTTTTTTTTTTT | 120 |
| chr11 | 70803935 | 70804055 | 788855_47188552_ENST00000601538.6,E | 1 | - | good | CGGGCCCCAGGGTGGCCCTGGGGTCCTGCTGATAACACTGCTGCTCCTGGCTCTTTCAGAAGGTTAATTTACATCTCCTCCGCTCGGTCCCATTTGGGAAGGGGTTGACGTTCCCAGTAG | 120 |
| chr11 | 70804055 | 70804175 | 788855_47188552_ENST00000601538.6,E | 1 | - | good | GGGCAGGAAACCCAGCTGCCTGGCTGGCAGGGGCAGGTCTGGGAGGGTTTCCAGAGAAACCAAACTCATTTTTTGCAGTGGAAGTCGGGTCTGGGCAGCAGGCAGGCAGGGTGGCTGTGA | 120 |
| chr11 | 70804175 | 70804295 | 788855_47188552_ENST00000601538.6,E | 1 | - | good | CATCCTTGCTGCCTGCATGGCAGTGCTTAGAGGCGGGCGGGTCGAGCCGATCCACTGCCTGCAGAGCCAAGGCCGTTTGGAGGAATTCTGTCCCCTTGATATCAAGGGAACAATCATTGT | 120 |
| chr11 | 70804295 | 70804415 | 788855_47188552_ENST00000601538.6,E | 1 | - | good | AGTGGGCCGCGCAGGCCATTTGCACACAATTGGCACGGAGCCCCTTCCTGCTGCCAGTGGCACCGCTCTTCCTTCCTGCCGAGCTGTCGGTGCCTCCTCCCCATCGTGGGGTGGCCCGCA | 120 |
| chr11 | 70804415 | 70804535 | 788855_47188552_ENST00000601538.6,E | 1 | - | good | GTGGGGCGTGCGGGGAGCTGGGCCTTTCTCCCAGGGAGAACAGAAATGACAGAGCTCGGTTCCAAAACTCACACTTCAAAGGATCGATGGCAAAGGACTGCTGTTCTGAAAAGAGATTCG | 120 |
| chr11 | 70804535 | 70804655 | 788855_47188552_ENST00000601538.6,E | 1 | - | good | TGACTCCTATCCAGGCCCCAGGACTGGGCCTCCTCAGCCTCTCTGCCCCTCCCCAGCACACTGGAACCCAGCAGCACCTCCTCAAACATGACCAGACTGGAACCCGCCCCTCGCAGGAGC | 120 |
| chr11 | 70804655 | 70804775 | 788855_47188552_ENST00000601538.6,E | 1 | - | good | GACATGGCAGGCGAATCCACATCTCACACCTTAATTCCCAGAAGCTCTGGGGGGGCCTTCCCCTGTCCAACTCCCTGGGTTTGATAGAATGATTCAACAAAGGAAAGAAGGGAGGTCACG | 120 |
| chr11 | 70804775 | 70804895 | 788855_47188552_ENST00000601538.6,E | 1 | - | good | GATGAGCAGGAGCTCAAGGGAGAGGGGGAGCCGGTGATGAGCCCCATCAGTGGATCCGAGTCTCCCACAGGCTGGCAGTGGTAGGCAGACTTGAGGACCCAGGACTCGCAGGGGTCCTGG | 120 |
| chr11 | 70804895 | 70805015 | 788855_47188552_ENST00000601538.6,E | 1 | - | good | AGGCTCCCATGCTGTGAACTTGGCACATGTCCCCAGTCCCTTAGACATAGAGCAGAGGCCCATGGGAGTGGGCACTAAACAGCTGAGAGTCCCTTAGAGGGGACCCTGAGTAGCAGCATG | 120 |
| chr11 | 70805015 | 70805135 | 788855_47188552_ENST00000601538.6,E | 1 | - | good | GTCCTGGACTTCCAAAGGGGCAGAGAGGCCTGTGAAGATGAACCTAGAGCCTGGGGCTGGGCATCTGTGGACTCACTCAGGCATACTGACCCATGCTCCGAGAGGGGCAGAGCTGCAGAC | 120 |
| chr11 | 70805135 | 70805255 | 788855_47188552_ENST00000601538.6,E | 1 | - | good | CCTGGGGCTCAGGGGTCGGAGGGTGGGAGCTGCTCCTCTGTGAGTCCTGCTTTGGAGCTCACTGATGCCCAAGGAGAAGGCTGGGGTGCGGCTGGCAGTTCCTGTAGGGCAGCCAGGTTA | 120 |
| chr11 | 70805255 | 70805375 | 788855_47188552_ENST00000601538.6,E | 1 | - | good | GCACACTCAGGGCCACCCCCTAGGCCCAAGTGGGAGCCAAGGAAGCTCACTGTCTCTCCAGAGCAGGAAGGGACACACGGTCTTCTTGTCCAGGCCTGTCCCTATTGTAAAGAGGAGAAG | 120 |
| chr11 | 70805375 | 70805495 | 788855_47188552_ENST00000601538.6,E | 1 | - | good | GTGGGGGAGCAAGAGGGGCCATCTGTGCAGGGATCAGGGCCGATGCCGGCCTTGTTAGCAACCCCGTTCTAGGAAGTGAGAGCTGAAGTTCCTAGTTAGAGAGTGTTTTCTGACCGGTCT | 120 |
| chr11 | 70805495 | 70805615 | 788855_47188552_ENST00000601538.6,E | 1 | - | good | TTCACACTTACCCCATCAACTTGGCTGTCTCTCTGAGGACTAGTGCTGAGAGCCACCTTCCCTCCTCCCCACTGAGGCCAGCCCTGAGATGTCGGGTGACAAGGATGTTAAGGCCCCGCA | 120 |
| chr11 | 70805615 | 70805735 | 788855_47188552_ENST00000601538.6,E | 1 | - | good | TGCCTTTTATTTGCCAAGATCTCCTATTACCTTGTTGAGATTTTTTTGTTCATTAAAAACTTAGTTTTTTGTTTTGTTTTTACAACACTGTAGGTTGGTGTAATTGCCACTTTAAAATCC | 120 |
| chr11 | 70805735 | 70805855 | 788855_47188552_ENST00000601538.6,E | 1 | - | good | TTGTTTAAGTTCACTTACCCTTTCCTTTGAAATGTCCCTGTAGCTGTTAAGTCCGTCTTTTGAGTTTTATAATTTCATAAATCATGTTTGCTAGCTTTTCTGTTTATCTGTTTTTATTGT | 120 |
| chr11 | 70805855 | 70805975 | 788855_47188552_ENST00000601538.6,E | 1 | - | good | TGATGGCCAGTGTTTCTGCTGCACACATATTTCATCATTTGATAGCATCAAACAAGTCCCTGGAGCTCTATTCATTTTTTTCAGTCATTTTTCTTTACATTAGATCATTTCTATTGGTTA | 120 |
| chr11 | 70805975 | 70806095 | 788855_47188552_ENST00000601538.6,E | 1 | - | good | GCCTCCTGGCCCCCAGCAGGTAATTGACTCAGTTCCTTTCTCTGCGCGACACACTGATCAGGTCAGCTTGACCTGGGTTACTGCAACATTCTAATGATGGCAGAGCCCCGCCCTCTTGCC | 120 |
| chr11 | 70806095 | 70806215 | 788855_47188552_ENST00000601538.6,E | 1 | - | good | GCCCTGCCATTGGAGGGCCTTGCCATTGGAGGGCAGGTTGCCATTGGAGTGGCATGGGTGGAAGGTGCGGCCCTTAGAGGTCAGTAGGCCCGGTTCAAATCCCGCCTTCCCCAGCTGGCT | 120 |
| chr11 | 70806215 | 70806335 | 788855_47188552_ENST00000601538.6,E | 1 | - | good | ATGGCAGGTGTGGGCTGCGGAGGTTGTGTGGGGTGGTAGGAACTCACTGGGTGGAGCAAGAAAGGCCTAGCACACGTGGCCAGCATGCCCGGGCCCCGCCATTGGAGGGCAGCATGCCGG | 120 |
| chr11 | 70806335 | 70806455 | 788855_47188552_ENST00000601538.6,E | 1 | - | good | GGAGCCAGTGATGGTTTCTGCCCAGGGGGAAGTCAGGTGTGTGAAGCACTGCAGGAAGATAAAAGTAGGGTTGGAGGGAAGTTGGCAGTGGTGTCATCTGAGGCACACAGCAGAAGCAAC | 120 |
| chr11 | 70806455 | 70806575 | 788855_47188552_ENST00000601538.6,E | 1 | - | good | CCAGGTTGGAAGGAAAGATTCTCAAAGCAGAATCCTGAGAACCAGCGTTGAGAAGCCTCTGGGGGGGCCATTGTCAACACCAGGTCAAGAAGGGGTTAGACATTGTTCCTGGGGTATTGG | 120 |
| chr11 | 70806575 | 70806695 | 788855_47188552_ENST00000601538.6,E | 1 | - | good | ATCCCCTAAGTCTGAAGTGTCGGGGTGGGCCCTGGCATGGGTCGGCTTTGGGGTCTCCAGGTAGATGGTCCGGTGTGACCTGAGGCCCGGGGCCTGCAGTGTTCAAAGGAAGGAATTCTA | 120 |
| chr11 | 70806695 | 70806815 | 788855_47188552_ENST00000601538.6,E | 1 | - | good | CGGCTTCCCCAGTGCTGGGTCTCAGGGATCGAAGGGGACACGTCCACTGGGAATGAGATTCCACTTGAGAAGGGCAGTGGGGGAGCCAGATGAGGGCATCGAAGAGCAGAGCTCCTTTCC | 120 |
| chr11 | 70806815 | 70806935 | 788855_47188552_ENST00000601538.6,E | 1 | - | good | CTGTGGGGACGATGTCCAAGGCTGTGCTTCTCCATGGAAAACACGGGCCCCTGGAAGGGCAGGAATGGGGAACAGAGTGACGTTCCCAGACCCTTTCCCTCACTCTCCAGGGAGCTGCTG | 120 |
| chr11 | 70806935 | 70807055 | 788855_47188552_ENST00000601538.6,E | 1 | - | good | ACCAACCCCAAGTGGACGGCGAGATCCCCCTTCACCGCGGTGACAGGGTCAAAGGTCAGTGTCTGCTGGTTGTCCTCCGCTCCTGGAGTGAGGTCAGGCTGGGGCCCTTCACCCGGCCTG | 120 |
| chr11 | 70807055 | 70807175 | 788855_47188552_ENST00000601538.6,E | 1 | - | good | CCAGGTCGCCTTTTGCTCTTGGTGCCAACAAGGACTCACTCTCGGCCTTCGAGTACCCGGGGCCCAAGCGGAAGCTCTACAGTGCCGTGCCCGGGAGGCTCTTCGTCGCTGTCAAGCCAT | 120 |

| | | | | | | | | |
|---|---|---|---|---|---|---|---|---|
| chr11 | 70807175 | 70807295 | 788855_47188552_ENST00000601538.6,E | 1 | - | good | AGGAGTTCTGATGCGGCCAGCCGAGGCGCAGCATGCGTTGAATGGCTTGGCTGTGGTTTGTGGCATGTGGCTTGTGACCTTGTGACTCTGCTCTCTCTCTCTCTGCCCCTCTCACCTGGT | 120 |
| chr11 | 70807295 | 70807415 | 788855_47188552_ENST00000601538.6,E | 1 | - | good | TCTCCCCGAGTGGCAGCTTTCTGTCTGTGCGGGGCAGACCTGCGAGTGGGATTGTTGGATCTGCCTGAGCCTCTGACCACAGTGCTGCCCCGGCCTGTTCCCATCTTCCTGTCTGGCATC | 120 |
| chr11 | 70807415 | 70807535 | 788855_47188552_ENST00000601538.6,E | 1 | - | good | TCCTGATGGAGGCTAAACAGCGTTCCATTGTATGTTTGGACCACCCTGTGTCTCTCCATTCTGTGGAGGGACCCTTGGGCTGCTGTGAACGGTGCCACTGTGAACGCAGGTGTGCAGACG | 120 |
| chr11 | 70808735 | 70808855 | 788855_47188552_ENST00000601538.6,E | 1 | - | good | CTTCCTCGTCCAGTGTGACTGACGCTCACAGTGAGGTGGATGAAGTGTCGGACACCCATGGCCTTCCTGGATTTCTGTCTCCACCTGAGTCTCTTTGGGATTTGTTTGTTTTTGTTTTTT | 120 |
| chr11 | 70808855 | 70808975 | 788855_47188552_ENST00000601538.6,E | 1 | - | good | AAAATCAATGACATACAGCAAGAACGGATATGAGTTCCTCTAGCGTGGTTCGCTGGGGCCGCTGGGAACTGCACTGGGAGAACCGAGGTCGTGACCCATTTTGATGGAAGGCCCTGGGTG | 120 |
| chr11 | 70808975 | 70809095 | 788855_47188552_ENST00000601538.6,E | 1 | - | good | GTCCTGGCACTGTGTAGGTGCATCTCAGAGTACTTTGTGTTGCGTGTCCTGTCTCTGGGCCCTGTTAAGGTGTGTCTTACTGTCGGGTGTAGAAGTCAGGTTGAAAACATAGCTCTGTGC | 120 |
| chr11 | 70809095 | 70809215 | 788855_47188552_ENST00000601538.6,E | 1 | - | good | TGGAGGATGCCAGCCGCAGGTGGACACTTTCCCTGTCTTTCTGAGCATGTGCCCGGGAGGGCCGTCCTGAATTTCTGGCCAACTTCCTCCGTGTAGATGCGTTTGTTTCTAAGGACGCCC | 120 |
| chr11 | 70809215 | 70809335 | 788855_47188552_ENST00000601538.6,E | 1 | - | good | CTCCCTGTCCTCATAGAAAGACTGATGCTGCCTCCATCGCCTCAGAGGCCACAGCACCTAAAGTTTGGGAGGGGAACCATGGTCAGAGACCACTCGCTTCCCTGCAGCCAGGCGCCCTGT | 120 |
| chr11 | 70809335 | 70809455 | 788855_47188552_ENST00000601538.6,E | 1 | - | good | GTGAAGGCAGGTGATCAGGGGGAGGCTTATGTTAAATGCCAAAAGAGGGTAGTCACATCCGCTTCTAGAACTTTCCCTGCAAGCCTGTTTTGCATGGGTCCCTGTGTGCATGGGCCCTTC | 120 |
| chr11 | 70809455 | 70809575 | 788855_47188552_ENST00000601538.6,E | 1 | - | good | TGAGAGAAGGAGTAAGAATCAGAACTAAATACAGCCCCATTGCCCTTCTGTAACTTTATACAAGCACAGAGCAAACTATAAGGCAATTATCAGGGACACATGAGTCGGGTGATTACAGTA | 120 |
| chr11 | 70809575 | 70809695 | 788855_47188552_ENST00000601538.6,E | 1 | - | good | AGACTGAGTATTCAGGCCCAATCTGCTCCCTACAGAGTAATCGGAGTACATGGAGCCCTGGCTCCTCAAACACGTTGACATGGGTAAGGTCCCCATGGCCAGATCTCACCATCCTAATGC | 120 |
| chr11 | 70809695 | 70809815 | 788855_47188552_ENST00000601538.6,E | 1 | - | good | TGTCCTCAGAGCCCAGGTAAGCATCCAAGACGCCCACCTGTTCCAGAAGAGGCCACTGTCCTGCCCTGTCTCTGCTCCCCATGGTTTCAGGACAAATCTGAGATTTGGAGTCAGGCAGAC | 120 |
| chr11 | 70809815 | 70809935 | 788855_47188552_ENST00000601538.6,E | 1 | - | good | CCTGTGCCCCCAGGAGTCTGTGACTGCCACATTTGTGGCTGTGACTTGGGGGAGGAATGGCATCCCTAGGGACAGGGGACAGGGAAGTCCTAGCTGCTTGCAGCCTCCCCCATCTCTCAA | 120 |
| chr11 | 70809935 | 70810055 | 788855_47188552_ENST00000601538.6,E | 1 | - | good | GGGGAACAGTGGTCCTTTGGAATTCTCCTGGCTGTGCACATCGATGCACTGCTGTGCCACACCGTGCCCGGTGGGCTCCATATTGTGCCCAGGCTGCATGGCAGTGACAGCTCAGAGACA | 120 |
| chr11 | 70810055 | 70810175 | 788855_47188552_ENST00000601538.6,E | 1 | - | good | CTCCCTGGTGGCTCACTGAGCTATGCCTTTGGACAAGAGGAGGCCACCAAACCTAGCTTTTGCAGGAGTTCCTGCTCCAGGAACTCCTGGGGTGTTTGGGGATAGATGATGTCCATCGAT | 120 |
| chr11 | 70810175 | 70810295 | 788855_47188552_ENST00000601538.6,E | 1 | - | good | GTGCAGATGGCTGGGCAGGCTGGCAGGGAAGCCTCTCAGGTGGGGGCCTGGCCCAGCGCTGCTGTTGCGTTCATCAGCTCCCCTTGCAGAGCAGGAAGTGCCCAGTGTCCTTTCCCTCTG | 120 |
| chr11 | 70810295 | 70810415 | 788855_47188552_ENST00000601538.6,E | 1 | - | good | ACAGCCGTGAGACGCTCAGTGCCTGCGGTACAGGCTCCCACCCCTCCCAGCTCTCGGTCCTGTGCCAGTCCAGCCCTGCCTGAGTCAGGCTGAGTTGAATGTGTCTGAGTGCTGGTACAC | 120 |
| chr11 | 70810415 | 70810535 | 788855_47188552_ENST00000601538.6,E | 1 | - | good | TTAAGGGAAGTTCAAGGAGAGGAGGGATTCTGACCCTGGGGTGCTGTATTCTCCTTGGAGTCCCTTGGTTGCCGGGATTCTCCTCCCCGCATCCCTGCTGGCCTGGGAGCCAGAGAGGAG | 120 |
| chr11 | 70810535 | 70810655 | 788855_47188552_ENST00000601538.6,E | 1 | - | good | AGGTCACACAGCTGGGGCAGGTCAGCCTGGGAGTGACGGGAGGTTCAGGCCGTCCACAGAGCCCATGATGGCCGCCAGCCTTGCTGGCTTCATCACTCGGGGCAGATGTGGGCAGGTGCA | 120 |
| chr11 | 70810655 | 70810775 | 788855_47188552_ENST00000601538.6,E | 1 | - | good | GCATTTTCTCATTAGTCTCCCAGCAACCACAGGTGGTGGGGATTCGCCCTACCCCCACCCTCTACCACCTATTTTACAAATAAGAAAATTAAGGCTCAGAGAGGTAAAGTGACTCACTGA | 120 |
| chr11 | 70810775 | 70810895 | 788855_47188552_ENST00000601538.6,E | 1 | - | good | AGATGATGTCGGCATAGCCCTGTGTTCTGGCTGCTGCCGGTCCAGCCTGGATAATGGTCAATGCAGAGACAGTGACTGCCCTCGGAGCTGCAGCGGGGCTGGGAACTGAGCTTCTCATCT | 120 |
| chr11 | 70810895 | 70811015 | 788855_47188552_ENST00000601538.6,E | 1 | - | good | AGGGGACACTGGGGCCACCTGTGACCTCATCACTAGGACTAACCTTCAGTCCTGATGGGGCCTAATGCATCCAGCTCCTCCAATCGCCCATGTCGCATCACCGGCCAGGAGAGGGGGTTG | 120 |
| chr11 | 70811015 | 70811135 | 788855_47188552_ENST00000601538.6,E | 1 | - | good | CCACAAGCCTCACCTTGCTCCCCCTGGAGTCAACCTATCCGGGTGCTTGGGGACTGATCATGGGGCTGGGCTGACCTGACCTGCAGCCACTCCACCTGCTGAAGAGCATCCATGCAGCCC | 120 |
| chr11 | 70811135 | 70811255 | 788855_47188552_ENST00000601538.6,E | 1 | - | good | GAACAGACCCTGTGACTCCTGCTCTGTGGCTCACCCTCTGCCCTGCCCCAGCTACTGTCAGCCTTGTTCATCTGATTTTTCCTCTAGACTTTGACCCACCCTTTCCCCTCGGGAGGAAGC | 120 |
| chr11 | 70811255 | 70811375 | 788855_47188552_ENST00000601538.6,E | 1 | - | good | GGGGCCGTGGGGTTCCTTGCAGGAGCTCTTTGGGTGTGGAAATCCTGAGAAGACGCACTCCTCATTTGGTTCCTAAGTTCTGGACTTAAGCAGCTGCTGTGTCAGACACATAACAGTTGT | 120 |
| chr11 | 70811375 | 70811495 | 788855_47188552_ENST00000601538.6,E | 1 | - | good | GAGGCCTGGATGGATAACAAAGAAGAGATGAGTGGATGGATTAATTCTGATGAAGTTAACTGACTTCTCAAGCAGGGCAGGCTCAGTGTCCCTTGCCGGCTGCTGTTGAGGATGCACAGT | 120 |
| chr11 | 70811855 | 70811975 | 788855_47188552_ENST00000601538.6,E | 1 | - | good | GAGAGGTAGCTAACTTGTCCAATGATGGATAGTAAGTGTTCAGCTGGGCTTCAGGCAAGATCTTTCTGAACTTTGGGTAGATGAGTGAATAGATGTTCAGAAAAATGATAAGATGGATGG | 120 |
| chr11 | 70811975 | 70812095 | 788855_47188552_ENST00000601538.6,E | 1 | - | good | CTGACTTTGCAGCTGCAGCAGGGCAGTGTGAAGGTGCTGATGTTTGCATAACACCTGGCAGAGTGCGTTATCACAGTTACTCCTGATCATCCGCTTTTTTGATGGGGAGGTTGAGGCTTT | 120 |
| chr11 | 70812095 | 70812215 | 788855_47188552_ENST00000601538.6,E | 1 | - | good | TGCAAAGAGAATCAAGAAACAGAGCTTCAAGGTTGGAGCTGTGTTCACTTTGACTTCTCTGTCAAGTGACACGCACTCTGGTGCCAGCTGGACCAGTGAGGTGAGACAGGCACTTTGTTA | 120 |
| chr11 | 70812215 | 70812335 | 788855_47188552_ENST00000601538.6,E | 1 | - | good | TGCCCGGTAATCACTGCCTGCAACCGTGTACACCACTCTGGTCTAGAAGAAATAAGGCTTTCCTGTTTTTAGATAAAGGTTCCCCAGGCTCTGGGGTTGATGGGTTAGGGAACTGATTCC | 120 |
| chr11 | 70812335 | 70812455 | 788855_47188552_ENST00000601538.6,E | 1 | - | good | TTTTCTCACCTGTAAAATGGGGACAAAAGTTTTCACCTTCTGCCTTTGTGGGACTTTTTGAGGTTCAAATTAACAGTGAATAAGAAGACATGTGAGCATTTGGCTGGAACCGTATGTCAC | 120 |
| chr11 | 70812455 | 70812575 | 788855_47188552_ENST00000601538.6,E | 1 | - | good | TGTAGCAGGTGAGATCTTGTTCACCTCATTCTGTCACCAGACAGAGCAGGGAACCCCGAGCTAGGATGTGAGGACCAGCTCTGTATTTCCATACCCAAGTCACCATTTATGGAGCCTCCA | 120 |
| chr11 | 70812575 | 70812695 | 788855_47188552_ENST00000601538.6,E | 1 | - | good | CCCTTCTCCCTGGGCTGCAGAGGTCTGGCTTCTGCTGCTGACTATGGCTGCTGTGGGAAGAGTGGGACCTGGCAGGAGGATGTAGTTCATCAAATGAGACATCACAAGGATAAGACACCT | 120 |
| chr11 | 70812695 | 70812815 | 788855_47188552_ENST00000601538.6,E | 1 | - | good | CATAAGGAAAACATGGACAGTTTGGGATGCAGGAACTTCAGACCAGTCCTCACTGTGCAGGCAGGACAGCAGGAGGGACCCCCAGCGTGGCCCGGGTGCCATGTCTCCTGAGCGGGGCAG | 120 |
| chr11 | 70812815 | 70812935 | 788855_47188552_ENST00000601538.6,E | 1 | - | good | CACTGATCCAAGTCACCCCACCGAGAAGATTACCAATTTTTTCATTTTAAGAATTCAGTGATTTGGCCCTTATACCAAAACAAGTTGGTTAGAATCCCTCCTGGCTGTTCTTGGCCAGAA | 120 |
| chr11 | 70812935 | 70813055 | 788855_47188552_ENST00000601538.6,E | 1 | - | good | GCAGAGTACTTGCTGGATGAATGAATGAATGTGTAACTGACTGGTGAATGAATGAATGGAGAAAGGTCAGTATGTCGGCCAGTTCCTGTTCACCCAGGCTGGCTGGGAAATGGCCCAACT | 120 |
| chr11 | 70813055 | 70813175 | 788855_47188552_ENST00000601538.6,E | 1 | - | good | AAATGCATCTGCTGACCTGTGCTTCTGCCCAGTTGACCGTCAGCTCCCTGAGGCCTGCATCCGTGTCTACCATTCACGGGGGTCTGCAGACCCACATGGCGATGGGCTCACAGCCCACCC | 120 |
| chr11 | 70813175 | 70813295 | 788855_47188552_ENST00000601538.6,E | 1 | - | good | CCTTCTCTCTGCACTACCCAGCTGTCCCCTCTTGTAGCCTTTTGTGGGTAGGCCCTGTGGACTCTAACCCTGACCTCAAGTCCTGGCCCTGTTCTCTGTGCTGACCCCTCAGATGGGCGG | 120 |
| chr11 | 70813295 | 70813415 | 788855_47188552_ENST00000601538.6,E | 1 | - | good | CTCCGGTCCAGCCTTGAGGCTGCATCCAGGGTGAGCTGGTGCTCGTTCACTGGCACCACTGCGCCGTGTGGCATCCATTTCTCCCCGAACATCCCTGCAGAGCCTTTCCCGTGCCCTTGC | 120 |
| chr11 | 70813415 | 70813535 | 788855_47188552_ENST00000601538.6,E | 1 | - | good | TGATTGTGCCTCTTGAAACCTCCAACTTTCTCACAATTCTACACATCAGGGGCTCCCCTAGCTCCCCTGGGATGGCCACAGGGGCCCTCAATGGGGCTCCCCACCTCCACCTTCTGTCTG | 120 |
| chr11 | 70813535 | 70813655 | 788855_47188552_ENST00000601538.6,E | 1 | - | good | CCCTCTGTGCCAGAGACCTGAGCTGACAGCTTTCTCTATTCCCCTACTGCCATCTTCTGCGTGAGCCGTAACCTGGAAGACACACCCACCTACCCACCTTGAGTAGGTCCCAGGCCTGGG | 120 |
| chr11 | 70813655 | 70813775 | 788855_47188552_ENST00000601538.6,E | 1 | - | good | GACCAGACAGGAGATGACACAGGACCTGAGCCTTCAACTTTAAAACAACTGTCAGCCAATAGCATTTAGTAACATTTGACAGACGCTAAGGGCTTGGGAACTCCTGCGTTTGCTGTGCTG | 120 |
| chr11 | 70813775 | 70813895 | 788855_47188552_ENST00000601538.6,E | 1 | - | good | CTCAGAGATGCCGTCACCAGAATCACAGAGGCAGCGTGAAGCCCTCAGGCCCGAGACGCACAGGGCTTGGGTGCGGGCTGACTTTCACAGTCCTGGGAAGCCAGGGAGGGAGGTGATATT | 120 |
| chr11 | 70813895 | 70814015 | 788855_47188552_ENST00000601538.6,E | 1 | - | good | TCTGGCAGACAGCCGGGGCCAGATGGAGGGGTAGCCGGTGTCCTGTGTCCGTCCCTGAAGCAGTCATTGCAGCCAGCAAGGTGAGACATTGTGAATAATCCTAAGTCCCTGTCGTGCATC | 120 |
| chr11 | 70814375 | 70814495 | 788855_47188552_ENST00000601538.6,E | 1 | - | good | GCCCGCAGAGGGGCTTCAAGTCAGTGCCAGGACTGCGGCTCCTGCTTGCCCTCCTGTGCCCCATCATCCTGACCTAGAAAGTGGGGGTGGCTGGGCCCTGCTGGCAGAGTAGGGGTGAGG | 120 |
| chr11 | 70814495 | 70814615 | 788855_47188552_ENST00000601538.6,E | 1 | - | good | CCAGTGGCCTCTGTTTTCCTGAAACTGGCATGGGAACTGAGGTGCAGCTGCGTTCAGGGTTTCAGGCGGATGAAATGGGCAGAAGGGGCCTTGGCCGCCCAGGGACACTGCTGGAGTGGG | 120 |
| chr11 | 70814615 | 70814735 | 788855_47188552_ENST00000601538.6,E | 1 | - | good | CTGCCGCCCTGTCCATGCGTCGTCCATTGCACGCATGAAAGAGCGGCACCCACCTGGCCACACAGGCAGGGGCTGGACCCCGGCAGGACCCAGTGGGGCTCTGGGAAGCCCGTCTTTGGC | 120 |
| chr11 | 70814735 | 70814855 | 788855_47188552_ENST00000601538.6,E | 1 | - | good | TGGCCTCACCCACCAGGGAATGTCCTGGCCACCCTCTCCTTCAAGAACGTTGCTTGGGTGGGGTCAGCTCAGCAGGAAAGAAGGACTCATGCTTTTGAGTTCAGTTGGTGGAGATGAATG | 120 |
| chr11 | 70814855 | 70814975 | 788855_47188552_ENST00000601538.6,E | 1 | - | good | GGGGGCATCCCTGTTCCTTCTGGGTCTCTCCCAGCTGCCAAGGGGCTGGTTGGCCTTCCCGGGTTCAGAACAACTCTGGATGTATTTCTACAAAGTGGCAGCCAGTCCCAAGGGCTGCCC | 120 |
| chr11 | 70814975 | 70815095 | 788855_47188552_ENST00000601538.6,E | 1 | - | good | GAGGCTCAGCCAGCTCCGCCCCTGGGGGCGCCCTCTGGTGGCGTCCACTGGCACTGCTGGACCACTGTCCTGAGAACATTCCTGAGACACCTGAGTCCCTGCTCTGGGAGTGTGTGGGGT | 120 |
| chr11 | 70815215 | 70815335 | 788855_47188552_ENST00000601538.6,E | 1 | - | good | TGCCCCCATCCTCCCCCACAGTCTCTCCTGCCTACCTGTCCACCTGTCCCTTGGCCTGTGTTCTTCCTTCCTCCCTGTGCCTCCTCCTTCTCATTTTCGTGTGTGTGTGTGTGTGTGTGT | 120 |
| chr11 | 70815335 | 70815455 | 788855_47188552_ENST00000601538.6,E | 1 | - | good | TGGAGACACCACAGGGTCCCTCGCTGCCCATCACTGCACAAGCCCCTCCCAGGTGAGCCAGAGCCTGTGGCAGGGCTGTCCCTCTGAGACAGAGAGAGGAACACGCCCCACACTTGTCCC | 120 |
| chr11 | 70815455 | 70815575 | 788855_47188552_ENST00000601538.6,E | 1 | - | good | CACCGGGCCCAGCCTGGGAGCTGGTGTTGCAGTGATGGTGTAACAGGAAGGGGGCTGCAAGGGGTGTGAGGGGGCCGATCTGTGGATGGTGAGTGTCCGCTGGGAGGTGAGGGCTGTTA | 120 |
| chr11 | 70815575 | 70815695 | 788855_47188552_ENST00000601538.6,E | 1 | - | good | CAGTGCCTCCTGGCCTGGGCTCAGAGCTGTAGGCAGGGTCTGCTTCCTGCATGGAGGTGCCAACCCTGGTTCGAAGCTGCCCTGAGCCAGCTTGCACAATGGCGAGCGCCCCAAGGAGCT | 120 |
| chr11 | 70815695 | 70815815 | 788855_47188552_ENST00000601538.6,E | 1 | - | good | TGGCAGTGTGGTCAGAACCAGCCTCAGGTCACACCCTCAAGTGCAGCCTTGATGGGGTTCAGGGAGGCCTCCCAGCCTCCCCAGGGCCCCCACCGGGACTCCTGCTGGCACCAAGCAACC | 120 |
| chr11 | 70815815 | 70815935 | 788855_47188552_ENST00000601538.6,E | 1 | - | good | GAGGCGACTGGCCAGGTCCCCTGTCCCTCTGCACATGCTCAGCACCCCTCTTGGGCCAGACCTATTTGCCTAAACGCTGACCCAAAGCCCCAGAAACCTTGGCCAGGACTGCAGTGGGCG | 120 |
| chr11 | 70815935 | 70816055 | 788855_47188552_ENST00000601538.6,E | 1 | - | good | CATGGGGCAGGGAGCACAGGGTATCAGGTGTTATTGTCCCGGTGGCAGAGGCCTGTGTGGGCTAAGAAGCAGGGGAAGCAGAGATGGCCAGGTAGGAGGAGGCCGACTGTCCTGGGCAG | 120 |
| chr11 | 70816055 | 70816175 | 788855_47188552_ENST00000601538.6,E | 1 | - | good | GCACTGACATCCACAGCTGCTCATAGACAGTCTTCGGTTCTGCAGAAACGTCGGCCACTCTTAGGGTCGTCTTCATCTTCATCCTTATTGTGGACAGTGTAGAAAGACCCCGCTGGCTGC | 120 |
| chr11 | 70816175 | 70816295 | 788855_47188552_ENST00000601538.6,E | 1 | - | good | CCTTGGGCAAGTCACTTGCTGGTTCTGTGCCTCTGTTTTCTCCTGTGTAAAGTGGGGCTGATGACTTTCACCCCCTCCCGAGTGTGCTGGGTGGATTAAGTGGCTTTCAGCACCTGGCCT | 120 |
| chr11 | 70816295 | 70816415 | 788855_47188552_ENST00000601538.6,E | 1 | - | good | TTCCTCCTGTCTGAAGGGTGGAGCAGCTGCCCATCAGCCCATGCTGAGCAACCGGTGCTGAACATCCCCCAAGCAGTGCTGACCGCCAGGGCTACCTGGCTCTACCCACCCGCCAGCTGA | 120 |
| chr11 | 70816415 | 70816535 | 788855_47188552_ENST00000601538.6,E | 1 | - | good | GGGAGGCTGCTAGCTCCCAGCCTCTGGGGGCCCTGGGTGACCCAGAGCTCTTTCTGGTTACCATCAGAAGCATAAGTGACAGCGTGCTCTCCCTGCCGTGGAGGGGAAAGCCCTCCCCAC | 120 |
| chr11 | 70816535 | 70816655 | 788855_47188552_ENST00000601538.6,E | 1 | - | good | TCATCCTCGAAGCTCAAAGCTCAGCCCCATGCCTTGTGCGCCTGCCTTACTATTCCTGAAAAACCAGCTCTTTCCTAGGGCAACGGGCCCTGGCCATGTGGCACTAACGGGCCACTTTCA | 120 |

| | | | | | | | | |
|---|---|---|---|---|---|---|---|---|
| chr11 | 70816655 | 70816775 | 788855_47188552_ENST00000601538.6,E | 1 | - | good | ATAGGGAAACGCTCATTCGGCCAGCCCTGGTGATGCTGGATGGTGGATCACAGGACTGTGCAACCCAACCCAGGGCCTTTTGGAGGCTCCAGGAGGTGGCTGAGCCCCTCTGCTCCTTGT | 120 |
| chr11 | 70816775 | 70816895 | 788855_47188552_ENST00000601538.6,E | 1 | - | good | GAGCACACTTGGTCTGAATGTCCTGCAAACAAGGTACAGCCAGGGGAGGGGGGCCTGGCTAGAGCTGTCTGCTTCCTGGCTGTGGACCGAGGCCCCATTGTGGAAGTGGCCTTTATTGAG | 120 |
| chr11 | 70816895 | 70817015 | 788855_47188552_ENST00000601538.6,E | 1 | - | good | GAGAGGCACAGAAGGGAGCCAGTTCACTGCACATTGTTTGCTTATGGAGCAGTTACCCAGGCCCATGCTGGGCGCAGGGACATTCAGGTGACATAGCCGGGGTGCAAAGGCTGGTGCCCT | 120 |
| chr11 | 70817015 | 70817135 | 788855_47188552_ENST00000601538.6,E | 1 | - | good | TTCTCCAGGCATCTTTTTGAGGTCCCCTGGGATGAGGGTGGGGCATGCAGGTGCCACTGTTGTGTTCTGTCCCTGCATAGGGCCATATGCATTCGGGGGGCAGAAGTGTCTTTGCACGGT | 120 |
| chr11 | 70817135 | 70817255 | 788855_47188552_ENST00000601538.6,E | 1 | - | good | AAGACCCGTGTCCTAACTAGGACCTGCCAGTGGCACTGACGTCCCCTCCTATGGGACAGCAGTGGTGAGCATTTGCCTTCGAGGCGACCAGGTGGGATGGAGAGTTGGAGGCAGGCATGC | 120 |
| chr11 | 70817255 | 70817375 | 788855_47188552_ENST00000601538.6,E | 1 | - | good | TTCACAGCCTCCCTGTCCATATCCATCTGTCCAGGATTCAGCATCCTCAATGGTAGCAATAGCTGTGCTGTAGGAGGAACCCTCTGGATGGGCACTGATGAGCTGAGAGGTCCTTCCCTT | 120 |
| chr11 | 70817375 | 70817495 | 788855_47188552_ENST00000601538.6,E | 1 | - | good | AGTCCCCCTAACTGTGGGCCACTGCCTGTCCCAAGTCGAGGGGCACTGCTCTTACCTGGTGGCCCTGACCCGTCAGTCCCTTCTTTCTTGGCCTTGTTCCATCTTCCCTTTCCCCACCCC | 120 |
| chr11 | 70817495 | 70817615 | 788855_47188552_ENST00000601538.6,E | 1 | - | good | ACCCAAGACCCAGAGCTCGGCCAAGACTGTCCCCTTTGTCACCTAGAACCGTTGGTAGAGTGACCAACCATTCAGGTTTGCCTGTGATGTGGGACTTTCAGTGCTAAAATCAGGACAGCC | 120 |
| chr11 | 70817615 | 70817735 | 788855_47188552_ENST00000601538.6,E | 1 | - | good | AGACCAACTTCAGGTTTTTCTCAAGATCAGCTGAAGGCGTTCAGGGTGGCCGCCTCCCACAGCACCTTGTCCAGCCAGTGCGCATTCCACATTGGTTCTTAGAGCCCAGGTCCATGTCCA | 120 |
| chr11 | 70818095 | 70818215 | 788855_47188552_ENST00000601538.6,E | 1 | - | good | ATGTTAGAGAGAGAATCATTGTGTATGTGTGTGTGTGTGGTCATCATGGTGGCCAGCCTGCCTCAAATGCTCATTGCCTGGCCTCCTGCATACACGCTTCTGTACTGTACCATTGTATAG | 120 |
| chr11 | 70818335 | 70818455 | 788855_47188552_ENST00000601538.6,E | 1 | - | good | AGACTAGATGTGCTAGATGTGGCTCGGGAAAGAGAGAAGCAGCCGGGTGTATTGCAGCTGCGGTCTGACCATCCCTGTTTCCCTGTTCATGTGTGAGGAACGGGGGACAGAATGCACATG | 120 |
| chr11 | 70818455 | 70818575 | 788855_47188552_ENST00000601538.6,E | 1 | - | good | CCACTTCACACACTTAGCAAGACTTCTGACTACAGTTTTAAAAAGAGTGTGTCCCCTTCACCCAGTACAAAATCTAATAAGACAGCAAAGGCAGCGTGGCCAGGGACTTGAGCTGGCTGC | 120 |
| chr11 | 70818575 | 70818695 | 788855_47188552_ENST00000601538.6,E | 1 | - | good | AACAAGGAGGTCCAGCTGCTTCTGGTCTAGATTCCTGCTTGGAAGGACCCCAAGGGAGGAGTCCCAGGTGCTGAGAACAATGGGAAGCTAAATTAAAGGGCTCCCCCGAGGCAGCCAATT | 120 |
| chr11 | 70818695 | 70818815 | 788855_47188552_ENST00000601538.6,E | 1 | - | good | TGAGAGCCCACTTCTCCCGGCCCCTAGACAGGGTCGAGGCCTGATCCTCTTAGAGCTGGCAGACCCCACGTCCTCTCAACGCTTTGGCCTTTAGCCCACAGACGCTTCCCACCAGAGCCA | 120 |
| chr11 | 70818815 | 70818935 | 788855_47188552_ENST00000601538.6,E | 1 | - | good | TGTGAGTTTGAGGAGGGTCAGTGCCTCACGTGGTCCCAGGAGGGGCTGATCACCCTGACTATGACCAGCAGCCCAGGCTTTGGGGCCACAGCCAAGGAGTTTGTGGGGAGAACATGCCGG | 120 |
| chr11 | 70818935 | 70819055 | 788855_47188552_ENST00000601538.6,E | 1 | - | good | GCCTGGCTGCGGGGAGTCTTGCAGGCCTGGGTCATGCTCTGTTGCCAGCTAAGGTCAGGGATGTCAGGGAGGAAGACCGCCACCACCCCTGCCCCCAGGGGCTCCACCTGGCAAGCTCCG | 120 |
| chr11 | 70819055 | 70819175 | 788855_47188552_ENST00000601538.6,E | 1 | - | good | TCCCCGGCCGTTGCCATGTTTTTGGTGATGTCGGCATTCAGTGCTGAGGCCCCTGACCCTCGCCCCCATCCTACATGCTTCCTGTTTCCTCCCTGGCCTCAAAGCCACTCGGGGTCCAGA | 120 |
| chr11 | 70819175 | 70819295 | 788855_47188552_ENST00000601538.6,E | 1 | - | good | TCTATGAGGAGCCCGAGGTCACACAGCCAGAACGTAGCAAGGGCAGGACTCGGCACAGGCCCCTGGCTCTGGCATCTGGTGACTTGGCACTCCCTCAGATTTGGGCCTTGGACCCGTGCT | 120 |
| chr11 | 70819295 | 70819415 | 788855_47188552_ENST00000601538.6,E | 1 | - | good | GGTTCCAGCCACGAAAGCTGCACCTGCAGGGGCAGTGGCCGTTGCCACCGATGCCTCCACATTCTCATGGGTACAGCTGCTCCAGGAGTTCACTGTGCTGGCTTCTTTGCTGCAGACGCC | 120 |
| chr11 | 70819415 | 70819535 | 788855_47188552_ENST00000601538.6,E | 1 | - | good | AGCTCCAACTCCCAAGCAGGTCTCTGCCTTTCTCACGCCACCTCCCAGAATTTTCAGTCGAGGAAGAGACTTCCCAGGGATCAGCTCATTGTCGGGGCCCCAGGTGCTAGGGTTGGTGTC | 120 |
| chr11 | 70819535 | 70819655 | 788855_47188552_ENST00000601538.6,E | 1 | - | good | GACTTTGGGAACTGCCTGTGGTCTCCTGCCCAGCACCCCATTCGGCGTCCAACTCCAGGTCTGGGTCTCTCCCCACCCTCCTGCCTCCCAAGCAGCCAGGGGAGCTGGATTTGCACTCAG | 120 |
| chr11 | 70819655 | 70819775 | 788855_47188552_ENST00000601538.6,E | 1 | - | good | CCTGGGTCCAGCATGGCAGGAGCACAGAGGGCATGTCTGTCAGTCTCTGAAGCAGCCCCAGGTGACAGGAGCCTGTGCCTCGTTCCATTTCCTAGACAAGGACATGGGCAGCATAGGGCA | 120 |
| chr11 | 70819775 | 70819895 | 788855_47188552_ENST00000601538.6,E | 1 | - | good | CCAGCCCAGAGGAGAATGCTGGGGGCCGGAGAGGCCGGCTCAGGGCTCTGGCGTCTGGCTGTCTCAGCAGCTGCAGGGATGGTCTCAGGTAGCGCTAACCCTCAGGGGAGCACTTGCAGG | 120 |
| chr11 | 70819895 | 70820015 | 788855_47188552_ENST00000601538.6,E | 1 | - | good | GGAGAGAGGCAGGGCTGCAAGCCTGGTCAGGGGTGTGTGCCCTCCCGAGGGCCATGTTTGCAAGCCTGCAAACACACCTTATCCAAATTGAGGGTGCACACGTGGGTGGCAGGGAGAAAC | 120 |
| chr11 | 70820015 | 70820135 | 788855_47188552_ENST00000601538.6,E | 1 | - | good | GGGGCAGGTGTGCAGGCTGAGCTGAGCATCCTGGGGTGGACTGGGGATACTGCTGGCCTCTCGGGTCCTGTTTCCTTTCCTTGTGTACTGACTTCTTGGTCCTAAGTGAAATGTCAAGAG | 120 |
| chr11 | 70820135 | 70820255 | 788855_47188552_ENST00000601538.6,E | 1 | - | good | ACCCAGATGTTGAGGAAACTTTCATGCCTGAGTTTTGCTCACTGTCCCTGCTGGCACGGTCTCCCCTTGGGGGGCTGGGGTCACCTTCCCGAGCCTCATGGAGGCACACGATGAAAGCTC | 120 |
| chr11 | 70820255 | 70820375 | 788855_47188552_ENST00000601538.6,E | 1 | - | good | CTGGCATGTCGGGTAAGGAGTGGCAGACGCCAAGGGAAGGAAGACCGCCACGTGCTGGGGGTCCCAGCTCCTCCAACTTGGGGTGGCCCAGAAGTTGAGCAGGAAGCCCATTTTATTCCA | 120 |
| chr11 | 70820375 | 70820495 | 788855_47188552_ENST00000601538.6,E | 1 | - | good | CAGCAAGCCCGAGGGGGCCGCGAAGACCATTGGGAGCTACGTGCCCGGGCCCCGCAGCCGGTCCCCATCGCTCAACAGGCTGGGCGGCGCAGGCGAGGACGGCAAGAGGCCGCAGCCTC | 120 |
| chr11 | 70820495 | 70820615 | 788855_47188552_ENST00000601538.6,E | 1 | - | good | CCGGGTCCTGCTGCGCTCCAACAGTGACAACAACCTCAATGCCAGCGCTCCCGACTGGGCCGTCTGCTCCACGGCCACCTCGCACCGCAGCCTGTCACCCCAGCTGCTGCAGCAGATGCC | 120 |
| chr11 | 70820615 | 70820735 | 788855_47188552_ENST00000601538.6,E | 1 | - | good | CTGACCACGACACAGATGCACTCACCGGCCTCTCTCCATGTCCTTCTCCCCAGTGCCCTTCCGAGAGGCCCCGGCGTACTCCAACCGCCGGCGGCGGCCCCCCAACACGCTGGCCGCCCC | 120 |
| chr11 | 70820735 | 70820855 | 788855_47188552_ENST00000601538.6,E | 1 | - | good | GTTCTGGGCCTGGTGGAGTCCAGCTGGGGTCTCGGCCCAGAACTCACAGCTCCCCTCTGCTGGGCTCGCATGGGGGCTTGGACGCACGCAGGCCTGCACCTCCCTAGGGTCCCCACCCAG | 120 |
| chr11 | 70820855 | 70820975 | 788855_47188552_ENST00000601538.6,E | 1 | - | good | CCCAGGCCAGATGGAAGCTCAGACCCCACGTCCCTTCCATGCGAAGTACTGGCTTGAGCTCCGTGCCGTGGAGATGTTCTCCGTCCCCAGCCTGTCTCTCTGCTTCCATGTCAGGGGTGA | 120 |
| chr11 | 70820975 | 70821095 | 788855_47188552_ENST00000601538.6,E | 1 | - | good | CCCAGAGGCTCAACTTAGGTTGTGGGGAGGAGGCATCTTGCGGCAGGCAGCTCAGTGCCCTCTCTGCTTTGTGTGATGGGGTGGCCCGTCCTTACACATGCCCAAGCCAGCCGGACGTCT | 120 |
| chr11 | 70821815 | 70821935 | 788855_47188552_ENST00000601538.6,E | 1 | - | good | GTGAGGGCCTGGCCCTGTCTAGAGCTTTGTAGGATGCCAGATAAGGGGTGCCCAGCCCCTGCTTCCTCACACAGTCCCCACTTGATGGGAGGGAGGAAGGCACCTTCTTAGTGAGGACTG | 120 |
| chr11 | 70821935 | 70822055 | 788855_47188552_ENST00000601538.6,E | 1 | - | good | GCTGTCTTAGGCTCTGGAGCTCTGGGTAAACCAAGGCAGCTGCGGCGTGGCTGCCACCCCTCAAAGCTCCCAGGGGCTGTTGCTGTCATTCTGAGCCCCTGGTCTCATTGAGTGAGGAGA | 120 |
| chr11 | 70822055 | 70822175 | 788855_47188552_ENST00000601538.6,E | 1 | - | good | TTATCACGGGCCGGGGAGAAGCCCGTGCCTTTGGCCCCCCAGTGATGGTGGTTTCTGCTGTTTCCTGCGGAGCTTGGAGGGCTGCCTTTCAGGAGGTCTTCTTTCTTGGAAAGGGCACTC | 120 |
| chr11 | 70822175 | 70822295 | 788855_47188552_ENST00000601538.6,E | 1 | - | good | CTTGCCTTGCAGTTTGTCAGATGGTAGCCCTGGTCCTGTGCAGTGGAAACTTCGGAGCCAAGCGTGGCTTCAGCCTAAGGGCTGAATCCTGTAGCCAGTGAAAAGCTGCTGCAGGCACGA | 120 |
| chr11 | 70822295 | 70822415 | 788855_47188552_ENST00000601538.6,E | 1 | - | good | CATCCATGTCCCCAAGAGCTTCGTCACTGCCACCTGTGTCCTAAACTACTTGTGACGTTCTTCTCCTCTGAGGCATGAAGGTTCCCCCAGACCCAATCCCAGGGAGGACACGTGTTTTCT | 120 |
| chr11 | 70824215 | 70824335 | 788855_47188552_ENST00000601538.6,E | 1 | - | good | CCACCTCCAGGTTCTTACCGACAGCAGGCACTGACGTCCAAGTGTGTCAAGTCCAAGTATCCAACCTCAGGACGCTTTGCGCAGCTCTGTGCGGCTCCCCAGGCTTGGCTCTAGCTCTGT | 120 |
| chr11 | 70824335 | 70824455 | 788855_47188552_ENST00000601538.6,E | 1 | - | good | AGTGGGATGAAAGTGCTGGCTGAGGAGCCCCGGCTGCTTAGCAGGGCATCTTCAATGACAGAAAGGCTTCACAGAGGAACCAGATCAGAGCAGGTGTGACACCGCGACCTCCCCAATGTA | 120 |
| chr11 | 70824455 | 70824575 | 788855_47188552_ENST00000601538.6,E | 1 | - | good | CCTCGAGGGGCTCTGGAAGGAGAGATGTCCACGGAGAGTGCCGTCAGCTTTTTCCGTTCCTTGTCTCTCCCCGGCCAGCCTGGGTGGCGTTCATGGATCCGTGTGCTTTGAGAAATTAGG | 120 |
| chr11 | 70824575 | 70824695 | 788855_47188552_ENST00000601538.6,E | 1 | - | good | CCGGTCCAGAGGGAGGGCACCAACATGGTTTTCCCTCAAGACCAACAAAACAGACTCGGTGACCTGTCCCAGGGATGAGGTTTTTCTCGTGGGTGGTGGCTGTGGATCCCAGGCCAAGCC | 120 |
| chr11 | 70824695 | 70824815 | 788855_47188552_ENST00000601538.6,E | 1 | - | good | GCCCAGCCCCCTCCAATGACAGGCAGTTCCCGGATCACCCGGAAGCCCTCTGGGCCAGTCGTATCTTTGTAAGCCTCCATGCAAACACTCAGTGATCTACAAGGTTTTTTTTAGTCTCTG | 120 |
| chr11 | 70824815 | 70824935 | 788855_47188552_ENST00000601538.6,E | 1 | - | good | TTTGAGATTTTGGCTCTCACACCAATACAGTCTTCCAACACGTAATCTCTGTAAATGAAAGATAATCGCTCTCTGAGACTGTGGGCTCATCAGCCCTTTGCAAGGTCTTTTAATCCACCC | 120 |
| chr11 | 70824935 | 70825055 | 788855_47188552_ENST00000601538.6,E | 1 | - | good | CGTTTCCGTGACTGGGGGCCCTGTGATGTCTCCTCCACATGGGTAGGGAGAGCCGTGGCCTCTCCTGGGGTGCCCCAGGAAGCTGGGGGCTCTGCGGCTGCTTTCTGCTGGCTGTGTCTG | 120 |
| chr11 | 70825055 | 70825175 | 788855_47188552_ENST00000601538.6,E | 1 | - | good | GCCATCACTTGCTATGAAGAGTATGCATACCTCCCCCAGTTACAAGGGTCAGAAAAAGAGAATTGGCTGCATTGTCGGGGCGGGGAAGCTGGGCGTGAGGCAGTCGGCTGTGGGAGGCAG | 120 |
| chr11 | 70825175 | 70825295 | 788855_47188552_ENST00000601538.6,E | 1 | - | good | AGCATGAAAATTTCAGTTGGGGGTTTTGGTGTTAGGTTAATTCTGTATCTGTTCTGTTCTTTTATAGACTTTTTTTTTCCTAATGAGTTTATCTTAGCAACTGCCAGGACTCCCCCAAAT | 120 |
| chr11 | 70825295 | 70825415 | 788855_47188552_ENST00000601538.6,E | 1 | - | good | CATACTCTTCTGACAGAGTCTTAGGAATGTATATAAATAACAACCTTTGTCTGTGCCCATTTTCTGACATTATTCACCAAAATTCACCTTCTGGGGTTTCAATTTTTATAATCCATGAAC | 120 |
| chr11 | 70825415 | 70825535 | 788855_47188552_ENST00000601538.6,E | 1 | - | good | TTCATTTGAGTAACTTGCGTGGTCTACAGAGTAAAAATGTATTTGAAAATGTCCCAGCTCTTTTATTCCTTAATCCTCCAAGGGTGGCCTCCCCATGTGAATTCTTCCCCCCTCTAAGTC | 120 |
| chr11 | 70825535 | 70825655 | 788855_47188552_ENST00000601538.6,E | 1 | - | good | CTCGTTGCAAAATAAACACCAAAGAAATTTCAGTACTCAGGAAACTAATTTAATATGAATCTTGGTAATTACGCGAGGTAATTTTCATTAAAATGTTCACACTGATGTCACCGGTGAGAT | 120 |
| chr11 | 70825655 | 70825775 | 788855_47188552_ENST00000601538.6,E | 1 | - | good | CATCATTTCCATGGATGGTTCCAGTTAGAGAGTTTTGTCAGAGAGCTCTCATTTTAGGAAATCAGATATCATAAAAAGGTATGACTTTAGAGAAATGAGGAAGGTACTTCCCAGCCCCAG | 120 |
| chr11 | 70825775 | 70825895 | 788855_47188552_ENST00000601538.6,E | 1 | - | good | CTAAGAAAATATCCACCTATTTAGAGTGCAGATTGCCTAAAAGAAGGAAACAGTTGGGTGGAATATTAATCATTTTCAGACAGAAAACAGTTATTTTTAACCCTTTTCACAAAAAAAAAT | 120 |
| chr11 | 70825895 | 70826015 | 788855_47188552_ENST00000601538.6,E | 1 | - | good | TTACATGGAATGGATGGGAAGGGAGCGTGGCCAACTTTAAAAGTGCACACTCTGTTTGTCATACCTTGTGTAATTTTTATTGTGGGGAAAAAAATGTATAAATAAATGCATGTAAGCCAG | 120 |
| chr11 | 70826015 | 70826135 | 788855_47188552_ENST00000601538.6,E | 1 | - | good | TCCTCAGTGACCAGAGGGCTGCTAGGCACTGGCCGAGGCAGGTCGGGGCACATGGCTGGCTGCACCAAATGGGTGGCTACCTTCCGAGCTGCTCCCGGCTCTCCCATCCCGGGGTTGCAG | 120 |
| chr11 | 70826135 | 70826255 | 788855_47188552_ENST00000601538.6,E | 1 | - | good | CTGCTTTCAAACCTTCAGCCCTTGTGGAACTCCATTCCGGAATGGCCATCCGTGGGGACCGTGCCACGGAGTGTGTTAGGAACTAAGTCCTAGGGAAAATAAAGGAAGTTTTGACTATAG | 120 |
| chr11 | 70826255 | 70826375 | 788855_47188552_ENST00000601538.6,E | 1 | - | good | CCTAAAATTGCCAGTTGTTAATCCCTAGCATGTCAGGGGATAATGACTTTGGTTTGGGATACGGTTTGGTTTGAGTTGGTCCGGAATATAACGCGGTGGAAGGTCTCGCCTTATCGTGTG | 120 |
| chr11 | 70826375 | 70826495 | 788855_47188552_ENST00000601538.6,E | 1 | - | good | TTTGTTAAATGCCTTCACCAAGAAGGAAGGTAAGGAAAAAGCTGTCGGGAGCCGGGTGCATGGGAATTAGCAGCTGCTTGCTGATTCTCGGGGTGGGGGAGAGGGAGGACTCCAGTTCCT | 120 |
| chr11 | 70826495 | 70826615 | 788855_47188552_ENST00000601538.6,E | 1 | - | good | TCTCCCCGGGTCCCAGCTGCTCTAGCACCGAGGGGCCGGGCTGCACGCCTGTCCTTCCTCCCATCCAGGAAGGTTATATAATGAGCGGACCGCTGCCAAAGTCAGAGGGAAAATGAAGTC | 120 |
| chr11 | 70826615 | 70826735 | 788855_47188552_ENST00000601538.6,E | 1 | - | good | TCTACGTGCAGTTGGTTCCCAGCTGACAGACAGAATATGGGGTGGGCTTTGTCACCAGCACGCTGGGTCTGGCATGCCCAGCTATGCTCGGAAGTGCACCGTGCCCAGACACCCGCCACC | 120 |
| chr11 | 70826735 | 70826855 | 788855_47188552_ENST00000601538.6,E | 1 | - | good | ACGCCTGATCCGCGTTAACCCTCCGCGGTGCCTCTTCCTCAATGCCCTGAGCACAAAAGCAGAGTAGCCCGGCTCACATGACGCGCGCCGCGAGAGGTAGCCGGGTTTGAGAGCAGCCCG | 120 |
| chr11 | 70826855 | 70826975 | 788855_47188552_ENST00000601538.6,E | 1 | - | good | ACAGTGCAGTATATTTAGGCACAGGCGCATGGAGCGCCCAGCACCAGCCCGAGTTTCCTTGGATAACTGGGGCTCGAGCTCACTCATGCCTCCCGGCGTCTCCCAGCTGCCAGCTCGAGG | 120 |
| chr11 | 70826975 | 70827095 | 788855_47188552_ENST00000601538.6,E | 1 | - | good | TGAAGCTTATCTGGTACTTGTACGCTTCCCTCCCCCTTGTCTGTCTTACCACATCTGCTAGCTGACAAACAAACTTGGCTTTTTCTTTAAAAAAATTTTTTTTTAAAAAAATAGAAAAGC | 120 |

| | | | | | | | | |
|---|---|---|---|---|---|---|---|---|
| chr11 | 70827095 | 70827215 | 788855_47188552_ENST00000601538.6,E | 1 | - | good | TGGGTGGGGAGCCAGAGAAGTCATGGGCACTTGTGTCTGATTCTTTTGAACAGAAATATAGGTCGTGTTCAGAAATTCCAGAATGCTGGCCATGTCTGCCCGGGGAACGTGGGTCTGGTG | 120 |
| chr11 | 70827215 | 70827335 | 788855_47188552_ENST00000601538.6,E | 1 | - | good | GAATCGGGGCCATCCTTGGCAGGGTCTCAGTTTAATTGTCTTTTTATCTCTCTCTTTAAAAAAAAAAAAAAAAAAAGTAGGAGGGGGTGGAGTCCTGTTTTTTCCAGGGTAATTACCTCGA | 120 |
| chr11 | 70827335 | 70827455 | 788855_47188552_ENST00000601538.6,E | 1 | - | good | ACTTTCTCCATGTAGGGCTTCCAAGGGGCAGCTTTCCTCTTTTCTCTTCTTCTGTTGGGTTGGAAATGCTGAGGAAATGCCTAAAGTGACCCTGAAGAGCTCGCTTTTATCAGCTGCCTG | 120 |
| chr11 | 70827455 | 70827575 | 788855_47188552_ENST00000601538.6,E | 1 | - | good | CCCCTGGATGCTCCACGTAGCTTCAGGAAGCTCCTTCTCCTGGTGTAGAAATAACTCACTGTCCACCTGCCACAGGGCACCAGCCTGCATCCACTTCTCTGTGTTTGCTTTATAAATCTG | 120 |
| chr11 | 70827575 | 70827695 | 788855_47188552_ENST00000601538.6,E | 1 | - | good | TGTGTTTTAAATTTCTGAAGCAGTTCTCTTAGCAACCAGAAAAAAAAAAAAAAAAAAACACACACACACACACAGCACACTAAATTACAGCCCAGCAGAGGCATCCGGCAACAGAACCTTG | 120 |
| chr11 | 70827815 | 70827935 | 788855_47188552_ENST00000601538.6,E | 1 | - | good | CTCTGCTCTCCCTAAAGCCAGAGGCCCTCACCGAGGCATGCCGCAGTCCCGTCCCTGGTACGCCTTCAGGGAAAGGCTCCTGGGGGGCCCCCAGCCTTCTTGTTCTCACACCGCCATTTT | 120 |
| chr11 | 70828295 | 70828415 | 788855_47188552_ENST00000601538.6,E | 1 | - | good | ATGAGTCGGTGATGCATGGTAGGGTGAGAATGTCACCCCGGCAGAGTTTTGACATGGCTCATGAGTTGTGTGAGTTAAATAAGAGATGAAGTTTATCGTTATTTCTGGGTTTTTGTGGGG | 120 |
| chr11 | 70828415 | 70828535 | 788855_47188552_ENST00000601538.6,E | 1 | - | good | GCTGTTCAGCCTTGCTCAGAGTGTCCGAGGAGGTGCATGAACTCTGTGGGACACACTGGTGGCTCTGACCTCAGAACCGTTTCTTAATGCAAGTCTTAGAAATGGCTTTGACTCAGTTTT | 120 |
| chr11 | 70828535 | 70828655 | 788855_47188552_ENST00000601538.6,E | 1 | - | good | ACTAGACACTCACCATCTGGAAAGCTTCTGGCCCCCCTGGCTGGTCCACCCGGAGGACACATCGTCAGGCCTCCCGAAGTGGGGGAAAGCTTTGTGCATCCGAGGAGGGTTCTTCTGGAA | 120 |
| chr11 | 70828655 | 70828775 | 788855_47188552_ENST00000601538.6,E | 1 | - | good | GGGTCCCCCAGCCTTGGAGGAGCTCCCTGTACACACCAATACGGACACGGTGGCATCCCCGTCTCAGCCTCCCAGCGCTCGCCAGCCAGGGCAGGCAGCCAGGCCAGCACATGGCAATGA | 120 |
| chr11 | 70828775 | 70828895 | 788855_47188552_ENST00000601538.6,E | 1 | - | good | TTTGGGAGGGAATCGATCAGTGACAGATGGAGGGGGTGTTGAGCGTCACTGAGTCTCCCCGAGAGTTCCCTGCGCTTTGATGAGTCCTCTGGGCTCACGTGGCTGAGCACACGACCAGCC | 120 |
| chr11 | 70828895 | 70829015 | 788855_47188552_ENST00000601538.6,E | 1 | - | good | ATCCTGGCCTCACCAGCCCAGAAACGTGCAGGCTCTCCCCACACCATGGCAGGCCGCAGAGGCAGCAGCGCCGGAGCAGTGATTGGCTCAGAAACTTGGCTTGGGATGAGGCTCTGTGCT | 120 |
| chr11 | 70829015 | 70829135 | 788855_47188552_ENST00000601538.6,E | 1 | - | good | GCAGGATTGTCACATGGTCCCAGGGAGGTGGCTTCTTCCTTACACCCTCCACCCAAGGTCTTCCGTGGCCGGCAAGGCAGCCAAAGATGACAGGACACCGCTGGCCTCAGGCCAAGGGCC | 120 |
| chr11 | 70829135 | 70829255 | 788855_47188552_ENST00000601538.6,E | 1 | - | good | CTCTGGTTTAGATGGGGCAGGGCCTCTTCTTGGTCCTTTGACATATTTGACATTGGAGGCCCCCGTGAGGGGAAAAAGGCTCCATTTTGTCCCCTGCCTGGTCCACCAGTCAGAACAGAT | 120 |
| chr11 | 70829255 | 70829375 | 788855_47188552_ENST00000601538.6,E | 1 | - | good | GGGCAAATGCACCTCCAGGAGCAGCTGCTGCACACGCACCGGACGGTGGCCGCTGAGGGTCCTTTCTGGGCCAGCTGTGAGTGCTGCTGCGTGGCTCCCGTCCAGAGGTTCTGGGTTTTT | 120 |
| chr11 | 70829375 | 70829495 | 788855_47188552_ENST00000601538.6,E | 1 | - | good | GGACCCCAGGGGTGGAAGCCAGCCGTATTCCTGGTGACGGGGCCTTGTCTTGGGAAGGCTCCTTCCCTTCTCCCTGGGCAGCCCAGGGTTCATGGAACCCCTGCCTGGGCTCCTGCTTGT | 120 |
| chr11 | 70829495 | 70829615 | 788855_47188552_ENST00000601538.6,E | 1 | - | good | CTCACCCCTGCACAGGCTGAGAAGCAGGCCTGGGTCTTCAGATCTGGCTGGCTGTCTTCTGCGCATGCCCCCAAACCAGATGGGACATGGGGTGCAGGGGTGGGGGTGGGCGGGGCAGAG | 120 |
| chr11 | 70829615 | 70829735 | 788855_47188552_ENST00000601538.6,E | 1 | - | good | GGAAAAGTTGTCTGCTTATTGAAAGAATTGAAGGTGTCTGGAAAAGGGATCCTTTAGCCTGGGGGTGTTCCCTGGATTCCTCAAATCGTCTTGTGTCCAAAGCCTCATCCTGCCCCGCCC | 120 |
| chr11 | 70829735 | 70829855 | 788855_47188552_ENST00000601538.6,E | 1 | - | good | CCGTGTGGCAGAGCCTGGCCCAGGACCGGGCACTCAGTAGGTGTGTGCATGAGACTGATGGAGCCACCACCGCTCCTGATCTTTGTGAATGGTTTACTCAGCCTCCTCCGAGTGTGGGCG | 120 |
| chr11 | 70829855 | 70829975 | 788855_47188552_ENST00000601538.6,E | 1 | - | good | AGGTAGTGAGTCCCCCGTCGCTGGAATTATCCACGGAGAGCCTACCACATGGCCAGGACAGTGGTGTCAAGAATGGCCCAACTCCCTTGGCAGGATTTCTGTGGGGTTGACCAGAAGACC | 120 |
| chr11 | 70829975 | 70830095 | 788855_47188552_ENST00000601538.6,E | 1 | - | good | ACTTCTTGGCAAATGAGGAAGTGGTCTGCACCCACCAGGGAAGGGGGCAAGTGGTGGGTGTGGGGGAAGGAGGCTCTTACAGGCTGCTGCCCGCGTCTGACAGTGAGAACCCCTGTGCAA | 120 |
| chr11 | 70830095 | 70830215 | 788855_47188552_ENST00000601538.6,E | 1 | - | good | AGGGCTGGAAGATAAATATTCCCCCAGAAAGGCAGGGATGGGGACACCAGATGAAAACTGCCCTGGAGGCTGCACAAAGGTTTGCGTTTGTCGGGGCAGAGTCTGTGCCTTCAGCGCCAG | 120 |
| chr11 | 70830215 | 70830335 | 788855_47188552_ENST00000601538.6,E | 1 | - | good | GGAACATTTGTCCACACAGAACTTCCCTCCCAGCCAGGCAGGCAGGGCAGGCTTCAGGCCACGGACATGCTCATAAAACGAGGGGCAGGAGGAATGCTGATGTTCCCATGGCCACATCCC | 120 |
| chr11 | 70830335 | 70830455 | 788855_47188552_ENST00000601538.6,E | 1 | - | good | GCCTGTGTGCAGATGGCAGGAGAATGCGAACCATCTACCTGGGGCTCAGGGGTGAGCTCGTCTCCTGCAATAACCCCAAGTGTGGATTTCTGGGAGGGCCCGGACCCCAAACACTCACAC | 120 |
| chr11 | 70830455 | 70830575 | 788855_47188552_ENST00000601538.6,E | 1 | - | good | TAGCGAATTCCCGGAGTTCTGCGGGAGGCTGTGCACCCGCCTCTGCTTCCCCAGCTCTCCAAGGTCACCTGGTCGGTGGCAGGTTTCTGGGTCTCCGTGGCTTGGAGGGTGAAGGGCCGG | 120 |
| chr11 | 70830575 | 70830695 | 788855_47188552_ENST00000601538.6,E | 1 | - | good | TGGAAGCCTCCAGGGAAAATGGGTTTTTAAGTTTCATCTTTTATCTTTGTCTTAATTTTCATCTCAACCACTAGCGCTCGGTCATCCTCGGAATACTTGTTTTAGAGATGGGAAGGGCGC | 120 |
| chr11 | 70830695 | 70830815 | 788855_47188552_ENST00000601538.6,E | 1 | - | good | AGCCTCTCTGAACCTCGTCTGTCTCATCTGACAGCCTTGCTGCTGGTCTCAGGGACCCAGGATGGCAGCTCCACGGGGACATTCCCAAAGCCTGAGGCTTTTCTCAGAGAAGAAGATCTC | 120 |
| chr11 | 70830815 | 70830935 | 788855_47188552_ENST00000601538.6,E | 1 | - | good | AAGGCAGAAGGAAGGGAGTGCTCAGCCATGCTGTGGGCATTTGGGCTCCAATGCGTGGAAACCTATGTGGGTGCAGCAGGGATGTCTGACCCTTCCCAGGCCCTGGGCAGCTTAGCGCTC | 120 |
| chr11 | 70830935 | 70831055 | 788855_47188552_ENST00000601538.6,E | 1 | - | good | CCAGAATGAAATCCATTTGAAATAAAACCTCATCTGCATTCAGTAGGAATGTGGCCGGCACAAGCCTCCAGGGTCGTTGTAGGCCCTCCACGCAGGACCCCTATAGGAACAGTGGGTCTA | 120 |
| chr11 | 70831055 | 70831175 | 788855_47188552_ENST00000601538.6,E | 1 | - | good | GGCAGGCAGGCAGGTGGATAGCTTCATCACCAACCACTCACATTGCCACCCTCTCTGCTACCCGCCTGCCCCCAGCCCAACTCACACCTGGAAGTTCTTCCCCTCCAATCTCTCCCTCTT | 120 |
| chr11 | 70831295 | 70831415 | 788855_47188552_ENST00000601538.6,E | 1 | - | good | CTGAAGAGCCAGGTTGGGTGGCTGAGAGTTTGGGGTTCATTTGGGGGTGACCCTTTTCCTTTCTGGAAACTCTGACTTCAGAGCCTCCCTCTCCTCTTTCCTCCTGGTTTTTCATCTGTC | 120 |
| chr11 | 70831415 | 70831535 | 788855_47188552_ENST00000601538.6,E | 1 | - | good | TGCCTGTGGGCAGGGGCTGCAGGGAGGGAGGCAAGGGGGCACGGGGCTTTGGATGTCACTGGCTCACGAAGCACCAAGGGCCAGGCACTGGAACTGGGCTGAGTGGAGCTGAGTTGGTG | 120 |
| chr11 | 70831535 | 70831655 | 788855_47188552_ENST00000601538.6,E | 1 | - | good | AAGAAGTGAAATTATTTTCAGTCCGTTGTCAGGCAGTTGTCAAAAGACATGGCAAATAAAGGGCTTAGTGAGTCTCAGGGTTTGAGGGGGTGGTGCAGTGTATGGGGTGAGTCTGTGGGA | 120 |
| chr11 | 70831655 | 70831775 | 788855_47188552_ENST00000601538.6,E | 1 | - | good | TGAGACCCGTGAGCACCTCTGAGTCTTGCTGGCATGTTTATAACACCTGGTACGGCGTCCCCGCCAGGACTTCAGAGCACCTTGAAATGTTCTGATGTGTGAAGAGCTTTTTTTTTATGC | 120 |
| chr11 | 70831775 | 70831895 | 788855_47188552_ENST00000601538.6,E | 1 | - | good | CAGAGTCAGCACCCAAACCTGCGTCTTGACTCCCACGTCCGGCTCTTTCCAAACCACATGGGCACTGAAGCCACAGCAAGTGGAGGCGGGCGTCTGATCGCCACTGAACGAGGATGTTTC | 120 |
| chr11 | 70831895 | 70832015 | 788855_47188552_ENST00000601538.6,E | 1 | - | good | TCAAACACGAGCTCGGGTGGCTCCACCCTGTGGGGTAAGTTGTGCTATTCCTACTTTACAGATGGGGAAACTGAGGGTCAGGGGGTGTCCAACCGGCCCATGGCTGCAGGGGGCATGATA | 120 |
| chr11 | 70832015 | 70832135 | 788855_47188552_ENST00000601538.6,E | 1 | - | good | ACAGACTAGGAAACAGAGGCACAGAGAGCTCATGGGACTTCTCTGGGCTCACAGAGCTGAGACTTGTCCCCAAAGGTGTCCCCACGGTGTTGCGTGCACCAAGCCCCTCTCCTTTGTGCT | 120 |
| chr11 | 70832135 | 70832255 | 788855_47188552_ENST00000601538.6,E | 1 | - | good | AACTTAAATGTATTCATTCAGGGATCACCGACACTGCCTCCAGGTACTACTGTTACAAATATTAACTTACTTTATCCTTATGACTCTCCTGTGAAGTGGCACCATTAATATCCCCATTAA | 120 |
| chr11 | 70832255 | 70832375 | 788855_47188552_ENST00000601538.6,E | 1 | - | good | AGGAGGCAGCACCCAGAAGGATAATGTGAATGCATCGTGAGCAATAGACCAGGACTTGTTTATTATAGAGATAGGGCAAAGCAGGAGGATTGCCTTGCTATCCACTTACAGATTTTTTAA | 120 |
| chr11 | 70832375 | 70832495 | 788855_47188552_ENST00000601538.6,E | 1 | - | good | CTGCCTTTGAGAAGCTGGGAGGCTCCTGGCTCCACAACATGCCCCAAGGGCTGACTGCTGCCACGGGAGCCCTTAGGAGTGGTCTGTCTGCTGCCTGGATCAGTCAGGGAGACTTCCTGG | 120 |
| chr11 | 70832495 | 70832615 | 788855_47188552_ENST00000601538.6,E | 1 | - | good | AGAAAAAAAGCTAGGACTCACCCCCTCTGTTAATTCACTCCATTCATTCATTCATTCATCTCCAGAGCACTAGGGATCACTGCCAGTGTGCAGGGCAGGGAGGAGGGCTGGACACGGAAC | 120 |
| chr11 | 70832615 | 70832735 | 788855_47188552_ENST00000601538.6,E | 1 | - | good | GGGCAGAAGAGGGGGTGGGGGTAGGGTGGGAGTTGGACAAGGAAGGTGGCTTCTGGCCTCAGGAGAGCAGCACAACTAGAGAGCAGCCTGGTGAACCTAGTAAAGTGCTCTGGAACACTC | 120 |
| chr11 | 70832735 | 70832855 | 788855_47188552_ENST00000601538.6,E | 1 | - | good | CACCTGCTCGTGGGGCAAGGCGGGGCAGTACAGGCCTTCTCAGGGTCTGGGGCAGCTGGTAGCGTGGCTGTGGCTTAGCCCAAGCCTCCCTGGGCCTCCACTGGGTACATGCAAGGCCCT | 120 |
| chr11 | 70832855 | 70832975 | 788855_47188552_ENST00000601538.6,E | 1 | - | good | TGCAGTGAGGATGCTGAGGACACTGGCTCAGCCACTCTGCAGATGCAGTGAGGATGCTGAGGACAACAGCCTAGCTGCTCTGCAGATGCAGTGAGGACGCTAAGGATGCTGTCCTGGCTG | 120 |
| chr11 | 70832975 | 70833095 | 788855_47188552_ENST00000601538.6,E | 1 | - | good | AACCACAGCAGCTGGGGAAGCTGAGGGATGAGCCCTCACTGTTTCCAGGCAGGACATGGGGTGCAGGGAACTCCAAATGCAGTGAGGACGCTGAGGACGCTGGCCAGGCTGCTCTGCAGA | 120 |
| chr11 | 70833095 | 70833215 | 788855_47188552_ENST00000601538.6,E | 1 | - | good | AGGCGCTGGCAGGAGCTTGACATTGTGCTGGAGCCCCGTGCCACACTCGCCGGCTTCTGTGAGTTGCTCCTTCTGCATCAGTGCCTTCCTGGCCCATGTGTCTGATGGGCTCGAACTCGA | 120 |
| chr11 | 70833215 | 70833335 | 788855_47188552_ENST00000601538.6,E | 1 | - | good | TCACTGTGCTCCAAGAGGGCTGGAGAGCCAGGCCTGGGAACTGCGGCCAGCCGGGGTGGCCAGTCGGGGCGTCTAGCCACAGGTGGGCTGCATGGAATACCTGGTGCTGTGCACAGTCAG | 120 |
| chr11 | 70833335 | 70833455 | 788855_47188552_ENST00000601538.6,E | 1 | - | good | GCTCTCTTGCCCGAGACCCAGCCTCACGGGCTCTACCGGTCAGGGTCTGGGTCACCAGCAGTAGAGTCCACCCTGGCTTGTTGATGCAGGAGGCAGAAGACTCTGACAGATGTCAGGCGG | 120 |
| chr11 | 70833455 | 70833575 | 788855_47188552_ENST00000601538.6,E | 1 | - | good | TTGTGACTCATGGACAGTTACCTTGTGGGATTTGAACCCAGAAAGTCCAGCTGCAGATGTTGCAGAGATGCTGTCAGGGTCCCTGGACCAGAGGCCTCCCTGAGAGATGTCGCATGCCCA | 120 |
| chr11 | 70833575 | 70833695 | 788855_47188552_ENST00000601538.6,E | 1 | - | good | GAGCACCCACTGTGCGTAAGTACTGCACCCAGTATGCACTTAACCCCCTGAGCAGCTCCTTGAAGTGACCATCATCACTGTGCTTGCTTTGTTTTTGGGGATCCAGAGTCCAGGGACATT | 120 |
| chr11 | 70833695 | 70833815 | 788855_47188552_ENST00000601538.6,E | 1 | - | good | TGAGAATGCCTGTCAGGACCTCCCTCGTCAACAAGCTGTTGTAATAATCACAATAATGAACAGTCAGCAATAATAAACAAACTTCAGTCGTGGTCGTGATGGCAACAGCTAACCTCTGTT | 120 |
| chr11 | 70833815 | 70833935 | 788855_47188552_ENST00000601538.6,E | 1 | - | good | TGAGTTACACAGAAGCAGTGTCCTGGGGAGACAACACTGTCCTGGCCTGGGTCAGGACCCTGTCTCTAGCGGCCTGATGGCCTTCCACTGCCTGCAGCCACCTGAGCTGCCCAGAGAGAC | 120 |
| chr11 | 70833935 | 70834055 | 788855_47188552_ENST00000601538.6,E | 1 | - | good | CTGGTTTGGTTGTGATCACACTCAGGGTTTTGCCCAGGGTTTAAGTTGAAGGTGACGCTCAAAGCTGGCCCTTGTGCCTTCATCAACTTGGAGCTCCTTCCTGGCAGAAGGGCCTGGGTC | 120 |
| chr11 | 70834055 | 70834175 | 788855_47188552_ENST00000601538.6,E | 1 | - | good | CCTGAAATATGGGGTGGAGCCTTAATAAATCTCCGGGATATCAGGGCATCCCCTGCCCCAGAACCCTGGGAATTTTCTTGGCATTGCTTCCTACATCCCAACATTTTATTTCTTCCCTT | 120 |
| chr11 | 70834295 | 70834415 | 788855_47188552_ENST00000601538.6,E | 1 | - | good | GCTTGGGCTGGTGGGCCACTGAGACCGTCTAGAGTTTGTTTTCTTTTTTCGGCTCCCCATGGGCCTTCAGTCTGACCAGAGTTAGAGCCACCTCCTGTGTATTTTGTGTTAAGGTGGAAA | 120 |
| chr11 | 70834415 | 70834535 | 788855_47188552_ENST00000601538.6,E | 1 | - | good | CTCTAGCTTGTACCTGCCCTGCACCTGCCACCCCTGCTCCTGGCTTGGCCCCCGCTGGCTGAGGCTCTTGCATACATACTTTCTCAGCATTCAGTCTTTGTTTCTGCAAGATAAAGTGGA | 120 |
| chr11 | 70834535 | 70834655 | 788855_47188552_ENST00000601538.6,E | 1 | - | good | TATTGTGAGATGGAGAAATGTTGCCTACTAATATTTAAATTGCAGTTATCTAAAGCTACGAGGGAAAGAAACAAGAGATGCTATTTTGGATCTGGTGGGGGATGGGCAGGAAGAATCCCG | 120 |
| chr11 | 70834655 | 70834775 | 788855_47188552_ENST00000601538.6,E | 1 | - | good | GACGGGAACGTTACAAACACCACAGACTCCTACCTCTTTGTGCAAAGATAACCCCGCATTATTGCTGAGATGCCTTTTTATATCTTGTGTTATTTATAAAAGCATGATTCATTTAACAAT | 120 |
| chr11 | 70834775 | 70834895 | 788855_47188552_ENST00000601538.6,E | 1 | - | good | CCAACCAGGGAGAGAGACCCTGTGGGAGGTTCCGACGCCCGCTCCTCAGTGCCCTCACCCCGATTTTCCACGGGCCTCCCTCGATCCTGCAGGAGCAAAGCTTCCCTCAATGACTCCAGT | 120 |
| chr11 | 70834895 | 70835015 | 788855_47188552_ENST00000601538.6,E | 1 | - | good | CAGAGCTCCTCCCCGCTGCCCCCCAAAATAGCCATGTTCTTGGCCCTAAAGCTCAATTCAGGGTCAGAAAACCGGTGAGTAGTATCTCTTGGAGTTACACCCCCAGCCCCTTTCAGCTCT | 120 |
| chr11 | 70835015 | 70835135 | 788855_47188552_ENST00000601538.6,E | 1 | - | good | CCAAGGCTGGAGCTGCGTGGAGGGAGGGCAGAGGAGGCAATGTCTGGAGGCTGTGATTCTTCTAGAATATACCTTGGGAACCCCTCACCTCTCTTTGTGGTCAGACCCACCTTCACCCTC | 120 |
| chr11 | 70835135 | 70835255 | 788855_47188552_ENST00000601538.6,E | 1 | - | good | TGGTTCATGCTGCCCGTAAGCCATTGCACGCCCCTTGGCTCCACCCCTCCTCCTTCAGGTGTGTCCCGCACACAGTAGTGTCCTGTGCACTGGGGCTGTGGCTCTGCAGGGAGCCTGTGG | 120 |

| | | | | | | | | |
|---|---|---|---|---|---|---|---|---|
| chr11 | 70835255 | 70835375 | 788855_47188552_ENST00000601538.6,E | 1 | - | good | CTGAGCGTCCTGTGTCCTCGAGAACTGGCGGAGCACGTCTCAGCACATAAGCCTCATTTTCCCCTGTGGGAGGCCCAGGTATCCACAGTCACTCAGCTCTCAGAAGGGCCACCACAGGGA | 120 |
| chr11 | 70835495 | 70835615 | 788855_47188552_ENST00000601538.6,E | 1 | - | good | CTCTAGTTCCTCCCAGGGACTGGACATCTCTTCTCATGGTGCAAGATCACCCTGGCCCCAGGGTCCTCCATAAGCCCCCAGCCCACTTGGCCTGCACCTCGGCCCCTGGTTGTGATTCCT | 120 |
| chr11 | 70835615 | 70835735 | 788855_47188552_ENST00000601538.6,E | 1 | - | good | ACTCACATCTTCCACCACCAAGCTCAGATTTCATTCTTTTTATAAGAAGCTGGAACAAAGGACCCCATCAGGTCCTGTGGAAAGAGAATGGCTCACTCCAAGAGGGGTCCTCCAGCTCAG | 120 |
| chr11 | 70835735 | 70835855 | 788855_47188552_ENST00000601538.6,E | 1 | - | good | CTAGCCTGTAGGATTCTCAGAGACCTCATGAGACCTCAGCAGAATGAGGCTGGTCACCCTATTCTACAGATGAGAAAACTGAGTCCCAGAACCCTGCCTTGAGTCAGAGACTGACCCAGG | 120 |
| chr11 | 70835855 | 70835975 | 788855_47188552_ENST00000601538.6,E | 1 | - | good | AGCTGCTGCCCTGCTCGGGGTGCAGTGGGCTCGGGGTGCCCACTCCCCCAGCCTCCACTGAGGTCCGAGAGGGAGCCCACTCTGCCCCTAGATCTCCAGGCTTCTCCTGCCTGGGTTTGT | 120 |
| chr11 | 70835975 | 70836095 | 788855_47188552_ENST00000601538.6,E | 1 | - | good | CTGGAAACGGAGCGACACTCCTACCCACATTTCAGGAGTCCCTCCTGCACCTTGGTGGGCTCTCTTGTATGGGAGAGGAGGTGGTGGGCAGCCCACCCCAGTGTGGCTTGAGACAGGGAG | 120 |
| chr11 | 70836095 | 70836215 | 788855_47188552_ENST00000601538.6,E | 1 | - | good | CTGAGTGTGCAGCCGCTCAGAAAGAGGAGGCAGCAGGCACGCTCCCTTAAGGACTGAGAGCTGGGCCTTCCCCTGGCCCCCCATGGGCAAGCAGGCCATGTTGTGAAAGGGTGTCTGTGG | 120 |
| chr11 | 70836215 | 70836335 | 788855_47188552_ENST00000601538.6,E | 1 | - | good | GCAGCAAGGGTGGAAGCGGGCACTGTCCTCACCCCCAGTCTCGCCCAAGCTTTCCCTCTGTGCCCAGTGGCAGCTGTGTGGAGCCGGCCTGAGAATCAGGGCTAAGGGGCCATGCTGCTT | 120 |
| chr11 | 70836335 | 70836455 | 788855_47188552_ENST00000601538.6,E | 1 | - | good | GGCACTTTGTAAACGTTGACTTGCACAACTGCTGGCCTTCAAGACAGCTGCTGTTATTATCCTTTCTCCAACTTGCAGGCTCAGGCTCGGACAGGCACAGCCACGTGCCTGAGGTCTCAA | 120 |
| chr11 | 70836455 | 70836575 | 788855_47188552_ENST00000601538.6,E | 1 | - | good | GAGGCAGGCAGGGCCCTCCAGACAGCGTCTACCCATGGGGGAGCAACAGGAACGATGTAGAGGTCCAGGCCTTGGGTGGCACCCGGGGCCACCCACAGGAAGTCCTTTCCCTCCATACCT | 120 |
| chr11 | 70836575 | 70836695 | 788855_47188552_ENST00000601538.6,E | 1 | - | good | TGAGACACCCAGGCGTTGGGCAGCTGCAGTGGTTCTTTCTGTGTGGTCCTGGCATCTGCAGTCTGGCACCAAATGCCCAGGCGGTGAGGAGCCTATGAACAGTGAGCAGAGGCTGTGATG | 120 |
| chr11 | 70836695 | 70836815 | 788855_47188552_ENST00000601538.6,E | 1 | - | good | TAAACCTTCAGGCCACAGGACTTGGTGACTGTAGTCAGCCTCAGATGGTCTCAGGTGCTGCTTTACGAAAGTCAGTCTGTGAGGGGAGCTGCTGGGTGGGCCTAGTGCTTGAATGTGTCC | 120 |
| chr11 | 70837175 | 70837295 | 788855_47188552_ENST00000601538.6,E | 1 | - | good | AGCATCTGGAGGAGCAGGAGCAGCAGAGACAATGGCGTAAGTCCCCTGGAAGCCTGTAAGGGGCAGTGGCCTGTAAGGGGGCAGTGGCATGCTCAGTCTGCCTGGTGTCTTCATTAGCTG | 120 |
| chr11 | 70837295 | 70837415 | 788855_47188552_ENST00000601538.6,E | 1 | - | good | GGAAGCAGCCAGACATGGGGAGAGCAGGCAGAGGAATGCCAGATGGAGGGTAGGCACAGCAGGGGCAGAGGTGTGTCCAGGCCAGTGGGAGGGAGCTGACAGGGCCCGTGTGGGTGAG | 120 |
| chr11 | 70837415 | 70837535 | 788855_47188552_ENST00000601538.6,E | 1 | - | good | GAAGGAAAAAGTCACAAGGGGCTAGCATAGAGATTAAGAGAGTAACCCTCTTGGACAGGATGGTGAAGGAAAGCCTCCTCATGCGTTGACCATGAGCGTGAATGCACAAGGCCTGGAGTT | 120 |
| chr11 | 70837535 | 70837655 | 788855_47188552_ENST00000601538.6,E | 1 | - | good | GGTGATGCAGTGGTGATTAAGACAGACTGCACTTGTCAGCAGTTTACAGTTTACTTGGGGGAGAAAGACCATATAAGCAAACTTAAAACTAACATAACTTCAATGTTCAATAAATACAAT | 120 |
| chr11 | 70838015 | 70838135 | 788855_47188552_ENST00000601538.6,E | 1 | - | good | GTGTTCATGGTGGGAAGACATTTCACACGGTGAGGCAAACACCATACAAGTCCAGGGTGGGAGGTAGGAACGTGCCTTCCTTTTATTTATTCATTTATTGGATGAAGTTACTCAGCATGC | 120 |
| chr11 | 70838135 | 70838255 | 788855_47188552_ENST00000601538.6,E | 1 | - | good | CATGTATTGATAAGCTCCCACTTTGTGCGGGGTCTTCTGTTGTGTCCTGAGCATAGAAAACTGTGTGAAGGCCCCTCCCCTGCCCCAGGTCTTCCAGAAGGCCCCAGTTGTAGGGAGGCA | 120 |
| chr11 | 70838255 | 70838375 | 788855_47188552_ENST00000601538.6,E | 1 | - | good | TTACTAGCTAGGAAGAGAAGGAAAATGCAATGTGCAACCAAAGGAAGCCACAATTGTTAGAAGCAGCATCAAACAACCCAAGTGTTGAAGCAGGCACACAAGGACCTAAGATAACCAGAA | 120 |
| chr11 | 70838375 | 70838495 | 788855_47188552_ENST00000601538.6,E | 1 | - | good | AAGGCATTGCACTGATAGTTTTACTACCTACTTGAAAAACTAAACACACCTCAGAGGAAGATAACAGAACCTAGAGTCTCTCCGACATAACAAACACAATGACCAGCATAGAATCAGAAA | 120 |
| chr11 | 70838495 | 70838615 | 788855_47188552_ENST00000601538.6,E | 1 | - | good | GGAGCTTGATAAACATCCAGAGCTGGAGCCACAGAGGGGCCCTAGGAGTCAGGACTCCATCCCAGGACCAAATCAACAGAGACTCCCCCAACAAAGCCTCAGATTGCACCTCCACGGGGC | 120 |
| chr11 | 70838615 | 70838735 | 788855_47188552_ENST00000601538.6,E | 1 | - | good | GAGTGAGAGTCCAGGGACTCAGTGGAAAACAGTAGCTAAAAGCCCAAGTGCCTTGGGGAGACTTCAGTAATTTCTTCCACAGGAGAGATAGAGTATAGAATTGAGTCCCAGCATGTTAGA | 120 |
| chr11 | 70838735 | 70838855 | 788855_47188552_ENST00000601538.6,E | 1 | - | good | GGGGTTGGAGGTTGGGGTCAGAGTTGCCTGAGAGACTGGAAGTCAATAGAAATCCTGGAAAGTGGAGGGCCCTGAAGTCTACACATAAATTCCTTTTGGATTGTTGAATCGGCCTGTGCA | 120 |
| chr11 | 70838855 | 70838975 | 788855_47188552_ENST00000601538.6,E | 1 | - | good | AGCAGACAGAACTAGAGATGCTGGGACCCTTGACAGAAGAGGGGCCCCCCACCCTTGTAAGGCCCACATTAATTAGATTTCCCCTGACATAAACTGGAGTCTTACTGTGTTGTGGATTCA | 120 |
| chr11 | 70838975 | 70839095 | 788855_47188552_ENST00000601538.6,E | 1 | - | good | CCATAGTGATGGAGCAGCTCATGTCAGACTAACCCTCCTGTTAATTACAGTGGCAAACGCGGGACAAATTCTATATAAACACCAACACCAGCTGCTTGTAGGCCTGGGCAAGTGGCCAAA | 120 |
| chr11 | 70839095 | 70839215 | 788855_47188552_ENST00000601538.6,E | 1 | - | good | TGAAAAAGTAATTTGAAGGAGCCTATTGGATGACTTCTCCCCCACTTTACTGAATATCCGCAAGTGTGTTGTATTAAATATAATTATAAATGTATTGATGAGCCCATGACTTCTGCTTTC | 120 |
| chr11 | 70839215 | 70839335 | 788855_47188552_ENST00000601538.6,E | 1 | - | good | CCCTTTCCCTGAGGGCTTTCTAGGCTGAGTTACCAAATAAAGCAACCAAGACACCTCCAAGCACATGGTCTGAGTTCTGGAGCCCACATGTTTTACCATTGTCCTTAGTTTTATTTAGAA | 120 |
| chr11 | 70839335 | 70839455 | 788855_47188552_ENST00000601538.6,E | 1 | - | good | GACGTACCCGCCCTCTCTACCAGGGGGGTGTGTGTTGCCTCCAGACCTGGGGAGACTGCGCTGGCTGCCGGATACCACGGGCTGTGGAGTGGGGTGGGGAGGGCCACTGGCCCAGACCCG | 120 |
| chr11 | 70839455 | 70839575 | 788855_47188552_ENST00000601538.6,E | 1 | - | good | TATAAGGAAGTCCAGGTCCAGGAGCCCAGTGGCACAGGAAGGCTGCACCTCCTGTCTGCTCAGGACTTTTTGGCTTACGTAGTCAGTGGTCCCAGCACATCCTGTTCCCCTCTCTCCTTT | 120 |
| chr11 | 70839575 | 70839695 | 788855_47188552_ENST00000601538.6,E | 1 | - | good | GGGACCTCCCCTTCAACCCACCTTCACTTCCAGCCTCCTGGAGGTACTCAAACCTTACAAAAGTTGGATTCTTGTATTGAAATAAACCCCCTCCAAGGCTGTGGTCCTTTCCTGATGGGG | 120 |
| chr11 | 70839695 | 70839815 | 788855_47188552_ENST00000601538.6,E | 1 | - | good | TTGGATTCGCTAACTAGGTGAAGAACTGAGGTGGAGAACCAGCCTCACTCAGCCGCCTGGTGAGTGCTTCCTTTTCCAGGGCGGCCTGGCAGCTTTACCGGGGACACCTCGATCTAAAAA | 120 |
| chr11 | 70839815 | 70839935 | 788855_47188552_ENST00000601538.6,E | 1 | - | good | TCTTGATGCAGATGTGGTGCCTGATATTCACTATGCTGCCAGCTCGGCCCTGGGAGGAATTTATTTGAAGGTTAAATCAGTCACTGAGCCTTAATTAGTTCCCAAAATATAAATGTGCAT | 120 |
| chr11 | 70839935 | 70840055 | 788855_47188552_ENST00000601538.6,E | 1 | - | good | TCTGCCTTCAGCCCCAGGGCTGCATAGCTGCTGGCCGGCCGAGCCCACCTCACGCTTGACATTCCCTCCTCTCCATGTCCCTTCCCTTTGGGTTTTGGTTCTTCCTAGACTTTGAGGTTT | 120 |
| chr11 | 70840055 | 70840175 | 788855_47188552_ENST00000601538.6,E | 1 | - | good | CTTTCACTGCCTGGGAAGCTGTGACCGTGACAATGCCTGAGTCCTCAAAGCCCCTCACACTGTTCTTCCCAGACTGTACGCCGTCACCATGTGGGGAGTCAGCTGGTGGGCCCCACGCCC | 120 |
| chr11 | 70840175 | 70840295 | 788855_47188552_ENST00000601538.6,E | 1 | - | good | TCTGTAGTGTGCCAGGTGAGCTCTCACCTGCCCTCCGAGGCACCCACCCAACTACCTCCCTGTCCAGGAGTCGGGACACCTCTGGTTGGCTTTGACATTGAGGTGGCGATCACAGTGAGG | 120 |
| chr11 | 70840295 | 70840415 | 788855_47188552_ENST00000601538.6,E | 1 | - | good | CCTTTTGATATATGGGGAAACTGAGGCTCCAGGGAGCTGGGACTCCCAGCCAGGCAGAGGTAGCACCCAGAACACGCAGGTGGGTCTGCGGACCTGCCCTTGGGCATCTCTGCCTCATGC | 120 |
| chr11 | 70840415 | 70840535 | 788855_47188552_ENST00000601538.6,E | 1 | - | good | CCAGGAGTCAGGCCCATGGACACTCTGCTCCTCTGAGGGCTCCTCCCAGACAAGGATGCTCCTCAGAGCTGAGAGGGGGGATCTTGCAGCCTGAAGGACTTTTGGGGTGCATTCTAGCCT | 120 |
| chr11 | 70840535 | 70840655 | 788855_47188552_ENST00000601538.6,E | 1 | - | good | TGGCCACTGAGCTACAGTATGTGGCTGCCGCTGTGTGCCCTTTTATTTGTGGACAGTTCAGAGGGAGAGGGCTCCCCTCCCAGGTCTCCACCTTTTGTGTCAGTTAATGCCCCTAGAATC | 120 |
| chr11 | 70840655 | 70840775 | 788855_47188552_ENST00000601538.6,E | 1 | - | good | CCAGGAGGGAGATCCTGAGTCTGCAGGCTCCAGCCTCCCGATATGGCCCCCCAGCCCTCCCCAGGCTCCCACTTCCTGGAAGAGCCCTGCCCTGATAGCTGAGGTCCCAAGAACTCTCCT | 120 |
| chr11 | 70840775 | 70840895 | 788855_47188552_ENST00000601538.6,E | 1 | - | good | AACACAGTCAAGTCTTGATGGCTGGGGGTGGCCTCATCAGCATCTCTTAGAAGTGCCCGGTGCTGGGATGGCTGCAGCATGAGAACCCCCCACTGCAGTTCCTTGTCCCTGTCCTCGGCAC | 120 |
| chr11 | 70841255 | 70841375 | 788855_47188552_ENST00000601538.6,E | 1 | - | good | GAGAGATGGAAGCCGTGAGCTGTTGCCCTCCATGCCTGGCGTGTCACAGGCATCACCTGCTGCCACTGCCCACCTTTCACTGCGGTGATAATTGCTATTTTTTTTAAGACGGAGTCTCGC | 120 |
| chr11 | 70841375 | 70841495 | 788855_47188552_ENST00000601538.6,E | 1 | - | good | CCCAAGCACATCTTGGCCTGGCTGTGTGACCTCCATCCAGTTGCTGACCTGCTCTGAACCTTGGACTCCTCTGCTGTGTAATGGGGAAAATGGCTGTGCCCTGGGTGGCGGTGGAGGAAG | 120 |
| chr11 | 70841495 | 70841615 | 788855_47188552_ENST00000601538.6,E | 1 | - | good | AGCCAGAATTCCTGAAGAACCAAGGTGGAGACCCTCCTGGAATGTGGCACCACAGGGCTTGAGTTTCACTCTCAGCAGCTGGGTCAGCGATGGGGCCAAAGGCAGACTCCACAGGCTAAG | 120 |
| chr11 | 70841615 | 70841735 | 788855_47188552_ENST00000601538.6,E | 1 | - | good | CCTGGCTTTGCGGGCTGGAGACTCACCCCGATTCTGCAGTAGCTGGCTCTAGTGCCTAAGGGAGAAGCAGACGGAAAGGGGCCCCGGACCTTCCAACAGTGATCGCAGGCACCCAGGTCA | 120 |
| chr11 | 70841735 | 70841855 | 788855_47188552_ENST00000601538.6,E | 1 | - | good | CAGACACGCACAGTCAGGGTTGGGGTTGAGGCCTGAGAAGGTCTGTAGATCAGAAGTGGCACTGCATGCTTGGGTCCTGAAGAGAAGCAAGAAGCCAGGCAGGAGGAGCAGTCACGTCAT | 120 |
| chr11 | 70841855 | 70841975 | 788855_47188552_ENST00000601538.6,E | 1 | - | good | GGGCAGTGACAGCACAAGGACCGTGTGTAAAGGATGGTTTGGGAGGCTGTGGGCTCTGGGGTTGTCTGCCCAGGTCGGTGACAGTAGCCAAGACTTGGAGAAGATTTGGGAGCTGACGAG | 120 |
| chr11 | 70841975 | 70842095 | 788855_47188552_ENST00000601538.6,E | 1 | - | good | GGGCCAAATAGCTGTGACACCCATCCAGGCTTCAGGGCGTGTGGAGTAGAGAGAGACAAGGTCTCTTGTCCTGGAGGAGCCCAGGGTTAAGGGAGAAGGGCAGTTAGGTCCCAGGAATAA | 120 |
| chr11 | 70842095 | 70842215 | 788855_47188552_ENST00000601538.6,E | 1 | - | good | GTAAAATGTTTAAATATTTTAATAACATAATGAGATTTCATATACTTGGAAGTAAATGCTTGGTGAGTTTTCAGATATCGCCAATATTGCATTTGTTCTGTTCGTGATTTGAACAAGCAT | 120 |
| chr11 | 70842215 | 70842335 | 788855_47188552_ENST00000601538.6,E | 1 | - | good | GGAGCCACCCGGCAGGGAATATGCTTTATTCCCTGGAGTGTCTGTCTCTTTCCAGCATGGGTTTTGGCCCCAAGTCAAGGGAAGGCAAAACCATCGTCAAATTTTACTAAAAAGAAATTA | 120 |
| chr11 | 70842335 | 70842455 | 788855_47188552_ENST00000601538.6,E | 1 | - | good | TGCAGCCTGGCCAAGGTGGCACAGCGAAACCCCAGTGCTGATTGAAGCAGAGTGGGCCATGGGCCTCCATGACTCATAGCTTGACCACCTCGGGGTCACCATGGTGCTGCAGAAAGGTCT | 120 |
| chr11 | 70842455 | 70842575 | 788855_47188552_ENST00000601538.6,E | 1 | - | good | CCCTCCCTGTGCCCAGTTCTCTTCTGCTTTTGAAGCTAAAGGAGCTCCTGAAAGGGACAAGTGCAGAGAGGGCCCTAGAACACCATTTGCACATGGAGAGACAGTGGCTCAGACAGGTTA | 120 |
| chr11 | 70842575 | 70842695 | 788855_47188552_ENST00000601538.6,E | 1 | - | good | TTAAGCTGCACGAGAATCCCAGGTGGCACAGCAGGGAGCAGGACCCACTCCTGGCCGAGGGACCTAGGCTGAGGCTCATGCTCGCTCCAGGGCAGGAGCTCACGGTGCAGGGCTGCTGAC | 120 |
| chr11 | 70842695 | 70842815 | 788855_47188552_ENST00000601538.6,E | 1 | - | good | GGCCGTCTCTTGGGCCTCTGGGTGGGGAGACCCTGCGTGCTGCCTCCCACCCCCCTGGGAGCCTCCAGCAGTGTCTCCTTTAGCTTCAGGACAGGGCTCAGCTTCCTTGCCGAGTTCCAT | 120 |
| chr11 | 70842815 | 70842935 | 788855_47188552_ENST00000601538.6,E | 1 | - | good | GCCGCCCAGCCCGGCCCCCTGAGCCTGTGGAGTGGAAGGTTCCACCGTCCGTGTTATTCTCACAGATCCCCGTGCAAGCTCAGATGCTGAGAGGAAACCGTGTGGAGGGAGCTTCACCTC | 120 |
| chr11 | 70842935 | 70843055 | 788855_47188552_ENST00000601538.6,E | 1 | - | good | CAGCCCGCTCTGGGAGTCTGAGGCCCAGGGTGTGGTGTGGGCAGCTCTGTGCAGATGCGAGCCACACCTGTGTTAGGGGTATGACACTGCGTCAGGGTAGTCAGATGACACGAGGGAATG | 120 |
| chr11 | 70843055 | 70843175 | 788855_47188552_ENST00000601538.6,E | 1 | - | good | CTGAGCATCTCCACTGCAGCCTCTGCTCCCTGACCCCATGACCTGAAACCTGTACCTTCCAGCCTCAGCTCTGGGCTCACACCCCAGCCTTCCACATGCTGTGAGAGTCCAGGGTCTGCC | 120 |
| chr11 | 70843415 | 70843535 | 788855_47188552_ENST00000601538.6,E | 1 | - | good | AATGGATTTGGGCCCACCCTAGTGGCCTCCCTTTAACTTAACCACTGTGAAGGACTCTGCTTCACATGCAGTCCCACTAGGAGTCAGGACCTCACAATACGAATTTCAGCTCTGTCTACA | 120 |
| chr11 | 70843775 | 70843895 | 788855_47188552_ENST00000601538.6,E | 1 | - | good | TTCCGACGGAAGCACCAGGAGATATTTGTGAACTCCTGGTCCCTGAGCACCAAATTGGGAGGGCTGGAGCAGTTCTGTTTTAGGGAAGTTGACTCGTGTGTCCTCTAAGAATCGGTTGAG | 120 |
| chr11 | 70843895 | 70844015 | 788855_47188552_ENST00000601538.6,E | 1 | - | good | TCCTACCCTAGGAAAACTGAGGGCTCACTGGAGAATCGTCCACGAAGGCCGTGTCCCCAAACTGAGGAGGAAGATGAAGATGGTATGGCCACCCACCCAAGGAGATGGCATTCACAGGCC | 120 |
| chr11 | 70844015 | 70844135 | 788855_47188552_ENST00000601538.6,E | 1 | - | good | GAAGCCAGGCGAGGGCTGGGTGTGTGTGGATGGATGGAGGCGCCCACTCATCCCCCAGGGAGCTCCCACCACCTTGGGGACTCAGATGGGACAGGGGATGGTGCACGCAGCACAGTGTGG | 120 |
| chr11 | 70844135 | 70844255 | 788855_47188552_ENST00000601538.6,E | 1 | - | good | GGGGCATTCATTAGATTTTTCTTTTTAAGCAGAGTATTAATGAGTTGAACTAATGTTGTAGGGGTGCCTGTGGCAGGGAGAACACACTGCCAGAAGAGTCCTCCTGTCCAGAGATGTGGG | 120 |
| chr11 | 70844255 | 70844375 | 788855_47188552_ENST00000601538.6,E | 1 | - | good | ATTTTCATGCCCCGCCCCCGGCCCCCCAACCCAAATGGAAAAACGCATGGCAATTTCAGGTAGAAACTGAACAGCTTGAATGCTATTCAAGGCATGCTCTGCTGACCCAATTAGGAAAAT | 120 |

| | | | | | | | | |
|---|---|---|---|---|---|---|---|---|
| chr11 | 70844375 | 70844495 | 788855_47188552_ENST00000601538.6,E | 1 | - | good | CCCCAAACCCCATGGTTCCAGTGAGTTTGTGCTATGTCGAGAAACGTGCCCAGCGTTTTAGATGATCCCCTTGTCGAGAATGCTGGCTGTGAAGGCTGGAGTAAGGATTTAGGGGGAGAG | 120 |
| chr11 | 70844495 | 70844615 | 788855_47188552_ENST00000601538.6,E | 1 | - | good | TATTACGTAACATCCCACGAGATTACTCAAAGGGCTGCAGCAGGAGGGCCAGGTGCTCAGTGTGGGGAGTGAAGGGCAGTGCCCGGCGTGGAGCCTGTGTTTCCAACTCCTCCACTGCGG | 120 |
| chr11 | 70844615 | 70844735 | 788855_47188552_ENST00000601538.6,E | 1 | - | good | GAAAATGTTCTCCGCCGTGTTATTGATCATGGGGAAAGTCGAAAAGCATCTCAGCTGGACTGAGATAATCCCTGTCTGAGGAATTTGTCATGTAAAAGAACAGCATGTATCCTCCAGGAA | 120 |
| chr11 | 70844735 | 70844855 | 788855_47188552_ENST00000601538.6,E | 1 | - | good | CTGGTGAAAGCAATTGGCCAGACACGTGCCCAGGGCCATGGGGGCGCTCAGACCTTTTGACCTGGCATTTCCGCTCCTGGGAATGTAAAATATTCAAATGAAGAAAGGTGCCACCTGCGT | 120 |
| chr11 | 70845215 | 70845335 | 788855_47188552_ENST00000601538.6,E | 1 | - | good | CCACTCCAGTGGAAAGTCCTCTCATCTTTGAAGTTTATTGTCCTCTGGTTGAAACCCATTTTCTCTATACCTCATTTTAGAAAAAGATTTTCTTTTCTTTTCTTTCTTTCTTTTTCTTTT | 120 |
| chr11 | 70845335 | 70845455 | 788855_47188552_ENST00000601538.6,E | 1 | - | good | AATGCCATGGACTTGGAGATTTGGAATAGGCCTGATGTTATCAGAGTCCATTTCTCTGTGAGAAGGAGAAACTATTCTGTTCTGATGAGTCCTTCAGCGTGAAATTATGTCTCCCCTAAT | 120 |
| chr11 | 70845455 | 70845575 | 788855_47188552_ENST00000601538.6,E | 1 | - | good | GGCATCCCCACCCATTTCTTCCAACACCGCCTCTCTATCCTCGGCACCTAAGATGGCCCCGTGCTGGCCCCGGGGCCAATAAGCACATTCCATTGGAAGTTCACGAAGCCGAGTAGGAAA | 120 |
| chr11 | 70845575 | 70845695 | 788855_47188552_ENST00000601538.6,E | 1 | - | good | CCCTGGCTTGGCTCGTCTTTCCATCAATCAATCCTGGGTTTTATTGGCATCTCAGTGTCACCAGGTTGTCTGCCATTCCTGCCTCTGGTCTCTGGGAACGGTCTTTGTGGGTGGTTTGGA | 120 |
| chr11 | 70845695 | 70845815 | 788855_47188552_ENST00000601538.6,E | 1 | - | good | CAAACAACTTGGAAGTGGGCTCTCCTTCTCGCTTTGTTCTGTGCTGCTTATTTCCATCTCTATGTAATTATGGGAAATGTGCATTCATTCACCCGGGGCCGTCTGTTTCATTCTGATTTT | 120 |
| chr11 | 70845815 | 70845935 | 788855_47188552_ENST00000601538.6,E | 1 | - | good | ACGGCTCCGTTTCCTGCCCCCAGTGCGTTGCTTGTTGTTTCATTGCAGGAAGCTCGGAGACAAACACTGGCGGTTTCTATCCAGCACCACCTCTCCGGTGTCTATTTTCAGCCTCTGCTG | 120 |
| chr11 | 70845935 | 70846055 | 788855_47188552_ENST00000601538.6,E | 1 | - | good | CCCAGTTCAGGTCTCTCTGGGCTCGGCACCAGCCTCTGGAAACCTTGGCGGCTGCCCTTGGCCACCTGTTTCTCCAGTGTCGGGAGACGGGGAACAGCGTCTCTGGGGTGCTTTCCTCTG | 120 |
| chr11 | 70846055 | 70846175 | 788855_47188552_ENST00000601538.6,E | 1 | - | good | CGGTAGAAAGACCCCACCACCGGCACCCTCTTTAGCTGACTTTGCCTGGGGGTGCTGTCCCACTGGAAGCGCACTCTAGACTCTCCAGTTCGCTTTTCTGCCCTTCCGCTTTCCCAGGGA | 120 |
| chr11 | 70846175 | 70846295 | 788855_47188552_ENST00000601538.6,E | 1 | - | good | CGAGACCGTGTTTCTAATAATAATAATAATACTAATTTACAGAAAAATAACCTGAGGCATTGAAGGCTCAGTTGTCCTCCTAGGAACCCCTAAGAAGAGGGGTGCCTCCGTTTTCTGCCA | 120 |
| chr11 | 70846535 | 70846655 | 788855_47188552_ENST00000601538.6,E | 1 | - | good | TTAGGCACCTGGGGTCTGTGGAGTGCTGAGAGCAGGGCTTGGGACACCAGCAGTTGACCTGTGCTTCCCAGTGTCATCCTTGTTATTGATTGACAGATGAGGCCGAGCGCAGCAGCTCAT | 120 |
| chr11 | 70846655 | 70846775 | 788855_47188552_ENST00000601538.6,E | 1 | - | good | ATTGGAACATAGACCTCGGAGCACTGGCTCCTTCCCAGCTGTGTGGCCGGGAGCAGGTGACTGGACTATCTGTGTCATGGGGATGATGACGCTGCCTTCTGTGCTGAGATGAGGTGGCCG | 120 |
| chr11 | 70846775 | 70846895 | 788855_47188552_ENST00000601538.6,E | 1 | - | good | GGCCTGGGAGGCACCGAAGGCTCTCCATTGGATGACCTCGTTCATGCTCACAGCACCCAGTGGTCCTGTTCTAGCCCCCATTTTTCAGAGAAGGGTGGGAGGGAGAGTGTGGTGCTTTAC | 120 |
| chr11 | 70846895 | 70847015 | 788855_47188552_ENST00000601538.6,E | 1 | - | good | CCAGCTGCAGTCCTGCCCATCCTCAGATCAGAGAGCAGCAGCCAGTCCCCGTGGCAGCAGGCTCTTTTATGAGCACAGAGAAACTCTTAAATAGCACCACGGGCCTTGAACCATGCTCTG | 120 |
| chr11 | 70847015 | 70847135 | 788855_47188552_ENST00000601538.6,E | 1 | - | good | TCAAGGACCCTCGCAAGATTGATGATGGGCCGGGGCAGAGGTCCCCTCCTCTTGATGGGGGCCTCCACCTCTCCTCTGTAAAGTGATGCCATCTTGCTCAGGGATGGGAATGCCCAGGCC | 120 |
| chr11 | 70847135 | 70847255 | 788855_47188552_ENST00000601538.6,E | 1 | - | good | ACGGCCTCCCGCGGGGTGCCATGGTGGGACTCCATAGCTGAGGCCTGGAACGTAGACCTGTGTCTCTGTCTCCCTAGCACACTCCCAGGACCCTGTACTGGGGTGTGGGTCGCAGCACAG | 120 |
| chr11 | 70847255 | 70847375 | 788855_47188552_ENST00000601538.6,E | 1 | - | good | AACCCCGTCACCACCCTGTGAGACAAGGCTGCATTTGGGGGACCCCGTCTCATTAGAACCTCAACAGAATTGCAGCGGTAGGAGGCGTGGACCTCATCAAGTACAGAGTGGGCAGGAGGG | 120 |
| chr11 | 70847375 | 70847495 | 788855_47188552_ENST00000601538.6,E | 1 | - | good | TGGCAGTCAGGCACATTTCCCCCTATTTTTAACGCTGACAGCAGATCCTGGGATTGACGGTTTTGCCTCCTGGCATCACTTTCTTGCTGAGAATCTCAGAGAGGAGGCCCTGCTGCTGCC | 120 |
| chr11 | 70847495 | 70847615 | 788855_47188552_ENST00000601538.6,E | 1 | - | good | CTGAGCGCAGAGAGATGCTCGGGCAGGTTTTCAGTGCTGGTCCCCCGTTCCAGAGCAATTTTAAATTAATTTAAAATTCTGAGTTGGGTAGGGGGGCAGCACCCTCCCTGGCAGGGGATC | 120 |
| chr11 | 70847615 | 70847735 | 788855_47188552_ENST00000601538.6,E | 1 | - | good | GGGGATGTGGGAGAGTTTGCGTGGTAGACTCTGCTTGTGAACTCGGAACCTGAGTGGAGTGAATTGAGAGGGAGCATGGGAAATTCAGGAGCAGAGATGAGACCCTGGATGGCGTCGCTG | 120 |
| chr11 | 70847735 | 70847855 | 788855_47188552_ENST00000601538.6,E | 1 | - | good | CTTTGGTCCAGGCCTTGCTGGGCAGGCCCTGGACTGGCTCTTGGCCCACCCCTGCAGCCAGATGATGAGGACTGGGTGGGTGGGGGCACCCCAGGCAAAGCCAGGCTGGTTCCTCTCTGT | 120 |
| chr11 | 70847855 | 70847975 | 788855_47188552_ENST00000601538.6,E | 1 | - | good | TCCAGCCACGGTCACCTCAGCAGCCCTGGGCTCCCCAGGTCTCCTGTAGCCGTGAAGCCTGGGAGGACGTGGTCTCCTGGTGGTCCTCCACTGGGAACCGTCTCCAAGCCCTGCCCCCCT | 120 |
| chr11 | 70847975 | 70848095 | 788855_47188552_ENST00000601538.6,E | 1 | - | good | AGCCCAGCGGACCTCAGACTCCTTTTGGGAACTGGGAATAGGTGTGGGGCCGCCCTTCCTACCGCAGGCATTGCCATGCCTTCTAGCAAGATGAAGCCACACCCCAGCTGGCTGGTCCCC | 120 |
| chr11 | 70848095 | 70848215 | 788855_47188552_ENST00000601538.6,E | 1 | - | good | ATCCAGAGTTCTGGCTTCTTTTTAGGCAGCAGTGATAAGATTGCTGTCGCTAAGCCTCCGTGGAGAGAAGGCTGTCCTGAGAGGGCCTTTCGTCTTCTGTTTCATTGGCTGTGAAATGA | 120 |
| chr11 | 70848215 | 70848335 | 788855_47188552_ENST00000601538.6,E | 1 | - | good | CCCCCACCCCACAATTCCTGGCGGCTCACGGGCAGATTCTTTTAGCCCATGCTGTAAGGCACATTTTGATAAAATAGGAACTCGTTGCTGGTATTTAAAATCAGGAGATTTCACATAAAA | 120 |
| chr11 | 70848335 | 70848455 | 788855_47188552_ENST00000601538.6,E | 1 | - | good | CCTATTTTGCACAGTCATCTTCAAGGGTCTCCAGGGTCCCGTGGAGCTCACTTTGGGAAGTGGCGGGAACGGGTGGGTTGATCACCCCCGCCAGGCCCTGACATTGCCCGTCTTTCAAAC | 120 |
| chr11 | 70848455 | 70848575 | 788855_47188552_ENST00000601538.6,E | 1 | - | good | CTTTAAGGTCCTGATAAGGCCTGCAGTGCACTTTAGGGCCTTTAAGGTCCTGGTAAGGCCTGCAATAAAAAAGCCCCTTAGCTTTGTTTGATCTGGCGTTTCCCAAACATGACTGTAGAA | 120 |
| chr11 | 70848575 | 70848695 | 788855_47188552_ENST00000601538.6,E | 1 | - | good | AGGTGCTGGATTCTAGTGAGTGCCGTGGCTGCTCTGCAAAGCACAGGTCTGTGGTCAAATCGCATTGGAAGCGCTACCTGCTTTACTTCCTTCCTGGACCCTCAGAATGTACTTAAGGGC | 120 |
| chr11 | 70848695 | 70848815 | 788855_47188552_ENST00000601538.6,E | 1 | - | good | TAATTAAGATCACAGCCAGACTGCACAGTTGAGGGTGGCTTGCATCACTGGACGTGGTACATCCCTGCCCAAGCTTCTCGGACCCTGAACTGCACACTTGCCATTTACCTTTCAAGGTAA | 120 |
| chr11 | 70848815 | 70848935 | 788855_47188552_ENST00000601538.6,E | 1 | - | good | TGTCTGTATGGTGCTCAATTTGTCTCCCAGCATCATGCACTTGTTGACATTCCTGGTGGTCCTGAAACAAGCTGGCATCAGGCTTGCTTGCTCGGGGAAAGGACTGGCTCTCCCATTCTG | 120 |
| chr11 | 70848935 | 70849055 | 788855_47188552_ENST00000601538.6,E | 1 | - | good | TCATATTTTCAGTGGCCCTGCAGCCAGCATCCCTGGGCACCACTCACATCCTTCTCTGACTCATTCACGCAGCATCAAGCCTCAGAGCAAGGTCCCCCTTCACCCACTCAGAGGCCCAGC | 120 |
| chr11 | 70849055 | 70849175 | 788855_47188552_ENST00000601538.6,E | 1 | - | good | CACCCTGCCTATGGGTGTATGTATCTTGTTGGTAATTGTGCCTTGTGTCCTGGTGGGGAGGGGGTGTGTGCCCCTATGTCTGAACATTCCGGAGTGTGAGGTCCAGGCCGTTTCTACCAT | 120 |
| chr11 | 70849295 | 70849415 | 788855_47188552_ENST00000601538.6,E | 1 | - | good | CATTTGTGCACCCGTAGACAGAATTCACACAGATGTGATCTAGGGAGAATAGATTGTGGATTGGTGTCTTTTTCTCTACCTCACCTTTTTAGACCAGCCATTGGCATGCATGCATGTGTA | 120 |
| chr11 | 70849415 | 70849535 | 788855_47188552_ENST00000601538.6,E | 1 | - | good | GTACAGTTGGCTCTGGGGTCCACCCTGAGCCCAGGCACCCACTTCCTACACTAGCCTGCCACAGCCTGCTGGCCTCCAGTGCAGAGAAGGACTTCCAGGTTCCATGTCCCTGTTCAGCCC | 120 |
| chr11 | 70849535 | 70849655 | 788855_47188552_ENST00000601538.6,E | 1 | - | good | GAGAGGAAGGAGTATGGCTACTGCATGCTAACATGTATATTAGGGTGTTATGTTCTCACACAGTGATCCCTTGTATTTTCATTTGACAAAGGGTAGGCACTTGCCCAGGGTTATGCTCTA | 120 |
| chr11 | 70849655 | 70849775 | 788855_47188552_ENST00000601538.6,E | 1 | - | good | CATCCAGGAGACTTGATAAATGGCAGCTATTAACAGATTTGACCCAACACTGCTGTTTGCATATTTTTTTTACTGTGGTTTCATTCTTTTGACATAAATGCCCTCATAAGAAAATTCAAA | 120 |
| chr11 | 70850135 | 70850255 | 788855_47188552_ENST00000601538.6,E | 1 | - | good | TTTGGCTTCTCTGATCCCATCTGGTCACCTGAGTACTGACCGCTGACACTGTTGTGAAGATTGAAGGAGAACCGTAGGAGGTTGTGATACAGGCTACAACATGGAGAGACCTTGAGGGCA | 120 |
| chr11 | 70850255 | 70850375 | 788855_47188552_ENST00000601538.6,E | 1 | - | good | TCAAAAACCACTGCTTCTCTGGCCTTGGGGGCAGGAGGTATAACACCTGGGGTGGAAGCTTGGAGACAGACATCTCTAGGTTCAAGTTTCAGGTGCAGCTGTGTGATCTTGGACATGTTG | 120 |
| chr11 | 70850375 | 70850495 | 788855_47188552_ENST00000601538.6,E | 1 | - | good | AGGTGCTTTAGGGACTGTCTGATGGGCAGGAGGGAGAGCTGGTAAACCACACCACCCTGGCCCCCAGTGAGTATGAGCATCTCATTTGGGTAAAGTGAGCCAGTTGCTTGAAAAAATGCA | 120 |
| chr11 | 70850495 | 70850615 | 788855_47188552_ENST00000601538.6,E | 1 | - | good | GAATTTGGGAGAGAGCTCAGCTCGGGCGTGCATGGTGAGGACACGTGGCTCTTGGCAGAGGGGAGGCTGAGGATCCATCCTTGGTATGGTTTCTACACACTGTCCCAGAGAGTTCCTCAC | 120 |
| chr11 | 70850615 | 70850735 | 788855_47188552_ENST00000601538.6,E | 1 | - | good | GATTCAGAACTGACTAGAAGGTCCGGGTGCCTCAGAGTCATAAGTTACTCTCCCCAGGAGCCCACGGGAGCTGCAGGCACAGTGACTCGAGGCCCAAGCCCCCACTACACTTAGATCTTG | 120 |
| chr11 | 70850735 | 70850855 | 788855_47188552_ENST00000601538.6,E | 1 | - | good | ACTCTCGGGTACTTGTTTCAGCTGGTCTGGACTCAGGGACCCCTGCCCAAAGCTGAGATCAGGAGCTGCCTTGTGAGGTCACCTTTCCCAGCTTCTCCCCATTCGCCAGCAGTCTGATGG | 120 |
| chr11 | 70850855 | 70850975 | 788855_47188552_ENST00000601538.6,E | 1 | - | good | ACTCTACCCCTGCAGCAAGCATGAAGAGTGGCCGTGAGCCCATTTTAAAAGTGGAAAATGTGTGGCTTGGAGGGCGAAGGGGTTGTCAGGGCTGCAGAGCCCAGCAGCACAATTGTCAGG | 120 |
| chr11 | 70850975 | 70851095 | 788855_47188552_ENST00000601538.6,E | 1 | - | good | AGGAATTCTCACCAGGGACTGACAGTCCCTACAGGGCTGGGTCCCAGCCTTCCTGTGAGAGCCACAGCCACTGTTAATGGGTCACCTTGGGGTACGGATGCTCCAAATTTGTGTTGGCTC | 120 |
| chr11 | 70851095 | 70851215 | 788855_47188552_ENST00000601538.6,E | 1 | - | good | AGATCTTTTCTGAAGGAAAGTCTGTTCTTGTTCTGCAAGGCTCTGGTATTTGGATCTGAGAGTTAAGGGCTCCATGAAGTGGAAAGTCCAGAGAGTAGGGAGGGATCCTGGAGAGGATTC | 120 |
| chr11 | 70851215 | 70851335 | 788855_47188552_ENST00000601538.6,E | 1 | - | good | CAGCCAGGTGACCAGATGAAGGGCCAGGTTACATTTCAGAGGAAGACAGGAGAACAGGAAGTGGGGGACAATTAGGAGCCCCAGCTACAACAGCCCTTTCTTGTGCCTTTGTTGGGGAGA | 120 |
| chr11 | 70851335 | 70851455 | 788855_47188552_ENST00000601538.6,E | 1 | - | good | ATAAGGACACTTTCCTTAAGTGGACCTGTTTCTTTTATTTATGTCCCAGTGGTCAGGATCTTATCACACAGCTCCCTTCTTAAGTGGACAGCCATGTTCTTAAACCTTCTTCTTTGTGGA | 120 |
| chr11 | 70851455 | 70851575 | 788855_47188552_ENST00000601538.6,E | 1 | - | good | GACTCCGTGCTCAATTGCCCTCATAGTCCAAGGCGGCTGCCTCACCTCCACCCATTACATCATCATTCCAATCAGCAGAAAGGAGAAAGGGGCAGGAGAAAGGCATTATTCCCTTTTCCT | 120 |
| chr11 | 70851575 | 70851695 | 788855_47188552_ENST00000601538.6,E | 1 | - | good | CTAAGTTTTTTAAAAAGTTTGTCTTTTCCATAAAAGTCCAGCTAGGTGGTCCATAACTGAGGCAGGGTTCTGTGGTGTCTGAGACGCATGTTCTTCTGACTTACTGCTTCATCATGTGTG | 120 |
| chr11 | 70851695 | 70851815 | 788855_47188552_ENST00000601538.6,E | 1 | - | good | AACTGGACCCCAAGCTCTGGGTCCTTAATAACGATGCTACTGCTCTCTCAAGTATTTCTAAATTGGCATTGCTAGCCTCTAATAGTCATGGTGAGATTTGGCTGTGAGTAACTAAAAATC | 120 |
| chr11 | 70851815 | 70851935 | 788855_47188552_ENST00000601538.6,E | 1 | - | good | CACATTCTATCACAGTCGTTTATGAGTCTGCCTCACTTTACCAGCCTGGGGGCCGGGAGGCTCTGACTTTTGCATCCACTCAGGATGATTAGCAAGAGTTTGAACCTCCACGCATGGTCC | 120 |
| chr11 | 70851935 | 70852055 | 788855_47188552_ENST00000601538.6,E | 1 | - | good | CCCAGAAGGTCTTGGTTTTAAAAGGGACTGTGTTTTATTTTCACGGCTTTCAGGAGTAGGTGGGAATGGGAACAGACTGCCTGGCAGAGTTTCATGGGGCAGTGATTCTTATTAATTGAT | 120 |
| chr11 | 70852055 | 70852175 | 788855_47188552_ENST00000601538.6,E | 1 | - | good | GGTGGGCCGTGTGTCATGTCTGAGTCCGAGCTCTCTCTCGCCTCTGCAGACCCTGCCCACGCTCCTCCCCGGCGGCCAGGGGACTCCTGCAGTAGTCCTGCTTCCCTGGGCACAGAGTGG | 120 |
| chr11 | 70852175 | 70852295 | 788855_47188552_ENST00000601538.6,E | 1 | - | good | TCTCCCAATATTCTTTCTTTAAAGCTTCTAAGCAGGAGCCAGTTTTCTTTGTGGTCTGTACAATCTGGCACTGTGCAAATGCTTTGGAAACAGCAGGATGAGGAGCTATATGCTCTAACA | 120 |
| chr11 | 70852295 | 70852415 | 788855_47188552_ENST00000601538.6,E | 1 | - | good | CCACAAACACAACACCCTTTGCAGGCCCCGTGGAGCCCCTGCATGGCCTTCTGAGGAAACTTCTTTCTGAAGGGCTTCTCTGGGATGGCCTTGAAGTTGACTGGGGTTTACGTTCCTCGC | 120 |
| chr11 | 70852415 | 70852535 | 788855_47188552_ENST00000601538.6,E | 1 | - | good | CCCCCAGAAACCTGGGGACCTTCGTGGAGTGCAGACTGTGGCAGTGTCCTGCTGGAAATGGACAGCCAGCCCGTTCTTCCTGCCCTTCCCCTGCAACCCTCCTGCCCTGGCCTCTGCATC | 120 |
| chr11 | 70852895 | 70853015 | 788855_47188552_ENST00000601538.6,E | 1 | - | good | GTTGTGGGGCTCTGCAGAACTTCCCACGTGGCCCACTGGTTCAGGGAACGTTGCCTTCCTCTGGCTGACAGTGATGTGGAGACTCAAGGCCTTGCCAAGGCAACCCCTCCCCTGGCCTCT | 120 |
| chr11 | 70853015 | 70853135 | 788855_47188552_ENST00000601538.6,E | 1 | - | good | GACCCTGCCAAGGATTGCTGTCTCCCGTAAACCCAATGCCAGGCACTGAAGGTGTCTCAGCAGGGGGCACCCCAGATGGCCTGTCGAGTGGGCCTGAGGGCTCTGCTTCCCCAGGGCAGG | 120 |
| chr11 | 70853135 | 70853255 | 788855_47188552_ENST00000601538.6,E | 1 | - | good | GAGTTTCCAGTGTCTGCTCCCTAGAAAGAGTCTGGCTCTGTTTCCAGAGCTCACGGAGTGTTGCTCATGTTTCATTTCCCCCAACTTCACTCTCCCCACAGGCTCTAAGGAGCCCAGGAG | 120 |

| | | | | | | | | |
|---|---|---|---|---|---|---|---|---|
| chr11 | 70853255 | 70853375 | 788855_47188552_ENST00000601538.6,E | 1 | - | good | GTCCCCACGGCCGCTGTGACTCACACCCTCTCCCGCACTTGGACTCTCCCAGGAAGCTGTGCTCTGAGCTCCAGGGCCCGTGGGAGGCGCCGGGACGCCTGGTCCAGGTCCACCTGGCTG | 120 |
| chr11 | 70853375 | 70853495 | 788855_47188552_ENST00000601538.6,E | 1 | - | good | CTGAGAAAGACGCTTAAGGCTGCACCCAGAACTGAGCACCAGTCCGCTTCAAACCTCTCTCGGTCACTGGGTGACAATGGCCGACGCACTCAGCCTCACTGCAGCCTCAGGGTCTTGTCT | 120 |
| chr11 | 70853495 | 70853615 | 788855_47188552_ENST00000601538.6,E | 1 | - | good | AGGCAGATGCCCAGGAAGCATCCTACACACATCACACGCTCCGGGCTGGTGGAGCTGTTGGGAGGAGACATTTGCCTTTGCCCTGCCTGCTGAGAGGCATCCCCCCTGGCCATAAAGAGA | 120 |
| chr11 | 70853615 | 70853735 | 788855_47188552_ENST00000601538.6,E | 1 | - | good | GAGAGAATGAATTTCTGTTGTCTGAAGTCACCCATTGTGTATTAAGCGGTCACGGCAGCTCCAGGAACATGGGACGCATCCTGTATTTTCACTCTGTCTCCCTCCCGTTACTGAGAGTGG | 120 |
| chr11 | 70853735 | 70853855 | 788855_47188552_ENST00000601538.6,E | 1 | - | good | CACACACAGACGAAGGCAGAGAATGGAGTGGAAGAGGCAGGGAGGGTCCTCTCCTAGAGGCCCTGGAGGGAGGAAGTGCTGCCACAGCCCTAATTTTGGACTCTTGGCCTCTAAGATTGT | 120 |
| chr11 | 70853855 | 70853975 | 788855_47188552_ENST00000601538.6,E | 1 | - | good | TTGGTTTGGTTAACATGAGGTCATGCTGTATTAGTATGACCAATGGCTGATGTCTTTATAAGAAGAAGGAGGGGGGATTCAGACACAGGAACCCACTGAAGCACACCAAGAGAGGAAGGC | 120 |
| chr11 | 70854095 | 70854215 | 788855_47188552_ENST00000601538.6,E | 1 | - | good | ATTTATTTGCCTCTGAATAGAGAGGTAATATGTGGTCTTCGTAGAAATTACACAAAAGCAAGGGGTAAATGCGTGTACATATGAAAATCATCCAAACCTTCTTCTCCCCCACTCCCCAAG | 120 |
| chr11 | 70854215 | 70854335 | 788855_47188552_ENST00000601538.6,E | 1 | - | good | CTGCTGGACTCACTAGATTCGATCTCAGCATTTTTAGCATTGGTAACTTTCCTACTCATGACAGTCAACAAGGACCTTTGAGATCTCAAGTAATTTTTTCTTAATCAAAAATATACTATT | 120 |
| chr11 | 70854335 | 70854455 | 788855_47188552_ENST00000601538.6,E | 1 | - | good | CACTTCACTTTCCTGTTGCTCAAGCTCTCAACTTTGTTGGCTGGGCTGAGTGCTAGTAACAGCACTAGCCACAGTTACAAGGCACTGGGTTTCTTGGGCCATGAGCCAGCAGCCTTCTCT | 120 |
| chr11 | 70854455 | 70854575 | 788855_47188552_ENST00000601538.6,E | 1 | - | good | CGTGTTTCACCCTCTGGACCTTGGTCCCTCTCCCTCAATCCAGCCATGCATGCGCTCAGTTGGATTTCAAGCTACAGGGCTGAATGTGGGCCACTGGGGGTGGAATCTGAGGCACCTGCT | 120 |
| chr11 | 70854575 | 70854695 | 788855_47188552_ENST00000601538.6,E | 1 | - | good | GTGCTTGTCAGCTTTGGCGGGACCTTTCACTGTGTGGGCCTGCATCCATCTTCCATTTGGGCCACTTCTGCATCTCCTTCCCCAGAAGGCCCTTCCTTGCCTGTGCCTGGATGGAAAGCA | 120 |
| chr11 | 70854695 | 70854815 | 788855_47188552_ENST00000601538.6,E | 1 | - | good | CATGGGGCAGTGTACTCACACCCCAGCAGCTTTGCCCCCTGTGGTGCCTGCTTCCAGCTGTGGGTGTTTCTCCCTGGGCTCCCCATTGAGAACAGAGAGGGACATTTCACTGGCCTTCAT | 120 |
| chr11 | 70854815 | 70854935 | 788855_47188552_ENST00000601538.6,E | 1 | - | good | AAAAACAAAAAGTACAGCTCTTGCTGTTCACTCCCTGCGTGCAACACAGGTGTAGTAGGGAGACCAGTGCCTAGTCCCTGCTCTGTGGTGGACAGAGGGCTTGTCCAGGACCCCAGAGGC | 120 |
| chr11 | 70854935 | 70855055 | 788855_47188552_ENST00000601538.6,E | 1 | - | good | AGAGTGTCCAGATTTTCAGGACTACGGGCCGGGCCCAGACCTCCTAGATTGATGGACAGTCATGAACATCTGGGGGTGGGTTTCTGATAGTCGTGGGTGAACCGTGATCTGGATTTCTGC | 120 |
| chr11 | 70855055 | 70855175 | 788855_47188552_ENST00000601538.6,E | 1 | - | good | TTATCCCTCATCATGAAGTCATTGAGGGGCAGCCGAGCCTGCTGTGCAAACAGCCAGAGCCACTGGGCCATATGCTGGCATCTAGCCAAATGGCTGTCCCTTCTTATTAGCAGGTCACTG | 120 |
| chr11 | 70855175 | 70855295 | 788855_47188552_ENST00000601538.6,E | 1 | - | good | CCTCACCTGCTAGAATGTCTGTCCTGTAGGGCTTAGAATACCTTGCTAATTTCCTCTGAGCTTGTTCATCATGTGACCAAATTGGATGAGGGAGGAGAACGTGCAGGGAAGCTGTTTTAA | 120 |
| chr11 | 70855295 | 70855415 | 788855_47188552_ENST00000601538.6,E | 1 | - | good | TTGGGGGAGCAAAGAGGAGGGATCTGAAACCTTCTGTGACTGAGTTGGGCCAGACTGATGGGCAATTTGCCCAGAACCCAACATTCTAGATATGGGAAGCTTCCAGAAACCTGGCTTACT | 120 |
| chr11 | 70855415 | 70855535 | 788855_47188552_ENST00000601538.6,E | 1 | - | good | CAGATGCCCTGGTCAGAGCTATAACATTGGGGTTCAAGAATCAGGAAGGGGAAGCTGGCTAAAGGGGGACCTGCCTGGAGGTTGTTGGAAGAGCTGATAGGATCAGATCTGAGCAAGGGG | 120 |
| chr11 | 70855535 | 70855655 | 788855_47188552_ENST00000601538.6,E | 1 | - | good | CCCTCAGGGTCTGCTGGTTTAGCGTTACCTCTTGTGAAGTTGGAGACCAGGATTTGAATGTAGGAATTCCTATTCCTTCACTTTTGTTCTAACCATGGGAGCTGTTGAAGATTTTTAAAG | 120 |
| chr11 | 70855655 | 70855775 | 788855_47188552_ENST00000601538.6,E | 1 | - | good | TTTAACTCCTTCCTTTCTTCCCACCTTTCTTTTTTCTTTCCTTCCTCTCTTCTGTTCATCTTACATTTTTTGTGAGCTTACCACATGCCAGGCCCTGTGCCATGGAAGGCAGCTTCTCTG | 120 |
| chr11 | 70856615 | 70856735 | 788855_47188552_ENST00000601538.6,E | 1 | - | good | CAGATGCCTCCCTCATAAGGCTGTTGGTGTTTTTAATGAAATAATGTATCAAACATGGTTTAAACTCACACCCCCTAGAGGCAAGGAAATATAATAGATGGGCAAAACAGTCCAAGAGGG | 120 |
| chr11 | 70856735 | 70856855 | 788855_47188552_ENST00000601538.6,E | 1 | - | good | ACTGGGCAAAGAGTCCTGGGCACAGATTGGCAGATGTTTGAATTCCAGCTGTCCCACCTGCTTATTGAGTGGGACTCTGTGTAAGATCCTTGAAATTCTCTGATTCAATAAAGCAGGAAT | 120 |
| chr11 | 70856855 | 70856975 | 788855_47188552_ENST00000601538.6,E | 1 | - | good | CAGACTAAGGATAGAGACGAGGTAGACAGAAGTTAAGGAAGGGCCAGCCAGGGGTGTCCATCTGGTGTGTATCCAGAGTTTCAAATGATGCTTGGGGCTGGGAAGAGCAGGGAGAACCAC | 120 |
| chr11 | 70856975 | 70857095 | 788855_47188552_ENST00000601538.6,E | 1 | - | good | TCAGGGAAATGTTTGCTTAATCCCTCTTGCAGGAACATTTGCACGCAGAGGCAGGCTGTTGCAGAGAGCATAAGTTAGCAGGAAGAGATGTAAGTCCAAAACAGGTATCACCAACCCTGT | 120 |
| chr11 | 70857095 | 70857215 | 788855_47188552_ENST00000601538.6,E | 1 | - | good | AGTGTCAGTCATCCTGTGGCCCGATTGGGCTCTCAGGCTCCAGCCCTTGGAGTCCACTGAAGTTTATGCAAAGCATGGAAACCAGCTATTGAGACTGCCTCCTGGCTGCTGGGGCCATCG | 120 |
| chr11 | 70857215 | 70857335 | 788855_47188552_ENST00000601538.6,E | 1 | - | good | TATTTACTGGGTGCTAAGTCCTGTGCATTATCCAATTTGATTCCTAAAACAACCCTTTGTGGGTGCTACTTGCATTATTCCCATTTTGTAGGTTGGGAGCTGAGGCCCGCTTACTTGTCC | 120 |
| chr11 | 70857335 | 70857455 | 788855_47188552_ENST00000601538.6,E | 1 | - | good | CCTGTGCTAGGGAAGGCCAGAGTCAGCCTGGGAGGGCTTAGCCTCCAGGTTGCTTGTGGCTAGGTTGAGAACGCTTTGCTTAGGAGCAGGTGTTTGAAAAGCGGGTCACCGTACATTGAG | 120 |
| chr11 | 70857455 | 70857575 | 788855_47188552_ENST00000601538.6,E | 1 | - | good | CTTGGCCCTCCCAAGTCCTGAGAAAGGCACCAAGGTCTTTATCAGGAAGTGAGGCCACTTGAGGAAAAGGAAATAGGATATCCAAGACCCAGCAGTCATCTGTTGGGCACATGACTGAGC | 120 |
| chr11 | 70857575 | 70857695 | 788855_47188552_ENST00000601538.6,E | 1 | - | good | CCGTGATGCCCCTAATAGCAAGAGGTGGGCCATCACGTCAACTAAAGTGCTGAGGATGATGGGGTGGGCAGCAGAACCCGTATGTCAACTGCTTGTGCTCTCTTGGGCACTGCCCAAGGG | 120 |
| chr11 | 70857695 | 70857815 | 788855_47188552_ENST00000601538.6,E | 1 | - | good | TGTTACCACCATTTTTGGCCTTCAAGGGTCATTGTGTCTTCGTGATGTCATGTGCTAAGGCTGCCGTCAGTTCTTCCTAGAGAGTTGCAAGTCTTGTGGGCTTAGATCTCCCCTTGTGCT | 120 |
| chr11 | 70857815 | 70857935 | 788855_47188552_ENST00000601538.6,E | 1 | - | good | AAGCACCCACACACTAGGTCAGATGTAGCCCCATCCTGGGTCTCAGCATCTTGAGAGGATGTGTGTGAGCCGTGGAACATGGTGTGTGACTCATGGAAGTGCCCCATCGCTGTGATCTGC | 120 |
| chr11 | 70857935 | 70858055 | 788855_47188552_ENST00000601538.6,E | 1 | - | good | GTGTGCTATGATCTGAATAATTCCCTCTCCAAATGGTGTCTTGCAACTTGCCACAGGCCCAGATAATCCATCATGCTACAGTACAAGGATTCCAAGGTGGGGTGGGTGGAGGCTGCAGTG | 120 |
| chr11 | 70858055 | 70858175 | 788855_47188552_ENST00000601538.6,E | 1 | - | good | TGGTTTCGCAGCAGCACTCGGCTCCTTCTTGACAGAGCACAGGCTGGGTACAACCCTCACCCTAAATTTAGCTCCTGGCACATTTGGGCTGGGATGAAAGCTGAGAAGCACAGGTCATGG | 120 |
| chr11 | 70858175 | 70858295 | 788855_47188552_ENST00000601538.6,E | 1 | - | good | CAGCGCAATACAAATTTAGTGGCAGTTGCTGCTGTTATTTTATATGAACATGGCAGTGAATTTTGATGACTTCATTTAAAATGGATTATGAACAGGGGACAGGAAGCAGATGCTCTGGGG | 120 |
| chr11 | 70858295 | 70858415 | 788855_47188552_ENST00000601538.6,E | 1 | - | good | TACAGGAGGCCATGGGCTTTTGCATGGCACACTGCCATGCCTGCAGCTTCAGACATGGGTATCACATGCACCCTGCCTACCTGACATGGCCATGGCTAAGAGAAGAGCAACCTGGCCGCA | 120 |
| chr11 | 70858415 | 70858535 | 788855_47188552_ENST00000601538.6,E | 1 | - | good | CAGCCATGGGCATGAGACTGGAGAGCCATGGTCAGGTTGGAATCTGCCTTGCTGCTAGGTGTTGGGTGTGCAGGTCATGCCTGGGTTTCCAGTCTAGCCCACTGAGTGGATGGCAAAGAC | 120 |
| chr11 | 70858535 | 70858655 | 788855_47188552_ENST00000601538.6,E | 1 | - | good | GAAGTTTCCATGATTGCTGTTGTTAAGAAGAATCCTTTGGCTGTTGTGTGGATAACTGTCTTTAGCAAGGAAGGAAGCAGGGCTCACACTGGAAGGCTGCTGCCATGTCCATGGGATACT | 120 |
| chr11 | 70858655 | 70858775 | 788855_47188552_ENST00000601538.6,E | 1 | - | good | AGGAGAAGAGGTAGTAAAGTTAGGCTGGAGATAGGAGCCAAGTCCCTTGAGCCTCATGTTCTAAATCAGGGGTCTGGGTTCTGTTCTAAATGGGGAGAACCTTTAAGGAGTTTTAACAAA | 120 |
| chr11 | 70858775 | 70858895 | 788855_47188552_ENST00000601538.6,E | 1 | - | good | TGGACAGCACATTCCTGAAAACAGCTTTGGCCAAGACCCTGCAGTAGGAAAGAACTTGCTTGGGTTCTCCCAGATGCCAGGGATAACCAGATGGAGGTGTGAGTGGGAGGTGCATAGTCC | 120 |
| chr11 | 70858895 | 70859015 | 788855_47188552_ENST00000601538.6,E | 1 | - | good | CCAACACCAGTGCTCCAGTGTGGAAAGGACAGCATGAGACTCAGGGCAAGGTGGACTGCCCAGTAGTAAAAACAAAGTGCTCAGGAATGCCTTTGGGAGGGACGAGCATGTACAGGTGTG | 120 |
| chr11 | 70859015 | 70859135 | 788855_47188552_ENST00000601538.6,E | 1 | - | good | CCACCGACCTATCCATTTATCTACTCACCCACCCACCATTCTGTTTATCCACTCACCTCTCCATTCATCTACCCACCCATCCAGAGCAGTGCTGTATGTAAACATTTTATGACTCTAGGC | 120 |
| chr11 | 70859615 | 70859735 | 788855_47188552_ENST00000601538.6,E | 1 | - | good | ACTTCGTGGAGGTCAAACCACATTCAAGGGCAGGGACTTGACCCTGTTTTGCCACTTGCTGGTTCTGTGTCTTCAGTTTCTCAGCTTCTCTGAAGAAGTTTCTCTGACTCAGTTGTATAG | 120 |
| chr11 | 70859735 | 70859855 | 788855_47188552_ENST00000601538.6,E | 1 | - | good | TGTCCTCGTGTTTAGCCCTGATAGGGAAAGAGGGCTTAGTGCATCTCTTCAGCTATGCTGGACACCAGCCTCGAGGGTAGAACCAATCTGTGGGTAACAAGCTTCCTGGCATGTTTTCTC | 120 |
| chr11 | 70859855 | 70859975 | 788855_47188552_ENST00000601538.6,E | 1 | - | good | TTGCCAAAGCTGAATTTAGAAGTCATTGGCATCCCATTGATCAATCCCAACTGTGAGCTGGACCCCGGGTCATGGTTTCTGACTGTCCCTGGAGGTGCGGTGCTCCAGAGTCTGGTCATC | 120 |
| chr11 | 70859975 | 70860095 | 788855_47188552_ENST00000601538.6,E | 1 | - | good | ACAGGGTCGGATGTTGGGGGAGGTTGGAGGTGGGGATGTTTGCTCCGCAGGGAGGGAGATAGTCGGGGTTCTTTGAGGAACACACGAGGATGACTGCCAAGCCTGGGCGCTGTGGGGTTG | 120 |
| chr11 | 70860095 | 70860215 | 788855_47188552_ENST00000601538.6,E | 1 | - | good | GGTTTGACCAGGTGTGATAGGGTATGGAGGTGGCGGGATCAGGTGGGCTGTGTTCACCTTGGTCTGGAGCAAAGAGGCTTCTGAGAGAAAGGGACTCCTACCATGTGAAGGGAGCAAGGG | 120 |
| chr11 | 70860215 | 70860335 | 788855_47188552_ENST00000601538.6,E | 1 | - | good | ATAACATGGCCTAAGGGGACATTCAGGCTGATGAGAGTCTGGGGGTGCCAAGTACAGAGCTGAGAAAATCCCCCGCACCCAAGTAACTCATAGCCATGGGGCTTTGGGGCGGGTAGAGAA | 120 |
| chr11 | 70860335 | 70860455 | 788855_47188552_ENST00000601538.6,E | 1 | - | good | TTCCTTCCTGCATTCCACAGACAATTTCCACCTTCAAAACGTTCTTTCTTGGGGTTACATAAAGAGACCCGTCACCTTAATGGGTGATGGGAGAGATCTGATGGGCAGGTGTGCACGAGG | 120 |
| chr11 | 70860455 | 70860575 | 788855_47188552_ENST00000601538.6,E | 1 | - | good | GGTAGAAAAGCAAGACTGAGTCCCAGTCCCTGGGGACCTCGCAGTCTCCCACTGCACACATCGGGATTGAAGATGGACAGACGAAATACACACTCTGTTGGTCTAATTCTTATTCCCTTA | 120 |
| chr11 | 70860575 | 70860695 | 788855_47188552_ENST00000601538.6,E | 1 | - | good | CATTTCATTTTATACTTTTATGACTTTGAGAGGTCAAAGGACAAGTATTCATCCATTTTCTTCAAGTGAGAGAGTAAGGGAGAAATGCTTAATCATTTGTCCTCGTTCATTCACCCAACT | 120 |
| chr11 | 70860935 | 70861055 | 788855_47188552_ENST00000601538.6,E | 1 | - | good | CCCACAAGGGGGCCCGGAAAGGCCAGCGCTGCTCCCCGTGTGCCCAGCAGGTCGAGGGATGTCCTTTCCCTGGCTCCTGGGAAGAGAGTATCCCCTGAGATGCACGCGGCTCCTGCAGAA | 120 |
| chr11 | 70861055 | 70861175 | 788855_47188552_ENST00000601538.6,E | 1 | - | good | GGTTAAAACCCTCAGGGAAGTGGCAGGATTTACTAACCTGGGCCTGATTAAAATGCCTGTTGGTGGCTGCCAGCTTGGACTGACAGTGCCTGCAGAGGGGGCAGGTCGGAATGCTTCCCT | 120 |
| chr11 | 70861175 | 70861295 | 788855_47188552_ENST00000601538.6,E | 1 | - | good | TTTTTAAGAGGACAGACTGGAGGAGCAAAACCATTACTAGGTGTGGGACTCATTTATTTTCACATGGAAACAGAGGCATAATAGTAGGTCTTGCTGTTAAACATGACCTTCCAACTGCAG | 120 |
| chr11 | 70861295 | 70861415 | 788855_47188552_ENST00000601538.6,E | 1 | - | good | ATTTCAATGCCATGTGGTGTAGGAAAACATAAATTTTCATCTTAATACACTTTTGAGGCATCAGAGCCATTACAGACCTTGTTTTATAGTGTTTTAAAAGCATTAGTCCTTCACGACCCA | 120 |
| chr11 | 70861415 | 70861535 | 788855_47188552_ENST00000601538.6,E | 1 | - | good | ATAAGACTTTTGATGGGAATTTTGATGGAAATCATGTTTTGATGCAAGTTCAGGTGCAGGTAGGGCTGGGGATTTTTTGTCAGAACACACAATAATTCTGCCTCTTATTTTAGAGCTTGT | 120 |
| chr11 | 70861535 | 70861655 | 788855_47188552_ENST00000601538.6,E | 1 | - | good | GGCCCTGAGCTACAGAGTTAACTGGCTCGTAGAGGAGGCCAACTCAGTGCAATTAACTCTGACATGGATGTTTGCATGGTTTAAAAAAAAAATTAACCCAAAAAGACTGCTTTATACCAA | 120 |
| chr11 | 70861655 | 70861775 | 788855_47188552_ENST00000601538.6,E | 1 | - | good | TTGTGGCCTGAGGTCAGCATCTGCTGGGATTGTCTCAAGGCACAAACCCTGCAGCCTGCAGAGGCAACTGTAGGTTGCTATATTTGCACCTGCCCAGAGGAGTCAAAGAAACAAACCCCA | 120 |
| chr11 | 70861775 | 70861895 | 788855_47188552_ENST00000601538.6,E | 1 | - | good | GATCACATCTCTGCTTTGTTCTTTTGGTTCTCCTCCCAGTTGGATCACTCATTTATTCCTGTGCTTATTTATTGTCTAGCTCTGAGCTCCCTGAGGTCTGGCTTATTCCCTGCCTTTCTT | 120 |
| chr11 | 70861895 | 70862015 | 788855_47188552_ENST00000601538.6,E | 1 | - | good | TCCCAGGTTTTCACTTTTTTTTTTTTGCCCCTGCCCCCTGCTCTCTGCATTTTGGCTGCACCTGTGTCCCACTATTCCGAGACCACCCTATCCAAATTGGGCTCTGCTCCTCTTCTAACT | 120 |
| chr11 | 70862255 | 70862375 | 788855_47188552_ENST00000601538.6,E | 1 | - | good | AGCCAGGCTCCACATTCTACTCCTTTCACCTACAACCCAGTCTCCACCCCAGGAGCTCGAAGAGGGATCTTTCAACAAAGACCATAAATCAGATTGTGTCACACACCCTCCCCCTGAGAC | 120 |
| chr11 | 70862375 | 70862495 | 788855_47188552_ENST00000601538.6,E | 1 | - | good | TCCATCAGCCAGTCCAGTCCAGGCTCCAAATACCAACTGACGTATTGCACTTCTTTCCATTTCCATTGTCTCTACTCAAGCCCATTCCAGAGACTACACAGACCAGTGTAGTAGCTTCCC | 120 |
| chr11 | 70862735 | 70862855 | 788855_47188552_ENST00000601538.6,E | 1 | - | good | AGACTCTAGACTGTGTCCTGCTGCCCAGTTGACTACTCCCGTGGGGGAATCCCAAAGGATCCAAAATGTGCAATGTGTCCAAAACCAAGCTCCCCACCCCCACCCCAAGCACATAGATTT | 120 |

| | | | | | | | | |
|---|---|---|---|---|---|---|---|---|
| chr11 | 70862855 | 70862975 | 788855_47188552_ENST00000601538.6,E | 1 | - | good | GAATCACTGAGGTAAGGCGGCTTGTCTCCTGCTGAGACCAAGAGCCCCAAGACGGGCACTTCAGCAAAATCAGCCCCACACCCATGTCTCCTCCTCCCAAATTCTTACCTCCAGTTTACT | 120 |
| chr11 | 70862975 | 70863095 | 788855_47188552_ENST00000601538.6,E | 1 | - | good | ATAAAAACCCCTTGATCCCTGCCATCTAAGATGGATGCTACCCCTGCTCCGTAAGTCCTGCGGAGCAGCCTCTGCCCTGCCCACCAGCTCACACCTTTTAGCCCATGTCCCTGAGTCCCT | 120 |
| chr11 | 70863095 | 70863215 | 788855_47188552_ENST00000601538.6,E | 1 | - | good | ACCCAGGCCAACCCCTCCTGACCAGGGACCTCTCTCACCTGCTGTCTGGCCTACAAATTATTTTTGAGATTTGTCCCTGGAATGTGACAGTGTTCACGGTTTCTCTAGACCTACCCAGGT | 120 |
| chr11 | 70863335 | 70863455 | 788855_47188552_ENST00000601538.6,E | 1 | - | good | CGTGGCCAGGAGTTCACAGGCGTCCCAGGGCACCGAGCAGAGGATGCTTGGGGCTCGTCCAGACAGGGCCAGCACTGGATGAACCAGGATGAGGTCGGGCTGTGTTCGGGGAGCCGGCAT | 120 |
| chr11 | 70863455 | 70863575 | 788855_47188552_ENST00000601538.6,E | 1 | - | good | ACTCTAAATTGTACCTTCACTGGGATCCAGAGTTAAAATGAGGATTTGCATGGGAGCAGGTGGGCGAGTCCTAGAGATAAGGAAAACCCCCCACAGAATCCGTGTCTTCCACCACATAGC | 120 |
| chr11 | 70863575 | 70863695 | 788855_47188552_ENST00000601538.6,E | 1 | - | good | TGCCAGTGCCTCTTCCCCATACCCCCAGCCCAGGATGCCAGAGAGAGTGGACCAGATGTTGACTGAAAGGAGGCACCAGGCCTAGCAAAGCTCCGTTTGCACCAGCCTGAGTTATTTTAG | 120 |
| chr11 | 70863695 | 70863815 | 788855_47188552_ENST00000601538.6,E | 1 | - | good | GGTTTCCTCTTCTTACAACACACCTCTCCTGTTGGATGAGGACCTACCTGAGTGACTCCACTTTAAAGACTCTCTGTAGCTGTAAAGATCTCATCTCCAAATAAAGAGGCCGAAACGGGG | 120 |
| chr11 | 70864055 | 70864175 | 788855_47188552_ENST00000601538.6,E | 1 | - | good | AAATGATGGGCCCCTTGGAAGCCTCTCTCCCCTTTGCCTCCTTGGTGACCAGAGCTCAAGGAAGAAAGCAAGGTGTGTGGGCACCCCGAACTGGAGGGTCCCTTTCTTGTGCTTGCAAGG | 120 |
| chr11 | 70864175 | 70864295 | 788855_47188552_ENST00000601538.6,E | 1 | - | good | TCTCCCTCAATTTTTCTTTCCTTTTTCCCTTCTCACTACCCACCTCCAATTTCAAAATGTGCAAATAGCAGAGCATCCCAGTGGAGTTGCCACGGCAACAGAAGCTGGTTTTCCCTTTCC | 120 |
| chr11 | 70864295 | 70864415 | 788855_47188552_ENST00000601538.6,E | 1 | - | good | CCCCGCGGCCCTGGGACTCATCTGAGGTCATTAAGCACTGTCAGGGTCACCTGTCCGAAGCTGGATTTTTCCTAAACTTCATTTGCTTTTCTCTCCTTTCTTTTGCCCTGGATACTCTCA | 120 |
| chr11 | 70864415 | 70864535 | 788855_47188552_ENST00000601538.6,E | 1 | - | good | CTCCAAGTCACAAGGTGACCTTGTGCAGGGCACTTCTCATCTTCGAGCCTCTGTTTCCTCTCTGACAGTGGGGGACCTGGGCTGACATTGGCTGGAGCCACCATGACTTAATGGGGAAGT | 120 |
| chr11 | 70864535 | 70864655 | 788855_47188552_ENST00000601538.6,E | 1 | - | good | AAAACCATCACGTTGGGAGTTAGGACTTCAACATATGAATTTGGGGGGTTGGCACAAACGTTTGGTTCATTACAAGAATAGAGGCCTTCCCCCAATACGGCTGCTTTTACCTGGAGAATC | 120 |
| chr11 | 70865135 | 70865255 | 788855_47188552_ENST00000601538.6,E | 1 | - | good | CCTGTGGTGGGATGCAGGTCTGACTCAAATCCTCTCACGGGTATCATAAGGACCCCTTGAACCCCACTCCATGGGATCTCCAAGGAGGAAGGTCTTAGCCCGCTCAGGCCACTGCAGCAA | 120 |
| chr11 | 70865255 | 70865375 | 788855_47188552_ENST00000601538.6,E | 1 | - | good | TTTGTGAGGCTAAATTTAGACGTCGCCTAGTTCTGCAGCATCTTTGCTTTTCTCTAGAATGGTTTGTCCAGTGAGAGAAGTCTGCCAGCCAGACCCTAGAAGTTGAGTCTCTCCTTAGAG | 120 |
| chr11 | 70865375 | 70865495 | 788855_47188552_ENST00000601538.6,E | 1 | - | good | GCACTTTCTCAGACACCCCCCGCGTGCCCTTACTCGTGGTGAATTCAGCCTCCTGGGCCCATTGTATTCTTCTCCTTGTCCCTTCAGAATAGTATTTCCCTGCTCCTGTGCTCTAGATTC | 120 |
| chr11 | 70865495 | 70865615 | 788855_47188552_ENST00000601538.6,E | 1 | - | good | GCTGCTGAGCCTGCCCGGCCACCCCACCCCACCCTGAGGCACACCTCCATCCAATGGGCGGTTGGGGAATGAGGCACGCGCAGCTGCATCTCAGCCTGGCCATTCCGGCACCTGACTTCT | 120 |
| chr11 | 70865615 | 70865735 | 788855_47188552_ENST00000601538.6,E | 1 | - | good | GAAGAAGAACACACTTCTTGTCCAGCTACATCAGAAGGCGGTGCGAGGCAGTGTCCTGTGGCTCCTTGCTAGCAGACTCAAGAGCAATTTTGGAGATGGATGCTTGTGGCCCCACCCTCT | 120 |
| chr11 | 70865735 | 70865855 | 788855_47188552_ENST00000601538.6,E | 1 | - | good | AGCACAAGGCAGGTGCTTGGTTTCTGCAGGGTGAGTGAAGGAATTAGTGAGTGAGTGAATGAATGAATGAATCTATCTATCTGAGGCTCATTTTGAGAGACAGGATCTGATTCAATGCTG | 120 |
| chr11 | 70865855 | 70865975 | 788855_47188552_ENST00000601538.6,E | 1 | - | good | AGCAGTGCCCGAGGGTGGGACTGCCAGGGAGGGGCCACGACCCAAGAAAGCCAGGTATTTGAAGTTGAGGAAGGGCAGCACTGTGGGCATATTCCTTGATGCTTGGGTAGAGCAGTACCT | 120 |
| chr11 | 70865975 | 70866095 | 788855_47188552_ENST00000601538.6,E | 1 | - | good | AGGCATCTGCGCGCGGAATTGGTGACCAAGATGCTGGACAGGCCATGGACCCTGGAGAACAGCCAGGCAGCCCCCCACTAAGAAACTGCAGAAAGCACCAGGGAGGAGAGGGCCCAGGGG | 120 |
| chr11 | 70866095 | 70866215 | 788855_47188552_ENST00000601538.6,E | 1 | - | good | CTGTCTGGATGGAGCCTAAGTCTCACCAGGAGGGTGTAGACTGCAGCAGACCGCTCCCTGGTGGCCTGGGGGTGAGAGTCTAATCCAGAAAGTGGAGACCACCCTAGGCCTTTCCGACAG | 120 |
| chr11 | 70866215 | 70866335 | 788855_47188552_ENST00000601538.6,E | 1 | - | good | GACTAGATCCCTCCCAAATGGTTCCCCCCATTTGGTGCCGCGGGGTGCGATGGGCTGTGGCTCCTCTGAGCTGCACTCTCTGCCCCTGCCTTCTGATGGCAGGGGAGCCACTCGGTGGTG | 120 |
| chr11 | 70866335 | 70866455 | 788855_47188552_ENST00000601538.6,E | 1 | - | good | CTTATCTGTTCCCTTCTTGTTTCAGCAGCATCTCGTTCTCAGAATCCTAATTAGCCGGGTGTGGCTCTTTTGACCATTTCTCTCAAAAATGAGTCTCGGTTCCCAACTTCCATTTTCTGT | 120 |
| chr11 | 70866455 | 70866575 | 788855_47188552_ENST00000601538.6,E | 1 | - | good | GGTGCTAGAGTTACTGACGTCTGCAGCAAAGTGCAATTTTGAGTTTCCGGAACATCAACACAAAAGAACACTTGGACTCCTTCTTTGTAGCCAAATGTTAGGTCTGGGATGTTTCTTGGT | 120 |
| chr11 | 70866575 | 70866695 | 788855_47188552_ENST00000601538.6,E | 1 | - | good | CGGGCCCCTGCGTGAACTGATAAGAAAAGTCATTCCTACTGAGATAGGACTTTAGATTTTATCCTTACTCCTTGTAAATTGCTCTCCTCGGATTCCCAAATGCAAGCTCCGTGAACCCTT | 120 |
| chr11 | 70866695 | 70866815 | 788855_47188552_ENST00000601538.6,E | 1 | - | good | AGAAATGTTTCCTTAGGGAAATGGGGAGATGAAAATTTGCCGGGTAGACATAGGAAAACTCCAGCAAAAGTGGACCTTTAGAAATCACAAAAATCTCCTATTTGTTGACTTGTGTGTCTC | 120 |
| chr11 | 70866815 | 70866935 | 788855_47188552_ENST00000601538.6,E | 1 | - | good | GGTTGTTGATCTGTCCTCCTGTGTTACCTAGAGATGCTCTCAAGACCAGCGAGGGCCGTCATGAAGCTCCTGCTGAGTCCATGTGCGGGCAGAGAGTGGCTGCCTCTAGGTCACCCCTTA | 120 |
| chr11 | 70866935 | 70867055 | 788855_47188552_ENST00000601538.6,E | 1 | - | good | GTCCATGGTGGTGTCCACGGCATATCATTCGTTTGGCTCCTGGCTTCATGGAATGTCAGCCCCAGGCCTTGTCTGCTGGGGTGAGAGCCACAGCGAGGGAGTGAAGGCAGAGGCTGGTGT | 120 |
| chr11 | 70867055 | 70867175 | 788855_47188552_ENST00000601538.6,E | 1 | - | good | AGTGGGGATCCACTACCCCAAGGGACCTCATTCTCTCTCTCTTTTTTTTTTTTTTTTTTAAATACACCGAATTGGATTCAAATCCCTCTGGGCTACGAGGTCCTGTTTCCCTTCTGTGCT | 120 |
| chr11 | 70867175 | 70867295 | 788855_47188552_ENST00000601538.6,E | 1 | - | good | ATTTACTGAGGACTCGGACAAGAGCGTCCTTCCACCAGCACCCTCCACCTACCCCTACCCGGATAGGAGACAGCTGACCTCAGGATGGCACCTTCGTATTCTAGAACTTGAGAGAGTCAG | 120 |
| chr11 | 70867295 | 70867415 | 788855_47188552_ENST00000601538.6,E | 1 | - | good | TCGTCCTCCGTTGGTGGAGTCCTGGCCTGATCCTTGGCCTCCGGACCTGCTCTCGGCCCTGCTGCCCGTGTGGAGCCAAGACGTGTGCACTCTCCTCATTGGTCCTCCCCTGTGTTCAGG | 120 |
| chr11 | 70867415 | 70867535 | 788855_47188552_ENST00000601538.6,E | 1 | - | good | GAAGGCATCCCTGGCTCAGGCAGCTTTGGCCCGTCCTGCCCCACCCCTCAGCCCTCATGCGCATGCCGTACTCCCCAGCAGTGGACATAGCTGTTTGCTGTTAACAGCGGCTCTTGCCTC | 120 |
| chr11 | 70867535 | 70867655 | 788855_47188552_ENST00000601538.6,E | 1 | - | good | CAGTGGGTAATGAATACAAATCTATAGCGTCGATTCTCAAGAGGGAACATGGCTGAGCTGACCCCACATTCACGAGCCGGTGGCCTGGGCAGACTTTTCTGGCTGGGGCAGCGAGGTTGT | 120 |
| chr11 | 70867655 | 70867775 | 788855_47188552_ENST00000601538.6,E | 1 | - | good | GAGTCCTAGGCAATAAAGGTCAGCAGTGGAATATTGTCACAGAATAATGACTCTGTGCCGCTGGGTAAATGTTAAAGCTTTTATTGGGGTGAGACGTGGATGTCTGGCTGGCTCTGAATT | 120 |
| chr11 | 70867775 | 70867895 | 788855_47188552_ENST00000601538.6,E | 1 | - | good | GCAAACAGCCCAGAGATGGATGGGCGGGAGGCGGGGGCCGGCCGGTGCAGGAGCAGTGGAGGCGGTGGACCTGGGTCCTGTGTGCTGAACGCTGAAATGCTTCCTTGGAAACTTCACAGA | 120 |
| chr11 | 70867895 | 70868015 | 788855_47188552_ENST00000601538.6,E | 1 | - | good | CTGGTAATTCCACCCAATTATTTGCCCAGAATTGACACTGATCAATTTAGAGGGAGGCACAGTCAGTGATGGAATTAAGTGGTAGCAAACACACTGTCAGTACATGAAGTCCAGACAGAG | 120 |
| chr11 | 70868015 | 70868135 | 788855_47188552_ENST00000601538.6,E | 1 | - | good | GGCTTCATATTTGTGTCTTCAATTATGCTGTCAAATGCTAAGGAGTTTAGACACTTACTGCCAGCGCCGGGCCCTTGGTGACGGCGTACATTATTTGCATAATCAGCACGGGTGGACCCG | 120 |
| chr11 | 70868135 | 70868255 | 788855_47188552_ENST00000601538.6,E | 1 | - | good | TATGCCGACCCCCTCAGCCTCTGTGTCTATGCGACCCACCTTCTTCACCTGCACCCTCTGGGCCAAGCCATTGAAAGTGAGAACGAGGGAGTCTTCCTCGAGAAAAGAAACTCTTCATTA | 120 |
| chr11 | 70868255 | 70868375 | 788855_47188552_ENST00000601538.6,E | 1 | - | good | AGAAACCGAGGCCTGGGAGTTTATCCAGCTTGACCAGGCTTATCATAGCAGGCACAGGTTATGACCCTGGGGTTGTCCAAGAGCAGATCACACCGCCCCATGTCCGTGGACAGGTGTCCG | 120 |
| chr11 | 70868375 | 70868495 | 788855_47188552_ENST00000601538.6,E | 1 | - | good | TCTGTATCACAGGGATAAATAATAGTCAATAGCCTCGTGCCCAGCTAGTAGCAGTGGTGGTAGAGGGACTCTTGACTGTTGTCATTGTTTATTATAGTATTATTTCCATTTTACACATGG | 120 |
| chr11 | 70868615 | 70868735 | 788855_47188552_ENST00000601538.6,E | 1 | - | good | ATGGGACTTAGGACTGGCCGTGGCAGCTCCAGTGACTGTGCTCCCTGTGTGGGTGGGGCCCTGGGCTGCCCCCGTGGGGTGTGAGGCTGACTGTGTAAGATGTGGACATGGCCCAGCTCA | 120 |
| chr11 | 70868735 | 70868855 | 788855_47188552_ENST00000601538.6,E | 1 | - | good | TCCTCTGGACTTTGGCACCCTGGGTGAGAATTGGGCAGGTTTGTGGAAGAGAATGGGTGAGGATGAGGAGGCCGCATGTTGTGTCCAAACAGGGCTTTCCGTGCCAGAGCCCTTTCCCGG | 120 |
| chr11 | 70868855 | 70868975 | 788855_47188552_ENST00000601538.6,E | 1 | - | good | TGCTGATATCCTGGGGGTGAAGCCTGCGTCACATGAATTTTGCAGGTGACACAGTTCAGCCTCTAGCACTCAGCGCCCACCTGGATGTGGCTTCTAACTGCAGGACGCCCTCCCCCAACA | 120 |
| chr11 | 70869215 | 70869335 | 788855_47188552_ENST00000601538.6,E | 1 | - | good | GCTGGTCTGGAGGCCTCTAGGACAGATTGTCGGGATCTATGGTGTATGAGTTTGCTCCAGCTGCCGTCATAAAGCACCACAGACAGGGAGGCTTACACAAGTCATTTATCGTCTCACCGT | 120 |
| chr11 | 70869335 | 70869455 | 788855_47188552_ENST00000601538.6,E | 1 | - | good | CACTCTGGTATGGGATGTGTGGAGTTCTGTTGGTCCCAGTGGGTTTAATAACGTGCCGATCGGTGCATCCCAAGCATTCTGTGGCCTGTGGACCGATACATTGTCCTCCCTCTGAAACCT | 120 |
| chr11 | 70869455 | 70869575 | 788855_47188552_ENST00000601538.6,E | 1 | - | good | TGGAGAGGCCATAGGCAGCAAGGTGGGACAGACAGCCTGCCATGAACCAACTGGTCTGCACAACTTTGCCGTGAAATATGAAAGCCTCCTACAGCACAGGCTGAGATTGGAAGCATTAAT | 120 |
| chr11 | 70869575 | 70869695 | 788855_47188552_ENST00000601538.6,E | 1 | - | good | ACCGAGCCCAGGGTGCCCAACCCAAGGAAGTATCTGCTGGTTATCAGGTTTCTAGCCCAGGACACAGAGCAGACAGAGTGCTGTGGACGGTCAGGAGCTCCATACCCGAGCCTGGCTTGG | 120 |
| chr11 | 70869695 | 70869815 | 788855_47188552_ENST00000601538.6,E | 1 | - | good | CACAGTGTCAGTTCTCATCCCATCTTCCAACAACACCACCATGGACTCCCCGTCGAGCCCCCTGTCTGGGGGAGGACAATCAAGGAGATGATGGGGACATGCCAAGATGACACGGTGGTC | 120 |
| chr11 | 70869815 | 70869935 | 788855_47188552_ENST00000601538.6,E | 1 | - | good | GGTGTGTGAGAAGGCAGAGCCTGGGGAGCTTCAGGCAGAGGGGAGGCATTCACATAATTGGTGGTGAGGTCCCTGCACTGACCCAGCGGCGACTGAGCACCCAGTGCGGCAGTAAGTGAC | 120 |
| chr11 | 70869935 | 70870055 | 788855_47188552_ENST00000601538.6,E | 1 | - | good | ACCTCACTGATCACATCTGCAAGGAGGCAATGACCCATTTCCAGTTAAGGCCCCATTCGTAGGTTCTGGGTGGACATGGATTTGGGGGGACACTCTTGTGGACTTTTGTGAGGGTTTCTG | 120 |
| chr11 | 70870175 | 70870295 | 788855_47188552_ENST00000601538.6,E | 1 | - | good | CTCATTTTGGCCCAGATGGACTGCAGGAGGGGACGGCTCTGGGTCTCACCCCGGGTTCATTTCTCTCAGTCCCCTGGAACGAAGTGGAGCACACTCTGCCTGGGTGAGCCTCCAGCACTC | 120 |
| chr11 | 70870295 | 70870415 | 788855_47188552_ENST00000601538.6,E | 1 | - | good | CCCAAAAACCTTCCTAGATCTTTCCTGGCGCTCACCTGCCCTTTTCCCATCTGTCCAGAACCAGCCCCTTCCACTTTTTCTATTTCCTGAATCTGCCCCTCCCTCCCCACTGGTTGGGCT | 120 |
| chr11 | 70870415 | 70870535 | 788855_47188552_ENST00000601538.6,E | 1 | - | good | TGTGGTTTCAGTCACCCAGGCTGTGGTTACAGGAGCTCCAGTGAACATACCCCATCTTTCCTGGCCAGCTGGCCCTTCCTCCTGGGCACCCCATCTCAGTTATGGCCCAGGAACTGAAAA | 120 |
| chr11 | 70871255 | 70871375 | 788855_47188552_ENST00000601538.6,E | 1 | - | good | CCCTGCTCCTATGGTGCCAGTGATCTTCGGGGCTGGTGGCTTGCAGAGCGCTCCCTGGACCTCCTCTTGGACCCTCAGACTCATCTCCTGCAGAGCCTGCCACCAGGCGAGCCCAGACCC | 120 |
| chr11 | 70871375 | 70871495 | 788855_47188552_ENST00000601538.6,E | 1 | - | good | GCCTCAGCACGTGAAGGAGGCTCAGTGCCAGGCTATGTGGCTCATCTCAGAAGGTGGTGTGGCCCCAGGTTCTGTGGTTGCCCTTGCTCTGCTCTCTGCTCCCCACCCCCGCCCATCCTG | 120 |
| chr11 | 70871495 | 70871615 | 788855_47188552_ENST00000601538.6,E | 1 | - | good | GACCTCACTTTCCTTTCTCTCCCTGCCCTGCTTGTTCAGCAAATGTGTTGGGGAATCCGCAGGGTCCCAGGCACAGCCTCTCCCTCCAGGGGCTCACAGGCTCCATGGCACACAGATGTG | 120 |
| chr11 | 70871615 | 70871735 | 788855_47188552_ENST00000601538.6,E | 1 | - | good | GTAAAACCCTGCAGGGCCCCTGAGTACCAGGTGTGTCTGGCCCGTGTTGCCTTTCAACATGAACTCTTTCTTGACTTTCTTCTCCATCAGGGATATGGTCATATTTATAATATTTGTCTC | 120 |
| chr11 | 70871735 | 70871855 | 788855_47188552_ENST00000601538.6,E | 1 | - | good | AATAGAAATCGTATATGGAAATCTCAGACGGCTTCAAAGAGTCTCTCAGTCAAATTCAGATGGGGACTGTGCCTCTTTCCTGGCTGTCCCCTCCCAGCCCCCATGTGGCCACAGTTTGCT | 120 |
| chr11 | 70871855 | 70871975 | 788855_47188552_ENST00000601538.6,E | 1 | - | good | GTTGAATGAACAAGAGTTCTTAATCCAAGAATAATGCTGGGTTTACCTTTCAAGTAGTTCAAAGGCTTCTTTCAAGACGGTGTTGTTATGGAGGATGGGATGCGTCTTGCCTTAAATATT | 120 |
| chr11 | 70871975 | 70872095 | 788855_47188552_ENST00000601538.6,E | 1 | - | good | TGGCGGTGTGACGTAACAGGCCACACACGCATCATCCTACTGCTGCAGAAGCAGGCAGCGTGCACATGCCAAGGCACCTCTGGCTCTGGTGTTCAGGGATGTCGTCGTGGGGCTGTATTT | 120 |
| chr11 | 70872215 | 70872335 | 788855_47188552_ENST00000601538.6,E | 1 | - | good | CTCATGAGTACCTTCCAACACGAGCTCTCAGGAGTAAAGACAGAGTTCCCTACCTTCAATGTGGATCTGCAAAAAGATCTGAAAAGCGAATGCCTTCTCCGTAGCTCGTTGGCAGTAGTA | 120 |
| chr11 | 70872335 | 70872455 | 788855_47188552_ENST00000601538.6,E | 1 | - | good | CACCTCCAAGCCTGCATCCTACTTGTCTTTCCTCTTCATGCGTGTGGGGTTTTTTTCCCCTCCTTTCCACTGTGCACCCCGTCTCCTCTGTAAACTTGAGGTAGGGAACTCTGTCTTCA | 120 |

| | | | | | | | | |
|---|---|---|---|---|---|---|---|---|
| chr11 | 70872455 | 70872575 | 788855_47188552_ENST00000601538.6,E | 1 | - | good | CCTCCATCTGAGGGTCTCTCTTCCCTCCATCCAGCCACTTGGTCGTTTGCACCCACTGTGTCCACCAAGCGCTTCTAGGGAAGGGTCTTGTCCTTTTTTCTCCCCGTCCCGCCGAGCCTC | 120 |
| chr11 | 70872575 | 70872695 | 788855_47188552_ENST00000601538.6,E | 1 | - | good | CACCAGGGAGCCGGAAGTAATGTGGTCCAGGGACCCATCTCTCATTCCACGTAGCTCACGGACCTGTGTGCCCCTGACTCTGGTCTGTGGGGGGCCCCTCTGTATGGGGGCCCCCTATTC | 120 |
| chr11 | 70872695 | 70872815 | 788855_47188552_ENST00000601538.6,E | 1 | - | good | GCAAAGCAGCCCCCACGACCCTTAGGACAATGAGGGGTGAAAATTGCCACCCATCAGTTTTCAAGTGGTCAGTGTGGAGGAAAACAGGGACCCGCAGGTGGCCAGAACGTGGAGGAGATA | 120 |
| chr11 | 70872815 | 70872935 | 788855_47188552_ENST00000601538.6,E | 1 | - | good | GCAGACAGTGGATATGTGCTCCTCCTGCCCCGAGGAGCTGGGCTCTACAGGGCATCGTGGGGTCCCTTCCTCTTAGCAATCAATGCCATGTTCCACTGTGGCCCCAGACTCCAGCACCAA | 120 |
| chr11 | 70872935 | 70873055 | 788855_47188552_ENST00000601538.6,E | 1 | - | good | GGCACCAGGATGGACGTGGGAACTACCTCCGCTCTTCCTTTCTCACAGGTGCCCAGGATGTCTTTAGCTTTTGTAATGTTTGGGGCTGGGGGAGGTGGAGGTCATGAGTGTGGGGAGAGG | 120 |
| chr11 | 70873055 | 70873175 | 788855_47188552_ENST00000601538.6,E | 1 | - | good | TGCCACGGTTGCTGTGGCAACGGCTCTTTGTTTTAGGGAACCACAAACATATCATTCTATAAACTGCAGGGCCCAAGCTGCAGGTTTCACCTTCCTGAGGGTCGCCACCCGGGAGGCAGA | 120 |
| chr11 | 70873175 | 70873295 | 788855_47188552_ENST00000601538.6,E | 1 | - | good | CTCCTCTTCCTCATGGGTTTTGTCAGAGGATGTTAATGCCTTCCAAGCAAATTGTATCAAATGCATTTGAAAGGCAAAACTCATTGTTTAGTGTCTGAGAATAGCCTCTGAGTGCTGGGA | 120 |
| chr11 | 70873295 | 70873415 | 788855_47188552_ENST00000601538.6,E | 1 | - | good | TGTTTCCCATTTCCTGTTTGCCTATTGAAAGTGTCTGGGGCCGTTGGGCCCTCCTGTCTCATGAATGGGTGTGATTCCTGGAAGCCCTGCAAGTCTCGACCTTGTCAGCTCTCACCTGCC | 120 |
| chr11 | 70873415 | 70873535 | 788855_47188552_ENST00000601538.6,E | 1 | - | good | CTCCTTCATGGGACCACTGCTAGAAGGTTAGGGATCCCAACTGCCTCACCTCTGGGCAGCCCCTCCTCCTCCACGGAACTGCTGTTGGAAGGTTAGGGATCCTGCCTGCCTCCCCTCTGC | 120 |
| chr11 | 70873535 | 70873655 | 788855_47188552_ENST00000601538.6,E | 1 | - | good | GCTGCGGGCTCTGGCCTGTGCTGCCATCTTTGCAGTGACCCTGAAGAAGAGAACTGCCCCAGATGGGACTGTGGAACGTCTCTCCTGAAGAAGCCCCGAAAGGGTGTGTGGCAGCCGCTC | 120 |
| chr11 | 70873655 | 70873775 | 788855_47188552_ENST00000601538.6,E | 1 | - | good | AAAGTTGTTCATGAACTGGCAAAAAAAAAAAAAAATGGTAAATGGAGGAAGCAATGGCTGGCTGTGAGCAGGGGACGCACTCAATGTCCTTTGCAGAGTGAGCAGGGTGGATTAAGTGGTT | 120 |
| chr11 | 70873775 | 70873895 | 788855_47188552_ENST00000601538.6,E | 1 | - | good | GGGCAGCTTGCCCAGCGTGCTAGTGACAGGAGGCTGGTGCTAGGAATGAAGGTTATTTGGTCTTCTCACTGGCATCACTGTTTGGTATTCTAATCCCATGAGATCGTTGTAAACAATAGC | 120 |
| chr11 | 70873895 | 70874015 | 788855_47188552_ENST00000601538.6,E | 1 | - | good | TGTTCTTTAGTCTGTTCAAAAATGTACTTTGCATTGGGATGCTGCCCCCAGCCTGCCTACATCATGCCTTTTTCCTGATGACACGTGTTCCTGTCACTGACGCCCGGGAGCTGTTTTTGA | 120 |
| chr11 | 70874615 | 70874735 | 788855_47188552_ENST00000601538.6,E | 1 | - | good | ATTTGAAACGGCAGTGGATGTGAACTACAAGGTGGAATTTTTTTTTTTTTTTTTTTTTTTTAGTAAATGCAGATATGCAGATTTCAATGCTTACATGAACTATTTTATTGGATTTGGATA | 120 |
| chr11 | 70874735 | 70874855 | 788855_47188552_ENST00000601538.6,E | 1 | - | good | TATTACACGGTAGATTTGAAACAGACATGAGGGCACAGTAACTATAAAGACAAAGTGAACTCGTGTTCCTTCAGTGCTGTCATTCTAGGTAGCCACTCGGAGTTTTCATCTGTGTTTAAA | 120 |
| chr11 | 70874855 | 70874975 | 788855_47188552_ENST00000601538.6,E | 1 | - | good | CTGTTATAGTTACATGTGGTTCTCAGGCATGAGTGGGCTTAGCTACAGCCTTTTGACTGGAGTGAAGTCTGCAGGGTCCGGAATGGCCTGGTCTCCATGCCCTGTGATCACCATTCCTGC | 120 |
| chr11 | 70875095 | 70875215 | 788855_47188552_ENST00000601538.6,E | 1 | - | good | GAGAGAATTCCAAATAACAATAAAGCCAGCTAAATGTCCCTTCCCTTTTAATGATCACCATGTCCCCCCGTTCTAAAAATTGCCAATGATAAAATACTCCTTCCCCCAATCGTTGGTCTA | 120 |
| chr11 | 70875215 | 70875335 | 788855_47188552_ENST00000601538.6,E | 1 | - | good | ACATGCAATATTAATGCTTTATCATCATGTTCCGAAAGCAGAGCTCTACTGGGGGCGGAGTGAGTGACGCTGTCTGCCCCAGGTACACGCAACAGGAGGGCACTTCTATTGCTTCATCTG | 120 |
| chr11 | 70876655 | 70876775 | 788855_47188552_ENST00000601538.6,E | 1 | - | good | CTGCGCTCAGCAGCAGCCCTGTTATTTCCTGCCTTCACTTGCACAGACCCTAGAAGTGTTGTCACCTGCTGCAGCCCACATGGATGAAACACTCCTCTGAACCTCAGCTTCCCCGTTAAT | 120 |
| chr11 | 70876775 | 70876895 | 788855_47188552_ENST00000601538.6,E | 1 | - | good | TCACGGGACCAGAGTATTGGAGTGGCCACTAGACAGGGCGCTCTGCCTCAGCATGCTGGCCCCTTCTGATGGAACATCCCCTACCCCACCGTTGCTCAGAGCATTTCACGAGACTCGACT | 120 |
| chr11 | 70876895 | 70877015 | 788855_47188552_ENST00000601538.6,E | 1 | - | good | CATGTGTGTGTTCCAGCTGTTAGAATCTCACCGGGTCTCAGTCCCTTGTGGAATTCTCCAAGGTCCAGGTGGTGAGGCCCTTTGTTGGCCAGTGTCCAGATGGCCCAGGCTGAGAGGCAG | 120 |
| chr11 | 70877015 | 70877135 | 788855_47188552_ENST00000601538.6,E | 1 | - | good | TAAATTATCCGTATTCTTTTTCTGTGTCCTGAAGAGAAATAAAAATATCAGTGAGGAAGAGATGGGAGGAAATTGTGTGGCTGGGACAGGAAGCAATAGTTACTCCTTTCTCACCAGGGC | 120 |
| chr11 | 70877135 | 70877255 | 788855_47188552_ENST00000601538.6,E | 1 | - | good | GGCCCTCGGAACACCTGCCTGCTCTCAGAGTTATCCAGCGAGCGGGGCACAAATTACATGTTGATGCGTTCCAAAAAGAGGGGAAAATAATTCAGAACACCATTGTTGCTCCTTTAGTTA | 120 |
| chr11 | 70877255 | 70877375 | 788855_47188552_ENST00000601538.6,E | 1 | - | good | AGTGGCCGGTTACTATAGAAATCCGGCCCAGGAAGCCTCTTGGCTTATGTGAGGTCACAGGCTGGCCATCGGCATAGGGTGAGGGCTCTGCCCAGGCCTCCTGGCCCTGACCCTGGCTGG | 120 |
| chr11 | 70877375 | 70877495 | 788855_47188552_ENST00000601538.6,E | 1 | - | good | GGGATCTCTCCCTGAAATCACGGTGGCTTCATTCAGTTCCAGGCAGCCAGCAGGCAGGGGAAGGGTGTTCTGCCAAAGAATAGAACAGCAGAAAGTCAGGCGACGTATGCCGGCAGACCC | 120 |
| chr11 | 70877495 | 70877615 | 788855_47188552_ENST00000601538.6,E | 1 | - | good | CCCTTGGTAGAGTCCTGGGGAGTCTCGGATGTGTACAACCAGAGCTGGTGAAGTAAGGAAAGGAAATGCATGACCCAAGAGGGCAGAGTTCTGCTGAAGCTGGTGGGGAGGGCAGGAGCC | 120 |
| chr11 | 70877615 | 70877735 | 788855_47188552_ENST00000601538.6,E | 1 | - | good | ACTCCTTGTTGCTCTGGCCAGAGCTTCAGGAGTCCAGCCAATGACATGGCTTTGGGTATCAACCATCCTTGAACCAATCACAGTGGCCAAAGGTGTTCATGTTCTATCCCCTCTATCAGG | 120 |
| chr11 | 70877735 | 70877855 | 788855_47188552_ENST00000601538.6,E | 1 | - | good | GGAGAGAAACAGCCACTTTCTCCGCAGGGAATATTTATAGAATGTGCCCGTCTATCCGTGTCTATATTTGGCTTCCCTCTGCTTGGCCTCAGCCTTACTCCTTGAGTAAGTAAGCTTCTT | 120 |
| chr11 | 70877855 | 70877975 | 788855_47188552_ENST00000601538.6,E | 1 | - | good | TTAACGTCTTCTCTTTTCCCCAGCACACCCAACCAATATGGCTTCTGCCACCTGGTAGGTGCTTGACAAATATTTGTTGAGTCAATTGCTGACTATTCAGACCTGTTCTTTCCTGGTGCA | 120 |
| chr11 | 70877975 | 70878095 | 788855_47188552_ENST00000601538.6,E | 1 | - | good | ACCCCCAGTGCCCTGCTCCTGAGGCCAAGTCAATGGCGGCTGAGCCATTCTCAGAACCGCTGACTCTCCCTCCTGCTTCTCTGCATTCTTCGCCGGACTCACGCTGCACAAAGGCAGGTG | 120 |
| chr11 | 70878095 | 70878215 | 788855_47188552_ENST00000601538.6,E | 1 | - | good | GCAGGAGGGAGCTGCGGCTTGGTTTTGTAAGAGAGATGTGGATGGGGCAGCCTTCTGCAGTGCTGACTTCATCCACAGAGCGAAGGCCATAAAGGCGACACCTTCCTTCCTGTGTCTGGG | 120 |
| chr11 | 70878215 | 70878335 | 788855_47188552_ENST00000601538.6,E | 1 | - | good | TGGCCCAGTGTTAGGGTCTAGACCTTGTCGAGGTCCTCCCAGGGTCCTGCCTTTGGAATGACTTGGTCATCAACTCCCAATTCCCAGTTCTCCATGAATATTCATCTGCATCCCAGCTCA | 120 |
| chr11 | 70878335 | 70878455 | 788855_47188552_ENST00000601538.6,E | 1 | - | good | AATATGGTAAATTGCTCCAGAAAGCAGGATGGGGAATTCCCGTGTGATGATGCCCTGTATGTTTGTTCCGAATTTTCTCCCTTCTATGTGGAGGGGAAGTTAGGGCTGCATTGAGAGGTG | 120 |
| chr11 | 70878455 | 70878575 | 788855_47188552_ENST00000601538.6,E | 1 | - | good | CAAGACCATTGGCTTAATAGGCAGGGTCCAAACCCACCATGCAGCCCGAGGGAGGTCCCTGGGTTCCTGTGGGTTGAAACGTGGCTTTCCATGATTTCTGAGCATCAGCCACAGCCAATT | 120 |
| chr11 | 70878575 | 70878695 | 788855_47188552_ENST00000601538.6,E | 1 | - | good | ATGGACTCATTTATCAAGTAGAGGCTGAAAAATAAGAACAGATCCCATTCCAGTCCTCTGCCAGTCCCCTGTCCAAGACCACCCCCGAGGAAGGCCGTCATGGTGGTGCCCTGAGTTGGG | 120 |
| chr11 | 70878695 | 70878815 | 788855_47188552_ENST00000601538.6,E | 1 | - | good | GAACAAGCTCGTAGGAGTTAGTGGGAAGGGAGCTGCTTTGAGCCCTGGGTAGCGCAGCAGAGAGCTTGTCAGGGTCCCTTTTGATGGAGCAGGAGGAAGTTCACCTCTCATGGCAAACTC | 120 |
| chr11 | 70878815 | 70878935 | 788855_47188552_ENST00000601538.6,E | 1 | - | good | ACATGACCTGGAGGGGCAGAGAGTCTGGCTGGAGCTGGAGGTGAGGGCAGGGAGCAGCCTCATTCCTGGCCTTAGACCAGATCCTTCTCCAAAATCGTTCAGCATTCATTCATGACCCTG | 120 |
| chr11 | 70878935 | 70879055 | 788855_47188552_ENST00000601538.6,E | 1 | - | good | CCAAGGAGTCCACCAAGATGGGGGCAGCTTCAGGAAGATACGTAGCCTCAGGAAACATTTTCCTTCCCCAGGACCAGACTTCAGGATGTCCCCTGGGGGAGAGATGGTAAAGGAGAAGTC | 120 |
| chr11 | 70879055 | 70879175 | 788855_47188552_ENST00000601538.6,E | 1 | - | good | CTAATGAAGCAGCAGCAGTTCATGGGATTCTGGTCCCCAATTCTCCGTGGATCCAGGTGCCAGATGGAGGCATCAGTATACCCTCGTCCTTAGGACCACCGGGCACATGGGCTGCCCTGA | 120 |
| chr11 | 70879175 | 70879295 | 788855_47188552_ENST00000601538.6,E | 1 | - | good | TTCTAATAAGCATAAGTGAAAAGACGTTATGGAGGCTTCAGACAGAAAAGGAAGAACTTGTACGCATATGTACCTAAGTGCTGTTGCTTTTAGCCAAGCCAGAAGGGACACATTAAACAG | 120 |
| chr11 | 70879295 | 70879415 | 788855_47188552_ENST00000601538.6,E | 1 | - | good | ATTCAGACAGAGAATCGAGAGTCCTTTCTTGAAGTGCCTTTTGGGATTTCAAATATCGGAGCCTCATAAGAATATGCTGGGTAGAATTAAATTGCACAGAGGAAAGGTTTCTAATTCTGG | 120 |
| chr11 | 70879415 | 70879535 | 788855_47188552_ENST00000601538.6,E | 1 | - | good | TTTTCCGTGTGGTCTTGGACCAGCCACGGCACCTCTCTGAGCCTCAGAAGCCACCACAGATAACAACCTCTCTCCCTCACCACTCCCTGCATTTGAGACCCTCTTTTGAGCTGAGCCCGA | 120 |
| chr11 | 70879535 | 70879655 | 788855_47188552_ENST00000601538.6,E | 1 | - | good | GGAGTCAGGAAATAGTCACAGAGATCTTAATTTTTGTTAATATGCAAAAGCACCTGGGTATTGGAGCTGAAGAAAAAGCGTTTGACCAAATCCCAGAGGGACGAGGGTTCCAATCCGGAA | 120 |
| chr11 | 70879655 | 70879775 | 788855_47188552_ENST00000601538.6,E | 1 | - | good | GGGTCCCACCGCCGGTGTCCTGCTGTCAGAGCCCATCGATCCTCCATAAGCCTGGGAGGGGGGCCCCGCATGGCCTTTGGGGCAAGGGATTCAACCCCTGAATTTCTAGCCACTGGCATT | 120 |
| chr11 | 70879775 | 70879895 | 788855_47188552_ENST00000601538.6,E | 1 | - | good | TGGACTCCGCTTTCCCTGACTCCTTGCTTGTGCCCTTCCCTGGGGAGTCCCTGGCTCTCCAGGCCTGGAAGCAACTGCAGGAAGCCAGCCGAGGCCCCATTCAGCTGGTGCCAGGAGAGA | 120 |
| chr11 | 70879895 | 70880015 | 788855_47188552_ENST00000601538.6,E | 1 | - | good | GAGTGGGCCACAGCACAGCACAGTTGCAACAGCAGGGTCCTGAAACTCGGAGAAACATCTGGCTACCCTCAAAAGCCAGGAAGCTCTGTCACCATTCCTGGGAGAGGCCTGGGATCCCTG | 120 |
| chr11 | 70880015 | 70880135 | 788855_47188552_ENST00000601538.6,E | 1 | - | good | GGATAGACATTGGGGTGCCTGGGCTAACTATCCCAAGGCTAGCCGGGGAGCTTGGAGCCCTGGCAGGCACAGCACCGGCCCCTGGGCCTCGCACTGGAGAGCTGCACAGCATCTACCACT | 120 |
| chr11 | 70880135 | 70880255 | 788855_47188552_ENST00000601538.6,E | 1 | - | good | TTATGGCTGGCTTCATAGACATCAGGGTGCATGTGCTGTTTATACCCAGGCTAACCAGGAAGCTCAGAGTAGACATCAGGCTGCCCAGGTTATCTATGCCCAGGCTAGCAGGTGAGCTCG | 120 |
| chr11 | 70880255 | 70880375 | 788855_47188552_ENST00000601538.6,E | 1 | - | good | CCCAACTTTCTTCGTAGACTCTTCCAGCAGCCCCCGAAGCCCGGTCTCCTCCCCAGCTGACCACAGCCCTTCTTTCGACCAGGATGCCCGATTAGTGCACCTGCCTTTTCACATTTAGAG | 120 |
| chr11 | 70880375 | 70880495 | 788855_47188552_ENST00000601538.6,E | 1 | - | good | TCTCCGGCTTTCTCTTTTCCAAGCCCCTTCCAGGCAGCTCACAGGCAGGAGCTGTCCCTGCCCCATCTTTCAGTCCTCTGTTCAGGGTGTTCCTTTTCTCAGCGCACCTCCATCCATCTC | 120 |
| chr11 | 70880495 | 70880615 | 788855_47188552_ENST00000601538.6,E | 1 | - | good | AGTCTTGACTTCTGTGTTCCATGGGCACATGCCCCTGGCCTCACAGAGGGGGCATGCCCGCTTCCAGGTGACTTGAATGGAAGTCTCAGAAGCTCCTAATTCTGAATGGCTCCACCTGCC | 120 |
| chr11 | 70880615 | 70880735 | 788855_47188552_ENST00000601538.6,E | 1 | - | good | GAGGTTCTGGTGAGGGTACACAGACTGATGGGAGCAGACATCGTGTTCCCCAGCCCCAGCCCAGGGTCTCCCCATCCCAGCATAGGGCACAAGCCCAGGGGTGGGCGAGGAGCAGAGAAG | 120 |
| chr11 | 70880735 | 70880855 | 788855_47188552_ENST00000601538.6,E | 1 | - | good | TGGCTGATGGCCCTGGGAGCTCTATCCTCTCAGGATGCCTATCCACAGGTGGCAGCTGGGACCCTAGCCCCTTCAGTCCTCCTTGTTCTATGGAGGGAAAATCTATGCGTCAGTGGGGCA | 120 |
| chr11 | 70880855 | 70880975 | 788855_47188552_ENST00000601538.6,E | 1 | - | good | CCAGGGAACACAGACTCAGAATGTGAAAGGGTCCTGGGAATCTTGGCAACCCCTGCTTAGTCTCATGACTCACAGACCCAGCGAGAAATCCCCCTTGGAGATTCTGCACCTGAGCAAATG | 120 |
| chr11 | 70880975 | 70881095 | 788855_47188552_ENST00000601538.6,E | 1 | - | good | ACTTTGCTTCAGCCTGGATGACAGACAGAGATTCTGTTTCTATTAACAAAAAGAAAGACTCTCAAGGTGCTTCCAGAAGTAATTGTCGTTCAGGGTTGGGATGCCCAAAAGTTGGTGACC | 120 |
| chr11 | 70881335 | 70881455 | 788855_47188552_ENST00000601538.6,E | 1 | - | good | GACATAAGACACTGTAACCAACCCCATTAGCCATTGTTTTAAGATTTGTGTCTATCTTGTACCCCTGATTCACCCACCTGCTTCATGGTGAAGCTGTAGAAAGACTCTTGGCCAGGCAAG | 120 |
| chr11 | 70881575 | 70881695 | 788855_47188552_ENST00000601538.6,E | 1 | - | good | TGGAATTCTAGAACTCTGTGCTCACCCATCCACCTGGGGAGAAAGATTATTGGGATCTACCCAGAAGAGGACAATCCAAGCCCCCTGGAGTTTACTATTATTATTATTATTATTATTATT | 120 |
| chr11 | 70882055 | 70882175 | 788855_47188552_ENST00000601538.6,E | 1 | - | good | TCCTCCTCCTCCATGCTGTTTGTCTCACCTGCCAGCATCATGCCAGTGAGTCACCCTTTAACCCTGGTCCTCAGTGTGTCCCCCTGGGGGCTTGGGGAATTCACACTGATGTTTTCCGCC | 120 |
| chr11 | 70882175 | 70882295 | 788855_47188552_ENST00000601538.6,E | 1 | - | good | CCCCGGAGCTCACCAGGCCTGGAAACACTTGGTGAATGGGCCGAGCAGAGCCACCCGTTCTTCTCTCCTCCACTTCCTTCTCCCTTGCTCATTCCTTCCCTTTCCCCACTCCCTCCCTCC | 120 |
| chr11 | 70882295 | 70882415 | 788855_47188552_ENST00000601538.6,E | 1 | - | good | GATATGTATTAAATCCCTTGCTTGTAGGCTATAGAACTGGTTCCCAGCTCCTAGCTACAGGCACAGCCCAGGGGCCTTTCTGCCTGCCTTCCACACTGCAGTGGCCCCTCTCCCGTCCCA | 120 |
| chr11 | 70882415 | 70882535 | 788855_47188552_ENST00000601538.6,E | 1 | - | good | ACATGTCAGCCTGCAGCAGTGACCAGATACTGCCAAGGTAAGGAATCCTTGGCAGCTTCTGTCATGATCTTTTTCACTGGGCCATTTTATCAAGGAAAATCAATTTCCCCATCACTCTGA | 120 |
| chr11 | 70882535 | 70882655 | 788855_47188552_ENST00000601538.6,E | 1 | - | good | TCCTGGCTGATTTTGCAAGCAAGTTTTGGGGTCTGCCCTACCTCTGACCTAAGCCTGCTTCTGCACAGACTAGCCCAGCTGGCCCCTGCTCCTGTTCTGTGACCTATTAATGACAAGGTC | 120 |

| | | | | | | | | |
|---|---|---|---|---|---|---|---|---|
| chr11 | 70882655 | 70882775 | 788855_47188552_ENST00000601538.6,E | 1 | - | good | GTGGACAGATGGCACAGAGGTGGCCTTTCCTGTGGACATGAGGATAAGCCACTGCCGCCTTTTAGCAAAGCCGTGGGGATCACCGTCCTCTCCCTGCACCGCCACCCCTCACCCCTGTGC | 120 |
| chr11 | 70882775 | 70882895 | 788855_47188552_ENST00000601538.6,E | 1 | - | good | GTTCACCTGCAGCTGTTTCCTCAGGACCTCCCCTCACTCCTGCAAAATCCACACAGTTCTGCCCTCCTGGAGGACCAGGCCAGCCGCTTGGCTTCCTGTGCTCGCAGATGCTGGGCCAGC | 120 |
| chr11 | 70882895 | 70883015 | 788855_47188552_ENST00000601538.6,E | 1 | - | good | CACTGCTTCTCTCTCTGCCACCTTGACCTCTGGGACCCTGGCTCCTGCCCTCCCCTCCCACAGGCCGCAGGCAAGCCCGCCCTCACCTCCACCTGCCTCCTGCCTGCCCTGCAGTTTGCT | 120 |
| chr11 | 70883015 | 70883135 | 788855_47188552_ENST00000601538.6,E | 1 | - | good | ATGAGGATCCTGGAGCCCAGACACTTGACAAGGGTCCCCAGGGTGGGGATGCAGCTGGGGCTGGAACCGACAGCCCTGACTCCGGAGCCGCCCCTTCACTCCTGCTAAGGCTGCCCATAA | 120 |
| chr11 | 70883135 | 70883255 | 788855_47188552_ENST00000601538.6,E | 1 | - | good | AAATAGCTCCTGTGCCTTCAGCCCTTTCAGAGCCAGGCCTGGGCTCAGTGCTCAGTGCTCAATCCTAATGACAGCCCTGTTAGGGAGGCGCCTTCACTGCCCCGTTTTCGGCGCAGGAGG | 120 |
| chr11 | 70883255 | 70883375 | 788855_47188552_ENST00000601538.6,E | 1 | - | good | CGTAGGTATCAGGGTGGGAAGGCTGAGGCCAGCATTAGGGGGTGCTAAATCCCAATGGGGCATAGAGTCAGGATGGCTTCCCAGAGGAGGAGGCAACTGCTGCCTTGATCTGAGCAGCAA | 120 |
| chr11 | 70883375 | 70883495 | 788855_47188552_ENST00000601538.6,E | 1 | - | good | ACATGGAAGCCCTGCACCAGGTCCAGGGGTCAGGTGGACCAGGACTTTGTGCCATAGCTGAGGAGCTGGACTTTTGGCTGTTGGTTTCCTACAGATGGAGGACATTTATGTTATACCTGC | 120 |
| chr11 | 70883495 | 70883615 | 788855_47188552_ENST00000601538.6,E | 1 | - | good | AGGTGGGCTCCCGGAGTCCTGTGTGGCCTCTGCCCCCTTTCCCAGCCCCCTTTGTTCTCTTGGTCTGGTCCTGCCTGCTCGGGTCAGGTTCTGGATTGTCCCTGCCCATGTCATTTCTCC | 120 |
| chr11 | 70883615 | 70883735 | 788855_47188552_ENST00000601538.6,E | 1 | - | good | TGCCCACCCCTTCTCTGTGCCTCCGGCGTGGGTCCAGGAGGGTGGTGAGGTGCACCCTGGGGGCTTTCCCGGGCCCCCTAGTTTTGCTCTGTCCTGCTCCACACCGCAGCACACCCTCTC | 120 |
| chr11 | 70883735 | 70883855 | 788855_47188552_ENST00000601538.6,E | 1 | - | good | CCGTGCCCTGCCAGCATCCTCAGCGCCTTCTCCCCTTGCCTGGGCCTCCCCAGCCCTGCCTGGCTTTCAGGGCACATGTCCTAGCCCAGGTGGAGGCCGGGTCTTGTTTCCACGTGGTCT | 120 |
| chr11 | 70883855 | 70883975 | 788855_47188552_ENST00000601538.6,E | 1 | - | good | CTGGCCCAGCAGGTTGGCCTCTGGTTCCTCAGCTCTGACTCCTGCAGAGGGCTGAAGACCACCAAGGTCATGACCAAGGCAGATTCCTCCTTCCTGGTCCCCTCTTGTAAGGTGTTCCCT | 120 |
| chr11 | 70883975 | 70884095 | 788855_47188552_ENST00000601538.6,E | 1 | - | good | TCTCCTCTTGCTCTCTGCATCCATTCTCCCAGAAAGTTCCTCCCTTGCCCCTGAGCGCCACCCACCCCTGCTCCTGGCTCCCTGCAGATCTCTGTTCCTACCCTTCTGAGACCGAGGACT | 120 |
| chr11 | 70884095 | 70884215 | 788855_47188552_ENST00000601538.6,E | 1 | - | good | ATTATCACAGCCCCGAGCCCTCTCTGGTGGGGTGGATGCCAGCCTGGTGGGAAACACTACTCGGGCATTCACAGGGTGTGCCCTGGTGCACTGGCCTGAGCCAAGCATGGGCTGGTGCAT | 120 |
| chr11 | 70884215 | 70884335 | 788855_47188552_ENST00000601538.6,E | 1 | - | good | CATTTGTCTGAACATCTGGTAGGTCACTCACAATCCCTAGGGGCCCGTTCCACGCATTGCAGGACTTTAGCCTGTCTGGCCCCTGCCCACCACGTGTGAGAAGCACCCCCTGACCCCGTT | 120 |
| chr11 | 70884335 | 70884455 | 788855_47188552_ENST00000601538.6,E | 1 | - | good | AACAAAGCCCATGCTGTCTTTCATTGCATTCCTGGAAAATTCGGGGAGTATTAAAGCTGTGCAGATTTGTGTGGTGTTTCTAAGTAACACGCAGGAGGTTGTGCAGATAAACAGATTTTG | 120 |
| chr11 | 70884455 | 70884575 | 788855_47188552_ENST00000601538.6,E | 1 | - | good | GTCACCATGAACCCCAGAGCTCCTCCAGCAGCTGGGAGATGCTCAGCCATGGGTCGGGTGTGGTCCAAGGCAGGGCTGGCTTCCAAAATAGTCAACAGCTCTTAGGAAAGTTCCAAACCA | 120 |
| chr11 | 70884575 | 70884695 | 788855_47188552_ENST00000601538.6,E | 1 | - | good | AGCAGGAACACTGTTCTATGAAGCAGGAACACTCGGTGGGGGGGACCGGACACCGTAGGCAGGGAGAGGTCTGTCCCAGAAGACCCAGTGCAGTTTCAATGGTCATTCCCTTCCAATCCG | 120 |
| chr11 | 70885055 | 70885175 | 788855_47188552_ENST00000601538.6,E | 1 | - | good | GCACTTACGGGAGATGACTTTGTTCCCTGGTTCCCAACCACACGAGACCCCAGTTCTCTTAGAAAAAGTGTTGTATAACATCTCCTTCATTATCCTCAAATGAAACTCTTAGATAACATA | 120 |
| chr11 | 70885175 | 70885295 | 788855_47188552_ENST00000601538.6,E | 1 | - | good | AGAGCCTTTCCCACTCCATCCCCCACGGAAGGGCAGAAGCTCAGAGAGTCTCCACTGTGCCCTAGCCTGACCCTCACCATGCGGTCCCTGCGCGGTTCCCCTCTATACAGCGCTCCCAGG | 120 |
| chr11 | 70885295 | 70885415 | 788855_47188552_ENST00000601538.6,E | 1 | - | good | GATCCCCCATGGTGCACACAGAGATCGACTCCTTCCGTCTTCTGGGTTGAACTCTTCCCCAAACCTGAGCTTCTCCCAGCTCTTTGCTCCCCGGTCCTGGGCAGTTCTCACCCTTAGAGA | 120 |
| chr11 | 70885415 | 70885535 | 788855_47188552_ENST00000601538.6,E | 1 | - | good | GTCCCCTCTTCAGCCAAATGGAGCACTGTTCAGGGGGACTGGGTGGAGGACACGGAGTCCCTGGCCGCACTGACTCCCACACAGTGGACACGTGGCTGCTGGGAAGTGTGTCCCACTGCG | 120 |
| chr11 | 70885535 | 70885655 | 788855_47188552_ENST00000601538.6,E | 1 | - | good | GCCCAGCATGCATCTCTGTGTGCCTTGTCTTTGTTGATGTTCGTTACCACGTTCTGTCCACACCCTTGCCCCATGCCTCCTGCGCAGCCCCTTTACTCACTGGAGCCCGCCCTGCGGCCT | 120 |
| chr11 | 70885655 | 70885775 | 788855_47188552_ENST00000601538.6,E | 1 | - | good | CCGCTGGAGGATCAGCCGTCTTGTGTGGCCTGCCCCGAATGCTGTGTACATGGTGCTCCCGGGGCAATTGTGGATTTTGTCCTGATGGGATGCTGCAGTGACAGTGAGGCCGGAGCTCCT | 120 |
| chr11 | 70885775 | 70885895 | 788855_47188552_ENST00000601538.6,E | 1 | - | good | CTGGCGGGGATGGGATCCAAGGTGACCGGCTTTGTGTAAGGGCCACAGGTGGCCAGCGTGTTGGGTTGGGAGGGTATTTTTACATTTGGCCGATTTCCAGCACCGACGCCCAGGGAGTGT | 120 |
| chr11 | 70885895 | 70886015 | 788855_47188552_ENST00000601538.6,E | 1 | - | good | TTGCTTGGTGTCATTTTCATGTCAGCTGAAGGGGTCGGTCACGTGCGTGTTGCTGTGGTCCTACTTTCTCCCGTGCGTATCTGTCTAGAAGTTTTGTTCCAAATTGTAAAGCGGAGTTGA | 120 |
| chr11 | 70886015 | 70886135 | 788855_47188552_ENST00000601538.6,E | 1 | - | good | AGCTGTGCAAGTCTGCGTGTGGTTTCCGGTCTGTGTGTGTGTGGTGCTGACTCCGCATGCCGGGCTGCTGGTTTCATCCGGACATAACCAGCTCTGTTGTTCGTCTGCCACTCTGGAGTC | 120 |
| chr11 | 70886135 | 70886255 | 788855_47188552_ENST00000601538.6,E | 1 | - | good | ACAGAAAACCCCCAACAGACACAGCAGTTTCAAGGCATATGGGAGCAGTCGGCCGCCTCTGTGACCTGTCTGTTGGGACTGACCTGCGTCTTGTCTCGTCCCTCTTCGCGGGACCGAGTG | 120 |
| chr11 | 70886255 | 70886375 | 788855_47188552_ENST00000601538.6,E | 1 | - | good | AGAATAAACCCTCCCAAGGAAAGTCAAGATTGTATTTTTACTTAGCGTTCTGCAGCAGTCACACAGCTGTTGACCAGATTCCCATAGAATATTTTCAAAACAAATTGCCGGAGAGGAAGG | 120 |
| chr11 | 70886375 | 70886495 | 788855_47188552_ENST00000601538.6,E | 1 | - | good | CTTTGTGTTTGCAAGTAGAACCAGCAGCTTCCAAGGAAAACCCCAAATCCTCACCTATTTCATTTCATTAGATCAGAGTGGTCGGGGTCCCCGTTATTGAGTCCCCTCCTGTGCATGTTA | 120 |
| chr11 | 70886495 | 70886615 | 788855_47188552_ENST00000601538.6,E | 1 | - | good | CTACCTGCTGTGAAATAAAATAGATCTGATTTGAAAAATTTCAGTTTACCTGGCATTTTGACTGTGAAATGTGCTCCTTCCAATGAGACACAAATGATGATCTGCAGTAATCACAATAGC | 120 |
| chr11 | 70886735 | 70886855 | 788855_47188552_ENST00000601538.6,E | 1 | - | good | CCGACGAAGGAATACTTGGATACTGTTGCTAGGTTTCAAGACCGGGTGCCTGTATAATTATGCTTTAGATATGTATAGATGATGAATAGAGGGTGGTGTTTTCAAACAGATTGCTTAATG | 120 |
| chr11 | 70886855 | 70886975 | 788855_47188552_ENST00000601538.6,E | 1 | - | good | CGAACAAGGTTTGTTTTCATACATCTTGTCCTTCCGCAGCTATGGAGAGAGAAGCGAGGGGTTTTTATATTTTAAAAGGAGGCACATTGAGCTGATGTGTGAGTTTGGGTGCCTGATTAG | 120 |
| chr11 | 70886975 | 70887095 | 788855_47188552_ENST00000601538.6,E | 1 | - | good | GATAAGCTTTGGACTGGGATAACCTATTTTTTACGGTCTTTGTTGTTTTTGCTTTTGCCCGGCTTTGTGGGGAGATGCACTGTCAGATGGATTAAATGACATGGGTTGTTCTTCCTCAAC | 120 |
| chr11 | 70887095 | 70887215 | 788855_47188552_ENST00000601538.6,E | 1 | - | good | TTTGCCATAGAGCCTGGACGATGGCTTCTGTGGGATGGAGATCCCAAATAGCAAGACTGTCCGGGTGCCGGGGAGCCCGCATCTGAAACAGTCGTGCACACATTAAGCATGTTGCCAGCT | 120 |
| chr11 | 70887215 | 70887335 | 788855_47188552_ENST00000601538.6,E | 1 | - | good | ATTTGCCAACTCGAAGAAAGTGATGGTTTAAATGTGGACGTGAATACCACAGTTAATCCTTGCTTGTTGTCGTCCAGTGTTTAGCCGGCCAAGTAAATGAGCCCAGCAGGTTATAAAAGC | 120 |
| chr11 | 70887335 | 70887455 | 788855_47188552_ENST00000601538.6,E | 1 | - | good | CGTTCTGTTCACAATCCGGCCAGCATGTATGTTGCTGTTGTCTGATGCTGTGATGCTGTTTCTGTCCCCTTCCTCTTAAAAAAAAAAAAAAAATTAAATGGGACCTCTGTTTGGTGATTTT | 120 |
| chr11 | 70887455 | 70887575 | 788855_47188552_ENST00000601538.6,E | 1 | - | good | GCTTTAGGGGAGAAGACAATGAACACTGTGAGGTGATGAATGGGTGCTCTGAGGTCTGTAGAACCCAGAAATGGATTCAAGTGTATCTCTTTAAAAGAGAGGTTTCAGCCTGCCGCGAAG | 120 |
| chr11 | 70887575 | 70887695 | 788855_47188552_ENST00000601538.6,E | 1 | - | good | TTTAAAAAGTCTCTTTAAAGGATCAAGTGGCCTCCCTGCTACTGAGGAGGGATCCTTTGTTTCCCAAACACAGAAGCAATCAGTGCCAGGAAATCAGTGCCCTTACGATGCTGGAATGTG | 120 |
| chr11 | 70887695 | 70887815 | 788855_47188552_ENST00000601538.6,E | 1 | - | good | AAGCAAATTCCAAGCTAGAGGTGGGTGTGGGGTGGATGTTGGTGCTTCAAAGTGGGTAAATTAAAATACCACACATTTTTCTGACTTGCCGTGTACAGCCTGGTCAATGGGATGTTTTCC | 120 |
| chr11 | 70887815 | 70887935 | 788855_47188552_ENST00000601538.6,E | 1 | - | good | TTGGCGTCAGGGCAAACCCTTGGTGCTCACAAGCAGAGCTTTCTCCTGTCAAGAAGTGGAGGTGAGACTCACCTTTGCCTCTATGGGGTTGGGTCACACTTCTCAATATGGGTCTGGTGG | 120 |
| chr11 | 70887935 | 70888055 | 788855_47188552_ENST00000601538.6,E | 1 | - | good | AGGAATCATGGCGTTTCTGCATTATGGTCTTCAGCAAGACTTGTCTTATACAACTTAGCTTAGGAGACATATGGATGAAGCGAGGCCTGTCCCTTAAGCCTGAGGCTGTTGGCTCATGCT | 120 |
| chr11 | 70888055 | 70888175 | 788855_47188552_ENST00000601538.6,E | 1 | - | good | TCTTCCTCAGGGTGAATCTTTTGTCCGTTCCTGGCCTCTTGGGTGGGGGTGGGTGGTGTCAGATGATCACCAGAGGACTCTAGATTCTTCACCTGCCATTTGTCTATCTCTGTCAACCTC | 120 |
| chr11 | 70888175 | 70888295 | 788855_47188552_ENST00000601538.6,E | 1 | - | good | GGGATTTCTGGATTTCTGCTTTAGATCGTGGGTGGCCATGTGGCTTCCTGATTCTAGGTTCTGTGGCTTGGCTTCCTCCACCCCATATCCTCCATAACCTCAGGGTGCTGCTGTAGTCCT | 120 |
| chr11 | 70888295 | 70888415 | 788855_47188552_ENST00000601538.6,E | 1 | - | good | GTTTTATCTGACTCGAACATTTTCAGAGTCCAGGAGCAGCCGAGCTCAGAGGCACAGAGGAGAGGAGGCAGTTGGAGGAAGTGGGCTGTGTTGGGTTTGATGTCGTGAGTCCTCAAGCCA | 120 |
| chr11 | 70888415 | 70888535 | 788855_47188552_ENST00000601538.6,E | 1 | - | good | GCCCTACTCTCTGTTCTTGACCTCAAACCTGCCTTCCCTGGTACCCGGTCTTGAGTGTCAAGGGTGGGACACATATTGGCAAATCAGTTGCTGCATTGAGTGTGTTTGCGAGAGGGCAAT | 120 |
| chr11 | 70888895 | 70889015 | 788855_47188552_ENST00000601538.6,E | 1 | - | good | TGCTTCATGACTTTTGTCTTCTCAGGCTGTTACCTCTTCAAGGACAGATGGTATCTGATTCTACCGTCCCCACTGTGCTAGGCACGTGGTGGGGCCTCAGGCAATGCATGTTGGATGGAT | 120 |
| chr11 | 70889495 | 70889615 | 788855_47188552_ENST00000601538.6,E | 1 | - | good | GGTTCCTCTGGAACTGCGGTTTCAAATCCCGCTGGTGGTTCTCATGCCGGACGTCCCAGCACCAGCTCCTCCTTTGCTTCTCAGGACCTGCCAGGCGTTTCTGCTCATCCACATCTTTCG | 120 |
| chr11 | 70889615 | 70889735 | 788855_47188552_ENST00000601538.6,E | 1 | - | good | CCACTGAGGCCCCCGCTGCCCAGCGGCCTGTATGATTTGGGGTGCATCTGAGTCCCTTGGGGAACTTGTGTGGAATGCAGATGCCCAGCGCCACCATCAGAGTCTGACTCAGTGGTCGAG | 120 |
| chr11 | 70889735 | 70889855 | 788855_47188552_ENST00000601538.6,E | 1 | - | good | CCCAGGAGCCTTCTATGTTGCTGGCATCCGCCCCTAAGACAGCTTCTGAACCACCCCACTCCTGGTCCTGGTCCTGGTCCTGACTGTCATCTTGACTCGCCTGCCCTAACCCTCTCCTAC | 120 |
| chr11 | 70889855 | 70889975 | 788855_47188552_ENST00000601538.6,E | 1 | - | good | AGGAAAACAGGCTTTTCCACACACGTTGCTTCTCTGGCTGTTGCAGCTCCTACTCCCCCATGTCCTGGCCCCAGTCTTCCAGCTGCGTCTGCCTCCTCTGCTCTCTCTCCATCCTCCATT | 120 |
| chr11 | 70889975 | 70890095 | 788855_47188552_ENST00000601538.6,E | 1 | - | good | CCTTCTACCCCCCAGATCTAAAAGTGATGTTTGAAGGATGGTGGATCCTGTGCACCCTTCCTAGGTGGAGTGAGCTACTAATCTGTTGCTGTTTAATGGGAGCACACACCACACCCGCTC | 120 |
| chr11 | 70890095 | 70890215 | 788855_47188552_ENST00000601538.6,E | 1 | - | good | CAAGTCTTCATTGAGCACCTTTTTCACAGACCCTGTGCTAATTTGGAGATTCAGAAATATTTGCATGTCCCAGGCCCTTTGGGGGAGGGGATGACCATGGAAGATAATGTCCCTCTGTCC | 120 |
| chr11 | 70891055 | 70891175 | 788855_47188552_ENST00000601538.6,E | 1 | - | good | ATGGCCCCAGTCTCTTCCCGTCTAGAGAGGAGCCCGCATACGCTGAGGAAGGCAGACTGCCTGTGCTAGAGGAGAGGCATCCTTTCTCCCAGGCTGTACTTGCACATGTCAGGTACCTGA | 120 |
| chr11 | 70891655 | 70891775 | 788855_47188552_ENST00000601538.6,E | 1 | - | good | CCAGGAGACCTTCAGAGGGGAGGTGCAGCCTGGCCCTCTACCCTGGGGGATGGAGACCGAGAGAGGGAGGGAGGAGGACACGCTTTTGGTGCTGCATAGTCACCTCTGTGGGATCACAGA | 120 |
| chr11 | 70891775 | 70891895 | 788855_47188552_ENST00000601538.6,E | 1 | - | good | TTTCAAAGAGGAATGTCTTCTGTGGCATTTCTCACCCACCATGAAATTTTCAATTGTCACTGGGATGCTCCTTCCCAGAATGCTCTTAGCAGTGTCAGGGTGAGGGTCCATCATGGTCAC | 120 |
| chr11 | 70891895 | 70892015 | 788855_47188552_ENST00000601538.6,E | 1 | - | good | TTTTAGTCAGAACTGGTCCTGGCCACCCCTGTCCTCAGCCGGGGCTCAGAGTAACTTCTCACCCACAGAATCTGTTGTTTTCTCTGCAAAGCCTAGATTCCTCCACCTTCCACATTAACA | 120 |
| chr11 | 70892015 | 70892135 | 788855_47188552_ENST00000601538.6,E | 1 | - | good | CCTCATGACGGTGTGACGGTGCTGCCTGGCAGCCCCCGAGGGAAGCCTTTGTAGCTCACAGTGGTCTCTCACGAGGGTGGCTTGAGGTGCTCATCTTCTGAGCTTTCGAGAGACATCATG | 120 |
| chr11 | 70892135 | 70892255 | 788855_47188552_ENST00000601538.6,E | 1 | - | good | GCTGTGCTTAGAAGGCAGGGTAGGGCTGGAGGAAGATGATGCTTAAGACCGCATGGGGTGGAGCCTGGCGGCTGGAGAGAGTTGTCTGAGAATGGATGGACTGATGGACTAAGGTGCCAG | 120 |
| chr11 | 70892255 | 70892375 | 788855_47188552_ENST00000601538.6,E | 1 | - | good | CAACAGGGCAGTCCCCCCCACCATAGTTAGAGGGGTATCTCCTCTCTGCTTCCTCATCCAGGCTCCCTCGCTCTAGGATTACTAGAGCTAAGCATTTGTCAGCTCCTAGCCAGCTCCCCA | 120 |
| chr11 | 70892375 | 70892495 | 788855_47188552_ENST00000601538.6,E | 1 | - | good | TGCAGGCATCAGTGTGGACTGGGATGGGGCAGGCTGGGGAGGCTTCATAGAGGCTGGACCTAGAACCAGAGCTGGGAAGGTGGCCAGAGGGAAGAGGAGGCTTCCTGGGCCACAGATCA[illegible] | 120 |
| chr11 | 70892495 | 70892615 | 788855_47188552_ENST00000601538.6,E | 1 | - | good | AGATTTCAACCCCATGTCCATCTGATGCTGAAGTACTTACCTTGAGAACCTGTCCTTTCAACCACCACATGCTGCTGCCTCCTTGTTGAAGCTGAGAAAAACCTTCCACCATACAAGAGC | 120 |
| chr11 | 70892615 | 70892735 | 788855_47188552_ENST00000601538.6,E | 1 | - | good | GGGGGTAAGCAGTGCACATACCTGGGAACAGTGAGTGCCAAGGAGATGCAGGCAGGGCACCGATGACATCTTAAATTCAGGACTTTTCCAGAGCCACATGCTGGAGGAATTGGGCAAACC | 120 |

| | | | | | | | | |
|---|---|---|---|---|---|---|---|---|
| chr11 | 70892735 | 70892855 | 788855_47188552_ENST00000601538.6,E | 1 | - | good | AGGGTGTGCCTGGTCATTATGAAGCAGTTCATGTATTTACCCACCAGTTCCAGCTCCTATGCGTGCTCCCTCTGGGTACCAAGACCCCCATCAGGGATTTGGGGAATTGCAGGGGCCTGT | 120 |
| chr11 | 70892855 | 70892975 | 788855_47188552_ENST00000601538.6,E | 1 | - | good | AGGGACTTAGGGAAGGAAGTCAGGGAAGGGCAGGGCACCAGCTGTACACATGTGTTCATGAGCAGTTGACTCTGTGGGCCCCGGAGGCTCAGGCCCCTGGGGTCCTCTGGGAACCTGCAG | 120 |
| chr11 | 70892975 | 70893095 | 788855_47188552_ENST00000601538.6,E | 1 | - | good | CCATGTGTAGAAGAGCCTAGGCGACATGGCATTTGGGGTGGGGCTCCCCTAGAAGCTGACCTTGAGGTGAGGATTTGAGGACAAGTTGTTTATTGTGGAGGAGATACAGGAAGCACCAGC | 120 |
| chr11 | 70893095 | 70893215 | 788855_47188552_ENST00000601538.6,E | 1 | - | good | GGGTGAATGAATGAGTAACCACAGTGAAAACAGTTAGCACTTACCAAGTACTACAGGCCAGCACTTCACGTACAGCATCACAGCCATGCAAACCTCCACGTCCAGATGCAGTTGTGGTCT | 120 |
| chr11 | 70893215 | 70893335 | 788855_47188552_ENST00000601538.6,E | 1 | - | good | GTGGTGGGGAGGGTCTCATACTCTCTTCCCCACTAGCTTGTGCGTTTTTTGCCTGCATGTGTCTTGCTAACTCCTGCATGCGGGTCTTTGCCCATTGTTTAAGAAAGCTGCCCAACTTCT | 120 |
| chr11 | 70893335 | 70893455 | 788855_47188552_ENST00000601538.6,E | 1 | - | good | AATCCTGAACTAATGCCTTCTTAACTAAACCTGTAGGTTGGAACCCAATTGCTTCTCCCCCTCCCCTGTGGTCACCCCCACTTTGCATCGCCCCTCTGGCAGGGGCCAGGTGCTGGGTAT | 120 |
| chr11 | 70893455 | 70893575 | 788855_47188552_ENST00000601538.6,E | 1 | - | good | TAGATCAATTGTGTTGAAAGTTTCCCGCAGAGAGAGATGAAGGGTGTGGAGGCTGAGTCCTCTAGGAACCTGACATCTCTCAATGTGTTGATTGATCCAAGCACAGTGAATCTGTTGTGA | 120 |
| chr11 | 70893575 | 70893695 | 788855_47188552_ENST00000601538.6,E | 1 | - | good | ATGCGGTTCCTTGTGCATCTACAAAGAAGAAGAAGAAAAAAAACCTCTTGTTTCATCTGAGTCATCTAAGGGTTCGTGATTATGAACTAGCAAACTGGCATCAACAAAAACTGTGCCATT | 120 |
| chr11 | 70893695 | 70893815 | 788855_47188552_ENST00000601538.6,E | 1 | - | good | GAAGCAGGACTGGCTTTTACAATCAGCTTACGCAGGGAGTCAGTGTCTGTTGTTCCCTGATTTTTGGGCTTTCCCAGGGCCCCAAAGGGAAAGGATAGTTTAGAAAATCTATACTTGCCC | 120 |
| chr11 | 70893815 | 70893935 | 788855_47188552_ENST00000601538.6,E | 1 | - | good | CCATTGAGAGAGGGGGGCACCCACCACAGGCTCACGTGACTAACATATTTTGAAGTGCCAGGATCGAGGTTTGAGTCTCTCATGTCCGTATGTTTGTAAAATCCCCAGTCCTGCAGGAAA | 120 |
| chr11 | 70893935 | 70894055 | 788855_47188552_ENST00000601538.6,E | 1 | - | good | ACTCTTTACGAACAGAGAGGTTTTTTTCATTATTATAAAAGGAATGCCCATTTGTTGGGCGAATACATTTTAATTACCCTTTACCTGTTCTGATCAATTGTTAACTGTAAACAGCACCCC | 120 |
| chr11 | 70894055 | 70894175 | 788855_47188552_ENST00000601538.6,E | 1 | - | good | CTGACTTTCCAAGTACGTTGAAGCACCGAGTTGATTTCACAATCCACCTTGGGAGACACTTGTTGTCTTTTCAGTTTTGGGATATAATTATGAATACCTTTCTTATTTATGATGCAGACA | 120 |
| chr11 | 70894535 | 70894655 | 788855_47188552_ENST00000601538.6,E | 1 | - | good | GAGGAAGGGGTGAAGTGGCTTCCTTATCCCAGCTCTCCACATGGAGCTGAGAGATATTGGGGCTGCTACCCCTGACCCTAAGTCTTGCCCCATGGAGTCACAGCCACCCTTCCTCCCTTC | 120 |
| chr11 | 70894655 | 70894775 | 788855_47188552_ENST00000601538.6,E | 1 | - | good | TGTTGCTGGCACCCCTGGAGGCAGAGCCCCCGCCAGCTCCACAGATCCTGAGTTGTCTGGGATGGTAACAAGCTAAGGGCTACCCCATTGGGCTACCCTCACTGGGGCCTGCACTGCAAA | 120 |
| chr11 | 70894775 | 70894895 | 788855_47188552_ENST00000601538.6,E | 1 | - | good | GTTTGAGAGCTAAAAATATCGGCTGCTGTGAAGGGAGTGGGACTCTCTTGACCAGGGGCACACAGTGAAGAAGGGAGGTAAGGGAGTCTGAGCACAGGTCCCGCCCATGACGACAGGGGT | 120 |
| chr11 | 70894895 | 70895015 | 788855_47188552_ENST00000601538.6,E | 1 | - | good | AGAGATTGATGAGTTAATGTCTGCAAAATGATTTAGAAAATGAACTGAGCTCTCAGAGTGCCAGAGACCGTCTTCCTGCTGCCTCCCTAATATGAAAGAGCAGCACACAGGATAGCTTTG | 120 |
| chr11 | 70895015 | 70895135 | 788855_47188552_ENST00000601538.6,E | 1 | - | good | CCAAAGGCTGCGCCTGACACTGACTTGCACATCCCTCCTGTGAGCCAGGGCAAATCACTTAATTGCTCTGAGATCCAGGGTTCTGCCAGGGCTGCCTCAGCACCTCCATGAGCGCGTTAT | 120 |
| chr11 | 70895135 | 70895255 | 788855_47188552_ENST00000601538.6,E | 1 | - | good | GCATGTGTGCCCTACGTCAACCACGGCCGGTACCTACCCCGCCTTGGAGAACCGCACGAGGATGCAGAGCCAGATGCATGGCTTGCGTCATCAAGCACCTGTGCAGAGCCTCGGGGCCTC | 120 |
| chr11 | 70895255 | 70895375 | 788855_47188552_ENST00000601538.6,E | 1 | - | good | CACCCCATCAAAGGGGTCAGGGGGGTGCTCCTGGGTGTCGTTTATTGCCTGTGCCCGTCACCTCATTTTTGGCATCAGGGAAAGCAGAACTGTGTTTGGCTGTCGAGGCTTTACCGTGTG | 120 |
| chr11 | 70895375 | 70895495 | 788855_47188552_ENST00000601538.6,E | 1 | - | good | CCCAGTCTTGCAACTTGGACGAGCCCACTGGCCTGTTTCCTCCAGCCCCGCGCTTCCACAGGACCTGCAGCGGGGAGCAGTCCCATTGGACCGGCAGTGGTGAGCTTTCGAAGATGGCAT | 120 |
| chr11 | 70895495 | 70895615 | 788855_47188552_ENST00000601538.6,E | 1 | - | good | CTGCTTCACTTCCTCCGCAGTTGTGCTGATGGCCATTCCCCTTTTTAGTACCCCTTGTGCCAACCAGGAGGAGAGTCTTTGGGGCTCCTCCTCTCTCCCATCGCCCTGCCATCGGCTGGG | 120 |
| chr11 | 70895615 | 70895735 | 788855_47188552_ENST00000601538.6,E | 1 | - | good | TTGGATGTGTTTATTGGAAGGTGCCGTAGCATCCACGCTTCTATTTAGGCAAAGAGCTGGTGACCCGATGAGGGTGGAGAGAGAGCCAGTGGCATCCTCTCGGCTCCCCTCTGGTTGCCT | 120 |
| chr11 | 70895735 | 70895855 | 788855_47188552_ENST00000601538.6,E | 1 | - | good | ATGACATCTTTTTTAAAAATCTAAAAAGAAAAGAAATCAAACTGGAGAATGTGCTAAGCATCAAGCAATGGATGTAGGGCACGGCTCTTGATTGCTGAAACAAATTAATTTGCAGCTGAT | 120 |
| chr11 | 70896095 | 70896215 | 788855_47188552_ENST00000601538.6,E | 1 | - | good | CCATGTGAACATCATTGATTTAGTAAAATGGGGAGAGGAACAGGAAGTGTTTAATTTATGCTAAGGGTTGGCTCTAGGATTCCCTATCTGATGTATGTATCTGATTTTCCCCATTAATGA | 120 |
| chr11 | 70896215 | 70896335 | 788855_47188552_ENST00000601538.6,E | 1 | - | good | TTTAACTTATAGTGGGGATGGGGCTAAGGAAGACCTTAGAGGTTAGGCTGTCTGAAACATGCCTCTCTAGAAGGATGATGTTCTGATCTATTCTGACAGTCCCTGTTGCAAGGTTTCTTT | 120 |
| chr11 | 70896335 | 70896455 | 788855_47188552_ENST00000601538.6,E | 1 | - | good | TGCTCTGAGGAATCAGTCACTCACCAGCTTTTTAAAATTCTCCTAAGATTGGTTTAGAGCCTGCTTTTAGCAAAACTTAACATAGTATTAGCCCATTTTCTGTCTCTGGTCTTAACTTTT | 120 |
| chr11 | 70896455 | 70896575 | 788855_47188552_ENST00000601538.6,E | 1 | - | good | CTTTGCAGGTGGCCATAATAGCAGGCAACTTTGAGCTGGCAGAATACATCAAGAACCACAAGGAAACAGACATTGGTGAGTAGGCACGTGGAGAAACTGTGTTGGGATTTGGTTTGGAGA | 120 |
| chr11 | 70896575 | 70896695 | 788855_47188552_ENST00000601538.6,E | 1 | - | good | TTGCGGCTGCCATTTTAAAGGTTCTGATTGGACTTCTTTTGGTGTTAGGCCTTCAATGATGTAATAGTATATGCAGAAATCATTGTTCTGCAGGTGACACTTAACTTAATGTGATTTTCA | 120 |
| chr11 | 70896695 | 70896815 | 788855_47188552_ENST00000601538.6,E | 1 | - | good | GACAGAGGCCCCCACTCATATAGGTTGTAGAGCAACGACAACCTTATCATCATAGAACAGCCAACTGCTTTGAGAGAAGTGAGTGTGGTGGTTGGCAAGGTGAGTGGATGAGTGTTGATA | 120 |
| chr11 | 70896815 | 70896935 | 788855_47188552_ENST00000601538.6,E | 1 | - | good | AGAGCTGGAACCAATCAAGCCCCGTGGGTACACAGGACATAGTCAATGGGGAGATTTCCAGACTCAGGCCTGGTTGAATCCTCAGCTGCTCTATGGGTAATTTGCCCCACATGCCTTCTG | 120 |
| chr11 | 70896935 | 70897055 | 788855_47188552_ENST00000601538.6,E | 1 | - | good | GAATTTTTCCCACATAACTGAAATCTGATGAGTTTTAAATGCTTTTTTTGAGGTTTCCCCTTCTTTCCAGTTATGTAGATTGGCATGCATCTAATCAGATAAACAGGGCTCCGGGGTGTC | 120 |
| chr11 | 70897055 | 70897175 | 788855_47188552_ENST00000601538.6,E | 1 | - | good | TCTCCCATATCAAAGATCATGCATCTTTAGGCAACAGATCTGTAGGGCCGGAGTTGGAACAATTTCCTGTAGGTGATGGTTTTGGTGCCACTCATCTGGCTGTGCATACATCTATGGGTA | 120 |
| chr11 | 70897175 | 70897295 | 788855_47188552_ENST00000601538.6,E | 1 | - | good | TGAGCCCAAGGTCTCTCCTTCTGAGGCACTGTGTGCCTTGCCCGTGTGGTCTCTGGTCCTGGCCTTCCTTATGGGTCTCCATGGCCATCCTAGGATACTTGGCTTTACCTGCCTGACTGC | 120 |
| chr11 | 70897295 | 70897415 | 788855_47188552_ENST00000601538.6,E | 1 | - | good | GCTGTTTGGATAAGGTCCACTTACATTATGACTTGACATCAGCGCATTGTAGATGTATGTGTATTTCTCCAGGCTGGCATCTGAATGCCTGGCTCCTTTTGTGGGTTATATGTATTGCCC | 120 |
| chr11 | 70897535 | 70897655 | 788855_47188552_ENST00000601538.6,E | 1 | - | good | AAATGAAGTAAGCCCATGTTGGGAAGATGTTATAGATATTTGTTATAGCCACACCCCACTCCTGGCACTAAAGTCTGCATTTGTTTATAATTTGTTGTAAAGAATTGGCTCATATAATTA | 120 |
| chr11 | 70898855 | 70898975 | 788855_47188552_ENST00000601538.6,E | 1 | - | good | CAGCTCTGTTACCCTCTTCTGTCAAATCCATTAAGTCCGATAGTCTTCTTTCATTTGTCTTATCTCCATCACCTTCCATTCAAACTGGCAGAGTTCCTGCTTTTATAATCCCATACATTA | 120 |
| chr11 | 70899455 | 70899575 | 788855_47188552_ENST00000601538.6,E | 1 | - | good | AATGTGGCCTGGAGACATGCTTATCTCTAAAGCCACCTTTTCTACCTTCATGCCCAAGGGAAGCCGCCACTCAGGAGGCTTCCCCCACATCTTAGTCCATTCTCACACTGCTATGAAGAA | 120 |
| chr11 | 70899575 | 70899695 | 788855_47188552_ENST00000601538.6,E | 1 | - | good | AGCAGACCAGGGCATGGGTCCACATCACCTAAGTCGAGTTCCCAAAGCTGAGTTGGAGGTGGGCTATAGGAAGAGATGGCCCCCTTTTGGGAGGACGGCACCTCAGCTTCTGACTCTCTG | 120 |
| chr11 | 70899695 | 70899815 | 788855_47188552_ENST00000601538.6,E | 1 | - | good | AAGGGCTGGTCTGTAGGAGTGCAGGAGAGATGCCTGGGCCTGTGGGCAGGGAGGCCTAGTGCCATCCTGAGCCAGCACAACCATCAGGGACCTGGAGAAGGACATGTCTGGCTCAGCCTG | 120 |
| chr11 | 70899815 | 70899935 | 788855_47188552_ENST00000601538.6,E | 1 | - | good | CCAGTGTCACATAGTACCAGTCTGGATTTACACCCAGGCAGCCCAGGCCAGAGCCCATGCCCTCCACCTCTGGGGCCGCATCCCTCCGCCAGTCCGCCAGTATCACCTCAGGCATTCGGC | 120 |
| chr11 | 70899935 | 70900055 | 788855_47188552_ENST00000601538.6,E | 1 | - | good | GGTTGATTGGCCATGTGCCACGCACTGCTCTAAACCTTTACTTGTGAGATCTCAATAAATCCACACACCAAACCTATTTCACAGGTGAGAAAACCTAGGTTTGTGGGTTACACGTAATGC | 120 |
| chr11 | 70900415 | 70900535 | 788855_47188552_ENST00000601538.6,E | 1 | - | good | TCAGCAGCTGTCCTAGGTGTGTGTTCCAGTGCAGATGTGGGTGTGCACTTGTGTGTATCTGACTGCGCATTTTTGTGGGGCGCTGGGTGACCTGGGGGTGGATTACCTTTACAAAGAGCA | 120 |
| chr11 | 70900535 | 70900655 | 788855_47188552_ENST00000601538.6,E | 1 | - | good | TTTGCACTGTGAGTTGAGTCCTTTAGACCAGCAGGGAGGTGCCTGGGCTTAAGCTCCTAGGGCTTTTCTTTGGAAGGAGAATCCCGGAATCACTAGGGGATGGTCCACGTACGTGTTATT | 120 |
| chr11 | 70900655 | 70900775 | 788855_47188552_ENST00000601538.6,E | 1 | - | good | GGCAGTGCCAGAGGACAGGCAGGAGGCTGCTTTCTGCCAAGAGGAATTTCTAGGCCTTGAAATCATCAAGGGGAGAAACAGTCAGGCAATTATCACAAGTAAATGTTTGCATTGTTCTCA | 120 |
| chr11 | 70900775 | 70900895 | 788855_47188552_ENST00000601538.6,E | 1 | - | good | TGCTGAGCCTGACACCAGCCTGACACTGAGCAATGCTGAGATGTCACATAAGACCCAGTTCCCTCTTCCTTCTCCCAGAAGACTTTTACTGGGAACCCATGATAAGCCAAGCATAGGGCG | 120 |
| chr11 | 70900895 | 70901015 | 788855_47188552_ENST00000601538.6,E | 1 | - | good | GCCTCGGCTGTGTCTCTGCTGTGAGTATTCTTTTCACTGCTGTGCACACCTGCAGCCACAGAGACAGACCCCACACTCTGCCTTCCTCATGCACCCAGACCAGCAGCCTGACACCAGCAA | 120 |
| chr11 | 70901135 | 70901255 | 788855_47188552_ENST00000601538.6,E | 1 | - | good | AGTCCATGAAAATGACAGCATATTATAAGCTAAGGTGATTAGATGAAGAGTTTGAACCCAAACGGCTTGAGTGAATGTCTTGACTTTGGTGCTGACTGTGATGTGGAGCCCGTTCCTTAT | 120 |
| chr11 | 70901255 | 70901375 | 788855_47188552_ENST00000601538.6,E | 1 | - | good | CCTGAGTGTGGCTTGTCATAACCCTCGAGAATAATTTGTCATTTGTTGTTTCTTATGCTTCTATTTTTTACAAAGGTGTTTGACCTGTGATTATATCATAACTTTTTAAAAAAATGCCAT | 120 |
| chr11 | 70901375 | 70901495 | 788855_47188552_ENST00000601538.6,E | 1 | - | good | GCTCCCCTCCAGGCCTCTAGGGCTCTGAAAAACCTGGACAAAGCAGCGTCCGTTCTTACAGGACTAGTCCTTCCCCTGCGCAGGGCACTTTTCACCATAGAAGATTCTGTGGGTGCCCAG | 120 |
| chr11 | 70901495 | 70901615 | 788855_47188552_ENST00000601538.6,E | 1 | - | good | CAGGTAGGCATCTGTCTGTATTACCGAAATGTTATTTGAAACAAGGAAGGAGGAGGACAGTTGAATTAATGGGGGAAAAACAGGCGAGCTGTAGAGAGCAGAGTTCTCATTTCTTCACTG | 120 |
| chr11 | 70901615 | 70901735 | 788855_47188552_ENST00000601538.6,E | 1 | - | good | TGCAGAGGGAGGCGACAGGAAGTCTCTCTACATCCTGTGGGGGTAAGGGGCTCCTTCCTGCTCTGGGTTTCTTTAAACCCTAACATTTCCCCCTCAAGGTTGCATTATGGCCGTACCCTG | 120 |
| chr11 | 70901735 | 70901855 | 788855_47188552_ENST00000601538.6,E | 1 | - | good | CCATCTTGCCTGTCTTGCTGCCACATTCACCACGTCTGACATCCATCCTCCAAACCTTTGCCTGAGCTGTAACCTCCACCAGGTGCTCCTTCTCCGGGTACTCCAAAGCTGGCGATGGCT | 120 |
| chr11 | 70901855 | 70901975 | 788855_47188552_ENST00000601538.6,E | 1 | - | good | TTTCTTGTCACCTCCTAGCCGCAGAAGCCAGGGTTCCCTTGGCTTCACATGTGGATTCCTCCCATGACTTACCAGTCGATGCCTTCTGTGCTGACCTCTGACCTTCCTCCCACCACACAT | 120 |
| chr11 | 70901975 | 70902095 | 788855_47188552_ENST00000601538.6,E | 1 | - | good | CCAGCCTCCTGTCTGTCCAAGAGTGCCAGCCGGGCCCCGGGCCATGGAGACATCTTGCCTGTTGCTGTTTTTCAGGGCTCCCTCGTATGGTCTGTCCCCAGATGCCTCCCAAGGCCATGG | 120 |
| chr11 | 70902095 | 70902215 | 788855_47188552_ENST00000601538.6,E | 1 | - | good | GCCAGTGAGGCGTCTCATACAAATGCCCAGTGTGGTGCAGTGACACTGCCCGGGTGGCCGCCTGCTGCTCTGCTGTCCAGTGGCATGGCACAGAACTGGCTGCCCATCGTCTGAATTCTC | 120 |
| chr11 | 70902215 | 70902335 | 788855_47188552_ENST00000601538.6,E | 1 | - | good | CACAGAAGAGAAACACAGACTGTTGCTTTGTGTTACATAACTCTTGTTTAAATGTCCATAAAAGGCCTTCCTCCTTCCCAGCCAGCAGATACGGAAGTGAGAATTAGGATGATTTAGAAT | 120 |
| chr11 | 70902335 | 70902455 | 788855_47188552_ENST00000601538.6,E | 1 | - | good | GACAGGTGGGGAGGCGGCAGGGGCTGGGACCTCGGCCGCACACTTGGCTGAGAGGCCAGCGAGGGAGGATGCCCTCTGCATCTTCCCTGGAGGCCCCGCCCCTCCCTTTCCTTTGAGATC | 120 |
| chr11 | 70902455 | 70902575 | 788855_47188552_ENST00000601538.6,E | 1 | - | good | CATAAGGTCACAGCAAAGGCTCCAGAAAGTGATGCTAGCTCTGTCTTGGGGCAAGAGCTAAGGGGAGCCCCCTGAGAGGCAGCCCCCAAGGGCACAGCGAGAAATCCCAGCTCTGAGGGA | 120 |
| chr11 | 70902575 | 70902695 | 788855_47188552_ENST00000601538.6,E | 1 | - | good | AGGACTTACAACTGGGGGTGATTCCCACAAAAAAAATCCTGTGGTGCCCTCATCACACAAAACTTCAGCCATCGATGCCGCCTGGAACTGTAGATCACACTTCAAATGCAGGATGTCAAT | 120 |
| chr11 | 70902695 | 70902815 | 788855_47188552_ENST00000601538.6,E | 1 | - | good | GCTTTCCTCTACTTGTTAATCCCAGTGCCAACTCTTGCAGGGGGCTTTGTCAGCACCAGCAAAAGGTTCCCATCTCAGATATGATTGCTTCATTCCAGGAAAACCCAGTTTAGGGCACTG | 120 |
| chr11 | 70902815 | 70902935 | 788855_47188552_ENST00000601538.6,E | 1 | - | good | GAGTGACTGCTGTTGAAAAAAAATGTGGGCTCCCTCCTGACATTCCTGCATCTGTAGGGGACCCTGTGGCTCTGAGTGCCAAGGGCAGAGCCCCCAGGGCCCCTGGCGGGGTGTCCATCT | 120 |
| chr11 | 70902935 | 70903055 | 788855_47188552_ENST00000601538.6,E | 1 | - | good | GGAGGAGATGGTTGTCAATTCTTGAGTTGGGAGAATAGATGGAAACTCCCCCTTCTCCCCCAGCTGGTAGATCAGTGGAGGGGGGGCCGTGCTGTCTCCTGGAGAGATGCACTGCCTTGCA | 120 |

| | | | | | | | | |
|---|---|---|---|---|---|---|---|---|
| chr11 | 70903535 | 70903655 | 788855_47188552_ENST00000601538.6,E | 1 | - | good | ACCAACCTCGAGCCCCGGGACCCCAGCCCCTCGTGGACAAGCCCTGGGTGCCTCTACTGTCTTTCCCAATGACGCTTGCCAATGATGAGTGTGATGGATCGAGTCCCCAGTCTTTCTGTG | 120 |
| chr11 | 70903655 | 70903775 | 788855_47188552_ENST00000601538.6,E | 1 | - | good | AAGAGAGGGGACAGGAAGTCTCTGCGGTGTTGCAATGCATGTGGCAGAAGAGACAACAGGGCTTCACAAGTCGGGGCTTCCAGCCAGTGGTCATGCACAGAGTATGTGGCAGGGGTCTGA | 120 |
| chr11 | 70903775 | 70903895 | 788855_47188552_ENST00000601538.6,E | 1 | - | good | TTTGGCAGAGCTCAGGGCTCTAGGAAGATGTGCAGGTGGCATTTTCACCAGCTTTGCGGGGCTTTGCATCTGGCACACAGCACCTTCTCCTGGTACTGCAAAGCTGGACAATGGCTTAAG | 120 |
| chr11 | 70903895 | 70904015 | 788855_47188552_ENST00000601538.6,E | 1 | - | good | TTGGAGGAAGCACCAGGTCGAGGGTCTGTGCTGGGAATTTGGGGTGCTGCTCCTGGAATGGCCTTACCTGCTTACACAGCATCAAAGTGGGGCATTTCGGGGAGAAGGAGGGTTTGGGG | 120 |
| chr11 | 70904015 | 70904135 | 788855_47188552_ENST00000601538.6,E | 1 | - | good | GGGCAGGCAACACCAAGCGAGGAACAAGTGCCCCCATGAAGTCTACAGGTGTCCTTGATGGGGTGGGTGCAAGGACTGGGTGGATAGCAGGGGCTGCCTAATTTAATACAGATTAGGGGT | 120 |
| chr11 | 70904135 | 70904255 | 788855_47188552_ENST00000601538.6,E | 1 | - | good | CCTGTGGGCACCGAGCACTGTCCTTTGGCATGAGACTCAGATAATTTCCAGGCACCCACCTTGTGGCAGTCTGATCCTTCTCACCACTTACTACTTGAGTGGTTTGTTTCTGAGTTTGTG | 120 |
| chr11 | 70904255 | 70904375 | 788855_47188552_ENST00000601538.6,E | 1 | - | good | AGCTTTTTAGGTGATGGCATGGGATTGCAGTCCACGCCACCTGCGGGACCTGAGCATGTAGCTTTACCTGCTGTGCTCACCAGCCCCCAGCCTGCCTGCACCTCCTGAACACGCTCGGGC | 120 |
| chr11 | 70905095 | 70905215 | 788855_47188552_ENST00000601538.6,E | 1 | - | good | GAGAACTGTGGCGAGGACCAACCAGACAGCCCCAGGTCCTCCTCACCTGCCTGTCTGGCTGGCTCATTTCCCACCATCAAGCCTGTCCAGGTACTGAGTCCCTGTGGGGCTGTCCTTGCT | 120 |
| chr11 | 70905215 | 70905335 | 788855_47188552_ENST00000601538.6,E | 1 | - | good | GTGTGCGGGTGGGCGTCTTGAGCCCAGGGAGGTATTTTGCACACAGTGCTGGAGCTGTCTAGGGAGCTGTCTTGAAAGGGGTTGCCTGATATAATAGGAGCAGTGGAATTCAGATCCTCA | 120 |
| chr11 | 70905335 | 70905455 | 788855_47188552_ENST00000601538.6,E | 1 | - | good | CAGGGAGAGTCATTTCGTCATTCAGAATGCAGGCTTACATGAAGCAGGAAGAGCATCACAGCAGGCCAAGTTCAGTGCTAGGTGGTATAGGATGCGGCACTTTGTGGTGTTCAGTGAAGG | 120 |
| chr11 | 70906055 | 70906175 | 788855_47188552_ENST00000601538.6,E | 1 | - | good | GTGACAGGTGCAACAGGCTGGGCGCTGTCTTTGTCTGTTCAGGCTGTCACATCACAAAATACCATAAAATGAGCTGGGTGCAATCCTGTGATCCCAGCACTTTGGGAGGCCTAGGCCGGC | 120 |
| chr11 | 70906175 | 70906295 | 788855_47188552_ENST00000601538.6,E | 1 | - | good | AGTCCCGCTCCATGGCGCCCTCACCACTGTGAAGAGCATTCGCATGTGCTGCCCATGTCCCAGCCCTGAGCAGGAAGGCAGGGCTTTGTGGCACCTTGTCGGAGCCTGGGGCTGGCCTGG | 120 |
| chr11 | 70906295 | 70906415 | 788855_47188552_ENST00000601538.6,E | 1 | - | good | CTCCACCCTCGCCTGGGCTCTACCTCTTGGCGTCAGGGAACAGGTCACACTGTGGGCACTGCAGCAAGTCCACAGCCTCTCCAGGGCCCTGGCCACCTGAGAGCCACACTCAAAGGCGAC | 120 |
| chr11 | 70906415 | 70906535 | 788855_47188552_ENST00000601538.6,E | 1 | - | good | GAGGCACATTTCGTTTTGGCAAGAGAAGGAGGCTTCGTCGGGGCGGTGGAGTGGCTTTCCTCCGAGTTCTGGCTGTGGATCCCAGGAGCTTGTTGAGCAAGTGATGAGGAGAGAGGTGGC | 120 |
| chr11 | 70906535 | 70906655 | 788855_47188552_ENST00000601538.6,E | 1 | - | good | CAAAGACGGGGACATCCCAAGTGCTCGGTTTCTCTCGTGCCTTTCCGAGAAGCCATTTTCCAGTTATGGCGAGGGGGTGTCAGGATCACAGAGCCGCCTCTAAATAGCTCTCTGTGTCGG | 120 |
| chr11 | 70906655 | 70906775 | 788855_47188552_ENST00000601538.6,E | 1 | - | good | GCTTGGAGGATGCCTGTAATTAAAACATTCTTACCAGGCCCTGGGGTCTTTGGGCATCAGTGGCTTCCTGAGTTGCCACAGGAGTCTGAAATAAATGAAGCACCATTATCTGGAAGGAAA | 120 |
| chr11 | 70906775 | 70906895 | 788855_47188552_ENST00000601538.6,E | 1 | - | good | CTGCCTCTCCAGATTCTGGGTTTTAGCCTGGGGATGGGCAGAGATTCCCTTGAAGTTTGCAGGGGAACTCGGCGACCCCTGACCTTGGTGAGCCAGGAGCCCAGGCCCACTCCCTGCCGT | 120 |
| chr11 | 70906895 | 70907015 | 788855_47188552_ENST00000601538.6,E | 1 | - | good | GTAAAGTCCCGGAGCAACCAGGGGCTTCTGGGTAATACACATCGATTTGCAAAGGCCCTGGCCCTGCATGGGGTTTTCGCAAAGCCCCAGGGAGCTGTTCTGGAACGACATAGGCCTCAA | 120 |
| chr11 | 70907015 | 70907135 | 788855_47188552_ENST00000601538.6,E | 1 | - | good | CCAGCTGCTTCCTCTGGCCGGGGGTCCTGGCCCACGACTCCCCTGAGCCGGCACTTCAGGAGTTGTGGACATTGCTTGAGTCAGCCCTGGCTCTATCTGGTCAGCCCTGTCAAGTTAGAT | 120 |
| chr11 | 70907135 | 70907255 | 788855_47188552_ENST00000601538.6,E | 1 | - | good | GGGGATACTGGCAGACGAGACCCAGGCAAGTCTTACTCAAATGCTCAGGGTCTTCCTCTAACCTGGGAGGCTCACTGGGTCACCCAGCATGGAACTGCTCCTACAATAGCTTCCCAGTCC | 120 |
| chr11 | 70907255 | 70907375 | 788855_47188552_ENST00000601538.6,E | 1 | - | good | CCTGAGTGAAATCTGGCCACAACCCAGGGGGTCCCCTTCACTCAGCCATGGGAGGGGTGCTGAGGGCTGGACCTGCTCCACCTTCTGTGGGACTGAGGGGCTGTTTCATCCCCAGCCTCT | 120 |
| chr11 | 70907375 | 70907495 | 788855_47188552_ENST00000601538.6,E | 1 | - | good | CTTATTCTCAGTTTCAGGAGCCTGAGAAGGGGGGAGGTTGTCTTTGGTATCGGTGCTAGAAGGCTCCCTCTGGGGCTGTCCAACGTCCAAGCCTGTGGTCCTGACCACCGCACAGTGCTA | 120 |
| chr11 | 70907495 | 70907615 | 788855_47188552_ENST00000601538.6,E | 1 | - | good | GGATCTCTGCATTCAATTTGGTGATACTGAATGGCGCTATTTTAAGTCATTTGCCAATTATTATTTGCTAATTAACTATTTTAATTCTCATGATAGTCATAGAAGAATCATACTTTTATT | 120 |
| chr11 | 70907615 | 70907735 | 788855_47188552_ENST00000601538.6,E | 1 | - | good | CTGTCAAAGATGATTATGTAGATTTCCCCTGCTTATCTGCTAATGCGATGAATTATATTTGTCAATTTTCTTATTTCTAAAATAAACTCACCTAGCCATGATATCATCTCCTTAATATAT | 120 |
| chr11 | 70907735 | 70907855 | 788855_47188552_ENST00000601538.6,E | 1 | - | good | TGTTGGGATTACAGGCATGAGCCTTCATGTCTGACCCAGTCTATTAAGTTTTATCATGAATCTATGTTGTCATATCCGATGTGTCTTGATAAAATATTTATCCAAAATCCGATCCTGCAA | 120 |
| chr11 | 70908095 | 70908215 | 788855_47188552_ENST00000601538.6,E | 1 | - | good | ACCCATGCAGCCCTTAGTTTTAGGGTGGTCTCTTAAGCAACAATCTGTTTCACTGCTGCACTAGCAGGCAAAGTTCACTTTTCTTCCCAGTTTATTAAGATTTTTTTTTTCTTTTTTTTT | 120 |
| chr11 | 70908215 | 70908335 | 788855_47188552_ENST00000601538.6,E | 1 | - | good | GAAAACAGATGTGTTCCAGCCAACAGCGAGGCCCATGTGAGGCCATCCTGCGTGTGCCAGCCAGGCTGACCCTCCACTTAAACGCAGTCACGTGAGGGAGTCCAGAGTGGCAGTCCTCAG | 120 |
| chr11 | 70908335 | 70908455 | 788855_47188552_ENST00000601538.6,E | 1 | - | good | ACAGGCGGAGTCTGTTTCCCTCCTTGGCAGCTTGACATGCTGTCTATGTGGAGCACTGTCCTGAGGACACCGTGTAAGGAAGCCAGCCTGGTCTGCTGGGGGATGGGAGGATGCATGGAA | 120 |
| chr11 | 70908455 | 70908575 | 788855_47188552_ENST00000601538.6,E | 1 | - | good | AGGTCAACCAGAAGGTCACCCAGAACACCTGCAGATCCCCATGACCATGTCTGAGCTGCAGTCATGGCAAAATCGCAGAATGAAACTGTGGCAGGAAGAAAAGTGCAGCCACTCCCATGC | 120 |
| chr11 | 70908575 | 70908695 | 788855_47188552_ENST00000601538.6,E | 1 | - | good | AACCCCAGCATCCCTGGGAGGTAACTTGCCAATGTCGGTGCAGCTCTGGAGGCCTGGTCCGAATCCCAGGTTGCTGTGCAGCCTTCGGAGCTTCTGCTCGATCCCAGGACCAAAGGACA | 120 |
| chr11 | 70908695 | 70908815 | 788855_47188552_ENST00000601538.6,E | 1 | - | good | TGATGTTGATTAGGTCAGAAACTTTCAAACGCTGAAAGTGGTTTTCCTCCTCATATATGATCCCCCACCGCCAACCCCTGGTTCAGCATTTCCCTTGTAATTACAGTCACCTTGGTGCTG | 120 |
| chr11 | 70908815 | 70908935 | 788855_47188552_ENST00000601538.6,E | 1 | - | good | TAAGGGGATGTGGCCTCCTCCAAGGAGGTGGCCAGTTTTTCCCCTGGCTGACGTTTCAACATCTTGGCTGGAAACTGAACGACTAAATGGAGGTAGATCAGTTGGGTCGCGGTTTGACGT | 120 |
| chr11 | 70908935 | 70909055 | 788855_47188552_ENST00000601538.6,E | 1 | - | good | GTGTGTTTGTTAAACTTACACGCTGATGTTTTTCGAGGTTTTTATTCCCTTGACTGCCATACGTATTTCACAGGACAGAGGATAATACTGCCCAGAAGGTCTCTTCTACCCTAAGAGGGG | 120 |
| chr11 | 70909055 | 70909175 | 788855_47188552_ENST00000601538.6,E | 1 | - | good | AAATGATGTCCTTCAATGCCATTGCTATTTCAAAGGAAAAATTTACATTTTATTGGCAAATCCTTCTGGGAGAGGGAGGTTGCATTTGGTCCACATGAAAGAATTATTTCCCATTCTTTT | 120 |
| chr11 | 70909175 | 70909295 | 788855_47188552_ENST00000601538.6,E | 1 | - | good | TCTAACTGCCCTGTGGGCTCACGCCTTTCTAGCCCCAGGGTCCCACCAGAGTCACTCTTGAAAATCATTCCTCTGCAAACATCAAAGAACTGTTGCTTTTCTATCAAGATTTCTGGGAGC | 120 |
| chr11 | 70909295 | 70909415 | 788855_47188552_ENST00000601538.6,E | 1 | - | good | CAGCACAGGAAAGCCAGCCTAGCAACATCTGTAATCCCCAGAAATCTCCTCTTCTCACTAGACTCAGCCTCCCTCATTCTGTGCCGTGAGTCTAGGGTTCCTGGAGGCCAAGTAGAAAAG | 120 |
| chr11 | 70909415 | 70909535 | 788855_47188552_ENST00000601538.6,E | 1 | - | good | CTCAAGGCATGCCGTGAGGGTGGAATTTAGAGACTGTATTTAGATGCTGGGCTGCGTGTCGTGTTGGGTAAACATGGGTAGTTATCATTATGATCAGCTTGATCACTCTCCTCATCATTA | 120 |
| chr11 | 70909535 | 70909655 | 788855_47188552_ENST00000601538.6,E | 1 | - | good | CCTAGGCTTTGGAATCGAATGGACAGGGCCTTCCTTTATGCAATGAAGAAATCTCACTCATTTGAGCTCTCGAAACCTGTTGTCTGATGTGCAGAACAAGGATCACGACAGCTCTACTTC | 120 |
| chr11 | 70909655 | 70909775 | 788855_47188552_ENST00000601538.6,E | 1 | - | good | ATCTCGTTGCTAATCTCCCTCTTGCCTTTTTTAAGCAGGGAGGCTGTGCCTGGGCTGCTGTGCAAGAGAACTTTCTGTAGTTAGAGCCACACAAAGCCATGTGTGTAATGCCTGCCTTCT | 120 |
| chr11 | 70909775 | 70909895 | 788855_47188552_ENST00000601538.6,E | 1 | - | good | GTGCTTAGTGATGGCAGCCTGATCAAACTCATACGCAAGCGTACCCCGTTGTCAGCCATGATTTCTTTCTTTGGTTACTGTCCTGCAGAAAATGGTGTTCAATTCTTGCTTTGGAACCAA | 120 |
| chr11 | 70910255 | 70910375 | 788855_47188552_ENST00000601538.6,E | 1 | - | good | CCACGTGAGGGGAGAGATTCTCAAGCCTTGTGCAGGCCTCAGACACCTCTGACCTGGGTGTTTGCGGGGGACCCTAGGTGTCCCTGAGGAAGGCAAAGAGATTCTGCGTACTCTTGTGCT | 120 |
| chr11 | 70910375 | 70910495 | 788855_47188552_ENST00000601538.6,E | 1 | - | good | ACTTTTCATAACCGAATCATGGGGATGGGAGAGGCAGTGATTGCTTCTGCTCCTGTGTGTCCATCTGTGACAATCGAGTGTGCGGTAGCCTCTTCCCACCCTCTGCTCCCTGAAAACAGA | 120 |
| chr11 | 70910855 | 70910975 | 788855_47188552_ENST00000601538.6,E | 1 | - | good | CATGGCTTAGCAAAAATCGAATTATGTGCACATGTGCCCCCAGCATGTGCACATGCACGCGCACACACACACACACGCACCACCAAGCAACATGTAAACAATTGTGTTATCCAGTTTGGC | 120 |
| chr11 | 70911095 | 70911215 | 788855_47188552_ENST00000601538.6,E | 1 | - | good | AACAACAAAAAAACTCATCCAACTCTGTGCTTAAGGCCTGTGTGTTTCGTGGCATGTACTTTTAGCTGAATTAAAACAGTACAAAAAATTAGGGGAAAGAGGCATGTGGAGACCTGTGCC | 120 |
| chr11 | 70911455 | 70911575 | 788855_47188552_ENST00000601538.6,E | 1 | - | good | AGCAATGGTGAGGTTGCAAAAAAGGAGAAGTTGGGGGGCAAATAAAGTTTAAAACCACCTACTTCACACACACACAAATAGCCAAAATTAAGAAGATGGACAGTGCCAAGTGTTGTCTGT | 120 |
| chr11 | 70911575 | 70911695 | 788855_47188552_ENST00000601538.6,E | 1 | - | good | AGCTGCAGGTGCACTCACAGCTTCTGTGAGGCTGATGTCTCCAGTTCTTGGCCTGAGTCCTGGAATCACTGCCTTTTTCCACCCTCTGGGAAAACGGATAATTAATTTTATTCTAATGAT | 120 |
| chr11 | 70911695 | 70911815 | 788855_47188552_ENST00000601538.6,E | 1 | - | good | GCCACCACGCCTGGCCACTGACCAGTTCTACCACCATTCTCACTCTTGTCATTTTGGGGTAGGAGATGATGTTCCTGAGCAAACAACAATAGCCATGAGTTGTACTCCCTGGCCCATAGA | 120 |
| chr11 | 70912175 | 70912295 | 788855_47188552_ENST00000601538.6,E | 1 | - | good | AAAAATGTAGTTGATACCCCATAAGCATGATTTTAGCTTATATAATTTGTTAAACTTCCTTCACATCATTGAATGAAGTAAAATATTTTCTGCTTGCATAAGGTCACGTGGTAAGTGTCA | 120 |
| chr11 | 70912775 | 70912895 | 788855_47188552_ENST00000601538.6,E | 1 | - | good | TGACACATGATAGTGACTTTAATGAAATTGTTAATTTGGATTTCTGTAGAAGCTGACCAAATAAAACCCCGTGTAAACACCTGTATTAGTTCATTTTCACACTGCTGATAGACATACCCA | 120 |
| chr11 | 70912895 | 70913015 | 788855_47188552_ENST00000601538.6,E | 1 | - | good | GTTCAGTTCCCTGAGACTGTAAATTGATAAGTCCAGAGAACTGAAGGAAATAAGACAGTTTTCCCGGGAGATCTTTTTGTGGTCAATTTTTGTTCCTGTCTCAGAAATTGTTCCCCGATG | 120 |
| chr11 | 70913015 | 70913135 | 788855_47188552_ENST00000601538.6,E | 1 | - | good | CATACCAGGAGGCTGTGAGAGGATGGGAGGGTAGGTTTTTTTTTTTTTTTTAATTTCCTTTTCTTTGCTGGAGAAGTGGAGGGGAAAGGAAATGAATGAGGAACTGGCTTAGAATCTGTA | 120 |
| chr11 | 70913135 | 70913255 | 788855_47188552_ENST00000601538.6,E | 1 | - | good | AGGAAAAGAAGGAAAACAAATCTCTTTATATAATAAGTTTCTTCTTTCTCACTGGGATTAATTATAAATTGAATGCCTTTTTTTCTAAACGACATGACCTAACCTGAATTTATTTATAAA | 120 |
| chr11 | 70913255 | 70913375 | 788855_47188552_ENST00000601538.6,E | 1 | - | good | GTGTTGAATTTAATTCTCATACCTCCCATCCATTTTAGCACATAGAGCCCCCTGGGTATTCCATATGGTCCCTGAAGGGTCCAAGTTGTCATAATAGGGTCTATCATTTAGGACAAATGA | 120 |
| chr11 | 70913375 | 70913495 | 788855_47188552_ENST00000601538.6,E | 1 | - | good | TTTACCTGCCAGTTACAAGGAACTTAATAATGATCTTTAGAATTTTCCTAATACAGTAAATGCCTTTTGATTTTTTTTTGTTCCCGGAGGAATTGAAATGCAATTTTATGTTATTTAATT | 120 |
| chr11 | 70913495 | 70913615 | 788855_47188552_ENST00000601538.6,E | 1 | - | good | AATTATTTGCAAGGATCTTAAAAACAACATTGAGGAAAACCATCCCTGGTTCTTTGCTGTTTCTTTCCCTCTTGGTCTCCCTCCCTTCTCCTTTCTCTGTTTGCTTCTTTCTCCTGAACT | 120 |
| chr11 | 70913615 | 70913735 | 788855_47188552_ENST00000601538.6,E | 1 | - | good | TCCGCATGAAGTCTGGGGCAGGGTTTAGAGCCAACGTCTAAACTCAGAATCTTCTCTTTTAGCCCCCAGATCCCAAAGTGGGAACTCTGAGGCCTTAGCTTATATTTTTAGTGGTTTTTA | 120 |
| chr11 | 70913735 | 70913855 | 788855_47188552_ENST00000601538.6,E | 1 | - | good | AAGACCGGCTCTGATTGAACCAGAAGCACAGAGAGCCTGGCCATCCGGCAATGACACCATGTAGGGTCCTAATGCCTGCTGCTCCTCGTCTCTGGAGAGGATGCCTTTTTAGGGTATGTT | 120 |
| chr11 | 70913855 | 70913975 | 788855_47188552_ENST00000601538.6,E | 1 | - | good | CTGCTCTAACGAGAACACCCTGATGCCAGCTGCTTTCCCTAATGAAGTGTGGCAGCATTACCTGAACAATGAGATGATTGTGTGGTTAAACGTGAGTGTTGTTTTCCAAATGGAAATGAA | 120 |
| chr11 | 70913975 | 70914095 | 788855_47188552_ENST00000601538.6,E | 1 | - | good | AGGGCCCGACAACTCCTTACAGAAATCCCCCACCGCCGCCCTGTTCTCTGCTTGCCGCCCCAGTGTAAATATGCTAAATCCAAAAGTGGATTAATTGAAAAGCAATAAACACTCACAGCA | 120 |
| chr11 | 70914095 | 70914215 | 788855_47188552_ENST00000601538.6,E | 1 | - | good | CAGAGCCCGATTTCCGGAAACAGGCTGTCCAATGCAGCTGAGCAGGGTGTGCCTTCTCAGGGCCCTGTAACAAGATGCTGCAGTGGCAGTGTGGCTGGGCGGGGCCCTGCACCTGCTTTG | 120 |
| chr11 | 70914215 | 70914335 | 788855_47188552_ENST00000601538.6,E | 1 | - | good | CATGCTCTCCTGGTGCCAGGTGGATTCTACCTGAGAGGAAGACCGAGAGGGTCGGGGGAGGGACTCTGTGTCTCAGCGGAACCCCATTGCACACTTGTTCCTGTGGCTCTGTATGAGCCC | 120 |
| chr11 | 70914335 | 70914455 | 788855_47188552_ENST00000601538.6,E | 1 | - | good | TTTTTAAGCTTCTCTCTGTATTCCCGGGAAGGCCCCAGCATGGACAGGTGGGAGGTGTGGTGTGATGACCTTTTTCTAGCCCCTGGTTTGAGGGGATGCCAGAAAAGCCTGGGTATTGCT | 120 |

| | | | | | | | | |
|---|---|---|---|---|---|---|---|---|
| chr11 | 70914455 | 70914575 | 788855_47188552_ENST00000601538.6,E | 1 | - | good | AAAGCACTGGGATTGCAGGTGTGAGCTACCGTGCCCAACCCTCCTCTTCATATACTCAGCCTCTTGTGTGTGTTTGCAAGGGCGTTGTGTTTTTGAAAGGCCTTTCTGTGCGAGGCTGGG | 120 |
| chr11 | 70914935 | 70915055 | 788855_47188552_ENST00000601538.6,E | 1 | - | good | ATTAGGGAAAGGAAGCTGAGATGCAGCTTGTGGAGAGGAAAGTGCATTTCCAAAGTGAATGGTGTTGTAGCCTACAAGATTCCTGCTGTTTCCAGTTAAATACTTTAACAACTGAATTTT | 120 |
| chr11 | 70915055 | 70915175 | 788855_47188552_ENST00000601538.6,E | 1 | - | good | AGGGATCATATTGGGAGCTGAGGATGAACGTGTCCCGCAAGCCAGCAGTACCCAGAAATCATCTGGAAAGTCATTGCCAGCCAATAGCTGTCGGCTTCACCCACAGATCACATTACTAAG | 120 |
| chr11 | 70915175 | 70915295 | 788855_47188552_ENST00000601538.6,E | 1 | - | good | TAACATGTGCTGGCGGATTTGGGAACAGCAGCCCTGCACATTTGAGATGAGATTCCCAGTGGCAGCAAAGTGTCTCCGAGGGCCCAACGTCTGTGTTTAAAGAAGGCATCCGTATTTTTG | 120 |
| chr11 | 70915295 | 70915415 | 788855_47188552_ENST00000601538.6,E | 1 | - | good | TTATTGTCTCTCATTATCTGTCCACAGATGGCACCATGAATCATAACTGAAAGAATTAAAATGTTAGGCGCCAGCGTGTAGTTGCTAGGGGATTTTTTTGGGGGTGTGTGTACAGTCTGC | 120 |
| chr11 | 70915415 | 70915535 | 788855_47188552_ENST00000601538.6,E | 1 | - | good | GAGTCCCATGTTAGCCATGGGAGTGGTTAGGCACTCCTGGGATGGCACCGTCCCTTCCCATACTTGGATGCAACTGATGCTGAAACTCTGCTGAACAAATGCCTGATAGCTATTCTTGGA | 120 |
| chr11 | 70915535 | 70915655 | 788855_47188552_ENST00000601538.6,E | 1 | - | good | CTTTAACTCCCTCCTGGACCTTGCAGCCATGCTCTCGGTATTCTCAGGAGTTTGAGCACAGTTGAGGGCATGATGTCACTCGGGTGGGCTCGGCGGCCAGACCTGCTGATCTGATTCACT | 120 |
| chr11 | 70915655 | 70915775 | 788855_47188552_ENST00000601538.6,E | 1 | - | good | AGCAAGCCTCAAGGGAGTCCCCTTTCTCATAGCCACAGCCCCTCCATCATCCCGTCCGTCCCGCGAGGCAGGCAGCCCTTTTGTGGGGTGGGCTCCTGTGAATTCTCTTATGCCTGTTGC | 120 |
| chr11 | 70915775 | 70915895 | 788855_47188552_ENST00000601538.6,E | 1 | - | good | TTTAGAATGAGAAAAAAATGATGTTGTGCCTGAGTGTCATAAGTCAAGTCATCCAGAAGTCTTTAAAGCATGGCCTTGATTTTATGACTGAGCAACGCATCTTTCTCCTTGGCAAGCTTA | 120 |
| chr11 | 70915895 | 70916015 | 788855_47188552_ENST00000601538.6,E | 1 | - | good | TGTGAATCGGGGAAGAGCACAACAAGGTGGAGGGTGTAGTGTCCAGAAACCCATGTCTGTGTCTGCTGACAGCCAGGCCTGCGCTCTGAAATAAATGGAGGTCAGGGAACACACACAGTG | 120 |
| chr11 | 70916015 | 70916135 | 788855_47188552_ENST00000601538.6,E | 1 | - | good | CTTTGGGCAAACCACTGAAAGAAGGTGGTAATTGCCCTTCATTCTTCTCTAATTCTTTTCTGCAGAAAATATGTAGGCCTCCCTAGACATGTCTGGCTCCCAAAAACAAGAGCAAGGTGA | 120 |
| chr11 | 70916135 | 70916255 | 788855_47188552_ENST00000601538.6,E | 1 | - | good | ACACGGGGCATGGGGCTCGACTTATCTGAACTCTACCCTCTCCCAAGAGGCAAGGGACGGCCAAGAATGTCCTTTATCCCTCAGGATTCACAGGTAGACACATTGCCCGGTTTAGAATGG | 120 |
| chr11 | 70916255 | 70916375 | 788855_47188552_ENST00000601538.6,E | 1 | - | good | GGGTTGGCGAGGGGTACTGTTCTATGCTGCCTCCCACTCTGCTGCAACTAGGCCATTTCCATGTCCCCCAACCTTTCCCCCAGACAGTGCCCGTGAGTATTAAACTCAGCATGAACTGTT | 120 |
| chr11 | 70916375 | 70916495 | 788855_47188552_ENST00000601538.6,E | 1 | - | good | CATACACATGAGTGACTGATGCTAGGATTTAGCTTTTCTGCTGTCTGCCAGCCTCGCATTTCAGAAGTGCCAAGAGCTTTCTGTACCTCTGTCCACTGTGTGGCCATGTGATGGCCAGCC | 120 |
| chr11 | 70916495 | 70916615 | 788855_47188552_ENST00000601538.6,E | 1 | - | good | TCATCATCCTCTTGCCGGTGAATCATGCTCCATTAGAATTCTGGAAATAATTCCTGAAGCTTGCTGTGCTACTGTGTGAAAGGGTGGCGGGGCAGCTGTGTGTGTGTGTACACGCACACA | 120 |
| chr11 | 70916615 | 70916735 | 788855_47188552_ENST00000601538.6,E | 1 | - | good | AGTCACATTAGCATTTTGCAAGCCTCTCTTCAGACAGCAGGAAGATGCTGAATGCTGTGCCAGGCGAAGGGAGGGCGAGGGGGCTTCTTGCCATTTAACTGACACAGGTGGATAATTAGG | 120 |
| chr11 | 70916735 | 70916855 | 788855_47188552_ENST00000601538.6,E | 1 | - | good | TTGTACATTAAGACACCAGAGCCTGCTGGGGTTCAGTTTCTTCTTGGACTCCATGTGCTCCTGGCTTCCCTTTCCCCAGGCTGCTTGGATACCTGCAAGCTTTCAGGATAGAGACCCTGG | 120 |
| chr11 | 70916855 | 70916975 | 788855_47188552_ENST00000601538.6,E | 1 | - | good | CATCCAAAAGCAATGGTTTCCTAACCTCCCTCCATACCCTCCCCACATGCACCTGGTGCTTAAGGTCGCAGGTACCGTGGAGATTGAGAAATGATACATTTGCTTATCATAGGAGGACAT | 120 |
| chr11 | 70916975 | 70917095 | 788855_47188552_ENST00000601538.6,E | 1 | - | good | GACTTGCTGTCTTCCTTCGAAGCAGCTCACTGCCCTGCTGCGTAGTGGCTTCTCTTGGTCGCTTTCCCTCTCCTCATGCTGACTTTTCATTTTCTGTGTCCTAATAAAAATTAAGCAAAG | 120 |
| chr11 | 70918055 | 70918175 | 788855_47188552_ENST00000601538.6,E | 1 | - | good | GTTGGCTCTCATGGCAGATCAGTGGGCCACCTGTTTTATAGCAATTGCTTTTCTTTCCTGGGTCAGGGTCCTCACAGTAAAATGACTTAAAATATGGGTTGCCTTGAGATGTTCATCTGT | 120 |
| chr11 | 70918175 | 70918295 | 788855_47188552_ENST00000601538.6,E | 1 | - | good | GACCTTGTAGCTGATGCAGGAGCCCTGTTAGGAGAGTCTAGTGACCTTTCTTCCCTAGGCCCCCGCCCCGCATGGAGCGGCTGCAAGCGTTCTGCCTTTGGGAGATGAATCTCTTGTTGA | 120 |
| chr11 | 70918295 | 70918415 | 788855_47188552_ENST00000601538.6,E | 1 | - | good | GGATGATGAAACTACAAGTGGGTGCTGGGTGACCTTGTGATTCTAGGACATAAATTGTATCAATAAAATTGATACCAATCGGCCTCTCTCCGTGGAGAGGGTACACGTCTGACTTGACCT | 120 |
| chr11 | 70918415 | 70918535 | 788855_47188552_ENST00000601538.6,E | 1 | - | good | ACAACACTCTGTCTCAAAAAAATAAAAAATCAATTAAAATAAGGATATCCATGTAGACTTTGGCACATTCTTCCCCAATTCCACTTCGACTGGCAGAGCCACCTGGGCTATACCTGGCAT | 120 |
| chr11 | 70919855 | 70919975 | 788855_47188552_ENST00000601538.6,E | 1 | - | good | CTATGTCTCTGCAGCTTTGGAAGCTGCCAGTTACCAAGCTTGGGTTGGTTGCTCTTGATTGATTATTTAAAAGACATCCATGGAATGAGCCCGGAGGACATAATGTTAAGCACCACAAGC | 120 |
| chr11 | 70920095 | 70920215 | 788855_47188552_ENST00000601538.6,E | 1 | - | good | AATTATGATTATACTAGTAGCTACTATTTATTAAGTGCTTGTGTATTGGGTAGTACTGATAATTATGCTGTATGCACTATCTCACTTTAATTGACAAAATGCCCTATGAGGTAGGGTCTG | 120 |
| chr11 | 70920575 | 70920695 | 788855_47188552_ENST00000601538.6,E | 1 | - | good | CCCCTGCAGTGGCGTGCTGTCACTGGAGTAGACCCCGTTAGGGACATGCAATTAAACTATTGTGCTTGAGAGACATTTGGAATAATTTCTCTCTATTTGTTTTTGCCAGCAGCTATGTTT | 120 |
| chr11 | 70920695 | 70920815 | 788855_47188552_ENST00000601538.6,E | 1 | - | good | TTTAACTTTTAATGAGCAACATCACTCTTCTTTTGGTTTATTCTACCATGCACAAAAAATTCTTCACATAATGCTGCCAGCACTTACCAGCAATACTCTGCTGTAGTGGTTTCAGATTCT | 120 |
| chr11 | 70920815 | 70920935 | 788855_47188552_ENST00000601538.6,E | 1 | - | good | TACTTTTTAGGTAATATGCATCATGAGTTATATTGATATTGCCTGTTGTGATTCAGGACAAGGTTTTGACTTAACCTCCTTGATCTTAACATCTGTATCGCCTTCAGCTATGCTAAAAAC | 120 |
| chr11 | 70920935 | 70921055 | 788855_47188552_ENST00000601538.6,E | 1 | - | good | TGGTATGGCGAGACATTTCAGATTTCTTGTCTCAGACATGGAATCAGCCATTTTTCAGGGAAGCTTGTTTCTTTTAGTCGAAACTGGTGTTTAAGGGCCACGTCTAAGTTTAAGGGGGCT | 120 |
| chr11 | 70921055 | 70921175 | 788855_47188552_ENST00000601538.6,E | 1 | - | good | AGCATTCATCTGCCTCTAGCTACTTCATCATATCTGTCTGTGCCCAGTGGAGGCTGATTCCGACCATTTCCTGAGTCTTTTGGACATGACCTCATAGTCATTGATAGCTTCCTGATTTTC | 120 |
| chr11 | 70921175 | 70921295 | 788855_47188552_ENST00000601538.6,E | 1 | - | good | TCAACAGCTGTCCTCTTTCCAAACCTTGGCACCTGTGGTCTTTGTAGCTTCAGTAGATCCCTGATGCTTTGAAAGATATGTTTACTGTTGTCTTTTGTTGTTTATTCTCTTTTTGGTGGG | 120 |
| chr11 | 70921295 | 70921415 | 788855_47188552_ENST00000601538.6,E | 1 | - | good | TTCTCTACAGCTAGCAGGAATCGGCTCTTGGCCCAAGGACCAATTCTCAGTCCTTTTCCTGAGTCTGGGATGTACAAATGATCCTAGGGGAGAAGCAGCTTTAGCACATCTGCTCACCTC | 120 |
| chr11 | 70921415 | 70921535 | 788855_47188552_ENST00000601538.6,E | 1 | - | good | GGAAAAGCAGCCGAATGCCTCATTCATTTTTCTACTCCTTCCTCTAGCCCAGACAACTCCCCTTCGGGTCTGCAGTTTTTGCCTTAGGAGCCCTACTTCACTGCTAAAAACTCCAGGGAT | 120 |
| chr11 | 70921535 | 70921655 | 788855_47188552_ENST00000601538.6,E | 1 | - | good | AGTGAGAGCCTCGTGTGCTTACCAGCATTTCTCTCCTGGGTGGGCCCTGCCCTCCATTTCTTATGCTTCGCTTTTTCACCTTCTGCTCCTCCCAATTTCAGAATTCGGCAAATATTGAAA | 120 |
| chr11 | 70921655 | 70921775 | 788855_47188552_ENST00000601538.6,E | 1 | - | good | GCCTTGTCACCTTGATGCAGTCAGAGACTGAGCTGAGTCAAGGCTGGGTTGTAGTTTTGAGAGGATTCATTCTAACTCCAGTTCTCCTTTCCTTCTAGATTGTAGCCCTCAGCACTTTCA | 120 |
| chr11 | 70921775 | 70921895 | 788855_47188552_ENST00000601538.6,E | 1 | - | good | TAGTAATTGTTTATAGAATGTCAGGCATTGCATATATATTTAAAAATTATGGAGACTCTAGGTAATGTTATTTTCTTTCAGAGAGGATTATCATTTCTTCTGTCAGGCAGATAGAGAGAA | 120 |
| chr11 | 70922015 | 70922135 | 788855_47188552_ENST00000601538.6,E | 1 | - | good | GTAGACGTTGGATTTTAAGGTCTAGAGTTTACGATTGATTCCCTTCTGCATTCAATTGTTTCTTGAAATTCTCCATCTTTTCAGCTCCTTTCTTCAGTTTTACTTTTACCTTCTTGATCA | 120 |
| chr11 | 70922135 | 70922255 | 788855_47188552_ENST00000601538.6,E | 1 | - | good | TTCCATTTGTCCTTCTCTCTGTGCTTCAGTTTGTGCATTTTTTATTGACCAATTTTCCAATTTCCTTATCCATCTCCTACATGTAATCCGATCTTAAGCCTACCTTTGAGGTCCTTAATT | 120 |
| chr11 | 70922255 | 70922375 | 788855_47188552_ENST00000601538.6,E | 1 | - | good | TTTGTCAGTTTAGGCTTAATGTATGTAGTACTTCTGCTCTATTTTTCATCTCTCCTTCTAGGATTTGATTCACATATATGTCAGAGCCTTTCAGTGTGTCTTATATGTGTCCGTTCTGTT | 120 |
| chr11 | 70922375 | 70922495 | 788855_47188552_ENST00000601538.6,E | 1 | - | good | GTTGCTTTAGAAATTTTCTTTGTCCTTGGTTTTCAGTATTTGTGTGCCTAGCTATGTTTTTTTCCTGTGTTTATCTCATTTGAGATTTATAGAGCTTTTTGAGTCTGTGGTTTTACATGT | 120 |
| chr11 | 70922495 | 70922615 | 788855_47188552_ENST00000601538.6,E | 1 | - | good | TTTTGTGAGGTACAGGTTCTAGATCAGTAGTTATTTTCTTTTTTCTTTGTCACATCCCTTATCTAGTTTCTTTTGAAAGACTGCTGTCGGTGTGATTGTAGATAATGGCCATTTGCTATG | 120 |
| chr11 | 70922615 | 70922735 | 788855_47188552_ENST00000601538.6,E | 1 | - | good | TTTTTTATGGGATCATCTTTCTTTTGCTAAAAGAAGTCCTTTTTAGAATTCATGCTGGTGATAGAATGCCTGCTTTGATTTTTCCAAAAATGTCATTATTTAGCCTTATTAATAGGATAT | 120 |
| chr11 | 70922735 | 70922855 | 788855_47188552_ENST00000601538.6,E | 1 | - | good | CTTCCTCCCCTGCATCTCCTAGCTGTGGTAACACGTTTAGAAACTCACCACTCCTACATCTTCCTGTTCTTTATTACTTGCTCACTTATTTTCCATTTGCTTTTTATTCCTTTCTGAATT | 120 |
| chr11 | 70922975 | 70923095 | 788855_47188552_ENST00000601538.6,E | 1 | - | good | CCCTGTCATAAAAAGTGTCAGTTGGATTGTGCCTGTTTGGGTTTTGGGTTTGGATTTCTGTGTTGTAGATTTATAGTAATATAATAGGTACTTTAACTCATTAATCGTAAGTTACAAATG | 120 |
| chr11 | 70923095 | 70923215 | 788855_47188552_ENST00000601538.6,E | 1 | - | good | GGCAGTCATATTGGAGGGAAACAGTGGGGACCCATCATCTATTATCACCCCCTCCTGACCATGCAACTGCAAATTTCTTTCTTCTTCCTTACAGTTGATGGGTTTTTACCATCAATGACT | 120 |
| chr11 | 70923815 | 70923935 | 788855_47188552_ENST00000601538.6,E | 1 | - | good | CATTCTTATAGGCTGGATGTCACTGGCTGGAAGAGAAGAGGAGCTGCCCAATTAGGCTAATCAGGGTTAAGACATTGATGCTTGGCCAGGTGTGGCGGCTCATGCCTGTCATCCCAGCGC | 120 |
| chr11 | 70923935 | 70924055 | 788855_47188552_ENST00000601538.6,E | 1 | - | good | TGGCCAGCCTCCTGCCCCTGTCTGCTCCCTCGTGCCTCCCCCTGAGCAGACACGGGGCTCACTGCAGTGTCTCACAAGGGCACTGGGGGAGCCATCACTGGGCAGTGGTTCACCGGTCTG | 120 |
| chr11 | 70924055 | 70924175 | 788855_47188552_ENST00000601538.6,E | 1 | - | good | GGGTGGTGGGGAGTGTGGCAGGAGGGATGCTGAGTCTTGTGCATTTGAAAGCAAAGGTTTTATGATGGCACCTAGGGCCCGGCAAGGCTTTCCCTGTGATGGTATTGGTTTCTGTGTAGT | 120 |
| chr11 | 70924175 | 70924295 | 788855_47188552_ENST00000601538.6,E | 1 | - | good | ACGGACTGAGCTCCTGTCCATGGAGAGACCTGCTCTGGGGCATGTCCTCAATATGGCAGGTCAGGCCGGTCAGTATCTGTACCGTCAGGAGGGACTCCTGGGGACTCTGGTAACGGTGCA | 120 |
| chr11 | 70924295 | 70924415 | 788855_47188552_ENST00000601538.6,E | 1 | - | good | ATTTCGCTGGCTGTTGGACAGAGCCCCTGGTGTTTGACTCTCACACCTGAGCCCGGCCCCGAGATCCTAGGAGCAGCGTGTCCCAGGTGAGACTGGGATGCAGCATCTTACCCAGGAGGC | 120 |
| chr11 | 70924775 | 70924895 | 788855_47188552_ENST00000601538.6,E | 1 | - | good | CGGGAGTGGCAGACCCTGTGCTGTCTGCGTAGCTTTTCCTTTATCTGTTCTCTGAAATAAGCCCCATTTCTCTTATAAGGTTTTGTGGTGCCTAATGAGATAATAAAAGTGTAAAACCCT | 120 |
| chr11 | 70924895 | 70925015 | 788855_47188552_ENST00000601538.6,E | 1 | - | good | ATCTCTGCTGGGCAAGTGGGTGCTCAATCTGCCCCCCTGTGAGCCCGGATGGTGATGGGAAGGGAGGGTCCTGGTGAACGCTTCCAATACAAAGTTAGCAGCTCCAGCCTTGCCTTGTTC | 120 |
| chr11 | 70925015 | 70925135 | 788855_47188552_ENST00000601538.6,E | 1 | - | good | ATAGTGGCCCACTGTGAAACCCCCACACATACTCCCCTCTGCAGTGTCGACCGCCATGGAGAGTTGTATCTGGCTTTGTCTATTTCCTGACCGCTTGGTGGAGGGAAGCTTAGAGTGCGC | 120 |
| chr11 | 70925135 | 70925255 | 788855_47188552_ENST00000601538.6,E | 1 | - | good | ATTTATTCCATAACGTTGCTGATGAAATAGGGCATATATTTAACCATTGCTTTCTTAAACATACAGGAACTGCTTAAGCTTTCAGAAACAGCTTAAGGCTGCTGATTCATGCAGATATAA | 120 |
| chr11 | 70925255 | 70925375 | 788855_47188552_ENST00000601538.6,E | 1 | - | good | CTATTGGGCACCTAATCTGCAAGAATTAACAGGGCTTTCTTCACTTTGGCGGGTCATCTTTAATAAAAATGCTTGTTTTTTTCTTAGCACAGAGTCAACATAGCAGGGTATCGTTGAAAT | 120 |
| chr11 | 70925375 | 70925495 | 788855_47188552_ENST00000601538.6,E | 1 | - | good | CATGTCTGCTTGGTGTTGATTATGGACACATGGGGTGGGGCACCAGGCACTGGAAGAGACATGTTCTGTGAACATGCTGTTTCCTTATTACACAGCTTGGTTTCCAGGTGATACCTACTC | 120 |
| chr11 | 70925495 | 70925615 | 788855_47188552_ENST00000601538.6,E | 1 | - | good | ATTTGTAGTGGATTTGCTTTTATATTTTAACCTAACACAAGCATGAAGCTCTACCATAGGAATGATTTGATTCATATGTTTTTTCTACCAGTTTTTAAGTATCTGACTTTTGCACAGGTG | 120 |
| chr11 | 70925615 | 70925735 | 788855_47188552_ENST00000601538.6,E | 1 | - | good | CTGCACACGGCAAGGCAAGCAGGGGAGGTCAAGCACCTCTCAAGTTCATGTTCATGTAACATCCAGTAGGCACGCTGTGCAAATCCGGCTTTTCTTCCCACTAGGGTGCTCCTACCAGTC | 120 |
| chr11 | 70925735 | 70925855 | 788855_47188552_ENST00000601538.6,E | 1 | - | good | CTAGGAGCAACAGCTCAGTATTGCTAATTTTGTATTTGTGGCAGGTTTATAGGATGTTAGCACCGCAGACCATGAGAATAATCTCTTTACACATCCTGGACAATAAGAAGTTAGTGCTGT | 120 |
| chr11 | 70926695 | 70926815 | 788855_47188552_ENST00000601538.6,E | 1 | - | good | ACTGAAATATAAAAGCTCATGCGCCTTCAGAGTTATTCACAGCCACACAGAGCAGTGGAGAGTCTGAGTCATCCAGCCCGCAGGTGCCCAGCTGAGGCTGAATGGGGCCACGCTCTGCCT | 120 |
| chr11 | 70926935 | 70927055 | 788855_47188552_ENST00000601538.6,E | 1 | - | good | CCCTGCCACGTCGTGCGGCAGAGATTCTGATCACCGGCTTTTGCCTGTTTTAAGCAGGTGGAAAGGGACCCAGCAGGAGGCAGAAGAGAGACAAGTGGTTATTGGTAAGTAAGGATCAGA | 120 |
| chr11 | 70927055 | 70927175 | 788855_47188552_ENST00000601538.6,E | 1 | - | good | GTGTTGCATTGGAGTCTACTTCTTATTACTCTAGGCCCTTATTTTTTTAAAACAAAGAAATTAACATATTGAGATGCCTTGGCAAGCACATGATGTGTGCCTGCCTGGTATTCTTTTTTT | 120 |

| | | | | | | | | |
|---|---|---|---|---|---|---|---|---|
| chr11 | 70927175 | 70927295 | 788855_47188552_ENST00000601538.6,E | 1 | - | good | AGAATCTTCATGAATATTTAAATGGAATTCAGAGTCAGCGTTCAGGGACTCTTATTTATTTTAACACTATGGGAAGTATTATTGTTTATAAAATAGCTACAAAGAATCTTTTGTCTCCGT | 120 |
| chr11 | 70927295 | 70927415 | 788855_47188552_ENST00000601538.6,E | 1 | - | good | TGTGCTTCCAACGGCTCTCAAGTCCGTCACACCAGGCAGTGAAATGACCCTGGGAGCATAGCAAGAGAGATGGGATAAACGGAATATTTTATTGTGAAATCTTTTCTGGAGATCGAGTTC | 120 |
| chr11 | 70927415 | 70927535 | 788855_47188552_ENST00000601538.6,E | 1 | - | good | GATGGCTGCTCTCTGGGGTTCTTTTCCTTGTGTTGACTCGCCTGCCCTCTCCCCATGTCGGGCCCCATCAGTCCCAACCTGGCCTAGATTCGCTGGTCACATAGAACAAGCTGAGTCTCC | 120 |
| chr11 | 70927535 | 70927655 | 788855_47188552_ENST00000601538.6,E | 1 | - | good | TCACTGGCCTTGAAGGTCACCAGGACCCTGCCTGTGTCTTAATGGGAATTTCCATCTCCTTAAGATCTTGCCTGCCTTTGCTTGAACCCCTGCAGAGATGGGGAACTCACTGCTTGTGGA | 120 |
| chr11 | 70927655 | 70927775 | 788855_47188552_ENST00000601538.6,E | 1 | - | good | ATGCTGATCCAGGAACATTCTCCCAGATATATCCAGGAACAGCCTCCCAAATAGACCCAGAAATAACAGTTTACTCTGAGGACCCTGGAACTCAGTCAAGGTGATGCATAAAATGAACTG | 120 |
| chr11 | 70928135 | 70928255 | 788855_47188552_ENST00000601538.6,E | 1 | - | good | TGGTGGAGGGGGCAGCTCCATGGCTGGGGTCTCCATCGACGGGAACAGGTATCGGCGACAATGCCCTTTTCCCTCTGGGTTTCAGGGCATGCAGTTTGTGGTGTCGACAGGCCTTGTATT | 120 |
| chr11 | 70928255 | 70928375 | 788855_47188552_ENST00000601538.6,E | 1 | - | good | TGTTCTGCCCTGAATAAGGCTTTGTAGCAAGTTGACCAGCTGTCTCAGATCTCTGCCTTCCAGTTTTCCCCCTTCAAGCATACAAGCCTTTCTTTCCCTTCTAGACACCTTTGGAATGCA | 120 |
| chr11 | 70928375 | 70928495 | 788855_47188552_ENST00000601538.6,E | 1 | - | good | TGCTGCATCTTCTGGGGGTGAGGTTGTTCTGGCAACCATGCTCTGTTTCTCATGTCAAAAAATGCATCTTAGGACTGTCCTGGGCAGCCCTTGATCCATGCCAGGTGCGCCACAAGAGCC | 120 |
| chr11 | 70928495 | 70928615 | 788855_47188552_ENST00000601538.6,E | 1 | - | good | TTCCCCCCCAGGGATCTGCTTTGAAACAAATGCCTTTTTCTGTTACCACAGGCTTTTCCTCCATTCTGAGGCCCTGTGATCCCATCTCGTGTTCCACATAAAAGCAAGCAGTAAACCCTT | 120 |
| chr11 | 70928615 | 70928735 | 788855_47188552_ENST00000601538.6,E | 1 | - | good | ATATCTTGGCCGGCTTGCAAGTTTTTCTCCCTTGGTCCTAATTCTGCCCCATGGATCCCATTGGAGGGGCTGGTTCTTCCTCCAGGGGACCACCTCCTATGCCCCCACCCAGTCTTCCTC | 120 |
| chr11 | 70928735 | 70928855 | 788855_47188552_ENST00000601538.6,E | 1 | - | good | CCTCTCTCCTCTGGTGTAGGCTTTTGGTGTGGGGGTTTGAAGGCAGCTCTACCACCTGTCCGGGTTCACGGTCGTAGGTGGGCTCCTAACTTCCAGCCCTAGGCACCTTTATCAGTGCTC | 120 |
| chr11 | 70928855 | 70928975 | 788855_47188552_ENST00000601538.6,E | 1 | - | good | CCCCAGGGTCTGCATTACCATTTGTCTGCTTCCTGACTCTGCCAGCCCCACCCTCCCAGGAGAGATGGGGAGTCACTTTCAGGTCACCCCCTACCGTTCTTCAGTGTGTGCATCTTCCTG | 120 |
| chr11 | 70928975 | 70929095 | 788855_47188552_ENST00000601538.6,E | 1 | - | good | GCCCTTGGGCGCATTGTTCTCCCTGTCCTGTGGGTGGTGTTGCTTCTTTTGCTGAAAAAGTAGTGCCCTCCCAGCTTCTCTCTCCACAAGACACGTGTGCACCTCCACAGGTCTGCACTC | 120 |
| chr11 | 70929095 | 70929215 | 788855_47188552_ENST00000601538.6,E | 1 | - | good | ATTCTACCCCTCAGATATTTCTGCAAAAGCAGAGGCTTCCTTATGATACACACACCACTAACAGATGGCAACCTCAGGTACAAGTTCTGGAAGTCACACTAGAGAGTAAGGAAGTGACGA | 120 |
| chr11 | 70929215 | 70929335 | 788855_47188552_ENST00000601538.6,E | 1 | - | good | CTTCCCAGCAGCTCTTTGAGGCACGTCCCATTTTACAGATGAGGAAATTGCCCAGACCGAGGTTGAGTAGTAAGTGCCTCGGGGCAGAGCCACATTTAAACCTCGGCATCCTGTCTCTCC | 120 |
| chr11 | 70929335 | 70929455 | 788855_47188552_ENST00000601538.6,E | 1 | - | good | GCTGACAGCACACCTTTATTTCAGGATTCTGACCTCCAGAACAGTAAGAGCATACATTCACATTGCATCATTTGTTACAGCAGCAACAGGGAACTGACACTCACGTGCAAGTCACTGAAT | 120 |
| chr11 | 70929815 | 70929935 | 788855_47188552_ENST00000601538.6,E | 1 | - | good | GTATCCTCATCAGCATCCTACTCACCACTGTCATTTCAGTGACGGTCTGTGGCACTTAAAGCTCACGTTTCTGTAGTCACCCAGGATCTGACAGCTTTCAATATTGCAGTTGCTAAAAGA | 120 |
| chr11 | 70930055 | 70930175 | 788855_47188552_ENST00000601538.6,E | 1 | - | good | GGAACCAAGAAAATATTCTGCATTGAGGTTTACACAGTAGGTCCTGCACTTGGCCCGTAGCCTTATCTTTGTAAATAAAAATTTTCTGGAGCATAGTCATGTCTGTTCATTGAAATGTTA | 120 |
| chr11 | 70930175 | 70930295 | 788855_47188552_ENST00000601538.6,E | 1 | - | good | CAGAGGGAGCAGAGCACCGGCAGAGCATGACGTCTGTGGTGTGGGAAGGACATGCTTCCTTCCACAGGGAAGCAGAAGGCAGGACTTTCTGATCCTCATCCAGGGAGAGATTGAATGATT | 120 |
| chr11 | 70930295 | 70930415 | 788855_47188552_ENST00000601538.6,E | 1 | - | good | CCCACATTCCCAGTGGCTTCAAGCCATCTGGTAACTGCGTCTAAAGGTTGACTATCAATTTGCCCCAGTTTCTTCTGTGTCCCAGGATCCAGACAAAAGGAGACATTGTGGTTCTCTGGG | 120 |
| chr11 | 70930415 | 70930535 | 788855_47188552_ENST00000601538.6,E | 1 | - | good | CAGGTGAGATAACAGAGAAGGTGCTGGAATAGCATTAAAACAATAAATACAATATTAATAACTAACACTTTGTATTTGCATCAGTCAGCTTGGACCCAGTTATGCCTCATAACAAACATC | 120 |
| chr11 | 70930535 | 70930655 | 788855_47188552_ENST00000601538.6,E | 1 | - | good | AAGAAATAAAAACAAGAGAAAAGGCATTTATGTTTGCAAGATCAACCTCTGAGACCAATGACCTCTACCCAGATGTGGCAGTGACTTGAGAGTGGCTTAGGTTCAGCAGCTCACATTGGC | 120 |
| chr11 | 70931375 | 70931495 | 788855_47188552_ENST00000601538.6,E | 1 | - | good | GACTTGTGGTCAAGGGTGTCCCAGCGCTGCCACCTTCTGAGGGCTGTTCGGAGGGGTAGACAGACATCTCTGGTGCTGGTTTCCCAGTGCTTTGTGCTATTCTTTTAAAAGACTGCTTTA | 120 |
| chr11 | 70931495 | 70931615 | 788855_47188552_ENST00000601538.6,E | 1 | - | good | GGGCTGGCCCTGGTGACCGGTGGACAGTGCCGCCACCAGGAAGGAGAACACAAGGACGGGTGATTCTATAGCCTCTTCGGTTTCCCTTTCCTGACTTGCCGTGGCCACGCCAGCAGCTCT | 120 |
| chr11 | 70931615 | 70931735 | 788855_47188552_ENST00000601538.6,E | 1 | - | good | CCTTGCTGTTGCCCTGTGTGCCACTAAGTGGTCTCTAATTATAGCTGTGTCCTTGGCATCCAGACACCTTCTCTTGCCCCTTTCAGAGGGGCGGGTGAGGGCTTAGAGATGACCTACGAA | 120 |
| chr11 | 70931735 | 70931855 | 788855_47188552_ENST00000601538.6,E | 1 | - | good | TGGAAAAATAATAAGCGTCGTTGAGAGGGTATCTGCTCAGAAGAGATTGGCCCTTCAGCAGCAGGCAGTGGCTTTCTAGATGTGAGACCTCATTGCTGTGATAGACGCTTGGCGCCAGCA | 120 |
| chr11 | 70931855 | 70931975 | 788855_47188552_ENST00000601538.6,E | 1 | - | good | TTTCAGTGTTTCTTAAAAAAATATTTGTTATGCTAATTTCGAATTGCTAGGATTTCTACAGGCTTGAAGATGGATGATGTTTTTCTTTCTTTATAAAAAACATTAGTATGATTTTCACTT | 120 |
| chr11 | 70931975 | 70932095 | 788855_47188552_ENST00000601538.6,E | 1 | - | good | TGAAAAAGACATTTCCACTCATTTCCGATGGAAGATACCGTATCATAGAGATTATGGTAATTCTAAGAAGTTTAGAGTGGGCTTATGGAAATTGGTAATAAATACTGCACGTAGAGGCTC | 120 |
| chr11 | 70932095 | 70932215 | 788855_47188552_ENST00000601538.6,E | 1 | - | good | GGCTGTTCAGATTTCCCCAAGGGGTGGATTTCCATGGCTAAGGCAGAAGCAACACTTTTGTCCAGAACCAGATGTCTACAAAATGTTTTCTTTTCTTCCCAGAACATCACAGTTTCAGCT | 120 |
| chr11 | 70932215 | 70932335 | 788855_47188552_ENST00000601538.6,E | 1 | - | good | TGAGGGCTTGCTTTATATAGGACAAGATTTTAAGATAAAAATGAGTGCATCTGGAAATTAATATTTCCCGTGGCTGTTATCAGAGACACTGCCTTTGTTTCCTCCACGGAACTTTGTTTC | 120 |
| chr11 | 70932335 | 70932455 | 788855_47188552_ENST00000601538.6,E | 1 | - | good | CCATGCATCTGCTTCCCTTAGCGGGGCCGCACGTGCGCGGACCCAGCAGAGTCCTGCTGCTTCTGCTGCGGAGGGGGTCTGAAGACCCCTTATTCCTTCTAGCTCCCGGAGTGTCTAAGA | 120 |
| chr11 | 70932455 | 70932575 | 788855_47188552_ENST00000601538.6,E | 1 | - | good | GAAGGTGTGGCCCAGATGCCTGACTTGGAAATTGGCTTTAATCTTTGTAAACAGAGGTAGTTTCCTGGGAATTGGTCTGAAGTCTTAGGGGCATGGATCTGCCTCTGGTCTGGACAGTTT | 120 |
| chr11 | 70932575 | 70932695 | 788855_47188552_ENST00000601538.6,E | 1 | - | good | CAGGTGGTCCCCACATGGGGTCTCTGAGGCCTGGCCACTTCTAAGGTACCCTGGCATTTGCTTTGGTCAGTGTGTGCAGAAGATTGAATGCTTGAGGGTGGATCTCTACTCACACAATGG | 120 |
| chr11 | 70932695 | 70932815 | 788855_47188552_ENST00000601538.6,E | 1 | - | good | CGTGAAGTGGTATGCTGTGCTTTTCAATAGTTTCTGTTTTCTGTGGCAACCACCAGGACAATATTGTATGGAAGGCATGAGCCTAGGCAGGGGGTGTAGTTAGGAGAGCCCCTGCCCCGC | 120 |
| chr11 | 70933535 | 70933655 | 788855_47188552_ENST00000601538.6,E | 1 | - | good | GAGCCACCACACCCAGCCTGTTGTTGCTTTTTTAAAAAATTACAATAAATACATATATACCCATTGTTGTCAGATCTTCCTGGTTTTCAAGAGAATTTAGAAATTCATATTTTTCTCAAC | 120 |
| chr11 | 70934255 | 70934375 | 788855_47188552_ENST00000601538.6,E | 1 | - | good | GGGAGGCCGGCCGGCTGCACGGGCCTCCATTACCTGGGGTGAGGAACCTGAGTGTGCGGCAGGAAGTGACTACCTGGTTTTAATGTGCAGTGGGCACTGTTGTTGCTGCTGTTTTTTTTT | 120 |
| chr11 | 70934375 | 70934495 | 788855_47188552_ENST00000601538.6,E | 1 | - | good | TACCCTGTAGCAGGGTGGGGACAGCCAAGGACTGCTTCCCAGGCAGAGCCCCTGTGTTTGAGGCTGCACCCTGGGCGGGGCTGGGCTGTTTAGGAAGCAGAGGCGGCCTGAGAAGCAGGA | 120 |
| chr11 | 70934495 | 70934615 | 788855_47188552_ENST00000601538.6,E | 1 | - | good | GAATTTGGTCTGACAATTGCAAAGAATGCCAGGAGAGGCTATCTGGGGGTGACTTAGTCTGAGACCCAGTGGCCTTCCTGGGGAGCATCATCTCATCTGGCCTCGGAAAGCCAAGGAGCT | 120 |
| chr11 | 70934615 | 70934735 | 788855_47188552_ENST00000601538.6,E | 1 | - | good | CCATTCGCCTCTTCTATATCATAATCAATAGTATTTGAAATATTCAGCAGCAGCATTTTCTCTTATCATTGGGACATAAAATAAGGAGCCTCCTTAATTGATGGTGGCTTTGATTTGATG | 120 |
| chr11 | 70934735 | 70934855 | 788855_47188552_ENST00000601538.6,E | 1 | - | good | TGGGGACAGGACATTCCCCAGCAAGGAGGTCCTCCTCTTTGAACGGCCACACTGGTCTCTGGGAGGAAAATCACGCCACTCGTGATTTACCCACACGTGGCATGTCAGCCTAAGGGGTTC | 120 |
| chr11 | 70934855 | 70934975 | 788855_47188552_ENST00000601538.6,E | 1 | - | good | CTCCAGACCTGCCCCGTCCCTTCTCTCCGTGAATGCTTCCACACGGCTGCTGCGTCGCTTTGGTTTGCATCGCGGTCCCCGGCAAAATGCAGGCAGTTCAACTCCCATTTATTGAAAACC | 120 |
| chr11 | 70934975 | 70935095 | 788855_47188552_ENST00000601538.6,E | 1 | - | good | TTTTTTTTGTTTCATATTTTCGGATTGCATTGGACTGCAGGTAACTGAAACCGCAGGTAAGGAGGGCTCCTGTCATAGGCACGGCCGTTTACCCGTGGAGAGAGGAAGCAGGGCGGCTAG | 120 |
| chr11 | 70935215 | 70935335 | 788855_47188552_ENST00000601538.6,E | 1 | - | good | CTGGGTGGATGCCAGAAATGGAGGCTAGTACAGACCCCTACATGTATGGGCTGTGTTTTTCATCTGAGAACTGAGAGAGCTCCTGAGTGACAAATGGGTGGGTGGCCTAGACAGCACGGA | 120 |
| chr11 | 70935335 | 70935455 | 788855_47188552_ENST00000601538.6,E | 1 | - | good | CCTTTCACTCCTGGCTAACCACGTAATAGGGGCTAATTACAAACTGAGGACTCGGTTCTTCAATCTGCAAAATGAGACTGTAGCAGACCCCTCTTACCTGTGTGGGCTGTGTTTCAGGAC | 120 |
| chr11 | 70935455 | 70935575 | 788855_47188552_ENST00000601538.6,E | 1 | - | good | TGTGGTTTTAGGTTTTAGGCTGTTGGAAAGTTCAGCATGACCCAGGGGGAGTGACCTCCTGAGATGGATTCCAGGTGGTTCAGAGGTGTCCTTGAACCCGTCTGCTTGGCTGTCACTCTC | 120 |
| chr11 | 70935575 | 70935695 | 788855_47188552_ENST00000601538.6,E | 1 | - | good | GCAGCTGTCCCACTCGGGGCCCGGCACATGGTAGATGTGTTTGCTGATTAAATAAATAAATAACAGGTTAGCATCCCAGTTTCAGAAAAGACCTCATGTCCAGGCCAGAATAGACTCGGT | 120 |
| chr11 | 70935695 | 70935815 | 788855_47188552_ENST00000601538.6,E | 1 | - | good | GCTTTTCAGAACATGGAGTTTTAAAACATTCCCAAGTTGCTGTCTCTTGGAGTAGGCCTTTCAGATCTTGGAAAGTGTCTAGTTAGGACAGAGTCTGTTTTTTTCATGTCTTCATTCAAG | 120 |
| chr11 | 70935815 | 70935935 | 788855_47188552_ENST00000601538.6,E | 1 | - | good | CCAAGGCTGTTCATGATGGGCACGTCCAAATCCCTGCTTAACCTGGAATGGTTCACCTTTGCGGGGGCGGTGTAAGAGGCCCGCTCACATGACGAGGAACTCAGGGCAAAGAGTCGCTAT | 120 |
| chr11 | 70935935 | 70936055 | 788855_47188552_ENST00000601538.6,E | 1 | - | good | TGGCCAAATTGATTTCAATCCGTGTGAGACCTCAGCCGCAGGCCTCCAGGAATGGTGCTCACTTCAGCCGGTGCAGCAGGGACAGGTAGGGGAAAGAGATGGTCCTTGGCGGGTGTGAAC | 120 |
| chr11 | 70936055 | 70936175 | 788855_47188552_ENST00000601538.6,E | 1 | - | good | ATTCTTCTTCATGGGTTTTTCCTTTTTCTGAACATTTTGAGAATAAGCTCCACAGAGCTGGCATTTAAGGAATGGTGTTTACTTGAGAAAGAGAAATCTCAGTCAGCTGAGACATTAATG | 120 |
| chr11 | 70936535 | 70936655 | 788855_47188552_ENST00000601538.6,E | 1 | - | good | GATCTTCAGAGACCAAAATGATTGTCAACAAGATAGAGAAATAAATAGCAAAGACAAACACACAAACACATGGTCTTGCTTCTTTTGCCTGTTGCCTTATTTTAGTTTTACTTTTTTATA | 120 |
| chr11 | 70936655 | 70936775 | 788855_47188552_ENST00000601538.6,E | 1 | - | good | TGCACATGGGCTGGAGTCCGGGAGGCGAGGAGGCCTGAACCCACCCGGCCTGCACCAGCTCTCTATGAACCTGTTCGTTAGTGGAGAACTGTTTCAGCTCTGGCTTTTGGTTGCTAACAT | 120 |
| chr11 | 70936775 | 70936895 | 788855_47188552_ENST00000601538.6,E | 1 | - | good | TGGCTGAATCACCTTGAGGACTTACAGTGGGTACCAGTGTGCGTTACCCTTCATTGGAGGTGTTTTCCTGCTCACAGTCTCTGGCTGTTCTTTAGGGAGGGCTGTTGGGAACCCTGAGGG | 120 |
| chr11 | 70936895 | 70937015 | 788855_47188552_ENST00000601538.6,E | 1 | - | good | AGCTAACAGTGACATTTCAAAAAAGGTCAGGCCAATCAAGACATCTCAGGGAGGGAAAGGAAGGTGGAAGAGTCAGTACCAAGGAGAAATGAAGGTTACTTGGAACTGGAAACCCAGTGC | 120 |
| chr11 | 70937015 | 70937135 | 788855_47188552_ENST00000601538.6,E | 1 | - | good | TCTTAGCTTGTACAGGGCTGGTAAGTGACATGTTCTGACTCCACGCAGAGGACTGGGGTTTGTCAGGGCAGAAGTATGTGACTGATGCTACAAATCTCGGTCTTCCAACTTAAAAGCGAC | 120 |
| chr11 | 70937135 | 70937255 | 788855_47188552_ENST00000601538.6,E | 1 | - | good | GTTTTCTTCCATCTCCTTTGAGCTTTGAGAGCATCTGTGCATCTAGTGTTTGGTGACTCTTAAACTCTTCTAGACACAGGCTCTATCGATCCTGAGGAACCCAGAGGTCCCTGCACAGCG | 120 |
| chr11 | 70937255 | 70937375 | 788855_47188552_ENST00000601538.6,E | 1 | - | good | AGGCAGCAGTGATCACTGAAGGCATTTGGGGCTTTTGTTTTGGACACGTCCCTGTTACCACATGACTTGGGGAGTTTCAAAAGATTTCTGTGCTCCCGTCACTTGTTCTCCCCAAGGGAT | 120 |
| chr11 | 70937375 | 70937495 | 788855_47188552_ENST00000601538.6,E | 1 | - | good | TGTAGGATCTGGGAGACCCCAGGGGGCTGACGATGTCACTTGTCCATCATCCGAAGGTCACATAGGAGGGGCCTCACCGGGCTCTCAGGGAAGCGAGGTCCTGGGGAGCCTCCATCAGGC | 120 |
| chr11 | 70937495 | 70937615 | 788855_47188552_ENST00000601538.6,E | 1 | - | good | AGAATGCACATGATCCAATGGTGGCCACCCCCATCTTTTGCACACTTGCCAAGGCCAGGCCCTCTGGGTACTCCAGGTGCATTGGCCACCAATTGATGAAGGTAAATGTCCTCAGGAAGT | 120 |
| chr11 | 70937615 | 70937735 | 788855_47188552_ENST00000601538.6,E | 1 | - | good | AGCTTTGCCTTTTAAAAAATGTTCTATTGCCCCTCAAGAAAAAAAAAAAAAGATGCACGCACACACACTATAAAGTACTCAAAAGATGGGAGTAGCCACTGTGACCGTGATCTTTGATAGC | 120 |
| chr11 | 70937735 | 70937855 | 788855_47188552_ENST00000601538.6,E | 1 | - | good | TTTTGTGAGATTCTGCCTCTGGGACTGAGCCCATCTGAGTGGATTTGTGTTCCTCAACAATAAGCGGTTCCTGAGTAAACAGGGACAACACCAGGACTTGACCTAGACCTTTTTTCTCCA | 120 |
| chr11 | 70937975 | 70938095 | 788855_47188552_ENST00000601538.6,E | 1 | - | good | CATTCCTGCTGTAATTATGAAGAGCCCTCTGCTCCTAGGGAGGCAGAAGCTGGAGAAGGACATGCCAGAAAAAAATAAAGTGAGAAGGTGATATAGAACTGGAGAGAATAAGATAGATCA | 120 |

| | | | | | | | | |
|---|---|---|---|---|---|---|---|---|
| chr11 | 70938095 | 70938215 | 788855_47188552_ENST00000601538.6,E | 1 | - | good | CCACCTGAAAACCCTCTCTGGGTTCCGTTCCGACCTTCACCCCACTCTGTCCCTAGCCAATTGGCTAATTTTGAGCCAATCGGATTTGTCCTTGGGGAAACTGGGATTGGGGCAGAGACT | 120 |
| chr11 | 70938215 | 70938335 | 788855_47188552_ENST00000601538.6,E | 1 | - | good | CCTTCGTTTGCTCCTTCTGTGTTATTTTATCCTCAGCACATCTCCTCTAGGCTCAAAGGGGTTAAAGTTGTGCCCAGGACACCCCCCTCCACCACACCCACACATACAGTGATGCTGGTC | 120 |
| chr11 | 70938335 | 70938455 | 788855_47188552_ENST00000601538.6,E | 1 | - | good | CTTGAGTAGAGCTGCATGTAAGTTTTCAGGAACATGCCTTTCCTTGTTGGGGAGGGAATCCTGTATTCAACAATATTCCAGAGTATTAACTAAGTGGATCACATTCATGAAGTGGTTCTA | 120 |
| chr11 | 70938455 | 70938575 | 788855_47188552_ENST00000601538.6,E | 1 | - | good | GGTGGCTGTGAGTAAATACTGTTTCTTTGGCTAAATCTTAGTGAGCAAAATGACACATATATGTATTTTTATAAGTCCCCTGATAGCCTCTGTGCCTCCCGTCTTTTGTTTCAGTAATCC | 120 |
| chr11 | 70938575 | 70938695 | 788855_47188552_ENST00000601538.6,E | 1 | - | good | TGTCCTTGGCTGGCTGCCCATTGCCTTCTGGGCTCTTTATCATCTCCTCGACTAGAAGCCGCGTTTCCAAAGGGCAGGGACTGCCTGTGCTCAGCACATTCCTCAGGATTGACCCGTACA | 120 |
| chr11 | 70938695 | 70938815 | 788855_47188552_ENST00000601538.6,E | 1 | - | good | TGCTCTAGGTGTGTGGAACTGGAGTCCACTAGCCTGACTCTGACCTCTGGCATTTTGCATATGCTGTTCCTCTGCCTGGAACTCTCCTCCTCCTCCTCAGCTCCTGGGCTTCCCTCTCCC | 120 |
| chr11 | 70938815 | 70938935 | 788855_47188552_ENST00000601538.6,E | 1 | - | good | CACCTTTCTAGATTACACATGGGGTCGTGTCACTCCCTGCAGCTGCCCCGCTAAGAAGGGCCCTCTGAGTGTGGCGGTTATCACGTCACCTCCCTTCTCTGCCGTGCTCACTGCATTCCA | 120 |
| chr11 | 70938935 | 70939055 | 788855_47188552_ENST00000601538.6,E | 1 | - | good | TCTTTCCAGCCTGTCTGCCTCCCTCTGTCCCTGCTGCCACCCATAGTTCCGGCGCCCTAGTGTCTCTCCTGGATTAGCCACCAGCTTCTCTCCCACTGCTGGTGGCCCAGACCCTCTCCT | 120 |
| chr11 | 70939055 | 70939175 | 788855_47188552_ENST00000601538.6,E | 1 | - | good | CAGCAAAACCCAAAGCCCCGGAGGGAAAGTCAGAAAGAACAGCCTGGTCCTCATCATCTTATTCATGAGAAAGAAAGTCACCCCGGGTTCCCTTAGGGTCTGTCATCTCCCTTTGGAAGG | 120 |
| chr11 | 70939535 | 70939655 | 788855_47188552_ENST00000601538.6,E | 1 | - | good | AGGGTGGACCAGGAGGGCCCAGGTCAGCAAATCCCAAAGTTGTTGAGCTGTTGGATGGAGAACGAAGCTCAGGAGAGGCCTTGGTGAGCAGGCTGCCCCAGACATGGAGCACCTTGCTAG | 120 |
| chr11 | 70939655 | 70939775 | 788855_47188552_ENST00000601538.6,E | 1 | - | good | CCGTTGTTCTGCCTACCTCTAAAGAGAGGAGCCACGCAGCGGGGAGGGCTGCGGGTAGGACGGGGCCTTTCTTGGTCCATCATGGGGAAGGGAATGAGGTGTGAGAAGTTCTGTAGGACC | 120 |
| chr11 | 70939775 | 70939895 | 788855_47188552_ENST00000601538.6,E | 1 | - | good | GGCCCACCTGGATGATCCAGGATAGTCTCTGCATCTGAAGATCGTTAACTTCATGACATCTATGAAGACAAAGTTCACTTATTTATGGGCTCCAGGGATTGGGACTTGATCTCTTCTAGG | 120 |
| chr11 | 70940495 | 70940615 | 788855_47188552_ENST00000601538.6,E | 1 | - | good | AGAGGAGGAGGGGCACTGGGTAATGGGAGAGATGCCTTTGTGTCCTGGGCCTGCTGTGATAAGTGACCACTGCCTGGGTGGGTAGAAACAGAAGAAATTCTTTGGGAGGCCAAGGTGGGT | 120 |
| chr11 | 70940615 | 70940735 | 788855_47188552_ENST00000601538.6,E | 1 | - | good | CTCCAGCAACCAATGTTCACCCTCTCGTCAGGGTGTTGCTGTCACCCCTTTTGACAATGGTTTATGGCAGTTATAATTTATCCTCTGAGAAAAATTGAGTCGGTTCTGGGGGGAATTATC | 120 |
| chr11 | 70940735 | 70940855 | 788855_47188552_ENST00000601538.6,E | 1 | - | good | TTAAGCTGGTGCTTTTTCAGCCATTTTTTCCTTTTTTCTTGCACCCTCCATGAAGTTTTAATGCCCCACATAGGCTGTAAATCTTTTTAGTTATCACTGCATTGCTGTGCTTCAAGCCCC | 120 |
| chr11 | 70940855 | 70940975 | 788855_47188552_ENST00000601538.6,E | 1 | - | good | AGGGCAGAGGCCAGTCCCGTGGTCATCATCTCCCTGCCACATAGCACACTTGGGAGCTCAGCAGATATTTTATGACTAAAGGAAGGAAGTGCCCATGACCCCTGTGTGGTGCTTGTTCAT | 120 |
| chr11 | 70940975 | 70941095 | 788855_47188552_ENST00000601538.6,E | 1 | - | good | GGAAGAATTGAAATTGGGGACTATCAGACACCAAACTTGTTTTCCATCAGCCCCTCTGCCTCCTGGGATTCAAACCCATAAATCTCTATCTTTGTCTCCCACCAGAAAGAATCCTTCAGA | 120 |
| chr11 | 70941095 | 70941215 | 788855_47188552_ENST00000601538.6,E | 1 | - | good | TACGTGTATTGAGTATCCACTGTATACACATAGGGGCCAGATACTTTATTGGTGTGACATGGTCCTTGAATGTTCTGCATGTTTAAATATTATGAACCAATTTTACCGACAAGCTTATGA | 120 |
| chr11 | 70941215 | 70941335 | 788855_47188552_ENST00000601538.6,E | 1 | - | good | ATGTAGTTCTGGGAGGCATCGTGTGTCTAAGTGTGTCAGCCTCCATTTTTGTGCAGTGTGTGGAGTTAGCTATCTGTCTAGTTCATGAATAGATAGAACTCAGGGGAGGGGAGAGAGGAC | 120 |
| chr11 | 70941335 | 70941455 | 788855_47188552_ENST00000601538.6,E | 1 | - | good | TTAGGAGATATGTTTCAGGTCAGGAGTATTAAGTAGTTTTCTCAGTTTACCGTAATGCTTCTGTTATCTGTAGTTCTGCAATATTTTGCTTTCAACACACGGCCTGCTGTCATCCTTCAG | 120 |
| chr11 | 70941815 | 70941935 | 788855_47188552_ENST00000601538.6,E | 1 | - | good | AGGAATGAGTCACCACACACAGCCCTCTTTTCTTTTGAAATGACCCAGTCTCCAGTATTATTATATTATTATTATTATATATTATATTATTGTTATAACACAAAACAGACAAAGACAGGC | 120 |
| chr11 | 70942655 | 70942775 | 788855_47188552_ENST00000601538.6,E | 1 | - | good | AGGAGGAAGGGCTTGTGAAATGCAAAAGGTGAGGGCAGAGTGCAACTGATGCTTGGTGAGATACTTATAATCCACTGTTTCTATCCTGAAGAAGTATCTGTCCCTGAATGTAAATCCTCA | 120 |
| chr11 | 70942775 | 70942895 | 788855_47188552_ENST00000601538.6,E | 1 | - | good | CCCTCAGTAACAAATGTAAGTGAACAGGCAAAGGGCCTGTCTGGTCTTATGGCTCCATGTAATTTCTTGAGCTGGCCACTGAGTGGCAGTGTTAACCCGGTACAGCTTGGTGCAGAGTTA | 120 |
| chr11 | 70942895 | 70943015 | 788855_47188552_ENST00000601538.6,E | 1 | - | good | TATTGTATTACAATGAATGGTTCATGCATAAGTCTCCCAAGCTAAACTTTAAAGTGCTTTGAGAGGGATGCACTTTCCCATCTATCTTTATTCTCCTGGCACCCAGCACCAGACCCAGAA | 120 |
| chr11 | 70943015 | 70943135 | 788855_47188552_ENST00000601538.6,E | 1 | - | good | CTTGCTCCATCGCCACCACCTTTGCCAGGTGAACTCCTTCCCACCCCTCAGAGCCGAAGTGCTTCCTCTGCAGGAAGCTTTTATTTTGATTCCCACAGTACTTGCTGGGTACCTTTTCTG | 120 |
| chr11 | 70943135 | 70943255 | 788855_47188552_ENST00000601538.6,E | 1 | - | good | CTGAGCCACCTCCAACAGGGGACATAATCTCCTCCAGACACAGACAGACCCCTTTGAACATGTCGGTGACTGAGCACCATCTTCTCAATGATGTGATGTCTCTCCAGCCAGGTGAGACTG | 120 |
| chr11 | 70943255 | 70943375 | 788855_47188552_ENST00000601538.6,E | 1 | - | good | TAGAGTGGGTTGTAGTCCTAGCCAGTTTCATGGGCAGGGCTCCATAGGTGCCCTCCCTAGCACGGCCACACTTTCTTTGTTGAGTTTATCTAAGAAGTGCCTTAGCCACCCTCTCTGCTG | 120 |
| chr11 | 70943375 | 70943495 | 788855_47188552_ENST00000601538.6,E | 1 | - | good | TTTGCATGGGAAAAGCTCCTGGTTCCTCCTGGGTTTGGCCTGCTTAGCAGCCTGGCCAGAGCCCTAGTTCTCAAGGTGCTTCAAGCCTGGGCATCCAATTGAGTTTCCACCTAAGGACAC | 120 |
| chr11 | 70943495 | 70943615 | 788855_47188552_ENST00000601538.6,E | 1 | - | good | CCTTCTGGCCCAGGCTCTTCTCCTGCCTCCTCAGCCCCTGTCCCTGCCGCCCAGGGGTATTGCCACCTGCTCACCCCATGAGTTTGCTGTGCCCTAAATCTGGACCCATTGCCATCTCTG | 120 |
| chr11 | 70943615 | 70943735 | 788855_47188552_ENST00000601538.6,E | 1 | - | good | TTCATTTGTTTAGAGAAAATCTTTAATATTACCTCTTCTCCGTCTCCTGATCTGGCACTGGGTGTTGCAAACAAGACTGCATTTCAGGGTCAGTGCTCCACAGCACTCAGGACTTGCCAT | 120 |
| chr11 | 70943735 | 70943855 | 788855_47188552_ENST00000601538.6,E | 1 | - | good | TGGCGTGGGATGTGGTTGGCCCTGTGGTGAGGAAGGGATTGGGTGGGAGGGGAGCAGCTCATACCCCATGTAGAACAAAGCATCCATCCAGACAGGACATTGTTGCTGTAAAAGTTAAAT | 120 |
| chr11 | 70943855 | 70943975 | 788855_47188552_ENST00000601538.6,E | 1 | - | good | TGAAACTCTCATCTTCCCTGGCAGGAAGTCAACAAGTAATATTTAAACAGAAGGAAATCAAGACGGATGATCAGATGTTTGCTATTTACAAATAGGCAGCAAGCACCTGAAAAAGCACTG | 120 |
| chr11 | 70943975 | 70944095 | 788855_47188552_ENST00000601538.6,E | 1 | - | good | ATTGCAGTCGACTGCCTACCTTGCTGTGGACAAGATGTAGCTAGCCATAATAGCATAAACACAGGAACCATGATTGAACCAAGTGGGCCTGTGGCTGTCTTGGGCTGTGTGATAAATGAG | 120 |
| chr11 | 70944095 | 70944215 | 788855_47188552_ENST00000601538.6,E | 1 | - | good | ACAGGAATAGACCTGCGCTCTGCAATTTACACCTCTGTCAGAACTTGGTGATAAATTCGTTACATGTAAATAGAAAGGCAGATTTCAAAACGTAGCTGTTATATACTCGAGGGAAATAGA | 120 |
| chr11 | 70944215 | 70944335 | 788855_47188552_ENST00000601538.6,E | 1 | - | good | GGGCCGGGCCTGGGATGAGTGTGGAGCTTCGGCCTAGGTGGGCACAGGAGGGTGGATGGCTCTGGGAGAGGGTGACTCAGAAACCCAGGCAACAGTGATTGCCAGATGTGTTTCCACCTT | 120 |
| chr11 | 70944335 | 70944455 | 788855_47188552_ENST00000601538.6,E | 1 | - | good | GGCCCCTCCTATGCTGCTTTCTCAGGAAGCTGCTGGAGAGGAGCTTCACCCAGAGAGGGAGTGAGCCAGGGGACAGGAAGACCTGGACTCCGGGGAGGGCCCTGGCACCGGGCAGAGGCC | 120 |
| chr11 | 70944455 | 70944575 | 788855_47188552_ENST00000601538.6,E | 1 | - | good | TGGGGTCTGACCTCTCAGCAGCCATATGGGAAGCGCAGTGGGAAAATCCTTTCAAACCAGTGTTCTAGCAAATCAGTGAGACGGTCCCAGAGGTGTTTCAGATGCACGTGTTCTCGAATC | 120 |
| chr11 | 70944575 | 70944695 | 788855_47188552_ENST00000601538.6,E | 1 | - | good | GTGTGAGAAGACACCCCCAGGCAGGATGGAGGTCCTAAAAGCTCAGAACTCAGGGAATAGAAACAGAAACGCAATCTTAAAAGTTTCAAGAGAGAAAAATCAAGTTGCGCATAGAGGATC | 120 |
| chr11 | 70944695 | 70944815 | 788855_47188552_ENST00000601538.6,E | 1 | - | good | GAATTGGAGGAGTCATCTTGTAGGTTCAGCATCTGAATAATGAGGGAGAAATAAACAAAGAGATCATCCAAGACAAAGCCACTGAGCAGATGGATGTGATTCCCAGTTGAAAGGGCCCTG | 120 |
| chr11 | 70944815 | 70944935 | 788855_47188552_ENST00000601538.6,E | 1 | - | good | TCTGTGAAACAGGGAGCATTTGAGAAACTCTTGAAGATCAATAATGATTGCAAAAAAATGGCATAGCAGTAATGTTGGAGATGAACACAAGGAGGCGGAAGAGAGCCCTGGAATCAGGTA | 120 |
| chr11 | 70944935 | 70945055 | 788855_47188552_ENST00000601538.6,E | 1 | - | good | ATGAAACAAGCAGAACTAAATAGTCCGGGCAGAAACCAAGACAATTTGGGGAACAGAAGGGAATTTCCAAAGGAGATACTGTACCTAATATCCTCAGTGCAGTAAGAGAATATAATAGCA | 120 |
| chr11 | 70945055 | 70945175 | 788855_47188552_ENST00000601538.6,E | 1 | - | good | TTGACAAGAGCCGCCCATACACAATTTCCATCCCCTGCATAAGTGCCTCAAGTAGAAGATGGAGATGGGCAGCTGATAATCACCTGTCATTAAAGGAAAGCTGTCATCTCAAAGGCAGAT | 120 |
| chr11 | 70945175 | 70945295 | 788855_47188552_ENST00000601538.6,E | 1 | - | good | CACAGGAAACACATGCTCACCCCAGGGCCTTTGCACTGCTGTGGACTCTGACTAGAATGCTTCCTCCAAGGATCTTATGGTTCATCTCCCCAGCAAAGCACCCCGAGGGAAGTCCTTCTT | 120 |
| chr11 | 70945295 | 70945415 | 788855_47188552_ENST00000601538.6,E | 1 | - | good | CAGCACCTTGGAGGAACCACGACCAGGCTGGTGGGGGCCTTGCTGATCTCACCAGCCTTCTGCCTTCACCTCCTCCCTCTGCTCAAGCCTCCCAGGCCTCCACTTTTGCTTTTAATCAAG | 120 |
| chr11 | 70945415 | 70945535 | 788855_47188552_ENST00000601538.6,E | 1 | - | good | GGTGAGGTGGCAGTCTGGAGAAAACCATCCCTGATGGCAGCAGGAAGGGCTGGAAGCCGTCGGAGCTCAGCGTATTGGAGGCCTGGCCCTTCCACATGAGGCCGTAGGTGTGGCTGATAA | 120 |
| chr11 | 70945535 | 70945655 | 788855_47188552_ENST00000601538.6,E | 1 | - | good | CACTCAGGCTGGGGCCTGCAAGGTACCTGCGGGTCAAGGCTCTCCATGAGCTTAGTGATGCTGAAGGCCTGGCAGGCACACAACCCTGGGCTCCGTTGTGTGAAATGAAGAAGTGGTTGT | 120 |
| chr11 | 70945655 | 70945775 | 788855_47188552_ENST00000601538.6,E | 1 | - | good | TGCTGCATCCCCAGGCACCTGCAGCTTTGGTGGTAACCTAGTGTCTCAGGAGTCTAAAGTCATTCTGTTGTCTCTCTAAGCAAAGCCACCTGAGGTCAACCCTGCCCACATGCCCTGATG | 120 |
| chr11 | 70945775 | 70945895 | 788855_47188552_ENST00000601538.6,E | 1 | - | good | GGACTTGAGGTCACGTTCATTCCCACCAAGGGCTGTAGATCCAATAATTAGGAGAGCCATTCCTTGTGCGCTGGAAGGGTGAGCCCTCGAGCTCTTCCCCCATAACCTCCCACTCAGGGG | 120 |
| chr11 | 70947215 | 70947335 | 788855_47188552_ENST00000601538.6,E | 1 | - | good | AAGCACATTTCCCATGTTTCAGGTGCTCTGCAGCCAGACCCTACTGCACAATGTTCTGGAAGTTGATGGGTTCTTAGTCCAGAATGCTGGCCTTGTTCCTTACATTTGGGTACTGCAGGG | 120 |
| chr11 | 70947695 | 70947815 | 788855_47188552_ENST00000601538.6,E | 1 | - | good | TGGGAAGCAACGTTGGTAAAATCTCATGAAGATGCTGAGTGAGGCATCTGGATTTGGCAGAATATTCTAGAGCTCCCTGTCCAGTCCAGTAACCACAGGACGCACATGGGTATTTATATG | 120 |
| chr11 | 70947815 | 70947935 | 788855_47188552_ENST00000601538.6,E | 1 | - | good | GCCACGTTGCTTGGCAGGGACTTTACTCTGGGGGAGATGTACAGTGCCAGCAGTGCAGAAGAGAACTCTAAAATGGCATTTAATCTGTGCATCCGTCACCCACCCAGATGCTATCCTGGA | 120 |
| chr11 | 70947935 | 70948055 | 788855_47188552_ENST00000601538.6,E | 1 | - | good | CAAAGGCCCTGGGGTCAGTGCATGCCTGGGGTGTGTGGGAAGCAGCCAGGAGACTGGCTAGCTGCAGAGGTGGGGGTGGGATGGTGCCCACACCAGTGGTGGGAAATGAGACCACAGAG | 120 |
| chr11 | 70948055 | 70948175 | 788855_47188552_ENST00000601538.6,E | 1 | - | good | GCTGGGAGCCGAGGGCGGCTTCTGTGACATTTGGGGCAAAGACCAGGAAAAGAGGAAGGAAAGAGTGGGCCCTCAGGATGTCAGAGGGAAAGGCATCTCAAAGAGTGGGGCTGACACGT | 120 |
| chr11 | 70948175 | 70948295 | 788855_47188552_ENST00000601538.6,E | 1 | - | good | TTAACAAGTAAGCAGAGAAACGTGCAACATGTCGGGGTTGGTAAATGACAGGAAGAAAAATACAGCAGGGTGAGGGTGGAATGGTACGGAGCTGGCAATGCGGAAGCGAGTTTCCTTCAC | 120 |
| chr11 | 70948295 | 70948415 | 788855_47188552_ENST00000601538.6,E | 1 | - | good | ATTCATTATTTCCACCAGTACATATGGAGTTTCTCGTAAGAGCCACGCTGTCTTCTAGGTGCTGGGGACACATCAGTGAGCCACACCACACTCTGCTCTCTGGAAGTGCACATTAGATAA | 120 |
| chr11 | 70948415 | 70948535 | 788855_47188552_ENST00000601538.6,E | 1 | - | good | GTTCCCATAGAAACAGGGAAGTGGAAGTTGTATTTCTAGATTAGGGCGTAGAAACCTGGGTAACCCAAAGTGTGGCTGATGTTCGGTGCATATCTATGATGAAAAGTAATCAGGGATCGA | 120 |
| chr11 | 70948535 | 70948655 | 788855_47188552_ENST00000601538.6,E | 1 | - | good | CACCTCCTCCACGTTGCAGCGTCGTGGCCCTGCTGAGGGCACGCAGTTGGCGGAGTCAGTCATTTAGAAGTAGCTTCTTTTAAAAGTTCCTCTGAGTCTTATTTGGGATTCAGACTCTGT | 120 |
| chr11 | 70948655 | 70948775 | 788855_47188552_ENST00000601538.6,E | 1 | - | good | ACCCAGGCAGCCTCTGTGCATGTGGACATTCGGCAGGGGTTGGGAGCGTGGCCCCTCTGGAAAGGCTGTGCTCTTCAGCCTGCCACACTCCCCACCGCTCCCTGCAGCCCGCCCCTGGTC | 120 |
| chr11 | 70948775 | 70948895 | 788855_47188552_ENST00000601538.6,E | 1 | - | good | AAAATTTAATTTTTTTAATTTAAAAAAAAATTGAATGTGTTGCTGGTATTTAAACTTCCAGGAATTTCATTAAAAAAATTCAGCTTTCCAGCTTCTCTTGAAAAATTAGGAAACGTGGCCAC | 120 |
| chr11 | 70948895 | 70949015 | 788855_47188552_ENST00000601538.6,E | 1 | - | good | TGGGCCCGGGGCCCAGCTCTCGTTTGGTTGATAGAATGAGCCGTAGTGTAAAGTTCACCGTCGGCCGTCTGGAGGTTGGATCTGACCCAAGAGAGCTTGATTGTAACCTGCCCAGATGTT | 120 |
| chr11 | 70949015 | 70949135 | 788855_47188552_ENST00000601538.6,E | 1 | - | good | TCTTTTGGTGTGTTTATCCAGTGTATTTAGAAGCAGCTGGTTGGCAGGGAGCTGGGCACTGGCTGGTACCTGCCATTCATCTTGTGATCAGTCAACCTCCTGCAGGGCAGGCAGTGAGTC | 120 |
| chr11 | 70949135 | 70949255 | 788855_47188552_ENST00000601538.6,E | 1 | - | good | GGAGGGGCGGTACTGAGGCTCTTGCAGACTGAATGAGGGAGTATCAGCTTCGATTTTAGCCTTTTCTCTTCCACATGTGAGTGACCAAGCAAATCATTATTTTTATTTCTTTTCCCTTCT | 120 |

| | | | | | | | | |
|---|---|---|---|---|---|---|---|---|
| chr11 | 70949255 | 70949375 | 788855_47188552_ENST00000601538.6,E | 1 | - | good | AAGATGATGCGCCCCCGGGGAGCTGCTTGCCGGGCAGCATGCCAGGCCTGGGGACGTTGCTGTGCACAGGGTTCTTCGCCCACTTTCACCCCCGCTGCCTGTGTTCTGTGGGCGGAGCCG | 120 |
| chr11 | 70949375 | 70949495 | 788855_47188552_ENST00000601538.6,E | 1 | - | good | GTTTCCTATTCTCAGTGTGAAGGGAGGGTGAGCAGGTGGAGCCTGAGAGTGAGGACGTCTCCCCACATGACGGGCGGAGTCTCTGAGCTTGTGCAGCCCAGGGTCCTGGGAGAGTTTGCT | 120 |
| chr11 | 70949495 | 70949615 | 788855_47188552_ENST00000601538.6,E | 1 | - | good | CAACCGTGAGGCTTGGTGCCCTAGTCTGGAGCCCAGGTCACCCTGTGGGTTCCTGGGTGGCTATGCAGCAGGGTGAAGAAACATGGCTGGTGCCACCTCCAGGATTCTGGTCCATCTCAG | 120 |
| chr11 | 70949615 | 70949735 | 788855_47188552_ENST00000601538.6,E | 1 | - | good | CAGGGAGGCCTGCAGCATGGGCCAGGGCAGTGCCAGCGAGTGTGTGGGATGGGGACTTCCTTCCATGGCAGGAGGAGGTGGTTGTGTCCAGCCGGCCCCCCATGATCCCTGCAAACACTC | 120 |
| chr11 | 70949735 | 70949855 | 788855_47188552_ENST00000601538.6,E | 1 | - | good | GCGCAGGTGGAGGGGTCAGACAGTGCCTCATGACTGTGGATACTGTCATCAGCATCAGCCCTACCGTAGGAACAGGCCAACCTTGGGAGCCCAGAGAGGCTGGCCCAGGCCACGGGGGCC | 120 |
| chr11 | 70949855 | 70949975 | 788855_47188552_ENST00000601538.6,E | 1 | - | good | CACCTACCGGCTCCTTGATCCTGGACTCCCGGCCACCAGCGCTGTGAGCCATAGATTTCTGCCATTTATAATCGACCCAGCCTCAGGTGTTTTGGCATAGAGGCCCCGACTCACTAAGAC | 120 |
| chr11 | 70949975 | 70950095 | 788855_47188552_ENST00000601538.6,E | 1 | - | good | ACCAGGGCGAAACTCCATCTCCAAAAAAAGGGCTGAAGGCAACTGCCTTGTCTTTCCGCCCATGAAGACACATTGAGAGCCACCACCTGTGCAGCAGGGAGCAGCCCTCCCCAGGCACCG | 120 |
| chr11 | 70950935 | 70951055 | 788855_47188552_ENST00000601538.6,E | 1 | - | good | ATCACTGCAGTTAAGGGAGTGTCCAGAAGTTTGCATAGCATCACATATAGGTGCTGTAGTCTTAATGTGCCTTCAAATTCATGTGTTGGCCAGGCGTAGTGGCTCATGCTTATAATTCCA | 120 |
| chr11 | 70952015 | 70952135 | 788855_47188552_ENST00000601538.6,E | 1 | - | good | GGTTTCTTCTCAGCTGGTGCCCCTGGGCAGGCCACTTAGGATTTACTTAAAAGTCTCCAAATGCTCCCTTACTGAGTCAAAGCGTTTGCCCCGTAAATTCATGTCTATCTAGAACCTGTG | 120 |
| chr11 | 70952135 | 70952255 | 788855_47188552_ENST00000601538.6,E | 1 | - | good | GACCTCGGCCAGGAGAGACGCCCCAAGGAAACCGGCCCACTCTCAGCCACAGATCTGGCCAGACGGCACAGCAGGGGCCTAGCATTCCCGTCTGCAGAACCTGGGTTTAAATCTGGCTTG | 120 |
| chr11 | 70952255 | 70952375 | 788855_47188552_ENST00000601538.6,E | 1 | - | good | TTGTGTTGTGTCCAACACCATTTCTATGAACAAGATGGAACTTGTTTCCAGAATTCTATTCCTGGTTCTCAGTCACCCAAGAAGGTGGTTTATTTTGGGTCTGCATTTACTGAGGCACAA | 120 |
| chr11 | 70952375 | 70952495 | 788855_47188552_ENST00000601538.6,E | 1 | - | good | TGAACAAGACAGCCAAGTAGCTAGACAGCTTGTGTCTATCCACTTTAAAAGAGGAGGGAGCAAATGCAGCTGGGAATTGGCAAAGGACTACGGTTTCCACAGCGCACCCTTGGAAAGGTT | 120 |
| chr11 | 70952495 | 70952615 | 788855_47188552_ENST00000601538.6,E | 1 | - | good | AAGCCACGGCATAGGAGCCAACTTGAGAACAGTTTCAGCACTGAGACGGGTACTAGCTCCATCGCCACCCAGAGACTGCATTTTGCACTTACTGGGTGATTTGATAAAATCCCATTTATT | 120 |
| chr11 | 70952615 | 70952735 | 788855_47188552_ENST00000601538.6,E | 1 | - | good | GGCCGCCCTGCCATCTGCGTCCCCCAGAGTGCTGCCCTGAAGCAGCCTCACCTTTGCATGAAAATCGTCAGCAGCTTGGGTGCGTCTCTGCTGTTTGCCACCAACGAAAGCCACACAGAA | 120 |
| chr11 | 70952735 | 70952855 | 788855_47188552_ENST00000601538.6,E | 1 | - | good | AAGCCACCTTGTTTGTGATCATTTGTTCCCACAGCCACAGAAAGCAGCTACCAGAGACCATAGGTGGGTGTCAGTTCCCACACTTCTGCTGCTCAGGCCCCCCCGGACACACGAAGCCAG | 120 |
| chr11 | 70952855 | 70952975 | 788855_47188552_ENST00000601538.6,E | 1 | - | good | GGGAAGGACCCTCCCCTAGAGCCTTCCTGGGGAACATGGCCCTGATGACACCTCGGGGTCAGCTTAGTGATGCCATTTTGGACACTGGCTTACAGGACTGCATGGAAGGTTTGTTGCTTG | 120 |
| chr11 | 70952975 | 70953095 | 788855_47188552_ENST00000601538.6,E | 1 | - | good | CCAGCGTAATCACCAAGGTCCTCGTAGGGCACACACAGGGAAGGAGGCTGCGTGAAGGTGGAGGCGGTGGGATTGGGACGTGGGATGGGGGCAGAGATCAGATGGTGCTGCCAGGAGGT | 120 |
| chr11 | 70953935 | 70954055 | 788855_47188552_ENST00000601538.6,E | 1 | - | good | TGCTGTCTGCATTGCAATTTCTATGAGCTGATGTGGGCAAATTGTAAAGTTCATGTAAGACTGTCTATTAGACACACACACACATACACACACGGCTCTCCTAGAGGTGCACGTGCCTGT | 120 |
| chr11 | 70954055 | 70954175 | 788855_47188552_ENST00000601538.6,E | 1 | - | good | CCTTCCCTGTTTTGTGGAAATCCACGTGGATGTGGGAACAAGGTCATGGCGTTTTCTCCCAGGACCTTCCTCTTCCCCGATGAGTGCCATGGGCAGGGTCCCAGGATGTCCCTGTTGTGC | 120 |
| chr11 | 70954175 | 70954295 | 788855_47188552_ENST00000601538.6,E | 1 | - | good | CTGGCGTTGCACTTTTCAGAAACCCACGAAATCTGCTCAGCGGACCCTTCTCTTCCCTCCAGGTGGCTTTCTGAGTTTCTTTCCTTGTTAAATGAGCCCCAACAAGACTGTTTGGTTGTG | 120 |
| chr11 | 70954295 | 70954415 | 788855_47188552_ENST00000601538.6,E | 1 | - | good | GCACTCTGCCAGTGGCTGTGCACTCTGCAGTTCCTCTGATCATGCTGGTGGAGGCAACTGTGTGGTTGAGGGACAGTCCGTGATGAAGGCTGCCTGACACCGTGGGTGATATTGTGGTTC | 120 |
| chr11 | 70954415 | 70954535 | 788855_47188552_ENST00000601538.6,E | 1 | - | good | CCAGACCCTTCGTCTGATTTCCTTTTGCACAGAGCTCCAAACTGAGTTCCCACATTTAGCAATTTAGCCAGAGGGGGGATGAGTACATTTTCTCATGCCACGCATGGTAGTGGCTGCCAGA | 120 |
| chr11 | 70954535 | 70954655 | 788855_47188552_ENST00000601538.6,E | 1 | - | good | AAGGATCCCGGGCAAGGCGTTTTTGGATGTGGCCTGTCCCCAGTGCATTCACTGGGGGAGGACTCAGCCCATGAAAGTGAGGGCGGATCTCCCGAAATGTCAGAGCCTGGCTTTTGCCCC | 120 |
| chr11 | 70954655 | 70954775 | 788855_47188552_ENST00000601538.6,E | 1 | - | good | ATTTCCAGTTCTCTGGCAATCAGGTGATGGACCCCAGCATCGCCACAGGCTAAGCCAGTGATGGCAGAGGCCCAGGAATGCCGTTGGCAATGGTTCTGTGTTGTATTGCGCTCTTCACTC | 120 |
| chr11 | 70954775 | 70954895 | 788855_47188552_ENST00000601538.6,E | 1 | - | good | CTCCTAAAGGGACAATGAACAGGGCATAGGCTGCTGCATGCCAGACCCCGTGTTCTTCCTTAACCCTTAACACTGCACATAGAAGGAGGAGAAAGGCCGAAGCCAGAAACAAAGAACTGA | 120 |
| chr11 | 70954895 | 70955015 | 788855_47188552_ENST00000601538.6,E | 1 | - | good | GCTTGGTGGTTTGGGTACCTTTCAGTGCATGGGGGCACCCTGCCTAGTCCTGTGCTGAGCCCTTTCCATGTGGTCCCACGAAGCAGGGCCTCCTGTGGATGAAGAGCCCAGGTCACACAG | 120 |
| chr11 | 70955015 | 70955135 | 788855_47188552_ENST00000601538.6,E | 1 | - | good | AGAATGGCTCCATTTTAATAAGGCCCAAATTTAATATGGGCAGGGAAGGACAAGGATGAAATGGGAGGGAAACGACAAAGAAGGCCTTGATGTGGATGTCTTTCTGATGAATGACACTGT | 120 |
| chr11 | 70955135 | 70955255 | 788855_47188552_ENST00000601538.6,E | 1 | - | good | CCACGTGCTGTCATTCTGTCGGTCTCCAGGCTGTATTCAACAGTTTTTAACCAGACATCCAGAAGCAAAGAAAGGGTGGAGAAAATCAGGAAAAAAAGGCAGTGGCAATGAGAGACGTTC | 120 |
| chr11 | 70955255 | 70955375 | 788855_47188552_ENST00000601538.6,E | 1 | - | good | CTGACTGGAGGAGTTGTTCCTGCCTCCCAGCAGCTGCATTCTGAAAGGGACTTCCACTGTTTTTGGATACAGGGACTTTAGCACCTGGCATTGTATATTTGGAGAGGGCAGCCTTCTCC | 120 |
| chr11 | 70955495 | 70955615 | 788855_47188552_ENST00000601538.6,E | 1 | - | good | AGCCTCCACAATCATTTAAACCAATCCCTTAAAATAAATCTCTTTGCATGTGTACACATACACACGTAAAGTACACACATACACAAATATATACATAGACACATATTACAAATATATGTG | 120 |
| chr11 | 71055935 | 71056055 | 788855_47188552_ENST00000601538.6,E | 1 | - | good | AACAACCCCAGCCAATCATGGGCCAGGGTGTTCTGTGGTGTAATGCACAAGCCCAGGTCCTAAACTTTTTAAAAATTGTGGCAAAATACATATAGCATCACATTTAACATTTGAACCAT | 120 |
| chr11 | 71056055 | 71056175 | 788855_47188552_ENST00000601538.6,E | 1 | - | good | GGGAGATGCTGGCTCTGCCGCCCCTCCATCCAGTGGGACAAGCTTCCTGAGGATCACACAGGAGGGGGCTTGCCTTCCATTGGCACCCCCATGGGGTCCGCTGTCCCAGAGTGGAGGGAG | 120 |
| chr11 | 71056175 | 71056295 | 788855_47188552_ENST00000601538.6,E | 1 | - | good | TGCTTGCTCACACTGGCTCCTTCCTATTAGAGCAAATGGCACAAATAATGTGGAAATGCATGCAGCAGTCAGCGCCGGAAAGGCAGCAAAGACGTAAGTGCTTCCAACTGTGCAGCTGTT | 120 |
| chr11 | 71056295 | 71056415 | 788855_47188552_ENST00000601538.6,E | 1 | - | good | CACGGTCGTGTTTGTATGTGTGACATGAGGTCGGCTCGTTTTGTTGCGGATGTTTTCTTATTCGGAGGCTGCTCTAGGTGCCTCCCTCTCTTAGTGCAAAGCTGCTGCCCCGTCTCAGTG | 120 |
| chr11 | 71056415 | 71056535 | 788855_47188552_ENST00000601538.6,E | 1 | - | good | GTTTCGAGGCGGAAATAAGGAGTTAAAAAACTACAACAGCCAGACTCCATTTCAGGTAAAAGAGATCTCAGCATCAGGAAGGATGTGTGTGGATCACAGTGGGTGATGGCTCTGCGTCTC | 120 |
| chr11 | 71056535 | 71056655 | 788855_47188552_ENST00000601538.6,E | 1 | - | good | TCCCTGGTTTCCCGTGAGCTTGAGACTCTGGACAGATTTCTTTGGAGAATGTCTGTGCTACCATTTTACAAACAGGTTTCTCCTTTTCTATTTGCAGGACAGCTGTGCAAGAGTGCTTCT | 120 |
| chr11 | 71056655 | 71056775 | 788855_47188552_ENST00000601538.6,E | 1 | - | good | GATATAGTCTTAAAAAGTGAAATTGTTTGGCCATAGTTGGCAGCAGGTGAGCCTCACCTGACAGCATACAGGAGCCAGATGAAAGCAACATGCTTTCCCTTTTAACTGTGCACTTCTGTT | 120 |
| chr11 | 71056895 | 71057015 | 788855_47188552_ENST00000601538.6,E | 1 | - | good | AGTTCCTGGATATGTTTTTCTCAGTTCATAGTCATCTGCCTTGTTTTGTCCTGAGACTGCCTAGGTGAAGAATGTTCTAGAATATATCCCCACTCTCCCCTATTTTAGAACATGTATGCT | 120 |
| chr11 | 71057375 | 71057495 | 788855_47188552_ENST00000601538.6,E | 1 | - | good | ATACCGTTAACTGGTGCATATCCTTCCAGACATAGTCCCTGCTTTTCTATAATATAGACTTTAAGCCTTTATTAGTTTCTCTTTAAGCCTTTAAAATGTAAATGAACTTAAACAATATTA | 120 |
| chr11 | 71058215 | 71058335 | 788855_47188552_ENST00000601538.6,E | 1 | - | good | TAGTAGTTATTTGCTGCTGCAGTCCTCACTGGGAGCTTCAGAGGGGTTGGCGAGAAGTTCTCCATCAGCTTCACGGGCAAATGGTAGCTGCCCAATCAATGGATGGATGCAGATTCATTT | 120 |
| chr11 | 71058335 | 71058455 | 788855_47188552_ENST00000601538.6,E | 1 | - | good | AGTATTCTACTGCTGCCTTTTTATTAGTAGTTATTTGCCGCTGCAGTCCCCACTGGTTACATCAACATTCTGCTGCTGCCTTTTATTGGCTACGTCAACATTCTACCACTGCCTTTTTAT | 120 |
| chr11 | 71058455 | 71058575 | 788855_47188552_ENST00000601538.6,E | 1 | - | good | ACAAGCACCAGCCCAGCCCCTGATCATTTAAATGCACTTCCAAAAACAGAGTTTGCTTATTCTAAACAGCCAACTGCATTTGCTAGGCAAAATTATTCTAATAGTCGGTACGGCTACATC | 120 |
| chr11 | 71058575 | 71058695 | 788855_47188552_ENST00000601538.6,E | 1 | - | good | GGTAAGGATGGAATTCCCAGGATCTGCCCAGTACAGGCTTCGGCTGTGACCTGCGCTGAGGCCCTGAGTCCTGTAGGAGAAGGTGCCCCCAGCCCGGTGTCAGGCCCCTGCTGCCGAGGC | 120 |
| chr11 | 71058695 | 71058815 | 788855_47188552_ENST00000601538.6,E | 1 | - | good | CTCTGACCTGTCGTGGGTGGGAAGCCCTGGTCATTTAATCCACTTACCCCTTTATTCGGTGTATATTTGTGCAGTGTTGAGTCCAGGTGGTCACACCTGTTGTCTGTACACTAGACTGTA | 120 |
| chr11 | 71058815 | 71058935 | 788855_47188552_ENST00000601538.6,E | 1 | - | good | ACCAATCAGCTCTCTTTCAAATCTGCTGATCTGCCAGGTCCCACCGCCTGTGCAGTGTCCACTCAGCTTGGCATTTTGCTGTTCCCAGGGATGGGGTTCCCCCGGGGCCCGGGCTGGGGT | 120 |
| chr11 | 71059535 | 71059655 | 788855_47188552_ENST00000601538.6,E | 1 | - | good | AGGCCTGGTGGGCTCCCCACCCTGAAATCAGCCGGATCCTGAGAGCTCAGTGTGCTCCTGGCCTGCCCGACCCACCCAGCCAGGGATGAAGACTCTCTGCCAGGGTGGCCTGCCTTCCTG | 120 |
| chr11 | 71059655 | 71059775 | 788855_47188552_ENST00000601538.6,E | 1 | - | good | GGCGGCCTTTACTACCCAACTCAGATCCGCCCACCTGTGTCTCCCAAGTGAGCGCAGCCCACACTCTTAGCTGCCGGAGAGGGTGTGGTTCCCTCATTAATAATCACCATCATGGGGTGG | 120 |
| chr11 | 71059775 | 71059895 | 788855_47188552_ENST00000601538.6,E | 1 | - | good | GCCTCAGTGGGGCTTGAGGCAGAGTGCACCCCCAAGCATGCAGGTGGCTTTTTACACAGTCGATGAACAAAGCTGGCCACAGTCACCGAGGCCCCAGGATGCTGGAGAACGCAGAGACGG | 120 |
| chr11 | 71059895 | 71060015 | 788855_47188552_ENST00000601538.6,E | 1 | - | good | CCTTCCTGTAGGGCAGTGGGACAGAACACTCAGCACGGTCTGTGGAGCGGTCCACACTCACTTCCACATGCCTGACTCATTGGCATCTGTGGCCAGAGCCCCCCACTCTGTACAGAGGCG | 120 |
| chr11 | 71060015 | 71060135 | 788855_47188552_ENST00000601538.6,E | 1 | - | good | AAGGCAAAATTACCCCCGGTTGAGAACCACTGTGTCAGAGGGTGGGGAAACCATGGTGGTTTTGCTTCTAATTGGAATCGGGGTGCAGGGGCTCTGGGGATCCCCCTCTAACCAACCTGA | 120 |
| chr11 | 71060255 | 71060375 | 788855_47188552_ENST00000601538.6,E | 1 | - | good | CTAGGGATGCAGCTCTTCACGACTCATATCCATCATCAGTAACATTTGGAGCCATGAAGACAGGGGTTGTTGACCTCAGCTTTATTGAGGTTTGGGGCTGGATGGTTTTTTGCCATGGGG | 120 |
| chr11 | 71060375 | 71060495 | 788855_47188552_ENST00000601538.6,E | 1 | - | good | AGACCCTGAATAGCATAGGGCTTGGCGTCCCCCCTTGAGGCCTGACCGGCTGGGGCCAGGGGCCAGTGGGCTTGGGGAGTGGCATGTGATTCATACAGTCCGTGTTGTTCTGAGACCGTT | 120 |
| chr11 | 71060495 | 71060615 | 788855_47188552_ENST00000601538.6,E | 1 | - | good | GGTCCTCATAGTGAGAGGTGACAGAAGCCGCCTCTGCCTTCCTGCTCCCCAGGAGTGGGTGAACATCTTCTGTGAAACCTGCTGGGCTGTCCCTGAGTCTGAGGACACGTTCCAGGGACG | 120 |
| chr11 | 71060615 | 71060735 | 788855_47188552_ENST00000601538.6,E | 1 | - | good | TCCATCTGCAGTGTTGTTCGAGCCTCACACGTGGCTTTGCCAGCTCTCCTTTCACCACTGCCCTGTGGGCCTCAGGCTGACAGCTCCATGGGCATCTGTCCCTCCCCTGCTGACATGCAT | 120 |
| chr11 | 71060735 | 71060855 | 788855_47188552_ENST00000601538.6,E | 1 | - | good | CTGTGCGGAGCGTCACCCTATATCGGCTGCTGCATGCTCTTGCAGACTCACACGTTGTTTTGAAACACCATGGGCTTCTAGCAGCTCGTTTCCTTGTGTTTGCTTTTTCTCAGCTCAGCC | 120 |
| chr11 | 71060855 | 71060975 | 788855_47188552_ENST00000601538.6,E | 1 | - | good | TTTTTTCCTGTACTCAAAGCAACATATATTTATGGCAATAATTAGTAGAGTATGGAAATCTCAGCGTAAAAACTGGAATCCACATTTGTGCTTTTTCCAGGGGTTTCCCACCCTGAAACC | 120 |
| chr11 | 71060975 | 71061095 | 788855_47188552_ENST00000601538.6,E | 1 | - | good | CCTGGGCACAGCAGAACTGACGCCAGGGCTTTCTGTTTTTGAAACTCTGGATTTAATTGTACCCCAGAAATGAAAGATTCGTTTGCTGAACATAGTTATGTCTGAAAATTCCCTTACATA | 120 |
| chr11 | 71061095 | 71061215 | 788855_47188552_ENST00000601538.6,E | 1 | - | good | CCAGTAAAACTTTATTTACAAAATTAGATGTGCAGGATTAGGCCCAAGGGCCGTAGTTTGCAACCCTAAAATGAAGTAAGCAGGCCTGCAGTCTTGACCTGCACCTCCCACAGTGTGGAC | 120 |
| chr11 | 71061215 | 71061335 | 788855_47188552_ENST00000601538.6,E | 1 | - | good | GTGCAGGACTGCTTTATTAAAATTGGGCTGGGTACCCTTTTTTCTGTAAAGACCCAGAGAGTAAATACCTCAGGCTTTGCAGCCACAGAAAATATGCAGGTGAGTAGATGTGGTCGCATC | 120 |
| chr11 | 71061335 | 71061455 | 788855_47188552_ENST00000601538.6,E | 1 | - | good | ATCACCCTGCACTCCCTGTCATCAGCCTAAGTGGGGCCTGAAGACCGTTGTTAATATGTCCACTGTTAGGGCATCCTGCCAGCGTGGTCATTTCTGTGAATGTGTGACCACGTTAGGGAG | 120 |
| chr11 | 71061455 | 71061575 | 788855_47188552_ENST00000601538.6,E | 1 | - | good | ATTGACAGTGCACTAAGCTCATTCTGCTGCCACTAAGCCCGGTATCCCCACTTTAAAAAGGAGGAAACTGAGGCTCAGAAGACAAGAGGCAGAACCGCTCTCTCAGTGCTGAAGCTGGTT | 120 |
| chr11 | 71061575 | 71061695 | 788855_47188552_ENST00000601538.6,E | 1 | - | good | CCATGTGCCTGGGACTTGGAGGGGCTGTACTAATATGAGTGGGCCCCTCTCCTGGGGATCAGGGGTTACTTTCCAGGGAAAGGGCATTGCTGGGGGTGGAAGGAGACCCCCACCTGTGTC | 120 |

| | | | | | | | | |
|---|---|---|---|---|---|---|---|---|
| chr11 | 71061695 | 71061815 | 788855_47188552_ENST00000601538.6,E | 1 | - | good | CAGTGCTCTCCACACTCTGTCTCATCCTGGTGGAGAAGGCTGCATTCTGTGCATCTCTTGAAATGCCGGTCACCTGCAGAAACTCACCATTCCCAGGTTCCCGGTGACACCCCGATGGGC | 120 |
| chr11 | 71061815 | 71061935 | 788855_47188552_ENST00000601538.6,E | 1 | - | good | CCCACTGGGGGTGATTTGACGGGAAATTTTGGGCTCATGATCAGGACACAGGGCAAGTCCTGCAGCTGGGGCAGAGCGGGGGAAGGGCTAGGTCGGGACTGGAAAAGCCAGGGACCTGCC | 120 |
| chr11 | 71062295 | 71062415 | 788855_47188552_ENST00000601538.6,E | 1 | - | good | CCCCACCAGGCCCCAGATGGGAACTGGCTGAGAGCCATCATCCCATTCTCCTGGCCCTGTGGTCTCTCCGCCTCCTCTTCGTTTTCTCTCTTACCAAAGCAAAAAACAAACTTTTTGCTC | 120 |
| chr11 | 71062415 | 71062535 | 788855_47188552_ENST00000601538.6,E | 1 | - | good | GCAGTTACAGCCCAGAGCAAATGGGTGAGCAAGACCCCCAGGGGCCAGCTTTCATGTCCCCACTGATGCCTACCCCCATGAGGTCCCTCCTCTCTTGCTGCCCTCTTCTCCCTTTGCTGA | 120 |
| chr11 | 71062535 | 71062655 | 788855_47188552_ENST00000601538.6,E | 1 | - | good | GTGTCCCAGGCTCCTCCCTGAGCAGGGCTCTCATATTCTGTATTCCAATTCGCCAGGCCACTCTCAGCCATCTGGGTTTTGCTGGTGCTGGGTGATTCTTCTAGGCCACTGGGCCAGGGT | 120 |
| chr11 | 71063135 | 71063255 | 788855_47188552_ENST00000601538.6,E | 1 | - | good | GTGACTGGAGGTAGTGGAGGATTTTTCTTGGATCCTAACTGTATCGCCCTGGAATGGCAGTGCCTGCTGAGGATTTATTCTTAACTTTCTATAGGAAGTACATGTCCTTTATAATAGGGA | 120 |
| chr11 | 71063255 | 71063375 | 788855_47188552_ENST00000601538.6,E | 1 | - | good | AAGTGAACCATGTACCCTTGAATTTCAAAATAACTAACTGGAACCTGAGCTCTTTTTGACGTCTGGGAAACACGTGGTAATGGGTCGGAGGTATGATATTGACTCCGTCAAAGTCATTGA | 120 |
| chr11 | 71063375 | 71063495 | 788855_47188552_ENST00000601538.6,E | 1 | - | good | TTAAACCAGAAGGAGCGGGTAGATGGGCATTTTGAACTGCAAACACACATGTTCACACACGTGTACACAGACACATGTGTGATATGCATACATACATGGAGGTATCATTGTTATTAAGCC | 120 |
| chr11 | 71063495 | 71063615 | 788855_47188552_ENST00000601538.6,E | 1 | - | good | ACAGTTTTTCCTTTCTCCTCCTTTCTGAAGCCGCAAGTTGTTGCAGTTTTTCACATTAACCTTCTCTGTAAAATGGGCCTCCTGTTCTCCCTGGCAGAGAGTTCTCCCATACCTTGAGAG | 120 |
| chr11 | 71063615 | 71063735 | 788855_47188552_ENST00000601538.6,E | 1 | - | good | CTTCACTGAATTATTTAAAAGGATAAAGACCTGTCAGATAAGATACACCTCTGAAAGGAATCCAGGGCAGCCAGCAGCCTCGCAAGGGAGCCAAAAGCTCCCACCAGCACCTCCTGGATC | 120 |
| chr11 | 71063735 | 71063855 | 788855_47188552_ENST00000601538.6,E | 1 | - | good | TACATTTATTAAGGGAAGTGTCTCGGAAAATTCATGCCAAGAGAAAAAGTGGCAGTGAAAATGATATTCTTCTTGATGCTCGGGTCTTCAAAAAGCATTCCAATTGCTTCTATATTTATA | 120 |
| chr11 | 71064095 | 71064215 | 788855_47188552_ENST00000601538.6,E | 1 | - | good | AGCCTGGCAGAGACCTTGACTTTCTGGTTTGCATCTGTGTCCCCAGGGACTAGGACAGTACCTGGCAGCTCCATGAGGACTCACTGCATGCATCGATGAGTGGCTTTCTCTGTCTGTCTC | 120 |
| chr11 | 71064215 | 71064335 | 788855_47188552_ENST00000601538.6,E | 1 | - | good | TCTCTCTCCTCCCATCCCAGGACTCAGCTTGCCCAGCCAGCCGCATCCTCTCCACTGTCTGCATCAGACCCACCCGTCCTGACGTTGTGTGGTTGTTTTCTGTGCCCCCTCTCCAGCCCA | 120 |
| chr11 | 71064335 | 71064455 | 788855_47188552_ENST00000601538.6,E | 1 | - | good | CAGCTTGCTGCTGCCTGGCCCCACCACTGCCTTCCTGCCTCCCGGCCCCTTGCCGGTTTCTACTTCGGTTCTTGGTGCTCTGCTGGGGTGGCCTCTGCCTCCGCCCAGCTGTCCCCAGGC | 120 |
| chr11 | 71064455 | 71064575 | 788855_47188552_ENST00000601538.6,E | 1 | - | good | CTTCCCTCAAATCCACTTTCTGTCACCCAAGTTCTTCATCCCTCCCCTGCTGAGAACCCCCTTTGCATAACCCAGGGCTTGTGTCCTGTGCCCTTCTCTCCTATCCAGCCCAGCTCATCC | 120 |
| chr11 | 71064575 | 71064695 | 788855_47188552_ENST00000601538.6,E | 1 | - | good | TACCCCTCTTGTCCCCTCCACCCCTCTTGTCCCCACCACCCTTCTCACCCTTGCCTGGGGCCCTGCTGTCTCTCCCTGGGACTATGTATTCACCATCCAGCCTCTTTCCTTCCAGTCTGT | 120 |
| chr11 | 71064695 | 71064815 | 788855_47188552_ENST00000601538.6,E | 1 | - | good | ACTCTGTAAATAACGCCCTTCCCCTGGAGCTCAGGCCCGAAACCTGGCCACCTCCTGTTTCTCACCGTCCCTCTCATCCCGGGACTTGTCCATTCCCTCTCCACCCTCTCTGGGCCCGTC | 120 |
| chr11 | 71064815 | 71064935 | 788855_47188552_ENST00000601538.6,E | 1 | - | good | AGTCCCCAGTCGGAGTCTCCCAGCTCACTTCTGCCCTGAGCTCTGGCAGTGATGTCTGCTCCTTGCCAATGGCCCCACGTGCCGACCTCTTAGGAGACTCCAGTGTGGCAGCTGCTCCCC | 120 |
| chr11 | 71064935 | 71065055 | 788855_47188552_ENST00000601538.6,E | 1 | - | good | GGCATCTCTCGGCGTCTGCACTGCACGCTCCTGTTACCCAGATTCCCTCCTGCAGTGACTCCCGAGGTGACCTCCGGCGGGCCTGTCTGTGTCCTTGTGCGCCTTCTAGAAAACCCCAAG | 120 |
| chr11 | 71065055 | 71065175 | 788855_47188552_ENST00000601538.6,E | 1 | - | good | GGCTGTTGTCTCACTGACACAGTGTGGTCAGACTTTCACCCCTGCTTGCTCAAACCCAGTCACACCCACATGGCACTTCCAAGGTCCCGTCTCTGTCCCCATCCAGCTCAACGTTTCTGT | 120 |
| chr11 | 71066375 | 71066495 | 788855_47188552_ENST00000601538.6,E | 1 | - | good | CTCTGCCTTCTCTTACGTCATGCCCATTTGCTTATGAAGACACTCCACCCACTTCTGTCTCCAGAAGTTCCTTGGCTTCACGGACCCCCTGCCCACCTTCCCTGCTCACTGCTCATCTCC | 120 |
| chr11 | 71066495 | 71066615 | 788855_47188552_ENST00000601538.6,E | 1 | - | good | AGCCTCAGTGCCGTTGAAAGGAAATGGCTTGACGGGGCCTGGAGCTTCCGAGTTAACATCCTGGTTGCAACGTTGTCTGTGCCTGTGGGCATGTCTGCCACCAGTGTTTCTACTTTTGAC | 120 |
| chr11 | 71066615 | 71066735 | 788855_47188552_ENST00000601538.6,E | 1 | - | good | TTTTACATTTATGGCTGCCCAAAAGGCCTTATTTCAAAAGATAGCTTAACCTGAAAACAAACTCCTGATTTACACTTTGGAGTGTGCAGTAATTTGTATCTGTGACATTTAAGTGAAGAC | 120 |
| chr11 | 71066735 | 71066855 | 788855_47188552_ENST00000601538.6,E | 1 | - | good | ACTTACAGGCACAGCTCTTCAAGGGGTGGGCTGGTCTGAGGGACCAGGTTCCAGCCCTGTCCCAATGACACAAGCGTCCCAGCCACTTCCCCGTTTTACTTATGCAGACATCTTAGTTGT | 120 |
| chr11 | 71066855 | 71066975 | 788855_47188552_ENST00000601538.6,E | 1 | - | good | CCTTGTGGATGCATTCCTGTAACCCACTGACTGGGGAAGGGGCCTCTACTGGGCACAGGTCAGGCAGAAGCACCTATGTCCACGGCAGTGAGGGGTGTGTTCGTACTTACCCTGGCCCTT | 120 |
| chr11 | 71066975 | 71067095 | 788855_47188552_ENST00000601538.6,E | 1 | - | good | CTCCCCCAGCTCTGGTGGGTGGCTGGCTTGGTTGGCTTTCCCTTTATACAGGCCAGAAGATGCAGGAGTGGTCAAAGGCCGCTCCGTTAGTAAATGAGGAAGCCAGGTTTCCAACCCATG | 120 |
| chr11 | 71067095 | 71067215 | 788855_47188552_ENST00000601538.6,E | 1 | - | good | CCTGTCCGAGCTGTGGCAGCTCCCATGGGGCCATCATGGGGTGATGGCCCCCTGAGCATTGGGCGTTGTGGGTGTGGGTCTCGCTGAATGGCGACGGTGTCCACACGCCCTTTCCTTCAC | 120 |
| chr11 | 71067215 | 71067335 | 788855_47188552_ENST00000601538.6,E | 1 | - | good | GTCTGGAAGGTAAAAATAATAAAAAACAGTCTGCCTGTCTTTGATACCAGCACGAGCCAGGGTAGGAAGACGTGGTGTTTCCTGTTTGCATCTGTGCCCTTGCTGGTGTCCCACGGGGAA | 120 |
| chr11 | 71067335 | 71067455 | 788855_47188552_ENST00000601538.6,E | 1 | - | good | ATCCTGGCTCTCCAAGAGCTCCAGAGGGAGGGGAGCAGCCTGGGGTTCATTTTTCTGCCCTGGATGATGAAGGAGCGAAGCAAGTTTTAAATATCAAAGGTTACTAGGCTAGGCCAATTT | 120 |
| chr11 | 71067455 | 71067575 | 788855_47188552_ENST00000601538.6,E | 1 | - | good | CTCCACACTTGTAATAAAATACACTATGATTAGAATGTTCTCTCTTCTTTTATGAAATCACCTAATTAGGGCTTTGATTGTAAGTATCAAATTGTGTCTAATCTGTCAACCCCTCACCTC | 120 |
| chr11 | 71067575 | 71067695 | 788855_47188552_ENST00000601538.6,E | 1 | - | good | ATGATGGTGATGCTGCTATTGACGATATTCCTGTAACAAGCCTCTCCTTGTTAGATCTAAAAGCACCCAGGTGTATGGATTGTTACATGTGCTTTATCAGTTTTAGGCAGGCCTTACCAC | 120 |
| chr11 | 71068175 | 71068295 | 788855_47188552_ENST00000601538.6,E | 1 | - | good | TTAGGATTCACAGTTTCTGGGCTCTAAACAGCAGGAGTGTCTGTCACTCAGTGAGCCATTGAGACTACTTGTGTACATCACCCCCTTCCTCTTTCTACTTGTAGATTAAGTCATGGTACT | 120 |
| chr11 | 71068295 | 71068415 | 788855_47188552_ENST00000601538.6,E | 1 | - | good | TTCAGATTCTTCCAGTCTCTTTGGATGAGTCTTACAGCCCTTGAGCTTTTCTGCTACAGACACTGGGGGTCAGGATGCAGTCTTCTGTTGTCTAGGACTGTGCCAAGCATTGCTGGGCAT | 120 |
| chr11 | 71068415 | 71068535 | 788855_47188552_ENST00000601538.6,E | 1 | - | good | GACAGCCCTTCTCCCTGAACATGGCCTGTGGTCGCCAAATACACAGTCATGGTTGCGGTGTTGCATTGTGCTGCTGGGGAACTTGTCAAGCAAAGGCACTTTATTTGCCTTCCCTAGATA | 120 |
| chr11 | 71068535 | 71068655 | 788855_47188552_ENST00000601538.6,E | 1 | - | good | CAGCAGCTGGCTCCACTGTGAGACAGGACCAGCATACCAGACTCCATGCCCATTAAGGTCTCTTTCCATTCTGCCCTCACAGAGCATGACCAAAGTCACTATGGCATCTAATGGTGGCTT | 120 |
| chr11 | 71068655 | 71068775 | 788855_47188552_ENST00000601538.6,E | 1 | - | good | AGGTTTCTAGCTGAGCTGTAACCTCCACATTGGAATTACCCAAAGAACCTGTGAATAGTACAGCTGCCAGGCTTCCAGCCCATAACCCAGAATCCCAGTGATGCTGAAGAATCTTCAGAG | 120 |
| chr11 | 71069495 | 71069615 | 788855_47188552_ENST00000601538.6,E | 1 | - | good | CTAAGAGGCTAAAAATAGAAAACAAATATATGGTAATATGGTATTCCTGGAACATAGTAGATGCTCAGTAAATAGTGGCTGTAATGATAGCAGTGGTGTTAGTGGTGATGGTGAAGGTGG | 120 |
| chr11 | 71069615 | 71069735 | 788855_47188552_ENST00000601538.6,E | 1 | - | good | CATCTGGACTAGCCCATTGCCTTGCCTTGCTATGCTGTCCTGGACAACCTATTGTCTCTCTAGGGCCCTGAATTTCCTTATCTGTAATGTGGGGATAATTAAACCGAATTCACAGGGTTG | 120 |
| chr11 | 71069735 | 71069855 | 788855_47188552_ENST00000601538.6,E | 1 | - | good | AGACTGTTCAGGGATGGGACAGAGGTGCCTTTGAATCAAGACAGGATTCAAGGAAAGGAGTCAGGAGGCAGCCCTGAGGAAGTGGAACAGTAATTCAAAGAGGCATGGTGGTCTGGGTTT | 120 |
| chr11 | 71069855 | 71069975 | 788855_47188552_ENST00000601538.6,E | 1 | - | good | TGTGGTTCATTAGAGCAGGCACTCATGCAGGGGGCAGAAACCACTCTAGCAACTTTAAGCAGAAAGGGTGCCATACAGGGCACACACACACAGGGCATGCAGGGAGTAGTGTTGAACAAC | 120 |
| chr11 | 71069975 | 71070095 | 788855_47188552_ENST00000601538.6,E | 1 | - | good | CCCTACCTTTGTCCTGCAGGCACTGCGTGTTGTACTTGACCCAAAGCTTCTTTCCTTCCTGAGTCACCATTTGAACACCTTGCTATACTCACTCCCCCTTGTCTACACATGTATGAGGCA | 120 |
| chr11 | 71070095 | 71070215 | 788855_47188552_ENST00000601538.6,E | 1 | - | good | TGTCTCACTGATGTAGTTAAAGAAGCTGGTCCAAACCACATCTCGTGTGATGCCAAGAGAGGCACAGCTTCATGTCTGCATCTTCCAGGTGCCAGTAAGTGACCAGGAGGAATGCAGAAC | 120 |
| chr11 | 71070215 | 71070335 | 788855_47188552_ENST00000601538.6,E | 1 | - | good | TGTCAGTCGGAATGTCTGCGAGTGGTCTAGGATCCAGCTTCGTGTATCCCTCATACTCCTCCTTAGGGTTCTAGGGTGTCTTTAATCTCCTCCCTCATGCCCTCCCCAAACTCTTTGGAG | 120 |
| chr11 | 71070335 | 71070455 | 788855_47188552_ENST00000601538.6,E | 1 | - | good | TGGGTAGAAATGCATTAGCTGGCTCCCCTGGTTTTGATTTCCAGCCTTGCCTAGCCTCCAAGAACCAGTGGGCAGGGTCATAGGCTCTGGACTGGTTGGCCCCTTGACCTCTGGCTACAG | 120 |
| chr11 | 71070455 | 71070575 | 788855_47188552_ENST00000601538.6,E | 1 | - | good | CTCCCCAGGATTGAGTGGCTTCTCCTGCAGCCAGGAAGCCTCTGGAGGTAGTATGGAGAGCTCTCCCATTCCAGGGCGGACAGCTGTTCCAACTCCTGCCTGTGTGCCCCCTCTTCTGTC | 120 |
| chr11 | 71070575 | 71070695 | 788855_47188552_ENST00000601538.6,E | 1 | - | good | AAAACTTTATTTACAAAACAGGGAGTGGGCTGATTTGGCCTGCAGGCCATTCTTCTCTAAGCCTTGGAATGGAAGGCTGCATCTCCTCCTCTCCCCCATTTGTCTTATTGATGGATCCCC | 120 |
| chr11 | 71070815 | 71070935 | 788855_47188552_ENST00000601538.6,E | 1 | - | good | CCTATGCCTCAGCTGCCGGCTCCAGGAGGGTCCCAAGCTGGCTGGGGCTCTCAGGACATCCTCTCCCCACCTCGGGCCTCAGGTTAGCACATGAGCCATCTGCTTTTGCTATAAAGAGCC | 120 |
| chr11 | 71070935 | 71071055 | 788855_47188552_ENST00000601538.6,E | 1 | - | good | GACAAGATGGTATTTGAAAGTGTTGATGTTTCTTTCTGATTAGAAAAGGTTAACACATTTTGATGATCTGTATTGAGCTTGACTTGGTATCACTCCACACTCCACATGCCCCATATCTGT | 120 |
| chr11 | 71071055 | 71071175 | 788855_47188552_ENST00000601538.6,E | 1 | - | good | GCTTCCTAATTACTGCTTTGGATGGAAAAAACATTCCTTAATTTTCCCCAGTAGATTGCAAGCACATTTTATCTTGAAAACTCAAGAAAAGTCTGTTTTTTGTCACTTGAAATAATTGTG | 120 |
| chr11 | 71071175 | 71071295 | 788855_47188552_ENST00000601538.6,E | 1 | - | good | TGCCACGGGGTGTGAATGCTGAAATGTCTGTGTGAGTGTGGAGATACATTACTCAGCTGCATTAGAAAAACACATCATGAGGCTTGGCTGTGGCTGGTAAACACATCTGCACCAACATCT | 120 |
| chr11 | 71071295 | 71071415 | 788855_47188552_ENST00000601538.6,E | 1 | - | good | CATAGCTGCGGTTGGGTGGCATTTCAGCTGAGCTGTGAGCAGCATTTTGCAGAGCTGCAGACTTCCCCCACTCAAAGAAAGCACATTGATGTAGTTTGTATGCAGCTGTGTCCCTAGGTA | 120 |
| chr11 | 71071415 | 71071535 | 788855_47188552_ENST00000601538.6,E | 1 | - | good | TGTCCAGGACCAGGCAGTTATTATGCCTTCCCTGTTTTCTCTGCTGATGCTCCCTTCCCCTTGGGTCCACCTGAGAGGAAGCACGTGTGGATATCATCAGAGGCATAAGCCGCCATCTTT | 120 |
| chr11 | 71071535 | 71071655 | 788855_47188552_ENST00000601538.6,E | 1 | - | good | TGCATCCTTCAAGATCCAATTCAGACTTCCTGACCCTCAGTCAAGCCTTCTGCTGATGACTCCCTCACATAGTTGTACATTTTTCTCTGCAGCCCTTTGTGAGACATAGCCAGGGCAAAA | 120 |
| chr11 | 71071655 | 71071775 | 788855_47188552_ENST00000601538.6,E | 1 | - | good | CCATTCCCCAGCCCTTCCATGGCTCCTCACCAGCCCAGGCCCCTCCAGCCACTCTGCTGGAGTCCCACTTTGCTTCTAGCTGTTCAATCTTCCTGGATTCTTTGCCCTGGAAAACTCCTA | 120 |
| chr11 | 71071775 | 71071895 | 788855_47188552_ENST00000601538.6,E | 1 | - | good | TGCCAGGGTCCATCCTCTGCAGGCTGTCAGCATTCTCTTCCCATCCCAAAGCAGTGCTCCCATCCTGTTGGCTCATGGTGGCCTCCGGGGTGGAATCTCAAGCCTCATCCTTTACCCAAG | 120 |
| chr11 | 71071895 | 71072015 | 788855_47188552_ENST00000601538.6,E | 1 | - | good | ATGTGGATGCTACACTAAGCGGCAGAGCAGCGTGGGCTGGGAGGGGATGTGGCTCTTTATCCTGGAGGAACAGAGGGAGATCTCAGAGGAGGAACTCTGCTGGATGCCCAGGGCTCTTTG | 120 |
| chr11 | 71072015 | 71072135 | 788855_47188552_ENST00000601538.6,E | 1 | - | good | TTCCTTGGTGCTCAAAGTCAGGTGAATTCCTGGACATCACTATTGAAGGACTTCTCTGTCCGTGCCTTCTTTTTTCTCTTTCTCTGATCTCAAGGAACTTCAAATCTGGAGGGTGTCGAG | 120 |
| chr11 | 71072135 | 71072255 | 788855_47188552_ENST00000601538.6,E | 1 | - | good | GGGGGGTTAAGTGGTGGGGGCGGCCTGTGGAGTGGAGCTGGTGAGAAGGGGCCCTGGGCTCACGGGGGCATAGGTGGGGTGCAGGCACTGCTTTAGGAGGACAGGGGGCTGTTTCTCTG | 120 |
| chr11 | 71072255 | 71072375 | 788855_47188552_ENST00000601538.6,E | 1 | - | good | AAGGCCAAGTGACACCCCGCCCAGTGTGTGCACCGTGGCCTCACGGGGAGGTGAAGCAGGGGCGTGGGGGGCTGTTGGTCCCGCAGGAGGCTGCTGGTCCTGTAGGAGCGGGGAAGCCA | 120 |
| chr11 | 71072375 | 71072495 | 788855_47188552_ENST00000601538.6,E | 1 | - | good | ACACTTCCCTGTAAGACCAAGTGTGAGAGCCGAGGAGAGCAGGACGTGAATGTATCTAGGAAACACTGAAGGCTTCTGGGATGTGAGCCCTGCCCTCTGGGAGGCTGCATCCTAAATGAC | 120 |
| chr11 | 71072495 | 71072615 | 788855_47188552_ENST00000601538.6,E | 1 | - | good | TGTCGTGGATAATGCAGCTATGAGCATAGATGCAACCATGTGTTCAAGACCCTGTTTTCATTTCTTTCGGGGAAAAATGCCCACAAGTGGAACCTTGATGTGAAAAAGCAAATCATCTTC | 120 |
| chr11 | 71072615 | 71072735 | 788855_47188552_ENST00000601538.6,E | 1 | - | good | GGAATTTTTTATCCTTCCTTTTTAAGCTGAATAATAGTTTTAAGGTAAATAATACTCCCTTGTGTGGATCATGTGCGTGGACATCTGTCAGGGGACCCTCGGTGGCTTCCACCTTTTGAG | 120 |

| | | | | | | | | |
|---|---|---|---|---|---|---|---|---|
| chr11 | 71073815 | 71073935 | 788855_47188552_ENST00000601538.6,E | 1 | - | good | TCAGTTCTGGAGTCAGCCATCTACCCCGTCAGCAGGTCTCTATATGGAACATGGAGAGGGGCAGATCAGAGTCATGAGATGGGGAAGGACGTGGCGGACAAAGCCCCAGGATAGTTCTGG | 120 |
| chr11 | 71073935 | 71074055 | 788855_47188552_ENST00000601538.6,E | 1 | - | good | GAACTGGGCTCAAGACTCAGGTCTGGTCAGTGCCACGCCTCCATGCTGTCCATGCTCCATAAGATGCGTGAGGAGCCGGGCTCTCTTGTCCACCCTCTGTAGCTGGGATGCAAACTCTGT | 120 |
| chr11 | 71074055 | 71074175 | 788855_47188552_ENST00000601538.6,E | 1 | - | good | GGGGGCATCTCAGGTCATCCTCACCTGCTGCCTCAGTGGGAACTGTTACCCTGTTGTAGAAGGGAGGAAACAGAGCCCCAGGAAAGCAAGGCGAGCATCCCACTCCCACAGCTGGTGCCA | 120 |
| chr11 | 71075255 | 71075375 | 788855_47188552_ENST00000601538.6,E | 1 | - | good | GGGTTCTGGTGGAGCAGCCGAGTGGCTGGTGTACTTTGTGTGTGCATGCGGCTGTCGTGGCGGAGGGTTCCTGATTCACAGCCTGGGTTCTGTGTTTCAGGCCTGCAGGTACGGGCACGT | 120 |
| chr11 | 71075375 | 71075495 | 788855_47188552_ENST00000601538.6,E | 1 | - | good | ATGAAGGTCTTTCTCCCTGATCCACAGCCAGGGGTCTGGTGGTGCATGCTGATGTCTTTAGAGCGCTTGCCCAGATGGTGGGTGGGTGGTTGGGGTCTGCACAGGCCCTGCCCTGCCTTG | 120 |
| chr11 | 71075495 | 71075615 | 788855_47188552_ENST00000601538.6,E | 1 | - | good | GATGGGAGAAAGACATCCTGCACCTCTGGGCTCTAGGCCCCGGGATCAGGATGGAAGGGACTTTGGGGCCCCTTCCCAGACCCTCTGTCTCACAACTGAGCACCAGTCCCCAATCCCACC | 120 |
| chr11 | 71075615 | 71075735 | 788855_47188552_ENST00000601538.6,E | 1 | - | good | GTGCAGGGCTGTGAAGGCCTCTCCTGGCCATTTTGTGCTGCCCCCTTGGAAGGGGGAGCTTGGGTAACTGAAGCCGGGTCTGAGCCTTCTGGTTTGCATCTTGTGGAGCTGGGCACTGAG | 120 |
| chr11 | 71075735 | 71075855 | 788855_47188552_ENST00000601538.6,E | 1 | - | good | CATAGTATGGGTCTGCCCTACCCCTGCGTCCTGCAACTTACTTAGGAGCTGGACTGCATTAAAGCAGTCGTGGGGCCGTCCGTCGCACGTCGAAGGACAGGGCTCACTCCTGTGGATGCA | 120 |
| chr11 | 71075855 | 71075975 | 788855_47188552_ENST00000601538.6,E | 1 | - | good | AGGGTGCACATAGGTACTCAGGCGAAAGTTCTGATAAATGGTTCAGGACTGGGTCTGTTTGACCCCTGGGCCCTGACGGTTCACAAGTCACACATGTTGCAGTTCTCAGAGAGGAGTTAT | 120 |
| chr11 | 71075975 | 71076095 | 788855_47188552_ENST00000601538.6,E | 1 | - | good | ACTGACTCACCAAGTGCTGACTCAGTGCCCAGCCTCGTGTGAGGCCCTGTGGTGTTCTTTGTCCCACGTGATTCTCATCTGAACCCTGTGAGGGTTAAGGGTTGTTTCTGCCTACAGATG | 120 |
| chr11 | 71076095 | 71076215 | 788855_47188552_ENST00000601538.6,E | 1 | - | good | TTTTCTCCAGGTCCAAGGTTGTGGCCACCATGTCGAGCAGTGTGTGGCTGGACAGAGGGGCCCTCATCACCTGTGCGGGGACCTCCCTGGCCCCTGTGAGCTTGGGTGGGGAGGGGCATC | 120 |
| chr11 | 71076215 | 71076335 | 788855_47188552_ENST00000601538.6,E | 1 | - | good | CCCCTTCCTGGGCTACCATGGCGCCTACTGCTCTCAACACCCCTAGGCTGCCTGCCCTTGGTGGGGCTCACCTCTTTCCCTCCCGACCCTGGGATTGAGATGTGTTAGGTCAGGACCGAC | 120 |
| chr11 | 71076335 | 71076455 | 788855_47188552_ENST00000601538.6,E | 1 | - | good | TCTTCCCTGTCATCTTTCCTCACTGTTAATTGCTGAGGTGTAGCTCTTTCTCGTGGACCCTGAGAAGGTGAAATTCACCTCTATCTGTCTCAATTAGCAGTAGAGCAGGATTCATGAGAA | 120 |
| chr11 | 71076575 | 71076695 | 788855_47188552_ENST00000601538.6,E | 1 | - | good | CAAACGCAGGTGGAGTTCCCAGAAGAAGGGGCCATGGGGGCCGGGCAGGTGACAGCAAGGAAACCTGCAACAGGGATCTACTTTATAAATTAATGAGATTTGTGGGAGACACAGCAATGG | 120 |
| chr11 | 71076695 | 71076815 | 788855_47188552_ENST00000601538.6,E | 1 | - | good | TTACTGTGCCTGGGGAGTTGGAACCTGATGGCTCCTGCCTATGTCCTGTGCCTCTGCAGGAGCAGGGTGGGGAGGCCTTCCCTGGACCTCTGAGAGTGGAGCTGGGGTAGCCCCCAAAGG | 120 |
| chr11 | 71076815 | 71076935 | 788855_47188552_ENST00000601538.6,E | 1 | - | good | CAGGCTCTGTCACTTTTCTGCTCCCCACTCTGGAAGAAGGAAAGAACTTTTTCCCCAGTAGTTGTGATGAATGTTCTAGCTTGGGACTTGGTAGCTTCTCTAAGGTCAGCTGGGCGTCCC | 120 |
| chr11 | 71076935 | 71077055 | 788855_47188552_ENST00000601538.6,E | 1 | - | good | CATCTTGGCTTTGTGCATGTCTCTTCCTGTCTCCTCTCTTTCTCTCTTTCTTTCCTTTCCTTTCTTCCTCTCCCTGCCCCTGTCACACATCCCTTCTGTGTTGAGGGTCCCAGCAGCTCC | 120 |
| chr11 | 71077055 | 71077175 | 788855_47188552_ENST00000601538.6,E | 1 | - | good | TTCAGTTCAAGGAACACAAATCCCAATGCAAACGACTGGGTGAGGGGAGGGAGTTATGCCTCATTGTTATGTCCCGGAAATTCCAGGGTCAGTTGCAGCTTAGAGGGGCTCAGACAGTGT | 120 |
| chr11 | 71077175 | 71077295 | 788855_47188552_ENST00000601538.6,E | 1 | - | good | TGTTCAGTGTTTTTATGCTAACATTAGTTGTCTTTATTGACCACAGAGCAATTAAATTAAGCTTTAACAAAGTGTTGCCTCCCAGAAGTCCTTTGGGGAGTGCCAAATAGCCAGGGATCC | 120 |
| chr11 | 71077295 | 71077415 | 788855_47188552_ENST00000601538.6,E | 1 | - | good | CCATTGAGCCTATTTTGCCCTTGCAAGGGAAGAGTGCTCTGGTGCCATCTGACTGATGGAACTTACAAGCTCATAAATTCAGATTAACCGGGGGAAGGGGAATAATGAGTGTGGATTAAT | 120 |
| chr11 | 71077415 | 71077535 | 788855_47188552_ENST00000601538.6,E | 1 | - | good | TTTCCCGGGTTTGTGTGTTATAAAACAATGGATGTGGGAAAAGTGGAAAATGGAAGCATGCTCTTGGGAACACGCCTACGAGACATCCACCCGGAGTCAGCGGTTCCGTTTCATCTCTTC | 120 |
| chr11 | 71077535 | 71077655 | 788855_47188552_ENST00000601538.6,E | 1 | - | good | AGGGGTTAAAACAGAAAGAAGAGACTAGCCGAAATGACACTTAAAAAAAAAAAAATTACTGCTAGTGCTAAGACTGTGGGATTCTGGGTGACTTTTGGTTTTTGTTCAGCCCTTTTCTGTA | 120 |
| chr11 | 71077655 | 71077775 | 788855_47188552_ENST00000601538.6,E | 1 | - | good | GAGCCCTGCGTGCAGCCGTCTGTGGACCCTTCTGCCCACCACCTGGCTTGGATCCCCCTCACCCCCCTGCATGGCAGCTGCATCTTCCTCAGCAGGTGTGAGCTCCGTGCCTCAACCACC | 120 |
| chr11 | 71077775 | 71077895 | 788855_47188552_ENST00000601538.6,E | 1 | - | good | CTCCCATTCCAAGCCAACACCACAAGGCATATTCTAGTTTTCTCCTGTACTGTATTTGTAACTTCCTTGACAGTGAGAGCCCAGCTCCCATTCCCTTTCCTGAGTGTCCTTGTGTGTGCA | 120 |
| chr11 | 71077895 | 71078015 | 788855_47188552_ENST00000601538.6,E | 1 | - | good | TGGGCTGTTGCTGGTCCTGCACCCTCTCAGTAGGCAGAGTTAGGGATGGGTAGACACACAGGCATCCACATTTTCATTTTTATTAGTCTCTATATTAGGAACCGTGAGTTTCTACTGATA | 120 |
| chr11 | 71078255 | 71078375 | 788855_47188552_ENST00000601538.6,E | 1 | - | good | ATTAGTTGGCTTGCTGTTGTAAAGAAAAGCTTTCTCTTCTTGTTTATTTATTCATATATTTACAACAAGGCAGACTTGCAGAGTTCTCTTTATTCAGGAGGTAATAATCTGTTATTATTA | 120 |
| chr11 | 71078375 | 71078495 | 788855_47188552_ENST00000601538.6,E | 1 | - | good | GTTTTCTCATACCCCTTTCACCTGACAGTTTTGGCATCCATTGGCCTTTCTTGCCAGAATTAATGCTATTATGGTTGCTAAATGGAGATTTTTTTTTAATCCTACCATTGCTTCTACATT | 120 |
| chr11 | 71078495 | 71078615 | 788855_47188552_ENST00000601538.6,E | 1 | - | good | CTAGTCCTCAGTTTGGTCCGGTGTCTGAGTCCCGCTTCCTGAGTCCCACCTCCTGCCTTACTTTGCTCCTTTGTAATTAATAGATCTTGTTTGGTGATCTTTGAGGCTATGCAAAAATCT | 120 |
| chr11 | 71078975 | 71079095 | 788855_47188552_ENST00000601538.6,E | 1 | - | good | GGTACTCATTGTGGAGGTCACACCGACCACCTAGTCAGGTGGGGTCTGTTGGGTTTCTCCTCTTTTGATGCAGGGCAGGTGAGCCCAAAAATGGGGGCTTAGCCCAGGAGGGCTCTTGGC | 120 |
| chr11 | 71079095 | 71079215 | 788855_47188552_ENST00000601538.6,E | 1 | - | good | GGTTTGCTCAGTGTTTGCTCGTGATGGGATCCAGCATATGCATTTGGCAGGAATGCCTTAGAAAGGATGCTGTGCTGTCAGCATCACACCTTCACGGGGATACATGGGGACCATTCGCCT | 120 |
| chr11 | 71079215 | 71079335 | 788855_47188552_ENST00000601538.6,E | 1 | - | good | ATAGAAGAGGATCCAGCCTGGGACCTGCCTGGAGCCTTTCCATCTGGGAGACTCCCTCGGGCTTCCCTTAATTCCTATGGCTGTGACTGTTTTGAAGATTTTGGGGCGGTCATTTTGTTG | 120 |
| chr11 | 71079335 | 71079455 | 788855_47188552_ENST00000601538.6,E | 1 | - | good | TAAATTACTGTTTCCATTTCCAGCTTATAGAATAGTTGCAAGAATAGTACAAGGAACTCCCATATGCCCTCCACGCGGAGAGCACATTTGAGTGGTGACGGTGTCCCAGTAACATCCCTG | 120 |
| chr11 | 71079935 | 71080055 | 788855_47188552_ENST00000601538.6,E | 1 | - | good | TGTTGCTTCTCTTGTGGGAGGATGAATTCCTTCTTTTAACACTTTGGTAGCTTTCAGTATCTTTGTGTCAAGCGAGTTCGTTCTTTCTTTCCTTCTTTCCCTCCTTCCCCACTTCCCTCT | 120 |
| chr11 | 71080055 | 71080175 | 788855_47188552_ENST00000601538.6,E | 1 | - | good | CAGGTGTCAAGGGCAGTGTTTTGCCTTTCATCCCATTACTCTCCACCCTCCTATCACCCTCTCAGCTGTCTCAACAGCCAGAGACCAGACACCCTTTTCAAAATAAGAAAAGGAGGTTAG | 120 |
| chr11 | 71080175 | 71080295 | 788855_47188552_ENST00000601538.6,E | 1 | - | good | GAGCCATCTCCAGGTGCCCCTGCCCAGGCCTCAGCAGGAGATTCAGAGGCAGCAGACATGGGTGGAGCCTCCCACACATGCATTTTCCAAAAAGCACCTCAGGAGAGTGGGTACGGGGTG | 120 |
| chr11 | 71080295 | 71080415 | 788855_47188552_ENST00000601538.6,E | 1 | - | good | AGCAGGTGGCGTTAAATGGCTTGAAGGAAGGAGCACCCTTTTTTCTAATTCACACCAAGTCTTGTGGTTGCACAGGGCGGGTGGGACGCTAGGCATCGCTGGGGCAAACAGTCTCGAGGG | 120 |
| chr11 | 71080415 | 71080535 | 788855_47188552_ENST00000601538.6,E | 1 | - | good | TCCCCTGCAGTGCAGTGACCTCCCCGCTGCTCTGGTTGCTGTGCTGGGCATTGCACAACCCCAGGTGGCGCCGTTCCTCTTGCCATAGTTTTAGGTCCTGTAGATGTATCACCAGTGTCC | 120 |
| chr11 | 71080535 | 71080655 | 788855_47188552_ENST00000601538.6,E | 1 | - | good | CCTCATGTCACAAGGCATATGTTTTGGGAGAAATGCCTAAGTGTGGTTGGGAACAATTTTGTTTGGGGCACAGAGCCAAGCAGAAGTGATAGGGGGTTCGTCTCAGCCGACCTGTGCTGC | 120 |
| chr11 | 71080655 | 71080775 | 788855_47188552_ENST00000601538.6,E | 1 | - | good | GAGGGGGATTTGCTTGTGTTTGGACGAGACGTTGTTTCTCTACTGACGGGCAGCCCACTTCCTGACCCAGTGAGAGCCGAAATGATGTGACTCTTTGTTGTGTGGGCTGTGGTGTCTCTA | 120 |
| chr11 | 71080775 | 71080895 | 788855_47188552_ENST00000601538.6,E | 1 | - | good | TTCCTGCATTCATTCAGGTGCATTGGCCTCCTTGACTACTGGGCCAGCAGGAAGACCACAGCCAGCCTTTGAAAACCGTGGTATGCGGGGCCACGGAGCAGCTGGTGGCTCAGGCCATGC | 120 |
| chr11 | 71080895 | 71081015 | 788855_47188552_ENST00000601538.6,E | 1 | - | good | AGGTACTGGGGGTGGGGACTCCAACGCATGAATGCTGCAGGGATGCAATTCAGCCCGCAACACCAACAAAACTGAACTAGCCACTTTCTGGTTTGCATCACAGCCCGTCCGCTGCCCGTG | 120 |
| chr11 | 71081255 | 71081375 | 788855_47188552_ENST00000601538.6,E | 1 | - | good | CTGTGATCATTTCAGGCTGAACAGCACACATTAGTAGACCCAAAGTGATTGTGTTTGCTGCATAGGGCTGCCACTGCAAAGTTCCTCAAACAATAGACATTCATTATCTCACAGTTCTGG | 120 |
| chr11 | 71081375 | 71081495 | 788855_47188552_ENST00000601538.6,E | 1 | - | good | CTTTCCAAAAATCATGTTATGAGAGTGATGGGCTCCATGGGGAAGGTGGGCAAGGGCCGCAGCTTTATGTGGGAATGCCAGTCATCTCTCCTTTTGGAATTATAGCATGATGTCTTTTGT | 120 |
| chr11 | 71081495 | 71081615 | 788855_47188552_ENST00000601538.6,E | 1 | - | good | CATGCGGGGAACAGGAGAGGGGATGGGTGAAGACCCCGAGACCAGCACCCCACCCTCTGCACTTTGGCAGGTCCTCTATCCTGCCCTCCTGACATAGGTGACATCTTTAAGAGACCTCAT | 120 |
| chr11 | 71081615 | 71081735 | 788855_47188552_ENST00000601538.6,E | 1 | - | good | CTCTGGCCTGGCAAGGTTCTGTTTGGAGAAGGAAGAGAGGCATCCTCTCTCTCTGAGCCTCCAGAGGGAATGAGGACCCTTGAGTACCTGCCTTCCCAGTCAGCCCTGACTCTTCAGGGA | 120 |
| chr11 | 71081735 | 71081855 | 788855_47188552_ENST00000601538.6,E | 1 | - | good | CCCCTGGGCACGAGCGGCCGAGCCTCTGTGGCCCCCGTGTCTGGAGCCGCAACAGTGAGCTGTGCTTTCAGGGGATCAGCTCTGAATTGACCTGGGAGGTCAGTTCCATATTCTGAGCCC | 120 |
| chr11 | 71081855 | 71081975 | 788855_47188552_ENST00000601538.6,E | 1 | - | good | TACTGCACCTTTCCAGATGAGCACCTCCAGGGTGTGAGAGTGTGGGGTCCCTGCCAGGTGGCAGGACCAGAGCACGGGAGCCCTTCCCCGTGTCACTCTGGACCTGGGGCTCAGGGTCTG | 120 |
| chr11 | 71081975 | 71082095 | 788855_47188552_ENST00000601538.6,E | 1 | - | good | CTGGCATGCATTGGCTCCGGGGGGACATTGTGAACAGGGTTCTGGCCCTTTGTGCTGGCATGTGCTTGGCAGCCTTGGAACTGGGACCGAGCCTCACTGTGGGACGTGGGTGCTGGCAGG | 120 |
| chr11 | 71082095 | 71082215 | 788855_47188552_ENST00000601538.6,E | 1 | - | good | AGGTGGAGAGCCCAGGAGCTCCACAGTGCCAGCCAGGAGTGGCTCATTCTGGCCAGGGGACGTGGGAGCAGTGTTGGAACCAGGCTGTGGTGCTGGCAGGTGACCCTCAGATCTGGGCAG | 120 |
| chr11 | 71082215 | 71082335 | 788855_47188552_ENST00000601538.6,E | 1 | - | good | GCCTAGGCACTGAGATGACCACTGTTCAGGCACCGCAGTGACCGCGGTGTCCAAGCCTGGGCTTGGTCACGGTAAGCACAATGCCAGGTAGCACAGGAAGGGCCAGCGTGAACAGCAGCC | 120 |
| chr11 | 71082335 | 71082455 | 788855_47188552_ENST00000601538.6,E | 1 | - | good | TTCCTTCAGATTGAGGTTTTTCTAGTGTACTGTGGTACCCACCGTGTCCAGGTGCTGAGGCGATCACTGTCCATCTAGGTATCGAGGTGATTGCTGTGTCCAGGCCCCAAGATGACCATT | 120 |
| chr11 | 71082455 | 71082575 | 788855_47188552_ENST00000601538.6,E | 1 | - | good | ACCTTCCTGCCAGGGTGGGACTGTGTACAGCCCAGCGTGAGCTGCCTGGTAGAGATGGGAGTCAATGCTTGACTGGAAAGTTAGCATTTAGAATGGGATGGAGCCACATGGGTTGTGGTC | 120 |
| chr11 | 71082575 | 71082695 | 788855_47188552_ENST00000601538.6,E | 1 | - | good | CAGCTCTTGAGCCTCTCTGGGCCCCACGGCGAGGGGACAGGCCCCTGATCCTCCCAGGCCTGTGGGAGGTTCTTTGCTGGAGCCCCACGTTATGTCGTTCTGGGTCCAAGCTTTCTGGTT | 120 |
| chr11 | 71082695 | 71082815 | 788855_47188552_ENST00000601538.6,E | 1 | - | good | TTGTTATTTATCGCATTTAAATGAAGAGTGTCCGCCTTAAAGAAATAGTTTCGGGCCACAGTGACTTCTCCTCTCCTCCTGCTCCAGGCTGCCCCCTGCCCATCCCTGGCCGGATGCCCT | 120 |
| chr11 | 71082815 | 71082935 | 788855_47188552_ENST00000601538.6,E | 1 | - | good | AAGAAACAATCCGTTGTATTCAGAGAGTGCCTACCACACCTGCTCCCAGCCCCTCGTTTGATTTTGCTCCCATTGTGCAATGGGTGTGACATCAGTCTGAATGTTGTTACTGGCTGCTAT | 120 |
| chr11 | 71083295 | 71083415 | 788855_47188552_ENST00000601538.6,E | 1 | - | good | GCTATGACAGAATTTACGTGAGGCATTTTCTGCAGTGCCCGGCATAGACCAGGTGCTTAATAAATGTCCCTATCATTATTGTCATCATCTGTCCCCCCATCATCATCCTCATTGTCATCA | 120 |
| chr11 | 71083415 | 71083535 | 788855_47188552_ENST00000601538.6,E | 1 | - | good | TGGGGTTGCAAACCCACCTCATCGCTCGCTGGCTCACCTGAGCAAGTGCCCCACCCTCTCTGAGCTTGTGTGCTAATCTCTCAGGTTGGCTGAGTAACAGTTTGTAGAGCTCTGGAGACA | 120 |
| chr11 | 71083535 | 71083655 | 788855_47188552_ENST00000601538.6,E | 1 | - | good | TTCCTTGGGGCAAAGCCTGGCTCTCTCCCTCCCTCTCTGAGTTTGACTCAGTATCTTCGGGATGAGGGACTCTCCTCATGCTGTCTGTTGGCCATATTCTGGCCTGGGTGTTATGCAGCC | 120 |
| chr11 | 71083655 | 71083775 | 788855_47188552_ENST00000601538.6,E | 1 | - | good | TACATCCACCGTGCAACTTCACAGCACGCTGGCCCTGGCCACAGGGGGTGTGGTCCTGGAGAGGATGCCGCTTGGTGAGGGATCCAGACCCTGCGTTTCCATGTGCTGAACCCCGGTACC | 120 |
| chr11 | 71083775 | 71083895 | 788855_47188552_ENST00000601538.6,E | 1 | - | good | AAATACAAAGAAGTATAAAGAAGCAAGTTAAGAAACACCCCAATGCATTCCCACAACCCAAGGACATCAGGGGTCAGATTTGGGGGTATTCATTCTGTATTTTTAATTGCTAAAGACTTT | 120 |
| chr11 | 71083895 | 71084015 | 788855_47188552_ENST00000601538.6,E | 1 | - | good | CCTGGGAGACAGAGCAAGACTCTGTCTCCCCCCCCCAAAAAAAAGTTGTTATTCAAATTACCCTGTTTTGTTTTGTTTTTGATTACCAAAAAATTACCCATGCATCATGGAAATCTTGGA | 120 |
| chr11 | 71084255 | 71084375 | 788855_47188552_ENST00000601538.6,E | 1 | - | good | CTCCACTTCACACAGTGGGGATATGCCCCATCCCCACAGGCTGATGTTAGAAGAGCCCCATTTGAGCTCAGGCTGTCTGCAAATTATCTTAAAAGTTGTTATTCGGGCCGGGTGCGGTGG | 120 |

| | | | | | | | | |
|---|---|---|---|---|---|---|---|---|
| chr11 | 71084375 | 71084495 | 788855_47188552_ENST00000601538.6,E | 1 | - | good | GCCACAGCCAGGCCTGCTCACAGATGCTGCAGGTGTACTCCAGGGTTCCCCAGGCCGCTGCTTTCCCATCATTGTCTCAGGCCAGGTGCACATGGCCTAAGGACACATTTCAGATCCACC | 120 |
| chr11 | 71084495 | 71084615 | 788855_47188552_ENST00000601538.6,E | 1 | - | good | CTCCGAGCTCTCAGGCTGATGGGCTTTGCATTTAAAGTCTGTGTCTCTGTTCTAGCACCTGGAGAATGGACAGCCCTGTCCCAGGGTTCTGGGCGTTCATCGTGTGGGAGAGCCTGCGCG | 120 |
| chr11 | 71084615 | 71084735 | 788855_47188552_ENST00000601538.6,E | 1 | - | good | GTAGGGTTCTCACATCCAAACCACTCACAGCACATTTGTGGCAGAAGAAATTACAGGGGCAGGTGGGTTTTCCTGGACCTCACTGGGCTGTGCCTGGCTCTGCCCTCTCGGGAGAAGCAC | 120 |
| chr11 | 71084735 | 71084855 | 788855_47188552_ENST00000601538.6,E | 1 | - | good | ATGAGTTCACAGGCCCAGAGAGGGCAGGTAACTTGCCCAAGGCCACGGAGCAAGCAGTTGGCTGAGCTGGATTCACTCCTGGCCTGTCTCTTACCAGAAGCCTGGTTCTCTATGCTGCCT | 120 |
| chr11 | 71084855 | 71084975 | 788855_47188552_ENST00000601538.6,E | 1 | - | good | AAATGGTCAAATGTGTATAGGACCCGTAGCAATTTGGGGGCATTCATTGTCCTGAGCTGCTTTAATTGCACTATGATTCTATATCCTCAGAGCTGGTTACTTCTCATTGCTGTTTCCCAT | 120 |
| chr11 | 71086655 | 71086775 | 788855_47188552_ENST00000601538.6,E | 1 | - | good | GGACAGTGCCGCCTGCCTCCCAGATCATCCTCATAAGTGCTCACATTTACCCTGCCAAGTGGCCCGGTAAATGCCACAGTGCGGGTCTCAAGTCCATCTCTAGCAGGATCTCATCCTTCT | 120 |
| chr11 | 71086775 | 71086895 | 788855_47188552_ENST00000601538.6,E | 1 | - | good | AACTTTGCCTTCGCGAGCCTCCTCTCATTCATCTGCAGGGCCCTTGTGCCTATCAGATGATAAGGGACGGAGCAGATAAGGGATGGGTGCGGGTGCCCTGCACGGGCCTGCAGGAGCAGC | 120 |
| chr11 | 71086895 | 71087015 | 788855_47188552_ENST00000601538.6,E | 1 | - | good | GCAGGGTTTGGAAAGCCAGCCAGGTGCGTTCACTGCCCTCTGGTGGTCTGCTGCCTGTTCATGAAAGGCCCGGAAAGCCATTTTCTCCCCTGCTAAAGATTCTCAGTGATAATGTGAAGC | 120 |
| chr11 | 71087015 | 71087135 | 788855_47188552_ENST00000601538.6,E | 1 | - | good | TCCCAGAGCCACCCGGCCACCTTGGTCCCGGGGCCTGAGCCTGTGCAGCTGCACTGGTACCTTGGAGGCTTGAGCTCTGGGGCAGTGCACCTGCTCCCACCTCAGCTTAGAAAATGTGGA | 120 |
| chr11 | 71087135 | 71087255 | 788855_47188552_ENST00000601538.6,E | 1 | - | good | GGAGGCTGGCAGGAGCCCTGACCATCAGTCACTCACTGCAGCTCTCAGCTGTGACTCCTGGCCTGCAAAGGGCCAGGTTCTGTCCCTGAGGCCCTGCGCTTTGCATTCAGAGCCTGGCTG | 120 |
| chr11 | 71087255 | 71087375 | 788855_47188552_ENST00000601538.6,E | 1 | - | good | TGGAGCTCCGTCCTTTCCCCTCCCACACCCCTCCTTTTTCCTTTTTCCTGCCCTCTCTCCCAACCAGGCAAATGTCTCCGGGCCCATCTCTCCAGCTGGCCCTGGGATGGAGGTAACGGT | 120 |
| chr11 | 71087375 | 71087495 | 788855_47188552_ENST00000601538.6,E | 1 | - | good | AGCTCTGAAGGTGTGATGCAGGCTAAAACCTCCTTAAACGTGGAGTAGTTCACAGAGTCCTGAATCCAGCCGCCAGCCCTTTTGCCAGAATAGCCCCTGTGTCTCAACTTTGGGGATTCC | 120 |
| chr11 | 71087495 | 71087615 | 788855_47188552_ENST00000601538.6,E | 1 | - | good | GCGTATGGGACCCGAAGCTCGCTTGTTCTCTGTGAGAATGCCTGCTTGAATTTGTGTTTTCCAGACATGTCTTCCCTTTAGACATCACAGTGCTCTGGTTGCATTGCTGTTTTCCTGTTG | 120 |
| chr11 | 71087975 | 71088095 | 788855_47188552_ENST00000601538.6,E | 1 | - | good | ACATTATGGAGAATCATCAATTTAGTCTACTGGTTTTGAGAGCAGTCACTGACTTCATGAAGACAGGAGCGCCCAAGGAAGAGGAAGGTTTAAATTCAGGTTTCCTTTAGTGAATGAGTC | 120 |
| chr11 | 71088335 | 71088455 | 788855_47188552_ENST00000601538.6,E | 1 | - | good | CGGTGATTCACGTGAACAGAACTGTCATTTGATGTCGCTGGATGACACCATAGAAACCACTCAATGTTGTTTGTGTGTTCAGGTGTGATGAGTTCTCCAGAAAAACAGAACCAATAGGGT | 120 |
| chr11 | 71088455 | 71088575 | 788855_47188552_ENST00000601538.6,E | 1 | - | good | CAGCTGCCGGCCCCCAGGGGACAAGGCTGAGTTGTGAGTCTCTAAGAGACATGGACGTTGGGGACGTCCCTCCCGCCTGGCCAAATGGATAATGAGTCTTGTTCTCAGTCAACTCAAAGT | 120 |
| chr11 | 71088575 | 71088695 | 788855_47188552_ENST00000601538.6,E | 1 | - | good | GTGGGGTGGTGCAGTCACCCACGAGACATTTCTACCAGCAAACATGGCTGGGCGAGGGGCGGGAGTCATGGAACCCTGAACACCCACATCTTTCTCGAGTCTCACAGGGTCTCCACCGAA | 120 |
| chr11 | 71089055 | 71089175 | 788855_47188552_ENST00000601538.6,E | 1 | - | good | ATCTCAGCTGTGTTGGGAGCTGTCCCAAGCTTCCAAGGTGACCTGCTTGGAAAAGGGCAGTTCTTGGGACTGCCAAAGCGCTAATGAGCTTGGTTTTTAAAACTCACTAGGCTGATTTGA | 120 |
| chr11 | 71089175 | 71089295 | 788855_47188552_ENST00000601538.6,E | 1 | - | good | TCCTCTCCTTGCTGCGCTTGGCTGTCAGCTCTGTGCTGGTGTTTTTGAGGCCAAGCCCGTCTGCATGGCGTGAGCTTCGGCTCCTGTTGAGGATGTGGAAGAGGCAGCTCCCGAAGGAGA | 120 |
| chr11 | 71089295 | 71089415 | 788855_47188552_ENST00000601538.6,E | 1 | - | good | TGTGCGTGGACATGCTTCTGAGCAGGCCTGACAGTGTTCTGTATTTCTGGGCACTGGTGCTCATGCAGTGGCCATGCAGACGCTCACGCCCTGAGACATGGCCCCGTTCCTCTGCCCGTC | 120 |
| chr11 | 71089415 | 71089535 | 788855_47188552_ENST00000601538.6,E | 1 | - | good | AAATGCCTGATGCAAGCCCCTGGTGTTCCCGTGGGGGGAAACCAGACATGAGATAGAAAGACGAACATAAATTACACACACACACACACTCGCACGCACGCATGTACATACAGGTCCACA | 120 |
| chr11 | 71089535 | 71089655 | 788855_47188552_ENST00000601538.6,E | 1 | - | good | TGCACCACAGACCCTACCCACTGCTAATGGGTGCTCATCAGCCACAAACGTCGTCTACTGGGTGTGAGCCCCAGCCATGTGTCTCGGTGTCCTGAATACACAGCCTGCAGCAGTGAACAG | 120 |
| chr11 | 71089655 | 71089775 | 788855_47188552_ENST00000601538.6,E | 1 | - | good | ACGCTCATTCCCTGGCCCCCTCCTCATTTGTGTCTGAGCTGTTTCAGGGGGTCCCCAAGAGGTCTCCTGGCCTGACCACGCTCGCCTCTTCACGCCTGACCCACTTCCAGCCATTCCGGG | 120 |
| chr11 | 71089775 | 71089895 | 788855_47188552_ENST00000601538.6,E | 1 | - | good | TGACCTCTTCCCCCTTTGTCAACCAACGGTTCCAGCCCCAGATGGCCCTGCCATCCCTCGGCTGTGTGTCAGTGTCAGGCCTGGCCCTTCTGAGCCTCTCCACAGCATCTGGCCCTGGCC | 120 |
| chr11 | 71089895 | 71090015 | 788855_47188552_ENST00000601538.6,E | 1 | - | good | TCGGAGGTTTTTGGGTTGGGGGAGCTGCTTCCCCTCCTCAGTCTCCTGCCCTGCCCTCGCTTTGTCCCTGGTGGAAACTCGCCAGTGTTTCTGTCCTTGTTCCCTCCTCCTTCCTCCCCT | 120 |
| chr11 | 71091095 | 71091215 | 788855_47188552_ENST00000601538.6,E | 1 | - | good | GGAGCCCGCAGGCAGGGAAGCAGTAATGACTGGTCTTTGGAAGTCCTCTTCCATGCACCCGCCATGAGATTTCTTGTCATTATTGTGCTTTTCTTTATGAAAAAGGAAAATAATTTTCCC | 120 |
| chr11 | 71091215 | 71091335 | 788855_47188552_ENST00000601538.6,E | 1 | - | good | GTGCTAGGTCAGTTCTGGTTCATCGTCGGCCCTGCAGCCTGGAGAAGCCCCTGTTCCTCCCGAGGGATGAGTAGGGGTCTGTGGGGCTGTTCATGGCAGCCCCCGGCCCACACCCAAGCT | 120 |
| chr11 | 71091335 | 71091455 | 788855_47188552_ENST00000601538.6,E | 1 | - | good | TTGTGTTCTCCTCTGCCATTACCACCTTACACACCTGTAAGGGAGGTTATGCACTCTGCCCACTGTGGTCAGACTGGGCAGTGGGGGTGCCAGCAGGCCCCGCCCTCCCTGGGCTTTCTT | 120 |
| chr11 | 71091455 | 71091575 | 788855_47188552_ENST00000601538.6,E | 1 | - | good | GTCTACCTAGGAGGTCAAGAGGAGGCTCTGGAACAGCAAGGGGAGGTCAAGGACAGAGGGCCCTGACTGGTTTCTTGCTTGGTTCTCTACACAGTGACTGCCATGGTTTCTCTGGCTACC | 120 |
| chr11 | 71091575 | 71091695 | 788855_47188552_ENST00000601538.6,E | 1 | - | good | TCTTCTCTTGGCTGAGCCCTGCCAGAGTTGGAGGACAGTGAGGGAGTGCTGTGGTGCAGGCTGCAGGCCAGGGTCTCCAGGCAGAGGGGTGAGGTGGGTCTGGGAGAGCTGATAGCACCT | 120 |
| chr11 | 71091695 | 71091815 | 788855_47188552_ENST00000601538.6,E | 1 | - | good | GCACCTGTGAGCTGATGGGCAGGGAGAGGAGATGATGCTCCCGGGAGCAGGAGAGCCAGGGCTGAGGAGGGGGCCCGCAGGCCCATGTCAGGGAGGACTCAGGGAAGGAATATCTGCCT | 120 |
| chr11 | 71091815 | 71091935 | 788855_47188552_ENST00000601538.6,E | 1 | - | good | TTGGCAAGAAACAGATTTCACCGGCTGTCTCCAGTGAAAAGTCTTTACCTGATCAGGTAGAGGTGTGGACGGGGTTAAAGGCTGAGGACCCCATAGAAGAGCAGTCTTGGGGAGCCATCA | 120 |
| chr11 | 71091935 | 71092055 | 788855_47188552_ENST00000601538.6,E | 1 | - | good | AAGCACCGGTGTTTTCTGAGCACAGGTGAGCCCCAGGAAGTCCTGATTTGCACGCACGAGGGGCGTGTTTTTCAGGCCCAACCTTGCAACAGTTGGGTTAGGAGGTAGGTCAGCCTGGTC | 120 |
| chr11 | 71092055 | 71092175 | 788855_47188552_ENST00000601538.6,E | 1 | - | good | AGGCAAGGCTGCTGAGTTTCTGCACAGCAGGAGTGCCCTGCAGTTATGGGGTGGACGCCCCCTTCCCAGGAGCCCTGCCCTCTCCAAGTTCCCACCGTCCTGACAGGGCAGGATACCAGA | 120 |
| chr11 | 71092175 | 71092295 | 788855_47188552_ENST00000601538.6,E | 1 | - | good | CAGAGTTCCTGAAAATATGTGAATTAAAAATGATTTTAAAATGTGAACGTCAGTATTTGAAAACAGACACATTTGCATAGAAATAGACATTTCTGACTTCTCTGGAGAAACTGGAAGATG | 120 |
| chr11 | 71092295 | 71092415 | 788855_47188552_ENST00000601538.6,E | 1 | - | good | GGGCCCGCTTCTCCACTCTGTTGTACCCACGAACTGACTTCCCCTTCTCCTCCGATCCCGCAGATAAGCAGGGTGGCCCAAAGGTTAGATCCTGCCTTCAGGTATTTTGTTTGGCCTGCC | 120 |
| chr11 | 71092415 | 71092535 | 788855_47188552_ENST00000601538.6,E | 1 | - | good | CCGCTGTATCACACAGCCATCGTCGGAGGTGATCCCTACTGCTGCGAGCTTCTCCTGCACGAACACGCCACTGTGTGCTGCAAAGATGAGAACGGCTGGCACGAGATCCACCAGGTACGT | 120 |
| chr11 | 71092535 | 71092655 | 788855_47188552_ENST00000601538.6,E | 1 | - | good | GATCACCTCACTCTGCAAAAGGGGTCATGAGACCAATAACGACGGCTTTCAATTTTTTTTTCCTAGACCCTTTTAGAGCTTGGTGCATCCCCAGATTATAAAGACAGTTACGGCCTCACC | 120 |
| chr11 | 71092655 | 71092775 | 788855_47188552_ENST00000601538.6,E | 1 | - | good | AGCCACCACACCCGGCCAGGTTATACCCTTCTTAATTCTGGGGAGCCCAGGAGAAGCACTGGGTACTCAGGGTGTGGGTTGCCTTGACCTGGGGGAGGGTGGTCCTGGTGCTGCTTTCTG | 120 |
| chr11 | 71093015 | 71093135 | 788855_47188552_ENST00000601538.6,E | 1 | - | good | AGCCTGCCCTGCATTTCTGGGCACTGTTTGCTATTTCAGAACTAGAAATGTAGTTGTCTAAAGTTATACCTGCCCCCCCCCCCCTTTATTTTTGAGACTGAGCTTTGCTCTTGTTGCCCA | 120 |
| chr11 | 71093135 | 71093255 | 788855_47188552_ENST00000601538.6,E | 1 | - | good | TCTCTCTTCCGCAAATGTTGCAGTCACTGTATTGCTGCCCATGCCACACCACTGGCACAAAATGTTGACCAGGCCGAGGAAGGGTCTGAGTGCTGCAGGTCTCCTCGTCTCCCAAATGTC | 120 |
| chr11 | 71093255 | 71093375 | 788855_47188552_ENST00000601538.6,E | 1 | - | good | GTTGGCTTCCTCGGAAATCAGGAGCATTCTTGAAGACGCGATGCTTGCTAGGTTCGCCACCCTCAGCGTGCGGAATGAGGTGGTTTTCCACTGGGCATGTAGGTTAAACCATGCACCTTT | 120 |
| chr11 | 71093735 | 71093855 | 788855_47188552_ENST00000601538.6,E | 1 | - | good | CAAGACGGGGGCCTCGCTGTCCCTAAAGTCTCAGGACTTTCACTTCATTTCAGCTGCACCCCTTCTGTTGGTGTGTGAGTCTGTTCCTGCGTTGCAATAAAGAAATGTGAGATCGGAGTA | 120 |
| chr11 | 71093855 | 71093975 | 788855_47188552_ENST00000601538.6,E | 1 | - | good | CTCCGTGCCCCTGCCTCACACTGCAGCCACGGCGCCTCTGGACAGGGTGGTGCACGCCCAGGCATCCTCCACCAGAGCCCACACCAGCAGGGATGGGACGTGCCGCTGCAGGTTGACCCC | 120 |
| chr11 | 71093975 | 71094095 | 788855_47188552_ENST00000601538.6,E | 1 | - | good | GAGATTTGGGAGGGGACACAGAGCCACATGGTAGATCCCCCCAGCACCCTGTGGCGGGTCCGTGGGAGCCTGCCCTGTTGTGGGTTCCGCGCCTCACCTTCCCCTGCTGCTGTCTTACTG | 120 |
| chr11 | 71094215 | 71094335 | 788855_47188552_ENST00000601538.6,E | 1 | - | good | ATGCTTTCAGAATTAATGGTTCCCAAGAAGTCATCTTTCACCTTCTATTCAGTCAAGGGACTTGTCTTTTCAACCCGTAAGGCCAGTGGTGAACCCCAACCCCATCCACAGGAAGACTCA | 120 |
| chr11 | 71094335 | 71094455 | 788855_47188552_ENST00000601538.6,E | 1 | - | good | ATCCTAGGGGGTCCGTGCACAGTGTTCCGGCCCACCCTAGGAGTCCGTGCACAGTGTTCCGGCTGGAGCTTGGAAGTGAAGCCAGGTTTCAGATTTTCCTGTTCTCTCTCCTGGTTGTCG | 120 |
| chr11 | 71094455 | 71094575 | 788855_47188552_ENST00000601538.6,E | 1 | - | good | CACAAAGCTGCCCGAGCGAGGAACCAAGTTGCCCTGAAGGTAAACTCGGGCCCATCTTACTTGGGTGCCACCCCGTGCTCTGATTCCGTGCGGCTGCCCCATCCCATCCCCGCAGTGCCC | 120 |
| chr11 | 71094575 | 71094695 | 788855_47188552_ENST00000601538.6,E | 1 | - | good | TCCTCAGAGACCCCCCTGACCTTAGCCGCTCAGCTGGACGACTCTGTGGAGGTCATCAAAGCTCTCAAAAATGGTGGAGCTCACCTGGACTTCCGTGCCAAAGATGGGATGACCGCCCTA | 120 |
| chr11 | 71094695 | 71094815 | 788855_47188552_ENST00000601538.6,E | 1 | - | good | GCTGTAAACACCCTGGAGGTGGAGGAGGGGAGTCAGTTCTTTCAGCTTAGTTTTGGGCATCTGAATATGGCTTTGTGAGCAGTGTGACAAATGTTGGTTCTAAAGTGTGTGTGTTTGGGT | 120 |
| chr11 | 71094815 | 71094935 | 788855_47188552_ENST00000601538.6,E | 1 | - | good | GTTGAATCCCTGCCTCTGCCACCGCTGTGTTGTACGCGTGGCCCTGCGTGTCACTGCATCGCACGGGGGTGCCTGAGCCACGCTGCCCGCCTGGACTCTTGGGGTTCCCAGGTGGTGGGA | 120 |
| chr11 | 71094935 | 71095055 | 788855_47188552_ENST00000601538.6,E | 1 | - | good | GGGCTTTAGTCCGTCACTCCTTCTGTACGGGAAGCACCCAGATGGTACCACAGGATTCTGTGTACTCAGAGGGAAGGGCAGCAAGGGAGCCTCTTGGACTGTGGCCCACACAGCAGTCCT | 120 |
| chr11 | 71095055 | 71095175 | 788855_47188552_ENST00000601538.6,E | 1 | - | good | GGACTTCATGTCTCATTTGTGTTTGTGTCCCTGCAGCCCATGGCTGGCATCTCCCAAGATGCCCATAAATGTGACATTGAATGAGGAAAGGTGGGTGTCTGCCTTTCCCATCTCTAATGA | 120 |
| chr11 | 71095175 | 71095295 | 788855_47188552_ENST00000601538.6,E | 1 | - | good | CTACAAAGGCAGACGGCATCCTTCTGTCTGCTGGGCCTCTGCCCTGCCTCCTGGGTCATCCTGCGCCGCGTTAATGAAGGAAGGTTTCCACTTACCCAGATTGGGAGCTCCAGGGGGCAC | 120 |
| chr11 | 71095295 | 71095415 | 788855_47188552_ENST00000601538.6,E | 1 | - | good | ATGGTGGGCATATTAGAAAGATGAAGAGTGGACGGAAAGATTTGTTGTAATAGACACTGACGTCCTTAGCCACGCTCATCTGCAGACTGGTCCAGGGTCATCTGCACGGGACCCTTGTCC | 120 |
| chr11 | 71095415 | 71095535 | 788855_47188552_ENST00000601538.6,E | 1 | - | good | TCATGGGAACTTGTGTGTTGTCCTGGTACTCTGTGCACTTTTCTGGAACGTTAAGAGCCTCTCAAATGAGGAACTCTCAAAAGAGGACCTCAAAGTAGTTCCTCTTGTTTAGCAGTGGCC | 120 |
| chr11 | 71095535 | 71095655 | 788855_47188552_ENST00000601538.6,E | 1 | - | good | GAGGGGGATGGGATGGGGTGAGGGGTCAGCAAGACACACGCCTTGTGAAAGGCATCCGAGTTTTCTTTCACTGGAATGTGCTTCGAGACCGAGTGAATATTCATTGTTGGTGTTTCTGGG | 120 |
| chr11 | 71095655 | 71095775 | 788855_47188552_ENST00000601538.6,E | 1 | - | good | CTAGGGTGAATGTTAAAAGTCCTCAGGAGGGTACCCATGTATGGATAAGTAAACATCACAATTAAAAATAGATGGGAATAGTCAGTGCACACTCACAAGGCAGGTGCCCTTTCTGGGATG | 120 |
| chr11 | 71095775 | 71095895 | 788855_47188552_ENST00000601538.6,E | 1 | - | good | GCTTTTCACATGGAGACAGGATGAATGAAGATGCTGGCATGTCTACACACAGCACACTCCTGACATGGAGGGGTTAAGCATGTGCATATCACCATGAAGTGCCAAGCTGCAGAGTGGTTC | 120 |
| chr11 | 71096135 | 71096255 | 788855_47188552_ENST00000601538.6,E | 1 | - | good | GTCCCAGAAAATTTCTATTTTTCTTTAAAACTTCTGCATCAGGAGGGTATGCAACAGAGTCATGTGCAATGGCATTCATTCATGAGAACATCGTTCCTAGTTGGGGAAGACAGGAAGCCT | 120 |
| chr11 | 71096255 | 71096375 | 788855_47188552_ENST00000601538.6,E | 1 | - | good | CATGTGGAACTCTGGCCATGTTCCAACTGGTCTCAATCCCCACACCCCTTGCTGAGCCTTCCTTGCCTTGTAGCTGTTTTCTATGCAAGAAACATCCCCCACTGCCACTGAAGTCCCTTA | 120 |
| chr11 | 71096375 | 71096495 | 788855_47188552_ENST00000601538.6,E | 1 | - | good | GGATGCCACCAGGGAAGAGACAGAGAATGAGGGGCTGACTGTTCCTCTCTCTAGCAGGCAGGGGTCACCCCTTGAGTTTGTTTTTCTCTGGCGTCTGCCCCATTTCCACCTCTCTGTGCA | 120 |

| | | | | | | | | |
|---|---|---|---|---|---|---|---|---|
| chr11 | 71096495 | 71096615 | 788855_47188552_ENST00000601538.6,E | 1 | - | good | CCAGTCTTTTCGTGCCCTTCGAGGATCAGTCACTTCCCTGACTGTCCACTCTCCCATGGTGGCCTTTATGCTCCAAGGAATAAACTCCCCAGATGCCCTCGGACACTCGTGCTTCAGCCT | 120 |
| chr11 | 71096615 | 71096735 | 788855_47188552_ENST00000601538.6,E | 1 | - | good | ATGGGAAAGAAAATCCTAACTGTTTTGAAGGCTAACCTTGCCCCGAACGAGTGTCACACGCTCATTAAGTCGCTGTAAGATTTGAATGTTCAGCGTCGATATTGGTGGCTCCCGACAGCC | 120 |
| chr11 | 71096735 | 71096855 | 788855_47188552_ENST00000601538.6,E | 1 | - | good | TCATTTATAAAATGGAACAGCTGAGGTTGTCAGCTATGAAAATTATATGTTAATTTAAAATATCTTTTAAAAAATCTGCTTTGGACTGCCTTTTTGCATTGGGCAAAAGGGAACAGTCAA | 120 |
| chr11 | 71096855 | 71096975 | 788855_47188552_ENST00000601538.6,E | 1 | - | good | GGCACTGACCAACAGGATGTGGTTTGGCTGCAGGCATCTGTAATTCTGAGTCAAAAGTCAGGATTCAAGCCTCTGATCTGATTTTACCACTTGCCACCAGCTCTCTCTGAGACAGCATTC | 120 |
| chr11 | 71096975 | 71097095 | 788855_47188552_ENST00000601538.6,E | 1 | - | good | GGCCCTGGAAGCCACCCAAGGACCACACATAGCTCCCCATCCTGCACGTGAGACACAGCCATGAACAGAGTGGGATCATTTTCTCCACCGGTGCGTTTATGCAGCCACCTTTAACTGCCT | 120 |
| chr11 | 71097095 | 71097215 | 788855_47188552_ENST00000601538.6,E | 1 | - | good | AAGAGGGGGTCTCCTGACCTCCCCAGGGTGGGTCCTGGCCAGAGGTGAACATGCAAGTCTGGAACATCACTTGAGGAGGGATTCCGGAACCCTTGAGAAGGCTGCCACTTGTCATTGGAT | 120 |
| chr11 | 71097215 | 71097335 | 788855_47188552_ENST00000601538.6,E | 1 | - | good | TATCTGTCACCGGTTTATAGCTCATCAACCAGCATCTTCTCGGGGGTGATTCCATCAGCCCCAAATGTGGAACCTCAGCGGACGTGCTCTTTGCAAATGTGGTGGCTAAGCCAGGGCGGG | 120 |
| chr11 | 71097335 | 71097455 | 788855_47188552_ENST00000601538.6,E | 1 | - | good | AAAATGCAAAACCTGAGGAATAGTCTTAAGAATCATGGTGCCAGATAACTGTGTGGTTCCCATGCTCATCAGGGGAGTGATAGTGCAGTCACTGGGTGTGCATCCCGGGCAGTGCTGTTA | 120 |
| chr11 | 71097455 | 71097575 | 788855_47188552_ENST00000601538.6,E | 1 | - | good | TATCTCAAAATATTACAGGTGTTTCTACTCCCACCCGTTGTTCACGTTGAGGGAGACTGGGGCCACTCTGAGGTTATGCTTGTTGAGGTTCCAGTCTATTTTTAAAACTAAAGAAATTTC | 120 |
| chr11 | 71097575 | 71097695 | 788855_47188552_ENST00000601538.6,E | 1 | - | good | CCTCCTGCTGCGCTTCCGGGAGCCCCCGCTTCTCACCCTTTCGAAGCACATCCTGCTTATCCCTGGAGCCAACAGTCTGCTGGGCAGCCTCTAGTTGGAGGGTCTGCGTGTCTCTGAGGT | 120 |
| chr11 | 71097695 | 71097815 | 788855_47188552_ENST00000601538.6,E | 1 | - | good | GTGCCCGCTGGGTAGCAGCAGCTGTCTTAAACACTTACACTCCTCCTGCCTCCTGCCCCCACCCCACCCCCCATTGCTGTGCCACCTCCCAGTGTCAGGACAAACAGCCGTGGCTAGTGC | 120 |
| chr11 | 71098295 | 71098415 | 788855_47188552_ENST00000601538.6,E | 1 | - | good | TCACTTCCCCTCATAGGAAGAATCGCTTGCCCAGTCGTTTGATTTTTGCAGCCCACTGATCTTGGACACTAGGATGCACCTAGGAGCACACACACGGGTGTGCACACATGCACACAGAGG | 120 |
| chr11 | 71098415 | 71098535 | 788855_47188552_ENST00000601538.6,E | 1 | - | good | CTTCTCTCACATCATGGTTTCCTTCTGAGCATCCCCAGCCTCCTGAACTATCATCTTCTAAAAAACCTGCTGGAGGCCACCCAGCCCTTCAGGGAAAGAGCTGGGTTTTGTTGACACCAG | 120 |
| chr11 | 71098655 | 71098775 | 788855_47188552_ENST00000601538.6,E | 1 | - | good | GAGTTTGTCTGCATTGCAAGATCGGCCCCTTCCTCCCGTTCACTTGTGAATCCCTTGCTTATTTCTATCAATGTGGTTGCACTCACTTTCTGGTTGACTTATAACCCAGCTTTATTTATT | 120 |
| chr11 | 71098775 | 71098895 | 788855_47188552_ENST00000601538.6,E | 1 | - | good | CCACACTATGCTCTCAGAAGGAAGCCACTGTGCACAGCCCACACATAAGGGGATCGGGGTGGGGGGTGGTGCGGGCTACGCATCACCCCCTGAGGACAGAGAAGTGATGCGAATTGCTTG | 120 |
| chr11 | 71099255 | 71099375 | 788855_47188552_ENST00000601538.6,E | 1 | - | good | CAGTATTCTTCTGAGCGGAAGTCGTATGGTACGCTTCTCACAGTTAGAGAACCGATATCGATGCATTATTATTAACCGAAGTCCACACCTTTTTCTTCTCTCATAAGCATTTACCTGCTT | 120 |
| chr11 | 71099495 | 71099615 | 788855_47188552_ENST00000601538.6,E | 1 | - | good | AAGGGTTCTCTGTATAATTTGGCTACAAGTACTTTATACGCTTTTCACAAACATTTTCTGCCTCTCTGAGCAGCAGCATTTTTTACATTAAATCTGTTAACCCGCACAATAGCCAGGAGA | 120 |
| chr11 | 71101055 | 71101175 | 788855_47188552_ENST00000601538.6,E | 1 | - | good | AGAAGGCATGGAGTTTTCCCCTATGCACTTTCACTTCCAGCTCGCCTGTTATTGACAGCATGCACTGGTTTAGGGCATTTGTTGCAATAACTGAAGTCCATAGTTTACATGAGGGTCTGC | 120 |
| chr11 | 71101175 | 71101295 | 788855_47188552_ENST00000601538.6,E | 1 | - | good | AAAATGCGGCAGAGGGAGTGGCGAGGGTAAGGGTGACACGGAGGCAGGGGCCAGGGTCCACAGGAGCCTGTCTTGGGGTCGGGGACAGAGCCCAGTTTTAGGTTGACAGAAAAACCAGG | 120 |
| chr11 | 71101295 | 71101415 | 788855_47188552_ENST00000601538.6,E | 1 | - | good | CCAATTATTTCAAATACAGTGTCTCATTCTTTCAGGGTTTTTCTATGTACACACTGCCATGTTCTGAAAGGGAATATAGGTGACTCTTAGATGGAGCGAGGTGCCGTGCCTTCAGTAAGG | 120 |
| chr11 | 71101415 | 71101535 | 788855_47188552_ENST00000601538.6,E | 1 | - | good | AGAAATTACCCAGTGCCAATGAAGGAGCTGGAAGACTTCGAGTAAATAACTCCCTTTGCTACCAGTGCCTTTGGTTTGTTCAGTACAGTTTCTCATAAAAAGAGACCTGTTTTTAAGGGT | 120 |
| chr11 | 71101535 | 71101655 | 788855_47188552_ENST00000601538.6,E | 1 | - | good | CCTGAATGAATCGCCAAGTGCGATTCCCCAGAACTAACGGCACCCGGGAATGTGAGGCAATTCAAGAGAATAATGGTTCTGGAACCTCGTAAACCTCCATCTGCAGAGGGTGAGAGGTGC | 120 |
| chr11 | 71101655 | 71101775 | 788855_47188552_ENST00000601538.6,E | 1 | - | good | CGAGGACATTCTGGGTCCCAGTGCACTGAGCACCTTCAGAGCATCGTCTTGAAGAGCAGAAGGGGCTTGTTTGATGTGTGCAGTGACTGACGGTTCATGCTCCGCAGTCTCACGTTGGCC | 120 |
| chr11 | 71101775 | 71101895 | 788855_47188552_ENST00000601538.6,E | 1 | - | good | CATACCAGGCTGAGCTCCTCCCATTCGGCCGCCCCCCAACTCCGGCTTCCCCTTCTTGCTGCTGTAGCTGTGTCACGGGGAGCCCCTTCCCAGCTCCCGCACCGCTCAGTTAGGAGGCCT | 120 |
| chr11 | 71101895 | 71102015 | 788855_47188552_ENST00000601538.6,E | 1 | - | good | GGTCAAAAAGAGATGGTTCCTTTACGGATCCAGATTCTGCAGAGCACCTGTGATGTGTCTGGACCTCACTGATGAGATTCAGTTACTGAATCTGTTTCCAGGGTCGGGGTGTACTCATTG | 120 |
| chr11 | 71102015 | 71102135 | 788855_47188552_ENST00000601538.6,E | 1 | - | good | GGTTGGCCCTTTTATCTTGCACTGTTGTCTCCAAATTTGAGAGAAGCCTCAGGCACCACAAAAGAGATCTGGCTAATAAAGCTAGAAATGCCTTTCGACTCTTTTAGTATCTGAATGAGA | 120 |
| chr11 | 71102135 | 71102255 | 788855_47188552_ENST00000601538.6,E | 1 | - | good | GACTTCCTCCAGCAGAGTCTCAGGCCTCTTCTCCTGCTTTGGTTGGAAACTCTGAGATGAGACCAGTTGTGTTTCCAAGCCCCAAACAGGAAGCGAACCTGGCAACTTGAACAAAATTTA | 120 |
| chr11 | 71102255 | 71102375 | 788855_47188552_ENST00000601538.6,E | 1 | - | good | AACGACCTGTCAATGAAAACAAAACAGCCTGTGTGATTGCAGGCGACCATCTGGTACCTGTTCTAAATGTAAATTTTATTAAGTAACAATGATAAAAGTAGTGTAAAAAAAAAAAAATAAA | 120 |
| chr11 | 71102375 | 71102495 | 788855_47188552_ENST00000601538.6,E | 1 | - | good | GCCGCCATGCTCAGTCCCGGGCCCTGCAGACCTGGGCCCCGCGTCCCTGTTTTGGGGCTGGACATGGGGCCGGAGCGGAGGGAAGCAGCCCTCTAGCCTGAGATAATTAAATCCCAGTGG | 120 |
| chr11 | 71102495 | 71102615 | 788855_47188552_ENST00000601538.6,E | 1 | - | good | TGTTCCCATCCCATATTTAGCCTCTCCACAGTGAGTTTGGTGACTTCTCAAAGCCACCCAGGGGACAGCAGAGTCTGTGCTTAGAGTTGAAAGCAGCCTTTTTGCTAAGCTCAGGAAAT | 120 |
| chr11 | 71102615 | 71102735 | 788855_47188552_ENST00000601538.6,E | 1 | - | good | AACAGGGGTCTCTTGGCATGAGCCGGCCTGGCTGGACGTGGATGACCCAGGCAGGCCCAGGTCGGGGACAAGCTGTTCACATTCCCACTTTCCACCTGGTTGTCGGCAGCAGCCTGGGCA | 120 |
| chr11 | 71102735 | 71102855 | 788855_47188552_ENST00000601538.6,E | 1 | - | good | TTACCACAGCCCAGTCTGGGCCAGTCACTGAGTCACCAGGGAGGGCAGAGGCAGGGCCTGGGCTCCACTTGCCCTGGGGGGTCCCTGCTCTCAACTCCTGGGGGAAAAGGCAGTGGGACG | 120 |
| chr11 | 71103095 | 71103215 | 788855_47188552_ENST00000601538.6,E | 1 | - | good | GGAGGAATCTGAAGCTCTGAGAGCCCAACTGGGTGGCCGCTAGGGCTGATGTCTGCATCCTCTATACTGTTGAACCCTAACCCCCTAGGTGATGGTATTAGGAGGATGGGGCCTTCAGCA | 120 |
| chr11 | 71103215 | 71103335 | 788855_47188552_ENST00000601538.6,E | 1 | - | good | GGCACTTAAAGAGTGATAGCTGCAGTCTATTGGTGACCTGCTGTGCTCCAGCAGCACACAGGGCCTTGCCAGGCCTCAGCTCTGTAATCAAGGGGGTCCCCTGCAGGCACCGCTGTCCTG | 120 |
| chr11 | 71103455 | 71103575 | 788855_47188552_ENST00000601538.6,E | 1 | - | good | ATGTTCCTGTCATCAGCATCAACTACGGCAGCTCAGTCAATGTTGTTTTCTCCTGGTTTTGGATGCAGAGGGAGAAACACAGAGTTGGCTTTGGCGGGCTGCCTCCAGGAAACAGGCAGG | 120 |
| chr11 | 71104175 | 71104295 | 788855_47188552_ENST00000601538.6,E | 1 | - | good | CCAGAGAGGAGACATTAACCCCTACTTCTCTGCAGCTGGCTGTGTGAAGACCAGGTTAATGAACCTCTCTCAGCTTCCATCCTCTTGTCTCCTAAATTGAATGATAACACCTACGGTGGT | 120 |
| chr11 | 71104295 | 71104415 | 788855_47188552_ENST00000601538.6,E | 1 | - | good | CGGATGACATCACTGTTGGAGACACTGGGCTTCAGAGATGCAGGATGGGGGGGGCCCTGTGTTGGGACTGCAGAGGCGACCCTTCCAATGCCCAGCTCAGTGCCAGTCCATGGAGAGCCC | 120 |
| chr11 | 71104415 | 71104535 | 788855_47188552_ENST00000601538.6,E | 1 | - | good | GAGAGGCCAGAGCCCAGTGCTGCCAGGGAAGACACCACCTTCACTTTGCTTCTGTCAAGGCAGAGCATAAGGGACGGATACGCATCCTCCTGACTGAAATGCAGACACTACCTGGCCACA | 120 |
| chr11 | 71104535 | 71104655 | 788855_47188552_ENST00000601538.6,E | 1 | - | good | AGGGTGGGTGCTCTCAGAAACAAAGGAAGCTGCGGGTAATTCCAGCACAATGGCTTATAGCAACCCTCAGGCACACATGGAGCAAGTATCAGTGGGGACAAGCTAAACGCAAGTTCCAAC | 120 |
| chr11 | 71104775 | 71104895 | 788855_47188552_ENST00000601538.6,E | 1 | - | good | AGGCCAAGTGCAGACCCACAGGTCGGCTGCAGACAGCGCAGATGGCCCTGGTCTCACACCACCCTGCTTCCCAGCTTCTGCGCCTGCCACCAGTGTCTCAAGAATTCTCATTTGTTTTTT | 120 |
| chr11 | 71104895 | 71105015 | 788855_47188552_ENST00000601538.6,E | 1 | - | good | TTGTACAGATCAAACTAACAGCGATCATACACAGTGGCATTTTTTTGTAAGGCCAAAAACATTACTTATGGCAAATAAAAAGGGAAAAGGGGTAACTAAGGCAGAAAGTTCTGAAACCGA | 120 |
| chr11 | 71105135 | 71105255 | 788855_47188552_ENST00000601538.6,E | 1 | - | good | TGGAATCTTAAAATCACACAGTGGGTTTCAGATCGGCTTCTTTCACATGGCAGCACGTGCGGAAGGTGCCTCCAGGCCCTTTCACAGCTTGGAAGCGCGTTTCCTTTTGGTGCTGGATAA | 120 |
| chr11 | 71105615 | 71105735 | 788855_47188552_ENST00000601538.6,E | 1 | - | good | TGTCAGTTTTGCCAGATGTCCTGTATCCACCGTCACAGTATCATAAAGATCAGCTTCACTGCCCGAGAGCTCCTCTGTGCCCCTCTTCATCCCTGAGCCCCTGGCAGCTGCTGACCCTTT | 120 |
| chr11 | 71105975 | 71106095 | 788855_47188552_ENST00000601538.6,E | 1 | - | good | CCCATATTGACTGCTCCCCCAGTTTCCTCTGTTAGTAATATCTTGCTTTAGTATAGGACATCTGTTACAATCAGTGAACCAGTTGTGATCCTCAAGTCCAAGCTTACATGAAGGTTCACG | 120 |
| chr11 | 71106095 | 71106215 | 788855_47188552_ENST00000601538.6,E | 1 | - | good | TGATATTTCTCTATAAAGCAGAACTTCCTCTTCCCCTGCCTCCCTTTTGTGAAATGAGCAGAGCTGATTTTTTAGAGCAGTTTTAGGCTGACAGAAAATTGAATAGAAGTACTGTGAGTT | 120 |
| chr11 | 71106215 | 71106335 | 788855_47188552_ENST00000601538.6,E | 1 | - | good | GCCACTGAGCCCAGCCTCTTATTTTCAATTTTAAAATTTAAGGCAGAATTATGGAATCAGTTCTCTCCTGGTTGTTGCCAAATGGCAATTTTATGAACCAGCTTTGCTTCTTGTAGTAGC | 120 |
| chr11 | 71107175 | 71107295 | 788855_47188552_ENST00000601538.6,E | 1 | - | good | GTGGGTTTGGTGCTCAAGGACAGAATGCGTGCAGGGAAGAATGCCTGCGTGCTTCCTTACGGGACACTTCCCTGCGTCACCCAGGGAGATGGTGACCACGCATAGAGGTTGGGGGTTTAT | 120 |
| chr11 | 71107295 | 71107415 | 788855_47188552_ENST00000601538.6,E | 1 | - | good | CTGGTGTCATCAGGTCCCCGATAAGTGGCTCCTGGGAAAGAGCACATAGTTAAGGACAGGCATTCTAGGGTCTCCAGCAAAGGCAGAGACCCACGGGGCCAGGGTCTGGGACAGGGCCCA | 120 |
| chr11 | 71107415 | 71107535 | 788855_47188552_ENST00000601538.6,E | 1 | - | good | ATCTGTGGAGATGGTACCACGCAGTTACCTTCCGAACGTTCTGAAATATTCCCACGATGACAAGCCTCTGGGCTGGTTTACTGATCACAAAGACCAAACCCAGAGAAGACAGAGGAACCC | 120 |
| chr11 | 71107655 | 71107775 | 788855_47188552_ENST00000601538.6,E | 1 | - | good | ATTCCAGAATAGACTCTCAATTCACTGCTAAAAACACATCAGTGCAGACTCTGAAACAGGTAAGCAGCGATTTCTCTGGGTCCTGCCTGGAGTTAAAACTGCTGCCACTCAAATCCTGGA | 120 |
| chr11 | 71107775 | 71107895 | 788855_47188552_ENST00000601538.6,E | 1 | - | good | CTCTGCACCTCTCTTCCTGATGTGCCCCCTGTAATTGGCCCCTGTCCTGAGCCAAGCCTCCCCCTGGCTTCCTCCACCAGACTGTTAATGTCTCGACTTTAAGGTTCTGAGATGACTGTT | 120 |
| chr11 | 71107895 | 71108015 | 788855_47188552_ENST00000601538.6,E | 1 | - | good | TCATCTTGGCCTCGTCGAGGGCTATTTCTGATGCTGCTGCGGGGGACTGGGCAGCGTGGCTCTCTGTCTCGCCTTGGCCTGGCCAGCAGTAGAAGGAGGCTGGATGTCCCCTCTGAGCTG | 120 |
| chr11 | 71108015 | 71108135 | 788855_47188552_ENST00000601538.6,E | 1 | - | good | TGACCTTGAGCCTTGGGGCACATCCAGGGGCAGCCTTCATTCTGAGCTGCTGACCCGGATTCTCCCTGGAGGCAGCCTTTGGCCGTGGCTCTCACCACCCTGGCAGAGCCACTTATGTCT | 120 |
| chr11 | 71108135 | 71108255 | 788855_47188552_ENST00000601538.6,E | 1 | - | good | TGCGCTGATGTGGCAGCTTCTCTTCAGTGATGGCCCCTGTGCCAGCTCCTGGCTCTGTGAGGGCATGAGCCCTTTCCATGTGAGGCTGTGAAGGTAAATGGACCCAGGTGCTCCTGAGCA | 120 |
| chr11 | 71108255 | 71108375 | 788855_47188552_ENST00000601538.6,E | 1 | - | good | GCTGGGAAGCAGAGAGGCCTGAGCTCAGGCATGGTGGTGCGATGGAGGGCGTGTACGTCTGTAGGAGGCTTGGGAGGGCCCTGGCCGACCATCTTTCCCTGACGTGGATGCGTCTCGTGC | 120 |
| chr11 | 71108375 | 71108495 | 788855_47188552_ENST00000601538.6,E | 1 | - | good | AGTGATTGGGACAAGGGCATGTGGTGGGCAGGGGTGGGTAGTAGGGCATCTGGAGGGCAGGTGGGCAGCTGCAGTGGCCCTCAGCTGTGGGGCAGGCAGTGGGAGGAGGAGAGGAGCT | 120 |
| chr11 | 71108495 | 71108615 | 788855_47188552_ENST00000601538.6,E | 1 | - | good | ACGTGGACAAGGCTGGGCTTCCGCTGGAGGTCTGGGCAGTCTCAGCCGGTGGCCTGGAGCAGGAGGCTGGGTTGGGCTTGTGTTGGTAGGGACATCTGGAGAAGGGGAGAGGGGAAAGG | 120 |
| chr11 | 71108615 | 71108735 | 788855_47188552_ENST00000601538.6,E | 1 | - | good | TGCCCCCGGGGATAGGGGGAAAGAAGGAAGGTGAGTGCCTGGCTACAGCTGAGCAGCCCCAGGCCCTGCGCCGATCCTCCATGAGGCAGGGAAGCCTGGGGGTCTGCTTGCAGGGCAGTG | 120 |
| chr11 | 71108735 | 71108855 | 788855_47188552_ENST00000601538.6,E | 1 | - | good | TCCCTTAGGCTTCGCTGTGAGCAGCAGGAGAAGCCGGCACGGGCTTGTGGCTGCTGGTAGCTTTGATGCCTGTGACGTTGGGAGGCAGGACTGCATGAGCGAGGCTCAGAGGGGACAGAG | 120 |
| chr11 | 71108855 | 71108975 | 788855_47188552_ENST00000601538.6,E | 1 | - | good | TGTGGGAGCCCCCTCGACCTCTGAACGCTGGGGGGGGGCCCGCTGGGTGTGGATTCAGCCTGCACAGCTCTGTCTGAGCACTGCCGGGGGACTGTCCTTGCCTTTTTGCAATTGTCTGTG | 120 |
| chr11 | 71108975 | 71109095 | 788855_47188552_ENST00000601538.6,E | 1 | - | good | GCCCTGCTTCTGGCCCCTGCATGGTATAGTGCCATCATCACATACTGGTGTCACACCTCAAAAATATAAAATACACAACAAATGTGTTTGTCAACCAAAGAATGCCACTTTCTCATCATT | 120 |
| chr11 | 71109095 | 71109215 | 788855_47188552_ENST00000601538.6,E | 1 | - | good | GAACCAAGAGGGGTGGAGGTGAGGGTCTCCATGCAAAGCAGAGTATGCCCCCTCAGGTGGGCTGTCCTGGGCTGGGAATACTCCTCCTGGACTCCCTCCCCTCTTGTCTTGAGTGTGCCA | 120 |

| | | | | | | | | |
|---|---|---|---|---|---|---|---|---|
| chr11 | 71109215 | 71109335 | 788855_47188552_ENST00000601538.6,E | 1 | - | good | CAACAGTGTCTCCAGACATCACCCACTGTCCCTGGGTGTGGGCACTGTCACCCCAGGTGACAGTCTCTTGCCAGCAGGGCACCAGCTGGAGTTGGCAGGCAGAGGTTAGGTCCTACTGGA | 120 |
| chr11 | 71109335 | 71109455 | 788855_47188552_ENST00000601538.6,E | 1 | - | good | GGCCCTTGGTGGTTCCACGAGTGTGTGAGTCTTTGCAGCGCGGCCATCCTATGTATCACAGGGTTCAGTGGCAGTACCCGCCTCCACTCAGCATCAATAGCAACCCCTAGTTGTGACAAC | 120 |
| chr11 | 71109455 | 71109575 | 788855_47188552_ENST00000601538.6,E | 1 | - | good | CCCGCTGGGCACGGGCAGCCCAGCAGATGGGCATCTGCACCCATGCATCTTATTCCCCTGCTGGAAACCACATTCCTCATGAGTGCGGGCCTCTGGGCATCGGTCGGGAAAGATGGCCCA | 120 |
| chr11 | 71109575 | 71109695 | 788855_47188552_ENST00000601538.6,E | 1 | - | good | TTGAGGAGGTGAATGAGTGAATGAATGAGCAGATGACACGCATGTGTTGTGCCAGCCTTGGGCACACAGATCTTTACAGACACGTATGTTTGTAAAGGTGGAGGATGTGTGTTTCTGCCT | 120 |
| chr11 | 71109695 | 71109815 | 788855_47188552_ENST00000601538.6,E | 1 | - | good | GGTGCATTCTCACTCTCCTCTGTCTTCTCAGTCAGAAGGGGCCGAGCCCGCCTTGTCCATGGGGCATCCCCAGCATCTTGCGTGGGGCCTGGAGCACGGCAGGTGTTCAGTGCAGAATTG | 120 |
| chr11 | 71109815 | 71109935 | 788855_47188552_ENST00000601538.6,E | 1 | - | good | CACTTGCTAGAGGGGACCTTACCAGGCTTTACGGAAGCGTAGAGAAGGCCAGCCCACTGCTCAGGCTACCTGTGAATAACGCTGCAGTTACCTTTTACTTGGTTTTGTTTGGTGTTTAAA | 120 |
| chr11 | 71109935 | 71110055 | 788855_47188552_ENST00000601538.6,E | 1 | - | good | TTCTAGACCAATCTGAAGAAATGCATGGATCACATTCAGCATCGCTTGGTGGAGAAGATCACCAAGATGCTGGACCGAGGCCTGGATCCCAATTTCCACGACCCGGAGACCGGAGGTGAG | 120 |
| chr11 | 71110655 | 71110775 | 788855_47188552_ENST00000601538.6,E | 1 | - | good | TTCATCCAGGATAGGCCGATTGTACTTGTCACGTGCCAGACACTGTGCTCACCCTGTGGTTCCAAAAGCCTCAGCTCCAGGAGCCTCAGGGCTTGTGGTGCACCCTGGCGAGGGTGGGTC | 120 |
| chr11 | 71111135 | 71111255 | 788855_47188552_ENST00000601538.6,E | 1 | - | good | AGGTGGGTCCTGGAAACTGGAGATTGGAAAAGTGCTCCAGCGGCCCAGCTGTCTCTTACGAAAGCAAAGGAGCATTCATAGCTCTCAGCAAGTGTCTGAGGCCATTTACGCTGCTTTTAC | 120 |
| chr11 | 71111255 | 71111375 | 788855_47188552_ENST00000601538.6,E | 1 | - | good | CTTCACCTTGAGCCACTCGGGGACCAGGCATCGCTATTTCTGCCTGTGTGGCCATGGGGTTCACCACAAAGTGGGTTAAGTACCAATCTGCGGGCCTCACTCCCTGGGGATGGAGGCAGG | 120 |
| chr11 | 71111375 | 71111495 | 788855_47188552_ENST00000601538.6,E | 1 | - | good | CTCTGGATGGGCAGAGAATCATGACTTTTCTTACAGCGAGGCTGGCAGAGGCTGGGAAAGTTCCCACCCCATCAGGTTGATCTGCGGACCATCATGACCTCACAGGAAATGAGTCCCGCA | 120 |
| chr11 | 71111495 | 71111615 | 788855_47188552_ENST00000601538.6,E | 1 | - | good | TTCTGTGTGTTTTGAGAGAGGTACGGAGTAGGGGCAAAGAATTAGGCTGGTGAGGAGTGTTAGACGCAGCAGGCGGAAGGGTGCCAAGTCCCACAGTCCTGCTCAGACAGCCCAGACTGG | 120 |
| chr11 | 71111615 | 71111735 | 788855_47188552_ENST00000601538.6,E | 1 | - | good | AGGGCATTTGATCAGTTCTAGTTCCGAAACATACAGAAAATGGGCAGGTGGAAATCTAGCACGTGCCAGGAGGTTGAAGGGCCCAGGAACCTGCTGTGTCAGAAGCCGCATTTGTCCCCC | 120 |
| chr11 | 71111735 | 71111855 | 788855_47188552_ENST00000601538.6,E | 1 | - | good | GCTTTTGTTTTTTATCACCCTCTGATGTGCAGATAAAAACGTATTTGTGAATTTTAAAAATGCTGATTTGTAAATAGCAATGCAGGTTTTCAGTGACACCATGTGTAAGTCATGTGTTTG | 120 |
| chr11 | 71111855 | 71111975 | 788855_47188552_ENST00000601538.6,E | 1 | - | good | CACCAGCCACTTTATGGCACAAGATGCTCTGGGCTCTTCCGATACTTCCCCTGCCTCTGCCCTGGAATTTGCCTGGAAAGCGGTATTCAGACGGCAAGATCTGGGTGCTTCTGGGTTTTG | 120 |
| chr11 | 71111975 | 71112095 | 788855_47188552_ENST00000601538.6,E | 1 | - | good | CACTGAGTTCTAATCTGTCATTATCATTATTGATTTTTGTGCTCAAGTTGTCCAGATGTGGGCGGCAGTTGCCCTTCCTGTGGTGTTCTGCCATGTGGTGCATCCCCACCATCCTTGACA | 120 |
| chr11 | 71112455 | 71112575 | 788855_47188552_ENST00000601538.6,E | 1 | - | good | GTGGAGAGTCCGGGCAGCTGCCTGAAGCGCTTCCCTTTGAGGGCCCTGATGCTCTCCAGGTGAGCTCCCCTGCAGCTGCGAGCTCACCAATGGCTTTCTCAGCCAGCAGGATCCTCTCGG | 120 |
| chr11 | 71112575 | 71112695 | 788855_47188552_ENST00000601538.6,E | 1 | - | good | AGCACTTTGCTTAACCAACAGGAGGAGGAATCTGTAATACCCAGGAGGCTTTTGTCCTGGCCCCTGGCCCCTGGTGCTGTTCCTAAGGAGAGCCCAATTGCACTCCCTGCAGAGCATGGG | 120 |
| chr11 | 71112695 | 71112815 | 788855_47188552_ENST00000601538.6,E | 1 | - | good | CTTGAGCTCAGAGTCCCAGATAAGACATCCACACCATGAACACCCTGCTGTCTTTCAGGGTGCTGTGCGAGTCGAGGAGGAGGCGTGGAGAGGGGGCTGAGGCATAGGGTGTGTGGGCGG | 120 |
| chr11 | 71112815 | 71112935 | 788855_47188552_ENST00000601538.6,E | 1 | - | good | TGTTGCTTCAGGAAAAACCTTGTTAAAGTGCTGTTCACATAAATTGCACTGAAGTTAGACTGCAGGTGAGCAGCAGGGGCGTTGCACAGAGTGTTCAGAACCTGGCGGAATGCACTATGC | 120 |
| chr11 | 71112935 | 71113055 | 788855_47188552_ENST00000601538.6,E | 1 | - | good | ATCCTACTGGCTGCTCCAGGGAGGGTCAAAAGCGGCATCTCTGAAGAGAGTGGGATCTGGACGCCTGGCGTGGTGGAGGGAAGCGGGTCCACAGTGAGCCCCGTATTAGTGTTCTGCTGT | 120 |
| chr11 | 71113055 | 71113175 | 788855_47188552_ENST00000601538.6,E | 1 | - | good | CGTGTGGCTGGGCTGGGATGTACAGGCAGACTGGCCCTTACGGTGCACACTGCGGAGACCCACCCGGGAGGCAGCAGGGGCTGCTTAGTGTGGCCGTGGGTGCTGGGGGAAGGGAGGCA | 120 |
| chr11 | 71113175 | 71113295 | 788855_47188552_ENST00000601538.6,E | 1 | - | good | AAGGTATGCAGGGAACGGGCTGTTATTTACACGTTAGGCAGGCGGCGTCCTCCTTGTTATCTTGTTGTGTTATCTTTAGACGCTCCTCTTCCAGGGGCACCTGCTGTGTACCTGGCATGG | 120 |
| chr11 | 71113295 | 71113415 | 788855_47188552_ENST00000601538.6,E | 1 | - | good | CTCTGAGAAGTATTGACTTGTTTTCTCTCTCTTTTTTGTTTTCTGTGCCAGTTTCGATACAAGAAGCGGGTGTATAAACAAGCCAGTCTCGATGAGAAACAGTTGGCCAAGCTCCACACG | 120 |
| chr11 | 71113415 | 71113535 | 788855_47188552_ENST00000601538.6,E | 1 | - | good | AGGATGTTGTAGGCTCTGGGGTGCAAATACCAGTCATTTATTTAAAACTGGAAGTTTGCTGTGGGGTTCCCCGTCTTCTGTGACATAGCCAAGGAAAAGGCCCGTTTTCCCTCTGAACAA | 120 |
| chr11 | 71113535 | 71113655 | 788855_47188552_ENST00000601538.6,E | 1 | - | good | GGTCACCGGGAGGGAGACCTCCCAGCTTCTCACCTGCAGGATCTTCCCTTCCCCTTGCTCCTGTTTCTTTATATGGCTTTTTAGCTAAACTGCTACAGAATCTTGACAAAAGTGATGCTG | 120 |
| chr11 | 71113655 | 71113775 | 788855_47188552_ENST00000601538.6,E | 1 | - | good | GGACAATGTAGGAGAAAAACCTGGTGTTTGAAGCAGCATCCTCATTTACCAAAAATGCCTTGCCAAGGGCTGATCCATTCCCCTGGGAGGAACCATAACCACTCAGTTAGCTGGGGAGCC | 120 |
| chr11 | 71113775 | 71113895 | 788855_47188552_ENST00000601538.6,E | 1 | - | good | AGGAGGCGCTGCTGTTTGGAGGGAAAAACAGTGGCTTCAGCGATCTGGGCAGAGGTTGCAGGAGTACAGGAGGTGGGGCCTTTCCCAGCCCAGGCTTTTAGGGTGTGTTTTAACAAAGGT | 120 |
| chr11 | 71113895 | 71114015 | 788855_47188552_ENST00000601538.6,E | 1 | - | good | CTGGAGGAGGGAGGAGGTATTTTAGAGTCTCACCAAGCCCACTCCGACAATGTTGCTTACTCCGTGCATCTGAGGAAGATGTGCTCCTTGGTTCATGCAGTGATGTAATTTCCAGTTCAA | 120 |
| chr11 | 71114015 | 71114135 | 788855_47188552_ENST00000601538.6,E | 1 | - | good | TGTGAGTTGAGAGCATTAGGGCTGGGATTGGAACAGAGGTCCCAGCTCCAAACTCCAGTGACTCCTAAATGCTCAGCATCCTTCCCAGACCTTCTGCAGCATCTCCAGCCAAGCGGGGGG | 120 |
| chr11 | 71114135 | 71114255 | 788855_47188552_ENST00000601538.6,E | 1 | - | good | CAGGTCTGTGCCAGGAAGCTCACATTCAATGCTCAGGCCACCCTGGGAGGTGGGTGCTATTATCCCCATGGGGCAGATAAGGAAACTGAGATGGGAAAGTGCCCAGCCCTGAGTTAACTC | 120 |
| chr11 | 71114255 | 71114375 | 788855_47188552_ENST00000601538.6,E | 1 | - | good | CCCCGCACCATCATTTTACCCAGGGGCGGTTTTTGCTGGCTCTGAGCACATGTGTAACCGTGCAGGGTGACGGCATCAAGGGCAGTAAGGAGGGAGAGATAAAGTGGACTTGTCTAGTGC | 120 |
| chr11 | 71114375 | 71114495 | 788855_47188552_ENST00000601538.6,E | 1 | - | good | GTCTTTTGTCAAATAGCCATTTTGTTTGGCGTGGTGGAGAAAGGAGACTAGGGTTGTTTATTTGGAAGCTTTTCTGTGAGATGCGACTTTCAGTTTCAGGATGAGCTCACCCGCCACCTG | 120 |
| chr11 | 71114495 | 71114615 | 788855_47188552_ENST00000601538.6,E | 1 | - | good | GCACTTTCTCCCCAAACTCTGTGCAGCCTGCGCTTCACAGAACTTTCCGATTTCACAGCTGGTTTATTTGGCCAGTTGAAAACAAAACCCCACAAGTATCAGTCTGTGCACAAGAATTTT | 120 |
| chr11 | 71114615 | 71114735 | 788855_47188552_ENST00000601538.6,E | 1 | - | good | ATGGACGCACAGAACCTGCTGTCCTGAAATCTAGAATCAGAAGTGGAAGTTCTAAAATGGGCCTGTAGCTCCTCCTGCCTGCTCTGTTCTGGGAAATGAATGGATGCCCTTCTGACCAGC | 120 |
| chr11 | 71114735 | 71114855 | 788855_47188552_ENST00000601538.6,E | 1 | - | good | TGGAATTGAATTCCCTGAGACTTTTAAGCTTAGCCACATGATCATGAGCCTGAGAAGGCACCTGGCATCATTCTCAGTGGCTTCTGAAATCTGGACCAGAGGTGTGATTGGTCCCGGGGC | 120 |
| chr11 | 71114855 | 71114975 | 788855_47188552_ENST00000601538.6,E | 1 | - | good | CTATGGGCTATAAATGTTTGTGACTCTGGAATAAACACTCCCTCCTTGAGTCCTTTACCCTCCTGCCCCAGAGATGCTGTGGGCCCAAGCGGATTGTTTTATGTCTGGAAAGGGGCCCCG | 120 |
| chr11 | 71114975 | 71115095 | 788855_47188552_ENST00000601538.6,E | 1 | - | good | GGGCATCAGTCTGGGCTCTTTGCCTCAGATAATTTCAGCACAATATGGTGGGCTGCTTATGAAATGTTGGAGAAATAGCAAGAAAAAGGGTCTGTGGTCCTGAGCCGGTGTCCTCTGTAG | 120 |
| chr11 | 71115455 | 71115575 | 788855_47188552_ENST00000601538.6,E | 1 | - | good | TGAGCCTGGCCATCCACAAGTCCCCAAATTGACTTTGCAGCTGCTGCTCTGGTTTCATTGTCATGAAATTGTTGCTGATACTTTGTTTTTTGTTTTGTTTTGTTTTTGAGACCGAGTCTC | 120 |
| chr11 | 71115575 | 71115695 | 788855_47188552_ENST00000601538.6,E | 1 | - | good | AGCCACCCATGCTCTCTCAGCCCCGGATTCCCATTCATGACATTGAGTGAATGTTAGTAGTGATGTCCGGGAGTGGTCGTGAGGCCCAGAGGAAGTGTTCATGGAGCAGTGTGTGCGTTG | 120 |
| chr11 | 71115695 | 71115815 | 788855_47188552_ENST00000601538.6,E | 1 | - | good | CAGTTCATTTCAGGGGCTGATCATTTCCATAACGGCAGGAAAACTGGGCGAGAAGACCAGAGAATTTGGGAGTCTCCTCTACATCTGGCCCCAGTACCTGCAGCTGTGCAACCTCGAGCC | 120 |
| chr11 | 71115815 | 71115935 | 788855_47188552_ENST00000601538.6,E | 1 | - | good | AATTGTCATTGAGGGCCAACTAAGGACCACACACTTTTCTGGCACAGAAAAGTCAAGGTTCTGATCCCACGGACCTTACACTCCAGTGGGAGGTGATGGACCAGGAAGAGTAAATCCATG | 120 |
| chr11 | 71115935 | 71116055 | 788855_47188552_ENST00000601538.6,E | 1 | - | good | AGGGGTTTCAGTGTAGAAAAAGATGAGATGCCACTTGGAAGTTGTGTGGTTTAGAGCAAATCACTCAGCTGTCCTGGCCTCAGTTTGCTTATCTGTAAAATGGGTATCCATTGGGAAAAA | 120 |
| chr11 | 71116055 | 71116175 | 788855_47188552_ENST00000601538.6,E | 1 | - | good | TGTCATGGACGCCCACCCAGGGCTAAGGTGATGCCGCGTCTGCACGTTTCTCAGTTCCGTTGTGACACTGAGGCTGAGGTTATTGGGATCCCCAGTGTGGACCATGTGTCAGATTCAGGG | 120 |
| chr11 | 71116175 | 71116295 | 788855_47188552_ENST00000601538.6,E | 1 | - | good | ATTGCACTCATTTTAGCTTAGTCAACAGCAAGAGGCTGCATGTGTGTAATTGCAGCTGGCACTTGAGCAACCTAAGTGGGTTCTCTACTCAGTGACTAGGGATTCCTCTGAAGCCCAAGA | 120 |
| chr11 | 71116295 | 71116415 | 788855_47188552_ENST00000601538.6,E | 1 | - | good | CTACGCCCCAAGTTTACAGGCATTGTTTCCTGCAGTGTATACGTGTTGGGTTACGTCAGTTAAGAGAAGCCAGAAACGGATCATTTCTACTCCTGCACAACATGTTGGCATCATTCAGCG | 120 |
| chr11 | 71116415 | 71116535 | 788855_47188552_ENST00000601538.6,E | 1 | - | good | TGTCAGCAGCCACAAGCTCCTGGGCCTCCCTCCAGAAGTGGCTGATCCGTAAGACGGGGCAGCCCTACACATTTCCACGTGATGCACGACCCCTGTCCCTACCTCTCTCACCCAGGCTTG | 120 |
| chr11 | 71116535 | 71116655 | 788855_47188552_ENST00000601538.6,E | 1 | - | good | CCATTCTCCAATTCATCTCTGAGTCCTTGAGTAACAGGGCGGCTGTGACCATCCAGCTGTTCTCAACAGCACAGCTTCACTTCTGCTTCCTGCCCGGGTGACCGGGGCTCTTTGAGCTGC | 120 |
| chr11 | 71116655 | 71116775 | 788855_47188552_ENST00000601538.6,E | 1 | - | good | GGCAAAGCAGGAGCAGATGTTAGGTGGCAAGAGGGGGTGTGAGAGCGAGCAGGGGAGGAGTAGTTCCTGGCTCCCCCGTGAACTAACAGAGCGAGAATTGACTGACTACCATGGGGAGGG | 120 |
| chr11 | 71140655 | 71140775 | 788855_47188552_ENST00000601538.6,E | 1 | - | good | ACTCTACACTTTTGGGCGAGGGTCTGAAGATCTGAGCCTCGGGGTTCCACCTGTCGGGAACAAACACCACTTGTGAAGTGAGCCATTGCGGGTTGGCGTTCAGCCCGGGGAGCAGCCACA | 120 |
| chr11 | 71140775 | 71140895 | 788855_47188552_ENST00000601538.6,E | 1 | - | good | GATTTGCACCTGCAAGTTGAATTCCCGGCAGGAAGGCAGCCAGCAGGCAGCGCTGCGTGGGAAGCAGCCCCGGCCCCAGTCGTCCTGGCATGAGTCAGAAGCACGTCTCCCTCGCCATTC | 120 |
| chr11 | 71140895 | 71141015 | 788855_47188552_ENST00000601538.6,E | 1 | - | good | GAGCCAGTGAGGGGCAGAGCCACTGTCCTGGGGACACCCAGCCTTGGGCCTTGAAACCTGCTGGATGGAAGCCCTGGGAGGCTGGCGTTGGGTGTGCCACCAACTGCTTGGTACTGGGCT | 120 |
| chr11 | 71141015 | 71141135 | 788855_47188552_ENST00000601538.6,E | 1 | - | good | CAGGTGGAGGAAGGCGTAGGACGCCAGCCAGCCCCGGGAGCCCCACTGCTCCCTCCTACTCATGGGGAGGGTTTATGCTTAATCCCACTTTACAGATGGGAACAGGAGGGTCAGACTTCA | 120 |
| chr11 | 71141135 | 71141255 | 788855_47188552_ENST00000601538.6,E | 1 | - | good | CTCCCAAACTGCTGTCAGCCTTGAGGTAGACAGCACTGTGAGCATCTTTGTTGAAGGATAACACACCCGTCCACAGGGCATGTGTCCCGTACACTCAAACGGAACGCAGCATGTAACCTG | 120 |
| chr11 | 71141615 | 71141735 | 788855_47188552_ENST00000601538.6,E | 1 | - | good | AGTGGAGTCTACGTGCAGGGCCTGGAATCCCTGTCTTTCCTGAGACATTGTATTAGCCCATTCTCATGCTGGTAATAACGACGCTGCTAACTACCGGAGACTGAGTAATTTGTAATAAAA | 120 |
| chr11 | 71141735 | 71141855 | 788855_47188552_ENST00000601538.6,E | 1 | - | good | CAGTAGTTTGGGCTTTCTGTTGGTTCGCATAGTGCATGGGTGAATTCCGTTTGGCTCCATCATGAAGTTAGATTTGGCCATCTGTTTGCCTTTGGAGAGGAGCAGGCCCAGGGCTGGGAG | 120 |
| chr11 | 71141855 | 71141975 | 788855_47188552_ENST00000601538.6,E | 1 | - | good | TGCAGCTCAGAGACCGTTGGAGGTTTGCATGCCGTCTTTCTTGCTGTGGTTCATTTTGTGTCTGTGCGGGGTCCCTGTGGCTTCCCTGACATTTCTGCCTGTCTCGTCTAGGTTGGCTGT | 120 |
| chr11 | 71141975 | 71142095 | 788855_47188552_ENST00000601538.6,E | 1 | - | good | TTTCTTTTTGAAAATACTCTCTTGAGTTGTTTGTTTGGTAGGTTTTCTGTCTTATTTTTGTCCTCCCTGGCATCAAGCATTCTCGGGGGCTTTTCTTCTTTTAATTGTGGTGAGTTTTCA | 120 |
| chr11 | 71142095 | 71142215 | 788855_47188552_ENST00000601538.6,E | 1 | - | good | GCCATGATTTTCTTGAGCTTTTAATTGTTGTGTTCTCAGTTAGACCCGTTCTAATTTTTGTGATTTCTAATTTACTGTAGTCATGCTCCTTTGAGTTTTTGTTTCCATAGCCCTGCACTC | 120 |
| chr11 | 71142575 | 71142695 | 788855_47188552_ENST00000601538.6,E | 1 | - | good | CATTTTAAGAATGTTTTCCTGTATTTTGTTACTTTTTCTGTGTTATCGGTTGGATCCTTTATTTCAGGGATAGTAATAATTCCTATACTGGATTTTCCTTGTTCATATCTGTTATCTTTT | 120 |
| chr11 | 71142695 | 71142815 | 788855_47188552_ENST00000601538.6,E | 1 | - | good | TTTGGACTAGATTCCTAAAGAAGTGGCTTCCGTTGCTACACCAGGAAGTGTTTGGTGTTGGGCATTCTTGGTTTTTCCTAGAAAAACTTTCTCTCTGCAGATTCTGTTTAAGGACATTTT | 120 |
| chr11 | 71142815 | 71142935 | 788855_47188552_ENST00000601538.6,E | 1 | - | good | GTCACAATTTCCCGTCTGTGGGGATGTTGTTTGATTGTCTTCCTGCCCTGAACATTGCTTTGAATACATTTGAGGCCAACAGATTTATTTTTTCCATCTCATAGGTGACTTTTTTTTTT | 120 |

| | | | | | | | | |
|---|---|---|---|---|---|---|---|---|
| chr11 | 71143415 | 71143535 | 788855_47188552_ENST00000601538.6,E | 1 | - | good | ATTCCATGGGCTGCAGTCAGACGCTTGTTTACAAGATGAGAGCTGAGACAAGAAGTCGGACCACCACCTTGATCAACCTCCCGTGGGAACCCGCGCATCGGGCAGGCTGAAACCTTTGGC | 120 |
| chr11 | 71143895 | 71144015 | 788855_47188552_ENST00000601538.6,E | 1 | - | good | TTTTAATTGGAGGGAATTCACTGTTCAGTGGGTTTTTTTTAGCAAGAGTTCCTAAGAGCAGTGCTTTCTCGGTGAATGGATGGTCGATGCTCGTTATCGGTGGTACTTGTGTTCTAAAAA | 120 |
| chr11 | 71144015 | 71144135 | 788855_47188552_ENST00000601538.6,E | 1 | - | good | ACTATCCACTGTGCCTCCAGGGCCCGGAGCCACTCTGTCTCTCTTGCCGGGGTCACTGCCATCATTTGCCTTTGCAATGCCCTTTTCATTTGCCAGATCCTTTTCTGGTTGTTATTTTGC | 120 |
| chr11 | 71144135 | 71144255 | 788855_47188552_ENST00000601538.6,E | 1 | - | good | ATGAGTAGCTAGTTATGTGTGCATGGGTGCACGCATGCGCACACTTTCCCTCCCCCACCTCCTGAGCCTTGTTGGGTGTGGACAGTGACTTGTTTCACACTGTCACTGTCCCTCCCACTT | 120 |
| chr11 | 71144255 | 71144375 | 788855_47188552_ENST00000601538.6,E | 1 | - | good | CTTGTAAAGTGCATCCATCACCTTCCCTGGTTTCATGAGGAGGTTGCACCTGACTGCAGAGTAGGCTGGGAAACGCAGTGGAGGTCTGAGCCCAGGAAGAAAAGGGCATCAGTTTGGTGA | 120 |
| chr11 | 71144375 | 71144495 | 788855_47188552_ENST00000601538.6,E | 1 | - | good | ATACAGACATTCAGGTACCCAGGCTCTTTCCATTAGATAGCTGGACCATTCCCTGGAGCTCTGGAATCCTCCACGGGGTCCTCTGTATCTGCCCAGAAGATGAGGGAGGAAGAGACAGAA | 120 |
| chr11 | 71144495 | 71144615 | 788855_47188552_ENST00000601538.6,E | 1 | - | good | CTCATTGCTATAATAAAAGGCCTCCAAATGTCAGGCACTTAACACATAGATGCTTTATTTTTTTTCTCTCTTCTCAGTCCACCTGGGTTGGCAGGATGGACAGGAGGGAAGTTCTGATCC | 120 |
| chr11 | 71144615 | 71144735 | 788855_47188552_ENST00000601538.6,E | 1 | - | good | GTGAAGGTAAAAGAAGTGATTATGAAACACTGGAAGGCTGGAATTTTTTTTCTTTCTTGTTACAGTTTTTTTTCCCTTTTCACTGTCCCCTCTTCCTATATCATTTAGGGGTAAACTAAG | 120 |
| chr11 | 71144735 | 71144855 | 788855_47188552_ENST00000601538.6,E | 1 | - | good | TTAAGAAATGAGCATCATCAATGTTATTTTATTTGAATATAAGAGGAGAAAAAGATGGTGAAGGAAATGCTTCTTTTATTTATTTATTTTTGTCACAAGAGACTTTTTCCAGCTAAATCT | 120 |
| chr11 | 71144855 | 71144975 | 788855_47188552_ENST00000601538.6,E | 1 | - | good | CACATCAATACTTGGCAGAGCCCTGTTCCGACACGTCCCTCTGCAGGATTACAGGCAGTGGGTGGAAGTGCATATGGATAGAAATACCTTGGTAAGCGTGGGCTCAAACCACACACGCTC | 120 |
| chr11 | 71144975 | 71145095 | 788855_47188552_ENST00000601538.6,E | 1 | - | good | TGGTCTTTTGGAGTTCCTTCTCCTATATTATTTAATGCCTCTTATAAAAGCTATAAATGATTGTTATGAAAAGTTCAAGTAATGCAGGAAAGCACAAATGCCAGACTTAGCAATGGTGAT | 120 |
| chr11 | 71145095 | 71145215 | 788855_47188552_ENST00000601538.6,E | 1 | - | good | AGATTAGACACCCGTTGTGGGGGAGGCTCAGGGTGGCCTGATGCCCTAGAGCTAATCTGGCCATGCCACTGGCCCCAGGGGCTTCCCTGGTAGTACAAAGGCTGTTCAGGAGGATAGGCG | 120 |
| chr11 | 71145455 | 71145575 | 788855_47188552_ENST00000601538.6,E | 1 | - | good | GCAGCTTGCGGTCCAGCAGGCACCCTCTGCTCAGCCCACCCCCAGCCCTGCTCCCTCTGAGAGAGACGCTAACTGGGCTGAGGGGACTAACGTGCTGACACGTGCCCTCTAGGGATGAAC | 120 |
| chr11 | 71145575 | 71145695 | 788855_47188552_ENST00000601538.6,E | 1 | - | good | CCATCCAGGGTCCCAGCTCCTGGTGCTGTGGAGGAGCTAAGAGATGCCCTGGGGTTAGCACCTCTCAGGGCTTGGTCCCCAGCGAGGGCTAAATGTGTGTCTCCTCTTCCCCCAGTGCCA | 120 |
| chr11 | 71145695 | 71145815 | 788855_47188552_ENST00000601538.6,E | 1 | - | good | ACCTCTATGAGTACAGTCTCCCGGCCCCAGCCACGATGGGGAAGGGGGGCCCAAGTGCTGGAGATCGGGGAAGCATCATGCTGCCTACTGGCTGAGTGTTGCTGGTGCCTCACTTGGTGC | 120 |
| chr11 | 71145815 | 71145935 | 788855_47188552_ENST00000601538.6,E | 1 | - | good | TGGGCTGGGAGGACCCTTCTCATTTTAGCTTTGGGTTAAAGGAATGGGGAGAGGTGGGCTGGTGGTGTAGGCTGCTGCAGGTGCTTGAGGGGGTAAGCCCAGTGCTCTGCATCCACCCCG | 120 |
| chr11 | 71145935 | 71146055 | 788855_47188552_ENST00000601538.6,E | 1 | - | good | TACGAAGCCCATTTGCTGTGGAGTAGACATGGTAGACATCCAAGAGTTAGGACTCAGAGGGGCATCTGGGTGAGGCTCGTCACAGTCTCCATCTGTGCTTTCTAGAAGGGTCTCCCTCAG | 120 |
| chr11 | 71146055 | 71146175 | 788855_47188552_ENST00000601538.6,E | 1 | - | good | GCGTGGAGCAGACATAATGGTTGCAAACAGGCGCCTAGTAGGGTTGGCCTGCGAACATTTCACACGCCGTCTGTCCACCTGGGGAATGTGGGTGGGGGTGGCAGTAGTAGGGGCAGTTGG | 120 |
| chr11 | 71146175 | 71146295 | 788855_47188552_ENST00000601538.6,E | 1 | - | good | CTTTTGCCTGAATCCACCAGAGCAAGCTGGGGGCCAGTCGCCCAGCCACGTTCCCTCTTGCTCACGGCGGGGACTGTGCCACATGTGGCCTCAAATCGGAGATGTGGGACAGAGCTGGAA | 120 |
| chr11 | 71146415 | 71146535 | 788855_47188552_ENST00000601538.6,E | 1 | - | good | CTGGCTGTGTCCTGGATGTCTGTCCTTATGCCTGGGGGTCCCTTTTGTTGGATTTGCCTTCCCCGAGGAGGTTGGCCCCTGGGAACTGGGGCTGTCTGTAAAGGGGAGCTTCGTGTGCAG | 120 |
| chr11 | 71146535 | 71146655 | 788855_47188552_ENST00000601538.6,E | 1 | - | good | GGGGTACAGCCCTGAGCACTCCTTCCCGCGTGGCTGCTGTCCGCCTCCTCCTGTCCCTCCAGCCCTGGGCCTAATGTCATTCCTGTGCCGCCCAGAGCCCTCAGGCTGCCCTGGCCCACG | 120 |
| chr11 | 71146655 | 71146775 | 788855_47188552_ENST00000601538.6,E | 1 | - | good | TGCACCCGCAACTGGCTTCTAGCACCAAGTGCAGGAGGCTCTCAGGCCTGGCCCTGCCTCTTCCTGGTGCCGGACTCTGCTGTGCCAGAGGCCCAGCATCCCATGCTCCCCCACTTCCAG | 120 |
| chr11 | 71146775 | 71146895 | 788855_47188552_ENST00000601538.6,E | 1 | - | good | TGTCCCCAACTCCATCCACAAACATTGACTGGGCACCTACTGTGTGCCAGACCTGAAGCAAGAGTCCCAGAGGTGGAGAGGAGTGTTTCACAGCCTCATAGAGAAAACCCGGGCGGCAGG | 120 |
| chr11 | 71146895 | 71147015 | 788855_47188552_ENST00000601538.6,E | 1 | - | good | CTCTGGGCACACCGTGCTCCTTCCTCGCTCCATACTTCTGCCCCCGTCCCCCCAGGCTCCCTGCCGTGCCCGCCTCCCCTCCCTGCTGCCGCTGGCCTCACATCATGCTGTTGCTGTCAT | 120 |
| chr11 | 71147015 | 71147135 | 788855_47188552_ENST00000601538.6,E | 1 | - | good | TGACCTGCAGCAGACGGTGAGCCCCGTGGTCTGCCCTTTGGTTCACATCTGGACAATGTCACCTGTGGCTTGAACGCCAGAGGCCTGCTCTGGTCCTCCCCGGACCCACGGTCCTGACTG | 120 |
| chr11 | 71147135 | 71147255 | 788855_47188552_ENST00000601538.6,E | 1 | - | good | GTCAGACAGCTCCAAAGAAGAGACCATCTATGACACGATCCGGGCCACTGCGGAGAAGCCGGGCGGTGCCAGGACGGAGGAGAGCCAGGGCAACACGCTGGTGATCCGCGTGGTCATCCA | 120 |
| chr11 | 71147255 | 71147375 | 788855_47188552_ENST00000601538.6,E | 1 | - | good | TCTCCCGTTCTTTGTCTTCCTTCTCCCCACCTCGAAGACACAGGCAGCCATGCCGCGCAGCCCAACATCCAGCGAGGACGAGATGGCCCAGAGCTTCTCCGACTACTCCGTGGGGTCGGA | 120 |
| chr11 | 71147375 | 71147495 | 788855_47188552_ENST00000601538.6,E | 1 | - | good | TGTGCATACCTGGGGCCCTCGGCTGGTGGTCACACCAACAAACACAGACTGTGAAACCGAGCCCACGTGTGTTCCAGCGCCAGTGTCCACCGTGCCCCTGGGTTTTCTTTCATTTTCACA | 120 |
| chr11 | 71147495 | 71147615 | 788855_47188552_ENST00000601538.6,E | 1 | - | good | TTCACAGCGCTGGTACCAGAGTTTGCAGGGTTTTCCTGGGTCCTGGCTGCCGCCTCTGGCTCAAGCTCCAGGGAGGAGCAGGCGGTGCCGACTTGTGTCCGCCCTCTGGGATCTGTACAT | 120 |
| chr11 | 71147615 | 71147735 | 788855_47188552_ENST00000601538.6,E | 1 | - | good | CAAAAGGCAGCCTTTTTTCTTACCTTGGGTAGCTCAGACAGCCCCAGTGGCCCTCTGCATCATCTTTGTAAGCTGTGGTCATCCCCGTGGCTAGCTCAGGCTCCTTTTGAGTGCCGGGGT | 120 |
| chr11 | 71147735 | 71147855 | 788855_47188552_ENST00000601538.6,E | 1 | - | good | ACGGTACCTATCATTGCTCTGTGTTTGAAGCAGAGAGGGTTATTATAGCGTGAACTTGCTGCCATTCTGACCTGAAGCCCATGTTTCTTCCTCTCCTGAAGCTCTGTCCTCCAAGGAAAA | 120 |
| chr11 | 71147855 | 71147975 | 788855_47188552_ENST00000601538.6,E | 1 | - | good | CTTGGGGACCCCCAAGTCTGATGGCTGCAGTCTGCTCATGACTCTTGGTTCCACTCCAGTCCTGGTCTGGGAGATTCTCTCATCTGTGGGCCTGGCCCTGGCCTTTTGTTCTGTTTTTAC | 120 |
| chr11 | 71147975 | 71148095 | 788855_47188552_ENST00000601538.6,E | 1 | - | good | AAAAAAAAAAACAAGAAGCCTTGGTCTAAGGTGGCCATTGGCAACCTGGCCTGACTCTTTGTGGGGTAATTTTTATTGCACTTGACCTTGAGGAAGCTGCACTCTCAGCTGCCCCTGCCT | 120 |
| chr11 | 71148455 | 71148575 | 788855_47188552_ENST00000601538.6,E | 1 | - | good | CACCCAGAAGTGGAATTGCTGGTTCAGCATGGCCCCTCCTCCCCCAGTCCCTGCCATCCCTGGGTCAAGCCAACTTTTATCCAACCCTTTCTGTACTCCTGCCCTGCAGAGGTGGGGGAT | 120 |
| chr11 | 71148575 | 71148695 | 788855_47188552_ENST00000601538.6,E | 1 | - | good | GCTTGATCAAGGGGTCCCTAAACAATCAGGGTGACCATGCATTTCCACATGCTGCTTTGGCAGGGAACAATCTTCTATGCCTTTCCATATGCACGTGGCCTCGTCCTTCCCCACAGAACG | 120 |
| chr11 | 71165975 | 71166095 | 788855_47188552_ENST00000601538.6,E | 1 | - | good | AGCCTCCTTCAAGGGAAGGGCTGACACCAGGCGAGCAGTAGTGTGTGCTTGTTGGTGTCGGTCCAGGCTGGGGGGCGTGTGGACACTCACATTCGTCTGTCTGTGCAGCACGCGGGAGGT | 120 |
| chr11 | 71166095 | 71166215 | 788855_47188552_ENST00000601538.6,E | 1 | - | good | CGTGAGAGGGTGAAGCGAGGTGACCCAGTACCTGGACGGCACCTGCTTTGGTGCTTGGCCTGTGTAGTTTTGATCTGTGGAACCCCCAGTCTGTCCTGTCATCCCTGCCTTTTTGCCAGG | 120 |
| chr11 | 71166215 | 71166335 | 788855_47188552_ENST00000601538.6,E | 1 | - | good | ACCTAGATCCTTCCAGTGCTCAAATTCTCTTCCCACTCAGAGGTCCTGTCTGCCCGTCTCTGGCGTCTTTGTGGCACAGCCCACAGACTATCTGTTGCAGACCTTTTGTTCCAGGACATC | 120 |
| chr11 | 71166335 | 71166455 | 788855_47188552_ENST00000601538.6,E | 1 | - | good | TAAACCAAGAAAAGTCATGTCTTGTAATACCGGGGTGTGCTGCATTGCTGTATGAGAATTCACTGCATTCTTGCTGTCTGGAAAGAAGTAGCTGTCCGGTTAATTTCTTGGCTTGTCCTT | 120 |
| chr11 | 71168855 | 71168975 | 788855_47188552_ENST00000601538.6,E | 1 | - | good | CCTGATTTCATAAATCACTGTTTGCTCAAACTAACTCTTTATGCATCTTATTCTGCCTCAGTTGACCTTTTATCAGTGTTGTCAAATGGTCCCTAGATGCTACCTGCCACTGTTGTGAAT | 120 |
| chr11 | 71169095 | 71169215 | 788855_47188552_ENST00000601538.6,E | 1 | - | good | ACTTCAACTAGGGACTTTCCAATGGGATAAGCCAGTACGGCGACCACATCATTGTAACCAACCAAGTGTCTCCTTTGCCTTGCGTCTGTGTTCACCCCATAAAGCTTTCCCCTTCCATCC | 120 |
| chr11 | 71169335 | 71169455 | 788855_47188552_ENST00000601538.6,E | 1 | - | good | CATTCTGTGTAGACTTATGTACATACATATTTTGGAGTCACATGAATTGTTTTATATTGTTAAATTAAAAAGGAAAACAGGCCTAAAGAATCCTAAGCAAAGCCAGTCAGACTTCATAAG | 120 |
| chr11 | 71169815 | 71169935 | 788855_47188552_ENST00000601538.6,E | 1 | - | good | TGGCCTCAAAATTTTAATCTAAATGATTTCTATAAAATGGAAACATTTTAATTTAGTTTTCTGAAAATTTACTCTCACTAAATATTGTTATGAATATAAAACAAATCACAGTTCTACCGC | 120 |
| chr11 | 71170055 | 71170175 | 788855_47188552_ENST00000601538.6,E | 1 | - | good | AAAATTTGGATGTATTTTGATAAAAGTATTAATTGAAATAAATTTAGCATTAAACATTTTTCATGATTCCAAAAATGATATCTGCTAAAATATTCCAAATTAACTGCTCAAAGGCAAAAT | 120 |
| chr11 | 71170175 | 71170295 | 788855_47188552_ENST00000601538.6,E | 1 | - | good | TAGTCTAGCATTTTGCACAATCTGTGTTCCATTGAGAGTAATATTCTAAATGATTAGAAATTTAATATATTCAATGTATACAAAATTTGGATGTCTATGAAAATGGACTAATACAGAAAA | 120 |
| chr11 | 71170295 | 71170415 | 788855_47188552_ENST00000601538.6,E | 1 | - | good | CTCATTGATGCATAGTGACATTTGGATAAGTGATAAATATAAATGTCATTCATTATTATATATGCCAGAAAATATATTTGTCCTGCTAGTCTGTTCAGCAAAATTACCACTACGTTGTTC | 120 |
| chr11 | 71170415 | 71170535 | 788855_47188552_ENST00000601538.6,E | 1 | - | good | AGACATCCCTTCATGATGGCAAAATGTAAATCATGTGGGAATTGACTATTTCATGAGATTGAAAGCTGGTGATACACCAGACACGGCTGAGTTTAATTGTTCTGTTACACACTTAGGCAT | 120 |
| chr11 | 71170535 | 71170655 | 788855_47188552_ENST00000601538.6,E | 1 | - | good | TCCCTACTATTTGGGGTCTGGGCAAGGAGCACCAGCGGAAGCCCACATGACGTGTTTCCTATTTGAAAGTTAGGTACACACCCAGCTAGCACATTAGATCATAGTTTCTTCTGCTTTCAC | 120 |
| chr11 | 71170655 | 71170775 | 788855_47188552_ENST00000601538.6,E | 1 | - | good | CACCTGGGAGCGTCCCAGCTCCTTGCCTCTTTGGAGACCCCTTTGAGCGTGGGCTCTGCCGTGACCTTGGAGGTCGGGAACAGGCAACACATCCCTCAGTGCCCACCTCTTGCCAGCCGG | 120 |
| chr11 | 71170775 | 71170895 | 788855_47188552_ENST00000601538.6,E | 1 | - | good | CTGCCAGCGACATCAGGAATCCCTGGAGGACGAGGACCATCGCATTTCCTAAAGCTCTCGTGATTTGATATGCATGAAGGTCTGACCCTTGTCCTGGGAGAAATAGCTGCTTCCTCCTGG | 120 |
| chr11 | 71171015 | 71171135 | 788855_47188552_ENST00000601538.6,E | 1 | - | good | TAGAAGAAGAGAAAACAGAAGTCTCACTCTGTCCCCGCGTGTGCAAAAAGACGAGGCCGTGCTGAGAGCAGCTAGAAGTTGGCCGTCTACGAGCCAGGAAGAACTCTCACTGGGAGCCTG | 120 |
| chr11 | 71171135 | 71171255 | 788855_47188552_ENST00000601538.6,E | 1 | - | good | TTTTCAGCCTCTGTCATCGCGGGGACTCTGTCAGCCCCCTGACAAGCCCACCCGCATGTGTCTGTATACATTTCCTGGTGATGCTGTCTCCTTGGAGAATCCGGACCAATACAGTGCACC | 120 |
| chr11 | 71171375 | 71171495 | 788855_47188552_ENST00000601538.6,E | 1 | - | good | CAGTAAGATGTAAAAGATGTCACCAATTTTCAACTTTGGCTGAGTCCCTGTTTCACTTGGGGGCATTTTAAGCTGAATGATCCCCAGATCCCCAGGCAGCCGCTTCCCCTCGGGTGCTCT | 120 |
| chr11 | 71171495 | 71171615 | 788855_47188552_ENST00000601538.6,E | 1 | - | good | GTCCAGAGAGAACAATGAAGAGAAGGGGCGAGGGCATGGAGCCGCTGGGCTGGACACACTCATCCTCTCTTGTCTTTGGACAACTCCAAAGCCTTTGGACTTCAGGACTTGACCAGCGGC | 120 |
| chr11 | 71171615 | 71171735 | 788855_47188552_ENST00000601538.6,E | 1 | - | good | TAAAGTGTTATCGTGGGCGTGTCTGTGTGGGGTTTCTGGAGGGGATTAGCACATAAGTCTGAGGGGACTGGAGGCAGGGATCTGCCCTGGGTATGGGTGGGCCCATCCAATATGCTGGGG | 120 |
| chr11 | 71171735 | 71171855 | 788855_47188552_ENST00000601538.6,E | 1 | - | good | GCAGTGATCCTGGTTAAACGAGGCCACAGGGATTAATCTGATTAATCCGATGGGATTAGTGCCCTGAGGGGTGATCTTTGGTGTCCTTTCGACTGGACTAAGGAACACCTGGGAACCCGG | 120 |
| chr11 | 71171855 | 71171975 | 788855_47188552_ENST00000601538.6,E | 1 | - | good | AGGGTCTCAGGACCAGAGCTGGGCCTTCCCAGGCGTGTGGACTAGATTGTGTCCCATCAAGATTCCTCCGTGGAAGCCGTGACCCCCATCGCAACTGTCTTTGAGAATGGGGCCTCTAAG | 120 |
| chr11 | 71171975 | 71172095 | 788855_47188552_ENST00000601538.6,E | 1 | - | good | GGATGGGGCTGAGTGATGAAGTCTGCCACCTTGTTCAACCCTTGGGCAGCCATGGGGGACTGCCCAAGGGGACATTCCCATCACCCTACTGCTGCTGCCCCAAGGAGGCCTCCCAGGCCC | 120 |
| chr11 | 71172095 | 71172215 | 788855_47188552_ENST00000601538.6,E | 1 | - | good | GGAAGAGAGCTGGCAGGACGGTGGCATGTTTTGAGGGCTTCCCTTGTGCCAGGTGGGAATCACTGTGTTGGAAATCCTAGCAATTGTTTGCTCCTTTTGGTTACATGGTCAGTGTGCCAC | 120 |
| chr11 | 71172215 | 71172335 | 788855_47188552_ENST00000601538.6,E | 1 | - | good | CCGGCGGCTGTTTCTTTTAAGTAAATAAAGTAACAAGAGAAAGCATGTGGCCCACCGTGGCCCCTGCATTCTTGAGGTTGACGCCTTTCTTGGACGCTCCTTCTCCTTGTGTGAGGTCAG | 120 |
| chr11 | 71172695 | 71172815 | 788855_47188552_ENST00000601538.6,E | 1 | - | good | TGCAGGAGGCAGGCATTGCCCTGGCGTTAACTATGCAGGCTTCCTGGGTGGCAGTCTGACTGTTCTCTCATGGGGGAGTGCCTTTAGGGGGTACATGTGCCCTGAGAGTTATTCCCCTCT | 120 |
| chr11 | 71172815 | 71172935 | 788855_47188552_ENST00000601538.6,E | 1 | - | good | TTTGAAGGAGCACGAATTTGCTGTGCTCGTGCATTTTTGATCACGTTCTCACATGGGACTCAGTCACTGTGTGTGTACCTGCCATTGCCTGCCCTGGCGTGCTGGGCTAGACCATGCACC | 120 |

| | | | | | | | | |
|---|---|---|---|---|---|---|---|---|
| chr11 | 71172935 | 71173055 | 788855_47188552_ENST00000601538.6,E | 1 | - | good | AAGAGACGACAGGGCACAGTGACAGGGTTGAGTGCTGGCTGGGATGCCCACTTCTCAGCCTCCAGCAGCCCAATGGCCCCGGAGGCCAGAGCACTTTGAAATGCAGCCCACGAGAGTGTC | 120 |
| chr11 | 71173055 | 71173175 | 788855_47188552_ENST00000601538.6,E | 1 | - | good | ATAGTCACAAAAGACCAGTAGCTTGAAAAAATAAATGTAATAGATTTATAAGACACTTCATCAAACAAGGCGTGATTTATAAGACACTTCATCAAACAAGGCGTTCTGCCTTTCCCTGCT | 120 |
| chr11 | 71173175 | 71173295 | 788855_47188552_ENST00000601538.6,E | 1 | - | good | CCAGCCAGAAGGAGAAACCTATACTGGAATCAATTTCCCTCTTCCGTTTCATCTTGTCCTCCATGAGGGGTGAGACAGATTTTCCTTTTATGGTTATTGACTCCAATAGTCCAGTATTTT | 120 |
| chr11 | 71173295 | 71173415 | 788855_47188552_ENST00000601538.6,E | 1 | - | good | GTGGTGCCAATCCTGAGGCCAGGCCAGCAGCCACCCTGCCTGGAGGCACCCAGGAGCTTCATCCTTTACATCTGCTCCAAAAGCAGGCAGGGTCACAGCAGACGCCTTCCAGCAGTGATC | 120 |
| chr11 | 71173415 | 71173535 | 788855_47188552_ENST00000601538.6,E | 1 | - | good | GTTTCCGAAGGGGACATTCGCCTGGTTCATCAAAGAAGAGGGTTCAAAACAGTATTAGCTTTGTGTGCTGGACAGTCAGGATATTTCCAGCAGGGTGAACTCACACCTGGACCACATGCA | 120 |
| chr11 | 71173535 | 71173655 | 788855_47188552_ENST00000601538.6,E | 1 | - | good | GACACTCTTCACGCCAGTGCACTTGGGTAGGTTTCAGCATGTTTTGGAATAGTTGACTCTTTGGGCAGAAGTTAAAATTGTTGCAGTGGAAGTCCTCCTAGCTTGTGGATGTGAAAGAAT | 120 |
| chr11 | 71174015 | 71174135 | 788855_47188552_ENST00000601538.6,E | 1 | - | good | GACTTGAAGCGTACATATCATCCAAAGAGAAATACATCGACAGCAGTTTTTAATGGTGCCACTTCCATAAGTGTGAAGCCTCGATGTATTGGGCTTTCATGGCTGCTGAAACAGATGGAA | 120 |
| chr11 | 71174135 | 71174255 | 788855_47188552_ENST00000601538.6,E | 1 | - | good | GTGTTCTTAATGACTGTTTCCAGTCACAGATACTGTGTTAAATCAGTGGCTTCTAAGCACCTTTGTTAATGGCAATGACTACCATGATTGTTTCCTCCTTCTGCAAAGATATCAGGATGA | 120 |
| chr11 | 71174855 | 71174975 | 788855_47188552_ENST00000601538.6,E | 1 | - | good | GAAGAAATCATGAACACAAAGCTGGGTATGTTCACTGTATCCAATCTACTCCCTGAGGGAGCCCCAGCATTGGAATCTAATCCTGTGTGCCCCGCACCAAAGCTTGTGTTCTTTTTTTTC | 120 |
| chr11 | 71174975 | 71175095 | 788855_47188552_ENST00000601538.6,E | 1 | - | good | ACATATGCGACCGTGTGTGTGTTTTGTAGCTGCAGAAAGGTAACCGGGGAAGGCTTATCTGAGAGCATGACATGTAAACTGGGAGAGGAGGAGGGGGAGACAAGGAGAGAATTCCAGGCA | 120 |
| chr11 | 71175095 | 71175215 | 788855_47188552_ENST00000601538.6,E | 1 | - | good | TGGACTAGGTGCCGAGGAAGCACTGGCGAAGACGATGGGTTTTGTCTCTTGCCCTCCTGGGAGCGACACACAAAATAGATAAGGCACAGATTTAACAAAATCCCTTGCTCTGGTAAATGC | 120 |
| chr11 | 71175215 | 71175335 | 788855_47188552_ENST00000601538.6,E | 1 | - | good | CTCAGGTTTCATCAGCTTTGCATTTCCTGGGGCGACCTGTCATGCTCAGCTGTGTTGTTCCCATGCCCATCCAGGAACCTGCCACTGCTCATTTATTGGTTCTTCGTACTTGTGCTGAGT | 120 |
| chr11 | 71175335 | 71175455 | 788855_47188552_ENST00000601538.6,E | 1 | - | good | CCTCCTCTCCCTCTTTTTCTTTCCTTCCTCCTGCTCTCTTTACAGACTCACTTTCCTGCTTCGGTGTCCACAGGACCACCCAGCAGCAGCCTTGGTTTCTCACTGACACCACCTCTGTGG | 120 |
| chr11 | 71175695 | 71175815 | 788855_47188552_ENST00000601538.6,E | 1 | - | good | AAACCCTGAGGAGGCAGAAAGCTGGGAGTCATCATAGGGCTCACCTCATTATTTCCTCTCTCCTGTGCATCCTGTCCTATACTTTGAAAATATATGTTATTTTTCTTTTTACATATTTTG | 120 |
| chr11 | 71175815 | 71175935 | 788855_47188552_ENST00000601538.6,E | 1 | - | good | TCTCTCTCCTCCTGGTTTTGTCCCACAAACTGGAGCTACCTTGGCCTTCCTGTGTTCTCAGCCCCATCGCCTCCACTCTGGGAGTTTGCCTGGGTGGTCCTTGCTTGCAGCATGGTCTGG | 120 |
| chr11 | 71175935 | 71176055 | 788855_47188552_ENST00000601538.6,E | 1 | - | good | ATGTATTATTTTTGCCAGTCCTTTTGGGTGGTTCTTTCTCCGGCCTAAGGTGGTTTGTTACATGCTTGCATCCATCATATTCTGAATAAGGGAACACATCTCTACTAGGCTCCGTTCCAT | 120 |
| chr11 | 71176055 | 71176175 | 788855_47188552_ENST00000601538.6,E | 1 | - | good | GGCAGTGTTCAGCCTATGTCTGATTATTTCCCATTAAGTGAGGCAAAACCCTTCTGTATGTTCACTGCCCCATGAATGGTGAGATGGGAACAGACACTGTTCCTGGCCTATGTAATTAGT | 120 |
| chr11 | 71176295 | 71176415 | 788855_47188552_ENST00000601538.6,E | 1 | - | good | TTATTTTCCTCATTATGGGTCATATTTTCTTCTTTGCATGTGTGATAATTTTTCATTGTTTTATTAATTGAATTTTCTATTGCTAAACATTATAAAATATGAATTTAATCTTGTTGGACC | 120 |
| chr11 | 71176655 | 71176775 | 788855_47188552_ENST00000601538.6,E | 1 | - | good | TTTTTTTAGAAACTCCAACTATGTACATATTAGGCTGCTTGACATTGTTTCATGGTTCACTGATACTCTCTTCTTTTATTATCATTTGTTGGTGTTATTATTATTTTCTTTGTGTGTTTC | 120 |
| chr11 | 71177255 | 71177375 | 788855_47188552_ENST00000601538.6,E | 1 | - | good | ATTGTTGTATATTTTTCTTCTATATGTCTTGTAAACCCCATGATACATTTTTTTCCTGCTTAGTCAGTTGTCTTAAAGGGGTTTAAATAAGAAAAACTTGCATATTTAGCCTCATGGTTA | 120 |
| chr11 | 71177615 | 71177735 | 788855_47188552_ENST00000601538.6,E | 1 | - | good | TTATTGGTTTAAAGTGCATTTCTTATAGAAAGCTTAGAGTTGGGTCACATTTAATGGAAAAATTGATAATGGTTTGATTTAAATCTGTCATCTTACTATTTATTTTCTTTTTGTTACATC | 120 |
| chr11 | 71178575 | 71178695 | 788855_47188552_ENST00000601538.6,E | 1 | - | good | ATACTAAGTATGCTTTTTTGTATCCAGTTACTTTTTATTTGTATTTGTCTTTATTTTTAAAGCATAGTTCTTTTTAAAGACTTTTTAAATTGTTGTAAAATGTGCATAGCATAATATCTA | 120 |
| chr11 | 71179055 | 71179175 | 788855_47188552_ENST00000601538.6,E | 1 | - | good | CTTTTTGCATCTGTGTTCATGAAGAATGTTGGTCTGTGTTTTCTTTTTCGTATCGGGGTAATATTGGCCTCACAGAAGGCATTGAGATGTTTCCTTTTATCCAGCTTTCTGGAAAAAAAT | 120 |
| chr11 | 71179415 | 71179535 | 788855_47188552_ENST00000601538.6,E | 1 | - | good | GGGAGGAAGTTCTCAGTCTTCCCATGTAAATACGGTGGACACTGTGGGTTTTTCAGAGATGCCTTTTCTCAGGCTGAGAAGACTCCCTTCCAGTTCTGCTTTGCTGAGGGATTTTATCCT | 120 |
| chr11 | 71179535 | 71179655 | 788855_47188552_ENST00000601538.6,E | 1 | - | good | GCAAACAAGGATGGTTTTATTTCTTCCATCTCACTCTGGATGCATTTTATGTATTTTTCTTGTCTTGCTGAGGAGAAGTGTGAAAGTGGACGCCTCAGCCTTGTAAGATCAAGGATCTTA | 120 |
| chr11 | 71179775 | 71179895 | 788855_47188552_ENST00000601538.6,E | 1 | - | good | TGTGTAAATCTTTCACATCTTTCGCCAAGATTTTCTCCTAAGCATGTCTTATTTTCTGTTGGCGTTTGTAAATAGAGTTGTTTTGACATTTCAACTTCCAATCATTCATTGCAATTATAT | 120 |
| chr11 | 71179895 | 71180015 | 788855_47188552_ENST00000601538.6,E | 1 | - | good | TAACCTTAATTGGCCAGCAGAGGTTTGCATGAAGGTGGACTTTGAGACTCACATTGCAAGGTGGCACCTCGGTTATCGAGTCATCTGTAGTTTCTTGCAGCAGTGTTTTGTTGTTTTCAG | 120 |
| chr11 | 71180015 | 71180135 | 788855_47188552_ENST00000601538.6,E | 1 | - | good | AGATAATGACATTTGTCCCCGGGAAATTCAGCAAGATGCTTGTCATTCCCAAGTGGCAGGCAGGTTACAGCGTGAACACTGGTTGTCAGCTCCCGGGATGTCCTGTGCCTGGAAGTCCTG | 120 |
| chr11 | 71180135 | 71180255 | 788855_47188552_ENST00000601538.6,E | 1 | - | good | GATAAATAACAAGGTGTTGGTCGGCACATCTGGAGTTAGATGTAATTGTGAGTACACCTGGAATAAGGATGCCCAGAGCTATTTGCTTCTGAGTCATGTTTTTCTGCAAGTGCAAAGTGA | 120 |
| chr11 | 71180255 | 71180375 | 788855_47188552_ENST00000601538.6,E | 1 | - | good | TTCTTTGGAAACCACTTCCTCTGTTCTCTTTCCTTCTATCCTGTACCTCTTGATAAAGAATGCCAGCAGGTAGACTCCATTCAAAGAAAAGCACATATTCTGTGCCCCGTGGTTTGTTCT | 120 |
| chr11 | 71180375 | 71180495 | 788855_47188552_ENST00000601538.6,E | 1 | - | good | TGGGCACAGGGTAAGGAACTTCGTGTTTTCTTGCTTTCCCTCTGCTCCTACCCTGGTAGGAAAAGCTGTTTTAACTGCCCAGACCATAAATTAACAAAGCTTGTCACCTCCTCCACATTT | 120 |
| chr11 | 71180495 | 71180615 | 788855_47188552_ENST00000601538.6,E | 1 | - | good | TTTTCTAGAAGCCTGTATGTATTTTCATTAGAAACTGAACTTTGTGTGTGGGGTTTTTTTTCTTTTTTTCTTTGCAACCTAGACTTTCACTGTCGACTTCTTGTACAAAGCGCTGTTAGA | 120 |
| chr11 | 71180615 | 71180735 | 788855_47188552_ENST00000601538.6,E | 1 | - | good | TGAAATTCATTTGATCCACATTGGTGTGTGGCCTAGTGTAGTGTGGCTTACGTGGATTGACAGGTTTGTTCAAACATGCCATTCCCGCCGCCCCCGCCAAAAAAATGCCATCTGTAGATG | 120 |
| chr11 | 71180735 | 71180855 | 788855_47188552_ENST00000601538.6,E | 1 | - | good | GACCTTGCCCCTGAATTTTTGCTTCCTCTCCAGTGGTATATGTAAGAGTGGTGGCTGGTAAAAATTCCGTCCTTCTTGGGTTGTAGCTTCAGCACTTCCTAAAAGTGTAAGTTCCTGAGC | 120 |
| chr11 | 71180855 | 71180975 | 788855_47188552_ENST00000601538.6,E | 1 | - | good | GGTAAGGCTGATTTGCCACATGGAAGTGGAAGGTTGCTTTATGGCCAGAAATACACTAGACAGCCGGACGGGGCTGTGCCTCGTGGGCGTTGGTGGAGGGGAGGGACTGTCCGATGGTCT | 120 |
| chr11 | 71180975 | 71181095 | 788855_47188552_ENST00000601538.6,E | 1 | - | good | AGGATCTTCCGCCATCTCTGGTTTCTTTCCCAGCTTCTCCTTTCCTGTTTGTCCTTGTGTTTATATGCATTAAAAATGAAATCCCTTCGCTGCTGTTTTTTGTGTGTGATTTCAAGCAGA | 120 |
| chr11 | 71181095 | 71181215 | 788855_47188552_ENST00000601538.6,E | 1 | - | good | AGGCTTCCTGCCTTCAGCTCAGCGGGAGGAACCTGCAATTCCGTTCCTGGGCCTTCTTGCACTTCTGTGCTGCAAGCTGGTCGGCTTCTCCAGACCTCCCCTCGTCAGCAGTTGCACTCA | 120 |
| chr11 | 71181215 | 71181335 | 788855_47188552_ENST00000601538.6,E | 1 | - | good | AAGGCCATGTGCCACTTGTTACCCAGTGGCTTCTTTGCACATGAGAGTGATGTAGTGGAGAAAGAGGAGCTGGCTGTGTTGAGGGGATCCAAGTGCTACCTTATTAACAAACTTTGAACC | 120 |
| chr11 | 71181455 | 71181575 | 788855_47188552_ENST00000601538.6,E | 1 | - | good | TTTCTTTGACACTGGGGTTGTCAAGATGAGATTCTGACCTTGGCTTCATTCTTCTGAGATCCTGAATTGTCATATGACCATTGCCTTTTTTGTTTGTTTGTTTGCTTTTAGAGCCTCATT | 120 |
| chr11 | 71181575 | 71181695 | 788855_47188552_ENST00000601538.6,E | 1 | - | good | GGTTGCCTTGGATGTAGGGTATTGCAGTGCTTCACCATGAGCGGGCGCCTTGCTTGGAGGCTTCTGCTGCTTCTTTTCAGCACCTGCCCTTTCCTCTTGCTGGTGGCCACTCCGATTGGT | 120 |
| chr11 | 71181695 | 71181815 | 788855_47188552_ENST00000601538.6,E | 1 | - | good | ACCTGAACATCTTATTGCTCCACAGGTGGCTCTGACGGCGTCCAGGATTGAGGATTCCTGGGCACGTCCCCACACCTGCGGCTCTTTGTTGTCCTGGTTCCGTTATGCATGCGGTGTCAC | 120 |
| chr11 | 71181815 | 71181935 | 788855_47188552_ENST00000601538.6,E | 1 | - | good | CATCCAGGTGGCCCCTTGGAATCGTAGGTTTCTTGCTAAAATACAAATTCTTCTTCCCACTTGTTCCCCTGGAGATTCGGATTCAGTAGATCTGGGTTGGCGGGGGAGGGGGGGCTGGGA | 120 |
| chr11 | 71181935 | 71182055 | 788855_47188552_ENST00000601538.6,E | 1 | - | good | AAAACACTTGTCATCACAGCCATCCTCTATGAGGGGGAGACAGGGATAGAGAGGGACCCCCTGCCAGGGTTACTGCAGGGCGCTGCCCCTCCGGCAAGGTTGGTCACACCCATGATGCTT | 120 |
| chr11 | 71182055 | 71182175 | 788855_47188552_ENST00000601538.6,E | 1 | - | good | AGGACCATGTTTGGAGAGATTTTTTTTTTTTGCCTAAAGAAGATGCTGCCTTGGGGAAGGAAAAGGCCCTAGCCTCTGGAACTTGCTGGGAAGCAGATTTCTGTTCCTCCTCAGGAGAG | 120 |
| chr11 | 71182175 | 71182295 | 788855_47188552_ENST00000601538.6,E | 1 | - | good | ATCATCTGAACAAAGACAGGCACTGTTTCTGCTCTGTGTGTAGAAACTCAGGAGCCCCGAAAACAATCTCCTCCCAGCAGATGGACAGCCGTGCTACTTACAGGCATGGGAACCTGGAAT | 120 |
| chr11 | 71182295 | 71182415 | 788855_47188552_ENST00000601538.6,E | 1 | - | good | TGTCAGATAATACATTTCTGCCGTTTTTGCTTCAATTCCAGTTTGTGGTACTTTGTCGTGGTAGTCCCAAGAAACAAATCCATCTTCAAGGCTGTTCCATCTACAAGCACATTTGAAAAG | 120 |
| chr11 | 71182775 | 71182895 | 788855_47188552_ENST00000601538.6,E | 1 | - | good | GCCAGCATGCATGGTGGGGCACGCTGTGGGAGAGAAACATTGAGCTTAGGGAAGCGAAATATTTTATGCTAGACAGTGAGCCTGTCTGCCGTTTGTACCAGAGGGAAACACCATATCTTC | 120 |
| chr11 | 71183015 | 71183135 | 788855_47188552_ENST00000601538.6,E | 1 | - | good | TCTGTGGGAGTATCCCATCCTCCTCATCGAAACTGTCAAGAGCCTTGAAGGCTTTGAGATTTCCCCTCCTGGCTGGCTAACAAGTGAGCTCACCAGAGTTTCATGAAGACACAAGACTCC | 120 |
| chr11 | 71183255 | 71183375 | 788855_47188552_ENST00000601538.6,E | 1 | - | good | TCATTGTCCTGATAGGAGCCCCTCTCTGCTCACCCGGGGGAGAGTGAGTTTCACCCACAGCATGAGTCGGGCAGCCTGTTCCATCCAGGGTCTCATCCTCAGCCTCGAAACTCCAGCACC | 120 |
| chr11 | 71183375 | 71183495 | 788855_47188552_ENST00000601538.6,E | 1 | - | good | TAGAGCTTTGCATTGAAAAGCAAGCACCTCCTCACTGACACCGTTCTAAAGCATGCCAGAGTCTTGGCCCAGCAGCTGCAGCCCTGTGGCCTCACAGCTGGGACTGCGGGTAGTACCACC | 120 |
| chr11 | 71183495 | 71183615 | 788855_47188552_ENST00000601538.6,E | 1 | - | good | TGGGAGCAGAGGGAGAGTGGCCTTGTGCCTGTGAGCCGCATGCATTGATGTAGTGATTTCTAGCAGTGTCGCAGTTGCTCAGGGCACAGCCAGATGAAGGGCTGCTGTTTTCTGTTTGCT | 120 |
| chr11 | 71183615 | 71183735 | 788855_47188552_ENST00000601538.6,E | 1 | - | good | CCCAGCATGCTGGGTGGGCCTGGGTGGTCTTTGGGGTTTCTTGGGGATGGAGCCACCTGCCCCCAGTCATCTCTCTCCTTACAGCAGCTGCAGTCAGGGTAGAGGGGTGTCAGGGATCCA | 120 |
| chr11 | 71183735 | 71183855 | 788855_47188552_ENST00000601538.6,E | 1 | - | good | TTGGCCTCATCAGCTGGGTCCAGGCACCATCTCCCACCATGAGACCGGGTGGGGTTGTGACTTGGGCACCTGTCTCCAACAGAGCCCTAAAAGCTGGAGCTCCTCCCCTCCCTGAAGTTT | 120 |
| chr11 | 71183855 | 71183975 | 788855_47188552_ENST00000601538.6,E | 1 | - | good | AGCGTAGAACACACCCCTTCCCATGTTACTTTTGGATGTGGAGGCCACGAAACCAGTTTGTTGACGTGGTCCAACCTCTCCCCCTACAAGGTTACATTAGCTTCCCTCTCCCCAATAAAC | 120 |
| chr11 | 71183975 | 71184095 | 788855_47188552_ENST00000601538.6,E | 1 | - | good | GTCGGTAATCTCAAGAGATGGGCAGCTTGATGGAGTCGTCAGGGTTGTAGCCCAGGGGCTCTAGTTAGTGCTCCCCAGTCAGGAGCGTCCTTCTTGGCCCGTGTGAAATGGGGAAGTACA | 120 |
| chr11 | 71184095 | 71184215 | 788855_47188552_ENST00000601538.6,E | 1 | - | good | AGGCCTCCCTCCATGGATCAGATGCTCAGTTTCAAAAAGAGCATAGTGACAGGTGGGTGCCTGGGAACCGTGCCAGGCAAGTCCAGGTGCCGCCGGGGAGGGGCTGTCTCCCTCTGCCCA | 120 |
| chr11 | 71184215 | 71184335 | 788855_47188552_ENST00000601538.6,E | 1 | - | good | CAGTGTGGAATAGACCATGGTTGTTTGACCAGCGAGGCTTGTGGGGCCTTTCCTATCTTGTGGGCCATTGAGTCAGAGCTGGCCCACCCAAAAATGGGAATATCAAGCCTGAAAATCACA | 120 |
| chr11 | 71184335 | 71184455 | 788855_47188552_ENST00000601538.6,E | 1 | - | good | CATTCAGGGGCAAACTCTGAATCAAGGGCCCCATGAAGCCAGCGACATTGTTTCAAGTGATGGACGCGAGAGCCATTGCTTTTCGTGGGGGCCAGGATGCCTCTGTGTGCAGCCTGGCCT | 120 |
| chr11 | 71184455 | 71184575 | 788855_47188552_ENST00000601538.6,E | 1 | - | good | TGGCATGGCCGTGGGCACCTGCAGTTCTACAGAGCCAGCCCTCGGGTTGAGCAGCTGCTGCAGCCCAGTGGGTATCATTGGGCTTCAGTATTGTTGAACCATCAGTGAACCCAGCCCCGA | 120 |
| chr11 | 71184575 | 71184695 | 788855_47188552_ENST00000601538.6,E | 1 | - | good | CAGACCAGACAGTGAGGTTGCTGCCAGCAGCTGCCCATGGGTCAGTAAAAACATAACCAAGTTCATCAGGAGGAGAGTTGGCCACAGGTTTGAGAAGGGCCCTGAGTCCTGCCCGCTAAG | 120 |
| chr11 | 71184695 | 71184815 | 788855_47188552_ENST00000601538.6,E | 1 | - | good | ACAAATGGACTCTGAGCTCAGGCATCTACATGCTGACCCAGAATGTTCAATCAGCCGCCGTGACTTGCCAGAATGTGTGGCCCCAAAGAAGGACGTCTTTAATCTGACTGTTCCAAGTGG | 120 |
| chr11 | 71184815 | 71184935 | 788855_47188552_ENST00000601538.6,E | 1 | - | good | GTGCCCAGGCAGTGGTGGATGGGGTGCTCTCTGGTATGATGGTGCATCTGGGTAGTAGTGGTGGCCTCTGGGCAGTGCAGGCCCAGTCAACAGCTTGGTCTTATTCAGTCTCATCAAAGA | 120 |

| | | | | | | | | |
|---|---|---|---|---|---|---|---|---|
| chr11 | 71184935 | 71185055 | 788855_47188552_ENST00000601538.6,E | 1 | - | good | GAGAGGAGGCAGGAGCAATGGGCCTGCCCTCACGGGGTACAGACGGGACAGCTGTGGGCAGGAGTCACAAGTCTGGTATGCAGAGATGGCCTCTGCGTCAGGAACAGAGTGTTTAGCTTT | 120 |
| chr11 | 71185055 | 71185175 | 788855_47188552_ENST00000601538.6,E | 1 | - | good | AGTGCTGGCTGATTGTATTGGGACAGCATCATGGTAGCCTATAATGGTCTCAGTGTTGCTGAGGCCCCCACCGGTACCCCTGGTGGGGAGAGCACCACATTTTGAGATGGTTGATCGGCT | 120 |
| chr11 | 71185175 | 71185295 | 788855_47188552_ENST00000601538.6,E | 1 | - | good | AACGCTCTGCCTGCTGGCAAGTGGCTTTTGTGATACAAGGTACACCATCATTGAGTGTAGATGGTCTGGAAAGTTGAAGTCATGCCACAGGTGAGTTTCAACAACCATAACAGTGTGGGC | 120 |
| chr11 | 71185295 | 71185415 | 788855_47188552_ENST00000601538.6,E | 1 | - | good | GAGATGATTCCTTCCTGGATTAGCCATATATAGTGATCAGACTTTTCTACTAAGGACCAGAAGGAGGTGAAGGTAGGAAGGGTGGAGAGTAGGGAGGGTAGACAGTCCCTCGAGTGTCTC | 120 |
| chr11 | 71185535 | 71185655 | 788855_47188552_ENST00000601538.6,E | 1 | - | good | GTCAGTCAACAGGAGAATCCAGGTGGCTGAACTGGAATGAGATTTGAACCCTCCAGGTCCCCGTTTGGCTGGGTAGTCACCTGGAGGCTGCACCAACCAGACCAGCCTGGGCTCGCTGCT | 120 |
| chr11 | 71185655 | 71185775 | 788855_47188552_ENST00000601538.6,E | 1 | - | good | CGTGCAAGAGGCATTTTTATAAAGCTGGAGGACCCATTGTATCTGCTTTGTCCCATGTGGAAGGACATGGTCAGCAGTGTACTCTCCCAAGTGTGTGTTGTCCTTGCAACCCAGGACCAT | 120 |
| chr11 | 71185775 | 71185895 | 788855_47188552_ENST00000601538.6,E | 1 | - | good | TGGGGTTGCTGTTCTTTCCCCACCCACCCCCCAAATTGGTGTCTCGTCACAATAGCTGCAACTTCACCTTCTTCCTTGATTGTCCCCTCCTTGCCCCCAGACTTTCATTCTGAAGGGTCA | 120 |
| chr11 | 71185895 | 71186015 | 788855_47188552_ENST00000601538.6,E | 1 | - | good | AGCAGCCAGTGAAATGTACCACCCTTTGGGGGTGTCACCGTACCATTGTATTTTCCATTGTTGGGAATGCGCCATGGGAGACAGGCAAGACTGTTAATGTCCCCCTCCCTCTCCAGGCTC | 120 |
| chr11 | 71186015 | 71186135 | 788855_47188552_ENST00000601538.6,E | 1 | - | good | AATGCCAGACCCAGCAGCCAGTGAAATTTATCACACTTTTGGGGCGTCACTGTACCAGTGTATTTTCCATCGTGGTGAATGAGCCATGGATGACTCAGAAAGGCAGAGAATGCTAGACCC | 120 |
| chr11 | 71186135 | 71186255 | 788855_47188552_ENST00000601538.6,E | 1 | - | good | ACATCAGCAGGTGTTGACTTGTGTTTTGCCTGGGACGTGCAGGACTGAAGTTCTCCCATGTGCTTTCTGACAGGTTTCTCAGTGACAGTAAGGTGAATGGGCATCAAATTGGGTTCAGAG | 120 |
| chr11 | 71186255 | 71186375 | 788855_47188552_ENST00000601538.6,E | 1 | - | good | ATTCAGTTCAGTCTGCGGGTCCAAACTGTTCAATGTGTCCATGCCAGACGGCATTTGTCCACTCCACCGAATGACTGGCCAGTGGTTCCAGGAGTAGGGGTGAGGCTGGGGCTGGGGAAG | 120 |
| chr11 | 71186375 | 71186495 | 788855_47188552_ENST00000601538.6,E | 1 | - | good | CAATTGCTGGAGCCCCTGCAAGAGGGACCTCCATAGGTTGGGGGATAGAGGTTGTCAGTGTCTCCAACATGGACTCCTGGATGACTGATAGACCTTAAGGCGCCTGGTTCAGTTAGATTA | 120 |
| chr11 | 71186495 | 71186615 | 788855_47188552_ENST00000601538.6,E | 1 | - | good | ATGACCCCTCTAAGGTACGAGTGTCCGTGTTCTCAGGAGGGAAGCAAGGTGACACTTGGCTTCCACACAGGAAGTAGAGAGCTGTGGGCACACGGTGCCCTGGAAAGTAAGGGTTGCTTA | 120 |
| chr11 | 71186615 | 71186735 | 788855_47188552_ENST00000601538.6,E | 1 | - | good | ACTAACCCGAAAAGTTAACCAAGTCACTGAGGGCAGCTCCACTTTGGAGGGAACTTCTCAGTGATCGAGGGCTATGTGACATGCAGGGAGTGCTTCTGCTTACATACCCGAAGGTCTCTT | 120 |
| chr11 | 71186735 | 71186855 | 788855_47188552_ENST00000601538.6,E | 1 | - | good | AGGGCCGAGGTAGTGGCATTGATCACCTTCCCTAAGTCCTGGGACACGTTTCCTACAACCTTTTCTAGTTAGATGTCTCCTATGGATAGCTGTCTGCAGCATTCACAGAAATAATGTGCC | 120 |
| chr11 | 71186855 | 71186975 | 788855_47188552_ENST00000601538.6,E | 1 | - | good | CTACGGCAGGAGGAATCGGCAATTGCACAGACTTCGCCTTGAACCACAAGGAAGAAGTCAAAGGCAATCCTGTCATCTGTGACACTTCTGGATGGTGAGGTGAGGCTGGGCTGAATGCCG | 120 |
| chr11 | 71186975 | 71187095 | 788855_47188552_ENST00000601538.6,E | 1 | - | good | TCCTGGATCCAGCCAGCTGAGCAAATCCCACAAGCCTTCAGGGTCCCCTTAGAAACCAAGCTAGTTTCTCTGTAAGTTTCTGTTGACCTTTCCGCTTGGCGGCATGTGTTAATAAACCAG | 120 |
| chr11 | 71187095 | 71187215 | 788855_47188552_ENST00000601538.6,E | 1 | - | good | TTGGCCATGCTGATTAACAGGCAGATTGGGACCAAATCCACCCCATTTGTCTCATGCAGTCTTTAATTAAGACAGCAGCCAACAGGACTGCAGTATTAACCTCAATTGTGACTTCCAGGG | 120 |
| chr11 | 71187215 | 71187335 | 788855_47188552_ENST00000601538.6,E | 1 | - | good | AAACCCGTGAAGAACTGTCCCTGAATAAAACATGAGAAACCCATGAAGAACTGTTCCTGAGCAATATCTACCCCTCATGTCTACATCATCCTGGCAAATTTTCCTGAGTGTGCCATTCTG | 120 |
| chr11 | 71187815 | 71187935 | 788855_47188552_ENST00000601538.6,E | 1 | - | good | GTGCAGGGAAGGCCACCCGGCACCACATGCACAGCACGTTGCGTTGTGGGAGGGAAGCGCTGAGTTTAGGGAAATAGAATCCTTGATAAAGGATGACTTCTGCACACTTGCCCTTTGCTC | 120 |
| chr11 | 71187935 | 71188055 | 788855_47188552_ENST00000601538.6,E | 1 | - | good | CGCCAGGTTAGCCTACCTCAGTTTCGTGATACCAGTAGAAGACACGAAGCCTCTGGATCAGAGATGAAGGTCAGTTCCTTCCTCACAGCAACACCAGTAGGAAGAGCATTTCCAAGGACA | 120 |
| chr11 | 71188055 | 71188175 | 788855_47188552_ENST00000601538.6,E | 1 | - | good | TCAGGAGTGGCTGGGAAAACGGGGCAGAGGCAGAGGGAAACGTCTCCGGTGATGTCTCAGTTTCTAATGGGAGGAGCAGATGTGTGGTGGCAAGGACTGTGTGGGGTCTGACCTTTGACC | 120 |
| chr11 | 71188175 | 71188295 | 788855_47188552_ENST00000601538.6,E | 1 | - | good | GGGGTAGGATGGTGAGTGGGCAGCTCCAGTGGTCCATGACCAGAGTGACACCAGCAGGGGATACTAAGGGGAGGTCTCAGCAGGATGTGAGAGAGATGGAGGTGGCAAGATCACCAGGTC | 120 |
| chr11 | 71188295 | 71188415 | 788855_47188552_ENST00000601538.6,E | 1 | - | good | GTCCTGGTTGGGAACGCACAACTTGCTGAGGGATGGTGTGTACTGAGGCCTGGGCATGGGCTGCTGAGGGCGTTTGGGGCTGACCTTCTCAGGCGTATGCTGGAAGGGGCTGGGCAGATG | 120 |
| chr11 | 71188415 | 71188535 | 788855_47188552_ENST00000601538.6,E | 1 | - | good | CAGTATTCCTGTGCCTTTTCGTACGTCCACTGTAAATATTTGCAGTGTTTTCCTGTTCCCTTGCGCCGAAGTCCTTCTCCCCAGAGGGAGGGGATTCTGTTTCTGGAGAGACCTCAGCAG | 120 |
| chr11 | 71188535 | 71188655 | 788855_47188552_ENST00000601538.6,E | 1 | - | good | CACACCTCAGTTTCATCTTGACTTTGGGTTATAGGCACAGTGAATTTCATTGTCTTCACGGCTGTACCTGGTTGCCAGGGTAATTCATGGGGCCTCGTTTATTTCCACCTTTCAATTACT | 120 |
| chr11 | 71188655 | 71188775 | 788855_47188552_ENST00000601538.6,E | 1 | - | good | GTGAGGCCAGCACCCAGCACAGTGCAGGGACGTGGCAGGCCCTCAGCAGATGAGGAGAGGTGTGGTGGGCGACAGATGAGTGAGCAGGGCACTCTAGCACCAGCACGGTGCATTTGGGCC | 120 |
| chr11 | 71188775 | 71188895 | 788855_47188552_ENST00000601538.6,E | 1 | - | good | CCACCCACCTTGACCCATCACTTGCCTCTGTGAGGGGTGACCCCTGCTAGAGAGAAGGTCAGGGTCCCTGAGAAACCACGTAGGCCTTAGCTCCATTTGGAGGTGGTCGTAGTTATCTCA | 120 |
| chr11 | 71188895 | 71189015 | 788855_47188552_ENST00000601538.6,E | 1 | - | good | CTGTGACCTGGCTGTTGCCCTGTGGTGGGGCAGCTTCCTGGGACTCCATCTAGGTGGGAGTGGGGGGCCGTGGAGCTGGGGCCCGCTCCTGCTGTATTGAGAGAACCCCAGGGTTTGCTT | 120 |
| chr11 | 71189015 | 71189135 | 788855_47188552_ENST00000601538.6,E | 1 | - | good | AGGGAATGAGTGGTCGGGGATTTCAGCCGAGAATGGCAGATGTGACTATTTCACAACCCAACCCGGAGAGCCTCATCACTGGGAGCTGGGAGCTCCGTCTTCCCTGGCCGTCTGTGGTTG | 120 |
| chr11 | 71189135 | 71189255 | 788855_47188552_ENST00000601538.6,E | 1 | - | good | AATGGAACAGAACGGGGTCTGGGAGCCCCTGGGGGAATCACACACCTGCTGGTGGATGGCGGGAAGCCGGGGTGGCCAGGAGGAGCCTCAGGAATGAGAACCTGGAACAAGGGGAGGGA | 120 |
| chr11 | 71189255 | 71189375 | 788855_47188552_ENST00000601538.6,E | 1 | - | good | TCTCTCTACTGAGAAACTCTCTGAGGTTTGGTAGAGTTTGTCAATTACACAGGCACTGATTGGTCACCAGCGAGGCACCAGCGACTGGAATAAGCCCTGAACAGCACCGCTGAGCCAGGA | 120 |
| chr11 | 71189735 | 71189855 | 788855_47188552_ENST00000601538.6,E | 1 | - | good | GCCTCAGCTGCGGCCTTTCCCGTGTTCATTCACCCCCAAGGGCAGCAGCACAGGCTTTGGGAGGGGAGCTCTGGACATGGAGTTTGTCCCTGAGACCTGCTGGCCTCAGATACAGAGCTT | 120 |
| chr11 | 71189855 | 71189975 | 788855_47188552_ENST00000601538.6,E | 1 | - | good | GGGCCCTCGTGCTCGGAGTGCTCAGCTTCCTTACTGTGCCCGTCTTGTCCTAACAGTCGTGGTGAAGCCTCCCCCAGATGCTGGGGCTTTGAAGCCAGGCTGGCAGAAGAGCTTCCCCAG | 120 |
| chr11 | 71189975 | 71190095 | 788855_47188552_ENST00000601538.6,E | 1 | - | good | GGACGTCTCTGCTGTTGGTCGACCTACGTGTGGATGACTTGGGGCTTCCTTGCAGATAAGGGCTGGGCTGCTGGGAGCAAGTCAGCTGTGCTCTTCACCATGCTACTGGACGCTTGGGCT | 120 |
| chr11 | 71190095 | 71190215 | 788855_47188552_ENST00000601538.6,E | 1 | - | good | GGCCAGCTTGGATAAGAGCGAGGACTGAATAATGGCAGCTGTAGCCACTGGGCAGTTTGGGCAGCAGTGCCTGTGCTTGGCTGGTTTCTTTTTCAGTCGTGGGGTTGGGGCCCTGCAGGT | 120 |
| chr11 | 71190215 | 71190335 | 788855_47188552_ENST00000601538.6,E | 1 | - | good | CTTGGAAGGAGAACACAAGAAGCGAAAGCCTTCTGAAGCTGCTGCCAACAACGCCCACCTTTGCAGCTGCAGAAGCATTTCCCCTCTGCTCCAAAGGGGCAAGGCCATCTTCTAGTCCAA | 120 |
| chr11 | 71190335 | 71190455 | 788855_47188552_ENST00000601538.6,E | 1 | - | good | GGGGTTAGTTCAGGAGTGCGGCAGACGGGTACCGAGATGCTTAAGGGTCTGGATGGAGCCTCACGCTGTCCCCTTGCTAAGTACTTAGCACCGCTGCCTCTAGGGCCTGGTTATTGATTA | 120 |
| chr11 | 71190455 | 71190575 | 788855_47188552_ENST00000601538.6,E | 1 | - | good | GTGACGCACTTTCTCTTGTTCCCCATCAACCATCTGTTGGTTCCAAGGCCTCATTTTCCAGACAATGGTGAGCCACACTTCTTCTGAGAAGTGGTATAGACGTCCCTTTCTCCCATTGCA | 120 |
| chr11 | 71190575 | 71190695 | 788855_47188552_ENST00000601538.6,E | 1 | - | good | CATGTCCTGGCATTCTGTCACGGCCTGCAGGACGTCACACACTTCTGCACATACGTTGTCAGAAAGGGGAGAGGCCTTAAGAGGCCGATTCCCTGCAGAGGCCTCATGCCTTGGTGGGTG | 120 |
| chr11 | 71190695 | 71190815 | 788855_47188552_ENST00000601538.6,E | 1 | - | good | GAGACAGAAATGATGGGTTCGTGCAGGCCACGCAGCCCAGGGTGGAGCCGTGGGATTGGGACTCGGCTCTGCCTGACCTGCAGAAGCCTCCTCTCAACGCCTGCGTGCAGCCCTTGTTCC | 120 |
| chr11 | 71190815 | 71190935 | 788855_47188552_ENST00000601538.6,E | 1 | - | good | TGCATGCATGCATATGTATGCACAAACACACCAGAGATACCTCACATTTAAGCTTCACAATGGCCTTCTCCAGCACCGCAGGGGCTGGACGTCCACCTTCCAGATGGGGAAAGTGAGGCC | 120 |
| chr11 | 71190935 | 71191055 | 788855_47188552_ENST00000601538.6,E | 1 | - | good | GGATTACAGGCGTGAGCCACTGTGCCTGGTATCTCTGGTGTGAGGTATTTCTTGGTGAGCCCCAATTCCTCTGAGAAGTGGCGTAGACGTCCCTTTCTCCCATTGCATGGGAGAAAGGGA | 120 |
| chr11 | 71191175 | 71191295 | 788855_47188552_ENST00000601538.6,E | 1 | - | good | CAGCGTCCTGTCTCCCTCATAACACACAGTTTATCAGTATTAAATTGTGAACAAAGACATAGCATGGATGGGGATACAGCTGAGTGTGCTGCGTTCATTTTTGCATTTTTTTTTTCTTTT | 120 |
| chr11 | 71191295 | 71191415 | 788855_47188552_ENST00000601538.6,E | 1 | - | good | GACCCAGCCCTGTGCGGACTCCAGCCTCTGGGATGTGAGATAATGAATGTCTGTTGTGTAAGCTGCCCTGTCTGCCGGGCTTTGTACGGTAGCCCAGGCGAAGACATCTGAGGTGATATG | 120 |
| chr11 | 71192255 | 71192375 | 788855_47188552_ENST00000601538.6,E | 1 | - | good | CCTCAAGGAGCCCCGTCACCACAGGTCATGAAGAACAGAGTGGAGAGTTGTCATGGGCTGCACTGTGTCCCCTCCCTGAATTCCTAAGTTGAACCTGTAACCCCCACTACCACAGAATGT | 120 |
| chr11 | 71192375 | 71192495 | 788855_47188552_ENST00000601538.6,E | 1 | - | good | AGGAAACAGGGTTCCAGGCAGTGACTGAGGTACAAGCATCACAAATGTGGATTCTGACATAAAGTAGCAGTTTTGTTTGACGTATTGGTCAATGGGCTCTCATAGTCATAGCCCAACAGT | 120 |
| chr11 | 71192495 | 71192615 | 788855_47188552_ENST00000601538.6,E | 1 | - | good | CCTTTCACCTGGACCTTGTGTTTCTTTTCATTTAAAATATCTTCATAAAGGCTCAGGTTCTGGTCTTTCTAATTACAGTGGGGACAGCCTGGGACCTTCCAATGTCATCCTGCCCCATTC | 120 |
| chr11 | 71192615 | 71192735 | 788855_47188552_ENST00000601538.6,E | 1 | - | good | ATTTGCATTTGTAAGTGAATATTCCAGTAATGACTGAAGACAGAGAGTATGAAATTGGATTGATTAGGTCATTTTCCTGGGAAAGGCGAGGCTTTTCCACCCAGGGTGGGGCACGTTGTT | 120 |
| chr11 | 71192735 | 71192855 | 788855_47188552_ENST00000601538.6,E | 1 | - | good | GGCATGATTGCATTCTCTGGGTGTCACCTTTTCCTATCACCTTTATGAGCTGATAGCCTCTGATAGCTTGATAAGCTCTCTAGGAAAAAGGAAGCCTTGATGAATATAGATCTCTTTGCA | 120 |
| chr11 | 71192855 | 71192975 | 788855_47188552_ENST00000601538.6,E | 1 | - | good | ACTCTTGTTTGAATTGGATGTTTGCCTTTGACAAGGAAGGTTCCCCCATTAAGGATGCAGTATTTTATACAAGGCGCTTCATCTTTTCCTCAAGCACCCTGCCTGCGATGCAATTAAGCA | 120 |
| chr11 | 71192975 | 71193095 | 788855_47188552_ENST00000601538.6,E | 1 | - | good | GTGTATCTAACATGGCAATAAAGAGAGGAAGGATTTTATTTCTCTCTTTTTCTAATACTTTCCAGACAGCCAGGTTTGGTTTTGCAGACTTCTTTAAAGGAGGAAGAGTAAGCTGTACCA | 120 |
| chr11 | 71193095 | 71193215 | 788855_47188552_ENST00000601538.6,E | 1 | - | good | AGTGTGACCATCACTGGGGCCCGTGCCTTTGGTTTGAGTTGATTTATTAGTATGATTTCGGGGAGGAATCTGGCTTTAGCAGAGTTATGTCTCTGCATTGCCTGGAGGCCATTTGTCTCT | 120 |
| chr11 | 71193215 | 71193335 | 788855_47188552_ENST00000601538.6,E | 1 | - | good | CTCCCACACACACTCACACACTCCTCTTGGAATTGCTGCAAGGCTCTGATGAGACGGCCCCAGAACCCATGGGCTGGGATTCCAGACCCACCACAGAAGCCCTCTCAGCCGGTTCTGGTG | 120 |
| chr11 | 71193335 | 71193455 | 788855_47188552_ENST00000601538.6,E | 1 | - | good | GCGCAAGGTCAAGCCCTGGGATATGGAACAGCTCTGCTGATCGTAGTGGAGCCCAGGAAATGAGGTATCTCTGGTGTGTTTGTGCACACATATACATGTATGTGTGCATGCACACACACA | 120 |
| chr11 | 71193455 | 71193575 | 788855_47188552_ENST00000601538.6,E | 1 | - | good | ACAGTCACGCCAGTACCCTTTGACCTCAGAAATTGTGGCTCTTTCCAGCCTGTTATCTCATGTTATGACAGAGACCTCAACCCTTCCTCTGGATGGGGTCATGAGATGTTTGCACGTCCA | 120 |
| chr11 | 71193575 | 71193695 | 788855_47188552_ENST00000601538.6,E | 1 | - | good | CACTTTGCCATAGCAGCCCTGGCCAACTCGTGCAGGTGCCCAGGGCCGGAGCAGGCTGGAGAGCTCTGGACGGGGAGCAGGCAGTGGCTAGGGGCACATGGGATGGGGACTCTGGTCCTA | 120 |
| chr11 | 71193695 | 71193815 | 788855_47188552_ENST00000601538.6,E | 1 | - | good | GGAGAGGGCCCTGGAACAATCTGTGCCTGCAGAGGTAACCCACCCTGTCTGGACCTGGGCCTCCTGGCCTCCGGGGGCGTGGGACAGCGTGTCTCTGTGGTTTCAGGTGCCTGGTCTGTG | 120 |
| chr11 | 71194055 | 71194175 | 788855_47188552_ENST00000601538.6,E | 1 | - | good | GGGTTACCCGAATACCTGCTGTCCACCCCATCCTGGAGTGGGTGGTGATGTGTTTGGTGTCAGTAAGGCTTGGGGATCTCTCTCTGCCTTGATTTCCATGGAAAATGGGAACGATCTTGC | 120 |
| chr11 | 71194175 | 71194295 | 788855_47188552_ENST00000601538.6,E | 1 | - | good | CCATCGGGCACCCACTGCTTAGTTGTCCCCGTGAGTAACTGTGGTCCCATCAGAACAGCCACCTACAGACCAGCATCCCCAGATGTGGCCTCAGGTCACTTTGAACACAAAACTTTCCCT | 120 |
| chr11 | 71194295 | 71194415 | 788855_47188552_ENST00000601538.6,E | 1 | - | good | AGAGAGTGAGGCTGAGGCAGTGGCAGCCTGCCCCGACTTCCTGCAGCCCCCACGCCCAGGTCCCCAGGTCTCAGCTGTCACACGCGGCACTGATGCTGGGGTCTCTTCATGACATCAACT | 120 |
| chr11 | 71194415 | 71194535 | 788855_47188552_ENST00000601538.6,E | 1 | - | good | GGCACCTGCTCTTTGGATCCAGCTGTTAACATCCTGTTTGGTGGCTTAATTCTCAGTTGACACAAAGGTGAGGTGATTTAAAGAAGTGCAGGTGAGTGGCGTTGGGGTTTGAGGCCACAG | 120 |

| | | | | | | | | |
|---|---|---|---|---|---|---|---|---|
| chr11 | 71194535 | 71194655 | 788855_47188552_ENST00000601538.6,E | 1 | - | good | CAGGTTGGCTCTGAAGACATGTGGCCATTTCCTCAGTCCCTTCACACCCGCATGTCACTTAGCATCCACTTCCTTGGTCACTGGTACTCTGTGTCTGTCTGGACCTGCCCCCCTCCCATT | 120 |
| chr11 | 71194655 | 71194775 | 788855_47188552_ENST00000601538.6,E | 1 | - | good | CAAGTTATGGAAGCCTGAATTATTCTGTCAAGTATACTTTGCCAGGAAGTTTGGAGCTCACTTTAATGGTCTGGTTCTCTTTTGCGTTTCTTTGTGGTCTTTAGACATTTTAATTACTTG | 120 |
| chr11 | 71194775 | 71194895 | 788855_47188552_ENST00000601538.6,E | 1 | - | good | GGTGGATGTAGCGCTGGGACCCAGGATTTGCTTCTTCCTCATGGTCCTTGCTCTCTGCAGCTGTAGCAGGTCTTCTGTGATTTGCAGAGGGCTGGGTCCTGCTGTCCACCAGTCCCTGCT | 120 |
| chr11 | 71194895 | 71195015 | 788855_47188552_ENST00000601538.6,E | 1 | - | good | TTGGTTTCTTTCTTCATGTCTGTTCTCAGGTTTAAGCCTTATTGGGATTGTAAACCATTAACCCACGAAGGAAGCAGACTGGACCTTCTGTCTAAGTAGCTGGAGTTGACCTTTGACTGG | 120 |
| chr11 | 71195015 | 71195135 | 788855_47188552_ENST00000601538.6,E | 1 | - | good | CAGGCGTCAGCCACCACACCTGGCCAGTTAGGCACTTTTTAATACTAAATATAAATAGGTGCCCTACGTAAAAAGTGAGCTCTATTTCTGGATTTTACATACAACACGCGTCAGGTGGCT | 120 |
| chr11 | 71195495 | 71195615 | 788855_47188552_ENST00000601538.6,E | 1 | - | good | CCTGAAATTCTATGAAAAAACTCTGTTTCGGGTATGTAAATAATCCTTATTAATTGCAATATACTGAAAATGGGAATCCTAAATTTTCTGAAACATCTAAGAAATGTTTTCACTTTTCTT | 120 |
| chr11 | 71195615 | 71195735 | 788855_47188552_ENST00000601538.6,E | 1 | - | good | CTTTTTTTAAGTTTGTAAATTTAATTTTGCAAGGATTTCTCTGTTTATGTCTAGTTGATTTTATGATCCTTTAAGAGTGGGCCTCTGATTACAGTGTGATCTCATTTACAAACCTGAATT | 120 |
| chr11 | 71195735 | 71195855 | 788855_47188552_ENST00000601538.6,E | 1 | - | good | GTACCAACACCAGGTCCTTTCTTGTTCCTCAAGGGAAGATTTCCCCTCAGAAAAGAGTATTTTTGTTATGTGTGATTTTAGCATTAAACTTTTAATTTATTAGAGTGATACATTCAAAAA | 120 |
| chr11 | 71196575 | 71196695 | 788855_47188552_ENST00000601538.6,E | 1 | - | good | ATGCGGGGGAGGGCATGCAGGGAGACCCCTGGTTTGTTGTGCCAGAGAGCCTTTGACATTTCTAGGACCTTGCTAGTCTCTCTCCCTTTTTAGCAGAAAGAACCAGAGGCAGGGCCACGT | 120 |
| chr11 | 71197055 | 71197175 | 788855_47188552_ENST00000601538.6,E | 1 | - | good | TTCTACCTTCAGTTCTGCTTTGACTCCCAGGACAACAGATGGTCTCCCAGGGCAGAGATTTATGTTGATTTTGTTCCATGTGAGGTCTCAAGCCCTTAGGGAGAGAGCGAGGCACGTGAG | 120 |
| chr11 | 71197175 | 71197295 | 788855_47188552_ENST00000601538.6,E | 1 | - | good | TCCACCCCACCCGGGAGGGCCCAGCCATGTGCTCAGGCCGGACACTGGGGGGAAGGAAATGAAGAAATGGTTTTGTGGATACAAACACGTTGGGTCACACTGAGTCACGGAACTAAGTTC | 120 |
| chr11 | 71197295 | 71197415 | 788855_47188552_ENST00000601538.6,E | 1 | - | good | TTATTTGCTTATTTATTAATCTGTTTTTGAACAGGCCTGAGCAGGTTTGTGAGCTGAGGTTGGCACAGGCTGTAGCACATGCTTGGAGTGGTAGCATCTTGGAACCTGCCCACACCCTCC | 120 |
| chr11 | 71197415 | 71197535 | 788855_47188552_ENST00000601538.6,E | 1 | - | good | GGAGAGACAGAGAAGAAGATGGCCTCATGGGGCAGGGTGTTGGCGGCCTCAGCTTGCCTGGCACCAGGGAGGCCTAACCCTAGATGAAAGAACTGAGCTAGAACTTAAGAGCCATCACAC | 120 |
| chr11 | 71197535 | 71197655 | 788855_47188552_ENST00000601538.6,E | 1 | - | good | TAAATAAAATGCCTGCCAGGTTCTCTTCAGAAGATGCAGACAAAATCTAAAATCTGAGACCCATGGGGTCCATCTCCCCAGGATTCACCATCTGATAGAGCAGATGGGATTTGCCCAGAT | 120 |
| chr11 | 71197775 | 71197895 | 788855_47188552_ENST00000601538.6,E | 1 | - | good | CTTTGCTTTAAATGCCCTTTCCGTGGGGCTGAGCATGGGTAAGCGATGTGGCTCAGAAATGGCCTTTCGGTTAATCAACAGTGTTTTATAAAATGCCTGCCAGTGTAGTCCCAGCTACTT | 120 |
| chr11 | 71197895 | 71198015 | 788855_47188552_ENST00000601538.6,E | 1 | - | good | CGCTTTCACCATCCTTAGCAGTGACTGCCCCTGTGTTTGAACTTGGACTTCTCTGACGCTGGAGTTTGTGACCAGGGTGCCCGACTCCTTCTTCCCTGGCTGGGGTTGGTGATTAAGACT | 120 |
| chr11 | 71198015 | 71198135 | 788855_47188552_ENST00000601538.6,E | 1 | - | good | CTCTAAAAAAAAAAATTAATTAAAAGCAAGGCAAAAATGCAAACTCAAGCCGCTGTCCTTGAAGGCTGGGTGTGGAGCTGACCGGAAACGGCACACAGAGTGAAGCTTCGGCAACTGCTG | 120 |
| chr11 | 71198255 | 71198375 | 788855_47188552_ENST00000601538.6,E | 1 | - | good | AGCTGCCGGCCCCTCCAAACTCCCCACAGGGGGAGATCCTAACTGGGGTCACTGTCCTGTTGGGACAGATGCTGGGCAGGAGGCCCCAGCACCAGCCACAACATTCATAACATACAAAGG | 120 |
| chr11 | 71198375 | 71198495 | 788855_47188552_ENST00000601538.6,E | 1 | - | good | GGGCTGTTTCTCAAGGGTGCAGAAACCCTTCTTGGAGGCCCCGCTAGAGACCTCTCCTCATATGGCATGAGCCTTCCTCAGCCATTCTCTGGCTGGGAGAACGAGATTGTTTAACCCGAG | 120 |
| chr11 | 71198495 | 71198615 | 788855_47188552_ENST00000601538.6,E | 1 | - | good | GGTAGGGGACGGCGTCTGCGGGGTCAGCTGTGGTCTCACCCCTCACAGCTCGTGGACTCACGATGTGGTAGTGGGATGTTCCTGACAGGGTCCTGGGGAAGAGAAGGCCTGTTTCCTGTG | 120 |
| chr11 | 71198615 | 71198735 | 788855_47188552_ENST00000601538.6,E | 1 | - | good | GGATTTTGAACCTCCAGTACATCTGCTGGACGGGAGGGGACCCCGGTGGGACTGACGTGGAAGGGAGTCTGGTCTTGGCCCATCCCCAGTGTGGTGTGGAGTCAGCCATGGATGGGGGTT | 120 |
| chr11 | 71198735 | 71198855 | 788855_47188552_ENST00000601538.6,E | 1 | - | good | GAGCCACGGCCACCAGTGCAGGATCCAGAACCAACTCTGCCTTTGAAGACCCCGCCGCCTCACAACCTCACTTGCCCACAGACACGAGACATCTGCCTCAGAGGCTCACTGCTGTCTTCT | 120 |
| chr11 | 71198855 | 71198975 | 788855_47188552_ENST00000601538.6,E | 1 | - | good | GAACGGGATGGAATTCGAGGAAATGGGCCTGGTCATGACCATCGATGCTGGGCCGAGCATCACAGGCAACTCTCAAAGCTGCAGTCTAAGGAGACTGCCACAGGATGCTCAGGCCTCCAT | 120 |
| chr11 | 71198975 | 71199095 | 788855_47188552_ENST00000601538.6,E | 1 | - | good | CTTCTTGGGGGGTCCCTCATGCCCCCTGGCCTGGAGTGCCTCACCTGTGGGCTCCGGGCCACATCATTGAATACGCCTCCGCCCCGTGTCCATTGCGGCATGGTGGCAGGTGACTGAGCA | 120 |
| chr11 | 71199095 | 71199215 | 788855_47188552_ENST00000601538.6,E | 1 | - | good | GAGGTCTGGTGACTTTCTCAAAGTGACACAGCAAGTGGCAGAGCTGAGAACGCTGGCCTACAGTTCCTGTGAGTAAGGGAAGTGACTGGTAACTGCTGAGACTGTCTCTGGTCTCTGGAC | 120 |
| chr11 | 71199215 | 71199335 | 788855_47188552_ENST00000601538.6,E | 1 | - | good | CTGAGTCTTCTATTTTAATATGCGGCTTGTTCACATCTGCAGAGGCTGACTTCTGTGGATTTGCCTTGTGAGGTGGGAAGTAAAGCTCCCATTTTACAGTTGGAGGGGCTAAGGTTCAGA | 120 |
| chr11 | 71199335 | 71199455 | 788855_47188552_ENST00000601538.6,E | 1 | - | good | GTCACGTGTAGCTGCAGGTATTGCCTCTACGGCAGGTCACATTTCATTCGGTCTGGCTCCTGCCTGCCCATTTGCCTGGAGCAAACCCTGGCTGCCTACAGGTAGTGACACAGTTTGTGG | 120 |
| chr11 | 71199455 | 71199575 | 788855_47188552_ENST00000601538.6,E | 1 | - | good | GGAGTGTGCACGTTTTAGCGAAGGCTTTGAGAGCCAAGGCAACCTTGCTGCTGTGTCTTCCTCCCATCCATGAAGTGCCTCTTCCTGGGGCGTGACTGCAACCTTCATTTTGACTGAGTG | 120 |
| chr11 | 71199575 | 71199695 | 788855_47188552_ENST00000601538.6,E | 1 | - | good | GACCCAGCTGCTTCAAGGGCATCCACTGGGCTCAGGTCCTAGGGGCAGTGGCCGGCTTGTCACATCTTCCCTGCATGCTCCCTTCGAGGCTGCAGCTGCTTTGCGTGGGCTGCGGCTTGG | 120 |
| chr11 | 71199695 | 71199815 | 788855_47188552_ENST00000601538.6,E | 1 | - | good | GGCTACAGGGTCTGCACGTCTGGCCAAGGTTGTCCTCGGTCTTTCCAGCGCAGGGGATTAGGATTCCCTATGGCAGGGTGCGGTGTCCGTGCGGCCAGCTTTCCTTCCACGGTTCCTGAC | 120 |
| chr11 | 71199815 | 71199935 | 788855_47188552_ENST00000601538.6,E | 1 | - | good | TTGAGAGATTAATCTATTTGGAAATGAAGCTTCATTCTCTTTGCCAGAGGGGATTCCTGCCTGGCATTTTGCTGTCTCCTTGAATCCTTAATACCCCCGACTCTGTCCCCAGCAGCTCAC | 120 |
| chr11 | 71199935 | 71200055 | 788855_47188552_ENST00000601538.6,E | 1 | - | good | ACAACATGGCTAAAAACCATTAAATACTTTAAAGTGGTAAATTCTATGTTATGTGAATTTTATCTGTCTAAAAAATTGCTAAGGAAACAAATGCCCTAAAAGACAAAACCATGTCCGCCT | 120 |
| chr11 | 71200415 | 71200535 | 788855_47188552_ENST00000601538.6,E | 1 | - | good | CAAATGTAGAGTGAAGTTAAACGTGGAGTTAGCACATGAGCCAGCAGTTCCACTCCTGGGTGTGTCCCCAAGAGAAATGGAATCATAGGTACGAGTGAAAATGTTTACATACATGCTCAT | 120 |
| chr11 | 71200655 | 71200775 | 788855_47188552_ENST00000601538.6,E | 1 | - | good | ACCATGGTGTGAAGCTCCTGACTGAGCATGCAGATAAAGGCTTGAAGACAAAATGACAGCAGTGGGAAGAGAATTACTGTCTGAGCCATACCCATGGTATTGGTTAACATGTGGCTGCCG | 120 |
| chr11 | 71200775 | 71200895 | 788855_47188552_ENST00000601538.6,E | 1 | - | good | ATATGACAGTCACTAAAATAAAGATCTGAAGTGTGTGTGTGTGTGTGTGTGTGTGTGTGTGTGTATTAATTGAGACTTGATACCTCTCTGTGGTCTGGTGACTGAGGACAAACACATCTG | 120 |
| chr11 | 71200895 | 71201015 | 788855_47188552_ENST00000601538.6,E | 1 | - | good | CTGATGCATCCTCGGGGCCCGTGGCAGGCCCTGCCACATTGCAGGTGCCCAATATTGGTTGAATGCACCACTGAACAAAGCTTTGAGCACATACCTTTATTTACACATAGGGTTTTCCCC | 120 |
| chr11 | 71201015 | 71201135 | 788855_47188552_ENST00000601538.6,E | 1 | - | good | AGATTAATTTTCCATCTTGGGAGGAATTTTTGGGCACTGCTAATTGTCTTAAACCATCACTGTCGTGTCTGTCTCTTTGATGGCTCGGGGGAGGTCGGTGTGGGGGCTTTATGTTGTTTA | 120 |
| chr11 | 71201135 | 71201255 | 788855_47188552_ENST00000601538.6,E | 1 | - | good | AGATTTCCACCGCTTTGTCAAGTTGGAATATGGGGGAGGGCACGTGGTTGCTGTGGAGAAGGAACAGTTGAGGGAAATGAATGCATGTCATCTGCAGGAGCTCCTCATGGCCTAATTGGA | 120 |
| chr11 | 71201255 | 71201375 | 788855_47188552_ENST00000601538.6,E | 1 | - | good | CAGCACAGCTCTCCTGAGGCGTGGAAAGACGTGCTTGTCCTCGGGATCACAACGGATCCCATATATCCATGAGACAAGAGGTGCTTCTCGTGGCCGAGACCTCTCTCTTTGCAGTGCCTC | 120 |
| chr11 | 71201375 | 71201495 | 788855_47188552_ENST00000601538.6,E | 1 | - | good | GTGAGTAGCAGCAGCAGCAGCTCCGACCATCATGGCATTGGAGTGCAGTGGACACAGCGTGGTGATCACCCTTTGTGGCACACGCCAGCTCGGCGCAATCTCCGGGCGGGCAGCGGTGGG | 120 |
| chr11 | 71201495 | 71201615 | 788855_47188552_ENST00000601538.6,E | 1 | - | good | TTATCTTAAAGCTGAGGTCACTGGTGTCTGGGCTTGAAAGAGAACCGGCTCATAGGGACCCTCACTCACGAGCCCCTGCTGTGCAGGTGACCGCCATCAGTGACCGCCGTCTGTGGCCAA | 120 |
| chr11 | 71201615 | 71201735 | 788855_47188552_ENST00000601538.6,E | 1 | - | good | GAGTCTGTGTGCTGGGCGTGGACAAGTCTCTGCTTTGTACCCACTCTGTGGCCGTGAACCTGTGACCTATACTTCTCAGCCTTGCTTTTCTTATCTGTAAAATGGGAATAGTCACTGGAT | 120 |
| chr11 | 71201735 | 71201855 | 788855_47188552_ENST00000601538.6,E | 1 | - | good | CGACTGGAGGAACTGGAGTAGGTGCCTCATCCCTGCCTCCTTCCCACAGATTCATCCCCCGTCTCCTGGTGGGAGACAACGTGCTCCTTCTGGACTTCAAAGCCAGCCCCACTACCCTCG | 120 |
| chr11 | 71201855 | 71201975 | 788855_47188552_ENST00000601538.6,E | 1 | - | good | ATCCTGTAGAACAGGGGACCCTAATCCATTTGGAGGGCTCATGGCTGTGCCTTTGAAATTTGGCAATGACAGAAGCCACTGGGCAGGTTCCGGGAGGCAGGTTTGTTCCTCGGGTCAGGG | 120 |
| chr11 | 71201975 | 71202095 | 788855_47188552_ENST00000601538.6,E | 1 | - | good | GTGATTAAGCACCTACTGTGTACCCGTGAGCCATGTGGGCACACCCCTCCTCCACACAGCTGTGAAGAATCAGAGAATCAGATAAACCAGGACTTGTCCAAATAACAATAATTGGTCGGT | 120 |
| chr11 | 71202095 | 71202215 | 788855_47188552_ENST00000601538.6,E | 1 | - | good | GGGCTGCCACCCGGCAGAAGGGTGTGTCCACCAGGCCCATGCAGATGCCTTAAAGGCAGGGAGTTGTTTGTGCACCTTGCACCCTCATCATCAAGCAGCCATTCATTCACTTGGCACACA | 120 |
| chr11 | 71202215 | 71202335 | 788855_47188552_ENST00000601538.6,E | 1 | - | good | CCTCTGTGCTGCTGTGCACATTCTGAGATGTGGTTCCATGACCTGTCTAGGCCCTGAGGGGAGGAATGAGGAGGGGATGGGGCTGGGGAACCAGTGGCCTCTGAGGTGAGTTCCTGATCT | 120 |
| chr11 | 71202335 | 71202455 | 788855_47188552_ENST00000601538.6,E | 1 | - | good | CTCACCTCTCTGACCTCAGTCTATGGATCTCTGAAATGGGTTCAGTCACCTGCCCACCCGCCAGGTAGGCTGTTCTTAAGGTACAAGGACCTGAGCTGCAGAGAGTGGCTGGTTGTGTGC | 120 |
| chr11 | 71202455 | 71202575 | 788855_47188552_ENST00000601538.6,E | 1 | - | good | AGCCACCTGGAAGCCAGCTCCTGAGGAGGGGTAGCCTGAGATGTGGGTATGATGTTGAGTGTGGAGTTGGAAGACCTGGGTCCTAGCTCTGCCTTGTCTCGCCGAGGGGCTGGGAGTACC | 120 |
| chr11 | 71202575 | 71202695 | 788855_47188552_ENST00000601538.6,E | 1 | - | good | TGCTCTGACTGGCCTATGACTCAGAGACTTCCCAAAGGTCACCTTTAGTGACTCTGTGTGCTGGCTGCTTTCCCGGTGGGTCATTCTGGCCAACATCGACCCCTTTTGACACCTGGGCAG | 120 |
| chr11 | 71202695 | 71202815 | 788855_47188552_ENST00000601538.6,E | 1 | - | good | GCAGGTGACTTTGGCACAGCAGGCAGTGGAGCCGGCCAGGTGGATGGAGACAGCCTTGCTGTGCCCTTTGGGTGGGACCTGGTCAGGAATCCCACATGGACAAGAGGATTGTCCATGGTG | 120 |
| chr11 | 71202815 | 71202935 | 788855_47188552_ENST00000601538.6,E | 1 | - | good | AGTGCCAACTATCCTGGTGGCATCTGGCTCCGTTTTTTCCAAGGCAGTTTCCCATGGCTTTGACCCTGGGTCAGTGCCGTGCTCATTGTTGCCCCTTGCATGCTCATTGACTTAACCAGG | 120 |
| chr11 | 71202935 | 71203055 | 788855_47188552_ENST00000601538.6,E | 1 | - | good | ATGCCTGTGTGTACAGTCCTGCGGACACACAGTGTTTTCCATATCTCAATTTACTTCTTATTTTGCTGCTCCACAGAAAATTTGGAAAACAGAGAAAAAGGGTCAAAGTCACCCAGCCCC | 120 |
| chr11 | 71203055 | 71203175 | 788855_47188552_ENST00000601538.6,E | 1 | - | good | GCGCCTCCCCCTCACAGATGGTCATTACGCTGGTGTGACCCTTGATTCCAAAAGCTGACCCACCCTTGCAGGTGGTACCTGAGATCACTGGAACCCCACAGGGCCACCTGCTCCATGAGC | 120 |
| chr11 | 71203175 | 71203295 | 788855_47188552_ENST00000601538.6,E | 1 | - | good | ATCTCTTAGGCATGTGGGAGATTGACTTGTTGGCAAATGCTGACCACTAAGGTCTTGGCAAAGGTTCGCTGCCTCCTCACCATCTCCAGCAGACAATTGGGGCACATCTCCCTAGACCCT | 120 |
| chr11 | 71203295 | 71203415 | 788855_47188552_ENST00000601538.6,E | 1 | - | good | TGGCATCTGGTCCAAGCTGCACAGGATCAAATCTTGGTTTGACCATTTATTAGCTCTCTGATTTAGATACTTCTTATATCTCATTGTTTTCTTGTTGAAATAGGGGTTATCATAGCACCT | 120 |
| chr11 | 71203655 | 71203775 | 788855_47188552_ENST00000601538.6,E | 1 | - | good | CTACCATGTGGCCTGGGGTCCCCACTGTAACACGTGTTGTTAGCACAGGCCATCTGATGTCACTCAAGGCCCCAGGTGAACTCAGGCACTCTTACCACACAGGATATTCCCAGGGCTCAG | 120 |
| chr11 | 71203775 | 71203895 | 788855_47188552_ENST00000601538.6,E | 1 | - | good | CACACAAGGAGTCCAGACCCCTGCACCCTGCTCCTGAGCCTCGGAGCCAGGGCTTTTCCTGGGGGTTCTTCACACAGGTATGACTGGCCTTAGTCCCCAGGCCCCTGAGAGGTCACGCTG | 120 |
| chr11 | 71204015 | 71204135 | 788855_47188552_ENST00000601538.6,E | 1 | - | good | GGACAGCTGTGTTCAGGGCTCCCCAAGGCCACCCCCATCCAGGTTCACTGATTCATGAGAAGGACTCACAGAGTCCTTAGTGAAGCTGTTTTAGCGACAGGCAGGGTTCATCACAGCGAA | 120 |
| chr11 | 71204135 | 71204255 | 788855_47188552_ENST00000601538.6,E | 1 | - | good | TGGAGTTTATGGGTTTTTTTTGGATTTACATCTTTGACTTTTATAAAAGTAACTTATGTTCTTGTAAATCCAGCCAGACCTCAGTTTCCCCACCCCTGGCCTGCAGTTTTCCTCGGAACA | 120 |
| chr11 | 71204255 | 71204375 | 788855_47188552_ENST00000601538.6,E | 1 | - | good | CCGTGGGACACTCCTACCAAGCCGCACCATCAGCCCAGCAGGGGCCCTCTGCCCGTGCCCTGAGCCGGGGGCCTGGTGCCGCAGTTACCATGTGAGCACGCCCCTTCCAGATCATTTCCA | 120 |
| chr11 | 71204375 | 71204495 | 788855_47188552_ENST00000601538.6,E | 1 | - | good | GTGGTGCCTCTTCCCGTCGGAGCCTGTCACCCGAGAACGTGTCCATGGGGTCCCTTCCTGGCACAGGCTCACTTTACAGAGGCAGCGCAGATGGGAGGTGCGCTGGGTCCTGTTACCTCC | 120 |

| | | | | | | | | |
|---|---|---|---|---|---|---|---|---|
| chr11 | 71204495 | 71204615 | 788855_47188552_ENST00000601538.6,E | 1 | - | good | AGTGCAAGGGCGTGGCTGGCAGGGTACAGGCGTCCAGGTGTCAGGGTGAAGTGCACCTGCCTGGGGAGTCCCTTGCCGCAGCCTTCACCCAGGGGAGAGGAGGTGAGCAAGCGGCACCCT | 120 |
| chr11 | 71204615 | 71204735 | 788855_47188552_ENST00000601538.6,E | 1 | - | good | TCTTGGTCTTGTTTTCTGTGTCTCTGGTCAGGGTTAGACTGGGGCTGTCTTGGAAGGAGACAGTGGCCCCAGCTGAAACCTCGAGGGGACCCTGTTCTCAGAAACCCTATCTTCCCCTAA | 120 |
| chr11 | 71204735 | 71204855 | 788855_47188552_ENST00000601538.6,E | 1 | - | good | TGGATGGGCCATTGGGAAGGTGGAGTGCTCCGAACCCCTGCCTCCTGCCCCCTTCCCAGGACTTGTCTGGCGTGGCTCCAGGGTGGCCCTCATCACTTAGCAGTCCCAGAATGAGTGCAT | 120 |
| chr11 | 71204855 | 71204975 | 788855_47188552_ENST00000601538.6,E | 1 | - | good | CACCCAAGGCTGACATCACCCGAGCTAGCTGTCCCCTGCCTCCCTCATCTCCATCTGCAGAGGGACCCTCCTACCTGGGACCAGGCGTCCCTTCCCTGCCCGGCCGTGGTCAGCATCCTG | 120 |
| chr11 | 71204975 | 71205095 | 788855_47188552_ENST00000601538.6,E | 1 | - | good | CCCTGGGCACTTGGGGCCTTCCTGCCTCTGGTGTTAACTGGGGCACCCTGCTGGGGCTGCCTTCATGGGGAAGAGGAGAGAGGGCAGGTGAGGGAGGGAGGAGGGAGGCCAGGCTGACAT | 120 |
| chr11 | 71205095 | 71205215 | 788855_47188552_ENST00000601538.6,E | 1 | - | good | GTGCTGAGAGTCCGTGGCCTGGGGTACTTCTGTGGTCAGGCCTCTGGCTGCAAAGGCCACGGGAACTTGGGTCTGAGTAGCTCATTACACAAGGGTGCTACGGAATCCAGGGCTGGGACT | 120 |
| chr11 | 71205215 | 71205335 | 788855_47188552_ENST00000601538.6,E | 1 | - | good | ACCCTCTGACCAGAGAGAGGCATGACCACCAGATTTCCAGATGACCCAAAGCTAGGAAGAATGTGGGCTGGATGCGGGGTGGTGACTGTGAGAAAAGGGGCTACCCCACAGGATGGGCCC | 120 |
| chr11 | 71205335 | 71205455 | 788855_47188552_ENST00000601538.6,E | 1 | - | good | GCACAGGCAGGAGCTGGAGGGTGGAGGTTCTCGCAGGAGCGCCCAGCAGGGGCACGAGGCAGAGCCACAGGACACTGCTGGATTTGCCACTGATTGAGGGTCAGGGTGGTGCTGGTCAGT | 120 |
| chr11 | 71205455 | 71205575 | 788855_47188552_ENST00000601538.6,E | 1 | - | good | AACAAACAGCTAGAGGGACTGGGGAAGGAGGCTGAAGGGGACTCTCCGTCCTGGCACGTGGGTCTCCTGGCACTGCCTCCTGAGGGGCTGTGCAGCCTCTACTGGAGCCCAGCAAGAGGG | 120 |
| chr11 | 71205575 | 71205695 | 788855_47188552_ENST00000601538.6,E | 1 | - | good | GCATTAGCAGACCTACCCCTCTCTGGAACTTTCCTCCTGGTAAAATTTGGAAGTGTGGTGCTTCCCTGACATCTTGGAAATCTCTGGGTGGCCGACTCGGACTTCCCAAAGTTGAATGCT | 120 |
| chr11 | 71205695 | 71205815 | 788855_47188552_ENST00000601538.6,E | 1 | - | good | TCCCACAGTGACCCTGTGGACTCTGCCACCCCCCTGCCACCAACAAGTGGATAAATAACACGCCTGACCCAACATGCCCAGATGCACATTAGGAGTGCCTTGGTAGGTGTTGGGGAGGGA | 120 |
| chr11 | 71205815 | 71205935 | 788855_47188552_ENST00000601538.6,E | 1 | - | good | TTTCACCTGCTGCAGCCCTGGGGGGACACAGGGCTGAGGGGCCTTTCCTTTCTCCCAGATCTCAGCAGCCATCCAGAAACGACTCTTGACATGGCCCCAAGAGACCATCCTTGGCCATCT | 120 |
| chr11 | 71205935 | 71206055 | 788855_47188552_ENST00000601538.6,E | 1 | - | good | ATGCAGGCTGGCGCGGGGCACACGGAGAGAAATTGGACCTGGCTCCTCTCTGGATGGGGATGGGGCCGATGAGAGCTCAGCCCTGATTGGAGCTTTTCTCCCCAGAGGTCCCCAGCCCCG | 120 |
| chr11 | 71206055 | 71206175 | 788855_47188552_ENST00000601538.6,E | 1 | - | good | TTGCACTAGCATCCTTCTTCTTCCATCAGCACGGGCAGATGTGTACTTTGCAGAGTGAGTTCGCGTATTCTTTCCCCTTGTGTTCATGGAGTAGCCACTTTAGGAAACGTGGAGGGTTCC | 120 |
| chr11 | 71206175 | 71206295 | 788855_47188552_ENST00000601538.6,E | 1 | - | good | CCGGGTGTGCGGATGCTGTTGATCACGTGCAGGCGTTGTTCCAGCCCTTCCTGGAGTCGCTCTGTGTGTGTGCCTGTCCGGAGAGGGAAGCAATGGGAGGGAATTCAGAAACAGAAGACT | 120 |
| chr11 | 71206295 | 71206415 | 788855_47188552_ENST00000601538.6,E | 1 | - | good | TCCGCATTCCAGTGTTAATTCTGCCTGGATGTTTTCAGAGGGAAGAGTTGGCTTCCAACAGCTGTTTAACAGTCACGATGTAACCTCTAAGGAATAGTTAACTAAACGCACAGAGGAGAT | 120 |
| chr11 | 71206415 | 71206535 | 788855_47188552_ENST00000601538.6,E | 1 | - | good | AGAATAGTGCGATCTCTCTGCGTGGATGTAGGCTGGCGGGGGTGCGGCGCTGCTGGCTCCCGGCTTTCTCTAAATATTTACGCTCCTGTGCTTCCTGACACTCTGAGTGGCAGTTGCAGA | 120 |
| chr11 | 71206535 | 71206655 | 788855_47188552_ENST00000601538.6,E | 1 | - | good | CCTTGACGTTTTTATTATATTACAAATTATCAACTGTTTACTGAGGTAGCAAGCTAGTTTTTTGGAGGTATGGGGACATGAGCGCTGTAATTTTGCTGTGGTTCTGTAACTGTGGATATG | 120 |
| chr11 | 71207495 | 71207615 | 788855_47188552_ENST00000601538.6,E | 1 | - | good | GAAACTTGCTAATTTCCCCTTTTCTTGGAAAGAAGAAGGGAAAAATCAGAGAGAATGTAATTCACCACAACAAGCAAACAAACCTAATGTAGGAGGCTTGGCCTATCTGAGACTGTTAGA | 120 |
| chr11 | 71207615 | 71207735 | 788855_47188552_ENST00000601538.6,E | 1 | - | good | GAGAGACGGGGTCACACGGGGCAGCTGGCCCTGGCGTCAGTGCTGTCCTGCTCTGTCGATTTGTTTTTGCTTCTTGGTTAAACTTTACCTGCATCTCAGCAAAGCACTGAAAATAGATTA | 120 |
| chr11 | 71207735 | 71207855 | 788855_47188552_ENST00000601538.6,E | 1 | - | good | CCTGAGTGTGCCACCTTCCAGAGAGGCCTGGTGCAGGGTCAGCCTCCAGTGACGGTGTCACACTGAGGACTAGTTAGAGCTGTTTGGATTCTGGAAGCGAGGACTGGAAGACCTGGCTGA | 120 |
| chr11 | 71207855 | 71207975 | 788855_47188552_ENST00000601538.6,E | 1 | - | good | CTTTTTTTTTGCAGTGGAATTTGTTGAAATTTAGGTAAGCCTGCCTGGGAAGCAGGCAGGAACACAGAGGAATTCTTTTCTCGGCATTTTGCTCTCCAGCCTGGAGAGGCAAGAGTGAC | 120 |
| chr11 | 71207975 | 71208095 | 788855_47188552_ENST00000601538.6,E | 1 | - | good | TGTTCAGGGCAGCTGTGGTTTTCTTCATGTGTGGGTTTTAGGTGTCTTAAGGCCCTTAAAAGCTTTTCTTGGGAGCCCTCAGACTGCCAAACCCAGTATACCTCTCGGACACCAATCTTT | 120 |
| chr11 | 71208095 | 71208215 | 788855_47188552_ENST00000601538.6,E | 1 | - | good | ATGCCCCTCCAGGGCACCCTCCTGCCACACTGGGTGCACAGTGGGGCCCCTGGGAACACCACTGCATGGCTTTCTGTGCCCCTCCTCGGGTTTAAGTACAGGGACGCTGGACTAGTTGGA | 120 |
| chr11 | 71208215 | 71208335 | 788855_47188552_ENST00000601538.6,E | 1 | - | good | CCTTCTCCCCCCAACTTCCCCATCTCCCCTGACACTTCTGCTGGGGCTGCCCTCCTTCCCAGGACAGCTTCTCCCTGCCCTGGCACATCCCACTCAACCAGCAATGCCCACTTCCAGCTC | 120 |
| chr11 | 71208335 | 71208455 | 788855_47188552_ENST00000601538.6,E | 1 | - | good | GATTTGGAAAGCAGCATCGGGCTAGGGTTTATGGGGTTGAGATTCTTTCTCCCCTTGTTCCATAGCCTCACCCAGTCCATCCTTGGAGCCTTCTGCACATACTGGGCTCTGGTGCGTGTC | 120 |
| chr11 | 71208455 | 71208575 | 788855_47188552_ENST00000601538.6,E | 1 | - | good | GTTTTATACCCACCTGTAATGTCCACAGCACCTGCCTAGGACTCGGGAAGGGAGGGTGGGGAGGAACACTTGGGTGCACAGCCCTGTCCTGGCAACGTAATGTCCCCTGGAGCTATCCGA | 120 |
| chr11 | 71208575 | 71208695 | 788855_47188552_ENST00000601538.6,E | 1 | - | good | TCACACCTAGGGGAATTGGAATCAGGATTCCAAGCCAGTGTTTCGTATTCCAGAGACTGCTCGTTGGTCTCCCTTGCTATGCCAGCCACTCTATCCTGGGAGCTCCCTTGAGGCAGGGCT | 120 |
| chr11 | 71208695 | 71208815 | 788855_47188552_ENST00000601538.6,E | 1 | - | good | TTTATTGAGCACTTCTGTCTCAATAAAAAGTCCAGTCTTCCTAGACTCTCTTGTTAAGCCTCCAACCTGCCTTGGACAGATAAGGAAACTGAGGCTGGAGATTCAGTAACTCACCAAGGA | 120 |
| chr11 | 71208815 | 71208935 | 788855_47188552_ENST00000601538.6,E | 1 | - | good | GTCATCTCAACAGGATCATGAAATTTCTTTTTCTTCTGTTTCCTGGCCTGTGATCTCCCAGAGTTTAGCAACCTCTGCTCTAAGTCAGAGAGTGTGTGTGTGTGCGTGCCTTAGATGCGA | 120 |
| chr11 | 71208935 | 71209055 | 788855_47188552_ENST00000601538.6,E | 1 | - | good | CTTTATTCTGCTCTGACCTTTCCCCTGACATTGGGGCCGAGTGTCCTCTCTGGTGAGGAACAGTGGCTGACGGTGCATCTTACTTCTGCAGATGCCTTTTCCTGGAGACATGTAACGTTG | 120 |
| chr11 | 71209055 | 71209175 | 788855_47188552_ENST00000601538.6,E | 1 | - | good | GTGATTCAGCCTGGCTGGTTCCTGACCAGCTCCTTCCATCTGTGGTGAAATGAGCAGAGCCAGGGCCTGAGAGGGACGTTCTGCTTTTGAAGCACAGTGGCTTAGACTCAGAGGACTGGG | 120 |
| chr11 | 71209175 | 71209295 | 788855_47188552_ENST00000601538.6,E | 1 | - | good | GGATTTCTTGATGAGAACAATCATGACAGTCCTTGCAAGGCCCTGTGGTTTTCCTCAAGTCCTCACCCAGAGCGACATTTATCCTGGCCTGGGGATGGGGCTGGAGGAGCCTCCTGCTTC | 120 |
| chr11 | 71209295 | 71209415 | 788855_47188552_ENST00000601538.6,E | 1 | - | good | TGCGAACCGAAGCTCCCTCCTGGGTCAGCCCTGCACGTGGGTTTTCTTGAGGCAGGGCGTGGGAGGGTGTGTCTGATTGGCCTGTGTCGGGAGTGGCCCACTCACTACAAATTGTGAAGG | 120 |
| chr11 | 71209415 | 71209535 | 788855_47188552_ENST00000601538.6,E | 1 | - | good | AGGGGGCACCATGGCTGTTCCTGATTCTGAGAGGAGGCACGACCTATGCAAAGGGGAGCACCCAGGTGTGTCCCTGGACAGGGAAGCCTGACCCCCTCCACACCTGCTTGGCTGGGCTCC | 120 |
| chr11 | 71209535 | 71209655 | 788855_47188552_ENST00000601538.6,E | 1 | - | good | TGTCTGCTGTGTGTTCTGCCACAAGGGATGGCTGCTGCCTGGCTTTGCAGCTCTCCCAGAAGTTCACTGAGTCAAGAGTGAGGAGCTGGGTCTGGGTATGACCATGTGTGTGCTGGTGGG | 120 |
| chr11 | 71209655 | 71209775 | 788855_47188552_ENST00000601538.6,E | 1 | - | good | ATGGGATGGGACAAAGACCCTCACCTCCAGAGCAGATGTGCTGTTTGGTAGTCCAGGGACACGCAGTATGGAGGGTCTCCTGCAGGAGCACCCAGCACATTGACACCAACACAGGGTTCC | 120 |
| chr11 | 71209775 | 71209895 | 788855_47188552_ENST00000601538.6,E | 1 | - | good | ATGGACACGTCCTGCTGTCCCAGAGAGCCCCCACGGGGTGCCAGGCAGGCCCCTGGGATGTAGTCTGGCAGAGACCCCCCAGTATATCTCTCTAGGCAGCTCACGGCAGGTGCTCTGTTG | 120 |
| chr11 | 71209895 | 71210015 | 788855_47188552_ENST00000601538.6,E | 1 | - | good | TCTTGTGAAGAATGGATGCCTCAGCACACAACCCACGTCCTGTGTCGAATCCTCGGTGGTTTATTGCCACTCCCAGAGATACCCCACTGGGCGCCAGGCATCTGATGAGTCTTGTGAAGA | 120 |
| chr11 | 71210015 | 71210135 | 788855_47188552_ENST00000601538.6,E | 1 | - | good | TTTTGCAGCCCCTGGAGACCCCTTGGGACTGAGGCTGCCTTCAGAAGTTCCCTTGCAGATGATCGAAAGCTTCTGTTCTTCACATTTTGGGGTCACGGACTCGTTTGAGCATCTGATGAG | 120 |
| chr11 | 71210615 | 71210735 | 788855_47188552_ENST00000601538.6,E | 1 | - | good | GACGCCTGGGTCTCTTGGTATCCACTGAGCCCCAGGTCAAGTGGGGGGAAGTGGGTGGATGGAGTGATCTGGTCTCCAGCTGCAGTGGATTTCCTGACACTTGTGGAACAGTCTGCCAGT | 120 |
| chr11 | 71210735 | 71210855 | 788855_47188552_ENST00000601538.6,E | 1 | - | good | TGTTTTTCTCTTCAGTCCTGAGGCTTCCGGGCTCATTCTTGGAGCAGGGAGAGTGCTGTCCAGGTGTGGAGTGTGTTTTATTTAGGACAATGGTGCTCTCTGTCCCCAGTTCCTTCAGCA | 120 |
| chr11 | 71210855 | 71210975 | 788855_47188552_ENST00000601538.6,E | 1 | - | good | ACTGGAATACACCCCTGCCCTCTCCTCATGCGTCTGTCTTGGACCTGTTATGTTTCCCCAAATCCGTGAGGATGAGGGGAGTGCCGGCGCTGCATCACACTCTCTTCACTGCCCCCAAGA | 120 |
| chr11 | 71210975 | 71211095 | 788855_47188552_ENST00000601538.6,E | 1 | - | good | CTGTCATCGTGAACTTGGTAAGGGGAGCAATTAGGGCTGCAGCAGCCCTGGGGGACCCCCCTGAAACAGGACCCTGAGGGGTCAAATGGCAGGATCCTGTCAGCGGTGCAGATGGACTGA | 120 |
| chr11 | 71211095 | 71211215 | 788855_47188552_ENST00000601538.6,E | 1 | - | good | CCAAAAAAAAAAAAAAAAAAAAAAGGAAATTGACCAATAGGACACAGTGTGCTGGACTGAGTCCAGGGTTGGGTGCCAGGGAATATGGAGTAGCCACGGGGCCTGTGGGTGTCTCAGGGAG | 120 |
| chr11 | 71211215 | 71211335 | 788855_47188552_ENST00000601538.6,E | 1 | - | good | AACACCTGAGCTCAAGCGATCTGACCTCCTCGGCCTCCCAAAAGGCTGGGAGCACTGAGTTACATGCTTTAAATGAGTGAATTCGATGATGTGTAAATTTTAAAATCTCAATGTTTTTTT | 120 |
| chr11 | 71211815 | 71211935 | 788855_47188552_ENST00000601538.6,E | 1 | - | good | TTCCAGTGTGTTCCCATCTCATATTTTGAAGTGATTCAGGCATGCAACTTAAAGAACCATGCACAGAGGAGATCTGGACAACTGCAAAGCTTTTCTCAAAAGAAGTGAAAAAAAGAATGA | 120 |
| chr11 | 71211935 | 71212055 | 788855_47188552_ENST00000601538.6,E | 1 | - | good | GCGACACGGTGTTTGCAGGTGGGGTTTTGCAAAAAACAAAATGGTTTCAAATATCTTTGACCTCAGTGAAAGCTGAAAACACAGTGAATAGGCTTTGAGGTGGACGTTAAGCCAGAGTCG | 120 |
| chr11 | 71212055 | 71212175 | 788855_47188552_ENST00000601538.6,E | 1 | - | good | AGCTTCCTGGCAGCCAGGGCAAGAAGAGAAAGATGGAATAACAGAATTCTTTTGTCTAATATCAGTTCATTTGGAAAGCAGACTTTTCTTGGTGCTCTCATCTAGGAAGAATTTTTTCAT | 120 |
| chr11 | 71212175 | 71212295 | 788855_47188552_ENST00000601538.6,E | 1 | - | good | CCAAATGAGTCTTCTGCATGCCACGCTGGTAAAACTCTGCTTGAACCATGAGGAGTCTCTAAGACCCAACTGTAGCATAAGAGAGCATTTGTACAGTTAACATCCGGTTTTTAATCTCTG | 120 |
| chr11 | 71212295 | 71212415 | 788855_47188552_ENST00000601538.6,E | 1 | - | good | TGCTTTGGCTTTAAAGCCAGGTCTGAGAACATTTTTAGGCATGTTATTTAAATTGCAGATTTTAAGACCTTATCTATAGGATATCAGATTCAGTAGGAATCTGTCTTTTTAGCTGTTTTC | 120 |
| chr11 | 71212415 | 71212535 | 788855_47188552_ENST00000601538.6,E | 1 | - | good | AACAGCCACATGATGCACATGGATGCTCACTGGGTGAGGACGAAGATGATACAAGAGGAGGAACGAGGCTTGTTTTCCCCAGAATGTCTTTTAAGTCCTAGGAAAAGGGCTGTGACATTT | 120 |
| chr11 | 71212535 | 71212655 | 788855_47188552_ENST00000601538.6,E | 1 | - | good | CAGCATGCTCATTTCTATAAATCCTTCCATCTCGGCCAAAGATGGTCTCATCATCTTAGAGCAATGACAGCCCTTTCTCGTGGAATGGGCACGTTTGCTCTGTTCACTGCAGCCCCTGGA | 120 |
| chr11 | 71212655 | 71212775 | 788855_47188552_ENST00000601538.6,E | 1 | - | good | AACGTCCTAAAGCTTTGCAAAGCTCTGGTGTTTGACGTTGAGGAAGTGGATTCATGGAAACTGCTGGAACCCTTTGAGGACTTGGATTGAGTTTATTCATCCAGGGAACAGGATTGTGTT | 120 |
| chr11 | 71212775 | 71212895 | 788855_47188552_ENST00000601538.6,E | 1 | - | good | CTCCGCCTGGGGTCCCTGCTCCTCTTCCTGTGCCTCCACCTGCTGTTCCGACCTGGTTACCCAGTGAGGGAAATGGGGAAAGAAACCTCTCCTACCAGAGCAGGGATTCCTGGTGGCAAG | 120 |
| chr11 | 71213015 | 71213135 | 788855_47188552_ENST00000601538.6,E | 1 | - | good | CAGCCCTTAAGCCCCCTGGACTTACCTGGGTGGCAGCAGCAGTCCTGCACCCTGCAACGCTGTAGAAATGGGCACAGTGTCTTTCGGTTCCTGCTCCTCTTCCTTTGCCTCCACCTGGGG | 120 |
| chr11 | 71213135 | 71213255 | 788855_47188552_ENST00000601538.6,E | 1 | - | good | GGGAAATATCTGCCTCTGTTGTGAGAGCCTCCCTGTGCCTGGCCCCTCAGCTGCTAGGAGTCCCTGCTGTCCCGGGAAGATTCTCCTTTTCCACCTTTGCTTGTTCTCTGTAAAGAACCT | 120 |
| chr11 | 71213255 | 71213375 | 788855_47188552_ENST00000601538.6,E | 1 | - | good | AAGGTTGGAGCAAACTGTGTTTTCCAGAATAGAGGGACGAAAGAGTATGGTAATCTTGGCGTCCTCACAAAATACCAGACAGTTGTTGGCAGGCACAGCTGTGCAGGCTGCTGGCCGTGT | 120 |
| chr11 | 71213375 | 71213495 | 788855_47188552_ENST00000601538.6,E | 1 | - | good | GTTTTACAGGAAAATGGACTTTGAGTTAAAAACCTTGCCCCTGGGGATTTATCCAGAGGGAAGAGTAAAGTGAAACTGTGATCATAAAGATGTCCAACTTGGCACTATTTAGGATAGTGA | 120 |
| chr11 | 71213495 | 71213615 | 788855_47188552_ENST00000601538.6,E | 1 | - | good | TCAGAGTTCAACAATGCTATTTAGGTCTAATGCCCCACATGGCTCGTGCAGTGATTAAAAACAATTCCAGAGCACAGAGGACAGAGTCTGCGCTCTCCTCTGTGTGGTGTTTGAACCTCT | 120 |
| chr11 | 71213615 | 71213735 | 788855_47188552_ENST00000601538.6,E | 1 | - | good | GAGCCAGAGAGGGAGATGACATGTTTGTAGAAGGAAGATCAGCCCGGTGCTCAGGTGTGGCCACCACCTTTTCCCACAGGGAGTCAGTTTCCGAGTGTGGGGAACATCAGAAGGAGACCC | 120 |
| chr11 | 71213735 | 71213855 | 788855_47188552_ENST00000601538.6,E | 1 | - | good | GTCTGCGAGGCTCCCCTAGATGTCAAAACGTCCTAAGACAGAAAATAGAAGACATGATTAGTCCAGTTTGCTTATCAGAAAGGCCGTTCCTCTGTGTGGATGGAGGCGGCAGGCTGGCAA | 120 |

| | | | | | | | | |
|---|---|---|---|---|---|---|---|---|
| chr11 | 71213855 | 71213975 | 788855_47188552_ENST00000601538.6,E | 1 | - | good | GAACAGGAGTTGAAAAAGGGGTGGTTGGCTTGTGCATGCGAGTGCATGGTAAATGGCCCAAGGAAGCCACTTACTGTCTTTGGGAACTAGCCAGCTCTGGAAGGGGCGGTCTCTCCCAGT | 120 |
| chr11 | 71213975 | 71214095 | 788855_47188552_ENST00000601538.6,E | 1 | - | good | GAGGCAGGGCCAGGCCAGGTGAGTCACTGCAGCAGGCCTGGGGCTGTAGGGGAGGTCTTTGCTTGCCTGGCACCTGGTCCTGGGGCGCGTTAGGGCAGGGGTGGTTCATATTGGCTGGGT | 120 |
| chr11 | 71214215 | 71214335 | 788855_47188552_ENST00000601538.6,E | 1 | - | good | CTCACCTGCGTGGAAACTGTCTTCCTTGCTGCGTCTGTCTCATTTGTGCTGCGGGATTTTAACTGATGGAGGTGGCTGAGTGAGGACAAAAAGATTTCCCAAAAGAAGGTTTTATGGCTT | 120 |
| chr11 | 71214335 | 71214455 | 788855_47188552_ENST00000601538.6,E | 1 | - | good | AAGAAGTCCCAGCTGTGTGTGTTTCGGCTGTGGGTGGAAAAGAAGGGAGGGTGTCCCTCTAGGGCCTGGGGAGCCCTGGTTTGTTGCTGGGAGGTCCAGAGGGGAGCCAGCACGCGGTTC | 120 |
| chr11 | 71214455 | 71214575 | 788855_47188552_ENST00000601538.6,E | 1 | - | good | TCTCCCAACCTAGCAGGCTGAGCAGAGATGACACTTCTTCGGAAACGTGTGGTTAGGGAAGGTCGTTTTCTCAAGCAAAGCTGATGGCGCTAAATCACGATGTCTTTTGGCTACAGAAGA | 120 |
| chr11 | 71214575 | 71214695 | 788855_47188552_ENST00000601538.6,E | 1 | - | good | GCACTTGGGTGGGCCAAGGGTCTGACAGTGATGCCCCCTTGGCCTGACTCCAGTTAGCTTCTGGCCAGACTTGCTGAGACAGCAGTGGGACCCTGAGCCAGACCCATCTTCACATTTTAG | 120 |
| chr11 | 71214695 | 71214815 | 788855_47188552_ENST00000601538.6,E | 1 | - | good | ACTTCACCTTCTTAGCAATTGCACGAGAGGACAGCTGCATGGGTCCCAGTCAGAACTGCGCGATACCCCACCCCAGAGGACAGGGCAGCCCGGGGTGGCACGCGACAGATGTGTGGGTGG | 120 |
| chr11 | 71214815 | 71214935 | 788855_47188552_ENST00000601538.6,E | 1 | - | good | ATGTGTGGAAGGCGGGATGCTTGGTGGCCTTCAATACCTGGAGACCAGAACGAAAGGCTGTGGCCACCCCCAGGCCTCCCATCGAGCACCTGCTGGTCTAGAATGGCCTTTTGTGACCCC | 120 |
| chr11 | 71214935 | 71215055 | 788855_47188552_ENST00000601538.6,E | 1 | - | good | CTGTCCGCTGAAGGGAAAGCCCCATGTACCTGCAGCACAGCCAGCTCCTCTGTGTATGTGGGATCCAGGAGGAAACCGCCACGCAGCCCTATTCTGTTCTCCCTCCCTGCCGAGAGGAGG | 120 |
| chr11 | 71215055 | 71215175 | 788855_47188552_ENST00000601538.6,E | 1 | - | good | ACTCGGATGGCATTAAGCTGCACTATCACGAGGGGTTTTGATGAAGGACATCGCACATCGGGGCATGCCTGCCTGATGGTGTGAGCGTGTGGGCTGTGGGAGAATGGCTGGTGGAGGGAG | 120 |
| chr11 | 71215175 | 71215295 | 788855_47188552_ENST00000601538.6,E | 1 | - | good | GGGACCAGATGTTACTTGGTGCTGCCCCTGCTGAGACTGTGGTGGACACCGAGGCTCAGAGAGAGCGTGGGCCCTGCTGCGGCTCCAGGGTGAGAGCTGGGGCCTGTGGTTCTTTCCAGC | 120 |
| chr11 | 71215295 | 71215415 | 788855_47188552_ENST00000601538.6,E | 1 | - | good | CAGGGCAGAACCTGCTGAAGGCTTCATGGGAGACACAAAGCCTTGAGATGACAGAGTGGCGTGTTCAGACAGGGTGAGGGGAGCCAGGGAGGCCTCAAAAGTACAGACGCAGGGGGCAG | 120 |
| chr11 | 71215415 | 71215535 | 788855_47188552_ENST00000601538.6,E | 1 | - | good | CTTTCTTTTCAGGCCTGACCTCACGTTCGCTCAGGGGGCCTGTCTTGGCAGGAGGCACCATCTGAAGCTCCCTCGGTGCCTCTGGGACCTGAGCCCTGCTGATGGGCAGCCAGGGGTGAC | 120 |
| chr11 | 71215535 | 71215655 | 788855_47188552_ENST00000601538.6,E | 1 | - | good | TGGGAGCCACTGTGCCACGGATGGCTATGTGGGGACACCTCCTCGGGGAACAGTTGGGGCTGTGGCCTTCTGCAGGTGCCTGCCTGGGAAAGGAGGTACATCCTCAGCAGGCCTTCCTTG | 120 |
| chr11 | 71215655 | 71215775 | 788855_47188552_ENST00000601538.6,E | 1 | - | good | GTCTGGGTTTTGCTAAATAATCGTGTTTGCAGAGGTTTGAGCTCCTGAGCTGGCACGCCTGCAGTGTGTGCAGGGAGAGAGGTGTAGATGGATGTCCGAGGACATTAGCAGGGCCCTGAG | 120 |
| chr11 | 71215775 | 71215895 | 788855_47188552_ENST00000601538.6,E | 1 | - | good | TTGAGCATCAGCAATGTGCCCAAGGTGGTGGTAGGTGCTGCGGGAGCCTGACTCCACTCCTTGTCCCTAAAAAGGTTGTGGTCTGACTACAGAGTGGGGGTGCCGCTTCTGGGGTGTCAT | 120 |
| chr11 | 71215895 | 71216015 | 788855_47188552_ENST00000601538.6,E | 1 | - | good | CGGATGAGGGCTCCACTTGCTCCACTCCCTTGCCTGCTTCTGTTCCCATCTCTCTTCTTGGTTATCTTCATCCTGGAGGGTGGGGGAGGCGCTGTCTCATTGTGGTTCGAAAGATTTGAT | 120 |
| chr11 | 71216015 | 71216135 | 788855_47188552_ENST00000601538.6,E | 1 | - | good | TCTCGGGTAGATTCCTAGGAGTGGAGCTGCTGGGCCGTGTGGAGTCGTGTGTCCCTCAGTGGAGGAGGGACAAACCATGTTCCAGAGAGTGGCTGCGCCGTTGGACCTCCTCTCCAGCGT | 120 |
| chr11 | 71216495 | 71216615 | 788855_47188552_ENST00000601538.6,E | 1 | - | good | TGTTGACAGCAGCACGTGGCTGTATTTTCAGAGTGGCGCAGTCATCACCCAAATCAATATTAACATTTGTGAAGAGTAATCTTTTTTTATATTCCCCCCCAAAACCTGTACCATTAGAAG | 120 |
| chr11 | 71218535 | 71218655 | 788855_47188552_ENST00000601538.6,E | 1 | - | good | GCTGGGTGGTGAGAAGGTAGCGATGAAGGTTCCATACCCTGCCCTCACGCTGCTCATCCCTGGGCACCACAGATTCAATTTAAAAATAGAACAGCCCGGTCCCGGGGGCTCACACCTGTA | 120 |
| chr11 | 71218655 | 71218775 | 788855_47188552_ENST00000601538.6,E | 1 | - | good | CTGCCACTGCAACTTGTGTCTGGTTCAGGGGTCATTGGATTTTCACGTGAGTCTTTCCCATCGAGAGACCTTGGAGTTGATGGAGATTGGGTGTGTAGGTGTTCTGTGCTGATGTGGACA | 120 |
| chr11 | 71218775 | 71218895 | 788855_47188552_ENST00000601538.6,E | 1 | - | good | AGAAGAGGGAGGCATTCAGGCACAGTCTCCAGCAGTGCTGGAAGAACGAGGCTGGGTTGAGTGGCAGGACAGCAAGTCTGGGGCTTCCTTCCCAGTTAGTTACGCCCAGTGCTGTTTATG | 120 |
| chr11 | 71218895 | 71219015 | 788855_47188552_ENST00000601538.6,E | 1 | - | good | CGTTACATTCTTCTGCGATGTCTGTGTAGGATGCTGACCACCTGTGGGGTGCACAGATGCCACTCTCTCTGGGTGCCCTACAGAGAAGGCTGCTTGGGACTCCTTCCTGCTGGGGGCTGC | 120 |
| chr11 | 71219015 | 71219135 | 788855_47188552_ENST00000601538.6,E | 1 | - | good | GATGTGATTCTGAGAAATGCTCTCTCCTCATCTGGCTTCTCTAAATGCTGGTATGTGGGACATTGCACGTGATGGTGTCCTCGTGGTCTGAAGCTTGCCCACAGAGCCTTGGGTAAGCTC | 120 |
| chr11 | 71219135 | 71219255 | 788855_47188552_ENST00000601538.6,E | 1 | - | good | TGACAGACTGCATGCTCTGCAAAGTTGAGAGTAGTTACCAGCTGGACTTTTACAGAAAATTTTGTGGGCCTTGTTCTAAATGATCTCAGCGCATTGCCTAGAACCATGCCACTTTTCTCT | 120 |
| chr11 | 71219255 | 71219375 | 788855_47188552_ENST00000601538.6,E | 1 | - | good | TCAGATGTGTAATTTGTGATATTTCCTGACAATCTGGGTTCTCTTTTGTGAATAAAGTTTTATTGGCACCCAGCCGTGCTCATCTCTTTCCACGTTGTGTGTGGCAGAGTTGAATCGTTG | 120 |
| chr11 | 71221295 | 71221415 | 788855_47188552_ENST00000601538.6,E | 1 | - | good | GGACCCTGTCCTCTGCACCTGCGCTCCGGCCACCGTCTCCTGCTGGGCACCCACTGCCTTGTGCCCAACCAGAAGCTGGCAGACTGTGGTGTGAGGGTCCAATCCTGCTCATTCCTTGTT | 120 |
| chr11 | 71221415 | 71221535 | 788855_47188552_ENST00000601538.6,E | 1 | - | good | GGTGTAATCAGGCACCCCCTAATGATTAATGACATGACCCAAGAGGATATGCTCCAGAGCACCCTCCTGGGTGGGGAAACAGGGGATCAAGCGGCATTTGCCAGGGAAGGACTGGGGTGA | 120 |
| chr11 | 71221535 | 71221655 | 788855_47188552_ENST00000601538.6,E | 1 | - | good | TTAATTGACAATGTTGCAATTCTCATTGACTGTTGAAGACACTAGATTAGATGAATTTGAAGATGTGGTTTTTCCTGAGGTGTTCTTTGCTATTGTCTCAAATTATATTTTCCACTTGTG | 120 |
| chr11 | 71221655 | 71221775 | 788855_47188552_ENST00000601538.6,E | 1 | - | good | CGATCACCTCCCCTTGCTGCTCAGAAAAAAACTGGAAAATTGCCCTCGAGGGGTAATTTAGTGAGCAGCCAGGGTATCGCATAATTAGTGGCAGAAGGGTTCCACTTAGGAGTAACGGAA | 120 |
| chr11 | 71221775 | 71221895 | 788855_47188552_ENST00000601538.6,E | 1 | - | good | TCATGAACATTGTCTTGTGGGCACCTCACGCCCCCGTGCCATGGCTGCAGCGTCATTTTACAGTGGAGACTCCCAGCGCCTAGGCACCTGCAGACATCCTGGGTTTCTAACCTTGTCTAC | 120 |
| chr11 | 71221895 | 71222015 | 788855_47188552_ENST00000601538.6,E | 1 | - | good | CTTCACCCTCCAGCGGCTTCTCCTGCTTCCTCAGGTGTCCCAGTTCCGCCCTAACAGGAGCCACCAAGGGACCCTACTCCTCAGTGGCTCTCCCAATGTCCTCAGACCCCCATTCATTGA | 120 |
| chr11 | 71222015 | 71222135 | 788855_47188552_ENST00000601538.6,E | 1 | - | good | GTGGGGTGGGTGGGGTTTTCCTCTTGGGGATGTCGATGATCTGGGAAGCCCAGGCGGTGCTGGTCCTGGTGCAGGGAAGGGTGAGCCAGAATCACCAACCCGGCCATGCTCACCTGGGCG | 120 |
| chr11 | 71222135 | 71222255 | 788855_47188552_ENST00000601538.6,E | 1 | - | good | GTTTTTCTGGCCAGGGCACTTTCAGCCCACACAGTTGGTGACCTTGGGACAGGGGAGCTGTGCTGCTTTCTGGGAAGATCAGTCCCCTCGGCTGGAGGCAGGATGGTGGTGCTGGATGGG | 120 |
| chr11 | 71222255 | 71222375 | 788855_47188552_ENST00000601538.6,E | 1 | - | good | GCTTCCAGAGCAGCAGCCCTGCAACCCTGCAACCCTGGCGGCTGCGATTAATGGGAAGGAGAGAAGGAAAAACTTGTCTGGGCAACTCACGAAGGCTGGGAGTGAATTTGTTTTAGTTTT | 120 |
| chr11 | 71222375 | 71222495 | 788855_47188552_ENST00000601538.6,E | 1 | - | good | TGAAACTCCCTTCTTAGCTCCTCTGGGAGCTTCGTAAGAAAGGAACACTCAGGCTGTTGGGTTGAGCCTTTGCTTGTCACTCGGGAGCTGCCGTTGGCTCACAAGTGATGGGGAGCGCTT | 120 |
| chr11 | 71222495 | 71222615 | 788855_47188552_ENST00000601538.6,E | 1 | - | good | TCATTCTCTCTCCCGGGGCTGGGGCAGAGCTAACAACCATGGCGTGCATGCGCTTCTGCCTTTGGAAGGCGCCTGCGTCCGGCCAGCTGGGCCTCGGTTCACAGCTGGGACCTGTCTCCC | 120 |
| chr11 | 71222615 | 71222735 | 788855_47188552_ENST00000601538.6,E | 1 | - | good | TCCCTGTCCACCCAGGAAGGAGCACGTGTCCCTGCAGGCTCGGGGTTGAGCCATAAGATGTTGTCAGCAGTCAGCTGCCTTTGGCTTCCTCGGAGCTTGGACTTGGGGCTTGCTATTTAT | 120 |
| chr11 | 71222735 | 71222855 | 788855_47188552_ENST00000601538.6,E | 1 | - | good | CACAGTTCCTCCTCGAAAGGTGTCACAAGGATGCATATGAGAACGTGTGACCCTGCTGGAGAAGGTGGTCCACCCCTTTATCCTAGAGCAGGAGCAGACTGAGAGAGGGAAGGACATTAG | 120 |
| chr11 | 71222855 | 71222975 | 788855_47188552_ENST00000601538.6,E | 1 | - | good | AGGGTGCTCCTGCCTTCCCTGGCTGTCTTGGCTGAGGGCTGGGTCTGTATCCTCCACAGTCCCCCTCGTCTGTGGCCGATGCTGCATACTTGTCCCTTGTGCAGAAGCACTGGCCCAGTA | 120 |
| chr11 | 71222975 | 71223095 | 788855_47188552_ENST00000601538.6,E | 1 | - | good | TCTCTGCACCTGCATGTCCCTCCAAATCTCTGAGCTCCCTCTCTAGCCTCTGCTGTCACTCACCTCCCAGATGTGGACAAGTGTCCTCCCAGATGGAGGGATGGTGACACGGCTCCGTGG | 120 |
| chr11 | 71223095 | 71223215 | 788855_47188552_ENST00000601538.6,E | 1 | - | good | TCCTGGCCACCATCGCAGTTGGGCGGCCTCACCTGCCTGGTTTTCCTGTCTCCAGCTCTACAGCTCACCAATCCCGTCTGAAATGCAGTCGGACTCTCTCCCTCCCCATGTTCAAGCCCT | 120 |
| chr11 | 71223215 | 71223335 | 788855_47188552_ENST00000601538.6,E | 1 | - | good | GCTGATCGTTTATCTCTGCTTCACCTTCCACCACCCCATGAGAATGCACACCCTGGGGTCCTAAACGCGGCAGGCAGGAGCAGAATGTCGGAGTCACGGCCCCATGCAGGACTACCCTGG | 120 |
| chr11 | 71223695 | 71223815 | 788855_47188552_ENST00000601538.6,E | 1 | - | good | TTTTCACGTGGATGGGGGTGTACCTGGTGGTGCCCCGTGCAGGGAAGGGCCTCTCAGCTGCACGGGCCTGCCCAAGAAGCTGCGTCAGATATACTTTCCTAATTGTGTGCACCATGAGAT | 120 |
| chr11 | 71223935 | 71224055 | 788855_47188552_ENST00000601538.6,E | 1 | - | good | ATTGAGTGGCTGAATCCAGCTTGGCTTGACCACACACTGGGCTCCTGCTGGTATAGGACCCTTGCCTCCTGGCATGGCCTTGTATACCAGGGACCATAAAGACAGATCTGTGTGGTCCAG | 120 |
| chr11 | 71224055 | 71224175 | 788855_47188552_ENST00000601538.6,E | 1 | - | good | AGTATCTTATCAACGTTGACGCTGACTACCACCAGCTCCATGCAGAAAGTGGAATTACCACCATTTTCAGTGAGGACACAGGCTCTCAGCAGTTAGATACTTGGTCACGGTCACGTGGGT | 120 |
| chr11 | 71224175 | 71224295 | 788855_47188552_ENST00000601538.6,E | 1 | - | good | CTGTGTGCTCTGAAGAGGGCAGATCGTGGGCTGGTGCTCAGTGGCTCTATTCTTGAGCGCCTCCTCTGTGCCTTGCTGGTATCACCTCACCTGTAATGTGCTTAAAGGCCCCGTCTGCCA | 120 |
| chr11 | 71224415 | 71224535 | 788855_47188552_ENST00000601538.6,E | 1 | - | good | ATGGCATTGTCCTTGGTGGCCCGCTTGTGTGACCTTGCGAGATATGAAAGGGACTATGCCTCTGGCTACTGGACCGTGCTCTGAGCTGTCCTCACCAGTGTGCCCAGGCTAGACCCAGTC | 120 |
| chr11 | 71224535 | 71224655 | 788855_47188552_ENST00000601538.6,E | 1 | - | good | CAGTGGCCACTGTCTGGGTTTGGTCCACCTCCCCACCTTTGCACTGTGGCCAGGGACTTTAAGTTTCTTGTTGTCAGGCCCTGTGTGCCCTGTGCTTGTGTTTGGGTGGGAAGAGGGTGG | 120 |
| chr11 | 71224655 | 71224775 | 788855_47188552_ENST00000601538.6,E | 1 | - | good | CCCTGCTTTGGCCCTCTAGAATACAGACAGTTCCTGCTCTAAGAGGCCATCTGTGAGGCTCAGTCTCCCAACACCTCTGGTGAGTTATAACATTTTCCCCTGTCCTTGTTACCTGCAAGC | 120 |
| chr11 | 71224775 | 71224895 | 788855_47188552_ENST00000601538.6,E | 1 | - | good | GTTTTTAGCACAGAAATATCAGAATCTACATGAAGTTTTAACATGTGCCTCTCAACAGTCTGTGTCTGACCATGCCCCGGTCTCATTATCTCTCCACAGGTCTGGAAATCTCTCCAGGCA | 120 |
| chr11 | 71225255 | 71225375 | 788855_47188552_ENST00000601538.6,E | 1 | - | good | TAGCTCCTTCCGTTGCTCTATTTGATGCCACGCTGGCAGGGATTTGGTGCTTCATTATAGCCACACAGGGTGGAAATCTAGACTCCCTACTCGGCCTTTGCTGCTGTGGGTGGGGCTGAG | 120 |
| chr11 | 71225375 | 71225495 | 788855_47188552_ENST00000601538.6,E | 1 | - | good | GTGGAAAGGGGTGTCACCTCATTACTGACAGGTGCCATAGAAATCCAGGCTCCCCATGCTGCCTATACTAATGCCGGAGTGAAAGGAGGATTTGTTACTGCCTGGTGGGGATGAATGTGC | 120 |
| chr11 | 71225735 | 71225855 | 788855_47188552_ENST00000601538.6,E | 1 | - | good | CACCGTGCCCGGCCAGTTCTATCATTTTCACTTGATTCTTTTTTATATCTTTGGTGAAAGGCACTGCTCTGGTGAGGCTTTCTGTTATTTCATCTGTTTCTAGCATGTTCCTAATTGTTC | 120 |
| chr11 | 71226935 | 71227055 | 788855_47188552_ENST00000601538.6,E | 1 | - | good | ATATCATGTATACTTAATTTTATCATCTATTGTAATACCAGTTTGAGTGAAATATACAATCTTCTTTGTCTCTTTACCCTTCTCCATTTATACTTGTTAAATATTTGCTCTATATATTTA | 120 |
| chr11 | 71229215 | 71229335 | 788855_47188552_ENST00000601538.6,E | 1 | - | good | GGGTTTTATGGTTGTTTCTTTGTGATTTTTCATGCTGTGTGCAAATAGGGACAATTTTGTTCCTTCTTTTCCAATCTTTGTTTGTTTGTTTTTTTGCCATATTACACTTGCTAGAACATC | 120 |
| chr11 | 71231375 | 71231495 | 788855_47188552_ENST00000601538.6,E | 1 | - | good | AATGTCTGTACCATTTTGTATTTCCACCAGCAACGTGTAAGATCCAGGTTTTCCACCGGCAATGTGATTCTTTATTTTCTTACTAGCAATTGGTGTCACCATTTTTCATTCTAGCTGTTT | 120 |
| chr11 | 71231495 | 71231615 | 788855_47188552_ENST00000601538.6,E | 1 | - | good | GCACCTGGCCCGTAAACATAAGTCTGAATTTCTGTGGGACGAATGCCTAGGAGTGTGGTTTTTGGGTCACATGGGAGTTACATATTTAGGTTTTTAAGGAACTGCTACACTGTTTTCTGT | 120 |
| chr11 | 71232215 | 71232335 | 788855_47188552_ENST00000601538.6,E | 1 | - | good | GTATTGGCCGGGATGCTGATGTTGACGCGGTCGAGATTCAAATGCTGCTGTCCACACCAGGACCCCTCATGGTGCCCTTTTATGGCCTGACACACTTCCTTCCACCCCTACTTGCTCTAT | 120 |
| chr11 | 71232335 | 71232455 | 788855_47188552_ENST00000601538.6,E | 1 | - | good | GAAGTTCAAACCTGTGTTGCTTAAGGGCCAACTATATTTTTAAGATGATTGCAGTTGTAAGAAATAGTTCAGAGATATACCCAGTTTACCCCCATAGTCTCGTCCTGGGTCAGTACGATA | 120 |
| chr11 | 71232815 | 71232935 | 788855_47188552_ENST00000601538.6,E | 1 | - | good | TTCTTTTGTTTTCCACAGTGTGCTTTTAAAAAGACTTAAAATTTTTACAGTTAACCTTTGAACAATGGAGAGGTTGGGTTCCAACTCCCACACAGTCAAAAGTCCAGGTATAACTTGTGA | 120 |
| chr11 | 71233415 | 71233535 | 788855_47188552_ENST00000601538.6,E | 1 | - | good | ACATGTTTATTCCCCCAATAAAGACGCCAAAACCATCAGCATTGCTCCCCACTCTGATCTTCCCTCCTCTTCCCAGCCCCTGGCAACGACTAATCTGTCTCCCGTCTTGGTGAGTTTGCC | 120 |

| | | | | | | | | |
|---|---|---|---|---|---|---|---|---|
| chr11 | 71233655 | 71233775 | 788855_47188552_ENST00000601538.6,E | 1 | - | good | GTGTCTGGCCTGCAGGCCAACATTTTAAATAGTTCAGTAATTCAGTTATTCGCTACTGGGGGAAGGAGCATCCCCCTTTAACAGATGCCGTTAAAATTTCCTAGAACCCAGTTGCGGATT | 120 |
| chr11 | 71234495 | 71234615 | 788855_47188552_ENST00000601538.6,E | 1 | - | good | GGGACCGAGGCTCGGGCAGGCTGGGTCTGGCTGCCTACAGACAGGCCCCCCGACTCGGAGTCCAGCACGTGGAGGACCAGTCTGGGGCGGGGGGCATTTCCTGTGGGTCCCAGAGCTGTT | 120 |
| chr11 | 71234615 | 71234735 | 788855_47188552_ENST00000601538.6,E | 1 | - | good | GAAGAGGACGCAGACTTAGCTCGAATGCGGGCAGTGGGGAAGGCATGGAAGGGGCTGGGGTCAGGCAAGGTGGGCTGCTCCAGAGACCAGGCCCCTGCTTTGTGGCACCTTCCAGAAAAG | 120 |
| chr11 | 71234735 | 71234855 | 788855_47188552_ENST00000601538.6,E | 1 | - | good | AGGGTGTGACATTTCCTCCCTCCGTTCATGGGGTGCCTGTGGGGCAGCAGGACCACCCGGCCATGCTCCTCCAGTGCCCGGGACTCTGGAATGAGTGTGGTCACTCATCCCTCCTTGACA | 120 |
| chr11 | 71234855 | 71234975 | 788855_47188552_ENST00000601538.6,E | 1 | - | good | GTAGCTCATGTTTCGTGTCTCCATTGTGTGTGGGTGGGTGTTTTGTTCTTTCTGGGTTTTGGCTGTTGTAAAATGTGCTGTTGTGAACATTTGTATACATGTTTCGAATATTAGAGGGGG | 120 |
| chr11 | 71234975 | 71235095 | 788855_47188552_ENST00000601538.6,E | 1 | - | good | CTCTGTGCTGGGAACTGGGGACAAAGCCCAGTGTATTTCTTATCCCACAGAGGATGATTCGTGTTGTAGCGTGGATCAGTGCCTCCTTCCTTTTTATGGCTGGAGATACTCCGTTGTACG | 120 |
| chr11 | 71235575 | 71235695 | 788855_47188552_ENST00000601538.6,E | 1 | - | good | TCCTGAACCCTTGTGAGCCGTCGAGAATGAGAGGTTCTTTTTAGTTCCTTTATGGAGGTAAAATACAGGTAACATAAAGTTAACGTTTTTTTTTTTTTCCCCGAGAGAGAGTTTCACTCT | 120 |
| chr11 | 71235695 | 71235815 | 788855_47188552_ENST00000601538.6,E | 1 | - | good | TAGACCTCAGGAGGTGGCTTCTGACAAAAACTGAAGATCGGAGGCTCTTCTGTCCCTGCTGCATGCCTGTACATGGTGGCCTTTTGCCTTCAGTGTTTTCCCCTTGCTCCATGAAGTGGA | 120 |
| chr11 | 71235815 | 71235935 | 788855_47188552_ENST00000601538.6,E | 1 | - | good | CATTGCTTCTGGCACAGACTTCAGCAAAGGCAGCCGGCAGGATGGGAGATGGTGTACACACACATGTGAGAGAGTGTGTGCCCATCGTCACGGCAGCCACTCCCCACAGGCTGTGTCACA | 120 |
| chr11 | 71235935 | 71236055 | 788855_47188552_ENST00000601538.6,E | 1 | - | good | GCTGAGCAGGAAGAACGGAATGGCCAGCCTGGCTTCCTCCTGGTCCTGGCTCTGCCGCCAGCACCTGCTGGTCTGGCCTCAGTTTTCTCAGCAGGAGAATGGGTGCATCTGATAAAGTCA | 120 |
| chr11 | 71236055 | 71236175 | 788855_47188552_ENST00000601538.6,E | 1 | - | good | GAGCTGGGGATGTGGAGAATGAGGAGTGTTCTATGGATGGTGAGACGGGTCCCGAGCAACATCACCCTCAGTCCGGTCCTGCCTTTGAAACAAACAGTGTCCATGGAGGGGGGCGGAGGAG | 120 |
| chr11 | 71236175 | 71236295 | 788855_47188552_ENST00000601538.6,E | 1 | - | good | CAGCATTTTGTGGTTCAACCAGGAGGCATAATGCATTTTTAAATTTTGTTCCTGAATAATTTTGCATGGTGTGTGTTGATAAGCTAATTTCAGTCTGCCCCGTTGCCTTCTCTGGCCATT | 120 |
| chr11 | 71236895 | 71237015 | 788855_47188552_ENST00000601538.6,E | 1 | - | good | GACTGATTGGAGCCATTACTGTGACAGCCTCCCAGGCCCGCAGCGCAGTCCCACCTGCGCCAAGTCCCTCCCGGCCGCTGCCTGCAGTGTTGCCCTCTGGGCACAGAAAGACCTCATAGG | 120 |
| chr11 | 71237015 | 71237135 | 788855_47188552_ENST00000601538.6,E | 1 | - | good | TTGCTTGCATCACATTTGCTGCAGTCCCCTCGGGTTTATTGATGTGGCCAGAGCGAACGGTGACCATTTCCAGGGTCACTGTGGGAGAAGACTTCCCAAAATGCTAGATCCAGGAGGTAT | 120 |
| chr11 | 71237135 | 71237255 | 788855_47188552_ENST00000601538.6,E | 1 | - | good | GCCCACTACAGCCGGGCTCAGTCCCTGGACTCAGTCCTTATTGCCCACTGGCTACTTGGTGGTTCTGGGCTCCCAAGGGCAGCAAGAGGGGGAGCTTCAGGATGCCATGGTCTGCAGTAT | 120 |
| chr11 | 71237375 | 71237495 | 788855_47188552_ENST00000601538.6,E | 1 | - | good | AACAGACCACCCCAGACAAAGGGTAAAACACCTGTTCATTTCAGGGCTGGATAGTAAATATGTCGTGCTTGCGTGCTGTGCAGTGTCTCTCCAAAACAGTGGTCAAAGGCAATACATGAG | 120 |
| chr11 | 71237495 | 71237615 | 788855_47188552_ENST00000601538.6,E | 1 | - | good | GCAAGTGACTGGATCCTTCGCTGCAGCCCTCGGCTTTGTGATCTTTAGGTTGGAGATAACGACAGCTATCTAATAAGCCACTGTTAAGTTGAGGGTCGTGTTAGTTAACCTTTGCTGCAT | 120 |
| chr11 | 71237615 | 71237735 | 788855_47188552_ENST00000601538.6,E | 1 | - | good | CAGTTAGGGGCCCTCCTATAAAGCCTGCATTGTGGTGGGCTTGGGGCCTTTGGTGTTAGACAAACAGGATTAACTCCTGCTTTGCCGGAACCTCCCTGTAAGTGCCACGATGTGATCTGG | 120 |
| chr11 | 71237735 | 71237855 | 788855_47188552_ENST00000601538.6,E | 1 | - | good | CCTAACATGGATGAACCTTTGTTCATCAGAAATCTCATCCTGCCGTCTTTTACCAAAGAGAGAGAGAGAGGAGGTGGGGAGCCTGGGAAGGGGAGATGGGGTAGGCTCTACAGCTAGTGATT | 120 |
| chr11 | 71237855 | 71237975 | 788855_47188552_ENST00000601538.6,E | 1 | - | good | TTGGGAGGGGGCCCCCTTATCCCATGGGAACTCCAGGTGGCTGGGCTGCTCTACCACACCCCAGCCTGGGTAGTGGTGTGTGTTCATCCTTTTCGGCCTTCGTGGCCCTCAGTGGCTTTT | 120 |
| chr11 | 71237975 | 71238095 | 788855_47188552_ENST00000601538.6,E | 1 | - | good | CAGCTCTTGCAAGGGACATCGGCAACATTGGGGGCCCCCTGAGTCCCACCCTCAGTCCGTGTGTGCATGTATGCAGGGCCTGGGGACAGAGGTGAGTCTGTGTGTCACAGGGAGAGCCCC | 120 |
| chr11 | 71238095 | 71238215 | 788855_47188552_ENST00000601538.6,E | 1 | - | good | TACAGGCAGAGCAGGGAATGCTGGGGTCTCCACCTGTTCTCTGCAGAGCCTCCCACGTGGGCATCATGAGTGCGTAGGGTTCAGGGATTCAGGATGATGGGGTGCCCAGCAGCAGCCTCC | 120 |
| chr11 | 71238215 | 71238335 | 788855_47188552_ENST00000601538.6,E | 1 | - | good | TACTGTCATTAGTGGTGATGTTTGCATTATCTGTGTTGGATTGTTGGGTCTATAAAGTGCCTGTCTTGTCAGGGCTGCCCACTGGCCTGTTGAGACCTCAGGTTTGCACACAGGTCCTGT | 120 |
| chr11 | 71238335 | 71238455 | 788855_47188552_ENST00000601538.6,E | 1 | - | good | TTGGTTTTCTTATCCTTGAAATGAGGGTACTAACATATCTCAGGTAGCTGTGAGCATTAAATAAGGTAATTTATGTAAAGCTGAGACTATGGTAAGCATCCACGTACCGCCAACTGTTCT | 120 |
| chr11 | 71238455 | 71238575 | 788855_47188552_ENST00000601538.6,E | 1 | - | good | TTGGGCTTTGGGGAGTGTTGGGTTCATTTTCCTTGCTTCAGTAAGTGGTACGTTGCGGTGCAGAGCAGCTCTGCTGTGTTGCTGTGTGACCTTGGACAGGCCAGTTGGCCTCTCTGGGCC | 120 |
| chr11 | 71238575 | 71238695 | 788855_47188552_ENST00000601538.6,E | 1 | - | good | TCAGCACACTGTTCTCTAGGCTGTGGTGATGGGTGATCTGCCTGGAGAATTTGGGCCACAGAGGGAGCATAGCACTGGCCAGGAGGGGCCTCTGGCCTCTCCGCTCTTACCTCGACATCT | 120 |
| chr11 | 71238695 | 71238815 | 788855_47188552_ENST00000601538.6,E | 1 | - | good | CAGCATAGGGACAGAGAGGACACAGAGGGCCCCAGCTCTTCAACGCTTCTGCTGATTGACTCGAGCAAGCTCTGGCCACAGACAGTGCATTTGCCCAGCAGAGTTGAGTGTCACTGTGTT | 120 |
| chr11 | 71238815 | 71238935 | 788855_47188552_ENST00000601538.6,E | 1 | - | good | ACACCCACAGGTGTGTAGGGGCAACCCACCCCTACATCTTCCCATGAATGAAGTCTGTCCCTGTGACTGTGTGTCTGGACAGCTCCTCTTCTGGGGAGAGTCCTCTGTGGCAGGTGCAGT | 120 |
| chr11 | 71239055 | 71239175 | 788855_47188552_ENST00000601538.6,E | 1 | - | good | AACAGAAAGGGATGGGAGCATTTAATCTCCTTCACTCCAGTGAGTCCTCCCTCTTGGCCTGCCTGGCGGCCCCTCCCGGTCATCTGTTGACAGCCAGTCCTTGCTCAAGACCCCTGAGCA | 120 |
| chr11 | 71239175 | 71239295 | 788855_47188552_ENST00000601538.6,E | 1 | - | good | ACCAGTTGCTCTCTGGGAGTATTTTAATGCCCATTTCACCGAAGGGGAAACTGAGCTCCAGAGAGGTGCATTGGCTGGGGCTTTGAGCCCATGTAGCATGTTGCGTGTTACAACCAGTAG | 120 |
| chr11 | 71239295 | 71239415 | 788855_47188552_ENST00000601538.6,E | 1 | - | good | TAGGTAGCTTAGTCAGTCATCATCATTATCATTATTAACGTATTATCTGCAGACATTGTTGGAGGACATTTCAGTACCTGCTGTGAGCCAGGAGCTTTGTGCATATGGTCTCATGGAATC | 120 |
| chr11 | 71239775 | 71239895 | 788855_47188552_ENST00000601538.6,E | 1 | - | good | GTGGGCTTAGTGGCCTGGGGTGGGTTGGGTGATTTTCTGAACGAAAGGAAATGCCTCTGAGGTCACAGACCATCCAGCCAGCTGCAGGACACTCCACATGTAGAGGCAAAAATGACTTCT | 120 |
| chr11 | 71239895 | 71240015 | 788855_47188552_ENST00000601538.6,E | 1 | - | good | TGGAGACCCTTGGATATTCAGACGGCTTTCTGGCCACTGCCTGGTGTTTGCCCAGCTCCCGGGGATGTGTGGTTGCCTGAGACACCTGGCAGACGCCCTGCGAGACGTAGCCTGTGCCTG | 120 |
| chr11 | 71240015 | 71240135 | 788855_47188552_ENST00000601538.6,E | 1 | - | good | CGTGGCAGGCTCTTTGTAGACCAAGCCTGGTTAAATAGCTTCTGGAGTTCATTCTCCCGAAGAACGAGGTTCTTACCAGAAGCAGTCTAGAATTAGTATCATATTACTATTGAACATTCA | 120 |
| chr11 | 71240135 | 71240255 | 788855_47188552_ENST00000601538.6,E | 1 | - | good | ACCTCTTGGAGAACAATGGACCTCGGGAGAAGATACACCGTGTCTGGGTCCCGCAGCGGGTTTTCAGCAGGTTTTATGCTGGATCCGAGAATCCCTATCCTGACAAGCCTCCAAGTACCC | 120 |
| chr11 | 71240255 | 71240375 | 788855_47188552_ENST00000601538.6,E | 1 | - | good | GGACTAGGACAGCTTGTCACGGCTTGTCACTCAGAGAGACCCGAGTGTGGGCAGGACCCCAGAGCTCATTGAGAATTAATCAGGGTTCTGCACTGCTGGCTCTGGAAAGTCCAGGTCCAA | 120 |
| chr11 | 71240375 | 71240495 | 788855_47188552_ENST00000601538.6,E | 1 | - | good | CCGCCTCCACCGCCACCTTCTCACCCTTGAAGGGAGCCCCCTTCCTGCTGATGTGCTTCCTTTGAGTTAGCAGCAAAGACTCTGTTTGGAGGAACAGGAGAGGGGCTAGGGAGAGAAGAT | 120 |
| chr11 | 71240495 | 71240615 | 788855_47188552_ENST00000601538.6,E | 1 | - | good | ATTTAAAATTACATTTGTGACTCCAATTATATTGCTATGGGGCAGCCTGGATTTGTAACCTTCATTCTCTAGGGGCTGGGGTCTGCCCCTTCTTGCTTGGCCCCCTGGCCTGCAGTGTCC | 120 |
| chr11 | 71240855 | 71240975 | 788855_47188552_ENST00000601538.6,E | 1 | - | good | TAATGCGTGGGAAATGGTGTCTCCTTTTAACTTGCATTTTCTCTGATTCATAGAAAGCTGCTCCCAGTTCCGGTTCATCGACCACTCAGCTTTCCATTTTCTGAATGGACCTCCGTGGCT | 120 |
| chr11 | 71240975 | 71241095 | 788855_47188552_ENST00000601538.6,E | 1 | - | good | AGTGTTCTCAAATTGCTTTCCCAGGGGTTGTACCAAGTTACAATCCCACCCACATGGTGTGAGAACTCCCTGGGCCCCTCCTCCTCAGCAAGACTTGGTATTTCCTGATATTTACCAGTC | 120 |
| chr11 | 71241095 | 71241215 | 788855_47188552_ENST00000601538.6,E | 1 | - | good | TCAGACTGTGCTGCAGGGAACATTCTTCTATGCAGGGCTGGCGGGGCTTCCCCGGGTACCTCCTTAGAGGAGAAGTTGCCTAGCCCGGGGCTTTCACCGTGCACAGAGCTAGCTTTCACC | 120 |
| chr11 | 71241215 | 71241335 | 788855_47188552_ENST00000601538.6,E | 1 | - | good | TAAAAAAATCCACCCTGAGTTGATCATGTAGCTCAGTTTATACATTCTATGGAATGGTTAACCGCATTTTATTCTTTCTCTTCCTGTTGACATTTGGGTTGTTTTCAGGATCTTGTTCTT | 120 |
| chr11 | 71241335 | 71241455 | 788855_47188552_ENST00000601538.6,E | 1 | - | good | AGGTACGTGTTTGTTTGTCCCATGGGGCCTTGGGTCGCCTGTGTGGGGCAGGGGAAGCGGGGCTTCCGGGCACTTCCCGCTCGCTGGAAGCTTTGCCTTGATGTTTCCTTAATAGTCACT | 120 |
| chr11 | 71241455 | 71241575 | 788855_47188552_ENST00000601538.6,E | 1 | - | good | GTGTCTCCTGCAAAATGTAGCTGATTAACTGATTAGATGCTTTCCGGTTGCCTCATGATCTTAAACGCAGGCTGATTGTGCAATGGGTCATTTGCTGATCTCATATGCAGTTGCTGGCGA | 120 |
| chr11 | 71241575 | 71241695 | 788855_47188552_ENST00000601538.6,E | 1 | - | good | GTGAGGCAAGGTGCTGCTCCCTGGGAACCTGGTGGGGGTGTCTCATCTTTGTGGGTTCAGCCCCAGCCTAGAGCTGGATCCCAAGAAAAACAGGGACCAGAGACAGTAGCAGGGGGGAGA | 120 |
| chr11 | 71241695 | 71241815 | 788855_47188552_ENST00000601538.6,E | 1 | - | good | GCTGCTTTCTCACAGAAGGAGGCCCCACCTTTCTTCATGGGTCTCAGGAGAAAGTAGGGGTGACCCTGAGTGTGGGAGAAATTGTGGGACATGACTCCTTCCCATGTCTGGTGTCTTCCA | 120 |
| chr11 | 71241815 | 71241935 | 788855_47188552_ENST00000601538.6,E | 1 | - | good | TGTGCAAGTATCTTGAGTCCTTGCTTTTAGAAAACACCTTCAAAAGCAGCGTCCAGCCCCAGGCCCGTCTGGTCTTGAGATACTAAAGCTGAGATCTTTTACCCACTCGCTCCAGCCTAG | 120 |
| chr11 | 71242415 | 71242535 | 788855_47188552_ENST00000601538.6,E | 1 | - | good | GGGTGAGCTGCATTGAGGTAGCGGTCCTTGAGCAAACACTTAAAGGAGGTAAAGGAAGCGGTGTGTGATACAGGAAGAGTTCAGTGGAAATTGAGAGCACTTTTTTTCCCAATGCACATT | 120 |
| chr11 | 71242535 | 71242655 | 788855_47188552_ENST00000601538.6,E | 1 | - | good | AGGGGAGCAAACAATAATTAAAGGAGATCATAAATAAGCAGTGATGTAGTGTGGGAGGAAAGGAGAGCTTGCTGGTGGGGGCTGGGAGCCAGGGGGGCTGCTGTTTTAGCAGGATGGTCG | 120 |
| chr11 | 71242655 | 71242775 | 788855_47188552_ENST00000601538.6,E | 1 | - | good | TATTCTGCCGTCTTTTGTGGCATGTGCTGCAGGTGTCTGCTAGGCTGCTCTAGACTCTGGGATACAACAGTGAACCAGACAGACAAATATTTCTGCCCTTGTAGGGTCGACATCCTCGCC | 120 |
| chr11 | 71244095 | 71244215 | 788855_47188552_ENST00000601538.6,E | 1 | - | good | GGCTGTGGGGTCAAGGAGGCTGGCAGAGCCTAAGGCAGTTTAGGAGACTCTGAGAGAGCAGCCTGGCCAATTTATAGAGTGAGGCAGGCTTGCAAGTGTAGACAGGAGAGTGGAGAGGAC | 120 |
| chr11 | 71244215 | 71244335 | 788855_47188552_ENST00000601538.6,E | 1 | - | good | AGGATATGTCTGTGGAGTTTAGTCAGGGGCTTCAGACAGCCACCAGCTGGTTCCGACCCCAGGGGCAGGCCCTCTGATTGTGGGGTCTAGGATGGTGGGAGCATGCAGAGGCACCTTCCA | 120 |
| chr11 | 71244335 | 71244455 | 788855_47188552_ENST00000601538.6,E | 1 | - | good | AGAGCCTCTGGGAGGAGCGTGTGATCCGCCTTGTCAGTGCATGGGTCCCGTTTGAAAGCTGGCTGTGCCATGTGGTCCAAAGGGTGCTTTGAATGATGCATGTACCTTTTCTTGTCGTTC | 120 |
| chr11 | 71244455 | 71244575 | 788855_47188552_ENST00000601538.6,E | 1 | - | good | TTCCGTTGTGCACTTGGGAGAGGATGAGCTTGAAAAAGCAGCTCCCATCTCAGTGTTATGATGAGAATGGTTTGACTTCATAGATCTCTCGAATAGGCTGGTTCCCGGGGCTGCACGCTG | 120 |
| chr11 | 71245415 | 71245535 | 788855_47188552_ENST00000601538.6,E | 1 | - | good | CAGGTAAAGCAGATGAGGTAGCACAGTGGTTGTCAGACTTCTAGCTTAAGGAATCTTTTTTGCTCTCTTAGCATTTTGAAGGACCCAGAAGCTTTTTATGTATGAGCACGAATCTATTGT | 120 |
| chr11 | 71245535 | 71245655 | 788855_47188552_ENST00000601538.6,E | 1 | - | good | CCAGCTGGGATCCACACCTGGCGGCTGCCCACTCCACCGTGCAGAAGAAGGCACCGGTCGCTTCCCCGGCCTCCAGATTGCCTCCTGGGTGGCACACCAGTTTATTTTTCTGTTCACTTT | 120 |
| chr11 | 71245655 | 71245775 | 788855_47188552_ENST00000601538.6,E | 1 | - | good | TTTGGCAGGGGAAGATACTGACCCGATGGCGTCGCTACTGCCCCTCGTCTTGGTTCTGGACTTTCCCTCCCTCGAGGGTCATTCAGCTCCAGTGGCTGAACTTGCCGAAGGTCGTGTGGC | 120 |
| chr11 | 71245775 | 71245895 | 788855_47188552_ENST00000601538.6,E | 1 | - | good | AGCCGGGCTCCTGTCTGACCCCTCCTCACGAGGTGGACAGGTCCCCCAGGAAAGGGACCACGCCTTGCTCCTGTCTGCATTTCAAGGTCCTACAGTGGCCCCAAGCATGGACCAGGGGAC | 120 |
| chr11 | 71245895 | 71246015 | 788855_47188552_ENST00000601538.6,E | 1 | - | good | CCGTTCCTGTCCATGTTTTTCCTGCCACCCCGCACAAATGTCCCTGGGATTTGAAGAATCTGCTCCACTCCCCTCTTCCCGTACAGGGGCCTGCAAACTAGAGGCATTTGCCCCTGGGCC | 120 |
| chr11 | 71246015 | 71246135 | 788855_47188552_ENST00000601538.6,E | 1 | - | good | GCCTGTCTTTCCTGAAAATCCTGCCCTGGATTCCTTCCCTGGGGAGAATGCAGGCTTCCTGGGGAGTGGAGGGGGATGCTTCTGTGGCCTCATTTGTACCCCTGAGAGCATAGGGCTCAG | 120 |
| chr11 | 71246135 | 71246255 | 788855_47188552_ENST00000601538.6,E | 1 | - | good | ACTGGGGTGGGTCCTGGGAGGCTTAGGGCCCAAGGAATGGGGTGCAGGGCATCCTGGGTTTCCAGACCTGGAGCTGACACAGACTTCCGGGCTCCTGGGAGGTGGGAAATGAGCTTCCAA | 120 |
| chr11 | 71246255 | 71246375 | 788855_47188552_ENST00000601538.6,E | 1 | - | good | CACTCCGGCTCATCAGTCCCCTCCTCTTCTCAAGGAGAATCCAGAGTCAGCACCTAGAGTCCTCGGTGGGGGTGGGGAATGGCCAAGAGTCTGGACAGCTGGCAGTCTGCTCCCTGGTCC | 120 |

| | | | | | | | | |
|---|---|---|---|---|---|---|---|---|
| chr11 | 71246375 | 71246495 | 788855_47188552_ENST00000601538.6,E | 1 | - | good | GGGGACAGTGATGATACTTACGTGAAGGCAGTGACATGTGATGCCAGGTGTGTGCTCTGCATACTGCCAGGCTTTGCTGCCTCCCTGCAGACTTGGGGGTGTCCTGCCCCACGTCTTCCT | 120 |
| chr11 | 71246495 | 71246615 | 788855_47188552_ENST00000601538.6,E | 1 | - | good | TAGACTGAGCTGGGTCTACGTGGAAAGGGTCCTTGCACCTCAGTTTGAAGCCCCCTGCAAGGGAGATCTTAGTTCACAAGATTACACCTGTACTCAAGGTCCTGCAGGTGATTCTTACCT | 120 |
| chr11 | 71246615 | 71246735 | 788855_47188552_ENST00000601538.6,E | 1 | - | good | CGGGGGATAGCTATGCCTTTCTTATGTAAAAGAGCCTCAGGGATGTTAGGGAGGGGAGGTGGGCAGGGGATCCAAGGCCTTTCTCCACTGAGCAATTTCAGGAAAGACTTGGCTACCCTT | 120 |
| chr11 | 71246735 | 71246855 | 788855_47188552_ENST00000601538.6,E | 1 | - | good | TGAGTGAGAAGGCAGTGGGTGTTTTTGGTGTTTTAATTGAAGCCCTAGTTGCATTAGCCCTTTTGTTTTGGTATCAGGAGGGCCCTTCACCTGGAGCTGCACCACCCCCTCACCAGCTTC | 120 |
| chr11 | 71246855 | 71246975 | 788855_47188552_ENST00000601538.6,E | 1 | - | good | AATTACTTCTGAATTGAAAAAGCTGTACAGTCATTGCTGGGGCGCAGTTGGTCAAAAACTATAAATATATTACAGATTTTTAGAAATGGAAAAGGGGAGCTGTTACTGGAGCTGCCTCTC | 120 |
| chr11 | 71246975 | 71247095 | 788855_47188552_ENST00000601538.6,E | 1 | - | good | AGACTTTTATGCTCATGGGTCTTTTATTAATCAACCTAGCATACTTCTCTGCACGAAAAATATACATAAATTGACTTTCATAAAATTTCCCATGACCTGGAGTTGCTAAGTATTTAAAAA | 120 |
| chr11 | 71247095 | 71247215 | 788855_47188552_ENST00000601538.6,E | 1 | - | good | CCATCGTGACTCACAGCCACTTTTTCTTGACGGATACGTAAGTGAGCTTTAGGACATTTTCATCGTTCTTTTCTTGAATTTATACAGCCACTTCCCCAAAGAATTTTATCCTAATGTAAT | 120 |
| chr11 | 71247215 | 71247335 | 788855_47188552_ENST00000601538.6,E | 1 | - | good | ACGTCTAAAATTTTTCTTTATAGGTTTAAATATTTGTAAAGGCTACGGTACCTCTGTGAGGTAAATATCGATATTTTAAAAGGAACTGTTGCCTGACTCTTTACCCACTGTGATATAAAC | 120 |
| chr11 | 71247335 | 71247455 | 788855_47188552_ENST00000601538.6,E | 1 | - | good | TTCTGCAACAATTTTTTATGAAAAATTCAAATATCCAGCAAAGCTGAAACAATTGCCAGTAAACTCCTTTCAGATTGTGCCATTCATCATTTACCAAGCTTCTCACACATTTTAAAGTCA | 120 |
| chr11 | 71247455 | 71247575 | 788855_47188552_ENST00000601538.6,E | 1 | - | good | AGCACTGCAGCTCTGCCATGGCCCTGCTGGGCTGGGGCCTGCCTCTTCCATTTTTTTTTTTTTGTCTGTCCCTGAAAGTGTTGTTTTTTTTAAAAAAAACAGCTTGTTCTCTCATACATT | 120 |
| chr11 | 71247575 | 71247695 | 788855_47188552_ENST00000601538.6,E | 1 | - | good | AGAAAGGCAGCACGGGAGCTCCCCCAGAGGGAGAGGAAGATCAGCTGGAGCAGCCCTGAGTTGCTGGACTTAGAAAGGCCTCACCAGAGACCCCCCAGGAGGCGTGCATTAGAGGTCATA | 120 |
| chr11 | 71247695 | 71247815 | 788855_47188552_ENST00000601538.6,E | 1 | - | good | AGCTCAGTGAGGGGCGTGTGAATGGAATTAGGCCACAAACTCCCAATCTGAGGCTGCAGATTCCAAATGGTGGAATGCAGAAGGGCTCCTGGCTCCCCGTCCCAGAAATTCTGCTGCAGC | 120 |
| chr11 | 71247815 | 71247935 | 788855_47188552_ENST00000601538.6,E | 1 | - | good | CATGCTGGGCTCCCCGATTGGCTGCCAGATGTCATTGCCAAACCCGGCACTGCACCAGAGCACGACTGGGGACAGTGTTTTAAAATTGTCCTTGCTGGGTATGGTAGCGACTCTGGGGTT | 120 |
| chr11 | 71247935 | 71248055 | 788855_47188552_ENST00000601538.6,E | 1 | - | good | TGCACATATGCCGGGGATGGGCCTTCAGGGAGGGACTGGCTGAAACGCAGCTCGCTCCCTGGGAGATTAAGTTCTTAAAAGCCTGTGGGGCTGCCAGGAGTTTGCTGGCAGCAGCAGGGA | 120 |
| chr11 | 71248055 | 71248175 | 788855_47188552_ENST00000601538.6,E | 1 | - | good | GATGCAAGCCTTCCCTGCATTCTAGGGTTTCTGTTCTTTGCAGGGCGGGGGACAATGGGCCCATGCCCACTACCCAGAGCCTGCCCTGGGTTTAGGGACCATAGCCTGGGTCTCTCTGGA | 120 |
| chr11 | 71248175 | 71248295 | 788855_47188552_ENST00000601538.6,E | 1 | - | good | GCCCCACGTTTTCCATTGTTGAAGGCAGATTCAAGGACTGGTTTGATGCCGCTGATGGACGGCTGTCTCCTTGGCCATGGCGCCAGTACTGATGCTGGGCTTTGGGGTACAGTCGCCTCT | 120 |
| chr11 | 71248295 | 71248415 | 788855_47188552_ENST00000601538.6,E | 1 | - | good | CTGCCCAGCTGGAGAAAGCCGCCAGGCAGCGGTTAGAAAATTAGTCTTCTGGTTCCCGTGCTGTGAAATCGCGCTGCGGTCAGCAGGTGAAAACCAGTTTCTCATCATACTTCTTGATGT | 120 |
| chr11 | 71248415 | 71248535 | 788855_47188552_ENST00000601538.6,E | 1 | - | good | GATTCCCATTTCACAGATCAGAAGCTTGAAGCTCAGAGGAGTGAAGCCGCTTGCCTTAGCCCACACAGCTAGTGGGTAGGGGAGCCCTTGACCCCACACCGCCCGGTGGTTAAATGCACG | 120 |
| chr11 | 71248535 | 71248655 | 788855_47188552_ENST00000601538.6,E | 1 | - | good | AGCGCAGGTAGTGAGACTCTCACCGAGCATTTTCCTTTTGATTGGCATTAGTGCCACACTCTTGCTATTTATTCCCTCAGTAAATCTCAGAAGTCCAATGAGCCAGCTCCGATTCATTAC | 120 |
| chr11 | 71248655 | 71248775 | 788855_47188552_ENST00000601538.6,E | 1 | - | good | GGGTTGTTTGATTGCATCCTCCTGGGACTGAACATGGGCCTTTGGCAGAATGACCCAGTCCTTGTTCCCAGGGTGGGGACAGAGCCACCCGCTAGTCACAATAATAACGTCAGGAGCTGC | 120 |
| chr11 | 71248775 | 71248895 | 788855_47188552_ENST00000601538.6,E | 1 | - | good | TTAGTCCGTGTTGTGACAGGGCTTGTGATGGAGGTGAGCATCGAGTGCTGCAGGGCATAGAAAGGGTCCCCAACCCAGCGTGGGGTGGGGTGGGGGGATACAAACCCTTGGAAGAGGTCT | 120 |
| chr11 | 71248895 | 71249015 | 788855_47188552_ENST00000601538.6,E | 1 | - | good | TTTCCGCGTGAATAATCATCTGAGGGCCTGCTGCCACATCCAAGCCTGGACTGGCCACAAGGGCTGCCTCCGGGAGCTTCTAGGATAATGGGGAGGAAGAGGCGTGACTGGACAGTCCAG | 120 |
| chr11 | 71249015 | 71249135 | 788855_47188552_ENST00000601538.6,E | 1 | - | good | CCTGCAGCCCCCCAGGGCTAAGTGACCTTTGTGACTTGGCCACTGCACCTAACCTCCCACCCAAAATGAGAGGCCTGTCATTCTCTCATGGTCAGAGGATATCTAGGGTGTATTTGTTTC | 120 |
| chr11 | 71249135 | 71249255 | 788855_47188552_ENST00000601538.6,E | 1 | - | good | CCAGACAAAGGCGGGTCTTTTCCATCTCTGGAGTCCCCCACCCCTTGCCATTCACCTGGGCACCGGGAGAATGAATCACTTGTTGAGAGGCTGTAAGCATATTGAGTGCAGGGCTTGGAG | 120 |
| chr11 | 71249255 | 71249375 | 788855_47188552_ENST00000601538.6,E | 1 | - | good | GACCCTTGAGAATCTTCTTGCCTCCACATGACCTCCAGCAAAGTTCTTGACCCTGTCCGTTTCTATAGACTGGGGAATTTGTGGAAGAAAGGGAAGGAGAGGGGAGAAGGGAGGATCGCT | 120 |
| chr11 | 71249375 | 71249495 | 788855_47188552_ENST00000601538.6,E | 1 | - | good | GGGCCCCTCCTTGGGGCAGCAGGGAGGGTGTGGCTGAGAAAAACCAGTGGAAATGGACGCGCACAGCCTCTTCAGGAGCCTCCACTTCTCCTGTAACTGGGACTGTGAGCCCCAGGGAGG | 120 |
| chr11 | 71249495 | 71249615 | 788855_47188552_ENST00000601538.6,E | 1 | - | good | CCTACTCTGAAAGAGTTCTGGGCCTGGACACCTGAGGTCTTACCTGCCTGCAATTCCAGCCTCCTGCTTTGTAAACTCAAGCCGGAGCTTTCACCTGGGGCTGTTGTACCTGGAGTCTGA | 120 |
| chr11 | 71249615 | 71249735 | 788855_47188552_ENST00000601538.6,E | 1 | - | good | CTTCCAAATACGTAACAGAGTTGCAGACGAGAGAGTTCTGGGATCCCCGTCAGTCCCGATGAGGACTTTTCTAAAGTGGGGCTCAGCTTGGGAGTTGCTGCAGGGCCTGAGGTTGTGCCT | 120 |
| chr11 | 71249735 | 71249855 | 788855_47188552_ENST00000601538.6,E | 1 | - | good | TCATTAGTGACGAGGAAAAGGATGGGAGAGAAAAACGGGCCGGGAATTAACCAGTATACAGAAACACACTCCTAAAAATGGCTCTTTATAGGAAGGCAGTGCTTTAGCATACTTGCCTTA | 120 |
| chr11 | 71249855 | 71249975 | 788855_47188552_ENST00000601538.6,E | 1 | - | good | GCCATGCTGGCTTCTAGGGGTCTTGGTCTAGGCCTGGGCTTGGGGAGCCAGGTGATATCAGATGTGAGTGCTGCCCAGAGGAAGGTAAAGGGATTCTGAGACCGCATTGCATTTTATATT | 120 |
| chr11 | 71249975 | 71250095 | 788855_47188552_ENST00000601538.6,E | 1 | - | good | ATCTTTCTGTGTACTGTCTCCAAGTACAGTCTGCTATAGAAAACCGTATTCCTATTTGGGTGAGTTTCAAGATTTTCATTCATTCATTTCTTCATTCAGGTGCTTAGGGTTGTATGCAGG | 120 |
| chr11 | 71250095 | 71250215 | 788855_47188552_ENST00000601538.6,E | 1 | - | good | TACAAGTAGGTGATTTCACCTATAAAGATGATCCCAGAGGGCTCGGTTGACAGCACTTGGCTGCTTTTAACAAATTAGATTCCCCCCCACCCCCGGGGAGGGGGGCGGTCTGTTGGGGGG | 120 |
| chr11 | 71250215 | 71250335 | 788855_47188552_ENST00000601538.6,E | 1 | - | good | TTCCATTATTACACCCAGGAATTCCTTGTTTTCCTGGAAGGAAGAATTGGGTGGAAAGCTTGGTGCTTCTTTGGGTCCCTTTGGAAGGAGGTTGGAATTCAAGTATCAAAGGTAACTCTT | 120 |
| chr11 | 71250335 | 71250455 | 788855_47188552_ENST00000601538.6,E | 1 | - | good | GGATGTCACTGTCCTCTGAGGGGGCTTGAGGGAGACTCTTGTCCTGATAGGAATCTGGGTGTCCCAGTAGTGCCTGAATGATTCAGCTCCTGGGAAGCACTGTCTATATAAGGCCTCTGA | 120 |
| chr11 | 71250455 | 71250575 | 788855_47188552_ENST00000601538.6,E | 1 | - | good | GGGGTGGAGTGAGGAGAGCGAGCTCGGCCCCCATCTGGGTTCAGTGTTAATTCTGAGGGAAAGCCGTCGTGTCTGCTCTAAGAGGGGGATTGGTAGGTGGGAGCTGTGGGTTTCTATGCC | 120 |
| chr11 | 71250575 | 71250695 | 788855_47188552_ENST00000601538.6,E | 1 | - | good | GCCTGGTGCCCCAGAGGGTTCAGCAGGTCCAGAACAGCCCCGGGCCGATTGCTTTTGGAGTTTTCCAACTACTTGTGGGTTGGAAACAAGGTTGAGGGTCCTAATAGGGTTGAGCCTGTG | 120 |
| chr11 | 71250695 | 71250815 | 788855_47188552_ENST00000601538.6,E | 1 | - | good | CCCAGCCACAAGGGAAGTTCTGCGCCCCTGGAAAAGGACCTGGCAGATTCAGCCTTAACCCCGGTGCCCTGCGGTCCGCATCACATAGTTAGTGACGCCCTTCTGTGCACTAGGAACCGA | 120 |
| chr11 | 71250815 | 71250935 | 788855_47188552_ENST00000601538.6,E | 1 | - | good | AACGTTAATGAGGCCAATCAGCTAATTAAGCTTCCAGATCCAGAGAGGTAATTTCGGAAGTCCCCCGGTGGGGGAGGGCGATTTGATCTCTATCTGGGATGTGGGCCTCGGTTTTGCCGA | 120 |
| chr11 | 71250935 | 71251055 | 788855_47188552_ENST00000601538.6,E | 1 | - | good | GCTGGCAGGGGCCTGGGCGTCCTCAGAGGCCATCCGAAAATAAAGTGGTTTTCGGGTGGAAATTCGAGGTTGTAAATTGGATAGTTGAGTGACCTCTTTTTCGTTTGAAACTCAAGTTCG | 120 |
| chr11 | 71251055 | 71251175 | 788855_47188552_ENST00000601538.6,E | 1 | - | good | CCGATCTTTTAAAGGGGCTCGGGGCGCCTGGAACTGAGAAGGCGCTCGAGCCAGGCAGGTGAGGGGTGGGTGGTGGGGGGGTCAGGCGAAAGTACGTAAGGCTCTCGGGGGCTGGCAGC | 120 |
| chr11 | 71251175 | 71251295 | 788855_47188552_ENST00000601538.6,E | 1 | - | good | GGGTCTGCGTGCGGACACCCTGCTCGAGGTGGGTGGGCGCGTGGGCGAGGCTTGCCTTTCCGTGGGGCTGGAGCACCCAGAGGGAGAAAGGGTGGTGCAGGTGACCGTGGGTTTTCGCAG | 120 |
| chr11 | 71251295 | 71251415 | 788855_47188552_ENST00000601538.6,E | 1 | - | good | AGGTGCGACCACCGCAGCGGGCCTCGCCGCCCGGGTTCGGGGGGATGCGGGGGCTGGGGTGGACCTCGCCGCCAGGGTTCTGGGGCTTGGCGTGGGGGTGCAGCGCGGCGCCCTCCAG | 120 |
| chr11 | 71251415 | 71251535 | 788855_47188552_ENST00000601538.6,E | 1 | - | good | CCCGCGGCCCAGGCGCCTCCACCGGGTGGCCCCGGCCCCGGGCGCCAGGCGCGGGGACCGCTCCCAGAGACGCCGACAGCGCACTCGTCACTTGGGCTGGAGCCGCCGGACTGAGCCAG | 120 |
| chr11 | 71251535 | 71251655 | 788855_47188552_ENST00000601538.6,E | 1 | - | good | CCACCGTCCGTGAATTATGCAGGAAGCTGGCACCGGCGCGCTCGAGTTGATCAAGTTTCCCTGGCAACTTTCGCCCGGGCATCCCTCCTCCCCGGCGGCCGGGCAGCTCCGCCCGCCCAG | 120 |
| chr11 | 71251655 | 71251775 | 788855_47188552_ENST00000601538.6,E | 1 | - | good | GGGGTGGCGGGTCGGGCGCCCAGGTCCCGGGGGTCGCGCTGAGCCACAGCCGGGTGCCCGACCCCGGCCCGCCGTGCTCACCAGCTGGGATAGGGAGACCCGGCGGGCGCAGCCCGGC | 120 |
| chr11 | 71251775 | 71251895 | 788855_47188552_ENST00000601538.6,E | 1 | - | good | TCCCCCGAGACCCCGCTGCGGTCACCCTGGCCCTGCACGCAGAGCTCCCATCCAGGCGCGCGCGGCGGCAGCAGCAGCTGTCGCGGAGAGGCCGGGGCCGGCGCGCCTGGTGCGGGCCC | 120 |
| chr11 | 71251895 | 71252015 | 788855_47188552_ENST00000601538.6,E | 1 | - | good | TCCTTGCCCCGGGTCACCGCCCGCCAGCCCTAGGACCCGGGCCAGCGAGCCCAGCGGAGGAGCCGGCGCGGGGCCGCCAGGTGGCGCGCGGGGGCGGAGACCCGAACCGTGGCCGCCA | 120 |
| chr11 | 71252015 | 71252135 | 788855_47188552_ENST00000601538.6,E | 1 | - | good | AGCGCAGGAAGGAGCCCCCGCTTTATCTCCTGCGCGCATGGACCACGTAGTCTCTGGCGCGTCCTGCCCCTTCCTCTGGAACAGCGCGTGTGGCGCGACCTCCCCCGCTCTCCCGGGGTC | 120 |
| chr11 | 71252135 | 71252255 | 788855_47188552_ENST00000601538.6,E | 1 | - | good | GGGGGAGTGAGCCGCGTGTCCAGGTGGGGGTCGATTTTCCGTAGAGAGCCCTTCCCCCTGCATGCCTTTCTTCCCGGGCAGGGGCCGGTGTCGAAAGCTTTGGAAGCGGCGTGGGGGCAG | 120 |
| chr11 | 71252255 | 71252375 | 788855_47188552_ENST00000601538.6,E | 1 | - | good | TGGCCCTTGGGCACCCGATCCTGCTCACTGACGGCATTTTGGGGGCATCTCGATTCCGGGAGCACCAACTACCGGGTAAGCGCAGTCCTGGACCCAGAGAGGCTGGAGAGGTGGGCGCTG | 120 |
| chr11 | 71252375 | 71252495 | 788855_47188552_ENST00000601538.6,E | 1 | - | good | GGCCGATCCAGGTGTGCGCGGACGCTGGCGCTCCCGGGGGGCTCGGACCCGCCCCCGACTCGCCGAGCCAGGTGAGGCGGCGGCCGGGGACGCAGGGCGCGGGCCGGGGAGGACGGG | 120 |
| chr11 | 71252495 | 71252615 | 788855_47188552_ENST00000601538.6,E | 1 | - | good | TCGCCGGAGCCAGCTCCCGCGCCGGGCGGGGCCGGGTCCTTCCCCACCCCCTGCACCTGAGCCGCGGCGGCGGCTCCTGAGCTCCGGCGGGGCTGGGCGCGGCGCCGCTCGGGTTCCCG | 120 |
| chr19 | 50659228 | 50659348 | 788855_47188554_ENST00000293441.6,E | 1 | - | good | CTTCCCCTGACTCCCCACCAAGATTTTGCAGAGAGAAATGGCTTTTGTACCAGGTCATTCTTTATAATTAAACTCACAGTATCCACCCCCTCGAGCCGGCCCCGCCTCCTTCCTGGGAAG | 120 |
| chr19 | 50659348 | 50659468 | 788855_47188554_ENST00000293441.6,E | 1 | - | good | CGCGGGCTTCAGTTCATCCCTCAAGGGATTGGGACCCAGGCGTCCACGTACAGGGACTGGAGGAAGATTAGGGACGCGGGATTATAACATGGGGGTTCATTCCTACCCCCCATCACCTCC | 120 |
| chr19 | 50659468 | 50659588 | 788855_47188554_ENST00000293441.6,E | 1 | - | good | AGAATTGTCTTCCAAGGGGGGGTGGGAGGGGAAGAGGAGGGGGGCGGATGAGGACGTCTGATTCTCGCTGAGACAGGGCTCGGCACGCGCTGAAAGCCCCGGTGCCCAGGTCTTCCCTTT | 120 |
| chr19 | 50659588 | 50659708 | 788855_47188554_ENST00000293441.6,E | 1 | - | good | GAGAAAAAGCAAAAAGAGGAACAATAACAAAAGAAGAAAATATAAAAAACAGAAAAAAAAAACACAAAACCCCTAGAACACACAAAAAAATCCAGAAAAAAAACCTGAAATAAAAGCCCG | 120 |
| chr19 | 50659708 | 50659828 | 788855_47188554_ENST00000293441.6,E | 1 | - | good | TCGTTCATTCTGGATGGGTGCCGCGGGGTTGGGGAGGGTGTAGGGGGTGGGGGGGGGAGGAGAGGTCGGGGTGGGGAGTGGGGAGGGACCGAATGCGGGGACAGCGCGGGGTGGAG | 120 |
| chr19 | 50659828 | 50659948 | 788855_47188554_ENST00000293441.6,E | 1 | - | good | GGAGGGGGGAGGGGTGGTGAGAGAGGGGCAGAGGCCGCGCTGGACCCCTACCCGCCCCCTCCCCCTTGTCGGATGCCCCCAGTCCGCAGGGGGCCTGCGCGGCGCCCGTCTATCAAGGG | 120 |
| chr19 | 50659948 | 50660068 | 788855_47188554_ENST00000293441.6,E | 1 | - | good | CCCCTCTGCTGCCCCCTACCCCCGAGAGGGGGGGGCGCTCCGTCCATGAGGGGAGGGGGCTGCAAGCCCCCTCCCCTCACCATGTCAGGGATCTGCGGGGAACCTCGTGGGAGGGTCGTG | 120 |
| chr19 | 50660068 | 50660188 | 788855_47188554_ENST00000293441.6,E | 1 | - | good | CTCCCCCTGAAAAGCCCCCTCCCCTGCTGTGGGTAAAGGGGGGAGGGCGTTGCCCTCAGGTCCCCCCCCCATGTTCAGCTGCCAAAGAAGGGAGCTCCCCCTCCTTCCCCAAGCCCATTC | 120 |
| chr19 | 50660188 | 50660308 | 788855_47188554_ENST00000293441.6,E | 1 | - | good | AGATTGCCCTCTCCTAACCTGTCCACTCCAAGAAGCTCCCTCTCCCTCCAGCATGGCTGTTCCTTCCCGCACGCTTCCTCTCTCCGCTTATTCCCGGGGCTTGGGAATGTTAAGCCACCC | 120 |
| chr19 | 50660308 | 50660428 | 788855_47188554_ENST00000293441.6,E | 1 | - | good | ACTGGCTTCCTGCTCACCATGCTTCCACACTCGAGCCTCTCCCTGCCTCCAGCCCTCACCCTTCCTCACATCCTCCCTCTTCTGTCTTTCCTGGGCCATCTGTCCATCCCTTTTGCACGA | 120 |

| | | | | | | | | |
|---|---|---|---|---|---|---|---|---|
| chr19 | 50660428 | 50660548 | 788855_47188554_ENST00000293441.6,E | 1 | - | good | CTTTTCCTGACACGCTTATTTTTTATTCCTGTTGACAAGCAGCTCTTGCGGCCATTCTCATTCGGATTCTCTTACCTCCTTAAGCTCTTTCTCTTCCTGGTGAGAGAAGACCCTATGCAC | 120 |
| chr19 | 50660548 | 50660668 | 788855_47188554_ENST00000293441.6,E | 1 | - | good | CGCACCATGACATGGCTGCACACCGCATCTTGTCTTTTCTACCTGGATGAGAGCTCTACCACTTCTTTCCTTTTGCACACTCATCTCATTCCATGCACTCTTGGGCTGTAACTGTCATTT | 120 |
| chr19 | 50660668 | 50660788 | 788855_47188554_ENST00000293441.6,E | 1 | - | good | TCTGCATTTTTGATTCTTTTGTGAACCCTCTTGCACACTTGCTTTTGCACACGCGCCCCCCCCCCAGCATTTTCAGTGCTCTCCGGAAGCCTTGTTTCTCCCCTTTAATAATGTTCTTG | 120 |
| chr19 | 50660788 | 50660908 | 788855_47188554_ENST00000293441.6,E | 1 | - | good | TTTTGGTCACGCCTTTTGCACACTTGCCTCCCAGAAACTGTTTCTCTCACACATGCTCATTCTCTTCTTCTTTGCCAGGCTCCCACTCCCTTGCCCCTTTCTGTTTGAAATCTTTGCCCT | 120 |
| chr19 | 50660908 | 50661028 | 788855_47188554_ENST00000293441.6,E | 1 | - | good | CACTTTCCCTTCTTATTCTTAAAAACCCCAGATTCTTCCACTCCCCAACTCTCCCCTCTCACCATGACACTCCTAGCCCCCCACCCCCCGCCTTCCCTCTTGCCCCTTTTGCACCGTCAC | 120 |
| chr19 | 50661028 | 50661148 | 788855_47188554_ENST00000293441.6,E | 1 | - | good | ACAGACTCCATGCACATCATTCATTCTCTGGAGTGCCCTGCTCTTTGCTCAGCCCTTGCACACTTGCTATCTTATGCACCCCTCCTGACACACTCTTACCTCCATTTTCTTACGTTGAAG | 120 |
| chr19 | 50661148 | 50661268 | 788855_47188554_ENST00000293441.6,E | 1 | - | good | GCTTACACACTTTTCCCCTACCAGGGCCCTCCTTTACACACTCCATTTCACATCTCCTCCTCCCCGCTCGCTCTGGACCCTTGTTGCAGACACACTTGGGACACATTTTCCCTCCCCCTC | 120 |
| chr19 | 50661268 | 50661388 | 788855_47188554_ENST00000293441.6,E | 1 | - | good | CTGTCTTGCACATTCTGCCTTTGCTTCTGGCTGGCGAGGCCCGCTCAAATACACTCTGCTCTCCTCGCACACGCTGCATTTCCTTTCTCACGCTCCATGCCTCCTTCTCTGCTTCTTGCT | 120 |
| chr19 | 50661388 | 50661508 | 788855_47188554_ENST00000293441.6,E | 1 | - | good | TCCCCTCTCCCCAGCACACACAGCTGCCCCCCTCAGCCTCTTGCACGCTCCGTTGCACGCCTGCTCACCTCTCTCCCCTCCCGTCCCCGGGCTGCAATTTTGCACAAATTTGCCCTCTCG | 120 |
| chr19 | 50661508 | 50661628 | 788855_47188554_ENST00000293441.6,E | 1 | - | good | CCCGATTGGATGGGGACTGTCCCCCAGGCCGCAGGGGAAGGGGGTGCCCTCGGATTCCACCACCTGCCACCCCTCTAACAACCCTTCCCACTCTCTCCAGGCCCCTCAGCTCTCCCCCCT | 120 |
| chr19 | 50661628 | 50661748 | 788855_47188554_ENST00000293441.6,E | 1 | - | good | GTGTGTGTGAAGCGGGGAGCGGAGGGGGGATAGGAGAAATTACAGAGGGGAATTGAGAAGGAGGGAAGGGCCTATTCTGGGGGGAGGGGATAGACAGAGCCATAGAGGGTCAGCCCTG | 120 |
| chr19 | 50661748 | 50661868 | 788855_47188554_ENST00000293441.6,E | 1 | - | good | GGACTCTGTTCAAACTGCGCCCTGCCCTCATCTCCCAAGGCCAGAGGTCACCAGGTGGCCCAATCCAGGCCGGACATTTGCACCTCACTGAAGGGGGGCAGCTGACACAAGAGTGTGTGT | 120 |
| chr19 | 50661868 | 50661988 | 788855_47188554_ENST00000293441.6,E | 1 | - | good | CTCAAATTCTTCCTGGAGAGGTGATGGCTGGCCTGGACGGACCAGCCCCGTCCACAGAACTCTTGAGCCTGCTGGCCTCTTGACCTCTGACCCCTGACTGTCATTCTCTCCCCGGGCCAG | 120 |
| chr19 | 50662108 | 50662228 | 788855_47188554_ENST00000293441.6,E | 1 | - | good | TCCCTCTCACCGACCCGCCTGCTCTCGCTGCCCCCGGACAAGCCGTTTGGCGCTAAACCTCTGGGGTTCTGGACCAAGTTCGACGTGGCTGATTGGCTGGAGTGGCTGGGTTTGGCGGAG | 120 |
| chr19 | 50662228 | 50662348 | 788855_47188554_ENST00000293441.6,E | 1 | - | good | ATCCGTGGCTCCCCAACTGGAGGGGCAGGAGGCTCGGCTGACCCCTTCGCCCCAGTCTTTGTGCCGCCACACCCGGGGATATCCGGGGGGCTCGGGGGAGCCTTGTCAGGGGCCTCGCGC | 120 |
| chr19 | 50662348 | 50662468 | 788855_47188554_ENST00000293441.6,E | 1 | - | good | CTCCGCCGGGCCCCCAGCCCCTCGCTGCTGCCCGCCTCGGAGCACAAGGTCAGCCCTGCGCCCAGGCCCTCGTCCCTGCCCATCCTGCCTTCCGGACCCCTCTACCCAGGCCTCTTTGAC | 120 |
| chr19 | 50662468 | 50662588 | 788855_47188554_ENST00000293441.6,E | 1 | - | good | GGAACAGGGGTCTCCCCTACAGCCGCTGCGGCCCCAGGGGCCACCTCACCCTCAGCCTCCTCCTCCTCCACGTCCACCCGCCACCTCCAGGGCGTGGAGTTCGAGATGCGGCCCCCTCTG | 120 |
| chr19 | 50662588 | 50662708 | 788855_47188554_ENST00000293441.6,E | 1 | - | good | CCCACTGCCCCCTTGTCACATTCCAGACTCTCCGACGACTCCCAGTCCTCACTCCTCTCCAAGCCTGTCAGCAGCCTGTTTCAGAACTGGCCCAAACCACCTCTGCCGCCACTCCCCACC | 120 |
| chr19 | 50662708 | 50662828 | 788855_47188554_ENST00000293441.6,E | 1 | - | good | ATTTCTTCCCGTCTCTCCTCCGTCTCTCCCCCTACCCTTATGTCTCTCCTCCTCTCTCCCTATATCTTGACCTCTGCCTCTTTCTCACCCCTGCCCTTGCCCCCTAAATCCCTGTCCTCC | 120 |
| chr19 | 50662828 | 50662948 | 788855_47188554_ENST00000293441.6,E | 1 | - | good | ATCACTTAATGTCTCTTTTTCTTTCATCTCTCTTAACTGTCGCCCTCTCCCTGTCCTCTCATCTTTCTCTCCCTTCATTTCTTTCCTCGACGGCCGGCCGTCTTACCCCTTGCTCCCTCC | 120 |
| chr19 | 50663308 | 50663428 | 788855_47188554_ENST00000293441.6,E | 1 | - | good | GAGTGAGCCAGGCAGGGGGAGCAGCCAGTTCAAAGGCCCTGAGGTGGGACAGTGCTGGAGGGGAGGGAACGAAAGGGATGAGGGGAACCTTTTTCTCCCCTTGTTTCTTCTCTCTTTCAT | 120 |
| chr19 | 50663428 | 50663548 | 788855_47188554_ENST00000293441.6,E | 1 | - | good | ATAATTGCAACCATCCTACAAAAAAAGCTACCACAATGGGATGGGTGGAGAAGGTCTCTCATGTAACAGGCAAGGTGGGCAGAGTAATGAAAGTGGAGAGGGAGCAGCTGCCCAGGGCAA | 120 |
| chr19 | 50663548 | 50663668 | 788855_47188554_ENST00000293441.6,E | 1 | - | good | GTGTGCAGAGTGCCCTTCTGGGTGATGGGATCCCCTGCTGGACAGGACAGGCAAAGACTGTGCAACTCCACAGGGCTCATAGTCTAGGGACATAGGAGTGGGGGCGGGGCAGTAAACAAA | 120 |
| chr19 | 50663668 | 50663788 | 788855_47188554_ENST00000293441.6,E | 1 | - | good | TGGTTAGAGAAGAGGATTTTTTTTTTCTTTTCTTTTTTTTTTTTTCCACTGCTTGTAGCAGCAACAGGAAGGATGCTTAATTCAGTCAACAATTGCATTAATTATTGACTCATGACCTAT | 120 |
| chr19 | 50663788 | 50663908 | 788855_47188554_ENST00000293441.6,E | 1 | - | good | AGTGTATATTATTAGAGCAACTTGAATCAATGCTCCTACGCTGTAAAAATAAACAAACAAATGTGGGTTTATTAGGCCTTTAAGAACTGAATGCAATTCCTTGACTATTCTTAAGTATTC | 120 |
| chr19 | 50664148 | 50664268 | 788855_47188554_ENST00000293441.6,E | 1 | - | good | AAATATCCTTGTCCATCTACAGAGTTCACTTGAATTCTGTTTGAAGGTCTGCTATCCAAATGTGTCTGACCTTCAAATGTGGGTTCAGGGAGTGTATATTTGGGGCAACTTAAATCTATG | 120 |
| chr19 | 50664268 | 50664388 | 788855_47188554_ENST00000293441.6,E | 1 | - | good | CCCAGAGCAGCCAGTCAGCTGCATCGTCGCAGAATTCGAGCCCATTCAAGACCTTCTAGTCCTAGAGAAAGTAGAGTTACATTTCTGCTTCAGATACACAGAGGGTTGGGAGCAGGTTTT | 120 |
| chr19 | 50664388 | 50664508 | 788855_47188554_ENST00000293441.6,E | 1 | - | good | AAGAGTTTTTGAGAGGATTTAGTGTGTGCCATTGTTGTAACATGGGATGGGGGAAGAGGGGGCCTCGAACAAGAGCCCAGCAACTTCTACGTGGGGAAATACGGGATGATGCAGATGCTT | 120 |
| chr19 | 50664508 | 50664628 | 788855_47188554_ENST00000293441.6,E | 1 | - | good | GGTGTGGTTCTAGAGCCCACCTTCTCAGCTTCCAGTCCCAACCCTGCACTTTCTACCTGGGTGATCTTAGGCAAATGAATCCACCTCTCTGTAGAAGTGGAATGGCCTAGTGCCAGCCCA | 120 |
| chr19 | 50664628 | 50664748 | 788855_47188554_ENST00000293441.6,E | 1 | - | good | GAGACTGGAGTGAGACCTGCTTATGTTTGTTTCCTTGGTGAGCTGCCACTTCTTTCTGACAGTCCCTGAAATCTGTGAGTCCCTTGAAGCCAGTGTGGTTCAAGACAGGATGCAGTTATA | 120 |
| chr19 | 50666068 | 50666188 | 788855_47188554_ENST00000293441.6,E | 1 | - | good | AGCGGGGGCTGGTGGCCAACAAGGGAGGCCAGAGGTGTTGATGGCATGGCTGTGTCCCTTTAGGGTCTCAGGGTGGCAGAAAGGAAACAGAAACCAGGTTGGCAGGAGTTTGCTGTCACA | 120 |
| chr19 | 50666188 | 50666308 | 788855_47188554_ENST00000293441.6,E | 1 | - | good | CTCGGCAGCTGGCGGCGCTCTGCCCTGGGCCCGAGGAGGCAGTGGGGGAGGCGGAGACAGCCACCACGGGGGAGCCAGCTATGTCCCCGAGAGGACCTCCTCCCTGCAGCGGCAGAGGT | 120 |
| chr19 | 50666308 | 50666428 | 788855_47188554_ENST00000293441.6,E | 1 | - | good | GGGCCCACCGCCACCACCTCTGCCCGGGCCCTTGGCCCAGCCTCAGGCCTCAGCCTTGGCCACAGTAAAAGCCAGCATCATCAGTGAACTCAGCTCCAAGCTTCAGCAGTTTGGGGGCTC | 120 |
| chr19 | 50666428 | 50666548 | 788855_47188554_ENST00000293441.6,E | 1 | - | good | AGGACCCCCCACGGCAGGCGTGGCGGGGGGTCCGGTGGCTGTAGAGCCAGAAGTCCCACCGGTGCCCTTGCCGACGGCCTCCTCTCTGCCCCGGAAGCTGCTGCCCTGGGAGGAGGGCCC | 120 |
| chr19 | 50666548 | 50666668 | 788855_47188554_ENST00000293441.6,E | 1 | - | good | CAGCGGAGGCCTGCGGCCTGGCCCCAGCGGGGGACTCCGAGACCCTGTTACCCCCACCAGCCCCACCGTCTCGGTGACAGGGGCTGGAACCGATGGGCTGCTGGCCCTGCGTGCTTGTTC | 120 |
| chr19 | 50666668 | 50666788 | 788855_47188554_ENST00000293441.6,E | 1 | - | good | TCTGGACACCTATGTGGCCTACCTGGACGGCCAGGCCTTTGGGGGCAGCAGTACTCCCGGCCCGCCATACCCTCCTCAGCTCATGACTCCCTCTAAGCTCCGGGGCCGGGCGCTAGGAGC | 120 |
| chr19 | 50666788 | 50666908 | 788855_47188554_ENST00000293441.6,E | 1 | - | good | TGACCACCCACTGGAGACCATCAGCAGCGCCTCCACGCTGAGCAGCCTATCTGCCGAAGGTGGTGGCAGCGCAGGGGGTGGGGGCGGGGCTGGGGCCGGTGTGGCCAGTGGGCCGGAGC | 120 |
| chr19 | 50666908 | 50667028 | 788855_47188554_ENST00000293441.6,E | 1 | - | good | CCCCCATCCACCAGGCCCGCCTGCCCCAGCAGCACCGGCTCCCGCTGCCCCACAGCCTGGCCCGGACCCTCCGCCTGGCACGGATTCTGGCATCGAGGAGGTGGACAGTCGGAGCAGCAG | 120 |
| chr19 | 50667028 | 50667148 | 788855_47188554_ENST00000293441.6,E | 1 | - | good | TGTGCCACCCCCGGCTGTGGCCGCAGCCCCTCCCACCCTGGACTCCACCGCATCCAGCCTGACATCCTATGACAGCGAGGTGGCCACCCTGACCCAGGGGGCCTCCGCCGCTCCTGGGGA | 120 |
| chr19 | 50667148 | 50667268 | 788855_47188554_ENST00000293441.6,E | 1 | - | good | GGAACCGCTGCCTCCGCCTCTGGAATTCTCCAACAGCTTCGAAAAGCCAGAGTCGCCCCTCACGCCTGGGCCTCCCCACCCGCTGCCCGACACACCTGCCCCTGCCACCCCGTTACCCCC | 120 |
| chr19 | 50667268 | 50667388 | 788855_47188554_ENST00000293441.6,E | 1 | - | good | CCGGTCCGTGCCCCCCTCCCCGACCTCCCCGAGGGCCAGCGAAGAGAACGGGCTGCCCCTGCTGGTCCTGCCGCCTCCCGCCCCCTCGGTGGATGTGGAAGATGGCGAATTCCTTTTCGT | 120 |
| chr19 | 50667388 | 50667508 | 788855_47188554_ENST00000293441.6,E | 1 | - | good | ACCCGAGCGGCTGCCGCTGCACGTGCGGTTCCTTGAAAACTGCCAGCCCCGGGCCCCTGTGACGAGCGGAAGGGGTCCCCCCTCGGAGGACGGGCCGGGGGTCCCGCCGCCCAGCCCACG | 120 |
| chr19 | 50667508 | 50667628 | 788855_47188554_ENST00000293441.6,E | 1 | - | good | GCTACACCGCCTGCCGCCCACCGCTCCCGGGGTGGGGCCCCTCCTGCTGCAGCTGGGGACGGAGCCCCCGGCCCCGCACCCCGGAGTAAGCAAGCCCTGGAGGTCCGCAGCCCCCGAAGA | 120 |
| chr19 | 50667628 | 50667748 | 788855_47188554_ENST00000293441.6,E | 1 | - | good | GCTGCGTCTCTGGGGGGCCTCCCCGCCGGACCCTGCGCGCCGGGAGCTGGGGTACAGGGCCGGGCTGGGCAGCCAGGAGAAGTCCCTTCCCGCCAGCCCGCCCGCCGCCCGGCGTTCCCT | 120 |
| chr19 | 50667748 | 50667868 | 788855_47188554_ENST00000293441.6,E | 1 | - | good | GAAGGAGTCCTCGGAGGGCGGCGGGGCCCCCCAGCCGCCTCCCAGGCCCCCATCGCCCCGCTACGAGGCCCCGCCGCCCACCCCGCACCACCACTCGCCCCACGCCCACCACGAGCCAGT | 120 |
| chr19 | 50667868 | 50667988 | 788855_47188554_ENST00000293441.6,E | 1 | - | good | GGGCAGCAGCGCCTTCACCAGCTTCCTGCCCCCGCGACCCCTGGTGCACCCGCTGACCGGCAAGGCCCTGGATCCCGCCTCCCCGCTGGGGCTGGCCCTGGCCGCCCGCGAGCGAGCGCT | 120 |
| chr19 | 50667988 | 50668108 | 788855_47188554_ENST00000293441.6,E | 1 | - | good | CTCCAAATCCATCGACGAGGGCATGTTCTCCGCCGAGCCCTACCTCCGACTGGAGTCTGCGGGCAGCGGCGCGGGCTACGGCGGCTACGGGGCCGGTAGCCGAGCCTACGGGGGTGGCGG | 120 |
| chr19 | 50668108 | 50668228 | 788855_47188554_ENST00000293441.6,E | 1 | - | good | GGGCTGGCAGAATGAGGCGCGCCGGCGCTCCACGCTGTTCCTGTCCACCGACGCGGGGGACGAGGACGGCGGGGACGGCGGGCTGGGCACAGGGGCGGCCCCGGGCCCGCGGCTGCG | 120 |
| chr19 | 50668228 | 50668348 | 788855_47188554_ENST00000293441.6,E | 1 | - | good | GTCACCCGTGCCCCCGTCCCCCTCGCCCGTGCCCACCCCCGCCTCGCCCAGCGGCCCGGCCACGCTGGACTTCACGAGCCAGTTCGGGGCCGCCCTGGTGGGGGCGGCCCGGAGGGAGGG | 120 |
| chr19 | 50668468 | 50668588 | 788855_47188554_ENST00000293441.6,E | 1 | - | good | CGGCCTCCCGCCCGCGCCGTCGCCCACGTCCCCGGCCTCCCCGCAGCCGCCGCCCGCCGTGGCCGCGCCCTCGGAGAAGAACTCCATCCCCATCCCCACCATCATCATCAAGGCCCCGTC | 120 |
| chr19 | 50668588 | 50668708 | 788855_47188554_ENST00000293441.6,E | 1 | - | good | TTTCCAGCTGCCCCCGCGGGCGGCCAGCGCAGCCATGTACGTGCCCGCCCGCTCGGGCCGCGGCCGCAAGGGCCCGCTGGTCAAGCAGACCAAGGTGGAAGGCGAGCCCCAGAAGGGCG | 120 |
| chr19 | 50668708 | 50668828 | 788855_47188554_ENST00000293441.6,E | 1 | - | good | GCCCAGCCTGCGAGGCTGGAGGGGCGGCGGGCCCAGCCCGACCCCGGGGGCCCCGTCCCCATCGCACCACGGCAGCGCGGGCGGGGGCGGCGGCTCCTCCCAGGGCCCGGCTCTACGC | 120 |
| chr19 | 50668828 | 50668948 | 788855_47188554_ENST00000293441.6,E | 1 | - | good | CCACCACCACGCCCCGCCCCCTCAGCCCCACCACCACCACGCCCACCCCCCTCATCCTCCCGAGATGGAGACAGGCGGCTCTCCCGACGACCCTCCACCCCGCCTGGCTCTGGGGCCCCA | 120 |
| chr19 | 50668948 | 50669068 | 788855_47188554_ENST00000293441.6,E | 1 | - | good | CTCCTCCCCTGCATCCTTTGACGGGCCCTCCCCTCCCGACACTCGCGTGGGGAGCCGCGAGAAGAGCCTGTACCACAGTGGGCCCCTGCCCCCGGCCCACCACCACCCGCCCCACCACCA | 120 |
| chr19 | 50669068 | 50669188 | 788855_47188554_ENST00000293441.6,E | 1 | - | good | CACCACGTCCCCACCGGAGCCTCCCTACAGCACACCTCCAGTCCCCTCCTCCTCAGGGCGCCTCACCCCCTCCCCTCGGGGAGGGCCCTTCAACCCTGGCTCTGGTGGCCCCCTCCCCGC | 120 |
| chr19 | 50669188 | 50669308 | 788855_47188554_ENST00000293441.6,E | 1 | - | good | CTTGAGCCTCTGTATCTCCCCAGGTGCGGCAGAAGATGACAGACCTTACCTAGCACCCCCAGCCATGAAATTCAGCCGCAGCCTGTCTGTGCCTGGTTCGGAGGACATTCCCCCGCCACC | 120 |
| chr19 | 50669308 | 50669428 | 788855_47188554_ENST00000293441.6,E | 1 | - | good | AGAGCTGGGCGAGTCCACGTGAAGTTCCTGGCATATACGGGGTGCCATTCTTGTCATTAGTATTGTTGCGGTTCTATGGGATAGTGGAGCAGGGAGACCCTCCCCGCCAGGGTCCCCCTC | 120 |
| chr19 | 50669428 | 50669548 | 788855_47188554_ENST00000293441.6,E | 1 | - | good | TGGCGTGGGGAGGAAGTTCTGCTGAATTTCAGTTTCCTCACCTTCCTTGCGGTTTGCAGTTGTGCAGTGCACAACCTGTACCTCTGTATGAAGCAGATTTGGGTTTACCTCAAATAGGGA | 120 |
| chr19 | 50669548 | 50669668 | 788855_47188554_ENST00000293441.6,E | 1 | - | good | AAGAAACACGCATCTATGCATAAACACGAGGGGACTATCCCTTCCCAGCTCCAGCCACTTTTGCTGGCCCCAAACCGTTCACTGAGCCTTTGAACGGAATTAGAAAATGGCGCCCCCTCC | 120 |
| chr19 | 50669668 | 50669788 | 788855_47188554_ENST00000293441.6,E | 1 | - | good | CCTGATTTTTTTTTTTATAGATCCAATCCCTTTTAAACTTTAGCCTCATCATAAGTGGTTCTCTCTGTAGAATCACAGCTTTGATGTGCTGGTTGCATTTCTTTCCCGCATATACGTGAA | 120 |
| chr19 | 50669788 | 50669908 | 788855_47188554_ENST00000293441.6,E | 1 | - | good | CTCCTGGAGTTGTTTTTCCCTGATTGGATTCAGCTTTCCCCAACTCCTGGCGTTCAAAGTCCACCAGTGTGAGTCTTACACTCCCACTGAGTGTGGGGTCCACCTGGCTCGCATCGTTTA | 120 |
| chr19 | 50669908 | 50670028 | 788855_47188554_ENST00000293441.6,E | 1 | - | good | GACAGCAGAAAGGCAGTGAATGAAGCTGGTCACTAGTTGAGAGGGAGTAGTCAGAGGTGAGAAGAGGTTCAAAGACCGAGCCTTGGCAGTGAAGTAAGCAAAGACTTAACTCCTGCACGC | 120 |

| | | | | | | | | |
|---|---|---|---|---|---|---|---|---|
| chr19 | 50670028 | 50670148 | 788855_47188554_ENST00000293441.6,E | 1 | - | good | TGGGTTGGAGCGTTTGGAGTCTGAGATGCCCGTGAGACAGCCAGCTGGGTGTGGGTATTTGAATATAGGGGCCCAGAGCTCAGGGCGGAGGATGGGGCCCCAGGCATAAACGTGGGCGTC | 120 |
| chr19 | 50670148 | 50670268 | 788855_47188554_ENST00000293441.6,E | 1 | - | good | GCTAAGAGTGGCTGCAAGGTTTTCAGTCTTAGCACCTGAAAGGACAGAGCTGCTGTTTGCTGCGGTGGTGGGGAGGGACCGTGGGCTGGGAGGAGTCTGTTGGTGAGGAGATCAGGAGCT | 120 |
| chr19 | 50670388 | 50670508 | 788855_47188554_ENST00000293441.6,E | 1 | - | good | GAGCTGACTTCGGTTTTAACAGAAACCCTCTGGCTGCTGTGAAGGGACTAGATTGAAGGGGATGAGCTGGGAAGCCGGGGGACCAAAGAGGAGGTGACAGCTATTGCTATCCAGATAGGA | 120 |
| chr19 | 50670508 | 50670628 | 788855_47188554_ENST00000293441.6,E | 1 | - | good | TGAGGGCAGTGGGGGGACAGGGTCCCGATCCTGCAGGGTCTGGGGAGTCATGGTGAGGACTTGAACCTTTGCCCTGAGTGAGGTGAGTGAATGGCAGGGCTCTAAGCAGAGGAGGGACAT | 120 |
| chr19 | 50670628 | 50670748 | 788855_47188554_ENST00000293441.6,E | 1 | - | good | GTTGAGGCATTCCAGGCAGAGCGAACTACCTCTGCAAAGGTCCTGAGGTCGGGCTGAGTCTGGTGCGTTTACAGAACAGGGAGGAGGCCTGTGTAGCAGGGGCCAAGTGAGCACTGGACA | 120 |
| chr19 | 50671468 | 50671588 | 788855_47188554_ENST00000293441.6,E | 1 | - | good | GCCTTATAGCAGGGGAAGACAGATAATGGATTATAAGGAAAATAAGCAAGAAAATTATCTAGTATGTTAGCAGTTGTACGTGCTATGGAAAACAAAAAAAAGAGTGGGCCGGGCGCAGCG | 120 |
| chr19 | 50671588 | 50671708 | 788855_47188554_ENST00000293441.6,E | 1 | - | good | TCCATCCATTCATTCATTTTACAAGTGTTTACCAAGCGCTCCCCACATGCCAGGAATTGTTCTAGAATGGGGATACGTCAGCGAATAAAACAGACTAAAGGTTCCTGCCCTCATGGGCTT | 120 |
| chr19 | 50671708 | 50671828 | 788855_47188554_ENST00000293441.6,E | 1 | - | good | TGTGTGAACTGGGGCAGGTGGCTTGACTTCTCTGGGCCTCTGGCTCTACATCTGGAGAATGGGGATGCAGTGAGCTAAAGCCCCCGCTACAGCTTTCTGGTTGTTTCTCTTATACCCGAT | 120 |
| chr19 | 50671828 | 50671948 | 788855_47188554_ENST00000293441.6,E | 1 | - | good | GCAGACTTGGAAATGTTCTGTGCTCCCAGAGAGACCATAGTTTCATCAAATTGGGTTAAGAGTTCAGGCTTTGGAAATGGGTGAACTTGGCCATGAATCCTGGCTCCCCTCCTTCCCAGC | 120 |
| chr19 | 50671948 | 50672068 | 788855_47188554_ENST00000293441.6,E | 1 | - | good | CTTCTTTCCTGCCTCCAGGACCTGGGTTGATGCTCCGGCAAAAATCTATCGGTATGCCACCCAGAAGAAGAGATGTGGGGGCTGGGGGAGGCCAGGAACGTGACAGTTCAGGAAAAGCAA | 120 |
| chr19 | 50672068 | 50672188 | 788855_47188554_ENST00000293441.6,E | 1 | - | good | GCCACAAGCCTTTCTGCGCTGACTGATCTCTCTCTATCTTTTCTCTCTCTCCCCCTCTGACTGCCGCTCCAAAGTCGAGCTTCGATCCCCACCACCGTGCCCAGCCAAGTTACGAGCGTC | 120 |
| chr19 | 50672668 | 50672788 | 788855_47188554_ENST00000293441.6,E | 1 | - | good | GTGGGGGATCAAGGATGGCAGCAGGGAGATGAGGGAAGAGGCTGCTCCATCCCGGAGAACACTGGTGGGGACCTGGATGTGGGCAGTGGCTGGAGGAAGGAAGCAGATCAAGAGTCGAT | 120 |
| chr19 | 50672788 | 50672908 | 788855_47188554_ENST00000293441.6,E | 1 | - | good | GAGTTGGGAGTTTGGTCCCCTCTGATCTCCTCCTAACTGACATGGGATTTCAAGCAGGGGCAATAGCATGCTTTTTGAAAGATGCCTCTGACTCTTGTGTGGAAAACAGACTGGAACAGG | 120 |
| chr19 | 50672908 | 50673028 | 788855_47188554_ENST00000293441.6,E | 1 | - | good | TCTTAAGGGCTGCAGGCAGGTGACCCTCCCAGCAGTGGGTGAGGGATGGGATGCTGCATCCAGGGGGAGGGCAGCTGCAGGGGCACCAGCAGTGCCTGCCATAGAGGGTCTTGTAGACCA | 120 |
| chr19 | 50673028 | 50673148 | 788855_47188554_ENST00000293441.6,E | 1 | - | good | GCACCCCGGCTTTAGTGCTGCCTCGGGCCTCCGTATCCGTCCATCGTGGGCTGTGGGCTGCTCCAGGAAGCAGGGACACATAACGGACTGGGCGAGTCCAGGACTCTGAAGGAGGGTGAT | 120 |
| chr19 | 50673148 | 50673268 | 788855_47188554_ENST00000293441.6,E | 1 | - | good | CTGTACCAAGGGTTTGCTACGGGGGAACCTTCAGGGAGTGTGGGAGACTGGCTGGGGTTGTGGAAGGAAGTCAAGCAGACACGGTTCACGCTGGAGTGGGAACCCATGGGGTCAGGCTGG | 120 |
| chr19 | 50673268 | 50673388 | 788855_47188554_ENST00000293441.6,E | 1 | - | good | AGAGGTGGGGGTGGAGGTGGACCAAGGGAGCAAGTGGGAAATGGGGGCAGGGCTGGGTCCCGCCTGCAGGACCTGTCTTCGTGTGGGCTTCCCTAGAACGGGAGCCGGAGGTGGGGATT | 120 |
| chr19 | 50673388 | 50673508 | 788855_47188554_ENST00000293441.6,E | 1 | - | good | GAGGAGAACTTCGCGCTGAAAAGGAACTGTGACAATAAGAGGGAAAGCTGTTGCCGAGAGAACCAGGGCTCGCAGGTGTTCTCCAATGAGGGTAGACCTGGGGCTGACAGAGGAGAAGGA | 120 |
| chr19 | 50673508 | 50673628 | 788855_47188554_ENST00000293441.6,E | 1 | - | good | AGGCTAGTGGAGGAAACAGACAGCAAGTAAACAAGATGGTTTCTGGGGAAGAGAGATGCTGTGAAGAAAATGGGACAGAGGGAAGTGCAGTCTGGGTAGGGGGCCGGGGAAGGTCTCGCC | 120 |
| chr19 | 50673628 | 50673748 | 788855_47188554_ENST00000293441.6,E | 1 | - | good | TTTTTTCTTTCACTTCTACTTTGAAAAGAAATGAAAACACTTCCCCTAAAAGCATCATGGGCCCTAGACACTGTAGGTGCGATGCCTGGGATACATCAGCCCCGCCTTCAGGGAGTTTCT | 120 |
| chr19 | 50674108 | 50674228 | 788855_47188554_ENST00000293441.6,E | 1 | - | good | GGCCCTATGCGGGGACCGGGGACTCAGTAGGAAAGGAGACTGATAAAGTCACTACCCTACCCTCAGGCAGGTTTCATAATGACTTCCATGAACCTTAGGCACTTTTGCCCTTCTGCATTA | 120 |
| chr19 | 50674228 | 50674348 | 788855_47188554_ENST00000293441.6,E | 1 | - | good | AAGATGGAGAAGGGAGACACATGGGGTGACCTGGCAGAAGCTGGGGAGTCAGGGCCCCGAAGTGCCTGACCTTGGCTGATTCTCCCTGTTACTCATCTAACACTTGCTTGTAGAACAGCA | 120 |
| chr19 | 50674348 | 50674468 | 788855_47188554_ENST00000293441.6,E | 1 | - | good | TGAATGGCTGGAGGAAAGAGGCAGGAGGGGGTATAGTGGGAAGTGCAGGAAGTAAGGGTTCATCCCAAAGGTGCGAACTTCATTCCGAGAGGGCTCCCTTTCTCTGCACTTCTGGTCTCA | 120 |
| chr19 | 50674468 | 50674588 | 788855_47188554_ENST00000293441.6,E | 1 | - | good | AGCCGGGCCAGGGTGGAGGGCTAAGATAACAGAGCTTTCCCAAGCAGAGCCCAGAGCCCAGGGAAAGGATGCTCCATGCAGAGTCTCGGAAGTGAGAAGCGAGGGAACTGGGAATGGTTG | 120 |
| chr19 | 50674588 | 50674708 | 788855_47188554_ENST00000293441.6,E | 1 | - | good | GTCAAGCGATCCTCCCTAATGTTGAGATTTCAGGCATGAGCCACCACGTTGGCCTGAAATGTACAGTTGAATGGCTAGTGGCTGCTCTAGGGGGGCACTGCAGGCCTAAAGGAAGCGGGT | 120 |
| chr19 | 50675428 | 50675548 | 788855_47188554_ENST00000293441.6,E | 1 | - | good | GGAAGTCCTCAATAAATATTTCTGGAATGAATGAATCAATGAGACAAACATAGAAACAGACAGTCCCAATTCAGGAAGGGAAGTGTAGTGAGAGGCATGTAATACAGACCCCAGGAAACT | 120 |
| chr19 | 50675548 | 50675668 | 788855_47188554_ENST00000293441.6,E | 1 | - | good | TCTGACCAATGTTATATTTTACGAATTTATGCTGTTTATGGCCTGCTTGCCCTTCCTGTGGGCTGGGACATTTGTCTAATTTGTCGTATCTGCAGTATCTGTCACAGTCCCTGGCATAAA | 120 |
| chr19 | 50676508 | 50676628 | 788855_47188554_ENST00000293441.6,E | 1 | - | good | AGAGTAACAGGAGGAAGCCTCTCCAGAGAGGAGACCTTGGAGCCGAAATCTGAAAGAAACCTGAAACTTGCAGGTCCTACCCACTCCCACCCTTCTATCACCTGTCTAACCTCAACTCCT | 120 |
| chr19 | 50676628 | 50676748 | 788855_47188554_ENST00000293441.6,E | 1 | - | good | CCAAGGAAATAGAGTAGAAAGAGTATTGGGGTGGGCTGCATGCCTAGGGAGACAGACAGTAAACAAGCAAGCAAATAATGATGATGACTCCAAACTGTCATCAGTTAAAGGAGTTATGAC | 120 |
| chr19 | 50676748 | 50676868 | 788855_47188554_ENST00000293441.6,E | 1 | - | good | CAACACTCCCACTTGGGCATCCCGGTTGGCCTTCCTTAGTCATATGGCCACACCTAGCTGCAAAGGAAGCTGGGAAGTGTAGTCTCTCAAAAGGTGCCAAATAAGATTGGCATTTTCTTA | 120 |
| chr19 | 50676868 | 50676988 | 788855_47188554_ENST00000293441.6,E | 1 | - | good | TCTAGCTATTGCACCAGCATTCCAGGTAGCAGGAACAAGGAAGGGCAGAAAGGCAGAAGGGCAAAAGAGGGCACCTTGCTGCCTCTTGAGTTACCTCCAAGACATCCCAGATGTAACACA | 120 |
| chr19 | 50676988 | 50677108 | 788855_47188554_ENST00000293441.6,E | 1 | - | good | AGGCAGGTTAGCAGCCTCCATGGTCTTAAAGACCCAGAATCCTTCTCAATTTCTGCTCCACCATCTTAGCCATGTTTCCAACTGCAGACTGGCCTCATAGTCCAAAATTGCTGCTGGAGC | 120 |
| chr19 | 50677468 | 50677588 | 788855_47188554_ENST00000293441.6,E | 1 | - | good | GAGTGAATCAGTCAAACAAAGTAACTTCCTTTAGAGAATTTACATTTGAAAGAGGAGACTAGCAGAATTGGGTTCTGCTGCACCTGAGGGAGAAAAAGCAAATAATGGTGGCTTGACCAA | 120 |
| chr19 | 50677588 | 50677708 | 788855_47188554_ENST00000293441.6,E | 1 | - | good | GAGTTGCTCGCCACATCTTGTCAGGGTGTCTTGTATTGCTTCCAATGAGTTCTTCCTTCTACAAATATTTATTGAGCACCTACTATATCCTGGACCTGCTGCTGGGCTCTCAAGGTATAA | 120 |
| chr19 | 50677708 | 50677828 | 788855_47188554_ENST00000293441.6,E | 1 | - | good | ATGGCAGGAAATCCGTCAAAGCAGGGATGCCTTCAGGCGACACCTCCTGAAGCCTGCTTGGAGCTGGCCCTGTGCTGTTTGATGGGGATGTTCAGAGGAACAGGCTGGGGTTTTGCCCTG | 120 |
| chr19 | 50677828 | 50677948 | 788855_47188554_ENST00000293441.6,E | 1 | - | good | GACAGAAAGCCAAGCAGAGTATTGGTCTGTGGCTGAAGGAGGTGGGGGAGGGGGTCAGAAAAACATTCAGCAAAGTGGAGAAGGATTGAGCAGGCTGAGAAGCGCTTAGAGAAGAGCAGC | 120 |
| chr19 | 50677948 | 50678068 | 788855_47188554_ENST00000293441.6,E | 1 | - | good | CATACGAAAAATAGCAGGTAAAATGGCTCACACAAGAGTGTCTCCTTCCACTTGCCACTTCTCTACTGCCTTTATCTTTTGCAAGGTTCTGCTCTCTGATACCACACAGCTGCAGGACGG | 120 |
| chr19 | 50678068 | 50678188 | 788855_47188554_ENST00000293441.6,E | 1 | - | good | CCTTACTCGCTGGCTGGTCTAGGATAAGGACCTTTTCTCTCTGATCCACGGCCTCTTTCATTATCATCCAAATGGGTATAATCCACCCTAACTCACAGGTGTGAGGATCCAGTGAGGACA | 120 |
| chr19 | 50678188 | 50678308 | 788855_47188554_ENST00000293441.6,E | 1 | - | good | TCCCCGTGTGCTTCCTGACTGGTCACACCCCCTTCTCCTGGCTCTTCGTACCCCTCCCATTTTATCTCAGGGTTTTTGAGCCACTCTGACCTCTTTTCTCCGATGCATCTTGAATCTCCT | 120 |
| chr19 | 50678308 | 50678428 | 788855_47188554_ENST00000293441.6,E | 1 | - | good | CCCACCACCTGCCCCCGTGCCCCATGCAGGTGTTTCCCTTGCCTTCCTCCCCTCCTCCCAGCTGTCCTCTCCTGCCTGGACTCTGTTCTTGCTGTCCTCTCCAGCCCTGCCCCTCAGCAG | 120 |
| chr19 | 50678428 | 50678548 | 788855_47188554_ENST00000293441.6,E | 1 | - | good | GTCCCTTCCCTGTCACTCTGACCCTTCCACTGTCACATCAACCCCTCCACTGTCACTCGGATCCCTGCCTCATCACCTTCACCCTCACCTGTCACCTGACCCTCGCCTCTCCCTCTGGCC | 120 |
| chr19 | 50678548 | 50678668 | 788855_47188554_ENST00000293441.6,E | 1 | - | good | CTCACCCTTCCCCATCACTCTGACTCCTACTCTGTCACCTGACCCCTTCTCTGTCACATTGACCTCTCCACTGTCCCCCTGACCCTTCCCTGTCATCCTGACCTCTCCCCCATCACCCTG | 120 |
| chr19 | 50679268 | 50679388 | 788855_47188554_ENST00000293441.6,E | 1 | - | good | AACCCATTTTTGTCCCTTCTTCTTCCACATGGTGCTGAAGTGTCTTTTGTCAGCTTGCAAATCTGACGAGGCGTCTGTCACTCTGCACCCTCCTCTGTCGGGGACACCCCTCCCCCTCAC | 120 |
| chr19 | 50679388 | 50679508 | 788855_47188554_ENST00000293441.6,E | 1 | - | good | CGTTGGTTGGTGGAGTTCAGGTGCTCCCTGGGGAGTCAGTGCGATGCCCCATCAGATTGTTATGGTTCCTCCGTCAGTCTGACATCTCTCCTGTCACCCTGACCCTTCTTATGTCAAGCT | 120 |
| chr19 | 50679508 | 50679628 | 788855_47188554_ENST00000293441.6,E | 1 | - | good | GTGACGGAGAACTGGGCCTTTGGGCGGCGGACTGAAGGGGAGTCTGCCCGGCAGCCAGTGACGTCATGCAAGGCTGCGCACTCTGATTGGCTGTCCCTTCTGCGCGGGAGCGAATGCTCC | 120 |
| chr19 | 50679628 | 50679748 | 788855_47188554_ENST00000293441.6,E | 1 | - | good | CAGTTGTCTCCCCGTCCATTCTCTCCCCTATCCCCTCTCAGGGCTGCCTCCTTAGCAGGAGTTCCCCAGCCACTGGTCAGCGGCAGGACATGCGCTGCCACTGAGGCGCCAGACCACGAG | 120 |
| chr19 | 50680228 | 50680348 | 788855_47188554_ENST00000293441.6,E | 1 | - | good | GTTCCCTCCTGGGGCTGCGCAGGCCCTCTCTGAGTCAGAGCTGCCCTCCCTCCTTCCTCTCTGGGCTCTTCCTGCTTCCCATGCTTCCCATCGCCCTCTCATTTCCCCTCTCATCTCTCA | 120 |
| chr19 | 50680348 | 50680468 | 788855_47188554_ENST00000293441.6,E | 1 | - | good | TGGCTTTCTCTGGTACTGTCTTTGTCACCCGCCACTGATGCCCGAGGGCTCCCTCGTGCCCCCTCATCTCTCAGTCTGAGACGTCATTGCTCCCTCTCAGTTGCACTCCCCACTTCTGGG | 120 |
| chr19 | 50680468 | 50680588 | 788855_47188554_ENST00000293441.6,E | 1 | - | good | AACCTAGAGGAGGTAGGGACAGCAGGGGCGCGGAGCCCAGCAGATCCACCCTAGGTCAGGGCACCTCTTGGGCTGTCTCCTCCTAGGGCTCTGCTGTCCATCCGCCCCTCGGTATGGCTT | 120 |
| chr19 | 50680948 | 50681068 | 788855_47188554_ENST00000293441.6,E | 1 | - | good | TTCCCTCTGGTCTAGATGCAGTGGGGTGGGGGGTGCAGTACCCCCCGAGGCTTCTTGGATCAGGTCTGCACAAAAGTGAACAAGAGCAAAGCATTGTATAAGACGCCACTATCAGCCAGG | 120 |
| chr19 | 50681068 | 50681188 | 788855_47188554_ENST00000293441.6,E | 1 | - | good | AATACAGATGCCCCTTCATGAGGTTTCTGCATCTTGGTGGCCTCTCTCGACGTCTGGTCTTGTTCTCGGTGTCTCGATTTCCCATTCATTTAGGCTCCTCTCCCCTCTTCACCCTGCATC | 120 |
| chr19 | 50681188 | 50681308 | 788855_47188554_ENST00000293441.6,E | 1 | - | good | CTCACCTCTTGGATCTGTCCCTGGGCCTTCTCCAGGTCCCCCTCTCATGTCGGAGGCTCTCTCTGCCGCCTTCTCTGCTCCTTCCCACCTGCCATCTTTATCTTTCCCGCTCTCTGAGTC | 120 |
| chr19 | 50681308 | 50681428 | 788855_47188554_ENST00000293441.6,E | 1 | - | good | CCCTGCCTCCTAGAAATGAAACTCACAAGCACTCACCTGAGTTCTGGGAGACCTCCTTTCCCTCTGTGTCCAGAGGACCCATAGTCATTTGCCCTGTACATCCCCCTCGCCTTTTCTTCT | 120 |
| chr19 | 50681428 | 50681548 | 788855_47188554_ENST00000293441.6,E | 1 | - | good | ACCCAGTTACAGGACAGTCAAGAGCCTAACCCTCAGGGAACTCACCCTCTGAGCTCCAAAGACCTCCACCCCCAGAGGAGCCCAGACACCAGCCAGTAATAAAACCAGCTGCCCACCTGC | 120 |
| chr19 | 50681548 | 50681668 | 788855_47188554_ENST00000293441.6,E | 1 | - | good | AGACTGAGTGTAAATCTCTGATAAACTAATTAGAGCAAGTCCTGTTTTCATTAATTCAACAAATGCGCAGAGCTCCTGTAAAGTGCCAGGCAGCCACAGTTCTAGGTCTACCCTGCACGA | 120 |
| chr19 | 50681668 | 50681788 | 788855_47188554_ENST00000293441.6,E | 1 | - | good | TAAATACATAAATTAATTCACTGAAATAAAATAAACTAGAAAGTCCACCATGGCTCACCAGTGATCTCTGTGTATGCCTCCAGAGACCCCGCAACACCAGAGACCCCACAATAGCGTATC | 120 |
| chr19 | 50682388 | 50682508 | 788855_47188554_ENST00000293441.6,E | 1 | - | good | GTTACTTAATTTCTCTGCATCTCAGTTTTCTGTTCTTTAAAATGGGGCTAAAGTGAAACTTGCCCAGTTCTTATGAGGGTTCCATGAGGTTAACTGTAACACAGCAAACTCCCAACAGAA | 120 |
| chr19 | 50682508 | 50682628 | 788855_47188554_ENST00000293441.6,E | 1 | - | good | GTCTCTAACCATACCCCAGGTGTTTCTGTTTGTTCCTGATCCCCCAGAAACTCTATATTAACAGACAAATGGGGATGCAAATCCTGACTTCCTCTAACTTAATTTGCCACCTTGGACCAT | 120 |
| chr19 | 50682628 | 50682748 | 788855_47188554_ENST00000293441.6,E | 1 | - | good | ACTCACTAAATTATTCAGTATTTATTGGAAGTCTGGAATGTACTCCCACTAATCCCATTTCTTCATGCCTCCCCCCAGTCTGTATTTTACACTCTCCTCCCTCTGAATTTCCCTCTCTGG | 120 |
| chr19 | 50682748 | 50682868 | 788855_47188554_ENST00000293441.6,E | 1 | - | good | GGGCAACAGAGAGAGACCACCCTGTCTCTAAAAAACAAACAAACGAAAAAGATTAAAAAGATTTTCAGCCCCAGGATTATCCTAAGTGGACTTAGATTCAGGATGCTACCATCCATATTC | 120 |
| chr19 | 50683108 | 50683228 | 788855_47188554_ENST00000293441.6,E | 1 | - | good | GCCAGCAGCAAATACTGCCCAACGTCCTCTGGGGGGCAAAACTACCTCTCATTGAGAACCACTGCCCTGTATCATTCTAAGGTTAAAAGATTTTCAGGGCCGGGCGCAGGGGCTCACACC | 120 |
| chr19 | 50683708 | 50683828 | 788855_47188554_ENST00000293441.6,E | 1 | - | good | AATTTATTAGAAGCCCAGAATCAATACGCGAAGGGCTTTTGTGGTTATTCTTGGACAGGCGCAAAGCAGAGAAAATCTTGAGTAGCCCTATGTGTGGGTTCCCAGACGAGGCTGAACGAG | 120 |

| | | | | | | | | |
|---|---|---|---|---|---|---|---|---|
| chr19 | 50683828 | 50683948 | 788855_47188554_ENST00000293441.6,E | 1 | - | good | AGACAGAACCAAAGTCCAAGACCTCTTCCTGTCCGTCATAGTTTCTAACATCTGCGCTGTTGACACAGTTGGCCCTCATGATTCCCAGATTCTGTGTCTGCGAATTTGCCTACTCGCTGA | 120 |
| chr19 | 50683948 | 50684068 | 788855_47188554_ENST00000293441.6,E | 1 | - | good | GGCGGTACAGAAACAGAAAGAAAGACAGAGCATGCACATTCCCATTTTACAGAAGATGAAACAGTCTTAGAGATTTGCATTCACTTGCCTGAGACCTTTCATCAAGTGTGTCTCTCATCA | 120 |
| chr19 | 50684068 | 50684188 | 788855_47188554_ENST00000293441.6,E | 1 | - | good | TCTTACCTTTTTTGACCCTGTTTCCTCCTCCTGTGATCTTCTTCTATCCTCAACTGCCCTGCAGCAAACTTCTCATAATTCCTCAACAAACTTCTTCACAACCAGCTTTAAGAAGGAAAG | 120 |
| chr19 | 50684548 | 50684668 | 788855_47188554_ENST00000293441.6,E | 1 | - | good | GAAAGGTTGAGTTTTTGTTGGAGGATGATGAAAATGAAGACATAGGATTTTTTTTCCTGTACAAGTGCACAGGCCCCTGAAGTCTATCCACAATCCCTCATGCTCTGGAGGGGCTCGTAC | 120 |
| chr19 | 50684668 | 50684788 | 788855_47188554_ENST00000293441.6,E | 1 | - | good | CTATTTTGGGAAATCTGCACCACTGTGATGGGATATGGAGATATCTGTGATTCCTACTGGGGATACAGGCATGGGTCTTGCTGAAACTAGCATGGTTTGCTGCTTACATATTTAATTGAG | 120 |
| chr19 | 50684788 | 50684908 | 788855_47188554_ENST00000293441.6,E | 1 | - | good | CAAAATATTTTTCAAAAGCAAAATTTGTGATATCATAATATCATCTTTTTTTGTCAACATGTTAGACATTGGCGGTGGATTTTAACAACTCTAATTTCAAAATCACGAGGCGCATAAACA | 120 |
| chr19 | 50684908 | 50685028 | 788855_47188554_ENST00000293441.6,E | 1 | - | good | TTGGCTTTTAGGGGGTCATGGATCTCTTTGGCAGGCTGCCGTGGTCCTTCTCAGAATAATGCTTATAAATGCATAACATGAAATAACAAAAGAGGCCAGTTACACTAAAATGTAGTTTCT | 120 |
| chr19 | 50685028 | 50685148 | 788855_47188554_ENST00000293441.6,E | 1 | - | good | AGAGGCCACGCTCTGCCTTGCTCCTCCCTTCCCGCTTCCTTCTTCTCTGCCCAGTCCCCATCCTCTGGCCGCATCCACCACGTAGTGGTCAGGGGTCCATTTACTCTAAAACTGGGTCCT | 120 |
| chr19 | 50685148 | 50685268 | 788855_47188554_ENST00000293441.6,E | 1 | - | good | AGGGGCCTGCTGCTCCCTTCCTGGATGTTGCCCCTACTCCACTGGGCTGTGTGCCTTCCCTCCAACACATCCCCCACTAGGCTTGCCAGTGCCACCCCTTACCCTCCAACTGTCTCCTCC | 120 |
| chr19 | 50685268 | 50685388 | 788855_47188554_ENST00000293441.6,E | 1 | - | good | ACGAGGTTGTTAGCCCATTTACAGAGGGGGAAACTGAGGCTCAAGGAAGGAGGGGGGATTTTAACCCAGGCAGGCAGCCCTCTGAACCCATGGTCCTGCCACAGCCCTGGCTATCCTGGT | 120 |
| chr19 | 50685748 | 50685868 | 788855_47188554_ENST00000293441.6,E | 1 | - | good | TATTAAAATATGAATGGTCATGTCTGTTGCCACTCTGTCTGCCTCTCCCCACAACCAGAGGCCAACCTGTAGGGGGTGTTGATAATTTGGGGGACACTGCATTTTATTTTATTTTTCAAT | 120 |
| chr19 | 50685868 | 50685988 | 788855_47188554_ENST00000293441.6,E | 1 | - | good | AGCTCTTGCCCCGCCTCTTTGTGTCCCTCTGCGTCCCATCTTAAGGGTCTCAAGGAAGGCAAAATGATGTCATGGATGGAGATTCCCCGGCATAATAATAGATACCCAACACGTAGTACT | 120 |
| chr19 | 50685988 | 50686108 | 788855_47188554_ENST00000293441.6,E | 1 | - | good | ATCCTAGGGTTTCCTCCTTTCTCCCAACTTTGGAGACCTCCTTTCCTCCTTATCCCCTTTCCCTCAAATCCCATCTCTGTCCTTTTCTCTCTCCACTTTCCACCCCTGCTATTATACTTG | 120 |
| chr19 | 50686108 | 50686228 | 788855_47188554_ENST00000293441.6,E | 1 | - | good | CCTGGTGTGGGGAGGAACTGCCAGGGGGGAGGCGGTTCACACACCAACCTGGAGCCTTTCTCTTTCCAAAACCCTCCTGACGCCTCTGTGCTTTAGGCTGGTCCTGGAGGGGCGACACAC | 120 |
| chr19 | 50686228 | 50686348 | 788855_47188554_ENST00000293441.6,E | 1 | - | good | GGACGAAATCCTGGCCGCAGCTCAACAGACCATCAGTGCAAGCGAAAGCCCTGGTCCCGGTGGCCTCGCGTCCCTGGGCAAACACCGACCCAAAGGTTTCTTTGCCACTGAGGTAGGCGG | 120 |
| chr19 | 50686348 | 50686468 | 788855_47188554_ENST00000293441.6,E | 1 | - | good | GGTTGATCCAGGTGCTGGATGAACTGCGGGCGGCAGCTCTGGTCTTTGCGGCCCGGGTCAAAGCAAACTTCATTTCATTGTCTTCCCACCTCTGTTCATTCATCTACCCCTAGACAAACT | 120 |
| chr19 | 50686468 | 50686588 | 788855_47188554_ENST00000293441.6,E | 1 | - | good | GGCTGCCCCGTTCCCTGCCCGCACCCCTTCCCACCCTGGGTGCTGCCCACTGGGCTGGGAGATTGTGTCTCCAAGCGGGACTGCCCGTCCACCCGGACCCAGGCAGGGCTGCCCTTTCCT | 120 |
| chr19 | 50686588 | 50686708 | 788855_47188554_ENST00000293441.6,E | 1 | - | good | GCCCCAGCCCCGGCACCCGCTGCATCCCTGTCCCACCATGAACCCCACGGCGCCCCTCACCTTCCTGCCCCAGAGAGGCTGCGATCCCACCTCCCCTCCCTCCGCCCGCCCTCTCCCGAC | 120 |
| chr19 | 50686708 | 50686828 | 788855_47188554_ENST00000293441.6,E | 1 | - | good | TGTTCTTTGCCCACAGAGTACGAGCAGCAGCCGGCGCCGGTGCCCAGCATGGAGAAAAAGCGGACCGTGTATCAGATGGCTCTCAGTAAGACGGCCCAGCCCTGCTCCTCGGGATGCCGG | 120 |
| chr19 | 50686828 | 50686948 | 788855_47188554_ENST00000293441.6,E | 1 | - | good | GACCCCGGCCCCTCACTGCCCCTCTCGCGCCCGCCACGCCCACTGGCCCGCACCTGCTGGCCACGCCCACAGGCCGAGCCCGCTCCGCCCCGCCCCCGGGGGCTCCCTGGGCGTCATCCG | 120 |
| chr19 | 50686948 | 50687068 | 788855_47188554_ENST00000293441.6,E | 1 | - | good | CTCCTCTACTGCCTCTCCCCCGGGCTAGTCTCCCCCTGGAAGAAGAAGAGTGGTGAGTGGGCACAGGACTGGCGGGAGGGCCCCCGTCTCCCGACCCCGCCGGCGGACTCGCCCCTCCCT | 120 |
| chr19 | 50687068 | 50687188 | 788855_47188554_ENST00000293441.6,E | 1 | - | good | GTCCCATTTCCCCTGACAGGGGGCTGGTCGGACAGCTGACCGCCCCCACCCCGGGGCCCCGAGGGAGGCGGGCTTCCAGACAGGGGGCCCCTCACCTTCCCCTGACGTCCTAACTCTTGG | 120 |
| chr19 | 50687188 | 50687308 | 788855_47188554_ENST00000293441.6,E | 1 | - | good | TTTTGTCCCACCTCACCCCCCAGCCCCATTCAACATCTCTCCCTCCACCAGCTCCCCCCAACCCCCAATAACAGAGGAAAGCGAGCCCCTCCCTCCAGACAAGGCCGAGTGGACCTGGGG | 120 |
| chr19 | 50687308 | 50687428 | 788855_47188554_ENST00000293441.6,E | 1 | - | good | TGTCTCTGGGAAGGGGTGTCTGAGAGACGACTTTCCTCTGTGCTCTTGTATCTGTTTCTCCGTTGGGGTCTCTCTCAGGGTCTGTATGTACCCTCTCTGCGGGTCTCTCTGTCTCCTTCC | 120 |
| chr19 | 50687428 | 50687548 | 788855_47188554_ENST00000293441.6,E | 1 | - | good | GGGGGGAGAGGGGAAGGAGGGCTGGACCATCCCTTGAGTTCGTTAGTGTTGTTAGTGACCAGGGGTCCTTGTCCTCATCCTGTCCCCAGTCCATGGGTCGGACTCTGGCCTGTGTGTGTG | 120 |
| chr19 | 50687548 | 50687668 | 788855_47188554_ENST00000293441.6,E | 1 | - | good | TTTCAGCACCCCAGCAGGCCAAGCGGCTGCCGCCCCCAACCATCTCCCTGCGTTCCAAATCTATGACCTCAGAGCTGGAGGAGATGGGTGAGCCTGCGGGGCTGCTGAGGCCAGGGGGCC | 120 |
| chr19 | 50687668 | 50687788 | 788855_47188554_ENST00000293441.6,E | 1 | - | good | CTGAGGCAGGGAGGGCTCCGCACGCTGGCAATGGCTATTCTCTGAGCCCTGGGAGCCCTGTGGTGGTAGAGGGTGGTGGGGGCATCTCCCCCATGATACTGATGCCTCCCCTCATCACCT | 120 |
| chr19 | 50687788 | 50687908 | 788855_47188554_ENST00000293441.6,E | 1 | - | good | ACTCTCGCAGCCACCCCACTGCCACCAGGCTCAGCCCTGCGGGAGCCCCAGGGCTGGTTGCTGCTGCCCAAGGCCTTCCCTTCCCTAGCACTGCTTCTCCGTGACCCTTGTGGGGAGAAA | 120 |
| chr19 | 50687908 | 50688028 | 788855_47188554_ENST00000293441.6,E | 1 | - | good | CACCGACAGGTGGTGAACATGATCCGCCAAGGGGGCAACACGCTGATGGTGAAGGTGGTGATGGTCACCAGGCACCCGGACATGGATGAGGCAGTGCACAAGAAAGGTGCTCCTGCGGCC | 120 |
| chr19 | 50688028 | 50688148 | 788855_47188554_ENST00000293441.6,E | 1 | - | good | GCGCAGGAGGCATCCTCCTTGGGGTCTCTGAAGCTGCAAGCACCCCTCCCCTCGTCTACTCTGTGTGATCTGGGGGTGTCTGTGCCACAGGTGAACGGGCAGAATGTGGTGAAGGTCGGC | 120 |
| chr19 | 50688148 | 50688268 | 788855_47188554_ENST00000293441.6,E | 1 | - | good | GCTGAGCCTCTCCCCCAACCCTGGGGCCAGGGAGGGTGGGAAAGGGGGCTTGCCCAGGGAGGCGGTGGTATGAAGGGTCGGAGGGAGAGGAACATAGCCCAGAGGGCCATGGACAGGG | 120 |
| chr19 | 50688628 | 50688748 | 788855_47188554_ENST00000293441.6,E | 1 | - | good | CCTTTGGCACCAGGATTCCCAGCCCCCTGCCCACCCCCCAGCCAGCCTCTGCTTTCCCCAGCGATTCCATCTAAAGGCCTCTGAGGCATCTCAAAAGAGTGACACAGAGCTGTCCCCAAC | 120 |
| chr19 | 50688748 | 50688868 | 788855_47188554_ENST00000293441.6,E | 1 | - | good | GCGAATGGGAGACTTCCTCATCGAGGTCAGTCCAGGCCCCCTCTCTTCCCTGGGACCCTGTGACAAGTTCCCAGACCCCCTCATGATTCACACCAGGATCTGGCTGCCCAGTTTTATTCT | 120 |
| chr19 | 50688988 | 50689108 | 788855_47188554_ENST00000293441.6,E | 1 | - | good | CCAGCCCGCCCACCACCCCACCCCGGGGTGCTGGTCCCTCAGCTGCGGTGAGGCCTCCGTGACTTGGGTCTGAGCCCCATCCCCTGCTCGGCCTCTCCACCCCCTCCCCTCCGACCCCGC | 120 |
| chr19 | 50689108 | 50689228 | 788855_47188554_ENST00000293441.6,E | 1 | - | good | GGGGTTTGGGTTCGTGCTCCGGGGGGCCAAGGGTGAGTGCCAGCCGGGGAGGGGATGGGAGAAGGGCTCTGTGACCCCAGAAGCAGGGCTGAAAATGGGGGCTGGAAAGGGGAAGGGA | 120 |
| chr19 | 50689228 | 50689348 | 788855_47188554_ENST00000293441.6,E | 1 | - | good | GCCTCTGTGTCTCATGATGTCTTCTCCACTCCCCCCACCACCCCCCCATTTCTCCTGCCTCCCTGCCAGCGATTACATCATTAAGGAGAAGACAGTCTTGCTGCAGAAGAAGGACAGTGA | 120 |
| chr19 | 50689348 | 50689468 | 788855_47188554_ENST00000293441.6,E | 1 | - | good | TCTGAGGGGGAAGCCCATCAGGGCTAGCATACACACACAGAGTGGGTGACAGAGATCCTAGGAGATGCAATTAATGGCCCTGGAGCAGAGGTTTGGGTCTGGGTTGCCGGGTGGGGGACA | 120 |
| chr19 | 50689468 | 50689588 | 788855_47188554_ENST00000293441.6,E | 1 | - | good | TGGTAATCTGGAAGCTCTTCTCCCCATCCTCCCCACCGTGCCTGGTGCTGAATGGATGGGTGGATGGATGCATGGATGGATGAATGAGGAGAGAGGGAGTGAGAGAGAGAGCATGCCGGC | 120 |
| chr19 | 50689588 | 50689708 | 788855_47188554_ENST00000293441.6,E | 1 | - | good | TCAGGTAGGATCTGTGATAGCCCCCCATTATAGATGAGGGAACTGAGGCACAAAAGGTTAAAGTTGTGGGCTCAAGGTCATAGGCAGGAAGTGACTAGCAGGGACCCAAGCCTGGCTGTC | 120 |
| chr19 | 50689708 | 50689828 | 788855_47188554_ENST00000293441.6,E | 1 | - | good | TTGACTTAACACTCTGACCTAAGGACATCGAGCAGATCATATTGAGCACCTACTATGTGCCAGGCACTGTGCGAAATACTAAATGTGTATTACCCCCACTGGATTCCCACTGCAACCCCA | 120 |
| chr19 | 50689828 | 50689948 | 788855_47188554_ENST00000293441.6,E | 1 | - | good | ACCTAGTGGCCACCATATTGGGCAGTGCAGAATTAGCAATTTTACCCTGTGTGGGGTTCCTGGGAGGTGACATATGTATAAAGGAATTTTGAGGGGTATCATCAAAGTACATGAGTCCTC | 120 |
| chr19 | 50690068 | 50690188 | 788855_47188554_ENST00000293441.6,E | 1 | - | good | CTATAATAGAATATGCTGACTGCATGCAGTAGCTCCATTTGAGATGCACCAGGGAATACAGAAGACAGTGCTGTCCACGGGGAGCTTTGCAGCAATGGAAATGTTCTATATCTGTACTGT | 120 |
| chr19 | 50690188 | 50690308 | 788855_47188554_ENST00000293441.6,E | 1 | - | good | GAACTGTATCTGAGTTTCATTGACATTTCCTGGGTACGTGAGCCTTACCTAAATGTGCTGGTGGCATTTGATGGCTTTACTGGAAACTACCACCCATATTCATGGGTGCTTTGCATTTTA | 120 |
| chr19 | 50690308 | 50690428 | 788855_47188554_ENST00000293441.6,E | 1 | - | good | AGTATTCCTGATGTAGACTGGGGTGTCGGATCCTCTGGGAATGGAATGGATGTATATTCCTGAGGTGTACAGTGGTCATGTCAGGAGTTTGGGGCTGTTTGAGTGAGCCTTGAGATCCAT | 120 |
| chr19 | 50690428 | 50690548 | 788855_47188554_ENST00000293441.6,E | 1 | - | good | GCCCTGGGACAGGCTTTAGACAGTGCTGAAGGAAGCTCAGCACAGCAGAGCACGCGTTCCAGTGACTCCTGGGTGTTCTTGGGTATTCTGAAGGCAGTGGGGTTGCAGCCAGGGTGTCTG | 120 |
| chr19 | 50690548 | 50690668 | 788855_47188554_ENST00000293441.6,E | 1 | - | good | AGGTAATTGGGGAGCATGCTTGGCTATGATTGGAGTGTATTAGGGCCACATGGATGTTACTGGGATCTGGGTATTATTGGAATACACTGGGATTGTGAGGGTGGGAGGTGCTACAAGTGT | 120 |
| chr19 | 50690668 | 50690788 | 788855_47188554_ENST00000293441.6,E | 1 | - | good | TGGGAATACACTGAGATCCGAGTGAGGGGGAGTTATTATTGGAACATACTGGGATGTGTGGGGGTTGCCGAAACATACTGGAGTATTCGAGGACGGCTGTTTTGCTGAATTTGTGGGGGC | 120 |
| chr19 | 50690788 | 50690908 | 788855_47188554_ENST00000293441.6,E | 1 | - | good | GGGAGCCATGGGGGTCCTGGGAGCATACAGGGGTGCTTGTGGGGGTGGTACCTAAATACAGTGGAGCACAGTAAGTATTATTGGCATTTGCTGGGATGTATGTGTTCAGAGGGCGTGTTA | 120 |
| chr19 | 50690908 | 50691028 | 788855_47188554_ENST00000293441.6,E | 1 | - | good | TGTGAACATGCGTGGTGTTAGGTGCACTGAGGTGCCACAAGGAGGGTGGCAGCAGGAAGGGAGGACACGTGCACATCCAGAACCCTGTCGCCATGGTGCAAATGTATGGATTTTTACAAG | 120 |
| chr19 | 50691028 | 50691148 | 788855_47188554_ENST00000293441.6,E | 1 | - | good | TTCCCTCCTTCTCTGCCAACACCTCTCTATCCAGGGGAGGAAGACATGGCGGGTATGGTGTGGATGAGTTTACTGGACTGTTACGAACTTTGAGCGAGGACTGTTGCTGGAGTATGTACA | 120 |
| chr19 | 50691148 | 50691268 | 788855_47188554_ENST00000293441.6,E | 1 | - | good | CTGACCTCTGAGCCAATTTCTGTGTCCAGTTTCCCCGTCCACTTGCTGTTCTGGCCTTCTCGGCCCCTTTCCTCCCTTCCCCTCTCCCTTTTCCTACCCTCATTCACCCCTTTCAGCCCC | 120 |
| chr19 | 50691268 | 50691388 | 788855_47188554_ENST00000293441.6,E | 1 | - | good | CAGTGAGGAAGGGTGCTGCTGGCTGCTGGGGGGCCTTGGAGACCTGGAAGAGATACAGGAGGAGGAGTAAGAAGGTGGGAATGGGGACATTACAGAATGCATGGTTAGAAACAGGCAGCT | 120 |
| chr19 | 50691388 | 50691508 | 788855_47188554_ENST00000293441.6,E | 1 | - | good | GGAAGGCTTATGTATGGGGTATGAGGCTGATGAGGGGGCAAGAGAGATTCCAGAGGGGAGGGAGAAAGATGGAGGAAGGGAGGGGTTTGTGGCATGAGTGAAAGAAAGTGAGGTTGCCT | 120 |
| chr19 | 50691508 | 50691628 | 788855_47188554_ENST00000293441.6,E | 1 | - | good | TGAGAAGACAATTTAATGATCTATCTGTGAAGGGCTTCAGATACTGCCCAGCCAATGGCAAGTGCTCCTTCTTGGCATATTGATATGGGAGTTTTATTGACTATATTGGCTGACATTTGG | 120 |
| chr19 | 50692108 | 50692228 | 788855_47188554_ENST00000293441.6,E | 1 | - | good | CATGGTGTTACTGCCACACCCTGGGATAAACTCCAGGTGCTGCAGGGCCGAGCTCAGGGTGGATTCCGAGCAGATGGATATATGTGCTCAAGGTATTTCGGTGCTGTGGAGTGGAGGGTA | 120 |
| chr19 | 50692228 | 50692348 | 788855_47188554_ENST00000293441.6,E | 1 | - | good | TCAACCCTTCCCTGGGTTGTTGTGAGGACTAACTGAATACATAAATGGAAATATATAAATGCAAAGTGCTTGGAATCATGGCTGGCATTCAGTAATAATTCGATAAGTGTTCCTCATTGT | 120 |
| chr19 | 50692468 | 50692588 | 788855_47188554_ENST00000293441.6,E | 1 | - | good | GGGAATGGAGCATGTGTAGGATAGGGGGTGGTTGAGAAGCAGAAGCAGGAACAGGGTGGGTCTGGATGTGGTTGGACCCCCCTCCCTGCCTCAGATCACCACTGGTGTGTGTGTGTATAT | 120 |
| chr19 | 50692588 | 50692708 | 788855_47188554_ENST00000293441.6,E | 1 | - | good | GTACATGGAGCTGTGGGAGCAAGAGAGCAGTTGGGAAGTTCAAGGAGACTGTGGTGGGATGAGGGCAACACCACGTATGTCCTAGCTGGGGATTGGGTGAGGGAGTCATGGAGCAGAGAG | 120 |
| chr19 | 50692708 | 50692828 | 788855_47188554_ENST00000293441.6,E | 1 | - | good | GGGAGTCAGGGATGAGGGGGGTGGGTTAGGGCGATGTGGAACCTCAACATGGCATGGAGGAGTAGGGAGTGAGGTGCAGAGAGGAGGGGTACACAGGAGTTGGGGAGAGGGAGTTTAG | 120 |
| chr19 | 50692828 | 50692948 | 788855_47188554_ENST00000293441.6,E | 1 | - | good | GGACAAGGGTGAGGAGGTCACAGAGGTCTGAGGAGGATGTTTAGGGGGAAAGGGGGGTAGGGTGTGTCAGGGGACACGGGAAATGGGGGGATGGCTCCATAGGGTGTGAGGAGGGCTT | 120 |
| chr19 | 50692948 | 50693068 | 788855_47188554_ENST00000293441.6,E | 1 | - | good | AGAGGGTACCCAGGAAGTATGAAAGGGACCAGAGGTACTTTAGGGGAGCCAGTGAGATTACTTACGGAGGTGCAGGAGGGTGTGGGAAGGGTGAGGCCAAAGGATGAGGGAGAGGTCGC | 120 |
| chr19 | 50693068 | 50693188 | 788855_47188554_ENST00000293441.6,E | 1 | - | good | AATACCAAGAGGACACAGGGAATGTAAATGAAACGCAGCTGCCAGTAAATGCAGGCGGGAGGGCATGTGGTGAAGTGAGTCGAGGGGGCAGAAATAGGTGGAGGTTTCCACGGATGCAAA | 120 |

| | | | | | | | | |
|---|---|---|---|---|---|---|---|---|
| chr19 | 50693188 | 50693308 | 788855_47188554_ENST00000293441.6,E | 1 | - | good | GAAGTGGGGGAGGGGTGGCGGCTGGAGCACCGAGGGTTCGGAGCAGAGAAATGGGGTGGGGAGGGAATTCAGGGAATGTGGGGGGGTATTTGGGGGTGGGGATTTGGGCAGTAGGGG | 120 |
| chr19 | 50693308 | 50693428 | 788855_47188554_ENST00000293441.6,E | 1 | - | good | AGAGGGAGCCTGAATGGGGTGCTGCAGGGGCAGCCAAGATGGGGGGCAAGGGGGAGGGACTAGAGGGGTCTGGGTAGGAGGTGCAGGGTTGAGAATGGAACAGGGGAGAGTGGGGAC | 120 |
| chr19 | 50693428 | 50693548 | 788855_47188554_ENST00000293441.6,E | 1 | - | good | CCTGTGTGTGGGATCCAGTGAACAAGAGCTGGCTGCTGAGCTCTGGAGGCCGCTCAGGGTCGCTTAGGGATGGCCTGGGGAAAAGCATTGTGTGGGCTCCTGCAGGGCAGGTAGTCAGCC | 120 |
| chr19 | 50693548 | 50693668 | 788855_47188554_ENST00000293441.6,E | 1 | - | good | AGGGGCTGAGTGCGGGGTTGGGGGTGCGCCCGTGCACAGCAGCAGGCTCTGGGCTGGCGGACCCTGGGTGTGTGAGAGGGACGTTGGGGTGGGGAGGGGAGTGCAGGCCGCTGCAGTG | 120 |
| chr19 | 50693668 | 50693788 | 788855_47188554_ENST00000293441.6,E | 1 | - | good | CCCGGCTGAGCTGCATGGGGAGCCGCATCCGCTTGTGTGTGCCACTTCTCAGCATGGGGAGGGGGGCTGGCGTCCCTCTGTGGTGGCCTCTGGGCAGGCCATGGACACGGGTACGTGGGA | 120 |
| chr19 | 50693788 | 50693908 | 788855_47188554_ENST00000293441.6,E | 1 | - | good | TGTGTCTCTGTCCCCATGTGCTCCTCTCCCCTGCCCCCTTGCCCCCGGGGGTCTGTGCATGTCAGGAGTACTTGAGTCACTGGCTCTGCCCTCTGCCCATGTGGGGCACGTGTGCTGAGC | 120 |
| chr19 | 50693908 | 50694028 | 788855_47188554_ENST00000293441.6,E | 1 | - | good | GTGTGTGTGGTGTGTGCTGGAGTGGGTCTGTCCCAGGGACTGCGGCTGTGCTTCCCAACACCCACCCTCCTGGGCCTGGCCTACCCCTCCCCCACAGGCCCCTACTTCAAGGGTCCATCC | 120 |
| chr19 | 50694268 | 50694388 | 788855_47188554_ENST00000293441.6,E | 1 | - | good | GGCAAGTCCCCCCATTCATACCCCTCTCTCCCATTCACAGTCATCCCCCCTCCATCACACACGCATCCCGCCCCCGGCTTCCCCTCCCCTCCTCTTTTTCCGCTGCGTGTCACTGTGCGA | 120 |
| chr19 | 50694388 | 50694508 | 788855_47188554_ENST00000293441.6,E | 1 | - | good | TTAGCGCCCCCATTCCCCGGAGCCCTCCCTGTGGGGAGCCACCGTGGCTTACCCGCATCCGCCTACATCCCAGGCCCCTGTCCCTCTGATCCCCACATCCCCACACCTCTTCACGGTGTT | 120 |
| chr19 | 50694508 | 50694628 | 788855_47188554_ENST00000293441.6,E | 1 | - | good | CCACCCCCTCACAGGCCCTTCCAGGCCCCCCTCTGCTCTTCCCCCAAGACCTTCCTCCCAGCAGGCCCCCTAAACCGACCCCCCTCCAGGATTTGGAGCCCACTCGCCTTTCTGCCGCCC | 120 |
| chr19 | 50694628 | 50694748 | 788855_47188554_ENST00000293441.6,E | 1 | - | good | AGCCCGAAATCCCTTTCGGGCCCCTGGTCTCAACCCTTCTCCTCTCCATCTTCCTCCCTGGAATCCTCTCCCCGACCCTCCCCCCTAATACGCACTCGTCAGCCCTGGGAATCTTTTCCT | 120 |
| chr19 | 50694748 | 50694868 | 788855_47188554_ENST00000293441.6,E | 1 | - | good | GGCCGGCAGGCCCCGGCCCGCGGTGGCGACCCTCCCCCGCCGCTAAACCCCAGCCCCAGGGCCTTTCCCCTTCCCCCAACCCGCCCCTTCCCCTCCGCTCCTCTTCCCTAACCTTCTCTC | 120 |
| chr19 | 50694868 | 50694988 | 788855_47188554_ENST00000293441.6,E | 1 | - | good | AGCTCCCTGCCGCAGCCGCCCCCCGCGCTGTCCTCCAGCTGGCCAGCCCTGGGGCCCCGGCGGCGCAGCGTCTGGTACATCTACAGGTAACGGAGCGCGCCGCCTCCTGCTTTGCGGGCC | 120 |
| chr19 | 50695108 | 50695228 | 788855_47188554_ENST00000293441.6,E | 1 | - | good | GCCCTCCCTCCCGCGGGAGGGGGTGCGCGCCCCCTCCCCCTCCCCCCTCCCGGCGCGGGTCAGCATGAGGCGGGGCCTGGGGGCCCGGACACGGGGGTTCGGCCGAGCGGGTACGGGG | 120 |
| chr19 | 50695228 | 50695348 | 788855_47188554_ENST00000293441.6,E | 1 | - | good | CCGCCTCCGGAGCTGTCCGCGGTGCTGAAGCCGGCGCCGTCCGCGGTCCTCGCGCTACCGCCGCGGCCCGACCCCGCGCCCCGCGCCCCCGCGCCCCCCGGCATGTGAGCCCCCGGCCCT | 120 |
| chr19 | 50695348 | 50695468 | 788855_47188554_ENST00000293441.6,E | 1 | - | good | CCCCTCCCCCTTCTCCCGCCAAGGGGGCCCCCGCGTTCCCCTCCCGGCCCCCTCCCCCCTGCCCGGCCCCCTCCGCCGCCCGGCGCAGGACGCCAAGGGGTTAAGCCTCGCGCCCGCCGC | 120 |
| chr19 | 50695468 | 50695588 | 788855_47188554_ENST00000293441.6,E | 1 | - | good | TCCCTCTCCCCGCGCTCTCTAGCTCTGCACCTGCGCGGGAGGGCTCTGGAGGGGGGAGGGGGATTAAGGGGGGGCCCAGACCGGAAGTGCCTCCCCTTCTCCTCCCCCCTCCAAGCCTCG | 120 |
| chr19 | 50695588 | 50695708 | 788855_47188554_ENST00000293441.6,E | 1 | - | good | TGGTGGGCTGCGTGTCCCCGCCTGGGGCCTCTCTCACCCGCGGTGTCCCTGCCGGGGAAGCGGGGCTCCGTCCGCGCGTCCGTGCACGCGGCTGCGAGGCATCTTGTGTGATGTTGTGCC | 120 |
| chr19 | 50695708 | 50695828 | 788855_47188554_ENST00000293441.6,E | 1 | - | good | GCCCCTCCGCCTGGGCTTCTCTGGTGGGGGAGGGGTGTTGCTCATGGCCTTGGGGCCCAGTCTGAATGTCCTGGCCTTTCTTGCCGACTGTGTCGAGGTGCGTTGCGTGTTGGTCTGTGG | 120 |
| chr19 | 50695828 | 50695948 | 788855_47188554_ENST00000293441.6,E | 1 | - | good | CCCAGCCTCCCTGCCGGAGCTGGGCGGTCCTTCCGTGTGTCTCCCCTCCTGGCGTGTGCCGCCCCCTCCCCGCGCTGGCTCGGGGTCCACGCTCTCTCGGGGAGCTGTGGTGGTGTGTTC | 120 |
| chr19 | 50695948 | 50696068 | 788855_47188554_ENST00000293441.6,E | 1 | - | good | GTGCTGCTGTGTGTGTGGTGCGTCTCAGCTTCCCCTCCTCCTGCCTCCCCGCTCTTCCCGGACCCTGTCTGCGCTGGCCTGGCCTGTGCTGTGCCGTGCGCCCGGCAGCCAGGGCCCAAC | 120 |
| chr19 | 50696068 | 50696188 | 788855_47188554_ENST00000293441.6,E | 1 | - | good | GGGTCACTTGTCCTACCAGCTGAATGCGGGTGGTCCCCGTCACCCTCAGCTTCCTCCTTCTACTCTGTCTTGTCAATGGGCTTCCTCCCCTACTCCTCCGTGCCTCTCGCTGGCTCCCAC | 120 |
| chr19 | 50696188 | 50696308 | 788855_47188554_ENST00000293441.6,E | 1 | - | good | CCTGCTCTGGGCACCGGGTGAGTATTGGGGTACACCCCCCCACACCCCGAGCTTCTATGTCGGCCTCTGCTTCTTGTCTACCAGTGTCTCTCTGGGTCTCTCCTTGGCCTCCGTCTGTGG | 120 |
| chr19 | 50696308 | 50696428 | 788855_47188554_ENST00000293441.6,E | 1 | - | good | GTCGGGGATGGGGGGGACAGAGAAATGGCAACATGGGGAGAGAGGGTGGGGCTGGGGTGCCACCCTGCGCCCCGAGTGTCTGAGGCGCTTGACCGTTGTGCGTGTTTGCCTGGCGCGTCT | 120 |
| chr19 | 50696428 | 50696548 | 788855_47188554_ENST00000293441.6,E | 1 | - | good | TAAAATATGACACTCTCCTCCACTGCTCCCCCCAATCCAGGGAAGAGTCCCCTTGTGCGTAGCATCCTTCCATTAAATGTGTCCAAACCTACGTGTCTGTGTTCTGGCCAGTGGGGATGG | 120 |
| chr19 | 50696548 | 50696668 | 788855_47188554_ENST00000293441.6,E | 1 | - | good | TGATCTGAGATTCAGGACCTGGGATGTTGGGGGAGGGAAGGGCAGAGGAGGGGAGGATTCCAGGGCTTGGGCCTCCAGAAGGACAGGGCTGTGGGGGAGGGAGGGTCATAATTTGGTGT | 120 |
| chr19 | 50696668 | 50696788 | 788855_47188554_ENST00000293441.6,E | 1 | - | good | TGAGTCTGCTCACACACATGGACAGGTGGGTCTCTGCACACATCCGTCCCTCTCTGTGCATGTCACTGTGCGTGTGTACTCGCACATTAACGTGTGTGCATGCATGTGCGTCTGTCTGAT | 120 |
| chr19 | 50696788 | 50696908 | 788855_47188554_ENST00000293441.6,E | 1 | - | good | GCTGGAGGGTCTGCCCTGCTCTGTATATACTGGTGCCTTTGTGGCTATGATGACTGTGGGGCTGGCTCTGCACATCTGTGCTTCTCTGAATCCGTGGGCAAGCACCACAGGGCATGTCAG | 120 |
| chr19 | 50696908 | 50697028 | 788855_47188554_ENST00000293441.6,E | 1 | - | good | GAACGTGTGTGCACGACCAGAGCTGAGGTTTCTTGGTGTGTGAACGTGTGTCTCTGTAGGTGTGCATGACACGCATCTGTGTGCATGTGTGGCCTGGATGTACCTGTACCTATGCCCGGG | 120 |
| chr19 | 50697028 | 50697148 | 788855_47188554_ENST00000293441.6,E | 1 | - | good | GGCTGCTGGCTGACTTGCTTCGGAAGCTTAATGGATGGGATTGGCCCAGGGAGGTGAGAGGCGAGGGGCAGGGAGGGGATGGGGTCAGGAGGGGAAGGACGGGGTGCCTGGGGGGCG | 120 |
| chr19 | 50697148 | 50697268 | 788855_47188554_ENST00000293441.6,E | 1 | - | good | TCTGTGCAGTCATGTGCAGTGGGAATGTGTGTGTCCCATGCACCGGTGTGGGGAGGGTGCAGTCAGGGCTGGGTGGGATTGGGGTGAGGAGATGGGTGCACTGACACCTTTCCCCTCCCT | 120 |
| chr19 | 50697268 | 50697388 | 788855_47188554_ENST00000293441.6,E | 1 | - | good | CCAGCTCTGGGGTAGCCAGAGACAACGGGGTGGTCAGGGCAGGGTGGGGATGCCTGGGGGGGCCGGGGCGCTCTTATGTCTGGCTGGCTGGAGGCCCTTGTCTCCTTCATGTGCCCCATC | 120 |
| chr19 | 50697388 | 50697508 | 788855_47188554_ENST00000293441.6,E | 1 | - | good | CTCTAAGCCCCCTTTCTAATCTGTTCCTTCTCAAGTTTGATCCAGGGGTGGGATAAGCCAGAATTAGCTTCAGACAGTACAGCCCAACCCCAAACCATCTTCGGCAGCCACTAAGACTGC | 120 |
| chr19 | 50697508 | 50697628 | 788855_47188554_ENST00000293441.6,E | 1 | - | good | ATTATACCGTGGGCTCCTACGACAGCTTTGATGCCCCAAGGTAAATGTCCTTACACCCTCGGGCAACCCCCCTCACCCCAAGTTCCCTTCACAATGTACACCTTTAAATATTTCCATCAC | 120 |
| chr19 | 50697628 | 50697748 | 788855_47188554_ENST00000293441.6,E | 1 | - | good | TGGGAAGACTTGGGCCAGGAGCTCTAGGGCTGGGATTGTGGTTTCCAAAACAAGAGTCCAAGGTTGTCTGTTGTCACCCTGCAGAAAGCCGCAGTGACAAGGCAAAGAGACTCTTCCGGC | 120 |
| chr19 | 50697748 | 50697868 | 788855_47188554_ENST00000293441.6,E | 1 | - | good | GAATCGCTCTCAGGAGAGCAAGCAAGGTAATGCGGAACCCCCAGCAGTGTTCAATGGACCCTAGGATTCCCACGTCCACTCCTTTCCCTTCCTAATGGGGGTGGGGTACTGGGAGGTAGA | 120 |
| chr19 | 50697868 | 50697988 | 788855_47188554_ENST00000293441.6,E | 1 | - | good | GAAATGTCTCTATGTCCCCGTTCCCCATCCAGTACTTAGCATCGGGGAAGGAGGCTTCTGGGAAGGCCAGGTCAAAGGTCGTGTTGGCTGGTTCCCCTCTGACTGCCTGGAAGAAGTGGC | 120 |
| chr19 | 50697988 | 50698108 | 788855_47188554_ENST00000293441.6,E | 1 | - | good | CCCAAGCTCCTCGCCCCTTCCTCAGATGTTTATTAAGGATGGCCTTGGGCCAGGAGCTGGTAACTGTGGGAATGCTCTAAGTTGAGTGTGTTGGCAGTTGAGGGGCAGCACAGAAACACA | 120 |
| chr19 | 50698108 | 50698228 | 788855_47188554_ENST00000293441.6,E | 1 | - | good | AATTGTGGACCTGCCTCTAACCTAGGAAGACCATGTTATGGAACAGTCGTCTTCCCCAATGCCTGTCATCCTTATTGGATCCAGAAACTAGAATGCAAGAGTGATATGATATGGGTCTCC | 120 |
| chr19 | 50698228 | 50698348 | 788855_47188554_ENST00000293441.6,E | 1 | - | good | GTGATGGGAGGGTGAGGTAGAACTTGTCGCCCCATCCGTGATCTAAGTCCACAGAGCCTAGGAGGGTGAAGGGGAATTACACAACTGAGCTGTGACCAGTTACCAGGGGAGACAAGGACC | 120 |
| chr19 | 50698348 | 50698468 | 788855_47188554_ENST00000293441.6,E | 1 | - | good | GTGATGCTAATCTCTTTTCCCTTCTAGAGCCAGGAGAGGCCGAGAGAGACAGAGACAGGGGTCTGGGGACAAAGGACTAAAGATTAAGTGGCAAGGAGACTTTGGGAGGTGGCAGGGACA | 120 |
| chr19 | 50698468 | 50698588 | 788855_47188554_ENST00000293441.6,E | 1 | - | good | TTCAGCAGCAGGCATGTAACAGGCAGTCTGTGATGTTTGCTAAATGGGGAAGCGGTGGGGCCTGGGAGGGTGGGCAGGGGGAAGCCGCTTCCATTAGGGGAGAGGGAGAGAAGGAAAGA | 120 |
| chr19 | 50698588 | 50698708 | 788855_47188554_ENST00000293441.6,E | 1 | - | good | CTCTCCAAGCCTCAGTGTTCCTATCAGTAAAAGGGGGATAATGAAAGTGGCCCCTCTTAGGGTTCTGGCCCAGGTGAGAGTTGGATACGCATTAGCTGTTCTTGTCATATATCTCTGTTT | 120 |
| chr19 | 50698708 | 50698828 | 788855_47188554_ENST00000293441.6,E | 1 | - | good | TTGCATGGAGAGGTAGCGCACATGAGTGACTGAAAGTATGCCCTCTGGAAGCAGACTGAGCTGTGTTCAAATCCAGATCTGTCAGTTACTAGCCTTGTAACCTTCAGCAAATGATCTTGG | 120 |
| chr19 | 50698828 | 50698948 | 788855_47188554_ENST00000293441.6,E | 1 | - | good | AAGGGTTGAACGAGGAAGGAAACTATGTGAAGGGCTGAATGGGGGCAGAGTCATGAAAAATGCCGGAGAAATGGAAGACCATGTGTTTGCCACTGGTGCCCTTGTCATGCACTGAAGCCG | 120 |
| chr19 | 50698948 | 50699068 | 788855_47188554_ENST00000293441.6,E | 1 | - | good | GGCCCCAACTCAACCCCTAGCATAGCTGGAGCCATCTCCAGGCAAGGGTTTTCTCCACATTCGCTTCAGCTTCTTTGTCTATAAAATGGAAAAACAGTTGTTACCTACCTGGGTAGGCAC | 120 |
| chr19 | 50699068 | 50699188 | 788855_47188554_ENST00000293441.6,E | 1 | - | good | AAGAACCCAGTCCCCTTGACAACCCATCTCCTGCTCTCTTCCATGACTTTCAGCTTCCCCTCATTTGGTCTCCCCAGATGCTCTCAAGGGAGGCAGTTTGTGGAGGGGAATACATCTGAT | 120 |
| chr19 | 50699188 | 50699308 | 788855_47188554_ENST00000293441.6,E | 1 | - | good | TGATTATATTCAACAGTGTCTACCCTGGTACCTGACTCATGACTGGCACTAACTAAATATTTTTACAGTGAACAAATGATTGATTACAAGACCACCAAGCCAGGCATGGCAGTACCAGCC | 120 |
| chr19 | 50699308 | 50699428 | 788855_47188554_ENST00000293441.6,E | 1 | - | good | TCCAAACACCCTCTTATTTTCCCTTGAGCTCTGACCCCAACTTCCAATCATACAGTCACTTGTATAGTTATGCCTCTATTGGCCACTGCTCTGGAAGCTCCTCAAAGGAAGGAACAACAT | 120 |
| chr19 | 50699428 | 50699548 | 788855_47188554_ENST00000293441.6,E | 1 | - | good | TGCTCTCTTTGCCTCCCCAGCTCCTAGGATCCTTCAGGTCTCAACCTATGTGTCACACCTCAGGGACACCTCCCTGGACCCCTTTGCCATTATAAGTTATATACTCCCATCTGGCTTCTC | 120 |
| chr19 | 50699548 | 50699668 | 788855_47188554_ENST00000293441.6,E | 1 | - | good | CTGAGCCCTCCAATTCTCCAGTCCTCCGATATACCAACCCCTCCATTCCTCCAATGTCCCAAGCTTTCTTGTACACCAAACTCTTCAAACATGCTGGTCCTTCTGCCTGGAACTCTTCCA | 120 |
| chr19 | 50700508 | 50700628 | 788855_47188554_ENST00000293441.6,E | 1 | - | good | ACATCCACGATGTCACTGAGCTTCACACAGCCTGATGAGGGAGCTGACTCCTGCTACTCTCAGAATCACAGCCCAGAGTCTTCACCTGGCCTCCAAGCTGCCTCTCTGCCCACCTTGTCA | 120 |
| chr19 | 50700628 | 50700748 | 788855_47188554_ENST00000293441.6,E | 1 | - | good | TGGGCAGTATCTTGCTGAGCTACAAGCTCAATGCTTCTACGCAGGGAGAGAACACACTGCCTTCTGTTTTTATATCCTCGTCTTCCATTAAGAGGGACTTAATTGCCTCAAAGCACACCC | 120 |
| chr19 | 50700748 | 50700868 | 788855_47188554_ENST00000293441.6,E | 1 | - | good | GGTGCAGCTCCTTCAGGGCCAGCTGCCTGGGATGTAGGCACGTGGGGTTGGGGGAGATGGGAGAAAAGAAAGGTGAGCTGAGGCCAGAGGTAAAGGACCTTCCGGGCAGGCAGAGGACC | 120 |
| chr19 | 50700868 | 50700988 | 788855_47188554_ENST00000293441.6,E | 1 | - | good | TCATAAGTCACTGGTCTCTGGGTCTCTAGGGCTCAGTTTAATGACCCAGCACACAGGAGGCTTTAATGCACATATGTTGAGTGAATGAATGATCAAATTCAGGAGCTACTGTGGGGGCCA | 120 |
| chr19 | 50700988 | 50701108 | 788855_47188554_ENST00000293441.6,E | 1 | - | good | TAGAAGGTCAACTTGAGTGAGGCCTTGAAGGGTATTCGGGGGTCAGAACATCAGGTGGGCTGTGTAACTGCATGTGTTTCCTAGTCTCTCTTCCCATAGCCTGAGTTCCTGTAGGTGAGG | 120 |
| chr19 | 50701108 | 50701228 | 788855_47188554_ENST00000293441.6,E | 1 | - | good | AGTGAAACTCCATCTCAAAAAAAAAAAAAAAAAAAAAGCCATCATTTTCCTAAGTGGCCCCTACTTCAGGCACTTAGAGGAGAGAGACCTGGGCCCCTGAATAAGCAGGGAGGATCCTACCA | 120 |
| chr19 | 50701468 | 50701588 | 788855_47188554_ENST00000293441.6,E | 1 | - | good | AAATGGGACTTTGATGAGGATTGGAGGAGTCAGATCATGTAAAGTACTTAGACCAATGTTTCCTACATAACAAGTGCCATATAAGAGCCATCATTTGGCCGGACAAGGTGGCTCACACCT | 120 |
| chr19 | 50701708 | 50701828 | 788855_47188554_ENST00000293441.6,E | 1 | - | good | TTGCAGGGCTGCTGACGTGTTCTTGGATAGAAGGATGCTTTAGAGACCACCACGTGCCGCAGGTGTGGCCGTTAATACGAGGCATTGGAACAATTCTTTGCCAGACTCTAGGCTGCCCAG | 120 |
| chr19 | 50701828 | 50701948 | 788855_47188554_ENST00000293441.6,E | 1 | - | good | ATAAGATGTTTGGCCGTGGCGAGGGTTCCCAAAATGGTGGTGCTGTTATCAGGGTTCACCTCCCTTTGGTTTTTCCACCTTCATCTGGAAAGTGGAGCCAGTCTCCCCTGCTTATAGGCT | 120 |
| chr19 | 50701948 | 50702068 | 788855_47188554_ENST00000293441.6,E | 1 | - | good | GCCACCTGGTGCAGAGCGTGAGGCCGAGCAGCCTTAACTTTGCCTCCTGCATCTGTCCAGGGGTCCTCAACTGAGAGACAGGCAGGGACCTACTTCTAGGGCCCCGACGAGCTGGCAGAT | 120 |
| chr19 | 50702068 | 50702188 | 788855_47188554_ENST00000293441.6,E | 1 | - | good | TCCTGATGGCCCTTTCTGGGGAGGGAGGTCATGGGCCCCATTGCATGGTTGAAGGAACAGAACCTCACACAGGTGAAGCCACTTGCCCAAGGTCCTCAGTCCAAGCCCACAGTCCTTCCT | 120 |

| | | | | | | | | |
|---|---|---|---|---|---|---|---|---|
| chr19 | 50702188 | 50702308 | 788855_47188554_ENST00000293441.6,E | 1 | - | good | ATAGAGTGTAGCTCAGATGACCCTGTCAGGAGGCAGTGAAGTGAGCCGTGTCCCACTGCGGGACACTGATGCCGTGTGCAGGGCATGGCTGGAACTTCTCACCAAATCGTCTCATTAGAC | 120 |
| chr19 | 50702308 | 50702428 | 788855_47188554_ENST00000293441.6,E | 1 | - | good | TGTGGGGGCGATCAGGGAAAATGTGGAGCAGAGAGCAGACCATTCTCGGGCACCTGGAGTATCTCATGCACTCTGGAGACCTCAGAGAGCCCCAGGACACCCCTTGAGGCTTGGAGGACC | 120 |
| chr19 | 50702428 | 50702548 | 788855_47188554_ENST00000293441.6,E | 1 | - | good | GGACGCTCCTTCATGGCTGTGAAGTCCTACCAGGCCCAAGCCGAGGGGGAGATCTCCCTGAGCAAGGGCGAGAAGATCAAAGGTACCCAGGGTGGAAGCAGGGCCTGGGCTGGGCTGGGC | 120 |
| chr19 | 50702548 | 50702668 | 788855_47188554_ENST00000293441.6,E | 1 | - | good | GTACGCAGCTCCAGCGGGACACCCCGGGAAGGGCCAGCCGGGGGCACGGGGGGCTCAGGGGGCCCCGGGGGCTCCCTGGGCAGCCGCGGGAGGCGGAGGAAGCTCTACTCAGCGGTA | 120 |
| chr19 | 50702668 | 50702788 | 788855_47188554_ENST00000293441.6,E | 1 | - | good | CGCCAGTGCTGGGAAACCCCCCCTCCCCCCTGATGCACCGTCCTCTTCCCCCACCCAAGGACCCCTGTCCTCCCAGAGGTGTCCCCTCCCTCCACCCTCCCCTCCTCTCATGCCCTCTGT | 120 |
| chr19 | 50702788 | 50702908 | 788855_47188554_ENST00000293441.6,E | 1 | - | good | GCGGTGCGCGCAGGACCGTGGGAGGGGGCAGAGCCGGAAGCCAGGAGGCAGGAGAGGGGGCTGGTTGGGGCAGCTGCGGGCAGGCAGAGGGAGCCCCCAAATGCCCTCTTCCTTGGA | 120 |
| chr19 | 50702908 | 50703028 | 788855_47188554_ENST00000293441.6,E | 1 | - | good | AGAGGCTGGCAACGGCGGGCAGAGGAGTTGGACCTGACCCAGGAGGAAGCTGGGAGTCGCGGAAGGCTCGTGAGCGGAGGGATGACACAGTCCCAGCTCAGCCTCAGGAAGCTGAATCA | 120 |
| chr19 | 50703028 | 50703148 | 788855_47188554_ENST00000293441.6,E | 1 | - | good | TCACTCTGGAGGGAGGATGAGGAAGGGGCGGGGGACATTCCGGGGTCAGGAAGCCCACCTGGCTAGAGGAGGGTGAGAAGGAGGCGAAGGCCAGCCAGGCGTGTCCCGAGGAAGTGGG | 120 |
| chr19 | 50703148 | 50703268 | 788855_47188554_ENST00000293441.6,E | 1 | - | good | TACCCCAAAGAAACATGGGCCCCAGCCCCTGGGGACAGACAGAATGACAGGGTGAGGTGGAGGGAAGCTGAACAGGAAGAGGATGGGGGGGGTGGCAGGGACTTGAGGGGGACAGAGA | 120 |
| chr19 | 50703268 | 50703388 | 788855_47188554_ENST00000293441.6,E | 1 | - | good | AAGTCCCTAGTGTCACCTTGGGGTAAAGCAAGGCCTTGGGAGCTACTGTCCCGACTCCACCCCCGGTGCACCACATGTTAGCTGGATGTCTGCTCCAGGACAGGGGCCCCAGGCAGAGGG | 120 |
| chr19 | 50703388 | 50703508 | 788855_47188554_ENST00000293441.6,E | 1 | - | good | CCGGCCCAGGTAGGGGAGGCAGCCCTGGCTCCACAGTCCCCAGCCCCGTCAAATCCCCTCTCCAGATCGGCGGCTTCCCCCGAGGCCAGCCCAGCTGCCTTCCTCTGTAGGCCCTCATCC | 120 |
| chr19 | 50703508 | 50703628 | 788855_47188554_ENST00000293441.6,E | 1 | - | good | GGCCCCCACCACCAAGCTCAGCAGCGGGACCCTCCGAAGTGCCAGCAGCCCCCGGGGTGCCAGGGCCCGCTCTCCATCCCGAGGGAGGCACCCTGAGGACGCCAAGAGGCAGCCCCGAG | 120 |
| chr19 | 50703628 | 50703748 | 788855_47188554_ENST00000293441.6,E | 1 | - | good | GCGGGCCAACAGTGACACCAGCATGGCGCTGCCCGACTGGATGGTGTTCTCCGCCCCGGGGGCCGCGTCCTCTGGGGCCCCTGGCCCTACCTCAGGGTCCCAGGGCCAGTCGCAGCCCTC | 120 |
| chr19 | 50703748 | 50703868 | 788855_47188554_ENST00000293441.6,E | 1 | - | good | CCTAACATCTGCCCCCCGCCGCCACTGTGCCCTCTCAGTGCCCTTCCAGGAGTCCCCCAAGTACGCGGCCCGGCGACGGGGGCCCCCAGGCACAGGGCTGACGGTGCCCCCGGCGCTGCT | 120 |
| chr19 | 50703868 | 50703988 | 788855_47188554_ENST00000293441.6,E | 1 | - | good | CTGGATGGTGGTGGGGGAGCAATTCCAGGCCCCCCTCCCTTTCTCATCTCTGTCTTTCTCCCTCATGCCTCTCTCCCACCATCTCTTGCCCCCGCATGGCCCCCTGCTTGCCTTTTCTCC | 120 |
| chr19 | 50703988 | 50704108 | 788855_47188554_ENST00000293441.6,E | 1 | - | good | GGAGCTCGGACTGCACAGAGAACCTGGGGCGGTTGAGGTTTCATGGTGGGATGGGGGAGTTGGGGGACCTGACTTGGGGGCCAGGATAGGATGGAGAAACCCTAGAGGACCAAAAATGT | 120 |
| chr19 | 50704108 | 50704228 | 788855_47188554_ENST00000293441.6,E | 1 | - | good | CTGGGCCCCCCTGGCCGACCTGCCTCTACCCCCTCCCTCCAGGTGGCAGTGATTGCTGGGAATTTTGAGCTGGGGGAGCTGATCCGAAACCACCGAGAACAGGATGTGGGTGAGTGACAG | 120 |
| chr19 | 50704228 | 50704348 | 788855_47188554_ENST00000293441.6,E | 1 | - | good | GAAAGCAAGGAGACAGGCCGTGGAGGGGGGAGCTGGAAGAAGACAGAGGGCGGCATTGTCGGAGTGGAAGGTTTGGGTCGGGCCAGGGACCTCGCCACGTTCCCTGCCCTCTCTCTGCTGC | 120 |
| chr19 | 50704348 | 50704468 | 788855_47188554_ENST00000293441.6,E | 1 | - | good | TGAAGAACAACAACGGACAGACCCCCTTCCAGGTCTGGGGCTGCAGGGGATGTGCTGGAGTTTCCAGGGCTAAGGAAAGGGTGACAGTGACACCCAGTGGATAAGGAAGGGGACAATGAG | 120 |
| chr19 | 50704468 | 50704588 | 788855_47188554_ENST00000293441.6,E | 1 | - | good | AACCTGCATCCACTCATGGTGATCTTAATTTTTTCTCTCTCTCTTTGTCCTGTGCCACTGGCCATTACCCGCAGGAGACCTGTGCCAGGATCCTCCTGTATCGAGGTGCCGACAAGGATG | 120 |
| chr19 | 50704588 | 50704708 | 788855_47188554_ENST00000293441.6,E | 1 | - | good | CTCCGTTTGGGCCGGCCACGTTTAAACTAAAGTGCTCCTGGTCCGTACCTGGCAGGTGGCTGCTGTCCTCAGTGGTGCGGGTTTAGGGTGGAGTATTTATTCCTGAGGCTCTTAGCACAG | 120 |
| chr19 | 50705428 | 50705548 | 788855_47188554_ENST00000293441.6,E | 1 | - | good | ACCAAAGGATTATCCGGCCCCAACATCCAGTGGAACATAGCATGAGCACATACCCTAAATGCCCTCAAATATCCTAGTGGCCATGTTAAAGAGAAGGGGAAAATAATTAGGTGACATTTA | 120 |
| chr19 | 50705668 | 50705788 | 788855_47188554_ENST00000293441.6,E | 1 | - | good | GAGCTTCTGCCTCTAAAGTGCCAAGCGCCTGGCACGTAACATGGTAAGCCTCCGTAAGGTCTAGACCTGTGGTTCTCAACCTGGGAGCTTCTGTCTCCCAGGGGACACTTGGCAATGTCT | 120 |
| chr19 | 50705788 | 50705908 | 788855_47188554_ENST00000293441.6,E | 1 | - | good | CTGGGATTACAGGAGTGAGCCACCACCCCTGGCCCACCTTGGGCTCTTTTGAAAGCCTTAGTTTCCTCCTCTGTAAAATAGAGACAATTTCCATGTCCTGGAATTGTTGGGCAATTCAAT | 120 |
| chr19 | 50706148 | 50706268 | 788855_47188554_ENST00000293441.6,E | 1 | - | good | AAGTGTGATTAAGATAGAAGACTTTGGAGTCAGAGAGATCTGGGTTCGAATCCAAGGCTGATAATCACCAGCTGTGTGACCTTGAGCCCCCCCAACCCCCCCTTTTTTTTGAGACAGGGT | 120 |
| chr19 | 50706268 | 50706388 | 788855_47188554_ENST00000293441.6,E | 1 | - | good | GTGATTGACACCGGTGAGAAATTGATCTGAATGTAGAAGGCATCCCTGTCCCCACTGGCCCAGTCCTGCCCCCACCCTTTCCTCCACAATGTGTGTCCCCTTCCTTCTGGGCCTGGCAGA | 120 |
| chr19 | 50706388 | 50706508 | 788855_47188554_ENST00000293441.6,E | 1 | - | good | GGTGAGGGTCAGGGTGGGGAGTGGCAGGGCAGGCAGCTCAGGTTAGCTCAGCTAACCCTAACCCTCCTAGGCCGTGGCATGGACTTTGTGCTTAGCCCAAAAAGTGATGATGGGTGCCAT | 120 |
| chr19 | 50706628 | 50706748 | 788855_47188554_ENST00000293441.6,E | 1 | - | good | GAGGACAAGGGTTTTTTTTTTAGACAGGATGGCCAGAGAAGACCTCTCTGAAGAGGCGGCATTTGTGCAAAAGCCTGAACGAAGTGAGGAAGCAAGCTATGCTGAGATCTAGCGGGAAAG | 120 |
| chr19 | 50707108 | 50707228 | 788855_47188554_ENST00000293441.6,E | 1 | - | good | TACAACTTATGGTGTAGAGGAGAAGAGACACTACTGAGACAAATAACAGGAAATGGAATTTCAGTGAGACGGGGTTACGAAGAAAATGAGTTAACATTGGCTTAGAGTGCCTGGCACATA | 120 |
| chr19 | 50707228 | 50707348 | 788855_47188554_ENST00000293441.6,E | 1 | - | good | GGCCAAGTTGGGTTTGGCCAGGCCAAGATAAAAGGGAGAGGACACTCCTTCTACCGAATAGTCCCTGTGGGTCCAGCATTAATCCAGGTGGTGGGTGCCACAGGCAAAATCATGCCCCTG | 120 |
| chr19 | 50707348 | 50707468 | 788855_47188554_ENST00000293441.6,E | 1 | - | good | ACATAGGCCCATACAAACCCCTTGGTGGATTTGTCATGTGGAAGGGGTAAGATCAAGGAGGTGCTTGGGAAATGGTGGTTAAATTAGCATCAGAGGGCGCTGGGGTAGAGTTGAGCTGGA | 120 |
| chr19 | 50707708 | 50707828 | 788855_47188554_ENST00000293441.6,E | 1 | - | good | GCACGGGCATACACAGACTCACATGCACACACACACTCCCAAACACACCCAGAACACACACTGGGCTAACTCTGCTGCCCGATGTGTGACTTTGGCAGTCAGAGTCGTCCATTCATAGGA | 120 |
| chr19 | 50708308 | 50708428 | 788855_47188554_ENST00000293441.6,E | 1 | - | good | GAGATAAGTGCTAGGATGCCGTTAAATGCGAGTTTCTCTTGGTGCCCAGGGGAGGGAGCCATCAACTCTGACTTGCACACCTTCTCAGCTTTGTAAGAGTAGGATGAGGAGCAGGTGACC | 120 |
| chr19 | 50708428 | 50708548 | 788855_47188554_ENST00000293441.6,E | 1 | - | good | GGATGTGAGATGAATCAGACACAGCATCTGGCCTGGGGAGGGCGAGGGCGGGGTTGGGGGGGGTGGGAGTTCACAGTCTGGTGAGGAGGCAGACGTGTAAGCAAATCATGGCTACGACA | 120 |
| chr19 | 50708548 | 50708668 | 788855_47188554_ENST00000293441.6,E | 1 | - | good | ACCAATGAATGAATGACTCGAACAAATGAATTTTGACTGACTCAACCCCACATCCTCCATCCTCATCCTTTCCATGCACGTTCCTTGAGCACCTACTATGTGCACAGCACCAGGCCTTGG | 120 |
| chr19 | 50708668 | 50708788 | 788855_47188554_ENST00000293441.6,E | 1 | - | good | ATGAAAATGAGTGAATGAATGATGGGGTCAGCCCTCTTGGATTTCCACCAAGACTGTGCACAGTGCCTGGCTCAGAGCATTCAACAGGTGTTTGCTTGTTTCAACAGACTGAATGACTCA | 120 |
| chr19 | 50708788 | 50708908 | 788855_47188554_ENST00000293441.6,E | 1 | - | good | ATCCGTGTCTAGGCTGTGAGCACCTCAAGGGTGGGGGTTGAGTCTGATTACTTGCTCTGTTCCAGGACCCCAGCAAACAGCAGGCCTAGTTCCATCTTTGTTGAGTGAGTGAATGAATGA | 120 |
| chr19 | 50708908 | 50709028 | 788855_47188554_ENST00000293441.6,E | 1 | - | good | TTGTCTTCCTCCCAGGCTTTAAGCTTTCCAAGAGCCAGGGCTGAGTCTGACAGTTTTTCCTCAGCCTCTGCCGCCTTGGCCTCCTGATATGCCCGTCTAGCATTTGGTGGTGAATGTCAG | 120 |
| chr19 | 50709028 | 50709148 | 788855_47188554_ENST00000293441.6,E | 1 | - | good | AAATAAATAAATATCTACCATGCAAAAGGATCAAGTGGGTTAATAGGTTTGCTTGCCTTTGCCATGGGCTTAACAAACCGATGCATAACACATACGAACTTGATTCCTCCATTTCCCCCC | 120 |
| chr19 | 50709868 | 50709988 | 788855_47188554_ENST00000293441.6,E | 1 | - | good | ACTAAGTGCTCATCCCTGGAAGGCAGCCATCAGAATAATAATTGTCATTGTTACCATCACCAGTGTCAGTCAGGGTCCAGTCGAGAGACAGAAACTGTACCGTTCAGATAAACAGAGAAA | 120 |
| chr19 | 50709988 | 50710108 | 788855_47188554_ENST00000293441.6,E | 1 | - | good | GCCCCACAGGCAGATGGGGGTGGGGAGCAAACAAGTCCCTCCCCCTCTTTGAGGCTCCATCCTTCTTCCTGAAATGGGAATAACGCCAGCTCTTATTGACGAGGGTGAGCCATCGTGTAT | 120 |
| chr19 | 50710108 | 50710228 | 788855_47188554_ENST00000293441.6,E | 1 | - | good | ACCAACATGGTGCCGGCACTTCTCTCCCCACGTGCTCCTGTCTCCTGCTCTGCGACACAGGCCTCACAACCAATCCTCTGACTCCCAGGCCCTCGTGCAGCTGATGGGGAAGAAGCCCTG | 120 |
| chr19 | 50710228 | 50710348 | 788855_47188554_ENST00000293441.6,E | 1 | - | good | AGCCCCCTCTGCCTCGGCAGAACTTGCAGTGCTGGGGATTTGGCGGGTGGCCACGTGCCGCCTTGTGGCAGCCAGCAGTATGCCTGGAGCCCAGGGAAGACTCAGTGTTTGTCAAACCGA | 120 |
| chr19 | 50710348 | 50710468 | 788855_47188554_ENST00000293441.6,E | 1 | - | good | TTCTTGATGCTTCCTGCCCTTGAAGGGCAGCTGGTGGGGGTACTTAGAGAGACAGGCGCTAGCATGACAGAGCTGGTCTGTCAGCCCCCTTCCTCCAGGAAGTCTTCCAAAACAGCCCTC | 120 |
| chr19 | 50710468 | 50710588 | 788855_47188554_ENST00000293441.6,E | 1 | - | good | GGCAGCTGGTTGGCAAGAGCCGGGTCTTTCAATGTCATGACCTCTGGTACTTTAGCCACCTGAGCTGCAGATGATCCTGGGCAGGTCAGCCCCATGGAAAGCCTCAATCTCCCCACCTGT | 120 |
| chr19 | 50710588 | 50710708 | 788855_47188554_ENST00000293441.6,E | 1 | - | good | GGGGGTCCTGACAACCTGGGTCCCTGAGCAGGAAGGGAGTGCTGAAGGGAGATGGGAAGAACACCCCCACTTCTGGAGTTCCTAGGAGAGGAGAGGTACGGAGCAGGCAGGGGGGCTCG | 120 |
| chr19 | 50710708 | 50710828 | 788855_47188554_ENST00000293441.6,E | 1 | - | good | GTGGTGTGTGCAGGTGCCACGTGGCGGCACGGCAGGGCACAGGGGAAAGGCCAACGTGCAGCCCCCGTGGCCCCTGAAGGGAATGACAGTGGGATGGGAGTTTTAGCAATTCAGGACCTT | 120 |
| chr19 | 50710828 | 50710948 | 788855_47188554_ENST00000293441.6,E | 1 | - | good | AGATGAGCAGGAGTAGAAAAGAGGAGAAAGGGGTGCTCCAGGGTGGGGGCACAGCCTGTGCAGAAAGCCAGAGGAAAGGGGGCTGTGGGTCCTCAGTTCCCCTGTGTCTGCATTCAGCCA | 120 |
| chr19 | 50710948 | 50711068 | 788855_47188554_ENST00000293441.6,E | 1 | - | good | GTGCTCAGAGCAAAGTCACAGGGAATCCAGATCCCTGCGGGAGCCCACATCTGGGACTGGGCTTGGGATGGAGTGGCTGGGGGTGAAGGTGGCTTCCTGGAGGAGGTGCCCCCAGTGTTA | 120 |
| chr19 | 50711068 | 50711188 | 788855_47188554_ENST00000293441.6,E | 1 | - | good | CATCCAACAGATATTTACTGAGCTCCTCTGTATCAGGCCTGGAGCTGGGTGCTGGGGACAAAGTGTTGAGCTCATGGCCTAGCAGAGGGCAGACAACATAGTTTGCTGCTACAGTCCAGA | 120 |
| chr19 | 50711308 | 50711428 | 788855_47188554_ENST00000293441.6,E | 1 | - | good | TGGAGCCCAGAACGCCTCGGGGAACACGGCTCTGCACATCTGCGCCCTCTACAACAAGGTCTGCCTAGCAGCGGCCACCCCAGGCCCCTCACATCCCCGATAGCCGCCTCCCCCAGGGCC | 120 |
| chr19 | 50711428 | 50711548 | 788855_47188554_ENST00000293441.6,E | 1 | - | good | TTGGCCCAGAGGGAGAACAGCCTGGGTCTGATCCATGCCCCTCGCTCCCTGTCCTGCCCAGGCCTGCCAGCGGGGTCACTCTCAGCACCTGGAGCATCTGCTTTTCTACGGGGCTGAGCC | 120 |
| chr19 | 50711548 | 50711668 | 788855_47188554_ENST00000293441.6,E | 1 | - | good | TTGGGTTTTGAGGCTGACAGCCTGGGTCCCAGAGCAGGAAGCGGGGCTAAAAGGTGATGGGGAGGTGGATGCGGTGAACGATAGGATGAGAAGGCCAGTGCACAGGTCATAGACAGACTC | 120 |
| chr19 | 50711668 | 50711788 | 788855_47188554_ENST00000293441.6,E | 1 | - | good | AGACCCTAAGGGGGCATAGGCTGGGGAACCTGATGCCTTGACCCCTGAGAGGGATGGGCTGTCCTCTTGACCCCCTTAAGGAGGCAGACCAGACGCCTGGGCCTTGGAATGACTGAACAT | 120 |
| chr19 | 50711788 | 50711908 | 788855_47188554_ENST00000293441.6,E | 1 | - | good | ACCCAGTCCCTGAGGCATCCAAGGGGCTGGGGCTGGGAACCAGATGCCAGGGTCCTCACAGAGCATGGGGGTGAGAATAACATGGTCCCCAAAGAGGAAATGGTGGGGGTGAAGGCAGAA | 120 |
| chr19 | 50711908 | 50712028 | 788855_47188554_ENST00000293441.6,E | 1 | - | good | CCCCCGATGCTGCGAGCTGCTCCTGTTCAACAGGGCCCAGCTGGGCATAGCTGATGAGAACGGCTGGCAGGAAATCCACCAGGTGCTCCTGATTCAGGCCATGAAGGGCTGGGGTGGGGG | 120 |
| chr19 | 50712028 | 50712148 | 788855_47188554_ENST00000293441.6,E | 1 | - | good | GTGTTTTTCTACTGGGTTCTCTCTCCGTTCCTAGGCGCTCCTGGACCTTGGGGGTTCCCCCAACTACAAGGACCGTCGGGGGCTGACCCCTCTGTTCCACACGGCCATGGTGGGTGGTGA | 120 |
| chr19 | 50712148 | 50712268 | 788855_47188554_ENST00000293441.6,E | 1 | - | good | CAAGGTTGCACAAGGGCATCACGGCAGAACTGAGACTGTCATTGGCCACTGTAGGTCACCCAGGACGGTCCAGTCCCCCTGGCCCTTCAGAATGAGCTGGTTAATGTGTGACTGTCATCT | 120 |
| chr19 | 50712268 | 50712388 | 788855_47188554_ENST00000293441.6,E | 1 | - | good | CGCCCCCAGGGGGAAGGCTGGGTGGGATGTCAGAGTGCAAGTGTCCTCTGGAGGCATCTACCCAAGTCCCACATTGTTCACACGGAATCAGAGTGGCCCAGGGAGGGAAAGGGCCAGGCT | 120 |
| chr19 | 50712388 | 50712508 | 788855_47188554_ENST00000293441.6,E | 1 | - | good | GGTCAGAGACCCTCCAGCATCCAGACTCTGTCCTCTCACTTGTCCCCAACAACCCACCTGTAGGAAGTGGAATGGGTGGGTACTGTGGCCCCTAATGTCTTTCAATGGCTCATCCAAGAT | 120 |
| chr19 | 50712508 | 50712628 | 788855_47188554_ENST00000293441.6,E | 1 | - | good | AAGTGGCCTCCAGCCCCAGTGAGGCCGGCCCGATCCTTGCCCTTACCCAAGCTTACCTGTCCCCTGGCTTCCTGCTGTGCCCCTTCTTTGAGATCCTGGACCTAATAACACCTCCCACTT | 120 |
| chr19 | 50712628 | 50712748 | 788855_47188554_ENST00000293441.6,E | 1 | - | good | TCTGTTGGGTGGCACTGCTACAGGCTGTCCTCACACGAGGTTGCTGCCAAGACTTTCTCCTATTAATGTTCATTATTGCCAGCCTTCCACACCCAGAGCTGTCCTTCAAGTGGGGGCAGG | 120 |

| | | | | | | | | |
|---|---|---|---|---|---|---|---|---|
| chr19 | 50712868 | 50712988 | 788855_47188554_ENST00000293441.6,E | 1 | - | good | CACTCACCCCCAAACCCACATCCTGCTCACACACGCCCCTCCCAGCCTCCAGACCTGCGCTATCCAAGCCAGTAGCCACTGGCCACCTGGGGCTGTTTAAATTTAAATTAAATAGAATGA | 120 |
| chr19 | 50712988 | 50713108 | 788855_47188554_ENST00000293441.6,E | 1 | - | good | CCCTCCCTTCTCTCCCTCAGATGCTTTCCTGAAGAGCCCCTAATGCAGTCAGATGCCAGCCACAAAGATCTGCAGTATACCCCTGGAGTTAACAGATCCTCTTCATGTATCTCCCATGCA | 120 |
| chr19 | 50713108 | 50713228 | 788855_47188554_ENST00000293441.6,E | 1 | - | good | TCTTCTCCCCCTGCCCCCGCCAGGACACCTGATTGCTCTTCACATGAGGGTCAGGATGGGCTGCCCCCAAAGACAGCCATTGAGAAGCACCCCGACGCACAGCTGCTCCAGCTAAACGGC | 120 |
| chr19 | 50713228 | 50713348 | 788855_47188554_ENST00000293441.6,E | 1 | - | good | CATGCAAATACAGCCCCTTCCCCACGTACACAGCCTCAAATGCACTTTCCCAAATAGACAAACCCATACACCCACAAAATACATACACAGCGGCAGCTGCACAGTCTAATTTTAAAATAC | 120 |
| chr19 | 50713348 | 50713468 | 788855_47188554_ENST00000293441.6,E | 1 | - | good | AATACGAAGTGAGCTCCCTGCTATGCCCCCAGATAGAGCCCCTCCACACACACCCTGAAAACAGCCCCCCAGCTACCCTTCCAAACCAGCACTGCACCAGCTCCCCTCCCTCCTCTCTCA | 120 |
| chr19 | 50713468 | 50713588 | 788855_47188554_ENST00000293441.6,E | 1 | - | good | ACACAGACCCGGGCACGGCCTCCTCCCCTAAACACCCAGAACGTCTCCATCCCCTGTGGCCCAGCTTAATACCAAATCTGCCTCTGTCTCCCCAAGGTGTAAACCCTTCTCATTCCTTAA | 120 |
| chr19 | 50713588 | 50713708 | 788855_47188554_ENST00000293441.6,E | 1 | - | good | TTAAAAATAGGCCCTCTCATGTCCCCAGTTCAAAGCCCCTTTCCATAGACTGAGACACAGCCCCTATTGTCACCCTTGTCAGGCCCCCCAGATGCACCCACCTGGGCTATAGCTGACATC | 120 |
| chr19 | 50713708 | 50713828 | 788855_47188554_ENST00000293441.6,E | 1 | - | good | CGCATGCGCCCGACACTGCCTGGCACTCACGGTGAGGCTTCGGGGACCCCCCATCCCCGCCATGGGGCCCTCTCTCCTTTTCCCTTTGTTCTTTCTCAAACCTCAGTTCTTTGGTTTCAG | 120 |
| chr19 | 50713828 | 50713948 | 788855_47188554_ENST00000293441.6,E | 1 | - | good | GACCCCCTTGACACTGGCGGCCCAGACCGAAGGCTCTGTAGAGGTGATTCGAACCCTGTGCCTGGGCGGGGCCCACATTGACTTCCGGGCCCGGGATGGCATGACCGCACTGCATAAGGC | 120 |
| chr19 | 50713948 | 50714068 | 788855_47188554_ENST00000293441.6,E | 1 | - | good | CCAAGGGGGAGGCCACGGGGCCAGAGGCTGTGGGGTGGAGAGCTGCAGAGCCTGGAAAATGGGAAGGGATGGAGGGGAGGAGAAGGGGCTTCATGTGACCAGCTTTTCCCGCCCTTCAG | 120 |
| chr19 | 50714068 | 50714188 | 788855_47188554_ENST00000293441.6,E | 1 | - | good | TCGGGAGGTAGGGAGGGGTGCGCATCAGGAGGACCTCAGGGCCAGGACAGAGCTGGGGAGCAGGGGTGCCATCTGCATTCACATCCATTCCTGACTGGTGTCTCCCAAACACATTGCTGT | 120 |
| chr19 | 50714308 | 50714428 | 788855_47188554_ENST00000293441.6,E | 1 | - | good | GTAAAGGAGGCTGCTAGAAGCTTTTCAATGGCGTATGTGACTCCACGTATATTTTCACTGGACGTCGCTGCTTGAGGACCCTGCTTCTCTGCCTTGCTCCTGACTGTCCTGTCTTCTCTC | 120 |
| chr19 | 50714908 | 50715028 | 788855_47188554_ENST00000293441.6,E | 1 | - | good | ACCACAGCATGCATTGAACTCTTTCCTCCCCTAAATGTTCCATCCCTCCAGACTTTCACGGTCCACTTTGGGAACCACTGGCCAAGTGTGGCTCCTGAGTGCTTGCAGTGTGTTTGGTCC | 120 |
| chr19 | 50715028 | 50715148 | 788855_47188554_ENST00000293441.6,E | 1 | - | good | TGTCCTGTGTCTCGATGGTGAGAGCTGGGAGGAAATCGGGAACTGCTTTGGTCAGAAATGAAGAACTCTTGATCCTTGAGAAGGAAAAACTCCCAGTCCCTCCTTGTCTACCCTCTCCCC | 120 |
| chr19 | 50715148 | 50715268 | 788855_47188554_ENST00000293441.6,E | 1 | - | good | GACAGGCTACATTTCTCCCAACCTCATAGCCTGAATCTCTGCAAAATCCAACCATTACACCCCCCTTTCCAGCCATATCTGTCCTCCTGGTACCTTGCCCGCCAGTCCTCAGCCCCTGTC | 120 |
| chr19 | 50715268 | 50715388 | 788855_47188554_ENST00000293441.6,E | 1 | - | good | CCTTTGAGTCTCCCTTCTCTTCCCCTGCCTCCCCCCATATCCATCCAAGGCCTCTTTACTTCCCCACCTTGGCCTAGGACCCTGACACTGAAGGTCCTTCTGCCTAGAGTCTATTTCAAA | 120 |
| chr19 | 50715388 | 50715508 | 788855_47188554_ENST00000293441.6,E | 1 | - | good | TGCTGTACCACATTTGAGACAACCACAAATACCCCCCATTATCCCCATCACAACCCCTTCTTACTTTCCTCTTTTGCCCTTCATCCCTCTGCCTGGGCCCACATCATTCCCACCTGGCCT | 120 |
| chr19 | 50715508 | 50715628 | 788855_47188554_ENST00000293441.6,E | 1 | - | good | ATAGCCCCCTACCACTGGAGCCCCCTACCCAACCACTTTCTCTCCCCACCTCTTCCCAATAGACCTCCACATTCCCCAAGCCAGCGATCCCTCTGGCCACTGCCCCCGCTTAACTTGGGA | 120 |
| chr19 | 50715628 | 50715748 | 788855_47188554_ENST00000293441.6,E | 1 | - | good | CCCTACCCTTGACCTCAGTTCCGATACAAGACCCGAGTTTACAAACAGACCAACCTGGATGAGAAGCAGCTGGCCAAGTTGCACACGAAGGTGAGAAAACCCTGCTGGCATCTATCCCCT | 120 |
| chr19 | 50715748 | 50715868 | 788855_47188554_ENST00000293441.6,E | 1 | - | good | CAGGGCTGTTTTCACTCCCACCCTGAGCTACCCCGGGGAATTAGAAGGGGATCAGACCTTCCCCTACCCCAAGCCTTCTCCCCAGGCCCCTGATTCCAAGTCTCTGTGTCTCTCTCACTA | 120 |
| chr19 | 50715868 | 50715988 | 788855_47188554_ENST00000293441.6,E | 1 | - | good | ATTCCCTGTGGGGTCCAGACTCCTCCTCCTTCAAGGCAGGGGTTGCAGTGCACATATACCATAATACCCAGCACTGTCGGAAAAGCCACTGCCACCCATCTACCGCCTAGACCAAGGTTC | 120 |
| chr19 | 50715988 | 50716108 | 788855_47188554_ENST00000293441.6,E | 1 | - | good | TACCATCTGTTCTGCTTCTGACCAGCTAAGTGGCCTTGGCCAAGTCACTTTCCTGCCCCAAGCTTCTCCTCACCCATCAAATTGGGGGAGGATTCCCAATTACAGTAGGCACTCTAGAAC | 120 |
| chr19 | 50716108 | 50716228 | 788855_47188554_ENST00000293441.6,E | 1 | - | good | AAACACATCTGCCCCCCACCCACTTTTTAACACACTCGAAACCTTAACGAGGGGGCACCCAGTGTGCGAGTCCAGACCTTCTAAGCATCACAGGGAAGTCAGAGAGTCCTGAGTTCAAAT | 120 |
| chr19 | 50716228 | 50716348 | 788855_47188554_ENST00000293441.6,E | 1 | - | good | CCAACTTCCTGGAGGAGGAGAGGCTGCTGCGGGAGTACCCCCAGTCCTTTGAGAAGGGGGTCCCCTACCTGGAGGTGAAGCTTCCCCAACTCCTTCACTGATAAGAGAAGGCATGCCCTA | 120 |
| chr19 | 50716348 | 50716468 | 788855_47188554_ENST00000293441.6,E | 1 | - | good | GCTTCAACCCCGATGCCACCATCTGGACGGCCAAGCAGCAGGTGCTCTGTGCCCTGAGCGAGAGCCTGCAGGATGTGCTCAACTATGGCCTGTTCCAACCGGCCACCTCCGGCCGCGATG | 120 |
| chr19 | 50716468 | 50716588 | 788855_47188554_ENST00000293441.6,E | 1 | - | good | TATCCTCCCAGGAATCACCCCCACACTCAGCTCCTGGCTCACTTCCCAACCCACCTCAGGCTCTCCTCTGGCCCCTGGCCCACCCTAGCCCCTATCTCTTCCCCAACCAGAAATGCCTTC | 120 |
| chr19 | 50716588 | 50716708 | 788855_47188554_ENST00000293441.6,E | 1 | - | good | GCATGATGGTCTTCAGGATTGGCATCCCGGACCTGCACCAGACAGTGAGTACCTGCCCACTGGCCAGCGGCAGGAGAGGGACAGTGCCCCAACCGACATGGGGAGTCCACTGGTGCTGGG | 120 |
| chr19 | 50716708 | 50716828 | 788855_47188554_ENST00000293441.6,E | 1 | - | good | ACGGGCCAGGTCGAGGCCCCCGGGGGACCCGGGGCCAGGGCAGTGGGGCACCTGGTAGCCTGGCCTCTGTTAGAGGCCTCCAGGGCCGCTCAATGTCCGTCCCAGACGACGCCCACTTCA | 120 |
| chr19 | 50716828 | 50716948 | 788855_47188554_ENST00000293441.6,E | 1 | - | good | GCCGTCCCCCGTCGCCCCGTGGCCCCACAATGACCCACAGCCCCGCGACAAGCGAGGACGAGGAACGCCACAGTGCCAGCGAGTGTCCCGAGGGGGGCTCAGAGTCCGACAGCTCCCCAG | 120 |
| chr19 | 50716948 | 50717068 | 788855_47188554_ENST00000293441.6,E | 1 | - | good | GGGCAGTGGGGGTAGGCTGCCCTTGACCGCTTGACCACCGAATAGCGCCTGCGGCCTCTGAGGGGTCCCGGGCTCCTAGTCTGATGGCCGCCCTTGGCCCCTGCAGCCGCCTGTGATGCT | 120 |
| chr19 | 50717068 | 50717188 | 788855_47188554_ENST00000293441.6,E | 1 | - | good | GAAGGAGACACATGCCAGAAGCATGTCAGGGACAGGGGTGTGTGCGTGCCAGCGTGGGAGGAGGCGGCAGGGGAGGCCGTGCAGGCCGAGGGTGTGTCCAGCCTCCTTCCTGCCTATCTT | 120 |
| chr19 | 50717188 | 50717308 | 788855_47188554_ENST00000293441.6,E | 1 | - | good | GGCCTGGCAAAGTGGGCCAGTAGCTCCCGCGGTGATTCTGCGGTTTCTGGGACACGGCCACGCTGGGTCACCAGCAAAGTGGGGCGCAGGGGGCGGGCTGCAAGCGGGGCCAAGGGCAG | 120 |
| chr19 | 50717308 | 50717428 | 788855_47188554_ENST00000293441.6,E | 1 | - | good | CTTCTCCCCAACCCCACCCCCCGGCCCTGGGATGAGGGGCCAGGCTGGGGCCAGGAGGGCAGGATGCCTGTCAGTTTGTCCACCTGTCTGTCCTTAGCTAATTTTAGCTGCATTTTCTGT | 120 |
| chr19 | 50717428 | 50717548 | 788855_47188554_ENST00000293441.6,E | 1 | - | good | CCAGCACTATTCGGCATGCCACGCGTTGTCAGCCACACTTGAGTGAAGTCACCAGCTCCACCCGCTCCAGGCTCACCACACCTCCCTCTCCCCCTTACTACATCTCAGTGTCAGCCTCAG | 120 |
| chr19 | 50717548 | 50717668 | 788855_47188554_ENST00000293441.6,E | 1 | - | good | AAGATGCCCCAGTCTCCAAAAGCCACCTCTTGTGGTTCCCTTGGGGACACCCTAGGCCAAATGCTCTGCCCATGACACCCCCATCTCAGTGCCCCTTGCTCCCCTGGGACGGCAGATTCC | 120 |
| chr19 | 50717668 | 50717788 | 788855_47188554_ENST00000293441.6,E | 1 | - | good | CAGTCCTAGAAAAGAGCTACAGGTCATCTGCAGCCACCCACATCTACCCGGGCCACCACACCACTGTCACCTGGACAGCACTCCCCAGGCCGACCTCCCCACCTACCCGCACACCCTCTC | 120 |
| chr19 | 50717788 | 50717908 | 788855_47188554_ENST00000293441.6,E | 1 | - | good | CTAGAGATGATCCCCCACTCAGTTCCCCAGACCCTCCTCTCACTTCTTAGCCTCCCTCCTAACACCCCCAGTCCTCAGAGTGGAGCCCCTAGTACCCAGCAAGCCAGCCCACCCACCCTG | 120 |
| chr19 | 50717908 | 50718028 | 788855_47188554_ENST00000293441.6,E | 1 | - | good | GACCAGACATTCTGGTTTCCGGTGCTGGGGTGGGAGTTGGGGGAGAACAGAAGGGAGGCGGACCCCATTTGGAAGGGGCTCCACCCTCATAAGGCCACCTCAAGGGAGAACAGCAAGGAT | 120 |
| chr19 | 50718028 | 50718148 | 788855_47188554_ENST00000293441.6,E | 1 | - | good | CTTCCCTTTCCCAACCCCCCAGCCCATCCCCACTCCCCCAAAGTCAGCAGTACCCTCCACTGCTTCCCATCTCCATGGCAACGGTGCAGGAACCGGGCCTGGTGTTGGGGGGAGGGACCA | 120 |
| chr19 | 50718148 | 50718268 | 788855_47188554_ENST00000293441.6,E | 1 | - | good | CCTGGAGGGGGAGGGGGCTTCGCAGCGCTGCCGCGCGTGGGGGAGGGGTCCGCCGGCCCTGGGTTAGAGGCTCGCTCGCCCACACCTCCTGGAACTCTCCTCCCACCACACACATTTAGT | 120 |
| chr19 | 50718268 | 50718388 | 788855_47188554_ENST00000293441.6,E | 1 | - | good | CCAGGGTCAGCCGAGGGGGAGTGTCTGCGGGCCCCAGATCGCGAGGTGGGCTGTGCGGCGGGGCAGGGAGGAGGCAGCCGGGGTCTCCGCAGGGACGAGGTGGGGCGGCGTGGCTGG | 120 |
| chr19 | 50718388 | 50718508 | 788855_47188554_ENST00000293441.6,E | 1 | - | good | GGGGAGGGGCCGGAGCCGGCCCGGGGGGGCTGGGGCAGCCGTACGCGCGGGGGGCCCTAGCCAGCTTTTCTTTCATCCCACCGGCCACCCCCTCCGAGCGGGGTTGAGGCTGGGGGTCA | 120 |
| chr19 | 50718508 | 50718628 | 788855_47188554_ENST00000293441.6,E | 1 | - | good | CCCCTCGCCGATTCCTCAGCCCGTCTCGGTACCAGCTGCGTGCCAGCTATTCTTAGCCGCGGAGCGTTTGATTCATGCCGCCCCGGCGGAGCGTGCCCGCTCCCTCCCGCGAGCCCTGAG | 120 |
| chr19 | 50718628 | 50718748 | 788855_47188554_ENST00000293441.6,E | 1 | - | good | CCCCTCCCCGCCTCCCTCCGCCTCCATTCATTCTCCCCGGCCCCGCTCTCGGCCACGCCAGCCGCCCCCTGTCCCCCCACAGCCTCCCAGGGCACCCCCTCTCTCGCGTCCTCCACCACC | 120 |
| chr19 | 50718748 | 50718868 | 788855_47188554_ENST00000293441.6,E | 1 | - | good | GGCCGGCGCGCCCCCTCGGCTCCCGCCCCCGACTCCTGCTCCCCGGTGCCCCCCGCCCCTCTCCCCGGCCGGGTTCTCCCCCACCAGTTTCCCAGCCTCGGCTCCGGCTCTCTCTCCCCG | 120 |
| chr19 | 50718868 | 50718988 | 788855_47188554_ENST00000293441.6,E | 1 | - | good | GCCTCCTCCGCCCCTTTCGACCCCTCCAGCTCCCGCTTCTCCTTCCAGGCTCCTCCAGTCCCTTTCCACCCCTGTCCAAGCTCCCTGCGTCCCCACCAGGTCTCCAGTCCCTACACCGCC | 120 |
| chr19 | 50718988 | 50719108 | 788855_47188554_ENST00000293441.6,E | 1 | - | good | CTGGTACCCGCCTCCCCGCCCAGCCCGGCCACTCCCCCGCCTCGCCCTCCCTGGCGTACCTGGGCCCCTCCGCCAGGTCACCCTGGTCCCCACTCCCCTCCAGCGAGGGTCTAGGCCCGG | 120 |
| chr19 | 50719108 | 50719228 | 788855_47188554_ENST00000293441.6,E | 1 | - | good | GGGTCCCCGGTGCCCCGTCCCAGATCGCATCCTTGATGGGAGGGCGCGCCCCCCACCCCCGGCCAGCAGAGCCCCTATCCCGAACCCGCCTCCCGGCTCTCCGGGGCCAAGGGCTCGGTC | 120 |
| chr19 | 50719228 | 50719348 | 788855_47188554_ENST00000293441.6,E | 1 | - | good | GGGGCGAGGGCGCCCCGGATGTCCTTGGCCCGGCGCCGGGCTCCTGGCCTGGGTGTGCGGCTGGGGGTACGGGTGAGGTGGGGAGCGTTGCCCTGGGCGACACCAAATTGGGTTACGCC | 120 |
| chr19 | 50719348 | 50719468 | 788855_47188554_ENST00000293441.6,E | 1 | - | good | GTGGGGGGAGGGGGGAGGGGCCGCGGGCGCCGCCGACCGGGACTCAGCAGCCCCGCCAGGCAGGTAAGGGGGGTCGAGAACCTGGGGGTTTGGGGCAGCTGCGGGGCCGGGGCCGCG | 120 |
| chr19 | 50719468 | 50719588 | 788855_47188554_ENST00000293441.6,E | 1 | - | good | GTGAGGTCCTCGGCCCTCCCGCCCTCCCCCCGGTCCCCGCCCGCCCCCTCCCCGGGCCCGCTCGCCCTCCGGGGCGGTGGGGCATGGCCTGAGGGTCCCCGTCTCCGGGGGGGTGGGGG | 120 |
| chr19 | 50719588 | 50719708 | 788855_47188554_ENST00000293441.6,E | 1 | - | good | AGGCGGAGACGCGGCGCCCGGAGCCGGCCGGACCGAGCCGGGGGCGGGCGAGGAGGGGCGCGGCGGCCGCGGCGGAAGGGGGGGCGCTCTGCGGATGGCCAGGCCCGGCCCGCGG | 120 |

**Supplementary Table 1c. The probe design for human TP53 capture**

| Chromosome | Start | Stop | Name | Score | Strand | Code | Seq | Len |
|---|---|---|---|---|---|---|---|---|
| chr17 | 7675318 | 7675438 | 788602_47142351_ENST00000413465.6,E | 48 | - | good | CTGAGTGACAGAGCAAGACCCTATCTCAAAAAAAAAAAAAAAAAAAGAAAAGCTCCTGAGGTGTAGACGCCAACTCTCTCTAGCTCGCTAGTGGGTTGCAGGAGGTGCTTACGCATGTTTG | 120 |
| chr17 | 7685158 | 7685278 | 788602_47142351_ENST00000413465.6,E | 24 | - | good | AATAAAGCTGCTATAAACATTCTTGTACAATTCTTTTTGTGATCATATGTTTTCGTGTTTCTTGGAGAAATACTTAGGAGGGGAATTGCGAGTTTGGAAGTAAAAAGTAGCTGTATTTTG | 120 |
| chr17 | 7666198 | 7666318 | 788602_47142351_ENST00000413465.6,E | 19 | - | good | CCAATTTCCTCATGGCTTCAGTCTAAACTTGTAATTCACAACCTTAAATCAATATGTTTCTATTTTTTTATTTAGAAAACATTTCCGGCCAGGCACGGTGGATCACACCTGTAATCCCAGC | 120 |
| chr17 | 7681078 | 7681198 | 788602_47142351_ENST00000413465.6,E | 16 | - | good | ATTCCATGTCAAAAAAAAAAAAAAAAAAAGGAAAGAAAAAAAATAACGATTAGAAAGGAAGAAATAAAACACATTCACAGCCAGTATGATTCTATACATACATGTCCTAATGGGGCCAGGC | 120 |
| chr17 | 7684798 | 7684918 | 788602_47142351_ENST00000413465.6,E | 15 | - | good | CTTTTACCTGTTATTTTGGTTTTTTGTTTGTTTTTATTGTTCAGTTGTGGGACTGCTTTATACATTCTGGATACAAGTCCTTTATCAGATCCATGTGTCGTGAATGTTTTCTTCTGATCT | 120 |
| chr17 | 7667038 | 7667158 | 788602_47142351_ENST00000413465.6,E | 8 | - | good | AAACCATTACCACAATCAAATTAATCAGCACGTCCATCATCACACACAGTTACCATTTGTGTGTGTGCACGTGTGTTCACCTACGACGAGGACACTTGGACCTACTCTGCAGATCTCAAG | 120 |
| chr17 | 7685038 | 7685158 | 788602_47142351_ENST00000413465.6,E | 6 | - | good | AACTTTTTCAGAAGCTCTGAGTTTTCCAGAGCGGTTGTACCATTTTACACTCCAACTAGCAAGGTATGGGAGTTATTATGGTTGTGCCACAGCCTTCCGGACATTAGGTATTGTCAGTCT | 120 |
| chr17 | 7670758 | 7670878 | 788602_47142351_ENST00000413465.6,E | 5 | - | good | ATAAATATTTTTAAAAGTCAGCTGTATAGGTACTTGAAGTGCAGTTTCTACTAAATGCATGTTGCTTTTGTACCGTCATAAAGTCAAACAATTGTAACTTGAACCATCTTTTAACTCAGG | 120 |
| chr17 | 7682638 | 7682758 | 788602_47142351_ENST00000413465.6,E | 3 | - | good | CTTTACTGGGGGGAAAAAAATCAGTCTGTGTAATTCACCACATTAACAAAGGGAAACATAAAAACCCTATGATCATTTCAACAGATGTAGCAAAAGCAGTTAATGATATTCAACACATAT | 120 |
| chr17 | 7667158 | 7667278 | 788602_47142351_ENST00000413465.6,E | 2 | - | good | CTTCCTTTTTTTTTTTTTAATTTTTTTTTATTTTAGGCTTTATTGGGGCATAATTGATCCCCCAAAATTGCATACATTCAAGGTATGCAGTGTGATGATTTGATATGGGGGTATATTGTG | 120 |
| chr17 | 7684558 | 7684678 | 788602_47142351_ENST00000413465.6,E | 2 | - | good | CCCAGGGTCATGAAGATATTATCTTACATTTTCTTTTAGGACCTTTATGGTTGTAAGTTTTACAGTAAGGTCCTTGAGCCATTAATTAATTCTTAAAATTAATTGTTTATGGTGTGAGGT | 120 |
| chr17 | 7662358 | 7662478 | 788602_47142351_ENST00000413465.6,E | 1 | - | good | GAAAACTAAAAAAATAAGAAAAATCAATTATCTCGAGATAAATATAATATAATGCAATTTCAAGTCATTAGAATCCCAAATTTTCATTGTGTGTGTGTGTGAGTTGGGTAAATTTATCATAAAT | 120 |
| chr17 | 7662478 | 7662598 | 788602_47142351_ENST00000413465.6,E | 1 | - | good | TGTGTAATAGGAATGGCACACACAAAGAAGGAATGGCACAGAGCCTGTATGCAGAAGACCACAAACCCTTATTTAACGACGTAAGCCAAGATCCAAAGAAAATGATAGATTCTCAGATGG | 120 |
| chr17 | 7662958 | 7663078 | 788602_47142351_ENST00000413465.6,E | 1 | - | good | GGATAGTATACCAAAACTCAGAAATATTATGGAATTAACAGAATTTAGTAAGGCAGATAAGTAGTAGGTAGAAAAATATTAATTTTATCTTCAGCAGAAGCACTGTGAAAAATTAGACA | 120 |
| chr17 | 7664878 | 7664998 | 788602_47142351_ENST00000413465.6,E | 1 | - | good | AATTTCAGTAGCCATTTCCTTAAAAAAGAAAACACTTTAACACAGTAATAGACTGATAATGGAATACCAATTTTCCTAATAAGTTAAACATTAAGATAATTTCAATTAAGGTCAAGAGCT | 120 |
| chr17 | 7664998 | 7665118 | 788602_47142351_ENST00000413465.6,E | 1 | - | good | CACCGTGCCCAGCCCAAAGTTGGTTTTAATAGCAGAAAATCTATCAACATAATTCAATATATTAAATTTAGAAAGAAAAATTATCTATCATATCAACAGATACTGAAAGGAATTTGATTA | 120 |
| chr17 | 7665958 | 7666078 | 788602_47142351_ENST00000413465.6,E | 1 | - | good | AAAAAAAAATGGAAGAAAAAAAAATTTCCCCCTCATTTTAGGAACACGAGGTCTCCAAATCTAAAATTCGTACTCTGAGGAGATTGAATAGCCTTAAATGCTTTCATCATTAAAAAGAAA | 120 |
| chr17 | 7666318 | 7666438 | 788602_47142351_ENST00000413465.6,E | 1 | - | good | TGGTACCTTACCCTGTGCCAGGAGCTGTTCTAGGCCCTGGAAACCTATGGCAGACATGTTCCCTACCCTCCCACTCAAAGAGCCCAGGCCTTATCCTAATGAGATCTGAAATCAAATCTC | 120 |
| chr17 | 7666438 | 7666558 | 788602_47142351_ENST00000413465.6,E | 1 | - | good | CTGAGACTTCAGGCCAGCTCTGCTACACTATATACTCTAACCTCTCTGGTCCTACTTGGTGACTTCTTCCCTCTGGTCGTGTTCAAGTTCCCGTCCCATCCAGTCAAGCAGGTACTCAT | 120 |
| chr17 | 7666918 | 7667038 | 788602_47142351_ENST00000413465.6,E | 1 | - | good | TAAACAGAAAATCTCCCTTTTTGACAACCATCCTCCACCCTTTCAATCCCAACCTTTTCCTAGATTATGTCCCTAGCTCTGTTTTTATTTCTGCTGTGCTGCTTCAGATCCATTCTGACT | 120 |
| chr17 | 7667278 | 7667398 | 788602_47142351_ENST00000413465.6,E | 1 | - | good | TCCAGACCCGCCTCTTTCAGAGACTTTTAACTTAAACATCTGTCCCTACCCAGCAGGCAAACTAGAGCTCCTGAAGCTCAGTCCCTGTCCTTGCCTCTGTAGACAGGTCACCTTGATGAG | 120 |
| chr17 | 7667398 | 7667518 | 788602_47142351_ENST00000413465.6,E | 1 | - | good | GCCCCTTGCCTTGCCCCAAGGATCCCAGACAGCTATGAAGCCACCAGCCTTCCTGACCTCAAGACCACCTTTTTTTTTTCTCTTTCTTACTAGGGAATGCCAAACACTCTCCCCAGGAGA | 120 |
| chr17 | 7667518 | 7667638 | 788602_47142351_ENST00000413465.6,E | 1 | - | good | CCCTTCCTCCTCCCTGGTGGTGGAGATGGGTGTTAGGGTCCAGTGGGTGCTACTGTCCAGGGGGGCTTCTGGGGCCACCAGCCTGTCAGCTCATCAACCAGGCTGAAGGTGCAAGCAGGA | 120 |
| chr17 | 7667638 | 7667758 | 788602_47142351_ENST00000413465.6,E | 1 | - | good | TGCCCATGCTCCTCAGGGCTGCTGGCTCCCAGGGAGGGGGGCTGGGAGCAGGCAGGCTCTTCCCCATCACCCACTGCTCTCTTGGAGCCAGTGCTTGAAGGGGCAGTCAGACATGGCTTG | 120 |
| chr17 | 7667758 | 7667878 | 788602_47142351_ENST00000413465.6,E | 1 | - | good | TCATCCTGGGCCTCATGCCAGGTAGCTGAATGAATTGAAGCTTTAAACTCTGCCAGGAAAACCTTTCAAAGGGCTTCTTGGGATAGGGAGGAGAGTCGGGTTGAGGAGCTCAGTACTGCC | 120 |
| chr17 | 7667878 | 7667998 | 788602_47142351_ENST00000413465.6,E | 1 | - | good | GGGGTGGGTGCTACTGGATGGAGAGAGGCCAGTGGGAGGCCTGCTAGCCAGGGTCCCAGGAAAGTGGGGGCAGCTAAGGTAAGAGTAGGGGTGTGGGGCTAGGTCCTTCCCAGCATCCC( | 120 |
| chr17 | 7667998 | 7668118 | 788602_47142351_ENST00000413465.6,E | 1 | - | good | TCGGCAGCCTCTCACCAAGGATTACGGATTTAAATGTCTGATTTAGCAAGGCTGAGCCTCCAGGGTGGCCATCTGCTCCATCAGAAAGTGGCAGGATACCTGGGTTCCCAAGGGGAACA | 120 |
| chr17 | 7668118 | 7668238 | 788602_47142351_ENST00000413465.6,E | 1 | - | good | CAACAAGGGAGGGGGTGGCTCCATTCATAACTCAGGAACCAACCGTCCCTCCTCCCCTCCGGCCACGGCTGGCACAAGGTTCTCTCCCTCCCCTGCTTCTAGGACTGGGCTGCTTCCCCC | 120 |
| chr17 | 7668238 | 7668358 | 788602_47142351_ENST00000413465.6,E | 1 | - | good | TGAGGAGGTGCTGGGAAGCAGCCACCTGAGTCTGCAATGAGTGTGGGCTGGGGGGCCCAGTGCCCGGGTTCCGGGAGGGGAACAAAGGCTGGAGACTGGGTCAGTCTGCGGGCTGCATG/ | 120 |
| chr17 | 7668358 | 7668478 | 788602_47142351_ENST00000413465.6,E | 1 | - | good | CCTTTTTATATCCCATTTTTATATCGATCTCTTATTTTACAATAAAACTTTGCTGCCACCTGTGTGTCTGAGGGGTGAACGCCAGTGCAGGCTACTGGGGTCAGCAGGTGCAGGGGTGAG | 120 |
| chr17 | 7668838 | 7668958 | 788602_47142351_ENST00000413465.6,E | 1 | - | good | AGGAAATCTCACCCCATCCCACACCCTGGAGGATTTCATCTCTTGTATATGATGATCTGGATCCACCAAGACTTGTTTTATGCTCAGGGTCAATTTCTTTTTCTTTTTTTTTTTTTTTT | 120 |
| chr17 | 7668958 | 7669078 | 788602_47142351_ENST00000413465.6,E | 1 | - | good | TGGTCGGTGGGTTGGTAGTTTCTACAGTTGGGCAGCTGGTTAGGTAGAGGGAGTTGTCAAGTCTCTGCTGGCCCAGCCAAACCCTGTCTGACAACCTCTTGGTGAACCTTAGTACCTAAA | 120 |
| chr17 | 7669078 | 7669198 | 788602_47142351_ENST00000413465.6,E | 1 | - | good | TACCTCACAGAGTGCATTGTGAGGGTTAATGAAATAATGTACATCTGGCCTTGAAACCACCTTTTATTACATGGGGTCTAGAACTTGACCCCCTTGAGGGTGCTTGTTCCCTCTCCCTGT | 120 |
| chr17 | 7669198 | 7669318 | 788602_47142351_ENST00000413465.6,E | 1 | - | good | ACAATCAGCCACATTCTAGGTAGGGGCCCACTTCACCGTACTAACCAGGGAAGCTGTCCCTCACTGTTGAATTTTCTCTAACTTCAAGGCCCATATCTGTGAAATGCTGGCATTTGCACC | 120 |
| chr17 | 7669318 | 7669438 | 788602_47142351_ENST00000413465.6,E | 1 | - | good | TGGTGTTTTGTTGTGGGGAGGAGGATGGGGAGTAGGACATACCAGCTTAGATTTTAAGGTTTTTACTGTGAGGGATGTTTGGGAGATGTAAGAAATGTTCTTGCAGTTAAGGGTTAGTTT | 120 |
| chr17 | 7669438 | 7669558 | 788602_47142351_ENST00000413465.6,E | 1 | - | good | CTCCCCTGCCATTTTGGGTTTTGGGTCTTTGAACCCTTGCTTGCAATAGGTGTGCGTCAGAAGCACCCAGGACTTCCATTTGCTTTGTCCCGGGGCTCCACTGAACAAGTTGGCCTGCAC | 120 |
| chr17 | 7669558 | 7669678 | 788602_47142351_ENST00000413465.6,E | 1 | - | good | CAAAAAGGGTCAGTCTACCTCCCGCCATAAAAAACTCATGTTCAAGACAGAAGGGCCTGACTCAGACTGACATTCTCCACTTCTTGTTCCCCACTGACAGCCTCCCACCCCCATCTCTCC | 120 |
| chr17 | 7669678 | 7669798 | 788602_47142351_ENST00000413465.6,E | 1 | - | good | TCCCGTTGTCCCAGCCTTAGGCCCTTCAAAGCATTGGTCAGGGAAAAGGGGCACAGACCCTCTCACTCATGTGATGTCATCTCTCCTCCCTGCTTCTGTCTCCTACAGCCACCTGAAGTC | 120 |
| chr17 | 7669798 | 7669918 | 788602_47142351_ENST00000413465.6,E | 1 | - | good | CAAAAAAGATAAATAAAGTAAAATGGGGTAAGGGAAGATTACGAGACTAATACACACTAATACTCTGAGGTGCTCAGTAAACATATTTGCATGGGGTGTGGCCACCATCTTGATTTGAAT | 120 |
| chr17 | 7670158 | 7670278 | 788602_47142351_ENST00000413465.6,E | 1 | - | good | TACTCAGCCCTGCCATGCACCGGCAGGTCCTAGGTGACCCCCGTCAAACTCAGTTTCCTTATATATAAAATGGGGTAAGGGGGCCGGGCGCAGTGGCTCACGAATCCCACACTCTGGGAG | 120 |
| chr17 | 7670278 | 7670398 | 788602_47142351_ENST00000413465.6,E | 1 | - | good | CAGGGACGAGTGTGGATACTTCTTTGCCATTCTCCGCAACTCCCAGCCCAGAGCTGGAGGGTCTCAAGGAGGGGCCTAATAATTGTGTAATACTGAATACAGCCAGAGTTTCAGGTCATA | 120 |
| chr17 | 7670398 | 7670518 | 788602_47142351_ENST00000413465.6,E | 1 | - | good | CAAAGGCAGCTGACCCCATCTCATTGGGTCCCAGCCCTGCACAGACATTTTTTTAGTCTTCCTCCGGTTGAATCCTATAACCACATTCTTGCCTCAGTGTATCCACAGAACATCCAAACC | 120 |
| chr17 | 7670518 | 7670638 | 788602_47142351_ENST00000413465.6,E | 1 | - | good | GGAGCCAGGGGGGGAGCAGGGCTCACTCCAGGTGAGTGACCTCAGCCCCTTCCTGGCCCTACTCCCCTGCCTTCCTAGGTTGGAAAGCCATAGGATTCCATTCTCATCCTGCCTTCATGGT | 120 |
| chr17 | 7670638 | 7670758 | 788602_47142351_ENST00000413465.6,E | 1 | - | good | TACTGTGTATATACTTACTTCTCCCCCTCCTCTGTTGCTGCAGATCCGTGGGCGTGAGCGCTTCGAGATGTTCCGAGAGCTGAATGAGGCCTTGGAACTCAAGGATGCCCAGGCTGGGAA | 120 |
| chr17 | 7671598 | 7671718 | 788602_47142351_ENST00000413465.6,E | 1 | - | good | TATCTCATGTGATTTATTGAATATTGTTACTGAAAGTGAGAAACAGCATGGTTGCATGAAAGGAGGCACAGTCGAGCCAGGCACAGCCTGGGCGCAGAGCGAGACTCAAAAAAAGAAAAG | 120 |
| chr17 | 7673278 | 7673398 | 788602_47142351_ENST00000413465.6,E | 1 | - | good | CCTGGTTGTAGCTAACTAACTTCAGAACACCAACTTATACCATAATATATATTTTAAAGGACCAGACCAGCTTTCAAAAAGAAAATTGTTAAAGAGAGCATGAAAATGGTTCTATGACTT | 120 |
| chr17 | 7673398 | 7673518 | 788602_47142351_ENST00000413465.6,E | 1 | - | good | CCTCTTATCAAGTGGAAAGTTTCCAGTCTAACACTCAAAATGCCGTTTTCTTCTTGACTGTTTTACCTGCAATTGGGGCATTTGCCATCAGGGGGCAGTGATGCCTCAAAGACAATGGCT | 120 |
| chr17 | 7673518 | 7673638 | 788602_47142351_ENST00000413465.6,E | 1 | - | good | TTTTATCACCTTTCCTTGCCTCTTTCCTAGCACTGCCCAACAACACCAGCTCCTCTCCCCAGCCAAAGAAGAAACCACTGGATGGAGAATATTTCACCCTTCAGGTACTAAGTCTTGGGA | 120 |
| chr17 | 7673638 | 7673758 | 788602_47142351_ENST00000413465.6,E | 1 | - | good | AATCTCCGCAAGAAAGGGGAGCCTCACCACGAGCTGCCCCCAGGGAGCACTAAGCGAGGTAAGCAAGCAGGACAAGAAGCGGTGGAGGAGACCAAGGGTGCAGTTATGCCTCAGATTCAC | 120 |
| chr17 | 7673758 | 7673878 | 788602_47142351_ENST00000413465.6,E | 1 | - | good | TTCCTTACTGCCTCTTGCTTCTCTTTTCCTATCCTGAGTAGTGGTAATCTACTGGGACGGAACAGCTTTGAGGTGCGTGTTTGTGCCTGTCCTGGGAGAGACCGGCGCACAGAGGAAGAG | 120 |
| chr17 | 7673878 | 7673998 | 788602_47142351_ENST00000413465.6,E | 1 | - | good | CCTCTTAACCTGTGGCTTCTCCTCCACCTACCTGGAGCTGGAGCTTAGGCTCCAGAAAGGACAAGGGTGGTTGGGAGTAGATGGAGCCTGGTTTTTTAAATGGGACAGGTAGGACCTGAT | 120 |
| chr17 | 7673998 | 7674118 | 788602_47142351_ENST00000413465.6,E | 1 | - | good | ACCCCTGGGCCCACCTCTTACCGATTTCTTCCATACTACTACCCATCCACCTCTCATCACATCCCCGGCGGGGAATCTCCTTACTGCTCCCACTCAGTTTTCTTTTCTCTGGCTTTGGGA | 120 |
| chr17 | 7674118 | 7674238 | 788602_47142351_ENST00000413465.6,E | 1 | - | good | GCATGGGCGGCATGAACCGGAGGCCCATCCTCACCATCATCACACTGGAAGACTCCAGGTCAGGAGCCACTTGCCACCCTGCACACTGGCCTGCTGTGCCCCAGCCTCTGCTTGCCTCTG | 120 |
| chr17 | 7674238 | 7674358 | 788602_47142351_ENST00000413465.6,E | 1 | - | good | CCTCCCCTGCTTGCCACAGGTCTCCCCAAGGCGCACTGGCCTCATCTTGGGCCTGTGTTATCTCCTAGGTTGGCTCTGACTGTACCACCATCCACTACAACTACATGTGTAACAGTTCCT | 120 |
| chr17 | 7674718 | 7674838 | 788602_47142351_ENST00000413465.6,E | 1 | - | good | CTCTGGGAGGAGGGGTTAAGGGTGGTTGTCAGTGGCCCTCCAGGTGAGCAGTAGGGGGGCTTTCTCCTGCTGCTTATTTGACCTCCCTATAACCCCATGAGATGTGCAAGTAAATGGGT | 120 |
| chr17 | 7674838 | 7674958 | 788602_47142351_ENST00000413465.6,E | 1 | - | good | TCAGCATCTTATCCGAGTGGAAGGAAATTTGCGTGTGGAGTATTTGGATGACAGAAACACTTTTCGACATAGTGTGGTGGTGCCCTATGAGCCGCCTGAGGTCTGGTTTGCAACTGGGGT | 120 |
| chr17 | 7674958 | 7675078 | 788602_47142351_ENST00000413465.6,E | 1 | - | good | CCATGAGCGCTGCTCAGATAGCGATGGTGAGCAGCTGGGGCTGGAGAGACGACAGGGCTGGTTGCCCAGGGTCCCCAGGCCTCTGATTCCTCACTGATTGCTCTTAGGTCTGGCCCCTCC | 120 |
| chr17 | 7675078 | 7675198 | 788602_47142351_ENST00000413465.6,E | 1 | - | good | CAAGACCTGCCCTGTGCAGCTGTGGGTTGATTCCACACCCCCGCCCGGCACCCGCGTCCGCGCCATGGCCATCTACAAGCAGTCACAGCACATGACGGAGGTTGTGAGGCGCTGCCCCCA | 120 |
| chr17 | 7675198 | 7675318 | 788602_47142351_ENST00000413465.6,E | 1 | - | good | TTTCTTTGCTGCCGTCTTCCAGTTGCTTTATCTGTTCACTTGTGCCCTGACTTTCAACTCTGTCTCCTTCCTCTTCCTACAGTACTCCCCTGCCCTCAACAAGATGTTTTGCCAACTGGC | 120 |

| | | | | | | | | |
|---|---|---|---|---|---|---|---|---|
| chr17 | 7675678 | 7675798 | 788602_47142351_ENST00000413465.6,E | 1 | - | **good** | TTTTATCCATCCCATCACACCCTCAGCATCTCTCCTGGGGATGCAGAACTTTTCTTTTTCTTCATCCACGTGTATTCCTTGGCTTTTGAAAATAAGCTCCTGACCAGGCTTGGTGGCTCA | 120 |
| chr17 | 7675798 | 7675918 | 788602_47142351_ENST00000413465.6,E | 1 | - | **good** | AACTTTGGGATTCCTCTTCACCCTTTGGCTTCCTGTCAGTGTTTTTTTATAGTTTACCCACTTAATGTGTGATCTCTGACTCCTGTCCCAAAGTTGAATATTCCCCCCTTGAATTTGGGC | 120 |
| chr17 | 7675918 | 7676038 | 788602_47142351_ENST00000413465.6,E | 1 | - | **good** | CTGGGCTTCTTGCATTCTGGGACAGCCAAGTCTGTGACTTGCACGGTCAGTTGCCCTGAGGGGCTGGCTTCCATGAGACTTCAATGCCTGGCCGTATCCCCCTGCATTTCTTTTGTTTGG | 120 |
| chr17 | 7676038 | 7676158 | 788602_47142351_ENST00000413465.6,E | 1 | - | **good** | CCCCCCGTGGCCCCTGCACCAGCAGCTCCTACACCGGCGGCCCCTGCACCAGCCCCCTCCTGGCCCCTGTCATCTTCTGTCCCTTCCCAGAAAACCTACCAGGGCAGCTACGGTTTCCGT | 120 |
| chr17 | 7676158 | 7676278 | 788602_47142351_ENST00000413465.6,E | 1 | - | **good** | CTACAGTCCCCCTTGCCGTCCCAAGCAATGGATGATTTGATGCTGTCCCCGGACGATATTGAACAATGGTTCACTGAAGACCCAGGTCCAGATGAAGCTCCCAGAATGCCAGAGGCTGCT | 120 |
| chr17 | 7676278 | 7676398 | 788602_47142351_ENST00000413465.6,E | 1 | - | **good** | CTGAAAACAACGTTCTGGTAAGGACAAGGGTTGGGCTGGGGACCTGGAGGGCTGGGGACCTGGAGGGCTGGGGGGCTGGGGGGCTGAGGACCTGGTCCTCTGACTGCTCTTTTCACCCAT | 120 |
| chr17 | 7676398 | 7676518 | 788602_47142351_ENST00000413465.6,E | 1 | - | **good** | GAGTGGATCCATTGGAAGGGCAGGCCCACCACCCCCACCCCAACCCCAGCCCCCTAGCAGAGACCTGTGGGAAGCGAAAATTCCATGGGACTGACTTTCTGCTCTTGTCTTTCAGACTTC | 120 |
| chr17 | 7676518 | 7676638 | 788602_47142351_ENST00000413465.6,E | 1 | - | **good** | ACTTTTCCTCTTGCAGCAGCCAGACTGCCTTCCGGGTCACTGCCATGGAGGAGCCGCAGTCAGATCCTAGCGTCGAGCCCCCTCTGAGTCAGGAAACATTTTCAGACCTATGGAAACTGT | 120 |
| chr17 | 7676638 | 7676758 | 788602_47142351_ENST00000413465.6,E | 1 | - | **good** | TCTTTTCCTGCTCCACAGGAAGCCGAGCTGTCTCAGACACTGGCATGGTGTTGGGGGAGGGGGTTCCTTCTCTGCAGGCCCAGGTGACCCAGGGTTGGAAGTGTCTCATGCTGGATCCCC | 120 |
| chr17 | 7676758 | 7676878 | 788602_47142351_ENST00000413465.6,E | 1 | - | **good** | GAAGGTTGGAAGTCCCTCTCTGATTGTCTTTTCCTCAAAGAAGTGCATGGCTGGTGAGGGGTGGGGCAGGAGTGCTTGGGTTGTGGTGAAACATTGGAAGAGAGAATGTGAAGCAGCCAT | 120 |
| chr17 | 7677238 | 7677358 | 788602_47142351_ENST00000413465.6,E | 1 | - | **good** | TGAAAGGAATCAAAGAAATGGAGCCGTGTATCAGGTGGGGAAGGGTGGGGGCCAAGGGGGTGTCCTTCCCCATACAGAGATTGCAGGCTGAGAATGACTATATCCTTGTTAACAGGAGGT | 120 |
| chr17 | 7677358 | 7677478 | 788602_47142351_ENST00000413465.6,E | 1 | - | **good** | ATGAGATTAAAGAAGCCGAGACGGGCCATTCGTGAGGGGTTTGTAATGCAGGGCTGAGGAGTGTCCGAAGAGAATGGGCAGGTGAGCGGTGAGACAGTTGTTCTTCCAGAAGCTTTGCAG | 120 |
| chr17 | 7677478 | 7677598 | 788602_47142351_ENST00000413465.6,E | 1 | - | **good** | CTGCACGGGAAGGAGCCTACCCCATGTTCCTGGCTAGCCAAGGAACCACCAGTTGATTAGCAGAGAAGGGCAGCCAGTCTAGCTAGAGCTTTTGGGGAAGAGGGAGTGGTTGTTAAGAG | 120 |
| chr17 | 7677598 | 7677718 | 788602_47142351_ENST00000413465.6,E | 1 | - | **good** | TAGGAAGTGTATATGCTATTAGAGGAAACAGATAAAATAAACAAGGAAAAGTATCAGACAATGTAAGTGCTATGAGAATGCAAATGAGGTGATGTGAATTAAAATAGGATGACTTAAAGT | 120 |
| chr17 | 7677718 | 7677838 | 788602_47142351_ENST00000413465.6,E | 1 | - | **good** | TTAACCTCTCATATAACAGGCTTGTGTGTTATTCACATTTATTGAGCCCAAGCAGGTGCAAGGCATTGTGATCTAATACTTTGGTCAGCAAGACAACAAGATAGATCACTGCCCTGCCCT | 120 |
| chr17 | 7677838 | 7677958 | 788602_47142351_ENST00000413465.6,E | 1 | - | **good** | CAAAAGAAAACATGGCAAAGCCTTTGAAAGCTTGTCTGGGAGAAGGTGCGATGATAGTTGCATAACTTCGTGCAAGATGCTGGTCCACACAGGGGCTGCCCCTTGCTCTTTCTCGCTCTC | 120 |
| chr17 | 7678798 | 7678918 | 788602_47142351_ENST00000413465.6,E | 1 | - | **good** | AAAGTGTAACAAACACTTACAGTAGGCATGTTTCTTAGCAAATCTGATGACAAATTTGGCATAAAGAAAGAGAGCATCCCTGAAAAAAAAAAAAAGAAAAAGAAAGAGAGCATCCTGCCT | 120 |
| chr17 | 7679638 | 7679758 | 788602_47142351_ENST00000413465.6,E | 1 | - | **good** | ACAATCACACAGTTCTCTCTAGAATAATAATATAGAACAAGTGAAATAGAACAATTGCAGAAAGAGCTAACCTTTGTTGAGCTCTTACTGTGTGCCCAGCACTTTCCTCAACTCTACATT | 120 |
| chr17 | 7679758 | 7679878 | 788602_47142351_ENST00000413465.6,E | 1 | - | **good** | AGACTTTGGGCCCCCCATTTCAGGACAGCACCTCCTGGCCTGTTGACTGAATAGATCCCTGAAGGAGGTGTACTTGCATTAATGGAGTGGGGGTGGGAGCAGTACCACAGATCCGCACTA | 120 |
| chr17 | 7679878 | 7679998 | 788602_47142351_ENST00000413465.6,E | 1 | - | **good** | GCTGGTACCCCTGGGAAGGTCCTGCTCTGAAAAACACGGAGATTTTAGTTGCTACTGAAGATTTGAGAGATAAAGACAGGGAGACCTGTCTGTAGACCTGTGTCCCTCCAAGTGGGATTG | 120 |
| chr17 | 7679998 | 7680118 | 788602_47142351_ENST00000413465.6,E | 1 | - | **good** | GGGGTCTCAGTTGATCTCATTGCCTTCCCACCCAGCCAAGGGCACCTGCATTTCTCTTGGCTCCCTGGCCATTTGGAAGGCCTAGTTCAGCCTGGCACATTTGTATCCTGGCCCACTGAT | 120 |
| chr17 | 7680118 | 7680238 | 788602_47142351_ENST00000413465.6,E | 1 | - | **good** | ACATGCAAGTATGTATGTAAGTACTTGTACTATATTGTTTAGGGAATCACTGGACATATAGGCCTTCAAGACTGATACCAGCAGCCACTGTTAAGATTCTGGTCAGGCCTGCCCCTGTTT | 120 |
| chr17 | 7680358 | 7680478 | 788602_47142351_ENST00000413465.6,E | 1 | - | **good** | TAAAAAGCTACCAAAACTAATAAATAAATATAGCAGGGTTGCAGGTTACAGGGCAATATAGTTATCCCTCTATCTGTAGGGGCTTGGTTCTGGGACTCCTCACACACCAAACCCACAGAT | 120 |
| chr17 | 7682518 | 7682638 | 788602_47142351_ENST00000413465.6,E | 1 | - | **good** | GCATGATTACAAACCAACCAACCTCCTAGCAAACTAGGGAAAGGAAACTTAACCTAGTTTGATAACAGGGCGTCCACAGTCGGAGTTCCACTAGCAGCATACATAATGGTAGAAAACTCA | 120 |
| chr17 | 7684438 | 7684558 | 788602_47142351_ENST00000413465.6,E | 1 | - | **good** | GTAGGAGTCAGTCTCTGGTATCTTTCCTGTATGGAAATCCAGTTATTCTGTCTCCACTTGTTGAAATAGGCTTCCTTTCTCTACTGAATGCTTTTAATTTTAATTATTTTACAGTTGGAG | 120 |
| chr17 | 7684678 | 7684798 | 788602_47142351_ENST00000413465.6,E | 1 | - | **good** | GTTGCTTGCCTATTTGTTTGCTTTACAGAGTTTACAGTATCTTAAGAGGAGTGGATTTATCTTTTTTATGTTCAGTATTTGCCTTGTCCTGTTTAGGACATCTTTTTTTTTTTTTTTAAC | 120 |
| chr17 | 7685278 | 7685398 | 788602_47142351_ENST00000413465.6,E | 1 | - | **good** | TCTGACATTCGTTCATGTTGTATATATCAGTATTTTGCTCCTTTTCATTTAGTATAGTCCATCGATTGTATATCCGTCCTTTTGATGGCCTTTTGAGTTGTTTCCCATTTGCGGTTATGA | 120 |
| chr17 | 7685758 | 7685878 | 788602_47142351_ENST00000413465.6,E | 1 | - | **good** | TGATGCCTACCAAGTCACAGACCCTTTTCATCGTCCCAGAAACGTTTCATCACGTCTCTTCCCAGTCGATTCCCGACCCCACCTTTATTTTGATCTCCATAACCATTTTGCCTGTTGGAG | 120 |
| chr17 | 7685878 | 7685998 | 788602_47142351_ENST00000413465.6,E | 1 | - | **good** | ACCGGCCACTTTGTGCCGTACTTACGTCATCTTTTTCCTAAATCGAGGTGGCATTTACACACAGCGCCAGTGCACACAGCAAGTGCACAGGAAGATGAGTTTTGGCCCCTAACCGCTCCG | 120 |
| chr17 | 7685998 | 7686118 | 788602_47142351_ENST00000413465.6,E | 1 | - | **good** | CTCCGGCCCGAGCCGGTTCTTCCTGGTAGGAGGCGGAACTCGAATTCATTTCTCCCGCTGCCCCATCTCTTAGCTCGCGGTTGTTTCATTCCGCAGTTTCTTCCCATGCACCTGCCGCGT | 120 |
| chr17 | 7686118 | 7686238 | 788602_47142351_ENST00000413465.6,E | 1 | - | **good** | GAAAAACAATCTACCTGTTATCTAGCTTTGGGCTAGGCCATTCCAGTTCCAGACGCAGGCTGAACGTCGTGAAGCGGAAGGGGCGGGCCCGCAGGCGTCCGTGTGGTCCTCCGTGCAGCC | 120 |
| chr17 | 7686238 | 7686358 | 788602_47142351_ENST00000413465.6,E | 1 | - | **good** | CCGGAGGTATTTTCAAGAATGAGTATATCTCATCTTCCCGGAGGAAAAAAAAAAAAGAATGGGTACGTCTGAGAATCAAATTTTGAAAGAGTGCAATGATGGGTCGTTTGATAATTTGTCG | 120 |
| chr17 | 7686358 | 7686478 | 788602_47142351_ENST00000413465.6,E | 1 | - | **good** | GCAAACCTCAATCCCTCCCCTTCTTTGAATGGTGTGCCCCACCCCGCGGGTCGCCTGCAACCTAGGCGGACGCTACCATGGCGTGAGACAGGGAGGGAAAGAAGTGTGCAGAAGGCAAGC | 120 |
| chr17 | 7686478 | 7686598 | 788602_47142351_ENST00000413465.6,E | 1 | - | **good** | AAATGTCCTTTAGGCCGGTTCCTCTTACTTGGCAGAGGGAGGCTGCTATTCTCCGCCTGCATTTCTTTTTCTGGATTACTTAGTTATGGCCTTTGCAAAGGCAGGGGTATTTGTTTTGAT | 120 |
| chr17 | 7686598 | 7686718 | 788602_47142351_ENST00000413465.6,E | 1 | - | **good** | GGCAGCCCTGTTATTGTTTGGCTCCACATTTACATTTCTGCCTCTTGCAGCAGCATTTCCGGTTTCTTTTTGCCGGAGCAGCTCACTATTCACCCGATGAGAGGGGAGGAGAGAGAGAGA | 120 |
| chr17 | 7686718 | 7686838 | 788602_47142351_ENST00000413465.6,E | 1 | - | **good** | GCCCCCGGCTCCGCTAGATGGAGAAAATCCAATTGAAGGCTGTCAGTCGTGGAAGTGAGAAGTGCTAAACCAGGGGTTTGCCCGCCAGGCCGAGGAGGACCGTCGCAATCTGAGAGGCCC | 120 |
| chr17 | 7686838 | 7686958 | 788602_47142351_ENST00000413465.6,E | 1 | - | **good** | CCTAGTGAAAACTGGGGCTCCATTCCGAAATGATCATTTGGGGGTGATCCGGGGAGCCCAAGCTGCTAAGGTCCCACAACTTCCGGACCTTTGTCCTTCCTGGAGCGATCTTTCCAGGCA | 120 |
| chr17 | 7686958 | 7687078 | 788602_47142351_ENST00000413465.6,E | 1 | - | **good** | CACCCGGTGCTGGGCGTAGGGAATCCCTGAAATAAAAGATGCACAAAGCATTGAGGTCTGAGACTTTTGGATCTCGAAACATTGAGAACTCATAGCTGTATATTTTAGAGCCCATGGCAT | 120 |
| chr17 | 7687078 | 7687198 | 788602_47142351_ENST00000413465.6,E | 1 | - | **good** | AGACCTGGGTGTAGATGATGGGGATGTTAGGACCATCCGAACTCAAAGTTGAACGCCTAGGCAGAGGAGTGGAGCTTTGGGGAACCTTGAGCCGGCCTAAAGCGTACTTCTTTGCACATC | 120 |
| chr17 | 7687198 | 7687318 | 788602_47142351_ENST00000413465.6,E | 1 | - | **good** | TATTTTCAGCTCGGGAAAATCGCTGGGGCTGGGGGTGGGGCAGTGGGGACTTAGCGAGTTTGGGGGTGAGTGGGATGGAAGCTTGGCTAGAGGGATCATCATAGGAGTTGCATTGTTGGG | 120 |
| chr17 | 7687318 | 7687438 | 788602_47142351_ENST00000413465.6,E | 1 | - | **good** | GACACTTTGCGTTCGGGCTGGGAGCGTGCTTTCCACGACGGTGACACGCTTCCCTGGATTGGGTAAGCTCCTGACTGAACTTGATGAGTCCTCTCTGAGTCACGGGCTCTCGGCTCCGTG | 120 |
| chr17 | 7687438 | 7687558 | 788602_47142351_ENST00000413465.6,E | 1 | - | **good** | CAGGGGTTGATGGGATTGGGGTTTTCCCCTCCCATGTGCTCAAGACTGGCGCTAAAAGTTTTGAGCTTCTCAAAAGTCTAGAGCCACCGTCCAGGGAGCAGGTAGCTGCTGGGCTCCGGG | 120 |
