## Supplementary Table 2 for "Transcriptional Determinism and Stochasticity Contribute to the Complexity of Autism Associated *SHANK* Family Genes"

**Supplemetary Table 2a-the probe design for mouse Shank3 gene**

| Chromosome | Start | Stop | Name | Score | Strand | Code | Seq | Len | GC |
|---|---|---|---|---|---|---|---|---|---|
| chr15 | 89552978 | 89553098 | 771629_41866724_SHANK3(58234)_444 | 21 | + | good | CACGCTTTATGGTATGTATGTGGAAGTCAGAGAACCTGTGGGAGTCAGTACTCTCTTTGCATTTTGTGGGTCCCGGGGATTGAACTCAGGCCGCCAAACTTGACTGCAAGTGCCTTCACC | 120 | 51 |
| chr15 | 89515658 | 89515778 | 771629_41866724_SHANK3(58234)_133 | 18 | + | good | AACCTGGGCTATGAAGTAAGACACACACACACACACACACACACACACACACACACAAACTGCCTATTGTGGTATGAAGTGAGACAGACAGACATACATACACACAAAACCTGCCCATTG | 120 | 45 |
| chr15 | 89514098 | 89514218 | 771629_41866724_SHANK3(58234)_120 | 17 | + | good | TTGGACTTGGTTAGCAGTCTCTCCAAACCACCGGGTATTTCTCCTTCCACCTCTCCCTCTTCTCTTTTTCTCTCCTCTTTTCTTCCTCTTCTCTCTCTCTTCTTCCCTCTCTCTCATTCG | 120 | 48 |
| chr15 | 89523818 | 89523938 | 771629_41866724_SHANK3(58234)_201 | 17 | + | good | CCGGGCGCGGCGGCGCGGCCCATGGAGCGGCCCCGGGCCGGGCCCCCGGCGGGCGGGCGGGCGGCGGGCGGGCGGTGGGCGCTGCGGCGGCGGCTGCGGTGAGGCCGGCGGCGG[illegible] | 120 | 93 |
| chr15 | 89520938 | 89521058 | 771629_41866724_SHANK3(58234)_177 | 10 | + | good | GGTACATACATATATGCACGTGATAACAATGAAAATGAGGGCATGAATGTGAAAGAGCAAGGAAAGGAATATGGATTGGTTTGGAGGGAGGGAAAAGAGAAAAGATGTAATTATTTAAAA | 120 | 35 |
| chr15 | 89540018 | 89540138 | 771629_41866724_SHANK3(58234)_336 | 9 | + | good | CATTGGCCAGTGTGGGCTGCATTGGCCAGTGTGGGCTGCATTGGCCAGTGTGGGCTGCATTGGCTAGCATGGCTGTGGGCTTCTTTGTGCTAATTTCCCCTAGACCTATATGGCATAGAC | 120 | 55 |
| chr15 | 89509178 | 89509298 | 771629_41866724_SHANK3(58234)_79 | 7 | + | good | CAGATACATCCATTTGTCCAAGAATTTCATGATCCCCAAATATTTTCATTGCTCCAAAGAATATCTCAGATATGTTAGGCAATTATTCTGTATTACCCTCTTCTGCCAACCTGTCAGCTG | 120 | 38 |
| chr15 | 89510498 | 89510618 | 771629_41866724_SHANK3(58234)_90 | 6 | + | good | TGGATTATAAGAATTCTTTATATATTTTGAATATACAAAGAATAGATGGCCAGCAAGATGGGTTAATGGTTAAAGGAGCTTGCTGCCAAGCCTGGCAACTAGATCAATCCCTGGGAAACC | 120 | 38 |
| chr15 | 89519618 | 89519738 | 771629_41866724_SHANK3(58234)_166 | 6 | + | good | CTCTTTTGGTTATTATTTACATGAGTTATCATTATTGTTATTATTGTTATTATTATGATGATGTTAAATATTCTAACTTTAGCCAGGCATAGTGTCACATGTCTTTAATTCCAGCACTTG | 120 | 28 |
| chr15 | 89539898 | 89540018 | 771629_41866724_SHANK3(58234)_335 | 6 | + | good | GTCCTGGCTGTGTCCTCAGCATCTGCAGTGGCCAGTGTGGTCTGCATTGGCCAGTGTGGTCTGCATTGGCCAGTGTGGGCTGCATTGGCCAGTGTGGTCTGCATTGGCCAGTGTGGTCTG | 120 | 60 |
| chr15 | 89510738 | 89510858 | 771629_41866724_SHANK3(58234)_92 | 5 | + | good | ATAGATGTAACAAATTAAAAAAATGTATCATATATATCTATTTTTTTGAGATAGGGTTTCACTATAGCCCAGGCTGTTGTTAAACTTGGTATGTAGCCCAGGCTGGGGTCAAACTCTTGA | 120 | 34 |
| chr15 | 89530058 | 89530178 | 771629_41866724_SHANK3(58234)_253 | 5 | + | good | TCTGTCTGTCTGTCTGTCTGTCTGTCTCTCTTTTAAATCCCCGTGCCTCCCTCTCCTCTTGCTGGCAGACTCCACACAGTTCCTGCCTGCTCCAGATTAATGCCTTTGCTGTGGTCTTTG | 120 | 52 |
| chr15 | 89507378 | 89507498 | 771629_41866724_SHANK3(58234)_64 | 4 | + | good | CAATCAATCAATCAATCAGTCATCACCACCACCACCACCACCATCATCACCATCATCGCCCCAACTGGCCTGTAGGTCAGTCTGATGGAGCTGTTTTCTTAACTGAAGCTTCCTCTTCCT | 120 | 48 |
| chr15 | 89509538 | 89509658 | 771629_41866724_SHANK3(58234)_82 | 4 | + | good | TTGTGTGTGTGTGTTCTCACATGCACAAGCCTGCTTATCTGTATGTGCATCACATGCATAATACATGTTCCCTTGGGGGCCAGAAGAGGGTATCTGATTTCCTGGAACTAGAGATACAGC | 120 | 46 |
| chr15 | 89519258 | 89519378 | 771629_41866724_SHANK3(58234)_163 | 4 | + | good | CGTGTGCGTGTGTGTGTGTGTGCATGTCTGTGTGCAGATGTGTTGTATGCCTATATAGGCCTGAGGTCAATTTCCAGTGTTATTCCTTGGTAGCCATCCTGAATCTTGCTGTTGAGACAT | 120 | 47 |
| chr15 | 89520578 | 89520698 | 771629_41866724_SHANK3(58234)_174 | 4 | + | good | GCTAGGCAGGCTGGCTGGAGAGCCCTAGGGATCTATCTGCCTCTCTCCACATCGCCTGCACTGGCATTTCAAGTGCATGCCAACACTCCTGGCTTTTTATGTGGGTGCTGCAGATTGAAT | 120 | 54 |
| chr15 | 89503778 | 89503898 | 771629_41866724_SHANK3(58234)_34 | 3 | + | good | CACTTCTCTTCTCCTCAGGCCTGTCGCTTTGGACACGTGCAGCACCTGGAGCACCTTTTGTTCTATGGGGCCAACATGGGTGCTCAGAATGCCTCGGGAAACACAGCCCTGCACATCTGT | 120 | 56 |
| chr15 | 89519498 | 89519618 | 771629_41866724_SHANK3(58234)_165 | 3 | + | good | TGTGTGGGTGCTGGGGATGGAACTCACGTATTCATTCATGCATATCAAGCACTTTACTACATGAGCTATCGTCCCAGGCTTACTGTATTTTCTTGTAAAGTTTACTTAGTCTCTACTCAG | 120 | 42 |
| chr15 | 89526218 | 89526338 | 771629_41866724_SHANK3(58234)_221 | 3 | + | good | CATGCGTCAAACTTGATGGATGCCTAGGTTATACAGGAACTCAGTTACTGACTTACTGCTGGCTAGGTTCCTTGCTTTCTTTTCCTTTCTTTCTCTCTCTCTCTTTCTTCTCTTTTCTTT | 120 | 41 |
| chr15 | 89532098 | 89532218 | 771629_41866724_SHANK3(58234)_270 | 3 | + | good | GACCCCCATTGAGGAGTTTACACCCACACCTGCCTTCCCTGCACTCCAATACCTTGAGTCTGTAGATGTGGAAGGTGTGGCCTGGAGGGCTGGACTTCGAACTGGGGACTTCCTCATTGA | 120 | 55 |
| chr15 | 89552858 | 89552978 | 771629_41866724_SHANK3(58234)_443 | 3 | + | good | GCAAATAGTCAACCACCGAGTTACATCTGCTTTGTTTTATTGAGATGTGAGACCTTTCTTTCCTTACATTTGTTGTATGTGTGCGTGTGTGTGCATGTGTGTGTGTGTGTGTGCATTATA | 120 | 40 |
| chr15 | 89503538 | 89503658 | 771629_41866724_SHANK3(58234)_32 | 2 | + | good | GACCCTGCTGGACCTGGGGGCTTCGCCTGACTACAAGGACAGCCGCGGCCTGACGCCCCTGTACCATAGTGCCCTAGGGGGCGGGGATGCCCTCTGTTGCGAGCTGCTTCTCCATGATCA | 120 | 65 |
| chr15 | 89509658 | 89509778 | 771629_41866724_SHANK3(58234)_83 | 2 | + | good | TGGTTGTGAGCCACCCAATTGGTACTGGGAACCAAACTTATGACCCTTTGCAAGAGAAGCATTCACTCTTTAATTTTCATTTTATGTGTATGAATACACTGTGGCTGTCCTCAGACACCC | 120 | 42 |
| chr15 | 89518058 | 89518178 | 771629_41866724_SHANK3(58234)_153 | 2 | + | good | TCTGAGTGCCGGCTGCCAGCAAGAGCCAGAGGGCAGCTTCGGGATTCCCTGGAACTGGAGTCTTGGATGGTTGTAAGTTGTCTGAGCTGTATCAGCTCAAATAGTGATGTTGGCTTCTCC | 120 | 53 |
| chr15 | 89518538 | 89518658 | 771629_41866724_SHANK3(58234)_157 | 2 | + | good | ACTATTGTTATGAAGTTCTGCCTGGTCTTGAACTTGTAGTGATCTTTCCGCCTATGCCTTGCCAGTGCTGGGATTATGGACATGGGCTAGCATGCCTGGTTCTTTCTACTTTTCTCTTTT | 120 | 44 |
| chr15 | 89537018 | 89537138 | 771629_41866724_SHANK3(58234)_311 | 2 | + | good | GTCTGGCACAAATCATTGGGAACACTCTGTGGTGTGGCACATACTTATTGCTCAGGAGGTTGAGGCAGGAGGATTGCTGTGAGTTCCAGGCCTGTCTGGATTACAGAGTGTGAGTTCAAG | 120 | 50 |
| chr15 | 89540978 | 89541098 | 771629_41866724_SHANK3(58234)_344 | 2 | + | good | GTCTGGAGGCTGGTCTCCTAGGTGACTGAATTCTATGTCAAGTCGACCACAGTAACTGTCATACAAGGTGTGCTGCCACACTGAACTCTGTACATTTTTTTGATAAGCCAAAGTAGCATT | 120 | 44 |
| chr15 | 89553218 | 89553338 | 771629_41866724_SHANK3(58234)_446 | 2 | + | good | TGCTGATATCACAGGTGTATACAGGCATGCCAGCTGACTCAGTCTTATACCTGATCGTCACTTTTCTTCCTTTTTTTTTTTTTTTTTTTTTTTTAAAGATAAAATGACTTTGGAAGTATTT | 120 | 32 |
| chr15 | 89499818 | 89499938 | 771629_41866724_SHANK3(58234)_1 | 1 | + | good | CCGCCGGGAACAGTCCGGGCCCGGGCGCTAGCACCGGGATGGACGGCCCCGGGGCCAGCGCCGTGGTCGTGCGCGTCGGCATCCCGGACCTGCAACAAACGGTGAGCCTTTGCGATCTCG | 120 | 73 |
| chr15 | 89499938 | 89500058 | 771629_41866724_SHANK3(58234)_2 | 1 | + | good | CCGTACCCCTGTGTGGCCCGCTAGCGGTGACCCCCGCCGGGCGCGCCCTGCGCCAGGACCCCCGGCCGTGGCCACTCGCCCCTCCCCCGCCGCCTCCGCGGGACCCTCCCCATCGCCGGG | 120 | 82 |
| chr15 | 89500058 | 89500178 | 771629_41866724_SHANK3(58234)_3 | 1 | + | good | GCGGGGCCCTGGAAGCCCAGCCCCGGGGCGGGGCCTCGGCCTCTGCGCGGTGGCGCGGGGTCCCTGCTCTCGCTCCCCAACTAGCCCTCCAAAGCGGGGCCCGGGGCGCGGGGCCCTGC | 120 | 82 |
| chr15 | 89500178 | 89500298 | 771629_41866724_SHANK3(58234)_4 | 1 | + | good | CCCGCAGGTCCGGGCTCACGACTCGCTCCGCTGCAGAAGTGCCTGCGTCTGGACCCAACCGCGCCCGTGTGGGCCGCCAAGCAGCGTGTGCTCTGCGCCCTCAATCATAGCCTTCAAGAC | 120 | 68 |
| chr15 | 89500298 | 89500418 | 771629_41866724_SHANK3(58234)_5 | 1 | + | good | GCGCTCAACTACGGGCTATTCCAGCCTCCCTCCCGGGGTCGCGCCGGCAAGTTCCTGGATGAAGAGCGGCTCTTACAGGACTACCCGCCTAACCTGGACACGCCCCTGCCCTATCTGGAG | 120 | 64 |
| chr15 | 89500418 | 89500538 | 771629_41866724_SHANK3(58234)_6 | 1 | + | good | GTGCTGGGCCGACTGGGGTGGGGGTGGAATGGAGCGCTCAGGGTGTGATCTTTGAGCTCCGTGGCTGGAGCAGTAGAAGGGCAGGCCGGGAGGAGTTGGCGACAGGCGCGGTGGGGGG | 120 | 68 |
| chr15 | 89500538 | 89500658 | 771629_41866724_SHANK3(58234)_7 | 1 | + | good | GTGTGCAGCCTGAGCAGTGTGCTGAGAGACCGAGGTGGGTACACCTCGAGTGCAGTGGGCAGGGTGAAGTGCAAGCTGCCTGTGTGGTAAGTGGTGTGTTGCAGCTTAAGCATGGAGAAA | 120 | 57 |
| chr15 | 89500658 | 89500778 | 771629_41866724_SHANK3(58234)_8 | 1 | + | good | CAGTGTGAGCGCCGTGTGTAGAGCGCTTGCAGCGAGCAATCTGGACAGCACACAATGAGCACATGCATGCCACACATGCACTGCACTCGGGGCGTGTGTGTGGTGCGAGCGGTTTAAGAG | 120 | 58 |
| chr15 | 89500778 | 89500898 | 771629_41866724_SHANK3(58234)_9 | 1 | + | good | TGAGTGCGTTAGGGGTCATGTGGTGCGAACAGTGTCACTTGTGTGCACATTTGCCTCCTCTTGCCACTCACTCCCATGTCTTCTGAGGTTAGCCCAAAATAATATTAACACCATTTGTGA | 120 | 47 |
| chr15 | 89500898 | 89501018 | 771629_41866724_SHANK3(58234)_10 | 1 | + | good | CATATAAAGGTGATTTGAGTGCCATTCCATAGGGACGGAGACCTAGTCTCAATAGGAAGGGCTCGTCCTGTGTGTGTGTGTGTGTGTGTGTGTGTATACACCAAGGTCAAAGTTGGTC | 120 | 48 |
| chr15 | 89501018 | 89501138 | 771629_41866724_SHANK3(58234)_11 | 1 | + | good | TGACCCTCATGGAATTTCCACCAGGAATTGGGTTATTTCTACTTGGAGTGGAAGTGTCTAGGAGAAGCTTAGTTTCAGATCTTGGGAGAGCAGTAGAGTTTTCCAGGTCCCAGTGGTGGA | 120 | 47 |
| chr15 | 89501138 | 89501258 | 771629_41866724_SHANK3(58234)_12 | 1 | + | good | AATGAAGCTGACTTGAGCTGTGGGCCCACCAGGCCATTAGTTGGCCAAGTACTTCTCTGGACTGCCCATCGTTTCTGGGTAGTAGGTGGTCAGTAGCTGTCACTTCATTTCTGTCCACTC | 120 | 52 |
| chr15 | 89501258 | 89501378 | 771629_41866724_SHANK3(58234)_13 | 1 | + | good | TGTACGTGTGTGTCTGGGACTTGCATGGCTGTGCGGGAGTGACACTGTGGGCAGGAGGGAGGCAAGCAGCCCGAGGTGGTAGCCACCTCTTGCTCACATGGGCCTGTGTGGTAGTCTTCT | 120 | 60 |
| chr15 | 89501378 | 89501498 | 771629_41866724_SHANK3(58234)_14 | 1 | + | good | CCCCACCTTTTCTTTATCTGAGCAGTTCCGATACAAGCGGAGAGTTTATGCCCAGAACCTCATAGATGACAAGCAGTTTGCAAAGCTACACACAAAGGTAAGGAGGCAGTCCCTGGAGGG | 120 | 49 |
| chr15 | 89501498 | 89501618 | 771629_41866724_SHANK3(58234)_15 | 1 | + | good | GACTCCCCATGGCCACTGACTGTCCCCATGCTGGCTTGACCCAGCCTGCTTTTCCCCCCAAGTTGATTTCAGGGCCTGAGCTGGAGCTGGGCATGCTGGGTGGGCACAGGCTCCATGCCC | 120 | 63 |
| chr15 | 89501618 | 89501738 | 771629_41866724_SHANK3(58234)_16 | 1 | + | good | ATGGCTGGTCCAGTGATAGGTTGCAGCTTAATCAGATATTGGCTGGTGTAGGGGTGAGAGCCTTGTTAGAAGCTTGGGGGCTGGGGGGGGGCAGGGCAGGGAAGCCAATAAGCATCCAA | 120 | 56 |
| chr15 | 89501738 | 89501858 | 771629_41866724_SHANK3(58234)_17 | 1 | + | good | TGTGTTGGCTCTGAGAGGGCAAGAAGTTCTCTATATAAGGAACACAAAATTAGGTAGCTCATGAATTCTCTTCCAGGAGGGATTTGCTACACATTTTACAGGTGAGGCAGAGGGCTTGAG | 120 | 44 |
| chr15 | 89501858 | 89501978 | 771629_41866724_SHANK3(58234)_18 | 1 | + | good | TCACCTGCTGTGACTGCCCGGTGAAGCTGGCTCAGATCTGTCTGACTAGGTAGGCCCTGTGGCCTGGGTACACTTAACCTGTATAGTGTCTTTGCATCTGGGACCTACTCCCAGATATAG | 120 | 53 |
| chr15 | 89501978 | 89502098 | 771629_41866724_SHANK3(58234)_19 | 1 | + | good | AGTATAGGGCCTGACTCTATTCACTTTCTTTTTGAGGGGGTACTAAACCCCTCCTGGGTGTCCAGGGGGCAAGGTGGGTGTCACAGCCCCTTTCCAACTCCCCCAGTTCTGTGGATCAAT | 120 | 53 |
| chr15 | 89502098 | 89502218 | 771629_41866724_SHANK3(58234)_20 | 1 | + | good | GGGAATTTGTTCATGGGATACTTTCTTGATGGGCTCTGGAGTATAGGGAAAGGTCTTTGTGACCCCATCTGTGGTCCTAGATCCTTACGCTTACTTGAAAGACTTTCCTGCTGCCCCAGG | 120 | 48 |
| chr15 | 89502218 | 89502338 | 771629_41866724_SHANK3(58234)_21 | 1 | + | good | ACAGGTGACAGTCAATGGCCATGGCTCTATGCTGGGTAGAGACCATGTCAGAGAGGTCTTTGTCTCTCCCTCCACCAGCCCTTGTGTGACCCTGGCACCCTGGGTTCTCCAGCTCTTTTC | 120 | 56 |
| chr15 | 89502338 | 89502458 | 771629_41866724_SHANK3(58234)_22 | 1 | + | good | TCAGGTGCCCGTAAGACATGAGAGCCACTTGCCTTTCAGAAAGTCATTTGGTTTTAGGATGGACCTGGACTTCTGACTGCAGCCCTGTCCCCTGGCTTGCCTGGGGTCCGGCTATCCCTA | 120 | 55 |
| chr15 | 89502458 | 89502578 | 771629_41866724_SHANK3(58234)_23 | 1 | + | good | GCTCTCTCTGTTACCTTGGGGCTGGCTAAGGTCTTCCTAGAGGATTTGCATAGCTTCTTGTAGGGTTTGAGCACCCTGACCCATGATGCTGGTCCATGAGGCACCCTTTTCTGTCTGTGG | 120 | 52 |
| chr15 | 89502578 | 89502698 | 771629_41866724_SHANK3(58234)_24 | 1 | + | good | TATAAGCTGCTTGTGAATGCCTGGGCTGGGACTCCTTGCCCTGTGTGGCTGGCACCTGTACCACCAGACAAGGCCAGTCAGTGAAACTTTGCCGGGTGAAGGACTGGAAGGCAAAGTCAT | 120 | 55 |
| chr15 | 89502698 | 89502818 | 771629_41866724_SHANK3(58234)_25 | 1 | + | good | GTCATAGGTGTGCCAGGGGTGTACCATGGGATGTAGGAGGAGCTAGTAAGATGGGGTGGAGGAATGAGACCAGAGTTGTTAGGATTTGACACCTGTCAGAATGGCCTCAGTGTTCTTTTA | 120 | 49 |
| chr15 | 89502818 | 89502938 | 771629_41866724_SHANK3(58234)_26 | 1 | + | good | GGCTGAGGGAAGGGACTCAGTTGGCCCGTGACCGGGGTGACCAGGCTGGGATGTGAGAGTGACCAGGCTCAGGATGACACTGTTGAGGGTCTGCAGGGAAGGGATTTGAAAGTACACGGG | 120 | 60 |
| chr15 | 89502938 | 89503058 | 771629_41866724_SHANK3(58234)_27 | 1 | + | good | TCCAGCCAAGCTGGAATGAGACCTGAGGTAAGAGTATAGGGTTGAGGATGAGCAAGCTAGGTGACAGGTTAGACACATTGGGGTTAGGGACAGGTGGCCTGGAGCCTGCTGGAGGATGTC | 120 | 54 |
| chr15 | 89503058 | 89503178 | 771629_41866724_SHANK3(58234)_28 | 1 | + | good | GGGTGATGCTAGGATGACCCTAGCTTTGGCACTCAGCTGACTTGGATTGCTCATGTTCTAGGCAAACCTGAAGAAGTTCATGGACTATGTCCAGCTACACAGCACAGATAAGGTGGCCCG | 120 | 51 |
| chr15 | 89503178 | 89503298 | 771629_41866724_SHANK3(58234)_29 | 1 | + | good | CCTGCTGGACAAGGGGCTGGACCCCAATTTCCATGACCCTGACTCAGGAGGTGAGGAGAAGAATGGGGAGGGGCTGTGTTTTTGACCGGCTGGGACCCTAACCTTCACTTATGTCCCATT | 120 | 56 |
| chr15 | 89503298 | 89503418 | 771629_41866724_SHANK3(58234)_30 | 1 | + | good | CCAGAGTGCCCTCTGAGCCTTGCGGCACAGTTGGACAACGCCACTGACCTCCTGAAGGTTCTCCGCAACGGCGGTGCTCATCTGGACTTCCGGACCCGAGATGGGCTGACAGCCGTCCAC | 120 | 63 |
| chr15 | 89503418 | 89503538 | 771629_41866724_SHANK3(58234)_31 | 1 | + | good | TGTGCTACCCGCCAGCGGAACGCAGGGGCATTGACGGTTAGTGAGGGCAGGACCCTGGTCTGGAGGGACTCAAGGCAGAAAGCTGGGATGGGCATGGATTTTGAGGTGCCTTAACCCTCA | 120 | 59 |
| chr15 | 89503658 | 89503778 | 771629_41866724_SHANK3(58234)_33 | 1 | + | good | TGCACAGCTGGGGACCACTGATGAGAATGGTTGGCAAGAGATCCATCAGGTGAGCAGGGGGACCTAGGCTGGGGTCCAGGCTGTATGTAGGTGGTCCTGGCCTCTGTCTGGGATACTTCA | 120 | 57 |
| chr15 | 89503898 | 89504018 | 771629_41866724_SHANK3(58234)_35 | 1 | + | good | GCCCTCTACAACCAGGTACATCTTGAGTCTTACCCCTCTGTGGGCTCACTTACATTTTCTGTATGTTGTAAAAGGGTGACTCCATGTGTGGATAGGATGCGCCTGCACCTAGGTGACTGA | 120 | 49 |
| chr15 | 89504018 | 89504138 | 771629_41866724_SHANK3(58234)_36 | 1 | + | good | ACATCCCAGCTCGTGTGACACTCCATGTGTAGGTTCTGTGCCCGGTCTGGTTACTGGGTAGTAGTTCCTGGGTGGCCCACTTAAAACACATCTGTTCTCTGAACACACGTATGTGGCAAT | 120 | 51 |
| chr15 | 89504138 | 89504258 | 771629_41866724_SHANK3(58234)_37 | 1 | + | good | ATATCCCCATGAATGTTCTTAGCATCTAGCTCTCAGATGAGAGCTACTTACATAGGTATGCGCTGACGAGTTTGTATGTGTGAGTGATTCTTGCCCCCTGTACACCACGTGTGTGTCCTG | 120 | 47 |

| | | | | | | | | | |
|---|---|---|---|---|---|---|---|---|---|
| chr15 | 89504258 | 89504378 | 771629_41866724_SHANK3(58234)_38 | 1 | + | good | CCATGGCCTAGCCCTCCCTTTTCTGCTGCCTGGTGTGATTGCTGCTGTCTCCTGACAGGAGAGTTGCGCGCGCGTCCTGCTTTTCCGTGGTGCCAACAAGGACGTCCGCAATTACAACAG | 120 | 58 |
| chr15 | 89504378 | 89504498 | 771629_41866724_SHANK3(58234)_39 | 1 | + | good | CCAGACAGCCTTCCAGGTACAGGCTGAGCCCGGGGCGGGTGGGGGGCGGTTGCTTATGAGTCCCTATTGCCACTTGACTTTGAACCTTGACTCTTCTTCAGAGGGCAAAAGGGAGTGCT | 120 | 57 |
| chr15 | 89504498 | 89504618 | 771629_41866724_SHANK3(58234)_40 | 1 | + | good | CACTCACTTTCAGGACTGCCACGGAAGGTGGTTTTATGAGTGGGCTAGTGTGGTGCCCAGAATGTGAGGGAGGGTTTGGTGTGGTGGTTCCTCCTGCCATGGAAAAACATCCGATGACTC | 120 | 53 |
| chr15 | 89504618 | 89504738 | 771629_41866724_SHANK3(58234)_41 | 1 | + | good | CAGAAATCTGGGAAAGAGCCAACTGGAGAGGGCAGAAAGACAGGTGTGGTGGGCCTGACTCCCCAGAAGCAGAAAGGGGTGCGGGGAACAAGGCTGAGATGGGTGTGGTCTGTACTACAC | 120 | 57 |
| chr15 | 89504738 | 89504858 | 771629_41866724_SHANK3(58234)_42 | 1 | + | good | CTGACTTGACCCAGGTTATAGTTCTGAGCTGTTTGTCAGCCAGGCTGGTTGGCACTGGCTAGCCAGGTAGCTATCATCTCTGCGGCCTGGAGTTTTGCAGGAGTTGAAGGTGTCGTGGGG | 120 | 55 |
| chr15 | 89504858 | 89504978 | 771629_41866724_SHANK3(58234)_43 | 1 | + | good | ATTCATGAACCTTGAAAGTAGCAGGGAAGGGGGTGTCTGTTATTCTTAGTTGGAGCAGAGGCCCCCTGAGGATGTCAGAAAGGCTATGTCTGGGAGTTCGAGGCAGGGGATGAAGGTTTG | 120 | 51 |
| chr15 | 89504978 | 89505098 | 771629_41866724_SHANK3(58234)_44 | 1 | + | good | AGCACACCCAGAAGTAGATACGCCCGGACCTATTAGCTGTCCGGATTTGGGGGACTCCGGAAGAATGAGGGCTGAAAGATATGGGGCTGAGACTGAGATAGGAACTGTGGCTGCATATAG | 120 | 52 |
| chr15 | 89505098 | 89505218 | 771629_41866724_SHANK3(58234)_45 | 1 | + | good | GGGCAGATAAAATGTTCTACCAGTGTGAGCAGATCTGTTCATGTTTTAACTCTAAGCCCTGTGTCCCAGAAGCACCCCAGGACTTAGGCGGATAACGGTAGTCGGCCAACCCTAGTTACT | 120 | 49 |
| chr15 | 89505218 | 89505338 | 771629_41866724_SHANK3(58234)_46 | 1 | + | good | GGCATCCTTGTCTCTTGTCTTTAAGAGGTGAGGTCCCAGTAGACATTTTGGCTTGACTTGAGAGGAGAGTGGCTTTGGGAGGATCCCAAAAGTAGGAGCAGCTGGGGTGGGATCAAATCT | 120 | 50 |
| chr15 | 89505338 | 89505458 | 771629_41866724_SHANK3(58234)_47 | 1 | + | good | GGGTTGCTGAGCCTCCTTCCTTGACCTCCATGTCTCCCCCCGCTCTCCCAGGTGGCCATTATTGCAGGGAACTTTGAGCTTGCCGAGGTAATCAAGACCCACAAAGACTCCGATGTCGGT | 120 | 57 |
| chr15 | 89505458 | 89505578 | 771629_41866724_SHANK3(58234)_48 | 1 | + | good | GAGTTCTGCCCCACTCAGCTCCTTCCTCGAGGTGCTGTGCTTGTCATAGGCAGTGGTGGTGTAACAGGACCCCAATATGTCAAGACCGTTGTCACCAAGTGACCTCTGCTGACACAGGAA | 120 | 54 |
| chr15 | 89505578 | 89505698 | 771629_41866724_SHANK3(58234)_49 | 1 | + | good | ATGAGATGAAAAAGAGCCTGTGTGGACTGAACAGAATTCTTGGTCAGGGCCTGGAGCAGGTCTGGGCTCTGGGCTGGAAGAGCCTTGGGCCCAGCAGGGCTTGGGCTTCCCATTAGGTGC | 120 | 57 |
| chr15 | 89505698 | 89505818 | 771629_41866724_SHANK3(58234)_50 | 1 | + | good | TAGGAAGGGTCTTCTGAGACCAGGGGGTGGTAGATCTCACTCCAGCCTGGAACATGGCTGGGGTTAGCTTCCTATGACTTTCTATGTGACAAGATCCCATGAACATAGCCGACGAGAAAA | 120 | 50 |
| chr15 | 89505818 | 89505938 | 771629_41866724_SHANK3(58234)_51 | 1 | + | good | AAAAATCACAGGGGTGTGGATGAAGACCGACGTGGGGATAGGCTTGTGGCTGGCGACTTGGCCAGCAGGTTTGAGCTCCCAGGAGGCAGCGCTTAAAACAAACTTGGCATATGTGGAAGT | 120 | 53 |
| chr15 | 89506058 | 89506178 | 771629_41866724_SHANK3(58234)_53 | 1 | + | good | TGTGTGTGAGTGAGTGAGAGAGAGAGAGAGAGAGAGCTTCTGGAGCCAGGCTGGTTCGCACTGGGTATAGGTGACTAAGTGTGCAGGGTACTGTGGGCCCCTCCTCCACTAAAGCAAAGC | 120 | 55 |
| chr15 | 89506178 | 89506298 | 771629_41866724_SHANK3(58234)_54 | 1 | + | good | CTTCTAGCCATAGGGATCTCTTAATGTTCTGGTAGGCTTTTCTGCCTGTGGCTCTTTCTGGAACACAGAGCCCCAGGCTTCCCTTTCTTCTGTGAGGATTACTTGAAGTGTCCCCATTTT | 120 | 48 |
| chr15 | 89506298 | 89506418 | 771629_41866724_SHANK3(58234)_55 | 1 | + | good | TTTTTTTCAGTGACATTGGGCAGCAAAAGGCTGAGGCTGGCATTAATCTTAGAACATTCCTCCGTTTTCTGTCAGTCTGGCCTGGGGGTTCATTTTTGGATGGGGTTAAGATTAAGGTCA | 120 | 44 |
| chr15 | 89506418 | 89506538 | 771629_41866724_SHANK3(58234)_56 | 1 | + | good | GTTTCATGTTTAGGGCTCTTCCTTGAGCTGCTCTCTGCTAATAAGGCTTGGTTACGAGCTCCTTGAGCCTTGCGAGGTGGGGCCGTAATGATGGTTGTACCGAGCGTGGGTGGCTTCCCC | 120 | 55 |
| chr15 | 89506538 | 89506658 | 771629_41866724_SHANK3(58234)_57 | 1 | + | good | AAGAGAGCTGAGTGACTTCATGATCCGTGGTACAAAGCTGGGTCCAGGTTCATGCAGAAAGAGTGGGGGGGGGGAATAGCTGGGTTGATATAGGGGCAAGAGAGAGGTGCAGTTTGAGTG | 120 | 52 |
| chr15 | 89506658 | 89506778 | 771629_41866724_SHANK3(58234)_58 | 1 | + | good | GGGGTGAGAGTATTAGTGACAGTGTTTTCTAGGACCACCCTGCTGTGGATCTTTGGGGGGTCTTGTTTCATCTGCTTGTCTAATCGTGAGAGAACTGTTAACAGTGCGGCACCTCACCAC | 120 | 50 |
| chr15 | 89506778 | 89506898 | 771629_41866724_SHANK3(58234)_59 | 1 | + | good | GTCATAGGTGAAATAGAGCATCTTAGAGTTATCGAGTCTTCTTTCACCTTGGTCTTTTATAGCACTTGATCCTGGCAGGGTTTGGGTTCTAGGAAGCGGCCTGTTTGTAACTGGGTCTTC | 120 | 45 |
| chr15 | 89506898 | 89507018 | 771629_41866724_SHANK3(58234)_60 | 1 | + | good | AAGGGATGAGGAGCTTGTATGGTCTCAAGCGTGAGGGGTAGAAATATTTCTCTACCCCTCATCCTAGGATTCTGGGGGTCACCATCTATTTTATCCCAGTGTGCTGGCTCAGTTGGGGTC | 120 | 50 |
| chr15 | 89507018 | 89507138 | 771629_41866724_SHANK3(58234)_61 | 1 | + | good | GGGCACATCATCCGCTGAGTAGAGTGAGCACGTGTGTCATCTGATCCTGATCTCTTTTAGGCATCTATTGTTAGGTTTCTATTGTTGTGGTAAAAACACCACGACCAGAAAGCAACTTGG | 120 | 45 |
| chr15 | 89507498 | 89507618 | 771629_41866724_SHANK3(58234)_65 | 1 | + | good | AGATGACCCTGGCTTGTATAAAGTTGTCAACAAATAAACAAACAAAAACCAAACCTGAGCAGAACATAACCAGTTGTCCTTGCGTAGGATGATTTATTTTGAACTTTCTTATTCCAGTTT | 120 | 35 |
| chr15 | 89507618 | 89507738 | 771629_41866724_SHANK3(58234)_66 | 1 | + | good | ATCTTCCCACTATCCTTTTGTGAAGTTGATTATTGGGCCTCTTAACAGAGAAGGAAACCAAATCTCCACAGGGTTCGTGACAAAACCCATCCTTAGATGTGGGGGCTCGGGTCTGTCATC | 120 | 47 |
| chr15 | 89507738 | 89507858 | 771629_41866724_SHANK3(58234)_67 | 1 | + | good | TGCACATCTTTACTCCAAGGACATTTCTTAAGTGATAGTTGTGCGTTCCTAGGTCTCACTCTGGTCTTTGGGCAGAGGCTTACACATGGAGAGCTATGTGGAACTCTCTGCCTATTCACC | 120 | 47 |
| chr15 | 89507858 | 89507978 | 771629_41866724_SHANK3(58234)_68 | 1 | + | good | CTCCTCTTCCTTCCTGAGGGTCGGGTAGGGTAGGCACTAGTGTCATGCTTCAAGAGACAGGACACGTCCTGTGTCTTGGCAGGCTTACTTCTCTTTAGGGACCAACAGACCTTGGGCAGG | 120 | 55 |
| chr15 | 89507978 | 89508098 | 771629_41866724_SHANK3(58234)_69 | 1 | + | good | ACTCTGTCCTGCCAGAGGCTTCCTCTCGCTTCCTGCCTCTGTCACGGGGCTGTCCTCTTTCAGTCAAGCGCCTGTGTGCATGAAAATACTCTATGCATATATGTTCTCATGGACCATCTA | 120 | 51 |
| chr15 | 89508098 | 89508218 | 771629_41866724_SHANK3(58234)_70 | 1 | + | good | TCTCATTTTTGGTAAGTGCCTTGGGAAACGCGCCTCCTATGGTGATGTCACCTGGAGAGCTTGTTCAGGGTAGGTTCTGAGTCTGGGCCTGGATGGAACCTGCGTTTCTACACCGGTCAT | 120 | 52 |
| chr15 | 89508218 | 89508338 | 771629_41866724_SHANK3(58234)_71 | 1 | + | good | CAGCTCAAGATGAAGCTTGCTCTGCTGGTCCCACCTCTACTTTTTATAGACAAAGCTAGCAGGTAGTGCTGCAGGCCCATGCCGGCCCATAGTGTAGTTTTAGTTTTTAAATTAAGGTGA | 120 | 46 |
| chr15 | 89508338 | 89508458 | 771629_41866724_SHANK3(58234)_72 | 1 | + | good | AATGTTCCTTACCTAAAATTGATGACTCTTAGTGGCAATCCAGGGCATCCCTCACATTTACAACATTGTATGCTGGTTTTATTGATCCCCAAATATTTTCTTTTTTTATTAGATATTTTC | 120 | 32 |
| chr15 | 89509298 | 89509418 | 771629_41866724_SHANK3(58234)_80 | 1 | + | good | AGAAAATAATTCCTCTTTGCATTTAGCTGTTATTGCTGTTTCATATCAATGGGATTATACTACCACTCTCTGTGTTTGACTTCTTTTACTCTTTTTCTTTTTGAGCTTCATCTGCCTTGT | 120 | 33 |
| chr15 | 89509418 | 89509538 | 771629_41866724_SHANK3(58234)_81 | 1 | + | good | AAGGTCACCACTTTATTCTACCTTAAGGATAAAAAACCATTCTACTGTCTTTATGGATGGTATTTGGCTTATCCATTTATTTGTGGGTGAATATATAGATTGTTTCTATGTTAAAATATA | 120 | 29 |
| chr15 | 89510018 | 89510138 | 771629_41866724_SHANK3(58234)_86 | 1 | + | good | GACACCACCGCTGGGCTGAGTGGCATACCCTCTTAACAGCTGGGCCATCTCTCTAGCTGGTTCCTATAACATATGAATATTGCTATCACCATCACAAACATGTCTGTACGTGTATTTATA | 120 | 45 |
| chr15 | 89510138 | 89510258 | 771629_41866724_SHANK3(58234)_87 | 1 | + | good | TGAATACTTGTTTACACTTCTTCTGGGTTTATCCTGAGGGGTGGAATTTCTAGGTGACATGACATTTCTTACAGTTTTAGAGGAACTTCCAAACTGTTTCCCACCAAACTTCTGTCAGCC | 120 | 41 |
| chr15 | 89510258 | 89510378 | 771629_41866724_SHANK3(58234)_88 | 1 | + | good | GTGCGTAAGTTTCTAACTTTTTAACTTCTTTACATTCTTCCTGAAGCCTGTCTGGTGCTTATTTATTTATTTTTTAAATTGTAGTTAATTCTGTTGGATATGGTGCGATATTTCATTGTG | 120 | 31 |
| chr15 | 89510378 | 89510498 | 771629_41866724_SHANK3(58234)_89 | 1 | + | good | GTTTGTGGTTTCCTAGTGACCAGTGATATTAAGCTTATTTACAAGAGTTTGTTGTTCACTTGTGTGTCTTTGGAGAAAAGTGAAGGCCTTTTCTAACTTAAAAACCGTTTTACTTATTGT | 120 | 34 |
| chr15 | 89510858 | 89510978 | 771629_41866724_SHANK3(58234)_93 | 1 | + | good | CAGTTCTTCTACCTTAGCTTCTGAGAATAGGGATTGCAGAAACCAGATCCTGTATACTGCTCTGTTTCTTGCCCAGATTCATTTTGAATTCTGGTCTCAGGTGATCTTCCTGCCTCACCA | 120 | 44 |
| chr15 | 89510978 | 89511098 | 771629_41866724_SHANK3(58234)_94 | 1 | + | good | CAGGCACTTGTCTTCGTAGACACAAAAACCTGCTCGTTTTTGTTTTTTATTTGCTCATGCTTTTGATGTAGCAAAGAACCCATTGTTGAATGAAAAACATAAGCAGTCCTAAGTTATTTT | 120 | 35 |
| chr15 | 89511338 | 89511458 | 771629_41866724_SHANK3(58234)_97 | 1 | + | good | AAGGAGCAAAAGGTCATATCTTTGTAGAGACAAGAAAAGATTTGTTTTTTGTAGTTAGTTGGTGGTCTCATTAGTGTGGCCTCTGGCTTCAGACGATGGTTTACTGTCACCCTGGCTTGT | 120 | 42 |
| chr15 | 89511458 | 89511578 | 771629_41866724_SHANK3(58234)_98 | 1 | + | good | TTGGTAGGAATTTAAAGAGTGGCCAGAGTCACTCTTATTAATGGCATTGCAATTAGATGATTTTGTAAAATTTATTATTAGTTTTTGAGACAGGGTCTCACTGTGGCTGGCCAGGAATTC | 120 | 37 |
| chr15 | 89511578 | 89511698 | 771629_41866724_SHANK3(58234)_99 | 1 | + | good | ACTACATAGACCAGGCTGGCCTCAAACTCCCAGGGAGATCCAAGTGCTGAGTTTAAAGACATGTTTCACCCTATGCCTGGCACTTAAAACAAGTTATCCACTTTGACCAGAACTCGGAGC | 120 | 48 |
| chr15 | 89511698 | 89511818 | 771629_41866724_SHANK3(58234)_100 | 1 | + | good | ATAATTGGAACCATGTGTCATGGGCACGATGTCTTTGACTCCAATCATACTGTTAAGATGTTGAAGTTTTTTTTTAAAAAAAAAAAATCTCATTTTTGGTGTGAGCACATTGTGGATGGCT | 120 | 34 |
| chr15 | 89511818 | 89511938 | 771629_41866724_SHANK3(58234)_101 | 1 | + | good | GTGGAGACACTTTTGAGACCCTAGAAAGAATGTAGAATGCTGGAGAATGAGAGGAGTATCAGGCAGATGCCCAGGAATAGAGTTTGTCCTTAGCTGAGGATTCTACCAACCAAACTTTTA | 120 | 43 |
| chr15 | 89512178 | 89512298 | 771629_41866724_SHANK3(58234)_104 | 1 | + | good | CATGTCTGGCTCAGGATAATATTTCTATAGGATAAAACCCTAGCAAAGGCAAATCATTAGAACAAACAAAATCAGTTGGGAGTCCAGAAGGGAGTTAGCCAAGATTTCTGGAGTTGAGCT | 120 | 40 |
| chr15 | 89512298 | 89512418 | 771629_41866724_SHANK3(58234)_105 | 1 | + | good | CTAAGCATGTTTCCTGGCTCGCATCTTGAACGCCCGTATTTCTGGCACAGAGATGCCTGGCCTCCCCTGGTGGCCAGCATCGGGGGTAGGGTTATCCTCTGGAATTACATCAATGTGTCT | 120 | 54 |
| chr15 | 89512418 | 89512538 | 771629_41866724_SHANK3(58234)_106 | 1 | + | good | CTCTCCAGCTTCGGAGTTTTCAGGAACCGAGCAGGTCTTGTTCTTAGTGATTCTAAGTTGGAAATGTGGGGAAAAAAATTCAAATGGAGGAAAATGGAAAATCACTATCCAGTTAGGTCT | 120 | 41 |
| chr15 | 89512538 | 89512658 | 771629_41866724_SHANK3(58234)_107 | 1 | + | good | ACAGGTAATAAATAGTAGTAATAGCATTCCAAAACAGTTAGCAGGCCCACAGTCTCAGAACAGTGTACTGCAGTTTTTCTCTGAAGCATAATTTTCTTCTCTGTAGCCACATCATTCTCT | 120 | 38 |
| chr15 | 89512658 | 89512778 | 771629_41866724_SHANK3(58234)_108 | 1 | + | good | TGAACCTTAGCCTATTTATACACATTTAGTAGGAAAGGCTTAGAACATAGTTAAATTGGTAGGCAGGGGCCCAGCTATGTAGAGCTCTGGGTTGTATCACCAGGACTGAATAAGGCAGTC | 120 | 43 |
| chr15 | 89512778 | 89512898 | 771629_41866724_SHANK3(58234)_109 | 1 | + | good | ACAGTAGAGCACACCTATAAGCCTCATGAAAGTGAATTTGAGGCCAGCCTGGACTACATGATACTATCTTCCAGCAACAGCAATAACAACACAACCACTCCTTTAGTAAGGACATAGTTA | 120 | 42 |
| chr15 | 89512898 | 89513018 | 771629_41866724_SHANK3(58234)_110 | 1 | + | good | ACGACATAGGCTCTTCTTAAGTCTGTTTAGCTGAAATGCTTGATAAGGGGCCTCCGTTTGGACTTTATATCAAGACAAACCTCCAACTTTCTCTGCAGTCGTTACTCTTTTTTTTTTTTT | 120 | 38 |
| chr15 | 89513018 | 89513138 | 771629_41866724_SHANK3(58234)_111 | 1 | + | good | TTAAGGTCTTTCAAATACCTTCACGTTTCTGGAGGAAACCTGGCAGGAAGTGAGATTTTACTCACAACAAGGCTTGGGACAGCTCTTGCAGGTTGCCATCCTATGCAGATTCTCAAAGCA | 120 | 45 |
| chr15 | 89513138 | 89513258 | 771629_41866724_SHANK3(58234)_112 | 1 | + | good | CATTCTAACCAAAGCCTTGGTGGTGGAATCGGTTTTTCCAATTGTACTCTATTAAAAAACAGAAGTTTCCTACGGAACTTGTGCAGACAATTATATTCCCACGTAATAAAAATATCCACG | 120 | 37 |
| chr15 | 89513258 | 89513378 | 771629_41866724_SHANK3(58234)_113 | 1 | + | good | ATCAGCTTTTCCAGTGCTGAAGGATCAGGTGGAGGGGGGGGTGGGGGGAAGGAGTGAATATTTCAGTTTTAATTTTCAAAAAGTAACCAATAGGCCACAGAGAGCCTAAAGGACAAGGGA | 120 | 47 |
| chr15 | 89513378 | 89513498 | 771629_41866724_SHANK3(58234)_114 | 1 | + | good | AGCATATTTCCTTAAATCTAGAACATTAAAATGAGAAGAAACTCAAATACACATACAACCACTCACCTCATCTTTTTTCTTAAAAAAAAAAGTTATTAAAACAGTTGAGCCAGGCAGTGG | 120 | 31 |
| chr15 | 89513858 | 89513978 | 771629_41866724_SHANK3(58234)_118 | 1 | + | good | CTGAAATTAACTGGACTCAGACACACTGAGATAACTCACCTGTTTTCCGGTCTGTTATTACTTCTTTTTCTGCTTGTTTATTATTTTGAGACAGAGTCTTATTCTGCATCCTAAAAGGCC | 120 | 38 |
| chr15 | 89513978 | 89514098 | 771629_41866724_SHANK3(58234)_119 | 1 | + | good | TGGGTTCCCCCTCTCCTCCAGGCTGGGCTTGCATTTATGATGATCTCTTACCTCATCTTTAAGTGCCAGGACTGCAGGTGTGAGCATTATGCCAGGCCCTTGTCCCAAGTCTTGTTTTGC | 120 | 52 |
| chr15 | 89514218 | 89514338 | 771629_41866724_SHANK3(58234)_121 | 1 | + | good | CTTTTCATTTACCCTTCTCTCCTCTTCCCGCCCCCACCCCTTCTCAGTGTTTCTATATTGCCCAGGCTGGCCCTGGACTCCTGGGTTCACCAAGTCCTCTGGACTCAGCCTCGGAACTAC | 120 | 56 |
| chr15 | 89514338 | 89514458 | 771629_41866724_SHANK3(58234)_122 | 1 | + | good | TTGTGAGCATGCGCCACCATCCCTGGCTTTCCCTTCTTCATTACAGTCCTGAAAATTCCTGTTCAAGCTAATGGTATGGTCGGAAAGTTGCTGGGAACCTGGATTCCACAGCACTTGTCT | 120 | 48 |
| chr15 | 89514458 | 89514578 | 771629_41866724_SHANK3(58234)_123 | 1 | + | good | AAGTGCAACACTGTCTTTGAAGGAGAGATAATTTTTGGCTGCAGCTTGTTGCACACACCTTTGGAAAAGACTGAGAAGAATAACTGTATACTTGACAAGATGGAAATCAGCCATGGCTGA | 120 | 41 |
| chr15 | 89514578 | 89514698 | 771629_41866724_SHANK3(58234)_124 | 1 | + | good | GATCTGATGATAAGGTCAGCCGACAAGCGACTGCGTTACTTCTTACCGATTACAAGATTTTATCACAACCAGACTTAACGACCAAAAAGAGGATATCAAGACAGGTCAAACTTCTAGGAG | 120 | 42 |
| chr15 | 89514698 | 89514818 | 771629_41866724_SHANK3(58234)_125 | 1 | + | good | TTCTGTACAATTTTAGAATGTTTGAGTCAGGAAAGTATGTGAAAATACAGTTTAAAATTTGGTACCTGGAGGGTGTATGCCTATGTTCCTAGGACTGTGGAGGCCAAGGCAGGATAGTAA | 120 | 39 |
| chr15 | 89514818 | 89514938 | 771629_41866724_SHANK3(58234)_126 | 1 | + | good | CAAGTTTGGGGCCCACCTGGGCTATATAGGAATTTCCTGTCCTTACCCCCCAGGCTCCACCCCAAAAGGAGGTTTAGTATTACATATGTAACACTGTTTTCAAACAATTTAATATATCAA | 120 | 42 |
| chr15 | 89514938 | 89515058 | 771629_41866724_SHANK3(58234)_127 | 1 | + | good | ACAAGCTTAATTAATTTAGCATCTCTTACTATTACAGGGGTCTTTCTGGATTGTTCCAAAGTTAGTTTAGGATAAATACAAGTCCGGGAGTCAGGCACGGTAGATACTCCAGGGCTTTAA | 120 | 39 |
| chr15 | 89515178 | 89515298 | 771629_41866724_SHANK3(58234)_129 | 1 | + | good | AAACCAAACCAAAAAACGAACAAAAAATCCCAAAACAACAAACAACCCCCCAAAAAACCCCACACCCCAAAATTACCTACTGTACTCTGAAATCAGTCACATAATTTTTCTTTTGTCCTT | 120 | 36 |

| | | | | | | | | | |
|---|---|---|---|---|---|---|---|---|---|
| chr15 | 89515298 | 89515418 | 771629_41866724_SHANK3(58234)_130 | 1 | + | good | AGGTCTGGCTGACCAGTAGCATGGGTGGGAAACACACTAGGAATGGCTTTCCATAGATTGCTAGCCTTTTCTTATTTATAGGTTGCCTGTGTGTGGACCCCATGGGGGCTTGTGGCCATG | 120 | 51 |
| chr15 | 89515418 | 89515538 | 771629_41866724_SHANK3(58234)_131 | 1 | + | good | ACTGTCATCCTTAGGGTGACATACCCAGGTTCCACTGGCATGGTGAGATGACACATGAGCTCAGGTTGGAACCTATGAGAAAGGCAAAAAGTATTTTTACTTAGACGGGAATACCTCTAT | 120 | 44 |
| chr15 | 89515538 | 89515658 | 771629_41866724_SHANK3(58234)_132 | 1 | + | good | TTATTTTTGAGCTCAAGAATGTGATCCATCTAACAGTGGTTAAAGAGCAGACTGAAAAGCTTGATTTTGGGATATGGTGTAATTCTGGATGTAGAGGTAGAGAGATAAGAGACTAAAGGA | 120 | 37 |
| chr15 | 89515898 | 89516018 | 771629_41866724_SHANK3(58234)_135 | 1 | + | good | TCCTGAGATACAGTCCCAGGTGCTGGGTTGAAATAATGTATTATTTCCTATCTTCCTGGAAATGTCCTACTGAGATCTTTTCATTAAAACCCAGTTAGATTCCCAAAGGCTCCCTACAGA | 120 | 41 |
| chr15 | 89516018 | 89516138 | 771629_41866724_SHANK3(58234)_136 | 1 | + | good | ACCCACGACAGTTTTTATTTCAACTTCAGAAGTCAGTATGACTTTAGGAGCCACAGTGAGTCAAAACACTGTCAAGTAAACCAAGAGAATCGCACACAAAATGTCGAGCGCAGGAAGGCT | 120 | 43 |
| chr15 | 89516138 | 89516258 | 771629_41866724_SHANK3(58234)_137 | 1 | + | good | CTCACTGGCTCCAAGAAGCTGCTCTTCTACTCCGTCTGACCAGAGAAATGTATGCATGTTTACATCTTATCGCATTCTGTAACTCATGAATAGGGGGGCGAGGATCACTCAGCCAGGAGG | 120 | 49 |
| chr15 | 89516258 | 89516378 | 771629_41866724_SHANK3(58234)_138 | 1 | + | good | AGAAGGGAGAAACACAACTTGTAGAAGCAAGCTGAGAAATACAATTTCAATTGGTTAGAAGTTCGTGAGTGTTTTCTAACTGGTCTCACTTCTAGTTTCAGTTCATGGTGTACACTAGCT | 120 | 38 |
| chr15 | 89516378 | 89516498 | 771629_41866724_SHANK3(58234)_139 | 1 | + | good | GTTGCTTTGTTCAGGTGGAATTTGAAGTTTGAGGGTCACCACATTAATGGCATTCAAATAACATTTTTTTTCTTCCCCCTTATCAGTCACCAGTCACTTCGTTTTCTTTTTAAGTTTCCT | 120 | 37 |
| chr15 | 89516498 | 89516618 | 771629_41866724_SHANK3(58234)_140 | 1 | + | good | GAGTTCAGCTTTTTATATTTAGGTCACGGATCAATTGTAACTTAATTTCTGTATGTTCTGTGAGTTCATAGTCAGCTTTCAGGCGGCCCTTTAGTTGTCTGGCGGTATTTGTTGGAGAAA | 120 | 39 |
| chr15 | 89516618 | 89516738 | 771629_41866724_SHANK3(58234)_141 | 1 | + | good | GTTTTCTTCTTTCGCCGGTTATTCGTTGCTGGTGTATAGAAACAAGCTGCCTTTTGTACAGAGGTCTTGGACCAACATTGCTGTATTTGCTTATTCGCTTTCGCGGCTTTCTTGGTGGGT | 120 | 45 |
| chr15 | 89516978 | 89517098 | 771629_41866724_SHANK3(58234)_144 | 1 | + | good | CCTCAGACTCCTGAGTGCTGTTATTAAAAGCATGGCCATCATGCTCAGCCTTATTTGGTTTTGCTTAAACTTCTCTATGTGGAACAGCGGTAAGGAGAGAGCAGGCATCCCGGTGCCGCT | 120 | 49 |
| chr15 | 89517098 | 89517218 | 771629_41866724_SHANK3(58234)_145 | 1 | + | good | CTTTATGCTAGGAAAGAGCCTTTAGTCTTTTCCCATAGATGTGATGTTAGCTGTTAGTTATAAAGACCCCTTAACATATCGAGCATGTGTCTTTTTCTGTTAGCAGTGTCCTGATGTTTT | 120 | 38 |
| chr15 | 89517218 | 89517338 | 771629_41866724_SHANK3(58234)_146 | 1 | + | good | ATCACTGAAGGATGTGCGATGATAGCAGATGCTTTTTCTGTATCAGCCCACCTAATGACGTGGGTTTTTTTCTCTCTGTTCCACTGATGCACCACGTTGGTTGCTTAAGCCTTTAGCTTG | 120 | 45 |
| chr15 | 89517338 | 89517458 | 771629_41866724_SHANK3(58234)_147 | 1 | + | good | GTCCTTGGAATGCGGTTATCTAGCTTGGGCATCAGTAAGACAGAATCAGCCAGGAATTGTTTTCTTCTTTCCCGACTCTTTGGAAGCATTTGAGAAGGATGGGTGTTGATTCCCAAGTGT | 120 | 45 |
| chr15 | 89517458 | 89517578 | 771629_41866724_SHANK3(58234)_148 | 1 | + | good | CTGTAGTAATACCATCTGTTATCACTTTCCGGTCGTTGGTAGGAAGATTGTGATTTCTCAGTTTCTTTGCCTGTTTCAGGTCTGTTTAGATGTTTCTGTTTCTTCTTGAGTCAGTTTTGG | 120 | 39 |
| chr15 | 89517578 | 89517698 | 771629_41866724_SHANK3(58234)_149 | 1 | + | good | TAATCTGTGGATGTTAAAAGAATTCCTACCTACCTAGTTTATCTAATTTGGTGGCAAGTGATTATTCACAGTGTCATCTTGTATTTTATTTCTGAATGGTTGGTTAGAATACCCTAAAGC | 120 | 33 |
| chr15 | 89517938 | 89518058 | 771629_41866724_SHANK3(58234)_152 | 1 | + | good | TTTCATTTTTTCCATATCCTTTATAAAAAGTCAGTCTGGCTAGAAGTGTGTTGGCTTCTCGGTTTACTTTATTTTATTTATGAATACAAAGATTTTGCCTGCAAGTGTGTCTGTGCATCA | 120 | 33 |
| chr15 | 89518178 | 89518298 | 771629_41866724_SHANK3(58234)_154 | 1 | + | good | TTTTTGTTCCACTGATACTTTATGTCCTTAAGACACAATCAACTTTTGTTTCCTTCAATCAGCTTTTGTTTCCTTCTTTCCTGTCTCTTTGGAAGAGTTTGAGATGTGGTGGTGGAAATC | 120 | 37 |
| chr15 | 89518298 | 89518418 | 771629_41866724_SHANK3(58234)_155 | 1 | + | good | GAACCCAGGTCTTCTGCAACAGCAACAAGTGCTACTAATGTTTGAGCCATCTCCCCCTATACACCTTTTTTTTTTCCTAGCCAGGGTTCTCTCTCTATGTAACTAACTGTCTTAGGATTC | 120 | 43 |
| chr15 | 89518658 | 89518778 | 771629_41866724_SHANK3(58234)_158 | 1 | + | good | CAATACTGGGACTGGAACCAGACACTTGGGGATGCCAGACAAGCTACCGCTGGTCCCTACTTTCTTGCTGCTTTTTAAATGTAGGTGCTTGCTGTTAAGTGAAATTTCCTAGGTGTGTTA | 120 | 46 |
| chr15 | 89518778 | 89518898 | 771629_41866724_SHANK3(58234)_159 | 1 | + | good | GTCAGCTCTGCTTGTCTTGTCTCCTGTGACCTCAACCATCTTCTTCTTCGATCTTCTAGTCATTACTGGAAATGAGTATTGAAGGTTTTGGGTGTTATTGTGGAACTGTCCATTTCTCTC | 120 | 42 |
| chr15 | 89519138 | 89519258 | 771629_41866724_SHANK3(58234)_162 | 1 | + | good | TTTATGGTCTTTGGTGTGTACATATTAATGTTTGTTAATCTTGATGAATTACTGTAACAAAAGTTAGAAAGATACATTATACGTAAGTATCTATCTATATTGTGTATGTGTGTGTGTGTG | 120 | 28 |
| chr15 | 89519378 | 89519498 | 771629_41866724_SHANK3(58234)_164 | 1 | + | good | GGTTTCTCACTGGACTGGCTTGCTGATTAGATTAGGCTGGCTGGCTGGCTAGTGAATCCTAATGGTCTTCTAGCTCTACCTCCTCGGCTCTGGGATAAGTGCAGGCAATGATGTTGACTT | 120 | 50 |
| chr15 | 89519858 | 89519978 | 771629_41866724_SHANK3(58234)_168 | 1 | + | good | AAATATTACAACTTTCAATATATTTATGACTTTGGATCTAAAGTGATCTCTTCTAAACAACGTATAGTTCAATATGTTCTATGTTTTTAAAATAAAGTTTTATTGGAACATAGGTGAAC | 120 | 24 |
| chr15 | 89519978 | 89520098 | 771629_41866724_SHANK3(58234)_169 | 1 | + | good | CCATTTGTTTACATACTATTTGTGGCAATTTTCACGCTACGGAGATAGAGTCTGAGATATGGCGTGGAGTCTCTTTGGTTGGCAGAGCCTGAAATAGTTCCTGCCTGGTTCTTTTTAGG | 120 | 44 |
| chr15 | 89520098 | 89520218 | 771629_41866724_SHANK3(58234)_170 | 1 | + | good | ATGTTTTCTGGCTCCAGTCTAGAATGCCTTTTTTTAGTTTCCCTGGCTATAGTCAATAGTATATAGTCTCACATAGCAGTAGTTGGATGAGACTTGTCACATGTAGCCTGCTTTATCCTG | 120 | 40 |
| chr15 | 89520218 | 89520338 | 771629_41866724_SHANK3(58234)_171 | 1 | + | good | AAGCCTGTAGAACGGGAAGTTCTGACCACAGAGCTGGGCATGCCGAGGCCTTCGCTGGCTGAGAAGAGAAGCAAACGTCGATCTGTAAAGTCAGGGTTGGAGGGAGCAGGACTGGGATGT | 120 | 56 |
| chr15 | 89520338 | 89520458 | 771629_41866724_SHANK3(58234)_172 | 1 | + | good | GGCCACCCAGTGGTGACCACATTAATGTGGTTAACTGTTCTGATTTCCTGGGATTGGGGCACTTCCTAGAGCATGTTTACATTCATTTATTCGTATGTGTGTGTACATTTATAGATATG | 120 | 41 |
| chr15 | 89520458 | 89520578 | 771629_41866724_SHANK3(58234)_173 | 1 | + | good | CAGGTAAGTGTACATGTGTATTGGAGTCAGAAGTCAACCTCGGGTATTTTTCTCAGGAGTTACCCCAAATTGATTTTGGAGACGGGGTCTCTTGGTAGGGCTTAGGGCTGACCGATTAT | 120 | 47 |
| chr15 | 89520698 | 89520818 | 771629_41866724_SHANK3(58234)_175 | 1 | + | good | TCAGGTCCCTAGTTTTTGCACACAAAACACTTTAATGACTGCACTATCTTCTAGGCCAGTATTTTAGTTTTAAAAGATCAGTTGCCCCACACAAAGAACTGCAGGCAACTGGGGCATACT | 120 | 42 |
| chr15 | 89521058 | 89521178 | 771629_41866724_SHANK3(58234)_178 | 1 | + | good | AAAATCAATGAATCTCGAGTCTTACTCCTGCCCCTCTACTCCAGTACCATTCAGGGAAACCCCCAGCTATGCAAAGCGACGGCGTCTGGCTGGCCCGAGTGGCCTGGCATCCCCACGGCC | 120 | 58 |
| chr15 | 89521178 | 89521298 | 771629_41866724_SHANK3(58234)_179 | 1 | + | good | CTTACAGCGCTCAGCCAGTGATATCAACCTGAAAGGTGATCAGCCCGCAGCTTCTCCAGGGCCCACTCTCCGAAGCCTCCCTCATCAACTCTTGCTCCAGAGGCTTCAGGAGGAGAAAGA | 120 | 56 |
| chr15 | 89521298 | 89521418 | 771629_41866724_SHANK3(58234)_180 | 1 | + | good | CCGTGACAGGGATGGTGAACTGGAGAATGACATCAGCGGCCCCTCAGCAGGCAGGGGTGGCCACAACAAGATCAGGTAGGAGGGCCTGGAAGGAAGGGTCAGGATGGGTCAGAGCTAT | 120 | 60 |
| chr15 | 89521418 | 89521538 | 771629_41866724_SHANK3(58234)_181 | 1 | + | good | GCAAGCCTAAGCTGGATAGTAGGGAGGAAGGGGAGTTCGTGGCCCCTGTGCAGTGATAGGCTGGGGCTAGGGGTGGAGTAAGTTCCTTGATGGTTTCACTCATGCTGATCTCTGGAAGAT | 120 | 54 |
| chr15 | 89521538 | 89521658 | 771629_41866724_SHANK3(58234)_182 | 1 | + | good | GCTAGGGCTCACTCTGAGGATAGGGGCAGCTGAGAGAGTTTTGGGTAACCAACTTGTGGTTAGGATGCTCACTACGGCATGAAAACTGGTGGCAAATTGGTTGGAAATATCTGTGTAGGC | 120 | 49 |
| chr15 | 89521658 | 89521778 | 771629_41866724_SHANK3(58234)_183 | 1 | + | good | TCAAAGCCCATAGGTGAGAGTTATGGCTGTCCGTCGTCCCCACATATCTGCCCGAGCCCTCTGTAGAGTAACAGTACTTTAAACCTCTCTCCTGAGTTCTGGAATCTTGAGAAACCAGGAT | 120 | 49 |
| chr15 | 89521778 | 89521898 | 771629_41866724_SHANK3(58234)_184 | 1 | + | good | ATTTGCAGTATCGACCTCCTTATAATCTAAGAGTTGGAAGCAAAGAACGGAGCCTCTTTCACTCTGTAGGCTTAAGGGGTGGTCACTACGCTGGTTTCTCTACCAGTACAGTGTAGGGTG | 120 | 47 |
| chr15 | 89521898 | 89522018 | 771629_41866724_SHANK3(58234)_185 | 1 | + | good | TTTCCTTTGAGAAGCCCCCTAGGGTGCCTGTGAGGGGGTCTTGGACCCTCCTCCAGCACTGCCTATGTTAACCACTGGTTGTACCAAATTCCCAGAGTCTTGAGGTTCGTAGCTCTCATT | 120 | 52 |
| chr15 | 89522018 | 89522138 | 771629_41866724_SHANK3(58234)_186 | 1 | + | good | TTTTGCTTCCTGCCGTTTCTCCAGAGCCAGGTATAAAGAAGGTGTTTGGAAGCCTGTTGGATACAGTGTTACATAAGCAGCTTGTTGTTTTCACAGTGTCGGCCAGCTCTGATAGCCAGC | 120 | 48 |
| chr15 | 89522138 | 89522258 | 771629_41866724_SHANK3(58234)_187 | 1 | + | good | CCTACGGGTTGATGTTCCCACTGGGCTGAGCAGCAAATGCATGCCACTTGGTTAACTGCTGTCGTTTAGTCTGGGCTAAACGACACTATTATACAGGCTGGTCCCCCAGCCCCATGCCCA | 120 | 54 |
| chr15 | 89522258 | 89522378 | 771629_41866724_SHANK3(58234)_188 | 1 | + | good | GGTTGGGTCAGAGAGAGTAGACATCGGCTTGTGTGGGTCCAGGGGTTTGGTAGAAGGCCTTTGATGTAGCATACACTAGGCTGTTCTTTCTGCAGCATCCCCTTGCCCGTGCAGCCTGAT | 120 | 54 |
| chr15 | 89522378 | 89522498 | 771629_41866724_SHANK3(58234)_189 | 1 | + | good | TTCCAGGCTGTAGACTTACTGGACCCCCAGGGTTGTCTGTCTGCTTGCAAAATCACATTCTCCTATGCAACTTTTGCACGGCCCTTTGGGTTTTCTAAGAATTGAACGGGGTGGTGGCT | 120 | 49 |
| chr15 | 89522498 | 89522618 | 771629_41866724_SHANK3(58234)_190 | 1 | + | good | GGAAGAATGCATTCAGGGATGTTGCGGTTTGCCAGGAGGACTGTGTGTGAGTGGAACTGTGTGGCTGTCACTGGGTCCCTTTCAGTTGTGTCTGTTGCAGTTGGCACAGAATAATCTCCT | 120 | 51 |
| chr15 | 89522618 | 89522738 | 771629_41866724_SHANK3(58234)_191 | 1 | + | good | CTAGTCTAGGCTTTCCAGTTCACCTTTGGGAGGGCAGAATTATTTCTTTTCCACATTTATTAGGCGACTGGAATGCCAGGTGTGTGTGTGTGCGCGTTCGCGTGTGCGTGCACGGCAC | 120 | 52 |
| chr15 | 89522738 | 89522858 | 771629_41866724_SHANK3(58234)_192 | 1 | + | good | ACGTGTGAGTTCAAGGCTCACCTCTGGCTCCCCACCTCTACCTACTCCCCATCACACAGCAGGGGTCCTGACTGGTTCTGAGGAGGTAGCCACCTGGGCTCCTGGCTGAGAAACACTTAG | 120 | 58 |
| chr15 | 89522858 | 89522978 | 771629_41866724_SHANK3(58234)_193 | 1 | + | good | ATGAGTTGTGATACAGGTTAAGGGGGCTTACCTGGTGTGACAGTGGTCTGCCAGCCAGGATGTGGCAGAGTGTGTTGGGGGTGCTGAGAAGGAAGAGACCCTCAGGCAGAGGTTGCCTGT | 120 | 56 |
| chr15 | 89522978 | 89523098 | 771629_41866724_SHANK3(58234)_194 | 1 | + | good | TAAAATAAAAACGAAAAAACAAAAACAAAAACAAAAAATCAGGATGTATTCATGGACCCTGGTGGGGCTGCAGTGGCTTCTATTGCTTCCTGTTGGCTGAGCCTGGCATAGAGCCCAGG | 120 | 43 |
| chr15 | 89523098 | 89523218 | 771629_41866724_SHANK3(58234)_195 | 1 | + | good | ACTATGGTCTGCAGGTTCTGGCCCCCAGCACCAGAGTGAAACCTACAAGGGGCAAATTTGAAGCCGAGGCAGTAACTTAATCTCTGGCCAAAAGGGCAGTGTCACAGTTGTTTAGGTGGT | 120 | 51 |
| chr15 | 89523218 | 89523338 | 771629_41866724_SHANK3(58234)_196 | 1 | + | good | GGGAACAGTGTAGCATATCTCTAAGTCTCAAAGAGCCTGGTCATTGGCAGAGCAGGTGAATTTGGCCTGTCGGGGTACCAGGGAGCCCAAGCAGGCTTTTGGGTAGAAAGGAAGACCAAG | 120 | 52 |
| chr15 | 89523338 | 89523458 | 771629_41866724_SHANK3(58234)_197 | 1 | + | good | ACCTCAGACTCTGTATATACCTAAGCCCGAGAAGTGGTCACCATAGATTGACATGGTGGTAGGGACAGCTGTCCAAGACCACAGGAGTCAGGCCCAGAGGCAGGAGAATTGGGGCTCCCA | 120 | 54 |
| chr15 | 89523458 | 89523578 | 771629_41866724_SHANK3(58234)_198 | 1 | + | good | GTGCATGTTGCTGGTCAGGGTGTGAGGAGGCCAGACCACTGCGCCCCGCCTGCCCTTCCCTGGTGCCCAAAGCCTGGGTGCCAGCCCTTCTCCAGCTCCTACCTGACAGTTCTGCCCGCC | 120 | 67 |
| chr15 | 89523578 | 89523698 | 771629_41866724_SHANK3(58234)_199 | 1 | + | good | GTTCCCCAGTGCGCCCACCTCTGGGCCCTCGGTGTTTCTTTCCAGCCGCCCGCGTGGGGTCTGCGGGGGGCACCGCGGACTCAGGGCGCGCCGGCCGCCGCCGCCCGCAGAGGGAGGAGC | 120 | 78 |
| chr15 | 89523698 | 89523818 | 771629_41866724_SHANK3(58234)_200 | 1 | + | good | TGGCGGAGGAGGCGGGGCGCGGAGCGGCCGTGGCATGGAGCGAGCCGGCGCGCCCGGGAGCGCAGCCCCGCGGCCCCGGAGCCCTGAGCGGGCGCGGCTCGTGCGCTGGCGGGAG | 120 | 85 |
| chr15 | 89523938 | 89524058 | 771629_41866724_SHANK3(58234)_202 | 1 | + | good | GCCGTGCGCCATGGAGGCCCCGGGCGCCGGCTTCGCGTGCCCGCTGCCCCCCGGCATCGCGTCCGTCCACCTACGTGTTCGTCTACAGCCCCAGTGGGCCCGGCGGATCCGGCCCCGCGCC | 120 | 78 |
| chr15 | 89524058 | 89524178 | 771629_41866724_SHANK3(58234)_203 | 1 | + | good | CGGCCCCGGCCCGGCGTCTCCCGCGCCCCCCGCGCCGCCGCCCCGGGGCCCGAAGCGGAAACTTTACAGTGCCGTCCCCGGCCGCAAGTTCATCGCTGTGAAGGCGCACAGCCCGCAGGG | 120 | 78 |
| chr15 | 89524178 | 89524298 | 771629_41866724_SHANK3(58234)_204 | 1 | + | good | CGAGGGCGAGATCCCGCTGCACCGCGGCGAGGCCGTGAAGGGTGAGGGGCGCGGGCGGGGTGAGGGGCGCGAGCCGGGTTGGGGGGCGGGCCCCGGGGCACGCGGCCTATTTCATTG | 120 | 78 |
| chr15 | 89524298 | 89524418 | 771629_41866724_SHANK3(58234)_205 | 1 | + | good | TTTGGGGGAGGGTCTCTGTGGTTGGGGTGTCCCTGGGTCATTCCAGCAAGCTGCTGTGCCGGGGTCTCGGCCCTGGTTTTATGGGCTCCTGGCATGAAGCTCTAGCGCTGTCA | 120 | 60 |
| chr15 | 89524418 | 89524538 | 771629_41866724_SHANK3(58234)_206 | 1 | + | good | TGGTAGAGGCTTGGCTCTTTTGGGGCGGTTCTTGTTACGATGGGGGCCAACGACAACTGGGTCCTTGCGTTGGAAGGACCCTAACCCGTTCCAGAGCTCCCACGCCTTGTGGGGTTCCTG | 120 | 59 |
| chr15 | 89524538 | 89524658 | 771629_41866724_SHANK3(58234)_207 | 1 | + | good | TGTCACTTCTGGGGCTTCTTCACTGGCAATCCTTGGGATCAGGTGAGGGGGGCTTGTCATCTTCGGGGGATCTTTAACAGTTGTCTGTGTACAGCACACATTTCTTCCATGGATGGGGA | 120 | 52 |
| chr15 | 89524658 | 89524778 | 771629_41866724_SHANK3(58234)_208 | 1 | + | good | AGACAGCTCCCCCTCCCATCATGAGGCTCTCTGCCGTCATTGGGGTCCTACCTTGGGCACTCTGTCAGTGAGGTGGAAATAGTTGTAAAGACCAGCCCTCCTGGTCTATAATGTGGCTTG | 120 | 54 |
| chr15 | 89524778 | 89524898 | 771629_41866724_SHANK3(58234)_209 | 1 | + | good | CTCCACTGGCTGCCTTGCCATGACCCTGAGCTCTGGACAGTGACTCTTGGGTGGATGTCAGTTACCATGGTTGGGGGGAACGTGATCCCTGTGGCAGGAGGAAGAGTATGTCTGGCCTC | 120 | 58 |
| chr15 | 89524898 | 89525018 | 771629_41866724_SHANK3(58234)_210 | 1 | + | good | TGTTTCTGCCTCTGCCCTCTCCCAGTGTGTCCATCTGTGTCTGTCTGTCTCCCACATGTCCATCCTTTGCTGAGGCCATCTGTTCAGTTCTCTTCTGTGTGGATCCTAGAATCTTCCCAG | 120 | 52 |
| chr15 | 89525018 | 89525138 | 771629_41866724_SHANK3(58234)_211 | 1 | + | good | GAAGAATGCTAGGGGGGAATGCTGGCTTGAAAAGAGAGCAGTGGTCAGGGTCGTGGGCAAAACCTGAGCCTCAGGTGGCAGGGAAGCAGGTACCCTGCTGTCCTCTTTTTCCCTGACAAC | 120 | 56 |
| chr15 | 89525138 | 89525258 | 771629_41866724_SHANK3(58234)_212 | 1 | + | good | CTGTCTGGCTTCTCTCCCTAGTGCTCAGCATTGGGGAGGGCGGTTTCTGGGAGGGAACCGTGAAGGGCCGAACAGGCTGGTTCCCAGCTGACTGTGTGGAAGAAGTGCAGATGCGACAGT | 120 | 59 |
| chr15 | 89525258 | 89525378 | 771629_41866724_SHANK3(58234)_213 | 1 | + | good | ATGACACCCGGCATGGTAAGTGACTCCAGTTTTTGCTCAGATACCTCCTAAAGGGCAGGCACTTCTTCTAGTCTCCCTATGGTCTTCTTGGTAACACATCTGGACATCCCCCCTAAGACT | 120 | 49 |

| | | | | | | | | | |
|---|---|---|---|---|---|---|---|---|---|
| chr15 | 89525378 | 89525498 | 771629_41866724_SHANK3(58234)_214 | 1 | + | good | GAGAAGCAAACTCAATAACACTCAAGATACATTCAGATTTGCACATATCCATGGTGGGCAAGCATGCATGTTCTTAGGAACAGGGGTATGTGAACCTGACAGTGGAGCCACCTTTGCCAT | 120 | 45 |
| chr15 | 89525498 | 89525618 | 771629_41866724_SHANK3(58234)_215 | 1 | + | good | GCCAGCTGGAGGAGTACACTCATGGTCATCCCTCTGTGTGCGTGTTGGCACACGAGTCTTCGGTGACAGTGACCACCCCTCTGGTACATTCCTCACGGAAATACCAGGTTCGTGGAACTC | 120 | 56 |
| chr15 | 89525618 | 89525738 | 771629_41866724_SHANK3(58234)_216 | 1 | + | good | CCAATTCATGAGTGGCTGGCACGTAAGGAAGGAACCAGTGTGAGAAAAGAGGCGTTTTTCCAAAGTGGACAGATTGTCCAAGTAGGCAGAGCAGGAGGCATGGGCCAGCTAAGAGGAAAT | 120 | 50 |
| chr15 | 89525738 | 89525858 | 771629_41866724_SHANK3(58234)_217 | 1 | + | good | GAGGCCAGTTGGGCCCCAAGGAGTGGCTCCGGGGTACCTGGGGTTGGTATGGTCTGAGGGCAGCTGATGAAGGATGGCTCCTGTTCTTACAGAAATGTGAGGTGAAGGTTGCAGCTCCTG | 120 | 58 |
| chr15 | 89525858 | 89525978 | 771629_41866724_SHANK3(58234)_218 | 1 | + | good | GGGCCCGCTGAGGAACTGTGGTCCAATCTAGCTGAATGCTTGGACTTCTAGGAAGGATCCCCTTAGATGCTGGTGAAGGGTCCTCTGGGGCCCTCAGTGTGGGCTGCTTTCATGTCTGGG | 120 | 57 |
| chr15 | 89525978 | 89526098 | 771629_41866724_SHANK3(58234)_219 | 1 | + | good | GTGTACTCTTTCATTGCGCCCTCGGGTATTCTGGGAAGCCACAAGATCGATTATCTTCTTTCCAGATCAGAGAGTAGGTAGGACAAATGAACTCTGGGGTGATTATCAGTGGTATGTCTG | 120 | 46 |
| chr15 | 89526098 | 89526218 | 771629_41866724_SHANK3(58234)_220 | 1 | + | good | GATCAGGGTACGAGTGTCCCTTTCTCTGTACATAGAAGGTTTGGGCCCAGGGTCGTGGTAACCTCTGTCAACCTCGACTTTAAGTTGGCAGCTTTCCAGGGGAGCCTACAATCATGTGGC | 120 | 52 |
| chr15 | 89526818 | 89526938 | 771629_41866724_SHANK3(58234)_226 | 1 | + | good | ACTTGGAGAATAGAAGGGGATCGTTGCACTCACTTGCTGGATGTGGCCTCCGGGTGGGAATGATTTCATAGGCCCGTGGCCCAGTTCCAACAGGACCGAGTAAAGAGCAAGGAGAAGGGG | 120 | 55 |
| chr15 | 89526938 | 89527058 | 771629_41866724_SHANK3(58234)_227 | 1 | + | good | CTACACACCTGTTGAATAGGAGCTAGTTGTATGCTTTTCCTACGGATGGCGGTACGGTTTTGAGAACTGTGGATCCAGGATCGTTTGGCCGTCATTCTGAAACCTAGGAAGGAGAGAGGT | 120 | 48 |
| chr15 | 89527058 | 89527178 | 771629_41866724_SHANK3(58234)_228 | 1 | + | good | CTGCGTTGGAGTCAGAATCCAAGAGGAGCAGATGTTGGTCAGTTTTGCTTCAAGCCGGACCTTGTGCTGGCTGTCAGTGGTTATGGCAAGTGTACCCCGTCCCGGGTGCACAGGCCACTG | 120 | 57 |
| chr15 | 89527178 | 89527298 | 771629_41866724_SHANK3(58234)_229 | 1 | + | good | GGAGGGGGCGGGATCCAGTTACCCCTGGATGCACAGGACACCGAGACCGAAGATCCGGGGTTAGCCTGAGGAAGCCAAAAAAAGATAAAAAAAGAAAAAAAAAGAGGCTCAAAGCAAAGG | 120 | 50 |
| chr15 | 89527298 | 89527418 | 771629_41866724_SHANK3(58234)_230 | 1 | + | good | TGTTGTTTCACTTGAATTCCCCGGGTAGTTTACAGATGGGTTCCTCATATTTGTGGAGATCCAGCACCGCATTCTAGAACTATGCCCGTTGCCAACTTCTGTATTTGAGCAGCGCCAGAC | 120 | 48 |
| chr15 | 89527418 | 89527538 | 771629_41866724_SHANK3(58234)_231 | 1 | + | good | ATGAGTTTTAGCATTTTTCTATGAATGGAGCTGTGGACCTATCCTAGCAATCCTGGGAGGTGGGGTGCCAGCGTTTAGTATGTTTCTGGGATGTCACTACCCACAGGGGGCGGGGATGTT | 120 | 50 |
| chr15 | 89527538 | 89527658 | 771629_41866724_SHANK3(58234)_232 | 1 | + | good | TGTTCTTTGTTTTGCTGCATAAGCAGTTAGCTTTGTGAGGAATGGCACGGCTGGAGGGGGGAGCATGAGTCTTAGTCGCCTGGTCCTTGGGGGTCAAGGTAAAGAACGAGATCTTGTAGT | 120 | 50 |
| chr15 | 89527658 | 89527778 | 771629_41866724_SHANK3(58234)_233 | 1 | + | good | AGGAGCTGACCGAGCAGATATTTTAGGGCCAAGATGAAGAAGATCTGGGCAGGGCAGAGTGGTGAGGACAATGTGTCAAGGTCAACAGGAGAGGACCACAGAGCAGCTGGTGTGGGGGTG | 120 | 55 |
| chr15 | 89527778 | 89527898 | 771629_41866724_SHANK3(58234)_234 | 1 | + | good | ATGGAGCTGCAGGCTTTCTAGCAACCACGGGACCTCTTAACAGGGCCATAGCTATCATAGATAACCTGAGGGAGTGGGGTTCAAAAGCACCTAGGTGATTGAGAGCTACGTGTATGACCA | 120 | 50 |
| chr15 | 89527898 | 89528018 | 771629_41866724_SHANK3(58234)_235 | 1 | + | good | TATGGTGAGGGCCCTGCTGGGACTGCCTTTGGGGAGGGTATAGGAGCTATAATGGCTTCCCTCTGACCCCAGCTACAATACCCATCCTGCTGCTCTGGGCAGAACTTCTTGTCTGGGACT | 120 | 56 |
| chr15 | 89528018 | 89528138 | 771629_41866724_SHANK3(58234)_236 | 1 | + | good | TAGGCTGAGTTGTGCCAGATTGCAGAGGGCATCCTAGCAACAGGAACTTGTTTCCAACAGTCCTCCCCCCACCCCCAGCCCTCCAGACTCTGATTCAGGAACCAGAGTGGGAATGAAGGA | 120 | 55 |
| chr15 | 89528138 | 89528258 | 771629_41866724_SHANK3(58234)_237 | 1 | + | good | GGACTATGAAAGACCAAGGATAGAAGCCGTGCTGGGGTAGAGGTTAGGTCAGTGTCTCCTAGGACCCAGGTCCTGGTCTTTACCTGAGCAAAGAGTCTGGCCTCCAGCAGCGCTCTAGCA | 120 | 55 |
| chr15 | 89528258 | 89528378 | 771629_41866724_SHANK3(58234)_238 | 1 | + | good | GGAATTGAGGAATGCAGCTAGTGCACACATCTCAAGGCCTGACATAACCTCCCATCCTGTTTGCCTTCCTCTTTCCACTCGGAGAGAATGTTCCATTTGGGAGCCAGTGCTGGGCACATT | 120 | 51 |
| chr15 | 89528378 | 89528498 | 771629_41866724_SHANK3(58234)_239 | 1 | + | good | CCTTTCAGCTGGCTTTCCTGTGTGTGATTAATAAAACGAGCACACGAATCTGTGTTATATAATAAATAGTTTGTCTGGTTGATAACAATTTAAAAAAAAAAAGAAAACAAGATGCTTTGA | 120 | 31 |
| chr15 | 89528498 | 89528618 | 771629_41866724_SHANK3(58234)_240 | 1 | + | good | AATAGGACCTGACCAGGGCGAGGCTGCTTAACTCTGCATAGCTCCTAGGGTTAAGATCTGGGGAGGGCAGGGCAGAGAGGTGAGTTGGGATGAGCTAAGGCCATGCTAGTGTGAGAGCGG | 120 | 57 |
| chr15 | 89528618 | 89528738 | 771629_41866724_SHANK3(58234)_241 | 1 | + | good | TGGAACCGGAGGCTTCCTGTCCACCGCTGGGTTCCCAGGCCTGGATCAGCTAGCGTAGTGTATCTGTGACCGGGTCAGCCTGAAGAAGGATGTGCGTGGGGTAAGGAGGGATTGAGGAGG | 120 | 60 |
| chr15 | 89528738 | 89528858 | 771629_41866724_SHANK3(58234)_242 | 1 | + | good | GCCGCACACGTGCTGGCGAGGCTTCATCTGCACTGGTCTCGGGACTGGGTGGCAGGTGTACATTTGTCTCTTAGAGTGGGACAGACGGGAAGGGGTAAGCAAGCGAGTCGATGGGGCCCG | 120 | 62 |
| chr15 | 89528858 | 89528978 | 771629_41866724_SHANK3(58234)_243 | 1 | + | good | AACGAGGCTGTTAGAACCTCGTCAAACAGGCTACTAGAGCCCTGTGGCCTCCCGACGTGACGGTTCCTACTGTGGAGAGAGAGAGGTTCAGATGTGACTGATGGGTTAACTTATCGAGAG | 120 | 52 |
| chr15 | 89528978 | 89529098 | 771629_41866724_SHANK3(58234)_244 | 1 | + | good | TCTCGCACAAAGCAAGAGGGACCTGAAGTCTGGACTGGGACTTTTCTTGATAGCCGTGTCATCTGATGGGAAGGTGCTCTAACCATTGCTCCACCCTGTCTCTTCCACCTGCCTTTTAAC | 120 | 51 |
| chr15 | 89529218 | 89529338 | 771629_41866724_SHANK3(58234)_246 | 1 | + | good | CACCCGTCCATCCATCCGTCCATCTATCCCCTCATGGACCTGTATATCCTTCGGCTCATCCGATATTCCCTCAAACCTTCAGCATCCCGCCCATTCATCTATGCATTTACTTTTTTGGGT | 120 | 49 |
| chr15 | 89529338 | 89529458 | 771629_41866724_SHANK3(58234)_247 | 1 | + | good | TATGTTCCTTCTCTTCCTACATTCATCCGTTTGCCCACGCATCCCCCATGCACACACTCACCTAGCCAGTTGTGTGCCTTCTGTTACGTACCAGATGCTCTGCTGGGCACTGGGACCACA | 120 | 53 |
| chr15 | 89529458 | 89529578 | 771629_41866724_SHANK3(58234)_248 | 1 | + | good | GGAGTCATGACATCTGGTCTTTCGCTTTCAAGGATCTCGTCGTTGCTGAGAAGACAGGCCATAAAAGAGCTGTATCTTTCTGCTGGGTATGGTGGCCTCGTGCCTGTAAGCCTAGCGTTC | 120 | 51 |
| chr15 | 89529578 | 89529698 | 771629_41866724_SHANK3(58234)_249 | 1 | + | good | AGAACACTGAGACAGATAATCCTAAACTCAAGGCTCAGACTGTATGCAGTGATACTTTTCTCAAAACAAAACAAAACAAAACACAAAAAAAAACCCCCCAAAACTGGGGGTGGTGGTGGT | 120 | 40 |
| chr15 | 89529818 | 89529938 | 771629_41866724_SHANK3(58234)_251 | 1 | + | good | CAAAGTCGTGACGGGTACTGTAGGACTCTGGTACCCAGCACCCACCTGAATCTCTATGACTCCCACTCCAGGGGATTCAGCTCCCCCATCAGGCCTGTGAATGCACTGGGCATACACGTG | 120 | 57 |
| chr15 | 89529938 | 89530058 | 771629_41866724_SHANK3(58234)_252 | 1 | + | good | GTACACATATATACGTGTAGGGAAACACAGGAAGTAAAAACCAAAATCTTTTAAAAACTCAAAACAGTACGGAACCCAACTTAAAAAAAAAAATCACAGGTTCCTCCTGTTGTCTGTCTG | 120 | 35 |
| chr15 | 89530178 | 89530298 | 771629_41866724_SHANK3(58234)_254 | 1 | + | good | CCTTGCCTGCCTCACCCCCAGAAGCCCTGCACCCTCACATCTACAGGCGGCTCCCCTGACGGTCGGGGTCTCTGCTCAGCTCTTCTTTCTGATGAGGCTGTTGCTATCCCTAGACTCGTC | 120 | 61 |
| chr15 | 89530298 | 89530418 | 771629_41866724_SHANK3(58234)_255 | 1 | + | good | CCCCTTTTCCTGCTTGCTTTTCTCCTGTTAGCTGGCGTTTTGTATTCATTGCTTGTTGATTCCATGTCTCAGAATGGCTTATTAACCCCTTAAAGGCGGAGACTTTGCTCTGTTCTAAAA | 120 | 42 |
| chr15 | 89530418 | 89530538 | 771629_41866724_SHANK3(58234)_256 | 1 | + | good | TAAACAGTGCCTGGTCGGTAAATGACTGTTGAAGTAAAAAGTAAGAAGGGCTATGCTATGGCAAGAGAGGGTGAAGGAGGGGAGAGATGCTGAGGAATGAAATTGAAAACTGTGGTGGCG | 120 | 46 |
| chr15 | 89530538 | 89530658 | 771629_41866724_SHANK3(58234)_257 | 1 | + | good | CCATCTTGAGGCCATCTGTGGTACTGCAAGTGCTGCTAGAGGAGCCATTCTGAAGCTTCAAGGGTCTAGTTGTAGATTTGAACCTTCTTCAGGAAATCTACAAGGCTAAGTGAGATTCAG | 120 | 46 |
| chr15 | 89530658 | 89530778 | 771629_41866724_SHANK3(58234)_258 | 1 | + | good | CAAGAGGTATGTCCTGGCCAGGTCCACAGTGTTCACGAAGGACTTTGAGATCAAAGGACAGATATGTATATAGTCCTGTGGAATACAGTCAGGGCAGGGGACTGAACACTTTATAATCCT | 120 | 46 |
| chr15 | 89530778 | 89530898 | 771629_41866724_SHANK3(58234)_259 | 1 | + | good | TTTTCTTCTTTCCTTGAATTTGTGCTGTTGTGAGGAAATCGGTGGACACCTTGGGATGCTTCCTTCCTGGTGCACACATAGGCTGGTAGCACAGCACACACATGTGTGTGAGCTCAACCC | 120 | 49 |
| chr15 | 89530898 | 89531018 | 771629_41866724_SHANK3(58234)_260 | 1 | + | good | CTGTGCACACAAACACATGTTTGTTACTAGATCCTGAGCTCATGGACGGAGAGCAGCCATGGCTCTCGTTTGACACACATAAGCACCAACCCAGCATTTATACCTGACTGTGAAGCACAC | 120 | 48 |
| chr15 | 89531018 | 89531138 | 771629_41866724_SHANK3(58234)_261 | 1 | + | good | ATTCATGTACTCTCATCTCTGAGGCTTGCAGACGGGACCATGCTGGCCTTGTATGGTGTCTAGATGCATGCTAAGTAAGGAGAGATGAGCACCATTGGTTTCCCCTTGGCTCTAACTCCC | 120 | 50 |
| chr15 | 89531138 | 89531258 | 771629_41866724_SHANK3(58234)_262 | 1 | + | good | AGAGGCCAGAATTCTATGTATGGTTATCTGAGCTCTACTCCCTTAGGACTTTAGGCCCCTGGGTCTTAGGATTTGCCCCCACCCCCATCCCCACTCACATCGAGGCCCAGAGAGTGTGGG | 120 | 55 |
| chr15 | 89531258 | 89531378 | 771629_41866724_SHANK3(58234)_263 | 1 | + | good | GGCAGAGACTGATCAGCGCAGTTGGGGGAAATTCAGCTAGGTGGAGGGTCACGACTGTTTCTTAGCCTTTGATGCTCCCCGCAGAAACCAGAGAGGACCGGACGAAGCGTCTCTTCCGCC | 120 | 57 |
| chr15 | 89531378 | 89531498 | 771629_41866724_SHANK3(58234)_264 | 1 | + | good | ACTACACTGTGGGTTCCTATGACAGCCTCACTTCACACAGGTACTGGCAGGGGTGTGGACACGCAGGGACAGTAGCAGGGCCTAGGGTGTCCAGTGAAAGGGGGAAGTCAGGGTATGGGG | 120 | 58 |
| chr15 | 89531498 | 89531618 | 771629_41866724_SHANK3(58234)_265 | 1 | + | good | AGTGGGTGGCTAGCTTCCTCTGCCTTTGTCATTCTGTGTGGTCTGGTGTGTATGGGACCCTGTGGGTTCAGCCTGACTGCCAGGCTCCCTGCAGACCTCCCAGCTCAACCTTGCTTTCCT | 120 | 57 |
| chr15 | 89531618 | 89531738 | 771629_41866724_SHANK3(58234)_266 | 1 | + | good | TTGCCAGCGATTATGTCATCGATGATAAGGTGGCTATCCTGCAGAAAAGGGACCATGAGGGGTTTGGCTTTGTTCTCCGGGGAGCCAAAGGTAAAGGGTGTGAATTTGGGGGGTATCGCA | 120 | 51 |
| chr15 | 89531738 | 89531858 | 771629_41866724_SHANK3(58234)_267 | 1 | + | good | TTGCTGTCAATGTCTTCTGTGGCCTGCCTCTGCCCCTTTTCCCCTTTTAAAGCCCTCTGCTTCCCCCCCACCCCCCGTGGTTATATGTCTTGGTTTGGAACATACTGGGGTACAAGACAG | 120 | 53 |
| chr15 | 89531858 | 89531978 | 771629_41866724_SHANK3(58234)_268 | 1 | + | good | TGGATCCCTCTCCTTTTAGCCTTTCATTTCCTTGTGGGGATCTCTGCTCCTGGTGAAACAGAGATCTTTTTGCTGAGGGTAAATGTGGAAGTTCACAGAGGGATGTACAGTCATTTGGAT | 120 | 45 |
| chr15 | 89531978 | 89532098 | 771629_41866724_SHANK3(58234)_269 | 1 | + | good | ATCCCTACCTGCTGCCTCTCCCACCCCATTGCTATGCCTTAGTAGTCAGGTGGCTTCTTGGGGCTCCCCGACATGATCTATGCTTGTGAATGCCTTTTATCTTGCTCTCCCTGCAGCAGA | 120 | 53 |
| chr15 | 89532218 | 89532338 | 771629_41866724_SHANK3(58234)_271 | 1 | + | good | GGTGAGGCCCGTCCCGACCCTGTCCAGCAGGAGGGGGCGAGAGCACTACTAACTGCCACCTGAAGAGGGGACAGCTGGCCTCTGACTACCTAGCTGCTTGTCTATCCCAGGTGAACGGAG | 120 | 62 |
| chr15 | 89532338 | 89532458 | 771629_41866724_SHANK3(58234)_272 | 1 | + | good | TGAATGTCGTGAAGGTTGGACACAAGCAAGTGGTGGGTCTCATCCGTCAGGGTGGCAACCGCCTGGTCATGAAGGTTGTGTCTGTGACCAGGAAACCCGAGGAGGATGGTGCTCGGCGCA | 120 | 57 |
| chr15 | 89532458 | 89532578 | 771629_41866724_SHANK3(58234)_273 | 1 | + | good | GAGGTGAGGGTTCTGATGGCTGGGCTTTCAGACCTTTGACTTCAAGCCCACATTCCTCTTCCATTTCTGATCCTACACAGCTGACATCACATTGCTGCCTGAGATGAGATAAAATGCCCT | 120 | 48 |
| chr15 | 89532578 | 89532698 | 771629_41866724_SHANK3(58234)_274 | 1 | + | good | TATTTGCTGCACAGGTCCTTGGTGTTAGATGCCTGACCTCTGACCAGGCCTCTGATCCCTGATCCTAGATTACCCCTCCTCAGTTCCCTCTCCCCTTCTCCTGTGAATCCCCCCAGCCCT | 120 | 56 |
| chr15 | 89532698 | 89532818 | 771629_41866724_SHANK3(58234)_275 | 1 | + | good | GGTGTGGCAGAGCCCTCCTTTCCGACGCCAGCAGCTGCCTGGAGGCCGGGTTGTCATGGCAACCGTGAGGTTGACGCTGCTGCTGCTGCTGCTTATTGTGCAGGGAAGGGGGAGGCGGCA | 120 | 65 |
| chr15 | 89532818 | 89532938 | 771629_41866724_SHANK3(58234)_276 | 1 | + | good | CCTGAGGGAAGAGGAAAAGCGTCTTCTCCCCGCGCCGACGCCAAAGACGCGGCACCGCCTGGGCCTAAGAGGCTTTGCCGCCGGCCCAGCCCAGGTGTGCCGCCTGCCGGGCCACACTTT | 120 | 69 |
| chr15 | 89532938 | 89533058 | 771629_41866724_SHANK3(58234)_277 | 1 | + | good | GGTCTTACAAAGAGCAGTGGTACCTGGGTCTGGGTGCTTTAGGCGGCAGGAGGTAGCCTGGCAGGAGATCTACCCAGCCCCCATGGGTGGTGCCCACTCACTGGGGCCTTGGAGCCCGCA | 120 | 63 |
| chr15 | 89533058 | 89533178 | 771629_41866724_SHANK3(58234)_278 | 1 | + | good | GGGTGGATCCTCAGAGCAGAACCAGCCAGCATCCTAGCCTTTGCTGGTGCTTTTATTACCATTTGAGGCCTTGGTCCTGGGAGTCTGAAGAGCCCTAGGGGCACAACTGCCCTGGCTCCA | 120 | 57 |
| chr15 | 89533178 | 89533298 | 771629_41866724_SHANK3(58234)_279 | 1 | + | good | CTGCTCCCTGAGTGCCTCACGCCATGAAGAAGTTTGCATCTAGCCGCAGCCTGAACAAGATCTTGGCACAGTGTGACTCCTCTTCCAGAGAGTACGAGGAGGTCCAGGCAGTGGAGCGCA | 120 | 57 |
| chr15 | 89533298 | 89533418 | 771629_41866724_SHANK3(58234)_280 | 1 | + | good | AGTGGCATTTGCACCTGGCCACTCCGCGCCGCCTGCTGCTGGACCGGAGGGCCAAGGCCTCCCTCTTCTTTGCAGCCCCACCACCCCCAAAGAGGGCTCCCAGCACCACGCTGACCCTGC | 120 | 68 |
| chr15 | 89533418 | 89533538 | 771629_41866724_SHANK3(58234)_281 | 1 | + | good | GGTCCAAGTCCATGACGGCTGAGCTCGAGGAACTTGGTGAGCATTTCGGGTGGGTGGACAGTGGGTGAGTGGCCAAAGTGGGGGTGGCTGCAGGTAGCATGGGAGGGCGGTGGGACTGA | 120 | 62 |
| chr15 | 89533538 | 89533658 | 771629_41866724_SHANK3(58234)_282 | 1 | + | good | CTGGGGGCTCAGACAGAAGGTAGAAAAGATGGGTTTGGAGACAGATCGGCAGTTAGGGGTGGGACTGGAGTGATGGGGAAAAGTGGGTACCTTGGGTGAGGAGTGTGACAGCAAGGGAG | 120 | 56 |
| chr15 | 89533658 | 89533778 | 771629_41866724_SHANK3(58234)_283 | 1 | + | good | AGATGAGGACTCGGCAGCATGAGGAGCCGGAGTTGGTCAGGGACAAGGGGGACAGATGGCTGGGGAACAAGGTCTGGCCCTGTAGCATGCCCGCATGCATGCACATGCATGTGTGTGTGA | 120 | 59 |
| chr15 | 89533778 | 89533898 | 771629_41866724_SHANK3(58234)_284 | 1 | + | good | GAGCTGTCCTCGGTTCTGTGCTTTTCCACTAAGACTTCGCATTTTGATTTAGCTGGCAGAGAGCAGTGGTAGAGCAACCTAGAGTCTGCTTCTGGATCTGCCTGGCACGAGCCGGGTCAC | 120 | 53 |
| chr15 | 89533898 | 89534018 | 771629_41866724_SHANK3(58234)_285 | 1 | + | good | CTTTGTGCTTATTTTTCTCGGTTGGTACCTGCTCACTTATCTGCCGAGTGAGTCTACTGACCTAGTAGAGGTATCTGTGAGCACCCCTAACACGGTGAGGGGGTGGGGCCTCGTGAACTG | 120 | 52 |
| chr15 | 89534018 | 89534138 | 771629_41866724_SHANK3(58234)_286 | 1 | + | good | CAGGAGGTGGACAAAAACGTATGTTAGTCTTTAATGGGTACCCCATCCTGACCTAGGGCGGCTGCCAGCTTTTATTATGGCCATATGCATAACACATATCTCGTATGCATGCGCACATGT | 120 | 47 |
| chr15 | 89534138 | 89534258 | 771629_41866724_SHANK3(58234)_287 | 1 | + | good | AGGACAGTTTCCAGCCTTGTTATAGGCATGTCTCATGTACATGACTATGATGCTTTGACATGTATAGAAGTGTCCAGCCTCATCATGGGACGTGTCCCATGTACATGAGCCGCGCATCTT | 120 | 47 |

| | | | | | | | | | |
|---|---|---|---|---|---|---|---|---|---|
| chr15 | 89534258 | 89534378 | 771629_41866724_SHANK3(58234)_288 | 1 | + | good | GACAAATATATGAGCTACCAGCCTTGTCGTGGGCACACGTCCCATATGCGTGACATATTTTGACACGTATGCCAGTTAACCTTGTCATGGGCATGTCTCATATTCACAGCCATGTCCTGA | 120 | 47 |
| chr15 | 89534378 | 89534498 | 771629_41866724_SHANK3(58234)_289 | 1 | + | good | TATGTGACATTTGCCAGCCTCGTCATAGGTATGTCCTACATGCATGATCACATCTTGACACATAGGACTGTTATTAGTCTCATCACAGGCATGTGCAGATATATGTGATTTTATTGTCAT | 120 | 39 |
| chr15 | 89534498 | 89534618 | 771629_41866724_SHANK3(58234)_290 | 1 | + | good | TGTGGGCATGTGCTAATGTGTTTGACTGTTAAGAGCCTTATTGTGGGTGCATCGTAACTGTCAACAGTCCCGTGGTGGGTATATTCCAATATGTGTGACAGATTGAAACACTCTTTGCTT | 120 | 43 |
| chr15 | 89534618 | 89534738 | 771629_41866724_SHANK3(58234)_291 | 1 | + | good | TTGTGGTTTGAGCCTAGTCAAATTCCCAGACTGTCTTCTGGCTTTTGTCCTAAAAGGAAACCCGTATCCTGGCTCTCAGGACTATCCCTGTCTTTTGCTTATGGACTCAGTGTTGACCGT | 120 | 47 |
| chr15 | 89534738 | 89534858 | 771629_41866724_SHANK3(58234)_292 | 1 | + | good | GTCCTGACCATAGGCTGAGTTGCTACATTCCAGGCAATTGCCTTGACTGAAGTTTGGTTTGGTTTGGAAAACAGTTACCAAGATAAAAGGGGCTGACGTGAGCAGGATTTTCTACGATAA | 120 | 44 |
| chr15 | 89534858 | 89534978 | 771629_41866724_SHANK3(58234)_293 | 1 | + | good | ACCTCCGAATAAAATAGAAATTACGGATAGTGTAGAGGAGGAGGAATCCTGTGGTTTCAGAAATAAAGCAGGAAACCAAAGCCAAAGGGATAGGAACCAGGTACTTCCAGTTCCTGATGG | 120 | 43 |
| chr15 | 89534978 | 89535098 | 771629_41866724_SHANK3(58234)_294 | 1 | + | good | CGGCTCTCAAGGGCGGTGCTGTGGTCTGAAGCCCCTGTGAGGCTTGAAGAGGTGCTATGGACTTGGAAAAGCCAGAAGCAACCATAGAATCTGGGATTAGCATAAAAAAAGGAGGGGCCG | 120 | 53 |
| chr15 | 89535218 | 89535338 | 771629_41866724_SHANK3(58234)_296 | 1 | + | good | TGAACATGGTTACGCACACATACATGCAGACAGAAACACTCAAAATAATCTAAAAAAAAATTTTTTTTGAAAAGGACTGGAGATAAAGAGCATGAGGGTAAAGTAAAAGGTTTCAGTTA | 120 | 32 |
| chr15 | 89535338 | 89535458 | 771629_41866724_SHANK3(58234)_297 | 1 | + | good | TATAAAGGCAGGAAACGTCACAGAATTAGTGAAATTAGCAGGATAGTGCGAACAACAGCAGCAAGGAAGTTTGCGATGGTCCTTAACTTATTGAAACAAACAAACAAACAAACAAACAAA | 120 | 37 |
| chr15 | 89535458 | 89535578 | 771629_41866724_SHANK3(58234)_298 | 1 | + | good | CAAGAAAGCTGCACACAGAAGAGTCTGTGGCTTGGAGACACAGAATCCAGGACTTACTAAAAAAAAAAAAAAAAGCAAAGCCATAGCAGTGAGCGAGGAGATCAGGCTGAGACCACCCAGA | 120 | 46 |
| chr15 | 89535578 | 89535698 | 771629_41866724_SHANK3(58234)_299 | 1 | + | good | ATCAGCTTAGGAAGCTGATGAGATGGCCATGAAGGACCAGTTAGGGGACCTGAGGACAAACACGGAGACCTCAGCATCTACTTGATAGGAATTCCAGAGGAAATGAATAGAAAGGCTAAG | 120 | 47 |
| chr15 | 89535698 | 89535818 | 771629_41866724_SHANK3(58234)_300 | 1 | + | good | AATCTCCCAAAAGCAAACAGATATTATGAGACTCCTTTCTGAAGAAGTATGTCACAAATAAATTATGATCTGTCAGGGCAGAAATTTAAAATGCCACCAACAAGCAGAAAGATGTTTAAA | 120 | 33 |
| chr15 | 89535818 | 89535938 | 771629_41866724_SHANK3(58234)_301 | 1 | + | good | AATACCAGCGCCGGTAGAGGGAGTCAGCGGCTCAGAGTGAGCGATGATGGTTTGACTTTAGGCTTCTTCAGCAGTGATAGAGACAGAAGACGATTAAAGTTATCCTAGCCAAACTGGCTG | 120 | 49 |
| chr15 | 89535938 | 89536058 | 771629_41866724_SHANK3(58234)_302 | 1 | + | good | ACTAGAGAGAGCAGAGACACCCACAGACTTCAGGCTCTAGGGTCTAATCTAACCCAGGTCAGTCTTAGAGATCAGACCTATGGCATAGATGGATAAACCAATATGGTACACAGCTCGGTT | 120 | 47 |
| chr15 | 89536058 | 89536178 | 771629_41866724_SHANK3(58234)_303 | 1 | + | good | TGACTGGACCTTGGTGCACAGTCCTGGAGTCTTGCTAGAAGCTGTTCTAATACCAGGTCAGAAGGGCTCTTCTTACTTCCTACTAAGAGGTGTTATCCATCCTGAACTCTGAACTCCAGG | 120 | 48 |
| chr15 | 89536178 | 89536298 | 771629_41866724_SHANK3(58234)_304 | 1 | + | good | ACCCTAGAGGCTTGCTGTGGGAATGGAAGAGCTCTGTCGGGCCGGGCTATGGCTTTTGCCTGTAGAAATGGGAGCAGAGGAGTTGAATCTTTTCCTCTGCTCCAGAACTACAACTGTAGT | 120 | 52 |
| chr15 | 89536298 | 89536418 | 771629_41866724_SHANK3(58234)_305 | 1 | + | good | TGTAACTACAGTTACAGATCTTGCTGGCCCAGAGGTATCATGGTATTTCTGAGTAGAGGCTAGAAGGGGTCTGGGAACAGCTCAGCTGGAGCCTCTGCCTATACTTCTGCGCCCAGGATC | 120 | 52 |
| chr15 | 89536418 | 89536538 | 771629_41866724_SHANK3(58234)_306 | 1 | + | good | ATAGCTATTGAAACTGAACCAAGGTTCTAAATGGGCCTGTATTTAGAGAAAAGCTACCCCGACACAAAGACAACTATCTTGGGAGGGGACCTGGTCCTTCTGGGAATGTCTGATTGGAGA | 120 | 45 |
| chr15 | 89536538 | 89536658 | 771629_41866724_SHANK3(58234)_307 | 1 | + | good | AAGGCAGACCAGCATAGCATTCTCCAAATCGCTGAGCATCCCACAATCCACTGTCTTGTTCTTTTTCGTTCGTTTGTTTGAGACAGGGTGATTCCGTAGTCTAGGCAGCTCTTGAACTGC | 120 | 48 |
| chr15 | 89536658 | 89536778 | 771629_41866724_SHANK3(58234)_308 | 1 | + | good | CTGTATTTTTTTGAGTTCTAGGATTGCAGGCATGCTCTATCATTCCTGGATTTGTTTGTGTTTTTGAAAGGTAGAAATAAGACATATTGAGTCACTACTGTTATAAACATGCCTATCCTG | 120 | 35 |
| chr15 | 89536778 | 89536898 | 771629_41866724_SHANK3(58234)_309 | 1 | + | good | CCAGTCACTGGCAGTCTCTTTAGGAGCCATGTTTGTCAAAGAGGTAACCTGAGCACTTGGGCAGAAGAAGCCAGTGTGTGGTGAGGGGGAAGTGGGTGCAGGGGTTGAGGTTCATGCTGT | 120 | 54 |
| chr15 | 89536898 | 89537018 | 771629_41866724_SHANK3(58234)_310 | 1 | + | good | GGAGAGGAGGAGGCAGGGCAAGGTAGGAGGAAGAAGAGAGTTGAGTTCCTCAGAACTAATGGAGTGAGCAGAGGCTGGTCCAAATTAAATTAATTAATTAAGTAAAATGGAAGCGGTGGC | 120 | 46 |
| chr15 | 89537138 | 89537258 | 771629_41866724_SHANK3(58234)_312 | 1 | + | good | GTCAATTTGGGTTCCACATCTCATATTAAGAAGGGTTGTTCATAAGAGGTGGGGGGAGAGGGTGGGGAAGAAGCACAGAGCACCCTGTCTCCATTCTTCTTGTCTGGGGCTCCCTTACCA | 120 | 51 |
| chr15 | 89537258 | 89537378 | 771629_41866724_SHANK3(58234)_313 | 1 | + | good | AATATGCCTATGGTCAGCTTGAGTCAGGGACATGGGTTAGCAGGGACCCACATCAGGGACCTAGTGTGGGCAAGCTGTCCCTGTTCATTTATTTAGGTGAAATTAAGACTTCCCTCAATT | 120 | 47 |
| chr15 | 89537378 | 89537498 | 771629_41866724_SHANK3(58234)_314 | 1 | + | good | CCTGGGTCTTTTCTTTTGTGATAAGAGGTTGGCCTAGGTCCCTGCCACTGCCTTGGTCTGGAGAAGATTTCTTTACCAAATATGGACCCTGCTTGCCGTTACACACCTGCTTTGGGATAT | 120 | 48 |
| chr15 | 89537498 | 89537618 | 771629_41866724_SHANK3(58234)_315 | 1 | + | good | GGAGGTGTGTATCCTTAGGGCGGCCCTTGCCCAGCCTGTTCAAAGGCCCTGGGTTCCATCTGCAGTACCGCAAAAACAAAACCCAATCCCCCCCCAAAAATCCCAGTGCGGAGTGTTCAA | 120 | 56 |
| chr15 | 89537618 | 89537738 | 771629_41866724_SHANK3(58234)_316 | 1 | + | good | AGGTGACCACCTTGGTGAATAGTGGACTTGAACCTTTGTTTTTGTAGTTCTATCTTTGTTCTGAACAGACATCTGGCTGCTTCAAGTACAGTTTAATTGTGCACATCTCCGTGTGTGCAT | 120 | 42 |
| chr15 | 89537738 | 89537858 | 771629_41866724_SHANK3(58234)_317 | 1 | + | good | GTGTGCACATGTGTGTGTGTGCATGCTTGTGAGTATTCGTGTGTACACGGCGTGTCTTCCTGCTTTTGCTTGTTCTCCTCTAGAGGTTTAGCGCATCCGTAGGCTTTGTTCTGCTGGTCT | 120 | 50 |
| chr15 | 89537858 | 89537978 | 771629_41866724_SHANK3(58234)_318 | 1 | + | good | CAGAAGGACGTACAGATGGGGACAGGTCTGCAGGACCTGGTTGTTGGGGAGGTGGGGAAATGGGGAACAGGATCCAGACATCATTGTGCAGCATGGCCCAGGGGCCGTGCCACGATCTTC | 120 | 59 |
| chr15 | 89537978 | 89538098 | 771629_41866724_SHANK3(58234)_319 | 1 | + | good | CTCTAAACTCCCTTTGCTCTGTTTCTCAATTCTCAGCTTCCATTCGGAGAAGAAAAGGGGGTAAGTCTGCTGCTGTGATTCTAGGCCTGGCTTGCCTGTCCCGACCGAGGGGTGCTGGTG | 120 | 53 |
| chr15 | 89538098 | 89538218 | 771629_41866724_SHANK3(58234)_320 | 1 | + | good | TGTGATGAATCCAAGTCAGAGTGGTCTGGAGTCTGGCATACTTATCTCGCTCCAGCTTGGGGATCTGGTAGAGACATGGTAAAGAAGTTCCCAGGGCCATCCCCAAGAGTTATAGGGAGC | 120 | 51 |
| chr15 | 89538218 | 89538338 | 771629_41866724_SHANK3(58234)_321 | 1 | + | good | ACAACCATAGTCCTTGTGGGTTTAGGTGAGCTGGGGGTGTAACCTCCGAAGATAGTGTGGTGGGGAGAAAGACCCAGAGCTGGGGAGGGGATGGCCTCTGGGGGTCAGGCCGTGGAGTTT | 120 | 58 |
| chr15 | 89538338 | 89538458 | 771629_41866724_SHANK3(58234)_322 | 1 | + | good | GGAGGAGTGAGGTGCTTGGGCTGTGTTGGGATGGGAGTGACAGGGTGTTCAACTCTGCTCTGTGCTGTATCTTTTATCTCCTCCTTGCCCTAAGCCAGTTTCCTGCTGGTGAGTTGTAGC | 120 | 53 |
| chr15 | 89538458 | 89538578 | 771629_41866724_SHANK3(58234)_323 | 1 | + | good | CCAGAGCAAAGCTGCTTTGGGTAGGAGTTTGAATCTGGTGGCTCTGGAAAAGGAGCTAAACTCTCCCGCTCCGAACCCCCACCCCACCCCTGGGCTGTTAAGTTCACCATCTCTTCTTCA | 120 | 54 |
| chr15 | 89538578 | 89538698 | 771629_41866724_SHANK3(58234)_324 | 1 | + | good | GGGTCCACCCCTGTGTCCTCTGAGGCCCTTTGTCATATGGTTTGTGTGTCCAGCATGGCTGCAGGGAACACAGTGTCTGTGTGGGGTCAGAGACACCAGGGCTGGGACTGGGTACTGAGG | 120 | 59 |
| chr15 | 89538698 | 89538818 | 771629_41866724_SHANK3(58234)_325 | 1 | + | good | TCCAGCATCTGGGGAAGTCTTACCTGGCTTCTGATGTCACCTCTGCCTCTTGAGTGGATGTGAGTCAGGCTCAGATATACCCAAAGCCTAGGCTGTGTGGTTTTGTGCATCCAGCATCCC | 120 | 52 |
| chr15 | 89538818 | 89538938 | 771629_41866724_SHANK3(58234)_326 | 1 | + | good | TCTCCCTCTCCCTCTTACTTCTGATGTCCTGGACTCTGCCTGGGTAGAGGGACCAGCTCTGGCTCATGATGGGCATGTGACCAGTAGGCTCCCCTCTAATCTCTTCTTGTTAAGATCGGT | 120 | 53 |
| chr15 | 89538938 | 89539058 | 771629_41866724_SHANK3(58234)_327 | 1 | + | good | ATCCTAACTCTGTGACAGTCGGGCTCCCAGGATGCTAGGCTATAGTCATGCACTTCTTCCAGATGTGGGAGAAATGGCTGATTTGTCACTTCCATGGACCCTTCTTTTCATCCCTGCTTG | 120 | 49 |
| chr15 | 89539058 | 89539178 | 771629_41866724_SHANK3(58234)_328 | 1 | + | good | GGTTCCCAACACACTCTGCCCTGCACCCAAGATGAAGTGGGCACGGAAGGCTGATGGGAATGCCCACACTTGGTTCCTAAAGGATTTTATTTAATTTTTAAATATCTAAAATCATTTAGG | 120 | 42 |
| chr15 | 89539178 | 89539298 | 771629_41866724_SHANK3(58234)_329 | 1 | + | good | ATTATTTTTTTCTAATTCTGTAGTGGATTTATGTTGGGGAAAAAAGTTCAACTTTTCATATGTAATTTCAGTTTTTCTAGTTTTTAAATTCTTCCTCTTGGAAGTAATTCTGTCTTGGCC | 120 | 28 |
| chr15 | 89539298 | 89539418 | 771629_41866724_SHANK3(58234)_330 | 1 | + | good | ACAGCATTATTGCACCAGAACCCTAAACGGTTGCTTTGTTAGACGTTGTGTGTAAACTGCTCGGATCTCTTGGATCTGTTAGGTTTCTTAGAGACACACAGTCTCTTAAGTCCACGGCTC | 120 | 45 |
| chr15 | 89539418 | 89539538 | 771629_41866724_SHANK3(58234)_331 | 1 | + | good | AGGTGACTGTGCCTTGGTAGGAAATATCACTGTGTCATGAACTGCCCTTTCTCCTTCTGTCTTTACTTCTGTGGTTTCTGGACTGGATGGTAGTAAGGGTCTGTGATCTGGGGTGATGCT | 120 | 48 |
| chr15 | 89539538 | 89539658 | 771629_41866724_SHANK3(58234)_332 | 1 | + | good | CTGTTGGCGACATTTCGTGCGTGGGAAACAGGAGAGCAACGATAGGGTATTGAGGTGGTGACAGCAGTAGCCATGGAAAACTTGGCACATACTGCACAGCACCAAGGACTTGGCCAGGGT | 120 | 53 |
| chr15 | 89539658 | 89539778 | 771629_41866724_SHANK3(58234)_333 | 1 | + | good | GGTGGTTTCACCTGTTCAAAAGGAAGTTAATCATTTCATTGCAAAGACAAATGAGAAGACCTTTTGCGTCTGTCTCTGCATTGTTACCTGGTGGTACTGAAAGCCATGCTTGGACCCTAC | 120 | 43 |
| chr15 | 89539778 | 89539898 | 771629_41866724_SHANK3(58234)_334 | 1 | + | good | CAATTTCTCTACAGTGGGAACTTGTTTTGCTGGCTAAGTCCTATATTGTTCAGTCCTCCAGTAGGGTAGAGCATTCTCTTTCTGAGGAACCCATGAATCATTGGTGAAGATAAGCTAAAT | 120 | 41 |
| chr15 | 89540138 | 89540258 | 771629_41866724_SHANK3(58234)_337 | 1 | + | good | ACTGCTCGTGACTATTTTACTAGATGAAGGCACTGGTTCTCAGGGGTATACATCTAAACAGAGTCAGTGTACAAGGTGCGAGTCTCAATACGCTATACAATAAGAGAGCCTCAGGAGTGA | 120 | 44 |
| chr15 | 89540258 | 89540378 | 771629_41866724_SHANK3(58234)_338 | 1 | + | good | GTTCACTTCTACACAACAACTGTTCGGTTAGAGACAGTAGGTGCTGGTGAGCAGCTGCGGTTTGGGTTTGAAAAGAGTAAGAGATGTCCTGGCTGACCATAGACGTTCTTCAGCAGTGGT | 120 | 48 |
| chr15 | 89540378 | 89540498 | 771629_41866724_SHANK3(58234)_339 | 1 | + | good | TCTTGGAGAATGGAAGGGTGTTTAGGACAAATGTCCCCCAATGGTAGACACAGCCTTCAAGAATTGGATATGGTCTGGGGACGGGTGGAGCCTGAGGCTTCTGAGGGAGAGGAAGAGGAA | 120 | 52 |
| chr15 | 89540498 | 89540618 | 771629_41866724_SHANK3(58234)_340 | 1 | + | good | GGCACTGCCCAACCTATAGAGCACGCCGGTGCCCTGACCCCAAGTGTACTCTGAGCAAGAGTTATAGAACCATATCTCAGACAGCCTTCAGACGCTTTAAAGGTCTCTGAAACCCTTTCC | 120 | 52 |
| chr15 | 89540618 | 89540738 | 771629_41866724_SHANK3(58234)_341 | 1 | + | good | CCTGCTGAAATGGGTTTCCCTTGTAGTTCAGTCTGATCTCAAACATGTCGTCTGGCCTCAGCCTCTCAAAGACTGGGATTGCAGGATTTCCTTTCCTTCTGCCATGAGAAAACACAGTGA | 120 | 48 |
| chr15 | 89540738 | 89540858 | 771629_41866724_SHANK3(58234)_342 | 1 | + | good | AATTACATGCTCAAGGCAGGAAGAGCTTACTTATATTTGGCTCTCAGTTTGAAGGCTTTTCCATCTCTAAATCAGGAAGAAGAGGGTGGTCAACATTCGGTGTTTTGTTTGTTCGGTTTT | 120 | 40 |
| chr15 | 89540858 | 89540978 | 771629_41866724_SHANK3(58234)_343 | 1 | + | good | TTTTTTTTTTTTTTTTTGAGCAGTTGTGAAGGAATGAGAGCTAAGGAAACGGCGCTGCTGGAGTTTGGGGTGGGCCTTCCCATATCAATCACCATCATCTTGAGAATCCCTCACAGGTGT | 120 | 44 |
| chr15 | 89541098 | 89541218 | 771629_41866724_SHANK3(58234)_345 | 1 | + | good | TTTCTTCCTCTCTTTTGAGGAGAGTGTGTGGTGTACCTATGTGCACGCATGTGCGTGCACCTGTACAGGCACGTGTGTAGGCCAGAGGTCTGCCTGCTGAGTGTCTTCCTCAGTCATTCA | 120 | 52 |
| chr15 | 89541218 | 89541338 | 771629_41866724_SHANK3(58234)_346 | 1 | + | good | TTCTCCATCTTACCCTTTGAGACAGGCTCTTTCACATCCACAGCTTGCCATGTGAACTAGACTGTCTGGCCAGGGAGTCCCTAGGATCTGTCTGTTACAGACATATACTACCACGCTCAG | 120 | 49 |
| chr15 | 89541578 | 89541698 | 771629_41866724_SHANK3(58234)_349 | 1 | + | good | GCAGTAGGTACTTTACTCACAGAGCCACCGCCACAGCCCCCAGAGTCCCATTTTAAAACCAGTAACTTTAATAAAAAAGTAAACCCGACTCCACAAAGGTTTTTATTTTATAGCGGATGT | 120 | 42 |
| chr15 | 89541698 | 89541818 | 771629_41866724_SHANK3(58234)_350 | 1 | + | good | GACAAGAATATGCCCGAGGCCGCCTGGCTCCTCTCAGCCAAGCCCTTCTTAGGCATCAGCCTTAAATAGCTCTGAAGTCTGAGGAGTTTGAGTAAGCTTTTGGAGAGGAGAGTCCTTTCC | 120 | 52 |
| chr15 | 89541938 | 89542058 | 771629_41866724_SHANK3(58234)_352 | 1 | + | good | CTGAGTACCGAGGCCATTTTAGATCTCCAGCCTGAGTTTTTTTTTTTTCCTTAGGAAATAGCTCCCCTCTGCCAGGCCAGCTCTTGGTTATTATCTTATTTACCCACTCAGAATATCTTG | 120 | 42 |
| chr15 | 89542058 | 89542178 | 771629_41866724_SHANK3(58234)_353 | 1 | + | good | CTTGAATGTTAGTGTCTGTCCCTGTGGCGGTGGTCCTACTAGAAGGAGCTTACCATGGCAGAGTGGTGGAGGATAGACCATGTGCTGAGAGAAGGCACTGGGTGAGATGAGACCAAGGGG | 120 | 54 |
| chr15 | 89542178 | 89542298 | 771629_41866724_SHANK3(58234)_354 | 1 | + | good | AATTGATCTAAGTGGTCTGCCGCCAGTCTGATATCCCTGTGGTTATCTCTGTGGTCACTGCAGGTGACTTGCAGTGTGTCTGGAGGTTGGTCTCCTAGGTGGTTGAATTCTTTTTTTTTT | 120 | 46 |
| chr15 | 89542538 | 89542658 | 771629_41866724_SHANK3(58234)_357 | 1 | + | good | TGATGCCACACTGAACTCTGTACATCTTTTTTTTTTTTTTTTTTTTTTTGATAAGCAAAAGTATCCCCTCTCCCCACCCCCAGCTTTTGGTTTTTAGAGACAGGGTTTCTCTGTATAGATA | 120 | 37 |
| chr15 | 89542898 | 89543018 | 771629_41866724_SHANK3(58234)_360 | 1 | + | good | TCTTCTTAAGGAGAGAGTTTGGCTGGTGAAGGAACTGGAACCTGGGGTCTGCAGAGTACAGATGGTGTCAGATCTAGGTCTCAGGCAGTGAGAGGCCTAGGGGTGGTGGGTAGGGCTCAT | 120 | 54 |
| chr15 | 89543018 | 89543138 | 771629_41866724_SHANK3(58234)_361 | 1 | + | good | TGGCTCCTGTTAGTGCTACGATGCCCGCGTAGGTGGGTGTGGCAGTCCTGCGGCTGAGGCGGCATCTCTTTGGTTGTGCAGAGAAGTTGGATGAGATCCTGGCAGTTGCCGCGGAGCCGA | 120 | 61 |
| chr15 | 89543138 | 89543258 | 771629_41866724_SHANK3(58234)_362 | 1 | + | good | CACTGAGGCCGGACATTGCAGATGCTGACTCGAGGGCGGCCACTGTCAAGCAGCGGCCCACCAGCCGGAGGATCACCCCTGCTGAGATCAGCGTAAGCCAGCCTTGGGCCAGGGCTCTGG | 120 | 66 |
| chr15 | 89543258 | 89543378 | 771629_41866724_SHANK3(58234)_363 | 1 | + | good | GATGGTCAGGTCCGGGCCTGGGGATCAGAGGGTCATGCGCCTCTCTTCTGAGGTAAGAGTTCTTTATCTGGATCCCTTCTCCAGAGGAGGTGGAACTCTCATGGCCTCTCTGGGAGACAT | 120 | 56 |
| chr15 | 89543378 | 89543498 | 771629_41866724_SHANK3(58234)_364 | 1 | + | good | TTGGGACCACGTTAGCCTGTAGGGTGGCCGTAGTTGAGTTCGCTTCTGTGGTTGTAGCTTTTAATGCTTCTGACCTTAGTCAGAGTCTGTGTTACTCTGGCTAACACAGAGGAGGGTCCA | 120 | 50 |

| | | | | | | | | | |
|---|---|---|---|---|---|---|---|---|---|
| chr15 | 89543498 | 89543618 | 771629_41866724_SHANK3(58234)_365 | 1 | + | good | CAGCTTTGCCGAGCCTATGGAACAGCATGTCCCTATTCAGATGGCATCTGTATCCCACCTGGCACCCTTGGATTCGTTCTCCCTCTCTGCTCAGACTATGCCCAGGGTCTGAACTTTCTA | 120 | 52 |
| chr15 | 89543618 | 89543738 | 771629_41866724_SHANK3(58234)_366 | 1 | + | good | TCGTGTGTTCATTTCACTACCCATAACGCTGCAGACCCCAAATGAACATGGGCACCGGTGCCTTTGGGGTGTCTGGTGTATGGGGTGTCTGGTCTAGCCTAGAATGGGTGGGGCTGGATG | 120 | 55 |
| chr15 | 89543738 | 89543858 | 771629_41866724_SHANK3(58234)_367 | 1 | + | good | GAAAACTGGGTGCTCCAGTGCCCTGGAACTTCCATGTTCCTCCCTGACCCCACAGTCATTGTTTGAGCGCCAGGGCCTCCCAGGCCCAGAGAAGCTGCCGGGCTCTCTGCGGAAGGGGAT | 120 | 62 |
| chr15 | 89543858 | 89543978 | 771629_41866724_SHANK3(58234)_368 | 1 | + | good | TCCACGGACCAAATCTGTAGGTACGGCTGGGCTGAGGGACTGCTTGGGGTTAGGGATGGCTGGGTTGGGGTGGGTAGATTCAGGGTTGGAGGACCCCATCTCTCTCTGCAAAAGTTCAAG | 120 | 57 |
| chr15 | 89543978 | 89544098 | 771629_41866724_SHANK3(58234)_369 | 1 | + | good | GATGCCACTTTGTTTATGGATCTGAGTTTGCTCTTCCACCCAGAGATGGGCCGTGGGAAGACACCTTTGGAATTCTTTTCCCGTCTGCCTCTCCCCATCCCCGGCCACCCTTTTAGTCCT | 120 | 53 |
| chr15 | 89544098 | 89544218 | 771629_41866724_SHANK3(58234)_370 | 1 | + | good | TCTTCTCTTCCTCTCTGTCATTTCTGCCATGCACGGGTGACAGCTGTTACCAGCACGGCCTTCTCCTGTCCTGTCAGCCAGAGCACAGGTCCAGTTTCCCAACCTAAAATTAACTCTTTT | 120 | 50 |
| chr15 | 89544218 | 89544338 | 771629_41866724_SHANK3(58234)_371 | 1 | + | good | CCTTGAGCCATCTCTCCAGGTAGCATCCTCTCCAGGTCCCTTAGTGGTCGGACCCAAAAGGTCACCCGTAATTGCTCTCTCTCGTCTCCATATGCTTATGAATTTGAAGCAGCTGGGTCT | 120 | 52 |
| chr15 | 89544338 | 89544458 | 771629_41866724_SHANK3(58234)_372 | 1 | + | good | CTCCGAAGCCAGATTGAAGCTGCTCCCACTAGGGTCAGCAGTAGCCTCGGGGCATCCCCCCTGGCTGGCACCATCCGCTTTCCTTCCACTGACCTTGGTTGACATTTGTGGCTTTGTTCC | 120 | 58 |
| chr15 | 89544458 | 89544578 | 771629_41866724_SHANK3(58234)_373 | 1 | + | good | CTCCTCTGCGAAACTCCTAGGATTGTGGGAATGAGTTTCTTGCTCTCCCCTCCTTTCTCACTGCATTTGTTCAGCATATCCAAAGAGTAGGTCAACTCAAGGCTGTGTCCTCGGGCTCCC | 120 | 51 |
| chr15 | 89544578 | 89544698 | 771629_41866724_SHANK3(58234)_374 | 1 | + | good | TTCCTTCTGTCTATCCTTCTGTCTATCCATCTGTGGAGCTTCCTTAGTCCCTGCCCGCCATCACACTCCACACCTGGCCGTGTCTTCTCCTAATCCCAGGCCAGGGTCCTTAACAAGCCC | 120 | 55 |
| chr15 | 89544698 | 89544818 | 771629_41866724_SHANK3(58234)_375 | 1 | + | good | TGCTGTCTGCACGTTTCTCAGGTATCTAGTCTCCTTGACCCACACATCCGTCCACACCTGTTCTTATACCCCTTGCCACACTCACCGTCTGTGCCACATCAGCGTAAACTCCTGTTCAGG | 120 | 52 |
| chr15 | 89544818 | 89544938 | 771629_41866724_SHANK3(58234)_376 | 1 | + | good | CCAATTACAGGCTAGACATGAGCTGCTTATATGCACGCTGGCTCATTTAATACCTACGGCAACCATGTGGGGGTGGGAAATCATTTATTTAAAGGGGAGTTAAAAATGGCTTGCCCCAA | 120 | 44 |
| chr15 | 89544938 | 89545058 | 771629_41866724_SHANK3(58234)_377 | 1 | + | good | AGTCACATAGCTTCCCTGCCTATACCTCATTAAGAATGGATACATGCTTAAGCCAAACAGCTATCCGAGTGTGCCTGTTGCCCATTCTGGGACTCCTGCTTACTATAAGCTGGCCCTAGG | 120 | 48 |
| chr15 | 89545058 | 89545178 | 771629_41866724_SHANK3(58234)_378 | 1 | + | good | CTGGATCTGGGTCTCAGTTGGAAGGTGACATGTCTAGAGTTATGTCAGTCCAACAGTTTATAATGTGGGTGTCAAGGTACACATATAAATGGAGAGTGACTGCGGAGATGCAGACACACA | 120 | 45 |
| chr15 | 89545178 | 89545298 | 771629_41866724_SHANK3(58234)_379 | 1 | + | good | GAAACAGCATTTATACTTAGAATAACAGGGCGAGCAGACCTTTCCTGGGGATTCTGGGGCAAACGTGTGATAGAGGCCAGTGTGTAAAAAGCACAGGCGTCTAGATGGTAGAGTCCTTCG | 120 | 48 |
| chr15 | 89545298 | 89545418 | 771629_41866724_SHANK3(58234)_380 | 1 | + | good | TGGGCTGGCAGTGATTCTTCCGTAAGTGGCTGGGTGGAGAAGGGCAGCTGGTGAAGACTGCAGTTCCTCTGGTGCCGAGAGACACTGCTTTATGCTCTTCCAAGCATCCTGGGCGTCCTC | 120 | 57 |
| chr15 | 89545418 | 89545538 | 771629_41866724_SHANK3(58234)_381 | 1 | + | good | CATGGTTTCTTTATTTCTTGCCTTACAAGTCTTACAGCAGGCTTGTCTTGTACCTCGTCCACAGCTTGGTCTTCCCCGTGTCCTCTTGCCACGGATCTGTTCTCTAGCCCGTTTCTCTGG | 120 | 50 |
| chr15 | 89545538 | 89545658 | 771629_41866724_SHANK3(58234)_382 | 1 | + | good | CAAAGGAAGCTTTCTGCAGTATATTTATAACCTGATATCCCTCCAGCCAGTCTCGGGTGGGCTCCTGTGGTCTCAGGTCCCCAGAGCAACAGCCCTGTCCTGCTTCATCTGGGTTATCCG | 120 | 53 |
| chr15 | 89545658 | 89545778 | 771629_41866724_SHANK3(58234)_383 | 1 | + | good | CTTCTGCCTTTCTGTGGCATGAGTCTTTCCTTGTTCCTTGTTTGCCTATCTGCCCTCTCTCTTCCTGGATGGTCTTTCCCAGAGCTGACTACTGTAGATCCAGACCTATGGGCTTGTTGT | 120 | 49 |
| chr15 | 89545778 | 89545898 | 771629_41866724_SHANK3(58234)_384 | 1 | + | good | ATAAGCCTCAAGCAAGCCCACCAGTTCCTCCCACTGAAATATAGCTCATCAGCGGGCTAGCAGGTGCTCAACTCTAGTTCCCCATGGCTGGAGCACTGCCGTGTGCCGGCAAGAGCATTG | 120 | 56 |
| chr15 | 89545898 | 89546018 | 771629_41866724_SHANK3(58234)_385 | 1 | + | good | GCCGAGTAGGCTCCGTGGACTCCCACCCCAGGTTCTGGCTGCACGTCCCTGGGCACGTAATACAACTGTTGGGATCCCAGGACTATAGTCTGCAAATATGGATATAGCCAGTAAGGACAG | 120 | 55 |
| chr15 | 89546018 | 89546138 | 771629_41866724_SHANK3(58234)_386 | 1 | + | good | CTCAAGCTTAAACCTTGTTGCAGTAGTTGCAGCAGCCCCTCCTTCCAGGCTGTGTCTGGAGTATACAGTCAGCACCCAGGAAACGACCCTGTTGTGGACTGGCAGGTAGAGAGCACCTTT | 120 | 53 |
| chr15 | 89546138 | 89546258 | 771629_41866724_SHANK3(58234)_387 | 1 | + | good | CTTTAGTCTGGTGTCAGCACAGACCGCTATTCCGTGCGAGGCAGGTGACTTTTTTTTTATCCTCAGGCACAACCTACCATGCCATTGGTATTAAATTCCACCCCCTCTCTCACTGTCAC | 120 | 48 |
| chr15 | 89546258 | 89546378 | 771629_41866724_SHANK3(58234)_388 | 1 | + | good | GCTTCATGCCTGTCTTCTGTAACCTTGAGGTCTCTGCGAGCTCTTCCTACATAGCTTCACCTAGCACACAGACCTGTGAGTGCTCAGTACGTGTCCCTGTGCTGTGTAGGAATGACATCA | 120 | 51 |
| chr15 | 89546378 | 89546498 | 771629_41866724_SHANK3(58234)_389 | 1 | + | good | GGAGGCAGACCTTAGAACACTGGTGATGACCTCAGCCATCTTTCGTGGCGTTGCTACCTGCGGCCTTGAGTTGTCAACTGTAAATAGCCCTGTGATGGGGGCACATAAGGTCCTGTGATG | 120 | 53 |
| chr15 | 89546498 | 89546618 | 771629_41866724_SHANK3(58234)_390 | 1 | + | good | AAACTGACTCCTAAAGGCGACTTGCATGAAGCTGGGATTCAAGCTCAAGTTTGCAACATACCTAGCCTTGTTAGTCCTGAGGCGACAGAGCCAGAGGCGTAACGAGTCTCCACCACGCCT | 120 | 52 |
| chr15 | 89546618 | 89546738 | 771629_41866724_SHANK3(58234)_391 | 1 | + | good | GCCTTCTACTCATTAGGATGGGCCACTTGGCTTTTTCAAGCCTCGCAGTGTAGGACTCTGCACCCGTGTCTGCATCCTGTCTGAAAAGACACAGGGAGGTTCTGTAGGGAACTAGTGACA | 120 | 52 |
| chr15 | 89546738 | 89546858 | 771629_41866724_SHANK3(58234)_392 | 1 | + | good | TCTCTAGACAAGAGGATGGGTTTCCTTTTCAGGGAAGTGTAGCTAATGAGAACATGCTTATATCCTGGCCTGGGGTTACATTAGAGAGTAGGGATGGACCTGGAACATGCCAGTTTTGTT | 120 | 45 |
| chr15 | 89546858 | 89546978 | 771629_41866724_SHANK3(58234)_393 | 1 | + | good | ACCCATCCATTCTTCCTTCCATCTTCCGGCCCAGGCTCTCTAACTTGTAGCCTCTGCCTAGGGTACCAACTTACTAGTTTCCTGCAGGCCCATTCCCGAAAGTCTCTCCAAGGAACCAAT | 120 | 52 |
| chr15 | 89546978 | 89547098 | 771629_41866724_SHANK3(58234)_394 | 1 | + | good | GAGGCTGCTTTCCCAGCCCCTACCCTGGTTGGCTGTTGGTGTCAGTTCTTGCAGATGCACGGGTGCACTCTCCTGTGTCCCCTCATTGATGTTGATGTTTTCCTTTTGATATGTGGAGGG | 120 | 53 |
| chr15 | 89547098 | 89547218 | 771629_41866724_SHANK3(58234)_395 | 1 | + | good | GGGGTCCTCCTTTCTGACCACCTTTCTTCCCGTCTCTGGTTACCATCCAGGTTTCCGATGACTCTGGCTTGTCCAACACTTTGTTTCTGAGTTGCTTCGCTTATCCTTGATGGGGAACCT | 120 | 51 |
| chr15 | 89547218 | 89547338 | 771629_41866724_SHANK3(58234)_396 | 1 | + | good | CCTGAGCAGCCCGTAGGTCCTCTCTGACAGTTTTAGGGGGCTGCCTGTACTCTTCTGGTGTCTTCTCATGGCCTGCTCCGTGCCCACAGGGGTTGGAGAGAGATCTTGGCTTCAGTTCAG | 120 | 57 |
| chr15 | 89547338 | 89547458 | 771629_41866724_SHANK3(58234)_397 | 1 | + | good | GCAGTCAGAGTTTGGGTCCTGTTCCAGATCCTCTGTAGGAGTGGGCCTCTACACTGTGGGTGGGGTCCCTCCTAAACCCTCCATCAGAGCTCTCCCTACCTGCAACAGGACATGGCATGG | 120 | 57 |
| chr15 | 89547458 | 89547578 | 771629_41866724_SHANK3(58234)_398 | 1 | + | good | AGTGTGGGTGTCCTGCTCAGAGCTCTTGCTGTTTCCCTTTCGCCTTTCCTTGGTGGGGCTGTGGCCATTATGGGGTGGAGGATAGGACTAGCCAGTGGCGCTGGGTCCAGAGAAGCCATT | 120 | 57 |
| chr15 | 89547578 | 89547698 | 771629_41866724_SHANK3(58234)_399 | 1 | + | good | CAATGGCCAGCATCCTTGCCAGACTTACCTGGCTCCTCCGTCTCCAGGGGAGGATGAGAAGCTGGCATCCCTACTGGAAGGGCGCTTCCCACGCAGCACGTCAATGCAGGACACAGTGCG | 120 | 61 |
| chr15 | 89547698 | 89547818 | 771629_41866724_SHANK3(58234)_400 | 1 | + | good | TGAAGGTCGAGGCATTCCACCCCCGCCGCAGACCGCCCCGCCACCCCCACCCGCGCCCTACTACTTCGACTCCGGGCCACCCCCCACCTTCTCACCGCCGCCACCACCGGGCCGGGCCTA | 120 | 74 |
| chr15 | 89547818 | 89547938 | 771629_41866724_SHANK3(58234)_401 | 1 | + | good | TGACACTGTGCGCTCCAGCTTCAAACCAGGCCTGGAGGCTCGTCTGGGTGCGGGGGCCGCCGGCCTGTATGATCCGAGCACGCCTCTGGGCCCGCTGCCCTACCCTGAGCGTCAGAAGCG | 120 | 68 |
| chr15 | 89547938 | 89548058 | 771629_41866724_SHANK3(58234)_402 | 1 | + | good | TGCGCGCTCCATGATCATACTGCAGGACTCTGCGCCAGAAGTGGGTGATGTCCCCCGGCCTGCGCCTGCCGCCACACCGCCTGAGCGCCCCAAGCGCCGGCCTCGGCCGTCAGGCCCTGA | 120 | 71 |
| chr15 | 89548058 | 89548178 | 771629_41866724_SHANK3(58234)_403 | 1 | + | good | TAGTCCCTATGCCAACCTGGGCGCCTTCAGTGCCAGCCTCTTCGCTCCGTCGAAACCACAGCGCCGCAAGAGCCCGCTGGTGAAGCAGCTTCAGGTGGAGGACGCTCAGGAGCGCGCGGC | 120 | 66 |
| chr15 | 89548178 | 89548298 | 771629_41866724_SHANK3(58234)_404 | 1 | + | good | CCTGGCCGTGGGTAGCCCGGGACCAGTGGGCGGAAGCTTTGCACGAGAACCCTCTCCGACGCACCGCGGGCCCCGGCCAGGCAGCCTTGACTACAGCTCTGGAGAAGGCCTAGGCCTTAC | 120 | 68 |
| chr15 | 89548298 | 89548418 | 771629_41866724_SHANK3(58234)_405 | 1 | + | good | CTTCGGCGGCCCTAGCCCTGGCCCAGTCAAGGAGCGGCGCCTGGAGGAGCGACGCCGTTCCACTGTGTTCCTCTCTGTGGGTGCCATCGAGGGCAGCCCTCCCAGCGCGGATCTGCCATC | 120 | 69 |
| chr15 | 89548418 | 89548538 | 771629_41866724_SHANK3(58234)_406 | 1 | + | good | CCTACAACCCTCCCGCTCCATTGATGAGCGCCTCCTGGGGACAGGCGCCACCACTGGCCGCGATTTGCTACTCCCCTCCCCTGTCTCTGCTCTGAAGCCATTGGTCGGTGGTCCCAGCCT | 120 | 64 |
| chr15 | 89548538 | 89548658 | 771629_41866724_SHANK3(58234)_407 | 1 | + | good | TGGGCCCTCAGGCTCCACCTTCATCCACCCTCTCACTGGCAAACCCTTGGATCCTAGCTCACCCTTAGCTCTTGCTCTGGCTGCCCGGGAGCGGGCTCTGGCCTCGCAAACACCTTCCCG | 120 | 63 |
| chr15 | 89548658 | 89548778 | 771629_41866724_SHANK3(58234)_408 | 1 | + | good | GTCCCCCACACCTGTGCACAGCCCCGATGCTGACCGCCCTGGACCCCTCTTTGTGGATGTGCAAACCCGAGACTCTGAGAGAGGACCGTTGGCTTCCCCAGCCTTCTCCCCTCGGAGTCC | 120 | 64 |
| chr15 | 89548778 | 89548898 | 771629_41866724_SHANK3(58234)_409 | 1 | + | good | AGCGTGGATTCCAGTGCCTGCTCGGAGAGAGGCAGAGAAGCCCCCTCGGGAAGAGCGGAAGTCACCAGAGGACAAGAAGTCCATGATCCTCAGCGTCTTGGACACGTCCTTGCAACGGCC | 120 | 60 |
| chr15 | 89548898 | 89549018 | 771629_41866724_SHANK3(58234)_410 | 1 | + | good | AGCTGGCCTCATTGTTGTGCATGCCACCAGCAATGGGCAGGAGCCCAGCAGGCTGGGGGCTGAAGAGGAGCGCCCCGGTACTCCGGAGCTGGCCCCAGCCCCCATGCAGGCAGCAGCTGT | 120 | 68 |
| chr15 | 89549018 | 89549138 | 771629_41866724_SHANK3(58234)_411 | 1 | + | good | GGCAGAGCCCATGCCAAGCCCCCGGGCCCAGCCCCCTGGCAGCATCCCAGCAGATCCCGGGCCAGGTCAAGGCAGCTCAGAGGAGGAGCCAGAGCTGGTATTCGCTGTGAACCTGCCACC | 120 | 68 |
| chr15 | 89549138 | 89549258 | 771629_41866724_SHANK3(58234)_412 | 1 | + | good | TGCTCAGCTGTCCTCCAGCGATGAGGAGACCAGAGAGGAGCTGGCCCGCATAGGGCTAGTGCCACCCCCTGAAGAGTTTGCCAATGGGATCCTGCTGACCACCCCGCCCCCAGGGCCGGG | 120 | 66 |
| chr15 | 89549258 | 89549378 | 771629_41866724_SHANK3(58234)_413 | 1 | + | good | CCCCTTGCCCACCACGGTACCCAGCCCGGCCTCAGGGAAGCCCAGCAGCGAGCTGCCCCCTGCCCCTGAGTCTGCAGCTGACTCTGGAGTAGAGGAGGCTGACACTCGAAGCTCCAGTGA | 120 | 67 |
| chr15 | 89549378 | 89549498 | 771629_41866724_SHANK3(58234)_414 | 1 | + | good | CCCCCACCTGGAGACCACAAGCACCATTTCCACAGTGTCCAGCATGTCCACCCTGAGCTCGGAGAGTGGGGAACTCACGGACACCCACACCTCCTTTGCCGATGGACACACTTTTCTACT | 120 | 57 |
| chr15 | 89549498 | 89549618 | 771629_41866724_SHANK3(58234)_415 | 1 | + | good | CGAGAAGCCACCAGTGCCTCCCAAGCCCAAGCTCAAGTCCCCGCTGGGGAAGGGGCCGGTGACCTTCAGGGACCCGCTGCTGAAGCAATCCTCGGACAGTGAGCTCATGGCCCAGCAGCA | 120 | 65 |
| chr15 | 89549618 | 89549738 | 771629_41866724_SHANK3(58234)_416 | 1 | + | good | CCATGCTGCCTCTACTGGGTTGGCTTCTGCTGCTGGGCCCGCCCGCCCTCGCTACCTCTTCCAGAGAAGGTCCAAGCTGTGGGGGGACCCCGTGGAGAGTCGGGGGCTCCCTGGGCCTGA | 120 | 68 |
| chr15 | 89549738 | 89549858 | 771629_41866724_SHANK3(58234)_417 | 1 | + | good | AGATGACAAACCAACTGTGATCAGTGAGCTCAGCTCCCGTCTGCAGCAGCTGAATAAAGACACACGCTCCTTGGGGGAGGAACCAGTTGGTGGCCTGGGCAGCCTGCTGGACCCTGCTAA | 120 | 57 |
| chr15 | 89549858 | 89549978 | 771629_41866724_SHANK3(58234)_418 | 1 | + | good | GAAGTCACCCATTGCAGCAGCTCGGTGAGCAGGGTGATGTGGGGAGGGGCCTGTCCCCCTGTCTTCTATTCCTTTTGTTCATTACAGGCTTCCTGCTGTCCTTCCTCATTCTGTCCCATC | 120 | 55 |
| chr15 | 89549978 | 89550098 | 771629_41866724_SHANK3(58234)_419 | 1 | + | good | TCTCATCTGTCCATGGCATGGGTTCTCCCTGTCCACCTTTTTATCTTTGGTTTTGCTTTGCAGTCTGAGATAGATGGCTCTTTTAGTACAGCTGCTCTGTGGACACCTGGTGCAGGCAGT | 120 | 48 |
| chr15 | 89550098 | 89550218 | 771629_41866724_SHANK3(58234)_420 | 1 | + | good | GGTGGGCCTGGGTACCTGTGAGACCCAGGGATGTTGTGAGACCTCCCAGGACCTGCTGGAGGCGGTGCTGCGGGCTTGCAACTTTGTATTCCAGCTTCTAACTCCAGCTGCTTTTGCCTT | 120 | 58 |
| chr15 | 89550218 | 89550338 | 771629_41866724_SHANK3(58234)_421 | 1 | + | good | ACCTGAGAATCTTTTCACCTTGTACTCTGGAGCCGCTGCTCTGTCTTGGCAGCTGTCTGACCCTGAGTCTACTGCCCCGTGGCTCCATTTTGGTTCTCTTTCTGATAGTATGGCTGCCCC | 120 | 53 |
| chr15 | 89550338 | 89550458 | 771629_41866724_SHANK3(58234)_422 | 1 | + | good | ATATGGCTGGTGTCCTGGAGCACCCCCCTCCCTTGATTCTTTTTCAGGAACCAGAGTACATTGGTTCCTTTCCTTAAGGGACTTGATACAAATTTCTTACCTTCTGCCCTAGAAACTTTT | 120 | 44 |
| chr15 | 89550458 | 89550578 | 771629_41866724_SHANK3(58234)_423 | 1 | + | good | TTTTTCCGTGTTGTGGCACCTCCTAGGGCCCTTCAACACAGAATCCTGGATTTGCGTTATAAAGCTTCATCAAGAAGAGCTGGAAATCACACCGTCTCTCTCACCTACTTTTTCCTTCTT | 120 | 45 |
| chr15 | 89550578 | 89550698 | 771629_41866724_SHANK3(58234)_424 | 1 | + | good | GTCATGTTTGCCCCATCCCCACGCCCGCGGTGATGGAGAAGTGGGAGGTGACGGAGGAGGTCTGGGTGCTACTAAGGTTGTTAGACTTTGTAGGCAGAAGGTGGCAGAGCCTTGGGTTGG | 120 | 58 |
| chr15 | 89550698 | 89550818 | 771629_41866724_SHANK3(58234)_425 | 1 | + | good | TCCCTCAAGCCCTTTCTTGGCAACCTTCCCCAGGGCTCCCATGTGGACTGCATGGGTGCTGGAGAGAGCCGACCAGTTTGTGGCCAGCCTCCTGTCTGTGTTTCTCTAGCTGCTTCTGCC | 120 | 59 |
| chr15 | 89550818 | 89550938 | 771629_41866724_SHANK3(58234)_426 | 1 | + | good | TCTGCCCCAGGGCTCTCTTATTACTGGGTTTGAGCGGGGTACCTACCACCCCCAATTCCCCACTCCTTGGTTCCTAGACACTGTTCACTGTCAGTTGTGGAAGACAGAGTTCACTGAGTA | 120 | 52 |
| chr15 | 89550938 | 89551058 | 771629_41866724_SHANK3(58234)_427 | 1 | + | good | GTGAGCAGTGCACAGCTTAAGGAAGGGCTTCAGGTCTTACTCAGAGGGCCTGCTCATTTTACAGTCTTCCAGTTTAGGGTCTAGTAGATGAACAGGAGCTTGGGAAGCTCACATTGCTCT | 120 | 49 |
| chr15 | 89551058 | 89551178 | 771629_41866724_SHANK3(58234)_428 | 1 | + | good | CTGAGAGGTTCAGGGACAGGGGCATTCCTGTCCTTCTCACGACCCGTTCTTAGACTTTCTCCCTTTTTGGCCTCTTCCATCTCTTAAGTCTGAATGCCCTTTTTTGTACTGCAGTGATGA | 120 | 48 |
| chr15 | 89551178 | 89551298 | 771629_41866724_SHANK3(58234)_429 | 1 | + | good | AGGCACCCTTAGAATGGGCAGTGGCTGTGAGCCAGTCAGGACACGTGGCTGGGTCCCAGGTTTGTCTACAGAACTGGGTTTTCAACTTTACTCGGACCATTTTCTTCCTTGTCTTTTCTT | 120 | 50 |

| | | | | | | | | | |
|---|---|---|---|---|---|---|---|---|---|
| chr15 | 89551298 | 89551418 | 771629_41866724_SHANK3(58234)_430 | 1 | + | good | CCACTATCAGCCCCTCTGTTTCTCTCTGCCTCTTTTTTTTTGATGAATGTTAGAAGTCACTTAAAGTACATAACATACTGCCTACCCTCAGTAATCACATATTAAGCCAATGAACAGAGG | 120 | 39 |
| chr15 | 89551418 | 89551538 | 771629_41866724_SHANK3(58234)_431 | 1 | + | good | AGGCTTGGTGCAAGTGCTTTGTCCTAAGGCAAAGTTCATGGAAGCACAGATGGGCAAAGATCTCTTGGGCATCTAGTAGCACCTTTTGGTAACACTCCATCTGTGCTCAGTGATTTCAGG | 120 | 48 |
| chr15 | 89551538 | 89551658 | 771629_41866724_SHANK3(58234)_432 | 1 | + | good | GTGGTTTGCAGAGGTAGACCTTTGCTGTGTAGCACACTTATGCATATCTTTTGTATATGAGTGATACATTTTATCATGTTCATCAGTCATTTTAACTTCACTGAAGCACCTTGCACCCTG | 120 | 39 |
| chr15 | 89551778 | 89551898 | 771629_41866724_SHANK3(58234)_434 | 1 | + | good | TTGGAGGCAGGGTCTTGCTATGCAGTCTGAACTGGCTTTGATCCTCCTACCTGTCCCCACAGTGTGAGCTGTCACATCCTCGCACCTGGATGAAACAATGGCCATCTGGAGTGTGTCCTG | 120 | 54 |
| chr15 | 89551898 | 89552018 | 771629_41866724_SHANK3(58234)_435 | 1 | + | good | CCACCTACCACAGTGCTTTCCTTTCTGGCTGCAGGCTTGGAGCACTTGTGGCTCTGTACACAGCTAGCTTATTTCAGACCTGGTCTTTTGTCTGGTCTTTGTGGACACCCTTACTTTGAT | 120 | 49 |
| chr15 | 89552018 | 89552138 | 771629_41866724_SHANK3(58234)_436 | 1 | + | good | GTCTGTGGGGGCTGTGCCTCAGCTCTGGCTTTGTAGCCAGTGGGGGCTGGAATCTGGCATTTGGGTCTTACTTTGGCATACAGATCACTAATGAGCCTTTCCCTTAGAGAAGTGTCCAAA | 120 | 52 |
| chr15 | 89552138 | 89552258 | 771629_41866724_SHANK3(58234)_437 | 1 | + | good | GCGTGTGAAGTTCTGATGTCTAGGTGTGACTGTGTCCTTCTCAGGACCTGTCCTGCTTTGGCAGAGCACACACCCTTGAGCCACATCTTTGGTCTGTGCATGGTTCCCGAGCCTTACCTA | 120 | 53 |
| chr15 | 89552258 | 89552378 | 771629_41866724_SHANK3(58234)_438 | 1 | + | good | CCAGGCTCATCTCATCTCCTCTTGATGGAGTCCACCAAGGACATGACCCTCTTCATTCATCATCTCTGAAGATGCCTCAGGCCAGGCAGCGATGCCTTCTAGCCACAGGTGCTCTGCAGC | 120 | 55 |
| chr15 | 89552378 | 89552498 | 771629_41866724_SHANK3(58234)_439 | 1 | + | good | CTGAGCTTTTTGTAGACAAGTTAAACTTGCCCGTGTTTCACAGGAAAAGCTCAGTAACTTGGAGATCAACTCGTTTGCACAGGGCCCAATGTTGGGTTTTAGGTTGAGGCTGGAAGGTAC | 120 | 46 |
| chr15 | 89552498 | 89552618 | 771629_41866724_SHANK3(58234)_440 | 1 | + | good | ATTGTTGTTGGTGCTACTGCTGAGGTCTGTGATCCCACCTGTCTCTCACTTCAGCTTGCAGGTCTCGCCCCTGCTGATCAGATCCCTGTCTACTGAGAGCCCTGGGCTGCATCTTACTTG | 120 | 54 |
| chr15 | 89552618 | 89552738 | 771629_41866724_SHANK3(58234)_441 | 1 | + | good | TTCATCACTTAGAGCCCGACCTTGGGCTCCACACCTGGTTGTGGAGCTCTGCCTGTCTGTCAGTCGTGAGACCTGCGCCTGTCTCCCGTTCCCCTTGTCTTCTGGGTCTCATGTTTGGAC | 120 | 57 |
| chr15 | 89552738 | 89552858 | 771629_41866724_SHANK3(58234)_442 | 1 | + | good | TTGAATCTTACATGTGGTTTGAAGCCTACAACAGGGCTCTTTTGCTTTGGCTTTCAGGGTCTTTCTGGGGAGGGGCTCTGGGGTGTGTGTGTGTCGAACTCAAGGCCCCGAGCATACTAA | 120 | 51 |
| chr15 | 89553098 | 89553218 | 771629_41866724_SHANK3(58234)_445 | 1 | + | good | GACTGAGCCATCTCACCAACCTGAGAGTTTTTCCATAGAGCTTAGGCTGGCATTGAACGCGATGTGTGGCTCAGCTTGGCCTTCAGCTTAAAATAAACCCTCCTGCCTATGCCTCCTTAG | 120 | 50 |
| chr15 | 89553338 | 89553458 | 771629_41866724_SHANK3(58234)_447 | 1 | + | good | GCTACAATACTTTACGGAGCCTTTAATTTTTTATTATTTTTTTTATTAGATGCTTGTGTGTGGGTGGTCAAAGTGTGCCCTGTTCTTTAGACCTATTGAGGTCACTGCTGAGATTTCACA | 120 | 37 |
| chr15 | 89553458 | 89553578 | 771629_41866724_SHANK3(58234)_448 | 1 | + | good | GGAGACCTACTTTCCAGTCAGACTGCTCCACAGAGCCCTGTCCTGACTGTTATCAGTCCTGCCTGCATGTTAAGGGGATGGACAGAGCCTAGGTTCTGGGGAAGTCATACATGAAGGTCC | 120 | 53 |
| chr15 | 89553578 | 89553698 | 771629_41866724_SHANK3(58234)_449 | 1 | + | good | CCAGTTGACCCCAGGAGACTGGGGACAGGATTCATTGTGTCCCCGCAGAGCAGTTGCATGAAGTTATAGGCAGATGCATTGCTAGGGAACAGAAAGAGACCATAGAAGAGGAGATGTAGG | 120 | 51 |
| chr15 | 89553698 | 89553818 | 771629_41866724_SHANK3(58234)_450 | 1 | + | good | AACACATGGCATGTCAGTGTTGAACGGGCCCGGCAGTGAGGAGTGAGGCCGAGGCAGGAGAGCTGCCGAGCCCTGCCTGTTCCTACTGGGAACTGTCCTGTAATGAGGCCTCTGAGTTGC | 120 | 60 |
| chr15 | 89553818 | 89553938 | 771629_41866724_SHANK3(58234)_451 | 1 | + | good | CAAGGGCCATAGGCCTAGGTCTGATCCACATGAGTGCAAAGGTGGGGATCGTGACATAGAAGCCAGGCCAGAGTACGAAGGAACCTTTTCTTGACTGTTGGTACCAGTGGGCTGACGGAG | 120 | 54 |
| chr15 | 89553938 | 89554058 | 771629_41866724_SHANK3(58234)_452 | 1 | + | good | GTAAGGCAATCCCAGGACGGTTTTTGTAGGACGACTTTTTTTTTTTTTGTCAGTGTGACTTATATCCCTGAGTTTGTGCCAGGCTTTGGGAGTCAAGGAAAGCCACTGAGAAGGAAGTGA | 120 | 44 |
| chr15 | 89554058 | 89554178 | 771629_41866724_SHANK3(58234)_453 | 1 | + | good | AGGGCCGGGATGTTGCTCAGTGGTTAGAGGGCTTGCCTTGTGTGCATGAGGCCTGGGCTTTGTCCCTAGCAGAGAAAAAGGGAGAGGGGGAGGGGGAGAAGAAGGAAGAGAAGAGGAGA | 120 | 57 |
| chr15 | 89554178 | 89554298 | 771629_41866724_SHANK3(58234)_454 | 1 | + | good | GGGAGAGAAGGGAGGAAAGGAGGGAAGATGGTGAAAGACCCTGAAACTCTGAGGGCCCTGGACTCAGGTGACGAGGAAGACCTGGGTTAATGGGAGGAATTGGTGACATTCCCCAAAGCT | 120 | 54 |
| chr15 | 89554298 | 89554418 | 771629_41866724_SHANK3(58234)_455 | 1 | + | good | GAAAAGGTAGAGGAGCCACAGTCCTCTCGTGGGTGCTGCTCACTGAGGCAGAGTTGGACATCCTGCATGTTAAAAGGTGTTAGGAGAGCCCAGTGGAAACCTTGAGCTCAGAATGCCACC | 120 | 53 |
| chr15 | 89554418 | 89554538 | 771629_41866724_SHANK3(58234)_456 | 1 | + | good | GATAGGCCCTCGCGCACGGCATTCCTGGGCTTGTTAGTCACATCAGACAATGGGTAGATGACATAGGCTGAGACCGTGCAGACCATGAAAGACCGCGGTAGGCTGGAGAAGCTGTCCAGG | 120 | 57 |
| chr15 | 89554538 | 89554658 | 771629_41866724_SHANK3(58234)_457 | 1 | + | good | AGGTAGGATGCCGAGGGAAGGGCTCCTCCAAAGAATCAGGTCATCCCAGGGGACCCCTGCGTCCAGCCTTGGTGAATGCCACAAACAAAGGCCAGTTCTCAGCGGCTCCCAGTTCCTGGG | 120 | 60 |
| chr15 | 89554658 | 89554778 | 771629_41866724_SHANK3(58234)_458 | 1 | + | good | GCAAGATCGAGTGTCTTAGAGTAGTGGATGCTGTGAGGTGACCATGCTGTGGATGCAGCCTGAGCAGAATGTGATTGGCCAGTCAGCTACTGGGTGGCAGTAGCCAGAGATGGGTGTGCA | 120 | 54 |
| chr15 | 89554778 | 89554898 | 771629_41866724_SHANK3(58234)_459 | 1 | + | good | GAGGTGCCGTAGACAGGTTAGAATCAATCAGATTTTCACCCTGCTGGGGTTTCTGTCCTGGGGAGGGGGGCTCAATGTGGGTTGAGCTCTTCCCCAAGATGGGGGCGGGGAACAGACACT | 120 | 57 |
| chr15 | 89554898 | 89555018 | 771629_41866724_SHANK3(58234)_460 | 1 | + | good | GATTCCTTGGCCACTCCTTCCTGTGCATACCCTTCCTTGGGCTTTTCCCCTAGGCCTCCCAACATTAGCAACACTGGTGGAGTTTTTGAGGTATATGACTGGCTCTCTCTCCACTCCCAG | 120 | 52 |
| chr15 | 89555018 | 89555138 | 771629_41866724_SHANK3(58234)_461 | 1 | + | good | TCTCCTGGGCACCTTAGAGGCTAGGTTGCCCTCAGCCCAGGCCTCCAGGTCACCCTGCCAGGAAGGCAGAATTTGGTGCCATTCAGACAGCTGTGGCTACTTGTCCTAGCTTAGGCCTTT | 120 | 57 |
| chr15 | 89555138 | 89555258 | 771629_41866724_SHANK3(58234)_462 | 1 | + | good | TGGACCCCAGAGCCTCCCCACTGCACTGTCTTATGGCTGTCTTGTCCCCTCCGGTTCAGCAGTAGACTGCTGAAACCTAACCTTTGGGGCAGGTCTTGCTTAGACATTTGTTCAGTGATA | 120 | 52 |
| chr15 | 89555258 | 89555378 | 771629_41866724_SHANK3(58234)_463 | 1 | + | good | CAGATGCCCATGCTCGGGTCAGCATGGATTCAGGGGGGAAAGCAGCGAAGGTGACATTGCCTGGCCTTGTTGGGCTCTGTAGGTGGCCAGAGTCTGACCTCACAACTGTGCTTCCAGCAC | 120 | 58 |
| chr15 | 89555378 | 89555498 | 771629_41866724_SHANK3(58234)_464 | 1 | + | good | TGCCTCCCCGAGGACCTCACCGCCTCACCTCTGCTGCCCACTGCACTTTGCCCCAAAGCCAGGATGGCTCCCCGAGGCCAGGTTCCATGGGACTGAAAGTTTCTTCCGGAGGCTTTGTCT | 120 | 62 |
| chr15 | 89555498 | 89555618 | 771629_41866724_SHANK3(58234)_465 | 1 | + | good | GCCTGTTCCCTTGTATAGCCCTCAGATGTTCCTGGAATCTTTGTGTTGGTGGCACTGGTTTGGCTTATCCGAGCAGCTCTGCCCTGGGTCTGAGCAGTTGGGAGGAAGCCGTAGGGATGG | 120 | 56 |
| chr15 | 89555618 | 89555738 | 771629_41866724_SHANK3(58234)_466 | 1 | + | good | ATGGCTGAGGAGAAAGCTGCTGCGAGTGATCGAAATGAGGTCTCTAGGGAAGGACTGATCTTGCACGGTGCAGGGGAAAAGTTGAAAGTTGCTTCAGGGAGCATGCTTGCTAGCTTGCTC | 120 | 52 |
| chr15 | 89555738 | 89555858 | 771629_41866724_SHANK3(58234)_467 | 1 | + | good | TTCTCAGTGATGGATTTCATCACCTCACAGCAACAGCTTGAGGAAGAAGGATGTACTGTGGTTCCTAGTTTCAGAGATTTCGGTCTCCGTGGTGGCTTGGCCCCCATTGCTGTTGAGCTA | 120 | 49 |
| chr15 | 89555858 | 89555978 | 771629_41866724_SHANK3(58234)_468 | 1 | + | good | GTCACAATAGGGAAGGCTCAGTGAATCATTTCCTGGCAGCCAGGAAGCAAGAAGGGAAAAGAGGGGGCAGGAATAAGACCATCAAGTCATGAAGTCATTGATGTTTGTTGTGATCATTAT | 120 | 44 |
| chr15 | 89556098 | 89556218 | 771629_41866724_SHANK3(58234)_470 | 1 | + | good | CTGGGATTAGAGGTGTGCGCCACCGCCACACCCGACCCCAGTCAGTGTATTCATTCATAGATTAGGGCAGCCCTCTCATGATCCAGTTACCTCTGTGATCCACCACTATACCTTCTGAGT | 120 | 52 |
| chr15 | 89556218 | 89556338 | 771629_41866724_SHANK3(58234)_471 | 1 | + | good | GCAGACCTTTTAGGGGGAATATTGCAGAGCCAAGCTATGCAGCAAGGAGAGGGGCACGTTGAGGGGACTCCTGCTGGTGTGGGGTTCTGGGCCAGGTGGGTAGAAACTTAGGCCATCCTA | 120 | 57 |
| chr15 | 89556338 | 89556458 | 771629_41866724_SHANK3(58234)_472 | 1 | + | good | AGGGAGAGGACGTTGCGGGGGTGAGATGGCCACTGGAGAGAAGTCGGTCAAACGTTAGAGTGACTATCAGACTGCAAGGCACTGCTAAGCAACACTTACGGTCTGCACCTTATGCTGGAG | 120 | 54 |
| chr15 | 89556458 | 89556578 | 771629_41866724_SHANK3(58234)_473 | 1 | + | good | GGAGCCATCGAGGCCACTCCGGGGCAGGCAAACCAGGTGCCTTCTGTCCAAGCTGTCCTGGGTATTTTCCGTTCCCCAGTACCTCTGCAGTAGACGGGCCTTTCCTTCTGAGTTCCGACC | 120 | 60 |
| chr15 | 89556578 | 89556698 | 771629_41866724_SHANK3(58234)_474 | 1 | + | good | TCCATAGAGCGTCCCACAGGGGTCCCTGAACCTGGCTGCCCATTGTGTGGTTGTGACTGAAGGCATTGTGGGCCTGGCCTGCTAACACTCTCTCTCATTGCTCTCTCCTGCCCATCTATC | 120 | 57 |
| chr15 | 89556698 | 89556818 | 771629_41866724_SHANK3(58234)_475 | 1 | + | good | TGCATGTGGCTGCTCTGTCTAGCTGCGCGGTGGTCCCGAGTGCCGGCTGTGTAAGTGAGCATGCCCATCCCATGCCTCTCGCAGTCCACATGCTGTGCCTGGCTGCGCCCCCTGGCTGCT | 120 | 65 |
| chr15 | 89556818 | 89556938 | 771629_41866724_SHANK3(58234)_476 | 1 | + | good | CTGCATGGCTGTGTGGGGCTCTGGTGCTACTGACAGCTACTGAGGCAATCCTTGGACACAGTGAGGGGTCGAGATGGGGGATTGGGTGGGGGTGCTATGGGAGTAGGCTTCCGGTAGAGC | 120 | 60 |
| chr15 | 89556938 | 89557058 | 771629_41866724_SHANK3(58234)_477 | 1 | + | good | AGCCGCGCTTCCGGTAGAGCAGCCGCCTCGGGCTGGCCTTCCCTCCAGCCAGGCAGGCTGGGGCAGGGCTGGAAGTCCAGGATCCTGGCAGGTGAGGCTGGGAAGCGAGGTACCATCTCT | 120 | 68 |
| chr15 | 89557058 | 89557178 | 771629_41866724_SHANK3(58234)_478 | 1 | + | good | CTGCCTTGTGTGTCTGCCTGTGTGCTACTGCGTTCCTGTGAGCACCGCTGCATGTCCACAGCACATGTAGTCTCCATTGCTGCTGTGGTGTGCTCTCCTGTGCAGGGAGACTCTCTTCCT | 120 | 57 |
| chr15 | 89557178 | 89557298 | 771629_41866724_SHANK3(58234)_479 | 1 | + | good | GTTACCGTGTTCTGTTGCCTTGCACCTCCTGTGGGTGTGAGCCCCGCCTGGTGTTTGTCTTCCGTGTTCATGGACCTGTCACGTCTTGTGCCACTGCCAGCCTTTACCATGACTCCGCCG | 120 | 58 |
| chr15 | 89557298 | 89557418 | 771629_41866724_SHANK3(58234)_480 | 1 | + | good | TGCCCGAAACTGGCCTGGGTTGCTCTGGGTCTCTTCAGACTTGAGAGCAGTATGCGGGGAGTGGTGAAGATATTGTTCAAATTTTATAGTCCAGAGCATCTGTGGATGGAACCCACGCCC | 120 | 52 |
| chr15 | 89557418 | 89557538 | 771629_41866724_SHANK3(58234)_481 | 1 | + | good | AGAGGCTGTGTTAGAGGCTGGCAGGGTGGGGCAAGCAGCTTCCGTCCATGAGAAAAGTTTCCTTTGGTGAACTGCCCTTGGGGCCTCCATGGAGCGCAGTGTGTGGGGACAGTCTTACTC | 120 | 57 |
| chr15 | 89557538 | 89557658 | 771629_41866724_SHANK3(58234)_482 | 1 | + | good | CTTAGAATCCAGTCAGTTTGAGAATCCCCCAGAGCAAGGCACTGGTCCTGACCTGACCCAGGATGTTTGTGTTTTCAAAAAGAATGACCACAGGATATTCCTCCAGCCCTAGACCTCTTG | 120 | 48 |
| chr15 | 89557658 | 89557778 | 771629_41866724_SHANK3(58234)_483 | 1 | + | good | GCTCTGCTGCCCTCTCTGTCCACTCCGACTTTATTTTCCATCCCTCACCTGGCCCTCACCCACTCTCCCTCCGCAGGCTCTTCAGCAGCCTCGGTGAGCTGAGCACCATCTCAGCGCAGC | 120 | 62 |
| chr15 | 89557778 | 89557898 | 771629_41866724_SHANK3(58234)_484 | 1 | + | good | GCAGCCCGGGGGGCCCGGGCGGAGGGGCCTCCTACTCGGTGCGGCCCAGCGGCCGGTACCCCGTGGCGAGACGAGCCCCGAGCCCAGTGAAACCCGCATCGCTGGAGCGGGTGGAGGG | 120 | 78 |
| chr15 | 89557898 | 89558018 | 771629_41866724_SHANK3(58234)_485 | 1 | + | good | TGGGGGCGGGCGTGGGAGGCGCGGGGCGGCCCTTCGGCCTCACGCCTCCCACCATCCTCAAGTCGTCCAGCCTCTCCATCCCGCACGAACCCAAGGAAGTGCGCTTCGTGGTGCGAAGTG | 120 | 70 |
| chr15 | 89558018 | 89558138 | 771629_41866724_SHANK3(58234)_486 | 1 | + | good | TGAGTGCGCGCAGCCGCTCCCCCTCACCATCTCCGCTGCCCTCGCCTTCTCCCGGCTCTGGCCCCAGTGCCGGCCCGCGTCGGCCATTTCAACAGAAGCCCCTGCAGCTCTGGAGCAAGT | 120 | 69 |
| chr15 | 89558138 | 89558258 | 771629_41866724_SHANK3(58234)_487 | 1 | + | good | TCGATGTGGGCGACTGGCTGGAGAGCATCCACTTAGGCGAGCACCGAGACCGCTTCGAGGACCATGAGATCGAAGGCGCACACCTGCCTGCGCTCACCAAGGAAGACTTCGTGGAGCTGG | 120 | 62 |
| chr15 | 89558258 | 89558378 | 771629_41866724_SHANK3(58234)_488 | 1 | + | good | GCGTCACACGCGTTGGCCACCGCATGAACATCGAGCGTGCGCTCAGGCAGCTGGATGGCAGCTGACGCCCCACTCCCTCTCCTGTTCCTGCTGCGCCCTGCCGGCAGGGCCCCCACCCCT | 120 | 70 |
| chr15 | 89558378 | 89558498 | 771629_41866724_SHANK3(58234)_489 | 1 | + | good | ACTCCAGGCCGCAGGCTCGGCTCGCCCCCTACCACGGCGCCCGGGCCAGGAATGTTGCATGAATCGTCCTGTTTGCTGTTGCTTGGAGACTTGCCCTGTACATTGCTTAGTGCCCTCCCC | 120 | 62 |
| chr15 | 89558498 | 89558618 | 771629_41866724_SHANK3(58234)_490 | 1 | + | good | TGCCGCTGAACCCCACCCAGCACACAGTAAGGGCGCGGACCAGGGGGGCTGGGTGGAAGGGGGTGGGGGCAGGGTGCTCTGGCCTGACCACTTCCTCCATAGCTCCTGGTGGCCATTCTT | 120 | 67 |
| chr15 | 89558618 | 89558738 | 771629_41866724_SHANK3(58234)_491 | 1 | + | good | CCAGAGGGGGAACCTAGTCCAGCATGCGAGGTCAGGACACGCCTTGGTGACTCGGGGGGAGGGGGGAGACATTGGGGTTCTCGATAGGGGCCAAGGAGCCCCCTGTTTTACATATTTTAA | 120 | 58 |
| chr15 | 89558738 | 89558858 | 771629_41866724_SHANK3(58234)_492 | 1 | + | good | TCCACTCTATATTTGGAAAGAGAAAAGGAACAAATATCTCTGTCCGTAATAGTTCCCGCCCTCTTCCCCTCTAGTCCTCTCGCTGGTCCCGCCACAGCTGCCCAGTCTTCCATCTCTGGC | 120 | 52 |
| chr15 | 89558858 | 89558978 | 771629_41866724_SHANK3(58234)_493 | 1 | + | good | CCCTCACTGCCACCCCATACAGGGCAGGGGACACTCCAGCTGGCCTGGGGTTAGCCAGGGTCCTGGCAGCCCACCCTGGGGACCCCGGCTCAGCCCCCCTTCCCTCGCTGAGCTATAGTA | 120 | 68 |
| chr15 | 89558978 | 89559098 | 771629_41866724_SHANK3(58234)_494 | 1 | + | good | TGCCCCACCCACCCTTTAGGTGCTGCTCAGGGGGACGGGTGGCAGGCATTGCCTGCTGGGCACTAGCAGGGCCAGGTGGCCTGGGAAATTATTGCCCTGGGGCTGGGCCCCGGTAACCCA | 120 | 67 |
| chr15 | 89559098 | 89559218 | 771629_41866724_SHANK3(58234)_495 | 1 | + | good | AACCCCAGCCATCATCTTCACAGGGTCTCTCCCACAGGAGGGGTCTAACCTTTCCCACTTCTTGGGCAACTACAGCAGAGAAGCCTCCCTGCCTCGCACCCCAAAGACTCCCCAATTCCT | 120 | 57 |
| chr15 | 89559218 | 89559338 | 771629_41866724_SHANK3(58234)_496 | 1 | + | good | GCCCTGTGTGTGTGTGTGCGCCACATAATGTGTGTGCACGCCTGTGTGCTTGTGAAAACTGGGTGTGGCTGAGCGCATGGGTGCCCTGTGTGTGCTTGACTGTGCAGTGGTCCCCAGGGG | 120 | 60 |
| chr15 | 89559338 | 89559458 | 771629_41866724_SHANK3(58234)_497 | 1 | + | good | CTGTTCTGGATGGGTGGGAGGTGGCGGCAGAAGCTTGCACAGGGTGCATGCATGGGTGTGTGCCCGGGAAGGGGCCCTGTCCTGCTGGTGGGTCCTGCTGTTCTCTCAGCCTGTCCCCTC | 120 | 65 |
| chr15 | 89559458 | 89559578 | 771629_41866724_SHANK3(58234)_498 | 1 | + | good | CTCCCCAAACCTCACCCAGCTTAAGCAGGGATTCTTCGTGAATCCTTTCAGCTTCGGGAGGCCTCAAGGGCTCCCCCCGCAGGCAGTACCCCTTCGGGCTTCTAAGGGAATTGTGGGGAC | 120 | 59 |
| chr15 | 89559578 | 89559698 | 771629_41866724_SHANK3(58234)_499 | 1 | + | good | CACTAAAATCAGGCCACAACAGCCCTTGGAGAGAGGCAAAGACTCCTGAGGGTCCCCTGGCCCCCTTACTGTGACTCCTCACACTCAGCAATGACCTGTGGGGTGGGGGGCCCTGGGGCA | 120 | 61 |

| | | | | | | | | | |
|---|---|---|---|---|---|---|---|---|---|
| chr15 | 89559698 | 89559818 | 771629_41866724_SHANK3(58234)_500 | 1 | + | good | TTTTTAAACATCGGGTTTGGAGTCTGGACTAAGCTCCATCCACGTCACTCACAAGTTTCTGTTTCTATTTCTAGCTTTTTTTAATAAAAAATAAATAAATAAATAAATATATAAATAAAT | 120 | 26 |
| chr15 | 89559818 | 89559938 | 771629_41866724_SHANK3(58234)_501 | 1 | + | good | ATATATATATATAAAAGACAGAAAACAGGTGTTTTCATGGCCCAGGGGCTTGGCACGCCGGTCTGTGCCCACCTGCCCCACCCTGGCCCATCGGCCCCATTCCTTAGACACAGAGTCACA | 120 | 53 |
| chr15 | 89559938 | 89560058 | 771629_41866724_SHANK3(58234)_502 | 1 | + | good | CCCACTAACCCTCTCACCAACAGAGCAGGTCACACACACAGCAGCAGTCACTGTAACAGACTGCCACATACACAGTCTCACATTTACCTGTGGGTTTTTGGTTCTGTTCAATTTGGGTTT | 120 | 48 |
| chr15 | 89560058 | 89560178 | 771629_41866724_SHANK3(58234)_503 | 1 | + | good | TTAACTTTACAGGGTCAGTTCCGCTTCTCCCCCCCCCCCTTTTGTATGGAGTTCCATCTGGGGGGCTTTCAACCCCCTGCTCCAGTCCTGAGGCCTCCTGACCCTGACGTTGTGATACAC | 120 | 57 |
| chr15 | 89560178 | 89560298 | 771629_41866724_SHANK3(58234)_504 | 1 | + | good | CCCACAGAGATCTATGTTTCTTATATTATTATTATTAATAATAATTATTATAATATTATGTAATAAATTTATAAGAAATGAAATCATGTCTCAGTTTGTGCTTGCTTAGGGTTAGGGAGT | 120 | 24 |

**Supplemetary Table2b-the probe design for mouse Shank1-2 genes**

| Chromosome | Start | Stop | Name | Score | Strand | Code | Seq | Len | GC |
|---|---|---|---|---|---|---|---|---|---|
| chr7 | 144215817 | 144215937 | 772751_42001748_Shank2(210274)_337 | 45 | + | good | GGGTCATGAGGAAAAGGCGCTTACCACTAAGCCTGATGACCTAAGTTCACTTCTGTGAACCCATATGGTAGGAGAGGACTGACTCACACAGTTTGTCCTCTGACCTACTCATTTAAGCTG | 120 | 48 |
| chr7 | 144341697 | 144341817 | 772751_42001748_Shank2(210274)_1386 | 44 | + | good | GTAATAGGTGGGTGGGTGGGTAAATGGATGTGTATGCATTCATGAATAAATGAGTGGATGACTAAATGAATGAGTCCATGAATGGATGGAAGGTGGATGGATGATGTGTAGATAGATAGG | 120 | 41 |
| chr7 | 44317328 | 44317448 | 772751_42001747_Shank1(243961)_60 | 42 | + | good | TTAGTATGCAGTGTGTGTGTGTGTGTGTGTGTGTGTGTATGTGTGTACATAACAGGCACATGCATGTGGAACCTATGAGCAGGCATGTACAAGTGTGGAAGCCAGAGGATAACCCCAG | 120 | 48 |
| chr7 | 144226377 | 144226497 | 772751_42001748_Shank2(210274)_425 | 40 | + | good | GATAGATAGATGAAGGAATGGACTAGAGAATAAATGAATGGATAGATGTGTGAGTGGGTAGATGGGTGGATGAGCGAGGAAGTGGGTGATGAGTAGATGGATAGATGTGTAGACAGATGA | 120 | 42 |
| chr7 | 144383337 | 144383457 | 772751_42001748_Shank2(210274)_1733 | 38 | + | good | CCTGAAGAGTCATGGATAGTTGATGGCTTCCAGGAGAGGGAGAGTCAGTTTTCTTTAAGGGTGTGACCCTGGTAGGTTGGTTATACACCAGTGGATAGTGCCACACGTAGGTATATATGG | 120 | 48 |
| chr7 | 144312417 | 144312537 | 772751_42001748_Shank2(210274)_1142 | 36 | + | good | AAAGTCAAAGCAGGAGCCTGTAGGCAGGCACTGCTGAGAGACCATGGAGGGAGCAAGCCAGTAACTCCCTGTGGCTTGTTCGCTCTGCTTCCTTTAGAACCCAGGACCCGCTAGCCCTGG | 120 | 57 |
| chr7 | 144276297 | 144276417 | 772751_42001748_Shank2(210274)_841 | 33 | + | good | TGTTTTATTTATTTGTGAATAGGAGTCTCTGCTCCATGACATTCTTGAGGTCAGAGGACAACTTGCAGGAGTCCATTCTCTCCTTTCACATGTGTGTTTTAGGGATCATACTCAAATTGT | 120 | 39 |
| chr7 | 144258537 | 144258657 | 772751_42001748_Shank2(210274)_693 | 32 | + | good | ATCTTTCATCTGAATCTGCTGAGTCAGGAAAGTGAGGCTTAATTAGAAGCGTTAATCATGTGTGTGCTTAGTTTAGTCTTTTTTTTTTTTTTTTTTTTTTTTTTTTTTTTTTTTTTTTTTTT | 120 | 25 |
| chr7 | 44321888 | 44322008 | 772751_42001747_Shank1(243961)_98 | 30 | + | good | CCTCTCCACCACCAGTCAATCTACCCATCCATTCATCTACACATTGACTTATCCATCCATTAATACACCCATCCACCTGTCCATCCATCTGTTTACATACCGGTCCACCTAGCCATCTAT | 120 | 46 |
| chr7 | 44347928 | 44348048 | 772751_42001747_Shank1(243961)_315 | 27 | + | good | ATTCCTGTATTTGTTTAAGATGAAAGAAAGAAAAAGAGGAGGCAGAAAAAGGAGGGAAAGAGAAAGAAAGGGCTGAAGAATGGAGGAAGGAGAGGAGGGAGGCAGGAAGAGAGAGAGAA | 120 | 42 |
| chr7 | 144236337 | 144236457 | 772751_42001748_Shank2(210274)_508 | 27 | + | good | TTGACCCCCCCCCCTCCCTTGGTGATGGCCTGTGACCTGGAAGTATAAACTGAGCAAATACTTTTCTCCCCTAATTTGCTTTTGGTCAGGGTGTTTGGCCACAGCAACATAAAGGAAACA | 120 | 48 |
| chr7 | 144226137 | 144226257 | 772751_42001748_Shank2(210274)_423 | 26 | + | good | GTTTAAGGTTTTGGGGATATCTCATTTGCATGGAGAAGCATTCTAGGAACTTAGACCTGTAATTTCACCAAGGATTACATGACATTTTTCAAGTTGAAGACTGGGAGTTAGACTCCCCAG | 120 | 40 |
| chr7 | 144229977 | 144230097 | 772751_42001748_Shank2(210274)_455 | 26 | + | good | TACCTCCAGGGCTGGAGGTTTGGGGGAGCCACCCTCTCCCAAGGACTCACTCCACTTCAAGGTTTCTCTGAACACGTGCATGCACACACACACACACACACACACACACACACACA | 120 | 55 |
| chr7 | 144282297 | 144282417 | 772751_42001748_Shank2(210274)_891 | 26 | + | good | CCATCCCTCAGCCACTAAAGCACACAGAGAGAATGGGGTGCTGGGCTGGGCTGGGCTGGGCTGGGCTGGGCTGGGCTGGGCTGGACTGAGCACCAAGGTCTCAGAGAAACCCTTGTGGCT | 120 | 63 |
| chr7 | 144347337 | 144347457 | 772751_42001748_Shank2(210274)_1433 | 26 | + | good | CCCCAGCTGTGTCACTAGTACCTACTAGGGCTGTGGGTCTCAACCCCTTTGGAAGTCAAACAACCCTTTCACATGGGTCACCTAAAACCATTAGAAAACATGGGTTTTTATATCATGATT | 120 | 44 |
| chr7 | 44332688 | 44332808 | 772751_42001747_Shank1(243961)_188 | 24 | + | good | TTTTTTGTTTTTATGTTGTGTGTATGAACCTTTTGCCTGTTTGTGTGTCTCTGCACCACTATAAGCTCCTGGGGCTGTAGTCACAGACAGTTGTGAGCCACCATGTGGTTGTTGGGAATC | 120 | 44 |
| chr7 | 144261537 | 144261657 | 772751_42001748_Shank2(210274)_718 | 23 | + | good | TCCATTTATCCATCTATTCATCCATTTTCTTTTCCATCCATCTATCCCTACATCTTTCTATTCTTTCATCCCTCTATCTAAATATCCATTCAGTCGTCATCTATCTTTCTACCTGTCCAT | 120 | 35 |
| chr7 | 144226497 | 144226617 | 772751_42001748_Shank2(210274)_426 | 21 | + | good | ATGGATGCATGCATAAATGGATAGGTAGATGGATGAATGAATTGATGAAAGGATGGGTGCATGGATGGTGTATAGACATATGAGTGGATATGTGGCTGGCTGGCTGGATGAATGAATGGG | 120 | 43 |
| chr7 | 144343257 | 144343377 | 772751_42001748_Shank2(210274)_1399 | 21 | + | good | GTTCAGTACAACTTAGCAGCTGCTGTTCGTTCAAACTTCAAGAGTCTGATGCTGGGTCGTATATCATAGGCATTTTTAAGCACAGCATAATTTGGGGGAAAAAAGTTTCCTGGTGATAAT | 120 | 40 |
| chr7 | 144205737 | 144205857 | 772751_42001748_Shank2(210274)_253 | 20 | + | good | CATGCACATGAGGAAGCCAGAGGAGGACATATGGTGTCCTGCTTTCTCACTCTACCTTATTCCTTTGAGATGGGGTCCCTCCATGAACCTGGAGCTTGACTGGTGGCCTGAAAGCCCCAG | 120 | 53 |
| chr7 | 144331737 | 144331857 | 772751_42001748_Shank2(210274)_1303 | 19 | + | good | GCCTGGAGCCAGACAGAGGGGAGCTTGTCTCCCAGAGACCTCCCAGAGTTCTGAGAACTTGTCAGATCACCCTTCAGGAAAGACTGGAGCTGAGACAGTGATGGGTCAGTGATGGGCAGA | 120 | 56 |
| chr7 | 144336417 | 144336537 | 772751_42001748_Shank2(210274)_1342 | 19 | + | good | GAGATGTTTTTCACCTAAATTGAAAATAGATTTTTTTTTAAATTTTGTTTGTTTGAGACAGGGTCTCATTTAGCCCAGGCTAGCTTCCAACTCACTGTAAGGCTGAGGATGACCTCCAAC | 120 | 38 |
| chr7 | 44315168 | 44315288 | 772751_42001747_Shank1(243961)_42 | 18 | + | good | TGTCTGGTAGGGGTGGGGTTGCATGTTCATGTCATGGCTCACATGTGGAGGTCAGAGGATAATTTCTAGAGCTCAGTTTTCTCCTTCTCTATGTGTGTTCTGGGGAATCAAATTCAGGTT | 120 | 46 |
| chr7 | 44349968 | 44350088 | 772751_42001747_Shank1(243961)_332 | 18 | + | good | TGATGAATAGAATGATGGACAGATAGAATAACAGATGAATAGCCAGACAAACGAAATGGTGGATGGAAGATACACGGACGGAATGATGGATGGATAGAATGATGGATGGATGCAATGACA | 120 | 40 |
| chr7 | 144192057 | 144192177 | 772751_42001748_Shank2(210274)_139 | 18 | + | good | ATAAATAAACTTTAAAAGAGTGATAAGGTGGATGATGTCACTGAGGATGAGACCCAAAGTTGTTCTCTGGCACATGCACATGCCTACACACACACACACACACACACACACACACACA | 120 | 42 |
| chr7 | 144262377 | 144262497 | 772751_42001748_Shank2(210274)_725 | 18 | + | good | TAGCATCAGCTTTGAGCCTCCACATGTGCATATACATGTGCATATTCAACCACACATGCACCAACACAGGTAAACATGCATGTACATGCATACACATATAAACATGCACACACACAATAC | 120 | 41 |
| chr7 | 144246177 | 144246297 | 772751_42001748_Shank2(210274)_590 | 16 | + | good | AATGATGTTGGATTTAGAACCTGCATAAAAGCTGGGCAGGCTGACTTATTCCTATACTCTCAGCACATTGGAGGCTGAGGCAGGAGGATTGTCATTAGTTTGAGGCCAATCTGGGCCATA | 120 | 46 |
| chr7 | 144255297 | 144255417 | 772751_42001748_Shank2(210274)_666 | 16 | + | good | GAGACACATCTTCACACAATGTAAAAGTGGTTACTCCAACAGATGATGGTGATGGTGGTAATGAAGATGAGGGTGATGGTGGTGGTGATGATGATGGTTGTGATGGCAGTGATGATGCTC | 120 | 45 |
| chr7 | 144348777 | 144348897 | 772751_42001748_Shank2(210274)_1445 | 15 | + | good | AACAACTGAAAACTATGCCCTGATCCTCAGAGAGGACAACACAAGGCCTAGTTTAAGCTTTTGAATCCTCCAAGTTCACCCCTAGTGACTCACTTCTCCCAATAAGGCCACACCTCCTAA | 120 | 46 |
| chr7 | 144348897 | 144349017 | 772751_42001748_Shank2(210274)_1446 | 15 | + | good | TCCTTCTCAAATAGTGCCACTCCCTGATGACTAAGCATGCAAGTATATGGGCCTTGGGGGCAGAGGGAGGGTAGCATTCTTATTTTAACTACTACACAGGGACACAAAGAGTCTGACATC | 120 | 47 |
| chr7 | 44343248 | 44343368 | 772751_42001747_Shank1(243961)_276 | 14 | + | good | TTCCAGTCCCAGAACTGCAGAAAAGGGCATTTTGGTACACACCTGCAATCCTACCGTCTAGGTAGAGGCAGGAGAATCAGAAGTCCAAGGTCATCCTGAGCTACTTAGTAAGTTTGAGCT | 120 | 48 |
| chr7 | 144218217 | 144218337 | 772751_42001748_Shank2(210274)_357 | 14 | + | good | ACATAGAGAGGCCAGAGGTCAATGTATCTTTCTTAATCCCTCCCCACCTCATGCTTTAAGACAGGGTCTCTCACTGAGCCCGGAATCACAGATTCCACTACACTAGCCGGCCAGCAAGCC | 120 | 52 |
| chr7 | 144308937 | 144309057 | 772751_42001748_Shank2(210274)_1113 | 13 | + | good | GGGAAGGTAGAAACCAGTGCATGCCACTGTAGTGCTTGCTCATGGTGTGAAAGGAAACATGTGCACATGCACACACACACACACACACACACACACACACTGAAGCATGCACAGGCACAC | 120 | 51 |
| chr7 | 144306177 | 144306297 | 772751_42001748_Shank2(210274)_1090 | 12 | + | good | TGAATATCTGAGCTTCCTGTCTCCACTTCCCAAGGGCAGAGATTACAGCCACATGCCACCATACCTGGCTTATGTGGTGCTATGAATTGAACCTAGGACCTTCTGCATGCTAAGTGAGCA | 120 | 48 |
| chr7 | 144372537 | 144372657 | 772751_42001748_Shank2(210274)_1643 | 12 | + | good | TGTGCATGCACATGTGTGTGTGCATGTGTGACTGTTTATGTCCATGTAAGTGGAGGCATCAGGACCCCTAGAGCTGGAGTTACAAGTGGTTGTGAGCCAGTTGACCAAGATCCTGGGAAG | 120 | 50 |
| chr7 | 44327408 | 44327528 | 772751_42001747_Shank1(243961)_144 | 10 | + | good | GAATGTTTGAATGCATGTGGGCACATGTGTGTGCTGGTGTTTGTGCATGTGTGTAGTGTGTATGGGGAGGCCGGAGGTTGACATCAGGTGTCTTCCTCAGTCAGCCTCCACATTTTCTTT | 120 | 49 |
| chr7 | 44330648 | 44330768 | 772751_42001747_Shank1(243961)_171 | 9 | + | good | TTTTCATGGTTCTAGGGATTGAAGTCAGATTCTCAGGCTTGTGAGGCAGGTACTTCATCTACTAAACTACATTCCCAGTTCTTCAGCCTCTTCTTTCTTCTTCTTCTTCTTCTTCTT | 120 | 40 |
| chr7 | 144257097 | 144257217 | 772751_42001748_Shank2(210274)_681 | 9 | + | good | AGCTGAGGTGGGGAGGGATGATTTTAAACTTCCACGACAGACACAGTCCATCATTGAGGGAAATCAGGGCAGGAACTCAAGGCAGAGCCCTGGAGCCAGGCCTGCTCACAATTCCATGCA | 120 | 53 |
| chr7 | 144345897 | 144346017 | 772751_42001748_Shank2(210274)_1421 | 9 | + | good | CACACACACACACACACTGCCACTTGCCCATAGGCCAATCTGATAGAAACATTTTCTCAACTGGGGTTCCCTCTTCACAGGTGCTGCTGAATTGTGTTGAGCTGACAAGAGCTAACCAGC | 120 | 49 |
| chr7 | 144362457 | 144362577 | 772751_42001748_Shank2(210274)_1559 | 9 | + | good | TCTTGGAGATCTGATGCCCTCTTCCGGCCTCCACAGGCACTGAAGGATGCAAAACCACACACAGGCACACACATATACACTAACTAAAAATAATTTTTCTAAAGAGCATTTTATATACAA | 120 | 40 |
| chr7 | 144384057 | 144384177 | 772751_42001748_Shank2(210274)_1739 | 9 | + | good | TCTAGTTTCCTTCCCTGGCTTTCCCACGGTGCTGGACTCTGACCTGGATGTCTAAGCCAAATAACCCTCTCTTCCTCGAGTTGCTTTTGGTTGGGGTGTTTTATTAGTCACAGAAAGGAA | 120 | 48 |
| chr7 | 144212577 | 144212697 | 772751_42001748_Shank2(210274)_310 | 8 | + | good | AATGGACCAATGGGCATACAAATGGATGGATGGATGATCAGGTGAATGGATGGATTGATTTATGGATGAATAGATGGGTGATCGACAGGTAGGTGGATGGGGAACTTAATAGTCTTCTTG | 120 | 42 |
| chr7 | 144266097 | 144266217 | 772751_42001748_Shank2(210274)_756 | 8 | + | good | GATTCTTGCATGAAGGGTTTTTTTTTGTTGTTGTTGTTGTTGTTATTTGTTTTTTGTTTTGGGATTCTACTTGCCTTTATTTTTCCCTTTGAAGCAGAAGAATTGGGTTTTAGTTAGCA | 120 | 32 |
| chr7 | 144266577 | 144266697 | 772751_42001748_Shank2(210274)_760 | 8 | + | good | CTGAGTTCCCTCAGACTCAGCAACCTTGCACATTCTCTCTTGCTCATGCTCTCTCAATCCCCCCTCTCGCCCTTGCTCTCTCCTGCACTCTCTTGCTCTCTCTCCCTCACTCATGCTCTC | 120 | 55 |
| chr7 | 144331857 | 144331977 | 772751_42001748_Shank2(210274)_1304 | 8 | + | good | ACCACACCCTGACCAAGACTCTTAGCTCTGCATTCCTCCTGCCCTCTGTTTAAGCCTGAGCCTCATGTCTAGATCCTGAAGACCGACTCTGGAGTCACTGGTGAACTCTTTGTCCCTTTT | 120 | 51 |
| chr7 | 144333657 | 144333777 | 772751_42001748_Shank2(210274)_1319 | 8 | + | good | TGTGACTGACCACCTGGTTGGGGGGGTCACCTGGTTATGTGCTCCGAGGCAGGCTCGGAAACAGAGAAGGAGCAACTTGTACTCCTGCCAGTCTCAAGTTTTGAGATTTCCAGGGTCTGA | 120 | 54 |
| chr7 | 144339057 | 144339177 | 772751_42001748_Shank2(210274)_1364 | 8 | + | good | ACATGTAAAGTGAATGTTCATGTGTGTGCACACACGTGTGAAGACCAGAGGTCAGTGTCAGGTATCTTCCTCTACTCTCCATCCTAATTTTTAAGACAAAGTATCTCACTGAACCTGGAT | 120 | 42 |
| chr7 | 144343377 | 144343497 | 772751_42001748_Shank2(210274)_1400 | 8 | + | good | TAAGTCAAGCCCATGTGCACAACAAAGTTTAAGATGTAATTACATTGAAACTCTATATAAAGTACATATGAGATCTTCTATAAAGATTGGTTCCATAGTAGGCTCAAGAGGTAAACAAAC | 120 | 32 |
| chr7 | 44332928 | 44333048 | 772751_42001747_Shank1(243961)_190 | 7 | + | good | TTGGGAGCAAAAGAGAAGCCCAGAAACCTCAAGAAGGGCCTGGGGTTCAGTTGTTAGAGGGATTAGCTAGCACACACAAAGCCCTGGGTTCAATTCCCAGCACCACATAAACTGGGTATG | 120 | 50 |
| chr7 | 144205617 | 144205737 | 772751_42001748_Shank2(210274)_252 | 7 | + | good | CAGGTTTATATGTAAGGGTAGAAATGTTGGGTTGTGTGATGGCTATGTTTAAGCACATGTGCAAGTGCATGTGTGCACATGTGTGTGTGTGTTATGTTTGCATATGTGTGTTGGTGTATG | 120 | 41 |
| chr7 | 144254817 | 144254937 | 772751_42001748_Shank2(210274)_662 | 7 | + | good | CAGCTATCATGCTTAACCTGACTAATCCTGGCACTGCGTCCTGCCACGTGGCTAGCTCTACTTCCCTACTCCACTTTACTTAGTCCATTCACCTCGGTGTCTGCTGGCAAATATCCCACA | 120 | 51 |
| chr7 | 144267777 | 144267897 | 772751_42001748_Shank2(210274)_770 | 7 | + | good | GGTATACTGCTTGCCTACCACTCAGCAAGCCGTGGCTTTGAGCCTCAGCACTGCATACACCTGCTGATGTAGCTCTTACCTGTATTCCTGGTACTCAGGAGGTGGAGGCAGGAAGATCAG | 120 | 53 |
| chr7 | 144280137 | 144280257 | 772751_42001748_Shank2(210274)_873 | 7 | + | good | CTCTCTCTGCCTTCTGGCTATGAGTGCGATGTGAGTGACCTTGTGTTCCTGCCACCGTGCTTTCCCCAGCATGATGGACTGTGCCCTTAAGCTGTGAGCCAAGGGAACCCTTCCTCCCTT | 120 | 56 |
| chr7 | 144297177 | 144297297 | 772751_42001748_Shank2(210274)_1015 | 7 | + | good | ACACACACACACACACACACACACACACACACACACATGAAAGGAAGCCAGTGAGGCAAGCAGGTGAAGGGACCGCCAGGCAGGGAAGTTGTGCAGCACATGAGGATCACAGGGTAGCTT | 120 | 54 |
| chr7 | 144302817 | 144302937 | 772751_42001748_Shank2(210274)_1062 | 7 | + | good | TGATCTTTTGGTTAATTGCAGACATGTCTATCATCCCTGCCAGGTCATCTCTGTCTTTCTCATTTTCTGGTGGATATGTGTGTGTGTGTGTGTCTGTGTCTGTGTCTGTCTATGTCTGGG | 120 | 44 |
| chr7 | 144327297 | 144327417 | 772751_42001748_Shank2(210274)_1266 | 7 | + | good | AGCCAGCCAACCTTGCAGAATCGGCAAGTTTCAGAATCAGAGAGAGAAAGACCCTGTCTCAAAAAATAAGGTGGGGGTGGCTGAGGAAACACTCAGTGGCAGCCTCGGGTCTCTACACAT | 120 | 51 |
| chr7 | 144374937 | 144375057 | 772751_42001748_Shank2(210274)_1663 | 7 | + | good | AGGTGGATCCCTGGGGCTCACTAGCCAGCCAGACTCGTCTAGTTGGTGAGTTTCAGACCAGTGAGAGACTGTTTCAAAGCTACTTCTAAGGGACAACACCAAGAGAGAGAGAGAGAGAGA | 120 | 51 |
| chr7 | 44332808 | 44332928 | 772751_42001747_Shank1(243961)_189 | 6 | + | good | AAATCCAGACTCTGGAAGAGCAACCCGTGCTCTTGGCTGCTAAGCCATCTCTTCAGCCCTCATTCTTCTTGAAGGATATCTTTGGGTCTTGGGACAACAGGACAGTGTGCACCAAGTGTA | 120 | 49 |
| chr7 | 44349488 | 44349608 | 772751_42001747_Shank1(243961)_328 | 6 | + | good | GGACTTATTTTGTTTTCTTGCATGTTTTGTTGTTTTTCTGAGACAGAGTCTTATGCCGTAGCTGACACTTCCTTGAATTCACTTGTAGACCGTGCTAGTCTGGAATTCATGGTAATCCTC | 120 | 40 |
| chr7 | 44357888 | 44358008 | 772751_42001747_Shank1(243961)_398 | 6 | + | good | CCTGCTGAGGCCCTCTCCGCTCCCCAACACGCTGCATTCTTTTTTTTTTTTTCTTTCTTTCTTTTTTTTTTTTTTTTTTTTTTTTTTCCTCAAGCGCCACCGCCTTCGCTAGCTTGCTCT | 120 | 42 |
| chr7 | 144229137 | 144229257 | 772751_42001748_Shank2(210274)_448 | 6 | + | good | ACCTGGCATTTTAATGTGGGTTCTGAGTATCAAACTTGGGTTCTTATGCTTAGAGGTCAAGCACATTACTGACTGAGCCATCTCCCCAGCCTCTAGCTGAGCACAGATGTGCCATGGCCT | 120 | 48 |
| chr7 | 144309177 | 144309297 | 772751_42001748_Shank2(210274)_1115 | 6 | + | good | GTGACACAACATTGGACAAATGCAACTTAAGAGAGAAAGGGTTTATTTTGGCTTGTAGCCTGAGGTTACAGCCATTATGTAGGGAAAGGCATGAAGGCAGGAGGTGAGGCGGCTGGTCAT | 120 | 47 |
| chr7 | 144365337 | 144365457 | 772751_42001748_Shank2(210274)_1583 | 6 | + | good | GCAATTTAAGGGAAGAGTGGTTTATTCTGGCTCACAGCTTGGGGGAATACAGTCCATCAGGACAGGGAAGGCAGGAGGAGCAGTCATGTTGTGTCCACAGTCAGGAAGGAGAAACTGGAC | 120 | 51 |

| | | | | | | | | | |
|---|---|---|---|---|---|---|---|---|---|
| chr7 | 144371937 | 144372057 | 772751_42001748_Shank2(210274)_1638 | 6 | + | good | TGTTAATTGTTAAGAATATTAGCATTGTAAGGAGAGGCCTGCTCTACCTGAAATTAAGGGAAGGGTGCCCTCAGGACAAGATCCCACACAGCTAAGCTTCCAACATCTTCCAACCTCAGC | 120 | 45 |
| chr7 | 44318768 | 44318888 | 772751_42001747_Shank1(243961)_72 | 5 | + | good | TGGATCCATGAGCTCTAGGTTCACTGAGAGGCTCTCTCAAAAAATAAAGTGAAGAGCAATTCAAGAAGACACATAGCATTGATTCCTATGTGCTAGCATTCACATGTGAGTGCATTCATG | 120 | 41 |
| chr7 | 44325008 | 44325128 | 772751_42001747_Shank1(243961)_124 | 5 | + | good | TTTTATCCTGAGACAGCACTTTGCTAAGTTGCCCATGCTGGCCTTGACCTTGTAATCTCCTGCCTCAGCTTCTGAAGTAAACTAGGATGACAGACGTGAGCCACCAGGCCTAGCTTCCCA | 120 | 50 |
| chr7 | 44327528 | 44327648 | 772751_42001747_Shank1(243961)_145 | 5 | + | good | TTTGTGAGAGAGAGGGTCTCTCACTGAACCTTCACTGACTCAACCAGGCTGGCTACCTGAACCAAGCAAGCTCCAAGAACCTACCTTTCTCTGCTTCCCTGCTGGGGGTTACAGATTCTC | 120 | 52 |
| chr7 | 44329448 | 44329568 | 772751_42001747_Shank1(243961)_161 | 5 | + | good | TCTTGATTGAAGAATGGCTCTCCTCTGTGAGGAAAGGCCATTCTCCTAGTCCGAGTGCATACTTCAAGAGAGATGCACATGGAGCAGTAAGGGAGGAAAGAGATACCTGCCTAGCCAGCC | 120 | 50 |
| chr7 | 44346728 | 44346848 | 772751_42001747_Shank1(243961)_305 | 5 | + | good | ATGACATGCCTCCAGCTCCTTCCCTGGGGGATTCTAGGCAGGGGCTCTACTCCTGCCCCTCACTGAGGAGATTCTAAGCAGTTCTCAACATGTGGGTCATAACCCTTCTCACAGGGGTCC | 120 | 56 |
| chr7 | 144226017 | 144226137 | 772751_42001748_Shank2(210274)_422 | 5 | + | good | TATTCACATACACTTTATTCAGGGTGAATCAAGGGATATATATATATATATATATATATATATATATATATGAACCTTGGAAAGGGGCTCTTTTTAGGGTGAAGTTTTACTGGCTGAG | 120 | 29 |
| chr7 | 144297657 | 144297777 | 772751_42001748_Shank2(210274)_1019 | 5 | + | good | GGGGGACTTTTGTACCGCATCATACATGTGGAAGTTAGAAGGCAACTTTTGGGAGTTGCTTCTCTCTTCCCACCATATGGGTTTCTGGATTCTAACTCAGATCATTAGCTTTACTAGCAA | 120 | 43 |
| chr7 | 144323937 | 144324057 | 772751_42001748_Shank2(210274)_1238 | 5 | + | good | CTTGAACTCCTGACCTTTCTGCTTCCACCTCCCTAGTGTTAGTATTAGCAGCATGTACCTCCAGACCTAGTTAATGCTACTGGGGATGGAACCCAAGATTTTATATATGCTAAACAAGCA | 120 | 43 |
| chr7 | 44332328 | 44332448 | 772751_42001747_Shank1(243961)_185 | 4 | + | good | CTCTGGGTATCCTGCTGTATCACTCTCTGACTTATCCCCCTTAAGACAGGGTCTCTCACGGGGCTTGGAGCTGGGCTAGGGGTCAGCAGACCCCAGGGATCTTTCTGTTTCCTAACAACA | 120 | 54 |
| chr7 | 144181497 | 144181617 | 772751_42001748_Shank2(210274)_51 | 4 | + | good | TGGTATCACAAAAAAAAAAAAAAAAAAAAAAAAAAAAAAAAAAAAAAAAGTCCTTTTAAAACAAGGACCTCACATTGCAGAGCAAAGTGCCTGAGACCGTAGAATATTCTGTATGTGCCA | 120 | 29 |
| chr7 | 144225297 | 144225417 | 772751_42001748_Shank2(210274)_416 | 4 | + | good | AGATGTGTGGATACGTGGAAAGAGATGAATAGATGGATTAGATAAATAATTAAATGGATAGATGAATGAGATAGATGGAGGGATGAAAGAGAGGATGGATGTGTAACTAGGAACAGAGGG | 120 | 38 |
| chr7 | 144225897 | 144226017 | 772751_42001748_Shank2(210274)_421 | 4 | + | good | TGGAGGGAAAGGATGAAGTAAAGTCCATAACAGGTAGTGGGTGGTGGGGGAACTCTTCTCCAACGATTTCAGTGTTACAGCTCTTTTAGGTGACATTCAATAAACAGCTCTCTCAGACGG | 120 | 46 |
| chr7 | 144244737 | 144244857 | 772751_42001748_Shank2(210274)_578 | 4 | + | good | GGTTGGCTTGGATATTTTATTTGTTTTGGTTTTGCTTTTGATGTGAATTCATGTTTCACTCTGTGTAGTACTAGCGTGTGCCTGGTACCTGTAGAGGCCTGAAGAGGGCATTGGATCCCC | 120 | 44 |
| chr7 | 144377337 | 144377457 | 772751_42001748_Shank2(210274)_1683 | 4 | + | good | CCCATTTAGTATTGCTCATTTGTAAATGTGTTTAGGGCTGATCACGCGGGATTAGATTACCTGTCAAGGGCTCATCCCTGAGTAAAACTGATTTTCCCTCATTTACCAGCAATTGGTTGC | 120 | 42 |
| chr7 | 44333048 | 44333168 | 772751_42001747_Shank1(243961)_191 | 3 | + | good | GGGTGCATGATGTCACTCCAGCATTGAGGGGGTGAAAGAAGGTGGATCCAAAGTTCAAGGTTGTGATCAACTATAAGTGAGTTTGAGGCTGACCTAGAGTACATGAGAACTTGTCTCAAA | 120 | 45 |
| chr7 | 144185337 | 144185457 | 772751_42001748_Shank2(210274)_83 | 3 | + | good | GCCGAACAAACCCTCCTTCTCCCATATTGCTTTCTTGTCAGGGCATTTTTATCACAGCAACAGAACTGAAACTAGGACAAAAATTCACCTGAACTAATCAAAGAGCATCCCACGGCTTCC | 120 | 44 |
| chr7 | 144189057 | 144189177 | 772751_42001748_Shank2(210274)_114 | 3 | + | good | GAAGAAGACACCCAATGTCAGCCACACAACCAGGGACACACATTCACACATTGCATGCATAGGTGCCTATCTACATGAACTCACTCTCTCTCTCTCTCTCTCTCTCTCTCTCTCTCTC | 120 | 48 |
| chr7 | 144193617 | 144193737 | 772751_42001748_Shank2(210274)_152 | 3 | + | good | ATAGGTCATGATAACCTTGAATTCCTGAGCCTCGTGTCTCCCCACCCAAGTGCTAGGGTTAAAGGCATAGACTACATGGTTCATATGGTGCTGGAGATGGAACTCGGAGCTTTGTGCATG | 120 | 48 |
| chr7 | 144205857 | 144205977 | 772751_42001748_Shank2(210274)_254 | 3 | + | good | TGAGCCCCCTGTCCCTGCCCCACACAGGGCTGGGTTACAGGTATGCATGTAGCTCCAGCCGGTTTTTTATGCAGCTCTGTGGTTTTGGATTCAGGTCCTCATGTTTGTGCAGCAAGCACC | 120 | 55 |
| chr7 | 144206217 | 144206337 | 772751_42001748_Shank2(210274)_257 | 3 | + | good | TCTCTCTCTCTCTCTCTCTCTCTCTCTCTCTCCCCCTCTCCCCCCCCCTCTCCTAGGGTGCTCAAGAGGTCAAGAGGAGTATGGCTTAGTTCTTTACAGCCTTATAAAAATGGGACCTTCCA | 120 | 52 |
| chr7 | 144208497 | 144208617 | 772751_42001748_Shank2(210274)_276 | 3 | + | good | GTTAGCCAGGTCTACCCAGAAATGTCATGCAGATCCTTGGTTAGCCAGGTCTACCCAGAAATGTCATGCAGATCCTTGCTGGAATGGTGACAGCTCTGTTTTGTGCAGCAGCTACTGCCA | 120 | 50 |
| chr7 | 144225777 | 144225897 | 772751_42001748_Shank2(210274)_420 | 3 | + | good | TTTTTTGTTTCTTCCCTTGGAGTGGAGTGGAGGTGCTGCTGCACACCCACCATGTGGGGAAGGATGAAGTAGAGCCAACTTCTGTGGTCTACTTCTTCCCTTGGGGTGTGGATACGTGCC | 120 | 52 |
| chr7 | 144237897 | 144238017 | 772751_42001748_Shank2(210274)_521 | 3 | + | good | ACACCAAGCTCCTCTACAGCTTTTTTTTTTTTTTTTTAAAGAGGGAAGCCAGGTGTGGTGACACATAGTTGACATCCAACCACTCAAGAGGCTTCTTAGTGAGGCCAGCCTGGGGTACA | 120 | 45 |
| chr7 | 144243177 | 144243297 | 772751_42001748_Shank2(210274)_565 | 3 | + | good | ACATCCATGCCTACCATGGGGGAATCAAAGGTTCTATGTGCGTGTGCATGTGTTTGTGTGTGTGTATGTGTGTGCATGCCAGAGCTCAACCTCAGAAGTCATTCCTCAGGTGTCGTTTAC | 120 | 48 |
| chr7 | 144252177 | 144252297 | 772751_42001748_Shank2(210274)_640 | 3 | + | good | GTCTCTCACTGAACTAGGAGCTCTCCATTGTCTGCTAGGCTGGTTGGTCAATGAATCTCAAGAATCTACCTGTCTCTTCCCCATTCACTGCTGAAGTTACAGCATGTGTGGCCACACCAG | 120 | 49 |
| chr7 | 144328977 | 144329097 | 772751_42001748_Shank2(210274)_1280 | 3 | + | good | CCAAGCCAGCAACACGGCATGCAGGCCTGTGATCCAGCACTGGGGAAGCAGAAATAGGAGAATCCTTGGGGCTTGCTGGCTGGCCAACCTAGTTGAATGGGTGGGTGAGTGCCATGCTCA | 120 | 57 |
| chr7 | 144361857 | 144361977 | 772751_42001748_Shank2(210274)_1554 | 3 | + | good | TAGCCTCTGTGGGTACTGCATACATGTAGTAAAGCTAAACACATGCAGCAAACACTCATGCACATTTAATAAAAATAAATAAAATGCCCAAGACAGGCTTACTAGAGGATGGGACATTGT | 120 | 38 |
| chr7 | 144364257 | 144364377 | 772751_42001748_Shank2(210274)_1574 | 3 | + | good | GGACATCTTTGAACATTTTGGCCTTCTGTTTTTGTTTCATTTTGATGTTTTTTATTGAAAAGAAAAACTTCCCTACAATATATTTTGATCATGTTTTTCCCTCCCCCATGATTTTCCCTT | 120 | 32 |
| chr7 | 144368217 | 144368337 | 772751_42001748_Shank2(210274)_1607 | 3 | + | good | CTCTCTGTGTGCCCAAATATCTATATGTGCCCTTGAAGGTCAGAAGTGGGTGTTGGGTCCCTCAGAACTGTCTAAGGTGGGCTGCTAGAAACTGAACCTGGGTCCTCTGCAAGAGCAAGT | 120 | 51 |
| chr7 | 144381177 | 144381297 | 772751_42001748_Shank2(210274)_1715 | 3 | + | good | GCCCAGTTTTTATTGTTTTTTAAAGACAGAGTCTTCTGCAGCACAGACTGGCCTGAAATTGGCTACATGGCTAAGGATGACATTGAACTCCTAATGCTCCTGTCTCCACAAGTGCTGGGG | 120 | 46 |
| chr7 | 144382137 | 144382257 | 772751_42001748_Shank2(210274)_1723 | 3 | + | good | CCATGTGGGTCCTGAGCCTTCATTTCATCAGGCTTAGCAAGCAAGCACCTTTATCCACTGAGCCATCTTGCCATCCCGGGAGCTTTCTCTAAAAACTGTTCCAAATGGAACAGGAGAGAT | 120 | 48 |
| chr7 | 144386097 | 144386217 | 772751_42001748_Shank2(210274)_1756 | 3 | + | good | CTGAATAAGGTGGTGAATGGTTGTAATCTCATCCCTGAAAAACAGAGACAGGCAGATCCCTGTGGCTCACCGACCAGCCAGTCCATTGTACATGGTGAGTTCTAAGCCAATGAAAGACCC | 120 | 48 |
| chr7 | 144390417 | 144390537 | 772751_42001748_Shank2(210274)_1792 | 3 | + | good | AGAAGTAGTAACACATACTTGAAATTCCAGCACATAAGAAGCAGAGGCAAGAGGGTCACCCATAGATAAAGTTTATCCCAAGCTATATAGTGAGTCCCAGGCCAGCCAGGGCTATATAGT | 120 | 43 |
| chr7 | 44316008 | 44316128 | 772751_42001747_Shank1(243961)_49 | 2 | + | good | TGTCTTTCTTTGTACATGATTTAGACTGGGTTGAAGAAGTTCCTGGAATATGTGCAGCTTGGGACGTCTGACAAGGTGGCAAGGCTGCTGGACAAGGGGCTGGACCCCAACTACCACGAC | 120 | 51 |
| chr7 | 44317088 | 44317208 | 772751_42001747_Shank1(243961)_58 | 2 | + | good | GATGTGGAGGCACAAGCCTGGCATCTCAGCACTGGGGAGGTAGAGGAAGGAGGGTGAGGGGTGTAAGGTCTTGATAGCCAAGATCTGGACACACACATACATACACACACACACACACTA | 120 | 52 |
| chr7 | 44321768 | 44321888 | 772751_42001747_Shank1(243961)_97 | 2 | + | good | TCACCATCTATCCGTCCTTCATTCACCTAATCACCCATTCATCCGTTACCAATGCATCTATCTATTCAGCCACACACCCACTCACCCATCTATATATGCTTCCACCCATCTGTCTGTCCA | 120 | 46 |
| chr7 | 44340848 | 44340968 | 772751_42001747_Shank1(243961)_256 | 2 | + | good | ATATCACATTCAGTGGGGACTGGTGCTTATTTTTTCCAGTGCCAGAGATGGAACCTAGGGCCTTAGTGATTGCTAGGCAGGAGCTTTACCATAGATCCAAACCCCAGCCCCTCACTGGGG | 120 | 51 |
| chr7 | 44342408 | 44342528 | 772751_42001747_Shank1(243961)_269 | 2 | + | good | CCCGTCTGCAGCGCAGACCCCCATCGAGGAATTCACCCCCACCCCGGCCTTCCCGGCGCTGCAGTACCTCGAGTCTGTGGACGAGGGTGGCGTGGCATGGCGGGCTGGACTACGAATGGG | 120 | 68 |
| chr7 | 144191697 | 144191817 | 772751_42001748_Shank2(210274)_136 | 2 | + | good | GTAAACAGGTAAATGGGGGTAGGAATTTGTAAGGGTGGGGATGGAAGGGGAACTGGATGTAAAGAGAATAAAAAATAAATTATTGAAAAAATAAAACAACCCAATACCCAGAGCTGGAGA | 120 | 38 |
| chr7 | 144195777 | 144195897 | 772751_42001748_Shank2(210274)_170 | 2 | + | good | GACTCCACATGCATCTACACACATGTGTACTCCTATGTATATATATATCTGTACATATAAATATTTACTTACACAAAATTAGTGGAAGAAAGGAAGGGAGGGAGGGAGGGAGAGAAGGAA | 120 | 38 |
| chr7 | 144208977 | 144209097 | 772751_42001748_Shank2(210274)_280 | 2 | + | good | CCCACACCTCTCTGCAGCCCAGACTCAGAGCACTGAGCGCTGTGTGTGCTGTTCCAGGTGCCAATGCTCTTCCCATTGGGCTGCCCTGGCTGACCCTTCCTAACTCAGCAGAGAAGGAAA | 120 | 58 |
| chr7 | 144209577 | 144209697 | 772751_42001748_Shank2(210274)_285 | 2 | + | good | ATGTTAAACTTTTTGAGAAATGGTAAAAATGTTTCTATGTGCTAGAAAATTGTGTTTGCATATGTGTGTATACATGTGGGGAAATCAACCCGTGTGTGTGTGTGTGTGTGTGTATAGG | 120 | 36 |
| chr7 | 144210057 | 144210177 | 772751_42001748_Shank2(210274)_289 | 2 | + | good | TGGCTGACCCTTCCTACCTCAGCAGAGAAGGGGAAGGCTAGAGGTCCCCTCCCCTGCAGGCTCAGCACCCCCAACCCCCAAGCCCTCTGGGATTTTATGTAAATCATGCACACACAAGAT | 120 | 57 |
| chr7 | 144215457 | 144215577 | 772751_42001748_Shank2(210274)_334 | 2 | + | good | CACTGGACTAAGTACCATGACTGTCCTCAGTGTTGATCCCTATGATGATCCACTGAGTCTTCTTGTGAACCTGTTGGGAGCTGGGAGGCTTATAGCTCCCCACCCCCAGATCCTGCAACG | 120 | 52 |
| chr7 | 144216417 | 144216537 | 772751_42001748_Shank2(210274)_342 | 2 | + | good | CTCTGAAGGACTTGGGGGACTCAGTGCAGAGTTGTATTGTTGTCTGAGATTGGTCACAGTGACACAGGTGGGACACCCATGTTGAGAACAGGCAGATGCTGCAAGATTGCACACCCCTCT | 120 | 52 |
| chr7 | 144216537 | 144216657 | 772751_42001748_Shank2(210274)_343 | 2 | + | good | AGATTAGCCTACACCAAGGACAGAGCACGTGCTGCCCTGGAAAGATCAGCCATGAAGGGAAGCCCTCAGGAGCCTCAGCAGTCAGTGCTGTGGTTTCTCCACTCAGCACAGTCAACAGCT | 120 | 55 |
| chr7 | 144220377 | 144220497 | 772751_42001748_Shank2(210274)_375 | 2 | + | good | TTATTAGTAGTAGTAGTATTAGTAGTAGTAATAGTAGTAGGAGTAGTATAGATGTACACTGTGACAGTCAAAGGGCAACTTTGTGGAAATATATTCTCTCCTCTACCGTTACCTGGTTCT | 120 | 36 |
| chr7 | 144221097 | 144221217 | 772751_42001748_Shank2(210274)_381 | 2 | + | good | TGTGGATAAGTCAGGCAGGAACTCAAGTAGCCTGTTGCATCATGTCCACAGTCAAAAGCAGAGAAAATAAAACCTTGCTTTCTTGGTTCTCTGAGCTAACCTTCTATACTCTTAGACAGT | 120 | 42 |
| chr7 | 144244137 | 144244257 | 772751_42001748_Shank2(210274)_573 | 2 | + | good | GGGGATTACAAGGGTGCATCACTATGCCTAGCATTTTTTAAGGTGGGCTTTGGGGATCAAACTCGGGTCCTCATGCTCGCACATCAGGCACTGTGTTGACTGACCTGTTCCCCTAGCCCA | 120 | 52 |
| chr7 | 144248337 | 144248457 | 772751_42001748_Shank2(210274)_608 | 2 | + | good | ATTGACCAGCTCAGAGCATCGTTGCCCCAATTATTGTGGGAAATATTTTTTAAAAAATGAACTGGCATGTTCCCACGCTTGAACCCCAGTGGCCTAGCCAGTCATGCACTGTTGAGCAAT | 120 | 45 |
| chr7 | 144249297 | 144249417 | 772751_42001748_Shank2(210274)_616 | 2 | + | good | TCCCAGAGTGCCTTTTTGGAAAAGGGCATGAGAGGAGGGACTTTCTCCAAGACCAGTATGAGTTAGCTCCAGGACCCCCCCTCCTCATCCCTAGATACTGAGCCCCCCACTCTAATCCCT | 120 | 54 |
| chr7 | 144250857 | 144250977 | 772751_42001748_Shank2(210274)_629 | 2 | + | good | GTCTTGGAGAAAGTCCCTCTTCTTGTTCCCTTTTCCAGAAAGGCACTCTGGGATGGGAGGCCAGAGTCTCCCAGGATGCCCCACTCTGGATTCCTTGCCAAGCTGCAGGCACCTCCTCCT | 120 | 57 |
| chr7 | 144252777 | 144252897 | 772751_42001748_Shank2(210274)_645 | 2 | + | good | TGGAATGCTACAATGACTTCGACATTGTCTGGTTCTAATGTCTGAGGTCCACCGAAGCCTAGAGAGATGGCTCAAAAGTTAAGAGCACTTGGTGCTCTTTGAGGAGACCCAAGTTTGAAT | 120 | 45 |
| chr7 | 144252897 | 144253017 | 772751_42001748_Shank2(210274)_646 | 2 | + | good | CCCAGCACCCATGTCCAGTGGCTCACAACTGCCTGCTCCAGTCACGAGGGTCTGGCATCTTCTGGCTTCCTCTGGCTCTTACACACATTTGGTGTGCACATAAATACCTTTTTTATACTA | 120 | 50 |
| chr7 | 144255777 | 144255897 | 772751_42001748_Shank2(210274)_670 | 2 | + | good | TGTATATGTATATGATGTATATACACATTCATATGCACTCAGACATCCATAAACATGTACATAGGCACTCATGAACAGGCACACAGGCATACAATATACACTTGTACATGTGCACATGTA | 120 | 36 |
| chr7 | 144257337 | 144257457 | 772751_42001748_Shank2(210274)_683 | 2 | + | good | ACTACATGACCAGGAAATGGTGTGCCCACGGTAGACTGGGTCCTCCTGCGCCAATGAAGACAATCAGGACAGTGCCTTGTAGACATCTTCACAGACCAACCTGATAAGGACAATCACTCA | 120 | 50 |
| chr7 | 144266697 | 144266817 | 772751_42001748_Shank2(210274)_761 | 2 | + | good | TCATGCACTCTCTCACTCTCCCATGCACTCTCTCGCTCTCTCGCTCTCTTCTCTCATCTCCCTGCATCTCCTCTACATCACTCCTTCCGTATTGCAGGCAACCCTCCCCACTTCCCACCT | 120 | 55 |
| chr7 | 144270177 | 144270297 | 772751_42001748_Shank2(210274)_790 | 2 | + | good | TGGCTATATGTTAGAGTGATGGATGAGTGATGGATGAGTGATGGATAGGTGAATAATGAATTTTGGATGTTAGATAGATAGGTTGATGATGAATCAATGAATGTTAGGTGGATGATGGGG | 120 | 38 |
| chr7 | 144270297 | 144270417 | 772751_42001748_Shank2(210274)_791 | 2 | + | good | GGTAGATGATTGCGTAGATGGATGGGTGAATGGGTATTAAATTGATCATAAAGGTAGACAGATGGACAGACAGACATAGACAGATGTTAGATGGATGTATGTCAGATAGATGAGGATAGA | 120 | 39 |
| chr7 | 144272097 | 144272217 | 772751_42001748_Shank2(210274)_806 | 2 | + | good | GGGTCACATGTGTGAAAAGAGAACAGCCAGCAGAGAGCCTTAGCCACCCTTCTCATCCATCAACCCACTCAATCGTCCTCTGTCCATCCACCCATCTATCCATTCATCCACCCATCCAAA | 120 | 51 |
| chr7 | 144272217 | 144272337 | 772751_42001748_Shank2(210274)_807 | 2 | + | good | AATTCATTTACTAGAAAAGAGAGGTGACTCAAAAGAGCACTTGCTGTTCTTGCAGAGGATCCAGGTTCAATCCCTGCCCCACTCTCATCAGGTGGCTCTCAGTGCCTAGACCTTATTTCC | 120 | 47 |
| chr7 | 144272337 | 144272457 | 772751_42001748_Shank2(210274)_808 | 2 | + | good | AGGGGATCTGACCCCTCTTATGGTCTCCAGAAGCACTTGCATATACATGGTACACATCCAGAGAAGCAGGCAAACACACAAACATACAAATACTAAGACTTTGTAATGCATGTACTAGGT | 120 | 42 |
| chr7 | 144280377 | 144280497 | 772751_42001748_Shank2(210274)_875 | 2 | + | good | TCACCAAGAGTTCAAGGCCAGCCTAGGCTACACAGTGAAACCTTATCTTCAAAAAGTAAAACATAAGCAAATGCCACTGTGGCTTGGCACCAGAAGGCAAGTCACAGGGGTAAGGTCCTA | 120 | 47 |
| chr7 | 144322137 | 144322257 | 772751_42001748_Shank2(210274)_1223 | 2 | + | good | CAGGGCATGGAGGTTGGAATGGCTCCGCACTGTGACTGCAGGAATGAGGCTGCATGCTCTCATCTGGATGGATCAGGAAGCAGAGACAGGACAGGAAGTAGGACAGTCTATAACCCTCAA | 120 | 53 |

| | | | | | | | | | |
|---|---|---|---|---|---|---|---|---|---|
| chr7 | 144338217 | 144338337 | 772751_42001748_Shank2(210274)_1357 | 2 | + | good | CGTACACTGGGAGGACCCAGCTCCGGGCATCATCCTCGACACTATAATATATGACATTGTCATCTACATCATTACACTTAAGAACCATGAAATCAAATTACACACACACACACACACACA | 120 | 42 |
| chr7 | 144361137 | 144361257 | 772751_42001748_Shank2(210274)_1548 | 2 | + | good | GGGCTTTGCAAGTTCAAGACCAGCCTGGGCAGCTTAGGGAGACACTGTCTCAAAATAAAAAGTTTTGTAAAAACAAGTAAAAAGAGACCTCAAGCTTGCCTAACAGACGAAGCTCCGAAT | 120 | 43 |
| chr7 | 144363057 | 144363177 | 772751_42001748_Shank2(210274)_1564 | 2 | + | good | TTGACCTTTTTGAGGCAAAAGCTGACTGTATAGCTCAGGTTAGCCTCCAATTTGTAATCCTTCTGCCTCAGCTTCCTGGGAGTTGGGATTACGGCCATATACACTGTACACAGTATCTGG | 120 | 46 |
| chr7 | 144365457 | 144365577 | 772751_42001748_Shank2(210274)_1584 | 2 | + | good | AAGAAGTGGGGTTTAAAACCTCAAGACCCACTCCAAGCCACATACATCCTCCAGCTAGGCTCTACTTCCTAAAGGTTCTAGAACTTTCCAGAACAGCACCTCCAGTTGGAGGCCAAGCAT | 120 | 48 |
| chr7 | 144371697 | 144371817 | 772751_42001748_Shank2(210274)_1636 | 2 | + | good | GTCCTAAGAACTTGCTTGAGGCTAGCTTCTAAAGTAATGGACTACTTTCTTTGGAAGAGGAAATTTCACAATAAGGTGACATTGAATGCATAATATGGTTGTTACTGATAACACTTACAC | 120 | 36 |
| chr7 | 144371817 | 144371937 | 772751_42001748_Shank2(210274)_1637 | 2 | + | good | GGTCTATGGTGAGAAAAAAAAAAGCAGATTAGGCAAAAAGAAACACAAAATATAGTGTAAAGAGAAAGAAGAGCTGTAAATGCAATGTCACAGCCAAGGCTTGTGCTGGAAGAGATCGAT | 120 | 38 |
| chr7 | 144375417 | 144375537 | 772751_42001748_Shank2(210274)_1667 | 2 | + | good | CAGCCCCATGAGTTACTCCCCTTCCATCTATATACACCACAGTCCCTACATCTAGATCCACCGTCTCTACCTCTGTTGCACTGTTACAAGCCTGCAGCGCCATGCCCATGCCATGCCTGT | 120 | 53 |
| chr7 | 144376617 | 144376737 | 772751_42001748_Shank2(210274)_1677 | 2 | + | good | TACATACATACATACATGCAACACATTCACACGTATATAATAAAAATGAAATAAATCTTTAAAGACCCAGCTCACGGGGTGTTTGAGCCCACATTCAGGTAGATCTTCCATCCTCAGTCA | 120 | 38 |
| chr7 | 144376977 | 144377097 | 772751_42001748_Shank2(210274)_1680 | 2 | + | good | TAGACATTTGCCACAGTCCCTACATCTAGATCCACCGTCTCTACCTCTGTTGCACTGTTACAAGCCTGCAGCGCCATGCCCATGCCATGCCTGTATCCTGGATAAAGGGTTTACATGCAG | 120 | 51 |
| chr7 | 144380697 | 144380817 | 772751_42001748_Shank2(210274)_1711 | 2 | + | good | TTTGCCATTCCGATGGATCTGAGGTATCAGCATCCTAGACATAGCTGTGCATGTCAACAGTATTCATTCACTCAGGACTTCATCCATTCTCCACAACAACGCATATCAGGGAGGAGGCAA | 120 | 46 |
| chr7 | 144386577 | 144386697 | 772751_42001748_Shank2(210274)_1760 | 2 | + | good | GAAGCCTAACCTGTTTTTTCGGCACCATCCCCTGGGCTGGGCGTGCTAGACTGAAGGGAAGGCAGAAAGTGAGCTGAGCGACAATGCTGTCCCCCTCTGCTTCCTAACAGCAGATGCAAT | 120 | 55 |
| chr7 | 144394017 | 144394137 | 772751_42001748_Shank2(210274)_1822 | 2 | + | good | ATGAGCACCAACATTCATCTCTAGCTGCCTCCTGAGTGTGGATACAACATAACTACCTGCCTCGCACTTCTGTCTCTAGGCGTTATTCTCCACAATGGACTGTGCCCTCAATAACTGGGA | 120 | 48 |
| chr7 | 144400137 | 144400257 | 772751_42001748_Shank2(210274)_1873 | 2 | + | good | GAAAAACCAAAAAAGAAAAAAAAAGAAAAAAGAAAAAGAAAGAAGCTCTTTGCCGATGCAGCCAAGACAAGGTCACAACCGGATAGGTGTACCCTAAAGGAACCATTGATTGGTATCCTT | 120 | 38 |
| chr7 | 144414777 | 144414897 | 772751_42001748_Shank2(210274)_1995 | 2 | + | good | GGGAAAACCTCAGCTTGCGTTCATCTCCCCCTAGAGCACAGGGTAGGGAGTGCTCCAGGGAGTCCCCACCGCTCCTCTCCGAGGGTGTGGCGTATGCAAACATGGATATGCAAACATTTC | 120 | 57 |
| chr7 | 144415857 | 144415977 | 772751_42001748_Shank2(210274)_2004 | 2 | + | good | AGACCAGCCATCACATCACTCAACAGGATGAAACCTGTTCTCTTCCAGCATCCTTGGAAAAACCTCAGCTTGCGTTCATCTCCCCCTAGAGCACAGGGTAGGGAGTGCTCCAGGGAGTCC | 120 | 53 |
| chr7 | 44310248 | 44310368 | 772751_42001747_Shank1(243961)_1 | 1 | + | good | GGGTGGCAGTGGAGGCGCCGACCGGGACTCAGCAGCCCCGCCAGGCAGGTAAGGGCACGGGACACTACAGGGTGGTTCTGGGGTAGGGTGCAATTGCGAGAACTGGGGTTGCGGGAGA | 120 | 68 |
| chr7 | 44310368 | 44310488 | 772751_42001747_Shank1(243961)_2 | 1 | + | good | GCGGAGGCGCCCCGGATGTCCTTGGCCTGGCGCCGGCTCCGGCCTGGGTGTACATTTAGGGTGTGTGTATGGGGGGGTGAGGGTGTGGAAGGTAGCATGGCCCCTAAGGGTGACATCAAA | 120 | 64 |
| chr7 | 44310488 | 44310608 | 772751_42001747_Shank1(243961)_3 | 1 | + | good | TTGGGTTGCGACCTGGTCCCGAGTGCCCCGTCCCAGATCGTATCCTTGCTAGGAGGGTGTGCCCCATCCCCCACGTGCTGCGCCCCCTAAGCCGAACTGGTCTCGGAGGCTCTCCGGGGC | 120 | 68 |
| chr7 | 44310608 | 44310728 | 772751_42001747_Shank1(243961)_4 | 1 | + | good | CAAGACTGGGTCTTGGTCCCCGCCTCCCAACCCGGCTCGGCCACTCCCCCGCCTCGCCCACCCCAGCACACCTGGGCCCCTCCCCTAGGTCACCCTGGTCTCCAACCACTCCAACCCAGG | 120 | 71 |
| chr7 | 44310728 | 44310848 | 772751_42001747_Shank1(243961)_5 | 1 | + | good | GTCTGGGCCAGAGCCTCCTACGACGCCTGGACCCGGCGCCCCTTCCAGGCGTCTGTAGTTCCTTTCCACCCGCGCCTCGCGTGCCCATCGCGCCTCAGGTTCCTTCTCCGCAGGCCGGCG | 120 | 71 |
| chr7 | 44310848 | 44310968 | 772751_42001747_Shank1(243961)_6 | 1 | + | good | CGCCCCCTCTGCTCCCGCCCCCAGTCCCAGCTCCCTCTCCCGACCCAGGGCTATTCTTTCACCAACCTCCCTGCTCCCGCCTTCCTCCACCTCCATTCATTCTCCCCGGGCCCGGCTCCT | 120 | 68 |
| chr7 | 44310968 | 44311088 | 772751_42001747_Shank1(243961)_7 | 1 | + | good | GGCCACGCCAGCCGTCCCCTGTCCCCCCGCAGCCTCTCAGGGCACCCCCTCTCTCCCCTTAGCCACCCCCCCCCCTCGCCGATTCCTCAGCCCGTCTCGGTACCAGCTGCGTGCCAGCTA | 120 | 72 |
| chr7 | 44311088 | 44311208 | 772751_42001747_Shank1(243961)_8 | 1 | + | good | TTCTTAGCCGCCGAGCTTTTGATTCATGCCGCCCCGGCGGAGTGTGCCCGCTCCCTCCCTCGAGGCCGGAGGGGGAGGGGCCCCGAGCTGTCCTGGGTGTTGGGGCAGTGGAGCCCGCGC | 120 | 72 |
| chr7 | 44311208 | 44311328 | 772751_42001747_Shank1(243961)_9 | 1 | + | good | TATAGGGGACCTAGCCTTTGCTGACCTTTTATCCCACCTGCCACCCCTTTCCTGCAGAGCTGGGGCTGCAAAGGGGATGTCAGCCAGGGTCAGCCGAGGGGGAGTGGCTGAGGTCACACT | 120 | 60 |
| chr7 | 44311328 | 44311448 | 772751_42001747_Shank1(243961)_10 | 1 | + | good | GCCAGGCGGGATGTGGTGTAGGGCGGTCAGCAGACTGAGCTGGAGGCCCGGGTCTCCTCCCCAGCCCAGCGGGTTGAGCCACAAGGCAGGGTACCTGGGATGGGGTGCTTCAGAGCTCGG | 120 | 68 |
| chr7 | 44311448 | 44311568 | 772751_42001747_Shank1(243961)_11 | 1 | + | good | TCCCCGCGAGGGGGAGGGTTCCTCAGCCCGTGGTTAGACGCCTGCTCGCCTACAGTTCTTGGAACTCTCTTCCTTAAACATTTCGTCTTTCCTTCCCCAACCCTCCCTCCCCCCAGCCCA | 120 | 60 |
| chr7 | 44311568 | 44311688 | 772751_42001747_Shank1(243961)_12 | 1 | + | good | TCCCCACTCCCCCAAAGTCAGCAGTACCCTCCACTGCTTCCCATCTCCATGGCAACGGTGCAGGAGCCGGGCCTGGTGTGGGGTGGGAGGGACCAGACCAGACATTCTGCTTTCCGGTTC | 120 | 62 |
| chr7 | 44311688 | 44311808 | 772751_42001747_Shank1(243961)_13 | 1 | + | good | TAGAGTGGGGAGTAGGGGGTGGCAAACAGGAGAGAAGAGGACCGCATTTGGAATGGAGCCACCCCTCAGTAGGCCACTTTGAGAGGGACGACCCTCCCCATCTCCTGGTCCCCATATTGT | 120 | 57 |
| chr7 | 44311808 | 44311928 | 772751_42001747_Shank1(243961)_14 | 1 | + | good | TGCAGCTCACTTCTTAGCTTCCTCCCCAGACACCCCACCCAATCCAAATTCTCCTCGGTGGAGCCTCCAGTACCAAGCAGTGCCAGCCCACCCACCCTGCTGTGCCATTCCTGAAACACC | 120 | 58 |
| chr7 | 44311928 | 44312048 | 772751_42001747_Shank1(243961)_15 | 1 | + | good | CACAGGTCACCTGCAGCTGCCCACACCCACCCAGGCCACCACAGCTTGTCACCTGGACTTTCCAGCCAGAGCCAGGGCCAGCCTCCCTATCCACCCACACATCCTCTTCACACGCCCCAG | 120 | 64 |
| chr7 | 44312048 | 44312168 | 772751_42001747_Shank1(243961)_16 | 1 | + | good | TAACCTCTCATCATGGGTCTCTTGGGCCCAGCTCAGGGGAACTTCTCTTTTCTGTTCATCCACCCCCACTCGTGTCCCTCCATCCCCTACAGGGACAGCTCATTCCCCAGCAGCAGCCCT | 120 | 57 |
| chr7 | 44312168 | 44312288 | 772751_42001747_Shank1(243961)_17 | 1 | + | good | CTAAACATGTGGTCAGCCACACTTCAGTGAAGTCACCAGCTCTCGGGCACCTCCTGCTGCCCCAGGCTCAAAACTCACACCTCCATCTCACCTTTACTATAAAAGTCAGTCTCCACCTCC | 120 | 52 |
| chr7 | 44312288 | 44312408 | 772751_42001747_Shank1(243961)_18 | 1 | + | good | CTGGGGTGAAGGGGCCAGGCTCCGGCCAGGAGGGCAGGATGCCGGGGTGGATTGGTCCACCTGTCTGTCCTTAGCTAATTTTAGCTTCATTTTTTTGTAGTTTGACAAAGTGGGCCACTA | 120 | 54 |
| chr7 | 44312408 | 44312528 | 772751_42001747_Shank1(243961)_19 | 1 | + | good | GCTCCCGCGGTGATTCTGTGATTTCTAGGACACACTGGGTCAGCCACAAAGTGGAGCGCAGGGGGGTGGGGGGCTGCAAGCAGGGCCAAGGACAGGGAGGAGAAAGGTGCCAGGAAGGC | 120 | 62 |
| chr7 | 44312528 | 44312648 | 772751_42001747_Shank1(243961)_20 | 1 | + | good | GCAGAGGTGGTGCAAACTTGTCTGTGCAGGGGGCTGAGGGCATGTCTAGTCTCCCTTGGCTGCCCCTGTGTCTGTCCACTGGTACTGGTGGCCTCTGAGTGGCTCAACAGTGAGGATCAC | 120 | 59 |
| chr7 | 44312648 | 44312768 | 772751_42001747_Shank1(243961)_21 | 1 | + | good | CCTTGCTTTTCTCAGCTGCCTGTGATGCTGCTGTCCCCCGTCGCCCCGTGGCCCCACGATGACCCACAGCCCCGCGACAAGCGAGGACGAGGAAAGACACAGTGCCAGCGAGTGTCCCGA | 120 | 65 |
| chr7 | 44312768 | 44312888 | 772751_42001747_Shank1(243961)_22 | 1 | + | good | GGGGGGCTCAGAATCGGACAGCTCCCCAGACGGGCCAGGAAGAGGCCCCCAGGGGACTCGGGGCCGAGGCAGCGGAGCACCTGGTAACCTGGCCTCTACCCGAGGCCTCCAGGGACGCT | 120 | 72 |
| chr7 | 44312888 | 44313008 | 772751_42001747_Shank1(243961)_23 | 1 | + | good | CATGTCTGTCCCTGACGACGCCCACTTCAGCATGATGGTTTTCAGGATTGGCATCCCTGACTTGCATCAGACGGTGAGTGGTCACTCACCTGCCAGCTGACAGACACATTGGCCAAGTCT | 120 | 54 |
| chr7 | 44313008 | 44313128 | 772751_42001747_Shank1(243961)_24 | 1 | + | good | TTCCCCACTCTCAATCCCAGATCTAGCTGGAATCTTATTCCTTTTCCAACCTACCCTTGACCCTCATACCCTTGACCCAACCTAGCCCCAGACATCCTGTCTCCCACCAGAAATGCCTAC | 120 | 51 |
| chr7 | 44313128 | 44313248 | 772751_42001747_Shank1(243961)_25 | 1 | + | good | GCTTCAACCCAGATGCCACCATCTGGACAGCCAAACAGCAAGTACTCTGTGCCCTGAGCGAGAGTCTACAGGACGTGCTGAACTATGGACTGTTCCAGCCAGCCACCTCAGGCCGGGATG | 120 | 57 |
| chr7 | 44313248 | 44313368 | 772751_42001747_Shank1(243961)_26 | 1 | + | good | CCAACTTCTTGGAGGAGGAGAGGCTTCTAAGAGAGTACCCTCAGTCCTTTGAGAAAGGGGTACCCTACCTGGAGGTAAGGGCACAGAACTCTTAACAGAAGTCATTCCCTGGAACTCATC | 120 | 50 |
| chr7 | 44313368 | 44313488 | 772751_42001747_Shank1(243961)_27 | 1 | + | good | CACTCCAACTCCAAAAGAACTCCCCAGTGGGCTCTTCAAGACTTCCCAGAGTCCAAATCACCAGCTGAGTTGCCTCTGATGGTGACCTTTCCTCTCCCTAAACTCCTTATGCCATCAAAT | 120 | 49 |
| chr7 | 44313488 | 44313608 | 772751_42001747_Shank1(243961)_28 | 1 | + | good | GAGGGAATCCCAACTACATAGTAGTTCTAGAACATTCCATAGATGGGGGGGGGGGCTGTCCCATGCTTATACTTATAAGGTAGGGATTATAGTGCCATAGACCTTAATGACCCAGTACTC | 120 | 46 |
| chr7 | 44313608 | 44313728 | 772751_42001747_Shank1(243961)_29 | 1 | + | good | TTGGAAAGGTCGCCCCCACCTACGGCCATTTCCATGCTAACCCTGAACTCATTCTGGGAAGACAAGAGGAGGGTCAGACATTGCTCCCCAACAAAGAATCTAATAAGTCTGTGTGTCCCC | 120 | 51 |
| chr7 | 44313728 | 44313848 | 772751_42001747_Shank1(243961)_30 | 1 | + | good | TCATATTCTACCCCTGGCCTCAGTTCCGCTACAAGACCCGAGTCTACAAACAGACCAACCTGGATGAGAAACAGCTGGCCAAGCTGCACACAAAGGTGAGAATGCCTGCTACCCCCCAGC | 120 | 54 |
| chr7 | 44313848 | 44313968 | 772751_42001747_Shank1(243961)_31 | 1 | + | good | ACCAGAGCACCCTGTCCGTTGCCTTCCTTCCCAGTTTCTGCCCAGCAACCATTCCCTCTGCCAACACCTCAATTATGCCAAGTCCCCGTTTAACTTTGAGGGTGCATCCTGTCTGCAGAC | 120 | 54 |
| chr7 | 44313968 | 44314088 | 772751_42001747_Shank1(243961)_32 | 1 | + | good | ACCCATTCATCTTCATCACACTCCCTTCCCACTGCCCTCTATGGCCTCACCTGCCCAGACTGATGCCATCTTTGCCTGGTATGGTCATGGGTGCTGGTGCTGGTGCTGGTGAGTGAATGG | 120 | 56 |
| chr7 | 44314208 | 44314328 | 772751_42001747_Shank1(243961)_34 | 1 | + | good | GCCATCTCACTGGTCTGACCTTGTTTAATACAGTCTTTTACTGCCTCACGGCCTAGGCCAGACTAGCTGACCAGGAGGACCTGGAGAAATGACTGTCTCCAACTCCCCAGCACTGTAATT | 120 | 51 |
| chr7 | 44314328 | 44314448 | 772751_42001747_Shank1(243961)_35 | 1 | + | good | ACAAGTGCACACAAGCACCCCTGGCTCTCTTACGTGGGCTCTGGTGAATCAAACTTGGGGCTCCATCCTTGCATAACAAGCTTTACCCCTAGCCCTCCGATCTGTTCTTTTAACTCTACC | 120 | 51 |
| chr7 | 44314448 | 44314568 | 772751_42001747_Shank1(243961)_36 | 1 | + | good | TTCTCTGCTCGGGAGCACTTAGGGCCCTTTTCTTACTTAGTGCATCAGCCAATTGTCTGCCTCGATAGTATTTCATAGACATCCACAGTTCCCCAGACCTGTGCCTGTATCTCCACAGGC | 120 | 50 |
| chr7 | 44314568 | 44314688 | 772751_42001747_Shank1(243961)_37 | 1 | + | good | TTTGCTCTTTTTTTTTTCCTCAGCCACATTTATTCTATTTGGTCCTTTTCCTCCTGACCTTGTCTGCTCTGGCCCTCTAGGAGTTGAGAGGAAGTTAGGAGGCTGATCTGGCCAGAGGG | 120 | 47 |
| chr7 | 44314688 | 44314808 | 772751_42001747_Shank1(243961)_38 | 1 | + | good | AAATAATTCTTGACATTTAATGGGAAGAAATTCCCTGATCGGTTTTTCCTCCAGGCTTCTCTCTCCCACACTATCCACCCCGATCTTCCCCGCCCCCTCCGTGCCTTCTCCTCCCTACCT | 120 | 51 |
| chr7 | 44314808 | 44314928 | 772751_42001747_Shank1(243961)_39 | 1 | + | good | GTGCCATCCACTGTGGGAGCCAGTGGCCATGTATGGCTCCTTTGTACATGTGACAGGTGTGAGCTGAGTCACACCGCAGAAAAAAACAGCACAAAAATTCCACTGCATCCGTTTCCAACA | 120 | 50 |
| chr7 | 44315288 | 44315408 | 772751_42001747_Shank1(243961)_43 | 1 | + | good | AGCAGGCTTGGCCACAAGCACCATTACCGCTGAGCCATCCCACCACCCCCAGGCCATGGGATTTTAATTATTTTCTTATGGATTGTGTATCAAAACAGTGACATTAAGCATATATTCCTT | 120 | 44 |
| chr7 | 44315408 | 44315528 | 772751_42001747_Shank1(243961)_44 | 1 | + | good | TGAGAACACCTGTCATCAGTATGGATTTCAACACAATTCACAGTCTCCCTTTGTCTTAGAGCTTTTATTGCTGTGAAGAGACTCTAATAAAGGACAACATTTTATAGGGTCTGGCTTACA | 120 | 38 |
| chr7 | 44315768 | 44315888 | 772751_42001747_Shank1(243961)_47 | 1 | + | good | AACTACCCTTTGTATTAGAATATTACTTTTTAGATGGGGTCTCCCTGTATATAGCCTAGGTGCGTCTAGAACTCACCATGCTCCTGTCTCTGCCCTGAGCACTGAGACAACTGACATGAC | 120 | 45 |
| chr7 | 44315888 | 44316008 | 772751_42001747_Shank1(243961)_48 | 1 | + | good | CCCACCACACCAGACTTCCTTTCTGCCTTTGCAAAGGGTTGGCTGTGTATTTCCACTGGGTAGTGCTGGCTCTCATTGCAGCTTTCTTCACTCAGTCTCCCTGATCTCCCTGACACCCCC | 120 | 54 |
| chr7 | 44316128 | 44316248 | 772751_42001747_Shank1(243961)_50 | 1 | + | good | TCAGATTCAGGAGGTAGGAAGGGGTATGGGCCATGAAGGACTGGGGTGAGGAGCAGGGTTGGGTAGAGAGGCCGGGGACACCACCTCAACCCCAGAGGGTGAGACCGAGGGACCAAAGG | 120 | 61 |
| chr7 | 44316248 | 44316368 | 772751_42001747_Shank1(243961)_51 | 1 | + | good | TGTCGGGGGGAGGGTGGCTGAGCTAGGAGGCCAGGGGTAGAGTCTGAAAGTGGAGGCAAAAGGTATGCCTCTGAAGGTCTTCTCTTGCCCTTCAGAGACACCTTTGACCCTGGCGGCCCA | 120 | 59 |
| chr7 | 44316368 | 44316488 | 772751_42001747_Shank1(243961)_52 | 1 | + | good | GACTGAAGGCTCAGTAGAGGTGATTCGGACGCTGTGCCTGGGCGGGGCGCATATAGACTTCAGGGCCCGGGATGGCATGACAGCGCTTCACAAAGCTGCATGTGCCCGGCACTGCCTGGC | 120 | 62 |
| chr7 | 44316488 | 44316608 | 772751_42001747_Shank1(243961)_53 | 1 | + | good | GCTCACGGTGAGGCTGCTGTAGTCCCCAGATGTGGGGCCCTTTCTCCTTTCCCTTCTATTTTTCCAACCCTACTCTCTTCAATTCTGATCAAAAGAATTCCCCTGATGTAGATACCTGTA | 120 | 48 |
| chr7 | 44316608 | 44316728 | 772751_42001747_Shank1(243961)_54 | 1 | + | good | ATCCTAGCACTGGGAAATTCAAGACTGGCCTAGGTTACCTGGCATCCCATCTCATTGCTCCCCAACCCAAACTCCCTCTATGTCCCTACCACAAAGTCTGTCCCACACACTGACAAGCAC | 120 | 52 |
| chr7 | 44316728 | 44316848 | 772751_42001747_Shank1(243961)_55 | 1 | + | good | ACAGTTCCCCAGTCAGTCTTCCAGATGCCCCAGCTTAGTCTACAAGTTGGTGGTGACTGAGCCCGGGTAAATCCCCTCCACTAAAGCTTCCTCCCTGAGCATTCAGAATGGCCCCCTTCC | 120 | 55 |
| chr7 | 44316848 | 44316968 | 772751_42001747_Shank1(243961)_56 | 1 | + | good | CTCTGCCCAGCCAATGCACACCCTATCTCCGCTTTCCCAAAGTACCTGCCGTTGAGGGCCCTAGAACACAGAAAGGGAGCCCCTGCTGCTTGGGGAGATCATCTCCAAGCACAGCTTTAA | 120 | 56 |
| chr7 | 44316968 | 44317088 | 772751_42001747_Shank1(243961)_57 | 1 | + | good | AATCGCCCCTCCCTTCATCCTTCCTAGCCTAGCACTTCACCTCCCAACACACACATTCTCTCGCGGACAAATCCAGACCCTTCTCACACCTAAGATGCAAAGCACACCTTCCAAAAACCA | 120 | 51 |

| | | | | | | | | | |
|---|---|---|---|---|---|---|---|---|---|
| chr7 | 44317208 | 44317328 | 772751_42001747_Shank1(243961)_59 | 1 | + | good | CAGAGTGAGTTTGAGGTTAACGGTGGCATCATGGCATCATGAGACTCTCCCATCAAAGAAACAAAAGGTGCTTTCCCAAACAAATTAGCCCTTTGTGCCCAGAAAAATGCACAGAACAAA | 120 | 43 |
| chr7 | 44317448 | 44317568 | 772751_42001747_Shank1(243961)_61 | 1 | + | good | CTACCAATTCTCAGGAACCAGTCACGGTATTTCTCCAGATGGGGCTTTCACTGGTACTCAGGGCTGCCCCATTAGGCTTGTCTGGCTGGCCAGCAAGCCCCAGAGATCCACCTGCCTCTG | 120 | 57 |
| chr7 | 44317568 | 44317688 | 772751_42001747_Shank1(243961)_62 | 1 | + | good | CCTCTCCAGCATTGAGACTGCAACCGTGTGCCACTTTTTGACATGGATTATTGAGACCAAGCTGGGGCCTTTATGTTTACATGACAAACACTTCACTAACTGAGCTGTCCACCCCCCACC | 120 | 49 |
| chr7 | 44317688 | 44317808 | 772751_42001747_Shank1(243961)_63 | 1 | + | good | CACCCACCCCCACCCCCAGCCCCTGCATACAGTCTTATTTAAAATATTCTTTTTCTTCTGTCTCTGTCAGGTCAACTGATCCCTCTTTAGGCATAGGTTAGGATAGGCTGATTCCAAA | 120 | 45 |
| chr7 | 44317808 | 44317928 | 772751_42001747_Shank1(243961)_64 | 1 | + | good | GACTGAGACACACCCATATTCACAGACGCCCCGGTTCACTGATGACTCCTCTTTTCACAAACACTTGGCACGAACAATTGCTGTTTATAGCCAGAACCAATGGATATTGTTTGCATTTTT | 120 | 43 |
| chr7 | 44317928 | 44318048 | 772751_42001747_Shank1(243961)_65 | 1 | + | good | GTCTGTCTGTCTGTTTGTCTGCGTGGTTTCAGATCTTTGCTATCAAGCCCCTTAACCACTGCTCACATCAGGCTGTCTAGTTTTAAATTAATTAGAATGGAATAAAATTAAAATCGAACT | 120 | 38 |
| chr7 | 44318048 | 44318168 | 772751_42001747_Shank1(243961)_66 | 1 | + | good | TCCTTGGACTCAACAGCCAGCCAGGTACTCATCGCCCCCCTGTGGCCACCATGTTGGATGACACCACTGGGGACAGTTCACTGTAGTATCCTGCTGCACAGAGCTGCTATGGGCTCTCTG | 120 | 57 |
| chr7 | 44318168 | 44318288 | 772751_42001747_Shank1(243961)_67 | 1 | + | good | CACAGGGACAACTGTCACACTTGTTCTTATACAAGCTTAGATTCCTCCCCTCTCTGCACCCTTCCTCTCTTTCCCCTAGAACTCCCACCCTGAAGGATGCTGGCTCGACATGGGTTCACC | 120 | 52 |
| chr7 | 44318288 | 44318408 | 772751_42001747_Shank1(243961)_68 | 1 | + | good | CCAGTCTTGCCTTTCCTAGTCTTCTGCCCTGGGTGTCTATATAGCGCTCCTGCACTTGACTGGGGACATTCTTAACACATAGACCTCCATCCTTCCATCTGTCTTCCCTACAACTCTGCC | 120 | 51 |
| chr7 | 44318408 | 44318528 | 772751_42001747_Shank1(243961)_69 | 1 | + | good | ATGAATTAAGACGCCTTTGGTCGTGGACATCCTCCAGTGGCTCATCGAGAGTCACCCACTAGAGAAAGGTTGAGCTGCATGTCAGAGCGGGAGGGTCCTGTGACGCCATCTGCCTTGTCT | 120 | 54 |
| chr7 | 44318528 | 44318648 | 772751_42001747_Shank1(243961)_70 | 1 | + | good | TCTCATTGTTCTTAACGGCTAGGACTAGTGCAGATTGGGCAAAGCCTGGCTCAGAGTCTACGGACTACCAGAGGGGCTGGAGAGATGGCGTAGTCAGTGGAGTCTGTGCAAGCATGAGAA | 120 | 52 |
| chr7 | 44318648 | 44318768 | 772751_42001747_Shank1(243961)_71 | 1 | + | good | CTTGAGTTCAACCTCCAGCACCCAAGCAGAGAAGCAGAGCGTGGTGGCAGGTGCCTGTAACCTCAGTGCCAGGGGGCAGAGATAGGAGGACCCCGGGTCTCACTAGTCAGCTGTTCTATC | 120 | 58 |
| chr7 | 44319008 | 44319128 | 772751_42001747_Shank1(243961)_74 | 1 | + | good | CCCTACCAGCCCCTTGAAACTAGAGCTGCTATCTGACCTTTCTTCCTGAGTTCTCTGTCCCTCCTTAGGCACTTCTGGACCTTGGGGGCTCCCCCAACTACAAGGACCGACGAGGGCTGA | 120 | 57 |
| chr7 | 44319128 | 44319248 | 772751_42001747_Shank1(243961)_75 | 1 | + | good | CCCCCCTGTTCCACACAGCTATGGTGGGGGGAGACCCCCGCTGCTGCGAGCTACTTCTGTACAACAGGGCCCAGCTGGGCATTGCAGATGAGAACGGCTGGCAGGAAATCCATCAGGTAT | 120 | 60 |
| chr7 | 44319248 | 44319368 | 772751_42001747_Shank1(243961)_76 | 1 | + | good | CCAGATTCAGTCCCAAGTACTGGGGAGAGCTTGGATACCTGGGCTCTGCAGCATCCAAAGGGATGGGATATCAAAGAGATGGCAGGTTTCCTTACAGAGAGGGGAGTGGAATTTCCTCAA | 120 | 51 |
| chr7 | 44319368 | 44319488 | 772751_42001747_Shank1(243961)_77 | 1 | + | good | AGAGGAAAAGGCAGAAAGAACTCCAGGAATAGATAAATGGATGAGGTTGGTTGCTTGATTCACCTGAAGGGCCAGCCTGGGGTATTTATACAGCGAAGTAACTGAATGCTTAGAGTCAGG | 120 | 45 |
| chr7 | 44319488 | 44319608 | 772751_42001747_Shank1(243961)_78 | 1 | + | good | AAGCTAAAGTCCTGGGTCTCTGAACAAGAATAGAGACAAAAAAAAAAAAAAAAAAAAAGAAGGGGACATGGATGCAGAGATCCCTGACCTCAAGGAAGAGGGCCTTGGATTCAGATGAGG | 120 | 42 |
| chr7 | 44319608 | 44319728 | 772751_42001747_Shank1(243961)_79 | 1 | + | good | TCTAACCAGGCCCATGCACTCCCTGTACCCTCAGGCCTGCCAACGAGGGCACTCTCAGCACCTGGAACACCTGCTCTTCTATGGGGCTGAGCCTGGAGCCCAGAATGCCTCAGGAAACAC | 120 | 60 |
| chr7 | 44319728 | 44319848 | 772751_42001747_Shank1(243961)_80 | 1 | + | good | GGCCCTGCACATCTGTGCTCTCTACAACAAGGTCAGCTCAAGCAATGTCTACCTGCCCCTTCAAAGGCACTTGCTTCATACCAGTTCAAGCCCTCTCCACTCCAACTTCCTATCTCTGCA | 120 | 52 |
| chr7 | 44320328 | 44320448 | 772751_42001747_Shank1(243961)_85 | 1 | + | good | ATCATCTACCCACCCATCATCTATTCAGCCATCATCCACCCATCTGCCCATCATGCTTAATGAGCTTCTTCCCTGTGTCCGGCTCCAAGCTTTATACTGAGCTTCTGATCTAATCCTTGG | 120 | 48 |
| chr7 | 44320448 | 44320568 | 772751_42001747_Shank1(243961)_86 | 1 | + | good | TGCTCAGTAAAGTTATGGAGAATCACGTTCCCTTGAGAGCCATGTTTGGACTTGGGTGGAGGTGGGTTTCTGAAGGATGTCAAGACAGCAGAGGCAGATGAGTGGAGCAGAGTGCTCCAG | 120 | 51 |
| chr7 | 44320568 | 44320688 | 772751_42001747_Shank1(243961)_87 | 1 | + | good | GAAGGGAGCACAAGCTGTGTACAAGACAAGAGGCCAGGCAAGTTCAGTCTGCAGAGGACAGAGCTGTGAACACAGTAAAGCCTGGAGGTGAGAGTGTTGTATATTTAGCAATTTAATGTT | 120 | 46 |
| chr7 | 44320688 | 44320808 | 772751_42001747_Shank1(243961)_88 | 1 | + | good | GTATGGAGGGTCCCTCTCCAGGAATTCTGGGGAAAGATGATCTGTAAAAAGGAGACAAAGCTCTCACCAAATGGGTCTGCTGCCAGCAAGAGCTGAGACCCCTTTGCCACCCAAGTACCA | 120 | 52 |
| chr7 | 44320808 | 44320928 | 772751_42001747_Shank1(243961)_89 | 1 | + | good | AGAGACCCCCAGGTTGGCCAAGCCTCTTGGAGCTACAGCCCCCACACAGCAAGATTCTTGCCCTTGAAGAGCTGCTTGGGGGACTTCAGGGTAGGGAGGGCAGCACCTGCTGGGATGCTG | 120 | 61 |
| chr7 | 44320928 | 44321048 | 772751_42001747_Shank1(243961)_90 | 1 | + | good | CTGATCTGCTCCCCCCCTTCCTCCAGGAAGTCTTCCAAAACAGCCCTAGCCCCTTCTGCCTCCTGAGAACTCCCAGTGCTGGGGATATGGCGGGTATCCAAGTGCCGCCTTGTGGCAGCG | 120 | 61 |
| chr7 | 44321048 | 44321168 | 772751_42001747_Shank1(243961)_91 | 1 | + | good | AGCAGGATGCTCAACGCCAAGGGAGGACTGGCGTTCCTTAAACCCAGAAACACAGTGCTGGCATTTCCTTCCCGCTTGCTGCTGCTGTCTCTACTGCTCCTCTGTGCCAGGTCCTCTGAC | 120 | 57 |
| chr7 | 44321168 | 44321288 | 772751_42001747_Shank1(243961)_92 | 1 | + | good | TCTGAGGCAGGTGCTTGACACAGAGAGAGAAAGCCCCAGCTCCACACACTCAGGAAGAGCCCGAACAAGTGCCTTCCACTCTCTGAGGCTTTGTCCTCCCTGAAATGGGAACCGTGTCAG | 120 | 56 |
| chr7 | 44321288 | 44321408 | 772751_42001747_Shank1(243961)_93 | 1 | + | good | CTGATTGAAAGGGATGAGTGGTCGCGGACATTAAGTATTCATCCCTAGAAGACAGCCGTGATAATAATTGTTATTATTAGTATCATCATCAGTGTCAATTGCCAGGATGCGGTCAGAGGC | 120 | 42 |
| chr7 | 44321408 | 44321528 | 772751_42001747_Shank1(243961)_94 | 1 | + | good | TGGCATCCATCTGAGCAGGGAAAATCGAGTGCAGAGAATTATACAGCAGCCACTAGTCCCCAGAGAGGGCCAGATGAACATGCTGCTTTGCCTTCTTCAGTTTTCCCCCACCAACCCCCA | 120 | 52 |
| chr7 | 44321528 | 44321648 | 772751_42001747_Shank1(243961)_95 | 1 | + | good | CCATCTGTGAAATGGGTCAGTCGTGAATATCCTTCATACAAAAGGTCAAACAGGTGAACATTCTGCTTGCTATCAGACCCAGTAGATGATTAGCAAATATGAGCTCATCACCCACTTCGC | 120 | 43 |
| chr7 | 44321648 | 44321768 | 772751_42001747_Shank1(243961)_96 | 1 | + | good | TCCCTGCCTCTGCTCATCCTGGCCATCCAGCTGCCTGTGGGTACATCAATAAGCCCATTCACCACCTATCCTCTACCCACTCACCCACCCAGCCGCCCATCCATTCAACCATCCACCTAC | 120 | 57 |
| chr7 | 44322008 | 44322128 | 772751_42001747_Shank1(243961)_99 | 1 | + | good | CCATTCATCTTTCTGCCCATCCATCCATTATCCACCCATCCATCCTCCTGGCAAAGGTCAGATTTTCTAGGCTATGGGCCTCTCAATGGTGGGAACAGGATCTGGTAGATTCTGGTTCTA | 120 | 48 |
| chr7 | 44322128 | 44322248 | 772751_42001747_Shank1(243961)_100 | 1 | + | good | TCCCAACCTCCCAGAACATGGCAATTATTTTACATCCATGCATCCATCCCTGAACCCATCCCTCCATCTATTTATATACTGTCTGTGCATTCATCCGCCTATCTGGCCCAGGTCTGACCT | 120 | 48 |
| chr7 | 44322248 | 44322368 | 772751_42001747_Shank1(243961)_101 | 1 | + | good | TCACCCTTTGCTGTCTGTCCCCTTAACCCTGCAATATGTACATGTAGCCCCTGCTGGTGGTCCAGTCTGTGTTTCTCTCTCAAGAGTGGGGAGAGATTCTCCTAGACTGCTCTGTCCCAA | 120 | 52 |
| chr7 | 44322368 | 44322488 | 772751_42001747_Shank1(243961)_102 | 1 | + | good | GCCGTCACCAAACAGTAATCCTCACTGCATTGTGTGCAGTGAGCAGATGAATGAATGCATGACAGATTGAATGAGTGATGGAACTGTGGATGGATCTCCAGCAGAACTCTGCACAGCTCT | 120 | 48 |
| chr7 | 44322488 | 44322608 | 772751_42001747_Shank1(243961)_103 | 1 | + | good | GGGCTCAGAACTCTTGACAGGTGGCTGCTGTTTTTGACAGATTGAATGACTCAACAAATGAATGAATAATTGAGTGGCTTGAGCAAATGAAGTTTGATTGGCAGCTCCACATCTTCCATC | 120 | 42 |
| chr7 | 44322608 | 44322728 | 772751_42001747_Shank1(243961)_104 | 1 | + | good | TTCATCTCTTCCAAGTGCATTTCCTGAGCACCTACTGTGTGCTTGAGCTACTCACTAGGACTTGGACATTTAAACTTGGTCAGGCACAGGAGAGACTAGGAAGCTTGAGCATCACACTAT | 120 | 46 |
| chr7 | 44322728 | 44322848 | 772751_42001747_Shank1(243961)_105 | 1 | + | good | GGCCCCATTGCCCCTCGAGGTGGACTGTGGGGAGTATGAAAGCATGAAAAAAACATACATCTGTGGTAGCATGGGAATATGCAGCTATACATGCACGAGGTTGTACATGCTACACACAAC | 120 | 48 |
| chr7 | 44322848 | 44322968 | 772751_42001747_Shank1(243961)_106 | 1 | + | good | ACAGTCAGGCACAAACTGGACTTGCTTTGTTATCCAGCAAGTGATGTTAGCAGTCAGACCCAAACAGACAGACACTAGGATCTGGAGACCAGGGTCACACCCTGCTTGCTCCAGTGACTG | 120 | 51 |
| chr7 | 44322968 | 44323088 | 772751_42001747_Shank1(243961)_107 | 1 | + | good | GCTGCAAGGGTTTAGGCACGAATGTTGCCTGCTCTGATCCCATCCTGTAGAATAGGCCTGAATTGAGTGCAGAGTATACACTTAACGGGGTTCCTATGAAGAGGGTATAATAAAGAGGCA | 120 | 47 |
| chr7 | 44323088 | 44323208 | 772751_42001747_Shank1(243961)_108 | 1 | + | good | CTTCAAAATTCACGTTCAAGGGAGCCAGAGTGGAGTGGAACTGCAGGCCGGTGTAGGTTTGTTCAGGCCAAGACAAAGACCAGATGATATTCTTCCACTGAGTAAATGTTCCCGGACAAT | 120 | 48 |
| chr7 | 44323208 | 44323328 | 772751_42001747_Shank1(243961)_109 | 1 | + | good | CCCTGTAGGCTGGGCACTGCTGAATGAGGTGTCACGCACCCATGCCTCTGTGGTACCTACCGTGGAAGAGAGCCGGTAACATAGAATATGGGATCTCTGGCATACAGTGTTCTGAAGGAA | 120 | 53 |
| chr7 | 44323448 | 44323568 | 772751_42001747_Shank1(243961)_111 | 1 | + | good | AGCATTCAGACTGAACAAGCAGATCTCCACGCATGTGAAGCCAGCATGGATTGCACAGCTAGTCCAGGCTATCCCAAGACCCTGTCTCAGGCAAAGAACAGAAGACGGATCAGTGAGGGA | 120 | 52 |
| chr7 | 44323568 | 44323688 | 772751_42001747_Shank1(243961)_112 | 1 | + | good | AAAGCTCTCTGAAAAGATGGTGCCTTAGTACAAAACCCAAGTGAAGCGAGAGTGTGTCGTGTAGAAATCGATGGATGGTGAATTCCAAGAAGCTCTTTAGATCAGTAGGTACAAAGACCC | 120 | 43 |
| chr7 | 44323688 | 44323808 | 772751_42001747_Shank1(243961)_113 | 1 | + | good | TGGAGCAAAGGTTTTCCCATTATGGGAAGAAGTAGCCAGCATGGTTGGAGCAAGGGAATGTGGTATTGAGGTTGAGATGGAATGTGGTATTGAGGTTGAGATGGGCTAGCCAAGTGGGTT | 120 | 48 |
| chr7 | 44323808 | 44323928 | 772751_42001747_Shank1(243961)_114 | 1 | + | good | GTTTGGGTGAGGCCTCGGGTCATGTCAGAGACCTAGTTCAGCTCAGAGTGATAGAGAGTGCTACGTGACAGATAGGAATGTATATGTCCCCCATTATCTCCCACGTGGCCCCCTACCCTT | 120 | 52 |
| chr7 | 44323928 | 44324048 | 772751_42001747_Shank1(243961)_115 | 1 | + | good | TGTGCTGTGTATACTATTCCTTCTAAGTCTGGCAGAGAGTGCAATCAAGACAGACCTAGGGCACTGGGGATGATGGCTTGGAGCTTAAAAGCACTTGCATGCAAGCATGAGGACCTAAGT | 120 | 48 |
| chr7 | 44324048 | 44324168 | 772751_42001747_Shank1(243961)_116 | 1 | + | good | TCAAATCTCAAGAACACACACACACACACACACACACACTGATGTAGTATGAATATGTGATCCCACTTCTCCTATAGGGAGATGGAAAGTAGAGACAAAGAACCTCCCCTCCGAAACAAA | 120 | 42 |
| chr7 | 44324168 | 44324288 | 772751_42001747_Shank1(243961)_117 | 1 | + | good | CAAACCTCATGGGCCAACTATGATGACATGGACACCATCAGACAAGAGTCTCAAACATTTCCTCCTGACCTCCACGTAGGCAGTGTCCCTCACAGAAACACATTCACACACCACACACAT | 120 | 48 |
| chr7 | 44324288 | 44324408 | 772751_42001747_Shank1(243961)_118 | 1 | + | good | ATCGTTCCCACACACAGACACACATGCACAGACACAGACCAGGAACGGAATCTAAGGTCAATCCTTTACCGGCGGCGTGACCTTGAACGATGCCGAGAACTTCGTTTCCCCATCTGTGAG | 120 | 52 |
| chr7 | 44324408 | 44324528 | 772751_42001747_Shank1(243961)_119 | 1 | + | good | ATCACCACGGTGATTTCTGCCGGGCAGGCGACTCGGTGAGCTCCTTTCTGAAGCTTGGTGACTGGCACACAAGCAAGAAGCCCTTCAAAGAGCTCAGAGGTGCTTCTGTCCACACAGGAC | 120 | 56 |
| chr7 | 44324528 | 44324648 | 772751_42001747_Shank1(243961)_120 | 1 | + | good | ACTTAGCCACGTCTGGAGCCCAGCCCCCTTAGCAGGAGACAACTTGTGGAGTCTGGTGGTAGAAAGGAGAGATACTCAGCCCCTCACAGGGTGGAGAGAACAGTATCTGCAACCACAGCT | 120 | 55 |
| chr7 | 44324648 | 44324768 | 772751_42001747_Shank1(243961)_121 | 1 | + | good | CAGACCTCTCCTGCCCTAAAATGTCCCTTAGAAAAAGGTTAGCTGCCTCTCCCCAACGCAGCAGACACAGTAAGGGAAAGGAAGCAGGCAATTTTGTGTCTGCAGCAGATTTTATTTAAC | 120 | 47 |
| chr7 | 44324768 | 44324888 | 772751_42001747_Shank1(243961)_122 | 1 | + | good | CTGAAGTTTCTACAAGTATTATCACTTCAATATGAAATCAACATAAAATTACTAGGGGTATATTTTTTGCATCCTAAGTCCTTGAAACCCCATATTTATTTACACACTGTATTGTGTGAC | 120 | 31 |
| chr7 | 44324888 | 44325008 | 772751_42001747_Shank1(243961)_123 | 1 | + | good | ACGCAATGCTACAGAACCTGCATGTGTTAGGCAGCCACCCTACACCGAGCCATTTCCCTCACTGGGGGACTCTAGGCAGAGGCTCTGCAACTGGTCTACATCGCAAGCTCACTTTTTACT | 120 | 53 |
| chr7 | 44325128 | 44325248 | 772751_42001747_Shank1(243961)_125 | 1 | + | good | GTGTGTTTTATGTACATAGTCCATCTCTCCTCCGACTGGCCGCATTTAAAGAATGACCTGCCTTGTCGAGGGGCTGCTGTCCCCGTGCTGGTTTAGGTCAGAGTATTTCTGAAGTCCCCA | 120 | 51 |
| chr7 | 44325248 | 44325368 | 772751_42001747_Shank1(243961)_126 | 1 | + | good | GCAACACAGCTCTGTTACTCACTCTAACTTACTATTCATTTTCCTCTCCTCCTGGACCACTGGCTGTCATGCTCAGGAGACCTGTGCCAGGATCCTCCTTTATCGTGGAGCAAACAAGGA | 120 | 49 |
| chr7 | 44325368 | 44325488 | 772751_42001747_Shank1(243961)_127 | 1 | + | good | TGTGAAAAATAACAATGGACAGACACCCTTCCAGGTCTGGGGTTCTGGGTTAATCGCCAGTGCAGGGACTGGGAGAGGGACACAGAAGACCAGGGGCCAAAAGCACTACCTGTGAGGGAT | 120 | 53 |
| chr7 | 44325488 | 44325608 | 772751_42001747_Shank1(243961)_128 | 1 | + | good | GCTGCATGGTCTGGGAGAAGGAAGAGGGTCGGTGGGGTCAGTGATGTGGATGGAGGGTTGATTGAGTGGGAGGCCGGGATAGAGGGGTTCAAGATGCCACCTGTCCGTGCTCTCTCCTGA | 120 | 59 |
| chr7 | 44325608 | 44325728 | 772751_42001747_Shank1(243961)_129 | 1 | + | good | GCATCCACCTCCCGGGACCCCTCCTGCAGGTGGCTGTGATTGCTGGGAATTTTGAGCTGGGGGAACTGATTCGTAACCATCGGGAACAGGATGTGGGTGAGTGAGAAGAAGTTAGGGTAA | 120 | 55 |
| chr7 | 44325728 | 44325848 | 772751_42001747_Shank1(243961)_130 | 1 | + | good | AGGAGAAACTGAGGGTTCATATTAGGGAGTACAGAGTTGGTGGCCTGATTGAAGGGTGTCAGGATGGCACATAGATGTTTGGGGGGGGATAACTCAGCGGGTCACCACCCTCACTGTCTC | 120 | 52 |
| chr7 | 44325848 | 44325968 | 772751_42001747_Shank1(243961)_131 | 1 | + | good | ATCTCTACCTTTCTCTCTCTCCCTCCACCGCTTGCCCTTACGGTTCCCACCTGCCTCTGTTTCCCCATCCTACCTGCCTGCCCCTGACCACCACTGTTCGCTCTCAGTACCTTTCCAGGA | 120 | 57 |
| chr7 | 44325968 | 44326088 | 772751_42001747_Shank1(243961)_132 | 1 | + | good | GTCCCCCAAGTATGCAGCCCGGCGTCGGGGACCCCCAGGCGCAGGGCTGACTGTACCCCCAGCGCTGCTGCGAGCTAACAGTGACACGAGCATGGCGCTGCCTGATTGGATGGTGTTTTC | 120 | 65 |
| chr7 | 44326088 | 44326208 | 772751_42001747_Shank1(243961)_133 | 1 | + | good | GGCCCCAGGGGCCTCATCCTCTGGGACCCCTGGCCCTACCTCAGGATCCCAGGGCCAGTCACAGCCCTCGGCCCCCAGCACCAAACTCAGTAGTGGGACCCTCAGAAGTGCCAGCAGTCC | 120 | 68 |
| chr7 | 44326208 | 44326328 | 772751_42001747_Shank1(243961)_134 | 1 | + | good | CCGAGGAGCCCGGGCACGCTCCCCGTCCCGGGGGAGACATCCTGAAGACGCCAAGAGACAACCTCGAGGCCGGCCCAGGTAGGCAGGACTAAGCCCCGCCCCCCAAACTTGCCATTCTAG | 120 | 68 |

| | | | | | | | | | |
|---|---|---|---|---|---|---|---|---|---|
| chr7 | 44326328 | 44326448 | 772751_42001747_Shank1(243961)_135 | 1 | + | good | AGCCCACACAGCCAACCCAGCTGTCCACCTTCTGTCCCAGGAGTGGTGGGTTCCATACCTACCTGCAGCTCTAATTGTCTCATGAGTAAAATGAGGGCGTTAGGGTAGGTACCTCACTCC | 120 | 54 |
| chr7 | 44326448 | 44326568 | 772751_42001747_Shank1(243961)_136 | 1 | + | good | TGGCATGGTAAGACAGAACCCTAGAAGAAAAAGGCCGCAACCTATGAGGACAGACAGATGGCAGGGTGAAGAGAGAGGAAAGAGAGCAGAAATAGGGGCTTGGGGTGGGCATGGACTTTG | 120 | 52 |
| chr7 | 44326568 | 44326688 | 772751_42001747_Shank1(243961)_137 | 1 | + | good | GAAGCAATATACAAGGCTGTGTACGAGTCCAGGGAGGGAGCTCTGGGAGCTGAGAGGACAGGAGAAGAGGCCGTTTCTGCGGATGTCAGGAAGAGGAGGCCCACCAGGTATGTCTCAGGT | 120 | 57 |
| chr7 | 44326688 | 44326808 | 772751_42001747_Shank1(243961)_138 | 1 | + | good | CCCAAGGAAGTGGAGAAGGCTGGGAACAGTTGGCTGTGGCCTTGGACCTGCCCCAGGAGGAAGCTAGGAGTCACTGAAGGCTCATGAGCTGTGCGATGTGCCAGAGCAGCTTCAGGAAGC | 120 | 59 |
| chr7 | 44326808 | 44326928 | 772751_42001747_Shank1(243961)_139 | 1 | + | good | TGAATCAGACTGTACTTGCAGACCGTGGGAGGGGTGGAGCTGGGAGCCTTAAAATGGGAGTGGGGCGTGCATCTGGTCTCCCAAGGACACTAGAGCTTGGAAAAGCACTCAATCCCTGTC | 120 | 55 |
| chr7 | 44326928 | 44327048 | 772751_42001747_Shank1(243961)_140 | 1 | + | good | CCCACCCTGATGCTCTGCCTTACTTCCATCCAAGGACGCCTGTCCCTCCAGAAGTGCCTTGTCCCTCCACTCTCCCCTGCCTCTCACAACTTCTGTGTATGCAGCTCCAGCGGGACACCC | 120 | 60 |
| chr7 | 44327048 | 44327168 | 772751_42001747_Shank1(243961)_141 | 1 | + | good | CGAGATGGGCCGGCTGGGGGAACTGGGGGCTCAGGGGGCCCTGGGGGCTCCCTGGGCAGCAGAGGCAGACGTAGGAAGCTCTATTCAGCGGTACCCGGTCGCTCCTTCATGGCGGTGAAG | 120 | 68 |
| chr7 | 44327168 | 44327288 | 772751_42001747_Shank1(243961)_142 | 1 | + | good | TCCTACCAGGCGCAAGGCGAGGGGGAGATCTCTCTGAGTAAAGGCGAAAAGATCAAAGGTGACAGAGTGGGGCTGGGTGGGGGCAGGGTGAGGGGAAGCTCAAAGACCGCCTGTGCCCTC | 120 | 61 |
| chr7 | 44327288 | 44327408 | 772751_42001747_Shank1(243961)_143 | 1 | + | good | TGCACTCTGGGGACTTTAAAGAGCCCTGAGAGGCCCCTTTTGGCTTGGAGGAGAGAGAGGCACTATATGGGACAGTGTGAAGAAGAGAGGAGATGAGCATGGATTGTCCCATGTGTGAAT | 120 | 51 |
| chr7 | 44327648 | 44327768 | 772751_42001747_Shank1(243961)_146 | 1 | + | good | ACAGCTGAATCTGACCTTTCACATGGGTTCTAGGGATTCAAGTTCAAGTCCTTAAGCTTGTGCACAAGGTCCTTCGTCTCCTGGGCCATTTTCTTGGCTCCCAGAGCTTTTCTTTTTGAG | 120 | 47 |
| chr7 | 44327768 | 44327888 | 772751_42001747_Shank1(243961)_147 | 1 | + | good | GCAGGGTCTTATGTTGCCCAAGCTCTTCTCAAACTCACTTAATAACACAAGTTCACCTTGAGCTTCTGCCTCCACCTCCTGACTGCTGGAGTTACAAGTTACCATGCCTGGCCTTCTCTC | 120 | 49 |
| chr7 | 44327888 | 44328008 | 772751_42001747_Shank1(243961)_148 | 1 | + | good | TGGAGGACAGTGACCATAGCACATAGACTGCAGCCTTCAAAGCATGAGCAGTGTGTGTTCCTGGATAAACGGGTACCTTGGAGATAGAGAGTCATGTGGAAGGCAATGTAATTCAGAGCA | 120 | 48 |
| chr7 | 44328008 | 44328128 | 772751_42001747_Shank1(243961)_149 | 1 | + | good | AACAGGGTAATTTTCTACCAGATTCTAGCCTGCCCAGGAGCTCTGCAGTGTGGTAGCTGAGCAGCAGGGCTGTTTGCATTCAAAGCTTGGCCCCACCACCTCTTAGAGGAAAGAGTTCTA | 120 | 51 |
| chr7 | 44328128 | 44328248 | 772751_42001747_Shank1(243961)_150 | 1 | + | good | GCAACCAGCTTCTCTGAACCTCAGTTTCACTGTCTTTAAAATAAGACTCTAGGGAGAATCATGACCGTCACTAAGTCTCTTAGGCCCAGGCTTTGATGGTGCATGAGAGCCGGCCTTCCT | 120 | 48 |
| chr7 | 44328248 | 44328368 | 772751_42001747_Shank1(243961)_151 | 1 | + | good | CCACATAAGCTCTCAGAGGCAGACTAGAGGTGGCTGGACAAGCAAGGCCAGTCCTTCCACGGGGCATAGATATGAGGAAGGCCTTCAAGGGTGTTTAGAGGTCCTCATCCAGCAAGATGG | 120 | 54 |
| chr7 | 44328368 | 44328488 | 772751_42001747_Shank1(243961)_152 | 1 | + | good | ATCTTTTGTGGGCTAGTTGTCTATCTTCCCATAAAAGACAAGGATAGGAAGGTCAACGGTGTGTGTGTGTGTGTGTGTGTGTGTATCTATTATCTATTATGTCTATATTGTCTGTCATGT | 120 | 38 |
| chr7 | 44328488 | 44328608 | 772751_42001747_Shank1(243961)_153 | 1 | + | good | ATGTATGTCTCATGTATCTAGCTTACAGTCTATCTATTAGATTTATCTATATTATGTACATATCAATCCATCATATATCTATCTCCTTATTTGTATATCTAGCAATCTATGCAGATGTCA | 120 | 28 |
| chr7 | 44328608 | 44328728 | 772751_42001747_Shank1(243961)_154 | 1 | + | good | GTCTTGAGGTAGTCTCACGCTGTGCCCAGCATAGCCTTGAACTCACTCTATAGCCAAGACTTGTCCTAACTGTCAACAGTCTTCTTGCCTTAGCGCCCTGAGTGCCAGGATGGTATAGAT | 120 | 50 |
| chr7 | 44328728 | 44328848 | 772751_42001747_Shank1(243961)_155 | 1 | + | good | GAACTAGCATGTTTTGGTTTTCGAACATTCTGTACCATTTATGTAGCAAACTCTTCCATGGTCTCTTTGTATCCTCAGCTACTAGGGTGCTTCAGGTTTTAGGGATATCTGGATATCTTA | 120 | 39 |
| chr7 | 44328848 | 44328968 | 772751_42001747_Shank1(243961)_156 | 1 | + | good | TCCATTCAGTTCACTTAAGGTCATCTGGCCTCTCTCAAAGCACCATTTTCTGTCAAGTCCTTCATTAAAAATGGTGGGAGTTTGTTTGGTTTACGTGTGTGCGTGTGCATGTGTGTGTGT | 120 | 43 |
| chr7 | 44329208 | 44329328 | 772751_42001747_Shank1(243961)_159 | 1 | + | good | CTTTGAGGCAGAGTCTAGGCTGAAAGCCCCAGCAATCCTGTCTCCACCCATCTTATCTTAGAACTGGGATTATAGGCATGCATGGGACACATAGCTTGTTACATGCATGCTGGGTCCTGA | 120 | 49 |
| chr7 | 44329328 | 44329448 | 772751_42001747_Shank1(243961)_160 | 1 | + | good | ACTCTGGTCCTAATGACTGTGCACAAATGTTCTTAACTACTAAGACACCTCTCCAGCCCTTCTTCCCTTTCTGAGAACTTGTCTGGAGGCTCTAGCAGGGGACTTCAAATATCCACATGC | 120 | 47 |
| chr7 | 44329568 | 44329688 | 772751_42001747_Shank1(243961)_162 | 1 | + | good | AAGCCAGCCAGTCCATCCAAATGAACCCTGGAAGTCAGTGGGATGAGACATGTCACAGTGGGGCTGCCTTCACATCCACTTCCACTACTTTTTCAATAATAGTTTCAGTTTATATGTATA | 120 | 43 |
| chr7 | 44329808 | 44329928 | 772751_42001747_Shank1(243961)_164 | 1 | + | good | GATAAGAATAGAACCTGGGTCCTTTACAAGAGCAGTCCGAGCCATCTCTCCAAGCCCCACCTCCCTATTTGTTTGAGACAAGGTATCTCACTATGTAGCTAACTGGGAACTCACTGTGTA | 120 | 47 |
| chr7 | 44329928 | 44330048 | 772751_42001747_Shank1(243961)_165 | 1 | + | good | GAGCAGGTTGTTTTTAGACTTGTAATAATCCTCCTGTATCAGCCTCTGGAATGCCAGGGTTAAAGACATATGCCCCCATGCCCAGCCTCTTCTGTTCAAGCCCCTTGATCGCTGTAGGTG | 120 | 49 |
| chr7 | 44330048 | 44330168 | 772751_42001747_Shank1(243961)_166 | 1 | + | good | TTCCTCTAAGGGCGGCTAATCTAGAAGCTCTTCTAGGGCAGGAACTTCATGGGTTGTGTCCAACTGTGTCTGTCCTGGTACTTGACACTTGATCAGCACTCAGTATGTGTTTTGTGAGCA | 120 | 48 |
| chr7 | 44330168 | 44330288 | 772751_42001747_Shank1(243961)_167 | 1 | + | good | AATGATTGATCTCAAGCAGTCAGGCATGGCAGAACCAACTGAGAGCCCAGATCCCTGGATGCCCTCTTGTCTACCCTCATCTATGACCTGGGGCTTTCACTCATGTCATTGCCCTGTATC | 120 | 51 |
| chr7 | 44330288 | 44330408 | 772751_42001747_Shank1(243961)_168 | 1 | + | good | TTCGGGAGCAGGGGGAGGTGTGTAGGGAGAAGTGTCAAGGAGCCACATGGCAATTTCTTTGTCATCTGAGCTATGGTGGAGACTTTTCAGTGTGGTCCTCAGTGTGTGTGTGTGTGTGTG | 120 | 52 |
| chr7 | 44330408 | 44330528 | 772751_42001747_Shank1(243961)_169 | 1 | + | good | TGTGTGTGTGTGTGTGTGTGTGTGTAAACTATATATACAGTTGTGTGCTTATGTATGTAGAAACTAGAAGTCAACCTTCAGTGATGTTCTTTAGGGCCAATCCTTCTTGCTTTATTGAGACA | 120 | 38 |
| chr7 | 44330528 | 44330648 | 772751_42001747_Shank1(243961)_170 | 1 | + | good | GGATCTCTCACGACCCTGAAACTTGTCAAGTAGGCTAGGTTGGATGACCTGCAAGCCCTAGCAATCTACTTGTCTCTACCTCCCCAGCCATCAGATTGCAGTGTGTACCACTGTGTCCAG | 120 | 52 |
| chr7 | 44331248 | 44331368 | 772751_42001747_Shank1(243961)_176 | 1 | + | good | AAAGATTGACTTTGAACTCACTCTGCAATACAGGCATCTCTTAAATTTTCAATCCTCCTGCCACAGCCTCTTGAGAAGCCTGTGCTACCAGCCCCAGCTTGCCTCCAGCTTTTTTACCCA | 120 | 47 |
| chr7 | 44331368 | 44331488 | 772751_42001747_Shank1(243961)_177 | 1 | + | good | GAAAGAGTTGGAGAAAATACTGAAGATACAGAAAAGACAGTGTGTCTGTTGCTATTAGCATCATTCTCTGAATGGGAATTCAGAGAACTACACAGGCTTGGGTTAAAATCCTGCCCTGCC | 120 | 42 |
| chr7 | 44331488 | 44331608 | 772751_42001747_Shank1(243961)_178 | 1 | + | good | AGTGACTACTAGCCTTCTAATCTGGGGCAGATGGCAAGCGGCATTAGCTCTCTGTGCTTCGGTGTCCTTGTCACTAAATATCCCTTCCCAGAGTTCTGGCCTGGAGTGAGCTCTCAGTGT | 120 | 52 |
| chr7 | 44331608 | 44331728 | 772751_42001747_Shank1(243961)_179 | 1 | + | good | CCTTGCCATCTATCTTTGCTTCAACACCAGATACGGAACAGAATGGTGATGGGTGGCAAGACAGGTCCACTAAAGGGCTGAAGTTGTCACTGCTTAGAGCAGTGGGAGGGGGGATGCCAA | 120 | 52 |
| chr7 | 44331728 | 44331848 | 772751_42001747_Shank1(243961)_180 | 1 | + | good | GAGAGCATCCTATGGCAAGAGGGGCAGAGTTGGGGAGGAGCTGGGCAGAGACTTCAGGGAGCAGCAGGGCTAAGAGTGGGCACAGCTGTCTTCAGAAGAGCCAGCGGAGCCACGGGGCT | 120 | 62 |
| chr7 | 44331848 | 44331968 | 772751_42001747_Shank1(243961)_181 | 1 | + | good | TGGGGGCTTAAGAAAACACTCCACTAGGTTGTAGGTGGTGACCAAAAGGGACAAGGACCAAGGGAGGACGGTGCCACAGAACCTAGTCTGCTCCCCAAATCTACCTGATATGCACCTCTT | 120 | 52 |
| chr7 | 44331968 | 44332088 | 772751_42001747_Shank1(243961)_182 | 1 | + | good | ATGTTAAGGAAGTTCACATTTAAGAATGTCCCAACTTAGCTCCCCATCATGGTCTCCCCCATTTCTCAGACCTTAGGGTTTTTTGGGGGGTTTTGGTTTTGGGTTTTTGTTTGTTTGTTT | 120 | 41 |
| chr7 | 44332208 | 44332328 | 772751_42001747_Shank1(243961)_184 | 1 | + | good | TGTCTCCAAGCCTGGGGGTATAGATGTGTACCACCACAACAGTTTTGTTTTGTTTGGAGTTTGGAGTGTGATATATGCATACTTTTGTACAGGTGCACATATTCAGAGGGCAGAGGAGGG | 120 | 45 |
| chr7 | 44332448 | 44332568 | 772751_42001747_Shank1(243961)_186 | 1 | + | good | GTGTTAAAGTTGTGTGTGGCCACACGCAGTTTCTGTTTGACCTTAGCTGCTGGCATCCAAACTCTGGTGCTTAGGGTTGTACAGTACATTCTTACCCACTGAGTCATCGTACCACTTCCT | 120 | 48 |
| chr7 | 44332568 | 44332688 | 772751_42001747_Shank1(243961)_187 | 1 | + | good | ACCCCTAAGTTTTTGTTTTGATTTTTGATAGGGTCTTGTGATATAGCCCAGATTAGCCTGGAAATCACATACACAAATCCTGCACCAGATTTTTCTTTTTAGGGTTTTTTTTTGTTTTGT | 120 | 34 |
| chr7 | 44333168 | 44333288 | 772751_42001747_Shank1(243961)_192 | 1 | + | good | AATATGGTGGGGCAGGGAGAGGAAGAAAAAGAAGAGGAGGAGGTGGGGGAAAGAGAGGAAAAAGAAAAAGAAAATTTAAAAAGGCAGCTTAAAACATTTCCTTGTTCCCACTGTCCCATC | 120 | 42 |
| chr7 | 44333288 | 44333408 | 772751_42001747_Shank1(243961)_193 | 1 | + | good | CAGTGCTCAGCATCGGGGAAGGAGGCTTCTGGGAAGGCCAGGTCAAAGGTCGAGTTGGCTGGTTCCCTTCTGACTGCCTAGAAGAGGTGGCAAACCGTTCCCAGGAGGGAAGACAAGGTA | 120 | 57 |
| chr7 | 44333408 | 44333528 | 772751_42001747_Shank1(243961)_194 | 1 | + | good | ACAAGAAACTTTCTCCCTCATGCCCTACAGTCTTCCTATCCCTCCTGCCCTGCCTGATAGGAGTGGGGCGCACAAGTCGGGGAGTAGGCTTTGCTGTTTACAAAGTAAAGGCTCAATGCT | 120 | 52 |
| chr7 | 44333528 | 44333648 | 772751_42001747_Shank1(243961)_195 | 1 | + | good | TTATGTCCCTGCAGAAAGCCGAAGTGACAAGGCAAAGAGACTCTTCAGGCATTACACAGTGGGCTCCTATGACAGCTTCGATGCCCCAAGGTAAATGTCCTCACACCCTTAGGTGCCCCA | 120 | 52 |
| chr7 | 44333648 | 44333768 | 772751_42001747_Shank1(243961)_196 | 1 | + | good | CCTCTCCCAGCTCCCTGCACACTGCACACTTGAAGCCTGCTTCTTTTCTTGGTCAACTTGGAAGCCAGGAGAAGGCAGAATTCTGTTAACGTAACAGCCCCATCTCAAATCTGCTTCTGG | 120 | 51 |
| chr7 | 44333768 | 44333888 | 772751_42001747_Shank1(243961)_197 | 1 | + | good | CAGCCACTGAGACCATCTAGTGTGGGCTGGGTGTGTCTGGGGCCCTGGGTTCTTGTGTGGGGACTGGTTAGATGCCCTGGTCTTCTTCCTGTTTCCCTGTCTGTGTGCAGTCTCATACTG | 120 | 57 |
| chr7 | 44333888 | 44334008 | 772751_42001747_Shank1(243961)_198 | 1 | + | good | TTGGACACATGTACATCCTGTATGTCACACCAGTGGGTGGTGGGGGAGGCAGGTGGAATCAATATGACCCCATGGGTGGGGACAGAGAGAGGGGCACACTGACACCTGTCCCCTCCTTGG | 120 | 57 |
| chr7 | 44334008 | 44334128 | 772751_42001747_Shank1(243961)_199 | 1 | + | good | CTGCTGGCTGACTTGCTTCTGAAGTCTGATCGATGGGATTGACTCAGGGAGGTGAGAGGGGCGGGGCAGGGTGGGGATTGGGCCAGGAGGGGATGCTTGTATATGGGGGGATGAGAATGT | 120 | 58 |
| chr7 | 44334128 | 44334248 | 772751_42001747_Shank1(243961)_200 | 1 | + | good | GTTTGTGTGCATGACCAGAGTTGAGGGGTCCCAGCGTGTGGGCGTGTCTATACCAGTGTGCATGACGCATCCGCGTGCATGCATGGGCAGAAGGCCCTTAGTCGGATCACTGTAAATAAC | 120 | 55 |
| chr7 | 44334248 | 44334368 | 772751_42001747_Shank1(243961)_201 | 1 | + | good | CCTAACGAAGGCTGTGTGCTACAGCCCTCGGTGCATATGGGTACATTTGTGTCGGTGTTGACTGCGTGGCTGGCTTTGCAACATGTGCGCTTCTCTGACTGTGGGCAAGCACTGCCAGAG | 120 | 56 |
| chr7 | 44334368 | 44334488 | 772751_42001747_Shank1(243961)_202 | 1 | + | good | GTGTCCATGTGTCTGCTCACACGTGAACAGGTGACAGATCTCTCTACACACCCCGTGTCACTGTGTGTTCTTTGGTGGCCCCAGTGTGCGTATGTCTTCTCTTCTATGATTGGTCTGAAA | 120 | 50 |
| chr7 | 44334488 | 44334608 | 772751_42001747_Shank1(243961)_203 | 1 | + | good | TTCAGAACCTGGAATGCTGAGGGAGGGAAGGGCAGAGGAGGGGCCTCAGGGCTGTGCTCTGGAGAAGGAATGTCATAATCCAAGATCTAACCGAGGACACCCTCCTCTGCCACCCCATCC | 120 | 57 |
| chr7 | 44334608 | 44334728 | 772751_42001747_Shank1(243961)_204 | 1 | + | good | CACCCCAATCCAGGGAAGGATCTTCCTGTTCACAGTATCCCTCCATTAAATGTATCCAAACCTGTGTCTGTGTTCTGGCCAGTGGGGGGAAGGTCTGGGTTGGGGGAGGAGAAAGGAAGG | 120 | 53 |
| chr7 | 44334728 | 44334848 | 772751_42001747_Shank1(243961)_205 | 1 | + | good | GGTGGTTGACAGGGTGGGGCTTGGGTGCCAGTCACCCTGTGCCCCAGGGTCTGAGGCAAATCATCTTGTGCCTGTTCACGGGGTATGTCTCTGGCTCTGGGTCCTGGGTGAGTATGGAGG | 120 | 61 |
| chr7 | 44334848 | 44334968 | 772751_42001747_Shank1(243961)_206 | 1 | + | good | CTCCCTCTACCCCAAGTCAGCCTCTGTCTGGGGATTCTGACCTGGGCCTGCCTGGTCTTGCAGAGGTGTCATTCTCTCCAAAGACTGAGTGGAGAGGGGGTCTCTTCAACCCCAGACTCC | 120 | 58 |
| chr7 | 44334968 | 44335088 | 772751_42001747_Shank1(243961)_207 | 1 | + | good | ATCCTCTTTTCCTAATAACGATCCTGTCTTGAAGCCTTGCAATTTTTGTCCTTTTCCCTCCTCTCCCGGGGCTGAGGTCCATCCAAGCCCCCTTCAGTCATATCTTCTTGCCTTTGCTCT | 120 | 48 |
| chr7 | 44335088 | 44335208 | 772751_42001747_Shank1(243961)_208 | 1 | + | good | ACCTGACCCAGTCTGCAGTTTCCTGGTTTGCCTGTTTACACCTTGCATCTTACTGTGGTCCCTAGCATCTCAACTGGAGCTGAGCCATGTTTGCTTGTGTACCTACCCCATGGTGTGTGC | 120 | 52 |
| chr7 | 44335208 | 44335328 | 772751_42001747_Shank1(243961)_209 | 1 | + | good | CACCATGTGCCACCACACTCCTCCTACCTGGGGGTGCGGTGGGAGGTCCATGGCTCTCTAATGAGTTGTGGTATATGTGTCCTCTCACTACTCTCTGGGTTCTGTGTTTTGGGTTCCAGT | 120 | 54 |
| chr7 | 44335328 | 44335448 | 772751_42001747_Shank1(243961)_210 | 1 | + | good | TGAATGTCTTGGCTCCCCTCAACCCACCCCTGCACCCTGCCTTTCCCTACTATTTTGGGGATGAGCTGTGTGTCTGTCTGCGCTGAGGGTGAGCTGTTTGCCCTCTCTTGGGGAGGGGTC | 120 | 58 |
| chr7 | 44335448 | 44335568 | 772751_42001747_Shank1(243961)_211 | 1 | + | good | TATCTCTCTCACCCTTGGGTGTCCCTTGTCCTGAAAGTGTGGTTCTGTCTGTCTGCGCGTCCCCGCACGCGATGGCGTGCCATCTTATGTGATGTGTGGCGCTGCGTCCTTCTCCCCGCG | 120 | 60 |
| chr7 | 44335568 | 44335688 | 772751_42001747_Shank1(243961)_212 | 1 | + | good | CTCCCGCTCTCTGGCTCGCTCTCTCTGCACCTGCGCGGAGAGGACTCTGGAAAGGGGGGAGTATGGATGGGGGGGACAGACCGGAAGTGACTCCCTCCCCCTCCCCAAGCCCTGCCCCTC | 120 | 68 |
| chr7 | 44335688 | 44335808 | 772751_42001747_Shank1(243961)_213 | 1 | + | good | CCCCTCCTCCCGCCAAAGGGGCCCCCGCGTTACCCTCCCCCGCCCGTCCCCCCTCCGTCCGGCTGCCCGGCGCAGGACGCCAAGGGGTTAAAGCCGGGCCGCTGCCGCCTCTGGAGCTGT | 120 | 77 |
| chr7 | 44335808 | 44335928 | 772751_42001747_Shank1(243961)_214 | 1 | + | good | CCGCGGTGCTGGAGCCTCAGCCATGCGCTGTCCCCCGCGCGCCCCCCGCGGCATGTGAGCCCCCCGGGGCTGCCCGCCCTCCCGAGGGGGGGTGCGCGCCCCCTCCCCCGGCCCCTCCCC | 120 | 84 |
| chr7 | 44335928 | 44336048 | 772751_42001747_Shank1(243961)_215 | 1 | + | good | CCTCCCGGCGCGGGTCAGCATGAGGCGGGGCCTGGGGGCCCGGACACGGGGGTTCAGCCGAGCGGGGACGCGGCGGCGGCGGCGACGGCGGCCGCGGTGGTGGCGGGGGCGGCCCG | 120 | 84 |
| chr7 | 44336048 | 44336168 | 772751_42001747_Shank1(243961)_216 | 1 | + | good | GGGACCCTGGCCGGGGCCGGGGCCGCGGTGGCGATGGCTTTGTCTGCGGTCGGGGGTGGCCCCGGCGGGGGCGCCCTGCCCCAACCGCCCCCTGCGCTATCCTCCAGCTGGCCCGCGCTC | 120 | 81 |

| | | | | | | | | | |
|---|---|---|---|---|---|---|---|---|---|
| chr7 | 44336168 | 44336288 | 772751_42001747_Shank1(243961)_217 | 1 | + | good | GGGCCCCGGAGGCGCAGCGTCTGGTACATTTACAGGTAACCGCGCACCGCCCGCTCGTCCCCCGCACGACCCTCCGTGCGTTTAGGGCTCCCCAAACCTGTCCCGTGTCCTCTTCGTAGC | 120 | 68 |
| chr7 | 44336288 | 44336408 | 772751_42001747_Shank1(243961)_218 | 1 | + | good | CTCCCTTCCCTAGGCCTGAGCCCTTCCAGACCCTCTCTCAGCAGCTCAGCAACCCGGTTCTCTTCCACCTCCCTCCTTCTTGTGCTTCAGGCGCTGCTCCAGGGCCTCCTCTGCTATTCA | 120 | 61 |
| chr7 | 44336408 | 44336528 | 772751_42001747_Shank1(243961)_219 | 1 | + | good | CCAAGACTTTTGCCTCCCCTCGAACCCCCACGAAAGGACCCCACTACAGCTATACTCTGCAACCCACTCTTGCCCTTTCTGGCACCCCTAGCTCCCCTACTCTAAGGAGTACTTCCCCAG | 120 | 57 |
| chr7 | 44336528 | 44336648 | 772751_42001747_Shank1(243961)_220 | 1 | + | good | GGCCCCAAACAGATACCCCGGGCCCCCATCCCTCCAATCCTCCGCATCCCCGCCCCTCTTCCTGGAGTCAGCAAGTCCCCCCATTCATACCCCTGTCTCCCATTCACAGTCTGTCCCCC | 120 | 64 |
| chr7 | 44336648 | 44336768 | 772751_42001747_Shank1(243961)_221 | 1 | + | good | CACACACACACCTCCATCACACAAGCATCCCGCCCCCGGCCTTCCCTCCCCTTCTCTTTTTCCGCTGCGTGTCACTGTGCGGCTGTGCGTGTGTCAGTGATGTGTGTGCGTATGACCGCC | 120 | 61 |
| chr7 | 44336768 | 44336888 | 772751_42001747_Shank1(243961)_222 | 1 | + | good | TGCGTGTGTGAGAGACAGACAGAGATGGGGAGGCTGTGGGCTCCCAGCCCACCCCCCCCCACCTATGTCCATCTGTGTGTCCCTGCGTGGGGGTGTCTGTCTCTGTGTGACTTGGAGCC | 120 | 63 |
| chr7 | 44336888 | 44337008 | 772751_42001747_Shank1(243961)_223 | 1 | + | good | ATAGCGAGGCTGTCTGGGGACTGTGCGTGTATGTGCGCCTGTGTGTGTCTGTGTGCGTGCGCGCACACGTGTGATAGAGTGCATCTGTCCCAGAGACTGCGGCTGTGTTTCCCAACACCC | 120 | 59 |
| chr7 | 44337008 | 44337128 | 772751_42001747_Shank1(243961)_224 | 1 | + | good | AGCCTCCTAGGCCTGGCCTGCCCCCTCTTCCCCCTCCCCCCTCCCCAGTTTGCCCTGTGTTTCTGTCCCCGCGTACCCTGCTCCGTGCCCTCCCCAGGAGTCTGTGCATGGGGAGGATCG | 120 | 68 |
| chr7 | 44337128 | 44337248 | 772751_42001747_Shank1(243961)_225 | 1 | + | good | ACCCTGGACCCCGAGGTTCTGCCCTCTGCGGGGATGGGGGCACGTGGGCTGAACCTCGGCAAGCTTCTGCATGGGGCTCTGGCACCTGCCTGTGTGCTGTTCTCAGCATGGGGACAGGGG | 120 | 67 |
| chr7 | 44337248 | 44337368 | 772751_42001747_Shank1(243961)_226 | 1 | + | good | CTGGCTTTCTTGCATGCGGGGGCCCTGTGCAGGCCGTGGAGAGGCCCGCCGAGATGGGAACACAGGGAGGGAACGAGGACTGGAGAATGCAGTGTGGATGGCAGCCGGCTCCAGGCTGT[illegible] | 120 | 66 |
| chr7 | 44337368 | 44337488 | 772751_42001747_Shank1(243961)_227 | 1 | + | good | GCATCCTGAGTGGGCAAGAGGGACCTGGGGTGGGGGTGGGGAGCGGTCGCGCCGCAGTCTCCATGAGGGAACCAGCAAGCCCTGGAGACCTTCTCTGGGACACGGCGGGATGGCCTGGG | 120 | 69 |
| chr7 | 44337488 | 44337608 | 772751_42001747_Shank1(243961)_228 | 1 | + | good | AAGGACATGGTGCCAGCTCCTCATGGGCAGGTAGTCCGCACCGCGAACCTTGGTGTTGTGAGTGGACTCAGAGGGAGCGGGGGTGGTGGTGCAAAGGGGATTGCTGGATAGAGGGGAAAT | 120 | 59 |
| chr7 | 44337608 | 44337728 | 772751_42001747_Shank1(243961)_229 | 1 | + | good | GGGACGCGTGGGTGCTGGAGAGGGGTTGGGTAATGGCGTTTGCGGGTTCTGGGTGCCAAGAAAGGTGCTGGGGTGAGAATGGAAGGCAGGCGGTAGGAAGAGGATTGGGGGGCGGGGA | 120 | 64 |
| chr7 | 44337728 | 44337848 | 772751_42001747_Shank1(243961)_230 | 1 | + | good | CAGCCGGCACTACCGCCTGGTGTATGGGAGGAATGTGAGCGGAAGAGGCTGCTAGGAAAACCAGCATGGACCAAGGAGAAAGAGCTTTCCGTAGTGGGCAGGCAGTGTGAAAGGAACGTG | 120 | 57 |
| chr7 | 44337848 | 44337968 | 772751_42001747_Shank1(243961)_231 | 1 | + | good | GGTGTTATTGGGGGACATCATGAGCTTCTCTGTACAGGAGAGAAGAGAATGTGGAAGGCTGAGGGTCAGGAATTTCGGAGAGGAGGCAGGCAGGGTGAGGAGGTGGCAGATGTACGAAGA | 120 | 53 |
| chr7 | 44337968 | 44338088 | 772751_42001747_Shank1(243961)_232 | 1 | + | good | GGTCACTTAGAAGAAGGGATGACTGGGTGCAGGGGTGGTCTCCTGGTATAGCATACATTATGGGCACCCTCAGACTGGCCGACTGATGGCATAGTTTTGGGAGCCATGGAAATACAGGAT | 120 | 52 |
| chr7 | 44338088 | 44338208 | 772751_42001747_Shank1(243961)_233 | 1 | + | good | GGTGTGCAGCGAGATGTAAATCAGAAGGATGGAGTCAATGGCAGGAAGCTGAGAGAGAGACAGGAGAGCAAGTAGGACATTTGGGGATGAGGGTACCACATCTGGCAGGGACATACACAG | 120 | 52 |
| chr7 | 44338208 | 44338328 | 772751_42001747_Shank1(243961)_234 | 1 | + | good | TGTAAACTGTGGAAATAGACAGTGCTGCAGCATGGTGGACTCCATATGAGAGGGTCCTGTGGTTATGGGGGCAGGTGGTGCTGCTGGGGTCTGTGGCTTGGGGGGGGGCAGCTTGTGGTC | 120 | 58 |
| chr7 | 44338328 | 44338448 | 772751_42001747_Shank1(243961)_235 | 1 | + | good | TTGACAATTCTAGGTGGTGTGAGATCTTGAGCAGGTCCTTGCTCTGTGCCTCACTTAACCTAGCATTAAAGGTGGGGACACTCACAGTCCCATCTAGGAGTGTTGTAATGACTTAAATAT | 120 | 44 |
| chr7 | 44338448 | 44338568 | 772751_42001747_Shank1(243961)_236 | 1 | + | good | TAAATGACCCAGAGAGTGATTAGTGTTGCCTGGATCTTGATGGGTTCCTTGTGTGACTGGCAGAAGTCAGGATAAACCCCAGAGGCTGGAGGGACAGTTGGGGGTTGTCTTAGTTAGGGT | 120 | 50 |
| chr7 | 44338808 | 44338928 | 772751_42001747_Shank1(243961)_239 | 1 | + | good | ATGCCCCCCCCCCCCACACACACACCATGGGCTGGGCCCTCCCTCTATCAATAAGTACTTATGAAACAGTACTTCAGATAACTCTAGTTTGTCAAGTTGACATAAGACTAGCCAGCACAG | 120 | 49 |
| chr7 | 44338928 | 44339048 | 772751_42001747_Shank1(243961)_240 | 1 | + | good | GGGTCTCCTGCTAAGCTTGGGGGTAAACCATGTGTCTGAGGTATTTAGGCAGTGTGACATGAAAGGAATTCTAGCTGCAGTTACCCAGATATTTTCTTGGGCATCAAAGAATGTTATTTG | 120 | 43 |
| chr7 | 44339168 | 44339288 | 772751_42001747_Shank1(243961)_242 | 1 | + | good | GCTGCACAGTGAGCTGTATGTGGAAAACAAACACAAAAGGATACTGCCTGTGTCTCTGGAGAGTTTCTTAACCTTTCTGTGCCTGACTTACTCACCTGTAAAATGGAAATAAATGGTATT | 120 | 41 |
| chr7 | 44339288 | 44339408 | 772751_42001747_Shank1(243961)_243 | 1 | + | good | TATGTCTTCTTCCAAGACTGGCTCTGAGGAGGGGGCTAACGCAGTGGGTCTGTAGTGAAGGGTTTCCCATGCTGCCAGGCCAGCAGCAAGCACTTGACCTCAGCATATGGACAAGAGGCC | 120 | 56 |
| chr7 | 44339408 | 44339528 | 772751_42001747_Shank1(243961)_244 | 1 | + | good | TGTTCATATCTGAAGACAAGACACTTGGGGTAACTGAGGCTTGTGTGTAGAGCTGGGGTTGAAAGAGATGGCAGAGTTTGATGTGGTGGCTCACAGCCATCACCCCAACACTAGGAAGGC | 120 | 50 |
| chr7 | 44339648 | 44339768 | 772751_42001747_Shank1(243961)_246 | 1 | + | good | TGGCAAGGAGGTATGAATGACTGAAGGTGACTTTGCCTTGGTGGGGTGTAAAGGCGGAGCCTGAAGGGCACTATGACCTCCTGGGAAGGTTCTTGGGAATGTGGAGAAGCTGTCAGAGTA | 120 | 52 |
| chr7 | 44339768 | 44339888 | 772751_42001747_Shank1(243961)_247 | 1 | + | good | GTAAGGACATAAGAGGTTGTCATAGTGTGTCAGGATGAGGCTACCACAGAATAGACAGTCAGAGCCAAGCTTCCTTGACCTCCCAGACTGTCCCTTCACCTTCCCTCCAGCCTGCTTAGA | 120 | 51 |
| chr7 | 44339888 | 44340008 | 772751_42001747_Shank1(243961)_248 | 1 | + | good | CCTATTTCTCCCATCTACCCTTAACCTTCCTTTCTTCAGCCCACCAGCCTTCTCCCAATATCTGCTCCATGCAGCTTCAAAGACTGCTGGGGGTGGGGCAATTGGAGAGGGCTGAGGGAG | 120 | 54 |
| chr7 | 44340008 | 44340128 | 772751_42001747_Shank1(243961)_249 | 1 | + | good | AGGAGGTAGAAACCTGTGGACAGGTTTATTGACCTCTGGATAAGTGCAGAGGCTGGAACATTCCTCTTCATGCATGCATGATGGGGTATTCATTAAGTTCCACAAGGAGGGAGGCAGCAC | 120 | 48 |
| chr7 | 44340128 | 44340248 | 772751_42001747_Shank1(243961)_250 | 1 | + | good | CTGGACTTGTGCAGAACACTCACTGCTATTCAGGTGCATGGGTTTTACCATGGAAGCTGTGGGGGGTCATGGGAGCTTACACAGACACTGATGGAAGGAGGTGGCACCACAATGCAGTAG | 120 | 52 |
| chr7 | 44340248 | 44340368 | 772751_42001747_Shank1(243961)_251 | 1 | + | good | GCACACAATTAGGTGTTATTCCCGTGTGCTGGGAGGTGCTCAGAGGGCGTGTGATGGGAATACACTGATATCAAAGTGAGGGGGAGTCAGGATTGGGGCACACTGGGATGTGTGGAGGTG | 120 | 54 |
| chr7 | 44340368 | 44340488 | 772751_42001747_Shank1(243961)_252 | 1 | + | good | TCACTGAAATATACTGGAGTACTTGTGGAGGGCTGTTAGCTGAGTACGTGGGGGTGGGTAATTGGGGGAGCCTGCATGGATATTATTGGAGTAGCTTAGAGCCATACAATCTAACTGAGA | 120 | 47 |
| chr7 | 44340488 | 44340608 | 772751_42001747_Shank1(243961)_253 | 1 | + | good | TTCGGTGGGGGTGAGGAGTGTCTGAGCATTGCTGCGATACAGAGAGTGCAGGGCACCAGCCCTGGGTTCTAGATGCTTTCTAGATGCTAGCAGAGACTGCAGTGAATTTGGTTCTGGTAA | 120 | 52 |
| chr7 | 44340608 | 44340728 | 772751_42001747_Shank1(243961)_254 | 1 | + | good | TACTGGGTATTCGGAAAGGAACTGGCTTGAGGTAGAGCAGTTTAGGTGTCCCTGTTCATGTTGGGAGGAACATGGCTGTGGCTTGCTGAGGGGTAGAGTGATCTAGGATCAGAGGCTGTG | 120 | 52 |
| chr7 | 44340728 | 44340848 | 772751_42001747_Shank1(243961)_255 | 1 | + | good | TACAACCAGCCTGAGCAAGCATGGGAGTGAGGGCTATGACTGGGGTTTTATCCGTTTCCTCTGTGTTTATGGTTTATGTCCTACTGAGATGTGCTGGTGAGGTACTGAGCTTTAGAGCTG | 120 | 48 |
| chr7 | 44341088 | 44341208 | 772751_42001747_Shank1(243961)_258 | 1 | + | good | TTACTGCTGAGCTACATCCCTAGCTCTCTTCTTACGTTATTTTATGGCATAGCATCCCTAAGTTGCTCAGGCTGGTCTTGAACTTTCAAACATACTTCCTCAGCACCCCAGCTCCAGGAG | 120 | 47 |
| chr7 | 44341208 | 44341328 | 772751_42001747_Shank1(243961)_259 | 1 | + | good | CTGAGAAGACAGGCTTGCATCATCAGGACTGGTCATTGGTGTGTGACTCCATGTGTGACTTATCACTTGAGTCTTATCGAGAGACGGCAGCAGTCGGCACGGTCCACCGGAACTTTCTAT | 120 | 51 |
| chr7 | 44341328 | 44341448 | 772751_42001747_Shank1(243961)_260 | 1 | + | good | ACGTGACGGGACTGTTCTGTGTCAGCACCATAGTTAGGAACGTTGTGTGGCCATCATAACCGAGGAACTGACACTTTAAATTTAATCCTGATTAATGTTGATAAAAATAGTCATGTGCAA | 120 | 40 |
| chr7 | 44341448 | 44341568 | 772751_42001747_Shank1(243961)_261 | 1 | + | good | CTAGCAGTTATCATACTGGGCAGGGTGATGCAAACATTCTTTTGGTTGGCTCTGATCCTCTGGTCCTCTGGAGCGTTATTGTAGTGAGGGAATGAGATGTGTTAGTACATTTGCTCTCTT | 120 | 45 |
| chr7 | 44341568 | 44341688 | 772751_42001747_Shank1(243961)_262 | 1 | + | good | GACCTAACACCTTGGCCTAGGGACACGGGGTGGGCTCTGTGCTGCTAAACCCTGTGTGTGTAGTGCCACACACAGGCACTGTAGTGTGTAGTGTTCCGACCTCCCCTACAGTCTCTTTAG | 120 | 56 |
| chr7 | 44341688 | 44341808 | 772751_42001747_Shank1(243961)_263 | 1 | + | good | GCAGGCCCCCATGACGGACTTGGAAATTGAAGCACAAAAAGGTTAAAGCCATAGGTTCAAGGTCACAGGCAGGAAGCTGCCAGCCAGGACCCAAGCAGGGCTGTGGTAATTCAGAGCCTC | 120 | 54 |
| chr7 | 44341808 | 44341928 | 772751_42001747_Shank1(243961)_264 | 1 | + | good | TTTCTGCCCACCCCACTGGGTGCCTGGTGCACCGTGGGCTCAGTAACAGCTCTGCTGAATGGATGGGTGGATGGATTCACGGAAGGATGAATGAGGAAAGAGTGGGTGGAAACAAGAGCA | 120 | 55 |
| chr7 | 44341928 | 44342048 | 772751_42001747_Shank1(243961)_265 | 1 | + | good | CCCCGGGCTTTGAGGGGGTGTCTTGAGCATAAGTACAGACTGGATGGGGCGAGGAACTGCAGTGAATGGTCTCTAAGCAGGGGCTGGGGTGCCCCGTGATGGGGTGGAGCACCAACCCTG | 120 | 62 |
| chr7 | 44342048 | 44342168 | 772751_42001747_Shank1(243961)_266 | 1 | + | good | TGCCTCATGAGCCCTTCTCCACCCCCACCCCCCTCCTGCCTCCTGCCAGTGATTACATCATCAAGGAGAAGACAGTCTTACTGCAGAAGAAGGACAGTGAGGGGTTCGGGTTCGTGCTCC | 120 | 57 |
| chr7 | 44342168 | 44342288 | 772751_42001747_Shank1(243961)_267 | 1 | + | good | GGGGGGCCAAGGGTGAGTGCTGGCCAGGGAGGGGATGGGAGATCCTCTGACCCAAAAAGCAGGGCCTGGATAGGGAGGATGGAAAAAGAGAAGGGACCCCTCACACGCCCTTAAGCCCA[illegible] | 120 | 62 |
| chr7 | 44342288 | 44342408 | 772751_42001747_Shank1(243961)_268 | 1 | + | good | CCATTGCCCCACCCTGGGGTGCTGGTCCCTCAGCTCCAGTGAGTCCTCTGTGACTTGGGTCTGAGCCCCATCCCCTGCTCGGCCTCTCCACCCCCTCCCCTCCGACCCCGCCGGCTCCTC | 120 | 70 |
| chr7 | 44342528 | 44342648 | 772751_42001747_Shank1(243961)_270 | 1 | + | good | AGACTTCCTCATCGAGGTGAGCCCTGGCCACACTGTCCTCCTGGGGCCCCTGATGATGACCCCTAAGTCCAAAAGCTATGTGCCACTTCTGGTGGCCCGTCCACCCCACTCAGTCTGTGG | 120 | 60 |
| chr7 | 44342648 | 44342768 | 772751_42001747_Shank1(243961)_271 | 1 | + | good | GACCACCCACACCTTTCCAGCAACTCTTCCTAGACCCTTGAAAGGTGCTCAGGCTGGGTGCGGGGGGCTCACACCTGTCATTCCAGCTTAGTCACCCCAGGACCACTGCTGCCACCACTA | 120 | 59 |
| chr7 | 44342768 | 44342888 | 772751_42001747_Shank1(243961)_272 | 1 | + | good | GTTCAAGACCAGCCAGGGCTACCTAGTGAGTACAGGCCAACCTGGGCTACAGAGTATAACTGTAGCTAAAGATGAACATTTTTTAAAAAGCATCACAGAATTGTCCCCCAACATATGCAA | 120 | 43 |
| chr7 | 44342888 | 44343008 | 772751_42001747_Shank1(243961)_273 | 1 | + | good | TACAAGGACATGATGGGTGTGGGGGTGCACCGGCTAGCCCAGTCAGTCCTGGGGATGTGGGGACTCCAGAGAACCATGGAGGGAAGGGGGTGGGGTAGGAGGTCACAGTTGGGGACTCTA | 120 | 61 |
| chr7 | 44343008 | 44343128 | 772751_42001747_Shank1(243961)_274 | 1 | + | good | GATCTAAAAAGCTTCTATCCTTCCTCCATGACACAGGTGAACGGACAGAATGTAGTGAAGGTTGGCCACCGTCAGGTGGTCAACATGATCCGACAGGGAGGCAACACACTGATGGTCAAG | 120 | 50 |
| chr7 | 44343128 | 44343248 | 772751_42001747_Shank1(243961)_275 | 1 | + | good | GTGGTGATGGTGACCAGACACCCAGACATGGATGAGGCTGTCCACAAGAAAGGTATCTCTTCCCAGCTATGGCCTTAGCTCAGTCAGTAGAGCACTGGCCTAGCACACACAGAGCCTGGG | 120 | 55 |
| chr7 | 44343368 | 44343488 | 772751_42001747_Shank1(243961)_277 | 1 | + | good | AGCCTGGGCTACAGGAGAGTTGGTCTTAAAATAAAGGAGCTTCCCCAGCTCTCTGCGGAGCCAGGCCTGGGTATCCCACTGCTTAGGGCTACTCTCCAAACTCCCACAAGGCCCTCCTTT | 120 | 56 |
| chr7 | 44343488 | 44343608 | 772751_42001747_Shank1(243961)_278 | 1 | + | good | CCCCTCAAAGCCTGCCTGTTCAGAACCTTTCTGGGGAGAAACTGAGGCAGAGAGGGCTCAGCATGCTGGGGTGCTGTTCTCAGAGTTCTGGAAGCACAGTGAAAGGAGTGGGTGTGGAGG | 120 | 57 |
| chr7 | 44343608 | 44343728 | 772751_42001747_Shank1(243961)_279 | 1 | + | good | GGTCTCCTCCCTGCAACACTGACACCTCCCACCCACCCCGCCTTTCCAGCATCCCAGCAGGCTAAGCGGCTCCCGCCCCCAGCCATCTCCCTGCGTTCCAAGTCTATGACCTCAGAGCTG | 120 | 64 |
| chr7 | 44343728 | 44343848 | 772751_42001747_Shank1(243961)_280 | 1 | + | good | GAGGAAATGGGTGAGCCTGTGGGGCCGGCTGGCCGAGGGCCAGGGGCGGCCAGGGGCAAGGGGCTGGACCACCCTTGACTTTGTCAGTGTTGTTAGTCACGAGGGGTCCTTGTCCTCCTC | 120 | 66 |
| chr7 | 44343848 | 44343968 | 772751_42001747_Shank1(243961)_281 | 1 | + | good | CTGTCCCGCCCCTCCCCCATGTCCTTGGGCCAGATTCCGTGGCCTGCCTGTGTGTCACTGGGAAGTTTTTGAGAGTCTCCTCTCTGCCCTCTTGTATCTCTGTTTCTCTGTCCTGGGTCT | 120 | 57 |
| chr7 | 44343968 | 44344088 | 772751_42001747_Shank1(243961)_282 | 1 | + | good | CTGGGGGTCTGTATGTGTCCTTCTACGGTGGGCCTCTCTCTCCTTTTGTCCCTTCCCCTCCTTCTACTGCCCCCACCCCATCCTCTTACTGGCCCCTTCCCCTCCTTCTACTTGCTCCTT | 120 | 58 |
| chr7 | 44344088 | 44344208 | 772751_42001747_Shank1(243961)_283 | 1 | + | good | CGACCCACAGAGGAAGGTTAGCTGCTCCTTGTAGGCAGATTCTCATGACCTGGGGTGGGTCTTATTTCCCCCTACCTGGGGCCACTTGGGCAGGTGACTGACTATTCCACACCGGGGGTC | 120 | 58 |
| chr7 | 44344208 | 44344328 | 772751_42001747_Shank1(243961)_284 | 1 | + | good | CCTAGGGAGGTGGGAATTCCAAAGAGGGGTCCTTCACGTCCCCCTGATGTCCTAACTCTTGGCTCCTCTGCTGTCTACCCCTGGGCTAGTCTCCCCCTGGAAGAAGAAAATTGGTGAGTG | 120 | 56 |
| chr7 | 44344328 | 44344448 | 772751_42001747_Shank1(243961)_285 | 1 | + | good | GGTTCAGGTCTGGTGGGAGGGTCCCAGTCTCCCAACCCCAGCGGCGGATTTACCCTTCCCTCGACCCCGGCCCCTCACTGCCCCCACCTCGCGCTGGCCACACCTGCTGGTCCACACCTG | 120 | 69 |
| chr7 | 44344448 | 44344568 | 772751_42001747_Shank1(243961)_286 | 1 | + | good | CTGGCCATGCCCTCTGCAGAGGTACCCTTGAGCTTCATCCCTCCCCTTTGCCCACAGAATACGAACAACAGCCTGCAGCAGTGCCCAGCATGGAGAAAAAGCGGACTGTGTATCAGATGG | 120 | 56 |
| chr7 | 44344568 | 44344688 | 772751_42001747_Shank1(243961)_287 | 1 | + | good | CACTCAGTAAGAAGCCGCAGCATGATGCCCTAGTCCCCCACCTCTGGGGCTGCTGCATGCATTCTGCCCTCCTCCTTCCCGTGTGCCCTTCAGTCACCCGTGAGCTCGGCACGCAGGGCC | 120 | 64 |
| chr7 | 44344688 | 44344808 | 772751_42001747_Shank1(243961)_288 | 1 | + | good | CTGGAACCCTGGGCACTACATACCGTCACGGTGCCACCTACTCTCCTGTGCTTTGTCATCCCTGGGCCGTGCCTGCTGGGCTGATGTCTCCACACAGAGACAGAGATAGGACCATGTGAG | 120 | 58 |
| chr7 | 44344808 | 44344928 | 772751_42001747_Shank1(243961)_289 | 1 | + | good | CTCTGTATCTTACCCCAGGGCAACCAGGTCTCTAGTGAGGCAAGGCGACTTTCACTTTCACTGTCTCTGACCGTTCTCACTTCCCCACAGACAAACTGGATGAGATCTTGGCTGCAGCTC | 120 | 52 |

| | | | | | | | | | |
|---|---|---|---|---|---|---|---|---|---|
| chr7 | 44344928 | 44345048 | 772751_42001747_Shank1(243961)_290 | 1 | + | good | AGCAGACCATCAGTGCAAGTGAGAGTCCTGGGCCTGGCGGCCTCGCATCCCTGGGCAAGCACCGACCCAAAGGATTCTTTGCCACTGAGGTAGGTGTGTGGGAAGGAAACTAAGGAAAGA | 120 | 57 |
| chr7 | 44345528 | 44345648 | 772751_42001747_Shank1(243961)_295 | 1 | + | good | CCCCTCTATTCTTGATGTCTCTGCTTCCCATGTTAAGTCTCATCCCCCTCTTCCCTTCCCACCTTAGAGATCTCGCATAAGACAAAAGGATGTTGCAGAGATGTGGCATGAAGGTCCACA | 120 | 48 |
| chr7 | 44345648 | 44345768 | 772751_42001747_Shank1(243961)_296 | 1 | + | good | ACCCCTATCTCAGACCCTGGGGAGCTGAGCTAGGAGAGGGGCTGTGAGCTTGAGATCAGCCTGGGCTATGTAAGGAGACCTTGTCTCCACACAAAATGTAATGTTGTCATAGTTACAGAC | 120 | 51 |
| chr7 | 44345768 | 44345888 | 772751_42001747_Shank1(243961)_297 | 1 | + | good | CCCCATCACGGTCCTAGATGCTCAGCAAAGGCTAGATGCTGTGAACACATTGATGCTTGGGCCCAAGACACAGCTCAGTGGTTGTGAGTATGCACTGCTTGTGTAGAAGATTGGATTTCA | 120 | 49 |
| chr7 | 44345888 | 44346008 | 772751_42001747_Shank1(243961)_298 | 1 | + | good | GTTCCCAGAATCCGCACTGGGCTGCTGACAACAGCCTAGAACTCTAGCTGCAGATTCTGTTTCCCTCTTCTGCCCGCTTAAGGCACAGCACAGTCCCATACACAGACACACACATAATTT | 120 | 51 |
| chr7 | 44346008 | 44346128 | 772751_42001747_Shank1(243961)_299 | 1 | + | good | TTCTTAAAATACAGTTGAAAATGATTTAAAAAGACACCTAAGGCCGACCTCTGACCTACACACACACACCACATACAGTGAACCATTTGGCCTGTGTGCCGTTTCCACCTGCCCTTTCCC | 120 | 45 |
| chr7 | 44346128 | 44346248 | 772751_42001747_Shank1(243961)_300 | 1 | + | good | ACAGGCAAAGACCTGATCTGTATCAAGAATTCAACATCTTTCCGGTACTGTGTTCCACACAGATTAGCTCATTCCTGTCTTTAGAACCTTCTTGTGAGGCACCACGCAGCAGGGGTTAGC | 120 | 47 |
| chr7 | 44346248 | 44346368 | 772751_42001747_Shank1(243961)_301 | 1 | + | good | TCATTTACAGGTATGGAAATGAAACTCAGGAAAGGCCCAAGCCCAGCCTGGGCCATGCCCTGCTGGCTCTGTCTGACCCATGAGTTTGATCCTTGGACACCAACACATCCACATCAGAAC | 120 | 52 |
| chr7 | 44346368 | 44346488 | 772751_42001747_Shank1(243961)_302 | 1 | + | good | TGCCTGTGCTCTTCTGTCTCCCAGCCGTCACCCCTGAAGGCGACCCACTGCCTGGCATCCCCCTTAGCCTCTGCCCTCTGACTGCCACACCTACAGCCTGGTGGCCAAGGGGCATTTACT | 120 | 62 |
| chr7 | 44346488 | 44346608 | 772751_42001747_Shank1(243961)_303 | 1 | + | good | CTAAGGGTCAAACCTTACTCAAGGGGAGCCCAGAATCTGCTTGACAGATTGGTTTGGGCTTTCTCAGAACTATCTGTGAGATAGTTAACTATGAGAACGAAGTTTCCCAGGACTTACTGC | 120 | 45 |
| chr7 | 44346608 | 44346728 | 772751_42001747_Shank1(243961)_304 | 1 | + | good | ATCTCAGCCTTCTCTCATAGATCAAGTACAGAGCACTACTTATAAAGACAAGCTGTGATACGGACAAGAGCCAGCCACACTAAGACACCCAAAATATAAGCTCTCATGCATTTTACCACC | 120 | 42 |
| chr7 | 44346968 | 44347088 | 772751_42001747_Shank1(243961)_307 | 1 | + | good | AATGGCACTGAGAACCACTGGCCTAGCCAGATGCTGTATCGATCAGTCATACTTCCCTTCCTAGGGGATTCTAGGCATTGACTCTTTACTAAGACACACACACACACACACGCACACACA | 120 | 48 |
| chr7 | 44347328 | 44347448 | 772751_42001747_Shank1(243961)_310 | 1 | + | good | GCCCAACATTCAAAATAATGTTTAGTGACTTTTGAAGTCAGAAGTGGTGGCCCACACCTATAATCTAAGCACTCCAGGGACTGAAGCAGGAGGATTTCCATGAGTTTTAAGGACCAACCT | 120 | 43 |
| chr7 | 44347448 | 44347568 | 772751_42001747_Shank1(243961)_311 | 1 | + | good | AGACTGAGCCTATCTCAGAAAACCAAAAGATAAAACAGATTTGTTGCTACAGCAATATCAGTGCCTCCTGGCCAGCACATTACACAGGGAGACTGAGCTGTGGCACCTAAGTGCTCTAGC | 120 | 47 |
| chr7 | 44347568 | 44347688 | 772751_42001747_Shank1(243961)_312 | 1 | + | good | TGCAGAGTCCACAGCAGCCTAGGGACTATCTGTAGGAATGAGCCGCTGAGATGGGGGTCAGGAATCTCCATGATTTCTGTGGGTGATGCAGCCACCAACTTCACTGAGACTAGCAGAGCT | 120 | 53 |
| chr7 | 44347688 | 44347808 | 772751_42001747_Shank1(243961)_313 | 1 | + | good | GATGCCAACAGCCTCAGGGGAGAAAGAGCTGAGTTCCCCACTGCATGATGTGGGGACAGGGGTGTCATTTATTCCCTCAGGACCACACACTCCATCCTGGAGTTCCCAATCTAGAGGCCA | 120 | 55 |
| chr7 | 44347808 | 44347928 | 772751_42001747_Shank1(243961)_314 | 1 | + | good | CTTCTCTGGCTGGTGGAGTGCCCTGCAGGCCCCGGCCCTCCCTTCCTGCCCATCCTTGGTTTCTCTAGATGTCTACATTGTTGGTTATCTGCTGTGCCATCCTGCACACAAGTACAGAGT | 120 | 56 |
| chr7 | 44348048 | 44348168 | 772751_42001747_Shank1(243961)_316 | 1 | + | good | CTAGGACAAAGAAGAAAGTGAGGCGGGCAGGGATGGGAATGGCAAGGGCACTTGGGCCTTGCAGCAAGCTGTGTGTTCAAATCCTGGTTCCTCTCCCTACGTGGTTCTGAGCTATGCTAC | 120 | 54 |
| chr7 | 44348168 | 44348288 | 772751_42001747_Shank1(243961)_317 | 1 | + | good | CTCATCCCCAGTGTGCTTCCTACTCTGGATAATTGGTTTGGATCTGGCAGCACTCAGTGGAGTCAAATGCGGTCAGCTCCAGTCCAACAGGGCTCTCAACAAACTGGCAGTCCTCTTCTG | 120 | 52 |
| chr7 | 44348288 | 44348408 | 772751_42001747_Shank1(243961)_318 | 1 | + | good | TGTGCGACTGTGCCAGCCCTCTGATCTCAGCCCAACACCATCCTCAGAGTCTGACAGCATGGCCCCCACACTGTCCTCCGACAGCCTCAGCTCTCATCCAACCCGTCTCTTATGCGTGTG | 120 | 59 |
| chr7 | 44348408 | 44348528 | 772751_42001747_Shank1(243961)_319 | 1 | + | good | TGCAAGTGTGTGTGCGAGTGTCTGTGCACATGTGTGCATGTGCCTGTGGAAAGCAGGGGCAGTCTTTGGTATCCATCCTCGGGCACCAATCAACTTGTTTGTTGAACTCTCTCAAAATCA | 120 | 49 |
| chr7 | 44348648 | 44348768 | 772751_42001747_Shank1(243961)_321 | 1 | + | good | ACCATGTATGGCTGTGGTACACGCAGTGTGATGCATTCTGAAACCAGACCAGGACAGGTATTTTCCTCCCTCTACCACTGTCTGGTCCCCATGAGACAGAGTCTCACTGAAGCACAGTTC | 120 | 51 |
| chr7 | 44348768 | 44348888 | 772751_42001747_Shank1(243961)_322 | 1 | + | good | ACAGTTTTATCTAAATTAGCTGTCTATCAAGCGCTAGGGACCCACCTCTTCCCCCTCACCACACAGGCACCCAATGCTAATAATACAGGCATGGACAGCCAGGCTTGGATTTTTATGTAG | 120 | 47 |
| chr7 | 44348888 | 44349008 | 772751_42001747_Shank1(243961)_323 | 1 | + | good | CTTCTGATGATTTGAACTCAGGACCTCATACCTGAAGAACTAACTTACCCACCTAACCATCTCCCCAGCCCCTCACCCAGTCAGAGTGTTTTCCCTTAGTCTGCAAACAGTGTCCTCCCC | 120 | 50 |
| chr7 | 44349008 | 44349128 | 772751_42001747_Shank1(243961)_324 | 1 | + | good | TCAGGCAGGGTTCCCCTCAAAACCTGCTCTTCTCCCTAGGAGCTTTTTCTGACATTCTCATTACCTCACCCTTCCTCATTCAGTTCCAGACACTCTGACACAACTTCTTTCACACTAATC | 120 | 47 |
| chr7 | 44349128 | 44349248 | 772751_42001747_Shank1(243961)_325 | 1 | + | good | TTTCTACTATGATGCTCGCCCCTCCCTTTTACTACTTCCTCTCTCCTGCCCACCCCCTCCTCCATCAACATGTTCCTCTCCATTACATGGCCCCTCCCCCATCTCCCTGGCCCCTCCTCC | 120 | 57 |
| chr7 | 44349248 | 44349368 | 772751_42001747_Shank1(243961)_326 | 1 | + | good | ATCGCATGGCCCCTCCTCTATCACCACAGTCCGTCCCCCATCTTTCTGGCCTCTCCTCCATCACATGGTCCCTCCCCCACCACCATGGCCCCTCCCCCATCAGCTTGATCCCTGTAAGGA | 120 | 61 |
| chr7 | 44349368 | 44349488 | 772751_42001747_Shank1(243961)_327 | 1 | + | good | GCCTGTCCCCAGACTCTCATACTCTGCTGCTCTAATGGGATAACAGGCACTTAAACTGTCCCTGGGTTACTTGGGACCTGTGAAAAGCACCCTCCACAGTGTCTCTGTCTTTCTAGGGTT | 120 | 51 |
| chr7 | 44349608 | 44349728 | 772751_42001747_Shank1(243961)_329 | 1 | + | good | CTGCCTCAGCTTCTCCAAATGACAGGTTCCATGTCATACAAGACTCCATGTTCTGTTCTCTAATATACTTATATCCTGCCTTTCATACAGCCATGCCTTGTGGAAGTGTCCTTGTCAGTG | 120 | 44 |
| chr7 | 44349728 | 44349848 | 772751_42001747_Shank1(243961)_330 | 1 | + | good | GAATGAACAAGACAAACACAGAAATAGACAGTTCTAAGCCAGGTAGGCCAGTGCAGCGACAGGGCTTTTAAGACCCAGGAAAGCCTTGAATTCCTCAGACTACCACTGCCCTCTAGAATA | 120 | 47 |
| chr7 | 44349848 | 44349968 | 772751_42001747_Shank1(243961)_331 | 1 | + | good | GTCCACACTTCTTGTCCTGTATGGAGTTACAGTCACTCTGGATTCAAACACAACCAGTGTTGTGGATGGATGGATGGATGGATGGAATGACAGACAGAAGGAATGGTCAGTGGAAGAAGA | 120 | 46 |
| chr7 | 44350088 | 44350208 | 772751_42001747_Shank1(243961)_333 | 1 | + | good | GACAGTAGAACAGTGGGTGGATGGTCAGACAGATGGGTGGAAGGATGGATGTGTGCAGTGATGGATGTGTGGAATGATGGACACGGGAGCAGACAGACAGATGGAATGATAAACAGGTAC | 120 | 50 |
| chr7 | 44350208 | 44350328 | 772751_42001747_Shank1(243961)_334 | 1 | + | good | AAATGATGAATTGACAGAACAATGGATTGGTGACCAGACAGACAATTTTTTTTTCTTGAATGAGTTGAAACTGGGGCTGGAAAGATGGTTCCATTGCTAAGAGGGTTTGCTGTGTCACCA | 120 | 40 |
| chr7 | 44350328 | 44350448 | 772751_42001747_Shank1(243961)_335 | 1 | + | good | TGAGGATGGGAGTTTGGGTTCCAGGCTTGGATAGCCCTGCAGCCCACCCACTTCCCCACATATGCAACTCTAGCTCTTGTGCCTTTCTCCGGCCTCTATGAGCCCCTCCAGCATGCGCG | 120 | 59 |
| chr7 | 44350448 | 44350568 | 772751_42001747_Shank1(243961)_336 | 1 | + | good | TGCGCGCGCGCACACACACACACACACTTTTAAGTCTGACGTGAGCTGCAGCTGCTGGGAATCAGAACTGGGTCCTCATGTCTGTATGTGTGGCAAAGCGTATTACTGACTGAGCAATCG | 120 | 52 |
| chr7 | 44350568 | 44350688 | 772751_42001747_Shank1(243961)_337 | 1 | + | good | CTCCAGCCCTGATTTCTTTTTGAAAGATTCCGCCAACTCTGGAAAAGAGACCAGAAAGGAGTGAAGGAGAGAGGAAACAACAAAGCCAGAGGGGTGGCTGCCACCACAGGCCAGAAGAG | 120 | 51 |
| chr7 | 44350688 | 44350808 | 772751_42001747_Shank1(243961)_338 | 1 | + | good | TGTCAATGGGACCTGGGTACAGGCCACGACTGCAGGTGGTACCTGCTTTAGTGGGACTGCTCGGCTGCCTGCTTTTCTTTTCTTTCTCCTGACTGCTTCCCCAGAGAATTCTCCCAGGAC | 120 | 55 |
| chr7 | 44350808 | 44350928 | 772751_42001747_Shank1(243961)_339 | 1 | + | good | TTGCTTTCTAGCTTCTGTGTGCCTCGTCTCTGCCATGAAAGAGCTCAGAGGAGAAAGTAATTTCCTTTTGGAGTGTGGAGAACTCTGACTGATCCCTCTTCGCTCTCTTTTCTCTCTTCC | 120 | 47 |
| chr7 | 44350928 | 44351048 | 772751_42001747_Shank1(243961)_340 | 1 | + | good | TCTAACTGCTGCCCCGAAGTCGAGCTTTGATCCTCACCACCGCTCCCAGCCAAGTTATGACCGGCCTTCTTTCCTGCCTCCCGGACCTGGCCTTATGCTCCGGCAGAAATCTATCGGTAT | 120 | 57 |
| chr7 | 44351048 | 44351168 | 772751_42001747_Shank1(243961)_341 | 1 | + | good | GTCCCCCAGAAAGGACCGTAGCCCAGGGGCAGGAGAGGACTGTCTAGGTTTGGAAATGTGCTGCAGCACCACATCCGACCCTGGTTAGAGGAGGTCGGGTTTGGATGAATTCCAACTTTG | 120 | 56 |
| chr7 | 44351168 | 44351288 | 772751_42001747_Shank1(243961)_342 | 1 | + | good | CTTCTCGCCTGCGGGACTGCGTGGGTGGTCTCTCGGCTTCCTGACCTTCATCTAGAAAACCGGGGTCTTTGGTGTCCAGAGGAAGGATTGTTGGTAGCATGCGGGTAGTAAGTGGAGGCA | 120 | 57 |
| chr7 | 44351288 | 44351408 | 772751_42001747_Shank1(243961)_343 | 1 | + | good | GATCCTTGTTGAGAGAGAAGCTAGCTACTTCCTAGGCGTCTCTGTGTAAAATAATAGCCTTTTCTTCTTCTCAAGCACGCTTTAAACTCGCTCCAAGTCACCATTATCATGGATTCAGGC | 120 | 43 |
| chr7 | 44351408 | 44351528 | 772751_42001747_Shank1(243961)_344 | 1 | + | good | CTTTGACTAATTAGGAAATGGCTCCCCCTCCTGGCCAGAGGGTGAAATTCCACTGAATTTCTCTTGCGCCTCTGCTGTTGTATAGCTTGTATGCAGAAGGAGCTGGGCCAGTCCCTAATT | 120 | 49 |
| chr7 | 44351528 | 44351648 | 772751_42001747_Shank1(243961)_345 | 1 | + | good | CCTCCATTGGTCTTACGTATCTCAACAGGGGCTGCGGAAGATGACAGACCCTACCTAGCACCCCCAGCCATGAAATTCAGTCGTAGCCTGTCAGTCCCGGGTTCAGAGGACATACCCCCA | 120 | 56 |
| chr7 | 44351648 | 44351768 | 772751_42001747_Shank1(243961)_346 | 1 | + | good | CCACCCACTACATCCCCACCAGAGCCGCCCTATAGCACACCTCCGGCCCCCTCCTCTTCCGGCCGCCTCACTCCCTCGCCTCGGGGAGGGCCCTTTAACCCTGGTTCCGGGGGTCCCCTT | 120 | 69 |
| chr7 | 44351768 | 44351888 | 772751_42001747_Shank1(243961)_347 | 1 | + | good | CCCGCCTCCTCACCTTCATCTTTCGATGGGCCATCTCCTCCTGACCCCCGTAGTGGGGGCCGAGAGAAGAGCCTATATCACAGTGGCGCCCTACCCCCTGCCCACCACCACCCACCTCAT | 120 | 63 |
| chr7 | 44351888 | 44352008 | 772751_42001747_Shank1(243961)_348 | 1 | + | good | CACCACCATCATCACGCCCCACCTCCTCAGCCCCACCATCACCATGCGCATCCTCCCCACCCACCGGAGATGGAGACAGGAGGTTCCCCCGACGATCCCCCACCTCGATTGGCTCTGGGA | 120 | 65 |
| chr7 | 44352008 | 44352128 | 772751_42001747_Shank1(243961)_349 | 1 | + | good | CCCCAGCCCAGCCTTCGAGGCTGGCGAGGTGGAGGACCCAGCCCGACCTCGGGGGCCCCATCACCTTCCCACCACAGTAGCAGCGGAGGCAGTAGCGGTCCTGCTCAGGCTCCCGCCCTG | 120 | 72 |
| chr7 | 44352128 | 44352248 | 772751_42001747_Shank1(243961)_350 | 1 | + | good | CGCTACTTCCAGCTGCCCCCGAGGGCGGCCAGTGCGGCTATGTACGTACCAGCCCGCTCCGGCCGCGGTCGCAAAGGCCCGCTGGTCAAGCAGACCAAAGTGGAAGGCGAGCCCCAAAAG | 120 | 68 |
| chr7 | 44352248 | 44352368 | 772751_42001747_Shank1(243961)_351 | 1 | + | good | GGCAGCCTCCCTCCCGCGTCCTCTCCAACATCCCCGGCGTTGCCTCGGTCCGAACCACCTCCGGCCGGGCCGTCGGAGAAAAATTCCATCCCTATCCCTACCATCATCATCAAAGCCCCA | 120 | 63 |
| chr7 | 44352368 | 44352488 | 772751_42001747_Shank1(243961)_352 | 1 | + | good | TCTACCAGTAGCAGCGGCCGCAGTAGCCAGGGAAGCAGCACAGAGGCTGAGCCCCCCACCCAGCCCGACGGCGCGGGCGGCGGAGGATCCTCGCCTAGCCCTGCACCTGCCACGTCCCCT | 120 | 72 |
| chr7 | 44352488 | 44352608 | 772751_42001747_Shank1(243961)_353 | 1 | + | good | GTACCGCCGTCCCCGTCGCCCGTGCCCACCCCTGCGTCACCCAGCGGCCCGGCCACCCTGGATTTCACTAGCCAGTTTGGAGCAGCCCTAGTGGGCGCGGCGCGGAGGGAGGGAGGCTGG | 120 | 73 |
| chr7 | 44352608 | 44352728 | 772751_42001747_Shank1(243961)_354 | 1 | + | good | CAAAACGAAGCCCGGCGGCGGTCCACGCTATTCCTGTCCACCGACGCAGGAGATGAGGATGGAGGCGACAGCGGGCTGGGCCCGGGGGCGCCCCCGGGTCCCCGGCTGCGCCATTCCAAA | 120 | 71 |
| chr7 | 44352728 | 44352848 | 772751_42001747_Shank1(243961)_355 | 1 | + | good | TCCATTGATGAAGGCATGTTCTCCGCCGAGCCGTACCTCCGGCTGGAGTCGGGGGGCAGCAGCGGGGGCTACGGGGCATACGCAGCGGGCAGCCGAGCCTACGGGGGCAGCGGGAGCAG | 120 | 71 |
| chr7 | 44352848 | 44352968 | 772751_42001747_Shank1(243961)_356 | 1 | + | good | AGCGCCTTCACCAGCTTCCTCCCCCCGCGGCCCCTGGTGCATCCGCTGACCGGCAAAGCCTTGGATCCCGCCTCTCCGCTGGGGCTGGCCTTGGCCGCCCGGGAGCGGGCCTTGAAGGAG | 120 | 72 |
| chr7 | 44352968 | 44353088 | 772751_42001747_Shank1(243961)_357 | 1 | + | good | TCCTCGGAGGGCGGGGTCACCCCTCAGCCGCCCCCGAGACCTCCATCACCACGCTACGACGCCCCACCGCCCACCCTTCACCACCACTCGCCCCATTCACCCCATTCGCCACATGCGCGT | 120 | 70 |
| chr7 | 44353088 | 44353208 | 772751_42001747_Shank1(243961)_358 | 1 | + | good | CATGAGCCAGTGTTGCGCCTCTGGGGAGACCCGGCGCGCCGGGAGCTGGGGTACCGGGCAGGCCTAGGGAGTCAGGAGAAGGCCCTTACCGCTAGTCCACCTGCAGCTCGCCGCTCGCTA | 120 | 69 |
| chr7 | 44353208 | 44353328 | 772751_42001747_Shank1(243961)_359 | 1 | + | good | CTGCACCGTCTTCCCCCGACGGCTCCCGGGGTCGGGCCGCTCCTGCTTCAACTGGGGCCGGAGCCCCCAACTCCCCATCCTGGGGTGAGCAAGGCCTGGCGCACAGCAGCCCCCGAGGAG | 120 | 73 |
| chr7 | 44353328 | 44353448 | 772751_42001747_Shank1(243961)_360 | 1 | + | good | CCAGAGCGGCTGCCTCTGCATGTGCGATTCCTTGAAAATTGCCAGGCCCGGCCTCCTCCAGCTGGCACGAGAGGGTCCTCCACTGAAGATGGGCCTGGGGTTCCGCCTCCCAGCCCTCGC | 120 | 66 |
| chr7 | 44353448 | 44353568 | 772751_42001747_Shank1(243961)_361 | 1 | + | good | AGAGTCTTGCCAACCTCTCCCACTTCCCCGCGAGGCAACGAGGAGAACGGCCTTCCGCTACTGGTCCTGCCACCGCCAGCCCCTTCGGTGGATGTGGACGACGGCGAGTTCTTGTTTGCG | 120 | 63 |
| chr7 | 44353568 | 44353688 | 772751_42001747_Shank1(243961)_362 | 1 | + | good | GAGCCGCTGCCTCCACCCCTGGAGTTTTCCAATAGCTTCGAAAAGCCAGAGTCGCCTCTCACACCCGGGCCTCCCCACCCGCTGCCTGACCCTCCCTCCCCAGCTACCCCTCTACCCGCA | 120 | 66 |
| chr7 | 44353688 | 44353808 | 772751_42001747_Shank1(243961)_363 | 1 | + | good | GCCCCACCTCCGGCTGTGGCCGCCGCCCCTCCCACCTTGGACTCCACAGCATCCAGCCTGACCTCTTATGACAGCGAAGTGGCCACGTTGACCCAGGGGGCTCCCGCGGCTCCTGGGGAC | 120 | 70 |
| chr7 | 44353808 | 44353928 | 772751_42001747_Shank1(243961)_364 | 1 | + | good | CCTCCTGCTCCAGGCCCTCCGGCCCCGGCAGCTCCAGCTCCTCCAGCACCGCAGCCAGGCCCCGACCCTCCACCTGGCACAGATTCTGGAATTGAGGAGGTGGACAGTCGGAGCAGCAGC | 120 | 69 |
| chr7 | 44353928 | 44354048 | 772751_42001747_Shank1(243961)_365 | 1 | + | good | GATCACCCGCTGGAGACCATCAGCAGTGCATCCACACTCAGCAGCTTGTCCGCCGAAGGAGGAGGCAACACAGGGGGCGTGGCTGGGGGCGGGGCCGGGGTGGCCAGCGGGACGGAGCT | 120 | 70 |
| chr7 | 44354048 | 44354168 | 772751_42001747_Shank1(243961)_366 | 1 | + | good | CTGGACACCTACGTTGCCTACCTGGATGGCCAGGCCTTCGGGGGCAGTGGGACTCCTGGCCCACCATATCCCCCTCAGCTCATGACTCCTTCTAAGCTCCGGGGCAGGGCGCTTGGGACT | 120 | 64 |

| | | | | | | | | | |
|---|---|---|---|---|---|---|---|---|---|
| chr7 | 44354168 | 44354288 | 772751_42001747_Shank1(243961)_367 | 1 | + | good | AGTGGAAATTTGAGACCTGGTCCCAGTGGGGACTTCGAGATCCTGTCACCCCCACCAGCCCCACCGTCTCTGTGACAGGAGCTGGAACAGATGGGCTACTGGCCTTGAGTGCTTGTTCT | 120 | 57 |
| chr7 | 44354288 | 44354408 | 772751_42001747_Shank1(243961)_368 | 1 | + | good | GGACCCTCAACGGCAGGTGTGGCAGGGGGTCCCGTGGCGGTTGAGCCAGAAGTCCCGCCTGTACCTTTGCCCACTGCCTCATCCCTGCCTCGGAAGCTACTACCATGGGAGGAGGGTCCG | 120 | 66 |
| chr7 | 44354408 | 44354528 | 772751_42001747_Shank1(243961)_369 | 1 | + | good | GGACCACCACCACCACCCCTGCCAGGGCCCCTGTCCCAGCCTCAGGCCTCTGCCTTGGCCACGGTCAAAGCCAGCATCATCAGCGAACTCAGCTCCAAGCTCCAGCAGTTTGGTGGGGCC | 120 | 66 |
| chr7 | 44354528 | 44354648 | 772751_42001747_Shank1(243961)_370 | 1 | + | good | TCTACCGCGGGAGGAGCTCTGCCTTGGGCTCGGGGAGGCAGCGGGGGCAGCACAGACAGCCACCACGGAGGAGCCAGTTACATCCCAGAGAGGACCTCCTCCCTGCAGCGGCAGAGGTGA | 120 | 68 |
| chr7 | 44354648 | 44354768 | 772751_42001747_Shank1(243961)_371 | 1 | + | good | GCAGAGGCCTGTGGCTGGTGGGTGGCTGTGGCAAGTGCTGGAGCCCAAGGTCCCCATAGAGCCTCGGAGTTACAGAAAACAAACTGGGCTGTCTTGCAGGAATCGGACCTCTAAGAAGCA | 120 | 57 |
| chr7 | 44354768 | 44354888 | 772751_42001747_Shank1(243961)_372 | 1 | + | good | CACTTTCAGCTCCATGGTCACCATAACCCATGGCCCAAGATGAATGGCTACATGTGTTTATTTTGTTTTTGATGGTCCTGGAGATCAAACCTAGGACTTCAGGCATGCTACACAAGTGCT | 120 | 45 |
| chr7 | 44355008 | 44355128 | 772751_42001747_Shank1(243961)_374 | 1 | + | good | GAGCTACACCCCCAGCTGTATTATGTTTTGAGTCAGGGCCTAAGTCACTTTGACTGGCCTTGAGCCTTTGTAGCCCAGGTTGGGTTTAGACTTCTAATTTTCCTGCCTCAACTTCCCCAG | 120 | 49 |
| chr7 | 44355128 | 44355248 | 772751_42001747_Shank1(243961)_375 | 1 | + | good | TAACTGGAATAACAGACCTACACCACCAAGTCTAGCTACAAAATATTAACAACAGCAACCAAATTGGAGTAACTGAAGGTGTGCGTGGTGCTTTACTGCAGCAGGTAGATGACATGCGCC | 120 | 44 |
| chr7 | 44355248 | 44355368 | 772751_42001747_Shank1(243961)_376 | 1 | + | good | CCTTCCTCCGTAGGAGTAGGGCACAGGGTGCCTAGAAAGGGATTGGGCAGTTGCTCCCAACAGGATAGAGGAGAGCAGAGCAGCTGTGGACCCACTAGATGGGAAGATATGGGTCTATGC | 120 | 56 |
| chr7 | 44355368 | 44355488 | 772751_42001747_Shank1(243961)_377 | 1 | + | good | CTAAGTTCCCCCAAACAGTTGTTCTCTCTGTGAAGCCATGACCCATGCTAACAATGGCTAGCCTGGATGGGGTCTGGGGAGGCAGTGAGGGGAGTTTGAAATTCTTCCTCTCCATCTGCA | 120 | 52 |
| chr7 | 44355488 | 44355608 | 772751_42001747_Shank1(243961)_378 | 1 | + | good | GAGCGCACTATAATTCTGTTCCCTGTGGATTCATATCTCCTTGGAGATATGAAACAACTCACTCATGTTATTCACGGGTGGGCGGAGGGGCGCCGCATTTAATCACCATAGGCACAGATT | 120 | 48 |
| chr7 | 44355608 | 44355728 | 772751_42001747_Shank1(243961)_379 | 1 | + | good | AGATGTTGAACACAGCAGTATGCCTTTATCCAGTGTCCACCTTGCTCCTAGTTGAGTAAGGTCCAGCTGCTACATCCCCAGCCCCTCGCTGGGGGGATCCCAAGCAGGGGCTCTACCACT | 120 | 56 |
| chr7 | 44355728 | 44355848 | 772751_42001747_Shank1(243961)_380 | 1 | + | good | GAGCTATACCTCAAGCTAAACAGTAGGCACAATAATTACAGATACCCCCACAAAACAACACAAGACCTGGCTGGTTAGAGTTATCTCAGATGAGGCCACCTGGCTGTGTAAGGGACCAGT | 120 | 47 |
| chr7 | 44355848 | 44355968 | 772751_42001747_Shank1(243961)_381 | 1 | + | good | GAGCCACCCAGTGGGAAGTTCAAAAGTCCTGTGGTGCAGCCACGCTGAAGGGAGCTCAAGGGATGAGGGGAACCTTTTCTCCCCTTGTTTCTTCTCTCTTTCATCTCCCTCTCTTCACCT | 120 | 53 |
| chr7 | 44355968 | 44356088 | 772751_42001747_Shank1(243961)_382 | 1 | + | good | TTCTCTCTCTCTGTTCTCTGTCCCCATCCCCTTCCATCTCTCTCCCCCTTGAATGCCCACCCTCTATTTGTTGTTGTTTTTGTTTGTTTCTGTCTCCCTGTCTTCCACTCTGCTATAACT | 120 | 47 |
| chr7 | 44356088 | 44356208 | 772751_42001747_Shank1(243961)_383 | 1 | + | good | GACCATAACAAAACAAAACAAAACAAAAGGCAGTATAGAATAGTGGTTAAGAGCTCAGACTTTGGTGACTTCTAACTGGGCTCTGCCACTCCCTGGCTGTGTGTCCTTAGGCAAGTTTCT | 120 | 43 |
| chr7 | 44356208 | 44356328 | 772751_42001747_Shank1(243961)_384 | 1 | + | good | TAAGCTCTCTGTGCTCTTGACGATAGTAGTAACTAACCTTCCACGGGGGCTGTTGTGGGGATTAAATTGAGATGTGATTCAGAAAGCCTTGAGAACGATGCCTGACACACATGAGCAGCA | 120 | 47 |
| chr7 | 44356328 | 44356448 | 772751_42001747_Shank1(243961)_385 | 1 | + | good | GCAAGTGGTGGCGGGTGTTATTGCTATCACTTAATGGTTCTTTTTCTTCCATCTCTCTTGGCTATCACCCTCTCCCTGTCCTTCTTCTCACCCTTCACTTCTGCCCTGACATTCATCTGA | 120 | 48 |
| chr7 | 44356448 | 44356568 | 772751_42001747_Shank1(243961)_386 | 1 | + | good | CCCTCCCTTACCACCACCACCCACCCCATCTTCTCCATCTCTCTGCCCCCACCCTTGTGTCTCTCCCTCTGAATCATGGCTTCTGCCTTTTTCACCCCGGCCCTGAACACTTCCTCCAAA | 120 | 57 |
| chr7 | 44356568 | 44356688 | 772751_42001747_Shank1(243961)_387 | 1 | + | good | CCCTGCCCCTCCCCCTGTCGCCCCACCCCCACACCCCTTGTCACATTCCAGGCTCTCCGAAGACTCCCAGACCTCTCTCCTCTCCAAACCCAGCAGCAGCATCTTTCAGAACTGGCCTAA | 120 | 62 |
| chr7 | 44356688 | 44356808 | 772751_42001747_Shank1(243961)_388 | 1 | + | good | ACCCCCTCTGCCACCACTCCCCACCGGATCTGGGGTCTCCTCTTCAACAGCTGCAGCCCCAGGAGCTACATCCCCTTCAGCCTCATCTGCCTCAGCTTCCACCAGACACCTGCAGGGTGT | 120 | 62 |
| chr7 | 44356808 | 44356928 | 772751_42001747_Shank1(243961)_389 | 1 | + | good | GGAATTTGAGATGCGGCCTCCGCTGCTCCGCCGGGCACCCAGCCCCTCGCTGCTGCCTGCCTCGGATCACAAGGTCAGCCCAGCGCCCAGGCCCTCTTCTTTGCCCATCCTGCCTTCAGG | 120 | 68 |
| chr7 | 44356928 | 44357048 | 772751_42001747_Shank1(243961)_390 | 1 | + | good | ACCCCTCTACCCGGGCCTCTTCGACATCCGTAGCTCTCCGACCGGAGGGGCAGGAGGCTCGGCTGACCCCTTCGCTCCCGTCTTTGTGCCACCACACCCCGGGATATCTGGGGGTCTTGG | 120 | 67 |
| chr7 | 44357048 | 44357168 | 772751_42001747_Shank1(243961)_391 | 1 | + | good | AGGAGCCTTGTCTGGGGCCTCTCGCTCCCTATCACCCACCCGCCTGCTCTCGCTGCCCCCGGACAAGCCGTTCGGCGCCAAGCCTCTAGGGTTCTGGACCAAGTTTGACGTCGCTGATTG | 120 | 64 |
| chr7 | 44357168 | 44357288 | 772751_42001747_Shank1(243961)_392 | 1 | + | good | GCTGGAGTGGCTGGGCTTATCAGAGCATCGAGCCCAGTTCCTGGACCACGAGATTGATGGCTCCCATCTGCCCGCCTTGACCAAGGAGGACTATGTGGACCTGGGCGTGACCAGGGTGGG | 120 | 62 |
| chr7 | 44357288 | 44357408 | 772751_42001747_Shank1(243961)_393 | 1 | + | good | CCACCGTATGAACATTGACCGGGCTCTCAAATTTTTCCTGGAGAGGTGATGGCTGGCCTGGACGGACCAGCCCGTCCATAGGACCCTTGAGCCTGCTGACCTCTTGACCTCTGACCCCTC | 120 | 58 |
| chr7 | 44357408 | 44357528 | 772751_42001747_Shank1(243961)_394 | 1 | + | good | TGACCATCATTCTCTCCCCTCTCCCCTCAGGGGGCCAGGGACTCTGCTAAAACTGTGCCTGGCCCCATCTCCCCAGGCCTGACTGAGGTCCCCAGGTGGACCCGATGCGGGCCAGACAAT | 120 | 63 |
| chr7 | 44357528 | 44357648 | 772751_42001747_Shank1(243961)_395 | 1 | + | good | GGCACCTCGCAAGAGGTGGGCAGCTGACACAAGACTGCAAGTGGAGTAGAGGGGGGGATTAGACAAAAATGACAAAGGGGTACTGAAGAAGGAGGGAAAGGCCGACTGGGGGTGGGGG( | 120 | 57 |
| chr7 | 44357648 | 44357768 | 772751_42001747_Shank1(243961)_396 | 1 | + | good | CAGACAGAGCCATAGAGGGTCAGCCCAGAGCCTGACCTGATGGGGTCCTGGGAAGGTGTGCCCTCCGATTCCATCATTCGCGCCCCTCAAATAACCCTTTTCACCTCTCCAGGCCCCTCA | 120 | 59 |
| chr7 | 44357768 | 44357888 | 772751_42001747_Shank1(243961)_397 | 1 | + | good | GTCTTCCCTCTCTCCGTGGCACACTCAGCTGCCCAGCCCCCGGCCTCTTGCACGCCTGTTCCCCTTCCTGGCTGCAGTTTTTGCACAAGCTTGCCCCCTGGCTGTCTTGCACACCTCTGG | 120 | 63 |
| chr7 | 44358008 | 44358128 | 772751_42001747_Shank1(243961)_399 | 1 | + | good | CTGCATACTTGCTTCTGGGGGAGAGGGGTTACTGCACCCTTTCTTTCTCTAGCCACCCTGGTCCCATGCTGCAGACACACATGGGACTTTCCTTCCCTCTCCTGCTCCCAGACCCAGGCA | 120 | 57 |
| chr7 | 44358128 | 44358248 | 772751_42001747_Shank1(243961)_400 | 1 | + | good | TACTCCCATAGCCCCTGGCCACCCGTGCTCTCTGCTCTCTCTGTACATGCTCTGAACACACTCACCTCCGCTCTCACGTCCAGGGCCACTTTCCTGTCTTGCCACCCCTAAGCCCTTACT | 120 | 58 |
| chr7 | 44358248 | 44358368 | 772751_42001747_Shank1(243961)_401 | 1 | + | good | CTCCCTCTCATGGTGGCACTCCTGTGCCCGGCCTTCTCTATTTTCTCTGACACTTAGCACACCCCTCACACTCATGGCCCTGAACTCTTTCCAAATGTTTAACCCCTTCTCCCCCGGCTC | 120 | 55 |
| chr7 | 144175497 | 144175617 | 772751_42001748_Shank2(210274)_1 | 1 | + | good | TGGATAACTGGGGCTCCGGCTCACAAATGCCTCCCGGTGTCTCCTGGCCGACAGCTCGCGCGCAATCGCCTGCTCCATGTTAACCCTGCGTGGTGCTGCCTCTCCGATGCCCTGCTTGCT | 120 | 63 |
| chr7 | 144175617 | 144175737 | 772751_42001748_Shank2(210274)_2 | 1 | + | good | CTCTCGGCAGCGTGGCACTGCTCACATGAGGATCACCGCCAGAGGTAGCCGGTTCTGAGAGCAGCCTCAGTCTGCGCACAGTTGCTTCCCAGCTGACAGACAGAATATGGGGTGGGCTTT | 120 | 59 |
| chr7 | 144175737 | 144175857 | 772751_42001748_Shank2(210274)_3 | 1 | + | good | GTCACCAGCACGCTGGGTCTGGCATGCCCAGCGATGCTCGGGGCTGCAGCGCGCCCAGACGCCCGCCACTGCTCGCCGGGTCCCAGCCGCGCCAGCTCCGAGGGGCCGGGCTGCGCGCCT | 120 | 78 |
| chr7 | 144175857 | 144175977 | 772751_42001748_Shank2(210274)_4 | 1 | + | good | AGCCTTCCTGCCCGCCGAGAGGGTTATATAATGAGCGGACCGCTGCCAAAGTCACAGGGAAAATGAAGTCTTTGTTAAATGCCTTCACCAAGAAGGAAGGTAAGGAGAAAGCTGTAGGAA | 120 | 48 |
| chr7 | 144175977 | 144176097 | 772751_42001748_Shank2(210274)_5 | 1 | + | good | GCGTGGGGTGCCTGGTGAGCAGCCGTGGGCTGCTCCTCGGGGGAGAGGGGACTCGGGTGCCTCCTCAAATTGCTGGTTACGGTTCATCGCTCAAGGAATAATTGCCCCAGCTTTGTGGTA | 120 | 61 |
| chr7 | 144176097 | 144176217 | 772751_42001748_Shank2(210274)_6 | 1 | + | good | TGGTTTGGCTCTAGTCAAGTCTGGAAGATTAACACAGATGAAGGTGCCTTTGTATTTTAGTGTGTTCCTGAGCTACCAGCCTCTGGCAGAACTGCATTCCAGAACATCTCATCCAGAGAG | 120 | 46 |
| chr7 | 144176217 | 144176337 | 772751_42001748_Shank2(210274)_7 | 1 | + | good | CTTATGCCAAGGGACACATTGGGAAACTAGGTCTTGGGGAAAATAAAAGAAGTTTTGGTTTTCTGTCTTCCAGTCACCTGGAGGCTGTGAGCACCTGCTAAGGAGGGTCAGGGCACATGG | 120 | 49 |
| chr7 | 144176337 | 144176457 | 772751_42001748_Shank2(210274)_8 | 1 | + | good | CTGGATGCACCCAAGGCATCCTCTTCTAGTGCCACTTGCCAGTTCTGCCTGGCTGTCCCTGGGTTGCGAAACAAACCATGGGGTGCATGAAGAATCCCTCTTTAACATACCTTGTATGAT | 120 | 51 |
| chr7 | 144176457 | 144176577 | 772751_42001748_Shank2(210274)_9 | 1 | + | good | TTTTATTGTGAGGAAGGAAAAAAACTGTATAAATAAATGCATGTAAGACAGCTAAGAAAATATCCACCCACCCAAGAGCCTGCAGATTGCTGAAAGAAGGAAACGTTGGGTGGCATATTA | 120 | 38 |
| chr7 | 144176577 | 144176697 | 772751_42001748_Shank2(210274)_10 | 1 | + | good | ATCATTTTCAGACAGGAAAGGAGTTATTTTTAACACTTTTCATGAAAAAAAAAAAACATCATTTGCGTGTATGGTCCCAGTCAGAGAGTTTTCTCAGATAGCAAGCTTTCTATCTTTCTG | 120 | 33 |
| chr7 | 144176697 | 144176817 | 772751_42001748_Shank2(210274)_11 | 1 | + | good | ATGAAACCAGGTTTCCAAAGAGGTGACTTTAAGGAAATTAGGAAGATCACTTTCTACCCGAGCTCACTGTGAACTAGACACCAAAGATACTTCAGCAGCCTTGAAACTAATTTGATATGA | 120 | 39 |
| chr7 | 144176817 | 144176937 | 772751_42001748_Shank2(210274)_12 | 1 | + | good | GGCTTGGTCGTTAAGCAAGACAATTTTCATTAAAATGTTCACTCTAATGTTACCTGTGTGGGTCCATTTGACTCACTTGCCTGGTCTTCGGGGTGAAATGTGCATAATGTGTCCCGGCTG | 120 | 45 |
| chr7 | 144176937 | 144177057 | 772751_42001748_Shank2(210274)_13 | 1 | + | good | TTTTGTTCTATGACCCTCCCTGTGTGGCCTCGCCTGCATGAGGACATCCTGCTTACTCCTTCCTTCTGCAGACTCAGGCTCTTCAAAAAGGGCATAGAATAGATCCTTTCTTTTTCTGAT | 120 | 47 |
| chr7 | 144177057 | 144177177 | 772751_42001748_Shank2(210274)_14 | 1 | + | good | GAGGCTTACCAAAATTTGCCATGAGGGACTTCGAATTTTTATAACCCACGGACAGCATGAATTGCAGCCATGGGTTAGAGTGTTAAGTAATGCTATCTCCATTCTGCTTCTATGAACATT | 120 | 41 |
| chr7 | 144177177 | 144177297 | 772751_42001748_Shank2(210274)_15 | 1 | + | good | TCTTTCTTACGGAGTCTGTCTTAGCAATGGACAGGGTATCAAGTGCCCTCACATCCCCTCACGTTGCCATTAAGGGTATTAATACCTTACTGGTTACATAGTCAGAGATGCCATCAACTG | 120 | 45 |
| chr7 | 144177297 | 144177417 | 772751_42001748_Shank2(210274)_16 | 1 | + | good | TACTCTTAGGGATGGGTGACTCCACAGATGCACAGTTGCAGGATGCCAGCATGTCCGGGTCTTGGGGCTGTTCGGGACCCTCCACATGCCTCACCCAGGGGATACTCCCAGCAGCCCCAA | 120 | 60 |
| chr7 | 144177417 | 144177537 | 772751_42001748_Shank2(210274)_17 | 1 | + | good | GGTATAGGCACTATTTATCCTCTCCATGTATGCTGACCTTTTGGCTTTCACACCAAGAACCCCGGATCTCAGTGAGTGGCAGATGAGACTGAGATTATGAGGTCACCTGTCCTTTACAAG | 120 | 47 |
| chr7 | 144177537 | 144177657 | 772751_42001748_Shank2(210274)_18 | 1 | + | good | GCCTTTTAAACCCCCCACCTCTGTAAGTCCCAAGATCACCCCTGACTCCCTGGTTGGCTCCATGGTGAATGCTCTGACTTCTCAAGGCCCTTCTCTGGCAGCGTCATCAGCAAGGCTTTC | 120 | 55 |
| chr7 | 144177657 | 144177777 | 772751_42001748_Shank2(210274)_19 | 1 | + | good | TTTGTGCCCAGCCAAACACACTTGGTGGCCTGACCTGGCTATAATTCTTTTGGAGGTGGCTGCTGGGGCTGTATAGGCCTAGCTCTTTAAGAGCACAGGCTCTTGGCTTCAGGGAAGACA | 120 | 52 |
| chr7 | 144177777 | 144177897 | 772751_42001748_Shank2(210274)_20 | 1 | + | good | CTACCAGCCTTTGGTGTTCCCAGTCTGTCCCTGACCAGCCCCCCGTGAGTCACATGCTGGAGAAACTGGGAGCGAGATCGCACATAGCCAGGAGGTTTGCTGACCATCTTTTCCCATCAA | 120 | 56 |
| chr7 | 144177897 | 144178017 | 772751_42001748_Shank2(210274)_21 | 1 | + | good | AAGCTTGCAGAGAAAGCAGGGCGCACGATACATGCTGCAGCCCACTGGAGAGATGAGGACTCCCTCCATATATTACTTGAGCCCCAAGCTTAGCAGCCTGTCACCCTGCTATATGGCTTC | 120 | 53 |
| chr7 | 144178017 | 144178137 | 772751_42001748_Shank2(210274)_22 | 1 | + | good | CCAGCTTGGCATCTAGCTCTGGTTCCTAAGAAAGAGGAAAATTAATTGTCCTGGGTCACAGCTGATCTGAAAGAGGATTGTCAGAACCACCTCTGCTATCTCTGTCCCTGTGAACTGTGC | 120 | 48 |
| chr7 | 144178137 | 144178257 | 772751_42001748_Shank2(210274)_23 | 1 | + | good | TGGGGCTCTGTCCCTAGAATGTCTGCCAAGGCCACTCTGTCCCCGTGAGCAGTTTGGATGCTAGCTCTATCCCCATGAGTTTCTCCAGGACCACTTCTATCCTTGTGGGACTTGCTAGGT | 120 | 54 |
| chr7 | 144178257 | 144178377 | 772751_42001748_Shank2(210274)_24 | 1 | + | good | CTACTTCTTGAGTTATGCCTGGGCTATCTCTGTCCCCCTGAGTTATTCGAGGGACACCTCTGTCCCTGTAAACTGTGCCTAGGCCACTTTTGTCCCCATGCGTTCCTCCAGGACCATCTC | 120 | 53 |
| chr7 | 144178377 | 144178497 | 772751_42001748_Shank2(210274)_25 | 1 | + | good | TGTCCCTGTGAGCTGTGCTGGAACACTTCTGACCCCATAAGATCTACCAGAGTCATTCTGTTCCCATGATCTGTTCATGAGACACCTATTTTAGTACCACTCATGTCCTCATCCTCCGGC | 120 | 48 |
| chr7 | 144178497 | 144178617 | 772751_42001748_Shank2(210274)_26 | 1 | + | good | TATGCAGATGAACAGAAAGCCATGGACTCTGATGGAGCAGGGATCTTAGCTCTCTATGAGGTGCCCACCTCTAACCCCTGACCTTGGGCAGCTGTTGCCATAGGTAGTTAACTGTAGTGC | 120 | 51 |
| chr7 | 144178617 | 144178737 | 772751_42001748_Shank2(210274)_27 | 1 | + | good | TCTGCTTCTTTGATGTCGGGGCCCTTACCCAGGTCAGTCCCCCTGGAAGCCCTGTTTTATTCTAGCTTCATAGGGTCCTGTTTATAGAATATAGAAATCTTGTAATAAGGTTTCACTGGG | 120 | 45 |
| chr7 | 144178737 | 144178857 | 772751_42001748_Shank2(210274)_28 | 1 | + | good | TACCAATTAAGAGGCTAGGGGTGTATAGTGTCTCTGGCCATCAAGGTGGGGGCATCTACTGTGGCTGCAATGTGAACACCATACCTGAAGGTACCCAGGTTTGCCTTATCTGAGTCCCCG | 120 | 52 |
| chr7 | 144178857 | 144178977 | 772751_42001748_Shank2(210274)_29 | 1 | + | good | GATATATTGGGCTGGAGTGTTTGAGGACCCAGCTCTGTCTAGACCTTTCTATGAAACACCCAGTGAAGTGGGGCCACCTGCTGCTGCCACTTGCTGAGAGGGAAAAGGGAAGATTCTGGG | 120 | 52 |
| chr7 | 144178977 | 144179097 | 772751_42001748_Shank2(210274)_30 | 1 | + | good | GTGTTGTGACCTCTTCTGCCTAAGAGATTAGAGGACAGAGGTGGCTAGCTTTCCCTGTGATCAAGGCTATAGGCCTGAGTGCATAAAGATGAAATACCCATTATTCTCCATACTGGAAGT | 120 | 45 |
| chr7 | 144179097 | 144179217 | 772751_42001748_Shank2(210274)_31 | 1 | + | good | TGACCATCCAAGATTAAAATGACCTGGGGCTGGTTCCCTGAGGTCTGTCTGTGACTTACAGATGATGTCTTCTCTGTATCCTCATGTCTGGGTCCTAAATCTCTGGATCCTGATTATTGC | 120 | 46 |
| chr7 | 144179217 | 144179337 | 772751_42001748_Shank2(210274)_32 | 1 | + | good | TATCACATGAGGTCATCCCAGTGACTCATTTTAATCTAGTTACCCCTGTAAAGGACCTCTCCCCACATGTGGTCACATCCTGAGGTCCTGGATTATATATACTAGTACTAGTGTAAGAAC | 120 | 42 |

| | | | | | | | | | |
|---|---|---|---|---|---|---|---|---|---|
| chr7 | 144179337 | 144179457 | 772751_42001748_Shank2(210274)_33 | 1 | + | good | TTTGGGTACATAGCTCTGCCCACAGAGGTCATGTTCAGATTTTTAGGGGATGCTCCTGAGCCGTGCAGTGTCCTAATGCTTGGTGACATGGGGAAGCCCATCCTTACGTGTGTTATGGCT | 120 | 51 |
| chr7 | 144179457 | 144179577 | 772751_42001748_Shank2(210274)_34 | 1 | + | good | GTGTCTCCCTGATGAAGGCATGGGCTGGCCCGATGGAAGGTCAGACCTTGAGTCCCTTCCATTGAGGTCCTGGTTTGAGCTTCCATGTGCTGGAGGCATTATCTGTCTCAGTGTTTCTCT | 120 | 53 |
| chr7 | 144179577 | 144179697 | 772751_42001748_Shank2(210274)_35 | 1 | + | good | GTGTTCATGTTGGGGGTGAGTCAGGCCTTGGGGATTCCAGCACTACGCAGCCTGCACCAAGCCCTCAGGGCCATCCCTGACTTGTACAGGGTGTGGATGCATGCAGCCCTGGCCTTCCTG | 120 | 61 |
| chr7 | 144179697 | 144179817 | 772751_42001748_Shank2(210274)_36 | 1 | + | good | TGTCCCTGTCTCCTGACATTATTCATCTCTCTCTCTCTCTTTTTCTCCCCAACCCAGTGCCCTTCAGAGAGGCCCCAGCTTATTCCAACCGCAGGCGGCGGCCCCCCAACACATTGGCTG | 120 | 57 |
| chr7 | 144179817 | 144179937 | 772751_42001748_Shank2(210274)_37 | 1 | + | good | CCCCCAGAGTTCTTCTGCGTTCTAATAGTGACAACAACCTCAATGCTGGTGCACCTGAATGGGCTGTCTGCTCTGCAGCCACCTCCCACCGAAGCCTCTCACCCCAACTGCTGCAGCAGA | 120 | 57 |
| chr7 | 144179937 | 144180057 | 772751_42001748_Shank2(210274)_38 | 1 | + | good | CACCCAGCAAACCTGATGGAGCCACCAAGAGCCTTGGAAGCTATACCCCTGGGCCCCGCAGCCGTTCCCCTTCACTCAACAGGCTGGGTGGTACTGCTGAGGATGGCAAGAGGACACAGC | 120 | 61 |
| chr7 | 144180057 | 144180177 | 772751_42001748_Shank2(210274)_39 | 1 | + | good | CACATTGGCATGTGGGGTAAGAAGAAACTGGGTAGTGGGCAGGGGCTTCCATGGGAGACGTACATGTTTCTCAATTCAAGCTGCCCCAGAAAACGCTGAATTCTGTCTGCTGAGCCCGGG | 120 | 53 |
| chr7 | 144180177 | 144180297 | 772751_42001748_Shank2(210274)_40 | 1 | + | good | GCCTCTCTGCTTAGCCCAGTACTCTTAATGCAGGGGCAAGCTGAGCTGTGCAGGGACCTTATGTCCTTTCCAGTGCCTTGACTCCTTGGTTCTCGAAATGAAAGGGCCGTAGGCAGTCAG | 120 | 54 |
| chr7 | 144180297 | 144180417 | 772751_42001748_Shank2(210274)_41 | 1 | + | good | AGCAGAGAGTACGGCCTGTGCCGAATACTCTCCCAGCAGTCTCTGTCATATACACACTATCTCCAAACCAGTGACCACATGCTCGCAGCAATGAGAAATATCAGCCTGGGGGGGGGGAGG | 120 | 54 |
| chr7 | 144180417 | 144180537 | 772751_42001748_Shank2(210274)_42 | 1 | + | good | GAGGAGGGAGACAAAGCTCAGAGGCGTGTTATAGCACAAAGAACTAGCCATATCTCCAGCAGCCTTGGCTTATGCTACAAATGCAAACTGTATCTGCCCAGCATGGCAGGGGCTCATAGC | 120 | 51 |
| chr7 | 144180537 | 144180657 | 772751_42001748_Shank2(210274)_43 | 1 | + | good | CTCTTCATCTACCTCTGAAGGAGACCCAAGGGACAGGCTAGAATTGGGCAGGCTGAGGGGACCACTCAGGGCATCATAGCAGGGACCTTCCCTAGGGAAGGGGCTTCCGACCTGCATAGC | 120 | 58 |
| chr7 | 144180657 | 144180777 | 772751_42001748_Shank2(210274)_44 | 1 | + | good | CAGGAGGGGTCTGCATTTCCATGTCCAGCTCCTGAGCACACCTCTGCCTTCCTAAGTTTCTTAGCCAAGAGAAAATACTCTCAAGGACCAGCTAATTATGGAAGCCTTGAGTCTATATTT | 120 | 46 |
| chr7 | 144180777 | 144180897 | 772751_42001748_Shank2(210274)_45 | 1 | + | good | GACAGCCATCAGAGAGATTTGGGGTAGCAGGGAATGAATGGTTTTACCGTCTTCCTGTGATCCCTGTGGCAGGCAGTAGGGGATGGTGGGATGTGGGCCTGCCATAGCAATGTCTTCACT | 120 | 53 |
| chr7 | 144180897 | 144181017 | 772751_42001748_Shank2(210274)_46 | 1 | + | good | GGGGCTTACTTCCATTCTCCTTGTCCTCGTATCTTCCTCAGTCTCCCTTTTGATTTTTCTTCTGCCCTGCCCTGCCTAACACAGGCTGGCCCAGAAAAGGTGGGAAGAAGTGTTCTCGCC | 120 | 52 |
| chr7 | 144181017 | 144181137 | 772751_42001748_Shank2(210274)_47 | 1 | + | good | CATCCACAGTCTCTGCCCTGTGCCTACATAGCACAGGTGTGAGTCCCTGTGTTCGCCTCTCTAGAGGCGGCGCCCTCTAGTGGCACTTAGAGGCACAGCTGAGTCACAGTCCTGGGAGAC | 120 | 59 |
| chr7 | 144181137 | 144181257 | 772751_42001748_Shank2(210274)_48 | 1 | + | good | TTGCATTTGCTCTGGGAGGTGGTCGGGGCATCCAGTATTCCTAGAGCCTCTCTCAGCTGTCCAGATGTAGAAGATTACTCAACATCACAGGTGGATCTGGCCCTACTCCTCATGGGGACA | 120 | 52 |
| chr7 | 144181257 | 144181377 | 772751_42001748_Shank2(210274)_49 | 1 | + | good | TTTCACTAGTGTCTCATTTGTCCTAAGTGACTTTAGATTTGGCCTGGAACAAAGGGGATTCCCTAGGAAGATCCCTAGAAAGACAGTGGTGCCATGGGCCTCTCTCCAGCTGAGTCCATA | 120 | 48 |
| chr7 | 144181377 | 144181497 | 772751_42001748_Shank2(210274)_50 | 1 | + | good | TGAGAAATGCTGCTCTTTCCAACGCCACCCATCTCTGCAGTTTCAGGTAGTATGGTGAGCATGGTGACATTGGCATCATCATCATCATCCATTATCCCAGGAAAAAGCCAGTTCCTCAGT | 120 | 47 |
| chr7 | 144181617 | 144181737 | 772751_42001748_Shank2(210274)_52 | 1 | + | good | CGGATTTCAGAAAGCACAGGCCACTCTAGCTTTTCCCCTGAAGGCTCTAGTAGATTCTGGGGTACCCACCTTGCCTATGCATCCCAACCTATCAGACATGGATCTGAGGGTGACAATTAG | 120 | 50 |
| chr7 | 144181737 | 144181857 | 772751_42001748_Shank2(210274)_53 | 1 | + | good | GTGTCTCCATGGCTCTATTCTTTGTCTACCTATATGTTCTCTCTCTGGGTCTATAGGAACATGGGCACCTGTACTCGGTGGAACAAGACATGGTTACCTCAACAAAGAGAGGAAAGGGCT | 120 | 47 |
| chr7 | 144181977 | 144182097 | 772751_42001748_Shank2(210274)_55 | 1 | + | good | GATTAGATCTCCCTAGACATCTGCTTGTCACCTACACCTAATGTCACCTGCATGTCACCAGGATACCAACAAAACAGTGTGGGTAAGACTACCAGGTTTATGGTTGGGGGAGGCAGGGCC | 120 | 50 |
| chr7 | 144182097 | 144182217 | 772751_42001748_Shank2(210274)_56 | 1 | + | good | AAGTGAAGGCTATTTTCTGTAGGAGCAGGTACACCCTGGCTACTTGTGGAATTCAGGTCCAGATAACTCTTCTCCTTGTGTAGTCTGCAGAGTGCAGTAGACTATGATTACTTTGGCAGG | 120 | 46 |
| chr7 | 144182217 | 144182337 | 772751_42001748_Shank2(210274)_57 | 1 | + | good | CCACAACACAGCACGGGTATTGTACTTCCTACCTGTGAATTGGATGGGCCCCTCCCTCTGTTTACTCTCAAAGATATAGTGGATCTGCCTACTGAAGGCTGATGACTGCACCTTGTCTCC | 120 | 50 |
| chr7 | 144182337 | 144182457 | 772751_42001748_Shank2(210274)_58 | 1 | + | good | TATGCCCCTCCTGAAGCAGTCTTTTTTGAGCCAGTGTGATGAGCCTATTGAGTCCTTGTCCCTGTCATAGCTCCTAGGGATGGGTAGTTAGGGGCAGGGTAAACTAAGGTGCCTTTTGTG | 120 | 50 |
| chr7 | 144182457 | 144182577 | 772751_42001748_Shank2(210274)_59 | 1 | + | good | CTTGCTAGTTTGGTGTTTCTTTCTCTAAATCTTCAATAGCCAGAAAAGAACAGGGTGACATGTGTCCAGTCTGTAGGCATAGCAATGATGAATGGTAATACACTAGAGTTCAAGGCCCAA | 120 | 41 |
| chr7 | 144182577 | 144182697 | 772751_42001748_Shank2(210274)_60 | 1 | + | good | TTGCTGGGTATGATATTGGCTATGCCCAAGCCCCAGCTAGGAGCTCTCTTGGCCTCCATTAATCCCAGGCCTTTCCCAGGGAGACATTGATTTCTGCAGGCGTCTGCTAGGTGACCCTTC | 120 | 54 |
| chr7 | 144182697 | 144182817 | 772751_42001748_Shank2(210274)_61 | 1 | + | good | CTGATCATACTTGGCATTCTTTGACTCTACACATCGGCCCCGCCCCATTTCTTCCCAAGACCAACCACAGCACCTCCTTATTGAATGTTTCACCTCTGTCTTGTGCCACTCCGGTCCCAC | 120 | 52 |
| chr7 | 144182817 | 144182937 | 772751_42001748_Shank2(210274)_62 | 1 | + | good | CCTACTGCCACAGCCACAGTGCCCTAGGCTCTACCTTGCTGTGTGTTCCTTCAGACAGCTTCTGTGGCCTCTTTTGCAGTGCCTGACCCTCACCCTAGCTCAGACAAGCCTGTGTGCACT | 120 | 57 |
| chr7 | 144182937 | 144183057 | 772751_42001748_Shank2(210274)_63 | 1 | + | good | CTTACTTTGACTTTGGTTGTCACCCCAGTCTCAGCAGAAACATTACCTGGTTGGAGTTCACTACAAAGCCTCTGCAGCTTCTCCCATCTTCTGTGCTCAGATGTGTGTGCTGCCCGGTGA | 120 | 50 |
| chr7 | 144183057 | 144183177 | 772751_42001748_Shank2(210274)_64 | 1 | + | good | TGGACCACCTGGGAAGCTATGAATAAATGAATGAGTAACTGATGGAAGAGGGAGTGAATGGAGGAAGCTCGATGTGTGGGTAGTTCCCGTTCATACAGGGTAGCTGGGAAAGGCCCAATT | 120 | 48 |
| chr7 | 144183177 | 144183297 | 772751_42001748_Shank2(210274)_65 | 1 | + | good | TATCCTTCCCAGTCACCCCACAGAACATTGGCTTTCTGGTACTGAATCTCCAGCAATCTAACCCTTGGACTAAAGCAAGTCGGCTGGTGCTCTGTTTGGCTGCTCTTGGCTAGAACATGA | 120 | 49 |
| chr7 | 144183297 | 144183417 | 772751_42001748_Shank2(210274)_66 | 1 | + | good | AGGAAATACAGATGCCAGGGCTGGGGAACTACTGAACCATTCACTCACTGTGTGTGTCCTCTCAGGTCTGCCTGTCTGAGGCTACCCCTTACCTGGGGTCATGGCTCTGTTGTTGCTCAA | 120 | 53 |
| chr7 | 144183417 | 144183537 | 772751_42001748_Shank2(210274)_67 | 1 | + | good | GGCTGCCATAAGGAGAGCATAGCCCCTGGGCAGGAGGTCACAGTTTATAAAACACCTTCCCTTGAGGTATAGCCTTTCCTATTTTTTTTTTAACTATACCACAGGATAGAGCATCAGGAT | 120 | 44 |
| chr7 | 144183537 | 144183657 | 772751_42001748_Shank2(210274)_68 | 1 | + | good | GCAAGGGTGGCTCTGACTCCCATGTGAGGCCATTATTCATGGAAACCCTCTGTCATCTGGGAAATGGGCTCAATTTTTCTACCTGCTGCCTTGGGGCACTTCATAAAGTTCAAGTTAGTA | 120 | 48 |
| chr7 | 144183657 | 144183777 | 772751_42001748_Shank2(210274)_69 | 1 | + | good | GTGATTATAATGATGCTTCAATACTCACACAGGACCGCTGAGCTTTACTGTGTGTGTGCCCTGCTTCATGACCAATGATCCCCCTACTCCTACCCTTGGAGGCTTCAGGCCCCCAGACTG | 120 | 52 |
| chr7 | 144183777 | 144183897 | 772751_42001748_Shank2(210274)_70 | 1 | + | good | ATGACTCTGAGCTAGATGTCTGTAGAGTGGGCTAGAAAACAGAGACCCAAGACTATGTTTACATTGACTTCTGTGTGGAGTAACATTCACTGGCTTGAATTAGCTAGAGTGCCCCAGCAT | 120 | 44 |
| chr7 | 144183897 | 144184017 | 772751_42001748_Shank2(210274)_71 | 1 | + | good | TTTGTTACCTGCTTTGCAGTTGACACTGCACTATGTGGAGTGGATTCCTATTGGTTTGTTCTCATGATTCATCATGATGGGGAAACTGATACTTTAAAAGATTCACTGATTTGTCCAGTA | 120 | 38 |
| chr7 | 144184017 | 144184137 | 772751_42001748_Shank2(210274)_72 | 1 | + | good | TCTGGTAATTAAGGGGTCAGCTGGGATTCAGGCAGTGTCTCTTGAAATGTGGATGTGTGAGTGAAGGGATAGTATTCAGTAAGGTGGGCATATGAGTGGATGGACAGATGGTTAGCAATG | 120 | 46 |
| chr7 | 144184377 | 144184497 | 772751_42001748_Shank2(210274)_75 | 1 | + | good | CCATGAAGACCCAAGATGTGACTGGCAAGGCAAGGATAGGCTATATAAAGAGGTGAGGCTGAGGCTGTGTTCTGTTCTCCATCTCATCTTACAATTGTCATCCTGGTGGTCACATAGCCT | 120 | 48 |
| chr7 | 144184497 | 144184617 | 772751_42001748_Shank2(210274)_76 | 1 | + | good | ACTCAGGCACCTGTGCCTCTGTAATTCTGTAACAACTGTATCCATGGTTATGATTTAGAAAACCTTGGGTATGTGTGGGTGTAGGTATACATTGCTATCCCCAGCTACAAGCAGCCTTCC | 120 | 45 |
| chr7 | 144184617 | 144184737 | 772751_42001748_Shank2(210274)_77 | 1 | + | good | ACCTCCCTCAGTGGCCAGAGCCCAGATATGTTTTCAGGCCCTACATTCATGCTAGAAGAAGCCAGGAGGCTTGGCCTTTTTCCCCTCTTTGTTTTACACGAATGAAGGGTAGATCCAGGC | 120 | 51 |
| chr7 | 144184737 | 144184857 | 772751_42001748_Shank2(210274)_78 | 1 | + | good | TTTGGGGGCAGGCAGGCCCTGTGTATTCAGTGCTCATCCATCCTCACCAACGTTATAAATGAGGGCCTAAATTTAACATGCCACTGCTCCAATTTCACATCTGTAGACTTGGCTCCTCAA | 120 | 48 |
| chr7 | 144184857 | 144184977 | 772751_42001748_Shank2(210274)_79 | 1 | + | good | AGACTCCAACCAGGGTCTAATGGCTAGATTGCACAGTCATGCTAGATAGAAAAGTAAGAACTTATGGGCCAATGATGTAATAGATAACACATCTCACTATTGGCCAGATAAAGTAAAAAC | 120 | 38 |
| chr7 | 144184977 | 144185097 | 772751_42001748_Shank2(210274)_80 | 1 | + | good | TACAGTCAAATATAACAGAGTGTACCTTTGCAGTTCTGTAAGAGCACCCCAAGCCACAGGGCCTTTACACACATGGGCAACATGGTACAAAAGGCCCTGGGAAGATAGTCTCAATGAAGG | 120 | 47 |
| chr7 | 144185097 | 144185217 | 772751_42001748_Shank2(210274)_81 | 1 | + | good | ATTGTCTAGGTAGATCTGGATAGGCTCGCCTGTGAGGGATTTCTTTTAACTGCCTTGAGTGTGTGAGGAGATACAGGTGGCACAATTCTCCAGGCTCGGGCTTCTGAGCTGAATGAGAGA | 120 | 49 |
| chr7 | 144185217 | 144185337 | 772751_42001748_Shank2(210274)_82 | 1 | + | good | GGAGAAAGCTAGCTAGCTGTACTCAGTTCTCTAAGCTCCTGACTAGATGGGTTGTGACTAGGTGTTTGAAGTCCCTGTCTTGACTTCCCTACAGTAATAGACTGTGACCTAGAATTGTGA | 120 | 45 |
| chr7 | 144185457 | 144185577 | 772751_42001748_Shank2(210274)_84 | 1 | + | good | AGAAATTCCCCTGCATGCCAGTTTCCCATTGGTCCTTGTGTTAAGTGGCTCCTTCATTTCCTTGCAGGAAGTCTGAGGCCACTTCTGTGGGCCTAAAGGCCATAGTTAGGTTTAGAAACC | 120 | 48 |
| chr7 | 144185577 | 144185697 | 772751_42001748_Shank2(210274)_85 | 1 | + | good | AGAGCTGTGGATTGTGGGCTCCTAGTCTAGGTTCAGACATGATTCCCATTCTGGAACCCTCTCCTATCCTTCCAATGCCCACTCCTTTCCCTTGGCGGTACCCTCTTGGAAGATTCCGGT | 120 | 52 |
| chr7 | 144185697 | 144185817 | 772751_42001748_Shank2(210274)_86 | 1 | + | good | GCTACTTTCTTGGCTAGGAAGCTCTGAGAAGTTGTCTTGACATCCTGACATGGCTTTTTGTTTTTGGGTTTTTTTTCCCAATGTGGTCATGTTTGTGTCTAAAGCTTCCCTTTCTGCAAA | 120 | 41 |
| chr7 | 144185817 | 144185937 | 772751_42001748_Shank2(210274)_87 | 1 | + | good | GAGAAGTGACAGGAGGTCCCCTTGAGGTTGTCCCTGATTAGGACTTGGGTGACAGTCACACTGGAATTCTGCTATGCCTCCTGATGAAGACACTGCAGACTGACACATGTGAGGACCTTC | 120 | 52 |
| chr7 | 144185937 | 144186057 | 772751_42001748_Shank2(210274)_88 | 1 | + | good | TCCCTACATGTCCTATTAATGGTCTTGATTAAAGGCTTCTCACTGTTGGGGGAAGATGGGCCTGGGACACACATTCAATGCAAAACTAGCATTGTAAAGAAAGTCCTTTTGGGAGTGACT | 120 | 43 |
| chr7 | 144186057 | 144186177 | 772751_42001748_Shank2(210274)_89 | 1 | + | good | GGCAGGCTGCCCATGGTCATGAGTACTTTGGTAACAGATTCCACATGTATCCTTACCAGCGTGACAAATATAGTGAATTGGAAAAAACATGACAAGGGCTTCCCGAATGACCTGGCCTTG | 120 | 47 |
| chr7 | 144186177 | 144186297 | 772751_42001748_Shank2(210274)_90 | 1 | + | good | ATTTGTTTTCATTATTGTAGTCAAGACACCCATGAGCTTTCCTGTCACCACCATTGTGTGTCCAGCTCAGTGGTCACTGAGCTAAGTGTCCTGAAGCTGTCCCCACTCTTCATCTCCCCA | 120 | 48 |
| chr7 | 144186297 | 144186417 | 772751_42001748_Shank2(210274)_91 | 1 | + | good | ACTTTGCAGCTTCCCAACTGAAACTCTGTCCCTGTGAACCACTGGCCCTCCATCTTACTTTCTGTCTCTGGATCTCGTAGTAGGGACCACATAGGGATGGAATCACTCAGCATATAGCCT | 120 | 49 |
| chr7 | 144186417 | 144186537 | 772751_42001748_Shank2(210274)_92 | 1 | + | good | TGTATGACTGTAATGTCTCCAGGGTTCATCCCTGTGGTGGCACACATCAGAATATCCTTTTCCTGGCTGGATGATGTTCTATTGCACGGGTGGTCATATGCTGTTCATCCTGTATCTATC | 120 | 46 |
| chr7 | 144186537 | 144186657 | 772751_42001748_Shank2(210274)_93 | 1 | + | good | TGTAGATACCTGGGCTGTGGATGATGCTGTAAGGAACATGATGTTGAAATGCCCCTTTGAGACCCTGCTTTATGTCCATGGGGATGTGCCCAACAGTGCAGCCACTGGGTCCTTTCAAGT | 120 | 50 |
| chr7 | 144186657 | 144186777 | 772751_42001748_Shank2(210274)_94 | 1 | + | good | CTTTTCAGCCACAGACACTGCCCCTTGACCCACCTATGCCCTTGCTGTCTGACTGCACACATATCAGTGAAAGCCACTCTCCACTCAGTGTTTGTAGACCACATTGGCCATGGTTGGCAA | 120 | 52 |
| chr7 | 144186777 | 144186897 | 772751_42001748_Shank2(210274)_95 | 1 | + | good | AGTGTAACTTTCTTCTCTCTCCTTCTGTGTTGCCTGATTCAGGTCGCCCTTCACTCCTGGTGCCAACAAGGACTCCCTCTCGACCTTTGAGTACCCAGGGCCCAGGAGGAAGCTATATAG | 120 | 52 |
| chr7 | 144186897 | 144187017 | 772751_42001748_Shank2(210274)_96 | 1 | + | good | TGCGGTGCCTGGGAGACTCTTCGTCGCCGTCAAGCCATACCAACCCCAAGTCGACGGCGAGATCCCCCTTCACCGAGGCGACAGGGTCAAAGGTCAGTCTTTGTCTTCTGCTTCTGGGGA | 120 | 60 |
| chr7 | 144187017 | 144187137 | 772751_42001748_Shank2(210274)_97 | 1 | + | good | AGGCAGGCCCTTTCGGTCTCCACAAAACACCTTGGTGAGTCTTCTGGAACTGGGGCAGAGTCCAAAGCATCAGTTCAGCTGGGCTCAGGACTCTTGAGTCTTGAATCATGACTCTTTGAC | 120 | 52 |
| chr7 | 144187137 | 144187257 | 772751_42001748_Shank2(210274)_98 | 1 | + | good | TCAGTGTCCCCATTGCTGAGCCTTTATGGATGGAAAATGCCCCCACACTCCTGGGTCCATTGTGGGAAGTACAGGAATGAGCCGAACTGGGTCTCAGTGAGTTAAGCCACTCTCCAGCCA | 120 | 53 |
| chr7 | 144187257 | 144187377 | 772751_42001748_Shank2(210274)_99 | 1 | + | good | GAAAACCCTATGAGAAAGTGGGCTTGGTGTAGCCCCTGGCATAGGTGTGCATTCGATGAGTAGAGGACGTCAGCCGGGTGACATAAATTCTCTAGGCGGAATCTGAGGGCTTTTGTGAAC | 120 | 51 |
| chr7 | 144187377 | 144187497 | 772751_42001748_Shank2(210274)_100 | 1 | + | good | AAGCTAAGAAGTACGATATTGGTCCAGGGGTCCTAGGGAGCCAGCAATCAGTCATGTGTGGGTGAGAGGAGTCTGTGAGGGATGCTGAGAGAACACGGATAATGACAGTGTGGAGAGAGG | 120 | 52 |
| chr7 | 144187497 | 144187617 | 772751_42001748_Shank2(210274)_101 | 1 | + | good | AGGTGGACAGTGACTTCAAAGCCTGTGGCTCAGGGCCACAGAGGGCATTGGAGGAGGGGTTGGTGAGGATCGCAGGAGCCACATAGCAAGAAATACAGCTCACAGGTGGAGCCAGAAGGA | 120 | 57 |
| chr7 | 144187617 | 144187737 | 772751_42001748_Shank2(210274)_102 | 1 | + | good | CACTTCATGTGCTCAGCATGCCCAGACCCCACCACTGGACACAAGGAAAAATAGTGGCTCTTCAGAGGTGGCAGATGGCTTCTGACCCTGCTCTCTTCATCTGGCCCTTGTTCAGCCCAC | 120 | 55 |

| | | | | | | | | | |
|---|---|---|---|---|---|---|---|---|---|
| chr7 | 144187737 | 144187857 | 772751_42001748_Shank2(210274)_103 | 1 | + | good | AGCAAGGGACACCTTTGTTTCTTTCTCTGTGCACTCACCTGAGGTCACTGTGAGGTTCTGCAGAGCGTGTTCCCCATCCTTTCCCAGCCCTGGCATCCGGCTTCCGTGCCGCTGCACTAA | 120 | 57 |
| chr7 | 144187857 | 144187977 | 772751_42001748_Shank2(210274)_104 | 1 | + | good | CAAGTCCCTGATGCCCTGTTCCCTTCTCTTTCAAGTTCACTTACTTCTTGCCTCTGAAAGCCCCTTTCCCCATCTCGGTCCTCCTGTGAGGTTCATAATTTCAAAAATTATGTTTTCCTCA | 120 | 45 |
| chr7 | 144187977 | 144188097 | 772751_42001748_Shank2(210274)_105 | 1 | + | good | CTTTCTGAGCGTGTGCTTTAAATTGCTCCTTCCTCATTGCGGCACCTTCTTAGGGCTCCACTGGGGCTCTATTCATTGGAAACATGTGCTATTTTACAACAGCAACACCGTGGCTTGGCG | 120 | 49 |
| chr7 | 144188097 | 144188217 | 772751_42001748_Shank2(210274)_106 | 1 | + | good | TATACACACCGCTTTAAAAATCCCCGTCTGATCATTCACCAGCAAGCGCTTGCAGGAGGTTTCCATCCTTCGCTTTTGTCTTCTCTCAAGGACACACTCTGCTTGCCTCATTCACACACC | 120 | 48 |
| chr7 | 144188217 | 144188337 | 772751_42001748_Shank2(210274)_107 | 1 | + | good | CTCTGACTTTTGTTTGTGTTCTGGACGTCAGCCTGGCATCCCTCAAGGTCTGTGGTTCCTAGTTTCAACTTTCTGATCCTTCCAGCTTCTCTCACTGTAGGCTTTTCTCTTTGCCCTCCT | 120 | 48 |
| chr7 | 144188337 | 144188457 | 772751_42001748_Shank2(210274)_108 | 1 | + | good | CCTTCGCTGCTGGCCTAGCAGTAGGGACTGTGCTCTGCCCAGGATAGACCAGGATATGGGGGGGGGGGGGCAGAGACTCTTGTAAAGGTCTCATCCTGCTCTTCTAGCAAGGGAGCGTCC | 120 | 60 |
| chr7 | 144188457 | 144188577 | 772751_42001748_Shank2(210274)_109 | 1 | + | good | CCCAGAGCCTGCCTGCTCTAGCCCCTCCCAACAGTGGCTGCCTGCAGCAGGCTTTGTAACTGTTTTCCCACTGGGGTCTAGTGCGATCCTCATTCACCATGCCCATAAGGTACCTGGACA | 120 | 57 |
| chr7 | 144188577 | 144188697 | 772751_42001748_Shank2(210274)_110 | 1 | + | good | GTGCAAGTGTGGTTTAATGTAGGAGTTTAAAATTTCACATAACCCATCCATGGGCCAGTCCCTTTCCCTATATAAGTAGATGATATTTGTTGATCTTACCACAGGTGTGTGGTGTGTCAG | 120 | 41 |
| chr7 | 144188697 | 144188817 | 772751_42001748_Shank2(210274)_111 | 1 | + | good | GCTCAGCAGTGATACACGGGATACAGACTTTTCAGGTGTGATGGGTGAGCGGTTGGATGGGTGCCTCCTTGACACCTTCATCTACAGTGCTCACTCAGACCCATCATAAATGGTGCTTAC | 120 | 51 |
| chr7 | 144188817 | 144188937 | 772751_42001748_Shank2(210274)_112 | 1 | + | good | AACATCCTTGCTCAGGCTGGAACTCCAGCTATAAAGTGCTTGTTGAGCAAGCATGAAGTGAGTTCAGGTTCCCAGTCCCTATGTGAAAAGCCAGGTGCAGCAGCTCGCATCTGGAATCCC | 120 | 51 |
| chr7 | 144188937 | 144189057 | 772751_42001748_Shank2(210274)_113 | 1 | + | good | AGCCCTTGAAGAACAGAGACAGGTTGATTCTGAAGCTCCTTGACCAGCCAGTCCAGTAGAATTGGTCAGCCCCAGGTTACATGGGTGACCTTGACTCAAAACGTAATACGGGAGGTGAAT | 120 | 49 |
| chr7 | 144189177 | 144189297 | 772751_42001748_Shank2(210274)_115 | 1 | + | good | TCTCTCTCTCTGTCTCTCTCTCTCTCTCTCTCCCCTTGTGGCCCTCCCTGGATCAGAGTCCATTTCAATTTCAAATCATATCAACAAAGCAGGCACATAGAAAGCCATAGCTCTGGATAG | 120 | 47 |
| chr7 | 144189297 | 144189417 | 772751_42001748_Shank2(210274)_116 | 1 | + | good | TTGCTGCTCACATCAGTGCTGTGGCCTCTGGAGACACGTCTCACATCTCCTGACTTTGCAGATTGCAGAAGGGCATTGTAGGGATGCAGTGGGGAGGCCTTCATCTGTCTGACCTGTTAG | 120 | 53 |
| chr7 | 144189417 | 144189537 | 772751_42001748_Shank2(210274)_117 | 1 | + | good | TTAGTGTTACTGTGTGGCTCTTAGACAGCATATGTGGAACCAGCATAGTCAGGCCAGAGGCAGAGGGGCCCGTGAAGCCCAGGTTCTGAAGACCTCCCAGGACCTTCTGTCCAGCAACCA | 120 | 55 |
| chr7 | 144189537 | 144189657 | 772751_42001748_Shank2(210274)_118 | 1 | + | good | TGCCCTGCCCAGACTGTGCCCCTTACTGCCATTGTCACAAGCAGCTGGAGCCATTGTCAGACAGGGACAGTGTTTGGAGTAAGCAGGACCCTGGCGCTTTCCCAACTTGTTCAGCATTAT | 120 | 54 |
| chr7 | 144189657 | 144189777 | 772751_42001748_Shank2(210274)_119 | 1 | + | good | CCAAGTAGCACTGGGAGCCAGCTGGATGCAGCCACTGCTAGTCCTGGGTAGCAGCCTCAAGGCAGAGGTCAGGGTGGCTGGACATCAGGCTACATCCAAGCTGAAGATGGCATACGTATG | 120 | 57 |
| chr7 | 144189777 | 144189897 | 772751_42001748_Shank2(210274)_120 | 1 | + | good | GGATGACCCGGGATGTGTCACAAGCAGGAAGACATTGTTCAGACCCTCACTCCAGGCTGCAGGCTCATTTTCACACCAAGACTGGGAGAGGAAATCCCACCCTGCACATCTGTAGAGGGG | 120 | 55 |
| chr7 | 144189897 | 144190017 | 772751_42001748_Shank2(210274)_121 | 1 | + | good | AGTCCCCTGTAGAGGCTATCACCTGACTAGTGAGACTGGACCTGTAGAAAGTAGGCAGGGAGCTCTTTGCTCCAGTCAATGACTTACAGTACTGTGCTGAGGTAGGTTAGGAAGGTCCTT | 120 | 50 |
| chr7 | 144190017 | 144190137 | 772751_42001748_Shank2(210274)_122 | 1 | + | good | CCTTCTAGAAGGTTCTGGGTGCTAAATGCACCCCTTCTCAGAGGGGCGCCTGGCATGTAGTGTCTTCACCCTGTCCTCCTTCTAGTCACCAGAAATACTGGCTCTTCCCTGACTCTCCCA | 120 | 54 |
| chr7 | 144190137 | 144190257 | 772751_42001748_Shank2(210274)_123 | 1 | + | good | GACCCTTTAGAGACAAATCTGCTCTCTGCCCCCAGGTCTGGCCTGGATGAGTTCCTCTGCCCAAGCTCATTACTCCTAGCCCAGTGCAGTTTTTCTTGTTCTGATGTGTGAAGGAGACTC | 120 | 52 |
| chr7 | 144190257 | 144190377 | 772751_42001748_Shank2(210274)_124 | 1 | + | good | CCTCATCTTTACACATCCATCACCTTAAACAAGGAGAACAATCTTAGCTATTCTGAATTAAAGCATGATTTACAAGGGAACCGAGCCTGACTCCCACTTTACTAGGCAAGGTGTACCCAC | 120 | 42 |
| chr7 | 144190377 | 144190497 | 772751_42001748_Shank2(210274)_125 | 1 | + | good | TCCCATACAGACACACCCACTTCTACAAGACACTCCCACAAGACACACCCACTTCCATAAGACACTCCCACAGACACATTTATGTGCATAAGACACCCCCACAAGACACACCCATTCTTA | 120 | 48 |
| chr7 | 144190497 | 144190617 | 772751_42001748_Shank2(210274)_126 | 1 | + | good | CACAGACATGTCTACTTTCACACTGGCATCCTGTTGTTTCTTTGTCCCCATGTCACTAAAATGCTCCAGAGAGCAGAGCTGGCAAATCCTAAAGTACCAGATGGCACCTCTCTGTGCACC | 120 | 48 |
| chr7 | 144190617 | 144190737 | 772751_42001748_Shank2(210274)_127 | 1 | + | good | TGGTGCATTCACAAGCTTTTTGCTTTTTCGGGGGAGGGGAGCTCATCCTGGTGACCCACCCAGGCTGTGTGCTTGAGGGGAATCAACTGCCGGGCTGCATGCAGAGCAGGCAGTGCGCTA | 120 | 58 |
| chr7 | 144190737 | 144190857 | 772751_42001748_Shank2(210274)_128 | 1 | + | good | AGATGTCCACAAGCAGGGATGGCGTTTCTGTGCCTCTGTTGACTCTGTGGCACCATGAGCTAAACGGTGTTCCTTTCTGACAAGAGAGGAAACTGAGGCACATAGTGATAAACAACTGCC | 120 | 49 |
| chr7 | 144190857 | 144190977 | 772751_42001748_Shank2(210274)_129 | 1 | + | good | TCACATGTGGCACAGGTAAGGTATGGCATGGTAGATCCCAGGCAGAAGAGGATGCCAGCCTGGCCTTACCATCTCACTTGCTGGGTGGTTTGAAACATCAGTGTTGACCAACTGGAGGCT | 120 | 52 |
| chr7 | 144190977 | 144191097 | 772751_42001748_Shank2(210274)_130 | 1 | + | good | CTGAAGAATGGGCATGGGGGTGTCTTCCTGGGCCTCATTGCACAACACAGACCCTCCACCCAGGACCCACACCTCAAAATGACCTTCTTAGGTCCAGAGCTGGGAGGCTGAGCCAACTGT | 120 | 57 |
| chr7 | 144191097 | 144191217 | 772751_42001748_Shank2(210274)_131 | 1 | + | good | CCTTCAGCTGAGTTGTGTGGAGGCAGGGGCTACGTCCATGAGTCCCTGCTTCAAGGAAATTGTTTCAAGCAGATATGATGGTATCAGCCAGTCGGGAAGTGGCTTGTCTCTTTATTTGGG | 120 | 50 |
| chr7 | 144191217 | 144191337 | 772751_42001748_Shank2(210274)_132 | 1 | + | good | TCATTTCCTCAGATTATTGGGCAAGAATTTACAAAGCTAGAGTTTGCTGTTCTGTCCTTTTCAAAACCACAGGCTGGGTAAATCTTTATCTGCTGTTTGGGGTCAGAATAATGGAGAACA | 120 | 39 |
| chr7 | 144191337 | 144191457 | 772751_42001748_Shank2(210274)_133 | 1 | + | good | AATACTTGTTCGAGGATCTCTTTTTCCCCACCACAGGGCCTCAATCCTCAAGTAATTTAAAGACAAATTGAGATGTGTGGACAGGATAGCAAGCTCCTTCATGTAACCATGTTCAGACAG | 120 | 42 |
| chr7 | 144191457 | 144191577 | 772751_42001748_Shank2(210274)_134 | 1 | + | good | TACACATCCCACTTTGAACAGGGCATCATAGCTTGGTCTTCATAACTGAAATCTTTATTCTAATGGAGCTAGAGTATACTGTGAGGTTCTTGGACCCTGATATATTTGGGCTCCTGCCCA | 120 | 42 |
| chr7 | 144192177 | 144192297 | 772751_42001748_Shank2(210274)_140 | 1 | + | good | CACACACGGATAAAAGCCTCTCTATCCACCCTCTGCCATCATCCACCAGGTAGGAGGGGTCCATGGGCTGGGGTGCCCTGCCCCTTCACACTGTTCTCCAAAGGCCTCATCTGCCAAGCT | 120 | 58 |
| chr7 | 144192297 | 144192417 | 772751_42001748_Shank2(210274)_141 | 1 | + | good | GCTGACAGCGCAGGCCTTGCTCCTTCTACCTGTGGTCTGGTGTCCAGTTGCTGGGGACCCTTTGCTTGGTCCCATCTCGCCCCCTGTACTGATGCCTTTCTCCTTTCTGTCACTTCTAGT | 120 | 57 |
| chr7 | 144192417 | 144192537 | 772751_42001748_Shank2(210274)_142 | 1 | + | good | TCTGAGCATCGGAGAGGGCGGCTTCTGGGAAGGCAGTGCCCGTGGCCACATCGGGTGGTTCCCAGCTGAGTGTGTGGAAGAGGTGCAGTGTAAACCCCGGGACAGCCAGGCAGGTAAGGT | 120 | 62 |
| chr7 | 144192537 | 144192657 | 772751_42001748_Shank2(210274)_143 | 1 | + | good | GACCTCACAGGGCCGGGTCGTGTCATGGTGGGGCTGCTTGGCCCAAGTGTGTGGCAAGTGTCGTGGCTATAGCCACACTGTGGAGGGAGGCCAGGTTACAAAACCCAGGCTTCTCTATTA | 120 | 58 |
| chr7 | 144192657 | 144192777 | 772751_42001748_Shank2(210274)_144 | 1 | + | good | CATTGCAAATGGCTTGGCTGGTTCTGTTCTGGTGGGCTCTGTCTAAACCAAAACCACTTGACAAGCTGGGGTGCAGGTTCTGTGAAAGGGCAGCTCTTTGCATGTCTGGTGCAAACCTAG | 120 | 51 |
| chr7 | 144192777 | 144192897 | 772751_42001748_Shank2(210274)_145 | 1 | + | good | AGCCGATTTAGGGATTTAGGACGTGTCCTTGTCATCCTCAGGAGAGTCTGGGGTGGAGTCAATGTCTTTTTTCTGCCTTCTGATTGGGGGTTAGAACACAGATGGCTTCCCGGTGAAAGG | 120 | 50 |
| chr7 | 144192897 | 144193017 | 772751_42001748_Shank2(210274)_146 | 1 | + | good | TCAAGCTCTGCAGACCCTCCCTAGGAAGTACCACACAGTGCTCATGGCCAAAGCTGTCCTGACTCCATGCTTTAGAGCTATTCTCCAAACCTGCCCCTGCTGAAAGCCTTCTGGTTAAAA | 120 | 51 |
| chr7 | 144193017 | 144193137 | 772751_42001748_Shank2(210274)_147 | 1 | + | good | CAGGTTGATCTACTATGATGTCAATGGCCACTTATATGTAATGGCCTGGGTAAGAAGCAGAGACCCCTGAGGCTTGGACAGAAGGGACTTCTTAGTGACCTATGTTTACCTGTGAGTTCT | 120 | 46 |
| chr7 | 144193137 | 144193257 | 772751_42001748_Shank2(210274)_148 | 1 | + | good | GTTTCCTTTTGTGTGAACTCCTGAGCTGTTTGTGTGCACCCCCAAATGCAGCCACATGCCTTCTGCAGATACTGGCTGTGTGGGTCTGTGACCACTGAACTTGGCTCAAATGTGCTCTTCAT | 120 | 50 |
| chr7 | 144193257 | 144193377 | 772751_42001748_Shank2(210274)_149 | 1 | + | good | GATATATTCTGAGGTTCTTTGTCTGATACTCCATCGAGAGTCCCTGGTCCTGGTCAGGTCACCTGGGCCAGGGAGGATCTAGGAACACATGCTGTGTGAAGCATCTCCCACTGTTGGGCT | 120 | 52 |
| chr7 | 144193377 | 144193497 | 772751_42001748_Shank2(210274)_150 | 1 | + | good | CCCTCAGGGCTGCCAAGCTTTGCAATCTGTTGGGCATTCAGAACACGTCCTCTCTGAGAACTTGACCTTGAGATATAGGGTCCATGTGTCCTAAGCCTCCCACCATGGGGCTGCTCAGGT | 120 | 54 |
| chr7 | 144193497 | 144193617 | 772751_42001748_Shank2(210274)_151 | 1 | + | good | GCCGCTGATAGGTTGCTATGTGTTATCTAATACCACACCCATGTCAGGCCCTTTGGTTTGTTTGCTAATGTTGTTGTTGAGATGGTCTTCTGTAGCCCGTGTTGGCTTCTAATTCCTCAT | 120 | 45 |
| chr7 | 144193737 | 144193857 | 772751_42001748_Shank2(210274)_153 | 1 | + | good | CTATCTCCCAAGCCCATCCAGACAGATCTTGAAAGACAGGGTTGGAAGTGGTGTGGTAGAGTCTGGCAGATGGGCAGTGGGTTAGTGGGTGCTAGAAGCCAGTTTTGCTGGATGAGGAAC | 120 | 52 |
| chr7 | 144193857 | 144193977 | 772751_42001748_Shank2(210274)_154 | 1 | + | good | ACTAGTCACCAGGTATCAACTGGTCTGTCCTAACACTTAACCCCCTTCCAGCTCTGGTCTGGCCCCAGCTTCCCTCTGGGGAGGGAAGATCTGGTTTCTGCTTTCTTGGCTTCCAGAGGC | 120 | 55 |
| chr7 | 144193977 | 144194097 | 772751_42001748_Shank2(210274)_155 | 1 | + | good | TTTATCTAAGGCTGCTCTGCTTCCTGTCTTCACCTTTGGAATGGGAATGTCACTTCCTCCCCTCCCATTCTTATCTTCAAAGCAACTGGGTCTCCCCAACCTCTGCTTTCCATTGGAAAT | 120 | 47 |
| chr7 | 144194097 | 144194217 | 772751_42001748_Shank2(210274)_156 | 1 | + | good | GCTCTGTAAGCCACACCCAAACAGGTCAGCCCCGGGGGTGATGCATGAGGGGTCAGTGGGTCATCCTGCTTTCTCCATCTTGTGGAATCGTGGCTTAGGTGAAGTGCAGCTCACCCTACT | 120 | 57 |
| chr7 | 144194217 | 144194337 | 772751_42001748_Shank2(210274)_157 | 1 | + | good | TTCCTGTAGATCTTGGGGGAGATTACAGCTTCTTTGCCCCCCCCCATTGGCAGTGGAGCCTGACAGACCAATACTCGCCCTGTGGCAGGCACCTTTAGGAGTAATGATGAGATTCCTTG | 120 | 53 |
| chr7 | 144194337 | 144194457 | 772751_42001748_Shank2(210274)_158 | 1 | + | good | TGGGGCTCCATGATCCCACTCCCATCCACACAGTGAATGTTGTGTACTTGAGTTGGTGCTGGCTCCAGGTTCTTCTGGGGCCCATCCTCACATTCTGAGCAGCCTGGAAGCACAGACTG | 120 | 55 |
| chr7 | 144194457 | 144194577 | 772751_42001748_Shank2(210274)_159 | 1 | + | good | GATCCTGACACTCTGCCCCAGCTGTCAAGAGCAGTGGCAGAGCAGTGATGTCTTATGTGTCACCTGTTCCTGAGCACCGCTGCCTCCAGGTCTTACTGAGCACAGGAGAAAGAGTTAAGT | 120 | 53 |
| chr7 | 144194577 | 144194697 | 772751_42001748_Shank2(210274)_160 | 1 | + | good | TACCTATCCTTGTGCCTCTTAAAATCCTAAAAGCTGACAAATTAGTGTCCATCCCCGTGCTGTATAGAATTCAGAAGACTCAGGGATCTAGCCTGGCTTTGATCTGGACATCAATGGGAT | 120 | 43 |
| chr7 | 144194697 | 144194817 | 772751_42001748_Shank2(210274)_161 | 1 | + | good | TGGCTCCATCTGGGATCTCTGGGGAGGGGGATCTGTTTCCTGCTTTCCTGGCTTCCAGAGGCTTTATCCTTGTCTGTAGTCCCTTCTCACTTCTGAGACGGCAGCTGTGAGTAGGGTCAC | 120 | 55 |
| chr7 | 144194817 | 144194937 | 772751_42001748_Shank2(210274)_162 | 1 | + | good | TCTCTGGTTGCATCTTTCAGAAAACCCCCACCCCTGCCTCCTCTTGTCACCACTTTTAGGGCCCAGCTAGGTGAGATCCAAGCTGCCAAAAACAGTTCCAATTTCATCTATGATCTTTTAAG | 120 | 48 |
| chr7 | 144194937 | 144195057 | 772751_42001748_Shank2(210274)_163 | 1 | + | good | CTTCCATGTGTCAATACGGAAACATATCAGGTCCTAGGAATGAGGGCATGAGCATCTTTAGGGTGGATGAGGAGCATCCTTTGCTCCAAGGTCATTGGCAAATGTGACCAATATCACCAT | 120 | 46 |
| chr7 | 144195057 | 144195177 | 772751_42001748_Shank2(210274)_164 | 1 | + | good | TTTGTAATATAATAGACCACACGTCCTGCTAGCATCCCTGGGAGAGTGGTAAACCTTTTCTCCTTATAAACTGATTAGCTCAGGAATTTTGTCACAGTAATGGAACATTTTCCCTGACGG | 120 | 41 |
| chr7 | 144195177 | 144195297 | 772751_42001748_Shank2(210274)_165 | 1 | + | good | CTGTCTAGCCATCCCTCCCATCTCACCAGAGACGAACAACTTTCTGAATTGTCACCACAGAGGAATTATGTCTGTTGTTGGATTCCAGGCCATCTTGACTTCTTCATTCAGCCAATGTCT | 120 | 46 |
| chr7 | 144195297 | 144195417 | 772751_42001748_Shank2(210274)_166 | 1 | + | good | ACCATGTTGCATTGTGGTCCTTTGTATGCTGCTGCGTGACATTCCCTTCTTTATAATTTTATGTGAGCAGTGTTCTCCACTTTCCTGAAGATCACTGACTAACTCCTCTGTGTGAGCTGA | 120 | 43 |
| chr7 | 144195417 | 144195537 | 772751_42001748_Shank2(210274)_167 | 1 | + | good | TGTAAACTAAGCTGCTGGGCTTGTATCCATACCTGGTTTCACTTCTCCTCACTTCCTTAGGTGAAAACCTGCTAGGTCACATGATAGGAACATGCTTAGTTTCATAAGAAATTGTTACAC | 120 | 41 |
| chr7 | 144195537 | 144195657 | 772751_42001748_Shank2(210274)_168 | 1 | + | good | GTGCCTGGGGAGATGACTTAGTAAAGTGTGCCTTTTGTGGACCCGAATTCTGATCCTAGAACTCACAAACCAAAGCCAGCTGTGCTGGTTCATGCCTGTAATGCCAGCACCAAGGAGGCA | 120 | 51 |
| chr7 | 144195657 | 144195777 | 772751_42001748_Shank2(210274)_169 | 1 | + | good | AAGAGAGAGGGTCACTGACTAGACAGCATAGTTCAGTGTGTGAGCTCCAGGCCAGTAAAAGACCCTGCCTCAAAATCAAGACAGATAGAGTTGCTGAGGATAATACCAGGTTGTCTTCTG | 120 | 47 |
| chr7 | 144196017 | 144196137 | 772751_42001748_Shank2(210274)_172 | 1 | + | good | GTCAGCCTCTGATACTGTCAGTCTCCCACGGCAGCCTGGTGGTTGTGACGTTGCTCCTTACTGTGAGGGACACCTCGTTTCTCGGATGGCTTTGGTGGTTAAGCCCCTCCTACAGGCATG | 120 | 57 |
| chr7 | 144196137 | 144196257 | 772751_42001748_Shank2(210274)_173 | 1 | + | good | GTTGCCACATGCAAAACTTCTGAGAGGTCTTGCTCTGTTCGGGGTTTCTGTTGCCTTTTATAACTGGAGCTCTCAACTCTTGTGCTGGGTGTGGAGTGATTTGCTTGTACGGACCTTGCT | 120 | 48 |
| chr7 | 144196257 | 144196377 | 772751_42001748_Shank2(210274)_174 | 1 | + | good | TGTCGATGCTCTGTCCCGGCTGCTGTAGATGCGCCTCCCAGTCTGCGACTTGCCTTTTGTTCTCTAATTGTGTTTTTTAACGAGTAGAAGGTTTTTAAATTTTAAATGAAGTCTAATTTA | 120 | 40 |
| chr7 | 144196377 | 144196497 | 772751_42001748_Shank2(210274)_175 | 1 | + | good | TCCTCCCCCCTTCATAGTTAGTACTTCTGTAGCTTTTTTTTTTTTTTTTTTTTTTTTTTTGGACACTGCAAATTTGAATATGCATAGAATCTAAAAGATGGTTCAAATATTGTTTCAGTGTT | 120 | 29 |
| chr7 | 144196497 | 144196617 | 772751_42001748_Shank2(210274)_176 | 1 | + | good | TCCATCTAGCCAGCTCTTTCCACACAAGGTCAGCTTTGTCTGAGCTGTGGAAGCCTTTGCCTCTGTATATGGCAAAGTAGAAACACTTTTGAATATTCCACAGAATAGGAAAGTAGGTCA | 120 | 43 |

| | | | | | | | | | |
|---|---|---|---|---|---|---|---|---|---|
| chr7 | 144196617 | 144196737 | 772751_42001748_Shank2(210274)_177 | 1 | + | good | AGGACTGAACCCCAACGAGAAAGGGCGCTTTCTACCTGTAGGTGGGACAGAGAATTTCTGTCACCTGAAGATGATTGTCAGAGACTGAGGACGAATAGCAAAATGAAGAGTGGCCCTGAC | 120 | 49 |
| chr7 | 144197097 | 144197217 | 772751_42001748_Shank2(210274)_181 | 1 | + | good | TGTGACTCCAGCACACATCAAGATGACAGAGTTAACCAGCATACTACTTCCTTGCAGAGGTTACTTGATGGCTCTGGGTAGATACCTTCATGTGTCTTATAATGCAGAAGGGTCTATGGA | 120 | 44 |
| chr7 | 144197217 | 144197337 | 772751_42001748_Shank2(210274)_182 | 1 | + | good | GATTACAACCATCTTCTTCACAGTTTCTGTGCCTGAGAGAGAACAGCATGGCCCAGTGTACAAACTCTTGCATGTCATCCCATGGCATTGGCAGCATAGAGTAGCCTGTGACCACAGTGC | 120 | 49 |
| chr7 | 144197337 | 144197457 | 772751_42001748_Shank2(210274)_183 | 1 | + | good | TATCTATGTGCATGGCAGGCAGCACTCTAGAAACTTTCGGTTAAACGTGTGCATGTCCCTCTCCCATCTTCCTGAAGTGCAGGCTCCTCAAGTCAGGAATAGCATCTTGCTCTTCTTAGT | 120 | 48 |
| chr7 | 144197457 | 144197577 | 772751_42001748_Shank2(210274)_184 | 1 | + | good | CCTACACGGTATATGCCAAGGGCAAAGTGACTGCCATTGGCTCTCTATTATCTATGGAAGGGTGCTGTTGCAATAGACCAGGAAACCGGGCAGCTGTTGTGTAGACTCTCAAGCCTCACA | 120 | 50 |
| chr7 | 144197577 | 144197697 | 772751_42001748_Shank2(210274)_185 | 1 | + | good | CCTGTATCAGGGAGCACCATGTCAGCTGTTGTAACAAATAGGCCTACCTCCTTAGCATATGAGGCAGCACTCCCCACCCCAGACCTCCTCCATCTATTGAGATGTCAGTTTCCTTGCTTG | 120 | 51 |
| chr7 | 144197697 | 144197817 | 772751_42001748_Shank2(210274)_186 | 1 | + | good | TTGACTAGAAGTGATAGACTAGTGATAGTGGGACCCTTCCTTCTTCAAATTCTAAGGTGGAGCTAAAAGCAGGGGTGCCCTAGAATTGGCTCTTGGATGAGAAACTAAGATAAAGACCCC | 120 | 44 |
| chr7 | 144197817 | 144197937 | 772751_42001748_Shank2(210274)_187 | 1 | + | good | AGTCTTAGTAGCACACACATATAATCCTGACACTCAGAAGGTGAGACAGGACTGTCACATATTCAGTGGCCAGCCAGGGTTGTGTAGTGGGTTCAGGGTCAGCCTGGATTGTATAGTGAA | 120 | 48 |
| chr7 | 144197937 | 144198057 | 772751_42001748_Shank2(210274)_188 | 1 | + | good | ACCTTGGCTCCCAAAGAATGAAAGCATTCCAAAATTGGCACTTGGTTTCTGGGAAAAAAAAAAAAAACAGTGTGGGAAGCCAGCACACATGTGTGTATTATTTCCCATGATGCATCTTGCA | 120 | 41 |
| chr7 | 144198057 | 144198177 | 772751_42001748_Shank2(210274)_189 | 1 | + | good | TGAAAATAGAAAACACTGTCTGTCAACCCTGGCTCTCCCCAGCAAAATTTAGTTGCTGAGACAAACATCCTGACCATTGATGAATATTGGTCATGTAAGACCAGAGGAGAAAATGACATC | 120 | 41 |
| chr7 | 144198177 | 144198297 | 772751_42001748_Shank2(210274)_190 | 1 | + | good | AGGAGTCCACTCCCCAGCCCCAAAGACACATAGAATCTGTTCACTTGAGAGCTATCATTAAAGAATAGTGGTCAGATGGCCAGATCTCGGATAAAAGGTACAACATTCTTAACCATGCCA | 120 | 44 |
| chr7 | 144198297 | 144198417 | 772751_42001748_Shank2(210274)_191 | 1 | + | good | CGATCATGGGTGCTCTGGTGGGACTGTGAGCTGTGTATCCTGCCAGGGCCTGTCAGGATCAGGACCTGTAAGAAGCTTCCAGAATCACATGGATTAAAGTGTCTGAGGTAGGATTTGAAC | 120 | 51 |
| chr7 | 144198417 | 144198537 | 772751_42001748_Shank2(210274)_192 | 1 | + | good | CTAGATGTCTGAGTCCTGAACTGTCTTCTATGGTCCCCTGCAGGGTCTTTTCTGGTGCCTTAGAGTGAGGCCCCTGAAGGAGCCGTCTCTTTCTCAGAATACTGCTTGACTGCCTGCCTC | 120 | 53 |
| chr7 | 144198537 | 144198657 | 772751_42001748_Shank2(210274)_193 | 1 | + | good | TGTGAACCATGCAGTGGTTGATGTTATACCCCAAGACCTCCCTGCAGGCTGCATGTACTTCTCACAGATGGAAAACTGGCCAGTGCCCTGTCCTTCCTGGATGTGACCTTAATAACAAAG | 120 | 49 |
| chr7 | 144198657 | 144198777 | 772751_42001748_Shank2(210274)_194 | 1 | + | good | GCAGTTCCTCAACTGTGATAAGCCTGTGCCAGGGGCCTGCCCACTGGTGACTATGCCTCACTGTCAGCCTTGGATCAGTCTCTAGGGTTTGTCCATTCTCTTGATGACTGTGTTCCTCTG | 120 | 53 |
| chr7 | 144198777 | 144198897 | 772751_42001748_Shank2(210274)_195 | 1 | + | good | TCCACAACAGTCACTAAGTATCTTCTGGTATCATTCACAGCCCTGAAGTGTGGGGCCTTTCCAGAGCTCATGGCTACAAGCCTTTATATTATAACAGCAGCCTTTCCGCCTTTAGTTTAG | 120 | 45 |
| chr7 | 144198897 | 144199017 | 772751_42001748_Shank2(210274)_196 | 1 | + | good | ACTTTCAAGTGACTCTTTTGATCTTCAAGTAAACTGGGGACTGGAGATAAATGAAGTAATGGTTCCCATTTTACAGATGAAGGCCAGAGAGGTGTATGTCTTATAAGCACTGAGCCAGCA | 120 | 41 |
| chr7 | 144199017 | 144199137 | 772751_42001748_Shank2(210274)_197 | 1 | + | good | CTTGAATATAGATAGACCTGAGTCAAAGATCTGTCCTTTCACCAAGAAGACTGTAGTGCCTACAGAACATCAGTCACCCTGGGTGATGACAAGAATTGGTAGCCATGGAGTTGGGTGGTC | 120 | 46 |
| chr7 | 144199497 | 144199617 | 772751_42001748_Shank2(210274)_201 | 1 | + | good | GTGATGGTGGTAATGATAGTACTGATGATGGTCATGATAACTGTGCTAATGATGATATTAATGATCGTTGTGGTGAGGTTAGTGGTGGTGCTGATCATGTAGGGACCAAAGTGTTAAAGC | 120 | 41 |
| chr7 | 144199617 | 144199737 | 772751_42001748_Shank2(210274)_202 | 1 | + | good | AATGACAGAAGACTAATACCAACAAAGTCTAGCCCTCCAGCTTGGGAGTGCTGCTGTATGAGCATAGATTGAGAGTTCTACTTCCTCCCTGCCTCTGCTCAGTGCTCCTCCCTTTTCCAG | 120 | 49 |
| chr7 | 144199737 | 144199857 | 772751_42001748_Shank2(210274)_203 | 1 | + | good | AAGTCCCTTAGACTGCTCTGTGCACTTCTGAACTGTCATCTGGTTCTCCATTTTTCCTCTTGGTATGGCAGGCTTCCTTTGAGAAGGTCACTTTGCCTTTGTCTCTGTGTATACTGACTT | 120 | 45 |
| chr7 | 144199857 | 144199977 | 772751_42001748_Shank2(210274)_204 | 1 | + | good | TTCCCCCTGGAGTGTGAGCTTTTCCCTCCCGATACATCTTATAATGGGTATAGACTCACCTGAGATGACTGCCGGGGCAGGGATGCCACAGCTGATGACCGATAATATCAGTAGCCTCTC | 120 | 52 |
| chr7 | 144199977 | 144200097 | 772751_42001748_Shank2(210274)_205 | 1 | + | good | TTCCTTCCATTTGAAATTTTGGGGCAGGGGTCATAGGTGGACATCACCATGCCCAGAAGCTGCTGTGCTCTCAGCTGCTGTGTACTTTATTTCCAAGATATATGCTAAGGGCTCATTTAA | 120 | 45 |
| chr7 | 144200097 | 144200217 | 772751_42001748_Shank2(210274)_206 | 1 | + | good | TAAGTCCATTCCTCTGCCCCGCATTAGTTCAAGGCATTGGCCCATTTATCTTTCAGGTTATCTGTCTATTTCTTGCTGATTCAAGGGTCCCTCTTCACACACCCTAGAAGCAAATCTTGG | 120 | 45 |
| chr7 | 144200217 | 144200337 | 772751_42001748_Shank2(210274)_207 | 1 | + | good | GCCAATGATGTTCACTGGAGCATGGGATACTCCATTCTCTGATGGGCTATTGATCGACTCTGTGACTCAGAAGGGGCAAGCGTGGGCCTCGTGTCTCCTGTAGGCAGTGGGACTGACTCC | 120 | 55 |
| chr7 | 144200337 | 144200457 | 772751_42001748_Shank2(210274)_208 | 1 | + | good | ACTGTAATCCCATTTCCTCCTTTCTGCTCAGATGCACACGGAGGCTCCATCCTGTCTGTGCCAGGGATGCCAGCTTCTTGTTTTAAACAAATTAGTGTTCCCTTCAGAAACACAAAAAGC | 120 | 46 |
| chr7 | 144200457 | 144200577 | 772751_42001748_Shank2(210274)_209 | 1 | + | good | TGGTTGGCTAAGGCTGGAAACCCAGCAGGGACTGCACCACACCCCTTCACTGGGACAAGCCCCTTCCCTGGGTTGAGATAACCAAGTGTCCCTGAGGCTTGCTCTCAGGGCTGCCTGAGT | 120 | 59 |
| chr7 | 144200577 | 144200697 | 772751_42001748_Shank2(210274)_210 | 1 | + | good | TGGGGAGTTTTGGATATGGGGGAAGGTTGAACAGTGAGGCCCTTTGTGTTCCTCTCTTTATAAAATTGTTCCCTTGTTGTGACAAAGATCCTGACCCAAATACGAACAGCAGCCTCTCCT | 120 | 46 |
| chr7 | 144200697 | 144200817 | 772751_42001748_Shank2(210274)_211 | 1 | + | good | AAAACAGCCTTATTAGGATGGCAGCCTGGCCCAAATGTCTTACCTGGCTTTCCTCTGATCCCTAAAGCAATCCCAGGCCTTATCACCCTATTGTGTACCCTGAGAAGGAGACCAAGAGAA | 120 | 48 |
| chr7 | 144200817 | 144200937 | 772751_42001748_Shank2(210274)_212 | 1 | + | good | CCTGAGCTGCACATTTGGCTTCCCAGAGCCCATGTTTGCAGACCAAAGGAAGGTCTCCCTCCTTACTTCCATTCAACAAGATCTAGCACAGGCTACAGTGAAGGCAAGAACCCGAGGGTG | 120 | 52 |
| chr7 | 144200937 | 144201057 | 772751_42001748_Shank2(210274)_213 | 1 | + | good | TGGAAACCCCAGAGGTCTGACTCTCTCTGTAGTAACACACAGATCTTTGTTTAAAAGGTAAAGCAGTTTTGTCTTTAAAAGTCAGCATGGTAGGGCTGGAGAAACGGCTCAGCAGTAAAG | 120 | 43 |
| chr7 | 144201177 | 144201297 | 772751_42001748_Shank2(210274)_215 | 1 | + | good | TACTCAGAATGAGTATATATTCTCTATATACTCAGATGTATGCACTCACACATTAGATGTATAAATAAATCATCTTAAAAGAAAAAGAAAAAAAACCAGCATAGCAGGCTAGGTAGGCAC | 120 | 32 |
| chr7 | 144201297 | 144201417 | 772751_42001748_Shank2(210274)_216 | 1 | + | good | AGTGATATACACCTTTACTCACAGCACTCCAGAGGTAGAGGCAGGCACATCTCTAAGTTTGAGCATAGCCTGGCCACCCATAGAGATAAGTCAGTGAAACCCTGATTCACCATATAGTGA | 120 | 46 |
| chr7 | 144201417 | 144201537 | 772751_42001748_Shank2(210274)_217 | 1 | + | good | AAGCCTATCTCAAAAAAAAAAAAAAAAAGTTCAGCAACAGATCCCTCATTTGGGTGATAGAGCCCGAGGGTCTTCCTAGGATTCGAGATACCTGAGGCTCCTGGATCCTCTGTGCTGTGGA | 120 | 46 |
| chr7 | 144201537 | 144201657 | 772751_42001748_Shank2(210274)_218 | 1 | + | good | GCCCAGGCCTGGTGCTCAGAGGGGTTAGGTGTGCTTCTGGGACTCTGGCTCCTTTTTGCTTTTGCTTCTCATGGCCCCATTCAAACTTAGTGGCAATTCAAGTCACCATCTTTTAATTCT | 120 | 49 |
| chr7 | 144201657 | 144201777 | 772751_42001748_Shank2(210274)_219 | 1 | + | good | TTCAACTGGAGTCAAAAGCCGTGAGCCATGTTCTGCCTGAGGAGAGTTGTTATCAGCTTCTAAAGTAGAAAACTCTTCCATTGATCATCTTCAGGCACAGCTGGGTCCAGGTGCTCACAC | 120 | 48 |
| chr7 | 144201777 | 144201897 | 772751_42001748_Shank2(210274)_220 | 1 | + | good | TGCTATGCTCAGGGAGGGCTTTCTAACTCCAGGCCCTTATCTCTCCATGGAGGCCTCTTCTGGTCAGTTTGGCCCTCACACGTTTCCTGGCAACCACAACTCTAATGTCCTTGTAGATTC | 120 | 52 |
| chr7 | 144201897 | 144202017 | 772751_42001748_Shank2(210274)_221 | 1 | + | good | CAAGTCCAGGACAAAGACATTTTGCTAAGCTTCGCCCTGGTCCTATGATGGCCTCAATGGGACTGGTTTGGCTCATAAGGGTACTTTCTGTAGAAGGGTTAGTGGAACGAGCTGGTATAT | 120 | 48 |
| chr7 | 144202017 | 144202137 | 772751_42001748_Shank2(210274)_222 | 1 | + | good | CAGGAAGGGTGGCTGGGCTAGACAATAGAGGGGGTGCTAAGTGCAGACAGTGGGGCTTCTGTGACCAGAGGGCAGAGTCCCTGTTGCAGATGTCTGAAGAGCAGAGAGGCCGTTGCCCTG | 120 | 59 |
| chr7 | 144202137 | 144202257 | 772751_42001748_Shank2(210274)_223 | 1 | + | good | CTCTTGTAGCTAGCAGGAATGTTTTAGCAGGGATGTAAATATTTGGGGAGTGTTCTTCACCTCCCTGGGGGTGGGGCTGGTGGTATACCTTGATAACTCTTGGTTAGTCAGTCACTGCAT | 120 | 48 |
| chr7 | 144202257 | 144202377 | 772751_42001748_Shank2(210274)_224 | 1 | + | good | GAGTAAACCCTCCATAATCTCAAAGTGGCCGGGTGGCCTATTCAAAAATAGAACGTGTCTGTTATCTTTAGAGCCGTTAAAAATACAAGCAAATGACTCTATTAGCACAATTGCAATCTA | 120 | 38 |
| chr7 | 144202377 | 144202497 | 772751_42001748_Shank2(210274)_225 | 1 | + | good | AAATATTAATAAGCAACCCCAGCCAGTTTCCAAAGGAGCCACTAACTGTTCCCTAATGGGACATAAAATAAATATGAGGCCTTGCCGTTCTGCCTGCTAACAACTGTGTGTTTGTTAAAC | 120 | 41 |
| chr7 | 144202497 | 144202617 | 772751_42001748_Shank2(210274)_226 | 1 | + | good | AACCTGGCACACACCCACACCTACTCGTACACACACATACAACCCCACCCTATCTTTTACACCATTTTATTAAAATATACTCAGAGCCACTTTGGGGAAGGGGGTTGGGACTGCTGTTTC | 120 | 47 |
| chr7 | 144202617 | 144202737 | 772751_42001748_Shank2(210274)_227 | 1 | + | good | TTAGCAGCTGCTCAGGGTTTTAAATACCAGTTTCCATGCCGGGGGTTTGCCCCCAAAATAGAGTTGTGCCCTCTCTGTGTGATGTTTTGGGTCACGTTGGAGGATTTAGTTGGAGGATTT | 120 | 48 |
| chr7 | 144202737 | 144202857 | 772751_42001748_Shank2(210274)_228 | 1 | + | good | GGCTTTCCTGTGAGATCCAGGCCTTCCTTGCCTCCTCCCAGAAGCAGTGGTGGACTGCTCTGGGTTACCATGACAACCTGGAGATACGGGGTACCAGCCCTGCCCTTTTTCCCTGTAGCC | 120 | 58 |
| chr7 | 144202857 | 144202977 | 772751_42001748_Shank2(210274)_229 | 1 | + | good | CTTCCTGCCTGCCAGGAAAGTCACCTTGAGTTCAGTCTCCTTGCACAGGCTTCTGAGGGAGAGGTACCAGTGTGGCCCAGATGCTCATAGGCAGGGAACATCTTCAGTGTTTCACCGGAC | 120 | 55 |
| chr7 | 144202977 | 144203097 | 772751_42001748_Shank2(210274)_230 | 1 | + | good | CACAGTAGAAAGGGGAGTAGGCTGGTGATATAGTTATGCTGTTTAGTTTGTCTATCCTGCAGCCTGGGTGTCTAACTCTTCACAGTTCTAGAGGTGGAAGTAGGGGGTCGAGGGGGGCGC | 120 | 52 |
| chr7 | 144203097 | 144203217 | 772751_42001748_Shank2(210274)_231 | 1 | + | good | ACAGGCTGGTCTAGGTTAAGGGTTTCCCTTGCTACACATGTATCCTAGTATCCTCACCTGCCCTCTGTTCCCTGTGTTCCAGCCCCACTCTCACAGGTATCTCTGACCCTCCTTAAGCAC | 120 | 52 |
| chr7 | 144203217 | 144203337 | 772751_42001748_Shank2(210274)_232 | 1 | + | good | CTAGTCTCTAAGCATAGCCTCACTAGGCTTGAGACCTTGACTGGGAGTGGTGCCGGTCAGGCACAGGATGAGGGGGCGAACTCAGAGGATGCAGAGGGTGAGTGTCTGGGTGATGGCTCA | 120 | 57 |
| chr7 | 144203337 | 144203457 | 772751_42001748_Shank2(210274)_233 | 1 | + | good | TAAGCACCTGCAGGTAAGGTGAGGTGATAGGGCTATGATGTGCCCTCCCTGACAGAAAGGACAAGGAGGGTGCAGCTGGCTTATGGCAGAACTTCCCATACCCCACATGGCCAAATAATG | 120 | 52 |
| chr7 | 144203457 | 144203577 | 772751_42001748_Shank2(210274)_234 | 1 | + | good | AGGTGCCAAGGAGTGGGAAGAGAATTGATGGGTGGATGGATAGTGGAATGGGGATTAGGTGGGTGATAAGGATAGATGGATAAATGTAGACAATGGGGAGATGAAAAGAGGTAGATGGAT | 120 | 45 |
| chr7 | 144203817 | 144203937 | 772751_42001748_Shank2(210274)_237 | 1 | + | good | TGCAGGTGGGTAGGCAGATGATTGGTTGAATGGACAAGCAGATAGGTGGGTGAGTGGGTGGGGTGGGCAGATAGGTAGGGGGTGGCTGGTTGTATACTTTGTCTCATTTGTACCTCATTC | 120 | 52 |
| chr7 | 144203937 | 144204057 | 772751_42001748_Shank2(210274)_238 | 1 | + | good | CAGTTTTGTTATCCCTCCCATCTTATACCCCTACAATGGACTCCATGAGATGTTTCAGAAGCCCTAGGATCTGTGAGCAAAGGCCAGCATGGGTGTATATGAGACCCCAGGCAAGGCTAT | 120 | 48 |
| chr7 | 144204057 | 144204177 | 772751_42001748_Shank2(210274)_239 | 1 | + | good | TAACCTCCCACTGGCTGGGCTTTGTTTAGGTACCACTTTCCATCAGGGATTTAGCACCTGCCTACCCCTCATGCCCTGCTAAATCATAGGGTCTGCAGCCTGCAGGCCTCACTGGTCCTT | 120 | 54 |
| chr7 | 144204177 | 144204297 | 772751_42001748_Shank2(210274)_240 | 1 | + | good | CTCATTACACCATGAGTGTCTGAGTGCTGTAGTGTTTCAGAAGGCAGGCAAGGTCATGCACACACAGGGAGGAATGGGCACTCGACATTTACTGGTGGCCCCTCTGCTCTGCCATTTGTC | 120 | 52 |
| chr7 | 144204297 | 144204417 | 772751_42001748_Shank2(210274)_241 | 1 | + | good | AGCAGGAGTGACAAGAGGCCTGAGGGCAGGCCCACTTTGGCACCAGTGCACCCAGCTCTGGGACAATGAGCCAGACCGGGAGTCCCCTGGAGCTGGGGCAAAAATGGATGCCTCATGCCA | 120 | 62 |
| chr7 | 144204417 | 144204537 | 772751_42001748_Shank2(210274)_242 | 1 | + | good | CCCCCAGAGCTGCATATCTTTCTGGAAAATGAAGGCTGTCATATGGCGCTCTGCTAGTGAATGGAGATGTTGTTTTTGGCTTCTCTGAAATAGACGATGGGAAACAGACCCAGAGATTAT | 120 | 45 |
| chr7 | 144204537 | 144204657 | 772751_42001748_Shank2(210274)_243 | 1 | + | good | GAATAAACATTGGCTTATGGGGGCTGCTCACTGCTAATGGCAAAACCTCCACACTTACAGATTAAATTTAAGACATGCATATGGAAAGGGACTCTGGGGAGGCCAGACTCCTGCTTGAGC | 120 | 47 |
| chr7 | 144204657 | 144204777 | 772751_42001748_Shank2(210274)_244 | 1 | + | good | AAATCTTACAGCCACTGTGGGAGGGTGCACTGTGTGGCCCCTGTCCCTAGCCAGGGTCAGGAGCACAGGCACGCTAGAGTCTAGGAGGAGCCCGTGTTCTCCAGGGGATCCAGGAAGATG | 120 | 60 |
| chr7 | 144204777 | 144204897 | 772751_42001748_Shank2(210274)_245 | 1 | + | good | CTGTAAGGGGCAGGGGGCAGGGGGGCAGGGGCCACCTTGGGACTGTTCACTGGCATGGACTAGACTAAGGTCAGTGTGTGACTTCAGCCAACGTGCCTTCAGGAATCTTTAGGGCAGTTA | 120 | 57 |
| chr7 | 144204897 | 144205017 | 772751_42001748_Shank2(210274)_246 | 1 | + | good | ATAAAATCAAGTCGCAGAATTAACTCTTGGTGGTCAGCTGCAGCCTGTCAGATGACCCCTGAAGAGCTCCGAGGGACATCTGGGGGTGGCTCGCCGGTGTTTTGTTAAACTGTAATATAA | 120 | 48 |
| chr7 | 144205257 | 144205377 | 772751_42001748_Shank2(210274)_249 | 1 | + | good | AAACCTCAGTGTATTGAATTCCAATATGCTCTAGTGCCTTAACTATAAAGCTGGCACCCAAAACAGTCTCTTGCTTCCCTCTCTCATCAGCCTTTGGCAGCCACCAACTCTCCTGCTGGC | 120 | 48 |
| chr7 | 144205377 | 144205497 | 772751_42001748_Shank2(210274)_250 | 1 | + | good | TCCGGGCAGTTCATGTCATGGAATTCATCAGAAGATAGTCTTTCACAAGTGACTATTTTAATTTAGCAAAAGGCTTATCATTCTTTTTTTGTGGCTAATTCCTTTGCATTTGCATATACA | 120 | 34 |
| chr7 | 144205497 | 144205617 | 772751_42001748_Shank2(210274)_251 | 1 | + | good | TGGCACATTTTATCAGTCTCTGGATGGGCACTTAAGTTGTTCCCACCCTTTGGCTGTTTGTGACTCATTTTTGGCTGTGAACATTCAGGTATAGGTTTTGGTTAGTCTATTCTGTTCTCT | 120 | 42 |
| chr7 | 144205977 | 144206097 | 772751_42001748_Shank2(210274)_255 | 1 | + | good | CTTACCTACTGAGCTATCTCATCAGCACTGATTAACATCTGAGGGCCTGCCAGGCTGTGGTTGGCTGTGACTGGCATTGGTGAGTAGCTTTGCTCTGTGGCTCTGTGAAAGTCACACACA | 120 | 51 |

| | | | | | | | | | |
|---|---|---|---|---|---|---|---|---|---|
| chr7 | 144206097 | 144206217 | 772751_42001748_Shank2(210274)_256 | 1 | + | good | CCCCCGTGCTGGAAAAAGTTCTTCGAACTGTAGGCACAATGCTACTTTGCTGAAGCAAAGTACAATATTATGTGGGCAGGTACCTAGGTTGGATGTAGCCTCTCTCTCTCTCTCTCTCTC | 120 | 48 |
| chr7 | 144206337 | 144206457 | 772751_42001748_Shank2(210274)_258 | 1 | + | good | TGGGTGCTGAAGCACAGTCGTGGGAGACAGAATATACCTGTTCTTGAGCCAGGAAAGCTTAGCTTCCCAGCCATCCCACACGCCACCTGCTTCATGCTTATTTGATGCTGTTGAAATTCC | 120 | 50 |
| chr7 | 144206457 | 144206577 | 772751_42001748_Shank2(210274)_259 | 1 | + | good | TACTGTTTTGGACAAAGGGCCCTGCTGTCTCCCTGCGCCAGGTCTCACAAACTTGTCATCAGGTTTTGCTGGCCCCAAGGTCCCTCCATGGATCCCTAGAGATCTTCACCAGTAGACCTC | 120 | 54 |
| chr7 | 144206577 | 144206697 | 772751_42001748_Shank2(210274)_260 | 1 | + | good | TGGGGAGGAACACACAGATTAAAGGTCATCTGTTCTAAGTGCCTCTAGAGCTGGCTACGACGCTGGTGGGAAAGGCAGGCGTGGACCGTGGAGGTGCCTGGCTGCTGGGGTATGTGTCCA | 120 | 57 |
| chr7 | 144206697 | 144206817 | 772751_42001748_Shank2(210274)_261 | 1 | + | good | GCTCATGCACTTAGTGAGTTGCTAATCTACTGTGGGGGAAAAGCAGCTTCATCCACTCTCAGCCAGAGCTGAGGTTGTGCACCAAGGCCTCAGCACTTGTAACAGCCCCATCCTTCAGGT | 120 | 52 |
| chr7 | 144206937 | 144207057 | 772751_42001748_Shank2(210274)_263 | 1 | + | good | GCCACTCAGTCTGGTGTAGCTGCTCCACACATGCTATTGACTTCCACAACTTAGTAGATTCCCTTGTCATCCGCACTTAGTGGGCCAAGATGTAAGTGTGCTCTTTGTTTGACTACAGCA | 120 | 48 |
| chr7 | 144207057 | 144207177 | 772751_42001748_Shank2(210274)_264 | 1 | + | good | GGTTCTGACACTTCATCTTACTGGAGGACTCCCAAGGCAAAGGAAGAAGAGAGGGCTGAGCAGCATAGACTAGAACTCTCTCCCTGAGGGTCACACACCAGACTCTCAGCTAGAGGACCT | 120 | 52 |
| chr7 | 144207177 | 144207297 | 772751_42001748_Shank2(210274)_265 | 1 | + | good | TTCCTGAGGGTCCAGATCCTCTGTCCAGCTGCTAGGAACCGTCCAGAAGCTCCATTGGCACACATGTTCTTCCACAGCTGGGTGTAAGTAGGCAGTGAGAGAGCTCATGGCAGCTTCGTG | 120 | 55 |
| chr7 | 144207297 | 144207417 | 772751_42001748_Shank2(210274)_266 | 1 | + | good | CGCCGGGCAAGCCAGCTGGCACCTCCCCATGAAATGCAGCACTGTAACCCTCACAGAACACCAGATGAGAGATGCCCGTGTACACATTCATGGCAGGGTGCAGAGGCATGCCATAGTGTG | 120 | 57 |
| chr7 | 144207417 | 144207537 | 772751_42001748_Shank2(210274)_267 | 1 | + | good | TGCATACACATGAGCACACGTGCATGTGACATACATGTTCATGGGGGCATGCATGCATAGACACATATCTGCACATGAGTGTGTACATGACTTATAACTGCAATATGTTCAGGTGTCCTC | 120 | 44 |
| chr7 | 144207537 | 144207657 | 772751_42001748_Shank2(210274)_268 | 1 | + | good | ACTGTAGTCAGACCTACAATAAGTATATGCATGATATCACTTATAACTGCATGTATGTACATTCGTAGTACTTAGATTTACATGAATTTGTGTGCACACACACAGTGCAGCTACACATAT | 120 | 35 |
| chr7 | 144207657 | 144207777 | 772751_42001748_Shank2(210274)_269 | 1 | + | good | GTTCATATACTTGGTGCTTGCACCTGTACATGTGTGCACACAATTTTCACCTATACACGCGTGTGCATGTCCTTTATATCTGGACACATGTGCATGTGTGTGCGTTCTGTTTTCTGCGCC | 120 | 46 |
| chr7 | 144207777 | 144207897 | 772751_42001748_Shank2(210274)_270 | 1 | + | good | ATTACCGAAGCCCGAGGCATGCTCCATGCAGGCTCACTGTGTCTGCCCAGCAGTCAGGTGGATACACGACTCCTGTACTATTTGGGACAGAGCTGTCCTTGAGTTCTGGCACAGATACAA | 120 | 53 |
| chr7 | 144207897 | 144208017 | 772751_42001748_Shank2(210274)_271 | 1 | + | good | AGCCCTGGTGACGACCTGACCCAGTTCCCATTCCTGAAACTCCGGCTTCCCTCGTGGACAGTCCTAGAAGGAGTCTGATTGCCAGGGTGACTGCCTCATTTGTCATCCAAACGGGGCCGC | 120 | 58 |
| chr7 | 144208017 | 144208137 | 772751_42001748_Shank2(210274)_272 | 1 | + | good | TTCCAAGAGCAAAGGAGAGCAGCAGAGCTGTGGCCGCTGAGGGAGTCGGCTGACCCACTTTCTCCAGGAGCTGAGGTTCCCTGGCCTGATGCAGGCAGTGCTCCCCCTCCAGGCAGAAAA | 120 | 61 |
| chr7 | 144208137 | 144208257 | 772751_42001748_Shank2(210274)_273 | 1 | + | good | GGCATGTAGTCACTTACTCTCCTACCCCCTACTGGCCAACAGAGGCTGCCTGATGCCTGGAGCTGGATTCCAGGCAGGAAGTCTGACATTGGGTGACATTTGGGGAAAGGCCATATTTTC | 120 | 53 |
| chr7 | 144208257 | 144208377 | 772751_42001748_Shank2(210274)_274 | 1 | + | good | CTGTGAATCTGGGCCTGCTGAGGAGGCTGGTACAGGCTGATGATAAGTGGCCATCACCTAGGAAGGGTCTGTAAGGGAGATATAGAGATATAGGGGTGCCCTCTACCTGCAGCAGAATGC | 120 | 53 |
| chr7 | 144208377 | 144208497 | 772751_42001748_Shank2(210274)_275 | 1 | + | good | TCACCCTATCCCAGAGGGTACCATGACACCAAGGGCAGATGCAGGGGATGAAGGGAGAGTTACCTCTCACACAGGGGCTTAAGCAAATACGAATACTCAGGACTCCCTTCCACCCATCTG | 120 | 53 |
| chr7 | 144208617 | 144208737 | 772751_42001748_Shank2(210274)_277 | 1 | + | good | CTGCCCCCTCCCACCCTGCCCTCTTTTCTTTTCCACACTCCTTGCTGCCCCAGGCTGTCCCCACCACTTTGTTTCTCCTACAAGTGAGACCTGCAGGGAAACCAGCTTACAAATGTTCAC | 120 | 55 |
| chr7 | 144208737 | 144208857 | 772751_42001748_Shank2(210274)_278 | 1 | + | good | TATCTCTTTCCACCACATGGGGCCCAGGGAGAGAACTCCAGTGGTCAAGCTTAGAGGCAAGTGCCTTTGTACCAGCTGAACTAGCTCACTGGGCCCAGTTGGAAACCATCTCCCTTGCTT | 120 | 53 |
| chr7 | 144208857 | 144208977 | 772751_42001748_Shank2(210274)_279 | 1 | + | good | TGGAAGACAATCAGAATGCAGGCTGCAAATTAAGATGCCAAACAAAGGCTGCAGGGACAGCCTGCCTCCTGAAGTTCAAGGCCCCCTTCTGTCTGAACAAGGAGCAAAGCATCAGGACAT | 120 | 50 |
| chr7 | 144209097 | 144209217 | 772751_42001748_Shank2(210274)_281 | 1 | + | good | GGGGAAGGATAGAGGTCCTTCCCCTCCCCTGCAGGCTCAGCACCCCTCCAGCCCTTCCCCTTTTCTTCACAAGAGGCCTCCTTGACTTTTATGGCTTCTTTGGGCCCCTCTTCTTTTGTC | 120 | 57 |
| chr7 | 144209217 | 144209337 | 772751_42001748_Shank2(210274)_282 | 1 | + | good | TCTCAGGAGGTAGTGGCACTTTATCTACCTTGATCATGACCTCTCACTCACTGCCTCAGAGCCCTGTGGGTATCCAGGCTCCATGAACATCTATCCTGTATGTGAGGTGGGACTCCTGTC | 120 | 52 |
| chr7 | 144209337 | 144209457 | 772751_42001748_Shank2(210274)_283 | 1 | + | good | CACACACCAGGAGCTATCTGCATTAGGCTGCTGGCTCTGGGACAGCTCAGACCCCCATGGTCAGGCTTTTGTTACTCAGCACCATGTCTGCTGAGCCCCTTCAGGTGTGAAACTCCTTCC | 120 | 56 |
| chr7 | 144209457 | 144209577 | 772751_42001748_Shank2(210274)_284 | 1 | + | good | TAAGGTCAGGACCCTCCCTCCATGCATGTCTGAGGACTCATCTGCCCAGCAGCAGAGGAGGACTTAGATTGCCTGCCTTTAATGTTTCAAGGTATATAATTTTATGGGAATAGAATTGCC | 120 | 46 |
| chr7 | 144209697 | 144209817 | 772751_42001748_Shank2(210274)_286 | 1 | + | good | GTCAACTGGTCAACCCCAGCTATTTCTCCTTAGGACATGTTCCACATTGTTATCTGAGACAGGGTCTTTCTCTCTGGCCTAAAACTCATCAAGTAATAGGCTAGACCCAGTAAGCCCCAG | 120 | 47 |
| chr7 | 144209817 | 144209937 | 772751_42001748_Shank2(210274)_287 | 1 | + | good | GAATACACCTATCTCTTCCTCTTTGGGTCTAGGCTTGCAGCTATACTATATATATAACCCAGGCTGCCTTAGAACTTACTATGCAGACCAGGCTGACATCTGCCTGTTCTTGCTTCCTGA | 120 | 45 |
| chr7 | 144209937 | 144210057 | 772751_42001748_Shank2(210274)_288 | 1 | + | good | GTTCTTTGGATTAAAGGCATGTGGCAATATGTCCAGCAACCTTTTTAAAAAATTAATTAATTAATTTACATGTTTGTTTATTTATTTGTTTGTGCCAATGCTCTTCCCATTGGGCTGCCC | 120 | 33 |
| chr7 | 144210297 | 144210417 | 772751_42001748_Shank2(210274)_291 | 1 | + | good | ATCCATCTCTCTAGTCATGCATTGACCCTTTTTCAAAAGGGTCCTTGGACTTAAAGTCGGGTCCTCTTGTTGGTGCAACAAACACATTAACAAATGAACTATCTCCCCAACCCCAAAGTT | 120 | 42 |
| chr7 | 144210417 | 144210537 | 772751_42001748_Shank2(210274)_292 | 1 | + | good | CTTGTATCTTCCTACAAAGGAAGCCACCTATGGCTGCTGTCTCTTGGGCAGAGCTTTCAGGCCTGGCCTGGCACTTATCTTCATATCCCTGGCATAAAGGCATAGCCACATCTGTCTGCT | 120 | 51 |
| chr7 | 144210537 | 144210657 | 772751_42001748_Shank2(210274)_293 | 1 | + | good | TGTCCTCATCTGTGCGCTCCCCAGGGCAGGACTTACCTGGCATCATCCCATGTCCCCACCTGGCACACAGTAGGCCCACTCTAGACTCCTGCCATGGAAACCAGAAGAATGACAGCGTGT | 120 | 57 |
| chr7 | 144210657 | 144210777 | 772751_42001748_Shank2(210274)_294 | 1 | + | good | CTTAGAGACAGCTAGTTATTTTCTAAGGAACGTGACACAGATTTAGAATTCAGTGTGGCTAAGAATCAGCCGGCTTCTGCCGCTGAAAACTTGCTTGTCGTTTTGTCTGCCAAGAATCCC | 120 | 44 |
| chr7 | 144210777 | 144210897 | 772751_42001748_Shank2(210274)_295 | 1 | + | good | AGAAGCTTCCCCAAGCGATCACTGATGGGAGCAGAAGCGTTGCCACTAAAATTAGGTGATGAAACAGTTGTTATGAAACACCAAAGAAACTGACAGGTACATTTAGTGAGCGGGGAACAC | 120 | 45 |
| chr7 | 144210897 | 144211017 | 772751_42001748_Shank2(210274)_296 | 1 | + | good | TCCTCCATGACATGGTGCTGTGGTCAGAGCTGTGTCCATGTGCTGGGGGCTGAGTTTAGTTGGCCCGGCTCTGTGGTTGCTTAAAGTAGAAACTACTTGATATTTGGGAGAATAGTGCTT | 120 | 49 |
| chr7 | 144211017 | 144211137 | 772751_42001748_Shank2(210274)_297 | 1 | + | good | TAGACCTCAAGATGCTTTAAAACTGATGTAAACCCCAAGTCTCCCTCCCTGTACAGGCCTGCTCTTCTCTTCTGCCCTAGCCCTTGCCCTCTGCTAAAGTACCTGTGACTTGCTGGCTGT | 120 | 51 |
| chr7 | 144211137 | 144211257 | 772751_42001748_Shank2(210274)_298 | 1 | + | good | GTATCCACAACGTCCATACTATATTGGACTGGCTTTACTCACAGCAGAAAAGCACCAAGGAGCTGTGGTTTGAATGAGATAGAGTTTGTCTTTTCTCTGTTCCAGAAGGAGACAGCCCAG | 120 | 45 |
| chr7 | 144211257 | 144211377 | 772751_42001748_Shank2(210274)_299 | 1 | + | good | AGATGGAACAGAGATGCTAAACTTACAGGGACCCAGTGTCTTCTTTTCTTGCTGTTCTGAAATCCTTAATAAGCTGCCCCATGGTTCAAGCTGGGTGCACAGATTCTAGCCATCACATTT | 120 | 44 |
| chr7 | 144211497 | 144211617 | 772751_42001748_Shank2(210274)_301 | 1 | + | good | AAAAAGTCCTATTAATGGCATGTCTAGCTATGTGATGTCAGCTGTAATATGATGTTATATGGGGTTTTGAGCCACTACTGGCAGCTGAATAGGAGACCAGATAGTAGAGAGAGCTTTAGT | 120 | 40 |
| chr7 | 144211617 | 144211737 | 772751_42001748_Shank2(210274)_302 | 1 | + | good | ACCTCTGACCAAACCCCTCTTCCCTTTTCCTCAGTGTTCCCTCCAGCCACATCTCATCCTCTCACACTTCAGAGCAGGTCAGGACATTTGGAGTAAGATATCACTATGGGATAAAGCAGT | 120 | 48 |
| chr7 | 144211737 | 144211857 | 772751_42001748_Shank2(210274)_303 | 1 | + | good | GTGATCCTTCAGGTGAGAACACAGCTGACTTCGCATCAGACAGCACTCAAGGTGGAGTCTGTCTGTAAGGGAGGATCAAATGGCCACCTGGGGGCCATAAGCACAGGTTCCAGATGCCTG | 120 | 54 |
| chr7 | 144211857 | 144211977 | 772751_42001748_Shank2(210274)_304 | 1 | + | good | CCCTTGCCTCAGTGGGACCCTGCCTGTCAGAGCGCACGTGTGCTTTAGTGAGTCCTCCTACTTAGAGCTGACCAGCAAGCAGCCCACTCAAGCTGAAGCTGCAGAGGAATTGAAGATCAA | 120 | 55 |
| chr7 | 144211977 | 144212097 | 772751_42001748_Shank2(210274)_305 | 1 | + | good | TAGCCTCGGGCATGGCTAGATCTAGGAGGACAACATTGTGTGGAGTCCTTTGGCAGTTGTGAGTGGCATAGAGCATCCTCTCTGGTCTCTGTGAGAGACAGCACAGGATTGGGTCTCATT | 120 | 52 |
| chr7 | 144212097 | 144212217 | 772751_42001748_Shank2(210274)_306 | 1 | + | good | GTGGGTTGGCTGATCTGATGAATCTTGACTCACTTGGTCCAATTGGCTAAGAGAGAAGGTGGGGCTGGCAGTGGTAATTCAGGAATGTGGGCTGAGTTTAGGACAGGGGTATCTCCTCCA | 120 | 51 |
| chr7 | 144212217 | 144212337 | 772751_42001748_Shank2(210274)_307 | 1 | + | good | GAAAGTAGTAAATGGGCTCTGGCAGGCCAAACTCATACTCCCTCTCTACCTACCCCTACACCTCAAACCCCCAGTCCCTGTCCTGATCCTACTCCTCTGCAGCAGGACAGAGCTGTCCCA | 120 | 55 |
| chr7 | 144212337 | 144212457 | 772751_42001748_Shank2(210274)_308 | 1 | + | good | GCCTTGGCCCCTAGATGCCCCCATTCAGTGGTTTGTCTTGTCTTTACCACTCTTCACTCTCTGGTGCCTGAGACAACCCCCAGAGCCGAGCTATGTAAACCTTAGTGAGGTCTCAGCACC | 120 | 54 |
| chr7 | 144212457 | 144212577 | 772751_42001748_Shank2(210274)_309 | 1 | + | good | TGCCAGCAAGGAGCATGGCATGGCACATTCTGAGGAAGTCACATGATGCTTGGGACAAAGATAGGGAAGAAGAGGCAAAGGGGAGGGGGAAATTGATGGATGAATGGTTGAGGGTAGGTA | 120 | 50 |
| chr7 | 144212697 | 144212817 | 772751_42001748_Shank2(210274)_311 | 1 | + | good | GTTGCCAACACAGGAAGAGTCACTGTGTCCTGCAGCTCTGGTGCAACACTAGCAGTTTAGTCTAGTGACATTTCATGTCAGTGCTGACAGATATGCTCTTACCCTACTGTAAAGTAGTAG | 120 | 46 |
| chr7 | 144212817 | 144212937 | 772751_42001748_Shank2(210274)_312 | 1 | + | good | TAGGCCCAGGTAGGGAAGGAGGTGGGAGGAGCTGAGGGGCTGATGGGAAGGCTCCAGGCAGCCCCAGGGTTGTGTGCAGGCTGTGTACTTTGGGCTTATGTTCTGACCACCAGCCATGAC | 120 | 61 |
| chr7 | 144212937 | 144213057 | 772751_42001748_Shank2(210274)_313 | 1 | + | good | TTTGGAATGACATGCTCTGCTGCCTGCATACATTCAGAGCATTTCAAACAAAGCCACCACCTGTTCCTGACTTCCCCTCCTCTTTCTTCTGCCTCTCCTCTTCTTCCTTCTCCTTCTCTG | 120 | 48 |
| chr7 | 144213057 | 144213177 | 772751_42001748_Shank2(210274)_314 | 1 | + | good | CCCCCTTGACCTCCACTTCCCTTCCTCCTACAGACATTGAGGGGTTCATAGCTGCTGAGGAGGGCAGGTGGCTGTGGTCCCTGCTGAGGTTTGCTGTGTCCCCTTGATGTTCATGTGTAG | 120 | 57 |
| chr7 | 144213177 | 144213297 | 772751_42001748_Shank2(210274)_315 | 1 | + | good | AAAATCTTAGCCCGAGGTAACAAGCCTGTGGCATTTGGATGTGGAGCCTCTGCTAGGTAGTGAAATCAGATGAGGTCACAAGGAAGACCTTGTGGTGGCTTTTGAAAAGACCCTGGCATA | 120 | 48 |
| chr7 | 144213297 | 144213417 | 772751_42001748_Shank2(210274)_316 | 1 | + | good | CTTGTTCTGTTCCCATGTGGTGCCCTTGGCTGTGTGATGACCTGGCAAGAGCCCCTTGCCAGGTGCCAGACAGATGCCAGTCAGTTTCCAGACTTTCAACCTCCAGACCTAAGGTCCATA | 120 | 54 |
| chr7 | 144213417 | 144213537 | 772751_42001748_Shank2(210274)_317 | 1 | + | good | TAAACCTCTGTGCTTTATAGACTCGTGAGGCCCAGATAATGTGTCATAGCAATAGAAAGTAGACTCAGACGCTTCCTGTGCTCTTTCGTGGGGAGCGTGGCACCTGTTCTGAAGAACACA | 120 | 48 |
| chr7 | 144213537 | 144213657 | 772751_42001748_Shank2(210274)_318 | 1 | + | good | GTGTTTCCCCTTGCTGGGCCCTGGCTTTGTCCCACAGTGACTGGATTTAGGAGGAAGAACACCCCTGCATAACCCGGACATCATGTTAAGCTCTGCCCCACAATACTTAGCCCAAGAATC | 120 | 52 |
| chr7 | 144213657 | 144213777 | 772751_42001748_Shank2(210274)_319 | 1 | + | good | TGCCATGGAAATTGCCACTGTCTGGAGACGCAAAGCAGATCTCTCTCTGCCCATTGCTCAGCTTCCTCGGGGGTCCCTCCATCTTGTGATTTACATCCTGATCCCAATGATGCTTGGAGA | 120 | 52 |
| chr7 | 144213777 | 144213897 | 772751_42001748_Shank2(210274)_320 | 1 | + | good | TATCAGGAGGAAAGATACTAACCAAGCTTGCTGGATGCAGTGGTGGCTCTCCAGGAGCGTTCTAGACCTAGGAACTCAGAAGAACTACAGACCCTAGGTCCCACCTAGTTGGGGGACTCA | 120 | 52 |
| chr7 | 144213897 | 144214017 | 772751_42001748_Shank2(210274)_321 | 1 | + | good | GATAGTCTTCTCAGGGCTTCCTGAGGCCAAGAGCTACCTGTAAAGGTGGACCATGGCAGATATGCTCAAAGGCCCCCTTAGCACACACAGTGTCCAGGCTCCAGCTTTCTTTTCAGGCTA | 120 | 52 |
| chr7 | 144214017 | 144214137 | 772751_42001748_Shank2(210274)_322 | 1 | + | good | TGGAGCCAGTGCTATGGGGCCTAGGTAGGCAGGGAGTGGTGCTGATGTTACCAGAGTGCACAGGCTCTGCCTCCACCCATCTGCCCACATGGAGAGGAATAGCCTCTCAGCCCAGCTGCT | 120 | 60 |
| chr7 | 144214137 | 144214257 | 772751_42001748_Shank2(210274)_323 | 1 | + | good | GTGACAAAGGCTGTAAGAACCAGATTGACAGAATACTGCTCACCAATCTGGAGACTGGATGATGGGAGGCTGAGCCTGGGAAGGGCTCACCCTCTGGGTCACAGATGGCCGCCTGCTCAC | 120 | 56 |
| chr7 | 144214617 | 144214737 | 772751_42001748_Shank2(210274)_327 | 1 | + | good | CATTCTTATTCAAACCATCATACAGACAAAAGGCAGGGAGACTGGCTACTGGTATTGTCCTGATTGGAGGGAAAGCACTTGACCATGGAGAACACAGCCTTGGCCACTTATTTATTTAAA | 120 | 42 |
| chr7 | 144214857 | 144214977 | 772751_42001748_Shank2(210274)_329 | 1 | + | good | CTCCTGGTACTGTTTTCCTCCGGAGACTTCTACCCTTTTCTATGAAACTGAGTCTCACTGTATAGCCCAGGTTGTTTTCAAACTTGAGATCTCCCAGCCTCAGCCTCCTAAGTGCTGGGA | 120 | 48 |
| chr7 | 144214977 | 144215097 | 772751_42001748_Shank2(210274)_330 | 1 | + | good | TGCAGACAGCACCTCACCTGGCTGGTGATATTTTTCATAAAGACACTCCCATTCTCATGAATTGCCCACTTTCATGGGCTGTCTAACCCAAGTCTTCCCCCAAAGGCTGGCCCACAAGGG | 120 | 51 |
| chr7 | 144215097 | 144215217 | 772751_42001748_Shank2(210274)_331 | 1 | + | good | CCAGGGTTTCAGTGTTTAAAGACCAGGGAACAGTTCAGGCCACACCCAGTAGCTTCATAAAAATTTCAAAAATACCTAACCCCTTGCCTGTTAACCACCTCACCTACCTGACACCTCAGC | 120 | 47 |
| chr7 | 144215217 | 144215337 | 772751_42001748_Shank2(210274)_332 | 1 | + | good | ATGTGAGAAGCACACCTCTGTCTCTGCCCCACACTTTGAATCCTATGCATTCAGCTTCCAGGGCCTGGTTGGCTCCTTCCCACCTGCAGGCCATAGGTATGATGATATAGACAGAAGTCC | 120 | 52 |
| chr7 | 144215337 | 144215457 | 772751_42001748_Shank2(210274)_333 | 1 | + | good | TCCAGACTATGGGCACCTCTGCCCAGCTCTCTGAGACAGATGCACAGTACCCTGGAGAACAGACCACTGGACTAAGTACCATGACTGCTCTCAGTGTTGACCCCTATAATGATCACAGGC | 120 | 52 |

| | | | | | | | | | |
|---|---|---|---|---|---|---|---|---|---|
| chr7 | 144215577 | 144215697 | 772751_42001748_Shank2(210274)_335 | 1 | + | good | ATTTCCCCATTAGAAAAAGAGGGTCAGGCATGGCTGTATACACCCATAATCCAGTCATTGGGAGACCAAGTTAGAAATGTACCTTGGCTACATAACAAATTCCTGTCTCGAAAGCAGGGG | 120 | 44 |
| chr7 | 144215697 | 144215817 | 772751_42001748_Shank2(210274)_336 | 1 | + | good | TAGAGGCTGGCATGTCCATGCCCAACACCACAAAAGACAGACATTGTTGCAGATCTAACTGCTGCCCCTAAACCTCAATAAGTACCAAGTATCTTTGCAGGAAGGTAGGGTCAGTGAGAT | 120 | 47 |
| chr7 | 144215937 | 144216057 | 772751_42001748_Shank2(210274)_338 | 1 | + | good | TGACAATACACCCACATACACACCAAATAAAATGTGATTTAAAAAAAATTAAAAAGAAATAAAAACTGGGTGTCATGGATATGTGTAAAGCCAGCTACCTAGAAGCTAGGAAAGCCCACA | 120 | 34 |
| chr7 | 144216057 | 144216177 | 772751_42001748_Shank2(210274)_339 | 1 | + | good | GGAACTGAAGCTGCCGGTCAGGGTTAGCAGACTGTGTCAAGGGAGAGGGACAGAGAAGAGATAAGCAAAGACACAGAGGTACATGGGAGTGGAGGCAGAGACTGGCATGGCTGGCCACCA | 120 | 56 |
| chr7 | 144216177 | 144216297 | 772751_42001748_Shank2(210274)_340 | 1 | + | good | GCCAGGGGACAGCAAGGGCTATTGGCAGCCCCCAGAAGCTGTGACAAAGCTGGAGCAGACTCTTTAGAATGTATAGAAGGGACGGGCCTGGCCATGCCCGAGTTTGGTGTCTAGTCTGGA | 120 | 57 |
| chr7 | 144216297 | 144216417 | 772751_42001748_Shank2(210274)_341 | 1 | + | good | ACATGAGATACTACATGTGAGGTGCTCTGTCTGTGGCAATCTGTTTCCATAACTTTAGGAAAGCCACTTAACCCACTGAGGTCGTAGGAGTCCCCAACACTCACCATACATCCCTAGGTT | 120 | 47 |
| chr7 | 144216657 | 144216777 | 772751_42001748_Shank2(210274)_344 | 1 | + | good | TAGACTCCAGGAAGCAAGGTGAGGGACCATGCCTGTGGTCTACAGTGTTAGAGCAGAGCCTGTGGGGGCTGCTGCAGCAGGGCTAGGTACCAGCTACTCAGAAAGAACATTTCAGCCCCT | 120 | 56 |
| chr7 | 144216777 | 144216897 | 772751_42001748_Shank2(210274)_345 | 1 | + | good | GAGCACAGGCCCAGGGCTCTCCAAGGCCCTCTTTGTTAGAGGGGCAGCCTTGAGAATATGGAATCCCCACTCTATAGTCTGAGTGACTATAACAACAGCCAAAGTTGGTGTGCTGAGCAC | 120 | 52 |
| chr7 | 144216897 | 144217017 | 772751_42001748_Shank2(210274)_346 | 1 | + | good | CTCATCTCCTGTAGCCATTTTAAGAGAGGCAGAATAGGAGCCACTTGATCCTTGAGGGCAGGTGAGGACCATAAACACTGTGCAAGAGGATTCCCACCCTGTCTCACTGGCATTGACTTC | 120 | 50 |
| chr7 | 144217017 | 144217137 | 772751_42001748_Shank2(210274)_347 | 1 | + | good | CACATTACATCACTACAGTGGGTTGGCACCTGGTTATGTTGGCCCCATGGAGCCTGCCCAGACACAGGGACCCAAGGATTAGTTGAAGGTAGCCAGGCCATTGGCCCATTTGGCTGGAGG | 120 | 56 |
| chr7 | 144217137 | 144217257 | 772751_42001748_Shank2(210274)_348 | 1 | + | good | TGAGATTTGAGTGGAACAGAGGCCTCACTGCCCTTCCTGTAAATATAATAATAATCCTTAACCCTGATAGCCAGAATTCATCTTTATGTGCCCAACAATGGCGGTAAAGCTCAGAGAAGG | 120 | 42 |
| chr7 | 144217257 | 144217377 | 772751_42001748_Shank2(210274)_349 | 1 | + | good | TGTGACTTCCCAAGGTTACACAGACACTTCCTCCCTCCTTAGGAGGTTGCCAAACTTTAATCCCTGTCCTTAAGGGATCCTACTTGGGAGCCAGCTCCCTGATGCATGTCCCCCTTCTCG | 120 | 52 |
| chr7 | 144217377 | 144217497 | 772751_42001748_Shank2(210274)_350 | 1 | + | good | CCACTAGATGGCAGGCTCTGGGTGCAGAAACCAAGAAGAGATGGTGGCCAGCCTGACTTCTGAGGCGCATCAGGGACCAGTCTAGGCTAAGAAGCCCAGTGGAAAGACATTTAGGGAGAA | 120 | 54 |
| chr7 | 144217497 | 144217617 | 772751_42001748_Shank2(210274)_351 | 1 | + | good | GATTCTAGCCTTGTTGGCTGCACTCAGACCTGTGATTATGCTGCTGCCTATGTAGACATAGACTGGGCTGGGGGAACACCATGAAAGACTCTGACTTGTTTCAGGGTGTGTCAGAGATGC | 120 | 50 |
| chr7 | 144217617 | 144217737 | 772751_42001748_Shank2(210274)_352 | 1 | + | good | TGAGAGGGCTTTCATCTCTTCTGGATGTTTCTGCAATATGGTGTATGTCTACAAAGCCCTTCCAGCCAGCATCCTCCCTGGAGCAACCTGGCTGGGGCCCAGCAGTACTGAGGGCCCGCC | 120 | 57 |
| chr7 | 144217737 | 144217857 | 772751_42001748_Shank2(210274)_353 | 1 | + | good | AACCTGCATGGGGAGCTGGACTCCTTTACCCAAATTCAAACCCATCTTCAACCTCAACCAGTGTGTGACCTCTCACAAGGACACAAGTCCTCTCAGTGCCTCATTTCTCCCTCCCCCAGA | 120 | 52 |
| chr7 | 144217857 | 144217977 | 772751_42001748_Shank2(210274)_354 | 1 | + | good | ACCATCAGGTTGACTTGTTAGCATCAACTGAGCAGGCAGAAGAGGTCACCAGCACCCCCCGCACTCCCCACTGGTGTGATGGTAATGGTTATGGATAGGGGTGGTTGAGCTTTGTTTGTG | 120 | 52 |
| chr7 | 144217977 | 144218097 | 772751_42001748_Shank2(210274)_355 | 1 | + | good | CATGCCACTGAACTTGGTGTTTACAAACATGGCATGGGCAGGGCATTCTCATGGGATAAGCTCCAGGACCCCAGAGAACCAAATTCCAGATGCCCAAGTCTCCTATGGGAAATGGCCTAC | 120 | 51 |
| chr7 | 144218097 | 144218217 | 772751_42001748_Shank2(210274)_356 | 1 | + | good | TGCTTGTGTGTGTATTACCCAGACTTCCCATACATTGTGAGTCAGCTCTTCATAGCTTGTAATACTTGACATTGTGTTGTATTGGTGTGAAAGTAGGGGGTGTGGGATGCACACCTGTGT | 120 | 44 |
| chr7 | 144218337 | 144218457 | 772751_42001748_Shank2(210274)_358 | 1 | + | good | CCCAAGATCCAGCCGTGTCTTCTTCCTAGGGCTGGGATTATAGGTGTGTGCCACCATGTCTGGCTTTTTATATGGACGATGGGGATATGGTTTTGAGTCCACAGGCTTGAACACCATGCA | 120 | 50 |
| chr7 | 144218457 | 144218577 | 772751_42001748_Shank2(210274)_359 | 1 | + | good | CTTTAACCCAGCCCTTCATCATGTCACTCCGGGAAAAATTATAAGGAAAGAAATCTCTTTGTTCAGTTTATTTGTCCACAGCATTTTGTTCAATATTTTCTATATGTAACTGGCTGGATC | 120 | 36 |
| chr7 | 144218577 | 144218697 | 772751_42001748_Shank2(210274)_360 | 1 | + | good | AGTGGATGCAGAACCTGAGAGTGCCCAGGGCTGACTGCACTTCAGATATGTCACTGCTGATATCAATGTTTTTTTTTATTATTATTGTATTATCTTGTTAGTAGCCCATCCTTCTAGTTA | 120 | 38 |
| chr7 | 144218697 | 144218817 | 772751_42001748_Shank2(210274)_361 | 1 | + | good | TTATCACTCTGGGATTAGTAGTGAGTCTTGGAGACCTAAGATGTCCCCATAGTCAGTGTCTCTCCTCATCCATGGATGTGTTCATTCTCATGGTACACATTGGTCCTGCCAGCCCCAGAA | 120 | 48 |
| chr7 | 144218817 | 144218937 | 772751_42001748_Shank2(210274)_362 | 1 | + | good | ACTGACTTGTGTGGGCAACAGGATGTGTCTTGGGGGTCCCATTGGCTGTGTGGACAACAGGATATGTCTTTGGGGTCCTATTGGCTGGGCACCTGGATCTTACCTCCACTGATATCATTG | 120 | 52 |
| chr7 | 144218937 | 144219057 | 772751_42001748_Shank2(210274)_363 | 1 | + | good | GTCCCTGTATTCAGGTCACCACTGGCAGGATGATTCAAAACCCATGCTCTGCCTGCTGAGCGGGTGTTCCACTGGCTTCGTAGAACCTACTTGTGGGATGAGGAAACCCAGGTCCTGGCT | 120 | 55 |
| chr7 | 144219057 | 144219177 | 772751_42001748_Shank2(210274)_364 | 1 | + | good | CTACATGACCACATCAGAAGGCCCGTGACCAGATGCACCACTGTGGTGCCAGGAAGTCCCTCAAGCATGTGGTCTGCAGCCCTGGGTAGCATCTCACTTTTCTTCATTTGTTTATAAAGT | 120 | 50 |
| chr7 | 144219177 | 144219297 | 772751_42001748_Shank2(210274)_365 | 1 | + | good | AGTTTATATCCACTGGACTTTCAGGCTAGACAGGTCTCAGTTAATGATATTTTTAAATTTTATTTTACCGATGACATGGAAGCAATAAGCACTCAGTCGACACAATAGGCACTCAGTCGA | 120 | 38 |
| chr7 | 144219297 | 144219417 | 772751_42001748_Shank2(210274)_366 | 1 | + | good | CACGGTATGCTGAACTGTAAGTCCTGTGCCAGTGACATGGGATATGTGTGAACTTCTGCACTGCTGAGCAGTGGTGGTGAATGCAGTTCGCAGTGGGTCTGTGGCTGCAGAGACAACAGC | 120 | 53 |
| chr7 | 144219417 | 144219537 | 772751_42001748_Shank2(210274)_367 | 1 | + | good | CAACCCTGTAGTATGGTGTGTTGTGTGACTTCGAAGGCAGATGCCAGAAGTCAGAGGTGCTCTGTGCACCTTTAGGTTAGTAAATTTTAATGTAAATAGATCGGTTGGTATGCAAAGCCC | 120 | 44 |
| chr7 | 144219537 | 144219657 | 772751_42001748_Shank2(210274)_368 | 1 | + | good | CTGAGCTGTTATTGACAGAACTGCTCAGAAGCAAATTGCACTCCTGGATAAACTGGACCTGATGGTGACACATTACCCCCTTCCTATACTGCTGAGTTTCACTTCCTAGGATTTTGGTGA | 120 | 45 |
| chr7 | 144219657 | 144219777 | 772751_42001748_Shank2(210274)_369 | 1 | + | good | GAGTCTTTGTATTTAGTTTCATAAAAAATGATAACTTTTTTTTTCTTATCTGTTTTTAAGACAGGCTCTTGCCATGTAGCCCAGGCTAGCTTCAGAAATGCAAACCTTCTTCCTTAGCCT | 120 | 35 |
| chr7 | 144219777 | 144219897 | 772751_42001748_Shank2(210274)_370 | 1 | + | good | CTCATGGTCTTTGCTTTTCTCATGCCTTTGTCTGCTTGGGTATCAAGTTCTGCCACGAGTTGTTACTCTTCTCATTGCTGTGACCAAATGTCTGGCAAGGTGTAATATAAAGGAAGAAGG | 120 | 43 |
| chr7 | 144219897 | 144220017 | 772751_42001748_Shank2(210274)_371 | 1 | + | good | GTTTAGTTTGCTCACAGTTCGGAGAACCCAGTCCATCAAGGCTGAAAAGGCAAAGGATGTATGGCAGAACCCACTAATAAACATGCAGCTGAAGAAGCTTTTTTAGAGGGTTCTTAAACT | 120 | 42 |
| chr7 | 144220017 | 144220137 | 772751_42001748_Shank2(210274)_372 | 1 | + | good | ACACCTTCAGTACCGACGGGGTGGGGATAAGACACTTTGTGCTATCTATTTCTTCAGTGGGAAGTTTTAAAGGTCTGTCACTTTCAAATATTCAATCTGTACAGCTGCAATGATTGAATT | 120 | 40 |
| chr7 | 144220137 | 144220257 | 772751_42001748_Shank2(210274)_373 | 1 | + | good | TATCCAAACAGAGTTGCTTGTAGTATGTTCTGTCTTAATTTTTTTAAATTTAATTATTTTGTGTATTTGCAAGTCAGGGGCCGAGGACAGCTTGTAGGAGTTGGTTCCACCGCATGGGTC | 120 | 39 |
| chr7 | 144220257 | 144220377 | 772751_42001748_Shank2(210274)_374 | 1 | + | good | CCGGTATTGTACACAGGGTGTCCAGCTTGGTGGCAAGCATTGTTACACACTGCCCCATCTCACCAGCTGGTCTCTTTGTTCTTTATCTTTTCTGATTTGTACATTTTATTATTGTTGTGT | 120 | 42 |
| chr7 | 144220497 | 144220617 | 772751_42001748_Shank2(210274)_376 | 1 | + | good | GCAGTTGTACTCAGGCCTTCAGACTTTTATCATCAGACAACAAGGAGTTACTCCAAGGTTATGCCCAGCACGTCTGCCACCTCCTCATTTCATTCCAGGACAGCCTCTAGGCAGCACAAT | 120 | 49 |
| chr7 | 144220617 | 144220737 | 772751_42001748_Shank2(210274)_377 | 1 | + | good | GAGAAAACTAGGCACGTAGACACTTACAATTTTCTCTTTGATCCTATGGTATGCGTCTCTCCCTGGTTCCACCAGCAGTTTCTAATGGAGCATCAGAGACCGTGAGGCATCTACACTGGC | 120 | 48 |
| chr7 | 144220737 | 144220857 | 772751_42001748_Shank2(210274)_378 | 1 | + | good | TGTGGAAAGGAGCAACACCTTGGCATTCACATGGCTGCCTGCTCTTGACCTACCCACTCTAGCCAAAGGGACATTGTTTTAGCTTCACTTCTGTGGTCATAATTAATTATCATAATTATT | 120 | 42 |
| chr7 | 144221217 | 144221337 | 772751_42001748_Shank2(210274)_382 | 1 | + | good | TCTGAAGGCTCCTCCTAAGGAATGATGCCATCCACAATGGGCTGTGTTAATTAACAGTCAAGACAATCCCCCATAGATATGCCTAAGGTTAATCTGTTCCAGTCTCCTCGATGGAGATTT | 120 | 44 |
| chr7 | 144221337 | 144221457 | 772751_42001748_Shank2(210274)_383 | 1 | + | good | CTTTCTCAGATGATTCAAGTTGTGTCAAGTTGACAGCTGACTAGCTCAGACCCCAGGTTCAGAGACTCTCACCCCTGGGAACTTGCTGTGCTTGTGGCCTTACCATTGTAGCATCTCATG | 120 | 49 |
| chr7 | 144221457 | 144221577 | 772751_42001748_Shank2(210274)_384 | 1 | + | good | GGGGAATGTATGTGGTATGAACATACTATATACAGCTATTCTGTCCCTCCTCTAAATGTGGCTGGGTCTGCTGATGGGGATCCTGGGAGCAACTTGAAGTTCTGGGGACCAAAAGAGCAT | 120 | 48 |
| chr7 | 144221577 | 144221697 | 772751_42001748_Shank2(210274)_385 | 1 | + | good | CTCAAAGCTAGAAGTTGGGCACCAAGAGCTAGGGTCTTTCTCCTTTGTGTAAACCTCAGACAAACTCTCTTTTTCTAACCAAGAGATGGAGTGGCTGGGTAAGGTGACACCAAGCTACTT | 120 | 46 |
| chr7 | 144221697 | 144221817 | 772751_42001748_Shank2(210274)_386 | 1 | + | good | CTACTCAGGATCCTCATAATGGTCCTGCCTCATCTCCCTCCCATTCTACATCCACCTCCAATGGCTATTGGCATCTACTTCTCCAGGCATAGTGCTTTGGGGCTGCGTGTCTAAGGTCCA | 120 | 52 |
| chr7 | 144221817 | 144221937 | 772751_42001748_Shank2(210274)_387 | 1 | + | good | CCTCAGGCTGGCAGGGTGTCGGGAAGAATGAGAGGTTGGCAGAGCAAATGCAATTATGGCAACCTTCTGGGAAGAGGCGCCCCATGGAGAGCCATGCATGGATGGGCACAGGGTGGGTTC | 120 | 58 |
| chr7 | 144221937 | 144222057 | 772751_42001748_Shank2(210274)_388 | 1 | + | good | CTTCTGGGCTCTGGGCTTATGCTCCAACTCTTGGGAAATGGCTGCTTGTTACCAGCTGTTCTAATTCAAGAAAGCCCTGGAAAGGATTTGGCACATGTCCTTGTGGCTTAGCCTGAGCCT | 120 | 50 |
| chr7 | 144222057 | 144222177 | 772751_42001748_Shank2(210274)_389 | 1 | + | good | ATGGCCACAACTCCCAGGCTTTGAAGATGTCTTCCATGGAACCTAGTGTTCTCTGAGTTGACATGATTGTTGTTGTATTTCTTGGAGTTATTCACAGGGAATCAGCTGTGTGGGCAGCTA | 120 | 45 |
| chr7 | 144222177 | 144222297 | 772751_42001748_Shank2(210274)_390 | 1 | + | good | GTTAGGAGGATAAGCCAACCTGAAAGCATCCTGCAGCCATCTCTTCTCTTACTGTGACCTACTGCCCAGGAGCATGGAGAGGGGACTGGCTGCACTCTGTTGACCCTCAGCGAAAGTCAG | 120 | 54 |
| chr7 | 144222297 | 144222417 | 772751_42001748_Shank2(210274)_391 | 1 | + | good | GTACCAGACAAAGCAAAGAAAAGGATCACTTCACACCAAGGAGCCTCCTGCAGCCAGGAAACTATAGACAGAGGAAAATTGCAGCTGGACGGTGCTGAACCATCTCACATGTCACCGTGT | 120 | 49 |
| chr7 | 144222417 | 144222537 | 772751_42001748_Shank2(210274)_392 | 1 | + | good | GTCACTGGTCTGGTATCTGAAGTGGAATTTTATAAATTCAAACTGGTTTTCGTCATTGTGAATTCAGAGCTCTAAGTCAGGGGCTGTTACAGACTCTTCAGGAGAGCAAAGCAGGGGGTT | 120 | 43 |
| chr7 | 144222537 | 144222657 | 772751_42001748_Shank2(210274)_393 | 1 | + | good | CCAAGGTCAAAGCCCCCTCTACTGCTGACCTTGGGCTGTGTGTGACCTTGAGCTCATCCCCTAGCCTTCCTCTGCTCAGCTGTAAAACAGATGAAGGACATGGATCTGTGGACATTAGAA | 120 | 52 |
| chr7 | 144222657 | 144222777 | 772751_42001748_Shank2(210274)_394 | 1 | + | good | TTCTGTAAAAGGCTCTTGGGAAGGTGATTTAAAATAGCATGCAAATGTGTTACTTGCTAGGGTTTCATGATATGGCTATAGAATAGTTGAGCTTCTGCATGGAGAAAGCCAGACTCAGGG | 120 | 41 |
| chr7 | 144222777 | 144222897 | 772751_42001748_Shank2(210274)_395 | 1 | + | good | CAGGTCCAGCTGTGGTCAGCTGTGGGTAGGGACCCTGCTCTAGAGCCTGGAAGATGCAACCAGGAGCCCATGTAGTGAAGGAGGTGAGTGCTCAGCTCAGAGTAGGGAGAACAGGAGGAT | 120 | 57 |
| chr7 | 144222897 | 144223017 | 772751_42001748_Shank2(210274)_396 | 1 | + | good | GAAAGGTGGCAAGGTTAAGGCTGGGTCTCTGATGCTGTGGAAGAGAAAGGATTGGCCAGAGAAGGAGGCACCCTTGGGAGTACAGGGCAAGGGATACAGGAGGGTGGACTGTGGAGAGG | 120 | 57 |
| chr7 | 144223017 | 144223137 | 772751_42001748_Shank2(210274)_397 | 1 | + | good | CCCCCAACTGGTGTAGGCATCTTTCCTGGACAGTGTGGTAGCCAGAAGTGTGGGCTCATGGATAGGGTGAGCCCTGAGTTTTCAAGTGCCTTCTGTCACTGATCTGTTCTCTAATCCTTG | 120 | 52 |
| chr7 | 144223137 | 144223257 | 772751_42001748_Shank2(210274)_398 | 1 | + | good | GATGACACAGTAGTCTCCAGGGACTGGTGAAGCAAATGACTGTGGATAAGGGGAAGGGCCCCGATGGCATAGATTTTGACCCCATATCCTGAGGGCCAGAGCTGTGAAAAGCCCACACCC | 120 | 54 |
| chr7 | 144223257 | 144223377 | 772751_42001748_Shank2(210274)_399 | 1 | + | good | CCTTGAGTGGTTCTTTCTGCCTCTTAGCTCCTGGAAGGTCCAAGTGGCTGCATTCCCCTGCCTCCAGCTTTGATGCTCATCTGACATTACTTTCTGTGTATGAGCCCCTCTTGCCCTCCT | 120 | 52 |
| chr7 | 144223377 | 144223497 | 772751_42001748_Shank2(210274)_400 | 1 | + | good | AGGAGGACACTTGCTATTGTCGTTAGTCAACCTGCTTAATCCTAAGTGACCTCACATCATGATACTTCACTTCCTCATGCCTGTTGAGAGACCTTTTCTCACACGTGATCCCGTGGGAAG | 120 | 47 |
| chr7 | 144223497 | 144223617 | 772751_42001748_Shank2(210274)_401 | 1 | + | good | CGTCTGAGGATGAAGATGGTACTACACATGAATATACTAACGGAGGCAAGAACCATGAAGGCATCCCTCAGCAGACCCTCATGTAACCCCATGAGGGCCCTGCAAAATGGAAGCACCATG | 120 | 50 |
| chr7 | 144223617 | 144223737 | 772751_42001748_Shank2(210274)_402 | 1 | + | good | TCAGAGCCCCAGGCTTCCAGAGCAGAGCACAGGAGGGAGGGTTCACAGGAGTCAGACTGCGTCCTGTGTTTCTACATGGGGACTCGGGGCGCCGCACTCTTGCATGTGTATGATGGGTTC | 120 | 59 |
| chr7 | 144223737 | 144223857 | 772751_42001748_Shank2(210274)_403 | 1 | + | good | CCACTGTGTTTATGGACAAGCCCTGCCCAGTTTGCCTCCTTGTCTACACATGGGTTTGGGGGACAGCAAGAGAGCCTTGAGACCTCAGACTGAAGATTCTGTTCATTCATGACAGGGTAA | 120 | 50 |
| chr7 | 144223857 | 144223977 | 772751_42001748_Shank2(210274)_404 | 1 | + | good | CCCTGGGTTTAGTTTGCTCTTGACCCTTCTTCCTCTGTGGTCCTGCTGAAGCAACACAGGCATAAGGAAAGGGTGGGTGTGGGATGAGGGTCTAGGCTATAGTTGAGCAGCTTTCAATAG | 120 | 51 |
| chr7 | 144223977 | 144224097 | 772751_42001748_Shank2(210274)_405 | 1 | + | good | GCAGAGGACCGTGCAGGCATGTGGACTGGGGCTTGGCAAGGTTCCCTGGGTTCAGAGAGAGCAACTGGGGCTATTGCTACAGCTCAGTCCAGATAAAAGGAAGATGCTGTAGAGATGCAT | 120 | 54 |
| chr7 | 144224097 | 144224217 | 772751_42001748_Shank2(210274)_406 | 1 | + | good | TCCTGAGCTAGCATGGCGTGGTGACAGGGAGTAGGGTAACTAGGCAGGGTGGGCCAGGATCTGTGATGATACCTGAAGGACAAGGGATGACACACTGCACCCTTGTGCCCATGTAATGGA | 120 | 55 |
| chr7 | 144224217 | 144224337 | 772751_42001748_Shank2(210274)_407 | 1 | + | good | TACTTTCCTTGTCACTATGAACAAATACTTGACCTGCCAAGGAGTGACTCAAGGAAGGGAGAGTTATTCCAGCTCATGTTTGAAAGTATATAGACCACTGAGACAGGGAAGGCATGGTGG | 120 | 44 |

| | | | | | | | | | |
|---|---|---|---|---|---|---|---|---|---|
| chr7 | 144224337 | 144224457 | 772751_42001748_Shank2(210274)_408 | 1 | + | good | CCGTGCACAGCTCCGTGACAGTGGGAGCATGAGGCTGCTTGTTCACACCTCAGTGTTCACATCTCAGATCAGGAAGCAGAGGGGAGAATGCTCAGCTTTTTTTACTTTCTGCTTTTCATC | 120 | 50 |
| chr7 | 144224457 | 144224577 | 772751_42001748_Shank2(210274)_409 | 1 | + | good | TTCAGACTTTAAGACTAGATGGTGAAAAGCACAGGGAGGGAGGGAGGGGAAGAGGGAGGGGAGGACAGAGCCTGGAGTTTGACTGCCTCAGTTCTGAAGAGTGCATTGGAGCTCTCCAGG | 120 | 54 |
| chr7 | 144224577 | 144224697 | 772751_42001748_Shank2(210274)_410 | 1 | + | good | GGAAGGGAACGAGTTGTGTGCTACACAGAGTACTGCAGACTATCTGGTTTCCTGGCTACCAAAAGGATGTGGTACAGGGGCAGGCAGCAGCCTCCAGTGGAAGGCTGTAAGAACCTGTAG | 120 | 53 |
| chr7 | 144224697 | 144224817 | 772751_42001748_Shank2(210274)_411 | 1 | + | good | CTGCCCGGTGGACAGATGGCCACTGCTAGCACAATTTCCTATTGAGGTAGTTGCCATGCTCACACACTACAGACAAGAAAGTAGAGACTCAGCAAGGTGCAGCAGGGCAAGGAGGAACCC | 120 | 53 |
| chr7 | 144224817 | 144224937 | 772751_42001748_Shank2(210274)_412 | 1 | + | good | AGGACAATGTGCACTAACACAGTCCTCCTGAACTTTTGCGGTACCTACCCTCAAAGTCCACACACATCCTGAAGCCAAGGGAGGAATCCTCAAGATGGAGCCTCTTTGCTTTCTGCTCCT | 120 | 50 |
| chr7 | 144224937 | 144225057 | 772751_42001748_Shank2(210274)_413 | 1 | + | good | TGTTCTATCTTTGTCAGTGTGTCTCCTCAGGACAACCAAGTGGCTCTTATGATGGCCACTTACTGAGAATTCACAGGCAGTAGGGCCTAACACACACAAACACACACACACACACACACA | 120 | 47 |
| chr7 | 144225057 | 144225177 | 772751_42001748_Shank2(210274)_414 | 1 | + | good | CACACACACACACACACACACACACACGGTTCTCTTAGTCTCTGACCTCCCAAGAGTGCTGAGGTCACAATGATGGGTGAGACAGAGAAAACATTAGAAATCTGGCAAGGGACAAGCCAACA | 120 | 48 |
| chr7 | 144225177 | 144225297 | 772751_42001748_Shank2(210274)_415 | 1 | + | good | AACAACATTGCAGGGGGATTTAGAAAGCATCTGGACAAATGGTAACAGTGGGGAGGCATCTCCTGGAAAGGGAGGGGCTTAATGTTAAACAGATGGATGTACAATGAAGAGTTGGTGGGT | 120 | 46 |
| chr7 | 144225537 | 144225657 | 772751_42001748_Shank2(210274)_418 | 1 | + | good | GAGAAATGGCCTAGATGGATACATAGATGGACACATGGACTGAGAAGATGGGTGGGTGAATGGGATGGGACAGATGAAGTAGGTAACAGAACTTGAATGAGCCTAATGGTGGAGAGAGGC | 120 | 48 |
| chr7 | 144227097 | 144227217 | 772751_42001748_Shank2(210274)_431 | 1 | + | good | AGATGTATGAGTGGATAGATAATGGTAACTTTCATTCCTTCCCTTTCATGTCCCTCAAAATGGTCTCAGTTGCCATCTGTCTCCGTGATCCACAGGAGTGGATGAACTTGACTACATCTA | 120 | 42 |
| chr7 | 144227217 | 144227337 | 772751_42001748_Shank2(210274)_432 | 1 | + | good | AAGCTCCTTAAGGGGCAGCTTTCCCGGGTGCCATAGCATTCCCCATCTTTCTCTGGAGACCAGGCTGGCTTTAGATGCCAGAGCTCAATCTGGAAGGGTCCAGCCCAAGGCCCTGTTCCT | 120 | 57 |
| chr7 | 144227337 | 144227457 | 772751_42001748_Shank2(210274)_433 | 1 | + | good | TGGCACTCTGCCGCTGGGTATGCTGGCTGCAGTTTGCCCCCACCACCCTGCCGTAGGAGGATGCAGCTCCCAATTAAACTGTTTCAGTGACAAAGACCCAGGGTAATCGTGTGATTAAAA | 120 | 53 |
| chr7 | 144227457 | 144227577 | 772751_42001748_Shank2(210274)_434 | 1 | + | good | ATAAACCAGGGCTGATGCTGCTCACTGGAGCCTAACAACTGAGGGTGGGGGCGAGGAGGAGGAGAGGGATGCTTCCATCCCTGCTCCTCTGACGCCATTGCCATAGGGGATGGGGCGGTC | 120 | 61 |
| chr7 | 144227577 | 144227697 | 772751_42001748_Shank2(210274)_435 | 1 | + | good | GCACTTTTGTTTACAGCAGAATGAAGGAGACATAGAAAGGAGCACTAGGTGGCAGGGACTTGAGGCAAAGTGGGAAATCCTTTCATGAAGTTAAAGTCCACTGTGGCACTGCAGAATGAG | 120 | 46 |
| chr7 | 144227697 | 144227817 | 772751_42001748_Shank2(210274)_436 | 1 | + | good | GGAGAATGAGCTGGGGTTCAGTGAGCAGCTACTGCATGAGTCCCTGGGACCTTTCTTTGGGTCATCATGGGAACCTGGTGCAAGGTATTTAGTGTTATTTACAGGAGTGTGGGCTGTGAT | 120 | 50 |
| chr7 | 144227817 | 144227937 | 772751_42001748_Shank2(210274)_437 | 1 | + | good | TCAGAGGTGCCTGTGCTGTACCCTTCACTCAGCATGTATTGAGTACCAACAGTATGCCAGGCACCGTTCTAGATGCAGAAAACAGGGAGAGTCAGACACAAGTCCTGCCACCATGTGATT | 120 | 50 |
| chr7 | 144227937 | 144228057 | 772751_42001748_Shank2(210274)_438 | 1 | + | good | TCCCAGAATCTCCACCAGCAGAGGGTTGAGCTGCCACTCTATCCCTCTTACTGTTTCCCACCTAGTGCAGGTTTGCTACCTGAGCCCAGGGTCCCTTCCCTGAATGAGAGCTGCTCTGAA | 120 | 55 |
| chr7 | 144228057 | 144228177 | 772751_42001748_Shank2(210274)_439 | 1 | + | good | GTGACAAAGCATGGCTGTGCTTGGGACATTTGGCAAATCCTTGTTTTTCTGGTGTCCTTCACACACTGAGGTTGTGTCTGTCACCAACTTCCTCTAGCCTGGAAATGCTCAGCTCCCTGA | 120 | 49 |
| chr7 | 144228177 | 144228297 | 772751_42001748_Shank2(210274)_440 | 1 | + | good | TGCTCAGGACTGGGTGGCTGAGTCCTACTGCACCACCCTTGTGGTCTGCCAGGTCACACAGTACCTTTATGTTAGTCTGAGGGACTTCAGTGACCTTTGGCATGCTCCTCTCAGGCCTTT | 120 | 54 |
| chr7 | 144228297 | 144228417 | 772751_42001748_Shank2(210274)_441 | 1 | + | good | AGCACTCTTGTCCTCACTGTAGAATCTTAGGGGCCCTTTAGGGTGATGAGCAAGGAACAGACTTCCAGCCCCCAACATCTGCATGACCCCGGACATGGCCGGAAGCCTTGAAACAGCATC | 120 | 54 |
| chr7 | 144228417 | 144228537 | 772751_42001748_Shank2(210274)_442 | 1 | + | good | GTGCACTTTTGTTATCACCCAGCTTCTCCCTCTGTCGAGGGTGTGGCTGGAGTCTTGCCTCCTGGAGGGACAGGTGCACCACCCCTGCAGAAGACTGGAGTCCAGAAGCAGCCATTCCCT | 120 | 58 |
| chr7 | 144228537 | 144228657 | 772751_42001748_Shank2(210274)_443 | 1 | + | good | GCAGCTCCATGGCTATGCTGAAACTGTAGGAGTTTCTGGTTTTGACTGACCCATGAATTAGAGCAGACACTAACTCTGAAGCTTCTGTGTGGAACTGCATGCTGCCCTTCCTGGAGGAGG | 120 | 50 |
| chr7 | 144228657 | 144228777 | 772751_42001748_Shank2(210274)_444 | 1 | + | good | CACCACGGTAGCACTCTGCCCTCCTTGCCCAGGGTCCTCACACAGCCAGTCATTTATGTATCAATTTGGCCCTGCTATATCCCTTCCTATGGCCCAGATGGGGCTTTGAACCTCTCTTGT | 120 | 53 |
| chr7 | 144228777 | 144228897 | 772751_42001748_Shank2(210274)_445 | 1 | + | good | GACCATAGGATGCGTCTTTGCCTTCTGTCTGGCAGTGCTGTCACCCAGAGGCTCATGCAGTGACTGTGCATCCTGTCACTGATCCAGAGCACTGAGAGCCTAGTCATCTCATGGGCTTCC | 120 | 55 |
| chr7 | 144228897 | 144229017 | 772751_42001748_Shank2(210274)_446 | 1 | + | good | CGCCTCCACAGCCACCATGTGCATTGCTGCTCACACTCGTGACCATGATCTGGATCAGATATGTGTACACATGAACCCGTGGAGATCAGCGTTCAACCTTGGGTTCCATCTTCAGGAATG | 120 | 52 |
| chr7 | 144229017 | 144229137 | 772751_42001748_Shank2(210274)_447 | 1 | + | good | TCATCCACCAAGGTGTCTCCCAGGCCTGGGGCTCACCTGTTAGGCTTGGCTGACTTGTAAGCCTCGGATTCTCTTGTCTCTGCTCCAGCACAACCCTGGACAATAAGCACATGCCACCAC | 120 | 56 |
| chr7 | 144229257 | 144229377 | 772751_42001748_Shank2(210274)_449 | 1 | + | good | GCAGCCATCTCCAGGGCAACACAGACTGCCCAACCCCCTACCTCATTCTCTGTGATGCTCAAAATACACCCGCCACCAGTGCACAGCCACATCTCCATTCTTATTGAGTTCTTCCTTCCC | 120 | 53 |
| chr7 | 144229377 | 144229497 | 772751_42001748_Shank2(210274)_450 | 1 | + | good | CTTGGGCCTAGGGAGGGCCTCTGCACATCTCCAGGGCCCCAACTATAGGCCATCATTCCCCTAAGTCCTGGGTAGAGCATCCTCTGTGGGCAGGAGGGAGGTGACAGCCAGAGGAGGACC | 120 | 62 |
| chr7 | 144229497 | 144229617 | 772751_42001748_Shank2(210274)_451 | 1 | + | good | ATGCCTGTGTCATTCCCAGTCCACCCACCTCAGCAAGGACATAGTAGAAGGCCTGCTCTGTGCCATGTGGGTGAAGCCCATCTGTGTCTCACCCCATGCTGGGCTGGGCAAGTAGCTCCA | 120 | 57 |
| chr7 | 144229617 | 144229737 | 772751_42001748_Shank2(210274)_452 | 1 | + | good | GACTGCCCCTCCTGTTCCCTGGGCAGGTGCTCCCCTCCACCCACATACTCTGGCTTCTCTTGTAACGTAAGGATTTGGGGTTTCTGACAACTGCTTGGAAGTATCAGATTCCAAGGCTGC | 120 | 54 |
| chr7 | 144229737 | 144229857 | 772751_42001748_Shank2(210274)_453 | 1 | + | good | CCCTTCTCATGTGGCAGACACAGCAGCAAAGACTCCACCCCTTGATCTCAGTGTCCTACCCTCCCATCCCCCCATTCCCATGTTCCATCCACCTGACCTCCACCCCTACCCCAACCAGCT | 120 | 58 |
| chr7 | 144229857 | 144229977 | 772751_42001748_Shank2(210274)_454 | 1 | + | good | AATCCAACTTTTACTAGCATATCTCCCTCTTCTTTCTGCCTGCTCCTGAACTGAGCCCTACCATGAGGTCCCCAGCCTGTGATGCAGCCTACTGTGCTGGGTCCAGCAAAGGCACTGAGT | 120 | 52 |
| chr7 | 144230097 | 144230217 | 772751_42001748_Shank2(210274)_456 | 1 | + | good | CACAGACAGACACACTTACCTTTGTGGCAGCTGAGTCCTACACCTACCCAGTTCTTTCTCCTTACCTCTCTCCCACATGCTCCATGCCCTTCTGTTCCAACACTCTTAGATGGGCTCAGC | 120 | 52 |
| chr7 | 144230217 | 144230337 | 772751_42001748_Shank2(210274)_457 | 1 | + | good | CCCACTTCACCTGTCCAGCTTCTACTGTCTCCTAGGCTAGAGCTGCCCAGACCAGACCTCATCCTTACAGGGCAGAGAGCATGGCTGCTTCAGCCTTTGTAGCCGTATGCACTCACAGCT | 120 | 56 |
| chr7 | 144230337 | 144230457 | 772751_42001748_Shank2(210274)_458 | 1 | + | good | GCTTCCTCTGAGCAGAGGACACACCAGGGAATGGGGCTTTCTATACCAATAAAACTGTATTGATGATGGTAGAAAGGGACGAAACACCTCACACTAATTATTAGTATGTATTGCCACTGT | 120 | 42 |
| chr7 | 144230457 | 144230577 | 772751_42001748_Shank2(210274)_459 | 1 | + | good | TGTGACAATTCCATCTTTGGGCCTTTCTTCTCCACATGCTAAACTCCTACTCATCCTACAAGACCCCACTATGTTAGCCTGCATCTCAGGCTTCCAGCAAAGCTCCTATGCTCCCTGTTC | 120 | 48 |
| chr7 | 144230577 | 144230697 | 772751_42001748_Shank2(210274)_460 | 1 | + | good | CCAGAACTGGAGAGCCTCACATCTCCCATCACAGATGAACCCACTCAGTTGTGATGAACTTCTCTGTGTATCTGTCTACCCCCTCTTGCCCTCCTCCTAGCTTAGTCATCTCTCTTTTGG | 120 | 50 |
| chr7 | 144230697 | 144230817 | 772751_42001748_Shank2(210274)_461 | 1 | + | good | TGGTGTCTGACATGCAGACAGCTCTGGAGTCACTCATTCAGCAAAAACTGCCAGACAGTGTACCTAGAAACACAATAGGGAGCCAGATGACACCTAGGGGCACAGGTCAATGCTGGGAGC | 120 | 52 |
| chr7 | 144230817 | 144230937 | 772751_42001748_Shank2(210274)_462 | 1 | + | good | ATGAAGCTGCTTCTGGGGAGGGGAGGGCCTATACATGGCACAGGCAAAGACCCAAAGGGAGTGAAAGGATCCAGGTGGATACGGGGATTAAGTATCCAGGCAGTAAGAACAGTGAGTCCA | 120 | 52 |
| chr7 | 144230937 | 144231057 | 772751_42001748_Shank2(210274)_463 | 1 | + | good | AAGGCTGGCAAGGGGACAGACAGCCTTATGGGGACAAGGGCTACATGAGACCCATTCGTGTTATGTCTGTGGGGACAGAGGTCACCATGTCAGGGAACATCACTGAAAAGGCACATGCAT | 120 | 52 |
| chr7 | 144231057 | 144231177 | 772751_42001748_Shank2(210274)_464 | 1 | + | good | AGGGACAAAAAGGCATGACAGCAGAGGCCTTGGCGAGGGGGGGGGAGTGTTTGCTGAACGGATTCATACTGTCCCATTGTGTGTATAGCGCTTTGGGGGCCCATTCTACCCCAATCCAGG | 120 | 56 |
| chr7 | 144231177 | 144231297 | 772751_42001748_Shank2(210274)_465 | 1 | + | good | GATCCTTTGCGTTATCAAATGAAAGCTAGCATGAACCTATTCCTCCCAGGAGCCTCTGTGGACATCAGCTGTTCAGGTCTATCTTGGGTGGGCACCTGTTGGGACAAGGAGACCAAAATG | 120 | 49 |
| chr7 | 144231297 | 144231417 | 772751_42001748_Shank2(210274)_466 | 1 | + | good | TGGAGAACACTGATCAGCTGGGGCCTGTGGCCACACAGGAGCAGCAGGCTCAGAGCCGCTTTGCTGTTCTGAGATATAGTTCTAATAGCAGCAGGATTAGCCTGCTGACATGCACAGGGG | 120 | 55 |
| chr7 | 144231417 | 144231537 | 772751_42001748_Shank2(210274)_467 | 1 | + | good | AGTGGCACCCTGCACACCCAGTCACTGTCTTCACTACTGCTTAGTTTCACAACCTCTTGGCATTATCAGAAGGAGATCCATGCCATCTGTCCATCCTCTGCCCAGCCGTCTGGTTGTCAC | 120 | 52 |
| chr7 | 144231537 | 144231657 | 772751_42001748_Shank2(210274)_468 | 1 | + | good | TCATCTGCTTCCTGTCTCCAAGGCTGTGTGTGTCTTGCAGGGGTCACAGTGAGTCCATCACTGTGCAGCCTTTATGCCTGGCTCCTTTTTCCAAATGATACATTTAAGGCTCACCTTTTG | 120 | 48 |
| chr7 | 144231657 | 144231777 | 772751_42001748_Shank2(210274)_469 | 1 | + | good | TTGCCTGTGACCCAATGGGTGTAATTAGGGATTCTCATCCCACCCAGAGAGTGCAGGGCTTCTTACCAACCTCCTCTACCCAGAGCCCCTTGTCCCTTGTCTCTGGAGACAGGTGCTCCT | 120 | 55 |
| chr7 | 144231777 | 144231897 | 772751_42001748_Shank2(210274)_470 | 1 | + | good | TCAGGAGCAACACATGTAGCCAGTGTGTGTAGTGAGAAGGACCAAGGACTGGTGTTTTGCAAATGCAGAGCAGGACCCCATCCTAGTGCTTCTGTCCTGAGAGGCCATGCCATGGCTGGA | 120 | 53 |
| chr7 | 144231897 | 144232017 | 772751_42001748_Shank2(210274)_471 | 1 | + | good | TGGGCAGTGTGTTGTTCTTCTGGGGTCGTCACCACTCACTTTCCTGGTTAGGCATGAGTTCTGTCCGGAGTATTGCTGGGCTCTTGTCTGGGGTGGTTGGGGTCCTGCCATTGGCTGAAG | 120 | 56 |
| chr7 | 144232017 | 144232137 | 772751_42001748_Shank2(210274)_472 | 1 | + | good | ATGCTTCTCAGTTTCCTGCTGATTGGCAGTGGCATTTACTATCCTGTGATCTGTAACAGATCAGAGCATGATTTGTCCCACTCAACCCAAATGAAGCTCCTGTAAGGGTCCTGCTTCTGA | 120 | 46 |
| chr7 | 144232137 | 144232257 | 772751_42001748_Shank2(210274)_473 | 1 | + | good | TGGAGAGACAGGACCCTATCCTGTTTTCCATAAACAGATTCGAGAACTGGGCATTGGCTGGGCTCCGCTAAAGCCTGTTGTTTCCCAAAGCTCTGCTCTGTCTCCATGGAGATCATAAAA | 120 | 48 |
| chr7 | 144232257 | 144232377 | 772751_42001748_Shank2(210274)_474 | 1 | + | good | CAGGGTAAGGGAGATGCAGGCCTGAACACCTGCCTGCCCCTGCCCTGCCCTGCCCTGCCCTGGCATAGGCTGCTTGGTTCATTATCCTTCATTTAAAGCATTTGTAGCATGGGGGCAGTA | 120 | 57 |
| chr7 | 144232377 | 144232497 | 772751_42001748_Shank2(210274)_475 | 1 | + | good | AACAGGAGAAAGACTTCTGCTGAGCCATGAATATGCATGGGAAATTCCAGCCTGCAACAGTCACAAAGCCCCTCTACTTCCTTGGCCTTAATGCACAGTGGTGTTCTTCAGTGCATGCAT | 120 | 48 |
| chr7 | 144232497 | 144232617 | 772751_42001748_Shank2(210274)_476 | 1 | + | good | AATGCGTGCACATGGGACCTAGAATAGCACTCAGCCCCCAGACTCCAGGAAGGCGGAGTTTGTGTTACTACCTACCTAGCACAGGACCTTGGGAGGCCACCTGTCCTCAGTGGCAGCCTG | 120 | 57 |
| chr7 | 144232617 | 144232737 | 772751_42001748_Shank2(210274)_477 | 1 | + | good | GGCCGCTAACATATTTGCACATGGTCACTGGCCTTGCCTGTTCCCAACATGGGAGGCAGAGCAGGGAGAGGTGGCTTCTCACCTACAAAGGCACAGTGGTGGCTAAGGGTGGGAACTCCT | 120 | 57 |
| chr7 | 144232737 | 144232857 | 772751_42001748_Shank2(210274)_478 | 1 | + | good | GAAGAGTGTGTTACTTCTCTGGCTGTTGGGCACAAACAACTTAAGTGATTTTAACTCACAGTTCCAGGTACTGTCACCTGTGAGTGATGTCACACACCACAGGCAAGTATGGCCACTCAA | 120 | 46 |
| chr7 | 144232857 | 144232977 | 772751_42001748_Shank2(210274)_479 | 1 | + | good | GAATTGTCACTGAACCAAGGACTGTCCTGTGAGGAGAAGGGTCATCCTTGCTGTTCTGGGTGTTTGTGAGGAAGGACAGTTTCTGAGGCTCAAGGACCACTGTTCTCTCCAGCACCCTCC | 120 | 52 |
| chr7 | 144232977 | 144233097 | 772751_42001748_Shank2(210274)_480 | 1 | + | good | TTACTCCCTACATTTGGTGTGTCCTCCCTTGTGCACTGGTGAACCACAAGGACTGAGGGGGAGAAATAGGTCCAACCCTGCTGTTGTCATGACCACTTGCCACATCTCCAGCCACCACCA | 120 | 53 |
| chr7 | 144233097 | 144233217 | 772751_42001748_Shank2(210274)_481 | 1 | + | good | CCCTAGAACCCCAAATTCCACATTGATGGACTCAGCCAATCAAGAGTGAAGAATATTCTTTTAACTGGGCTCAGTAGTACACACCTGTAGTCCTAGCACTCGGGAGACTGAGGCAGTAGG | 120 | 48 |
| chr7 | 144233217 | 144233337 | 772751_42001748_Shank2(210274)_482 | 1 | + | good | ATTGTGCCAATACTATGCCATCTGTTTATTTCGGAGTCCTGGGTCCCTGCAGATATGGGGACAACTGTGTGTTAATTATCTTGCTTTAATGTCTTCTTCAGACAGTGGCCTCTGGTGGTG | 120 | 45 |
| chr7 | 144233337 | 144233457 | 772751_42001748_Shank2(210274)_483 | 1 | + | good | TGAGGCAGGTGTGGACAATGCTCACTGTCATAGTTGTTCCAAAATGAACTTAATGGCACTCTGGGCTTTGCAGTTTCTGTGTGACACAGGCTGATTCCCCCCCCCCCCCCCCGAGCTATC | 120 | 53 |
| chr7 | 144233457 | 144233577 | 772751_42001748_Shank2(210274)_484 | 1 | + | good | TCTCTGGCAAGCCCGGAATCTGTGTGGATTCCAACCCTGATCTCCCAGGTGAGTCGGTGAATCTGGCTGCCACACTCAGCATTTAGGAGTCCCCAAGTATTTCCAGCTGACTCATGGAAG | 120 | 53 |
| chr7 | 144233577 | 144233697 | 772751_42001748_Shank2(210274)_485 | 1 | + | good | CGTGACACACAGCCACTGCTGATAGGTTAAGGTTGCTGGCTCTGGGTGTCAGTGGCATTTATGAATATCATAGGGCAACATTTACTCATGCAGCTTCCGGAAACACAGCGAAGGAAATCA | 120 | 48 |
| chr7 | 144233697 | 144233817 | 772751_42001748_Shank2(210274)_486 | 1 | + | good | CCCCAGCGAGTACTCGCTGCCTGACTTATCTTTGCCTGTTACAGTTGCCTAATCACATAACTATGTATGAGGAAACACATCCACACTAAGTCTCAGAGCGAGCTCTGCTGAACTAGGAGA | 120 | 48 |
| chr7 | 144233817 | 144233937 | 772751_42001748_Shank2(210274)_487 | 1 | + | good | GATGTGGTATCACTGTATTCCAGAAGAGGCTCATGTAACCTGGGTTAACCTTGAACGTGCTAGGTAACCAAAGCTGGTCTTGAACTCTTGAACTCTGGCCTCTGCTTTTACAGTGCTGGG | 120 | 48 |
| chr7 | 144233937 | 144234057 | 772751_42001748_Shank2(210274)_488 | 1 | + | good | ATGACAGGCATGAGCCATCACACTTGGCTCTTAGCTTCGTCTTTGTATGAAGGATGGTGCATAGCCTTATAGCTGTTTCTATACTTCTGCATTGAAGTGTGTGTGTGTATGTGTGTGTGG | 120 | 44 |

| | | | | | | | | | |
|---|---|---|---|---|---|---|---|---|---|
| chr7 | 144234417 | 144234537 | 772751_42001748_Shank2(210274)_492 | 1 | + | good | CCAAGTTCATACTCAAATGTTCCTCTGGTGCTGTCCGTGTTTCTGTTTCTTTCTTGAGACAGTGTCTCTCATTTGGCCTGGAACATTCCAAGTAGTATGGATTGTTTGCCCAGCCAGCAC | 120 | 46 |
| chr7 | 144234537 | 144234657 | 772751_42001748_Shank2(210274)_493 | 1 | + | good | CAGGGAGCCCCCTGTTTCCACAGACCCAGCATGAGCCACCGTGCCTGGCTCTTTTTTATCTAGGGATCAAACTCCTATCCGTGTGCAAGCGCAGGGGCTTTATTGACTGAGCCTCCTATC | 120 | 55 |
| chr7 | 144234657 | 144234777 | 772751_42001748_Shank2(210274)_494 | 1 | + | good | AGTCCAGCAGTGACGTCTTGATATGCACTTGCCTTCTGGATTCTGAAACTTGGGAAATTCTCTCTTATCTCATCATCTCTCACATCCCCCTCTGCCTCATGATGTGGACACAAGCTTGCC | 120 | 48 |
| chr7 | 144234777 | 144234897 | 772751_42001748_Shank2(210274)_495 | 1 | + | good | ACTTACCTCCTGGGTCCTCGTCTTTGCTGTTTAACTGGTTCCAGACCTGAGGGAAGCATCCATTCCTAATGCAGGCAGTGGCAGCACTGACTGCTTAGTCAGGGTGAGTTCTGCTGGGCA | 120 | 53 |
| chr7 | 144234897 | 144235017 | 772751_42001748_Shank2(210274)_496 | 1 | + | good | CAAAGTGGCCTGGCTTTTATTGTCCACCAGGCCAAGAGGCCCCTGTATGCTGGATACCAGCCAGGGGTCATCCCCTGCTGGGTAGCCAGAGAAGCCACAGCGTACACACTGCCATCTAGC | 120 | 58 |
| chr7 | 144235017 | 144235137 | 772751_42001748_Shank2(210274)_497 | 1 | + | good | TGTTGGAACCCTACACCACACACCTTCACACTGTGCGTGCACTCCACAAGAGACAGAATGTGAACTGAGCTAGCAGGATACAAGGAAGGAAGAACCGGGTGAGCTAGGGCGCATCTTCAC | 120 | 52 |
| chr7 | 144235137 | 144235257 | 772751_42001748_Shank2(210274)_498 | 1 | + | good | CATGGAAAGAGACCCAGGTGACTAGGGGTGCAGATGAGGAACAGGGGGACCCAGATGAGCTGAAGCCTTGGCTGGGTTGGCAAATATGTAAGGACAGTCCCCCTTCCCCCAGTGCTGGGA | 120 | 57 |
| chr7 | 144235257 | 144235377 | 772751_42001748_Shank2(210274)_499 | 1 | + | good | GAGGAAAACAATAAGTCTAAGACATTTCTCCAGAGAAGTTCTAAATCACACCCTCTCATCCATTCCTGACTGGGCATGTGTGTGTGTGTGTGTGTGTGTGTGTACAGGTTCATGTGGGGG | 120 | 46 |
| chr7 | 144235377 | 144235497 | 772751_42001748_Shank2(210274)_500 | 1 | + | good | TGCATATCATTGCATTTGTAAGAATCTGTGTGTATGCACATTGCCCGAGTGTGTGTACATGGGTGTGTCTACAAATACATTCATGTGAGTGTGTGTATCTTGGCATTTGCAAGAGTATGC | 120 | 41 |
| chr7 | 144235617 | 144235737 | 772751_42001748_Shank2(210274)_502 | 1 | + | good | ATCCACACACTTCTTGGCACATGCTGTCCTATCTGTGTCTGAGCGATTAGAGACACCAACTGCTCATACCTTTGCCAGCCACCCTCCTCAGGAGCTCTCTGAGTCCGCCTATATTGTAGG | 120 | 52 |
| chr7 | 144235737 | 144235857 | 772751_42001748_Shank2(210274)_503 | 1 | + | good | CGTGATGGATGGGTGGAGTTCAGCTGTTCTGTTCCCAGCATATGTGTGCCTTGCCCAGCTCAGCAAGCTCATTGTGCCTTTCACACACCCTAGCTGGTTGTCCTCCCAAACTCTGGGGAG | 120 | 55 |
| chr7 | 144235857 | 144235977 | 772751_42001748_Shank2(210274)_504 | 1 | + | good | TAGCCTTTTCCTCGTCCCTACTTTGTTGCTGATGGAGGAGTCCTGGGGGCCAACAACTCCAATAATTACCCTCAGGTCCCATAAACAACCAACCCCAGAGCAGGAGTTCAAATATATGTG | 120 | 48 |
| chr7 | 144235977 | 144236097 | 772751_42001748_Shank2(210274)_505 | 1 | + | good | GCCATGAACATCGAGCTCTTTGCTGTTCCTAGAAGTGTCTATCTGAGTGTGAAGGAAAAACAGACAGAATAAAGACAGTCAGTCAGATAGACACTCCTGGGACATGTCCACCTTTGTGAT | 120 | 44 |
| chr7 | 144236097 | 144236217 | 772751_42001748_Shank2(210274)_506 | 1 | + | good | AAATAGCCTGAGCCAGTTAGTTTTTGTCAACTTGCCACAACCTATCACTTGAGGAATTGTCTAGATCAGGCTGGTCTCTGGGCATCTCTGAGAGATTGTCTTGACTAATTAACCTAGCCC | 120 | 44 |
| chr7 | 144236217 | 144236337 | 772751_42001748_Shank2(210274)_507 | 1 | + | good | ACTATGGGTAGCTCCATTCCTTAGGAAGGTGTCCTGAACTATGTACAAATGAAAAAGTCAAGCTGCATACCTATCTCTCTGCTTTGACTGTGGATGTGATTGCTTGCTTGAGTTCTCACC | 120 | 43 |
| chr7 | 144236457 | 144236577 | 772751_42001748_Shank2(210274)_509 | 1 | + | good | TAAAGGAATACAGTCCTGTACTAGCAATGACAGTAGCACTTTGCCCAGTACCCTATAGTTTATGCAGCCTGTTGGGACAGGTGGGTGTTTTTAGGTGTTATCAAGGCTGTAGCAGAGTCC | 120 | 46 |
| chr7 | 144236577 | 144236697 | 772751_42001748_Shank2(210274)_510 | 1 | + | good | TTGTGTGCTGTGGTTCCTGCCATGTGGACCTGGCACTGATACCCTTGGTCCACTGGCCCTGCAGCAGTGTTCATACTTACCGACCCATTACTGGTAGATGGGGAAACTGAGTCCTTAGAG | 120 | 53 |
| chr7 | 144236697 | 144236817 | 772751_42001748_Shank2(210274)_511 | 1 | + | good | AGAGTGGCTGAGACACAATTGTCCAGGAACACCAGCCAGGGACCAGGCAGCAGCCTGTCATCTTTGATACTCCTCAATCACTACTCCTGAACTCCTGACCAGCCTCACTCCTCAGACATA | 120 | 52 |
| chr7 | 144236817 | 144236937 | 772751_42001748_Shank2(210274)_512 | 1 | + | good | AAGATAATCGAGGAAGGTAGAGACTATCATGTCCACCCCTGAGCCCCTCACCTCTTTGATGAGGAAGGGCGTATCCAAGGACACGAGCTTTAAGATGTATGCTCCAAAGACTTCGAGGAT | 120 | 48 |
| chr7 | 144236937 | 144237057 | 772751_42001748_Shank2(210274)_513 | 1 | + | good | CACATGTGCCCGCTGTGTGGGGCGGGCCTGGAGAGGGAAGGCAGAGCTTCCGTGGAAACCCTGGCTATTTTTACTCCTGTCTCTCCTGCTTCTGCCTCCCACTGGATTCTCGGTGAGCTG | 120 | 60 |
| chr7 | 144237057 | 144237177 | 772751_42001748_Shank2(210274)_514 | 1 | + | good | CGGCGGCAGCAGCCCCCTGGGTCCCTCGCGTGCCAGGCACAGGCTATTTTTATGACCCTCTAATTTGCCCATAGATCATGTCAGACCACCAGGGGCAAGGTGGGGAAGAAAGGGGTACAG | 120 | 59 |
| chr7 | 144237177 | 144237297 | 772751_42001748_Shank2(210274)_515 | 1 | + | good | AGAAGCAAAACGTCTGTGGGAGGGGGCCTGCACCCACTATTAAAGTGCAGGGGATAAGATTAAGGTATAGCTGGAGATAAACCAGGCCTCCCAAGTAGGAGGGAACCCACTGCCTTTTAC | 120 | 51 |
| chr7 | 144237297 | 144237417 | 772751_42001748_Shank2(210274)_516 | 1 | + | good | AGCCACAGCAACCAGCATGGCCTCACTGTGCCCAGCATCCCCCGGAATAGACCAGGAGACCAGGGTTCTAGTCCTGAGTCCAGCCTTAACTGGCTGGGTGACTTTGGGACATTCACCTAA | 120 | 57 |
| chr7 | 144237417 | 144237537 | 772751_42001748_Shank2(210274)_517 | 1 | + | good | CTCCTTTGGGCTCGGTTTTCTCTGTCTTTGCCATTTATAGAGCATCTAGGTCTGCATGCTAGATAAGCAAGAAGTAAATTCTGACACCAACTCTACAAAGTCCCCACAGGCCATCTGAGT | 120 | 45 |
| chr7 | 144237537 | 144237657 | 772751_42001748_Shank2(210274)_518 | 1 | + | good | CTGGCAAGGGTAGGGAGCAGCTCTAAGACCTCTGCACAGGCATTGAAGACCTAGATAATACCAGAGTTAGCCTTTCCCTGAGACCTACAGCATCTCTGGTGCCCTTTCACGTGCTTTATA | 120 | 50 |
| chr7 | 144237657 | 144237777 | 772751_42001748_Shank2(210274)_519 | 1 | + | good | CATCTTATGCCTACAGTATCCCTCTTATGGATAATTTTATGTATGCCCGTCAAACAAGTCAAGACAGCTGGAGGTGTGCCCAAGGTCACACAGCTGGAGGCTGGGACGTGACCCTCCCCA | 120 | 51 |
| chr7 | 144237777 | 144237897 | 772751_42001748_Shank2(210274)_520 | 1 | + | good | CCCACTCAGCCTCCCCCCCCCTCCCCCCGGGGCAGAAGGACATTTGATGAACATTCCTTTCCCACTCTTTCCTAACTCATCTGCAGCTCAGAAAGGAGAGGGCTCTTGTCTAAGGCCAT | 120 | 57 |
| chr7 | 144238017 | 144238137 | 772751_42001748_Shank2(210274)_522 | 1 | + | good | TAGCAAGATCTTCTGTCTCTAAAAATAAGCAGCTTAAGATTTGTTTAATTAATTAAGTTAAAAGAAAAAAATGACAGATGGCAAACAGAAGAAGAGTCCCTGCCCTCTTATTTGACTTCA | 120 | 32 |
| chr7 | 144238137 | 144238257 | 772751_42001748_Shank2(210274)_523 | 1 | + | good | ATCGCTGGGGATGAAGAGAAAGGGGAGGCACCATAGGGAAGCACTCGGAAGGTGGTCAGCTCACCCCACACCCTGAACCCCAGCTCTCAGCTCTGCCTCTGAATTTCAATCTGGTGTTCG | 120 | 57 |
| chr7 | 144238257 | 144238377 | 772751_42001748_Shank2(210274)_524 | 1 | + | good | CCCAGCAAGATCCTTCAGGCAGAAGTTGCAGCATGTCTGAAGAATGTGCCCTAGTCTGCCTCCCATGCACTACCAGGAGTCCCAATCACCTGGAAGGGTGCACTGTCCCTGAGCCCCGGG | 120 | 58 |
| chr7 | 144238377 | 144238497 | 772751_42001748_Shank2(210274)_525 | 1 | + | good | CCTCCCCTTTGGCATTAGTTCCTTGAAGTGATGGGACTACCTTGCTGTCCACTATACAGAAAGCAGGGAAACTGCCCAGGCGGAGAAGATGCCCTGTTCCAAAATGAGAAAGCAATTACT | 120 | 48 |
| chr7 | 144238497 | 144238617 | 772751_42001748_Shank2(210274)_526 | 1 | + | good | CAGGACAAGGACGGTCACTGCCTACAAGTGACACCTCTGGCCCCTGCTAATAGTGGTTGAATCATTTTCCTCTCCCTGCTAATGGCGTGCGTAATTGCACTGGTTTTTCAGGCGTCCTTC | 120 | 52 |
| chr7 | 144238617 | 144238737 | 772751_42001748_Shank2(210274)_527 | 1 | + | good | ACCCGTACTTATTTAGACTAGTGACTGATGCGTTGCCATGGAAACCGGCACTAGGAGGAAAAAAAAAAAGTCCCCGGTACAGTGAGGAAGTGGAAGGCTGCAGCTCCAGTCTGCAGCTAC | 120 | 50 |
| chr7 | 144238737 | 144238857 | 772751_42001748_Shank2(210274)_528 | 1 | + | good | AACTTGTGTCAAAGAACCGAATTTTCAGCCGATCCTGATTTTTCCATCTTCAAAAGTCAAATTCGTGAACCTCCTCGTGGATTAAAGTCTCTGGGGTTGTCTTTCTTAGTCCTGTGATCT | 120 | 41 |
| chr7 | 144238857 | 144238977 | 772751_42001748_Shank2(210274)_529 | 1 | + | good | ATTCTTTTTTTCTTCTCTTCTCCTTAAATTCTGACTGTAGAAGACTGCAACTTTTTTTTTTTAACTCCTTGTTGGATTAACAGAGAATATAGTAAGTGAGGCCAGCCTTGAGGACCCAGT | 120 | 36 |
| chr7 | 144238977 | 144239097 | 772751_42001748_Shank2(210274)_530 | 1 | + | good | CTCCACATGGACCCAGAGGAGGAAGAGGCAAGGGACCGGTCTGAGTGAGATGGTCATACCCAGGTTAGGAACGATGGCTGAGAGCTCCCGGTTCAAGAAGAGAGTTCATTTTGGAGGTAT | 120 | 53 |
| chr7 | 144239097 | 144239217 | 772751_42001748_Shank2(210274)_531 | 1 | + | good | GGTGCTACCTCCCCAGCCTGCTTGGGGTTGGCCGATGTGAGTCTTTGTTCTCTGCTGGAGCAGTGGAGACAGTAGCCCCGGAACTCTGATGCTACTGGCGCTCGCTCGCTCAGTGATTAT | 120 | 58 |
| chr7 | 144239217 | 144239337 | 772751_42001748_Shank2(210274)_532 | 1 | + | good | GGTGGCATGCTCGCTCGGTGTAGTCAGCAGGCTGATGGGTTCCTCCAGGCTATTGTTTCCATCATGGCTTTCTCCTACTTCTGACTTAGACGAAAACAGAGGTAGCTGACGGATCACCCC | 120 | 53 |
| chr7 | 144239337 | 144239457 | 772751_42001748_Shank2(210274)_533 | 1 | + | good | AGGATGCAGGCATCTCTGTGGCAGAGAATCCCCCACAGTCTTATTTGTATTCTTGAGTGGCTCTCTGAGCACTGGTGGGTTTCTGAGTCCAACTAGAACATCTGTTCCTAGTGAAATACT | 120 | 47 |
| chr7 | 144239457 | 144239577 | 772751_42001748_Shank2(210274)_534 | 1 | + | good | GTGTCGAGGGGGCTGACTGTATGGGGAGGGGGTGACTGAGGCACCAGTGTCTCAGGATCCCTGCAAAGTGCATGGGAAGTTCCCCAAGCTGGGACGGGGATGCCTGCTTCCCTCAGTGCA | 120 | 62 |
| chr7 | 144239577 | 144239697 | 772751_42001748_Shank2(210274)_535 | 1 | + | good | GGCTGCCCTTCTTCAGAGAAAGCATGGCCAGCTCAGCTTCTGTTATTGTTCTTTAATCAAGCAGACTCCCTGCTACTCTGGTGGTGTGGCAGTAGAAAGAAAGAAAGAAAGAGAAGCTAA | 120 | 46 |
| chr7 | 144239697 | 144239817 | 772751_42001748_Shank2(210274)_536 | 1 | + | good | CAGGAGCTGCTGTCTTCCCAGGGTTCAGACTGGCTCAAGGACGTAGTGAGCAAGAGGTTATCTCACCCCACCCCTGGATGTCACTTTGAGGTCTGCAAAGATAAATGTGAGTTGCCAAGA | 120 | 52 |
| chr7 | 144239817 | 144239937 | 772751_42001748_Shank2(210274)_537 | 1 | + | good | TTCTAGTGACGGGGGCCCTAGTTCTCACTGCACGGCAACCCTGTGTTCTGTTTGTCACCTGTGGAGTTTGTGCTTCTGGACAGGGGCTGCAGTTATTGACTGGCAGGGAATGAGAGCTCA | 120 | 54 |
| chr7 | 144239937 | 144240057 | 772751_42001748_Shank2(210274)_538 | 1 | + | good | GGGGTCAGATGTCTGTCTGTAAAGGTCATGGCCAGGCTGAGTCTGTGCCTTCTGTTTCCATTTGGACTGCCCAGCCCTCACCCAGAGCCACCCTGATAAGGGGTGTGCTCTGATCCCCCG | 120 | 58 |
| chr7 | 144240057 | 144240177 | 772751_42001748_Shank2(210274)_539 | 1 | + | good | TGGTCCTGAGCTCTGGTTAGCCAAGGTGTGGATGCTGAAGCTCCAGTCTGCAGAGTCCTGGCTCCCTCCCACAGGAGGCAGCAGCTTCCTCCTGCTCAACAGGTCTCTCTTCTCAGTGCC | 120 | 59 |
| chr7 | 144240177 | 144240297 | 772751_42001748_Shank2(210274)_540 | 1 | + | good | CTAGTTCCCAAGCCTCAGCTGCTGGTGTCCTTAAAAAAGATTAAAAAGCTAAAAGGATCCTCCCGCCTCCAGTCCGGATCCCAAATTTAAATCCAGGCTACACTCCTCCAACCTGTGAGC | 120 | 48 |
| chr7 | 144240297 | 144240417 | 772751_42001748_Shank2(210274)_541 | 1 | + | good | CTTGGACTTCACCTCCAGGAGCCTTCTCTAGGAGGGACTGACTCTGTCTTTTGGAAGGGGTGGGGGTGGGGTGTTGTTCACCAGGCTGCTCCACTAAGGGCAGTGCAAATTGGCAGCTGT | 120 | 57 |
| chr7 | 144240417 | 144240537 | 772751_42001748_Shank2(210274)_542 | 1 | + | good | TTTAGTTGAGAAATTCTGAGTCCCTGGGGGGAGGATGCCAGTTGTGAAGTCGGACCCAAGACCAGATGTTAGAAGTGGAGATATCGGATAGATAGAGGCTGCATGCTTGGGCACACAAAA | 120 | 48 |
| chr7 | 144240537 | 144240657 | 772751_42001748_Shank2(210274)_543 | 1 | + | good | TGGGGCCCACCCAGGCCACATGACTTGGCTCAGCAGAGCATGGAAGCCTTTGCTGCCTTGGGGGAAGCAGCTAGGTTGACTCTGGGACAGGAAATTTTGTCTACCTGTTCTTATGGGAGT | 120 | 55 |
| chr7 | 144240657 | 144240777 | 772751_42001748_Shank2(210274)_544 | 1 | + | good | GCCCCTCCCCTGCCATTGTCAGTGAAGCCCTGGGCTGCTGGAGCCTGAGGCACCTGTCCTCTTGGCCTCTTCTTTTCACCGAGTGCTCCTAGAACACCAGAGGCCACCTTGGCAGGGCAC | 120 | 62 |
| chr7 | 144240777 | 144240897 | 772751_42001748_Shank2(210274)_545 | 1 | + | good | TCCCAGCCAAGCTAATCAGCCACATGCAAATCAGACCCAATTCATCCATTTAGTGTAATTTCCATTCAAAGTGCTGGTTTGGGTCTGTGGATTGTGCAGGCTTGTGGAGTCTCTTGGGCT | 120 | 47 |
| chr7 | 144240897 | 144241017 | 772751_42001748_Shank2(210274)_546 | 1 | + | good | GATTTTTCCTTTTTTTTTTTTTTTTTTTTTTTTTATTCAAACATTTTAATTACTACAGAACAGAAACCACCTCACTGACAGCCAGTGGCTCTATTCATTTGTTCAACCAATTTGAGAGGG | 120 | 30 |
| chr7 | 144241017 | 144241137 | 772751_42001748_Shank2(210274)_547 | 1 | + | good | TGGTCACTGGGCAACGTCCCCAGAAACCGCTACTACCCCCCTCCTCAATGTCCTCAGTTCCCCTTGGCCATAGAACAGCAAAGAGGGCAGAGGTGGGTAAGAGAGATGCTGATAGTCACC | 120 | 56 |
| chr7 | 144241137 | 144241257 | 772751_42001748_Shank2(210274)_548 | 1 | + | good | ACCAGGAGCACCCGGCACTCACAAGAGAGTTATCAGCTGAGCTCAGAAGGGCAAAGGGAGACAGAGGCAAAGAAGCCATGCACCCTTAGATGAGCACTTCCAAAGTGAAGAAAGTGAGGG | 120 | 52 |
| chr7 | 144241257 | 144241377 | 772751_42001748_Shank2(210274)_549 | 1 | + | good | TTGTTGTGGACCCCTCTCCCCAAAGCCAAGGGTGCCCACCCCTTCCTGTATTCCTAGTAAATCTCATACAGCATTGTGGGAGTGTTACTTAACGGGAACAGCTCACTGTTCAGATTTACT | 120 | 48 |
| chr7 | 144241377 | 144241497 | 772751_42001748_Shank2(210274)_550 | 1 | + | good | CTCTTCAGAAGCTCTGTGTGCCTTGTGGTGACCTTGATTCATTCAGTCAGGCTGACCTGGGTTCGGATCCCAGGACCTCTACTCGGGAGGCAGATGACTCTGGAGGCAATCGAGTCCTGC | 120 | 56 |
| chr7 | 144241497 | 144241617 | 772751_42001748_Shank2(210274)_551 | 1 | + | good | CCAGCTGCCTCCCGACAGAACAAGCTTGAGACAGCTGTGTGAGTTCAGTAGCATGCCCTAGGGAGCTCGGTCATACCCCCCCCCCGCCCCCACTGTAGGTCATGGGGTTATTTCTGGGCC | 120 | 61 |
| chr7 | 144241617 | 144241737 | 772751_42001748_Shank2(210274)_552 | 1 | + | good | TCAGTTTTCTCATTGGAGTTTTTACAATAAAAGCATGACTCTCTTCTGGGTGTTGGTGACTGAAGAAGCAGGGTGTGGCCATGCAGGCAGAAGTGCCTGTGACTGTGAGAAAAGGGGTCT | 120 | 47 |
| chr7 | 144241737 | 144241857 | 772751_42001748_Shank2(210274)_553 | 1 | + | good | TGGAGTCATACCTGGAAGTCACTGGGTCAGGGGTCACATGTGGGTAGGCATGGAGACCTGTGAAGCCACTTGGGATTCTGGCTTTCTGTGTCTTGCTTGTCCTCAGCATTCCCAGGATGT | 120 | 53 |
| chr7 | 144241857 | 144241977 | 772751_42001748_Shank2(210274)_554 | 1 | + | good | TGATGTGGCTGCCAAACCCTGTAGCCAGTGAAGGGGTTGTTTCAAAAACTACTAGGACCTAGGAGGTATAGTGGGATGAAGGGAAGCCACACCCAGCAGACCCTGCTCCATCTCATTGGC | 120 | 52 |
| chr7 | 144241977 | 144242097 | 772751_42001748_Shank2(210274)_555 | 1 | + | good | CAGGACCGTAGCTACTCCTGGGAAAGGAGGGAGCATGGATTTTTAGCTTCAAAGCCATGCGTGAAAGCGGTTGCTGACTACCACAATATAGGTGTGCCCTGGAAGAAAGGCCTTGGCACA | 120 | 52 |
| chr7 | 144242097 | 144242217 | 772751_42001748_Shank2(210274)_556 | 1 | + | good | GAGTGAGGTGGGACCTCTGATTCTGTGAGGGGGAAGCTCTGTAGAAGATCAGTTAAGGGCTTGATGCTAGAGTTTTAGTGTCTGGCTTTGGGTGCCCTCAGTAGCTGGCTTGATCTAGTC | 120 | 51 |
| chr7 | 144242217 | 144242337 | 772751_42001748_Shank2(210274)_557 | 1 | + | good | AGGCTTGAGGCAGAGAAGGATCTCATGTATTCTGCACTAGTTCAGTGTGTGTAACACGTGCTGACTGAGGCCAGGTGATGACTTTGAGCAGATCCCTGTCATGGTATTTCTTCCCAGCCC | 120 | 50 |
| chr7 | 144242337 | 144242457 | 772751_42001748_Shank2(210274)_558 | 1 | + | good | TCCCTCCCTGTCCGCCTTCTGGAGCCTCATTTCCAATGTGCGAGTATTCTGTGTCATTATAGCGAGGAAAAGCGACACAAGCTGAGACACCCAGATAGGAAGCCGGTGCCAACTTAGAAC | 120 | 52 |
| chr7 | 144242457 | 144242577 | 772751_42001748_Shank2(210274)_559 | 1 | + | good | ACACTCATAATGACCTCAGCAGAGTTAACCTGGGATCCCTTGCAGCCACCTTCATCAGTCTGAGGGCAGAATCCTCAGGACCTGAGGATTTCCCCTTGTTCCATCTGTCTCGGTCTCAAC | 120 | 52 |

| | | | | | | | | | |
|---|---|---|---|---|---|---|---|---|---|
| chr7 | 144242577 | 144242697 | 772751_42001748_Shank2(210274)_560 | 1 | + | good | ATACTCACAGAGCGATGCTTCCAGTATATGATCTTAGGAAGCCACAGCACATGCCCACCATGAGGGAGCTGACCGTCCAGTTCCCCATGGGCTCTCCAACATGTGAGTGTCCTAAGCGTC | 120 | 53 |
| chr7 | 144242697 | 144242817 | 772751_42001748_Shank2(210274)_561 | 1 | + | good | CAGGAGAGCCCGAACACTGGCCTGAGGGGGGGGCCTTCACTGAGGAATCATGACCCTCTCTCCCTCCCTGGTTTGGTTCGGAGCTGGTGTTCTGTGACCCCTCTTGTCTGTGCATGAGCC | 120 | 61 |
| chr7 | 144242817 | 144242937 | 772751_42001748_Shank2(210274)_562 | 1 | + | good | CTGATGTCTCGGGGGGTCTTTGATGGGGCAAGGAACACAGATGCACCTGGTTAAAAAAAAAAAAAACCACTGTGTAAAGTGACAGCCGCCTGACAGTCCCTTAAGTCGGTATTCAGCTCAT | 120 | 48 |
| chr7 | 144242937 | 144243057 | 772751_42001748_Shank2(210274)_563 | 1 | + | good | GTTTCATAAAAGGTTTTCAGTTGCTAGCAGATTCCCAGGAGAACTGAGAAGAAAACCACTTTGCAGGGGCAGGGAGGAGCACCTCTGTGAAGTGGTTTGTGTTTATTTGGGAAGGGAAAG | 120 | 46 |
| chr7 | 144243057 | 144243177 | 772751_42001748_Shank2(210274)_564 | 1 | + | good | CAGTTCGTGTGCTGGCAACCTTCCTCTTCTGACCTGGTTTTAGATGCTGAACTTGGCTCTGGCACAGGGCCTTAGTGTTTGGTCCCCATGGATGCAAGTCCTGGGACGCTGGTGGCCCCT | 120 | 57 |
| chr7 | 144243297 | 144243417 | 772751_42001748_Shank2(210274)_566 | 1 | + | good | TTGACTTACTTTCTTTCAGTCTTACTTTTATTATATTTAATCATATATACATGTGGGGGCAGGATATTCACATTAATGTTGGTGCCCTTAGAGGTCAGAGGTGCTGAATCCCCTGAATCT | 120 | 37 |
| chr7 | 144243777 | 144243897 | 772751_42001748_Shank2(210274)_570 | 1 | + | good | TGTGGTATATTTGTGTGATATGTGTGTGTGGTATATGTATGTGTGTAGTATGCTCATGGACTTCAGAGGCCAACTTTTAGAAGTTGGAGGAAGGGTCTCTCTTGGTGTATCTGCTGTACT | 120 | 42 |
| chr7 | 144243897 | 144244017 | 772751_42001748_Shank2(210274)_571 | 1 | + | good | CCATATTCCAGGCTAGCTGGTTTGTGAACTTCAGACAGTTCTCTTGTCCCACCTCCCATCTTACCATAGGATTACAAATGGCTGCCATGACATCTGGATCTTCAGGTCTGTCAGGCCATC | 120 | 48 |
| chr7 | 144244017 | 144244137 | 772751_42001748_Shank2(210274)_572 | 1 | + | good | TTGTCACCTGCTGAGCCATCTTGCTGGCGCCTACCTTGTTTTGTTTTGGAGCTAGTCAATTAGACTAGACTAGCTGGCCGGAAAGTTCTAGAGCTCTGCCTGTCTCTGCCTCACCACTTA | 120 | 50 |
| chr7 | 144244257 | 144244377 | 772751_42001748_Shank2(210274)_574 | 1 | + | good | AAGCGTGAGGCTCTTCAAGAAGGCATCTCTCCTCTGCTTACTGCTGGGTTCATAGCCAGATGCTGAGACACAGAGCTGGCCCAGGGAGGAGGGAATGAGCAGACATCACACACCACAGAT | 120 | 54 |
| chr7 | 144244377 | 144244497 | 772751_42001748_Shank2(210274)_575 | 1 | + | good | GCATGCACACACCAGGGCAGCATGCCATGCTCCAGGGAGCTGGGGATAAACAAAAGTGATGTCTCAGCTGTCTGTTGGCGCCATATCTCACCACAGAGGCAGCCGTGCCTCCTAGGAACC | 120 | 57 |
| chr7 | 144244497 | 144244617 | 772751_42001748_Shank2(210274)_576 | 1 | + | good | AGTGAGATTCAGGGAGGTGAATGTCAGCCTTGATCAGTAGACGGAAAGCCTCCAGGCCCTGGGTTGCTGGTACTGAGCGAGCTCTGTCAGCTAAAGGGATCATCCCACAAAGGCATGGAG | 120 | 54 |
| chr7 | 144244617 | 144244737 | 772751_42001748_Shank2(210274)_577 | 1 | + | good | CAGAGCGAAGATGCTGTCCTCAGCCCTATTGAATGAGGAAGGGGAGGATCCAGAGCGTGGAGCCCAAGCCCTGGTCTCACTGAGGTAGCACAGGCACTGTCTGGTTTTAAATTGTCATTT | 120 | 52 |
| chr7 | 144244977 | 144245097 | 772751_42001748_Shank2(210274)_580 | 1 | + | good | TGAGATGGGATCCTCTGCAGCCCAGGCTGGCCTCACATTCACTGTGTAGCTGAGGATGGCAATGAACCACTGCTCTGTGGTCAAGTGCTTCTAACAGAGTTCTGAGTCCCAGAGTACTGA | 120 | 52 |
| chr7 | 144245097 | 144245217 | 772751_42001748_Shank2(210274)_581 | 1 | + | good | GTCTCAAAGCACTAAGTCCCAGGTTCTGAGTCTGATGAACAAGACTCCCACTTCTGGCCATGCTGCCCTATAAAGATGGCCTCAGCCTGTGGCAATGGTGGACAGAGGAGTGGCCTTTGA | 120 | 52 |
| chr7 | 144245217 | 144245337 | 772751_42001748_Shank2(210274)_582 | 1 | + | good | ACATAGCCATTTCTTGGTCCCCAGTATCTATGGGTGTCCCACTCCAAATGGCCAAGGGGAATTAAAGTCACTAATGGAATCCAGGTTGCTGTCTGTCAAGTCTAAGGCCCCTAAATATCT | 120 | 46 |
| chr7 | 144245337 | 144245457 | 772751_42001748_Shank2(210274)_583 | 1 | + | good | GGGTGAGCTTTCTGTGACCTCAACCGTCCTTAAAGAGAGAGGGTAGAGTTCTGAGTCACAGTGATGAGATGTGAGAGAGACTGACATGCCACAGCTCTCTCTGATTGGAGGATGGCTGGC | 120 | 52 |
| chr7 | 144245457 | 144245577 | 772751_42001748_Shank2(210274)_584 | 1 | + | good | TCCAAGGAACACAGTCTGTTTAAGTAGGTAGGGAATGGGAGGGAACACCTGCAGAAATGCCTTCAAAAAGGGCCACAACTTAACTACAATCTTGCGGGTCACCCAGGAAGACTGAGCCAG | 120 | 49 |
| chr7 | 144245577 | 144245697 | 772751_42001748_Shank2(210274)_585 | 1 | + | good | GATTCTGTAGGAAGACAGGTCTGTGTGGCCACAGGCCCCCAAATCTGCCCTCTGTCATTAGGTCAGTGTAGTGCACAGTGCATTTGTGCAGTCTGACGATGCCTCCTTTGTGTCTGCTCC | 120 | 53 |
| chr7 | 144245697 | 144245817 | 772751_42001748_Shank2(210274)_586 | 1 | + | good | CTTGCTAGATCAGTTCCTGGGACCCAAAGGTACTCAAAAAATGTCCTGCAACTCATCTAAGAGATATACTTGTGTGCAGAGTGGCTCCCCAGATCCTACTTTTGTTTTATATGCTCTTCT | 120 | 42 |
| chr7 | 144245817 | 144245937 | 772751_42001748_Shank2(210274)_587 | 1 | + | good | TTTCAAATCAGAGTCCTCTACCTGCTGACTCTGGGTGCTGAGGGCCTTAGGCCTTTGTCCCCTCATGGTAGGTCACTGTGACTGTCTGATGGCCAGAGCAGAGAATATACAGACTGTGAA | 120 | 51 |
| chr7 | 144245937 | 144246057 | 772751_42001748_Shank2(210274)_588 | 1 | + | good | GGTGAACTTCAGTCTGTAAAATGGGGTACGAGCTCCCAGCCTAGGACCCCTCCAGACCTGCTCAGCCCTAAAGAGCCCCCAGAGTTCTAGCCTGCCCTGGTAGCTGAGAGGGTGTGGCCC | 120 | 60 |
| chr7 | 144246057 | 144246177 | 772751_42001748_Shank2(210274)_589 | 1 | + | good | GGCAATAGGCCTTGCTGCTTCCATAGCTGTGTGTCTTCCCTACTTCACAGAGAGCCCAGCACGGTCTTTGGTTTTCGTCTCTGGCCTCAGTTGTCCCCTCAGCCCACTTGGAGTTGGGTG | 120 | 56 |
| chr7 | 144246297 | 144246417 | 772751_42001748_Shank2(210274)_591 | 1 | + | good | TAGTGAGTTGCATGCCATCCTGAGCTACCGAATAAGACCCTGACTCAAGAAAACAAGATAAATGAATGACTAAAACTTACAGGAGACACAGGAGCCTGGGCAAGGAGCCGGGTGCAGAGC | 120 | 48 |
| chr7 | 144246417 | 144246537 | 772751_42001748_Shank2(210274)_592 | 1 | + | good | ACCAGTGAGCCAGAGCCAAACAGCCTGGCCATGTTGTCAGCCAGAGACCCTGAGCCCTTTGCAGAAAGTTGGAACAGTTCCCCAAAGCTGAAAAAAAATCCCTGGCATCACAGCCCATGT | 120 | 52 |
| chr7 | 144246537 | 144246657 | 772751_42001748_Shank2(210274)_593 | 1 | + | good | GAGTGGGTCTCTGTCCGGCATTGGGGGCTGGACATAGCACCCTCTTGCTCTTCTGCCACACCAGGAAGCCAGCATCTTAGTCTTCAGCTCCCCAGCCTCCAGAATGGCAAAGAGATTAAT | 120 | 55 |
| chr7 | 144246657 | 144246777 | 772751_42001748_Shank2(210274)_594 | 1 | + | good | TTCCATTCTTTATAGACAACCTGGCCTTGGGTATTGTTACAGCATCAGAAAGGGACTGAGCTAAGAATGAAAACTAAGCAAGGGACTGGATTCAATAGAAAGAAGGGCAGAAATAGCCAA | 120 | 41 |
| chr7 | 144246777 | 144246897 | 772751_42001748_Shank2(210274)_595 | 1 | + | good | AGTGTTCGTCTCAGAAGGAGTGTGTCTAGGGCACAGACACTGACATGTTCTGCAGGAGCAGTGTCTCAGCTCAGAAAATGGGCTCACCACTGCCCCTCTGGCCCTGTCTCTGAGTCACTA | 120 | 54 |
| chr7 | 144246897 | 144247017 | 772751_42001748_Shank2(210274)_596 | 1 | + | good | GCACGATCCTTTTCCACTACAAGAGGAAGGCAGTTCTTGTGTTCCAGGCCCCCAGGGACATCCATCCTGGAGTATAAACGCCTCAGAGTCTTCTGAGTCTTTGAAGCCATTGTCTTAGAG | 120 | 50 |
| chr7 | 144247017 | 144247137 | 772751_42001748_Shank2(210274)_597 | 1 | + | good | GGATTCCTGCTCCATCAGGGGCCCTCTCCCTTCTCCCTCCCAGAGATCCACACTCAGCCACTACCAGCTTTACCTCAACCTTAAGCCCCCAGGCTACAAGTGGGCGTTCCCCCATGTTAC | 120 | 58 |
| chr7 | 144247137 | 144247257 | 772751_42001748_Shank2(210274)_598 | 1 | + | good | TTATAGCATCCCATCTGTCAGCTCATGCAGGTCCTTGGGGATCAACCCTCTTTCTGTCATGGCTCTCTCCTCTACATCCCCATTTGTGGAGTGGGACAGCAATATCCCCACTGCCCCTTG | 120 | 52 |
| chr7 | 144247257 | 144247377 | 772751_42001748_Shank2(210274)_599 | 1 | + | good | GCTCTCAGTGGGATTCCACCATGATTGTCAACCAGCAGTTGAAACCTGATAAGATACAGAGGTTTGTCAAGAATACTAACAGCTTGTCAGGATCTAGACTCACCACAGAGACCAGCTTCT | 120 | 45 |
| chr7 | 144247377 | 144247497 | 772751_42001748_Shank2(210274)_600 | 1 | + | good | AGGCACGTCTGTAAAGGAGCTTCTAGATTAGGTCTGAGGGGGCAAGAACTGCCCTGAATGTAGGGTCAATCTCTTTCTGATTCCCCACTATGGATGCAGTGTGATATGTGTGCCACCACA | 120 | 49 |
| chr7 | 144247497 | 144247617 | 772751_42001748_Shank2(210274)_601 | 1 | + | good | CCTTCCCACCATGATTGACTGTGTGGTAGAATAAACGCTTCCTTTTCTAAGTTGCTCTGTTGGTCATTTGTTATCTAGATGAGACAAGTCACTAGTACTGATAACTAGTATCTCTGACCT | 120 | 40 |
| chr7 | 144247617 | 144247737 | 772751_42001748_Shank2(210274)_602 | 1 | + | good | ACAGTGCTCCATAGCTCAGCCATAGCGTTCGTTACCCCTGCTCGAAATTCACACTAACCCCTGCTGGGCAAGATCATCCAACACTCTGCCAGAGAGAATGAAGTGCCAATGGCCTCATGG | 120 | 52 |
| chr7 | 144247737 | 144247857 | 772751_42001748_Shank2(210274)_603 | 1 | + | good | CATAGATTAAGACTGTAGAAGCCACAGCTGATCTCAGAAGCTAAGCAGGGCCAGGCCTGGTTAGTATGGGAAAGGAAAATAGCCTGGTGGTCCTGGGGTCATCTCATGCCATTGTAAGCC | 120 | 50 |
| chr7 | 144247857 | 144247977 | 772751_42001748_Shank2(210274)_604 | 1 | + | good | AGGGAGCACCTGTATCCATCCAACTTGGGTCTGATGTCAATGTACACGGCACCCAGGGAAGGCGCTCAGGCAGCCTTCAGCTTACACCTGACCATTGTCCTCTCTGATGTTTTATCCTTC | 120 | 52 |
| chr7 | 144247977 | 144248097 | 772751_42001748_Shank2(210274)_605 | 1 | + | good | TGTGACTTCCTGGTGTGTCCTGGCTGGGTCAAACACGACCTCGGGAGGATATTTCTGGACCCTCCTCCCACAAAGTACTCAGCAACATGCCATTGAGACCTACTTCCCTCTGAGGGAGAT | 120 | 53 |
| chr7 | 144248097 | 144248217 | 772751_42001748_Shank2(210274)_606 | 1 | + | good | GGGTAGAAGATGAAAGCTAACTGGCCAGGGTCCAGGGCATCTTTCTTGTGTTCTTGCTTAGACTTAGACTAACACCCCTGTGTTCCATGTTCCTCAGGCACAGTGCATGCATTTTGAGTT | 120 | 48 |
| chr7 | 144248217 | 144248337 | 772751_42001748_Shank2(210274)_607 | 1 | + | good | GAGCCTTTGTATTAAACTGTCTTCTGCATGCCTAGCATTGCTCTCCATCTCTTCCATAGTTTCTATGCTTCTTGTCAATCAAAACAAGGGACATTTACCACTCAGCTCTTCTAGAAAGGA | 120 | 41 |
| chr7 | 144248937 | 144249057 | 772751_42001748_Shank2(210274)_613 | 1 | + | good | GCATCAACATACAAATGGCACCAGGCCCATCTACAACACCTGACCCTCCAGTGGTTAAGAGGGTTAGTGCCCTCCATCAGGGCAAGGGAATTGTCTGCCTTTGAAGTTAGTCTTCAGCCA | 120 | 51 |
| chr7 | 144249057 | 144249177 | 772751_42001748_Shank2(210274)_614 | 1 | + | good | GCATCCAGGGCAGAATCCCAGGGTTTCCCTGTCATTAACACAGACACTGGAGATACTTGACTCTTATGTGCCCAATGTTCTCTGTCTGGCCACCTCTTCAGCAAAAAGTAACCTAAAATT | 120 | 46 |
| chr7 | 144249177 | 144249297 | 772751_42001748_Shank2(210274)_615 | 1 | + | good | ATCAATCACAAGGCCTCCACAGGAATTCTGGTTGAGGGGTATGCTGCCCCCTGGGGGCATCAGGAAATGGATGCAGGGCTACTGATTGTCACAGGAAATATGGCATTCCTTTGTATCCCC | 120 | 52 |
| chr7 | 144249417 | 144249537 | 772751_42001748_Shank2(210274)_617 | 1 | + | good | TCTCTAGCATGCCAAAAGCTTCTCATGTAAGTCTCGCAATCCTTCCACATACTTTAAGTCAGTCCCAAACATAACACCCAGTGCAGTCTGTGTGCTCCGCAAACATTATTCTACTGAGTT | 120 | 43 |
| chr7 | 144249537 | 144249657 | 772751_42001748_Shank2(210274)_618 | 1 | + | good | GCTTAGGGGAGTAAACACCGGACAAACCTGTTCACACTCAGTGCACATACAGTAATGTCCAAATATGGTCTTTTAAAAAGTTTAACATCATTAATTTTAATTATATCTCTGTGTGGGCAT | 120 | 36 |
| chr7 | 144249777 | 144249897 | 772751_42001748_Shank2(210274)_620 | 1 | + | good | CAAATGCAGCAAGCACCTGTAACTACCGAGCCGCTCTCCAGCCCCTGTTTTTCAAATATTTTCAGACACACCTTAGGTTGATAGAATCCATGGGTGTGAAAGCCATAGCTACATGAGGCC | 120 | 48 |
| chr7 | 144249897 | 144250017 | 772751_42001748_Shank2(210274)_621 | 1 | + | good | TGGTGCAGTTAGGGATGGCGCAGGGGAATTCAAAGCTATGCCATGTCCAGAACAACCCATTCAGCAGCATGGTCCGGCAAGCACAACCTGACTCACCCTAGAGTGTTTCTACCTAAACTG | 120 | 52 |
| chr7 | 144250017 | 144250137 | 772751_42001748_Shank2(210274)_622 | 1 | + | good | ATACGGGATGGAATGGAATATCAATGTCGCGTGGGACAGCTAAAAAAAACCAAAATGTACTCCAGTCTGTGGGGCTCAGCTTGGGCCTTTTAGGCTCGTCACTCAGGGCAGAACCCAGGG | 120 | 51 |
| chr7 | 144250137 | 144250257 | 772751_42001748_Shank2(210274)_623 | 1 | + | good | GCAAGTGGGTCCTTCCAGGAACTGAGACCGTATGGCTTTATCATCGATGCCAAGAGTCTCATGCGCTCTGTCACTGTGTGACCTCAGTCCACCCCTGTCCTCCACTTGTTGCAGTGCAGT | 120 | 54 |
| chr7 | 144250257 | 144250377 | 772751_42001748_Shank2(210274)_624 | 1 | + | good | AATGGGTCCTCATTTGAACAGGATGTCCAGGAGCAGAAGGTGTCTTGTTTCAGGGCCCTGGGATGGTGTCCAGTTGGCTTGGATGATATAGACCCTTCTAAGCTCAGGCAGCAGCAAGCA | 120 | 52 |
| chr7 | 144250377 | 144250497 | 772751_42001748_Shank2(210274)_625 | 1 | + | good | GGGTATCATATGGGGGCATAAATTCCTCTTGTATGTAAGCCAGACACCAAAGTGGGTGTTAGGGAGGGATGTGTGCAGAATGTCTTTACCCTACTCTCTCTGTATCTCTATGTTGTGGGC | 120 | 47 |
| chr7 | 144250497 | 144250617 | 772751_42001748_Shank2(210274)_626 | 1 | + | good | TCTGCTGTAGACAACATTGTAGGTGGGAACCTGAGGTTGCTGAGAGGACCTTTCCTGCTTTGATTAGAAGACCAGGAAAGAACTGAGGCTCAGGTTTGTCTTGTCTGTCTTAGGGGCAGG | 120 | 49 |
| chr7 | 144250617 | 144250737 | 772751_42001748_Shank2(210274)_627 | 1 | + | good | GGCATGGCTCAGTGATGGAAGGTTCATTTAACATGCCTGAGACCTTGGAAACCATCCCCATTCATGGAACAAAAAGGAAAGAGGAGGAGAAGGGAGGATGGAGAAGGAGAGAATGCGGCA | 120 | 49 |
| chr7 | 144250737 | 144250857 | 772751_42001748_Shank2(210274)_628 | 1 | + | good | CTGTTTTATGTGCTTCATAGTCTATCTATCCACTGGTATCCCAAGAGAGGAGGGGGTTACCCCAGGGAATATATCTGTGGCTTGAAGGGCTTTACATTCTTCTGGTCTTTACTCATACTG | 120 | 44 |
| chr7 | 144250977 | 144251097 | 772751_42001748_Shank2(210274)_630 | 1 | + | good | GGTTGCTTTGCATAAGGGCTATTTTAACTGCTGACAATCGGAGACTGTCATGAAATCTGTCGCTGCCTTCCACACATCCCAAGGGGTGTATGAAATAGAATTTCCAAAGATTCACACCCC | 120 | 45 |
| chr7 | 144251097 | 144251217 | 772751_42001748_Shank2(210274)_631 | 1 | + | good | CCATTTTTCCATGTAGATTTCTCAGAGTGGCGTTTTTAGGGCATATCAGCTTAAATGCTGCCTTGTGGGGAGGGCTCATAGCTGCCTCCCGGGAAAAGGAGATCCCCAATTAACCTGGGA | 120 | 49 |
| chr7 | 144251217 | 144251337 | 772751_42001748_Shank2(210274)_632 | 1 | + | good | TATGTGTCATGAGGGAGCACCCACCCAGGGCATCCATTAAGGCGAGCATCCTGTTCTCTAGGTGAGGCAGGAACACCCAGAGGCCCATTGAGAAGGTTTCTCCTGGTCACTGCCTTGTCC | 120 | 56 |
| chr7 | 144251337 | 144251457 | 772751_42001748_Shank2(210274)_633 | 1 | + | good | AGAGCTGGAACTGGGGGTTCCCCAGTGCAGATGGGCTCTTTGCACTCTGCTAGCAACCTCCAGGAGGCACTAACGTATTCTCTGTAGAGCCTTAAGCCCATCACAGCCAGACCAAGCAGA | 120 | 55 |
| chr7 | 144251457 | 144251577 | 772751_42001748_Shank2(210274)_634 | 1 | + | good | TGGAGGTACCAATTTTCTGACGTTCACTCATTTACCAGAGATCCTGGGTGCCCACGTGGCCAGGTTGTCATAGGAGCCAAGGATATGTTGGCACAGTGAAATCCGAACTGGGGAAAGAAG | 120 | 50 |
| chr7 | 144251577 | 144251697 | 772751_42001748_Shank2(210274)_635 | 1 | + | good | AATCTAGCAAATGCAAAGGCCCTGAGATGCAAATCAGCTCGTCTTCTAGGAGCAAAGAGCAAGGGGCACACTGGTGGGGATGAGGATGCAGGGATGGGCAGGGCTGATAGCATCACTGGA | 120 | 53 |
| chr7 | 144251697 | 144251817 | 772751_42001748_Shank2(210274)_636 | 1 | + | good | TTCACATTCCTTCAAAGTCACTAGGGGCCATGGTTGGATAGGTGGAACAAATCACCAGGAAACCTGGAGACAGGAAGCTACTTCACAAGTCCAAGTCAGAGATGGTGATCCCTTGGACCA | 120 | 48 |
| chr7 | 144251817 | 144251937 | 772751_42001748_Shank2(210274)_637 | 1 | + | good | GGATGATAAAGAGGCCTGAACTTTGGGATGCTGCAGACATTGCAGTGTAGCAGCTGTGTGAGGAACTCTGATGCTAAGTGAGAAAGGAGACTTGGTTGTGAACTAGGTCTTCTAGAAACT | 120 | 46 |
| chr7 | 144251937 | 144252057 | 772751_42001748_Shank2(210274)_638 | 1 | + | good | TGTAGAACTTTCTATACAGTAAAGGAAGCAATAGTTCTCTCTTGATTGGCTGGCTTGTCTGTCCCTTTATCATCCACAAGGGGGCCTTCTTTTCAGTCAAAGGGATGTGTTCTACACTGT | 120 | 42 |
| chr7 | 144252057 | 144252177 | 772751_42001748_Shank2(210274)_639 | 1 | + | good | GCATGTAGAATACCCAGCTAGATACATGTATGTGTGCATTCACCCATGTGTGCACACGTGGAAGCCAAAGGAAACTCTGAGTATCCTCTATCACTCTCCACCTTCCCGATTTGAGATAGG | 120 | 47 |
| chr7 | 144252297 | 144252417 | 772751_42001748_Shank2(210274)_641 | 1 | + | good | CCATTCATATGAATGCTTAGGATTTGAACTCAGGTTTGCTGCCTGCTATGTCAGGACGCATTCCTTTCCTCTACTAGGCTGTTTGCTCCTGATGAGTTTGGGGTTTATCCTGATGAAATC | 120 | 44 |

| | | | | | | | | | |
|---|---|---|---|---|---|---|---|---|---|
| chr7 | 144252417 | 144252537 | 772751_42001748_Shank2(210274)_642 | 1 | + | good | CCCACTCAGCCTTTGTCTTGTTATCAGGTCTCTGCTATATACAACTGTCATGACTCTTAGTGCTGCTGACAGAGATGTAGCTGTACAGGGCCAGAGCCCAAGTGGAAGAATAGCCTGCCT | 120 | 49 |
| chr7 | 144252537 | 144252657 | 772751_42001748_Shank2(210274)_643 | 1 | + | good | TGTACATTTGAAAAGGCCACACATGTCTGTCACAGCCAGCTTTCCCTGAGTGATGCTGTGTGACAAGCATCCCTCAGTGTCGGGCAAGGAACAGCAAAGACAGTTAACATTCTTGCCACA | 120 | 48 |
| chr7 | 144252657 | 144252777 | 772751_42001748_Shank2(210274)_644 | 1 | + | good | CATCTGTGCCATGTGCACACTGGTTCCTTGGAGTCTCCTTGAGAGCAGTGATGGCCCTAATGTGGGCTCTGTGCTCATGGGCAGCTATCCCATGAAGCTCTTGGTATGTTGCTCATGTCA | 120 | 52 |
| chr7 | 144253017 | 144253137 | 772751_42001748_Shank2(210274)_647 | 1 | + | good | TTAGAAAGAGGTCTATTGGATCAGAGAGCTAAGGAATGAGATGGTAAGAGATACAAAAAGCATATACAATGGCTCTCATGACCTCTCAGGCCACAGCCCAGGGCCTCTGTAGCAGGGCCA | 120 | 48 |
| chr7 | 144253137 | 144253257 | 772751_42001748_Shank2(210274)_648 | 1 | + | good | TGTGCCTTCTTCCAATACTCACTATCCCCACCTCACATAGAGCAAGCTTACCCAGCACCCAGGCTGCAGAATGTTTCAGAGAAAGGCTGCATTCCATACACAGGCTGAGGGTCACTGGAT | 120 | 51 |
| chr7 | 144253257 | 144253377 | 772751_42001748_Shank2(210274)_649 | 1 | + | good | AGGACTGGTGGAGTCAAAGGCCATCATAATGCTTGGACCAAGTTGGACCACATCTTGACGTGCTCACAGGTTGGAGGTATGCCCTGGCATCTGAGACACCATCTTGCTAGGAACCTCATA | 120 | 51 |
| chr7 | 144253377 | 144253497 | 772751_42001748_Shank2(210274)_650 | 1 | + | good | TGGCAAAAAGAAAACACTAACTTTCCTGTGAGGATGTGTGGGCACAGACTCTATGACAGTGCCATCTCGTGGCTGGCTGCTCCCTCGAGGTCCTGTCCTCTAGCCATTGATGGGCCAGTA | 120 | 52 |
| chr7 | 144253497 | 144253617 | 772751_42001748_Shank2(210274)_651 | 1 | + | good | TACATGGCTGATACATACATGTATTGTCCAAGCTCTTGTGCTGTCAGTAATTGGAGGCTAAATCACATCGGAGCAGAAATGGCAGTAAGTGGGTTTGTGTGTTAGAAGTAAGAACTTTCT | 120 | 41 |
| chr7 | 144253617 | 144253737 | 772751_42001748_Shank2(210274)_652 | 1 | + | good | GGAGCTCCAACTGTGTTCTGTCTGTCTGTTCCTTTCACTGTCTCTCTCTCTGGATCCTGTGCTGTCCTGCCTGTACATCATCTCACAAGGCAGGCCACCCTTGGAGGTCCCATAAGTGGG | 120 | 54 |
| chr7 | 144253737 | 144253857 | 772751_42001748_Shank2(210274)_653 | 1 | + | good | AATCATGGCAGCGGGGCAAGAACCTTGCACCAGATGAGGTCACAGGGGTGGCCTTATCAGAGTCCTATGCCAACTACTAGTGCCCAGAACTGGAAATCTCTACTGAGATCTCTACTGGGT | 120 | 52 |
| chr7 | 144253857 | 144253977 | 772751_42001748_Shank2(210274)_654 | 1 | + | good | CTATTGAGGAGAGACAGCAGGGCCTCCTGCCTCTTTAGCAAGCATAGAGAAGTGGTGGAATAGCCACAGCCTGGGGGTGATGTGTCAGGTTCAAATGAGCTGGTGCTTCCCTGAAGAACC | 120 | 53 |
| chr7 | 144253977 | 144254097 | 772751_42001748_Shank2(210274)_655 | 1 | + | good | AAGTGGGATTCATCTGTCCTGGCCATGGGCCTCCAGCCTCCTGCTCCCCCTGCCTGTTCTGGGGTTTCTTACTTGTATCCTGTGTACCCTATGGGGTTCCAGAAGGGGTTAGCACATAGT | 120 | 55 |
| chr7 | 144254097 | 144254217 | 772751_42001748_Shank2(210274)_656 | 1 | + | good | ATATAGTGTAAAGCTGGTGACAATAAACACTGGACGTAACATAAGGCAGCTCTGTCACAGTGCAGGAAGGGTCCCAAGAACCACCCAGCCCGCCCATCTATCCCAGGTCAAAACAGAAGC | 120 | 50 |
| chr7 | 144254217 | 144254337 | 772751_42001748_Shank2(210274)_657 | 1 | + | good | TGGGTGAGGGCATATCCACAGTATGGCCATCTGTATTCAGATGATTGAATTGAAGGCCCACACTGCTTCTCATGAGCTTTGTGACTTCGGAAGGCTTGGGAAGACAGAAGCTCTGAACCT | 120 | 48 |
| chr7 | 144254337 | 144254457 | 772751_42001748_Shank2(210274)_658 | 1 | + | good | AAGGTGTGACATGTGATAGCTATAGCCAGCATAACACACGCTTCCCATGTGCTACGTTGCCCAACACCTCTAATCACTCAGGTGCCTCTGTGAGAGCAGGGCTGGGGCTCTCCCCACATC | 120 | 54 |
| chr7 | 144254457 | 144254577 | 772751_42001748_Shank2(210274)_659 | 1 | + | good | ACAAGTGAGAAGACGCGGCATAGCCAGTTGGTGACAGAGCCAAACTTTGAACTCCACTAGTAGCTTTAGAACCTCATCCCTAATCACCACACCCTGCCCCTTCTGTAAGGTGGAGACAGG | 120 | 51 |
| chr7 | 144254577 | 144254697 | 772751_42001748_Shank2(210274)_660 | 1 | + | good | AAATGGGCTGCCTGCTTGGTGCTATGTGAGGACTGGAGCAGGTGGAGGACGAGCGATGATGTGGATCACGAGTAAAGCGCCTTGTCAGCAGAGAGCAGCGCAGGGAATCTGCTGCAGTGA | 120 | 57 |
| chr7 | 144255417 | 144255537 | 772751_42001748_Shank2(210274)_667 | 1 | + | good | ATCCACATGTATGTTCACATACACGCATGTACACATTTACCTGTGCTCACATGCTCAAATACACATCCACACATGTACCTGTTTACACATTCCTTTCTTACCTATATGCACATACGTACA | 120 | 39 |
| chr7 | 144255537 | 144255657 | 772751_42001748_Shank2(210274)_668 | 1 | + | good | TTTGTGTGCACATGTATATAATCACATGTATAAAGACATACACATGTATACATGCACTTTCACATATGCATGTGTGTACACACTTCCATGGACCTGCATACACATGCTTTGCCACACAAT | 120 | 38 |
| chr7 | 144255657 | 144255777 | 772751_42001748_Shank2(210274)_669 | 1 | + | good | CATGTGCTTGCATGTATACTCATATGCATCTGCACTCACAGACATGCTTATATGAATGTATGCACACCCTTACATAATACAGTCACCATGAGCCTGTACATACAAGGTCTCCCATACACA | 120 | 42 |
| chr7 | 144255897 | 144256017 | 772751_42001748_Shank2(210274)_671 | 1 | + | good | TTCTACTCCACATCTGCTTCTCTGCTCTTGGCCCTGTGCCCTTTGGCCCTGCTGAGCCCTGCCCACTCTGAGCACCCACTGGTTTCCTAGACAACCAGTACAGAACAGCCACATGAGATG | 120 | 56 |
| chr7 | 144256017 | 144256137 | 772751_42001748_Shank2(210274)_672 | 1 | + | good | GAGGCAACCAGAGCAACTCAGAGCAATACTAAGAAGGAGCTGTCCTTGGGCCCTTCGATGGGCAGCTATGCTTCATGTCTCACTTACCCTTCCTAGCCCTGGCACATGCGTGACCTGAGC | 120 | 55 |
| chr7 | 144256137 | 144256257 | 772751_42001748_Shank2(210274)_673 | 1 | + | good | AGGGATGTCCCAGGGAAAGTCAGACCTCTGTGTTTCCCTCCATGGTGATCCGATCCCTGGGGACCTCATCCTGTCATAGAAGGCCAGTATACCTGGGCTCCTTCACCCACCCCTGGTGTG | 120 | 57 |
| chr7 | 144256257 | 144256377 | 772751_42001748_Shank2(210274)_674 | 1 | + | good | AAGTGTGAGGCAGGTGTGCACAGCTTGGCTAGATCGTCCCCAGCCTGGCTAGTGTTCCTTGTCCACAAGTTCTCACTCCAGTGCCCTGGACTGCAGTGATTTGCTCCTTGTGTGATGGGG | 120 | 56 |
| chr7 | 144256377 | 144256497 | 772751_42001748_Shank2(210274)_675 | 1 | + | good | CCTTTGGTTAGTTCAGCAATCTATGATGAGCAGGGAAGCCATGGTCAGAACTACCCATGTAGACTCGGTGATGTTTAGAGGACAAAGTCATTCCACTTGTGGAGCCAGAAGTCTAAGATC | 120 | 46 |
| chr7 | 144256497 | 144256617 | 772751_42001748_Shank2(210274)_676 | 1 | + | good | AAGAACTGTGTCTGGTGCACACCCTTCCTACTGCTGGCGACTTCTGCCTTACCCACACATGGCACAGGCTATCAAGTGGTAAGCCTGGGAAAGCTGCTAGTTGAGGTCTCTCTTTTTATC | 120 | 51 |
| chr7 | 144256617 | 144256737 | 772751_42001748_Shank2(210274)_677 | 1 | + | good | CTTAGTAAGCCACTTATTAGCTCATGAGGGTCCCCCTTCCTCATGGCCTCTGCCTCAAACCATAAGTCAGGAGGATGAGTCAGAGAAACTAGGTAATGCCCCATCTTGGAGCCCTGAGGA | 120 | 51 |
| chr7 | 144256737 | 144256857 | 772751_42001748_Shank2(210274)_678 | 1 | + | good | CAGCCCAGGATGTAGGTCCCTGACCCAACCTACAGAGCGAAGGAAGGCACTGTCTGGGGTGCTGATCTGAACAACCTGAGTGACATGGGGTAGGTGTGTGATTTGGTTTCGGTGATGCCA | 120 | 55 |
| chr7 | 144256857 | 144256977 | 772751_42001748_Shank2(210274)_679 | 1 | + | good | TCCGTCAAATGTACCCAGTCACCCCTGCCCTCCCTGTGAATAGATGCCCTCAGGTTATGTATGAGGACAGGGCTGTTATAATTCAGATGGTGACAGCATGGGAGTGACAGGCATAGACAG | 120 | 51 |
| chr7 | 144256977 | 144257097 | 772751_42001748_Shank2(210274)_680 | 1 | + | good | AGGATGAGAGCCCCTTTACTATCTTCTGTCTGTAGTCCAGCATCTTGCTGCTCAGTGCCCCCTTTTTGTATCCATGTTCTAGTTGCATTCTGTCGCTGCAATAAAACTGTTACCAACATC | 120 | 45 |
| chr7 | 144257217 | 144257337 | 772751_42001748_Shank2(210274)_682 | 1 | + | good | GCATTACCCCTGACCAAAGAATTCATTCTCAGCCAACACCATGCAGCAGAAACCCTGGGGAGCTTGCCTGCTTGCTCGCTCTCAAGTTAGTGCTCAGCTAGCTTTCTTAGACAGACTAGG | 120 | 51 |
| chr7 | 144257457 | 144257577 | 772751_42001748_Shank2(210274)_684 | 1 | + | good | GTTGAGACTCTTGCAAGTGTTTGAGGCTGTGCCACATGTGTGGTTAATGTTAAGTCTGAGTGACACATTTATAATCCTGACATCTAGCTTTGAGCCATCCGTTTACTATTTGAAAGGGTA | 120 | 41 |
| chr7 | 144257577 | 144257697 | 772751_42001748_Shank2(210274)_685 | 1 | + | good | CATTTTCTAAAGCTGAAATTGCTTGTTTTGTTTGCAAATGCCATTGCTTTCCACTAGCATTGTGAGGGGATAGTTCTGGCAATTGTTGGCACATTTTACCAATGGCCTTCATGCTGTGTG | 120 | 41 |
| chr7 | 144257697 | 144257817 | 772751_42001748_Shank2(210274)_686 | 1 | + | good | CCTTCAATACGAGGCCAAGGAGTGGTTGATTAGAGTTGTAAAGCACACAGCCTGACATTTAAACTACCAAGATGCCTTAACTTTAACTTAAATAAGGCACTGCCCTTGTGGAGGGGACCC | 120 | 45 |
| chr7 | 144257817 | 144257937 | 772751_42001748_Shank2(210274)_687 | 1 | + | good | ACAGTCTGGACTGGCCTTCAAGGACAGTGAACACGAAGAGGGGTCGCTGAGCTGTATGTATGTCCCTCACTGGGTTTCCATGGTAGCATTTGACTGGGTACAGCAGCGGTGCTCCCAGTA | 120 | 54 |
| chr7 | 144257937 | 144258057 | 772751_42001748_Shank2(210274)_688 | 1 | + | good | GGACCTGGGTACACAGAAGCACTCAGTGAGTGTCAGTAGAGGGAGGGGTAGCATTGTCATGGCTGAGGAAGGCATTGGAAGCCTGGAGCTGGCACAGAGAAAGGTGAGAGACAGCGGAGC | 120 | 57 |
| chr7 | 144258057 | 144258177 | 772751_42001748_Shank2(210274)_689 | 1 | + | good | CCGTCTGTCGAAGAGAACTAATGATGTACCAGGGCAGCTGGGAAATGCAAGTCTAGCTCCCATCTCAGGAATGGCTCCTTCCACCCCAGGAGGGGACTCCCCACTCCATAAGAAGTGCGG | 120 | 56 |
| chr7 | 144258177 | 144258297 | 772751_42001748_Shank2(210274)_690 | 1 | + | good | TGTTCTGTCTCTGCCTCTTAGCCTGTAAAGACCTGGGGCCTCTCTCATTAACCTGCACTGGGAAGGGAAAGTTAAACAAATAATCACGTTGACCTTGGAATGAGAGGAGAAACTGAACTG | 120 | 46 |
| chr7 | 144258297 | 144258417 | 772751_42001748_Shank2(210274)_691 | 1 | + | good | AGGCTCACCTGACTCTCAGAGCTCAGGCTTTGCTTTAAGCCATTTGCTTAATTTCTCCCAAAGTTCCACAGGACCTCAGAAGGAGGCAGGGCAGGGACTCTTGTCGGTGAGGGGAAATAC | 120 | 52 |
| chr7 | 144258417 | 144258537 | 772751_42001748_Shank2(210274)_692 | 1 | + | good | CTTGAAGCTCTCTGTGTATTGTGGTGCTTCCAGCAGATGCATCTTCACGTGTTCCCCTCCCCTCCCCAGCCCTGTTACCATCTGCTAATCTAATTACAACAGTCATATCATTTCCCACCG | 120 | 49 |
| chr7 | 144258657 | 144258777 | 772751_42001748_Shank2(210274)_694 | 1 | + | good | TTTGTGGAAAGCAACCATCATCAGGATATTCATCCTGATCATGTCTCTCTGGGATAAAGCCATCATTGGAGGTTCCTGAGATGTCCAGAGGCCTGCCAGCCTGGGTAGCATCCTTTGGGG | 120 | 50 |
| chr7 | 144258777 | 144258897 | 772751_42001748_Shank2(210274)_695 | 1 | + | good | CTAGCGGGGCTAGCAGATTCCACAGGAGGTGGCTGTCACTGTTGTGACTGGGAGGCTTATAGCCCCCTGTTCTGCAGTACCTCATTTGCCCCTCTGTGAACAGGCACAAGATTTACAGCA | 120 | 54 |
| chr7 | 144258897 | 144259017 | 772751_42001748_Shank2(210274)_696 | 1 | + | good | TCCCCACATTTTGCAGACTTGGGCCCCCAAATCAGCCCCATCTCCCATGCTTTTCCCTATGTAGAGGGTGTCACCCCTGTCTGGGGCCAAAAAGGAAGGTCTAGCCTTTTGCTCATGCAA | 120 | 53 |
| chr7 | 144259017 | 144259137 | 772751_42001748_Shank2(210274)_697 | 1 | + | good | GATATGGGATTTGGCTTTGATGAACGGTTTCTCTCTTCCCTCTGCAGAAACCCGCGCGGACCGCAGCAAGAAGCTCTTCCGGCATTACACGGTGGGCTCCTATGACAGCTTTGATGCTGC | 120 | 54 |
| chr7 | 144259137 | 144259257 | 772751_42001748_Shank2(210274)_698 | 1 | + | good | CAGGTAAGGCATGCTTCTGCCTTCTGGAGACCTCCGGCCTTGGGTGCTCCTGCATGCCTGGGTAGAGGGGCAGTGCCTTCTGTGCTTTGTTATTCGTGTGCCACCTCTGCTGGGATTGTG | 120 | 58 |
| chr7 | 144259257 | 144259377 | 772751_42001748_Shank2(210274)_699 | 1 | + | good | GCAACCATACCATACAGGAGAACCTGTCGCAGACAGGGCCTGTGGAAGAAGCCCCTGAGAGAGGGCAGCATGCTTGAAGGAGGAGGGAAAGATGGCATCCTTGAGAGAAAACAATAGGGA | 120 | 53 |
| chr7 | 144259377 | 144259497 | 772751_42001748_Shank2(210274)_700 | 1 | + | good | TGTGAGAAGGGGCATGAACAGGGACCCTAAAGCAGAGTGATTGGCTTCCTCCAGCAGGTATTTAGGCCTTCCTGGCTCTGGGCTTACACTTCACCCCGAGTCCCGCATTCTTGTTCAGGG | 120 | 55 |
| chr7 | 144259497 | 144259617 | 772751_42001748_Shank2(210274)_701 | 1 | + | good | AATCTTGCTCTCCTTGATGAGCAGGGCCCCAGGGAAGCAGACCTTCTAGGCTCATGGGTTGGGAGTTTGTAGAAAGTTGTGTGTTTCCTGTGGTATGGCTGTCACAGGATCCAGGATAGA | 120 | 51 |
| chr7 | 144259617 | 144259737 | 772751_42001748_Shank2(210274)_702 | 1 | + | good | ATCTTCCCACAGAAGCCTTGCTCGAGTGGCTACTGGGAAACCTGGCCAGAATGGACCATTAACAGGTGGCCAGAGTCAGTAGTAGCCTTAACCCATATGTCCATCTAACAAGGAAGATCC | 120 | 50 |
| chr7 | 144259737 | 144259857 | 772751_42001748_Shank2(210274)_703 | 1 | + | good | AGGCAGAGCCAGTACGCCATAAGTTCCTGGGAGGTCTGGGAAAAGCTGTTACTGTGGCTACTTGAAGAGCTGTTCAGGTAGGGAACAGTGGGCCTTGTGGGGTGGGGGTGGGGAGAACCT | 120 | 57 |
| chr7 | 144259857 | 144259977 | 772751_42001748_Shank2(210274)_704 | 1 | + | good | AAGGTAGAAGGCTTCAAGCAGTGGTCACTCTCAGACTGTGCCTTTGATGAAGCATATCTGGGGATTAGGGCCAAACTGGGGCCAACCAGCAACTTCTGAGCAGAGTGTAGAAGAGTTTAG | 120 | 49 |
| chr7 | 144259977 | 144260097 | 772751_42001748_Shank2(210274)_705 | 1 | + | good | ATGAATAGTTAGGAGTCAGCCATAGCTCTCCCATGATGAAGAGTTGGACAAGCTGTGGGCCTCTCTTGACCTGCCATTTCAGTTGGAGCAAACAAGGGTCACATGTCTATCTGTCCATCT | 120 | 48 |
| chr7 | 144260097 | 144260217 | 772751_42001748_Shank2(210274)_706 | 1 | + | good | CACCAGCACCACTACCATGCCCTGGACACAGCCTCAGCAACCTAGTAAAGCAAGGTCACTGTCAGTGTGATAGAAAAACATGGAGACTGGAGAAGAGGCCAAGGGGGGTAGGGTATGAGT | 120 | 52 |
| chr7 | 144260217 | 144260337 | 772751_42001748_Shank2(210274)_707 | 1 | + | good | CCTCAGCCTCCATGTATTTACACCTGCGTGTGAGGCCGTGCCTGGGACCACCTGAGGTCAACATGTTTCTGTTCACAATCACCTGGGGTTAGGTGTGTGAGGAGAAGTCTTGGTGAATAT | 120 | 52 |
| chr7 | 144260337 | 144260457 | 772751_42001748_Shank2(210274)_708 | 1 | + | good | CAAATTCTTGCTGCCCAGAGCACCATTGGGACAGAAAACACTGTTTCTGTGACTGACAAAAGACTGCTCATACAGATGTAGGGATTAGTAAGGGAGACAGCTTTGACCGTGGCAGGGATC | 120 | 48 |
| chr7 | 144260457 | 144260577 | 772751_42001748_Shank2(210274)_709 | 1 | + | good | TGCAAGGCAGGTAGAACCAGGAATTCATGGGTGGAGAGACAGAATGGTCTCCCAAACCATAAGAACCCAATGGTCATGTCGCCCTGGAGGGGACATCGGAAGGAGAGGCTAGATGTCTCA | 120 | 52 |
| chr7 | 144260577 | 144260697 | 772751_42001748_Shank2(210274)_710 | 1 | + | good | CATCTGACTGAGCAGGCAGCAGCCATCCTTGGGGGGCAGGCATCTGCAGTTTAATGCACTCATCAGCTCATTAGGAAGGGCCAAAAGCACAGGGAGACCCAGGAGTCGCATCACCCCACT | 120 | 57 |
| chr7 | 144260697 | 144260817 | 772751_42001748_Shank2(210274)_711 | 1 | + | good | GGTCAATTAGCCACTCTGGGTGGTGTGACTGTGGTCTGTGCAAGCGCCAACAGCTGAGGCTGTCTTGCTGCAGATGCTAATGTAATTAAAGAAAATGCAATGGGAAGTAAGGCTCAATGG | 120 | 48 |
| chr7 | 144260817 | 144260937 | 772751_42001748_Shank2(210274)_712 | 1 | + | good | AGGCATTTTCCGAATCAGTTGAAAGCTTTCTCCCCTTCGCACCCCTTCATGTGTCAAGTTAGCCCAGAGGCTCAGTGCTGAAGAGGTCAGCAGTTAAAACAAACCCAGGCATTAGCATGT | 120 | 48 |
| chr7 | 144260937 | 144261057 | 772751_42001748_Shank2(210274)_713 | 1 | + | good | GTGCCCAGTTGCTCCACCTGATTCTTGCCTTAGGATTCTCTTCCATCTGCAAGGGGAACATGAGAGGGATTTAGAAATAGATACGATAGTCAGAATCAGACAGCAGAGGTCCTGGGGCCA | 120 | 49 |
| chr7 | 144261057 | 144261177 | 772751_42001748_Shank2(210274)_714 | 1 | + | good | CCACTGTACAAGGCTTTGCAGGTACCTGAGGGTCAGTGTTAAAAGCACATCTCAACATGAGGTGACAGCAGTGAGTCCTCATTCCTCATTAATTGGGTCCCTGTCTCATCAGCATGGAGC | 120 | 49 |
| chr7 | 144261177 | 144261297 | 772751_42001748_Shank2(210274)_715 | 1 | + | good | ACCAGGAGGAGCTACATGGGTGGGCAGAGGAACCTTGGCGGTTGCTATGTGCTGAACAGGCTCCATTCATCCACCCATTTTTTTTTACCACCCATCTACTCAGGTATTCATCTGCATACA | 120 | 49 |
| chr7 | 144261417 | 144261537 | 772751_42001748_Shank2(210274)_717 | 1 | + | good | TCAGTACTCATATATCCATCTGTCTCTGTCTCCCCACCCACCCATCCATCAGTCTAATGACCTGGTTGTTCATCCATTCATCTACCCATTCATATATACACCCCTATATCCATCTTTCTG | 120 | 43 |
| chr7 | 144261777 | 144261897 | 772751_42001748_Shank2(210274)_720 | 1 | + | good | TCTGTCCATCTATCTATCTAATCATAACTAGTCATCATCTACCCACTCACATCCATCTTTTTGATCTACCCCACTCATCTCTCCATCAGTCTAACCAGTAAGCCACCAACACTTGCTTAT | 120 | 41 |
| chr7 | 144261897 | 144262017 | 772751_42001748_Shank2(210274)_721 | 1 | + | good | TCTGCGCGTTAAAAATCATGAGGTCATACTTGATGGCTCATTGGATAAAGTGCCTGCCATGTAGTGTGAAGCCTGGAGTTCTGGTCCCCAGAACCTGCATAAAGAGCTTGACAGGCACCA | 120 | 48 |

| | | | | | | | | | |
|---|---|---|---|---|---|---|---|---|---|
| chr7 | 144262257 | 144262377 | 772751_42001748_Shank2(210274)_724 | 1 | + | good | GAAACCCACAAGTTGAGAATCTGTCCTAGAGCACGCTGGTTGGCTAGATTAGCCGGAACTGGAGGACTCTGGGCTCAGCGAGAGACCCTGCATTAGCAGAGAATAATTGAAGAAAGCATC | 120 | 50 |
| chr7 | 144262497 | 144262617 | 772751_42001748_Shank2(210274)_726 | 1 | + | good | ACACACAAAGATTTTGAGGTCAACAAGAGAAATCTCATCAGTCTCGTGAAGATGGTAGTGTTATTAGTGAAGACAAACAGTAAAGTCACAGCAGAGATTTGCCCTGGGTGGTCAGTCAGA | 120 | 42 |
| chr7 | 144262617 | 144262737 | 772751_42001748_Shank2(210274)_727 | 1 | + | good | GAGACACAGAGGAGACCGGCAGCCTTTGAGCCGCAGAGCAAGCTGTTGTGAGGTCAGAGCAAATGTCCTTGAGAGCAAACTTGAGGTGACTCCCAGAGCCGAGAGAGAGCCAGACAGGAC | 120 | 57 |
| chr7 | 144262737 | 144262857 | 772751_42001748_Shank2(210274)_728 | 1 | + | good | TCAGTAACTGCAAAGAGATTGGAGCAGGCCAAGAATGTGAGATTGTGGGGAACAAGGGATGCACAGAGTGGAGAGGAACTGGAGCCCATGCGACCTGATCATGAGACAAGCAGACAAATG | 120 | 50 |
| chr7 | 144262857 | 144262977 | 772751_42001748_Shank2(210274)_729 | 1 | + | good | AATGTAGGTCAGTGAGGGGACTGCCTCTGTCATCTAGGCCGAGCATGAACTGTGTGCACTGGCACGGGAGAGAGATGGATCTGAGGTGACATTGGAATGAACAGGGCCCTGGGCATGGAC | 120 | 56 |
| chr7 | 144262977 | 144263097 | 772751_42001748_Shank2(210274)_730 | 1 | + | good | GAGGGGAAGCAAGGTGGATCGGCTCTCCACCCTCTCTGAGGTTTACCAGGGAGGACTTGGTGAGACAGTGGGTGGGTACCAGGCCCAGGACTAACTAGCAGCAAGGTCCTCCTGGTCCCC | 120 | 61 |
| chr7 | 144263097 | 144263217 | 772751_42001748_Shank2(210274)_731 | 1 | + | good | ACTACACACAGCTGCATAGCCACTGCACAGCCTGCTTGGAAGGAGCACTTCCTGAATGCCTGCCAAGGCAAGCCACCACTATTGCTCCTCCTCCTCAGGAAACCTAATTTCCTCCTGGAG | 120 | 54 |
| chr7 | 144263217 | 144263337 | 772751_42001748_Shank2(210274)_732 | 1 | + | good | ATTGATGGAGAAGGGCAACAGATTGTGCCACTTCAACAGGTGACAACCTGCCACCCGCAGCTACTTCTTCTCAAGACTTTCAAGTACCATTTGTACATTTTGAATTGGTAATAGAAAATA | 120 | 41 |
| chr7 | 144263337 | 144263457 | 772751_42001748_Shank2(210274)_733 | 1 | + | good | GGTACTTGGTAAAAGATTCAAAATGTTTAGGACATAGTAGGGTGTGAAGAACAGCCCCCTCCTTACTCTGTCCCACCCAGCCCCTGAATTCTTCCCAGATAGAACCTCTCTTGATAAATT | 120 | 44 |
| chr7 | 144263457 | 144263577 | 772751_42001748_Shank2(210274)_734 | 1 | + | good | CCTGAGGCTTTCTAAGAGAAAGACTCTGTCTATGTGATACCGTGAGAAGCCTCTATATGGAGGGAGGGGTATGCTCTGTGTACACATTCATGTATGTGCATATGGGTGTGTAGGAGTTGG | 120 | 47 |
| chr7 | 144263577 | 144263697 | 772751_42001748_Shank2(210274)_735 | 1 | + | good | GACACTAGTACATGTGTGCTCATACATGCAGAAGCCAAAGATCAATTTGAGTATCTTCCTCTGGCACCATTCATCTCATTGTAAAATTATTTTTAAGCTTTATTTATTTAAGTATGTAT | 120 | 32 |
| chr7 | 144263697 | 144263817 | 772751_42001748_Shank2(210274)_736 | 1 | + | good | GTGTTTCTATGTGTATTATTATTATTTGCAACAGAGCAGCTCACAAGCCTGAGATTCACAAATTTAGCAAGGTCTGATGTCCAGTGTGCTCCAGGGAGTCTTGTGAGGCACTCTCCCCTC | 120 | 45 |
| chr7 | 144263817 | 144263937 | 772751_42001748_Shank2(210274)_737 | 1 | + | good | ACTCCAGCCCTGGGATTACAGGTGTGTGTTACCACACCTAGCTTTGTATGGGTATTGGAGATCTGGACTCAGGGCATCATGATTGTGTGGTGAATATTGGCTGACCCATTTTCCCAGTCG | 120 | 49 |
| chr7 | 144263937 | 144264057 | 772751_42001748_Shank2(210274)_738 | 1 | + | good | CTACAAACCCTTTTAAAACTCTTGAAAAAAGATAAATAAAAACTGCCCGCGTTCCCTCCCAGACTGTCTAGCTGAATCACCCTGTGACCTTGGGCAGCTCACCTCCTGTCTCCGAGTCTT | 120 | 47 |
| chr7 | 144264057 | 144264177 | 772751_42001748_Shank2(210274)_739 | 1 | + | good | CAAATTGCTATCTGCAAGACAGAAACACACTTAAACCTGTTCCCTGTCTCCAGTTGGGTTCCTGCTAAGAACAAATGAGAAAATGTGTTAAGGGCTTGCAGCACCAAGAGGCCTGCACTG | 120 | 46 |
| chr7 | 144264177 | 144264297 | 772751_42001748_Shank2(210274)_740 | 1 | + | good | TATAGATCCAGTGGGCTGCAAGCTGTAAGTGCTGCCGGTGGGCACCTGGCTTTCCCCTGACACCTCTCGCCGTGTGTTCTAGAGCACCGCCCTTGGAAAGGAGGAATGGTAAGAAAGAGC | 120 | 57 |
| chr7 | 144264297 | 144264417 | 772751_42001748_Shank2(210274)_741 | 1 | + | good | GACTGCCCCACTCTTCTTCCAACATTATTTTGTGTTCGTTCGTTGCTATTGTTTTTAGTCAGACTTGAAGAAAAGAGCACCAAGACTGAGTGTGTACAGGGCAGATACAGAAGGGAGCAT | 120 | 43 |
| chr7 | 144264417 | 144264537 | 772751_42001748_Shank2(210274)_742 | 1 | + | good | CCTCTGCTTGGCTGATGACTGGTTCAGGTAATTAATGGTGGCACTTGCAAAGCTGCTGCCTAGGGACATAGTTTGGGACTACAAAAGAGACCGTGCGTGCGTGCATGCGTGCGTGCGTGT | 120 | 53 |
| chr7 | 144264657 | 144264777 | 772751_42001748_Shank2(210274)_744 | 1 | + | good | TGTGTGATCAAGTGTTTCTTCAGTCTCTCTTTATCTCCTTTTTAGATGTATAGTCTCATTGGCTGGCCATGGCACCCCTTTCTCTGCTTCCCAGTGCTGGAGCTACTGGGCACGCATTGG | 120 | 48 |
| chr7 | 144264777 | 144264897 | 772751_42001748_Shank2(210274)_745 | 1 | + | good | CGTGCCCAACTTTTATGTTTGTGATTAGGAATCTGAATTCAGGTCCTCCTGAGTAGATAGTGAGCACCCTGCCTGCTGAGCCAATTCCATAGGCAGTGGTTTCACTTTTGAAGTTGTGTG | 120 | 46 |
| chr7 | 144264897 | 144265017 | 772751_42001748_Shank2(210274)_746 | 1 | + | good | TTTCTCTATTTGTATTTTTCTGACAGGGTTTGGTGCTCTGGTAAATGTTTGTAACTAAGTGGGACAGGAACCCTGAGGGCATTGATCAAGGAGAAGCTGGCACTGTCACCTGGCTGGCTC | 120 | 47 |
| chr7 | 144265017 | 144265137 | 772751_42001748_Shank2(210274)_747 | 1 | + | good | TTCTTCCAGTCTCCCAGTGTGATGTATATGAAGGGACAGGCCTCCATATGGGTGAGCCATTTTCCTGGGGAACACACCCCAGACTATGACTATCATGGTCTCCCTTCAAGCTACTGTGCT | 120 | 50 |
| chr7 | 144265137 | 144265257 | 772751_42001748_Shank2(210274)_748 | 1 | + | good | CTGGGGCTTCGTCCTGATGGACACCATGGAAGGCCTGGCTGCTCCTGTTCAGCCTCTTTCCCTCACTCAGTTCACTACAAGAATTGCGAAGGTCCTCTCCCTCCCACCCTCGCCTCCATG | 120 | 58 |
| chr7 | 144265257 | 144265377 | 772751_42001748_Shank2(210274)_749 | 1 | + | good | TTACCCAACACCCAACTCTTCATAGCTTTTCTAGCAGCTGGCTTCCTGGGACCAAGGACTCAGTTTAAGTTGCTTTCCCCCCTTCAGAAATTCTAAGCTGTGTTTTTAAGAAAGGCAACC | 120 | 45 |
| chr7 | 144265377 | 144265497 | 772751_42001748_Shank2(210274)_750 | 1 | + | good | AACAACCCATTAAAGCAGATCACACAGACAGTGCTAAGAATGGAAAGGCCACCCTTCCCTCCCCAGCCTCATCTGCAGGGCCCTGCTGCCCACCCAGGTACTCCATTGACTTTCCCTCCT | 120 | 55 |
| chr7 | 144265497 | 144265617 | 772751_42001748_Shank2(210274)_751 | 1 | + | good | TAGTCCTCCCCTGGTTACTCCAGCTCTACAATAACCTCATCTCGAGTCTTCATTTCGCTATGAAATTACCGTGACTCTGCTGGAGAGAACCTGGATCATAGTTCCGCCTATCCGCAAGAC | 120 | 48 |
| chr7 | 144265617 | 144265737 | 772751_42001748_Shank2(210274)_752 | 1 | + | good | TGGGTCGTTGTTGTTATTTCTAATTGTTCTAGACAAAGACGAAATTCAACCATGACCATAACAGCAGCAGAAGCAGAGAAGGAAGGGTGTGGGGAAGGACAGGAAGAGAGAGGGGGAGAC | 120 | 47 |
| chr7 | 144265737 | 144265857 | 772751_42001748_Shank2(210274)_753 | 1 | + | good | GAAAGGCAGCTAGAATCCACGGGAAACAAAACACACACAAGCACCGCAGCCCTAATAGTAAACATGGCACTACTTCCTGTTTGCATTAATATTTATATTTCATTAATATAGTATTTTATA | 120 | 35 |
| chr7 | 144265857 | 144265977 | 772751_42001748_Shank2(210274)_754 | 1 | + | good | TTTTTATTCATATTTACATCTTTTAGTTTCTCTCAGTCCAACTAGACTAATAGCCAAAAAAAAAAAAAAAACCAAGCCAGTGGAAAGCATTAAGAGAACATGACCGTTCACATCCACACCA | 120 | 33 |
| chr7 | 144265977 | 144266097 | 772751_42001748_Shank2(210274)_755 | 1 | + | good | AGAGCAGGGCTCCAGTTCCTCGTATGGGCATGAGAAGGGTCCTCATGGAACCACAAGCCAACAGACTCTTTTTCTTTTCTTCTTATTCAAAGTCAGCCCACATTTACGCGTGATACAACG | 120 | 48 |
| chr7 | 144266217 | 144266337 | 772751_42001748_Shank2(210274)_757 | 1 | + | good | AGCCATTAGCCATCCCTCCTTCCTGCCTGTGTGGTACATCCTGAGAATGTGAGTCACTCCTGAGAGACTCCTTTGCTGCTTCACTCAGTTTTCCCCTAACAATTTCAACAAAAGGGACCT | 120 | 48 |
| chr7 | 144266337 | 144266457 | 772751_42001748_Shank2(210274)_758 | 1 | + | good | GCAGTGTAAATGACCCTGTCTGCAGTCTCTACCTTTAAAATTCCAGGCTTAGAGTCACTCACTGGCCTCTTCATGGTGTGGAAGTGCGATGCTCTCAGAGCCTCGTTCCAAGTCTCTGCC | 120 | 51 |
| chr7 | 144266457 | 144266577 | 772751_42001748_Shank2(210274)_759 | 1 | + | good | CCTCTGTTCTCCTCTCCAGAAGTTTGTAGGAGTGTCTGCATTTCTTTGTCTCCATAAGATCTTGATGCTTTGACATACTGACCTTAGTCTGACTCCCCTCAAACATTATGTTTACTGCTC | 120 | 42 |
| chr7 | 144266817 | 144266937 | 772751_42001748_Shank2(210274)_762 | 1 | + | good | GTATGAATTGTCTATACTCCCACTTTGTCACTGTCCAAGGGCTTTTCATTTCACGGCAGGGTGAGCTTCTGACACTTCACATAAGTGCCTCTGAGAGCAGACCGGCCAGTGCCAGTGTTC | 120 | 50 |
| chr7 | 144266937 | 144267057 | 772751_42001748_Shank2(210274)_763 | 1 | + | good | ACAGAGGCCGCCGTGTGTGGCTGCCCTCCAGGCACAGCCTAGGTAGAATGTAGGTTCTTTCTAGTCAGAACCAGAAAGAATGATGGAAGACGTGCTCCTTGCATGCTCCCACAAGAGATC | 120 | 53 |
| chr7 | 144267057 | 144267177 | 772751_42001748_Shank2(210274)_764 | 1 | + | good | ATAAACAAAGATAAATCTAGTTCCTCTTACACACCTGCTAGGCTAGTTTTGTTGCTATGAATAAACACCCTGACTACGCTGACTAAAAGCAACTTGGGGCAGAAAGGATTTATAACATAT | 120 | 37 |
| chr7 | 144267417 | 144267537 | 772751_42001748_Shank2(210274)_767 | 1 | + | good | ATGGAGATAATTCTTCAAATGAGGTTCCTGCTCCCCAAGGTTTGTGTCAAGTTGACAATACATAACTGACACAGTCCCACCACAAAGGGACTACGTAGTTTGCTCTGGAGTGAGCACACG | 120 | 46 |
| chr7 | 144267537 | 144267657 | 772751_42001748_Shank2(210274)_768 | 1 | + | good | GGTGGCCAGGTGATTGACAAGGATGAAGAAGTAGGTTCCAGGAATGCCACATGCAACTGAGTCCCTGTCAGCCACCAGGAAATGGAGGGGGCGGGAATGCTCATGCCTGGCTGGCTTTCT | 120 | 56 |
| chr7 | 144267657 | 144267777 | 772751_42001748_Shank2(210274)_769 | 1 | + | good | CCTTTCCCCTTTTCATTCAGTCAGGGACCCCAGCCCATCTGGTGATGCTCCCCGCATGAAGGGAGTGTCTTCTTGGTTAAGCCTCTCTAGCGGAGGCTACAGATGTGGTTCAGTGGGGTA | 120 | 55 |
| chr7 | 144267897 | 144268017 | 772751_42001748_Shank2(210274)_771 | 1 | + | good | GAGTTTAAGGTCATTCTAAGCTACTGAGAGAGTTGCAGGCCAGCCTGAGTTTGTAAAGTCTCTGTCCAATCAAAATAAGCCAACAAACACTCCCTGGAAACACCCTCATAGATATCCCCA | 120 | 44 |
| chr7 | 144268017 | 144268137 | 772751_42001748_Shank2(210274)_772 | 1 | + | good | AGGTGTGCCTCATAAGAAAATCCTGAATCCTAGCAGGCTGATAATAGAGATGAGCCCTTGTGACCCCCAGCCCTTCCTGGTGTTTAGGACCTGACTTAGGGACTCTAGAACTTCAGGGTT | 120 | 49 |
| chr7 | 144268137 | 144268257 | 772751_42001748_Shank2(210274)_773 | 1 | + | good | TCCTTTCCCAGAGCACATGAGGTGGCAGAAGGGGTGGGGGCTTCGTTAGAAGCCAGTTTAAAGATCTGAAGGAGCTGAGTTATTTCTTGGGGGAGAGAAGGCGGGAGACCCCTTGTATCT | 120 | 52 |
| chr7 | 144268257 | 144268377 | 772751_42001748_Shank2(210274)_774 | 1 | + | good | GGCCTGAAAACCAAGCTCAGAAACCACCCTGTGCATAGCTCGTCACCACTGACTTCTGGGTTGGGGCTTTTCCCTGAAGCTACTGAGTGTGTCCCACTGGAGCCTGGAGCAGAGAAGCCC | 120 | 57 |
| chr7 | 144268377 | 144268497 | 772751_42001748_Shank2(210274)_775 | 1 | + | good | AGGCCCTTGCTGTAGGATTGCTTGCGTTGTGGGATCATGCAGGACAATCCATGAACTTCCGTCCATGATGGATTCCATCCTGGCTTCCATCCCTGCAGGGAGGCGCCTGGCTCAACACTC | 120 | 56 |
| chr7 | 144268497 | 144268617 | 772751_42001748_Shank2(210274)_776 | 1 | + | good | TCAGGCCACATTTAGCACTTTTCCCAGGTGGCGAGTGGGTGTCCTGATGGGGGAGAGCAGGCAGATGGGTGGGGCGATGGAGAAGCCGCAGAGCAGTTGCTAGGGGAGGCTGGCATTCAG | 120 | 61 |
| chr7 | 144268617 | 144268737 | 772751_42001748_Shank2(210274)_777 | 1 | + | good | GCACCCCTCCAGGCCGCACCCAGCAGACTGCCCTTGGCCCTTGGGTTATGTTGAGAGATGGGCCAAGGTAGGCAGGTGCACCATGAGCTTCCTGTCTCTCCACCTCCTTAGCCACACCAG | 120 | 62 |
| chr7 | 144268737 | 144268857 | 772751_42001748_Shank2(210274)_778 | 1 | + | good | CAGCAGAAGGAGGGTGTTGACCCTGGAATTTTCCATGGTCACATCTGCCCAAGGCATCCTCAGTGTGCTAAGCTCAGTGCTGTATAAGGTGCCCCCCCCCACCTCCCCTCTGTACAGCTG | 120 | 57 |
| chr7 | 144268857 | 144268977 | 772751_42001748_Shank2(210274)_779 | 1 | + | good | CCCTGTAATTCGCATTTACTAATTTCCATACTGCGAGAACTCTGCCCCACTAATCTCAAGCTGCTGATGTGTTTACTGATCCTGGAGCTGAGGAAACATGCCCACAGCTAGCCCTGGCAA | 120 | 48 |
| chr7 | 144268977 | 144269097 | 772751_42001748_Shank2(210274)_780 | 1 | + | good | GCTGGCACGCTGCAGCACGCCCCCAGCCAGCACTATGTCCCTCTTCTGCCTTCTTCTCTAGCTCCACCATGGTAGGCTCTCAATCCTGGGACAGAATGAATCCTACAGGGTGGTCTTCAA | 120 | 57 |
| chr7 | 144269097 | 144269217 | 772751_42001748_Shank2(210274)_781 | 1 | + | good | ACCTAATGGACCAGTCTGGGAGTTCTGATACCAAGGCTGTCTTTCCTGTCCTGATACAAACTCCTTGGGTGGGCCTCAGTTTCCTAACCAGTAAATGGCTCTATTCTCTTCATGTTCTTC | 120 | 47 |
| chr7 | 144269217 | 144269337 | 772751_42001748_Shank2(210274)_782 | 1 | + | good | TTTTCTACCACCATTTTTGGGGGGGGGGATATTTGGCCAGAAGCCAGTGTAACTATTTGTCATAATTTGCTGTAAGGGAATACCACTGATGAATATGTCAGATGCAGAAGCATGCATCTG | 120 | 42 |
| chr7 | 144269337 | 144269457 | 772751_42001748_Shank2(210274)_783 | 1 | + | good | TTAATCTAGCACATCTGTGATGAGAGGAGAGGCAGAAACAGGATTCCTGAAAGCTCACATGCCAGCTAGCCTCATGTTTGTAATGGTGGGGTGTTAGAAAACCTCTCTCAAACAAAAGAA | 120 | 43 |
| chr7 | 144269457 | 144269577 | 772751_42001748_Shank2(210274)_784 | 1 | + | good | AGGCAAAAAACTGACTGATGGTTGAGGTTCTCCTCTGAGCTCACGTGTGCTACATGGCACAAGCTCCCCACTAGACACACACACACATACACACAGGAAAAATTGCCAGCATTAATGGGC | 120 | 48 |
| chr7 | 144269577 | 144269697 | 772751_42001748_Shank2(210274)_785 | 1 | + | good | ATAAGCAGTAGTATCCTGAACAAATATGTCACTCACTGTATGTGGACATAGCCAGTGGTCACCATGAGCATCCACACATGTTGACCCTAAAGCAGGCCTGTCAGGACCAGGTTTCAGAGC | 120 | 48 |
| chr7 | 144269697 | 144269817 | 772751_42001748_Shank2(210274)_786 | 1 | + | good | CACAGAGGTGGTCACAGAACAGCCAGGTGGACAAGAATGCTCAAAACTGGTACCTAAGCTTGTGCAGCCCATCCAGGTGCCAGCCAGGGAGAGGGGGGCGAGGCTGCCAGGTCTGCAGGT | 120 | 60 |
| chr7 | 144269817 | 144269937 | 772751_42001748_Shank2(210274)_787 | 1 | + | good | GTTCTTAGGACTGAAGTGGGCACACAAACTCTCAGCAAATGTCTGTGTTAATCCTCACCTTTGGCAAGCTCCAGGTGTTCCATCCTCCTCTAGGGAATAGCAGCCTTTGGGGTCTGACTC | 120 | 50 |
| chr7 | 144269937 | 144270057 | 772751_42001748_Shank2(210274)_788 | 1 | + | good | TGGGACCCACCTGACTCACTGTTGCCTAGGCCAAGGACAGCAGCCAAGTGTTTGATCTGCATTTGCTTGTGTCTCGTCTACCGTGTATCAGTGCTGGCTCTTGCTTGTCTCCTTTCCTTA | 120 | 52 |
| chr7 | 144270057 | 144270177 | 772751_42001748_Shank2(210274)_789 | 1 | + | good | TCACCTGGGGTAGGGCCTGGCAAATGTTCAACCTCAGAAAGTCGAGGAAGAAAGAGGATTAATGGATGAATGAATGATCAATGGGTGGATGGATAAAGGGTGGATTGGTGGCTAGATAAA | 120 | 45 |
| chr7 | 144270897 | 144271017 | 772751_42001748_Shank2(210274)_796 | 1 | + | good | TTCTGAGGCTTCTTTATAGGCAGAACTGTGCACACTGTTGGCTCTCAATAAATCCTATAAGGAGAGCAAGGGACAGTGTGGCCCCGAGTAGAAGACGAGGTACCCAGTGCCAAGCATCAC | 120 | 50 |
| chr7 | 144271017 | 144271137 | 772751_42001748_Shank2(210274)_797 | 1 | + | good | ACCACAGAAAGAGTTTATGCCATCAGAGGAAGTTCAGCTGACCCAGCCCCACACAGCTCTCCTGCTTCCATTCCCTCAAAGGCAACACCGGGCACCCTGATTCATTGCCAGAGTTAGAAA | 120 | 52 |
| chr7 | 144271137 | 144271257 | 772751_42001748_Shank2(210274)_798 | 1 | + | good | GCAGCACAACATGGTCAGGTCTGCAGGAGGGCTCAGCCATGAAAGCCTGCTGCTGTCTCTGAGATTTCCTCATGGTGAGTTTGGGCTAGGTGGCCAGCCTGTCCATGATCCTTGGCAGCC | 120 | 57 |
| chr7 | 144271257 | 144271377 | 772751_42001748_Shank2(210274)_799 | 1 | + | good | GCCAGCCAGTCAGCAGCACTTAGCTCTTCAGGACACATGCTGATTAAAACCTGACTCTGCTAACCAAGGTCAAGGCTGAGGCTGGCATGGTGGCCAGAAGCCTGGCATAGCAGATGGCAC | 120 | 55 |
| chr7 | 144271377 | 144271497 | 772751_42001748_Shank2(210274)_800 | 1 | + | good | TCTTGTGCCTCAGGTTCCCTGGCCTGTTCCTGGCCCCACAGAACTGTAATGCTTGTAGAAGAGTAGGAGTCAGTGATAAACAAAAAGGCCAAAAGCCAGGGGAGGGCAGGGCTAGGAGGG | 120 | 54 |
| chr7 | 144271497 | 144271617 | 772751_42001748_Shank2(210274)_801 | 1 | + | good | ACCTGGGACAGGGGAGAGGACAGTTGATAGGAAGGCTTGGAGCATGCCCCCTTGGAAGGGCTCAGGAGTGTGGCTCCCTCTGGCACTGGCAGATAGCATGGCTAGGACTGAAGTCAGTTG | 120 | 58 |
| chr7 | 144271617 | 144271737 | 772751_42001748_Shank2(210274)_802 | 1 | + | good | TCTGTCAGGGACTGGGAAAGGTTGTGGGCAGAAGTGACCTCAGGCTTTGGCTGCATCTTTCTACACCAGTTCCCACCCATCCGCTGCAGAACTTGGGGGCTTGCTCATCTCTCCTTAGTG | 120 | 55 |

| | | | | | | | | | |
|---|---|---|---|---|---|---|---|---|---|
| chr7 | 144271737 | 144271857 | 772751_42001748_Shank2(210274)_803 | 1 | + | good | CCAGGGTAGAGGCAGATGCACAGGCCCTTCATTCCTTGGATGAAGATGAGGAGGTATGGGATAGAAAGGTCACAGGAATGACAGGGACAGGCTGGGACCTCAGAGCACTGGGAGGCTAGG | 120 | 56 |
| chr7 | 144271857 | 144271977 | 772751_42001748_Shank2(210274)_804 | 1 | + | good | TGGGGCATTCATGCATCCTGCTTAGTCCTCTGTCTCCACCCCTCTGTGGTGACTGGTGGACATAGGGCCCCCTTCCCATCAGTGTGGCTTGTGCTGTCCTGGTATGAGGAGAGTGATACG | 120 | 57 |
| chr7 | 144271977 | 144272097 | 772751_42001748_Shank2(210274)_805 | 1 | + | good | CAGGGCAAGCTTCCCTCAGCAAAGCCTTAAATATTGTTGGTTTGTGGTTCTGAGGGCATTTCCCCCTTTTCAAGTGCAGTGAGTGGCTAGGTTAATGCAGTAATGACAGCCCTAGAGGAG | 120 | 48 |
| chr7 | 144272457 | 144272577 | 772751_42001748_Shank2(210274)_809 | 1 | + | good | ACATAGCTCTATACCACATGACTGGGCACATATGGTCAGCTGAGTGGCCTTCAAAGATGTCCATTTCAAAATTCTGTCAACTTTTAAGTCAAAGGAGGACTTGGCAGATTTGGCTATTAC | 120 | 42 |
| chr7 | 144272577 | 144272697 | 772751_42001748_Shank2(210274)_810 | 1 | + | good | ATTTGACTTGTACATTAAGGCTATTGCAATAAAGGATCATCTTGGGGTACCCAGGAAATGTATAGATGCCATTTCATGTCTCCTCCTAAGAGGACAGAGCTAGAGGAGACCCTGTGACCA | 120 | 43 |
| chr7 | 144272697 | 144272817 | 772751_42001748_Shank2(210274)_811 | 1 | + | good | CAGAGGCAGGAAGTAGGGAGATGCAGCTGTGGCCCTAGTAATATCTAGAGCCACAAGAAGCTGGGGAAGGGCAATGGACCCTCACCTAGAACTGACTCTCAGGCTTGAGTTTTGTCATCA | 120 | 52 |
| chr7 | 144272817 | 144272937 | 772751_42001748_Shank2(210274)_812 | 1 | + | good | CAGACTTCTGCTGATTGGAGTTACCATTGTTTGTCACAGCAGCTGCAGGAGCCTCATACTGTCTGTGTGGTTTAACAGCCTTAACAAGGTACATGTCCCACAAACAATGACCTTCCTTAG | 120 | 46 |
| chr7 | 144272937 | 144273057 | 772751_42001748_Shank2(210274)_813 | 1 | + | good | ACAGAGGCCACTCAGGAAGCTAGGCTCATTGTCACCTTGTAGGGCCACATGTCTGGTCTCCAAGTCATGACCATAAGTGACATCAGAAGTCAGGAGCGATGTGGAATTTTACTTGAGTGC | 120 | 49 |
| chr7 | 144273057 | 144273177 | 772751_42001748_Shank2(210274)_814 | 1 | + | good | GTAAATGTCTGAGAAGCAGGTAGCTAACCAGAGTTGGTGCCCAACAAGCCCTCAGATTGCTGGGTGATAGCAAGGACTAATACTGGCTATGCACTGGACACAGAGGAGGAAGCTGGCAAC | 120 | 51 |
| chr7 | 144273177 | 144273297 | 772751_42001748_Shank2(210274)_815 | 1 | + | good | TGGCAGGACAGGCTGAGTCTTGGAGGTTTCTCAGGGCATGATATGTGCCCTGAGATGTGGCTGAGCAACCATTGGTTGGAGTTCTGCATGACAGGATGGAAGCAGCATACAACTAGCAGC | 120 | 52 |
| chr7 | 144273297 | 144273417 | 772751_42001748_Shank2(210274)_816 | 1 | + | good | AGTCAGACCCAGGCCCAGGCCCCACCTTGCTTTCCCTGTCCACATTATGGCTTAGGTCTCATATGTCTATCATTCTTTCCTGACATTAGCTTCCTGAAGATGTCATCACTCTGGTTGGCC | 120 | 50 |
| chr7 | 144273417 | 144273537 | 772751_42001748_Shank2(210274)_817 | 1 | + | good | TTGTCCTCAACCATCTATATGCAAGCCATTATATCAGCCTAGCTGGAATGCCCCAGAAACCTGGACCACTCTGAGAACTCGCAAAACTCCAGAGCAACTAAGCCAAGGAAAAGTGTGTGT | 120 | 47 |
| chr7 | 144273537 | 144273657 | 772751_42001748_Shank2(210274)_818 | 1 | + | good | CCTTAGCTCACTGCCTCTTGAACTAAGCCCATGCAAGAGCAGGCACTGTGGCCAGATGTTCCCTGGAGAGCTAAGCACAACATAGGACTTTGCTCTACCTCCTTGTTTCCAGCCAGTGTC | 120 | 52 |
| chr7 | 144273657 | 144273777 | 772751_42001748_Shank2(210274)_819 | 1 | + | good | CTCTAGGAAGCATTTGGCAGAGACTGCGACCTTGGTGGCAGGCCTGAAGAGGCCAGAACCAGTATTTGTCCCATAGGTCAATTGCAACTCTCCAATCTCTTGGCTTTTTCTTCATGGTTA | 120 | 48 |
| chr7 | 144273777 | 144273897 | 772751_42001748_Shank2(210274)_820 | 1 | + | good | CAGAGTTGCAGCTAGCACTCTGGTTATATCTCCACAGTGCATCCTTTGCAGGGATGTTGAGGAAGGCCATGCTTCTCCAGCATACTGAGTCGTTGAATAATTCCCAGAGGCTCTGGTAGA | 120 | 49 |
| chr7 | 144273897 | 144274017 | 772751_42001748_Shank2(210274)_821 | 1 | + | good | TTTCTACTCTACTCTCGTTGGCTAAAGCCATGTTCTATGGCACAGCTGTCCCTGGTGTGACTGGGAACTGCAGGGGAGACTGGGAAGCAAGAATGTAGGTTTTCTAGAGTCTGTTGTGGG | 120 | 50 |
| chr7 | 144274017 | 144274137 | 772751_42001748_Shank2(210274)_822 | 1 | + | good | AAATAAACCAGGGGGTGGAACAGAGACTGCTGGAATCTGGATGTGAGGGGAAATGAGATGAGGTTTGGCTTCATCACTAAGGGCACAGCAAAAGAGCCTCCTAGATATGATGACTATTTC | 120 | 46 |
| chr7 | 144274137 | 144274257 | 772751_42001748_Shank2(210274)_823 | 1 | + | good | AAATGGGCTTTGAGGGATGAATAGGAGTTGAGCAGCTAGAGTTAGCAATCAGGGAAAGTACTCTGGGCTGGGAATATAGCTTGATGGTAGAGGACTGGCTTTGCACCCCTGGGTGCATCT | 120 | 49 |
| chr7 | 144274257 | 144274377 | 772751_42001748_Shank2(210274)_824 | 1 | + | good | GTCATGCTTGATGTCTGCTATGTACTTGTTTCTATGTGGGAACCCCATCCTTCTTTTCATTGAACACCCCATGAAGTGGAACTGCTGCCTTCTTTGGAGATTTTCTGTTTGGCTTGGGGT | 120 | 45 |
| chr7 | 144274377 | 144274497 | 772751_42001748_Shank2(210274)_825 | 1 | + | good | GTAATCTACTGGCAGGCTGTCGGAGTATTGTAGGTGCTATCTGGCAGCTCAGCCACTTCCCAGAGCTGCTGGAGCCTCCTCTGCCTCCAAGGAGACTCCCATCCCTGTCATCCTGGGGTA | 120 | 57 |
| chr7 | 144274497 | 144274617 | 772751_42001748_Shank2(210274)_826 | 1 | + | good | GGAGCTGTGAGCAGTCAGGGCCCCCTAACTGGGCAGCATCATAGGTCACAGAACTGTCTCCTGCCCCCTGCCCCTGGGCAGCTGGTCTCCTTCTCTGGATTTGTTCTCTTGTGCAAAGTT | 120 | 57 |
| chr7 | 144274617 | 144274737 | 772751_42001748_Shank2(210274)_827 | 1 | + | good | CTCCACCTCTCCGTACATGCCTTTGTTTCCCTCATCTATACATGAGTCAGTACCTTAGAAAGCTGAGGCAGGAGGATTATAGTAAGTGCATGGCCAGCCTGAGCTATATAGTGAGTTCCA | 120 | 47 |
| chr7 | 144274737 | 144274857 | 772751_42001748_Shank2(210274)_828 | 1 | + | good | GGCCAGATTGGGCCACATGATAAGATCCTGTCTCAAATAAAACCAAAAGCGGAAGGCGTCTCAGGGAAACAGAAGGTTCAGTCAGTTACGCTGAGAGCACTCACAGCCCGTGTACACAGT | 120 | 50 |
| chr7 | 144274857 | 144274977 | 772751_42001748_Shank2(210274)_829 | 1 | + | good | CATTATGATAATTGACATATTAATTACCACTTGATAATTATTGCTCAGGGAGGCTGAGTCCAGGCAGACCTCAACTGTGAGCCAAGTTAGGGGGCGAGAGGGCGCGCTTCGCTTTGTGCA | 120 | 48 |
| chr7 | 144274977 | 144275097 | 772751_42001748_Shank2(210274)_830 | 1 | + | good | AAGGGTTCTCCGTCTCGGCCTCCAAACCTTCTAGGGCAGGTCACGGGTGACCTGGTTTTTCCACAGCACTCTGTGGTATATTAAGCCAGGCACACCTGTACCGGGCAGTGTCACAGCTA | 120 | 55 |
| chr7 | 144275097 | 144275217 | 772751_42001748_Shank2(210274)_831 | 1 | + | good | CTGAGGAAAATTAATTCTTTTCTTAGCTTCTGCTATAGGCAGTGATGAACCACATAGGCATGTTATTTGTGTCCATTCTAAGTCCTGTAAGGTTACTGAAGACATGTCCTGAGAGAACCC | 120 | 40 |
| chr7 | 144275217 | 144275337 | 772751_42001748_Shank2(210274)_832 | 1 | + | good | AGCCTGAGGCCCACTTCATATCTGTCAATAAAGTTTTATTGGAGCAGAGCAGAGCCCAAGCTTTCTCAGGTTGGCAGGGCTGCTAGTGTCACACCTAACAAGCAAGGCTCCCTCAGAGCC | 120 | 52 |
| chr7 | 144275337 | 144275457 | 772751_42001748_Shank2(210274)_833 | 1 | + | good | TGGACTGTTGATTCTAGGTCTTTATAGGAAAACTTGTCAGCTGCTGGCCTGGAATATGGAAATTTTAAAGCACTCCCTCTGCCTGGGTGTGGAGGTGGCTCTGGAGAGGTGGAGGTGAGG | 120 | 50 |
| chr7 | 144275457 | 144275577 | 772751_42001748_Shank2(210274)_834 | 1 | + | good | AGTTCAAAGCCAGCCTCCCCATACAGTAAATTTGAGGCCACCCTAGGCTATATTAGGCCCTGTGTTTAAAGAGAAAAAAATCCTAAATAGTCAGGAGTGTGTCTCAGTGGGTAAATGCTT | 120 | 42 |
| chr7 | 144275577 | 144275697 | 772751_42001748_Shank2(210274)_835 | 1 | + | good | GCCCGACGTGCAGGAAGCTCTAGGCCAATCCCCAGTGCCTCAAAATACATGTGTGAAGGTGCATGTCTATGATCTTATCACTATGGAAGTGAAGGTAGGAGGGTCAGGAGTTCAAGGCTT | 120 | 50 |
| chr7 | 144275697 | 144275817 | 772751_42001748_Shank2(210274)_836 | 1 | + | good | TCCTTGGCTACATAGTGAGTTTGAAGCCAATGTCAGTTGAATGGGACCCTGTTTGTAAAAAAAGAGAGAAAGAGGCCTCTAGGACTTGTAACTTCTTCTGTAAGCAGCACTGTGGTGACA | 120 | 43 |
| chr7 | 144275817 | 144275937 | 772751_42001748_Shank2(210274)_837 | 1 | + | good | GTTTCCTCTCTCTGAGCATCAACACCTGTTGGAGAGGCAAAATAGTCGCACGCTACAGGAATTCCATCCACAGTGTACCACAAATGCTTCCTGTTGCCCCATCAGATGGCACTGTGACTG | 120 | 50 |
| chr7 | 144275937 | 144276057 | 772751_42001748_Shank2(210274)_838 | 1 | + | good | TGGAGTTCTTTTCAAAGTTCAGTGTTGAGACATACCCATACTCAGTATAGCGTAGCTGTGCAGGCCTGTGGTCTACGCCAGGCAGCCTACGTGTGCCAGAGGCATCCCTGTCCCCACTCA | 120 | 53 |
| chr7 | 144276057 | 144276177 | 772751_42001748_Shank2(210274)_839 | 1 | + | good | GGCAAACACACACTACCATGTTCACACATCAGAGCACCCTTATGGAGGAGTACCTGGTGACCTTATCCTGTGGTGCTCCCAAGCAGCAGCAGGCAGGTATGGGGTCAGCTGTGACCTGGT | 120 | 55 |
| chr7 | 144276177 | 144276297 | 772751_42001748_Shank2(210274)_840 | 1 | + | good | CTTGTAGTTTGGGCCTCAGCCTGGGCTTCTGATGGTATCCAGGGCAGGAAAGCCCAGTGAAGCCTCCTCACCAGAATCTGAGGTCCCAACAGACATTCACACTTTTTTTTAATTGGCACT | 120 | 50 |
| chr7 | 144276417 | 144276537 | 772751_42001748_Shank2(210274)_842 | 1 | + | good | CAGGCTCAGTGGCAGGTGCCTTTACCTGTTGAACCATCTCACTAATTTAATAGTTAAAGCTAGGATGTCTTCCCCATCCCACAGACGACCAGAGTGGGCTAGCCACATTAAAAGAGCC | 120 | 48 |
| chr7 | 144276537 | 144276657 | 772751_42001748_Shank2(210274)_843 | 1 | + | good | AATGCCTGTTTCTCAGCCTGTTGGGCTAGCTTTGAGACAAACAGATGGGCCAGGCCTAGCCTGTCATTCTAATTAGCAGTTAGAAGGAAAGGCACAAACAGCCCCTGCCTATGCCTGTAG | 120 | 50 |
| chr7 | 144276657 | 144276777 | 772751_42001748_Shank2(210274)_844 | 1 | + | good | CAAAATGCCAGGCAGCTGGATTGCCCTGTGATATTCTAGAGAATATTCCACACCGAATGGAGGCTGGGATTCCAGAGCAGCCCAAGCACACAAGTCACCAGGCCACAGAATCAGGAGCCA | 120 | 52 |
| chr7 | 144276777 | 144276897 | 772751_42001748_Shank2(210274)_845 | 1 | + | good | AGGGGAACAGGATTTTCAGTGGAACCCAGGTTGAACTCCTGTAGGAGTGGCCTAGGAAGGAGATACTCCTGATTCTAAAAATAAGTCTCATGTCCCAGCAGAACAAAGCCAGAGGGTTAC | 120 | 48 |
| chr7 | 144276897 | 144277017 | 772751_42001748_Shank2(210274)_846 | 1 | + | good | CCATGCCTCATGGGCTACACAAGGATGTCACCCAGGAACTGGGAACCAGTAACTGCTGAGATCCTAGAGGCCAGTGGATGGCCATGTGTGTGTGTGTGTGTGTGGAGTATTCTCCCC | 120 | 55 |
| chr7 | 144277017 | 144277137 | 772751_42001748_Shank2(210274)_847 | 1 | + | good | ATAACTAGCTACCCAGCCCTGCCAGAGCTGGTACCCACCAAACACACAAGTCACTGAGGGACCTACACAGTGACAACAACATGGCTCCTGAAGACCATGGCAGCTTTGTGCAGAGAACAT | 120 | 52 |
| chr7 | 144277137 | 144277257 | 772751_42001748_Shank2(210274)_848 | 1 | + | good | GGGCTGTGCAAAGACACTATAGTTCATGAAGGGTATCTATGCCCAGAAGAGGAGACAGCAGTCAGGCTGGAGCATGGGGAGTTGGGAGAGGCATTACAAAGGGTCTTAGCTATAGTGAGG | 120 | 51 |
| chr7 | 144277257 | 144277377 | 772751_42001748_Shank2(210274)_849 | 1 | + | good | AGTATAGATTTTATTCCCAAAATGATGAGACTCTGCAGAAGGAATATTTTAAATGGGAGAAACAGTCTGACTTAAGGGATTTTTAAAATATATGTTTATGTGTATATGCATGCATACATA | 120 | 29 |
| chr7 | 144277377 | 144277497 | 772751_42001748_Shank2(210274)_850 | 1 | + | good | TACAATATATATGTACAATGAAGAGAAAAGTAAGGGGAGACCCAGCATCCGGTCTTGGAGGATTCTGAGTCCTGAGAACATGTCAGTTTTTTCTCTGTCTTGGTCTCCAGAACCATGTTA | 120 | 42 |
| chr7 | 144277497 | 144277617 | 772751_42001748_Shank2(210274)_851 | 1 | + | good | GTTCCAGAGTCAGGGTGGAAGGCCTTCATGACAGTTCCTAGAGAATGGTGCCCAAATGTTTTCTGTTTTCCTTGGTACTAGACCCTTTTCTTCTGTTGTTTTCTTGAATCAGCTTCTCC | 120 | 44 |
| chr7 | 144277617 | 144277737 | 772751_42001748_Shank2(210274)_852 | 1 | + | good | CAGAGCGCTATGAGCCGCCATGCTGGGCTAAGCAATTCCCCTTCTAGCCAAGGAGGGAGGCCTCAGGCAGGATGCTTAGGCTTCTGCAAGCTCCAGGAGAGAGGCTGCTGCCCCTCCCTC | 120 | 62 |
| chr7 | 144277737 | 144277857 | 772751_42001748_Shank2(210274)_853 | 1 | + | good | ATGGGCCCCTGACCTTCATCTGCCTGCTCCGTGGGAACATCTTGTGTCTAACAGAGAGCTATGTCCTGTTGCTTGCTGCAGGCAGATCACCCAAGCCTGTCTTGCTAATGCCCACACGAG | 120 | 55 |
| chr7 | 144277857 | 144277977 | 772751_42001748_Shank2(210274)_854 | 1 | + | good | GGTGGGGTACACATGGCAGTATCTACATTTGGGCATTGCATTCCCAGAATGCAACAGGCTCTGAAGCACAGGGCTTTCTCCTAAGCTCATAGGTTTCCAGCCCTGCTATGTGCCTGGTCC | 120 | 52 |
| chr7 | 144277977 | 144278097 | 772751_42001748_Shank2(210274)_855 | 1 | + | good | CTCTCAAATGATCTTTTCCTCACCAGCCTGAGTATGTCAAGGTTTTGGCCAAGACCAACATGGCAAGCAAGCAAGATGACTCTCTTAAGGACCCTGTGTGAGTTCAGGATACAGGGACCC | 120 | 48 |
| chr7 | 144278097 | 144278217 | 772751_42001748_Shank2(210274)_856 | 1 | + | good | AGACCCTACTCTGTGCATGGTTTTCCTCATTTTAGTTGGGAACAGTGATCATAGCTTGTCACCTAGGCAAGGTACTTCTGGGGCCATTGTCTTGAGTATGCTTTGGCAGGTAGCACCGGA | 120 | 48 |
| chr7 | 144278217 | 144278337 | 772751_42001748_Shank2(210274)_857 | 1 | + | good | TGGTGCTCAGGCAATACTCGTGGGTGACTTATTGCAAGGTAATGAGGTGATTTAATTCGAATCAAGTTCTCTGCTCATGATTGTTTCCATCGGGAAGAAGTTGTATTGCTTATCACGTTA | 120 | 41 |
| chr7 | 144278337 | 144278457 | 772751_42001748_Shank2(210274)_858 | 1 | + | good | GCAACGAGGAGGGGAAATGGCCGTAATAAACGCAGGCAAGGCTTTTGCATAATTGCCCATCTGAGAAATCACTTCTGTATCTAAGGAGGGGGGTGTGCAGTTTCCACAGATAAGGGTGACA | 120 | 48 |
| chr7 | 144278457 | 144278577 | 772751_42001748_Shank2(210274)_859 | 1 | + | good | CAGTGACACCTCCATTCATCCCAAGAAAGAGTGGCTTCCAGGAGCCAGGAGGAGTGCAGTCCATGACTGTGTCTATCCCTTCCCGCAGGTGACACAATTCTGTCTCCCAGTAACATCCTT | 120 | 52 |
| chr7 | 144278577 | 144278697 | 772751_42001748_Shank2(210274)_860 | 1 | + | good | GCTTGACTATCGCTGAAAGGTGGCACCCAGCGTGAGACTTATTAGCCCAGAAGCAAGAGAGCCTTATGAATGCTCCAATCAGGAAGCAGGATGTTAGAAGATGTTATCATGCGTGGTTAA | 120 | 47 |
| chr7 | 144278697 | 144278817 | 772751_42001748_Shank2(210274)_861 | 1 | + | good | AGCCAAGAGGACCCCCACTTTGGTGGCAAGTGACAAATAATGAGGGACCCCTCCTGTCTACCAGCAGGTTTCCAAGCCTTCCCGTGAACCAGGCTATTCTGCCTTTGCCCACCCATAGAT | 120 | 54 |
| chr7 | 144278817 | 144278937 | 772751_42001748_Shank2(210274)_862 | 1 | + | good | AGGTCCTCTCTGCCAACCTACAGCCAGCTTACCCTCCATGAGCTGGGCTGACTGCCCACACCAGCAGCAAGAGGGCCTGGCTGGCATCTCTGCCGGCTGAGCTGGAACACCTGTTAAGAC | 120 | 61 |
| chr7 | 144278937 | 144279057 | 772751_42001748_Shank2(210274)_863 | 1 | + | good | TTGGCCTGTGCCCCAGCACCTCTCTACCCACCCCCTCCATGGAGGAGCTGACAGCATCCTGCATGGCCCCATCCTTCTGATTGGCAGAGTCCAGAAATGTCTAGGGCATCCACTGTCGC | 120 | 59 |
| chr7 | 144279057 | 144279177 | 772751_42001748_Shank2(210274)_864 | 1 | + | good | TGCTGCTCCCTGGAGACACGTCTTTCTAATAGTGCTCTAGCTCCTGTGTGGACAGGCCACCAAGATACAGTCACAAGAACAGTGGGGTTGCCGATTTCAGCCTTCTGCTCCCAGCACTGG | 120 | 54 |
| chr7 | 144279177 | 144279297 | 772751_42001748_Shank2(210274)_865 | 1 | + | good | AAATGCTTTGCAGAGATAATCTTGGGTTTCCGCCGCCTCTGGACAGGGAGCAGGTACCACCTCCCCACCACCAGTGGATTTTCCCAGTGTGCCCATGGCCCTTACTCAGCATACCTTTTG | 120 | 54 |
| chr7 | 144279297 | 144279417 | 772751_42001748_Shank2(210274)_866 | 1 | + | good | GCCCATTGGAGTTGTCTAAAAATTTTCCCTTGAACCCATTCATTTGAGTCACTGTGGCCACTGCCTGGCTGCGCCCTCACACACTTAGCAATGAATAGCACTGGAGGATTCTCCCGAGGT | 120 | 50 |
| chr7 | 144279417 | 144279537 | 772751_42001748_Shank2(210274)_867 | 1 | + | good | GAGAAAAATCCAACTTACAAGTGATTTTTCACACCTGAAGGAATGATGTTTACAGATACCAGATTTAACAAGAAATGGAGTTGACCCCAGGTGAGATTCCAGAGACACTTAATCAAATGG | 120 | 38 |
| chr7 | 144279537 | 144279657 | 772751_42001748_Shank2(210274)_868 | 1 | + | good | CAGGCTGGCTTCTGTGTTGCTGCTTAAGCCCTGAGGACTTGAGAGGCGCTGAATTTATTATTTCCCAGCTTGCAAATAGGCTTGAGAGTTAAAGCCAAAACCTGGGGGGGTCCCCCCTTG | 120 | 52 |
| chr7 | 144279657 | 144279777 | 772751_42001748_Shank2(210274)_869 | 1 | + | good | TCTTCAAGACTGTGGATAACACCCTGTCACCTCTTGATGGTGGCCAACACACCTGCCTGGTCTCAGCTGATCAAAGTCCATCCTCAGTAGGGTCTATGACATCCCTTTCTGTAGGCCTAG | 120 | 51 |
| chr7 | 144279777 | 144279897 | 772751_42001748_Shank2(210274)_870 | 1 | + | good | CTATACAGGGGTTCCCTGTTGTCTGGGTCTGTGCAACAGCCAGCTATTATATGGAATCAGCCTGGCTTTAGTATCTACCTTGTAGAATGTAGTGACTTCTTGTAGAATGAACCCCAATCC | 120 | 45 |
| chr7 | 144279897 | 144280017 | 772751_42001748_Shank2(210274)_871 | 1 | + | good | AGAGAGCCATAGATTATGTGTCCTCAGTCACACCCACAGGCCACTCCCACCACCCAAGACATCAGTCATAGCCTCCAGGGAGCAAGACTAGGCTACTTGTGCTGACACCAAGGAGAGTTG | 120 | 53 |

| | | | | | | | | | |
|---|---|---|---|---|---|---|---|---|---|
| chr7 | 144280017 | 144280137 | 772751_42001748_Shank2(210274)_872 | 1 | + | good | TGCAAGGAAGAGGAGCTGACTTTCATGGTAGGACTATGTAGGAATTCCAGCCACTTGTGTGTCAACCCTGCTGGGTCCCAGACTGAATAAAGAGAAAGTGAACCGAGCACCAGCGTTCAC | 120 | 50 |
| chr7 | 144280257 | 144280377 | 772751_42001748_Shank2(210274)_874 | 1 | + | good | AAGTTGCTTCTGTCAGGTGTTTTGTCACAGCAACAGGAAAAGTGCTACACCGGGTGTGATGGCTCACATATGTAATTCCAGCAACTTAGGAGCCTGAAAGCAGAAGCTGGGAGGGGGGGA | 120 | 50 |
| chr7 | 144280497 | 144280617 | 772751_42001748_Shank2(210274)_876 | 1 | + | good | GGACCCACTCTGTGGCAGACTTCAAGTCCCCACAATACTCCACATGTACTTGATCAGTTAACGCAAGATAGATTGGGGGATGGGAACAAACATGGCTGCCTTCTGCATGTAGTTCCCAGA | 120 | 49 |
| chr7 | 144280617 | 144280737 | 772751_42001748_Shank2(210274)_877 | 1 | + | good | AGTCAACGTGCTTCCTGTAGAAACAGCTCTTCCATCTTCCATGGCAATCACAGATACGAAGTATGGACTTGTACTTCTCTGACTTGTACTGTGCATGACGTACACAGCTGTACTTAGCA | 120 | 45 |
| chr7 | 144280737 | 144280857 | 772751_42001748_Shank2(210274)_878 | 1 | + | good | TATCTGTATCCCTCTCCTCCACACACACACACATCTGCTGCAGGTAGATATTGTCATAGGAGGGTCCATTTGAGGTATGCTGGGAACTCTTCTTCCCTGTGCAGGAAACACATCTGAGGA | 120 | 48 |
| chr7 | 144280857 | 144280977 | 772751_42001748_Shank2(210274)_879 | 1 | + | good | ACATGTGCTGGCTTGTATTTGTGTACAGTCAATGAACACTGGGAGGTTTTTTTTAGAACAGGGAATATTGGGGGTTTTGCTTCCAGGATACCCACACTGAATGCTTTATGTGCACGGTCA | 120 | 43 |
| chr7 | 144280977 | 144281097 | 772751_42001748_Shank2(210274)_880 | 1 | + | good | GTTATGGGTAAGGCAACAGGTCTCCAGAAGAAGCCTGAATGGGCTCGCCCAGCAGCAGTGTGCGGACTATGGGAGCTTCAGATGTGTCCCAGTGAAATGTAAACAATGGCCTTCGGGTCG | 120 | 54 |
| chr7 | 144281097 | 144281217 | 772751_42001748_Shank2(210274)_881 | 1 | + | good | AATCAGGGTATATCTGCTGGGAAGCCAGGCAGAGGCTACCAGGGACTTAGGCAGCGTTAGGTCTACTTATGGTGAAGTGCCAGGGCAGCCAGTAAGGCTGAACTGGTCTCACCCAGGGTC | 120 | 56 |
| chr7 | 144281217 | 144281337 | 772751_42001748_Shank2(210274)_882 | 1 | + | good | AGGTCAACAAGAGGAAAAGGCTACTGAGAGAGAGTCCAAGAGAGACCTCCCCTCTGCAGTCATCTGCAGCATCTCCAACCCACCTCCCCTCTGCCATGTATTGCACCCCTGCCTCAGGTT | 120 | 55 |
| chr7 | 144281337 | 144281457 | 772751_42001748_Shank2(210274)_883 | 1 | + | good | CTGCCAGCCCCTGGGCATTCAGGGTGATCTGTAACAACAACTCACTGTCTCTCAGGAGATCAAGCCCCTTGAGGAGAAGGGGGCATTTGCTTCTGGGTAAAGCACAAGAGTGTGCACAGA | 120 | 53 |
| chr7 | 144281457 | 144281577 | 772751_42001748_Shank2(210274)_884 | 1 | + | good | AGCTTGAGGGCCGTGTTTCTGGCACTGGGTCTGCCCACAGTGATATGTCTTATCAGTGGAAACTCAGTTTCCACGCCTAAGTCTTTACCCACTCACTTCAGACTGGTGAATGTCTGAATT | 120 | 48 |
| chr7 | 144281577 | 144281697 | 772751_42001748_Shank2(210274)_885 | 1 | + | good | TGGGGTTTAGGCCCAAGGCTTGTTGGTGGTTAAGGGTGGCGGGCACATGGTGGGGCTTGGCCCCTGAAAAACATCATAGAGAAGAACTAGAAAACATCTTCTAGGAATGGGAGGGGTTTG | 120 | 52 |
| chr7 | 144281697 | 144281817 | 772751_42001748_Shank2(210274)_886 | 1 | + | good | GGGAGAGGAGGCCTGTCTACACACACAGGGCTGTCACAGCAGAAAGCTGTTAGGAAGGTCTTGGAAGATCAGAATCTGGGAGAAGCCTCGGGCACGGGGCTGTGGAAGAGGTGAGTAATA | 120 | 55 |
| chr7 | 144281817 | 144281937 | 772751_42001748_Shank2(210274)_887 | 1 | + | good | CAGTCATTGCCTCTAATAGTAGCTCTTTAACTGAAAATAATTCCTGTGTGTTAGAGTTGACTGCCCACTAGGGGTTGTCCTGAGAGTATAGGGTCCTTCTGTTCCCCCCTTCCATGCCTC | 120 | 47 |
| chr7 | 144281937 | 144282057 | 772751_42001748_Shank2(210274)_888 | 1 | + | good | ACCCCATGATCATGCCTGCAGTTGTGATTCTCAGTGTATAGGGCAAGATCTGAGGCTTAAGAGGTTAGGGTGACCTTGGGCTCAGCATCACACTTCACTTTGAACCCCTGCTCAGTGACA | 120 | 50 |
| chr7 | 144282057 | 144282177 | 772751_42001748_Shank2(210274)_889 | 1 | + | good | CCCCTGTCCCTTACAAGCAGTCAGAAGGAGGCAGCCAGTGTCCAGTTAGCATAGCTGGCCTGAGGAGGTTCCTGTGACTGGTTTGAGGAAGTGTCTGGTTCATCAGTCAGGTTCTGCTGA | 120 | 54 |
| chr7 | 144282177 | 144282297 | 772751_42001748_Shank2(210274)_890 | 1 | + | good | CTGATGGAATACATTCCAGAAGGCCTCTGTGGCTTCTTCCCTGAGCTGAGGCCAAAGTGGGAGCTGTTGCTCACCTGTGTCCAGAGGATCATGCAGACATTAGACTCTCATGTTGGGGCA | 120 | 52 |
| chr7 | 144282417 | 144282537 | 772751_42001748_Shank2(210274)_892 | 1 | + | good | AAGAAGGGACATAAAGAAGCCACAGGATTTCTACTGTTTAACCACTGAGGGAGCACTGGGGCTGAGACAGGTAGCTTCCTATGTGAAAGTTGGAGGAGAAAGGAAAAGGGTGTCCAATAA | 120 | 46 |
| chr7 | 144282537 | 144282657 | 772751_42001748_Shank2(210274)_893 | 1 | + | good | GGGACCTGAGTCTATATTCCAGATACCTAGGGTGGGCACGAAAGTTCAAAGATTGAAGGGTCTGGCATAAACTTGCTTCACCCAGTGTCTTGTAAAGTGCCCGAATCCACAGCAACAAGT | 120 | 48 |
| chr7 | 144282657 | 144282777 | 772751_42001748_Shank2(210274)_894 | 1 | + | good | CCCTCCCTACCCTAGCTCTCCTTCGAACCATTAATGTGATGGAGGTCCTGGCTGCTAGCCAGAGCCTCTCCAGTTACTGACCACACTTGAAAGGGGTTCCCTGGAGCCTTGTCACTAAAC | 120 | 54 |
| chr7 | 144282777 | 144282897 | 772751_42001748_Shank2(210274)_895 | 1 | + | good | CACACTATCATGGGTATAATGGCATCCAACCACAGCAGCCCCAAGTGACTGTGAGCTGTCCTTACAGCCTGTCAATATGGGACGCAGCGGACCAAAGGGTCTGTCACTTCCTACAGGCCA | 120 | 53 |
| chr7 | 144282897 | 144283017 | 772751_42001748_Shank2(210274)_896 | 1 | + | good | GCCCAATGACTCACCCATAGGCTCTCATCACCCCTGTGGGATCTGCTGGCACAGGGGGCTTTGAGAACTGAGTAGAAAGGGTCTCACATAGCAGGATCCAGGAGGCTGGCAGGATATGTC | 120 | 56 |
| chr7 | 144283017 | 144283137 | 772751_42001748_Shank2(210274)_897 | 1 | + | good | AGTAGACAGAGCAAGGTTTTAGGTGAGCTTCTCAGCCACCTTCTCAGACTTCTGTGACTAGTCATCCTACCTAGACCACTGTGTTTGCAGATTATGTCACTGCCCAGCCTCTGTTACCTC | 120 | 48 |
| chr7 | 144283137 | 144283257 | 772751_42001748_Shank2(210274)_898 | 1 | + | good | CCCCCATAGCAAATTCCCAGAACACAGATTCTCCCTGGGACATTATTAGGATAACCCGGTGATGGGTAACTGTTGAGTGACTTCTGTGAACTCGACACTATCAGTCTTGCTTGCCCATTG | 120 | 48 |
| chr7 | 144283257 | 144283377 | 772751_42001748_Shank2(210274)_899 | 1 | + | good | GGAACCTTGAAAAGCCACCACTCTGTCCTCACCTGTTTCCAGGTGTGTCTAACCAGCAGACCACTGGCCACATGCACCCCAAGGCAGCTTTGAATAGGGCCAACCCAAAATTTTATACTT | 120 | 50 |
| chr7 | 144283377 | 144283497 | 772751_42001748_Shank2(210274)_900 | 1 | + | good | ACTCAAACATTGTAAGATTGTTTTGCCATATTTTTTTTTTCTTGTAACTTCATTGTGTGAACTTTGTAATTGAGAATGTCAACATAATGTCACAGTGACAAAAGGTTGGCCAGGCCTGCT | 120 | 34 |
| chr7 | 144283497 | 144283617 | 772751_42001748_Shank2(210274)_901 | 1 | + | good | AGACTGGTTAGAAGGGGCTTTCCTGCTCCTACAGCTCCAGGGTGCCCTGTTCTAGTTGACTGCTATCTGCAGGCCAGACCCTACCTAAGTCCCATGCCTCCTGTAGACCCAAAGCCCTAG | 120 | 56 |
| chr7 | 144283617 | 144283737 | 772751_42001748_Shank2(210274)_902 | 1 | + | good | AGAACTGACTGGCACCACTCTGGGACTCAGAGCCGTCACTGTTCTTCACTCCCAACCCATGTTCTATCCACAGCCCTCTTCTTCAGTGTCTGGGGCTCAGCTACTCCTGTTGTGATTTCT | 120 | 52 |
| chr7 | 144283737 | 144283857 | 772751_42001748_Shank2(210274)_903 | 1 | + | good | GCCCACCATGTCCTGTGTCCTCCTACTGGGACTCTACAGAAGAAAGCCCTCTCCCTCCTTCTCCAGCCCTCAACCCCCAGACTTCCTGCCTTCTGACTTACTTCCGGGGGGCGGGGGCAA | 120 | 61 |
| chr7 | 144283857 | 144283977 | 772751_42001748_Shank2(210274)_904 | 1 | + | good | TGGCTGAATCTGGGATACAGACAGCACTGTCTCCTGCCCCAGAGCAGCCACCAAAAAACGGCCGACAAGGGCCAGAGTTGGTGGCTCTCAATTTATTAGGCAGCATCTTTTCATCCTGTG | 120 | 52 |
| chr7 | 144283977 | 144284097 | 772751_42001748_Shank2(210274)_905 | 1 | + | good | TCCCCTGTGCGCACCTCTAGGTGGACAGTCAGAGTTCTGTTAATGTTGGTGACAGGCAGCTTTGGCCGGAACAGCAAAGTTGGGGCTTTTTAGTCTGTTTCCTTTAAAAATGGAAATGCT | 120 | 48 |
| chr7 | 144284097 | 144284217 | 772751_42001748_Shank2(210274)_906 | 1 | + | good | ATACAAAGAGGTGAGCCCGGTGGTTCTTGGTCTTCCTTGCAGTGGGGATGCTGCTGCCCTCGCTGTCCTCAGCAAGGCATCTTCCTCAGGAGGGAGCCGCTGCCCTCCTTGGCTCTGGTA | 120 | 60 |
| chr7 | 144284217 | 144284337 | 772751_42001748_Shank2(210274)_907 | 1 | + | good | GGACTGAAATGGCCTGAGCCCCTCACTCCCTCACCCCTCAGCGTTGGCCTCCATGGCGGGGAGAGACACAGAGCTGGCGGCTCCCCCTCCCGCCGGCCACCAGCCCAGCCTGCGGCTCCT | 120 | 71 |
| chr7 | 144284337 | 144284457 | 772751_42001748_Shank2(210274)_908 | 1 | + | good | CTGCTCTGCGCGGGATGACGCCCAGGCAGCCGCGCCGGTCGCACTGTGGGCTCATCAGATGACTGTGGGGGCTCCTGCTGCACACTACATTCCCTAGGACGGGGGAGAGGGAGGGAGGGG | 120 | 68 |
| chr7 | 144284457 | 144284577 | 772751_42001748_Shank2(210274)_909 | 1 | + | good | GCGCTGGCTGCAAAGCACAGGTCTCTGCATCCCCAGTGCCTGCTGCTGGCTCGTGCACAGAACAAACGGAGAGAACCCGATCTCCTCTTTAAGCCCCTTCAAGAAATAGCAAAGCAGCTG | 120 | 56 |
| chr7 | 144284577 | 144284697 | 772751_42001748_Shank2(210274)_910 | 1 | + | good | TCTTGGTTATTCCTCCGGCTGATTTCTTCTGCTTCCCTGGAAGATGCGTCTGGCCGAGACAGGCCTGGGGACCTCCTTCACCACGCTGCTGCCTCTGCCGCCGCCGCCACCACCCGGGCC | 120 | 65 |
| chr7 | 144284697 | 144284817 | 772751_42001748_Shank2(210274)_911 | 1 | + | good | ACTTACTTTCTTTGCTCTGGGCGGCGGTGTCCTGGCCGGCCCCCGGGTCCCTTGAGCTGGGCCCCCTGCTCCCCCAGCGGCCTGGGCAGCCCCTCCAGCATCCATGACTCCTTCCCCTGG | 120 | 70 |
| chr7 | 144284817 | 144284937 | 772751_42001748_Shank2(210274)_912 | 1 | + | good | CTGGCTCCCGCGCCCGCCGCCCCCCTCCACATCCCACTGTGCCCCGCTCGCCCATCCTGCTACCCCCCCACCCCCAGCTCCTCCTGCACCGAGCTGCTGCCCAGGCTCCTCCATGCTGCC | 120 | 75 |
| chr7 | 144284937 | 144285057 | 772751_42001748_Shank2(210274)_913 | 1 | + | good | TGGATCTGAGCTGGGTGATTAATTGGCTGTGATGATGAGCGTCCCCGGCGGTGGAGCAGCCACCGTGATGATGACCGGTTACAATAATGGTCGCTATCCCCGGAATTCTCTCTACAGTGA | 120 | 53 |
| chr7 | 144285057 | 144285177 | 772751_42001748_Shank2(210274)_914 | 1 | + | good | CTGCATTATTGAGGACAAGACGGTGGTCCTGCAAAAGAAGGACAACGAGGGCTTTGGATTTGTGCTCCGAGGGGCAAAAGGTGAGTGTTCTGGGATCCCTTTCCACAGTGTGTGTGTGTG | 120 | 52 |
| chr7 | 144285177 | 144285297 | 772751_42001748_Shank2(210274)_915 | 1 | + | good | TGTGTATGTGTGTGTGTTGTACACAGGCTTGTATGAAGAACCTTCCCCAGCTTGGAAATAAGCATGAAACAAGCATTTTCCTGGGGGCACTTGGTTGCTAGACAACGATGCTCTTTAACA | 120 | 44 |
| chr7 | 144285297 | 144285417 | 772751_42001748_Shank2(210274)_916 | 1 | + | good | TTCTGCCTTCCTCAGAAAACATCTAGGCTTCCTGCAGGCATGTTTTTAATTTTCCATCTACAGTACTGCTCCAAATGAGCTGCTCTCCCTCCCTCCCCCCCCCACCTCTTTCTCCCAATC | 120 | 50 |
| chr7 | 144285417 | 144285537 | 772751_42001748_Shank2(210274)_917 | 1 | + | good | CTGGGGAGACCCTGCAGCTTGGGCTTCAGTTTAGTCACAGAGTGCTGACTCTCAAAGCTCCTGGTGTCTGATACTCAGAATGGCCCAGCCCAAGTTTGAGGCACCCGGAAGCTGGCCTGT | 120 | 57 |
| chr7 | 144285537 | 144285657 | 772751_42001748_Shank2(210274)_918 | 1 | + | good | GCTGGGGATGGGTTTCAGCTCTAAGGGTGGCAGGTGTCTGCCAACTCACCTCTAAGGTGACTTTGCAAAGGCAAGGAGAAAAGAAGGTCCCCTCTACTGTCAACTAAGCTGGCAGCACCC | 120 | 54 |
| chr7 | 144285657 | 144285777 | 772751_42001748_Shank2(210274)_919 | 1 | + | good | CACACCCTGTGGCCTGGGAGTGGGAGGCTCTGGGATACATCTCAAAGCAGTGCCCCACGCTGGCCCCTAGTCTGGGGCTTCCCTTCCAAACTACAGCAAAATGGCATCAGCTTGCTCTAG | 120 | 58 |
| chr7 | 144285777 | 144285897 | 772751_42001748_Shank2(210274)_920 | 1 | + | good | AAGAGGCGTCTGCCTGAGGCAGGTGGAGGTCAGCCATGTGGCAGTTACCTGTGCAGGAGGCATCCTTCCCTGGCTAGCTTGCCCACCTGCCACGGGCTCTGACCGCCCACAGCAATCCAT | 120 | 62 |
| chr7 | 144285897 | 144286017 | 772751_42001748_Shank2(210274)_921 | 1 | + | good | CCAAAAAGAGAAGAGCAGAGTCTTCTCCACCCCCAGGGTACAGGTCCATGCCTGTGTGATGCGCCTTCCCCTCCCCCAATCTGGGCATCACACCTTCCCACCCACCCACCCACCCACCCC | 120 | 62 |
| chr7 | 144286017 | 144286137 | 772751_42001748_Shank2(210274)_922 | 1 | + | good | CAGCCTCTCAGGAGGACCAAGAGAGTCCTTCAGCAGCCTCATGGGCCCCAGAAGCCCAGCCTTGCTCCGCTGAGAGGCATGGTGGTCATGTGATGGTCATAGAAAGCCAGGTTCCCCCCT | 120 | 60 |
| chr7 | 144286137 | 144286257 | 772751_42001748_Shank2(210274)_923 | 1 | + | good | GGAGAGAGTTCAGAGCAGATTCCAAATCAGCAGCCACCCACTGGGTCAACATTGTTCTCTGTCCCTGTTCCAAGTTGAATCCCCTGGGTTGCAAGTCTGTATAGCTGTGAAACTGGCAGG | 120 | 51 |
| chr7 | 144286257 | 144286377 | 772751_42001748_Shank2(210274)_924 | 1 | + | good | CGGTTTCGGTATGGTCTTTCCCTAGCTCTGTGTGGCTGGGGAAGGAACTTTTCCATTTTTAGAACTGAAAATGTTTAGTCATTGATACTTTTAAAGAAAATGACAGACATATAGACATGT | 120 | 38 |
| chr7 | 144286377 | 144286497 | 772751_42001748_Shank2(210274)_925 | 1 | + | good | TCCGCAGACCACTTTATTCCATGGTTTCGGCAGAGCGGCCAGGAGGCGTGAGGGGACACTGGCATGCACCTGGCTTCATAGCTTACATTAGTACCAATGTGCATGTGAGTAAGGAGCCCC | 120 | 55 |
| chr7 | 144286497 | 144286617 | 772751_42001748_Shank2(210274)_926 | 1 | + | good | TCTCTGGGACAAGCCAGCCAGGACCACTGCCTCCCAATAGGGAATGCTGTGACCCTCTTCCTCCCAGAGTCCCGTGGGCAGGTGGGATGACCAACTTGGAAAACATCCCCCAGACTCAAA | 120 | 57 |
| chr7 | 144286617 | 144286737 | 772751_42001748_Shank2(210274)_927 | 1 | + | good | CACAGTATTTCTTTGTCTTCCTAGCCGATACCCCCATTGAGGAATTCACACCCACGCCAGCATTTCCAGCCCTGCAGTACCTGGAGTCCGTGGATGAAGGTGGGGTGGCATGGCAAGCCG | 120 | 56 |
| chr7 | 144286737 | 144286857 | 772751_42001748_Shank2(210274)_928 | 1 | + | good | GACTAAGGACCGGGGACTTCTTGATTGAGGTAGGGCACAGGTGTTTGTATCACACTACTTGGGGAGAGCGCCGACTCTCATGTGAGAGAGGATTGTGGTTTTGAGCACAGCATCCGGATG | 120 | 52 |
| chr7 | 144286857 | 144286977 | 772751_42001748_Shank2(210274)_929 | 1 | + | good | GTGGATGACTTAGGCCACATGGCTTGTATTCTCTCGAGCATACCCTACCCACCGAGGATACTTTTCCTCTGTCCCTGATGTTCAAGTGTTTTCGTAGCTCTTTGGGAAGAGAAAGCATAC | 120 | 48 |
| chr7 | 144286977 | 144287097 | 772751_42001748_Shank2(210274)_930 | 1 | + | good | ATGGAACTTGGCCAGTGGTCTTCAAGAGGAAGATGGAGACATTGTCATGATGCTGGAGCCACATGCAGATCTGTGGGTGGCTCAGCTCTGACTCTAATAATAGGAGGGGAGTCCTTGCCC | 120 | 52 |
| chr7 | 144287097 | 144287217 | 772751_42001748_Shank2(210274)_931 | 1 | + | good | ACTCCTGTTGTGTCCTAAGGACAAGTCACCAAGCCTTTGTGTGGTGTTGGCTAGCATGACGTGTGTTGTGGGTGGCTATGAGTTGTTTGGTCTATGTTCATTGCATGGTTGTTGGTCTTT | 120 | 47 |
| chr7 | 144287217 | 144287337 | 772751_42001748_Shank2(210274)_932 | 1 | + | good | AAGGACCACAGAGCATGGGGCGGGGGTGGGGCTTGGGAGGGGCTAGAGTAGGCCCAGAGAGGAGACTGCAGTGTGCAGAAGCTTTCAAAGCTAATTTTCATTCAAACCCAAAAGCAGGAT | 120 | 54 |
| chr7 | 144287337 | 144287457 | 772751_42001748_Shank2(210274)_933 | 1 | + | good | AATGTTTGGTCATTTGAGGAGTCTTCTTGTGTGCAAGTGTTGGTGTCTGTGAACGTCACCTGCGGGTTGTATGTTGGTGCCAGTGAGGAAGTTGTTGCATCCTGGTGCTTCAGGGCATTC | 120 | 49 |
| chr7 | 144287457 | 144287577 | 772751_42001748_Shank2(210274)_934 | 1 | + | good | TGCCTCTTTGCGGGTTTAGGTGTTACTTTGGATCTCACTATATGAGACAGCTTGCTACTCTTTGGAAGCAGTGTGACTGAACGAATATGCACTAGGTTGGAAAGATGGCCTCTTTTATGA | 120 | 43 |
| chr7 | 144287577 | 144287697 | 772751_42001748_Shank2(210274)_935 | 1 | + | good | TCTGATAGACCCAAACGGGCCTGGTCAGTAGGTTTTTGTCTCCCAGTGTTCACTCAGAGTCTTGCCTCAAGTTTGTCAGAAATCACACACAGGTGGAGATAGTAAACAGGTGTATCTCAC | 120 | 47 |
| chr7 | 144287697 | 144287817 | 772751_42001748_Shank2(210274)_936 | 1 | + | good | CCAGCCAATTGGGTGACCTCAATGCACATCCATTCAATAAGAGAGGATACCCTAGCCTGTCCCAGTAAGATGGCACATGAAGCCTTTGCTCCGTGTTATATGTTACCTGCCCACAATAGT | 120 | 48 |
| chr7 | 144287817 | 144287937 | 772751_42001748_Shank2(210274)_937 | 1 | + | good | TTTTCTCTGCCCATCATGGTTTTGTATATCTCTGGAGTCCGTTCACACTAGTAAGGGTGACAGGAGTAGAGGGAGAAGAGAACACATCTGTTAGGGAAAGAAGCAGGTAAAAAGGAGTCA | 120 | 44 |
| chr7 | 144287937 | 144288057 | 772751_42001748_Shank2(210274)_938 | 1 | + | good | CATATGGCTCTCTTGCCATTGCTGAAAGATGCTTTAAACTACTCTTGACAGCCCAGAGCCTGGTCTGAGGGAGGCACGAGCTAATTCAGGTTTCTGTGGGTCAGAGCAGTTAGAATCTCT | 120 | 48 |
| chr7 | 144288057 | 144288177 | 772751_42001748_Shank2(210274)_939 | 1 | + | good | TCCAGAATGGAGAAGCCGGAGCTGGCTGCAACTCCTTGAGTTCATGGGGTTTGTAACCTCCTGGAACTGCAGACTGGGTTGAATCATCTTCTGTAGGATGCGGTTGTGACGCAAACCCTC | 120 | 52 |

| | | | | | | | | | |
|---|---|---|---|---|---|---|---|---|---|
| chr7 | 144288177 | 144288297 | 772751_42001748_Shank2(210274)_940 | 1 | + | good | TTCCTGTGTATGAGAGACACTAGAGACAGAGGGGGTGTGGCCTGGGACTGGGGCTATAAGTGGTGTGTGGGTGGCCGTGTGATGGCAGAGAAGGGGGCTCCAGGAGCGTGAGATGGCTT | 120 | 59 |
| chr7 | 144288297 | 144288417 | 772751_42001748_Shank2(210274)_941 | 1 | + | good | CAATAATAAATGAGGGCTGTAGATGGATGCACACCCCCTGATGTTCAGCTCACTTGACTGTTCCATGATAAACTAGTCAATAAAACAAGCCTGGGCCTCTGAAGTGGAAATGCCTTGGG | 120 | 45 |
| chr7 | 144288417 | 144288537 | 772751_42001748_Shank2(210274)_942 | 1 | + | good | GTGACACTTTCCTGGGGATGTCCTGCATACCACCCCACAATTTTCTGTCCTTGTAGCACTTCCTTGACTTTTTTTTAAACCTCTGATATACTAAACTGTCTTGGGGAACAGGCATT | 120 | 42 |
| chr7 | 144288537 | 144288657 | 772751_42001748_Shank2(210274)_943 | 1 | + | good | TTGTTCCAGTGGCCTCTGAGTTCCAGGATTCTAGGCTTGTCTGACTTGGTCCCCAAGGTCTGTTAGGGCTATATGTTGTTCTGTAATTTTGTACATGGGCTATGGAAGGACTTAGCACAT | 120 | 46 |
| chr7 | 144288657 | 144288777 | 772751_42001748_Shank2(210274)_944 | 1 | + | good | CAGAGCCATCGTCTCACAAAGCTGATTGTCACATTCATATACTGATGATGGGACTGTGGGATGCAGACCCCCGGAGGTGGCAAACAGGCTCCTGTCTTCTACTTCCCAGGCTGCCTGAGG | 120 | 54 |
| chr7 | 144288777 | 144288897 | 772751_42001748_Shank2(210274)_945 | 1 | + | good | TGAAACTCAAGATTTCTCTATGTTAGGTGCTTAGATTTAAGTCTGTACCTCCCATGAGTTGTTTCCTTGGCTGCTGGCTTTTGCCATCAGAATATTAAAATCATTTATAAGCCCTTACAT | 120 | 37 |
| chr7 | 144288897 | 144289017 | 772751_42001748_Shank2(210274)_946 | 1 | + | good | CGGCGTGTGCTTCCCAGGGGACAACTGCTCTGTATTTTCAGTGAACGGTTTTCCACCTTGAAGGGATTGTTTTTAAAAAGCTTTATGTGTTCCCTTATGAATATGTGTGGGTTGAAGCTG | 120 | 44 |
| chr7 | 144289017 | 144289137 | 772751_42001748_Shank2(210274)_947 | 1 | + | good | TCCCCTACAGCACTCCACGAAGCATGGGTACCCATGGACAAAGCTACCTCAAGTGCCAGCGTGTGGTCCTCATCCCCGTGCCCCTTGGCAACCCTTTTCTCTACTGCCTGAAAAACAAAA | 120 | 54 |
| chr7 | 144289137 | 144289257 | 772751_42001748_Shank2(210274)_948 | 1 | + | good | ACCCTGCAAAGTATATCAGACTTCGTATTTGCTTTCTAAAATTGCATGCTGTGGTTTGAACTTCGGGACCAAGGGGCCACTTTAGAAGTGGGGGTTGACTTTTAACATCTCAAGTATCCA | 120 | 42 |
| chr7 | 144289257 | 144289377 | 772751_42001748_Shank2(210274)_949 | 1 | + | good | CAAAGCTCCATCCCTCTTTCTTTAACTCGGTCAGATTGCCCTGACATTGTCATCGTGCCTGGCGCTGGAACCCATAATGAAGAGCTTCCTGTTGTACGGGATTTTCTCTAGTCTACCATC | 120 | 48 |
| chr7 | 144289377 | 144289497 | 772751_42001748_Shank2(210274)_950 | 1 | + | good | TAAAGTTTCTTTCTCTCCTGATAAAGTGTTAAGGAGAGCGTTTTTTTTTTTTCCCCTCTGTTGTTTTTTTCCTTCCTTGGTGACTACACTTCACTCCCAAAACACTCTTAAGATTTGAGT | 120 | 36 |
| chr7 | 144289497 | 144289617 | 772751_42001748_Shank2(210274)_951 | 1 | + | good | AAAGCGCTGTCAACCTTTTAAGTCTTTCACTACTTACTTCGTCTTTGTATGCACAAGTGCACACTTGTACGTGTATGTGTGCATACATGTGTGCATGTAGACATGCACACCACAGTGCTC | 120 | 43 |
| chr7 | 144289617 | 144289737 | 772751_42001748_Shank2(210274)_952 | 1 | + | good | ACCAAGTAGGGTCCAGTAATCAAACTCAGGTTCTAAGGTTTGGTGACAACTGTTCTTATCGATTGAGCCTTCTTGCCAGCCCCTTTGATAAGCTATCTTGAAGTCAGGGATGAGGAAAAA | 120 | 43 |
| chr7 | 144289737 | 144289857 | 772751_42001748_Shank2(210274)_953 | 1 | + | good | AAGCCCCCCGGGGGTCAGCATGAGAACTCAGCTCTGAGTGAGGGTGTATGTGTGCACCCCATGTTTGAGGGCATTTGACTGCAGTCACTCTGATGTCCACACACTGGTCAGAGTAGTGTG | 120 | 55 |
| chr7 | 144289857 | 144289977 | 772751_42001748_Shank2(210274)_954 | 1 | + | good | GCTACATGACCCCGATGTACCAAGGTTAAAAGATATTCTTGCTTCTTGTAGAAAGTGGCCCAGTGCTGATGGAGAACCCATTCAGCTCCTTTGAGACTTGAGAGCAGCACTCCATGCTTA | 120 | 47 |
| chr7 | 144289977 | 144290097 | 772751_42001748_Shank2(210274)_955 | 1 | + | good | CACTATATAGTAAAGGCTTTGTCCATGGCTGTTAGACTAAATCGCTTAGGGAATCGTGACAAGGGGAAACATCTGGGCGTGCTTAGTATAGGCAGTGGGTTGTTTTTGTTTCTGCTTTAA | 120 | 42 |
| chr7 | 144290097 | 144290217 | 772751_42001748_Shank2(210274)_956 | 1 | + | good | AGCGTTTTCTGTCCATGATGCTAGAGTTTGTAGAAACAGAGCCCGGAGCTCGAACACTGGTGACTGTGTCTCTATGTTTTCCTCATTAAGAGACTTGACAACTGGCCCAGTAACAATAGC | 120 | 46 |
| chr7 | 144290217 | 144290337 | 772751_42001748_Shank2(210274)_957 | 1 | + | good | TAGAAAGGTCTTCAGAGGACATGGAGTGGCTGGGAAAGTGCGCTCTCTGTGCAGCTTGGGTGAGAGACTGGGGATTCCAATAGCCAAGAAAGCCATAGAAAGATGTACATGACTGGCACA | 120 | 50 |
| chr7 | 144290337 | 144290457 | 772751_42001748_Shank2(210274)_958 | 1 | + | good | GGAAAAGGAGAGGAGAGCTGGCTGTCACTGTGATGTCCTGTAAGCTCGTAGGCTCATCCTCTGAGACTTGGCTGGCATTGCCGTCACCAAAAATGCCTATTAACAATGCAAGGCCTGGTC | 120 | 52 |
| chr7 | 144290457 | 144290577 | 772751_42001748_Shank2(210274)_959 | 1 | + | good | CCAGGCCCTGTCTGCTAGGCCGGAACCACAGTGGGAGTTTGGGCACAGTGCTTTCACAAAACTGAGGTATGAGGCCAGGGAGATGGGTCTCCCTGGAGAGTGCTGCACAAACTTGTGGAC | 120 | 57 |
| chr7 | 144290577 | 144290697 | 772751_42001748_Shank2(210274)_960 | 1 | + | good | CTGAGGTCAGATCCCAGTACCTATGTAAAAAGTGGGGTGCAGAGGCATGCATCTATTCCAGCACTGGGAAGTTGAGGCAAGAGGATTCAGGCCTCACTGGCCAGAACTGAAATACCCCA | 120 | 52 |
| chr7 | 144290697 | 144290817 | 772751_42001748_Shank2(210274)_961 | 1 | + | good | TATGTTATTAGTAGCCCTTAAGCCAACTCACCAGAAAATGCACAGTAAGCAAACAGCATCTGATGCAAGTGCCACCGAAGGCCCATGTCCTTCCTGACTGTCATTAAGGTATACCAGTCA | 120 | 45 |
| chr7 | 144290817 | 144290937 | 772751_42001748_Shank2(210274)_962 | 1 | + | good | CAGTCATTTCTTCCTCTGATTTGCACAGAACTTTAAAAAAAAGACATCTAATGTCATTATTTTATTTTAGAATAGACATGGAGACCTTCAGTGACTTCCCACAAGGTCACACACTTGATT | 120 | 34 |
| chr7 | 144290937 | 144291057 | 772751_42001748_Shank2(210274)_963 | 1 | + | good | GTGTTAGACGCTCAGCTCTGACCAGACTGCCTAACTGGATGCCCTCCACTTCCTAGAGCTAAGACCTACACCCTTCCCCCACCTGTGGGTAATGCAGGAATTAAGGCAAAGGCATGTGC | 120 | 54 |
| chr7 | 144291057 | 144291177 | 772751_42001748_Shank2(210274)_964 | 1 | + | good | TAGGTTTTTGTTTTGTTTTGTTTTGTTTTTTCTTTTAAATTAGCAACCATTTTCCAGAACCATCGTACCTGTTGCTGGTGACGACATATTTGTGCTGTTAGAGCAAAACACAGAGTGCC | 120 | 36 |
| chr7 | 144291177 | 144291297 | 772751_42001748_Shank2(210274)_965 | 1 | + | good | AGGAATTCTGGGGAGGTCAACAAGACAGCATGTTACGGGAAGCAGAGAGTGGGTAATTTCGCATCTAAGATTCCATCCTTAGGGAACTGCATCGAAAGCAAGCAGCATTCACGGCACACA | 120 | 48 |
| chr7 | 144291297 | 144291417 | 772751_42001748_Shank2(210274)_966 | 1 | + | good | AATAAAATAATAAGAGAATGGGAAGGAGAGACAGCCAACATCTGCTCTGGCTCCACACAGAAGCCTTTCACAAAGCCCTACAAGGAGTCCTCCATGCCCATGGGGTATAATGCACTTCTA | 120 | 46 |
| chr7 | 144291417 | 144291537 | 772751_42001748_Shank2(210274)_967 | 1 | + | good | TTTAGTGAGCAAGGCAGAATCAAAATTATCTGCATAAGCCCATCTACAGGCAGGGAGGAAATCCCTGCCCTACCCCCACCCTGGGGGATGAAAGAGCACCTTCCACTTGCTTCAGACTTT | 120 | 50 |
| chr7 | 144291537 | 144291657 | 772751_42001748_Shank2(210274)_968 | 1 | + | good | TCTGTTTCTCAGCTTTTCTGCAGTGGTGTGTGTGTGTTCATTAATTTTACAAATAGAACATTCCTTCGAAGGTTATGGAGCACTGTGTTGTGGAATGGTGTCCTGGCCTCCTGGGCCAAG | 120 | 45 |
| chr7 | 144291657 | 144291777 | 772751_42001748_Shank2(210274)_969 | 1 | + | good | GTGGCATGTATCCAGAAAAGGGTTTTCTGGTGTTAGTAATTGTCCAGGGCCTTCCCTCACCAGCCAGCCCGTGCTGGGCTGCGAAAAATTCCTGTGGTTTCTGGGATTGTTGTCTTAGCC | 120 | 52 |
| chr7 | 144291777 | 144291897 | 772751_42001748_Shank2(210274)_970 | 1 | + | good | CTGAAGTGGCCGATCTCGTTTTTTGTGTTAATTTCCCTAAATAGGTTTTTCTTTGCAATTAAAGCAGGCGGCAATATCCTTCATCCTCCAACAGCCACGCTGCAGATCGCTAATTACCTT | 120 | 42 |
| chr7 | 144291897 | 144292017 | 772751_42001748_Shank2(210274)_971 | 1 | + | good | TTCCTCATATTAAACACTTGCACCAGCGATGCCTGGAACCGTTTTTTATAATCTCAGTTTATATTGAAGCGAGGACGCTAAGAGGGATGTCCTGGGGTTCTCCTCAGAGACTGAGTGTGT | 120 | 45 |
| chr7 | 144292017 | 144292137 | 772751_42001748_Shank2(210274)_972 | 1 | + | good | TGTAAAAGCTTTCTTAGGTGGGTTCTTAAAGATGGTACTTGTCTCACACACACACACCTGCCTAAACCATTGCTTCCAATACTACTGCTCCCTCCAAGACTGCATCCCCCAGCCTCAGAC | 120 | 48 |
| chr7 | 144292137 | 144292257 | 772751_42001748_Shank2(210274)_973 | 1 | + | good | CTCAGCAAGGCTTGTGGGAAAAGCAGCCAAAACTGGTCAGTCCGAGTATTCTATTGAGAGCATTATTCATAAACATATGAGGGGGGCTTGCCACCTCATTGGTAGCGTGTGGTGCTGAGA | 120 | 48 |
| chr7 | 144292257 | 144292377 | 772751_42001748_Shank2(210274)_974 | 1 | + | good | ATTAAGTCACCTGAGATAAACTACGTGTTACCTCAGGCTCTCCAAGGAAGGTTGGAGGTCAGAAGTTCTAGGCACAAAGCTAGCACTTGTCAGTATCTGTTGATGGGTGTCTTGTCTTTC | 120 | 45 |
| chr7 | 144292377 | 144292497 | 772751_42001748_Shank2(210274)_975 | 1 | + | good | TGCCTCATTAACATGCTTGACTCAGAACAGAGAACATTTGTCAAATAAATAATAATAAGTTATAGTGATGGTGAGTGATTTTGTGGTGGCAGTGACCCCCTTCAGATGTGTACCATTGTA | 120 | 38 |
| chr7 | 144292497 | 144292617 | 772751_42001748_Shank2(210274)_976 | 1 | + | good | CAGTGACAAATACAAGTCAGACTAGGTGTTGTTCAATACAAGACCCTGTTAAAGCCTCCCCTAAGCCCCTGGCAGACAAACTTGACTTATATCTCTGCTCCCTGAGCATTCACCACCCTG | 120 | 48 |
| chr7 | 144292617 | 144292737 | 772751_42001748_Shank2(210274)_977 | 1 | + | good | TACAGTCCTGCCCTGGCACAGGCCTGGTATTCACATAATCATGGAGTTTCCATTTGCTATGCATATGCTAACCCCTAGGGACAAATCCCAGGGTTGAGGAAAGGCCTTTCTGTGGCTTCC | 120 | 50 |
| chr7 | 144293097 | 144293217 | 772751_42001748_Shank2(210274)_981 | 1 | + | good | GACTATTCCCACTGGTGACCCCAATGGCCAATGTAGGAATTCTAGCCTCTGCCTCCTCCTAAATCTTTATATCTTATTCAAATAGTGCCCGTGGGCATTCAATTCAGTATGACAAGCCAG | 120 | 45 |
| chr7 | 144293217 | 144293337 | 772751_42001748_Shank2(210274)_982 | 1 | + | good | GCACCAGCAGACATTAGCAGGGCTGGAGCTGAATTCCTAGCCTGGTGTTGTGTTTGTTCTTGGGTTGTCAGCTTTGGTTCGCATCAAAGATGAGCAGTGGCTGCCTTAGTCCCCCTCTTG | 120 | 52 |
| chr7 | 144293337 | 144293457 | 772751_42001748_Shank2(210274)_983 | 1 | + | good | AGTGTGGAGTGTATCTGTTCCCATAGAGCTGTTACTTCTGGCCAAGCCATTGCAGTTTGCTTGCAGAGGATCACCGCATTGGAAATCTTTGCCATTCTAGGACCACAGCCATGAGAGTGA | 120 | 48 |
| chr7 | 144293457 | 144293577 | 772751_42001748_Shank2(210274)_984 | 1 | + | good | GATCCCCAAGGCTGTTGACATTGGGTTCTTTACTTTTAAGGGTTTAAAGTGGCTTTTCTCTTAGGAAAGCCCCTCGTCATCTGGAAAGCCAGATTCTTTTTCTTCAAACATCCCTGGACC | 120 | 44 |
| chr7 | 144293577 | 144293697 | 772751_42001748_Shank2(210274)_985 | 1 | + | good | CGGAAAACTGATAGCCTCACCTTCCATACTGACTAAGAACAGGTGGTTTTTTGGCTCCAGAGTAGCAAAGAGATTCTGCTGAAAAGGTAGATGCAGGCTGTTTCACCATAGCAGCTCCGA | 120 | 47 |
| chr7 | 144293697 | 144293817 | 772751_42001748_Shank2(210274)_986 | 1 | + | good | AGGCTTGGCTCTTTCATTGTGGGGTTCTAATTTGAAGCATCAGTGATTTCAATTTCAAGCTAAAAAAAGATCCCGAATCACTTCAGGGGACCATGGCTGCTGTGCTCAGACTCTGTGGGA | 120 | 45 |
| chr7 | 144293817 | 144293937 | 772751_42001748_Shank2(210274)_987 | 1 | + | good | TTGGCAGATGGCAGTGCAGTGGGTTGGAGAGAGAAAGGACCCATGGTCAAGGGCACAGGTTGTTCAGCCAAGCCAGGTCTGAGTCTGGGCTCTGCCTCCTCTTAGCTGTGCGAGCATGGG | 120 | 58 |
| chr7 | 144293937 | 144294057 | 772751_42001748_Shank2(210274)_988 | 1 | + | good | CACATCACTCACTTCAAGTCTCTCAGTATCTTACCTGTAAATAAAGACACATGTGGAACTTACAGCACAGATCCTGACATCATGGACGATGAGCAAATGTTGGGGGATTAGTGACATCTG | 120 | 42 |
| chr7 | 144294057 | 144294177 | 772751_42001748_Shank2(210274)_989 | 1 | + | good | AAAACCCAATTCATTGGTTTATTTCTGCAGCTGATAAATGGGTACCCAGGAGGCACTTGGCTGGAGGAAAATCAAGAATTAGCTACAGCTTCTTGGGAGCTCACAGTCTAGAGGAGAAAC | 120 | 44 |
| chr7 | 144294177 | 144294297 | 772751_42001748_Shank2(210274)_990 | 1 | + | good | AAAAAGAAAGTATATAGCAGGGAAAAAGATAGTATTGCCACCAGGCTTTAGTGGATAAGAAAGGTCCCCACAACTGGTGTGGAGAAGGGAGGGCTTGAGGGTCAATAACATTCCAGGTGA | 120 | 44 |
| chr7 | 144294297 | 144294417 | 772751_42001748_Shank2(210274)_991 | 1 | + | good | ATGTGGGCAGTAGACAGATGTAATGGTGCTGACAATGGCTGAGGCTGCTGCATTAAGGCCTTGAATGGTTTTTTTTTTTTGAGCAGGTGCAGCTATGTGTGGCTGGTCAGTGTGCGGAG | 120 | 48 |
| chr7 | 144294417 | 144294537 | 772751_42001748_Shank2(210274)_992 | 1 | + | good | AGCGGTCTGTGGTCCCAAGAAACCGCTCTGTGCCGTTTCCTCTGTCCATGGTGCTGAAACGGGCTCTCTATATTTCAGAAATACCCAACTCTCCTCCCTGTATCTTCTATCATATCCCCT | 120 | 50 |
| chr7 | 144294537 | 144294657 | 772751_42001748_Shank2(210274)_993 | 1 | + | good | CCCTGTGTCCTTGGCCTGTCCGGGTATGGAAGCTCATTACTTGAGACAAGGCTCATCTGCTTGGTAGAGAATTCTCTTCCACGAGGGTCTGTTGTGGCAGAGCATCTGTGGTATCTCTC | 120 | 52 |
| chr7 | 144294657 | 144294777 | 772751_42001748_Shank2(210274)_994 | 1 | + | good | CTGTGTTTTGTCCCCGCCTTCGCCTTGTTGCTAAGGTCTCAGCTCACGCGCCAAACTCCCAAACAGGCCTTTCCAGGACCATCTGGCTGCCCCTGACTCTCATGCTCATGCTGCATGATT | 120 | 56 |
| chr7 | 144294777 | 144294897 | 772751_42001748_Shank2(210274)_995 | 1 | + | good | CTGTCACTGTCAGAATCATCTCTGGTGTCATGTGATTGACTTAACATCCCTTTCCCTTCTTATACAGTGTCTCTCCCTATGGGTTATACACACTGTCAACACAGCACCTGTCACATAGCAGAG | 120 | 46 |
| chr7 | 144294897 | 144295017 | 772751_42001748_Shank2(210274)_996 | 1 | + | good | GTCTTAAAGTGTGAATCTGGTGGTGGCAAGCCCTGGGACTCTTGCATTACCATCTTTCAGATAATAAAACTGAGCCTCAGAAAATTGCCTGCCTGGGACCTCGTGGAGGTTAGAGTCTT | 120 | 48 |
| chr7 | 144295017 | 144295137 | 772751_42001748_Shank2(210274)_997 | 1 | + | good | GGCAGCCTGCAAAGCATGTCTCTCTTTTTGTGTGTCTGGTCAGTGCCCTGGCTAAGTCCTGAGTCATGTGGTTGTTGTGTCTTATGCCCATCCATCTTTACCTAATTTATCTCTACCCGG | 120 | 48 |
| chr7 | 144295137 | 144295257 | 772751_42001748_Shank2(210274)_998 | 1 | + | good | CACAGACTGTAAAGATGAGGTCCTAAGGAGGCAGATTGGAGGTTTGGGGTATAGGGTCCTCTGATGGACCCATTGGTTAACCCTGGAAGGAGAACACAGGGACCTTTTTGCCTTCCACCC | 120 | 52 |
| chr7 | 144295257 | 144295377 | 772751_42001748_Shank2(210274)_999 | 1 | + | good | ATCCTATCAAGCATCCTGTCTCTTAGAGTGCCTGGCAGGAATCTTGTTAGCATTGATTTGTTTCTTTCTGTGGTGCCCTTTTGTGCTTTGGGAGTTTTATGATCCAAGTGGTTAGGTTGGT | 120 | 43 |
| chr7 | 144295377 | 144295497 | 772751_42001748_Shank2(210274)_1000 | 1 | + | good | AAATGAACTGAAGGAAGGGTGTGGGCCAAATGCCCTGTAGGACTCTAATTGATAGGGCTCGTCAGCCCTGTAAGTTAGTATGTGTGCACATCAACACTTGCAGAGCTGAGATGAGAAGG | 120 | 48 |
| chr7 | 144295497 | 144295617 | 772751_42001748_Shank2(210274)_1001 | 1 | + | good | AAGAGGAGATGGAGGTGTGGAAGCCACAGTGTAAGGCTGTGGGAGTCAGAAAATGTCCTCTGCAATTTAGTCTCACAGGATCAACCACTAACTAGCAAATACACTACTTTCTCCCAACTA | 120 | 45 |
| chr7 | 144295617 | 144295737 | 772751_42001748_Shank2(210274)_1002 | 1 | + | good | GACTGCCCTGGGTGGGGCAAGATGAGAGAGGGTGGGGGCATAGGAACCTCAATCAGTATGATCAAGGCACTCAGGGCTAGTGTTTTATACAAAAAGGGTTATTACTTTTATGGATGGTGG | 120 | 48 |
| chr7 | 144295737 | 144295857 | 772751_42001748_Shank2(210274)_1003 | 1 | + | good | CTGTTGTTTGGCCATGGAAAAGGCAGAAACTGCCTAGCAAGGCCAGGTACATCTCCCAAGGGAGAGTTTCCCCATGTGCTCTGCCATGGAGTGTGGTCTCTCTCATAAACTGTGAGAATC | 120 | 51 |
| chr7 | 144295857 | 144295977 | 772751_42001748_Shank2(210274)_1004 | 1 | + | good | ACAAGGGCAGAAGCTGACTTGTTTATGACTGCCCTGTGATAAGTCCTAGGCACTAGAAATGGGATCAAAGCCAAAGAGAAAGGTAATTACAGTTAGCCAGAACTAAGCTGACTAACTAGG | 120 | 43 |
| chr7 | 144295977 | 144296097 | 772751_42001748_Shank2(210274)_1005 | 1 | + | good | ACTAAGTCCTGACTAAACAGGACTCACTAGGCCTGGGCATGGCAGTGCTAGGCAAGCCAGTGCCTGCTGGATGCCTGGCTCCCAATACAGTAACTTGGGTTTTGGGTTTTTCTAGACATA | 120 | 51 |
| chr7 | 144296097 | 144296217 | 772751_42001748_Shank2(210274)_1006 | 1 | + | good | CATCCCTTTCCAACACCATTCTGGCTCGTTCTGACATGTCCCTCGCCATTGCTAACTCTTTAACCCTCACCAGAATGCAATCCCCAATTCCCAAAGATACTGGAGCATAGTAGGGTTAAC | 120 | 48 |
| chr7 | 144296217 | 144296337 | 772751_42001748_Shank2(210274)_1007 | 1 | + | good | ATAGGGAATAGCAGCCTCAGGTCACAGAGCTTCAGAGTAGCAGAACTGGGATTTGAGCTGACCTGGTCCCCTTATCAGGCTCTAACCATTGCACACTCCAGTCTCTCTGCCCTTGTGGTT | 120 | 52 |

| | | | | | | | | | |
|---|---|---|---|---|---|---|---|---|---|
| chr7 | 144296337 | 144296457 | 772751_42001748_Shank2(210274)_1008 | 1 | + | good | TAGCAGCTGAGACGTTAGTCCTGGGTGGTACCGTTGCGATTTCCTGTCTCCTGGTGGGGGAGGGCTGTTACCAGGTTCTCACCTGCTACCATCTGTGCTCTGTCCACAGCTCCAGGGAGA | 120 | 57 |
| chr7 | 144296457 | 144296577 | 772751_42001748_Shank2(210274)_1009 | 1 | + | good | AGCTAGCCCTTACCTTAGCTTATAGGGCCCCAGGGTTTCAGCCTCAAAAGGGGTAACTCCTCATTCTACCCTTCACCTACCTTGACTTGATCACTTGACTTTTTGCTGGTCCAATGATAC | 120 | 48 |
| chr7 | 144296577 | 144296697 | 772751_42001748_Shank2(210274)_1010 | 1 | + | good | AGCTCTGCCTCTGGCTTTTCCCATTCCTTCACTTCCGCTGGGACACTCCCCTCTTTTTCTTACCATTGTTCCTCTAAATTTCTCCTAGCCTCCTTACCTAAATTGCATGAACTCCCTC | 120 | 48 |
| chr7 | 144296697 | 144296817 | 772751_42001748_Shank2(210274)_1011 | 1 | + | good | AGTCTCCTGTCTCTGCTAGGGATATGTCTCATCTCTGCATTTTCACGTGTTGCTTTTGCATGGCATGTAGAATTAATGGTGAGTATCTCTCATCAGAACCTGAGCCCATCAGAAGCTGAT | 120 | 44 |
| chr7 | 144296817 | 144296937 | 772751_42001748_Shank2(210274)_1012 | 1 | + | good | GGGTTTAGTCACAGAATGTCAGTCACACAGGTGACACAGCAGACAGCCATGATCAGGACCTCGGGAGACTAGATTCTTTCCTGTCAAAGAGATAGGTGTTTCTTCTCAGACTCACCCCAC | 120 | 49 |
| chr7 | 144296937 | 144297057 | 772751_42001748_Shank2(210274)_1013 | 1 | + | good | TCCCCACCCCCGAGAGTACCAGGAACCTCAGGGATCAACAAACAGCAGTCCAGCCCTGTCTGCCAGTGCCTAGCTCTCTACTCCACACCAATTGCAACCTTGGAGTTTTCAGAGGATGTG | 120 | 55 |
| chr7 | 144297057 | 144297177 | 772751_42001748_Shank2(210274)_1014 | 1 | + | good | TGTACTGATCTCAAAAGTATTAGAGATTTTGTTAACATTCCACCCCCACTCTGACTGGTAGTCGTCTTTCATTAACTCTTCTTCATCCTGGCTTGAGTAAGTCGTAATACACACACACAC | 120 | 40 |
| chr7 | 144297297 | 144297417 | 772751_42001748_Shank2(210274)_1016 | 1 | + | good | TATGGCTTTGTCTATCTTGTAGCAATGCATGTTATCAGACATCCTCATTTGAGTATATCTTTTTATCCTTGTGCTAAAATATACACTCTAGAGCTGGAGAGTGCTCGCTGCCCTTGCAGA | 120 | 40 |
| chr7 | 144297417 | 144297537 | 772751_42001748_Shank2(210274)_1017 | 1 | + | good | GGACCTGGGCTCACTTGTCAGCACCCACGTCAAGTGAGTTACAGCTGCCTCTGACACCAGCTTCAGAGGATCTAATGCTTCAGGCACCTATACTCACCCATGCACAAAGCTCCACACACG | 120 | 54 |
| chr7 | 144297537 | 144297657 | 772751_42001748_Shank2(210274)_1018 | 1 | + | good | TGTACACGTAAATAGTTTTTTAACTGAAAAACACACATGCTGTGAACATTATTTTATGCATTTAAAAGTATACAATGCTTTCAATCTTTAAATTTTATATTTATTTGTGTTTGTACATGG | 120 | 24 |
| chr7 | 144297777 | 144297897 | 772751_42001748_Shank2(210274)_1020 | 1 | + | good | GTCCCTGTACCCAGTGGGCCATCTTCCCAGCCTTCAATGGCTTTCTCATTCAGCATGGTACACTGCCATCACACCTGTTGCTCTGGAACATTGTTATTGACCCAAAGGTGAATCCTGTTG | 120 | 50 |
| chr7 | 144297897 | 144298017 | 772751_42001748_Shank2(210274)_1021 | 1 | + | good | CTGTTGGTCATTACCCCTCATCTTCACGTGAGCCCTCAAAATGGCCACTTCCTGCTTCCTGTTTCTGCAGGCGGGGGGGGGGGGGTTGCCTGCTCTAGACATTTCAGACATAGGACAAGG | 120 | 56 |
| chr7 | 144298017 | 144298137 | 772751_42001748_Shank2(210274)_1022 | 1 | + | good | TACTGTGGAGCCTTCAGGAATGCTTTGTGTTATCAACTTGATAGAATCCAGGATGACCTGGGACTCTGGGCAAGACTGTGGGAGGTTATCTTCATTCTGTTCATTGATATGGGAAGGCCT | 120 | 46 |
| chr7 | 144298137 | 144298257 | 772751_42001748_Shank2(210274)_1023 | 1 | + | good | ATCTCAACCACGGAAGGAACCATTTCCTGGGCAGGGGATCCTGGCCTATATGAGACAGGGAAAACAAACCGAGCACTAGCAAGCATGAATTCCTTGCTTTCCATTTCTTGAGTGTTCGTG | 120 | 48 |
| chr7 | 144298257 | 144298377 | 772751_42001748_Shank2(210274)_1024 | 1 | + | good | GGATGTGGACCATCTCCCTCGGACTCTTAGGAAATGCCCACCACTGCCCTGCACCCGTGGCTACCCAGCCATGATGGCTTTACATAAACCTGAAACTATGAATCAAAATGAACCCTTTCT | 120 | 50 |
| chr7 | 144298377 | 144298497 | 772751_42001748_Shank2(210274)_1025 | 1 | + | good | CCTTGGGCTGATTCTATGGAGGTATTTCATCACAGCAACACAAGAAACTCAGACCCATCCCAGCACCACTGTTCATTGGTTCCATCCATGCCATACATGGATTGGTTCTCTGTGCCTTTC | 120 | 48 |
| chr7 | 144298497 | 144298617 | 772751_42001748_Shank2(210274)_1026 | 1 | + | good | ATAGGTAAATGGACTTCATGGTGTGGCCTGACCACATCAAGGTGCCTATTCATCATTCACGGGAATTTTGGTTTGTCTCCACTTAGGCCTTTTATACATAAAGCTGCCACTGTGGTTTAT | 120 | 42 |
| chr7 | 144298617 | 144298737 | 772751_42001748_Shank2(210274)_1027 | 1 | + | good | AACTTTTCCTCAAAACTCAGATGCTGGAGACCCGGGAGATGACTCGGTGGGTGTGAGCACTTGGTGAGGACCTGAATTTGGATCTCAACACTCATCTAAAAATCCAGGTGTGGCCTGTAA | 120 | 48 |
| chr7 | 144298737 | 144298857 | 772751_42001748_Shank2(210274)_1028 | 1 | + | good | CTTCAGCACTGGATGAGATGGAGAGGGGAGATCTTTGAGGCTCGCTGGTCACCAGCCTAAGTTCAGTGACAGACCCTCTCTCAGGGGAATAAATTAAAAATAAATAAATAAATAAATAAA | 120 | 41 |
| chr7 | 144298977 | 144299097 | 772751_42001748_Shank2(210274)_1030 | 1 | + | good | GGAAAGGATTCCAGGATCATGTTTTCCATTAGAAACACTATTTTGGAATCCATACTGTTATGTTGCTGTTTGGGGTGTGAAATATGGATGATACTTGTCACCTAGCTAGTAGTGATGCTGTTT | 120 | 38 |
| chr7 | 144299097 | 144299217 | 772751_42001748_Shank2(210274)_1031 | 1 | + | good | CAGAAGGCTGTGGAGCCCTTAAAAGGTAGAAACTCAGTGGAAGAAGCAGGACTCCGGAGAAAGGGTCTTAAGGGTTATTTCCTGGTCTCTCTTGCTGTCCCATCTCTGTGTCCTAGTTAA | 120 | 48 |
| chr7 | 144299217 | 144299337 | 772751_42001748_Shank2(210274)_1032 | 1 | + | good | GTGAGATGTAAAGCAGCTTCTGTGTGCTCTGATACCCACAGAGCCACCAATCTGTCCCTGTCATGATGGACAGGAATCATCTCGAGCCATGAGCCCAATCCCTCAGCTCCCTAAAGTTGC | 120 | 52 |
| chr7 | 144299337 | 144299457 | 772751_42001748_Shank2(210274)_1033 | 1 | + | good | TGCCATAGTGATGAGAAAAGTAGTAAGCACATGTGTGCTGATGGTCTTAAGAGACAGGCCTCTGGGAAGTCATTAGGGTTAAATGAGGGTGGTCCATGACAGCCTTTGTGGCTTTATATA | 120 | 44 |
| chr7 | 144299457 | 144299577 | 772751_42001748_Shank2(210274)_1034 | 1 | + | good | AAGAGGGAGAAAGATCTGAGCTGGCACACTTGTCCTGCCTCACCTCACCATGACCCAGCACGGGCCTTCAGCCAGATGTGGAGCAGATGCTGGCTCCATGTGCCTGGAATCCCCAGCTCTC | 120 | 58 |
| chr7 | 144299577 | 144299697 | 772751_42001748_Shank2(210274)_1035 | 1 | + | good | CAGAACCATGAGTCACATAAACATCCATTCACTATGAATTACTTGGGCCCAGGCATTTTGTTACAACATCTAAAAATGTAGCAAAGATGGCTTCTGGGAACATTTGTATTCAAGCTTTTA | 120 | 37 |
| chr7 | 144299697 | 144299817 | 772751_42001748_Shank2(210274)_1036 | 1 | + | good | TCTGAGCGCTTGTTTTCAGTCTGTTCACACCTGGTGAGTGGAGTTGCTGGATCACATGGCCAGTCTGTGCTCATCTGTGGATCAGTGCTCACTGTCCACTTAGCAAATATCTACCACGGA | 120 | 50 |
| chr7 | 144299817 | 144299937 | 772751_42001748_Shank2(210274)_1037 | 1 | + | good | CCCTCACTGTGCAGAAGCAATGGTAACTGGAGAGGAAACAAAGCCCGACCCACAGAGGCCAAGCTCAACACACAAGTCTTGCAACACAACACTTACTGCAGGAAACGTATGGACTGCAAG | 120 | 51 |
| chr7 | 144299937 | 144300057 | 772751_42001748_Shank2(210274)_1038 | 1 | + | good | AACACAGCATTAGGCGAAATGTCTGTGTCTCCCGAGCACCCTCTCTGCAGCAAGCAGCTTTCTTGGAATTTCTCCTTCATACACTCACATGCTCAGTTTAGTTAAAATAGTTCTGTCTTC | 120 | 44 |
| chr7 | 144300297 | 144300417 | 772751_42001748_Shank2(210274)_1041 | 1 | + | good | CAGAGCAAGGGGAGGTGCTGACTTGGCAGGTTGAGCATCCTTGCAAAGTGAAGCTCAGCTCTGGCCCCAAGAACAGGCAAGATGAGGAGCCACCATGGCTTCTTGCAGCCTTCCGCTCTG | 120 | 57 |
| chr7 | 144300417 | 144300537 | 772751_42001748_Shank2(210274)_1042 | 1 | + | good | GGTCTGGCTGAGTACACATCCGTGTCTGAGGGTTGGGTGCTTCTCTCAGCCCACAAGGCTTGCTGGCTGCTGTTGGGTACAATGATCCTCGAAGTCCTGGCTCACTCAGCAAGTAAAGCT | 120 | 55 |
| chr7 | 144300537 | 144300657 | 772751_42001748_Shank2(210274)_1043 | 1 | + | good | CCTCAGTCACGGTCCAGGATCAGCCAACATCCTTCTTGGCCCTAATGATAAAAGGGGCACATGTCTGAGGTCCAAAAGGAGGTCGTGAATCTCCATGAGGCATGTGCAGAACATGGAGGG | 120 | 52 |
| chr7 | 144300657 | 144300777 | 772751_42001748_Shank2(210274)_1044 | 1 | + | good | AAGTGCTCAGTGAGAATGTGCTGAAAGATGCATCCATTCACATGTCTGTCCACCTCCCAGCTCCCTTCCACCCTCCCATTCACCCATCCTTCCACCCATTCACTCATCTGTTCCTCCATC | 120 | 52 |
| chr7 | 144301017 | 144301137 | 772751_42001748_Shank2(210274)_1047 | 1 | + | good | CCATAAATTAATCATCCTATCCACTTAACCACCCATCAGTCTACCCAGAATTTATATCTCCATCCCTCCCATCATTCATTCATCTATCCACTCACCCACCAAGGGTCCATTCAGTCATTC | 120 | 42 |
| chr7 | 144301137 | 144301257 | 772751_42001748_Shank2(210274)_1048 | 1 | + | good | AATTGGGGGTGTAGCTTATTGGTTCCCATCAAACCCTTCTTGTTCACATGATAAAGAAATGGGAGCCTTAGCTGATATGGTATATTTGTGGTGCAGATACTAGATGATAAGGAAAGACAG | 120 | 40 |
| chr7 | 144301257 | 144301377 | 772751_42001748_Shank2(210274)_1049 | 1 | + | good | AAACAAGAAAGATGGGGATTTGGCAGAAATGATAAAGTAACAGCAGTAAAAAGCCCGAAGACAGATCTATTATGTACCAGTGTGTAGAGTCAATCCTCAAAGAGGTGGTGCCATGTGCTC | 120 | 42 |
| chr7 | 144301377 | 144301497 | 772751_42001748_Shank2(210274)_1050 | 1 | + | good | CTGCTCAGATTCCTATCAGTAGCTGGAAATACCACTAGAGATGGGAAACCTGAGAACCTCTCCTTGGCATGTGGACCTTAATTCTCTGGAACTCCTTCTCAGCAGCGACTGGGGCCACA | 120 | 50 |
| chr7 | 144301497 | 144301617 | 772751_42001748_Shank2(210274)_1051 | 1 | + | good | GTGTGCATCTGAGTTACATATCTGGATCTGGATCAGCTTTGATATGTGGGCTGAGGGTTTGGGTTAGTGTCTGGTGACGTAGGTATGGGTGAGACCTGTAGCCCATGTCTCTGGCTCTGT | 120 | 50 |
| chr7 | 144301617 | 144301737 | 772751_42001748_Shank2(210274)_1052 | 1 | + | good | CACACTCAGGGCTCAAGGACTTCCAAAGTAAAGCAACTGTAAAAGAAGAAACTGGGGGTGTAGAGGGACACTGGTGACATCATCACATGGGGAAGGCCCAGAGAGTTCTATGCACTGAGA | 120 | 49 |
| chr7 | 144301737 | 144301857 | 772751_42001748_Shank2(210274)_1053 | 1 | + | good | GATGAGCTGTAGCCACAGGGTGCACCAGTGTTGATGGTGGACAACCTGCTCGTGCACCTGGGCTGGGACAGACTTGGGGGGAAGGGCTGGCAGTGCCTTTGACCAGCTGGCCATGTAGAG | 120 | 61 |
| chr7 | 144301857 | 144301977 | 772751_42001748_Shank2(210274)_1054 | 1 | + | good | CCAGGCACGTCAGACAGCCAGATGGACCTTCCCATCCTGCCCGGAAGAGCAAATCATTCCAGCCCACGGCACATGATGGACCACTGACAGAAATAAGCAGAGAGCATAAAGGCCCCAAG | 120 | 56 |
| chr7 | 144301977 | 144302097 | 772751_42001748_Shank2(210274)_1055 | 1 | + | good | ACATAGAAAACCAGAGCATTTCTAGACCTTTTGGGCACCATTCTGCAAGATGTTGGGTCTACATAGTGCTGAGGTCAATGGCAGTGAATGTTTCTGTCATCGTTCTGTGGGCGTGCTGTA | 120 | 46 |
| chr7 | 144302097 | 144302217 | 772751_42001748_Shank2(210274)_1056 | 1 | + | good | GTATGCTGATGGTGCCAGCATCTCCTCACCCTCAAAGGCTCCAGGAGGGAAGCTAAGCACATGGGGAAACTGAGGGTGAGCATGATGTAGTCAGGAGATGTTATGGTAAGGCAGTGATGG | 120 | 52 |
| chr7 | 144302217 | 144302337 | 772751_42001748_Shank2(210274)_1057 | 1 | + | good | GGCAGAACTCAAGCCCACATCCAACACTGGATCCACAGTTCCCAAACACTGTTATTCCAGGGCAGCAAAAACACTGGGGACTCTTGGGGATCCGGCTACCCCGAGGTGGGGTCAGCCCA | 120 | 57 |
| chr7 | 144302337 | 144302457 | 772751_42001748_Shank2(210274)_1058 | 1 | + | good | GCTCGCTCTGCCTAGCCTCAGCCTTCCAAGCTTGTCTCCACAAACCAGAAGTGGTCATTCGGTTGGTCACAGCACATGAGTTAACTCAACCTCCAGGCCTGGTGGCAGGCAGTGCACCCT | 120 | 57 |
| chr7 | 144302457 | 144302577 | 772751_42001748_Shank2(210274)_1059 | 1 | + | good | CCAAACCATGGTATACGCCTCATTACCTCCTTGCAGTTATAATAGCAGTTCGGCCACTGTGGCACCCACACGGGGTGGGCAGTAATTTGTTTTAATTCCTGTGTTTGCCTGACCTCATCT | 120 | 48 |
| chr7 | 144302577 | 144302697 | 772751_42001748_Shank2(210274)_1060 | 1 | + | good | GCTCCGGGAAGAAATCTTATTTATTGTCAGTTAACGTTCCCTGAGTGGTTTGATGAGCCTTCCAGATAAAATGGCCTGAGCTGTCCTTCTGTTGGGTCCTTTTCCCCAGAGAGCAGGCT | 120 | 48 |
| chr7 | 144302697 | 144302817 | 772751_42001748_Shank2(210274)_1061 | 1 | + | good | GAAGGCCCTGTGGCCATTCCTATGCCAAGCTTGTTCTCTCTTCTTGCCCCAGAATTCTGGCGTCTGGCTGCAGGGGCTTTCATAGACCCTGCCTCCCCCACCCCCATTCATGATGCTCCT | 120 | 57 |
| chr7 | 144302937 | 144303057 | 772751_42001748_Shank2(210274)_1063 | 1 | + | good | ACACAGATGTGGGTCAAAAAAGCAGGAGCTGTTGTCTGTTGAGGACATGAACAAAGAAGGCATTAATGAGACTGGAAAGACAGGTTGATGGGTAAAATGGGTGCCATGCAAACCGAGGAT | 120 | 45 |
| chr7 | 144303057 | 144303177 | 772751_42001748_Shank2(210274)_1064 | 1 | + | good | CTGAGCTCGGATCCCCAGAACTCATGTAAAAATCTGGCTCTGGCTGTGTGTGTCTATTATCCCAGCACTCCCCACAGAAAGACTGAAGGTGGAGACAGGAGGATCCCCATAAGTCTCACCA | 120 | 51 |
| chr7 | 144303177 | 144303297 | 772751_42001748_Shank2(210274)_1065 | 1 | + | good | TAGACTTGTGTTCATAGCAGTGCATGAGAAACCTTGTTTCAAGAAAGGAGAAGACACCAGGGGTTGTCCTTCACATATGACATGGCCTACATGTGCACATACATGCACCATACACACACA | 120 | 44 |
| chr7 | 144303297 | 144303417 | 772751_42001748_Shank2(210274)_1066 | 1 | + | good | CACAGAGGATTTTTGTTTGTTTTTTTAATTTTGTTTTTAAGTAAAAATAGATGGGAATAAAATGACCTGAGCAAACTTGGGCTCTGAGACCAGGTGAGATTTGACAATCAGGGGTGTGT | 120 | 35 |
| chr7 | 144303777 | 144303897 | 772751_42001748_Shank2(210274)_1070 | 1 | + | good | ACTTGGCCCATGAGCTTCCAAGGATTCTCTTGCTTCCACCCCACATCCTGTTGTGTTGCACTAGGATTAGAGACACACACCATGTCCAACTCTCTGTTGGTTCTGAGGATCCAAACTCAG | 120 | 49 |
| chr7 | 144303897 | 144304017 | 772751_42001748_Shank2(210274)_1071 | 1 | + | good | TTCCTCACACATGTTTGACAAGTACTCCCCTTCCCCCCAAGCCATTTTCCAGGCTCACACAAAAGTTTCTTAAGATGGGAGAGAGTGGAGAACAGAGAAGAGATGGTTCACAGGCAGCTGT | 120 | 48 |
| chr7 | 144304017 | 144304137 | 772751_42001748_Shank2(210274)_1072 | 1 | + | good | GGTGAAGCTGCTGTGTTGGGATGGCATTCGTGCTTTCTTCAAAAGTTTGCACAAAATACTGCCCACACCCTCATGTGGCCAAGGATAGCTCTGATGTAGTCCAACACAAAACTGTAAGCT | 120 | 47 |
| chr7 | 144304137 | 144304257 | 772751_42001748_Shank2(210274)_1073 | 1 | + | good | TAAAACACTATGAAGAGACTTGGTGATTTTTTTTGTTTCAGAGTGAACACTCAATGTCAAATGGCTGAAGAGATCATAAATACCAGGAAGAAATCATGAAAGGCAGCTGGGGTGGCAGCT | 120 | 39 |
| chr7 | 144304377 | 144304497 | 772751_42001748_Shank2(210274)_1075 | 1 | + | good | AGAAAGGAAAGTTCTGGAAGAACTGAGACATGCCTTCTGATCATGGGGTTACCTTAAAACAGGGCAGGAAGGGAGTGAGGGAACAGGGACCCAGCTCTCTCCCTCCCCCTCCCCCTCTCC | 120 | 55 |
| chr7 | 144304617 | 144304737 | 772751_42001748_Shank2(210274)_1077 | 1 | + | good | TCCCTCTCTCCTTTCCCCTTCCCCCTCCCCTCCCCTCCCCCTCAAGTTGCCATTTTCTTCTTATATATCTTTTATGTATATATTGTATATCTAAGAGCAAGCAAGCCCAGGAAATCACAG | 120 | 45 |
| chr7 | 144304737 | 144304857 | 772751_42001748_Shank2(210274)_1078 | 1 | + | good | CTCTCAGCCCAGGGGTCTCTACTGCTTATTAGTCATGCAGAGCTTGCTTCTGTCATTCTCCACCTCTGTCTGGTGCTACAGGAAGTCCAGTGTACACTCCCTTGATGTTATTTCTCTAGG | 120 | 49 |
| chr7 | 144304857 | 144304977 | 772751_42001748_Shank2(210274)_1079 | 1 | + | good | CTGAAGGATGGTGATGGCTGGTGAGGTCCATGTCAGTGGGCAGGAAGAGCACCCAGTAAAAACACAGGCCGTGGTACATCTTCAGCTAGCATGGACATGAGAGGCACATCTTGAGCTTGT | 120 | 52 |
| chr7 | 144304977 | 144305097 | 772751_42001748_Shank2(210274)_1080 | 1 | + | good | GTGGACTTGGGAGGCATGTCCTCCTCTAGCAAATACACTGAAAGTACACCTTACTGCACCCCTAATGCCATTAGCCAACCTCAGCCACCTCCTCATAAGTAAGCTGAGCCACCGTTCCAC | 120 | 52 |
| chr7 | 144305097 | 144305217 | 772751_42001748_Shank2(210274)_1081 | 1 | + | good | CCACAAAAGCAGCTTCCTCTGGGATAGAATCAGTGTGGGCCCTGTTCATCCACAGTCACTGCTAAGTGGACCTTTGTCACTGCCTAGCTGCTATATCCTCTGGGACCCAATCTCCAGCC | 120 | 53 |
| chr7 | 144305217 | 144305337 | 772751_42001748_Shank2(210274)_1082 | 1 | + | good | AGCTCTGCAAACATAGATTTAAATTCAGATGGAATGGGAAATCAGCTACTGGGCCCAGGGGCACTGTTAAGGAAAGAGTTTCTCTCTTTCCAGCCCAATGGCCCAGTGCTAAGCAAAGGC | 120 | 48 |
| chr7 | 144305337 | 144305457 | 772751_42001748_Shank2(210274)_1083 | 1 | + | good | TGAGGTCAGGGCACTGGGCAAGAAGTCCTGGGATGCTTGGATCTGGGTTCCTTAGACCCTGCTGACAACTGGAGAAATGTGCAACAGGGGGCCTCCCTGAAGAAAGCTGTGGCATGTGGT | 120 | 56 |
| chr7 | 144305457 | 144305577 | 772751_42001748_Shank2(210274)_1084 | 1 | + | good | GTGTCCCAGCCTCTGTCCTATGAGCCTTGCTTGCCCTAGCACATGGAGCCCTGTGCTTATCTCCCTCCCAGAAGCATCTGAAATGAGCCACCTGAAAGAGTATAACCAGTGAGAACAGAC | 120 | 52 |
| chr7 | 144305577 | 144305697 | 772751_42001748_Shank2(210274)_1085 | 1 | + | good | AGATCAGCCCCTTTACCCAGCAAAACCCTGACCCGAGGCACAGCATCTCATGCAATCCCTGTCCCTGGGTACCACCTCTGGCTACCCTTTGCAGAACTTAATCCATGTCAGACTTTGTAT | 120 | 52 |

| | | | | | | | | | |
|---|---|---|---|---|---|---|---|---|---|
| chr7 | 144305697 | 144305817 | 772751_42001748_Shank2(210274)_1086 | 1 | + | good | CAGTTAGGGAACTGCCAGACTCTATAACAAACATGCTCTTCCCCAGAAGGATGTGGCTATTCATTTGAACAGAGTCAGCTCCCCTCTGTGCCCTGATACAAGCTGTTTTCATTCAACTTC | 120 | 46 |
| chr7 | 144305817 | 144305937 | 772751_42001748_Shank2(210274)_1087 | 1 | + | good | AGCATATGGGGACCCAGGTGGCCAGCGTCTACTACAGCCACTTCTAGTCATTAGGGAGTGAAATCCCACAATATCCCATTCCCATCAGGACCAGTCAACTGCAATTGCACTTAGAGCCCT | 120 | 51 |
| chr7 | 144305937 | 144306057 | 772751_42001748_Shank2(210274)_1088 | 1 | + | good | TCCCCCATTGTCAGGAAGCTGAGGCACAGGAAGGGACCCCACAGTGGGCCTGGAAGGAATACAGACCCCACCCTGTGCTGCTGATGAAACAGGCCTTATATTCAGACCTTGGTTGAAGCC | 120 | 56 |
| chr7 | 144306057 | 144306177 | 772751_42001748_Shank2(210274)_1089 | 1 | + | good | CTTGTGGGTAAAACACATAGCAGAAGACATGTGGGGGGAGTTTTGATGCTGTTTTTAAGACCAGGTCTCATATAGGCCAAGGCAGCCTTGAACTCACTATGTATCCACCAAAGATGGCCT | 120 | 46 |
| chr7 | 144306297 | 144306417 | 772751_42001748_Shank2(210274)_1091 | 1 | + | good | CCCTACCAGCTGAGTCGAATCTGTCTTATTGTTTGGTTTTACTTAGGTGTGGTTTGAGACAGCACTGTGAAAGACATCAGGGAGGGTCCCTAACCTAGGTGTCTGCTCACAGCGCTCACA | 120 | 49 |
| chr7 | 144306417 | 144306537 | 772751_42001748_Shank2(210274)_1092 | 1 | + | good | GCTGTGACATGCCATGGCAGGGCATAGGAACAATCACAAACTGGTGGCTCGAACAAGCAGGCAGCAGTCCTGCAGGGTTCTTCCTGCTTCTTCCAGCTGGCGGTCTTCAGGGTTCTCTTG | 120 | 56 |
| chr7 | 144306537 | 144306657 | 772751_42001748_Shank2(210274)_1093 | 1 | + | good | GCTTGTGGCTACATTATTCTTCTGTCCATCTCTGTCTTCATGGGCTGTTCTGTGTAATGGCCTCATCTCCCTGTGCCCTTCTAAGCAGTGCACTTGGTAGCAGACTTAGAGACTAGGCAG | 120 | 49 |
| chr7 | 144306657 | 144306777 | 772751_42001748_Shank2(210274)_1094 | 1 | + | good | GAAAACCAGGGTGACCTTCCACATCTAGGTCTCCAACTCATTTGCATATTCTAAGATGCCCTGGTCCTAGGCTGCTATTCTTCATTCCCCAACTCAGAATCATGACAGATACTGCTTAGG | 120 | 46 |
| chr7 | 144306777 | 144306897 | 772751_42001748_Shank2(210274)_1095 | 1 | + | good | GTGCCCATTCCTACAGTTTAGGTCTCAAGACAATGGGTTATAAGAAGAAACAGTTGGAAAAGAATTTAGGATGAAACAAGATCCAGACCCTCCATGTTGCAGACCAATGCCTATACCCCG | 120 | 43 |
| chr7 | 144306897 | 144307017 | 772751_42001748_Shank2(210274)_1096 | 1 | + | good | CCCCCCAGACCCATGTGGGCAAAGACCGCTTTCTCTAGTGTGGTAGACAGTCTTGGAGCGTATCTGGACAAGGTAGCATCTCTCCAGATTGGGATTGCTAAATTTATTTTAAAAGGGACT | 120 | 48 |
| chr7 | 144307257 | 144307377 | 772751_42001748_Shank2(210274)_1099 | 1 | + | good | GCTGCATGAGGATAGAGGGAGGGACACCGCTGTGAAGGACAATCAGTCCTCATCAGCCTCTAGATCACTCTAGACTCACAGCCTGTGAGACAAACCTCTTTCTTTAGCCACTCAGAACTG | 120 | 51 |
| chr7 | 144307377 | 144307497 | 772751_42001748_Shank2(210274)_1100 | 1 | + | good | GAGTATTCAGTAACAGCCAATGTGGACCACAGTTTCTTCTCAAGTCTGACCAAAGCCTGAAACTGTCCTCTGGGTCCCCAACACAGAGACATATGTGTGTCCTGAGAAGCCCCAAGAGCA | 120 | 49 |
| chr7 | 144307497 | 144307617 | 772751_42001748_Shank2(210274)_1101 | 1 | + | good | AGCTTCTTGCCTCCTCATGCCTGCTCTGGGCACCTCACCATGCCAGTCATTGTGTGCAGCTATTTAAATGATGTCAGAATAGGAATTTTTCTAGGAAATTTTCCTGCTGGTAATTATTAA | 120 | 42 |
| chr7 | 144307617 | 144307737 | 772751_42001748_Shank2(210274)_1102 | 1 | + | good | CATATATATATATATTACAAGTAACTTGTAAGCAGACTTATTGAAGCTGGGCCACATTGGCCCCTGATCTAATGGCTCATCTGAGCTAAGTCACAGCCCTGCTTGTCTGTGACAATGACT | 120 | 42 |
| chr7 | 144307737 | 144307857 | 772751_42001748_Shank2(210274)_1103 | 1 | + | good | CATCTTTTGCTTTAATACAAAGGAGGCAGGATTGGGAGGAGGAGGAGGAACATCTGGTGGGGCTGGGGTCCCAGCACCCTAGGGTGATCCACCACCGAACAATGTTCTCCCTTGCCTATC | 120 | 54 |
| chr7 | 144307857 | 144307977 | 772751_42001748_Shank2(210274)_1104 | 1 | + | good | ACTGAGGTGTGTAGGTGGCCGTGTCAATGCAGTGCCCGATACCCAAGTGGGCTCTCAGTGACTGGAAATGATTCCCTCCTCGCCAGCTCTGAGAGTTGCATTTCCTGGCAGTGTGTGCTT | 120 | 55 |
| chr7 | 144307977 | 144308097 | 772751_42001748_Shank2(210274)_1105 | 1 | + | good | CCGCCTGCAGCAGTGGTCTTATTCTGCGCCAGGCCCGTTCCCCCCTGGGTCTGAGCACAGAAGCTCGGGCTCGGAGCAGTGCAGCGAAGATTAGTGTATTTATCAGATTAGAGGATCAAC | 120 | 57 |
| chr7 | 144308097 | 144308217 | 772751_42001748_Shank2(210274)_1106 | 1 | + | good | TTCTTTGCAGGAGAAAATTAGCCCCAATTTGCCATGTTTCTCACAACCCACTGTGGGCTGCTGGCTGTCCTGAGGATGCCTAGCCGTGCCCTTCCCCCAGGGCTGGCAGCAGATCCTAGG | 120 | 57 |
| chr7 | 144308217 | 144308337 | 772751_42001748_Shank2(210274)_1107 | 1 | + | good | GCAGGGCCTACCTCACACAGGCCTGGAGGATGAGTCTTGGAGAGGCAGAGTGGATCCTGTGTGAATGACCAGGAGTTGCAGACCCCAAGCAGGGCTCAAGCCAGAACACAGCAGACTGGC | 120 | 59 |
| chr7 | 144308337 | 144308457 | 772751_42001748_Shank2(210274)_1108 | 1 | + | good | TGGCTGGTCAGCTAGAGATCAGCTGAACACCACATCAGTAATTTACTGGGACCACCATGGGCTCTGTAGAGTTTGGGATTCATAATGAAGGGGGTCTAAGGCTCATGGTTTTACAAGGCT | 120 | 48 |
| chr7 | 144308457 | 144308577 | 772751_42001748_Shank2(210274)_1109 | 1 | + | good | GCTGAGAACTGGGTTGCAAGGGACCTAGGGCTGTTGGGATCTGGAATTGTAGGGCTCTCCAGGGTCTGTGGCTTCCCAGGTAGATGGTACAGCACACTTATATGCCCTGTGGCCCTTCTC | 120 | 56 |
| chr7 | 144308577 | 144308697 | 772751_42001748_Shank2(210274)_1110 | 1 | + | good | CAGGACCATACTTACTAGACACAATAGAGTCTTCACTATGGTGGCTGTTCTCTAGCAGGCTGGCTGAATCCTCCAGGATGTGGCTTTTTTGAGCTCTCTCCTGAATGTGCTTACCCTATG | 120 | 48 |
| chr7 | 144308697 | 144308817 | 772751_42001748_Shank2(210274)_1111 | 1 | + | good | TCAGCTCTTGGACTACATACACAGCCCCTGTGTTCTGGGGCTACTGTGCTCTGGCTAGGAATGTGGGAGACCTCACAAGGACACTGGCAACTTATCATGCCACCTGCAAACTCTCATGGT | 120 | 52 |
| chr7 | 144308817 | 144308937 | 772751_42001748_Shank2(210274)_1112 | 1 | + | good | TGCCTGGCTTATGGACTTTCTATACAGACCATCCACTGGATATGTGCATTCAATAGGTCTGGCTCCTTTTAAGAAACTCAACTAAAAGTGAGATAAGACCTAGGCTGCTAGAACTGTGCT | 120 | 42 |
| chr7 | 144309057 | 144309177 | 772751_42001748_Shank2(210274)_1114 | 1 | + | good | AGTTATGTATGTATACACATTGCATGTGTGCCCAGGTGTGGGGAGCCCACAGCGTCTGCCTACTCTGTTTTATTCTGTTCTCCCCACTCAACAACCTGCTATTTGCTTGTATGCTTGGCT | 120 | 48 |
| chr7 | 144309297 | 144309417 | 772751_42001748_Shank2(210274)_1116 | 1 | + | good | ATTGTATCCACATTGGGGAAACAGAGAGACATGAATGCTTTAGTACTCAAATTGTTTCCTCCCTTTTCTCCAGGCTGGGACCCCAGCCCATGAGTGCCTTCCATATTTATGGTTGATCTT | 120 | 44 |
| chr7 | 144309417 | 144309537 | 772751_42001748_Shank2(210274)_1117 | 1 | + | good | CCACCTCAGTTAACCCTGTCTAGAGCCTTACTCACAGACATGCCCAGAGGTCTCTTCCCACGGTGATACTAAATCAAAGTAGCAATCACAACTAACCACAACACTTTCCAAATTAACACG | 120 | 45 |
| chr7 | 144309777 | 144309897 | 772751_42001748_Shank2(210274)_1120 | 1 | + | good | ACCTTCCCCCAGGTTGGTCATTATAGTAACAGTGTTGGCACATCCACCATATACAGAGTGATGCCGGTATGTGGCTATTTGCCCACCTCTCTGGCACACCAGGGTTGCCTGGCAGCATCC | 120 | 53 |
| chr7 | 144309897 | 144310017 | 772751_42001748_Shank2(210274)_1121 | 1 | + | good | AGTGTAGATAGGGTAGTGAGATAATTTCATTATAGAAGACGGTTTTCTACAAAGTATCAAAATGTGGAATTTATCAAGGGAAGAAAGGAGTTTGGGGTATCAGAAATGTGGCCTGACTCA | 120 | 37 |
| chr7 | 144310017 | 144310137 | 772751_42001748_Shank2(210274)_1122 | 1 | + | good | AGGGTTCCCAACCTGCCTCCCAAAACTGTGCTTGAGCCTGGACAGAGCCACAACACTAGACTTGGAAGTCTCTAGGACACTTGTCCCGCGCTCATCCTGAAAAGGATAAGTTGTACCCAG | 120 | 52 |
| chr7 | 144310977 | 144311097 | 772751_42001748_Shank2(210274)_1130 | 1 | + | good | TCCCTCCCTCCCTCTCCTTTCAAACTTAAGGAAAAATATCTTCTGATTTTCCATGTGAGTCTGGCAAACATGTAGGGAATTGCCTCTGGAGGCCTCCCACCTACTCTGAGCCTATGTTGC | 120 | 48 |
| chr7 | 144311097 | 144311217 | 772751_42001748_Shank2(210274)_1131 | 1 | + | good | CCACACTGGCCTGTGAGTATGGCCACTCCAAGGTGCTTGTTAGACACTCAGGCCCATTCCTTCCATCTGGATGCTGTCCTGCCAGGGGACCATTATGACTAACCCAACTATCCATCTCAG | 120 | 53 |
| chr7 | 144311217 | 144311337 | 772751_42001748_Shank2(210274)_1132 | 1 | + | good | AAGTGACCTACATTCTCCTAGCTAAATATCCATCCGTCTGTCCGTCTGCCCCCCACTTTGAGGGCAGTTGTGGTTATGTTCAAACACCTTTATATAAAGACACCTGCTGTGTTGGCCACC | 120 | 48 |
| chr7 | 144311337 | 144311457 | 772751_42001748_Shank2(210274)_1133 | 1 | + | good | TTGGGCTGGAAACTACAAGCTCTGACCTGAGGTGGGTGGGCTGCCACCTGCCCAGTACATTGTTTGGAGCTGAGCCCCCAGGGTTGCTGCCTCCATGGTGAAGTTGAACGCTGTGGAAGG | 120 | 58 |
| chr7 | 144311457 | 144311577 | 772751_42001748_Shank2(210274)_1134 | 1 | + | good | GAGACCCAGAGCATAGCTGATTTGGGAGAGGAAGGACTAGAGCGAAGTGGGTATTTAAGTGATCCAGGATCTGAAGGCACTGGACAGCACAGACCCATCTTCCATTTCTGCAAGTATGCC | 120 | 50 |
| chr7 | 144311577 | 144311697 | 772751_42001748_Shank2(210274)_1135 | 1 | + | good | CTGTGCATGGATGGAAGTTGCCATGATGCCAGGGAAGGCCTGCCCAGCCTGTCCTGCCTCAGGCACCCTGAGGTGTAGTCATTGCTTGACAGTCAACAGTCATTGCTCTGTTTCTGCAGC | 120 | 56 |
| chr7 | 144311697 | 144311817 | 772751_42001748_Shank2(210274)_1136 | 1 | + | good | CAGCTCTGCGCCTACTTAAGTCCCCTCCTATGAACCATCAAGTGTTCCTCCCATCAACACCATAGCCCACAGAGGGCACCAGCTCTGAATGCAGGCCCCTGCCTCCGGCTGTAGCTAGAC | 120 | 57 |
| chr7 | 144311817 | 144311937 | 772751_42001748_Shank2(210274)_1137 | 1 | + | good | CTGCGCTATCCCTCCTGAAAGCAGCCTCAGGATGTCTTTTCAAGAGATGAAGAAATGTACAGGCAGGATCTGTGATCTCACCACACCCTTTAGGTATTTTCTGGCCCCTCGCTAAGAAAA | 120 | 48 |
| chr7 | 144311937 | 144312057 | 772751_42001748_Shank2(210274)_1138 | 1 | + | good | AAGCCAAAGACAATGACAGAGGAGGTCTCACACAGCAGAAGACTCTCAGCCTTCAGAGAGGCCAGAATCTAAAGTCCAATTGTCAGGAGGACCTGGCTTGTATCTCTGTCGCCTTCATTC | 120 | 49 |
| chr7 | 144312057 | 144312177 | 772751_42001748_Shank2(210274)_1139 | 1 | + | good | CTGCCACCATCTTCACAGAATACTCAGCCTGTGTCTCCATGTCACTATTATTCCCTTATAAGAGACTCCAGCCACAACCGGGGAGATGGTTGAGCTCGTAAAGTGCTTGCTGCATAAGAT | 120 | 48 |
| chr7 | 144312177 | 144312297 | 772751_42001748_Shank2(210274)_1140 | 1 | + | good | GAGGATGAGTTCATATCCATGGCAGAGCAGCTCATGCCTGTAACCGTAATGCTAGGTGAGGCAGGAGGCAGAGACAGGAGGATTCCTGGACTCTGACCAGCCAGTCTCGCCAAGTCAGTG | 120 | 55 |
| chr7 | 144312657 | 144312777 | 772751_42001748_Shank2(210274)_1144 | 1 | + | good | CTTGCATCAACTAGCCAGCCAGACACAGACAGAGAAAAGAATTTCAGAAGGGATGCAATGTTGCCCTCCACACACAGGTGCTGGGATTAAGGGTGTGTGCTGCCACACCCAGTTAGTGGT | 120 | 51 |
| chr7 | 144312777 | 144312897 | 772751_42001748_Shank2(210274)_1145 | 1 | + | good | TGGTTTTAATTACCAACTTCATGGGACTGAAAGGTGCTGAGCTGATTAGTAAATCACACTTATGGGTGTGCCTGAGAGGACAGTTCCAAAGATGATTGGCATGTGGGACAGCAACTGAGA | 120 | 44 |
| chr7 | 144312897 | 144313017 | 772751_42001748_Shank2(210274)_1146 | 1 | + | good | GAGAAAGACTTGCCCCTTTCCTGAATGTAGGTGGGCTCCATCCAGTAGAGAGGCCCTTGGGAGTCATGGAGAAACAGGAATCCTCCCTCTCCTCTTCCTCTCATTTCTCCCCCTCTTCTC | 120 | 53 |
| chr7 | 144313017 | 144313137 | 772751_42001748_Shank2(210274)_1147 | 1 | + | good | TCTTCCTCATCCCCATTCTGACCCCCCCATGTCCCCCCACCCCCATCTTTCTGCTTTCCACACACCTGAAGTGAGCTGCTCTATTCCACCATACCACACCCAGCCCAGAAACACAAAGT | 120 | 55 |
| chr7 | 144313137 | 144313257 | 772751_42001748_Shank2(210274)_1148 | 1 | + | good | CAGCCCACCATAGCCTGAAGCCATGAGCCAAAATAACTAGTTCCTCATTTAAATCCTTTCCCTGATGTGTTTGTGTCGTGTCTACACTCTACTAATGTGAAGTCCAAAGGACTCCAGATA | 120 | 43 |
| chr7 | 144313257 | 144313377 | 772751_42001748_Shank2(210274)_1149 | 1 | + | good | CAAGTTTTTCTGCTCACAGATTCCCCCCTTTCCCATCAGTCAGGGTAGCCTGGGAGTAGCCTTCTGTTTCCAGCTTTGCTGTCCTGTCCTTGATCTGCATCCTATCCTGAGGGATGAGTG | 120 | 52 |
| chr7 | 144313377 | 144313497 | 772751_42001748_Shank2(210274)_1150 | 1 | + | good | GTCTTCTGGGGCCTCTCTACTTCTTGTTGCCTGTGCGGTCAGGGCTAGGCCCAGAGGTAGGATTGTTAGCTTGATCTACCCTGGAGTACCACTGTTACTGTTGGAGAGCCCTGGCGAGAA | 120 | 55 |
| chr7 | 144313617 | 144313737 | 772751_42001748_Shank2(210274)_1152 | 1 | + | good | CTTAGTGGAGAGTGAAAACCTTTCAGCCATGCCTCTTGCATCTCACTGCCAAGCATGTGTTGTGTCCAAGTTGGCCTCGGCTGAGGAAATGGTTCAACCAAGTATCCTAAGGTATCTACA | 120 | 48 |
| chr7 | 144313737 | 144313857 | 772751_42001748_Shank2(210274)_1153 | 1 | + | good | GCCACACTCCCCACAACGAGGAAAAAGGAGATATTAAGAGTAGCTTATCAGTAGACAGGGTCTTCTCTGCATCCCCCACTGAATACTTGAGGCCCAGAGATCAATTAGGGTTGCCATAGA | 120 | 48 |
| chr7 | 144313857 | 144313977 | 772751_42001748_Shank2(210274)_1154 | 1 | + | good | CAGTGGGTCTGGATGTAGATGAGGTATGAACTCAGACACTAGAGGATACGTAAACTCTGGTGTCCTAAACCCTACAGGGAAAGCTCTAGTGGACAGTGGCGAGTCCATAATGTTAGGAGC | 120 | 48 |
| chr7 | 144313977 | 144314097 | 772751_42001748_Shank2(210274)_1155 | 1 | + | good | TATTAAAGTAACTGGGACTGGTACTTAGAAGTCTAAAGGAGTGACAAGATAGGCCCTAAAAAGGACCTGATCAGGAGCCTGGAAGCTTCCAGGGACCAGGAGACACAGCCAGTTGAAGGT | 120 | 48 |
| chr7 | 144314097 | 144314217 | 772751_42001748_Shank2(210274)_1156 | 1 | + | good | CCAGGACTATCTTGACATGTTGTCTCCCACCCCCAAACACTTATCCCATTCCCCAGTTCCACCCCTTCCTCAACAGCGGGTGTGTGCAATTCTAAGGGTCTTCAGTGATAGAAGATAGTG | 120 | 50 |
| chr7 | 144314217 | 144314337 | 772751_42001748_Shank2(210274)_1157 | 1 | + | good | GTGACTCTTGTCACCCTGTGTGGCCTCAAAATTGAAGTTGTCTAGACATGTGTGCTTCTGTCTATCTGTCAGAAATGAGATCCCTGAGGCCAGCATGGTCACCAAAGGACTAAGCTAAGG | 120 | 48 |
| chr7 | 144314337 | 144314457 | 772751_42001748_Shank2(210274)_1158 | 1 | + | good | GACAGAGCACTGCACATATGCACATACAGGTTGCTTCCAAGAAGCAGTTCCCTTCAAACCTCTGCCCTTTCTCTATCCCATCAGACCTGGTCACTGACAACCAGGGACAGCCTGGATGTG | 120 | 52 |
| chr7 | 144314457 | 144314577 | 772751_42001748_Shank2(210274)_1159 | 1 | + | good | GTCCAGTGGAACACACATAACCTAGGCCTTCCAGACAGGATGAGGGGCCAAGACTTCTTTAGGTGCTGGGGCTGGGGAGTAGCTTAGAATGCTAGTAAGGACAGGTCTCCGTGATACCTC | 120 | 53 |
| chr7 | 144314577 | 144314697 | 772751_42001748_Shank2(210274)_1160 | 1 | + | good | CGCCCCTAAATTCAGGCTGCAGCAGCCATGTGCTTGCATGCCCAAGCTCCACCCCTTCCGAATGTTGGCTTCTTTTCTCCCAGGCTAGAATCGACTCCAGGCTTAAAATTAATTTCTCTT | 120 | 50 |
| chr7 | 144314697 | 144314817 | 772751_42001748_Shank2(210274)_1161 | 1 | + | good | CCGCTTCATAGCGATGCTGCGGTCTCTCCCAAACAGCCTGTCTCCGATTGTGCAGAATGTGCTTTGCCTTGTTGGGCCAGGCCTTTGAGCCCCAAAACTCAGATTATTTATCTCAGCATC | 120 | 52 |
| chr7 | 144314817 | 144314937 | 772751_42001748_Shank2(210274)_1162 | 1 | + | good | ATCCACTCTGCCAGGTGTGCACCTGCCCTGCAGAGAGTGCAGACAAAATATGGGGACTGCTGCAGCCCCGCCCCACTCAGGGCACTGATGCCAACAGGAAGCGGATCTGGAAACCAGGAT | 120 | 59 |
| chr7 | 144314937 | 144315057 | 772751_42001748_Shank2(210274)_1163 | 1 | + | good | CTGGAAATTGGGGACAGATGGCTGCCTGGTGCTGGGATGCAGGGAGGGGAGCAACTCCACCTTCTCAGGGTAAGGTGACCTTGGACCAGTCACCTGACTTCTAGGCTTTCTGAAGCTGGA | 120 | 57 |
| chr7 | 144315057 | 144315177 | 772751_42001748_Shank2(210274)_1164 | 1 | + | good | TCCGTGGTTAAAGAGCTGCTCTGAGACCTGAGATTCCTGTAGATAGAGCTCTGTGCAATGAGGTGCTGCCTCAGCAGACAGGGCTGGCCCTCAAGAACATTCCCATCTGCTCTGACCTAA | 120 | 52 |
| chr7 | 144315177 | 144315297 | 772751_42001748_Shank2(210274)_1165 | 1 | + | good | CCAGGGAGGGACTTGAGGCCCCAGGAAAGGAAATTGGCTTGACAAGGGTAGGACTGGAAAGACGTCCCTCCACCCACTCATCCCTCTGCCTTGTGTTATTCCTTCTTCCCACCAGGACTG | 120 | 56 |
| chr7 | 144315297 | 144315417 | 772751_42001748_Shank2(210274)_1166 | 1 | + | good | CAGTCACAGGCTGAGTGCCCACCTGGTCCAACATGCCTTGCACCATACAAACCTCCCTGCTGACCGCCACCATCTCTCCAGCTTGCAGTAACAGTTGTTGCTTTCTTTCTTTCTTTCTTT | 120 | 51 |
| chr7 | 144315657 | 144315777 | 772751_42001748_Shank2(210274)_1169 | 1 | + | good | CCTGTCTATGCTGCCTCCTCCCAGGGACACCAGTCTCTTGCAAGCACACACATACATATACACAGATACACATTGCCATACACCACTAGCTATAGAGTTTTCCATGTGTGTGCATCTGTG | 120 | 48 |
| chr7 | 144315777 | 144315897 | 772751_42001748_Shank2(210274)_1170 | 1 | + | good | GAGGCCAGAGGACATCAGGTGATCTTCCCTATCAGTCTTGGCCTCTTCCTTTGAAGCAAGATCTCCTGGATCCAAACTTCACATTTTCTCAACTAGGCTGGAAGACAGCAACACTCAATG | 120 | 48 |

| | | | | | | | | | |
|---|---|---|---|---|---|---|---|---|---|
| chr7 | 144315897 | 144316017 | 772751_42001748_Shank2(210274)_1171 | 1 | + | good | ATCCTGCTTATGTCCCGCCCCTCCTTCCCCTTCCCCTGCCCCGCCCCCACCGAGTTACAGCTGTGTTATGGGAGGCCTGGCTTGTTACATGGGTATTGAGATCTGAACTCTGGTCCTCAT | 120 | 57 |
| chr7 | 144316017 | 144316137 | 772751_42001748_Shank2(210274)_1172 | 1 | + | good | GCTTGAGCAGCAGGTACTCTTAACCACCAAGTCATCTTTCCAGTCCCCAGTCTTCTGTTTTGAGACTTGTCTCCCCTGCTGGTGATGTTGGCATTGTGAAGGGCACCTTGTAAGGTTTGG | 120 | 50 |
| chr7 | 144316137 | 144316257 | 772751_42001748_Shank2(210274)_1173 | 1 | + | good | CATCACCTAATCTGTACTAGTCAGCAAATCTTGGGTTAGGCTGCAGTCATAAACTTATAGTGGCTTACAAACCTAGCTTGCCCACTTGCATGACCCTATCCCTGGCAGTTCACACTGAGG | 120 | 47 |
| chr7 | 144316257 | 144316377 | 772751_42001748_Shank2(210274)_1174 | 1 | + | good | GCGCGAGGGACCCAACCACACAGACTTTCCCCAGCTTCCAGTATCAGAAGGTGAGGTGGAGACATGATAACAGTCATGTTGCTCCTGGCTTTTCCTAAGCGCAGCTTTGCCAGCCTGTCC | 120 | 55 |
| chr7 | 144316377 | 144316497 | 772751_42001748_Shank2(210274)_1175 | 1 | + | good | CACTGGCCACACCCAGCTTCCAGAGTGCGGATGAGTACAATCCCCCACGTAGTCAGAAGTAGGAAGTGAGAAATGTAGGAAGCAGAGCCCTAAAACACAGCCCCGGAAGCAAGTGCCTCT | 120 | 54 |
| chr7 | 144316497 | 144316617 | 772751_42001748_Shank2(210274)_1176 | 1 | + | good | TGCACTGGGAGAAAGGCATCCCCACACATGCCTAACTGTTCCCAAACCAGCCAAAGGCCTCCCCAAGGTCTCTGGGTTCAGGGCCAACAAGTGAATGGAAAAGTGAGTAGCCTCTGTCCA | 120 | 54 |
| chr7 | 144316617 | 144316737 | 772751_42001748_Shank2(210274)_1177 | 1 | + | good | TGTAGCATCTTCCCTCTTCCTACCGCCCACCCACCATGAACTCAGACCTAGCCTCTGTCCTGGGGCTAGGGCCCCTCCTCTGTGGTGAGCACCAGGCCCACCTGGGATCAGAAACAGGGT | 120 | 61 |
| chr7 | 144316737 | 144316857 | 772751_42001748_Shank2(210274)_1178 | 1 | + | good | TCCCCTACAGCAGCAGGCGTCCCTGCAGGCTTCACATCCTTCCACCGAGAAGTGGAGGAGATGCATCTGTTTCATAATGCTTACTAAATAGAGAGTGTGTCAAAGTGCTAGCGAGGAGCC | 120 | 52 |
| chr7 | 144316857 | 144316977 | 772751_42001748_Shank2(210274)_1179 | 1 | + | good | AGGTAAAGGAAAATGCTGGCAATGTGCAGGGCCGAGGCGGAGTGGACAAGAAAACAGGCCTCCCTAAATGGGCTGGCAAACTGTGTGGCTGGGGGTGTTTTCCGTTGCCAGCTGAGTTGT | 120 | 55 |
| chr7 | 144316977 | 144317097 | 772751_42001748_Shank2(210274)_1180 | 1 | + | good | GACTCTCAGCAACCGTCTCCTAGCAACCATACATGCGGGGTACCTAGCTCAGCCGGGATGGGTGGCTGGGACAGAGGCTGCCGGGAGAGAACAAGGCAGCGGAAGTGGAGAGGACTGCCC | 120 | 62 |
| chr7 | 144317097 | 144317217 | 772751_42001748_Shank2(210274)_1181 | 1 | + | good | CAGAGGCCTGCAGCTCCAGGCTATCTAAGGAGGAAGGGGGCCACTGAGACGCCAGGTCCCCACTCACCCACTCATATTGTCCCAAGGGTGTCCTTCATGCCATCCATGGTTCCTGCCTGT | 120 | 59 |
| chr7 | 144317217 | 144317337 | 772751_42001748_Shank2(210274)_1182 | 1 | + | good | GTTGTACACACAAGTGTGTTCCATTTCCCATTTTCATGCCACAGTTTAAAGATCTCTCTGTTCCACAGTTTGCTAAACAACTAGACTGGGGTTGCTCCTGGCATGGGCTTGCTGACCACC | 120 | 47 |
| chr7 | 144317337 | 144317457 | 772751_42001748_Shank2(210274)_1183 | 1 | + | good | AGAAAGTGTGGGGTGTGAGGAGTTAAATATCCCAGTGTGAAAGAGCAGGAGGCAGAGTCTCCTCGCCAATGCTCAAAACCTGGACAGGCTGCCTACAGTCTGTGCCTCAGTTTCCCAATT | 120 | 51 |
| chr7 | 144317457 | 144317577 | 772751_42001748_Shank2(210274)_1184 | 1 | + | good | TGCAAAAAAGAGGCAGGATTGCATGAGGGTAGGGATATGGGGGTAGGGACTAGCCTGACTCAAAAATGGGCCTGGACTTAGTCTTTGGCTCTTGGGAGGAGCTCTGGTCCCAAATATTAA | 120 | 49 |
| chr7 | 144317577 | 144317697 | 772751_42001748_Shank2(210274)_1185 | 1 | + | good | ATTTTACAGTGATGGTCTAGGCCACATCCTCTCATTCTTGACTTGAGTTTCTAAAGGATTAGCTCCGTGGACCTGTGGACTAAAACAGAACCTCACAACCCAAGGGAAACCACTCTGATC | 120 | 45 |
| chr7 | 144317697 | 144317817 | 772751_42001748_Shank2(210274)_1186 | 1 | + | good | CCAGCTCTGAGCATCAAGGGGGCTTTCCTGGCCAGCAATGTCCTATGCATCTTGTTGTGTAGGTCAGTCCTGAAAGGGTCCTCATTTCTAACACATAGAAAGAGTCCTTTGTCTGACTCT | 120 | 48 |
| chr7 | 144317817 | 144317937 | 772751_42001748_Shank2(210274)_1187 | 1 | + | good | CAGTCTGTGGTGGATTTTAACCCTTTCCTATAACCAGCATTGAGGGCACAGGAGCATCCAGTGAGGACCGAGAGTCTTTCTTACAAATCCTTCAGGTCATGGGGACCCCAGAACCTGTAA | 120 | 49 |
| chr7 | 144317937 | 144318057 | 772751_42001748_Shank2(210274)_1188 | 1 | + | good | CTGGTGTCTGATGTAAGTGTGGGTCAGCCTTTCCTTTTGATTAAAGGCTCCTCAAGCCTCTCCCATGAGCCATCAGGAGAGCTGGCAGGTAGAGGTGGCACTCCAGGTGGTGACATTTGC | 120 | 53 |
| chr7 | 144318057 | 144318177 | 772751_42001748_Shank2(210274)_1189 | 1 | + | good | AGAACTAACCTTGAAGCATGGGAAGGGGGTCAGCAATTTCAGAGGCACTTGGGCCAGTCCCCCGCTAGGGCCTGCATTGTGTATGACAGCATCCCCCGGTTGTGACAGTCAAAATGTCTC | 120 | 54 |
| chr7 | 144318177 | 144318297 | 772751_42001748_Shank2(210274)_1190 | 1 | + | good | CTGGTATTGAGAACTTTCTCCTGGATGTAGAAGGGGGCAAGAGCAGCCTTGATGGAGGACCAACCTCAAGAAAAGAGATGTCTCCTGCTGTTTCAGAGACAGGCTGACCAGTCACAGAGT | 120 | 50 |
| chr7 | 144318297 | 144318417 | 772751_42001748_Shank2(210274)_1191 | 1 | + | good | CCTCTATGGGAGAGCATCTCAGACTGACCATCTCAATCTGAACAGGGCATGTGGTCTGCTGTCCCTCCAAGAGAGCCAGGATGTCTCCAGAGAATGCCAGGCCTGGAGAACCACATACCC | 120 | 55 |
| chr7 | 144318417 | 144318537 | 772751_42001748_Shank2(210274)_1192 | 1 | + | good | CCTTCCACAAACAAGAGGATCAGGGCTCCCTATCCTCTGGCTGGGTCAGAGCTGGTATCCAACTCCATCTCTTCCCTTCCCTGGAGAGAGGAATAGAAAGAAGCCCTGCTGTGAGGTCTG | 120 | 54 |
| chr7 | 144318537 | 144318657 | 772751_42001748_Shank2(210274)_1193 | 1 | + | good | CCAGGAGCGTGACTTGAGTCAAAGGATCAGTTTTTCATCTCTAAGTGGGCAGAGATGGTATTTCAGAACCCCACACCCTTCTCTAGACCTGCCCTCCCTAGCCCAGACCTTGAACTGGCC | 120 | 52 |
| chr7 | 144318657 | 144318777 | 772751_42001748_Shank2(210274)_1194 | 1 | + | good | TCCATGGCCAGTCTTAGAGGGCTGTCCCTAAGGCTGTGGGTCACAGGCTGAGCTCAGCCCTCCTGGAGGACCCAGCAGGGTGGTGACCCGGGGTTAGCTCTACTCCTTGATGGCACTTGG | 120 | 62 |
| chr7 | 144318777 | 144318897 | 772751_42001748_Shank2(210274)_1195 | 1 | + | good | TCCCACCACATTTCAGCTCTAGTACTAAGGCAGAACGCCTGGCAAACACCCTCACCCACATCCCCAGTTGTATGAGCATGACTCTGCCTACCTCCAGCTCGTACCTCTAGCTCCACTCAG | 120 | 54 |
| chr7 | 144318897 | 144319017 | 772751_42001748_Shank2(210274)_1196 | 1 | + | good | CCGCAGCGTCTGTTTCCCAAGAATAGATGAGAAGTGCTCACAGCACTGTGGCCACCTCTAATTCTCTCATCGTTCTGGAGCCACTCTCCAGTAGACATTTCTAGAGATGACCAGTATGCT | 120 | 49 |
| chr7 | 144319017 | 144319137 | 772751_42001748_Shank2(210274)_1197 | 1 | + | good | TGATAAAGTACCAAGGGCCACATATAAGCACTGGAAATACAGCCAGTGCGATAAAAGAGTGAATCTGCTTTCACAGTAATGAATGAATCCGTTAGAAATGACGGGCTGCCATGTTGACCT | 120 | 42 |
| chr7 | 144319137 | 144319257 | 772751_42001748_Shank2(210274)_1198 | 1 | + | good | CCACCTGGTGAGGCTGGTTTGTTCCTGGGAAACAAAGGAACTGAGAAAAAATATTCTTTTCCTTCCCTCCATCTTCTCTTCCCATCAGCCCCTCCTCCCTATACTTCCAGGCACCCACCT | 120 | 50 |
| chr7 | 144319257 | 144319377 | 772751_42001748_Shank2(210274)_1199 | 1 | + | good | ATCCTTTCCCTGTGCTGGACCACTGCCGTTGTTCTGGGAGCCGGGCAGGTGATGTGTTTGGAGGTGGGTAGTAAATAGATGAACCAGGTTATTCCCCAAGAGAGCCATCGGTGCTCAGAG | 120 | 54 |
| chr7 | 144319377 | 144319497 | 772751_42001748_Shank2(210274)_1200 | 1 | + | good | CATAACGCAGTGCCGTGTGGGCTGGTGAGGAAGGCAGGGACCTTCTTCCTAAGGTGGCTCAGAGCTCAGCTCAGAACCTGGAACTGGAAGCAGAGAAACCGCTGTCTCTTCCAAGGCTGG | 120 | 57 |
| chr7 | 144319497 | 144319617 | 772751_42001748_Shank2(210274)_1201 | 1 | + | good | GAGCAGCACTGTGAAGCCCAGAGGCCCAGCTGAAGGCCAGCAAGGCTCATAGTGACCCATGTGCACCCAAGTGTCGGGAGCAGAGCAGGAGCCCTCAGAGTTCTGATTGTACTTCTACAA | 120 | 57 |
| chr7 | 144319617 | 144319737 | 772751_42001748_Shank2(210274)_1202 | 1 | + | good | TGAAGAAGGTGTGCAGTGGCCCAGACACATGGGGGCCTATTGCATTGGTTTGTTTGATGTCCATTTAAATGGGGAAAGTATCCCCCAAACTCATGCCACTGGGAACCTCCTTTAGAAGTG | 120 | 48 |
| chr7 | 144319737 | 144319857 | 772751_42001748_Shank2(210274)_1203 | 1 | + | good | GGACCTTTGCAGAGGTTATCAGCTGAGGATGCTATCCCTGAGTGGGGTCCTTCCTCAGTTGTAGTGAAGAGAAGGGACTTGAAGGAATAGAAAAAGGGCCCAAGCCAGAAATGGGGGCTC | 120 | 52 |
| chr7 | 144319857 | 144319977 | 772751_42001748_Shank2(210274)_1204 | 1 | + | good | GGGATGGGTGTTAGCCCCACCCAACAGCTGGAAGAAGGAAGAGCAAATGCTTCCCGCCTCCCAGGTCCCACTGTAGACCTCTCAGGCAGTGTCCGCATCCTCAGCTTGTGATGCTTTGCT | 120 | 57 |
| chr7 | 144319977 | 144320097 | 772751_42001748_Shank2(210274)_1205 | 1 | + | good | CAGCATCCTTGGAATGCTGACAGACAGAAGCCGGAGGGCTCTGACACACGGACAGTCGCACTCAGCACTGACACCATGACACTGTAATGCAAAAGAGTCAAGCCAGTCCTCCCTCCCTCC | 120 | 56 |
| chr7 | 144320217 | 144320337 | 772751_42001748_Shank2(210274)_1207 | 1 | + | good | TGGTCCTGCTGGCCCCTACTGGGATGAAGAGGTCTGGAGTAGGTTGGACACAGACAGGACACAGAGGCTCCTGTACTGGGACATCAATCCCCTGACATGTCCTACCAGGGCCAGATGGTC | 120 | 57 |
| chr7 | 144320337 | 144320457 | 772751_42001748_Shank2(210274)_1208 | 1 | + | good | CTGAAGGTGCCAAGAAGGGACACACACACACACACACACCAAGATAGATTTCCCTATTTCCCTGTGCCTATCCTCCAGCAGTGTCCCCTTCTCTGTCAGAGTCGTCTGGCTTGGCTCAGC | 120 | 53 |
| chr7 | 144320457 | 144320577 | 772751_42001748_Shank2(210274)_1209 | 1 | + | good | CAGAGGGCTCAAGTCCTGTTCAGCTCCTGGGTTCCTACTATAACTTTAGACAGTTCCCTGGGCAAGTGTTGCCCATCTGTCCTGTAAGATGGGGATCCGTATGGAGGGTGTTCTGAGTAC | 120 | 52 |
| chr7 | 144320577 | 144320697 | 772751_42001748_Shank2(210274)_1210 | 1 | + | good | TATCCAGGACCACCATGGGGTATATATACCCCAGAAATGGGATCACCAGAAAGGTAGGGGAGGGCTGGTAGGTGTGGAACTGTGAGTCAGGAGGATGTCTCCCCACCCTGCTAGCTCTAT | 120 | 53 |
| chr7 | 144320697 | 144320817 | 772751_42001748_Shank2(210274)_1211 | 1 | + | good | CAAATCCCCACAACCTGTGGAATGGGTGTTGTCACCACCCTACCATAGTTGGGGAAACTGAGGCCTGACAGGTGTTCCACAGATGTCTTGTTACACCTGAGGTCACCCAGGTGTCCTTTG | 120 | 52 |
| chr7 | 144320817 | 144320937 | 772751_42001748_Shank2(210274)_1212 | 1 | + | good | GAGAAGGTAGAAAGAACATAAGGTGGGGATGGGTGACACCATTCCCACAGGACACCAGTGTCTCTGCTCACTTGGCAGGCTACCTGTGCCACTTAGTTAACTGTTAGCCACCATCCTTGA | 120 | 51 |
| chr7 | 144320937 | 144321057 | 772751_42001748_Shank2(210274)_1213 | 1 | + | good | AGCCCAGAAGTGCCTTAAAGGCAAAACCATATAGTGAGTTACAGCACTTCTGCATGGGAAAGGATAGAGTCTCAATGTCTGTGTGACTGCCTGGATCTCAGGGAGCCTGGCTGGCTGCTG | 120 | 51 |
| chr7 | 144321657 | 144321777 | 772751_42001748_Shank2(210274)_1219 | 1 | + | good | CACCACTTGTGCACTGGAGCACTCTTCTGCTCTCCTGTGCAGCAGGGCTTTAGAGGGATTTGGGGCTCAGATGAGGCCAGAATGGTGGGCCTCACAGCGGGATTAGTGCCCTCTGGGGAG | 120 | 59 |
| chr7 | 144321777 | 144321897 | 772751_42001748_Shank2(210274)_1220 | 1 | + | good | ATGCAGGGCTCTGCCTTCCAGAAGGACATGCTATTGCACTGGCTAGCACAGTCCCCAAATGTGAAATGTTCTCACCTCACCCCCTAGTTATGTCATGATTCCTTCAGAATTCTCCCCTCT | 120 | 49 |
| chr7 | 144321897 | 144322017 | 772751_42001748_Shank2(210274)_1221 | 1 | + | good | GGTCCCCAAGGCCTCAGGTAACGAACAAGGGCATGTCTGAGTCACAGAGAAGTTTGACCTGCTTCTGAGCCCCAGATCAGACTTCTCACAGGTGAGCTCTGAGCTAAGCACCTTCTGAAG | 120 | 53 |
| chr7 | 144322017 | 144322137 | 772751_42001748_Shank2(210274)_1222 | 1 | + | good | GTGAAGCATTCAGGTGGGAGCCAGGTGTCATTACAGGGCAACTAGTATTCTTGTCACTGTGATCTTAGAAAGGTTGTATCCTGACACATGGTTTGGGGGTACTGTCCATGGTGTATGGGG | 120 | 48 |
| chr7 | 144322257 | 144322377 | 772751_42001748_Shank2(210274)_1224 | 1 | + | good | GGCCCACCCCACTTCCTCTTACTGGTTGAGCCACCTGCTCTTACCTCTAAATGTTAAAATACACGTGCCTGTTGGGGAGAGCTCACACCCAATCCTCACTCGGCACGTGTGTAGGAAGAC | 120 | 53 |
| chr7 | 144322377 | 144322497 | 772751_42001748_Shank2(210274)_1225 | 1 | + | good | AGACCAAGAGCAATGCCAGACCAGATGAGGTGCATAAAGTGGGGGATCCTTGCGCAAGAACCAGAAGGATATGGGGAGAAGGACGTGGGGAGGATGTGGGTGAACACATACATCCAAGGA | 120 | 52 |
| chr7 | 144322497 | 144322617 | 772751_42001748_Shank2(210274)_1226 | 1 | + | good | CAGAACCTGCCGTGACTAACCTCATCTACCTTCTACTGGTCACTGGGGTCAGAGGCCCACTGGGTATGATCCCTGCTCCCACCTTGAGGCTCGGTCCCCTTGACACGACCTCCAAGTTAG | 120 | 57 |
| chr7 | 144322617 | 144322737 | 772751_42001748_Shank2(210274)_1227 | 1 | + | good | TAAAATAGTTCTAACATAAGTGTCCCTGTGAGCACTGGGGACAGTCCATGTCTCTGTGGAGTCTTCCACTAGGTTTCCAGAGCCATGTTTTTATTACATAGGAAGCATTCTGAGTCTTGT | 120 | 42 |
| chr7 | 144322737 | 144322857 | 772751_42001748_Shank2(210274)_1228 | 1 | + | good | GATGGGAGATGGCCATGCTTTGACGTGGGCTGATTTAACTCAAACATCGCACACTGCACAGGAGCCTTAGTAATTGTGCCGAAGCATCCTCATAAGCCTCTCACATCTTGGTCTTCTTCT | 120 | 48 |
| chr7 | 144322857 | 144322977 | 772751_42001748_Shank2(210274)_1229 | 1 | + | good | CCAAACACCAAGCTTGTGTTTTGTAGCTTGATTACCGAGTCCCAGCCATGCAAAAGTGTCAAGCAATACAGACCTGAGTTGGGGACTGGCCAGGACCCTACCACTTGATAGAGCCCCTTG | 120 | 51 |
| chr7 | 144322977 | 144323097 | 772751_42001748_Shank2(210274)_1230 | 1 | + | good | GGAATTAGCTAAGGGTGCAGGCCTCCCGCTCCACTGGACACGAACATTGGTTTATCCTACAGACTTTGGTCTTGATTTGAACCTTTTTTCTCTGAGTAGGCTCATTGTTAAGCTTTAAAG | 120 | 44 |
| chr7 | 144323097 | 144323217 | 772751_42001748_Shank2(210274)_1231 | 1 | + | good | TCACCTGGACCTAGACATGCCTTCTGTCAGACTGACCACCTCTTAACACTGTGCTGAAGGAGGCTTTATACCGAAAGTGGTTCACAACTGAACCTATGTGGTACCTGTATGTCCCCTCTT | 120 | 48 |
| chr7 | 144323217 | 144323337 | 772751_42001748_Shank2(210274)_1232 | 1 | + | good | CCTGCTCAATTGTGCCTTTATCAAGTTGCCTACCTGGCAGCGGCCTCCCTGGGCATGTATACTCAGAAACCCAGCACCAATGTCCACCTGAGCTGCAAGAACCTTTGAGCTAAAAAGCTT | 120 | 51 |
| chr7 | 144323337 | 144323457 | 772751_42001748_Shank2(210274)_1233 | 1 | + | good | GCTCAGACCCCATAGCTGGACACTGGAGACCTGGAGGAGTCAGGGCCCCTGGTCCACCCTTATGGCCAGTCCCATCCCCTCTGTGCTGGCTGCTTAGCCTCTCAGGCACTGCACAGCTTT | 120 | 62 |
| chr7 | 144323457 | 144323577 | 772751_42001748_Shank2(210274)_1234 | 1 | + | good | GACTCCCATCCTGCCCCTGCCTTCTCCCCCGCAGGAATCACCCACATGCCCCTCCTGGACTGAAAGCTTCCCCATGGGGTCCTGGGAGTTGATTTTTGTATCACTGACAATTCTTGACTT | 120 | 55 |
| chr7 | 144323577 | 144323697 | 772751_42001748_Shank2(210274)_1235 | 1 | + | good | GTGAAGCCACCCAAGATCTGTGATTCCCATTCTCTGAGTCTCTGGGAGCTCTTGGAGGGAAGAAAAATGCTTATTGGAATCTGGGTGCTGATCAAGGAGCCTGTCAGGGTGTGTCACATC | 120 | 50 |
| chr7 | 144323697 | 144323817 | 772751_42001748_Shank2(210274)_1236 | 1 | + | good | CAACTGGCAGCATTTAACAGTCTTTGACCTACAAAAGTGGCTATTTCAGGTGGCTTGTACCTCAGGAGGAAAACTGAGACTTGAATCGGGAAGGGAGTGCTGATGCAGGATACAGTGAGA | 120 | 47 |
| chr7 | 144323817 | 144323937 | 772751_42001748_Shank2(210274)_1237 | 1 | + | good | GCCAGGCTTGCACAACCAGAAGTGCCTCTAATACCAGGTTTCTATATACAAGCATTTTTTAAATATAGGATCTCATATAGCCCAGGCTATCTGAGACTCAATAGATATCTGAGGATGAC | 120 | 40 |
| chr7 | 144324057 | 144324177 | 772751_42001748_Shank2(210274)_1239 | 1 | + | good | CTCTGCTTCCTGAGCCCTATCCCTAACTCCATCTGTAATACATGCAGGATGGCAGATGCCAGTTAGACACACTTGCTCTTCCTCACCCTGACCCAGTCAAGAAGAAACCAGACCAAGGCA | 120 | 51 |
| chr7 | 144324177 | 144324297 | 772751_42001748_Shank2(210274)_1240 | 1 | + | good | GGGAGAGGGGCAGCAAGCAAGTATCATTGCCCACAGGCCTGATACCATCAAAGCAGCCCCAAATCAGGCTGCGGCTTGCTCCAAGAGCCAATTCCCAGAACCAGCCTGCCAGGCCAGATG | 120 | 58 |
| chr7 | 144324297 | 144324417 | 772751_42001748_Shank2(210274)_1241 | 1 | + | good | GCTGTGGCCCTGGCCATGGAACCTGTGCTCAGGAACCTTGTCACTGATGTGTCTAAGTCCTATAGGAGCCACATCATACACTTGGCATTAGCGGGTTGTCAAGAGAAGCCCAAGGGTTTG | 120 | 52 |
| chr7 | 144324417 | 144324537 | 772751_42001748_Shank2(210274)_1242 | 1 | + | good | TGCTAGGCCACCCAAATGCTCAGAACATTCTGGGGTCCCACAGCACATGCCCTCTTGCACTAAAGCTGAGAGACCTAGAAACCAGCTGCATTCGTCAGCACTTGAGTCCTCCCCAGCAAC | 120 | 54 |
| chr7 | 144324537 | 144324657 | 772751_42001748_Shank2(210274)_1243 | 1 | + | good | TAAAGAAGCCAAAGAAGGGGGCTCAGAATCCCTGTATGGTCCATGTATGGGATTCACATCACAGCCTGAGACTGATTGATGCCTGAGAATGGGAGAGGGATCTTGTGGGGCTGAGCCCCA | 120 | 52 |

| | | | | | | | | | |
|---|---|---|---|---|---|---|---|---|---|
| chr7 | 144324657 | 144324777 | 772751_42001748_Shank2(210274)_1244 | 1 | + | good | ACCTATAGGATCTGACACTGCTCCGAGGTCCATAATGGCAGAACTAGGTTGGGAACACCTGGTGGGCCTGGGAAACGGCTGGCTCACGTGGCAGGTGGGGACAGCCCCACGTTTATTACA | 120 | 57 |
| chr7 | 144324777 | 144324897 | 772751_42001748_Shank2(210274)_1245 | 1 | + | good | GACATCTCACAGTAGACAGTGGCTTCACTTTTCCTGTCAGTTCAAGGCAGGTAGAGAGACATGCATGGGCAGCAACAGAGCAAGAAAAGAAAGGAGTGTGTTGAAAATAGACACAGGCCC | 120 | 48 |
| chr7 | 144324897 | 144325017 | 772751_42001748_Shank2(210274)_1246 | 1 | + | good | GAGGTCCCTGGGAGTGGCCAAGAGGGGCTCTAGCCCAGCCCTTAGCACAGGTTTGTCCTCTCTTGTTTGTCATGAGCCTTTCTGACTGTTCTCTTTGGAATTATGAGTTGCTCTAAATAT | 120 | 49 |
| chr7 | 144325017 | 144325137 | 772751_42001748_Shank2(210274)_1247 | 1 | + | good | ATTTGTAGCTGGAGAAGAAAGACCGTTATAGGAATCACGTGTGTGCACGTGTGTGTCTTTTCGGCTCACTACACTGCTGTGAGACACCTGAGCCTGCCAGCTTTATGGGGACAAAAGACT | 120 | 48 |
| chr7 | 144325137 | 144325257 | 772751_42001748_Shank2(210274)_1248 | 1 | + | good | GCTCAGCTCAGGTCCTGGAGGTTTCAAGGCTACAGTCGGGCTCAGTCAAAGCCTGGATGACTCATAGATTTGATATCTGGGGAGGGAAGGAGACATCAATGTTCAATAAGTCACCTTTCC | 120 | 49 |
| chr7 | 144325257 | 144325377 | 772751_42001748_Shank2(210274)_1249 | 1 | + | good | AACCAGAGCAGAAGGACAGTCAGACTCCAGTTTTTGTAACAATTCTCTCTCCCAAACCTGCCATCTGAAGACACCTCCCCAGAATCTAAAACCTCCCTTGGGTCACCTCCAGAAGGTTCC | 120 | 49 |
| chr7 | 144325377 | 144325497 | 772751_42001748_Shank2(210274)_1250 | 1 | + | good | ATGGGACTTTCTAGGAGAGCTGCCCTGGGGACCAAGCATTTAGCTCTGACCTGTAAGACACAGTGACCTTCAGGACACAGTGAACCTGGCGGCTTAGATATAAAGGAGATGCACCATCAG | 120 | 51 |
| chr7 | 144325497 | 144325617 | 772751_42001748_Shank2(210274)_1251 | 1 | + | good | TCTTGGTAGGTACCTCTGTCCGTAGCTGCCTGTCACGTCATACCTGTGTTCTGTGAGGAAGGCCCTGGTGCATTTGCTTCTTCCTAGGATCTATGCACTAGGATCAGAGAGGCTCCCTGC | 120 | 53 |
| chr7 | 144325617 | 144325737 | 772751_42001748_Shank2(210274)_1252 | 1 | + | good | AGGGCCTCCTGTGCCCCAACAGCCTAGCCCTGGTGGCTGGCTCTCTTACTCTGTGGCAGCCGCTGCACAGAGCTAACCCATCCAGTCCTTCAAACCCTGCAACACTGCCCCTCTCCCCTT | 120 | 62 |
| chr7 | 144325737 | 144325857 | 772751_42001748_Shank2(210274)_1253 | 1 | + | good | TTCCGGGATCTCAACAGCAAGCTTCCCTCTGGCCTGCCTTCCATGTCCATATCCCTGTCCCCACCGTGGGCGCCATCGGCAGCTGGCAGGTGGCAGGGAAGGAGGCGTTTGCTTCTCTTC | 120 | 62 |
| chr7 | 144325857 | 144325977 | 772751_42001748_Shank2(210274)_1254 | 1 | + | good | TAGCAGCCGCCAGTTTCTCGCTCCCAAGTGTTGCCAGGAGAACAGAGAGTTGCCTTTATCCATCCTGCTCAGAGGAAGTGAAGCCGATTTCCCCAGCACTCCCAGTGCTCCCGCATGGCC | 120 | 57 |
| chr7 | 144325977 | 144326097 | 772751_42001748_Shank2(210274)_1255 | 1 | + | good | CATCCTGAGGGTTTGGCTACTGAGAAGTCCTCCCCTTTATGACAGCACCCTAAGATGCCCTCTCAGCTGCTCAGACATTGGAGTTGACGTATATACACTGTTTTTTAATCTTTCCAAAAT | 120 | 44 |
| chr7 | 144326097 | 144326217 | 772751_42001748_Shank2(210274)_1256 | 1 | + | good | CATGGCTCAATTTAGGGTAGAAAGAACCCAAAAGATGAGTGAATTCCAAGGCAGCCTTGAGGTTGGACACCACCTGTCCCTGTGATAAGAAAGTCTCCAAAGGTCAGCTTCCAAGTTCCT | 120 | 47 |
| chr7 | 144326217 | 144326337 | 772751_42001748_Shank2(210274)_1257 | 1 | + | good | GAAGGTTCTGTGGCTAGGAAATCCCAGGAGGGACAGATGGGGAGCAGCCAGAGGCCTCTTTGTGATGAGATGACAGGTTGTGGTGACAGTCCAGCAATGCCAGAGCCAATAGATGACTGT | 120 | 53 |
| chr7 | 144326337 | 144326457 | 772751_42001748_Shank2(210274)_1258 | 1 | + | good | GAACTTGCCTGTGCCAGCTGTTGGGGCTTCCATAATGACCCCCTTTCCATACTGCCACACTCCTGGAACCTCAAAGTGCCTTCCTAGAGTGGCTGCCACCGCCACAAGCACAATTCTTCC | 120 | 55 |
| chr7 | 144326457 | 144326577 | 772751_42001748_Shank2(210274)_1259 | 1 | + | good | CAGTGAGGGCAGGCAAAACTGGAATGCAGGCACCCAGAGCCACACATCTGCCTCTTCAACCCTGGCTATGGGTCACCGGCTCTCTGGGTGCTCGCAATCCCAGGAGCTTCATGCTCTGGC | 120 | 60 |
| chr7 | 144326577 | 144326697 | 772751_42001748_Shank2(210274)_1260 | 1 | + | good | CTTAGGGAAGCAGCAACCTAATGCCTCCCCACTGTGAGTAAACTGCATTTACTACCAGATCTGCTCCAGTACACCAAGCAACCATTCTTCCTAGTAGCTCACTTGTCTTTAATTAATATG | 120 | 43 |
| chr7 | 144326697 | 144326817 | 772751_42001748_Shank2(210274)_1261 | 1 | + | good | AAATAAAGACAGTGATTGCAAGTGTGGCCCACTCAGAAAACTGCCACCCTGAGTCGGGGACAGCAGAGAACAAAGGGCCCTTTGGAGCCTCTACCTGTCTCCGGGTCCTGCTGTTGTCTG | 120 | 54 |
| chr7 | 144326817 | 144326937 | 772751_42001748_Shank2(210274)_1262 | 1 | + | good | GAGGGAGCTGAGGGACAAGGGGGATTGGTCTGGGGTTGTAGGCATCACTGTCTCACCCAACAGACCCCGAGGACTTGTGCTGGGTCTGGAGAGCCAGACTGGTGAAAAAACGGGTATGCC | 120 | 58 |
| chr7 | 144326937 | 144327057 | 772751_42001748_Shank2(210274)_1263 | 1 | + | good | TGGATCCTGGGTAGCACGAGAGGCCTGGCTGCTGCATCTGTGAGATGAGGGATGAGGACTGAAAAGATGGCTAAGATGGTAAAGTGCTTGCTGGATAAGCATGAGGTCCTGAGTTTAGAT | 120 | 50 |
| chr7 | 144327417 | 144327537 | 772751_42001748_Shank2(210274)_1267 | 1 | + | good | AACACAAACATACACCACAAACATTTACCACACATATACACACACACTCATAGAAAGAAGTGACCTATGCAGAGGCCACCCTGGCAAACATAAGTACTGCAAACGCCTACTCTCAGGCCC | 120 | 45 |
| chr7 | 144327537 | 144327657 | 772751_42001748_Shank2(210274)_1268 | 1 | + | good | CGAGGTTTTCACTATAATAGAAACTGTGTGTCCTCTTTTCCAAGTAAAGTTGTTTAAAAGTAAATGTTTGTGACCTCCTAGGGTAGGGTAACCTTCATAGGTGTATGGCCCTCACTCTCC | 120 | 41 |
| chr7 | 144327657 | 144327777 | 772751_42001748_Shank2(210274)_1269 | 1 | + | good | CTAGTAGCCACTCCCCTAACACTCTCTAGCAGTGCACAGCTGGCTCTTTGTATCTTCTCCACGGATTCACGTGTTCTGGGTGTTTCTAATCATGAGTCACACTGTGTAGCCTTTTGTCTG | 120 | 48 |
| chr7 | 144327777 | 144327897 | 772751_42001748_Shank2(210274)_1270 | 1 | + | good | TGTGGTGGTGGGTATCTATGTTTTGTTCCTGTTCATAACTGAGATGTGTTTTGTGTTTTATGTTACATATTTTCATTCTATATCTACTATTGGGGGAAGCTGTCCTCCAAGGTCCCTGTG | 120 | 39 |
| chr7 | 144327897 | 144328017 | 772751_42001748_Shank2(210274)_1271 | 1 | + | good | TTATAGTCTTAGTCAACAGGGCCCAGATACTTTTTAGAAGATGCTATGGACCCTATGGGGAGGTCCTTTGGGAGGCCCTCAACTCATTGGGATGTGTCCTTGAAGGGACCTTGTCCTCTG | 120 | 49 |
| chr7 | 144328017 | 144328137 | 772751_42001748_Shank2(210274)_1272 | 1 | + | good | CTGTCAACCTCTTGCCACAGTACACCGTGCCCACCAGCAATGAAAAGCAAAGGGGCTTTGTAGTCCTCCTAGACTGGAGCCTCCAGGCCTGACAGCTAATAAACCTTTTGAGTTCATTAC | 120 | 50 |
| chr7 | 144328137 | 144328257 | 772751_42001748_Shank2(210274)_1273 | 1 | + | good | AGTAGTGGGAAGCTGGCTAGTGCTGTGTGTGTGTGTGTGTGTACAGGCATATGTGCATAACATATCCAAAGCGGGTGTATGTCATGGGGTTTTGTATGTAATGGGGCGTTCCCACCACCTCT | 120 | 49 |
| chr7 | 144328257 | 144328377 | 772751_42001748_Shank2(210274)_1274 | 1 | + | good | GAATCTGTGAATGGAGCTACAAAACACAAGTGTGTGTTCATGTGTCTTCTACTGCATTTTCAGTCCTCTGCAGCAGTGGAGCCTCTGGGTCACATCCAGAAGCCAGACTTGAAAAGCCAT | 120 | 47 |
| chr7 | 144328377 | 144328497 | 772751_42001748_Shank2(210274)_1275 | 1 | + | good | AGTGTAACCACGTGACTGACCACAGCACCTCTGGGTGCTCCTCCTGAACATCAGCCAGCCACTGTGCCTGCCCACAGACCCAAGCTTGACTGCATGGTACAGATGAAGAGGCTGCAGCTT | 120 | 57 |
| chr7 | 144328497 | 144328617 | 772751_42001748_Shank2(210274)_1276 | 1 | + | good | TGAGAAAGACAGATGGCCCTGGTGAAGGTTCAAACCTGCCCCTCACCAGGAGCTCAGTATCTCAGAGAATTAGTGTCTACTTCTGGGGACCCCTTAGGGCATATAGACCCCCTACATGTC | 120 | 52 |
| chr7 | 144328617 | 144328737 | 772751_42001748_Shank2(210274)_1277 | 1 | + | good | CGACTAAGCCTCAGTCTGTGGCTCTCTTCACTGAGTGGCAAAGGCAGGTGCTCATGTGTAGAACTGAAATGGCCAGCGCGGTGTTCAGCCGGCTTCCTTCTGTTTCTATGGAGAATGTGC | 120 | 53 |
| chr7 | 144328737 | 144328857 | 772751_42001748_Shank2(210274)_1278 | 1 | + | good | CTGCCTCAGACATGGGGGTGCAGTCAGGAGAAAAGAGGCTGTCTTTCCTGTTTGACCTTGGATTGCTCACTCAACATCTCTGGTCAGAGTCAAGTGGAGAAACCAGCAGTTGCTCATCGA | 120 | 51 |
| chr7 | 144328857 | 144328977 | 772751_42001748_Shank2(210274)_1279 | 1 | + | good | CCCTGTGAGGAACACAGGCAGAGCTGGGGACTCTAAAGAGACTTTAGGGCTAGAGAGGAAAAGGCTCCTTGGGAAAAGGTCTTGCTGCATGAGCAGTGGAGCCTAAGTCTGTCCCTAGCA | 120 | 53 |
| chr7 | 144329097 | 144329217 | 772751_42001748_Shank2(210274)_1281 | 1 | + | good | GAGAGAAACCCTATCTCACTGAGCAATGTCAGGGGAGGTGGAAGAAGATGCCTGACATCATCCATGCACACACATGTGCACCTGCACTCTTACACGGGCACACATGGACGCACACATGTT | 120 | 52 |
| chr7 | 144329217 | 144329337 | 772751_42001748_Shank2(210274)_1282 | 1 | + | good | GCTCATAGAGTCATATTTTGTGGCCTCTCCTGGCCCTCCACTGACTTCATAGCAGACCATCAGTCTGTGGTTGCTTTTGAGCACAGTGGAACAGCCAGTGTGGAGTGGTGGAATACCCAG | 120 | 52 |
| chr7 | 144329337 | 144329457 | 772751_42001748_Shank2(210274)_1283 | 1 | + | good | GCTCTGGTGTCTGGAAGCAGTTGTGACCTTTCTTCCCCTTGGCAAGGTCAGAGAGGTGGATGGGGACCCAGCCCCTCCTCTCTGGTGTTCTTGGGTGTCTTCCTCGAAGTCTCTGGCCCA | 120 | 58 |
| chr7 | 144329457 | 144329577 | 772751_42001748_Shank2(210274)_1284 | 1 | + | good | CTGTGCCTAAGGTGGCCTTGTCAGGAGAGGACAATGACCAGGGCCCGGTGTCAGCCAGTGTCTCCTTGGGCCTCTGCACAGGCCTTGTCAGGGGTTTTGCTTCTAGGCATATCTGCCCAG | 120 | 59 |
| chr7 | 144329577 | 144329697 | 772751_42001748_Shank2(210274)_1285 | 1 | + | good | GCCCCAGCATGCATCCTGGGAGAGCAGATGGTTGGTACCCAAAGCTGGGAGAGCCTTCTTGTGGGGACTGGCATCCCTCAGTGCCCAGGGAACAGATCAATGCTTCCAGGAAAGTCTCCT | 120 | 57 |
| chr7 | 144329697 | 144329817 | 772751_42001748_Shank2(210274)_1286 | 1 | + | good | CCTGAGAGGCCCAGACCTTAGAGTACTGTTAGCTTGCATTTTCACTGACCCAGTTCTATGCAGAGGCAGAGGCAAGGTCCAGATGGCAATAGGACTTTGCCTATCTCCAGGGCTGGGGGC | 120 | 54 |
| chr7 | 144329937 | 144330057 | 772751_42001748_Shank2(210274)_1288 | 1 | + | good | ACACACACACCTCCTGCTACTTGACCCAGCTTCAGATCAGAATTGAGTGGAAGAGTGCCCCAGGCAGGGAGCAGACAGGACTCTATCCCAAGCCACACCTGGCATGTTATAGCCCTGGGT | 120 | 55 |
| chr7 | 144330057 | 144330177 | 772751_42001748_Shank2(210274)_1289 | 1 | + | good | GTGGTCCTGGGGTGCCAAGATGGCCAGGCCATGGGATGGTCCTCAAGGACAAGGACTGAAAGGAGTGCAGGGTCATGGTAGAGGATTGTAGTGGCCAGAGTTCTGATTCCTGGCTTACCA | 120 | 56 |
| chr7 | 144330177 | 144330297 | 772751_42001748_Shank2(210274)_1290 | 1 | + | good | GCATGGCTCAAAACCCAAGTCCTGCTAGGACTGGTTGACTTGGCCAAAGGAGTCAGGATAGGCTATGCAAAGGCCCTGAGGACACTTCACAGTCCCAGGGAGAAAGCCAAAAACCCAGGA | 120 | 53 |
| chr7 | 144330297 | 144330417 | 772751_42001748_Shank2(210274)_1291 | 1 | + | good | ACAGTGAGTTTCCAAATATGGGGCAGGCTGGTTTTGTATAAGACTGAGGTAGCTCAGTCTAGTCTTAGCCACCTCCCAGCCAGTGTGACCCTGGACAGTGAGACTCTGAGGCAGCCCTGT | 120 | 52 |
| chr7 | 144330417 | 144330537 | 772751_42001748_Shank2(210274)_1292 | 1 | + | good | CTGTATTCATTTGCAGGTCCCTCTGTCCTCCAATGGGCAGGAAGCAGGAACTCTGTGACCCATGCAATGTCTAGTCAGCTTCTATGCTCCTTGGAAGTTGGTAGCCACCTCTGTCCATGG | 120 | 52 |
| chr7 | 144330537 | 144330657 | 772751_42001748_Shank2(210274)_1293 | 1 | + | good | TTGCCACAGAAGCTGGACCTTGTTAGATGCCACCCAGAGCCACACTCCTCTTTGTGGCTGGATTTGGGAGAGCCACCGCCTGTCCAGAGACATTCTTGCCCATATCCCTTTCTAGAAGCC | 120 | 54 |
| chr7 | 144331017 | 144331137 | 772751_42001748_Shank2(210274)_1297 | 1 | + | good | ATATGGTCTTCTTCTGAAACATGCAGGTCCCAGCAACTCAAAGTGTGAATGGATGAAGGGCAGGCGCTCAGATCCTTAGGAAAGCTCAGGTCTTACCTGAAGCAAGGGGCAGCAGGGGCT | 120 | 52 |
| chr7 | 144331137 | 144331257 | 772751_42001748_Shank2(210274)_1298 | 1 | + | good | TTCTGGGGAGAGGCCCTTTCTGAGCCAGCATTCTTGTTGATCACAGCACACAGCCCTGTCATCTGTGTCTCCTGAGCCCTGCTCTTTTGTGCTGGCGCCCACCACAGCGTCACAGCATAG | 120 | 57 |
| chr7 | 144331257 | 144331377 | 772751_42001748_Shank2(210274)_1299 | 1 | + | good | GGGCAGACATAGAAGCTAGGACAGCTGACTTGTGGGATCAAGTATCCTGAGGGTGGGCACAGGTATTCAGGTTTGAGCTGCCCTTTTGCCTATGAAGCTTGCCCTGTAGAAGCTGCAGTC | 120 | 52 |
| chr7 | 144331497 | 144331617 | 772751_42001748_Shank2(210274)_1301 | 1 | + | good | ATGGAAGACTCAGGCCTTCTCTGAGCCCAGGGTCTTCCCTGTCTATGGATACAGCCACAGTGCCTAGCCCAGGAGCTAAGCATCACTTTGCCCAGGCTAGCCCTACTACCTCAGGCCCTT | 120 | 57 |
| chr7 | 144331617 | 144331737 | 772751_42001748_Shank2(210274)_1302 | 1 | + | good | CCAGAAAGCCCCCAGGAGATGGGCTTGTAATCATTCTTGCAGAGGTGAAGATGTTCTGAGGAGACACCTGTACCCTGCCCATTGCCCCAGGAGTCTCACACTGGGGAAGGAAAGGCTGAA | 120 | 54 |
| chr7 | 144331977 | 144332097 | 772751_42001748_Shank2(210274)_1305 | 1 | + | good | CCCAATGTGGCCGCACTGAATGGCTCTCCTCTCTTCTTTTCGTTGCATATGCAGGTGGCTGTCACCTCTGCATATCTACAGGCTGGGACGGGTGCCTGTGAGAGAATGCCACAGGCATTT | 120 | 54 |
| chr7 | 144332097 | 144332217 | 772751_42001748_Shank2(210274)_1306 | 1 | + | good | ATAGCCTGCCATGGAGGCTGAAAATGGGGAGAATGAGGGACATCTGGTGACCTGCGTCCTCTCTGTGCCCCTTGGGTCTTCGCAGAATTACAGAACTTGGGGGGAGGTGTGCATGGAAAA | 120 | 54 |
| chr7 | 144332217 | 144332337 | 772751_42001748_Shank2(210274)_1307 | 1 | + | good | CCTGCTTCCCCTGCATCTCAGGCTCAGCACCTTACTGCTGTGTGACCTTAGGCTGGTGACACAGCCTTTCTGTTTCTTACCATCATTATGGTGTTTGTTAGTAGTTCACCTGCTAAGAAT | 120 | 48 |
| chr7 | 144332337 | 144332457 | 772751_42001748_Shank2(210274)_1308 | 1 | + | good | GATTTGTCCAACAAGGGTCCTGCTGGACAGAATTGACAATGACAAAAATATTTGCCACAGTCAATGAGCTTGCTGTCTAAGGAGGCAAAACGTGAATGGCAGCTGATGGGTGACAGCACT | 120 | 46 |
| chr7 | 144332457 | 144332577 | 772751_42001748_Shank2(210274)_1309 | 1 | + | good | GGGGACCCTCTCCTGTCATGTGACAGATCTTAGAGGGCTTCAGAGGACTGTAACCTGCATGAACCTAGGCAGGACACATGTCAGGTGTAGTAGGCTGAGCTGAGCACTGTGGCAGGGGAG | 120 | 56 |
| chr7 | 144332577 | 144332697 | 772751_42001748_Shank2(210274)_1310 | 1 | + | good | GGGAGAGTTTGGAAAGGCCCTGCTTTGTCCTGACCTGAGTCTCATTCAGGTGAGTGACCTGTCAGATGGTCACCACAGCTTTGTGCTAGCTACCGGGCAGGCTCAAACAAAATAAGAGGG | 120 | 52 |
| chr7 | 144332697 | 144332817 | 772751_42001748_Shank2(210274)_1311 | 1 | + | good | ATCTAGGACCCACTTCTGTGCTCTGAGTAGCTTGGGACTGGGCTGCTGCAGGGAGCAGACAGGAGGGGCAGCTCTTATGACAATGTCACAGACCTTCCAGGCCCAGCCCTGTATGATTCT | 120 | 56 |
| chr7 | 144332817 | 144332937 | 772751_42001748_Shank2(210274)_1312 | 1 | + | good | TGTGGCCTAGACACCAGACACTCACTGTCCCAGCCTTGGTGTGGCCCACATGTAATTATGTTCCCATCCTAGAGGTACCCGAGAGGGGCAAGCAGAGCCAAGGGCAGATAAGGAACACGG | 120 | 56 |
| chr7 | 144332937 | 144333057 | 772751_42001748_Shank2(210274)_1313 | 1 | + | good | TTAGGACTGCACAGACCTGCTGTGATGGACCTCTGCCTCTCCAGGCCTACCCTGGGATCTGCCACCAGTAGAGGCCCTCCAGTGTACCTCTAGCTACCTTTCGGCTTCCCCTCCTGAGTT | 120 | 58 |
| chr7 | 144333057 | 144333177 | 772751_42001748_Shank2(210274)_1314 | 1 | + | good | TAGCAATGCCCATGCTCACAGACTGGCTCTGATGGCCTGGGATCCTCTGAGAACTGTGTTGTTACTCAGTTCCAGGCCTATAAGATTCTGTCAAAGGATGGCACAGGTCCTAAGTCCCTC | 120 | 51 |
| chr7 | 144333177 | 144333297 | 772751_42001748_Shank2(210274)_1315 | 1 | + | good | ACCCAGAACCCAGAGCCCAGAGGCCTTACAAAACAAAGATAAAGCACAAACGTATACTTTATATCACAAAAAAGTTCTGCCTCTCCTTTTAGAGTAACCAGATGTCAATGGTACACCAGC | 120 | 42 |
| chr7 | 144333297 | 144333417 | 772751_42001748_Shank2(210274)_1316 | 1 | + | good | TTGGAGAGGACTGGTGTCCCACCTCTCCCTCCCTGGGTGTCAGATGCTGTTGCCCACCAGGCCGGGAAGGGGTGAGACAGTCAGGGGTGGAGTGTAGAGGGGACTTCTTCAGATTCTTAG | 120 | 59 |
| chr7 | 144333777 | 144333897 | 772751_42001748_Shank2(210274)_1320 | 1 | + | good | CCATGGTCCACTTCACCAGTTCTGTGCCAGTTCTTCTGGTGGTGTGGTCCACGTGCCCCATGCCTGACACTTAAGCACAAGAACCTGGGTTCAGTCCCCAGAACCATGTGAAACAGCCAG | 120 | 55 |
| chr7 | 144334017 | 144334137 | 772751_42001748_Shank2(210274)_1322 | 1 | + | good | CCAACACCTTCTTCCTCTAAGGCACTGTATATATGCAGTCCACTTATAGACGAGCAGGAGTGTGTGTGTGTGTGTGCAGAAATAATAATTTTGTAAGTGAATAGTGTGTACCTGTATTAT | 120 | 39 |
| chr7 | 144334137 | 144334257 | 772751_42001748_Shank2(210274)_1323 | 1 | + | good | CAGCATTGAGGAGGGAGAGACACCAGCATCCCCCGGGCAGTCTAGCTGAATCTGAATTCCAGACCAATGAGGGGTGTGTCTCCAAATATCAAGTGGAAGATGCCTGAGGACCAGAACCCA | 120 | 53 |

| | | | | | | | | | |
|---|---|---|---|---|---|---|---|---|---|
| chr7 | 144334257 | 144334377 | 772751_42001748_Shank2(210274)_1324 | 1 | + | good | AGCTGGCCTCTTGCCTCCACAGATATGCCCACACAAACACCCTCCTGCTCACACTTGGGTATGTGGTGCTTTTACTACAAAGTGTCATCTTACAAGCTGTGAAGCCCTGTCCATCTTGTT | 120 | 49 |
| chr7 | 144334377 | 144334497 | 772751_42001748_Shank2(210274)_1325 | 1 | + | good | TGTATGGCCCCAGTTGGGTAGCATCACATGACTAAACCCAGGAAGATTGGGCAGAGCCAGGAGAGAAATCAAGGCACTCATGTCCGCCAGGGTCTCTGCTCACCAGCTGACTCACAGGGC | 120 | 56 |
| chr7 | 144334497 | 144334617 | 772751_42001748_Shank2(210274)_1326 | 1 | + | good | ACATAGCAAGGAAACACCTCTTGTTTCCAGCCTGGTCCAGGTGGGAACGTGGGGAGTAACACTGACTTTGTGACCGACCAGATATGCTTGCAGCCCTCTGGTGTGTACCTCTGGACGTCA | 120 | 53 |
| chr7 | 144334617 | 144334737 | 772751_42001748_Shank2(210274)_1327 | 1 | + | good | CCCATGACATCAGAACAGTCACTAGGTCAGCAGGAGCCACGTATTCAGCTTGGCCAGCAAGTGGCTAGTGGCTTTCCATGTGGCACTCCAGAAATCTGGATGGCATTTCCCCAGCACCTC | 120 | 54 |
| chr7 | 144334737 | 144334857 | 772751_42001748_Shank2(210274)_1328 | 1 | + | good | CACACTTGCAGTGCCCTGCGCTGGTCGTTCTTACCATAGGAAAGGTTTCTGTGGGTTTTCATAGGCATCTTTGACATTTCCTGACTATAGGCAGACTAGAGAACATCTTTTCTGTTTCTT | 120 | 44 |
| chr7 | 144334857 | 144334977 | 772751_42001748_Shank2(210274)_1329 | 1 | + | good | TGGGGTTTGGAGCTAAGGATAGACCCAGGACCAGTCACATGCTAAACAAATACTCCATCACCAACCTACACCCCAGGCCTCGCTATTGGAGGAACAAGATCTCTTTAGACCTACACCTTT | 120 | 48 |
| chr7 | 144334977 | 144335097 | 772751_42001748_Shank2(210274)_1330 | 1 | + | good | CAATCCTGCTGCCTCAGACCCCTGAGTATGAGGTTTAATGGTGTGCACCATGGAGCCCAGGTTTTTTCCTTATGTTTCTGGGCCACATTTTATCTCCCATGTTTATTCTTGTTTTCCCTT | 120 | 45 |
| chr7 | 144335097 | 144335217 | 772751_42001748_Shank2(210274)_1331 | 1 | + | good | TGCTGCTCCTTTTACTCTGTGTTCTTTATGTATTCTGCTTATCAGTCCCTAGTTTGTGCCGTCTCTCTTTTATGGCAGCCAATTGTTTCTCGTTTAGTGTGAAAGAATTTGTCAAACAAA | 120 | 38 |
| chr7 | 144335217 | 144335337 | 772751_42001748_Shank2(210274)_1332 | 1 | + | good | CTCCGAGCTTGTGACTGCTGGCTTCCTCTGAAACATTCTTCCTTACCACAGTGTCTCAGGCTACTTTCCACCATATTTTCCAGAAATTTCTTATGCTTGCTTCTTGTTTGTAAGCATTTG | 120 | 42 |
| chr7 | 144335337 | 144335457 | 772751_42001748_Shank2(210274)_1333 | 1 | + | good | TGTTTCTTATGTGTGCTGTGAGGCCAGGAACCAAGTTCATCTTTTCTGCTTGCTGTGGCTAACACAGTCTTCCTGTAACATTTCTGAGTATCACATTGTATCAGTGACCTCTCACTGCTG | 120 | 44 |
| chr7 | 144335457 | 144335577 | 772751_42001748_Shank2(210274)_1334 | 1 | + | good | GGACAAACACCTGACCAAAACAGCTTAAGAGGAGAAAGGTTTATTTGTTTCACAGTTGATGGCTCACAGTTTGAGGACTTACAGTTTGAGGGCACAGTCCATAAAGGGAAGGCAAGAAGG | 120 | 44 |
| chr7 | 144335697 | 144335817 | 772751_42001748_Shank2(210274)_1336 | 1 | + | good | GGATGGTGCCGTCTACATTTGAGAGGGTCTTTCCCCAGTTAAACCTCTCAGGTGACTTTCTCACTGCTAAGATTACAGGTGTGTCTCACAGGTGATTCCAAATCCCCTCTATTGGACCTT | 120 | 48 |
| chr7 | 144335817 | 144335937 | 772751_42001748_Shank2(210274)_1337 | 1 | + | good | GAAGATGGACCCTCATACTCTCCTTTCTTGGCGCTCCCCATCAGCAACTCAGTAGTCACTATGTATCCACTTGCTGTGTGTCCATGTTTCAAAGTCCTATCTTTGAGGTGTCTAGAGGGT | 120 | 48 |
| chr7 | 144335937 | 144336057 | 772751_42001748_Shank2(210274)_1338 | 1 | + | good | CTTCACTACACTTGCACTGACCTGGAATAACTGGCTTTGGGAAAATGAATGTCTTTGTTACAGTGCCTGCTGAACCGGGAATGGGTGAATCTCTCCACTTGATTCAGCCTTCCTTAATGT | 120 | 45 |
| chr7 | 144336057 | 144336177 | 772751_42001748_Shank2(210274)_1339 | 1 | + | good | CTTCCAAGAAGACCTTGTGATCATGCCTAAAGCCTCCACAGCAATTTTTTATATTATTATTCATTTCTCAGGACATTAGCTGACGCTTCATGGATATTTTTTGAACTTGCTCTTGCTATT | 120 | 36 |
| chr7 | 144336177 | 144336297 | 772751_42001748_Shank2(210274)_1340 | 1 | + | good | TGGCTGAAATATCAAGAAGTGGCTGGCTGTGGAGTATCGATTCTGGATTCTACCAGCACCTTGCTGAATCCTCTAATTAATTCTCCTATTTTATGAGCTGCTTGTGCCTTTTATGGCCCA | 120 | 43 |
| chr7 | 144336297 | 144336417 | 772751_42001748_Shank2(210274)_1341 | 1 | + | good | CTTCTTCCTCTCTCTGCCTGCTGGGAAGGGCGATGAAGAGGGCAGGTGTCCCTGGTTTCCCAATCTTAAGATTTGATCTTCAGATTTCTGCCACTTTGGGTCATCTTTTTCTAGTATGCA | 120 | 48 |
| chr7 | 144336537 | 144336657 | 772751_42001748_Shank2(210274)_1343 | 1 | + | good | CTCCACCTCCTGAGTACTGGGATGACAGCAGTGCCCTGCTGTGCCCAGATTACAAATGGATATTAAGTTTTCATCTACTGCTTGTTTTTATCATTTGGGAGGAGCACATGGATGATTAAT | 120 | 43 |
| chr7 | 144336657 | 144336777 | 772751_42001748_Shank2(210274)_1344 | 1 | + | good | AACAGCAATTGATTTTCCACCAGCTCACTCACTGTAGTACCAGCTGCTAATCTTCTATTCTAACAGTGACTGAGACCTCTGCTGCCTCCTTCCCTGCTAGTGCCACAAGGCCTCAGCTGG | 120 | 50 |
| chr7 | 144336777 | 144336897 | 772751_42001748_Shank2(210274)_1345 | 1 | + | good | ATTGCTCATCCTAGGAAGAAGAGCTTGGGGTGGAGCCTAGAGTGGGTGGAGCATGTGGTGGGTGGATTCACAGTGGGTAGAGTCTGTAGTGGTGGTGACCAGGGGTGGAGCCTGTGAAGG | 120 | 57 |
| chr7 | 144336897 | 144337017 | 772751_42001748_Shank2(210274)_1346 | 1 | + | good | TGGAGCCTCTGTAGGCAGCTGGCTCAGAAAGCTGCTATCTGCAAGGGCTCTTTTGTGAGTTTTGTGGCATCTAGTTAGTGCTGATTCTTCAGTTATGATTCCTCCAAGTCAGGGCTGGAA | 120 | 48 |
| chr7 | 144337017 | 144337137 | 772751_42001748_Shank2(210274)_1347 | 1 | + | good | GCAGAGCCTTATTTGGCTCCTTCCAGTTGCCCAAAGACAACTATGCCCATGTCTGCCCCCGCCCCCTTTCCTTCCTGTCCTCACAAGACTCAGCCAATATTCAGTGTGTATGTATGTGTG | 120 | 52 |
| chr7 | 144337137 | 144337257 | 772751_42001748_Shank2(210274)_1348 | 1 | + | good | TGTGTGCAGGGGCTGGGGAGTGTCCCCACCTCTCACTCCCCTATCACAGCTGGAAGTGACCAGTCTGCCCTCTCAAATAGTGTCCCCAGTGGCTCTGGACTCTACAGGCCTTGCCCACCC | 120 | 61 |
| chr7 | 144337257 | 144337377 | 772751_42001748_Shank2(210274)_1349 | 1 | + | good | TAGCAACAGCTGGAGTTTAGGAGTTTAGAAGGCCTCCATCAGCATAGGTCAGGGGCCATCTGTGGACAGCATGGACCCATAGTTGCAGGACCTGCTTTTCCCTAGACCGCCTGGTCTGGG | 120 | 55 |
| chr7 | 144337377 | 144337497 | 772751_42001748_Shank2(210274)_1350 | 1 | + | good | ACCCTTGAAAACATCTCTGAAGAAACATTTACAGTGAGACCAGATATCATGGTTTGGGTATGGAATGTCCCCACAAGTGCGTGTGTTTGAACACTTGACCCCCAGCAGATGGGAGTTGTG | 120 | 46 |
| chr7 | 144337497 | 144337617 | 772751_42001748_Shank2(210274)_1351 | 1 | + | good | GAACATTTAAGAGGCGGGGTCTTGGTGGAGGAAATGGGTCACGGAGGGATAGGCCCTGAGGGGTAGAGTGGAGCTCCATGTCTTGTCGAATTTCTGTTTTTTGATCTGCCCAGATGTAAG | 120 | 51 |
| chr7 | 144337617 | 144337737 | 772751_42001748_Shank2(210274)_1352 | 1 | + | good | CAGGCAATCTCCTGTTCCCATTTGTCATGCCTTCCCAGAATGACATAAGATCCTTCAGACTGTGAGCCAAAACCATCCTCTTTTTTAAAACTCCTGGTCAGGCATTTGACCACCATGAGG | 120 | 46 |
| chr7 | 144337737 | 144337857 | 772751_42001748_Shank2(210274)_1353 | 1 | + | good | AGAAACATAACTAATAGACCAGGTAGGCTCTCTCCCTACAGGGAGCCTGGCTGGACACCATGTCTCATGGGGTCAAATGATGACTCCCAAGACAGCCCTGGCAGCCCTGAGGAGCCACAG | 120 | 55 |
| chr7 | 144337857 | 144337977 | 772751_42001748_Shank2(210274)_1354 | 1 | + | good | TGGTTGTGTAGGGAGAGAGTAAGTCGATGCCAGCCAAGGGCAGATTGCAGAGAGAGCATGAAAACCTGGCCCGAGTAGGCACCCAGTGGTACCTTGTGCCAGAACAGACTGCATTTGTCA | 120 | 53 |
| chr7 | 144337977 | 144338097 | 772751_42001748_Shank2(210274)_1355 | 1 | + | good | CCATGTGGTCTTATGCAGCCACCTTGCTACTGCCAGGGGTGGGGATGGAATCTCACACCCAAATACCAGGAGACTAGGCTCAGCCTTTGCACACGGCCAGCCAGGAGCTCAGGACACCCA | 120 | 58 |
| chr7 | 144338097 | 144338217 | 772751_42001748_Shank2(210274)_1356 | 1 | + | good | AATATGGTGTGAACCAAGTGCTTTCCCTCAGAGCCTCTGCCCAGGCTAGACTTCAGCCATGTACCATCTCCATCCCCACCTATCACATATTTCACTTCCTCTACTGGAAGATGCTGAGTT | 120 | 48 |
| chr7 | 144338337 | 144338457 | 772751_42001748_Shank2(210274)_1358 | 1 | + | good | CACACACACACACACATACATTGTAAAAGAAAAAATTTAAGTTTTTAAAAAATATTTTAAGAAACTTTCCCCTTTTGTGCTGTTCCTGTTCATAGCTGTCATCAGCCATGCACACACA | 120 | 33 |
| chr7 | 144338457 | 144338577 | 772751_42001748_Shank2(210274)_1359 | 1 | + | good | GCTTGGGCACATCTGCAGCAAAGTGAGCTCTGCTGAGGCCCACGAGGCCACGTTGTCGTTTCTCCTGTGATGCGGAGGGACAAAGGTGAGAGGTCTCAGCTGACACCTAGGATATTCTTT | 120 | 55 |
| chr7 | 144338577 | 144338697 | 772751_42001748_Shank2(210274)_1360 | 1 | + | good | GTCCTGCACCTCGTCTACTGGGCCATCTATCCCCCCACTTGTGTCCTTGGAATCCTAGACCAAGACTTAAGAACAAGTCTCTGTTTCTGGGTGCTCAGGATTCCGATGAGCAAGGGGCCC | 120 | 54 |
| chr7 | 144338697 | 144338817 | 772751_42001748_Shank2(210274)_1361 | 1 | + | good | CAGGACCCCTCTTGTTGTCCCTCTTGACCCTCAGAAGTATCCTGGACCACACCCTACTGTGATGGAAAACTCCTTCTGGAACTGACCTAACAGTCAGTGTCCCCCTTCTTTCTTCACATC | 120 | 51 |
| chr7 | 144338817 | 144338937 | 772751_42001748_Shank2(210274)_1362 | 1 | + | good | CAAATACTGGGTCTGTCCTTCAGGAAACAAGGAAGTCAGTTAGTCCAAGGAAATGTTGGGATTCTGCCAGATACCCCTCTGGCCTTCTGCAGACTTGCCGCACATGCTACCATGTGCAAG | 120 | 50 |
| chr7 | 144338937 | 144339057 | 772751_42001748_Shank2(210274)_1363 | 1 | + | good | GTAGGAAGCTGGTCAGGGAGAAGATTCCCCATGGCCACCGTGACAAAGGCTGGCATGGCAGTCTTGCTCGGAGGTCTGTATTCTTTTAATCTGCGTGTGAATGTTTGTGTGTGTGCACAT | 120 | 51 |
| chr7 | 144339177 | 144339297 | 772751_42001748_Shank2(210274)_1365 | 1 | + | good | GGAGCTCTCTCATTCAACTAGGCTGGCAGGCCAGTGAGCCCAGGGATCCTCCAGTCTCACCTTGATGCTGGGATCATAGGTGCATCACCAAGCCTGGCTTTGTACACCAGTGCTGAGAAT | 120 | 55 |
| chr7 | 144339297 | 144339417 | 772751_42001748_Shank2(210274)_1366 | 1 | + | good | TTGGACTCAAATCTTCAGGCACTTTATGAACTGAGCCATCTTCTGAGTCTCTCATTCATTGCTCTGTAGGCCTGAAGTCCAAAGTCACAGGGTTAGATACTCCACAGGCTCCCATGGAAG | 120 | 47 |
| chr7 | 144339417 | 144339537 | 772751_42001748_Shank2(210274)_1367 | 1 | + | good | GTCCCACCTCACCCTTCCTGCTGCTGGCAAATCCTAGCATCCTCAGCACTCCTCGGCTTTTGGCCCCATCACTGGCATCTCTCTGGGTGTGTGTCTCAGATAATCCCTGGTATGGAGTGG | 120 | 57 |
| chr7 | 144339537 | 144339657 | 772751_42001748_Shank2(210274)_1368 | 1 | + | good | TAAGGACCCACATTCCAGCATGAACTCATCTTTGTAGAACATCTTTGCCATTTCCAATGCAGGGCCCACTGTGAGAGATTGCAGTAGACACAAAGTTAATGTTGACACCATTCACTGAAC | 120 | 43 |
| chr7 | 144339657 | 144339777 | 772751_42001748_Shank2(210274)_1369 | 1 | + | good | AGGGAAACCTCACCAGTCCTGGACTGAGCCTCTGCAGGGAACCTGTGAGCTGTGTGGTTTTTATTGCAGGGTACGTGGCCCCAGACTGAGGACACTTGGGTCTCAGAATCATCATCTCAG | 120 | 54 |
| chr7 | 144339777 | 144339897 | 772751_42001748_Shank2(210274)_1370 | 1 | + | good | AGATATAAGCATTGTCTTCAGAACAAGCCCCCATTATGTTATAAATAAATTTATTCAATTATAAGTTGCCACAGATGGCACCGGTATTTTTAACAGATGCAGGCAGACAACTGGCTGTAG | 120 | 37 |
| chr7 | 144339897 | 144340017 | 772751_42001748_Shank2(210274)_1371 | 1 | + | good | TATGCCTACAGGAGAACTGGCACAAGGCTTCCTGGGGTGGTCTTCAGGGCCACCCTCACTCCTGAGGATGGGCAGGAGAGACCATGGAGCTTCCATATATTAGCAAGGAGGTGGCATCTG | 120 | 56 |
| chr7 | 144340017 | 144340137 | 772751_42001748_Shank2(210274)_1372 | 1 | + | good | TGTGAGCTGATGATCCAATGAGTGAGAGGAAGCAGCAACAGGTCACAGATACCAGTAATGGTGGAGGTTGGGGGTGGCAGGTCCTCCTTCTAGGTCCTCTGTCTGCTCTGGGAGACTCCA | 120 | 54 |
| chr7 | 144340137 | 144340257 | 772751_42001748_Shank2(210274)_1373 | 1 | + | good | GACCTCCTCTGTCCTGACACTTGTTCAAGTGCATGCTAAAACACCTCCAAGATAAAAGTAAGCCCTTTAGGAGTGAGGAAGGGAACCCAGCCTGGCCTCAAGCTGAGTATCCTCCTGCCT | 120 | 51 |
| chr7 | 144340257 | 144340377 | 772751_42001748_Shank2(210274)_1374 | 1 | + | good | CAGCTTTGCAGACACTTGGGATTGCAGGTGTGCACTGCTAGGCGTAGTAAAACTTCACATTTAAAACCTGGAGTTTAGCTACCCTTGCACTATGTTTGGCCACCAGCTTCATCTGTTTTC | 120 | 46 |
| chr7 | 144340377 | 144340497 | 772751_42001748_Shank2(210274)_1375 | 1 | + | good | AGAACATTCTCTCACATACATACACAGTGACTCCCTGGATCCACCCATCTGAGCCTTCCCCCAACCTCTCCTCCTGCCCGGCATCACTAAGTAGCGGTCTCTGCGGACTTTCCCGACTCA | 120 | 55 |
| chr7 | 144340497 | 144340617 | 772751_42001748_Shank2(210274)_1376 | 1 | + | good | GACAACTCAGAGGAACAGACTCATCACAACATGTGCTGTTTTACAAGCGGCTTCTGCCATTACTCATGCCATTACTCATGACCACAGAGTGCCATCCATATTGCAATACAGACAAGGTCC | 120 | 46 |
| chr7 | 144340617 | 144340737 | 772751_42001748_Shank2(210274)_1377 | 1 | + | good | CTGTTCCTTTCTGTGGTTGAGTAATGGTTTTCTGTATGGACACACCATGTCTTGCTTATCTGTTGATCTGCAATGGACGTTATTAGAAGGAAAGCCAGAGTGATAGCATTTTCAAGACTT | 120 | 40 |
| chr7 | 144340737 | 144340857 | 772751_42001748_Shank2(210274)_1378 | 1 | + | good | TTTTTTTTATATAACTTTTGAGCAAATAGTAATTCCATGAGTCAGGCAGCCAGTGCGGAGTGGCTCTATAATAAAGCTAAGAATGAGGGAGAGACAGCCTTCATAGACTAAACGTGGACA | 120 | 39 |
| chr7 | 144340857 | 144340977 | 772751_42001748_Shank2(210274)_1379 | 1 | + | good | CGAATGAAAAGGACCCCTGCCTTGTGCTTTCACTAGTCACCATGTGGTCAGCCCCATCTCAGAGCTGCTGGTTTAATTGTGAGCTATGGTTGTTCCATACTGGCTGGTGTGAGGTCTGCT | 120 | 51 |
| chr7 | 144340977 | 144341097 | 772751_42001748_Shank2(210274)_1380 | 1 | + | good | CTTTTTCAGCCTAGACAACTGCCCAAGCCGCCTCTCACGCAAGATCAAGATCGGGGACAAAGCCCTGCCCTCCCTGCTTGGGGTTCTTTGAGTCTGTTCTGCAGTTTAAGTCTGTCTCAC | 120 | 53 |
| chr7 | 144341097 | 144341217 | 772751_42001748_Shank2(210274)_1381 | 1 | + | good | ATCAAGTTTGCTGGTTGAGAGACCTGCTTTCTGTGGGTACCGGGGGCTCATGGACCTGGGCTTACTTTGTTGCAGGACCACAACACGCCCCTGAGGATGCAGGAGCCAGTCGCTTACCAT | 120 | 56 |
| chr7 | 144341217 | 144341337 | 772751_42001748_Shank2(210274)_1382 | 1 | + | good | CACAGGAATCTGGCCATTCCTGCCTGGCTTTGACCCTTGGACAGCAAGAGACTAGGTGTGACCCCCGGGTATCAGAGCTGTGGACACAGCTAAGACCACATAGGAAGGCTGGGGTGTGGC | 120 | 57 |
| chr7 | 144341337 | 144341457 | 772751_42001748_Shank2(210274)_1383 | 1 | + | good | TGCAGCCTTGGGCTGACAACTGTAAGAGAGTCTCCTCTTATGCTTCAACTTTGCCCAGTGGCTGTGGCTGGTGGAAGGCAGCCATAGGCAGCTTGCTTTTGTCAGGGAAGTGGAGAGCTG | 120 | 54 |
| chr7 | 144341457 | 144341577 | 772751_42001748_Shank2(210274)_1384 | 1 | + | good | GACCACACACACCTTCCTGTGAGCCCTTCCCCCATACCTGCAGCTGGCAACCTCTATGAATGTGTAACTCATTGTTGTATCATTGCAACCTGGAACAGAGACTGAAGTGTGCCCACTGGA | 120 | 51 |
| chr7 | 144341937 | 144342057 | 772751_42001748_Shank2(210274)_1388 | 1 | + | good | TGCACAGACAGTTGGTTAGATGGACACATATGTATGGATGCATGAATGATTGGGTACATGAGTGAATTAATGCATGCATGAATGGATCAATGAATATATAGGCATACACACATAGATGGG | 120 | 38 |
| chr7 | 144342057 | 144342177 | 772751_42001748_Shank2(210274)_1389 | 1 | + | good | TGGATGAATACGTCGTACATACATGTGTGCATGCATGCATTAAGGAACTAATGTGACAACATCTATCCAGTCTTCCAGCTGGGGCCTGCCCTCTTACAGCTCCTCTGGCTACCGGTTTGG | 120 | 49 |
| chr7 | 144342177 | 144342297 | 772751_42001748_Shank2(210274)_1390 | 1 | + | good | GAATCTACCCTCTTCTTGTAACAAGGGCCTCTGGCTGTGAATTCTGGGGAGCTTGACCAGGACCACAAGGGAGACAGTTGTCCTTGAGGGATACAAACAGCACCATTACACTTTGGTTTC | 120 | 49 |
| chr7 | 144342297 | 144342417 | 772751_42001748_Shank2(210274)_1391 | 1 | + | good | TCATCCTTCTGTTCATGCTGTGAACTGGGAAAGTGCACCTGGCTTTGGATGCCACCTCAGGCTGCCAGCCTGCCCAACATCTATCACTAAGAAGGAACCTTATGTTGTCAGAGCCATTCA | 120 | 50 |
| chr7 | 144342417 | 144342537 | 772751_42001748_Shank2(210274)_1392 | 1 | + | good | GCCTATGCAGGACTCAAGTGAGTCTCCTTGGGCTCCGCCCACCCTCTCAGAGTCTCATTTCTAGAATCAGGAGCCTTATTTAGCCTCCTGCATTCAAGGCTGCTGCATGGCCAAGATCAC | 120 | 53 |
| chr7 | 144342537 | 144342657 | 772751_42001748_Shank2(210274)_1393 | 1 | + | good | CCCCGAGCTCCACAACACCTGGCTCACTGATGTTCCTCACTGGCACAATCAGGGGCAGTAACCCAATCCTGCCACACTTTCCTGTCCTCCCCCTCACTGTTTGTCATCAGGGTCCCTTCC | 120 | 57 |
| chr7 | 144342657 | 144342777 | 772751_42001748_Shank2(210274)_1394 | 1 | + | good | CTGCCCCTCACCCTCTCTGTCTCCCTTCTTGTTCTCTCTGACCTTAGAAGGCTGTTCCGATAACATTGTGAGCCCCTGCAGGCAGGGCAGCCACCACTGCTTGAGTGGGAAACTGAGGCA | 120 | 57 |
| chr7 | 144342777 | 144342897 | 772751_42001748_Shank2(210274)_1395 | 1 | + | good | CGGGTTGGTGGACTAGCAGACCCTTCTCAGGACATCTTTGAGCTAAGAAGTCTAAAAAAGGCCTTTCAAGGCAGAAGAACAATACAAAACACAATAGAGCTAGTGGGTAAATACTTCATT | 120 | 42 |

| | | | | | | | | | |
|---|---|---|---|---|---|---|---|---|---|
| chr7 | 144342897 | 144343017 | 772751_42001748_Shank2(210274)_1396 | 1 | + | good | ATGCTGTGTAAGCCTAGATATGCTGAGCTGCACATTTCTCTACAAGCCTGCTAAGTTAGATTCTCAGATAGGAAGAGTTATCAGAGATCAGAGTCACTGAGAGGCCACACCTTGAACAGA | 120 | 44 |
| chr7 | 144343017 | 144343137 | 772751_42001748_Shank2(210274)_1397 | 1 | + | good | AGCCATGCCCTCCCACATTCACAGAAAGAAATGGAACCAATGCCATCAAAATGCAAATAGGCCAAAATGACCTATATGAACATGAGTGTATTTTTCTTAAAAATGACAGGGGAGGGGGTA | 120 | 41 |
| chr7 | 144343137 | 144343257 | 772751_42001748_Shank2(210274)_1398 | 1 | + | good | CTTTGAGAGTCAGCAGTGGTGAGCTCAGGACTGAGGTATGGGAGCAAAAAAGATAGGCAGTTGTGGGAGATCAATGCAAACATCACAGCACAGCGTGTCATGCCGAAGGAAGGAAGTTCG | 120 | 50 |
| chr7 | 144343497 | 144343617 | 772751_42001748_Shank2(210274)_1401 | 1 | + | good | TCAAGGGTCTGGTAGAGGATCGAGCTATAGATTGTCTGAGACAAACAGGCTCAAGAGCAGAGTCTGGAGGATCTGGTGGGGTCTGTCTGCATTCTCTGTCATAAAGATAGCACAGAGGTC | 120 | 49 |
| chr7 | 144343737 | 144343857 | 772751_42001748_Shank2(210274)_1403 | 1 | + | good | AGCCTAAAGATCTATATGTCTTCTATACCCAGTCTTGTCCAGGAGTTATACATCTTGGGGTGAGGCCCCCAGAAAGCCTCTAGCTTCCTGCACAGGGTCATTCTGCTTTCTGTGAGCTGA | 120 | 48 |
| chr7 | 144343857 | 144343977 | 772751_42001748_Shank2(210274)_1404 | 1 | + | good | CAGACTCAAGTCACAATCAATAAATGGTCATCTGGGTTATAAGCAGAGGATGAATGTACCAAAGAGGATGTGATGGGCAGGGGATGCTAAGTGGATGCAGAGTCCACACAGTGGTCAATC | 120 | 46 |
| chr7 | 144343977 | 144344097 | 772751_42001748_Shank2(210274)_1405 | 1 | + | good | AGACCTTAGGGCTCTGACAAAAGAGAGAAGGAAGCCACAGGTCTAGAGAAGACTTTCAAGTCTAGAGTGGCTTTTCTGGAAAGTGAGGAATGCAAACATGGCTTATGGCTGGGATGAGCA | 120 | 47 |
| chr7 | 144344097 | 144344217 | 772751_42001748_Shank2(210274)_1406 | 1 | + | good | TGTGAGAAGCAGATGGGAAGATAAGAGGCCTAGCCAGGTAACTCAGACTCCAGGCACTGTGAGGTGTTGGCCATCCTCTGAGTGGCACACAGAATTGCATGGTGGGTGAGACCTTTGATT | 120 | 52 |
| chr7 | 144344217 | 144344337 | 772751_42001748_Shank2(210274)_1407 | 1 | + | good | TCCAAGATGGTTTTGACTGCTGAGTTGGGAGCTGTCCACAGAGGATGGGTACAGCCTTAGAGTCTACAAGAGGCTCCTAGCTGTGGGCCACAGCATGTACAGGATTCAGGCCTATTGAAA | 120 | 51 |
| chr7 | 144344337 | 144344457 | 772751_42001748_Shank2(210274)_1408 | 1 | + | good | AAGTGGGGAGCTAGGCTGGTCAGCAGATGGCCTGAGGTCAAAATGATCAAGGAGGGTTCCAGAGTACTCTAGACTTTTCTCCTAAGCAGCTCAAGACAGAGGGCTCAACCCCTTTAAACT | 120 | 50 |
| chr7 | 144344457 | 144344577 | 772751_42001748_Shank2(210274)_1409 | 1 | + | good | TGGGCTTCACGGACTGGGAGCATCTATTCTTTCCACTATCTTGGAAACTGATGCTCCTGGCACTGCCCCTCAGCATCTTACAGATGCTGGTATTACATGTTGGCCCTTGTCTCACCCCTG | 120 | 52 |
| chr7 | 144344577 | 144344697 | 772751_42001748_Shank2(210274)_1410 | 1 | + | good | AGCCTGGTCTCCCTCTCTCCTTGGTATTCACTTCTCAGCCTAAGCTTTCCCCAAAAGACTTCTCCTGGCCCAGAACTAGGTAGACACTCTCAGGCTAAACTGGACTACCATTGGAGTCTC | 120 | 51 |
| chr7 | 144344697 | 144344817 | 772751_42001748_Shank2(210274)_1411 | 1 | + | good | AGAAACCAGTAACTCATTAAGAAGAGAGATAGTCTCTTAGTGCTTCTGGAGCTTCCAGCACCCACCAGCCTGGATGGCCTCTGCAATTAAGGCTCCAGACCCCACACTGGCAGAAAGACA | 120 | 50 |
| chr7 | 144344817 | 144344937 | 772751_42001748_Shank2(210274)_1412 | 1 | + | good | TGGCCTGTGCACTGAAGGGTTGGCCCTACAGTTCACCCTACCCAAGAGTCCTAGGTCTCTGTAGCCACCACCACAACAACAAAGCCTGGGCCAGGCCAAGGCTGAAGATGGCCTCAGGCC | 120 | 59 |
| chr7 | 144345057 | 144345177 | 772751_42001748_Shank2(210274)_1414 | 1 | + | good | TGCCCCATATGTCCCTCTTGATACCTCTCATCATCCACATGTTGATGGCAGCAGACATTGTCTGGGATGGAGCCATGCCACCTGTGCCCAGGATCCCAGCCCACACTGTATGTATGTGAT | 120 | 52 |
| chr7 | 144345177 | 144345297 | 772751_42001748_Shank2(210274)_1415 | 1 | + | good | TCAGTTTCCTGTTGCTGATGGGAGATGGCTGTGGCTGACTGGCTAATGACAAGAGCTGCAGCACTGTGACAGATATTGCTGGTGTCCCTTGGGCACCTAAGCCATTCATGGTATACAGCT | 120 | 51 |
| chr7 | 144345297 | 144345417 | 772751_42001748_Shank2(210274)_1416 | 1 | + | good | GAGCCTTTACCCAGTACCTTGTTCATAAAAGATTCAATGGCAAGATGCATGCAACACAGCCATGCCATGCCTCGGCGCTGGGTTTTGTTCCCATCAGCACCCATGCTACTGATGAACAAT | 120 | 48 |
| chr7 | 144345417 | 144345537 | 772751_42001748_Shank2(210274)_1417 | 1 | + | good | ACACACTTAATTCTCTTGTGGTCCTGGAATCTAATTGGGTCCCATGAATTTAAAGCAAAGTGTGATGGGGCTGGTCCATTCCCAAGGCTCCAGAGAAGGTCCACTTCCTCTGCATCCAGG | 120 | 48 |
| chr7 | 144346017 | 144346137 | 772751_42001748_Shank2(210274)_1422 | 1 | + | good | ACACTTGCCTTCCCCAGACCTAGAGGGACTTGTGAGATCTGTGTTTCCTCTCTCTACCTCTACTGCCATGGTCACTGGTGTGTGAGCATTGGCATGCCCACCACCCAGGGAACCACATGA | 120 | 54 |
| chr7 | 144346137 | 144346257 | 772751_42001748_Shank2(210274)_1423 | 1 | + | good | TCTAGATTATTTCTGAACTGCAGCCTGGGGCTTGTAACACTCATTTATGGGTAAAAGTAGTAACTTCTGTGCCTAACCTGCAGCTCACAGGCCACATGCCACCCAGGATAGCTATGAATG | 120 | 46 |
| chr7 | 144346257 | 144346377 | 772751_42001748_Shank2(210274)_1424 | 1 | + | good | TATATCCCAACACAAAACAGAAACTTACTTAAAACACTGAGAATTCTTCACAGTTTGTTTTTGTGGCTCTTCAGTATGAACTTCATAATGTAATGATGTGGCTTAGAGAACCCAATCAAG | 120 | 34 |
| chr7 | 144346377 | 144346497 | 772751_42001748_Shank2(210274)_1425 | 1 | + | good | GATTATACATCCCTGAATGGAAAGGACCTAGGGACTTATAACTTCTTTGAGTCTTGGTGGTGTGCTCAGGATCTGGACCCAACTTGACCCACTCACAGTCTTCAGAGGACTGTCCCCCAC | 120 | 49 |
| chr7 | 144346497 | 144346617 | 772751_42001748_Shank2(210274)_1426 | 1 | + | good | AGGCTTCCAGAGGGATGAGAGCCATGAAGTTGCTTACCTACAGATAGACACATCTGCCTCTGGGGTCTGCACCATTCCACATAAGTCTTCCAGATGACAGCTGCCTAACTGTAGCTCAGG | 120 | 51 |
| chr7 | 144346617 | 144346737 | 772751_42001748_Shank2(210274)_1427 | 1 | + | good | TTCTGACATCAGCACCCCTTGCTAGGTGTCCCAGAGTGGGTACTGAGTCACTCTTCCATCCTCCATTGACTGGGGATTCTGAGTTCCCTCAGAGGATGTCTGGGCTCCACGGTTCCTCGT | 120 | 55 |
| chr7 | 144346737 | 144346857 | 772751_42001748_Shank2(210274)_1428 | 1 | + | good | GTGGCACAGTCTCCAGATGGAGAGAAGAGGCTGTGCCCTCATGATACCCAGCCAGGTGTGCTGCAGGCTTCAGGTTTACCACCCACGCTGATCCACAGAGGTTAGAGAAGAGGCCTCAGA | 120 | 57 |
| chr7 | 144346857 | 144346977 | 772751_42001748_Shank2(210274)_1429 | 1 | + | good | ATGGCAGATGAGGGGCAGTGCTTTCAAGACTCCATGGTATCCAAAGTTGTCAAGGGAACAGAGTAACGATGTCTGCTGTCCTTCCTACCTGCTCGCATGAGTGCTTCATTTGTCCCATCA | 120 | 49 |
| chr7 | 144346977 | 144347097 | 772751_42001748_Shank2(210274)_1430 | 1 | + | good | TCCTCTGGGCCCTCAGTCCTCAGAACACATATCAACCCAATTTTCCCCAAGTGGATAACACCACTGACAGAAAAAAATGAGGCTGAAGAGAGGGACAGAGACTGACTGGTATGGAGACTT | 120 | 48 |
| chr7 | 144347097 | 144347217 | 772751_42001748_Shank2(210274)_1431 | 1 | + | good | CTTTGAAGCTGCACAGAGCTCTTGACTATCTTCTAAGCAGGGCAACTGAGTCACTGACCCCTGTTCGGTAGACCAGTGGGATACAAATGAGAATGTGGCTTCCTAGCTCCAGCCCTGTGC | 120 | 52 |
| chr7 | 144347217 | 144347337 | 772751_42001748_Shank2(210274)_1432 | 1 | + | good | TTCCTGCTGACACTGACATTCATCCAGACGCAGAAGGCCCCACCTGCCACCCCTCCCTCATGAGTGGGACTCCCACTAAGTGTCTGTCACAGGTTCAGTGCCCCAGCCCATCTTCTCTGT | 120 | 57 |
| chr7 | 144347577 | 144347697 | 772751_42001748_Shank2(210274)_1435 | 1 | + | good | GCTCTAGGATAAAGACCATGAGGAGGTGGGTGATGTGGACCCCAGAATCCAGCACTCAGACTCCCAGGCTTATTTCCCAGAATTCTAACCCCATCTGACATTCCATAAACCCCATTCTAA | 120 | 48 |
| chr7 | 144347697 | 144347817 | 772751_42001748_Shank2(210274)_1436 | 1 | + | good | CTCTGGTGACACTCAGAACAAATGAACAAGCATTCTCTCTCTGCTCTTGGCTGGGATGGAATGGGAGCAGTTGTTTGAAGTACCATGCCTTCCCTGCCTTCAGGCCTTCCCTGCTTTCAG | 120 | 51 |
| chr7 | 144348057 | 144348177 | 772751_42001748_Shank2(210274)_1439 | 1 | + | good | CTTTCTTTCCCTGGAACTCCAGCTCATGGAATTGTACCACTCATATTCAGGGGAGGACTTCCCTTCTCAGTAAAACCAATCTAAAAACTTTTTTACTGGTGTGCCTGGTATTTTGTTTCT | 120 | 41 |
| chr7 | 144348177 | 144348297 | 772751_42001748_Shank2(210274)_1440 | 1 | + | good | ATGGTGATCTCCCCCCAATCCCCATTGACAATCAAAATTAACCATTGCAGGAAAGGTGCTGGCCTTCATGCCAGATAGCCCTTGTGTGCAAAAAGGATAACCAGTTGCTGGGTATGGTGG | 120 | 48 |
| chr7 | 144348417 | 144348537 | 772751_42001748_Shank2(210274)_1442 | 1 | + | good | TTTCCAAAAAGAAAAGGGTGGAGGGGAACAACTGCATAGACTCTTTATCAGTGGCCAAGAAGGAAACATTCACTGAACTAATGAATGGAAGGGAGATAATCACCCAAAAATTGTATTATA | 120 | 38 |
| chr7 | 144348537 | 144348657 | 772751_42001748_Shank2(210274)_1443 | 1 | + | good | AGGAAAAAGGAGCCTGTCCCAACCCATGTGCCTGTCTCCAAAGCCCCTCACTGGAGCCTCCACTGTGGGGGACACAGGATGTCTTAGTCACTGTTCTATTGCTGTGAAGAGATACCATGA | 120 | 52 |
| chr7 | 144349017 | 144349137 | 772751_42001748_Shank2(210274)_1447 | 1 | + | good | TCAGGACACTGTCTCTAGCAGGATGCCCCAAGTGTTCTTGCCTTAAGCCACCTTCCATGACTCTGACAGAACATCTCAGTTTGAGTAAATGACAAAGACATTTTTTGGTTCACTGTTCTG | 120 | 44 |
| chr7 | 144349137 | 144349257 | 772751_42001748_Shank2(210274)_1448 | 1 | + | good | AAGGAAGTCTGGTGAAATGTAGGGCCACAGCTTCCTGAGACTTGGCTCCTGTGTTTGTCTTGGGTTTGAGTCCTTTGAAGTGTTATAATGGCTGTTACTTTATTTTCTATTGTTTTGAGA | 120 | 41 |
| chr7 | 144349257 | 144349377 | 772751_42001748_Shank2(210274)_1449 | 1 | + | good | CAGAGCCTTACTAATGCAGCACAGAGTAACATCAAGCTTTCAATCCTCTGAATCCATCTCTCAAGTACTGCAATGTCAGGTGTGTGGGACCACACTGCACTTAATAGTTACATTTAGACA | 120 | 42 |
| chr7 | 144349377 | 144349497 | 772751_42001748_Shank2(210274)_1450 | 1 | + | good | AGTGCTTGGGGTGGACATTCAGACCACAGCTGCCCAAGCATACTATGCCAGGCAGCATGGGTAGATAGATGGTAGTGCTCCCCCACAACTGAAGACCCAAAACCTATGTTGCCCTGAGAT | 120 | 52 |
| chr7 | 144354657 | 144354777 | 772751_42001748_Shank2(210274)_1494 | 1 | + | good | TCTCCATCACCACCTCATCCTACCTCAGCCACAGCTAGAAGACTAGGCACCCATTAGAATAGTCTATGTGAAATGAGAGGCTTGTGTAGAGCTTTGGGCAAGAAGGGTGGGAATTGTACT | 120 | 48 |
| chr7 | 144354777 | 144354897 | 772751_42001748_Shank2(210274)_1495 | 1 | + | good | ATGTGATATCCCTCTGGGAGGGGAGGGATAAAACTGAGGGTACAGTATCACTCAGGTCCCCTACTTTTACCAGAGTGACAGCTATGCCCCCTAATCTTCAGAATGGGAGGTGTACTATAG | 120 | 48 |
| chr7 | 144355257 | 144355377 | 772751_42001748_Shank2(210274)_1499 | 1 | + | good | CTTACTCAAACCACAACAGACTCAGAATATAAAGGTGGCCCCTCCCAGATGCGCCATCAAGTGTAGGGCTCAGAGATCAGTCACCTCACCCATCCTGCCCCCTCACCCCAAAGACAAACA | 120 | 52 |
| chr7 | 144355377 | 144355497 | 772751_42001748_Shank2(210274)_1500 | 1 | + | good | ACAAGAAGCAGTGACTCAGGCACTGTGCCCAACCTGAATCTCGATCCAGGCAATATTTCAGGGGCAATCTGGGAAGTCATTATCCTGTGTTTGTCCCAGTACCTTTTCCTCCATATAGAA | 120 | 47 |
| chr7 | 144355497 | 144355617 | 772751_42001748_Shank2(210274)_1501 | 1 | + | good | TGCCATCATCCCCAACACACAAATATACAGAGACAGAGACAGAGAGACAGAGAGAGAGAGGTGGCTTGTTCCTTGGAGCCCTGTGATGGGAAGACATTGGGTTTCCCAAGAGTCTCACTG | 120 | 50 |
| chr7 | 144355617 | 144355737 | 772751_42001748_Shank2(210274)_1502 | 1 | + | good | AGATATCACATCAGCACACAGTAAACACTTCAAACTTTCAGTGTGAAGCTTTATGAACATCAACATCTATGTGATTGGCAAGGCACCAACCAGGTCAAGGTCTAACGAATGACAGATGAC | 120 | 41 |
| chr7 | 144355737 | 144355857 | 772751_42001748_Shank2(210274)_1503 | 1 | + | good | AATCCTCTGCAATGGAGGAGGGCCAACCTCACCTCCCTTTCTCCTATCCCCAGCTACAGCAAAATCAGGCATGTGACTTGAAAACTTTTCCATGAGTAGAAGTGTTACCCACAGCAGTGC | 120 | 49 |
| chr7 | 144355857 | 144355977 | 772751_42001748_Shank2(210274)_1504 | 1 | + | good | CTTCTTGTAGTCCACACCACTATTCCAAGATCTTTGAAAATCAGTACTAATCCTTCTTACCCATGAGTGACATCACAACTTGCTCAACCATGTGTCCACCCATACCCTTACTGATGATTT | 120 | 41 |
| chr7 | 144355977 | 144356097 | 772751_42001748_Shank2(210274)_1505 | 1 | + | good | AGTTTCTGAAGGCTGTGAATGTTGATGTATTTGCCCTATACAGACACTTCATTTCAAAGAAATATTCCTGTTGGGAACAGTAGTGTGTGTGTGTGGGGTGGGGTGGGGTGTCTCACAGTA | 120 | 44 |
| chr7 | 144356097 | 144356217 | 772751_42001748_Shank2(210274)_1506 | 1 | + | good | TGATCATTTGGTGAAGAAACCTTACCAAGGTACCTCCAAGCCAGCCTTATCATCTTCATCTGCATTCCTAGCTCCCTGCCCTGAGCTGAGCTACACACACACACACACACACACACACAC | 120 | 49 |
| chr7 | 144356217 | 144356337 | 772751_42001748_Shank2(210274)_1507 | 1 | + | good | ACACACACACACACACACACAGGGGTCCCCACTTTCCCCCTTTATTCCTTTGGATTATGATGAGAGACAGTTGGGCCCTGTGGCTACCATGTGCATCTCCCATGTGGCCACGATGCCTGGCA | 120 | 54 |
| chr7 | 144356337 | 144356457 | 772751_42001748_Shank2(210274)_1508 | 1 | + | good | TCTGCCCATCTGTTGGCACCTGTTGGGACTTTACAAAGAGAGGGGTTCCCTCTTAATTTCTTCACTCTCATTCTTTCCTTGTCCTCTCCCCTTCCCCCCCATATGAGCTCAATCCTGGGA | 120 | 50 |
| chr7 | 144356457 | 144356577 | 772751_42001748_Shank2(210274)_1509 | 1 | + | good | CCAAAGAGCAACAGGCATTTCTTCTCAGGGTTCTGGATCTGGAGGTCTATGAGTCTGGGTTTCTAAAGACTGCAGGGCCTATAGGGGATGAGGTAGAAGGCAGCTGCCCAGAGCCCTCTT | 120 | 52 |
| chr7 | 144356577 | 144356697 | 772751_42001748_Shank2(210274)_1510 | 1 | + | good | TCATAAGGGCTTCAGTCCTGTCAGGAGGTCTCTATGCCAGTGATCAACTCTCTTCCCATAGGCTCTGCCTCCCAACACCATCTCCTTCAGGTAGGTTGCAGCACATGACCTAGGCAACAC | 120 | 52 |
| chr7 | 144356697 | 144356817 | 772751_42001748_Shank2(210274)_1511 | 1 | + | good | AGAAAAGTTACCCACAGCCCCTCCTCAGTGCCCACTCTCACTAGGCCTGAACTCTTGCCCTTGCCCTCCTCCTGCCCTCCCCTTTACGGCCTCTGGTCCAGGCTCAGCACCTGCCTGGCT | 120 | 62 |
| chr7 | 144356817 | 144356937 | 772751_42001748_Shank2(210274)_1512 | 1 | + | good | ACATCTTTCCTCCCTACCCTGATGTGCTGGTCATCCTGCTGGGTCAGGTGGCCACTTGTGTTCAGTTGGCATATAAGGTCATGGTTTGCCTAAGGTGCTGGGGCCCTCCGCCAAAAAGAT | 120 | 53 |
| chr7 | 144356937 | 144357057 | 772751_42001748_Shank2(210274)_1513 | 1 | + | good | GGAGATGTTTGGACTTCTTTCTTAGAAAATGAACAAAAGAGAAGCCAGCCTGGTCAGGTGGTACTGCCATCCGTTCTTCCCCTTTGGGGTTCTGTTCCTCCTGGGTAGCCCCATTGTCCA | 120 | 50 |
| chr7 | 144357057 | 144357177 | 772751_42001748_Shank2(210274)_1514 | 1 | + | good | CCCAACACACAGGGTGGAAATGACCCTCCTTCACTCATATCTAACTCTGTGTGCACCTCCTCTGTGGTGGGCTGAGTGAGAAGTGGCCCCTCTAGCTTGGCCATTTGGCACGTGTTCCCC | 120 | 56 |
| chr7 | 144357177 | 144357297 | 772751_42001748_Shank2(210274)_1515 | 1 | + | good | AGTTGGTGGCTCTATTTGAGGGGTTAGTGGAGAGGTGCAGTCTTGCTGAAGGACTAGGGATGGGCTTCACCTTACTCTTGGTTCACTTTCTCTGCTTCATGGTTTTGGGCAAAGATGTGC | 120 | 49 |
| chr7 | 144357297 | 144357417 | 772751_42001748_Shank2(210274)_1516 | 1 | + | good | GCTCCCAGCTTCCTCTTCTAGCTGCCACTTGCTGCCAATGACAGAGTCTAGAGCCATCAGCCAAAGTAAACTCTCCTGTAAGTGCCTTCATCACAGTTTTGTATCACAGCAACAACAACA | 120 | 48 |
| chr7 | 144357417 | 144357537 | 772751_42001748_Shank2(210274)_1517 | 1 | + | good | ACAACAACAAAAAGTAACTAAGACATCCTCCTAGCATTCCTAGACATGGGACCCTAGCTGTGCTATTCCTGGAGCCAGGTGTCTCCTGCCAGGCAGCAACAGGTCTAGTCACTTGTCATC | 120 | 49 |
| chr7 | 144357537 | 144357657 | 772751_42001748_Shank2(210274)_1518 | 1 | + | good | AGGGAATCCAGAAACCTATCAAGTGGCCCCAGAGTTCTATGTCATACAGTCTAGAAGGAAGGGAGTTGGAGTCCAGGACTACCTGGTCTTAGGATCAGGTAGGATTCAGGCTGTGTTAAG | 120 | 48 |
| chr7 | 144357657 | 144357777 | 772751_42001748_Shank2(210274)_1519 | 1 | + | good | AGAGATCTCAAAATATCCTCCTGCTGTTGCCCCAAAGTCAAGAGGAAGAAATAGCACCCACAGTGCATGGCTGAAGCTGAGGACCAGGCAGGTGGCTGACCTTCACTGACTCCAGAGCCC | 120 | 53 |
| chr7 | 144357777 | 144357897 | 772751_42001748_Shank2(210274)_1520 | 1 | + | good | ATGAGCCAGCTCTATGCACCCTGATCATGCCTTGAGCTGCCTCTGTTGCCCGAGTCTGGATGCCTGCCTTTGATACTGCCTGGCTGTTGGTGCCTTGTCATAGTGTCAGTGATTTGAGCC | 120 | 54 |
| chr7 | 144357897 | 144358017 | 772751_42001748_Shank2(210274)_1521 | 1 | + | good | ACTCCCAAGACTTCTCTCTGCCTGGTGGTTTCTCTTCAGTACCCAGCTCTTTACCCAAGCAGAACATGTCTGATCCACCCAGCTTTCTCCACCCAGTCCATGACTGCCCACTCTGAAGAG | 120 | 52 |
| chr7 | 144358017 | 144358137 | 772751_42001748_Shank2(210274)_1522 | 1 | + | good | AACACATCATTGACCTAGGGAGAGATGGGCAGGTGACCACAGAGGGTGCTGCTGAGAACAGGAGGCAGGAGCATCCATTACCCAGAGGTTTAGCAGCTCCTCCGTAGAGGTGGTGGGGTG | 120 | 57 |

| | | | | | | | | | |
|---|---|---|---|---|---|---|---|---|---|
| chr7 | 144358137 | 144358257 | 772751_42001748_Shank2(210274)_1523 | 1 | + | good | CCGGTGGGATGCACAGCTCTGAGGAAAAGACAGACTTTTGTCTCATTCACATACTGTTTCAACTAAGGAACCCCAGCTTCTTGGAGCTCCACTCTTCCTCACTTGGTGCCCATCTGTTTA | 120 | 48 |
| chr7 | 144358257 | 144358377 | 772751_42001748_Shank2(210274)_1524 | 1 | + | good | CTGTTGGATCAAGCAGGCCAGTCAATATGAAGTACCCGGGTTATACAGGCCCTGCCCTCCCTCCAAACAGCACCTGTTTCAGCTCAAGCCACTGAAGCTATGACAAGTCACCAAAGGCCA | 120 | 52 |
| chr7 | 144358377 | 144358497 | 772751_42001748_Shank2(210274)_1525 | 1 | + | good | GGCAGGCATCCATCATCCCTGTCCTGGAGGTGGGAGGTCCAAGTGATAGACCTGGCTCCTTCCAAGGTTCTACTCCTTGGCTTGCAGATGCATCTCCTTCCTGTGTCCTCATGTGCATGT | 120 | 55 |
| chr7 | 144358497 | 144358617 | 772751_42001748_Shank2(210274)_1526 | 1 | + | good | TCTCTGTGTGTCTGTGCCCCAGTGTGCTCTTACAAAGACAGCAGGAACAAGGAATAACCTACTGGCCTCATTTTAGCATATGAACCTCCTGAAACACCCTGACTTCAAACAGTGACGCTC | 120 | 48 |
| chr7 | 144358617 | 144358737 | 772751_42001748_Shank2(210274)_1527 | 1 | + | good | TGAGAGACCAGGAGTTAGGAGGTCTTCAGATGATCTTGGATCCCACGTAAGGACATGGCAGTCTCCCCTGCCACAAGCTCTGCAGACCAGCTCCCACAGGTTAGAGCCAAGTGTTTTTGA | 120 | 52 |
| chr7 | 144358737 | 144358857 | 772751_42001748_Shank2(210274)_1528 | 1 | + | good | CTTAAGGTGGGAGGCCACCTATCTCTGATGATAGCAATGGCACTTAGAGGTGTCTCAGAGAGCGTTCTAGACCATTTTGCCATCTGTGACAGAGAGGGGCACATCTGAGGACCCAAGGGA | 120 | 52 |
| chr7 | 144358857 | 144358977 | 772751_42001748_Shank2(210274)_1529 | 1 | + | good | TGATCAGAGCCACCAGGAGGTCTCTCTGATCTTGTTTGGCAAATGTGCCACTGACAGAGAGCCCTCCTGCTTCCTGCTTCCTGCTTCCTGCCTAGGTGATTAATGATGGAGTTCAATGGC | 120 | 52 |
| chr7 | 144358977 | 144359097 | 772751_42001748_Shank2(210274)_1530 | 1 | + | good | TCAGGGCGGTCCTGTGCACATCAATTATTCAGATCGACACCTCTAAAGGTCTGGAACAGGAAATAGAGAGGAAGAGATCGCTAAGCAGTTTTGAAGGGGTCTAAATGGATTGCTCGTTAA | 120 | 45 |
| chr7 | 144359097 | 144359217 | 772751_42001748_Shank2(210274)_1531 | 1 | + | good | CCGCCATGACAGATCGGGTCTGGTGAGAGTCTGTAAGTTGGCCTAGCTGCCCAGCAATAGAACCATCCTATCTGGAGGGGAGGCTGGGAGGCTGTTCTGAACTTTGGAACAGCAGGAGCT | 120 | 56 |
| chr7 | 144359217 | 144359337 | 772751_42001748_Shank2(210274)_1532 | 1 | + | good | TTTTTCAAGCTCATTGCTCACCCGGGCCTGTCTGTCTATTATAATAATGAACCTCAGATTGAGTCAGGCAGGTGTGCTAGAATTAATCTGGAAACCAGGCTGTGGAGCCACTCCAGGGTG | 120 | 48 |
| chr7 | 144359337 | 144359457 | 772751_42001748_Shank2(210274)_1533 | 1 | + | good | AGCTGGGTCTGATGGTTACCATGGCCTTGGGATCTGAAGGGGCTAAATGTTCACTGTGTGGGGAGGGCTGAGGCACTCATTACTCACTGTGACATCAAACTGGGCCAGCTCGTGATGTGT | 120 | 53 |
| chr7 | 144359457 | 144359577 | 772751_42001748_Shank2(210274)_1534 | 1 | + | good | CCAATAGAAATTATTTAAGGCAGTAGTATGATCTGCTTTATTGATCGAGATGCTGCAAACCCAGTTAGGGAAGCTTCAGGTAAGGCAGAGTGGTGCCCCCCCCCCCCCGCCACCTTATCT | 120 | 49 |
| chr7 | 144360177 | 144360297 | 772751_42001748_Shank2(210274)_1540 | 1 | + | good | ATGTAGAGATACAGATTAGGAGTCACCGTTCCCTATGAAGCATCTAGACACTCAGCTTTAAAGGAGACATGGTGAAGGGAATCTCAGATGGGCTGTCAGGTGACAAAGGACAGCAGTTCT | 120 | 46 |
| chr7 | 144360297 | 144360417 | 772751_42001748_Shank2(210274)_1541 | 1 | + | good | TGAGGGATGAGATACAGGAAGCCAGCATGGGTTCTGTGCTTTGCTGCAGTAGTATGTGAGGAGATGGTCCCACGCATGGGAAGGAACCAAGCTACAGAGCAGGAACTGAGGGTCTGGAGA | 120 | 53 |
| chr7 | 144360417 | 144360537 | 772751_42001748_Shank2(210274)_1542 | 1 | + | good | GACCAAGGCAGTTCAAGTGTGTGAGGAGAACCAGGGGGAACATGTCAGCCAAAGAGGCCCTGGAGAACTGCAGAGTTCCCAAGGGTCAGCTGAGTGACTAGAACGTAGGGGAGAAGAAAT | 120 | 53 |
| chr7 | 144360537 | 144360657 | 772751_42001748_Shank2(210274)_1543 | 1 | + | good | TTCTAGAAGCTGGATAGAGTGTTAACCCAAAGATGAGAGATTGTACCCCACTCATGCAGTACCAAGCAGTGTGCCACCGAGTGGTATGTCACTGAGCGGTGTGCCTCTGTCTCCACCAGA | 120 | 51 |
| chr7 | 144360657 | 144360777 | 772751_42001748_Shank2(210274)_1544 | 1 | + | good | CCTCAGACTGCTCAATGCTTGTGGGGTTCTTAGAAGGCCTGAAACATGAAGGAGGGTCAGCCTGTGACTGGCAGAGATTTTTATCCCCCTAACACATTTTTAAATAAAGCACTCAGGAGA | 120 | 46 |
| chr7 | 144360777 | 144360897 | 772751_42001748_Shank2(210274)_1545 | 1 | + | good | AACTCAAGAACATTCATAAGAAGATAGAAGTTGTCAGTTGCCAGCTAGACAAAATCCACAATATCTCACACTCTCTTGGGAGTCCCAGCTCAGGAGATGAGGTCTTGCGTCAGGTCATGA | 120 | 44 |
| chr7 | 144360897 | 144361017 | 772751_42001748_Shank2(210274)_1546 | 1 | + | good | CGAAAGGAGACATGACCTGTGAGAAAAGGAGGCTCAGTTAACCATAAGCAAGGAGACATGGCCCATGAGAAAAAGAGGCTCAGTTAACCACAAGCAACTTCAATCTACACAGATGTCCCA | 120 | 46 |
| chr7 | 144361017 | 144361137 | 772751_42001748_Shank2(210274)_1547 | 1 | + | good | ATTGCAGACAGAGGCTTTGAAATAGTCTTCGTAACTATATTCCCCATAGCCAAAGAGTTAGGTAAAGCCAGCCCTGGTGCCACAGTCCTGTAATCCCGAGTACCAGTGAAGCTATAGCGG | 120 | 48 |
| chr7 | 144361257 | 144361377 | 772751_42001748_Shank2(210274)_1549 | 1 | + | good | CTAGTGTGTATCACAGCATAAACTGCATGGTGAGTGCACTTGTAATTCCAGCATGAAGTTCAAGGCCATCCTCAGCTATGTAGAAAGTTTAAGAAAAGCCTAGACGGTGAAAAACAGCCT | 120 | 42 |
| chr7 | 144361497 | 144361617 | 772751_42001748_Shank2(210274)_1551 | 1 | + | good | TCTAGGCTAGACTGATCTATATAGCGAGACCTTGTCGGAAGGAAGGAAGAAAAAGAAAACAAGAAAAGGTTGTCTAGATTCTCAAAAAAAAAAAAAAAAAAAAACCTGAAAATTCAGAGAA | 120 | 32 |
| chr7 | 144361617 | 144361737 | 772751_42001748_Shank2(210274)_1552 | 1 | + | good | ACGTACATCTTGAAATGTTTTGTGTATTTTGTCATAATTAAAATTTAATCAAGTCAGAATTTCAGAGATGGAAACTGTACTCTCTGAGGTAAAGAATACTCAGGATTAGACACTGGAGAG | 120 | 32 |
| chr7 | 144361977 | 144362097 | 772751_42001748_Shank2(210274)_1555 | 1 | + | good | GGAAGAAAAGTGATGGACGTGGCAAGGCAGCTGTGTGAAAGAAGCCATCTTAAACGAAGCAGAGGAAGACTTGCTGTAGCCTCCCCCCATAACCCCCCCCTTTGAAGAATGGTCTTCTGA | 120 | 51 |
| chr7 | 144362097 | 144362217 | 772751_42001748_Shank2(210274)_1556 | 1 | + | good | GCCATGAGACCCTGGCACCTGACAGATGCCAGCAGAGTCTTGAGAGAGGAGGAACTGATGTGTGGAAGTCAGAACACAGAGTTGAAGAAATAATTATGTGAAATTTCCCAACTTGAAGTA | 120 | 45 |
| chr7 | 144362217 | 144362337 | 772751_42001748_Shank2(210274)_1557 | 1 | + | good | AACTGTGAATTCACAGATTCAAGCAGCCTGTGATCCCCAAACTCAGGACATGTTTAAGAGAAGGAAAACTACAGAAACACCGGAGCACCCAGCTTCCTAATTGCTCCAGATGAAAAAGCA | 120 | 44 |
| chr7 | 144362337 | 144362457 | 772751_42001748_Shank2(210274)_1558 | 1 | + | good | CCTTGAGGTGGTGAGGGGGCTCAGCGGCTAAATGTGTGCTGCTCTTGCAGCTTCTGAGTTCAGTTCCCAGAACCCACGCTACCATTTGCCCAGTGGCTCACAGCCACCCAGGACCCCAGC | 120 | 60 |
| chr7 | 144362577 | 144362697 | 772751_42001748_Shank2(210274)_1560 | 1 | + | good | AAGAGAAGAGTCAGGCCAAGGGTGGATGTATGGCGGGAAAGGACACAGGGAGCAGGAAGTGGAGAGACAGCTTTCCATTGTGAGACAGAAAATGACATTTTATAAAAAGAGAGTAAAATA | 120 | 43 |
| chr7 | 144362697 | 144362817 | 772751_42001748_Shank2(210274)_1561 | 1 | + | good | AAGAGACATGGCACAAACACAAAAAGCTTGGCCAATTCATCCCAGAAGACCATAATATTGGGCACCTCCTGGGTAACAAGAAACAACTGTACATGGAAACCCATGCAGGGACAAATGCAG | 120 | 45 |
| chr7 | 144362817 | 144362937 | 772751_42001748_Shank2(210274)_1562 | 1 | + | good | GAAATATTTTCTGGTTATTTAAATCTCTTCAAGGCAGACGGGGTGGCTCAGTGGGTCAGGCACCTTCTGCTCAGCCTAATGATGTGACCTCAATCCCTGGAACCCCTATGGTAAATGGAG | 120 | 48 |
| chr7 | 144362937 | 144363057 | 772751_42001748_Shank2(210274)_1563 | 1 | + | good | AGAACCACCTCCATCTACACACACACACATTAAATGCACATATATACAAATAAAATGTAATGAAAATGAAATTTAAACCTGTTCAAATGTCAACTAGATTTATTTATTCACTTATTGATT | 120 | 27 |
| chr7 | 144363177 | 144363297 | 772751_42001748_Shank2(210274)_1565 | 1 | + | good | GATGATGAGACAAGGATATGGAAAGGTGATTATATAGTAATGAAGACTGAAAGAGGAAGAACTGCCCATTAAGATCCCCTGTGATGTGGCAGGGCATCAGGAAAACATGAACAGAGGGCA | 120 | 43 |
| chr7 | 144363297 | 144363417 | 772751_42001748_Shank2(210274)_1566 | 1 | + | good | GTGCAGATGTTGGAAGTGAATGACAACAGCTAAAATAACAAACACTGGAGTTCACAGGCTGATGGAAGAAACAAGATGGATCATAAGAAGGTTTTATCCTGGTGAGCTGGTTTTCTGGTA | 120 | 41 |
| chr7 | 144363417 | 144363537 | 772751_42001748_Shank2(210274)_1567 | 1 | + | good | TCTAGCACTCAAATAGCTGTAGCAGGAGGATGTCTAAATTGAAGGCTGCCCTGGCTACATAGCTAGAATCTTTTTCAAAAAACAAAAAGCATAAGAATAGAGGATCGGGACACAAAATAT | 120 | 38 |
| chr7 | 144363537 | 144363657 | 772751_42001748_Shank2(210274)_1568 | 1 | + | good | ACATGAGAAAAATATAAAGCAATCAGGGGGCACAAGCATGGCTCAGTAGCAAGGTACACAAAAGTCCTGGGTTTGATCCTCAGCATCCCACAAAAACAAACAAACAAACAAAACAAAAAA | 120 | 39 |
| chr7 | 144363657 | 144363777 | 772751_42001748_Shank2(210274)_1569 | 1 | + | good | CAAAAAAAACCCCACAAAACTATTCAGATCAATAATTACACTAAATATAAATGATACAAATACCCAACTCAAAGGCAGGGTTTTGGGGTTAAACACTTGTCCTCAAAGTATTTTAATTAT | 120 | 30 |
| chr7 | 144363777 | 144363897 | 772751_42001748_Shank2(210274)_1570 | 1 | + | good | AAAGACACAAGTATATTCAGATGGGGAGAAATTATATACCACATTGACATTACTCCAAAGAAAATTATCTATATTATCATGAAGGTGGACTCCAGAGTAGAGGGCACAAACTGATAGGAT | 120 | 35 |
| chr7 | 144363897 | 144364017 | 772751_42001748_Shank2(210274)_1571 | 1 | + | good | AAAGGTCATTTTATAATGGTGAGGGGCCAGCTCTGTGAAAGGGCAGAAAACACAATGACATCACCACTGACATCACGCTTGGGGTCCCCATGCATACATACCCAGCACACCACACAAATG | 120 | 48 |
| chr7 | 144364017 | 144364137 | 772751_42001748_Shank2(210274)_1572 | 1 | + | good | TGTAAAAATTAACAAATAAGCATCTCAGTGGGAAATGATGAAACAGGAAGTTGGTGCTCAAGAAAAATGCAAATTAGTAGCAGTGGAGTGAGGGTGGGCCATCCTGCAGGTCAGCCCACT | 120 | 43 |
| chr7 | 144364137 | 144364257 | 772751_42001748_Shank2(210274)_1573 | 1 | + | good | GTGTGCTTTGTCAGGGTTGCTCACACTTGTGTGCAAGAATGTCTTTCTCTTCTCTGCTTTAGTCCCCCATACATCCATTCCTCCATCTGTCTATCCATCCATCCATCTGTCTATCTCACA | 120 | 46 |
| chr7 | 144364377 | 144364497 | 772751_42001748_Shank2(210274)_1575 | 1 | + | good | CAAGGTCCTTTCCACCTCCCTACCCATGGTCACCACATCTTTGTAACCTGTCCTGGACTGGGTTGGGGGTAGCCCAGCAGCTCTGTTCTTTCAAGCATTTACTGTGTCTGGTTGTACTTT | 120 | 51 |
| chr7 | 144364497 | 144364617 | 772751_42001748_Shank2(210274)_1576 | 1 | + | good | TTTCATTTCAATATGAATTTTATGATCAGCTTGTCATTTTTTTTAAACCAAAGCCATGTGGCCCTTTGATTGGAAACATATTCACTCTGTAATCTGTCTGGATATGACCACACTATAACT | 120 | 32 |
| chr7 | 144364617 | 144364737 | 772751_42001748_Shank2(210274)_1577 | 1 | + | good | AGACAGCTTTGAACTGCCTTCTGCACCTTGGGGGACAGAGGTGCTCTCATTCCTGGGTCCCATCCACTCCTCTCAGCCATGCCTTCTTGGATTTGGTGTCTAAATCTTGCAACTTTTGTC | 120 | 51 |
| chr7 | 144364737 | 144364857 | 772751_42001748_Shank2(210274)_1578 | 1 | + | good | AGATTTATCTCTGCTTATTGTTTTTTTGGAAATACTATAGACATATCTTTAAATTTCTATTGATGGCCATGAAGTGCTCTGATATTAAATTACAGTTTACTTGGGGCCTATGATACTGTG | 120 | 31 |
| chr7 | 144364857 | 144364977 | 772751_42001748_Shank2(210274)_1579 | 1 | + | good | AATTTTGGTCCACTCTACACATAGGCCCAAGGTGGGAAAAGAGAAGGAAATTTGGTTTTTAAGCCCTTGCTTATTATCAGAGTCATCTTCACCAATTTGTGCAGAGTTGCTGGGCTAGAC | 120 | 42 |
| chr7 | 144364977 | 144365097 | 772751_42001748_Shank2(210274)_1580 | 1 | + | good | AGGAAGTCATGTAGACACACAAACCCTGAAGAAGGCAACAGTGCCATTGCTACAGGAACACAACGTGCCCGTGTCCTCCGACCACTGTGCACCCTGTGACCATGTCTTCCACCTCTGCCT | 120 | 54 |
| chr7 | 144365097 | 144365217 | 772751_42001748_Shank2(210274)_1581 | 1 | + | good | ACGTGGATGCAAACCCTGAGCTGCCCCAACTTCCCAGAGGGCAAGTTTCCAGGAAGCCCCTTGCAGGGGAGAGCTGTACCATGCCACCTTCAGAAAGGTGATATTTAACATCAAGGGCTT | 120 | 53 |
| chr7 | 144365217 | 144365337 | 772751_42001748_Shank2(210274)_1582 | 1 | + | good | CAGGGGAAAGGGCAAAGTACAGGAGTGTGTGCTACTCCGCTACTTTCCACACTGAAGAGAGAAAGAGATGTGTCTTGATGTATTTCTTTTCTTGTTGCCATGACAACATGCCTAACAAAA | 120 | 43 |
| chr7 | 144365577 | 144365697 | 772751_42001748_Shank2(210274)_1585 | 1 | + | good | CCAAACACCTGAGCCAGTGGGCAACATTTCACTTTCAAACCATGGCACATGGGAAGTCTGGACTTCCTCCATGATTTTTCTGTATACCTAAAACTATTCTAAAACCAGCCTGTTAGATGA | 120 | 42 |
| chr7 | 144365697 | 144365817 | 772751_42001748_Shank2(210274)_1586 | 1 | + | good | AAGCCTTTAAAGAGGCGGGTGTGTTGCCTGAGTTCAGAGCCCATTTGCAGTTGGTCATGTACCACCCACAAGGGATAGCAGTACACAGCCTATGTTGAGTGGGCTTACATGGTACTGAGC | 120 | 51 |
| chr7 | 144365817 | 144365937 | 772751_42001748_Shank2(210274)_1587 | 1 | + | good | ATAGAACATAGCTGTGCAGAACCTTGGACCATGAAGAGGCAGGGCCCAGGTGTATGCTGAGTTCTACCTCCCTGCAAAGCTTCAGATACCTAGCTTTTGGTGGTGGGGGGGACATGGATC | 120 | 52 |
| chr7 | 144365937 | 144366057 | 772751_42001748_Shank2(210274)_1588 | 1 | + | good | ATGTCCCTGTGTGACAGATGCAGAGACTGTGGCATGGAGAAAGTAACTCTGTCTAAGACTGCCCAGCTGAGGTGTAGCAAACTTGGAATTTGAACCCAGGCAGGCTGGTTCTAACCAGGG | 120 | 51 |
| chr7 | 144366057 | 144366177 | 772751_42001748_Shank2(210274)_1589 | 1 | + | good | AGGAAGCCAACTCCTCCTTCTCTTTGAGCACCCCTACAGATGTTACTACCATCATGTGTTACAAAGTTGATGAACCCCCAACTTTCTGCTTCCACATGGGCAAGTGCTTCCACCTTCCTG | 120 | 48 |
| chr7 | 144366177 | 144366297 | 772751_42001748_Shank2(210274)_1590 | 1 | + | good | GAGGGTGGGTGCTGAGTGAGTCCCAGTAAGCCTGCAGCCTGGGGACCTGGGTGAATAAGACGTATCTGTTTCAGAATCCTCAGGAGAGCCAGACCTCCTGGGTGGATCACTGGCACCTTG | 120 | 57 |
| chr7 | 144366297 | 144366417 | 772751_42001748_Shank2(210274)_1591 | 1 | + | good | CCATGTGCCAGCTGGGGAGAGTGGGCCTAGGGCCCCTTCTGCAGTTTCCCCTCCTCCCCCAGCTCCTGGTGGCATTTGTGTGCTGAAAAGATGGATGGTGGAGCTCCTGGTGGCATTTAT | 120 | 59 |
| chr7 | 144366417 | 144366537 | 772751_42001748_Shank2(210274)_1592 | 1 | + | good | GCTCAACTGCTGCTTAATACATTTGCCAAAGGCTGTCCTTTTAAAGGGGAGCATGTCCAAACAGCAGCAGTGACTCAGGCCCCAGATCGGTCCCCTGGGAACTATGGCAGCATAGAGCCT | 120 | 52 |
| chr7 | 144366537 | 144366657 | 772751_42001748_Shank2(210274)_1593 | 1 | + | good | GGCTGTGGGGTCAAGGGTGGCCTCTGCTGCCTTGGGAGCTCCAGAATGCTAGAGAGCCGCTCCTGGAACCCAGAGCACTGCAGCCCTATCCCTGCCCGCCATACTTCACTGCCACCCTCC | 120 | 64 |
| chr7 | 144366657 | 144366777 | 772751_42001748_Shank2(210274)_1594 | 1 | + | good | TGCCCTGCCAGCCTCCCCAGGCTACCATCCCACATTCACCTCTCAGCTTCAGCAGCAAAGCTAACCATGGGATGTTCCAGGGCCTCCTATCTCATCATAGCTCTTTCTCTCACTACTTCT | 120 | 53 |
| chr7 | 144366777 | 144366897 | 772751_42001748_Shank2(210274)_1595 | 1 | + | good | CCAAAGCATGATTCTGGAAACTTCTATAAAGGGGTAGAAGGTAAAGAGTTGGAGAAATAGCTCAGCCAATGAGAGTGCTTGCTTCTCTTCACAGTATACACCTCAGGTGGCTCACAGCCA | 120 | 45 |
| chr7 | 144366897 | 144367017 | 772751_42001748_Shank2(210274)_1596 | 1 | + | good | CCCATACTTCTAGTTTCAGGGGTTAGACGCCCCCTTCTGGGCCCTGAAGGGACTACACCCACATGCATACACACAAAGACATAGGGGGGTGGTAATCTTTTTTAAAAGAAAGAAAGATG | 120 | 47 |
| chr7 | 144367017 | 144367137 | 772751_42001748_Shank2(210274)_1597 | 1 | + | good | GTGACTTGGGGAAACTCTGTCATCTTAGTCCTCCTTTTAGATTTCTGGGTGAGCAGCCTGGGCTTGCGCTCTCCTTGGGAACTCACTGTATGTTATGTTTACTTACCTTCATTTTGCCTA | 120 | 46 |
| chr7 | 144367137 | 144367257 | 772751_42001748_Shank2(210274)_1598 | 1 | + | good | ATTTGTATCGAGGCCAGGTGATGGCTTGGACTCTGGCATCTAGAATAGGGCAAGCTTCTAGACTGGGCTCCAAGAAGCTGATGCACATGTGAATGAGCAGAGGAGCCCTTCCATGCAACA | 120 | 51 |
| chr7 | 144367257 | 144367377 | 772751_42001748_Shank2(210274)_1599 | 1 | + | good | CTGCTGTGCAGTGAGTGTGAGCATGTGAAGCATGGAAGCCACCAGGGCCCAAGTCACAGCAGTGATGTCAAGACACAGGTTGCCATGGCTTTGGAAAATTAAGACCAAGCTGTCTGGATG | 120 | 52 |
| chr7 | 144367377 | 144367497 | 772751_42001748_Shank2(210274)_1600 | 1 | + | good | CATACAGCAGCTCTGATCAGAAATGCCCAAAGCCAGATGCAAGCATCAAGTCCTGTGGGGGATGGGCAGACAGGCCACATCTCCAGACAGTGGAGAATCAATGGGCAGCAATGTGAAATG | 120 | 52 |
| chr7 | 144367497 | 144367617 | 772751_42001748_Shank2(210274)_1601 | 1 | + | good | AGCTGTGGAGCCACAGAAGACAGAAAAACTGTGAATATGGCCAAGCAAAAGAAGGGAGACTGGAAAGGCAGCATGCAGTGTGATTCCAGCTCTGTGACACTCTGGAGAAGGCAAAATGGT | 120 | 48 |

| | | | | | | | | | |
|---|---|---|---|---|---|---|---|---|---|
| chr7 | 144367617 | 144367737 | 772751_42001748_Shank2(210274)_1602 | 1 | + | good | GGAGACATGCAGGGGTCAGTGATTGTGGGTGAGGCAGAGGACATGTGGCCAGGGTGACAGCATACCTGTGGTAGTGTGATGGTGGACATGGGACATGTGACCTTTGCTGCAGCCCAGAGG | 120 | 57 |
| chr7 | 144367737 | 144367857 | 772751_42001748_Shank2(210274)_1603 | 1 | + | good | CTGAGCACCCAGACTGAGCCATGTTGTGAATTGATACTTGGTAAATAAAATGTTTCCCTGTTAGTTTCTTAAAAGCCACATGTGTCCCACATTAGGAAAAGACATTAGCAATATAAGACT | 120 | 38 |
| chr7 | 144367857 | 144367977 | 772751_42001748_Shank2(210274)_1604 | 1 | + | good | CCTTGGAGTAGGAGGGGAGCCAGAGGTGGGGACTTGACTCTTCCTCAGCAATTTGTCCACAAATCCAAAATGCCCCACTCCATACACATAAAAGTCTTAGTTAAAATGAACTGTTCTCAC | 120 | 46 |
| chr7 | 144367977 | 144368097 | 772751_42001748_Shank2(210274)_1605 | 1 | + | good | TTCCAACTTCCAAGTTCTCACTTAGAGGTACATGGCAAGATCCATCTCCCAAACAAAGGACACAAAAGACAGATGATTCTAGAATGTTTTAGAAAGCTGCATAAGAACAGATCTTCAGGC | 120 | 40 |
| chr7 | 144368337 | 144368457 | 772751_42001748_Shank2(210274)_1608 | 1 | + | good | GCTTAACCTCCAGCCCTGTTCGCTTGGTTTCTTAGGCAGGGCTTGCTATATAGTGCAGGCTGGATTCATGCTACTTCCTTAGCCCTCCAGAGTGGGGTTGTTAGCCTGGGGCCCCATACC | 120 | 55 |
| chr7 | 144368457 | 144368577 | 772751_42001748_Shank2(210274)_1609 | 1 | + | good | CAGACCACAGGCTGTGGGTCTAACTAACAAGCTATAAGTCCTTGTGACTGCCTTAAGTAGCCTGTGGCTTGTCTCTAGCCAGCCTTCACTGTGCAGGTGTGGGTGGGAGGTGACAAATGG | 120 | 52 |
| chr7 | 144368577 | 144368697 | 772751_42001748_Shank2(210274)_1610 | 1 | + | good | CCTTCGAAGATGAAAGCCCACAGATCAGAGCCTCGGAAATGAGGCGTCTCATCCCTTAGAGAGGGGATTAATTCTTTGCTGTTGTCTGAGCTGGAGCACTTGATAAGTGGTTTGCGATAA | 120 | 48 |
| chr7 | 144368697 | 144368817 | 772751_42001748_Shank2(210274)_1611 | 1 | + | good | GGTGGAACACAGAGACGCCAAATGGAGCCATAAATCTCCATGGGCTGGATGGTTGGCCACAGTCTGAACTTGCTGCAAGAGGGGAGATGGGAGCCTGGTGATACCTCGGGTTGTGTGAAC | 120 | 55 |
| chr7 | 144368817 | 144368937 | 772751_42001748_Shank2(210274)_1612 | 1 | + | good | TCTTTGGGAGGGTGCCTCCAGAGTTTGGCCTATGCAGAGTTCAAAATGCCTGTGCTCTGTCATGTCCCTGTATCCGGAAAGGCCAGCTGCTGCTTGCTTTTTAGTGTCAGCTTGTGGTCC | 120 | 52 |
| chr7 | 144368937 | 144369057 | 772751_42001748_Shank2(210274)_1613 | 1 | + | good | TGATGTCTCTACCCTGGCAAGCAATCTAGCCCCTCCTTTTTCAGCTTGCTACCTCCTCCCAGAAGCCCTCTGTGCAGCTAGTGACCAACTTGGTTAGTCACCTAGCAAGTGTTGTTTCAC | 120 | 51 |
| chr7 | 144369057 | 144369177 | 772751_42001748_Shank2(210274)_1614 | 1 | + | good | CCACCCTCTTCAAACAAAACTGTGTCTGTACCAGAAGAAAGATTGTGGGATGGGAAGGTGACCAGCACAGGTTTGGTGAGGGGACTGTCTCTTGGGGTCATTTGCACTCCACCCCCATCT | 120 | 52 |
| chr7 | 144369177 | 144369297 | 772751_42001748_Shank2(210274)_1615 | 1 | + | good | GCTTCAGATGGACTCACCTGGGTTCCTGTCGGTCCATGCTTGAAGCACAGTGCTGTCAGAGCTCCAGACACACCAGTCAGTCCACTGCAGTTGTAACTGAGGGACCAGCTAGTCACAACC | 120 | 55 |
| chr7 | 144369297 | 144369417 | 772751_42001748_Shank2(210274)_1616 | 1 | + | good | CCCATGAGCTGTTGACCATGAGCGAACTAAAGCCCTCAGTGTCTATGTCTGTTGCTATTGCCCCAGGGCTGACTGCTCACCCCCAAGCAAGAGGAGTTTTTGCCCAAGGACCCAAGCTAA | 120 | 53 |
| chr7 | 144369417 | 144369537 | 772751_42001748_Shank2(210274)_1617 | 1 | + | good | GGATGGGCATGAGAGCCAGTAACAGAAGCCACCCCATGTCCTCATCCTCAGGGCTGTTTGCAGTGTGATTTAAGGATATCTGGGGGCTTGTGGACCTGTTGTCAGGTCAGTCTTGAGCAT | 120 | 52 |
| chr7 | 144369537 | 144369657 | 772751_42001748_Shank2(210274)_1618 | 1 | + | good | GCCTCCTGAGCACTCCTCAGACGCTGGTTGACATTTGTGCCAGAGCTGTTTGAACGAAGGGGCTCAGTTCTCTGTGTCTCCTCGGCTGCTCCGAGAGGCCTTGTCGTGTGTCCTAGCTTC | 120 | 57 |
| chr7 | 144369657 | 144369777 | 772751_42001748_Shank2(210274)_1619 | 1 | + | good | ATTTCTGTTTCTGGGATTTAAACAAATTGGACGAAAGCATCTTGGAGGAGAGAGGGCTTATCTGGCTCACACCTCCTGGTTGCAGTCTGTCACAGCAGAGACGTCACGGTGGCAGGAGGG | 120 | 51 |
| chr7 | 144369777 | 144369897 | 772751_42001748_Shank2(210274)_1620 | 1 | + | good | TAAAGCCTCCGGTCACAGCACATCTGTAGTGAAACAGAGAGAACTCAGTGCAGGCAGACAGGCTTGCTTGCCTTCTCCTCCTTCATCAGGGCCCTGCCTAAGAAATGGTGCTACCCACAG | 120 | 53 |
| chr7 | 144369897 | 144370017 | 772751_42001748_Shank2(210274)_1621 | 1 | + | good | TGACTGGGCCTTCCCAAAATGACCAACAATCAGGAGAATCTCCGCAGACATGCAAGGCAGGCTGATCTAGACAATCCCTCAAGACTCTCTAAACCAGGGTTCTTAACCTTCCTGAGGCTG | 120 | 50 |
| chr7 | 144370137 | 144370257 | 772751_42001748_Shank2(210274)_1623 | 1 | + | good | TGTGGCCCCTGTGCAAGGGTCGATTGACTACCAAAATCAAGGGATCACAAACCACAGACTGAGAACCGCTGCTCTAGGCGATCCTAGGCTGAGTCAAGACGCAGTTAAAGATGAACCATC | 120 | 51 |
| chr7 | 144370257 | 144370377 | 772751_42001748_Shank2(210274)_1624 | 1 | + | good | ACATCTTGACAGTAACATGCAGGCTGCAGATTCTGTGGCCTACGTGTGCTGGTGTTGACTCTCCCTGGCCAACTGCCATTGCCCAGAGCCACAGCTTCCTGGGGAATTCCAGAAAGACCT | 120 | 54 |
| chr7 | 144370377 | 144370497 | 772751_42001748_Shank2(210274)_1625 | 1 | + | good | CCAAAGCAGAACTGCTCCCTCTTTCCCTGAGGATTCCCAATGAAGGAGCCACGGCTCGGGGACAAAGGTCCCAAGACTCTCTGGCCACTAGCATTGACAGGTTACATCCTGTTCAACCTT | 120 | 53 |
| chr7 | 144370497 | 144370617 | 772751_42001748_Shank2(210274)_1626 | 1 | + | good | CAGATCCGGAGTGAGGAGGAACAGCTCCTGCCTGCCTTACCGTGCTGCCTTGATAGCCAGAGCACAGCAGAAGGCAAAGATCACTTGGCCAAGTCATTGAGGCCACCCTTGATAGATGAT | 120 | 54 |
| chr7 | 144370617 | 144370737 | 772751_42001748_Shank2(210274)_1627 | 1 | + | good | GTCCCATCTCCTTGGAGGAGAGGAGCAACAGCCTCCATCTCCCACGCTGGGCCTTGTATATACTGGGCAACTGCTTTACCATTGAGCTACATCCACAGCCCGTAGCCCCATTCCTATGGA | 120 | 55 |
| chr7 | 144370737 | 144370857 | 772751_42001748_Shank2(210274)_1628 | 1 | + | good | CATCCACAAGAGGACAAATGTTATTGTAGAATTGAGTGAGAGTATACCAGGCACATCCATCTTTCTGAGATTGTAGCAAGATACTGTCATCTACAGAGTCCATAGTTGAGTCGAGATTCA | 120 | 40 |
| chr7 | 144370857 | 144370977 | 772751_42001748_Shank2(210274)_1629 | 1 | + | good | CAAACAGTAGCTTGTTATTATCCATAAGATGGAGAGCCTGTCACAGACAGGTATTATAGTGTGAGCCTGAGCCTCGTCTGGAGTCTAATGGTCACCAGAAATGAGATTAGTGAGGTTGTC | 120 | 44 |
| chr7 | 144370977 | 144371097 | 772751_42001748_Shank2(210274)_1630 | 1 | + | good | TTCCCTAACTTACTTTGACTTAATTACATTACATCAACAATGAACTGTCACAGGTTCAAGGCCCATGGGTGCCCATTTCTGAGGTTGTTATTTTGAGTGTGTTGTTGTTGTTGTTGTTGT | 120 | 38 |
| chr7 | 144371097 | 144371217 | 772751_42001748_Shank2(210274)_1631 | 1 | + | good | TGTTGTTGTTGTTGTTGTTTTTGGTTTGGCATGTTTTTGGTTTTTGTTGTGAGATGACCTTGAATTTGCAATCCCCTATCTTCATCTTTCAGGTAGCAGAGATTCAGGTACACTCATCTG | 120 | 38 |
| chr7 | 144371217 | 144371337 | 772751_42001748_Shank2(210274)_1632 | 1 | + | good | AGAGACACTTTTCAAGCACCACAGAATGCTGTGACCCTAAAATGAGGTTGCAGCCCAGGTACCAGTCAGCTGCCCCAGAGGTCTGACCATGGGGACAGAGGGAACTGTACCTCAACTTCT | 120 | 52 |
| chr7 | 144371337 | 144371457 | 772751_42001748_Shank2(210274)_1633 | 1 | + | good | GTTTTTGAACATCTGGGAACATCAGGAGATGTGCAGACCCTGGCAGAGCACAGGCTGTCAGCCAGCCTCTACACAGTCCCAGATGCTCCCACCAGAGATGCTACCTGAGAGCAGGGAAGG | 120 | 56 |
| chr7 | 144371457 | 144371577 | 772751_42001748_Shank2(210274)_1634 | 1 | + | good | GGGGAGGAGAACAAGAAGGGACAGTCTCCAAAAGAAAGTCCCCTGTCTGGGGCAGAGCAGTCCCAGGAGTAAGAAAGGGGTGCTCCAAGCATCTGGCATGCATACACAAAGGTGGGAGAC | 120 | 55 |
| chr7 | 144371577 | 144371697 | 772751_42001748_Shank2(210274)_1635 | 1 | + | good | ACCACTCCAGAAGGTGGCAGCCATGAGAGAAATACAGACTGGGCTCTCAGGCAGGAAGAGGCTGGGTTAGAAGTTATTTGTGTGAGGTCTGGGCAAAGAATGTGACTGTTTTCCACCCTT | 120 | 50 |
| chr7 | 144372057 | 144372177 | 772751_42001748_Shank2(210274)_1639 | 1 | + | good | TCCACAGGCCACGTGTATTGACTGATGCTTTCAAAGCTGTGAGCTAGATAAGCCCCCTCCCCCTACTGTTCTATTCATAGTGATGAGAAACAATCAACCATGCCTGTGCCGGGCCTCAGG | 120 | 51 |
| chr7 | 144372177 | 144372297 | 772751_42001748_Shank2(210274)_1640 | 1 | + | good | TATGTAGGCAAACTCTCCATCCTACCCGGGCAAGCGCAGGAATAGATATACAAGATTGTGCCCAGGACCTTCCAGAGTTGGCAGAATCTCAGGGGCAGCAATGAAATAATGGACCCCACA | 120 | 50 |
| chr7 | 144372297 | 144372417 | 772751_42001748_Shank2(210274)_1641 | 1 | + | good | CTACAAGACCAGACCTGGGGTAAGCCTAGGCTACCCAGCTGCTGCACAGACCAGCAGAGCACACTTCCAGGGACCTCCAAGGGTCTGACTCAGCCCAGCTGGCAGACAGGCAGCAAGGAC | 120 | 61 |
| chr7 | 144372417 | 144372537 | 772751_42001748_Shank2(210274)_1642 | 1 | + | good | TTCACAGAGTCGGGAAACCCCAGTATGAAGGGCAGAAAGCAAAAGGGTCTAGTTCTTTTTTTTGTAATATGTATTTAATTATATTTTACTTTGATGGTAGTGGTGGCGTTTGTGTATGTG | 120 | 37 |
| chr7 | 144372657 | 144372777 | 772751_42001748_Shank2(210274)_1644 | 1 | + | good | CAAACTCGGGTCCTCTAGAAGAAAAGTGTGCTCTCTTCACTACTAAGCCACCTCTCCAGCCCCAGGGTCTGGTCCGTAAGGAAGGCCTACAGAACAAGGCTGGGCATCTGGGTGGGAGAC | 120 | 56 |
| chr7 | 144372777 | 144372897 | 772751_42001748_Shank2(210274)_1645 | 1 | + | good | AATGATTCTATGACCAAAAATAAGATTCTTCTTTGTCCCCACTTCTTGGCATAGAACCAACACCCAAGGTATTTTCCTGTGTAGTCTGTTTATAAGATGTTTTAGGGGAAGGCCCTCCAT | 120 | 39 |
| chr7 | 144372897 | 144373017 | 772751_42001748_Shank2(210274)_1646 | 1 | + | good | AAAATCAAAATGTTGGAGTCTCCTACCCACCCCTACTCAGCCGCCTAGAGATGTCATCAATCATGACCAGCCTGGGATTTAGCCATGGATGTCCAGGCCAGCCTAAATGAGAGCAGAGAC | 120 | 50 |
| chr7 | 144373017 | 144373137 | 772751_42001748_Shank2(210274)_1647 | 1 | + | good | TTGTTTAATTAGAAGTGAAGTCCCACCATAGAACCTTAAAGATGCAGTAAACTACAGGTGTGGACAGGGAGTCAGAGACTCAGAAGAATACACAAGTTATTTCAAAAGCAATAACCTTTG | 120 | 38 |
| chr7 | 144373137 | 144373257 | 772751_42001748_Shank2(210274)_1648 | 1 | + | good | GTGTCAAGAATCTGTAAAACTCCATCTGCCCAAGACGGGGTACGCAGCTGCTGAGCGCACTGCTCACTGAAGTGTGCAAGGCTGTAGCTCTGGGAGGCTGCAATAGCCCTTGGCACCAGG | 120 | 57 |
| chr7 | 144373257 | 144373377 | 772751_42001748_Shank2(210274)_1649 | 1 | + | good | AACAAGAAGTGAGCATTCCCCAGTCTCTCGCAGATTTAGCATTTGCCGCTGAGTCTCCATTTGGACCTAGACAACCTTCACAGCCACAGCAAGAACTCTGAAGCAGGGCCGGGGAAGCGC | 120 | 53 |
| chr7 | 144373377 | 144373497 | 772751_42001748_Shank2(210274)_1650 | 1 | + | good | CTGGGTTCGTGGCACCCAGAGGGGACATGCAAACCCTGGGCTTTGTCATGGGCATTGCCAGGTGCTGTTTCTGTTATCACATGGGGAGTGCCGTCTCAAGTCTTGTGAAGAAGAATAGTG | 120 | 53 |
| chr7 | 144373497 | 144373617 | 772751_42001748_Shank2(210274)_1651 | 1 | + | good | GGGGATTTGTAAGCTATGAGAGCCTATAGGCCCAAGCCCTTTGAACCTGAAGGGTCTTGGGAACTCTGCAGTGGGTGTCAGAGCAGGACTTGATTGTGCAGATTCAGCAGGTGTCACCCC | 120 | 53 |
| chr7 | 144373617 | 144373737 | 772751_42001748_Shank2(210274)_1652 | 1 | + | good | TGTGGTGTCACCAGGAGCTCTTGTAGACACAGACTTGGTATGAACAAACTCAATAGCATAGTAGGTTCCGTGCCCAGGGGTCCTGTGTCCCCTGTCTCATTATCCCAGCACTCTTGAGGG | 120 | 52 |
| chr7 | 144373737 | 144373857 | 772751_42001748_Shank2(210274)_1653 | 1 | + | good | GTTGACATTATAATTTGCCAGCTCTGTGGAGGAAGCTGAGGCCCAGAGGCCAGGACTCTCAGCTCTGTCCTAGCCTCTGAAGCAGTCACTACAGGAGAGCAAGCACATAGGGCAACCTGG | 120 | 54 |
| chr7 | 144373857 | 144373977 | 772751_42001748_Shank2(210274)_1654 | 1 | + | good | CTCACCTACATAAATGAGGACTCACCTCCATATATCCCAGTGTCCATATGAGCAAAGGCTGAAGTGCCAAAGGTGCCATTGCTAGAGGCCTAACCAGAGACCTAGGGCGTATTGTAAATA | 120 | 47 |
| chr7 | 144373977 | 144374097 | 772751_42001748_Shank2(210274)_1655 | 1 | + | good | TGAAATACCACGGGTGATATCAATGGCCAGGTTACAAGGACAGAGATGCATAGTCAGTGGTTTCTGGAGCTGATTCTGTCCTTTCTCAGGTAGAAAGTGGCCAGCTGTGTGATGAGGGAT | 120 | 48 |
| chr7 | 144374097 | 144374217 | 772751_42001748_Shank2(210274)_1656 | 1 | + | good | TCCAATCCAAGGTACACCCATGTGCTCCCATTCATCCAGAACCTACCAGCCTCTGCCTGCCTGGCTCTCACACTGGGCATTGGCCTTTCTTCTATTATACAGACACAGGTTTTCAAAACA | 120 | 49 |
| chr7 | 144374217 | 144374337 | 772751_42001748_Shank2(210274)_1657 | 1 | + | good | GACTGACACAGAGCCCTGGCTCCTGGGGCTGAGACATCACTGTTGATTCCTGGGTTGGTCCAGAGACTTTTCAGGCAGAAACTCTGACAGGTGCTCAGGGTCCCTGGTGACATTTGCTTC | 120 | 55 |
| chr7 | 144374337 | 144374457 | 772751_42001748_Shank2(210274)_1658 | 1 | + | good | CAAGGTCAACTGAGGTCTCCCCTCTGTGGTAGTGCTGAGACAGGGGCTCCAGGTACTAGTATAGATGGCAGAAACACTCTATTTCCTGTCAACAGAAGGTCACATCATCATAGAGGTGAC | 120 | 49 |
| chr7 | 144374457 | 144374577 | 772751_42001748_Shank2(210274)_1659 | 1 | + | good | TTGAAGACACAGCCTAAGAAGGTGTTCACTCCCAGGGGAGCTCTCCATGTGGATTCCAGTTGGTCATGTCCATGGAAGGCCTTTTGGAACCCAGAGGTAACACCTTGCTTGGGCACCTTC | 120 | 52 |
| chr7 | 144374577 | 144374697 | 772751_42001748_Shank2(210274)_1660 | 1 | + | good | AGACCTCAGTTCCAAGTAAGAGGCTCTCTAATCCCCAGTCCATCCCACCATTGACTCACTATTGGGCTCCTGCCCCAAGTTTTCTAGACACAAGCCAGATGGTGGAGTGTGTGGGGTGGC | 120 | 53 |
| chr7 | 144374697 | 144374817 | 772751_42001748_Shank2(210274)_1661 | 1 | + | good | CCTGGTCTCTGCAGGTCCAGCTGTGGGAACAGTCTGGCAAATGGAAATCTCTCACCTGCCCTCCCAGCATGAAGCCCACGGAGAAAGTGTTCCCTTACAGGGCCTCTAGAGAGCTGGGGA | 120 | 57 |
| chr7 | 144374817 | 144374937 | 772751_42001748_Shank2(210274)_1662 | 1 | + | good | GATTATCCCAGTAAAGTGCTTTGCTTGGCAAGCATGAGAACCTGAGTCCAATCTTAGGTACCCACATATAACGCAGGGCATGGCACTGCAGGCTTGTAACAGTGCAACAGAGATACAGAC | 120 | 48 |
| chr7 | 144375057 | 144375177 | 772751_42001748_Shank2(210274)_1664 | 1 | + | good | GAGAGAGAGAGAGAGAGAGAGAGAGAGAGAGAGTTAGTTAGTTAGTTGGGGGAAGACTGGTCTATGCTATAACTCCCCAAGAGGGGCAGCTGCATAGCCCAGGGGCTGTCATCACCAGGTCA | 120 | 52 |
| chr7 | 144375177 | 144375297 | 772751_42001748_Shank2(210274)_1665 | 1 | + | good | AGAGTTGTCCTCAGGAGCTGAGCTACTCCCTCCAGTTTCTGAATCACACACAGATACTTAGAGCATCTCCGCTCAAACATCAGGCTTGGGGAATCTGAGGACTGATGTCCATTGGGCCCC | 120 | 52 |
| chr7 | 144375297 | 144375417 | 772751_42001748_Shank2(210274)_1666 | 1 | + | good | AGGCAGTTCCTCCTTCAGTTACTCCCAGATATAAGCATTTTTAAACCATGGCCCTAGAATCCTGGTCCCTTGGAGCTCCAAGAAAGGGAAGAGGGTTGAGCCAGTTCTACCCCAGCCCTC | 120 | 51 |
| chr7 | 144375537 | 144375657 | 772751_42001748_Shank2(210274)_1668 | 1 | + | good | ATCCTGGATAAAGGGTTTACATGCAGTGGCCTGGGCCATCACCACCATAAAAAGCACCAGCATGCCTTCAGGAGGTTCTCAGGAGGGGGTGGCTCAGCTCTTAACCCTCTGGTGACAGCC | 120 | 55 |
| chr7 | 144375657 | 144375777 | 772751_42001748_Shank2(210274)_1669 | 1 | + | good | CCTGGGCTATGCAGCTGCCCCTCAGCTGCTGCAGAGCCATCCTCCAGCACTAGTCCACGTGGTGATGCAACTGGGGTTTATTTTAGACCAAAGCCTTCTCCTCAGTCCAACACACTTGAA | 120 | 54 |
| chr7 | 144375777 | 144375897 | 772751_42001748_Shank2(210274)_1670 | 1 | + | good | CTTTAATAAAGCAAAGAGTGACAAACCCAGCTTGCCTTTACAGACATTTCCAACATACAGGTAAGACTTGGCCTGTATTAGGGAGAGGCTGGGAAAAGGGCCTCAGGGCCAGTGCCTCCA | 120 | 48 |
| chr7 | 144375897 | 144376017 | 772751_42001748_Shank2(210274)_1671 | 1 | + | good | ACAGGCAATGGGCTTGGCTGGATGAGGGGTTTGGACTGGGATCACTGGTCCAGCAATCGGGACAGGCCCTACTAGTGGCCTGTGCCTGGTACTAGAGGCATTTGTGTGACCCAGTGGTCA | 120 | 57 |
| chr7 | 144376017 | 144376137 | 772751_42001748_Shank2(210274)_1672 | 1 | + | good | CGTGGACACCTACAGAGTGTGAGCTATTTCTATGCTGGTCTACCCACATCAATGTAGCCCAGCATGACGTAACAGGGAAGCCCTACCTACTGCAGGTACATGGAGCTCACACAGGATGGG | 120 | 52 |
| chr7 | 144376137 | 144376257 | 772751_42001748_Shank2(210274)_1673 | 1 | + | good | AGCTGGGCCTTGTGAGGCTGACATGCTTCTGAAAACACCTGTTATCACATAAACCTGCATCCGGCAAGCCCTGGTTCTCACAGATGGTGGGCCTGGAAAGGTGGCATTGCCCCTCTGGCC | 120 | 56 |
| chr7 | 144376257 | 144376377 | 772751_42001748_Shank2(210274)_1674 | 1 | + | good | AAGAGTCAGAGGGAGGAGGACCAGGTTAAAAGAAAACAGTCAGAACCCTCCAGCAGCTCTGTATTCTATAGTGAGGTATGCAGCCCAGGGCTGCAGTCTAGGTGCATGATGGTACGCCTG | 120 | 52 |
| chr7 | 144376377 | 144376497 | 772751_42001748_Shank2(210274)_1675 | 1 | + | good | CTTGCCTGTCCAGCCTCAGTGTTGAGGTTAGGGCTAGAGAACTACAAGGTGGAAAGACAGATGGTGCAGCAGGCTGAGAGGTCACCAGCCCATCAGCATTAAGCAAGTCACCAAATGAAC | 120 | 52 |

| | | | | | | | | | |
|---|---|---|---|---|---|---|---|---|---|
| chr7 | 144376497 | 144376617 | 772751_42001748_Shank2(210274)_1676 | 1 | + | good | AAACTGTGCCCAGTCTCAGATCTGTCACCCGGGAGCTCAGTTGGTCCTCAGTCAGAAAAGTAGATTCACCAGAGGGACAGAAGGAAGAAGGCATGTGGGCATACATGCAGTACACATACA | 120 | 50 |
| chr7 | 144376737 | 144376857 | 772751_42001748_Shank2(210274)_1678 | 1 | + | good | CACCTAAATGTCCTTCTAGGCATACTCAGGGGTGTGTCCCCCAGGTGTTTGCAAAGCCATCAGGCTCACAGTGAGAGTAAACCGTTATACAATGTTCCAGTGTATCCTGCGTTGGAGTGT | 120 | 49 |
| chr7 | 144377097 | 144377217 | 772751_42001748_Shank2(210274)_1681 | 1 | + | good | TGGCCTGGGCCATCACCACCATAAAAAGCAAATCACACATTTGCACAGTGAATTTAATTTTTTGAATTAAATTTAAATTGCACAGTGAATTTAATTTTTTGAGAATTTCATATGAGTACT | 120 | 30 |
| chr7 | 144377217 | 144377337 | 772751_42001748_Shank2(210274)_1682 | 1 | + | good | GACTATATTTATATCATTTCTGCCCTTCCTTGTCCCCCTCCAACTCCTCCATCCCCAGCTCCCTCTCCTAACCTCTTTTTCTATAATGATGATTGTTTTATACACACACAACCTACTGAG | 120 | 42 |
| chr7 | 144377697 | 144377817 | 772751_42001748_Shank2(210274)_1686 | 1 | + | good | ACACACACACACACACACACACAATCTTTTCTCTTCCTCTTCTGAGATATTTCCTGAGCCTTGAGTATTCTTTGTGTGGTGGTGGTGGGTGGTGGGGAGACCCAGAGCATCCCACATGCT | 120 | 49 |
| chr7 | 144377817 | 144377937 | 772751_42001748_Shank2(210274)_1687 | 1 | + | good | AAGCAAGCTCTGTGCAGGCAGAGGTCAGCGATCAGCCCCACTGCACAAGACTCACACTGTTCCACCACATGGATGACATTACTCCCAGTGGCAGTGTTAATCTGACAAGCAGATTAACAC | 120 | 51 |
| chr7 | 144377937 | 144378057 | 772751_42001748_Shank2(210274)_1688 | 1 | + | good | TAAACTAATGAGATTTCATGGGACTATGTCACTAGGTACCCACAGGCACCCTAGGACTGAGGCAGCAATTTTCCCCATTCTACAGAAAAGGAATTTAAGGTAGCACGTGGGGTGTAGACC | 120 | 46 |
| chr7 | 144378057 | 144378177 | 772751_42001748_Shank2(210274)_1689 | 1 | + | good | ATCCTTCTTGTGCCTAGAAAGTCGTGCAGAGGAAGTCCTGCAACCTAGGATGAGCACACAGACAGACAGATGCATCATAGAGAAGCAGGACCACTGTGCCTGCCACTCTCCACCTACCTT | 120 | 52 |
| chr7 | 144378177 | 144378297 | 772751_42001748_Shank2(210274)_1690 | 1 | + | good | CCCAGTGTGAGGATGCTAAAGATGTCAGAGAGGGGTGAGGATAATCCTACACTCAAGACAGCCACAGCCTCCGCCCCTGATGAGGTGTGTTTGCTAGAGAGAGTGGCCACAGCCCTCAGC | 120 | 56 |
| chr7 | 144378297 | 144378417 | 772751_42001748_Shank2(210274)_1691 | 1 | + | good | AGCTCCTGTGAAGCTATGACCTTGTGATTTGCAAGGAGCAAGCAGCTGGCTCACACCCGATGATCAGTAGCCTGCCAAATGCTCTTAGAGCCACGCTCTGTCTGCCTTAGGGCCTTCACG | 120 | 54 |
| chr7 | 144378417 | 144378537 | 772751_42001748_Shank2(210274)_1692 | 1 | + | good | GCTCTTGGAGAGGTCTGGTCAGCTCTACTGCACAAGTCTTCAGACTGGCCCACTGTACAGATGATGTCATGGTCAGCCCACCAAGGGCTATAAATGCACAGCAGCAGATGGACCTGACT | 120 | 53 |
| chr7 | 144378537 | 144378657 | 772751_42001748_Shank2(210274)_1693 | 1 | + | good | CAGGAGACAGCTGGGCAAGCTGCTATGGACAGCCACACTCTAGGTCCTCAGCTCATGCTGGCTCTATGTTGAAGGGCTAGTCAGGTCCCTGTAGCGGTTCACCTCTCCAGCCATACCTGG | 120 | 57 |
| chr7 | 144378657 | 144378777 | 772751_42001748_Shank2(210274)_1694 | 1 | + | good | GGCCTTAAACCCGGCACTCTGTGGCCTTACTGAGTTGGAGTGATGGGAATGGCTCAGCTTTGGCAGTGAGGGAATGAAGTCTGCTGTAGAGTGATGTCACCCTGGCTGTACAACACCATC | 120 | 53 |
| chr7 | 144378777 | 144378897 | 772751_42001748_Shank2(210274)_1695 | 1 | + | good | CCTACTAGACATGTTTCCACTCCTTTGGACTGAGATGCTAGAGCTCATTGGGAGAAGAGGCAAGGCCCCTCAGGGTCCTGATCCACACACAGCTCATCCCCAGCTAGGCAGCTCAGCCTT | 120 | 55 |
| chr7 | 144378897 | 144379017 | 772751_42001748_Shank2(210274)_1696 | 1 | + | good | TGTCAGAGGATACGGGAAAGAGGGCTTAGACAAGTGGGACAAAGGGATAACTAAGGATGAGCAGAGTCTGATGGGTCAGGCAGGCCTGTCTTCCCAGCTACTCAAGAGACTTAGGCAAGA | 120 | 51 |
| chr7 | 144379017 | 144379137 | 772751_42001748_Shank2(210274)_1697 | 1 | + | good | GGATAGCAAGTTCAAGGCTGGCCTGAATTCTAGAGTGAGAGCAAGGATAACCAAGATAACTTAGTGAGTGAGACCTTATCTCAGAATCAAAAATGAAAAAGAATCTGCTAGGATATAGCT | 120 | 39 |
| chr7 | 144379257 | 144379377 | 772751_42001748_Shank2(210274)_1699 | 1 | + | good | ATGGTGATGGAGGTGATTATAGCAGTAATAGCAACCAGGAATGAAGGCAGCTCAGATTTGACAAGGCAGCCATTGCTCCAACACAGCCTGACAGCATGTGGGGAAAGCTGCTTAAAAGCA | 120 | 48 |
| chr7 | 144379377 | 144379497 | 772751_42001748_Shank2(210274)_1700 | 1 | + | good | TACCAGAGGGAGGTATTAGGTGAAGATGGATGGCCACAGCCCCCAGCAGCTCGTTTCTAGGGGCAACAATGGTTGCTAAGAGAGGAGCTGTTGCTCGGGTTTGGGGGGGGTGGCATTTTG | 120 | 56 |
| chr7 | 144379497 | 144379617 | 772751_42001748_Shank2(210274)_1701 | 1 | + | good | GACCCAAGGAAAGAGCTGTTAGTGGATCCCTAGATCTCTGACAAAAACTCCTGAGCCCACCCCACCCCCACCCCCGTAGCAGTGAGCAGGGGAATGGGCGGGGGTTGGGGGGGGGGGGCT | 120 | 63 |
| chr7 | 144379617 | 144379737 | 772751_42001748_Shank2(210274)_1702 | 1 | + | good | TTGCTCTGTTCCTATATTCTTTGTGCTCTCCACAGTGGCTGAGGGACTCACACCCACTCTGGGTGGATGATAGGGAAGACGGGAGGCAGCTTTTAGAGAACATTTGTGAACAACCTATAC | 120 | 48 |
| chr7 | 144379737 | 144379857 | 772751_42001748_Shank2(210274)_1703 | 1 | + | good | CAGCCAGCCATCGGGCAGGGCAGGGCAGGGCCCAGGGCTGAGGATGAGCATCATGACCAGGCATGGAGGATCTTAGAGACTTTAGTGGGATATGAGGTATTGTGTGGAGACTCTCAGAGA | 120 | 57 |
| chr7 | 144379857 | 144379977 | 772751_42001748_Shank2(210274)_1704 | 1 | + | good | GATTTCAGGTGAGTAGAAGGAATAAGATCAGGCTTCAGAATGGGGTTAGCCTTTGCATGTTCAACTGATTTCAAAACTCAACGGTGAAGAGGGTAGCAACCCCCAGGCCATCAGGAAACT | 120 | 46 |
| chr7 | 144379977 | 144380097 | 772751_42001748_Shank2(210274)_1705 | 1 | + | good | GCCCAGTTAGTGCCTTATTGTGCCTTTGGAATTCTGCATCTTTGCAAGAATAAATGATTGGGGGGGGGAGTGTCTGTGAGAAAGATGGTAAATCTAATGACTTAAAAACCATGCATCCAG | 120 | 42 |
| chr7 | 144380097 | 144380217 | 772751_42001748_Shank2(210274)_1706 | 1 | + | good | ACTGGCCTGCTGGAATCCGCTCAGACAGCAGGGAAGGCATCTTTAAGGAGAAGACCTGGTGTGCAGAAACAGATGGCCCGCACTCGACTCTGGATGCCACTAAATGGACTAAACAGTCCC | 120 | 54 |
| chr7 | 144380217 | 144380337 | 772751_42001748_Shank2(210274)_1707 | 1 | + | good | AGGGAGCCAGATGCTGAGCACTTCAGAGGAGTATGTGGAGAGGGAACTGCCAAGTGGACCGAAAGTAAGCTTTGCAGGACCAGGTCCCCGAGCAGAAAGCCTGGTGCCTAGAACCTGAGG | 120 | 57 |
| chr7 | 144380337 | 144380457 | 772751_42001748_Shank2(210274)_1708 | 1 | + | good | GATGGTGGGGAGTTGGAGCAGTATCTCCAGCGTGCCCACCTAGACCAGTGTGCTGCATCCTGCATGGTCTGGTCCTGGAACTCAGGGATGGAGAAATGGTAGACACTGAAGAAACCACAG | 120 | 55 |
| chr7 | 144380457 | 144380577 | 772751_42001748_Shank2(210274)_1709 | 1 | + | good | ATACATATACTGAAAGTATTGGATCCAGTGGGCTGTGCTTGTACTGGTGGTGGAGGGCACCTACTACGCACTCCAGGAAAATCAGGATGTGTGTTATGAACAACACAGGAGGAGGGTTTA | 120 | 47 |
| chr7 | 144380577 | 144380697 | 772751_42001748_Shank2(210274)_1710 | 1 | + | good | GCCTGTCTGTGTGGAAGGTCTTATAGGGCCAATCTCAGGTGGGAGGAATACAGGGACCAGCTGTGCTGCTGACTTGTACACACTGTCCACATCTGTATACAGACCAAATTGTATACAGGC | 120 | 50 |
| chr7 | 144380817 | 144380937 | 772751_42001748_Shank2(210274)_1712 | 1 | + | good | GAGGATGAAGAAACTGGACTGTTTGCTTCAGCTGTTTGTGAGACCAAAGGACTGAAGTCACCACCAGATCCACTGGTGGGGGCCGGAGTGGGCAGGCTGTGTGCCACTGACTCCAGGGCA | 120 | 57 |
| chr7 | 144380937 | 144381057 | 772751_42001748_Shank2(210274)_1713 | 1 | + | good | AGGAAGGTGTGGCCAGCCTGCTGCCCTCCTCCCAGGTGGGGAGGGGATGCACACGGGCTCAAATTCCTTCTCCATTACGTTGTTCTTTTGTGTTGTTGCTCCTGTTGTTTTGCGGGGAGG | 120 | 57 |
| chr7 | 144381297 | 144381417 | 772751_42001748_Shank2(210274)_1716 | 1 | + | good | TGACAGGTGTGTATCAGCGTGACTGGTTTATGAAGTACTGGGGATCAAACCTAGAGCATCGTGCATGCTAGACAAACCCACTACCAGCTAAGCCGTATCCCCACCCCACCCCCAATCTCC | 120 | 52 |
| chr7 | 144381417 | 144381537 | 772751_42001748_Shank2(210274)_1717 | 1 | + | good | TTTCTCCTTTTTAACAAAAAGATCTAGGTATTTGGTTCTAAAGTTGCCAAGGATGGGACTTGTGTGAGGAAAGATAATCAGACATTCAAGGACATTCCTAGGTACATAGGAAGTTTGAGG | 120 | 38 |
| chr7 | 144381537 | 144381657 | 772751_42001748_Shank2(210274)_1718 | 1 | + | good | CCAGCCTGAGCTACATGAGACATTGTCTAGAGAAATGGTGGGGAAGGGAGGAAGTGGGGAGAGGGAGAGGAACCCATCCCAGTTTTAATGGTTTGCTGTCCTGTTCTGAACAGGACAGGA | 120 | 52 |
| chr7 | 144381657 | 144381777 | 772751_42001748_Shank2(210274)_1719 | 1 | + | good | GGATCCTGAGCCGGGAGACAAACACTGGATAGAAATTAAAGCCGTGCACTCCATGGTCTTGTTGAATAATCATGTGTGCATACTAGCTCGTGAATTGCGAGAAAACACCACGCTCCTGCA | 120 | 48 |
| chr7 | 144381777 | 144381897 | 772751_42001748_Shank2(210274)_1720 | 1 | + | good | AGACTCGGGAACAGGGGAGGAGGTGGGAGCACGTGGAGACGCCCTCAACTGTCTCCACTGTGTTCCTAGCAGCCTAATGCTAGTCTGAAGCAACACGGCTTGTTTACAGAGGACAGTGAA | 120 | 55 |
| chr7 | 144381897 | 144382017 | 772751_42001748_Shank2(210274)_1721 | 1 | + | good | TGCAAGCTTATCTGATGCTTTAAAACAACAACAAAAGTCTTTCTGAGTTCATTATGATGAAGGCACCAGATCGTGAACCAGAAATAATAAACGAAAGAAGAACTACACTGTTCCCTTGTG | 120 | 37 |
| chr7 | 144382017 | 144382137 | 772751_42001748_Shank2(210274)_1722 | 1 | + | good | CTTCTCTTTGCAAGGTCCCCAGACAGGGCGAGTTAAGAAGTGCTCATGCTCCCCGCCCCCTCCCTCTCCTCTCTTAGTTTTCTTTCTATGATGTGTGTGTATGTGTAATCTCTCCTTCTA | 120 | 49 |
| chr7 | 144382377 | 144382497 | 772751_42001748_Shank2(210274)_1725 | 1 | + | good | GGTATTCTAGCTGCCATGGGCAACTGTATTTATGTGCACATACCTCTACAGACATTTTATTTTGTAATTAGTTTTGCTTCATAGGGGGCAAAGTGAGGGCCAGGGAGATGGCTCAGTGGG | 120 | 45 |
| chr7 | 144382497 | 144382617 | 772751_42001748_Shank2(210274)_1726 | 1 | + | good | CAACGGTGCTTACCGCCAAGACTAACCTGCATCCCAACCCTGGACTCGGTAGAAGGAGAATACCAACCCCTGCAGGTTGTCCACTGACTTTCACACATGCAATTATAAAGTCTAAATTAA | 120 | 47 |
| chr7 | 144382617 | 144382737 | 772751_42001748_Shank2(210274)_1727 | 1 | + | good | AAAAAAATAATAATAAAAAGTACAAAGGCCCAGATTTCCTCCCCGTTTCTAATTGGTTATGACCTGTTGGTCATGTGACAGCTGCCCTCTTCATTTCTCCCAGTTGGAATCTGTTATGTG | 120 | 39 |
| chr7 | 144382737 | 144382857 | 772751_42001748_Shank2(210274)_1728 | 1 | + | good | ACTGCTGACAAAGTGCGTGTTAGTAACTGAGCATCCTCTGCGCTGTACATCCTGTGGGTTCTGAGCTATGTGTGCTGACAATTGTCCACCATTACAGTACCACACAGATGGTCTCATTAG | 120 | 48 |
| chr7 | 144382857 | 144382977 | 772751_42001748_Shank2(210274)_1729 | 1 | + | good | GCAGAAAGTCCACTGGGTTCCGTCTGCTCTCCCACAAACTAGAACCACTGACAGCCAATGCTATCTCTGCAGTTTTCTGTAGTCTTGAACTTGGATTCAGGGCACAACCTTTTTATTAAG | 120 | 46 |
| chr7 | 144383097 | 144383217 | 772751_42001748_Shank2(210274)_1731 | 1 | + | good | AGTGGTACCACCAAGAATGTGTTCGTCCACTAGTCTACCAAAAGACATCATAATTGCTTCTAAGTCTTAGCAATTAAGAGTAACATTTCTAGAAATATCCATATTCACGTGTTAGTGTGG | 120 | 37 |
| chr7 | 144383217 | 144383337 | 772751_42001748_Shank2(210274)_1732 | 1 | + | good | ACCTGAGTTTCTAGCTTCCTTGGGTAAATACCAATAAAATGGGTGGCTGGGTTATACAACAAGAATATGTTCCATTATATAAGAAACTGGCAGACTGGGGGCTGTGAGCCCCCACCCCTA | 120 | 45 |
| chr7 | 144383457 | 144383577 | 772751_42001748_Shank2(210274)_1734 | 1 | + | good | ACAGTACAAATCCAACTCATGGGTTATAAAAATCACATTTTTAGTTTTTTTTAAATGAGGAGATAAAGTTGGGAGAGGGTGGGAAGGATGGATGGGAGAGGAGTATGATCCAAATACACT | 120 | 37 |
| chr7 | 144383577 | 144383697 | 772751_42001748_Shank2(210274)_1735 | 1 | + | good | GGATAAAATGAACAAATTATTACACTATTACATTTTTTTCAAAGAAGCTGCTAGTATATTCCTACAGAGGCTGGACCATGTACAGGATGGGGGGTTCCTCAGGGCACAATGTTTTCATGA | 120 | 38 |
| chr7 | 144383697 | 144383817 | 772751_42001748_Shank2(210274)_1736 | 1 | + | good | GGTTGTCCTAATGGGTCCATAACAGTACCCCATGTCCTATGAATCTACAACTTCTTAGTACCACATTGTTCAGGTTCAGCTAAGAAATGTTTCTCTCTCTCTCCCCCCCCCCCTTCTCTC | 120 | 47 |
| chr7 | 144383817 | 144383937 | 772751_42001748_Shank2(210274)_1737 | 1 | + | good | TCTCTCTGTTTTTTCCTGGCTGGTGGTGCTGTTTGGGGGAGAATATGTATCCTAGTTACCTTTGTCAAGACACAGTGTAAAGTCATCTGAGAAAAGAGCCTCAATTGACATAAAAGGGCC | 120 | 43 |
| chr7 | 144384177 | 144384297 | 772751_42001748_Shank2(210274)_1740 | 1 | + | good | ACTAGAACAGTGTGGAATTTTTGAGGTCATGGGGCCAAGCAGAAGATAGCGGACTACTGGTTGGTAAACCTTGAGAATAACTCTGCTTCCTGCCCTGCTTCCTAGTCTGCAGAGATGTAA | 120 | 47 |
| chr7 | 144384297 | 144384417 | 772751_42001748_Shank2(210274)_1741 | 1 | + | good | GCCAGCAAGCACACCTAGCCCTTCCTGCCCTTGTGCCTTCCCTGCCAACAGGAGCCCTTCCTTCCTTGGCACACTTATTGGGCTACTCGTGCATGGAGCCGGCCTTTGCATACTTATCAG | 120 | 57 |
| chr7 | 144384417 | 144384537 | 772751_42001748_Shank2(210274)_1742 | 1 | + | good | CCCTGTGTCTGTCTTTGTTGGTGGTTATCTGAGCAGAGAGCTCACCCATTTCTAATTCAGTTGTTTGCTTCCTTACTGTTGAAGAGTTCTTTGGCATTTTAAACACCAGCATATATCTGA | 120 | 41 |
| chr7 | 144384537 | 144384657 | 772751_42001748_Shank2(210274)_1743 | 1 | + | good | TGTGTACTTTGCACAGACTTTCTTCCCCAGTCTATGGCACTTTTATTCTTAACAGTGGGTCTTTCTTCATGCAAACCTTCTGCTTGATGCAACCCACCATACCCACAAGCCCTTCCTAGA | 120 | 45 |
| chr7 | 144384657 | 144384777 | 772751_42001748_Shank2(210274)_1744 | 1 | + | good | CCCTGCTTTGGATGTTTCATTTGTAGACTCAAGGTCACCTGGATTTTCTCTAGCATTATCTCCCAGGAGTATCATAGTGTGGAGTGTTATATTTAGAATTGTGATCCATTTTGGGTTAAT | 120 | 38 |
| chr7 | 144384777 | 144384897 | 772751_42001748_Shank2(210274)_1745 | 1 | + | good | TTCTAAAGGTTTAAGGTCTGTGTCAATGTTCAGTTTCTGCACGTGGGTGTCCGATTGTTTGGCAGACTGTGAGAAAGCTTGTTCTTTCTCCACGGCCTCGCCAGTGAGCCTTTGTCAAAA | 120 | 47 |
| chr7 | 144384897 | 144385017 | 772751_42001748_Shank2(210274)_1746 | 1 | + | good | GATCAGTTGGCCAAACTATGGGGTCTATTCTGGGCTCTGTGTTACGCTCTGCTGACCTTCTGTCAGCTCTCCCACTGGCACCATGCTGTCCCACATCCAGCTGCTGGATAGTGCATCTCT | 120 | 54 |
| chr7 | 144385017 | 144385137 | 772751_42001748_Shank2(210274)_1747 | 1 | + | good | GAGTCTAGTCATCTTAGGTCCGGGCTTTGTCCTGCTCCTGTGTCCCATTTGCTAACTTGATCCGTAACCTTTCCAGGTAAATTTTAGGTCAGTTTCCCAGCATTCACTTACAATGTGTCT | 120 | 45 |
| chr7 | 144385137 | 144385257 | 772751_42001748_Shank2(210274)_1748 | 1 | + | good | GGCTAAAAGTGACTCTGTTAGAAGTCTCTCCTGAATATGAGATGTCTATACGGAGCCCCAGGTATTTGGTGTAAAAATTAACATTAAAGACAAAGCCCAAAACAAACACACTCCAAAAGG | 120 | 39 |
| chr7 | 144385377 | 144385497 | 772751_42001748_Shank2(210274)_1750 | 1 | + | good | GACCTCTCCACAAGGTGTACTGTGGCACATGCACTCACATAGACAATCACAAAATAAGTAAGTTAGTAAAGAACTAAAAGCACATGCCATGAAAAAAATGGGAGGGGTCGCTGTGTGGGG | 120 | 44 |
| chr7 | 144385497 | 144385617 | 772751_42001748_Shank2(210274)_1751 | 1 | + | good | AGATGACGACTATTAAACAAATGTGTGACTCGTGTGCTCAGCAAGATCAAAGAAGATGCCGCCTCCCTGAAGTAAGAACAAAACGCCCTTCTGAAAGGGAACATTCAGAGAACAAAATAA | 120 | 42 |
| chr7 | 144385617 | 144385737 | 772751_42001748_Shank2(210274)_1752 | 1 | + | good | GAGTTCTAGAAAATTTTAAAAACACGATGGCAGATGTGAGTATTTTGATAGATGGACTAGGAGATGACATCGAAGAGAACCTTTGAGAACAGAAGAGATGGTGAGGGATACAGGGAGGAT | 120 | 40 |
| chr7 | 144385737 | 144385857 | 772751_42001748_Shank2(210274)_1753 | 1 | + | good | GGGAAGACACGGGCAGGGGATGATTAACAAATCACAGGGGAAAACAGTCACACTATACACACCAGACACACCAGACACCCGTGAAGCTCGGGATGTGGACAGAAGGGACTCAGAGAATGT | 120 | 52 |
| chr7 | 144385857 | 144385977 | 772751_42001748_Shank2(210274)_1754 | 1 | + | good | CACACAGTAGTAAACCGTCGCAGTGGGGCTCAGGAGATAGCTCGGCCAGAGACACACTTGCTTTGTTAGCACAGGAAGCTGAATCTAATGCCCAGAACCACATCAAAAACCCTGAATGAA | 120 | 49 |
| chr7 | 144386337 | 144386457 | 772751_42001748_Shank2(210274)_1758 | 1 | + | good | ACACACACAAAACTATCATATCTCTTAGTCCCTCTGTAGCAGGCCTCACATCTGAAATAAATGGAAGCCAGCCTTCAGAACCCTCAAGATCACTTGATATCTTCATTGATTTGTCTAACT | 120 | 40 |
| chr7 | 144386457 | 144386577 | 772751_42001748_Shank2(210274)_1759 | 1 | + | good | ATCCTATCAGTTTGAGAAAGGGGTGTGTGATGGTTAATATTGTCAACTGGGTCTGGAATCACCCAGGAGCACTCTGGACACATCTTTAAGGGATTATCGAGATTAAGTTCCCAGGAGCCA | 120 | 45 |
| chr7 | 144386697 | 144386817 | 772751_42001748_Shank2(210274)_1761 | 1 | + | good | GTGCCCAGCTGCCTCTTGCTTCTGCAGCCAGGCCTTCTCTACAGTGGTGGGCTGTACCCCTAAACCATGACCAAGATAAACGCTTCCTCCTTTATGCTGCTTCTGTAGGGCATTCAGTCT | 120 | 53 |

| | | | | | | | | | |
|---|---|---|---|---|---|---|---|---|---|
| chr7 | 144386817 | 144386937 | 772751_42001748_Shank2(210274)_1762 | 1 | + | good | CAGCTTTAGAAGAAAGGATTAGTGGGGGCAGTGGTGTCATTTCCCACTCTGTCTGTGGGTTGCTGAGCTCTGTGTGTATTTATGCAAAAGTTTACACATGTTTACACATGTAGGGGGTCT | 120 | 45 |
| chr7 | 144386937 | 144387057 | 772751_42001748_Shank2(210274)_1763 | 1 | + | good | TTGATATTTGACACATGTTTATCACTGCCTTGATATCTTCCCTCTTTACTTCCCATAATGCTCTTTCTTTCGTTTTAAATTTTATTACAGGTATATGATGTTTTGCATGTACATCTGTAC | 120 | 32 |
| chr7 | 144387177 | 144387297 | 772751_42001748_Shank2(210274)_1765 | 1 | + | good | GCAGCCTGTACTTTTAACGCTCACTCGCCTCTCCAGCCCCATTAACACAATTTTGAGGTAGAAACCACAAAGCATAAACCTTGACCCTTTAAAGTGTACAGCCCAATGGGTTTTAAATTT | 120 | 42 |
| chr7 | 144387297 | 144387417 | 772751_42001748_Shank2(210274)_1766 | 1 | + | good | CAGCTGTGCAGCCATTAGCACAATAAGACTGTAGAACAAGATGTTGCCCCAAGAAGCAATCTCACACCCATGATCCTCCTGCCCATCAGGAGCCTCTGCGCATAGTCTTCATGTGTATTC | 120 | 49 |
| chr7 | 144387417 | 144387537 | 772751_42001748_Shank2(210274)_1767 | 1 | + | good | AGCTGCTTTTAATATTCCTGGGCATGTACCTTGGTGTAGAGTTCCTGGGGTCATGTAGTGGCATATTTGACAGGTTCCCAACAGAACACCGTGCTACTGCTTGGAGCAGGCTGGCTTGTG | 120 | 50 |
| chr7 | 144387537 | 144387657 | 772751_42001748_Shank2(210274)_1768 | 1 | + | good | GATACTGATGGAGCTTCCACCCTTACACCGACTTTCACACTATCCCTTAGTGGCTTCGCACTGAAGACACATAAACTGGCTGGAGAGATGGTTCTGCATTTAAAAGCACTTGCTGCTCTT | 120 | 47 |
| chr7 | 144387657 | 144387777 | 772751_42001748_Shank2(210274)_1769 | 1 | + | good | CCAAAGGAGCTAAGTTGGGTTTCCAGCACCCTTGTCAGGCAGCACACAACTATGTCCCTCCAGATCCAGGAGACGTGGCGCCCTCTTCTGGAACTTGCAGTCAATGCATGCACAAAGGGT | 120 | 54 |
| chr7 | 144387777 | 144387897 | 772751_42001748_Shank2(210274)_1770 | 1 | + | good | ACATACATAGCTCCCCATCCCCCCGCAATACACACATAATTAAAAAAATAAAATAAGCCCTTTTAAAAAGAAAATCTGTGAATAGCATTTTAGATCTTTCTTTTTAATCCATTTTTATAT | 120 | 28 |
| chr7 | 144387897 | 144388017 | 772751_42001748_Shank2(210274)_1771 | 1 | + | good | CTCTGGTGTTTCGTTTTGTGGTTGTTTTTGTTTTTGAGACAGGCCTGGGAGTTCTGTGTGCTGGGATTAAGGGTGTATGCCGCCATGTCTGGCTAGGCCTGCTTGTTAATTGGAGTTTTA | 120 | 47 |
| chr7 | 144388017 | 144388137 | 772751_42001748_Shank2(210274)_1772 | 1 | + | good | TCCTGCTAACATTTAAGTCATTACTGACATTTAGTCTGGATCTGTGGCTTTGCTTTCTGCCTATCTGCTCCTTTCATCCTGTCTCCCCTTGTGGCTTATTTGTTCTTGGACTCTGTTTTG | 120 | 42 |
| chr7 | 144388137 | 144388257 | 772751_42001748_Shank2(210274)_1773 | 1 | + | good | GTTTGCCAATGGGCTTTAGTGTCCCTCTCTGCATTCCACATGGCTCAGGTCACAGGCACTCCCCAGTGCTCGGGGCCTACTTGGAGTTGGTAGTACAGATGTGACATAAACTTTTTATAT | 120 | 49 |
| chr7 | 144388257 | 144388377 | 772751_42001748_Shank2(210274)_1774 | 1 | + | good | GTTGTATTCTAGTCTTTCGTGTCCCTGCATCTGAATCTGCATGATCACAGATAGCCCCAATACCGTTAGTTTTGCCTCAGGGGACACACAGTTCTGTGGACAGCAAAAGAGTCCATTCCT | 120 | 47 |
| chr7 | 144388377 | 144388497 | 772751_42001748_Shank2(210274)_1775 | 1 | + | good | CAGCCAATCTTCATCCTCTGCTAATGCCTTCCAAGTCTTTCGAGTGTTGGAAACGTTGCCCCGACTTCTGGCCTGTGGGGCTTCCGACATGAAGTCCTTTGAAGTATCCCTCTTCTCTGG | 120 | 52 |
| chr7 | 144388497 | 144388617 | 772751_42001748_Shank2(210274)_1776 | 1 | + | good | TTCAAGGTATCCACATTGCCTTCCACTTCCAGGGTCTAGTTGTAATGTTCCTGCCTGAGATTCACAATATAATTTGAGGCTCACAGTATTTCTTGAACCTGTGGCTTTACATCCTTCACC | 120 | 42 |
| chr7 | 144389337 | 144389457 | 772751_42001748_Shank2(210274)_1783 | 1 | + | good | TCTATGACTCTAGCTAGTCTCCTCTCTTTCAGAACCCCATATTACCTTTACATTAAGCCACTGGCTGATGCCCCGTACATCCTTAGGTCTCTGCTCATTTTTAAAATCTGTCTTCTTCTG | 120 | 42 |
| chr7 | 144389457 | 144389577 | 772751_42001748_Shank2(210274)_1784 | 1 | + | good | ACTCACTGCTCTGTCATCTTTGCTCTGCTGCCAAGCCCACCCAGGAGAGCACTATAAGAACTGTGCTAACACCCCTGTATGACACTTCCAATGCCATGCCCGGTCATCTGTGTTTCTCAA | 120 | 51 |
| chr7 | 144389577 | 144389697 | 772751_42001748_Shank2(210274)_1785 | 1 | + | good | GCAGGCTTGAATTGACCCTGAACGTTGTGACAGTTGGGGCGTGGCAACTTTGGACTCTGTTACAGTTCCCGGATGAAAATTCATGTTTTCCGTGGAGCTGACTGCCACCTTAATTAGACT | 120 | 48 |
| chr7 | 144389697 | 144389817 | 772751_42001748_Shank2(210274)_1786 | 1 | + | good | CCGAGGCCCCACTCTGTCAGTGTTGCTTGTAGTAGCCGAAGCTCACGGTTCCATCCCAATTCTTTTGGCCCTTTTGGTAGTTTGCTCTGAGACTGTCTCACACACACAGCTCACACAGTC | 120 | 52 |
| chr7 | 144389817 | 144389937 | 772751_42001748_Shank2(210274)_1787 | 1 | + | good | AGCCTGAGATAGGGGGAAAGTTCACCCAGAGGACATGCAGCTGCCCTTTCACTCCTTCCTGCTGGGACCTGGCCACCCTGAGCTCTGCCCTATGGTCCTCCAAACTGGTGTGGCTGCAGT | 120 | 59 |
| chr7 | 144389937 | 144390057 | 772751_42001748_Shank2(210274)_1788 | 1 | + | good | TTTCTCTCAGAGCTGCAGATGAGCCACAGGGCACTGCTCCCAGTGCCCAACCAAAGCCTGCCTGCCTTTGTTTGCACTCCAACACTGCCATCTCTGCTTCCATTCTGGTTTGGGTAGGGA | 120 | 55 |
| chr7 | 144390057 | 144390177 | 772751_42001748_Shank2(210274)_1789 | 1 | + | good | TTGTGGCTGTCCAGTCTTCACAACTGTTACCTGAAGAAGAGTTAGAGTCAGGGGCTTACTTTACCATGCTCTCCACCTTCAAGTTTTATACCCAGCTAAGCTACTGATTACATTTGAGGG | 120 | 44 |
| chr7 | 144390177 | 144390297 | 772751_42001748_Shank2(210274)_1790 | 1 | + | good | TCATAGGGCAGTGTATCGAAAATAGTCTCCAAATTGCTATTAGGAAACTCCACTCCCCCTATCACACCACCAAAAAAAAGAAAGTTTTAAACCAAATAAACCATTCACACACGGAAAGTC | 120 | 38 |
| chr7 | 144390297 | 144390417 | 772751_42001748_Shank2(210274)_1791 | 1 | + | good | TGAGATTCAATTTATCCACCATGAGTATCTGAGGACACCCAGGGATGGTGGCATAGGGGCCTCAGACTAGCCTAGAAAACTTGACTATGCAGTACCAATCAGAAGTCCTCAAGAGAGACC | 120 | 48 |
| chr7 | 144390537 | 144390657 | 772751_42001748_Shank2(210274)_1793 | 1 | + | good | GAGACCCGGTGTCTAAAACAAACCAAGAAAATCTCAGAAGACAGTTATTCACAACAAAGTTAAAACCTCATGGACTGTGAAAAGGGAGTCTCAGAATTAGAGAAGTGTTTGGGTTTGAAT | 120 | 38 |
| chr7 | 144390657 | 144390777 | 772751_42001748_Shank2(210274)_1794 | 1 | + | good | TTCTGTTAAGTACACAGAAAATTAAACCAATAAAAAACAAAAAGCTAAACAAAATTATTTTGCTCCGGGGAGGAAAACCATACATGAGATGATAAAGTAACCAAATCAGAGTATATGGTT | 120 | 30 |
| chr7 | 144390777 | 144390897 | 772751_42001748_Shank2(210274)_1795 | 1 | + | good | TAAGTATGGGAGGATTATCGCACAGTGGAGAGGAACAGTGGATGTTGGTCTAGTTAAATGTGATGTAAATGCATCCCGAGAGTCAGAGCTGGAAGCACAGCAGGAGGGCTGAAGGAAAAG | 120 | 48 |
| chr7 | 144390897 | 144391017 | 772751_42001748_Shank2(210274)_1796 | 1 | + | good | AAGTACCCGCCCCCAAACTCCAGTCTTCCTTTCCATAGTGGCTCGTCCACGGATGAAAGCTCAAGTGTAGTTCACAATAAACAGTCAAGAAAGTCAACAGCAGGGTCAGAATTGGCTAAA | 120 | 47 |
| chr7 | 144391017 | 144391137 | 772751_42001748_Shank2(210274)_1797 | 1 | + | good | GGCCTTGGCTGAGAACGGGCATTGAGCAGGAAGTCTCCTCTGTCCGCTGTCCCCACACAAAACTCTGACATTTAAGAAACACTCAGTTTGAAGACCCAAATTCATTATGAAGGACTATAT | 120 | 46 |
| chr7 | 144391137 | 144391257 | 772751_42001748_Shank2(210274)_1798 | 1 | + | good | AGGACCCTGTTCTTCACAAAACAACTAAATCCCATTTAGCCTGGTTTATCCACTCTCCATAGGTCCACAGCTATGTGTAAGGATGTATGGAAAGGCCCATTGCTTCCTTAAGGGTAACTG | 120 | 44 |
| chr7 | 144391257 | 144391377 | 772751_42001748_Shank2(210274)_1799 | 1 | + | good | GATCAGCTTTTAATACAAAAACAAAAACAAAAACAAGCAAGCAAACAAAAGAATCTCTATATTGGGTAAACTGCAGGGGCATCAGTATGCACACGGGAGCTCTAATGAGGCACATGATAC | 120 | 38 |
| chr7 | 144391377 | 144391497 | 772751_42001748_Shank2(210274)_1800 | 1 | + | good | AAGTGGGATATGTCATGGAACCTAGGGAGGACTGGGCAAAGGCACATGGTCTGTGCAACACAGAGACTCCTCACCAAGACCAGCTTCCAAAGGCGACAAGCCAGAAAAACCCGGGTACAT | 120 | 52 |
| chr7 | 144391497 | 144391617 | 772751_42001748_Shank2(210274)_1801 | 1 | + | good | TTTCCTTACCACAGCAGGGATCAAGTTCTAGGCTTTGTAGTTTCCCAGCCTACAAAATGGCTACAAAAAACAGTTTTTTATCCCTTCTTGCTGATATGACATGTGGCACTCGTATCCCCC | 120 | 43 |
| chr7 | 144391617 | 144391737 | 772751_42001748_Shank2(210274)_1802 | 1 | + | good | CACCCCCACCCCGTGGTAACAGACACAAATTGCAGGTTCAGGCCCTTTGTGCGTTAACCCCATGAACGTTCAAAGTGGTACTGAGAGTCAATGAGGGAGGGGCAAGCCTTCTTGCATACT | 120 | 52 |
| chr7 | 144391737 | 144391857 | 772751_42001748_Shank2(210274)_1803 | 1 | + | good | CTGGCTCCCATCAGCACACGAAGCATCTATTTTCAGGACTGTGTCTGGCTCTAAGCATCCGTTGCACACTAAACTGGTTTTCGTAGCCATTTTGTAGGCTATAAAACGACATATCACACT | 120 | 45 |
| chr7 | 144391857 | 144391977 | 772751_42001748_Shank2(210274)_1804 | 1 | + | good | TGTATCATGTGCCTCATTAGAGCTCCTGTGTGCATACTGGAAGCTGATATGCAAAGCCATATGGCCTTCAGCTCTGACCCGAAGGTTAGGGCAGAGTTTGTGCAGAGTTCAGTACTTTTG | 120 | 48 |
| chr7 | 144391977 | 144392097 | 772751_42001748_Shank2(210274)_1805 | 1 | + | good | AGTTTTGCTTTCAGTTTTGTTTCTGCAAATCAGGAAGGGTGTTGGAGTCCCACATGGTTGCAGGGAGGGAGAGGAGGAAGGGTGAAGACCACAGCAATGGGGCCCAGATCAGCGTGCCTC | 120 | 53 |
| chr7 | 144392097 | 144392217 | 772751_42001748_Shank2(210274)_1806 | 1 | + | good | CATTGTGGGCACTGCTGTGTCAGCTACTTTTCATGGTAAAATGTCATAAGCAAAAGCAAGCTGTGGGGGAGTTTATTCTGGCTTACTGTGCCCAAGAGTTAAGAGTCCATCATGGCAGAA | 120 | 45 |
| chr7 | 144392217 | 144392337 | 772751_42001748_Shank2(210274)_1807 | 1 | + | good | GATATGGCCGCCAGGGGCTAGCATCATGGCAGGAGTGGAAAGGTGAGAGCTCACATCCTTAACAGCAAGCACAAAGCAGGCAGCAAACAGGAAGTGGCATGAGGCTTCCTGTCTCAAAGC | 120 | 54 |
| chr7 | 144392337 | 144392457 | 772751_42001748_Shank2(210274)_1808 | 1 | + | good | TCAGTCCCAATGACATACTTCATTTAGCAAGGCCACACCACTTAAACTTTCTCACGTTGTAACCACCAACTCAAATGCTCAAATCTCCGAAGGACATTTCTCACTCAAACCACAACATTT | 120 | 41 |
| chr7 | 144392697 | 144392817 | 772751_42001748_Shank2(210274)_1811 | 1 | + | good | ATGGGGCATCATGGGGAAACACATGCCAAGGCAAGATGAAAAGCCCAGCGGGACAAACAGCAAAACCCATAGCTCCAATATCAGGTGTCCCTGCAGTTGGAAAAGCCAACCTTCTGTCTG | 120 | 51 |
| chr7 | 144392817 | 144392937 | 772751_42001748_Shank2(210274)_1812 | 1 | + | good | GGAGAAGGACTGTGCCTCATCTTCCCCTGCAAGGAGAGGTTGGAGGTGTGGGGCAGGCTCTGGGACTCCACAGCCCAGCCTGGGATGGGGACTTTGTGCATGGCACACCCAGCACTGTGG | 120 | 62 |
| chr7 | 144392937 | 144393057 | 772751_42001748_Shank2(210274)_1813 | 1 | + | good | AGGATGACAGCAGCGTGGACACACCTGCCCTATTCCTTCTAGAAGGTTCTCTTGTCCTCTGTGTGGGAATGGGAGGAGGCACAGTGGCTCTGAAGCTTTGGGGAGAACAGAGTCCAATGG | 120 | 54 |
| chr7 | 144393057 | 144393177 | 772751_42001748_Shank2(210274)_1814 | 1 | + | good | CAGAAAAGGAAGGCGCCTGGAGAGACAACTCCTAGGAAAGAGTCTCTCATCAGGGGTCCCCATCCCTGTGTCCCTAGAAAGCAGTAACACTGCTCTCTGGACATGGCATGTTCTTGCCCT | 120 | 53 |
| chr7 | 144393177 | 144393297 | 772751_42001748_Shank2(210274)_1815 | 1 | + | good | CTGGACCTGTAGGCAGCTGTGATTACCTGCACAAGTCTGGAACCATTCCTTTTCTGTCATGGGAGGGGAGAGGGGCTCATGGCTCCCTGTCCATCCTCAAGAATTAATAGACCTTTAAGG | 120 | 51 |
| chr7 | 144393297 | 144393417 | 772751_42001748_Shank2(210274)_1816 | 1 | + | good | ATTGCTAGAAAGAGGAGATCACAAGGCCACGCATCCCATGGATATTTGTGGGCAGCTAACGGTTATCAGGATGGCTAGTCACTTAATCAGTGCCAGGGTTTCAGCTGGCTATAAGTTGCC | 120 | 48 |
| chr7 | 144393417 | 144393537 | 772751_42001748_Shank2(210274)_1817 | 1 | + | good | CAAGCTCTCATAACACCATTGCAAAGCAGGGAAGTGTAGCTACAGGACCTGCCCACAAGGCCTGCTCCTTCCTGAGATAGCATTTCCCAGTGTCCCCTTCTCTCTGAGTTCCCTCCATAG | 120 | 52 |
| chr7 | 144393537 | 144393657 | 772751_42001748_Shank2(210274)_1818 | 1 | + | good | CAGCACTGGCCCTTGCAGGCTGTGTCAAAGCTACTCTGTCACCTCTGTAGGACTTGGGGACTATAGCTCCTTCCCTGCCTCCCAGCTTCTCTTCAGCCAGGGAAGCAGCTGGCAGGGGTG | 120 | 59 |
| chr7 | 144393657 | 144393777 | 772751_42001748_Shank2(210274)_1819 | 1 | + | good | CTGCCTTCCCCACCAAGGCCTGTGTGTAACTGCGGCATAGCAGGTAGAATGCTGACTCTTAAGGGCCTCAAATGAAGGGTAGTAGGAAACCCAGCTCCTGAGCAGTGCAGCATGGCTGGT | 120 | 55 |
| chr7 | 144393777 | 144393897 | 772751_42001748_Shank2(210274)_1820 | 1 | + | good | CACTGGTTTCTGTCCAAGGCTGTTCAGTTGCTGTTCTATAATAAAGTGCACAGCCCAGGGAGACAGAGTTATTTTCTCCCTGCTTTGGAGGCTGGGACCGGCTATCCTGGCTTACAGGCT | 120 | 51 |
| chr7 | 144393897 | 144394017 | 772751_42001748_Shank2(210274)_1821 | 1 | + | good | GTTGTGCCCTTGATGGTCCCTCACAGGGGGACAGGCCTTGGTACCCTTTGAAGGAAGCACCCATCAATTTATGAGTCACCCTTGAGGGAGCCCCTGGACAGAATGAAAAGGAGCATGGAG | 120 | 54 |
| chr7 | 144394137 | 144394257 | 772751_42001748_Shank2(210274)_1823 | 1 | + | good | ACCAAATAAGCCCTCTCCTTAAACTGCCTGAGTCAGGTGTTTGTCCCAAAGGCCTCAGTCACCTCTGGGGTAAGGTCTGTGCTGTGACAGTTGTGACTGCTGGGATGGCAAGCTTACTGA | 120 | 52 |
| chr7 | 144394257 | 144394377 | 772751_42001748_Shank2(210274)_1824 | 1 | + | good | ACCCCTGAACACTCAGTGAAGCCATAGCTCCACTCCTGTAGCCTCAGCCCAGCTCGGCTTCTTTGGGAAACTCAGTATTTAGTTTCTCAGTATTTAGTAGCCTTGACATCCTTTGTCTAA | 120 | 47 |
| chr7 | 144394377 | 144394497 | 772751_42001748_Shank2(210274)_1825 | 1 | + | good | TGACAACTGTGTGTCACCGGGGCTGCGAGCTTCAGGCTCCAGCTGTTTTGTGACACTTGGAGCCCAGTCTTCTGACATTTCCCAAATTTGGCCCAAGGCGAGCTTGGAGGTCTCAGTATC | 120 | 54 |
| chr7 | 144394497 | 144394617 | 772751_42001748_Shank2(210274)_1826 | 1 | + | good | CATGCCAGCCCTTGCATCCATGTCTGTCTGATAGGCACACACATGGACTCCATGGGAAGACAGTCAGCAAGACAGGGGCTCCTGCACAGCCACCCACGTCACTGTGTAGGAATGAGGTTT | 120 | 55 |
| chr7 | 144394617 | 144394737 | 772751_42001748_Shank2(210274)_1827 | 1 | + | good | GGGGATTTGTATGAAAGTGCCGGAGTTCTTAAGTTCCAATAGCTAAGTGGTGTCCAACACTGTTGGGCAAGATTTGGCCCCTGGCAATCTTGATTGGAAGACACAGTGCTCAGATGGACA | 120 | 48 |
| chr7 | 144394737 | 144394857 | 772751_42001748_Shank2(210274)_1828 | 1 | + | good | TGTTCACAATGCCCAGCTTCCAACCTGCCCATCCTCAAGACTCTGGCATTGGGGACAAGGCCTAGGGGTCAGGGGGATCTACTCCATGAGGTCCAAGAGCTCTTGGCAAAGGGTCGAGGT | 120 | 57 |
| chr7 | 144394857 | 144394977 | 772751_42001748_Shank2(210274)_1829 | 1 | + | good | AATGCCCACCTCTGCTCACCTCTATGTCTGCTTCTTCCTTGGGGGTGGCAGGAGAGCGCCTGATTCTGAAAACCCAGGTGTTCTGAGCCTGGGGTTGTGTTCACCGCAACAGTGTTGGGA | 120 | 56 |
| chr7 | 144394977 | 144395097 | 772751_42001748_Shank2(210274)_1830 | 1 | + | good | TCCAAGCGTGCACCGCCACTATCCCGGGAACACAGAGATCATTTTCCCTCCTCCCTGCAGTGCCAGGGCAGAAGCGTCATCTCTCTTAGCACCTTGTAATGGCAAGCTTTTCTGCCTATG | 120 | 54 |
| chr7 | 144395097 | 144395217 | 772751_42001748_Shank2(210274)_1831 | 1 | + | good | TGTAGAATAAAAGAGAAAGCCTAGAGGCAGTGAGCCAAAGCACAGGAGAAGCAACTAGGGACCAGGTCTCCACCTCACTGTAAGAGGGGAGAAGCTGTTCCTATATCCAGCCCTGGGCAA | 120 | 51 |
| chr7 | 144395217 | 144395337 | 772751_42001748_Shank2(210274)_1832 | 1 | + | good | CTCTTGAGAATTCTGATAGGTGGGACCCACAGGTCCTGGCAGAGGATACTGGATGAAGCATCCCCTTTCTCCCAGTGTGGTAGCCCCTGCAGCCACAGCAACACACATGCTGACCTCCCC | 120 | 57 |
| chr7 | 144395337 | 144395457 | 772751_42001748_Shank2(210274)_1833 | 1 | + | good | AAAGCAACTACTCATCCATAGAGACCAATGTCCCCCGGGTCTCCTGTAGGATAGCAGCAGCACGTGCCTCACCACCTCTGCCTCCTCTCACATGTTGCTGACCTGCCATGCCTCTCTCCC | 120 | 57 |
| chr7 | 144395457 | 144395577 | 772751_42001748_Shank2(210274)_1834 | 1 | + | good | TGCAGGTTAACAATGAAAATGTCGTCAAGGTGGGCCACAGGCAGGTGGTGAACATGATCCGCCAGGGAGGGAATCACCTCATCCTTAAGGTCGTCACGGTGACCAGGAATCTAGACCCTG | 120 | 53 |
| chr7 | 144395577 | 144395697 | 772751_42001748_Shank2(210274)_1835 | 1 | + | good | ATGATACAGCCAGAAAGAAAGGTGTGTGTCCCTCAAGGTGTTCTTCAAGGGGGAAGGCAGGGGCCGTGGGGAAGAGAGCAGGATTCTTTTCAGCTGAGAGCCAAGGCAACGGTGTGTGGC | 120 | 55 |
| chr7 | 144395697 | 144395817 | 772751_42001748_Shank2(210274)_1836 | 1 | + | good | TTAAGTTTGTGTCCGAGGCCATGATATTGAACCTGAGTCACCATTCTGCTAAGGAAGGCCATTGCTGGGTCAGCAAACAGGTACCAAGTGCAGAACAGGTGCCTCTGCCCCATTGGTAGG | 120 | 51 |
| chr7 | 144395817 | 144395937 | 772751_42001748_Shank2(210274)_1837 | 1 | + | good | GCAGATGTCAGGGATAAAAGAGTCCTTCTAGCCAGCTGTCCCTAAAGCAGCTCTGTCATGGAGCCAGGGACAGCTAGGAGAGGATGTCGAGGTGCCCATACAGAGTTTACTCATTAGCAG | 120 | 52 |

| | | | | | | | | | |
|---|---|---|---|---|---|---|---|---|---|
| chr7 | 144395937 | 144396057 | 772751_42001748_Shank2(210274)_1838 | 1 | + | good | GGCAGTCTAGGAGAGAGGGCTGGAAACAACTAAGACCACAGTGAGAAGGTAGAGGAATGAAATTTCCTGGATACCAACAGGCCTCTTTCCCCAGCTCCTGTGAAGCCTGTGCCCCTGCCC | 120 | 54 |
| chr7 | 144396057 | 144396177 | 772751_42001748_Shank2(210274)_1839 | 1 | + | good | TGACAATCTGGCAGGTGTCACAGTCAGAGGTGCACAGGCAGAACAGAGCCTGAGGTCCCCTCCCAGCCCAGAGCACCTGAGAACCTGAGAAGTTGCTACCAGTCATGGCAGCCCAACTGT | 120 | 57 |
| chr7 | 144396177 | 144396297 | 772751_42001748_Shank2(210274)_1840 | 1 | + | good | GACAACCCTACCCCCAACCACCTGTCAGCATCCCCTACAAAACACAGATTGGAGGGCCTCTGCCTCTCAGATTCTTCCTGGGTCCAAAAGTTAAGGACATGGGGTGGGATATAGGATGGG | 120 | 53 |
| chr7 | 144396297 | 144396417 | 772751_42001748_Shank2(210274)_1841 | 1 | + | good | GTGAGGCCTTCTGGTGATTCTCATACTTGCTCTGCCTCTGCCTACCCTTCAAGCTCCCCCACCTCCAAAGCGGGCTCCGACCACGGCCCTCACCCTGCGTTCCAAGTCCATGACAGCGGA | 120 | 61 |
| chr7 | 144396417 | 144396537 | 772751_42001748_Shank2(210274)_1842 | 1 | + | good | GTTGGAGGAACTCGGTAAGGCTACCTCGCCCTGCTGCCCTCAGCCAATGCATGACCTCCTGCTGGCTGGCTTGTCCGTGCCCCGCCCCTGCATGCCCTACAGCACCACACTTCGCTTGGC | 120 | 64 |
| chr7 | 144396537 | 144396657 | 772751_42001748_Shank2(210274)_1843 | 1 | + | good | ATCTCTAGTGGGGACCGAGGTGGGACCTATAGGGATGGGGCTGTTTCTCTTTCTGCTTATCTAACATTGTTTTGTTTTGACTTTTTGGGCCTCTCACTAGTGGACAAAGGTAAGCAGCCC | 120 | 48 |
| chr7 | 144396657 | 144396777 | 772751_42001748_Shank2(210274)_1844 | 1 | + | good | CGCTGGCCCCGCAGCAGCATCCCTGCTGCCTCGGGCACCCTCACCAACCCCACCTCAGCCTGGCCTCTAGCTTGCTCCAACCACCGAAGGCCCTGCCTTCAGGGACTCTTCAGACAATCT | 120 | 65 |
| chr7 | 144397257 | 144397377 | 772751_42001748_Shank2(210274)_1849 | 1 | + | good | CATTAAGAAAAGGGATGACTCTGACACGACTAAGAACCAAATGCTGTGTGTCACCCATGTGCCACCAGGTAGCCAGGGAACCTCTGCAAAGAGGGTGACTCTGAGCCAGAGATTGTGAGT | 120 | 50 |
| chr7 | 144397377 | 144397497 | 772751_42001748_Shank2(210274)_1850 | 1 | + | good | GGAAGCACTGACCCCAAGAAAGGCTGGTGAGAGTGGACTCCTGGGCACAGAGGCAACTGCTCAGATGCAAAGCTAGTAGCTCTTAGTGGTGTCCTGCCAGGATGTCACTAACAACCAACA | 120 | 53 |
| chr7 | 144397497 | 144397617 | 772751_42001748_Shank2(210274)_1851 | 1 | + | good | ATATTATGTGGGAAACCTGCATGTCTCCACCAGCATGGAGCAGCCTCCTACCTCCTTGCTGATGATAAACATTGTCTTTCTTGCTGCTTTCTTGGACTCTGAAGCTTGCCTCAAATTGTC | 120 | 46 |
| chr7 | 144397617 | 144397737 | 772751_42001748_Shank2(210274)_1852 | 1 | + | good | AGTTTTCTAGAAGAGGGTACCCTGACACAGGAATGCTTCTCCACCCAGAAACTGGAGTACCTCTCTTTTCAGCCATGCCCCTGACCCTATCACATTCCTTCAAACATTCCACTGGGAACA | 120 | 48 |
| chr7 | 144397737 | 144397857 | 772751_42001748_Shank2(210274)_1853 | 1 | + | good | TTCTAATTGTGTCCCCCATGACAGAATATCTGCCCTACTTACGATGGGCCAAGGCCAGCCTGCCCTACTGAAGGGGCTACTACCATGTTCCCTAGGAGTTCATTAACCAAGGGCTGTCTC | 120 | 52 |
| chr7 | 144397857 | 144397977 | 772751_42001748_Shank2(210274)_1854 | 1 | + | good | CAGCTCCCAGGGCCTCTAGCTCCTGGCATCTCCGGCTCCTGGGACCTCCAGCTCCTTGGGTCTCCATGCCCTTTTAAAGGGCCCCTCCTGTCCCACTTCATGAAGCTCAATCCCACCTGT | 120 | 61 |
| chr7 | 144397977 | 144398097 | 772751_42001748_Shank2(210274)_1855 | 1 | + | good | GGAGTGCTGGTGAGATGCAGGGGCCAAAGTAAGAACGAAAGCCTTATAACTACCACACAGGAGACAGGGCAAGAAAAAAAGAACAACCAAGAACAAACTGTGAGGCCACTCTGCACCCCT | 120 | 49 |
| chr7 | 144398097 | 144398217 | 772751_42001748_Shank2(210274)_1856 | 1 | + | good | GCCTCTCACTCCCAGTAGTGGTCTCCATGCTGCTTCCCCTGTCTGTGTCCCATGATCACTGCGTTGAGTAAGACCCCGGGTGTGGGCCACTATGGGAGGGTGCCACCGTGGGTGGCTCAC | 120 | 62 |
| chr7 | 144398217 | 144398337 | 772751_42001748_Shank2(210274)_1857 | 1 | + | good | GCGCTTCCCTTTCTGTTTGCAAGCCTCAGTCCGGAAGAAGAAGGGTAAGGGAAGCATCCGTCCCCATGCTCTTGCCCAGTAGAAGAGAAATCCCCTCGTGACCATCGTGTAAAGCCCAGA | 120 | 54 |
| chr7 | 144398337 | 144398457 | 772751_42001748_Shank2(210274)_1858 | 1 | + | good | GTGTGCTGTGACCCTTGTGTCGTCTTCCCTATTTGGGGGGGGGTCATACCTAAGAGCTCCTCCCTGGGATGTATCCTGTTCTGGTCTGTCTGATCTGCGTGGCTTCCGTCTGTGACCCCC | 120 | 57 |
| chr7 | 144398457 | 144398577 | 772751_42001748_Shank2(210274)_1859 | 1 | + | good | TTGACCTATTTGTAAGTCCCCCAATCTGTGCCCACATCAACACACTGACAAAGTGGCCAATCCTGGATAATTCTTGCTGTGCCTGCCTGGCTGCAGGTGCATTGACTTTTCTTCTGTGGA | 120 | 48 |
| chr7 | 144398577 | 144398697 | 772751_42001748_Shank2(210274)_1860 | 1 | + | good | GCTGGTCAGAGACATCCATGTGTGCTGCCTAGCCACTGAACAGACTGGCATCCAGGCAAGGGGGAGCCCAGGCTTTTTTCCTCTTCAGCTCCCCTCACACTGGGGAAGAGGGAGCAGTGT | 120 | 58 |
| chr7 | 144398697 | 144398817 | 772751_42001748_Shank2(210274)_1861 | 1 | + | good | GGGACTTTTGGCTGCCTGTCCCCAGGCTCCCATGGGAGGAGCAGATATGACACAAGTCTCATGAGTAGCCAGGCAGAGCCCTGCCTCCTTCAGAGCACCCTGTTATAGATGTATACCCCT | 120 | 56 |
| chr7 | 144398817 | 144398937 | 772751_42001748_Shank2(210274)_1862 | 1 | + | good | GACTGCTCATGACTCCCCAATACTGAGCCAAATACAACGTGTGACCCAGAAGTACAATCCTAAAACATCGTTCTCCAAGGCTGAGATAGATGAACCCTAATTAGGCTAAGGAAGTCCCTA | 120 | 45 |
| chr7 | 144398937 | 144399057 | 772751_42001748_Shank2(210274)_1863 | 1 | + | good | ATGATTGCTGAGCTCAGTCTGAGGCCTGGTGTGGACCTAGCTTCCTAGATGCTCACATGGGGGATCAGGAATGCCTGAGTCTAGCAGGATCAGCCTTACAGCAGGGCAGGTACCCTGGAG | 120 | 56 |
| chr7 | 144399057 | 144399177 | 772751_42001748_Shank2(210274)_1864 | 1 | + | good | CTGCCAGGGTAGTCAACTCAGGACTAGCAGCCATCTTTACAGCAATCCATGCCTTTACCACATTGGACTCCAGCAGGCTCTGTAGGGTCCTCCCCTTGCCTAGGGTATGGCTGATGACCA | 120 | 54 |
| chr7 | 144399177 | 144399297 | 772751_42001748_Shank2(210274)_1865 | 1 | + | good | GTCAAAGCCTACCCTTCTTCTTAGGAATCCTATATTTATATTACTATATGAGCACCCCACCCCTATCAAAGGCCAGACGCAACTCTAGCCTGTCAAACAAGGCATGCCACTACTGTGGGC | 120 | 47 |
| chr7 | 144399297 | 144399417 | 772751_42001748_Shank2(210274)_1866 | 1 | + | good | TCCCAAGGTCCAGGGTGGCCCATCAGAGCTGGTGAAGAAAAGCTCATTGGCAAGAAGATGGGATCTGGGCAGTCAAGTCCCTGTCCGAGGGCACCTCCCACTGAAATGCTGCAACAGCTG | 120 | 57 |
| chr7 | 144399417 | 144399537 | 772751_42001748_Shank2(210274)_1867 | 1 | + | good | GTGACAGAGGGATTTTTAGCCAGCACGAGGTGAATCCATCAGAGATACATGAGTCTCAGACAGTGGAAAATGCCTTCCATGTTAGCAGCAGAAAAAACAAAAAACAAAAAACAAAAAAAA | 120 | 38 |
| chr7 | 144399537 | 144399657 | 772751_42001748_Shank2(210274)_1868 | 1 | + | good | AACAGTTGAGAAGCAATGCCTCAACCTGGAACTAGAGCAGGGGTTAGGGGTGAGGGTGGAGGCAGATGCCAAAGGAACCAGTAGCCACAGCCAAGGAGAAGGGACTAGGGGTGCCTGTTC | 120 | 56 |
| chr7 | 144399657 | 144399777 | 772751_42001748_Shank2(210274)_1869 | 1 | + | good | CCATAGCTACTGCATCTCTCACCTTACTTTGCAGAAGCCCTTGCCAACTTGGTTCCTTCACTCAGTTGGTACACTAGCACCTGCCATGCATGCCTTGGGGGCAGTGTGGGCACATGGTGC | 120 | 54 |
| chr7 | 144399777 | 144399897 | 772751_42001748_Shank2(210274)_1870 | 1 | + | good | CTGAGTGTGGAAGGGTGCTCAGCCATGGCTCCAGCGCCTCCCACAGCATGTCCAGGCTCCCTGGGAGGGCTGCCTACCTGGAAGGGAATACACTCCAGGGAGGCAGGCTAGAGTGAAAGA | 120 | 62 |
| chr7 | 144399897 | 144400017 | 772751_42001748_Shank2(210274)_1871 | 1 | + | good | ACTTGCTCTTCCCAGCTGCCCTGAGCTGAAAAGTGGCCCCACCTCCTGCAAGTCTTTGTCTACCCAGAGACCATGAACTAAACCACATTTAGAAATAAGCTCTTTGGCTGGGTGTGGTGG | 120 | 51 |
| chr7 | 144400257 | 144400377 | 772751_42001748_Shank2(210274)_1874 | 1 | + | good | ATGAGAAGAGGAGGAGACTTAAGTGGTTTTAGAGACTCTTGGGACCTCCAGAAACAGGGCTGTCCCCAACTATCTACAGATAGAACCTGGTCCTGCTAACCCCAATCGTGGATGTCCATC | 120 | 49 |
| chr7 | 144400377 | 144400497 | 772751_42001748_Shank2(210274)_1875 | 1 | + | good | ACCCAGAACAAAGCTATTGTCCTTTTAAGCTATTGTTTATGGTGCCAGGAGATTGCTGAAAGTTAGCAGCCTCAGCTAAGTGTGGTAGTGTATGCCTGTAATCACAGCACTATGGACATG | 120 | 43 |
| chr7 | 144400497 | 144400617 | 772751_42001748_Shank2(210274)_1876 | 1 | + | good | GGAGACAGAAAGACCATGAGTTTAAGACCAGCCTGAGCTACATGAGACCTTTCTCAAATGGAGTAAGGCAGCCTTAACCCTGGATCCACCCCACCAGCCACATTGGCTTAGAGATACTGG | 120 | 50 |
| chr7 | 144400617 | 144400737 | 772751_42001748_Shank2(210274)_1877 | 1 | + | good | GTTTTCAATGAGCTTTTTGCCCTAGTGCTCATGGTGAAAATAAAATCACCATGGCCAACTACCATGTGCACGCCAGCCCAGGTCTGCTGATGTGAGAATAGAGACAGAGGTGACCAGGCA | 120 | 48 |
| chr7 | 144400737 | 144400857 | 772751_42001748_Shank2(210274)_1878 | 1 | + | good | GAGTGAGTGGTGGAACAGGTTGTTTGACCATGACTTCCAGCCTTGCTGGTCACCTCTAGATAAGCACACAGGCCTTCCCAGGCACAAACCTTGGAATCTACAGATCCACAGGGTCCAGGC | 120 | 53 |
| chr7 | 144400857 | 144400977 | 772751_42001748_Shank2(210274)_1879 | 1 | + | good | TGCATACCCACACTTCACTGCCTGGGAGAGCAGACAGGAGGCAGCCCTCTGGTATAGCTCAGATGACAATGTGACATTTTGGCTTTTACTCCTGGTATTAGAGTTGCAGAGGAAGCCTAC | 120 | 50 |
| chr7 | 144400977 | 144401097 | 772751_42001748_Shank2(210274)_1880 | 1 | + | good | ATGGTGCGTCCTCACCCTGAGGTGGGTGTAGTGGCACTGTCCTGCAGCGGGAGTGGGCTATTGGCTCTGGGACACTAAGGGAAAAGCATGGCGGCTGTGGGCAGATACACTTCTAGTTAT | 120 | 57 |
| chr7 | 144401097 | 144401217 | 772751_42001748_Shank2(210274)_1881 | 1 | + | good | CTCAGATGCCATGGGCACCCAGGAATCTTGCCGCTGGCTTTGTGGATGATTCTAGAACACTTGATCTTGCTCTCTCACAAAGAGCCAATGTAGTATGTGAGAAACAACTAGGATCCTGCT | 120 | 48 |
| chr7 | 144401217 | 144401337 | 772751_42001748_Shank2(210274)_1882 | 1 | + | good | CTAAGGCTTGAGTGCTAGGCTCTGACTGTATTATCACCCCGTGTGTGGCCAAGGGACAGGGCACCTTAGCAGGGCTTCCACTCACTGCCTGGGGAATTTGAGGTGCCTGGAAGACTCCCT | 120 | 57 |
| chr7 | 144401337 | 144401457 | 772751_42001748_Shank2(210274)_1883 | 1 | + | good | CAGGACTGTGCCACGGGCTACAATACCAGATGGGTCCAGCTTTTCCTTTGCTTTAGCCATTGGATTGTTTCCCTTCTCCTAGCATTTGTGGCTTCTTCCCACAGAGCCATGACATCTGGG | 120 | 51 |
| chr7 | 144401457 | 144401577 | 772751_42001748_Shank2(210274)_1884 | 1 | + | good | GAACTGTCGCATGGAAAGCAACAGGTGCCTCATGTTGAGGACACTGGACATACTGTCCAGTGCTTGCTTCCCAGGCAAACTCACAGACTGGGAGAAGCAGTGATGGGCAGGGCAGGCTAA | 120 | 54 |
| chr7 | 144401697 | 144401817 | 772751_42001748_Shank2(210274)_1886 | 1 | + | good | CTCTCTCTCTCTCTCTTTCCTAGAAGTCTACTATTTATAGCCCATGGGCAACACGTCTGCCTTGTGAGAGGCTATACGCTTGCTTAGAGACTGAGAACTCAGCTCATGGCTGGGCTGTTT | 120 | 48 |
| chr7 | 144401817 | 144401937 | 772751_42001748_Shank2(210274)_1887 | 1 | + | good | AGTGACCCACAACAGCCCTCATGTATGTCTATAGTATATCCAGTCATCAAGACCCCATGTTAGCCCTGGAAGGGCCTGGCCCCCTCAGCGATCTGTACAACTCCAGTCCTGGGCCTTTAA | 120 | 52 |
| chr7 | 144401937 | 144402057 | 772751_42001748_Shank2(210274)_1888 | 1 | + | good | GTTTAAACCTCATTCTCAGAAGCCAGCCATACAACAGTTTCCTTTAAAACCTTTATTTCCTGCCAATAAGATCTCAGAGAAGAAGCCTGTGTGGCCAGGTCTCAAGTTCCCATTGGCCAG | 120 | 44 |
| chr7 | 144402057 | 144402177 | 772751_42001748_Shank2(210274)_1889 | 1 | + | good | CAAATCTAAGGATCAAGTTGAAGAGCCCTCTGTCACCTGACTCACACCAACAGCACTAACACTAAAAGATCTGGAGCCCTGGGTCTAACAGGCTTATAATGCAGCTGCTAAGGGAGATGG | 120 | 48 |
| chr7 | 144402177 | 144402297 | 772751_42001748_Shank2(210274)_1890 | 1 | + | good | GAGAGGCAGAGCTCAAAAGAGCAGGCCTCAAGACATAACCATACTAGCACTGTCCTATTACAGACTCCACAAACCCCCTCTGTTCTGCAGTCAGCTAAGTGAGGTTGGTGACTCTGGTCA | 120 | 50 |
| chr7 | 144402297 | 144402417 | 772751_42001748_Shank2(210274)_1891 | 1 | + | good | CCCTTACAGGAGCACTATCAAGCTGTGTTCTCCCATGAAAGACTGTTCTTCCTATAGCCACGTGGAGCAGGCTGAAGCCTCAAAGCCCCATATGCTTAGAACCATCATGGGAGACCCTTA | 120 | 50 |
| chr7 | 144402417 | 144402537 | 772751_42001748_Shank2(210274)_1892 | 1 | + | good | TTCCAAGGCCCTGAAATCTCAAATCACAGCCTATTGCCCACTCCTGCTGGACATGTGAAGATGCCACAGGCACCCACGTGCCAGGCAGTAGACCCACATGCCACCGGGGAGTGGGGGAGG | 120 | 59 |
| chr7 | 144402537 | 144402657 | 772751_42001748_Shank2(210274)_1893 | 1 | + | good | GGAGAATGCACACACTGCCAGCTTCTGCCTGGGAGACTCTCAGTGATTCCACAACTCTTCAGCACGTGTGTTCATTCTTCTAGTGTCCTTTGTTTTCCATGTGCCCAGACCCAGGCCCTA | 120 | 52 |
| chr7 | 144402657 | 144402777 | 772751_42001748_Shank2(210274)_1894 | 1 | + | good | CACAGCAGACTCTGGCCAATCACCTCCTTATTCTGAGCCTGTGCCCAAGGCCCTCCTAGTAGGACTATGACCCTGTGTCTCTTATCTGATCCTCCAAGTTGTCTGGCCCACAACCTTCTC | 120 | 53 |
| chr7 | 144402777 | 144402897 | 772751_42001748_Shank2(210274)_1895 | 1 | + | good | TTGCCCAGCTGCTACCATTAGCCATTGAATTCGACAAAGACCAGTTTTATGGGGTAGGAAAAACAGGAGCTTCCTGGAGCTCAACCACACCTCTCCTTGATAGTCCTACCCAACTACTCA | 120 | 48 |
| chr7 | 144402897 | 144403017 | 772751_42001748_Shank2(210274)_1896 | 1 | + | good | CAGGGCAAGCAAGGTAGGCTACCCACACAGCCAGCAGGGGCCTTCCCATCTCAAGGTTCGCTCAGGTTAGAGGTGGCTAGATTCTTGCAGGTCTCCAGAGCTCTCCATATGAAATACCCA | 120 | 55 |
| chr7 | 144403017 | 144403137 | 772751_42001748_Shank2(210274)_1897 | 1 | + | good | GACAGGCCTCATAACTCAGTGGCTTCCACTTGACCCACATTTGAGATTCAGGATAGTGTGTCCCCATGCTAAGCCTCTGTGGGTTCAGAGCAAGTATCACTGAGGAGAGGAGAGTAGTTT | 120 | 49 |
| chr7 | 144403137 | 144403257 | 772751_42001748_Shank2(210274)_1898 | 1 | + | good | CTAGAGTGAACGCAGTGTTTTCTGCCCAGTGCCCACTGCAGTGCTGGGACATAGCTCTCCATTGCTAGAGAGAGCTCAACTCCTTCCTCGAAAGAACATACCATGGGCACACCTCAGCTA | 120 | 52 |
| chr7 | 144403257 | 144403377 | 772751_42001748_Shank2(210274)_1899 | 1 | + | good | AAGCTAGGCCAAATCACCAGTGCTAAGAGACCTAGAGTATGGGGTTGGGATGTAGATCAAGCTAGACCAGTGTGTTCTGGCTGGCTCACCAAAGTAGGCTGGCAAGTGGACAGTGGGGAC | 120 | 52 |
| chr7 | 144403377 | 144403497 | 772751_42001748_Shank2(210274)_1900 | 1 | + | good | TACCCAGAAAGAAGTTTGATCTCAGCCGTAAGAGCCAATCTGGACCCCTGGAGAAAGATGGCAGAGATGGAGGCAGGTGGGCATGGCCTATATCACATGGCACAGGACGAGGTCCACCAG | 120 | 54 |
| chr7 | 144403497 | 144403617 | 772751_42001748_Shank2(210274)_1901 | 1 | + | good | TTCTGTTAGTAAGGAAGGTTAGTAAGAAGATGATAGACACTTCAGCAGAAGTGCCAAAGCCTAGGAGAGTGAAGGTTTGGTTTTGAGACTCAGGGGCCAAGGACATGGCTGATTTGAGCT | 120 | 45 |
| chr7 | 144403617 | 144403737 | 772751_42001748_Shank2(210274)_1902 | 1 | + | good | CCAAGTGTCCTCACTGCTGCTGGAGTTGGGAGCTGAGCAATTGCCTGACCAATCTCTGGAGGCTGAATTCTGGCCCAGAAAGAGGCCCGTGGCTGCAGGTCCCAGATATCAGAGTCAGCC | 120 | 57 |
| chr7 | 144403737 | 144403857 | 772751_42001748_Shank2(210274)_1903 | 1 | + | good | ATGAAGGGACAGGCCACCTGAGACTGCATGGATTCTGCCGGCCTTTTAAAAATTCTCCACACATGTGAAAGGCTCCAAGGCCATGAAGGACCATGTGGCATGGGCTACATCTGGACTGCA | 120 | 52 |
| chr7 | 144403857 | 144403977 | 772751_42001748_Shank2(210274)_1904 | 1 | + | good | TCCATCATCTGTGGACTGCTAGAATCAACACTGGTCAAGAGTTAGCCATTTTCCCAGGCTGTCAGAGGATCCATGATGGATACTTCCATCGGATGGATATTCCCCCTAATCATAGCACAC | 120 | 47 |
| chr7 | 144403977 | 144404097 | 772751_42001748_Shank2(210274)_1905 | 1 | + | good | TCTCTCCCTCGAGGGGATCCTGGCTCTGCCCTGGCTTCTTACACAGTGTTGCTTCATTCCAATGCTGAGCACTAACGCAGCTCTGAGGGCTTAGCCAATGGGTTAGCTCCTCCTGATAAA | 120 | 53 |
| chr7 | 144404097 | 144404217 | 772751_42001748_Shank2(210274)_1906 | 1 | + | good | CCACCATTGCCAAGTAGTATTTGACTTGGTAGCTCAGTCAGTGAACCCAGTGTTCACTAAACAGGGAGCCACCAGACCCCAGAAATAGGCAAAGCTTGCCTGGAAGGACTTGGGCATCTA | 120 | 50 |
| chr7 | 144404217 | 144404337 | 772751_42001748_Shank2(210274)_1907 | 1 | + | good | CAGCAAAAGTTCTAAAAGCTAAGCATCTCCTGCACCTCAAAAGGATCCAGCGTGGGAGGTGTCCACAGGCATGGCCCCCCATAAGCTTCATCCTGACCCAGGGCATCTTATAAAAAGCAC | 120 | 51 |
| chr7 | 144404337 | 144404457 | 772751_42001748_Shank2(210274)_1908 | 1 | + | good | AGCCCGGGACAGCTAGGGCCTTCTCCTGAGTGCCTCTGGGGACTGGGTGACTCTGACTGCAGATCTCAGGAAATGGGGTCCCCCTGTGTTCTGGCAACCCTAGACAAAGCCTTATATCTG | 120 | 57 |
| chr7 | 144404457 | 144404577 | 772751_42001748_Shank2(210274)_1909 | 1 | + | good | GTCTCAGGATCTCAGCTGTCTTAGAGCTGAACCCGGTGTCAAGGTCATTCTAGAACAAGTTTTGGTTTTCAAGTTTAAACAAAAAGCTTTTGGTGTTTTATTTTGTTTTGTTTTTGTTTG | 120 | 36 |

| | | | | | | | | | |
|---|---|---|---|---|---|---|---|---|---|
| chr7 | 144404577 | 144404697 | 772751_42001748_Shank2(210274)_1910 | 1 | + | good | TTTCGTTTCTTCCATTTTCCAGGGCTCCTTGAAAATTGGTTTTTATTTTCTAGAATGTTCTAAAATGGGGAGAATAAATAGAAAGTTCCCTGTGTAGCATATGCTCAGAGGTCTGGGGTT | 120 | 38 |
| chr7 | 144404697 | 144404817 | 772751_42001748_Shank2(210274)_1911 | 1 | + | good | CCTGCCATCCTGTCTGGCTTACTAATCACAGTCAACATCCCAGCCTTGGTACAGGGGAGGCTTGCCTTGAAAACTATGTTTAATTACAAAAAGGACCACATTTGTGTGTGTGTGTGTGTG | 120 | 46 |
| chr7 | 144404937 | 144405057 | 772751_42001748_Shank2(210274)_1913 | 1 | + | good | CCAGCCTGAGCTACACAATGAGACCCTGGCTCTAATAGCAATTCCAAAATTAACACTTATCAATAACAATACCTGTCATAGCTGAAGATGGCTGAGTCACAGGTAATGGGTGGTATAGCA | 120 | 42 |
| chr7 | 144405057 | 144405177 | 772751_42001748_Shank2(210274)_1914 | 1 | + | good | CTCTCCAAGGGGGCAGGCTGAGATGTGGACGTCTCCTGCTGACTTTGTCCAGATAGTTAGTTACACACCTAAGGCAGGCAAGGGATGCATGGTGGACCCCCTAGTGGGGCATGTGGAGGG | 120 | 57 |
| chr7 | 144405177 | 144405297 | 772751_42001748_Shank2(210274)_1915 | 1 | + | good | GGCATAGATGCATGTTTGCTCCCCACACACAGCCTGCTCGATCGTCCAAGCACATGAACAGTGACTGCATGCCTGAAGTCTCCTGACTGAGCCCATCCACCCTCCAGGCTTCCATAAGAC | 120 | 55 |
| chr7 | 144405297 | 144405417 | 772751_42001748_Shank2(210274)_1916 | 1 | + | good | TAAACCACTCCAAAACCTACTTTGTCTTTTAGATAAACCGGAAGAGATAGTCCCAGCCTCCAAGCCCTCCAGGACTGCAGAGAACGTGGCCATCGAATCCAGGGTGGCGACCATCAAGCA | 120 | 51 |
| chr7 | 144405417 | 144405537 | 772751_42001748_Shank2(210274)_1917 | 1 | + | good | GCGGCCCACAAGCCGGTGCTTCCCAGCTGCCTCTGATGTGAATGTGAGTGAGTGGAACCGGGTAGGGTTTGGGCTGGCCTGACTGATGGGCGGGCAGGCAGAGGGTTCAGCATGCAGGGA | 120 | 63 |
| chr7 | 144405537 | 144405657 | 772751_42001748_Shank2(210274)_1918 | 1 | + | good | GCGGCAGGCCAGTTCAGCAGCTGACCTTCACAAGCCCTCCCTGCACGTGTTCAGAGCCCCATCTCCAGAGGCTAGTACTCTGATCCGGCCCAGAGCTACAGCAAGTCAGTTACTGAGCTG | 120 | 59 |
| chr7 | 144405657 | 144405777 | 772751_42001748_Shank2(210274)_1919 | 1 | + | good | AGGTGCAAATCTCACCACTCCTGCATGCCACCCTTACCTGGGTTTATCCAAACCTTGAGGGTGTCCCTCCACGTGTCCAACCAAGAGACTTTGGACAAAGGGGATGTGGGAGTGACATCA | 120 | 52 |
| chr7 | 144405777 | 144405897 | 772751_42001748_Shank2(210274)_1920 | 1 | + | good | CTCCTCAGACAGTGCCCCTTGCCTGAGCACAGCCTCATGCACTGTGGTCCTTTGCACAAACACTTGGCACGTGCTCTGAGGTGGGGTGACAGCATCCTTAGGAGGTCAGTGACTTGATGT | 120 | 56 |
| chr7 | 144405897 | 144406017 | 772751_42001748_Shank2(210274)_1921 | 1 | + | good | GTTGGTCTTCATTGTGCATGGGAAGCGCCGGATGCCAGCTCACTGGTGCTGGTGCTGGTGCTGTGCCACCCACCCCAAGTAAAAGGCACTCTCAGTAAAGAGGTGATGAATTTCCTAGGA | 120 | 54 |
| chr7 | 144406017 | 144406137 | 772751_42001748_Shank2(210274)_1922 | 1 | + | good | GCCAGGAGTCAGTCCTGGACAACAGTAAGATCTAAGAACCCCTTAGCACCTCCTTCTGGTTTCTGGTCACCATCACCGTCCAGTGCCAGGTCTAAGCACTAACTGTCCCATCACATAGCT | 120 | 52 |
| chr7 | 144406137 | 144406257 | 772751_42001748_Shank2(210274)_1923 | 1 | + | good | GAGTATTTTCTCTGACTCACTTTCAGAGTGACTTGGTTTAATCCAGCCCTTTTACAAAGTGACACAGTTGCCACGAATTTGATAAGAGTTTCATAAGATAAGCAATAAAATAGACACTCC | 120 | 37 |
| chr7 | 144406257 | 144406377 | 772751_42001748_Shank2(210274)_1924 | 1 | + | good | CCGAGGTTCAGATACTCACTCCTCCCCTTCAGAATCCCATAGGGAGGAACCCTGGCTTTCAACACACCCACACTCCACCCTGATACTCTCCAACCTGATAGCCCCCATGGGACAGGTTCC | 120 | 56 |
| chr7 | 144406377 | 144406497 | 772751_42001748_Shank2(210274)_1925 | 1 | + | good | TTTGAGCTCCCAGGGGCCAGCATCCTACCTTCCCAGCCTCAGATTCCAGGGTCAGGTGGGTAGAGGCTCTGGAGCCTAGATGAGCCTTTCAGAGTATACCCACAGACATGTTCTTGGTGA | 120 | 55 |
| chr7 | 144406497 | 144406617 | 772751_42001748_Shank2(210274)_1926 | 1 | + | good | GCCTTAGTGGCCAATCTTTCCCAAGTCCCTCCCAGGGCTGTGGAGCATAGCTGGCCAGCCATCTCAGACAAAACAGCATAGGTGTGTACCAGCTGTGAGGGCGTAGGCAGGTACTCTCTG | 120 | 57 |
| chr7 | 144406617 | 144406737 | 772751_42001748_Shank2(210274)_1927 | 1 | + | good | AGAAGTCCGCCTTCAGGCCAGGGAAGGGACAGCCCGCTGCAGTCTCTGTGATACTTTCCAGGGTCACTTGGCAGTTCTCACAGGTCAGGCACCCTACCCACCTGCACAGGTTTGGTACCT | 120 | 58 |
| chr7 | 144406737 | 144406857 | 772751_42001748_Shank2(210274)_1928 | 1 | + | good | GGACACCTCCATAGTATCTGGAAGAGGAAAACAGAATCCAAACTGAGGACTTGACAGAGGTCACAGGGCATAAGGGCATGGCCTGGATCAGACCCCACTGCCTACCACGCTGTGCCTGTC | 120 | 54 |
| chr7 | 144406857 | 144406977 | 772751_42001748_Shank2(210274)_1929 | 1 | + | good | CCTGTGGGCCAGCCATCCTGGCCCTCTTCTTCATGAGCTCTTCTGCTGACTGGCCGGATGAATGTCTCAGCAGAGTGTGGCTGACTAAATCTGGACACGCTGGCATCAGATACATGAGAG | 120 | 55 |
| chr7 | 144406977 | 144407097 | 772751_42001748_Shank2(210274)_1930 | 1 | + | good | CTAAGAGGACTATGGGGAGGGCATGGCCTGAGGAAGCTGTGGTGATTGATACCCGAAGCTCCCAGAGCTTCCTGACGCCTCTAGCCCTGTCACCCTGGGTGGGGCTCCCAGGGATGCACC | 120 | 62 |
| chr7 | 144407097 | 144407217 | 772751_42001748_Shank2(210274)_1931 | 1 | + | good | TCACCTGGCTCTCTGGTTTGCAGTCCGTGTACGAGCGCCAAGGGATTGCTGTAATGACGCCCACGGTCCCTGGGAGCCCGAAAGGCCCATTTCTGGGCCTCCCTCGAGGTACGATGCGAA | 120 | 61 |
| chr7 | 144407217 | 144407337 | 772751_42001748_Shank2(210274)_1932 | 1 | + | good | GGCAGAAATCGATAGGTAAGAAGCGTGACCCTCCACCCGAAGACGTTGCCCCTAAGCTTTCCAACATAGCCCTTAGCTCCTCAGGGATTCTGGCCTCCTGACTCCAGGCTCCATGATGGC | 120 | 56 |
| chr7 | 144407337 | 144407457 | 772751_42001748_Shank2(210274)_1933 | 1 | + | good | CTTGGAACCTAGACCCTAGACACTCCCCACCGGGACTGCCTATCAGGGGGCAAGTCCCCAGCTTCTCTCCAACAGGCAGCCCCTTGAACACGGTGGGGGTGTGAAACCTTCCTTTAGAAT | 120 | 57 |
| chr7 | 144407457 | 144407577 | 772751_42001748_Shank2(210274)_1934 | 1 | + | good | GGTTTCTCTAATCTCCTGACAGTAGAGAGAGCAGGGAGGAGGGCGGGCAGGGTGGGCAGACCCCCTCCAGTCAATGTTTCATAGGCTGAGCTCAAGGGACACACTTCTCCGGGTGGCTGG | 120 | 59 |
| chr7 | 144407577 | 144407697 | 772751_42001748_Shank2(210274)_1935 | 1 | + | good | GTCAGGTCTGAAGCGCTGTGTGTAAGAAAGCAGCTTGCCTTCCTCGCTAACATCCCATTTCCTTCACTGGCTCGGACAGGGCAAGGGAAACCTCACCCCTCAGGACAAGAAAGACCTGAG | 120 | 54 |
| chr7 | 144407697 | 144407817 | 772751_42001748_Shank2(210274)_1936 | 1 | + | good | TGTGGTCCCTACCACATAAACTTTCTGGAAGTCTAGGCTACTGCTGGACACAGTGGAGCCTTAGCATGTAAAGAGAAGGGGTATGCAGCCTCCAGCCTCAGCAGAGCACTGCCAGGAACA | 120 | 52 |
| chr7 | 144407817 | 144407937 | 772751_42001748_Shank2(210274)_1937 | 1 | + | good | CTGCTGCTCTGTGTGTTCCGGAAGCTCCTCCTGCGGCTGCTGCCAGTGACATGGCATGTCTGTTTTTAGTCAGAGGAGGAGGCAGGAGGCAGCATCTAGGCAGTAATGACCATGGTGGCG | 120 | 57 |
| chr7 | 144407937 | 144408057 | 772751_42001748_Shank2(210274)_1938 | 1 | + | good | TCTGCAGCTTGGGTGGCCCACCCACCCCTGGCACAGTCAGGGCCGGGCTTAGCTGTACACACACCTTATTGTATTTCTGGATCTTTCTGGCTAATAACAGTTGATTTCCTCCTGTCAGAC | 120 | 52 |
| chr7 | 144408057 | 144408177 | 772751_42001748_Shank2(210274)_1939 | 1 | + | good | AGCAGAATCTTTCTATCAGGTCAGTGAACCAACCCCCCACACCCCAGTTTGGAATCTGTAACTTGAATGTATCCCAGTTTACTTTATAAGGGGTATTCTTAAATTATTACTCCTTGGGAA | 120 | 40 |
| chr7 | 144408177 | 144408297 | 772751_42001748_Shank2(210274)_1940 | 1 | + | good | TGTCACCATTAGGCGACAACTCACAGTGAATGGTACACAGAGATCTCCTTCAGCAGCAATCCTCTGAGATGCCAAATTAGCCTCCCACAGGAGTATGTTCAATTTGGAATACCCTTATCA | 120 | 45 |
| chr7 | 144408297 | 144408417 | 772751_42001748_Shank2(210274)_1941 | 1 | + | good | AATGGAAGTTCTCATGAAATAAGTGACTGTAGTTTGTCCTGGGATTTTTTTTTTATTTCAAGTAATTCATTTTGTTGCCTCTGAAAACCTGCATGCCCCTGTGTTGAGAATTCATCCCTG | 120 | 37 |
| chr7 | 144408417 | 144408537 | 772751_42001748_Shank2(210274)_1942 | 1 | + | good | GTGTCTGCGCTGTGTACGCTGCTTCTGTCTGTGTGTCTTAGGCTGTACTGTCACACCAGCTGGTACATACACCCAGGGAAAGCTCAGGTCACACTGTGGCTTCAGCCACACCCCCACTCA | 120 | 56 |
| chr7 | 144408537 | 144408657 | 772751_42001748_Shank2(210274)_1943 | 1 | + | good | TCACACACACACACACAAAATCACCAGTCAAGTCTCAGAACATGGTTCTGCCTTTGTTCCAAGTCATTGAGGCTTGGTTAACCATGACAGCTAACCTCAGCTCTCCAGAGAGTCCCACAG | 120 | 48 |
| chr7 | 144408657 | 144408777 | 772751_42001748_Shank2(210274)_1944 | 1 | + | good | GCCCACATCAGAGCTTCCGTGGACATGGCAGCTGGCATGGAGTAGCTTACTGGCATCTTATAGCTGCTGTGGCCCAGGACTAGGTGGAGAAATCAAGTCTCCCTCCCTTCATGGACAGAG | 120 | 55 |
| chr7 | 144408777 | 144408897 | 772751_42001748_Shank2(210274)_1945 | 1 | + | good | TCCATGCCAGAATGGCCCTGTCTTTCAAGCCAGCTACCCGCTGAGGGCTGAGGAATTCAACACACACAGAGCCTTCAGTGTGACCCAGCAGTAGCAGGTGACAAAACTGGTCAATGTCCC | 120 | 54 |
| chr7 | 144408897 | 144409017 | 772751_42001748_Shank2(210274)_1946 | 1 | + | good | TTTGGAGAAGACCACACAGTGACAGGAGCAGCTCCTCCTTGCACATTTGGGACACTACTAGGGAGACAGCTACTCTAGCAAGGTAGCAGGTCTACCTGCTAACTGGACAGGAGGGACAGC | 120 | 53 |
| chr7 | 144409017 | 144409137 | 772751_42001748_Shank2(210274)_1947 | 1 | + | good | AACCTTGACCTTTATGCCATTCATATAGCACCTCAGGGTCAGAAGAGAGCCTCAGTCCCTTAGAAACCCTCACTAGGTCTGAAAGACGCAAAGACGGATATCACCTGAACCTCACACTTG | 120 | 48 |
| chr7 | 144409137 | 144409257 | 772751_42001748_Shank2(210274)_1948 | 1 | + | good | TACCCACAACCACAACTTTGGATTCAGGTTGCCAGGGCTTGGCCCTGTTTCAGGCACTTACCACCTTTGCCTCACTCTAGACACCACACCATGCCCCAGAGAAGCAAATCCCTACATGCG | 120 | 53 |
| chr7 | 144409257 | 144409377 | 772751_42001748_Shank2(210274)_1949 | 1 | + | good | TTCGTTCATCATAAATACCTACCTGTCTTTCTGTTTCTGGCTCCTAGAAGGTGTGTGACTAACGGCAGCATCCTGGCCTGTGCTGTGCTTCCTAACAAGTGTCTCCTGTCTGAGTCTGTG | 120 | 48 |
| chr7 | 144409377 | 144409497 | 772751_42001748_Shank2(210274)_1950 | 1 | + | good | GAAACACAGTGTATTACACAGTCCTGAGGACTCCAGTGCACTTTGTGGTTGCTGGGCTTAAAACACCGTGTTCTCTGCTTGCAGGGATAACAGAGGAAGAGCGGCAGTTTCTGGCTCCCC | 120 | 52 |
| chr7 | 144409497 | 144409617 | 772751_42001748_Shank2(210274)_1951 | 1 | + | good | CAATGCTGAAGTTCACCCGAAGCTTGTCCATGCCAGACACTTCTGAGGACATCCCCCCTCCGCCACAGTCTGTGCCCCCCTCTCCCCCTCCTCCTTCCCCCACCACATACAACTGTCCCA | 120 | 60 |
| chr7 | 144409617 | 144409737 | 772751_42001748_Shank2(210274)_1952 | 1 | + | good | GGTCCCCGACTCCAAGAGTCTATGGGACAATTAAGCCCGCGTTCAATCAGAACCCCGTCGTCGCCAAGGTGCCCCCAGCCACCAGGTCTGACACTGTGGCCACCATGATGCGGGAAAAGG | 120 | 60 |
| chr7 | 144409737 | 144409857 | 772751_42001748_Shank2(210274)_1953 | 1 | + | good | GGATGTTCTACAGGAGAGAGCTGGACAGATTTTCCCTGGACTCAGAAGACGTCTACAGCCGCAGCCCCGCCCCACAGGCCGCCTTCCGCACCAAGCGGGGACAGATGCCTGAGAACCCGT | 120 | 62 |
| chr7 | 144409857 | 144409977 | 772751_42001748_Shank2(210274)_1954 | 1 | + | good | ACTCAGAGGTGGGAAAGATAGCCAGCAAGGCCGTCTATGTCCCTGCCAAGCCAGCCAGGCGGAAGGGCGTGCTGGTAAAGCAGTCCAACGTGGAGGATAGCCCCGAGAAGACGTGCTCCA | 120 | 60 |
| chr7 | 144409977 | 144410097 | 772751_42001748_Shank2(210274)_1955 | 1 | + | good | TACCCATCCCAACCATCATCGTCAAGGAACCCTCCACCAGCAGCAGCGGCAAGAGCAGCCAGGGGAGCAGCATGGAGATTGACCCCCAGGCCACCGAGCCCGGCCAGCTGCGGCCAGATG | 120 | 65 |
| chr7 | 144410097 | 144410217 | 772751_42001748_Shank2(210274)_1956 | 1 | + | good | ACAGCCTCACCGTCAGCAGCCCCTTTGCTGCGGCCATCGCTGGGGCTGTGCGTGACCGGGAGAAGCGTCTGGAAGCCAGGAGGAATTCCCCAGCCTTCCTCTCCACCGACCTGGGAGATG | 120 | 65 |
| chr7 | 144410217 | 144410337 | 772751_42001748_Shank2(210274)_1957 | 1 | + | good | AGGACGTGGGTCTGGGGCCGCCTGCTCCCCGGATGCAGGCCTCCAAGTTCCCAGAGGAGGGTGGGTTTGGTGACGAGGATGAGACGGAACAACCGCTGTTGCCTACCCCCGGGGCAGCAC | 120 | 67 |
| chr7 | 144410337 | 144410457 | 772751_42001748_Shank2(210274)_1958 | 1 | + | good | CCAGGGAGCTGGAGAATCACTTCCTAGGTGGTGGTGAGGCTGGTGCTCAGGGGGAGGCTGGGGGACCCCTGAGTTCCACATCCAAAGCCAAGGGGCCTGAGAGTGGCCCAGCAGCCCCCC | 120 | 66 |
| chr7 | 144410457 | 144410577 | 772751_42001748_Shank2(210274)_1959 | 1 | + | good | TCAAGAGCAGCAGCCCAGCCGGCCCTGAGAATTACGTGCACCCACTCACAGGGCGGCTGCTTGACCCCAGCTCCCCACTGGCCCTGGCACTCTCAGCCAGGGACCGAGCCATGCAGGAGT | 120 | 66 |
| chr7 | 144410577 | 144410697 | 772751_42001748_Shank2(210274)_1960 | 1 | + | good | CCCAGCAGGGGCACAAAGGAGAGGCCCCCAAGGCTGACCTTAACAAGCCTCTCTACATCGATACCAAAATGCGGCCCAGCGTGGAGTCCGGCTTTCCTCCAGTCACCAGACAGAACACCA | 120 | 58 |
| chr7 | 144410697 | 144410817 | 772751_42001748_Shank2(210274)_1961 | 1 | + | good | GGGGTCCCCTGCGACGGCAAGAGACAGAGAACAAGTACGAGACAGACCTGGGCAAGGACCGGAGGGCGGACGACAAGAAGAACATGCTGATCAACATCGTGGACACTGCCCAGCAGAAGT | 120 | 58 |
| chr7 | 144410817 | 144410937 | 772751_42001748_Shank2(210274)_1962 | 1 | + | good | CAGCTGGCCTACTGATGGTGCACACAGTGGACGTCCCCATGGCCGGGCCACCCCTGGAGGAAGAGGAGGACAGAGAGGATGGGGATACAAAGCCAGACCACTCACCCTCCACAGTGCCAG | 120 | 62 |
| chr7 | 144410937 | 144411057 | 772751_42001748_Shank2(210274)_1963 | 1 | + | good | AAGGCGTTCCCAAAACCGAAGGTGCTTTACAGATCTCCGCTGCCCCGGAGCCCGCCGTCGCGCCCGGCAGGACCATCGTGGCGGCGGGCTCCGTGGAAGAGGCGGTGATTCTGCCATTCC | 120 | 67 |
| chr7 | 144411057 | 144411177 | 772751_42001748_Shank2(210274)_1964 | 1 | + | good | GCATCCCCCCTCCCCCTCTGGCATCCGTGGACTTGGATGAGGACTTTCTTTTCACAGAACCGTTGCCTCCCCCCCTGGAATTCGCCAATAGTTTTGATATCCCCGATGACCGGGCAGCTT | 120 | 57 |
| chr7 | 144411177 | 144411297 | 772751_42001748_Shank2(210274)_1965 | 1 | + | good | CAGTTCCCGCTCTGGCTGACCTGGTCAAGCAGAAGAAAAACGACACCTCTCAGCCCCCTACGTTGAACTCCAGCCAACCAGCCAACTCCACAGACAGTAAGAAGCCAGCCGGTATCTCGA | 120 | 56 |
| chr7 | 144411297 | 144411417 | 772751_42001748_Shank2(210274)_1966 | 1 | + | good | ACTGTCTGCCCTCCTCGTTCCTGCCACCCCCCGAAAGTTTCGATGCAGTCACCGACTCGGGGATTGAGGAGGTGGACAGCCGGAGTAGCAGCGACCACCACCTGGAGACTACCAGCACCA | 120 | 62 |
| chr7 | 144411417 | 144411537 | 772751_42001748_Shank2(210274)_1967 | 1 | + | good | TCTCCACGGTGTCCAGCATCTCCACGCTGTCCTCAGAGGGCGGCGAGAGCATGGACACCTGCACAGTCTATGCAGACGGGCAAGCCTTTGTGGTTGACAAGCCCCCAGTACCTCCAAAGC | 120 | 60 |
| chr7 | 144411537 | 144411657 | 772751_42001748_Shank2(210274)_1968 | 1 | + | good | CAAAAATGAAGCCCATCGTTCACAAGAGCAACGCACTTTACCAAGACACGCTCCCAGAAGAGGACACGGATGGCTTTGTGATCCCCCCACCTGCACCCCCGCCCCCGCCGGGCAGCGCCC | 120 | 62 |
| chr7 | 144411657 | 144411777 | 772751_42001748_Shank2(210274)_1969 | 1 | + | good | AGGCCGGTGTGGCGAAGGTCATCCAGCCAAGGACCTCCAAGTTGTGGGGTGACGTTCCAGAGGTCAAAAGCCCAATTCTCTCAGGCCCAAAGGCAAATGTCATTAGTGAGCTAAACTCCA | 120 | 53 |
| chr7 | 144411777 | 144411897 | 772751_42001748_Shank2(210274)_1970 | 1 | + | good | TTCTGCAGCAGATGAACAGGGGGAAATCGGTCAAGCCCGGGGAAGGGCTGGAGCTGCCGGTGGGAGCCAAGTCGGCCAACCTCGCTCCAAGGTAAGTTCCCAATCAGCTTGGGGAATGTG | 120 | 60 |
| chr7 | 144411897 | 144412017 | 772751_42001748_Shank2(210274)_1971 | 1 | + | good | TGGTCCAGTGCTGGCCAAAAGAAACCATTGGGGAGCCACCTATGCAGCCGAGAAGAGGGGACCTGCATGGCCTCTGCCACACCCATTAGGCTATAACCATGTTGCCACCATCCTCCTGTC | 120 | 57 |
| chr7 | 144412017 | 144412137 | 772751_42001748_Shank2(210274)_1972 | 1 | + | good | ACCGGACTACGGCAGAGCCAAGGCCATGGCTGCCATGCAGCCTATGACACACCTTCCTAAAGACACATAAAGGAACCAGATTGCACGGGACCAGGCCCACGGTCGCTTCCAGAAGGAAGC | 120 | 57 |
| chr7 | 144412137 | 144412257 | 772751_42001748_Shank2(210274)_1973 | 1 | + | good | TTTGGTCTCTGCTCCGTTTCCTCTGGCCCTTTGCCAGGACTACCTCAAAGGCAGGCAGAGCTCCATTAAAACCCAAACTGGTCAGCAGAGGTGGCCAGTGTCTGAACATGGGTGTACCAT | 120 | 53 |
| chr7 | 144412257 | 144412377 | 772751_42001748_Shank2(210274)_1974 | 1 | + | good | TCCCTCTCCCTTAATCCCCCTTGGGGTTGAGGATTTGGAAGGAGCTGGGTCATTGCTCTGGATGAACACCTGCCCCTCTGCTGCATCACCCCACCCAGGGGCAGACGTCACAAAGAAATC | 120 | 57 |
| chr7 | 144412377 | 144412497 | 772751_42001748_Shank2(210274)_1975 | 1 | + | good | ACTGCCAGCTGCGCCCGATGGGCCGTTTCCAGCCTCACGTAGGGCTTAACAAGCTACCTTCATTGGGATGGGGGGGGGGGTAGAATAAACACATTCTTCCTAAGTCCAAACAGAAGGTTC | 120 | 54 |

| | | | | | | | | | |
|---|---|---|---|---|---|---|---|---|---|
| chr7 | 144412497 | 144412617 | 772751_42001748_Shank2(210274)_1976 | 1 | + | good | TAGAACAGCCCCCTGTGTGGTTTCAGGATGGTACTCGGCAGCCTTGCAGGCCCCACCCTCACTCATGGCCTTTCTGTTTTTGTTCCTTGTGTGATGTACACTCGTGCTGTCTTTCCGCTT | 120 | 53 |
| chr7 | 144412617 | 144412737 | 772751_42001748_Shank2(210274)_1977 | 1 | + | good | CCGTGTGTGTAGTGCTCAGCCTCTCTGGCTGACAGTTGAGCTCAGGTTTCATCGGTTTGACTAATGAAGTCTTTCTTTATTCTCCCGCTGGGCAGGCATCTGATTCAAACAACCGAGCCC | 120 | 51 |
| chr7 | 144412737 | 144412857 | 772751_42001748_Shank2(210274)_1978 | 1 | + | good | ATTAATCAGGGACATCACACTGCATCTGAGTTCTTGAGGTTTGGGCTGTGAGCCACAATTCTTCCTGTTACCTTTTGGTGTGGTTGTAAATGGAGGACTTGGACGGCTCCTTAACTCCTTT | 120 | 46 |
| chr7 | 144412857 | 144412977 | 772751_42001748_Shank2(210274)_1979 | 1 | + | good | GAGAAATACTGATTGGCGGCTGTTGTAAAGAAGTGTGCGGATGCCACCGCTTGGTAATGGGAAACCTGCTACAGGTGGTGTGCAGAAGCAAGCTCCCGTGGCTCCCTGCTAGCTTGGGCG | 120 | 56 |
| chr7 | 144412977 | 144413097 | 772751_42001748_Shank2(210274)_1980 | 1 | + | good | GGCCTGCAGAAACCCTCCACTCTGGTGGGTGGGAGGTGGGGTAACCACACACCTGCATATTTTAAACATGTCCAGGAGTCTCACAGCAACCTTGGAATATGCTTGGTTCATATTTTTCCT | 120 | 49 |
| chr7 | 144413097 | 144413217 | 772751_42001748_Shank2(210274)_1981 | 1 | + | good | CTATAGAGTCTTTTATGCCACAGCCTCTTGTGAAAGTAATATTTGCAAGCCCTTGGGGTGGCGTCCCTCACGACTGGGCAGCATGATCCTGTGTGTTAAGTCCATTCTTCAGAGAGGCCG | 120 | 50 |
| chr7 | 144413217 | 144413337 | 772751_42001748_Shank2(210274)_1982 | 1 | + | good | AGCAGGTATTGCAGCCTCTCTCTAAAAGTGAGGTTACAATAGCTGCCCAGTCTTCAACAGGAGAAAAGCTGAGGTGGCCGCTGGCATTGGGAAGGTGCGGGGGGGGGGGGGGGGGTTGCA | 120 | 58 |
| chr7 | 144413337 | 144413457 | 772751_42001748_Shank2(210274)_1983 | 1 | + | good | GGGGAATTCTGAATCGCAGCAGATGACCCTTGCTGCCTTTCTTTTTCTGGTTAGACAATGTCTGGTGGCTCCCAGGAGAGGGAAGGAGACTTACCCTCCACTGGGGGCTGGGACTATTGT | 120 | 53 |
| chr7 | 144413457 | 144413577 | 772751_42001748_Shank2(210274)_1984 | 1 | + | good | CTTTCCCTACTCCAGGTAGGTGCCAGATGTGGGCTGGGTGCTCAGAGAGAGCATCACTAGCACTCAAAAACTGGCTCCACCCAGCCAGTCTATTCCTGGCGCAGACCTCTCAGAGCAAGG | 120 | 57 |
| chr7 | 144413577 | 144413697 | 772751_42001748_Shank2(210274)_1985 | 1 | + | good | ACTTTAGCCAACGCCTACAAAGGACCAACAAACATTGGCCATTCTGTCACATAGGTACCAGCAACGTGGCCATCAACCAGCAGAGAGAGAAATCTGAGGCAAAGAGTCAAAGGGACTGTA | 120 | 48 |
| chr7 | 144413697 | 144413817 | 772751_42001748_Shank2(210274)_1986 | 1 | + | good | TGGGAAGGTTCTAGAAGCTGAGGCTGGGGGTTGGCTTCGGAAAAAGGGATCACCACTGAAGACCATCTACAAGACTGAGCTCCTGGGAAGAAAGGAACCGCCATTAGACCTAGAACTCCC | 120 | 52 |
| chr7 | 144413817 | 144413937 | 772751_42001748_Shank2(210274)_1987 | 1 | + | good | CTGCTAATGAGGGGAGAGATGACTGTTGTTGACCTCAAGCAGTAGAGGAAACACAGGCCTCTGACCCTCCTTTTCAAATCTAAGGATTGCACTGGGTTGGGCATGGGCTTAGGGTCCTGG | 120 | 52 |
| chr7 | 144413937 | 144414057 | 772751_42001748_Shank2(210274)_1988 | 1 | + | good | GATGCCAGCAGCAACTGCGAGACACAGAGGGCCACAGGCTGGGGTAGCCGTTCACAGAACCATGCATGCATCCTGTATGGGATATGCAGGTGACCGAGCTCTGCCTCTTGCCTTGGTGCA | 120 | 58 |
| chr7 | 144414057 | 144414177 | 772751_42001748_Shank2(210274)_1989 | 1 | + | good | CTGGCCTCTTGCCTACATTGCTCAGACACCTCTCCGTTGCCTTCACTGCCAGGCCCAGGATGACAACTAGAGATTCACTCAGTCTGAGCATCTTGCTAATGCCACTTGTAAATGCTTCTA | 120 | 50 |
| chr7 | 144414177 | 144414297 | 772751_42001748_Shank2(210274)_1990 | 1 | + | good | AAAGTTAAGTTGTGGACCACCCTGGGCCAAATGAGGTTCAGAATCAAAAGGAGAGAAAGAAAATGAAGGTTTCAAGAGATAACTGAAAGCTGGCCATGCTCTCTGACGTGACACACATTA | 120 | 42 |
| chr7 | 144414297 | 144414417 | 772751_42001748_Shank2(210274)_1991 | 1 | + | good | AGGCAGAGAGACCCTGTTTCCTCAAGCATGACTGCTCACCAAGGAAAGCCTGGCTCTCAGCGTGTTCCTCAGTGGTTCCACCACATGGGCAAGAGCTTTGCTGTGCTTGTAAATTGAAAT | 120 | 51 |
| chr7 | 144414417 | 144414537 | 772751_42001748_Shank2(210274)_1992 | 1 | + | good | AATGGGCCTTTTGCTACTTGGCAGCTCCAAGAGACCTCGTCTTATGCTGGCACCAGGACAGGATCCAGGAAGAGCAGAGAGTTAGAAGGCCTGGCATTTGCTTCCTGGGGTGTAAGCAAT | 120 | 52 |
| chr7 | 144414537 | 144414657 | 772751_42001748_Shank2(210274)_1993 | 1 | + | good | CTAACTTTAGGATACCCGGGGTTTAGTATACCCAGTAGACTGACCCCAAGATCAAGACAGTGTGGAATTTAGACATATTTAACTACAAATATATTGAAAAAGTCAGCTGGAACTCTCTTT | 120 | 38 |
| chr7 | 144414657 | 144414777 | 772751_42001748_Shank2(210274)_1994 | 1 | + | good | CAATATATTTGTGTATATGTTAGTGTATGATATCAGAGCCACTGAAAGACCATGGTGCAAAGCCGTGAGCAATTCTGGCTATAGGCCTGATGCTGAAACCTGTTCTCTTCCAGCATCCTT | 120 | 42 |
| chr7 | 144414897 | 144415017 | 772751_42001748_Shank2(210274)_1996 | 1 | + | good | ATGGGTAACCTTGGCATGTCCTTCAGTTGGACAGGGCTTTTGGCTTTCAGAGCCTCCTGTCATCTGAATCTGGCCCACAGTGGAGGCAGCAGGCCTGACACACATTTTTACTGCTTCATG | 120 | 52 |
| chr7 | 144415017 | 144415137 | 772751_42001748_Shank2(210274)_1997 | 1 | + | good | GATCTAAGCAGAGCACTTAGGTGTCCTGGTTAAAGTGCATAGGGCCAATGACGAGCCATGGTAGGGAGTTTACTAAGCTGCTACCACTGTCTACAGCCCAGTACCAAGCCCAAGAAAGAA | 120 | 49 |
| chr7 | 144415137 | 144415257 | 772751_42001748_Shank2(210274)_1998 | 1 | + | good | GAGACAAGAGGACATCACCACAGAGGATTCTGCAGTACAGTCCTCCACAGCTGGGAAACGTAGTCATTGTCATCCACTCAGGCAGCTTGTTGATGGCCCAGGACACCAGGCGCCATATGG | 120 | 54 |
| chr7 | 144415257 | 144415377 | 772751_42001748_Shank2(210274)_1999 | 1 | + | good | CACTCAGAATTTGTCATTTAAGAGAAGCACGTCCCTGTTAAGAAAGCATAGCCAGTGTTCCCTTGGGGAATGCCAGCTCTTGAGGTGAGAAATGGCCAAAACGAAACAACAGAACTATAT | 120 | 43 |
| chr7 | 144415377 | 144415497 | 772751_42001748_Shank2(210274)_2000 | 1 | + | good | GTAACAAGTACCACGGAGAAGACCAGGCTCCAAAAGGGCAGAACTAGGATGGAGCTGGCACTCAGAGGAAGGTATTTCAGTGACCTAAAGTTCACACCGGAACCAGTTATGCAGAAACC | 120 | 50 |
| chr7 | 144415497 | 144415617 | 772751_42001748_Shank2(210274)_2001 | 1 | + | good | AGAAGCATGGAGCCCAGGACATGATGGGTGGAGCATGTGAGGCTAGGCCAGATCAGACTCTCCATGCCCGTGGGGGGAGTCAGCGTCAACCCACCAGGCAGGGGAGTGGGCAGGCTGTGG | 120 | 63 |
| chr7 | 144415617 | 144415737 | 772751_42001748_Shank2(210274)_2002 | 1 | + | good | GGACACAGGATTGCATCCAAGGGATGAAAACCACGCACAGACTTATCATGGCTTACCAGGAGTGGACTCCAGTTTGTTGACACTCCCGGGATGCGTGTCACATGAGGAAGATGCTTTCAA | 120 | 50 |
| chr7 | 144415737 | 144415857 | 772751_42001748_Shank2(210274)_2003 | 1 | + | good | GGTCTGTGCTGAAGACAGGTGGTTTCTGTTGCTAGAGACAAGAGGTACCTTGGAGACACTGGAGAGCTAGACCCGGCTGCCTGTCTATAAGCACTGACTCCTCAAGCTGGCTGTGACCAG | 120 | 54 |
| chr7 | 144415977 | 144416097 | 772751_42001748_Shank2(210274)_2005 | 1 | + | good | CCATGGCTCCTCTCCAAGGGTGTGGTGTACACTCTACTGCCAGCACTTTGTGGTGAGTCCACAAGGGGCAGAGTTCGGCCTTCACCTTTCTGAAGCACACAGGAAAGAAACAGTGGTACT | 120 | 53 |
| chr7 | 144416097 | 144416217 | 772751_42001748_Shank2(210274)_2006 | 1 | + | good | GAGGAGGTACAGACACCCAAGGTCCCTGGGCAGGCAGCGCTCCAGGTGCTGATGGGGCTTCACTCAATCCCTGCCACCCCTGCTGTCCTCCCAGGGCAGTCCTCCGACCCTGGCCATTGT | 120 | 65 |
| chr7 | 144416217 | 144416337 | 772751_42001748_Shank2(210274)_2007 | 1 | + | good | CCTCTGGCTCTCAGCTGGGTCACCAGTATTAATCAATACGATTGTTAAACTGCTGTGACCACCTGGTCCTAGAGCATGGGCTTCTGTTTCATGACGTCTCAAGCTCTCTGCACTGCCACA | 120 | 50 |
| chr7 | 144416337 | 144416457 | 772751_42001748_Shank2(210274)_2008 | 1 | + | good | CAGCTCTGGTACCTCAGTCACCAAGCAAAGCAGGAGGGTGATGGCACTGCCAGGGAGCAGCACTCAGTAGTGAGGTGAGCCCTGCTGGGGACAGCCTGGCACTTCCTGCCACTATCCACA | 120 | 60 |
| chr7 | 144416457 | 144416577 | 772751_42001748_Shank2(210274)_2009 | 1 | + | good | TATTAAGGTGACTAGTCTGCCAGAGTCTCTTCCTTTGACCACCAAACAAACGGCAGGTCTCCTTTTTAGAACATATTTATCTATTTGTATGTGCAGTGGATGTTTTGCTTGCTTGTGTGT | 120 | 40 |
| chr7 | 144416817 | 144416937 | 772751_42001748_Shank2(210274)_2012 | 1 | + | good | GATCCTATCACAGGTAAACAGTTCTTCCTGAAAGTAGGTGCCTTCCTGTCCTGGCAACTTGTGAGCAGGGTGGGAACCCAGACAAGTGTGCTGAGCCTGAGACAGCTTCCTCGGCAGGAA | 120 | 53 |
| chr7 | 144416937 | 144417057 | 772751_42001748_Shank2(210274)_2013 | 1 | + | good | AGCAGGCAGTGTCCCACAGCCATTCCTTCTGTGGTTATGTATTCTGTTCCCATTCTGGAGAGAGGCGTGGAGTGAGCAGAGTCCTCACAGTTCTCTGAGCCTGTGGTGACTGCAAGGCAC | 120 | 54 |
| chr7 | 144417057 | 144417177 | 772751_42001748_Shank2(210274)_2014 | 1 | + | good | CAGGATCAGAAGTGCTGAGGGGGTCCCAACTTCTGAGACTCAGGCAAACACATGACCTGGCTCTCCTTCCCGTGCCCCATTTGTGGAAGCCTAGATAGGCAGAGCAGCTTCATAAGAAAG | 120 | 53 |
| chr7 | 144417177 | 144417297 | 772751_42001748_Shank2(210274)_2015 | 1 | + | good | CAGGCTGATGCCCAATGGGGTTCTATACCTGTAATGCCAACGCTAGGGAGGCTGAGGCAGGGGGATTTGATAAGTTCTAAAATAACCTGGGGCTACAGTGAGACTCTGTCTCAAGTCAC | 120 | 50 |
| chr7 | 144417297 | 144417417 | 772751_42001748_Shank2(210274)_2016 | 1 | + | good | TAAAAAGAGGAGAGAGAGGGGGGGAGGAAGGAGGGAAGACAAATAGCTTAGAATTTAGTGGCTCATACAGAGGGGAGCCTGGGACTGAGGGTCCATGCCTCAGAACCCTAGCATTAGTGCT | 120 | 51 |
| chr7 | 144417417 | 144417537 | 772751_42001748_Shank2(210274)_2017 | 1 | + | good | GGCCCCTGAGGAGCCCTCTGGCCCACACCTCTGCTCAGGTTCAAACTGAGTAACCACAGTGTGAGTCTGGTTGCCCTTCCATCTCCCCAGCTCATCAGCTCCTGCCAGGAAAGACAGGTGA | 120 | 58 |
| chr7 | 144417537 | 144417657 | 772751_42001748_Shank2(210274)_2018 | 1 | + | good | AGAGCGTCTGTGAGCAGCTGGATTCGCTCTTGGGCTGGTTTTGTTTTGTTTCTTAATTCTCACATTTTTTAAATGAATTTGATTTTAAAAGCAGGCTAGAGATGATCCTCCCAGAGGCGG | 120 | 42 |
| chr7 | 144417657 | 144417777 | 772751_42001748_Shank2(210274)_2019 | 1 | + | good | GTGTTTGTGAGTGTGTATCTATTCTAAATCAGGGTAACAGACTTCTGTATATGCCGCCCAGCAGCCCCGCGTTGCCTAGCAACATAATGATCTGCGCACACGGGAGGATCCTGTGAGTC | 120 | 51 |
| chr7 | 144417777 | 144417897 | 772751_42001748_Shank2(210274)_2020 | 1 | + | good | ATCACTGTGTGATTCACCAGGAAGGAGGCAGGCTACCCAGAAGGTGCAGAGCAAAGCCGGCAGGCAGACCAGTCCAACTTAACTCAGCAGTTGTCATGCTGGGTGGGGTACCGGTAAGCC | 120 | 57 |
| chr7 | 144417897 | 144418017 | 772751_42001748_Shank2(210274)_2021 | 1 | + | good | CCAAGCATGTGGCCGGACCAGATCCATGCCATTAGCCACTAGGCTGGTCACCATGTTGGGGTGTAAACGTGATACAGAGCCCTAGCCTGGGCAGGTTGGCCTGGGTCCACTGGGAACAAG | 120 | 58 |
| chr7 | 144418017 | 144418137 | 772751_42001748_Shank2(210274)_2022 | 1 | + | good | ATTCATTCTTTCCAAGTCAAAGCCAAGTGAGGGAAATAGCTGTGGGCCACTTGTCAGAGATCCTTTCTTTCTTCCGCAGTGCCTCATGGCACGCCAGAGAAGGCAAAGAGTAGTTCCCTT | 120 | 48 |
| chr7 | 144418137 | 144418257 | 772751_42001748_Shank2(210274)_2023 | 1 | + | good | GCTCGTATGAGTGCCTGCCCCAAGCCCCCTGGGCTGCCTGCCAAGAGGGCTCTGAGGTGCATTGTTCAGCCAGGGTTCAACCTACCATGTTTCCACCCGTGAGGAGGCTGCTGCCTCTCC | 120 | 62 |
| chr7 | 144418257 | 144418377 | 772751_42001748_Shank2(210274)_2024 | 1 | + | good | TTGCTTTCCAGTCAGTTCTGCAGTGTGGTCTCAACCGTTCCCTGTGTCGTTTCCATTTGCTCCCATGGTGCTGTTTGCAGAAGAGGGTCATGCTGAGCAGAAGTAAGTGCGGCGGCCTGA | 120 | 53 |
| chr7 | 144418377 | 144418497 | 772751_42001748_Shank2(210274)_2025 | 1 | + | good | CCAGATTTGTGTGTCCGTCCCGTCCCCCCCCCCCCCCCCCCCGTGTGTCACTGTCTCTGCTCCTACCTCTGCTTCCCAGATTCAGCTGCCTGCTGGCTGAATCCGTCCTGACCAGAAGG | 120 | 64 |
| chr7 | 144418497 | 144418617 | 772751_42001748_Shank2(210274)_2026 | 1 | + | good | CCAGCCCAGTCCAGGGTGCTGGCTTAGAGACCCTTCAGAGCAGTGCCTGGGTATCTGCTGGCCCTCAGGGGACTGGCATGCCAACCACATGGCTCCAAAATAAGGGTCAGGCCAGTGCTG | 120 | 61 |
| chr7 | 144418617 | 144418737 | 772751_42001748_Shank2(210274)_2027 | 1 | + | good | GGCACAGAACTTCACAGCTCCCTCCCAATCCTTCACTGCCTTACTGGCACTCAGGAGCTCCTGTAGGACACCCCCTGCCTCCCCCATCCTGAAGTATCACATGCTAGAAATGTGCCCCCC | 120 | 57 |
| chr7 | 144418737 | 144418857 | 772751_42001748_Shank2(210274)_2028 | 1 | + | good | TCCCCAGTTGCAAAGAGCCCCTGGGCAACTTCAATGCTTATCCAGGCTGAACCTGGAATGGCTTACCCTACAACCAGGAAATCGTTGGGTGACATAGCATGTTCTGCAGACCCTTAACAT | 120 | 50 |
| chr7 | 144418857 | 144418977 | 772751_42001748_Shank2(210274)_2029 | 1 | + | good | GTCCCTGGCCCACCTCTCCATGACTAACTAAAGCTGAATCTTTAGAGTTCCCCCGAAAGCTGCAAAGCCCAGATGGGCTTCAGGACCTTAACTTCATGAGCTGCCTCAGTCCTGTTAAGA | 120 | 51 |
| chr7 | 144418977 | 144419097 | 772751_42001748_Shank2(210274)_2030 | 1 | + | good | TGACCCGCACACTGCAGGGACAGGCTCTCTGAGCAAACTTTGGTGGGAGGGCACTCTTGGACTTACAGGTACCTACAGTTCCCAGAAGCCACTGCTGCAGGTCCAGTGCTCCCACCCACA | 120 | 58 |
| chr7 | 144419097 | 144419217 | 772751_42001748_Shank2(210274)_2031 | 1 | + | good | GCAGAGTCCTAGTCCACTGAACTCACACCTGCAATGTCCAGGTCTCCCCTGCAATGTCCAATATTTGAAGTAATCATTGCTAGCCCTAGTAGGAATAGTCCATAGAACAGAGAGGTCACC | 120 | 48 |
| chr7 | 144419217 | 144419337 | 772751_42001748_Shank2(210274)_2032 | 1 | + | good | TGCTAATGGCTTCATCCAGACACACAGCAGGCTAAGAAAACAAACAACATCCCAGGAAGAGAAAGAGGCAGTGGAAGGTCCAGGGACAACTGTATGTTGCCTCAGACTCTTTCCCCTCAC | 120 | 49 |
| chr7 | 144419337 | 144419457 | 772751_42001748_Shank2(210274)_2033 | 1 | + | good | TTATGATCTGAGGAGGTCCTAGTCTGCTAGTATGTATGTGCCCTGCAAGGGAACTAAGTAAGGGGTCCCTAAAGTTCAGTGGACTGACACCATAGTAAGGAGTTTAGCTCAAAGCAGGCC | 120 | 48 |
| chr7 | 144419457 | 144419577 | 772751_42001748_Shank2(210274)_2034 | 1 | + | good | CTGTGACATCACACCCAGAATCCCCACCCTGACAGTTATGAGGTGGACTGGGGAGCCCTCAGTTCTTACCAGTTTTATGCAGGGCATGAAGGCGGGGCTAAACTCTCAGCCATTTCAGTC | 120 | 53 |
| chr7 | 144419577 | 144419697 | 772751_42001748_Shank2(210274)_2035 | 1 | + | good | TAGCTTGTGCCCATGGCAGGATGGACCTTTCCAGAAATGAGAGGTGTGTGGTGTGTCTACATTCACGTGGGTGAGCCCAGGTCAAGAGACACACCTACTTTCTTTAACCTGGTGGTTGTC | 120 | 51 |
| chr7 | 144419697 | 144419817 | 772751_42001748_Shank2(210274)_2036 | 1 | + | good | TGCAGTTTGGCAATAGAGCAGGAAGCAGTGCTTAGAGAGAGGCTTCACCTTCCTAGGTGTGTGACTAGAAAGTCTTCAGCAGCCCTGACTAACATGGTCAGAGAACACATTTCATAGAAT | 120 | 46 |
| chr7 | 144419817 | 144419937 | 772751_42001748_Shank2(210274)_2037 | 1 | + | good | GACCACCCCCAGATAGATGGGTGAGAAGAACTTCCTAGTCACTCAGGACCTCCTCCCAGTGCAAGTCACCCACCTACAGCAGGCCTGATAGGCTCTGTGAGAGTTGAACCTGTCTCTCCC | 120 | 56 |
| chr7 | 144419937 | 144420057 | 772751_42001748_Shank2(210274)_2038 | 1 | + | good | TATTCTGAGGACTTCACTTACCCAAATGTAGCAGGAAGGACACCACTGGGAAAAGGACCTTCTGTCTGCTATGGTAATAGGGACAGACGACCCTGCCAGCAAGTCTCTGCCTGCAGGGAG | 120 | 52 |
| chr7 | 144420057 | 144420177 | 772751_42001748_Shank2(210274)_2039 | 1 | + | good | TGCACTGCTATACCCTGAAGCCTGCTGTCAAACCTTCCAGTCAGTGGTAGGCAAGGTTCCCTATGAAGAGGCCACCTGCAGAACTTACCCTGGGACACACTAGCAACCTCTGTGCCCTGA | 120 | 54 |
| chr7 | 144420177 | 144420297 | 772751_42001748_Shank2(210274)_2040 | 1 | + | good | GCATCCAGGGAAAATCTCTTCCAACCCTACGTCCCAATTCTGAAGTAGGGCTGGCCCTCGCATTTGTCTGAAACCCAAGAAATTTAACCCTGAGCAGACCCCTGACAGGCTGCTGAGAAAG | 120 | 51 |
| chr7 | 144420297 | 144420417 | 772751_42001748_Shank2(210274)_2041 | 1 | + | good | GGCCCCTGGGAGGTGTATGTTGTTGCAGAGGGCCTGAGCTCATTCTGGCCTGCACAGAGGGTCTCTCCAGGCGCCTCCCAGCCTGGTCCAGGTGAGTGGTTGGCTTGCTGTGCTGTGCTC | 120 | 63 |
| chr7 | 144420417 | 144420537 | 772751_42001748_Shank2(210274)_2042 | 1 | + | good | CACCTACATCCCTCTGACTGGCATGGTTGTCTGTATCACTGCCAGTGCAGACCCTCTGCTGTGGTGTGGGTGCCTCCCCCTCCCATCCCATGAGCCTCAGTGACCCAGAAGTGATGTTCT | 120 | 57 |
| chr7 | 144420537 | 144420657 | 772751_42001748_Shank2(210274)_2043 | 1 | + | good | TCTCTGTCTTTGCTGTTTCAGAAGCCCGGAGGTCATGAGCACCGTCTCAGGTACACGGAGCACGACGGTCACCTTCACTGTCCGCCCTGGCACCTCCCAGCCCATCACCCTGCAGAGCCG | 120 | 62 |
| chr7 | 144420657 | 144420777 | 772751_42001748_Shank2(210274)_2044 | 1 | + | good | GCCCCCCGACTATGAAAGCAGAACCTCAGGACCTAGACGCGCCCCAAGCCCTGTGGTTTCACCAACGGAATTGAGCAAAGAGATCCTGCCCACCCCTCCCCCTCCGTCCGCCACTGCAGC | 120 | 62 |

| | | | | | | | | | |
|---|---|---|---|---|---|---|---|---|---|
| chr7 | 144420777 | 144420897 | 772751_42001748_Shank2(210274)_2045 | 1 | + | good | CTCTCCCTCCCCCACACTCTCAGATGTCTTTAGCCTTCCGAGCCAGTCCCCTGCAGGGGACCTCTTTGGCTTGAACCCAGCAGGACGGAGCAGGTCACCATCTCCTTCAATATTGCAACA | 120 | 57 |
| chr7 | 144420897 | 144421017 | 772751_42001748_Shank2(210274)_2046 | 1 | + | good | GCCAATCTCAAATAAGCCTTTTACAACTAAGCCTGTCCACCTGTGGACGAAACCAGATGTGGCAGACTGGCTGGAAAGTCTGAACTTGGGTGAACACAAGGAGACGTTCATGGACAATGA | 120 | 48 |
| chr7 | 144421017 | 144421137 | 772751_42001748_Shank2(210274)_2047 | 1 | + | good | GATTGACGGCAGCCACCTGCCAAACCTTCAGAAGGAAGACTTGATAGATCTTGGGGTGACTCGAGTTGGGCATAGGATGAACATAGAAAGGGCTTTGAAACAGCTGCTGGACAGATAAGG | 120 | 49 |
| chr7 | 144421137 | 144421257 | 772751_42001748_Shank2(210274)_2048 | 1 | + | good | GTGGCTGCCCTGGGCCTTCACAGACTCCTTTCTTATAAGTAGAGATGGGCTTATGTTGAAATGTGTGGTGGCCAAGCAGAGGCTGACAGCACCAACCTCCAGTCTGTGGGTTTGTCTCTG | 120 | 52 |
| chr7 | 144421257 | 144421377 | 772751_42001748_Shank2(210274)_2049 | 1 | + | good | GGTACCCAAAGATGTGCTGTGGGTGTCCCTTGGTGTGGCCAAACTGGGGCCACAGATATCTTGGGTGGCTTCTGTCCAAAGGGACAGTAAGGGGGTTTCTGTCCTGGTCTCAGAGGCTGC | 120 | 57 |
| chr7 | 144421377 | 144421497 | 772751_42001748_Shank2(210274)_2050 | 1 | + | good | CATAGACGGTATGTGGGACCCTCTCCTCCTGTGGCAGACACCAGTGTACTCCAGATACCTCCTGAGGCCTCCCTCCTCTGCCTTCTGGGCAACTCTCACCCCTCAGCCCTGGAACACAGC | 120 | 60 |
| chr7 | 144421497 | 144421617 | 772751_42001748_Shank2(210274)_2051 | 1 | + | good | CTTTCTTCAGGCCTTTGGCCTCTGGGTGGGCTCATTGGCTCTGTCCCTCCCCTGTGCTTGGCTCTCAGTGAGCAGTTCCAGGTGGAGCCCTGCCTGAGCCAGACTGTAGAGAACACTGTC | 120 | 59 |
| chr7 | 144421617 | 144421737 | 772751_42001748_Shank2(210274)_2052 | 1 | + | good | CCATCCACCTCTTTGTCCGACTCCTGTGACGTGCATCGGAGCTTCTTTGGCCCCTAGGTCCTTGGCAGTGGTGGCTGTGGACAGGATCGTGCCCTGCTTGCTTAACTTGCTGTCTTTATT | 120 | 55 |
| chr7 | 144421737 | 144421857 | 772751_42001748_Shank2(210274)_2053 | 1 | + | good | TCTGCAAACAGATTAGCATAATTCCAGGGTGGCCAAAATGAAGTCACACGGAGTTAGTCAAAGCACAAAGTCACGATCCTGAGGAGGAGGGAGGCCTGGCTGTAGAGCCAGCCAATGTGT | 120 | 50 |
| chr7 | 144421857 | 144421977 | 772751_42001748_Shank2(210274)_2054 | 1 | + | good | GGCTGCCCAAGCCCACAGCCTCCTCCAGGGCAAGGACAGGTGTCTGCCCAGAGCATCTGCCCAGGAGGGAGAGGTTGGGGCTCTTGAAATTTACTCTAATTGGATGTCTGCAGAGCTCCA | 120 | 58 |
| chr7 | 144421977 | 144422097 | 772751_42001748_Shank2(210274)_2055 | 1 | + | good | CAGAGGTCAGAACAGGTGAGCAGCTACTTTTACAAGGACAAGAGGACCTGAGGCTTGTGGTAAGCATGGCTGCCATCGGGAAGCTGCTGTCCTCCCTGGTAGCAGATGGAATGCCATACC | 120 | 54 |
| chr7 | 144422097 | 144422217 | 772751_42001748_Shank2(210274)_2056 | 1 | + | good | CTTCATCTGCACCAGGCCTTGGCTGAGCATCCTGCCGAGGCAGCCCTTTGGCTCTCAGAGGCCAAGAAAGGTGGAACTCAGGGGCCTTTCTCCCTTCTCTCACAAAGTGGGGATCCCACC | 120 | 59 |
| chr7 | 144422217 | 144422337 | 772751_42001748_Shank2(210274)_2057 | 1 | + | good | AGCTTGTCTTAACATCTGGCTCTCAGAGAAGCCCGTGGTCAGGACCTGGTGGAGGACAGAAAGTGGGGCACAGGCTGTGGCAAAGTTTAGGTGTGTGTGGAAAAGAAGGTGGGGAGCATC | 120 | 54 |
| chr7 | 144422337 | 144422457 | 772751_42001748_Shank2(210274)_2058 | 1 | + | good | GTGGACCAAGCCTCACCTCAGGCCCAGCCACCTGAGCCCACCCTTCTCGTCCCCAGATGTCTCCCTCCCAAAACAATACCTCTAATGTCCCAGGGAGGTCACTGACTGCACACAGCCACC | 120 | 60 |
| chr7 | 144422457 | 144422577 | 772751_42001748_Shank2(210274)_2059 | 1 | + | good | ATGCAGCCAGCCAGGCTCCCGAAACCCCCTCTCCCCCACGCTCAGAGGAGACTCGAATGTCAGGTGACGGTTTTATCATTTCAGAACTAGCCCAGCCCATCTCTAATTATAAAACATCTT | 120 | 51 |
| chr7 | 144422577 | 144422697 | 772751_42001748_Shank2(210274)_2060 | 1 | + | good | TTTCCTTTTTTTTTTTTTCTCTCTTCACTTATGCTCACAACCACGTGTCTGACGAGGCCTGGAACAGCGCTACAATGAACACATAAAATTTAGCAATTAAAAAAAATCTTTCTTTACTGC | 120 | 35 |

**Supplemetary Table2c-the probe design for mouse Shank2 gene bost**

| Chromosome | Start | Stop | Name | Score | Strand | Code | Seq | len | GC |
|---|---|---|---|---|---|---|---|---|---|
| | | | | | | **good** | **CATTCTTCCTTCTATCCAGCATATATACAGCCAAGCTAGTCATCCATCTATCCATCCAATCTAATCCATATATCCAGCCAGCCAACTGGCCAGCCTCCTATCCATCTATCTACCAGTCCT** | **120** | **45** |
| chr7 | 143709026 | 143709146 | Target=1;ProbeIdx=15349;Target #1 | 50 | + | good | TTCCCTATTCTCTCCATGATGGGCTGTAAGCTGTAAGATGAAACTTTCCTCCCCAAGTGCTTTTGGTCACCTTGTTTATCACAGCGATAAAAGCAAACCAGAACAGAAGGGGTGCTAGC | 120 | 45 |
| chr7 | 143563106 | 143563226 | Target=1;ProbeIdx=757;Target #1 | 48 | + | good | GAGGTCAGAAGAGGGCAACCTTGGAGCTGGACTTGCATGCAGTTGTAAACTGTCTGACTTGGGAGCTGGGAACTGAATTCAGGTTTTCTGCAAGAACAGTACAATCTCTCTAGCCCTGAA | 120 | 48.33 |
| chr7 | 143588906 | 143589026 | Target=1;ProbeIdx=3337;Target #1 | 48 | + | good | TGTGAATAACAGAGTGACTTTAACTTTAAAATCTTTATTTATGTGCACATGTGTTAGTATGTATGTGCATGCATGCGTGTGTGTGTGTGTGTGTGTGTGTGCGCGAGCGCGCACGCAT | 120 | 42.5 |
| chr7 | 143619146 | 143619266 | Target=1;ProbeIdx=6361;Target #1 | 44 | + | good | TCTGTGGCTTTGGAAACTGCCAGTCACCAGGTTTGAGTTGGTAACTCTTGATAAATTGCATAAAAGATAGTCATGGGGCTGGAGAGGTGGCTCAGCAGTTAAGATCACTGGGTGCTCTTC | 120 | 46.67 |
| chr7 | 143663546 | 143663666 | Target=1;ProbeIdx=10801;Target #1 | 42 | + | good | ATTACTTACCTTTCTATTGCTCTGGCCTGACCAAAGCAACTTACAGAAGAAAGTGTCTATTTGAGCTTATGGTTCCAGAGGGATCCAAATCTGTGATGGCAGAACGGAGGCATAGCAGCT | 120 | 44.17 |
| chr7 | 143602586 | 143602706 | Target=1;ProbeIdx=4705;Target #1 | 34 | + | good | CAGAGAGAGAGAGAGAGAGAGAGAGAGAGAGAATGTCAATGAAATGTATGGTCTTTTCTGAGTCAGGTTCTCATTGCATAGTCCAGGCTGGCCTTAAATTCAAGAGATCCACCTGCCTCTGC | 120 | 45.83 |
| chr7 | 143679266 | 143679386 | Target=1;ProbeIdx=12373;Target #1 | 34 | + | good | ACATCTAAGATGCAGGGAGAGACCCTGTCTCAAAAGCAAACTCTATGGCTCCTAAGGAACAACAAAATTGTCCTCTGGCCTCTACATATACACATGAATGCACACACACACACACATACA | 120 | 43.33 |
| chr7 | 143655386 | 143655506 | Target=1;ProbeIdx=9985;Target #1 | 31 | + | good | GCATGGAACCCAGGGCTTTGTGCATGCTAGGCAAGCACTTTACCAGCTGAGCTATACTCCCTGCCCTACTTTATTGGCTTTGATTGACAGACATAGTAAACGACGAACTCTTCTTTCCAT | 120 | 46.67 |
| chr7 | 143718986 | 143719106 | Target=1;ProbeIdx=16345;Target #1 | 29 | + | good | TGAGGCAGGGTCTCTCAATCAAACCCAGAGCTCATCTGTCCTACGGCTAGGCTTGCTAGTAGCAGCTTGCTCTGGGGATCCTTTTTCTGAGGATGGATTACGGGGTTCTGGGGATCTGAA | 120 | 52.5 |
| chr7 | 143595266 | 143595386 | Target=1;ProbeIdx=3973;Target #1 | 27 | + | good | GTACCAGCATTCCTCTCTCTCTGCTTCCTGATTGTGGATGGGAAGTGAGTGACTGCTTCATGCTCCTGCTGCCAAGCCTTTCCCGCCATGATGGACTGCACCATAACCCAGGGAGCCCAG | 120 | 55.83 |
| chr7 | 143720066 | 143720186 | Target=1;ProbeIdx=16453;Target #1 | 27 | + | good | CATATGCATGTACACATGTGCACACACAGACACATGCAGGCACTCACACATGCATGCACATGCGTGTGCACACACACACATAAGCACACATTACCTTTTGGTAATATGCCTCTTGTGTGT | 120 | 45.83 |
| chr7 | 143639666 | 143639786 | Target=1;ProbeIdx=8413;Target #1 | 26 | + | good | AAGAGAGGAACTATCTAGATCAGGTTGGCCTGTGGGCATTATCTTGATTCTTAATTGATGTGGGAAGACGTATAGCCTACGGTGGGCGGCACCATTCTCTAGTCAGGGAGTCCTGAACAG | 120 | 48.33 |
| chr7 | 143681306 | 143681426 | Target=1;ProbeIdx=12577;Target #1 | 26 | + | good | AATGTGGGTGGTACCATCCTTTGCTCTGGGGTCCCAGACTGAATAAAAAGAAGAAAGCAATTTAAGCACCAGTGTTTATCTCTCTGCCTCCTGACTGTGGATTTGATGTGACCTCACACT | 120 | 45 |
| chr7 | 143684906 | 143685026 | Target=1;ProbeIdx=12937;Target #1 | 25 | + | good | AGGGGATCTAATGCCCTCTTCTGGCCTCTATATTTAACAGTCATATATATGGTATACTGATATATAGGCAGGCAAAACACAAGTAGACTCAAAGTAATAAAATGTTTTTTGAGACAGCAT | 120 | 35 |
| chr7 | 143607266 | 143607386 | Target=1;ProbeIdx=5173;Target #1 | 23 | + | good | TAGCTTTTTTATTTTGTTTATGTGTGTGTGCATACATGTGTGGGTACCTGTGAAGGTCAGAAGAAAGCATTGGGTGCCTGGAGCTGGAGTTATAGGCAGTTATTAGTGTCCTGAAGTGGG | 120 | 43.33 |
| chr7 | 143643746 | 143643866 | Target=1;ProbeIdx=8821;Target #1 | 23 | + | good | GATGTAGGATGTAGAGGATGTCGTTGGGCGAGGGTTGGGTAGTAGGGAAGCCCGGGGACACCGCTCCAGGGGTGGGGAAGCACAGACTGAGGAGCAGGACAGCCAGAGCTGGCTCAGGG | 120 | 63.33 |
| chr7 | 143558186 | 143558306 | Target=1;ProbeIdx=265;Target #1 | 22 | + | good | GGTGACCCTTCTGTGTGAACATGGACTGTCACTGAGGTGACCCTGTAGCTTGTCTTTGCTACTTTGTGGGTTATTTTTTCTTCTACCTTTCTGTACCCCATTCTTCCACCCTTTCAAACC | 120 | 45.83 |
| chr7 | 143567546 | 143567666 | Target=1;ProbeIdx=1201;Target #1 | 22 | + | good | TCTCTCTCTCTCTCTCTCTTTCTCTCTCTCCCCTTTCTTACTTCACTCCCCCTTTCTATTATAGGGTATACTATAGCCCAAGGTGGCCTCAAACTCACAAAGTAGTCAAGGATGACCT | 120 | 45 |
| chr7 | 143680346 | 143680466 | Target=1;ProbeIdx=12481;Target #1 | 22 | + | good | TGGGAGGCAAGAGGATCCGGAGTTCAAGGCCATCCTTAGCTACATAGCAAGTCGGAAACTAGCCTAGGATACATGAGACCATGTTTCAAAATAATTGATTGATTAAGTCTAGCCCCGAGT | 120 | 45 |
| chr7 | 143617106 | 143617226 | Target=1;ProbeIdx=6157;Target #1 | 21 | + | good | GCCTCCCAAGTGCTGAAATTAAAGGCATGTGCCACCACAACTGACCCCTACCCACCTATGTTTATCAGTCCAGTCTCATCTGAGGTCCAGGTTGGTCTTCAGTTTGTCTGTGCTTTCTCT | 120 | 49.17 |
| chr7 | 143689826 | 143689946 | Target=1;ProbeIdx=13429;Target #1 | 21 | + | good | GGCAGACCCATTCCATAGGCTGGGATCCTGGACTGAATAAAAAGGAGAGAGTGGGCTCTCTTTTACTTCCCTCTGTCTTTGCTTCCTGACTGTGGATATAAGTCCTGCCGCCATGCCTTC | 120 | 50.83 |
| chr7 | 143596466 | 143596586 | Target=1;ProbeIdx=4093;Target #1 | 20 | + | good | AAATGCCCTATGGATGGAGCTCATGGAACGTCTTCTCAGCTGAGATTCCCTCCTTGCAGATAACTCTAGCTTGTATCAACTTGACATAAAATTAGCTTCTATGCTCCCTAGGCATGTAAC | 120 | 43.33 |
| chr7 | 143611946 | 143612066 | Target=1;ProbeIdx=5641;Target #1 | 20 | + | good | ATAGAGCTGTGTGTCCTGGTCAGGGTGTGGTCCTGTCCAGCACTGACCCATCACTATCTCAGCTCCAGTCTCTCCTGCAAGGTGCACTGACAAGTTCTCAGAACTCTGGAGGCTCTCTAG | 120 | 54.17 |
| chr7 | 143715506 | 143715626 | Target=1;ProbeIdx=15997;Target #1 | 20 | + | good | GTATACAGATATGCAAGCAGGCGCACACTCATACAAATGAAATAATAAATAAACAAACAAACAAATAAATAAATAATAAATAAAAGAGTTAAAGGTAGTAGCTGAGGCATATCAACCCCA | 120 | 30 |
| chr7 | 143566586 | 143566706 | Target=1;ProbeIdx=1105;Target #1 | 19 | + | good | AGTGAGAGAGCCTAGCAAGCAAAACAAACAAACAAACAAACAAACAAACAAACAAACCACTTAGGCTCTGGCAAGCGTCCAAAACAAAGAGAAAGAGAAGAAGAAAACGGGAAAGCCGTG | 120 | 41.67 |
| chr7 | 143641826 | 143641946 | Target=1;ProbeIdx=8629;Target #1 | 19 | + | good | TTTTTTGATTTAATAAAGAAAAATTAATTAATTAAGGGAGCTTGCCTGCATGTATGTGTGTGCACCACTTCTGTGCCTGGTATCATAGAGAACGGAAGAGGGCACTGGGTCCCCTGGAGC | 120 | 43.33 |
| chr7 | 143579186 | 143579306 | Target=1;ProbeIdx=2365;Target #1 | 17 | + | good | CACAACAATCCTCCTACCCACCCACCCACCTATCTACCTGGCCTTTATCATCCATCCTCACATCTATCCATTCATTGTCTATCCATTCTCCCATCCATCTCTCCTCACAGCATCTATCCA | 120 | 47.5 |
| chr7 | 143582186 | 143582306 | Target=1;ProbeIdx=2665;Target #1 | 16 | + | good | TTTGAGGGTCTCTCAATCAAACCCAGAACTCATCAATTGGCTAGTCCTGCTAGTCATCTTGCTCTGGGGATGCCCAGGCTCTGCCCTCTGGAGGCTGTAATTACAGGGGGGGGGCTGCCA | 120 | 55 |
| chr7 | 143564186 | 143564306 | Target=1;ProbeIdx=865;Target #1 | 15 | + | good | CACTTTTATGTGGGGGTCTGGGGATTGAACTCAGATTCTCAGGTTTTCTCTGCAAGCACTTTACCTACTGAGCCATCTCAGTTGGCCCAGATGAATTCTTCAGAGTTCTAAATAACATTC | 120 | 43.33 |
| chr7 | 143617706 | 143617826 | Target=1;ProbeIdx=6217;Target #1 | 15 | + | good | AAAGGCACAGCAGCAGGCTCAGCTGAGAGGTCACATCTAGATCTAAAATCAGGTGTCAGAGAGAGGTAGGGGGAGGAAAGAGAGAGACAGAGGGAGACAGAGACACAGAAAGAGACAAAG | 120 | 50 |
| chr7 | 143631146 | 143631266 | Target=1;ProbeIdx=7561;Target #1 | 15 | + | good | CACACACACATACACACACACACACACACACCTACACCCCTACAAATAGGAACATGTACACATACCTACACGGGCTCAGAAATTGTTCCTTCAAAGTTAAAGAACACCCATAACAACTCC | 120 | 43.33 |
| chr7 | 143655506 | 143655626 | Target=1;ProbeIdx=9997;Target #1 | 15 | + | good | CAGACACAGTGCTGATATTGCCCAAGACAAAACCATAAACCTGCCCGAATCATTGTTAGTTTCTGTGATTTGTGTGTGTGTCGTGTGTGTGTGTGTGTGTGCATGTGTGTCTGTGTTTGT | 120 | 44.17 |
| chr7 | 143555786 | 143555906 | Target=1;ProbeIdx=25;Target #1 | 14 | + | good | GAGGATGACCTTGAACTCCTGATCTTCTTGCCTCCAAATCCTAAATGCTGGGATTACAGGCATGCACAGCCATACTATATATATATATATATGTCACTCTCTTCAGACACACCAGAAGAGGC | 120 | 43.33 |
| chr7 | 143588306 | 143588426 | Target=1;ProbeIdx=3277;Target #1 | 14 | + | good | GGGCTTTGATATGTTACCACAGGTGCAATTGAGAGCCCTGTCCCTTCTGCCAGAGGTAAGGGAGATAGCTGCTTTTAGGGACAGAAAACACATTTCAGATTTCTGAGGAATGCTGTCT | 120 | 46.67 |
| chr7 | 143598626 | 143598746 | Target=1;ProbeIdx=4309;Target #1 | 14 | + | good | AAGGAAGATTTATGGGGTTTTTTTTTGTTTGTTTGTTTGTTTGGTTGGTTGCTTTTTGTTTGTTTTGTCCATTTGCCTTGTGGGTCCTGCAACTGCCCAAAGTTCTACAGAACACTTAGGG | 120 | 39.17 |
| chr7 | 143644226 | 143644346 | Target=1;ProbeIdx=8869;Target #1 | 14 | + | good | CATGAAACACCAGGAGTGGTGGCTCATACATTATAATCCCAGTGCTAGGGAGGTAGAGACAGAAGGAGCCCTGTGGCTTCCTGGCCAGCCAGCCTTGCCTAATCAGTGAGGCCCACGTCC | 120 | 55.83 |
| chr7 | 143654906 | 143655026 | Target=1;ProbeIdx=9937;Target #1 | 14 | + | good | TTGAACAATGGCAAAACACATCCATATTCTTCACAAAAATAGTCTGTAAACCACATACTAATATTCTTTTCCATTGAAACCTCTGGGGTCAGCAGGCCTACATAGTTAAAGTCACCTTCA | 120 | 36.67 |
| chr7 | 143678306 | 143678426 | Target=1;ProbeIdx=12277;Target #1 | 14 | + | good | GCAATAATGTAAGGAGAGAAGCCTCCATTATGGCTCACAGTTTGAGTGCCCACTCCTCCATGGTGGGAAGGCATGACATCAGGGCCATGAGACAGCTGGTCACATTGCATCTGTAGTCAG | 120 | 50.83 |
| chr7 | 143680706 | 143680826 | Target=1;ProbeIdx=12517;Target #1 | 14 | + | good | GGCTGGTCCTTCCTATATTAATTAACAATGAAGACACTCACCCACAGGTTTTGCCACAGACCAATCTGATCTGGGTAGTCTTTCAACTGTGACTTCCTTCTCAGCTGACTCTAGGCTGTG | 120 | 45.83 |
| chr7 | 143640146 | 143640266 | Target=1;ProbeIdx=8461;Target #1 | 12 | + | good | GATGCCACCCCAGTTGAGGGAGGGTCTTTCCCACTCAGCTCAACATTTCTAAAATATCCTCACAGGCATACCCAGAGGTGTGTCTCCTAATTAAGTGATTCCAAATCCAGTCAAGTTGAA | 120 | 45.83 |
| chr7 | 143657306 | 143657426 | Target=1;ProbeIdx=10177;Target #1 | 12 | + | good | GAAACAGATGCATGCTGGTATTCAGTTTATTTCCTCTCTCTTTATTCAGTCTGGGAATCCAGCCCACAGGATGGTGCCCTCCACACTCAGGATGGGTCTTCCCTCCTCAATGAACTCAAT | 120 | 47.5 |
| chr7 | 143680826 | 143680946 | Target=1;ProbeIdx=12529;Target #1 | 12 | + | good | AGGTGAGATGGAAGGTGGAGACAGGAGAATCCCTGAGAGCTTTTGGGCCTGCTAGCCTGGTGCACAGATCAGTGAACAAGAGATCCTGCTGCCTCCAAGAAGGTGGAAGGTGAGGCTTGA | 120 | 54.17 |
| chr7 | 143695226 | 143695346 | Target=1;ProbeIdx=13969;Target #1 | 12 | + | good | TCCCTCTCATCTCTACATCCCCAGTGCTGGGATCACAGGTACATGGAATCATGTCCAGTTGTTTATGTGGGTGCTGGGGATATGAACTCAGAGCCTTATACTTACATGCCAAACACGTTT | 120 | 46.67 |
| chr7 | 143571506 | 143571626 | Target=1;ProbeIdx=1597;Target #1 | 11 | + | good | GTGTTATATGATGGTCCAGGTGTGTGATGAAGAAGGCAAGGGAAGAGCAAAGTACACTGTAGCTCTGGAACTCTTTTTCTTATTTTAAAGATTTATTTATTATATTTATATGAGTACACT | 120 | 33.33 |
| chr7 | 143572586 | 143572706 | Target=1;ProbeIdx=1705;Target #1 | 11 | + | good | AGGATCAGGAGTTCAAGGCTGTCTTCTCATATAAAGAGAGTTTGGGGCCAGCCTGGGCACCACCTGAGACCCTATCTCAGACCAATCAAAAGAAAACAAAAATAGACATTCGAGTAACAG | 120 | 45 |
| chr7 | 143656826 | 143656946 | Target=1;ProbeIdx=10129;Target #1 | 11 | + | good | CCACACCTCTAAATCCTTCCCAAACAGTTCTAGCAACTGGGAACTGAACATTCAAACACACAAGCCACATCATTCTCAGTGATGACCATCACCAGGGTCATCACCAATTACTATTACCAT | 120 | 43.33 |
| chr7 | 143602826 | 143602946 | Target=1;ProbeIdx=4729;Target #1 | 10 | + | good | TGCTAATGTAGATGGAAAAAAAATTAAATCCATTGCTTCAAAACTATTTGATCAAAAAGCTGGGGGTGGGGTGGCCTGCACTTGTAATCCAAACACTGAGGAGGCAGAGGCAGGAAGATT | 120 | 41.67 |
| chr7 | 143653706 | 143653826 | Target=1;ProbeIdx=9817;Target #1 | 10 | + | good | TGGTTTTACTTTGTTCAGACAGAGACGGTTTCCCGGTTCTTTCTCTCTTGGAATGAGAAAGTTCTTAGATTGACTTGGATTTTTTTCTTACAGGAGCTCACAGTTGAGAGACTTTGGACTA | 120 | 40 |
| chr7 | 143562266 | 143562386 | Target=1;ProbeIdx=673;Target #1 | 9 | + | good | TTTCTTTTTTGAGGCTGGGTCTCAATCTGTGGCCTGGGGTATCCTAGAGTCACTTCATAGAACAAGCTAGCTTTGAACTTTAGGGCATCTGCCTGCCTCAGCCTCCAAGGGTTGGGACTA | 120 | 49.17 |
| chr7 | 143624786 | 143624906 | Target=1;ProbeIdx=6925;Target #1 | 9 | + | good | TCCCTCCTTCCTAGGGGTCATCCCTCCTTCCTAGGGGTCACCCCTCCTTCCTAGGGGTCATCCCTCCTTCCTAGGGGTCATCCCTCCTTCCTAGGGGTCACCCCTCCTTTCTAGGAGTTG | 120 | 60 |
| chr7 | 143642186 | 143642306 | Target=1;ProbeIdx=8665;Target #1 | 9 | + | good | CTTTCATCTCCTGAGTGCTGGGTGGGATTGCAGGTGTAAATTACATCACGTGTGCCTGGCTTATGCTGTGCTGGGGATTGAACTAGGGGCTTTGTGCATGCTAAGCAAACACTGTGTGA | 120 | 50 |
| chr7 | 143724026 | 143724146 | Target=1;ProbeIdx=16849;Target #1 | 9 | + | good | AAGGCTCAGGGAACAGCACAGAAGGAAGAGAGGAAAGAATGCGAAGCTGGAGGATATGTGGAGCCTTGTGAAATGCCATCTTGGGGACATGACATGGCTGTTGTTCCCATGAACTCACAG | 120 | 50 |
| chr7 | 143726426 | 143726546 | Target=1;ProbeIdx=17089;Target #1 | 9 | + | good | ATACACACACACATATACACAGAGATTGCTTTGTGGTTTAGAGCCCTTGTTGCTGTTGCAGAGGAACTGGGCTCAATTCCCAGCACTTGGTAGCTTACCACCATCTTTAACTCTAGTTGC | 120 | 45 |
| chr7 | 143607146 | 143607266 | Target=1;ProbeIdx=5161;Target #1 | 8 | + | good | AACCTGGAGTTACAGGAAATTGTAAGCCACCCAGCATGAATGCTGGGAACTGAACTTGGGCCCTTTGGAAGAGCAGGGAGCTTTCTTCAAATTTTGTTTTAATTGTTGGATATTTTTATT | 120 | 40 |
| chr7 | 143627066 | 143627186 | Target=1;ProbeIdx=7153;Target #1 | 8 | + | good | GGATTTAGTGGGAAAAATTGGAAGGGTTATAGGAGCAGAACATAAGCATGACACCTAACACCTCCTGACAGGAAGACATAATTGCAATGTGGTCTCAACAACAGGTAATCATAGCAACCA | 120 | 41.67 |
| chr7 | 143632226 | 143632346 | Target=1;ProbeIdx=7669;Target #1 | 8 | + | good | TGAGACGGAGTGCACTCTGTGACTCAGGCTGTCATGGGACTCACTATGTAGCTCATGCTGGCCTCACACTCCCAGCAGTCCTCCTGCCTTAGCCTCCCAGTGCCGGGCTGGACCTTATGA | 120 | 58.33 |
| chr7 | 143662586 | 143662706 | Target=1;ProbeIdx=10705;Target #1 | 8 | + | good | TTTTTCTATTTCTTCAGTCCAGGATGCCAGCTCATGGGATGGTGCTGTCTATGCTCAGGGTATGTCTTCCCCCCTCAGTTGAACCTACCTGGAAACACCCTCACAGACTTGCACAGAGAT | 120 | 49.17 |
| chr7 | 143669306 | 143669426 | Target=1;ProbeIdx=11377;Target #1 | 8 | + | good | GGGACAAGCTCCCCTTTGGCACCAGACTTCAGCCTTTCCCCCCCTAGCATGAGACTCCTGGGGCAATCCAATGCCTTGGGGTCCACTGTTTCCTTGGTCTGGTAGGTAAAAAGAAGGGTA | 120 | 55 |
| chr7 | 143715626 | 143715746 | Target=1;ProbeIdx=16009;Target #1 | 8 | + | good | CATTGACTGTCAACTTGAGGAGATCTAGAATCACCAAGGAGACGGTTGTGTCTGGCTATGACCATGAGGGATTAGCTAAACGAGGTTGACTGAGGAGTGAAGACTCATCCTGAGTGTGG | 120 | 48.33 |
| chr7 | 143569826 | 143569946 | Target=1;ProbeIdx=1429;Target #1 | 7 | + | good | GACGTTGAAAGCTCAGAGCTACCATCTTGAATTCCAAGAAGCAGAGGGAATTAACTGGAAATGTAGTGATTCTTTTAACTCCCAGAGCCCGCCCCCAGTAACATACTTCCTCCAGCAAGG | 120 | 46.67 |
| chr7 | 143602706 | 143602826 | Target=1;ProbeIdx=4717;Target #1 | 7 | + | good | TATCATTTTGTCCCATTTTCAGTATGAGAAGCACTTAGAGCTCACAAGAGAGTCACAGAGAAGCAGGAGGCACCGGGTCCCCGTCTTGCCTAGCTGGCAGCTGTTGTCTGCTTGTCTAGG | 120 | 50.83 |
| chr7 | 143568746 | 143568866 | Target=1;ProbeIdx=1321;Target #1 | 6 | + | good | CCTCCTGAGTGCTGGGATTAAAGGCGCGCGCCACCACAATCCTGTATGTTTTATGTGATGCTGTGAGCAGAACCCAGGCATGTGCTGGTCCCTAGCTCCTAGTGTGTGTGTGTGTGTGTG | 120 | 54.17 |
| chr7 | 143588666 | 143588786 | Target=1;ProbeIdx=3313;Target #1 | 6 | + | good | AACTCGTGGGAGTTAGTTCTCTCTTCTACTATGTAGGTTACAGGGATTGAACTCCAGTCATCAGGCTTGGAAACTCAGCCCTACTCACTGGTCCAGAATGGACTCTCTTTTCTACTTGCT | 120 | 45.83 |

| | | | | | | | | | |
|---|---|---|---|---|---|---|---|---|---|
| chr7 | 143596106 | 143596226 | Target=1;ProbeIdx=4057;Target #1 | 6 | + | good | CATTGTGCCAGCTGGGGAGACAGTTCAGTTGGTAAAACGCTTGTCCTGCAAGCTCAAGGATCTGAGTTCAATCCCCATCACCACATTGAAGAGCCAGGCATAGTGGTACACTCTTGTAAT | 120 | 48.33 |
| chr7 | 143600786 | 143600906 | Target=1;ProbeIdx=4525;Target #1 | 6 | + | good | TTGCCACACCTGTCTCAAAAAATAAGGTGGAGAGAGGTAGAGAAAGACACCTTAATTCTATCAACCTACACACACATACACACAAATTTGCACACACATAAAGCACATTCCTCTATCATG | 120 | 39.17 |
| chr7 | 143663186 | 143663306 | Target=1;ProbeIdx=10765;Target #1 | 6 | + | good | AGGGGAGATGTCTCAGTGGGTAAAGTACTTGCCATACAAGTACAAGGACCTGAATTCCCACCCACAGAATCCACCTAAAGCCAGATATACTGGAGCATAACAGTCATCCCTGTGTTTCCA | 120 | 46.67 |
| chr7 | 143695106 | 143695226 | Target=1;ProbeIdx=13957;Target #1 | 6 | + | good | GTGTCCAAACTCCAGCACCTGTTAGAGACCCTGTTTCAGCTATTCTTGACTGTCAACTTGATGGGATTTAGAATCAGGATAGAAACACTTGTGGGCATGTCTGTGAGGGAGCTTCTAGAT | 120 | 45 |
| chr7 | 143574506 | 143574626 | Target=1;ProbeIdx=1897;Target #1 | 5 | + | good | GCTTACAGCATGGTGTGTGTGTGTCTGTGTGTGTGTGTGTGTGTGTGTAAAACAACTCAGATAAATTTTCCTGCTTTGTTCATATCTGTAAATGAAAGTATTTAGGTATTTAAAGTTG | 120 | 35.83 |
| chr7 | 143587946 | 143588066 | Target=1;ProbeIdx=3241;Target #1 | 5 | + | good | TTCCATTAGGGGGAAGGTTTTTAGTAGGAACAAGAGAGGGAAACTATCCAGAGGCCTCTGGATGAGTTCAGAGCAGAGAGTAGACTGGACATGTCCAGGGCCAGGGAAGCAGGAGACTAG | 120 | 50.83 |
| chr7 | 143598266 | 143598386 | Target=1;ProbeIdx=4273;Target #1 | 5 | + | good | AAATCCTTCCCAAATGCTCCCACCAACTAGGGGCCAAGCATGTAAACATACAGGCCTATGGGGAGCACTCTCATTCAAATCACTACATTGTCCCAACATAAAATAACACAAACACACATT | 120 | 42.5 |
| chr7 | 143631386 | 143631506 | Target=1;ProbeIdx=7585;Target #1 | 5 | + | good | TGTTACATAGTGAGAATCTAGCTCCAGTTGTCAATCAAAAGAGAATTGTGTTCATGGTAAGACTCCATACCAGGGGTTGGAGAGATGGTTCAGTAGCTAAGAGCCCTGGCTGCTCTTCCA | 120 | 45 |
| chr7 | 143631866 | 143631986 | Target=1;ProbeIdx=7633;Target #1 | 5 | + | good | GAGGATCTGGGTTCAATTGCCAGCAACCACATGGTGGCTCACAAATGTCTATAACGCCAGTTATTAATCTCAAAGACAGAAGGAAAGCACTGGCCCAAATTATTGTGGCCAAATTAATTA | 120 | 41.67 |
| chr7 | 143631986 | 143632106 | Target=1;ProbeIdx=7645;Target #1 | 5 | + | good | TCAAATGCCCTCTGCTGGCCTCATCAGGTATTGCATGCATGTAGTGCATAGACCTACATAAAAACACTCATGCACATAAAATTAATTAATTAATCCCCATACCAAGCCAGCATCACAAGG | 120 | 40.83 |
| chr7 | 143632586 | 143632706 | Target=1;ProbeIdx=7705;Target #1 | 5 | + | good | TAGTCAGCAGGGGGTGCTGGAGTCTCTGTCTCAGACCTAGCCAGCAGAGGGTGCTGGAGTCTCTGTCTCAGACCTAGCCAGCAGAGGGTGCTGGAGTCTCTGTCTCAGACCTAGTCAGCA | 120 | 58.33 |
| chr7 | 143635226 | 143635346 | Target=1;ProbeIdx=7969;Target #1 | 5 | + | good | GGTGGTGATGCTGGTGATGCTGTAGCTGTTGATGAGGTGATGGTAGTGATGGTGATTATAGTGATGAAGAAGCTATTATGGTGAAGTTAATGGGAATGATGTTGATAATGGTGATAGTAA | 120 | 40 |
| chr7 | 143638946 | 143639066 | Target=1;ProbeIdx=8341;Target #1 | 5 | + | good | TAAAATTTTATAGCTTCTGCCAAGGTGTTTTCAAAAGCATTGTGGTGATGAAGCCTCAGCAGAATTTATCTTCTCTTTCTTGGTGTGTGTGTGTGTGTGTGTGTGTGTGAGAGAGAGA | 120 | 40 |
| chr7 | 143644586 | 143644706 | Target=1;ProbeIdx=8905;Target #1 | 5 | + | good | CTGGATGTGGTAGTTTGAATGGGAAATATCTCCCCAAACAGCTCAGGCTTGGAACACTTGGTTCCCAGCTGGTAGTGCTGTTAGGGGAGGTTTAGGAGGAGCAGCCTTGCACATCCTTAG | 120 | 50.83 |
| chr7 | 143655026 | 143655146 | Target=1;ProbeIdx=9949;Target #1 | 5 | + | good | CTATAGTTATAGAGATTGTCAACTTGGGGTTTAGAATCTCCATGAAAACAAACCTCTGGGCGTGTCTGTGAGTAAGAGTCTACTTAGTTTGGGTTAACTGATATATGAAGACCCACACTG | 120 | 40 |
| chr7 | 143684786 | 143684906 | Target=1;ProbeIdx=12925;Target #1 | 5 | + | good | TTATTATTTAGTAATTATGTATTTGTGTGTCAATTTGGCAAGGAGTCAGTTTCCTGGTTAGTGACTTGACATAAGCTAGCGTTGCCTGGGAGGAAGAACCTTAGTTGAGGAATTGCCTCT | 120 | 39.17 |
| chr7 | 143562746 | 143562866 | Target=1;ProbeIdx=721;Target #1 | 4 | + | good | TACCGCCGTGCCTTCCCCACCGTGATGGGCTGCATCCCCTTGGACTACAAGCCAGAGAAAGCCCTTTCTCCTTTAAGTTGCTTCTGCCAGAGTGTCCACCCCAGCAACGAGAGAGCACCA | 120 | 57.5 |
| chr7 | 143570066 | 143570186 | Target=1;ProbeIdx=1453;Target #1 | 4 | + | good | GATACAATGTGACTGGCTACCTTACTCACACTCCTGCTGTCCTGCCTTCCCCACCACGATGGACTACATCCTTAACTCTTATCCAAAACAAACCCTTCCTTCGTTGAGTTGCTTTTGGCA | 120 | 47.5 |
| chr7 | 143574746 | 143574866 | Target=1;ProbeIdx=1921;Target #1 | 4 | + | good | TAAGGAGCAGCAAGGGGAGAGAAGCCTGTGAGCTGGAGGGGATTTAGAGGGAGGAGTGGAGGCTCTGAAATGTATAACAGGTACTTGTGAATGGGCACTGAGGAGCCTGGCAGCCAGTAT | 120 | 53.33 |
| chr7 | 143598506 | 143598626 | Target=1;ProbeIdx=4297;Target #1 | 4 | + | good | AACTGTATTAAAGGGTCACAGCATCAGGAAGATTGAGGACTGCTACTGTGGAGTGTTGTCACTCTGGGTCACAGAAACTTGTAGGTTGTCCTGGTGATCTGTGGTCACTTGAATGTGGCA | 120 | 46.67 |
| chr7 | 143621906 | 143622026 | Target=1;ProbeIdx=6637;Target #1 | 4 | + | good | CAAACTAATATCTTTTAATGAACACTTATGTTTTTTATTTATATCTGTGTATACTTGTGTGGGCATGAGTTTGTGGGTTCAGGTACCCATGGGAGCCAGAAGAGGGTGTCAGATACCCTG | 120 | 39.17 |
| chr7 | 143626946 | 143627066 | Target=1;ProbeIdx=7141;Target #1 | 4 | + | good | GGGGGTGCTGGAGTCTGTCTCAGACCTAGCCAGCAGGGGGTGCTGAGGTCTCATACATCCTCTGTGTCTAGAATCTAGCCAGCAGAGAGCACTGGGGTCTCAGTCTCAGAACTGGCCAGC | 120 | 58.33 |
| chr7 | 143635346 | 143635466 | Target=1;ProbeIdx=7981;Target #1 | 4 | + | good | ATAATTATTCCTCCTATCTCTTTAGCATGGCTCAAGATTAACCCATCTCACTAGGAGTGCTTTCCTTTTCCTAGAGGTCATCCCTCCTTCCTAGGGGTCATCCCTCCTTCCTAGGGGTCA | 120 | 46.67 |
| chr7 | 143642066 | 143642186 | Target=1;ProbeIdx=8653;Target #1 | 4 | + | good | TGCTGGGCACTGAACTTGGATCCTCTGCAAGAGCAGGAAGCATTCTTAGCCACTGAGCCGTCTCCCAGTGGTCTACCCTTTGTCCTTTTGTTGATAGCCACTCGTCCACACGGTCCTCCA | 120 | 54.17 |
| chr7 | 143643866 | 143643986 | Target=1;ProbeIdx=8833;Target #1 | 4 | + | good | TAACTTTGAGGCATCTCTCCAGTCCCGGCACCTGACATCTTTACATGGATTCTGGGGTCCATACTCAGGTCCTCATGCTTGCATGGCAAAAGCTTTACCAACTTCCCCATCCCCTGTCTG | 120 | 50.83 |
| chr7 | 143645186 | 143645306 | Target=1;ProbeIdx=8965;Target #1 | 4 | + | good | GCCTCTATTAACCAGATCTAGAAACTTCCTGACAGGTGTGCGAGGTCTCTCTCCATGTGATTATTCTAGACCCTGTCAAGTTGACAAGCAGTAAGCATCAAAGAGACACCTTGAGTCTTT | 120 | 44.17 |
| chr7 | 143652746 | 143652866 | Target=1;ProbeIdx=9721;Target #1 | 4 | + | good | TTGAATACAGGTTCTCTGTGAGAGCAGCCAGGGTTCTTCTGCTCGTAACCACTGAGCCGTCTCTCCAGCCCCTCAGCCCCTTTGGACACAGAGTCTGTCCCTGTCACCATTGGCTAACCA | 120 | 55 |
| chr7 | 143670746 | 143670866 | Target=1;ProbeIdx=11521;Target #1 | 4 | + | good | AAAATGCTTAAGATTTTGAGGTCACTTTCTTGAGACCTGTCATAGTTAGGGTTACCATTGCCATGATGAAGCATAATGACCAAACACATTGGGAAAGAAATGGGTTTTTTTGGCTTACAC | 120 | 37.5 |
| chr7 | 143679746 | 143679866 | Target=1;ProbeIdx=12421;Target #1 | 4 | + | good | AGCCACCCAGACAAGCACAAGGACTTGAGTTTGATCCTCAGGAACCACATAAAAAAATGTAGGTATGAGGCATGCACTTTTCACCTCAGCACTGGGGGAGGGAGGACAGAAAACTAGTAC | 120 | 47.5 |
| chr7 | 143706266 | 143706386 | Target=1;ProbeIdx=15073;Target #1 | 4 | + | good | ATCACCAGCTTTGCAGATGTGTGCCGCCTTGTATGCACAGCTTTCTGTGGGTACTGGGGATCTGAACTCAGGTTCTCAGACCTAGGCAGCAGGCACTTTATCTTCTGAGCCAGCTCTCAG | 120 | 52.5 |
| chr7 | 143556146 | 143556266 | Target=1;ProbeIdx=61;Target #1 | 3 | + | good | AGTCTTGAGCTTGACTTGGTGGCAACCATGAATGGGAGCACAGACTGTCTTCTATGGAAGATTAGACAGGGGCTCCATGGTTCCCTACTGCAGATCCCAGCGAAAAGAGACAGTCCGTGG | 120 | 51.67 |
| chr7 | 143558546 | 143558666 | Target=1;ProbeIdx=301;Target #1 | 3 | + | good | CTCACAGTGCCCTTCCTACTCCCCCCCCCCCCACACCACACATTTAAATCCAGGTTGTTCATATGAGAGCAAACATGCAATATTTGTCTTTTTGAGTCTGGCTTATTTTGCTTAATAAA | 120 | 43.33 |
| chr7 | 143591066 | 143591186 | Target=1;ProbeIdx=3553;Target #1 | 3 | + | good | TTCTTTGAGACAGGGTCTCTCAGTGTCTCACTGGCCTGGTGTTCCCAAGCTAGGCTAGGTAGGCTAGCCGGCTGGTGAGCCCCAGGATTCCATCTGTGTCTGCGTCCCCTGCACCGGGGG | 120 | 60.83 |
| chr7 | 143614946 | 143615066 | Target=1;ProbeIdx=5941;Target #1 | 3 | + | good | TATAAGTGGATGCCCCCATGCCTGGCTTTTTATGTAGGTTCAGGGGATCTGAACTCAATTCCTTAGGCATGCACAGAAAACACTTTAGCAATGGAGCCATCTAACGCCCCAGCCCCACTC | 120 | 49.17 |
| chr7 | 143615066 | 143615186 | Target=1;ProbeIdx=5953;Target #1 | 3 | + | good | TGTTGTGTGATGCTTTTCCTGGGGAAACATGTCACAAAAATATAATCATCCCTGACTGATAGGAACCAATGACAGAATAAAGTAAGGATAACACTGAAGTCCAACTTAGTGAGCCAGTAA | 120 | 38.33 |
| chr7 | 143630066 | 143630186 | Target=1;ProbeIdx=7453;Target #1 | 3 | + | good | ACTGTACATTCATCCATCCAGGCATGACAGTTCACTCTCTAGCACTTGGAAACCAGAGGCAAGACAGTCACCCCAGGTTGACACCAGCCTAGTCTACATAATGAGTTCCAGGTCATCGGG | 120 | 50 |
| chr7 | 143631746 | 143631866 | Target=1;ProbeIdx=7621;Target #1 | 3 | + | good | TCTTTTTGAGGCCTGACTCTCAAAGGCAAGCAGACTCAACAGAGTCTCATGGAGGATCATAGTGACTATGCTAGGCAGAGGGAATTCTGGGAAGGAAAAGTACAGTATCTCATTAAAGGG | 120 | 45 |
| chr7 | 143641946 | 143642066 | Target=1;ProbeIdx=8641;Target #1 | 3 | + | good | GAGTAGACTTTGAGGGTTTAGAGCCTTCAAGTTCACTCTCCCTGCTTCAGGCTTACAGTTAAAGATGTGAACTCTCGGCTTCCTGCCCCTGCCACCATGCTTGCTGCTTTCTGCCGTGCC | 120 | 52.5 |
| chr7 | 143655146 | 143655266 | Target=1;ProbeIdx=9961;Target #1 | 3 | + | good | CCCTCCTGTCCCTCTCATGTCTGCTTGGAGGTGGCCCTCCTGTCCCTCTCATGTCTGCTTGGAGGTGGCTTTGGCTTGGTGGCACTCCAGTGCCCACCCCAACCCCTGGCAAAACCACAT | 120 | 60.83 |
| chr7 | 143661746 | 143661866 | Target=1;ProbeIdx=10621;Target #1 | 3 | + | good | TTGTGTCACCCTGTACTCTGCTCTGAGCATCAGCATTGTTAGAAGAGGGATGTGTGTAGAGCTCAAACCAAGCGAAGTCCCAGGTTCCCAAGCTATTCCCTATGCTTGCCTGCCTCAGCC | 120 | 51.67 |
| chr7 | 143683226 | 143683346 | Target=1;ProbeIdx=12769;Target #1 | 3 | + | good | GAGAGAGGAGTCATTTGATATGGCCACATTAGGAAGAGGGACAAAGAGGTGACGTGACTCCCAAGTCTATCTTTGGGCTCCTGGTGTGAAATTTAGGTTTTAACAGAAGACAGGAGGAGC | 120 | 46.67 |
| chr7 | 143715386 | 143715506 | Target=1;ProbeIdx=15985;Target #1 | 3 | + | good | GTTAAGTCTACACTGTGGCTTTCTCTCTTTTTAAAAGAATAAACTGCATGTGTATGGAGTGTGTGTATGTGTACAGGTACACATCTGTATGTGTATGGAGTGTGTGTGTGTGCATAGGTG | 120 | 40 |
| chr7 | 143555906 | 143556026 | Target=1;ProbeIdx=37;Target #1 | 2 | + | good | TCCCTGGGCCTACAACGTGGCCGGGGCCATCTGGTTTTTAGGTTCTCAGGGTCATCTGGTTCTTAGGCTCTTAGGGTCATCTGGTTCTTAGGCTCTTGGACACCCAGGCACCACTGAAAG | 120 | 55 |
| chr7 | 143558306 | 143558426 | Target=1;ProbeIdx=277;Target #1 | 2 | + | good | AGCTGAGCAGGCTCTGGAGAGCAAGCCCGTAAGCAGCGTAAGTTACTGAGGAACACTGCTTATGAGCTTGCACTCCATAGCCTCTGCTTCCACTCCTGCCTCTGGGTTCCTGCCTTAGCC | 120 | 55.83 |
| chr7 | 143562986 | 143563106 | Target=1;ProbeIdx=745;Target #1 | 2 | + | good | CCATGTACATGCATGTATGCATGTTTTGCATGTGTATAGGCACACATCTGTAGGTGTGCATGCATGTGGAGGCCCCTAGGGTTGATGTAATCTTCTTTGATCACCTTTACCAGCTTAGTC | 120 | 45 |
| chr7 | 143564066 | 143564186 | Target=1;ProbeIdx=853;Target #1 | 2 | + | good | CTCTCCGTTCATCCTGGACTTGAAAAGTCACAGGGTGTTTGCTATGTAACTGAGTCCACAGATCAAAAAGGAGGGACTCTTGAGTCTCAGGACTTATGTTAGATAAGACTTTGTGTTCTG | 120 | 43.33 |
| chr7 | 143568866 | 143568986 | Target=1;ProbeIdx=1333;Target #1 | 2 | + | good | AACTAATGGGGTGGGCAACTGGAAAGGGGGCAAGACTAGATATTGGGAAGATCGTGCCTCACTGAAGAACTCAGCCATCGAGGGATTCAGGGGGCAGAGAGGTGCAATTAGGAATTTAAC | 120 | 50 |
| chr7 | 143568986 | 143569106 | Target=1;ProbeIdx=1345;Target #1 | 2 | + | good | CAGCACCACCATGGGCCGCCATTCAGACCGAATACAAAGGAGAAATGGATCTGAGTGCCAACATTCATATCTCTCTGCTTCCTGAGAGAGAGCGCAATGTGGCCAGCCACCTCATGCTCT | 120 | 52.5 |
| chr7 | 143569946 | 143570066 | Target=1;ProbeIdx=1441;Target #1 | 2 | + | good | ATATTTTAGGGACAGAGACATGGCTCAGTAGATAAAGTATCTGCTGTGCAAGCCGAGGGCATGAGTTTGCATCCATTTAAAAAGCTAGGCATGGTGGTACACATCTGTGGTCCCAGTGCT | 120 | 45.83 |
| chr7 | 143573306 | 143573426 | Target=1;ProbeIdx=1777;Target #1 | 2 | + | good | TAAAGCTAGATGGTTATGAGCTGTCATTTGGATTCTGGAAACTGAACGTGGGCCAGGGCTCTTCACCACTGAGACATCTCTCCAACCCCTGTTGATCCTTCCATTTACTGATAACCACAG | 120 | 46.67 |
| chr7 | 143579306 | 143579426 | Target=1;ProbeIdx=2377;Target #1 | 2 | + | good | ACAGCCTCCAGTTTCATTCATTTTCCTGCAAATGTCACGACTTTATTCCTTATGGCTATTTAAGATTCCATTGCAGGCATACACTGTATTTTCTCTATTCATGCATCAGACATCTGTGTG | 120 | 38.33 |
| chr7 | 143591186 | 143591306 | Target=1;ProbeIdx=3565;Target #1 | 2 | + | good | TCTGGTCTTGCTGTGTGCGCGCGCGTGCGTGTGCGTGTGTGTGTGTGCGCGAGCACGCATGTGCATGCATGCCTATTGTGGGAGGTCTATTGAGTCATCCCAGCTGCCATAGTGTTCCTT | 120 | 57.5 |
| chr7 | 143591546 | 143591666 | Target=1;ProbeIdx=3601;Target #1 | 2 | + | good | GTTGCTTCATCTCCATTGCCAACCTGACTGGCTTTGGAATCACCCGGGAGACTCACCTCTCAGTGTCTGTGAGGGCATTCTAGAGGCTCAACTGGGGTGGGAATACCCACCCTGGATTCA | 120 | 54.17 |
| chr7 | 143596346 | 143596466 | Target=1;ProbeIdx=4081;Target #1 | 2 | + | good | TTTATCTAACAGCCAGAAATCCGTCTATAGTCTAGCTGGAGCTCATTCCTGGATATGTGGACTGCTTGCCACCTAAGAGTGTTATCAGAAATTTCACGTTGCCAGATTTGACTGGGGAAG | 120 | 44.17 |
| chr7 | 143598746 | 143598866 | Target=1;ProbeIdx=4321;Target #1 | 2 | + | good | CTATTGTTCTTCTCCCAACAGTAGAACCAGCCAGCCAATCAAAGCTCATTTGCCCTGATAAAGAGATGGGAGGAATAATTGTCCTTAGAATGAGATAATGTGATTTTTTTTCTCCCAGAA | 120 | 39.17 |
| chr7 | 143599106 | 143599226 | Target=1;ProbeIdx=4357;Target #1 | 2 | + | good | CCCCTGTTCTCTGCTGTTCTTAGGTTCTCTTAGGTTCTTTGATTAACATGACTAAGAGCCCTTCGGGGACACCCCCCTCCCCCAACTTCCCCAGTTAAATCTGGCAATCTGAAATTTCTG | 120 | 48.33 |
| chr7 | 143599226 | 143599346 | Target=1;ProbeIdx=4369;Target #1 | 2 | + | good | TCACATTATCTCATTCTGAGGACAATTATTCCTCCCATCCCTTTATCAGGGCAAATGAGCTTTGATTGGCTGGCTGGTTCTACGGTTGGGAGAAGAATAATAGCATCTCTCTACCTGGGG | 120 | 45 |
| chr7 | 143599586 | 143599706 | Target=1;ProbeIdx=4405;Target #1 | 2 | + | good | ATATACATATGCATGTTTGGCTACATGATGTCCACAGTGGATGTCTAGAATCATCCTTCACTATTCTTTCACCTGTTTGCTGAGGCAGGATTTTAAGGCTCTATCAGTCAAACCCAGAGC | 120 | 41.67 |
| chr7 | 143604746 | 143604866 | Target=1;ProbeIdx=4921;Target #1 | 2 | + | good | TGGACCCACTCTTGTGCAGACCCAGTGCAGGGGTCCACAGCTGCTGAGTTCAAGGCTGCAATGTCTGTAGTTTGCCAGGAGATGGCACTGTGCAGCCCTTCTCCCTTTCTTCCAGCTCTT | 120 | 55.83 |
| chr7 | 143609546 | 143609666 | Target=1;ProbeIdx=5401;Target #1 | 2 | + | good | GAAAGCTAAGACAGGAGGAGGATTGCAAGTTCACTTCCAGCTTGGGTCACATAGAAAGACCCTGTCTCAGATTTAAAGGTAAGAGAAGGGCTATGGGTGTAGTTCGGTGGTAGAGCCCTT | 120 | 47.5 |
| chr7 | 143613386 | 143613506 | Target=1;ProbeIdx=5785;Target #1 | 2 | + | good | GGGTGCTCGACAGAAATCGTTATTTGTAATTTTGTTGTTTGTTATAGGTGTGTGTGTGTGTGTGTGTGTGTAGGGCAGAGGGCAACATTGTGTTTCTCTAGAGCTGTCCATGTTCCCT | 120 | 44.17 |
| chr7 | 143614826 | 143614946 | Target=1;ProbeIdx=5929;Target #1 | 2 | + | good | TCGGTTTAAGGAAAGAATGTTTTATTTGTCTCACGCTTTGAGGGCGCTCATGGTGTCACGGTGGGGAAGGAAGGCATGGTGGCAGGTGTGTGAGGGAGCTGGTCACACTGCAGTCACTGT | 120 | 52.5 |
| chr7 | 143629706 | 143629826 | Target=1;ProbeIdx=7417;Target #1 | 2 | + | good | GACAGAGAGAGACACAAGCACTGGTGCTCAGCCTATTGCTTCGTTTTGTTCAATTCGGGACCCTATCCCATGGGATGGTGCCATCCTCATTTATACCTCAATGAACCTAATGTAGATACT | 120 | 45.83 |
| chr7 | 143629826 | 143629946 | Target=1;ProbeIdx=7429;Target #1 | 2 | + | good | GCTTGCAAGCAGCTCGAAAGGTCAGACAGTTACTTCTAGGCAGCTCATCTGGTTTAAGAGCATCTTAGCTTATATGACTTTAGAGAGGCAGGGGCCTGTGCATCTGTTCAGTTTCAGGGA | 120 | 47.5 |
| chr7 | 143630306 | 143630426 | Target=1;ProbeIdx=7477;Target #1 | 2 | + | good | CATCCTGAGAATTATGAATTGTCTACTTCCTGACTCTGAATGAACCTCCCTTTAGAACCTCCTCGGTTTTAAGGACTTTCACCTTTAATATGGAATGTCTCAACTCAGGAAATTGCCTCA | 120 | 40 |

| | | | | | | | | | |
|---|---|---|---|---|---|---|---|---|---|
| chr7 | 143630426 | 143630546 | Target=1;ProbeIdx=7489;Target #1 | 2 | + | good | TTCAGTCTCTGTGTCTTAGGGCCAGGCAACAGGGGGTGCTGGTGTCTCATGCAGGCTCTGTGTCTCAGATCTAGCCAGCAGGGGGTGCTGGGGTCTCATGAAGTCTCTGTGTCTTAGACC | 120 | 56.67 |
| chr7 | 143635106 | 143635226 | Target=1;ProbeIdx=7957;Target #1 | 2 | + | good | AACAATCTCCAGAACAGGCAAGCTGTGAATCTCAGCAGTTGCCTGAGGGAGAGTGGAGGAGAGAAGGGGAAAAGCCATGCATGGAGGACAGGCTGCTCCATGGTGGGAGCAGAGGGTAAG | 120 | 55 |
| chr7 | 143641706 | 143641826 | Target=1;ProbeIdx=8617;Target #1 | 2 | + | good | CCTAGGGCTCTATGTTTTGCCCAGCTTAACATACTTAGTGAGTTCCAGGCCATGAGAGGCCTTTAAAAAGAGGGCTGGGGCAGAGAGATGGCTCACTGATTAAGAACACCAGCTGCCCTTG | 120 | 50 |
| chr7 | 143653826 | 143653946 | Target=1;ProbeIdx=9829;Target #1 | 2 | + | good | CTTTTGACAAAGAACATTGACTCTGCTGTATAGATTTCATTTATGTGTATGTTTTTATACATTTGTGCACATGCACTGAGCTGCACATGGTGGTCAGAAGAGGGCGTTGGATCCCTCTGG | 120 | 41.67 |
| chr7 | 143659106 | 143659226 | Target=1;ProbeIdx=10357;Target #1 | 2 | + | good | GATCCTGGAGCTGACAGATTCCCTGAGACAGGGTGGCCAGCACACTCCAGGAATCCTCCTGTCTCTGGTGTACCCATCTGACTTTTTACCTGGATCCTAGGGATGCAAACTCCAGTCCTC | 120 | 54.17 |
| chr7 | 143667146 | 143667266 | Target=1;ProbeIdx=11161;Target #1 | 2 | + | good | GTTCTTATTCCTCCTGCTCCATAAGTGGACACCTTCTTCGGGGCCACAATTGTCAGAGTCAGTCTGGAAAACAGCTCTGTCCCTGACTTAGCCTCTCTGAATCCTTATTCCTTCTGCTCT | 120 | 48.33 |
| chr7 | 143670026 | 143670146 | Target=1;ProbeIdx=11449;Target #1 | 2 | + | good | GTAAGTGGGAACTGCCACATTTACCTCAAGGCCAAATGGGAACACCCTAGTGTCCTGTGGATGCACAGAGCATTCGCTAGCTGAGCTTCTGTCAAGTCTCCTCTTAGAGACCCTATCTGC | 120 | 50.83 |
| chr7 | 143670146 | 143670266 | Target=1;ProbeIdx=11461;Target #1 | 2 | + | good | AAACACTGGGGAGTTGAGGGATTTCGGTGGAAACCCTGGTGAATAGAAGGTTCCCTTCAAAGCCATTGCAGTGTCGGAGGAGAAATTCACATTTTATTGCTTAGTATGTAACCACATTTG | 120 | 43.33 |
| chr7 | 143670266 | 143670386 | Target=1;ProbeIdx=11473;Target #1 | 2 | + | good | GCTGTGTGGAAAATATAAGCAAAATGTATTTTCTTTTCTATCAAATAAGACAAAACTACATTTAAAAAATTGTAAAGAAGCCAATTGGTAGAAATTAACATTCTGCCATAGTTGGCTTGC | 120 | 29.17 |
| chr7 | 143681546 | 143681666 | Target=1;ProbeIdx=12601;Target #1 | 2 | + | good | AGTTTTTGGAATCAAAATATATGTTCAAGTTTTATTTCTTCCTAGAAACTGCATGGCTATTAGCCAGCCAGGCAAGTAAGGATGTGAGGCTGAGCTCTGGTCACCTGGGTAAAACACAAG | 120 | 40.83 |
| chr7 | 143681666 | 143681786 | Target=1;ProbeIdx=12613;Target #1 | 2 | + | good | CCTAGGAGAGGGGTGAGTATATCAGAAATACTCATCATGAGCAAACAAGATGGCTCAGTAAAGCAAGATGGCTCAGTAAAGCAAGATGGCTCAGTAAAGCAAGATGGCTCAGTAAAGGAG | 120 | 44.17 |
| chr7 | 143694506 | 143694626 | Target=1;ProbeIdx=13897;Target #1 | 2 | + | good | TCACAAACACATAGCATATCACACCTCACAGACCACACATACATCACAGCACACATGCTAAAACTCTTATGTGCCTACCACACCTCACCCAGACTACAGATACCACATCCTATCTACACA | 120 | 44.17 |
| chr7 | 143704586 | 143704706 | Target=1;ProbeIdx=14905;Target #1 | 2 | + | good | GGACCTGGTACTGTCTGATTAGGCTAGGCTGGCTTGCCAGAGAAACCTCAGAATCCTCCTGTCCTCCCTCCCCCAGTGCTGAGGTGAAAAGTGCATGCCTTATACCTACATTTTTTATG | 120 | 50 |
| chr7 | 143706866 | 143706986 | Target=1;ProbeIdx=15133;Target #1 | 2 | + | good | AGAGGAGGAGGCAGCAAGCTCTGAGGGGCTTACAGTTTAGGTACCACGATCATGTCTGCCACCTACCCGGCATAACTCTTTTTTTGTTGTTGTTTTTTTTTGTTTTTTTTTGTTTTTTT | 120 | 40 |
| chr7 | 143707226 | 143707346 | Target=1;ProbeIdx=15169;Target #1 | 2 | + | good | GGCATCGTTACTGGAGCTCTTAGGGTTGGAACCTGGACTCTCTCTAATGGTCTGAACAGGCCAGGCTTGGTAACTCATCCTCAGTAGGCCAATTAAAGAGACAGCTAATAGTGAAAAGCA | 120 | 47.5 |
| chr7 | 143715266 | 143715386 | Target=1;ProbeIdx=15973;Target #1 | 2 | + | good | CTGAAATTGGATGTACCTAAGAGGAGGTGGAGGGGCCATTGGGATGAATATGACCAAAATACATTGTACACATGTTAAAAAAAACAAAAAACAAACAAACAAACAAACAAATGGTGTCTG | 120 | 35.83 |
| chr7 | 143726786 | 143726906 | Target=1;ProbeIdx=17125;Target #1 | 2 | + | good | AAAGTCCAACTGTTGGTTTGTTTTAAGCTCAGAAAAACCAAGGTCTGTGGAGAGTTGGACCCTGTGCCCCTTGCAGTCCCACCTGACAGTGCCCTGGCTCCTGTCTTTACAGGTCCAGA | 120 | 50.83 |
| chr7 | 143555546 | 143555666 | Target=1;ProbeIdx=1;Target #1 | 1 | + | good | AGCTTCTCCACTTTCCTCGCCATCCAGAGTGCAGGCTGTGGCTGCTGTAGGGAGCCTTCCTGTGGGACCTGGTGTGCTGACCCCTCTGGTAAGTTGGAACCTGGTTCCCTTGTCCTGTCA | 120 | 58.33 |
| chr7 | 143555666 | 143555786 | Target=1;ProbeIdx=13;Target #1 | 1 | + | good | CACATCTGTATATACACATGTGGTCTGGGTTGGACATCTTCCTCTCTTGTTTTCCAGCATACATTTAGAAGTAGGGCCTCTCACTAAGCCTGGAGCACAGCCTGGCTTTCTCTGCCCTCC | 120 | 49.17 |
| chr7 | 143556026 | 143556146 | Target=1;ProbeIdx=49;Target #1 | 1 | + | good | CCCCTATTGTATGGCTTTCGGTTGTCTTCTTTGTAGTGACTGTTGTGTCACAATGTCGATAGATCAGCTATGCCCCTGAGAGATGATCTTGAATATTTCGCTTTCCCAAAGCATTCTGGG | 120 | 44.17 |
| chr7 | 143556266 | 143556386 | Target=1;ProbeIdx=73;Target #1 | 1 | + | good | TCACTTCAAGCCATGTTCTCTACCGGTGGGTGCCTCAAACACCTGCCAGTGCTTCTTCAGCCTAGAGGGTTCGCCCAAACTTGGGATGTGCTTTATCAGCCTTCCTGCTTGGGTTGGCCT | 120 | 54.17 |
| chr7 | 143556386 | 143556506 | Target=1;ProbeIdx=85;Target #1 | 1 | + | good | TTTCAGGAGGGGGAGAGAAATAGATGCGGATGAGCGGGGCTGGTAGTGCATATGTGTCTTGGAGGGTGGGTATGGGGTCCAGTGTCTCATTTCCATTCAGGGGCTGACTGTGGCTCCAGG | 120 | 55.83 |
| chr7 | 143556506 | 143556626 | Target=1;ProbeIdx=97;Target #1 | 1 | + | good | CTAGTGCCTGCCTACTGCCTTTTCCAAAAATAACATTTTAGTCTGTTGATTGTGCTAGAATACACACAGCACAAACTTCCCACGTAGCAGTTTCCAGTGGATAGTTTGATGACTTTTTCT | 120 | 40 |
| chr7 | 143556626 | 143556746 | Target=1;ProbeIdx=109;Target #1 | 1 | + | good | CTGTCATGTGGTCATTCCCAGTGGCTTTCCAGAAGGAGGCTCTGCCCTCTTGGACACTCCCTCCCTCCACCCCCAGACTTTCCGTGTATATGGGTCTGACTGGGGACTGACACAATGGA | 120 | 56.67 |
| chr7 | 143556746 | 143556866 | Target=1;ProbeIdx=121;Target #1 | 1 | + | good | ATGAAATCTGACCAGGGTGTGATGTCTGTTCCATGTTGGAGTGTGTGTCCGAGCCTCAGTTTTTCTTATAGGGGCCATTTTGCATTTGGGGAGGCGTATTGCATCATATGGCTGTGTCAC | 120 | 47.5 |
| chr7 | 143556866 | 143556986 | Target=1;ProbeIdx=133;Target #1 | 1 | + | good | GTGGGTAGATGTTTTCCTTTCTCTTGGATAGATGCTTTGCAGTGGGTGCTGTGGGACTCTTGAATTTCCAAGAGCTGGGTGATGACTGCCATCCTAGTGGATATGCAGAGCTGTCCCATC | 120 | 49.17 |
| chr7 | 143556986 | 143557106 | Target=1;ProbeIdx=145;Target #1 | 1 | + | good | GTGGTGTGGAAGGACTTCATTTGGCCATCAGCAGCAGGCCCTTTGTGGTATCTGTGCTGTGGGAGGTCAACCCACCTCTTTATCCTGAGAAATTTGAGGTCTGGGAGCCTAAGGGTTGGG | 120 | 53.33 |
| chr7 | 143557106 | 143557226 | Target=1;ProbeIdx=157;Target #1 | 1 | + | good | TTTTTGCCTACAGAACCGTACAGTGGTTTCTGCTGAATAACTTCGTGTGCACAGGGCTGGGTGTGTACAATCAGATGTGGCTGGGCTGTACGTGGGGAGACAGCAGCTGGATGTCACCAT | 120 | 51.67 |
| chr7 | 143557226 | 143557346 | Target=1;ProbeIdx=169;Target #1 | 1 | + | good | CATTTTAGTCAGGGCCTTGGGTCCTTGGCTGTCTTGCTGGGGAGAGAATGAGTCACCCCAAGGTTCTGTGGGGCCTAGCCTCATGCTCAGCAGTTCCTCACATATGTGCCAGAGAACCAT | 120 | 54.17 |
| chr7 | 143557346 | 143557466 | Target=1;ProbeIdx=181;Target #1 | 1 | + | good | AATTGAGAAAACTGGTCCTTGTTGACCTGACATGTCAGCCCCAGGCCTGTGATCATAGGTAGGATCTATGCCAGGAGATGAAGCCTGGAGGACCCAAGATTGGGTAGAGTGTGGACAGGG | 120 | 51.67 |
| chr7 | 143557466 | 143557586 | Target=1;ProbeIdx=193;Target #1 | 1 | + | good | AATGTGGACCTGTCTTCCTAGGAGTTTGGCTTTATCAAATAGGTGTTACTCTTCCATCTTCATGCTGTCTATGCAATGTGAGCAAAATCAGGTCACTCTGTCCCCTTGCTGACCAGAGCC | 120 | 45.83 |
| chr7 | 143557586 | 143557706 | Target=1;ProbeIdx=205;Target #1 | 1 | + | good | TAGTCTGTAATGGGGTACCATGGGAACCAATGACCCTCTCTTGGTGAGTGCTACTTAGCTGGATGGCAATGGCTGAGCTCAAGGACCTACCCAAAGCTGATGAACCATCTTTATCATCTC | 120 | 48.33 |
| chr7 | 143557706 | 143557826 | Target=1;ProbeIdx=217;Target #1 | 1 | + | good | TGAGATTGCTTTGGGGGACCTGCCACCAGAAGCATGTGACAGGTTGGCAGATGGCCCTAAACCCACAGAATCCTGATCCCTTGATCTTGTGTGGCCCCAAGGGAACTGGCAACAAGACTG | 120 | 54.17 |
| chr7 | 143557826 | 143557946 | Target=1;ProbeIdx=229;Target #1 | 1 | + | good | AGTTGAGGGAGGGCCTTGTGTATGGGAGATGGCCTGGATAATCTTGCTAGTCTAATGTCATTAAAGCCCTTGTATAGGGGCTGAGAGAAGCTTGGAGAGAGAGAAGATGATGCCTCAAGA | 120 | 47.5 |
| chr7 | 143557946 | 143558066 | Target=1;ProbeIdx=241;Target #1 | 1 | + | good | AGCCAAGGCAGAACCTGGAGAGATGCAGCCATGGGGGCAGGTGGGTAAAGTGCTTATTGTGAGAGCCTGGGGACCTGCCTGGGTTTGATCCTCAGAACCCACAGAAATGTAGAAGGAGAG | 120 | 55 |
| chr7 | 143558066 | 143558186 | Target=1;ProbeIdx=253;Target #1 | 1 | + | good | CCTTTCATGGTCCCACAAGAGGACAGACTCCATCAAACAGTTCTCTGATCTTGATATGTGTGACATACATACATCTTATATACACACAATGATGATGATGATGACAACAATGAAGAAGAA | 120 | 37.5 |
| chr7 | 143559026 | 143559146 | Target=1;ProbeIdx=349;Target #1 | 1 | + | good | TGTCTAAAGCCACAGGAACTGAGAGAAGCAGGGGGATGCTCTTCTGGGCTTTGAGGCTCTGGGAGAGTGCAGTATTGCTTACATAGATCCAGCTGGGGGTTCTGAGTTGTTGACATTGGC | 120 | 51.67 |
| chr7 | 143559266 | 143559386 | Target=1;ProbeIdx=373;Target #1 | 1 | + | good | TTCCAAGAGCAAAATCTCTGCTGCTTGAGACCAGGCTTTTGTACTTTGTGGAATGGTTGTAGGAGACCGCCACAGGATGTGATGGGCTCCTTTTCAAGCCTCTGGCTTTCACTCAGCAAG | 120 | 49.17 |
| chr7 | 143559386 | 143559506 | Target=1;ProbeIdx=385;Target #1 | 1 | + | good | AAGACAGGTGTGAGCATCCATTTCTGAGGGCTCAGGTTCCCTTGACAGCACACTGGTTACTGTGGGAACACCTGGTGGCAGTTTCCTAGGTGGACAGCAAGGATGGCAGGCAGCACTGTG | 120 | 55 |
| chr7 | 143559506 | 143559626 | Target=1;ProbeIdx=397;Target #1 | 1 | + | good | ATCTTTGGGATCCTGAGGCTGTGGGGAGGATTAGCTTTTGTGACCATAGTGGATGAGGTAGAGGTTGATAAAAGGACCTGGCTTGAGGTAATACATTGGAGATACAGGAAGATACTTGAT | 120 | 44.17 |
| chr7 | 143559626 | 143559746 | Target=1;ProbeIdx=409;Target #1 | 1 | + | good | ATCCTGCTGTGAATTCTGCTTGAGCATACACTGGAGTACTCGTGTGCATGTAACACACACACACACACACACACACACAGACATCAGGCCCAGAAAAGTTGGCCTGGTCTATGACATG | 120 | 48.33 |
| chr7 | 143559746 | 143559866 | Target=1;ProbeIdx=421;Target #1 | 1 | + | good | ATGCACATCAGATCTTCAATATGAGGGACTCTCAGGGAACGTTGCAGGAATCAAAGGTGAGGCAGGAAGGTGGATAAGAGACCACCAGGACATCACCTACCCCAGGAACTGTGGGCCAGT | 120 | 51.67 |
| chr7 | 143559866 | 143559986 | Target=1;ProbeIdx=433;Target #1 | 1 | + | good | TTTGGCTGTCACTCCATGTGTGGTCCGTGAGTCTAGAACTCTCTGGACCTGCACAATAGCGGCTTCATGTTTTCACCTGAGAGAGTAAAATGCAGCTAGAGAGTCAACATCAGCACCGTG | 120 | 49.17 |
| chr7 | 143559986 | 143560106 | Target=1;ProbeIdx=445;Target #1 | 1 | + | good | GGCTTTGTAAACTGCAGCTTTGAGAAAGTGTGTCTTGTTAAACCTGAAATGTCACAGTGAGGATCAGGCTCAAGTGCGCGCGAAGGTGGCAACAGTGATTGAAGGCAGGCAGAAGAGAGC | 120 | 49.17 |
| chr7 | 143560106 | 143560226 | Target=1;ProbeIdx=457;Target #1 | 1 | + | good | CGGTGCCGGGGATGTGCAGGCTGCAATATGAATTGAGCCCTTGGCTTTGAGAGCCTGTGGAGGTAAAGTGGGGCTCACAGCCACTTCCTTAGAAGTCCTGCCCTAGGAGAGGCACACCAG | 120 | 57.5 |
| chr7 | 143560226 | 143560346 | Target=1;ProbeIdx=469;Target #1 | 1 | + | good | AGATGGACAGGATCTAACAGCAAAGGTGGCTTGGCTTTCAGACCTTTGGCAAACAGTGTATGCTGTCCAGCAACTATGCCTCCAGGAGATGGGGTACAGGAGAGATGTGAGCAGGCACCA | 120 | 51.67 |
| chr7 | 143560346 | 143560466 | Target=1;ProbeIdx=481;Target #1 | 1 | + | good | TACATTGGACCAGAAGATGCCGCTGGAACATCATAGGTCACTGTAGAAGGCAAGACGGCCACATGGATACCATGCCTCATCAGGTAGGATTCTGATCACCTGAACTGTACATGATGGGCA | 120 | 49.17 |
| chr7 | 143560466 | 143560586 | Target=1;ProbeIdx=493;Target #1 | 1 | + | good | GCCACCCTGTCTAGGATGAGCAGAGCCATGGCCAAAGCTGTAGGCATCCACACTGCTGTGAGCAGAATCAACAACAGGGATTCCAGAAGTCAGCAGAAGTCTAAGCGGATAGCCTCTGGC | 120 | 54.17 |
| chr7 | 143560586 | 143560706 | Target=1;ProbeIdx=505;Target #1 | 1 | + | good | TAGGAAAAGAGGACTTTGCTTTAATGGGTAGGGGCCCAGTTTTATCTTTTAGGGTTATTTTTCTTGGCAATTTACAAATTGATGTAAGATTCACGTAAAACTTACTCTGGTGGCCGACTG | 120 | 38.33 |
| chr7 | 143560706 | 143560826 | Target=1;ProbeIdx=517;Target #1 | 1 | + | good | GATCTGGAATCAACTGGAACAGTCCTCTGGGTGCTCCCGTGAAGGATTTTTTTTTTCTCAGATTATTTGAAGTAGGAAGACAGACCTACACTGATGTCAGCAGCAGCCCAGATAAAAGA | 120 | 43.33 |
| chr7 | 143560826 | 143560946 | Target=1;ProbeIdx=529;Target #1 | 1 | + | good | ATGGAATGAAGGAAACGTTGCTTGCTTACCATTGCTCTCAGTGGCAAGTTCAGAGGCCGACCACTGAGCATATCACCAATACCTGGACCAATTTTATCAGCCTTCTGCTGTAGACAGAGT | 120 | 46.67 |
| chr7 | 143560946 | 143561066 | Target=1;ProbeIdx=541;Target #1 | 1 | + | good | CCCAGCAGCCCTTCAGGACTGAGCAGCTGCCAAACTCTCCATCACTCTGGATGAGCTAGCTAGAGTTGAACTACCCAGACAGCATTGTGCAGGCCAGCATGACACATCCCCTTTTAAAAT | 120 | 51.67 |
| chr7 | 143561066 | 143561186 | Target=1;ProbeIdx=553;Target #1 | 1 | + | good | GCCCCCATCCTATTGGTGCTCTTCCACTGGAGAGTGCTGTCTCCTATGCTTACCATGTAAAGTGTCCAAGTCAGTGGCTCCTGACTATGCTGTACAACCATACCACTGTCTCTTCTAGA | 120 | 50 |
| chr7 | 143561186 | 143561306 | Target=1;ProbeIdx=565;Target #1 | 1 | + | good | ATGTTCTTACCTCCCCAAGAAGAAACCCCTAGGTCAACAGCAATCATGCCTTACCTTCCCTTGGCTCTGAGTAACTACACTTCTGGACACTGTATAGAAATGGTTAGACAGGATATGGCC | 120 | 45.83 |
| chr7 | 143561306 | 143561426 | Target=1;ProbeIdx=577;Target #1 | 1 | + | good | TTCTGTGCTGTGTGATTTCACTTGGCATAGGGTCCTCAAGGCTCATCCATACTGTAGCCTGTACCAGTGCCTTGCTCTTTTTCACAGCTGGAGAATATTCCATCGGCCTGGTTCCTTGTA | 120 | 48.33 |
| chr7 | 143561426 | 143561546 | Target=1;ProbeIdx=589;Target #1 | 1 | + | good | GATTGATTCCTACTCCTTGATGATTGCCAGTGATGTGGTGATGAGGGGGAGGATGGCAGATAGACAAATGCTTTTACTGCTCATGGGTTTGGAGGAAGGAGGGAAACTGCCCACAAAGTG | 120 | 48.33 |
| chr7 | 143561546 | 143561666 | Target=1;ProbeIdx=601;Target #1 | 1 | + | good | ACAAGCCTCTCTTGGCTGGCAGACTGGCCAAATAGAGTGGCTTACTCCTGATTGCACACTGAGGAGGGGAGGGACATTCCTGCCACTCTCTTAAGGTGCAGCAACCCTTCCTGGATACCA | 120 | 54.17 |
| chr7 | 143561666 | 143561786 | Target=1;ProbeIdx=613;Target #1 | 1 | + | good | AGGACACTTCTCAGACCAAGTAACCAGTTTCCAGCTGGAAAAATAGGAGTTTTTCCCCCCACAGTGATGGCCAGCCTTGCTCCAGGTTGTAGGGTGGGGATACTCCTGAAACCAGGGTTC | 120 | 51.67 |
| chr7 | 143561786 | 143561906 | Target=1;ProbeIdx=625;Target #1 | 1 | + | good | TGTTGCAAGGTGTCACAGAGACTCTAGGGCTCAAGGGGTCATGGAGAGGGGCTGAAGCTTGGCACGGTGTAGCAGGTCAGAGTCCCTGAAGAGAGGCCAGAAGGTCACTGGTGAAGGGAC | 120 | 57.5 |
| chr7 | 143561906 | 143562026 | Target=1;ProbeIdx=637;Target #1 | 1 | + | good | AGAGTGTGGTGACGCCAGGACCATGGAGATCAAGGACACTGGAATACGAGGACAATGGGAGGATCACCAAGGACAGCAGCAGCTGTGGTGTGGAGTCTGCCTGAGCCTATGTGTTATGTG | 120 | 54.17 |
| chr7 | 143562026 | 143562146 | Target=1;ProbeIdx=649;Target #1 | 1 | + | good | TGCTGTGCAGGACAGAGCTGGAGAGGTGGAGCTGCCCAGGCCTGTAGGAGCCCAGAAGATGGTGAGCGAGGGCCAGATGCAAGACTCTGAGTTACAGGATGTAAGACGCGTACTGCTGTGCT | 120 | 58.33 |
| chr7 | 143562146 | 143562266 | Target=1;ProbeIdx=661;Target #1 | 1 | + | good | AAGAGACATTGGGTATTTTAAAGAGACTGGTCTTTTTTAAAGTGTTGAAATTTTTAAAAAATTGTATAGGTATGATGTTTGCCTGGTTGTATGTCTGTGAACCATGTGCATGCAGTACT | 120 | 32.5 |
| chr7 | 143562386 | 143562506 | Target=1;ProbeIdx=685;Target #1 | 1 | + | good | TCCCTGAGCTAACTCTCCAGCCCCAAGTGCTGAAATTGTTAAAGACTACAGGAATTTAAGTCACACTTAGTTTTTCATTGTGATATTAACATAAGATCTTGGAGATAAACAGAGAGGATG | 120 | 37.5 |
| chr7 | 143562626 | 143562746 | Target=1;ProbeIdx=709;Target #1 | 1 | + | good | CAGGGGGTCCCAGCTTCACTGCACTGAGGCCACCCACAGAATTCACAGCCCCCCCAAAGGGACTATTCAAACTTGAGCCCTTCCCTGTTCCTGAAGGCTTGGCAGGAGGTGGAATGAGTT | 120 | 56.67 |
| chr7 | 143563226 | 143563346 | Target=1;ProbeIdx=769;Target #1 | 1 | + | good | GGCGCAGGCTGGCCCTCTGAAGGATTGCCTCTTCTCCATGTCATTAGTGGCTCTTGTATTGCTTTGGAACGTGGCCTCCCGTGTGTGGGCACTCTCAGTCAGGTGTCGGGCCTGCCTTCC | 120 | 59.17 |

| | | | | | | | | | |
|---|---|---|---|---|---|---|---|---|---|
| chr7 | 143563346 | 143563466 | Target=1;ProbeIdx=781;Target #1 | 1 | + | good | TCTGCTTTAGGCGCCGAATGGAGTAGGTGGCTGAGTTCTTCCCTTCTCTTAGCCAGCAGCTGCTGAGCCGCACACTGAGCGCTCTGCGGGCTGAGGCTAGTGTGTCCTGAGCTTCTGCTT | 120 | 58.33 |
| chr7 | 143563466 | 143563586 | Target=1;ProbeIdx=793;Target #1 | 1 | + | good | CAGTAGCTTGGGCTGCTCCCATGGAGAAAGAGCAAGATGGACTTGGGAGCTTTGCAGGCTGGGTCCCTGTGCCAAAATGGAGGCACTGGCCAGGATGGGTGGCAGGCTGAGGCTGCTCCA | 120 | 60 |
| chr7 | 143563586 | 143563706 | Target=1;ProbeIdx=805;Target #1 | 1 | + | good | GGCATGGAGGTTGAGGATTATAGAATCTCAGAGCATTCCCTCACCTTTCTCTGCCCCTCCTTGCGAGATCAGGTGTTTTGCTCAAGTGGGACAGAGTTGGGGGTGGGGAGGTGGATAGAG | 120 | 53.33 |
| chr7 | 143563706 | 143563826 | Target=1;ProbeIdx=817;Target #1 | 1 | + | good | GGACCTCCAGACCGCTGCTCTTCAGCTGGTGGCCTCCATTACCAGTGTGCTTCATTTATAAATCCAGGCAAGCAGTGCCTTCTCACCTACAGACCCAAAGCCCCTTAGAGTCTATCAAAA | 120 | 50.83 |
| chr7 | 143563826 | 143563946 | Target=1;ProbeIdx=829;Target #1 | 1 | + | good | CATTCTTTCTTGGGGATTTGAGTTTCTCTCGTCTACCTCATTGCACAGAAGATAAAGAAAAAATTTAAGTTCTCTGAAATATTCTAGAATAATTTTAAGCTTTCTTCCCTCCTCCCCTTT | 120 | 34.17 |
| chr7 | 143563946 | 143564066 | Target=1;ProbeIdx=841;Target #1 | 1 | + | good | TATCTACCTGATATTTGTATAGGTTCTGAGGATCTGATCATGGTCCTTACATTTGTGAGGCAAGTGCTTTAACTGAGAGCTATTTCCAAATCCAGTTTTCAGTTTTCTCAACAGTAGCTC | 120 | 38.33 |
| chr7 | 143564306 | 143564426 | Target=1;ProbeIdx=877;Target #1 | 1 | + | good | TCAGTGCAATAATCTCTCATGTACACCAATCCCTTGGGCGTATATTTGTCTGGGATAAAAAGGCTTGCTGGTTAGCACCCAATACCAACGTGAGCAGGTGCAGTGTATAAAACAAAATCA | 120 | 42.5 |
| chr7 | 143564426 | 143564546 | Target=1;ProbeIdx=889;Target #1 | 1 | + | good | CTTCCTGGCAGTTCTGAAGATGGCTTCACAACCAACCTTAGGCCTGAACCTCCATGGCAGCAAGAACAGCATTCCCCAAGGGAAATGGCTAGATGGCCCCCACAGTCTGATTGTCCATCT | 120 | 52.5 |
| chr7 | 143564546 | 143564666 | Target=1;ProbeIdx=901;Target #1 | 1 | + | good | CAGCAGCTAGGGTTAGCCCTTGACCAGGGCATGCAGCCACTCCTATAGAATCCAGGATTTGCCTACCTCAAGAACTTCAATCCATGTGGGCCTGCCATGGGTTTACATATCTCCTGACAT | 120 | 50.83 |
| chr7 | 143564666 | 143564786 | Target=1;ProbeIdx=913;Target #1 | 1 | + | good | CCAGTGGGTGTGGCCTTTGTGCTCATCATCTGTCCAGAGTTCATCTGTCACAGGGGTTGGCATATTCTACATGGCAGGATAGCTTGGTAGAAAGCTTGTGCCCTGTATGTGGTATTTGGG | 120 | 50 |
| chr7 | 143564786 | 143564906 | Target=1;ProbeIdx=925;Target #1 | 1 | + | good | AGTCACCTGCAGAGAACCATTCTCAGATTTGGCCTTGGTGGCCACCACAAGCATAAAGCTTGTCCCTCAGTTATCTGAGTCTCTGACTTTTCCCATTCCTAGGGCTACTGGGGACACACT | 120 | 50 |
| chr7 | 143564906 | 143565026 | Target=1;ProbeIdx=937;Target #1 | 1 | + | good | GAACATTGCTAAAGCTTTGGGTCTCCCTGTATCCAGTTTCCCAGTGTGGCAGGCATGATCTTGCTCTTGTCCTAGATGTACCAGATGATGGCCTGTGAAGACCTGAACCATGTCCGCCTA | 120 | 50 |
| chr7 | 143565026 | 143565146 | Target=1;ProbeIdx=949;Target #1 | 1 | + | good | TGTTGGCCTTGTTTTCAGATTGATTCTTTAATGAGGTGACACAGTACAATGAGGTCAGTAATCTGGGTCCTGTTAAACAGGGAAGGATTATAGGAGGTAACTGGGACATAGGCCCACACA | 120 | 43.33 |
| chr7 | 143565146 | 143565266 | Target=1;ProbeIdx=961;Target #1 | 1 | + | good | GAGAGGCCATGTAAGGACAGGGGGATGTGGGCATGTTTGCATGGAGATGTTGCTTACCTGCTGAATATGTTGCTCACATGTGGTGCTGAATATCTGCTTTCTCTATGAGGGGCTTGACCA | 120 | 49.17 |
| chr7 | 143565266 | 143565386 | Target=1;ProbeIdx=973;Target #1 | 1 | + | good | TGGGAACAGCTGAGACCCTCAGGAGACCCTCAGGAGACCCTTAGAGATCATACCTGTTAACAGTCTCCAGTGGGACAGGGTGGACTTCTGACATCAAGACAAGTACTCCGCACCCCCAAA | 120 | 53.33 |
| chr7 | 143565386 | 143565506 | Target=1;ProbeIdx=985;Target #1 | 1 | + | good | GTTCTATCAAAGGTAGATCTTTGTCTCGGGAGGGTGCACTACTTAGTAGTATAGTTTCTTGTATTACCTAGGGGGTCTCTTTGTTGGCACAGCTTTTATGAGTTCTTGCAATGGCTGAGA | 120 | 42.5 |
| chr7 | 143565506 | 143565626 | Target=1;ProbeIdx=997;Target #1 | 1 | + | good | CTCTTCAGAGGTATAGCTGCTTCTGAGTTATACCCATGCACCTGAAAGTGACCCCAGCAAGCTCACTGTTCACCTAACTGCATCAGACAGAATTCTTTCTTTGGTCTGCTGGTGACTTCT | 120 | 46.67 |
| chr7 | 143565626 | 143565746 | Target=1;ProbeIdx=1009;Target #1 | 1 | + | good | CTGTCTAGGATGAATGATGTTTCTTTACATCTCCCTAGGAAAAGTCCCATGACACATGGGGCAAAGCCTGGTCTAGTGGCTCTCTGGAGAGACCTACCCTGGTCATGCCTGCATTCTACA | 120 | 49.17 |
| chr7 | 143565746 | 143565866 | Target=1;ProbeIdx=1021;Target #1 | 1 | + | good | CTCCTGCCTCTGAAATGTGAGCTGATACATGTTGGCTGTGTGGCTGCACCCTTGAGAAATGTGCTCAGAGATGCCAGGCAGGGAGCTGCAGAATAGAAAGTGGCACTGTCTGCTGTCTCT | 120 | 52.5 |
| chr7 | 143565866 | 143565986 | Target=1;ProbeIdx=1033;Target #1 | 1 | + | good | GCTGTGTGGATTAGGGGGATGGAGACCCTTCCCAGGCCTTAGTTTCCTCCCTGGATTGGGGTTGGATTTAAGGTCTCCTCTGTGTGGGTTGAAGAAGCATCAATGCCAAGTACAGTTCCA | 120 | 51.67 |
| chr7 | 143565986 | 143566106 | Target=1;ProbeIdx=1045;Target #1 | 1 | + | good | GCATGGACCCTGGAATGGCTCTGCTGGCTGCAGATGTGTTTAAAGCATGGATGCCTCGAGCCAGTAACCTTGTTCCCACTGACTTGGCTTTCCTAGAGACCAGGGTGAGTGACTTGGAAG | 120 | 53.33 |
| chr7 | 143566106 | 143566226 | Target=1;ProbeIdx=1057;Target #1 | 1 | + | good | CAGGATAGAGATGCATAGAGGATGGCTGTGTGAAGATACTGGGAAGAAGACACCATCTGCAGGGAAAGGACACAAGCCTTGACACACATCTGTCTTAGCTCTCCAGCATCCACGATGACA | 120 | 49.17 |
| chr7 | 143566226 | 143566346 | Target=1;ProbeIdx=1069;Target #1 | 1 | + | good | GGCAGTGGGTGCAACTGACCACAGACTATAGACTAGAGCTGTGGGTGGCCATGAGGGGCTCCCATGCTGTTCCCTTAAGATTCAAGATTATCCCAGGCCATCTGCTCCTAGAGAGATAGT | 120 | 52.5 |
| chr7 | 143566346 | 143566466 | Target=1;ProbeIdx=1081;Target #1 | 1 | + | good | CTGGGTTGTTCTCGCACTTTCTAAAAGTTTATGTTCCTGGGCTGAAATTAATTCGATCTACGTTGGCGTGTGGGCAAGTGTGGTGTGGTTTATTTATTGTGGATTGACAGGTTTTTTTCA | 120 | 40.83 |
| chr7 | 143566706 | 143566826 | Target=1;ProbeIdx=1117;Target #1 | 1 | + | good | AACATACCATCTGTAGGTGTTTTATAGAAGTTTGTGTCTATTTTAATTAGCAATTGTACTGGGTTTTTTTTTTTTTTTGGTACAGTTTTATTTTGTGTGTTTGTGTGTGTCTGTATGGGT | 120 | 30 |
| chr7 | 143566826 | 143566946 | Target=1;ProbeIdx=1129;Target #1 | 1 | + | good | GACCTGAACTTCAACTGCGGACCTCTTGTCCACGGCATTGTTAGTGGGGGAACTTTCTGATTTTTCTGTTTTGTTGGTTTCCGAGTTACCGCCCTCATGAGAAGAAAAGCTATTTTAATT | 120 | 43.33 |
| chr7 | 143566946 | 143567066 | Target=1;ProbeIdx=1141;Target #1 | 1 | + | good | GTCCAGAGCACACCACCTCCTCCACAGCTTTCTGTGGAAACTGCGTCGTCGTCCTGTTCTCATCGCCCCTACCCTGTCCCTATTGGTGCAGAATGCCAGCAGGTAGCATCTCTTAAATAA | 120 | 53.33 |
| chr7 | 143567066 | 143567186 | Target=1;ProbeIdx=1153;Target #1 | 1 | + | good | AGCAAGGCACAGGCTGGGTGCCCATGGCTTGTTTGTATAAATACCATGGTGTTGGCGAAGCTGAGCAGCACAGTAACCGCACACCTGGAATAAGACAAGCCACAGCCACCAGCCGCCGAG | 120 | 55.83 |
| chr7 | 143567186 | 143567306 | Target=1;ProbeIdx=1165;Target #1 | 1 | + | good | GCCTACACGTGCAACTAGAGATGACATCTGTCCCCAGGACAGTCAGGGTTACTCTGTTATCCTCATGATGCAGTCAGGCCACAGTGGACTCTGGTTGTCAGCTCTGGGGGACATTCTAGG | 120 | 53.33 |
| chr7 | 143567306 | 143567426 | Target=1;ProbeIdx=1177;Target #1 | 1 | + | good | CCTGGTGGATTCCCCAGGGCAAGCTATGGTTCATGTGGACATGAGCTGTTATCAAGTGAACCATCTGTGTGAACATGGGCTGTTACCAAGGTGACCCTTCTGAATGTGGGTAGGCATTGA | 120 | 50 |
| chr7 | 143567426 | 143567546 | Target=1;ProbeIdx=1189;Target #1 | 1 | + | good | TGAACACGGTATGTTTTGTTATTAATTAAAATTGCCTTTCCCAGTTTCCTGTGTGTATATGATGTGTCCTGATGGTTCTCACCCATCCTCCCATGCCTTACCTTCCTCTGTCTCTGTTCA | 120 | 41.67 |
| chr7 | 143568146 | 143568266 | Target=1;ProbeIdx=1261;Target #1 | 1 | + | good | CTGCTGATCTCTTTCCCACATTCCTCTCCTTTATTTTGTTTCCATCCCTCTGGGTTTGTCCAGGCCTCCAGGGTGGCCTTGGGGCTGGGTCTCTCCAGGAGCTGGCAGACTCGGCAGCTG | 120 | 57.5 |
| chr7 | 143568266 | 143568386 | Target=1;ProbeIdx=1273;Target #1 | 1 | + | good | GTGCACAGCGAGGGCAATGGCACACTTTCCCAGGGTCACCTGGGTGTCCCTGGGCTCCCAGCCTACCACCTCCACAGTGGCATGGAAACTGTTCCAGGTAAAAGCTGGAATGATCCAGAA | 120 | 57.5 |
| chr7 | 143568386 | 143568506 | Target=1;ProbeIdx=1285;Target #1 | 1 | + | good | CTTACCTTGTTTCCTGCATTACTGCCCTGCATTTGAAAATAGGTGGTTTTTTTTCAATCTATTTCAGTGTATTGTTGGTATCAGGATGGAGGCTAAACCCAGAACACCCAATGTCTCCAT | 120 | 40 |
| chr7 | 143568506 | 143568626 | Target=1;ProbeIdx=1297;Target #1 | 1 | + | good | TTGGTGGAGGAGGGGATCCCCCCAAGTCTTTCCTACCCAACCCCCAGGCTTCTTAGCCCCTGTCTCCATTCTCTCCTCTGTTCCTCCCTTCTCCTCTTTCTTACCGCCCCAAAAGAAATG | 120 | 55 |
| chr7 | 143568626 | 143568746 | Target=1;ProbeIdx=1309;Target #1 | 1 | + | good | AACTGTTTTGTGCTAAGCAGCAATGTACCAGTGGCTTCTGCCTTCTCTTTCTTCTGTCCGCAAGGGCGACAACAAACTGCTCCACTCTCTGACTCCCCCAAACTCTTGTTGTTAATGGAG | 120 | 48.33 |
| chr7 | 143569106 | 143569226 | Target=1;ProbeIdx=1357;Target #1 | 1 | + | good | GCTTTTCACAGCACCCCAGCTCTGCCCTCTAGGACCTTAAGTCCCTCTTCAGTATCCTTTCATTAGCCTCGATGGCCAAGTGCCATGCTCATGTGCCCACCTCAGCTGTTCCTTCACTGG | 120 | 54.17 |
| chr7 | 143569226 | 143569346 | Target=1;ProbeIdx=1369;Target #1 | 1 | + | good | TCCTGCCAAGTCCTCAGAGATAGACCTGCGAAAGACCAGTGAGATCCCAGTTCTTGCTCTTGATCCCTGGAGGTGACATGGGAAGTTTGGAACACATTGTTGCCTCAGGGCTCCAGGGTC | 120 | 53.33 |
| chr7 | 143569346 | 143569466 | Target=1;ProbeIdx=1381;Target #1 | 1 | + | good | TTTCTAGTGCAAGTTCTAGTACTGGAGTTTGTGACCCTCAAAGCCTATTCCTATCAACTTTTCCTGGAACTTTACAGCAGTATCCACAGGCTGCCCAGACACACTTTCATCTGATACTTA | 120 | 42.5 |
| chr7 | 143569466 | 143569586 | Target=1;ProbeIdx=1393;Target #1 | 1 | + | good | AGGACCTAGGGAGAGGGTGACATGAGACACAAATGGCCATGTTCTTAACGAGAGTATTGAGTTCCTAGACTGTGTTACTGTCACCTGGGATCTCCTCCGAGAGAAGAGCAGATCTCATGA | 120 | 48.33 |
| chr7 | 143569586 | 143569706 | Target=1;ProbeIdx=1405;Target #1 | 1 | + | good | AAATCACTTGCAGGTCTAATATGTCCTCCCCCACACTGGAGTGGAGGAGGCAGGAGAGCTGACATCCACCTGAGTCTAGTGGAGGACAGGAGGACAGCCAGCAGCTCCTGTGGCAGTTAG | 120 | 55.83 |
| chr7 | 143569706 | 143569826 | Target=1;ProbeIdx=1417;Target #1 | 1 | + | good | CACACAGCTCCAGCCCATGGGCTTACAGATGCTCCAGGAACATGGGCTAGGCTGGTCCCGAGCAGTGGTTGGATTGGCTTATGAAGGATATGTCATAGCTCTACTCATAAGACAATGATG | 120 | 50.83 |
| chr7 | 143570186 | 143570306 | Target=1;ProbeIdx=1465;Target #1 | 1 | + | good | CCAGTGAACAAGCTGTCCCCAGGTCTTATTTAGCCTAGCTTCTCTCTCTCCTGCAAGCCTTTGGTTGGTGAGTGAGACCCACATCCCAAGGTAGGATGAGCAGGATCTCCGTGGGGCTGA | 120 | 54.17 |
| chr7 | 143570306 | 143570426 | Target=1;ProbeIdx=1477;Target #1 | 1 | + | good | GAGCCCTGGAGCAGACAAAGTGCAGAGGCATTTCAGAAGACAGGTCTCTGGGTAGGCATCTGACTGTGGTCTGATTGGGCTTTGGAGGGGAGAAGTGAGGGCAGGCTCAGGCTGGGTTTT | 120 | 55.83 |
| chr7 | 143570426 | 143570546 | Target=1;ProbeIdx=1489;Target #1 | 1 | + | good | GACCCCTCTTAAGGCTTTGGGCACTGAACATGAAGGCTGCTAAGTGTGGAGGCCGCTGGGCGATATCACATCCTAGGACCTGATCCCAAGAACCCAGCACATGGCTGGCTAGAGCTGGAT | 120 | 55.83 |
| chr7 | 143570546 | 143570666 | Target=1;ProbeIdx=1501;Target #1 | 1 | + | good | TGTTTTAAACTTTCCAGACATATTCTCTGAAGAATACCAAAACATTGGTATTCCAATTGCAGAGGACCCTGGTTCCCACAGCTCTCTGAATAATAGTAATCAACCAAGTCCAGTCTGGCC | 120 | 40.83 |
| chr7 | 143570666 | 143570786 | Target=1;ProbeIdx=1513;Target #1 | 1 | + | good | TAAATACTGTCATCAAACATGGATGATGGCAGGTTGGATAGCCACAGCCCCACTGAGGGCTGTCCCAGGTAGGCCTGTGCTTGGCAGTCAGGGCTCGTGGGACTGCAGGCTTTCTTCAGT | 120 | 55 |
| chr7 | 143570786 | 143570906 | Target=1;ProbeIdx=1525;Target #1 | 1 | + | good | CTTGGCCAGAGTTGGTGTCCCTTCAAGGGTCATCCCACTCTGTGCCAGGACTGGCGCTGGCTATTGAGGGCATTAAGAACTCTCTACAGAGCTGTGTGAGTCCAAATACTTGCTTCAATA | 120 | 50.83 |
| chr7 | 143570906 | 143571026 | Target=1;ProbeIdx=1537;Target #1 | 1 | + | good | GAGAGACTGGGGTCCAGAGGTCTTTGTTGGGTAGTCTACTCTATCTCTTTCAATCTTAGTTTTCCAGACATAGTCTCCTACTGAATCACTTTGCCTAGATTAGTGGTTCTCAACCTGTGA | 120 | 43.33 |
| chr7 | 143571146 | 143571266 | Target=1;ProbeIdx=1561;Target #1 | 1 | + | good | CTAACTGAGCCACCTCCCCACCCCATAGCTCTATAAGTCTGTTGTTTGTCACACCTACATCTCAGGGACTTTACCCAGTTCTGGTGGACATTGTGGTTTAGACTGTGCATCGGCCTCAGG | 120 | 50.83 |
| chr7 | 143571626 | 143571746 | Target=1;ProbeIdx=1609;Target #1 | 1 | + | good | CCCCAACACTGACAATGACACTGGAGATGATTGTAGCAGAGGTCTCATGGGAGGAGATCCTTCTGCTTGGGGTCTTGGCTGATGAGTCTTGCTGTACTGGTCCAGCCTAGAGTAGAACCC | 120 | 52.5 |
| chr7 | 143571746 | 143571866 | Target=1;ProbeIdx=1621;Target #1 | 1 | + | good | AGCAAGACCGCCTTTTTCTCTGTTGTAGCCAAGGTCCCTGATTAATAATGTCAGTGTATTTTAACCAGGTCAGATGCTGCAGAAATGTAACCAGGATGCTGACTTTTCCAATGATGGATT | 120 | 41.67 |
| chr7 | 143571866 | 143571986 | Target=1;ProbeIdx=1633;Target #1 | 1 | + | good | AGTCCATGTGTTCAACCTTGTGTCTGTGTATGTACACTTTATACCCAAAATGTTAATGGCACCTTTAGTGACTCCTGGATTGTAGGGCTAGCTGGATGTTTCTCCATCATTTTGTAGGCA | 120 | 41.67 |
| chr7 | 143571986 | 143572106 | Target=1;ProbeIdx=1645;Target #1 | 1 | + | good | ATGTGAGGGATTCGGTTATCCTTTGTACATTGATCTCCTACCTGTCTGACCTGTGACTTTCTTTGCTAGTTCAGTTTATCTTTAGCTTCTCTTGGGCTTTCAGTGCAGTGTAGGTAATTA | 120 | 40.83 |
| chr7 | 143572106 | 143572226 | Target=1;ProbeIdx=1657;Target #1 | 1 | + | good | TACAAAAATACTGATTACCTCCATTACTTCCGATCCTTGGGCAGGGGTTCACTTTCCTGCCTGCTTGCGTTAGCTGGAGATGCTATGACTCAGTGCTTCTGTCCAAATTAGGGAATGTAT | 120 | 45 |
| chr7 | 143572226 | 143572346 | Target=1;ProbeIdx=1669;Target #1 | 1 | + | good | TGCTTCCCAGCATTACAAGGGGTTGATCCTGATGGAGGTGTCCCTTGTTCCTTAGTGTCTATGGAAGACTGTCCATGGCCTGGATTTTTTATTCCTGCTGCACACAGGGATTTACTGGAC | 120 | 48.33 |
| chr7 | 143572346 | 143572466 | Target=1;ProbeIdx=1681;Target #1 | 1 | + | good | ACAGGGCGGAGAGCTACAGACAGGGCAGCATAGCACAAGAATTCATGGCCTGTGGCTCTTTACGTCATCAAACCTTTGCCTTGCTTGTCCTTCCTATGCATGATGGGCAGCTCTGCAGAA | 120 | 51.67 |
| chr7 | 143572466 | 143572586 | Target=1;ProbeIdx=1693;Target #1 | 1 | + | good | TCTCTTTAGCCCTGCTTTGGAACTCTTGAATTCTACTATAGAGAAGCTAACGTGCAAATCTAAACTAAAACCAGGTAGCACAGGGTCTGCTTGCAGTAGGAGCCAGGGCAAAAGCTGGCT | 120 | 45.83 |
| chr7 | 143572826 | 143572946 | Target=1;ProbeIdx=1729;Target #1 | 1 | + | good | GGAGCCTGAGGAGACACACTATGTGTGCTCTCCATTATCTGGGTGGAGAGCTGAGTGAAGCAATTGTCTGAAATTGCCCCAATGCTAATCAAGGAAGGATGGGAGCCACCACATGGGTTG | 120 | 50.83 |
| chr7 | 143572946 | 143573066 | Target=1;ProbeIdx=1741;Target #1 | 1 | + | good | TCTCATTACCAATATCAACCAGGAATGGCACATGCTTCCAGGACAATGACAGCACCCTCTCAGCAACTTTTTTTTTTTCCAACCTGCCTAGAATCTTCCTCAGGAGACAACTCTCCAAAC | 120 | 44.17 |
| chr7 | 143573066 | 143573186 | Target=1;ProbeIdx=1753;Target #1 | 1 | + | good | CTTTGGTAGAATGAATCAGCCCAGCATGGCAGATCCCTTCAGCAGTGCTGGTTAGGGGAAATACTTTAACCTGGTAGCATCTGTTTTCTGATTTAGACCCTTGAGTCTGACATGATGAAG | 120 | 45 |
| chr7 | 143573186 | 143573306 | Target=1;ProbeIdx=1765;Target #1 | 1 | + | good | AGAGGGAAAGAGACAGCAGGATCGCTGGGGCTTGCTGGTCATTCAGTCTAGCCATCAGTGAAATAAGGGTCAGTGAAATAATGTAGAGAGCTAAAGGTGGGATGTATGTATGTATGTATG | 120 | 45 |
| chr7 | 143573426 | 143573546 | Target=1;ProbeIdx=1789;Target #1 | 1 | + | good | ACACGGACACAAACACACAAAGTTATTGAGTGTATATCATGTCCCCACCATCCTTTTCTCCTGAATAAAGAGCCACTTCAGATTTCTGGTCACCACTGTCGTGTGAGCACAGTCACTGCC | 120 | 45.83 |
| chr7 | 143573666 | 143573786 | Target=1;ProbeIdx=1813;Target #1 | 1 | + | good | TTCAAAGATGCCGGGTGTAGCCAGGCAGCTTCCATTCCTGCTGGACAGGTTGTCTGTGGCCTCCAGGGAGAAGCTCGCTCAGTCCAAGAGACCTTGGTAGAGTCTGTCTTAGCTGTCCTC | 120 | 55.83 |
| chr7 | 143573786 | 143573906 | Target=1;ProbeIdx=1825;Target #1 | 1 | + | good | TGGGAAGCTTCCTCGCCAAACACTATGGTGGCTTTGGCTCTAGTGGGACTCAGGCATTTTTACTCAATTCAAGCGTGTTTCCTAAATCCGCTTGAGCTGAGGAGGCCAAGAGAGCATGCA | 120 | 50 |

| | | | | | | | | | |
|---|---|---|---|---|---|---|---|---|---|
| chr7 | 143573906 | 143574026 | Target=1;ProbeIdx=1837;Target #1 | 1 | + | good | TGAATGATCTGCATGCAGTTTTCTAGCGCCGGGTGGCCATGGCAGCTTGAGAGGGTGCTGAAATCAGCAGCTCAGTCATTGGGGAGAAGGGCAGGCTGAAGGGAAGGGCATCAGAGAAGG | 120 | 55.83 |
| chr7 | 143574026 | 143574146 | Target=1;ProbeIdx=1849;Target #1 | 1 | + | good | GGGTGCTCATGGGACCTTTTACTATGGTTTATGCAAAGACACACATAGGGCCAGACTTGGGGCCTGGTCCTTAGAAAGCATTTGGGGAAATTTCCAGTGAACCAAGACAGATGGAGAATC | 120 | 47.5 |
| chr7 | 143574146 | 143574266 | Target=1;ProbeIdx=1861;Target #1 | 1 | + | good | TGACCCTGAGTTCACAGGATGTTTCACCAGTGTTCCAGCTGTGACAGCGTGGGTGCTATCCCAGAGCACCCTGGACAGGTGGTATAAACAGTAGGAATCTACTGTCTCACATCCTGGAGG | 120 | 52.5 |
| chr7 | 143574266 | 143574386 | Target=1;ProbeIdx=1873;Target #1 | 1 | + | good | GTGGAGGTGGGAGGAGACAGAGATGTTCAGGGCAGGCTCCTCCAGAGCCTCTCTCCTTTTCTTGCAATGGAGTTTTCTAGCTGCTCTTCACAAGGTCACGCCTTTGCCTGGTATGTGACT | 120 | 53.33 |
| chr7 | 143574386 | 143574506 | Target=1;ProbeIdx=1885;Target #1 | 1 | + | good | TAGGTTCACTGAGGTGGAAAGATCCTGAGTGTGGGTGGCTCCATGTTCTGATCTGGGATTTTACAATGTTTAAAGGGGGAAACTTGGTCAGAGCATCCTTCCCTCAGCTTTGGGAGTATG | 120 | 47.5 |
| chr7 | 143574626 | 143574746 | Target=1;ProbeIdx=1909;Target #1 | 1 | + | good | AGGAGAAAAAGAATTCACCATTATTGAATCAGGACTGACTGTACTATATTTGATTTACTTGGGTTTTGTGGTGCTGGGGCTAGAACCCAGACAGGCACTCTACCACTGAGCCTTCTTGCG | 120 | 44.17 |
| chr7 | 143574866 | 143574986 | Target=1;ProbeIdx=1933;Target #1 | 1 | + | good | TCTGTCCAAAGTCCTTTAACCACCCATCGAAAGAGTTTTATCCAAGTCATGTGACCAGGCCCAGGGAGTAGGACTCCCACAAATGAATCTGAAAAGGCACACCGTTCAGCCTATTCACTC | 120 | 47.5 |
| chr7 | 143574986 | 143575106 | Target=1;ProbeIdx=1945;Target #1 | 1 | + | good | CATAGACCACTGGACATCAGCTTATGGACCTCCTAAGATTTTAGAGTTGATCTCCAAAACTTTTAGCTGTTGCCTGGCAGTTTTTGTCTGTCTTGAATGATGTCCTGGATGAGTGAGCAG | 120 | 43.33 |
| chr7 | 143575106 | 143575226 | Target=1;ProbeIdx=1957;Target #1 | 1 | + | good | GTGGGGTGGGGTTGGCATGCTTAGGCAGCAGGGCAAATATCTCACTAGAGTCTTGCTCAGTGGGGGGGACACATCCTGTGTTGGTTATTTTCACCTTTGCTGGGACCAAATGCCTGATGA | 120 | 52.5 |
| chr7 | 143575226 | 143575346 | Target=1;ProbeIdx=1969;Target #1 | 1 | + | good | GAAGCATTTTAATAAAGGAAAGGTTTGCTTAGGTTTGTAAGGATGCTATCTGTCATAGCACGGCGGGCAGTCAGCAGTGGGAGACCACATGGCAGTCCAGAACCTAAGCGCATGGATGGT | 120 | 48.33 |
| chr7 | 143575346 | 143575466 | Target=1;ProbeIdx=1981;Target #1 | 1 | + | good | TCTGCCCACGGTTAGGGTGTGTCTTTCCATCTCAGTAAACCTAATCTAGGAAGTCCCTCTTACACTTGCTCAGAAGTTTGTCTCCATCAAGTTGGCCCTCAAGATGGACCATGGCACAGC | 120 | 49.17 |
| chr7 | 143575466 | 143575586 | Target=1;ProbeIdx=1993;Target #1 | 1 | + | good | CACACACAGGGTTTCATTCAGCTGTACCCATGGTGCTCTGGTGGGGCAGGTAGAGCGGGCTGCTTCTCTGGAGAATTCGGGGGGCAGTGGGGTGGAGGGGTGGAGCTGGTCCCTGAAGCT | 120 | 61.67 |
| chr7 | 143575586 | 143575706 | Target=1;ProbeIdx=2005;Target #1 | 1 | + | good | GTCTGGCCTCTGTGTCACTGACCCTTGCCTGCTCTACTCAGTTGAGTTCCCAGGAGGCTGGTGCTCTTCTTGGCAAATGTGACCTCTGTTCTCCAGGTCCCCCCGGAAGCAGCCCTATCC | 120 | 58.33 |
| chr7 | 143575706 | 143575826 | Target=1;ProbeIdx=2017;Target #1 | 1 | + | good | TGCCCTGGGACCTTGACCTGGCTTTAAAGATCTAAACAATGCCCCAAGGGCAGCTTGGGAAGGGAAGGTCTTGCTGCTTTAAGGACCAACAAATTTCATACATTTTATGCACAAAGTAAA | 120 | 44.17 |
| chr7 | 143575826 | 143575946 | Target=1;ProbeIdx=2029;Target #1 | 1 | + | good | CTACAAAATAAAATCTCTCATTGTGTTGGTTTGAGATTCCCTCAGGAAGGAGACCTCTGTGCATTTCAAACTAAAAATCTGTCTGTGAACCCCTGAGAGTTTTGAATTGATCCCCAAAGG | 120 | 40 |
| chr7 | 143575946 | 143576066 | Target=1;ProbeIdx=2041;Target #1 | 1 | + | good | CATTGCAGCCATTGCCTGGACTGTTTTATCTGTATCTTCCATTGAAGGTCAGTTTTTATGCCCCAGTACAAGAATGAATATGAAACTGTGCGTCTGCCATAAGGGGTGGAAATTTGCTTC | 120 | 42.5 |
| chr7 | 143576066 | 143576186 | Target=1;ProbeIdx=2053;Target #1 | 1 | + | good | TCTGTGGTGATTTTTCTGTTTCCTTATTAGTAGCTGTCCAGCAGGGCTCATGCAAGAGGCCTGCACTGAGGGATGACCTTCCTGGCCTGGGAGACTGCAGCTCTGCATGTACCCCTGTGA | 120 | 53.33 |
| chr7 | 143576186 | 143576306 | Target=1;ProbeIdx=2065;Target #1 | 1 | + | good | ATAGCTACACCAAAAGCTACTGCCCAGGGTGGCACAAAACCACCCTTCCTCTTTGCCTCACCCTACTTGCTTTCTGTTTCTGGGCAACCTCTTTTGCAGTCTGATCCTGAGGGGATCCAG | 120 | 51.67 |
| chr7 | 143576306 | 143576426 | Target=1;ProbeIdx=2077;Target #1 | 1 | + | good | GGACCAAAGACAGTTCCTGCTCTCCAGAATGAAGCCAGACAAAAAGCAAGGCCTCCTATTCTGTGTGAGCATGGGATAAATGACCAGTGGTTGTTTAGTGTTATGTGTGCCCCAGGTAGC | 120 | 48.33 |
| chr7 | 143576426 | 143576546 | Target=1;ProbeIdx=2089;Target #1 | 1 | + | good | CAGCAGGATGTTCACAGAGCAACGTTGGTTTCCTGCTTACCTGAAAGTGAATGTGCCCAACTGCCCTCTGCCCTCGCAGGAGTTAATCATTAATGGCCCCTGTCAGGTGGCAGGGTGTCT | 120 | 53.33 |
| chr7 | 143576546 | 143576666 | Target=1;ProbeIdx=2101;Target #1 | 1 | + | good | GCTGTTCCCAAGGCGCTGCACCACATTCCTTATTAGGGTCACGACCCTCCAGGTCAGGTCTCTGGCACTAGGATCTCAGGACCCTCCTGCAGTGACCCTTGGAGCTGATTCTTCCTTCAA | 120 | 55.83 |
| chr7 | 143576666 | 143576786 | Target=1;ProbeIdx=2113;Target #1 | 1 | + | good | TCACCCCACTTCCAGACCCAGCCTCGGGCCTCTTTGGATCTGTCTGGTGGAGCCCTTCTTAGGTACCTACCTGTTTAGGCCTGTCTTCTCCAAGTCAGCAATAGGGTACATTATGGGGTG | 120 | 54.17 |
| chr7 | 143576786 | 143576906 | Target=1;ProbeIdx=2125;Target #1 | 1 | + | good | TCCTTCTGGATGAGGAGGTTACTGGTAGTGGCTTGGACTGGGGAGCCTCGCCTCATTATTCTCTTTATTGGTTTGTGAGCTCTGTTTTTGGAGTCAGTCTCTGGCTTTGGCCTGTTATGA | 120 | 48.33 |
| chr7 | 143576906 | 143577026 | Target=1;ProbeIdx=2137;Target #1 | 1 | + | good | ACACCCCCATCACCCTCCTTTATTATGGAAAGAAGACTGTGTGTTAAGCTTACCTCAGAGTTCTGGTTTACTTCCAGCCTGTAGGCACATCTACATGTGACCTGAAAAAAAAAACCAAGC | 120 | 43.33 |
| chr7 | 143577026 | 143577146 | Target=1;ProbeIdx=2149;Target #1 | 1 | + | good | CTTTAAAATGGTACATTTCCAGTATTGATACTTGTTACATGCCTGCCCACATCTGACCTAAACCATCCCCACAGGACTGGGACTCTGGGCTCTGTTCCCTCTTGCTGGTGGTAGTTTGAC | 120 | 47.5 |
| chr7 | 143577146 | 143577266 | Target=1;ProbeIdx=2161;Target #1 | 1 | + | good | ACTCTACTGGGGGCAGATTGTGGGCCCATGCTCTGCCCATCTCTCCCTCACTGCATCCCGCCAGCTCCATAGCCTTGCTCCTGTCTGGCTTGTGTCCACTGAATGCCCACTGTATCCCAG | 120 | 59.17 |
| chr7 | 143577266 | 143577386 | Target=1;ProbeIdx=2173;Target #1 | 1 | + | good | ATGATGGATGGACAGCAGCAAATGAGACACACTAAAGAACCAGTAGTCACAGGCATGCAGGTTCATCCGGGAGCTGTAATGATGCAGTCACTAAGAAAACTTTTGAGAGAGCTGGTCAGA | 120 | 45.83 |
| chr7 | 143577386 | 143577506 | Target=1;ProbeIdx=2185;Target #1 | 1 | + | good | GGTCAGGTTGGGTCTGATGGGGGAGGGAATCAGGGTTGTGGGGAGGAGTTGAGGTAGATGTGGGTGTTTGCTTCAAGCGATTGGGTGCAGTTTCAGGCAGCTGGGAGTAGAATGCACATA | 120 | 54.17 |
| chr7 | 143577506 | 143577626 | Target=1;ProbeIdx=2197;Target #1 | 1 | + | good | AGAGGGCCTTCAGGTGCGCTGGCTGCCTGCCTCTCTACCCTCCAGGGCAGGAATTCTGAGGACTGGAAAATCTCATTTGCTCCAGACTTGGACAAAGCTGGTCTGGGCTATACAGGCATC | 120 | 55.83 |
| chr7 | 143577626 | 143577746 | Target=1;ProbeIdx=2209;Target #1 | 1 | + | good | GCTGTGTGCTGTGTGCTGTGTGCTGTATGCTGGGGCCAAAGACAAGTGATCCCTTCATGGGGAGCCAATGGAACATCGTCTCTTCTCACTCTTCAGCGTATGCTTCTGCCTTTGACCCGT | 120 | 53.33 |
| chr7 | 143577866 | 143577986 | Target=1;ProbeIdx=2233;Target #1 | 1 | + | good | TCCTGGACTCAGGAACCCCACAGCAGCCATTGCTCATGCACCACATCTATCTATGCCAGACCTGGCTTCTTTCTCTGTCCCTGCTGGAGTGGAAAGTGGGGCAGCCTTAGAAGCGGGAAC | 120 | 55.83 |
| chr7 | 143577986 | 143578106 | Target=1;ProbeIdx=2245;Target #1 | 1 | + | good | CACTGGGGCCAAACTAACATCTGTCAGTACAAAGGACTCTGTAAGATGGAACATGAAAGGCCCGACCTCCCCCAGACATGGTAGTTTTCTTTTTTCAGCCCAACACAGGTTGTTTTAGAG | 120 | 46.67 |
| chr7 | 143578106 | 143578226 | Target=1;ProbeIdx=2257;Target #1 | 1 | + | good | ATAGCTTGCCTACTTGCTCATCCCCAGAGAGGCCCCGGTGTCTATCCCTGCAGGTTGCGTGGCCGTGGTGAGGGACCCTTGGTGGTAGTGGCTGAGCAGGCTGGCTGCATGGTGAACTAC | 120 | 60.83 |
| chr7 | 143578226 | 143578346 | Target=1;ProbeIdx=2269;Target #1 | 1 | + | good | TCAGCTGCAGCCCGTGCAAGCTGGCAGAGCTGCAGCAGGGCCATGCTGCCTTCAGCCTACTATGGTGCTATGGCACCAGTGTGTGAGCATCTTCATTCCTTTCTTGCAGGCTGCCAGCAG | 120 | 58.33 |
| chr7 | 143578346 | 143578466 | Target=1;ProbeIdx=2281;Target #1 | 1 | + | good | GTCTTGACCACCCTTAATGCTACCTCATTCACAGTAATACGGGGTCAACAATCCCGTGTCCTATTGGCCTACATGCAGCTGGGACACGTGCCCTGTATGCTCTGGTGTCAAGCCTGCAGT | 120 | 52.5 |
| chr7 | 143578466 | 143578586 | Target=1;ProbeIdx=2293;Target #1 | 1 | + | good | ATTGGGAAGCCCTCGCAGCTCCTGGGCCCAGCTGAGCTGTTGCTACTTCTGCTGCTGCCACAGCTGTGGTCACTACAACCACTGTCACCTCTGCTGCTGCTGTCACAGCTGAAGCTGCTG | 120 | 59.17 |
| chr7 | 143578586 | 143578706 | Target=1;ProbeIdx=2305;Target #1 | 1 | + | good | TTACTGCTGCCAGCACCACTGGCAAGATTTAGCAAGCAGCAGATTATGCCCCCTCTTTGTAGCCCCCCCAACATCTCTTGACAGCCCCACACAGGAGAACCCAATGAGCATGAACATTTC | 120 | 51.67 |
| chr7 | 143578706 | 143578826 | Target=1;ProbeIdx=2317;Target #1 | 1 | + | good | TGGTGTTGAATGCAAGAGTGAGAATGGCTTGCTTATCTGCCAAGAGGTTCTGTGCACAAGTCCAGCCATTAGACAACCACTGGGCAAGTGCCTGCACTATCCCACCTCTTCCTTTATCCA | 120 | 49.17 |
| chr7 | 143578826 | 143578946 | Target=1;ProbeIdx=2329;Target #1 | 1 | + | good | TGTATTGGCTCTTGGATCATCACTGAGCCAGGCTGGGAGCCTTGTCCTGACCACTGCTCCCCTGTGCTCCAGCACAAGCTTCTGGCTAGGGTGAGTGTTTGCCCTTCTCTGCTCCCAAAC | 120 | 56.67 |
| chr7 | 143578946 | 143579066 | Target=1;ProbeIdx=2341;Target #1 | 1 | + | good | ACAGCAGCAGAGGCTAAAGCTGAGCCAGAGATACTAAGAGTAGAAGATGCTGGAGGTACCAGGTGGGCCACACTTGGACTTTTATGTTTATGCCCCAAAATTAGCGAAAGATTTCCCAAT | 120 | 45.83 |
| chr7 | 143579066 | 143579186 | Target=1;ProbeIdx=2353;Target #1 | 1 | + | good | CCCCCAAGCAGCAAACCCCATGCCAGGGGGGCCGCCCCATGTCCTCTGGGTCTGTATGGGATGGAGGACTCTGTCCTCCTTCCATTATCCCAAACTTTGTTGGATAATGTCTCTCACTCA | 120 | 55.83 |
| chr7 | 143579426 | 143579546 | Target=1;ProbeIdx=2389;Target #1 | 1 | + | good | GTGGAGAAGCTGGTCTGTGTCCATCTAGTCTGTGAAGCTACAACTGTTTTGACATTAAAGAGTATAAGGTTGGCTCGAGGTTAATGAGGACCCATGGCTCCAACCCGTGGCTGTCAACCT | 120 | 48.33 |
| chr7 | 143579546 | 143579666 | Target=1;ProbeIdx=2401;Target #1 | 1 | + | good | GCCTGCTTGTTAGGCCACCTGAGAGCATTGAAAGTGCAATAGCCGTAGTGAGGCAGTTAAATATCAGTCTCTGGCCTGGAGCTGGTGTGTTCGAAAGAACTGCACAGCCGGGATGAGAGC | 120 | 52.5 |
| chr7 | 143579666 | 143579786 | Target=1;ProbeIdx=2413;Target #1 | 1 | + | good | ATCAATAGCTTCCTGAGAACCAGGGGCCTGGGTGGAAATCATGACACCGTGCATAATAATCTCATGAGTCAAGCCATACACCCCCCCCCGCCCTCCAAAACCCATAAATCCAAAGCACTT | 120 | 50 |
| chr7 | 143579786 | 143579906 | Target=1;ProbeIdx=2425;Target #1 | 1 | + | good | TGATATCCAAACCTTCCAGACCAGAAACATAAAGATGAATGGAGAATTCCACTCCTAACCTCAGGGCCTGGGTTGTAGCTGAAACGTGCGTGCAGAGACAGTGGTGTCGCAAGCCGCCAG | 120 | 50.83 |
| chr7 | 143579906 | 143580026 | Target=1;ProbeIdx=2437;Target #1 | 1 | + | good | CAGGCTACATATGTATGACACACAAAGGAATTCATCGTCAGACCTGAGTCCCCTCCACAAGATGGTTCATTATGTATGTGCACATATTTTCAAACTGCAAGCCGTCTGGCATCCCTGGGG | 120 | 46.67 |
| chr7 | 143580026 | 143580146 | Target=1;ProbeIdx=2449;Target #1 | 1 | + | good | ATGCAGGATTCTTATCTTGTGATACTGTCACATCAGGTTTGAATACAGACCCTTGTTAACTCCACACGGGTGAATGAGGGGCTGGTGAGCTGGCTTAGCAGTTAAGAGCATGCTCTTCCA | 120 | 46.67 |
| chr7 | 143580146 | 143580266 | Target=1;ProbeIdx=2461;Target #1 | 1 | + | good | CAAGTGATAAAGATAAGTCTAAAAAGAGGATAGTGCATGCATGAACAAGTGAATGCTAAAAGTTACTTTGAAGCCGGTGCCTAACCAAGATGGACAGGGAGGGCTGAGTCCTTCAACCCC | 120 | 44.17 |
| chr7 | 143580386 | 143580506 | Target=1;ProbeIdx=2485;Target #1 | 1 | + | good | AAACCTTGGGTGCCTTGCTGGATCTTTGTGTGGCTGAGAAGATGCTGATAGAAATCATGAATGTGTTCGGCTAGTGAATACAGGAAGGCTCACTAGTCCTGAGATAGCAGATTAATTAAT | 120 | 42.5 |
| chr7 | 143580506 | 143580626 | Target=1;ProbeIdx=2497;Target #1 | 1 | + | good | CCCTGAGATTTTAACTGAGTACCACATAGAGTAAATTTTTACTTAACAAAACTTCCTGGCTGTAAGGGGGGTCATTGCTGGGGCTGTGCCTGCAGGTGGGGCTGGACAAAAAGAAACAGT | 120 | 45.83 |
| chr7 | 143580746 | 143580866 | Target=1;ProbeIdx=2521;Target #1 | 1 | + | good | GCCAGGTCCATTGGGAAGATCCTACTGAAGAACTTCATGGGGAGGGGTGTCATCAGGTGACCTGGGATTTCTGAGGGTCTCTGAGGACCCTGTGGGGCTTCTGGGCGCTAGGAAGAGGGG | 120 | 58.33 |
| chr7 | 143580866 | 143580986 | Target=1;ProbeIdx=2533;Target #1 | 1 | + | good | CCACTAGCTTCTACTCTTTGATGTCCTAGGTCAGGGATGGATCCCTTACAATTGGGATAAAGCCAGGTGTCTGGGGTTTTGTTATGATATCTCTGGATCAGTGGGCACTTCTTTTCATTC | 120 | 45 |
| chr7 | 143580986 | 143581106 | Target=1;ProbeIdx=2545;Target #1 | 1 | + | good | CTCTAACTCTGGCCCATGCCCAAGGCTATGGGGGCAGCAAACTGGTGCAGTTATCCCAGGACACCTGATGGTGAGGGCTAATGGCCTTGATGTGCACAGAGGTGATAAAGGGTGGAGCGA | 120 | 55.83 |
| chr7 | 143581226 | 143581346 | Target=1;ProbeIdx=2569;Target #1 | 1 | + | good | GGTGTACCTTGTCGATTCTGCCTCTTCTATAGGGCAGCTGCAACTTAGTGCTGTTTCAAAGTGGCCAAATGGGCCAGGATTGGGTCTGGTGTAGCTGTAGCCTGAGGCCTTGTAGGCTAA | 120 | 51.67 |
| chr7 | 143581346 | 143581466 | Target=1;ProbeIdx=2581;Target #1 | 1 | + | good | CATTGAGGTATGTCTACATGGAGACAACCATGTTTTTCTGAAACTTACTGCCAGATTAGGGACTTGGGAGTGAGAAAGGAGCCTGGGTACCCTTATCTCCACAGGTACAATCTTGGCATA | 120 | 45 |
| chr7 | 143581466 | 143581586 | Target=1;ProbeIdx=2593;Target #1 | 1 | + | good | TATGTAGTGTTTGAATCAGACAGTGCTTGTCCTCCTGAGCCTTGTGGAGGATACAGAGCAGTGGTCATGTTAGAGAAGAGGACCCCCGAGGAGCACCCTGACTTAGGAAGAGTGGCTGGG | 120 | 52.5 |
| chr7 | 143581586 | 143581706 | Target=1;ProbeIdx=2605;Target #1 | 1 | + | good | TGATTGTGCTGGTACCCACTGTGGTCCCTCTGAGGAAGAAGACTACTGTGTGTCAATATGGTGTGAGGCCAGAAGGAGACCTAGCTTTGCACGTGGTGGCCCAGTATTAGCTTTGTACAT | 120 | 50 |
| chr7 | 143581706 | 143581826 | Target=1;ProbeIdx=2617;Target #1 | 1 | + | good | GGTGGCCCAGCATACAAGGTAGGCAGAAGGTGCATCTGCTAGAGCCTGATGCTGCCTTCTCGACCTGTGCAGCCAAGTAGTACACCATTGGAGTGTGGGGGTGTGTTTTCATTCATCGAC | 120 | 54.17 |
| chr7 | 143581826 | 143581946 | Target=1;ProbeIdx=2629;Target #1 | 1 | + | good | AGACATGAATTAGCTGTGGGCAATTGGTTTAGATTCACATGCACTGGCATTCCCCACAGCCACGGTGTGAGTGAGTAACTGGCAGTGAACTTACCTTGTGTTATGAAGAGCCTTGTTAGG | 120 | 46.67 |
| chr7 | 143581946 | 143582066 | Target=1;ProbeIdx=2641;Target #1 | 1 | + | good | TATCTGTTGCCATCCCTCCTCCCTTTCTCTCTTACATCCCTACTATTTTTTAATCCATTTAAGTCACTCAAACAACTACACACTCACTTACCCCAGCCATTCATATATTCATCCATGTA | 120 | 38.33 |
| chr7 | 143582066 | 143582186 | Target=1;ProbeIdx=2653;Target #1 | 1 | + | good | TCGATCATCTTCTCATTCATATTTACCTCTCACCCATGCACATATTCACTCAGGCAGCAATTCATGTCCCATCCAACCACCTCTCCATCCACCTACCAGACCCACATATCCCTCCATCTG | 120 | 47.5 |
| chr7 | 143582666 | 143582786 | Target=1;ProbeIdx=2713;Target #1 | 1 | + | good | CCATCCATCTATCACCCTTCACTCCTTTCATAAGGAGGGAAACCCCTTTATTGGTATTCTAGAGTCTCTGCTTTGTGGCTCTGTGAATTCAGAGACAAAAGGCAGATCCACCAAAGAAAA | 120 | 43.33 |
| chr7 | 143582786 | 143582906 | Target=1;ProbeIdx=2725;Target #1 | 1 | + | good | GAAGGTTGCTACCACAGGCAGCCCACACCTCCCTCCCCTTCCTGGGCTAAGCCTTTGGGAAGAATAACTCAAAGGCTTGTTTAGAGCTCTGGCCTGGGGTAGCATCTCTGCAGGAGAACA | 120 | 55 |
| chr7 | 143582906 | 143583026 | Target=1;ProbeIdx=2737;Target #1 | 1 | + | good | ATGAAGATGAGAGAAGACTTGATCTCAGGTGACAGTCTGCTGGATGGGCGGCACAGGGAGGCAGTGCTGGAGAGAGGCCACTCGGTCCAGCCTGTCATGTGGGTTCCCCTGGGCTGACTG | 120 | 59.17 |
| chr7 | 143583026 | 143583146 | Target=1;ProbeIdx=2749;Target #1 | 1 | + | good | TCCTCTAGATGATGCGTATCAAAGTTGTTTCTAGAAGTAGACCTTTGTCCTCCCTGGCAGCCAGAGGGGAGCAACTTTGTAAAGCGAGGCTCTACTCTTAGGCCAGCCAGGAGACAGGAG | 120 | 50.83 |

| | | | | | | | | | |
|---|---|---|---|---|---|---|---|---|---|
| chr7 | 143583146 | 143583266 | Target=1;ProbeIdx=2761;Target #1 | 1 | + | good | TGTCCTTGCCTCTGCTTCTGCTTAGTGGCTCCCAGCATAAAATATTCTCTATGCACGAAAGTAACCTGAAATGTGTATATGTGTGCCTGTGTGTGTGCCTGTGTGCATGTGTGTGTATGT | 120 | 45 |
| chr7 | 143583266 | 143583386 | Target=1;ProbeIdx=2773;Target #1 | 1 | + | good | GTGCCTGTGTGCATGTATATGTATGTGTGTGAGGGGATTGGAGGTAACATTCCTTCTGTCCCTCTCTCCCTTTCTCCATCCCTCTTTCTTGCTTCCTTCTTCCATTAACTATTTCCCACC | 120 | 46.67 |
| chr7 | 143583386 | 143583506 | Target=1;ProbeIdx=2785;Target #1 | 1 | + | good | TGGAGACAGTAAACTACAACTCTCTGCACTGACAACCCCCCCCACCCCCACAAGTGCTTCCAGCTTTACCAGAGCTTAACACTCCTCCTCTTACTCTGACCTTTCCAGCAAGCACATAG | 120 | 51.67 |
| chr7 | 143583506 | 143583626 | Target=1;ProbeIdx=2797;Target #1 | 1 | + | good | CCCTGTGGGGCTCAGCATTCCTAACTCTACCTTCCTGAGAGGTGCCTTCTTCATAGATGTCTCCCTGCTGTGATGCCCCTTGAATCTGTCTGGAAGGACTGTCCCTGTCCCATAGCTTTC | 120 | 53.33 |
| chr7 | 143583626 | 143583746 | Target=1;ProbeIdx=2809;Target #1 | 1 | + | good | CAACCATAGAACTAGGAGAATCTTCCTCAGGTTTCTTCCTCTGCTTGGAGTCATGTTTGAGATCCTCCATACAAAGCTCAGGTTTATATTTGTTCATAGCCCACGGATAGAGGAGACCAT | 120 | 43.33 |
| chr7 | 143583746 | 143583866 | Target=1;ProbeIdx=2821;Target #1 | 1 | + | good | AACTTGGCCAGGTAGTGCCTGAGTGACGTAGGCCTAGGGTGCCTGAGACTACATGGCCATCACCTTCTTGACACAATTTCTATTCAGGTGGAATTGCTAGTATGGTGTGGCACACTCCCT | 120 | 50.83 |
| chr7 | 143583866 | 143583986 | Target=1;ProbeIdx=2833;Target #1 | 1 | + | good | GCCAGGTCCTCAGAAACCCAGGTTTAAGCATCTATTCAGCCCTTTCTGGACTCCTAGCAGGGTGAGGGTGGGGCTGGGGATGGGAGAGTCACATCTGCAGTTGCCCATCGCTGGATGACT | 120 | 56.67 |
| chr7 | 143583986 | 143584106 | Target=1;ProbeIdx=2845;Target #1 | 1 | + | good | TTCCTGGGTTCTCTTGATATTTTGCTTTTATTCCTCCCTCTGGTAGCCCAGGCTGGCCTTAAACTCATTTTTTTGAGCTTGACTGAACTTATAACCCTCTTATCTCCACCTCCTAAATGC | 120 | 42.5 |
| chr7 | 143584106 | 143584226 | Target=1;ProbeIdx=2857;Target #1 | 1 | + | good | TACCATTACAGGTGTGTGCTGCCACAACTGGCCATGCTTGTCTTCCAGGTGCTCCCTGAAGACCCTGCTCTGAGGTAGCCACCAGCAACCTGCTGGGAGCTTTTGTGCTGGAGCCGTCCC | 120 | 58.33 |
| chr7 | 143584226 | 143584346 | Target=1;ProbeIdx=2869;Target #1 | 1 | + | good | TACGTTGGCTGTAAGGAAGGTGCTTCCCCAGCTTCTACATCTGAGGATCCTTGCCTCTCCTGACTTCAGCTTGCTCAAGACTTGGGTCTCTGTTCTGCTCTTAGACCTGGAGATGATCTT | 120 | 50 |
| chr7 | 143584346 | 143584466 | Target=1;ProbeIdx=2881;Target #1 | 1 | + | good | TCTAAGGCTCTCCTCGTCTTTCTGTACTATTTTTACTCTGTGCTGCTAGCCACTCTGTGTTTGAAGTAGGGGGTTATTACAGCATAGCTCGCCGCCTTCTGACCCCGTGCCTGCGTTCTT | 120 | 50 |
| chr7 | 143584466 | 143584586 | Target=1;ProbeIdx=2893;Target #1 | 1 | + | good | TCCTCTCCTGCTGCCACTCTGGCCTCAAAGGAAAAAAAAAAAAAAAAAAAAACAAAGGTAGCTGTTCTTCCTATGTGGGCAGTTCCAAACTGCCAATGCAGCCATCCGGTAAATTCCTATA | 120 | 42.5 |
| chr7 | 143584586 | 143584706 | Target=1;ProbeIdx=2905;Target #1 | 1 | + | good | GTCGTGGCCATTGCTGTAGTCAGCTGGGACTCCTTCAGGGCCGCTTTGCCACAGTGTGAGGGACATTCCTGGCCCTGTTCCTACCTGCTGCCTCTGTCCAGGATCAAGGAAGCAGGGGGG | 120 | 60.83 |
| chr7 | 143584706 | 143584826 | Target=1;ProbeIdx=2917;Target #1 | 1 | + | good | GGGGGTGCCCAGTTCTTCCCACCTTTAGGGATCTGACATGCAAACACGTGGATGCCGTGAACTGGCAGTCACGTTGGCAGACTCAGGAACAAGTCTCCATGCGCTCCTGGCTTCTCGCTG | 120 | 57.5 |
| chr7 | 143584826 | 143584946 | Target=1;ProbeIdx=2929;Target #1 | 1 | + | good | ATATTCATATTTATTTATTCTCCTTTCTCTTTCCTTCACAGAGATCCAGGCACCATGCCACGCAGCCCCACATCCAGTGAGGACGAGATGGCCCAGAGTTTCTCTGACTATTCTGTGGGC | 120 | 47.5 |
| chr7 | 143584946 | 143585066 | Target=1;ProbeIdx=2941;Target #1 | 1 | + | good | TCGGAGTCAGACAGCTCCAAAGAAGAGACCATCTATGACACGATCCGGGCCACCACAGAGAAGCCAGGTGGGGTGAAGATGGAGGAGCTCCAGGGTAACACCCTGGTGATTCGAGTGGTC | 120 | 56.67 |
| chr7 | 143585066 | 143585186 | Target=1;ProbeIdx=2953;Target #1 | 1 | + | good | ATCCAGGATCTGCAGCAGACGGTGAGCGAGCTCCTGCATACAGCATGTGCTTCCACACCCGATCATATCTATGTCTAGAAGCACTGACTGTGGCTGTGGGTGGCTCTGGTCCACCCTGAG | 120 | 55.83 |
| chr7 | 143585186 | 143585306 | Target=1;ProbeIdx=2965;Target #1 | 1 | + | good | CCTACAGGCTGGTGCTCCAGACTCAGGGCTCCTGGGACATTTCTTGCCCCTAGCATCTCCTTTCTATTGCAGCCTACAATATTCATTGAGCATTCTCCATTCATTCCACATTGCTATAGT | 120 | 47.5 |
| chr7 | 143585306 | 143585426 | Target=1;ProbeIdx=2977;Target #1 | 1 | + | good | AAAACCTTGTGGCAGGGGAATGTATAAACTAAAGAGGTTTGCTCAGGCTCATTGGTTCTACAGGGCCAAGGTCGAAGGCTGTCTATCCTCGGTGGTGGCCTTCTTCTTAGCAGAGTTCCA | 120 | 49.17 |
| chr7 | 143585426 | 143585546 | Target=1;ProbeIdx=2989;Target #1 | 1 | + | good | AGGTGAGTCAGGACGTCACAGGCAAAAAGTAGGGGGTTGTGTGCATGCCTCTGTCTGATATTTCTCCTTATTCTCCTGTAGCTACCAGGATTTGAGAGTTGGATTCCCCACTTGACTTCA | 120 | 47.5 |
| chr7 | 143585546 | 143585666 | Target=1;ProbeIdx=3001;Target #1 | 1 | + | good | CCTAATTTTAACTGTTCCTTGTTGAGTTCAGTTTCCACCCATTTCATACTTCATGATAGAGATTAGATTTGAAGCCTAGAACCCTAGAGAAATACCCAACCCATGTCCAGGCCATAGTAA | 120 | 39.17 |
| chr7 | 143585666 | 143585786 | Target=1;ProbeIdx=3013;Target #1 | 1 | + | good | GCCCCTTCTGCTGGGTCAGGCCTCCATGCCTTGGGACCCAGAGCAAAGGACCCACAGGAGCCTTTCCTGGTCTTACAGATGAAGTTCAATGGTGAACACATCCACCAGTAGGCAGTGTCT | 120 | 55.83 |
| chr7 | 143585786 | 143585906 | Target=1;ProbeIdx=3025;Target #1 | 1 | + | good | ATGATACCTCAGCAGGGAAGGAAGCCCACTGCCCCTGCCTCCTCTCATCTGCTTACAGCCTGTTTTCGCCCACTTCTGTAGGAAGGGCTTCTGGTTTGAGTGTACCTTGGTTCCCCACGT | 120 | 55 |
| chr7 | 143585906 | 143586026 | Target=1;ProbeIdx=3037;Target #1 | 1 | + | good | GGCTCTGGTCACTTACTACCTGGTTGTCCCTGAACCCTGGGCCCAGTATCATTTCCTGGGATGTCCCCAAAGTCCCCAGGCCACAGCTTGCCCATGCTGATCACCCTGCTCTGTCCTTGT | 120 | 57.5 |
| chr7 | 143586026 | 143586146 | Target=1;ProbeIdx=3049;Target #1 | 1 | + | good | GGAGGGGCTCTTCCCCATGGATGTTGGCTTCTGCTGTTTGTGTTGGTGGGGGTGGGGGGTGGGGATGCAGAGGCAATCTGGATGTGATAGATGGCCGTGGTTGCTGTGTTCAGGCGTCAC | 120 | 59.17 |
| chr7 | 143586146 | 143586266 | Target=1;ProbeIdx=3061;Target #1 | 1 | + | good | TTCCCAGGAGGTTCCCAGACTGGTGTTCTGCCATTCCTCTCAGCTTTTCTCCTTCCTTGTTCTTTAACTCCTTGGGGCTTTTCTGATGAAAGTGGAAGCTCAAAGCAGAGATAGGCCTGA | 120 | 48.33 |
| chr7 | 143586266 | 143586386 | Target=1;ProbeIdx=3073;Target #1 | 1 | + | good | CCTCTTCCGTCCCCATGAGGGACAGGTTGTGAATTACTGGTGCATGGCTGGTCTCGGGTATCAGGCTCTGGTTTCTCTGGGGCATTGCACGGGGTCATGGCAGGGCCTTTTGAGAATCTG | 120 | 56.67 |
| chr7 | 143586386 | 143586506 | Target=1;ProbeIdx=3085;Target #1 | 1 | + | good | CTCCACCTCTGATGTCTGTGACAAGAGACCATGCCAGTTAATGACAGGCCAGGAGAAAGAGTGAGGGATGTGCTCTGAGACAACAGGATGAAGAAGGTGGAGTGGGATGATAGTGGGGTG | 120 | 51.67 |
| chr7 | 143586506 | 143586626 | Target=1;ProbeIdx=3097;Target #1 | 1 | + | good | GCAGGGTGAGAGGTGAGTGTGGTAATCAGGTGGAGGGATGAGGGATGTGAGGGGTTGGGTAGTGAGGTAATAGAGTGGTGTTGGGGGATGGGATAGAGGGGTGATGGGATGGTAGAGTG | 120 | 55.83 |
| chr7 | 143586626 | 143586746 | Target=1;ProbeIdx=3109;Target #1 | 1 | + | good | TGAGGTGAAGAATAGAAGTAGTGGGTGATGGGGTGAGGGAGGTGGCAGGGATAGGGAGGTGGGGGAGATGACAGGAAGAGGCAGGTGAGGCAGGGTAGCATCTCTCTAAGCTCCCATTA | 120 | 55.83 |
| chr7 | 143586746 | 143586866 | Target=1;ProbeIdx=3121;Target #1 | 1 | + | good | CACACACACACACACACACTCACACACGTACACTCATGCACTATTCATTAATGAACATCCCTGACATTATTCATTTGAGTTTAAAAGGAAGAAAAGGCAGTGAGGAAACAATGAGCATTC | 120 | 39.17 |
| chr7 | 143586986 | 143587106 | Target=1;ProbeIdx=3145;Target #1 | 1 | + | good | ATATTTTTTTGTTGTTGCTACAAGAAAGATGGTTTCCAAGGACATCTAGAGGGAAGTATTATTAATCCCATGAAGAGGGCTAGATTTTCTGGTTTCTGTTAGGCCCTCCCCCTCTGGCTG | 120 | 41.67 |
| chr7 | 143587106 | 143587226 | Target=1;ProbeIdx=3157;Target #1 | 1 | + | good | TGCTTTTCCCAGGTCACTTTTGGATGAATGGATCAGATTGCTAAAGTGAGCTGCCTTTCCAACTCCCAGGCTCTTGGAACAGGTTCTCATATGTGCAGTCACCTGGGGATTCAGGCTCCT | 120 | 49.17 |
| chr7 | 143587226 | 143587346 | Target=1;ProbeIdx=3169;Target #1 | 1 | + | good | TCACTTGGTAGCCAGGTCTGTCCCAGGGTCTGGAGTCCTCCACATCCTCCTATGTATCTGTCCTAAAGGTGACAGATGGAAAGGGTGACCTGGGAAGTATCTCACTTCCCTGTCTTCCCC | 120 | 53.33 |
| chr7 | 143587346 | 143587466 | Target=1;ProbeIdx=3181;Target #1 | 1 | + | good | AGCTTCCTTAAGGAGGTCGCTTAATTGCAGGTGACCTTGGAAGTTTTGGTGAGCAGCAGCTAGCTATTTGTACTTCCCTCCCTCCACCCCCATCCCCATCCCAATCGTCCCTCTTCCGTT | 120 | 52.5 |
| chr7 | 143587466 | 143587586 | Target=1;ProbeIdx=3193;Target #1 | 1 | + | good | GGTAAGGACTCTATGCAGACTTTGCTTGTCTTACTTAATGCTCATCCCAGCCTGCACCTACCCCAGAGTCACCCTGCCTCTGTTTCTACCTGGACGCTCTTTGCAAGGGCAGAGACCTCT | 120 | 52.5 |
| chr7 | 143587586 | 143587706 | Target=1;ProbeIdx=3205;Target #1 | 1 | + | good | GTAGTAGCCTTTTCTTTCCCAGGTCCTGGTTTTATTTGCCCCTGCACTGGGCAGAGCCTCCCGATGACTGTGAACATGCTGCTGATCCTCACCGCGTGTGGTTTCTCTGCTCTATAAAGA | 120 | 51.67 |
| chr7 | 143587706 | 143587826 | Target=1;ProbeIdx=3217;Target #1 | 1 | + | good | TACCACAGTTCCTGTTGAGGCTGAGTTGTGGATGCTGAGCTCAGGGCCAGTGGCACTGGAAGCCACACCCATGGGCCTTTGTGTCTTCCTAAGGAGCAGCTCAACCTTCCTCCAGTCCTT | 120 | 55.83 |
| chr7 | 143587826 | 143587946 | Target=1;ProbeIdx=3229;Target #1 | 1 | + | good | GCTTGTTCCTATGCAGGGATCCCACAAACACTGAGAGGGGTGACTTCATTTTAAAATATTCCTTTATCACCTTTGGCTCAGACATGCATTTTCCTCTCTATCATAGTTTGGGGCCAAAAC | 120 | 42.5 |
| chr7 | 143588066 | 143588186 | Target=1;ProbeIdx=3253;Target #1 | 1 | + | good | TGTTGCTCCTGATTCAGCCACTAGACTATAACTGCTTTTGTTGTTATTGTTAAATATTTAGAGTTAGGGTCTACAGTTTCATGTAGCCCAGGCTGGCCTCAACAAATTCATTAAATAGAT | 120 | 36.67 |
| chr7 | 143588186 | 143588306 | Target=1;ProbeIdx=3265;Target #1 | 1 | + | good | AGGGGTTATTGTTGGTTCTTCCTGGAAGTGTGGCCTGAGAGTTTGTCTTTTGTTCTTGCTAGGTAGTTTCCTGGTATTCTGTGGCATTTCCCATGTTGCTCTCCTGGCTCCTGTCTATCA | 120 | 46.67 |
| chr7 | 143589026 | 143589146 | Target=1;ProbeIdx=3349;Target #1 | 1 | + | good | GGGGAACCAGCTAGTGTCACTCTGGGAGTTTCTGCTCCGTGTCTGCATCTCAGTCCCTCTCTGTTGAGTTCTCCAGTGTCTTTGTATTAGGCATTTGGCTGGAGCTGTTTGTCCCCAGGA | 120 | 52.5 |
| chr7 | 143589146 | 143589266 | Target=1;ProbeIdx=3361;Target #1 | 1 | + | good | TTCTGATCTGGAGTACGATCTGTTTTCATAGCTTCGTACTCTCTGCTTCTGACAAGTATTCTCTGGAGCTGTTGTTTGGGATATGTTCTGATGTCTGATTATTTCCCTCCTGAGTATAAT | 120 | 40 |
| chr7 | 143589266 | 143589386 | Target=1;ProbeIdx=3373;Target #1 | 1 | + | good | GTTTCTTAAGACAGTCCTTCCTCTGTGGGGCCAGTTTGAGAAGTTAGATATATCAGTCTGCCAGCCTGCCTGCCTGTCTGTCTGGTGTGTGTGTGTGTGTGTGTGTGTGTGTGAATCTTT | 120 | 48.33 |
| chr7 | 143589386 | 143589506 | Target=1;ProbeIdx=3385;Target #1 | 1 | + | good | CTAGAGGGCTGGGCAGCTATCTCTTTATCTTTTTTCTTCCAAACGTTATCTTAAGTGCTTTTCCTTGGGAAGTCCCCTGAACAGAGACTAAGTGTCTTCATCACCGATGGACCCCCTCTG | 120 | 46.67 |
| chr7 | 143589506 | 143589626 | Target=1;ProbeIdx=3397;Target #1 | 1 | + | good | CCCCCATCACTCAGTGGAGAGCACCGCCTAAAGGGTCTCCCAAGAGCTGGTTCCATAACCTTATCTCTCACCCAAGGAGGGGGAGAGAGCGGCAATGAGATGGAAGAGGCACTCCTCCTA | 120 | 56.67 |
| chr7 | 143589626 | 143589746 | Target=1;ProbeIdx=3409;Target #1 | 1 | + | good | CCACAGCCTTCTGTATCTCACAGATCATTGGATCGGGTCGGTTCTGACCGTGTGTAATGCCTTGGGGATTTTCTAGGCTGGCTTTTGTATTTTACAACATTTAGTTTGACTTACTTGATT | 120 | 42.5 |
| chr7 | 143589746 | 143589866 | Target=1;ProbeIdx=3421;Target #1 | 1 | + | good | CCCCGAGACTGTCTCCCTCAGTGTTGGTCTCCAAAGTAGAGAGCCCATGCCCCATCATCCATTTGAGCTATCTTTCTGTTACAGTGTTTTATCTGGTAAATCTTTATTTCCCAGAACTGC | 120 | 45.83 |
| chr7 | 143589866 | 143589986 | Target=1;ProbeIdx=3433;Target #1 | 1 | + | good | CTATTGACGTAGGTGCACATGCTACAGACTGTAAGAGCACTCTAGGGTGGCAGACCTAGAACCTGAACATCAAAGCTGGGAGGGCAGAGGGAGTGGTGAGGGTTTCGCTGCCGGGAGCAG | 120 | 55.83 |
| chr7 | 143589986 | 143590106 | Target=1;ProbeIdx=3445;Target #1 | 1 | + | good | GTACAGACCATAGTACACTTCTTCTCTGCTGGAGTCTATCCCCAGGATGGGTGGGACTCACTGCACTCAGGCAGGATGCAGCCTTGCCTCAAGCTGATCCTGACTCAACACATCCCTGTG | 120 | 54.17 |
| chr7 | 143590106 | 143590226 | Target=1;ProbeIdx=3457;Target #1 | 1 | + | good | GGTGTGTCCCATGCAGGAGGAATCCCAGTGTGACATAATTATACCAAAAGGTGTTAAATGGTTTCCTGAATGTTGACCTCATGACAGACACCATATTAAGGTCTTGAATGTGTTCTGTAG | 120 | 41.67 |
| chr7 | 143590226 | 143590346 | Target=1;ProbeIdx=3469;Target #1 | 1 | + | good | AGTCCTTGTAACATCCTTGACCTTGGTAGTGGGTCTGAGCATCTTTGTTGAAGGCTAACACACCTATGGAGAGGACCAGAATCTTAGGTGTTCAAATAGAACCTTGGTGATGTACAGCTG | 120 | 45 |
| chr7 | 143590346 | 143590466 | Target=1;ProbeIdx=3481;Target #1 | 1 | + | good | GGAGGAGACAGAACCATACAGCAGCCTGGTCACCAGGGAGGCTCAGCATGTGCTGAAGGTTTTAGTAAGGCTCTTTGATAACACATGAAGTGCGTTTGCTTGTGACATTTTCGTACATAC | 120 | 47.5 |
| chr7 | 143590466 | 143590586 | Target=1;ProbeIdx=3493;Target #1 | 1 | + | good | ATGCAATTAAACTTTGTTCTTGTTCATCTTCTTCCCCACTGTCCTTCCCTGCCCCCTCCATCTCCTACCAGTTGTTCTTCCCTGTCACCTCCATGTCCCCCTCCCTCTCTCTCCACTGTC | 120 | 51.67 |
| chr7 | 143590586 | 143590706 | Target=1;ProbeIdx=3505;Target #1 | 1 | + | good | TCTACCTCCCCTCACTCTCCTTCCCTGTCCTCACTCTACCTTCCTCTCAACTGCCCTTCCCTGTCTGTCTACCTCTTTGTTGGTCTGCTTCCTCCTCCCAAATAGCCTGTCTTCTCCTTT | 120 | 52.5 |
| chr7 | 143590946 | 143591066 | Target=1;ProbeIdx=3541;Target #1 | 1 | + | good | AGCTGCTGTGAATAGTGTAGCAGCAAACACCCCATTTTGTAGAAGGAGCATGGAGGATCAGAAGGGAGCCGGTGAGAGACAGAGCAGATGGTCTAAGGTCTGAAAACCCGCTGAGCAAAA | 120 | 50 |
| chr7 | 143591306 | 143591426 | Target=1;ProbeIdx=3577;Target #1 | 1 | + | good | GGCCCATGATGTGCCCACTGACTGCCAGGTACCCAGGCCTAGTTTATACATGCAACTTGTATTCAGTGTGGGAAGGTGGCTACTGAGCAGCTTGTGTGGTAAACAGTCCTGGGACCACAG | 120 | 53.33 |
| chr7 | 143591426 | 143591546 | Target=1;ProbeIdx=3589;Target #1 | 1 | + | good | AGCATCCAGGATGGGTTAGGTGACTAAGGAGGGTTCACAGGACTCTGAGAAGAAGCCTAGTGTGGTATAGCATGAGCAACCACAGAACTGACATCTGGCAAGCCCAAGCTCCATGTTCCC | 120 | 51.67 |
| chr7 | 143591666 | 143591786 | Target=1;ProbeIdx=3613;Target #1 | 1 | + | good | TGCCTGCTTTGGACAGTTCCTGGCTGGTCACCTTTACTGTGGTACAGGCCCATTTCTCCCACAGGTCCTGCATCTGCTTGTTTGCACTGGGTTACTGGGAATGCTGGGAAAGGCTGCCCT | 120 | 55 |
| chr7 | 143591786 | 143591906 | Target=1;ProbeIdx=3625;Target #1 | 1 | + | good | GGTCACCTAGTACTGAAGACCAGAAGGGCAATCGCACTTTGTAACCGGAAGGGCTCTCTCAGGAGTGTAATCTGTTCTGGAACTTTCTAAAGACCATTTCACTTCATTCTGTTCAAGACA | 120 | 45 |
| chr7 | 143591906 | 143592026 | Target=1;ProbeIdx=3637;Target #1 | 1 | + | good | AACAAAGCCACTGGAGGCAGAGACAGGCAATCCCACGGGTATGAGTTGTCAGCAACAGAGCGGACATTGTTCCCGCATGCCGTCCCTTGCGGGCTAAATCCTCTGTCTAGCTGGGGACAT | 120 | 55.83 |
| chr7 | 143592026 | 143592146 | Target=1;ProbeIdx=3649;Target #1 | 1 | + | good | GACACAATTCCTGGAATCCTGCCCGGAGGAGCTGAGTGGAGCTGCCAGCCAGCTTGCCCCAGGGGCCCCCAGTGTCTACACACTGGGTGGCGGTGCCTGGAGAAGCTGTGGGGCTGTCCT | 120 | 65 |
| chr7 | 143592146 | 143592266 | Target=1;ProbeIdx=3661;Target #1 | 1 | + | good | TCACTGCCCCATGAGGCACAACCCTGCTGTCTTCTGCTGCCTGCTATTTCTGAGCTCACAACTCCTGTGAGCCCAGGCCAGGAAAGCCTTGGGCAGTACTGACAAGGCACCCTTGGAGCC | 120 | 58.33 |
| chr7 | 143592266 | 143592386 | Target=1;ProbeIdx=3673;Target #1 | 1 | + | good | CTGGCTGAGCAGGGGCTGGGTGGTGGTTTTAGAGGACTGGGAAGAGTTAATAAAGGGTGCCAGGAGTAAGTTGAAGCACATTTCTGCATCATTCCGAATGTGGTATACAACTGGCAGTGG | 120 | 50 |
| chr7 | 143592386 | 143592506 | Target=1;ProbeIdx=3685;Target #1 | 1 | + | good | CCATTCAATGTGGCCTTTTAAAAAGTGGCATGTGACCAAATCAGTACCAAGCAAAGTGGGCTCTTCTGTTATATTTTTGGAGAACAGTGAATAAGGGAGTGGGTACAGCGCTGGTTGTGG | 120 | 44.17 |

| | | | | | | | | | |
|---|---|---|---|---|---|---|---|---|---|
| chr7 | 143592506 | 143592626 | Target=1;ProbeIdx=3697;Target #1 | 1 | + | good | CCGAGCTGCGCAGCAGCAATGATCGAACCCACATTTATTCCCGGGAGGTGGCAGAGGAGTACTGCGGAGTGGGGGGAGGGGCTCTCTAGGGTTCCCAGCAGCAGCCCCTCAACCTCCAGT | 120 | 62.5 |
| chr7 | 143592626 | 143592746 | Target=1;ProbeIdx=3709;Target #1 | 1 | + | good | CAGGCTTGGCACCAGGAGCTCAGCCAAAGCTATCAGTCCAAGCCCCATCCCCTCCACTCCACCTCCCTGGCTCATCTCCTGGCAGCCGTTATAGGGAAACAAGGCTGGCCCCTTTTTCCT | 120 | 59.17 |
| chr7 | 143592746 | 143592866 | Target=1;ProbeIdx=3721;Target #1 | 1 | + | good | TAATTAGTACTTCCATTTATGACATAGATGTATCTTCCTATGGGTTATTTCATTTAGCAAAAAGCTTTTCATCTACCACCCTGACCCCCTTAAAGACAATTTATTGCCATTCATCCTTCC | 120 | 35 |
| chr7 | 143592866 | 143592986 | Target=1;ProbeIdx=3733;Target #1 | 1 | + | good | TCTCTCTGTGTTTGACTCTCTCTCCCTCTCTTGCTCATGTACCTCCCAGAAGCCACTGCTCCAGATCTGGCACTCCTAGAAGTCATTTCCTCTCCATGTCACTCTCCCTGCCTCCCGCAT | 120 | 53.33 |
| chr7 | 143592986 | 143593106 | Target=1;ProbeIdx=3745;Target #1 | 1 | + | good | CAGCAACCATCCATCACCTGCGTCTGAATGGTCTACCTAATTTTCCTGGCTCAGAAGCACTCTTGACCATTACATTTGTCATTGAGATGGAACTGCATCTCCGGGCACAGGGTGGTCATC | 120 | 49.17 |
| chr7 | 143593106 | 143593226 | Target=1;ProbeIdx=3757;Target #1 | 1 | + | good | TCCCTCTTGTGCTTTTGGTCACGGGCAGGTGACTGTCCAGTTGGCCTTACATGTATGAAATGTCCTGTGTTTTTGAGTACCAGGGGAACAAAGACACTTGAACCCATGTCCCTGTCTAGT | 120 | 48.33 |
| chr7 | 143593226 | 143593346 | Target=1;ProbeIdx=3769;Target #1 | 1 | + | good | AGCTGAGAAAGGACCATAGTGACCATCCATGCTGAGAGAGTGCAGGAAGGCTGGCATGCAACGTATCACCCTGATCAGCTGTTTGAACATTGTTCTTGGCACCATGGAAGGAGAGAATGT | 120 | 49.17 |
| chr7 | 143593346 | 143593466 | Target=1;ProbeIdx=3781;Target #1 | 1 | + | good | GATATGTCTTCCACAGCTCATGGTGTCTATGTTTTTAGAAAGCAGATATGTTTTTAGAAAGCAGGTGAAGGTTTTTAGGCATGGTGACACCTTCATTCCACATCAGGAAGATTTGTTCTG | 120 | 39.17 |
| chr7 | 143593466 | 143593586 | Target=1;ProbeIdx=3793;Target #1 | 1 | + | good | GATTCTTTAATATGCACTGCCTAGAGGATGCTGGCTCCATATAGTTCACCTGCCATCTTGTTCCCAACAGCATCATCTCACTAGTCACTAAATTTTAGACTAATGATTAAGTAAGTGTCT | 120 | 39.17 |
| chr7 | 143593586 | 143593706 | Target=1;ProbeIdx=3805;Target #1 | 1 | + | good | TAGCCAAGTCACTAACAGGCCTCCAGCAGAGGGCTAGTTCTCCAGTGCCTTGCTCCCTAGTCAAAGCTCCTAGTCCCCACATCTAGCTTAGATCTCAGCTTGCAGCTAGCTCACACCTTG | 120 | 53.33 |
| chr7 | 143593706 | 143593826 | Target=1;ProbeIdx=3817;Target #1 | 1 | + | good | TAGCTCTCAGCTCACCATTCCCAGTCTCTGATTCCAGCCAGAGGCTCCTGTCTGCAGAATTCTTTTCCCTGGAATCTCAGCTCCCCACTGTGCATGTCATTATCTTATCTTCCCTCCCCA | 120 | 50.83 |
| chr7 | 143593826 | 143593946 | Target=1;ProbeIdx=3829;Target #1 | 1 | + | good | GCTCAGGGACAGCTGGCTCCTTATACTATCCATCCAGCAGGTATCCCAGCTCATTGTCTGCCTGTAACCCTGGGCCCACCATCACCACATCCCTAACATAACAACTTCTGTCAACTGTGA | 120 | 51.67 |
| chr7 | 143593946 | 143594066 | Target=1;ProbeIdx=3841;Target #1 | 1 | + | good | GCTCGCTCCTCTTGGATCCTTTGTGGAAGCCCAGAGGCCTCTGACTGAGTCTGGGTGCAGCCTGGAACATAGTTCTTCCCTTCCCCCACACCTGGCCCAGCTCTGAGTATCTTCCTCAGC | 120 | 58.33 |
| chr7 | 143594066 | 143594186 | Target=1;ProbeIdx=3853;Target #1 | 1 | + | good | TAGGTTTACCTTTAAAAAGTAGTGTCCTCTCTTCTAAACAGTCAAAAGAGCTTTATTGATTTGTAGGTTTTTATAGATCCCAGACTCTGTTTGTTTTAAGCATTCGGGTCAACAATTTCT | 120 | 33.33 |
| chr7 | 143594186 | 143594306 | Target=1;ProbeIdx=3865;Target #1 | 1 | + | good | AGTGAACTTAGCAAGCTGTTTGCCCATTGACTTCATCCAGGTTTGACACACTCCAACATCCCAGAAAGATCCCTCACGCCCACTGCCAGCCTGACAAATGTGCTTCTACCCTCTGTCTCT | 120 | 50 |
| chr7 | 143594306 | 143594426 | Target=1;ProbeIdx=3877;Target #1 | 1 | + | good | GTTTTTCTTGTTCTGGACATTTCCAAGTAGATCAGGCCACAAGTCAGTAGCTTCAGTGACTGGCATCTCACTCAGCATCATGGTGGTCACTGCCTTATTTGGTTCTTATAGGACTTCTAG | 120 | 44.17 |
| chr7 | 143594426 | 143594546 | Target=1;ProbeIdx=3889;Target #1 | 1 | + | good | CTGCCCCAAAGGGAGCTCAGACACTGCTCTATCCAACAGGAGCTCTGCCTGACTTTCTCAGGCCTTTGGCCGTCTTGGAGTTGTAATTTCTCTCACAGCAGAATGTATGGTCATAGGGGA | 120 | 51.67 |
| chr7 | 143594546 | 143594666 | Target=1;ProbeIdx=3901;Target #1 | 1 | + | good | GGACTGAGACACGGCCAGGGCATGGTGGTTTTGGTGGCCGAGTGAGTTCTGTTCAGAGCATGTGATTTCACTGAATTTGTGCTCATGGCCTTGGCTGCTCTGGCTGGAGGAAGGCTGGAA | 120 | 55 |
| chr7 | 143594666 | 143594786 | Target=1;ProbeIdx=3913;Target #1 | 1 | + | good | AAAGAGGGAGACTCCTGGGGAGAAGGAGAAACAGCAGGGTGGGGAGCCAGGTACAGAGCTGGGAAGGAGAGGAGGGACTGCATAAGTCCCTGGTGACTGGGCCAAGGACAGGAATGATA | 120 | 56.67 |
| chr7 | 143594786 | 143594906 | Target=1;ProbeIdx=3925;Target #1 | 1 | + | good | GATCAGAGCCACAGGCTCTTACAGAGTCCTAGGTGGAGGGGAGGAGTTTACAGAGCCCTTAACAGTGACCTGTCAATCAATTTGAAATAGAGTGCCCATGTCATGGATGTTTCTGAGACT | 120 | 47.5 |
| chr7 | 143594906 | 143595026 | Target=1;ProbeIdx=3937;Target #1 | 1 | + | good | AAGACCAAAACCCCATTTTCTTTCTTTTTCTTTCTCATTTTGTATATGTGTGGTATACATGTGTGTATACGCATGTTTGCATGTATGTGTGAGTTGGTGTGGTTATGTGCTGGAATACAT | 120 | 35.83 |
| chr7 | 143595026 | 143595146 | Target=1;ProbeIdx=3949;Target #1 | 1 | + | good | GTGTGTATATGCATGTTTGCATATGTGCAGGAGTTGGTATGCTTGTGTGACTTGTGGGTGTGGTGGCCGGAGGTGAATGTAGAGACTCATCCTTGATTGTGGCTCTCCCACCTTGTCAGC | 120 | 50 |
| chr7 | 143595146 | 143595266 | Target=1;ProbeIdx=3961;Target #1 | 1 | + | good | CTCTTGGCCTCATGCTAGCACAGCAAGTTTTAGCGGCCAAATCATTTCCTCAGCCTCCATTTTCTTTCTTTAAAAAAAAAAAAAAAAAAAACCTCATCGTAATTCATATAACCGCACTGTT | 120 | 36.67 |
| chr7 | 143595386 | 143595506 | Target=1;ProbeIdx=3985;Target #1 | 1 | + | good | CACTCATGGACAGTGAGTGCTTCAGTGAACCTTAGAGTATTCACAGGTGGGGGGCACAGCCCCCCAGTCATCGAGTTCTCTAGTCCCTGAAGGGATTCATGCACCGACTAGCCAGCTGTC | 120 | 55 |
| chr7 | 143595506 | 143595626 | Target=1;ProbeIdx=3997;Target #1 | 1 | + | good | CTCTCATGGATCGCTCTGTCTCTCCCTGTTCTGCACTCGCATGGTAAGTGCACTCACAGGGATATGGTCTGAGTCCTGTCTCTACTATTGGAGACATATCTGTCACCTCAAATCTATGCA | 120 | 48.33 |
| chr7 | 143595626 | 143595746 | Target=1;ProbeIdx=4009;Target #1 | 1 | + | good | CACACCAACACTCGCAAATCCACTGGGGTCCTGGTTGTTTCTTGTTTTTGAATTTTTTCACACCGTATCTTTCAATGTAGATCAGATTGGCCCTTGAGCTTCTGGCAATTCCCCTGCCTC | 120 | 45.83 |
| chr7 | 143595746 | 143595866 | Target=1;ProbeIdx=4021;Target #1 | 1 | + | good | AGCCTCGTCAGTGCTAGGATTGAAAGGGTAAGCAGCCATACTGACTGACTTCTTAAACACCATAAGACAGTCTCTCTGATCAAGTCAGAAGTGCAGAAGTTCAAAAGGTTCTTCTGAACT | 120 | 43.33 |
| chr7 | 143595866 | 143595986 | Target=1;ProbeIdx=4033;Target #1 | 1 | + | good | GAGGGCTGACTTCACTTGGCTCATCTCTCTCAGAATGAACTATCCTTGTATGGCTTGAAAATAAGATTTTATTTGTAGGTGCGTATGCGAGCCATGGCTCATGTGTGGAGGTCAGAGAAC | 120 | 45 |
| chr7 | 143595986 | 143596106 | Target=1;ProbeIdx=4045;Target #1 | 1 | + | good | CAGTGCTCACCCAAAGGAGGTTTGTTCAGGACTAAAGGCCTGGCTACAGATTACCAGCCTGTGAGTTTGATAGGGCACATTTTCTGCTCTGATAGAAAAGCCACCCAACAGGACTGACAT | 120 | 48.33 |
| chr7 | 143596226 | 143596346 | Target=1;ProbeIdx=4069;Target #1 | 1 | + | good | TCCGCCATGGTGGCCTGTGCCTTCATACCGCGAGCCAGATAAACTCTCCCTTCCTTGAGTTGCTTTTGTCTCATCAATGAGGAAAGTCACAAATACAGCTGTAGAGTAAATGACATCAAT | 120 | 45.83 |
| chr7 | 143596586 | 143596706 | Target=1;ProbeIdx=4105;Target #1 | 1 | + | good | AATGGTCCTTGTTCAGAGGCTTGGAGTTCAAAGAACCAGTGGTCAGAACTAATACCTATCCATCTAGCCAGGCTTCGGGAGGCTGAGTAGACCTGAGGGCCTCAGAAGTATATTTTATTT | 120 | 45.83 |
| chr7 | 143596706 | 143596826 | Target=1;ProbeIdx=4117;Target #1 | 1 | + | good | CTCATTTAGGGAAAATGTCAAGAATAATAAATATCATATGAGCTCTAGGGTGTCTCTTACACAGATCTACGACCACAATCTTCAACTTTTTAGTGATAAGATTAAGCCTCACTGGTACAG | 120 | 35.83 |
| chr7 | 143596826 | 143596946 | Target=1;ProbeIdx=4129;Target #1 | 1 | + | good | TCTTAGTCCCCAAAGTGCTGAGATCACAGGACGATGTCACCATGTTCCCTTACATGTCCTCTTACAATCCATTGCCCACATATCTATCTACCCACCAACTCAATCATCCAGCTTTTATAT | 120 | 43.33 |
| chr7 | 143597186 | 143597306 | Target=1;ProbeIdx=4165;Target #1 | 1 | + | good | TCCTGTCCATCCATTCTTCTACCCACTCACTCATTCCTTCCTTCAAGCAAACCTTCAAGCATCTGTCTGTCCATCTATGCTTTTGCAATATAGGAACTGGTGCTTAGGATTTTGTGTCTA | 120 | 42.5 |
| chr7 | 143597306 | 143597426 | Target=1;ProbeIdx=4177;Target #1 | 1 | + | good | CTCTGAGTGACAACTACCCAGTGTGCTTTGTAGTGCATAGCTGGATTGCTGTTGGAGAGATTGAAGGCAAAGAGTCTGGTTTTCTTTGTGGTTTCATTGAGTCATCATGAAAACCCAAGG | 120 | 44.17 |
| chr7 | 143597426 | 143597546 | Target=1;ProbeIdx=4189;Target #1 | 1 | + | good | AAACTCCCATCCCTGTCTTCTCTGCAATGCTACCATGATATCCTGCTTCCCTGTGCTTCAGCCGAGAAATTCCACTTTGAAGCTTTCATCAGGAATTTTGTGCATCATAAGTTGCTTGAA | 120 | 43.33 |
| chr7 | 143597546 | 143597666 | Target=1;ProbeIdx=4201;Target #1 | 1 | + | good | AGAGAGCAAAGGCCTGGGGCCTTTCCATGTAGGAAACCAAGGAAAGGCAACCCTCATCCTGCCGGTATTAATCCTCTCATGGCTTGATTGACAGCTGGAGAGGGACTTGACATTTATATC | 120 | 49.17 |
| chr7 | 143597666 | 143597786 | Target=1;ProbeIdx=4213;Target #1 | 1 | + | good | TAACACTTTCAAACAACTAAAACTGGCCAAGCCACATCTCTTCTCTGTGTTAGGATCCTGTGCCGCCCCTTCCTCTGTTGAATATCGAGATGGAGTTTTATATGCAAGGTCAGGTGACCT | 120 | 45 |
| chr7 | 143597786 | 143597906 | Target=1;ProbeIdx=4225;Target #1 | 1 | + | good | AGCTTGAAGAGATCTCTGAGCCCAGACATAATCAGCTAGCAATAGTTTGCTTTTCTGCTTAAGCTTCATGGTTAATTCCCACTTAATTCTATTTGAGAAGATAACTTCTGGGGTTAGAAG | 120 | 38.33 |
| chr7 | 143597906 | 143598026 | Target=1;ProbeIdx=4237;Target #1 | 1 | + | good | CCTTGGTTTGGAATTACCAAAATTCTATACAGAAACTCAGTAAAATCCCACATTGAGGTGGAATGCTGGGGCGCAGAGGTCAGGCTAGAGAAGAGCATGTAGCACCTTTTTTGCTGGTGA | 120 | 45 |
| chr7 | 143598026 | 143598146 | Target=1;ProbeIdx=4249;Target #1 | 1 | + | good | TTTATGATAAGACACCAGGAAGGTGTCCCCTCATTCATGTTAAACGCAAGGGCAGGACTGTCTTTGCTCTGAGTATGTTTGTTATGCAAAGGTGCGGTCAAGAAACTGCTGAAGTGGCCA | 120 | 45.83 |
| chr7 | 143598146 | 143598266 | Target=1;ProbeIdx=4261;Target #1 | 1 | + | good | CGGAGATAGCACGAGACAGAGACTAACAAGAATGCAAGAAAATATTGGGTGATAGAGTAGTATTGATATCAAGGAAGCAAGAGGACAGCAGGTGAGAGGCCAGCAGGAACAGCTGGGTTA | 120 | 45.83 |
| chr7 | 143598386 | 143598506 | Target=1;ProbeIdx=4285;Target #1 | 1 | + | good | TTGGGGGAAGGGGGCCCCACAGGGTTCTTAGTCATGTTAATCAAAGAATTTAAGAACAGACTCAGAAAGAAAAAAAAGACTGGAGGTAGATCCTATTAGTACCAATAAGAAAAAACCCCA | 120 | 40 |
| chr7 | 143598866 | 143598986 | Target=1;ProbeIdx=4333;Target #1 | 1 | + | good | GGGCAGGCAAGCTGTAAATCTGGGTACTTGCCAGGAGGGAGAGATGGAGAAGAGGAAGGCAGTTTGTGCTGTTTGTAAGCAGACATGTGTAGGAAAAGCAGCTATGTGGTGGCAAGGAAG | 120 | 50.83 |
| chr7 | 143598986 | 143599106 | Target=1;ProbeIdx=4345;Target #1 | 1 | + | good | ATAACACTCTTAAGAGGTCTCAGGCAGTCCACATATCCAGGAATGAGCTCAGAAAGACCAAAGTAAAAAGGCCCAGGATCTTGGAGACAGCTCACTGAGGCTCTGGCCATAGTCCAAAAG | 120 | 47.5 |
| chr7 | 143599346 | 143599466 | Target=1;ProbeIdx=4381;Target #1 | 1 | + | good | AAAGAGACAGAAGGGAAGAAAAGAAAGAAAAATTATATTTTTCTGAGTCAGGGCTCAGGTGAGCAGACCTAAGAGAACGTAGGAACAGCACAGAGAACAGGGGTTCTGGGAGGAAAAAAA | 120 | 41.67 |
| chr7 | 143599466 | 143599586 | Target=1;ProbeIdx=4393;Target #1 | 1 | + | good | TTAGATTCCATATTTATGGCCAGAAGGAAACAGCTCCCTTTTAATGAGTTCCCAGTGCCTCTCTAAGAATTGGATTTTCAGGATCCGCTCATCTTGTCTCTGGCTGGGCTTCCCAACTCA | 120 | 45 |
| chr7 | 143599706 | 143599826 | Target=1;ProbeIdx=4417;Target #1 | 1 | + | good | GTTGTTCCTACTTCCTCTCTGGTGGATAAACTACCTTTCTGCCTCCCTGCCCTGCCCCCACTCTCCCCTTTCCACAGCCTTCTGCCTGTCTCCTCCTGATTAGCTTTGGCTACTTCCCCT | 120 | 55 |
| chr7 | 143599826 | 143599946 | Target=1;ProbeIdx=4429;Target #1 | 1 | + | good | CCACTCTGTGTGACCGGGAGGGACTTGCGACTCATGAGCGAGACTTCAGGGTTCCCCACATTGCTGCTCCTGGCAGTTGGGTGGCTCTGGAAATGTGAAAGTTGAGGTGTCCCACCTTGG | 120 | 57.5 |
| chr7 | 143599946 | 143600066 | Target=1;ProbeIdx=4441;Target #1 | 1 | + | good | GTGCAGGGGTGTTAATTGTTACTCTTAGAACAGCGAGGGGGAGCAGGATGCATCCTTAGTGCTGCTGACAGTCTAGCTAAGGGCCATCTCTCCCCTTCACTTGATGGGTTTTTCACTGAA | 120 | 50 |
| chr7 | 143600066 | 143600186 | Target=1;ProbeIdx=4453;Target #1 | 1 | + | good | GACAGTGGCCAGCGTCTGACCGACTACTTTGTGATATCTCTGCCTGAGAGCGGCTAAAGAGGAAAATTAACTCATCTTCAAGGAGCACACGAGCCCTGGCGGGTCTGAGCAGTGCCTCCC | 120 | 55 |
| chr7 | 143600186 | 143600306 | Target=1;ProbeIdx=4465;Target #1 | 1 | + | good | AGTTCAGTCTCACCTTGGCAATAGCGGAGGCTTCACCACAGCATCAGATGGCTGTGGTTAAGCTTCTGCAGCTGGAAATGTTCTAATTGGAATAGGGAATCGACAGCCACGATCTCCTTC | 120 | 49.17 |
| chr7 | 143600306 | 143600426 | Target=1;ProbeIdx=4477;Target #1 | 1 | + | good | ATTAAATTCTCAGAAGCATGAGCCTGGCCCTTGGACTTCAGCATCATGGAGTTAGCCTGCCTTTCTAGGTCAAGGTCACTAGTTCCCTATAAAGCCCTTGTTTAAAGCTGCAGGTGTTGC | 120 | 46.67 |
| chr7 | 143600426 | 143600546 | Target=1;ProbeIdx=4489;Target #1 | 1 | + | good | CCTAGCTGTGTCTATAGGTCTCAGTATGATTGATGGGCTTTTGCTGAAGACAAAACCCATTTCATTTGGACAGTGGCATTTGTGACCCAGTCATAGCATGGGTTGACTTTAGGTTGCTGT | 120 | 44.17 |
| chr7 | 143600546 | 143600666 | Target=1;ProbeIdx=4501;Target #1 | 1 | + | good | TTTTCAAAGGTGCAGACTGTACAGAGTGCAGACAGACAGACATCCAGGAAAAAAATTTCTCCCCAGTGTACCTTGCCTCAGCTTCCACCAAGCCTGTGCTCACCACAGGTCACCCAAATG | 120 | 49.17 |
| chr7 | 143600666 | 143600786 | Target=1;ProbeIdx=4513;Target #1 | 1 | + | good | CCCAGCGCTTGATAGGGAATGCCAGGTAGATCCCTGGGGCTCACTGGACAGTCAGCATTGCCCAGTCAGTGAATTCTAGGCTAGTGAGAGGAAAACAAGGTGAATGGCACCTGAGAAATG | 120 | 52.5 |
| chr7 | 143600906 | 143601026 | Target=1;ProbeIdx=4537;Target #1 | 1 | + | good | ACACCTGAGATTGTCCTCTGGTCTCTGTGCACACATATGCATCTGCACACAAGAACGCCTGCACACACATGAACACAGACAGAGCATCATGAAGAAATTCATGATGATGCGGATGGGTAG | 120 | 47.5 |
| chr7 | 143601026 | 143601146 | Target=1;ProbeIdx=4549;Target #1 | 1 | + | good | TGTGGAGAGGTTTTTGGTGTGAGATGCAGTTCTTGTCAGGAGTATGTTAGGAGCACAGTGGGAGTGCATGATCCACAGGCGTGCTGTGGTGTGATCTGTGCCAGGGACTGGGCTTTCAGC | 120 | 53.33 |
| chr7 | 143601146 | 143601266 | Target=1;ProbeIdx=4561;Target #1 | 1 | + | good | TCTGGAGATGGCCCCAGATACCTTAGGACCCCTCCCTGAGTGCTTAGCCTTAGTTCTCCTGGCTCCTCAGGTGTAAGTGGATGATTTAAGTCTCTCTCAGGGTTAACAGAGGTTGAGGAG | 120 | 51.67 |
| chr7 | 143601266 | 143601386 | Target=1;ProbeIdx=4573;Target #1 | 1 | + | good | GCAGTCATCTTAGAGTGTATGTGTAATGGCAGTCTGGGGTTGTTGATCTTTCTGTCCAGGCTGCCCACTGTAACGAGACATATTTATGCAGTGTAGTATTTACTATTCTCCCTCGCTTTG | 120 | 44.17 |
| chr7 | 143601386 | 143601506 | Target=1;ProbeIdx=4585;Target #1 | 1 | + | good | TCTAGACGTTCCCTCACCTCACTGCAGTCTCATGGAGCAGTCCTTATACTCATCGCAGGCTCCAGACAGACATGGCCACACTCTAACTGGCTTCATCCACAGTGCTGCTGAGACTCGGAG | 120 | 54.17 |
| chr7 | 143601506 | 143601626 | Target=1;ProbeIdx=4597;Target #1 | 1 | + | good | CACTTGAGGATAGGAAATGAGGGGCAGCTGCCCTTCTGCAGTGGTGGACATGTGATCCCAGCACTGCTGTGTGCCCCGCTGAGGTTCACATGGACTGACAGTGCAGACGTGCCCCCCAGC | 120 | 60 |
| chr7 | 143601626 | 143601746 | Target=1;ProbeIdx=4609;Target #1 | 1 | + | good | CTTTACTATGGGCACATGGCCCTCTGTGGGTTATAGATATTTGCTCCTGACCATCCAGCCATACACGTAACATTCATATAGCAATAGGGCAGGTGTGACCTCTGTTCTTGTGGTGTAGCC | 120 | 47.5 |
| chr7 | 143601746 | 143601866 | Target=1;ProbeIdx=4621;Target #1 | 1 | + | good | CTGACCCCAGGACCTCACAATGTGACTGTTTGGAGCTAGGGCTTTCAAGAGGGGATTTGGGTAAATTAAGGCCAAGAGGGTGACTCTGATTCACGTGATTTGAGACCTAATAAGAAGTGG | 120 | 47.5 |
| chr7 | 143601866 | 143601986 | Target=1;ProbeIdx=4633;Target #1 | 1 | + | good | GGATCAGACACAGGATAGCCTGTGAGGACAGGGAGAAGACACCAGCTGTAAGCCAAGGAGACAGGTCTCAGGAGGAGCCGGCACTGAGACTCCATCTGGGACTTCTGATGCTGTGGGAAC | 120 | 56.67 |

| | | | | | | | | | |
|---|---|---|---|---|---|---|---|---|---|
| chr7 | 143601986 | 143602106 | Target=1;ProbeIdx=4645;Target #1 | 1 | + | good | AAATGGATGCATAGTGCAAGAGTGAAACATACAGGTTGGGGCTTTTCTGGTTGGATTTGATGAAGATCAAGACCCTCTGAACTTTACCCTTGTAAACTAGCTTGGCAAACTTTGGCCAAT | 120 | 41.67 |
| chr7 | 143602106 | 143602226 | Target=1;ProbeIdx=4657;Target #1 | 1 | + | good | GAACATGGTGTGGAAGATTTGTTCTGGCGGCCTCTCAAGCTCCTGCCCTGAGTTGAAGACACAGCATGGCTGTTTTCTCTACCTGTCCACGGTGTATGCTAATGCTGTCTGCGCCTGAAT | 120 | 51.67 |
| chr7 | 143602226 | 143602346 | Target=1;ProbeIdx=4669;Target #1 | 1 | + | good | ATTTTCAGTTCTTGGCTCACATCTTGCAGAACACTTTCAAATGCATGATCACCAAATGCTGTCGGGGATGTCTGTGGCTGGGGTGGTGGTGCTCTGACTGTTCTTTGCATGAAAACACCA | 120 | 46.67 |
| chr7 | 143602346 | 143602466 | Target=1;ProbeIdx=4681;Target #1 | 1 | + | good | TTTGTCTTGGATTTGACTTTGACAGTGAAGTTCACTGGTGGATGTATCAGTGTCCTACACTATTTAGCATTATGAACAGCCCCAGCTCATGCTTGCAGCTCTGTAGGTTTGAGTGGTTAT | 120 | 42.5 |
| chr7 | 143602466 | 143602586 | Target=1;ProbeIdx=4693;Target #1 | 1 | + | good | TACCAGCCCTCCACCACCATCACCGTTATTCCTAATGGATAATTCACCATTATCTAATTGATATGCCCAGATGTGTGATTCCAAATTTAATCAAGTTGGCAATCGAGATTGAAGGTCAGG | 120 | 40.83 |
| chr7 | 143602946 | 143603066 | Target=1;ProbeIdx=4741;Target #1 | 1 | + | good | GGCAGTGTTTTACAGATCTGTTGTAACTGACCCCAGTCAGGCTTCCTTATACCACTGAGTTTAGATGTCCAAGATCAAAGTGCTATCTAGGATTCCCCAGGATGCCCTGAGGAGCCTCTG | 120 | 48.33 |
| chr7 | 143603066 | 143603186 | Target=1;ProbeIdx=4753;Target #1 | 1 | + | good | TCCCACTCTACAGTTCTAGGCCTTTGTAGGAGAACCTCAGCATTTTTCCCCTATAGGACCAATCCCACGATAATCCTCCTGAAGCCTCCATCCTATGCACCTTCTGTGACATCCCCATTT | 120 | 48.33 |
| chr7 | 143603186 | 143603306 | Target=1;ProbeIdx=4765;Target #1 | 1 | + | good | ACCATCGCTGCAATGCCACGGTATTGGGTCAGAGTCATACTTCTCCAGTGCAGCCTCATCTTAATTGACCACACCTACAAAAACCCCATTTCTAATTATATGTCTGTCTTCTTGAGGGTA | 120 | 44.17 |
| chr7 | 143603306 | 143603426 | Target=1;ProbeIdx=4777;Target #1 | 1 | + | good | TATATTGCTTATGGTCACCTGTATCTTTCTGGCTGTCATGTGTGTGGAGTGAGGCCCGCTTTACTGAAGAATCAACATGGTAGCCAGTGGCCACTTAGTAGGTCTGCCCAGCAGGTGGAT | 120 | 49.17 |
| chr7 | 143603426 | 143603546 | Target=1;ProbeIdx=4789;Target #1 | 1 | + | good | GGCAGACCCAGGCCTTCCTGACCTGTGAATTCTAGCTCTTCCCAGAAGGGACTCCATGTGGCACCCCTGGTGCTTTCCAAAGCTGCTGTTTTATTTCTCAAGGTTGGTTCTTACACTAGG | 120 | 51.67 |
| chr7 | 143603546 | 143603666 | Target=1;ProbeIdx=4801;Target #1 | 1 | + | good | GAGGCACAGACTGGGCGTCCCTGCTCTGCCCACCAAGTGGCAGTGAGTGCCAGTCTTTGTGTAGCCTACAATGGCAAACAAGTCCTCTGGGAGGCATCATACAGCCTTGGGCCTGTCTAG | 120 | 57.5 |
| chr7 | 143603666 | 143603786 | Target=1;ProbeIdx=4813;Target #1 | 1 | + | good | GCACTTGCCAGTTGGCAGGTTCTTGTCAGCTGAGGGGAGAGAGAGGCTGATAAAATGCTCTGCAGTGTGGCTTGTGAAGGAACCTGAGTTGGGACGTCCCATCTGTGGACTGTAACCCAA | 120 | 53.33 |
| chr7 | 143603786 | 143603906 | Target=1;ProbeIdx=4825;Target #1 | 1 | + | good | ATCCCGCAATGATTTCTGAACTGGGAGTGGTGGGGTTGTGTGCAGCCCCAGGAGGCTATATCTCTTATTGTACTTTTGTTATGTCAAGATCACAGAGAGTTCTCACCTAGCCCCGGGTGG | 120 | 50 |
| chr7 | 143603906 | 143604026 | Target=1;ProbeIdx=4837;Target #1 | 1 | + | good | ATGGAGGAGGGGGGGCATGTGTCTCCTCCCTAGAAGGAACAACTGATCCCACCAGGTCTTGTAACCCAGAAAGGTCAGAGGGTCACCCTGAACTCAGCCAATGACATCCGAGAGGGGCCA | 120 | 56.67 |
| chr7 | 143604026 | 143604146 | Target=1;ProbeIdx=4849;Target #1 | 1 | + | good | ACAGGAGCGTTCATTCATGTGTAGGTGTCAGCTTCCCTGGCTGGGGTTGTAGCCGCTGGACTGGTGTTCAGGGTGGACAGCAGCTGAGCCATTCTGCAGCCATGAGAACCTACTGGTGCT | 120 | 56.67 |
| chr7 | 143604146 | 143604266 | Target=1;ProbeIdx=4861;Target #1 | 1 | + | good | CAGGAACAGGATCCGGCTCTACCCTTAGGCACACACAGAATCCCTTGTGAAGCACTAATTCCAGCCTCTTTCAGCAATCCCTTGGCTCTTCTGAGCTGTCTCCACAGCAGGGAAGCACAT | 120 | 52.5 |
| chr7 | 143604266 | 143604386 | Target=1;ProbeIdx=4873;Target #1 | 1 | + | good | CCCTGCACTCCTGCTATGGCTGAGCACTAGAAACTAATTTTAAGCCAAGACAAATGAAAAGTCTTAGTTAGACATGAGCTTGGTCGGGCTTTGGCTTCCAAGGGAGTGCCATAGAAGTCT | 120 | 45.83 |
| chr7 | 143604386 | 143604506 | Target=1;ProbeIdx=4885;Target #1 | 1 | + | good | CCTGGGAACACAACTGAGTGTCGCTATTCCAGTTCTCAGAGGCAGGGATGCATCTTTAGGAGTGTGGGCGATGGAGGCTCTGCTGATGCTGTTTAACCTTTTAAGAAATTACCTCTTGCA | 120 | 48.33 |
| chr7 | 143604506 | 143604626 | Target=1;ProbeIdx=4897;Target #1 | 1 | + | good | GTAATTGGCTTGATGTTCCCCAACCCTGCTCAAATAGTTTCTCGTGATTGCCACCCCTCACCCTTCAAGTACAGGTTGGGTTTTGGTTTTTGCATTTTGACATAGGTGTACGTGGTGTGA | 120 | 45.83 |
| chr7 | 143604626 | 143604746 | Target=1;ProbeIdx=4909;Target #1 | 1 | + | good | TTGCCTTTACATCTGGTGCCATCAGCCAGCTCTGCTGGATGCACTGTCCCTGCTTTCTAAGGCTGGAATTACAGATGCACGCCACACCCACCTGGCATTTGTAAGAGCTCTAAGGATTTG | 120 | 50.83 |
| chr7 | 143604866 | 143604986 | Target=1;ProbeIdx=4933;Target #1 | 1 | + | good | AATTCCAATTCTCACACTTGCACACAAGCGCTTTTAACCGTAGAGCTTTGTCACTAGCTGTTTAGTGCATACGGTTCAGAGACAGTGCTAGTGATATGAAGGCTCTAGTCTCTCAGGTTT | 120 | 43.33 |
| chr7 | 143604986 | 143605106 | Target=1;ProbeIdx=4945;Target #1 | 1 | + | good | CCCATGGTGAACTCCTGGGGCAAGGTGTCAGAGTTGCCAGAGAGCAAGCCGTGGACACCTCAAGTGGCACTGTCCCACAGTCAGCTGCTGCACCAGGGGTTCCATGTAGTGGATTGAGAA | 120 | 57.5 |
| chr7 | 143605106 | 143605226 | Target=1;ProbeIdx=4957;Target #1 | 1 | + | good | TCATTCACAAGACTCATTTTCTATCTTCGTCAGGAAAGAGAGTGACTTTTACACAGGGGTACACCGGATCTATTAGTTGTTACAACCAAATAGGAAATCATGCAGTATATCCAACCATGT | 120 | 38.33 |
| chr7 | 143605226 | 143605346 | Target=1;ProbeIdx=4969;Target #1 | 1 | + | good | GTCCCAGGGCAGCTGTAAATGCCTCAAACAAGAGAGATTTCTTTTTCTAACTCAATCACATAGTTCTCAAACATGAACTTTGTAGATGGCAATGTTGTACTCCAATATGAAATGCACTTG | 120 | 38.33 |
| chr7 | 143605346 | 143605466 | Target=1;ProbeIdx=4981;Target #1 | 1 | + | good | AGAGGACTTTTTAAAAACATGTCGTCATTGCGTGTGAGGAACCTGAGTGTCTGTAAGTCTGAAGGTTCAGAGGGGGTTCTGTAACCCCTAGGATGTCTTAGGGACAGCTGTATTTAACTA | 120 | 44.17 |
| chr7 | 143605466 | 143605586 | Target=1;ProbeIdx=4993;Target #1 | 1 | + | good | TGACTCCCAGTGTACGCCCCCACCACAGTGCACTTGTGCACTCTGGTGTGCACATATAAGCATATGAGAATCTTGGTTTCTGTCCCTAGGCTGGCTGGTCAGGAAGTCTCAGGGATTAGC | 120 | 52.5 |
| chr7 | 143605586 | 143605706 | Target=1;ProbeIdx=5005;Target #1 | 1 | + | good | CTGTCTCTCCCTCCCTAAGGCTGAGGGTTCTGGGGCTCAAGCTCACGTCCCATGGTCCCATGGTTAGAGGGCAGGCACTTCATTGATTTCCAGCCTGGACTGAGTTCTTTTGCTTTCCCC | 120 | 55.83 |
| chr7 | 143605706 | 143605826 | Target=1;ProbeIdx=5017;Target #1 | 1 | + | good | AGCCTTTTTGTTTGAGAGACAGAGGCAAACTTTACTTTGTAGTTAGTAAGCCATATCTCTTGGCTTCTCTGTTCTCAACACGTAGCAGTGGGGCAGAGGGGACAGGAACAGAAGGAACAG | 120 | 45.83 |
| chr7 | 143605826 | 143605946 | Target=1;ProbeIdx=5029;Target #1 | 1 | + | good | GAATCCTTTGATCCTGTGTCCGGGTTAGTAGCTCTGATGACCTTTGGCATCCTGCCTCTTTTCTCTGTTCTTCCTGCAGAAGTGCATTCGATTTAACCCAGACGCCACCGTGTGGGTGGC | 120 | 51.67 |
| chr7 | 143605946 | 143606066 | Target=1;ProbeIdx=5041;Target #1 | 1 | + | good | CAAGCAACGGATACTGTGCACGCTGAATCAAGGCCTGAAGGATGTACTCAACTATGGGCTCTTTCAGCCCGCCAGCAATGGGCGTGACGGCAAGTTCCTGGATGAGGAAAGGCTCCTGCG | 120 | 55.83 |
| chr7 | 143606066 | 143606186 | Target=1;ProbeIdx=5053;Target #1 | 1 | + | good | GGAATACCCACAGCCCATGGGCCCGGGTGTGCCTTCCTTGGAGGTAACTAGCCGTGTGTGAGTGTGTGAGGGAGGGATGGTGCTGTGCAGCCTGGGTGCCACCGAGGGCATTGTTTTAAC | 120 | 60 |
| chr7 | 143606186 | 143606306 | Target=1;ProbeIdx=5065;Target #1 | 1 | + | good | CAGGCACCTGTTGGTTTGCGATTAGAAAATTACCCACCCTGCAGGTAAAGTCTGGCTCCTCTGGACCTTCCATGTTAGGCATGTTGTGACAGGTACCCAGTGGTCCTTCTTGGTGGATTT | 120 | 50 |
| chr7 | 143606306 | 143606426 | Target=1;ProbeIdx=5077;Target #1 | 1 | + | good | GGGAATTTGGATCAGTGGCATGATTAGTTAAAGACAAAGTGAGAGCTGAGTTTATGGTGGCATATGGATGTGGTCTGGGGCAGGATGCAGGGCTAAAGCATTGTAGTCTAGCAGCAGGCA | 120 | 47.5 |
| chr7 | 143606426 | 143606546 | Target=1;ProbeIdx=5089;Target #1 | 1 | + | good | CAGACATCTTCCAATTCCATCCAGAAGCTATAGGCTGAGCTTATAAGATATACACTCCCCAGGCAGTCCCGTCAATCCAGTTCTTCAGGACTGACTTGTTTTGACACTTAAAAATCAAGA | 120 | 42.5 |
| chr7 | 143606546 | 143606666 | Target=1;ProbeIdx=5101;Target #1 | 1 | + | good | AAGTTCCCATCTGAGTCATGGGCATACCTGTAGACTTGTCATCTTGTGTGGGGCAGAGGGAGAAGAGAACAAAGTTGTCTCTTTCAGATATTATGGGATTGGTGATGGTGCTAGAGAGAT | 120 | 45 |
| chr7 | 143606666 | 143606786 | Target=1;ProbeIdx=5113;Target #1 | 1 | + | good | TTGTGTATGCTTCAAATATGTGAAGCACCCACAGAAGCTACATCTATGTATGCTATGACATCATCAGATCCTCTGCAAATGGAATTACGGATGGTTGTGAGCCAGCGCATGGGGTACTAG | 120 | 44.17 |
| chr7 | 143607386 | 143607506 | Target=1;ProbeIdx=5185;Target #1 | 1 | + | good | GAACAGAACCAGGGTCCTCTAGAAGAACAGCTCTTGCTCCTAAGCAAAAGCCCTCTCTCCAGTTTCCTAAGTTTAAAAAAAAGTCAAAAAAGGGAAAATAATACAAGGTCAGATTTCAGT | 120 | 39.17 |
| chr7 | 143607506 | 143607626 | Target=1;ProbeIdx=5197;Target #1 | 1 | + | good | TGGGGGACTAGCGTATGCCTCCACCTGGAATGTTCCCTTGTTCCCATGTTCCCGCCCTTTATTCCTGGGAGTGTGACAGTCTCAGGAAAAGATCAGGAAGGAGACACAAAGTGACCTGGG | 120 | 53.33 |
| chr7 | 143607626 | 143607746 | Target=1;ProbeIdx=5209;Target #1 | 1 | + | good | TTCAGGAAGGTCATTCTACTCTGGGGCCTTTGGGTGTTGTCTTTATGAGACAGATAAGGAAACTGAGACTGAGGTGGGAGAGTGACCAGTTCAGAGTCAATCAACAAATAGAAGACCTCA | 120 | 45 |
| chr7 | 143607746 | 143607866 | Target=1;ProbeIdx=5221;Target #1 | 1 | + | good | GGTTTCCTCTCAAAATCTCCATCATACACCTGCCCTTAGTGTTCTTCGAGAACCTTTTGCAGTGCCTCTAACAGTCAGCTACACAAAGGCCTGGGTGTCTGGGAGGAGAGGGGTGTGTGG | 120 | 51.67 |
| chr7 | 143607866 | 143607986 | Target=1;ProbeIdx=5233;Target #1 | 1 | + | good | TAAAGCTGACCACTGGGTCACCGCTTGCCCGGGAGTGTTGCCATTAACGCCAAGTCCATTCCCCCACTCTGCAATGAAATTTCCAGTCCCAAGTAAGCAGCACTGTCCTGAGGGGCCAGC | 120 | 55.83 |
| chr7 | 143607986 | 143608106 | Target=1;ProbeIdx=5245;Target #1 | 1 | + | good | GGGGACTTGAGTCCTGGGCAGGGGGATGAAGAGATGAAGAGATGGCTCATGCAGCTCTGACAGAGGAGAGTGACAGGGAAGAAAAATGTCACATCCACAGCACCCTCCAGACAAGCTCAG | 120 | 54.17 |
| chr7 | 143608106 | 143608226 | Target=1;ProbeIdx=5257;Target #1 | 1 | + | good | GGCAGGTAGATTCCCACAGACCGTGAGCCAGCATGATGAAATCATGAACTCTGGTGTGTCAGGCCATCTTATTCTCTGTACTAACCGAGGACTGGCTTCCATAGCTGAGAACCTTCCCTG | 120 | 50.83 |
| chr7 | 143608226 | 143608346 | Target=1;ProbeIdx=5269;Target #1 | 1 | + | good | TGTCCAGATGGTTGACAGATGGTGGGACTCTTAAGGGGATGGGCTGTGCAGTATGGACACCAGTGTCTAGGTTAACACTGAGGTTCCTTGGCCTTTGCCCTGATCGCTTTTCCTTTGAGC | 120 | 51.67 |
| chr7 | 143608346 | 143608466 | Target=1;ProbeIdx=5281;Target #1 | 1 | + | good | ATCCATGTCTGGCATTTAGTCATCGGCAACCTTTTTCTTGTTTCCATGGCAGTTTCGCTACAAGAAACGTGTGTACAAGCAGTCCAACCTGGATGAGAAGCAGTTGGCCAGGCTCCACAC | 120 | 48.33 |
| chr7 | 143608466 | 143608586 | Target=1;ProbeIdx=5293;Target #1 | 1 | + | good | AAAGGTATGGGGTGTGTGTGGGGATAACGCTTTCTCCACTCATCAGGTCACCTCTGATCCTGTTGTGTTGTCTCTGGAGGGCTCATTCCCCTGAGAGTGGCCCCTGTGTACCCTGTGGGA | 120 | 55 |
| chr7 | 143608586 | 143608706 | Target=1;ProbeIdx=5305;Target #1 | 1 | + | good | TGAAGCCAGACTGGGGCATACAGGATGCCTCACCATGCACTTTGCTGGAGACCCAGCAAGATAGCAGGCAGCAAGGTGGCCAGGGAAGCTGGTGGCGGTTGAACAGGGGATTGCTGTCCC | 120 | 59.17 |
| chr7 | 143608706 | 143608826 | Target=1;ProbeIdx=5317;Target #1 | 1 | + | good | CCTGGTGACCAAGGGAGGTTAAAAGGCAGCAGGAGTGGCATCGTGGGATTTGGATATATGGTGAGGTTGAGGGATGCCTGCTAAGCACGGTCTCTGGTTGCTGTTGCTGCCGAATCAGAA | 120 | 53.33 |
| chr7 | 143608826 | 143608946 | Target=1;ProbeIdx=5329;Target #1 | 1 | + | good | CAGACCTTATAAAGGCAGGAAGTTCTGGTATAATGGGAGAAGATTTCTGTGCTAGCTGGAGGGCTCTGTGCCCACCGAAGGCATGATGAGTCCCTGTGTCACATTCCAGAAATGCCAGTG | 120 | 50 |
| chr7 | 143608946 | 143609066 | Target=1;ProbeIdx=5341;Target #1 | 1 | + | good | AGGCCCTGGGAGCCAGGAGCATGCCTGTAGCAGGAGACAGAGATCCTGAGGAATGTGGGTGGAGCCTGGGGGACATGGGGTGGACACTGCCTCTCCTTAGGCACCCTGAGGTGCCTGGTG | 120 | 63.33 |
| chr7 | 143609066 | 143609186 | Target=1;ProbeIdx=5353;Target #1 | 1 | + | good | CTGGCCGGTGAGTTTTCCTGCTACAGGGAGTACACCTGGGGCTTTCACAACACGCAAGTTCCTTGTGTCTCAGATGGAGCGGGGTTGTCACTGTGGGTTTCATTCTTGTCTTCGGTTAAT | 120 | 51.67 |
| chr7 | 143609186 | 143609306 | Target=1;ProbeIdx=5365;Target #1 | 1 | + | good | TTTTTTTTAATTAAGTTAGTTCTTTGACAGCCTCATACTGTATGGTCCATACTATATAAAAGCATTTTGATTGATCTACAATTCTCTCTTACCTCCCTCTCATCCCTATCAGTCCTCATA | 120 | 33.33 |
| chr7 | 143609306 | 143609426 | Target=1;ProbeIdx=5377;Target #1 | 1 | + | good | CTTCCCATCACTCTTGTTCCACCAAAGCCAAAGACTCCTCAGTTTTCTTAATGTATCCACAAATAGTTCAGCTATGAGGAATAGGTCAGAGCCCCTCATCCATCCACGCCTGAGTGCTGA | 120 | 46.67 |
| chr7 | 143609426 | 143609546 | Target=1;ProbeIdx=5389;Target #1 | 1 | + | good | GTGGTCTTTCCTCCCTTTCTTCCTTAATGTTCTCTGAGCCTTAGAGGGGATAGTCTCAATGTCTCATTTCAGGCTGGGCAGTCACAGGTTACTTATTGTCAGTGCCTGTGCAACCACCAG | 120 | 48.33 |
| chr7 | 143609666 | 143609786 | Target=1;ProbeIdx=5413;Target #1 | 1 | + | good | GCTCTGCAAATACTAGGAAGAGAGGTTTCTCTGATTTGGGCCAAGAGTAACATTGTTTAGGGAGTAAGCAGATATTTAGTAGGCAGCTTGGCACCATGTCAATTTAGTGTTTCCTCACAA | 120 | 41.67 |
| chr7 | 143609786 | 143609906 | Target=1;ProbeIdx=5425;Target #1 | 1 | + | good | GGGCCTGTGACTTTCCCTACCATGCGTTTTTTGGACCCAGTTTATAGCACTGGGTTTGGATTTGTGCTGTGGAGTGAGCCTCAAAGGCAGTCAGGGAGTGGTTGGCTATCCTCATAACTA | 120 | 50 |
| chr7 | 143609906 | 143610026 | Target=1;ProbeIdx=5437;Target #1 | 1 | + | good | TTCCGTTGCTATTATGCTATTGCCTAGCAGGTTGCTATCATGGATTGTAGGATTTACAGCTGGGCAAGACCGTTGGTGCCTTATCTCCCACAAGATAGGGCTATTTCATAACTTCTTAGA | 120 | 43.33 |
| chr7 | 143610026 | 143610146 | Target=1;ProbeIdx=5449;Target #1 | 1 | + | good | ATTACGAAAGCTAGCCGCCCATGGGGAAGTGTCCAGCTCAGGTCCATTTTGAGTTCTCTATATCACGAAATAAGCTGTGCCCTATCTTCAGCTCTTGGTCTCATTATCTAGTTCTCATGA | 120 | 45 |
| chr7 | 143610146 | 143610266 | Target=1;ProbeIdx=5461;Target #1 | 1 | + | good | GCAACCCAGACAGACAGCAACAGCCTGTGCTGTTTTAGGGGCCTCTGTGGCCTTTCTGACCAACATCGTTTTATGTTTGTGCCTAAGTTATCCAGAGGTGACTGGCAGTTGCCCTTGGTC | 120 | 51.67 |
| chr7 | 143610266 | 143610386 | Target=1;ProbeIdx=5473;Target #1 | 1 | + | good | CCCTGCCTTTGGGTATACCACCACTTTCCTTGACACTGTGACACCTCAGCCATGTCCTGGCCAAGTATGGTCCAGGCTCTCCTTATTTTTATTATTATTGTTGTTATTTAATGTGTAGGC | 120 | 44.17 |
| chr7 | 143610386 | 143610506 | Target=1;ProbeIdx=5485;Target #1 | 1 | + | good | ATAAATACTTGTGGCTTTAAAGAAAGGTATCAATGATGTTTGGGCGGCACTTGACTCTCTGACAGCACAACTTGAATTTGGACTTCTGGGTGCCCAGGTTATAGCTTCAGGAGTGAACTC | 120 | 43.33 |
| chr7 | 143610506 | 143610626 | Target=1;ProbeIdx=5497;Target #1 | 1 | + | good | TCTGGGACCTCTTTAGGTGATTATGGACTCCACTCATCAGGGTTCCACCCTGATGACCCAGAAACCCCTAGTGTCGTTATATAGGAATTACAGTCCATTGTGAGAATTCTGGAGGGACAG | 120 | 47.5 |
| chr7 | 143610626 | 143610746 | Target=1;ProbeIdx=5509;Target #1 | 1 | + | good | ACATTGAGATGTTGGTACAAAAGTGGCAAACAGCTAGGGTTCTGAGACATTCCTCAGGACAAGCCCATGGTGTTGGCTGTGTGCTGACCTCACATGTGGCCTCCTGCCTGTGAGGCCTCT | 120 | 52.5 |
| chr7 | 143610746 | 143610866 | Target=1;ProbeIdx=5521;Target #1 | 1 | + | good | TGGGGGGGGGGGGGGGCGTCAGCAGTGCTCTGTGCAGAGGAGGCCAGTGCCTCCTAGTGGACAATCTTCTTAGTCTATAAAGATCTTCCCTCCCTCCATCCCTTCCTCCTCTCCCATCTA | 120 | 58.33 |
| chr7 | 143610866 | 143610986 | Target=1;ProbeIdx=5533;Target #1 | 1 | + | good | CCAACTAGCTTTGAAACCATTATGTAGCTAAGGATGACCTTGAACCACCAATCCTCCTGTCTCCACCAGGAGCTGGGAGTTGCAGCAAACTCAGGGCGTCATTCATGGTAGGCAAGTTCT | 120 | 49.17 |

| | | | | | | | | | |
|---|---|---|---|---|---|---|---|---|---|
| chr7 | 143611226 | 143611346 | Target=1;ProbeIdx=5569;Target #1 | 1 | + | good | CTGCCCACTGAGCCTCACCCCTGCCCCTGTAAATACATTTTTCACAGACACTTCTGTCTTGTTTGTTTCGTCTCCAGACCAATCTGAAGAAGTTTATGGACCACGCTCAGCATCGTTCAG | 120 | 48.33 |
| chr7 | 143611346 | 143611466 | Target=1;ProbeIdx=5581;Target #1 | 1 | + | good | TGGAGAAGCTGGCCAAGCTGCTGGACAGAGGCCTGGATCCCAATTTCCATGACCTGGAGACTGGAGGTAGGTGTTCATGAGGGAGGCCTTTTCTGTATACTCTAGCGGGTAAAGCATTTG | 120 | 52.5 |
| chr7 | 143611466 | 143611586 | Target=1;ProbeIdx=5593;Target #1 | 1 | + | good | TCTATTCCTTGGTTCAGGTTACTCTGCATCATTCATCTCCCATGCTTGGCTTGGTTTTACCTGGGCTTTGCCGTGGCTCCTTGTCTTAGTTAGGATTTCTACTGCTGTGATGAAACACCA | 120 | 45.83 |
| chr7 | 143611586 | 143611706 | Target=1;ProbeIdx=5605;Target #1 | 1 | + | good | TTCTGATATGAAGGCCTGCATTTTTCTCCTTGCTGTTGCCTCTGGCTCAGTACATGCACACAGTAGGTGCTCCTTGCATAGGTGCTAGGAAATAAGAAAGCAAATAGATGCCATAGATGT | 120 | 44.17 |
| chr7 | 143612066 | 143612186 | Target=1;ProbeIdx=5653;Target #1 | 1 | + | good | GTGGGGCCATTGCCTGCTGACTGGGGCCTTCTGCGAGGCCCTGGGGTGCATGTGCTAAGCAGCCAGCATTGTAGCACGCTTGGGTTTTTCCCCCTGCCTGGGTGCAGACTTCCTCACAGA | 120 | 61.67 |
| chr7 | 143612186 | 143612306 | Target=1;ProbeIdx=5665;Target #1 | 1 | + | good | CAGACAGCTGCATGCACATATCTTGTGCTCCCTGCTGGGTGTTGAGTTCCTTAGGCTTGAAATATCCTTGGGAGAAAGCTGCACTGGTTGGGGAGGGGGCTCTTGGCCCTGCACTGCTAA | 120 | 54.17 |
| chr7 | 143612306 | 143612426 | Target=1;ProbeIdx=5677;Target #1 | 1 | + | good | CACCTGGTCTGACTGGTCTCTGCTGTGGGGTAGGTCTAGGGTGTGTGGGGCGCTGCTGAGTCTCTCACTAGATGCCAGTAGGGCAATCCAGAGTGCCTTCCCGTGTTGCTAGTATCTTCT | 120 | 56.67 |
| chr7 | 143612426 | 143612546 | Target=1;ProbeIdx=5689;Target #1 | 1 | + | good | GCTTTTACTGGGTAGAGCTCAACAGCAACCCAGGTGAACACTGGCCGAGGGATGATATGATGGCCAAGGATGGCAGGCAGAGATCCAGGGACTTTCCCCAGCAGCTTGGATCTGCTTGGC | 120 | 55.83 |
| chr7 | 143612546 | 143612666 | Target=1;ProbeIdx=5701;Target #1 | 1 | + | good | CTGGTACGTGATCTCGGGTCAGGAGAGACTTGACCTTAGCAGTAGGGTCCTTTTTCCTGCCTTCTTTTCTTCTCTCCCTCCCTCTCCTCCCAGGATACCCCAGGTCCACTGCTAGCTCCC | 120 | 55.83 |
| chr7 | 143612666 | 143612786 | Target=1;ProbeIdx=5713;Target #1 | 1 | + | good | ACATGGCTTCCTGTCACACCTCGAAATTAATATGTGCACAAGATATGGAGGTGTCTTATAAAGTGTGGATCTTGTCTGTCATTTGGGGCAGCCTGGCTGACCTGTCAAGCTGAGGTGGGC | 120 | 48.33 |
| chr7 | 143612786 | 143612906 | Target=1;ProbeIdx=5725;Target #1 | 1 | + | good | CCCACTGGGTACTGATCTAGCTAGCATGACCTAGTCTGAGCCCTGATGCAGGCTGGGTTGCCTCTTTGCTCCTGCACTTCTTTGGGTTTAATTGTAAGCAGCGTGAGAAACTGGGAGAGA | 120 | 51.67 |
| chr7 | 143612906 | 143613026 | Target=1;ProbeIdx=5737;Target #1 | 1 | + | good | GCTTGTTTCTGAGGATGGTGGTCACATCGAACATATCGGGAGAGATTAGGTGTCCTGGCGAGGGGATGTGAGAGGAGGTCCTGACCAGAATAAGCAGGCACCAGCCCAAGGAAGAAGCCC | 120 | 55 |
| chr7 | 143613026 | 143613146 | Target=1;ProbeIdx=5749;Target #1 | 1 | + | good | CCTGAAGGAAGAAAGTGATGTTTCTGTGCCTTGTGGTGACCAGGTGCCTCCTGTGGCCTCAGGGCTCCTGACTTTGTACTATCTGGTCCCTGCTTTCCCCATGGCTCTATGTTGATGCTC | 120 | 53.33 |
| chr7 | 143613146 | 143613266 | Target=1;ProbeIdx=5761;Target #1 | 1 | + | good | CCAAGTATTGACCTTGGGCTAGCTCTGGCTTGTCCCATTCAATGACATGCAGCCCCTGGACTCTTGATATGGAAATGAAGTATAGCCAGGAATGACACATGTCCACAACCCCAGCACTCA | 120 | 49.17 |
| chr7 | 143613266 | 143613386 | Target=1;ProbeIdx=5773;Target #1 | 1 | + | good | TCAGGTTCTTTCCCCATCACACACACACATACACCCAAAAAAGAGGGAAAAGGTCTATGGGCCTGTGAGTTCCTAAGGATGAACTGGAGCCTTCAAATCTCCACAGATGACCTTCATCCA | 120 | 46.67 |
| chr7 | 143613506 | 143613626 | Target=1;ProbeIdx=5797;Target #1 | 1 | + | good | GGCCCGTGGCTCTGTCCTCAAGGTCTGTGGCCTGTATTTTCTTCTAATTATGAGTCAGTGGACTTGGTGGTAAGCTACTCAGGTCTGGCTCTCTCTCCTTGTTTCATGGTGACTGTTTCT | 120 | 49.17 |
| chr7 | 143613626 | 143613746 | Target=1;ProbeIdx=5809;Target #1 | 1 | + | good | TGAAGATTCTACAAGGGACCAGACCAGGCAGTATGAGCCTCTCAACTTGCCGTGTTCATTAGAACAGGCTAGTGTCCTCAGTGGGAGTTTTCTTCCTGATGGGAGCTGCGGCTCCTTTGT | 120 | 50.83 |
| chr7 | 143613746 | 143613866 | Target=1;ProbeIdx=5821;Target #1 | 1 | + | good | TCCCATCCCTTTTTACATCAGGGTCTTTGGTTCTGAGGTTTGAAGCAGCTGTGATCCTGGAGCAGACCCCAGTTGACACTGACAGTTTTGAAGCAGAGGTGTACATGTGGTGGCCTCTCT | 120 | 50 |
| chr7 | 143613866 | 143613986 | Target=1;ProbeIdx=5833;Target #1 | 1 | + | good | TAGCCCCATCTGGAGCTAGAACAGAGGTAGTAATGATGGGCAAGGAGCTGGCTCTTAGAGAAATTCCAAGAGATTCTTCCCTCTCAGCTCCTGCTATGAGGGACCTGGAAGCTCAGAGCA | 120 | 50.83 |
| chr7 | 143613986 | 143614106 | Target=1;ProbeIdx=5845;Target #1 | 1 | + | good | TGGACTCCACATGTGATGATTATTCTGTGTTCTCTGAGATGCAGCCAGTGTTCAGTGAAACACACTGATCTAGCTAAGGAGGGATTGACAAAGAAAGCCAAAGAACATGTCTGTCCTCTC | 120 | 44.17 |
| chr7 | 143614106 | 143614226 | Target=1;ProbeIdx=5857;Target #1 | 1 | + | good | TCCTGTCATTGCCCTGTAAGAAACAGGCTGTGGTATCTACAGACCTCTTAATGGAATCCATTGGTCTTAGAATAATCTAGAGATGACTAAAAGTCAAATATAACATAGCTTTATAGAAGG | 120 | 36.67 |
| chr7 | 143614226 | 143614346 | Target=1;ProbeIdx=5869;Target #1 | 1 | + | good | ATAGTCAAGGCTATACAGTAAAACCCTGTCTGGAGAGGAGGGATGGGATAGGAGGCTTTTAGGTGTCGCAGGGCCCCTAGACTTTGTCCTCCATGGGCACTGAGAGACAGCTGTGTAATT | 120 | 50.83 |
| chr7 | 143614466 | 143614586 | Target=1;ProbeIdx=5893;Target #1 | 1 | + | good | GTGGCTACAACGTAGGCTGTTAGCTGATAAAAATGAGCAAACGCTGGAGAGAAGCAACGGGAATGCTAAGTGGGTCCTGATACAGCACTCCTGGTCAAGGGTGGCTGTTCTGGAGTCATC | 120 | 50.83 |
| chr7 | 143614586 | 143614706 | Target=1;ProbeIdx=5905;Target #1 | 1 | + | good | AGCAAAGGCAGGCCTAGCCAGATGTGGCTGCAGAGCACAGCAGACATATGGGGTTGGAGAGGGACACACCATTCTAGAAACCCTCCCCTACCGTACCCATGGAGGGCTTGGGGTTTTTA | 120 | 55 |
| chr7 | 143614706 | 143614826 | Target=1;ProbeIdx=5917;Target #1 | 1 | + | good | TTGTTTCCAGGGCGCTTGCCTCCTCTAGCTCTAACAGACTTTGGTTATTCCTTCCACATTTTGGAAGCTCCAAGCTGGGTTTGGTGACTTTCTGAGTCACCCTCTGAATAGCAGACTCGC | 120 | 49.17 |
| chr7 | 143615186 | 143615306 | Target=1;ProbeIdx=5965;Target #1 | 1 | + | good | TGCACACCAGATACCCTATGGAGAGTCAAACTCAGCTTCCTTCTGAAGGCCAAGGCATGCCACCAGCCTTGGCCTCAGCTCTGCATCTCAGACTCAGCTGCACTGGGGAGCTGTAGCTGA | 120 | 55.83 |
| chr7 | 143615306 | 143615426 | Target=1;ProbeIdx=5977;Target #1 | 1 | + | good | GACAGCAGTCATTCTGGGTTGACTGACAAGCAAGTGACATGGGGATCCTGGTGGGGTGACCTGCCACTTAAAGCCACATGGTCCGTCAACAATAGTAGATGTCCATAGCTGCTGGAAATG | 120 | 50.83 |
| chr7 | 143615426 | 143615546 | Target=1;ProbeIdx=5989;Target #1 | 1 | + | good | TCAATCTCATTAAATAATATGAAGAGAAAACTGAAGACCTGTCCTAGTGGAGGGGTCCTTGGGTTCCTCTCTGCCATGGTGGGACCCCTGGGATGTCAGATGGATGACAGCTCCAAGCCC | 120 | 50 |
| chr7 | 143615546 | 143615666 | Target=1;ProbeIdx=6001;Target #1 | 1 | + | good | TGGGAGAGGCAGTCACAGTGACAGCCCTGGGGGACTTAGTCCATCTCTTTCCCACTCTGGGTTCCTGAGCTTTGAGGCTTGCTCAGATGAGAGAAAGGCATCTGATTAATGAACCAGAAA | 120 | 50.83 |
| chr7 | 143615666 | 143615786 | Target=1;ProbeIdx=6013;Target #1 | 1 | + | good | CTTGCTGTACCTCCAAAAACTCTGAGAAAGGCTTGGAATAGCACAGTCCCCTGCCTGGATGCTCCTGCAGAGCCACTTTGCTCATCTGGACTCACTTCTAAGACCTGGTCACCACTGGTC | 120 | 52.5 |
| chr7 | 143615786 | 143615906 | Target=1;ProbeIdx=6025;Target #1 | 1 | + | good | CTTATGTCAAGTTGACAAAAATCCCAGGCAGCACACCAAACGAAAGGTTCCGGGCAGTAATGCTCCAGAGCTGTCGGTGGGGGTGGGGCTGCTTCTAGCTAAGTTCTTTATCATCCTTGA | 120 | 50 |
| chr7 | 143616266 | 143616386 | Target=1;ProbeIdx=6073;Target #1 | 1 | + | good | GAGGCGCTGTTTCCACCCACATATGGCTGTGGTGGGCTCCTGGGGAGCCCGGTCCAGCTCCCACACGGCCCAATTAGGAGCTTTTGTGGACATTCTGTTTCCCCTTACTTGGAGCTCTGT | 120 | 58.33 |
| chr7 | 143616386 | 143616506 | Target=1;ProbeIdx=6085;Target #1 | 1 | + | good | GACTTGGGGAGTGGAGGGGCCTGTCTGATGACTTATGGTCTGTGTCTGTCACACAATGAATCACAGTGCATGATCGATACCCCAGGAATTGCTGTGTCTGGAGGGAGCAGGGTTCTGAGA | 120 | 52.5 |
| chr7 | 143616506 | 143616626 | Target=1;ProbeIdx=6097;Target #1 | 1 | + | good | CCTCATAAACATGAGTCTGCAGACAATGAGAGAGATGTGCAAACGAAGGCAGCCTGCCGCCAGTTCATAAATACTCCCCCCTCCAGTGAGTCGGGTGCGTCTGGTTGGTCCTTGGAGTTC | 120 | 53.33 |
| chr7 | 143616626 | 143616746 | Target=1;ProbeIdx=6109;Target #1 | 1 | + | good | TAATAAAGAGGCCTATTGTTAGGCGAGAACTCAGATGTGGTACCTAGTTATTTTGCAATCCTTCTTTATGCACCTGCTCCCATAGGAGCATGGGAAGCTCATGAGCTAGAAGATGAGGTT | 120 | 43.33 |
| chr7 | 143616746 | 143616866 | Target=1;ProbeIdx=6121;Target #1 | 1 | + | good | CATCAAATATGGAAGGTGAACCACGAAAGGCAGCCTGGGAAGAAAATGAACCTGGGATTGGGTCTAGGGTCCATAAAGAACCAGTTCTAGGGTCACAGAAAAATCCCACTGGGCAGCTGG | 120 | 49.17 |
| chr7 | 143616866 | 143616986 | Target=1;ProbeIdx=6133;Target #1 | 1 | + | good | GATGTAGCTCAGAGCCAGACCATACAGAGCCTGTATGCCTGAGGTCTTGGGTTCCAACCCAACAACAACAAACAAGAAACGGATGTGGTGGCATACACTTTGCAATTCCAACACTCAGGG | 120 | 49.17 |
| chr7 | 143616986 | 143617106 | Target=1;ProbeIdx=6145;Target #1 | 1 | + | good | GTGGACATTCCCCCATGCTCCTGCACCAGTCTCTCTGTCACTAACAGTCAAAACTGGTATACTGTGTTTGTCACAACTGAGGCTCAGGGATTACACTAGGGCCACTCTTGGGGGTGGGGT | 120 | 52.5 |
| chr7 | 143617226 | 143617346 | Target=1;ProbeIdx=6169;Target #1 | 1 | + | good | GGGGGATGTCACAGATTCTGTGGGGTTTGTCAAGCTTGTGACTCCTGTACCCATCATCCAGCATAATCGGGGACCCCATCACTGTCTACACACCCCACGTGCTTGGCTGTCTTCTCCTCT | 120 | 54.17 |
| chr7 | 143617346 | 143617466 | Target=1;ProbeIdx=6181;Target #1 | 1 | + | good | TAGCTACAGTCACCTGTGTCCCTGTTGTCACCTCCTTTCCTACAATGTCATTGTGGGAACTGGGCAGTATGTTGCAGCCATACTTGGCTTCATTCAATCAGTGGCCTGTGCCCAAGTTTC | 120 | 49.17 |
| chr7 | 143617466 | 143617586 | Target=1;ProbeIdx=6193;Target #1 | 1 | + | good | ATCTGTGTCTGTTTGTGTCTCTTTAACGACATATTTACTTAATTGTGTGTTGGGAAGAGTATGTGCACTGTGAACATATGTGGAGGTCAGAGGAACGCTGTGGAGTAATTTATCACCGTT | 120 | 40 |
| chr7 | 143617586 | 143617706 | Target=1;ProbeIdx=6205;Target #1 | 1 | + | good | TTCCCTTTCAGTAGGTTTCTTTTTAATTTTTAATAAAATATTTATTGACTTAGAATAGAAAAATGATTCCATGCAATCTTTAACAATTATGAACATAGGTTTTCCTTTCCTCTTTCTTTT | 120 | 24.17 |
| chr7 | 143617826 | 143617946 | Target=1;ProbeIdx=6229;Target #1 | 1 | + | good | AATTTTTACATTTTTTCAGATAGTTCAGGCTGACCTTAGACTCAAATCAATTCTCCTTCCTCAACTTCTCAGAGTGCTCAGAGAAATAGGTCTATGCTACCATACTCAGCTTAGACATAT | 120 | 36.67 |
| chr7 | 143617946 | 143618066 | Target=1;ProbeIdx=6241;Target #1 | 1 | + | good | ATTTTCAGCTTTCTTGTGCAAACTCCTAGAGCACAAGTGTGGCTGGTAGAATGTGCTTGATTTAAAAAAAAGAAAACTATAATTATTCCAGGTATATGGCTATTTTGCCTGCCTGCATGT | 120 | 36.67 |
| chr7 | 143618066 | 143618186 | Target=1;ProbeIdx=6253;Target #1 | 1 | + | good | CTGTGTATCACATTGTGCAGTGCCAATGGTGGCTAAAAGAGGTCATCAGATCTGGAACAGGAGTTGCAGACAGTTATAAACCTTCACGCGAGTCTTAGAATTTCTACTGCTGTGATAAAA | 120 | 42.5 |
| chr7 | 143618186 | 143618306 | Target=1;ProbeIdx=6265;Target #1 | 1 | + | good | TATGCTGAGCTGTCCTTCAGGACACCCTAGGTGGCTGCACCCAGTCTCATTTGTGTCAGTGATGAGGGAGCAGGGCCCTCTGTCCTCAAGGGTTAAGGGCTCAGTGTTTTGTATTTGGCT | 120 | 53.33 |
| chr7 | 143618786 | 143618906 | Target=1;ProbeIdx=6325;Target #1 | 1 | + | good | GTTCTGAGTGGCTGCAGCGTGACCCCTGTGGGGTGTCGCCTGGAGGTGTGAGTGCATTTCCTAGGGACCTGGGGCACAGCACACAGTGTTCCACCATCTGTCTGTCAGTTCTCTTTGCTG | 120 | 58.33 |
| chr7 | 143618906 | 143619026 | Target=1;ProbeIdx=6337;Target #1 | 1 | + | good | AGGATTTTGTTCAGGTGTCTCATGCACTTGGTGGGGGAGGTTATATAAAGTTTTCTGTGCATATCGAATGCCGGCATCTCCTGCATTTGTATTTGATGTTGGAAATATTTTCTCTAAGCC | 120 | 40.83 |
| chr7 | 143619026 | 143619146 | Target=1;ProbeIdx=6349;Target #1 | 1 | + | good | CCTGTGTCTTAATTGCAGAGTCATCTCTCTGACCCAAAGGTTTTTGTTGTTGTTGTTGTTCATTTGCTAAAGAAACCTTTCTTAGAAAAAGCTTTGCCTTAGGGAGTTTCTGCAGACTGC | 120 | 40 |
| chr7 | 143619386 | 143619506 | Target=1;ProbeIdx=6385;Target #1 | 1 | + | good | TCACCCATTGCTACCCTTTGACATCAGTGTGGCATATTTTGCCACCATTGGCCCTCAGATCCACATGCGAGTGTTATAAAACCCATGCAGTATTCTGATTTCCTCAGTTTCTCTAGATGT | 120 | 44.17 |
| chr7 | 143619506 | 143619626 | Target=1;ProbeIdx=6397;Target #1 | 1 | + | good | TTTCTTCCTGTCAGAGGATCTCAGGGCCCTATGGCAATTTTACTTGCCATCTCTCTGTGGGCTGCATGGCTTTCATGGTTGCTTGGACTTGATGACCCTGGCAGCTTTGAGGACTGGCCT | 120 | 51.67 |
| chr7 | 143619626 | 143619746 | Target=1;ProbeIdx=6409;Target #1 | 1 | + | good | GGTATTTATGCAGATGCCCCTCCGTGCACATTTGTCCAGTGCTTTCTTTCTGACTGGACCTAGATCCTGTGAACAGGAGGAAGTGTGGAGCTGGGAGGAAGGCGGCAGAAGTGAAGCAGC | 120 | 53.33 |
| chr7 | 143619746 | 143619866 | Target=1;ProbeIdx=6421;Target #1 | 1 | + | good | TCCTTGGGTGGGGGCTCCTGCTGTCGGCCAAACTGACTGCTGTGACGCTGACCTCCCCTAGCATGGCCCTGACTGCAGCTTTTCTTCCCTGAGAGCCACCCGCTGAGCCTGTCCCATACA | 120 | 61.67 |
| chr7 | 143619866 | 143619986 | Target=1;ProbeIdx=6433;Target #1 | 1 | + | good | GTGTGCTTTGGAAGGGGCTCAGTTGGCACAGCTCACTATTAAGGAGTCTGTCTGTGAGATTGTCCCCCTCTGATGCCACATACTGGTCACCCAGTCTCTGTGTGGGTTATGCTTTATGCA | 120 | 50.83 |
| chr7 | 143619986 | 143620106 | Target=1;ProbeIdx=6445;Target #1 | 1 | + | good | GATGTCCCTGAGGGTTGGGCTTCAGGATCCCTCAGGTCTGAGCTCACATCACACAGAACCTGTGCACATCCCCCGGCAGCCTGCAGCATCTCTAGATTGCTTATAAAACCTGATTCATGG | 120 | 53.33 |
| chr7 | 143620106 | 143620226 | Target=1;ProbeIdx=6457;Target #1 | 1 | + | good | ACGCTGTGAAAACCACTCTGACATTGTGTTATTTCCTGGTTATATTTTATTGTTGTTATTTTTCAACCTGTGCATTGCATTAACTTTACCCGGGAAGATGTGAGTGAACTTCTGTTCACT | 120 | 36.67 |
| chr7 | 143620226 | 143620346 | Target=1;ProbeIdx=6469;Target #1 | 1 | + | good | CTGGATAGAGCAGAACAAAGAAACAACAACAAGCCCAGACTAATGACTTAATCAGGCTGAGTGACAGGAACAGTGAGGGGTCAAGAGTTACTTATAGGAGTGCCGGTGACCCTAAAACAG | 120 | 45.83 |
| chr7 | 143620346 | 143620466 | Target=1;ProbeIdx=6481;Target #1 | 1 | + | good | CTGCATCACTGCAAAGCCCCACCCCCAATTTGGATAATGTCCCCCTATCTGCAAAATGGAGCCCTGCTTCATTAACCTTCAGTTTGTATCCTATAGCACCTTGGAAGAGACCACATGTAG | 120 | 47.5 |
| chr7 | 143620466 | 143620586 | Target=1;ProbeIdx=6493;Target #1 | 1 | + | good | CGGAGCAGTTTCATACACCTGGGAGGGAAGGCCGGGTGGGAGCAGCTGGATCTCAGGTCTGGGTCTGATGACCTTCCCTTCCCTCCCCTTGTCCTATGAGAGAGCCTTGGGGCTCTCTTA | 120 | 59.17 |
| chr7 | 143620586 | 143620706 | Target=1;ProbeIdx=6505;Target #1 | 1 | + | good | CAGGCATCAGCTGCTCTGGTGAAGATGGGGCCCAGCCAGTATGAGAGAGGAAAGTGTTTCCTGATGCTGGGGTTGGCTCGGCTCACCAGTGAGTGCCTGGTTGACACTGTGGCCTCTGCT | 120 | 58.33 |
| chr7 | 143620706 | 143620826 | Target=1;ProbeIdx=6517;Target #1 | 1 | + | good | GGGAAGCAGTTCCCGAAGGCGCATTTGTTTTGCATCTTGCATTTCACATTTAGGGTGGCTCAGGTCAGCCTGTGTGCCCCACCTCCAGTCCCCACATCTGTGATACACTCATTATTCCTC | 120 | 51.67 |
| chr7 | 143620826 | 143620946 | Target=1;ProbeIdx=6529;Target #1 | 1 | + | good | CCTGTGTTCAAGTTCCCATCCCGGGAACCCTCAGTCTCCTAAGTCAGTGTTTCAAGAACCCTGCTAGAGGTCACCCAGCCCTTCAGAGGAAAACTGATTTTCACTGCCATAGTTGTTGCC | 120 | 50.83 |
| chr7 | 143620946 | 143621066 | Target=1;ProbeIdx=6541;Target #1 | 1 | + | good | AGAGCTACCCTCCAGCTGCTGCTCTGAGTCCTTTGCATCCCCTCCTTGCCCTTCCTGGTGTGGTGTGCTCTATCCCTGTAGCCTGGCTGGAGCTCTCAGCTGGCCTGCCAGGGTCCTGCC | 120 | 62.5 |
| chr7 | 143621186 | 143621306 | Target=1;ProbeIdx=6565;Target #1 | 1 | + | good | TCTTTCCCTTCCTCCTGGTGTCTCTTGGAATACACCTGGTTACCTCCCAGCGACAATGCTCCCAGGATATTTCTAGGGTGATGGTGTGCCTATAAGGTAAAGTGACCCAAGTTCAGGTAA | 120 | 48.33 |
| chr7 | 143621306 | 143621426 | Target=1;ProbeIdx=6577;Target #1 | 1 | + | good | CAGCCTCTGCTCAGGTGTACTTGCTAAATTCTAACGTGTTTTAAAAACCAAGGAGATTTCAACAAGTGAAAAATGAGGACTTTTTTTTCAAGACACCGTGTTGCACATACCTGTGTGGGG | 120 | 40.83 |

| | | | | | | | | | |
|---|---|---|---|---|---|---|---|---|---|
| chr7 | 143621426 | 143621546 | Target=1;ProbeIdx=6589;Target #1 | 1 | + | good | CCAGCTCTCATGTAACCAGAGGCCCACTGTTTTGACTGGACGTGGCCAGCAAGCCCTCAGGATCCACCTGTCTCTGTTGAGGTGACAGACAGGCACACTTGCACTCCGCTTTTATGTGGG | 120 | 55.83 |
| chr7 | 143621546 | 143621666 | Target=1;ProbeIdx=6601;Target #1 | 1 | + | good | TGCTGGGGATCATTGTGCAGCAGGTACTCTACCCACTGAGCTTCCTCCTCTGCCTCAAGATGGAGACTCAAGACCAACAGTGCTCAACCTGTGGGTCGCAACTCCTTTAGGGGGTTGAAC | 120 | 54.17 |
| chr7 | 143621666 | 143621786 | Target=1;ProbeIdx=6613;Target #1 | 1 | + | good | CCTAGGGGACCCCAGTTTTTAATATCTGACAGCCACAAAGAGTCTGCATGCATCACAGAATGGAAAGAGAAAGGGTGCTAACTTTCCAGTGTCCAATAAGAGTTCTCTCCAGTTGTGCCT | 120 | 45.83 |
| chr7 | 143622026 | 143622146 | Target=1;ProbeIdx=6649;Target #1 | 1 | + | good | ATGAGGGATGTGTCAAGATGGGGCCCAGGCTTCTGTCCTCAGTGAGTGGGCTGGAACTCTCCAGGACTCATACCACCAAGTCCCACATTGTGGAAACCCTGCTGCTGCTATTTTGATTTA | 120 | 51.67 |
| chr7 | 143622146 | 143622266 | Target=1;ProbeIdx=6661;Target #1 | 1 | + | good | CATGTGTGACCCCTTGTGTCTCTGTCATTGTCATGAGGTTGTCTGAATCACCTGACAGTGTGAAAAAAATGGGAAGCGTAGGATCACAGAAAGAGCTGTCCAGTTTCAAGCGCTCTAACA | 120 | 45.83 |
| chr7 | 143622266 | 143622386 | Target=1;ProbeIdx=6673;Target #1 | 1 | + | good | GGGCAGAGTGCACAGTAGGGCTTGGCAGCCCTTTCTGCACTTGTTCTAATGCCTCTTTGTTTGGTCCCCTCAGAGACGCCCCTGACTCTGGCTGCCCAGCTTGATGATTCCATGGAGGTC | 120 | 56.67 |
| chr7 | 143622386 | 143622506 | Target=1;ProbeIdx=6685;Target #1 | 1 | + | good | ATCAAAGCTCTCAGAAACGGGGGTGCACATTTGGACTTCCGCTCCAGGGATGGGATGACTGCCCTTCACAAGGCTGCCCGCATGAGGAACCAGGTTGCCCTGAAGGTATGTCCTGAGCCT | 120 | 56.67 |
| chr7 | 143622506 | 143622626 | Target=1;ProbeIdx=6697;Target #1 | 1 | + | good | GCCTTGTGTCAGCAGCAGGGCCCTTTATCTGGTCCCTGAGCTATCGTCATGATGGTCTTCTTTCTAGGGGACCATGAATCTCTTCCCACTGGAGCTCAGAAGGGCAGCTGAGTGTCAGGC | 120 | 55 |
| chr7 | 143622626 | 143622746 | Target=1;ProbeIdx=6709;Target #1 | 1 | + | good | TCTCCTGCCCCCTCTGTGCTCTAGCTGCTTCTAGAACTACTCACCTTAAGCTACCTTGATCCTTCTAGTCCACAGAAGGGCTCATGTTTCCAATGCATGAGTTGGTGCTAAACCCTGTCG | 120 | 50 |
| chr7 | 143622746 | 143622866 | Target=1;ProbeIdx=6721;Target #1 | 1 | + | good | TGGTCCCATAGAGGAGGCCTCAAATCCAGTAGAAGTACTGACAGCATGGTTGCCCCCTGTATTGCAACTCTGGGAGCCAGACGGCTGTACCCACACCCATCTTCTCCTGAACTTCCATGT | 120 | 53.33 |
| chr7 | 143622866 | 143622986 | Target=1;ProbeIdx=6733;Target #1 | 1 | + | good | GGCATATAGTACACCAGTCTTCCCCGTGGCCATCTTCCCCACCCCCTTTCCCATGCCTGCTATTCAGGTCCATTCCCCATACATACCTGTCTTTCGCACAGCCATCTCCCCCCACCCAC | 120 | 57.5 |
| chr7 | 143622986 | 143623106 | Target=1;ProbeIdx=6745;Target #1 | 1 | + | good | TTTTCCCATGCCTGTCTTCTCAGGCCCATCTTCTTTGTCCTCCCCACACCCTCCTCCCCGCACAGTCCTCCCCATGCCCATCTTTTCCCCACTAACAGGCAGCATTCAGTCTCTTCCTTT | 120 | 55 |
| chr7 | 143623106 | 143623226 | Target=1;ProbeIdx=6757;Target #1 | 1 | + | good | CTTCCTTTCTCCCACATCTTTGCTCCTTGCCCTTGTCTGGCAGCACATGCAGCCTTGTCCAGAGAGAAGGTGGCTCCTGATGAGGAGGCCAGGTCCTCTGGCTGCATCTTGATCCAAGAT | 120 | 55 |
| chr7 | 143623226 | 143623346 | Target=1;ProbeIdx=6769;Target #1 | 1 | + | good | GGAGGTTTTCCTCTCATTCCTGCCATCCTTACCTCCCTCAACTGCACCTTCCTACATGTTCCTGCCCCTTCCCCAGGGCTCACTACCCCTGACTCACTGGGGTCCCGCTGTCCTGTGTGT | 120 | 58.33 |
| chr7 | 143623346 | 143623466 | Target=1;ProbeIdx=6781;Target #1 | 1 | + | good | AAGTTCAAATGTATGGCTCTCTGTAGCTGTCTTCTGGGATACAGGATTCTAGGTGCCAGCAGGGACCTATCAAGGAGCCACAGCTGCCAACTGTCTCCCACCTGGGTTCAGGTGCTGTTC | 120 | 52.5 |
| chr7 | 143623466 | 143623586 | Target=1;ProbeIdx=6793;Target #1 | 1 | + | good | CTGCATCCCCAAATATCACCTGTCACCTTGATTGGCAGTTAGGACTCTCCAGTTTTGTGTGTGTCAGGGGAAGGCAATCAGGTGTCCAGGCATGTCTCCCGCCTGTGGGAAAGCAGTCTAC | 120 | 52.5 |
| chr7 | 143623586 | 143623706 | Target=1;ProbeIdx=6805;Target #1 | 1 | + | good | ATGTGTATTCATTCCGTGTACGAGACCAGTGACAACTTTAAATTCTTCTCCTAGACCCTCCTGGAGCTTGGCGCCTCCCCAGATTACAAGGACAGCTACGGGCTCACACCGCTGTATCAC | 120 | 50.83 |
| chr7 | 143623706 | 143623826 | Target=1;ProbeIdx=6817;Target #1 | 1 | + | good | ACAGCCATTGTGGGGGGTGACCCTTACTGCTGTGAGCTCCTTCTCCACGAGCATGCCTCCGTGTGCTGCAAGGATGAGAATGGCTGGCACGAGATCCACCAGGTATGGCTCTGCTCGCCA | 120 | 59.17 |
| chr7 | 143623826 | 143623946 | Target=1;ProbeIdx=6829;Target #1 | 1 | + | good | TGCCCACTGCTGAGATCCTAGCCCTGCTGACACCAGACCCAGCCCAAAGAGCACCTGAGTGCCGGGCCCTGAAAAGTTTACATTAATCATCAGCATATACATGCTGACACATTTATGGCTA | 120 | 50.83 |
| chr7 | 143623946 | 143624066 | Target=1;ProbeIdx=6841;Target #1 | 1 | + | good | TTTCCCACTGTGCAGGAGAAATTCAGTGGCCACGTGAGGCTGCTGTGTATCTACAACAACCATTATTTAGAGTAGGGATGGGGCTGAAGGGGTGTATCCCTGTTCATGATATCTAGGCCC | 120 | 49.17 |
| chr7 | 143624066 | 143624186 | Target=1;ProbeIdx=6853;Target #1 | 1 | + | good | TGACATTCTAGGAACATGGCAAAGAAGGTACTAGGACCCAGGATGAAGCCTGATAGTTTTTCACAAGGCCTGAAAATTTCAGACATTATCTTCTAATCATGAGACTGGGAGATATGTCAC | 120 | 40.83 |
| chr7 | 143624186 | 143624306 | Target=1;ProbeIdx=6865;Target #1 | 1 | + | good | CTATAACCTCGTAACCATGAAACTGGAGGGTGTGTCACCTATGACCTTGTAACCATAGGGCCAGGAAGTGTGATAACTAGGACCCTGTCAAGAAACGAAATTATCCTGATGGTTCTAGAA | 120 | 44.17 |
| chr7 | 143624306 | 143624426 | Target=1;ProbeIdx=6877;Target #1 | 1 | + | good | AAGCTGTCTCTCATGAGTCAGGTGCAGGAAGAAAGGGTTCAAGTTCAAACTGAGCCCTGTATACCAGCCAAGCAGCTGGGGGAGCCTCTTACCAGGCTGCTGCCTCCCTGTGGACTTCAC | 120 | 55 |
| chr7 | 143624426 | 143624546 | Target=1;ProbeIdx=6889;Target #1 | 1 | + | good | CCTGGGGGCCTCATCACTGTGAAAAACAACTCTTTAAAACATGATTTCTGATCTCTCTGTGCCTCCATATACATTCAAAGTCCAGCCATCTCTGAGGACCTTCAGGGGTCCCATT | 120 | 45 |
| chr7 | 143624546 | 143624666 | Target=1;ProbeIdx=6901;Target #1 | 1 | + | good | CCAGGACTTCCCAGTGTTACCAAGATTCGAGACGCTCCAATCCCTTCTGTAATACTGTGTGGCATTTATTGCACACATTTACCAAAGCCTGTTCAGCCATCTTCAGGTTACTCTCTTGAT | 120 | 45 |
| chr7 | 143624666 | 143624786 | Target=1;ProbeIdx=6913;Target #1 | 1 | + | good | CAGGCATGATATACAGTAACCTGAATGATTTAAGTTTTTTGAGAGTGATGGTTCTACCATGTTTAAAGACAAGAAGAAAAAGAAGTCTGTGTGTGTTTAGCACAGATGCAACAATTCCCC | 120 | 36.67 |
| chr7 | 143624906 | 143625026 | Target=1;ProbeIdx=6937;Target #1 | 1 | + | good | AAGTATTTCTTCTCGCCATGGCTGTAGACTCTGTAGGTGACAGATTCACAGATACAGAGCGCAGCCAGCGTACACATTTCTACCCAAGCTGTTCTGGCTCTTTGGAGAACCCTAACGTTT | 120 | 47.5 |
| chr7 | 143625026 | 143625146 | Target=1;ProbeIdx=6949;Target #1 | 1 | + | good | CCAGCCCTGAGAACTGGGGTAGCATCTTGCCCACCTGCTTCCCTCTTTCCTCTCAATCCCTGCTGCTTCCCCTTCCTTCCCTGAACCAAGGTTGTACCCAGCCACCAGGAGAGCCAGTCC | 120 | 59.17 |
| chr7 | 143625146 | 143625266 | Target=1;ProbeIdx=6961;Target #1 | 1 | + | good | TTCCTGTGGTGTGGGGCAGCGTTGAGAGTAGCTTCCCAGTTAGTTGTTCAGCTGTCTCCCTGGTCTGGCCCTCACTCGTGCACGGTACCTGCTTGTTTGTATCTGCAGAGGCCTCATGGG | 120 | 56.67 |
| chr7 | 143625266 | 143625386 | Target=1;ProbeIdx=6973;Target #1 | 1 | + | good | GCTCCTGGCCTGTGGTCCCTCCCCCCACCCCATCTTCCCAGCACTGGCTGGGGACTCCACTGCCCATCTGTTATTCACACAGCCGTGAATGTTGCTGACCGATGGTCAGTGAAACCTGCC | 120 | 60.83 |
| chr7 | 143625386 | 143625506 | Target=1;ProbeIdx=6985;Target #1 | 1 | + | good | ACCCTCTCAGTGTTCTGACGCCAGGATGAAGTGGTGGACAGACACCCTGATCCAGGAGGGAAACAAGCTAGATGGGAAACAAACAGATATTTAAAAGATCACAAGTGTGTGTGCACACAC | 120 | 47.5 |
| chr7 | 143625506 | 143625626 | Target=1;ProbeIdx=6997;Target #1 | 1 | + | good | ATATGTGGGCATTTAGGAAGCAGGTATATGTACTCTGTGTCCCTATATGGTAGTAGTTTCTGGGCTCATATAGTGACAACAGGCACTGGTGTCCTGGAACAGAAAATGGGTGACTGTGGA | 120 | 45 |
| chr7 | 143625626 | 143625746 | Target=1;ProbeIdx=7009;Target #1 | 1 | + | good | TCTTCTACTCTGCATATTTATAGCTTGTTCACCCATGTCTGTCTGTCTGTCTGTCTCCCTGCCTCACCTCCAGCCTTCTTGCCTGTCCCTTCACCAGGGCAGCCCCTCTTCACAGTCCAG | 120 | 53.33 |
| chr7 | 143625746 | 143625866 | Target=1;ProbeIdx=7021;Target #1 | 1 | + | good | TGACACCACACTTACAGGTCTGGCCCTTTTCATGTAAACCATGTGACCCTGGTTCTTACCTTGGACAGCTCTTGGCTGTGGGACCGTGGCTGTCTCTGATTCCTGGGCTCAACAGAAGCC | 120 | 53.33 |
| chr7 | 143625866 | 143625986 | Target=1;ProbeIdx=7033;Target #1 | 1 | + | good | AAGTTCCGTCAAGGGTCTGGATTGAGAACTTTACCACTTCAGTGTCATCTAGACCTGTCAGTCCCTCAGCCTCTCAGCCTCTCAGCCCACAGACCATGGTTCTATGGAGACCTTGGAAGA | 120 | 50.83 |
| chr7 | 143625986 | 143626106 | Target=1;ProbeIdx=7045;Target #1 | 1 | + | good | CTTTACACTGTTGACTCTGTAGCCAGTCTCTTGCCTTTTCTTGCAGGGTTTGAACTGCCTCTAGTGGTCAAGTTTCAATCTAGCCTGGGAACCTTGCTTTTTGTCTCTCTTAGCTGCA | 120 | 45.83 |
| chr7 | 143626106 | 143626226 | Target=1;ProbeIdx=7057;Target #1 | 1 | + | good | GGTTGCTGTGTTTACTAAACAGGGTGACCCTGGGCAGACCTGGCAGGAGTAGGAGGCACAGCCAACACCTTCAAGGTTATCCTTCCAGCCACAGTTCACACTTACCCAGCTAAACACTGC | 120 | 53.33 |
| chr7 | 143626226 | 143626346 | Target=1;ProbeIdx=7069;Target #1 | 1 | + | good | TCTACCCACTACCTACATATACCCTCAAAATAACCCCAGCAGAGTGTTTCCTGTTCCTGTTAACAGGTGAATGAACAGGCCCAGAGAGGGTAACTAACTTGGCTTAAGGTCACAGAGTGA | 120 | 45.83 |
| chr7 | 143626346 | 143626466 | Target=1;ProbeIdx=7081;Target #1 | 1 | + | good | GCAGTTGGCTGAATGAAATTGATCCCTACCTTGACTTATCTCATTGCCTTCTCCTTCATCTTACCCGCAGACCTCTCAAGAATAAATCATTCTTAACATAACTGTATTTGCGGAGGAAGA | 120 | 40.83 |
| chr7 | 143626466 | 143626586 | Target=1;ProbeIdx=7093;Target #1 | 1 | + | good | AGTGGCATGGGATGGGTGGGTTTTGTAGGGGTCCACCTAGACTAGTTCTGGCCCTCTTAGAACAGTCCCAGAGCCCCAGATTTCTGGGCTCAGGATTTAAAGTTGGTTCCTCGGTGGGTA | 120 | 53.33 |
| chr7 | 143626586 | 143626706 | Target=1;ProbeIdx=7105;Target #1 | 1 | + | good | GCACCACCATCTGATGTAGCATCCTTCTATGGAGATGTCTGCTTTCTCCTGCTGGCCCGTGGAGTCACCTTCTGGTAGCACACATGGCAAGTGTCTGGCCAGCTCCCTGGGGTTCTCAGA | 120 | 55.83 |
| chr7 | 143626706 | 143626826 | Target=1;ProbeIdx=7117;Target #1 | 1 | + | good | GCTGTCCCTTGTATGTCTGTTCTGAAGCCAGGTTTACATCACAGGACATTCTGCTGAGGTGTGCACAGGCAGGTATTGCTGCTGGCTATATCTAGAGCTCTTGTTTGAATTCACATCTG | 120 | 46.67 |
| chr7 | 143626826 | 143626946 | Target=1;ProbeIdx=7129;Target #1 | 1 | + | good | CCCACCACTGTGGGACCTGGGAGTTTTCAGTTCTGTTCCTCCCCCCTGCCCCAGACACACATACAGCTATAGTGCAGCCTGCAGTGTATTGGGTCCAACACAGACAAGCTTCTCAGGGAG | 120 | 55.83 |
| chr7 | 143627546 | 143627666 | Target=1;ProbeIdx=7201;Target #1 | 1 | + | good | AGGAGAGCCACCTGGCTAAGCGGCTAGGTCATGATGGCTCTATGGACTCGACTTACTCCCATGCTCTATGAGCCCACATCATCAGTGTGGGGTGACCTCCTCATACCACCCATGGAATG | 120 | 55 |
| chr7 | 143627666 | 143627786 | Target=1;ProbeIdx=7213;Target #1 | 1 | + | good | ATTTTTGTCCAGTATGGGCATACTGAGATGGAAATCCCCTCATACTTCCTAGGTGGCCAGGGCAAGGACAGTGTCTCCTCAGGTATTTCAGAGCTCTTGAAAAGGCTGAATGTAAGAGCA | 120 | 46.67 |
| chr7 | 143627786 | 143627906 | Target=1;ProbeIdx=7225;Target #1 | 1 | + | good | CCTGCGATATGCTTTCTACCATGCCTGTGTTCTAGAGCTTAGTTTGTGTTCTCAGCATACACATGACCCTCTCCACAGTCATCACCCTGCCTGTGACCTCACATGTCTGACTTCTGTGTG | 120 | 49.17 |
| chr7 | 143627906 | 143628026 | Target=1;ProbeIdx=7237;Target #1 | 1 | + | good | AGGTCCTCTGGAAGAACAACAAATGCTCTTAATTGCTGAGGCATCTAGCCCCATTGGCTAGTCTACTTTTGCTCACTCCAGTGATCTCATTCTCACTCACCGCCTTCCTTTTAGACCAAA | 120 | 45.83 |
| chr7 | 143628386 | 143628506 | Target=1;ProbeIdx=7285;Target #1 | 1 | + | good | GAGTCCAAAGAGTGACATTGTAACACCCTGGTTAGAACTCTGCTGGTGTCCAGGAGCTTTTCAGCCCCTTCCTTCTGCAGTATGACTTTGGAGATTTACCACAAGCGCCCAGCAGGTGGC | 120 | 51.67 |
| chr7 | 143628506 | 143628626 | Target=1;ProbeIdx=7297;Target #1 | 1 | + | good | AGTGCAGTGCTCAGAGGAAGGTTCCTTGTTTCCTCATTCATACCACATTTGTGGTTTGGCAAGTGGCAGATAGGCTGCTGGGTACAGCTGGGGTGAACCCAGCACTCTGAGGGCACCTTA | 120 | 52.5 |
| chr7 | 143628626 | 143628746 | Target=1;ProbeIdx=7309;Target #1 | 1 | + | good | TTCTAAATACTCACGTCCAGTCCTCCCCAAGGCTGTCTGAGACAATAGATTTGGGTGACCCCCAAGGAAAAAAAATTTTTTTTGAAGGCAGGGAAGTGGGAAGACAGGGTCTGAGTCAGT | 120 | 45.83 |
| chr7 | 143628746 | 143628866 | Target=1;ProbeIdx=7321;Target #1 | 1 | + | good | GTCGTGTCCTGAATTTGGTTTGCTCCCTACCTGTGTCCTGCTTGGAAGCTGTGTCCGGTGGGCTGTGCAGGGGACAGGGACTGGCTTTTCCCTTTGGTAACATTCAGGTTCCTTGCTTTA | 120 | 53.33 |
| chr7 | 143628866 | 143628986 | Target=1;ProbeIdx=7333;Target #1 | 1 | + | good | GAACTGTCTTCAGGGAGTTTGCAAGGTAGGGCATCATACAGCAGATGCCTCCTGGTGGGCTTCACTCCAGAGCCCGCTCATTTTTACCAACTGCCCCAGCAATGTCAGAGGGTACATCCC | 120 | 54.17 |
| chr7 | 143628986 | 143629106 | Target=1;ProbeIdx=7345;Target #1 | 1 | + | good | GAGTCTACCCTGGACCCCTTTTGTGGGCAAGCCTGCTGTGCCCTCTGGTCTCTGGCTTCCCAGGGTTTCCTTCGGACTCGGGGACAGTTGTTTTCTAGATTGTCTTTCTGCCTGCGCT | 120 | 56.67 |
| chr7 | 143629106 | 143629226 | Target=1;ProbeIdx=7357;Target #1 | 1 | + | good | CCTCAGCCGTGTCTTGAGGACACACAGGTCTGTATTTGTCTCCAAGCTGCTCCCACTGTGCTCTTGGCCTTGTGTCCTGTCAGGTGGCCTGTGGTGACCACTTGACTTGACTAGGGCGTC | 120 | 56.67 |
| chr7 | 143629226 | 143629346 | Target=1;ProbeIdx=7369;Target #1 | 1 | + | good | CCTGTGAATACTACTTCTTCCCTGTTGCAGTGAGCTAGTGATCCCCAAGGCTGCCAGTGCATGCACGATTTCACCTGCAGCTGTGGCATTGCGATTTCTTGTCTGTATTGATTTTTGCCT | 120 | 48.33 |
| chr7 | 143629346 | 143629466 | Target=1;ProbeIdx=7381;Target #1 | 1 | + | good | TTGCCCAGTAGAGATTCTCCTCCATTTTTGCCTCTAGCTGGGTATCCATCAGACAGAAAAGCTACTTTTGTTTATCTTTTATTTATAGCAAGGAAGACTCATAGGTTTCTCCTTTCTGGC | 120 | 40 |
| chr7 | 143629466 | 143629586 | Target=1;ProbeIdx=7393;Target #1 | 1 | + | good | AGATATTATTGGATAACACCTGGTGTTAGCATCTGTCTGGATCTGTGGGTGCCCCTTTGGAATGGATCTGAACATTGTTTTAGTTGTTTTTCTTATTGCTTGACAAAATGCTGGACAGAA | 120 | 39.17 |
| chr7 | 143629586 | 143629706 | Target=1;ProbeIdx=7405;Target #1 | 1 | + | good | CCCTCACAGGCATGCCCAGAGGGGTGATTCCATGGTGATTGTAAATACCCATAAGTCAATATGAACCATACCAGGCATGTTCCTGTCATTTTTTTTTCTGACACATGGTTAACTCAGGC | 120 | 44.17 |
| chr7 | 143629946 | 143630066 | Target=1;ProbeIdx=7441;Target #1 | 1 | + | good | GTTTGTTAGCATTCCTTATAGGGGTGTGGGGTGGGATAGTTATAGGAGCAGGGATGACTGTAAGTCCACCCAAACACACTGATGCCTCATGACAGCTAGAACCCTAGAGCTCCCAGCACA | 120 | 50 |
| chr7 | 143630186 | 143630306 | Target=1;ProbeIdx=7465;Target #1 | 1 | + | good | GCCATGGAATGACCATTTCTTCAAGACCTTACCTCTGTCTAGTGTAGCATGGTGCATGGCTGCTGATAACATGTTTTCCTTAGCTAGAGTTAGAGACACATGCATATGTGTTCTCTCTCT | 120 | 43.33 |
| chr7 | 143630546 | 143630666 | Target=1;ProbeIdx=7501;Target #1 | 1 | + | good | TGAGAAGGTACAGCCAGATATGGAGACTCACACCTGTGATCTCACATTTAAGAAGCTGAGGCCAGGGGTTGACTTGAGTTCTAAGTCAGCCTAGGCCACAGAGCTACAGAGTAAAGCC | 120 | 48.33 |
| chr7 | 143630906 | 143631026 | Target=1;ProbeIdx=7537;Target #1 | 1 | + | good | TGCACCCCTCTCATACTCTGAACCACCCCTGTCATCTATGACCCTGCACAGTGGACATGCTGTACAAATACTTGTTCTGTTCTTTTGTTTGGGAAACAATGACAAGAAAAAGATGTCTGT | 120 | 43.33 |
| chr7 | 143631506 | 143631626 | Target=1;ProbeIdx=7597;Target #1 | 1 | + | good | GTACCATCACTACAAATATGGTGTTTATCCTGAATATTTCTGATTGATCACTGATGGTCTCATAGCTCAGAGGGACAGCTGTCTCTGCTTCCATCTTTGTTTCTCTGTATATATGTCAAA | 120 | 38.33 |
| chr7 | 143631626 | 143631746 | Target=1;ProbeIdx=7609;Target #1 | 1 | + | good | CCTTGGCTTTTCTTTAAATTGAATCTTAGCTAATTTTTCTTTAAATTGAATTTAGTCTCTTCAGCTTAGAGAAACCACCCTGAGTTCTTCACCTGAGCACCTGTTTGTTGTTAAGCTCTGGCT | 120 | 36.67 |
| chr7 | 143632706 | 143632826 | Target=1;ProbeIdx=7717;Target #1 | 1 | + | good | GCAGATGGTCTGCTCAGCCATCCAGTGTCTGATCCCTGGGCTCTCCCTACCCACACTCCTGCATCTCCACCAGCAGGGATTAAGAAGGGAAGAGGCTAAAGGCCTAATCTATGTGCCTGG | 120 | 55.83 |

| | | | | | | | | | |
|---|---|---|---|---|---|---|---|---|---|
| chr7 | 143632826 | 143632946 | Target=1;ProbeIdx=7729;Target #1 | 1 | + | good | CTCAGTGTCCATCACATTTTGACCCTCTCTGAGAAAGTGGTGGCAGACTCCTGGGAATGCAACTTCAACCTTCATGAATTCAAATTATGTGGGGGAGGGGGTAATGAGTCTGGATTAATT | 120 | 45 |
| chr7 | 143633186 | 143633306 | Target=1;ProbeIdx=7765;Target #1 | 1 | + | good | GCTCAGTGAGCCTGTAGTAACCAGTCCTCTTAACTGACTAGAGAGCAATTAAATTAAGCTGCTACAAAATGCTGTCTCCACAAAGCCTTCAAGAAGAGCCGATTAGCCAAATTGTATCTT | 120 | 41.67 |
| chr7 | 143633306 | 143633426 | Target=1;ProbeIdx=7777;Target #1 | 1 | + | good | CAGTTTAAGGAGCAGAATTCCCATCTCAAACTGGTTGAAGCAAGCGGAGGATTAGGCCTTATTTTAATGTCTCTGAAAAGCCCCAGGGGTAGAGTAGCTCAGAGGAGTCCCAGCAGGGT | 120 | 49.17 |
| chr7 | 143633426 | 143633546 | Target=1;ProbeIdx=7789;Target #1 | 1 | + | good | CTCCTCTCCCCTCCCCTCCCCTTTCTGTGGTGTGCTAGCCCCAGCAGCTGTAAAGAATGTTACTTTTCCAGAAGTCCTGAGCCATGCTGGCCTTTGCTCCCTGAGTCTGCTGGGCTCCAA | 120 | 56.67 |
| chr7 | 143633666 | 143633786 | Target=1;ProbeIdx=7813;Target #1 | 1 | + | good | GGTCCCTCCCTGAGCCTTGCCTGCAATGATGGAGCTGCCAGCTGCATAGCTGTGACCTGTCCTTCTGCAGGATCAGGTGTACCTGGGCCATCTGAACCAACAGGAACTGGGCAGGCAATC | 120 | 58.33 |
| chr7 | 143633786 | 143633906 | Target=1;ProbeIdx=7825;Target #1 | 1 | + | good | TCAGGAATGAAGCAGGTGCTTAAGTGAAGTGGGGCCAGCAGGCAGGCAGGTAGGAGAGCGGGATCTCCTGGCAGAGCCATCTAGTCAACAATGAGAGAGCATCACCGGGGGCATGGCACG | 120 | 58.33 |
| chr7 | 143633906 | 143634026 | Target=1;ProbeIdx=7837;Target #1 | 1 | + | good | ATAAATGGATGTTTCTTGGATGCTTTTAACAGCGATAGTCTACTTCTCCCTCTTTGCAAAGTGACTCTTACTCTCACTGACCCTGCTGAGGTAAAGTGTCCTCTGCACACTGATCAGAGG | 120 | 44.17 |
| chr7 | 143634026 | 143634146 | Target=1;ProbeIdx=7849;Target #1 | 1 | + | good | CATAGTCGTTGACCCAGGACCAATGTCATCATCTAGGAGAGTTTTCTTTCTTTGTCCTTGGTGTGGCAGACCTGCCTCCTCCTGTGAGGTCACTGCCTGGCGGAGGTGCCCAGTCACCTG | 120 | 55 |
| chr7 | 143634146 | 143634266 | Target=1;ProbeIdx=7861;Target #1 | 1 | + | good | TGCATGGACCTTCCTGCCCCCTTTGAACTGGGATGGGGTGCTGTGCCTATTTTTAGGGCCCAGTGTCATGTCATCTTTGATGTACTTGTCCTCTTAACTCAAGAGAGGAACACTGCTTCC | 120 | 50 |
| chr7 | 143634266 | 143634386 | Target=1;ProbeIdx=7873;Target #1 | 1 | + | good | ACTTAATGACAAGGACATCCATAAGCTTGAAAGTGAAGGCCAAGCTAATGGATGGTCTGGGACCCAGTCCAAGTGTGTTGGCTTTCAAACTACCTGGGACATAGGCACTGTGTCAACCCC | 120 | 48.33 |
| chr7 | 143634386 | 143634506 | Target=1;ProbeIdx=7885;Target #1 | 1 | + | good | TAGAAGGACCTGAAAGGAGCAGATCACATAGTGGGGCTGTCCTACCCTGGGTCCCACCTCCTGTGGAAGGTTGGATTTCATTTAATGCAGTGGAAGCTGGGTTTGCATGTGGAAGGACA | 120 | 51.67 |
| chr7 | 143634506 | 143634626 | Target=1;ProbeIdx=7897;Target #1 | 1 | + | good | AGTGTAGGCCTCTTTGTACACACAGGTATAGGGCTCAGAGCATCTCCATGCCCATACATAACAGCATCCCACCTGGGAGCATGGACCAGTAATACAGAGCTGAGTATCCTGATTGGTATG | 120 | 49.17 |
| chr7 | 143634626 | 143634746 | Target=1;ProbeIdx=7909;Target #1 | 1 | + | good | TCACAGAGTTCAGCACCCATATCAGGGAAAGATCTATTTGTCTTTTGAACAATGTGTCTTTTTGGTTCTGTGTGGAAAGAACCCTGGGGCCCCTTCCAGACCCCTCGTCCAAGATCAGAG | 120 | 47.5 |
| chr7 | 143634746 | 143634866 | Target=1;ProbeIdx=7921;Target #1 | 1 | + | good | TACAGGTTTCCAACTAGTCAGGGGTTCCCTGTTTCCACTCTTAGCCAGTCCTTGGTGTGTGATGTCTGTGGGCCTAAGTCTACAAAGCTAGGTGGGGTGAAGTCTTCTAAGAGCTCTGTC | 120 | 50 |
| chr7 | 143634866 | 143634986 | Target=1;ProbeIdx=7933;Target #1 | 1 | + | good | ATGCTTTTAAATGAGTGGTAGGATGCTGGGCATAGGCCTGTTGCCTGGGCAACCTTAGGTACCAAGTCTCACTCAACATCTGTGTCTTAGGCCCAGCCAGTAGAGGGTGCTGGGGTTTTA | 120 | 50.83 |
| chr7 | 143634986 | 143635106 | Target=1;ProbeIdx=7945;Target #1 | 1 | + | good | AGGGGGCATTGGGGTCTCTGTCTCAGGACTGACCAGCAGGAGCACCAAGGTCTCACTCAGTCTCTTATCTCAGGCCTGCAGGTACGGCCATGTCCAGCATCTGGAACATCTGCTTTTCTA | 120 | 55 |
| chr7 | 143635466 | 143635586 | Target=1;ProbeIdx=7993;Target #1 | 1 | + | good | TGGCGCAGACATGAGCGCACAGAACGCCTCAGGGAACACAGCCCTGCATATCTGCGCCCTCTACAATCAGGTCAGTGGCATGCACATCCCTCACCCTAACCCTGGGGCGCATTGGACACT | 120 | 59.17 |
| chr7 | 143635586 | 143635706 | Target=1;ProbeIdx=8005;Target #1 | 1 | + | good | GGGTGGAGAGACTGGCCAGGGTATCTCTGTGACCCTCACCTGCTGCCTTACTGAGAACCAGAGGCTGGTGAAGCTGTGAGCAGCCCCAGGTCACATTCCAGATCTACAGTCAGCCCTGCA | 120 | 58.33 |
| chr7 | 143635706 | 143635826 | Target=1;ProbeIdx=8017;Target #1 | 1 | + | good | CAGTGCCAGTGACCCAGGCTGCTCATTGTCTAAAAGACACAGGAGAAGCTGGAACCTCCTGTTCCCCTGGCTGAGTGCCAATTTTGCTTGGTCCCAGAGCCTGGCACTTGGGTAACCCTT | 120 | 55 |
| chr7 | 143635826 | 143635946 | Target=1;ProbeIdx=8029;Target #1 | 1 | + | good | CCGCTCTCCTTTTCACTTGTGGTTAAATGTGAATAACAAACATGACTCTCATAAACACGTGGGCACACAGTCAGCACATTCACAACACTGTATGGACATCCCCACTCCC | 120 | 45.83 |
| chr7 | 143635946 | 143636066 | Target=1;ProbeIdx=8041;Target #1 | 1 | + | good | CACCTCCAGAATTTTTCATCTTCCCAGACTGACTCTCTGTCTCCCCAGGAAACACTGGCTCCCGACCCCCCACCCCCACAACCCCTAGTTCCCCATCCTGCTCTTTGCTCTAAGGACCTC | 120 | 56.67 |
| chr7 | 143636066 | 143636186 | Target=1;ProbeIdx=8053;Target #1 | 1 | + | good | TGAAGACTAGGATTTTGCAGGAGATAGCCACCTATGCTAAGCTTCTTCCACTCAGCATCATGTTCTCATGGTCTAACCTGTACTAACGAGTGCCAGACACTCTGCCTTCTGAAGTTTGTT | 120 | 45 |
| chr7 | 143636186 | 143636306 | Target=1;ProbeIdx=8065;Target #1 | 1 | + | good | TCCACACCACTGAGGGACGGCACCTTTGCTTACCTGCTTTCCCCTGGGTGGACATGTGGCTTGTTCTGCCTTTGGCTGTTGTGGTTTCATGTTGCCATGAGGCCTGGGTGTGCAGATGC | 120 | 56.67 |
| chr7 | 143636306 | 143636426 | Target=1;ProbeIdx=8077;Target #1 | 1 | + | good | AGGGTCTAGCCTCAGCTATCTTTTGACCTCCATGGGGACAAACTCCTAAGCAGATTTCAAGGTACATGACACACTGGTGCCCACACCCCTCCCCCAAGCTTCCCTGGCAGACCCTGGTGG | 120 | 55.83 |
| chr7 | 143636426 | 143636546 | Target=1;ProbeIdx=8089;Target #1 | 1 | + | good | GCAGTGTTGTTCCAGAATCCCTATGAGGAACATGGTCCCTGCTCTCTGGAAGGGAGGCACCTTAGTAGCTGGGACTTGGATCCTGGACCATGAGTAGGTTGAGCTGGATGTGGAGGGGAA | 120 | 54.17 |
| chr7 | 143636546 | 143636666 | Target=1;ProbeIdx=8101;Target #1 | 1 | + | good | GTAAAGATGGATGTCCCGGGGTGGGGGTGGGGTGGGGTAGAGCATAGTGCTGCTCAAAGGATGTGGGTGGACTTCAGGGCCAGCTTCAGCAAGCAGGAAAGTCACCAATGTCCTTGGTGC | 120 | 57.5 |
| chr7 | 143636666 | 143636786 | Target=1;ProbeIdx=8113;Target #1 | 1 | + | good | TCAAGGTCAGGTTGACAGGGCAACCCCACTGAAGACGCTGCTTTCTTGTTTCTCTTCTCACTTGGTGAAACCTCAAGTCTGGGGCTAATGACCCACGGACTTGACCCCAGGCAGCCCAGC | 120 | 55 |
| chr7 | 143636786 | 143636906 | Target=1;ProbeIdx=8125;Target #1 | 1 | + | good | AGAGGGACTGGGGAAAGACTCTCCCTCTTGAAGGGCAGAGGGATCTCAGTGGTGAGGCTCTGCTGAGCCTGGGATGTGGTCTGGTGAACACTGAGTTCTCTTCCAGTTTCAGAGCAATA | 120 | 53.33 |
| chr7 | 143636906 | 143637026 | Target=1;ProbeIdx=8137;Target #1 | 1 | + | good | ATCTCATCACACCACTGACAGTGACCCTAGATGACAATTCAAGCCTTAGCTTACATCCAAGTTTGTTACCAGGCCCTAATGGCTGGCCCCAAGTCCACTCTACTTCTAGCCAGTAACTTT | 120 | 46.67 |
| chr7 | 143637026 | 143637146 | Target=1;ProbeIdx=8149;Target #1 | 1 | + | good | CCTGGATCATTCACTCTTGAGAACTCCTACAAACCCCTTTAAGGCCCCACTCACATGGTCCACACCCAGTCAAGCCTTCTGACCATAGAATCTGTGAGTGTCGTTGCTCATGGTTAGGCTG | 120 | 50 |
| chr7 | 143637146 | 143637266 | Target=1;ProbeIdx=8161;Target #1 | 1 | + | good | ATGGAATCGGTGTTTCAGCTGGAGTGTCTTGCAGGGCTCCTGGCTTGCACGCAGAGTATATTCACAGTGCAGATGCCCCTGGGGTCCACCGTGGAGCTGGAATGGGGGAACCCTTGTTAA | 120 | 55.83 |
| chr7 | 143637266 | 143637386 | Target=1;ProbeIdx=8173;Target #1 | 1 | + | good | GTGTGGGAATATTCAGCTGCCTTAGAAAACACATCATGAGGCTTGGCTGGGGCTAGGAAAACATCTGCACCAACATCTGCTTTGTAATGATTGCCTGCTATGGGAAAACAAGGCTTAATT | 120 | 44.17 |
| chr7 | 143637386 | 143637506 | Target=1;ProbeIdx=8185;Target #1 | 1 | + | good | TCACTCAGTATATTTCAAGCATATTTTAATCATCAAAAAATATAGGAAATCACAGTTTCTGTCACTTAAAATAATTGTGGGCATGATGATGGTCTTTGAAAAGGAGGCTGCAGAGTTTTA | 120 | 31.67 |
| chr7 | 143637506 | 143637626 | Target=1;ProbeIdx=8197;Target #1 | 1 | + | good | TTAGGCAGCCTCTGTTCTTGACTATATAGGAGACTGAACGTGCCGGGGTTACTCCACATACCCCAGAGCTGTCTTCATCCCTAGGAGGATCCCTAGATGGCCAGAGCTCCCAGGAAAAC | 120 | 52.5 |
| chr7 | 143637626 | 143637746 | Target=1;ProbeIdx=8209;Target #1 | 1 | + | good | CTTCTCCCCATATAATGTCAGTTAGAATAGAGATAGCTGACATTACAGGGCCCATTGGTAAACATTTGATGCTTTGGTCACTGCCTGGACCCTCTGGTCACATGTCAGCTGAGCAGCCAT | 120 | 46.67 |
| chr7 | 143637746 | 143637866 | Target=1;ProbeIdx=8221;Target #1 | 1 | + | good | GGTATAGCGGTCAGAGTGACAGGTCATGGAAAGTGGCCATGCTGCAAGAAAGTTATTTGTTCAGTAAACAGGGAGGAGGGGTGGGGTTGGAGGAAGGGGGTGATCTGGCCCGCAGGCTA | 120 | 54.17 |
| chr7 | 143637866 | 143637986 | Target=1;ProbeIdx=8233;Target #1 | 1 | + | good | TTAGCCATCTTTGGAGTAGAAGGCTACACTATCTTTTCCTCGCACCTCATTTATAAATTTGTGCCTCCCCAGGACTGGGTGACTTCCATCCAACAGGCCAGTGAGTCTGAGCAAGGGTGG | 120 | 49.17 |
| chr7 | 143637986 | 143638106 | Target=1;ProbeIdx=8245;Target #1 | 1 | + | good | GGAGAGCTGTCCACTCCACAAAGCTGCCCTCTCATGAGGGCAACCTGCTGCCACCCAGCTTCAGACAGTATGCCTGTTCTCTGAAGCCTGGGACAAATTGAGCCAAGGTGACTAATCTTG | 120 | 54.17 |
| chr7 | 143638106 | 143638226 | Target=1;ProbeIdx=8257;Target #1 | 1 | + | good | TGGCACCTCCACCGGATACTGTGTTGGTTTATAGGCTCTGGAGTTACTGACCACACAGCCCCTTGACACCATGACAAGTTGGGATATCAGTGAGTGTACCTGAATCTGTCCTCCCCTGGA | 120 | 51.67 |
| chr7 | 143638226 | 143638346 | Target=1;ProbeIdx=8269;Target #1 | 1 | + | good | TGGTAGGGGGTCTTCAGTCTGTCTCAGTCCATAGCTGTGTCTGAATTGTGTGAGTGAAGTTGCCTTAGCACTGCTCTGAGGGTCACAGTCTTGCATAGCTTCCTCATGGGAGCTGTCTCA | 120 | 50.83 |
| chr7 | 143638346 | 143638466 | Target=1;ProbeIdx=8281;Target #1 | 1 | + | good | AGAGTGAAGGATAAGTCAGCCTTTGGCACTGCACATAGAGAGTCCTCAGTCTTTTCTCCCCTTATCACCGCCTTGCTGGCATCTGTGCCCATGGCACTTTGTCTTCAGATATGTGTGGCT | 120 | 50 |
| chr7 | 143638466 | 143638586 | Target=1;ProbeIdx=8293;Target #1 | 1 | + | good | TCACAGGCTCGGTTGTTGATAACAGAGACCATTGCATAGGAAGTGGTGTCATACTGGGTATGCAGAGGATGTACCCCAGGGAGGGAGAGAATTCAGAAGGGCAAGATTCAGGGCAAGGAG | 120 | 50.83 |
| chr7 | 143638586 | 143638706 | Target=1;ProbeIdx=8305;Target #1 | 1 | + | good | AGACATGGCCTCAGGGACATGGGATAGTGATGAGGAGGCCTGGCCCACTGGGCTCTGATGGGCATAAACTGGGGGATGTGCAATTAGGTGACCTGAATGCACTTTGGATCTAGGGTTGCT | 120 | 54.17 |
| chr7 | 143638706 | 143638826 | Target=1;ProbeIdx=8317;Target #1 | 1 | + | good | AAGAGGCTGAAGTAGAAATCAAACATCTGTGATACCTGGCATGGGGTGAGGACTCAGCAACTCGAAGCTGTAATGGTGGTGTTGAAGATAGTTGAAGATGATGATGTTCATGGAGATTAT | 120 | 42.5 |
| chr7 | 143638826 | 143638946 | Target=1;ProbeIdx=8329;Target #1 | 1 | + | good | TGATATTAAACTTGGTGATGATGATGGTGATGGTGTGGTGATACAACTGATAATGATCATCCTTAATGAGCTACTCCTCAGATCTGGAAGCCCCAGGTGTGTAGAATGCTATTCATGCTT | 120 | 40.83 |
| chr7 | 143639186 | 143639306 | Target=1;ProbeIdx=8365;Target #1 | 1 | + | good | TATCAATTTCATCAGGGCACTTTTAATAAAACACACTGTGATCAGAAGATTCTCTTTCCTCTCATGCAATCAACTAATTAGGACTCTGATTGTAAGCAGCAAATTGATTCTAATCTTTGG | 120 | 34.17 |
| chr7 | 143639306 | 143639426 | Target=1;ProbeIdx=8377;Target #1 | 1 | + | good | GCTCTCTCGCCTCATGCTGGCTTCCCAAGAGCTCCAGAGGGAGAAAGCACCACCCTGGGTAATGAGAGATCAGAGGAAGCTTAAATATCAAAGGTTACTAGAGCAGCCTGGTTTGTCTAG | 120 | 50.83 |
| chr7 | 143639426 | 143639546 | Target=1;ProbeIdx=8389;Target #1 | 1 | + | good | TGTGATGGGAATGTTTTCAATGTGTATCTCTGTGCTTTGCTGGTTTCTTATGGGGCATGTGTCCACTTCCCTGGAGCCATCTGAGGGCCATCATCCCTATAGGCATTATGGCTCTGGGTC | 120 | 49.17 |
| chr7 | 143639786 | 143639906 | Target=1;ProbeIdx=8425;Target #1 | 1 | + | good | TCAAGTTGACAGTTAAAGCTAACTAAGGTAGATGGCATCCATACAGTCTTTCCGAGAGCCTTGTGGATACCACTGACAGGTATGTGGATCATACTGTTCAGCAGATCCAGGTTAAGAAGT | 120 | 43.33 |
| chr7 | 143640266 | 143640386 | Target=1;ProbeIdx=8473;Target #1 | 1 | + | good | GATTCCACTCCCCTCAGCCTTTGGGGTCTGGATGGAACTCGGGGGGTACAGGTGAAGCAGAAGCCTGGTGACCTACAGCATGTGAAATAAGTGGTCTCAATTACCCAGGTCCTTTTTGGG | 120 | 53.33 |
| chr7 | 143640386 | 143640506 | Target=1;ProbeIdx=8485;Target #1 | 1 | + | good | GGACACACTATTAAAGGTTGTAGAGTTATCCTGGTGATGCCTCACAGTGGTCACACCTGCTCACCTGATGTCTCCACCTGGTGGCCTCTTAGGAGACCCAGTCCCTATGCTCCTCCCTGC | 120 | 54.17 |
| chr7 | 143640506 | 143640626 | Target=1;ProbeIdx=8497;Target #1 | 1 | + | good | TCTCTAACTAAGCATCCTTTTCTCTGGAGTTAGAGGCTGGGATTTAGATGACCTTCCCACCAACTTGCGACCATAGCTTCTCTTCTTGTCATACCAGGGCTCAGACTACTGATCAGCTGC | 120 | 47.5 |
| chr7 | 143640626 | 143640746 | Target=1;ProbeIdx=8509;Target #1 | 1 | + | good | CTCATATCCATATGATTGGCCCTGTTCTCCTTTGAAGGCCACACAAAACAGCTCACCTTAAGCTGACTGCTTCTTGTCCCCCGACTCCCACCCATCCCAAGTTCACAGGGACCGTGACCC | 120 | 53.33 |
| chr7 | 143640746 | 143640866 | Target=1;ProbeIdx=8521;Target #1 | 1 | + | good | CCATGCTTCAATTCTGTGTCCCCAGGGACTGGCGTGGCCATTGGCTAGCCAACTCTCCACAGGCATGGATGACTGAGATTTCTCAGTCTGTCCCTTCACTTGCAATGTGGACAGGATGGT | 120 | 53.33 |
| chr7 | 143640866 | 143640986 | Target=1;ProbeIdx=8533;Target #1 | 1 | + | good | GGGGCCCACAGCTCCTGGAGTCAGAGGCACATAGCCCATAGCTGAGTGCAGAGTGGCCTCAGAATGACTCCCCTCAGGGAAAAGACATATTTTTTTAAGTGTCTTTGAAAATTTGCACCA | 120 | 49.17 |
| chr7 | 143640986 | 143641106 | Target=1;ProbeIdx=8545;Target #1 | 1 | + | good | AGAGAAAAACTGGCAGGGAAATGGCCCTTAGAAGATGAATACAAATTGCTTATGTATTCATACTCCATTGATTTTTTTTTTTAAAGGATGGAAGCCTGCCAGCTGAGACTCGAGCCATCC | 120 | 40 |
| chr7 | 143641106 | 143641226 | Target=1;ProbeIdx=8557;Target #1 | 1 | + | good | AAGGGCGTGCAGGGGAATGGCTAAGCTCTGCATTAATGCCTTGGGGACTTCCTCACCCTCCTGACACTGCAGATTGCCCTGTTTCTCACCAACCCTTCATTTCTCTTCTTATCCAGTTCA | 120 | 50.83 |
| chr7 | 143641226 | 143641346 | Target=1;ProbeIdx=8569;Target #1 | 1 | + | good | AGCAGCTCTCCCACATGCTTGCCCTTCCTTTTTGTGGCATGGATCTCTTGCCACCCCATCAAGGTGCTGTGCCAGGCATGTGAGATGGAGCAGCCGGGTATAAAACACATGCCTGCACAA | 120 | 54.17 |
| chr7 | 143641346 | 143641466 | Target=1;ProbeIdx=8581;Target #1 | 1 | + | good | GTGTGTGCACACAGGTGCATATGCATACATACATATGCACACACGTGTTGCATATTATTAAGCCACAAAACCTCTGGCCCTTTAGTTCTGAAGTAGCTGACTGACTGGAACTGTTAACCA | 120 | 43.33 |
| chr7 | 143641466 | 143641586 | Target=1;ProbeIdx=8593;Target #1 | 1 | + | good | AGAATTTGGGGATCCTGCTAGGGTTGCCCTATGAAATCAGAGAAGTCAGGGCTCCACTCTTGAGGTCCTTAGTTTCATTAATAAAAAAAAAAAAAATGGAAGCTTTTATTAGAGTTCTGAA | 120 | 37.5 |
| chr7 | 143641586 | 143641706 | Target=1;ProbeIdx=8605;Target #1 | 1 | + | good | TCTCTCCTTCCTTGGGGTGTTCTCTCCTTCATAGAAGGGGGATATTTCTTTCTTAGGGGTAGCCCTTCCTTGTCCCCCTCTATAGGGGTTTGTTCCTTAGCCAGACCACTGATCTGCCCAACTGGAACGT | 120 | 50 |
| chr7 | 143642306 | 143642426 | Target=1;ProbeIdx=8677;Target #1 | 1 | + | good | TAGGTCCCCACTCAGGCCTGAGCTGCTGTGTCAGAATGCACTCTGCCTTTGTAGCCTGGCCATCATTGCAAGAATGGGTTGGGCTCTGGGGGGTAGTTCATATGGGTTCTGAAAACAGCC | 120 | 54.17 |
| chr7 | 143642426 | 143642546 | Target=1;ProbeIdx=8689;Target #1 | 1 | + | good | CCTTTCGGGTGAGTAGAAGGGTTTTCTGAGCCCCTTCCCATGTCCCACTGGATGAGTGGGGATGACAATACCAGGGGAGGGTTTATTCCTGGCCTTTTATGAAAATTTCAGTGACAATGT | 120 | 48.33 |
| chr7 | 143642546 | 143642666 | Target=1;ProbeIdx=8701;Target #1 | 1 | + | good | AGGTGCTCTTGAGGGGTTACAGGGGCCAGGGAGGGGTTGCCCAGGCTTCCCCATGGGGCTCTCAGTCTGGAGTTTTGATTGGGAAACAGGGGGCTCTGAGTTGCTTTTGGAGCTGAGGGG | 120 | 60 |
| chr7 | 143642666 | 143642786 | Target=1;ProbeIdx=8713;Target #1 | 1 | + | good | GTATGGTGCAAAGAGCCCTACACAGGTTCAATTCCTGGGGAAACCTAGAGCCAGGCTTCCTGTCTGATTCCTTCTAGCTGTTGTGTGCTGTTCCCAGGTAGGGTCCACCTGCTTCCTCGC | 120 | 54.17 |

| | | | | | | | | | |
|---|---|---|---|---|---|---|---|---|---|
| chr7 | 143642786 | 143642906 | Target=1;ProbeIdx=8725;Target #1 | 1 | + | good | CTGAGCTGACACAGTTTTCTTTTCCTTTGTTCTGTTTCCTTTCCTTTTCTCTCACCCCAACATTCCCTTCACTCAAGCTGTATTGTGATTGGTGTCATTCATTGCCCTAAAGATTTAGGT | 120 | 40.83 |
| chr7 | 143642906 | 143643026 | Target=1;ProbeIdx=8737;Target #1 | 1 | + | good | TGTAGATGACAGGGCCACAATGGGGCAGGTTGAGGCTGGCTGAGGCTTGGGCTCAGGATGCCTGGGGAGCCCCTGGGTCAGGCCAGGGCAGCCCTTCTGTGTCCTCTCATCCACGCAGCC | 120 | 65 |
| chr7 | 143643026 | 143643146 | Target=1;ProbeIdx=8749;Target #1 | 1 | + | good | CTCATCCTCCTTCTCTTGACACAACCTCACAGCCCCCAGCTCCTGTTAAGGGAGCATCAGCATGTGCATACCCTTTTCAGTTCGGGTGGCTGCAGGAGGACCTCCTCCCAGCTCCCAGGG | 120 | 58.33 |
| chr7 | 143643146 | 143643266 | Target=1;ProbeIdx=8761;Target #1 | 1 | + | good | TCCTGATGGTCAATGCATCTGGGGCACAGAGGGAAAACACCTGCTTCTCCGCAGGGTGTAATCACAGACTGAGCCTCTACCCTTGCTGCACACGTGCCTGCTGGGCATCCCTGAGTGCGG | 120 | 58.33 |
| chr7 | 143643266 | 143643386 | Target=1;ProbeIdx=8773;Target #1 | 1 | + | good | ATATGCAGGAATCTGCTTGGCAAGGGGCCTGACCAGCTGCAGTCCAATGACAAACACTGGCTTAAACTAATCATGCAAATATTTATTCCAAACTAGAATTCAGCAATCTGAATCCTAACA | 120 | 40.83 |
| chr7 | 143643386 | 143643506 | Target=1;ProbeIdx=8785;Target #1 | 1 | + | good | GGGTCCGTTTACTCAGGTAGCCGTGCACCACACAGTGAAGCTTTGCTCAGCAAGGACCAAGCAACACCACCAAGCAACTCCTCACCTAGGGATGTGGTAGGCATCCTAGTTTGCTAAGTT | 120 | 52.5 |
| chr7 | 143643506 | 143643626 | Target=1;ProbeIdx=8797;Target #1 | 1 | + | good | CAGTTCGTGAGGGTCAAATGGTAACTATGTTGTCTCCTAGGAACACATTAAGTGACCATGCGACTGCAGTCAGTCCTTCTGTCAGTCCTTCTATATCAACACCATAATCTCAGTCTTTTG | 120 | 43.33 |
| chr7 | 143643626 | 143643746 | Target=1;ProbeIdx=8809;Target #1 | 1 | + | good | GGAAGCTTCTAAAGGTCAGTGAGACATCTCTGTTGGCGCTAAGTGCAAATAAATGGCCAGGGGTAAAGGCTCCCTTTCACTATAGAAACACTGAGATGCCCTCCTTCACTTTGCCAGTGC | 120 | 48.33 |
| chr7 | 143643986 | 143644106 | Target=1;ProbeIdx=8845;Target #1 | 1 | + | good | TGAGACATGTAACCCTGAGCTATGGGCAAGGACCCATGTATTTGCCTAGATCCCATTCCTTCTCTGGACTTTGGAAATGTGATGAAACCTGGTTCCTCTGCTGGCCAGATCTTTGCCTTC | 120 | 48.33 |
| chr7 | 143644106 | 143644226 | Target=1;ProbeIdx=8857;Target #1 | 1 | + | good | AACATTGTGTCCCAGGAGCTTGAGTTATAGGCAGTTGTGGAGTTATAGGCAAACCCTTGTGGGTCCTGCAATTGCAGAATGTTCTAGAACACCCTTACCCTGTCTCAGAATGTTTACACA | 120 | 45 |
| chr7 | 143644346 | 143644466 | Target=1;ProbeIdx=8881;Target #1 | 1 | + | good | ATTCCCGCAGTTTCTTTCTTTCTGTGACTTGCCTCCAGTCACTTGTCACATGTTTAATAGATGTGTGTGGAACATCTGTATTGTCAGGAGTAGCCTGGTTTGCCATGGACAGGTGTGGTT | 120 | 45 |
| chr7 | 143644466 | 143644586 | Target=1;ProbeIdx=8893;Target #1 | 1 | + | good | GAGAGAGAGAGAGAGAGAGAGAGAGGTCCTCTCTATCCTGAAACTTACTAATTTGGACAGGCTCCAAGGATCTGCTGCCTTAGCGCTGAGATTGCACACAGGTGCCACACACAGCTCACTTG | 120 | 50.83 |
| chr7 | 143644706 | 143644826 | Target=1;ProbeIdx=8917;Target #1 | 1 | + | good | TCTTTGCTCTCTCTGGAAGTGCCACACCCATTTAACCTTTCAGGCACAGGCCCCAGACTTCCCATGAGTCTGAATTTGGCAGCACGTGTCTTGAGGTACAGCCCATGCCACCATCTCACA | 120 | 52.5 |
| chr7 | 143645306 | 143645426 | Target=1;ProbeIdx=8977;Target #1 | 1 | + | good | AGTGGACTTCTTCCATTTGCAGGACAGCTGTGCCAGAGTGCTGCTGTTTCGGGGTGGAAACAAGGAATTGAAAAACTACAACAGCCAGACACCATTTCAGGTAAAGAGTCACTGACGGCA | 120 | 48.33 |
| chr7 | 143645426 | 143645546 | Target=1;ProbeIdx=8989;Target #1 | 1 | + | good | CCGTGGCAGGCTATTGCTTAGAGGCAGCGCAGGGCCAATGACTGCAGCCACAATACCTCCTGGGAGGAGGCTGCACTGCATCCCTGTGGTACAGCTGAGGTGGTTCCATTTGCCTATCTA | 120 | 57.5 |
| chr7 | 143645546 | 143645666 | Target=1;ProbeIdx=9001;Target #1 | 1 | + | good | CATAGCCTGCTTCTTTTAGACCAAAAAGTACAAATAATGCAGAGACTTATCTCTGGAGAAGGTTCTAGAAGGGCAGTGAGGTAAGACCACTTCAAATTCTGCAGCTGTTGGAAAGTGCTG | 120 | 42.5 |
| chr7 | 143645666 | 143645786 | Target=1;ProbeIdx=9013;Target #1 | 1 | + | good | ACTAAACTGTATCTCAGAGTGCAAACTTGATGGGAAGGAGGCTTTGCTGCTCTCTGTGTCCTCCTTCTCTGCCATCCCAGAGTCCAGGAGAACAACCCCCGCTGGACCTTGGCCTGCCAC | 120 | 55 |
| chr7 | 143645786 | 143645906 | Target=1;ProbeIdx=9025;Target #1 | 1 | + | good | CTTCTGGGAACATGTAAATCAGCTCATAATTGCTCATAAATTAGGGAAAATGCATAGGACATAGAAGTTAACAACCTAACCATTTAAAAAGATTTGATAAAACCAATTTAGTGACTTTAA | 120 | 30 |
| chr7 | 143645906 | 143646026 | Target=1;ProbeIdx=9037;Target #1 | 1 | + | good | GGGCATTCAGAATGTTGTTTGGCCATTACTACTCTCTGGCTAAGAACTCTACCATCACCCCAAAAGGAGGTCCTGTCCCCTCTAACTATTACTCTCAGTCCCTCCCTCAACCCTAGCAGC | 120 | 50 |
| chr7 | 143646026 | 143646146 | Target=1;ProbeIdx=9049;Target #1 | 1 | + | good | CTCTCATCCCCTTTCTGCCTGAATGGAGTCCTCTCTCCTGAGCAGTTCATACAAAGGGCGCACATGGGGTATGACTGTCATCTTGACCTCTTCAGTTGAGGGTGACATCTGGTGGTCAGC | 120 | 52.5 |
| chr7 | 143646146 | 143646266 | Target=1;ProbeIdx=9061;Target #1 | 1 | + | good | TTGACAAGGCACGTCTCCTGATTGTATTCCACCGTGGTGGGTTTCATGTGAGGCTCCAAGTCAGGTTCCCATGTTGAGCAGTGTAGGAAGACAGATGAAAGGGTTTTCTTGCTATATTGA | 120 | 46.67 |
| chr7 | 143646266 | 143646386 | Target=1;ProbeIdx=9073;Target #1 | 1 | + | good | TATATAGGATTAGAAGGTAGAGCCCTTTGTTGGTGACCAGGAACTCTGTAATCTCTGCATTCAACCTGTTCTAAAGCCACTGCTCCGTTGAGGGACAAGTCAGTGATGACACCACAAGGC | 120 | 46.67 |
| chr7 | 143646386 | 143646506 | Target=1;ProbeIdx=9085;Target #1 | 1 | + | good | ACTGGGTGATATCATGGTTCTAGAAACTGTGGTTTGGTGGAAAATGTATAAAGTCCTTTAAGCCATTCTAGTCCCCCTTTCTCCGGGGTTTTGGAATGTCCTTCCTGCTTAAGGGGTTCA | 120 | 44.17 |
| chr7 | 143646506 | 143646626 | Target=1;ProbeIdx=9097;Target #1 | 1 | + | good | CTTTGTGGGGTGTCATAAGCAGGGTTCTGAGACACCCCACTGTAGTCTCCCCAGGTGTGTCACACATGATATAAGCTAGCTTTGGCAAAGTCCACAGGGACTGAGCCTGGACACACACAT | 120 | 51.67 |
| chr7 | 143647106 | 143647226 | Target=1;ProbeIdx=9157;Target #1 | 1 | + | good | GCCCATGTGTGGCTGTTCTCTGGGAGTTACTATGTGTAAAGCTAGCCAGCAAGGTTTCTGATTAGCCAGTCTCCACTAGGAGCGGGTTCCCCTTCTGGTCATCTCAGATGTAGCCCCAGC | 120 | 53.33 |
| chr7 | 143647226 | 143647346 | Target=1;ProbeIdx=9169;Target #1 | 1 | + | good | ATAGATGACTATATCACTGGTAATTGTGATCAGTCCAGCCTCCAGCCCTGAGAAGGAAGACTGAAAACCCCAGCCCTGATCTGGGGTTTTGGGCTTATGTCCCTACACCAGCCTCAGCCT | 120 | 50.83 |
| chr7 | 143647346 | 143647466 | Target=1;ProbeIdx=9181;Target #1 | 1 | + | good | TGAGGGAGACCATGAACTAAGTTGGTCTGTCTCATTATCATATGAAAGAAGCCCAAGAGGTGTTAAGAGCTCTGTTCTGTGAATAGAGGTAAAGGCCAAATATGAGATGTGTGATGTGGT | 120 | 41.67 |
| chr7 | 143647466 | 143647586 | Target=1;ProbeIdx=9193;Target #1 | 1 | + | good | GGTGGTGGTGGTGGTGTGTGTGTGTATATACATGTTCATGTACAGATTCAGTACATATTATCTGCCTTGTATTTTGAGACAGGGTCTGTCACTTGATCTTGAACCTGTAAATTAAGCTGG | 120 | 40.83 |
| chr7 | 143647586 | 143647706 | Target=1;ProbeIdx=9205;Target #1 | 1 | + | good | GCTCCTATCTACCCCTTAGTGCTGGGCCTGCAGACACATGCCACCCCACCCCAGCTTTTTTAAAAGTGGGATCTGTAGATCAAACTAGGACCCTCACATGAGAGTCATCATCTCCCAAGT | 120 | 50.83 |
| chr7 | 143647706 | 143647826 | Target=1;ProbeIdx=9217;Target #1 | 1 | + | good | CCTGAAATATGTATTTCTTGCTTTACATCAATCACAACAGCACAGGGTCTGGAGTCAGGCAAGACTGTGGTCGTCGGAGGGAAAGTATCGCCGGCCCTGCTGGCTGACAGCCTGCCGTCC | 120 | 54.17 |
| chr7 | 143647826 | 143647946 | Target=1;ProbeIdx=9229;Target #1 | 1 | + | good | CCGCCAAGGGCATGTGGCTGGGGAAGCCACTCATTTCTAAGTCTCTGCTCTCCAAATTTCTGTGAGTCTTATTTGGGATTCGGGCTCTGCGTTCCCATAGAAACAGGGGGGTTGGAGTTG | 120 | 52.5 |
| chr7 | 143647946 | 143648066 | Target=1;ProbeIdx=9241;Target #1 | 1 | + | good | TATTTCTAGACGAAGGACAGGAGCCCGAACTACCTAAAATGTGACTGGCAGACGATACATATTTATGATGGAAAGAAATCCAGGTTGAATCTGTCATCTTGGCCAGGCCAATTCACACTA | 120 | 42.5 |
| chr7 | 143648066 | 143648186 | Target=1;ProbeIdx=9253;Target #1 | 1 | + | good | CGCTTCAAGTGACAGCCACTTGTGGGTGATGCTGCTTGCAGGTGTTAGGGGAGATGAGACCCAGCCAGGCTATACCCTCTCCCACTAACAGCACATCAGGCAGACAGGTAGCCAGCAGAT | 120 | 55.83 |
| chr7 | 143648186 | 143648306 | Target=1;ProbeIdx=9265;Target #1 | 1 | + | good | AAGCAGACACATGCTGTACTGTGTCAGGTCTGCTAAACACAAGAGAAGCAAAGCACAGTGGGAAAGCAGAGTGAAATGGAGGGTTTCATGACATCAGAGCAGAGGCCAGGAGAGAGGGAG | 120 | 50 |
| chr7 | 143648306 | 143648426 | Target=1;ProbeIdx=9277;Target #1 | 1 | + | good | GGAGGGAGAGAAGAAGGGAGCGGCTGATTTGTGCAAAGGCCCTGGGGTCAACATGTGCCTGGTGGGCTTTTGGAGCAGCAAGACTGGCCAGTTGGGGAGGGTGCAGAGACCAGCTCCTGT | 120 | 60 |
| chr7 | 143648426 | 143648546 | Target=1;ProbeIdx=9289;Target #1 | 1 | + | good | GATTGGAAATGAGTGGGTCCAGAAGGCTTGCAAGGGCTTTGGGTGTGATTCTGTGTACATGGCCAGCAGAGCAGAAGAAGGTTCTAGATGGAAATCAATCATTCATTGTGAGATCCAGTC | 120 | 46.67 |
| chr7 | 143648546 | 143648666 | Target=1;ProbeIdx=9301;Target #1 | 1 | + | good | ACCATGCCATGTGATCTGCAGTATGGATGACATCTTTGACCACAGATGAGTTGTCTGGGCTCTGAGACACAGCGAACACCCTGCTGTCAGAGTATGGGACACGAGACAGTGGAATCAGAA | 120 | 50 |
| chr7 | 143648666 | 143648786 | Target=1;ProbeIdx=9313;Target #1 | 1 | + | good | TTTTAGGAAGTGAGACTCCAGCTCCGAGTCCCATTACCTTGTATCTCAAGTGCTTGGTGCATCTGGGAGTCATTCTAGAAGTTGGCACCTGTGCTGCTTAATCATGATCAGCAACTGGAT | 120 | 46.67 |
| chr7 | 143648786 | 143648906 | Target=1;ProbeIdx=9325;Target #1 | 1 | + | good | CAGGGTCTCACATGGGACACTACATTTGTACTCACAGGTGTCCCCCTCCCTACTCTCTTTGCCTCCTCTCTCACTTCAGCCACCCAGGCCTTCTCCCAAGCACACCAAATGCATCCACGC | 120 | 55.83 |
| chr7 | 143648906 | 143649026 | Target=1;ProbeIdx=9337;Target #1 | 1 | + | good | CCTGGAATCTATATGTAATCTGTAAATATGATATGTAAATATGTATATGTAAATATGTACGCGTAACTAGAACCCACCTGCAGTATCCTACCCCAGCAGGAGCATCGAGGCAGTCCCATG | 120 | 40.83 |
| chr7 | 143649026 | 143649146 | Target=1;ProbeIdx=9349;Target #1 | 1 | + | good | TTGAGAAATAGTTGACACCATATACAGTTTCCTGTTTGCATATTAATATTTCACACTGAAAATAGGGACAGATAGCTGAGGATAGCCCATTGGGGAGGGCAGAAATCGAGAAGACTGGAC | 120 | 40.83 |
| chr7 | 143649146 | 143649266 | Target=1;ProbeIdx=9361;Target #1 | 1 | + | good | AGCAGAAGCAAAGATGCTGGTGAGAACAGAGACAACTTGGGACATGGAAGGGACCTTCCAAATTGTCATAACCAGATGACACATGATCATCATGTCACCATGATCTGATGTCATGATCCC | 120 | 45 |
| chr7 | 143649266 | 143649386 | Target=1;ProbeIdx=9373;Target #1 | 1 | + | good | TAGTGTGGCGGGAATGAGTATGGCGTCCATGAGAGGTGGCATTCAAGACTCTCCAGGGGAGTTGTAGCAAAGAGACAACAGCAACTGGAGGGAGGTCAAGATCAGTGCAATGCTGACTGA | 120 | 51.67 |
| chr7 | 143649386 | 143649506 | Target=1;ProbeIdx=9385;Target #1 | 1 | + | good | GGACTAACTGGGTTTAGGGTCTGAATGTAGATTTCCAGAGGAAGAAGAAACATGCAGACCTCTCAGGACAGTGTCCAAGAGGAAGGGCATGAGGCCCGCAGCTGGAAGGGTCTTCAGAAC | 120 | 51.67 |
| chr7 | 143649506 | 143649626 | Target=1;ProbeIdx=9397;Target #1 | 1 | + | good | ACCAGTTAGAAACACATTCTCTAAGGGCCCAGAGCTCATAAAGATGGTGTCCTAAAGGCTTGCAGGGAAACCTCAGTTTCAGAGGAGAGTCAGGGGTCAGACCCTTGGAACAGCAGGGAA | 120 | 50 |
| chr7 | 143649626 | 143649746 | Target=1;ProbeIdx=9409;Target #1 | 1 | + | good | GGCTGGCATCCTTCCTACTTGTACATAGGAGCTATCCAGTATGTGATTAGCACGCATCCTCTCAAGGACCGCACCTGGGGCTGCTGCAGGCCACTGCCCGCAATGAGGGACTAGATTGTT | 120 | 55 |
| chr7 | 143649746 | 143649866 | Target=1;ProbeIdx=9421;Target #1 | 1 | + | good | ACATAGGGAAACCCTGAGCTAGGGAAGACCCAGCAGCAGGGAAGCCAGATGGACTTAGGAGCAGCAAGGAAGGAGGTGATCTATCACGAATGACACCGGACAGAAGGAGCTTGTCTGTGG | 120 | 54.17 |
| chr7 | 143649866 | 143649986 | Target=1;ProbeIdx=9433;Target #1 | 1 | + | good | GTTTGCACTGGAGGAGCTTATACTGCCCAGCCAACACCTGTCAGAACCCAGCAGTAACTGTAGGCAAGCAGGCAGATGTTAAAATATGACCACTACTCCCTGGAGGGATGCCTGCTCAAT | 120 | 50.83 |
| chr7 | 143649986 | 143650106 | Target=1;ProbeIdx=9445;Target #1 | 1 | + | good | CTACTTACAGGGGACCATCTGCCCGGCTATCGCAGATGTAGGTAACATCCCATGGTGATTAACCGAAGCTGGACATACTTATTCGTGGCTGTGACAAATGAGCGAAGCTCATGTCTTCCT | 120 | 49.17 |
| chr7 | 143650106 | 143650226 | Target=1;ProbeIdx=9457;Target #1 | 1 | + | good | AAGTTATGTTTAACCAGACATAGCCAAGGAGGACCACCAGCCACTTTTTCTGTACACTTGGGCAAAAGCCACGTATGAAACGTGGTCAGCACAGGCATGGTGTGTGGGACATGGCTTATA | 120 | 47.5 |
| chr7 | 143650226 | 143650346 | Target=1;ProbeIdx=9469;Target #1 | 1 | + | good | ACTGGTGTGGAAGATTGACTTATTTCAACCAGCTCCCAGGATGTGTGAAAGTCAAAATTTGATTTAGGGTTTTGAAAAATAATCTCTCATGTTTCCTCTTCCCACTGGTACATAATCTCA | 120 | 37.5 |
| chr7 | 143650346 | 143650466 | Target=1;ProbeIdx=9481;Target #1 | 1 | + | good | TCTGCATTGCTGAAGAGAAGAGAGCTTTAGGATCTTCCCAGGGGTCTGGGATGCCACCTCTCTGTCCCAGTCATCCCTGCTGTTCCTTAGCCTTTCCCCAGCATTACAGAAGGCCGGGGC | 120 | 55.83 |
| chr7 | 143650466 | 143650586 | Target=1;ProbeIdx=9493;Target #1 | 1 | + | good | TCCTGCACACTCCTCAGGTGTCTCTTGCACCCTGAGAGGAACACCTCAGCCTCTTCAGGTTTCCGGGGTCTCAGAGTCACCATGTGCTGTCTATGCTATAGTGTTACCTACATCAGCCCC | 120 | 54.17 |
| chr7 | 143650586 | 143650706 | Target=1;ProbeIdx=9505;Target #1 | 1 | + | good | AACCCAAAGGACAACAGCAAGCCTAGCCCCTAGCCACATTTCTCCTGGACCTGATACATCAGAGTGGGCCACCAGCTTCTTTTGCCCAGTTCCCTGGGCATGGCTCAGATATCCTCACAG | 120 | 54.17 |
| chr7 | 143650706 | 143650826 | Target=1;ProbeIdx=9517;Target #1 | 1 | + | good | CTAAGCCCCTGTTTATCAACGAGCTTCAATCACTCTCCCTGGTACCACTATCTCTAGGGATAAGAACTCTTCGAGTCAGGTGACTGCTTGAGCGCTGAGGGAGGCAGGACTCTGCAGCCA | 120 | 52.5 |
| chr7 | 143650826 | 143650946 | Target=1;ProbeIdx=9529;Target #1 | 1 | + | good | AGTAAGACTCCACGCCCCACTCCCAGCTGGCCAGGGAAATTCCTGTCTCCCTTCAGACTTACCTCCGCAGGAGCCTACAGCTTGTTATGCACATTTTGGGGGTTATAAATAGTCCTCGAC | 120 | 52.5 |
| chr7 | 143650946 | 143651066 | Target=1;ProbeIdx=9541;Target #1 | 1 | + | good | TCAGCCTCTTCATCTGATGCACAGCTGTGTCCTCCATCTCTGTGATCCTGGGGTCTAGCCCAAGACCTTGACCCTCAGGAAAGGGAACTGGGCAGAGAGTTACTGGGCTGCTGACAACTT | 120 | 54.17 |
| chr7 | 143651066 | 143651186 | Target=1;ProbeIdx=9553;Target #1 | 1 | + | good | CATGTAATTTTTCATGCATAGCACTAGGGGGCGGTGTGAGCATAGAGCACCTTGGTACCAGGTGTACAAGAGGAGGAGCTTGTAAAATGCATTGTTGAGAGTCCTGGTTGGTAGGATCTG | 120 | 47.5 |
| chr7 | 143651186 | 143651306 | Target=1;ProbeIdx=9565;Target #1 | 1 | + | good | CTAGGAAACACCTCTGGGTGTGTCTAAGGGGCATTTCAAGAGGGTTTTTTTTTTTTTCTTGTTTTTTAAATTTATTTTTGGTGTGTGTATGTGTGTGTGTGAATGGGTGCCTACAGAGCA | 120 | 38.33 |
| chr7 | 143651306 | 143651426 | Target=1;ProbeIdx=9577;Target #1 | 1 | + | good | CTGTAAGTAAGGAGGCAGAAGTTAGAGTCAGTTCTCTAATTGTAGTCTCTCAATTTATTTCTCGACACAAGTTCTCACTAAATCTGGTGCTCACAGATCTGGGGAGCCTGGCTGACTAGG | 120 | 44.17 |
| chr7 | 143651426 | 143651546 | Target=1;ProbeIdx=9589;Target #1 | 1 | + | good | GAGCCTCAGGGACTGTCCATCTCCACTACCCAGCACTGTGATTAAATGTGTGCATCCCTGCCCTGTGCCTGGCTTTTTACATGGGTGGTGGTGGGGACTCAGGCATGCGCAATAGGTTCT | 120 | 55 |
| chr7 | 143651546 | 143651666 | Target=1;ProbeIdx=9601;Target #1 | 1 | + | good | TCACCCACTGAGCCTTCTCCTCAGCCTCTCTAGAGAGGTTCAACTGCAGAAGGAAGATCCTGAAAGAGGGCAGCACCATCTCACAGGCTGGCATCCAGGGCTGAATAATGAAGAGAATGT | 120 | 51.67 |
| chr7 | 143651666 | 143651786 | Target=1;ProbeIdx=9613;Target #1 | 1 | + | good | GAGCTAAGTACGAGCATTCATTCATCTCTCTGCTTCCCACTATGGGTGTCACGTGACCAGATGCCCACCATGGTGAATGGTACCCTTGAGTTGTGAGCCTAAATTAAGCCTTCCCTCATC | 120 | 49.17 |
| chr7 | 143651786 | 143651906 | Target=1;ProbeIdx=9625;Target #1 | 1 | + | good | TGTGGGTTCCCAGGGTGGTGGACGTCCGTCTTGGTGGTGGGCAACTTCATTCACCGAGCCATCTCTCTGGTCTTTAGTAGCTTTTGTGAATTATTTTGTCCCAACATTGAGAAAAATCAT | 120 | 46.67 |
| chr7 | 143652026 | 143652146 | Target=1;ProbeIdx=9649;Target #1 | 1 | + | good | TAATATACCCCCATAGTAGGAGATATTAGAATAAGGCTCTCACTCAGACATCAGATATGTGCCCAGGATCCTGAGATACTGCACAATTTGACTGTCACAGGGATGCTGAACACTCCATCT | 120 | 43.33 |

| | | | | | | | | | |
|---|---|---|---|---|---|---|---|---|---|
| chr7 | 143652146 | 143652266 | Target=1;ProbeIdx=9661;Target #1 | 1 | + | good | CTACTACAACCAGAGTGTACCACAACCAGAGAGCTTCAGTCTCTCTCCAGAGGCACAGGGGCCTGTGAGAGACACAATGTGGCACTTGGGGCTCCATTTCATTCCTGCCTCACAGGCTGA | 120 | 54.17 |
| chr7 | 143652266 | 143652386 | Target=1;ProbeIdx=9673;Target #1 | 1 | + | good | CCTGGGATATTGCTTATGAAGGTTAAACATGCAGAGGTCCCGGGTTACACCAGAGTAGCCAGCCCTTGGCTCCCCTTGTGGTCCACAGAGGACACTCAGTGAATCTCAGCCTCCTCCTCC | 120 | 55 |
| chr7 | 143652386 | 143652506 | Target=1;ProbeIdx=9685;Target #1 | 1 | + | good | TCCTCCCCTGCTTCTCATACTTTGCAGAATGCTCCTGCATGACTTACAGACTGGCAGGGGTGTGCTGTGCACCACAGTGAGTGCTGCAGGTTCACAGCACCAACCACCGCACCCAGAACT | 120 | 55.83 |
| chr7 | 143652506 | 143652626 | Target=1;ProbeIdx=9697;Target #1 | 1 | + | good | GGTCGTGTGGCATCCAGCCAGGAAGCAGAGTGCTTGCTAGTATCACTTTCTCCTTTTTACTCACGGGTGGTGAGTAAAACCCCTGGGAAGGCACTGCCCACAGGTAGGGTGGACCTTCTC | 120 | 55 |
| chr7 | 143652626 | 143652746 | Target=1;ProbeIdx=9709;Target #1 | 1 | + | good | GCACAGGCATAGCTCTCGGTTAGCCTATGTCCCCAAGACTAGATACTTTTGGGATTGTGGTAGGCAGTGACACTTGCAGTGAGCAGCCAGTTCCTCTGCTTGCTTTGTTCAGAGCAGGTT | 120 | 50.83 |
| chr7 | 143652866 | 143652986 | Target=1;ProbeIdx=9733;Target #1 | 1 | + | good | GTATGCACCAAGGTATACTGAACCCGATGCTAGCACAGCTGTCCATTTGCGTCCCTTCTAGGGTTCGTCTTGGGAAGACTTTTCATGCACTCTTAGCAAGACCACTCACTGGGATCTCTG | 120 | 50 |
| chr7 | 143652986 | 143653106 | Target=1;ProbeIdx=9745;Target #1 | 1 | + | good | TGGAGTGAGGGAGGCTTTAGGGGTCATGTGGAGGATGTGGCACTGGTTCTTTTAATAGATTTTCATGTGCCCCCTTCACCATTGCTAGCCTTCCTGGAAGTGCAGAGAGTGGCCACAGGT | 120 | 51.67 |
| chr7 | 143653106 | 143653226 | Target=1;ProbeIdx=9757;Target #1 | 1 | + | good | CCACGGCCATTACTCCCACTTCTGCAGACACACACAGCATGGATCCTGTCAGTCACTCTGTTAGACCTAGAACCCTCTGGGAGAAGTCTCAGTGTGCCCGGGATGGTGAGCGACAATGCC | 120 | 55.83 |
| chr7 | 143653226 | 143653346 | Target=1;ProbeIdx=9769;Target #1 | 1 | + | good | TGGAGCAGGGCTCCAGTGGAAGGAGAGATGCTGGGGCTATGGATCCAAGTCTTCCTAGAGGAGAGCCACAGCTGACAATGACCCTGCAGGAAAGGTCATGCCAATCAGGAAATCCCAAGA | 120 | 54.17 |
| chr7 | 143653346 | 143653466 | Target=1;ProbeIdx=9781;Target #1 | 1 | + | good | CAGCAAGGAACGGTGGCAGCGTCTTCTGAGAGAGAATGCAGTTCGTACACAAGTCCAGCGTTGATGAAGGTTTAGAAAGCATATAATTCGCCCCGTGCAGGGCCATGTGCCACCGCCATT | 120 | 52.5 |
| chr7 | 143653466 | 143653586 | Target=1;ProbeIdx=9793;Target #1 | 1 | + | good | GCTGTCTCCACACAGGGGGCACCCTGTGATGTGGGGGCCCTGGTTAGTGCTTGCTGCTCTTCCAATCTGTACCAGCTTAGAAGGAGCTCTTCACACTGGGGAGAGTACTTGCATGATGGC | 120 | 56.67 |
| chr7 | 143653586 | 143653706 | Target=1;ProbeIdx=9805;Target #1 | 1 | + | good | ACATGTACATGTACCCTACACATCTGCACCCCCACACAAGCATACACACATAGGTGTTCAATAGTGCATGAGATGGTCTTAGCCTGTGAGTCTTCCCCATTTTTATTAACATTTTGAGAA | 120 | 41.67 |
| chr7 | 143654186 | 143654306 | Target=1;ProbeIdx=9865;Target #1 | 1 | + | good | TATGCCTCAGAGGTTTGGTATTTAAGAAATGGTATTTATTCAAGAGAGAGAAATCTCAGCCAGCCATGACATTAATAAGGACAAATTGATCTCAATTAGTGAGAGATAGCAACACTGGGA | 120 | 36.67 |
| chr7 | 143654306 | 143654426 | Target=1;ProbeIdx=9877;Target #1 | 1 | + | good | TTAGGAGGGGTCTGTTCCCTGCTGGCTTCAGGGACAAAAGATAGTCCCTAGAGGATATGAATCTCAGGACATCTGTGACCAATACCTTCAAGTAATTCTGTGTTTTATTTGAGGCTCTTC | 120 | 44.17 |
| chr7 | 143654426 | 143654546 | Target=1;ProbeIdx=9889;Target #1 | 1 | + | good | ACAGAACACGAGCAGCAGCTGACAATACAAGTTAGCACCATGGGAGAACTCCTACCAGGAGCAGGGTACACACTTGTTATCCCAGCCTTCAGGAGGCAGGGCAGGGTGCAGGGACAGGGT | 120 | 55.83 |
| chr7 | 143654546 | 143654666 | Target=1;ProbeIdx=9901;Target #1 | 1 | + | good | ATATAGCCGGTTCCAGGCTGGCAAGACTGTCACAGATCAAGTTTCCAAGGTGAGGAGATGGCTCAGTTAATACAGTACTTGTCTCAAGCATGAGGACCTGCATTGAGCCTCAGCACCGA | 120 | 49.17 |
| chr7 | 143654786 | 143654906 | Target=1;ProbeIdx=9925;Target #1 | 1 | + | good | TTCTGCCCCCAATTTTATGGGGATTGACACACAATCCAACTGCTTTCTGGGAAGGCGTCATTCAGGGTGAATCCGATTATTCATCTGTGAAGTATAGTTGGCTCCTCAGCTCGGACCTGG | 120 | 48.33 |
| chr7 | 143655626 | 143655746 | Target=1;ProbeIdx=10009;Target #1 | 1 | + | good | CACAGACTATGTGTGTTTGTTGGCTACACGAGTGGAAAATAGTAAGTTGGCATCTAAGCTCTGAAGATGACTTGATTTCAAGGCAACAGTTTTGGTCTGTGCTTTTCACTGGAGTGGTAC | 120 | 42.5 |
| chr7 | 143655746 | 143655866 | Target=1;ProbeIdx=10021;Target #1 | 1 | + | good | TGCAGGGACTGGAAGCATTGCAAGGAGGCCCCTGGCCCCTGGTAATGCACCCCTAAGGATCCTCAGGGCCCGGCTGCCTCTTAGCTGCCTCTGCCATGGTCTTTACTGCATGTGTAGAGA | 120 | 59.17 |
| chr7 | 143655866 | 143655986 | Target=1;ProbeIdx=10033;Target #1 | 1 | + | good | GTTTCTCCTGAGCCTGCCCTTGCCCCCTGATCCGGGGCTGCTCTTCTGAGGTCTGACTAGGAGGCATGGTTCAGGTCAGTGTGAAGCAGACCCCTGGATTCCTAGGAATTGTATGGAATG | 120 | 55.83 |
| chr7 | 143655986 | 143656106 | Target=1;ProbeIdx=10045;Target #1 | 1 | + | good | TTGGTCACATGGGTCTCCTGAGGCCTGGATGGCTTCTTTGAGGGGCCTGCCAGGTGTTACTGTCAGCCCCGAAGACCCCATAGCTCATTTGCTTCCTCTCTGTCAGAGAGGACTCAACAG | 120 | 55.83 |
| chr7 | 143656106 | 143656226 | Target=1;ProbeIdx=10057;Target #1 | 1 | + | good | AGGAGCCAGGGTAAGGTGGGGGTGTGTGTCAAGATAAAAGCATTTGGTCTGGAAGTTGATACTTCCTGCAGCTCAAGTTAGGGGTGACACAGACCAGTGCTTCCTCCAGACCTCTGATGG | 120 | 51.67 |
| chr7 | 143656226 | 143656346 | Target=1;ProbeIdx=10069;Target #1 | 1 | + | good | ATGGCTTTAGGATCTGTAGAGCCAGTCCCTGGCAAGGGAAGAGCAGATGTTTCCAGGGAGACATAGGCACAACAATACCTCTTCTAAGAACACCCCAGTTTGGGTTGAATAAGATGTCTC | 120 | 47.5 |
| chr7 | 143656346 | 143656466 | Target=1;ProbeIdx=10081;Target #1 | 1 | + | good | CACCCATTGCCAGGGGAAGATGCAGAGCCATGGGGAAGGGCTTGTGGAGATTGCTGTTGAAGACTGCTCAGAGTCTCTTTTAGGAGTTCCTTCAGAACATTTGTTATGCTAATTTTGGAC | 120 | 47.5 |
| chr7 | 143656466 | 143656586 | Target=1;ProbeIdx=10093;Target #1 | 1 | + | good | TGCCAGGGATGATGCAGGCGTGAAGTGGATGGTGTTTTTCATTCTTTTTGTAAAACATTATAAGGGTGATTTTTCACTCCAGAAATGTAACAAACATAACGGGGTTGTTGATAACAAGAA | 120 | 37.5 |
| chr7 | 143656586 | 143656706 | Target=1;ProbeIdx=10105;Target #1 | 1 | + | good | TTCATGCAAGGAACATGGCCTGGGGTTGGAATGTGGCTCATGCTTGCCCGGGATTCCATAGCACAAACTGGGTATGCTGTCGCATGCCTGCAACCCCCAAAGTCAGGAGGCAAAGGTAGG | 120 | 55 |
| chr7 | 143656706 | 143656826 | Target=1;ProbeIdx=10117;Target #1 | 1 | + | good | GTAAGCCCTTGGAAAGCAGCAGACCTCACTGGAGAGAAGAGCCAGGAAGGCCGACTCCTGCCATCAGCTTAGGGGCAAGGTAGCCTTTCAGAGTTGTGCCTTTCCTTGGTAACATGTCTT | 120 | 53.33 |
| chr7 | 143656946 | 143657066 | Target=1;ProbeIdx=10141;Target #1 | 1 | + | good | TGCCATTGAGACACCTTCTCTCTCAGCAGAGGTTGGTGGCTGCAGCAGCAACGGCAGAGATAGACAAATCCAGAAACAAAAGTAGAGAAGAGAGAGAGGAAAAGAAAAGGAGAGGAAGG | 120 | 47.5 |
| chr7 | 143657066 | 143657186 | Target=1;ProbeIdx=10153;Target #1 | 1 | + | good | GAAGGGAGGGGAGGGAAAGAGAGGAAAGGGAAGGAGAGTGAAGAGAAGAGAGAGGCATAGGGCACCTGCCTTTTCCTTAGACCTCATAATATTAGGACTTCCTGGTGCCTCCTGTTGGAG | 120 | 51.67 |
| chr7 | 143657186 | 143657306 | Target=1;ProbeIdx=10165;Target #1 | 1 | + | good | ATGAAGAGTAACCATCACAGTGAGGAAACAGCTGAAGCTGGGGTGGTGGTGTAGCCCGCACCCCTAACACTTGGAGGAGGTGGGGGCAGTTGTTCAGAGTTAGTCTCTGATAAACAGTGAAG | 120 | 51.67 |
| chr7 | 143657426 | 143657546 | Target=1;ProbeIdx=10189;Target #1 | 1 | + | good | CCAGGTGGCCAGGGGTGGGGGTGGGGGTGTAGCTGGAATGACACCGAGGCAGTGTGTTTTTATAATAATCAGTGACGACACCACTTGCATTAGTCTGCACAGACTTGCATGGTAACAGGT | 120 | 52.5 |
| chr7 | 143657546 | 143657666 | Target=1;ProbeIdx=10201;Target #1 | 1 | + | good | TGTGGGCATGATGAAAGAGATGCTAAGGCCTCGCTCATACCAGTTCCATGGAACTCAAGGCTCCTCCAGCCCTCTGCTCCTGTATATCCTGAGCCTCGGGGAACACTGTAGAATGGCATC | 120 | 53.33 |
| chr7 | 143658146 | 143658266 | Target=1;ProbeIdx=10261;Target #1 | 1 | + | good | ACTGGGTTATGATCTGTTCACTGTGGTGTCTGCCCACCCTTCCAGGGGAGTCCTGCATAACTTTAGAGTCACTCTGTCCCATTCTTCATGGTGTACTTGTGACTGACTCCACCCCACTCC | 120 | 51.67 |
| chr7 | 143658266 | 143658386 | Target=1;ProbeIdx=10273;Target #1 | 1 | + | good | GTTGTCATGAGTTAGGTCAGGGACTCATTCTTGTCCATGAGGACTGCCTACTCTTGTCAGTGCGTGAGCCTGACACATCTGCCTAGTCTTGTTCCATCTTTCTTTCCTTGGTTCTGGCTC | 120 | 49.17 |
| chr7 | 143658506 | 143658626 | Target=1;ProbeIdx=10297;Target #1 | 1 | + | good | TGCCCCTGGGTCCTATTGGAGGGTACTGACTCTCCCCCGAGGGAGACCCAGCCCCCAGCCATCTTTCTTTCTCTTCTCTAGGCCTCTGGAAACTAGATGTAAGAAAGAGGCTCTTGCTGC | 120 | 55.83 |
| chr7 | 143658626 | 143658746 | Target=1;ProbeIdx=10309;Target #1 | 1 | + | good | ATCATTCCCCAGGAGGCCCTTTCTGGGGGGTGAGGTTGCCTTGGCAACCACACTCTGCAGCCCTCCTTTGGTGCTGTGGGCCCCTGGAGTCTATTTCTTCCTGAATGAACCTTCGCAG | 120 | 57.5 |
| chr7 | 143658746 | 143658866 | Target=1;ProbeIdx=10321;Target #1 | 1 | + | good | GTTGACCTGAGGTGGCCTCCTATCAGTCAGAGAGTCTCTCCTTTCGGACACTGGGAATGCTGTTCTAGTGGACACCTGTGGCCCCAGATTCTCTCCATAGACAGAAGGGCCTGATAGCAA | 120 | 53.33 |
| chr7 | 143658866 | 143658986 | Target=1;ProbeIdx=10333;Target #1 | 1 | + | good | GGCCATCACTTAACCAGGTTTCAGGCCGGTTGTTTAAAATGGGCCTTCTGTTAACTCAAATTCTCTAGAAAAGCAGCAAGGCTTAAAATATGTCAATATGCATTTATAAGCACGTGGACA | 120 | 40 |
| chr7 | 143658986 | 143659106 | Target=1;ProbeIdx=10345;Target #1 | 1 | + | good | AAATGAAAACAAAACAAAACAATAACAGCAAACCTGTCTATTCTGAATTAAGTATAAGTGATAAGGCCTTGGAACTTATCACTTTGGGATTTGGGTCTCCTTGAATGACACACTCTACTG | 120 | 35 |
| chr7 | 143659346 | 143659466 | Target=1;ProbeIdx=10381;Target #1 | 1 | + | good | TGGAAACAGTTACATGATCTTCATAGCCACAGAACAAGGGAGTTGTAAGTCAATATATTTGACCAGTGTGTCAGTAGAGATGTCCTCTGAAGCTCAGATCTTATCTTATGACATTCCATC | 120 | 39.17 |
| chr7 | 143659466 | 143659586 | Target=1;ProbeIdx=10393;Target #1 | 1 | + | good | TTTCAGGTGGTGGAGGAAAAACTCTGCTCTGCTCCGTGATGGATTTACCTCATAGCCATGAGAACACTGAGTTGGATACAGAGGTTAAAAGTAAATGGCTGTTAAAGAGCCCACACAGCC | 120 | 45.83 |
| chr7 | 143659586 | 143659706 | Target=1;ProbeIdx=10405;Target #1 | 1 | + | good | AAGGTGATTTTGACTCTCAGAGGACAACATTCCCACTCTCCATAGTCCTTGTCCTCTGACCAAGGTCAAGAGTCAGGATTTTGCTATAGATATGATGCAATTTCTTCAGAGAAGGCCAAG | 120 | 43.33 |
| chr7 | 143659706 | 143659826 | Target=1;ProbeIdx=10417;Target #1 | 1 | + | good | GTGATCTTGGGTGCTGGGGAGTTTACCTCCCACAATTCTAAGCTGAAACCTCAAGAAAGCTTAGAGGGTGGGACTTCCAGGCCAAGGTCAGGAGAGAAGCAGTCCACTTGGGCTAAAGCT | 120 | 51.67 |
| chr7 | 143659946 | 143660066 | Target=1;ProbeIdx=10441;Target #1 | 1 | + | good | ACCAGGCTGAATGGACCCCTAACCATTCTTCAGTCTTGTTCTGTCAGGGCCCCCTGGGATTGGATGCTTCCCCCACCCCACCCCACCCGCACTTAGGGAGAGCCATCTGCTTTAGAGT | 120 | 58.33 |
| chr7 | 143660066 | 143660186 | Target=1;ProbeIdx=10453;Target #1 | 1 | + | good | TACATTGAAATGCTAATCTGGTCCAAGCTTGTAGAGCATCCATGGATAATCTTGGCATCTCAATTAAGCTATGGTGGCAGGCCTGGCAGATCACATCCCCCTCCCCCTCCCCCCCCACCA | 120 | 51.67 |
| chr7 | 143660186 | 143660306 | Target=1;ProbeIdx=10465;Target #1 | 1 | + | good | AGGCCTTTGTGGTGCTGGTCTCCACATCCACGTGATCTGAATGCTACCCTTTGCTTGACCCCTCAGAAAAAGGGCGCTGTTGCTCCTCCAGGGACCTAACTATCTCGGATTCTCTTCCCA | 120 | 53.33 |
| chr7 | 143660306 | 143660426 | Target=1;ProbeIdx=10477;Target #1 | 1 | + | good | ACATTGGCCAGTGGGCTTCCTCACCTGTCAGCTGCGTACACCCCCTCCCCTGGTCAATGACTTGGCCTCTGAGAACAGGGCCTCCCACACCTCCCCCAGTTCTCAAATCCACAGCATCAG | 120 | 59.17 |
| chr7 | 143660426 | 143660546 | Target=1;ProbeIdx=10489;Target #1 | 1 | + | good | GCAGAGAAGTGACCCTAAGGTCAGGGCAGGAGGGACAGTATCAGTGTCATCTTCTTACGTGTGAGTCTACTCTGGAGCAGGAATGCCTTAGAAACCTCCATGAATATTCAAAGGCATTCAG | 120 | 48.33 |
| chr7 | 143660546 | 143660666 | Target=1;ProbeIdx=10501;Target #1 | 1 | + | good | CGTCAACTCCGCTCAGGGATTATTTATTTGAACACTGTAGGAAGCGTCATTGTTTATAAAATAGATAGGAGAGAAGCCTTTGTCTTTGTGCTTCTTTTGAGCCTCCTTCTTATAGGACTG | 120 | 40 |
| chr7 | 143660666 | 143660786 | Target=1;ProbeIdx=10513;Target #1 | 1 | + | good | AATTCTTTTTCCTTTCAACACAGAAGGCAGCAGAGAGCCTTGAGCTCTGCAGGTGGGAAGGGAGAAGCCAGGAGATGGTGGGAACTGAGGGTGGCCTGCTGCCTTGGGGACTTTATGAGG | 120 | 54.17 |
| chr7 | 143660786 | 143660906 | Target=1;ProbeIdx=10525;Target #1 | 1 | + | good | ACACCCTCCCACCCCACAGGCTTCCTAGATTATAGATTGGGAGCTGGTGAATCTGCCTTCTCACCTTTTCTCAAAAACAAAAAGAAAAATACAAAACAAAGCACATTTAAGGCTAATGGA | 120 | 40.83 |
| chr7 | 143660906 | 143661026 | Target=1;ProbeIdx=10537;Target #1 | 1 | + | good | GGTTAGAGACTCAGGAATGCAGGCCTGTGACTATACCTGCTTGGTGCAGCTTCCCTTTGATAGAAGGTGAATGAGTGGGGGCAAGAAGTACTAGGGGAACATTCTAGAACATGGTATTTC | 120 | 47.5 |
| chr7 | 143661026 | 143661146 | Target=1;ProbeIdx=10549;Target #1 | 1 | + | good | TTTGAAAATAGTTTGAAATTAAAATTCAAGTTTGACTTTCAGGGGGCCTCGTCAGCTATCAGGTACACCATTTTGTAAGCATGACAAGCCCCTTCCTTACTTTGGGTCAACATTAATAAAA | 120 | 36.67 |
| chr7 | 143661146 | 143661266 | Target=1;ProbeIdx=10561;Target #1 | 1 | + | good | ATGCATGTGTTTCAGCTCACAGTCAGCAGAACAGGGCTTGCTTAAAACAGAACTCTGGCAACACAGGGTATATTTGGAACCACACAGGACCCATGCTGAGGGTTCTGGAAACAGCCTAGT | 120 | 48.33 |
| chr7 | 143661266 | 143661386 | Target=1;ProbeIdx=10573;Target #1 | 1 | + | good | AGGTGTAACATCACCTTGGCTCCCTCATCCACCTATCACTCCTATGGTGCAGCCCCCCCCCCCACCTGAGTTGGGATGGTGGGAGGCAGGGAGGGGCCCGCTGACCTTGTAGGCTCTAA | 120 | 61.67 |
| chr7 | 143661386 | 143661506 | Target=1;ProbeIdx=10585;Target #1 | 1 | + | good | TGGAGGACACTATCTTCCTTAGAATGTTGAGTCTGGGATAGGGGGTACTGAGGTGGACCCCAGCTTATTGTCAGGGATCCCCTTTGCAAAGAGCAAAGAAGACTTCCCCATGACAACAGC | 120 | 50 |
| chr7 | 143661506 | 143661626 | Target=1;ProbeIdx=10597;Target #1 | 1 | + | good | TCATGGAGCCCAAAGTCAGAAAGAAGAGCACACTGTGTCAGATGCCTGACATCACATCCAACCTCTGCAGGCCAGAACCTCATGGAGAAAAAAGTGGTCCTCAGGGATCGGGCAAGCCA | 120 | 52.5 |
| chr7 | 143661626 | 143661746 | Target=1;ProbeIdx=10609;Target #1 | 1 | + | good | AATGAAACTATTTCAGACATAGGATCAGTGCTGAAACCTTCCCCTTCCCCCTCCCCCTCTTCCCTTTTCCCCCCTCTATTCCTGGCTAGAAGTGGTCAACCCTGGCCTCTGTCTTAATGG | 120 | 50.83 |
| chr7 | 143661866 | 143661986 | Target=1;ProbeIdx=10633;Target #1 | 1 | + | good | ATAAAACAAGGCCAATCAGGAGGGTTTGTGGGGCTTATGTGAAACAGCTGTAGATGGTAACAGCCACCCTTGGCCTTGGCATCTGGTGAGAGCCTTGGCACCCTTATCACTGGGACAAAG | 120 | 50.83 |
| chr7 | 143661986 | 143662106 | Target=1;ProbeIdx=10645;Target #1 | 1 | + | good | TGTGAAAGGAGAGGAGCTCAGAGCCCTGGGATTCCGTAGCCTCTCATTAGGCCCCATCTCCTCAAACATGCTTGGGGATTAAGCTTCTGACATGATCTTTGGGGACCATGTCGGGACAGA | 120 | 52.5 |
| chr7 | 143662106 | 143662226 | Target=1;ProbeIdx=10657;Target #1 | 1 | + | good | GACGGTGGCTTGTGTCACAGGTCACATCATATGACAATTCGGGGTCTGGCGTAATCCCAAATTTCTCCTACTGTGTAGACCCTCTGAACTTTTGGTTGTATTCAGTTTTTGCTTTTT | 120 | 42.5 |
| chr7 | 143662226 | 143662346 | Target=1;ProbeIdx=10669;Target #1 | 1 | + | good | ATTTTCAAATGTTTTTTTTTTCCCAAAAATTGTAACACATATTTAGTATGTGTGTATACACACACGTGCACGTGCACACACAAGCAACTGTGTACACATGCACAGGTATGGGGAACAGAG | 120 | 37.5 |
| chr7 | 143662346 | 143662466 | Target=1;ProbeIdx=10681;Target #1 | 1 | + | good | GCTGCCATACCCCACTGAAAGGTTTTTAACAAAGGGTTGTCCTTGCCTCGGCTTGTCCTAACCCAGGCTGGAAAGCTACTCCTGACAACAAAGCCAACGCCGAGCTGCTGAGCTGGCCTC | 120 | 55 |
| chr7 | 143662706 | 143662826 | Target=1;ProbeIdx=10717;Target #1 | 1 | + | good | ACTGTTACACTTCCTGTGGCCTCCTCACTGTTGTGGACCCTGTGGGCTGTGCATTCGATGAGCGATGAGCGAGATTAAAAAGGGCATTTGCAGCGTGGCCGTATTTGTTTATGCCGGCAG | 120 | 52.5 |
| chr7 | 143662826 | 143662946 | Target=1;ProbeIdx=10729;Target #1 | 1 | + | good | CCATGGGCACACAGAGTGTCCTTTGTTTAATCTTATTTGATTTTTAATGGCTGCTCTTTAAGTTTTGAGTTTGGGCTTATCAATATGTGGTATATTTGCATCGTTTTGAAGTTTAAGCTA | 120 | 34.17 |

| | | | | | | | | | |
|---|---|---|---|---|---|---|---|---|---|
| chr7 | 143662946 | 143663066 | Target=1;ProbeIdx=10741;Target #1 | 1 | + | good | TGAAGCAGAATAGGGACACAAAAGCCCAGCTTTCTCCCTCTCCCCTCCACTGTCTTCTCCTCTTCACAGGTGGTTCTAACTCCTTCCCTGGGCAGGGAGATGGCTCTGTAGGCAAAGTGC | 120 | 54.17 |
| chr7 | 143663066 | 143663186 | Target=1;ProbeIdx=10753;Target #1 | 1 | + | good | CATAAGAGTGTTTTAAAACATGTGGCAGGCACTGTAGTTTATTGAAGGTCTTTCCTGTTCAGGAGTAAATACAGCAATTACTATGTCACTGGCAAGACAGGGAATGTTTCTGCCTCTTTT | 120 | 39.17 |
| chr7 | 143663306 | 143663426 | Target=1;ProbeIdx=10777;Target #1 | 1 | + | good | TATAGTGAGGAAATAGGCTCCTAATAAGTTATAGGGCCAAAGCCATACCGTCGACAAACTTGAATCCGGGACTAGAAGTTGGGCCTTGTCACACCTGGACCCCTTACATAAACCATAGCT | 120 | 45.83 |
| chr7 | 143663426 | 143663546 | Target=1;ProbeIdx=10789;Target #1 | 1 | + | good | AGACATTCATACACAATATGTAATATATCCACATTAGGTGTTACAGCCCCACTTGTAATTCCATTACTCTGAGGAGTGGGAGTCACCGTGAGCCCTAGGCCTTGGCCAGACACACCTGAA | 120 | 45.83 |
| chr7 | 143663786 | 143663906 | Target=1;ProbeIdx=10825;Target #1 | 1 | + | good | GGACATTAATTTGCATCTCATAATCAATAAAAGAGGAGACCTGAATGGTGGTTGTACCTGGACCCGCTGCCAGTACTAAGCGTGTCTGGTGACTTTGTGGTCCAAGGACTTGATCATCTT | 120 | 45 |
| chr7 | 143663906 | 143664026 | Target=1;ProbeIdx=10837;Target #1 | 1 | + | good | TAAGATGGGCAAGCTCATCTGCTTCTGGTTCCCTCTGTAGGAAGAGGAGCTGAGAGTACTCACTGTGGTCTTGGCTTGCAGTGCAAACTTGAGTGACCTGAGGTTTCTATGCAGAAAGTT | 120 | 48.33 |
| chr7 | 143664026 | 143664146 | Target=1;ProbeIdx=10849;Target #1 | 1 | + | good | CCCTGCTGGGAGCTGGAGGCTCCCTGCTGCGGTGCCCATCCTGGTGGCCTCTGTGTCTGTGGAACACCCGCAGTTTAGCAATTGCTTTATTCTCTCAGGTGCCTCAGCAAGTGAGAACAT | 120 | 57.5 |
| chr7 | 143664146 | 143664266 | Target=1;ProbeIdx=10861;Target #1 | 1 | + | good | TTAAAACACAGGCTGGCAGCCAGTCTCTTGCGCCTGGGTGTGGGCTTTCAGTGTCCATGGGGTAAGAGGTGACTCATCCTCTCCCCACAAAACAGCTCCAACGCACAGCAGCTCTTCCCG | 120 | 56.67 |
| chr7 | 143664266 | 143664386 | Target=1;ProbeIdx=10873;Target #1 | 1 | + | good | GATATCTTCACCTTTTCCTTTGCATTGGAAGAGTCAAATAGTGCACACCATGGTGTACCATGGCCCGCTGGCATGCACACTCTAAACTCCAAGCGCAGTGGGGATGCTGTGTCTCTTTGT | 120 | 49.17 |
| chr7 | 143664386 | 143664506 | Target=1;ProbeIdx=10885;Target #1 | 1 | + | good | AGAGAGAGATGTCACAGTGCAGGGACCTGCTGGGCTCCATGTCTTCACAGCTCCAGGAACTCATCTTCTGGGGAAGCACCTGCAGGCCCCCCAGGTCAAGTGAATTTTTGCTCTCAGTGC | 120 | 55.83 |
| chr7 | 143664506 | 143664626 | Target=1;ProbeIdx=10897;Target #1 | 1 | + | good | GCATGCCAGAAATGTGGATGGATGGCAGCACTGCTTGTGGTGAAGGGGGACCTTCTCGCCATTCAGTGACACAGGTGGATAATTGTAATAGCACCTTCGCTCTAGTCAATCATGCTCCAT | 120 | 50 |
| chr7 | 143664626 | 143664746 | Target=1;ProbeIdx=10909;Target #1 | 1 | + | good | GCGTGCATGTGAATGTGCTACAGATAACCAGGACTTGGCTAGCTTTGCAGTCTAGAAATGCCACATGCTTTTCTTCTGTGTGACCACGTGACAGGTCAACAGTGCATGTGAAGAGGAGGG | 120 | 49.17 |
| chr7 | 143664866 | 143664986 | Target=1;ProbeIdx=10933;Target #1 | 1 | + | good | GCTCTTGGGTCTGGGCTCCCACTTTGTTCCTCCTGGGTTATTCCCACACCCACCTGACCTCTCCCTGACTCCCAGATTAAACCTGGCATAAAATGCCACAGGGGCTTGGTTATGTGGTTT | 120 | 53.33 |
| chr7 | 143664986 | 143665106 | Target=1;ProbeIdx=10945;Target #1 | 1 | + | good | CTGCTATCCATACACCAAGGCAGAGGACTGAGGTTGTTTATCCCTCTGATTCACAGAGGGTTTGCAGTGGCTTTGCATAAACCTCTGGTAAGAGGGGATTAGCACTCCTGCACACTCCTC | 120 | 50 |
| chr7 | 143665106 | 143665226 | Target=1;ProbeIdx=10957;Target #1 | 1 | + | good | TGATCCTTTACTGCAGGACTTCCTAGCTGAACCTGGTATAGCAGCTGACTGATGGGCACAATGTGAATGGGAAAGAACATGCAAGAGCATGCACACACGTGCCTGCTGGTCTCAGGGTTA | 120 | 50 |
| chr7 | 143665226 | 143665346 | Target=1;ProbeIdx=10969;Target #1 | 1 | + | good | CATCCCCAAATCAGAGATGGGAGAATACGCAAAATGAAAAAGGATGTCTGATATCTGACGGCCACTAGCCAAATAGCCAATCCATTTTAAAGTGTGACCTTGACCACAGTTTTAATTTGC | 120 | 40.83 |
| chr7 | 143665346 | 143665466 | Target=1;ProbeIdx=10981;Target #1 | 1 | + | good | TGGGCTTACAAAGGCTGCTGGGTGGGGTAGGATGGGGTGGGGTGGGGTGGGCACCTCCTTTGTCACAGCCCAGAGGACTGTCCTCTCACTCAGCTTGGCAGGCAGTGGGTGAGTGCTCTC | 120 | 62.5 |
| chr7 | 143665466 | 143665586 | Target=1;ProbeIdx=10993;Target #1 | 1 | + | good | TTTGGGCCCAGCAGGTTTCCTGCATTTTGCTATGAGGCCTGTTGCCTGTGACTCTTTCCTGGATTGTGTGGACACAATCCAGAGTCTCCTTTCAAGTAATTAAGAACAACTCAACTGGTT | 120 | 45.83 |
| chr7 | 143665586 | 143665706 | Target=1;ProbeIdx=11005;Target #1 | 1 | + | good | TGAGGTGATGTGGTGTCAAGTGAGCCTAATGTCTTTCTCATCAAGTCTGCTGGGTCCAGTCTGAGCAGTGGATGGGGGCACAAGGCCTGTCCTGGTGTGGCTCCTATTCCTGCCAACAGC | 120 | 55 |
| chr7 | 143665706 | 143665826 | Target=1;ProbeIdx=11017;Target #1 | 1 | + | good | TCCCAGGCACTGACACTGAAACTCTGCTGCATACACGCCTTGATGGCTATTCTCGGATCATTGTCCCTCATTATCTGTCCACAGATGGCACTGTGAATCATAACCAAAGGAATTAAAATG | 120 | 45 |
| chr7 | 143665826 | 143665946 | Target=1;ProbeIdx=11029;Target #1 | 1 | + | good | TGAGGTGCCAGTATGTAGTTGCTAAGGTGTGTGTGTGTGTGTGTGTGTGTATACAGAGCGCTGACATACACCAGCAGATCTGGAAATGGTTGGTCTGTGTACTGGAGATGAGAGTCCTAG | 120 | 48.33 |
| chr7 | 143665946 | 143666066 | Target=1;ProbeIdx=11041;Target #1 | 1 | + | good | CCAGCAGCTAAGTATCCCCAGGAGACTAAGCACTGGATTTAAAAGCAGACTTTAGTGTTTGGATCAGCCCTATCAGCTCTCAGAGATGGTTTTGTCACGTGGACAAGTAGTACCCAGAAA | 120 | 45.83 |
| chr7 | 143666066 | 143666186 | Target=1;ProbeIdx=11053;Target #1 | 1 | + | good | GCGCCAGGAATGCCATGCAAAGCCGGCAGCTGAGACCAGCAGGTGACCCTCAGATCCCCACATGAACTCTTAAAAGTCTAGAGAAACTACCAAACTGTGATTTTTACCGGGAGGGCATCT | 120 | 51.67 |
| chr7 | 143666186 | 143666306 | Target=1;ProbeIdx=11065;Target #1 | 1 | + | good | TGGGAGTGAAGGTCACACTGCCTAGGAGGTGCCTGCAGCTGTCAGTCACTGAGATGAGGTCCTTCCTCAATCTCCAGTGCTTGCTTTCAAGGCCTTTGCAGTACCAATGGGCTTAGCTGT | 120 | 53.33 |
| chr7 | 143666306 | 143666426 | Target=1;ProbeIdx=11077;Target #1 | 1 | + | good | GGCAGCTCAGGGTCCCTGGCCTTCCTCTGCAATCCAGAAGGGCTCTGGGATGGGGGTAGCTGTGAGTGAGTAGCATCCCTCAGCCTCTGACCAGGAAGCTGGTAATAAACATTGTCCTGG | 120 | 57.5 |
| chr7 | 143666426 | 143666546 | Target=1;ProbeIdx=11089;Target #1 | 1 | + | good | CCTCACACAGATTGTGTCTAAGAGACAAGGGAGCCTGAGAGGATTGGCCTCAAAATGCTTGCATCCTCAGCCATGTGGCTGGGCTGACCCTAGCCCTAGGGCAGCAACAGGTGTGGAATA | 120 | 54.17 |
| chr7 | 143666546 | 143666666 | Target=1;ProbeIdx=11101;Target #1 | 1 | + | good | AAGGTCTCTGGGGGAGATGCTACAGAGCCACTGTGTCAGGCTGGAGTCTGTGCTCTATCCCTAGCAGCCCCCAACCCAAACCTCTACTCTGTAGTGTTCTGTGCCTGCTCCACAGTGTCG | 120 | 56.67 |
| chr7 | 143666666 | 143666786 | Target=1;ProbeIdx=11113;Target #1 | 1 | + | good | ATGGCTCCACTAAAACGTGATCAGGCGTGATGACTGCCCATGTGGCTGTCTGGAATATTTACTGGTGCCCAGGGCTTGCTCTGCTGACGTGTGGTGCATGCTGCCTAAGCAATCAGACAG | 120 | 53.33 |
| chr7 | 143666786 | 143666906 | Target=1;ProbeIdx=11125;Target #1 | 1 | + | good | CAGTTGGGGCAGAGTAGTGGCCCTTGATGGAAATTGAGGGACACAGATCCAAATGGAAGAACTACTTGAACCCTAACTCCGCTAAGCCACACTGTTCCTAATGTATATGCTTTGACACTC | 120 | 46.67 |
| chr7 | 143666906 | 143667026 | Target=1;ProbeIdx=11137;Target #1 | 1 | + | good | CTAACACTCATGCCGGGTGCCGCCATATGCACAGGTGCTTGTGATGATCGTGTTCAATGGCAGTGTCTTCCTCACTCACTCTCCATCCTTTTTTTCTGAGGTTTTTCCAGGTCCCTCCCT | 120 | 50.83 |
| chr7 | 143667026 | 143667146 | Target=1;ProbeIdx=11149;Target #1 | 1 | + | good | ATGAAAACACTTTATCTACCAAGCCATCTCCCTAGTTACCATCCCCAAACCCCACACAAAGGATGCCTATTTAAGGGGCTTCTGGATGAAGTTGCAGGTTTTAGAACCAATGTCTAACTC | 120 | 43.33 |
| chr7 | 143667266 | 143667386 | Target=1;ProbeIdx=11173;Target #1 | 1 | + | good | CCCGAATCATCTTTTAGCCACCAAATGTCAAAGCTGGAAACTGTGAATCCTTGACTGACATTTTTAGTGGTTTTAACTTATTTGCAAGGACATTTTTAAAAGCCAGTGATACCAGTTGTC | 120 | 37.5 |
| chr7 | 143667386 | 143667506 | Target=1;ProbeIdx=11185;Target #1 | 1 | + | good | CCTACAAATGGGTGAAGGAACTCGGGCAGCTGCTCTTGGGATGGTTTTAGCAAAGAAAACACCTTTTGATTTTTAAAAAAATATCCTTTTAGGTGTCAAAGTGAAATTTTCAGTTATTTA | 120 | 35 |
| chr7 | 143667506 | 143667626 | Target=1;ProbeIdx=11197;Target #1 | 1 | + | good | ATTGTGTGGCATTTACATTACATATATTCATTCCATGTGTTTTAGGGTAGGTGGACCCTGGGGATCCAAGTGTCTTCAGAGCAGACCAACTCACAGTGACAAGGGAAATGAGGGAAGGGG | 120 | 45.83 |
| chr7 | 143667626 | 143667746 | Target=1;ProbeIdx=11209;Target #1 | 1 | + | good | TGGGAGCAGGTCTGATACCTAAGGAGTTTCTACTCTGGCTCTGGGACTCATTAAAAGTGTGATGTCTTTTCCCCCAAAGGACCCCATCTCTCTGGGATTTATTTATAAACACTGGGAGTT | 120 | 45 |
| chr7 | 143667746 | 143667866 | Target=1;ProbeIdx=11221;Target #1 | 1 | + | good | TGTAGGGATGCTGGTGATGCAGGAGGAGGCAGGAGTGAGTTTTGCTTTATTTTCTTTCTCTTTGCTGGAGAGTGTGGAAGACATTCTACTCACAGAATCAGGGAGACTTAGAGTGGTGAG | 120 | 46.67 |
| chr7 | 143667866 | 143667986 | Target=1;ProbeIdx=11233;Target #1 | 1 | + | good | CCCGTCAAACCTGCTGTAGGGAGGCTGAGGCAGGAGGATGACAAGAATTTGAGGCCTACCTGGGTGACATGCAAGACTCTGTAAAGCAAAGACAAAGAGATACAGAGTCAGCCAGGCTTA | 120 | 50.83 |
| chr7 | 143667986 | 143668106 | Target=1;ProbeIdx=11245;Target #1 | 1 | + | good | GTCGTCTCTGGTTCCAGGACCTGCAGCTGCCAAACAGCCGTGGTTTCCAGGGGAAGCCTTGGCTTCTGTGGTCCAGCACAACTTTGATTTATAGTTTAGAGGCTGTTCCCCAGTGTGGCT | 120 | 54.17 |
| chr7 | 143668106 | 143668226 | Target=1;ProbeIdx=11257;Target #1 | 1 | + | good | TGTGAGTTATGGTGATGTTTATGAAACTGTTTTTTTTTCCCCAGTTACTTATAGAAGAAGTTGAGCCAGTAACATCCCCTCAAAACTCCAGTTTGTATGTCTCCAGGTAGACCTGATTTT | 120 | 37.5 |
| chr7 | 143668226 | 143668346 | Target=1;ProbeIdx=11269;Target #1 | 1 | + | good | AGGCTGTGTGACTTTCCTCCATCGCTAAATGATGTGAACTAGTCCGTGTATGTGTAGTGTGAATGGAAAATAGTATGAATCACCCCCTGTAGGTTATTTAGTCACCTGTAGCCAATCTC | 120 | 42.5 |
| chr7 | 143668346 | 143668466 | Target=1;ProbeIdx=11281;Target #1 | 1 | + | good | GCCACACTGTGTGGAGGTGGAGCCAGCACTGAACTGTTCCGTCCCACCCCGGACACAGCCATTTTCCTTGTCCAGAGTATACCTGTGGCATACGATTCCTGCCTATTAGTCACTCAGGAA | 120 | 54.17 |
| chr7 | 143668706 | 143668826 | Target=1;ProbeIdx=11317;Target #1 | 1 | + | good | CATCTTGGTTCATGGCAGAGCTTGAGTTCACATGGTTCCTGCTTCATTTAACCATAGCTGCCAAAGCACAATGGTAGTGAGGGATGCTGGCTGTTTGGGTATGCCAGAGAGAAGCAGGAA | 120 | 48.33 |
| chr7 | 143668826 | 143668946 | Target=1;ProbeIdx=11329;Target #1 | 1 | + | good | TGTTCTTTTGAGCAAAGAGGTAAATTTCTCAGTGTGTATGTGGTTTTGCACTGTCTGAGTTCTAAGGCACCCATTAGGCCTTGGGGTCCCTAACAATGAGCAGGAAGGCTGCAGCTTCTC | 120 | 46.67 |
| chr7 | 143668946 | 143669066 | Target=1;ProbeIdx=11341;Target #1 | 1 | + | good | TATCTTTGTGTCCTGAGCACCATGACATCTGGGCGGGAAGAGCTGTTGAGCAAACAGTGGTGGTGTACCAAAATGCCTGACAGAAGTAAATCCAGGAAGGGGTCTTATTTTGGCTCCTCA | 120 | 48.33 |
| chr7 | 143669066 | 143669186 | Target=1;ProbeIdx=11353;Target #1 | 1 | + | good | TGCACAGTCTATCACAAGAAGACATGCCTATGGAGGATGTGAAGACATCTAGTGGGAATGGCAGCTGCTCACATCCCACCCTGGGAGCAGAGAAAGATGCACGCTGGTCCTTAGCTTGTG | 120 | 51.67 |
| chr7 | 143669186 | 143669306 | Target=1;ProbeIdx=11365;Target #1 | 1 | + | good | GTATCTCCTAAGTGATTCCAAAGCCAATCAGGTTGATAATGAGCATAAGCTATAACATTCTCAGCTATGGTATTCTATCTTGTGGCTCAGCAAGCTAACACATTGTGAATACATGCGTGC | 120 | 40 |
| chr7 | 143669426 | 143669546 | Target=1;ProbeIdx=11389;Target #1 | 1 | + | good | AAAATTGGGCTGTTTTCTACCTCTGCAGCTGATCTCATCAGGTGCTTCTTCTGGCTTCCTTGAGCAGAAGGCTGAGCCTTGTCCCAAACACAATTTTTTTCTGCAGTTAGAACCCAGGGA | 120 | 46.67 |
| chr7 | 143669666 | 143669786 | Target=1;ProbeIdx=11413;Target #1 | 1 | + | good | GGCTATTCAGATGATGGCATCTCTCTTACCCCCTCTTAATGTTCAAATGGATGAGGTCTAGTGTTAGGTAATGGAAATCCTACTTTCTCACCTGTCAATCAATCAGCCTTCTCATGCTGT | 120 | 42.5 |
| chr7 | 143669786 | 143669906 | Target=1;ProbeIdx=11425;Target #1 | 1 | + | good | GAGACCCAGTCAGATGAGAGGAGTACATATGAGTGGCCACTGCTGTGACCAAGACCTTCTTCGAGGCCACAGTGTCATAGCCAGTCTGGAAAACAGCTGTCCCTGACTTAGTCTCTCTGG | 120 | 52.5 |
| chr7 | 143669906 | 143670026 | Target=1;ProbeIdx=11437;Target #1 | 1 | + | good | GATAATGTAAGATATGGTGATAATTAGAATAAGAACCACAAGGAGGCTTCCACATTTCTCTGATGCTCCCCCCCCCCCCCGCCAATCTGTATGTGTGTGTGTGTGTGTGTGTGTGTGTGT | 120 | 46.67 |
| chr7 | 143670386 | 143670506 | Target=1;ProbeIdx=11485;Target #1 | 1 | + | good | GTTCATGAGTAAACGTTCACACATGTGTTCACATGTTAGAGTACAATGGCCGAGCATGTGTTATGGCTCATCCTCATTATTAGCCTGACTGGATTTGGGATCACCTAGGAGACAGCTCTG | 120 | 45 |
| chr7 | 143670506 | 143670626 | Target=1;ProbeIdx=11497;Target #1 | 1 | + | good | CTGAGCTTCAGGGATCTGCCTGCCTCTGCCTCCTCAACTCTAGGATCATAAACACAGACCAGCATTTTTTAATGGTTCTAGGGAACAAGTTCAGGGACTCACATTTTCATTTCATTTCAA | 120 | 43.33 |
| chr7 | 143670866 | 143670986 | Target=1;ProbeIdx=11533;Target #1 | 1 | + | good | TGACCAAGACAGACTTGGATGTACCTCTGATGTCTCTGACAGCATTTTACTTTGCCACAGGTATTCTTTTATGACAGGACACTGTATGCTATTGAATGGAATCCCCCCTATGCAGAATGG | 120 | 43.33 |
| chr7 | 143670986 | 143671106 | Target=1;ProbeIdx=11545;Target #1 | 1 | + | good | ACCAGGCAGGTGGCCACCCTCCAAGGAGATGGCAGATGTTCTACTTGGCCCAGCTTTCCATGTCTCTGCTGGAAACTGGACCCTGGAATGAAGGTAGACCAGTTAGGCCATCGTGTAGCT | 120 | 55 |
| chr7 | 143671106 | 143671226 | Target=1;ProbeIdx=11557;Target #1 | 1 | + | good | TTGCTTGCTGGTTAGGTCAATGTAGAAGATTGCTTTTCTCCTCAAATGGGATCTCCTAGCCCCAACTACTGGCCCAACAACATGGCTCCCTTGTAATTAGGAGCCTGGCATGGTGCCCAG | 120 | 50 |
| chr7 | 143671226 | 143671346 | Target=1;ProbeIdx=11569;Target #1 | 1 | + | good | AATCCCTGGGAAAGACATCCACAGCACCACTGCTTACTGTAAAGCTGTCTTAACCAGATCGTGCTGTGCCATCTGAGAAGTTTGGGCTTGGACTCCAGCTCCAGACTCCAGGGTACATGC | 120 | 51.67 |
| chr7 | 143671346 | 143671466 | Target=1;ProbeIdx=11581;Target #1 | 1 | + | good | AACTGTGAGCCTCTTTCTTGTCTTAAAAAAGGATGACTTTTAGCTTTAATCATGTGTACATGGGTGTGGAGGGTCCAAAGCCAGACTCAGGAAGAGTAGGGGATTTGGGTTTTGCCACAC | 120 | 44.17 |
| chr7 | 143671466 | 143671586 | Target=1;ProbeIdx=11593;Target #1 | 1 | + | good | GAAAGTCTACCCTGGGGTTACCATGTAGGAAAATCAGTTTGGCCTCCTGGAGGATAGAGGATATGAGGAAGAAGTTAGGTGAGCACAGGCCACTTTGAGACCTACATTGACACTGGCTTG | 120 | 48.33 |
| chr7 | 143671586 | 143671706 | Target=1;ProbeIdx=11605;Target #1 | 1 | + | good | CACCTGATAGCTCTGCCACCCTGCAACAGTCACATGGGCAAGCCTAGAGTGGCAATGCTCAGAACCTGTTATAACTTAGTGTGATAGCATATTGTTCAGGAGTGTCGTTATCCCTCAATG | 120 | 47.5 |
| chr7 | 143671706 | 143671826 | Target=1;ProbeIdx=11617;Target #1 | 1 | + | good | TCCATGGAGATCAGTTCTAAGATCCTGCCATATGCCAGACCCATGGATATTTGTATTTGGGTTCTGGAAAGGGTTGGAATGGATAGAGCAGATACGTATTAGATTGGTTTACATGATACT | 120 | 40.83 |
| chr7 | 143671826 | 143671946 | Target=1;ProbeIdx=11629;Target #1 | 1 | + | good | GTCTGGGTAGTCCAGCAGTGGCTGTCTTACACCGTGAAGGCTGGGAGCTCTGTAGCTAGCTGCACATCCACAATGTTGGGCACCTCAGCATCCCAATCTGGTGCTGAAGGCTCAGAAGAT | 120 | 55 |
| chr7 | 143671946 | 143672066 | Target=1;ProbeIdx=11641;Target #1 | 1 | + | good | TCCTAGAGAGCTGTCTCCAGTCCATACTGGAGACAAAGAAGTTGGGTTCTGACAGCTCAAAGGATGGCAGCAGCAGGAACAGGATAGACACATTCACCAGCCAGACACCAAGGCAAGCAG | 120 | 51.67 |
| chr7 | 143672066 | 143672186 | Target=1;ProbeIdx=11653;Target #1 | 1 | + | good | GTAGCAGGTCTTTCCCCAAGATCTCTTTCTATCTGAGCTGCCATGGGAAGCCACTGCCCACTTAGGGGGAGGTTCTCCTCTCTTCAGTTAGTCCTTCCAGGAAATGCCCTCACAATCCCA | 120 | 52.5 |
| chr7 | 143672186 | 143672306 | Target=1;ProbeIdx=11665;Target #1 | 1 | + | good | CCCAGAGTATAGCTCTTAATTCTTTCCAGATTCAGTAAGTTGACAAACAGATCAACCACCCCAATGCTTACATCCTTCTAGAGAAGGAAGTGATGTTGAACAGGACTTGTATGATCAGGG | 120 | 41.67 |
| chr7 | 143672306 | 143672426 | Target=1;ProbeIdx=11677;Target #1 | 1 | + | good | TGCTTCAGATCACCTCTAGCTGACCACTGATGGTATCAATATGTAAATGACATGCAAATGGAGTGGTTAGGGACGGTACCATGATTGCTCCTTTGCAGATGCATTTTCTTTTGAATTTTA | 120 | 40 |

| | | | | | | | | | |
|---|---|---|---|---|---|---|---|---|---|
| chr7 | 143672426 | 143672546 | Target=1;ProbeIdx=11689;Target #1 | 1 | + | good | GCCTAGGGTGGGATCTCTCCTCAGCCCTGTCACAGTTGTCAGTCCCAGGACATTGTTGTCTACAGACCTCACAGGCTTTATCCTGGAGATGGGCAAAGCCTCCTGTGGAGTTTGTCTGAG | 120 | 54.17 |
| chr7 | 143672666 | 143672786 | Target=1;ProbeIdx=11713;Target #1 | 1 | + | good | GTGGTAAGCAATCTCTGAACTTACTGGGCCAGGCACCACTCAACCTCCCCACTCACTGAAGAAGTTTGTGTTTAAACATCCGAACCCAGCTCTGATAATCCGCTGTGTCCCCAGGCTTCT | 120 | 50.83 |
| chr7 | 143672786 | 143672906 | Target=1;ProbeIdx=11725;Target #1 | 1 | + | good | GTGTCAATCAGAACAAGAGAGGAGATGTGGCTGCCTTGAGCCTCCCTCTTCCTTTCTGCAGAGGCTTTTTCTCTATGATGGGAGGGGAAGTGGGGATGGCAAAGCTGCCCACCTGCTAGC | 120 | 54.17 |
| chr7 | 143672906 | 143673026 | Target=1;ProbeIdx=11737;Target #1 | 1 | + | good | TGAGGCACAGCCTCATAAGCTGGGGCAGGGTAGAGTATAGTGGCTTTCCTCTGTGCTCTGGCTGTGGATCTAAGAAGAGACTGTTGAGAGGGCAATGAGAAGGGGAGTCATTGGCCAAGC | 120 | 53.33 |
| chr7 | 143673026 | 143673146 | Target=1;ProbeIdx=11749;Target #1 | 1 | + | good | TTGGGCAAGCTCAGCTCCACAATGCAGAGTGAATGCCCCTTGGTACACAGTAGGGAGGCAGTGACCCCAGGGTGCCAACTGCCCTGCAGCAGGAGACCAAGGGAATGAAGGGGCAGGGCC | 120 | 60.83 |
| chr7 | 143673146 | 143673266 | Target=1;ProbeIdx=11761;Target #1 | 1 | + | good | AAAGGTGGCAGTCTCTCTTACATCTTCCTTTGAGAAGAAATACTCAGAGAAAATACTTTGCTGCCCCTCCCCTTCAGCCAGGACCAGGTTTGCTTGTTGGGTTTCTTGGAGCTCTTCATT | 120 | 45.83 |
| chr7 | 143673266 | 143673386 | Target=1;ProbeIdx=11773;Target #1 | 1 | + | good | CTTTTAGGTAGCCTGTGGGGCAGGAGGGATGTTACCCAGGAGACTCCCAGGCTTATGAACCACACACAGTGAACTCTTGCAGCTATTCCAGGCTGAAGTCCAAGGTCCAAGTGTGACTTT | 120 | 51.67 |
| chr7 | 143673386 | 143673506 | Target=1;ProbeIdx=11785;Target #1 | 1 | + | good | GATAAGCACACTCCTTAAGGGTAGCTCAGCATGCGCACTAGTCTAGGACAAAACAAGTTCACAGTGACCCTAGCTGGGAAACAGGCAGGTGCCCTGGGGTGGGCTGAGGTGCCAGCTAAC | 120 | 55 |
| chr7 | 143673506 | 143673626 | Target=1;ProbeIdx=11797;Target #1 | 1 | + | good | ATATCTGTAGGGACTGGAAAGAATAGCTCTTCTGTCTGCTGTGCTGCCCAGCCCCTGGCCCACAACATACCTGGCTCCTATGCATCATCCAGCAAGGCCTCCTGTGCAGGACCTGCACAA | 120 | 55 |
| chr7 | 143673626 | 143673746 | Target=1;ProbeIdx=11809;Target #1 | 1 | + | good | TCCTCAGCCTCTCTCCATGCTGCCCTGACTCCTTCCACTGCCTTGTCCTTGAGTCTTTGGTTCCAGGCTTAGTCGGGTACACTGTATCAGAGACATCAAGAAGAGCAGACCTTATACCCC | 120 | 52.5 |
| chr7 | 143673746 | 143673866 | Target=1;ProbeIdx=11821;Target #1 | 1 | + | good | TCCACACACAAACTCTTCTGATGGAATGGGCACAGGACCCAGAAGGTGAAGGGATTCTTTGAAACTATAATGAAGAGAAAATTGAAGTAAGCATCAGCCAGGAGGGAGAAGGAGGGAGGG | 120 | 46.67 |
| chr7 | 143673866 | 143673986 | Target=1;ProbeIdx=11833;Target #1 | 1 | + | good | AGGGAGGGAGGGAGGGAGGGAGGGAGACAGAGACAGAGAGAGCGAGCTGGGACTTCTGGGTGCTGCTTCGGGGCACCTCCTGTTCCATCAGGTACAGTGTTGAAGTGCTGCTCTGGTATG | 120 | 60 |
| chr7 | 143673986 | 143674106 | Target=1;ProbeIdx=11845;Target #1 | 1 | + | good | TAAGGAGGTCTTGAGGCTTGGCAGGATGCAGGGCTCTGACCTGCCTGTGGCATTTCCCTGAGGTCTATTGAACAGGCTGGGATGCCTGTGTGCTGCCTGTGACAGCAATGCCATCCTTCC | 120 | 56.67 |
| chr7 | 143674106 | 143674226 | Target=1;ProbeIdx=11857;Target #1 | 1 | + | good | CAGGCATGGTCTGAGCTTGGTGGGTGAGGTTGAGTCTCTTCCCGTGAGCCCGACTTGTAGGGTGTGATTGGCAGGTTATGGATTTAGAAGCCACGCTTCAAAGAGCGTGAGATGAAATTA | 120 | 50.83 |
| chr7 | 143674226 | 143674346 | Target=1;ProbeIdx=11869;Target #1 | 1 | + | good | ATGTTAAGTGTAAGTGTAAGTGTTTGAGGCGCCACAGAGAAGACGCTTGTCAGTTTCTGAAAACAGATGACACGTCTCTCCCTCCCCAGAGCTAGAGGATCCGGTGACATGGGAACTGTGCCTCCT | 120 | 50.83 |
| chr7 | 143674346 | 143674466 | Target=1;ProbeIdx=11881;Target #1 | 1 | + | good | CAGGTGATTCATTTCCTGGATGCTCTGCTGGCCAGTCAGCTCTGTGGACAGGAGCCAGATAAGCACCTTTTCCTGCCTCAACAGGGGATCATGCTGGAGAAGGGCATGTCCCTGTCTACT | 120 | 54.17 |
| chr7 | 143674466 | 143674586 | Target=1;ProbeIdx=11893;Target #1 | 1 | + | good | GAGTGTCCTGAGTCCTCAAGGTTCCAAGAGCCATGTGAGGGTAGAGCTCCCTGGGATCAGGTCAGATCTACCCTGACTCTCCATAAAGGAGTCCCACTGAGGGTCTCTTGGGAGGCACCT | 120 | 55.83 |
| chr7 | 143674586 | 143674706 | Target=1;ProbeIdx=11905;Target #1 | 1 | + | good | GGAGTCTCCTCAAAGGGTCACCACCCACACATCACATCCATAGGCCTCATGGAAAAGACAGGGCAAGGAGGGCTAGTGACTGTGAGTGAGTACCAGCCTGGTGCTATACATCCTGACATG | 120 | 53.33 |
| chr7 | 143674706 | 143674826 | Target=1;ProbeIdx=11917;Target #1 | 1 | + | good | TTAAACGTGCAGTTGGTGAAATGTCTTAACACATCCTCTGCCCTATCAGGGCCAGGGGAGTATCTGACCCTTCTTGGCTTGAGCCAATACCTTGGCTTTGACACTGTCTTAGATTCTCTG | 120 | 47.5 |
| chr7 | 143674826 | 143674946 | Target=1;ProbeIdx=11929;Target #1 | 1 | + | good | AAGTTGACAAAAATTAACTGGCACAGGCACATCCTGCTCTGTGATGGACAGGTCCCATGACACCATTGTCTTACCTTCCTCTTCTGCCCAGCAGCTCAAAGCAGTGCATGGAACAGATGC | 120 | 49.17 |
| chr7 | 143675306 | 143675426 | Target=1;ProbeIdx=11977;Target #1 | 1 | + | good | CTATGACATTTGCGCTGTTACTGAGTTATTCTTACTGACAAACATGGCTGAAATCAAAGAAGACAGCATCCCCCTGCCTTCTCTCACATCTAGCCCTACAGCCTCAGTGGACTATGGAGC | 120 | 46.67 |
| chr7 | 143675426 | 143675546 | Target=1;ProbeIdx=11989;Target #1 | 1 | + | good | TCTGGCCCTTCCCTCCCTCTGAAGACATCTTTGGAAGCATGGACAGGCAGTGGTGGAGGGGGGTTGGGTGGCCACTGCTGCCTGGAGGTTCCCCAGAGAGAAGCAAGCAAGCAAGTCATA | 120 | 58.33 |
| chr7 | 143675546 | 143675666 | Target=1;ProbeIdx=12001;Target #1 | 1 | + | good | AATATTTACATCAACTTCATTTGCATAGCAAACATAATTACTTCACACAGAACCATGTGGAGACTTCATCCTTGGTAGCTTATATGGCAGCAGGTATATCTCTTCTGTCCCTGCCACCTG | 120 | 40 |
| chr7 | 143675666 | 143675786 | Target=1;ProbeIdx=12013;Target #1 | 1 | + | good | AGTCCTTTAGGAAGTGGCTTTCCAGCCATGTACTGACAGTTCCTTACTTCAGGGTGAATGGGCCTCTTGTTCTACCCACCAGGACAGGCTATGACACTGCTAAACCTTCCATTCTGGCTG | 120 | 50 |
| chr7 | 143675786 | 143675906 | Target=1;ProbeIdx=12025;Target #1 | 1 | + | good | TTTTCTGCCTTGTGGTCCTGTCCTGTTCTTTGCTTGTCACCATGAGCAGTGGGAATGATGGTTGCTCCCTCAATGCCATGCCGAGAAATGTCCTCAGCTTTAATAGTTTCACCTGCTACT | 120 | 47.5 |
| chr7 | 143675906 | 143676026 | Target=1;ProbeIdx=12037;Target #1 | 1 | + | good | GGGTTTACCTTTTACAGAGTGCTAGAACCCATTTCTGGCAAGTTCTATGCCACTTTAGAGTGGGGATGGGTTTTCCAAGCTCCCCTAGCATGCCCTCTCCTCTCCAGCTGAGAGCATGCC | 120 | 51.67 |
| chr7 | 143676026 | 143676146 | Target=1;ProbeIdx=12049;Target #1 | 1 | + | good | TCTGTCAGCATCTTCATCGGTGTGACTTAGTTGTACAAAGTGATGGGTTTGCTGTGGTGTTTTCACAAAAATCCATGTTGTAGCTTTGGTCCTCTTCACCCCACTCCACAACCTTGTTCC | 120 | 45 |
| chr7 | 143676146 | 143676266 | Target=1;ProbeIdx=12061;Target #1 | 1 | + | good | GCCAAGCGTATTCAAACCACCACACCTACACTTAAGCCATGTGTATATCTATACATAGACAAAGACGTGCCAATGTTATATCTAAATATACATATACATATACATATGTTTAAAGTGTATTTAA | 120 | 32.5 |
| chr7 | 143678906 | 143679026 | Target=1;ProbeIdx=12337;Target #1 | 1 | + | good | TATATAATGCACATATTGAGTACGTGTTTATTTAATACATTGTTTAGGTAATACAGATGAGAGAAAATGAATTCCTTTAAAATTAGTATTTCCAGTGTACACTCCAAAGCCTTTGCAGTG | 120 | 30 |
| chr7 | 143679026 | 143679146 | Target=1;ProbeIdx=12349;Target #1 | 1 | + | good | TGTAATGGTTTCAAAGACATCTATCTACACTACATTCCTAGTCTTACCATCTGTGTTCACTGAGGTTCTTAAAAGGGAGAACAATACACACACACACACACACACACACACACACACA | 120 | 40 |
| chr7 | 143679146 | 143679266 | Target=1;ProbeIdx=12361;Target #1 | 1 | + | good | CTTCTTAGTAAAAGAAAGGACTCCATCTGGCTCTTGCTCTCTGTTCTATCAAAAGGCCAGGCAAAACAAAACTCCACTTAAAATAATATAAAGAAGCAACTTGGAAGAAACTCAGGTTCC | 120 | 38.33 |
| chr7 | 143679386 | 143679506 | Target=1;ProbeIdx=12385;Target #1 | 1 | + | good | TCTAGGGTCAGATGTTCTTGAATCAAAGTACTGCCCAAGGTTTCACTCTCTGCTGAAAATCACACTCTACTCAGCAGCCTTACTGCATTAAATCACCCAGTCTTTTAGGAAGGAGAGCAG | 120 | 44.17 |
| chr7 | 143679506 | 143679626 | Target=1;ProbeIdx=12397;Target #1 | 1 | + | good | AGCTCTGACCACCCAGAGCAAGGCAGAAGATCCATTTGCATTAGGAATAAAATCCCAGGGGGGACCTTGCTTAAGAATAGTTACACTCCCCTCTGCAGAAATAAAGCTTTCTGCCTTATC | 120 | 45.83 |
| chr7 | 143679626 | 143679746 | Target=1;ProbeIdx=12409;Target #1 | 1 | + | good | CCAAGGAAGCTGGAAACAGAGGTCAGTGCTGCAGCCTCGAGGATTGCTTCTTCCTGGGGAGACCCACCCCCACAGGGTTCCTTTACGTTGCTATGTGATGTCAGTTCAGTGTAGGTGTGT | 120 | 54.17 |
| chr7 | 143679986 | 143680106 | Target=1;ProbeIdx=12445;Target #1 | 1 | + | good | ATGACTCTGAGCCTGCATCTTATGGTTTGTTGTGGGTGTACTTATAATGGCTTCATTCCGGGGGCTCTAGAACAGTATAGGCATTGGCTTAGGGTCTCTATTTGGGGCTCCAGGACCATT | 120 | 47.5 |
| chr7 | 143680106 | 143680226 | Target=1;ProbeIdx=12457;Target #1 | 1 | + | good | CCCTGGTTACTTGATGTCACTTGCTGAGTGCTTCTGACTGTCAGGTGGCAGGCAGCAGAGCTACAGCACTATGGTGAGATGCTTTTTAGCTGGATCTCTCTCTCTCTCTCTCTCTCTC | 120 | 50.83 |
| chr7 | 143680226 | 143680346 | Target=1;ProbeIdx=12469;Target #1 | 1 | + | good | TGAATTTGAGATCCTCTTGCCTTTACCTCCTAAGTGCTGAGATTATAGCCTGTTCCACCACATCTGCTTTAGAACGAATGGTGCTTTAGAAGGTGGGGCTCAGGGATTCTTGCTGTTAGG | 120 | 45.83 |
| chr7 | 143680466 | 143680586 | Target=1;ProbeIdx=12493;Target #1 | 1 | + | good | CAGGCTCTCAAGCTCTCTATCATTGAGCGGCATCCCCAACCCTCAGGCTGTGTGCATGGGTGGAACAGTTTGTGCCCAGGGAGCCTTAAGAGCATTTGAAGGACAGAATACCCAACAAAG | 120 | 52.5 |
| chr7 | 143680586 | 143680706 | Target=1;ProbeIdx=12505;Target #1 | 1 | + | good | CTAGAAACTCCCTCAGACATGTCCAGAGGCCTGTCTTCATGGTGGTTCTAAACTCCCCCTGTCAAGTTGACAATCATAATGAGTCCTCACAGAAGAACTCCTGGGAGTGGGTGGGATAGT | 120 | 49.17 |
| chr7 | 143680946 | 143681066 | Target=1;ProbeIdx=12541;Target #1 | 1 | + | good | AGCCCAGCCTTCTGATACATAGAATACAGTCCTCAGTGAGGAGTTCACAGACCCTATGAACCCTTGCTGCCACTGTAGACTGTCCCCATGCCTCTGGACAGACAGTTGTCCCCACACAAA | 120 | 51.67 |
| chr7 | 143681066 | 143681186 | Target=1;ProbeIdx=12553;Target #1 | 1 | + | good | GATTGCAGTGGAACAAAAATGTCATGATGTGGTTGCTGAGTATTGATGGCCAGGCATCACTTGGCTCGGGCCTTTGTGGTGGTTCACTTTGTCAATTTGATACAATCTAGAATCACCTGG | 120 | 45 |
| chr7 | 143681186 | 143681306 | Target=1;ProbeIdx=12565;Target #1 | 1 | + | good | TGTAATAGTCAAGAAGTTGACTCAGTAGAAGCAAGCAAGCATCCATGCGTTTATTCTCCCTCTGCTCCTGACGGTGGATTGATGCTTTAAGTCCCTGCCTCAGTTTCCCCTCAGTAATTG | 120 | 45.83 |
| chr7 | 143681426 | 143681546 | Target=1;ProbeIdx=12589;Target #1 | 1 | + | good | ATGTAAAGGGTTTCATTAGGAATAAAAATCCAGTGTGACTCCTGCTGCCTGCTGGCTTCAGTGGGTGCAACATTTTTTTCCTGCTGAAGTGCTTTGGATTGTGTGGTCATTTTCTTGAGA | 120 | 42.5 |
| chr7 | 143681786 | 143681906 | Target=1;ProbeIdx=12625;Target #1 | 1 | + | good | CCCATTTCTTACAACTCTAACCTGGAGTTGTAAGGCAAATAAACCTTTCCTTCTCTAAGTATCTGTCACAGCAACAGACATGAAATGAGGACAGCGTTTGACGATGTAAGCGCAAAGAGA | 120 | 41.67 |
| chr7 | 143681906 | 143682026 | Target=1;ProbeIdx=12637;Target #1 | 1 | + | good | CCCGTGGTTGTGCAGCTGACGTTGAATTCTGTGATTTTTCACTCTGCAGGTGGCTATAATAGCGGGCAACTTTGAACTTGCAGAGTACATCAAGAACCACAAGGAGACAGACATTGGTGA | 120 | 46.67 |
| chr7 | 143682026 | 143682146 | Target=1;ProbeIdx=12649;Target #1 | 1 | + | good | GTAGGCTTAGGGAGAAACCACAGCCATTGTCATCTGAGATCTTCCTGGGACTACTTGTGGAGCTCTCAGCCTTGGGAGTCCTTCCCTTATAGTGGGGAGGGTTGTGAAGACCTTGGGGGA | 120 | 53.33 |
| chr7 | 143682146 | 143682266 | Target=1;ProbeIdx=12661;Target #1 | 1 | + | good | TGAGCAATACACTTCCCTTGAAGAGTACCACTCTCCTCTCTCCTGACAGTGCCTGCTACAGGCACATGTGAAGTGTGCAGGGCCTCAGGAATGGGGATGTTTGAATTCTGCTGGGGTTTG | 120 | 51.67 |
| chr7 | 143682266 | 143682386 | Target=1;ProbeIdx=12673;Target #1 | 1 | + | good | GTCTAGGGTTCCCACCTGATTCAACTCTCTTGTTTCCCAATATTGATTTAAAATGTATGCCTGTAGCTGTGCATTTCTGCAGATGGTTAGGGTCTAGCAACATTGGCAGGTGACGCTGAT | 120 | 44.17 |
| chr7 | 143682386 | 143682506 | Target=1;ProbeIdx=12685;Target #1 | 1 | + | good | CCTAGGGGAGGTGGGGGGTCGGGAGGGCTGGGGGATATCTCTATACCTTCCTCTCAAGTTTGCTATGAGCCCCAAGCTTTCTCTAAACAGTCTTTAAAAACACTACAGAAACAAAATC | 120 | 48.33 |
| chr7 | 143682506 | 143682626 | Target=1;ProbeIdx=12697;Target #1 | 1 | + | good | AGAGAAACTGTGTGGGTACCAAGGAGGAAATGCACTGGGGAGCTGGCTTAGATGTGCTTATTGGAAGGCACAGACTCATGTGTGGATGGGTGGCAACTGAGAGGGAAGAAGAATGTGAGG | 120 | 50.83 |
| chr7 | 143682626 | 143682746 | Target=1;ProbeIdx=12709;Target #1 | 1 | + | good | TCCTCTCTTCTGCCCTCCCGTCACCTCTACTTGGTGTCCCCCTCCCCTTTCTAGTACCACTTGCACCAACCAGGAGGCCCATCTCGGGGGCTCCTCCCTGCTCCCACCGTCCTACCATGG | 120 | 62.5 |
| chr7 | 143682746 | 143682866 | Target=1;ProbeIdx=12721;Target #1 | 1 | + | good | GCTGGGCCCAGTCTTGCACAGTGGATGAGCCAGCCGGCCTGCGCCCTCCAGCGCTGGGCTTCCACAGGACCTGCAGCGGGGAACAGCCCCACTGGACCGGCAGTGGTGAGCCCTTGAAGG | 120 | 69.17 |
| chr7 | 143682866 | 143682986 | Target=1;ProbeIdx=12733;Target #1 | 1 | + | good | TGGAGTTGTCCCATGGAGGGGCTCTGGGATGGAGCTTCTTGGGGCCTTGTGGAAGAAGCAGATGGCATCTGCCTCTCACTGCCTATGCCCTCACCCTGCTTTGGGTGTCAGAAGTGAGCG | 120 | 57.5 |
| chr7 | 143682986 | 143683106 | Target=1;ProbeIdx=12745;Target #1 | 1 | + | good | AAACTACTCTGTCTTGCTTTTAAAACGTCTTCACCATGTTGAATGTGTGTCTTGCATCGAATGTGGCTGGGATCTGTCTTGCTGGGAAGTTGGATCGTGATGTAGAGCCAGACGATGGGC | 120 | 46.67 |
| chr7 | 143683106 | 143683226 | Target=1;ProbeIdx=12757;Target #1 | 1 | + | good | TCATTCCAGCTGTGAGCAGATTGCAAGTGCCGAACCTAATGTTGATATCTTGTCCTCTCACATGCATTCTTCACCTTGACCAAGTGAGTTCTTACCTGCAGAAACAGTTGCACCATCCAC | 120 | 45.83 |
| chr7 | 143683346 | 143683466 | Target=1;ProbeIdx=12781;Target #1 | 1 | + | good | AGGTCCTATATTAATTTATAACCTTCCCTTATATTTTACCACTTCAGAGGGAAATGGCTTCCCTTTGAGGGATGGCAGGCCTGGGAACTTCTTCCCAGAGTATCCTATGAGGTTGGAGTC | 120 | 44.17 |
| chr7 | 143683466 | 143683586 | Target=1;ProbeIdx=12793;Target #1 | 1 | + | good | AGTGCCAGCAGGACACCTCATATGGGAATGCAAGTGATACTGCCCTGTGGAGGCTGAGAGGGAAGATACATTTGAGGGAGACAGTCACTGTTCAGGGCTCCTAAAGTGGGGAGTCTGTA | 120 | 51.67 |
| chr7 | 143683586 | 143683706 | Target=1;ProbeIdx=12805;Target #1 | 1 | + | good | TCCTAAGACTCCAGTCTGGTGTAGGGAGCTGCAGACACCATGTCTCCTTCAGCACATAAGAGAATGTCTACAGGGACCAGAAGTATGAGGGGTTAGAGATGTCTGGGTCAGAGCTCCCCA | 120 | 51.67 |
| chr7 | 143683706 | 143683826 | Target=1;ProbeIdx=12817;Target #1 | 1 | + | good | TGACCAATGTGATCTCTGACCACCTCCTCCGGGGAAGACCCTGGCAGATTAGTGGGTCCAGAAATCCCCCCCCCCGTGCCCTTTAGAAGGTGGGGGGGGGCTCAGAGACTGATAGCTT | 120 | 60 |
| chr7 | 143683826 | 143683946 | Target=1;ProbeIdx=12829;Target #1 | 1 | + | good | CTGTAGCTCCTCCCTCACCCCAGGCTCTAAGGGTGATGTCAAAAGTACTGAAGATGTTGGGTCCCCTTAGAGGTTATGTGACTTAAAGATCAGTGGCTGCTTACTTTGAAAAAGGCACCT | 120 | 47.5 |
| chr7 | 143683946 | 143684066 | Target=1;ProbeIdx=12841;Target #1 | 1 | + | good | GACTTTTCAGAAGTTCAGCCTTTGGTACACATTGTCTCATCTTTGCTGTTACAGTGCCGCCTCCTCCTGAGACCTGGTCATCATTTGCTCTAGATGTTTTTTTGTTTGTTTGTTTTTTT | 120 | 40 |
| chr7 | 143684066 | 143684186 | Target=1;ProbeIdx=12853;Target #1 | 1 | + | good | TTTTTTTTTTTTTTTGCTATCCCACTACTCCCTAACTCTACCGTTTTCTGCTAGCCTGGCAATTGATCTGCAGAGATAGCTTGGGAGCCACTCCTTCCTCTCTGTAATGGACTATGCTC | 120 | 43.33 |
| chr7 | 143684186 | 143684306 | Target=1;ProbeIdx=12865;Target #1 | 1 | + | good | ACCTGAATGAGAGATCTCTTGGTCTCCTCAGGCAGTGTTGCACAGTGTGTATTCGATAGTTGGGTTTTCTGAGAGTTTGTGTGGCATGCAGATTCCCTGGCAACCACTTTGTGTGACTCC | 120 | 48.33 |
| chr7 | 143684306 | 143684426 | Target=1;ProbeIdx=12877;Target #1 | 1 | + | good | ACGAGGTGCCACTGAAATCACTATGCTAACGTCCTCTGGCAGTTCAGAGGCTGGGCACCCACCTTTGTGTCTCAGGGCCTACCTGGGCATCCCCATCTCTCATGAGCTCATTCAGATAGG | 120 | 55 |
| chr7 | 143684426 | 143684546 | Target=1;ProbeIdx=12889;Target #1 | 1 | + | good | ATTGGCCACACATCAGTTCCCAAGAATGACAAATGACAGCAGGCTGAGCCGTTTAAAGTAATGAGTTTACTCTGCCATTCTAGGGGACACGAGGTCAGAATCAAGATACTGGGTTGTTCC | 120 | 45.83 |

| | | | | | | | | | |
|---|---|---|---|---|---|---|---|---|---|
| chr7 | 143684546 | 143684666 | Target=1;ProbeIdx=12901;Target #1 | 1 | + | good | GCAAGACTCTGGAAGGGTTTACTTCAGACTCTTTCCTGTGTTAGCTGCTGCTGCTGGTCTTTGGCCTCTATGGGTTTGTGACACGATTGCAGTCTCTGGTCTGTGATAACCTGACCTTCT | 120 | 49.17 |
| chr7 | 143684666 | 143684786 | Target=1;ProbeIdx=12913;Target #1 | 1 | + | good | CCTGATGTCTCACACCCCACGCCATCCTCCGCCATGATGGACTATGTTCTCAGGCTGTCTATCAAATAAACTTGACCTTTTTGTTTGTTTGTTTGTTTTTGTTTTTTCATTTTTTTGTTT | 120 | 38.33 |
| chr7 | 143685026 | 143685146 | Target=1;ProbeIdx=12949;Target #1 | 1 | + | good | GCACCACCACGCCCAGCTCTTAAATTGCTTCTTACCAGGCATGTGGTCCCAGCAGTGAGACAATTAACTGTTCAGCGTGTCTCAACTTCCTGCCTGATCTTAGGACACCAGTTTGGTCCT | 120 | 50.83 |
| chr7 | 143685266 | 143685386 | Target=1;ProbeIdx=12973;Target #1 | 1 | + | good | GTATCTAAGATGACCTTCCACTCAATATCCTTAAACTGACCATCTGTGAGAAGACCCATTTCCAAGGCCATAATTGTAGGGCCCAGAGTTTAGTGTTGCCTGTCTCTGGGCCATCGCCTC | 120 | 47.5 |
| chr7 | 143685386 | 143685506 | Target=1;ProbeIdx=12985;Target #1 | 1 | + | good | TTCCAGGGCAACAGTATTGGACTCTGCACACCCCTCAGTACCTGCTGCTCAGGGAGGCCAGGAAGTATGTGCTAAGTGAGTGACCCTCCTCCATGCTGTTTGCCATTCAAGGCTGGCCTT | 120 | 55 |
| chr7 | 143685506 | 143685626 | Target=1;ProbeIdx=12997;Target #1 | 1 | + | good | TCTGTCACTTTCCTTCCAGCATGTGTATTTGTGCTCATTCATTTAGAACTCTGGGGACTGTTTTGGATGGTGTTCCTCAGGAGCTGTCTACCTTTGGTTTGGAACGGCCTCTCTCTGGCC | 120 | 48.33 |
| chr7 | 143685626 | 143685746 | Target=1;ProbeIdx=13009;Target #1 | 1 | + | good | TGAAGGTTTATTTGCTGGGTAAGCTAGCTGATTAGAGAGTCCCAAGAAGCCTCCTGTCTCAGCCTCCTGTGCATAAACCTCTGTGGAGGCTTTTTACATGGCTTCGGGGATCCAAACTCA | 120 | 48.33 |
| chr7 | 143685746 | 143685866 | Target=1;ProbeIdx=13021;Target #1 | 1 | + | good | GGTGCTCACACGGTGGACTAAGCTAGTATTCCATGTTGTAGTCTCATAGTTTAAGCTTTGCTACCAATCTGTCCCTAAGTGTGTCCCCCATTTGTGAGTCTGGCACCCCTCACTGCTCAT | 120 | 48.33 |
| chr7 | 143685866 | 143685986 | Target=1;ProbeIdx=13033;Target #1 | 1 | + | good | GAGCAGAGAGTCCCGTGTATCCTAAGCTGAGGAAGGCTGAGAGAGGCTGAGCTGCTGTCCTTGAATCTCAAGCAGGAGGAAATGAAGGTAGGAACCCCCCACCCCCAACACACACCTCTG | 120 | 55.83 |
| chr7 | 143685986 | 143686106 | Target=1;ProbeIdx=13045;Target #1 | 1 | + | good | TCAGGGTGGGTCAGGTTAGGGCACATGTCTGTATGTAGGGTAGTGGAAAGCTCTCCACATCCCAAGTTGGGGATGGTTAATCAGGGAAAAAACTTCCTACCGAAAAATCAAACTGTCCTT | 120 | 46.67 |
| chr7 | 143686106 | 143686226 | Target=1;ProbeIdx=13057;Target #1 | 1 | + | good | GCTTTTGAATGGGCAAATCTTTGTTTTCTATACTTGCTAGTGTGATAGAAAAACCAGAGCACTTCAGAAGTGCAGAGGTGGTCTTGGGGAAGGCAGCCATAAGACCCTGCGGTCAGAGCC | 120 | 47.5 |
| chr7 | 143686226 | 143686346 | Target=1;ProbeIdx=13069;Target #1 | 1 | + | good | ATTTGGCAAATGGACACCTGGAGTTCATGTGGCTGCAGCTCTCCTCCCCAATAGGCACAGCATCATCATAGCGTGCACATTTATGTGCGAGCCGGTCACAGTGTGCGTGCTGCTCTTGTC | 120 | 53.33 |
| chr7 | 143686346 | 143686466 | Target=1;ProbeIdx=13081;Target #1 | 1 | + | good | TGATTCTGCGCTGCTGTTTTTTTTTTTAAAGCACCCCCCCTTAAAATATTAAACGTGCCTTCCATCAGATTATTTTATCTGCCAGCTGGAGGAAAGTGCTTGTTGGGAAAGGCTGTGAAG | 120 | 41.67 |
| chr7 | 143686466 | 143686586 | Target=1;ProbeIdx=13093;Target #1 | 1 | + | good | AGCGAAGTTAATCCCTGCTTGCTGTCACCCACTGTAAGCTGTGGCTGGGCAGATGAGCCGAACCGGCTTGAGGTGCCTTGACCACAGACTACTTAGGATGACTTCCATAGGATGGCTTCC | 120 | 54.17 |
| chr7 | 143686586 | 143686706 | Target=1;ProbeIdx=13105;Target #1 | 1 | + | good | ATTAAAGCCAAGGCCAGGGGCGAGCCGGGGAGCCCACGTCCGAAATAAGCATGCAAGTATTAAGCATGTGGTACGCTGATAAGCTGCGGACCCGGCCAACCTGTTTTCCCCAATCTTTGG | 120 | 55 |
| chr7 | 143686706 | 143686826 | Target=1;ProbeIdx=13117;Target #1 | 1 | + | good | TGGTTTTGTTCTTGCCTGGCTCTGAGGGGAGATGCGCTGTCAGATGGATTAAATGACATGGGTTGCGCTTCTTCAGCGGAACAAGGTTTGTTTTCCTGTCTTGTCCTTCCGCAGCTATGG | 120 | 50 |
| chr7 | 143686826 | 143686946 | Target=1;ProbeIdx=13129;Target #1 | 1 | + | good | GCTGACTAAGGAGCACTTGGATACGGTTGCTAGGTTTCAAGACCGGGAGCCTGTATCATTACACTGTGGATTTGCATAGATGATGGATAGAGGGAGATGTTTTCAGATTTCTTAGTGTAA | 120 | 43.33 |
| chr7 | 143687066 | 143687186 | Target=1;ProbeIdx=13153;Target #1 | 1 | + | good | TATAGTGTGTGTGTGGGGGGGGGGTGTGAGAATGTGAAATTTGAGCTTGTTTTGACTGATGGTTCTTTGAGAATATACCTGTTTGGAAGATACCACCTGCTGGGAAATTAAATAATTCCG | 120 | 41.67 |
| chr7 | 143687186 | 143687306 | Target=1;ProbeIdx=13165;Target #1 | 1 | + | good | ACCTGAGAGATTTCTGTTTTCCTGGCATTTTCACTATGAAACGAGCTACTTCCGATGAAGCAGGCAAGGCTCTCAGAAGTGAGCCGTGGGAGCCTTTATATTCGCCAGAGAGACTCTCCA | 120 | 48.33 |
| chr7 | 143687306 | 143687426 | Target=1;ProbeIdx=13177;Target #1 | 1 | + | good | GCAGGTCCTCGGGAAAACCCAAACGGTGACCTATTTCCTTAAATTTAACCAAATGCATCAGGGCCCCCACTGTGGTGTCTCCCCTCTGTGGTTCTCCCCAAGGAAAGGGAGAAATGATAC | 120 | 50.83 |
| chr7 | 143687426 | 143687546 | Target=1;ProbeIdx=13189;Target #1 | 1 | + | good | TACAGTCGGGCAGGCATTACACAGTTGTGGATAGGATTCCCACAGCAGATTTTCAGAGCAAGTAGCTTAGCAGAGGAGGGGGACAGAAAGCTTCTCAGACACTGCAGTTTCTAAACAGAT | 120 | 47.5 |
| chr7 | 143687546 | 143687666 | Target=1;ProbeIdx=13201;Target #1 | 1 | + | good | GGTCTGCTACCTGGCTGCCGGGGGCTCACCCTATTCTTATTGCCCTTTGAGGAGATTAGAAAGACATGTAGAATTTTATCAGTCTAGGGTGTGTGTGTGTGTGTGTATATGTGTGTGT | 120 | 46.67 |
| chr7 | 143687666 | 143687786 | Target=1;ProbeIdx=13213;Target #1 | 1 | + | good | TGTGTGTGTGTGTGTGTGTGTCCGTCCTTGCATTGGGTTGATTGTTCAGATCTGTCTGTGCCTGGGGTCTGGGTTGCTGTCTGCCGTTGAAGGCATTGGTCTAGTGGTTTATTGTGGTTC | 120 | 50.83 |
| chr7 | 143688146 | 143688266 | Target=1;ProbeIdx=13261;Target #1 | 1 | + | good | TGTGTTCGCTGCCTCTTCTATTCAGGAGTTTTATCCCCAGTCATAAAGCATAGTTGTGCCACCTGAGTGATATGGTTTTGTTTATTTGTTTGCTTGTTTTTCCCAGTGACATCCTTGTGC | 120 | 41.67 |
| chr7 | 143688266 | 143688386 | Target=1;ProbeIdx=13273;Target #1 | 1 | + | good | TGCAGCTGTGTTAGGTGGGGGTGGTGTCTTGAATTGGGGATTTCTAACACAAGACCTAGGTGAATATCCACTTCAGATCTACTGCCATCCATGGCCCATCCTATGCGTGCGCTCGGCGCC | 120 | 53.33 |
| chr7 | 143688386 | 143688506 | Target=1;ProbeIdx=13285;Target #1 | 1 | + | good | TAAGGCACTGTGGTTTTGTTCCTAGTAGGCTATGGCAACAGGCCGTGGCCTTCTTTCTGTCTTGTGAGCATCTCCATGTGATTTGTCTTCGTGATCTCCGTCTGTCCATGTGCACTCCAA | 120 | 49.17 |
| chr7 | 143688506 | 143688626 | Target=1;ProbeIdx=13297;Target #1 | 1 | + | good | CCACGTGCACTCCCACGTGCCCCCCTAAGCAGCTCTCCATCTGTTCTAGCTTGCTGCATCCGCTCTCCCTTGAACCAATCGGAGTGCTGCTTGAGAGAGGGGTGGGTGTGAACACAATGT | 120 | 57.5 |
| chr7 | 143688626 | 143688746 | Target=1;ProbeIdx=13309;Target #1 | 1 | + | good | GTCCTGCTTTGGTCCCTCCTTTCCAGGTCACCCTGCTTCCCCTGCAGTGGACATACCTGGCTGCTGGCAAGTATATATAGCACCCCTGCTCTCTCCTGCACACATACAAACACCTTTGAG | 120 | 54.17 |
| chr7 | 143688746 | 143688866 | Target=1;ProbeIdx=13321;Target #1 | 1 | + | good | TTCTAGGTCGATCTGAAACCTTTCCTGGCCTATGACTTCCAGGTTCTGCACACCCCGAATGCAGGAAGCTCCAGAGGTATGTCTATGTCACTGTCCTCTACCCTCATCCCTGCAGTCTCT | 120 | 51.67 |
| chr7 | 143688866 | 143688986 | Target=1;ProbeIdx=13333;Target #1 | 1 | + | good | AGCAGTTTCCTTCAACATGAAACACCTGTCTCCCTGGCTGCAGCCTATAAGAGTGCAACTCCCCTCATGCCAGTCAGCATATCCATCTTATCCTCAAACTCAGAACCGACTGGATGTGGT | 120 | 49.17 |
| chr7 | 143688986 | 143689106 | Target=1;ProbeIdx=13345;Target #1 | 1 | + | good | TGACTAGCCATTTAAAGGTTCCCGTGGCCCCTCCCACACTGTCCTGGGTACCCACTGAAGCTGTCACAGCAGGGTATCCTGACCTGGAGACCCAGCACTGTCATCCGCTGCTTGCTCCCT | 120 | 58.33 |
| chr7 | 143689106 | 143689226 | Target=1;ProbeIdx=13357;Target #1 | 1 | + | good | CCTCTGTGATCACTATCCACTCCAAAGTCTCTCTTGCTGTTATGAGGCAAGAGGGCTAGGCCAGAGATGGTGGGGGGGGGGGTTACAGAAGTGATAGACAGCTCTTAGTAAAGGTCTGAA | 120 | 50.83 |
| chr7 | 143689226 | 143689346 | Target=1;ProbeIdx=13369;Target #1 | 1 | + | good | CCAACTAACAAGCCTTCTGTCATTTATGTGGAAGAGCGTCAAGTTTCTGGGGAATTCCTGAATAATGAAACACCTGGGAAAGAAGGTGGTGGCTGTGGGACACGGCCACACACCCGAGCA | 120 | 50 |
| chr7 | 143689346 | 143689466 | Target=1;ProbeIdx=13381;Target #1 | 1 | + | good | GAGGGCTGCAGGTCTACAAGGGTCTGATGCCTCATCCAGACTTCGTAAGGGATAATCTGCACAGAGCTATACTGTTCCTGTTCCTGAATACCTCAATCTCATACATCTACTGGGACTTGG | 120 | 48.33 |
| chr7 | 143689466 | 143689586 | Target=1;ProbeIdx=13393;Target #1 | 1 | + | good | AAAATTCATGGAAGTCTTGAAAGCCAGGATTGTCCTTGTTCTTGCCTATGTAGCGTACCACCACCATCACTTGAGCCAATCCATCTTTTCCCAGCCCTACCTACCCACTTCTTTTGTTTT | 120 | 44.17 |
| chr7 | 143689586 | 143689706 | Target=1;ProbeIdx=13405;Target #1 | 1 | + | good | GCCTGCAGGCTCTTCTTCTGGCTTGGGTCCTCCATAGGGGAAGACATCGTCCTCCCAGGCCTTGTGCTCTTACCACAGAGTCCTCCCACCTGATTTTGCTTTCCCTTGTGTGGGCTCCTG | 120 | 57.5 |
| chr7 | 143689946 | 143690066 | Target=1;ProbeIdx=13441;Target #1 | 1 | + | good | GGTCCTTTCTGACCTCCTTCATCCTTTCTTGTTCTCTTAGACCGCCCTGAGCCCTGTATCAGCAGGGAAGGTAAGGGCAGTCAGCAGGTACTAGAACTTTCCATTACATCTTAAAAACTA | 120 | 46.67 |
| chr7 | 143690066 | 143690186 | Target=1;ProbeIdx=13453;Target #1 | 1 | + | good | TAACGTTAGAACTTTGTAGACAGTTGCTTCATGGTTGCCATAGGCAGCCAGGGAGAAAGACTGAAGTCTACATAAGAAGTATGGAGTGTAGGCAACCTCAGAAAGGAGGAACAGTGTGTC | 120 | 45 |
| chr7 | 143690186 | 143690306 | Target=1;ProbeIdx=13465;Target #1 | 1 | + | good | TCACTCATCTAACTGACTTGAAGTGGGCTCCATCCATGCTCTATCCATCCATACATACATGGAAAGTGGGTCTCACTCTACAGCCCAGACTGATCTCGAACTCCCTATATACCCCAGACT | 120 | 47.5 |
| chr7 | 143690306 | 143690426 | Target=1;ProbeIdx=13477;Target #1 | 1 | + | good | GCCTTCAAACTAGTGGCAGTTTTCCAGCATCAGCCTCCTGAGTACTGGGACTTGATTGCTTTTTACCCAACTGGGATGGAACTGTACTGTCAGTCATGATCCCCACCCTACCCTGCATTC | 120 | 50 |
| chr7 | 143690426 | 143690546 | Target=1;ProbeIdx=13489;Target #1 | 1 | + | good | TACCAGTGACCAGAGCAGAGGGAGACTCCAAGAATGCAGACCCTAGACTCAGCTATGGGTGAGAGCCTGATTAGGCTGTGGACAGAGGTCTGTCAGAACAAGTGGCCCTAAACCCTTGCC | 120 | 54.17 |
| chr7 | 143690546 | 143690666 | Target=1;ProbeIdx=13501;Target #1 | 1 | + | good | CTGCCCCTGCAGACCTCAGACCTGCCTCACCACAGCCTACCTAACACCCGTGGGCCTCAGTGGCTGGCAGTGAATATAGCAATGGCTGCCCACCCACCTGGAATTGCTGGCCTCCTCCCT | 120 | 61.67 |
| chr7 | 143690666 | 143690786 | Target=1;ProbeIdx=13513;Target #1 | 1 | + | good | CTCAGGAGCTCATTCCTGTGCAGTTTCCCCTCATGCCAACCTGGGAAGGTCACACAGAGTGGCCTTTCTGTGGTTTCTACACCCTGGGGTCCCTGTCACTCATAACTGCATCCCTACTGG | 120 | 55 |
| chr7 | 143690786 | 143690906 | Target=1;ProbeIdx=13525;Target #1 | 1 | + | good | CTGAGTTTACCAGGACCCTCTTCCTGCTTTGTGCTTTTCCTCTGCACACTCATGCTTCTCCTAGTCCTCGGGTCCTCAGGCTCCACTGCAGACACTGTCAGCTGCTCAACTCTCAGAGCC | 120 | 55 |
| chr7 | 143690906 | 143691026 | Target=1;ProbeIdx=13537;Target #1 | 1 | + | good | TAGGGCAGGAACAGGTCTTCTAGTCAGAGCTTTCTCAGGTCAGTCTTTAGCCGCTTTTCCAGGTCTCTGGGATGTTAGGTTCTGAAGCTCAGCCTGCCTGTTGAGAGTGTATAGAAGGCT | 120 | 50 |
| chr7 | 143691026 | 143691146 | Target=1;ProbeIdx=13549;Target #1 | 1 | + | good | TGCTGGCTACCAGCCCGAATCCCAGTCTCCAACCACAGCCTTGGGCTGCTTAAGGGTCTTCAGGGGCTCAGGGCAAGTCAGTGAGGCAGGGCGGCCACCCTGCTTCTCTCCTTTCCCGTC | 120 | 62.5 |
| chr7 | 143691146 | 143691266 | Target=1;ProbeIdx=13561;Target #1 | 1 | + | good | CATTCCCTTCACCTTCCTTTTGTTTTCTCTTCCTTCTTTCACTCCTCTTTCCTGTAGTATTTGACCTTCACGCTGGCACTGGTGTCCTACCCTAAGTCCGTACTTGCTTCTGCCCCTCTG | 120 | 48.33 |
| chr7 | 143691266 | 143691386 | Target=1;ProbeIdx=13573;Target #1 | 1 | + | good | GAGGGTCCAAGAGGCTCCTTCCCAGGTTCCCACTGTTCTTGCGTAGCTCACCCCTGTAATCTCAGCATTTAGGAGAATAAGCAAGGAGGATTGCTGTGGATTCTAGGTTAGCCTGGCTAT | 120 | 50.83 |
| chr7 | 143691386 | 143691506 | Target=1;ProbeIdx=13585;Target #1 | 1 | + | good | ACAGTGAATTTTAGGTAAACCTTCACTACTGCAGTGAAACTCTGTCTTTCAAAAAAAGTATGTGTGGGGGGCATGTATGAAAGTGAACATGTAGTTGGAGGAGTTCTGGTCCGGGTCCTT | 120 | 42.5 |
| chr7 | 143691506 | 143691626 | Target=1;ProbeIdx=13597;Target #1 | 1 | + | good | GCTGCTCACTGATCCTGAGAAATGTCTCTCCTGAGCATACATGGCACATGGAGACAAGCCATGGCTGTAAGACTTGCCCTACCCTAGCAACCATGGGGAAAGCTGTGGTGATGACCTCAG | 120 | 52.5 |
| chr7 | 143691626 | 143691746 | Target=1;ProbeIdx=13609;Target #1 | 1 | + | good | CATGACCTTTGTGTCCTGCCCGACAGGTTGTTCTGCCCAGGATCTCTCTTTGGTTTGCATTCAGTTTCCTGGTCCTTATGATATCAAATGCAGAAAGGACTGACTCAGACCACATAGAGA | 120 | 46.67 |
| chr7 | 143691746 | 143691866 | Target=1;ProbeIdx=13621;Target #1 | 1 | + | good | GCATAGTGTAGCCCCACTCAGGGATGGAAGTATGATTAGATCCCTTCCTGACCAATCAGAGGCCTGGAATTTCAGCTCCTGCTGATCATAAGAGGACTCATCTTCAGTGACCTTCCAGCC | 120 | 50 |
| chr7 | 143691866 | 143691986 | Target=1;ProbeIdx=13633;Target #1 | 1 | + | good | CACGGGTATGAAGGTTCAGTTTGACAGGAGTATGAAGCAGTTGGTCCTGTAGGATCCACACTCAGGAAGCAGAGAATGACTAGAACCCTGCTCCCTGCAACCCCACTTTCAGCAGAATTA | 120 | 49.17 |
| chr7 | 143691986 | 143692106 | Target=1;ProbeIdx=13645;Target #1 | 1 | + | good | GCTTATTTCAAAGACATCAGCCATTCAGTTTATCCAAGGCCAAATGAGACTAGGGATTAGGCTCTGATTGTTTGCTTGCTTCAACTCCTGGGCATTGCGTTAATGAACAGCTGCTTTGAA | 120 | 42.5 |
| chr7 | 143692106 | 143692226 | Target=1;ProbeIdx=13657;Target #1 | 1 | + | good | AGTAGGTATCCCATTTAACCCTGGGCTAGACGTGTACTGGGGACCAGCTCTGGTTATTCACTGATCTCAGCTCCTGGGCTTTGCCCTGAAGAACTGCTTGGCAAATTACTAGGCATGTAA | 120 | 49.17 |
| chr7 | 143692226 | 143692346 | Target=1;ProbeIdx=13669;Target #1 | 1 | + | good | GACCAGTCCTGACTCAAAGCTGCCAGTAGCATACCAAGCATGGTTTAGGATCAGTCCTGCCAAAATCCCAAGGTCCTTGTAAGCTCTGTTTCACCCTGACCCCCTCACCTATGCCTTCCC | 120 | 52.5 |
| chr7 | 143692346 | 143692466 | Target=1;ProbeIdx=13681;Target #1 | 1 | + | good | TGGCAATGAGTTCAAGTACAGTTCTGGAAGTGATCACAGCGCCCATATTGGCTCCAATTGAGCTTTCTGTGAAGCTCTCTGAGGACTCTCAGCCCTCAGTGAGCCAGGGAATACCTATGT | 120 | 49.17 |
| chr7 | 143692466 | 143692586 | Target=1;ProbeIdx=13693;Target #1 | 1 | + | good | TCAACACCAGAACTTGAAGACCTTCCTTTAAAAAGCCTCATACCCACAATGGCTTCTGGGTAGTAGAACTGAAGAAAAATCATTAGGCCAAGGACCAGAAAGGGTGAGATTACATCTGTA | 120 | 41.67 |
| chr7 | 143692586 | 143692706 | Target=1;ProbeIdx=13705;Target #1 | 1 | + | good | ATTATTTTGGTGATCTTGGACTAGCCATGTTCTCTTTGGACCTCAGCAGAGGGTCAGCTCTTCCCCTACACCCCTTGGAGCTATATATATCAGATTCTCTCTGGGTCAGAGCTGAGCCCA | 120 | 48.33 |
| chr7 | 143692706 | 143692826 | Target=1;ProbeIdx=13717;Target #1 | 1 | + | good | AATGCAAGAAGGATTTAAACAGGCAGCAGCAGAGTTGGAGGCTCTTTCTTGTGGCTCTTTAAGGCGGTCCAGCTATTCCAGCCACGTGGTAGGGGCTGCAGACTTGCAGGATGCGTGTGG | 120 | 53.33 |
| chr7 | 143692826 | 143692946 | Target=1;ProbeIdx=13729;Target #1 | 1 | + | good | TGGAATTAAACGGAGCAGAGGAAAGGTTCCTAATTCTGCTCTAATAAGCAGCAATAGCAGGACTTCACCAGGGCTCCAGTGAGAAGAGGGAGGGTTCCCCCATGCGTGTACCTCCATGGG | 120 | 51.67 |
| chr7 | 143692946 | 143693066 | Target=1;ProbeIdx=13741;Target #1 | 1 | + | good | GCTCACTCCTACCCAGCTGGAAGGGACAAGTCAGAGCGCAGGGAAATGGCAGGAATTATCAGAAGAAAAACAGAGGTGAGCGATCACAGCTTGGTTTTGTGGCTCTGCTGGTCCTAGCTT | 120 | 51.67 |
| chr7 | 143693066 | 143693186 | Target=1;ProbeIdx=13753;Target #1 | 1 | + | good | CTGAGGCAGCCAGGACAGGCATGAGAGACCCTCACATCACTATACTGCCATGCCCCGAGGCCATTGCTGTATGGGCTTTACTGTCCAAGAGATGCCAATGTAGCTGATAAAGAGCCCTAA | 120 | 52.5 |
| chr7 | 143693186 | 143693306 | Target=1;ProbeIdx=13765;Target #1 | 1 | + | good | GGACATTTTGCTTCTTCTAGACCAAATCCGCGGGATGTCCCTCATGATTCTTAGAAAGAAGCAGTGTCAGGTGGTCTGAAGGAGCAGAAAGCTCAGCTGGAACAGCAGGGAGGGCATGAA | 120 | 50 |
| chr7 | 143693306 | 143693426 | Target=1;ProbeIdx=13777;Target #1 | 1 | + | good | TGAATCAGGAGCCTTTTGCAAAGGGTCTTTGTCTAGTGGCCCTGCCCAGGCTCCTGGGAGCCAGATGGAAAGAACCACTTCTAGCTATGGGAAGTTTCCAGAATTGGGAGGCTGCTCCTG | 120 | 53.33 |

| | | | | | | | | | |
|---|---|---|---|---|---|---|---|---|---|
| chr7 | 143693426 | 143693546 | Target=1;ProbeIdx=13789;Target #1 | 1 | + | good | ACCGAGAAAGAGAAACTTATTTGCTATGCCAGAGGAACCTTCATCAAAGGAGAGGCTGATCCCCTGCTCAAGGCTACCCTTGACCAAGGCCACTGTGGTCATGTGGTGTGACGCTGCACT | 120 | 51.67 |
| chr7 | 143693546 | 143693666 | Target=1;ProbeIdx=13801;Target #1 | 1 | + | good | CCTGTGGTAGGAAGGTAGCTTTCTGTGATTTCCAGAGTGAGGTCTTACTGAGAGAGTAAATTCTCTAGAAAGTGGGTGGAGGCAGCCAACTGATGTACAGACACTGAATCCTCACATGCA | 120 | 46.67 |
| chr7 | 143693666 | 143693786 | Target=1;ProbeIdx=13813;Target #1 | 1 | + | good | CCCCCCCCCAGTCATGTCCTTCTCCCAGCAAGGGATGGGGTGGGGGGGAAAGTTGTTACAGGAGAAGAAAGACTGAGCTCTGGGGTCTGGCTTTGTTCTAGTCCCTCCCAGAGATCCACT | 120 | 57.5 |
| chr7 | 143693786 | 143693906 | Target=1;ProbeIdx=13825;Target #1 | 1 | + | good | TGGTCTCCAGACCATTAATTCCATTCTTCCTGAACTTCCATGAATCTCCTGGCTAAATACTGGCTGCTGTGTTTTTTGTGTAACATGAAATACATGCTGCTGGCTGAAATACAGCTTGTG | 120 | 41.67 |
| chr7 | 143693906 | 143694026 | Target=1;ProbeIdx=13837;Target #1 | 1 | + | good | GCTTTAGAGAGGTATATATAGGGTGGCCTTTCCGATGTCTCTTTCAATCCAAGCCAGCAGGACTTTCCTCTGCTGCTGGGGCAGGTGCTCAGCCTGGCTTCCCTACTCCCCTACACCACT | 120 | 54.17 |
| chr7 | 143694026 | 143694146 | Target=1;ProbeIdx=13849;Target #1 | 1 | + | good | ATGTCCACCAACTCTCTTTATTTGCCTGTCTGTCTGCCCTAAGACTCCAGGCTCATCAGCCTGTGATCTGTACTATTCCCAGCCCAACACCCTGAAGGCTCTCGGGCAATATTAATTAAG | 120 | 48.33 |
| chr7 | 143694146 | 143694266 | Target=1;ProbeIdx=13861;Target #1 | 1 | + | good | TTGCCAGCTCATGTCAGGGAAGAGGACCTGTCTTTCTCCGTAGTGACTGTTCATAGAACATAAATCTTTTCTTCCATGTCAATGCAAGTGACTTTACTCAAGGAGAACCCTGGTCCTCGC | 120 | 45.83 |
| chr7 | 143694266 | 143694386 | Target=1;ProbeIdx=13873;Target #1 | 1 | + | good | TGCCAAGACTGGCATCTGAATGCCTTCTGGGGAAATGGCTTTGGCATCAGCCCATCTAAGCAGTCACAGTGGCTACTAACATGCATTTTTCTTTCTTCTCATTCTCAGATCTGAGAAAGG | 120 | 45.83 |
| chr7 | 143694386 | 143694506 | Target=1;ProbeIdx=13885;Target #1 | 1 | + | good | CTTTCAGCTAAGCCCAGATCCCACATGATAAAAGCAGAACCGTCTCCCCAAAGTTGTCTTCTAACCTTTACATATGTACGTAGCACAGATATGTAATTAAAAAAAAAATCCCAGCTAGAT | 120 | 38.33 |
| chr7 | 143694626 | 143694746 | Target=1;ProbeIdx=13909;Target #1 | 1 | + | good | CTGGAAGAATGAAGAGGGGCACATGGCAAGACTGGAGGGTAGAGGCCTGTGAAGCTGGGACTGGGCCAGGCCTAGCTCATGATGGCTTTCTTCCGTTTTAGGTCAATGGCTAGTACTAGG | 120 | 54.17 |
| chr7 | 143694746 | 143694866 | Target=1;ProbeIdx=13921;Target #1 | 1 | + | good | CCACATGGTCACCCTATAGCCTGCCAGGCAGAGCAGAGTGCTATGCCCAAGAACAGGACAGAATGGGTCAGGAGGAGGGCAGGTGACAACTGTCAGCAATCACAGCAGATCTGAGGAATA | 120 | 54.17 |
| chr7 | 143694866 | 143694986 | Target=1;ProbeIdx=13933;Target #1 | 1 | + | good | GTGAGCTACTATCCCAGGAGCTCTGTGGGTTCTAAGAGGAGCTCAGGGTTCAGGCCCAACTCTTGGCTCTGATCTCGGCTACTGTGTCTTCTGGTTGCATGAAGAGGCTGATGGGAAGGA | 120 | 54.17 |
| chr7 | 143694986 | 143695106 | Target=1;ProbeIdx=13945;Target #1 | 1 | + | good | CACACCAGGCTGCCCTCTAACCTTTATTTGGCACATGAACATGTGTACACACACACACAAATGTATACAGAGGGATAGCCTGTCAATAAAATGTGTTCTGTCACTGAGCACCACGTGTTA | 120 | 44.17 |
| chr7 | 143695346 | 143695466 | Target=1;ProbeIdx=13981;Target #1 | 1 | + | good | GTGGTTTCTAGGGACCTCAGTTACCCAGTCAGCCACATGTGACCCTCAGGCACCTGTAATCTCAGCTATAGCCCTGATAGAGGTTTAGCAGGATCCATGCCATAACCCCGTGGCAACCCT | 120 | 53.33 |
| chr7 | 143695466 | 143695586 | Target=1;ProbeIdx=13993;Target #1 | 1 | + | good | CCCATGCTCCTGCTGTGGAAAGAAGCAGGCCTGCTTTCGTCAATTCCACCATCCTGTGGCACCAGGTTAACCAGCAGCCAGTGCAGACGGCAGGGTAGGAAGTGTTCAGTGAGTGTGAAT | 120 | 55 |
| chr7 | 143695586 | 143695706 | Target=1;ProbeIdx=14005;Target #1 | 1 | + | good | TTCAGTGCATGTGGATTTCAGCGCTTTTGAAAACCCCATTACTGAATTTGGATAAGACCTTTAACATTGACATTTATTCTTTTCTAATGTGTCATTTAAATACAAATACATGAATCAAGA | 120 | 30.83 |
| chr7 | 143695706 | 143695826 | Target=1;ProbeIdx=14017;Target #1 | 1 | + | good | AAATAACACTTATTACATGACTGTTACTGAAAGTAGTCTTGCTGTGTGAAGGAGCAATGTGAGACATGTCCTTCATGTGTGGCCCCATGCAGGGAATGCCTGTGGTTGGAAATGAGGGCA | 120 | 45 |
| chr7 | 143695826 | 143695946 | Target=1;ProbeIdx=14029;Target #1 | 1 | + | good | TAAACAGAGGGTGTCTAGTGGTGCCTGGCTCAGTGCCAGGAGATCAGAGATGCAGAGATGCTTGTGTCGATGGTTGCTGACTTACGATGGTGCAACTTGGAATGGTTTCACTTTCCGATG | 120 | 50 |
| chr7 | 143695946 | 143696066 | Target=1;ProbeIdx=14041;Target #1 | 1 | + | good | GTAGGAACCGGAGGCCCTTGATTAGAAAGCCAGGCTTGGATTTGGGGTGTTCTCCTGGACTGACAATCTCATCATCTCTTGGGGTGCTGGGTCATAGCAGAACTCTTCAGCACCCACCAG | 120 | 53.33 |
| chr7 | 143696066 | 143696186 | Target=1;ProbeIdx=14053;Target #1 | 1 | + | good | CCCTGAGGTGATGAAGAACACTTGGTACTACGCTGTGTGGCTAAACTGTGGCATGAGGTCAGCCATGTGCAGTTTCTTTTCTTTTTGACAGTGTTGAAAATAGAACTTAGGATTTTGTGC | 120 | 43.33 |
| chr7 | 143696186 | 143696306 | Target=1;ProbeIdx=14065;Target #1 | 1 | + | good | CCAATAGATAGTCCCTCTACTGCTGAGTTATCCTCCCTACCCCATAGCCCTCAATGAATTTTCTACTGAACAGTAATTTCAACTTGCAATGGGTTTACCAGGCTCAGATCTCACTGTGAT | 120 | 43.33 |
| chr7 | 143696306 | 143696426 | Target=1;ProbeIdx=14077;Target #1 | 1 | + | good | CTAGTTAGAGCCTTGCCCTTCAGACTATCTCCACTCCACTTCCTAGCCTCTCATGCCCATGTTACTTACCCCAGGCACATCCGTGGAGGGACTGACCCAGCATGCTGGTAGCATACCCCT | 120 | 55 |
| chr7 | 143696426 | 143696546 | Target=1;ProbeIdx=14089;Target #1 | 1 | + | good | GAGCTAAGAGCAGAGCCATCTCTGCCCAGTGTCTGCCTCCTGTGTGTAGGGGATCATCCACAGCAGTAAAGGTTCCGTGAACCTGTAAGAGTCTGGGGAAGGAAAAGTAATACTGTGGG | 120 | 51.67 |
| chr7 | 143696546 | 143696666 | Target=1;ProbeIdx=14101;Target #1 | 1 | + | good | CAGGCCTGGCCCTGAGTCTTTCCAGCGGGATCAGATTAGGTGGTTGCTGTGGGCTCCAGTCTTTTCGGAAAGACTTTGGAGAGGTGAGACCTTCCATGGGGACTGCAGGAGGTTCTCCCG | 120 | 57.5 |
| chr7 | 143696666 | 143696786 | Target=1;ProbeIdx=14113;Target #1 | 1 | + | good | TGAGGCTGCCCCTTATGGTGTCTCTGTGCCTTTTACCCTCCTGGTGGTCAAAGGTAGCTCCAAGCCATTTCCTTCTTTCTCCTTCCCTCCCTCTTTCATTCTCCCTTTTCCCTTCCCGCC | 120 | 53.33 |
| chr7 | 143696786 | 143696906 | Target=1;ProbeIdx=14125;Target #1 | 1 | + | good | TGTGTGGTCATGGAGCACAGATAACAGTGTGAAAGGTATTGGTGTTTTCTTGTGATTGCATATACTTCTCATATTCATAACTCTTCAGTGGGATGTCGTCTCCCACAAAACTATCATGAT | 120 | 39.17 |
| chr7 | 143697146 | 143697266 | Target=1;ProbeIdx=14161;Target #1 | 1 | + | good | AGTCACTTTATAGTTAATGCTTTTGGTCTATGCCTCTTTGTGTTAATCAAGATTTTTTTGATAAGGCAAGATCATAATATAAATATTCCATTTGCTTTCATCACAGTCAAGTCTGTAACA | 120 | 30 |
| chr7 | 143697266 | 143697386 | Target=1;ProbeIdx=14173;Target #1 | 1 | + | good | TACTGTTCTAGTGATGACAATGATTATAATAATAATTCTAAAGCCTTTAAATTACCGGTATTTATTCATCTATGTATAGTGTATGTAGATATATATGTGGATATGCACATGGATATGTGC | 120 | 28.33 |
| chr7 | 143697386 | 143697506 | Target=1;ProbeIdx=14185;Target #1 | 1 | + | good | ATGGATATATATGTGGCTATGGCGTGGCTATATACGTGGCACAGTTTCCTGTAGAGGTCAGTGGGAATTATTTCTCTTTCTCACCACCTCTTCTCTGTGAGTTGAACTCAGGTCTTCATA | 120 | 43.33 |
| chr7 | 143697506 | 143697626 | Target=1;ProbeIdx=14197;Target #1 | 1 | + | good | CTTGCTGGTAAGTGCTTTTACCTGCTGAGCCATCTCTTGGGCCTAATAATGACTTATTTTAAAACTAGGGATTCACTTTGCATCTCAGGAACAGCTGTGACTCTGGGGAATGGCGGGTGC | 120 | 46.67 |
| chr7 | 143697626 | 143697746 | Target=1;ProbeIdx=14209;Target #1 | 1 | + | good | CTCCCTCCTGAGCCTTCCCACTCCTCCCACTCCTGGGATTTCATCGAAGGTTAAGGCATTCAAAGCAGAGCCCACCCAAGTCATTTGAGAAACAAAACTTACTGTCTACTGTCTCAGAAT | 120 | 48.33 |
| chr7 | 143697746 | 143697866 | Target=1;ProbeIdx=14221;Target #1 | 1 | + | good | AGGCTCTAGCTCTGGATGTGGAGGTTGCCATGGCAACACTTCTTTGTTTTTGGGAAACCACAGATACTAGCCTAAGAACTACAACAGGCCAGATGTGGATTTTGCCTCTGGGGTAAGGGG | 120 | 49.17 |
| chr7 | 143697866 | 143697986 | Target=1;ProbeIdx=14233;Target #1 | 1 | + | good | TAGAGAGCAGGGACCATCTAGCACAACTGGTCCCTGATGCACATCCATCAAAGTTATGACATCATTTGTATTCCCACATACAATCCACACAGATCATGGACAGTGAGAGCCTGCACCTCA | 120 | 46.67 |
| chr7 | 143697986 | 143698106 | Target=1;ProbeIdx=14245;Target #1 | 1 | + | good | AGTCCCCATGAGCTGCTTTTCTAGGTATCACAGATATCTGTCTTCCTAGCCAGCCAGACCAAGCCATGGGCAGGTGACCCCAGAACATGGGAGGTAGGATCATTTACTAAGATTGGCTAG | 120 | 50 |
| chr7 | 143698106 | 143698226 | Target=1;ProbeIdx=14257;Target #1 | 1 | + | good | ACTGGCCTGTCAGTGAGCTCTGGGGGTCTGCCTGTCTCCATCTCCATAGTACTAGGGTTATAGATGTGTGTGCACCATCATCTCTGGCTGACTTGCTCCTTTTCCCTCTCTTTTCTCTTC | 120 | 50.83 |
| chr7 | 143698226 | 143698346 | Target=1;ProbeIdx=14269;Target #1 | 1 | + | good | TTTTTTCTTTTCTTTCTCTTTTTGGCTCATTCTGAAGATTCCAACTCAGGACCTCATGCTCACGTAGCTGGCGTTTTACCAACTGAGCTCTCCCCCATCAACCCCTACAAATCGGTTTTC | 120 | 44.17 |
| chr7 | 143698346 | 143698466 | Target=1;ProbeIdx=14281;Target #1 | 1 | + | good | ACTTAGCTAAGATGGAAGAGCAGAGAGCTCAGCAGATACCCAGCACTGGGAGGGGGCTCAACAGAGTTGCAGCATCCTGTTGTGTGTGCCTCGTGGATCCCACATATCTGCATTCACTGT | 120 | 52.5 |
| chr7 | 143698466 | 143698586 | Target=1;ProbeIdx=14293;Target #1 | 1 | + | good | CTGTGTAGGCACCTCTGATTGGACAGGCATTTCTTCCATGCCTCCAACCACGGTGAACATTTCTACTGCCATGCTTGGGATGTGGTTTTAACTTTTCTTTCAAGGCCTCATGTATCTGGA | 120 | 46.67 |
| chr7 | 143698586 | 143698706 | Target=1;ProbeIdx=14305;Target #1 | 1 | + | good | GCTTGGAACCCCATAGTAGCAGTGTTTGAAGGTGGTCAGGCCTTTAACAAAGTGGATTCCTACCTTCTCCAGCATCCTGCCTTGTGATGCAGTCTTTTCTTCTCTTGCCTGCCCACCATT | 120 | 49.17 |
| chr7 | 143698706 | 143698826 | Target=1;ProbeIdx=14317;Target #1 | 1 | + | good | GTGATACCATCTACCATGAGGCCCTTGCCAGTACCTCAGAACTTGCTAAATGCTGGACTAACAGAAACCTTTTTTTGTTTATGAAGTTGCCCAGCTTCAGGCATTTCATCATGGTCATAT | 120 | 42.5 |
| chr7 | 143698826 | 143698946 | Target=1;ProbeIdx=14329;Target #1 | 1 | + | good | CGAAGGAACCAATACATCTATCTATTGTATTTCTTGTTACCAAAGAAGGGTCTGGGTCCTTTCCCTACCTACCTCCCAGTCAATCCAACTTCCCTTTGATCTTCATGGATGTGTCTACCT | 120 | 43.33 |
| chr7 | 143698946 | 143699066 | Target=1;ProbeIdx=14341;Target #1 | 1 | + | good | GGACCCTTCCATGAACTTTAAAGTCTATACTCTTGTCTTCACCGCCCTGTGCCACCCCGCCCCAGTCCCCATGCTGAGCAGAAGCCAGTATAACAATCATCAGAGGGCTTCAGGCATCTC | 120 | 53.33 |
| chr7 | 143699066 | 143699186 | Target=1;ProbeIdx=14353;Target #1 | 1 | + | good | ACAGCTAAAGAAAGATCTGGAAAGTAGAAAAAAATTTCCATACCTGTCAGCCTTGACCTGAATCAATTTTCATTTATAGTTTTTATTTGGCATTTCTAGTGTAGTTTTATCTCTGTCTTC | 120 | 31.67 |
| chr7 | 143699186 | 143699306 | Target=1;ProbeIdx=14365;Target #1 | 1 | + | good | TGTGGAAGATCCCTCATGTGATCCGTGATGGCATGTTGCTCAGACCCACTGGCTGTGTGACATGCTAGGGCAACATGTTACTGTCACAGAACCTCGGTGATACACAGATTCTCAATGCAC | 120 | 50 |
| chr7 | 143699306 | 143699426 | Target=1;ProbeIdx=14377;Target #1 | 1 | + | good | CTCTGGTCTCCAGGGTTGGCAGCATGGGCCTGTGTATGTTGACTGAACAAAAGAGTTCTTTAAAAATAATCCTGGGTTGTCTTCTAGGCCATGCAGAAGTGCCTTTCAGGATGGTATTGC | 120 | 46.67 |
| chr7 | 143699426 | 143699546 | Target=1;ProbeIdx=14389;Target #1 | 1 | + | good | AGGAGAGTGGGTCAGGCTGTGCTTTAAATGCTAATAGAAAGCCAGAGTGGGAGTGGCAAGTGCCTTCAAAGACCCTCTCAGCCAAGGTCTGACAGGATCTGTGGTGATTTATGGCTCCCC | 120 | 51.67 |
| chr7 | 143699546 | 143699666 | Target=1;ProbeIdx=14401;Target #1 | 1 | + | good | AAGTCTTCCCCAACGAGTAACACATGGGCCCTGTGAGTGACAGGCATGACTGGCCCTCACTAACTTGTCATGAATGCTCATCTTTTCCCCCCAAGTCCATGGTCACTTCCATCCCGCTCT | 120 | 52.5 |
| chr7 | 143699666 | 143699786 | Target=1;ProbeIdx=14413;Target #1 | 1 | + | good | TCTATTACATTCCTTTCTTGTCCAGCTTGTTTAGCAAGTGTGCTAGTGGAGCCTTCAGGGTCCCCTCACAGCCCCAGTGCAGGGGCTCATGGCTTTTGTGAGACTGAAGTGCCCAGAGAC | 120 | 52.5 |
| chr7 | 143699786 | 143699906 | Target=1;ProbeIdx=14425;Target #1 | 1 | + | good | CCCTGAGACTCTGCTGTGCTACCCCTTGGTCTCTCTGCCCTGTATGATCCTCATCTGGTAGTCTCAGCACAAGTCTTTCCTAATCTCCATACAGAGTCTGGGATTCTCAGCCTCAGCTCC | 120 | 52.5 |
| chr7 | 143699906 | 143700026 | Target=1;ProbeIdx=14437;Target #1 | 1 | + | good | AACAGGACCCCACCTGCCCAGCCAACTGAGAGACAATCATTTCGATGCCTGTCATGAGCTGAATGGTGTCCCCTAAACTGTGTTGAAGTCTAAGCCCTGACTCCTCAGAACGGGCCTTTC | 120 | 52.5 |
| chr7 | 143700026 | 143700146 | Target=1;ProbeIdx=14449;Target #1 | 1 | + | good | TTTGGAAACAGGGTCTTTCCTGAGGCTATCAGACTAAAGTGAAGAATTAGAGTGTCCTACAAATTGGGCCTTGTGAACACAGACACGTGGGTGGAAGTCACCTGCAGATGAAGGCAGAGA | 120 | 46.67 |
| chr7 | 143700146 | 143700266 | Target=1;ProbeIdx=14461;Target #1 | 1 | + | good | TCAGGCAAGGTTCCCATGAGCCAAGAATGGCAGCAGATCTGATCCCAGAGGCCACTTCAGCCAACTTCAGCCCTGAGTTCACCCTTCTGCTTGAGGAGGGATGTAGAGCCCCCCACCTCG | 120 | 57.5 |
| chr7 | 143700266 | 143700386 | Target=1;ProbeIdx=14473;Target #1 | 1 | + | good | GGACCTGGAGGTCAGGTGCAAGAGGCTCTGTGCCAACTGCCATTTTGACTTTACTTCCTCAGAAGTCAATGAGGCAGACCAGTGCTGTTTCCTTTGAAGGCAGACACTTTCTCTGCTGTC | 120 | 50.83 |
| chr7 | 143700386 | 143700506 | Target=1;ProbeIdx=14485;Target #1 | 1 | + | good | TGCACCCTCCCTGACCTATGTGGGGCTGGAACCCTGACCTGAGGACATTTTGTTTGTATCATGTCCTGTAGCAGCCCACATCCCCCTGCCCTTGCCATAGATCTTGGTCCTTCCTAAGCA | 120 | 54.17 |
| chr7 | 143700506 | 143700626 | Target=1;ProbeIdx=14497;Target #1 | 1 | + | good | GCCCCTCTCTGAGTTCTCCAGAGGGTGACCATCCATGTGCTTTGGCTCCCCAGTTCCACTCTTACACTTATCCTCTGAAGGCATTGGGAGAATCTTTCAGGAGCATGGGGATTCATGCAT | 120 | 51.67 |
| chr7 | 143700626 | 143700746 | Target=1;ProbeIdx=14509;Target #1 | 1 | + | good | CCCTGCCTAGTGGTATAAGAGTTCTAGTCTTACCCAGAAAGTGGCCACCATCTGAGGCCTTCTAGACTAGGGGACTAACACATACGGGAGGGGTGCTTAGTCTGAATGATGGGCAAGTGG | 120 | 51.67 |
| chr7 | 143700746 | 143700866 | Target=1;ProbeIdx=14521;Target #1 | 1 | + | good | GGTCTCTTAGTGAAGAATGCCAGGCACAAGACCCTCGGGCCAGTGGATCTGTGACTAGACATGTGAGTGGTCTCTCAGAAAGGGTAGCTGCTGGCCTTGAGGGGAGAGTTTCAAGAGTCC | 120 | 54.17 |
| chr7 | 143700866 | 143700986 | Target=1;ProbeIdx=14533;Target #1 | 1 | + | good | TCCTGACTCTGCTTCCCCAGTACCAGGATAGTAAGCATACCCTGCTACACCCAGGTTTTTGTATGAGAGATCAAACTCGGGTCCTCATGCTTGTATGGAAAGTGTTTTTTGTGATGGAGC | 120 | 46.67 |
| chr7 | 143700986 | 143701106 | Target=1;ProbeIdx=14545;Target #1 | 1 | + | good | TGTCTCTCCAGCCCCAGAATTTGTGTCTTCTGAACAAATGAATCTCAAGGTGAAGCTCTCTCACACTGGGTGTCAGGGGGCATGTCCATTCATGTGTAGCCCCAGGCCTGCTGTGTTCTT | 120 | 50.83 |
| chr7 | 143701106 | 143701226 | Target=1;ProbeIdx=14557;Target #1 | 1 | + | good | TTCCAAGTTCCCTGGGCATTGTTACTCTCACAGATGCTCTGGGTTTTCGCACAAACTAGGTGTGGACCCTGCCTGCCCTTTCTTTCTTGCACCGTTGCTTTGTCTGGTGAGGGCAACCAA | 120 | 51.67 |
| chr7 | 143701226 | 143701346 | Target=1;ProbeIdx=14569;Target #1 | 1 | + | good | GCACTGGGAGCCATAGTTGGCCATTAGGAATAGTAGCTCTCCTACTGCCTAGGTGGGCTCTGGGTCTGATGTAAACATTCCTCAACAAGTGTTTCAGGGGCATCTTGGATACCCACATGA | 120 | 50 |
| chr7 | 143701346 | 143701466 | Target=1;ProbeIdx=14581;Target #1 | 1 | + | good | ATAATGTGCTCTCTAGGTCTGCGTGGATGAGAAAGGGTCTTAGTACATTCAGATCATGCTAACAAAGCGCCATAAACCCACAGTTCACTGACAACTCAGACCCCTCTCTTATAGCACTGG | 120 | 45.83 |
| chr7 | 143701466 | 143701586 | Target=1;ProbeIdx=14593;Target #1 | 1 | + | good | AGGCTGGGAACAGGATTGGGAGCCAGCAAGTTCCATAGGCTACCATCCTAAGGAAACTGACACTACTCAGGAGGCCTCACCTCAAAACCTAAGTGTTTCCCAAAGGCATTAAGATTTTGG | 120 | 48.33 |
| chr7 | 143701586 | 143701706 | Target=1;ProbeIdx=14605;Target #1 | 1 | + | good | GGAACATGGCATGGTGCCTGATCTATCACAAGGAAAGACCCCCCCCATGCACCCCAAGGAATTTCCCAGCTCACGGCATGGCCTTGCTAGGCCACTTCTCTTCAAACCTCAGTTTTCTGT | 120 | 53.33 |
| chr7 | 143701706 | 143701826 | Target=1;ProbeIdx=14617;Target #1 | 1 | + | good | CTGATGGCCGACCTGGGCATCAGCTTGGCCCCACATGTCAGAAATTCACGAGCATGGGAGACTCTCTCTAGATCAACTAGCACAGACCCAGCAGCCTGAATATCTTCCTTACTAATACCT | 120 | 50.83 |

| | | | | | | | | | |
|---|---|---|---|---|---|---|---|---|---|
| chr7 | 143701826 | 143701946 | Target=1;ProbeIdx=14629;Target #1 | 1 | + | good | TTGCCTTTTTCTCTGAAGCTCCTGTCTCCCCATCCCTTCTCACTCCCCCTCCAATTTCCAGATGTGCTAAACAGGAAGATGCCCAGTGGCGTTGCCATGGCAACAAAAGCAGACCTGGCT | 120 | 52.5 |
| chr7 | 143701946 | 143702066 | Target=1;ProbeIdx=14641;Target #1 | 1 | + | good | ATCTTAGTGTTGCCCCTCTGAGGCACCTCTCTCCTTCAGCCTTCTGGATGACAGCCTGGGAATAAAGTGGGAAAGAGTAGGTGGGATACCGAGCTGAGGTGTCTCTTTGTTTCAGTAAGT | 120 | 50 |
| chr7 | 143702066 | 143702186 | Target=1;ProbeIdx=14653;Target #1 | 1 | + | good | GACTGGGGGTGGCGACCCTGCTCAAGCCCCCTTACCTAGATCTGCTTGCACTGCCAAGTCAAAGTACCGTAAATTGGGGAGTCTTATAGCCTCCTAGTTCTGGAGGTCAGAAACACAAGT | 120 | 52.5 |
| chr7 | 143702186 | 143702306 | Target=1;ProbeIdx=14665;Target #1 | 1 | + | good | GTGGCACGGTGGCTCCCTTTGTATGCCTGTCCTTAGCTGTGGATGGTGACTTCATCCCCCACCACCTAATTCTCTCATAGGGCCTGCATCTCCACCCTTTGTACTACATTAGGGCCTCAGT | 120 | 53.33 |
| chr7 | 143702306 | 143702426 | Target=1;ProbeIdx=14677;Target #1 | 1 | + | good | TTAACCCATTTGATGACCTTTTCCATAAATAAAGTCACATTATAAAACATGAAGGTGAGGACTCAACACAGGAGGGATTTGGGGGTTTGGTTTGGTTTTTGGGTTGTTCGTTTGTTTGT | 120 | 38.33 |
| chr7 | 143702426 | 143702546 | Target=1;ProbeIdx=14689;Target #1 | 1 | + | good | TTTTCAGGGAGAGACAGTTTAGCCTGTAGTGCTTCTGATAATTGAGGGGATCCCACAACTCATCTTTATATGGTGACAACTCTGGCTGCCAGAGACAGAGTGTCCATATGACACCTGTGA | 120 | 45.83 |
| chr7 | 143702546 | 143702666 | Target=1;ProbeIdx=14701;Target #1 | 1 | + | good | GGGGAAGATAGACTGTGCATTCCCAGCCCCAGTGACTCTTGATACATAGCAGGTTAGAGGCTCCTGGGGACATTCCAGGGGACAAAGCAGCATACCAGTGGTGGCCCATTTGGAGGGCCT | 120 | 55.83 |
| chr7 | 143702666 | 143702786 | Target=1;ProbeIdx=14713;Target #1 | 1 | + | good | GGCTCATCCCAGATCCTCAGCTTCATGCTTGGGGCCAGCTACCTACTTACTGCTATGATTACTTACACCCTAACTTTGCCCTCCTTAGAGGTCAGGAGCCATATCTGGCTTGTCCCCTGC | 120 | 52.5 |
| chr7 | 143702786 | 143702906 | Target=1;ProbeIdx=14725;Target #1 | 1 | + | good | CACACCATCTCCTGGGCCTGGCCCATAGTAAGCCTCAGCTAAACTCGTCTTTTTGGATAGAAGGACACATGGCATCCCCCATGCAGACCACTTGCAGCCTTGATTATTCTCCTGCTAGGA | 120 | 51.67 |
| chr7 | 143702906 | 143703026 | Target=1;ProbeIdx=14737;Target #1 | 1 | + | good | CATGGCAAAACACAGGCTGCACAGATCCCAAACCTACCGGGTGAGCATGGGAGCTGGGTTTGCACACTGTGGCTCTGAGCCACAGAGGGTTAACTGGCTTGGAAAGGAGGCCGGCTTGGT | 120 | 57.5 |
| chr7 | 143703026 | 143703146 | Target=1;ProbeIdx=14749;Target #1 | 1 | + | good | GCAATTAAGTCTGTGATGCTTATTTGCAGATTTTTTTAACACCAATTTCAGTCTAAAAAGCCTGCTTCTCACCAAATAAGATTTCTGATGAGAATTTTGATTGAATTCATGTGTTGTGAC | 120 | 32.5 |
| chr7 | 143703146 | 143703266 | Target=1;ProbeIdx=14761;Target #1 | 1 | + | good | GCAAGTTCAGGTACAGGTCGGGCTAGGAACACACAATTATCCCACCTCTTGTTTCAGAGCTGGTGATGGAAGGTTGGGTGGTGGGGAGGCATCTAACTCTCTCTTGTAATGTCCTTCTGC | 120 | 50.83 |
| chr7 | 143703266 | 143703386 | Target=1;ProbeIdx=14773;Target #1 | 1 | + | good | AGCTGGGGCTGTGCCAGGCCTCATTTGCTAGTGTTTTAAAAACCACTCATCTTTTACAGTCTATTTTTAAGAGGACAGGTTGGAGGAACAAAACCATTAGCAGATTTGGGACTCATTCAT | 120 | 41.67 |
| chr7 | 143703386 | 143703506 | Target=1;ProbeIdx=14785;Target #1 | 1 | + | good | TTTCATGTAGAAATGGGCATAATAGGGCTCTGGGTTAAACATGACATGTGGCTGGGGGCACAGGGCTTACAGACCCAGGGCCTGATTAGTGCCTTGAACAATCTGCAACCTTGAGCTGAC | 120 | 50 |
| chr7 | 143703506 | 143703626 | Target=1;ProbeIdx=14797;Target #1 | 1 | + | good | CATGCTTGTGGAGTGGCTGGGCCCTGCAGGGTTCCCACTGCCTGGGACCTGAAAGGGCTTCATCCTTATTTGCAACCCCAGGGTTGAGGGATGGATACGCTGAGCTGGATATAACATCCA | 120 | 55.83 |
| chr7 | 143703626 | 143703746 | Target=1;ProbeIdx=14809;Target #1 | 1 | + | good | GGAGCTGCCTTCTGAGGGAAAACTTGTATGGCACACACAGCCACTTTTCTTACTATAGTATCTTCCCACAAGCAGTGATCTCTGGGAATGCTGGCTTGTGCGTAGCCTGTCCCCATCACC | 120 | 50.83 |
| chr7 | 143703746 | 143703866 | Target=1;ProbeIdx=14821;Target #1 | 1 | + | good | TCTTCAGGACCCTGTCCCATAGGGTGCTCGTTTAGCCATCCTGGATGAATGAAGGAAAGAATGGGTGATATATTATTAGGTTTCTACTGCCGAGATAAAACACTGACCGAAGCCAACTTG | 120 | 45 |
| chr7 | 143703866 | 143703986 | Target=1;ProbeIdx=14833;Target #1 | 1 | + | good | CGGCTTACTGGATGTGCCTTAAGGCTTGCTCAGCTTGCTTTCTTATAGAACCTAGGATCACTTGTGAAGGGAGGTGTGGATGAGGGGAGGAAGAAGAGAAGGACAGATTGGACACAGGCA | 120 | 50 |
| chr7 | 143704106 | 143704226 | Target=1;ProbeIdx=14857;Target #1 | 1 | + | good | GGGAGGAAGGGTTCACACAGCTTCTGGGGACACAGCTTCTGTCCACGGTGGTGATGGGGACAAGATAAGGGCCAGGCTGGAGGAGAAATGCAGTTAGAGAATACGATGGAGACCTCATTA | 120 | 53.33 |
| chr7 | 143704226 | 143704346 | Target=1;ProbeIdx=14869;Target #1 | 1 | + | good | TGTATGAATCACATGCATGCATGCACTACATAACATCACATGCCACACACTGCCAATACATGTCATTATGACTGGAGTTGAGCATGCAGGACCATGACTTAAAGCCCTTGCTGTGGTGTC | 120 | 45 |
| chr7 | 143704706 | 143704826 | Target=1;ProbeIdx=14917;Target #1 | 1 | + | good | TTTTTAAGATGGCAAAGCCACTGTTTTACAATCACTTTATGTTTCTCCATTCACTGCACTGAGACGTCAAAGGGCAAATGTTCAGCCCTTTTTCCGGAAGACCCAATGAAGGTGGACTAT | 120 | 41.67 |
| chr7 | 143704826 | 143704946 | Target=1;ProbeIdx=14929;Target #1 | 1 | + | good | TAGGAGTGTGTCCTGATCCCAGGGGACTTTGTGCTTCCCTTTAGGTGGGGGTGTGGGACTGAAGGGGTTGGGGATAGACAGACGGGGACACTACTGGTGTCTTGCTGGGCACAATGAGCT | 120 | 56.67 |
| chr7 | 143704946 | 143705066 | Target=1;ProbeIdx=14941;Target #1 | 1 | + | good | GAGAGTTCTCCTTATTTCCTGCATTCAACTCTAGTTCAGCACATTTTTAGGTCTTGGGTCTGCCACATGGTAGCTGATGGGAAAAGTCTGAAAGGTGCACCTACTGAATCATGGGCTCCA | 120 | 45.83 |
| chr7 | 143705066 | 143705186 | Target=1;ProbeIdx=14953;Target #1 | 1 | + | good | AGGCAAAGGGACTGGGAGGGGACAGTCAGGCTGCTGAAAGTTTGAGATTCCTCACTACAGAGTTGAAGAACCCCCTCCCCGCAAATCAGGGACTCCCAGGCCCCGTGGTGGGGTAGCCAG | 120 | 59.17 |
| chr7 | 143705186 | 143705306 | Target=1;ProbeIdx=14965;Target #1 | 1 | + | good | AAGTAGGTTAGCCAGCTGGGATGTCTGTGCTGAGCTTGGTCAGTGTCAGGGAGGTTTCAGGACAAGATGATCCTTAAACCTTAGGAAGGGCTTGGACCAGGGTGATATGAGAGGTAGAG | 120 | 50.83 |
| chr7 | 143705306 | 143705426 | Target=1;ProbeIdx=14977;Target #1 | 1 | + | good | CAGGCAATACGGTCAGGGACTAAAGGAGATGCCTGACTGTCATACACAGAGCCAGCTGTGGACAGTCACCTGCAGGCTGAGGGAGAGGGGTATCCTCCTCTGATGCCCTTGGCTCTGCTG | 120 | 57.5 |
| chr7 | 143705426 | 143705546 | Target=1;ProbeIdx=14989;Target #1 | 1 | + | good | CCCTCAGCTCACCATCACTGTTAGCTGCCTATGGAGCTGGGTTCTATCCAGCTTCTGTCCTTGCCCATGCAGCTCCCTGACTGTGCTGGGTGCTCATTCTGGGGTAGGTCCCTATATGGT | 120 | 55.83 |
| chr7 | 143705546 | 143705666 | Target=1;ProbeIdx=15001;Target #1 | 1 | + | good | AATGGTTGTTTAACAGGTCCTTTCTTCCCAGAGAGGCTGATTGACTGCTCTCTTGATTTAAAATCTGAAGGTGTTTGTTATCTTCTGCTGGAAGTTTCCTGGACCGTAAGTTGGCAGTGA | 120 | 42.5 |
| chr7 | 143705666 | 143705786 | Target=1;ProbeIdx=15013;Target #1 | 1 | + | good | TGGTATCTATCCAGCCACCTATTCTTTACCCATCTTTTCAGAGGCTCATGCACCCACCCACCTACTCACCGGCCCACTTATCTAACTCCTTACCCATTCTCCCACTCACCCATCCATATC | 120 | 50 |
| chr7 | 143705786 | 143705906 | Target=1;ProbeIdx=15025;Target #1 | 1 | + | good | CCATTTACCCATTAACCAACCCACGGATTCTACCTATTCACATATCTGTCTGTCTACCTTGTATCTGCCCGTCCACCCATCTCCATCTACTCACCCTGCTGTCCTTCATCCACTCATTC | 120 | 47.5 |
| chr7 | 143706026 | 143706146 | Target=1;ProbeIdx=15049;Target #1 | 1 | + | good | ATCCGCATACAGACACATCTACCATCTTATCCACTCAGTCACCCACCACATACTCCCTCCCCACTCCTGTCCATTTATCTACCACTATACCTACTTGCCCACTGGCTCATCACCTCATCC | 120 | 50 |
| chr7 | 143706146 | 143706266 | Target=1;ProbeIdx=15061;Target #1 | 1 | + | good | AGTGGTGAGAAGGGACAGCAAGGGTCACAGAAGACGAAATCTCCACCAACAGAGAACTCAGGGAAGGAAGGCCTTACTAAGGGCTAAGTCTGGGCTGTGAGCTGTGCACTGGGAAGAACA | 120 | 52.5 |
| chr7 | 143706386 | 143706506 | Target=1;ProbeIdx=15085;Target #1 | 1 | + | good | AGCCATCTGAAGACAGTTGGATGGCCTATTCCTGGATGCAGACTGTCAAGCCACTATAGAGGGGAGAACTTTCTAGAGATCTCCTTGGCCCAGGTTGCATAGCCCATGAGAAGAGGCAGG | 120 | 51.67 |
| chr7 | 143706506 | 143706626 | Target=1;ProbeIdx=15097;Target #1 | 1 | + | good | AGTGTCAGGCTCCAGCAGGAAGCTAGCTTCTTTGAGCCTAATAGTAAAAAAATCAAGAGGTGGACTCTTTCATGATTTAGTAGTCCTTAGGGAACCGGTTTTAACAAAGATACCACTGCA | 120 | 41.67 |
| chr7 | 143706626 | 143706746 | Target=1;ProbeIdx=15109;Target #1 | 1 | + | good | GTTTACTTTTGTGTGTGTGTCGAGGGGTACACACGTGTGTACATGCACGACACACAGATGAACCATGGGTGTCATTCCTCAGGATCCATCCAATTTGCTTTTTGAGAAAGAATCTGTCACTG | 120 | 45 |
| chr7 | 143706746 | 143706866 | Target=1;ProbeIdx=15121;Target #1 | 1 | + | good | TGGTTCTTGTGAGAGTCAAACTCAAGTCCTTGTGCTTGTGTGGCAAGCCAAACCATTTTATCCCCCAGCCCTGTCATTTACATTCCTAAGATAAGTGCTTTGCCTGTGGTATGAGCCTTGTC | 120 | 45.83 |
| chr7 | 143706986 | 143707106 | Target=1;ProbeIdx=15145;Target #1 | 1 | + | good | TATAGCGGCGGTGGCGGGGGGGGGGGGATATGTGAGGGAGGCAGGGCTCCTGATGCTGGGGATAGAATATGCTGGATCATATCTGGGTATCTGATGTGGACTACTGACTCATTGGCCAAG | 120 | 56.67 |
| chr7 | 143707106 | 143707226 | Target=1;ProbeIdx=15157;Target #1 | 1 | + | good | ACAGGTGATGACAGTTGCCGCTGGTACTGCACATGCTCATGGCAGTGAATTTTGATGGCCCCCATCTGAGATGGATTATGAACGGGACAAGGAGCAGATGCTCATGGTGACCTTTGTACA | 120 | 50 |
| chr7 | 143707466 | 143707586 | Target=1;ProbeIdx=15193;Target #1 | 1 | + | good | GCCCGACACTCTTTCTCAACAGAGCGCAGGCGGGGTGCAGCCTTCCCCCTCATTTTAACTCCTAGCACATTTGGGCTGGGATAGAAGCCGTGAAACTGAAGTCACAGTGCGCTGTGACTT | 120 | 55 |
| chr7 | 143707586 | 143707706 | Target=1;ProbeIdx=15205;Target #1 | 1 | + | good | GAATAATTCCCTTTCTGAATAGTGCTTTGCAGCCAGTCACAGTCTGGGAAATCCACCATGGGCAGAACAAGGGTTTCTGGATGGGGTGACAGGCCTTTCTGGCAACTCCAGACCCTAGTT | 120 | 50 |
| chr7 | 143707706 | 143707826 | Target=1;ProbeIdx=15217;Target #1 | 1 | + | good | AACTTAGATCTCAGATTAGACTCCATGAGGCTGAGGCTACTGAAGTAATTCATGTGCAGCTCCCAGAAATGACTAATTGCCGTCACCTCCTGGGGCTCCTGTCACTGGCCTTCTGAGGAA | 120 | 49.17 |
| chr7 | 143707826 | 143707946 | Target=1;ProbeIdx=15229;Target #1 | 1 | + | good | TTGGCTATTTGAAGCAGTGCCATTGTCCTCCTATGGACAAGTCCCTCTCACAGGAACGTCATCAATTCCTGTTGGAGGGTGCCATATCATGTGTAATTAGGTCCCCTTTGCCCAGTGTGC | 120 | 49.17 |
| chr7 | 143707946 | 143708066 | Target=1;ProbeIdx=15241;Target #1 | 1 | + | good | CCTGAGCAGTGGGAGGTGGCCATTGTGTCCTGTGATGTGGAAGCCAGGACCACAGTCACCTGCTGGTGAGTTCTCAGGCTCTGAGCAGGTCAGCTTTCCTGAGGTCTGTGAAAGACATCA | 120 | 55.83 |
| chr7 | 143708066 | 143708186 | Target=1;ProbeIdx=15253;Target #1 | 1 | + | good | GGGTCCTTGTGGAGGAAATGAAGCTCCTGAGAGGAAGGTAGGACTCAGCAAGACTCAGTGTTGTCACTGAGTGTGAAGCTCACAACTTTCTTGTGCCCAGGAAGACAGGGATAAACAGTGA | 120 | 49.17 |
| chr7 | 143708186 | 143708306 | Target=1;ProbeIdx=15265;Target #1 | 1 | + | good | GCTCTGCCTCCAGCCTGCTAGTCAAGTCCAGATGCTTCTCTGGCCAGTGGCAAGTGACCACAGCAAGGTGTTCATCCCCCAAACAACTGTGTGTGGGTGCTGCTCACACTGTCTATAATC | 120 | 54.17 |
| chr7 | 143708306 | 143708426 | Target=1;ProbeIdx=15277;Target #1 | 1 | + | good | TATGGGAATCTAGCCCATCCACTGTCAGTCCAGTTCCAAGCCCCCAGAATCTGCTGAGGATTTTACAGAGTATGGCTGCCAGCTCTTGAGAGTGGACCCTGCCTCTTTGCCCCACTGCTG | 120 | 54.17 |
| chr7 | 143708426 | 143708546 | Target=1;ProbeIdx=15289;Target #1 | 1 | + | good | GGGAATGTTCTCTTCTGTACATCCTGTGGCAAGAACTGACCATAGGCACAGGTTACTGCAAAGAGCCTAAGCTGTCAGTGATGACTATAACTCCAAAACATATCTGACTTCCCCATCAGA | 120 | 45 |
| chr7 | 143708546 | 143708666 | Target=1;ProbeIdx=15301;Target #1 | 1 | + | good | CAGAGTCCAGTGACAAGGTAGCCAGCAGGTAGAGCAACCTGTGAATAATTCAGAATACTGAGTGGTGCTGGAGGCTGGGAAGGCAGGCAGAGCAGGGCTGGGCTCAGATGGACCAGCTAT | 120 | 55 |
| chr7 | 143708666 | 143708786 | Target=1;ProbeIdx=15313;Target #1 | 1 | + | good | GTTTGGATCTCAACCTCTGGATGGGGCTCTGTTGTAAGGTGCTTACAGCCCTCTCCTTCAGCAAAACAGGATTTGGTACCTTTCTTCAGGACTGTTGAAGTGTTCTATAAACTATTTCCTA | 120 | 44.17 |
| chr7 | 143708786 | 143708906 | Target=1;ProbeIdx=15325;Target #1 | 1 | + | good | AAAGGAGGTAGAGACTCAGACCAGGCAGAAGCATGTAGAAATGGGCTACACAGGTCAAAAGGGTTTATAAACAACTGGTCTATGTGTCAGACAATTCATCTTACCCATCCCTTATCCATC | 120 | 43.33 |
| chr7 | 143708906 | 143709026 | Target=1;ProbeIdx=15337;Target #1 | 1 | + | good | TCAATCTGATCCATGTGTCCAGTCATCCTTTCTTTCTCTTTACTATTTTAAGTCTTTTGGGAGGTGATCTCGTCAGATCCTAGACACCTCACCACTTGAGGAAGCCTCCCTGCCTTCAGA | 120 | 45 |
| chr7 | 143709266 | 143709386 | Target=1;ProbeIdx=15373;Target #1 | 1 | + | good | AACTGCCTTTTATCTACATCCACTGCCCACTGCCAGGCTGGGAGCAGAAAGGACTCGGGTGTGATTGTTCACACACTGCCCTCCCTGGAATCTGCCAGAGGCTCACAAGGGCCTGAGCTC | 120 | 56.67 |
| chr7 | 143709386 | 143709506 | Target=1;ProbeIdx=15385;Target #1 | 1 | + | good | CATTTACTCACCCCTGGAAGCTTACTTCCTTTCCAGCCAGACTTGAGTTTTATTGACATTCAACATGTTGGGGACATGATTCCTAAGGTCTTTCCAATCCCTGAGACCTCTGTGTTGA | 120 | 44.17 |
| chr7 | 143709506 | 143709626 | Target=1;ProbeIdx=15397;Target #1 | 1 | + | good | TCATAAGCTTGTCTGCCAGCCTGCTGAGCTGTCCCTTTCCCTGCCTTGGGTTTGCAGCCTCTGGTACAGATGAGTACCAGGTTTGCTCCCATCCTGTTCTGCATATTGGCAACAACAGGT | 120 | 52.5 |
| chr7 | 143709626 | 143709746 | Target=1;ProbeIdx=15409;Target #1 | 1 | + | good | AAAGGGACAAGATGAACCTCTGCCATGGAGGAGGAGCAGGTATGGGTGGAGGGAATCTGTTGACACTTTTGTAACACACCAGGGACACCCTCGTGGGGTCTGCAGCATCCTGGGAAGGTG | 120 | 55 |
| chr7 | 143709746 | 143709866 | Target=1;ProbeIdx=15421;Target #1 | 1 | + | good | AACTCTGGGTCCTGTGTGAGTATAAGTTTCAGCTTTCTGCTAGGACTCCTAAGCCTTGAAAACTGGAGGAGTGGTCCAGGGTTGTCGTGAGACTGGGGTGGCCAGCATCTGGGGCTGGAG | 120 | 54.17 |
| chr7 | 143709866 | 143709986 | Target=1;ProbeIdx=15433;Target #1 | 1 | + | good | GAAAATTGCCTCAAACCCTATTGGTGACTGCCACCATCCTGGTGTATAAATCTTGGATTGCCACATCTTGAGAAAGGGAACGCAAACTTTCTTTTTTTTTTTTTCTTTTCCATTTTTCAT | 120 | 36.67 |
| chr7 | 143709986 | 143710106 | Target=1;ProbeIdx=15445;Target #1 | 1 | + | good | TTTCTAGCAGCAGAGAAAGTGACTGTAGCTTGTAGAGAGACTCAGGTGGGACAGTGTGTAGAGAAGACTCATCGGAAGGGGCACCAGCTTAAGGCTAGAGAACCTTAAAGTAAATATGTA | 120 | 45 |
| chr7 | 143711426 | 143711546 | Target=1;ProbeIdx=15589;Target #1 | 1 | + | good | GGCAGAAGCAGAGGTCAAGGGTAAAGTATATATAACTGGCAACAGGAGACGCCATGTGGAGAAGGTAGGAGAACAGAGAAGGGAAGAGCTAAGGTTTTCATGTCTTCTGTGACCTAGATG | 120 | 46.67 |
| chr7 | 143711546 | 143711666 | Target=1;ProbeIdx=15601;Target #1 | 1 | + | good | AAACTGAGGCCCTTGGCCACTTATTCAGAACACCAGCCTTTCTGACCTGGAAGCTTATAGAAGCAAAGCAAACTCCTCCATTCTGGGCCAGTTCTTTCCACGAGGAACCAGATTGAGTTT | 120 | 47.5 |
| chr7 | 143711666 | 143711786 | Target=1;ProbeIdx=15613;Target #1 | 1 | + | good | GGGAGGGACATACCGCAGTGTGCGTTCTTTTTAATTAGCCCTTGTGGTAAAGTCGTGAGGAGCTATGGAGCCTGCCAGAGCCTTCGGGGCACCTGTTGGTGGCCGGCCAGTTGGCTGTCC | 120 | 58.33 |
| chr7 | 143711786 | 143711906 | Target=1;ProbeIdx=15625;Target #1 | 1 | + | good | TCTCATATCACTGGGTCACCAGGAATTCTATGGAGCCAGGGCTCAGATCTCCTCGTTCAAATGACAAGCTATGCCTGCTTCTGAAGGCTGTGGGTTGAATAGCCTTCTCTATGGTAGATG | 120 | 48.33 |
| chr7 | 143711906 | 143712026 | Target=1;ProbeIdx=15637;Target #1 | 1 | + | good | CAGGGTTTGTCCAGGCAAGGAGGTCACGGCCAGGGCCCTAATGTCCTGGCAGCACTGTATGCTCCAGGTCTGACATTGCGAAGGCCCAGGCTAGGTCCTCAAGAGTACACCAACTCCTTC | 120 | 58.33 |
| chr7 | 143712026 | 143712146 | Target=1;ProbeIdx=15649;Target #1 | 1 | + | good | GTTCTCCCAGAGTCCTATGCCACTCAATAGCCAACCAGGTGCAAAAGGCCTGCATTTAGCTCATGTTCCAGACTCTGAGGCTCTGGTTCCGAGTTCGAGCTTGTGCCAACTACCATTCAA | 120 | 50.83 |
| chr7 | 143712266 | 143712386 | Target=1;ProbeIdx=15673;Target #1 | 1 | + | good | AGCTGGTAAATCATTCATGATTGGGGTGCATTGCATGTATCTCTGACAGAGCCCCCCAAGTTAGGTTCAGACCTACTTGCTCTCGGCATTCTAGGTTCTAGTAAGTTTCTCAGCTATTCT | 120 | 45 |

| | | | | | | | | | |
|---|---|---|---|---|---|---|---|---|---|
| chr7 | 143712386 | 143712506 | Target=1;ProbeIdx=15685;Target #1 | 1 | + | good | GAGAAACACTAGTCAGGGGTAAGTTCTGCCCAACCCTATTCCAGTGCTTCTAGGTGCATCATGGTGTGCTTGACAATACTTAAGATCCCGATAAGAAACCTAACCTTGTATGCCTCATGG | 120 | 45.83 |
| chr7 | 143712506 | 143712626 | Target=1;ProbeIdx=15697;Target #1 | 1 | + | good | AACCTGAATTACACCATGCCTCCAGTGTCCAGGGTCCCATGGAACTAACTTTGAAAAACTGCACTGAGGCCACCTACGGGCTGCCTGGGGAACACTGAGCTTGACAGAAATGCCCTGATA | 120 | 51.67 |
| chr7 | 143712626 | 143712746 | Target=1;ProbeIdx=15709;Target #1 | 1 | + | good | GACAGGGATACTTATTTGTCCCGCTAGGCACATCAGAAAGAAACAAGTCCTAAAGATGCCTAACTTTCAAACACCCAGTTCCGATTTCTGGTGGCTTGTGGGCAGATCCTAAACTATAAG | 120 | 44.17 |
| chr7 | 143712746 | 143712866 | Target=1;ProbeIdx=15721;Target #1 | 1 | + | good | TAAGGCATATTTGGATTTTAGGGGGGGAAAAAAAGGAAATGATTTACTAGTATTTAAAAATCGAGAAATTTCACATGATGATCCAAGCTTCTGCCTGCCTTTTTTTAGGAAGCGGGAAGG | 120 | 36.67 |
| chr7 | 143712866 | 143712986 | Target=1;ProbeIdx=15733;Target #1 | 1 | + | good | ATATTGCTGTCACCAAGACTGTGCTGTCCCAAGAGAACCTTCTCTTGTCTCCTGTGGGCAGACTAGTGAGCTTCGAACTCTCTTAGCCACAGCAACGGGCATGGACCCACCCTTCCTGCA | 120 | 53.33 |
| chr7 | 143712986 | 143713106 | Target=1;ProbeIdx=15745;Target #1 | 1 | + | good | CAGCCCCATCTTCTAGCATGACAAAACAGCTAAACCTGACCCAGCTGGCTAGAATTTCAAACGGGCCACCCTGGGGTCACCTCTGTCCAACCTGGATCTTTCTACTAAGGACAGTCCCTG | 120 | 52.5 |
| chr7 | 143713106 | 143713226 | Target=1;ProbeIdx=15757;Target #1 | 1 | + | good | AGCCTTAGTAGAGTCCTGGCCCCTCCTGGAGGCCAGATTTCAGGAGGACCTAGGCAGTGCTCTTCTGAAGCCCATGGTGACTGCAACATTGTGAGTTGGAGTTGTGAGCTGAATTAATCA | 120 | 51.67 |
| chr7 | 143713226 | 143713346 | Target=1;ProbeIdx=15769;Target #1 | 1 | + | good | GAGGTTGCAGCTGGGGGCTGGGGATTTTCGGGGCACAGAGGAGACGTCTGGTAGGGGCTGCTGCTGAGCATGAGGGAACACTCTGGCAGGTTCTCAGTGCTGTCCCCTTTCCAGAGCAAT | 120 | 60 |
| chr7 | 143713346 | 143713466 | Target=1;ProbeIdx=15781;Target #1 | 1 | + | good | TTAAAATTAATTAAAAATGCTGGTGTGAGTGGGGCTCCTCACAGGCAGCCTGGCAGCCTGGCACAGTCCCCCCTATTTTTAACACTGACAGCAGATTCGAGGATTGACGGCTGTGCTTCC | 120 | 50 |
| chr7 | 143713466 | 143713586 | Target=1;ProbeIdx=15793;Target #1 | 1 | + | good | TGGCATTTCTTTTCTTGCTTAAAAACATAGCAGCGGCCCTTCTGCCGCCAAACCTTCTACCACCGTGCTCTGCGAGGCCACATTTAGGGAACCCTTCCTTAGGGTTTCATCAGAGTCCAC | 120 | 50.83 |
| chr7 | 143713586 | 143713706 | Target=1;ProbeIdx=15805;Target #1 | 1 | + | good | TGGAAGGTGTGAACATGACTGTACTTATATTGGTTAGAGGGACAATTGGCCTTGGCCCAGGCTGAGACCCTAAGAACAGGCTCTGTGTGTGCTTCCATCTCCCCAGGATCCTTATAGGCC | 120 | 50.83 |
| chr7 | 143713706 | 143713826 | Target=1;ProbeIdx=15817;Target #1 | 1 | + | good | CATGCCAAGCAGAAGGGTACAGGGCACAAAGTGGTCCAGAGCAAGTCTAGTGAGCTTGTGTGGGAGCCTCAAGTAAAGCCATTAAGTTCTGGGAAGAGGTGTGTCTACCCAGCTGAGCAG | 120 | 52.5 |
| chr7 | 143713826 | 143713946 | Target=1;ProbeIdx=15829;Target #1 | 1 | + | good | GTTGGACTCCATGCAAGAGGCGAATTCTTGGTCAGTTTCATGTGTATAGAGCTGTCCGTGGTACTTAGGAGCTTCTTGTTCTGAGTGTGTAACAAAGGCCCTCCAAGGGTGGGGCTCATT | 120 | 49.17 |
| chr7 | 143713946 | 143714066 | Target=1;ProbeIdx=15841;Target #1 | 1 | + | good | CCAACAGTCTTACAGATGGTAGGAAGGATGGTGTCATGGTGTCCCCAAGAGCAGACTTTGGAGCTTCAGGTCTCAGCCAAACTGTTTAACCTCTCTAAGCTAAGGTCCATGACCCCCTGA | 120 | 50 |
| chr7 | 143714066 | 143714186 | Target=1;ProbeIdx=15853;Target #1 | 1 | + | good | AGTCAATGGCACAGTATTGTGGCCTCCCCTGTGTTGATATAGGCATTAGACAGGTCAACACTGGACATGTTGCATGTGTCATTCCCAAGACAAGCAGTTCATAAATGGCTGGTCTTTGTT | 120 | 45 |
| chr7 | 143714186 | 143714306 | Target=1;ProbeIdx=15865;Target #1 | 1 | + | good | GGCCAGAACATGTGGCCAATATTTTCACTATTCACCTACAGATTAATAACCAGAGACATTGGGGTTTCAGAGTCCCCTCCCCACAGAGTACCAGCAAGGATGGAGCCTTTGCTTTCCACC | 120 | 48.33 |
| chr7 | 143714306 | 143714426 | Target=1;ProbeIdx=15877;Target #1 | 1 | + | good | TTCTCTAAAAGCCCCCACCATCTTCTTTGCCACCTTTGTGGACGGTGGCTCTGTCAACCTGAGACTGCGTTGCAGACTCTAGCTTGCTGTCCTGCTCTGCCTCCAAATGGCAACGGCCCC | 120 | 55.83 |
| chr7 | 143714426 | 143714546 | Target=1;ProbeIdx=15889;Target #1 | 1 | + | good | AGTGCTGGTCTCCGTCTCAGTCATTCTGGATCTGGCATCCCGGAGGCCACCCGCTGCCACAGGCTTTTCCAGGGCGTGCGGACTATCCCAGTGTCTTCTGAGGTGAGGTGTCCCCTCCTC | 120 | 61.67 |
| chr7 | 143714546 | 143714666 | Target=1;ProbeIdx=15901;Target #1 | 1 | + | good | AGTGGCCCTCCCGGCCTCTGTTCTTGATGCTGTCTGACTGCAGGAAGCTCACAGACGACAACTCCGCCCTGTGGCGTCCAGTTTTCTTCAGTGGGTATCTATTTTCAGCCCTGCAGCAAG | 120 | 55.83 |
| chr7 | 143714666 | 143714786 | Target=1;ProbeIdx=15913;Target #1 | 1 | + | good | CAGCTGGGATGCTTGTCTCTCCGCCTTTGTTCTGCTCCTCCTGTTTCCCCGCTCTGTAATTATGGGAGATGTTTATTCATTCACTAAGGGCCTGCCTGTGATTCTTAATTCTGTGATGCC | 120 | 48.33 |
| chr7 | 143714786 | 143714906 | Target=1;ProbeIdx=15925;Target #1 | 1 | + | good | CCTCTCTATTACCGAGCACATAAATCTTATTCTGTATGGGGAACCCAACATCATTGGCCCTACTGTGCTACCTGCCATCCTAGTTTGAGCCTCAGGGAAGAGACTCTATGAATGGTAGGA | 120 | 46.67 |
| chr7 | 143714906 | 143715026 | Target=1;ProbeIdx=15937;Target #1 | 1 | + | good | GGTTACTGGATCCCAGTCTTGTTCACGGTTGCCAAGACAACATCACCCACCAAAAACCCATCACAGTTCTTCTGAGGTCATCCTAGGGGACCAGAGCCAGCAGCCAGCCAGTAGATAGCA | 120 | 52.5 |
| chr7 | 143715026 | 143715146 | Target=1;ProbeIdx=15949;Target #1 | 1 | + | good | CCTTCTAAGCTGGCAGCTTGCTGCCCACTGGGGAAGATCCAGGAGCAGGGAAGAAGAACGCTTGCCTGATATCACAAGGGTTATGGTCCATTTTCTGCTGGAAGGGTGACCTCTGGCTTT | 120 | 53.33 |
| chr7 | 143715146 | 143715266 | Target=1;ProbeIdx=15961;Target #1 | 1 | + | good | CAAATGTCTCTCTATAATATCCTATTTTTTAAGAGAAAATTTACTCCTAGTGGCTAATTAGTTCTTCCTGGAGAGCGATTTGCACTAACCATAAGCACATTCAGGCCTTTTGCCCTGTGG | 120 | 38.33 |
| chr7 | 143715746 | 143715866 | Target=1;ProbeIdx=16021;Target #1 | 1 | + | good | CTTCCTTCTTTGAGAAGAAAGAACTCAAAGAAGCAAGCACCACAGGCACAAAGATGTCCGGGCCATGCTACTGATCAAGCAGAGAGTCCAATTGCATCTCACCCAGCTCTGAAGAACCTG | 120 | 48.33 |
| chr7 | 143715866 | 143715986 | Target=1;ProbeIdx=16033;Target #1 | 1 | + | good | GCCTGAAAAGGACAGCGAGAAGCCTCCAGGGACATTATATAATGTCACACGGGACAGCACAGAGCTGTGGCACAGGGCCGAAAGCTGTGAGGGAAGGAAAGGCAGGCTTCAAGGTCTGTC | 120 | 55 |
| chr7 | 143715986 | 143716106 | Target=1;ProbeIdx=16045;Target #1 | 1 | + | good | CCTCTGCAGAAGCTGAAGTTGTGCCCTGCTGCACGTGGGGCCATGTCAGCCATAGAAGCTCACACAGCATTTTAACCACCTCCTGGGTAATACCAGAAGGATCAGGTTCAGGGGCAGGAA | 120 | 54.17 |
| chr7 | 143716106 | 143716226 | Target=1;ProbeIdx=16057;Target #1 | 1 | + | good | ACTTCATTTCCCTCCCCCACCCTCCTGGGACAAATGTGTGGCCGACCACATCAGGTAGCAGCGAGCAGGGAGAGTACACTTCCTGGATTGTGCTGCTGACCCAGATAGAAAAATGGAGCG | 120 | 55 |
| chr7 | 143716226 | 143716346 | Target=1;ProbeIdx=16069;Target #1 | 1 | + | good | CTCCTTGGGTGTTGGTAAAAGTAGACTATAAAGGGACCGATGCGGTAGAGGCCCATCAGTCAGAACAGGAGGGCTGGAAAGAGGGCTGGCTACAGACAGGTGAGGTGTGCAGGCTTACAG | 120 | 54.17 |
| chr7 | 143716346 | 143716466 | Target=1;ProbeIdx=16081;Target #1 | 1 | + | good | CCTTGGGAGCTTGTACCCCATGGGACCTGTGCTGTTGGTGCAGAAGCACTGGATTATGTCTTCAAGATGAGGTGGAATGGGATGAGGCCCTCTACCCAAGATAGATTGAAGACCCCAAAA | 120 | 50.83 |
| chr7 | 143716466 | 143716586 | Target=1;ProbeIdx=16093;Target #1 | 1 | + | good | TAGCTCATGAAGAGGAAGTCCAACTTCCCAGTTTCAGCGCTTAGTCCAGGACCAGAGTTTCACCTCAAAGGTAATGGACCACACCCCCTGTAGGGCTGTGTGCACTCCTCATTCTGCCTG | 120 | 52.5 |
| chr7 | 143716586 | 143716706 | Target=1;ProbeIdx=16105;Target #1 | 1 | + | good | GGCTGTTGGAGAAGCACTTAGCTGGGTCTTAAGTGGCACATGTCCATCATTCCGTATCCCAGAAGTCCAAGATGGAGGCTTGCAGGACTGGCTCCTTCTGAGGCCTCTCCCAATTTGCCC | 120 | 54.17 |
| chr7 | 143716706 | 143716826 | Target=1;ProbeIdx=16117;Target #1 | 1 | + | good | ATAGTGTTTTCTCCCCATGTCCTCATACAGCTGTCCTCTGTGGGCTCCTATCCTGACCTCCTCTTCATAATAAGGACCCAAACCATATGTGACAAGGGCTCAAGCCAGTGACCTATTCTA | 120 | 47.5 |
| chr7 | 143716826 | 143716946 | Target=1;ProbeIdx=16129;Target #1 | 1 | + | good | ACCTAATCACACCTGGAAAGGGTCTTCTTCCAAAGGTAGAAGCCACAGATGGAAAAATAAGAATTTGGAAGTGTTGATAATTCTGTGTATAATATCACTCCTTCCACATACCCACCCTAG | 120 | 40 |
| chr7 | 143716946 | 143717066 | Target=1;ProbeIdx=16141;Target #1 | 1 | + | good | ACCACCAGAAGCATGATTAGCTTCCTCTGAGGGACTCTCCAGTCCCCATGAAATGTCACCTTCAACCCCAACAGTGAATAACCACTCGAATATCCCATGTGCCTCATTACCTGAGAGCCT | 120 | 49.17 |
| chr7 | 143717066 | 143717186 | Target=1;ProbeIdx=16153;Target #1 | 1 | + | good | GAGGCTCTGGCCCTGCTCTGAGCATAGACTGGTAATCTACACCATGTTAGATCCAGCCTTCCATGTTCAACATCTGCTGGAGCAGACTGGGGATCATGGAATGGACACCTTACCAGGAGA | 120 | 51.67 |
| chr7 | 143717186 | 143717306 | Target=1;ProbeIdx=16165;Target #1 | 1 | + | good | AGTGGCAGAGGTCATGTTAGTCCAGCTATTCTCTAAAACCCTCAATGGAGGGTAAAATGCAGCTGGGGAGACAGAAAGGTCCCTCCTGCCTCCCCACACCTCCAACTACTGTCAGAGGGA | 120 | 52.5 |
| chr7 | 143717306 | 143717426 | Target=1;ProbeIdx=16177;Target #1 | 1 | + | good | GACAAGATGAGTGCAGCCTCTATCTCTGTTACTTTCCATGCCAAGATACTCTCCCCATCTCATCCTACCCCTGTGCAAGTGAGTAGGCTCATGTTAGGGTCATCTCACTCAGATCCCATC | 120 | 49.17 |
| chr7 | 143717426 | 143717546 | Target=1;ProbeIdx=16189;Target #1 | 1 | + | good | AAAGTGACCTGGTCTAGATTCAGTGGCAGTGCACCAGTGAGTCCAGGACAGCCTCTAAAGATGCTCCATGTGGCACACCCTCCTAGACTGTTTGCAGCGTGTCCCTGGATAGCACACATC | 120 | 53.33 |
| chr7 | 143717546 | 143717666 | Target=1;ProbeIdx=16201;Target #1 | 1 | + | good | TTCCTTGTAACAACTGCCTGTGCGATGCTTCTGTCTGGCTGGAGGCCTTCCTCCTGGACTCTTCTTAGAGGGCTGGTGAGGACTGTTTGTGGCTGAAGGGAGAACTTCCAGTAGCATCAT | 120 | 52.5 |
| chr7 | 143717666 | 143717786 | Target=1;ProbeIdx=16213;Target #1 | 1 | + | good | GTGGAAGACACCCTGTCCTGACTGTTGGATCTTGCTGGCTGCCTCTCTCCTAGGCCTCCCAGCCCAGCAGTGTCTCCTTCAGTGTTAAGAATGGCTTAGCTCCTTCACCTAGCTGAATGG | 120 | 55 |
| chr7 | 143717786 | 143717906 | Target=1;ProbeIdx=16225;Target #1 | 1 | + | good | AGGAGCCAGATTCTATAAAAACTCAAGTTATCTGCTCTGCAACCTTAGGATTCTATTAGCTGGTTGGGAAGAGGTGCCCATCGCCCACCAAGGAATGTGAGCAAGCTCCTGACACAGCTC | 120 | 48.33 |
| chr7 | 143717906 | 143718026 | Target=1;ProbeIdx=16237;Target #1 | 1 | + | good | TGCAAACCCACTGCTGTGCCCAGTTCTCCATGCTTCTTAAAACCAGGACTTTACAGAGCATGTCACTCGAGTCTCATTTACCCATGAAGATGCAGAGACTCACAGGGAATAAATAACTTG | 120 | 45 |
| chr7 | 143718026 | 143718146 | Target=1;ProbeIdx=16249;Target #1 | 1 | + | good | GTTCAAGAACATAGGTAATCCTTGACACACAAAACCTAGATAGGTATCCTGCAAGACCTCCTCAGTAACTGTCTTGGTCTTGAGCCTCCAGACCATCTCTGGGGCTCTTCTAAGAAATAT | 120 | 44.17 |
| chr7 | 143718146 | 143718266 | Target=1;ProbeIdx=16261;Target #1 | 1 | + | good | AAGCCCTTGTTTGAGCCAGGGCTTCCACAAAGCAGTGGATCTATCAGGAGCACACACAAATGTCTGTCAACTTGCCTGGGAACCATAAGGAGTCTACCTAACAGTCTCTCTACTCAATGG | 120 | 48.33 |
| chr7 | 143718266 | 143718386 | Target=1;ProbeIdx=16273;Target #1 | 1 | + | good | TTGGGACATTTCTTGGCTTAACATACCATTTTAATACAAGTGAGTCCAACTCTGTTGTGGGAAGCCGTGGCAGGGAAATTGGGGCCAGCATTTTAATAATAAAATCCATCCCCAGTTCAT | 120 | 41.67 |
| chr7 | 143718386 | 143718506 | Target=1;ProbeIdx=16285;Target #1 | 1 | + | good | GGCAAGGGCTCACAACACCACCCAGTTTGGGTCCTTCCATGAGCATGTGGGCATGCGCCCTCCTTGTCTTCAGTCCCTGGCTGTATACACTGCTCATCAGTCCTTCCAGTATTCCCTGTT | 120 | 55 |
| chr7 | 143718506 | 143718626 | Target=1;ProbeIdx=16297;Target #1 | 1 | + | good | TCTCTCTGGGCTTCAGGGCTTTCCTGATCCCAGTTGCATCTAATAGTAACCAGACCATGAGTACAGCCCTGGAATTGATTTGCTTTAAGGTTAAATCAGTCACGGCGGCTCTTTAGTTCC | 120 | 46.67 |
| chr7 | 143718626 | 143718746 | Target=1;ProbeIdx=16309;Target #1 | 1 | + | good | CAAAATAAAACTGGGCACTTGGATCCAGTGAGGAAAAAGTTGATTTGAGTTCCCGAACCAGCAACCTCAAGAGCATTTCATGTTTTGGGTATCTCTTTGTCTCAGACTCATTTATCCAAG | 120 | 40.83 |
| chr7 | 143718746 | 143718866 | Target=1;ProbeIdx=16321;Target #1 | 1 | + | good | TAGCCTTCACCTCTGGTCTCCCAGGGCCCTTGTATAGGAAAGAGTTGGATTTGTTTATTGCAATAAAATCCCAACTCATGACAAGAGCAGAGGTCCTGGGAAGGCTGTTTCCCCTCCTGC | 120 | 49.17 |
| chr7 | 143719106 | 143719226 | Target=1;ProbeIdx=16357;Target #1 | 1 | + | good | ATCTGTGGTTGGAGACCAAGTTGCTTGTGTCAGATAACTGTCCCACTAGGAACAGTGAGGAAGTATGTATAAAAGCCACTCCTGAGTGGCCCAAAGCAGATAGAAAAGGAAAGGGCCCTG | 120 | 47.5 |
| chr7 | 143719226 | 143719346 | Target=1;ProbeIdx=16369;Target #1 | 1 | + | good | AGGAAAGGGAAGCCCCAGGAAAAGCCCTCACAGATAGTGGGATATCACTGGCTAGAAGATTCCATTTAGACATTGGGGTCAGAGTTGTCTAGATGGCTAAAAGAGAGAGGGAAGTTCTTA | 120 | 45.83 |
| chr7 | 143719346 | 143719466 | Target=1;ProbeIdx=16381;Target #1 | 1 | + | good | CAAAAAGTAACACCTATAAAATCTTCACCCAAATCTTCCTCAAATCCCTGACAGGCTCCTGAGCTACCATATATGTGCAAGGGAAGAACTGAAGGGTCCAATGGAAAACAGTAACTTAAA | 120 | 40 |
| chr7 | 143719466 | 143719586 | Target=1;ProbeIdx=16393;Target #1 | 1 | + | good | CTTGGAGAGTCCAGGGGAGATGTCAGGAGTTTCCATCTGCAGAGGCAACGGTGGAGACATAAGTCCCACCTTGGTGGAACTTAGCAAGCACCTTGGCCTGTCTATGGGATCTCTGGAAGG | 120 | 54.17 |
| chr7 | 143719586 | 143719706 | Target=1;ProbeIdx=16405;Target #1 | 1 | + | good | GAGATTAGAGTCAGACCTCATTCTAAAACCAAAGCAGTAGCAATCCCTCTAAAAGAACCCTGTAACTGGCTTCATGTTCATGTTCTGCCAGTGATTTTATTTACTGCTAGAAAACTCAAC | 120 | 39.17 |
| chr7 | 143719706 | 143719826 | Target=1;ProbeIdx=16417;Target #1 | 1 | + | good | ACTCATCAGATAACAGATACAGTTGTGCAGTGTCTGAATTGGACAACAACAGATACACAAAGGAGAACATTTTGCCATATGTGGGCATGAATCTTGATAATCAGCATGAATAGAACCATC | 120 | 38.33 |
| chr7 | 143719826 | 143719946 | Target=1;ProbeIdx=16429;Target #1 | 1 | + | good | TAGGAGACATCCCCATATGTGTCTGGGAAGTTTTTTTCAGAGAGGTTTAACTGAGGAGTGAAGACCCACTCTCAATGTGGCTGTGGTCCAGAAGTGAATGAAAAGGAGAAAGTAAACTGA | 120 | 43.33 |
| chr7 | 143719946 | 143720066 | Target=1;ProbeIdx=16441;Target #1 | 1 | + | good | ATAGGCCTTTCCTCCCTTGTATTGCTTTTGTTAGGCCCCTACACTCATTGTTAAGCTCTTGCTATGCCAGGGAGTGGGTTTGCCAAAGCGAGAATGACTGTGACTCCGTCTTCTGCCACA | 120 | 49.17 |
| chr7 | 143720186 | 143720306 | Target=1;ProbeIdx=16465;Target #1 | 1 | + | good | GACCTCCCAGAAGGTGCAGGAAGGAGTATTCATGGTGTGAGGAATTCTAGAAACAGTTGTCTTATATAATGAATAACACCCAGGGTGAAGCAGGGAGAGAACCATTTGAAGGCATCTTTG | 120 | 44.17 |
| chr7 | 143720306 | 143720426 | Target=1;ProbeIdx=16477;Target #1 | 1 | + | good | AAGGGAGCTGGGCTCCTTACTGTAGTGAAGTAGTGACTAAGACAGATATTTTGTTCAGGAAGAAACATAGTATGAATTCTTAAATAATACGACTTCAGTTTCTGATATATGCAATGAGGA | 120 | 35.83 |
| chr7 | 143720426 | 143720546 | Target=1;ProbeIdx=16489;Target #1 | 1 | + | good | CAAATAATGAACGGCAAGGACAAGACCATTTCTGGAGTATCTCATCTGTGTTACTCCCATGAGAGAACCTCTCCTGTGTCTTGTTTTAGAAGACCATGAATCATGAGCCCCTTCCTTCAC | 120 | 44.17 |
| chr7 | 143720546 | 143720666 | Target=1;ProbeIdx=16501;Target #1 | 1 | + | good | AGGCTCACCACCTAGCCCTAATCTTTGCCAATATCCCTGTCTACACATACCATCACATGAAGCACCAGGAACACAAGTACACACACTGGGGAATGCTGGCCTTCAGTATGGGGTCCTGGT | 120 | 50.83 |
| chr7 | 143720666 | 143720786 | Target=1;ProbeIdx=16513;Target #1 | 1 | + | good | TGCCATGGTTGTGCTTTCTGGTCTCAGCTTCTGTATATGAAGTCATGGTGGCCATACAGTCACTGTGGTCCCTCATCATGGCCCTGGCTCCTGCAGTCCACCTTTATCCTCCAGGGTGTG | 120 | 53.33 |
| chr7 | 143720786 | 143720906 | Target=1;ProbeIdx=16525;Target #1 | 1 | + | good | GTGAGGTGCTATGCGGAGGAAGCAGTGGCTGTGTAGGTCAGGACAACTCTAAAAGCAGAGTCCTCCCTAGAGCTCCCAGAGAGTCCAGCCAAAGCAATGGTGGCTGGGAGGAGTTGATTC | 120 | 55 |
| chr7 | 143720906 | 143721026 | Target=1;ProbeIdx=16537;Target #1 | 1 | + | good | TCCTTAATAGCTGAGAGCTGGGCATGTGTATCCACTATGGAGAAGCAGCCAGGCCATGTGAGGATCAGCTCTCTACCATGCATGCAGCCAAGGCAGGATGCAGGCCTTGTTATATGTTGG | 120 | 51.67 |

| | | | | | | | | | |
|---|---|---|---|---|---|---|---|---|---|
| chr7 | 143721026 | 143721146 | Target=1;ProbeIdx=16549;Target #1 | 1 | + | good | CCATTCTGTCTGAGTCTCTGGGTTACCTCATAAGGCTTCCAGCTAAAGAGGGTGGAGGCAGGCAGCCCATGCCACATGCTTGTGATAGGAAGATGCTGAGCTACATACTTTGTGGGGTAG | 120 | 51.67 |
| chr7 | 143721146 | 143721266 | Target=1;ProbeIdx=16561;Target #1 | 1 | + | good | GGCTTAGCACATGCTCCCATCTCCAGAGAGGCTGAGAGTGAGTTCAGTCTCTCTGGGTCTTAAAGCCCTTCCTCAGGATGCTCAAGGAACTTTTGAGGCAGGCTAGGGCTTGTTTCTCTT | 120 | 51.67 |
| chr7 | 143721266 | 143721386 | Target=1;ProbeIdx=16573;Target #1 | 1 | + | good | TAGATGAGGAAGCTGTATCTTGGAATCCCACCTTGGGCCACTCACTGACCCAGGATGCCCATCTGCAAGCTCCAGGCTCAAATTTCATGAATTCCTAGGAGCTGGACCAAAGGACCCAAC | 120 | 51.67 |
| chr7 | 143721386 | 143721506 | Target=1;ProbeIdx=16585;Target #1 | 1 | + | good | AGGCCTGGAGGCTGGGGAGGGAGGGGGTGCTGCACGGACTTGGCTCCCAGGCCGAGGACTGTACTCTGCTAATCAGCACTGCTGGTAATTGGCTTACTCCCTGTAGACGGCTATCTTTGT | 120 | 59.17 |
| chr7 | 143721506 | 143721626 | Target=1;ProbeIdx=16597;Target #1 | 1 | + | good | CTCTCTGAAGTCTACAGGGGACCTTCGCATGTCCTTCCATCTTGGGGGAAGGGTTCCTGTAGGAAGGTAGGACTCTAGGGCTCTCTTGGGAGAGCCAGACAGGTGGGGTCTTCAGGTCCA | 120 | 56.67 |
| chr7 | 143721626 | 143721746 | Target=1;ProbeIdx=16609;Target #1 | 1 | + | good | GGTAGAAGCCAAGAACCTGATCCATGCCCTTCTGTCCACCAGAAGTTTCCAGAGCTCAGAGACAGGCTTATGTGGGAAAAGACCAGTTCTGATACAAGGGTGCTGGCTGAGTCTAAGGGG | 120 | 51.67 |
| chr7 | 143721746 | 143721866 | Target=1;ProbeIdx=16621;Target #1 | 1 | + | good | AAATAGAGGTGATGTTTGAGCCAGAATGGGATCAGGACATGGGTTATAACAGCCTTAGGGTCCGAGTATAGAGCAGGCAGTCCTGAGCAGGCAGGGCAAGGTGGGCCTGGCTGGTGAGAC | 120 | 54.17 |
| chr7 | 143721866 | 143721986 | Target=1;ProbeIdx=16633;Target #1 | 1 | + | good | AGCATTTGACACTTAAGATCTTTCCAGAATATATTCCAGTCTCTCTTACCTCTCTCAGGCCTCCAGAAACCCCGCACTGGCTACGGTCCTGGACAGGTATCCAAGGTTACTTACAAGAGG | 120 | 47.5 |
| chr7 | 143721986 | 143722106 | Target=1;ProbeIdx=16645;Target #1 | 1 | + | good | CTCTTACCCAGTTCTTACCCAGCCCCTCAGCACCAGTCCAGTCCAGCCAGGCTCAGGCCAGTGCTGAAGCCTGATCCTCGGCAGTGCCACTCACCCTGGGTACCATACAACTTCCAAAAG | 120 | 58.33 |
| chr7 | 143722106 | 143722226 | Target=1;ProbeIdx=16657;Target #1 | 1 | + | good | TTGAGCCCCTCAGTGGCTCCTGCCAGGAATGGTGCAGACGCCGCAGATGCAGACGACACCCACATCTTTGTGCAAAGCTAACCCCATGTTACTGCCGAGACGCCTTTTAATATCTGGCCT | 120 | 55 |
| chr7 | 143722226 | 143722346 | Target=1;ProbeIdx=16669;Target #1 | 1 | + | good | TATTTATATGGGTAATGACTCATTCACCAATTACTGTGAGATGGAGATACTTTGCCTACTAATATTTAAATTGCAGTTATCTGAAGCTACTAGGGAAGGAAGTGAGGTCTGCTGGTTTGA | 120 | 36.67 |
| chr7 | 143722346 | 143722466 | Target=1;ProbeIdx=16681;Target #1 | 1 | + | good | GTTTGGTGGGAATGGGCTCCAGGAATAGAGCTCTTGTCTCTCTCTACCTGCCCAGTTGCACCCGCCTCTGTCTCTGCCTCCCTGCTCCAGTCAGATACCACTGGCCAGGGGCCCTACACC | 120 | 60 |
| chr7 | 143722466 | 143722586 | Target=1;ProbeIdx=16693;Target #1 | 1 | + | good | TGGTCCATTTCCTTTCTGACATTTGTTGGAGTTAGGGTCAGGTGAGCCACTGAAAACCTCTGGAGCCTCTCTTTCTCCTCTCAGCTCCCCATGGACCTCTGTTAGGCCAGGGGACAGAGC | 120 | 53.33 |
| chr7 | 143722586 | 143722706 | Target=1;ProbeIdx=16705;Target #1 | 1 | + | good | AGCCACTGAGCCAGTGTTGAAAGCGGCTCAGGTACTCCCCTGCCTCAGCAGCCCTGTCTCTCTCTCCCCTGTCCTCCTCCACTGATATAAGGGCCAGTGGAGCAGAAGCTCAGGAGGCTA | 120 | 59.17 |
| chr7 | 143722706 | 143722826 | Target=1;ProbeIdx=16717;Target #1 | 1 | + | good | AGGTGAGAAAATAAGGATTGCCTCAAGGCATCAAGGTGTGCTATGCCACAAAGCCTGGGAAGTTGGACTAGCTGCTTCTAGAATCTCCCATTTTCTTCTCTTCCACTGTGGTGGTACCAC | 120 | 47.5 |
| chr7 | 143722826 | 143722946 | Target=1;ProbeIdx=16729;Target #1 | 1 | + | good | ATGGGGTGGGGAGGGGGGGTGGCTTTGTGCCTTGAGGTATCTAGCCACAGCTGAAGGGAAGCATGTGACTTCATGCACATACCTAGCATGCCCTCAGTACCCATTCTCAGCGAATCCTCA | 120 | 55 |
| chr7 | 143722946 | 143723066 | Target=1;ProbeIdx=16741;Target #1 | 1 | + | good | GAACATAGCCTTGTGCTGGCTGAGGCAGGTCTCTTCTCTAGGAGAGGCTCCTGCAGGAGTCTAATGTGAACTCCCAAATATACAGTGATGGAGGGAGCACCATCCTGGCCTAGGTGAGTG | 120 | 53.33 |
| chr7 | 143723066 | 143723186 | Target=1;ProbeIdx=16753;Target #1 | 1 | + | good | CCTAGTTAAACTGCTACCTGCCATTTACATATGCAGTTTTGCTTCTCTTTTTGCTGCCGCCTCACAAGGTACTCCCTTGTATCAAAATGCCCTTCGCTGTGCAAATCAGATGGGTGAGTT | 120 | 45 |
| chr7 | 143723186 | 143723306 | Target=1;ProbeIdx=16765;Target #1 | 1 | + | good | CAAGCAATGTTAGAAGCTAAGGACAGGCTTTAGCTTTCACCCAGCAAAATAAGGCTGCAGGGAATGCTGAAGACACCAAAAGGATTCTGGGAACTGAGTTCTGGTGCTCCTGTTCAAATG | 120 | 45.83 |
| chr7 | 143723306 | 143723426 | Target=1;ProbeIdx=16777;Target #1 | 1 | + | good | ACAATCCTGTCAGAGCCCTCTTGTACAGGCCTTACCAGGTCCCTAGACAACTGCTTGGTGTGGTGTGGTACCCCTATAAGCCCTTGATTGGAGTGGAACAGCCCTGGGTAGGGGTGGGGC | 120 | 56.67 |
| chr7 | 143723426 | 143723546 | Target=1;ProbeIdx=16789;Target #1 | 1 | + | good | TGGGCAATGGGCTGCCTGTTGCCTCATGTCACTTAAATAGGACTGGAGACAGACCAGCCTGCTGCGCCTGGAACCCAACGGGTGAGGACTGGATTCTTTTACTTGTTCATTTAGTTTCCA | 120 | 50.83 |
| chr7 | 143723546 | 143723666 | Target=1;ProbeIdx=16801;Target #1 | 1 | + | good | GCATGTTGGGAATTTCCAGCTCCCAGTTTTCAGGGGAAAGAGGAAAGACTGGCATATCCAGGAAAAGCCCAAAGGACTGTCTGCTGCACTAGGAATCTTGATGAACAGTGTCTCCAATGA | 120 | 47.5 |
| chr7 | 143723666 | 143723786 | Target=1;ProbeIdx=16813;Target #1 | 1 | + | good | CAAGGTCAGCCAGGGAGGTGTCTTAGAGGAGGTAACACCCTGAAAGATGACAGAGGCATCTGGACATGCTAGGGAGGGTCTATGTAGGGGAAGGGATTGTCCACCACCTTTCCACCTGAT | 120 | 53.33 |
| chr7 | 143723786 | 143723906 | Target=1;ProbeIdx=16825;Target #1 | 1 | + | good | GACTTGTAGTTTGGTGTGGTTGGGATCTTCTTGCTGTAGTTGTGAAGCAAGGTGTTCTGTGGCCCAGTCTGGCCTCAGACTTGCTGTGTAGTTGAGGATCTTAAAATCGTGATCTACCTG | 120 | 47.5 |
| chr7 | 143723906 | 143724026 | Target=1;ProbeIdx=16837;Target #1 | 1 | + | good | ACTGAGCCGTGTCCCCGGATTCCATCACACATGTTCTAATGGCACGTATTACTCAGCCAGTGCTGACAGTACTGTTGCAAATAGAAACACTTGAACTCTTGCCACCTCTCATGAGGCAGC | 120 | 49.17 |
| chr7 | 143724146 | 143724266 | Target=1;ProbeIdx=16861;Target #1 | 1 | + | good | TCCAGGGACATCCTTTTCTACAGAAACAGAAGTACACAGAGAAGGGGTAGCTTTTCTAGAGTTACAGAGCCTCATTCCCCACTGGGTAGTTCTTCTCCAGAGTGCTGTGTGTGGGAACAC | 120 | 48.33 |
| chr7 | 143724266 | 143724386 | Target=1;ProbeIdx=16873;Target #1 | 1 | + | good | TTTGTTATACCCTGAACACATGAACTCAGGTAACCCCACAGCCAGTACTGGGAATTGCCAGGCACCCCACTTCTGCTGGTGGGGGTTTCCCGAAGGAAAAAGGGGCTGCCCGAACCGCTA | 120 | 55 |
| chr7 | 143724386 | 143724506 | Target=1;ProbeIdx=16885;Target #1 | 1 | + | good | GCCCATGGCTACAGAGGACAGTGCCTCAGCCCACACTGTCCAGCTCCCGCACGCACACCCTCCAACACTGAAAGCACAGCAAGGGGAGACACGCTTGTCATCAGCAGCAAGCACTTCCGA | 120 | 60 |
| chr7 | 143724506 | 143724626 | Target=1;ProbeIdx=16897;Target #1 | 1 | + | good | GTCTCACCACGTGCTGGGAGCACCTGCCACGTCCTCCCTGCCAGGACTTGAGAGCCTCAGAATGTTCTGATGCATAGAGAGCCTCTTTTTGTACAAGAAGTAAAATTATTTTCAGCCTGT | 120 | 49.17 |
| chr7 | 143724626 | 143724746 | Target=1;ProbeIdx=16909;Target #1 | 1 | + | good | TGTCAGGCAGCTGTCAAAAGACATGGAAATGAAGGGCTTAGTGAGCTCTGAGAGGAAGTCCCATGGCCAGGGTGGGTGTAGGGACACAGGCTGTTGGTCATGCCGGTTGTGGAGTGGAGA | 120 | 55 |
| chr7 | 143724746 | 143724866 | Target=1;ProbeIdx=16921;Target #1 | 1 | + | good | GAAAATGGTGGTGTGGGGCTCATTTCGGGGGTCACTCTCCTTCTGGCAACTGATCCCGGGGGCATGATCTGATTTTTTTCCGAGCATGCATGGGCATTGAGTGGTCCCAGATGCTGATGG | 120 | 54.17 |
| chr7 | 143724866 | 143724986 | Target=1;ProbeIdx=16933;Target #1 | 1 | + | good | AGTTCACCTCCCTGCAGTGCTGGGGGTCCATGGTTACATGTCTAGACTGGAGCTCTTTGCAGTCTGTACATCATATACTGCCCATGGCCTGTCATCCAGAATGAAATCAGGTTGGTGGGG | 120 | 51.67 |
| chr7 | 143724986 | 143725106 | Target=1;ProbeIdx=16945;Target #1 | 1 | + | good | GGGAAAGCTTGTGGATATGGGAGTGATCACCCAGGAGGCTGAGGCAGGAGGATGGCAAGTTCCGGGCCAGGCTTAGCTTCATAGTGAGACCCTCTCTCTCAAAGCAAAAGAGAAAAAGGA | 120 | 52.5 |
| chr7 | 143725106 | 143725226 | Target=1;ProbeIdx=16957;Target #1 | 1 | + | good | AATGAGGAAATTGGTGGTGTTGGCTTTTGGACCATGGCAGCTTCACCCCCATAATGTCAGGGCAGGGCACCCCAGCACCTCCAGGACCCAAGTACCTTACACACCTCTGCAAATCTCTAG | 120 | 53.33 |
| chr7 | 143725226 | 143725346 | Target=1;ProbeIdx=16969;Target #1 | 1 | + | good | AGTGTTCTGTGGGTACTCTCAGGGAACCCAGGATGGGGATGCTCCCAGAACATTCCAGAAGTCTCAAACCCTCTCATGAGAAAGCACTGTGAATAAAGGATCTCTAGCAGTCTTGAAGAG | 120 | 48.33 |
| chr7 | 143725346 | 143725466 | Target=1;ProbeIdx=16981;Target #1 | 1 | + | good | AAACAGCTTTTGGATTTTATCTCTTCCTTAATTTTAATTTTTAATCCCTGCTCTATAGTTTCTGGTGTCCTTAGAGATGCTACCAGTGTTTCCCAGGGATCTGCGGGCTTGCTGTCAGCC | 120 | 41.67 |
| chr7 | 143725466 | 143725586 | Target=1;ProbeIdx=16993;Target #1 | 1 | + | good | TCAGTCAACACCATCCAGAGAAGCCTGAGTTGGTAGTAGGGACCTGGGTCCCCACGGTCACTGGGTGGAAGACCTCAGGGGATGACGTAGACAGTTCTGTGCTAGGCGAGGAGAAGTCGG | 120 | 57.5 |
| chr7 | 143725586 | 143725706 | Target=1;ProbeIdx=17005;Target #1 | 1 | + | good | GCCAATGTCTGTTGACAGACCCGTGAGCATAGAACCACAGCAGTGTAAGCCAAAAGTCAGCTTCTTGCATGGGACCGTTGTTCACACACAACTGATGTTCTGCCAGCCATGTATGCTGTA | 120 | 49.17 |
| chr7 | 143725706 | 143725826 | Target=1;ProbeIdx=17017;Target #1 | 1 | + | good | TGTAACATAAGATGAGAGATAAAATAAACACCCCTGTGTTTGTTTTCTGAAGCCAGGAAGAGAGAGTCATTGGTGTGGGGTAGGGTACAGCGCTAACATCTCCTTGGGGTAGGGTGCCAT | 120 | 45.83 |
| chr7 | 143725826 | 143725946 | Target=1;ProbeIdx=17029;Target #1 | 1 | + | good | GACCCGCCCTTGACAACAAGAATCACTGCTTGCAGGTAGAGACCCCTTTGTTGCTGGGAGTGTTCAAGGACAGCATGTTTGCGGGGATGTATCAGAAGCAGCCCATTCCCCCAGCAGGAA | 120 | 54.17 |
| chr7 | 143725946 | 143726066 | Target=1;ProbeIdx=17041;Target #1 | 1 | + | good | TGCTCAGGGTATAGCATAAGGCACCTCCACCTCTCCATAGCAGAGCCTATCCCAGGCTGACTATACAGGATTCTGACTGTCCAGATCACAATAGATGCTCACACAGTCTCTAGCCCACCT | 120 | 50.83 |
| chr7 | 143726066 | 143726186 | Target=1;ProbeIdx=17053;Target #1 | 1 | + | good | GCCCACAGTCCTGCTTGCCACAGCCACTCCACAGGGTCACCCCAGCTCCAAGCTGAGCTGCTCTCAGCCTTGTCAAAGAAGCTTCATTTAGCAGTGAACAGTGGCAAAGGCAAACTCATA | 120 | 54.17 |
| chr7 | 143726186 | 143726306 | Target=1;ProbeIdx=17065;Target #1 | 1 | + | good | ACTGGTCAGTTCTGAGAATTAAGGGACTGTAATGAACAATCCAACCCTGTGTTGGACCTCTGTAGCAATCCAGCCCTGCGTGGGACCTCTGTAATAACCCCTCTGCACTCCCTGACCACC | 120 | 51.67 |
| chr7 | 143726306 | 143726426 | Target=1;ProbeIdx=17077;Target #1 | 1 | + | good | CGCTGTGGTTGCCTTGTATAAGACCTCCTCTACAAGGCCAAGCCAACCAGAATCCCAGTATGGATGGCAAGGAGCTCATGAGCACCACCTTCCAGCTGAAGGTCTATTGTCAGTGGCTGA | 120 | 52.5 |
| chr7 | 143726546 | 143726666 | Target=1;ProbeIdx=17101;Target #1 | 1 | + | good | CTGCTGGAGGAGGGAGCAGTCACTGGTAACTAGCCCATGTTCCAGTGGATGGCTCCATACCCCTGCACACACAGGCAACACTGACTAGACTCAGTGAGCTTTAGCAAGAGAAAGCAGGAC | 120 | 54.17 |
| chr7 | 143726666 | 143726786 | Target=1;ProbeIdx=17113;Target #1 | 1 | + | good | AAGAGAGTGTCCTCAGGTATGGGATTATACTGTGGAGTCCTTTGTCACATCATCTTGTATCTATAGCCTACCTAGACATGCCACCTGTGCCCCATACCTGTTGCTCTCCCACCCTACCCC | 120 | 50 |
| chr7 | 143726906 | 143727026 | Target=1;ProbeIdx=17137;Target #1 | 1 | + | good | ACCCTACCCAGCCAGCACAGCTCACTCTCCAAGGCCTGATGGGACATTGGATTCGGGGTTGGGTCTAGTTGGCACTACGGGAACTGGGTGGAGACATGATTCTGCTTGGGGATATTCTCA | 120 | 55 |
| chr7 | 143727026 | 143727146 | Target=1;ProbeIdx=17149;Target #1 | 1 | + | good | GCTCTTCTCATTGGAGAGCTCAAGGCTCTGTACACTTCCACAGCAATTTCTACTTGTGAGCAAATGCATCCGAAGGAGCAGCTACCTAATATATGGCTGGTTATGGTGGCAGCTGGGGGC | 120 | 50 |
| chr7 | 143727146 | 143727266 | Target=1;ProbeIdx=17161;Target #1 | 1 | + | good | TTTGTATCTGGAACGTGTGACGCTGCTGCCTTGCTGGAGTTTGGGAGTTTGGATTTCTGCTCTAGGGAAGATGGGACAGGGACCTGCTTGATCCTCACTCTTTTGACATATCTTAGGTCA | 120 | 48.33 |
| chr7 | 143727266 | 143727386 | Target=1;ProbeIdx=17173;Target #1 | 1 | + | good | GAGACCCAGTGTGGAAAAGAAGGTTCTGTCATTTCCTTCATTTAGAAAAGGTATGGGGTGTCACACTGCCCTCCTGGGAAGAGGTCTCCTCCCACAACCTCCACCGAAGGTCAGTGTCAG | 120 | 51.67 |
| chr7 | 143727386 | 143727506 | Target=1;ProbeIdx=17185;Target #1 | 1 | + | good | ATGTAGGATCTTCTGGTGGGAAGGGTGACTGTGGCACAAAGCCTGGCTTCTAGCTTTACCACCCAGTTGTGGGCACACATTCTCTGCACCACACAGTGGCCAAAGAGATGCCAAAGCAGT | 120 | 52.5 |
| chr7 | 143727506 | 143727626 | Target=1;ProbeIdx=17197;Target #1 | 1 | + | good | GGGTGGCAGGGAAACTAGGCTTGGGTGCAGCTTTGTGCTTTCTGGAAGATATTGATCCCCAACAGATGGAGCAAGCACTGAGCGGCACTGGTGTCTCAGAGAGTCCCCAGAACTCTGGCA | 120 | 55 |
| chr7 | 143727626 | 143727746 | Target=1;ProbeIdx=17209;Target #1 | 1 | + | good | AGTCCCCCGAAGCCCATGTGGTTGTGGGTCTGACTGGCGGGGGTCCCTGGCCTCTGGAGTGGGGCTTCCTGTATTTGTGGACACTGGTGACACCATCCCAATATGGCCTCCAGCCAAGTG | 120 | 60.83 |
| chr7 | 143727746 | 143727866 | Target=1;ProbeIdx=17221;Target #1 | 1 | + | good | CTTCATGATGATCTAGAAACCTCCGTGTTGACGCCTACCCAGGAAAGGTATCTCTAGCTCATCCCTGGAAGATGGATGCTCATGGATGATGCCACAGTCAAAGGGTCTCCCGAAAGCTGC | 120 | 50.83 |
| chr7 | 143727866 | 143727986 | Target=1;ProbeIdx=17233;Target #1 | 1 | + | good | TCAACCTTGCCTACAATGTCCAAAGGACCACAGAGGCCACATGCATGGCGCGGCAGCGAGTGTTAGACAGCCACTCCATCTCTAAGCCAGCCGTTTTGATGCCACCGGCTGTGATACTAC | 120 | 55 |
| chr7 | 143727986 | 143728106 | Target=1;ProbeIdx=17245;Target #1 | 1 | + | good | AGGGGAGGATGTCACACGGGCGGTTTTGACATTGCTGACGAACCGTGTGGGCCCAATGGAGATGATGTTTAAATGCTCAGCCGACACGAAGCTCACACACAGACACACATAGTGATTCAG | 120 | 51.67 |
| chr7 | 143728106 | 143728226 | Target=1;ProbeIdx=17257;Target #1 | 1 | + | good | GCACAGCCTTTTCCTTCCCAGACAACGCCTGGCACAGCAGCGACCATTCTGAGCGTGCCTGGGATCCCAAGTGTGCTGAGGAAGGAAGGTGCCACTGCTCAGGCATGCCAGCCTCCTCTT | 120 | 60 |
| chr7 | 143728226 | 143728346 | Target=1;ProbeIdx=17269;Target #1 | 1 | + | good | CCTTACACCGCCATTTTTGTTGACATAAACTGTGTCGACCATATGGGATTGAATTCAGTGTTCCAGTTGCTTTTTCCTGTCTCTTCCTTTTCCTTTTCTCCCCCCTCCCCCGCCCCCTCC | 120 | 49.17 |
| chr7 | 143728346 | 143728466 | Target=1;ProbeIdx=17281;Target #1 | 1 | + | good | CCGGGTTTGGGAAGCTGTTCTCTTGGCAACCACAGAAGCAAAGCACACTCTTAATTGCTGCCCCAGTGGTAGATGCCCAGTGCCGAGCCTGACCCTTGCACTCCCTCCTCCTTTGATAGC | 120 | 55.83 |
| chr7 | 143728466 | 143728586 | Target=1;ProbeIdx=17293;Target #1 | 1 | + | good | ATCTTCACCCTGGTGTGGGAATCATTCCTTGTGTGCCCCCACCACCACCACTGCAGGGCACAGCAGTGAGCAAGCACCCCCACCCACACTCTGCACTCTCTTTGTTTGCTATACAGACAG | 120 | 55.83 |
| chr7 | 143728586 | 143728706 | Target=1;ProbeIdx=17305;Target #1 | 1 | + | good | AACTTTTCTCAAGGTGATGTTTGGAGAGAGCTTCCCTTGCTTGACTTCTGTTGGGGGTGTGAATGCCAGAGACACGTGGGGAAACAGCTCTGAAGGGCCCATTTTTGCCGGCTGCCCTGT | 120 | 52.5 |
| chr7 | 143728706 | 143728826 | Target=1;ProbeIdx=17317;Target #1 | 1 | + | good | TTTGGGGTGCTTTACCTCTGACTACTTTGACAGTTATTATTTCCCATTCCTTACACAAAGCGAATGTAAGCGGCAGGGGGGCGGAGTCCTGCTTTCCCAGGGTCATTATCAAGATGGGGAG | 120 | 48.33 |
| chr7 | 143728826 | 143728946 | Target=1;ProbeIdx=17329;Target #1 | 1 | + | good | GGCCAGAGAAGTGATGCCACTTGTGTCTGATGCTTCCAAAGGGAAATAGGTCACTGTCAGAATTCCTGAATCCTGGCATGCCTACCGGGGTGTCAGGGATCTTCTGGTGACTTGTCTGAA | 120 | 50.83 |
| chr7 | 143728946 | 143729066 | Target=1;ProbeIdx=17341;Target #1 | 1 | + | good | GCGTGGATATTTCTCCCTTGACTTTCTTACCACATCTGCTAGCTGACAAATAGACTTGGCTTCTTCTTTTAAAAAAAAATGTAAAAAAAAAAAAAAAAAATAGAAAAAGCACAGTACTTT | 120 | 30 |
| chr7 | 143729066 | 143729186 | Target=1;ProbeIdx=17353;Target #1 | 1 | + | good | ATATTTAGCCACAGGCGCACGGAGCGCCCAGCAGCAGCCGGGTTTCCTTGGATAACTGGGGCTCCGGCTCACAAATGCCTCCCGGTGTCTCCTGGCCGACAGCTCGCGCGCAATCGCCTG | 120 | 64.17 |

| | | | | | | | | | |
|---|---|---|---|---|---|---|---|---|---|
| chr7 | 143729186 | 143729306 | Target=1;ProbeIdx=17365;Target #1 | 1 | + | **good** | CTCCATGTTAACCCTGCGTGGTGCTGCCTCTCCGATGCCCTGCTTGCTCTCTCGGCAGCGTGGCACTGCTCACATGAGGATCACCGCCAGAGGTAGCCGGTTCTGAGAGCAGCCTCAGTC | 120 | 61.67 |
| chr7 | 143729306 | 143729426 | Target=1;ProbeIdx=17377;Target #1 | 1 | + | **good** | TGCGCACAGTTGCTTCCCAGCTGACAGACAGAATATGGGGTGGGCTTTGTCACCAGCACGCTGGGTCTGGCATGCCCAGCGATGCTCGGGGCTGCAGCGCGCCCAGACGCCCGCCACTGC | 120 | 65.83 |
| chr7 | 143729426 | 143729546 | Target=1;ProbeIdx=17389;Target #1 | 1 | + | **good** | TCGCCGGGTCCCAGCCGCGCCAGCTCCGAGGGGCCGGGCTGCGCGCCTAGCCTTCCTGCCCGCCGAGAGGGTTATATAATGAGCGGACCGCTGCCAAAGTCACAGGGAAAATGAAGTCTT | 120 | 65 |
| chr7 | 143729546 | 143729666 | Target=1;ProbeIdx=17401;Target #1 | 1 | + | | | | |

### Supplemetary Table2d-the probe design for mouse Trp53 gene

| Chromosome | Start | Stop | Name | Score | Strand | Code | Seq | Len |
|---|---|---|---|---|---|---|---|---|
| chr11 | 69586596 | 69586716 | 788425_47129966_NM_011640.3,XM_006 | 30 | + | good | TACAAACACCTCGGCTGTAGCTTTGGTTTGTTCTCTGACTTGCTTATAACTTAATATCCCTCTTATTCTAACCTAAGTTCTGCCACGTGGTTGGTTACCTCTGCTCAGCCCCCGGCTTCT | 120 |
| chr11 | 69588156 | 69588276 | 788425_47129966_NM_011640.3,XM_006 | 15 | + | good | AGCCAGGGCTACATAGAGAAACCCTGTCTCGAAAAACACAGAAAGGGAGGAAGAAGGAAAGGTCCCAGTCCTCTCTTTGCTGGCTGCCCTCTGCCACTGCGAGGGCGTCCAATGGTGCTT | 120 |
| chr11 | 69586716 | 69586836 | 788425_47129966_NM_011640.3,XM_006 | 14 | + | good | GTCCTCCATGTTCCTGGGGGAATCCCTACACTTTCAGAATTTAATTTCCCTACTGGATGTCCCACCTTCTTTTTATTCTACCCTTTCCTATAAGCCATAGGGGTTTGTTTGTTTGTATGT | 120 |
| chr11 | 69586836 | 69586956 | 788425_47129966_NM_011640.3,XM_006 | 3 | + | good | TTTTTAATTGACAAGTTATGCATCCATACAGTACACAATCTCTTCTCTCTACAGATGACTGCCATGGAGGAGTCACAGTCGGATATCAGCCTCGAGCTCCCTCTGAGCCAGGAGACATTT | 120 |
| chr11 | 69583236 | 69583356 | 788425_47129966_NM_011640.3,XM_006 | 2 | + | good | TTGGGACAAGCCTCGAAGTAAGTTGGATGCCTAGGATCCCAGCACTACTGTGGTTAAGACAGTAATGAATTTAAGGCCAGCCTGGGATACATAGTGAGACCCTGTCTCAAGAAAGGGGGA | 120 |
| chr11 | 69584916 | 69585036 | 788425_47129966_NM_011640.3,XM_006 | 2 | + | good | ACGACCTGGAAGATAGAGCAGGAAGACCTCCTATTGTCAGTGTGGCTATGTCTCCAAGACTGAGACATTGGGCCGCCATCCCAGCATGTGCGCGCGCGCGCGCACACACACACACACA | 120 |
| chr11 | 69587316 | 69587436 | 788425_47129966_NM_011640.3,XM_006 | 2 | + | good | TGGGCTGGTAGGCTGAGAACACAGTCCTGAGGGTTCTTCTTTGTCCCATCCACAGCCATCACCTCACTGCATGGACGATCTGTTGCTGCCCCAGGATGTTGAGGAGTTTTTGAAGGCCC | 120 |
| chr11 | 69587436 | 69587556 | 788425_47129966_NM_011640.3,XM_006 | 2 | + | good | AAGTGAAGCCCTCCGAGTGTCAGGAGCTCCTGCAGCACAGGACCCTGTCACCGAGACCCCTGGGCCAGTGGCCCCTGCCCCAGCCACTCCATGGCCCCTGTCATCTTTTGTCCCTTCTCA | 120 |
| chr11 | 69587556 | 69587676 | 788425_47129966_NM_011640.3,XM_006 | 2 | + | good | AAAAACTTACCAGGGCAACTATGGCTTCCACCTGGGCTTCCTGCAGTCTGGGACAGCCAAGTCTGTTATGTGCACGGTGAGTGGGCCCCGGGGAGTTGTCTTTCGTGTGACCTTTCAATG | 120 |
| chr11 | 69588396 | 69588516 | 788425_47129966_NM_011640.3,XM_006 | 2 | + | good | CTGGCGAAGACGTGCCCTGTGCAGTTGTGGGTCAGCGCCACACCTCCAGCTGGGAGCCGTGTCCGCGCCATGGCCATCTACAAGAAGTCACAGCACATGACGGAGGTCGTGAGACGCTGC | 120 |
| chr11 | 69588636 | 69588756 | 788425_47129966_NM_011640.3,XM_006 | 2 | + | good | CCCCAGCATCTTATCCGGGTGGAAGGAAATTTGTATCCCGAGTATCTGGAAGACAGGCAGACTTTTCGCCACAGCGTGGTGGTACCTTATGAGCCACCCGAGGTCTGTAATTTTGTTTTG | 120 |
| chr11 | 69589116 | 69589236 | 788425_47129966_NM_011640.3,XM_006 | 2 | + | good | CCTGGATCCTGTGTCTTCCCCCAGGCCGGCTCTGAGTATACCACCATCCACTACAAGTACATGTGTAATAGCTCCTGCATGGGGGGCATGAACCGCCGACCTATCCTTACCATCATCACA | 120 |
| chr11 | 69589596 | 69589716 | 788425_47129966_NM_011640.3,XM_006 | 2 | + | good | CTTTGAGGTTCGTGTTTGTGCCTGCCCTGGGAGAGACCGCCGTACAGAAGAAGAAAATTTCCGCAAAAAGGAAGTCCTTTGCCCTGAACTGCCCCCAGGGAGCGCAAAGAGAGGTACGCA | 120 |
| chr11 | 69590676 | 69590796 | 788425_47129966_NM_011640.3,XM_006 | 2 | + | good | GAGATGTTCCGGGAGCTGAATGAGGCCTTAGAGTTAAAGGATGCCCATGCTACAGAGGAGTCTGGAGACAGCAGGGCTCACTCCAGGTAAGTGGCCTGGGGCAGCGCCTGCCTGTGGTGC | 120 |
| chr11 | 69591396 | 69591516 | 788425_47129966_NM_011640.3,XM_006 | 2 | + | good | CAATGGTCAAGAAAGTGGGGCCTGACTCAGACTGACTGCCTCTGCATCCCGTCCCCATCACCAGCCTCCCCCTCTCCTTGCTGTCTTATGACTTCAGGGCTGAGACACAATCCTCCCGGT | 120 |
| chr11 | 69591516 | 69591636 | 788425_47129966_NM_011640.3,XM_006 | 2 | + | good | CCCTTCTGCTGCCTTTTTTACCTTGTAGCTAGGGCTCAGCCCCCTCTCTGAGTAGTGGTTCCTGGCCCAAGTTGGGGAATAGGTTGATAGTTGTCAGGTCTCTGCTGGCCCAGCGAAATT | 120 |
| chr11 | 69591636 | 69591756 | 788425_47129966_NM_011640.3,XM_006 | 2 | + | good | CTATCCAGCCAGTTGTTGGACCCTGGCACCTACAATGAAATCTCACCCTACCCCACACCCTGTAAGATTCTATCTTGGGCCCTCATAGGGTCCATATCCTCCAGGGCCTACTTTCCTTCC | 120 |
| chr11 | 69591756 | 69591876 | 788425_47129966_NM_011640.3,XM_006 | 2 | + | good | ATTCTGCAAAGCCTGTCTGCATTTATCCACCCCCCACCCTGTCTCCCTCTTTTTTTTTTTTTTACCCCTTTTTATATATCAATTTCCTATTTTACAATAAAATTTTGTTATCACTTATAT | 120 |
| chr11 | 69580356 | 69580476 | 788425_47129966_NM_011640.3,XM_006 | 1 | + | good | ACTTTCCCCTCCCACGTGCTCACCCTGGCTAAAGTTCTGTAGCTTCAGTTCATTGGGACCATCCTGGCTGTAGGTAGCGACTACAGTTAGGGGGCACCTAGCATTCAGGCCCTCATCCTC | 120 |
| chr11 | 69580476 | 69580596 | 788425_47129966_NM_011640.3,XM_006 | 1 | + | good | CTCCTTCCCAGCAGGGTGTCACGCTTCTCCGAAGACTGGGTAAGTAATTGATGAGCGTGACGAGACCTCTCGGTCACTGGCTCTCTCCGTTTGCATCCATAAAACTAGAGAAAACCGTGG | 120 |
| chr11 | 69580596 | 69580716 | 788425_47129966_NM_011640.3,XM_006 | 1 | + | good | GGTTTGGGGGTGGGGCAGTGGGGGGACTCAGCGCGATGGAGATGGGCGGAATGGAAGCTTGGCGGGCGGGATGAACGGGAGTGTATATGTCAGATGCTGTAGTGAGGGTAGCTGATGAT | 120 |
| chr11 | 69580716 | 69580836 | 788425_47129966_NM_011640.3,XM_006 | 1 | + | good | ATGATGTTAGGACCGACGAGCCTCACTGTCATGCACCTGCAAAGTAGAGCATATAGGGACCACTGAGATGGCCTAAGGGGTTTTCTCTCCGCTACGCGTTGTACACACTTATCTGCCCGC | 120 |
| chr11 | 69580836 | 69580956 | 788425_47129966_NM_011640.3,XM_006 | 1 | + | good | TGCTAGGTGATGGAAGCTCCGGAAATAACATGCACAAAGCACCAGGATTTAAGATTTTTCGAGATTCATAGCTTAAGACTTAAGACCCCCCATAGCATCCTAATGAAACCCTGGGTTCCG | 120 |
| chr11 | 69580956 | 69581076 | 788425_47129966_NM_011640.3,XM_006 | 1 | + | good | TTCCTGGATGAGATCGGGGTGATCCGGGGAGCCTTAGCTGCTAAGGTCCCGCAACTTCCGGACCTTTGTCCCTGGAGTGATTTCTTTTTTTTTCCAGCCGCTTCTCGACCCTGCTAGATG | 120 |
| chr11 | 69581076 | 69581196 | 788425_47129966_NM_011640.3,XM_006 | 1 | + | good | AAGAAAATCCAAGAAAAGCCTGAAGCACTAGCGGTGCTAGCCAGAAGTATTTGCCCTCGGGGCCCGACTCAGCCTCTTGGTCTGAAAGGCCCGCCGGCCCTGTTATTGTTTGGCTCCTTT | 120 |
| chr11 | 69581196 | 69581316 | 788425_47129966_NM_011640.3,XM_006 | 1 | + | good | ACGTTTCTGCCGCTTGCAGGAGCATTTCCGGTTTCTTGTTTTCGGAGCAGATCACTGCTCGCCCGGCGACGGGGGAGTAGCGAAAGGGGAGAAATGGATTCTAGGCTGGTTCTGTGGTTT | 120 |
| chr11 | 69581316 | 69581436 | 788425_47129966_NM_011640.3,XM_006 | 1 | + | good | GAGGAGGAAAACTGCTGTCCTCGACATCTTATTTTTCTGGATTACTTGGTTATTGCTTTTGCAAAGGAGGAGGTGTTTATTTAAAAGAGTGCGCCGATAGGTCGTTTCTTCCTGCCGGAA | 120 |
| chr11 | 69581436 | 69581556 | 788425_47129966_NM_011640.3,XM_006 | 1 | + | good | AAGCAAATTACCGAGTATCCGGTTTTAGGGTGAGCCATTCCCTTGCTTAACGCATTCCGCGCGTCCTGAAAGCGGAAGGGGCGGGCTTGGGCCCCGGGCGCCCGGCGGTCCTAAGTGCAG | 120 |
| chr11 | 69581556 | 69581676 | 788425_47129966_NM_011640.3,XM_006 | 1 | + | good | TCAGCCCGCACCATCCGGTTCTTCCCAGCGGGAGGTGGCGGGACTCTGCCTGGGCGTGGGCTATGCCTGTCCCCGCCACTTCAATTTTTAGAACAGGTCATTTCATTACAGTTTCTTATC | 120 |
| chr11 | 69581676 | 69581796 | 788425_47129966_NM_011640.3,XM_006 | 1 | + | good | GCGCACCAGACATCGTGCGTGCCTTTTTTTCTTTTGTAAAAATACTATTTCCACACTGCACTTGGATCGGGAAGTAAATTTTTTTGTCCATCTGAAGATTTAGTTTCGGTCTAAAGAATA | 120 |
| chr11 | 69581796 | 69581916 | 788425_47129966_NM_011640.3,XM_006 | 1 | + | good | AATGATAGGTAGACCGGGAGGCGGGGGGACTTTGTTTTCGTGGCTTCAGCTGTAAAGCCGTAAGGAAGACTAGATCCTCCAGTTCATTGAGAAAGTATACAAATGACTTCACTTAAACTT | 120 |
| chr11 | 69581916 | 69582036 | 788425_47129966_NM_011640.3,XM_006 | 1 | + | good | TGACGCGCGAAGGGAACCCTAGCTGTGAAGGTTAGACTGGACAGCCACAGCGGTCCAGGGATGCCTTCTTATGCAAATAGCCTCCCTGCTCTTGCAATCTCTTTGACTGGATTTAGTGAA | 120 |
| chr11 | 69582036 | 69582156 | 788425_47129966_NM_011640.3,XM_006 | 1 | + | good | GGACTCTCTTGTGAATGTTAAAAGATCCCAACCCGAAGTCAAAAGGAGGTGCCCCTAGGCCCCATCTTGTTTACAAATGTAGATATATGTAAATTGGTTTTTTTGTTTTTTTTTTTTTTA | 120 |
| chr11 | 69582156 | 69582276 | 788425_47129966_NM_011640.3,XM_006 | 1 | + | good | CATAAAGACAACTTTGGAAGATATTTACCCTGCTCTTCCAGTTTCTCTTCTCTGAATAATTTACTGCAGAGGGGTTTGGGAATGTCTGGGATGGGTAGTATATGCACTTAAATTATGCTT | 120 |
| chr11 | 69582276 | 69582396 | 788425_47129966_NM_011640.3,XM_006 | 1 | + | good | AAATCATCTGTAGCTTACATATGACCCTGCACTGTAAATGTTATTCAAATAAATGTTACCGTGTTGACTGAAAAAAGGAGCCAGGCTGGACTGGGACCGGGTGCTGCAAGCCTTTAACCC | 120 |
| chr11 | 69582396 | 69582516 | 788425_47129966_NM_011640.3,XM_006 | 1 | + | good | CAAAGCTAGGGAGGCCGAGGCAGGTCAATATCAGAGATTCAGGCCAGCCAAAGATATGCAAGGAGACCCTATCTCAAAAAGAAAAATATGTCACGTATGTTAATATTCACACTGACACAA | 120 |
| chr11 | 69582516 | 69582636 | 788425_47129966_NM_011640.3,XM_006 | 1 | + | good | GATAACCTGCTGCTACAGTTGAGTAGCAGTGGGAAGCCTCGACAGTCTCGAGTTCCAGACCACAGTGTCCAGACCATACATAGAGGGAGTTCAAGGTCATCCTTCCCTGGGTAGTTGTAA | 120 |
| chr11 | 69582636 | 69582756 | 788425_47129966_NM_011640.3,XM_006 | 1 | + | good | GCTGACTTGAGATAAATGAGAGTGTCTTCAAAAACAGAGAAAGAGTACAAAAGTTAACATTGGCAAACCAGAGCCTACTGGCCTTTCTCCTGATGGGTAAATGAGGACGTCTCGATGGCC | 120 |
| chr11 | 69582756 | 69582876 | 788425_47129966_NM_011640.3,XM_006 | 1 | + | good | CGTGCCTTTAATGCTAACACTCTTGGGATGCTGAGGTAACTTGGGCTACTTAGCAAGACCTTGTCTCAGAAAGGCAAACAGCTGGAGATATGGCTTGGAGTAGATTATTTCACCTGGCAA | 120 |
| chr11 | 69582876 | 69582996 | 788425_47129966_NM_011640.3,XM_006 | 1 | + | good | ACATGATACCCTTGGTTTAAATCCTGAGACCACAAAAAACAGGTTAAACCCAGCTTGACCAAGTGCCATTGGTCCATGGATTGCTGTATTGGAATCAAACAGAAATCTATGTCATTCACA | 120 |
| chr11 | 69582996 | 69583116 | 788425_47129966_NM_011640.3,XM_006 | 1 | + | good | GCAGTAACCTCCTGGGAATACTTCAAGAGACGGAGAAAGGGCGACTGACTGTGCCCTCCGTCCTTTTTCGCAATCCTTTATTCTGTTCGAATCATAATTTGTTTTCCTGAGACAGGGTTT | 120 |
| chr11 | 69583356 | 69583476 | 788425_47129966_NM_011640.3,XM_006 | 1 | + | good | AATAAGTTCCAAAAGTGACATTTTCCAAGGTGCGAAAGCGATGGTTCCTTAGAGAAAGTGACGTTGGAAGGGACAGAAGTAAATTTGACTCCTTAGTTATTAATTTGGTATCGATTTGAG | 120 |
| chr11 | 69583476 | 69583596 | 788425_47129966_NM_011640.3,XM_006 | 1 | + | good | ACCTGGATAAAAAGCTGGTAAGCTTTAGGAAGGAAGGTAACGGTGGGCACGCATCTTTACGTCTTGGTTGGTTGGTTTTTTTTTGTCTCTGGCTGATAAAAGTAAACCCACTGTCTTTGT | 120 |
| chr11 | 69583596 | 69583716 | 788425_47129966_NM_011640.3,XM_006 | 1 | + | good | CTTGGTGCAAGAGAGGGCTACTACTAGGCTATTTCAAAGTTTGCCAAATAAACACTTACCATGTTCTCTTGTATCCACGTTTTAGTTACATAAACAGTCTTAAACCAGATGTGGTGGCTC | 120 |
| chr11 | 69583716 | 69583836 | 788425_47129966_NM_011640.3,XM_006 | 1 | + | good | ACACCTGCAAGCCCAATGGTCAGGAAGCTGCAACAGAAGAAGTGCCATTGTAGCCGAGCCACAGAGTGAGGCGACTGTCTCAGAGTACAAAATAAAACTAAATCAAACATAATGGTCATC | 120 |
| chr11 | 69583956 | 69584076 | 788425_47129966_NM_011640.3,XM_006 | 1 | + | good | TGTGTGTCAGAGGTCAGCTGACAACTTAATAGAAATTAGTTTTGGGGTACTCAGTCTTGGTACCAGGTGCCTATTCTGGCCACACAGACCCACCTATTTTCTGATTTGTATAAACTATTT | 120 |
| chr11 | 69584076 | 69584196 | 788425_47129966_NM_011640.3,XM_006 | 1 | + | good | TGAGTTTTTCTACCAATCGTATTCCTCACCCAGTGTGTTTCTAAACGTTTTCCAAAGCCGTGTTTTATGTGTCTGTTCCCCACCAGTGTGTTGTGATTTCTTCTTGCTCCACACCCGGAA | 120 |
| chr11 | 69584196 | 69584316 | 788425_47129966_NM_011640.3,XM_006 | 1 | + | good | TAATGCTGACAGGAGCAGTTTTTGTGCAAGGCTCCTGCTGTCCGTGTTCTAGGAACTGCACCTAGATGTGGTTTTGCTGAGTGAGATGTGTGCTTTAAGTGATACCAGACACAGCCTTCT | 120 |
| chr11 | 69584316 | 69584436 | 788425_47129966_NM_011640.3,XM_006 | 1 | + | good | TATTTTTTCTTAGAACCAACAATATATCGGTCCTTGGCCCCCACCCCCATGCCCCTTTTTTCCAATGCTGGGTGTTGAACCCGGAGCCTTAGACATAACACACGAACTCTGCAGCTGAGC | 120 |
| chr11 | 69584436 | 69584556 | 788425_47129966_NM_011640.3,XM_006 | 1 | + | good | TACCCTGCCAGCTCCGAAAGATTTGTATACTAAGTTCTATGAAACATTAAGGTCGTTTTATTCAGGGTAAGGTGTGGCTTCTGGCTTCCCTGGGTCCAGTCCTTTGATGGTGGTGGGTAG | 120 |
| chr11 | 69584556 | 69584676 | 788425_47129966_NM_011640.3,XM_006 | 1 | + | good | TACTGTTCGTTCCATTCCGTTTGGGGTTTTGATTGACAAGCCTTGCACCTTTCCAACTGTTCTACCTCAAGAGCCAAAGATAAAGGGTAAAAGATTCCCTTCCCTTATCCCTGTCAATAG | 120 |
| chr11 | 69584676 | 69584796 | 788425_47129966_NM_011640.3,XM_006 | 1 | + | good | CAGCCTGCCTAGCTTCCTCAGGATCAAATGAGATGAGCCCCTGAGAAGAGCAAGGCCCGCTGGGCCTGGAAGGCCAGCCCTGGTTGTACTCAAACCTCTCGAGTCTATTGCCTTTCCCAG | 120 |
| chr11 | 69584796 | 69584916 | 788425_47129966_NM_011640.3,XM_006 | 1 | + | good | CCAACATTTTCTTACACATCCAGCCTCTGTGGATACTGTGACCCTCCTGATCTGGTTCTTGTGAAAAGTTTCATATTGGCAACTGTTCTTAAGGAAACACTGAAAACCTAATTACTACTA | 120 |
| chr11 | 69585036 | 69585156 | 788425_47129966_NM_011640.3,XM_006 | 1 | + | good | CACACAGTATTCACTAAGCAAGAGTTCCTTGACAGGGTGAACTGGCATTAACGGGTAGTGGTTAGCGCCACAGAAGCACTGCTAGCTCAGAACAAATGAGGTGCTCATACCCACAGGGCT | 120 |
| chr11 | 69585156 | 69585276 | 788425_47129966_NM_011640.3,XM_006 | 1 | + | good | GAGCTCTTACATGAGTGTGTACTGTTGGGCGTGGGGTGGGGCTGGACCCAGGGCTTCACAGATGAGCTCCGCCCAAGGCCAAATCTCAAAAAATGAAAAGGTCAAAAGGTAACCCTTTCA | 120 |
| chr11 | 69585276 | 69585396 | 788425_47129966_NM_011640.3,XM_006 | 1 | + | good | GATAGATACCGGAGATATGATATTTACAAAAAATATGATAATTAAATGCTACAGTTTTAATTATGAATCTGAAGCAGCTCTGGGTCTCCCTGACCCCTTTCCAGGGCCAGCCACCTCATT | 120 |
| chr11 | 69585396 | 69585516 | 788425_47129966_NM_011640.3,XM_006 | 1 | + | good | ACTCAGATGTAGGGGCTATATAATACCCAGTCCTTTGCTTTCTTCATCTAGCTGATTGCAAGAGAACTGTGCCTAAGAGCCTGTGACGCACTAGATTGTTTGGGTGGCCATCTTTAATAA | 120 |
| chr11 | 69585516 | 69585636 | 788425_47129966_NM_011640.3,XM_006 | 1 | + | good | AGAGTCATTCACCTGCCCAGGGGCAGATGTGACCATCGAGACAGATGAAATAGCCCATATGGAAAGGTTTCTCAAACTCTTAAATAAATGAGAACTTGATTGTTCTACATACAAGTGAAT | 120 |
| chr11 | 69585636 | 69585756 | 788425_47129966_NM_011640.3,XM_006 | 1 | + | good | ATATTAAATGTTGAGATTATAGTCAGTATTAACAGTAATTATTCTGAAATGCTTGCCTTTGCTGTTAGAAAGGAAACATGATGTCCCAGGGGGGTTGGCCGAACTCAATCCCCAGAACCCA | 120 |

| | | | | | | | | |
|---|---|---|---|---|---|---|---|---|
| chr11 | 69585756 | 69585876 | 788425_47129966_NM_011640.3,XM_006 | 1 | + | good | CTTCTTATAAGTTGTCATCTGACCTCCATACGCATGTCTTGGCACCTTTAAATAAGGCACAATAAATAAGTAAATGTCTAAGTAAGGAAGATAATAGCTCCGTGGTTAGTAACAAGAGAA | 120 |
| chr11 | 69585876 | 69585996 | 788425_47129966_NM_011640.3,XM_006 | 1 | + | good | AATGAGTGAGAGAGTATCATACGGCTTAACCTCTGAAGGAGGTGGTGGTGGTGTGGCTGGCTGGAGATTGGCTGGCTGTGACTGTCTCCGAGGAGCTATGGCATACAAGATAAGGAAGAG | 120 |
| chr11 | 69585996 | 69586116 | 788425_47129966_NM_011640.3,XM_006 | 1 | + | good | CCTACCTCATAGCCTAGGGAGAACAGAAACTGTGACTTTGCTCTTGTAGAGGTAACCACACTGATTCAGCCAGGAGGAAGTAAGTGCTCCCTAGAGCCCTTGGGGAAGAAGGCAGTAGGA | 120 |
| chr11 | 69586116 | 69586236 | 788425_47129966_NM_011640.3,XM_006 | 1 | + | good | AACCTGCTGAATCTTCAGGAATTTGTAAGGCGCTGGGGACCTGTCCCTAGGGGGCAGATGAGACACTGATGGGCGTACTTAGAGATTTGCCATGAAGTGGGTTTGAAGAATGGAGCTGTG | 120 |
| chr11 | 69586236 | 69586356 | 788425_47129966_NM_011640.3,XM_006 | 1 | + | good | TGTGAAATGGTGGATGGGGGGGGGGAGTCCCTCCCCAGAGGGAAGGGAAGAGAGATGAGATGTAGGGTGCAGATGTAGGGGGGCTTGGGGCTAGAAGTACCTCCCTGATTACCTGTTCCT | 120 |
| chr11 | 69586356 | 69586476 | 788425_47129966_NM_011640.3,XM_006 | 1 | + | good | TGAAGCAGTGTGTGGTTCGAGAAGCTGATAGGAAGCCAGGCCAAGGCTTGAGCAGCCTGAATAAAAGACGGAAGAGCTGCCCCATTCCTGCTTCTCTGGAAATGGTGTCCCTCACGGCCA | 120 |
| chr11 | 69586476 | 69586596 | 788425_47129966_NM_011640.3,XM_006 | 1 | + | good | TCTTGGGTCCTGACTTCTTCTCAAAGGAGCCTGGCCGACTTCTTGGATACTTGTAACTTTGCGCTTTCCCACCCTCGCATAAGTTTCCTGAAATAATGACTCTGAAACTCAAAATATATT | 120 |
| chr11 | 69586956 | 69587076 | 788425_47129966_NM_011640.3,XM_006 | 1 | + | good | TCAGGCTTATGGAAACTGTGAGTGGATCTTTTTGGGGCCCTTAAGATACATCCCGCCATACCTGTATCCTCCCCTTGCCTGAGAGAAACAAAAACAGTAGTGTTCAAACATGGTATGGTG | 120 |
| chr11 | 69587076 | 69587196 | 788425_47129966_NM_011640.3,XM_006 | 1 | + | good | TTGGGTGTCTGTAAATCCTGCGGGGCGGGGTGGCGGGGGGGGGGGGGGACTGCAGGGTCTCAGAAGTTTGAGGTCATCATTGACTACATAGCAAGTTGGAGGCCAGCCTGGGATAAGTGA | 120 |
| chr11 | 69587196 | 69587316 | 788425_47129966_NM_011640.3,XM_006 | 1 | + | good | GATTCTGTCTTCAAAAAATGGAAGGAAATCAGGAACTAACTCTCTGCTCTTGTTTTCCAGACTTCCTCCAGAAGATATCCTGGTAAGGCCCAGAGCAGAAAGGGACTTGGGCTTTGGTGT | 120 |
| chr11 | 69587676 | 69587796 | 788425_47129966_NM_011640.3,XM_006 | 1 | + | good | CCTCTTTTTCTCTGTCTAGATTTGGGGGTTCCTCTTCAGCCTGTAGACTGTGCCTCAGAGTTTAGTTTTGCCCTGAACCTTTTTAGCCTCTTCTCCTTTCTCTCATATTCTCTGCATCTC | 120 |
| chr11 | 69587796 | 69587916 | 788425_47129966_NM_011640.3,XM_006 | 1 | + | good | TCCAGGGGACGTGGAACTCTCTTCCCTCACATTCCTTTCTTGGCTTTTGAAAATAATCTTCTGAAGCCAAGCACAGAGGCGTGTGCTGTAGGCCCAGCTGCTAGAGACAGTTGAGGCAGG | 120 |
| chr11 | 69587916 | 69588036 | 788425_47129966_NM_011640.3,XM_006 | 1 | + | good | ATTGCTTAAGGCCAGCCTGAGCATGGAAGTAAGACCCCTTCTCACCAAAAACAAAAACAGCCCAGTGGAGTGACACACACCTGTAGCTCCAGCACTGGGGAGGCCAAAGTGGGAGGGACA | 120 |
| chr11 | 69588276 | 69588396 | 788425_47129966_NM_011640.3,XM_006 | 1 | + | good | GGACAATGTGTTTCATTAGTTCCCCACCTTGACACCTGATCGTTACTCGGCTTGTCCCCGACCTCCGTTCTCTCTCCTCTCTTCCAGTACTCTCCTCCCCTCAATAAGCTATTCTGCCAG | 120 |
| chr11 | 69588516 | 69588636 | 788425_47129966_NM_011640.3,XM_006 | 1 | + | good | CCCCACCATGAGCGCTGCTCCGATGGTGATGGTAAGCCCTCAACACCGCCTGTGGGGTTAGGACTGGCAGCCTCCCATCTCCCGGCTTCTGACTTATTCTTGCTCTTAGGCCTGGCTCCT | 120 |
| chr11 | 69588756 | 69588876 | 788425_47129966_NM_011640.3,XM_006 | 1 | + | good | GTTTGTGCGTCTTAGAGACAGTTGACTCCAGCCTAGACTGATGTTGACTTTCTAGCAACCCGTTTGCCTCACCCTCCTGAGTGCTAGCTAGGCTTAGAGGTGCAAGCTGCCGTGCCCAGC | 120 |
| chr11 | 69588876 | 69588996 | 788425_47129966_NM_011640.3,XM_006 | 1 | + | good | CCAGGGTCTACTTTAACAGCAGTCTCTGGGAGAAGGGGCTTCCCCATCCAGGCGGGAAATAGAGACGCTGAGTCCGGTTCCCTCCCATGCTAAGCAAGTGTTGGGCCCCACAGCTCCAGC | 120 |
| chr11 | 69588996 | 69589116 | 788425_47129966_NM_011640.3,XM_006 | 1 | + | good | AGGTGTGCCGAACAGGTGGAATATCCCTACTCTACAACTAAAACTGAAACTTATTAGAGGCTATAGCCAGCCATTCCCGGCTGCTGCAGGTCACCTGTAGTGAGGTAGGGAGCGACTTCA | 120 |
| chr11 | 69589236 | 69589356 | 788425_47129966_NM_011640.3,XM_006 | 1 | + | good | CTGGAAGACTCCAGGTAGGAAGGCGCGTGGTAGGTTAGGTTAGCCTGTTTCTTCCCCAGCTTCTGCCTGTTTCTGTTCCACGAGTCCCGCCCCCTACCACATGCCCAACGCTCTTTGGTT | 120 |
| chr11 | 69589356 | 69589476 | 788425_47129966_NM_011640.3,XM_006 | 1 | + | good | CCTACCCTATCTACCTAAATGAAGTCTCCTCCTCTGTTTCCTCTTGGGCTTAGGGACGTCTCTTATCTGTGGCTTCTCGGGGTTCCTGTAACTGGACCTTTGGCTGCAGATATGACAAGA | 120 |
| chr11 | 69589476 | 69589596 | 788425_47129966_NM_011640.3,XM_006 | 1 | + | good | GGGGTTGGGAACAGGTGGGGGCCTAGTTTACACACAGTCAGGATGGGGCCCAGCTTTCTTACTGCCTTGTGCTGGTCCTTTTCTTGTCCCGGATAGTGGGAACCTTCTGGGACGGGACAG | 120 |
| chr11 | 69589716 | 69589836 | 788425_47129966_NM_011640.3,XM_006 | 1 | + | good | GGCGGGACCAAGGAGGCGGAGGAGCCTGTTGAGCTTCACCCCAAAGTCACCTCTTGCTCTCTCCTTCCACAGCGCTGCCCACCTGCACAAGCGCCTCTCCCCCGCAAAAGAAAAAACCAC | 120 |
| chr11 | 69589836 | 69589956 | 788425_47129966_NM_011640.3,XM_006 | 1 | + | good | TTGATGGAGAGTATTTCACCCTCAAGGTACCAAGGCTGGAGAGTCGCATGCCAGAGACAAGCTGTACCCATTATTGCCTCTGTCTCTCGCATGTATAAAATAGTGTTATTAGCAGGTTGC | 120 |
| chr11 | 69589956 | 69590076 | 788425_47129966_NM_011640.3,XM_006 | 1 | + | good | CAGGTCTTTTTCAGTGGCTTTATCTCTAGCCGTGACATTAGCTTGAGAGCTCATGCTCTGAGGCTGTGCCTCCTCCGACAGTGGTTCTCAGTGTAACTAACTTGACAACACCAACTTACA | 120 |
| chr11 | 69590076 | 69590196 | 788425_47129966_NM_011640.3,XM_006 | 1 | + | good | CCATAAGACAGGTGCTCCTCCACTGGGGATGGGAACATGTCCTAGGAAACTCACCATAAATTGAAAATAACACACGACAAAAATGTGTTTAGAGGCAGGCCTGGTGGCACCTGCCTGTAA | 120 |
| chr11 | 69590196 | 69590316 | 788425_47129966_NM_011640.3,XM_006 | 1 | + | good | TTACAGCACTGAAGAGGGTGGTTCAAAGCTGGCAAGAAAAAGTAACCCAAGGAAAGGACATTGGGGTTTAAGGGTATAACTCAGTTCCAGGATACTTGCCAAGTGTGTGCCAGGCCCCGG | 120 |
| chr11 | 69590316 | 69590436 | 788425_47129966_NM_011640.3,XM_006 | 1 | + | good | ATTAGGTCCCCAGCGGCTCACGGTGCCTTTAGTCCACCTAACCCACAACTGGACTCTACAGCTCAGCTCCCGGTTGCCCTGTTAAGCGTCTGTTCCCTCTAACTGAAAAGATCCAAGCTT | 120 |
| chr11 | 69590436 | 69590556 | 788425_47129966_NM_011640.3,XM_006 | 1 | + | good | GTTGTACACGTTCTACTGAATGCCTGTTACTTTCACACCATCTTATTAAAGATGATCTCCCCCCCCCCAAAAAAAAAAACAAAACAAAACAAAACAAAAACCTGTAAGTGGAGCCAGCT | 120 |
| chr11 | 69590556 | 69590676 | 788425_47129966_NM_011640.3,XM_006 | 1 | + | good | TAAGTTGGGAACCAACTTTCAGAAAGAAAGTTGTTAAAATCGTGAAAGTGGTTGTGTGACCTTGTCCAGTGCTTCCATCTCACTTCATCTCTGCTGCAGATCCGCGGGCGTAAACGCTTC | 120 |
| chr11 | 69590796 | 69590916 | 788425_47129966_NM_011640.3,XM_006 | 1 | + | good | TCTACCCAGACCTCCCTCCAGCTCAGCCTTTGTAGTGAAAGATAAAAACCCCACCCTGCTAGATGCTTAGGGCTGCACCCTACGAGAACTGACTTCTTGACTTTTTAGGCTCTGTTAAGG | 120 |
| chr11 | 69590916 | 69591036 | 788425_47129966_NM_011640.3,XM_006 | 1 | + | good | GGTATGAGGGACAAGGTATGGTGTCATGCTCCTATAATCTCAGCAGTAAGGAAGACAAAGTCAGGAGGATTTGGGGAAGTTTGAAGCCTTCATAAACTATATAAAACTTTAGGCCAGCTA | 120 |
| chr11 | 69591156 | 69591276 | 788425_47129966_NM_011640.3,XM_006 | 1 | + | good | GTGAAAGGGAGGATAAACTGATTCTCAGAAGTATTCCAGTGTGTTCTGTGAATATCCCTACCCATAGTAGAAGCCATCTTAAATTCCTTTTTTTCAGCCTCCAGCCTAGAGCCTTCCAAG | 120 |
| chr11 | 69591276 | 69591396 | 788425_47129966_NM_011640.3,XM_006 | 1 | + | good | CCTTGATCAAGGAGGAAAGCCCAAACTGCTAGCTCCCATCACTTCATCCCTCCCCTTTTCTGTCTTCCTATAGCTACCTGAAGACCAAGAAGGGCCAGTCTACTTCCCGCCATAAAAAAA | 120 |
