## Supplementary Table 3 for "Transcriptional Determinism and Stochasticity Contribute to the Complexity of Autism Associated *SHANK* Family Genes"

Supplementary Table 3a The numer of isoforms of top 100 genes from SIS in human brain

| Gene_name | Transcripts_per_gene | Gene_abundance |
|---|---|---|
| SEPTIN4 | 692 | 829 |
| MADD | 660 | 1156 |
| DST | 600 | 1568 |
| BAZ2B | 564 | 797 |
| SORBS1 | 491 | 1094 |
| TTC3 | 464 | 1450 |
| MACF1 | 462 | 1418 |
| ACADVL | 441 | 640 |
| MAP4K4 | 423 | 796 |
| MAP4 | 373 | 1140 |
| ANK2 | 372 | 1291 |
| PDE4DIP | 369 | 912 |
| RHOT2 | 368 | 421 |
| ZBTB20 | 361 | 665 |
| SYNE1 | 360 | 912 |
| NRCAM | 359 | 1508 |
| ANKRD36C | 357 | 404 |
| NDRG2 | 345 | 619 |
| RAP1GAP | 340 | 790 |
| GTF2I | 339 | 947 |
| DTNA | 325 | 1177 |
| PCM1 | 314 | 609 |
| SEMA3B | 314 | 514 |
| SGSM3 | 313 | 553 |
| KTN1 | 310 | 832 |
| PUM1 | 308 | 572 |
| SPTAN1 | 306 | 936 |
| SEC31B | 303 | 219 |
| MYCBP2 | 300 | 1021 |
| NBPF20 | 300 | 325 |
| MICAL2 | 291 | 658 |
| GFAP | 287 | 499 |
| ANK3 | 284 | 511 |
| NBPF10 | 276 | 295 |
| UBR4 | 276 | 687 |
| SNHG14 | 265 | 390 |
| HERC1 | 263 | 704 |
| CAMK2G | 258 | 525 |
| MINK1 | 255 | 455 |
| GRIA2 | 252 | 1010 |
| NT5C2 | 249 | 659 |
| USP54 | 247 | 367 |
| EPB41L2 | 245 | 601 |

| | | |
|---|---|---|
| SMARCC2 | 245 | 553 |
| EPB41L1 | 241 | 522 |
| SRSF11 | 241 | 642 |
| BBS2 | 239 | 598 |
| RASSF4 | 238 | 574 |
| ARPP21 | 236 | 793 |
| SPECC1 | 236 | 554 |
| SEPTIN2 | 234 | 686 |
| BAG6 | 228 | 476 |
| KANK1 | 227 | 405 |
| FN1 | 225 | 538 |
| PHLDB1 | 225 | 376 |
| MEG3 | 223 | 1106 |
| PAM | 221 | 583 |
| KIDINS220 | 220 | 670 |
| PPFIA2 | 215 | 617 |
| EXOSC10 | 214 | 355 |
| ATXN2 | 212 | 389 |
| MATR3 | 211 | 1501 |
| HERC2P2 | 210 | 370 |
| LIMCH1 | 209 | 657 |
| NCOR1 | 209 | 583 |
| CELF1 | 208 | 388 |
| CROCCP2 | 208 | 297 |
| KIAA1109 | 208 | 556 |
| SCAPER | 208 | 446 |
| ASPH | 207 | 653 |
| SIPA1L1 | 207 | 514 |
| SORL1 | 207 | 690 |
| CHD4 | 206 | 696 |
| R3HDM1 | 206 | 509 |
| SRRM1 | 206 | 530 |
| HECTD1 | 204 | 488 |
| PAN2 | 204 | 297 |
| UBAP2L | 204 | 467 |
| RYR2 | 203 | 473 |
| EPB41L3 | 202 | 515 |
| MBD1 | 202 | 500 |
| GGT7 | 201 | 318 |
| HDAC6 | 201 | 348 |
| ADD3 | 200 | 766 |
| CLIP1 | 200 | 475 |
| MYT1L | 200 | 619 |
| SNRPN | 200 | 607 |
| CAMK2B | 198 | 420 |

| | | |
|---|---|---|
| TRIP12 | 198 | 610 |
| SUN1 | 197 | 549 |
| DDX5 | 196 | 871 |
| HID1 | 196 | 289 |
| RBM39 | 196 | 731 |
| GIGYF2 | 195 | 528 |
| PLXNB1 | 195 | 391 |
| TRO | 195 | 515 |
| HIP1R | 194 | 460 |
| JMJD1C | 194 | 585 |
| CAMKK2 | 193 | 632 |
| SEC31A | 193 | 541 |

**Supplementary Table 3b The numer of isoforms of top 100 genes from SIS in mouse brains**

| Gene_name | Transcripts_per_gene | Gene_abundance |
|---|---|---|
| Sorbs1 | 158 | 218 |
| Rap1gap | 147 | 215 |
| Sorbs2 | 147 | 210 |
| Arpp21 | 145 | 286 |
| Camk2b | 126 | 227 |
| Rnf112 | 124 | 233 |
| Pde4dip | 120 | 190 |
| Ttc3 | 120 | 251 |
| Atp2b1 | 118 | 375 |
| Sez6 | 116 | 153 |
| Mcf2l | 112 | 208 |
| Cplx2 | 110 | 161 |
| Mink1 | 109 | 168 |
| Syne1 | 109 | 207 |
| Ank2 | 106 | 229 |
| Bag6 | 106 | 148 |
| Map4 | 102 | 252 |
| Synj1 | 102 | 164 |
| Madd | 101 | 140 |
| Gtf2i | 98 | 192 |
| Pde2a | 98 | 166 |
| Chd4 | 97 | 201 |
| Add1 | 94 | 188 |
| Epb41l3 | 94 | 212 |
| Myt1l | 94 | 143 |
| Mast3 | 93 | 142 |
| Sptan1 | 92 | 202 |
| Meg3 | 91 | 386 |
| Srcin1 | 91 | 145 |
| Kidins220 | 90 | 168 |
| Trim9 | 89 | 197 |
| Kifc2 | 88 | 172 |
| Sbf1 | 87 | 127 |
| Mycbp2 | 86 | 133 |
| Grin1 | 85 | 129 |
| R3hdm1 | 84 | 157 |
| Trip12 | 84 | 181 |
| Kif1a | 83 | 167 |
| Abca2 | 82 | 144 |
| Cbarp | 82 | 149 |
| Chd3 | 81 | 136 |
| Smarca2 | 81 | 182 |
| Trak1 | 81 | 157 |

| | | |
|---|---|---|
| Gria2 | 80 | 185 |
| Kdm4b | 79 | 121 |
| Mical2 | 79 | 123 |
| Phldb1 | 79 | 88 |
| Ptpn5 | 79 | 125 |
| Atp2b2 | 78 | 129 |
| Kalrn | 78 | 106 |
| Zmynd11 | 78 | 188 |
| Aatk | 77 | 113 |
| Dgkz | 77 | 98 |
| Arhgap33 | 75 | 115 |
| Bcan | 75 | 124 |
| Entr1 | 75 | 143 |
| Slc4a3 | 75 | 147 |
| Snap91 | 75 | 137 |
| Dlgap1 | 74 | 213 |
| Myh10 | 74 | 219 |
| Sptbn1 | 74 | 155 |
| Tsc2 | 74 | 115 |
| Dclk1 | 73 | 391 |
| Ogdh | 73 | 204 |
| Ptk2 | 73 | 156 |
| Smarcc2 | 73 | 161 |
| Akap8l | 72 | 132 |
| Celf3 | 71 | 99 |
| Pum1 | 71 | 96 |
| R3hdm2 | 71 | 166 |
| Adgrb1 | 70 | 108 |
| Zfr | 69 | 198 |
| Zmynd8 | 69 | 120 |
| Ndrg4 | 67 | 136 |
| Tcf4 | 67 | 105 |
| Aplp2 | 66 | 140 |
| Arhgef2 | 66 | 161 |
| Dctn1 | 66 | 100 |
| Epb41l2 | 66 | 106 |
| Ncald | 66 | 204 |
| Clasp2 | 65 | 125 |
| Kif1b | 65 | 258 |
| Pde1b | 65 | 88 |
| Pi4ka | 65 | 137 |
| Ptprn | 65 | 141 |
| Rims1 | 65 | 93 |
| Sgip1 | 65 | 216 |
| Sh3glb2 | 65 | 161 |

| | | |
|---|---|---|
| Eif4g3 | 64 | 102 |
| Celf1 | 63 | 96 |
| Cltc | 63 | 328 |
| Dbn1 | 63 | 137 |
| Hdac5 | 63 | 104 |
| Herc2 | 63 | 119 |
| Huwe1 | 63 | 149 |
| Inf2 | 63 | 81 |
| Dnm1l | 62 | 161 |
| Lrrc45 | 62 | 105 |
| Pum2 | 61 | 111 |
| Ubap2l | 61 | 118 |
