## Supplementary Table 4 for "Transcriptional Determinism and Stochasticity Contribute to the Complexity of Autism Associated *SHANK* Family Genes"

**Supplementary Table 4-Top ASD risk genes from recent large genomics studies**

| ENSEMBL_GENE_ID | GENE_SYMBOL | CLASS | SUB_CLASS | FOUNCTION |
|---|---|---|---|---|
| ENSG00000005339 | CREBBP | ASD, non-synaptic | Gene expression regulation | CREB binding protein(CREBBP) |
| ENSG00000005483 | KMT2E | ASD, non-synaptic | Gene expression regulation | lysine methyltransferase 2E (inactive)(KMT2E) |
| ENSG00000011485 | PPP5C | ASD, non-synaptic | Other | protein phosphatase 5 catalytic subunit(PPP5C) |
| ENSG00000015171 | ZMYND11 | ASD, non-synaptic | Gene expression regulation | zinc finger MYND-type containing 11(ZMYND11) |
| ENSG00000021574 | SPAST | ASD, non-synaptic | Cytoskeleton | spastin(SPAST) |
| ENSG00000021645 | NRXN3 | ASD, synaptic | Neuronal communication | neurexin 3(NRXN3) |
| ENSG00000022355 | GABRA1 | ASD, synaptic | Neuronal communication | gamma-aminobutyric acid type A receptor subunit alpha1(GABRA1) |
| ENSG00000036257 | CUL3 | ASD, synaptic | Neuronal communication | cullin 3(CUL3) |
| ENSG00000042753 | AP2S1 | ASD, synaptic | Neuronal communication | adaptor related protein complex 2 subunit sigma 1(AP2S1) |
| ENSG00000048740 | CELF2 | ASD, non-synaptic | Gene expression regulation | CUGBP Elav-like family member 2(CELF2) |
| ENSG00000048828 | FAM120A | ASD, synaptic | Neuronal communication | family with sequence similarity 120A(FAM120A) |
| ENSG00000049618 | ARID1B | ASD, non-synaptic | Gene expression regulation | AT-rich interaction domain 1B(ARID1B) |
| ENSG00000054523 | KIF1B | ASD, non-synaptic | Other | kinesin family member 1B(KIF1B) |
| ENSG00000055163 | CYFIP2 | ASD, synaptic | Neuronal communication | cytoplasmic FMR1 interacting protein 2(CYFIP2) |
| ENSG00000055609 | KMT2C | ASD, non-synaptic | Gene expression regulation | lysine methyltransferase 2C(KMT2C) |
| ENSG00000061676 | NCKAP1 | ASD, synaptic | Neuronal communication | NCK associated protein 1(NCKAP1) |
| ENSG00000064419 | TNPO3 | ASD, non-synaptic | Gene expression regulation | transportin 3(TNPO3) |
| ENSG00000065526 | SPEN | ASD, synaptic | Neuronal communication | spen family transcriptional repressor(SPEN) |
| ENSG00000065883 | CDK13 | ASD, non-synaptic | Other | cyclin dependent kinase 13(CDK13) |
| ENSG00000066032 | CTNNA2 | ASD, non-synaptic | Cytoskeleton | catenin alpha 2(CTNNA2) |
| ENSG00000067715 | SYT1 | ASD, synaptic | Neuronal communication | synaptotagmin 1(SYT1) |
| ENSG00000067798 | NAV3 | ASD, synaptic | Neuronal communication | neuron navigator 3(NAV3) |
| ENSG00000070808 | CAMK2A | ASD, synaptic | Neuronal communication | calcium/calmodulin dependent protein kinase II alpha(CAMK2A) |
| ENSG00000072518 | MARK2 | ASD, non-synaptic | Other | microtubule affinity regulating kinase 2(MARK2) |
| ENSG00000080298 | RFX3 | ASD, non-synaptic | Gene expression regulation | regulatory factor X3(RFX3) |
| ENSG00000080503 | SMARCA2 | ASD, non-synaptic | Gene expression regulation | SWI/SNF related, matrix associated, actin dependent regulator of chromatin, subfamily a, member 2(SMARCA2) |
| ENSG00000080603 | SRCAP | ASD, non-synaptic | Gene expression regulation | Snf2 related CREBBP activator protein(SRCAP) |
| ENSG00000081189 | MEF2C | ASD, synaptic | Neuronal communication | myocyte enhancer factor 2C(MEF2C) |
| ENSG00000082898 | XPO1 | ASD, non-synaptic | Differentiation/Proliferation/Cell cycl | exportin 1(XPO1) |
| ENSG00000083799 | CYLD | ASD, non-synaptic | Gene expression regulation | CYLD lysine 63 deubiquitinase(CYLD) |
| ENSG00000084676 | NCOA1 | ASD, non-synaptic | Gene expression regulation | nuclear receptor coactivator 1(NCOA1) |
| ENSG00000088038 | CNOT3 | ASD, non-synaptic | Gene expression regulation | CCR4-NOT transcription complex subunit 3(CNOT3) |
| ENSG00000092964 | DPYSL2 | ASD, non-synaptic | Cytoskeleton | dihydropyrimidinase like 2(DPYSL2) |
| ENSG00000095787 | WAC | ASD, non-synaptic | Gene expression regulation | WW domain containing adaptor with coiled-coil(WAC) |
| ENSG00000096433 | ITPR3 | ASD, non-synaptic | Other | inositol 1,4,5-trisphosphate receptor type 3(ITPR3) |
| ENSG00000100207 | TCF20 | ASD, non-synaptic | Gene expression regulation | transcription factor 20(TCF20) |
| ENSG00000100393 | EP300 | ASD, non-synaptic | Differentiation/Proliferation/Cell cycl | E1A binding protein p300(EP300) |
| ENSG00000100888 | CHD8 | ASD, non-synaptic | Gene expression regulation | chromodomain helicase DNA binding protein 8(CHD8) |
| ENSG00000101040 | ZMYND8 | ASD, non-synaptic | Gene expression regulation | zinc finger MYND-type containing 8(ZMYND8) |
| ENSG00000101126 | ADNP | ASD, non-synaptic | Gene expression regulation | activity dependent neuroprotector homeobox(ADNP) |
| ENSG00000101266 | CSNK2A1 | ASD, non-synaptic | Gene expression regulation | casein kinase 2 alpha 1(CSNK2A1) |
| ENSG00000101337 | TM9SF4 | ASD, non-synaptic | Other | transmembrane 9 superfamily member 4(TM9SF4) |
| ENSG00000101489 | CELF4 | ASD, non-synaptic | Gene expression regulation | CUGBP Elav-like family member 4(CELF4) |
| ENSG00000102879 | CORO1A | ASD, non-synaptic | Cytoskeleton | coronin 1A(CORO1A) |
| ENSG00000102974 | CTCF | ASD, non-synaptic | Gene expression regulation | CCCTC-binding factor(CTCF) |
| ENSG00000103197 | TSC2 | ASD, non-synaptic | Other | TSC complex subunit 2(TSC2) |
| ENSG00000104517 | UBR5 | ASD, non-synaptic | Other | ubiquitin protein ligase E3 component n-recognin 5(UBR5) |
| ENSG00000105409 | ATP1A3 | ASD, synaptic | Neuronal communication | ATPase Na+/K+ transporting subunit alpha 3(ATP1A3) |

| | | | | |
|---|---|---|---|---|
| ENSG00000106829 | TLE4 | ASD, non-synaptic | Gene expression regulation | TLE family member 4, transcriptional corepressor(TLE4) |
| ENSG00000108509 | CAMTA2 | ASD, non-synaptic | Gene expression regulation | calmodulin binding transcription activator 2(CAMTA2) |
| ENSG00000108510 | MED13 | ASD, non-synaptic | Gene expression regulation | mediator complex subunit 13(MED13) |
| ENSG00000108557 | RAI1 | ASD, non-synaptic | Gene expression regulation | retinoic acid induced 1(RAI1) |
| ENSG00000108671 | PSMD11 | ASD, non-synaptic | Cytoskeleton | proteasome 26S subunit, non-ATPase 11(PSMD11) |
| ENSG00000108819 | PPP1R9B | ASD, synaptic | Neuronal communication | protein phosphatase 1 regulatory subunit 9B(PPP1R9B) |
| ENSG00000108883 | EFTUD2 | ASD, non-synaptic | Gene expression regulation | elongation factor Tu GTP binding domain containing 2(EFTUD2) |
| ENSG00000109118 | PHF12 | ASD, non-synaptic | Gene expression regulation | PHD finger protein 12(PHF12) |
| ENSG00000110066 | KMT5B | ASD, non-synaptic | Gene expression regulation | lysine methyltransferase 5B(KMT5B) |
| ENSG00000111300 | NAA25 | ASD, non-synaptic | Differentiation/Proliferation/Cell cycl | N-alpha-acetyltransferase 25, NatB auxiliary subunit(NAA25) |
| ENSG00000112282 | MED23 | ASD, non-synaptic | Gene expression regulation | mediator complex subunit 23(MED23) |
| ENSG00000112640 | PPP2R5D | ASD, non-synaptic | Gene expression regulation | protein phosphatase 2 regulatory subunit B'delta(PPP2R5D) |
| ENSG00000112655 | PTK7 | ASD, non-synaptic | Cytoskeleton | protein tyrosine kinase 7 (inactive)(PTK7) |
| ENSG00000113569 | NUP155 | ASD, non-synaptic | Other | nucleoporin 155(NUP155) |
| ENSG00000113595 | TRIM23 | ASD, non-synaptic | Other | tripartite motif containing 23(TRIM23) |
| ENSG00000114030 | KPNA1 | ASD, non-synaptic | Gene expression regulation | karyopherin subunit alpha 1(KPNA1) |
| ENSG00000114554 | PLXNA1 | ASD, synaptic | Neuronal communication | plexin A1(PLXNA1) |
| ENSG00000114861 | FOXP1 | ASD, non-synaptic | Gene expression regulation | forkhead box P1(FOXP1) |
| ENSG00000114933 | INO80D | ASD, non-synaptic | Gene expression regulation | INO80 complex subunit D(INO80D) |
| ENSG00000115306 | SPTBN1 | ASD, synaptic | Neuronal communication | spectrin beta, non-erythrocytic 1(SPTBN1) |
| ENSG00000115677 | HDLBP | ASD, non-synaptic | Gene expression regulation | high density lipoprotein binding protein(HDLBP) |
| ENSG00000116539 | ASH1L | ASD, non-synaptic | Gene expression regulation | ASH1 like histone lysine methyltransferase(ASH1L) |
| ENSG00000117139 | KDM5B | ASD, non-synaptic | Gene expression regulation | lysine demethylase 5B(KDM5B) |
| ENSG00000118058 | KMT2A | ASD, non-synaptic | Gene expression regulation | lysine methyltransferase 2A(KMT2A) |
| ENSG00000119042 | SATB2 | ASD, non-synaptic | Gene expression regulation | SATB homeobox 2(SATB2) |
| ENSG00000119669 | IRF2BPL | ASD, non-synaptic | Gene expression regulation | interferon regulatory factor 2 binding protein like(IRF2BPL) |
| ENSG00000119772 | DNMT3A | ASD, non-synaptic | Gene expression regulation | DNA methyltransferase 3 alpha(DNMT3A) |
| ENSG00000119866 | BCL11A | ASD, non-synaptic | Gene expression regulation | BCL11 transcription factor A(BCL11A) |
| ENSG00000120156 | TEK | ASD, non-synaptic | Other | TEK receptor tyrosine kinase(TEK) |
| ENSG00000120251 | GRIA2 | ASD, synaptic | Neuronal communication | glutamate ionotropic receptor AMPA type subunit 2(GRIA2) |
| ENSG00000120733 | KDM3B | ASD, non-synaptic | Gene expression regulation | lysine demethylase 3B(KDM3B) |
| ENSG00000121741 | ZMYM2 | ASD, non-synaptic | Gene expression regulation | zinc finger MYM-type containing 2(ZMYM2) |
| ENSG00000121879 | PIK3CA | ASD, non-synaptic | Differentiation/Proliferation/Cell cycl | phosphatidylinositol-4,5-bisphosphate 3-kinase catalytic subunit alpha(PIK3CA) |
| ENSG00000121989 | ACVR2A | ASD, non-synaptic | Other | activin A receptor type 2A(ACVR2A) |
| ENSG00000123066 | MED13L | ASD, non-synaptic | Gene expression regulation | mediator complex subunit 13L(MED13L) |
| ENSG00000126461 | SCAF1 | ASD, non-synaptic | Gene expression regulation | SR-related CTD associated factor 1(SCAF1) |
| ENSG00000126464 | PRR12 | ASD, synaptic | Neuronal communication | proline rich 12(PRR12) |
| ENSG00000126705 | AHDC1 | ASD, non-synaptic | Gene expression regulation | AT-hook DNA binding motif containing 1(AHDC1) |
| ENSG00000127191 | TRAF2 | ASD, non-synaptic | Differentiation/Proliferation/Cell cycl | TNF receptor associated factor 2(TRAF2) |
| ENSG00000127955 | GNAI1 | ASD, non-synaptic | Other | G protein subunit alpha i1(GNAI1) |
| ENSG00000128573 | FOXP2 | ASD, non-synaptic | Gene expression regulation | forkhead box P2(FOXP2) |
| ENSG00000130699 | TAF4 | ASD, non-synaptic | Gene expression regulation | TATA-box binding protein associated factor 4(TAF4) |
| ENSG00000130811 | EIF3G | ASD, non-synaptic | Gene expression regulation | eukaryotic translation initiation factor 3 subunit G(EIF3G) |
| ENSG00000131095 | GFAP | ASD, non-synaptic | Cytoskeleton | glial fibrillary acidic protein(GFAP) |
| ENSG00000131242 | RAB11FIP4 | ASD, non-synaptic | Differentiation/Proliferation/Cell cycl | RAB11 family interacting protein 4(RAB11FIP4) |
| ENSG00000131473 | ACLY | ASD, non-synaptic | Gene expression regulation | ATP citrate lyase(ACLY) |
| ENSG00000131653 | TRAF7 | ASD, non-synaptic | Gene expression regulation | TNF receptor associated factor 7(TRAF7) |
| ENSG00000132510 | KDM6B | ASD, non-synaptic | Gene expression regulation | lysine demethylase 6B(KDM6B) |
| ENSG00000132535 | DLG4 | ASD, synaptic | Neuronal communication | discs large MAGUK scaffold protein 4(DLG4) |
| ENSG00000134138 | MEIS2 | ASD, non-synaptic | Gene expression regulation | Meis homeobox 2(MEIS2) |

| | | | | |
|---|---|---|---|---|
| ENSG00000134698 | AGO4 | ASD, non-synaptic | Differentiation/Proliferation/Cell cycl | argonaute RISC component 4(AGO4) |
| ENSG00000135316 | SYNCRIP | ASD, non-synaptic | Gene expression regulation | synaptotagmin binding cytoplasmic RNA interacting protein(SYNCRIP) |
| ENSG00000135365 | PHF21A | ASD, non-synaptic | Gene expression regulation | PHD finger protein 21A(PHF21A) |
| ENSG00000135387 | CAPRIN1 | ASD, non-synaptic | Differentiation/Proliferation/Cell cycl | cell cycle associated protein 1(CAPRIN1) |
| ENSG00000136451 | VEZF1 | ASD, non-synaptic | Gene expression regulation | vascular endothelial zinc finger 1(VEZF1) |
| ENSG00000136531 | SCN2A | ASD, synaptic | Neuronal communication | sodium voltage-gated channel alpha subunit 2(SCN2A) |
| ENSG00000136535 | TBR1 | ASD, non-synaptic | Gene expression regulation | T-box brain transcription factor 1(TBR1) |
| ENSG00000136854 | STXBP1 | ASD, synaptic | Neuronal communication | syntaxin binding protein 1(STXBP1) |
| ENSG00000136928 | GABBR2 | ASD, synaptic | Neuronal communication | gamma-aminobutyric acid type B receptor subunit 2(GABBR2) |
| ENSG00000138028 | CGREF1 | ASD, non-synaptic | Differentiation/Proliferation/Cell cycl | cell growth regulator with EF-hand domain 1(CGREF1) |
| ENSG00000138081 | FBXO11 | ASD, synaptic | Neuronal communication | F-box protein 11(FBXO11) |
| ENSG00000138668 | HNRNPD | ASD, non-synaptic | Differentiation/Proliferation/Cell cycl | heterogeneous nuclear ribonucleoprotein D(HNRNPD) |
| ENSG00000138834 | MAPK8IP3 | ASD, non-synaptic | Differentiation/Proliferation/Cell cycl | mitogen-activated protein kinase 8 interacting protein 3(MAPK8IP3) |
| ENSG00000139613 | SMARCC2 | ASD, non-synaptic | Gene expression regulation | SWI/SNF related, matrix associated, actin dependent regulator of chromatin subfamily c member 2(SMARCC2) |
| ENSG00000140332 | TLE3 | ASD, non-synaptic | Other | TLE family member 3, transcriptional corepressor(TLE3) |
| ENSG00000141431 | ASXL3 | ASD, non-synaptic | Gene expression regulation | ASXL transcriptional regulator 3(ASXL3) |
| ENSG00000142611 | PRDM16 | ASD, non-synaptic | Other | PR/SET domain 16(PRDM16) |
| ENSG00000142949 | PTPRF | ASD, non-synaptic | Other | protein tyrosine phosphatase receptor type F(PTPRF) |
| ENSG00000143442 | POGZ | ASD, non-synaptic | Gene expression regulation | pogo transposable element derived with ZNF domain(POGZ) |
| ENSG00000143624 | INTS3 | ASD, non-synaptic | Gene expression regulation | integrator complex subunit 3(INTS3) |
| ENSG00000144285 | SCN1A | ASD, synaptic | Neuronal communication | sodium voltage-gated channel alpha subunit 1(SCN1A) |
| ENSG00000145362 | ANK2 | ASD, synaptic | Neuronal communication | ankyrin 2(ANK2) |
| ENSG00000145864 | GABRB2 | ASD, synaptic | Neuronal communication | gamma-aminobutyric acid type A receptor subunit beta2(GABRB2) |
| ENSG00000146830 | GIGYF1 | ASD, non-synaptic | Other | GRB10 interacting GYF protein 1(GIGYF1) |
| ENSG00000146872 | TLK2 | ASD, non-synaptic | Gene expression regulation | tousled like kinase 2(TLK2) |
| ENSG00000148143 | ZNF462 | ASD, non-synaptic | Gene expression regulation | zinc finger protein 462(ZNF462) |
| ENSG00000148737 | TCF7L2 | ASD, non-synaptic | Gene expression regulation | transcription factor 7 like 2(TCF7L2) |
| ENSG00000148948 | LRRC4C | ASD, synaptic | Neuronal communication | leucine rich repeat containing 4C(LRRC4C) |
| ENSG00000149187 | CELF1 | ASD, non-synaptic | Gene expression regulation | CUGBP Elav-like family member 1(CELF1) |
| ENSG00000150051 | MKX | ASD, non-synaptic | Gene expression regulation | mohawk homeobox(MKX) |
| ENSG00000151240 | DIP2C | ASD, synaptic | Neuronal communication | disco interacting protein 2 homolog C(DIP2C) |
| ENSG00000151623 | NR3C2 | ASD, non-synaptic | Gene expression regulation | nuclear receptor subfamily 3 group C member 2(NR3C2) |
| ENSG00000153827 | TRIP12 | ASD, non-synaptic | Gene expression regulation | thyroid hormone receptor interactor 12(TRIP12) |
| ENSG00000154122 | ANKH | ASD, non-synaptic | Other | ANKH inorganic pyrophosphate transport regulator(ANKH) |
| ENSG00000154358 | OBSCN | ASD, synaptic | Neuronal communication | obscurin, cytoskeletal calmodulin and titin-interacting RhoGEF(OBSCN) |
| ENSG00000154640 | BTG3 | ASD, non-synaptic | Differentiation/Proliferation/Cell cycl | BTG anti-proliferation factor 3(BTG3) |
| ENSG00000156113 | KCNMA1 | ASD, synaptic | Neuronal communication | potassium calcium-activated channel subfamily M alpha 1(KCNMA1) |
| ENSG00000157087 | ATP2B2 | ASD, synaptic | Neuronal communication | ATPase plasma membrane Ca2+ transporting 2(ATP2B2) |
| ENSG00000157103 | SLC6A1 | ASD, synaptic | Neuronal communication | solute carrier family 6 member 1(SLC6A1) |
| ENSG00000157388 | CACNA1D | ASD, synaptic | Neuronal communication | calcium voltage-gated channel subunit alpha1 D(CACNA1D) |
| ENSG00000157445 | CACNA2D3 | ASD, synaptic | Neuronal communication | calcium voltage-gated channel auxiliary subunit alpha2delta 3(CACNA2D3) |
| ENSG00000157540 | DYRK1A | ASD, non-synaptic | Cytoskeleton | dual specificity tyrosine phosphorylation regulated kinase 1A(DYRK1A) |
| ENSG00000157933 | SKI | ASD, non-synaptic | Gene expression regulation | SKI proto-oncogene(SKI) |
| ENSG00000158321 | AUTS2 | ASD, synaptic | Neuronal communication | activator of transcription and developmental regulator AUTS2(AUTS2) |
| ENSG00000158445 | KCNB1 | ASD, synaptic | Neuronal communication | potassium voltage-gated channel subfamily B member 1(KCNB1) |
| ENSG00000158796 | DEDD | ASD, non-synaptic | Differentiation/Proliferation/Cell cycl | death effector domain containing(DEDD) |
| ENSG00000159459 | UBR1 | ASD, non-synaptic | Other | ubiquitin protein ligase E3 component n-recognin 1(UBR1) |
| ENSG00000160305 | DIP2A | ASD, synaptic | Neuronal communication | disco interacting protein 2 homolog A(DIP2A) |
| ENSG00000160551 | TAOK1 | ASD, non-synaptic | Cytoskeleton | TAO kinase 1(TAOK1) |
| ENSG00000160877 | NACC1 | ASD, non-synaptic | Gene expression regulation | nucleus accumbens associated 1(NACC1) |

| | | | | |
|---|---|---|---|---|
| ENSG00000162105 | SHANK2 | ASD, synaptic | Neuronal communication | SH3 and multiple ankyrin repeat domains 2(SHANK2) |
| ENSG00000163041 | H3-3A | ASD, non-synaptic | Gene expression regulation | H3.3 histone A(H3-3A) |
| ENSG00000163625 | WDFY3 | ASD, synaptic | Neuronal communication | WD repeat and FYVE domain containing 3(WDFY3) |
| ENSG00000164134 | NAA15 | ASD, synaptic | Neuronal communication | N-alpha-acetyltransferase 15, NatA auxiliary subunit(NAA15) |
| ENSG00000164190 | NIPBL | ASD, non-synaptic | Gene expression regulation | NIPBL cohesin loading factor(NIPBL) |
| ENSG00000165025 | SYK | ASD, non-synaptic | Other | spleen associated tyrosine kinase(SYK) |
| ENSG00000165671 | NSD1 | ASD, non-synaptic | Gene expression regulation | nuclear receptor binding SET domain protein 1(NSD1) |
| ENSG00000165699 | TSC1 | ASD, non-synaptic | Differentiation/Proliferation/Cell cycl | TSC complex subunit 1(TSC1) |
| ENSG00000166206 | GABRB3 | ASD, synaptic | Neuronal communication | gamma-aminobutyric acid type A receptor subunit beta3(GABRB3) |
| ENSG00000166313 | APBB1 | ASD, synaptic | Neuronal communication | amyloid beta precursor protein binding family B member 1(APBB1) |
| ENSG00000166963 | MAP1A | ASD, non-synaptic | Cytoskeleton | microtubule associated protein 1A(MAP1A) |
| ENSG00000167522 | ANKRD11 | ASD, non-synaptic | Gene expression regulation | ankyrin repeat domain containing 11(ANKRD11) |
| ENSG00000167971 | CASKIN1 | ASD, synaptic | Neuronal communication | CASK interacting protein 1(CASKIN1) |
| ENSG00000168036 | CTNNB1 | ASD, non-synaptic | Gene expression regulation | catenin beta 1(CTNNB1) |
| ENSG00000168137 | SETD5 | ASD, non-synaptic | Gene expression regulation | SET domain containing 5(SETD5) |
| ENSG00000169375 | SIN3A | ASD, non-synaptic | Gene expression regulation | SIN3 transcription regulator family member A(SIN3A) |
| ENSG00000170004 | CHD3 | ASD, non-synaptic | Gene expression regulation | chromodomain helicase DNA binding protein 3(CHD3) |
| ENSG00000170027 | YWHAG | ASD, non-synaptic | Differentiation/Proliferation/Cell cycl | tyrosine 3-monooxygenase/tryptophan 5-monooxygenase activation protein gamma(YWHAG) |
| ENSG00000170471 | RALGAPB | ASD, non-synaptic | Differentiation/Proliferation/Cell cycl | Ral GTPase activating protein non-catalytic subunit beta(RALGAPB) |
| ENSG00000170871 | KIAA0232 | ASD, non-synaptic | Other | KIAA0232(KIAA0232) |
| ENSG00000171587 | DSCAM | ASD, synaptic | Neuronal communication | DS cell adhesion molecule(DSCAM) |
| ENSG00000171862 | PTEN | ASD, synaptic | Neuronal communication | phosphatase and tensin homolog(PTEN) |
| ENSG00000173064 | HECTD4 | ASD, non-synaptic | Other | HECT domain E3 ubiquitin protein ligase 4(HECTD4) |
| ENSG00000173175 | ADCY5 | ASD, synaptic | Neuronal communication | adenylate cyclase 5(ADCY5) |
| ENSG00000173575 | CHD2 | ASD, non-synaptic | Gene expression regulation | chromodomain helicase DNA binding protein 2(CHD2) |
| ENSG00000174672 | BRSK2 | ASD, synaptic | Neuronal communication | BR serine/threonine kinase 2(BRSK2) |
| ENSG00000175115 | PACS1 | ASD, non-synaptic | Cytoskeleton | phosphofurin acidic cluster sorting protein 1(PACS1) |
| ENSG00000175224 | ATG13 | ASD, non-synaptic | Differentiation/Proliferation/Cell cycl | autophagy related 13(ATG13) |
| ENSG00000175727 | MLXIP | ASD, non-synaptic | Gene expression regulation | MLX interacting protein(MLXIP) |
| ENSG00000177030 | DEAF1 | ASD, non-synaptic | Gene expression regulation | DEAF1 transcription factor(DEAF1) |
| ENSG00000177565 | TBL1XR1 | ASD, non-synaptic | Gene expression regulation | TBL1X/Y related 1(TBL1XR1) |
| ENSG00000179010 | MRFAP1 | ASD, non-synaptic | Other | Morf4 family associated protein 1(MRFAP1) |
| ENSG00000179915 | NRXN1 | ASD, synaptic | Neuronal communication | neurexin 1(NRXN1) |
| ENSG00000181090 | EHMT1 | ASD, non-synaptic | Gene expression regulation | euchromatic histone lysine methyltransferase 1(EHMT1) |
| ENSG00000181652 | ATG9B | ASD, non-synaptic | Differentiation/Proliferation/Cell cycl | autophagy related 9B(ATG9B) |
| ENSG00000181722 | ZBTB20 | ASD, non-synaptic | Gene expression regulation | zinc finger and BTB domain containing 20(ZBTB20) |
| ENSG00000182568 | SATB1 | ASD, non-synaptic | Gene expression regulation | SATB homeobox 1(SATB1) |
| ENSG00000182831 | HAPSTR1 | ASD, non-synaptic | Differentiation/Proliferation/Cell cycl | HUWE1 associated protein modifying stress responses(HAPSTR1) |
| ENSG00000182934 | SRPRA | ASD, non-synaptic | Other | SRP receptor subunit alpha(SRPRA) |
| ENSG00000183134 | PTGDR2 | ASD, synaptic | Neuronal communication | prostaglandin D2 receptor 2(PTGDR2) |
| ENSG00000184156 | KCNQ3 | ASD, synaptic | Neuronal communication | potassium voltage-gated channel subfamily Q member 3(KCNQ3) |
| ENSG00000185630 | PBX1 | ASD, non-synaptic | Gene expression regulation | PBX homeobox 1(PBX1) |
| ENSG00000186487 | MYT1L | ASD, non-synaptic | Gene expression regulation | myelin transcription factor 1 like(MYT1L) |
| ENSG00000188994 | ZNF292 | ASD, non-synaptic | Gene expression regulation | zinc finger protein 292(ZNF292) |
| ENSG00000196092 | PAX5 | ASD, non-synaptic | Gene expression regulation | paired box 5(PAX5) |
| ENSG00000196361 | ELAVL3 | ASD, non-synaptic | Gene expression regulation | ELAV like RNA binding protein 3(ELAVL3) |
| ENSG00000196535 | MYO18A | ASD, non-synaptic | Cytoskeleton | myosin XVIIIA(MYO18A) |
| ENSG00000196628 | TCF4 | ASD, non-synaptic | Gene expression regulation | transcription factor 4(TCF4) |
| ENSG00000196712 | NF1 | ASD, non-synaptic | Gene expression regulation | neurofibromin 1(NF1) |
| ENSG00000197102 | DYNC1H1 | ASD, non-synaptic | Cytoskeleton | dynein cytoplasmic 1 heavy chain 1(DYNC1H1) |

| | | | | |
|---|---|---|---|---|
| ENSG00000197170 | PSMD12 | ASD, non-synaptic | Cytoskeleton | proteasome 26S subunit, non-ATPase 12(PSMD12) |
| ENSG00000197283 | SYNGAP1 | ASD, synaptic | Neuronal communication | synaptic Ras GTPase activating protein 1(SYNGAP1) |
| ENSG00000197724 | PHF2 | ASD, non-synaptic | Gene expression regulation | PHD finger protein 2(PHF2) |
| ENSG00000198216 | CACNA1E | ASD, synaptic | Neuronal communication | calcium voltage-gated channel subunit alpha1 E(CACNA1E) |
| ENSG00000198719 | DLL1 | ASD, synaptic | Neuronal communication | delta like canonical Notch ligand 1(DLL1) |
| ENSG00000198728 | LDB1 | ASD, non-synaptic | Gene expression regulation | LIM domain binding 1(LDB1) |
| ENSG00000198963 | RORB | ASD, non-synaptic | Gene expression regulation | RAR related orphan receptor B(RORB) |
| ENSG00000204406 | MBD5 | ASD, non-synaptic | Gene expression regulation | methyl-CpG binding domain protein 5(MBD5) |
| ENSG00000204435 | CSNK2B | ASD, synaptic | Neuronal communication | casein kinase 2 beta(CSNK2B) |
| ENSG00000205726 | ITSN1 | ASD, non-synaptic | Differentiation/Proliferation/Cell cycl | intersectin 1(ITSN1) |
| ENSG00000213923 | CSNK1E | ASD, non-synaptic | Gene expression regulation | casein kinase 1 epsilon(CSNK1E) |
| ENSG00000214753 | HNRNPUL2 | ASD, non-synaptic | Gene expression regulation | heterogeneous nuclear ribonucleoprotein U like 2(HNRNPUL2) |
| ENSG00000231887 | PRH1 | ASD, non-synaptic | Other | proline rich protein HaeIII subfamily 1(PRH1) |
| ENSG00000251322 | SHANK3 | ASD, synaptic | Neuronal communication | SH3 and multiple ankyrin repeat domains 3(SHANK3) |
| ENSG00000273079 | GRIN2B | ASD, synaptic | Neuronal communication | glutamate ionotropic receptor NMDA type subunit 2B(GRIN2B) |
