## Supplementary Table 5 for "Transcriptional Determinism and Stochasticity Contribute to the Complexity of Autism Associated *SHANK* Family Genes"

**Supplmentary Table 5a. The RT-PCR human primers used for the validation**

| Transcripts | Features | Forward | | Reverse | |
|---|---|---|---|---|---|
| TALONT000460240 | Fusion: SH3exon20-ACRexon2 | F1 | TGGTGTTTGCTGTGAACCT | R1 | GTGGCTGTTGTACGTGAAGA |
| | | F2 | GAGACCACAAGCACCATCTC | R2 | GTGAGCACCCATCGTGAAT |
| TALONT000456172 | new exon1 | F | CCTGCAGCAGACGAAGTG | R | GAGTCAGGGTCATGGAAGTTG |
| | | F2 | GACCTGCAGCAGACGAAG | R2 | GCTCCTCATCCAGGAACTTG |
| | | F3 | TCATCGCAGGGAACTTTGAG | R3 | TTGGGTCCACATTGTCTCTTAC |
| | | F4 | GAGGTTATCAAGACCCACAAAGA | R4 | CTGTTGCCTGAAGGGTTACA |
| TALONT000460246 | Fusion: SH3exon20-ACRexon4 | F | TCGCTACCTCTTCCAGAGAA | R | GTCGATCTTGCCTACAGGATAC |
| TALONT000456151 | new exon1 | F1 | GGAGGAGGAAGCCAGACT | R1 | TAGGGAGTTAGTGGCAGATCC |
| | | F2 | ACTGGTCCCAGGGAAGT | R2 | TGTTTCTGGGATGGGAAATAGG |
| TALONT000459573 | exon19-ACRexon2 | F | TCCCAGGCCCAGAGAAG | R | GTGAGCACCCATCGTGAAT |
| TALONT000458917 | intron14 retention | F1 | CATTGATGACAAAGTGGCTGTC | R1 | CCTCTTCTGGCTTCCTTGTC |
| | | F2 | CCACGAGGGCTTTGGTTT | R2 | CCTTGTCACAGACACAACCT |
| TALONT000456776 | exon16 forward extention | F | CTCGTCATGAAGGTTGTGTCT | R | ACTGTGCCAGGATCTTGTT |
| TALONT000457890 | novel exon on intron17 | F | GCCTCCATTCGGAGAAGAAA | R | GTCTGCGTCTGCGATGT |
| TALONT000457195 | nove exon on intron16 | F | CCCAGCACCACACTGA | R | TGTGATCCTCCGACTGGT |
| TALONT000456173 | 2 novel exons on intron8 | F1 | TCATCGCAGGGAACTTTGAG | R1 | GAAGTCAGGAGTTTGAGACCAG |
| | | F2 | GAGCTTGCAGAGGTTATCAAGA | R2 | CAGCACTTTGGGAGGTCAA |
| | | F3 | GAGGTTATCAAGACCCACAAAGA | R3 | AAATGGTGGCTCAAGCCTATAA |

**Supplmentary Table 5b. The RT-PCR mouse primers used for the validation**

| Transcripts | features | Forward Primer | | Reverse Primer | |
|---|---|---|---|---|---|
| PB.13554.151 | novel exon1, intron1 retention | F1 | TGAGGGTAGCAGGGAACA | R1 | ACGGTTTAGCTGCATCGG |
| | | F2 | TAGCAGGGAACAGACGTACA | R2 | GTTGCAGGTCCGGGATG |
| | | F3 | GCTAGCACCGGGATGGA | R3 | CGTCTTGAAGGCTATGATTGAGG |
| PB.13554.190 | split exon1 | F | CCTAAGCCTTTCCTCGTAGGT | R | GGCACGGTTTAGCTGCAT |
| PB.13554.659 | exon21-Acr exon2 fusion | F | CACCATGCTGCCTCTACTG | R | GGTACCTGCGGCTGTTATG |
| PB.13554.478 | intron19-Acr exon4 fusion, intron2, 11 retention | F1 | GCTGAGATCAGCTCATTGTTTG | R1 | TTTGGGCAATGTGTTCTGTATG |
| PB.13554.637 | exon9-exon19 splicing | F1 | CGAGGTAATCAAGACCCACAAAG | R1 | GTGATCCTCCGGCTGGT |
| | | F2 | GCCATTATTGCAGGGAACTTTG | R2 | GCCGCTGCTTGACAGTG |
| PB.13554.606 | exon5-exon21 tail splicing | F1 | GTGCCCTCTGAGCCTTG | R1 | GCTGAAGAGCCGAGCTG |
| | | F2 | AGTGCCCTCTGAGCCTT | R2 | TGACTTCTTAGCAGGGTCCA |
| PB.10584.110 | novel exon9a | F1 | CTTTGAGCTTGCCGAGGTAAT | R1 | GGCTTTGGTTAGAATGTGCTTTG |
| | | F2 | GAGGTAATCAAGACCCACAAAGA | R2 | AACCGATTCCACCACCAAG |
| PB.10584.116 | novel exon11a | F1 | GCCCGAAGCGGAAACTTTA | R1 | CCAAGGATTGCCAGTGAAGAA |
| | | F2 | CAAGTTCATCGCTGTGAAGG | R2 | GCACTGGATCCCAAGGATT |
| PB.10584.1123 | exon18-22 splicing | F1 | TCTTCCAGAGAGTACGAGGAG | R1 | GACAGCAGGGTCCAATGAA |
| | | F2 | CTCACGCCATGAAGAAGTTTG | R2 | GTCATTTCCCTCCGTCACTAC |
