## Supplementary Table 6 for "Transcriptional Determinism and Stochasticity Contribute to the Complexity of Autism Associated *SHANK* Family Genes"

### Supplementary Table 6a. Mouse GERP score

**Comparison of Shank3_known_exons vs. Shank3_PFC_novel_exon**
D: 0.15003064348112852
p-value: 1.4165964969182617e-142
--------------------
**Comparison of Shank3_known_exons vs. Shank3_ST_novel_exon**
D: 0.09604900498226407
p-value: 8.391330023440328e-59
--------------------
**Comparison of Shank3_known_exons vs. non_transcribed_regions**
D: 0.6984571863104513
p-value: 0.0
--------------------
**Comparison of Shank3_PFC_novel_exon vs. Shank3_ST_novel_exon**
D: 0.05406483031568332
p-value: 0.0
--------------------
**Comparison of Shank3_PFC_novel_exon vs. non_transcribed_regions**
D: 0.5484265428293228
p-value: 0.0
--------------------
**Comparison of Shank3_ST_novel_exon vs. non_transcribed_regions**
D: 0.6024081813281872
p-value: 0.0
--------------------

**Supplementary Table 6b. Mouse PhyloP score**

**Comparison of Shank3_known_exons vs. Shank3_PFC_novel_exon**

D: 0.18361353523931612

p-value: 6.760053055365219e-214

--------------------

**Comparison of Shank3_known_exons vs. Shank3_ST_novel_exon**

D: 0.12821643547280087

p-value: 1.2285234431441975e-104

--------------------

**Comparison of Shank3_known_exons vs. non_transcribed_regions**

D: 0.562087557362343

p-value: 0.0

--------------------

**Comparison of Shank3_PFC_novel_exon vs. Shank3_ST_novel_exon**

D: 0.05558535822918709

p-value: 0.0

--------------------

**Comparison of Shank3_PFC_novel_exon vs. non_transcribed_regions**

D: 0.38521758745925183

p-value: 0.0

--------------------

**Comparison of Shank3_ST_novel_exon vs. non_transcribed_regions**

D: 0.4388526416435776

p-value: 0.0

--------------------

**Supplementary Table 6c. Human GERP score**

**Comparison of novel_exon vs. known_exon**

D: 0.29911039761846003

p-value: 0.0

--------------------

**Comparison of novel_exon vs. non_transcribed_regions**

D: 0.09669690399342157

p-value: 2.807809940773249e-264

--------------------

**Comparison of known_exon vs. non_transcribed_regions**

D: 0.37095974538375465

p-value: 0.0

--------------------

### Supplementary Table 6d. Human PhyloP score

**Comparison of novel_exon vs. known_exon**

D: 0.2962083760306952

p-value: 0.0

--------------------

**Comparison of novel_exon vs. non_transcribed_regions**

D: 0.13357773142368867

p-value: 0.0

--------------------

**Comparison of known_exon vs. non_transcribed_regions**

D: 0.39900906321663954

p-value: 0.0

--------------------
